# In vitro culture of aberrant basal-like cells from fibrotic lung tissue

**DOI:** 10.1101/2020.08.16.247866

**Authors:** Petra Khan, Julien Roux, Sabrina Blumer, Lei Fang, Spasenija Savic, Lars Knudsen, Danny Jonigk, Mark P. Kuehnel, Amiq Gazdhar, Thomas Geiser, Michael Tamm, Katrin E. Hostettler

## Abstract

**Rationale:** In idiopathic pulmonary fibrosis (IPF) atypical epithelial cells are present in the alveolar compartment. Their origin and contribution to IPF pathogenesis is unknown. We recently cultured a distinct population of cells, which readily grew from fibrotic lung tissue, but only rarely from non-fibrotic tissue. Here we aimed to characterize these fibrosis-enriched cells and determine transcriptomic differences between cells derived from IPF and patients with other interstitial lung diseases (ILD).

**Methods:** Cells were cultured from peripheral lung tissue of ILD patients and analysed by bulk or single cell RNA sequencing (scRNA-seq), TaqMan-PCR, immunofluorescence (IF), immunoblotting or electron microscopy (EM).

**Results:** scRNA-seq demonstrated an overall homogeneity and epithelial origin of the cells. The majority of cells expressed basal cell markers (Cytokeratin (KRT) 5 and 17, TP63), of which a fraction co-expressed mesenchymal cell markers (VIM, FN1, CDH2), alveolar (SLC34A2, ABCA3, LPCAT1, EMP2, HOPX) and/or secretory epithelial cell markers (SCGB1A1, MUC4). Interestingly, most of the cells showed closest transcriptomic similarity to recently described aberrant basal-like cells. Cells derived from IPF versus other ILD patients revealed significant transcriptomic differences with an up-regulation of fibrosis-associated and a down-regulation of inflammatory pathways in IPF cells.

**Conclusion:** We here confirm the presence of aberrant basal-like cells in fibrotic lung tissue and, importantly, are the first to describe their *in vitro* characteristics and a way of culturing these cells *in vitro*. Cultured basal-like cells co-express epithelial and mesenchymal markers, suggesting a partial epithelial to mesenchymal transition (EMT). A subset of cells co-express alveolar, ciliated or secretory epithelial cell markers, possibly indicating differentiation towards these cell linages. Furthermore, cultured basal-like cells display a disease-specific transcriptome, possibly induced by their specific microenvironment. Our findings will contribute to a better understanding of the cells origin and their potential contribution to IPF pathogenesis.

## Introduction

Idiopathic pulmonary fibrosis (IPF) is a rare, chronic, and irreversible interstitial lung disease characterized by a progressive destruction of the lung parenchyma and the respective loss of lung function [1]. The initial events leading to the development and progression of IPF are still not elucidated, however, evidence suggests that repeated injury to the alveolar epithelial cells plays an important role [2]. In the healthy lung, the alveoli are lined with alveolar epithelial cells type 1 and 2 (AT-1 and 2). AT-2 cells maintain the alveolar epithelium by their ability to self-renew and they serve as progenitors for AT-1 cells [3]. The conducting airway is lined with p63+/KRT5+ basal cells and different secretory and ciliated cells. Basal cells have the ability to differentiate into all other types of airway epithelial cells, and therefore serve as progenitors to regenerate the tracheobronchial epithelium after injury [4]. During normal lung homeostasis there is no overlap of epithelial cell types of the alveolar and the conducting regions. Interestingly, severe lung injury in mice through H1N1 influenza infection or bleomycin resulted in an accumulation of p63+/KRT5+ basal-like cells in the alveolar region [5, 6], and it was suggested that these cells could contribute to alveolar epithelium regeneration [7, 8]. However, the presence of basal-like cells in bronchoalveolar lavage (BAL) fluid or in the alveolar region of lung tissue derived from IPF patients [9-11] was associated with increase mortality [11] and pathological bronchiolization and honeycomb formation [11, 12]. Two recent studies performing single-cell RNA-seq analysis of cells derived from dissociated human fibrotic and control lung tissue described a novel fibrosis-enriched aberrant basal-like cell population, named “aberrant basaloid” by Adams et al. [13] and “KRT5-/KRT17+” by Habermann et al. [14]. These aberrant basal-like cells were shown to express some but not all canonical basal cell markers along with mesenchymal cell markers, senescence markers, and IPF-associated molecules [13, 14]. Interestingly, these unique cells were shown to localize on the surface of IPF fibroblastic foci (FF).

Recently, we observed a fibrosis-enriched outgrowth of a distinct cell population from peripheral lung tissue of ILD patients *in vitro*: Similar to mesenchymal stem cells (MSC), these cells stained positive for CD44, CD90 and CD105 and had the capacity to differentiate into adipocytes, osteocytes and chondrocytes [15]. However, when cultured in an epithelial cell specific medium, these cells acquired cobblestone morphology typical of epithelial cells [15]. A potential for epithelial differentiation of mesenchymal progenitor cells was described earlier [16-18], but is controversial. On the other hand, epithelial cells were shown to acquire MSC-like features when cultured under specific conditions [19]. Therefore, we here aimed to characterize our fibrosis-specific cell population in greater detail.

## Methods

### Cell culture

Fibrosis-enriched cells or fibroblasts were cultured from transbronchial or surgical lung biopsies derived from IPF or non-IPF (other interstitial lung diseases or non-fibrotic diseases) patients and from lung-explants derived from patients undergoing lung transplantation due to terminal fibrotic lung disease. Cells were cultured and conditioned medium collected as previously described [15, 20]. The culture of fibrosis-enriched cells is described in the methods of the supplement document. IPF was diagnosed based on ATS/ERS guidelines [1, 21]. The local ethical committee of the University Hospital, Basel, Switzerland (EKBB05/06) and of the Medical University of Hannover, Germany (2699-2015) approved the culture of human primary lung cells. All materials and instruments used in this study are listed in supplement table 1.

### TaqMan RT-PCR, Immunoblotting, Immunofluorescence & Electron microscopy

TaqMan® PCR, immunoblotting and IF stainings were performed as previously described [15, 22]. Details for primer and antibodies are listed in supplement table 1. For electron microscopy (EM) imaging, cells were fixed in 4% paraformaldehyde/0.1% glutaraldehyde/ 0.2M HEPES and embedded in epoxy resin (Epon®) and subjected to EM imaging as previously described [23]. Semi-thin sections were cut and stained with toluidine blue for light microscopic investigation. Afterwards, ultrathin sections were used for transmission electron microscopy (Morgagni, FEI, Eindhoven, The Netherlands) to characterize the ultrastructure of isolated stem-like cells.

### RNA sequencing

Bulk and scRNA-seq datasets were generated at the Genomics Facility Basel of the ETH Zurich, Basel, and data were analysed by the Bioinformatics Core Facility, Department of Biomedicine, University of Basel. The processed bulk and scRNA-seq data can be obtained from the gene expression omnibus database (GEO) under accession GSE145441. A detailed description of the analysis methods used for these datasets and other public scRNA-seq datasets can be found in the supplement document.

## Results

### Fibrosis-enriched cells with distinct morphology

We here confirmed our previous findings [15], showing an enhanced outgrowth of cells with distinct morphology from fibrotic lung tissue: peripheral lung tissue pieces obtained from explants derived from all 15 IPF patients that were put in culture showed outgrowth of the cells. In contrast, peripheral lung tissue pieces obtained from surgical lung biopsies of non-fibrotic donors (e.g., emphysema) showed outgrowth of the cells for only 3 out of 62 donors (5%).

The morphology of these cultured fibrosis-enriched cells impeded a clear classification to any known plastic adherent lung cell type as they neither displayed the morphology of cubical epithelial cells nor that of spindle-shaped mesenchymal cells (Figure 1 A, B). They were long and flat cells showing a certain polarity. Microvilli-like structures were observed on their surface and the cells were joined by desmosomes and tight junctions. Multi-lamellate bodies were occasionally seen. Bundles of intermediate filaments were often found, oriented along the axis of the cell and connected to the plaques of the desmosomes. Occasionally, late endosomes and autophagosomes were detected in the cells (Figure 1 C).

**Figure 1:**
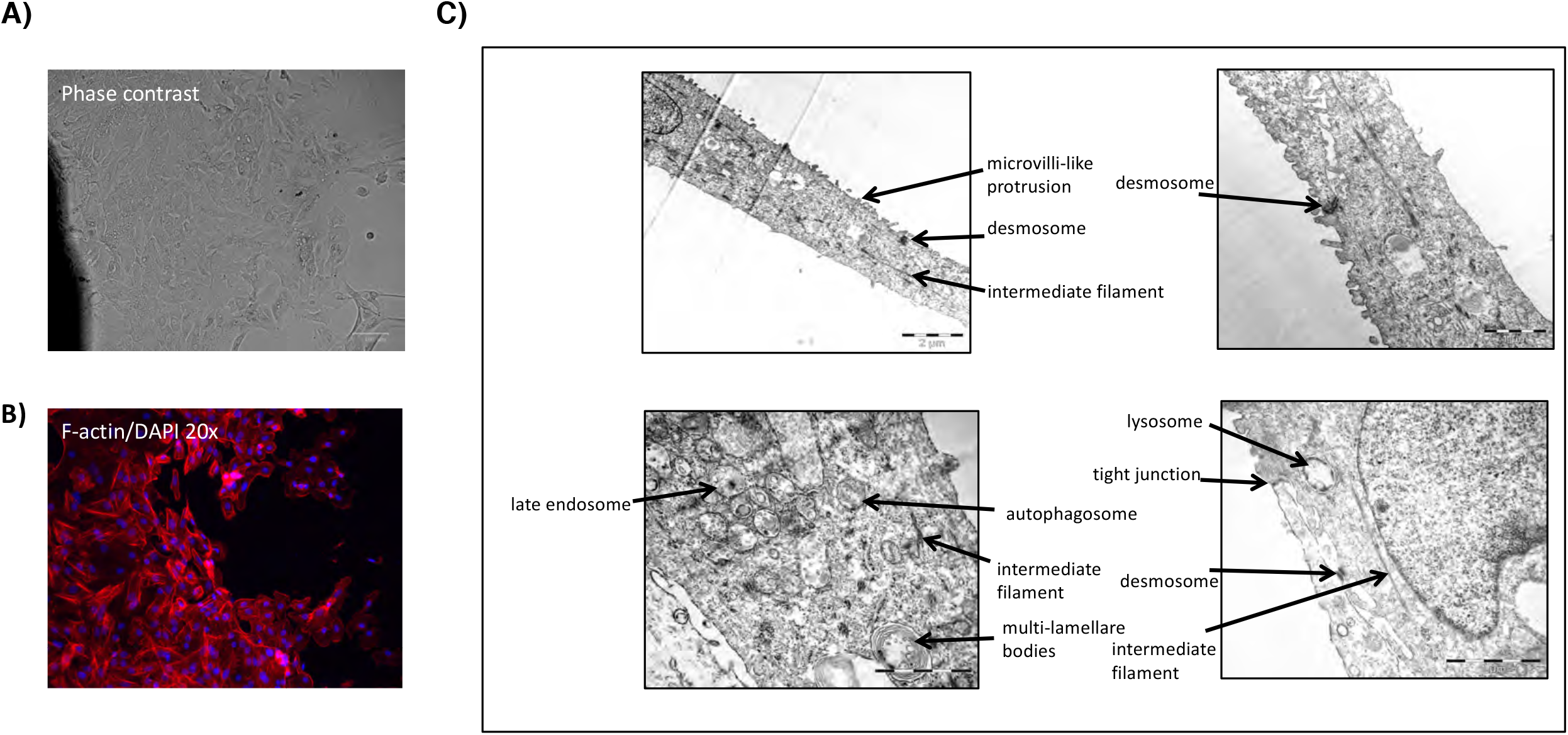
Morphology of fibrosis-enriched cells. Fibrosis-enriched cells were cultured in DMEM/10%FCS for 5 days and then analysed by phase contrast microscopy A), immunofluorescence for F-actin B), or electron microscopy C).

### Fibrosis-enriched cells express mesenchymal- and epithelial marker

We performed bulk RNA-seq on the cultured cells derived from eight different ILD patients (patients details see supplement table 2). We observed expression of mesenchymal stem cell markers, as described previously [15]: CD44 was highly expressed, whereas CD90 (THY1) and CD105 (ENG) were expressed at lower levels (Figure 2 A). Furthermore, other mesenchymal markers such as vimentin (VIM), N-cadherin (CDH2), or fibronectin (FN1) were also highly expressed. We observed moderate to high expression of the typical epithelial cell markers E-cadherin (CDH1), EPCAM, NKX2-1, SOX2, SOX9, basal cell markers p63 (TP63), cytokeratin 5 (KRT5) and 14 (KRT14), AT-1 markers EMP2, HOPX, AT2 markers SLC34A2, ABCA3, LPCAT1, and the club cell marker SCGB1A1 (Figure 2 A). The RNA expression of CDH1, CDH2, FN1, SCGB1A1 (n=5), TP63, KRT5 and SOX9 (n=11) was confirmed in cells derived from independent ILD patients by TaqMan PCR and compared to the expression levels in primary human lung fibroblasts (Figure 2 B). The expression levels of CDH1, KRT5, SCGB1A1, and TP63 and were much lower in fibroblasts compared to the fibrosis-enriched cells, whereas the mesenchymal markers FN1 and CDH2 were expressed at similar levels (Figure 2 B). Protein expression of mesenchymal (FN1, CDH2), epithelial (CDH1, NKX2.1, SOX2, SOX9) and basal cell markers (TP63, KRT5 and KRT17) in fibrosis-enriched cells was confirmed by IF stainings (Figure 2 C).

**Figure 2:**
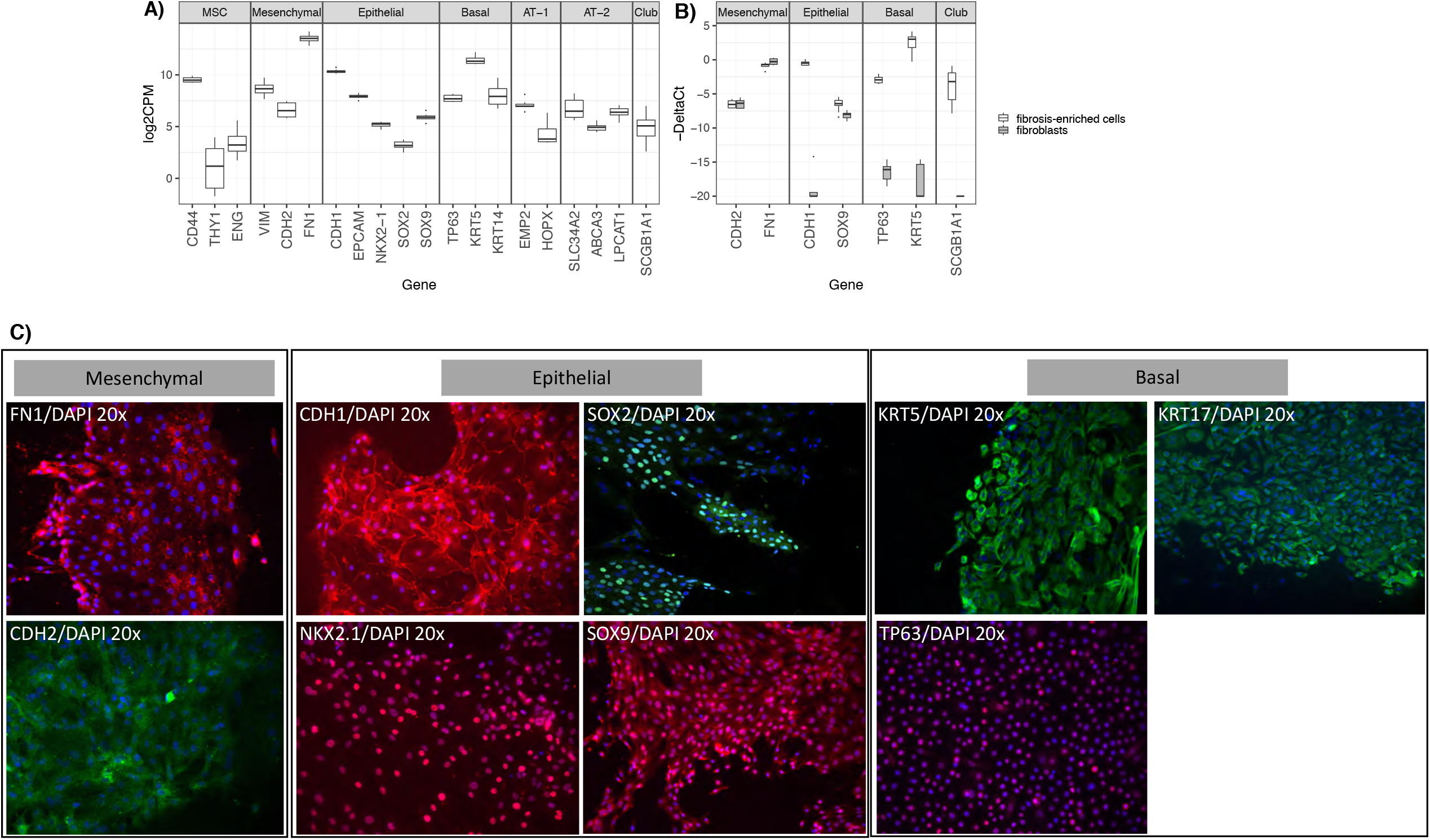
Expression levels (normalized log-transformed CPM) of mesenchymal stem cell (MSC), mesenchymal, epithelial, basal, AT-1 and 2 and club cell markers in fibrosis-enriched cells are shown across all bulk RNA-seq samples A). RNA expression (-delta Ct) of CDH2, FN1, CDH1, SOX9, TP63, KRT5, and SCGB1A1 was confirmed in fibrosis-enriched cells of 5-11 additional patients and compared to expression levels in human lung fibroblasts B). Protein expression of FN1, CDH2, CDH1, NKX2.1, SOX2, SOX9, KRT5, KRT17 and TP63 was detected in fibrosis-enriched cells by IF C).

### The transcriptome of fibrosis-enriched cells shows closest similarity to basal epithelial cells

In order to characterize more specifically the cell-type composition of cultured fibrosis-specific cells, we reanalysed a publicly available scRNA-seq dataset of the whole lung (GEO accession GSE122960)[24], where each of the main cell types of the lung could be identified based on the expression of established cell markers (Supplement figure 1). After aggregating cells into “pseudo-bulk” samples [25, 26] (see methods in the supplement document), a principal component analysis (PCA) allowed us to visualize the similarities of the transcriptome of our fibrosis-enriched cultured cells samples to the main lung cell types. This revealed that they lied in-between epithelial and mesenchymal samples and did not closely match any of the cell types (Figure 3). They however showed closest distance to epithelial cells, notably basal epithelial cells (Figure 3, in orange).

**Figure 3:**
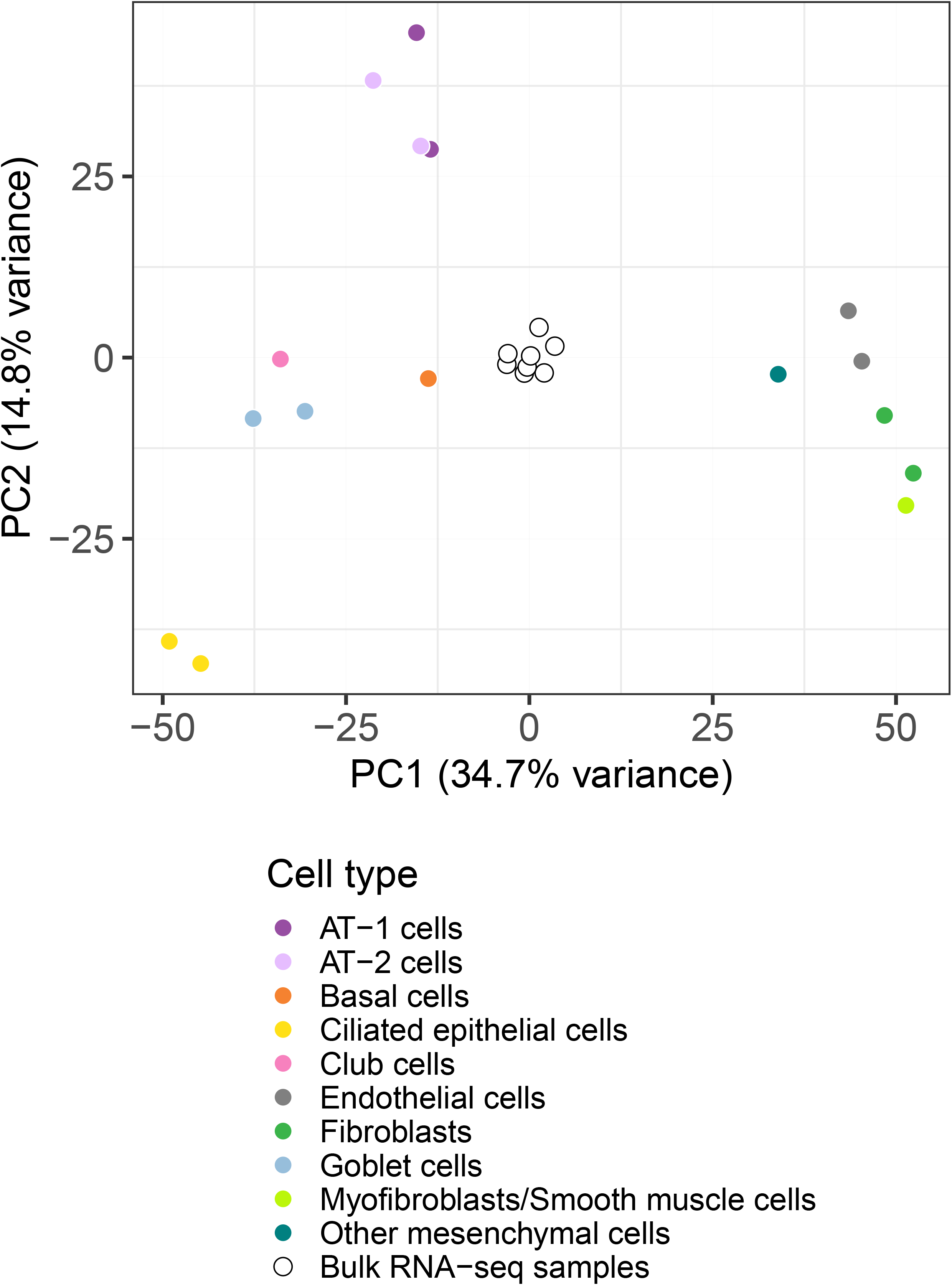
Principal component analysis (PCA) was used to visualize the similarities of the transcriptome of fibrosis-enriched RNA-seq cell samples to pseudo-bulk samples generated from main lung cell types isolated from a publicly available scRNA-seq dataset of whole lung (GEO accession GSE122960; see methods). White circles represent the 8 bulk RNA-seq samples derived from fibrosis-enriched cells. Filled and colored circles represent pseudo-bulk samples corresponding to reference cell types.

### scRNA-seq analysis of fibrosis-enriched cells

The high expression of both epithelial and mesenchymal markers in our samples could be due to the joint presence of both mesenchymal and epithelial cells within the samples, or to co-expression of these markers by the same cells. In order to disentangle these two possibilities, we performed scRNA-seq using the 10X Genomics technology on cultured cells obtained from IPF patient 6 (Patients details in supplement table 2). After quality control and filtering, a total of 7,498 cells were further analyzed (Methods in the supplement document).

Based on their expression profile, cells were grouped into seven clusters, each encompassing between 2% and 32% of the cells, visualized on the tSNE reduced dimensionality projection (Figure 4 A; PCA in supplement figure 2). The relative expression of cluster-specific genes (Supplement figure 3 A), as well as unbiased annotation of cells based on the above-defined pure cell types used as reference, revealed close transcriptomic proximity of the majority of cells to basal cells (60%, orange), followed by AT-1 cells (19 %, purple), goblet cells (14 %, blue), and fibroblasts or other mesenchymal cell-types (7 %, light, medium and dark green) (Figure 4 B). Clusters 1, 2 and 4 were composed of a large majority of cells matching basal cells; Clusters 3, 5 and 6 were composed of cells matching more differentiated epithelial cell types (Figure 4 C). Cluster 7 mainly matched mesenchymal cells. The expression across cells of different clusters of the basal cell markers KRT5, KRT14, KRT17, ITGB4, and TP63; markers of transitional epithelial cells KRT8 and KRT18 [27]; the general epithelial marker KRT7 [28], mesenchymal markers FN1 and VIM; the transcription factors SOX4 and SOX9, IPF-related matrix metalloprotease (MMP)7 [29], and secretory cell markers MUC4 and SCGB1A1 can be seen in Figure 4 D. Interestingly, most of the cells from the more differentiated clusters 3, 5 and 6 retained a strong basal cell signature (i.e., they had high transcriptional similarity scores to basal cells). A large proportion of cells from clusters 1, 2 and 4 displayed a strong mesenchymal cell signature. This was mirrored by the atypical co-expression of the mesenchymal markers FN1 and VIM in the majority of the KRT17 positive cells (Figure 4 E).

**Figure 4:**
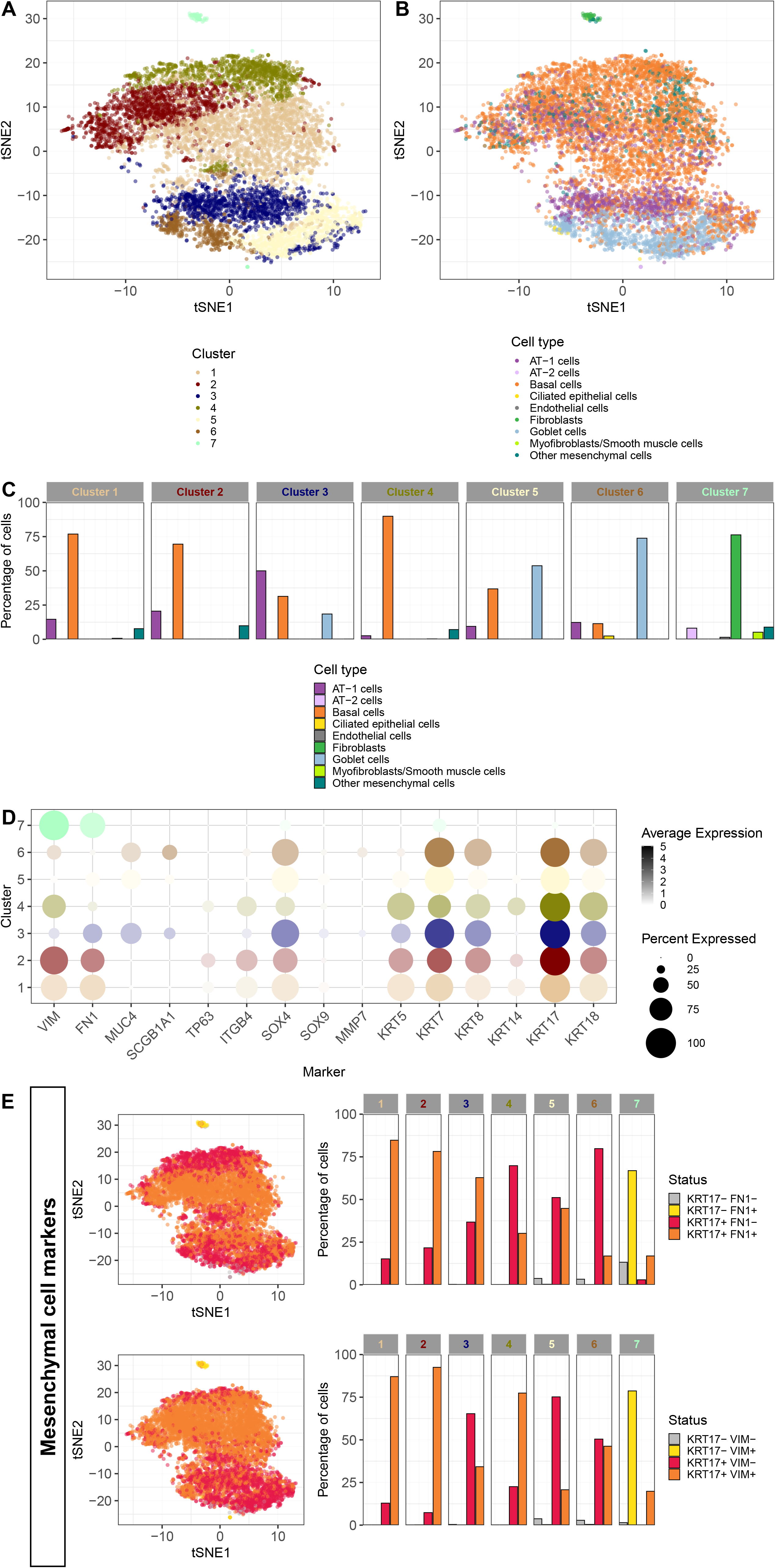
Single-cell RNA-seq of fibrosis-enriched cells from patient 6 was performed using the 10X technology. tSNE showing the separation of the cells into 7 clusters A). tSNE showing the best matching cell type for each cell, using as reference the cell types obtained from reanalysis of a publicly available scRNA-seq dataset of whole lung (GEO accession GSE122960) B). Relative proportions of cells with different cell type annotations across clusters C). Average expression of selected markers across cells from different clusters, shown by color intensity, and percentage of cells with detected expression of these markers across clusters, shown by diameter of the circles (D). tSNE showing co-expression of basal cell marker KRT17 and mesenchymal markers (FN1, VIM). The proportion of cells expressing/co-expressing the different markers across clusters is shown in the respective barplots E).

We replicated these findings in a second batch of scRNA-seq, including two additional patients (Supplement figures 3-6). The majority of cells showed again closest transcriptomic proximity to basal cells (54 to 74%, orange) and other more differentiated epithelial cell types such as AT-1 cells (8 to 15%, purple) or goblet cells (7 to 9%, blue)(Supplement figure 4 C), and the majority of the KRT17 positive cells co-expressed mesenchymal markers (Supplement figure 6 A).

**Figure 5:**
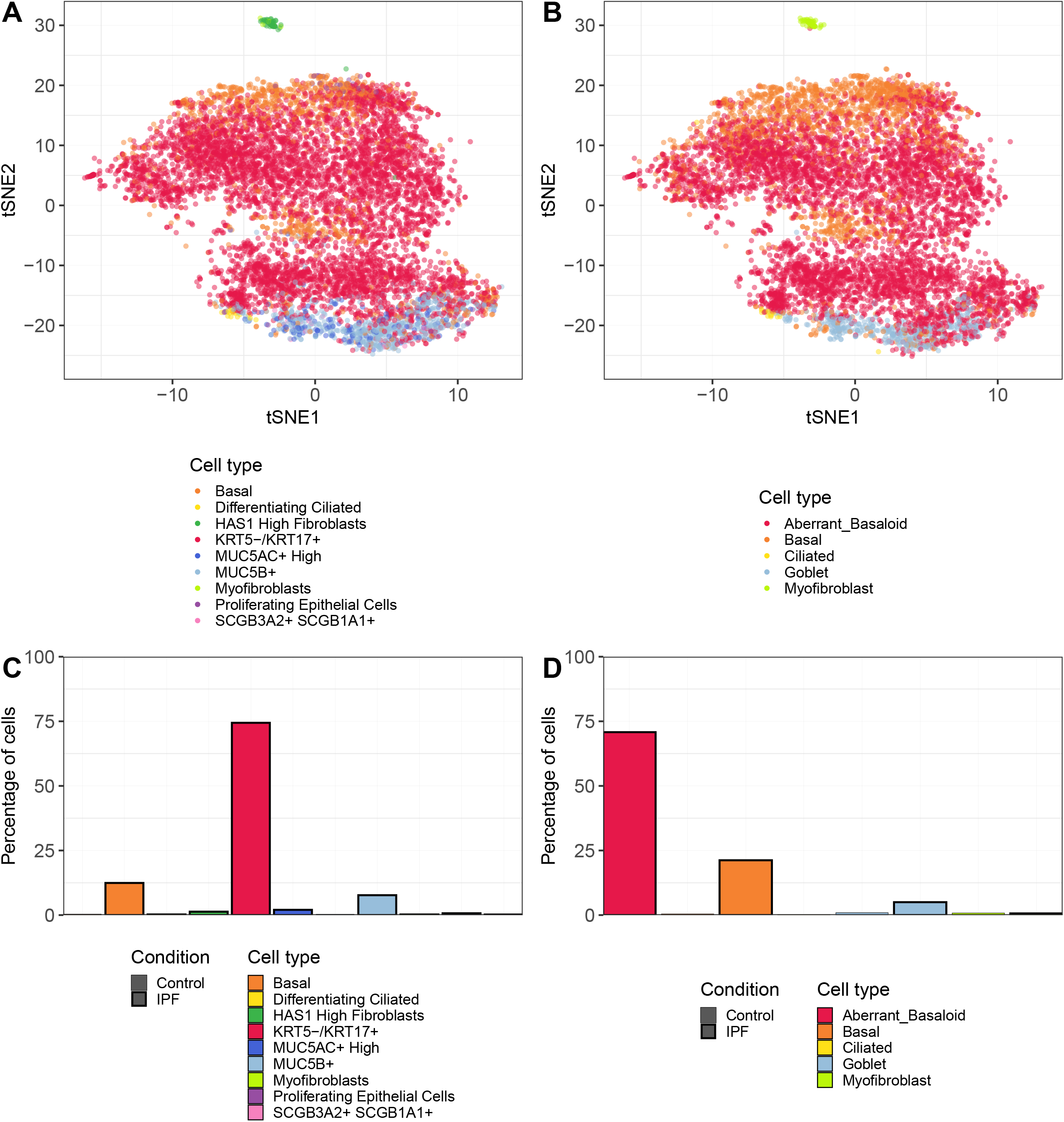
tSNE showing the best matching cell type for each cell after reannotation of cultured cells using as reference the cell-type annotation of clusters from the authors of the publicly available scRNA-seq datasets GSE135893 A) and GSE136831 B).The relative proportion of the different annotated cell types is shown in respective barplots (C, D).

**Figure 6:**
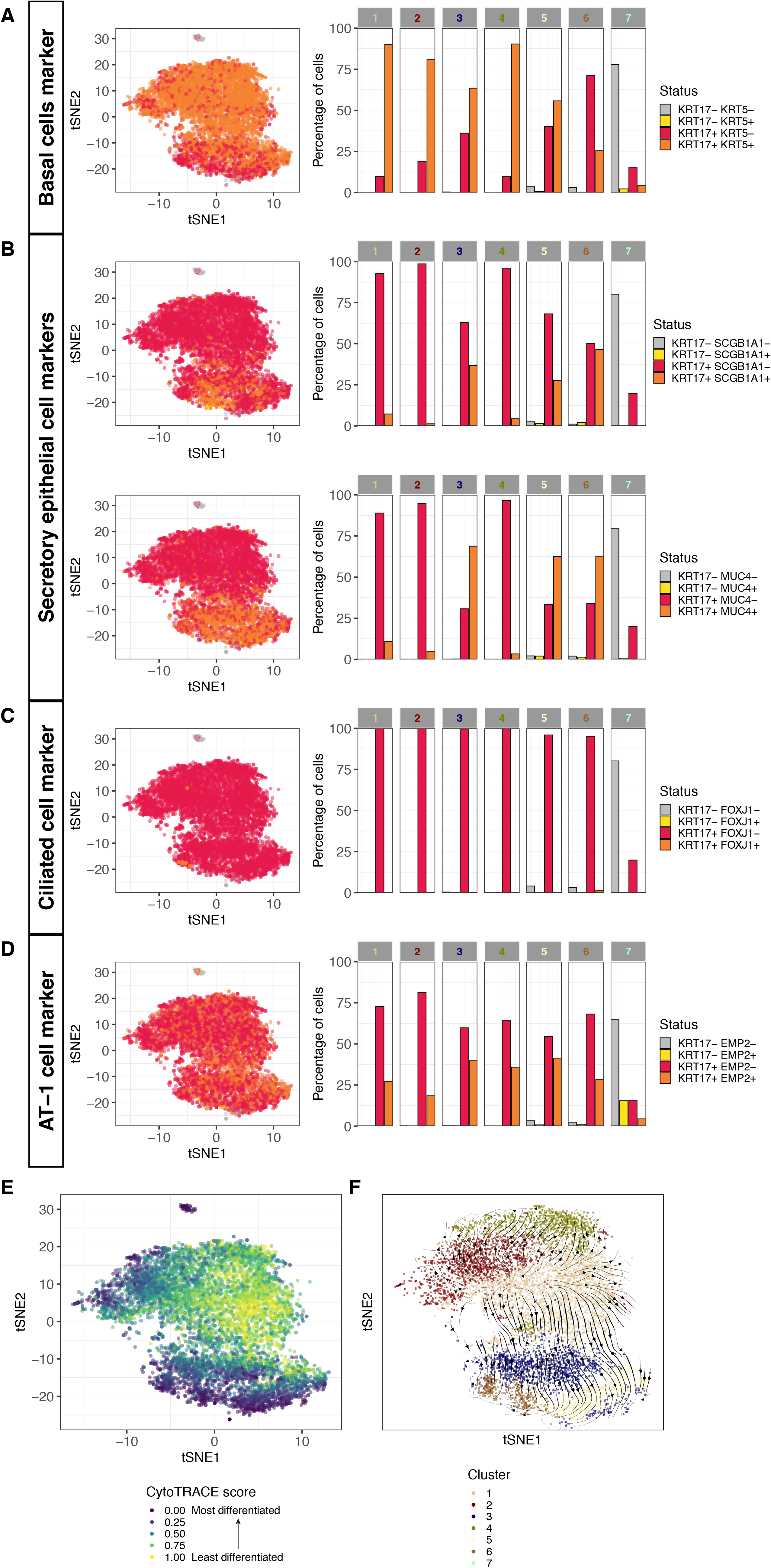
tSNE showing co-expression of basal cell markers KRT17 and KRT5 (A). Co-expression of KRT17 and secretory cell markers (SCGB1A1, MUC4) B), ciliated epithelial marker FOXJ1 C) and AT cell marker EMP2 D). The proportion of cells expressing/co-expressing the different markers across clusters is shown in respective barplots. tSNE showing the CytoTRACE differentiation state scores across cells E). Streamplot of the RNA velocity vector fields showing the differentiation trajectories inferred from comparison of exonic and intronic signal F).

The high proportion of cells co-expressing basal epithelial cell and mesenchymal markers is unlikely explained by cell-doublets formation in the droplet-based scRNA-seq experiment, especially that in this second replication batch, a much lower number of cells was loaded into the 10X Chromium controller, further reducing the chance of doublets. The possibility of droplets contamination with the ambient RNA solution was also excluded (Supplement table 3).

Thus, our cultured basal-like cells displayed obvious similarities to recently described aberrant basal-like cells also co-expressing mesenchymal markers [13, 14]. We compared our dataset to the two scRNA-seq datasets used in the studies describing these aberrant basal-like cells (GEO accessions GSE136831 and GSE135893). We notably reannotated our cells by using this time as reference the annotation of cell clusters from the authors of the two respective datasets. Surprisingly this revealed that 71-74% of the cultured basal-like cells matched best to this newly described aberrant basal-like cell population (Figure 5 A, B). From the remaining cells, the annotation was unchanged, with 13-22% of cells matching best basal cells and 6-8% secretory epithelial cells (goblet/MUC5B+ cells) (Figure 5 A, B). The relative expression across cells of specific markers (KRT5, KRT14, FN1, ITGB4, VIM, TP63, SOX4, SOX9, MUC4, SCGB1A1, MMP7, KRT7, KRT8, KRT17, and KRT18) supported these findings (Supplement Figure 5). Similar to the aberrant basaloid cells virtually all of our cultured basal-like cells expressed KRT17. In contrast, they also co-expressed varying levels of KRT5, which was detected above background levels in 77-79% of cells matching the aberrant basal-like cells (Figure 6 A). This might indicate a differentiation path towards airway basal cells *in vitro*.

When visualizing co-expression of KRT17 and the secretory cell markers SCGB1A1 or MUC4 (Figure 6 B) a second differentiation gradient from aberrant basal-like cells (KRT17+/SCGB1A1-, KRT17+/MUC4-) to secretory cells (KRT17+/SCGB1A1+; KRT17+/MUC4+) was apparent (Figure 6 B). Similarly, the co-expression of KRT17 and FOXJ1, a marker for ciliated epithelial cells, or EMP2, a marker for alveolar epithelial cells was detectable in a fraction of cultured cells (Figure 6 C, D). These differentiation gradients, all starting from a “core” of aberrant basal-like cells and pointing to cells at the edges of the tSNE plot, were confirmed by the analysis of differentiation state across cells with CytoTRACE [30] (Figure 6 E) and by RNA velocity trajectories [31, 32] (Figure 6 F).

### Basal-like cells do not undergo a full EMT in vitro and do not induce α-SMA expression in cultured fibroblasts

Co-expression of epithelial and mesenchymal markers in basal-like cells suggests that the cells underwent a partial EMT. In order to determine whether the cells become fully mesenchymal after a prolonged culture period, we observed cultured basal-like cells over a period of ten days. Interestingly, after ten days of culture the cells did not undergo dramatic morphological changes, and still highly expressed the epithelial cell markers E-cadherin and Nkx2. (Figure 7 A, B).

**Figure 7:**
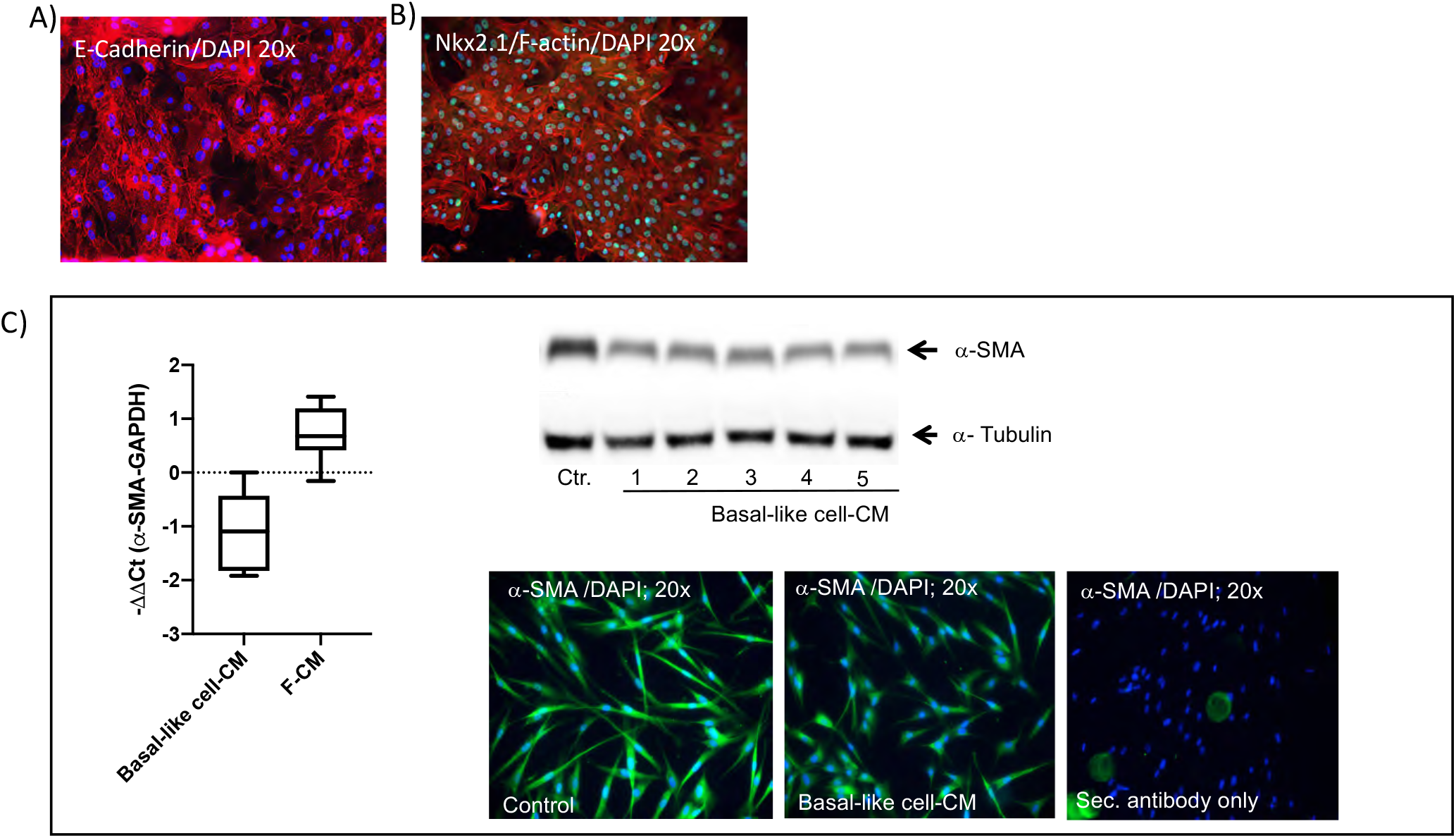
E-cadherin/DAPI A) Nkx2.1/F-actin/DAPI (B) expression in basal-like cells cultured for 10 days. IPF fibroblasts were treated with control medium (corresponds to -ΔΔCt=0), basal-like cell-conditioned medium (CM) or fibroblasts-CM (F-CM) for 24 h. α-SMA RNA expression was measured by TaqMan RT-PCR and data expressed as -ΔΔCt (C). Changes in α-SMA protein in IPF fibroblasts treated with basal-like cell-CM of five different patients (1-5) was determined by immunoblotting and confirmed by immunofluorescence. Fibroblasts incubated with secondary antibody only served as negative control C).

In order to determine whether the cells may induce differentiation of neighboring fibroblasts to myofibroblasts by paracrine mechanisms, we investigated the effects of basal-like cell conditioned medium (CM) on α-SMA expression in cultured human IPF lung fibroblast (Figure 7 C). IPF fibroblasts in control medium (DMEM/10%FCS) expressed α-SMA RNA, which was significantly down-regulated after incubation of the cells with basal-like cell-CM. This effect seemed basal-like cell-CM-specific, as CM derived from fibroblasts (F-CM) up-regulated α-SMA RNA. α-SMA protein expression in control medium (Ctr.) was inhibited by basal-like cell-CM derived from five different patients (1-5; Figure 7 C). α-Tubulin served as a control for equal protein loading. Immunofluorescence analysis confirmed reduced α-SMA in fibroblasts incubated with basal-like cell-CM (Figure 7 C) when compared to control medium. Fibroblasts incubated with secondary antibody served as negative control.

### Bulk RNA seq demonstrates a clear separation of cultured basal-like cells derived from IPF versus other ILD patients

Among the bulk RNA-seq samples of cultured basal-like cells used above, four were derived from IPF patients and four of patients with other ILDs (non-IPF) (Patients details can be viewed in supplement table 2). As we could observe a disease-specific separation of the transcriptome on the first component of a PCA (Figure 8 A), we aimed at characterizing the transcriptomic signature associated to IPF. At a FDR of 1%, 101 genes were up-regulated in IPF compared to other ILDs, and 87 were down-regulated. The differentially expressed genes are shown in a volcano plot (Figure 8 B), heatmap (Supplement figure 7) and are listed in supplement table 7. Functional enrichment analysis using the MSigDB Hallmark pathways showed that pathways related to epithelial to mesenchymal transition (EMT), angiogenesis or myogenesis were up-regulated, whereas pathways overall related to inflammation were down-regulated in IPF cells (Figure 8 C; Functional enrichment analysis results on c2 and c5 collections in the supplement table 4 and 5). An enrichment analysis on transcription factors “regulons” [33] revealed a disease-dependent involvement of particular transcription factors, for example TGF-β1 signaling-associated SMAD7 (whose targets are overall up-regulated in IPF) or pro-inflammatory NFκB signaling-associated transcription factors (RELA, NFKB1, NFKB2, whose targets are down-regulated in IPF; Figure 8 D, supplement table 6).

**Figure 8:**
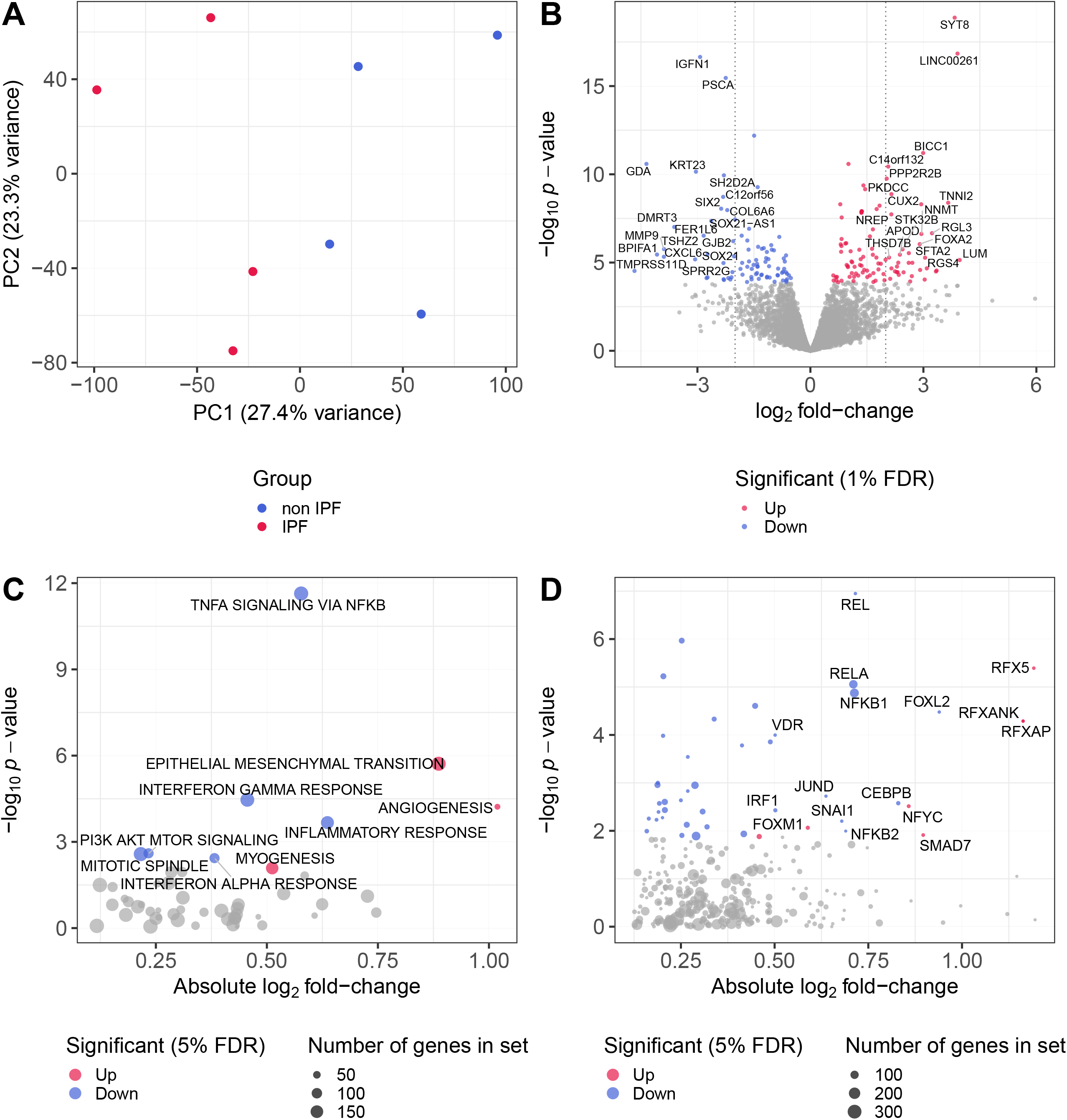
Principal component analysis (PCA) based on normalized log-transformed CPM values of the bulk RNA-seq IPF and non-IPF samples a). Volcano plot showing the differential expression analysis results for IPF versus non-IPF stem-like cells (genes colored are significant at an FDR threshold of 1%; genes labelled are significant, with an absolute log2 fold-change larger than 2) b). Functional enrichment results on gene sets from the “hallmark” collection of the MSigDB v7.0 c) or on DoRothEA regulons to show involvement of key transcription factors d). Only regulons with average absolute log2 fold-change larger than 0.5 are labelled.

## Discussion

In this study we confirmed our previous findings of a fibrosis-enriched outgrowth of a distinct cell population from peripheral lung tissue derived from ILD patients. Although the cells morphology impeded a clear classification to any known lung cell type, the cells displayed some characteristic features of epithelial cells on EM images. Bulk RNA-seq highlighted the expression of epithelial and mesenchymal markers and disease-specific transcriptome differences. scRNA-seq showed an overall homogeneity and epithelial origin of the cells. The majority of cells displayed close transcriptomic similarities to the recently described fibrosis-enriched aberrant basal-like cells. In this study, we therefore confirm the presence of aberrant basal-like cells in fibrotic lung tissue and importantly, demonstrate for the first time that these cells can be cultured and characterized *in vitro*.

Our previous observation of a disease-enriched outgrowth of a distinct cell population from peripheral lung tissue derived from IPF and other ILD patients [15] was confirmed in this study. Based on the expression of the mesenchymal stem cell markers CD44, CD90 and CD105, their differentiation potential and the lack of the current advanced transcriptomic tools we earlier named these cells mesenchymal stem cells [15]. Using RNA-seq in this study, we confirmed the expression of mesenchymal cell markers, but however surprisingly, observed closest transcriptional similarity of these cells to epithelial cells. Epithelial origin of the cells was corroborated by EM images, showing polarity to some extent, microvilli-like structures on the cell surface and occasionally the presence of multi-lamellate bodies in the cells. Furthermore, the cells were joined by desmosomes and tight junctions. In line with our findings, epithelial cells with similar characteristics reported here were detected in the alveolar compartment of fibrotic lung tissue and were suggested to be bronchial basal cell-derived [34, 35]. The disease-specific presence of atypical epithelial cells in the peripheral IPF lung was observed before: ΔNP63 [12], KRT5 [10], or ΔNP63 and KRT5/6 [11] expressing cells as well as recently described KRT17+/KRT5-aberrant basal-like cells [13, 14] or KRT8+ alveolar epithelial progenitor cells [27, 36] were detected in the alveolar compartment of lung tissue derived from IPF patients, but were absent in tissue from control donors. In addition, KRT5/6+ and KRT5+ cells were present in BAL of IPF patients, but were only rarely observed in BAL from control donors [11].

scRNA-seq data demonstrated an overall homogeneity and epithelial origin of the cultured cell population. Importantly, comparison of our dataset to those describing the aberrant basal-like cell population [13, 14] showed a high transcriptomic similarity of the majority of our cultured cells to aberrant basal-like cells. In all likelihood, the *in vitro* culture process resulted in a strong and specific enrichment of these cells in our samples. Therefore in addition to confirming the presence of aberrant basal-like cells in fibrotic lung tissue, we show for the first time that these cells can be cultured *in vitro*. Cultured basal-like cells partially co-expressed alveolar, ciliated -or secretory epithelial cell markers and showed typical characteristics of alveolar epithelial cells (microvilli-like structures and multi-lamellar bodies) on EM images. This may indicate either recruitment of basal-like cells from the conducting airways that may subsequently acquire the phenotype of alveolar or secretory epithelial cells or a disturbed differentiation program in resident alveolar epithelial cells. Studies in mice suggested migration of airway-derived progenitor cells into the alveolar compartment after severe lung injury with the ability to differentiate into alveolar epithelial cells [5-7, 37]. However, more recent studies suggest that basal-like cells are derived from resident AT2 cells rather than from phenotypically similar airway basal cells in the human lung: Xu et al. identified several different types of atypical epithelial cells in the peripheral IPF lung, proposing the initiation of an abnormal differentiation program in resident alveolar epithelial cells through the IPF microenvironment [38]. Habermann et al. suggested that human aberrant basal-like cells might derive from transitional AT2 cells [14]. This is in line with a recent study showing that cultured human AT2 cells trans-differentiate into KRT5 positive basal cells via a KRT17+/KRT8+ alveolar-basal intermediate when co-cultured with adult human lung mesenchyme in a 3D organoid system [39]. Similarly RNA velocity analysis of IPF-derived epithelial cells showed transcriptional trajectory of AT2 cells, through a alveolar-basal intermediate cell, to more mature KRT5+ basal cells [39]. Interestingly, alveolar-basal intermediate cells show similarities to the recently described KRT5-/KRT17+ aberrant basal-like cell [13, 14] and to our cultured basal-like cells, expressing KRT17 along with varying levels of KRT5, seemingly differentiating towards a more mature airway basal-like cell *in vitro*. Airway basal cells have the capacity to differentiate into secretory or ciliated epithelial cells [4], which likely explains the presence of secretory or ciliated epithelial cell markers in the most differentiated subset of our cultured cells.

The high expression of mesenchymal markers in aberrant basal-like cells *in vivo* [13, 14] and in the here described cultured cells points to a partial EMT within these cells. Interestingly, others have demonstrated that epithelial cells that underwent a partial EMT acquire MSC-like properties [19], which may explain our previously observed similarities to MSCs [15]. More importantly, aberrant basaloid cells localize on the surface of fibroblastic foci (FF) in IPF lung tissue [13, 14]. Earlier studies suggested that epithelial cells covering the surface of FF may contribute to an increased number of myofibroblasts by either undergoing an epithelial to myofibroblast transition [40] or by secreting factors inducing fibroblasts to myofibroblasts differentiation [41]. However, we here show that cultured basal-like cells do not undergo a full transition into mesenchymal cells *in vitro*. Furthermore, mediators secreted by cultured basal-like cells rather reduced α-SMA levels in cultured fibroblasts, suggesting a different functional role of the cells in fibrotic lung diseases.

After determining an overall homogeneity of the cultured cell population using scRNA seq, we went back to analyse our bulk RNA-seq samples. Interestingly, we revealed a significant difference in the transcriptome of cultured basal-like cells derived from IPF compared to patients with other ILDs. IPF derived basal-like cells were enriched with fibrosis-associated genes mediating EMT, myogenesis or angiogenesis, whereas pathways involved in inflammation, which is known to play a minor role in IPF pathogenesis [2], were down-regulated in IPF cells. This suggests epigenetic changes of the cells, potentially induced by the disease-specific microenvironment that persists *in vitro*. Basal-like cells accumulating within the alveolar region in mice after severe lung injury were suggested to help regenerating the alveolar epithelium by their ability to differentiate into alveolar epithelial cells [5, 6]. Furthermore, we previously observed anti-fibrotic effects of basal-like cell-derived conditioned medium (CM) *in vitro*: IPF fibroblast proliferation was inhibited and epithelial wound repair increased after treatment with basal-like cell-CM [15]. However, in IPF the presence of basal-like cells in the alveolar region was associated with pathological bronchiolization [12]. Furthermore, enrichment of genes that are highly expressed in airway basal cells in BAL from IPF patients was associated with higher mortality [11]. Interestingly, it was shown that bronchial epithelial stem cells are able to contribute to pathological bronchiolization and alveolar regeneration, depending on the activation of specific signaling pathways [8]. Therefore, it can be speculated that the fibrotic microenvironment directs basal-like cells to contribute to pathological bronchiolization rather than alveolar epithelial regeneration. Studying the differentiation potential of fibrosis-specific basal-like cells exposed to different pro-fibrotic factors/microenvironments would therefore be of uttermost interest in future studies.

It is important to mention that our results are limited by the low number of patients included in the study. Furthermore, we had to exclude two outlier samples from our bulk RNA-seq analysis, both coming from our only female patients, leaving the possibility that the reported differential expression patterns are only male-specific. An increase of the patient numbers in future studies would corroborate our findings. Regarding the scRNA-seq analysis of basal-like cells, it would have been interesting to compare our results to basal-like cells grown from non-fibrotic tissue. However, we clearly showed in this and our previous study [15] that under standardized cell culture conditions the basal-like cell population readily grows from fibrotic, but only rarely from non-fibrotic tissue, making it difficult to provide a control group of cultured non-fibrotic basal-like cells.

To the best of our knowledge, this is the first study describing the culture and *in vitro* characteristics of aberrant basal-like cells from peripheral lung tissue of IPF patients. Culturing these cells *in vitro* opens up the opportunity to study their role, function and origin in the IPF lung. Understanding the cells origin, differentiation potential, and how they are influenced by pro-fibrotic factors/different microenvironments may lead to the development of novel therapeutic targets in IPF.

## Acknowledgments

Genomics facility Basel, ETH Zürich for bulk and single-cell RNA-sequencing. Calculations were performed at sciCORE (http://scicore.unibas.ch/) scientific computing center at the University of Basel.

## Author’s contribution

**PK:** Conception and design of the study, cell culture of human primary lung cells, immunofluorescence, PCR, data analysis and interpretation, drafting the manuscript; **JR:** Data analysis and interpretation of bulk and scRNA-seq, revision and editing of the manuscript for important intellectual content; **SB:** Acquisition of immunofluorescence images, cell culture of human primary lung cells; **LF:** Acquisition of immunofluorescence images; **SS:** Provided lung biopsies of non-IPF donors, revision of the manuscript for important intellectual content. **LK:** Provided samples of IPF lung explants, acquisition of electron microscope images, revision of the manuscript for important intellectual content; **DJ:** Provided samples of IPF lung explants, revision of the manuscript for important intellectual content. **MK:** Provided samples of IPF lung explants, revision of the manuscript for important intellectual content. **AG:** Revision and editing of the manuscript for important intellectual content; **TG:** Revision and editing of the manuscript for important intellectual content; **MT:** Conception and design of the study, revision of the manuscript for important intellectual content; **KEH:** Conception and design of the study, cell culture of human primary lung cells, data analysis and interpretation, revision and editing of the manuscript for important intellectual content.

## Figure legends

**Supplement figure 1:** Heatmap showing the average centered and scaled normalized log-transformed expression of relevant cell-type markers across cells of each cluster of cell-types from the public dataset GSE122960, separating cells from fibrosis or control patients.

**Supplement figure 2**: PCA on scRNA-seq normalized log-transformed expression levels of cells from patient 6. The four first principal components are shown, and the cluster membership of cells is indicated with colors. Blue contour lines represent the cell density in the 2D space.

**Supplement figure 3**: Relative expression across cells of the strongest cluster-specific genes in the dataset of patient 6 (A) and the dataset of patients 7 and 8 (B). Cluster membership and cell-type annotation based on the pure cell types of the dataset GSE122960 are shown as heatmap metadata.

**Supplement figure 4**: tSNE showing the cell distribution of patients 7 (blue) and patient 8 (pink) A); the separation of the cells into 6 clusters B); the best matching cell type for each cell using the reference dataset GSE122960 (analogous to Figure 5) C), using the reference dataset GSE135893 (analogous to Figure 5) D), and using the reference dataset GSE136831 E). The respective relative proportion of annotated cell types are shown in barplots F-H).

**Supplement figure 5**: Heatmap showing the centered and scaled normalized log-transformed expression of selected markers across cells from patient 6 (A) and patients 7 and 8 (B). Cluster membership and cell-type annotation based on the three reference datasets are shown as heatmap metadata.

**Supplement figure 6**: tSNE showing co-expression of basal cell marker KRT17 and/or mesenchymal markers (FN1, VIM) (A), basal cell marker KRT5 (B), secretory epithelial cell markers (SCGB1A1, MUC4) (C), ciliated epithelial cell marker FOXJ1 (D), and AT cell marker EMP2 (E) in cells derived from patients 7 and 8.

**Supplement figure 7:** Heatmap showing the centered and scaled normalized log-transformed CPM values for differentially expressed genes between IPF and non-IPF bulk RNA-seq samples.

