## Supplement Figures for "In vitro culture of aberrant basal-like cells from fibrotic lung tissue"

Supplement Figure 1

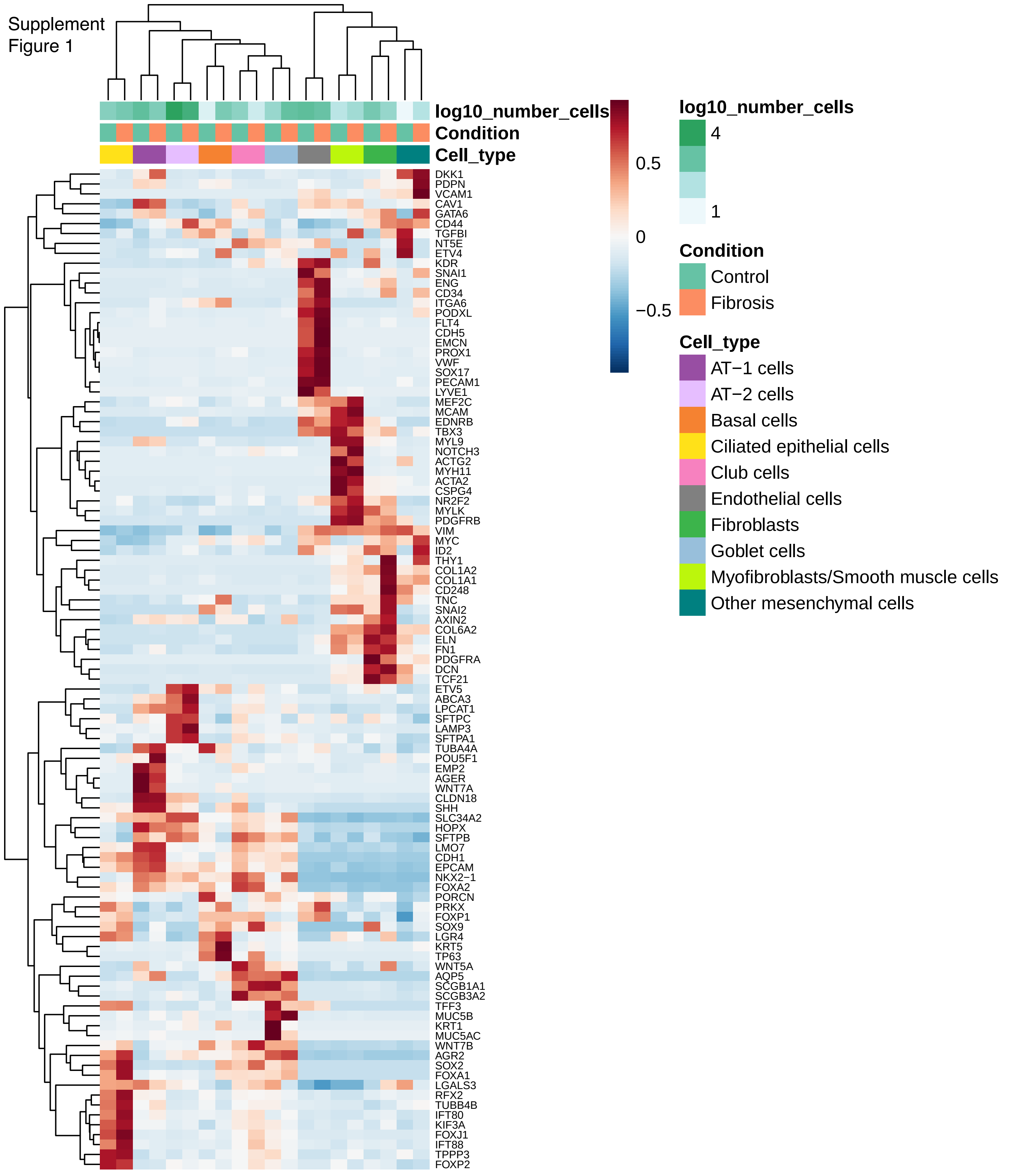

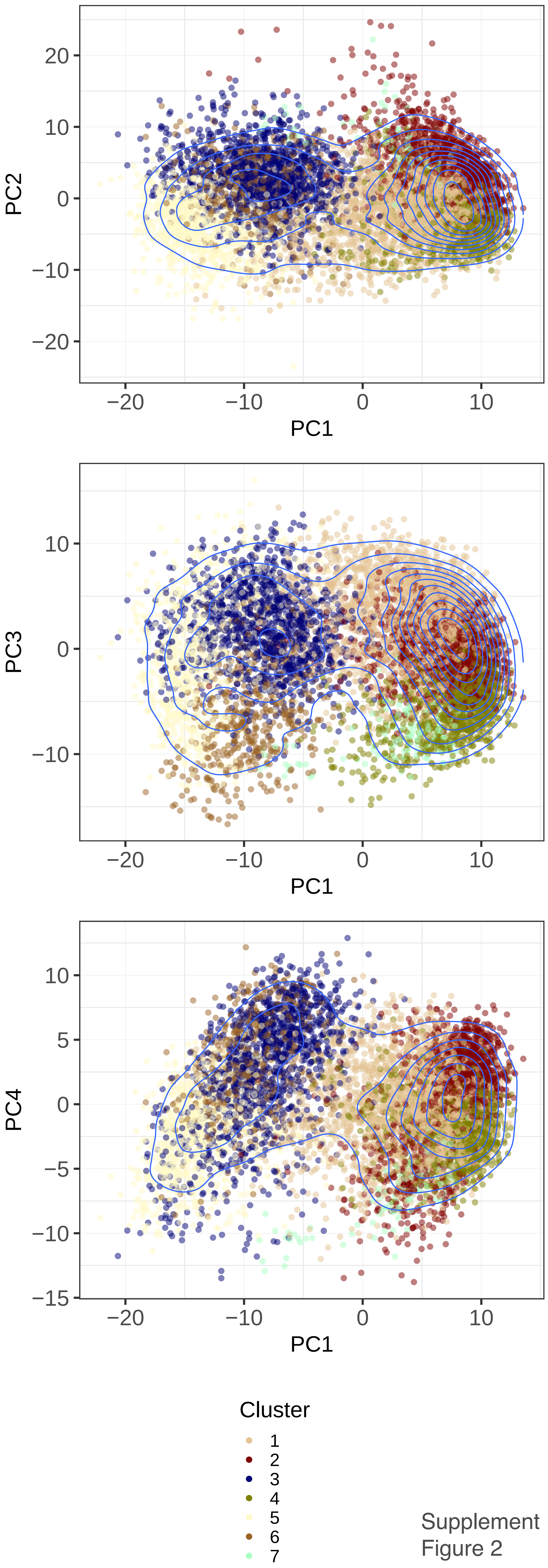


Supplement Figure 2

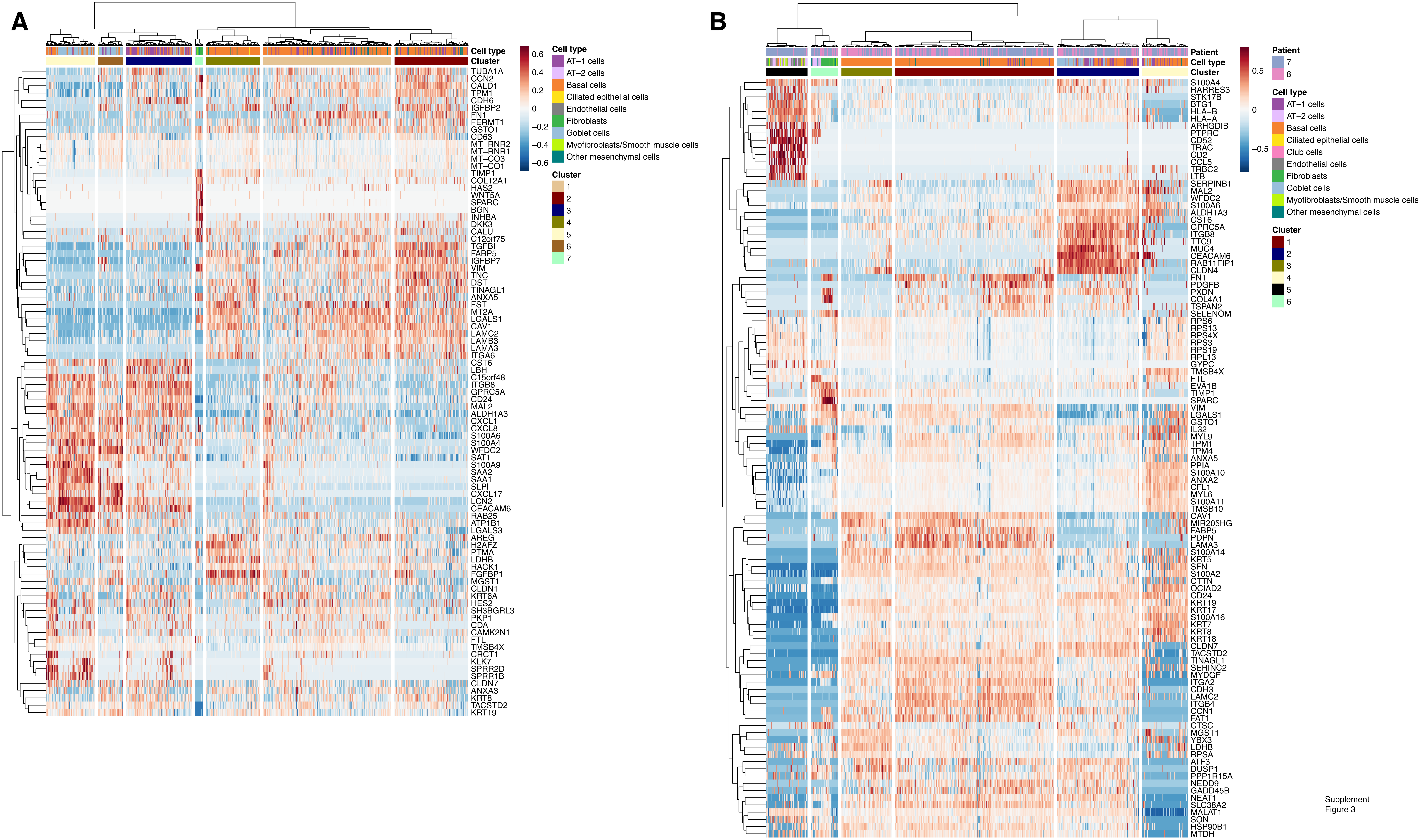


Supplement Figure 3

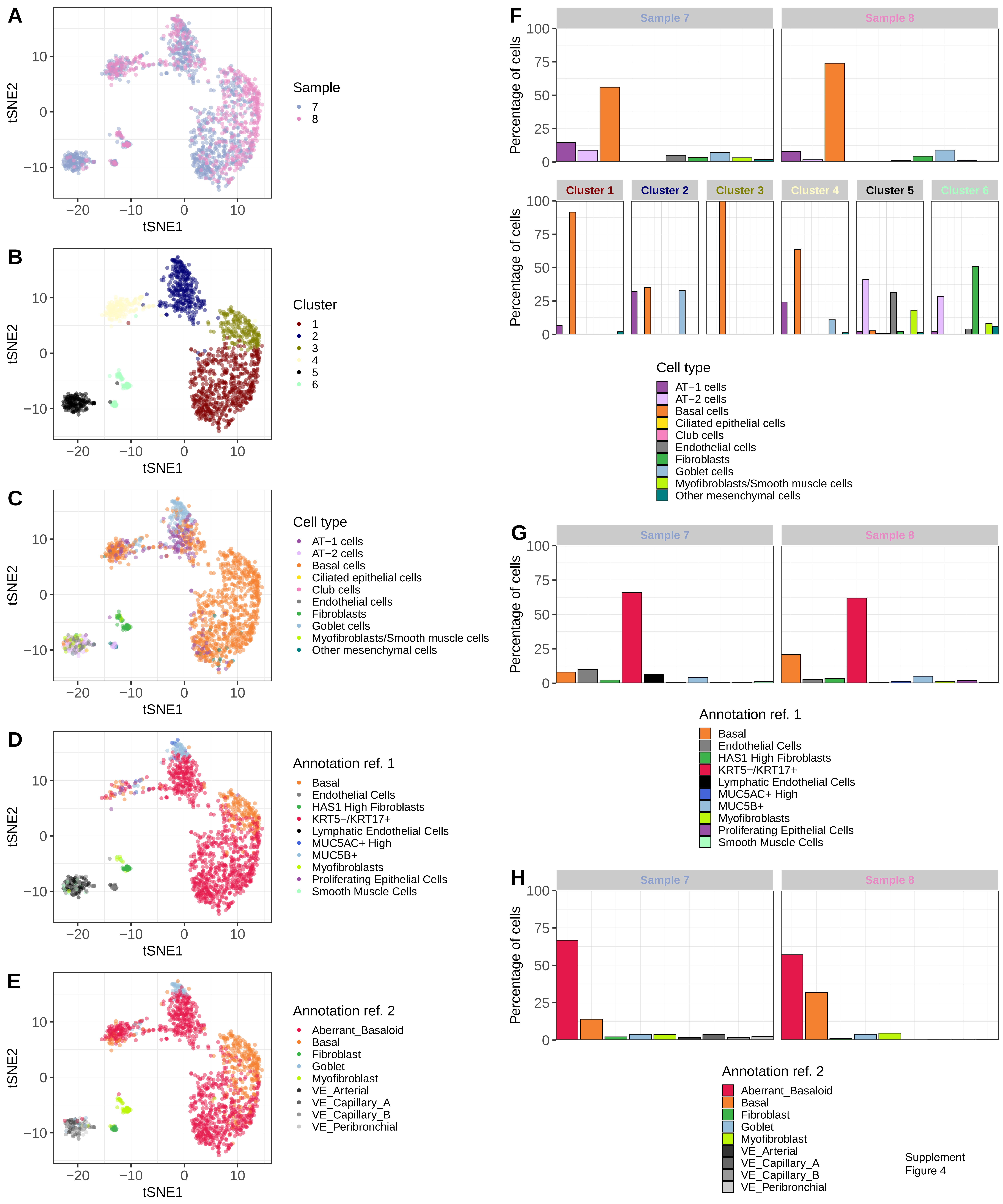


Supplement Figure 4

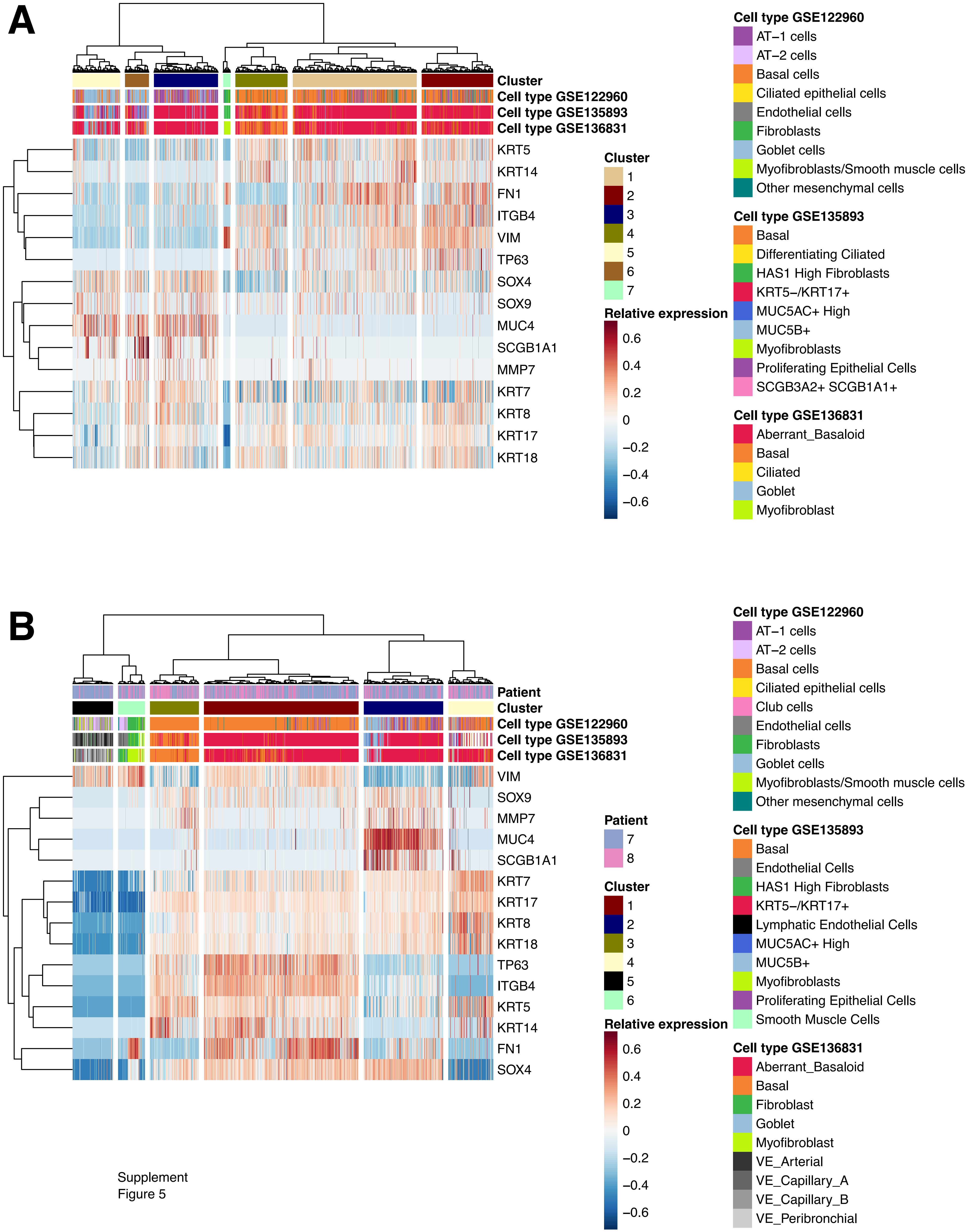


Supplement
Figure 5

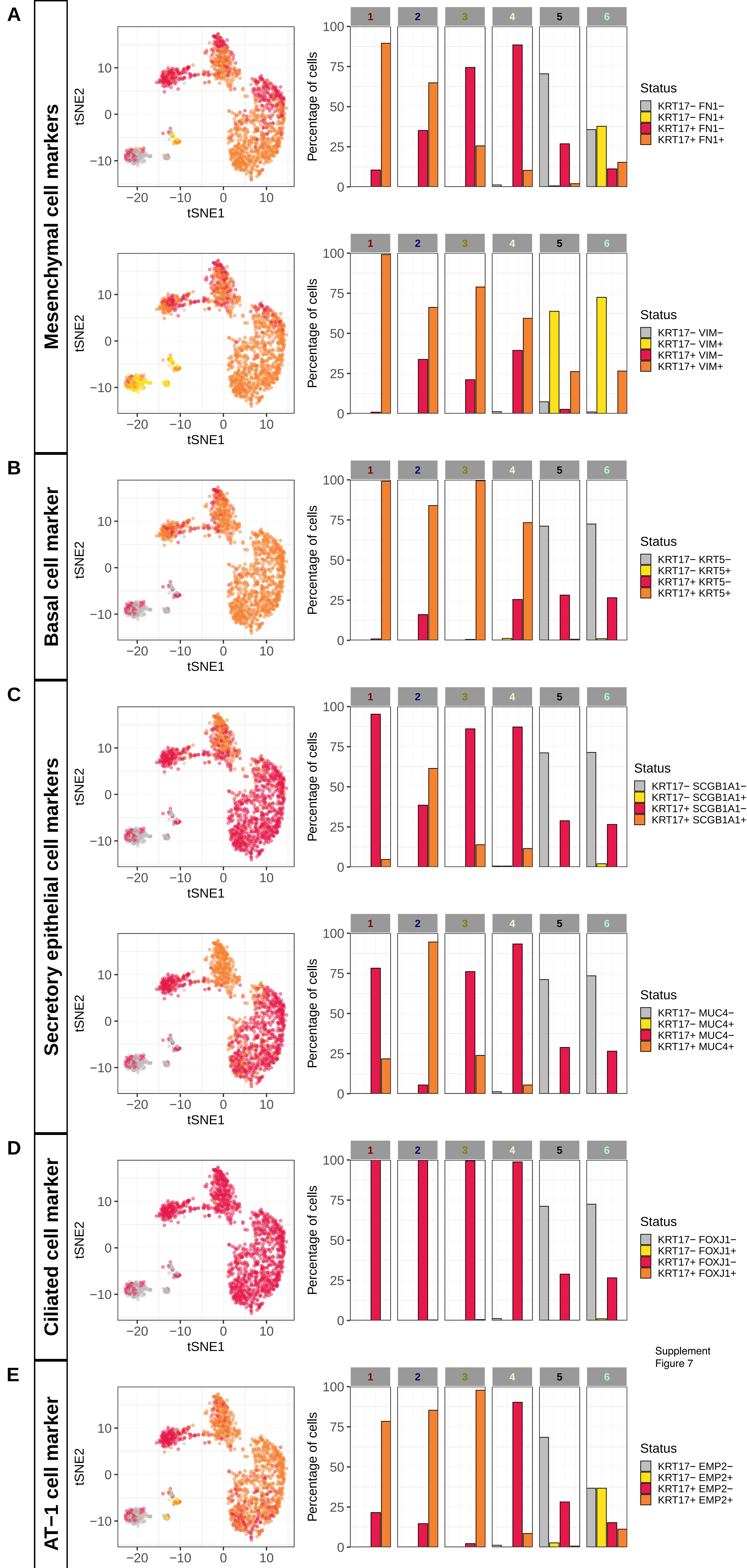


Supplement Figure 6

Supplement
Figure 7

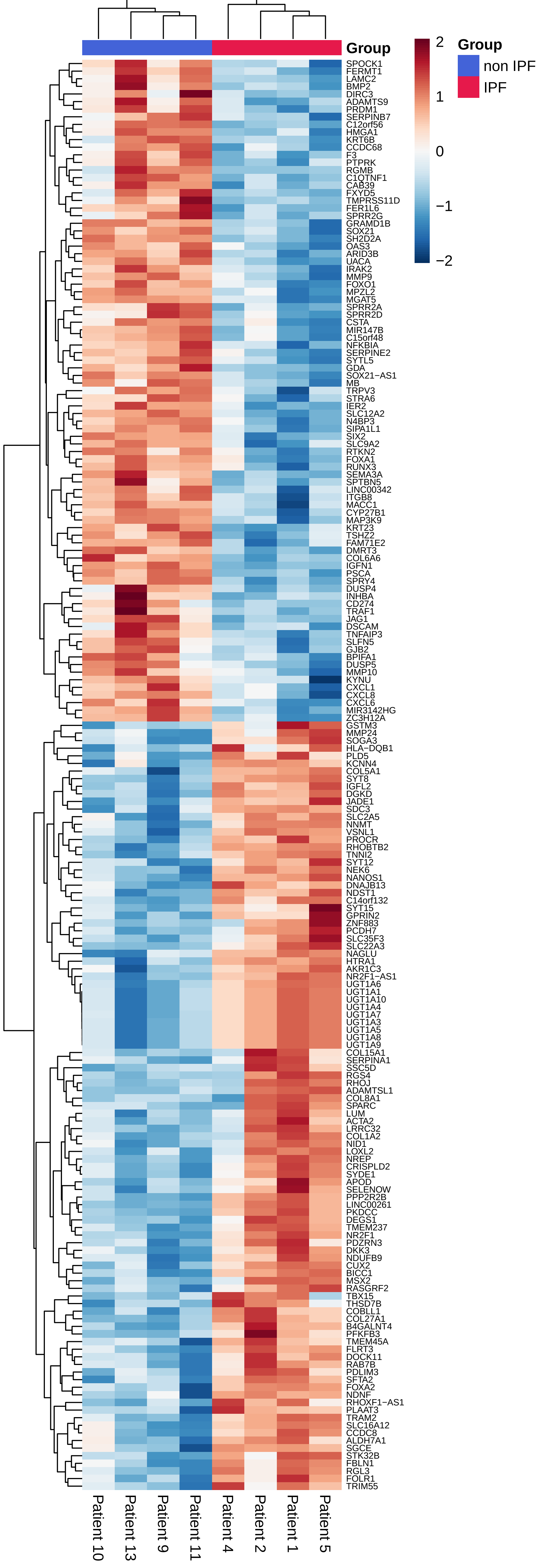
