## Supplement Document for "In vitro culture of aberrant basal-like cells from fibrotic lung tissue"

*Materials & methods*

*Cell culture and treatment*

For the culture of basal-like cells, lung tissue was cut into small pieces and placed into flasks containing DMEM with 10% FCS, 25 mM HEPES and 2 mM l-glutamine, 1× (vol/vol) MEM vitamin-mix, 100 U/l penicillin, 100 µg/ml streptomycin, and 0.25 µg/ml amphotericin B and maintained in a humified atmosphere of 5% $CO_2$ at 37°C. After five days, tissue pieces were removed and cells were harvested for down-stream analysis. Cells derived from several tissue pieces of the same patient were pooled.

*TaqMan RT-PCR*

Total RNA was isolated using a Quick-RNA MiniPrep Kit and expression levels determined by TaqMan® Gene Expression Assays. Samples were run at 50°C for 2 min, 1 cycle; 95°C for 10 min, 1 cycle; 95°C for 15 s, 60°C for 1 min, 40 cycle and expression levels were quantified using the -ΔCt value relative to GAPDH. Missing -ΔCt values (because the target genes was expressed at too low level to be detected after 40 cycles) were arbitrarily set to -20.

*Immunofluorescence*

Basal-like cells were fixed with 4% PFA/PBS (15 min) and permeabilized in a 0.5% Triton/PBS solution (15 min). After blocking with 3% BSA/PBS (30 min), the cells were incubated (4°C, overnight) with a primary antibody, which was detected by an Alexa488 or Alexa594 conjugated antibody (1h). Nuclei were visualized by DAPI and slides examined on an OLYMPUS BX63 upright fluorescence microscope.

*Bulk RNA-seq*

Total RNA was extracted using the MicroElute Total RNA Kit. RNA was quantified using the QuantiFluor RNA System and quality-controlled on a Bioanalyzer instrument using the RNA 6000 Pico Kit. When all 10 samples had been collected for the study (Nov. 2015 to Nov. 2016), RNA-seq library preparation (Illumina Truseq stranded kit) was performed at the Genomics Facility Basel of the ETH Zurich, Basel, starting from 50 ng of total RNA. Sequencing was performed on the Illumina NextSeq 500 platform to produce single-end 81nt reads. Data analysis was performed by the Bioinformatics Core Facility, Department of Biomedicine, University of Basel. Reads were mapped to the human hg38 genome assembly with STAR (version 2.5.2a_modified)[1] with default parameters, except filtering out multimapping reads with more than 10 alignment locations (outFilterMultimapNmax =10) and filtering reads without evidence in the spliced junction table (outFilterType="BySJout"). All subsequent RNA-seq data analysis was performed using the R software (version 3.6) and Bioconductor 3.10 packages [2, 3]. Gene expression was quantified by the qCount function of the Bioconductor package QuasR (version 1.18)[4] as the number of reads that started within any annotated exon of a gene from the UCSC knownGenes annotation (downloaded on 2017-04-03). Read and alignment quality was evaluated using the qQCReport function of QuasR.

The R Bioconductor package edgeR (version 3.28)[5] was used for differential gene expression analysis. Between samples normalization was done using the TMM method [6]. Based on a principal component analysis (PCA) performed on the normalized log-transformed CPM (counts per million reads mapped) values, two female patients (IPF patient 3 and non-IPF patient 12) were clear outliers, so they were excluded and only male samples were kept for further analysis. Genes with CPM values more than 1 in at least 3 samples were retained for the differential expression analysis. A generalized linear model was fitted to the raw counts (edgeR function glmFit) to test for differences between the IPF and non-IPF groups using a likelihood ratio test (edgeR function glmLRT). P-values were adjusted by controlling the false-discovery rate (FDR; Benjamini-Hochberg method) and genes with a FDR lower than 1% were considered significant. Gene set enrichment analysis was performed with the function camera [7] from the edgeR package (using the default parameter value of 0.01 for the correlations of genes within gene sets) using gene sets from the c2, c5 and hallmark collections of the MSigDB Molecular Signature Database (MSigDB v7.0)[8], or DoRothEA v2 regulons [9]. We used the "TOP10score" regulons containing the targets with the highest quality score possible and at least 10 targets per transcription factor, retaining only regulons with confidence levels A to D. We filtered out sets containing less than 10 genes, and gene sets with a FDR lower than 5% were considered significant.

*Reanalysis of public scRNA-seq datasets and comparison to bulk RNA-seq samples*

The dataset of Human pulmonary fibrosis and control patients [10] was downloaded from GEO (accession GSE122960; HDF5 processed data files produced by CellRanger v2.0). Processing of the UMI counts matrix was performed using the Bioconductor packages DropletUtils (version 1.5.4)[11, 12], scran (version 1.12)[13, 14] and scater (version 1.12)[15], following mostly the steps illustrated in the simpleSingleCell Bioconductor workflow [16]. The resulting filtered and normalized expression matrix included 14,983 genes and 108,509 cells from 17 patients. Clustering of cells was done on normalized log-count values using a shared nearest-neighbours graph approach (scran function buildSNNGraph), resulting in 93 clusters. Annotation of cell type and identification of clusters of cell doublets was done based on the expression of known marker genes and following the annotation of the original study [10](Supplement figure 1). Clusters of cell doublets and clusters of immune cells were filtered out for further analyses. Cells of each condition (control or fibrosis) and each remaining cell type (AT−1, AT−2, basal, ciliated, club, endothelial, fibroblast, goblet, myofibroblasts/smooth muscle, or other mesenchymal cells) were aggregated using a "pseudo-bulk" approach [17]. Four groups of cells were not aggregated because the number of cells was too low (Other mesenchymal cells-Control, Basal cells-Control, Myofibroblasts/Smooth muscle cells-Control and Club cells-Fibrosis). The UMI counts of grouped cells were summed to produce 16 pseudo-bulk samples which were used as reference set for comparison to the bulk RNA-seq samples, and for annotation of the scRNA-seq dataset (see below).

For comparison to our bulk RNA-seq samples, after merging the bulk and pseudo-bulk datasets to keep 11,065 genes in common, a PCA was performed on the normalized log-transformed CPM. Since the first component (40% of variance explained) was separating bulk and pseudo-bulk samples, we used the function removeBatchEffect from the Bioconductor limma package (version 3.42) to subtract this effect from the data, and used the resulting log-CPM values for PCA (Figure 3).

The IPF Cell Atlas dataset [18] was downloaded from GEO (accession GSE136831; R ".rds" files including exonic UMI count matrices for each patient, and the "GSE136831_AllCells.Samples.CellType.MetadataTable.txt.gz" file including the annotation for cells). Only non-immune cells from control and IPF patients were retained. We additionally excluded cells from patients with less than 50 cells left. The final dataset was composed of 30,617 cells, 7,898 from control patients, and 22,719 from IPF patients.

The Habermann et al. dataset [19] was downloaded from GEO (accession GSE135893; Full Seurat object downloaded as ".rds" file). Only non-immune cells from control and IPF patients were retained. The final dataset was composed of 44,137 cells, 12,151 from control patients, and 31,986 from IPF patients.
For both datasets, cells of each condition (control or IPF) and each cluster (i.e., cell type annotated by the respective authors) were aggregated, keeping only samples where at least 65 cells could be aggregated. The pseudo-bulk samples were inspected for quality issue using PCA, and then used as reference set for reference-based unbiased annotation of the scRNA-seq dataset (see below).

*Single cell RNA-sequencing*

Single-cell suspensions were generated from stem-like cells grown from cultured peripheral lung tissue biopsies from 3 IPF patients (IPF patient 6-8; see patient details in supplement document 2), controlled for viability, and loaded on separate wells of a 10X Genomics Chromium Single Cell Controller in 2 batches (IPF patient 6 on April 8$^{th}$ 2019, and patients 7 and 8 on May 27$^{th}$ 2019) for an expected recovery of 10,000 cells.
Single cell capture and cDNA and library preparation were performed at the Genomics Facility Basel of the ETH Zurich, Basel with a Single Cell 3' v3 Reagent Kit (10X Genomics) according to the manufacturer's instructions. Sequencing was performed on one flow-cell of the Illumina NexSeq 500 platform (with 56nt-long R2 reads). Read quality was assessed with the FastQC tool (version 0.11.5). Sequencing files were processed with Kallisto (version 0.46.0) and BUStools (version 0.39.2) to perform sample and cell demultiplexing, and pseudo-alignment of reads to human transcripts (Ensembl release 96) and their intronic regions (extended by 50nt on each side). The commands "bustools correct", "bustools capture" and "bustools count" were used to generate gene-level spliced and unspliced UMI counts table [20, 21]. For the main analysis, the spliced counts table was used and processed as indicated above (reanalysis of dataset GSE122960). The *emptyDrops* function of the Bioconductor package DropletUtils (version 1.5.4) was used to estimate the expression profile of the ambient RNA solution [12].

After quality filtering, the resulting dataset consisted of 17,869 genes and 7,498 cells derived from patient 6, 905 cells from patient 7, and 550 cells from patient 8. The 2D *t*-distributed stochastic neighbor embedding (tSNE) used for visualization of cells was calculated using the 500 most variable genes and the denoised principal components as input, using a perplexity of 200. Clustering of cells was performed on each batch separately using normalized and denoised log-count values with hierarchical clustering on the Euclidean distances between cells (with Ward's criterion to minimize the total variance within each cluster; package cluster version 2.1.0). The number of clusters used for following analyses was identified by applying a dynamic tree cut (package dynamicTreeCut, version 1.63-1). No cluster of cell doublets could be identified, consistent with the expectations from the number of loaded cells in the 10X Genomics Chromium wells. The scran function findMarkers was used to find markers up-regulated in any of the clusters. The top 20 markers for each clusters were extracted and pooled to form a list of 89 unique genes for patient 6, and 105 unique genes for patients 7 and 8 (Supplement figure 4 A) and B). The Bioconductor package SingleR (version 1.0.5) was used for unbiased cell-type annotation of the cells [22] using as reference the pseudo-bulk samples resulting from the reanalysis of the 3 public datasets (see above). The pruned scores from SingleR were used subsequently to remove low-quality assignments, and for clarity, only the cell types matching at least 10 cells in the dataset were considered for plotting. Cell type annotation was double-checked by direct integration of our dataset with the reference public datasets, using the Seurat functions FindIntegrationAnchors and IntegrateData (not shown). For co-expression plots of KRT17 with other cell-type specific markers, we used a cutoff of 0.5 on the normalized log-counts to account for potential background noise when determining the present/absent status. A cutoff of 0 was only used for FOXJ1 given the small number of cells where it was detected. Differentiation states across cells were inferred using the R package CytoTRACE (version 0.1, available at https://cytotrace.stanford.edu/) [23]. RNA velocity calculations were performed based on the spliced and unspliced UMI count tables produced by Kallisto and BUStools (see above), normalized separately, and filtered to keep only genes expressed at sufficient levels in both tables, and only epithelial cells. RNA velocity was calculated using the python package scVelo (v0.1.25)[24] called from R using the package reticulate (v1.14). A minimum of 20 cells with expression in unspliced and spliced matrices was required for every gene, and the top 500 genes were retained for further analysis. The first- and second-order moments for velocity estimation of cells were calculated using the 30 nearest neighbours calculated from the neighbour graph obtained from euclidean distances in PCA space (10 first principal components retained). The likelihood-based dynamical velocities were calculated using default parameters, and visualized on the tSNE embedding using a stream plot.


1. Dobin A, Davis CA, Schlesinger F, Drenkow J, Zaleski C, Jha S, Batut P, Chaisson M, Gingeras TR. STAR: ultrafast universal RNA-seq aligner. *Bioinformatics* 2012; 29: 15-21.
2. Amezquita RA, Lun ATL, Becht E, Carey VJ, Carpp LN, Geistlinger L, Martini F, Rue-Albrecht K, Risso D, Soneson C, Waldron L, Pages H, Smith ML, Huber W, Morgan M, Gottardo R, Hicks SC. Orchestrating single-cell analysis with Bioconductor. *Nature methods* 2019.
3. Huber W, Carey VJ, Gentleman R, Anders S, Carlson M, Carvalho BS, Bravo HC, Davis S, Gatto L, Girke T, Gottardo R, Hahne F, Hansen KD, Irizarry RA, Lawrence M, Love MI, MacDonald J, Obenchain V, Oles AK, Pages H, Reyes A, Shannon P, Smyth GK, Tenenbaum D, Waldron L, Morgan M. Orchestrating high-throughput genomic analysis with Bioconductor. *Nat Meth* 2015; 12: 115-121.
4. Gaidatzis D, Lerch A, Hahne F, Stadler MB. QuasR: quantification and annotation of short reads in R. *Bioinformatics* 2015; 31: 1130-1132.
5. Robinson MD, McCarthy DJ, Smyth GK. edgeR: a Bioconductor package for differential expression analysis of digital gene expression data. *Bioinformatics* 2010; 26: 139-140.
6. Robinson MD, Oshlack A. A scaling normalization method for differential expression analysis of RNA-seq data. *Genome Biol* 2010; 11: R25.
7. Wu D, Smyth GK. Camera: a competitive gene set test accounting for inter-gene correlation. *Nucleic acids research* 2012; 40: e133.
8. Liberzon A, Birger C, Thorvaldsdottir H, Ghandi M, Mesirov JP, Tamayo P. The Molecular Signatures Database (MSigDB) hallmark gene set collection. *Cell systems* 2015; 1: 417-425.
9. Garcia-Alonso L, Holland CH, Ibrahim MM, Turei D, Saez-Rodriguez J. Benchmark and integration of resources for the estimation of human transcription factor activities. *Genome research* 2019; 29: 1363-1375.
10. Reyfman PA, Walter JM, Joshi N, Anekalla KR, McQuattie-Pimentel AC, Chiu S, Fernandez R, Akbarpour M, Chen C-I, Ren Z, Verma R, Abdala-Valencia H, Nam K, Chi M, Han S, Gonzalez-Gonzalez FJ, Soberanes S, Watanabe S, Williams KJN, Flozak AS, Nicholson TT, Morgan VK, Winter DR, Hinchcliff M, Hrusch CL, Guzy RD, Bonham CA, Sperling AI, Bag R, Hamanaka RB, Mutlu GM, Yeldandi AV, Marshall SA, Shilatifard A, Amaral LAN, Perlman H, Sznajder JI, Argento AC, Gillespie CT, Dematte J, Jain M, Singer BD, Ridge KM, Lam AP, Bharat A, Bhorade SM, Gottardi CJ, Budinger GRS, Misharin AV. Single-Cell Transcriptomic Analysis of Human Lung Provides Insights into the Pathobiology of Pulmonary Fibrosis. *American Journal of Respiratory and Critical Care Medicine* 2018.
11. Griffiths JA, Richard AC, Bach K, Lun ATL, Marioni JC. Detection and removal of barcode swapping in single-cell RNA-seq data. *Nat Commun* 2018; 9: 2667.

12. Lun ATL, Riesenfeld S, Andrews T, Dao TP, Gomes T, participants in the 1st Human Cell Atlas J, Marioni JC. EmptyDrops: distinguishing cells from empty droplets in droplet-based single-cell RNA sequencing data. *Genome Biol* 2019; 20: 63.
13. Lun AT, Bach K, Marioni JC. Pooling across cells to normalize single-cell RNA sequencing data with many zero counts. *Genome Biol* 2016; 17: 75.
14. Vallejos CA, Risso D, Scialdone A, Dudoit S, Marioni JC. Normalizing single-cell RNA sequencing data: challenges and opportunities. *Nat Meth* 2017; 14: 565-571.
15. McCarthy DJ, Campbell KR, Wills QF, Lun ATL. Scater: pre-processing, quality control, normalization and visualization of single-cell RNA-seq data in R. *Bioinformatics* 2017; 33: 1179-1186.
16. Lun A, McCarthy D, Marioni J. A step-by-step workflow for low-level analysis of single-cell RNA-seq data with Bioconductor [version 2; referees: 3 approved, 2 approved with reservations]. *F1000Research* 2016; 5.
17. Lun AT, Marioni JC. Overcoming confounding plate effects in differential expression analyses of single-cell RNA-seq data. *Biostatistics* 2017.
18. Adams TS, Schupp JC, Poli S, Ayaub EA, Neumark N, Ahangari F, Chu SG, Raby BA, DeIuliis G, Januszyk M, Duan Q, Arnett HA, Siddiqui A, Washko GR, Homer R, Yan X, Rosas IO, Kaminski N. Single-cell RNA-seq reveals ectopic and aberrant lung-resident cell populations in idiopathic pulmonary fibrosis. *Science Advances* 2020; 6: eaba1983.
19. Habermann AC, Gutierrez AJ, Bui LT, Yahn SL, Winters NI, Calvi CL, Peter L, Chung M-I, Taylor CJ, Jetter C, Raju L, Roberson J, Ding G, Wood L, Sucre JMS, Richmond BW, Serezani AP, McDonnell WJ, Mallal SB, Bacchetta MJ, Loyd JE, Shaver CM, Ware LB, Bremner R, Walia R, Blackwell TS, Banovich NE, Kropski JA. Single-cell RNA sequencing reveals profibrotic roles of distinct epithelial and mesenchymal lineages in pulmonary fibrosis. *Science Advances* 2020; 6: eaba1972.
20. Melsted P, Ntranos V, Pachter L. The barcode, UMI, set format and BUStools. *Bioinformatics* 2019; 35: 4472-4473.
21. Melsted P, Booeshaghi AS, Gao F, Beltrame E, Lu L, Hjorleifsson KE, Gehring J, Pachter L. Modular and efficient pre-processing of single-cell RNA-seq. *bioRxiv* 2019: 673285.
22. Aran D, Looney AP, Liu L, Wu E, Fong V, Hsu A, Chak S, Naikawadi RP, Wolters PJ, Abate AR, Butte AJ, Bhattacharya M. Reference-based analysis of lung single-cell sequencing reveals a transitional profibrotic macrophage. *Nature immunology* 2019; 20: 163-172.
23. Gulati GS, Sikandar SS, Wesche DJ, Manjunath A, Bharadwaj A, Berger MJ, Ilagan F, Kuo Angera H, Hsieh RW, Cai S, Zabala M, Scheeren FA, Lobo NA, Qian D, Yu FB, Dirbas FM, Clarke MF, Newman AM. Single-cell transcriptional diversity is a hallmark of developmental potential. *Science* 2020; 367: 405-411.
24. Bergen V, Lange M, Peidli S, Wolf FA, Theis FJ. Generalizing RNA velocity to transient cell states through dynamical modeling. *bioRxiv* 2019: 820936.

**Supplement Table 1: Chemical, reagents, instruments**

| Product | Company | Catalog Number | Country |
|---|---|---|---|
| DMEM | Thermo Fisher Scientific | 32430-027 | Waltham, MA, USA |
| FCS | Thermo Fisher Scientific | 16140063 | Waltham, MA, USA |
| HEPES | Thermo Fisher Scientific | 15630080 | Waltham, MA, USA |
| L-Glutamine | Thermo Fisher Scientific | 25030081 | Waltham, MA, USA |
| MEM-vitamin mix | Thermo Fisher Scientific | 11120052 | Waltham, MA, USA |
| Penicillin/Streptomycin | Thermo Fisher Scientific | 10378016 | Waltham, MA, USA |
| Amphotercin B | Thermo Fisher Scientific | 15290018 | Waltham, MA, USA |
| Quick-RNA MiniPrep Kit | ZymoResearch | R1050 | Orange, CA, USA |
| GAPDH TaqMan® Gene Expression Assay | Thermo Fisher Scientific | Hs03929097_g1 | Waltham, MA, USA |
| KRT5 TaqMan® Gene Expression Assay | Thermo Fisher Scientific | Hs00361185_m1 | Waltham, MA, USA |
| CDH1 TaqMan® Gene Expression Assay | Thermo Fisher Scientific | Hs01023894_m1 | Waltham, MA, USA |
| TP63 TaqMan® Gene Expression Assay | Thermo Fisher Scientific | Hs00978340_m1 | Waltham, MA, USA |
| SCGB1A1 TaqMan® Gene Expression Assay | Thermo Fisher Scientific | Hs00171092_m1 | Waltham, MA, USA |
| SOX9 TaqMan® Gene Expression Assay | Thermo Fisher Scientific | Hs00165814_m1 | Waltham, MA, USA |
| FN1 TaqMan® Gene Expression Assay | Thermo Fisher Scientific | Hs01549976_m1 | Waltham, MA, USA |
| CDH2 TaqMan® Gene Expression Assay | Thermo Fisher Scientific | Hs00983056_m1 | Waltham, MA, USA |
| alpha-smooth muscle TaqMan® Gene Expression Assay | Thermo Fisher Scientific | Hs00426835_g1 | Waltham, MA, USA |
| Rabbit Polyclonal Fibronectin antibody | Abcam | ab2413 | Cambridge, MA, USA |
| Rabbit Polyclonal E-cadherin antibody | Abcam | ab15148 | Cambridge, MA, USA |
| Mouse KRT5 Monoclonal Antibody (XM26) | Thermo Fisher Scientific | MA5-12596 | Waltham, MA, USA |
| N-cadherin (D4R1H) Rabbit mAb | Cell signaling technology | 13116 | Danvers, MA, USA |
| Monoclonal Rabbit NKX2.1 antibody | RnD systems | MAB94581 | Abingdon, UK |
| Polyclonal Goat p63 antibody | RnD systems | AF1916 | Abingdon, UK |
| SOX2 Rat Monoclonal Antibody (Btjce) | Thermo Fisher Scientific | 14-9811-82 | Waltham, MA, USA |
| Polyclonal goat SOX9 antibody | RnD systems | AF3075 | Abingdon, UK |
| Ki67 FITC-confugated antibody | Thermo Fisher Scientific | 11-5698-82 | Waltham, MA, USA |
| alpha-smooth muscle actin antibody | Abcam | ab5694 | Cambridge, MA, USA |
| alpha-tubulin antibody | Cell Signaling Technology | 2125 | Danvers, MA, USA |
| Picro Sirius Red | Sigma-Aldrich | 365548-5G | St Louis, MO, USA |
| Alexa488 Goat anti Rabbit IgG antibody | Thermo Fisher Scientific | A11008 | Waltham, MA, USA |
| Alexa594 Goat anti Rabbit IgG antibody | Thermo Fisher Scientific | A11012 | Waltham, MA, USA |
| Alexa594 Donkey anti Goat IgG antibody | Thermo Fisher Scientific | A11058 | Waltham, MA, USA |
| Alexa488 Donkey anti Mouse IgG antibody | Thermo Fisher Scientific | A21202 | Waltham, MA, USA |
| Alexa488 Donkey anti Rat IgG antibody | Thermo Fisher Scientific | A21208 | Waltham, MA, USA |
| DAPI | Thermo Fisher Scientific | D1306 | Waltham, MA, USA |
| OLYMPUS BX63 | | | |
| paraformaldehyde | Merck | 1.04005.1000 | Darmstadt, Germany |
| glutaraldehyde | Merck | 1.04239.0250 | Darmstadt, Germany |
| Electron microscope | Morgagni, FEI | | Eindhoven, The Netherlands |
| MicroElute Total RNA Kit | Omega Bio-Tek | | Norcross, GA |
| QuantiFluor RNA System | Promega | | Madison, WI |
| RNA 6000 Pico Kit | Agilent | | Santa Clara, CA |

**Supplement Table 2: Patient details**

| IPF<br>Patient ID | Sex | Age (years) | Clinical diagnosis |
|---|---|---|---|
| 1 | Male | 65 | IPF |
| 2 | Male | 63 | IPF |
| 3 | Female | 49 | IPF |
| 4 | Male | 77 | IPF |
| 5 | Male | 60 | IPF |
| 6 | Male | 60 | IPF |
| 7 | Female | 62 | IPF |
| 8 | Male | 68 | IPF |
| **Non-IPF<br>Patient ID** | **Sex** | **Age (years)** | **Clinical diagnosis** |
| 9 | Male | 37 | Non-classifiable ILD |
| 10 | Male | 67 | Vasculitis-associated ILD |
| 11 | Male | 73 | Non-classifiable ILD |
| 12 | Female | 75 | Carcinoma |
| 13 | Male | 72 | Chronic eosinophilic pneumonia |

Abbreviations used: IPF, idiopathic pulmonary fibrosis; ILD, interstitial lung disease.

**Supplement table 3:** Ambient_RNA_top_50_genes

| | |
|---|---|
| MT-RNR2 | 0.159517772374171 |
| MT-CO2 | 0.0477388337582393 |
| MT-CYB | 0.0472479251207663 |
| MT-RNR1 | 0.0362529243176831 |
| MT-CO3 | 0.0321255719045492 |
| MT-ATP6 | 0.0304290908499241 |
| FGF23 | 0.0224240674558514 |
| MT-CO1 | 0.0210167208803602 |
| MT-ND4 | 0.018772969692355 |
| APOOL | 0.0147611583505042 |
| S100A2 | 0.0114138490578216 |
| MALAT1 | 0.0107640010005182 |
| TMSB4X | 0.00894465188708348 |
| MT-ND1 | 0.0084390892620596 |
| MT-ND2 | 0.00656957842015833 |
| KRT17 | 0.00628833455260871 |
| S100A10 | 0.00595523810848544 |
| MT-ND5 | 0.00568864823038249 |
| AC010970.1 | 0.00467019599201318 |
| TPT1 | 0.00390706133523167 |
| MT-ND3 | 0.00354127522734618 |
| S100A11 | 0.00351534893932046 |
| TMSB10 | 0.00339698979843498 |
| KRT19 | 0.00329384826152357 |
| FTL | 0.00325326798474621 |
| ACTG1 | 0.00317267104621892 |
| KRT7 | 0.00303345815264456 |
| MYL6 | 0.00279279456825102 |
| AL135790.1 | 0.00223256131217869 |
| EEF1A1 | 0.00209560288219747 |
| CD24 | 0.00205276814709606 |
| ANXA2 | 0.00202796908998037 |
| S100A6 | 0.00196653506228803 |
| MT-ND6 | 0.00192144586780884 |
| RPS19 | 0.00185888461070968 |
| RPL8 | 0.00181661349121955 |
| MT2A | 0.00178505105543063 |
| MTATP6P1 | 0.00173545294222761 |
| RPL13 | 0.00172474425872548 |
| EIF1 | 0.00168190952483132 |
| RPLP1 | 0.00163231141214426 |
| AD000090.1 | 0.00152691542366233 |
| MYL12B | 0.00143673703903146 |
| SAT1 | 0.00136572156209339 |
| KRT6A | 0.00130992368804706 |
| C15orf48 | 0.00130485115407603 |
| RACK1 | 0.00121636361802482 |
| S100A16 | 0.00121241831391721 |
| MARCH6 | 0.00120960023955731 |
| DYNLL1 | 0.0011876192596286 |

**Supplement table 4:**
edgeR_DE_GSEA_results_c2.CATEGORY.v7.0.entrez.hsa_IPF_effect

| | NGenes | Correlation | Direction | absLog2FC | PValue | FDR |
|---|---|---|---|---|---|---|
| KEGG_ASCORBATE_AND_ALDARATE_METABOLISM | 15 | 0.01 | Up | 0.958564859030419 | 1.2576347739059e-27 | 5.65306830870701e-24 |
| REACTOME_GLUCURONIDATION | 13 | 0.01 | Up | 0.99181144989429 | 3.96415100879193e-25 | 8.90942939225986e-22 |
| ZHANG_RESPONSE_TO_IKK_INHIBITOR_AND_TNF_UP | 214 | 0.01 | Down | 0.602417896519383 | 1.02677151484743e-20 | 1.53844598641307e-17 |
| BOSCO_EPITHELIAL_DIFFERENTIATION_MODULE | 54 | 0.01 | Down | 1.07529536129393 | 1.53611875749025e-18 | 1.72621345372967e-15 |
| KEGG_PENTOSE_AND_GLUCURONATE_INTERCONVERSIONS | 17 | 0.01 | Up | 0.787240126903798 | 2.87938165460172e-18 | 2.58856410748695e-15 |
| KEGG_METABOLISM_OF_XENOBIOTICS_BY_CYTOCHROME_P450 | 40 | 0.01 | Up | 0.831452055344611 | 5.79718831282077e-18 | 4.3430602443549e-15 |
| PHONG_TNF_TARGETS_UP | 62 | 0.01 | Down | 0.708013085396942 | 4.41413767910867e-17 | 2.83450698108478e-14 |
| TIAN_TNF_SIGNALING_VIA_NFKB | 26 | 0.01 | Down | 0.967468349921907 | 9.90113201308045e-15 | 5.56319854984958e-12 |
| LIN_SILENCED_BY_TUMOR_MICROENVIRONMENT | 104 | 0.01 | Down | 0.714335019312897 | 3.93151764517049e-14 | 1.96357464611570e-11 |
| ONDER_CDH1_TARGETS_3_DN | 58 | 0.01 | Down | 0.878679855336258 | 2.58018096432612e-13 | 1.15979134346459e-10 |
| KEGG_PORPHYRIN_AND_CHLOROPHYLL_METABOLISM | 30 | 0.01 | Up | 0.537419230453302 | 3.93241549866123e-13 | 1.6069279696802e-10 |
| ANASTASSIOU_MULTICANCER_INVASIVENESS_SIGNATURE | 56 | 0.01 | Up | 1.51580534238146 | 4.47799712306763e-13 | 1.67738308901575e-10 |
| FARMER_BREAST_CANCER_CLUSTER_4 | 19 | 0.01 | Up | 1.55448651801345 | 1.52732758272125e-12 | 5.28102883410155e-10 |
| REACTOME_COLLAGEN_BIOSYNTHESIS_AND_MODIFYING_ENZYMES | 49 | 0.01 | Up | 1.00807082117032 | 2.61907781590219e-12 | 8.40911055891452e-10 |
| KEGG_STEROID_HORMONE_BIOSYNTHESIS | 30 | 0.01 | Up | 0.950862297846994 | 3.7670487393389e-12 | 1.08363324088346e-09 |
| KEGG_DRUG_METABOLISM_CYTOCHROME_P450 | 39 | 0.01 | Up | 0.793611318827354 | 3.85720397199895e-12 | 1.08363324088346e-09 |
| ZWANG_CLASS_3_TRANSIENTLY_INDUCED_BY_EGF | 212 | 0.01 | Down | 0.46378345334927 | 5.50724373032596e-12 | 1.45618003340089e-09 |
| REACTOME_KERATINIZATION | 63 | 0.01 | Down | 0.907177716165867 | 6.86707014510059e-12 | 1.7148600167904e-09 |
| KEGG_STARCH_AND_SUCROSE_METABOLISM | 30 | 0.01 | Up | 0.593047056597582 | 9.31477538818039e-12 | 2.20367975630899e-09 |
| KEGG_RETINOL_METABOLISM | 29 | 0.01 | Up | 0.823154631659858 | 1.29570645291555e-11 | 2.89127512522319e-09 |
| REACTOME_FORMATION_OF_THE_CORNIFIED_ENVELOPE | 61 | 0.01 | Down | 0.891833256935227 | 1.35076257240683e-11 | 2.89127512522319e-09 |
| REACTOME_PHASE_II_CONJUGATION_OF_COMPOUNDS | 70 | 0.01 | Up | 0.540343967414742 | 5.4140416526191e-11 | |

1.10618714675104e-08
ONDER_CDH1_TARGETS_2_UP 224 0.01 Up 0.705713151232459
6.2523715481103e-11 1.22193087429373e-08
NABA_CORE_MATRISOME 154 0.01 Up 0.938916708850495
1.05122903561438e-10 1.9688643812861e-08
SCHUETZ_BREAST_CANCER_DUCTAL_INVASIVE_UP 309 0.01 Up
0.865603231579515 1.30497900345665e-10
2.34635224821506e-08
NABA_COLLAGENS 30 0.01 Up 1.28506989046779
1.45656968708572e-10 2.42491879387049e-08
REACTOME_COLLAGEN_CHAIN_TRIMERIZATION 30 0.01 Up
1.28506989046779 1.45656968708572e-10 2.42491879387049e-08
KOBAYASHI_EGFR_SIGNALING_6HR_DN 18 0.01 Down
0.589573796841963 5.14817099215253e-10 8.26465307490201e-08
HOLLERN_EMT_BREAST_TUMOR_DN 111 0.01 Down
0.367361173488558 7.10759753214022e-10 1.10167761748173e-07
SEKI_INFLAMMATORY_RESPONSE_LPS_UP 67 0.01 Down
0.661280645569614 1.57119089596799e-09 2.3541676924587e-07
PID_CD40_PATHWAY 27 0.01 Down 0.468292330224766
2.88840773531204e-09 4.13138072606666e-07
REACTOME_ECM_PROTEOGLYCANS 52 0.01 Up
1.07850194226203 2.94113867039228e-09 4.13138072606666e-07
GHANDHI_BYSTANDER_IRRADIATION_UP 70 0.01 Down
0.772849250229913 4.50858502297733e-09 6.14123929644943e-07
THUM_MIR21_TARGETS_HEART_DISEASE_UP 15 0.01 Up
1.46210894733469 4.8717268821201e-09 6.44070951033231e-07
HINATA_NFKB_TARGETS_KERATINOCYTE_UP 81 0.01 Down
0.690993957181174 5.27747114781927e-09 6.77778080269931e-07
ONDER_CDH1_TARGETS_2_DN 451 0.01 Down 0.528064725465505
7.39225929629739e-09 9.19740661311774e-07
LINDSTEDT_DENDRITIC_CELL_MATURATION_B 46 0.01 Down
0.527090459368152 7.57072401969647e-09 9.19740661311774e-07
LIANG_SILENCED_BY_METHYLATION_2 46 0.01 Down
0.814285779455829 1.0676479466584e-08 1.26291513690251e-06
SENESE_HDAC2_TARGETS_UP 109 0.01 Down 0.404271182550095
1.38528087561234e-08 1.59662500919935e-06
SENESE_HDAC1_TARGETS_UP 417 0.01 Down 0.360376822537735
1.94748241382153e-08 2.18848336253195e-06
REACTOME_SCAVENGING_BY_CLASS_A_RECEPTORS 13 0.01 Up
1.51306890128243 2.38231836818762e-08
2.61183440609838e-06
SESTO_RESPONSE_TO_UV_C1 73 0.01 Down 0.513186070464607
2.5683600797251e-08 2.74875679961055e-06
CHEN_HOXA5_TARGETS_9HR_UP 216 0.01 Down
0.263054431634624 3.06529552512107e-08 3.20430311288819e-06
AMIT_EGF_RESPONSE_60_MCF10A 39 0.01 Down
0.516737193208124 3.18375167163289e-08 3.25249176454314e-06
SESTO_RESPONSE_TO_UV_C2 54 0.01 Down 0.360240579137045
3.32621941384416e-08 3.32252361449545e-06
BILD_HRAS_ONCOGENIC_SIGNATURE 239 0.01 Down
0.480338202484268 3.68239801200032e-08 3.59834327476988e-06
AMIT_SERUM_RESPONSE_60_MCF10A 58 0.01 Down
0.523510395278958 3.9420178980574e-08 3.77007881952511e-06
SANA_TNF_SIGNALING_UP 76 0.01 Down 0.686265362373307

5.04735744840674e-08 4.72663994387256e-06
REACTOME_COLLAGEN_FORMATION 71 0.01 Up
0.938216707464169 5.26512080736172e-08 4.82994245491652e-06
ONDER_CDH1_TARGETS_1_DN 165 0.01 Down 0.47089651187
5.49083091257435e-08 4.8396568307476e-06
AMIT_EGF_RESPONSE_120_HELA 66 0.01 Down
0.440910987157818 5.49104556992497e-08 4.8396568307476e-06
YAO_TEMPORAL_RESPONSE_TO_PROGESTERONE_CLUSTER_16 65 0.01
Up 0.845799877825494 5.62046894886499e-08
4.85846306252848e-06
RASHI_NFKB1_TARGETS 16 0.01 Down 0.681467981125494
6.8581654149365e-08 5.81650066795086e-06
REACTOME_BINDING_AND_UPTAKE_OF_LIGANDS_BY_SCAVENGER_RECEPTORS 26
0.01 Up 1.29516917627458 7.42547967723795e-08
6.18102428688604e-06
KEGG_DRUG_METABOLISM_OTHER_ENZYMES 31 0.01 Up
0.640120892935911 9.04332395475933e-08 7.39086203211694e-06
BOQUEST_STEM_CELL_UP 195 0.01 Up 0.814343031943684
9.71554311728049e-08 7.79845827003139e-06
AMIT_SERUM_RESPONSE_240_MCF10A 54 0.01 Down
0.432792911911316 1.08062636258839e-07 8.52178157865755e-06
ZWANG_CLASS_1_TRANSIENTLY_INDUCED_BY_EGF 440 0.01 Down
0.343682980576616 1.1810341291377e-07
9.15301450081716e-06
PEDERSEN_METASTASIS_BY_ERBB2_ISOFORM_1 43 0.01 Down
0.553578153038284 1.24469938794171e-07 9.48292160813219e-06
ROZANOV_MMP14_TARGETS_SUBSET 31 0.01 Up
0.82118790470737 1.38158830074797e-07 1.01967436278198e-05
GILMORE_CORE_NFKB_PATHWAY 13 0.01 Down
0.428455031715683 1.38376276150613e-07 1.01967436278198e-05
PID_INTEGRIN1_PATHWAY 56 0.01 Up 0.963883473240334
1.58348457729852e-07 1.14802631854142e-05
REACTOME_CROSSLINKING_OF_COLLAGEN_FIBRILS 18 0.01 Up
0.774402027262086 1.69719584768776e-07
1.2109357675169e-05
JAEGER_METASTASIS_DN 221 0.01 Down 0.558660701404633
1.75145488802379e-07 1.23012339401046e-05
UZONYI_RESPONSE_TO_LEUKOTRIENE_AND_THROMBIN 34 0.01 Down
0.606900146565268 2.01719982351984e-07
1.39497126257257e-05
RASHI_RESPONSE_TO_IONIZING_RADIATION_2 110 0.01 Down
0.395986433256175 2.44063026435436e-07 1.66221712701104e-05
DIRMEIER_LMP1_RESPONSE_EARLY 56 0.01 Down
0.478564918995447 2.99104383315171e-07 2.0066779149279e-05
NAGASHIMA_EGF_SIGNALING_UP 55 0.01 Down
0.440255224182171 3.07831848102603e-07 2.03485905473706e-05
NAGASHIMA_NRG1_SIGNALING_UP 165 0.01 Down
0.38363239367955 3.22080119066977e-07 2.09818860174792e-05
ANDERSEN_CHOLANGIOCARCINOMA_CLASS2 151 0.01 Down
0.572463703679325 3.53751032736274e-07 2.27158698878507e-05
BIOCARTA_NTHI_PATHWAY 22 0.01 Down 0.43092594144598
3.59339365367505e-07 2.27497246102385e-05
COLDREN_GEFITINIB_RESISTANCE_DN 220 0.01 Down
0.356250783432735 3.8435035460738e-07 2.39952061661135e-05

BIOCARTA_IL1R_PATHWAY 28 0.01 Down 0.463618769857485
5.32984214234697e-07 3.26942286697582e-05
GARGALOVIC_RESPONSE_TO_OXIDIZED_PHOSPHOLIPIDS_BLUE_UP 135 0.01
Down 0.308016132537063 5.38236467533283e-07
3.26942286697582e-05
REACTOME_BIOLOGICAL_OXIDATIONS 134 0.01 Up
0.519435999244741 6.88199245685577e-07 4.12460747914222e-05
NABA_ECM_GLYCOPROTEINS 108 0.01 Up 0.775569258866001
7.3991948892574e-07 4.37623434568579e-05
DORN_ADENOVIRUS_INFECTION_32HR_DN 38 0.01 Down
0.288484727074426 8.75987259123929e-07 5.11371783085982e-05
WAMUNYOKOLI_OVARIAN_CANCER_GRADES_1_2_UP 138 0.01 Down
0.335598651617923 8.91184327880045e-07
5.13573532541128e-05
AMIT_SERUM_RESPONSE_120_MCF10A 65 0.01 Down
0.432481252089242 9.19630695736045e-07 5.19561171801586e-05
REACTOME_DISEASES_ASSOCIATED_WITH_GLYCOSAMINOGLYCAN_METABOLISM 34
0.01 Up 0.919397244033627 9.24691740692478e-07
5.19561171801586e-05
KEGG_NOD_LIKE_RECEPTOR_SIGNALING_PATHWAY 52 0.01 Down
0.495402238713484 1.1797589029663e-06
6.54693366522658e-05
LIEN_BREAST_CARCINOMA_METAPLASTIC 27 0.01 Up
1.20559034189529 1.45979985128694e-06 8.00219552626193e-05
DEBIASI_APOPTOSIS_BY_REOVIRUS_INFECTION_UP 277 0.01 Down
0.27909504989153 1.50329704317485e-06
8.10466527704622e-05
BIOCARTA_CD40_PATHWAY 14 0.01 Down 0.396167867113994
1.51455368914768e-06 8.10466527704622e-05
PICCALUGA_ANGIOIMMUNOBLASTIC_LYMPHOMA_UP 175 0.01 Up
0.838239682063361 1.78394049045854e-06
9.43389706424837e-05
BIOCARTA_TNFR2_PATHWAY 17 0.01 Down 0.427297944821957
2.35254250962748e-06 0.000122961378846227
ZHOU_INFLAMMATORY_RESPONSE_LIVE_UP 313 0.01 Down
0.467178994622328 2.8013329216042e-06 0.000144735534282884
BIOCARTA_INTRINSIC_PATHWAY 13 0.01 Up
0.744343045763485 3.19507240273013e-06 0.000163202846025817
LEE_SP4_THYMOCYTE 14 0.01 Down 0.748859456621776
3.88901382338447e-06 0.000196417046473182
WANG_TNF_TARGETS 16 0.01 Down 0.90941646096457
4.068833095927e-06 0.000203215608513243
GARGALOVIC_RESPONSE_TO_OXIDIZED_PHOSPHOLIPIDS_GREY_UP 17 0.01
Down 0.606321448051247 4.36589775253855e-06
0.000215656158216052
GALINDO_IMMUNE_RESPONSE_TO_ENTEROTOXIN 70 0.01 Down
0.5240183570281 4.73102221988879e-06 0.000231151574765218
DORN_ADENOVIRUS_INFECTION_48HR_DN 39 0.01 Down
0.307432031812019 4.93165706530623e-06 0.000236020259588399
TURASHVILI_BREAST_LOBULAR_CARCINOMA_VS_LOBULAR_NORMAL_DN 64
0.01 Up 0.994857530565027 4.93568507259388e-06
0.000236020259588399
PICCALUGA_ANGIOIMMUNOBLASTIC_LYMPHOMA_DN 130 0.01 Down
0.294049682907123 5.12722565270801e-06

0.000236039634774122
ZHOU_INFLAMMATORY_RESPONSE_LPS_UP 248 0.01 Down
0.519824569391725 5.15290828138526e-06 0.000236039634774122
REACTOME_ANTIMICROBIAL_PEPTIDES 23 0.01 Down
1.51689079625682 5.22834236316861e-06 0.000236039634774122
KEGG_ECM_RECEPTOR_INTERACTION 60 0.01 Up
0.957882027681829 5.23546547923979e-06 0.000236039634774122
REACTOME_ASSEMBLY_OF_COLLAGEN_FIBRILS_AND_OTHER_MULTIMERIC_STRUCTURE
S 52 0.01 Up 1.03653269075071
5.23802480212894e-06 0.000236039634774122
LANDIS_ERBB2_BREAST_TUMORS_65_DN 35 0.01 Up
0.72356022332596 5.25115983924632e-06 0.000236039634774122
SENESE_HDAC1_AND_HDAC2_TARGETS_UP 209 0.01 Down
0.514697122323876 5.38014765454264e-06 0.000239443205021477
REACTOME_RAF_INDEPENDENT_MAPK1_3_ACTIVATION 21 0.01 Down
0.382916830799805 5.46810201407088e-06
0.000240971750522045
ZHU_CMV_8_HR_UP 38 0.01 Down 0.580005343942038
5.61798106639402e-06 0.000245173057217875
DAZARD_RESPONSE_TO_UV_NHEK_DN 294 0.01 Down
0.243159095978187 5.92060378059757e-06 0.000255895326863328
LIM_MAMMARY_STEM_CELL_UP 402 0.01 Up
0.526760539926095 6.64471025370837e-06 0.000284456881813516
REACTOME_NON_INTEGRIN_MEMBRANE_ECM_INTERACTIONS 52 0.01 Up
0.694544635305357 7.54799137231595e-06
0.000320077558665662
REACTOME_COLLAGEN_DEGRADATION 50 0.01 Up
1.11389229851519 8.15357623543922e-06 0.000342526403535508
REACTOME_INTEGRIN_CELL_SURFACE_INTERACTIONS 66 0.01 Up
0.915979101064372 8.42932684803544e-06
0.000350831705388142
DURCHDEWALD_SKIN_CARCINOGENESIS_DN 238 0.01 Down
0.324665596476488 8.97684668214039e-06 0.000370191980148817
GRAHAM_CML_QUIESCENT_VS_CML_DIVIDING_UP 17 0.01 Down
1.14847903862383 9.27801536076962e-06 0.000379133445878722
APPEL_IMATINIB_RESPONSE 32 0.01 Up 0.822500470444935
1.04570054153819e-05 0.000423461615694968
GROSS_HYPOXIA_VIA_ELK3_DN 141 0.01 Down
0.428545742535806 1.14761549486534e-05 0.000460583182983901
BIOCARTA_AMI_PATHWAY 11 0.01 Up 0.798874179298572
1.2034320912181e-05 0.000478710376108439
PID_IL23_PATHWAY 24 0.01 Down 0.796643938076091
1.25374013010097e-05 0.000494347533754726
HOLLERN_ADENOMYOEPITHELIAL_BREAST_TUMOR 16 0.01 Up
1.3290946750272 1.35299932477896e-05 0.000528846257815774
GHANDHI_DIRECT_IRRADIATION_UP 86 0.01 Down
0.683041674112105 1.42055585036767e-05 0.000550465392017471
REACTOME_EXTRACELLULAR_MATRIX_ORGANIZATION 231 0.01 Up
0.770704047861096 1.44662357179912e-05
0.000555775466259577
PAPASPYRIDONOS_UNSTABLE_ATEROSCLEROTIC_PLAQUE_DN 36 0.01
Up 0.637638267478494 1.46782946230029e-05
0.00055914351127456
PHONG_TNF_RESPONSE_VIA_P38_PARTIAL 155 0.01 Down

0.457801268642031	1.49972614978902e-05	0.00056461258092874
GARGALOVIC_RESPONSE_TO_OXIDIZED_PHOSPHOLIPIDS_TURQUOISE_UP	78
	0.01	Down	0.294869146380404	1.50730833618351e-05
	0.00056461258092874
PID_NFKAPPAB_CANONICAL_PATHWAY	22	0.01	Down
0.40601082346006	1.62683013528295e-05	0.000604347227941891
REACTOME_TRIF_TICAM1_MEDIATED_TLR4_SIGNALING	93	0.01	Down
	0.300643972938414	1.72975952207754e-05
0.000637317135388404
WESTON_VEGFA_TARGETS_6HR	44	0.01	Up
0.974437034340311	1.79789372505356e-05	0.000657035145863071
REACTOME_TRAF6_MEDIATED_NF_KB_ACTIVATION	22	0.01	Down
	0.304095005310491	1.88704294429183e-05
0.000684053067305789
WINZEN_DEGRADED_VIA_KHSRP	91	0.01	Down
0.685170224841889	1.92509685002439e-05	0.000692264827268769
GRAHAM_NORMAL_QUIESCENT_VS_NORMAL_DIVIDING_UP	56	0.01	Down
	0.716377871139364	1.95669391681604e-05
0.000698042790165724
AMIT_EGF_RESPONSE_40_HELA	41	0.01	Down
0.401847316231371	2.08474044580403e-05	0.000737866795581819
REACTOME_SIGNALING_BY_HIPPO	20	0.01	Down
0.274773790262441	2.2059479118478e-05	0.000774666864355928
REACTOME_SYNDECAN_INTERACTIONS	26	0.01	Up
0.734155626846581	2.28902929029255e-05	0.000797611368981783
BUYTAERT_PHOTODYNAMIC_THERAPY_STRESS_UP	803	0.01	Down
0.270551396735539	2.50480942133811e-05	0.000866086026839602
DOANE_BREAST_CANCER_ESR1_DN	33	0.01	Down
0.772496336649511	2.54331149013396e-05	0.00087268588917192
BROWNE_HCMV_INFECTION_12HR_UP	101	0.01	Down
0.325850317116116	2.57645371472487e-05	0.000877360564218809
HINATA_NFKB_TARGETS_FIBROBLAST_UP	75	0.01	Down
0.634001471367114	2.63411712420335e-05	0.000888204675889107
REACTOME_A_TETRASACCHARIDE_LINKER_SEQUENCE_IS_REQUIRED_FOR_GAG_SYNTHESIS	21	0.01	Up	0.757448174915337
2.65314683551744e-05	0.000888204675889107
DORN_ADENOVIRUS_INFECTION_24HR_DN	42	0.01	Down
0.316932465420259	2.66757800322646e-05	0.000888204675889107
RHEIN_ALL_GLUCOCORTICOID_THERAPY_UP	61	0.01	Down
0.478989867145575	2.7456976873828e-05	0.000907493463587184
SCHOEN_NFKB_SIGNALING	34	0.01	Down	0.892277032216759
	2.94271114338204e-05	0.000965509970036661
REACTOME_RIP_MEDIATED_NFKB_ACTIVATION_VIA_ZBP1	16	0.01	Down
	0.277706310740737	2.97928156549546e-05
0.000970425408471167
REACTOME_REGULATION_OF_INSULIN_LIKE_GROWTH_FACTOR_IGF_TRANSPORT_AND_UPTAKE_BY_INSULIN_LIKE_GROWTH_FACTOR_BINDING_PROTEINS_IGFBPS	81
	0.01	Up	0.579583595942958	3.02183908203371e-05
	0.000977206235520973
BIOCARTA_VITCB_PATHWAY	10	0.01	Up	0.628027785231512
	3.15541058299816e-05	0.00101311218361262
CREIGHTON_ENDOCRINE_THERAPY_RESISTANCE_3	662	0.01	Down
	0.365109931910636	3.32671250250423e-05
0.0010605370708338

GU_PDEF_TARGETS_DN	33	0.01	Down	0.382201063473003
	3.38374347990588e-05	0.00107112161564626
ROY_WOUND_BLOOD_VESSEL_DN	21	0.01	Down
0.516125681147911	3.45882890043996e-05	0.00108723328024319
REACTOME_DEFECTIVE_B4GALT7_CAUSES_EDS_PROGEROID_TYPE	17	0.01
	Up	0.874921635696109	3.49139676855377e-05
0.00108984919962842
REACTOME_INTERLEUKIN_1_FAMILY_SIGNALING	124	0.01	Down
0.298265696507947	3.54536229442218e-05	0.00109906231127088
REACTOME_CHEMOKINE_RECEPTORS_BIND_CHEMOKINES	22	0.01	Down
	1.37492701251537	3.81072199746904e-05
0.00117323256017968
DACOSTA_UV_RESPONSE_VIA_ERCC3_DN	840	0.01	Down
0.247868678152022	3.92840116963248e-05	0.00120123559574816
PHONG_TNF_RESPONSE_NOT_VIA_P38	329	0.01	Down
0.389810692838076	4.00676042626299e-05	0.00121691811594947
AMIT_DELAYED_EARLY_GENES	18	0.01	Down
0.301516894087959	4.15944228243828e-05	0.00125481161473558
KIM_WT1_TARGETS_UP	209	0.01	Down	0.402625421629059
	4.27217028439167e-05	0.00128022702855604
KEGG_RIG_I_LIKE_RECEPTOR_SIGNALING_PATHWAY	53	0.01	Down
	0.322322502642082	4.4032098359191e-05
0.00131075683526201
BIOCARTA_NFKB_PATHWAY	21	0.01	Down	0.384389244558911
	4.7128859596482e-05	0.00139371199925123
PLASARI_TGFB1_TARGETS_10HR_UP	166	0.01	Down
0.532106276800972	4.76130966882848e-05	0.00139882921316235
SESTO_RESPONSE_TO_UV_C3	21	0.01	Down	0.440624026512512
	4.82314081761308e-05	0.00140779337501109
REACTOME_INTERLEUKIN_1_SIGNALING	97	0.01	Down
0.23319630281794	4.88876670650054e-05	0.00141774234488516
DACOSTA_UV_RESPONSE_VIA_ERCC3_COMMON_DN	458	0.01	Down
0.242581789239296	5.03131128208798e-05	0.00144972719314009
KEGG_TOLL_LIKE_RECEPTOR_SIGNALING_PATHWAY	78	0.01	Down
	0.506461225921203	5.28442283858323e-05
0.00151296055155615
REACTOME_ZBP1_DAI_MEDIATED_INDUCTION_OF_TYPE_I_IFNS	19	0.01
	Down	0.24592336251646	5.46750739685797e-05
0.0015554712499289
CHARAFE_BREAST_CANCER_BASAL_VS_MESENCHYMAL_UP	122	0.01	Down
	0.37632630007468	5.68778964142627e-05
0.00160516912928568
MUELLER_METHYLATED_IN_GLIOBLASTOMA	27	0.01	Down
0.694706997953358	5.71361647799128e-05	0.00160516912928568
VECCHI_GASTRIC_CANCER_ADVANCED_VS_EARLY_UP	151	0.01	Up
	0.896839705275309	5.81477559098248e-05
0.00162344200506002
BROWNE_HCMV_INFECTION_18HR_DN	140	0.01	Up
0.469485791160688	5.92228241027723e-05	0.00164325058235779
IGLESIAS_E2F_TARGETS_UP	128	0.01	Up	0.711545713413399
	6.21062948582975e-05	0.00171268586127636
KEGG_FATTY_ACID_METABOLISM	34	0.01	Up
0.314995582843419	6.30223637582295e-05	0.00172735076276367
REACTOME_TOLL_LIKE_RECEPTOR_4_TLR4_CASCADE	114	0.01	Down

0.348941375840157 6.44395809235641e-05
0.00175549040152376
CLASPER_LYMPHATIC_VESSELS_DURING_METASTASIS_DN 32 0.01 Up
1.09621464009093 6.56224075142063e-05
0.00177694410708649
BILD_CTNNB1_ONCOGENIC_SIGNATURE 75 0.01 Down
0.354287362666284 8.69934393608968e-05 0.00234152999956426
REACTOME_MYD88_CASCADE_INITIATED_ON_PLASMA_MEMBRANE 80 0.01
Down 0.278570044145668 8.89374626005113e-05
0.00237960651422201
REACTOME_APOPTOTIC_CLEAVAGE_OF_CELL_ADHESION_PROTEINS 10 0.01
Down 0.411317080866444 9.06019144690723e-05
0.00239912938123428
PETROVA_PROX1_TARGETS_DN 52 0.01 Up
0.5610389698559 9.07345928386714e-05 0.00239912938123428
DAZARD_UV_RESPONSE_CLUSTER_G2 25 0.01 Down
0.534297928628487 9.1740533876182e-05 0.00241154210393823
GARGALOVIC_RESPONSE_TO_OXIDIZED_PHOSPHOLIPIDS_MAGENTA_UP 26
0.01 Down 0.305500001792176 9.4542739437646e-05
0.00245774577228176
MOSERLE_IFNA_RESPONSE 31 0.01 Down 0.512430868116595
9.48023226249407e-05 0.00245774577228176
LINDGREN_BLADDER_CANCER_HIGH_RECURRENCE 45 0.01 Up
0.830889782500288 9.51385460238099e-05 0.00245774577228176
VILIMAS_NOTCH1_TARGETS_UP 35 0.01 Down
0.799126574071735 9.65797300552715e-05 0.00246800714165421
KIM_GLIS2_TARGETS_UP 61 0.01 Up 0.974970850793609
9.66338725097086e-05 0.00246800714165421
REACTOME_TOLL_LIKE_RECEPTOR_9_TLR9_CASCADE 90 0.01 Down
0.303797372127544 0.000101287450153925
0.00257224343752483
GABRIELY_MIR21_TARGETS 275 0.01 Down 0.266282690739961
0.000103830984290044 0.00262202401339185
SERVITJA_ISLET_HNF1A_TARGETS_UP 119 0.01 Up
0.735758047790871 0.000105214786634849 0.00264212550795333
FINETTI_BREAST_CANCERS_KINOME_GRAY 15 0.01 Down
0.216293448936811 0.00011141658617571 0.00278231974922119
DORN_ADENOVIRUS_INFECTION_12HR_UP 26 0.01 Down
0.358155602137041 0.000118096844940165 0.00293284706080686
REACTOME_NCAM1_INTERACTIONS 23 0.01 Up
1.09907349176317 0.000122137296206716 0.00301652278268784
REACTOME_TAK1_ACTIVATES_NFKB_BY_PHOSPHORYLATION_AND_ACTIVATION_OF_IK
KS_COMPLEX 30 0.01 Down 0.289353872740819
0.000123823853215099 0.00304146568416323
ITO_PTTG1_TARGETS_UP 11 0.01 Up 0.620651178719379
0.000132676873342106 0.00324120948735199
HOLLERN_EMT_BREAST_TUMOR_UP 119 0.01 Up
0.57678832488246 0.000134431723494741 0.00326632755193978
REACTOME_CHONDROITIN_SULFATE_DERMATAN_SULFATE_METABOLISM 43
0.01 Up 0.550653665540779 0.000141281329549779
0.00341429879745299
PID_ECADHERIN_STABILIZATION_PATHWAY 39 0.01 Down
0.245848102204874 0.000146886011062564 0.00353076267233275
REACTOME_SIGNALING_BY_INTERLEUKINS 488 0.01 Down

0.401204649505357	0.000148780828911195	0.00355728630827565
ZHU_CMV_24_HR_DN	73	0.01	Up	0.704826890471312
	0.000167457622732202	0.00398265615968915
RICKMAN_TUMOR_DIFFERENTIATED_WELL_VS_POORLY_DN	361	0.01	Down
	0.368572951094349	0.000172900399407451
0.00409045944913943
TURASHVILI_BREAST_LOBULAR_CARCINOMA_VS_DUCTAL_NORMAL_UP	57	0.01
	Up	1.19319948649462	0.000181231123480656
0.00426509895311806
ALTEMEIER_RESPONSE_TO_LPS_WITH_MECHANICAL_VENTILATION	96	0.01
	Down	0.799087300259894	0.000184083564031676
0.00429659700887411
IKEDA_MIR30_TARGETS_UP	105	0.01	Down	0.26619623206869
	0.000185320572355134	0.00429659700887411
RIGGI_EWING_SARCOMA_PROGENITOR_DN	144	0.01	Up
0.658069353830784	0.000185437112285112	0.00429659700887411
CHANG_IMMORTALIZED_BY_HPV31_DN	53	0.01	Down
0.569908559590225	0.000190110995704198	0.00438230218302755
SENESE_HDAC1_TARGETS_DN	230	0.01	Up	0.404649208265293
	0.000191875915679489	0.00440041959683318
KRIEG_HYPOXIA_VIA_KDM3A	48	0.01	Down	0.447659222568717
	0.000199393736749173	0.00454961851110422
REACTOME_CYTOKINE_SIGNALING_IN_IMMUNE_SYSTEM	673	0.01	Down
	0.388999495443087	0.00020811972338424
0.00472473816470788
SENESE_HDAC3_TARGETS_UP	455	0.01	Down	0.384936612934246
	0.000210221525089021	0.00474847113203593
HAHTOLA_MYCOSIS_FUNGOIDES_CD4_UP	55	0.01	Down
0.602442381219861	0.000223706023965873	0.00501459326900236
KEGG_JAK_STAT_SIGNALING_PATHWAY	97	0.01	Down
0.3949419626883	0.000224234315254611	0.00501459326900236
GRAHAM_CML_QUIESCENT_VS_NORMAL_DIVIDING_UP	45	0.01	Down
	0.823578807530156	0.000225370323466965
0.00501504754447528
REACTOME_MET_ACTIVATES_PTK2_SIGNALING	25	0.01	Up
0.894004465780724	0.000227685413188596	0.00504160557774748
SENESE_HDAC2_TARGETS_DN	108	0.01	Up	0.593481800822802
	0.000234922069287225	0.00517634657571606
JI_CARCINOGENESIS_BY_KRAS_AND_STK11_DN	14	0.01	Up
1.33811902627748	0.000236243466041996	0.00518007014565255
REACTOME_INTERLEUKIN_10_SIGNALING	34	0.01	Down
0.873537427154279	0.000242998559030919	0.00530232292642708
PID_SYNDECAN_1_PATHWAY	39	0.01	Up	0.978573887628075
	0.000246718490287272	0.00535748605720428
GENTILE_UV_RESPONSE_CLUSTER_D2	36	0.01	Down
0.364620897945342	0.000253353678039157	0.00547511914800966
PID_NFKAPPAB_ATYPICAL_PATHWAY	16	0.01	Down
0.337590061427793	0.000255511410010911	0.00549532912918204
REACTOME_LAMININ_INTERACTIONS	29	0.01	Up
0.790227289742602	0.000257053090466229	0.00550216019831285
REACTOME_TOLL_LIKE_RECEPTOR_TLR1:TLR2_CASCADE	88	0.01	Down
	0.368285139580326	0.000263893971601392
0.00562181707274055
REACTOME_DDX58_IFIH1_MEDIATED_INDUCTION_OF_INTERFERON_ALPHA_BETA

62	0.01	Down	0.236751300933091
0.000271397832457577	0.00575440215517362
LINDSTEDT_DENDRITIC_CELL_MATURATION_A	53	0.01	Down
0.736833872125024	0.000277053037570806	0.00584672959568438
REACTOME_ANTIVIRAL_MECHANISM_BY_IFN_STIMULATED_GENES	81	0.01
Down	0.201339540758914	0.000280398689938416
0.00588968276295878
NAKAYAMA_SOFT_TISSUE_TUMORS_PCA1_UP	66	0.01	Up
1.3084063339869 0.00028665407729002	0.00598197158210321
BIOCARTA_STRESS_PATHWAY 23	0.01	Down	0.255584178244987
0.000287557954041732	0.00598197158210321
MAINA_VHL_TARGETS_UP	10	0.01	Up	0.672347464770928
0.000289574434160318	0.00598197158210321
MACAEVA_PBMC_RESPONSE_TO_IR	90	0.01	Up
0.292063634474107	0.000291304764784874	0.00598197158210321
CONCANNON_APOPTOSIS_BY_EPOXOMICIN_DN	129	0.01	Up
0.401348182868535	0.000291446446380557	0.00598197158210321
REACTOME_HEPARAN_SULFATE_HEPARIN_HS_GAG_METABOLISM	42	0.01
Up	0.539600075097029	0.000300889493248851
0.00614771941887994
BROWNE_HCMV_INFECTION_16HR_DN	69	0.01	Up
0.568621173387624	0.000306839596059582	0.00624092300582725
REACTOME_MAPK_FAMILY_SIGNALING_CASCADES 232	0.01	Down
0.30485868431744	0.000312245389235106	0.00632226587662973
NAKAYAMA_SOFT_TISSUE_TUMORS_PCA1_DN	45	0.01	Up
0.549742056555447	0.000318753819920328	0.00641957742949582
SIG_CD40PATHWAYMAP	33	0.01	Down	0.297481911611406
0.000319907751770203	0.00641957742949582
LANDIS_BREAST_CANCER_PROGRESSION_DN	65	0.01	Up
0.600979687413678	0.00032170963598411	0.00642704361666032
KEGG_T_CELL_RECEPTOR_SIGNALING_PATHWAY	89	0.01	Down
0.391589465071248	0.000329138867308049	0.00654636817942336
REACTOME_FATTY_ACID_METABOLISM	136	0.01	Up
0.315585622646845	0.000332176720552293	0.00657768440036369
PID_ANGIOPOIETIN_RECEPTOR_PATHWAY	41	0.01	Down
0.387346377156737	0.000335024783853482	0.00658577478635473
RICKMAN_TUMOR_DIFFERENTIATED_WELL_VS_MODERATELY_DN	96	0.01
Down	0.487354415850319	0.000335515556412732
0.00658577478635473
HANSON_HRAS_SIGNALING_VIA_NFKB	16	0.01	Down
0.578778576645964	0.000352416373567768	0.00688744173559616
YAGUE_PRETUMOR_DRUG_RESISTANCE_DN	11	0.01	Up
0.712928914757502	0.000369733142902285	0.00717070422814567
BIOCARTA_MAPK_PATHWAY	79	0.01	Down	0.226957714820128
0.000370100863388164	0.00717070422814567
WENDT_COHESIN_TARGETS_UP	33	0.01	Down
0.194690546573735	0.000374031906444792	0.00719231972035361
REACTOME_TOLL_LIKE_RECEPTOR_CASCADES	133	0.01	Down
0.386526918637659	0.000374416643951667	0.00719231972035361
PID_BCR_5PATHWAY	57	0.01	Down	0.291047421855223
0.000383814314872681	0.00730473794207754
WONG_ENDMETRIUM_CANCER_DN	47	0.01	Up
0.856521158692381	0.000384135068458885	0.00730473794207754
KEGG_GLUTATHIONE_METABOLISM	42	0.01	Up

0.45193049057846	0.000385144136211875	0.00730473794207754
ZHENG_FOXP3_TARGETS_IN_THYMUS_UP	175	0.01	Down
0.244192546422183	0.000388074965658257	0.00732939903627674
BERENJENO_TRANSFORMED_BY_RHOA_DN	365	0.01	Up
0.39757851961146	0.000390285481190714	0.00734030643494669
DEBOSSCHER_NFKB_TARGETS_REPRESSED_BY_GLUCOCORTICOIDS	11	0.01
Down	0.920835139847936	0.000410655989541785
0.00769124447079301
LANDIS_ERBB2_BREAST_TUMORS_324_DN	136	0.01	Up
0.510737237958521	0.000417279720748939	0.00778287279986092
DAZARD_UV_RESPONSE_CLUSTER_G28	18	0.01	Down
0.716927107943971	0.000422083321006225	0.00783993606579745
DELPUECH_FOXO3_TARGETS_UP	64	0.01	Down
0.41255287023569	0.000457132418140725	0.00845600913391999
REACTOME_CS_DS_DEGRADATION	11	0.01	Up
0.874615760361958	0.000466786928633019	0.00859229044706961
REACTOME_NITRIC_OXIDE_STIMULATES_GUANYLATE_CYCLASE	14	0.01
Up	0.833858493196057	0.000468322838605574
0.00859229044706961
REACTOME_NOD1_2_SIGNALING_PATHWAY	34	0.01	Down
0.332092700387773	0.000492962300193006	0.00900758349336407
FOSTER_TOLERANT_MACROPHAGE_DN	376	0.01	Down
0.367483025826227	0.000504459158636382	0.00918033974927343
REACTOME_SIGNALING_BY_TGF_BETA_FAMILY_MEMBERS	92	0.01	Down
0.214345215425549	0.000515671403556477
0.00934654418946115
WARTERS_RESPONSE_TO_IR_SKIN	73	0.01	Up
0.286077130280011	0.000522001027723634	0.00942327156472987
CHEN_HOXA5_TARGETS_9HR_DN	41	0.01	Up
0.289027234638515	0.000526552676896163	0.00946741713059301
REACTOME_DISEASES_OF_GLYCOSYLATION	115	0.01	Up
0.697534385239999	0.000530302554547174	0.00949685252067548
ZHAN_MULTIPLE_MYELOMA_DN	28	0.01	Up
1.03319187720854	0.00054384398176231	0.00970070911913327
KEGG_EPITHELIAL_CELL_SIGNALING_IN_HELICOBACTER_PYLORI_INFECTION	60
0.01	Down	0.303444864446131	0.000551391456944188
0.00979646086547085
REACTOME_SYNTHESIS_OF_BILE_ACIDS_AND_BILE_SALTS_VIA_7ALPHA_HYDROXYCH
OLESTEROL	17	0.01	Up	0.642114702598474
0.000564378420364038	0.00998772047061556
KEGG_ERBB_SIGNALING_PATHWAY	77	0.01	Down
0.249496542067331	0.000572984149505744	0.0101002500079542
REACTOME_INTERLEUKIN_6_FAMILY_SIGNALING	20	0.01	Down
0.504268544010022	0.000590695529616441	0.0103539637821698
REACTOME_INTERLEUKIN_6_SIGNALING	11	0.01	Down
0.348610853992223	0.000591984136155202	0.0103539637821698
REACTOME_TRAF6_MEDIATED_IRF7_ACTIVATION	15	0.01	Down
0.271317507198518	0.000595097935724492	0.0103680822522542
BIOCARTA_DEATH_PATHWAY	28	0.01	Down	0.315369053368412
0.000605403146255021	0.0105069001637696
GEISS_RESPONSE_TO_DSRNA_UP	37	0.01	Down
0.571180557262613	0.000608132517742973	0.0105136756432872
GENTILE_UV_HIGH_DOSE_DN	281	0.01	Down	0.314154840291338
0.000639744878050695	0.0110178284553175

REACTOME_HS_GAG_DEGRADATION 20 0.01 Up
0.45234497877213 0.000654859786324669 0.011230423439299
ROY_WOUND_BLOOD_VESSEL_UP 44 0.01 Up
0.874552847623547 0.000662990535347509 0.011230423439299
VANHARANTA_UTERINE_FIBROID_UP 36 0.01 Up
0.669599958318905 0.000663637833573075 0.011230423439299
LIU_PROSTATE_CANCER_DN 406 0.01 Up 0.518428238061293
0.000664664766110804 0.011230423439299
ZHOU_INFLAMMATORY_RESPONSE_FIMA_UP 332 0.01 Down
0.503718407040449 0.000666279844421668 0.011230423439299
KEGG_LYSOSOME 116 0.01 Up 0.295867771421691
0.000669576628434783 0.011230423439299
REACTOME_TRAF3_DEPENDENT_IRF_ACTIVATION_PATHWAY 13 0.01 Down
0.291084142341108 0.000669674935977307
0.011230423439299
NABA_BASEMENT_MEMBRANES 33 0.01 Up 1.07563000570581
0.000672076508380738 0.011230423439299
RODRIGUES_THYROID_CARCINOMA_ANAPLASTIC_UP 671 0.01 Down
0.247433880802414 0.000694977444374226
0.0115700874535635
MIKKELSEN_MCV6_ICP_WITH_H3K4ME3_AND_H3K27ME3 15 0.01 Up
0.870014061669615 0.000701650552848335
0.0116380783581301
DUTTA_APOPTOSIS_VIA_NFKB 28 0.01 Down
0.560032277830026 0.000707639977489057 0.0116942709515195
SENESE_HDAC1_AND_HDAC2_TARGETS_DN 206 0.01 Up
0.492448618655745 0.000721764818107981 0.0118840031406424
PID_IL2_PI3K_PATHWAY 30 0.01 Down 0.293015640039718
0.000727943556785489 0.0119419937509152
GARGALOVIC_RESPONSE_TO_OXIDIZED_PHOSPHOLIPIDS_YELLOW_UP 32 0.01
Down 0.267902935071892 0.000749980924644453
0.0122587791137339
SESTO_RESPONSE_TO_UV_C0 107 0.01 Down 0.267179927441
0.000759780568095363 0.0123739625130024
KEGG_CHEMOKINE_SIGNALING_PATHWAY 131 0.01 Down
0.532397354072528 0.000789839482925182 0.0128170703095621
LANDIS_BREAST_CANCER_PROGRESSION_UP 42 0.01 Down
0.326952894243329 0.000811346566455339 0.0131187151662473
KORKOLA_YOLK_SAC_TUMOR 42 0.01 Up 0.762544954513693
0.000831876019906996 0.0134024469873905
PID_AR_TF_PATHWAY 45 0.01 Down 0.228067248601411
0.000836197281453609 0.0134239527861928
ZHENG_GLIOBLASTOMA_PLASTICITY_DN 44 0.01 Up
0.571742353466507 0.000872411052012019 0.0139554721665268
BIOCARTA_AKT_PATHWAY 20 0.01 Down 0.296501317900237
0.000876544551464948 0.0139652548453898
FERRARI_RESPONSE_TO_FENRETINIDE_UP 21 0.01 Down
0.602608353368506 0.000879236289487279 0.0139652548453898
PID_MET_PATHWAY 75 0.01 Down 0.191718181000593
0.00091688382076977 0.014511946388592
KEGG_GLYCOSPHINGOLIPID_BIOSYNTHESIS_LACTO_AND_NEOLACTO_SERIES 22
0.01 Down 0.465018380017714 0.000922978490949507
0.0145571519888352
REACTOME_TNFR1_INDUCED_NFKAPPAB_SIGNALING_PATHWAY 30 0.01

Down 0.346541552056135 0.000967310536427953
0.0152030100043484
VANTVEER_BREAST_CANCER_POOR_PROGNOSIS 47 0.01 Up
0.543019508333756 0.000989909095502246 0.0155039769487199
NAKAYAMA_SOFT_TISSUE_TUMORS_PCA2_UP 56 0.01 Up
0.585649647295328 0.000999483683007225 0.0155995803997135
THEILGAARD_NEUTROPHIL_AT_SKIN_WOUND_UP 71 0.01 Down
0.457992499085284 0.00102679587942836 0.0159704064983754
DAUER_STAT3_TARGETS_UP 41 0.01 Down 0.47765438553866
0.00104022733670758 0.0160554554664961
CHARAFE_BREAST_CANCER_LUMINAL_VS_MESENCHYMAL_UP 435 0.01 Down
0.314045393382397 0.00104132178910143
0.0160554554664961
REACTOME_DISEASES_OF_IMMUNE_SYSTEM 20 0.01 Down
0.495135367555853 0.00104570882208538 0.0160554554664961
HUPER_BREAST_BASAL_VS_LUMINAL_DN 57 0.01 Down
0.505768481672527 0.00104655137968484 0.0160554554664961
MCGARVEY_SILENCED_BY_METHYLATION_IN_COLON_CANCER 24 0.01
Up 0.667420857602285 0.00106073497395368
0.0162176996868088
PID_LYSOPHOSPHOLIPID_PATHWAY 58 0.01 Down
0.41106978259546 0.0010685030440241 0.0162810887555536
ZHU_CMV_ALL_UP 102 0.01 Down 0.37505752353856
0.00107380934680292 0.0163066655874295
MILI_PSEUDOPODIA_HAPTOTAXIS_UP 493 0.01 Down
0.179895185980101 0.00110245485307792 0.0166853015642601
REACTOME_GLYCOSAMINOGLYCAN_METABOLISM 99 0.01 Up
0.558228424080693 0.00111064129549491 0.0167527940377504
REACTOME_NUCLEOTIDE_BINDING_DOMAIN_LEUCINE_RICH_REPEAT_CONTAINING_RE
CEPTOR_NLR_SIGNALING_PATHWAYS 49 0.01 Down
0.344056547892225 0.00114804638480675 0.0172590919722621
REACTOME_ESTROGEN_DEPENDENT_NUCLEAR_EVENTS_DOWNSTREAM_OF_ESR_MEMBRAN
E_SIGNALING 23 0.01 Down 0.343387647016589
0.00116057940759824 0.0173893481238469
ZHU_CMV_ALL_DN 106 0.01 Up 0.615755086301466
0.00119773521395803 0.0178402762322205
REACTOME_INTERLEUKIN_17_SIGNALING 67 0.01 Down
0.302939166992761 0.00119861255219813 0.0178402762322205
BROCKE_APOPTOSIS_REVERSED_BY_IL6 135 0.01 Down
0.368412628132095 0.00122112947350559 0.0180604578793117
LANDIS_ERBB2_BREAST_TUMORS_324_UP 142 0.01 Down
0.292669665001432 0.00122144142276101 0.0180604578793117
REACTOME_SIGNALING_BY_EGFR 46 0.01 Down
0.217537917290248 0.00123799789380823 0.0182308306532362
CHICAS_RB1_TARGETS_CONFLUENT 494 0.01 Up
0.497525330087012 0.00124107545715023 0.0182308306532362
CHANGOLKAR_H2AFY_TARGETS_UP 41 0.01 Down
0.326028404881599 0.00125545440113022 0.018345683636126
SEITZ_NEOPLASTIC_TRANSFORMATION_BY_8P_DELETION_UP 65 0.01
Down 0.704607626234125 0.00125705685426625
0.018345683636126
RICKMAN_HEAD_AND_NECK_CANCER_C 71 0.01 Down
0.854277584488404 0.00126800370669259 0.0184455555391042
REACTOME_ACTIVATION_OF_NF_KAPPAB_IN_B_CELLS 64 0.01 Down

0.158781223384266 0.0012931260922405
0.0187389820171066
ROZANOV_MMP14_TARGETS_UP 220 0.01 Up
0.495328345097014 0.00129651243766855 0.0187389820171066
AMIT_EGF_RESPONSE_240_MCF10A 21 0.01 Down
0.487018957612558 0.00130363363481019 0.0187815166297173
ZHENG_IL22_SIGNALING_UP 41 0.01 Down 0.934066966433332
0.00130982179144304 0.0188103800400525
HOLLERN_SQUAMOUS_BREAST_TUMOR 99 0.01 Down
0.659616158586134 0.00131795988310056 0.0188322000662425
STOSSI_RESPONSE_TO_ESTRADIOL 28 0.01 Down
0.772608434541931 0.00131972036059319 0.0188322000662425
FARMER_BREAST_CANCER_CLUSTER_1 26 0.01 Down
0.702296973917677 0.00134658023217143 0.0191546776696537
REACTOME_ARACHIDONIC_ACID_METABOLISM 37 0.01 Up
0.55511318007252 0.001356709574328 0.0192378849735153
PLASARI_TGFB1_TARGETS_1HR_UP 30 0.01 Down
0.498652482748491 0.00136286339855969 0.0192643741400183
SHAFFER_IRF4_MULTIPLE_MYELOMA_PROGRAM 33 0.01 Down
0.433013856858708 0.00137300438342509 0.0193468799482626
REACTOME_DISEASES_OF_SIGNAL_TRANSDUCTION 333 0.01 Down
0.24064583151112 0.00138291672746915
0.0194256584061682
AMIT_EGF_RESPONSE_120_MCF10A 41 0.01 Down
0.404336603913655 0.0013886895158996 0.019445979358158
REACTOME_DOWNSTREAM_SIGNALING_EVENTS_OF_B_CELL_RECEPTOR_BCR 78
0.01 Down 0.175691829023719 0.0014357834467547
0.0200430018421192
PYEON_CANCER_HEAD_AND_NECK_VS_CERVICAL_UP 169 0.01 Down
0.258004300616824 0.00144678353698123
0.0201340309558223
REACTOME_MITOCHONDRIAL_FATTY_ACID_BETA_OXIDATION 32 0.01
Up 0.225758671138407 0.00145725275288985
0.0202023487355861
PID_REG_GR_PATHWAY 65 0.01 Down 0.378277020903208
0.00146068149923593 0.0202023487355861
STEGER_ADIPOGENESIS_DN 20 0.01 Up 0.813770287089334
0.00148230855051225 0.0204385795538422
PID_PI3K_PLC_TRK_PATHWAY 32 0.01 Down
0.204928470260355 0.00149190773779582 0.0205080283834624
SIMBULAN_PARP1_TARGETS_DN 16 0.01 Up
0.498186396699243 0.00150089150471291 0.0205686198587943
PID_HIV_NEF_PATHWAY 33 0.01 Down 0.357105425261344
0.00152063891821594 0.0207759025452299
KOINUMA_TARGETS_OF_SMAD2_OR_SMAD3 821 0.01 Down
0.262410423176558 0.00153214926784263 0.0208548334363383
MISSIAGLIA_REGULATED_BY_METHYLATION_UP 117 0.01 Down
0.408510368762767 0.00153569518741446 0.0208548334363383
REACTOME_DISEASES_OF_METABOLISM 79 0.01 Up
0.424224398550718 0.00156782849199261 0.0212014238570075
REACTOME_SIGNALING_BY_THE_B_CELL_RECEPTOR_BCR 101 0.01 Down
0.211285588008481 0.00157065053267709
0.0212014238570075
NIKOLSKY_BREAST_CANCER_21Q22_AMPLICON 14 0.01 Up

0.471069078927273	0.00161305250998685	0.0217085959053619
PETROVA_ENDOTHELIUM_LYMPHATIC_VS_BLOOD_DN	148	0.01	Up
	0.507611678560523	0.00161982344511744
0.0217346459277698
GARGALOVIC_RESPONSE_TO_OXIDIZED_PHOSPHOLIPIDS_BLACK_UP	33	0.01
	Down	0.32286351419739	0.00163282635338587
0.0217786023512677
WAMUNYOKOLI_OVARIAN_CANCER_LMP_DN	181	0.01	Up
0.433436972641361	0.00163713853937771	0.0217786023512677
HERNANDEZ_MITOTIC_ARREST_BY_DOCETAXEL_1_UP	20	0.01	Up
	0.608993113879393	0.00164597578919824
0.0217786023512677
PID_AVB3_INTEGRIN_PATHWAY	66	0.01	Up
0.688792514258781	0.00164717472681369	0.0217786023512677
SIG_IL4RECEPTOR_IN_B_LYPHOCYTES	25	0.01	Down
0.227048846355031	0.0016473247607188	0.0217786023512677
REACTOME_CONSTITUTIVE_SIGNALING_BY_AKT1_E17K_IN_CANCER	25	0.01
	Down	0.242383959383637	0.00166375774959502
0.0218880665711008
REACTOME_CLEC7A_DECTIN_1_SIGNALING	97	0.01	Down
0.186452088190094	0.00166534344100478	0.0218880665711008
MUELLER_COMMON_TARGETS_OF_AML_FUSIONS_DN	26	0.01	Down
	0.44076515997615	0.00167172310848062
0.021907858229214
REACTOME_GAB1_SIGNALOSOME	16	0.01	Down
0.299718536762596	0.00172339396321463	0.0225193484437493
BURTON_ADIPOGENESIS_PEAK_AT_0HR	51	0.01	Up
0.687460354186022	0.0017642514228680 1	0.0229864062196861
AMIT_EGF_RESPONSE_60_HELA	45	0.01	Down
0.346851602843817	0.00179085729911188	0.0232656172240113
ELVIDGE_HIF1A_TARGETS_DN	89	0.01	Up
0.436817080773776	0.00180311683922047	0.0233290261386329
LA_MEN1_TARGETS	18	0.01	Up	0.682476876303999
0.00180611815266835	0.0233290261386329
REACTOME_OVARIAN_TUMOR_DOMAIN_PROTEASES	35	0.01	Down
0.265264167178286	0.00185024437916021	0.0238077494875744
SHIN_B_CELL_LYMPHOMA_CLUSTER_7	24	0.01	Down
0.245855898392309	0.00185398378240665	0.0238077494875744
BIOCARTA_GCR_PATHWAY	14	0.01	Down	0.231939277512439
	0.00185907009346799	0.0238077494875744
ROSS_AML_WITH_MLL_FUSIONS	67	0.01	Up
0.43101860090115	0.00188077192595748	0.0240172437703946
REACTOME_INTERLEUKIN_20_FAMILY_SIGNALING	18	0.01	Down
	0.437648941230595	0.00192359639492099
0.0244945206662035
CREIGHTON_ENDOCRINE_THERAPY_RESISTANCE_5	454	0.01	Down
	0.321379486700503	0.00193565552581195
0.0245784508150416
JI_CARCINOGENESIS_BY_KRAS_AND_STK11_UP	12	0.01	Down
0.504422542050866	0.00194470633067733	0.0246031412827528
BIOCARTA_TOLL_PATHWAY	22	0.01	Down	0.271623334233163
	0.00195204100137232	0.0246031412827528
FLORIO_NEOCORTEX_BASAL_RADIAL_GLIA_DN	126	0.01	Up
0.445687688527196	0.00195402034214522	0.0246031412827528

MARTINEZ_RESPONSE_TO_TRABECTEDIN_DN 264 0.01 Down
0.202422925295793 0.00196973447163184 0.0247317219273328
WAMUNYOKOLI_OVARIAN_CANCER_GRADES_1_2_DN 61 0.01 Up
0.49167946044114 0.00199470857580432
0.0249755293822853
PID_IL6_7_PATHWAY 42 0.01 Down 0.361060793183823
0.00200298522919578 0.0250094961256529
BIOCARTA_RELA_PATHWAY 15 0.01 Down 0.305188376737348
0.0020112316834799 0.0250165600767469
WEINMANN_ADAPTATION_TO_HYPOXIA_UP 25 0.01 Up
0.717473629787672 0.00201468181263234 0.0250165600767469
BRUECKNER_TARGETS_OF_MIRLET7A3_UP 95 0.01 Down
0.391111154288557 0.00202865988035759 0.0251207332292214
PID_IL5_PATHWAY 12 0.01 Down 0.26147937394204
0.00204012947928643 0.0251906287144128
REACTOME_DEFECTIVE_EXT2_CAUSES_EXOSTOSES_2 13 0.01 Up
0.536465334768756 0.00204551267647624
0.0251906287144128
BURTON_ADIPOGENESIS_1 31 0.01 Down 0.436113800063187
0.00207666499889566 0.0254583587036714
RICKMAN_HEAD_AND_NECK_CANCER_E 58 0.01 Down
0.868104413662139 0.00208041041114451 0.0254583587036714
CHIANG_LIVER_CANCER_SUBCLASS_UNANNOTATED_UP 68 0.01 Down
0.345464032463195 0.00208424382713038
0.0254583587036714
PID_ERBB1_DOWNSTREAM_PATHWAY 103 0.01 Down
0.24069344359675 0.00209496934979699 0.0255200195862804
HECKER_IFNB1_TARGETS 75 0.01 Down 0.699734275506628
0.00210491703148848 0.02557189745011
DORN_ADENOVIRUS_INFECTION_12HR_DN 31 0.01 Down
0.323631276473875 0.00211462192920219 0.0255823438116256
PID_IL2_1PATHWAY 50 0.01 Down 0.376258254292499
0.00211715948785867 0.0255823438116256
BIOCARTA_HCMV_PATHWAY 16 0.01 Down 0.215876210991585
0.00213156490008943 0.0256873571739999
ALONSO_METASTASIS_DN 17 0.01 Down 0.513605566168924
0.00215298719978751 0.0258761429493178
REACTOME_NEGATIVE_REGULATION_OF_MAPK_PATHWAY 39 0.01 Down
0.231363597280604 0.00217958296988282
0.0261259345323288
SESTO_RESPONSE_TO_UV_C5 46 0.01 Down 0.213819267589643
0.0022039299856962 0.0263475140577245
ABBUD_LIF_SIGNALING_2_UP 10 0.01 Up
1.01646644931863 0.00221817741491975 0.02644749994712
PID_ECADHERIN_NASCENT_AJ_PATHWAY 39 0.01 Down
0.206991580492144 0.00222791545322504 0.0264933332334564
REACTOME_TRANSPORT_OF_NUCLEOSIDES_AND_FREE_PURINE_AND_PYRIMIDINE_BAS
ES_ACROSS_THE_PLASMA_MEMBRANE 10 0.01 Up
0.450239132978605 0.00226937502562048 0.026915147071673
GARGALOVIC_RESPONSE_TO_OXIDIZED_PHOSPHOLIPIDS_GREEN_UP 26 0.01
Down 0.354023415560621 0.00228852518687098
0.0270126473409912
REACTOME_FCERI_MEDIATED_NF_KB_ACTIVATION 75 0.01 Down
0.173448783479085 0.00228961482467579

0.0270126473409912
NADLER_OBESITY_UP 53 0.01 Up 0.596709802716945
0.00232031527558169 0.0273031862925123
BILD_SRC_ONCOGENIC_SIGNATURE 58 0.01 Down
0.339896404972105 0.00234419518162769 0.0275121601603563
REACTOME_SIGNALING_BY_ACTIVIN 12 0.01 Down
0.34735391072282 0.00235717383982922 0.0275924385677925
KEGG_CYTOKINE_CYTOKINE_RECEPTOR_INTERACTION 140 0.01 Down
0.646387237198879 0.00242285456303551
0.0282876136645315
HINATA_NFKB_TARGETS_KERATINOCYTE_DN 21 0.01 Down
0.673818909969904 0.00249973803353595 0.0291096436806842
LIM_MAMMARY_LUMINAL_MATURE_DN 83 0.01 Up
0.60786552587474 0.00251719899771581 0.0292372338365182
KEGG_VALINE_LEUCINE_AND_ISOLEUCINE_DEGRADATION 42 0.01 Up
0.211349802157874 0.0025286915054013
0.029295021434997
CUI_TCF21_TARGETS_2_UP 336 0.01 Up 0.409614067642523
0.00257684388367008 0.029776126624928
BROWNE_HCMV_INFECTION_16HR_UP 210 0.01 Down
0.287747280747664 0.00258786410823492 0.0297880121582235
IIZUKA_LIVER_CANCER_PROGRESSION_G1_G2_DN 25 0.01 Down
0.188218327824954 0.00259112630786772
0.0297880121582235
SCHEIDEREIT_IKK_TARGETS 16 0.01 Down 0.384697893382352
0.0026403216776945 0.0302558593778557
HOSHIDA_LIVER_CANCER_SUBCLASS_S1 225 0.01 Up
0.570301791886213 0.0026452842570628 0.0302558593778557
JECHLINGER_EPITHELIAL_TO_MESENCHYMAL_TRANSITION_UP 58 0.01
Up 0.861162710911368 0.00265454935053414
0.0302847698747486
REACTOME_O_GLYCOSYLATION_OF_TSR_DOMAIN_CONTAINING_PROTEINS 28
0.01 Up 1.17113018645917 0.00267797596659002
0.0304746885311953
DAZARD_RESPONSE_TO_UV_NHEK_UP 188 0.01 Down
0.384827490358288 0.00269332643045942 0.0305168017584982
PELLICCIOTTA_HDAC_IN_ANTIGEN_PRESENTATION_DN 49 0.01 Down
0.191314410143042 0.0026952547937984
0.0305168017584982
NEWMAN_ERCC6_TARGETS_UP 22 0.01 Down 0.614932391314771
0.00274009001147509 0.0309464939738205
PID_IL2_STAT5_PATHWAY 23 0.01 Down 0.377512521909755
0.00279950289807214 0.03153825946575
REACTOME_MISCELLANEOUS_TRANSPORT_AND_BINDING_EVENTS 22 0.01
Down 0.383425178100004 0.00282539327858228
0.0316922761509392
REACTOME_ONCOGENIC_MAPK_SIGNALING 75 0.01 Down
0.221096915292995 0.00282727535851538 0.0316922761509392
DAZARD_UV_RESPONSE_CLUSTER_G24 18 0.01 Down
0.798562160661242 0.0028534115455631 0.0319056838241446
ROSS_LEUKEMIA_WITH_MLL_FUSIONS 64 0.01 Up
0.36920687015574 0.00287740631196472 0.0320493141222998
XU_HGF_TARGETS_INDUCED_BY_AKT1_6HR 18 0.01 Down
0.339500815093652 0.00288051677539692 0.0320493141222998

KIM_WT1_TARGETS_8HR_UP	162	0.01	Down	0.362645746039563
	0.0029048811395254	0.0322405943757202
REACTOME_SIGNALING_BY_RAS_MUTANTS	52	0.01	Down
0.248922206672381	0.0029455893767835	0.0325487561036747
PID_TOLL_ENDOGENOUS_PATHWAY	21	0.01	Down
0.780425363078301	0.0029471287506553	0.0325487561036747
OSWALD_HEMATOPOIETIC_STEM_CELL_IN_COLLAGEN_GEL_UP	188	0.01
	Down	0.493294472772747	0.00295889713200355
0.0325986338440097
NABA_PROTEOGLYCANS	16	0.01	Up	1.3924747807134
0.00296688321753248	0.0326066994200697
LUI_THYROID_CANCER_CLUSTER_4	12	0.01	Up
0.870422474995277	0.00299988070804184	0.0328889360552392
REACTOME_REGULATION_OF_MECP2_EXPRESSION_AND_ACTIVITY	27	0.01
	Down	0.223314877043938	0.00302712144415766
0.0329892435750622
KANG_DOXORUBICIN_RESISTANCE_DN	19	0.01	Down
0.228600811134015	0.00302994493108406	0.0329892435750622
BIOCARTA_PLATELETAPP_PATHWAY	10	0.01	Up
0.599777638441014	0.00303104729621818	0.0329892435750622
REACTOME_GLUTATHIONE_CONJUGATION	28	0.01	Up
0.452364824943262	0.0030576046805146	0.0331979058910945
SHEN_SMARCA2_TARGETS_UP	428	0.01	Down	0.143743420171449
	0.00309370680930623	0.0335089448381483
BURTON_ADIPOGENESIS_11	49	0.01	Down	0.345577381278162
	0.0031014693902113	0.0335122714158649
BIOCARTA_41BB_PATHWAY	15	0.01	Down	0.272055476309005
	0.00312253257335482	0.0336056257503516
REACTOME_INTERLEUKIN_2_SIGNALING	10	0.01	Down
0.556565329287001	0.00312506152695149	0.0336056257503516
PID_BMP_PATHWAY	35	0.01	Down	0.458519756048996
0.00318642430835555	0.0341837166254372
DER_IFN_BETA_RESPONSE_UP	102	0.01	Down
0.285531020055515	0.00322216739640355	0.0344848629686523
REACTOME_ANTIGEN_PRESENTATION:_FOLDING_ASSEMBLY_AND_PEPTIDE_LOADING_
OF_CLASS_I_MHC	25	0.01	Down	0.255695405074499
0.00332561998652562	0.035507510307441
IZADPANAH_STEM_CELL_ADIPOSE_VS_BONE_DN	91	0.01	Up
0.577634294119104	0.00333492757294202	0.0355225105222142
PID_EPHB_FWD_PATHWAY	38	0.01	Down	0.23069829852871
	0.00337070221767337	0.0358186914147561
WINTER_HYPOXIA_DN	36	0.01	Up	0.57409186165028
	0.00339090340769316	0.0358988530570041
HOFFMANN_SMALL_PRE_BII_TO_IMMATURE_B_LYMPHOCYTE_UP	43	0.01
	Up	0.411227611321977	0.00339421945470664
0.0358988530570041
GENTILE_UV_RESPONSE_CLUSTER_D4	53	0.01	Down
0.24701243165412	0.00340220498382286	0.0358988530570041
REACTOME_OTHER_INTERLEUKIN_SIGNALING	224	0.01	Down
0.303154163972779	0.00342238921504983	0.0360272588329016
KEGG_GLYCOSAMINOGLYCAN_DEGRADATION	16	0.01	Up
0.342025936509863	0.0034583258192433	0.0363205013025669
DACOSTA_ERCC3_ALLELE_XPCS_VS_TTD_UP	22	0.01	Up
0.816301323686145	0.00347498910961296	0.0363449786287439

DIRMEIER_LMP1_RESPONSE_LATE_UP 56 0.01 Down
0.373470700226294 0.00347682776648718 0.0363449786287439
WANG_CLIM2_TARGETS_DN 173 0.01 Down 0.184700794050394
0.00352291864579116 0.0366585277519232
OSMAN_BLADDER_CANCER_UP 383 0.01 Down 0.253220981529479
0.00352313325669207 0.0366585277519232
REACTOME_INTERFERON_SIGNALING 172 0.01 Down
0.323906988189258 0.00357515036531539 0.0371138588731933
REACTOME_SIGNALING_BY_CYTOSOLIC_FGFR1_FUSION_MUTANTS 18 0.01
Down 0.172477748554628 0.00362977730057088
0.0375889401957133
GU_PDEF_TARGETS_UP 68 0.01 Up 0.608828021764305
0.00364256450987455 0.0375889401957133
PID_TGFBR_PATHWAY 53 0.01 Down 0.256153266110701
0.00364600176314372 0.0375889401957133
BERTUCCI_MEDULLARY_VS_DUCTAL_BREAST_CANCER_DN 155 0.01 Up
0.498991919504169 0.00370928936202669
0.0381197231257051
BIOCARTA_RNA_PATHWAY 10 0.01 Down 0.28857667208566
0.00371444688076949 0.0381197231257051
CHIBA_RESPONSE_TO_TSA_UP 47 0.01 Up
0.6190147052462 0.00375693117576836 0.0383852646667362
KEGG_PRIMARY_BILE_ACID_BIOSYNTHESIS 10 0.01 Up
0.711533299304179 0.00375740076826784 0.0383852646667362
MATZUK_MALE_REPRODUCTION_SERTOLI 15 0.01 Down
0.405031325249401 0.00377466746353895 0.0384018590313674
DAZARD_RESPONSE_TO_UV_SCC_DN 113 0.01 Down
0.240165414928561 0.0037761116110933 0.0384018590313674
WESTON_VEGFA_TARGETS_12HR 27 0.01 Up
1.21126311037271 0.00381008409212619 0.0386598826052082
DACOSTA_UV_RESPONSE_VIA_ERCC3_XPCS_DN 84 0.01 Down
0.315142299525181 0.00383519415541444 0.038827021911234
AIGNER_ZEB1_TARGETS 35 0.01 Down 0.368089145080401
0.00386989019655197 0.039090239176407
SCHAEFFER_PROSTATE_DEVELOPMENT_AND_CANCER_BOX4_DN 32 0.01
Down 0.202745596306855 0.00388740395608116
0.0391791048936879
PEDERSEN_METASTASIS_BY_ERBB2_ISOFORM_3 19 0.01 Down
0.334916945683828 0.00391134089037956 0.0392651940610542
SANDERSON_PPARA_TARGETS 13 0.01 Up 0.481013389408063
0.00391611538688994 0.0392651940610542
PHONG_TNF_RESPONSE_VIA_P38_COMPLETE 217 0.01 Down
0.332518126611857 0.00392272195282314 0.0392651940610542
REACTOME_SIGNALING_BY_NODAL 13 0.01 Down
0.302226397921151 0.00393088705839252 0.0392651940610542
PID_IFNG_PATHWAY 36 0.01 Down 0.214326885406358
0.00396793282454193 0.0395473570871751
RODWELL_AGING_KIDNEY_NO_BLOOD_UP 189 0.01 Up
0.630837046727133 0.00397752095064363 0.0395552138786352
BIOCARTA_VIP_PATHWAY 22 0.01 Down 0.331211588569941
0.00399086742463738 0.0396003290811149
REACTOME_METABOLISM_OF_PORPHYRINS 15 0.01 Up
0.26854457581561 0.00401771396020032 0.0397301279222845
REACTOME_SIGNALING_BY_BMP 20 0.01 Down

0.279597666308918	0.00402162585197763	0.0397301279222845
SHETH_LIVER_CANCER_VS_TXNIP_LOSS_PAM6	44	0.01	Down
0.336354209035433	0.00403070379096122	0.0397324858341462
KEGG_NEUROTROPHIN_SIGNALING_PATHWAY	116	0.01	Down
0.278650554952174	0.00404654327077566	0.0398013391731654
SCHLINGEMANN_SKIN_CARCINOGENESIS_TPA_UP 30	0.01	Down
0.682472449868275	0.00406004365861355	0.0398469350337727
TSENG_IRS1_TARGETS_DN	107	0.01	Up	0.50883730720504
0.00413716352896349	0.0405153596137057
CHARAFE_BREAST_CANCER_LUMINAL_VS_MESENCHYMAL_DN 438	0.01	Up
0.457399345460167	0.00416239675584799
0.0406086187368794
LANDIS_ERBB2_BREAST_PRENEOPLASTIC_DN	53	0.01	Up
0.419258399194952	0.00416475489159097	0.0406086187368794
CROONQUIST_STROMAL_STIMULATION_UP	50	0.01	Up
1.09387602274982	0.00418762651950517	0.0407432493618523
KHETCHOUMIAN_TRIM24_TARGETS_UP	43	0.01	Up
0.674515534794305	0.0042107448057128	0.0408796930921793
DING_LUNG_CANCER_EXPRESSION_BY_COPY_NUMBER	100	0.01	Down
0.175457482566255	0.00426668282953283
0.0412893349185931
WAMUNYOKOLI_OVARIAN_CANCER_LMP_UP	267	0.01	Down
0.239937962391443	0.00427131050881998	0.0412893349185931
PID_IL1_PATHWAY 33	0.01	Down	0.290875681477825
0.00429965980478349	0.0414741863143815
URS_ADIPOCYTE_DIFFERENTIATION_DN	28	0.01	Up
1.12765804173177	0.00435656824038593	0.0418839554870566
REACTOME_PROGRAMMED_CELL_DEATH	162	0.01	Down
0.216580707395548	0.00436077667807397	0.0418839554870566
FRIDMAN_IMMORTALIZATION_DN	33	0.01	Up
0.517178007478661	0.00439852768030114	0.0421564646544854
SENGUPTA_NASOPHARYNGEAL_CARCINOMA_DN	249	0.01	Down
0.50797089558733	0.00446105291720434	0.0425779904582048
REACTOME_MAPK6_MAPK4_SIGNALING	85	0.01	Down
0.206898796910732	0.00446145350518676	0.0425779904582048
KEGG_LONG_TERM_POTENTIATION	51	0.01	Down
0.284654487510019	0.00449623785575941	0.0427969963192892
MCCLUNG_DELTA_FOSB_TARGETS_2WK	35	0.01	Up
0.488961574077867	0.00450344366162932	0.0427969963192892
TSENG_ADIPOGENIC_POTENTIAL_DN	33	0.01	Up
0.722274436247841	0.00452652287527854	0.0429255703045929
GRANDVAUX_IRF3_TARGETS_UP	13	0.01	Down
0.347444789951987	0.00454370299693384	0.0429478968309438
MORI_PRE_BI_LYMPHOCYTE_DN	66	0.01	Down
0.323887320901334	0.00455473465877027	0.0429478968309438
LIM_MAMMARY_LUMINAL_PROGENITOR_UP	44	0.01	Down
0.759332620540138	0.00456680826007765	0.0429478968309438
PID_IL3_PATHWAY 22	0.01	Down	0.205495654319771
0.00456709559181116	0.0429478968309438
KEGG_CHRONIC_MYELOID_LEUKEMIA	71	0.01	Down
0.255918088577677	0.00461761302962588	0.0433322976370947
KEGG_DORSO_VENTRAL_AXIS_FORMATION	20	0.01	Down
0.270865242895727	0.00465106475043733	0.0434931230679929
REACTOME_SIGNALING_BY_TGF_BETA_RECEPTOR_COMPLEX 72	0.01	Down

0.178522734303949	0.00465410282440591
0.0434931230679929
ZWANG_CLASS_2_TRANSIENTLY_INDUCED_BY_EGF	40	0.01	Down
0.467513576211982	0.00474299035889892
0.0442318291768685
REACTOME_C_TYPE_LECTIN_RECEPTORS_CLRS	120	0.01	Down
0.245380295642193	0.00475746489388896	0.0442749579669376
WHITFIELD_CELL_CYCLE_LITERATURE	34	0.01	Up
0.283196986729231	0.00477152971292799	0.044314103428949
KEGG_ASTHMA	16	0.01	Up	1.12082965820949
0.00478957039046852	0.0443899358869195
BLUM_RESPONSE_TO_SALIRASIB_UP	241	0.01	Down
0.227746117045644	0.00483624004104059	0.0446476458186772
PID_ERBB_NETWORK_PATHWAY	11	0.01	Down
0.344824938745618	0.00483724216100017	0.0446476458186772
SIMBULAN_UV_RESPONSE_IMMORTALIZED_DN	31	0.01	Down
0.266636449942912	0.00484901660975186	0.0446646099607267
BROWNE_INTERFERON_RESPONSIVE_GENES	63	0.01	Down
0.433179969279215	0.0048652515529228	0.0447225066061104
LINDGREN_BLADDER_CANCER_CLUSTER_2B	326	0.01	Up
0.678853607456959	0.00488990630695539	0.0448574058158459
REACTOME_MUCOPOLYSACCHARIDOSES	10	0.01	Up
0.237531391632422	0.00491372785570031	0.044984127721737
REACTOME_METABOLISM_OF_FAT_SOLUBLE_VITAMINS	30	0.01	Up
0.515783813342139	0.00497708530935621
0.0454715415966589
JI_RESPONSE_TO_FSH_DN	53	0.01	Down	0.261689230996155
0.00500534221157064	0.0456369436937323
NABA_SECRETED_FACTORS	146	0.01	Down	0.807988240474156
0.00503093108275746	0.0457529416005382
MOOTHA_FFA_OXYDATION	23	0.01	Up	0.228766445173908
0.00503842182252868	0.0457529416005382
KEGG_GNRH_SIGNALING_PATHWAY	77	0.01	Down
0.307240521067439	0.00516594622021289	0.0468163876206793
REACTOME_INTERFERON_ALPHA_BETA_SIGNALING	53	0.01	Down
0.358181181786727	0.00517684819120179
0.0468207899787768
SMID_BREAST_CANCER_RELAPSE_IN_BONE_UP	67	0.01	Down
0.572710650762278	0.00519392016261644	0.0468808657248211
REACTOME_SYNTHESIS_OF_BILE_ACIDS_AND_BILE_SALTS_VIA_27_HYDROXYCHOLES
TEROL	10	0.01	Up	0.656855295695844
0.00520874942294878	0.0469038124376174
PID_CD8_TCR_PATHWAY	42	0.01	Down	0.412976891024029
0.00521733175056924	0.0469038124376174
WONG_ENDMETRIUM_CANCER_UP	19	0.01	Down
0.72962053654023	0.00524301742753909	0.0469969721560045
BOYAULT_LIVER_CANCER_SUBCLASS_G123_DN	36	0.01	Up
0.348719661433758	0.00524860512176068	0.0469969721560045
PID_ARF6_PATHWAY	28	0.01	Down	0.21435316565402
0.00531082806335204	0.0474103367301911
REACTOME_SIGNALING_BY_FGFR1_IN_DISEASE	32	0.01	Down
0.27186678918425	0.00531586422959206	0.0474103367301911
PRAMOONJAGO_SOX4_TARGETS_UP	52	0.01	Down
0.332037608844466	0.00534524031359105	0.0475295744939191

SIMBULAN_PARP1_TARGETS_UP	28	0.01	Up
0.647411994301825	0.00535038146694618	0.0475295744939191
SIMBULAN_UV_RESPONSE_NORMAL_DN	33	0.01	Down
0.286897617630502	0.00542195778280068	0.0480704146621086
REACTOME_EGFR_DOWNREGULATION	28	0.01	Down
0.239680201439666	0.00543476023845227	0.0480890694327617
PID_FOXO_PATHWAY	47	0.01	Down	0.24268675212496
0.005552567751927	0.0488803140639362
REACTOME_DISEASES_OF_CARBOHYDRATE_METABOLISM	26	0.01	Up
0.190306410892331	0.00556064139542609
0.0488803140639362
REACTOME_FC_EPSILON_RECEPTOR_FCERI_SIGNALING	121	0.01	Down
0.211260345085473	0.00556367975247539
0.0488803140639362
GROSS_HYPOXIA_VIA_ELK3_ONLY_UP	30	0.01	Down
0.530650712819233	0.00556767982218806	0.0488803140639362
CUI_TCF21_TARGETS_2_DN	740	0.01	Down	0.336729417144291
0.00571059630629216	0.0500372912217997
HIRSCH_CELLULAR_TRANSFORMATION_SIGNATURE_UP	233	0.01	Down
0.359684635817837	0.00579903551959431
0.0507133553707712
CHIANG_LIVER_CANCER_SUBCLASS_PROLIFERATION_DN	99	0.01	Up
0.474585070957857	0.00581216461701615
0.05072947563784
GROSS_HYPOXIA_VIA_ELK3_AND_HIF1A_UP	135	0.01	Down
0.320456202845354	0.00593320355396182	0.0516135829742229
REACTOME_BILE_ACID_AND_BILE_SALT_METABOLISM	27	0.01	Up
0.484265571595025	0.00593642322528882
0.0516135829742229
BIOCARTA_HER2_PATHWAY	21	0.01	Down	0.227938364951472
0.00599650066483879	0.0520352712132246
PID_VEGFR1_2_PATHWAY	64	0.01	Down	0.239512486635269
0.00605766512681126	0.0524647490270069
CROONQUIST_NRAS_VS_STROMAL_STIMULATION_DN	79	0.01	Up
0.646509761874923	0.00608838923715837
0.0526294415788978
GAZIN_EPIGENETIC_SILENCING_BY_KRAS	22	0.01	Down
0.224536705050586	0.00614807041132231	0.0530057770981041
SA_PTEN_PATHWAY	16	0.01	Down	0.213228799670396
0.0061555095984895	0.0530057770981041
REACTOME_REGULATION_OF_LOCALIZATION_OF_FOXO_TRANSCRIPTION_FACTORS
11	0.01	Down	0.262904702582319
0.00618759613272094	0.0531802000317029
GESERICK_TERT_TARGETS_DN	21	0.01	Down
0.559007718717955	0.00623214159437227	0.0534179234966948
REACTOME_CYTOCHROME_C_MEDIATED_APOPTOTIC_RESPONSE	13	0.01
Down	0.226042867740783	0.00623902332275079
0.0534179234966948
ANDERSEN_LIVER_CANCER_KRT19_DN	49	0.01	Up
0.416844539756629	0.0062589719840635	0.0534327076496479
RODRIGUES_THYROID_CARCINOMA_DN	75	0.01	Down
0.346086808222918	0.0062679407665348	0.0534327076496479
LOPES_METHYLATED_IN_COLON_CANCER_UP	16	0.01	Up
0.777925381480567	0.00627641148810102	0.0534327076496479

BENNETT_SYSTEMIC_LUPUS_ERYTHEMATOSUS 24 0.01 Down
0.494328917167211 0.00630870537705983 0.0535093749883475
LIM_MAMMARY_STEM_CELL_DN 397 0.01 Down
0.295693461964973 0.00630922552699092 0.0535093749883475
REACTOME_SHC1_EVENTS_IN_EGFR_SIGNALING 13 0.01 Down
0.315752452394373 0.00640551007691714 0.0542236681652402
KEGG_GLYCOLYSIS_GLUCONEOGENESIS 44 0.01 Up
0.343290833167628 0.00653705808951006 0.0552332257750897
JAATINEN_HEMATOPOIETIC_STEM_CELL_UP 268 0.01 Up
0.362151685798348 0.00655965526642479 0.0553201696483667
RHEIN_ALL_GLUCOCORTICOID_THERAPY_DN 350 0.01 Up
0.264217389052928 0.00659128756098703 0.0554828419225406
DELYS_THYROID_CANCER_DN 155 0.01 Up 0.579406198283365
0.00677832829690336 0.056850210679585
DAZARD_UV_RESPONSE_CLUSTER_G6 140 0.01 Down
0.23500104172324 0.00677902400984596 0.056850210679585
PID_GMCSF_PATHWAY 34 0.01 Down 0.325427630446679
0.00683251052616598 0.0571920573838288
UROSEVIC_RESPONSE_TO_IMIQUIMOD 19 0.01 Down
0.555913836782517 0.00690930865134987 0.0577274022078396
REACTOME_DEGRADATION_OF_THE_EXTRACELLULAR_MATRIX 110 0.01
Up 0.862708206850497 0.00694396786728929
0.0579093424183031
MOROSETTI_FACIOSCAPULOHUMERAL_MUSCULAR_DISTROPHY_UP 16 0.01
Up 1.2361143791219 0.00697026723716992
0.0580210207982941
SCHRAETS_MLL_TARGETS_UP 27 0.01 Up 0.675927435406201
0.00699468528977088 0.0581006198685412
PID_ER_NONGENOMIC_PATHWAY 34 0.01 Down
0.390336191866096 0.00700568097191308 0.0581006198685412
REACTOME_SYNTHESIS_OF_BILE_ACIDS_AND_BILE_SALTS 25 0.01 Up
0.504676570679983 0.00704649255724546
0.0583314623293155
NEMETH_INFLAMMATORY_RESPONSE_LPS_UP 75 0.01 Down
0.471609123621557 0.00709080830903219 0.0585339356692012
HILLION_HMGA1B_TARGETS 64 0.01 Up 0.310108569652897
0.00709699553720015 0.0585339356692012
WANG_LMO4_TARGETS_DN 337 0.01 Down 0.209292623636775
0.00716377510530004 0.05897650018008
PEDERSEN_TARGETS_OF_611CTF_ISOFORM_OF_ERBB2 74 0.01 Down
0.388827757190528 0.00723486258958165
0.0593720580979959
BIOCARTA_KERATINOCYTE_PATHWAY 41 0.01 Down
0.290061768492424 0.00723823978591808 0.0593720580979959
GOTZMANN_EPITHELIAL_TO_MESENCHYMAL_TRANSITION_UP 59 0.01
Up 0.473782548685661 0.00726591423122144
0.0594713511998345
TSUDA_ALVEOLAR_SOFT_PART_SARCOMA 10 0.01 Down
0.256248631734156 0.00727680604224894 0.0594713511998345
REACTOME_SIGNALING_BY_BRAF_AND_RAF_FUSIONS 59 0.01 Down
0.212942933741001 0.00733669342876068
0.0598519727083108
BIOCARTA_HIVNEF_PATHWAY 55 0.01 Down 0.291611209295016
0.00740888152978997 0.0603313812978368

| | | | | | | |
|---|---|---|---|---|---|---|
| REACTOME_INTERLEUKIN_15_SIGNALING | 14 | 0.01 | Down | 0.327225765074247 | 0.007503691662476 | 0.0609346597931739 |
| KEGG_ENDOCYTOSIS | 164 | 0.01 | Down | 0.249932542283204 | 0.00751007820365258 | 0.0609346597931739 |
| REACTOME_GLYCOSPHINGOLIPID_METABOLISM | 37 | 0.01 | Up | 0.250073777005092 | 0.00752426562158726 | 0.0609397729171797 |
| SMIRNOV_RESPONSE_TO_IR_2HR_UP | 51 | 0.01 | Down | 0.300865944003848 | 0.00756287856744687 | 0.0610328481328081 |
| PID_FAS_PATHWAY | 34 | 0.01 | Down | 0.274664563188621 | 0.00756958487257586 | 0.0610328481328081 |
| REACTOME_DISSOLUTION_OF_FIBRIN_CLOT | 11 | 0.01 | Down | 0.40572202848112 | 0.0075764914923486 | 0.0610328481328081 |
| REACTOME_NEGATIVE_REGULATORS_OF_DDX58_IFIH1_SIGNALING | 34 | 0.01 | Down | 0.223903981232667 | 0.00760596182598902 | 0.0611606411588921 |
| KIM_MYC_AMPLIFICATION_TARGETS_DN | 86 | 0.01 | Down | 0.280593913401551 | 0.00781908165545396 | 0.0627620929308313 |
| HOEBEKE_LYMPHOID_STEM_CELL_UP | 82 | 0.01 | Down | 0.399643151364205 | 0.00785400948034696 | 0.0629215129773551 |
| STAMBOLSKY_TARGETS_OF_MUTATED_TP53_DN | 45 | 0.01 | Down | 0.317966044640022 | 0.00786693888615653 | 0.0629215129773551 |
| TANG_SENESCENCE_TP53_TARGETS_UP | 22 | 0.01 | Up | 0.484734998510403 | 0.00791014545240169 | 0.0631547136919105 |
| KEGG_RIBOSOME | 85 | 0.01 | Up | 0.117213154699309 | 0.00793963534143489 | 0.063253578153999 |
| MOOTHA_GLYCOLYSIS | 16 | 0.01 | Up | 0.341094207399316 | 0.00795369982711317 | 0.063253578153999 |
| NAKAYAMA_SOFT_TISSUE_TUMORS_PCA2_DN | 42 | 0.01 | Up | 0.845881865764468 | 0.00796474421249464 | 0.063253578153999 |
| CHICAS_RB1_TARGETS_GROWING | 202 | 0.01 | Up | 0.488932095387538 | 0.00801914209763395 | 0.0635732693630769 |
| MITSIADES_RESPONSE_TO_APLIDIN_UP | 409 | 0.01 | Down | 0.272974852133635 | 0.00804206091035939 | 0.0636427179437772 |
| SETLUR_PROSTATE_CANCER_TMPRSS2_ERG_FUSION_DN | 11 | 0.01 | Up | 0.659604600601425 | 0.0080828356681327 | 0.063852981244739 |
| SCIAN_INVERSED_TARGETS_OF_TP53_AND_TP73_DN | 29 | 0.01 | Down | 0.326805834516529 | 0.00813612899594037 | 0.0641612277837754 |
| ZHAN_VARIABLE_EARLY_DIFFERENTIATION_GENES_UP | 10 | 0.01 | Up | 0.744517774020744 | 0.0081987508863214 | 0.064541830532425 |
| PID_CD8_TCR_DOWNSTREAM_PATHWAY | 39 | 0.01 | Down | 0.491021209527138 | 0.00829264254344352 | 0.0651668325747878 |
| REACTOME_SMOOTH_MUSCLE_CONTRACTION | 33 | 0.01 | Up | 0.512507915570587 | 0.00831590672996341 | 0.0652356034052104 |
| BONCI_TARGETS_OF_MIR15A_AND_MIR16_1 | 83 | 0.01 | Down | 0.353554195936211 | 0.00848482526405856 | 0.0663695315527674 |
| REACTOME_DEUBIQUITINATION | 242 | 0.01 | Down | 0.192346332555055 | 0.00848998457015378 | 0.0663695315527674 |
| REACTOME_TIGHT_JUNCTION_INTERACTIONS | 22 | 0.01 | Down | 0.445473308573671 | 0.00851375865249945 | 0.0664398353176824 |
| GRANDVAUX_IFN_RESPONSE_NOT_VIA_IRF3 | 14 | 0.01 | Down | 0.268806111537189 | 0.00862661387785532 | 0.067203863745164 |

PID_INTEGRIN3_PATHWAY	37	0.01	Up	0.804980496619065
	0.00865736989105857	0.0673267779590108
AMIT_EGF_RESPONSE_240_HELA	59	0.01	Down
0.354860931763735	0.00868137477443118	0.0673968559776652
SHEN_SMARCA2_TARGETS_DN	109	0.01	Down	0.602510357798248
	0.00872263106158285	0.0676003907272671
KEGG_BETA_ALANINE_METABOLISM	18	0.01	Up
0.3917983277478 0.00874911777762945	0.0676889576771848
DUTERTRE_ESTRADIOL_RESPONSE_24HR_DN	490	0.01	Down
0.358937373246067	0.00880878220562045	0.0680334639420342
REACTOME_DEATH_RECEPTOR_SIGNALLING	132	0.01	Down
0.275354334449171	0.00884121908855836	0.0681668607256772
XU_AKT1_TARGETS_6HR	23	0.01	Down	0.280713036437266
	0.00885838039687888	0.0681822258287167
REACTOME_SIGNALING_BY_EGFR_IN_CANCER	23	0.01	Down
0.259232505807501	0.00894022597270297	0.0686945568329912
REACTOME_SELENOAMINO_ACID_METABOLISM	112	0.01	Up
0.153532037793002	0.00905652760186383	0.0694694395398941
HUMMERICH_SKIN_CANCER_PROGRESSION_DN	88	0.01	Up
0.49655208194042	0.00911832274057044	0.069824294239973
REACTOME_CELL_SURFACE_INTERACTIONS_AT_THE_VASCULAR_WALL 108	0.01
	Up	0.529555300829231	0.00915780462823485
0.0698512735613759
REACTOME_ACTIVATED_TAK1_MEDIATES_P38_MAPK_ACTIVATION	22	0.01
	Down	0.254066596134896	0.0091756098319692
0.0698512735613759
CHIBA_RESPONSE_TO_TSA_DN	21	0.01	Down
0.443498526968321	0.00918653212689734	0.0698512735613759
BERENJENO_TRANSFORMED_BY_RHOA_FOREVER_DN	28	0.01	Down
	0.488527940189916	0.00920275894318011
0.0698512735613759
HUMMEL_BURKITTS_LYMPHOMA_DN	15	0.01	Down
0.357937063777103	0.00920504245720398	0.0698512735613759
SAGIV_CD24_TARGETS_DN	44	0.01	Down	0.411227484748065
	0.00921508458774103	0.0698512735613759
REACTOME_CELL_CELL_JUNCTION_ORGANIZATION	47	0.01	Down
	0.387303492097308	0.0092672508921874
0.0701284389905427
REACTOME_RAF_ACTIVATION 28	0.01	Down	0.145549214812872
	0.00929522880403733	0.0702219386120131
PID_PI3KCI_AKT_PATHWAY	34	0.01	Down	0.182090211070795
	0.00931090853473141	0.0702223722543921
NOJIMA_SFRP2_TARGETS_UP 29	0.01	Down	0.582595750438217
	0.00934429034695087	0.0702560002220145
MODY_HIPPOCAMPUS_PRENATAL	35	0.01	Up
0.182802421698192	0.00934662694833474	0.0702560002220145
IRITANI_MAD1_TARGETS_UP 13	0.01	Down	0.500651037870251
	0.00937127337198803	0.0703236624492257
KEGG_ACUTE_MYELOID_LEUKEMIA	53	0.01	Down
0.268392145464886	0.00941735749093776	0.070551703202942
PID_TCR_JNK_PATHWAY	11	0.01	Down	0.314429737664273
	0.00944568641350751	0.0706461903971984
KEGG_CYTOSOLIC_DNA_SENSING_PATHWAY	37	0.01	Down
0.385150642991367	0.0094642666860662	0.0706675725504763

BRIDEAU_IMPRINTED_GENES 42 0.01 Up 0.418423825089018
0.00957724216509048 0.0713925431709481
LEE_AGING_CEREBELLUM_UP 59 0.01 Up 0.589796921683076
0.00960048369552903 0.0714473082970248
XU_HGF_SIGNALING_NOT_VIA_AKT1_6HR 25 0.01 Down
0.361332761970947 0.0096026076520008 0.0717733423794618
REACTOME_TRANSLOCATION_OF_SLC2A4_GLUT4_TO_THE_PLASMA_MEMBRANE 62
0.01 Down 0.164938418346549 0.00977742899604889
0.0724122995952158
REACTOME_FOXO_MEDIATED_TRANSCRIPTION 50 0.01 Down
0.244962401579774 0.00977847961163426 0.0724122995952158
JAEGER_METASTASIS_UP 33 0.01 Up 0.513379074946545
0.00993316941956406 0.0734368364160204
WU_HBX_TARGETS_2_DN 13 0.01 Down 0.32130209375109
0.00997162102695259 0.0735094806388314
CORRE_MULTIPLE_MYELOMA_DN 42 0.01 Up
0.659793875755483 0.00997570260059781 0.0735094806388314
BUYTAERT_PHOTODYNAMIC_THERAPY_STRESS_DN 613 0.01 Up
0.285600205192083 0.010013706011615 0.0735850428218653
KERLEY_RESPONSE_TO_CISPLATIN_UP 42 0.01 Up
0.363696137324 0.0100186977101183 0.0735850428218653
HIRSCH_CELLULAR_TRANSFORMATION_SIGNATURE_DN 100 0.01 Up
0.288434261603719 0.0100748166228234
0.0738253117266451
BROWNE_HCMV_INFECTION_18HR_UP 157 0.01 Down
0.284268669839004 0.0100852312702425 0.0738253117266451
LIN_NPAS4_TARGETS_DN 43 0.01 Down 0.497137279829229
0.0101006822495855 0.0738253117266451
KATSANOU_ELAVL1_TARGETS_UP 131 0.01 Up
0.524577276794554 0.0102217842400889 0.074589156102597
MARZEC_IL2_SIGNALING_UP 98 0.01 Down 0.445524715367104
0.0102545803735409 0.0747071941313879
KEGG_APOPTOSIS 80 0.01 Down 0.3050681166044
0.0103402648656357 0.0752095316683373
AMIT_EGF_RESPONSE_480_HELA 161 0.01 Down
0.301099875085792 0.010497436925444 0.0762293683035072
REACTOME_PI3K_EVENTS_IN_ERBB2_SIGNALING 12 0.01 Down
0.27438992990786 0.0105876637926209 0.0767605624965017
GARGALOVIC_RESPONSE_TO_OXIDIZED_PHOSPHOLIPIDS_LIGHTYELLOW_UP 11
0.01 Down 0.235807209448443 0.0106230110655955
0.0768928095649788
PID_WNT_SIGNALING_PATHWAY 22 0.01 Up
0.310493154642069 0.0107450675656781 0.0776512519416773
ZHU_CMV_24_HR_UP 81 0.01 Down 0.279632644529152
0.0108559177032693 0.0782936277048456
BLUM_RESPONSE_TO_SALIRASIB_DN 308 0.01 Up
0.294268672281606 0.0108687928115292 0.0782936277048456
SHAFFER_IRF4_TARGETS_IN_ACTIVATED_DENDRITIC_CELL 56 0.01
Down 0.296559537551271 0.0110298491716565
0.0793266752425538
WHITEFORD_PEDIATRIC_CANCER_MARKERS 92 0.01 Up
0.235386677468315 0.0110939512456882 0.0796055963069388
WANG_LSD1_TARGETS_UP 21 0.01 Up 0.602373318676867
0.0111040509197888 0.0796055963069388

ZHOU_CELL_CYCLE_GENES_IN_IR_RESPONSE_24HR 103 0.01 Up
0.302575033507692 0.0111418578162424
0.079740181675176
HSIAO_LIVER_SPECIFIC_GENES 125 0.01 Up
0.534162250359504 0.0111686211771401 0.079740181675176
REACTOME_JNK_C_JUN_KINASES_PHOSPHORYLATION_AND_ACTIVATION_MEDIATED_B
Y_ACTIVATED_HUMAN_TAK1 22 0.01 Down 0.259826496166291
0.0111760432603695 0.079740181675176
LEE_BMP2_TARGETS_UP 633 0.01 Up 0.455312739029691
0.0114158023313504 0.0811713815842377
FALVELLA_SMOKERS_WITH_LUNG_CANCER 70 0.01 Down
0.390294067985838 0.0114261776118435 0.0811713815842377
BIOCARTA_MET_PATHWAY 31 0.01 Down 0.219208623216252
0.011430808574599 0.0811713815842377
JOHNSTONE_PARVB_TARGETS_2_DN 331 0.01 Down
0.237743947043025 0.0115088518127736 0.0813063876532751
BANDRES_RESPONSE_TO_CARMUSTIN_WITHOUT_MGMT_48HR_DN 23 0.01
Up 0.321671297794746 0.0115096256193406
0.0813063876532751
KEGG_OTHER_GLYCAN_DEGRADATION 14 0.01 Up
0.274253086035732 0.0115153138075369 0.0813063876532751
WATANABE_ULCERATIVE_COLITIS_WITH_CANCER_UP 18 0.01 Up
0.434954254077038 0.0115221732892405
0.0813063876532751
HAHTOLA_MYCOSIS_FUNGOIDES_UP 15 0.01 Down
0.653943068035582 0.0116286667750771 0.0818031287165667
REACTOME_DECTIN_1_MEDIATED_NONCANONICAL_NF_KB_SIGNALING 59 0.01
Down 0.147409547326331 0.011628965350364
0.0818031287165667
SCHAEFFER_PROSTATE_DEVELOPMENT_12HR_UP 84 0.01 Up
0.489994641236189 0.0116587376722297 0.0818844153698006
BIOCARTA_GLEEVEC_PATHWAY 23 0.01 Down
0.237484722609717 0.0116795842384121 0.0819028567108615
GENTILE_UV_RESPONSE_CLUSTER_D1 17 0.01 Down
0.630257843077129 0.0117024624824219 0.0819354655116612
CHIARADONNA_NEOPLASTIC_TRANSFORMATION_KRAS_DN 131 0.01 Up
0.448487119071859 0.0117498165424027
0.0821390752069988
SCHUETZ_BREAST_CANCER_DUCTAL_INVASIVE_DN 78 0.01 Down
0.370857454090729 0.011801366893884
0.0823713419068455
REACTOME_FOXO_MEDIATED_TRANSCRIPTION_OF_OXIDATIVE_STRESS_METABOLIC_A
ND_NEURONAL_GENES 17 0.01 Down 0.325056355852087
0.0118461021763443 0.0825553942366937
REACTOME_TRANSPORT_OF_VITAMINS_NUCLEOSIDES_AND_RELATED_MOLECULES
32 0.01 Up 0.500127938032043
0.0119354597467086 0.0828765981996974
REACTOME_FGFR1_MUTANT_RECEPTOR_ACTIVATION 25 0.01 Down
0.291560618762499 0.0119441467085207
0.0828765981996974
BROWNE_HCMV_INFECTION_24HR_DN 121 0.01 Up
0.472719326761911 0.0119736878910328 0.0828765981996974
VECCHI_GASTRIC_CANCER_EARLY_DN 218 0.01 Up
0.533096250931017 0.0119772440606403 0.0828765981996974

REACTOME_P75NTR_SIGNALS_VIA_NF_KB 16 0.01 Down
0.254202396753382 0.0119888421262951 0.0828765981996974
XU_RESPONSE_TO_TRETINOIN_AND_NSC682994_UP 16 0.01 Down
0.43046164587595 0.012002817670301
0.0828765981996974
REACTOME_MET_PROMOTES_CELL_MOTILITY 36 0.01 Up
0.67760200895248 0.0120552141000916 0.083063887085624
REACTOME_BMAL1:CLOCK_NPAS2_ACTIVATES_CIRCADIAN_GENE_EXPRESSION 24
0.01 Down 0.286234904155407 0.0120669006155534
0.083063887085624
KEGG_PROSTATE_CANCER 80 0.01 Down 0.292404676764105
0.0120966554693865 0.0831094796411724
HOFFMANN_IMMATURE_TO_MATURE_B_LYMPHOCYTE_UP 28 0.01 Up
0.659331853729561 0.0121105025950985
0.0831094796411724
DEMAGALHAES_AGING_DN 14 0.01 Up 0.600146018779846
0.012139851322503 0.0831838897784314
WANG_ADIPOGENIC_GENES_REPRESSED_BY_SIRT1 20 0.01 Up
0.44722755650468 0.0122506200130858
0.0836629698655768
REACTOME_DOWNSTREAM_SIGNAL_TRANSDUCTION 29 0.01 Down
0.351037404463007 0.0122623570368718 0.0836629698655768
REACTOME_CYTOSOLIC_SENSORS_OF_PATHOGEN_ASSOCIATED_DNA 61 0.01
Down 0.202107646944743 0.0122656055931958
0.0836629698655768
LINDVALL_IMMORTALIZED_BY_TERT_DN 69 0.01 Up
0.668285943812067 0.0123366419864206 0.0840200086802433
SMID_BREAST_CANCER_RELAPSE_IN_BRAIN_DN 67 0.01 Down
0.411815362977533 0.0124388755964053 0.0845881177092918
MOOTHA_HUMAN_MITODB_6_2002 399 0.01 Up
0.170855069110134 0.0124982131837533 0.084863245107207
REACTOME_RESPIRATORY_ELECTRON_TRANSPORT_ATP_SYNTHESIS_BY_CHEMIOSMOTI
C_COUPLING_AND_HEAT_PRODUCTION_BY_UNCOUPLING_PROTEINS 108 0.01
Up 0.120800588811481 0.0126033314927066
0.0854383722318874
REACTOME_DEADENYLATION_OF_MRNA 25 0.01 Down
0.129926689373653 0.0126209297357004 0.0854383722318874
REACTOME_TRANSCRIPTIONAL_ACTIVITY_OF_SMAD2_SMAD3:SMAD4_HETEROTRIMER
43 0.01 Down 0.17040121757057
0.0127202414031318 0.0859811806121466
DAVICIONI_PAX_FOXO1_SIGNATURE_IN_ARMS_DN 13 0.01 Down
0.564584307355575 0.0127429357394816
0.0860052494729278
MCBRYAN_PUBERTAL_BREAST_5_6WK_UP 101 0.01 Down
0.418615760849067 0.0127996087050611 0.0862582325775855
DER_IFN_ALPHA_RESPONSE_UP 72 0.01 Down
0.281564216815639 0.0128342597916915 0.0863238271289516
BROWNE_HCMV_INFECTION_2HR_UP 27 0.01 Down
0.724322008853132 0.012884097326419 0.0863238271289516
PID_TCPTP_PATHWAY 40 0.01 Down 0.385403037107925
0.0129033526350953 0.0863238271289516
REACTOME_TICAM1_RIP1_MEDIATED_IKK_COMPLEX_RECRUITMENT 19 0.01
Down 0.242291356465798 0.0129039971349862
0.0863238271289516

PID_ARF6_DOWNSTREAM_PATHWAY 15 0.01 Down
0.213601814007317 0.0129053641447509 0.0863238271289516
REACTOME_SIGNALING_BY_MODERATE_KINASE_ACTIVITY_BRAF_MUTANTS 39
0.01 Down 0.232936879218922 0.0129692952839532
0.086622342338505
DAZARD_UV_RESPONSE_CLUSTER_G3 13 0.01 Down
0.664401724005097 0.0129885336454177 0.086622342338505
NIKOLSKY_BREAST_CANCER_20Q11_AMPLICON 27 0.01 Up
0.253011678517549 0.0130214273379207 0.0867130605688202
NUYTTEN_NIPP1_TARGETS_DN 800 0.01 Down
0.267975907203011 0.0130508667940437 0.0867805417740037
KUROZUMI_RESPONSE_TO_ONCOCYTIC_VIRUS 29 0.01 Down
0.813597763571297 0.0130798744288975 0.0868449565109218
BROWNE_HCMV_INFECTION_20HR_UP 207 0.01 Down
0.284288652782493 0.0131006131533951 0.0868543600656507
REACTOME_REGULATION_OF_RAS_BY_GAPS 66 0.01 Down
0.156893890233909 0.0131692803576236 0.0869582767080899
REACTOME_SIGNALING_BY_SCF_KIT 40 0.01 Down
0.399653404472787 0.0131711400107913 0.0869582767080899
BOWIE_RESPONSE_TO_TAMOXIFEN 18 0.01 Down
0.485744380186422 0.0131743240129498 0.0869582767080899
REACTOME_TRANSCRIPTIONAL_ACTIVATION_OF_MITOCHONDRIAL_BIOGENESIS 54
0.01 Down 0.259555121116173 0.0131999915259769
0.0869999441484843
WU_SILENCED_BY_METHYLATION_IN_BLADDER_CANCER 42 0.01 Up
0.684096569286902 0.0132238428037192
0.0870295364607876
PID_ERBB2_ERBB3_PATHWAY 41 0.01 Down 0.198581532621951
0.0132938881814019 0.087362613121932
WORSCHECH_TUMOR_REJECTION_UP 33 0.01 Down
0.76063383854412 0.0133742709707635 0.0877625518446453
NUNODA_RESPONSE_TO_DASATINIB_IMATINIB_DN 12 0.01 Down
0.421773031062715 0.0134288698312452
0.0878711344778546
KEGG_UBIQUITIN_MEDIATED_PROTEOLYSIS 127 0.01 Down
0.166320959388485 0.0134299153250915 0.0878711344778546
BIOCARTA_GSK3_PATHWAY 17 0.01 Down 0.255972580924916
0.0134529570523423 0.0878718471205591
REACTOME_REGULATION_OF_TNFR1_SIGNALING 33 0.01 Down
0.348171809975497 0.0134691218389467 0.0878718471205591
REACTOME_COHESIN_LOADING_ONTO_CHROMATIN 10 0.01 Down
0.164965204085412 0.0134984377678643 0.0879103511631031
PID_NFAT_TFPATHWAY 31 0.01 Down 0.533824385674914
0.0135141385214025 0.0879103511631031
BIOCARTA_IL7_PATHWAY 15 0.01 Down 0.461945764010319
0.0136996655537248 0.0889884344855388
CHEN_METABOLIC_SYNDROM_NETWORK 1000 0.01 Up
0.467142207360414 0.013743640392735 0.0891452576700486
REACTOME_SURFACTANT_METABOLISM 20 0.01 Up
0.809490332392404 0.0138027034671389 0.0893993545890338
KANG_FLUOROURACIL_RESISTANCE_DN 14 0.01 Down
0.38856675567824 0.0138595408739704 0.0896383255086286
REACTOME_MET_ACTIVATES_RAS_SIGNALING 10 0.01 Down
0.211217132273486 0.014002412742646l 0.0902856497887625

SCHLOSSER_SERUM_RESPONSE_DN 637 0.01 Down
0.231202964777438 0.0140086745713733 0.0902856497887625
REACTOME_BIOSYNTHESIS_OF_SPECIALIZED_PRORESOLVING_MEDIATORS_SPMS
13 0.01 Up 0.598531928899267
0.0140198851062416 0.0902856497887625
AMIT_EGF_RESPONSE_480_MCF10A 42 0.01 Down
0.389388939949994 0.0141700538221241 0.09102165329077
REACTOME_INSULIN_PROCESSING 21 0.01 Down
0.234401126455029 0.0141786398909698 0.09102165329077
TENEDINI_MEGAKARYOCYTE_MARKERS 47 0.01 Down
0.535648905003146 0.0142120659963918 0.09102165329077
SMID_BREAST_CANCER_ERBB2_UP 110 0.01 Down
0.434327067732252 0.0142151725495263 0.09102165329077
REACTOME_TNF_RECEPTOR_SUPERFAMILY_TNFSF_MEMBERS_MEDIATING_NON_CANONI
CAL_NF_KB_PATHWAY 12 0.01 Down 0.540360979644816
0.0142822408804275 0.091221729381847
COULOUARN_TEMPORAL_TGFB1_SIGNATURE_UP 106 0.01 Down
0.327319789260524 0.0142870072268788 0.091221729381847
BURTON_ADIPOGENESIS_6 166 0.01 Up 0.377633456526179
0.0143689357506008 0.091614703828299
NADERI_BREAST_CANCER_PROGNOSIS_DN 13 0.01 Up
0.546588640875308 0.014403087657307 0.0917023782147237
NAKAMURA_ADIPOGENESIS_LATE_DN 33 0.01 Up
0.822845558940684 0.0144312919368627 0.0917156235576864
GRAHAM_CML_DIVIDING_VS_NORMAL_QUIESCENT_DN 79 0.01 Down
0.605110515401108 0.0144652859214071
0.0917156235576864
KEGG_ADHERENS_JUNCTION 67 0.01 Down 0.241750548175134
0.0144663797780644 0.0917156235576864
DACOSTA_ERCC3_ALLELE_XPCS_VS_TTD_DN 29 0.01 Up
0.521783692150608 0.014513857202361 0.0918870255276237
MOOTHA_VOXPHOS 85 0.01 Up 0.126752245535098
0.0148232711782839 0.0937139296011055
YANG_BREAST_CANCER_ESR1_LASER_UP 30 0.01 Up
0.382928401558169 0.0148861639034949 0.0939793634075977
BOWIE_RESPONSE_TO_EXTRACELLULAR_MATRIX 17 0.01 Down
0.393309308594695 0.0150675494382846 0.0949910725457073
DISTECHE_ESCAPED_FROM_X_INACTIVATION 13 0.01 Down
0.241122811653815 0.0151539595258908 0.0954020281076738
REACTOME_ERKS_ARE_INACTIVATED 13 0.01 Down
0.259707515557879 0.015260908843168 0.0959409583916648
KOBAYASHI_EGFR_SIGNALING_24HR_UP 97 0.01 Up
0.406752625979934 0.0153296417887224 0.0962384634641162
BIOCARTA_EGF_PATHWAY 26 0.01 Down 0.232734103188903
0.01540124690452 5 0.0965531448198602
ZHENG_RESPONSE_TO_ARSENITE_DN 13 0.01 Down
0.543738723675145 0.0154798651622689 0.0969108550200539
REACTOME_IL_6_TYPE_CYTOKINE_RECEPTOR_LIGAND_INTERACTIONS 13
0.01 Down 0.556736403888939 0.0155780332961026
0.0973897909123522
WARTERS_IR_RESPONSE_5GY 44 0.01 Up 0.363171829919401
0.0156027496625275 0.097408832962585
ABRAHAM_ALPC_VS_MULTIPLE_MYELOMA_UP 26 0.01 Up
0.853172393940725 0.0156300276349705 0.097443792259629

REACTOME_NONSENSE_MEDIATED_DECAY_NMD_INDEPENDENT_OF_THE_EXON_JUNCTIO
N_COMPLEX_EJC 93 0.01 Up 0.120478124423366
0.015757628755274 0.098103242735397
PID_SMAD2_3PATHWAY 16 0.01 Down 0.148156388933595
0.0158784254465095 0.0986123877311205
GAURNIER_PSMD4_TARGETS 53 0.01 Down 1.08362694415633
0.015890143546898 0.0986123877311205
MASSARWEH_TAMOXIFEN_RESISTANCE_UP 530 0.01 Down
0.354642589989674 0.0159276058080503 0.0986123877311205
REACTOME_ACTIVATION_OF_RAC1 12 0.01 Down
0.310415230623214 0.0159565812663089 0.0986123877311205
GARGALOVIC_RESPONSE_TO_OXIDIZED_PHOSPHOLIPIDS_CYAN_UP 17 0.01
Down 0.194142506331441 0.0159704088207883
0.0986123877311205
ZHENG_BOUND_BY_FOXP3 391 0.01 Down 0.310910410191681
0.0159710385468867 0.0986123877311205
REACTOME_REGULATION_OF_RUNX1_EXPRESSION_AND_ACTIVITY 17 0.01
Down 0.26106860879718 0.0159983973902925
0.0986458110690876
IKEDA_MIR1_TARGETS_UP 52 0.01 Down 0.262791314085393
0.016040071146313 0.0987672874009272
DURAND_STROMA_S_UP 233 0.01 Up 0.562534652515775
0.016102886301858 0.098797509561786
MATZUK_SPERMATOGONIA 14 0.01 Up 0.562221945404945
0.0161038769020342 0.098797509561786
REACTOME_NOTCH_HLH_TRANSCRIPTION_PATHWAY 28 0.01 Down
0.230236802782245 0.0161109175770387
0.098797509561786
KEGG_B_CELL_RECEPTOR_SIGNALING_PATHWAY 66 0.01 Down
0.35767085728474 0.0163743989343259 0.0999379107743539
BIOCARTA_IL6_PATHWAY 21 0.01 Down 0.27018331933174
0.0163843512216484 0.0999379107743539
PID_KIT_PATHWAY 45 0.01 Down 0.302718511280603
0.0163887802971725 0.0999379107743539
KAAB_HEART_ATRIUM_VS_VENTRICLE_UP 203 0.01 Up
0.381567903098965 0.0164039526719193 0.0999379107743539
PID_AP1_PATHWAY 54 0.01 Down 0.534856355840747
0.0164080485320296 0.0999379107743539
AMIT_EGF_RESPONSE_40_MCF10A 16 0.01 Down
0.444683562857057 0.0164635465619121 0.100140246002429
DAZARD_UV_RESPONSE_CLUSTER_G4 17 0.01 Down
0.545311024848292 0.0165282368957735 0.100397871414192
LIU_VAV3_PROSTATE_CARCINOGENESIS_UP 63 0.01 Up
0.706936125796701 0.0171992074045207 0.104243124553561
SANA_RESPONSE_TO_IFNG_UP 70 0.01 Down
0.622254321582302 0.0172076525959382 0.104243124553561
PID_TCR_PATHWAY 53 0.01 Down 0.427448031741733
0.0173881703724958 0.105194920355813
PODAR_RESPONSE_TO_ADAPHOSTIN_UP 142 0.01 Down
0.301525740672793 0.0175400299234019 0.105971014120553
GAUSSMANN_MLL_AF4_FUSION_TARGETS_B_UP 16 0.01 Down
0.464026745700076 0.0175742004250833 0.106034940819798
REACTOME_APOPTOTIC_CLEAVAGE_OF_CELLULAR_PROTEINS 34 0.01
Down 0.241718097597785 0.0178234336028003

0.107280577164454
HATADA_METHYLATED_IN_LUNG_CANCER_UP 228 0.01 Up
0.412441916342218 0.0178588208472846 0.107280577164454
DAUER_STAT3_TARGETS_DN 47 0.01 Down 0.383591011652484
0.0178731692897833 0.107280577164454
SASAI_RESISTANCE_TO_NEOPLASTIC_TRANSFROMATION 49 0.01 Up
0.676479372757777 0.0178761184196165
0.107280577164454
REACTOME_INTRINSIC_PATHWAY_FOR_APOPTOSIS 49 0.01 Down
0.236606121616324 0.0181451007551214
0.108749637192361
REACTOME_REGULATION_OF_CHOLESTEROL_BIOSYNTHESIS_BY_SREBP_SREBF 55
0.01 Down 0.22868094120546 0.0182180787598329
0.108916540620176
ISSAEVA_MLL2_TARGETS 50 0.01 Up 0.518669618468789
0.0182214101326746 0.108916540620176
REACTOME_RUNX3_REGULATES_NOTCH_SIGNALING 14 0.01 Down
0.309003768862882 0.0183432226134154
0.109499051324438
REACTOME_INTRACELLULAR_SIGNALING_BY_SECOND_MESSENGERS 258 0.01
Down 0.265875353009794 0.0184566878736701
0.109906809338619
REACTOME_PI_METABOLISM 76 0.01 Down 0.239288917023325
0.0184604318243954 0.109906809338619
PAL_PRMT5_TARGETS_UP 187 0.01 Up 0.272413238276319
0.018654138127352 0.110913162542919
PID_IL12_2PATHWAY 41 0.01 Down 0.616598720034751
0.0187196092897864 0.11115540787
LEIN_ASTROCYTE_MARKERS 24 0.01 Up 0.605134213837632
0.0187597810872325 0.111246986790383
REACTOME_CLASS_I_MHC_MEDIATED_ANTIGEN_PROCESSING_PRESENTATION 332
0.01 Down 0.20165487543038 0.0188520706025759
0.111552021522237
VALK_AML_WITH_FLT3_ITD 27 0.01 Up 0.639208681694241
0.0188608534720578 0.111552021522237
SMID_BREAST_CANCER_NORMAL_LIKE_UP 293 0.01 Up
0.661230008507418 0.0189545370481683 0.111939029368775
KEGG_ARACHIDONIC_ACID_METABOLISM 34 0.01 Up
0.620989088409156 0.0189760935214698 0.111939029368775
REACTOME_ANTIGEN_PROCESSING:_UBIQUITINATION_PROTEASOME_DEGRADATION
276 0.01 Down 0.162214685159337
0.0190054696021388 0.111965381207882
REACTOME_SIGNALING_BY_EGFRVIII_IN_CANCER 14 0.01 Down
0.190238750100167 0.0190913018990365
0.112323824654672
REACTOME_INTERLEUKIN_4_AND_INTERLEUKIN_13_SIGNALING 81 0.01
Down 0.601771238433997 0.0192499864871499
0.112962808011458
ZHAN_MULTIPLE_MYELOMA_PR_UP 29 0.01 Up
0.291543848125183 0.0192501692851562 0.112962808011458
REACTOME_PI3K_AKT_SIGNALING_IN_CANCER 74 0.01 Down
0.414717992297593 0.0192781591863913 0.112979563941107
NAKAMURA_ADIPOGENESIS_EARLY_DN 33 0.01 Up
0.854163163584869 0.01930858990341 3 0.112990462799934

TSAI_RESPONSE_TO_IONIZING_RADIATION 128 0.01 Down
0.408095746859606 0.0193302927459732 0.112990462799934
KEGG_PYRUVATE_METABOLISM 32 0.01 Up
0.230020190387425 0.0193811594743681 0.113032399118257
REACTOME_DETOXIFICATION_OF_REACTIVE_OXYGEN_SPECIES 33 0.01
Up 0.339576332437498 0.0193877596707844
0.113032399118257
REACTOME_SUMOYLATION_OF_IMMUNE_RESPONSE_PROTEINS 11 0.01
Down 0.241955142541572 0.0194379493212742
0.113178215283844
ZHANG_INTERFERON_RESPONSE 22 0.01 Down
0.461975148421767 0.0196898474933179 0.114496590533588
REACTOME_CREB1_PHOSPHORYLATION_THROUGH_NMDA_RECEPTOR_MEDIATED_ACTIVA
TION_OF_RAS_SIGNALING 18 0.01 Down 0.278264201042018
0.0197340167981682 0.114605175074633
REACTOME_WNT5A_DEPENDENT_INTERNALIZATION_OF_FZD2_FZD5_AND_ROR2 13
0.01 Up 0.334430891986968 0.019825156412084
0.114985907190087
YAN_ESCAPE_FROM_ANOIKIS 19 0.01 Down 0.587647467122505
0.0199020080433278 0.115282894529328
PEREZ_TP63_TARGETS 286 0.01 Down 0.406399750843889
0.0200641828866894 0.116009121418302
REACTOME_COMPLEX_I_BIOGENESIS 49 0.01 Up
0.128248568527356 0.0200789981008763 0.116009121418302
REACTOME_TRANSPORT_OF_THE_SLBP_DEPENDANT_MATURE_MRNA 36 0.01
Down 0.156280221884052 0.0201705785128531
0.116388639814216
GINESTIER_BREAST_CANCER_20Q13_AMPLIFICATION_DN 174 0.01 Down
0.202778490601308 0.0202381597586843
0.116628882199085
REACTOME_TRANSPORT_OF_MATURE_MRNAS_DERIVED_FROM_INTRONLESS_TRANSCRIP
TS 43 0.01 Down 0.142939971733994
0.0203857661599247 0.117329089486379
BERTUCCI_MEDULLARY_VS_DUCTAL_BREAST_CANCER_UP 185 0.01 Down
0.331996366796582 0.0204179094108147
0.117355523312368
SIG_PIP3_SIGNALING_IN_CARDIAC_MYOCTES 60 0.01 Down
0.259714122711794 0.020442575028606 0.117355523312368
WILSON_PROTEASES_AT_TUMOR_BONE_INTERFACE_UP 19 0.01 Down
0.554626802185589 0.0205120561387836
0.117556929595076
REACTOME_FOXO_MEDIATED_TRANSCRIPTION_OF_CELL_CYCLE_GENES 15
0.01 Down 0.310758952682379 0.0205299643453025
0.117556929595076
YAO_TEMPORAL_RESPONSE_TO_PROGESTERONE_CLUSTER_5 25 0.01 Down
0.636334592107737 0.0206661270591376
0.118186057418351
REACTOME_METHYLATION 13 0.01 Up 0.448146195780986
0.0206989693315569 0.118223465241866
REACTOME_NUCLEAR_IMPORT_OF_REV_PROTEIN 34 0.01 Down
0.164689293074104 0.0207270345678272 0.11823352840404
WORSCHECH_TUMOR_EVASION_AND_TOLEROGENICITY_DN 12 0.01 Down
0.342169149057034 0.0208223477506168
0.11862668331942

KEGG_PENTOSE_PHOSPHATE_PATHWAY 22 0.01 Up
0.251238306760929 0.020859122192827 0.118685764881971
REACTOME_RESPIRATORY_ELECTRON_TRANSPORT 89 0.01 Up
0.115662834383019 0.0209469552460615 0.118907173115509
REACTOME_REGULATION_OF_MRNA_STABILITY_BY_PROTEINS_THAT_BIND_AU_RICH_ELEMENTS 85 0.01 Down 0.142587344438629
0.0209509412919874 0.118907173115509
HUPER_BREAST_BASAL_VS_LUMINAL_UP 51 0.01 Down
0.81290783274703 0.0209794652025055 0.118918910574101
DASU_IL6_SIGNALING_SCAR_DN 15 0.01 Up
0.483407932625037 0.0210248252953687 0.11893771204737
SHETH_LIVER_CANCER_VS_TXNIP_LOSS_PAM4 195 0.01 Up
0.36416992665752 0.0210357021307361 0.11893771204737
PID_ERBB1_RECEPTOR_PROXIMAL_PATHWAY 35 0.01 Down
0.214569284067998 0.0212020436422917 0.119727620819223
LEE_NEURAL_CREST_STEM_CELL_UP 93 0.01 Up
0.698127614752134 0.021385153654132 0.120610120044321
REACTOME_MITOCHONDRIAL_IRON_SULFUR_CLUSTER_BIOGENESIS 13 0.01
Up 0.169946168663523 0.0215252631896098
0.121248193029193
YIH_RESPONSE_TO_ARSENITE_C4 17 0.01 Down
0.280191854326927 0.0215873830119383 0.121359734791366
REACTOME_SIGNALING_BY_ERYTHROPOIETIN 24 0.01 Down
0.33153353057627 0.0215990629217114 0.121359734791366
CHANDRAN_METASTASIS_UP 181 0.01 Down 0.288261244688861
0.021772854009668 0.122157125904665
HAMAI_APOPTOSIS_VIA_TRAIL_UP 607 0.01 Down
0.245724538259502 0.0217953314739802 0.122157125904665
RIGGINS_TAMOXIFEN_RESISTANCE_UP 50 0.01 Up
0.630561769819122 0.021958313482951 0.122847880579073
CROMER_TUMORIGENESIS_DN 38 0.01 Down 0.805739479353208
0.0219732360368353 0.122847880579073
KRIGE_RESPONSE_TO_TOSEDOSTAT_24HR_UP 714 0.01 Down
0.285215555754898 0.0220588282304039 0.123057187378463
ZHOU_TNF_SIGNALING_4HR 51 0.01 Down 0.317576726853714
0.0220654267023451 0.123057187378463
BIOCARTA_PYK2_PATHWAY 26 0.01 Down 0.181002891673626
0.0220953480131395 0.123062189405581
MAHAJAN_RESPONSE_TO_IL1A_UP 64 0.01 Down
0.617119981675732 0.0221329374263619 0.123062189405581
WANG_BARRETTS_ESOPHAGUS_AND_ESOPHAGUS_CANCER_DN 30 0.01 Down
0.599493882972614 0.0221484563357318
0.123062189405581
NUYTTEN_EZH2_TARGETS_UP 966 0.01 Down 0.347124625889633
0.0222815317066283 0.123648746939869
REACTOME_SIGNALING_BY_FGFR3 33 0.01 Down
0.257799036981005 0.0224221226251229 0.124275513193499
GRAHAM_NORMAL_QUIESCENT_VS_NORMAL_DIVIDING_DN 66 0.01 Up
0.335511280790789 0.022465115271802
0.124360459540332
REACTOME_GLUCONEOGENESIS 26 0.01 Up
0.198385617558964 0.0225210548826179 0.124516779455556
THUM_SYSTOLIC_HEART_FAILURE_DN 167 0.01 Down
0.244011427977686 0.0227173752808979 0.12541221804851

NGUYEN_NOTCH1_TARGETS_DN 79 0.01 Down
0.321278793174653 0.0227388115037899 0.12541221804851
REACTOME_GAMMA_CARBOXYLATION_HYPUSINE_FORMATION_AND_ARYLSULFATASE_AC
TIVATION 30 0.01 Up 0.243774603809299
0.0228450992348391 0.125601222552596
REACTOME_REGULATION_OF_PTEN_STABILITY_AND_ACTIVITY 66 0.01
Down 0.130593672268874 0.0228641598760916
0.125601222552596
GYORFFY_DOXORUBICIN_RESISTANCE 32 0.01 Down
0.59336386482314 0.0228901617698265 0.125601222552596
HUMMEL_BURKITTS_LYMPHOMA_UP 37 0.01 Up
0.377130569143536 0.0228922728277326 0.125601222552596
REACTOME_LISTERIA_MONOCYTOGENES_ENTRY_INTO_HOST_CELLS 18 0.01
Down 0.171623199913012 0.022930838900134
0.125601222552596
BURTON_ADIPOGENESIS_7 44 0.01 Up 0.583872674505757
0.0229407349756799 0.125601222552596
HOLLEMAN_VINCRISTINE_RESISTANCE_B_ALL_UP 37 0.01 Down
0.234848814986464 0.0230433152608396
0.125876893774385
KEGG_PROPANOATE_METABOLISM 29 0.01 Up
0.250170452622544 0.0230731438651613 0.125876893774385
GOERING_BLOOD_HDL_CHOLESTEROL_QTL_CIS 10 0.01 Up
0.475983642451091 0.023075096878775 0.125876893774385
BROWNE_HCMV_INFECTION_30MIN_DN 95 0.01 Down
0.331004501936868 0.0232347089489307 0.126479002631966
XU_HGF_SIGNALING_NOT_VIA_AKT1_48HR_UP 33 0.01 Down
0.447340125798833 0.0232551176661907 0.126479002631966
STAMBOLSKY_TARGETS_OF_MUTATED_TP53_UP 41 0.01 Down
0.327696017432962 0.0232698854675496 0.126479002631966
WILLERT_WNT_SIGNALING 20 0.01 Up 0.393654883784358
0.0233468244474194 0.126743932235689
TSAI_DNAJB4_TARGETS_UP 13 0.01 Down 0.295756483912663
0.023476499504828 0.127294168002656
KIM_LIVER_CANCER_POOR_SURVIVAL_DN 22 0.01 Up
0.55848599081608 0.0235216777351325 0.127385471589664
ONDER_CDH1_TARGETS_3_UP 17 0.01 Up 0.538052743123279
0.0235887340619813 0.127559049795791
DE_YY1_TARGETS_UP 11 0.01 Down 0.614657941728051
0.023610484856529 0.127559049795791
REACTOME_DEFECTIVE_TPR_MAY_CONFER_SUSCEPTIBILITY_TOWARDS_THYROID_PAP
ILLARY_CARCINOMA_TPC 30 0.01 Down 0.1711892480621
0.0236459204372443 0.127597133691973
BIOCARTA_PTEN_PATHWAY 17 0.01 Down 0.198077012333178
0.0237185064773173 0.127672756304976
REACTOME_ERBB2_REGULATES_CELL_MOTILITY 11 0.01 Down
0.267383229171748 0.0237680036289931 0.127672756304976
SANA_TNF_SIGNALING_DN 71 0.01 Up 0.511125264751376
0.0237799780928008 0.127672756304976
CROONQUIST_NRAS_SIGNALING_DN 58 0.01 Up
0.252643066778214 0.0237999405075064 0.127672756304976
GRANDVAUX_IRF3_TARGETS_DN 15 0.01 Down
0.365565556251002 0.0238019510085807 0.127672756304976
KOKKINAKIS_METHIONINE_DEPRIVATION_96HR_DN 65 0.01 Down

0.257096401353603 0.0240235375563089
0.128707748886304
STONER_ESOPHAGEAL_CARCINOGENESIS_UP 33 0.01 Down
0.232195627909596 0.0240676368413768 0.128790509049987
REACTOME_CGMP_EFFECTS 10 0.01 Up 0.583602551055557
0.0241018157448558 0.128820049670781
PID_FAK_PATHWAY 57 0.01 Down 0.199796062905517
0.0241388393446213 0.128864706477521
IVANOV_MUTATED_IN_COLON_CANCER 12 0.01 Down
0.166857102211762 0.0241797334686968 0.128929895541865
COWLING_MYCN_TARGETS 33 0.01 Up 0.802164236920307
0.0243537810724333 0.129599963577636
KEGG_GLYCOSPHINGOLIPID_BIOSYNTHESIS_GANGLIO_SERIES 11 0.01
Up 0.441268977420804 0.0243630632309461
0.129599963577636
TAKAO_RESPONSE_TO_UVB_RADIATION_DN 92 0.01 Down
0.279562763642253 0.0245170992095362 0.130179810469261
LEE_LIVER_CANCER_SURVIVAL_UP 105 0.01 Up
0.465360255342145 0.0245299887580565 0.130179810469261
YAO_TEMPORAL_RESPONSE_TO_PROGESTERONE_CLUSTER_12 70 0.01
Up 0.260441730311882 0.0246992391996284
0.130923443634823
FARMER_BREAST_CANCER_CLUSTER_3 16 0.01 Down
0.34279252424378 0.0247814284743227 0.13120438279397
REACTOME_THE_CITRIC_ACID_TCA_CYCLE_AND_RESPIRATORY_ELECTRON_TRANSPOR
T 154 0.01 Up 0.124791002460947
0.0248538747369526 0.131433137579532
REACTOME_EXPORT_OF_VIRAL_RIBONUCLEOPROTEINS_FROM_NUCLEUS 33
0.01 Down 0.167778216976183 0.0249434095047292
0.131642996461158
REACTOME_ACTIVATION_OF_GENE_EXPRESSION_BY_SREBF_SREBP 42 0.01
Down 0.245157776424218 0.0249521319210025
0.131642996461158
PID_ERBB4_PATHWAY 32 0.01 Down 0.305961616820955
0.0250077340198914 0.1317816698938
FLECHNER_PBL_KIDNEY_TRANSPLANT_OK_VS_DONOR_DN 39 0.01 Down
0.285310892285335 0.025064367038696
0.131925444776275
BIOCARTA_TID_PATHWAY 15 0.01 Down 0.30155012575093
0.0253395012813414 0.133217611999567
REACTOME_INTERACTIONS_OF_REV_WITH_HOST_CELLULAR_PROTEINS 37
0.01 Down 0.162687203403482 0.0254072935926308
0.133309255101033
REACTOME_NOTCH2_INTRACELLULAR_DOMAIN_REGULATES_TRANSCRIPTION 10
0.01 Down 0.429788593052854 0.0254930240739071
0.133309255101033
AFFAR_YY1_TARGETS_DN 191 0.01 Up 0.51810848441551
0.0254962562797368 0.133309255101033
KEGG_PROTEASOME 41 0.01 Down 0.119535637718637
0.0254977252140693 0.133309255101033
DE_YY1_TARGETS_DN 84 0.01 Down 0.235949767864686
0.0255052189959707 0.133309255101033
EINAV_INTERFERON_SIGNATURE_IN_CANCER 27 0.01 Down
0.358547516878714 0.0257276018706455 0.134315412785774

INGRAM_SHH_TARGETS_DN	52	0.01	Up	0.632346158751284
	0.0258659726055833	0.134881144851621
REACTOME_HOST_INTERACTIONS_WITH_INFLUENZA_FACTORS	43	0.01
	Down	0.162525564538312	0.0259074514060611
0.134940896952775
SCHEIDEREIT_IKK_INTERACTING_PROTEINS	51	0.01	Down
0.378739149672431	0.0259445991191275	0.134977978056109
LEE_AGING_MUSCLE_UP	29	0.01	Up	0.340191929251139
	0.0262511351129338	0.136263202749265
HILLION_HMGA1_TARGETS	63	0.01	Up	0.337845615294442
	0.0262522655352 31	0.136263202749265
WORSCHECH_TUMOR_EVASION_AND_TOLEROGENICITY_UP	19	0.01	Down
	0.67751900084649	0.0263447826861878
0.1365856957029
KEGG_LIMONENE_AND_PINENE_DEGRADATION	10	0.01	Up
0.297823849849556	0.0265995772089221	0.137735363807784
PEDERSEN_METASTASIS_BY_ERBB2_ISOFORM_4	92	0.01	Down
0.467554121843055	0.0266278156059988	0.137735363807784
REACTOME_TRANSCRIPTIONAL_REGULATION_BY_RUNX1	171	0.01	Down
	0.208083477070018	0.0267897360808627
0.138220704113871
LIU_SOX4_TARGETS_UP	131	0.01	Down	0.237789187235006
	0.0268011693949637	0.138220704113871
BIOCARTA_TALL1_PATHWAY	12	0.01	Down	0.261033296431044
	0.0268138941017342	0.138220704113871
REACTOME_DOWNSTREAM_TCR_SIGNALING	88	0.01	Down
0.302213726868344	0.0268852740839708	0.138429904934076
SCHLINGEMANN_SKIN_CARCINOGENESIS_TPA_DN	16	0.01	Up
0.790504572259644	0.0270294383697774	0.139012958206121
REACTOME_IKK_COMPLEX_RECRUITMENT_MEDIATED_BY_RIP1	22	0.01
	Down	0.305991928771516	0.0277011115550571
0.142115050028509
BOYERINAS_ONCOFETAL_TARGETS_OF_LET7A1	10	0.01	Down
0.262188157727834	0.0277180070553313	0.142115050028509
PID_TELOMERASE_PATHWAY	62	0.01	Down	0.176503948226834
	0.0277274524749727	0.142115050028509
REACTOME_UB_SPECIFIC_PROCESSING_PROTEASES	170	0.01	Down
	0.186434500484802	0.027855148048578
0.142493397933475
REACTOME_TRANSCRIPTIONAL_REGULATION_OF_WHITE_ADIPOCYTE_DIFFERENTIATI
ON	74	0.01	Down	0.28009376386349
0.0278646711420522	0.142493397933475
CHEN_HOXA5_TARGETS_6HR_UP	10	0.01	Down
0.496051091807963	0.0279435921442979	0.142734598509795
BIOCARTA_PCAF_PATHWAY	12	0.01	Down	0.24486216348469
	0.0282479859803238	0.144125649241266
RODWELL_AGING_KIDNEY_UP	423	0.01	Up	0.535845984743135
	0.0283842164630545	0.144656522677358
SHAFFER_IRF4_TARGETS_IN_PLASMA_CELL_VS_MATURE_B_LYMPHOCYTE	57
	0.01	Down	0.346490078952835	0.0284951532493124
	0.145057433585118
LEE_LIVER_CANCER_ACOX1_DN	34	0.01	Up
0.559669233313769	0.0285724664934254	0.145286467067814
ONDER_CDH1_TARGETS_1_UP	135	0.01	Up	0.357346500203756

0.0286551872582837	0.145542448277949
PID_SMAD2_3NUCLEAR_PATHWAY	69	0.01	Down
0.271550729944785	0.0287119092446589	0.145665950400386
STEARMAN_TUMOR_FIELD_EFFECT_UP	28	0.01	Up
0.836390682230201	0.0288081583397088	0.145989483356247
REACTOME_APOPTOTIC_FACTOR_MEDIATED_RESPONSE	15	0.01	Down
0.2170971956348 0.0288566535571968	0.146070560517567
DACOSTA_UV_RESPONSE_VIA_ERCC3_TTD_DN	82	0.01	Down
0.27074256469804	0.0289151164482115	0.146201854257267
GARGALOVIC_RESPONSE_TO_OXIDIZED_PHOSPHOLIPIDS_BLUE_DN	56	0.01
Up	0.331077618063136	0.0289945242800195
0.146438636672683
CAVARD_LIVER_CANCER_MALIGNANT_VS_BENIGN 25	0.01	Up
0.502878830822971	0.0290577027563891	0.146593012222187
BEGUM_TARGETS_OF_PAX3_FOXO1_FUSION_DN	42	0.01	Up
0.420429328222111	0.0291278652425201	0.146704420563346
VERRECCHIA_EARLY_RESPONSE_TO_TGFB1	53	0.01	Up
0.671202797552495	0.029145060636945	0.146704420563346
REACTOME_NUCLEOBASE_BIOSYNTHESIS	15	0.01	Up
0.192228118068013	0.0292341386624043	0.146988202782447
BIOCARTA_TCR_PATHWAY	39	0.01	Down	0.341251723875943
0.0292992588443772	0.147151026263101
CHEBOTAEV_GR_TARGETS_DN 90	0.01	Up	0.582910832729044
0.0293429923792517	0.147206195027608
SMITH_TERT_TARGETS_DN	82	0.01	Up	0.434197126381698
0.0294992719232079	0.147825225523768
REACTOME_HDMS_DEMETHYLATE_HISTONES	28	0.01	Down
0.213007729006043	0.0295904852600292	0.148003021328057
PID_CXCR3_PATHWAY	36	0.01	Down	0.404910629825698
0.0296006042656113	0.148003021328057
SIG_BCR_SIGNALING_PATHWAY	40	0.01	Down
0.248507841671957	0.0298314719947273	0.148991629573666
MINGUEZ_LIVER_CANCER_VASCULAR_INVASION_DN	10	0.01	Up
0.386663923281826	0.0298673249962996
0.149005134138032
WONG_EMBRYONIC_STEM_CELL_CORE	313	0.01	Up
0.167003819302206	0.0299041226520887	0.149023316320553
LEE_LIVER_CANCER_DENA_DN	37	0.01	Up
0.555273519870095	0.0299528260325963	0.149100723163367
REACTOME_CELL_CELL_COMMUNICATION	103	0.01	Down
0.393543409322355	0.0299909534771686	0.149125371548532
LIU_LIVER_CANCER	28	0.01	Up	0.552786921832435
0.0300582063098791	0.149294626920339
PID_CDC42_PATHWAY	63	0.01	Down	0.185646381807956
0.0301639066840471	0.149649358902787
MOROSETTI_FACIOSCAPULOHUMERAL_MUSCULAR_DISTROPHY_DN	10	0.01
Up	0.86264795109546	0.0301962110177593
0.149649358902787
PID_RET_PATHWAY 35	0.01	Down	0.224867476275416
0.0302412289292617	0.149687716720475
REACTOME_TNF_SIGNALING	42	0.01	Down	0.301807983651826
0.0302705527250082	0.149687716720475
PID_ERBB1_INTERNALIZATION_PATHWAY	39	0.01	Down
0.179028168642703	0.0303248608387715	0.149791482934371

REACTOME_HS_GAG_BIOSYNTHESIS 23 0.01 Up
0.490291219300798 0.0303651103520307 0.149825654261666
REACTOME_SRP_DEPENDENT_COTRANSLATIONAL_PROTEIN_TARGETING_TO_MEMBRANE
110 0.01 Up 0.123750479714261
0.0306805064567077 0.151215873380374
TORCHIA_TARGETS_OF_EWSR1_FLI1_FUSION_TOP20_UP 14 0.01 Down
0.97029641574029 0.0308202061362606
0.151622609395009
REACTOME_REGULATION_OF_FOXO_TRANSCRIPTIONAL_ACTIVITY_BY_ACETYLATION
10 0.01 Down 0.30219408304508
0.0308304927668606 0.151622609395009
DODD_NASOPHARYNGEAL_CARCINOMA_UP 1295 0.01 Down
0.385635646149134 0.0309521198730665 0.151902902073445
BASSO_CD40_SIGNALING_UP 90 0.01 Down 0.583772970898348
0.0309550741488934 0.151902902073445
MOOTHA_MITOCHONDRIA 416 0.01 Up 0.168078970936561
0.0311165517852215 0.152528789830502
BIOCARTA_RANKL_PATHWAY 12 0.01 Down 0.390564539881694
0.0312252753850626 0.152830214127541
SENGUPTA_NASOPHARYNGEAL_CARCINOMA_WITH_LMP1_UP 340 0.01 Down
0.386360473276877 0.0312460437782448
0.152830214127541
HAHTOLA_CTCL_PATHOGENESIS 10 0.01 Down
1.56655457174218 0.0313267103382821 0.153054447104887
REACTOME_SUMOYLATION_OF_UBIQUITINYLATION_PROTEINS 39 0.01
Down 0.17105181445431 0.0313805747591958
0.153054447104887
REACTOME_DEGRADATION_OF_BETA_CATENIN_BY_THE_DESTRUCTION_COMPLEX 81
0.01 Down 0.129826731424047 0.0313940378711247
0.153054447104887
KEGG_PPAR_SIGNALING_PATHWAY 45 0.01 Up
0.424272133996651 0.0314904829125204 0.15331472148810 5
PID_ATF2_PATHWAY 49 0.01 Down 0.396303045375771
0.031532671746684 0.15331472148810 5
COATES_MACROPHAGE_M1_VS_M2_UP 71 0.01 Up
0.525951483491832 0.0315504161414787 0.153314721488105
BIOCARTA_LAIR_PATHWAY 11 0.01 Down 0.942100995452708
0.0315838558616207 0.153314721488105
VALK_AML_CLUSTER_6 31 0.01 Up 0.604954665787569
0.0316398383501905 0.153420791137116
BIOCARTA_AT1R_PATHWAY 24 0.01 Down 0.185568815984043
0.0316766357541919 0.153433704434367
BIOCARTA_SPRY_PATHWAY 17 0.01 Down 0.20134696890229
0.0318831957769201 0.154240586743636
PID_AVB3_OPN_PATHWAY 31 0.01 Down 0.457703077680805
0.0319385880492327 0.154240586743636
CHANDRAN_METASTASIS_TOP50_UP 36 0.01 Down
0.194700300590539 0.0319461593455672 0.154240586743636
REACTOME_PEROXISOMAL_PROTEIN_IMPORT 55 0.01 Up
0.248338351093031 0.0320072147238966 0.154326051765087
SMID_BREAST_CANCER_RELAPSE_IN_PLEURA_DN 19 0.01 Down
1.0286326253714 0.0320325264286598 0.154326051765087
REACTOME_MAP2K_AND_MAPK_ACTIVATION 36 0.01 Down
0.23439076560807 3 0.032185164559678 4 0.154895411879823

BIOCARTA_RACCYCD_PATHWAY 26 0.01 Down
0.195560107707342 0.0322571935204708 0.155076026603761
KEGG_AUTOIMMUNE_THYROID_DISEASE 23 0.01 Up
0.887394598073384 0.0323503457213479 0.155357696599849
GENTILE_RESPONSE_CLUSTER_D3 51 0.01 Down
0.19083089567283 0.0324239771034536 0.15554511961582
XU_HGF_SIGNALING_NOT_VIA_AKT1_48HR_DN 14 0.01 Up
0.584498183573107 0.0324800311677397 0.155647910553294
REACTOME_MUSCLE_CONTRACTION 109 0.01 Up
0.499143654791024 0.0331797752714286 0.158831831570896
VANTVEER_BREAST_CANCER_METASTASIS_UP 57 0.01 Up
0.413886240023093 0.0332631793801537 0.158955715001259
RAY_ALZHEIMERS_DISEASE 11 0.01 Down 1.28740905801729
0.0332763799368598 0.158955715001259
REACTOME_SYNTHESIS_OF_PROSTAGLANDINS_PG_AND_THROMBOXANES_TX 11
0.01 Up 0.602735998482811 0.0339018653330169
0.161771639779099
PARK_TRETINOIN_RESPONSE 11 0.01 Up 0.857707750846269
0.0340030684661836 0.162082494968712
WANG_SMARCE1_TARGETS_UP 255 0.01 Up 0.61115126581969
0.0340958429780064 0.162265376678292
REACTOME_INTERFERON_GAMMA_SIGNALING 78 0.01 Down
0.473900872776064 0.0341701071824851 0.162265376678292
PID_SHP2_PATHWAY 48 0.01 Down 0.425441418138524
0.0341881757915005 0.162265376678292
ENK_UV_RESPONSE_EPIDERMIS_DN 483 0.01 Down
0.348421169012224 0.0342520700130208 0.162265376678292
HUANG_FOXA2_TARGETS_DN 36 0.01 Up 0.692987358339445
0.0343044254412497 0.162265376678292
BIOCARTA_IGF1MTOR_PATHWAY 18 0.01 Down
0.197460982265308 0.0343476739071902 0.162265376678292
BURTON_ADIPOGENESIS_12 31 0.01 Down 0.201081087399686
0.0343517948344637 0.162265376678292
REACTOME_PHOSPHORYLATION_SITE_MUTANTS_OF_CTNNB1_ARE_NOT_TARGETED_TO_
THE_PROTEASOME_BY_THE_DESTRUCTION_COMPLEX 15 0.01 Down
0.163646433280531 0.0343631170252024
0.162265376678292
REACTOME_ANCHORING_FIBRIL_FORMATION 15 0.01 Up
0.766058089589354 0.0343663267180721 0.162265376678292
ALONSO_METASTASIS_UP 177 0.01 Up 0.247060350973078
0.0344626504912774 0.162549437521817
RUIZ_TNC_TARGETS_UP 147 0.01 Up 0.471995932707412
0.0345845239268236 0.162581768824764
TIAN_TNF_SIGNALING_NOT_VIA_NFKB 19 0.01 Down
0.437062574545244 0.0345952480286787 0.162581768824764
ALONSO_METASTASIS_EMT_UP 30 0.01 Up
0.51618859839567 0.0346184117900117 0.162581768824764
DAVICIONI_TARGETS_OF_PAX_FOXO1_FUSIONS_DN 42 0.01 Down
0.448839701839091 0.0346491515022255
0.162581768824764
GAUSSMANN_MLL_AF4_FUSION_TARGETS_C_UP 142 0.01 Down
0.291656485657292 0.0346503525103723 0.162581768824764
BIOCARTA_PPARA_PATHWAY 43 0.01 Down 0.286208743086992
0.0353221963060304 0.165400470348627

REACTOME_NETRIN_1_SIGNALING 38 0.01 Down
0.363939996971252 0.0353246833225099 0.165400470348627
AMIT_SERUM_RESPONSE_40_MCF10A 31 0.01 Down
0.573643250960398 0.03541944012054 0.165671574757365
KEGG_MAPK_SIGNALING_PATHWAY 208 0.01 Down
0.350267292383255 0.0355023559692709 0.165886788026895
REACTOME_SHC1_EVENTS_IN_ERBB4_SIGNALING 10 0.01 Down
0.29021309053122 0.0358330985525815 0.167258336442216
REACTOME_DOWNREGULATION_OF_SMAD2_3:SMAD4_TRANSCRIPTIONAL_ACTIVITY
23 0.01 Down 0.178972218612281
0.0360729734192288 0.168157855480011
POMEROY_MEDULLOBLASTOMA_DESMOPLASIC_VS_CLASSIC_DN 52 0.01
Up 0.487728952366567 0.0361006297081669
0.168157855480011
RIGGINS_TAMOXIFEN_RESISTANCE_DN 201 0.01 Down
0.369297659076789 0.0361826433648625 0.16836540571952
WIKMAN_ASBESTOS_LUNG_CANCER_UP 16 0.01 Down
0.338115382060802 0.0362831068196243 0.168658288680674
BIOCARTA_MTOR_PATHWAY 21 0.01 Down 0.152316515910859
0.0364485275337547 0.169252201719243
REACTOME_P75_NTR_RECEPTOR_MEDIATED_SIGNALLING 90 0.01 Down
0.257545730026958 0.0366227025030529
0.169692110781728
BENPORATH_ES_CORE_NINE_CORRELATED 92 0.01 Down
0.244183475593851 0.036628272641442 0.169692110781728
REACTOME_SIGNALING_BY_ERBB2 44 0.01 Down
0.247745552564938 0.0366565160331608 0.169692110781728
BIOCARTA_IL2_PATHWAY 20 0.01 Down 0.359455274497388
0.0367211249090442 0.169816313236784
NIKOLSKY_BREAST_CANCER_19Q13.1_AMPLICON 20 0.01 Down
0.20687054916105 0.036842083134926 0.170200579333497
BAELDE_DIABETIC_NEPHROPATHY_UP 60 0.01 Up
0.6312888369498 0.0369115212400024 0.170346291554221
GARY_CD5_TARGETS_DN 417 0.01 Down 0.183628692522676
0.0370605236868638 0.170858516894824
PID_HDAC_CLASSI_PATHWAY 62 0.01 Down 0.206066178950296
0.0372458936852527 0.171438876025231
RICKMAN_HEAD_AND_NECK_CANCER_F 22 0.01 Up
0.695993767960749 0.0372626878479756 0.171438876025231
BIOCARTA_HSP27_PATHWAY 12 0.01 Down 0.37132010974217
0.0374975638278339 0.172316468358702
ZWANG_EGF_INTERVAL_UP 60 0.01 Down 0.36320028802244
0.0375301051219509 0.172316468358702
PRAMOONJAGO_SOX4_TARGETS_DN 45 0.01 Up
0.240540361715672 0.037696754268914 0.172905010651804
KORKOLA_TERATOMA 30 0.01 Down 0.608874505860469
0.0378217707209665 0.173135111218605
TIEN_INTESTINE_PROBIOTICS_24HR_DN 211 0.01 Down
0.224922883940417 0.037823955331851 0.173135111218605
REACTOME_METABOLISM_OF_VITAMINS_AND_COFACTORS 152 0.01 Up
0.336463038947614 0.0378661692522332
0.173152015044546
CAFFAREL_RESPONSE_TO_THC_UP 31 0.01 Down
0.216949225753114 0.0379955320503132 0.173553458107666

ZHANG_ANTIVIRAL_RESPONSE_TO_RIBAVIRIN_DN 39 0.01 Down
0.312610741086487 0.038031180475206
0.173553458107666
WANG_TARGETS_OF_MLL_CBP_FUSION_DN 42 0.01 Up
0.219830455876249 0.0381928542014951 0.174095008229538
REACTOME_TNFR2_NON_CANONICAL_NF_KB_PATHWAY 85 0.01 Down
0.24416922098341 0.0383070334476166
0.174095008229538
NAGASHIMA_NRG1_SIGNALING_DN 47 0.01 Down
0.216373156115188 0.03831851133209 0.174095008229538
ISHIDA_E2F_TARGETS 46 0.01 Up 0.224838472779301
0.0383212384213475 0.174095008229538
REACTOME_SUMOYLATION_OF_SUMOYLATION_PROTEINS 35 0.01 Down
0.156878100269014 0.0383648928943757
0.174095008229538
REACTOME_SIGNALING_BY_FGFR3_FUSIONS_IN_CANCER 10 0.01 Down
0.173118482036928 0.0384107460709771
0.174095008229538
WEST_ADRENOCORTICAL_TUMOR_MARKERS_DN 16 0.01 Up
1.0352596030739 0.0384209673334154 0.174095008229538
ABDULRAHMAN_KIDNEY_CANCER_VHL_DN 12 0.01 Up
0.417406231445259 0.0384707296216757 0.174144944259247
BILANGES_SERUM_SENSITIVE_VIA_TSC1 12 0.01 Down
0.36809267246159 0.0386866276468022 0.174946067678447
DAVICIONI_MOLECULAR_ARMS_VS_ERMS_DN 151 0.01 Up
0.444853402700368 0.0388657530979105 0.17557945746242
KEGG_INTESTINAL_IMMUNE_NETWORK_FOR_IGA_PRODUCTION 28 0.01
Up 0.91860433379271 0.0390454870488398
0.176214321570818
KONG_E2F3_TARGETS 71 0.01 Up 0.259598594859255
0.0391164485473238 0.176252618239622
REACTOME_APOPTOTIC_EXECUTION_PHASE 44 0.01 Down
0.230336619503956 0.0391868318976776 0.176252618239622
REACTOME_SMAD2_SMAD3:SMAD4_HETEROTRIMER_REGULATES_TRANSCRIPTION 31
0.01 Down 0.162419802717213 0.0392039828709703
0.176252618239622
REACTOME_NEGATIVE_REGULATION_OF_THE_PI3K_AKT_NETWORK 82 0.01
Down 0.407000333782431 0.0392457326376837
0.176252618239622
GENTILE_UV_LOW_DOSE_DN 61 0.01 Down 0.362426997666851
0.0392500268871772 0.176252618239622
RASHI_RESPONSE_TO_IONIZING_RADIATION_1 35 0.01 Down
0.465784951490022 0.0393626744782806 0.176582057664542
XU_GH1_AUTOCRINE_TARGETS_DN 116 0.01 Down
0.357523412446024 0.0394046826780971 0.176594265840525
TURASHVILI_BREAST_DUCTAL_CARCINOMA_VS_DUCTAL_NORMAL_UP 39 0.01
Up 0.814960662447984 0.0394640552211365
0.176684191453196
LIAN_LIPA_TARGETS_6M 56 0.01 Up 1.03763254982252
0.039565919067998 0.176944318015587
HUMMERICH_BENIGN_SKIN_TUMOR_UP 11 0.01 Down
0.747521455640028 0.0396008863011526 0.176944318015587
NELSON_RESPONSE_TO_ANDROGEN_UP 79 0.01 Down
0.339983132626394 0.0397266995353002 0.17733020299024З

REACTOME_NF_KB_IS_ACTIVATED_AND_SIGNALS_SURVIVAL 13 0.01
Down 0.25166260792489 0.039800524031681
0.177483487621435
KORKOLA_SEMINOMA_UP 38 0.01 Up 0.251914256586576
0.039845321376404 0.177507155190224
REACTOME_BLOOD_GROUP_SYSTEMS_BIOSYNTHESIS 16 0.01 Down
0.403373179588301 0.0398990336052668
0.177570451540272
GOZGIT_ESR1_TARGETS_UP 121 0.01 Up 0.456150661517557
0.0400794658188568 0.178197031509161
MOLENAAR_TARGETS_OF_CCND1_AND_CDK4_DN 45 0.01 Up
0.337411729494261 0.0402984263367682 0.178993504331792
WANG_RECURRENT_LIVER_CANCER_DN 14 0.01 Down
0.287279479465896 0.0406622309024623 0.180136318524568
KEGG_VEGF_SIGNALING_PATHWAY 65 0.01 Down
0.30933412490446 0.0407117198199944 0.180136318524568
SATO_SILENCED_EPIGENETICALLY_IN_PANCREATIC_CANCER 30 0.01
Up 0.706979372403779 0.0407182806281169
0.180136318524568
REACTOME_EFFECTS_OF_PIP2_HYDROLYSIS 21 0.01 Up
0.311440548927137 0.0407255263003188 0.180136318524568
KEGG_PEROXISOME 68 0.01 Up 0.238901421345148
0.0407560925338122 0.180136318524568
REACTOME_VXPX_CARGO_TARGETING_TO_CILIUM 17 0.01 Down
0.196926371008636 0.0408899462553156 0.180550401196114
REACTOME_RORA_ACTIVATES_GENE_EXPRESSION 18 0.01 Down
0.276226926860372 0.0410332216673538 0.181005231987002
REACTOME_DOWNREGULATION_OF_ERBB2_SIGNALING 23 0.01 Down
0.225377383356875 0.0413503563447583
0.181925646037388
PID_NFAT_3PATHWAY 51 0.01 Down 0.269097418838796
0.0414458630293807 0.181925646037388
SCHAEFFER_PROSTATE_DEVELOPMENT_6HR_DN 421 0.01 Down
0.22274947566152 0.0414559266366762 0.181925646037388
IYENGAR_RESPONSE_TO_ADIPOCYTE_FACTORS 10 0.01 Down
0.792421905999353 0.0414850735153996 0.181925646037388
HONRADO_BREAST_CANCER_BRCA1_VS_BRCA2 17 0.01 Up
0.429945498406764 0.0415205304835945 0.181925646037388
YIH_RESPONSE_TO_ARSENITE_C3 26 0.01 Up
0.360117587915179 0.0415238370778275 0.181925646037388
IGARASHI_ATF4_TARGETS_DN 92 0.01 Down
0.321248430964792 0.0415251863925162 0.181925646037388
REACTOME_TRANSPORT_TO_THE_GOLGI_AND_SUBSEQUENT_MODIFICATION 161
0.01 Down 0.225250328318136 0.0416179753791498
0.182064910875418
REACTOME_INTERLEUKIN_2_FAMILY_SIGNALING 34 0.01 Down
0.444316260491193 0.041637981842031 0.182064910875418
LIU_TARGETS_OF_VMYB_VS_CMYB_DN 37 0.01 Down
0.401393485345831 0.0417333828236052 0.182304718942765
BOYAULT_LIVER_CANCER_SUBCLASS_G3_DN 36 0.01 Up
0.46771845553886 0.0418073273262377 0.182450423622756
REACTOME_MITOCHONDRIAL_TRANSLATION 94 0.01 Up
0.110354921729002 0.0418664509211776 0.182531228797957
NIKOLSKY_BREAST_CANCER_15Q26_AMPLICON 18 0.01 Down

0.274776165389656	0.0419719685970064	0.182813952367775
REACTOME_MAP3K8_TPL2_DEPENDENT_MAPK1_3_ACTIVATION	16	0.01
	Down	0.215346942131786	0.0420370830093889
0.182920317644921
REACTOME_ESR_MEDIATED_SIGNALING	163	0.01	Down
0.278672597427239	0.0421168661578885	0.183090245048074
SA_TRKA_RECEPTOR	16	0.01	Down	0.183597546117971
	0.0422606975381288	0.183538005250134
ELVIDGE_HIF1A_AND_HIF2A_TARGETS_DN	102	0.01	Up
0.438318375913622	0.0423815687397814	0.183885281356484
REACTOME_TRANSCRIPTIONAL_REGULATION_BY_RUNX3	95	0.01	Down
	0.229665396265952	0.0425298082681764
0.184350518963793
REACTOME_REGULATION_OF_RUNX2_EXPRESSION_AND_ACTIVITY	66	0.01
	Down	0.194537386603833	0.0426070673703604
0.184363932573381
VILLANUEVA_LIVER_CANCER_KRT19_DN	41	0.01	Up
0.332490301904034	0.0426149334691307	0.184363932573381
KEGG_ENDOMETRIAL_CANCER	48	0.01	Down	0.199070654340677
	0.0426862030054464	0.184494694720655
FERREIRA_EWINGS_SARCOMA_UNSTABLE_VS_STABLE_DN	76	0.01	Up
	0.261367958704671	0.042924414638861
0.185030480980078
REACTOME_METABOLISM_OF_POLYAMINES	56	0.01	Down
0.156596953168677	0.0429379111741254	0.185030480980078
REACTOME_PURINE_RIBONUCLEOSIDE_MONOPHOSPHATE_BIOSYNTHESIS	12
	0.01	Up	0.179754081639167	0.0430108853090481
	0.185030480980078
ROSS_AML_OF_FAB_M7_TYPE	52	0.01	Up	0.421871995410087
	0.0430285614240976	0.185030480980078
PAPASPYRIDONOS_UNSTABLE_ATEROSCLEROTIC_PLAQUE_UP	52	0.01
	Up	0.616615256590402	0.0430345433976606
0.185030480980078
BIOCARTA_P38MAPK_PATHWAY	34	0.01	Down
0.167525971206384	0.0430571486329614	0.185030480980078
KEGG_ONE_CARBON_POOL_BY_FOLATE	16	0.01	Up
0.227476573532532	0.0431382319465253	0.185201864947117
REACTOME_TGF_BETA_RECEPTOR_SIGNALING_ACTIVATES_SMADS	32	0.01
	Down	0.187570361107343	0.0432117588261108
0.185340511377259
KIM_BIPOLAR_DISORDER_OLIGODENDROCYTE_DENSITY_CORR_DN	39	0.01
	Down	0.719015002593127	0.0434582427944597
0.186220020363295
SEIDEN_ONCOGENESIS_BY_MET	86	0.01	Down
0.205185884375681	0.0436061605285795	0.186675896739014
WANG_SMARCE1_TARGETS_DN	339	0.01	Down	0.377795683627836
	0.0437325580914445	0.187038866432962
BIOCARTA_IL2RB_PATHWAY	34	0.01	Down	0.320519436991433
	0.0438682682561939	0.187440937083262
PIONTEK_PKD1_TARGETS_DN	12	0.01	Up	0.594087195046564
	0.0441323956710937	0.1883904259654
PILON_KLF1_TARGETS_DN	1900	0.01	Down	0.215645998413663
	0.0442436787427695	0.188680423775172
CASORELLI_APL_SECONDARY_VS_DE_NOVO_UP	36	0.01	Up

0.498437935244274	0.0442842818871649	0.188680423775172
TURASHVILI_BREAST_LOBULAR_CARCINOMA_VS_DUCTAL_NORMAL_DN	75	0.01
	Down	0.501452353296959	0.0443507597585778
0.188784720752658
WESTON_VEGFA_TARGETS	80	0.01	Up	0.801101143383834
	0.0444028985206301	0.188827841863985
REACTOME_N_GLYCAN_ANTENNAE_ELONGATION	12	0.01	Down
0.384863027491557	0.0446806139879051	0.18982926264237б
REACTOME_ACYL_CHAIN_REMODELLING_OF_PI	10	0.01	Up
0.626178035413811	0.044732302806318	0.189869406151463
REACTOME_DOWNSTREAM_SIGNALING_OF_ACTIVATED_FGFR3	18	0.01
	Down	0.313578803941877	0.0449785826323363
0.190729109056564
REACTOME_ENDOGENOUS_STEROLS	16	0.01	Up
0.463305136333518	0.0450197073879899	0.190729109056564
REACTOME_INTERACTIONS_OF_VPR_WITH_HOST_CELLULAR_PROTEINS	37
	0.01	Down	0.182735610551541	0.0451713770774991
	0.191191468892051
REACTOME_HYALURONAN_METABOLISM	12	0.01	Up
0.463454787463155	0.0459110602701865	0.194139431716358
REACTOME_N_GLYCAN_ANTENNAE_ELONGATION_IN_THE_MEDIAL_TRANS_GOLGI	20
	0.01	Down	0.459484720893017	0.0461453285293553
	0.194857268621988
DUTERTRE_ESTRADIOL_RESPONSE_24HR_UP	269	0.01	Up
0.292609021648073	0.0461675174821841	0.194857268621988
NIKOLSKY_BREAST_CANCER_8P12_P11_AMPLICON	41	0.01	Down
	0.345686446109801	0.0462523811804925
0.195032320268587
BIOCARTA_TGFB_PATHWAY	19	0.01	Down	0.196922323695676
	0.0463635079062696	0.195317683260246
KEGG_ALDOSTERONE_REGULATED_SODIUM_REABSORPTION	28	0.01	Down
	0.417962041840013	0.046421122293349
0.195377289053
KEGG_TGF_BETA_SIGNALING_PATHWAY	68	0.01	Down
0.374150353063768	0.0466437800796183	0.196130768435813
KEGG_PANCREATIC_CANCER	68	0.01	Down	0.264269159305898
	0.0467969265726978	0.196590827050726
REACTOME_ACTIVATION_OF_NMDA_RECEPTORS_AND_POSTSYNAPTIC_EVENTS	57
	0.01	Down	0.207301861380091	0.0468621704503844
	0.196681098202127
BIOCARTA_LIS1_PATHWAY	15	0.01	Down	0.21026188789954
	0.0469426136396206	0.196834933125088
REACTOME_INTERLEUKIN_37_SIGNALING	19	0.01	Down
0.283204187513384	0.0469915688348971	0.196856572146191
LI_INDUCED_T_TO_NATURAL_KILLER_UP	255	0.01	Down
0.425795576743248	0.0471015464695551	0.197034243362966
LE_NEURONAL_DIFFERENTIATION_DN	15	0.01	Up
0.407488660582799	0.0471216488576615	0.197034243362966
PID_RHOA_REG_PATHWAY	42	0.01	Down	0.327601728549082
	0.0472256569936095	0.197285620990962
REACTOME_UPTAKE_AND_ACTIONS_OF_BACTERIAL_TOXINS	20	0.01	Down
	0.253911039715961	0.047272041805592
0.197296033348316
GINESTIER_BREAST_CANCER_ZNF217_AMPLIFIED_UP	66	0.01	Down

0.175033351427647 0.0473549250677326
0.197458616121946
BIOCARTA_NFAT_PATHWAY 38 0.01 Down 0.325485900024144
0.0474937095977467 0.197853776313134
TURASHVILI_BREAST_NORMAL_DUCTAL_VS_LOBULAR_UP 65 0.01 Down
0.4208046300461 0.0475523830057556 0.197887733927651
NIKOLSKY_BREAST_CANCER_20Q12_Q13_AMPLICON 107 0.01 Down
0.350736384529099 0.0475899088711436
0.197887733927651
RIZ_ERYTHROID_DIFFERENTIATION_HEMGN 16 0.01 Down
0.418199736389777 0.0476833615844564 0.19809307793173
REACTOME_METABOLIC_DISORDERS_OF_BIOLOGICAL_OXIDATION_ENZYMES 25
0.01 Up 0.577845825565349 0.048166713326442
0.1997778234366
AZARE_NEOPLASTIC_TRANSFORMATION_BY_STAT3_DN 17 0.01 Down
0.594658615872951 0.0481777887887152
0.1997778234366
RUTELLA_RESPONSE_TO_HGF_VS_CSF2RB_AND_IL4_DN 225 0.01 Down
0.337259923287007 0.0485107279233504
0.200973015682451
KIM_BIPOLAR_DISORDER_OLIGODENDROCYTE_DENSITY_CORR_UP 627 0.01
Up 0.201427934376543 0.0488951827842735
0.202379232610782
LIU_IL13_PRIMING_MODEL 14 0.01 Down 0.374594710376681
0.0489916436579501 0.202591939505507
BURTON_ADIPOGENESIS_8 70 0.01 Up 0.336966718874279
0.049103180883822 0.202866542346305
TAKEDA_TARGETS_OF_NUP98_HOXA9_FUSION_3D_UP 139 0.01 Down
0.489866171898128 0.0491784960373343
0.202991129189915
SHIN_B_CELL_LYMPHOMA_CLUSTER_3 24 0.01 Up
0.620734262913256 0.0492478443319369 0.203090880983538
BIOCARTA_FMLP_PATHWAY 31 0.01 Down 0.27512099493352
0.0493125546352752 0.203171341050011
HOLLEMAN_ASPARAGINASE_RESISTANCE_B_ALL_DN 14 0.01 Down
0.173328624182597 0.0493683600372279
0.20321499850489
BIOCARTA_PITX2_PATHWAY 13 0.01 Down 0.238720388510265
0.0495730466182768 0.203870855031248
BAE_BRCA1_TARGETS_DN 28 0.01 Down 0.336673016506086
0.0496193391705615 0.203874707103907
KEGG_ADIPOCYTOKINE_SIGNALING_PATHWAY 54 0.01 Down
0.3080745396139 0.0497568449199062 0.204252984397241
LIU_CDX2_TARGETS_UP 15 0.01 Down 0.888913904097969
0.0498958024918809 0.204466256710905
REACTOME_METABOLISM_OF_ANGIOTENSINOGEN_TO_ANGIOTENSINS 10 0.01
Up 0.594110810120542 0.0498997738847303
0.204466256710905
ENK_UV_RESPONSE_KERATINOCYTE_DN 474 0.01 Down
0.19356949442576 0.0500594013763803 0.204933523849571
REACTOME_AUF1_HNRNP_D0_BINDS_AND_DESTABILIZES_MRNA 53 0.01
Down 0.118474342492537 0.0501769078878947
0.205079356461868
ROVERSI_GLIOMA_LOH_REGIONS 28 0.01 Down

0.344925556239295	0.0501862718816585	0.205079356461868
LY_AGING_MIDDLE_DN	12	0.01	Up	0.354067704583694
	0.0503235493422809	0.205453546134017
GARGALOVIC_RESPONSE_TO_OXIDIZED_PHOSPHOLIPIDS_PINK_DN	27	0.01
	Up	0.47310298047525	0.0503887524335272
0.205533069136756
HOFMANN_MYELODYSPLASTIC_SYNDROM_HIGH_RISK_DN	11	0.01	Down
	0.271431391107635	0.0506081529113389
0.206240840740225
REACTOME_TRAFFICKING_AND_PROCESSING_OF_ENDOSOMAL_TLR	12	0.01
	Up	0.464763496187917	0.0507838260289062
0.206769291666606
KEGG_RENAL_CELL_CARCINOMA	67	0.01	Down
0.260415261674963	0.050904191133427	0.207071800131
BROWNE_HCMV_INFECTION_14HR_UP	139	0.01	Down
0.242265905814893	0.0509544891046417	0.207088995050058
PID_FGF_PATHWAY	43	0.01	Down	0.476724752335165
0.0512067151814699	0.207749086878658
REACTOME_PERVASIVE_DEVELOPMENTAL_DISORDERS	13	0.01	Down
	0.269046365417949	0.0512606868780889
0.207749086878658
NIKOLSKY_BREAST_CANCER_5P15_AMPLICON	18	0.01	Up
0.336602819243882	0.0513232643212862	0.207749086878658
BIOCARTA_VDR_PATHWAY	24	0.01	Down	0.142201175139615
	0.051403642772508	0.207749086878658
SCHMIDT_POR_TARGETS_IN_LIMB_BUD_UP	23	0.01	Down
0.244803576890197	0.051470997238593	0.207749086878658
OKUMURA_INFLAMMATORY_RESPONSE_LPS	151	0.01	Down
0.360656077864416	0.0514934867537309	0.207749086878658
DAZARD_RESPONSE_TO_UV_SCC_UP	105	0.01	Down
0.369311851877327	0.0515789149296017	0.207749086878658
NABA_MATRISOME	534	0.01	Up	0.829223248985927
0.0516078577625368	0.207749086878658
SABATES_COLORECTAL_ADENOMA_UP	99	0.01	Down
0.725225494225808	0.0516240047839788	0.207749086878658
GRAHAM_CML_DIVIDING_VS_NORMAL_QUIESCENT_UP	136	0.01	Up
	0.294779377670249	0.0516506041910206
0.207749086878658
VANLOO_SP3_TARGETS_DN	71	0.01	Up	0.422897273961663
	0.0516694072407047	0.207749086878658
LEE_EARLY_T_LYMPHOCYTE_UP	70	0.01	Up
0.351252519089193	0.0516715192726005	0.207749086878658
REACTOME_SYNTHESIS_OF_PIPS_AT_THE_PLASMA_MEMBRANE	50	0.01
	Down	0.246819797513443	0.0517493271969421
0.207875983691023
KEGG_BUTANOATE_METABOLISM	25	0.01	Up
0.289539177192857	0.05188599164505	0.208070437714787
PID_RHOA_PATHWAY	40	0.01	Down	0.203344461748144
	0.0519003207729304	0.208070437714787
KANG_AR_TARGETS_UP	14	0.01	Up	0.829256138758971
	0.0519996493154665	0.208070437714787
MARTORIATI_MDM4_TARGETS_NEUROEPITHELIUM_DN	94	0.01	Up
	0.56749139790705	0.0520039271626827
0.208070437714787

REACTOME_TRANSLOCATION_OF_ZAP_70_TO_IMMUNOLOGICAL_SYNAPSE 11 0.01 Up 1.24688294882978 0.0520291817555996 0.208070437714787
TURASHVILI_BREAST_CARCINOMA_DUCTAL_VS_LOBULAR_UP 17 0.01 Down 0.440661106420431 0.0522940294547777 0.20878795095363
REACTOME_NUCLEAR_EVENTS_KINASE_AND_TRANSCRIPTION_FACTOR_ACTIVATION 25 0.01 Down 0.273977238211296 0.0523014978362153 0.20878795095363
LU_EZH2_TARGETS_DN 360 0.01 Down 0.288072663892253 0.0524453949228001 0.209176619501319
LIAN_LIPA_TARGETS_3M 45 0.01 Up 1.13034529852953 0.0525113388020141 0.209253960917601
BROWNE_HCMV_INFECTION_1HR_UP 35 0.01 Down 0.561120206334652 0.0528059494719459 0.210077606866186
REACTOME_CTLA4_INHIBITORY_SIGNALING 19 0.01 Down 0.288864437917583 0.0528193978356454 0.210077606866186
XU_HGF_TARGETS_INDUCED_BY_AKT1_48HR_UP 13 0.01 Down 0.538425950586035 0.0528582365663307 0.210077606866186
REACTOME_REGULATION_OF_RUNX3_EXPRESSION_AND_ACTIVITY 55 0.01 Down 0.187367491916965 0.053345156212139 0.211729866899918
STAMBOLSKY_RESPONSE_TO_VITAMIN_D3_UP 70 0.01 Down 0.442302948216518 0.0533684293318355 0.211729866899918
CAMPS_COLON_CANCER_COPY_NUMBER_UP 58 0.01 Down 0.260402971811551 0.0534152767662974 0.211729866899918
BURTON_ADIPOGENESIS_4 41 0.01 Up 0.276380874744076 0.0535215864926194 0.211964344743898
SCHWAB_TARGETS_OF_BMYB_POLYMORPHIC_VARIANTS_UP 12 0.01 Up 0.451251619743485 0.0535687825867791 0.211964505041877
CAIRO_LIVER_DEVELOPMENT_UP 142 0.01 Up 0.460450561363689 0.0537540569155275 0.212510541631747
BIOCARTA_HDAC_PATHWAY 20 0.01 Down 0.230410418995029 0.0538520551549712 0.212710885695602
YORDY_RECIPROCAL_REGULATION_BY_ETS1_AND_SP100_DN 54 0.01 Down 0.324719985939773 0.0540782626103472 0.213416848492986
REACTOME_RAS_ACTIVATION_UPON_CA2PLUS_INFLUX_THROUGH_NMDA_RECEPTOR 10 0.01 Down 0.336620724284263 0.0542386727061454 0.213696804265698
WOOD_EBV_EBNA1_TARGETS_DN 44 0.01 Down 0.404800681654606 0.0542442833519825 0.213696804265698
BIOCARTA_NPC_PATHWAY 11 0.01 Down 0.218685710128252 0.0543372051134183 0.213770744416248
REACTOME_EXTRA_NUCLEAR_ESTROGEN_SIGNALING 63 0.01 Down 0.403008403445471 0.0543581670451104 0.213770744416248
RICKMAN_TUMOR_DIFFERENTIATED_WELL_VS_MODERATELY_UP 101 0.01 Up 0.276837882490318 0.0548735864105394 0.215609065485467
REACTOME_PEROXISOMAL_LIPID_METABOLISM 25 0.01 Up 0.228262921025858 0.0550753076820263 0.215864289910914
REACTOME_EPHB_MEDIATED_FORWARD_SIGNALING 39 0.01 Down

0.245448784097435	0.0551047823573947
0.215864289910914
FARMER_BREAST_CANCER_CLUSTER_6	13	0.01	Down
0.301312838083042	0.0551381945062379	0.215864289910914
BEIER_GLIOMA_STEM_CELL_UP	29	0.01	Down
0.439662784585242	0.0551493094189683	0.215864289910914
GINESTIER_BREAST_CANCER_20Q13_AMPLIFICATION_UP	97	0.01	Down
0.233500540182683	0.0551786583109321
0.215864289910914
REACTOME_METABOLISM_OF_CARBOHYDRATES	242	0.01	Up
0.359206840790369	0.0554476176744227	0.216727862127417
PID_EPO_PATHWAY	31	0.01	Down	0.31078818032893
0.0556221437550814	0.217221143509202
GAUSSMANN_MLL_AF4_FUSION_TARGETS_E_UP	79	0.01	Up
0.65058302365555	0.0557191042426449	0.21741091455789
SENGUPTA_NASOPHARYNGEAL_CARCINOMA_WITH_LMP1_DN	113	0.01	Down
0.916820299030384	0.0558698733220212
0.217778596304166
PID_PDGFRB_PATHWAY	126	0.01	Down	0.337577390332482
0.0559102336229159	0.217778596304166
REACTOME_GRB2_EVENTS_IN_ERBB2_SIGNALING	12	0.01	Down
0.25374986841215	0.0559983462229455	0.217932957811377
BEGUM_TARGETS_OF_PAX3_FOXO1_FUSION_UP	49	0.01	Up
0.391347112987086	0.0562525154071614	0.218732748058123
REACTOME_NEURODEGENERATIVE_DISEASES	20	0.01	Down
0.273719680053236	0.056680578123346	0.220198647321612
BIOCARTA_TPO_PATHWAY	21	0.01	Down	0.260184566585014
0.0567654972475252	0.220198647321612
KRIGE_AMINO_ACID_DEPRIVATION	28	0.01	Down
0.343401478069854	0.0567764699100663	0.220198647321612
REACTOME_IMMUNOREGULATORY_INTERACTIONS_BETWEEN_A_LYMPHOID_AND_A_NON_
LYMPHOID_CELL	62	0.01	Up	0.739501943313333
0.05692550687669	0.220397071772248
MIKKELSEN_IPS_HCP_WITH_H3_UNMETHYLATED	34	0.01	Up
0.5952762707734 0.0569256952897842	0.220397071772248
REACTOME_MET_ACTIVATES_RAP1_AND_RAC1	10	0.01	Down
0.156236874744876	0.0571640506376283	0.221129438568106
CHARAFE_BREAST_CANCER_BASAL_VS_MESENCHYMAL_DN	46	0.01	Up
0.633105730974107	0.0572673979209983
0.221338739170153
REACTOME_UPTAKE_AND_FUNCTION_OF_ANTHRAX_TOXINS	10	0.01	Down
0.209967110090312	0.0573493850825762
0.221465194111839
ZHAN_MULTIPLE_MYELOMA_PR_DN	42	0.01	Down
0.284133798493376	0.0575203741167161	0.221934834038317
ZHOU_TNF_SIGNALING_30MIN	51	0.01	Down
0.324492663867049	0.0577997241483738	0.222821406558268
REACTOME_COMPLEMENT_CASCADE	21	0.01	Up
0.993723039971059	0.0578980137972011	0.223009059141747
REACTOME_RHO_GTPASES_ACTIVATE_IQGAPS	23	0.01	Down
0.209293412637351	0.058019525439675	0.22328575929053
WATANABE_ULCERATIVE_COLITIS_WITH_CANCER_DN	13	0.01	Down
0.661136904430581	0.0583484965136363
0.224219041388074

REACTOME_PI_3K_CASCADE:FGFR3 11 0.01 Down
0.396368276396316 0.058373089127902 0.224219041388074
TAVOR_CEBPA_TARGETS_UP 43 0.01 Down 0.544608435168291
0.058461887543473 0.224219041388074
MARIADASON_RESPONSE_TO_CURCUMIN_SULINDAC_7 13 0.01 Down
0.550192572129484 0.0584846074737381
0.224219041388074
REACTOME_SIGNALING_BY_FGFR4_IN_DISEASE 10 0.01 Down
0.184274216455868 0.0585414040686203 0.224219041388074
REACTOME_BETA_CATENIN_PHOSPHORYLATION_CASCADE 17 0.01 Down
0.151512010314209 0.058591678551876
0.224219041388074
ALCALAY_AML_BY_NPM1_LOCALIZATION_DN 147 0.01 Up
0.374150927347438 0.0586226563457278 0.224219041388074
BIOCARTA_IL10_PATHWAY 12 0.01 Down 0.525908734491292
0.0586610884699388 0.224219041388074
KEGG_PATHOGENIC_ESCHERICHIA_COLI_INFECTION 47 0.01 Down
0.247094882769163 0.0588745820403865
0.224843879584994
LEE_AGING_MUSCLE_DN 33 0.01 Up 0.49935130071276
0.0589601209582585 0.224979408919671
BUCKANOVICH_T_LYMPHOCYTE_HOMING_ON_TUMOR_UP 14 0.01 Down
0.423865057310965 0.0590191022652384
0.225013456049403
REACTOME_SIGNALING_BY_NOTCH 183 0.01 Down
0.232020939705289 0.0593666960489684 0.226146863339079
HOLLEMAN_ASPARAGINASE_RESISTANCE_B_ALL_UP 27 0.01 Up
0.203041066250563 0.059560880904845
0.226694462038339
HUANG_GATA2_TARGETS_DN 63 0.01 Down 0.287246105724057
0.0598329181996958 0.227537197383784
REACTOME_SIGNALLING_TO_ERKS 30 0.01 Down
0.195652113333639 0.0599766337797664 0.227890928858876
BIOCARTA_CDMAC_PATHWAY 14 0.01 Down 0.427749471291887
0.060151088508169 0.228150095616937
YUAN_ZNF143_PARTNERS 21 0.01 Down 0.181051779166202
0.0601742661118347 0.228150095616937
BILANGES_RAPAMYCIN_SENSITIVE_VIA_TSC1_AND_TSC2 63 0.01 Up
0.215079779330981 0.0601971108791294
0.228150095616937
RHODES_UNDIFFERENTIATED_CANCER 58 0.01 Up
0.29766548495798 0.0604440877532306 0.22886312246038
REACTOME_CELL_JUNCTION_ORGANIZATION 74 0.01 Down
0.361772300599074 0.0604870721875265 0.22886312246038
MATZUK_SPERMATID_DIFFERENTIATION 23 0.01 Down
0.387748271281282 0.0606280408727021 0.229203569152898
SOTIRIOU_BREAST_CANCER_GRADE_1_VS_3_UP 120 0.01 Up
0.204992174946638 0.060686061771351 0.229230124085902
BROWNE_HCMV_INFECTION_4HR_UP 40 0.01 Down
0.44360740798855 0.0608965800781455 0.229832180899466
GARGALOVIC_RESPONSE_TO_OXIDIZED_PHOSPHOLIPIDS_YELLOW_DN 23 0.01
Up 0.404474119145125 0.0609980857827191
0.23002214395413
SANSOM_APC_TARGETS_REQUIRE_MYC 203 0.01 Up

0.193511898842709 0.0610843085618763 0.230154205352585
PEREZ_TP53_AND_TP63_TARGETS 173 0.01 Down
0.373845213651714 0.0612147576198623 0.23045254229588
REACTOME_CLATHRIN_MEDIATED_ENDOCYTOSIS 125 0.01 Down
0.258511395880004 0.0613144501408823 0.230634689023653
REACTOME_CELLULAR_RESPONSE_TO_HEAT_STRESS 94 0.01 Down
0.177314536382275 0.0614131675983625
0.230812866517257
REACTOME_GAP_JUNCTION_ASSEMBLY 20 0.01 Down
0.44608094581221 0.0614870817093156 0.230897604246761
JOHANSSON_BRAIN_CANCER_EARLY_VS_LATE_DN 39 0.01 Up
0.569595603583503 0.061585442320709 0.231070756939167
SIG_INSULIN_RECEPTOR_PATHWAY_IN_CARDIAC_MYOCYTES 46 0.01
Down 0.268458042138352 0.0616646364525684
0.231070756939167
WANG_METHYLATED_IN_BREAST_CANCER 34 0.01 Down
0.391363010630136 0.0617521845586884 0.231070756939167
PETROVA_ENDOTHELIUM_LYMPHATIC_VS_BLOOD_UP 101 0.01 Up
0.490689589183267 0.0617628471111424
0.231070756939167
GERHOLD_RESPONSE_TO_TZD_DN 11 0.01 Up
0.624254124129595 0.0617902224340108 0.231070756939167
REACTOME_THE_ACTIVATION_OF_ARYLSULFATASES 10 0.01 Up
0.260268985244225 0.0623642063019592
0.233023364361851
REACTOME_MAPK_TARGETS_NUCLEAR_EVENTS_MEDIATED_BY_MAP_KINASES 31
0.01 Down 0.302021274679185 0.0624844741375495
0.233278829940436
WU_HBX_TARGETS_3_UP 16 0.01 Up 0.437241855613114
0.0626819113627988 0.233782564274293
BIOCARTA_RAS_PATHWAY 22 0.01 Down 0.163681602370591
0.0627587625407405 0.233782564274293
REACTOME_PHASE_I_FUNCTIONALIZATION_OF_COMPOUNDS 63 0.01 Up
0.522342125493675 0.0628149322795611
0.233782564274293
BURTON_ADIPOGENESIS_PEAK_AT_24HR 35 0.01 Up
0.466092610811906 0.0628274388528021 0.233782564274293
HOSHIDA_LIVER_CANCER_SUBCLASS_S3 196 0.01 Up
0.394795869933471 0.0630311014721779 0.234346402909379
DEURIG_T_CELL_PROLYMPHOCYTIC_LEUKEMIA_DN 272 0.01 Down
0.330907016458156 0.0630847699966157
0.234352100111395
KANG_IMMORTALIZED_BY_TERT_UP 74 0.01 Up
0.627085968274531 0.063251837015501 0.234778701391145
PID_NECTIN_PATHWAY 30 0.01 Down 0.265184024365754
0.0634671140955101 0.235315432003743
LEE_BMP2_TARGETS_DN 827 0.01 Down 0.208406601488568
0.0635011388254818 0.235315432003743
KEGG_INSULIN_SIGNALING_PATHWAY 115 0.01 Down
0.268565723382107 0.063749092819951 0.235871843368718
REACTOME_SIGNALING_BY_LEPTIN 10 0.01 Down
0.314821631181168 0.0637562379739694 0.235871843368718
NADLER_HYPERGLYCEMIA_AT_OBESITY 49 0.01 Up
0.323825916717404 0.0638136030028853 0.235889922284514

| | | | | | | |
|---|---|---|---|---|---|---|
| FARMER_BREAST_CANCER_CLUSTER_2 | 25 | 0.01 | Up | 0.276162231431677 | 0.0639139297924993 | 0.235979246316816 |
| XU_GH1_EXOGENOUS_TARGETS_UP | 28 | 0.01 | Up | 0.502658038017721 | 0.0639427635181051 | 0.235979246316816 |
| CROONQUIST_IL6_DEPRIVATION_DN | 77 | 0.01 | Up | 0.255797112048729 | 0.0641448125404069 | 0.236530707439811 |
| LIU_COMMON_CANCER_GENES | 56 | 0.01 | Down | 0.273338129219245 | 0.0642903734249564 | 0.236873138151786 |
| KIM_ALL_DISORDERS_OLIGODENDROCYTE_NUMBER_CORR_UP | 680 | 0.01 | Up | 0.207092281912601 | 0.0645235698884763 | 0.237537630342917 |
| BIOCARTA_ARAP_PATHWAY | 17 | 0.01 | Down | 0.224319026447451 | 0.0646056845240926 | 0.237579314947774 |
| PID_SYNDECAN_2_PATHWAY | 31 | 0.01 | Down | 0.357872961459807 | 0.0646839582356164 | 0.237579314947774 |
| RADAEVA_RESPONSE_TO_IFNA1_UP | 44 | 0.01 | Down | 0.35113320786153 | 0.0647576237519406 | 0.237579314947774 |
| CHESLER_BRAIN_HIGHEST_GENETIC_VARIANCE | 29 | 0.01 | Up | 0.32794816383209 | 0.0648361177463315 | 0.237579314947774 |
| GOBERT_OLIGODENDROCYTE_DIFFERENTIATION_UP | 484 | 0.01 | Up | 0.253326630860568 | 0.0648653616550187 | 0.237579314947774 |
| REACTOME_BASIGIN_INTERACTIONS | 20 | 0.01 | Up | 0.350091209211831 | 0.0648974077623039 | 0.237579314947774 |
| SHEDDEN_LUNG_CANCER_GOOD_SURVIVAL_A4 | 170 | 0.01 | Up | 0.431056764687639 | 0.0649048718033073 | 0.237579314947774 |
| LINSLEY_MIR16_TARGETS | 205 | 0.01 | Down | 0.186528289481124 | 0.0649670729657264 | 0.237613501204996 |
| POOLA_INVASIVE_BREAST_CANCER_UP | 195 | 0.01 | Up | 0.834052236421122 | 0.0650202707447314 | 0.237614729266315 |
| REACTOME_PLATELET_ADHESION_TO_EXPOSED_COLLAGEN | 10 | 0.01 | Up | 0.987481711137433 | 0.0651420434249982 | 0.23786635677934 |
| MYLLYKANGAS_AMPLIFICATION_HOT_SPOT_8 | 11 | 0.01 | Down | 0.264189561391661 | 0.0652611516889228 | 0.237895339054373 |
| SPIELMAN_LYMPHOBLAST_EUROPEAN_VS_ASIAN_DN | 571 | 0.01 | Down | 0.196967872702289 | 0.0652889638176872 | 0.237895339054373 |
| REACTOME_DEGRADATION_OF_AXIN | 53 | 0.01 | Down | 0.126459471501714 | 0.0653087538138145 | 0.237895339054373 |
| FEVR_CTNNB1_TARGETS_UP | 573 | 0.01 | Down | 0.378791837844229 | 0.0653908075388218 | 0.238001360232392 |
| FORTSCHEGGER_PHF8_TARGETS_UP | 243 | 0.01 | Down | 0.294375566070053 | 0.0655356835595333 | 0.238335677669986 |
| REACTOME_ELASTIC_FIBRE_FORMATION | 38 | 0.01 | Up | 0.6542864710239 | 0.0657787707558535 | 0.239026333506517 |
| BIOCARTA_BIOPEPTIDES_PATHWAY | 24 | 0.01 | Down | 0.2298267008731 | 0.0658962079333057 | 0.239259656429894 |
| REACTOME_NRAGE_SIGNALS_DEATH_THROUGH_JNK | 55 | 0.01 | Down | 0.297948962525638 | 0.0660183392244011 | 0.239509632617985 |
| CASORELLI_ACUTE_PROMYELOCYTIC_LEUKEMIA_UP | 145 | 0.01 | Down | 0.342787440341719 | 0.0662376627688077 | 0.24011527536928 |

IRITANI_MAD1_TARGETS_DN 40 0.01 Up 0.158920904336158
0.066302193868466 0.240151781981269
KEGG_PRIMARY_IMMUNODEFICIENCY 21 0.01 Down
0.494548051864689 0.0664378454355739 0.240449368142436
XU_HGF_TARGETS_REPRESSED_BY_AKT1_UP 10 0.01 Down
0.243489551326109 0.0666315836567498 0.240956531405543
HAHTOLA_CTCL_CUTANEOUS 20 0.01 Down 0.551952665983792
0.0667793208451988 0.241231083237807
COLINA_TARGETS_OF_4EBP1_AND_4EBP2 304 0.01 Up
0.367017005846434 0.0668148384051323 0.241231083237807
LE_EGR2_TARGETS_UP 95 0.01 Up 0.233370852422317
0.0669798568683744 0.241574413332974
NEWMAN_ERCC6_TARGETS_DN 31 0.01 Up 0.675620307512189
0.0670192293574192 0.241574413332974
BILANGES_RAPAMYCIN_SENSITIVE_GENES 32 0.01 Down
0.272824386047154 0.067071160809689 0.241574413332974
CHOW_RASSF1_TARGETS_UP 26 0.01 Down 0.224650227399147
0.0671545703037543 0.241681179756105
REACTOME_VIF_MEDIATED_DEGRADATION_OF_APOBEC3G 52 0.01 Down
0.125240944211274 0.067426097976219
0.242464248322484
REACTOME_HOST_INTERACTIONS_OF_HIV_FACTORS 125 0.01 Down
0.18698785565057 0.0679865774023064
0.244284304894778
GEORGES_TARGETS_OF_MIR192_AND_MIR215 811 0.01 Down
0.262252637644302 0.0680542475606358 0.244332142799567
KEGG_AXON_GUIDANCE 109 0.01 Down 0.364275826449476
0.0681107479261116 0.24433748389597
REACTOME_RAB_GEFS_EXCHANGE_GTP_FOR_GDP_ON_RABS 86 0.01 Down
0.206328363686624 0.0681644504572961
0.24433748389597
MAHAJAN_RESPONSE_TO_IL1A_DN 66 0.01 Up
0.309342023859082 0.0682746423556002 0.244413718863325
REACTOME_REGULATION_OF_HSF1_MEDIATED_HEAT_SHOCK_RESPONSE 77
0.01 Down 0.183649830952783 0.0682944673842795
0.244413718863325
WANG_IMMORTALIZED_BY_HOXA9_AND_MEIS1_DN 15 0.01 Down
0.411869291490009 0.0685877449937144 0.245268030029233
CLASPER_LYMPHATIC_VESSELS_DURING_METASTASIS_UP 16 0.01 Down
0.509026859883381 0.0688513658242232
0.246015015405313
DITTMER_PTHLH_TARGETS_DN 66 0.01 Down
0.254181295066828 0.0690679596967427 0.246434622427904
MARKEY_RB1_CHRONIC_LOF_DN 98 0.01 Up
0.652268360076239 0.0690784481110476 0.246434622427904
BARRIER_CANCER_RELAPSE_TUMOR_SAMPLE_UP 16 0.01 Down
0.316366461417683 0.0695359275298469 0.247869939925981
REACTOME_TRISTETRAPROLIN_TTP_ZFP36_BINDS_AND_DESTABILIZES_MRNA 17
0.01 Down 0.203312962662925 0.0696809928312485
0.248190224070097
REACTOME_TCR_SIGNALING 105 0.01 Down 0.301046188117844
0.0698135421379378 0.248465456777538
VANASSE_BCL2_TARGETS_UP 21 0.01 Down 0.527962282084456
0.0700013073324806 0.248811064060859

| | | | | | | |
|---|---|---|---|---|---|---|
| REACTOME_PEPTIDE_LIGAND_BINDING_RECEPTORS | 54 | 0.01 | Down | 0.942100767673092 | 0.0700213561817545 | 0.248811064060859 |
| NICK_RESPONSE_TO_PROC_TREATMENT_DN | 26 | 0.01 | Down | 0.27193006675089 | 0.0701026842953405 | 0.248903290606284 |
| REACTOME_TNFR1_INDUCED_PROAPOPTOTIC_SIGNALING | 13 | 0.01 | Down | 0.350228288110298 | 0.0703093651814729 | 0.249440091942163 |
| HEDVAT_ELF4_TARGETS_UP | 10 | 0.01 | Down | 0.746646193845328 | 0.0705653600988698 | 0.250150862495599 |
| REACTOME_UCH_PROTEINASES | 87 | 0.01 | Down | 0.137685571465325 | 0.0706491624718573 | 0.250250579441291 |
| PID_FOXM1_PATHWAY | 32 | 0.01 | Up | 0.464170155394095 | 0.0707783503460326 | 0.250510775437336 |
| REACTOME_GENE_EXPRESSION_TRANSCRIPTION | 1279 | 0.01 | Down | 0.20779209105972 | 0.0708828989378966 | 0.250683423073049 |
| MIKKELSEN_NPC_ICP_WITH_H3K4ME3 | 343 | 0.01 | Up | 0.334124496803545 | 0.0710498461425011 | 0.250729451713537 |
| YIH_RESPONSE_TO_ARSENITE_C2 | 14 | 0.01 | Down | 0.329187607515051 | 0.0710562795824413 | 0.250729451713537 |
| REACTOME_RUNX3_REGULATES_P14_ARF | 10 | 0.01 | Down | 0.530753535806029 | 0.0710632528327134 | 0.250729451713537 |
| BIOCARTA_PDZS_PATHWAY | 14 | 0.01 | Down | 0.223429702946869 | 0.0716983572256722 | 0.252771855474037 |
| REACTOME_EUKARYOTIC_TRANSLATION_INITIATION | 117 | 0.01 | Up | 0.11345658112912 | 0.0717805539537565 | 0.25286331506437 |
| KATSANOU_ELAVL1_TARGETS_DN | 113 | 0.01 | Up | 0.531852308670569 | 0.072028627167314 | 0.253371314824859 |
| SHIN_B_CELL_LYMPHOMA_CLUSTER_2 | 24 | 0.01 | Down | 0.334393559361556 | 0.0720655851943499 | 0.253371314824859 |
| SARRIO_EPITHELIAL_MESENCHYMAL_TRANSITION_DN | 142 | 0.01 | Down | 0.475158542572785 | 0.0720938624384859 | 0.253371314824859 |
| WANG_BARRETTS_ESOPHAGUS_DN | 24 | 0.01 | Down | 0.617018552040111 | 0.0723097639273686 | 0.253931553791814 |
| LOPEZ_TRANSLATION_VIA_FN1_SIGNALING | 31 | 0.01 | Down | 0.148710125492956 | 0.0724203850965366 | 0.254121491810252 |
| ZHONG_SECRETOME_OF_LUNG_CANCER_AND_ENDOTHELIUM | 53 | 0.01 | Up | 0.436558987678871 | 0.0725971350192879 | 0.254542996810998 |
| REACTOME_NOTCH1_INTRACELLULAR_DOMAIN_REGULATES_TRANSCRIPTION | 44 | 0.01 | Down | 0.2026387362206 | 0.072808399185792 | 0.254891026443458 |
| FERRANDO_HOX11_NEIGHBORS | 16 | 0.01 | Up | 0.315258256331145 | 0.0728098059963071 | 0.254891026443458 |
| KIM_ALL_DISORDERS_DURATION_CORR_DN | 131 | 0.01 | Up | 0.182398411141093 | 0.0729414569575114 | 0.255065137346304 |
| REACTOME_ESTROGEN_DEPENDENT_GENE_EXPRESSION | 103 | 0.01 | Down | 0.213512372181627 | 0.0729730292830582 | 0.255065137346304 |
| KEGG_GLYCOSAMINOGLYCAN_BIOSYNTHESIS_HEPARAN_SULFATE | 18 | 0.01 | Up | 0.363936817987835 | 0.073280032542542 | 0.255939196797767 |

RORIE_TARGETS_OF_EWSR1_FLI1_FUSION_DN 12 0.01 Up
0.583815573014271 0.0735120375268968 0.256550162021274
FOSTER_TOLERANT_MACROPHAGE_UP 129 0.01 Down
0.456230510039355 0.0736437126850888 0.256644468254235
MOOTHA_GLUCONEOGENESIS 23 0.01 Up 0.280341942747555
0.0736532511786346 0.256644468254235
PETRETTO_HEART_MASS_QTL_CIS_DN 24 0.01 Up
0.361246434314816 0.0737347399517627 0.256729400529181
ROSTY_CERVICAL_CANCER_PROLIFERATION_CLUSTER 111 0.01 Up
0.258643560589833 0.0738488003939509
0.256927521494434
SCHLOSSER_MYC_AND_SERUM_RESPONSE_SYNERGY 33 0.01 Up
0.143653086038914 0.074139387948806
0.257497401250478
REACTOME_ER_TO_GOLGI_ANTEROGRADE_TRANSPORT 136 0.01 Down
0.191830779848183 0.0742114635839998
0.257497401250478
ONGUSAHA_BRCA1_TARGETS_DN 10 0.01 Down
0.274313038372667 0.0742500971559719 0.257497401250478
REACTOME_SYNTHESIS_OF_ACTIVE_UBIQUITIN:_ROLES_OF_E1_AND_E2_ENZYMES
29 0.01 Down 0.122302546057089
0.0742656252837673 0.257497401250478
CASTELLANO_NRAS_TARGETS_DN 12 0.01 Up
0.389285487798395 0.07429902768006 0.257497401250478
BIOCARTA_BCELLSURVIVAL_PATHWAY 14 0.01 Down
0.281662939573357 0.0743716139242766 0.257550388743932
BURTON_ADIPOGENESIS_3 84 0.01 Up 0.299019249152301
0.074435606127816 0.257573556231357
REACTOME_GENERATION_OF_SECOND_MESSENGER_MOLECULES 24 0.01
Up 0.736944255610897 0.0747386739244984
0.258423337915862
REACTOME_DEACTIVATION_OF_THE_BETA_CATENIN_TRANSACTIVATING_COMPLEX
38 0.01 Down 0.223810386513802
0.0747965483694703 0.258424661737716
BROWNE_HCMV_INFECTION_8HR_UP 88 0.01 Down
0.312680569131717 0.0751149650290411 0.259291014719942
REACTOME_SIGNALING_BY_NOTCH4 79 0.01 Down
0.180836339887317 0.0751626678932334 0.259291014719942
REACTOME_ACTIVATION_OF_BAD_AND_TRANSLOCATION_TO_MITOCHONDRIA 15
0.01 Down 0.178286171517602 0.0755346935020221
0.260374576143857
HOEGERKORP_CD44_TARGETS_DIRECT_UP 20 0.01 Down
0.526841740445624 0.0757534227258834 0.260671945491332
MARTIN_INTERACT_WITH_HDAC 36 0.01 Down
0.230794123897619 0.0757658439916524 0.260671945491332
GROSS_ELK3_TARGETS_UP 25 0.01 Up 0.468575502964765
0.0757949349849102 0.260671945491332
REACTOME_PKMTS_METHYLATE_HISTONE_LYSINES 49 0.01 Down
0.21942878327157 0.0758740374100751
0.260744494004807
VANDESLUIS_COMMD1_TARGETS_GROUP_3_DN 19 0.01 Up
0.535833396819994 0.0759383567598045 0.260766167788633
YAO_TEMPORAL_RESPONSE_TO_PROGESTERONE_CLUSTER_13 166 0.01
Up 0.153261601116612 0.0761686423596732

0.261357288096741
VETTER_TARGETS_OF_PRKCA_AND_ETS1_UP 16 0.01 Down
0.239133823972394 0.0765646279135176 0.262406811396375
REACTOME_CHONDROITIN_SULFATE_BIOSYNTHESIS 17 0.01 Up
0.747999159837986 0.0765912650838807
0.262406811396375
REACTOME_INTERLEUKIN_3_INTERLEUKIN_5_AND_GM_CSF_SIGNALING 42
0.01 Down 0.353073435106318 0.0767294329536658
0.262679970393548
REACTOME_GLYCOGEN_STORAGE_DISEASES 12 0.01 Up
0.176834368634788 0.0769610170942205 0.263272276893852
SCHRAETS_MLL_TARGETS_DN 25 0.01 Up 1.24350055944553
0.0770701947799719 0.263445266567281
BURTON_ADIPOGENESIS_PEAK_AT_2HR 44 0.01 Down
0.427284403981274 0.0773262484540283 0.26411967082132
KANG_FLUOROURACIL_RESISTANCE_UP 21 0.01 Up
0.335034771801279 0.0774653951986342 0.264254469657472
SWEET_LUNG_CANCER_KRAS_DN 299 0.01 Up
0.533032712822478 0.0775310493664446 0.264254469657472
GEORGES_CELL_CYCLE_MIR192_TARGETS 56 0.01 Down
0.262858267165656 0.0775420790830267 0.264254469657472
BIOCARTA_TOB1_PATHWAY 11 0.01 Down 0.317969973114878
0.0777442110329145 0.264552188008699
ELVIDGE_HYPOXIA_DN 144 0.01 Down 0.192648722170977
0.0777839294994974 0.264552188008699
WHITFIELD_CELL_CYCLE_M_G1 122 0.01 Down
0.228250582967413 0.0778060050161289 0.264552188008699
SWEET_KRAS_TARGETS_DN 52 0.01 Down 0.335147223204636
0.0778700794098774 0.264569922106877
CHICAS_RB1_TARGETS_LOW_SERUM 84 0.01 Up
0.26636330118573 0.0783482057270064 0.265993341950826
KYNG_RESPONSE_TO_H2O2 64 0.01 Up 0.284172743630487
0.0786163492911354 0.266702256651814
PID_AR_PATHWAY 57 0.01 Down 0.216485695952831
0.0788177938629528 0.267183999558049
FIGUEROA_AML_METHYLATION_CLUSTER_5_UP 10 0.01 Down
0.359704570713316 0.0789447531988641 0.267268645733852
WONG_PROTEASOME_GENE_MODULE 49 0.01 Down
0.11913596898305 0.0789938308792527 0.267268645733852
PID_ECADHERIN_KERATINOCYTE_PATHWAY 20 0.01 Down
0.181972910324813 0.0790211413081843 0.267268645733852
REACTOME_SYNTHESIS_OF_PIPS_AT_THE_LATE_ENDOSOME_MEMBRANE 10
0.01 Down 0.220355869806678 0.0791172241892073
0.267392423105629
KANG_DOXORUBICIN_RESISTANCE_UP 39 0.01 Up
0.211057782155659 0.0794978287071736 0.268476889585834
BIOCARTA_PDGF_PATHWAY 27 0.01 Down 0.255313651690216
0.0796787726554666 0.268885948263005
LIM_MAMMARY_LUMINAL_MATURE_UP 91 0.01 Down
0.42060633159608 0.0798807238478668 0.269365231580016
YOKOE_CANCER_TESTIS_ANTIGENS 25 0.01 Up
0.371588990141864 0.0801800570604663 0.270171931399397
NAM_FXYD5_TARGETS_DN 18 0.01 Down 0.298622091396814
0.080366869906596 0.270469670636098

KEGG_TYPE_I_DIABETES_MELLITUS 29 0.01 Up
0.825794788023623 0.0803887608386711 0.270469670636098
MORI_IMMATURE_B_LYMPHOCYTE_DN 72 0.01 Up
0.192965352365692 0.0805715640134495 0.270702762141652
MACLACHLAN_BRCA1_TARGETS_UP 20 0.01 Up
0.345459367003399 0.0805784862615196 0.270702762141652
REACTOME_ESTABLISHMENT_OF_SISTER_CHROMATID_COHESION 10 0.01
Down 0.173686019447355 0.0806835528493923
0.270853301014203
WONG_MITOCHONDRIA_GENE_MODULE 211 0.01 Up
0.161560026082559 0.0808054702833034 0.271060140987648
WANG_THOC1_TARGETS_DN 10 0.01 Up 0.485275100884475
0.081037440555236 0.271589811530229
MONNIER_POSTRADIATION_TUMOR_ESCAPE_UP 373 0.01 Down
0.269255695658587 0.0810842106948981 0.271589811530229
REACTOME_ADHERENS_JUNCTIONS_INTERACTIONS 23 0.01 Down
0.321702042460918 0.0812306378749777
0.27180702213218
SHAFFER_IRF4_TARGETS_IN_MYELOMA_VS_MATURE_B_LYMPHOCYTE 94 0.01
Down 0.295213959268612 0.0812699972737819
0.27180702213218
ZHANG_RESPONSE_TO_CANTHARIDIN_UP 14 0.01 Down
0.306578479617728 0.0814320956920055 0.272146669245773
REACTOME_RETROGRADE_TRANSPORT_AT_THE_TRANS_GOLGI_NETWORK 49
0.01 Down 0.169390531674103 0.0822969858455724
0.274679567258681
REACTOME_SUMOYLATION_OF_DNA_REPLICATION_PROTEINS 44 0.01
Down 0.155689656341384 0.0823122084755158
0.274679567258681
REACTOME_FORMATION_OF_FIBRIN_CLOT_CLOTTING_CASCADE 18 0.01
Up 0.572614274757907 0.0828128846298261
0.276145338583878
REACTOME_CIRCADIAN_CLOCK 66 0.01 Down
0.231323730716823 0.0829183357965508 0.276264593796662
HUANG_FOXA2_TARGETS_UP 42 0.01 Down 0.377715526915163
0.0830222385411948 0.276264593796662
KYNG_ENVIRONMENTAL_STRESS_RESPONSE_DN 16 0.01 Down
0.216091812748656 0.0830330291922781 0.276264593796662
YU_MYC_TARGETS_UP 36 0.01 Up 0.244224410936555
0.0832105427050752 0.276650435990616
MARCHINI_TRABECTEDIN_RESISTANCE_DN 43 0.01 Up
0.461680446806012 0.0833865965647815 0.277030858506055
WALLACE_PROSTATE_CANCER_UP 19 0.01 Down
0.233401855271842 0.0834910650032643 0.277173070302565
REACTOME_ANTIGEN_ACTIVATES_B_CELL_RECEPTOR_BCR_LEADING_TO_GENERATION
_OF_SECOND_MESSENGERS 24 0.01 Down 0.336328503090956
0.0836418491752341 0.277456686043096
MATZUK_OVULATION 12 0.01 Down 0.664470802867343
0.0836999480032121 0.277456686043096
SARTIPY_BLUNTED_BY_INSULIN_RESISTANCE_DN 17 0.01 Up
0.20437794379688 0.0838506885715352
0.277737054850599
REACTOME_TCF_DEPENDENT_SIGNALING_IN_RESPONSE_TO_WNT 172 0.01
Down 0.221347399695816 0.0839081024442968

0.277737054850599
ABE_INNER_EAR 32 0.01 Up 0.395868719930915
0.0841311653736443 0.278159516545345
REACTOME_HIV_INFECTION 223 0.01 Down 0.171471824922274
0.0841594977756772 0.278159516545345
REACTOME_DEADENYLATION_DEPENDENT_MRNA_DECAY 56 0.01 Down
0.138761481366536 0.0845590379356628
0.27927470648112
REACTOME_TRANSLATION 289 0.01 Up 0.11440340166498
0.0846947882143486 0.279517674760277
GAL_LEUKEMIC_STEM_CELL_DN 162 0.01 Up
0.649459743883337 0.0848400057326057 0.279791508267104
SHETH_LIVER_CANCER_VS_TXNIP_LOSS_PAM5 64 0.01 Down
0.335416534958488 0.0850649081615128 0.280261760143377
LINDGREN_BLADDER_CANCER_CLUSTER_2A_DN 137 0.01 Up
0.333889068768987 0.0851072975741291 0.280261760143377
REACTOME_PKA_MEDIATED_PHOSPHORYLATION_OF_CREB 15 0.01 Down
0.327355878732759 0.0852132748064706
0.280297083191602
SHAFFER_IRF4_TARGETS_IN_ACTIVATED_B_LYMPHOCYTE 76 0.01 Down
0.234204682533458 0.0852427392042091
0.280297083191602
BHAT_ESR1_TARGETS_NOT_VIA_AKT1_UP 180 0.01 Down
0.385391360192688 0.0856043789904401 0.281117184634472
REACTOME_PROLACTIN_RECEPTOR_SIGNALING 11 0.01 Down
0.315540301825674 0.0856172248642028 0.281117184634472
REACTOME_SIGNALING_BY_NUCLEAR_RECEPTORS 196 0.01 Down
0.298766402125562 0.085703487799696 0.281121884886269
NADLER_OBESITY_DN 35 0.01 Up 0.358324393882842
0.0857437384158121 0.281121884886269
REACTOME_SIGNALING_BY_LIGAND_RESPONSIVE_EGFR_VARIANTS_IN_CANCER 18
0.01 Down 0.171442611578488 0.0858793680252884
0.281361340578478
HAHTOLA_MYCOSIS_FUNGOIDES_CD4_DN 110 0.01 Down
0.163906323557315 0.0859770399547607 0.28147617960426
BIOCARTA_INSULIN_PATHWAY 20 0.01 Down
0.270631078092989 0.0860719903095707 0.281581947919593
DOUGLAS_BMI1_TARGETS_DN 299 0.01 Down 0.270517902208645
0.0862621382933966 0.281919627810712
MARTENS_TRETINOIN_RESPONSE_DN 704 0.01 Up
0.196821068963756 0.0863006469115773 0.281919627810712
LEE_METASTASIS_AND_ALTERNATIVE_SPLICING_UP 61 0.01 Up
0.369463395684293 0.0871645396227709
0.284534935079416
REACTOME_REGULATED_NECROSIS 19 0.01 Down
0.272881210571629 0.0874309093744384 0.285197342262773
REACTOME_VESICLE_MEDIATED_TRANSPORT 582 0.01 Down
0.263061691823569 0.0875109170668014 0.285251321403388
CHIANG_LIVER_CANCER_SUBCLASS_CTNNB1_UP 120 0.01 Up
0.395269918209728 0.0875883400629715 0.285296803321056
CHEOK_RESPONSE_TO_MERCAPTOPURINE_DN 17 0.01 Up
0.379224563446439 0.0876993858339067 0.285451657728755
REACTOME_BETA_OXIDATION_OF_VERY_LONG_CHAIN_FATTY_ACIDS 10 0.01
Up 0.233931561461732 0.0878409114719239

0.285705424794716
REACTOME_NEGATIVE_REGULATION_OF_NOTCH4_SIGNALING 54 0.01
Down 0.127482123926939 0.0882076988541762
0.286305597673738
REACTOME_GASTRIN_CREB_SIGNALLING_PATHWAY_VIA_PKC_AND_MAPK 16
0.01 Down 0.419676358463709 0.0882923760596913
0.286305597673738
BROWNE_HCMV_INFECTION_6HR_DN 146 0.01 Down
0.271990178631624 0.0883124771846223 0.286305597673738
MARSHALL_VIRAL_INFECTION_RESPONSE_DN 19 0.01 Down
0.412979486789772 0.0883274901065734 0.286305597673738
REACTOME_TRANSPORT_OF_MATURE_TRANSCRIPT_TO_CYTOPLASM 81 0.01
Down 0.125401058907787 0.0883439074468242
0.286305597673738
KEGG_GLYCOSAMINOGLYCAN_BIOSYNTHESIS_CHONDROITIN_SULFATE 19 0.01
Up 0.341453433581584 0.0884242827245695
0.286359618765807
GROSS_ELK3_TARGETS_DN 28 0.01 Down 0.558572075045184
0.0885248272688464 0.286478832666281
PID_TNF_PATHWAY 46 0.01 Down 0.335012489174802
0.0886390793164382 0.286642202537691
CHIANG_LIVER_CANCER_SUBCLASS_UNANNOTATED_DN 188 0.01 Up
0.206309077551965 0.0887138599030503
0.28667778595558
STARK_PREFRONTAL_CORTEX_22Q11_DELETION_DN 487 0.01 Up
0.162485227181879 0.0887879741697515
0.286711166589823
REACTOME_VEGFR2_MEDIATED_CELL_PROLIFERATION 18 0.01 Down
0.248072978018328 0.0889359019933102
0.286982684465132
REACTOME_NOREPINEPHRINE_NEUROTRANSMITTER_RELEASE_CYCLE 11 0.01
Down 0.300273996579494 0.0890901532431589
0.287274202889526
WEST_ADRENOCORTICAL_TUMOR_DN 438 0.01 Up
0.453072135294299 0.0893441724049524 0.287886777749291
GARCIA_TARGETS_OF_FLI1_AND_DAX1_UP 49 0.01 Up
0.280653382505327 0.08981703039425 0.289057502091045
REACTOME_RAB_GERANYLGERANYLATION 57 0.01 Down
0.298530566591504 0.0898415938818106 0.289057502091045
PROVENZANI_METASTASIS_UP 188 0.01 Up
0.305963943924638 0.0900777445303415 0.289057502091045
KOHOUTEK_CCNT1_TARGETS 40 0.01 Up 0.31909328766299
0.0900926426733881 0.289057502091045
REACTOME_METAL_ION_SLC_TRANSPORTERS 20 0.01 Up
0.438359463121582 0.0902363633416939 0.289057502091045
REACTOME_VISUAL_PHOTOTRANSDUCTION 54 0.01 Up
0.475693965454264 0.090276646496907 0.289057502091045
KEGG_SYSTEMIC_LUPUS_ERYTHEMATOSUS 61 0.01 Up
0.618315960815244 0.0902785089524696 0.289057502091045
WAKABAYASHI_ADIPOGENESIS_PPARG_RXRA_BOUND_36HR 127 0.01 Up
0.263304493799786 0.0902859957944793
0.289057502091045
VANASSE_BCL2_TARGETS_DN 45 0.01 Up 0.437900052370591
0.0902862587176479 0.289057502091045

KEGG_TIGHT_JUNCTION	103	0.01	Down	0.29725676782535
	0.090352686162935	0.28906428775971
WANG_NEOPLASTIC_TRANSFORMATION_BY_CCND1_MYC	18	0.01	Up
	0.76096803582781	0.0906061616102867
0.289669058633171
REACTOME_GENERIC_TRANSCRIPTION_PATHWAY	1031	0.01	Down
0.221892796832799	0.0909293433741173	0.290495663444675
WILLIAMS_ESR1_TARGETS_UP	20	0.01	Up
0.396638852541643	0.0910178085468107	0.290571768052496
SCHAEFFER_PROSTATE_DEVELOPMENT_AND_CANCER_BOX1_UP	14	0.01
	Down	0.160571080125594	0.0910918545832319
0.290601764621453
HOUSTIS_ROS	30	0.01	Up	0.319871095681654
0.0917438006237971	0.292449202738387
GENTILE_UV_RESPONSE_CLUSTER_D6	35	0.01	Down
0.392978783226755	0.0918010734291132	0.292449202738387
CREIGHTON_ENDOCRINE_THERAPY_RESISTANCE_2	342	0.01	Up
	0.268858770354996	0.0919080339239857
0.292460513120941
REACTOME_FRS_MEDIATED_FGFR3_SIGNALING	13	0.01	Down
0.367306153186626	0.0919487673635307	0.292460513120941
CHUNG_BLISTER_CYTOTOXICITY_UP	129	0.01	Up
0.336754310944074	0.0919998143610702	0.292460513120941
REACTOME_BRANCHED_CHAIN_AMINO_ACID_CATABOLISM	19	0.01	Up
	0.184627950250186	0.0921659568485862
0.292482032527594
MIKKELSEN_ES_HCP_WITH_H3_UNMETHYLATED	27	0.01	Up
0.466459109595281	0.0921821651963218	0.292482032527594
REACTOME_SYNTHESIS_OF_PIPS_AT_THE_EARLY_ENDOSOME_MEMBRANE	16
	0.01	Down	0.226464797831463	0.0922017886744384
	0.292482032527594
REACTOME_DEGRADATION_OF_GLI1_BY_THE_PROTEASOME	56	0.01	Down
	0.122752864056122	0.0925560083509746
0.293398630139373
REACTOME_GOLGI_ASSOCIATED_VESICLE_BIOGENESIS	55	0.01	Down
	0.180031004241171	0.0926982922533212
0.293642581873629
CERIBELLI_PROMOTERS_INACTIVE_AND_BOUND_BY_NFY	15	0.01	Down
	0.437258734540247	0.092779365932101
0.293692429482249
KEGG_OXIDATIVE_PHOSPHORYLATION	105	0.01	Up
0.147551174387132	0.0930106449690494	0.294217346330667
SOTIRIOU_BREAST_CANCER_GRADE_1_VS_3_DN	52	0.01	Up
0.216870340648562	0.0930907332111333	0.294230880657814
REACTOME_FBXL7_DOWN_REGULATES_AURKA_DURING_MITOTIC_ENTRY_AND_IN_EARL
Y_MITOSIS	53	0.01	Down	0.123221373468294
0.0931458383039088	0.294230880657814
PID_HIF1A_PATHWAY	18	0.01	Up	0.246789564861567
	0.0936060445111059	0.295104171276408
WHITESIDE_CISPLATIN_RESISTANCE_UP	11	0.01	Down
0.269263041987727	0.0936173059486194	0.295104171276408
VIETOR_IFRD1_TARGETS	19	0.01	Up	0.382954080084908
	0.0936192543359638	0.295104171276408
FOURNIER_ACINAR_DEVELOPMENT_LATE_DN	19	0.01	Down

0.367432621680238	0.0938227635926096	0.295538417903841
MANALO_HYPOXIA_UP	183	0.01	Up	0.463190825557234
	0.0939576821091286	0.295565850385203
SIG_CHEMOTAXIS	39	0.01	Down	0.266128927400179
0.0939629811346951	0.295565850385203
PID_RAC1_PATHWAY	50	0.01	Down	0.2316060112369
0.0940682523580273	0.295690065978555
WINNEPENNINCKX_MELANOMA_METASTASIS_UP	137	0.01	Up
0.197466915179625	0.0942982187125437	0.29607790789079
CUI_GLUCOSE_DEPRIVATION	48	0.01	Down	0.189710039719896
	0.0943233735483007	0.29607790789079
MULLIGHAN_NPM1_MUTATED_SIGNATURE_1_DN	106	0.01	Down
0.276712473702895	0.0945052278931453	0.296441730202155
BOYLAN_MULTIPLE_MYELOMA_C_DN	50	0.01	Up
0.335052802467374	0.0946448709062157	0.296655582493369
BIOCARTA_MEF2D_PATHWAY	17	0.01	Down	0.282070538394164
	0.0947053973032223	0.296655582493369
REACTOME_DISEASES_ASSOCIATED_WITH_O_GLYCOSYLATION_OF_PROTEINS	47
	0.01	Up	0.89825226528542	0.0950730246080541
	0.297599753212537
PUIFFE_INVASION_INHIBITED_BY_ASCITES_DN	143	0.01	Down
0.268199464664583	0.0956702450452837	0.299260787389388
BIOCARTA_TFF_PATHWAY	20	0.01	Down	0.185866896397037
	0.095840658807529	0.299585369499195
VANHARANTA_UTERINE_FIBROID_WITH_7Q_DELETION_DN	36	0.01	Up
	0.225933476706171	0.0959708269222793
0.29978378527842
ZHAN_LATE_DIFFERENTIATION_GENES_UP	27	0.01	Up
0.456568492378653	0.0961945634102343	0.300274001756252
REACTOME_TIE2_SIGNALING	15	0.01	Down	0.423083394990522
	0.0962940795583567	0.300376049697997
PID_FCER1_PATHWAY	52	0.01	Down	0.33069542776368
	0.0963894629429803	0.300465073459567
SEITZ_NEOPLASTIC_TRANSFORMATION_BY_8P_DELETION_DN	30	0.01
	Down	0.437716242585467	0.0966367648050597
0.301027205681735
BIOCARTA_RANBP2_PATHWAY	11	0.01	Down	0.266224106598106
	0.0967521951363257	0.301178058959684
POMEROY_MEDULLOBLASTOMA_PROGNOSIS_DN	41	0.01	Up
0.209153732974001	0.0971254187300467	0.302130627814228
BILANGES_SERUM_RESPONSE_TRANSLATION	23	0.01	Up
0.12607306639188	0.0976360838429444	0.303493545259633
REACTOME_PHASE_0_RAPID_DEPOLARISATION	18	0.01	Up
0.462968101419805	0.0976985895418661	0.303493545259633
VERRECCHIA_RESPONSE_TO_TGFB1_C2	21	0.01	Up
0.814788325685563	0.0978039047845848	0.303610878457672
HOFMANN_MYELODYSPLASTIC_SYNDROM_RISK_DN	13	0.01	Down
0.340926966939421	0.0980175114083396	0.304063984665622
REACTOME_DEFECTIVE_CFTR_CAUSES_CYSTIC_FIBROSIS	58	0.01	Down
	0.128355751510927	0.0982599305388191
0.304605784670339
WOO_LIVER_CANCER_RECURRENCE_DN	50	0.01	Up
0.337079452637312	0.0984176650848126	0.304717703733643
MCCLUNG_COCAIN_REWARD_4WK	55	0.01	Down

0.494510320293087	0.0984316141982757	0.304717703733643
BYSTROEM_CORRELATED_WITH_IL5_DN 55	0.01	Down
0.233989302259964	0.0991373429723086	0.3066912296356
BOSCO_INTERFERON_INDUCED_ANTIVIRAL_MODULE	69	0.01	Down
	0.423478008773557	0.0993087787148312
0.307010289080582
REACTOME_TRANSCRIPTIONAL_REGULATION_BY_RUNX2	104	0.01	Down
	0.212713247197983	0.0995021314881654
0.307396619270999
GRABARCZYK_BCL11B_TARGETS_UP	69	0.01	Down
0.310287070672214	0.0997242881566377	0.307791197776486
BOYAULT_LIVER_CANCER_SUBCLASS_G56_DN	16	0.01	Down
0.49988357667964	0.0997668020378954	0.307791197776486
REACTOME_SHC1_EVENTS_IN_ERBB2_SIGNALING 18	0.01	Down
0.299384481307036	0.0999424698233737	0.308121674798398
REACTOME_TICAM1_DEPENDENT_ACTIVATION_OF_IRF3_IRF7	12	0.01
	Down	0.204955565485087	0.10011881547369
0.308453787220177
REACTOME_NCAM_SIGNALING_FOR_NEURITE_OUT_GROWTH	41	0.01	Up
	0.748502684055856	0.100490013328925
0.309288156256124
INGRAM_SHH_TARGETS_UP	94	0.01	Up	0.461440267094343
	0.100527251677463	0.309288156256124
SESTO_RESPONSE_TO_UV_C4 20	0.01	Down	0.426113597572617
	0.100759257951555	0.309789920993323
KIM_MYCL1_AMPLIFICATION_TARGETS_DN	17	0.01	Down
0.297690243504064	0.101052192386943	0.310478198755509
REACTOME_PERK_REGULATES_GENE_EXPRESSION 30	0.01	Down
0.229491151120666	0.101674609692542	0.312177165688508
LI_WILMS_TUMOR_VS_FETAL_KIDNEY_2_DN	43	0.01	Up
0.642841698572524	0.102032042310467	0.313060771457714
PID_IL27_PATHWAY	18	0.01	Down	0.544054772550467
	0.102274678138432	0.313591185697306
REACTOME_TRANSCRIPTIONAL_REGULATION_BY_SMALL_RNAS	67	0.01
	Down	0.158336363026043	0.102348551165216
0.313603774701872
REACTOME_REGULATION_OF_APOPTOSIS	51	0.01	Down
0.118337745859056	0.10256038647191 6	0.314038785552631
REACTOME_METABOLISM_OF_WATER_SOLUBLE_VITAMINS_AND_COFACTORS	105
	0.01	Up	0.303861042861785	0.102689378845146
	0.314219712667753
REACTOME_ACTIVATION_OF_IRF3_IRF7_MEDIATED_BY_TBK1_IKK_EPSILON	16
	0.01	Down	0.30141116172198	0.102842633891861
	0.314474584587697
STARK_HYPPOCAMPUS_22Q11_DELETION_UP	43	0.01	Down
0.25683116992442	0.102958793911334	0.314615757057406
VANDESLUIS_NORMAL_EMBRYOS_DN	10	0.01	Up
0.315049634925048	0.10313875801981	0.314865848156763
NATSUME_RESPONSE_TO_INTERFERON_BETA_UP	60	0.01	Down
0.415500247754247	0.103180732888746	0.314865848156763
SUZUKI_RESPONSE_TO_TSA	12	0.01	Down	0.189284543066671
	0.103369400485203	0.315227581533913
REACTOME_INTERLEUKIN_27_SIGNALING	10	0.01	Down
0.601314467947909	0.103753852619731	0.316185469508943

| | | | | | | |
|---|---|---|---|---|---|---|
| WANG_RESPONSE_TO_ANDROGEN_UP | 25 | 0.01 | Down | 0.176707086621396 | 0.103894563311583 | 0.316399771060683 |
| PID_FRA_PATHWAY | 34 | 0.01 | Down | 0.72861690395414 | 0.103971680828768 | 0.316420247342797 |
| XU_GH1_AUTOCRINE_TARGETS_UP | 140 | 0.01 | Up | 0.396566089787274 | 0.104142562534484 | 0.316725858316987 |
| HAN_JNK_SINGALING_UP | 27 | 0.01 | Down | 0.398897459453862 | 0.104315763126636 | 0.317038103620168 |
| NAKAMURA_TUMOR_ZONE_PERIPHERAL_VS_CENTRAL_DN | 597 | 0.01 | Down | 0.381514707965371 | 0.104465448084245 | 0.317278506174785 |
| PID_ERA_GENOMIC_PATHWAY | 53 | 0.01 | Down | 0.330516730490177 | 0.10497972639063 | 0.318625165513761 |
| REACTOME_ACTIVATION_OF_BH3_ONLY_PROTEINS | 29 | 0.01 | Down | 0.192604193961724 | 0.105238156855421 | 0.319194004767286 |
| CHEN_ETV5_TARGETS_SERTOLI | 20 | 0.01 | Up | 1.18236142625731 | 0.105483568715081 | 0.31972261724497 |
| LUI_THYROID_CANCER_CLUSTER_5 | 11 | 0.01 | Up | 0.520361532404542 | 0.105696069263989 | 0.320150829745033 |
| REACTOME_ADRENALINE_NORADRENALINE_INHIBITS_INSULIN_SECRETION | 15 | 0.01 | Up | 0.276746097502134 | 0.106088309029777 | 0.321005585775977 |
| MARKS_ACETYLATED_NON_HISTONE_PROTEINS | 12 | 0.01 | Down | 0.177021324634035 | 0.106165317396468 | 0.321005585775977 |
| LU_TUMOR_VASCULATURE_UP | 25 | 0.01 | Up | 0.903448286570668 | 0.106261264707431 | 0.321005585775977 |
| CHICAS_RB1_TARGETS_SENESCENT | 514 | 0.01 | Down | 0.453293162397293 | 0.106281591877912 | 0.321005585775977 |
| HATADA_METHYLATED_IN_LUNG_CANCER_DN | 14 | 0.01 | Down | 0.560352617787972 | 0.106335331973399 | 0.321005585775977 |
| MORI_PRE_BI_LYMPHOCYTE_UP | 63 | 0.01 | Up | 0.309789509303858 | 0.106542051690175 | 0.321413773387474 |
| PID_RETINOIC_ACID_PATHWAY | 27 | 0.01 | Down | 0.201047416625507 | 0.106815545125267 | 0.322022719877984 |
| REACTOME_SIGNALING_BY_FGFR4 | 31 | 0.01 | Down | 0.251243711280306 | 0.107527666270331 | 0.323786692210059 |
| GILDEA_METASTASIS | 29 | 0.01 | Up | 0.713190055405346 | 0.107668099713104 | 0.323786692210059 |
| KEGG_PARKINSONS_DISEASE | 101 | 0.01 | Up | 0.141128860369037 | 0.107673598805416 | 0.323786692210059 |
| WANG_PROSTATE_CANCER_ANDROGEN_INDEPENDENT | 58 | 0.01 | Up | 0.361657932463738 | 0.107688788621588 | 0.323786692210059 |
| KYNG_WERNER_SYNDROM_UP | 15 | 0.01 | Down | 0.304412647897141 | 0.107816319693173 | 0.323822772732036 |
| KEGG_FOLATE_BIOSYNTHESIS | 10 | 0.01 | Up | 0.737238596548539 | 0.107863361731195 | 0.323822772732036 |
| CUI_TCF21_TARGETS_UP | 26 | 0.01 | Up | 0.92513013107109 | 0.107943564542765 | 0.323822772732036 |
| BROWNE_HCMV_INFECTION_20HR_DN | 91 | 0.01 | Up | 0.400056062537767 | 0.107988951351573 | 0.323822772732036 |
| REACTOME_ADAPTIVE_IMMUNE_SYSTEM | 607 | 0.01 | Down | 0.307064545398158 | 0.108232940554382 | 0.324338045194631 |

KEGG_TRYPTOPHAN_METABOLISM 28 0.01 Up
0.544263451037716 0.108359379585785 0.324500607087346
REACTOME_ACYL_CHAIN_REMODELLING_OF_PC 20 0.01 Up
0.496784450386647 0.108503622834938 0.32471623478232
CHEOK_RESPONSE_TO_HD_MTX_DN 21 0.01 Down
0.139727521103493 0.108853548134417 0.325218774718384
PARK_TRETINOIN_RESPONSE_AND_RARA_PLZF_FUSION 21 0.01 Up
0.748849867988295 0.1088734664957 0.325218774718384
MUELLER_COMMON_TARGETS_OF_AML_FUSIONS_UP 14 0.01 Up
0.175770049313572 0.108957581914488
0.325218774718384
REACTOME_COPI_MEDIATED_ANTEROGRADE_TRANSPORT 88 0.01 Down
0.176466033505996 0.108960950995748
0.325218774718384
SESTO_RESPONSE_TO_UV_C6 38 0.01 Up 0.335547719750937
0.109077419406921 0.325350365118852
CHEOK_RESPONSE_TO_MERCAPTOPURINE_UP 12 0.01 Up
0.255874060422204 0.109642977944091 0.326820415025655
MCMURRAY_TP53_HRAS_COOPERATION_RESPONSE_DN 53 0.01 Up
0.487658611783168 0.109888999716058
0.327336682388124
REACTOME_PLASMA_LIPOPROTEIN_ASSEMBLY 10 0.01 Up
0.652399566195462 0.110008468487662 0.327475540299364
XU_CREBBP_TARGETS_DN 33 0.01 Down 0.370188645070267
0.110183174524355 0.327778537052929
FRASOR_RESPONSE_TO_ESTRADIOL_DN 69 0.01 Down
0.504487137160509 0.110457552954676 0.328377447441315
KEGG_NON_SMALL_CELL_LUNG_CANCER 50 0.01 Down
0.271768345811574 0.110562984176031 0.328473637720595
BIOCARTA_MAL_PATHWAY 17 0.01 Down 0.131021766097445
0.110783178594864 0.32891042786256
YAMASHITA_LIVER_CANCER_STEM_CELL_DN 41 0.01 Up
0.51152845399242 0.110944293124945 0.329052022267242
KEGG_MTOR_SIGNALING_PATHWAY 48 0.01 Down
0.233570007503972 0.110977278255203 0.329052022267242
LEIN_CHOROID_PLEXUS_MARKERS 71 0.01 Up
0.538680420749507 0.111400767496388 0.330089947195955
TCGA_GLIOBLASTOMA_COPY_NUMBER_UP 65 0.01 Down
0.283259423725046 0.111546078572917 0.330302782071978
REACTOME_ERYTHROPOIETIN_ACTIVATES_PHOSPHOINOSITIDE_3_KINASE_PI3K
11 0.01 Down 0.48367976891663
0.111711396713509 0.330527885139676
LIU_TOPBP1_TARGETS 10 0.01 Down 0.399118377122317
0.111769342833892 0.330527885139676
CHEMNITZ_RESPONSE_TO_PROSTAGLANDIN_E2_UP 126 0.01 Up
0.224599984367923 0.111842694838142
0.330527885139676
REACTOME_NONSENSE_MEDIATED_DECAY_NMD 113 0.01 Up
0.125920209568951 0.112009494678274 0.33080333677979
BIOCARTA_AMAN_PATHWAY 12 0.01 Down 0.199931723064231
0.11241198365124 0.331774042358714
GENTILE_UV_RESPONSE_CLUSTER_D5 36 0.01 Down
0.259021623044849 0.112511185344213 0.331775517785591
VALK_AML_CLUSTER_8 20 0.01 Up 0.339533059290084

	0.112560103364411	0.331775517785591
PID_HDAC_CLASSII_PATHWAY	31	0.01	Down
0.232542391707929	0.112714748469765	0.332013626717951
POMEROY_MEDULLOBLASTOMA_PROGNOSIS_UP	41	0.01	Up
0.375296176613361	0.112909601599151	0.332369783358339
BIOCARTA_CXCR4_PATHWAY	17	0.01	Down	0.306953983926252
	0.11300170542241	0.33242321065035
KEGG_GALACTOSE_METABOLISM	21	0.01	Up
0.309911954970475	0.113498865239029	0.333261182743975
MASRI_RESISTANCE_TO_TAMOXIFEN_AND_AROMATASE_INHIBITORS_DN	17
	0.01	Up	0.4637584608491	0.113503127935098
0.333261182743975
KAAB_HEART_ATRIUM_VS_VENTRICLE_DN	227	0.01	Up
0.307332454153373	0.113508981263855	0.333261182743975
RICKMAN_METASTASIS_DN	250	0.01	Down	0.307292154393288
	0.113656529014213	0.333415265821494
BIOCARTA_RAB_PATHWAY	13	0.01	Down	0.144333738946347
	0.113709811458142	0.333415265821494
VALK_AML_CLUSTER_9	29	0.01	Up	0.66833463966694
	0.113855037617864	0.333623464206193
CADWELL_ATG16L1_TARGETS_UP	53	0.01	Down
0.48845976230899	0.114130523006946	0.334077700957929
REACTOME_RRNA_PROCESSING_IN_THE_NUCLEUS_AND_CYTOSOL	190	0.01
	Up	0.11160458380876	0.114158698258371
0.334077700957929
REACTOME_HIV_LIFE_CYCLE	147	0.01	Down	0.162940584937358
	0.114960773018737	0.336057819943669
EGUCHI_CELL_CYCLE_RB1_TARGETS	16	0.01	Up
0.195442805648387	0.114984855856143	0.336057819943669
BOYAULT_LIVER_CANCER_SUBCLASS_G1_UP	113	0.01	Down
0.228398829480486	0.115378899614305	0.336832552310643
PID_CERAMIDE_PATHWAY	43	0.01	Down	0.267166245611929
	0.115399806575838	0.336832552310643
LANDEMAINE_LUNG_METASTASIS	18	0.01	Down
0.495522596038082	0.11565842781685	0.33722834754443
RUTELLA_RESPONSE_TO_HGF_VS_CSF2RB_AND_IL4_UP	380	0.01	Up
	0.485887485376032	0.115732296357728
0.33722834754443
PACHER_TARGETS_OF_IGF1_AND_IGF2_UP	31	0.01	Down
0.414594638252927	0.115787208772881	0.33722834754443
REACTOME_ERYTHROPOIETIN_ACTIVATES_RAS	13	0.01	Down
0.332129700804277	0.115860253715296	0.33722834754443
SMID_BREAST_CANCER_RELAPSE_IN_BONE_DN	240	0.01	Down
0.545022948984772	0.11591052212595	0.33722834754443
KEGG_SNARE_INTERACTIONS_IN_VESICULAR_TRANSPORT	36	0.01	Down
	0.12274924925395	0.116009101535567
0.337296837905804
WANG_RECURRENT_LIVER_CANCER_UP	19	0.01	Down
0.273516817022338	0.116399294195499	0.338212558118143
JOHNSTONE_PARVB_TARGETS_3_DN	798	0.01	Down
0.192566787675368	0.116812747452667	0.339194638113525
KIM_GERMINAL_CENTER_T_HELPER_DN	10	0.01	Down
0.179552559354031	0.117115909200166	0.339764512658387
ONGUSAHA_BRCA1_TARGETS_UP	10	0.01	Up

0.547549684078118	0.117168536325125	0.339764512658387
SIG_PIP3_SIGNALING_IN_B_LYMPHOCYTES	31	0.01	Down
0.362947443550682	0.117235763989579	0.339764512658387
APPIERTO_RESPONSE_TO_FENRETINIDE_DN	47	0.01	Up
0.24998361215287	0.11752417476743	0.340318093674079
VECCHI_GASTRIC_CANCER_ADVANCED_VS_EARLY_DN	103	0.01	Down
	0.382479310661929	0.117578197881167
0.340318093674079
ASTON_MAJOR_DEPRESSIVE_DISORDER_UP	38	0.01	Up
0.339036272578288	0.117773684007041	0.340529701737571
REACTOME_PROTEIN_LOCALIZATION	152	0.01	Up
0.164962100382548	0.117906539842874	0.340529701737571
REACTOME_SIGNALING_BY_NOTCH1_IN_CANCER	54	0.01	Down
0.220053069261413	0.117957206218449	0.340529701737571
KRASNOSELSKAYA_ILF3_TARGETS_UP	33	0.01	Down
0.618134984431012	0.11800192186032	0.340529701737571
CREIGHTON_ENDOCRINE_THERAPY_RESISTANCE_1	472	0.01	Up
	0.314690568821739	0.118030094617828
0.340529701737571
REACTOME_CLASS_A_1_RHODOPSIN_LIKE_RECEPTORS	91	0.01	Down
	0.7394466339687 0.118415824391617	0.341237117221918
KINSEY_TARGETS_OF_EWSR1_FLII_FUSION_DN	292	0.01	Up
0.377586520689002	0.118427119658775	0.341237117221918
BAUS_TFF2_TARGETS_UP	16	0.01	Down	0.280312920708462
	0.118621951936726	0.341579547697362
REACTOME_SIGNALING_BY_FGFR_IN_DISEASE	51	0.01	Down
0.227083614822523	0.118912264506576	0.342196305350229
STAEGE_EWING_FAMILY_TUMOR	16	0.01	Up
0.432575583285332	0.119007813753372	0.342252157915168
REACTOME_TP53_REGULATES_TRANSCRIPTION_OF_ADDITIONAL_CELL_CYCLE_GENES
_WHOSE_EXACT_ROLE_IN_THE_P53_PATHWAY_REMAIN_UNCERTAIN	20	0.01
	Down	0.228300255459342	0.119158036449938
0.342465072789304
HOFFMANN_PRE_BI_TO_LARGE_PRE_BII_LYMPHOCYTE_UP	22	0.01	Up
	0.376634316139093	0.119712729169426
0.343730811332454
FRASOR_RESPONSE_TO_SERM_OR_FULVESTRANT_UP	21	0.01	Down
	0.278897876459683	0.119806096954025
0.343730811332454
GUENTHER_GROWTH_SPHERICAL_VS_ADHERENT_UP	16	0.01	Up
	0.500996589952451	0.119879508245587
0.343730811332454
LENAOUR_DENDRITIC_CELL_MATURATION_UP	97	0.01	Up
0.614974528172835	0.119980541386864	0.343730811332454
REACTOME_DEVELOPMENTAL_BIOLOGY	734	0.01	Down
0.352192317620961	0.12007846673563	0.343730811332454
LU_AGING_BRAIN_DN	116	0.01	Down	0.295405047948625
	0.12011679398127	0.343730811332454
UDAYAKUMAR_MED1_TARGETS_DN	228	0.01	Down
0.276974992686671	0.120133727386715	0.343730811332454
PID_PI3KCI_PATHWAY	43	0.01	Down	0.363741595851361
	0.12030296079840	0.343851663412974
MILI_PSEUDOPODIA_HAPTOTAXIS_DN	647	0.01	Up
0.266889890370809	0.12032895807533	0.343851663412974

HOFFMANN_LARGE_TO_SMALL_PRE_BII_LYMPHOCYTE_UP 127 0.01 Up
0.192961128379698 0.120659498425869
0.344577157194588
REACTOME_CHROMATIN_ORGANIZATION 221 0.01 Down
0.17830292402042 0.121029236980453 0.345323381586474
TUOMISTO_TUMOR_SUPPRESSION_BY_COL13A1_UP 12 0.01 Up
0.994185355076433 0.121074449250341
0.345323381586474
REACTOME_PROLONGED_ERK_ACTIVATION_EVENTS 11 0.01 Down
0.135822213964478 0.121266604075358
0.345652114977003
SU_THYMUS 10 0.01 Down 0.607021214311836
0.121670500285675 0.346583586048232
BIOCARTA_HSWI_SNF_PATHWAY 12 0.01 Down
0.16994357418435 0.121862449108862 0.346781590954395
REACTOME_THE_CANONICAL_RETINOID_CYCLE_IN_RODS_TWILIGHT_VISION 13
0.01 Down 0.728763335596435 0.12189430783269
0.346781590954395
HUTTMANN_B_CLL_POOR_SURVIVAL_DN 51 0.01 Down
0.396078121757326 0.122539453129047 0.348375989952504
OSWALD_HEMATOPOIETIC_STEM_CELL_IN_COLLAGEN_GEL_DN 245 0.01
Down 0.278818207854535 0.122609747743017
0.348375989952504
REACTOME_RESPONSE_TO_ELEVATED_PLATELET_CYTOSOLIC_CA2PLUS 100
0.01 Up 0.440285019013692 0.122750538588299
0.348555698644602
LIEN_BREAST_CARCINOMA_METAPLASTIC_VS_DUCTAL_DN 89 0.01 Down
0.42018923031881 0.122983139754102
0.348995715400687
REACTOME_SPHINGOLIPID_METABOLISM 78 0.01 Up
0.25201281606954 0.123314077791642 0.349598440547148
RAMALHO_STEMNESS_DN 50 0.01 Up 0.494022708059317
0.123351084918304 0.349598440547148
KEGG_GLYCEROLIPID_METABOLISM 38 0.01 Up
0.274486524975202 0.123577381770113 0.34992469927264
PID_NOTCH_PATHWAY 51 0.01 Down 0.25141274896914
0.123621895983304 0.34992469927264
KANNAN_TP53_TARGETS_UP 52 0.01 Up 0.373380856833987
0.123857467274882 0.350370871869475
RIZ_ERYTHROID_DIFFERENTIATION_HBZ 24 0.01 Down
0.406183852435169 0.124295938654578 0.351390090724734
HANN_RESISTANCE_TO_BCL2_INHIBITOR_UP 23 0.01 Up
0.586779225597251 0.124427939934182 0.351542168450126
HEDENFALK_BREAST_CANCER_BRACX_DN 18 0.01 Down
0.155974369154091 0.12451984073410 4 0.35158083172098
REACTOME_GAP_JUNCTION_TRAFFICKING_AND_REGULATION 33 0.01
Down 0.38025355960663 0.124665870624294
0.351708788878734
KEGG_WNT_SIGNALING_PATHWAY 121 0.01 Down
0.288882983544511 0.124721648381024 0.351708788878734
REACTOME_P38MAPK_EVENTS 13 0.01 Down 0.20946952677666
0.124866615701752 0.351896826068575
KORKOLA_EMBRYONAL_CARCINOMA_UP 38 0.01 Up
0.272177360181597 0.125254785779124 0.352769587767645

HESS_TARGETS_OF_HOXA9_AND_MEIS1_DN 68 0.01 Up
0.874363346042795 0.125541943627514 0.353356942145069
JOHNSTONE_PARVB_TARGETS_1_DN 60 0.01 Down
0.413021539902325 0.125714117594063 0.353620124271159
REACTOME_CROSS_PRESENTATION_OF_SOLUBLE_EXOGENOUS_ANTIGENS_ENDOSOMES
45 0.01 Down 0.187487648649124
0.125807076769458 0.353660293982936
BIOCARTA_IGF1_PATHWAY 20 0.01 Down 0.25113202840444
0.126069306399766 0.353919112365256
REACTOME_SIALIC_ACID_METABOLISM 26 0.01 Up
0.38524969320657 0.126141688253478 0.353919112365256
GARGALOVIC_RESPONSE_TO_OXIDIZED_PHOSPHOLIPIDS_TAN_DN 11 0.01
Down 0.215436009241245 0.126173593105933
0.353919112365256
REACTOME_SHC_MEDIATED_CASCADE:FGFR3 11 0.01 Down
0.396866429542535 0.126214090572081 0.353919112365256
REACTOME_RUNX1_REGULATES_GENES_INVOLVED_IN_MEGAKARYOCYTE_DIFFERENTIA
TION_AND_PLATELET_FUNCTION 52 0.01 Down
0.188382436989228 0.126511245014026 0.354531200958883
TURASHVILI_BREAST_DUCTAL_CARCINOMA_VS_DUCTAL_NORMAL_DN 168 0.01
Down 0.532155781344075 0.126643630989143
0.354681072458691
PID_INTEGRIN_A9B1_PATHWAY 20 0.01 Down
0.385661243542691 0.127166647202333 0.355924084168424
WILCOX_RESPONSE_TO_PROGESTERONE_DN 49 0.01 Up
0.668875103524599 0.12732185996322 0.356136752043978
RICKMAN_HEAD_AND_NECK_CANCER_A 65 0.01 Up
0.645872430024005 0.127771092165286 0.357171056768011
HORTON_SREBF_TARGETS 24 0.01 Down 0.239180455929817
0.127885973604222 0.35727001326972
SARTIPY_NORMAL_AT_INSULIN_RESISTANCE_UP 29 0.01 Down
0.238939082354231 0.128269745699558 0.357973416046429
REICHERT_MITOSIS_LIN9_TARGETS 23 0.01 Up
0.309166338582195 0.128340265110609 0.357973416046429
REACTOME_MOLECULES_ASSOCIATED_WITH_ELASTIC_FIBRES 31 0.01
Up 0.651704583855137 0.128376673340788
0.357973416046429
MORI_LARGE_PRE_BII_LYMPHOCYTE_UP 72 0.01 Up
0.192925464489447 0.12851774574038 0.358067147390379
REACTOME_SIGNALING_BY_FGFR2 60 0.01 Down
0.206186770325974 0.128640843101527 0.358067147390379
BECKER_TAMOXIFEN_RESISTANCE_DN 48 0.01 Down
0.468526291476182 0.128726715270826 0.358067147390379
HOWLIN_CITED1_TARGETS_2_UP 13 0.01 Down
0.463443390122306 0.128728923288733 0.358067147390379
VERRECCHIA_RESPONSE_TO_TGFB1_C3 14 0.01 Up
0.571258340458886 0.128868794337889 0.35823452724107
REACTOME_PTEN_REGULATION 133 0.01 Down
0.171194344125775 0.129132290262839 0.358745145075067
REACTOME_PTK6_REGULATES_RHO_GTPASES_RAS_GTPASE_AND_MAP_KINASES 13
0.01 Down 0.195497965152317 0.129230795342664
0.358797050688866
REACTOME_PI_3K_CASCADE:FGFR1 11 0.01 Down
0.46636772183971 0.129673092760038 0.359347895559104

SENGUPTA_NASOPHARYNGEAL_CARCINOMA_UP 269 0.01 Up
0.506946633760288 0.129707769187063 0.359347895559104
KRIGE_RESPONSE_TO_TOSEDOSTAT_24HR_DN 952 0.01 Up
0.210842502830902 0.12974620775566 0.359347895559104
VERRECCHIA_RESPONSE_TO_TGFB1_C4 11 0.01 Up
0.59292115842876 0.129821353496862 0.359347895559104
PID_NEPHRIN_NEPH1_PATHWAY 28 0.01 Down
0.290575124168008 0.129906637143784 0.359347895559104
LI_INDUCED_T_TO_NATURAL_KILLER_DN 114 0.01 Up
0.487887727147961 0.130036338297325 0.359347895559104
ICHIBA_GRAFT_VERSUS_HOST_DISEASE_35D_UP 110 0.01 Up
0.721544494674751 0.130044067745418 0.359347895559104
BIOCARTA_CALCINEURIN_PATHWAY 17 0.01 Down
0.237336804627969 0.130068748848646 0.359347895559104
REACTOME_FOXO_MEDIATED_TRANSCRIPTION_OF_CELL_DEATH_GENES 15
0.01 Down 0.263717165720687 0.130309482383587
0.359791844787606
BILANGES_SERUM_AND_RAPAMYCIN_SENSITIVE_GENES 48 0.01 Up
0.10306560354435 0.130389740521069
0.359792439313816
GAVIN_IL2_RESPONSIVE_FOXP3_TARGETS_UP 15 0.01 Down
0.450083843489658 0.130600396535072 0.359953070778032
KAUFFMANN_MELANOMA_RELAPSE_UP 51 0.01 Up
0.201743704794504 0.130620138735788 0.359953070778032
REACTOME_RRNA_PROCESSING 200 0.01 Up
0.110692822631677 0.130688189434872 0.359953070778032
BASSO_HAIRY_CELL_LEUKEMIA_UP 67 0.01 Up
0.46769789714401 0.13079587716202 0.360029067877085
WANG_RESPONSE_TO_FORSKOLIN_UP 18 0.01 Down
0.176733308218301 0.13089033575142 0.360068579683375
MARKS_HDAC_TARGETS_DN 13 0.01 Down 0.57597912257392
0.13128174317137 0.36092442541609
SHIN_B_CELL_LYMPHOMA_CLUSTER_5 10 0.01 Down
0.554212516136558 0.131405119830054 0.361042795621085
SAGIV_CD24_TARGETS_UP 20 0.01 Down 0.29606288625629
0.131890942131109 0.362156252217063
KEGG_COLORECTAL_CANCER 58 0.01 Down 0.242125622495758
0.132209889047381 0.362810409809511
UDAYAKUMAR_MED1_TARGETS_UP 133 0.01 Down
0.208835344291833 0.132731957773548 0.363488683784651
KAMMINGA_EZH2_TARGETS 36 0.01 Up 0.17166037761595
0.132732898607816 0.363488683784651
SPIRA_SMOKERS_LUNG_CANCER_UP 35 0.01 Down
0.569507418056306 0.132863465134384 0.363488683784651
REACTOME_DOWNREGULATION_OF_TGF_BETA_RECEPTOR_SIGNALING 26 0.01
Down 0.178409273594467 0.132881919095664
0.363488683784651
TIEN_INTESTINE_PROBIOTICS_2HR_UP 30 0.01 Up
0.264114934495346 0.132884131595354 0.363488683784651
MCDOWELL_ACUTE_LUNG_INJURY_UP 41 0.01 Down
0.667656519668874 0.132942246082751 0.363488683784651
MARTENS_TRETINOIN_RESPONSE_UP 379 0.01 Up
0.614014355285589 0.133244659570412 0.36409406976839
LEE_LIVER_CANCER_CIPROFIBRATE_DN 34 0.01 Up

0.648674440496707	0.133430657008571	0.364380803920734
LIANG_HEMATOPOIESIS_STEM_CELL_NUMBER_QTL	12	0.01	Up
	0.156937647055928	0.133819047192534
0.365219561099235
MIKKELSEN_ES_ICP_WITH_H3K4ME3	516	0.01	Up
0.373111910376451	0.133948002062402	0.365349677955398
KRASNOSELSKAYA_ILF3_TARGETS_DN	38	0.01	Up
0.387709328341535	0.134101583102604	0.36554676534033
ODONNELL_TARGETS_OF_MYC_AND_TFRC_DN	35	0.01	Up
0.238087608015558	0.134201971618447	0.365598704499951
KORKOLA_YOLK_SAC_TUMOR_UP	20	0.01	Up
0.192986014277271	0.134492400039587	0.366167981936974
CHUNG_BLISTER_CYTOTOXICITY_DN	36	0.01	Down
0.304910542934994	0.134644687233215	0.366360695589166
REACTOME_DOWNSTREAM_SIGNALING_OF_ACTIVATED_FGFR4	17	0.01
	Down	0.332587858843321	0.13492903831159
0.366912297163095
BIOCARTA_WNT_PATHWAY	21	0.01	Down	0.175143212271436
	0.135241199568674	0.367538810194189
REACTOME_CRISTAE_FORMATION	29	0.01	Up
0.108857808452517	0.135480861459809	0.367821150644691
REN_ALVEOLAR_RHABDOMYOSARCOMA_DN	406	0.01	Up
0.359978118095357	0.135598533494511	0.367821150644691
REACTOME_INTEGRIN_SIGNALING	21	0.01	Down
0.286229454808779	0.135649499118516	0.367821150644691
REACTOME_SIGNALING_BY_WNT_IN_CANCER	31	0.01	Down
0.15753495415325	0.135781480408912	0.367821150644691
VILIMAS_NOTCH1_TARGETS_DN	12	0.01	Up
0.663455216064056	0.13582876719771	0.367821150644691
KEGG_VIRAL_MYOCARDITIS	52	0.01	Up	0.544212997766167
	0.135836064531744	0.367821150644691
SCIBETTA_KDM5B_TARGETS_DN	69	0.01	Up
0.265384096548384	0.135946510543914	0.367898594157069
REACTOME_COPII_MEDIATED_VESICLE_TRANSPORT	63	0.01	Down
	0.226405795276949	0.136112868167753
0.367956469780125
YAO_TEMPORAL_RESPONSE_TO_PROGESTERONE_CLUSTER_8	41	0.01	Down
	0.34073445422844	0.136244857275435
0.367956469780125
OHM_METHYLATED_IN_ADULT_CANCERS	21	0.01	Up
0.52428336382062	0.136288030193742	0.367956469780125
MORI_MATURE_B_LYMPHOCYTE_DN	60	0.01	Up
0.264834252963513	0.136295333077621	0.367956469780125
WU_HBX_TARGETS_1_UP	13	0.01	Down	0.448585986339886
	0.136507237768072	0.368307343197769
SASSON_RESPONSE_TO_FORSKOLIN_UP	79	0.01	Down
0.327033831329886	0.136683181074191	0.368560827191656
REN_BOUND_BY_E2F	54	0.01	Up	0.16853738514423
	0.137117999525895	0.369511635413008
ZHAN_MULTIPLE_MYELOMA_CD1_VS_CD2_UP	56	0.01	Down
0.490185981684057	0.137349699271671	0.369914258973134
ZHOU_CELL_CYCLE_GENES_IN_IR_RESPONSE_6HR	73	0.01	Up
	0.264453858113167	0.137554444850145
0.370243850060718

REACTOME_REGULATION_OF_IFNA_SIGNALING 11 0.01 Down
0.254991164290077 0.138289618786513 0.371630537919232
REACTOME_ASSEMBLY_AND_CELL_SURFACE_PRESENTATION_OF_NMDA_RECEPTORS
20 0.01 Down 0.219178700853432
0.138302049532807 0.371630537919232
SATO_SILENCED_BY_METHYLATION_IN_PANCREATIC_CANCER_2 32 0.01
Down 0.535261836809153 0.138317661832898
0.371630537919232
PLASARI_TGFB1_TARGETS_10HR_DN 191 0.01 Up
0.506865514390276 0.138449557692335 0.371705846866982
WINNEPENNINCKX_MELANOMA_METASTASIS_DN 28 0.01 Up
0.287444675998159 0.138511077531078 0.371705846866982
REACTOME_SIGNALING_BY_NOTCH1 67 0.01 Down
0.237354055545673 0.138885319630681 0.372487775501141
TAKEDA_TARGETS_OF_NUP98_HOXA9_FUSION_16D_UP 118 0.01 Down
0.580206501326396 0.139173479067624
0.372671056404368
GAVIN_FOXP3_TARGETS_CLUSTER_P6 64 0.01 Up
0.402402267885841 0.139185158337963 0.372671056404368
ELLWOOD_MYC_TARGETS_UP 12 0.01 Down 0.469096604489447
0.139214297741776 0.372671056404368
HAHTOLA_SEZARY_SYNDROM_DN 21 0.01 Down
0.730837012695502 0.139310206745495 0.372671056404368
BIOCARTA_CDK5_PATHWAY 12 0.01 Down 0.255657820419946
0.139368197066906 0.372671056404368
ZHAN_EARLY_DIFFERENTIATION_GENES_DN 37 0.01 Down
0.367724096642919 0.139548450077055 0.372931202792129
KEGG_DNA_REPLICATION 34 0.01 Up 0.124034669129386
0.139649784210577 0.372980261453681
RAO_BOUND_BY_SALL4 180 0.01 Down 0.286540971698895
0.139852917566957 0.373300988398736
HALMOS_CEBPA_TARGETS_UP 45 0.01 Down 0.75259655659372
0.139937978673183 0.373306358537659
REACTOME_TRANSLESION_SYNTHESIS_BY_POLK 17 0.01 Up
0.139385932819339 0.140023942204392 0.373314128237689
JOHANSSON_GLIOMAGENESIS_BY_PDGFB_UP 53 0.01 Up
0.348731462985498 0.140204080744861 0.373430287446322
STREICHER_LSM1_TARGETS_UP 42 0.01 Down
0.340640242342831 0.140233665230121 0.373430287446322
DACOSTA_UV_RESPONSE_VIA_ERCC3_UP 299 0.01 Up
0.201151441785293 0.140492769287869 0.373787813301502
REACTOME_IRAK2_MEDIATED_ACTIVATION_OF_TAK1_COMPLEX 10 0.01
Down 0.254818929433222 0.140534239038829
0.373787813301502
PID_AURORA_B_PATHWAY 29 0.01 Up 0.238228158169841
0.140666078455029 0.373917222149826
KALMA_E2F1_TARGETS 11 0.01 Up 0.240201742578015
0.140959638752811 0.374471254133281
REACTOME_CA_DEPENDENT_EVENTS 26 0.01 Down
0.272375754564747 0.141041119743636 0.374471254133281
REACTOME_PLATELET_HOMEOSTASIS 57 0.01 Up
0.506433256089606 0.141667399493192 0.375806334446435
YAMAZAKI_TCEB3_TARGETS_UP 143 0.01 Up
0.399548713536844 0.141718942714087 0.375806334446435

REACTOME_INLB_MEDIATED_ENTRY_OF_LISTERIA_MONOCYTOGENES_INTO_HOST_CEL
L	13	0.01	Down	0.159107799120098
0.141828257131641	0.375806334446435
REACTOME_VIRAL_MESSENGER_RNA_SYNTHESIS	44	0.01	Down
0.143230223288355	0.141878386997909	0.375806334446435
GAUSSMANN_MLL_AF4_FUSION_TARGETS_A_UP	162	0.01	Down
0.393341357796452	0.142844819050563	0.378143381408881
RAMASWAMY_METASTASIS_DN	38	0.01	Up	0.531473535124812
	0.143195706101189	0.378660544487127
REACTOME_MITOTIC_METAPHASE_AND_ANAPHASE	175	0.01	Down
0.156401894545481	0.143208659761538	0.378660544487127
REACTOME_GROWTH_HORMONE_RECEPTOR_SIGNALING	19	0.01	Down
	0.355038258723001	0.14429838749734
0.381074871198144
NIKOLSKY_BREAST_CANCER_16P13_AMPLICON	87	0.01	Up
0.228390887738704	0.144445340725112	0.381074871198144
REACTOME_RHO_GTPASES_ACTIVATE_WASPS_AND_WAVES	34	0.01	Down
	0.258464549191853	0.144460857926703
0.381074871198144
PID_NETRIN_PATHWAY	26	0.01	Down	0.233763022227539
	0.144535111149593	0.381074871198144
NIKOLSKY_BREAST_CANCER_6P24_P22_AMPLICON	20	0.01	Down
	0.253618528640604	0.144545640799296
0.381074871198144
LINDVALL_IMMORTALIZED_BY_TERT_UP	63	0.01	Up
0.467922089281114	0.14463656262916	0.381091060385741
BOYAULT_LIVER_CANCER_SUBCLASS_G6_UP	55	0.01	Up
0.259457277474356	0.14479230576332	0.38116875557981
TERAMOTO_OPN_TARGETS_CLUSTER_6	23	0.01	Up
0.526291437151284	0.144835647281494	0.38116875557981
GENTILE_UV_LOW_DOSE_UP	24	0.01	Down	0.20166818560894
	0.144944002574045	0.381230714786619
KAAB_FAILED_HEART_ATRIUM_UP	26	0.01	Up
0.420717892903145	0.145103153418199	0.381426125505733
REACTOME_ADVANCED_GLYCOSYLATION_ENDPRODUCT_RECEPTOR_SIGNALING	11
	0.01	Down	0.185202533906782	0.145323461028687
	0.381781973888922
REACTOME_INTRA_GOLGI_AND_RETROGRADE_GOLGI_TO_ER_TRAFFIC	181	0.01
	Down	0.17741741329877	0.145492343579232
0.381786194414013
SA_PROGRAMMED_CELL_DEATH	12	0.01	Down
0.315504647197212	0.145494939050324	0.381786194414013
REACTOME_BUDDING_AND_MATURATION_OF_HIV_VIRION	28	0.01	Down
	0.166045287341524	0.145789368434631
0.382335595748929
PID_P38_MK2_PATHWAY	20	0.01	Down	0.207232941096542
	0.146042606334243	0.38277639386147
GAUSSMANN_MLL_AF4_FUSION_TARGETS_D_UP	26	0.01	Up
0.555449830947514	0.146424342920333	0.383553275889801
MARTORIATI_MDM4_TARGETS_FETAL_LIVER_DN	446	0.01	Down
0.252157901791851	0.146555439660094	0.383673093344277
SHEPARD_BMYB_MORPHOLINO_DN	138	0.01	Up
0.458296346671485	0.147109083702143	0.384898330175282
KEGG_COMPLEMENT_AND_COAGULATION_CASCADES	37	0.01	Up

0.825098591966199 0.147482177552718
0.385650022163739
ROSS_AML_WITH_PML_RARA_FUSION 64 0.01 Up
0.320543195359702 0.147856336785452 0.386403624331748
ENK_UV_RESPONSE_EPIDERMIS_UP 283 0.01 Down
0.365246816473713 0.147992816210174 0.38653556587143
RAY_TUMORIGENESIS_BY_ERBB2_CDC25A_DN 227 0.01 Down
0.279129598144919 0.148107073294829 0.386547575898605
REACTOME_INTERLEUKIN_12_FAMILY_SIGNALING 47 0.01 Down
0.336225059056214 0.148169404510188
0.386547575898605
HASINA_NOL7_TARGETS_DN 12 0.01 Down 0.545431927941294
0.148256739245032 0.386551068971242
DEN_INTERACT_WITH_LCA5 26 0.01 Down 0.128508657959945
0.148783350201315 0.387669450801792
KEGG_GLIOMA 57 0.01 Down 0.33828936614425
0.148858169540354 0.387669450801792
NAKAMURA_CANCER_MICROENVIRONMENT_UP 17 0.01 Up
0.860447574257139 0.149266024217054 0.388506530894995
TARTE_PLASMA_CELL_VS_B_LYMPHOCYTE_DN 32 0.01 Down
0.328985995706462 0.149899107690451 0.389928523766539
GAZDA_DIAMOND_BLACKFAN_ANEMIA_ERYTHROID_UP 25 0.01 Up
0.216192698654982 0.150014082099057
0.390001908059723
PID_IL4_2PATHWAY 42 0.01 Down 0.512441042970173
0.150364416375606 0.390540129862257
ZHAN_MULTIPLE_MYELOMA_LB_UP 36 0.01 Down
0.369230081360702 0.150418293619329 0.390540129862257
KEGG_ALLOGRAFT_REJECTION 23 0.01 Up
0.926788683313594 0.150622577680973 0.390540129862257
DER_IFN_GAMMA_RESPONSE_UP 69 0.01 Down
0.254141184836647 0.150732237600154 0.390540129862257
NOUSHMEHR_GBM_SILENCED_BY_METHYLATION 42 0.01 Down
0.638360041830733 0.150751377936972 0.390540129862257
SMITH_TERT_TARGETS_UP 142 0.01 Down 0.238726257939847
0.150825509081319 0.390540129862257
KENNY_CTNNB1_TARGETS_DN 48 0.01 Down 0.252881748612772
0.15082929153301 0.390540129862257
REACTOME_SUMOYLATION_OF_DNA_METHYLATION_PROTEINS 16 0.01
Up 0.193990659818917 0.150956524924227
0.390586813814755
REACTOME_INTRA_GOLGI_TRAFFIC 42 0.01 Down
0.192977578164046 0.151021108433825 0.390586813814755
KRIGE_RESPONSE_TO_TOSEDOSTAT_6HR_UP 870 0.01 Down
0.262522645001672 0.151371977705394 0.391269143062534
REACTOME_AKT_PHOSPHORYLATES_TARGETS_IN_THE_CYTOSOL 14 0.01
Down 0.12565051539839 0.151638863159157
0.391590834161712
SARTIPY_NORMAL_AT_INSULIN_RESISTANCE_DN 16 0.01 Down
0.204056102345166 0.151670665689776 0.391590834161712
VANTVEER_BREAST_CANCER_ESR1_DN 241 0.01 Down
0.425322368807049 0.151962369798281 0.392118744112097
REACTOME_HEME_BIOSYNTHESIS 10 0.01 Up
0.178621620845698 0.152380921286815 0.392774400987734

ROYLANCE_BREAST_CANCER_16Q_COPY_NUMBER_DN 17 0.01 Up
0.375521630981231 0.152391224765875
0.392774400987734
REACTOME_ANTIGEN_PROCESSING_CROSS_PRESENTATION 91 0.01 Down
0.273467556278556 0.15258495765351
0.393048357967063
REACTOME_INTERLEUKIN_7_SIGNALING 21 0.01 Down
0.321417470371356 0.152698895711159 0.39311657286464
REACTOME_REGULATION_OF_IFNG_SIGNALING 12 0.01 Down
0.223678022125091 0.152858050680369 0.393301051979541
RICKMAN_HEAD_AND_NECK_CANCER_B 21 0.01 Up
0.832579363934779 0.153003702484289 0.39345059649135
PLASARI_NFIC_TARGETS_BASAL_UP 17 0.01 Up
0.790067038136234 0.153535346107779 0.394591984422221
REACTOME_INTERLEUKIN_RECEPTOR_SHC_SIGNALING 21 0.01 Down
0.393162881526542 0.153667776873013
0.39470666116811
TANG_SENESCENCE_TP53_TARGETS_DN 42 0.01 Up
0.328990542539713 0.154210746849326 0.395787899609342
PYEON_CANCER_HEAD_AND_NECK_VS_CERVICAL_DN 19 0.01 Down
0.646524807891783 0.154314804488124
0.395787899609342
PID_ARF6_TRAFFICKING_PATHWAY 40 0.01 Down
0.372532793288953 0.154375016776369 0.395787899609342
DORSAM_HOXA9_TARGETS_DN 29 0.01 Down 0.322986763495989
0.154449762967438 0.395787899609342
BIOCARTA_TEL_PATHWAY 16 0.01 Down 0.207932541589016
0.154571323940751 0.395787899609342
ABE_VEGFA_TARGETS_30MIN 20 0.01 Down 0.583406270158752
0.154617030414684 0.395787899609342
EBAUER_MYOGENIC_TARGETS_OF_PAX3_FOXO1_FUSION 31 0.01 Up
0.824020737172604 0.154882626409229
0.396242120494868
NIKOLSKY_BREAST_CANCER_22Q13_AMPLICON 14 0.01 Down
0.275941546571622 0.155088598970821 0.396543374501614
VANHARANTA_UTERINE_FIBROID_DN 55 0.01 Down
0.431788440865177 0.155329615626374 0.396837135385546
REACTOME_MITOCHONDRIAL_TRNA_AMINOACYLATION 21 0.01 Up
0.115017889636107 0.155380057459079
0.396837135385546
LU_TUMOR_ENDOTHELIAL_MARKERS_UP 19 0.01 Up
0.981365688788896 0.155659185272326 0.397316402302396
LIU_IL13_MEMORY_MODEL_UP 12 0.01 Down
0.151950755124959 0.155827639328842 0.397316402302396
MISSIAGLIA_REGULATED_BY_METHYLATION_DN 108 0.01 Up
0.215533891856262 0.155890673307789 0.397316402302396
PID_CASPASE_PATHWAY 49 0.01 Down 0.328110106909168
0.155921275564277 0.397316402302396
ZHENG_FOXP3_TARGETS_IN_T_LYMPHOCYTE_UP 10 0.01 Up
0.341690536171733 0.156434614117087 0.398261063927434
GERY_CEBP_TARGETS 105 0.01 Down 0.520096428191117
0.156494495677371 0.398261063927434
PID_TCR_CALCIUM_PATHWAY 20 0.01 Down 0.392725577366258
0.156557797543888 0.398261063927434

CARDOSO_RESPONSE_TO_GAMMA_RADIATION_AND_3AB	13	0.01	Down
	0.165596798036891	0.156911999553416
0.398936333706224
FONTAINE_PAPILLARY_THYROID_CARCINOMA_DN	47	0.01	Down
0.428627337783468	0.157116414907312	0.399155151735122
LEIN_LOCALIZED_TO_PROXIMAL_DENDRITES	23	0.01	Up
0.308637039203265	0.15717566597801	0.399155151735122
SINGH_NFE2L2_TARGETS	13	0.01	Up	0.239693580463148
	0.15752800365011	0.399662358506878
REACTOME_SIGNALING_BY_NTRK1_TRKA	72	0.01	Down
0.215715205220114	0.15755321452151	0.399662358506878
REACTOME_STRIATED_MUSCLE_CONTRACTION	18	0.01	Up
0.586870686255327	0.157909893661231	0.400229708903838
REACTOME_TFAP2_AP_2_FAMILY_REGULATES_TRANSCRIPTION_OF_GROWTH_FACTORS
_AND_THEIR_RECEPTORS	10	0.01	Down	0.474464718239265
	0.157978755997292	0.400229708903838
DORSAM_HOXA9_TARGETS_UP	32	0.01	Down	0.217125012746434
	0.158043989611638	0.400229708903838
MULLIGHAN_NPM1_SIGNATURE_3_DN	134	0.01	Down
0.314760751548868	0.158815325436934	0.401910748328819
SU_LIVER	22	0.01	Up	0.362217928065832
0.158886629539558	0.401910748328819
COATES_MACROPHAGE_M1_VS_M2_DN	62	0.01	Up
0.478761879303709	0.159159378328611	0.402374243862265
BASAKI_YBX1_TARGETS_DN	355	0.01	Down	0.349795992923561
	0.159298978904644	0.402500792679243
YIH_RESPONSE_TO_ARSENITE_C1	20	0.01	Down
0.233255696151279	0.15982174011983	0.403594787549795
LOPEZ_EPITHELIOID_MESOTHELIOMA	13	0.01	Down
0.408847707290239	0.159982266271372	0.403773322228982
SNIJDERS_AMPLIFIED_IN_HEAD_AND_NECK_TUMORS	35	0.01	Down
	0.282173851081051	0.160239421364178
0.40419539788551
PID_REELIN_PATHWAY	24	0.01	Down	0.250547168885371
	0.160329484641901	0.404195756290155
BRUECKNER_TARGETS_OF_MIRLET7A3_DN	70	0.01	Up
0.509807159245341	0.1604396912725	0.404199983054205
HASLINGER_B_CLL_WITH_6Q21_DELETION	14	0.01	Down
0.129685578135563	0.160511005506509	0.404199983054205
STEIN_ESRRA_TARGETS_UP	376	0.01	Up	0.230843913445957
	0.16065077811487	0.404325446599296
REACTOME_G_PROTEIN_MEDIATED_EVENTS	38	0.01	Down
0.305153384116532	0.160811050079001	0.404502333578686
BIOCARTA_HIF_PATHWAY	13	0.01	Down	0.260816163839136
	0.161144315228218	0.40507994997905
REACTOME_DARPP_32_EVENTS	20	0.01	Down
0.213099454709758	0.161220918912685	0.40507994997905
REACTOME_PEPTIDE_HORMONE_METABOLISM	45	0.01	Down
0.380110656656716	0.161438024271793	0.405187382077195
LEE_AGING_NEOCORTEX_UP	66	0.01	Up	0.435174402564501
	0.161443960244773	0.405187382077195
RODRIGUES_NTN1_TARGETS_DN	137	0.01	Down
0.390193476778185	0.161654625951534	0.405264702905706
TONKS_TARGETS_OF_RUNX1_RUNX1T1_FUSION_GRANULOCYTE_UP	46	0.01

Up	0.391129558072004	0.161655086164612
0.405264702905706
REACTOME_CLEC7A_DECTIN_1_INDUCES_NFAT_ACTIVATION	11	0.01
Down	0.244955937148402	0.161980900664287
0.405855155231868
OHGUCHI_LIVER_HNF4A_TARGETS_UP	34	0.01	Up
0.460358080101663	0.162145884667013	0.405965337462843
REACTOME_NEGATIVE_REGULATION_OF_FGFR3_SIGNALING 22	0.01	Down
0.26486744095526	0.162205505246555
0.405965337462843
FOURNIER_ACINAR_DEVELOPMENT_LATE_2	265	0.01	Down
0.207635967175355	0.162422684598625	0.406282675164618
REACTOME_ATF4_ACTIVATES_GENES_IN_RESPONSE_TO_ENDOPLASMIC_RETICULUM_S
TRESS	25	0.01	Down	0.249742310958728
0.162635720011534	0.406520179851605
REACTOME_DEGRADATION_OF_DVL	54	0.01	Down
0.125748847053321	0.162698510245392	0.406520179851605
LAU_APOPTOSIS_CDKN2A_UP 54	0.01	Down	0.232771132620925
0.162826799512552	0.406614702116066
REACTOME_MISMATCH_REPAIR	14	0.01	Up
0.150502309831993	0.162978199004711	0.406766798737466
REACTOME_RAB_REGULATION_OF_TRAFFICKING	116	0.01	Down
0.190844898370675	0.163124622141764	0.406906313278153
NEMETH_INFLAMMATORY_RESPONSE_LPS_DN	28	0.01	Up
0.304339586076649	0.16331589509154	0.407157486653616
REACTOME_NUCLEOBASE_CATABOLISM	28	0.01	Down
0.399863540380413	0.163632040829518	0.407657779883138
REACTOME_DUAL_INCISION_IN_GG_NER	40	0.01	Up
0.127499876026989	0.163829144101897	0.407657779883138
ZHONG_SECRETOME_OF_LUNG_CANCER_AND_MACROPHAGE	66	0.01	Up
0.359599771457039	0.163833421554403
0.407657779883138
HOSHIDA_LIVER_CANCER_SURVIVAL_DN	82	0.01	Up
0.302278456598382	0.164017934813662	0.407657779883138
DACOSTA_UV_RESPONSE_VIA_ERCC3_COMMON_UP 67	0.01	Up
0.212725554922565	0.164043867500054	0.407657779883138
KEGG_PATHWAYS_IN_CANCER 271	0.01	Down	0.378628513234968
0.16409047637917	0.407657779883138
TAKAO_RESPONSE_TO_UVB_RADIATION_UP	80	0.01	Up
0.155929408701881	0.164151408584756	0.407657779883138
HOLLMANN_APOPTOSIS_VIA_CD40_DN	234	0.01	Up
0.257848144820509	0.164420165447236	0.407941656234129
BUCKANOVICH_T_LYMPHOCYTE_HOMING_ON_TUMOR_DN	17	0.01	Up
0.301223887538413	0.164447226050332
0.407941656234129
REACTOME_RUNX1_REGULATES_TRANSCRIPTION_OF_GENES_INVOLVED_IN_DIFFEREN
TIATION_OF_HSCS 82	0.01	Down	0.153195471939468
0.164554894925251	0.40798359221677
ZHENG_RESPONSE_TO_ARSENITE_UP	12	0.01	Down
0.39313986291034	0.164743202722997	0.408225301124515
HOFFMANN_PRE_BI_TO_LARGE_PRE_BII_LYMPHOCYTE_DN	59	0.01	Up
0.420413964518283	0.16561310213916
0.410154762598085
NIKOLSKY_BREAST_CANCER_1Q21_AMPLICON	30	0.01	Up

0.205380430273259 0.165833193144205 0.410473680166962
REACTOME_METABOLISM_OF_FOLATE_AND_PTERINES 16 0.01 Up
0.219202832763987 0.166494541885412
0.411652751084602
INAMURA_LUNG_CANCER_SCC_SUBTYPES_UP 14 0.01 Down
0.198674742915658 0.166583259402107 0.411652751084602
MASSARWEH_TAMOXIFEN_RESISTANCE_DN 211 0.01 Up
0.399242693941273 0.166584283475615 0.411652751084602
REACTOME_PHOSPHORYLATION_OF_CD3_AND_TCR_ZETA_CHAINS 15 0.01
Up 0.983986968503533 0.167025187686359
0.412515504752849
KYNG_DNA_DAMAGE_BY_UV 47 0.01 Down 0.287326510111299
0.167743002972581 0.414060844789541
GOTTWEIN_TARGETS_OF_KSHV_MIR_K12_11 60 0.01 Down
0.309847708030579 0.168126985802373 0.414780900758324
ABDELMOHSEN_ELAVL4_TARGETS 14 0.01 Down
0.345675251538829 0.168646941648875 0.415667421021387
REACTOME_CELLULAR_SENESCENCE 144 0.01 Down
0.230753935814241 0.168793184331829 0.415667421021387
CERVERA_SDHB_TARGETS_1_UP 95 0.01 Up
0.56539495860264 0.168794143059162 0.415667421021387
BIOCARTA_MITOCHONDRIA_PATHWAY 19 0.01 Down
0.31557656924886 0.168947387418236 0.415667421021387
DEBIASI_APOPTOSIS_BY_REOVIRUS_INFECTION_DN 261 0.01 Up
0.257590534359441 0.168948693705467
0.415667421021387
REACTOME_CELLULAR_RESPONSE_TO_HYPOXIA 71 0.01 Down
0.156553099146247 0.169293936829794 0.416197084425077
ZHAN_MULTIPLE_MYELOMA_CD1_VS_CD2_DN 43 0.01 Up
0.382820244176126 0.169349158490204 0.416197084425077
WEST_ADRENOCORTICAL_CARCINOMA_VS_ADENOMA_DN 18 0.01 Up
0.777975599106726 0.169585660693445
0.416550570938271
GRESHOCK_CANCER_COPY_NUMBER_UP 274 0.01 Down
0.28763702316404 0.169837620381057 0.416941618576106
REACTOME_PLASMA_LIPOPROTEIN_CLEARANCE 26 0.01 Up
0.385693315963155 0.169935271834692 0.416953628218855
REACTOME_INNATE_IMMUNE_SYSTEM 795 0.01 Down
0.392385993844111 0.170192924703059 0.417357990474768
PODAR_RESPONSE_TO_ADAPHOSTIN_DN 17 0.01 Down
0.27298558062838 0.170546004017303 0.417995795015145
HERNANDEZ_MITOTIC_ARREST_BY_DOCETAXEL_1_DN 30 0.01 Up
0.296400443903333 0.170902609789386
0.418463125941376
EPPERT_LSC_R 40 0.01 Down 0.221516235818427
0.17092286968373 0.418463125941376
MATTHEWS_SKIN_CARCINOGENESIS_VIA_JUN 13 0.01 Down
0.414536858732081 0.171323253402028 0.419077675142854
MURAKAMI_UV_RESPONSE_6HR_DN 18 0.01 Down
0.396391921346545 0.171398353757569 0.419077675142854
CUI_TCF21_TARGETS_DN 25 0.01 Down 0.427146758932268
0.171538443133399 0.419077675142854
REACTOME_PROTEIN_UBIQUITINATION 68 0.01 Down
0.143677688216883 0.171546812516763 0.419077675142854

REACTOME_SIGNALING_BY_WNT 247 0.01 Down
0.23654332703236 0.171863106490015 0.419447159622619
REACTOME_WNT5A_DEPENDENT_INTERNALIZATION_OF_FZD4 13 0.01
Up 0.156635184432795 0.171979691501071
0.419447159622619
REACTOME_PLATELET_ACTIVATION_SIGNALING_AND_AGGREGATION 194 0.01
Up 0.391368967616206 0.172065361540049
0.419447159622619
REACTOME_TRAF6_MEDIATED_INDUCTION_OF_TAK1_COMPLEX_WITHIN_TLR4_COMPLE
X 15 0.01 Down 0.348093441457593
0.172071315315708 0.419447159622619
REACTOME_RNA_POLYMERASE_II_TRANSCRIBES_SNRNA_GENES 74 0.01
Down 0.13022140414543 0.173111928969287
0.421554694866698
BAELDE_DIABETIC_NEPHROPATHY_DN 405 0.01 Down
0.324063476605359 0.173123463119894 0.421554694866698
MAHADEVAN_IMATINIB_RESISTANCE_UP 16 0.01 Down
1.22335007884735 0.173549718174199 0.422363824143489
BIOCARTA_SODD_PATHWAY 10 0.01 Down 0.418729635795018
0.173832141519493 0.422496002165543
ZHOU_INFLAMMATORY_RESPONSE_FIMA_DN 219 0.01 Up
0.295143265007701 0.173836050267973 0.422496002165543
XU_RESPONSE_TO_TRETINOIN_UP 15 0.01 Down
0.452636540609987 0.173886007565351 0.422496002165543
STARK_HYPPOCAMPUS_22Q11_DELETION_DN 17 0.01 Up
0.109176442993679 0.174446604016616 0.42362911132074
VALK_AML_WITH_11Q23_REARRANGED 16 0.01 Up
0.471575454577717 0.174699456036657 0.424014068512296
RAO_BOUND_BY_SALL4_ISOFORM_A 109 0.01 Down
0.279892325752633 0.174953090765079 0.424337273227035
CAIRO_HEPATOBLASTOMA_CLASSES_DN 159 0.01 Up
0.45384616440781 0.175021424819338 0.424337273227035
MARSON_FOXP3_TARGETS_DN 40 0.01 Down 0.315037629933496
0.175168273772375 0.424464361513113
MIKKELSEN_IPS_WITH_HCP_H3K27ME3 28 0.01 Up
0.721417734233457 0.175559951971367 0.425184258680655
MARTINEZ_RB1_TARGETS_UP 567 0.01 Down 0.352420127636825
0.176212962934182 0.426535954975308
DAVICIONI_TARGETS_OF_PAX_FOXO1_FUSIONS_UP 228 0.01 Up
0.422948466500983 0.176313883127079
0.426550540719171
REACTOME_BUTYRATE_RESPONSE_FACTOR_1_BRF1_BINDS_AND_DESTABILIZES_MRNA
17 0.01 Down 0.171142297130193
0.176804539837537 0.42750748067226
BOSCO_ALLERGEN_INDUCED_TH2_ASSOCIATED_MODULE 119 0.01 Down
0.404550878047636 0.177016866250817
0.42779076010614
BROWNE_HCMV_INFECTION_48HR_DN 363 0.01 Up
0.401645393014201 0.177231338805661 0.42807891882399
WEST_ADRENOCORTICAL_TUMOR_UP 276 0.01 Down
0.234380099271181 0.177621096644612 0.428789919128643
ASTON_MAJOR_DEPRESSIVE_DISORDER_DN 130 0.01 Down
0.312665096317448 0.177948659594715 0.429350093869159
SLEBOS_HEAD_AND_NECK_CANCER_WITH_HPV_UP 61 0.01 Down

0.287408559038922	0.178268993085463	0.429892233862208
REACTOME_TYPE_I_HEMIDESMOSOME_ASSEMBLY	11	0.01	Down
0.397312309462276	0.178599734798038	0.430458878239776
PID_INSULIN_PATHWAY	42	0.01	Down	0.252935822803487
0.179424456397078	0.4320294888789
REACTOME_INTRAFLAGELLAR_TRANSPORT	46	0.01	Up
0.173717557648315	0.179443616404206	0.4320294888789
COULOUARN_TEMPORAL_TGFB1_SIGNATURE_DN	115	0.01	Down
0.328189357885735	0.179733745789625	0.432349087796597
REACTOME_HEDGEHOG_LIGAND_BIOGENESIS	58	0.01	Down
0.161981649131828	0.179824708266691	0.432349087796597
REACTOME_FRS_MEDIATED_FGFR1_SIGNALING	13	0.01	Down
0.42653645317719	0.179901006178016	0.432349087796597
PID_IGF1_PATHWAY	29	0.01	Down	0.174004410449286
0.180128910550117	0.432349087796597
REACTOME_RESOLUTION_OF_ABASIC_SITES_AP_SITES	37	0.01	Up
0.144509276831101	0.180140113490396
0.432349087796597
KIM_HYPOXIA	18	0.01	Down	0.332227901665857
0.180196580914941	0.432349087796597
ZHOU_INFLAMMATORY_RESPONSE_LIVE_DN	315	0.01	Up
0.271275180206362	0.180249653065811	0.432349087796597
SMID_BREAST_CANCER_RELAPSE_IN_BRAIN_UP	27	0.01	Up
0.585686378502729	0.180350325771235	0.432359847648908
MIKKELSEN_MEF_ICP_WITH_H3K4ME3_AND_H3K27ME3	19	0.01	Down
0.548308381001176	0.180537661607285
0.432578245695494
FURUKAWA_DUSP6_TARGETS_PCI35_DN	59	0.01	Up
0.316690277830548	0.18093672245229	0.433100809619347
BIOCARTA_INFLAM_PATHWAY	16	0.01	Down	0.901343957146302
0.181084640458447	0.433100809619347
GRAESSMANN_RESPONSE_TO_MC_AND_SERUM_DEPRIVATION_DN	76	0.01
Down	0.306604718391129	0.181089377834528
0.433100809619347
REACTOME_DOWNSTREAM_SIGNALING_OF_ACTIVATED_FGFR2	19	0.01
Down	0.377408517447127	0.181141161754032
0.433100809619347
BOQUEST_STEM_CELL_DN	167	0.01	Down	0.546205521112154
0.18155312350921	0.433855018699573
BASAKI_YBX1_TARGETS_UP	260	0.01	Up	0.284668315016522
0.181729204976127	0.434045045891442
MARTINEZ_RESPONSE_TO_TRABECTEDIN_UP	65	0.01	Up
0.292833673625553	0.182169316284015	0.434865149599918
SONG_TARGETS_OF_IE86_CMV_PROTEIN	52	0.01	Up
0.202650329137225	0.183037860390556	0.436706572428635
REACTOME_NUCLEAR_SIGNALING_BY_ERBB4	24	0.01	Up
0.574692195956927	0.183413096807624	0.436743643890125
REACTOME_REGULATION_OF_TP53_ACTIVITY	151	0.01	Down
0.183417282838824	0.183416448947976	0.436743643890125
SCHAEFFER_PROSTATE_DEVELOPMENT_48HR_UP	343	0.01	Down
0.410848461247072	0.183677283428139	0.436743643890125
CHARAFE_BREAST_CANCER_LUMINAL_VS_BASAL_UP	347	0.01	Down
0.289559997749282	0.183731898384246
0.436743643890125

CHNG_MULTIPLE_MYELOMA_HYPERPLOID_UP 52 0.01 Up
0.108026026147142 0.183791003811706 0.436743643890125
WANG_METASTASIS_OF_BREAST_CANCER_ESR1_DN 26 0.01 Up
0.390770198416387 0.183853856340901
0.436743643890125
HOLLEMAN_ASPARAGINASE_RESISTANCE_ALL_DN 26 0.01 Down
0.130529595269767 0.183903034968726 0.436743643890125
REACTOME_INTERLEUKIN_35_SIGNALLING 11 0.01 Down
0.509105082219071 0.183924932784501 0.436743643890125
SUZUKI_RESPONSE_TO_TSA_AND_DECITABINE_1B 11 0.01 Down
0.560815991177725 0.183927857148833
0.436743643890125
ZHAN_MULTIPLE_MYELOMA_HP_DN 44 0.01 Up
0.350660538183436 0.184046026597848 0.436793500294259
BIOCARTA_SPPA_PATHWAY 13 0.01 Up 0.431346692126727
0.184517542906109 0.437681454017393
REACTOME_CELLULAR_RESPONSES_TO_STRESS 367 0.01 Down
0.215186595211798 0.184784415848533 0.438083306560736
KEGG_ARGININE_AND_PROLINE_METABOLISM 41 0.01 Up
0.360904284401997 0.185165346028301 0.438754997573648
BERENJENO_TRANSFORMED_BY_RHOA_FOREVER_UP 18 0.01 Up
0.974085594335342 0.185369762123558
0.439007945598205
RODRIGUES_THYROID_CARCINOMA_ANAPLASTIC_DN 484 0.01 Up
0.326950280140225 0.185897886865626
0.440026857009472
WU_HBX_TARGETS_1_DN 21 0.01 Down 0.22489541880089
0.18600404112051 0.440046402545626
BIOCARTA_ERK_PATHWAY 26 0.01 Down 0.152450115947842
0.186323255861264 0.440452469772635
POOLA_INVASIVE_BREAST_CANCER_DN 92 0.01 Up
0.4558056962517 0.186371656842614 0.440452469772635
CAIRO_PML_TARGETS_BOUND_BY_MYC_UP 19 0.01 Up
0.219703225691107 0.186581480735225 0.440487120952897
REACTOME_NEGATIVE_REGULATION_OF_MET_ACTIVITY 18 0.01 Down
0.154976883806246 0.186582308853018
0.440487120952897
YAMASHITA_LIVER_CANCER_STEM_CELL_UP 39 0.01 Down
0.39332340922176 0.187262187642311 0.44186012254708
BURTON_ADIPOGENESIS_PEAK_AT_8HR 36 0.01 Down
0.265916456189574 0.187541307733244 0.442190008563477
MONTERO_THYROID_CANCER_POOR_SURVIVAL_DN 10 0.01 Up
0.244589866937422 0.187809036151646 0.442190008563477
KEGG_LYSINE_DEGRADATION 41 0.01 Up 0.233944540825461
0.187866189095604 0.442190008563477
MEDINA_SMARCA4_TARGETS 31 0.01 Down 0.197751103645187
0.187888984152923 0.442190008563477
DAVICIONI_PAX_FOXO1_SIGNATURE_IN_ARMS_UP 44 0.01 Up
0.401826868269966 0.188017895253675
0.442190008563477
KEGG_GLYOXYLATE_AND_DICARBOXYLATE_METABOLISM 14 0.01 Up
0.143799733649124 0.1882638581302 0.442190008563477
MIYAGAWA_TARGETS_OF_EWSR1_ETS_FUSIONS_DN 200 0.01 Up
0.624368317856121 0.188289499463696

0.442190008563477
MALONEY_RESPONSE_TO_17AAG_UP 35 0.01 Up
0.274069311546838 0.188339161684587 0.442190008563477
HOEBEKE_LYMPHOID_STEM_CELL_DN 80 0.01 Up
0.265169161439888 0.188373625067074 0.442190008563477
MYLLYKANGAS_AMPLIFICATION_HOT_SPOT_12 11 0.01 Down
0.160183987315131 0.188385732235608 0.442190008563477
TAKEDA_TARGETS_OF_NUP98_HOXA9_FUSION_10D_DN 108 0.01 Up
0.728625106164961 0.188605066981684
0.442473787099515
PID_TRAIL_PATHWAY 28 0.01 Down 0.214319295407591
0.188969908620647 0.443098455529373
PID_TCR_RAS_PATHWAY 13 0.01 Down 0.360539484313865
0.189184111021338 0.443147659688768
REACTOME_ACTIVATION_OF_AMPK_DOWNSTREAM_OF_NMDARS 19 0.01
Down 0.163593515572365 0.189188066505617
0.443147659688768
HEIDENBLAD_AMPLIFIED_IN_PANCREATIC_CANCER 51 0.01 Down
0.174118428746983 0.189376657339676
0.443358372261376
MULLIGHAN_MLL_SIGNATURE_2_DN 232 0.01 Down
0.349953210772197 0.189501270559004 0.443419162500115
NIKOLSKY_BREAST_CANCER_12Q24_AMPLICON 13 0.01 Down
0.159128327315729 0.189804671228327 0.443715927802754
YANG_BREAST_CANCER_ESR1_LASER_DN 45 0.01 Down
0.281748950610652 0.189825523729632 0.443715927802754
ZHU_SKIL_TARGETS_UP 16 0.01 Down 0.389165831552734
0.190183495975139 0.444277569234791
PID_MAPK_TRK_PATHWAY 32 0.01 Down 0.311119261146407
0.190318682314032 0.444277569234791
DELACROIX_RAR_TARGETS_UP 41 0.01 Down
0.371786376441153 0.190362313313951 0.444277569234791
SERVITJA_ISLET_HNF1A_TARGETS_DN 59 0.01 Up
0.459288868370406 0.190777770334707 0.444902362289431
GALLUZZI_PERMEABILIZE_MITOCHONDRIA 40 0.01 Down
0.299295335544992 0.190899423509403 0.444902362289431
MEISSNER_NPC_HCP_WITH_H3K27ME3 26 0.01 Up
0.608110992922039 0.190926953694396 0.444902362289431
POS_RESPONSE_TO_HISTAMINE_UP 11 0.01 Up
0.280316809000024 0.191500271661864 0.445686892737768
HOFMANN_CELL_LYMPHOMA_UP 42 0.01 Down
0.301604928173116 0.191537231519054 0.445686892737768
VERHAAK_GLIOBLASTOMA_MESENCHYMAL 207 0.01 Up
0.495446770785316 0.191561084932006 0.445686892737768
WANG_NFKB_TARGETS 10 0.01 Up 0.697375679561344
0.191817667938603 0.445775792390654
RICKMAN_TUMOR_DIFFERENTIATED_WELL_VS_POORLY_UP 227 0.01 Up
0.240337009721463 0.191893861413286
0.445775792390654
REACTOME_DEGRADATION_OF_CYSTEINE_AND_HOMOCYSTEINE 12 0.01
Up 0.165947022395051 0.191995571301319
0.445775792390654
XU_HGF_TARGETS_REPRESSED_BY_AKT1_DN 55 0.01 Up
0.541797596556809 0.191995980882827 0.445775792390654

VERRECCHIA_DELAYED_RESPONSE_TO_TGFB1	36	0.01	Up
0.493665769313378	0.192204527638891	0.446029608537332
REACTOME_P75NTR_RECRUITS_SIGNALLING_COMPLEXES	13	0.01	Down
	0.204877097706945	0.192657555058718
0.446664692191924
REACTOME_DAG_AND_IP3_SIGNALING	30	0.01	Down
0.28549291390406	0.1926769384116 0.446664692191924
RAGHAVACHARI_PLATELET_SPECIFIC_GENES	61	0.01	Up
0.4598374918779 0.193154457156357	0.447505427046729
CHEMNITZ_RESPONSE_TO_PROSTAGLANDIN_E2_DN	336	0.01	Down
	0.294113098118311	0.193238717218621
0.447505427046729
REACTOME_SULFUR_AMINO_ACID_METABOLISM	22	0.01	Up
0.133338426079379	0.193471269806656	0.447700431426266
FRIDMAN_SENESCENCE_DN	11	0.01	Up	0.824177908781597
	0.193556124406529	0.447700431426266
AMIT_EGF_RESPONSE_20_HELA	11	0.01	Down
0.359906250390393	0.193621721622394	0.447700431426266
BIOCARTA_IL4_PATHWAY	10	0.01	Down	0.239118091696345
	0.19405587978582	0.448473614209388
CARD_MIR302A_TARGETS	65	0.01	Down	0.284701713020284
	0.194603438120271	0.449334081913977
BHAT_ESR1_TARGETS_VIA_AKT1_UP	243	0.01	Down
0.355684358166466	0.194640718473555	0.449334081913977
REACTOME_PROCESSING_OF_CAPPED_INTRON_CONTAINING_PRE_MRNA	238
	0.01	Down	0.102342373177794	0.194728096010774
	0.449334081913977
KANNAN_TP53_TARGETS_DN	22	0.01	Up	0.586666046219745
	0.195149483981646	0.449624555920709
SANA_RESPONSE_TO_IFNG_DN	83	0.01	Up
0.320408322316332	0.195244322959816	0.449624555920709
REACTOME_ROS_AND_RNS_PRODUCTION_IN_PHAGOCYTES	28	0.01	Up
	0.428281965030806	0.195281315583351
0.449624555920709
REACTOME_CYTOSOLIC_TRNA_AMINOACYLATION	24	0.01	Down
0.109761589624897	0.195312827570451	0.449624555920709
REACTOME_CELL_DEATH_SIGNALLING_VIA_NRAGE_NRIF_AND_NADE	72	0.01
	Down	0.269318019990444	0.195354117400032
0.449624555920709
CHEN_LIVER_METABOLISM_QTL_CIS	72	0.01	Up
0.295869945788574	0.195683105648121	0.450151258898825
REACTOME_SUMOYLATION	167	0.01	Down	0.225828656410365
	0.196269313933074	0.451268831779626
QI_HYPOXIA_TARGETS_OF_HIF1A_AND_FOXA2	33	0.01	Up
0.265068736961292	0.196883623674814	0.452395127352749
REACTOME_PIWI_INTERACTING_RNA_PIRNA_BIOGENESIS	20	0.01	Up
	0.29256673472065	0.196960459227882
0.452395127352749
MCBRYAN_TERMINAL_END_BUD_UP	11	0.01	Up
0.678312854045888	0.197103363521765	0.452492144550732
FARMER_BREAST_CANCER_BASAL_VS_LULMINAL	288	0.01	Down
0.342733476625253	0.197328138689022	0.452776918533514
REACTOME_METABOLISM_OF_AMINO_ACIDS_AND_DERIVATIVES	296	0.01
	Up	0.203210989607305	0.197537388002537

0.453025795444593
HOLLEMAN_VINCRISTINE_RESISTANCE_B_ALL_DN 16 0.01 Up
0.187758475942032 0.198022850624503
0.453907554083193
KRIGE_RESPONSE_TO_TOSEDOSTAT_6HR_DN 864 0.01 Up
0.2183757906275 0.198566423659822 0.454921546560091
SU_PLACENTA 20 0.01 Down 0.618980585350637
0.198957840755009 0.455586089757395
WEIGEL_OXIDATIVE_STRESS_BY_HNE_AND_TBH 55 0.01 Down
0.285265491238699 0.199335437139052 0.456218324816721
PID_THROMBIN_PAR4_PATHWAY 10 0.01 Down
0.270965989087014 0.200091423712403 0.457600143917601
MORI_SMALL_PRE_BII_LYMPHOCYTE_DN 65 0.01 Up
0.390417232943571 0.200142799319689 0.457600143917601
TAKEDA_TARGETS_OF_NUP98_HOXA9_FUSION_10D_UP 135 0.01 Down
0.583099442744261 0.200650922822392
0.458528672133528
ROETH_TERT_TARGETS_UP 14 0.01 Down 0.321735827789503
0.201149490082723 0.459434429838334
KEGG_VIBRIO_CHOLERAE_INFECTION 46 0.01 Down
0.234530788242492 0.201302919565513 0.459493242225504
ACEVEDO_LIVER_CANCER_WITH_H3K27ME3_UP 161 0.01 Up
0.51790193655551 0.201716260736393 0.459493242225504
WANG_BARRETTS_ESOPHAGUS_AND_ESOPHAGUS_CANCER_UP 21 0.01 Up
0.685115482646454 0.201842791894406
0.459493242225504
WILENSKY_RESPONSE_TO_DARAPLADIB 28 0.01 Up
0.93895606353935 0.201852342603987 0.459493242225504
REACTOME_NOTCH4_INTRACELLULAR_DOMAIN_REGULATES_TRANSCRIPTION 17
0.01 Down 0.378931737735146 0.201961488565004
0.459493242225504
KONDO_PROSTATE_CANCER_WITH_H3K27ME3 44 0.01 Up
0.707031926655235 0.202080580406778 0.459493242225504
REACTOME_CDK_MEDIATED_PHOSPHORYLATION_AND_REMOVAL_OF_CDC6 68
0.01 Down 0.115323720427312 0.202149099524971
0.459493242225504
COLLER_MYC_TARGETS_UP 24 0.01 Down 0.294526655505744
0.202166561467421 0.459493242225504
KIM_MYC_AMPLIFICATION_TARGETS_UP 177 0.01 Up
0.275441696896831 0.202188060982196 0.459493242225504
RUTELLA_RESPONSE_TO_CSF2RB_AND_IL4_DN 291 0.01 Up
0.487805985923693 0.202247660745217 0.459493242225504
REACTOME_CELLULAR_RESPONSES_TO_EXTERNAL_STIMULI 452 0.01 Down
0.202765869009381 0.202352214684215
0.459493242225504
REACTOME_HEDGEHOG_ON_STATE 80 0.01 Down
0.19381150173838 0.202443570453451 0.459493242225504
REACTOME_EXTENSION_OF_TELOMERES 28 0.01 Up
0.12131514009995 0.202504140789483 0.459493242225504
XU_CREBBP_TARGETS_UP 21 0.01 Up 0.245098765474099
0.203061154123057 0.460524665884532
IKEDA_MIR133_TARGETS_UP 42 0.01 Down 0.233731543091133
0.2035660676185 0.461436951056559
AMIT_SERUM_RESPONSE_480_MCF10A 34 0.01 Down

0.448684666407159	0.203712297417619	0.461535673836793
REACTOME_SIGNALLING_TO_RAS	19	0.01	Down
0.231635891241925	0.203846288454449	0.461606582671411
REACTOME_OPIOID_SIGNALLING	62	0.01	Down
0.279483496271813	0.203972288603921	0.461659333975139
VANDESLUIS_COMMD1_TARGETS_GROUP_4_UP	10	0.01	Up
0.325220027100956	0.204229052513102	0.462007846525614
BIOCARTA_GRANULOCYTES_PATHWAY	10	0.01	Down
0.935508257201257	0.2044118671216 0.4620544813701
VALK_AML_CLUSTER_16	20	0.01	Up	0.295811720135495
0.204455253269217	0.4620544813701
MULLIGHAN_MLL_SIGNATURE_1_UP	347	0.01	Up
0.326747756712716	0.204850721520806	0.462715574490465
REACTOME_REGULATION_OF_TP53_ACTIVITY_THROUGH_ACETYLATION	29
0.01	Down	0.147298629336652	0.205025327718623
0.462877372222606
RAO_BOUND_BY_SALL4_ISOFORM_B	393	0.01	Down
0.308159158872572	0.205446133939146	0.463371963135198
WALLACE_PROSTATE_CANCER_RACE_UP 218	0.01	Up
0.725535389142114	0.205450572308888	0.463371963135198
BIOCARTA_BCR_PATHWAY	29	0.01	Down	0.246718851764741
0.205920565908976	0.464002000968574
REACTOME_PROCESSIVE_SYNTHESIS_ON_THE_C_STRAND_OF_THE_TELOMERE	11
0.01	Up	0.149971164284371	0.205936371953794
0.464002000968574
HASLINGER_B_CLL_WITH_MUTATED_VH_GENES	11	0.01	Up
0.307065204556505	0.206314305488421	0.464443771522618
REACTOME_DEFECTS_IN_VITAMIN_AND_COFACTOR_METABOLISM	20	0.01
Up	0.168509011158703	0.206339090485132
0.464443771522618
DELASERNA_MYOD_TARGETS_DN	50	0.01	Up
0.420219335748287	0.206876579393696	0.465420532720052
LANG_MYB_FAMILY_TARGETS 28	0.01	Down	0.217047823962734
0.207266848167237	0.466065273892811
AKL_HTLV1_INFECTION_UP	26	0.01	Up	0.320549359455167
0.207372816018546	0.466070404001681
REACTOME_METABOLISM_OF_NON_CODING_RNA	54	0.01	Down
0.131352220327849	0.207664250247114	0.466327917272485
REACTOME_CD28_CO_STIMULATION	30	0.01	Down
0.352408911681908	0.207694881063296	0.466327917272485
KEGG_RNA_DEGRADATION	56	0.01	Down	0.150400961450051
0.207961851789465	0.466694220565974
PID_TRKR_PATHWAY	54	0.01	Down	0.236739922907893
0.208402200173151	0.467449046795565
REACTOME_GENE_SILENCING_BY_RNA	88	0.01	Down
0.194100065682998	0.208746723205555	0.467988289680284
BURTON_ADIPOGENESIS_10	28	0.01	Down	0.325248970489697
0.209148629940566	0.468317811187179
JAERVINEN_AMPLIFIED_IN_LARYNGEAL_CANCER 38	0.01	Down
0.212616619901973	0.209203402435941	0.468317811187179
SASSON_RESPONSE_TO_FORSKOLIN_DN 83	0.01	Up
0.372036367186136	0.20930895500776	0.468317811187179
SABATES_COLORECTAL_ADENOMA_DN	120	0.01	Up
0.685614766085618	0.209310452208018	0.468317811187179

REACTOME_CYTOSOLIC_SULFONATION_OF_SMALL_MOLECULES 16 0.01 Up 0.396415395473698 0.209458874978479 0.468416737825006
RORIE_TARGETS_OF_EWSR1_FLI1_FUSION_UP 26 0.01 Up 0.488170281557373 0.210060984560494 0.469529649726215
REACTOME_PRE_NOTCH_EXPRESSION_AND_PROCESSING 68 0.01 Down 0.21341030508896 7 0.210480905928811 0.470234429498014
MCBRYAN_PUBERTAL_TGFB1_TARGETS_UP 154 0.01 Down 0.351796047562323 0.211050545739218 0.471272828165815
VALK_AML_CLUSTER_7 16 0.01 Down 0.398069943060791 0.211762753872449 0.472628390594169
REACTOME_TP53_REGULATES_TRANSCRIPTION_OF_GENES_INVOLVED_IN_G2_CELL_CYCLE_ARREST 16 0.01 Up 0.187444853236324 0.212479346627784 0.47399238863121
PEDERSEN_METASTASIS_BY_ERBB2_ISOFORM_6 25 0.01 Down 0.442576874341211 0.212779289577885 0.474426044966564
WUNDER_INFLAMMATORY_RESPONSE_AND_CHOLESTEROL_UP 47 0.01 Down 0.644696947363975 0.213212913661378 0.475157187361376
BIOCARTA_RAC1_PATHWAY 20 0.01 Down 0.270613913146278 0.213400492342614 0.475171562220112
RIZ_ERYTHROID_DIFFERENTIATION_CCNE1 37 0.01 Up 0.247937055866499 0.213523476721914 0.475171562220112
WENG_POR_TARGETS_GLOBAL_DN 19 0.01 Up 0.409775143090676 0.213677492958265 0.475171562220112
HADDAD_T_LYMPHOCYTE_AND_NK_PROGENITOR_UP 71 0.01 Up 0.35421240834286 0.213717326860769 0.475171562220112
SETLUR_PROSTATE_CANCER_TMPRSS2_ERG_FUSION_UP 61 0.01 Down 0.228146320541417 0.213747919646066 0.475171562220112
BHATI_G2M_ARREST_BY_2METHOXYESTRADIOL_UP 94 0.01 Down 0.423543394422126 0.213909085400477 0.475294779473626
NGO_MALIGNANT_GLIOMA_1P_LOH 15 0.01 Up 0.217938891948157 0.214197769517972 0.475634129479578
TING_SILENCED_BY_DICER 27 0.01 Up 0.761339939847658 0.214527173901234 0.475634129479578
GREENBAUM_E2A_TARGETS_UP 30 0.01 Up 0.403337323788133 0.214542574228432 0.475634129479578
CHEN_LVAD_SUPPORT_OF_FAILING_HEART_DN 25 0.01 Down 0.563878717857406 0.214563807668238 0.475634129479578
REACTOME_ABC_TRANSPORTER_DISORDERS 66 0.01 Down 0.157200611083789 0.214590881998795 0.475634129479578
REACTOME_SYNTHESIS_OF_VERY_LONG_CHAIN_FATTY_ACYL_COAS 17 0.01 Up 0.272205301129104 0.214807841569782 0.475880358726551
LU_IL4_SIGNALING 77 0.01 Down 0.372327884593088 0.215030217799843 0.475932825946982
HOWLIN_CITED1_TARGETS_1_UP 27 0.01 Down 0.387440305315574 0.215043285761584 0.475932825946982
OUELLET_CULTURED_OVARIAN_CANCER_INVASIVE_VS_LMP_DN 22 0.01 Up 0.48841156521804 5 0.21525091828976

0.476076034702849
SAKAI_TUMOR_INFILTRATING_MONOCYTES_UP	24	0.01	Down
0.332767979669256	0.215319817252701	0.476076034702849
WOOD_EBV_EBNA1_TARGETS_UP	102	0.01	Up
0.373943008317774	0.215544773362251	0.476339113207138
REACTOME_AMYLOID_FIBER_FORMATION	54	0.01	Up
0.233259203142573	0.215797726664502	0.476663774622573
LEE_LIVER_CANCER_CIPROFIBRATE_UP	49	0.01	Down
0.626523337582051	0.216045128868263	0.476726857846336
MACLACHLAN_BRCA1_TARGETS_DN	16	0.01	Up
0.381914155687411	0.216107374315831	0.476726857846336
WANG_ESOPHAGUS_CANCER_VS_NORMAL_DN	98	0.01	Down
0.34484384608407	0.216173062032188	0.476726857846336
REACTOME_NEUREXINS_AND_NEUROLIGINS	30	0.01	Down
0.337060246520358	0.216250514604823	0.476726857846336
SMID_BREAST_CANCER_BASAL_UP	517	0.01	Down
0.523304166655794	0.216443227309216	0.476917797428885
YAO_TEMPORAL_RESPONSE_TO_PROGESTERONE_CLUSTER_14	136	0.01
Up	0.217636042179373	0.216573714454117
0.476971507335256
BIOCARTA_EIF4_PATHWAY	23	0.01	Down	0.214055819851812
0.216700975805895	0.47701806378428
VALK_AML_CLUSTER_12	23	0.01	Down	0.315292567317657
0.217039606091883	0.477529627696043
REACTOME_NF_KB_ACTIVATION_THROUGH_FADD_RIP_1_PATHWAY_MEDIATED_BY_CASPASE_8_AND_10	12	0.01	Down	0.24135282440287
0.217288264515752	0.477842832190951
PECE_MAMMARY_STEM_CELL_DN	133	0.01	Down
0.34348795516987	0.217576832313484	0.47824345293355
REACTOME_O_LINKED_GLYCOSYLATION	78	0.01	Up
0.670937877295501	0.217754528467479	0.478400100420976
RODRIGUES_THYROID_CARCINOMA_UP	14	0.01	Up
0.24780106281718	0.218519661292378	0.479632347206623
FISCHER_DIRECT_P53_TARGETS_META_ANALYSIS	292	0.01	Up
0.295702908636325	0.218528820262328
0.479632347206623
HOSHIDA_LIVER_CANCER_LATE_RECURRENCE_DN	47	0.01	Up
0.411466058975389	0.21873393078364	0.479783287972484
REACTOME_PRE_NOTCH_PROCESSING_IN_GOLGI	17	0.01	Down
0.336634158115181	0.218887285878994	0.479783287972484
YAO_TEMPORAL_RESPONSE_TO_PROGESTERONE_CLUSTER_17	179	0.01
Up	0.170539587422178	0.218917802810137
0.479783287972484
TONKS_TARGETS_OF_RUNX1_RUNX1T1_FUSION_ERYTHROCYTE_UP	128	0.01
Up	0.543444129143196	0.21927320733159
0.48029013099159
REACTOME_SIGNALING_BY_NTRKS	90	0.01	Down
0.262719637593095	0.219362767280475	0.48029013099159
WANG_RESPONSE_TO_GSK3_INHIBITOR_SB216763_DN	310	0.01	Up
0.270573725237825	0.219551154240975
0.480468567825307
HSIAO_HOUSEKEEPING_GENES	384	0.01	Up
0.188626330210036	0.219747640276392	0.480638996083281
GINESTIER_BREAST_CANCER_ZNF217_AMPLIFIED_DN	309	0.01	Down

0.185639435216201	0.219987284912234
0.480638996083281
BOYLAN_MULTIPLE_MYELOMA_D_CLUSTER_DN	31	0.01	Down
0.25673163392534	0.220008004406389	0.480638996083281
HELLER_SILENCED_BY_METHYLATION_DN	87	0.01	Down
0.335051775249625	0.220056741699531	0.480638996083281
BIOCARTA_P53_PATHWAY	16	0.01	Up	0.249512103363556
0.220319008556187	0.480978117270549
REACTOME_SUMOYLATION_OF_CHROMATIN_ORGANIZATION_PROTEINS 60	0.01
Down	0.181718403021003	0.220547009908911
0.481242140553667
ODONNELL_TFRC_TARGETS_UP	266	0.01	Down
0.419218668519715	0.220697721528634	0.481337340257744
SUZUKI_AMPLIFIED_IN_ORAL_CANCER 13	0.01	Up
0.474043194613758	0.22100347269565	0.481770421807442
REACTOME_ACTIVATION_OF_THE_AP_1_FAMILY_OF_TRANSCRIPTION_FACTORS 10
0.01	Down	0.356795220946284	0.221410141972706
0.482338334203225
REACTOME_VLDLR_INTERNALISATION_AND_DEGRADATION	12	0.01	Up
0.164460396821773	0.221478603291536
0.482338334203225
MATTIOLI_MULTIPLE_MYELOMA_SUBGROUPS	11	0.01	Up
0.213227646222356	0.22167004026964	0.482521467802437
SHI_SPARC_TARGETS_DN	10	0.01	Up	0.782831257152434
0.221901929697242	0.482792436587174
REACTOME_BIOSYNTHESIS_OF_THE_N_GLYCAN_PRECURSOR_DOLICHOL_LIPID_LINKE
D_OLIGOSACCHARIDE_LLO_AND_TRANSFER_TO_A_NASCENT_PROTEIN 71	0.01
Up	0.276971683891992	0.222048945154696
0.482878572070806
REACTOME_SPRY_REGULATION_OF_FGF_SIGNALING	16	0.01	Down
0.151129225225991	0.222282196660138
0.483143361116202
REACTOME_THE_ROLE_OF_GTSE1_IN_G2_M_PROGRESSION_AFTER_G2_CHECKPOINT
67	0.01	Down	0.146177429600719
0.222531241257073	0.483143361116202
GAZDA_DIAMOND_BLACKFAN_ANEMIA_MYELOID_DN	35	0.01	Up
0.3004491052141 0.222693273160485	0.483143361116202
KEGG_FOCAL_ADHESION	165	0.01	Up	0.530237869837938
0.222723938198233	0.483143361116202
SUMI_HNF4A_TARGETS	12	0.01	Down	0.653377599852715
0.222741475333459	0.483143361116202
BROWNE_HCMV_INFECTION_4HR_DN	205	0.01	Down
0.288480113393053	0.222841945613003	0.483143361116202
REACTOME_PI_3K_CASCADE:FGFR4	10	0.01	Down
0.436962616974214	0.222952570669779	0.483143361116202
PID_AMB2_NEUTROPHILS_PATHWAY	34	0.01	Down
0.672046520168747	0.22303058383006	0.483143361116202
REACTOME_ACTIVATION_OF_THE_PRE_REPLICATIVE_COMPLEX	29	0.01
Up	0.154311769567132	0.223880088593172
0.484749999145621
HANN_RESISTANCE_TO_BCL2_INHIBITOR_DN	33	0.01	Down
0.544430047557051	0.22408758637118	0.484819331860672
WELCH_GATA1_TARGETS	15	0.01	Up	0.315705931484177
0.224190083645466	0.484819331860672

SERVITJA_LIVER_HNF1A_TARGETS_UP 100 0.01 Up
0.408118573173468 0.224235682077494 0.484819331860672
PID_ERB_GENOMIC_PATHWAY 12 0.01 Down 0.215172063189653
0.224374082015598 0.484885335894285
EHRLICH_ICF_SYNDROM_UP 11 0.01 Up 0.32009636139481
0.224591957491633 0.484951461479419
WHITESIDE_CISPLATIN_RESISTANCE_DN 12 0.01 Up
0.292580076736958 0.224636697652369 0.484951461479419
CHIARADONNA_NEOPLASTIC_TRANSFORMATION_CDC25_DN 139 0.01 Up
0.369837493002431 0.22472834132628
0.484951461479419
WIELAND_UP_BY_HBV_INFECTION 90 0.01 Up
0.671423750413809 0.224861088198036 0.485005082269755
RUAN_RESPONSE_TO_TNF_DN 62 0.01 Up 0.421745026628889
0.225190194989888 0.485401413192879
KEGG_TYROSINE_METABOLISM 26 0.01 Up
0.369542016853038 0.225349317810201 0.485401413192879
BOYAULT_LIVER_CANCER_SUBCLASS_G6_DN 18 0.01 Down
0.280278820491765 0.225368798516916 0.485401413192879
KANG_IMMORTALIZED_BY_TERT_DN 65 0.01 Up
0.712417030702169 0.225495495960439 0.485441692692612
CROMER_METASTASIS_DN 77 0.01 Down 0.269463171363843
0.226668466501232 0.487733248886088
KORKOLA_EMBRYONIC_CARCINOMA_VS_SEMINOMA_DN 12 0.01 Up
0.567979058392095 0.227717814063118
0.48953711922841
HOOI_ST7_TARGETS_UP 83 0.01 Down 0.406356780434745
0.227724608744517 0.48953711922841
FISCHER_G2_M_CELL_CYCLE 201 0.01 Down 0.281113115417419
0.227885533850219 0.489648888459242
DOANE_RESPONSE_TO_ANDROGEN_UP 143 0.01 Down
0.309860049827012 0.228260225572152 0.489794716665903
SASSON_RESPONSE_TO_GONADOTROPHINS_UP 77 0.01 Down
0.331013338465844 0.228264935627375 0.489794716665903
SHIPP_DLBCL_VS_FOLLICULAR_LYMPHOMA_UP 42 0.01 Up
0.258462406132843 0.228337235648272 0.489794716665903
JECHLINGER_EPITHELIAL_TO_MESENCHYMAL_TRANSITION_DN 56 0.01
Down 0.330236430746592 0.228389260540986
0.489794716665903
RICKMAN_METASTASIS_UP 322 0.01 Up 0.218774325843325
0.228605190971475 0.489913306812428
ULE_SPLICING_VIA_NOVA2 35 0.01 Down 0.251044637665766
0.228662540087314 0.489913306812428
ODONNELL_TFRC_TARGETS_DN 89 0.01 Up
0.326815541205488 0.228773491528801 0.489917505679829
FERRANDO_T_ALL_WITH_MLL_ENL_FUSION_UP 75 0.01 Up
0.354285065146278 0.229077675345698 0.490335309847101
REACTOME_REGULATION_OF_ACTIN_DYNAMICS_FOR_PHAGOCYTIC_CUP_FORMATION
54 0.01 Down 0.307432623301021
0.229267897790806 0.490508900794705
KEGG_MISMATCH_REPAIR 21 0.01 Up 0.125059465158026
0.229475931888643 0.490615160018862
MARSON_BOUND_BY_FOXP3_STIMULATED 864 0.01 Down
0.254928796661375 0.229535857957656 0.490615160018862

KYNG_ENVIRONMENTAL_STRESS_RESPONSE_NOT_BY_4NQO_IN_OLD 11 0.01
Up 0.321084454485031 0.229960993197873
0.491290239745455
PID_MTOR_4PATHWAY 68 0.01 Down 0.185108918110734
0.230122833054017 0.491402439229361
LUCAS_HNF4A_TARGETS_UP 50 0.01 Down 0.358927155536295
0.230682787214354 0.492364258560551
BIOCARTA_FCER1_PATHWAY 35 0.01 Down 0.276007604550178
0.23110719521711 0.492918028423487
NIKOLSKY_BREAST_CANCER_11Q12_Q14_AMPLICON 117 0.01 Up
0.292509929696557 0.231161558157221
0.492918028423487
TERAMOTO_OPN_TARGETS_CLUSTER_4 13 0.01 Up
0.421739793487603 0.231411163657983 0.493121833554324
ZHONG_SECRETOME_OF_LUNG_CANCER_AND_FIBROBLAST 110 0.01 Up
0.35085818133971 0.231704038632164
0.493121833554324
OLSSON_E2F3_TARGETS_UP 24 0.01 Down 0.620503301937625
0.231743249531907 0.493121833554324
CHARAFE_BREAST_CANCER_LUMINAL_VS_BASAL_DN 443 0.01 Down
0.417625105030494 0.231754326896656
0.493121833554324
HOSHIDA_LIVER_CANCER_LATE_RECURRENCE_UP 54 0.01 Down
0.462594013520377 0.23180565835379 0.493121833554324
BIOCARTA_CTCF_PATHWAY 22 0.01 Down 0.252997637292256
0.232093424342422 0.493500445799047
MARKEY_RB1_ACUTE_LOF_UP 202 0.01 Up 0.229195108716667
0.23241885080462 0.493816329220788
REACTOME_INTERACTION_BETWEEN_L1_AND_ANKYRINS 17 0.01 Up
0.566177890979244 0.232461702476349
0.493816329220788
LIN_APC_TARGETS 68 0.01 Up 0.223355267854862
0.232571989732369 0.493817238472839
FONTAINE_FOLLICULAR_THYROID_ADENOMA_DN 56 0.01 Down
0.345690483934828 0.233139505444526 0.493889379880378
MCCLUNG_CREB1_TARGETS_UP 74 0.01 Up
0.332491279050954 0.233140302461991 0.493889379880378
REACTOME_STABILIZATION_OF_P53 55 0.01 Down
0.156466421386697 0.233273744566023 0.493889379880378
REACTOME_FORMATION_OF_THE_BETA_CATENIN:TCF_TRANSACTIVATING_COMPLEX
49 0.01 Down 0.215408171659914
0.233317853751342 0.493889379880378
REACTOME_SIGNAL_REGULATORY_PROTEIN_FAMILY_INTERACTIONS 13 0.01
Down 0.73626286574758 0.233375684088167
0.493889379880378
WANG_HCP_PROSTATE_CANCER 98 0.01 Down
0.376366890975745 0.233376625392552 0.493889379880378
BENPORATH_ES_WITH_H3K27ME3 489 0.01 Up
0.558565573682175 0.233596364370225 0.493889379880378
KEGG_NUCLEOTIDE_EXCISION_REPAIR 42 0.01 Up
0.135142286867456 0.233608690751702 0.493889379880378
EHLERS_ANEUPLOIDY_UP 40 0.01 Up 0.269941885967083
0.233679861117477 0.493889379880378
REACTOME_G_ALPHA_12_13_SIGNALLING_EVENTS 65 0.01 Down

0.291619568951682 0.233704718799903
0.493889379880378
GOLDRATH_IMMUNE_MEMORY 59 0.01 Up 0.383034357130355
0.23412890298519 0.494553298363922
PID_CXCR4_PATHWAY 91 0.01 Down 0.486345450682459
0.234449617036292 0.49499813460692
REACTOME_IRON_UPTAKE_AND_TRANSPORT 51 0.01 Up
0.337756940731212 0.234741964991086 0.495382691377903
RASHI_RESPONSE_TO_IONIZING_RADIATION_3 41 0.01 Down
0.201957136682763 0.234953616607458 0.495596671351724
REACTOME_RET_SIGNALING 31 0.01 Down 0.347548787117521
0.235120362114074 0.495715772843698
MOLENAAR_TARGETS_OF_CCND1_AND_CDK4_UP 56 0.01 Up
0.232315549200475 0.235466915403278 0.496213682483701
REACTOME_LYSINE_CATABOLISM 11 0.01 Up
0.349387559888984 0.236034603126615 0.497056009061817
JIANG_AGING_HYPOTHALAMUS_DN 31 0.01 Up
0.279156018598922 0.236087781834701 0.497056009061817
MISHRA_CARCINOMA_ASSOCIATED_FIBROBLAST_UP 23 0.01 Up
1.01083003208616 0.23651606734017
0.497582707233923
CHIANG_LIVER_CANCER_SUBCLASS_INTERFERON_UP 17 0.01 Up
0.917730734649325 0.236559342682735
0.497582707233923
KEGG_SMALL_CELL_LUNG_CANCER 77 0.01 Down
0.405756965697822 0.237420989498806 0.499161528436452
REACTOME_TP53_REGULATES_TRANSCRIPTION_OF_CASPASE_ACTIVATORS_AND_CASP
ASES 10 0.01 Up 0.260496761252274
0.238027123853467 0.500003718104363
BIDUS_METASTASIS_UP 191 0.01 Down 0.186108743240581
0.238214499862791 0.500003718104363
KEGG_LEUKOCYTE_TRANSENDOTHELIAL_MIGRATION 90 0.01 Down
0.456586363956567 0.238218570515426
0.500003718104363
KEGG_FC_EPSILON_RI_SIGNALING_PATHWAY 63 0.01 Down
0.37789592946325 0.238266510384771 0.500003718104363
GUO_TARGETS_OF_IRS1_AND_IRS2 77 0.01 Down
0.278759331936869 0.238523381070482 0.50020137065105
EBAUER_TARGETS_OF_PAX3_FOXO1_FUSION_UP 144 0.01 Up
0.506308308345902 0.238671248117635 0.50020137065105
REACTOME_PROCESSING_OF_INTRONLESS_PRE_MRNAS 19 0.01 Down
0.0963439548802781 0.238694536161625
0.50020137065105
TERAMOTO_OPN_TARGETS_CLUSTER_7 16 0.01 Down
0.36119623915776 0.239134728726574 0.500672207711515
CAIRO_HEPATOBLASTOMA_CLASSES_UP 567 0.01 Up
0.150015006890678 0.239254848473151 0.500672207711515
IVANOVSKA_MIR106B_TARGETS 87 0.01 Down
0.234831905881656 0.239406035424709 0.500672207711515
REACTOME_NEPHRIN_FAMILY_INTERACTIONS 17 0.01 Down
0.248197304863873 0.239457053817144 0.500672207711515
LY_AGING_MIDDLE_UP 11 0.01 Up 1.77546164804181
0.239616844859248 0.500672207711515
REACTOME_SIGNALING_BY_NTRK2_TRKB 23 0.01 Down

0.33722430898293	0.239678229184011	0.500672207711515
REACTOME_CYTOCHROME_P450_ARRANGED_BY_SUBSTRATE_TYPE	33	0.01
	Up	0.564264606118447	0.239698907896592
0.500672207711515
SASAKI_TARGETS_OF_TP73_AND_TP63	11	0.01	Down
0.313989671729263	0.240076242941671	0.501153650777321
SAKAI_CHRONIC_HEPATITIS_VS_LIVER_CANCER_UP	81	0.01	Down
	0.248657868168844	0.240152383487063
0.501153650777321
KEGG_GRAFT_VERSUS_HOST_DISEASE	25	0.01	Up
0.922641398328367	0.240462360146212	0.501567660722609
REACTOME_ATTENUATION_PHASE	24	0.01	Down
0.258784396901806	0.240768462144453	0.501609484511633
MIKKELSEN_ES_ICP_WITH_H3K4ME3_AND_H3K27ME3	60	0.01	Up
	0.719516882082789	0.240804496699782
0.501609484511633
BOYLAN_MULTIPLE_MYELOMA_C_D_DN	195	0.01	Down
0.55561667548603	0.240875465288611	0.501609484511633
TCGA_GLIOBLASTOMA_COPY_NUMBER_DN	27	0.01	Up
0.304701778088113	0.240928782438402	0.501609484511633
CHASSOT_SKIN_WOUND	11	0.01	Down	0.567648140100446
	0.241120652589283	0.501776543235567
GAVIN_FOXP3_TARGETS_CLUSTER_P4	83	0.01	Down
0.419821833848732	0.241467189476768	0.502109246876107
HADDAD_B_LYMPHOCYTE_PROGENITOR	229	0.01	Down
0.310716731318127	0.241503935872334	0.502109246876107
LIU_BREAST_CANCER	22	0.01	Down	0.148774733836098
	0.241887870612324	0.502583412222581
JI_METASTASIS_REPRESSED_BY_STK11	21	0.01	Down
0.338755260910754	0.241955618253541	0.502583412222581
JAZAERI_BREAST_CANCER_BRCA1_VS_BRCA2_UP	46	0.01	Down
0.197612801790005	0.242312164066207	0.502706907475173
YAUCH_HEDGEHOG_SIGNALING_PARACRINE_UP	84	0.01	Down
0.533331095004457	0.242342796853983	0.502706907475173
SMIRNOV_RESPONSE_TO_IR_6HR_UP	156	0.01	Up
0.318657605709929	0.242350582535862	0.502706907475173
PID_EPHRINB_REV_PATHWAY	24	0.01	Down	0.462808522832775
	0.242487133889711	0.502758148908787
KIM_ALL_DISORDERS_CALB1_CORR_DN	19	0.01	Down
0.598053191795351	0.242640394677819	0.502843971450806
ZWANG_EGF_PERSISTENTLY_DN	57	0.01	Up
0.27861847914002	0.24316120661232	0.503691070839806
SHIPP_DLBCL_VS_FOLLICULAR_LYMPHOMA_DN	26	0.01	Up
0.45251928432004	0.243838592653383	0.504861572536598
CADWELL_ATG16L1_TARGETS_DN	51	0.01	Down
0.241007544031815	0.24408967627475	0.505148754537294
VERNELL_RETINOBLASTOMA_PATHWAY_UP	68	0.01	Up
0.256070735251676	0.244237213080879	0.505221478508306
REACTOME_SIGNALING_BY_PDGF	52	0.01	Up
0.771906482943649	0.244602835203347	0.505561249635802
BIOCARTA_NKCELLS_PATHWAY	15	0.01	Down
0.2862128037577 0.244714747964936	0.505561249635802
LUI_THYROID_CANCER_CLUSTER_1	50	0.01	Up
0.294864352797629	0.244738883027254	0.505561249635802

FLECHNER_PBL_KIDNEY_TRANSPLANT_REJECTED_VS_OK_UP	61	0.01	Down	0.197632091795512	0.245000956268774	0.505835862225162
REACTOME_BIOTIN_TRANSPORT_AND_METABOLISM	11	0.01	Up	0.216612112317812	0.245113378999637	0.505835862225162
REACTOME_FATTY_ACYL_COA_BIOSYNTHESIS	28	0.01	Up	0.223310594174441	0.245268496674272	0.505835862225162
REACTOME_RHO_GTPASE_EFFECTORS	246	0.01	Down	0.229838101153469	0.24532195320375	0.505835862225162
REACTOME_BBSOME_MEDIATED_CARGO_TARGETING_TO_CILIUM	21	0.01	Up	0.155599361327863	0.245680196683934	0.506113131059034
QI_PLASMACYTOMA_UP	199	0.01	Down	0.472628425079296	0.245681613341671	0.506113131059034
HEIDENBLAD_AMPLICON_12P11_12_DN	21	0.01	Down	0.267752935880883	0.246094103814703	0.506341900588374
SILIGAN_TARGETS_OF_EWS_FLI1_FUSION_UP	12	0.01	Down	0.209948309581163	0.246094900532227	0.506341900588374
FAELT_B_CLL_WITH_VH3_21_UP	42	0.01	Down	0.185443184530778	0.246130601287118	0.506341900588374
BIOCARTA_MPR_PATHWAY	16	0.01	Up	0.338922317410833	0.246257574928393	0.506371362901704
YU_MYC_TARGETS_DN	45	0.01	Up	0.437350296073119	0.246444947613382	0.50652493805311
REACTOME_EPHA_MEDIATED_GROWTH_CONE_COLLAPSE	22	0.01	Down	0.343695381147823	0.247009393910389	0.507453028165996
NABA_ECM_REGULATORS	145	0.01	Up	0.75825064251347	0.247191693417807	0.507503691178392
ZAMORA_NOS2_TARGETS_DN	68	0.01	Up	0.258504587148321	0.247259862887804	0.507503691178392
KEGG_CARDIAC_MUSCLE_CONTRACTION	42	0.01	Up	0.24893123354582	0.24749703382944	0.507592921939892
REACTOME_CYCLIN_A:CDK2_ASSOCIATED_EVENTS_AT_S_PHASE_ENTRY	84	0.01	Down	0.142625145030268	0.247529184625638	0.507592921939892
ROSS_ACUTE_MYELOID_LEUKEMIA_CBF	67	0.01	Up	0.45535153151186	0.248110841292901	0.508267921264856
WOTTON_RUNX_TARGETS_DN	25	0.01	Up	0.421974629571883	0.248124071395549	0.508267921264856
SANSOM_APC_TARGETS_DN	262	0.01	Up	0.45394148372322	0.248197572230558	0.508267921264856
DANG_MYC_TARGETS_UP	122	0.01	Up	0.176248683862772	0.248598537698839	0.508857207174991
REACTOME_NUCLEAR_PORE_COMPLEX_NPC_DISASSEMBLY	35	0.01	Down	0.232602973146546	0.249217826823208	0.509861128462658
BROWNE_HCMV_INFECTION_10HR_UP	80	0.01	Down	0.336555651224201	0.249403416686028	0.509861128462658
REACTOME_SHC_MEDIATED_CASCADE:FGFR1	11	0.01	Down	0.466865874985929	0.249429281755147	0.509861128462658
WAKASUGI_HAVE_ZNF143_BINDING_SITES	57	0.01	Up	0.175082373454019	0.249590975451465	0.509959743024698

| | | | | | | |
|---|---|---|---|---|---|---|
| REACTOME_ENDOSOMAL_SORTING_COMPLEX_REQUIRED_FOR_TRANSPORT_ESCRT | 31 | 0.01 | Down | 0.153753434961604 | 0.249716776952015 | 0.509966829184893 |
| LEE_DOUBLE_POLAR_THYMOCYTE | 17 | 0.01 | Down | 0.378142503237002 | 0.249821347689685 | 0.509966829184893 |
| SCHAVOLT_TARGETS_OF_TP53_AND_TP63 | 15 | 0.01 | Up | 0.38584399773326 | 0.249952520238259 | 0.510002986142067 |
| VANTVEER_BREAST_CANCER_METASTASIS_DN | 108 | 0.01 | Up | 0.334476030866268 | 0.250418840657948 | 0.510722635552393 |
| REACTOME_FORMATION_OF_ATP_BY_CHEMIOSMOTIC_COUPLING | 16 | 0.01 | Up | 0.0915513132313944 | 0.251062166997822 | 0.511802467417328 |
| IWANAGA_CARCINOGENESIS_BY_KRAS_PTEN_UP | 144 | 0.01 | Down | 0.31381623032067 | 0.251443376067164 | 0.512347223672666 |
| BIOCARTA_CARM_ER_PATHWAY | 23 | 0.01 | Down | 0.260470423869998 | 0.252154301673952 | 0.513167374020174 |
| BENPORATH_SUZ12_TARGETS | 494 | 0.01 | Up | 0.57937863292955 | 0.252288408204926 | 0.513167374020174 |
| MURAKAMI_UV_RESPONSE_24HR | 16 | 0.01 | Down | 0.39999425009368 | 0.252332071710893 | 0.513167374020174 |
| PID_BETA_CATENIN_DEG_PATHWAY | 16 | 0.01 | Down | 0.133322388498098 | 0.252457194328333 | 0.513167374020174 |
| REACTOME_ZINC_TRANSPORTERS | 12 | 0.01 | Up | 0.163773724368315 | 0.252746944470053 | 0.513167374020174 |
| RUAN_RESPONSE_TO_TNF_TROGLITAZONE_UP | 10 | 0.01 | Down | 0.539261408010166 | 0.252763378515087 | 0.513167374020174 |
| REACTOME_ACTIVATION_OF_KAINATE_RECEPTORS_UPON_GLUTAMATE_BINDING | 19 | 0.01 | Down | 0.400773709635365 | 0.252944200263326 | 0.513167374020174 |
| KEGG_HEDGEHOG_SIGNALING_PATHWAY | 36 | 0.01 | Down | 0.348338187166128 | 0.252989805708418 | 0.513167374020174 |
| HOFFMANN_SMALL_PRE_BII_TO_IMMATURE_B_LYMPHOCYTE_DN | 28 | 0.01 | Up | 0.34684729792163 | 0.253097720387172 | 0.513167374020174 |
| ALONSO_METASTASIS_NEURAL_UP | 16 | 0.01 | Up | 0.490311773922301 | 0.253118731028426 | 0.513167374020174 |
| REACTOME_MEIOTIC_RECOMBINATION | 41 | 0.01 | Up | 0.160727433727713 | 0.253192973084294 | 0.513167374020174 |
| IVANOVA_HEMATOPOIESIS_STEM_CELL_SHORT_TERM | 16 | 0.01 | Down | 0.329839416818468 | 0.253215847736762 | 0.513167374020174 |
| COLDREN_GEFITINIB_RESISTANCE_UP | 84 | 0.01 | Up | 0.272338957154248 | 0.253467466373261 | 0.513261178794031 |
| VART_KSHV_INFECTION_ANGIOGENIC_MARKERS_UP | 112 | 0.01 | Down | 0.774908299513294 | 0.253667493508326 | 0.513261178794031 |
| REACTOME_MITOCHONDRIAL_BIOGENESIS | 90 | 0.01 | Down | 0.208369547201277 | 0.253836067306102 | 0.513261178794031 |
| ZHAN_MULTIPLE_MYELOMA_CD1_DN | 42 | 0.01 | Down | 0.289482598302622 | 0.253926100426332 | 0.513261178794031 |
| ALCALAY_AML_BY_NPM1_LOCALIZATION_UP | 115 | 0.01 | Up | 0.442500709725554 | 0.253958340699276 | 0.513261178794031 |
| BIOCARTA_ARENRF2_PATHWAY | 17 | 0.01 | Down | 0.276779746539709 | 0.253986728788167 | 0.513261178794031 |

YAMASHITA_METHYLATED_IN_PROSTATE_CANCER 42 0.01 Up
0.914798771440534 0.254061428880249 0.513261178794031
MCBRYAN_PUBERTAL_TGFB1_TARGETS_DN 52 0.01 Down
0.68058201157147 0.25493809150496 0.51472407534997
MITSIADES_RESPONSE_TO_APLIDIN_DN 231 0.01 Down
0.227423025469856 0.255014575262377 0.51472407534997
PUJANA_ATM_PCC_NETWORK 1173 0.01 Down 0.236211720581699
0.255361162439793 0.515192291367536
REACTOME_MITOTIC_PROPHASE 100 0.01 Down
0.179919835650063 0.255713529757976 0.51566417169937
REACTOME_GLUCAGON_TYPE_LIGAND_RECEPTORS 14 0.01 Up
0.360988310301208 0.255824494524938 0.51566417169937
MARTIN_VIRAL_GPCR_SIGNALING_DN 37 0.01 Down
0.265293718413287 0.256421589727114 0.516349380925282
LU_TUMOR_ANGIOGENESIS_UP 23 0.01 Up
0.689211387544228 0.256538524323444 0.516349380925282
KIM_MYCN_AMPLIFICATION_TARGETS_DN 103 0.01 Down
0.363120680738379 0.256619009806175 0.516349380925282
DARWICHE_PAPILLOMA_RISK_LOW_UP 117 0.01 Down
0.357024435147875 0.256671778648366 0.516349380925282
REACTOME_O_LINKED_GLYCOSYLATION_OF_MUCINS 40 0.01 Down
0.442254179956626 0.256738791183094
0.516349380925282
REACTOME_SIGNALING_BY_VEGF 98 0.01 Down
0.317065893885285 0.257227175180901 0.516975813689206
SEIDEN_MET_SIGNALING 20 0.01 Down 0.153496051101998
0.257329298892341 0.516975813689206
KANG_AR_TARGETS_DN 15 0.01 Up 0.87814002116195
0.257395299451934 0.516975813689206
REACTOME_CILIUM_ASSEMBLY 180 0.01 Up
0.159211812121169 0.257531783212827 0.517018921635399
KEGG_NATURAL_KILLER_CELL_MEDIATED_CYTOTOXICITY 90 0.01 Down
0.433382080481911 0.257695628826698
0.517032800263092
ZHANG_RESPONSE_TO_CANTHARIDIN_DN 65 0.01 Up
0.147850919223429 0.25776874424685 0.517032800263092
VERRECCHIA_RESPONSE_TO_TGFB1_C1 18 0.01 Up
0.58142092580339 0.257935802146896 0.517137123394423
KEGG_CIRCADIAN_RHYTHM_MAMMAL 13 0.01 Down
0.192381782234203 0.258238040635424 0.517512257091498
BARRIER_COLON_CANCER_RECURRENCE_UP 36 0.01 Down
0.235493902361485 0.258413398394214 0.517632899189835
GUTIERREZ_CHRONIC_LYMPHOCYTIC_LEUKEMIA_DN 45 0.01 Up
0.455940809589984 0.258562812195958
0.517701488116183
BRUINS_UVC_RESPONSE_EARLY_LATE 274 0.01 Down
0.239406054388272 0.259076859675598 0.518441393016541
ABBUD_LIF_SIGNALING_1_UP 36 0.01 Down
0.530853115211096 0.25916302783274 0.518441393016541
REACTOME_HEMOSTASIS 444 0.01 Up 0.393506485137058
0.259485416258187 0.518665038227668
REACTOME_TRANSPORT_OF_CONNEXONS_TO_THE_PLASMA_MEMBRANE 13 0.01
Down 0.358551602071572 0.259545564844043
0.518665038227668

ENK_UV_RESPONSE_KERATINOCYTE_UP 477 0.01 Down
0.279394879069887 0.259646444659694 0.518665038227668
LIANG_HEMATOPOIESIS_STEM_CELL_NUMBER_SMALL_VS_HUGE_DN 27 0.01
Up 0.256177867302262 0.259736373982309
0.518665038227668
HELLER_HDAC_TARGETS_UP 264 0.01 Up 0.31166995340833
0.259876918311047 0.51871525213506
HARRIS_BRAIN_CANCER_PROGENITORS 24 0.01 Up
0.725264353782047 0.260206991193032 0.519143553223558
IVANOVA_HEMATOPOIESIS_INTERMEDIATE_PROGENITOR 124 0.01 Up
0.191934479153203 0.260692056348232
0.519734274748549
REACTOME_RAP1_SIGNALLING 13 0.01 Down
0.455134134650699 0.260734324707003 0.519734274748549
TERAO_AOX4_TARGETS_SKIN_UP 26 0.01 Down
0.134733268141565 0.261311640431342 0.520089598092148
FOSTER_KDM1A_TARGETS_DN 181 0.01 Up 0.232930601714442
0.261336189262468 0.520089598092148
REACTOME_RESOLUTION_OF_AP_SITES_VIA_THE_MULTIPLE_NUCLEOTIDE_PATCH_RE
PLACEMENT_PATHWAY 24 0.01 Up 0.159600820859023
0.261347469538035 0.520089598092148
BIOCARTA_EIF_PATHWAY 16 0.01 Down 0.126843748072264
0.261375395348201 0.520089598092148
JAZAG_TGFB1_SIGNALING_VIA_SMAD4_DN 48 0.01 Up
0.529059064806567 0.261562410528243 0.520177471284588
HAHTOLA_MYCOSIS_FUNGOIDES_DN 12 0.01 Up
0.395343655748912 0.261750433176143 0.520177471284588
GRADE_COLON_AND_RECTAL_CANCER_DN 78 0.01 Up
0.310407577361317 0.261766727485148 0.520177471284588
REACTOME_COSTIMULATION_BY_THE_CD28_FAMILY 56 0.01 Down
0.497168559246298 0.262120013994653
0.52052756445466
DIAZ_CHRONIC_MEYLOGENOUS_LEUKEMIA_DN 93 0.01 Up
0.4584764742495 0.26221674963251 0.52052756445466
BROWNE_HCMV_INFECTION_48HR_UP 146 0.01 Down
0.309502423909287 0.262290307784161 0.52052756445466
ANDERSEN_LIVER_CANCER_KRT19_UP 35 0.01 Up
0.152853181323519 0.262408874880667 0.520533050568666
REACTOME_SCF_SKP2_MEDIATED_DEGRADATION_OF_P27_P21 59 0.01
Down 0.141794587246788 0.262636722319387
0.520626834009607
ZHENG_IL22_SIGNALING_DN 18 0.01 Up 0.58492592567453
0.2627950767621 0.520626834009607
REACTOME_SYNTHESIS_OF_PIPS_AT_THE_GOLGI_MEMBRANE 15 0.01
Down 0.150375567538833 0.262803623218642
0.520626834009607
GRAHAM_CML_QUIESCENT_VS_NORMAL_QUIESCENT_UP 69 0.01 Up
0.484505900898848 0.262923952427089
0.520635756017518
DAIRKEE_CANCER_PRONE_RESPONSE_BPA_E2 100 0.01 Down
0.224445893199664 0.263450961779501 0.521449613914071
PEDERSEN_METASTASIS_BY_ERBB2_ISOFORM_5 11 0.01 Down
0.675262944939254 0.263848052667043 0.522005720395405
STREICHER_LSM1_TARGETS_DN 19 0.01 Down

0.552446449157947	0.264600730140045	0.523100945073231
REACTOME_TRANS_GOLGI_NETWORK_VESICLE_BUDDING	69	0.01	Down
	0.175969402298856	0.264634382446391
0.523100945073231
LINDGREN_BLADDER_CANCER_CLUSTER_3_UP	294	0.01	Up
0.226316791601842	0.264779990950766	0.52315870739503
REACTOME_PYRUVATE_METABOLISM	26	0.01	Up
0.150457628917521	0.265056206004719	0.523474361156068
TIEN_INTESTINE_PROBIOTICS_6HR_UP	51	0.01	Up
0.212232883969512	0.265554939624862	0.524229009053032
REACTOME_PLASMA_LIPOPROTEIN_ASSEMBLY_REMODELING_AND_CLEARANCE	46
	0.01	Up	0.324317119645904	0.266007575406437
	0.524892033122008
MEISSNER_BRAIN_HCP_WITH_H3_UNMETHYLATED	14	0.01	Down
0.432621647482373	0.266148570505629	0.524939808873542
VICENT_METASTASIS_UP	11	0.01	Down	0.343721454342584
	0.266288788143928	0.524986009959192
LU_EZH2_TARGETS_UP	260	0.01	Up	0.24399313148663
	0.266449723762067	0.5250729979441
PURBEY_TARGETS_OF_CTBP1_NOT_SATB1_UP	243	0.01	Down
0.329144517176866	0.267216648888304	0.526250336407933
RODRIGUES_DCC_TARGETS_DN	116	0.01	Up
0.309197763688891	0.267281316578267	0.526250336407933
REACTOME_APC_CDC20_MEDIATED_DEGRADATION_OF_NEK2A	23	0.01
	Up	0.115788643771286	0.267620814182437
0.526688073445734
PID_P38_ALPHA_BETA_DOWNSTREAM_PATHWAY	36	0.01	Down
0.209615005241501	0.26823633326987	0.527557013086491
HOLLERN_MICROACINAR_BREAST_TUMOR_UP	35	0.01	Down
0.457157567204523	0.26835457265237	0.527557013086491
REACTOME_DNA_STRAND_ELONGATION	30	0.01	Up
0.123011116927855	0.268521893292757	0.527557013086491
BROWNE_HCMV_INFECTION_24HR_UP	124	0.01	Down
0.260300112009095	0.26853180109942	0.527557013086491
SHIN_B_CELL_LYMPHOMA_CLUSTER_8	28	0.01	Down
0.398307635583416	0.268985820354783	0.528218113802861
PID_AJDISS_2PATHWAY	43	0.01	Down	0.328484436672582
	0.269770576513041	0.529527834683895
REACTOME_HDR_THROUGH_HOMOLOGOUS_RECOMBINATION_HRR	62	0.01
	Up	0.155128522606483	0.27005377260005
0.529852338645668
PID_LKB1_PATHWAY	45	0.01	Down	0.207799265977432
	0.27035924154888	0.53022024029765
ZHONG_RESPONSE_TO_AZACITIDINE_AND_TSA_DN	66	0.01	Up
	0.187349998740882	0.270861898850503
0.53087698023333
MIKKELSEN_IPS_ICP_WITH_H3K4ME3_AND_H327ME3	54	0.01	Up
	0.581240369375073	0.270930320946665
0.53087698023333
KEGG_SPLICEOSOME	126	0.01	Down	0.106691356598752
	0.271552965368234	0.531531210974387
KYNG_WERNER_SYNDROM_AND_NORMAL_AGING_UP	70	0.01	Down
0.226824028392831	0.27161558796207	0.531531210974387
PID_BETA_CATENIN_NUC_PATHWAY	61	0.01	Down

0.368305524406256	0.271618952526845	0.531531210974387
CEBALLOS_TARGETS_OF_TP53_AND_MYC_DN	28	0.01	Up
0.486201708673329	0.271898268126509	0.531846264242237
FLECHNER_BIOPSY_KIDNEY_TRANSPLANT_OK_VS_DONOR_UP	533	0.01
	Down	0.272764278710714	0.272056500325732
0.531919597998245
REACTOME_CHOLESTEROL_BIOSYNTHESIS	24	0.01	Down
0.201730409658665	0.272172430566399	0.531919597998245
REACTOME_SYNTHESIS_OF_PC	25	0.01	Up
0.337745444522185	0.272348688218267	0.532032748170844
REACTOME_HSF1_DEPENDENT_TRANSACTIVATION	32	0.01	Down
0.233102270883802	0.272769108530263	0.532448226276029
KORKOLA_EMBRYONIC_CARCINOMA_VS_SEMINOMA_UP	15	0.01	Up
	0.339057836627064	0.272798279224404
0.532448226276029
NABA_ECM_AFFILIATED	89	0.01	Up	0.789881073278972
	0.273249831691817	0.532880459937151
PID_S1P_S1P3_PATHWAY	24	0.01	Down	0.342314661700674
	0.273256832070107	0.532880459937151
CEBALLOS_TARGETS_OF_TP53_AND_MYC_UP	20	0.01	Down
0.271363887663078	0.27361226780194	0.533342213256599
PID_PRL_SIGNALING_EVENTS_PATHWAY	22	0.01	Down
0.257583316925867	0.274194658078881	0.534245768558548
REACTOME_GRB2:SOS_PROVIDES_LINKAGE_TO_MAPK_SIGNALING_FOR_INTEGRINS
	11	0.01	Down	0.316319087996553
0.274765181588129	0.535125429479481
CHOW_RASSF1_TARGETS_DN	30	0.01	Down	0.214889137965299
	0.274936577512363	0.535227334741478
KYNG_DNA_DAMAGE_BY_4NQO_OR_UV	55	0.01	Down
0.373041193432656	0.27524036433917	0.535496543966169
PEDRIOLI_MIR31_TARGETS_DN	320	0.01	Down
0.434405175922217	0.27531312861086	0.535496543966169
WIERENGA_STAT5A_TARGETS_GROUP2	45	0.01	Down
0.597942812784628	0.275517245913936	0.535582726823631
REACTOME_G1_S_SPECIFIC_TRANSCRIPTION	27	0.01	Up
0.185144890999296	0.275595739075208	0.535582726823631
REACTOME_HEDGEHOG_OFF_STATE	101	0.01	Down
0.172729632087497	0.275843497356122	0.5358325499636
TAKEDA_TARGETS_OF_NUP98_HOXA9_FUSION_8D_UP	109	0.01	Down
	0.540696507950827	0.276141948932955
0.536180587668956
VART_KSHV_INFECTION_ANGIOGENIC_MARKERS_DN	89	0.01	Up
	0.454562646383087	0.276492145472055
0.536586425277778
REACTOME_GPVI_MEDIATED_ACTIVATION_CASCADE	31	0.01	Up
	0.480547876775473	0.276589710204363
0.536586425277778
REACTOME_SUMOYLATION_OF_RNA_BINDING_PROTEINS	47	0.01	Down
	0.177534048207329	0.276824121331499
0.536606403883892
HOWLIN_CITED1_TARGETS_2_DN	15	0.01	Down
0.320127006562757	0.276838765429755	0.536606403883892
HERNANDEZ_ABERRANT_MITOSIS_BY_DOCETACEL_4NM_UP	15	0.01	Up
	0.434539915102563	0.277172182764942

0.536677705262214
STEIN_ESRRA_TARGETS 511 0.01 Up 0.222642336199577
0.277212958556819 0.536677705262214
REACTOME_TNFS_BIND_THEIR_PHYSIOLOGICAL_RECEPTORS 17 0.01
Up 0.406065653748703 0.277233733396854
0.536677705262214
WU_APOPTOSIS_BY_CDKN1A_VIA_TP53 41 0.01 Up
0.214495892090357 0.277413265652641 0.536794071936556
ZHENG_FOXP3_TARGETS_IN_T_LYMPHOCYTE_DN 33 0.01 Down
0.406478021814689 0.277558234740224 0.536843487589204
FIRESTEIN_PROLIFERATION 132 0.01 Down 0.343251538814882
0.278016177556133 0.53730377228419
SENESE_HDAC3_TARGETS_DN 496 0.01 Up 0.306563731703487
0.278035277938382 0.53730377228419
COLIN_PILOCYTIC_ASTROCYTOMA_VS_GLIOBLASTOMA_DN 26 0.01 Up
0.570265682308758 0.278388100562866
0.537325484016178
CHNG_MULTIPLE_MYELOMA_HYPERPLOID_DN 30 0.01 Down
0.135861791526021 0.278456026274719 0.537325484016178
REACTOME_PROCESSIVE_SYNTHESIS_ON_THE_LAGGING_STRAND 15 0.01
Up 0.125511055634981 0.278485566384467
0.537325484016178
GRABARCZYK_BCL11B_TARGETS_DN 48 0.01 Up
0.283945226439002 0.278524666909387 0.537325484016178
SMID_BREAST_CANCER_LUMINAL_A_DN 14 0.01 Up
0.314488441640296 0.279001640270246 0.537819418544634
REACTOME_HSP90_CHAPERONE_CYCLE_FOR_STEROID_HORMONE_RECEPTORS_SHR
48 0.01 Down 0.213584313911148
0.279019996450742 0.537819418544634
BIOCARTA_EIF2_PATHWAY 10 0.01 Down 0.122910962631151
0.279319503658209 0.538165953254886
REACTOME_CHROMOSOME_MAINTENANCE 71 0.01 Up
0.150702745367643 0.279662222683001 0.538595411722404
REACTOME_DCC_MEDIATED_ATTRACTIVE_SIGNALING 13 0.01 Down
0.287805224679224 0.280199058913957
0.538603951131497
HOLLEMAN_VINCRISTINE_RESISTANCE_ALL_DN 19 0.01 Up
0.183403651207597 0.280388264167032 0.538603951131497
HAN_SATB1_TARGETS_UP 389 0.01 Down 0.391041259384831
0.280427383010071 0.538603951131497
REACTOME_REGULATION_OF_LIPID_METABOLISM_BY_PEROXISOME_PROLIFERATOR_A
CTIVATED_RECEPTOR_ALPHA_PPARALPHA 108 0.01 Down
0.279088554164538 0.280447246143483 0.538603951131497
TURJANSKI_MAPK1_AND_MAPK2_TARGETS 10 0.01 Down
0.298077132001698 0.280535316299948 0.538603951131497
REACTOME_ASYMMETRIC_LOCALIZATION_OF_PCP_PROTEINS 62 0.01
Down 0.177708135465142 0.280595798211921
0.538603951131497
SCIBETTA_KDM5B_TARGETS_UP 19 0.01 Down
0.210370980963059 0.280595957554492 0.538603951131497
KAYO_AGING_MUSCLE_UP 162 0.01 Down 0.348902679700546
0.280645183264733 0.538603951131497
BOYAULT_LIVER_CANCER_SUBCLASS_G1_DN 39 0.01 Up
0.215184321907066 0.280745062847852 0.538603951131497

SCHAEFFER_PROSTATE_DEVELOPMENT_12HR_DN 28 0.01 Up
0.564136362964059 0.280898501458765 0.538668414700148
WANG_ESOPHAGUS_CANCER_VS_NORMAL_UP 106 0.01 Up
0.646781206833462 0.281158526589061 0.538896493160368
HOFMANN_MYELODYSPLASTIC_SYNDROM_LOW_RISK_DN 24 0.01 Down
0.177195002519291 0.281257213115512
0.538896493160368
ANDERSEN_CHOLANGIOCARCINOMA_CLASS1 51 0.01 Up
0.619837363752312 0.281555490136935 0.539238145788463
GAUSSMANN_MLL_AF4_FUSION_TARGETS_F_UP 158 0.01 Up
0.512071918911556 0.281942595738159 0.539433206133733
SAKAI_CHRONIC_HEPATITIS_VS_LIVER_CANCER_DN 24 0.01 Down
0.376004460806897 0.281962482866583
0.539433206133733
THEILGAARD_NEUTROPHIL_AT_SKIN_WOUND_DN 213 0.01 Down
0.260938101861655 0.282017360270139 0.539433206133733
OZEN_MIR125B1_TARGETS 22 0.01 Down 0.203406281294318
0.282297260886971 0.539533142722964
BIOCARTA_PGC1A_PATHWAY 13 0.01 Down 0.237207378138627
0.282309666670614 0.539533142722964
VERHAAK_GLIOBLASTOMA_NEURAL 93 0.01 Up
0.321940289444511 0.282758909359847 0.539760957579811
JIANG_TIP30_TARGETS_UP 42 0.01 Down 0.45311492237057
0.282799636049367 0.539760957579811
REACTOME_NOTCH3_INTRACELLULAR_DOMAIN_REGULATES_TRANSCRIPTION 19
0.01 Down 0.299673885143105 0.282849627662916
0.539760957579811
UEDA_PERIFERAL_CLOCK 146 0.01 Down 0.26969509369059
0.282963494052321 0.539760957579811
TRAYNOR_RETT_SYNDROM_DN 11 0.01 Up 0.469931272440888
0.283029271861094 0.539760957579811
REACTOME_DNA_DAMAGE_RECOGNITION_IN_GG_NER 38 0.01 Up
0.127024197384363 0.283323022718356
0.54009202167897
REACTOME_G_ALPHA_Z_SIGNALLING_EVENTS 29 0.01 Up
0.394429878799651 0.283686769195104 0.540099159936124
REACTOME_ACETYLCHOLINE_NEUROTRANSMITTER_RELEASE_CYCLE 10 0.01
Down 0.229727427522903 0.283687233950876
0.540099159936124
REACTOME_SEROTONIN_NEUROTRANSMITTER_RELEASE_CYCLE 10 0.01
Down 0.229727427522903 0.283687233950876
0.540099159936124
WANG_MLL_TARGETS 193 0.01 Up 0.62029553024072
0.284139666854513 0.540731499792987
PID_INSULIN_GLUCOSE_PATHWAY 22 0.01 Down
0.171429308911063 0.284330510760314 0.540865698632083
NUYTTEN_NIPP1_TARGETS_UP 689 0.01 Down
0.324581428432562 0.284527273962071 0.540924852284683
CASORELLI_ACUTE_PROMYELOCYTIC_LEUKEMIA_DN 609 0.01 Up
0.236800366896884 0.284614078631505
0.540924852284683
LAIHO_COLORECTAL_CANCER_SERRATED_DN 83 0.01 Down
0.20912006608939 0.284810923921904 0.540924852284683
LEE_DIFFERENTIATING_T_LYMPHOCYTE 164 0.01 Down

0.403695235981942	0.284911353953971	0.540924852284683
MCBRYAN_PUBERTAL_BREAST_5_6WK_DN	111	0.01	Down
0.295240455794094	0.28499538303029	0.540924852284683
CAIRO_LIVER_DEVELOPMENT_DN	141	0.01	Up
0.516200243632457	0.285222694178265	0.540924852284683
SHARMA_PILOCYTIC_ASTROCYTOMA_LOCATION_UP	20	0.01	Down
	0.700571860740878	0.285237728953076
0.540924852284683
BEIER_GLIOMA_STEM_CELL_DN	64	0.01	Up
0.471709086799659	0.285420414540523	0.540924852284683
FLECHNER_PBL_KIDNEY_TRANSPLANT_OK_VS_DONOR_UP	146	0.01	Down
	0.206981436482797	0.285444660649448
0.540924852284683
PID_RANBP2_PATHWAY	11	0.01	Down	0.20348733660268
	0.285740667573584	0.541137583511613
REACTOME_TRNA_PROCESSING_IN_THE_NUCLEUS 58	0.01	Down
0.144185222292042	0.285797691492007	0.541137583511613
NUNODA_RESPONSE_TO_DASATINIB_IMATINIB_UP	28	0.01	Down
	0.310651310643928	0.285988118086039
0.5411771735595
VALK_AML_CLUSTER_10	24	0.01	Up	0.396789271506383
	0.286059391407646	0.5411771735595
MISHRA_CARCINOMA_ASSOCIATED_FIBROBLAST_DN	22	0.01	Up
	0.527070528595326	0.286210986882402
0.541236174184433
KIM_GERMINAL_CENTER_T_HELPER_UP 59	0.01	Down
0.272839339673327	0.286573851998193	0.541694476338047
SA_MMP_CYTOKINE_CONNECTION	12	0.01	Up
0.727310173905447	0.286856703657057	0.542001211827856
SCHLOSSER_SERUM_RESPONSE_UP	107	0.01	Down
0.260284665213176	0.287096600291316	0.542226562314901
REACTOME_NOTCH4_ACTIVATION_AND_TRANSMISSION_OF_SIGNAL_TO_THE_NUCLEUS
	11	0.01	Down	0.249521093539429
0.287561055474	0.542600618901595
LIU_NASOPHARYNGEAL_CARCINOMA	61	0.01	Down
0.258225309029419	0.287748806672718	0.542600618901595
BOHN_PRIMARY_IMMUNODEFICIENCY_SYNDROM_DN	34	0.01	Up
	0.427481551704195	0.287753843972716
0.542600618901595
ACOSTA_PROLIFERATION_INDEPENDENT_MYC_TARGETS_UP 77	0.01	Up
	0.239240190277634	0.287777502883515
0.542600618901595
KARLSSON_TGFB1_TARGETS_DN	180	0.01	Up
0.323361188169121	0.288270792104372	0.543302813630672
REACTOME_COPI_DEPENDENT_GOLGI_TO_ER_RETROGRADE_TRAFFIC	81	0.01
	Down	0.178150873808595	0.288437932182567
0.543389985398423
REACTOME_FRS_MEDIATED_FGFR4_SIGNALING	12	0.01	Down
0.398712926734067	0.289051694694229	0.54431812637225
BRUINS_UVC_RESPONSE_VIA_TP53_GROUP_C	65	0.01	Down
0.41567331574121	0.289488097484827	0.544542701034476
BIOCARTA_NGF_PATHWAY	19	0.01	Down	0.226876101440146
	0.289531128635245	0.544542701034476
DARWICHE_PAPILLOMA_RISK_HIGH_UP 110	0.01	Down

0.374139996642229	0.289773311735581	0.544542701034476
REACTOME_MHC_CLASS_II_ANTIGEN_PRESENTATION	105	0.01	Up
	0.331867569068849	0.28985252219555
0.544542701034476
OUELLET_OVARIAN_CANCER_INVASIVE_VS_LMP_UP	114	0.01	Down
	0.196233578994865	0.290083991631873
0.544542701034476
BIOCARTA_CREB_PATHWAY	19	0.01	Down	0.206376261560241
	0.290198602811702	0.544542701034476
REACTOME_SIGNALING_BY_NON_RECEPTOR_TYROSINE_KINASES	46	0.01
	Down	0.26705963379947	0.290225960096499
0.544542701034476
CHIANG_LIVER_CANCER_SUBCLASS_PROLIFERATION_UP	149	0.01	Up
	0.333922653693416	0.290254284049942
0.544542701034476
APRELIKOVA_BRCA1_TARGETS	45	0.01	Down
0.278549322512247	0.290379633975557	0.544542701034476
ACEVEDO_LIVER_CANCER_WITH_H3K27ME3_DN	121	0.01	Up
0.395335967290587	0.290454283831806	0.544542701034476
REACTOME_MITOTIC_G2_G2_M_PHASES	183	0.01	Down
0.150804079733981	0.290503536614165	0.544542701034476
SCHWAB_TARGETS_OF_BMYB_POLYMORPHIC_VARIANTS_DN	13	0.01	Down
	0.284996921676682	0.290708900174963
0.54470050282887
LEI_HOXC8_TARGETS_DN	13	0.01	Up	0.828222759520552
	0.290973356852128	0.544878680114809
BIOCARTA_CTLA4_PATHWAY	11	0.01	Down	0.76311768384794
	0.291046431803261	0.544878680114809
THEODOROU_MAMMARY_TUMORIGENESIS	16	0.01	Down
0.784332544884105	0.291529154741691	0.545555183415447
MCBRYAN_PUBERTAL_BREAST_3_4WK_DN	28	0.01	Down
0.374625644213146	0.292404040349742	0.546964694703324
MCGOWAN_RSP6_TARGETS_UP	10	0.01	Up	0.287375705941126
	0.29299287910837	0.547797990271361
BOYAULT_LIVER_CANCER_SUBCLASS_G5_DN	27	0.01	Up
0.525723675500072	0.293093251746969	0.547797990271361
HOLLEMAN_DAUNORUBICIN_B_ALL_DN	12	0.01	Down
0.101482050101717	0.293892449177425	0.548981605138985
MARSON_BOUND_BY_FOXP3_UNSTIMULATED	1037	0.01	Down
0.234313777482899	0.293970795009908	0.548981605138985
GUTIERREZ_MULTIPLE_MYELOMA_UP	35	0.01	Down
0.164072509819265	0.294579137812168	0.549889212817979
LI_WILMS_TUMOR_VS_FETAL_KIDNEY_1_UP	170	0.01	Up
0.404970825597238	0.294742918451565	0.549966549788205
RAMALHO_STEMNESS_UP	191	0.01	Down	0.167138917692606
	0.29536530624177	0.550669848043939
REACTOME_FCERI_MEDIATED_MAPK_ACTIVATION	29	0.01	Down
0.321781882181054	0.295454091996675	0.550669848043939
REACTOME_PENTOSE_PHOSPHATE_PATHWAY	14	0.01	Up
0.186424531883272	0.295487357838038	0.550669848043939
REACTOME_M_PHASE	321	0.01	Down	0.159876129887634
	0.296291122166581	0.551938911785654
PID_P38_MKK3_6PATHWAY	24	0.01	Down	0.213032069481955
	0.296615359356262	0.552314018353934

BIOCARTA_STATHMIN_PATHWAY 14 0.01 Down
0.536530251545512 0.296827517952166 0.552394330009515
BIOCARTA_ECM_PATHWAY 18 0.01 Down 0.198825845157749
0.296904271702556 0.552394330009515
KIM_WT1_TARGETS_12HR_DN 203 0.01 Down 0.303960607302506
0.297030106321073 0.552399804680688
CHANDRAN_METASTASIS_DN 270 0.01 Up 0.40333810307279
0.297165244448867 0.552422569808791
PANGAS_TUMOR_SUPPRESSION_BY_SMAD1_AND_SMAD5_UP 109 0.01 Up
0.421921523729046 0.297623207978641
0.553045192171968
REACTOME_SEMA4D_INDUCED_CELL_MIGRATION_AND_GROWTH_CONE_COLLAPSE 19
0.01 Down 0.222131874719316 0.297842436769049
0.553166084252089
CREIGHTON_ENDOCRINE_THERAPY_RESISTANCE_4 267 0.01 Up
0.371359642592132 0.298018112567257
0.553166084252089
TARTE_PLASMA_CELL_VS_B_LYMPHOCYTE_UP 72 0.01 Up
0.227628070238056 0.298057454073095 0.553166084252089
REACTOME_CARGO_CONCENTRATION_IN_THE_ER 28 0.01 Down
0.3476262699005 0.29849075530566 0.55374161993353
SCHAEFFER_SOX9_TARGETS_IN_PROSTATE_DEVELOPMENT_DN 35 0.01
Down 0.315034036909814 0.299037981723368
0.554508958988619
CHOI_ATL_CHRONIC_VS_ACUTE_DN 18 0.01 Down
0.117670792000404 0.299151106907097 0.554508958988619
BIOCARTA_IL3_PATHWAY 13 0.01 Down 0.257519959915736
0.300042063790156 0.555881645746715
ZIRN_TRETINOIN_RESPONSE_WT1_UP 22 0.01 Up
0.678595144791222 0.300147584958068 0.555881645746715
HAMAI_APOPTOSIS_VIA_TRAIL_DN 149 0.01 Up
0.364276848738818 0.300317223972616 0.555881645746715
JAZAG_TGFB1_SIGNALING_UP 88 0.01 Down
0.302914972726776 0.300387755982651 0.555881645746715
KEGG_GLYCEROPHOSPHOLIPID_METABOLISM 64 0.01 Up
0.296621313043196 0.30050998869066 0.555881645746715
REACTOME_TELOMERE_C_STRAND_LAGGING_STRAND_SYNTHESIS 23 0.01
Up 0.127145204957235 0.301116418601982
0.556774291080177
COLIN_PILOCYTIC_ASTROCYTOMA_VS_GLIOBLASTOMA_UP 26 0.01 Up
0.59184148909969 0.301605569218022
0.557449438172289
REACTOME_CD28_DEPENDENT_PI3K_AKT_SIGNALING 20 0.01 Down
0.415462284610632 0.302005868686219
0.557852820434209
GAVIN_FOXP3_TARGETS_CLUSTER_T4 88 0.01 Down
0.374599215894412 0.30207202779463 0.557852820434209
SAMOLS_TARGETS_OF_KHSV_MIRNAS_DN 54 0.01 Up
0.471240844133068 0.302461949046361 0.558343515796055
ZAMORA_NOS2_TARGETS_UP 50 0.01 Down 0.216074491915211
0.303081908454209 0.559258283457171
SCHAEFFER_PROSTATE_DEVELOPMENT_48HR_DN 282 0.01 Up
0.509165492913055 0.303399611615418 0.559437940487557
LEE_LIVER_CANCER_MYC_UP 46 0.01 Up 0.433789771238245

0.303444088564652 0.559437940487557
MANALO_HYPOXIA_DN 277 0.01 Up 0.193945750449447
0.30367517663206 0.559437940487557
REACTOME_TRANSCRIPTIONAL_REGULATION_BY_MECP2 45 0.01 Down
0.307283536690089 0.30367710228913
0.559437940487557
BIOCARTA_CASPASE_PATHWAY 21 0.01 Down
0.420586027906533 0.30421646377823 0.560201968325746
SA_CASPASE_CASCADE 17 0.01 Down 0.422743659607321
0.305591044269637 0.562429930388316
TARTE_PLASMA_CELL_VS_PLASMABLAST_UP 265 0.01 Up
0.574033414469632 0.305692636470641 0.562429930388316
YANG_BREAST_CANCER_ESR1_BULK_DN 20 0.01 Down
0.429930739859495 0.305801724108798 0.562429930388316
REACTOME_GAP_FILLING_DNA_REPAIR_SYNTHESIS_AND_LIGATION_IN_GG_NER
24 0.01 Up 0.12924493673836
0.306378839413028 0.563260892908614
KUMAR_AUTOPHAGY_NETWORK 59 0.01 Up 0.356919143461383
0.307183890499996 0.564392605946197
WEIGEL_OXIDATIVE_STRESS_BY_TBH_AND_H2O2 32 0.01 Up
0.192995338745072 0.3073360802303 0.564392605946197
BIOCARTA_ERK5_PATHWAY 13 0.01 Down 0.17504883215832
0.307371101080376 0.564392605946197
ROESSLER_LIVER_CANCER_METASTASIS_UP 89 0.01 Down
0.321535311914861 0.307592982913052 0.564569399017627
QUINTENS_EMBRYONIC_BRAIN_RESPONSE_TO_IR 68 0.01 Down
0.264102541884878 0.308150565301899 0.56536195552328
REACTOME_PI_3K_CASCADE:FGFR2 12 0.01 Down
0.490532866741758 0.308504647630813 0.565780657323747
REACTOME_ACYL_CHAIN_REMODELLING_OF_PE 19 0.01 Up
0.494674662203294 0.308995168680925 0.566172477861129
REACTOME_LAGGING_STRAND_SYNTHESIS 20 0.01 Up
0.117198467815384 0.309100623082427 0.566172477861129
BIOCARTA_CCR5_PATHWAY 15 0.01 Down 0.420570773282472
0.309320783689614 0.566172477861129
BHATI_G2M_ARREST_BY_2METHOXYESTRADIOL_DN 106 0.01 Up
0.295760898450351 0.309331097062101
0.566172477861129
REACTOME_APC_C:CDH1_MEDIATED_DEGRADATION_OF_CDC20_AND_OTHER_APC_C:CD
H1_TARGETED_PROTEINS_IN_LATE_MITOSIS_EARLY_G1 68 0.01 Down
0.11847362815893 0.30934807689142
0.566172477861129
REACTOME_PCNA_DEPENDENT_LONG_PATCH_BASE_EXCISION_REPAIR 20 0.01
Up 0.166255123163658 0.30957910494222
0.566182621190448
DANG_REGULATED_BY_MYC_DN 227 0.01 Up
0.455325434357341 0.309636899819443 0.566182621190448
PID_MYC_REPRESS_PATHWAY 60 0.01 Down 0.474639106869348
0.309731493994952 0.566182621190448
REACTOME_GPCR_LIGAND_BINDING 135 0.01 Down
0.63985333522189 0.310137970324972 0.566661477615143
REACTOME_PYRIMIDINE_SALVAGE 10 0.01 Down
0.238195597167548 0.31024558318373 0.566661477615143
CREIGHTON_AKT1_SIGNALING_VIA_MTOR_UP 34 0.01 Down

0.161991414592823	0.310647967062103	0.567165967483409
SUNG_METASTASIS_STROMA_UP	108	0.01	Up
0.4842862754522 0.311594356354715	0.568662863099652
PID_INTEGRIN_CS_PATHWAY 22	0.01	Up	0.639860123700042
	0.311959861672652	0.569098854796497
PID_HDAC_CLASSIII_PATHWAY	22	0.01	Down
0.29266097699793	0.312132653640623	0.569158803500053
FIRESTEIN_CTNNB1_PATHWAY	29	0.01	Down
0.291093239843582	0.312245964278338	0.569158803500053
HE_PTEN_TARGETS_UP	17	0.01	Down	0.295890005638181
	0.31243338841552	0.569269590971935
INAMURA_LUNG_CANCER_SCC_UP	12	0.01	Down
0.294380408364389	0.312590877641202	0.569285602857616
DEMAGALHAES_AGING_UP	48	0.01	Up	0.667073426540803
	0.31269547351623	0.569285602857616
ZHANG_RESPONSE_TO_IKK_INHIBITOR_AND_TNF_DN	91	0.01	Up
	0.306968711366541	0.313184845424651
0.569945700479274
REACTOME_UNFOLDED_PROTEIN_RESPONSE_UPR	86	0.01	Down
0.189168038157718	0.313612828461942	0.570204002512726
FERRANDO_LYL1_NEIGHBORS 12	0.01	Up	0.464874497633453
	0.313777554844965	0.570204002512726
LEE_RECENT_THYMIC_EMIGRANT	201	0.01	Down
0.267735591581503	0.313952594096208	0.570204002512726
REACTOME_NUCLEOSOME_ASSEMBLY	38	0.01	Up
0.184348111390788	0.314017826844682	0.570204002512726
BURTON_ADIPOGENESIS_PEAK_AT_16HR	38	0.01	Up
0.22560641226626	0.314054927252837	0.570204002512726
KEGG_SPHINGOLIPID_METABOLISM	33	0.01	Up
0.24818127310272	0.314087899938044	0.570204002512726
MATZUK_POSTIMPLANTATION_AND_POSTPARTUM	11	0.01	Down
0.266903854922306	0.314645804197065	0.570986229255474
TOMIDA_METASTASIS_UP	23	0.01	Down	0.324285684933522
	0.315149294384809	0.571669119555979
KEGG_PROXIMAL_TUBULE_BICARBONATE_RECLAMATION	13	0.01	Up
	0.52748650423483	0.315558574283052
0.572078768627791
REACTOME_CONSTITUTIVE_SIGNALING_BY_ABERRANT_PI3K_IN_CANCER	48
	0.01	Down	0.511373140178716	0.315629665449816
	0.572078768627791
WHITFIELD_CELL_CYCLE_G2 147	0.01	Down	0.300687593128725
	0.31821425153695	0.576499711383949
MARIADASON_REGULATED_BY_HISTONE_ACETYLATION_DN	35	0.01	Up
	0.275618615396741	0.318443086231166
0.576499711383949
WAGNER_APO2_SENSITIVITY 20	0.01	Down	0.331794941333988
	0.318453566933558	0.576499711383949
REACTOME_MRNA_DECAY_BY_5_TO_3_EXORIBONUCLEASE	15	0.01	Down
	0.157865541096351	0.318976562952881
0.577026785959692
SASSON_RESPONSE_TO_GONADOTROPHINS_DN	84	0.01	Up
0.370537766072799	0.319089374476612	0.577026785959692
MODY_HIPPOCAMPUS_POSTNATAL	41	0.01	Up
0.262378399311356	0.31916959107749	0.577026785959692

NIKOLSKY_BREAST_CANCER_7P22_AMPLICON 31 0.01 Up
0.240133168582494 0.319461967581321 0.577026785959692
LINDGREN_BLADDER_CANCER_CLUSTER_1_UP 115 0.01 Up
0.205176830556557 0.319497012464358 0.577026785959692
REACTOME_EPH_EPHRIN_SIGNALING 82 0.01 Down
0.299241644080399 0.319514943326735 0.577026785959692
WU_HBX_TARGETS_3_DN 11 0.01 Up 0.608871407528816
0.320038982843152 0.577741055373481
OUYANG_PROSTATE_CANCER_MARKERS 19 0.01 Down
0.359170408895057 0.320203435482834 0.577805878159509
BORCZUK_MALIGNANT_MESOTHELIOMA_DN 64 0.01 Down
0.463952477869274 0.320722317658424 0.578294554416417
LINDGREN_BLADDER_CANCER_WITH_LOH_IN_CHR9Q 113 0.01 Down
0.171728201158797 0.320731551537292
0.578294554416417
ZHANG_BREAST_CANCER_PROGENITORS_UP 394 0.01 Down
0.183002089030819 0.320988768753854 0.578526269265668
JACKSON_DNMT1_TARGETS_UP 72 0.01 Down
0.384735830129607 0.321590569825464 0.579321032675562
IIZUKA_LIVER_CANCER_EARLY_RECURRENCE 11 0.01 Up
0.768228031636704 0.321866191476853 0.579321032675562
REACTOME_CELLULAR_HEXOSE_TRANSPORT 12 0.01 Up
0.253705868208593 0.32189479718731 0.579321032675562
GALE_APL_WITH_FLT3_MUTATED_DN 15 0.01 Up
0.267983917865577 0.321945259093115 0.579321032675562
REACTOME_G1_S_DNA_DAMAGE_CHECKPOINTS 65 0.01 Down
0.162328854401841 0.322268052368642 0.579669826089254
BERNARD_PPAPDC1B_TARGETS_UP 35 0.01 Down
0.20359816610069 0.323093529207058 0.58092216551429
REACTOME_REGULATION_OF_TP53_EXPRESSION_AND_DEGRADATION 36 0.01
Down 0.25898948391504 0.323439326944678
0.581311385292414
CHO_NR4A1_TARGETS 24 0.01 Down 0.535190738901857
0.32389827155761 0.581795264277699
WIERENGA_PML_INTERACTOME 39 0.01 Down
0.202829648717149 0.323982676214511 0.581795264277699
CHEN_ETV5_TARGETS_TESTIS 15 0.01 Up
0.195528876818027 0.324096850222772 0.581795264277699
BIOCARTA_PRC2_PATHWAY 14 0.01 Down 0.133812769396553
0.324339381132584 0.581998210854677
ZHENG_GLIOBLASTOMA_PLASTICITY_UP 193 0.01 Up
0.362430022542509 0.32455114036763 0.582099701718298
CHESLER_BRAIN_HIGHEST_EXPRESSION 33 0.01 Down
0.262014501192166 0.324654939312074 0.582099701718298
SA_REG_CASCADE_OF_CYCLIN_EXPR 10 0.01 Up
0.400669088251354 0.32516199197185 0.582704962874415
REACTOME_RESOLUTION_OF_SISTER_CHROMATID_COHESION 101 0.01
Down 0.185057434898137 0.325341968033845
0.582704962874415
REACTOME_MRNA_SPLICING 187 0.01 Down 0.0895452115888942
0.325381414196837 0.582704962874415
KEGG_MELANOGENESIS 73 0.01 Down 0.315425477024467
0.326151187351381 0.583850890937657
BIOCARTA_NKT_PATHWAY 13 0.01 Down 0.633979858880721

0.326364720087853	0.583902792625328
LEE_AGING_NEOCORTEX_DN	42	0.01	Up	0.43628030859606
0.326665205014056	0.583902792625328
IIZUKA_LIVER_CANCER_PROGRESSION_L0_L1_UP	15	0.01	Down
0.280962733608953	0.326668220199393
0.583902792625328
KOKKINAKIS_METHIONINE_DEPRIVATION_48HR_UP	123	0.01	Down
0.469987762741333	0.326699782748098
0.583902792625328
REACTOME_FACTORS_INVOLVED_IN_MEGAKARYOCYTE_DEVELOPMENT_AND_PLATELET_PRODUCTION	113	0.01	Down	0.245794781735196
0.327062772679811	0.584319222255863
REACTOME_PINK_PARKIN_MEDIATED_MITOPHAGY	21	0.01	Up
0.0875528646088845	0.328516175731198	0.5866826419991
WANG_LMO4_TARGETS_UP	331	0.01	Down	0.2201525051254
0.328932087399632	0.58688532867321
GROSS_HYPOXIA_VIA_HIF1A_UP	72	0.01	Up
0.220421293490613	0.329007072714979	0.58688532867321
ELVIDGE_HYPOXIA_UP	166	0.01	Up	0.422338062156732
0.329130466045365	0.58688532867321
ABE_VEGFA_TARGETS	15	0.01	Down	0.589986835603761
0.329151927382683	0.58688532867321
MOHANKUMAR_HOXA1_TARGETS_DN	122	0.01	Down
0.553845026866059	0.329663891995169	0.587565104884332
GRUETZMANN_PANCREATIC_CANCER_UP	331	0.01	Up
0.465416522435328	0.329817986844972	0.587606758172077
TOOKER_GEMCITABINE_RESISTANCE_UP	75	0.01	Up
0.210485226801485	0.330501836818871	0.58803082870874
PUJANA_BRCA1_PCC_NETWORK	1413	0.01	Down
0.182831612875999	0.330531586344599	0.58803082870874
MIKKELSEN_MEF_LCP_WITH_H3K27ME3	20	0.01	Down
0.679617922760943	0.330571813389547	0.58803082870874
SCHLOSSER_MYC_TARGETS_AND_SERUM_RESPONSE_UP	48	0.01	Up
0.175948128732118	0.330579289020464
0.58803082870874
DIRMEIER_LMP1_RESPONSE_LATE_DN	25	0.01	Up
0.280156471586427	0.331181624810727	0.588791000325978
IVANOVA_HEMATOPOIESIS_STEM_CELL_AND_PROGENITOR	441	0.01	Up
0.275618592302139	0.331268618425895
0.588791000325978
NAKAMURA_METASTASIS	38	0.01	Up	0.337533453970339
0.331685253553239	0.589298503842613
REACTOME_GLOBAL_GENOME_NUCLEOTIDE_EXCISION_REPAIR_GG_NER	83
0.01	Up	0.127975293631112	0.331999434617724
0.589307289266965
REACTOME_KERATAN_SULFATE_KERATIN_METABOLISM	29	0.01	Up
0.696927223231512	0.332291064684983
0.589307289266965
KEGG_NICOTINATE_AND_NICOTINAMIDE_METABOLISM	17	0.01	Up
0.469878001774734	0.332433823490984
0.589307289266965
PID_P53_DOWNSTREAM_PATHWAY	127	0.01	Down
0.305869137664137	0.332440356670703	0.589307289266965
LEE_LIVER_CANCER_ACOX1_UP	53	0.01	Down

0.528997121787579	0.332461526172991	0.589307289266965
KEGG_LINOLEIC_ACID_METABOLISM	14	0.01	Down
0.567189530910521	0.332476815479649	0.589307289266965
RODRIGUES_NTN1_AND_DCC_TARGETS	32	0.01	Down
0.393067814197864	0.332746910776387	0.589553552991666
MATTIOLI_MGUS_VS_MULTIPLE_MYELOMA	15	0.01	Down
0.404361563942684	0.333320395896037	0.590261024776553
HOELZEL_NF1_TARGETS_DN	83	0.01	Up	0.47781061512747
0.333408841358769	0.590261024776553
FRASOR_RESPONSE_TO_ESTRADIOL_UP	28	0.01	Up
0.544726360537206	0.333809410907126	0.590554187243664
TAGHAVI_NEOPLASTIC_TRANSFORMATION	12	0.01	Up
0.387734939546598	0.333837194613159	0.590554187243664
SPIELMAN_LYMPHOBLAST_EUROPEAN_VS_ASIAN_UP	464	0.01	Up
0.189582312966429	0.33400798942165
0.590623883733405
IWANAGA_E2F1_TARGETS_INDUCED_BY_SERUM	24	0.01	Down
0.224170778335437	0.334479213156517	0.591224562775675
REACTOME_NUCLEOTIDE_SALVAGE	20	0.01	Down
0.261121080785775	0.334698372727372	0.591379396780479
POS_RESPONSE_TO_HISTAMINE_DN	11	0.01	Down
0.124848913859056	0.335012169266531	0.59170088356428
VALK_AML_CLUSTER_1	23	0.01	Up	0.403423089367918
0.335259389705141	0.59170088356428
TSENG_IRS1_TARGETS_UP	94	0.01	Up	0.359776193427675
0.335308777405708	0.59170088356428
KORKOLA_TERATOMA_UP	16	0.01	Up	0.446838360543299
0.33540686347537	0.59170088356428
REACTOME_RNA_POLYMERASE_I_TRANSCRIPTION_TERMINATION	31	0.01
Up	0.154759014422305	0.336065174417834
0.592629642608146
REACTOME_DEFECTS_IN_COBALAMIN_B12_METABOLISM	12	0.01	Up
0.15962532998876	0.33637445890164
0.592857523719508
REACTOME_APC_C:CDC20_MEDIATED_DEGRADATION_OF_CYCLIN_B	21	0.01
Up	0.10262259737244	0.336678715584507
0.592857523719508
TOMIDA_METASTASIS_DN	17	0.01	Down	0.287425132651304
0.336807702685426	0.592857523719508
BASSO_B_LYMPHOCYTE_NETWORK	136	0.01	Up
0.227409097223728	0.336922039438886	0.592857523719508
SPIELMAN_LYMPHOBLAST_EUROPEAN_VS_ASIAN_2FC_UP	12	0.01	Down
0.172219015345649	0.336984023215349
0.592857523719508
BIOCARTA_GPCR_PATHWAY	27	0.01	Down	0.262573813065903
0.337004474754691	0.592857523719508
KEGG_THYROID_CANCER	26	0.01	Down	0.188768127679725
0.337117648637833	0.592857523719508
REACTOME_RHO_GTPASES_ACTIVATE_KTN1	11	0.01	Down
0.111314005194727	0.337636154834462	0.593537159163436
MCCABE_HOXC6_TARGETS_CANCER_DN	20	0.01	Down
0.202063898967406	0.338150319706776	0.594208634512102
REACTOME_RA_BIOSYNTHESIS_PATHWAY	14	0.01	Down
0.607593357098881	0.338343256995972	0.59431533419183

MARTENS_BOUND_BY_PML_RARA_FUSION	396	0.01	Down
0.339790329168544	0.338723468343402	0.594750777423278
REACTOME_ACTIVATED_NTRK2_SIGNALS_THROUGH_FRS2_AND_FRS3	10	0.01
	Down	0.270742190313369	0.340557456326601
0.596998364571056
REACTOME_SYNTHESIS_OF_SUBSTRATES_IN_N_GLYCAN_BIOSYTHESIS	56
	0.01	Up	0.310517046900499	0.340564680220145
	0.596998364571056
REACTOME_IRAK1_RECRUITS_IKK_COMPLEX	14	0.01	Down
0.164033086385175	0.340590787643748	0.596998364571056
FUJII_YBX1_TARGETS_DN	174	0.01	Up	0.228268671561946
	0.340712924852207	0.596998364571056
BIOCARTA_PTDINS_PATHWAY	22	0.01	Down	0.280241917352199
	0.340874062980356	0.596998364571056
GENTILE_UV_HIGH_DOSE_UP	19	0.01	Down	0.591428697165497
	0.340874064514386	0.596998364571056
LIANG_SILENCED_BY_METHYLATION_UP	28	0.01	Up
0.848886669223158	0.340936608089969	0.596998364571056
HOFFMANN_LARGE_TO_SMALL_PRE_BII_LYMPHOCYTE_DN	47	0.01	Down
	0.383363329770493	0.341066028969627
0.596998364571056
VALK_AML_CLUSTER_13	23	0.01	Up	0.431422403910987
	0.341202206996129	0.59700425085543
CAFFAREL_RESPONSE_TO_THC_8HR_5_DN	11	0.01	Down
0.128723545722598	0.34170110784642	0.597441926768051
BRUINS_UVC_RESPONSE_LATE	992	0.01	Up
0.278559888301253	0.341990913116883	0.597441926768051
REACTOME_VEGFR2_MEDIATED_VASCULAR_PERMEABILITY	26	0.01	Down
	0.299626534030259	0.342009614155682
0.597441926768051
REACTOME_ATF6_ATF6_ALPHA_ACTIVATES_CHAPERONE_GENES	10	0.01
	Down	0.167165210627464	0.342034329261434
0.597441926768051
NIKOLSKY_BREAST_CANCER_17Q11_Q21_AMPLICON	87	0.01	Down
	0.26672044180158	0.342116912013562
0.597441926768051
MOREIRA_RESPONSE_TO_TSA_DN	13	0.01	Down
0.475486884062953	0.342650384834515	0.598012606845408
JOSEPH_RESPONSE_TO_SODIUM_BUTYRATE_UP	29	0.01	Down
0.445191301333847	0.342709783144332	0.598012606845408
MCBRYAN_PUBERTAL_BREAST_6_7WK_UP	165	0.01	Down
0.342983531957607	0.343025494864681	0.598309373447949
MCBRYAN_PUBERTAL_BREAST_3_4WK_UP	179	0.01	Down
0.458459489786935	0.343146065572594	0.598309373447949
TIEN_INTESTINE_PROBIOTICS_2HR_DN	88	0.01	Down
0.253214704987468	0.343596819308871	0.5987036590269
MCDOWELL_ACUTE_LUNG_INJURY_DN	32	0.01	Up
0.592049699142258	0.343750212041519	0.5987036590269
PID_P75_NTR_PATHWAY	63	0.01	Down	0.325778494003442
	0.343771778408994	0.5987036590269
KOBAYASHI_RESPONSE_TO_ROMIDEPSIN	17	0.01	Down
0.501264867596966	0.344164599358265	0.598971752079347
LI_CYTIDINE_ANALOGS_CYCTOTOXICITY	14	0.01	Down
0.230924273652138	0.344192221495206	0.598971752079347

REACTOME_RUNX1_INTERACTS_WITH_CO_FACTORS_WHOSE_PRECISE_EFFECT_ON_RUNX1_TARGETS_IS_NOT_KNOWN 36 0.01 Down 0.210182741182145 0.34506291242475 0.600093096982077
LEIN_NEURON_MARKERS 36 0.01 Up 0.347547865134864 0.345103594148758 0.600093096982077
KINSEY_TARGETS_OF_EWSR1_FLII_FUSION_UP 1138 0.01 Down 0.216377930646293 0.345274124668243 0.600157459545147
REACTOME_BASE_EXCISION_REPAIR 58 0.01 Up 0.138005088784445 0.34560028660858 0.600492187207409
PARK_APL_PATHOGENESIS_UP 12 0.01 Down 0.378170759071962 0.345815910541353 0.600634666879204
BIOCARTA_PRION_PATHWAY 11 0.01 Up 0.297888873310995 0.34602132096582 0.600759303878471
YAGI_AML_FAB_MARKERS 160 0.01 Up 0.352016143287237 0.347152936238624 0.602491292815682
KEGG_GLYCOSYLPHOSPHATIDYLINOSITOL_GPI_ANCHOR_BIOSYNTHESIS 25 0.01 Up 0.12142035861559 0.347371664972177 0.602638222327262
REACTOME_DEPOLYMERISATION_OF_THE_NUCLEAR_LAMINA 13 0.01 Up 0.189239937117911 0.348179499592023 0.603752178919341
REACTOME_INTERLEUKIN_12_SIGNALING 39 0.01 Down 0.242451433281299 0.348305206065206 0.603752178919341
MAGRANGEAS_MULTIPLE_MYELOMA_IGG_VS_IGA_DN 20 0.01 Up 0.341617124334058 0.348433001477475 0.603752178919341
MURAKAMI_UV_RESPONSE_1HR_UP 13 0.01 Up 0.280905940749402 0.348637080181012 0.603752178919341
LOPEZ_MBD_TARGETS_IMPRINTED_AND_X_LINKED 15 0.01 Down 0.372246839491559 0.34868786119939 0.603752178919341
RAY_TARGETS_OF_P210_BCR_ABL_FUSION_UP 11 0.01 Down 0.477879866630562 0.348932514484303 0.603752178919341
REACTOME_RHO_GTPASES_ACTIVATE_FORMINS 116 0.01 Down 0.180845929937634 0.348953984612335 0.603752178919341
REACTOME_INHIBITION_OF_THE_PROTEOLYTIC_ACTIVITY_OF_APC_C_REQUIRED_FOR_THE_ONSET_OF_ANAPHASE_BY_MITOTIC_SPINDLE_CHECKPOINT_COMPONENTS 19 0.01 Up 0.117921805248902 0.349503247478249 0.604141182527297
MIKKELSEN_MEF_ICP_WITH_H3K27ME3 71 0.01 Up 0.688546966098633 0.349528055408312 0.604141182527297
BIOCARTA_ETC_PATHWAY 10 0.01 Up 0.11639385218874 0.349582027976307 0.604141182527297
HOQUE_METHYLATED_IN_CANCER 36 0.01 Up 0.613008940561857 0.349855241576338 0.604380980355741
MARZEC_IL2_SIGNALING_DN 30 0.01 Up 0.333129790927186 0.350383660116954 0.605061295515062
PID_IL8_CXCR1_PATHWAY 23 0.01 Down 0.613343132156586 0.350643615702387 0.605228573334949
ZHENG_FOXP3_TARGETS_UP 21 0.01 Down 0.43263844089813 0.350749818362078 0.605228573334949
WHITFIELD_CELL_CYCLE_G2_M 174 0.01 Down 0.270278629637795 0.35153350195558 0.60618326333323
GHO_ATF5_TARGETS_UP 12 0.01 Down 0.253944070990091

0.351572807009951 0.60618326333323
REACTOME_PROCESSING_OF_CAPPED_INTRONLESS_PRE_MRNA 28 0.01
Down 0.0799782889978126 0.352224028798759
0.606797776548787
MULLIGAN_NTF3_SIGNALING_VIA_INSR_AND_IGF1R_UP 21 0.01 Up
0.210618314362423 0.352240574088411
0.606797776548787
REACTOME_RNA_POLYMERASE_II_TRANSCRIPTION_TERMINATION 65 0.01
Down 0.0943427721919817 0.35233419283478
0.606797776548787
REACTOME_ACTIVATION_OF_THE_MRNA_UPON_BINDING_OF_THE_CAP_BINDING_COMPLEX_AND_EIFS_AND_SUBSEQUENT_BINDING_TO_43S 59 0.01 Up
0.127081319315108 0.352997106142612
0.607299387074682
MULLIGHAN_MLL_SIGNATURE_2_UP 381 0.01 Up
0.324625832374398 0.353047197397343 0.607299387074682
TAVOR_CEBPA_TARGETS_DN 24 0.01 Up 0.410577453648652
0.353161997879987 0.607299387074682
REACTOME_G_PROTEIN_ACTIVATION 13 0.01 Up
0.279539794273498 0.353381584193011 0.607299387074682
REACTOME_ADP_SIGNALLING_THROUGH_P2Y_PURINOCEPTOR_12 13 0.01
Up 0.279539794273498 0.353381584193011
0.607299387074682
KIM_ALL_DISORDERS_OLIGODENDROCYTE_NUMBER_CORR_DN 22 0.01
Down 0.370761468807861 0.35343608377917
0.607299387074682
FINETTI_BREAST_CANCER_KINOME_RED 11 0.01 Up
0.269168347499759 0.353748732357029 0.607604337770288
REACTOME_DISORDERS_OF_TRANSMEMBRANE_TRANSPORTERS 133 0.01
Down 0.251553727735697 0.354040761127809
0.60765762526833
MOREAUX_B_LYMPHOCYTE_MATURATION_BY_TACI_DN 75 0.01 Up
0.127172986780149 0.354050126936097
0.60765762526833
BOYAULT_LIVER_CANCER_SUBCLASS_G3_UP 183 0.01 Down
0.163076301270485 0.35420160749118 0.607685582317883
REACTOME_NEGATIVE_REGULATION_OF_FGFR4_SIGNALING 21 0.01 Down
0.277936134780876 0.354358087543151
0.607722092142871
DEURIG_T_CELL_PROLYMPHOCYTIC_LEUKEMIA_UP 305 0.01 Up
0.354288183940336 0.35484635971979
0.607942988497624
YAO_TEMPORAL_RESPONSE_TO_PROGESTERONE_CLUSTER_0 61 0.01 Up
0.3892219378413 0.354895295221051 0.607942988497624
EPPERT_CE_HSC_LSC 30 0.01 Up 0.516469085286392
0.354976878302994 0.607942988497624
REACTOME_CLASS_I_PEROXISOMAL_MEMBRANE_PROTEIN_IMPORT 19 0.01
Up 0.137989534260998 0.355110731471073
0.607942988497624
CAMPS_COLON_CANCER_COPY_NUMBER_DN 31 0.01 Up
0.356002963098418 0.355163134103395 0.607942988497624
LEIN_CEREBELLUM_MARKERS 53 0.01 Down 0.321749673270542
0.355499410766991 0.608286962846451
REACTOME_DISEASES_ASSOCIATED_WITH_N_GLYCOSYLATION_OF_PROTEINS 17

0.01	Up	0.17579667143911	0.355732941803223
0.608329974062079
MILI_PSEUDOPODIA_CHEMOTAXIS_DN	413	0.01	Up
0.273131008319029	0.355795217310168	0.608329974062079
AZARE_NEOPLASTIC_TRANSFORMATION_BY_STAT3_UP	117	0.01	Down
0.658939811900253	0.356553162141069
0.609345097497595
KEGG_OOCYTE_MEIOSIS	91	0.01	Down	0.237150202772318
0.356809265113954	0.609345097497595
BIOCARTA_CERAMIDE_PATHWAY	21	0.01	Down
0.199178112375649	0.356833229807523	0.609345097497595
REACTOME_MECP2_REGULATES_NEURONAL_RECEPTORS_AND_CHANNELS	12
0.01	Down	0.268929297501848	0.356953670734483
0.609345097497595
VILLANUEVA_LIVER_CANCER_KRT19_UP	154	0.01	Down
0.233982717967459	0.357200776811959	0.609345097497595
RIZKI_TUMOR_INVASIVENESS_3D_UP	161	0.01	Up
0.348798185646274	0.357262461370325	0.609345097497595
REACTOME_NUCLEAR_ENVELOPE_REASSEMBLY	13	0.01	Down
0.166545387318814	0.357337859177678	0.609345097497595
MATTIOLI_MULTIPLE_MYELOMA_WITH_14Q32_TRANSLOCATIONS	32	0.01
Up	0.395743343073524	0.358057877128143
0.610225182194269
GRYDER_PAX3FOXO1_ENHANCERS_IN_TADS	942	0.01	Down
0.252095460374726	0.358200605463647	0.610225182194269
KONDO_PROSTATE_CANCER_HCP_WITH_H3K27ME3	53	0.01	Up
0.348146224625588	0.358319409924459	0.610225182194269
KEGG_NOTCH_SIGNALING_PATHWAY	42	0.01	Down
0.231746338354777	0.358512844769235	0.610225182194269
REACTOME_STING_MEDIATED_INDUCTION_OF_HOST_IMMUNE_RESPONSES	15
0.01	Down	0.23267386541865	0.358532748870982
0.610225182194269
BIOCARTA_RHO_PATHWAY	19	0.01	Down	0.166936234253173
0.358723662345859	0.610319024316667
MARSON_FOXP3_TARGETS_STIMULATED_UP	20	0.01	Down
0.486771084321201	0.358984439525866	0.610531613949591
WATANABE_RECTAL_CANCER_RADIOTHERAPY_RESPONSIVE_DN	88	0.01
Up	0.416884849914308	0.359332256506844
0.610873208060009
HERNANDEZ_ABERRANT_MITOSIS_BY_DOCETACEL_2NM_UP	58	0.01	Up
0.468909992280146	0.359462881752321
0.610873208060009
VALK_AML_CLUSTER_11	34	0.01	Up	0.54247517685124
0.359592994110519	0.610873208060009
REACTOME_TRAF6_MEDIATED_IRF7_ACTIVATION_IN_TLR7_8_OR_9_SIGNALING
13	0.01	Down	0.244443532673451
0.359767635346644	0.610938995422428
SCHLOSSER_MYC_TARGETS_REPRESSED_BY_SERUM	158	0.01	Down
0.125776017435794	0.360232926083707
0.611281397860709
REACTOME_INFECTIOUS_DISEASE	356	0.01	Down
0.162914641203291	0.360326382669327	0.611281397860709
CROMER_TUMORIGENESIS_UP	51	0.01	Down	0.998112842425317
0.360464526672551	0.611281397860709

RIZKI_TUMOR_INVASIVENESS_2D_DN 52 0.01 Down
0.364980884921249 0.360513233755003 0.611281397860709
REACTOME_PYRUVATE_METABOLISM_AND_CITRIC_ACID_TCA_CYCLE 50 0.01
Up 0.128665850764652 0.361814679055089
0.613256780675951
REACTOME_PHOSPHORYLATION_OF_THE_APC_C 17 0.01 Up
0.115989645667433 0.362512052743758 0.614207190758837
BILBAN_B_CLL_LPL_DN 37 0.01 Down 0.35682854529227
0.363201424303304 0.614974424449223
LIANG_HEMATOPOIESIS_STEM_CELL_NUMBER_LARGE_VS_TINY_UP 33 0.01
Up 0.393213187822314 0.363238508768117
0.614974424449223
NIKOLSKY_MUTATED_AND_AMPLIFIED_IN_BREAST_CANCER 67 0.01 Down
0.322956898921296 0.363533288743976
0.615241766906692
WAKABAYASHI_ADIPOGENESIS_PPARG_BOUND_36HR 29 0.01 Up
0.293738416960985 0.363790150194296
0.615444759173264
PID_HEDGEHOG_2PATHWAY 18 0.01 Down 0.415877076468359
0.364039394439223 0.615634717082131
SATO_SILENCED_BY_METHYLATION_IN_PANCREATIC_CANCER_1 288 0.01
Down 0.551340202445536 0.365118224131256
0.617226933986459
STARK_PREFRONTAL_CORTEX_22Q11_DELETION_UP 166 0.01 Down
0.278369584093649 0.365605301874414
0.617817981926877
REACTOME_GLYCOGEN_SYNTHESIS 13 0.01 Up
0.16253040891549 0.365882386930876 0.618053862929083
HAHTOLA_MYCOSIS_FUNGOIDES_SKIN_UP 160 0.01 Up
0.259572902465954 0.366065054209099 0.618130134737001
KEGG_TYPE_II_DIABETES_MELLITUS 30 0.01 Down
0.424488524884854 0.366331419130809 0.618347626358613
KEGG_GLYCOSPHINGOLIPID_BIOSYNTHESIS_GLOBO_SERIES 12 0.01
Up 0.46171289605193 0.366738542469417
0.618770588569403
WONG_ADULT_TISSUE_STEM_MODULE 584 0.01 Up
0.425199837165214 0.366857312244151 0.618770588569403
HORIUCHI_WTAP_TARGETS_UP 277 0.01 Down
0.42813545892767 0.367033104997771 0.618834886333451
REACTOME_SIGNALING_BY_NOTCH2 30 0.01 Down
0.330580183390488 0.367252798952688 0.618973127593675
KEGG_BASE_EXCISION_REPAIR 31 0.01 Up
0.157853006839014 0.367533250492561 0.619085307264169
OSADA_ASCL1_TARGETS_DN 21 0.01 Up 0.523950958449393
0.367594813145287 0.619085307264169
PECE_MAMMARY_STEM_CELL_UP 134 0.01 Up
0.212916950373332 0.367873958941333 0.619323387805728
BOQUEST_STEM_CELL_CULTURED_VS_FRESH_DN 14 0.01 Up
1.00501654777859 0.36841530465813 0.619949847517996
KOHOUTEK_CCNT2_TARGETS 37 0.01 Up 0.311583547275148
0.368646464455327 0.619949847517996
PID_HIF2PATHWAY 29 0.01 Down 0.203815403772493
0.368659831460646 0.619949847517996
NIKOLSKY_BREAST_CANCER_12Q13_Q21_AMPLICON 39 0.01 Down

0.371620768473693	0.369285632367314
0.620769976623438
HASLINGER_B_CLL_WITH_11Q23_DELETION	22	0.01	Down
0.297324775449692	0.370150193866757	0.621488682260389
HENDRICKS_SMARCA4_TARGETS_UP	51	0.01	Up
0.405936257567337	0.370227984373468	0.621488682260389
REACTOME_ROLE_OF_PHOSPHOLIPIDS_IN_PHAGOCYTOSIS	21	0.01	Up
0.50461206324235	0.370279942598928
0.621488682260389
REACTOME_FRS_MEDIATED_FGFR2_SIGNALING	14	0.01	Down
0.450094525140554	0.370433004471872	0.621488682260389
CHAUHAN_RESPONSE_TO_METHOXYESTRADIOL_UP 46	0.01	Up
0.189894595685269	0.370651266800562	0.621488682260389
REACTOME_SEMA4D_IN_SEMAPHORIN_SIGNALING 23	0.01	Down
0.213009141821895	0.370697047031706	0.621488682260389
TAKEDA_TARGETS_OF_NUP98_HOXA9_FUSION_16D_DN	96	0.01	Down
0.576047650810925	0.370802901308222
0.621488682260389
FINETTI_BREAST_CANCERS_KINOME_BLUE	19	0.01	Up
0.42679010633548	0.37081927604502	0.621488682260389
BENPORATH_PRC2_TARGETS	247	0.01	Up	0.637538870137399
0.372318222969945	0.623768323611593
REACTOME_GOLGI_TO_ER_RETROGRADE_TRANSPORT	114	0.01	Down
0.174962455293564	0.372555953945401
0.623934058489038
DELPUECH_FOXO3_TARGETS_DN	35	0.01	Down
0.265311603203421	0.372802811026797	0.62401491299431
BIOCARTA_PAR1_PATHWAY	16	0.01	Down	0.296730045433838
0.372909063834865	0.62401491299431
MORI_LARGE_PRE_BII_LYMPHOCYTE_DN	46	0.01	Up
0.397078544581391	0.37302070549849	0.62401491299431
WATANABE_COLON_CANCER_MSI_VS_MSS_UP	23	0.01	Down
0.401204659484451	0.373805516329718	0.62496632913391
REACTOME_NEGATIVE_REGULATION_OF_FGFR1_SIGNALING 23	0.01	Down
0.304788006641981	0.373867510353968
0.62496632913391
SESTO_RESPONSE_TO_UV_C7 67	0.01	Up	0.330505722967527
0.374675319636334	0.625964076178117
REACTOME_POST_TRANSLATIONAL_PROTEIN_MODIFICATION	1171	0.01
Down	0.276729819678757	0.374742898552906
0.625964076178117
PAL_PRMT5_TARGETS_DN	18	0.01	Down	0.351041801607349
0.375151189389762	0.626413297290855
BORCZUK_MALIGNANT_MESOTHELIOMA_UP	305	0.01	Down
0.186122825558842	0.375442615724502	0.626641268930351
GAJATE_RESPONSE_TO_TRABECTEDIN_UP	61	0.01	Up
0.493423978687513	0.375603359850428	0.626641268930351
REACTOME_FCERI_MEDIATED_CAPLUS2_MOBILIZATION	25	0.01	Down
0.326900810143754	0.375705944330878
0.626641268930351
REACTOME_GABA_B_RECEPTOR_ACTIVATION	23	0.01	Down
0.282114448732428	0.37659612136549	0.627893013923545
ZHANG_ANTIVIRAL_RESPONSE_TO_RIBAVIRIN_UP	25	0.01	Up
0.385404362485061	0.377435646481307

0.629059410802178
MALONEY_RESPONSE_TO_17AAG_DN 75 0.01 Up
0.13322723419045 0.37799370334213 0.629756003158959
WANG_TUMOR_INVASIVENESS_DN 196 0.01 Down
0.199038485463545 0.37818760612422 0.629845605605175
PID_NCADHERIN_PATHWAY 30 0.01 Down 0.215900461470685
0.378415706289992 0.629913598679612
REACTOME_TP53_REGULATES_TRANSCRIPTION_OF_DNA_REPAIR_GENES 61
0.01 Up 0.155078393704605 0.378508705235513
0.629913598679612
LUI_THYROID_CANCER_CLUSTER_3 28 0.01 Up
0.100719269231219 0.379188590126135 0.630531048954778
FIGUEROA_AML_METHYLATION_CLUSTER_3_DN 30 0.01 Up
0.45323981530354 0.379199862923018 0.630531048954778
KEGG_INOSITOL_PHOSPHATE_METABOLISM 49 0.01 Down
0.221399670154212 0.379401499294042 0.630531048954778
BURTON_ADIPOGENESIS_2 64 0.01 Up 0.302319869390064
0.379440820338749 0.630531048954778
IVANOVA_HEMATOPOIESIS_EARLY_PROGENITOR 391 0.01 Up
0.221374696597213 0.379969989861315 0.631056077281981
CASTELLANO_NRAS_TARGETS_UP 57 0.01 Down
0.47265481529402 0.380037553103965 0.631056077281981
REACTOME_SLC_MEDIATED_TRANSMEMBRANE_TRANSPORT 154 0.01 Up
0.459117514699158 0.380246331245667
0.631169593408151
REACTOME_NUCLEAR_RECEPTOR_TRANSCRIPTION_PATHWAY 37 0.01 Down
0.395970199380729 0.380579346812884
0.631291095329211
REACTOME_ORC1_REMOVAL_FROM_CHROMATIN 69 0.01 Down
0.124162924604089 0.380600415648979 0.631291095329211
KUMAR_PATHOGEN_LOAD_BY_MACROPHAGES 205 0.01 Down
0.318566229402542 0.381021286141357 0.631756060938916
MATZUK_IMPLANTATION_AND_UTERINE 15 0.01 Down
0.434356553951651 0.381777839749235 0.632539896073322
VANHARANTA_UTERINE_FIBROID_WITH_7Q_DELETION_UP 64 0.01 Down
0.168094498522319 0.381886023078927
0.632539896073322
GAVIN_PDE3B_TARGETS 19 0.01 Up 0.570453629275959
0.381916190866072 0.632539896073322
REACTOME_TRAFFICKING_OF_GLUR2_CONTAINING_AMPA_RECEPTORS 11 0.01
Up 0.327963427531164 0.382712615080276
0.633625489792206
LIU_VAV3_PROSTATE_CARCINOGENESIS_DN 13 0.01 Down
0.364307159725728 0.382869729401434 0.633652221524097
REACTOME_ER_QUALITY_CONTROL_COMPARTMENT_ERQC 21 0.01 Up
0.153798640156539 0.383356721653374
0.634224683044504
BIOCARTA_EICOSANOID_PATHWAY 12 0.01 Up
0.616716081418133 0.383736053724845 0.634545948655575
BYSTRYKH_HEMATOPOIESIS_STEM_CELL_AND_BRAIN_QTL_TRANS 129 0.01
Down 0.220335076716843 0.383833244581648
0.634545948655575
BIOCARTA_EFP_PATHWAY 14 0.01 Down 0.316648054794427
0.3842377692743 0.634981166502933

DAVICIONI_MOLECULAR_ARMS_VS_ERMS_UP 276 0.01 Down
0.280822207627178 0.384468386239721 0.635128774769402
REACTOME_NEUROTRANSMITTER_RECEPTORS_AND_POSTSYNAPTIC_SIGNAL_TRANSMIS
SION 108 0.01 Down 0.280071354071107
0.38504249883624 0.635744492391946
NAKAMURA_METASTASIS_MODEL_UP 37 0.01 Up
0.307677921159464 0.38525556879374 0.635744492391946
WELCSH_BRCA1_TARGETS_UP 187 0.01 Up 0.281823685099263
0.385279626574694 0.635744492391946
SANCHEZ_MDM2_TARGETS 13 0.01 Up 0.602411484780056
0.385406839103015 0.635744492391946
DUAN_PRDM5_TARGETS 57 0.01 Up 0.45248806213435
0.385829351019161 0.636152385228423
SHEPARD_BMYB_TARGETS 46 0.01 Up 0.392064412654402
0.385937164520113 0.636152385228423
ONKEN_UVEAL_MELANOMA_DN 489 0.01 Up 0.267942363272287
0.386840575862718 0.637407767046524
REACTOME_MITOTIC_TELOPHASE_CYTOKINESIS 14 0.01 Down
0.249547141295325 0.387105669979215 0.637610841537768
REACTOME_REGULATION_OF_TP53_ACTIVITY_THROUGH_METHYLATION 19
0.01 Up 0.175539625721981 0.387277841149651
0.637651394964884
IVANOVA_HEMATOPOIESIS_MATURE_CELL 253 0.01 Down
0.327480673984961 0.387414006596017 0.637651394964884
KYNG_DNA_DAMAGE_BY_GAMMA_AND_UV_RADIATION 69 0.01 Up
0.398659951890572 0.387626048361898
0.637761010844008
REACTOME_INSULIN_RECEPTOR_SIGNALLING_CASCADE 38 0.01 Down
0.365737196641853 0.388066308941643
0.637761010844008
RIZ_ERYTHROID_DIFFERENTIATION 70 0.01 Down
0.163427523212622 0.388126754712271 0.637761010844008
REACTOME_ABC_FAMILY_PROTEINS_MEDIATED_TRANSPORT 85 0.01 Down
0.155611600547249 0.388172565888345
0.637761010844008
SANSOM_WNT_PATHWAY_REQUIRE_MYC 50 0.01 Up
0.359133903480741 0.388190016834084 0.637761010844008
MOOTHA_PGC 398 0.01 Up 0.214804213770354
0.388366431226262 0.637817723186718
KEGG_HEMATOPOIETIC_CELL_LINEAGE 44 0.01 Up
0.833177152777692 0.388608643608629 0.637875581476874
REACTOME_NEF_MEDIATED_DOWNREGULATION_OF_MHC_CLASS_I_COMPLEX_CELL_SUR
FACE_EXPRESSION 10 0.01 Down 0.181713242129438
0.388747919357027 0.637875581476874
KLEIN_PRIMARY_EFFUSION_LYMPHOMA_DN 48 0.01 Up
0.463954593707082 0.388918270793069 0.637875581476874
KEGG_PANTOTHENATE_AND_COA_BIOSYNTHESIS 12 0.01 Down
0.457642798825671 0.38912622853221 0.637875581476874
MARCHINI_TRABECTEDIN_RESISTANCE_UP 19 0.01 Up
0.402954312496461 0.389204666003853 0.637875581476874
KEGG_AMINO_SUGAR_AND_NUCLEOTIDE_SUGAR_METABOLISM 41 0.01
Up 0.323317330433715 0.389253107895676
0.637875581476874
REACTOME_RESOLUTION_OF_D_LOOP_STRUCTURES_THROUGH_SYNTHESIS_DEPENDENT

_STRAND_ANNEALING_SDSA	24	0.01	Up	0.194095083158751
	0.390956317641723	0.640428792132676
OHGUCHI_LIVER_HNF4A_TARGETS_DN	75	0.01	Up
0.500557511728841	0.391096114439198	0.640428792132676
BOYLAN_MULTIPLE_MYELOMA_PCA1_UP	75	0.01	Down
0.754682113205392	0.391287097721609	0.640508195287194
CHIARETTI_T_ALL_RELAPSE_PROGNOSIS	18	0.01	Up
0.498592091813469	0.391461580849166	0.640560540923554
FORTSCHEGGER_PHF8_TARGETS_DN	738	0.01	Down
0.304624685167417	0.392328732243157	0.641745870244902
HUMMERICH_MALIGNANT_SKIN_TUMOR_UP	12	0.01	Down
0.939821226408646	0.392733951808575	0.642175013961274
REACTOME_SIGNALING_BY_HEDGEHOG	132	0.01	Down
0.208236003765803	0.393694358565934	0.643511324274137
VALK_AML_WITH_CEBPA	25	0.01	Up	0.44024763185984
	0.394732447837287	0.644973592522211
BIOCARTA_MCM_PATHWAY	18	0.01	Up	0.148355647459031
	0.395164868670527	0.645042487083794
BOYLAN_MULTIPLE_MYELOMA_PCA3_UP	62	0.01	Down
0.29571831664819	0.395207729203101	0.645042487083794
MMS_MOUSE_LYMPH_HIGH_4HRS_UP	35	0.01	Down
0.1680385200209	0.395412809558497	0.645042487083794
BILANGES_SERUM_SENSITIVE_GENES	56	0.01	Up
0.470025923256409	0.39557832203941	0.645042487083794
REACTOME_ASPARAGINE_N_LINKED_GLYCOSYLATION	272	0.01	Down
	0.220842997597296	0.395617667016835
0.645042487083794
PID_ATR_PATHWAY	37	0.01	Up	0.203996354624281
0.39574780986961	0.645042487083794
YAO_TEMPORAL_RESPONSE_TO_PROGESTERONE_CLUSTER_11	90	0.01
	Down	0.182034989824972	0.395779127781336
0.645042487083794
BIOCARTA_ETS_PATHWAY	17	0.01	Down	0.348009137171184
	0.396369621022895	0.645770730879997
MAHADEVAN_GIST_MORPHOLOGICAL_SWITCH	14	0.01	Up
0.708804581119567	0.39687173351313	0.646354508022289
MAGRANGEAS_MULTIPLE_MYELOMA_IGLL_VS_IGLK_UP	30	0.01	Down
	0.371809299478841	0.397031003803081
0.646379703764886
REACTOME_SIGNALING_BY_RHO_GTPASES	364	0.01	Down
0.269147171249153	0.398833545567192	0.648844326651043
DARWICHE_PAPILLOMA_RISK_LOW_DN	128	0.01	Up
0.299091297198223	0.398936193324657	0.648844326651043
TONKS_TARGETS_OF_RUNX1_RUNX1T1_FUSION_MONOCYTE_UP	190	0.01
	Up	0.353553152141893	0.39897791298409
0.648844326651043
KIM_TIAL1_TARGETS	28	0.01	Up	0.191052568977329
	0.399229595108327	0.64901881736417
REACTOME_G_BETA:GAMMA_SIGNALLING_THROUGH_CDC42	13	0.01	Up
	0.257058985913293	0.39966148469784
0.649433420396002
MATTHEWS_AP1_TARGETS	15	0.01	Up	0.655191701118164
	0.399773587149219	0.649433420396002
GRATIAS_RETINOBLASTOMA_16Q24	18	0.01	Up

0.163723569964459	0.400616157983235	0.650421592587874
NIKOLSKY_BREAST_CANCER_17Q21_Q25_AMPLICON	252	0.01	Up
	0.26263226819714	0.40067127694679
0.650421592587874
REACTOME_KSRP_KHSRP_BINDS_AND_DESTABILIZES_MRNA	17	0.01	Down
	0.173380541653268	0.401274748088751
0.651141430384499
IZADPANAH_STEM_CELL_ADIPOSE_VS_BONE_UP	116	0.01	Down
0.39457309233528	0.401404427941145	0.651141430384499
CHIARADONNA_NEOPLASTIC_TRANSFORMATION_KRAS_CDC25_UP	53	0.01
	Up	0.385905575802065	0.401773091859808
0.65150434628782
YORDY_RECIPROCAL_REGULATION_BY_ETS1_AND_SP100_UP	23	0.01
	Down	0.290534858816974	0.402326060222876
0.652165755752552
PID_SYNDECAN_3_PATHWAY	12	0.01	Up	0.420035950993809
	0.403296191488405	0.653404647127431
REACTOME_SHC_MEDIATED_CASCADE:FGFR4	10	0.01	Down
0.437510585435054	0.403381066914043	0.653404647127431
MEISSNER_NPC_HCP_WITH_H3_UNMETHYLATED	326	0.01	Up
0.407153980985387	0.403568715774879	0.653467297412265
FLECHNER_BIOPSY_KIDNEY_TRANSPLANT_REJECTED_VS_OK_DN	474	0.01
	Up	0.283275948415276	0.403764604405333
0.653467297412265
BARRIER_COLON_CANCER_RECURRENCE_DN	17	0.01	Up
0.256979984582675	0.403855873684376	0.653467297412265
MIKKELSEN_MEF_HCP_WITH_H3K27ME3	187	0.01	Down
0.624868352333453	0.40457520833696	0.653995775101316
DOANE_BREAST_CANCER_ESR1_UP	85	0.01	Down
0.510180978294347	0.404588499483185	0.653995775101316
LEE_TARGETS_OF_PTCH1_AND_SUFU_UP	43	0.01	Up
0.527029920982477	0.404618965641103	0.653995775101316
YANG_BREAST_CANCER_ESR1_DN	25	0.01	Down
0.353338399704931	0.404823480280132	0.654091137260674
REACTOME_TELOMERE_MAINTENANCE	49	0.01	Up
0.123556649582564	0.405098584588665	0.654300444745257
VERNELL_RETINOBLASTOMA_PATHWAY_DN	23	0.01	Down
0.255848739597071	0.407032914327961	0.656688092445468
RIZ_ERYTHROID_DIFFERENTIATION_12HR	30	0.01	Up
0.402890160502978	0.407204421686595	0.656688092445468
IWANAGA_CARCINOGENESIS_BY_KRAS_UP	134	0.01	Down
0.27291624320898	0.407258356520292	0.656688092445468
RUTELLA_RESPONSE_TO_CSF2RB_AND_IL4_UP	314	0.01	Down
0.377608870314249	0.407413671774589	0.656688092445468
LEIN_LOCALIZED_TO_DISTAL_AND_PROXIMAL_DENDRITES	12	0.01	Up
	0.249479090094924	0.407553922207208
0.656688092445468
KEGG_HOMOLOGOUS_RECOMBINATION	24	0.01	Up
0.147064595492144	0.407587443077211	0.656688092445468
REACTOME_EPIGENETIC_REGULATION_OF_GENE_EXPRESSION	106	0.01
	Down	0.147868957763232	0.407651222740353
0.656688092445468
REACTOME_PRESYNAPTIC_FUNCTION_OF_KAINATE_RECEPTORS	13	0.01
	Up	0.39427923212036	0.407745598668587

0.656688092445468
DORN_ADENOVIRUS_INFECTION_24HR_UP 10 0.01 Down
0.307632027724325 0.408009808092135 0.656878254790168
KYNG_RESPONSE_TO_H2O2_VIA_ERCC6_UP 37 0.01 Down
0.29311910534642 0.408344360234926 0.657181489171498
LEE_LIVER_CANCER_E2F1_DN 39 0.01 Up
0.420215142708248 0.408527814982605 0.657241420310239
BIOCARTA_ATRBRCA_PATHWAY 23 0.01 Up
0.166515771461113 0.408899760541223 0.657604444949122
FAELT_B_CLL_WITH_VH_REARRANGEMENTS_UP 46 0.01 Down
0.312989248119211 0.409678127439945 0.658393740833055
BIOCARTA_CARM1_PATHWAY 10 0.01 Down 0.241269423307687
0.409741435177651 0.658393740833055
RASHI_RESPONSE_TO_IONIZING_RADIATION_4 37 0.01 Down
0.269054579647746 0.409829963704313 0.658393740833055
REACTOME_TRANSCRIPTIONAL_REGULATION_BY_THE_AP_2_TFAP2_FAMILY_OF_TRAN
SCRIPTION_FACTORS 28 0.01 Down 0.346262137480901
0.41057999627342 0.659072242865025
BROWNE_HCMV_INFECTION_12HR_DN 84 0.01 Up
0.379002048825953 0.410641156662002 0.659072242865025
STEARMAN_LUNG_CANCER_EARLY_VS_LATE_UP 118 0.01 Down
0.199010159053616 0.410692180704101 0.659072242865025
YAO_TEMPORAL_RESPONSE_TO_PROGESTERONE_CLUSTER_1 63 0.01 Down
0.416761851895723 0.410858004398688
0.659103044172771
NAKAJIMA_MAST_CELL 30 0.01 Up 0.374038227712207
0.411428462657822 0.659388853407342
PUJANA_BRCA_CENTERED_NETWORK 112 0.01 Up
0.154939341155768 0.411433019658086 0.659388853407342
REACTOME_HOMOLOGY_DIRECTED_REPAIR 108 0.01 Up
0.153131273300263 0.411747819921159 0.659388853407342
WENG_POR_DOSAGE 15 0.01 Up 0.593409011005805
0.411873205107083 0.659388853407342
MORI_EMU_MYC_LYMPHOMA_BY_ONSET_TIME_UP 99 0.01 Up
0.204078074871403 0.411936055547951 0.659388853407342
SANSOM_APC_MYC_TARGETS 227 0.01 Up 0.205567358179436
0.412065446521969 0.659388853407342
ACEVEDO_LIVER_CANCER_WITH_H3K9ME3_UP 73 0.01 Up
0.457232316778964 0.41221696678810 2 0.659388853407342
SUZUKI_RESPONSE_TO_TSA_AND_DECITABINE_1A 16 0.01 Up
0.909896230365425 0.412329855403298
0.659388853407342
REACTOME_NEGATIVE_REGULATION_OF_FGFR2_SIGNALING 23 0.01 Down
0.31971422006771 0.412356410884992
0.659388853407342
REACTOME_RESOLUTION_OF_D_LOOP_STRUCTURES 30 0.01 Up
0.186210665330171 0.413163073226456
0.660419903619201
REACTOME_VOLTAGE_GATED_POTASSIUM_CHANNELS 12 0.01 Down
0.702237621222853 0.413377481813344
0.660419903619201
REACTOME_PHOSPHOLIPID_METABOLISM 179 0.01 Down
0.258686315811794 0.413441959685079 0.660419903619201
GRAESSMANN_APOPTOSIS_BY_DOXORUBICIN_DN 1642 0.01 Down

0.223814856501526	0.414531051371743	0.661924360893778
CHEOK_RESPONSE_TO_MERCAPTOPURINE_AND_HD_MTX_DN	23	0.01	Down
	0.200903533884916	0.414913560006539
0.662122542454973
PID_IL8_CXCR2_PATHWAY	28	0.01	Down	0.506534936126552
	0.414949766873339	0.662122542454973
TONKS_TARGETS_OF_RUNX1_RUNX1T1_FUSION_SUSTAINED_IN_GRANULOCYTE_UP
	13	0.01	Up	0.47826552471664
0.415134158653605	0.662181704452787
MARSON_BOUND_BY_E2F4_UNSTIMULATED	600	0.01	Up
0.181025042121999	0.415621124273505	0.662693781067242
LEE_TARGETS_OF_PTCH1_AND_SUFU_DN	55	0.01	Up
0.451752219157995	0.416022073995048	0.662693781067242
REACTOME_FORMATION_OF_SENESCENCE_ASSOCIATED_HETEROCHROMATIN_FOCI_SAH
F	13	0.01	Down	0.192475448798249
0.41602375619153	0.662693781067242
KAPOSI_LIVER_CANCER_MET_UP	18	0.01	Down
0.26296778881776	0.416545709203204	0.662693781067242
REACTOME_DOPAMINE_NEUROTRANSMITTER_RELEASE_CYCLE	14	0.01
	Down	0.248117438965697	0.416598711277449
0.662693781067242
LI_WILMS_TUMOR	15	0.01	Up	0.663727576175851
0.416739028455385	0.662693781067242
KRIEG_KDM3A_TARGETS_NOT_HYPOXIA	185	0.01	Down
0.264186278920504	0.417051285148322	0.662693781067242
REACTOME_INFLUENZA_INFECTION	153	0.01	Up
0.129566112883171	0.417135504204262	0.662693781067242
JIANG_TIP30_TARGETS_DN	22	0.01	Up	0.284953495841948
	0.417144845249428	0.662693781067242
MAHADEVAN_IMATINIB_RESISTANCE_DN	10	0.01	Up
0.565404173399008	0.417247974907439	0.662693781067242
KEGG_HUNTINGTONS_DISEASE	158	0.01	Up
0.191194189126792	0.417329245332864	0.662693781067242
LIU_SOX4_TARGETS_DN	299	0.01	Down	0.182789937389459
	0.417554407320534	0.662693781067242
GARGALOVIC_RESPONSE_TO_OXIDIZED_PHOSPHOLIPIDS_RED_UP	15	0.01
	Down	0.328349429555215	0.417682031112961
0.662693781067242
REACTOME_SIGNALING_BY_FGFR1	38	0.01	Down
0.298496660435221	0.417960063191802	0.662693781067242
NUTT_GBM_VS_AO_GLIOMA_UP	43	0.01	Up
0.295832868768842	0.418198286427068	0.662693781067242
LEE_METASTASIS_AND_RNA_PROCESSING_UP	16	0.01	Down
0.10068188968298	0.418304267463036	0.662693781067242
GRAESSMANN_RESPONSE_TO_MC_AND_DOXORUBICIN_UP	536	0.01	Up
	0.272208490692388	0.418371412344121
0.662693781067242
KYNG_RESPONSE_TO_H2O2_VIA_ERCC6	15	0.01	Up
0.161766896626511	0.418465330306053	0.662693781067242
PIONTEK_PKD1_TARGETS_UP	28	0.01	Down	0.338998115422094
	0.418470923927291	0.662693781067242
FLECHNER_BIOPSY_KIDNEY_TRANSPLANT_OK_VS_DONOR_DN	19	0.01
	Down	0.273720970040374	0.418631651471773
0.662693781067242

AUNG_GASTRIC_CANCER 39 0.01 Down 0.272640138818424
0.418651292092744 0.662693781067242
REACTOME_GLYCOGEN_BREAKDOWN_GLYCOGENOLYSIS 12 0.01 Down
0.189929262419813 0.418725873221266
0.662693781067242
REACTOME_TRANSPORT_OF_INORGANIC_CATIONS_ANIONS_AND_AMINO_ACIDS_OLIGO
PEPTIDES 64 0.01 Down 0.457954917049733
0.418846058289663 0.662693781067242
VISALA_RESPONSE_TO_HEAT_SHOCK_AND_AGING_UP 14 0.01 Up
0.166826880973881 0.419572631795196
0.663545653776578
ZHAN_MULTIPLE_MYELOMA_HP_UP 39 0.01 Up
0.34406465883882 0.419679709385275 0.663545653776578
REACTOME_INFLAMMASOMES 16 0.01 Down 0.374636328210768
0.419852319631161 0.663585153566128
HUANG_DASATINIB_RESISTANCE_DN 61 0.01 Down
0.307380451535446 0.4201388320499 0.663624921884246
MCBRYAN_PUBERTAL_BREAST_6_7WK_DN 65 0.01 Up
0.548632777307995 0.420172753655743 0.663624921884246
KEGG_BIOSYNTHESIS_OF_UNSATURATED_FATTY_ACIDS 21 0.01 Up
0.313704384257402 0.420648789761996
0.663712241083561
SU_PANCREAS 23 0.01 Down 0.29675705324417
0.420738390033374 0.663712241083561
REACTOME_REGULATED_PROTEOLYSIS_OF_P75NTR 11 0.01 Down
0.230357096461046 0.420811927535943
0.663712241083561
TOMLINS_METASTASIS_DN 20 0.01 Down 0.235718433209489
0.420870989011824 0.663712241083561
REACTOME_TP53_REGULATES_TRANSCRIPTION_OF_SEVERAL_ADDITIONAL_CELL_DEA
TH_GENES_WHOSE_SPECIFIC_ROLES_IN_P53_DEPENDENT_APOPTOSIS_REMAIN_UNCE
RTAIN 11 0.01 Down 0.223190633319841
0.421044546938245 0.663712241083561
BOYAULT_LIVER_CANCER_SUBCLASS_G2 26 0.01 Up
0.227946277785908 0.421113973653018 0.663712241083561
REACTOME_TICAM1_TRAF6_DEPENDENT_INDUCTION_OF_TAK1_COMPLEX 11
0.01 Down 0.174315310291538 0.421342438355873
0.663839558503207
HOWLIN_CITED1_TARGETS_1_DN 34 0.01 Down
0.381614026394024 0.421831519054284 0.664372222764562
MALIK_REPRESSED_BY_ESTROGEN 12 0.01 Down
0.402793233239623 0.422065397557818 0.664372222764562
ZAIDI_OSTEOBLAST_TRANSCRIPTION_FACTORS 11 0.01 Up
0.407558024737691 0.422123930637506 0.664372222764562
MIKKELSEN_ES_HCP_WITH_H3K27ME3 12 0.01 Up
0.577403388443124 0.423434121054392 0.666007921283946
MIKKELSEN_MEF_HCP_WITH_H3_UNMETHYLATED 84 0.01 Up
0.437707690673007 0.423459541497112 0.666007921283946
PANGAS_TUMOR_SUPPRESSION_BY_SMAD1_AND_SMAD5_DN 122 0.01 Down
0.273053423483801 0.424151042654459
0.666862167447288
AZARE_STAT3_TARGETS 24 0.01 Down 0.580533207433071
0.424523871766765 0.667214966290773
YAO_TEMPORAL_RESPONSE_TO_PROGESTERONE_CLUSTER_15 27 0.01

Up 0.587338710812425 0.42488404258545
0.667547630696119
SIG_REGULATION_OF_THE_ACTIN_CYTOSKELETON_BY_RHO_GTPASES 31 0.01
Down 0.238062315676481 0.425270336937803
0.667626144168754
ROVERSI_GLIOMA_COPY_NUMBER_UP 69 0.01 Up
0.282164145736612 0.425376705466597 0.667626144168754
PETRETTO_CARDIAC_HYPERTROPHY 30 0.01 Up
0.560957077020439 0.425379594415865 0.667626144168754
CAIRO_HEPATOBLASTOMA_DN 182 0.01 Up 0.439183545186912
0.425773332655989 0.668010865720305
STEARMAN_LUNG_CANCER_EARLY_VS_LATE_DN 50 0.01 Up
0.554026519609909 0.426111182143023 0.668307663549508
HADDAD_T_LYMPHOCYTE_AND_NK_PROGENITOR_DN 39 0.01 Up
0.760896793854423 0.426442889312089
0.668594624156902
BIOCARTA_CBL_PATHWAY 11 0.01 Down 0.302678220763418
0.426666844153839 0.668712505045852
KIM_GASTRIC_CANCER_CHEMOSENSITIVITY 68 0.01 Up
0.297112507081496 0.426994834999047 0.668791807735872
MARIADASON_RESPONSE_TO_BUTYRATE_SULINDAC_4 15 0.01 Up
0.327285912661273 0.427015014060501
0.668791807735872
LEE_CALORIE_RESTRICTION_MUSCLE_DN 42 0.01 Down
0.345054720476831 0.427286418217102 0.668854722712461
KEGG_GLYCOSAMINOGLYCAN_BIOSYNTHESIS_KERATAN_SULFATE 13 0.01
Down 0.487527345499443 0.42737124460676
0.668854722712461
HESS_TARGETS_OF_HOXA9_AND_MEIS1_UP 59 0.01 Up
0.275159634666987 0.427501583615773 0.668854722712461
REACTOME_HDACS_DEACETYLATE_HISTONES 49 0.01 Down
0.15442652511907 0.427664083331852 0.668876149817911
RODWELL_AGING_KIDNEY_NO_BLOOD_DN 131 0.01 Up
0.269246395139509 0.427949920956783 0.66909039815678
BYSTRYKH_HEMATOPOIESIS_STEM_CELL_SCP2_QTL_TRANS 16 0.01 Up
0.478033821264863 0.428426296936636
0.669602296498671
NOUZOVA_TRETINOIN_AND_H4_ACETYLATION 131 0.01 Down
0.158213010720238 0.428887375821376 0.669889241449454
GARY_CD5_TARGETS_UP 443 0.01 Down 0.252716035251053
0.42890795036519 0.669889241449454
KARLSSON_TGFB1_TARGETS_UP 116 0.01 Down
0.199801653117782 0.429132705165633 0.670007471246794
ZWANG_EGF_PERSISTENTLY_UP 29 0.01 Up
0.234416330541264 0.429334895966661 0.670090401864632
TIAN_BHLHA15_TARGETS 14 0.01 Down 0.203428997671374
0.429588925501136 0.670254154851652
REACTOME_SIGNALING_BY_FGFR 70 0.01 Down
0.248548437596551 0.429814374881204 0.670373218282794
WANG_CLASSIC_ADIPOGENIC_TARGETS_OF_PPARG 18 0.01 Up
0.489331731454523 0.430484281133453
0.671185169509147
CAIRO_PML_TARGETS_BOUND_BY_MYC_DN 11 0.01 Up
0.432204143866419 0.430795165474823 0.67143698641101

| | | | | | | |
|---|---|---|---|---|---|---|
| TONKS_TARGETS_OF_RUNX1_RUNX1T1_FUSION_HSC_DN | 148 | 0.01 | Up | 0.49656383904904 | 0.431047033316964 | 0.671596677559706 |
| TIEN_INTESTINE_PROBIOTICS_24HR_UP | 536 | 0.01 | Up | 0.191942211249345 | 0.431246434631209 | 0.671674540425254 |
| MARTINEZ_TP53_TARGETS_DN | 480 | 0.01 | Down | 0.31528013145085 | 0.431490352647365 | 0.671821660945586 |
| KEGG_P53_SIGNALING_PATHWAY | 61 | 0.01 | Up | 0.286177593483335 | 0.433025234393035 | 0.673977987741237 |
| PARENT_MTOR_SIGNALING_DN | 35 | 0.01 | Down | 0.565341020372442 | 0.433986726846786 | 0.675148956865428 |
| NIKOLSKY_BREAST_CANCER_16Q24_AMPLICON | 42 | 0.01 | Up | 0.234222629078621 | 0.434149897995459 | 0.675148956865428 |
| REACTOME_TRNA_MODIFICATION_IN_THE_NUCLEUS_AND_CYTOSOL | 43 | 0.01 | Up | 0.162984243856639 | 0.434228172257609 | 0.675148956865428 |
| FLECHNER_PBL_KIDNEY_TRANSPLANT_REJECTED_VS_OK_DN | 47 | 0.01 | Down | 0.130259884469214 | 0.434389788921408 | 0.675166701660349 |
| MARIADASON_RESPONSE_TO_BUTYRATE_SULINDAC_6 | 44 | 0.01 | Down | 0.262065922172481 | 0.434735396559279 | 0.675470310243332 |
| STAMBOLSKY_BOUND_BY_MUTATED_TP53 | 14 | 0.01 | Up | 0.222946544858 | 0.435142012892662 | 0.675765123280346 |
| REACTOME_ASSEMBLY_OF_THE_PRE_REPLICATIVE_COMPLEX | 65 | 0.01 | Down | 0.132501032874281 | 0.435225813547631 | 0.675765123280346 |
| GARGALOVIC_RESPONSE_TO_OXIDIZED_PHOSPHOLIPIDS_BLACK_DN | 11 | 0.01 | Up | 0.465389875683414 | 0.435547334848076 | 0.676013688322791 |
| LUI_THYROID_CANCER_PAX8_PPARG_DN | 46 | 0.01 | Up | 0.191360699717874 | 0.43582079046205 | 0.676013688322791 |
| REACTOME_FORMATION_OF_TC_NER_PRE_INCISION_COMPLEX | 53 | 0.01 | Up | 0.101692730556658 | 0.435978656115551 | 0.676013688322791 |
| IIZUKA_LIVER_CANCER_PROGRESSION_L1_G1_UP | 17 | 0.01 | Up | 0.330343593680346 | 0.435987471067357 | 0.676013688322791 |
| HEIDENBLAD_AMPLICON_12P11_12_UP | 31 | 0.01 | Down | 0.273163542391886 | 0.436267617759446 | 0.676214807527141 |
| REACTOME_INTERCONVERSION_OF_NUCLEOTIDE_DI_AND_TRIPHOSPHATES | 26 | 0.01 | Up | 0.191329892451248 | 0.436582492934144 | 0.676335356655457 |
| REACTOME_FORMATION_OF_INCISION_COMPLEX_IN_GG_NER | 43 | 0.01 | Up | 0.125463127004572 | 0.436788652910978 | 0.676335356655457 |
| HOEGERKORP_CD44_TARGETS_TEMPORAL_DN | 20 | 0.01 | Down | 0.310416745559212 | 0.436821272961215 | 0.676335356655457 |
| PID_LYMPH_ANGIOGENESIS_PATHWAY | 23 | 0.01 | Down | 0.472189745182008 | 0.437051606930192 | 0.676335356655457 |
| BOYLAN_MULTIPLE_MYELOMA_D_CLUSTER_UP | 23 | 0.01 | Down | 0.259082993562041 | 0.437097711030946 | 0.676335356655457 |
| REACTOME_POTASSIUM_CHANNELS | 35 | 0.01 | Down | 0.454947904660513 | 0.437294160837695 | 0.676389712042246 |

RAMPON_ENRICHED_LEARNING_ENVIRONMENT_EARLY_UP 12 0.01 Down 0.0990590660485651 0.437573774645505 0.676389712042246
REACTOME_FORMATION_OF_THE_EARLY_ELONGATION_COMPLEX 33 0.01 Up 0.10276507219029 0.4376087462587 0.676389712042246
GOLDRATH_ANTIGEN_RESPONSE 285 0.01 Up 0.323297280042901 0.437734743566384 0.676389712042246
CHIANG_LIVER_CANCER_SUBCLASS_INTERFERON_DN 43 0.01 Up 0.414464523940041 0.437981256101379 0.676538057105052
PID_AURORA_A_PATHWAY 27 0.01 Down 0.153309215213076 0.438355916210818 0.676884178415536
KEGG_CYSTEINE_AND_METHIONINE_METABOLISM 27 0.01 Up 0.161431176572269 0.438886321599916 0.676890607487293
REACTOME_P130CAS_LINKAGE_TO_MAPK_SIGNALING_FOR_INTEGRINS 11 0.01 Down 0.316237967627002 0.438925392881957 0.676890607487293
TOYOTA_TARGETS_OF_MIR34B_AND_MIR34C 396 0.01 Down 0.210827843086807 0.439102339562099 0.676890607487293
RIGGI_EWING_SARCOMA_PROGENITOR_UP 313 0.01 Up 0.514377052084638 0.439193082183818 0.676890607487293
REACTOME_TRANSMISSION_ACROSS_CHEMICAL_SYNAPSES 143 0.01 Down 0.294158515930872 0.439252184124335 0.676890607487293
BIOCARTA_GH_PATHWAY 22 0.01 Down 0.262525231797943 0.439263604458384 0.676890607487293
GHO_ATF5_TARGETS_DN 16 0.01 Down 0.253696927054756 0.439994670058223 0.677537072537983
REACTOME_AFLATOXIN_ACTIVATION_AND_DETOXIFICATION 11 0.01 Up 0.540397102371208 0.440055869206427 0.677537072537983
AGUIRRE_PANCREATIC_CANCER_COPY_NUMBER_UP 282 0.01 Up 0.1996908320107 0.440135317421782 0.677537072537983
KEGG_ARRHYTHMOGENIC_RIGHT_VENTRICULAR_CARDIOMYOPATHY_ARVC 49 0.01 Up 0.443149730660082 0.440362794611642 0.677655173495148
DIAZ_CHRONIC_MEYLOGENOUS_LEUKEMIA_UP 1346 0.01 Down 0.189036863244909 0.440994466413195 0.678394978277656
REACTOME_G_PROTEIN_GATED_POTASSIUM_CHANNELS 12 0.01 Down 0.209820279011124 0.441385842356613 0.67864485962728
HOLLEMAN_VINCRISTINE_RESISTANCE_ALL_UP 25 0.01 Down 0.279781515277249 0.44145885863185 0.67864485962728
GAJATE_RESPONSE_TO_TRABECTEDIN_DN 17 0.01 Down 0.20987190954479 0.441899065115806 0.679089332545486
KEGG_VALINE_LEUCINE_AND_ISOLEUCINE_BIOSYNTHESIS 10 0.01 Up 0.0874057851024065 0.442238017573584 0.679107479664922
FARMER_BREAST_CANCER_APOCRINE_VS_BASAL 295 0.01 Down 0.351843557497687 0.442379939699363 0.679107479664922
REACTOME_HOMOLOGOUS_DNA_PAIRING_AND_STRAND_EXCHANGE 40 0.01 Up 0.160211850884993 0.442387369541789 0.679107479664922

ZHAN_VARIABLE_EARLY_DIFFERENTIATION_GENES_DN	30	0.01	Up
	0.154518213985806	0.44251519642682
0.679107479664922
MATZUK_EMBRYONIC_GERM_CELL	16	0.01	Down
0.484592404402454	0.443264487511565	0.679548306672166
MCLACHLAN_DENTAL_CARIES_DN	60	0.01	Down
0.505794968498108	0.443300525038605	0.679548306672166
RAY_TARGETS_OF_P210_BCR_ABL_FUSION_DN	14	0.01	Up
0.490331400552161	0.443406883531678	0.679548306672166
KAUFFMANN_DNA_REPAIR_GENES	209	0.01	Up
0.158332983738307	0.443407159837478	0.679548306672166
REACTOME_PRC2_METHYLATES_HISTONES_AND_DNA	33	0.01	Down
	0.147304364929331	0.44388529659847
0.679931248024804
GARGALOVIC_RESPONSE_TO_OXIDIZED_PHOSPHOLIPIDS_SALMON_DN 12	0.01
	Down	0.249034883526414	0.443959557942781
0.679931248024804
REACTOME_TRANSCRIPTION_COUPLED_NUCLEOTIDE_EXCISION_REPAIR_TC_NER
	77	0.01	Up	0.111646785543087
0.444426397734909	0.680207133662866
PID_LPA4_PATHWAY	12	0.01	Down	0.398176711205816
	0.444442347401076	0.680207133662866
MIDORIKAWA_AMPLIFIED_IN_LIVER_CANCER	45	0.01	Down
0.243354501208187	0.444918410762103	0.680703967452571
ABE_VEGFA_TARGETS_2HR	27	0.01	Down	0.386463846397516
	0.445514278702306	0.681142420714724
KEGG_RIBOFLAVIN_METABOLISM	11	0.01	Down
0.356905701250033	0.445629396748995	0.681142420714724
CHANG_IMMORTALIZED_BY_HPV31_UP	55	0.01	Up
0.422103692814223	0.445659590505451	0.681142420714724
KEGG_BASAL_CELL_CARCINOMA	40	0.01	Up
0.362882468421478	0.446695219553354	0.682493205945726
REACTOME_GLUTAMATE_AND_GLUTAMINE_METABOLISM	13	0.01	Up
	0.16561846637907	0.447035351226343
0.682653894002144
DUNNE_TARGETS_OF_AML1_MTG8_FUSION_DN	16	0.01	Up
0.393745104837206	0.447254509059295	0.682653894002144
JOHNSTONE_PARVB_TARGETS_2_UP	137	0.01	Down
0.305465868005801	0.447306541142687	0.682653894002144
SCHUHMACHER_MYC_TARGETS_UP	78	0.01	Up
0.186267649944589	0.447661446416288	0.682653894002144
BANDRES_RESPONSE_TO_CARMUSTIN_MGMT_48HR_UP	15	0.01	Up
	0.256927475891401	0.447691408682721
0.682653894002144
REACTOME_INWARDLY_RECTIFYING_KPLUS_CHANNELS	14	0.01	Down
	0.230698834747744	0.447842578119174
0.682653894002144
MCBRYAN_PUBERTAL_BREAST_4_5WK_UP	232	0.01	Down
0.454391064717446	0.447863477956023	0.682653894002144
XU_GH1_EXOGENOUS_TARGETS_DN	67	0.01	Up
0.355696472118139	0.448444405796558	0.683288475640379
XU_HGF_TARGETS_INDUCED_BY_AKT1_48HR_DN	17	0.01	Up
0.315719112748649	0.4485838246084 0.683288475640379
PROVENZANI_METASTASIS_DN	132	0.01	Down

0.320395062309019	0.449206057727316	0.684004481532616
YAMASHITA_LIVER_CANCER_WITH_EPCAM_UP	49	0.01	Up
0.264167956679164	0.449701109107683	0.684526408885552
KOINUMA_COLON_CANCER_MSI_UP	14	0.01	Up
0.613183931473367	0.45019496105167	0.685046157727575
KEGG_TASTE_TRANSDUCTION	12	0.01	Down	0.354791027643694
	0.450902806231815	0.685685059639619
OSMAN_BLADDER_CANCER_DN	390	0.01	Down	0.2051015404165
0.450919919086699	0.685685059639619
MIZUSHIMA_AUTOPHAGOSOME_FORMATION	22	0.01	Down
0.118231420545416	0.451865384754388	0.686708290823482
APPIERTO_RESPONSE_TO_FENRETINIDE_UP	35	0.01	Down
0.439386193347018	0.452149142181866	0.686708290823482
MILI_PSEUDOPODIA_CHEMOTAXIS_UP	78	0.01	Down
0.235957512678079	0.452171937429326	0.686708290823482
LANDIS_ERBB2_BREAST_PRENEOPLASTIC_UP	18	0.01	Up
0.497862119357417	0.452203902299779	0.686708290823482
REACTOME_TGF_BETA_RECEPTOR_SIGNALING_IN_EMT_EPITHELIAL_TO_MESENCHYMA
L_TRANSITION	16	0.01	Down	0.169087994758559
0.452387560777791	0.686755179228696
FRIDMAN_SENESCENCE_UP	75	0.01	Up	0.42311656849491
	0.452702936378465	0.68700192404497
REACTOME_VITAMIN_B5_PANTOTHENATE_METABOLISM	14	0.01	Down
	0.37869812985819	0.453346648837846
0.687746603620019
MIYAGAWA_TARGETS_OF_EWSR1_ETS_FUSIONS_UP	224	0.01	Down
	0.456436461489421	0.453524105873624
0.687783689575553
FAELT_B_CLL_WITH_VH3_21_DN	49	0.01	Down
0.178298243554297	0.453712863809005	0.687837882907749
KIM_WT1_TARGETS_DN	429	0.01	Up	0.247806034921332
	0.454052723421001	0.688121035663317
DELLA_RESPONSE_TO_TSA_AND_BUTYRATE	19	0.01	Down
0.203012866577963	0.454955519473092	0.689256845308914
PUJANA_BREAST_CANCER_LIT_INT_NETWORK	98	0.01	Down
0.192577104390764	0.455200108825759	0.689307777853879
TONG_INTERACT_WITH_PTTG1	52	0.01	Up
0.154737085441329	0.455295838141973	0.689307777853879
GRAESSMANN_RESPONSE_TO_MC_AND_DOXORUBICIN_DN	721	0.01	Down
	0.181399095661363	0.455668918923032
0.689347179471618
DING_LUNG_CANCER_MUTATED_SIGNIFICANTLY	17	0.01	Down
0.419643963217605	0.455681784328748	0.689347179471618
MANN_RESPONSE_TO_AMIFOSTINE_UP	19	0.01	Up
0.161992323385103	0.455781939352536	0.689347179471618
HU_GENOTOXIC_DAMAGE_4HR	31	0.01	Up	0.181238407178568
	0.456105501302515	0.689427201518187
CAFFAREL_RESPONSE_TO_THC_24HR_3_DN	12	0.01	Up
0.213993779016707	0.456141601182445	0.689427201518187
ACEVEDO_LIVER_CANCER_DN	433	0.01	Up	0.322939089446474
	0.456475043571917	0.689699267514544
SA_G1_AND_S_PHASES	13	0.01	Up	0.295432866692607
	0.457044410002655	0.690327494274844
REACTOME_METABOLISM_OF_LIPIDS	591	0.01	Up

0.293337276170182	0.457765122143256	0.691183817277103
REACTOME_BETA_CATENIN_INDEPENDENT_WNT_SIGNALING 126	0.01	Down
	0.229522828980999	0.458235918862322
0.691241632037533
STEIN_ESTROGEN_RESPONSE_NOT_VIA_ESRRA	16	0.01	Down
0.260330860646365	0.458315718092513	0.691241632037533
REACTOME_CARGO_RECOGNITION_FOR_CLATHRIN_MEDIATED_ENDOCYTOSIS	87
	0.01	Down	0.288171553810379	0.458423057399955
	0.691241632037533
REACTOME_INTRINSIC_PATHWAY_OF_FIBRIN_CLOT_FORMATION	10	0.01
	Up	0.597652127149035	0.458460512031037
0.691241632037533
REACTOME_ENDOSOMAL_VACUOLAR_PATHWAY	11	0.01	Down
0.352734409762156	0.458572312955712	0.691241632037533
REACTOME_ION_TRANSPORT_BY_P_TYPE_ATPASES	37	0.01	Down
	0.408837376374887	0.458773474529689
0.691313029839407
WIERENGA_STAT5A_TARGETS_DN	139	0.01	Up
0.501518733467898	0.459044220232601	0.691412747959527
CERVERA_SDHB_TARGETS_1_DN	17	0.01	Down
0.529986664739922	0.45914728646478	0.691412747959527
REACTOME_OXIDATIVE_STRESS_INDUCED_SENESCENCE	81	0.01	Down
	0.234852627856562	0.459798577223692
0.69193620650897
ZHAN_MULTIPLE_MYELOMA_CD2_UP	33	0.01	Up
0.385227064467773	0.459802769486606	0.69193620650897
BIOCARTA_TNFR1_PATHWAY	29	0.01	Down	0.211296044329031
	0.460885741515435	0.693333804588983
REACTOME_SYNAPTIC_ADHESION_LIKE_MOLECULES	11	0.01	Up
	0.301447373758117	0.461487361169673
0.694006586971455
TIMOFEEVA_GROWTH_STRESS_VIA_STAT1_DN	15	0.01	Up
0.22389641035769	0.461776244449019	0.694098699815923
CHIARADONNA_NEOPLASTIC_TRANSFORMATION_KRAS_CDC25_DN	44	0.01
	Up	0.662744446637709	0.462065815179827
0.694098699815923
KIM_MYCN_AMPLIFICATION_TARGETS_UP	70	0.01	Up
0.382647823761018	0.462195411882423	0.694098699815923
OISHI_CHOLANGIOMA_STEM_CELL_LIKE_DN	247	0.01	Up
0.620833931744861	0.462319330884499	0.694098699815923
ROZANOV_MMP14_TARGETS_DN	25	0.01	Down
0.345953608870263	0.462320691267825	0.694098699815923
PID_IL12_STAT4_PATHWAY	21	0.01	Up	0.608447865661036
	0.462489808640823	0.694108722743024
GAVIN_FOXP3_TARGETS_CLUSTER_T7	90	0.01	Up
0.231813611210172	0.462693860527596	0.694108722743024
REACTOME_RNA_POLYMERASE_I_PROMOTER_ESCAPE	51	0.01	Up
	0.145968267741652	0.462790621148129
0.694108722743024
GERHOLD_ADIPOGENESIS_UP	40	0.01	Up	0.384342125235756
	0.463555487671139	0.695023988352825
BLALOCK_ALZHEIMERS_DISEASE_UP	1460	0.01	Down
0.291425019080633	0.464012922822189	0.695477855313684
RODRIGUES_NTN1_TARGETS_UP	15	0.01	Up

0.378685971799239	0.46442005183442	0.695816466546473
PID_ALK1_PATHWAY	23	0.01	Down	0.291949291974144
	0.464616663013781	0.695816466546473
THILLAINADESAN_ZNF217_TARGETS_UP	38	0.01	Up
0.239404878245369	0.464707049564728	0.695816466546473
NIKOLSKY_BREAST_CANCER_8Q12_Q22_AMPLICON	95	0.01	Down
	0.282831366657885	0.464858030931937
0.695816466546473
LEI_HOXC8_TARGETS_UP	11	0.01	Up	0.550936735815911
	0.465225373991194	0.69613450602211
REACTOME_RETROGRADE_NEUROTROPHIN_SIGNALLING	12	0.01	Up
	0.144671045951639	0.465629268737438
0.696404008725366
REACTOME_NEDDYLATION	209	0.01	Down	0.173537271033519
	0.465805470606371	0.696404008725366
KEGG_ALANINE_ASPARTATE_AND_GLUTAMATE_METABOLISM	24	0.01	Up
	0.310478214771033	0.465870267905935
0.696404008725366
PEART_HDAC_PROLIFERATION_CLUSTER_UP	50	0.01	Up
0.184729495159321	0.466132777115575	0.69656477165376
SMID_BREAST_CANCER_LUMINAL_A_UP	41	0.01	Up
0.609776000879623	0.466407825125905	0.69660248563388
NIKOLSKY_BREAST_CANCER_8Q23_Q24_AMPLICON	126	0.01	Down
	0.287131697699694	0.466467960346603
0.69660248563388
REACTOME_RHO_GTPASES_ACTIVATE_PKNS	49	0.01	Down
0.168099430160448	0.466828255006844	0.696909002409752
MARTIN_NFKB_TARGETS_UP	29	0.01	Down	0.552052770399393
	0.468159760199481	0.698664715171537
KEGG_PHOSPHATIDYLINOSITOL_SIGNALING_SYSTEM	67	0.01	Down
	0.266743625800203	0.468587294434176
0.698784148093001
REACTOME_PECAM1_INTERACTIONS	12	0.01	Up
0.626611389458898	0.46860270359929	0.698784148093001
REACTOME_ACYL_CHAIN_REMODELLING_OF_PG	11	0.01	Up
0.307109417342474	0.46875125026640	1	0.698784148093001
ROYLANCE_BREAST_CANCER_16Q_COPY_NUMBER_UP	46	0.01	Down
	0.23698730594015	0.468861621946272
0.698784148093001
DASU_IL6_SIGNALING_UP	57	0.01	Down	0.575451965961738
	0.469480849443896	0.699475113772063
ELVIDGE_HIF1A_TARGETS_UP	65	0.01	Down
0.214894608569281	0.469850589464628	0.699586298329229
REACTOME_REGULATION_OF_MITOTIC_CELL_CYCLE	80	0.01	Down
	0.142407043594364	0.469955312703702
0.699586298329229
BOUDOUKHA_BOUND_BY_IGF2BP2	93	0.01	Up
0.171517987901135	0.470022385084376	0.699586298329229
LEE_INTRATHYMIC_T_PROGENITOR	16	0.01	Up
0.457291248792569	0.470746708176216	0.700229208229079
MEISSNER_BRAIN_HCP_WITH_H3K27ME3	99	0.01	Down
0.60589521784748	0.47076588815757	0.700229208229079
CHEN_LUNG_CANCER_SURVIVAL	26	0.01	Up
0.352013874074069	0.47099213226521	0.700333984297756

REACTOME_THE_PHOTOTRANSDUCTION_CASCADE 16 0.01 Up
0.171140680422263 0.471335544155093 0.700612854159109
REACTOME_CELL_CYCLE_MITOTIC 453 0.01 Down
0.175447648457189 0.471513342322423 0.700645445864228
MELLMAN_TUT1_TARGETS_DN 44 0.01 Up 0.335297693094174
0.471828458621096 0.700760306322614
REACTOME_REGULATION_OF_SIGNALING_BY_CBL 22 0.01 Down
0.307605616773636 0.471902435425707 0.700760306322614
GALE_APL_WITH_FLT3_MUTATED_UP 58 0.01 Down
0.191982843433633 0.472133454398905 0.700871612467675
BIOCARTA_LONGEVITY_PATHWAY 13 0.01 Down
0.361680540945483 0.472368696582472 0.700871612467675
REACTOME_CHK1_CHK2_CDS1_MEDIATED_INACTIVATION_OF_CYCLIN_B:CDK1_COMPL
EX 11 0.01 Down 0.228781910107401
0.472445158126152 0.700871612467675
PEDERSEN_METASTASIS_BY_ERBB2_ISOFORM_7 365 0.01 Down
0.334572671782398 0.473020231885193 0.70149321752687
TOOKER_GEMCITABINE_RESISTANCE_DN 122 0.01 Down
0.246572009467071 0.47360203398776 0.702010595408943
WOO_LIVER_CANCER_RECURRENCE_UP 97 0.01 Up
0.585930230701375 0.47368145403233 0.702010595408943
GROSS_HYPOXIA_VIA_HIF1A_DN 100 0.01 Down
0.368726074590562 0.474227676894027 0.702588466591513
BIOCARTA_TRKA_PATHWAY 11 0.01 Down 0.150597029625975
0.475350013118165 0.703875745278175
REACTOME_GENE_AND_PROTEIN_EXPRESSION_BY_JAK_STAT_SIGNALING_AFTER_INT
ERLEUKIN_12_STIMULATION 33 0.01 Down 0.228725487933433
0.475537216426223 0.703875745278175
GROSS_HYPOXIA_VIA_ELK3_AND_HIF1A_DN 89 0.01 Up
0.304549251990008 0.475566326676267 0.703875745278175
DOANE_BREAST_CANCER_CLASSES_UP 51 0.01 Down
0.493419125173247 0.475918837189383 0.703965272775432
ACEVEDO_NORMAL_TISSUE_ADJACENT_TO_LIVER_TUMOR_DN 312 0.01
Up 0.253589524591303 0.476135861003134
0.703965272775432
CAFFAREL_RESPONSE_TO_THC_24HR_5_UP 31 0.01 Up
0.193561675787162 0.47619194003824 0.703965272775432
BOCHKIS_FOXA2_TARGETS 337 0.01 Down 0.299750978613834
0.476253257955526 0.703965272775432
DING_LUNG_CANCER_BY_MUTATION_RATE 13 0.01 Down
0.339360588124684 0.476954317973844 0.704698377604648
MEINHOLD_OVARIAN_CANCER_LOW_GRADE_DN 17 0.01 Down
0.148555145838059 0.477062772647596 0.704698377604648
SAKAI_TUMOR_INFILTRATING_MONOCYTES_DN 74 0.01 Down
0.190825267710679 0.4774521666557 0.705041882101634
HEIDENBLAD_AMPLIFIED_IN_SOFT_TISSUE_CANCER 10 0.01 Down
0.224389337892563 0.477830479158663
0.705368802567551
IIZUKA_LIVER_CANCER_PROGRESSION_L0_L1_DN 17 0.01 Up
0.241480706043029 0.478096774125319
0.705530203444947
HOLLEMAN_ASPARAGINASE_RESISTANCE_ALL_UP 21 0.01 Up
0.167280206457548 0.478620590526406 0.705894036890171
OUILLETTE_CLL_13Q14_DELETION_UP 65 0.01 Down

0.225978619395335	0.478718112452722	0.705894036890171
GRAESSMANN_APOPTOSIS_BY_DOXORUBICIN_UP	1024	0.01	Down
0.283178416385707	0.478814442375557	0.705894036890171
REACTOME_PKA_ACTIVATION_IN_GLUCAGON_SIGNALLING	13	0.01	Down
	0.300259683083646	0.480520690623251
0.70817721454148
MARKEY_RB1_ACUTE_LOF_DN	198	0.01	Up	0.547051111104351
	0.481596979525998	0.709530784322963
FIGUEROA_AML_METHYLATION_CLUSTER_4_UP	87	0.01	Down
0.284145769779438	0.482136365311545	0.710092713655109
WOTTON_RUNX_TARGETS_UP	17	0.01	Up	0.381368094257993
	0.482466744994666	0.710282865505572
PID_RXR_VDR_PATHWAY	24	0.01	Down	0.318291651351001
	0.482627530183044	0.710282865505572
BOYAULT_LIVER_CANCER_SUBCLASS_G12_UP	37	0.01	Down
0.155717218114001	0.482739522607235	0.710282865505572
FIGUEROA_AML_METHYLATION_CLUSTER_7_UP	92	0.01	Down
0.237705589501949	0.483276777759162	0.710840679328349
DARWICHE_PAPILLOMA_PROGRESSION_RISK	48	0.01	Up
0.383529170784035	0.48394416517197	0.711408947594719
SPIRA_SMOKERS_LUNG_CANCER_DN	13	0.01	Down
0.250346772300483	0.483979657785239	0.711408947594719
REACTOME_CD209_DC_SIGN_SIGNALING	20	0.01	Down
0.308618054858433	0.484171586445485	0.711458411596095
REACTOME_RNA_POLYMERASE_III_TRANSCRIPTION_TERMINATION	23	0.01
	Up	0.138458785360605	0.484597075722915
0.7115996099975
BARRIER_CANCER_RELAPSE_TUMOR_SAMPLE_DN	10	0.01	Up
0.174361781146158	0.484664567200798	0.7115996099975
KEGG_ALZHEIMERS_DISEASE	138	0.01	Up	0.23507112780419
	0.484829181709814	0.7115996099975
ICHIBA_GRAFT_VERSUS_HOST_DISEASE_35D_DN	29	0.01	Down
0.511903313120373	0.485038876134136	0.7115996099975
DARWICHE_PAPILLOMA_RISK_HIGH_DN	135	0.01	Up
0.288149224615027	0.485059222476605	0.7115996099975
PID_INTEGRIN4_PATHWAY	11	0.01	Up	0.555924017565686
	0.485459940025419	0.711955115958976
LEE_LIVER_CANCER_MYC_TGFA_UP	54	0.01	Down
0.414874942414825	0.486049253754148	0.712586887027036
BYSTRYKH_HEMATOPOIESIS_STEM_CELL_AND_BRAIN_QTL_CIS	61	0.01
	Down	0.164845363001779	0.486590775711797
0.712792356204487
TOMLINS_METASTASIS_UP	11	0.01	Up	0.216594775180903
	0.486605935740123	0.712792356204487
REACTOME_DOWNREGULATION_OF_ERBB2:ERBB3_SIGNALING	12	0.01
	Down	0.146156591882579	0.486821024242828
0.712792356204487
CROMER_METASTASIS_UP	71	0.01	Down	0.256081331609073
	0.487015721342039	0.712792356204487
NAKAMURA_CANCER_MICROENVIRONMENT_DN	42	0.01	Down
0.174249370226916	0.487044138744877	0.712792356204487
KYNG_NORMAL_AGING_DN	16	0.01	Down	0.358877420420565
	0.487252718961198	0.712792356204487
SHEDDEN_LUNG_CANCER_POOR_SURVIVAL_A6	387	0.01	Down

0.242264087932979	0.487299423941355	0.712792356204487
LI_DCP2_BOUND_MRNA	85	0.01	Up	0.138769112254839
	0.487803846845871	0.713298077935001
GUO_HEX_TARGETS_UP	70	0.01	Down	0.317337945390289
	0.488487006756591	0.714064746462074
REACTOME_REGULATION_OF_FZD_BY_UBIQUITINATION	16	0.01	Up
	0.198100698570492	0.488864194427351
0.714383795172608
SMID_BREAST_CANCER_LUMINAL_B_UP	114	0.01	Up
0.453506553606652	0.489472086607154	0.714850164730063
REACTOME_RECEPTOR_TYPE_TYROSINE_PROTEIN_PHOSPHATASES	10	0.01
	Up	0.301069469298052	0.489501403123278
0.714850164730063
ZHAN_LATE_DIFFERENTIATION_GENES_DN	15	0.01	Up
0.385961727532381	0.489884936378274	0.715177911341455
BROWNE_HCMV_INFECTION_6HR_UP	41	0.01	Down
0.351120391224559	0.490679104081796	0.715778111634951
RIZKI_TUMOR_INVASIVENESS_2D_UP	58	0.01	Up
0.249161602543104	0.490716388426441	0.715778111634951
BIOCARTA_DNAFRAGMENT_PATHWAY	10	0.01	Down
0.420220928408593	0.490773779768392	0.715778111634951
REACTOME_ANCHORING_OF_THE_BASAL_BODY_TO_THE_PLASMA_MEMBRANE	93
	0.01	Up	0.154091300756528	0.4910588114594
0.715961517194292
YAO_TEMPORAL_RESPONSE_TO_PROGESTERONE_CLUSTER_6	63	0.01	Down
	0.375994983751875	0.491596820094773
0.716360272954972
BIOCARTA_EPO_PATHWAY	18	0.01	Down	0.290753816965508
	0.491651043841176	0.716360272954972
REACTOME_ATF6_ATF6_ALPHA_ACTIVATES_CHAPERONES	12	0.01	Down
	0.160169804381732	0.492324333174613
0.717108839150967
REACTOME_RNA_POLYMERASE_II_PRE_TRANSCRIPTION_EVENTS	79	0.01
	Down	0.126143693482911	0.492867413311255
0.717285209874654
REACTOME_DECTIN_2_FAMILY	10	0.01	Down
0.782260164579036	0.493035893460034	0.717285209874654
FLECHNER_BIOPSY_KIDNEY_TRANSPLANT_REJECTED_VS_OK_UP	74	0.01
	Down	0.918918132368577	0.493217898872079
0.717285209874654
PID_P53_REGULATION_PATHWAY	59	0.01	Down
0.181820738860199	0.493291626844758	0.717285209874654
SESTO_RESPONSE_TO_UV_C8	70	0.01	Up	0.333637348435984
	0.493332057242835	0.717285209874654
DARWICHE_SKIN_TUMOR_PROMOTER_DN	139	0.01	Down
0.277048324770432	0.493402862943811	0.717285209874654
BENPORATH_MYC_TARGETS_WITH_EBOX	197	0.01	Up
0.23399299328776	0.494475756892189	0.718612520928027
HEDENFALK_BREAST_CANCER_BRACX_UP	17	0.01	Up
0.293025505005847	0.495351380231423	0.719426127597992
ALFANO_MYC_TARGETS	211	0.01	Up	0.32193580196352
	0.49548141117699	0.719426127597992
OSADA_ASCL1_TARGETS_UP	33	0.01	Up	0.583454354284858
	0.495515748841687	0.719426127597992

CORRE_MULTIPLE_MYELOMA_UP	59	0.01	Up
0.563116028328281	0.49575905413186	0.719546964263065
MARCINIAK_ER_STRESS_RESPONSE_VIA_CHOP	19	0.01	Down
0.340641000686315	0.496211868758407	0.71959220971599
REACTOME_POLO_LIKE_KINASE_MEDIATED_EVENTS	14	0.01	Up
0.30698670420827	0.496425049347316
0.71959220971599
KEGG_SULFUR_METABOLISM	10	0.01	Up	0.473916963571839
0.496456399943037	0.71959220971599
BIOCARTA_UCALPAIN_PATHWAY	12	0.01	Down
0.240618180329618	0.496478907000424	0.71959220971599
KAAB_FAILED_HEART_ATRIUM_DN	143	0.01	Up
0.249735374032486	0.496590663968632	0.71959220971599
KIM_LRRC3B_TARGETS	29	0.01	Down	0.770483724070558
0.496851337596592	0.719737918948334
HU_GENOTOXIC_DAMAGE_24HR	31	0.01	Up
0.272446725025876	0.497838970071041	0.720801833914681
ONO_AML1_TARGETS_DN	23	0.01	Down	0.509976164806286
0.497906494839841	0.720801833914681
REACTOME_NUCLEOTIDE_EXCISION_REPAIR	109	0.01	Up
0.121478520822968	0.498564437753874	0.721521940664412
GARGALOVIC_RESPONSE_TO_OXIDIZED_PHOSPHOLIPIDS_GREEN_DN	22	0.01
Down	0.22400400042194	0.498913908563687
0.721795307046596
LINDSTEDT_DENDRITIC_CELL_MATURATION_D	57	0.01	Up
0.430971148214735	0.499079112978205	0.721801998982314
REACTOME_RHO_GTPASES_ACTIVATE_CIT	17	0.01	Down
0.189908576140128	0.499552047139921	0.722253603053697
DUTERTRE_ESTRADIOL_RESPONSE_6HR_DN	86	0.01	Down
0.403255689971013	0.50059610003209	0.723320965084385
REACTOME_PREFOLDIN_MEDIATED_TRANSFER_OF_SUBSTRATE_TO_CCT_TRIC	24
0.01	Down	0.128755361320604	0.500612129561184
0.723320965084385
TAKEDA_TARGETS_OF_NUP98_HOXA9_FUSION_6HR_UP	55	0.01	Up
0.582577876790033	0.501469567577107
0.724236385930989
REACTOME_COBALAMIN_CBL_VITAMIN_B12_TRANSPORT_AND_METABOLISM	16
0.01	Up	0.254350542873095	0.501577576399671
0.724236385930989
REACTOME_SHC_MEDIATED_CASCADE:FGFR2	12	0.01	Down
0.490989507125791	0.501729055792903	0.724236385930989
REACTOME_SWITCHING_OF_ORIGINS_TO_A_POST_REPLICATIVE_STATE	86
0.01	Down	0.114515744855816	0.502003568959837
0.724400013635464
REACTOME_REGULATION_OF_EXPRESSION_OF_SLITS_AND_ROBOS	159	0.01
Up	0.12848576447302	0.502753883677019
0.725034274665713
IIZUKA_LIVER_CANCER_PROGRESSION_G2_G3_UP	23	0.01	Up
0.54738993608878	0.502978466408265
0.725034274665713
REACTOME_SIGNALING_BY_ROBO_RECEPTORS	201	0.01	Up
0.169583014664802	0.503069172263556	0.725034274665713
MATZUK_EARLY_ANTRAL_FOLLICLE	10	0.01	Up
0.626808834133535	0.503088298705753	0.725034274665713

VERHAAK_GLIOBLASTOMA_PRONEURAL 108 0.01 Down
0.362405978222082 0.503434819770981 0.725301126561077
BIOCARTA_MHC_PATHWAY 12 0.01 Down 0.564826789774704
0.503850819475415 0.725667873611661
PID_EPHA2_FWD_PATHWAY 19 0.01 Down 0.20216184062976 8
0.504442161145282 0.726286839957732
YAGI_AML_WITH_11Q23_REARRANGED 299 0.01 Up
0.297979162033155 0.504929532006582 0.726504310178234
JIANG_HYPOXIA_VIA_VHL 19 0.01 Up 0.208900995467309
0.505219373292416 0.726504310178234
GRYDER_PAX3FOXO1_TOP_ENHANCERS 414 0.01 Down
0.243378174162498 0.505308015050539 0.726504310178234
ZHAN_MULTIPLE_MYELOMA_CD1_AND_CD2_DN 49 0.01 Down
0.243430701428729 0.505528683213285 0.726504310178234
KOKKINAKIS_METHIONINE_DEPRIVATION_96HR_UP 113 0.01 Up
0.529411192995832 0.505622827860812
0.726504310178234
REACTOME_GABA_RECEPTOR_ACTIVATION 26 0.01 Down
0.282503827007315 0.50567857192064 7 0.726504310178234
REACTOME_DOWNSTREAM_SIGNALING_OF_ACTIVATED_FGFR1 20 0.01
Down 0.420157423920845 0.50576762028041 9
0.726504310178234
HUTTMANN_B_CLL_POOR_SURVIVAL_UP 236 0.01 Up
0.342535429324612 0.505886204862708 0.726504310178234
CHIN_BREAST_CANCER_COPY_NUMBER_DN 11 0.01 Up
0.216798064508406 0.506517202918039 0.72698011440279
KEGG_CALCIUM_SIGNALING_PATHWAY 88 0.01 Down
0.390045214590274 0.506540982938719 0.72698011440279
LI_WILMS_TUMOR_VS_FETAL_KIDNEY_1_DN 148 0.01 Up
0.221240026793953 0.506733076365039 0.727023676431807
REACTOME_CALNEXIN_CALRETICULIN_CYCLE 26 0.01 Up
0.142548887370002 0.507282630836915 0.727579906066348
NIKOLSKY_BREAST_CANCER_7Q21_Q22_AMPLICON 57 0.01 Down
0.355651868115754 0.507481535861781
0.727633015533877
GUENTHER_GROWTH_SPHERICAL_VS_ADHERENT_DN 26 0.01 Up
0.321059554486954 0.508146359190099
0.728353917270246
MULLIGHAN_NPM1_MUTATED_SIGNATURE_2_UP 105 0.01 Down
0.279543682047852 0.50888113398393 0.729084616349279
MCCABE_HOXC6_TARGETS_DN 14 0.01 Up 0.567479577784083
0.508980539733935 0.729084616349279
TSENG_ADIPOGENIC_POTENTIAL_UP 23 0.01 Down
0.357083129514375 0.510274152139041 0.730704783008916
BORLAK_LIVER_CANCER_EGF_UP 48 0.01 Down
0.450216363497841 0.510490376435467 0.730781605757141
REACTOME_COOPERATION_OF_PREFOLDIN_AND_TRIC_CCT_IN_ACTIN_AND_TUBULIN_
FOLDING 27 0.01 Down 0.127162705477064
0.510936292117998 0.73118708470882
POS_HISTAMINE_RESPONSE_NETWORK 29 0.01 Up
0.464586945063593 0.511272252397236 0.731435001440349
REACTOME_GAP_JUNCTION_DEGRADATION 12 0.01 Up
0.316118965438313 0.511534316591118 0.731577077021023
NAKAMURA_METASTASIS_MODEL_DN 39 0.01 Down

0.392653311262825	0.512146135693295	0.732205669834709
BHAT_ESR1_TARGETS_VIA_AKT1_DN	79	0.01	Down
0.329182741127634	0.512299628838746	0.732205669834709
MEISSNER_NPC_ICP_WITH_H3_UNMETHYLATED	10	0.01	Up
0.557316554497946	0.512690497330809	0.732506567473286
MORI_SMALL_PRE_BII_LYMPHOCYTE_UP	59	0.01	Down
0.302487490746535	0.512836077383411	0.732506567473286
REACTOME_RECYCLING_PATHWAY_OF_L1	38	0.01	Down
0.181809161127814	0.513017252421887	0.732532576123375
IWANAGA_CARCINOGENESIS_BY_KRAS_DN	87	0.01	Down
0.336758953099237	0.513499484989622	0.732688615932743
REACTOME_MTORC1_MEDIATED_SIGNALLING	22	0.01	Up
0.14212539642395	0.513544514447887	0.732688615932743
BIOCARTA_NO1_PATHWAY	16	0.01	Down	0.304475605794805
0.513615534772875	0.732688615932743
NIKOLSKY_BREAST_CANCER_14Q22_AMPLICON	12	0.01	Down
0.41169325755154	0.514227632895278	0.732755560182288
KEGG_BLADDER_CANCER	40	0.01	Down	0.383883672357906
0.514325005584674	0.732755560182288
LINDSTEDT_DENDRITIC_CELL_MATURATION_C	53	0.01	Down
0.417113291626253	0.514571428866464	0.732755560182288
HAHTOLA_MYCOSIS_FUNGOIDES_SKIN_DN	27	0.01	Down
0.297966857757137	0.514572042621123	0.732755560182288
RODRIGUES_THYROID_CARCINOMA_POORLY_DIFFERENTIATED_DN	741	0.01
Down	0.301238980450951	0.514631489854969
0.732755560182288
GARGALOVIC_RESPONSE_TO_OXIDIZED_PHOSPHOLIPIDS_RED_DN	21	0.01
Up	0.201330524408524	0.514640556951164
0.732755560182288
MIKKELSEN_MCV6_HCP_WITH_H3K27ME3	150	0.01	Up
0.655460162063971	0.514976326303019	0.733001452416742
YAGI_AML_WITH_T_9_11_TRANSLOCATION	108	0.01	Down
0.366518539966853	0.516481977828522	0.734776560493154
BENPORATH_CYCLING_GENES	583	0.01	Down	0.258649100163496
0.51655037400631	0.734776560493154
HOELZEL_NF1_TARGETS_UP	101	0.01	Down	0.388531781349009
0.516740649534727	0.73481468511819
OUELLET_CULTURED_OVARIAN_CANCER_INVASIVE_VS_LMP_UP	66	0.01
Up	0.22693643904066	0.517564161123926
0.735323898077732
SABATES_COLORECTAL_ADENOMA_SIZE_UP	20	0.01	Down
0.255048508478137	0.517596407637101	0.735323898077732
KLEIN_PRIMARY_EFFUSION_LYMPHOMA_UP	41	0.01	Up
0.306698955873813	0.517736797751029	0.735323898077732
LOPEZ_MESOTELIOMA_SURVIVAL_TIME_UP	10	0.01	Down
0.185405826207625	0.517753089525255	0.735323898077732
REACTOME_REPRESSION_OF_WNT_TARGET_GENES	13	0.01	Down
0.170557600658931	0.517918126608687	0.735325956761229
ASGHARZADEH_NEUROBLASTOMA_POOR_SURVIVAL_DN	35	0.01	Down
0.226361343682049	0.518161910223355
0.735439781008519
SHEPARD_CRUSH_AND_BURN_MUTANT_DN	127	0.01	Up
0.319326837645065	0.518559968744738	0.735552360170717
MAYBURD_RESPONSE_TO_L663536_UP	26	0.01	Down

0.195114207919495	0.518568504867854	0.735552360170717
CHIANG_LIVER_CANCER_SUBCLASS_POLYSOMY7_DN	24	0.01	Down
	0.312930356956477	0.519612844898927
0.736659707105615
REACTOME_FCGAMMA_RECEPTOR_FCGR_DEPENDENT_PHAGOCYTOSIS	78	0.01
	Down	0.360481834591296	0.519676959117221
0.736659707105615
MARTINEZ_RB1_AND_TP53_TARGETS_DN	484	0.01	Down
0.318126029936697	0.520293149275132	0.737137666173782
BIOCARTA_CELLCYCLE_PATHWAY	22	0.01	Up
0.307460051251966	0.520342116745141	0.737137666173782
LI_CISPLATIN_RESISTANCE_DN	25	0.01	Up
0.779860245924176	0.520672965622864	0.737342381005551
JISON_SICKLE_CELL_DISEASE_UP	160	0.01	Up
0.371005553314964	0.520907821495163	0.737342381005551
SCIAN_CELL_CYCLE_TARGETS_OF_TP53_AND_TP73_DN	18	0.01	Up
	0.246352555547552	0.520978732385679
0.737342381005551
HUANG_DASATINIB_RESISTANCE_UP	78	0.01	Down
0.37425743036833	0.52157572249269	0.737954948884055
REACTOME_GABA_SYNTHESIS_RELEASE_REUPTAKE_AND_DEGRADATION	12
	0.01	Down	0.280753667498628	0.521778225491862
	0.738009164123953
ZHONG_RESPONSE_TO_AZACITIDINE_AND_TSA_UP	149	0.01	Down
	0.329984297435342	0.522164524205825
0.738323226267751
ENGELMANN_CANCER_PROGENITORS_UP	44	0.01	Up
0.526102611443566	0.523038672072927	0.739027460681695
KEGG_DILATED_CARDIOMYOPATHY	56	0.01	Up
0.483662563899911	0.523078769351483	0.739027460681695
REACTOME_PLATELET_SENSITIZATION_BY_LDL	16	0.01	Down
0.44678741932102	0.523155813101035	0.739027460681695
PLASARI_TGFB1_SIGNALING_VIA_NFIC_1HR_DN	89	0.01	Down
0.459704483747383	0.523630294407069	0.739323295413208
KEGG_AMYOTROPHIC_LATERAL_SCLEROSIS_ALS	38	0.01	Down
0.327692376663836	0.523787758628297	0.739323295413208
REACTOME_DNA_REPLICATION_PRE_INITIATION	78	0.01	Down
0.129540741131243	0.523948708593899	0.739323295413208
KAMMINGA_SENESCENCE	32	0.01	Down	0.251848208889476
	0.524023141087092	0.739323295413208
ZHAN_MULTIPLE_MYELOMA_SPIKED	18	0.01	Up
0.458782662798668	0.524336376481273	0.73950801835889
KAN_RESPONSE_TO_ARSENIC_TRIOXIDE	103	0.01	Up
0.485537683955605	0.524483106235404	0.73950801835889
RATTENBACHER_BOUND_BY_CELF1	332	0.01	Down
0.272313355083092	0.525171585763965	0.740246559425845
VERHAAK_GLIOBLASTOMA_CLASSICAL	135	0.01	Up
0.302400958428124	0.525607565299711	0.740628842013229
REACTOME_MTOR_SIGNALLING	38	0.01	Down
0.150269055228746	0.526008857072133	0.740962022105684
REACTOME_GOLGI_CISTERNAE_PERICENTRIOLAR_STACK_REORGANIZATION	13
	0.01	Down	0.214793083088824	0.526439975764641
	0.740967871487003
SU_TESTIS	46	0.01	Down	0.226248060647201

0.526620698987817	0.740967871487003
JI_RESPONSE_TO_FSH_UP	66	0.01	Up	0.34071271183001
	0.526632076000019	0.740967871487003
REACTOME_S_PHASE	153	0.01	Down	0.135832079893218
	0.526672380289427	0.740967871487003
SHI_SPARC_TARGETS_UP	19	0.01	Down	0.337724014222562
	0.527558128828327	0.741981786321442
SEMENZA_HIF1_TARGETS	32	0.01	Up	0.347994896136557
	0.527970008946946	0.742209959769533
REACTOME_ACTIVATION_OF_ATR_IN_RESPONSE_TO_REPLICATION_STRESS	34
	0.01	Up	0.125431779427908	0.528050600966177
	0.742209959769533
JIANG_AGING_HYPOTHALAMUS_UP	41	0.01	Up
0.253038299170839	0.528501118667587	0.742451622006019
VARELA_ZMPSTE24_TARGETS_UP	38	0.01	Down
0.355545457777881	0.528552878847444	0.742451622006019
REACTOME_ONCOGENE_INDUCED_SENESCENCE	32	0.01	Down
0.261800001810272	0.529781649864186	0.743688809938614
SMID_BREAST_CANCER_BASAL_DN	521	0.01	Up
0.412339703363317	0.529831183783993	0.743688809938614
YAGI_AML_WITH_T_8_21_TRANSLOCATION	311	0.01	Up
0.328848869483864	0.529929979584734	0.743688809938614
WACKER_HYPOXIA_TARGETS_OF_VHL	13	0.01	Down
0.513128977364343	0.530688622818364	0.743879403917891
TONKS_TARGETS_OF_RUNX1_RUNX1T1_FUSION_HSC_UP	152	0.01	Up
	0.390749989655634	0.530855342482837
0.743879403917891
KYNG_ENVIRONMENTAL_STRESS_RESPONSE_NOT_BY_GAMMA_IN_WS	30	0.01
	Down	0.298118060005784	0.530891270978293
0.743879403917891
PID_ALPHA_SYNUCLEIN_PATHWAY	27	0.01	Down
0.492039116576897	0.531099727811732	0.743879403917891
ACEVEDO_LIVER_CANCER_UP	912	0.01	Down	0.237428560971646
	0.531107005770232	0.743879403917891
DACOSTA_UV_RESPONSE_VIA_ERCC3_TTD_UP	57	0.01	Up
0.229964719401431	0.531117785709903	0.743879403917891
REACTOME_REGULATION_OF_PTEN_GENE_TRANSCRIPTION	58	0.01	Up
	0.200635446419389	0.531224223932465
0.743879403917891
REACTOME_CELL_EXTRACELLULAR_MATRIX_INTERACTIONS	18	0.01	Up
	0.271826818905777	0.531869326870947
0.744225029993224
KYNG_NORMAL_AGING_UP	15	0.01	Down	0.290097291509468
	0.531959222720801	0.744225029993224
REACTOME_KINESINS	42	0.01	Down	0.226998256832412
	0.531967746689261	0.744225029993224
MORI_IMMATURE_B_LYMPHOCYTE_UP	43	0.01	Down
0.458230703558482	0.532446030092523	0.744662384961385
NATSUME_RESPONSE_TO_INTERFERON_BETA_DN	47	0.01	Up
0.240010038739135	0.532682049613716	0.74476075054857
REACTOME_PURINE_CATABOLISM	16	0.01	Down
0.503713029240637	0.533098844216249	0.74511172411444
KEGG_PURINE_METABOLISM	124	0.01	Up	0.273216981545046
	0.533347530591804	0.745227587817892

REACTOME_CHAPERONE_MEDIATED_AUTOPHAGY 13 0.01 Up
0.260548642862043 0.534253365842483 0.746261304991287
YAO_TEMPORAL_RESPONSE_TO_PROGESTERONE_CLUSTER_7 66 0.01 Down
0.218999841837803 0.534942541143169
0.746991836731452
KEGG_ALPHA_LINOLENIC_ACID_METABOLISM 12 0.01 Down
0.355585626116896 0.535119843420917 0.747007359061187
GYORFFY_MITOXANTRONE_RESISTANCE 28 0.01 Down
0.486570095803667 0.535376804103462 0.747114940814856
HERNANDEZ_MITOTIC_ARREST_BY_DOCETAXEL_2_UP 49 0.01 Up
0.592672973240927 0.535529330212562
0.747114940814856
HENDRICKS_SMARCA4_TARGETS_DN 39 0.01 Up
0.625577112890535 0.535756641744612 0.74720015657525
REACTOME_PROSTACYCLIN_SIGNALLING_THROUGH_PROSTACYCLIN_RECEPTOR 11
0.01 Up 0.264928051587354 0.535980630221707
0.747280686366803
REACTOME_TP53_REGULATES_TRANSCRIPTION_OF_CELL_CYCLE_GENES 46
0.01 Down 0.22711323895811 0.536924792596482
0.747876232950803
BIOCARTA_NDKDYNAMIN_PATHWAY 17 0.01 Down
0.210863883857852 0.536982907428166 0.747876232950803
REACTOME_SIGNALING_BY_INSULIN_RECEPTOR 58 0.01 Down
0.292016319531501 0.53700724030052 0.747876232950803
AFFAR_YY1_TARGETS_UP 135 0.01 Up 0.393279511901639
0.537272211407725 0.747876232950803
BIOCARTA_MYOSIN_PATHWAY 10 0.01 Down 0.240749170272987
0.537335987849659 0.747876232950803
REACTOME_MITOCHONDRIAL_PROTEIN_IMPORT 63 0.01 Up
0.0989568564580295 0.537510871445076 0.747876232950803
OUYANG_PROSTATE_CANCER_PROGRESSION_UP 19 0.01 Down
0.336958190007255 0.537718788024378 0.747876232950803
MIZUKAMI_HYPOXIA_UP 10 0.01 Up 0.735006928519403
0.537854328115186 0.747876232950803
URS_ADIPOCYTE_DIFFERENTIATION_UP 43 0.01 Up
0.351396678654423 0.537905197136807 0.747876232950803
KUNINGER_IGF1_VS_PDGFB_TARGETS_UP 46 0.01 Up
0.403897365517649 0.538312828264106 0.748024710362915
LIAO_HAVE_SOX4_BINDING_SITES 39 0.01 Up
0.356853453338498 0.538344813798449 0.748024710362915
REACTOME_DUAL_INCISION_IN_TC_NER 64 0.01 Up
0.109444226898253 0.539040337108725 0.748759677164314
VANDESLUIS_COMMD1_TARGETS_GROUP_2_UP 12 0.01 Up
0.393524632312017 0.539431384267458 0.748955582129626
KEGG_FRUCTOSE_AND_MANNOSE_METABOLISM 30 0.01 Up
0.300881399578549 0.539514610664233 0.748955582129626
VERHAAK_AML_WITH_NPM1_MUTATED_UP 154 0.01 Up
0.764691055909908 0.540029728949761 0.748998840980757
MIKKELSEN_NPC_LCP_WITH_H3K4ME3 41 0.01 Down
0.393566531848797 0.540054825513698 0.748998840980757
REACTOME_REGULATION_OF_TP53_ACTIVITY_THROUGH_PHOSPHORYLATION 88
0.01 Down 0.168176851731186 0.540149344992571
0.748998840980757
PID_VEGFR1_PATHWAY 23 0.01 Down 0.317411623486293

0.540303770221087 0.748998840980757
GRYDER_PAX3FOXO1_ENHANCERS_KO_DOWN 405 0.01 Down
0.276392872863595 0.540378919088008 0.748998840980757
HELLER_HDAC_TARGETS_SILENCED_BY_METHYLATION_UP 353 0.01 Up
0.393359112392957 0.540838425757506
0.749296752609299
GREENBAUM_E2A_TARGETS_DN 15 0.01 Down
0.412540056869683 0.540927244097258 0.749296752609299
BIOCARTA_NUCLEARRS_PATHWAY 17 0.01 Up
0.396918351562237 0.541267030545159 0.749536445563922
YOSHIMURA_MAPK8_TARGETS_DN 309 0.01 Down
0.285601487261609 0.541634901335334 0.749814869572629
ADDYA_ERYTHROID_DIFFERENTIATION_BY_HEMIN 62 0.01 Down
0.418512765460472 0.541981238643362
0.749846813239767
BIOCARTA_ACTINY_PATHWAY 11 0.01 Down 0.26234455407516
0.541991612061403 0.749846813239767
HOSHIDA_LIVER_CANCER_SUBCLASS_S2 107 0.01 Down
0.2159182093257 0.542373142231284 0.750143776716807
RODWELL_AGING_KIDNEY_DN 124 0.01 Up 0.271078461008786
0.542947920437549 0.750498615071934
REACTOME_SIGNALING_BY_GPCR 368 0.01 Down
0.459883523071361 0.543255096108977 0.750498615071934
MOOTHA_GLYCOGEN_METABOLISM 15 0.01 Down
0.13763145336253 0.543268375681371 0.750498615071934
GROSS_HYPOXIA_VIA_ELK3_ONLY_DN 41 0.01 Up
0.258527456843078 0.543297551377992 0.750498615071934
REACTOME_AMINO_ACID_TRANSPORT_ACROSS_THE_PLASMA_MEMBRANE 23
0.01 Up 0.469390415017285 0.543879206158288
0.751071284694779
BIOCARTA_D4GDI_PATHWAY 11 0.01 Down 0.544255233283384
0.54422306150193 0.751156735219567
MYLLYKANGAS_AMPLIFICATION_HOT_SPOT_23 18 0.01 Down
0.397767210386424 0.544432030137733 0.751156735219567
REACTOME_ENOS_ACTIVATION 10 0.01 Up
0.415678969687279 0.544471980908063 0.751156735219567
REACTOME_G_ALPHA_I_SIGNALLING_EVENTS 178 0.01 Down
0.538279748269069 0.54475455380777 0.751156735219567
SHIPP_DLBCL_CURED_VS_FATAL_DN 37 0.01 Up
0.277088447035542 0.544776631104736 0.751156735219567
SUH_COEXPRESSED_WITH_ID1_AND_ID2_UP 17 0.01 Up
0.277023598065156 0.5450121345843 0.751251010412888
MILI_PSEUDOPODIA 47 0.01 Down 0.221864080258648
0.546585257203976 0.752958133473569
ZWANG_TRANSIENTLY_UP_BY_2ND_EGF_PULSE_ONLY 756 0.01 Up
0.49115897007656 0.546730342627125
0.752958133473569
REACTOME_SUMOYLATION_OF_DNA_DAMAGE_RESPONSE_AND_REPAIR_PROTEINS 76
0.01 Down 0.20611113847109 0.546753136297604
0.752958133473569
MARTINEZ_RB1_AND_TP53_TARGETS_UP 500 0.01 Down
0.406187529487496 0.547792608763151 0.754158583886789
LIANG_HEMATOPOIESIS_STEM_CELL_NUMBER_SMALL_VS_HUGE_UP 33 0.01
Down 0.248325821043321 0.548711989448432

0.754602361776506
KEGG_HYPERTROPHIC_CARDIOMYOPATHY_HCM 53 0.01 Up
0.497606013800446 0.548834664516138 0.754602361776506
NADERI_BREAST_CANCER_PROGNOSIS_UP 36 0.01 Up
0.330292765798665 0.548897735402551 0.754602361776506
REACTOME_MEIOTIC_SYNAPSIS 39 0.01 Down
0.139790542716393 0.54894984867222 0.754602361776506
REACTOME_TP53_REGULATES_METABOLIC_GENES 80 0.01 Up
0.153953084918871 0.548954332148871 0.754602361776506
LE_NEURONAL_DIFFERENTIATION_UP 10 0.01 Up
0.516411347001894 0.549634171982948 0.755250184343641
BIOCARTA_CPSF_PATHWAY 10 0.01 Down 0.090741768630703
0.549872638924094 0.755250184343641
VERHAAK_AML_WITH_NPM1_MUTATED_DN 189 0.01 Up
0.486794276804717 0.549959507746438 0.755250184343641
BIOCARTA_ERAD_PATHWAY 19 0.01 Down 0.100808339691265
0.550097687105914 0.755250184343641
SMIRNOV_CIRCULATING_ENDOTHELIOCYTES_IN_CANCER_UP 140 0.01
Up 0.589919263364298 0.55098346892619
0.756235326052893
PID_CIRCADIAN_PATHWAY 16 0.01 Down 0.186304617780136
0.551283432531467 0.756265726275541
REACTOME_E2F_MEDIATED_REGULATION_OF_DNA_REPLICATION 21 0.01
Up 0.145094545709236 0.551401647621944
0.756265726275541
FARMER_BREAST_CANCER_CLUSTER_5 19 0.01 Down
0.134709528383747 0.551563023728739 0.756265726275541
IVANOVA_HEMATOPOIESIS_STEM_CELL_LONG_TERM 164 0.01 Up
0.405610909378521 0.551678602104004
0.756265726275541
DARWICHE_SKIN_TUMOR_PROMOTER_UP 106 0.01 Up
0.391941771797961 0.551876942198985 0.756304651566624
REACTOME_REGULATION_OF_BETA_CELL_DEVELOPMENT 19 0.01 Down
0.440738798482193 0.55204350651615
0.756304651566624
SHEDDEN_LUNG_CANCER_GOOD_SURVIVAL_A5 65 0.01 Down
0.220131114800175 0.552634598134733 0.756883765574535
KOKKINAKIS_METHIONINE_DEPRIVATION_48HR_DN 56 0.01 Up
0.367632370731056 0.553089212524373
0.757195542247641
PID_WNT_NONCANONICAL_PATHWAY 31 0.01 Down
0.313776304456321 0.553229054216234 0.757195542247641
AMUNDSON_RESPONSE_TO_ARSENITE 204 0.01 Down
0.281553453903497 0.553367598728254 0.757195542247641
ZHAN_MULTIPLE_MYELOMA_SUBGROUPS 30 0.01 Up
0.166329425837418 0.55395942861997 0.757774690093355
ZHAN_MULTIPLE_MYELOMA_LB_DN 37 0.01 Down
0.309983391113109 0.554484779858334 0.758262575437545
BIOCARTA_P53HYPOXIA_PATHWAY 20 0.01 Down
0.265503543154335 0.554830743573166 0.758359321191901
REACTOME_CASPASE_ACTIVATION_VIA_EXTRINSIC_APOPTOTIC_SIGNALLING_PATHW
AY 22 0.01 Down 0.301178893081349
0.554892949365998 0.758359321191901
ZHAN_MULTIPLE_MYELOMA_MF_UP 37 0.01 Up

0.507964003936164	0.556144471509872	0.759719742305993
PID_INTEGRIN2_PATHWAY	20	0.01	Up	0.640287418594448
	0.556292716864549	0.759719742305993
LEIN_PONS_MARKERS	65	0.01	Up	0.467165027861537
	0.55639541527727	0.759719742305993
REACTOME_THE_ROLE_OF_NEF_IN_HIV_1_REPLICATION_AND_DISEASE_PATHOGENES
IS	24	0.01	Up	0.362974617774903
0.556745686605208	0.75986899744368
DORSEY_GAB2_TARGETS	25	0.01	Up	0.829384165564383
	0.556854586074295	0.75986899744368
DARWICHE_SQUAMOUS_CELL_CARCINOMA_UP	112	0.01	Down
0.354248404411218	0.557270680034454	0.75986899744368
LIEN_BREAST_CARCINOMA_METAPLASTIC_VS_DUCTAL_UP	65	0.01	Down
	0.515361548092369	0.55728441553567
0.75986899744368
REACTOME_FORMATION_OF_TUBULIN_FOLDING_INTERMEDIATES_BY_CCT_TRIC 20
	0.01	Down	0.136305614192978	0.557452176438969
	0.75986899744368
REACTOME_CARDIAC_CONDUCTION	63	0.01	Up
0.451352218642231	0.557519010805174	0.75986899744368
BIOCARTA_EGFR_SMRTE_PATHWAY	10	0.01	Down
0.219784331771366	0.557920493622925	0.760079440143338
LAIHO_COLORECTAL_CANCER_SERRATED_UP	112	0.01	Down
0.299436037001472	0.558011602329926	0.760079440143338
REACTOME_MITOTIC_SPINDLE_CHECKPOINT	93	0.01	Down
0.162708720650487	0.558668406100406	0.760743558140359
BIOCARTA_IGF1R_PATHWAY	21	0.01	Down	0.192965499822953
	0.559050216701094	0.760920931111429
BRUINS_UVC_RESPONSE_VIA_TP53_GROUP_D	194	0.01	Down
0.356690435594318	0.55915631638741	0.760920931111429
TAKEDA_TARGETS_OF_NUP98_HOXA9_FUSION_6HR_DN	38	0.01	Down
	0.631519206708859	0.559306508652316
0.760920931111429
BROWN_MYELOID_CELL_DEVELOPMENT_UP	126	0.01	Up
0.548164084437758	0.560606745732139	0.762146853346365
KAMIKUBO_MYELOID_MN1_NETWORK	18	0.01	Up
0.148913209586745	0.560684799918961	0.762146853346365
KIM_WT1_TARGETS_8HR_DN	100	0.01	Up	0.353671909639171
	0.560716272306213	0.762146853346365
YANAGIHARA_ESX1_TARGETS	26	0.01	Down	0.303327102911372
	0.561187883567009	0.762451128361253
REACTOME_YAP1_AND_WWTR1_TAZ_STIMULATED_GENE_EXPRESSION	11	0.01
	Up	0.198393238703539	0.561304617167479
0.762451128361253
REACTOME_UNWINDING_OF_DNA	10	0.01	Up
0.134636415152797	0.561478505954985	0.762451128361253
DORN_ADENOVIRUS_INFECTION_48HR_UP	12	0.01	Down
0.360704840406514	0.561618617575997	0.762451128361253
PUIFFE_INVASION_INHIBITED_BY_ASCITES_UP	72	0.01	Up
0.302875029841569	0.562320321021446	0.763173261772766
PETROVA_PROX1_TARGETS_UP	18	0.01	Up
0.395994565635798	0.562583040885096	0.76329935670948
WEI_MYCN_TARGETS_WITH_E_BOX	731	0.01	Down
0.1940142504211 0.562902546471029	0.763409136283332

MCLACHLAN_DENTAL_CARIES_UP	206	0.01	Up
0.912663965808979	0.563003623310177	0.763409136283332
SEMBA_FHIT_TARGETS_UP	10	0.01	Down	0.595235170880705
	0.563264526727859	0.763532583727903
REACTOME_MICRORNA_MIRNA_BIOGENESIS	24	0.01	Down
0.127068339688909	0.563773197728866	0.763768920711984
BANDRES_RESPONSE_TO_CARMUSTIN_WITHOUT_MGMT_48HR_UP	12	0.01
	Up	0.31069524282621	0.563867563273492
0.763768920711984
KASLER_HDAC7_TARGETS_1_DN	17	0.01	Up
0.550345286719603	0.563948620209805	0.763768920711984
GROSS_HYPOXIA_VIA_ELK3_UP	185	0.01	Up
0.269790944036137	0.5642005206629 0.763879921801125
BIOCARTA_BARRESTIN_SRC_PATHWAY	14	0.01	Up
0.46625144719591	0.565594276843809	0.765405504681571
REACTOME_FGFR2_MUTANT_RECEPTOR_ACTIVATION	22	0.01	Up
	0.280936550446649	0.565727194241525
0.765405504681571
BROWNE_HCMV_INFECTION_1HR_DN	157	0.01	Down
0.295721506427533	0.565838151736788	0.765405504681571
REACTOME_CDC6_ASSOCIATION_WITH_THE_ORC:ORIGIN_COMPLEX	10	0.01
	Up	0.140016749705191	0.566159934558302
0.765486124423401
DODD_NASOPHARYNGEAL_CARCINOMA_DN	1267	0.01	Up
0.282121835160389	0.56643730053079	0.765486124423401
BAKKER_FOXO3_TARGETS_DN 139	0.01	Down	0.210121182461513
	0.566510653619418	0.765486124423401
LEE_CALORIE_RESTRICTION_NEOCORTEX_UP	52	0.01	Down
0.350658941285997	0.566681816138592	0.765486124423401
REACTOME_NICOTINATE_METABOLISM	24	0.01	Up
0.408643595109969	0.566749237392898	0.765486124423401
BIOCARTA_CHREBP_PATHWAY 13	0.01	Down	0.158639416466625
	0.567176130271502	0.765832594043377
FRASOR_RESPONSE_TO_SERM_OR_FULVESTRANT_DN	43	0.01	Up
	0.219481460914441	0.567368350109583
0.765862082204978
SATO_SILENCED_BY_DEACETYLATION_IN_PANCREATIC_CANCER	34	0.01
	Down	0.499770093233542	0.567663160151212
0.76602999245863
THUM_SYSTOLIC_HEART_FAILURE_UP	370	0.01	Up
0.479033532811989	0.567952668851129	0.766173018635439
NUTT_GBM_VS_AO_GLIOMA_DN	37	0.01	Up
0.153314022521561	0.568273498786534	0.766173018635439
REACTOME_PCP_CE_PATHWAY 86	0.01	Down	0.190531634213985
	0.568280499250401	0.766173018635439
KARAKAS_TGFB1_SIGNALING 17	0.01	Up	0.562772629663096
	0.568728004564842	0.766546440935222
CREIGHTON_AKT1_SIGNALING_VIA_MTOR_DN	23	0.01	Up
0.120881496801426	0.569228780806941	0.766991417783933
REACTOME_TRANSCRIPTIONAL_REGULATION_OF_GRANULOPOIESIS	44	0.01
	Down	0.252830889149532	0.569578546868687
0.767099141912473
KEGG_PHENYLALANINE_METABOLISM	14	0.01	Up
0.504060541345396	0.570101706160423	0.767099141912473

| | | | | | | |
|---|---|---|---|---|---|---|
| REACTOME_SENESCENCE_ASSOCIATED_SECRETORY_PHENOTYPE_SASP | 69 | 0.01 | Down | 0.257257807518155 | 0.570131244351074 | 0.767099141912473 |
| EPPERT_PROGENITOR | 121 | 0.01 | Down | 0.205193159184673 | 0.570172563932615 | 0.767099141912473 |
| SASAKI_ADULT_T_CELL_LEUKEMIA | 144 | 0.01 | Up | 0.280177014655539 | 0.57019061638686 | 0.767099141912473 |
| SCHAEFFER_PROSTATE_DEVELOPMENT_6HR_UP | 120 | 0.01 | Down | 0.388126620295393 | 0.570333586016037 | 0.767099141912473 |
| CHEBOTAEV_GR_TARGETS_UP | 56 | 0.01 | Up | 0.492950559380627 | 0.570503321782735 | 0.767099141912473 |
| KYNG_WERNER_SYNDROM_AND_NORMAL_AGING_DN | 170 | 0.01 | Down | 0.297555508236431 | 0.57145950535624 | 0.76812260335951 |
| REACTOME_FOLDING_OF_ACTIN_BY_CCT_TRIC | 10 | 0.01 | Down | 0.116913753885411 | 0.571695184716405 | 0.76812260335951 |
| MIKKELSEN_IPS_ICP_WITH_H3K27ME3 | 20 | 0.01 | Up | 0.918154400901228 | 0.57181010518354 | 0.76812260335951 |
| YAO_TEMPORAL_RESPONSE_TO_PROGESTERONE_CLUSTER_2 | 36 | 0.01 | Down | 0.41032136403263 | 0.571948020788494 | 0.76812260335951 |
| LOPEZ_MESOTHELIOMA_SURVIVAL_WORST_VS_BEST_UP | 12 | 0.01 | Up | 0.27601518032326 | 0.572387650355283 | 0.768353900069438 |
| LENAOUR_DENDRITIC_CELL_MATURATION_DN | 119 | 0.01 | Down | 0.714518999270201 | 0.572462115980545 | 0.768353900069438 |
| ZHANG_PROLIFERATING_VS_QUIESCENT | 48 | 0.01 | Up | 0.31994496427974 | 0.572656602466474 | 0.768385500921433 |
| ODONNELL_TARGETS_OF_MYC_AND_TFRC_UP | 68 | 0.01 | Up | 0.395654459356489 | 0.573447112502727 | 0.76914845325656 |
| BREUHAHN_GROWTH_FACTOR_SIGNALING_IN_LIVER_CANCER | 21 | 0.01 | Down | 0.384801286226876 | 0.573856126175777 | 0.76914845325656 |
| MARSON_FOXP3_TARGETS_STIMULATED_DN | 10 | 0.01 | Down | 0.311364549788486 | 0.573866849448564 | 0.76914845325656 |
| SMID_BREAST_CANCER_LUMINAL_B_DN | 399 | 0.01 | Down | 0.715594544697652 | 0.573909657891547 | 0.76914845325656 |
| REACTOME_CLASS_B_2_SECRETIN_FAMILY_RECEPTORS | 43 | 0.01 | Up | 0.423256228395825 | 0.574144729769818 | 0.769234146144659 |
| REACTOME_SIGNALING_BY_NOTCH1_HD_DOMAIN_MUTANTS_IN_CANCER | 14 | 0.01 | Up | 0.289116299777305 | 0.574843742861138 | 0.769941187175451 |
| WENG_POR_TARGETS_GLOBAL_UP | 11 | 0.01 | Down | 0.387986487760415 | 0.575871807593154 | 0.770950553686085 |
| REACTOME_SYNTHESIS_OF_LEUKOTRIENES_LT_AND_EOXINS_EX | 11 | 0.01 | Up | 0.588272238614975 | 0.576131468520807 | 0.770950553686085 |
| REACTOME_SIGNALING_BY_ERBB4 | 44 | 0.01 | Down | 0.401391230970611 | 0.576319229218468 | 0.770950553686085 |
| REACTOME_RNA_POLYMERASE_I_TRANSCRIPTION_INITIATION | 47 | 0.01 | Up | 0.126685758857923 | 0.576399879334214 | 0.770950553686085 |
| IVANOVA_HEMATOPOIESIS_LATE_PROGENITOR | 430 | 0.01 | Down | 0.27880794540438 | 0.57645490788407 8 | 0.770950553686085 |

REACTOME_SYNTHESIS_OF_GLYCOSYLPHOSPHATIDYLINOSITOL_GPI 18 0.01 Up 0.13618373776762 0.576640988274371 0.770970030426323
FONTAINE_THYROID_TUMOR_UNCERTAIN_MALIGNANCY_DN 18 0.01 Down 0.538478275188613 0.577005958211782 0.771086903876193
TAKEDA_TARGETS_OF_NUP98_HOXA9_FUSION_3D_DN 21 0.01 Down 0.831358378480758 0.57711566008513 5 0.771086903876193
LASTOWSKA_COAMPLIFIED_WITH_MYCN 32 0.01 Down 0.220435064457311 0.577465546748237 0.771086903876193
DOUGLAS_BMI1_TARGETS_UP 489 0.01 Up 0.306676436697433 0.577633800354117 0.771086903876193
VALK_AML_WITH_EVI1 17 0.01 Up 0.248743794898894 0.577712827449724 0.771086903876193
MOREAUX_MULTIPLE_MYELOMA_BY_TACI_UP 243 0.01 Down 0.432384764174508 0.57776590969529 0.771086903876193
NAKAMURA_ADIPOGENESIS_EARLY_UP 54 0.01 Up 0.546912610854413 0.578068884501308 0.771086903876193
COLLIS_PRKDC_SUBSTRATES 21 0.01 Down 0.238916733767603 0.578100748846 0.771086903876193
STANELLE_E2F1_TARGETS 24 0.01 Up 0.238621939867562 0.578924034450223 0.771955958129264
CHIBA_RESPONSE_TO_TSA 42 0.01 Up 0.232190147480141 0.579142293124269 0.772017973782203
REACTOME_TRANSCRIPTION_OF_THE_HIV_GENOME 68 0.01 Down 0.12232854716376 0.579667125486696 0.772434824429937
FONTAINE_PAPILLARY_THYROID_CARCINOMA_UP 56 0.01 Down 0.333049455731175 0.579798686902471 0.772434824429937
PID_EPHA_FWDPATHWAY 25 0.01 Down 0.552300444562624 0.580266928467769 0.772829583248185
BIOCARTA_ALK_PATHWAY 28 0.01 Down 0.32554209660249 0.580439333818496 0.772830214903478
KEGG_REGULATION_OF_AUTOPHAGY 19 0.01 Down 0.117001241930301 0.581319292319626 0.773772644055884
HAHTOLA_SEZARY_SYNDROM_UP 78 0.01 Down 0.395881225476198 0.582403414395628 0.774986189374881
MCCLUNG_COCAINE_REWARD_5D 54 0.01 Down 0.265352764331619 0.582765494292525 0.77501480965294
FURUKAWA_DUSP6_TARGETS_PCI35_UP 63 0.01 Up 0.474676080490838 0.582996066170412 0.77501480965294
REACTOME_FERTILIZATION 11 0.01 Down 0.319500076097062 0.583036725193083 0.77501480965294
WILLIAMS_ESR2_TARGETS_DN 11 0.01 Down 0.322296784144634 0.583114590933535 0.77501480965294
WANG_IMMORTALIZED_BY_HOXA9_AND_MEIS1_UP 24 0.01 Down 0.740649470216565 0.583377170539724 0.775134608801672
WIERENGA_STAT5A_TARGETS_UP 164 0.01 Down 0.432015689061213 0.584079204343213 0.775838068416886
REACTOME_GLUCOSE_METABOLISM 79 0.01 Up 0.213661497537224 0.584282424975956 0.775878729768663
REACTOME_CASPASE_ACTIVATION_VIA_DEATH_RECEPTORS_IN_THE_PRESENCE_OF_LIGAND 14 0.01 Down 0.322285411329615

0.585324747735614	0.777033296240869
ROSS_AML_WITH_AML1_ETO_FUSION	58	0.01	Up
0.340380628348404	0.585677102812943	0.77726631322555
HELLER_HDAC_TARGETS_SILENCED_BY_METHYLATION_DN	248	0.01	Down
	0.371853672644721	0.585851347791703
0.77726631322555
PARK_HSC_AND_MULTIPOTENT_PROGENITORS	39	0.01	Up
0.140649448282517	0.586080665591898	0.77726631322555
WEBER_METHYLATED_IN_COLON_CANCER	10	0.01	Up
0.376807881769943	0.586228475101467	0.77726631322555
MINGUEZ_LIVER_CANCER_VASCULAR_INVASION_UP	12	0.01	Down
	0.17327293202541	0.586364864993957
0.77726631322555
WANG_TARGETS_OF_MLL_CBP_FUSION_UP	32	0.01	Down
0.426478846760814	0.58761351234403	0.77869184492524
NAKAYAMA_FRA2_TARGETS	40	0.01	Up	0.299894317744147
	0.587846861004644	0.778771482527519
REACTOME_DNA_DOUBLE_STRAND_BREAK_RESPONSE	53	0.01	Up
	0.188857769267058	0.588570784105035
0.779283025410967
MURATA_VIRULENCE_OF_H_PILORI	18	0.01	Down
0.73247157958203	0.588579726645213	0.779283025410967
BOHN_PRIMARY_IMMUNODEFICIENCY_SYNDROM_UP	40	0.01	Down
	0.19340117582593	0.588983596389496
0.779588123018487
DELASERNA_MYOD_TARGETS_UP	68	0.01	Up
0.399606439210036	0.589396828146976	0.779905429061129
WEINMANN_ADAPTATION_TO_HYPOXIA_DN	37	0.01	Down
0.739993738506078	0.589660643988929	0.780024895447392
LEE_LIVER_CANCER_MYC_DN	37	0.01	Up	0.404420903592646
	0.590291858517815	0.780630157116086
REACTOME_ADENYLATE_CYCLASE_INHIBITORY_PATHWAY	11	0.01	Down
	0.360980815701123	0.590562144205346
0.780757893589126
REACTOME_ROLE_OF_LAT2_NTAL_LAB_ON_CALCIUM_MOBILIZATION	14	0.01
	Down	0.394672935944175	0.591245663080413
0.781431712892225
REACTOME_RNA_POLYMERASE_I_TRANSCRIPTION	71	0.01	Up
0.135958874956238	0.591510732590047	0.781552246617361
WU_APOPTOSIS_BY_CDKN1A_NOT_VIA_TP53	10	0.01	Up
0.194313135420096	0.591807346441464	0.781714376213453
REACTOME_TP53_REGULATES_TRANSCRIPTION_OF_GENES_INVOLVED_IN_CYTOCHROM
E_C_RELEASE	19	0.01	Up	0.242986240445994
0.593304682056233	0.783461969989062
CHIANG_LIVER_CANCER_SUBCLASS_CTNNB1_DN	136	0.01	Down
0.817197710560828	0.594111811406666	0.784297383927448
REACTOME_RRNA_MODIFICATION_IN_THE_NUCLEUS_AND_CYTOSOL	60	0.01
	Up	0.101875352047039	0.594516536181154
0.78460124196544
XU_RESPONSE_TO_TRETINOIN_DN	11	0.01	Up
0.213363563218301	0.594803882773768	0.784684305661936
REACTOME_LEADING_STRAND_SYNTHESIS	14	0.01	Up
0.107445411678555	0.594928612613099	0.784684305661936
VALK_AML_CLUSTER_5	28	0.01	Up	0.612079658624272

0.595180674938737	0.784786486902207
GUILLAUMOND_KLF10_TARGETS_UP	33	0.01	Up
0.488384213263086	0.595440776130943	0.784899204899879
KYNG_DNA_DAMAGE_DN	168	0.01	Down	0.387883059406174
0.595742133063954	0.785066223430804
PID_RAC1_REG_PATHWAY	34	0.01	Down	0.360088504686659
0.596003655620989	0.785180665889902
WEIGEL_OXIDATIVE_STRESS_RESPONSE	31	0.01	Up
0.328163687408049	0.596581022935545	0.78571101614277
MYLLYKANGAS_AMPLIFICATION_HOT_SPOT_24	12	0.01	Up
0.420179798951669	0.597102728575175	0.786167769462627
REACTOME_GLUTATHIONE_SYNTHESIS_AND_RECYCLING	12	0.01	Up
0.324224142012066	0.597388984649123
0.786176499218336
BERTUCCI_INVASIVE_CARCINOMA_DUCTAL_VS_LOBULAR_UP	22	0.01
Down	0.28205835051808	0.597459159361476
0.786176499218336
REACTOME_CELL_CYCLE	541	0.01	Down	0.171229159690869
0.598657068717683	0.78747978737551
LI_WILMS_TUMOR_ANAPLASTIC_UP	13	0.01	Up
0.209058166626921	0.598832436790157	0.78747978737551
TAVAZOIE_METASTASIS	68	0.01	Down	0.568736077371187
0.599122136156496	0.78747978737551
REACTOME_RHO_GTPASES_ACTIVATE_ROCKS	18	0.01	Down
0.17031411162775	0.599150361028753	0.78747978737551
TERAO_AOX4_TARGETS_HG_UP	25	0.01	Down
0.143644829493046	0.600097349255793	0.78826158783755
CHIN_BREAST_CANCER_COPY_NUMBER_UP	23	0.01	Down
0.388749256449568	0.600276976094728	0.78826158783755
REACTOME_FGFR2_ALTERNATIVE_SPLICING	27	0.01	Down
0.112454030947831	0.600315018172957	0.78826158783755
ABRAHAM_ALPC_VS_MULTIPLE_MYELOMA_DN	20	0.01	Up
0.212460841410419	0.600446646664243	0.78826158783755
REACTOME_ACTIVATION_OF_HOX_GENES_DURING_DIFFERENTIATION 72	0.01
Down	0.243971164896448	0.601146716986079
0.788950216891219
RASHI_RESPONSE_TO_IONIZING_RADIATION_6	65	0.01	Down
0.547127194740285	0.601466915706393	0.789132983489795
REACTOME_DISEASE	897	0.01	Down	0.311460360257153
0.601637093308015	0.789132983489795
REACTOME_GLYCEROPHOSPHOLIPID_BIOSYNTHESIS	104	0.01	Up
0.272796039916192	0.601961540893067
0.789268515415807
BIOCARTA_CK1_PATHWAY	10	0.01	Down	0.274350217583032
0.602091599412859	0.789268515415807
HO_LIVER_CANCER_VASCULAR_INVASION	10	0.01	Down
0.166977757088738	0.602777862283315	0.789880144746047
BAKER_HEMATOPOESIS_STAT1_TARGETS	12	0.01	Down
0.702229314172384	0.602909627724958	0.789880144746047
LEE_LIVER_CANCER_DENA_UP	51	0.01	Up
0.532563055337258	0.603455997372629	0.790365590964443
SILIGAN_BOUND_BY_EWS_FLT1_FUSION	33	0.01	Up
0.338997027083105	0.603915276919367	0.790736722910736
JIANG_AGING_CEREBRAL_CORTEX_UP	26	0.01	Up

0.304244489007118	0.604342949917715	0.790824246775549
PID_GLYPICAN_1PATHWAY	22	0.01	Up	0.612670203979465
	0.604429128414305	0.790824246775549
GEISS_RESPONSE_TO_DSRNA_DN	14	0.01	Down
0.320276945719444	0.604509924787717	0.790824246775549
CERVERA_SDHB_TARGETS_2	88	0.01	Down	0.522639813568081
	0.605222259836437	0.791318952983457
PHESSE_TARGETS_OF_APC_AND_MBD2_UP	10	0.01	Up
0.390220212188442	0.605240169156201	0.791318952983457
REACTOME_TRANSCRIPTIONAL_REGULATION_OF_PLURIPOTENT_STEM_CELLS	16
	0.01	Up	0.385937002271105	0.606056676112628
	0.792156080001822
HERNANDEZ_MITOTIC_ARREST_BY_DOCETAXEL_2_DN	17	0.01	Up
	0.324064762029082	0.606484118588491
0.792484335190485
LASTOWSKA_NEUROBLASTOMA_COPY_NUMBER_DN	755	0.01	Down
0.187873785934254	0.60701300869372	0.792944921266571
AGUIRRE_PANCREATIC_CANCER_COPY_NUMBER_DN	221	0.01	Down
	0.177394330592465	0.607208019332536
0.792969217576917
VECCHI_GASTRIC_CANCER_EARLY_UP	371	0.01	Down
0.321598624733674	0.607598011005628	0.793248056773249
REACTOME_DNA_REPAIR	291	0.01	Up	0.155649685262853
	0.608293143291403	0.793909217067493
WEIGEL_OXIDATIVE_STRESS_BY_HNE_AND_H2O2	33	0.01	Up
0.259002257167581	0.608457675817022	0.793909217067493
KRIEG_HYPOXIA_NOT_VIA_KDM3A	699	0.01	Up
0.288873038082914	0.60870611767217	0.794002901606617
REACTOME_METABOLISM_OF_STEROIDS	113	0.01	Down
0.332242471606607	0.609141570371443	0.794340400005697
BOQUEST_STEM_CELL_CULTURED_VS_FRESH_UP	344	0.01	Up
0.547797658353373	0.609320160222821	0.794342842285841
NELSON_RESPONSE_TO_ANDROGEN_DN	13	0.01	Down
0.225626181968141	0.610019547733561	0.794689651647964
KEGG_MELANOMA	52	0.01	Down	0.400602443440085
0.610113005423461	0.794689651647964
EHRLICH_ICF_SYNDROM_DN	10	0.01	Up	0.415309940213513
	0.610116571265211	0.794689651647964
REACTOME_NEUROTRANSMITTER_RELEASE_CYCLE	29	0.01	Down
0.362346032691225	0.611616436080871	0.796348188152404
REACTOME_SIGNALING_BY_FGFR2_IN_DISEASE	32	0.01	Down
0.25072957107453	0.611744225515072	0.796348188152404
STEIN_ESRRA_TARGETS_DN	99	0.01	Up	0.196731963087082
	0.611986914399803	0.796433462717752
REACTOME_OTHER_SEMAPHORIN_INTERACTIONS	16	0.01	Up
0.661612465470764	0.612464574367892	0.796666694070947
DANG_MYC_TARGETS_DN	29	0.01	Up	0.210580571551327
	0.612549731380309	0.796666694070947
REACTOME_WNT_LIGAND_BIOGENESIS_AND_TRAFFICKING	16	0.01	Down
	0.239910733057564	0.612729752524323
0.796666694070947
HELLER_SILENCED_BY_METHYLATION_UP	220	0.01	Down
0.583675862253797	0.613249384364861	0.796666694070947
GRUETZMANN_PANCREATIC_CANCER_DN	157	0.01	Up

0.307905468409857	0.613336464786379	0.796666694070947
ABRAMSON_INTERACT_WITH_AIRE	42	0.01	Down
0.130971977803719	0.613657189568944	0.796666694070947
REACTOME_RNA_POLYMERASE_III_TRANSCRIPTION	41	0.01	Up
	0.141831894304003	0.613693492463429
0.796666694070947
REACTOME_METALLOPROTEASE_DUBS	24	0.01	Down
0.18119271238459	0.613705623469264	0.796666694070947
TERAMOTO_OPN_TARGETS_CLUSTER_1	11	0.01	Down
0.234767430898752	0.613761237278685	0.796666694070947
REACTOME_NUCLEAR_ENVELOPE_BREAKDOWN	51	0.01	Down
0.225895104062075	0.614350233058787	0.797201009699552
PASINI_SUZ12_TARGETS_UP	78	0.01	Down	0.367534818423848
	0.614917345725706	0.797706628870721
GARCIA_TARGETS_OF_FLI1_AND_DAX1_DN	140	0.01	Down
0.216175102138073	0.615642855961075	0.798187932877556
WANG_RESPONSE_TO_BEXAROTENE_DN	23	0.01	Down
0.348928175672513	0.615805095335829	0.798187932877556
WEST_ADRENOCORTICAL_TUMOR_MARKERS_UP	15	0.01	Down
0.452263291217637	0.616093959954632	0.798187932877556
KONDO_COLON_CANCER_HCP_WITH_H3K27ME1	19	0.01	Up
0.246391190015724	0.616132124478546	0.798187932877556
KEGG_PROTEIN_EXPORT	23	0.01	Down	0.152787106868126
	0.616176224045633	0.798187932877556
ELVIDGE_HYPOXIA_BY_DMOG_UP	131	0.01	Up
0.429317381763599	0.616479302989794	0.798291623869368
KYNG_DNA_DAMAGE_BY_4NQO	31	0.01	Up	0.369840736764248
	0.61661146119565	0.798291623869368
KEGG_VASOPRESSIN_REGULATED_WATER_REABSORPTION	38	0.01	Down
	0.182840403044232	0.616955144153871
0.798420321733465
REACTOME_G_ALPHA_S_SIGNALLING_EVENTS	63	0.01	Down
0.31920739649822	0.617282706250806	0.798420321733465
REACTOME_PHASE_2_PLATEAU_PHASE	11	0.01	Up
0.601101747978126	0.617318886317059	0.798420321733465
KANG_GLIS3_TARGETS	12	0.01	Down	0.372811981815492
	0.61742136559411	0.798420321733465
GRADE_COLON_VS_RECTAL_CANCER_UP	35	0.01	Up
0.384741313806141	0.617891486190826	0.798798455688169
GOLUB_ALL_VS_AML_DN	17	0.01	Down	0.66182380930676
	0.618895412140063	0.799639006125934
KUNINGER_IGF1_VS_PDGFB_TARGETS_DN	38	0.01	Up
0.75917157678388	0.618897464363098	0.799639006125934
BANDRES_RESPONSE_TO_CARMUSTIN_WITHOUT_MGMT_24HR_UP	10	0.01
	Up	0.173936906880213	0.619192762272051
0.799790651268066
BASSO_CD40_SIGNALING_DN	58	0.01	Up	0.379869194294303
	0.619763187479528	0.800208182965851
BENPORATH_EED_TARGETS	498	0.01	Up	0.516891249246264
	0.619872056304137	0.800208182965851
WU_HBX_TARGETS_2_UP	21	0.01	Down	0.303690209451098
	0.620282278133808	0.800507849615695
MARKEY_RB1_CHRONIC_LOF_UP	97	0.01	Up
0.416966709967312	0.620533122865335	0.800601718507371

MASRI_RESISTANCE_TO_TAMOXIFEN_AND_AROMATASE_INHIBITORS_UP 13
0.01 Up 0.889857166680558 0.621008713227328
0.800985413473985
GAUSSMANN_MLL_AF4_FUSION_TARGETS_G_UP 165 0.01 Down
0.396728016156014 0.621898799401277 0.801903357231423
REACTOME_IMPORT_OF_PALMITOYL_COA_INTO_THE_MITOCHONDRIAL_MATRIX 13
0.01 Up 0.245512625803059 0.622342219090801
0.802244988475237
WENG_POR_TARGETS_LIVER_UP 29 0.01 Up
0.508844492570574 0.622695388539801 0.80247011797202
PATIL_LIVER_CANCER 606 0.01 Up 0.26968744472446
0.623015110751197 0.802652027178741
HOLLMANN_APOPTOSIS_VIA_CD40_UP 184 0.01 Up
0.263523417642494 0.623757810457369 0.803154176335832
REACTOME_MITOPHAGY 28 0.01 Up 0.0924565599606888
0.623910076764214 0.803154176335832
IKEDA_MIR30_TARGETS_DN 17 0.01 Up 0.201564024469338
0.623940908512731 0.803154176335832
REACTOME_METABOLISM_OF_RNA 654 0.01 Down
0.123778955126612 0.624655368796992 0.803772603142153
KEGG_GAP_JUNCTION 67 0.01 Down 0.3722362063324
0.625032736580676 0.803772603142153
LY_AGING_OLD_DN 51 0.01 Up 0.329858738924999
0.625060194654937 0.803772603142153
CAFFAREL_RESPONSE_TO_THC_8HR_3_DN 10 0.01 Up
0.20865064823906 0.625136600797546 0.803772603142153
WIEDERSCHAIN_TARGETS_OF_BMI1_AND_PCGF2 52 0.01 Up
0.482256625828884 0.625832429645681 0.804437166501955
YU_BAP1_TARGETS 25 0.01 Up 0.254155018538066
0.627064169345385 0.805790006062752
LI_LUNG_CANCER 38 0.01 Down 0.409644913612516
0.627266211142848 0.805819268101486
REACTOME_SUMOYLATION_OF_TRANSCRIPTION_FACTORS 19 0.01 Up
0.32871388341263 0.627918973130869
0.806413655908603
REACTOME_HSF1_ACTIVATION 27 0.01 Down
0.217725337058443 0.62808769951858 0.806413655908603
REACTOME_MITOTIC_PROMETAPHASE 170 0.01 Down
0.159593253871857 0.628326815247266 0.80649030112406
PID_ENDOTHELIN_PATHWAY 51 0.01 Down 0.435401181700893
0.628857169741141 0.80688866171214
REACTOME_INTEGRATION_OF_ENERGY_METABOLISM 70 0.01 Down
0.307352263221379 0.629053322114851
0.80688866171214
MULLIGHAN_NPM1_MUTATED_SIGNATURE_2_DN 67 0.01 Down
0.303811555716696 0.629175697286107 0.80688866171214
BIOCARTA_ARF_PATHWAY 15 0.01 Up 0.244454336871386
0.629365873481528 0.806902339218331
REACTOME_E3_UBIQUITIN_LIGASES_UBIQUITINATE_TARGET_PROTEINS 49
0.01 Down 0.143851234331415 0.629820365798972
0.807212548629705
JAZAG_TGFB1_SIGNALING_DN 23 0.01 Down
0.179292962556941 0.629966990120802 0.807212548629705
BLALOCK_ALZHEIMERS_DISEASE_DN 1081 0.01 Up

0.217773131809289	0.63045460595529	0.807607139860082
REACTOME_N_GLYCAN_TRIMMING_IN_THE_ER_AND_CALNEXIN_CALRETICULIN_CYCLE
	35	0.01	Up	0.127507473735456
0.631044323138084	0.808048451070861
MONNIER_POSTRADIATION_TUMOR_ESCAPE_DN	340	0.01	Down
0.275276698058426	0.631158645541667	0.808048451070861
HUNSBERGER_EXERCISE_REGULATED_GENES	19	0.01	Up
0.328386802017091	0.631640809670684	0.808245394789074
HOOI_ST7_TARGETS_DN	102	0.01	Down	0.479789254261098
	0.631672096083207	0.808245394789074
PURBEY_TARGETS_OF_CTBP1_AND_SATB1_DN	135	0.01	Up
0.227452884074502	0.633041321982497	0.809766858938908
MARTINEZ_RESPONSE_TO_TRABECTEDIN	50	0.01	Down
0.255277415137928	0.633706852527421	0.810387568168067
REACTOME_ACTIVATION_OF_APC_C_AND_APC_C:CDC20_MEDIATED_DEGRADATION_OF
_MITOTIC_PROTEINS	71	0.01	Down	0.139477128639961
	0.634327567677605	0.810950630463832
PID_S1P_META_PATHWAY	16	0.01	Up	0.170963111362146
	0.634961264174795	0.81152996373776
MAEKAWA_ATF2_TARGETS	12	0.01	Up	0.326092701897477
	0.635386782887889	0.811842975861587
REACTOME_ACYL_CHAIN_REMODELLING_OF_PS	14	0.01	Up
0.427269983159722	0.635916311747929	0.812288667606405
MIKKELSEN_MEF_LCP_WITH_H3K4ME3	97	0.01	Up
0.597296403142993	0.636369670340791	0.812636837551663
YANG_BREAST_CANCER_ESR1_UP	29	0.01	Up
0.356478874700983	0.636628446293385	0.812736400479627
BILD_E2F3_ONCOGENIC_SIGNATURE	221	0.01	Up
0.301064647841729	0.637002072255999	0.812982485744098
BIOCARTA_CCR3_PATHWAY	13	0.01	Down	0.180060030500494
	0.637641071763047	0.813488170622287
VANDESLUIS_COMMD1_TARGETS_GROUP_3_UP	64	0.01	Down
0.451445421967356	0.637760247669174	0.813488170622287
BRUINS_UVC_RESPONSE_MIDDLE	85	0.01	Up
0.184142061021064	0.638300411217602	0.81370791972681
BONOME_OVARIAN_CANCER_SURVIVAL_OPTIMAL_DEBULKING	201	0.01
	Down	0.294726211212798	0.638400762643325
0.81370791972681
REACTOME_IRF3_MEDIATED_INDUCTION_OF_TYPE_I_IFN	12	0.01	Down
	0.246624284424522	0.638476155235188
0.81370791972681
MARTINEZ_TP53_TARGETS_UP	506	0.01	Down
0.405071716900903	0.638656627540864	0.81370791972681
GOUYER_TATI_TARGETS_DN	16	0.01	Up	0.431421960342265
	0.639155587612144	0.814112883626123
TAKEDA_TARGETS_OF_NUP98_HOXA9_FUSION_8D_DN	145	0.01	Up
	0.714876250429749	0.639594693926755
0.814441402039876
GENTLES_LEUKEMIC_STEM_CELL_UP	20	0.01	Down
0.693624430151914	0.639987297991791	0.814500959094886
REACTOME_DNA_DOUBLE_STRAND_BREAK_REPAIR	136	0.01	Up
0.176450794428095	0.640003868192021	0.814500959094886
LEIN_MIDBRAIN_MARKERS	47	0.01	Up	0.364105870573964
	0.640220321573506	0.814545809644186

REACTOME_ENERGY_DEPENDENT_REGULATION_OF_MTOR_BY_LKB1_AMPK 27
0.01 Down 0.153648412255108 0.641930129406312
0.816490076876449
LEE_LIVER_CANCER_MYC_TGFA_DN 34 0.01 Up
0.509083003449812 0.642200789345657 0.816603266791719
KEGG_TERPENOID_BACKBONE_BIOSYNTHESIS 13 0.01 Down
0.220715726688314 0.643083965606042 0.817495029807455
PID_AR_NONGENOMIC_PATHWAY 27 0.01 Down
0.265991189390609 0.643297558307401 0.817535347636915
BOYAULT_LIVER_CANCER_SUBCLASS_G23_UP 44 0.01 Up
0.22789259045931 0.644072961325966 0.818032398701949
HU_ANGIOGENESIS_UP 19 0.01 Down 0.365158635777613
0.644169743438796 0.818032398701949
BURTON_ADIPOGENESIS_9 75 0.01 Up 0.489711848101322
0.644234636575061 0.818032398701949
MORI_PLASMA_CELL_UP 44 0.01 Down 0.187484689906487
0.644675103604299 0.818360517001221
REACTOME_RECOGNITION_OF_DNA_DAMAGE_BY_PCNA_CONTAINING_REPLICATION_CO
MPLEX 28 0.01 Up 0.111169671110093
0.645003219413627 0.81854586992215
VALK_AML_CLUSTER_3 28 0.01 Up 0.420233610454081
0.645779331929388 0.819299491115609
PID_PLK1_PATHWAY 38 0.01 Down 0.22636451013402
0.646060684089552 0.819364073863802
LIN_NPAS4_TARGETS_UP 142 0.01 Down 0.202240575034748
0.646377760349828 0.819364073863802
REACTOME_MRNA_DECAY_BY_3_TO_5_EXORIBONUCLEASE 16 0.01 Up
0.134655787858716 0.646721148677917
0.819364073863802
BIOCARTA_EDG1_PATHWAY 19 0.01 Up 0.220006704867738
0.646830399965215 0.819364073863802
SARTIPY_BLUNTED_BY_INSULIN_RESISTANCE_UP 16 0.01 Down
0.348965461536849 0.646901274937665
0.819364073863802
MOREIRA_RESPONSE_TO_TSA_UP 27 0.01 Up
0.32278589456677 0.646923937295358 0.819364073863802
KYNG_ENVIRONMENTAL_STRESS_RESPONSE_NOT_BY_GAMMA_IN_OLD 27 0.01
Down 0.3796039713252 0.647726800819782
0.819954254855599
REACTOME_REGULATION_OF_KIT_SIGNALING 15 0.01 Down
0.402385414317326 0.647883338265548 0.819954254855599
BILD_MYC_ONCOGENIC_SIGNATURE 201 0.01 Down
0.299507728532101 0.647937155338618 0.819954254855599
YEGNASUBRAMANIAN_PROSTATE_CANCER 88 0.01 Down
0.37870609057307 0.648346776154016 0.820241699637574
PID_PS1_PATHWAY 40 0.01 Down 0.219468688297933
0.648991985760122 0.820826948787773
WAESCH_ANAPHASE_PROMOTING_COMPLEX 11 0.01 Up
0.082033653049017 0.649327663315775 0.820869870195267
PARK_HSC_VS_MULTIPOTENT_PROGENITORS_UP 14 0.01 Down
0.19139693410303 0.649391158712874 0.820869870195267
BILBAN_B_CLL_LPL_UP 51 0.01 Down 0.306429372008918
0.649736739323528 0.821075806370328
OKAWA_NEUROBLASTOMA_1P36_31_DELETION 18 0.01 Up

0.196247440388969	0.650108087475132	0.821203798556864
RUAN_RESPONSE_TO_TNF_TROGLITAZONE_DN	27	0.01	Up
0.331365343413798	0.65020340802311	0.821203798556864
BRUNO_HEMATOPOIESIS	54	0.01	Down	0.45002070874229
0.65070909293544	0.821611621557529
TAKADA_GASTRIC_CANCER_COPY_NUMBER_DN	19	0.01	Up
0.544777949521679	0.650968896826083	0.821708843367942
REACTOME_TRANSLESION_SYNTHESIS_BY_POLH	19	0.01	Up
0.112426591036128	0.651384251932962	0.8219429208287l
BERTUCCI_INVASIVE_CARCINOMA_DUCTAL_VS_LOBULAR_DN	27	0.01
Up	0.679991992584063	0.651520050481133
0.82194292082871
BIOCARTA_G1_PATHWAY	27	0.01	Down	0.276342301835345
0.651836069949842	0.822055171047216
PEPPER_CHRONIC_LYMPHOCYTIC_LEUKEMIA_UP	19	0.01	Up
0.72285994575196	0.652137809187421	0.822055171047216
NOJIMA_SFRP2_TARGETS_DN	23	0.01	Down	0.33101879123088
0.652319847578453	0.822055171047216
REACTOME_MRNA_CAPPING	29	0.01	Up	0.116241854837901
0.652340555089081	0.822055171047216
MAYBURD_RESPONSE_TO_L663536_DN	51	0.01	Down
0.187165911747209	0.652937501914872	0.822337468008285
MOREAUX_MULTIPLE_MYELOMA_BY_TACI_DN	172	0.01	Down
0.139065683193123	0.65308143074366	0.822337468008285
GOTZMANN_EPITHELIAL_TO_MESENCHYMAL_TRANSITION_DN	180	0.01
Down	0.374616861612367	0.653113406182331
0.822337468008285
BIOCARTA_RB_PATHWAY	10	0.01	Up	0.264216885214756
0.653332919288226	0.82238349823595
GRASEMANN_RETINOBLASTOMA_WITH_6P_AMPLIFICATION	14	0.01	Up
0.134879750104114	0.653906107517559
0.822844574459459
JAZAERI_BREAST_CANCER_BRCA1_VS_BRCA2_DN	34	0.01	Down
0.383745175327106	0.654065331377897	0.822844574459459
XU_RESPONSE_TO_TRETINOIN_AND_NSC682994_DN	15	0.01	Down
0.110206748103296	0.654482566081622
0.823139097520115
AMBROSINI_FLAVOPIRIDOL_TREATMENT_TP53	95	0.01	Down
0.364224907937354	0.655602143860264	0.824316541720807
GRAESSMANN_RESPONSE_TO_MC_AND_SERUM_DEPRIVATION_UP	170	0.01
Up	0.483998865761565	0.65582467297343
0.824365745250438
ACOSTA_PROLIFERATION_INDEPENDENT_MYC_TARGETS_DN	98	0.01	Up
0.397913795981478	0.656967651976308
0.825491427745888
PID_HEDGEHOG_GLI_PATHWAY	44	0.01	Down
0.245895223814626	0.657087503553902	0.825491427745888
KEGG_PROGESTERONE_MEDIATED_OOCYTE_MATURATION	73	0.01	Down
0.268738170837651	0.65751120984818
0.825792927708178
ZHU_CMV_8_HR_DN	46	0.01	Up	0.416256752730881
0.65794955960124	0.826112645365244
GROSS_HIF1A_TARGETS_DN	23	0.01	Up	0.435265311875689
0.658568785441441	0.826504511477765

KANG_CISPLATIN_RESISTANCE_UP 21 0.01 Down
0.294561399566284 0.658629401582503 0.826504511477765
CHENG_RESPONSE_TO_NICKEL_ACETATE 43 0.01 Down
0.245032326820396 0.659977717642658 0.827965347698506
REACTOME_INSERTION_OF_TAIL_ANCHORED_PROTEINS_INTO_THE_ENDOPLASMIC_RE
TICULUM_MEMBRANE 21 0.01 Up 0.174385943506366
0.660814363798706 0.828783639864727
FINETTI_BREAST_CANCER_KINOME_GREEN 11 0.01 Down
0.662961947219242 0.661686350329119 0.829645786535394
WALLACE_JAK2_TARGETS_UP 22 0.01 Up 0.158141549976599
0.661931909943725 0.829722235135819
GRADE_COLON_AND_RECTAL_CANCER_UP 282 0.01 Up
0.23490329928679 0.662291151140078 0.829941099630513
REACTOME_CYTOSOLIC_IRON_SULFUR_CLUSTER_ASSEMBLY 13 0.01 Up
0.138881048311206 0.662616352910617
0.830117197974701
OISHI_CHOLANGIOMA_STEM_CELL_LIKE_UP 322 0.01 Down
0.18960718811259 0.663403348302422 0.830871566068372
REACTOME_VASOPRESSIN_REGULATES_RENAL_WATER_HOMEOSTASIS_VIA_AQUAPORIN
S 28 0.01 Down 0.407200463297898
0.666164541659808 0.833989139282315
DALESSIO_TSA_RESPONSE 13 0.01 Down 0.443437504688636
0.666320824358875 0.833989139282315
BOSCO_TH1_CYTOTOXIC_MODULE 59 0.01 Down
0.687247257741702 0.66644916313728 0.833989139282315
LEE_NAIVE_T_LYMPHOCYTE 16 0.01 Down 0.391590563338842
0.667124967337829 0.8346024848827
RIZ_ERYTHROID_DIFFERENTIATION_6HR 22 0.01 Up
0.515859319785286 0.667823604082737 0.835244045729522
WILCOX_RESPONSE_TO_PROGESTERONE_UP 122 0.01 Down
0.32763029797361 0.668038650124589 0.835280593132136
CHEN_LVAD_SUPPORT_OF_FAILING_HEART_UP 90 0.01 Up
0.390310827670953 0.668315505731703 0.835281454847768
BIOCARTA_MCALPAIN_PATHWAY 15 0.01 Down
0.245879230784918 0.668679421749715 0.835281454847768
MELLMAN_TUT1_TARGETS_UP 19 0.01 Down 0.181170331616547
0.668738798048096 0.835281454847768
LEE_LIVER_CANCER_MYC_E2F1_UP 50 0.01 Up
0.701432032085674 0.668782637596689 0.835281454847768
RODRIGUES_THYROID_CARCINOMA_POORLY_DIFFERENTIATED_UP 611 0.01
Down 0.198130176176646 0.669011572243258
0.835335282564846
REACTOME_PD_1_SIGNALING 16 0.01 Up 1.04961668228245
0.669357116416641 0.835534639903582
RUTELLA_RESPONSE_TO_HGF_UP 389 0.01 Up
0.443747374917438 0.670426567699446 0.836637263134094
MARSON_FOXP3_TARGETS_UP 57 0.01 Down 0.373435600924799
0.670824217460092 0.836901153894841
VISALA_RESPONSE_TO_HEAT_SHOCK_AND_AGING_DN 13 0.01 Down
0.248474229124399 0.671164988869717
0.83693517756293
FIGUEROA_AML_METHYLATION_CLUSTER_1_DN 35 0.01 Up
0.354155893480729 0.671223874330226 0.83693517756293
REACTOME_GLYOXYLATE_METABOLISM_AND_GLYCINE_DEGRADATION 21 0.01

Up 0.132210953207627 0.67338541294479
0.839397512808329
SAFFORD_T_LYMPHOCYTE_ANERGY 68 0.01 Down
0.365849915944678 0.673727673976266 0.839591320910263
OUELLET_OVARIAN_CANCER_INVASIVE_VS_LMP_DN 12 0.01 Down
0.267368214602352 0.674048540057054
0.839758366839373
GAL_LEUKEMIC_STEM_CELL_UP 104 0.01 Down
0.415925769210164 0.674604908537918 0.839770772318817
BIOCARTA_AKAPCENTROSOME_PATHWAY 11 0.01 Up
0.240488366797486 0.674610366580183 0.839770772318817
MODY_HIPPOCAMPUS_NEONATAL 29 0.01 Up
0.278294910703362 0.674618967484593 0.839770772318817
SCHAEFFER_SOX9_TARGETS_IN_PROSTATE_DEVELOPMENT_UP 16 0.01
Down 0.429670300345816 0.675315720338372
0.840164836108227
WU_ALZHEIMER_DISEASE_UP 10 0.01 Up 0.314195773537216
0.675617641982586 0.840164836108227
ACEVEDO_LIVER_CANCER_WITH_H3K9ME3_DN 71 0.01 Down
0.29673332532051 0.675658956158221 0.840164836108227
FONTAINE_FOLLICULAR_THYROID_ADENOMA_UP 65 0.01 Up
0.234127056108446 0.675834548066881 0.840164836108227
KEGG_HISTIDINE_METABOLISM 18 0.01 Up
0.34330525588484 0.67587008840208 0.840164836108227
RUAN_RESPONSE_TO_TROGLITAZONE_UP 20 0.01 Up
0.376033207577286 0.676396914409656 0.84051924326317
REACTOME_ION_HOMEOSTASIS 30 0.01 Up
0.352969337804752 0.676632793807873 0.84051924326317
PLASARI_TGFB1_SIGNALING_VIA_NFIC_10HR_UP 39 0.01 Down
0.622126381453878 0.676844253813629
0.84051924326317
REACTOME_SIGNALING_BY_FGFR2_IIIA_TM 19 0.01 Up
0.221901660187745 0.676903150303153 0.84051924326317
NAGY_STAGA_COMPONENTS_HUMAN 13 0.01 Up
0.135833988354471 0.677843430150022 0.841331274888355
REACTOME_CELL_CYCLE_CHECKPOINTS 241 0.01 Down
0.159041088375639 0.678062053072486 0.841331274888355
SA_B_CELL_RECEPTOR_COMPLEXES 24 0.01 Down
0.277322824601813 0.678118622674196 0.841331274888355
ZHAN_MULTIPLE_MYELOMA_MF_DN 34 0.01 Down
0.244081009693282 0.678313548173393 0.841340893774669
REACTOME_ALPHA_LINOLENIC_OMEGA3_AND_LINOLEIC_OMEGA6_ACID_METABOLISM
11 0.01 Up 0.124626662857752
0.678593023602938 0.841455349267643
PARK_HSC_VS_MULTIPOTENT_PROGENITORS_DN 13 0.01 Down
0.229590022956222 0.678812527669126 0.841495397648296
REACTOME_GLUTAMATE_BINDING_ACTIVATION_OF_AMPA_RECEPTORS_AND_SYNAPTIC
_PLASTICITY 20 0.01 Up 0.295374380293727
0.679866933997562 0.84235261410724
PID_LIS1_PATHWAY 25 0.01 Down 0.190736814574612
0.679878817348402 0.84235261410724
PUJANA_CHEK2_PCC_NETWORK 701 0.01 Down
0.164430814499484 0.680123365655834 0.842423402761911
SMITH_LIVER_CANCER 42 0.01 Up 0.356169279580826

0.6804450223567 0.842538114065408
RHODES_CANCER_META_SIGNATURE 63 0.01 Down
0.269618903733137 0.680685949915746 0.842538114065408
PID_PDGFRA_PATHWAY 20 0.01 Down 0.217481847685578
0.680778293723151 0.842538114065408
HOFFMANN_IMMATURE_TO_MATURE_B_LYMPHOCYTE_DN 29 0.01 Up
0.295298379214248 0.681128052903396
0.842738947922039
REACTOME_REGULATION_BY_C_FLIP 10 0.01 Down
0.186618253875626 0.681406851942215 0.842851898591155
BIOCARTA_TH1TH2_PATHWAY 11 0.01 Up 0.753224081318494
0.68175878200515 0.842945122877509
BIOCARTA_BTG2_PATHWAY 10 0.01 Down 0.18829356788009
0.681877155374766 0.842945122877509
YAGI_AML_SURVIVAL 108 0.01 Down 0.244881549584463
0.682044807987875 0.842945122877509
UEDA_CENTRAL_CLOCK 74 0.01 Up 0.297548017871741
0.682431543444337 0.842961853789348
REACTOME_RUNX2_REGULATES_BONE_DEVELOPMENT 25 0.01 Down
0.212557007662129 0.682438333353724
0.842961853789348
REACTOME_PLATELET_AGGREGATION_PLUG_FORMATION 23 0.01 Down
0.409174009935663 0.682620945004055
0.842961853789348
ZWANG_EGF_INTERVAL_DN 194 0.01 Down 0.337214112776105
0.682817200354976 0.842972621696132
PID_CDC42_REG_PATHWAY 29 0.01 Down 0.338372771714554
0.683573913648471 0.843675107591949
REACTOME_COPI_INDEPENDENT_GOLGI_TO_ER_RETROGRADE_TRAFFIC 44
0.01 Down 0.169626657329653 0.684280663073553
0.844315558747082
LI_WILMS_TUMOR_VS_FETAL_KIDNEY_2_UP 22 0.01 Up
0.394658828768162 0.684647089849205 0.844352252530643
BIOCARTA_CARDIACEGF_PATHWAY 14 0.01 Down
0.333637202572367 0.684689168622353 0.844352252530643
NABA_MATRISOME_ASSOCIATED 380 0.01 Down
0.784768531040812 0.684873929416401 0.844352252530643
GOLDRATH_HOMEOSTATIC_PROLIFERATION 138 0.01 Down
0.216170254038958 0.685103426471346 0.844403592538717
DITTMER_PTHLH_TARGETS_UP 109 0.01 Up
0.216333948062549 0.685465725727971 0.84452499291456
LASTOWSKA_NEUROBLASTOMA_COPY_NUMBER_UP 171 0.01 Up
0.202357203586818 0.685577686127971 0.84452499291456
QI_PLASMACYTOMA_DN 86 0.01 Up 0.393342590820406
0.685782568374182 0.844545930093685
REACTOME_TRANSCRIPTIONAL_REGULATION_BY_TP53 342 0.01 Down
0.185343684999385 0.686029371119524
0.844618467045264
ONDER_CDH1_SIGNALING_VIA_CTNNB1 79 0.01 Up
0.769521481741166 0.686219260782639 0.844620913805576
GOLUB_ALL_VS_AML_UP 22 0.01 Up 0.222022610226729
0.686623826177418 0.844887516744455
ROESSLER_LIVER_CANCER_METASTASIS_DN 48 0.01 Up
0.267533616581771 0.687009719722634 0.845131004420701

MEISSNER_ES_ICP_WITH_H3K4ME3	24	0.01	Up
0.269604853723746	0.687336007929586	0.845301054895619
HOFMANN_MYELODYSPLASTIC_SYNDROM_LOW_RISK_UP	14	0.01	Down
	0.481297337841274	0.687756530750328
0.845586872462452
PID_HES_HEY_PATHWAY	38	0.01	Up	0.254032925404894
	0.68835746890473	0.846094291147597
DURCHDEWALD_SKIN_CARCINOGENESIS_UP	61	0.01	Up
0.328621573638548	0.688908899908882	0.84631050062726
REACTOME_ABORTIVE_ELONGATION_OF_HIV_1_TRANSCRIPT_IN_THE_ABSENCE_OF_T
AT	23	0.01	Up	0.0759189423196672
0.68890992698446	0.84631050062726
REACTOME_IRS_MEDIATED_SIGNALLING	33	0.01	Down
0.386586072317483	0.689231965971527	0.84646018497726
VALK_AML_CLUSTER_2	23	0.01	Down	0.476289882961247
	0.689408395373025	0.84646018497726
LOCKWOOD_AMPLIFIED_IN_LUNG_CANCER	210	0.01	Up
0.177655304384399	0.689612545322803	0.846479626222282
CHIARETTI_T_ALL_REFRACTORY_TO_THERAPY	21	0.01	Up
0.559308708909967	0.689990440824227	0.846712266313104
DING_LUNG_CANCER_MUTATED_FREQUENTLY	11	0.01	Down
0.39877321196737	0.690402414296826	0.846986586316657
BIOCARTA_SHH_PATHWAY	13	0.01	Up	0.309860271738471
	0.691106588683197	0.847563082952108
REACTOME_SIGNALING_BY_TYPE_1_INSULIN_LIKE_GROWTH_FACTOR_1_RECEPTOR_I
GF1R	37	0.01	Down	0.39918215822159
0.691404923526628	0.847563082952108
KYNG_ENVIRONMENTAL_STRESS_RESPONSE_NOT_BY_UV_IN_OLD	21	0.01
	Up	0.314287986516472	0.691704863019637
0.847563082952108
HELLER_HDAC_TARGETS_DN	251	0.01	Down	0.379816208793326
	0.691720371787761	0.847563082952108
KYNG_ENVIRONMENTAL_STRESS_RESPONSE_NOT_BY_UV_IN_WS	11	0.01
	Up	0.454132040383508	0.691981185882215
0.847563082952108
KEGG_GLYCINE_SERINE_AND_THREONINE_METABOLISM	20	0.01	Up
	0.390593625990889	0.692003673956448
0.847563082952108
LI_AMPLIFIED_IN_LUNG_CANCER	168	0.01	Up
0.332394780733661	0.692463756995951	0.847797690339864
REACTOME_ABC_TRANSPORTERS_IN_LIPID_HOMEOSTASIS	11	0.01	Down
	0.15617547298272	0.692982283180088
0.847797690339864
LEE_LIVER_CANCER_HEPATOBLAST	16	0.01	Down
0.709858278210839	0.693016577043616	0.847797690339864
WATANABE_COLON_CANCER_MSI_VS_MSS_DN	38	0.01	Down
0.425565047889789	0.693040850608583	0.847797690339864
LEE_LIVER_CANCER_SURVIVAL_DN	169	0.01	Down
0.190558889296682	0.693138267408009	0.847797690339864
REACTOME_HATS_ACETYLATE_HISTONES	96	0.01	Down
0.154166643669829	0.694026584430436	0.848627253573192
GAUSSMANN_MLL_AF4_FUSION_TARGETS_A_DN	64	0.01	Up
0.433626847081506	0.694194084847303	0.848627253573192
PID_ANTHRAX_PATHWAY	15	0.01	Down	0.432051511852387

0.694726032321951	0.849046632758883
BRUINS_UVC_RESPONSE_VIA_TP53_GROUP_A	577	0.01	Down
0.388604647147952	0.69604376058477	0.850425850456249
REACTOME_G_PROTEIN_BETA:GAMMA_SIGNALLING	23	0.01	Up
0.347040530064099	0.696784553292362
0.851099610611186
REACTOME_PLASMA_LIPOPROTEIN_REMODELING	13	0.01	Up
0.36109635860469	0.697967460116736	0.852142743541687
BAKER_HEMATOPOIESIS_STAT3_TARGETS	16	0.01	Down
0.591084939973407	0.698017704498441	0.852142743541687
KAYO_CALORIE_RESTRICTION_MUSCLE_UP	76	0.01	Up
0.592850526991253	0.698750563710575	0.852477535866192
MCBRYAN_PUBERTAL_BREAST_4_5WK_DN	161	0.01	Up
0.306349764410063	0.698939029689451	0.852477535866192
WANG_CISPLATIN_RESPONSE_AND_XPC_DN	201	0.01	Down
0.278488584974847	0.699027782192799	0.852477535866192
MATZUK_MEIOTIC_AND_DNA_REPAIR	27	0.01	Up
0.186858542614158	0.699050544427761	0.852477535866192
MEISSNER_NPC_HCP_WITH_H3K4ME3_AND_H3K27ME3	82	0.01	Up
0.498449354008939	0.699797778748033
0.852847301146591
REACTOME_NEURONAL_SYSTEM	202	0.01	Down
0.336828417454406	0.700161475940915	0.852847301146591
KEGG_SELENOAMINO_ACID_METABOLISM	25	0.01	Up
0.222493548110638	0.700233726759509	0.852847301146591
TONKS_TARGETS_OF_RUNX1_RUNX1T1_FUSION_MONOCYTE_DN	41	0.01
Up	0.347847194262342	0.700290705701428
0.852847301146591
AMUNDSON_DNA_DAMAGE_RESPONSE_TP53	15	0.01	Up
0.28421942559534	0.700334766548761	0.852847301146591
CHIARADONNA_NEOPLASTIC_TRANSFORMATION_KRAS_UP	114	0.01	Down
0.384047072636949	0.700757428268927
0.852847301146591
HOLLERN_MICROACINAR_BREAST_TUMOR_DN	15	0.01	Up
0.457857565805766	0.700777639020791	0.852847301146591
BERENJENO_ROCK_SIGNALING_NOT_VIA_RHOA_UP	21	0.01	Up
0.48775065058347	0.700871619674195
0.852847301146591
KEGG_NEUROACTIVE_LIGAND_RECEPTOR_INTERACTION	63	0.01	Up
0.52270373097517	0.701487509953676
0.853299361315414
TOMLINS_PROSTATE_CANCER_UP	40	0.01	Down
0.258359485214133	0.701622789637769	0.853299361315414
PILON_KLF1_TARGETS_UP	380	0.01	Down	0.417052859306338
0.702146098502296	0.853704818168196
LUND_SILENCED_BY_METHYLATION	15	0.01	Up
0.305248291454985	0.702975425208699	0.854482027126312
KEGG_ABC_TRANSPORTERS	27	0.01	Up	0.294755738238053
0.703627290419655	0.854799477304745
BARIS_THYROID_CANCER_UP	22	0.01	Down	0.198713142786994
0.703730933149926	0.854799477304745
BHAT_ESR1_TARGETS_NOT_VIA_AKT1_DN	84	0.01	Up
0.361095482948524	0.70380708910008	0.854799477304745
FERRANDO_TAL1_NEIGHBORS	13	0.01	Down	0.83903332043303

0.704015975283773 0.854822206618195
DAWSON_METHYLATED_IN_LYMPHOMA_TCL1 33 0.01 Up
0.404634532584979 0.704890691681216 0.855653162059699
ACEVEDO_LIVER_TUMOR_VS_NORMAL_ADJACENT_TISSUE_UP 821 0.01
Up 0.213576768126156 0.705789255788113
0.856512609278501
PLASARI_NFIC_TARGETS_BASAL_DN 12 0.01 Up
0.581501864064668 0.706492956150483 0.857135178919412
VARELA_ZMPSTE24_TARGETS_DN 28 0.01 Down
0.492307696454731 0.706873774529605 0.85736578966826
GAVIN_FOXP3_TARGETS_CLUSTER_P2 62 0.01 Up
0.524287416921309 0.707167815363956 0.857491052080114
BACOLOD_RESISTANCE_TO_ALKYLATING_AGENTS_DN 48 0.01 Up
0.256899085948158 0.707419906468349
0.857565393628703
ELVIDGE_HIF1A_AND_HIF2A_TARGETS_UP 39 0.01 Down
0.196643537186161 0.707744921148995 0.857605940614601
WNT_SIGNALING 71 0.01 Down 0.307719726147083
0.707834936525065 0.857605940614601
VALK_AML_CLUSTER_15 26 0.01 Up 0.377923548971535
0.708483009473541 0.858159829583285
PID_INTEGRIN_A4B1_PATHWAY 30 0.01 Down
0.456476819719967 0.709359131254851 0.858576635579556
BROWNE_HCMV_INFECTION_8HR_DN 36 0.01 Up
0.323477256992642 0.709448993741556 0.858576635579556
REACTOME_SIGNALING_BY_RECEPTOR_TYROSINE_KINASES 396 0.01 Down
0.39577357921899 0.709543957418591
0.858576635579556
REACTOME_GLUTAMATE_NEUROTRANSMITTER_RELEASE_CYCLE 16 0.01
Down 0.333632884056592 0.709591145979544
0.858576635579556
LOPEZ_MESOTHELIOMA_SURVIVAL_OVERALL_DN 11 0.01 Down
0.385342493319448 0.709826309069931 0.858630048242556
TANAKA_METHYLATED_IN_ESOPHAGEAL_CARCINOMA 65 0.01 Down
0.355591762730966 0.710496884673297
0.85904628330771
REACTOME_CYCLIN_A_B1_B2_ASSOCIATED_EVENTS_DURING_G2_M_TRANSITION
23 0.01 Down 0.259863585042178
0.710552632110805 0.85904628330771
KYNG_ENVIRONMENTAL_STRESS_RESPONSE_NOT_BY_4NQO_IN_WS 35 0.01
Down 0.316265065243105 0.710901899761456
0.859237440018216
REACTOME_G2_M_CHECKPOINTS 136 0.01 Down
0.142681649050227 0.711130141523012 0.859282254340306
REACTOME_SIGNALING_BY_NTRK3_TRKC 15 0.01 Up
0.326067349894764 0.711337764727133 0.859302137180452
GRAESSMANN_APOPTOSIS_BY_SERUM_DEPRIVATION_UP 456 0.01 Down
0.362581287696508 0.712076945992826
0.859783568590013
PID_BARD1_PATHWAY 29 0.01 Up 0.189803375377139
0.712118848912262 0.859783568590013
JEPSEN_SMRT_TARGETS 29 0.01 Down 0.620819334121118
0.712349135270941 0.859830656026552
REACTOME_FANCONI_ANEMIA_PATHWAY 36 0.01 Up

0.152445773622677	0.71274452350793	0.860076948501516
SILIGAN_TARGETS_OF_EWS_FLI1_FUSION_DN	16	0.01	Up
0.272709212143054	0.714113476801507	0.861497605534829
BIOCARTA_ATM_PATHWAY	19	0.01	Down	0.188712318011446
0.714321608442565	0.861517475167515
TORCHIA_TARGETS_OF_EWSR1_FLI1_FUSION_UP 216	0.01	Down
0.355419444541541	0.714652326611696	0.861685141662976
FARMER_BREAST_CANCER_APOCRINE_VS_LUMINAL	288	0.01	Down
0.383982188683631	0.714933113555735
0.861792530285071
REACTOME_PROTEIN_METHYLATION	17	0.01	Down
0.131460683246627	0.715262337648676	0.861896404457278
REACTOME_REGULATION_OF_GLYCOLYSIS_BY_FRUCTOSE_2_6_BISPHOSPHATE_METAB
OLISM	10	0.01	Up	0.2283977341686 0.715620329207028
0.861896404457278
NUYTTEN_EZH2_TARGETS_DN 913	0.01	Up	0.217672039311647
0.715766276812016	0.861896404457278
REACTOME_EARLY_PHASE_OF_HIV_LIFE_CYCLE	14	0.01	Up
0.35383635285218	0.71590480652984	0.861896404457278
MA_MYELOID_DIFFERENTIATION_DN	30	0.01	Down
0.395845760476082	0.715978014292208	0.861896404457278
CONCANNON_APOPTOSIS_BY_EPOXOMICIN_UP	221	0.01	Down
0.311415182659203	0.716715452660777	0.862375588600181
MARKS_HDAC_TARGETS_UP	20	0.01	Up	0.172406153315396
0.716759777310406	0.862375588600181
RAHMAN_TP53_TARGETS_PHOSPHORYLATED	21	0.01	Down
0.15649208569173	0.717221572941292	0.862610937940503
MATHEW_FANCONI_ANEMIA_GENES	10	0.01	Down
0.256712618508031	0.717339196000356	0.862610937940503
SUBTIL_PROGESTIN_TARGETS	34	0.01	Down
0.262053062817722	0.717633506478091	0.862734049644028
REACTOME_ACTIVATION_OF_MATRIX_METALLOPROTEINASES	25	0.01
Up	0.896108636239131	0.718407778181153
0.863412274727701
REACTOME_IRE1ALPHA_ACTIVATES_CHAPERONES 48	0.01	Down
0.164939878401801	0.718942830263308	0.863412274727701
OUYANG_PROSTATE_CANCER_PROGRESSION_DN	20	0.01	Up
0.205753437653809	0.718958168793702	0.863412274727701
REACTOME_CD28_DEPENDENT_VAV1_PATHWAY	10	0.01	Up
0.43556877740268	0.719013192236878	0.863412274727701
LEE_CALORIE_RESTRICTION_MUSCLE_UP	26	0.01	Down
0.269477742256152	0.719158077103562	0.863412274727701
ONGUSAHA_TP53_TARGETS	30	0.01	Up	0.263652111416926
0.720686780857458	0.865016576756816
REACTOME_METABOLISM_OF_NUCLEOTIDES	86	0.01	Up
0.264948861032201	0.72096113318433	0.865114867502285
REACTOME_PROCESSING_OF_SMDT1	16	0.01	Up
0.0859160421800522	0.721643146403855	0.865438632077171
MEINHOLD_OVARIAN_CANCER_LOW_GRADE_UP	12	0.01	Down
0.253744366174183	0.721728362062404	0.865438632077171
KEGG_N_GLYCAN_BIOSYNTHESIS	46	0.01	Down
0.186161784596971	0.721894687775337	0.865438632077171
CHIARETTI_ACUTE_LYMPHOBLASTIC_LEUKEMIA_ZAP70	55	0.01	Up
0.256684872764631	0.722001083490409

0.865438632077171
BYSTRYKH_HEMATOPOIESIS_STEM_CELL_QTL_TRANS 596 0.01 Down
0.264230817930677 0.722449157381171
0.865744858018759
KAYO_AGING_MUSCLE_DN 108 0.01 Down 0.227268888333329
0.722659657010487 0.865766300176476
REACTOME_SIGNALING_BY_NOTCH3 42 0.01 Down
0.266560284750208 0.723354526900105 0.86605131707818
FIGUEROA_AML_METHYLATION_CLUSTER_3_UP 130 0.01 Down
0.233495375371138 0.723436216070916 0.86605131707818
TAYLOR_METHYLATED_IN_ACUTE_LYMPHOBLASTIC_LEUKEMIA 65 0.01
Up 0.264081074055287 0.723475571886222
0.86605131707818
RAY_TUMORIGENESIS_BY_ERBB2_CDC25A_UP 133 0.01 Up
0.380124088465771 0.724266523784028 0.866171800195689
STEIN_ESRRA_TARGETS_RESPONSIVE_TO_ESTROGEN_DN 35 0.01 Up
0.22486157040429 0.724301664247534
0.866171800195689
ZHANG_GATA6_TARGETS_DN 58 0.01 Up 0.486447274320201
0.724316573949505 0.866171800195689
HU_GENOTOXIN_ACTION_DIRECT_VS_INDIRECT_24HR 46 0.01 Down
0.193793230865239 0.724347007104693
0.866171800195689
REACTOME_L1CAM_INTERACTIONS 88 0.01 Down
0.308791996516231 0.724630121644448 0.866279892763775
REACTOME_BASE_EXCISION_REPAIR_AP_SITE_FORMATION 31 0.01 Up
0.122772626836177 0.724843768840034
0.866291931977233
ONKEN_UVEAL_MELANOMA_UP 761 0.01 Down 0.27540082746773
0.725025639176496 0.866291931977233
PID_A6B1_A6B4_INTEGRIN_PATHWAY 44 0.01 Down
0.371455652448152 0.725476078093528 0.866599779705132
BROWNE_HCMV_INFECTION_30MIN_UP 33 0.01 Up
0.594921148675884 0.72570379524586 0.866641487680696
REACTOME_SUMOYLATION_OF_INTRACELLULAR_RECEPTORS 25 0.01 Up
0.330718087314156 0.726758999013422
0.867671102407791
KEGG_BASAL_TRANSCRIPTION_FACTORS 32 0.01 Down
0.161477197898226 0.728040655637557 0.868970458600855
BENPORATH_SOX2_TARGETS 683 0.01 Down 0.254514388173832
0.728552966284568 0.869351097278772
REACTOME_TRP_CHANNELS 17 0.01 Up 0.589341447964027
0.729511747848103 0.870264147180792
DELACROIX_RAR_TARGETS_DN 21 0.01 Up
0.397969158282548 0.730060499958271 0.870687701595232
WALLACE_PROSTATE_CANCER_RACE_DN 79 0.01 Down
0.258386558029086 0.730268495850295 0.870704745052274
CHEOK_RESPONSE_TO_HD_MTX_UP 15 0.01 Up
0.939311440855877 0.730656827283073 0.870936737904379
BIOCARTA_G2_PATHWAY 21 0.01 Up 0.20631103520725
0.731472459579132 0.871602813861408
REACTOME_GLUCAGON_LIKE_PEPTIDE_1_GLP1_REGULATES_INSULIN_SECRETION
25 0.01 Down 0.284991528611208
0.731603429743959 0.871602813861408

BERNARD_PPAPDC1B_TARGETS_DN	50	0.01	Down
0.431289660751643	0.732075711045229	0.871934372323344
DAZARD_UV_RESPONSE_CLUSTER_G5	10	0.01	Up
0.100028220235743	0.732331456541739	0.87200791977619
LIU_PROSTATE_CANCER_UP	82	0.01	Up	0.277050958426694
0.732843555734251	0.872386595080894
TURASHVILI_BREAST_LOBULAR_CARCINOMA_VS_LOBULAR_NORMAL_UP	80
0.01	Down	0.475183426473994	0.733516872859597
0.872956935002353
MARIADASON_RESPONSE_TO_CURCUMIN_SULINDAC_5	22	0.01	Down
0.218612027016189	0.733721777071657
0.872969663297274
PURBEY_TARGETS_OF_CTBP1_NOT_SATB1_DN	320	0.01	Down
0.344918231141033	0.734804249001314	0.874026223673169
REACTOME_PROCESSING_AND_ACTIVATION_OF_SUMO	10	0.01	Down
0.0811415309105412	0.735956768038788
0.875165521781574
FARMER_BREAST_CANCER_CLUSTER_7	19	0.01	Down
0.463568450216077	0.73710761505161	0.876147741931738
DURAND_STROMA_NS_UP	144	0.01	Up	0.444209714127366
0.737297592721415	0.876147741931738
RASHI_RESPONSE_TO_IONIZING_RADIATION_5	110	0.01	Down
0.276773404966417	0.737367498938324	0.876147741931738
BLALOCK_ALZHEIMERS_DISEASE_INCIPIENT_DN	150	0.01	Up
0.243522949080726	0.737846598114663	0.876485322020458
GRAHAM_CML_QUIESCENT_VS_NORMAL_QUIESCENT_DN	36	0.01	Up
0.754719973454625	0.738111626669411
0.876568497193924
YOSHIMURA_MAPK8_TARGETS_UP	657	0.01	Down
0.4396356712019	0.738496412161011	0.87679381211403 7
REACTOME_MICROAUTOPHAGY	26	0.01	Down	0.23412769849688
0.738705666475595	0.876810660366464
ZIRN_TRETINOIN_RESPONSE_UP	18	0.01	Up
0.657493717561091	0.739103689210967	0.877051500264862
LEE_LIVER_CANCER_MYC_E2F1_DN	37	0.01	Up
0.50738970623536	0.739512841167941	0.877284371084621
VETTER_TARGETS_OF_PRKCA_AND_ETS1_DN	13	0.01	Down
0.565954952934426	0.739823251875878	0.877284371084621
REACTOME_INSULIN_RECEPTOR_RECYCLING	21	0.01	Down
0.177027866697421	0.740104306326731	0.877284371084621
REACTOME_ORGANELLE_BIOGENESIS_AND_MAINTENANCE	270	0.01	Up
0.175597723814539	0.740190947306398
0.877284371084621
CHEN_NEUROBLASTOMA_COPY_NUMBER_GAINS	40	0.01	Down
0.342144033738337	0.740441778263761	0.877284371084621
KEGG_CELL_ADHESION_MOLECULES_CAMS	91	0.01	Up
0.612780464895246	0.740512482202652	0.877284371084621
NAKAMURA_ADIPOGENESIS_LATE_UP	90	0.01	Down
0.429333595182486	0.740677507957965	0.877284371084621
SMID_BREAST_CANCER_RELAPSE_IN_LUNG_DN	15	0.01	Up
0.620434895559403	0.741058073822884	0.877284371084621
REACTOME_RECEPTOR_MEDIATED_MITOPHAGY	11	0.01	Down
0.107408338543416	0.741421522950247	0.877284371084621
REACTOME_GLYCOGEN_METABOLISM	22	0.01	Up

0.172615149767654	0.741556244753433	0.877284371084621
RICKMAN_TUMOR_DIFFERENTIATED_MODERATELY_VS_POORLY_DN	14	0.01
	Up	0.183755490959815	0.741772188583841
0.877284371084621
BIOCARTA_P27_PATHWAY	13	0.01	Up	0.105284417408721
	0.742023742753411	0.877284371084621
PID_P38_ALPHA_BETA_PATHWAY	28	0.01	Down
0.411950090508158	0.742129008857138	0.877284371084621
KEGG_LONG_TERM_DEPRESSION	48	0.01	Down
0.42367752076873	0.742301061686806	0.877284371084621
KEGG_ETHER_LIPID_METABOLISM	24	0.01	Up
0.377819665085314	0.742353225542978	0.877284371084621
REACTOME_GLUCAGON_SIGNALING_IN_METABOLIC_REGULATION	21	0.01
	Down	0.306377423739001	0.742700669921704
0.877284371084621
KEGG_CELL_CYCLE 114	0.01	Up	0.210834111625466
0.742910891928826	0.877284371084621
ROZANOV_MMP14_CORRELATED	10	0.01	Down
0.439338608991921	0.7429697489649 0.877284371084621
REACTOME_PURINE_SALVAGE 11	0.01	Down	0.261872286138669
	0.743008142540412	0.877284371084621
REACTOME_MACROAUTOPHAGY 87	0.01	Down	0.116054230956793
	0.743237205804354	0.877299291084095
RUAN_RESPONSE_TO_TROGLITAZONE_DN	13	0.01	Up
0.492441058646412	0.743411123412529	0.877299291084095
PID_ATM_PATHWAY 33	0.01	Up	0.148371414619476
0.744509859990792	0.878365307259478
SMIRNOV_RESPONSE_TO_IR_2HR_DN	49	0.01	Down
0.271353035086062	0.744925522492007	0.878625091472466
LOPEZ_MBD_TARGETS	858	0.01	Up	0.245281020777565
	0.745594472908791	0.879183409161862
REACTOME_NOTCH3_ACTIVATION_AND_TRANSMISSION_OF_SIGNAL_TO_THE_NUCLEUS
	24	0.01	Up	0.23812518576025
0.746054280137008	0.879494883088343
OZANNE_AP1_TARGETS_UP	15	0.01	Down	0.760888521224987
	0.746904864403859	0.880266745017134
KIM_LIVER_CANCER_POOR_SURVIVAL_UP	22	0.01	Down
0.381735179810719	0.747231378370945	0.880420719731952
MEISSNER_BRAIN_ICP_WITH_H3K4ME3 24	0.01	Up
0.251424294972082	0.747992081022826	0.880935563820675
LOPES_METHYLATED_IN_COLON_CANCER_DN	21	0.01	Down
0.402898285176016	0.748060299689325	0.880935563820675
LEIN_MEDULLA_MARKERS	61	0.01	Up	0.324610148633414
	0.748518375500109	0.881070654336323
TONKS_TARGETS_OF_RUNX1_RUNX1T1_FUSION_SUSTAINED_IN_MONOCYTE_UP	16
	0.01	Up	0.270588732124233	0.748665223118218
	0.881070654336323
FUJII_YBX1_TARGETS_UP	38	0.01	Up	0.527272924614345
	0.748763047734095	0.881070654336323
REACTOME_REGULATION_OF_TP53_ACTIVITY_THROUGH_ASSOCIATION_WITH_CO_FAC
TORS	11	0.01	Down	0.183201039928977
0.749795146948137	0.881904516852299
STEGMEIER_PREMITOTIC_CELL_CYCLE_REGULATORS	10	0.01	Down
	0.19467558120329	0.749950339579289

0.881904516852299
SANSOM_APC_TARGETS_UP 104 0.01 Up 0.277928224098924
0.75013227544453 0.881904516852299
REACTOME_TERMINATION_OF_TRANSLESION_DNA_SYNTHESIS 31 0.01
Up 0.121290388646604 0.750256478852768
0.881904516852299
FISCHER_DREAM_TARGETS 866 0.01 Down 0.185373511375425
0.75071565179825 0.88221355681912
HUMMERICH_SKIN_CANCER_PROGRESSION_UP 73 0.01 Up
0.556456896055128 0.751799738020162 0.88325661850513
IVANOVA_HEMATOPOIESIS_STEM_CELL 136 0.01 Down
0.409349515227524 0.752579375935219 0.883941545552342
LI_CISPLATIN_RESISTANCE_UP 19 0.01 Down
0.471299260341585 0.753492107728151 0.88466645114223
REACTOME_POST_TRANSLATIONAL_MODIFICATION:_SYNTHESIS_OF_GPI_ANCHORED_
PROTEINS 57 0.01 Down 0.575033190607916
0.753642472127692 0.88466645114223
BYSTROEM_CORRELATED_WITH_IL5_UP 29 0.01 Down
0.22563272053263 0.753786987291377 0.88466645114223
RUIZ_TNC_TARGETS_DN 125 0.01 Up 0.316647573242789
0.754657525245281 0.885456950137702
TORCHIA_TARGETS_OF_EWSR1_FLI1_FUSION_TOP20_DN 15 0.01 Up
0.239109434548461 0.756155726810691
0.886871436601444
AMIT_SERUM_RESPONSE_20_MCF10A 15 0.01 Down
0.431922115511452 0.756275162243635 0.886871436601444
PID_RB_1PATHWAY 57 0.01 Down 0.288069860399684
0.756454969506103 0.886871436601444
WANG_RESPONSE_TO_BEXAROTENE_UP 24 0.01 Down
0.240034326194613 0.756870076939948 0.887010363151483
DUTERTRE_ESTRADIOL_RESPONSE_6HR_UP 201 0.01 Down
0.300598445056821 0.756997105190653 0.887010363151483
REACTOME_SLC_TRANSPORTER_DISORDERS 67 0.01 Down
0.34449858891519 0.757166785079932 0.887010363151483
REACTOME_RHO_GTPASES_ACTIVATE_NADPH_OXIDASES 20 0.01 Down
0.588742060694747 0.757362797280399
0.887010363151483
REACTOME_INOSITOL_PHOSPHATE_METABOLISM 41 0.01 Up
0.271524779713495 0.758168450604263 0.887449154471441
TERAO_AOX4_TARGETS_SKIN_DN 15 0.01 Down
0.242582428155393 0.758291656084575 0.887449154471441
JOSEPH_RESPONSE_TO_SODIUM_BUTYRATE_DN 46 0.01 Up
0.45629253629465 0.75833361881904 0.887449154471441
REACTOME_ION_CHANNEL_TRANSPORT 121 0.01 Down
0.328304299331886 0.758527174967581 0.887449154471441
BOYLAN_MULTIPLE_MYELOMA_C_D_UP 115 0.01 Up
0.274291270193566 0.758966470579235 0.887732054450602
GUILLAUMOND_KLF10_TARGETS_DN 23 0.01 Down
0.4812423344911 0.759592412100104 0.887914039321695
YAO_HOXA10_TARGETS_VIA_PROGESTERONE_UP 57 0.01 Up
0.53234312560747 0.759680217589512 0.887914039321695
DARWICHE_SQUAMOUS_CELL_CARCINOMA_DN 133 0.01 Up
0.303606944809154 0.759934497997185 0.887914039321695
BIOCARTA_AGR_PATHWAY 26 0.01 Down 0.289852341874088

0.760076295233025	0.887914039321695
PID_CMYB_PATHWAY	69	0.01	Down	0.420067611246518
0.760163348142772	0.887914039321695
KEGG_REGULATION_OF_ACTIN_CYTOSKELETON	170	0.01	Down
0.340664286626842	0.760381670666992	0.887914039321695
AMIT_EGF_RESPONSE_20_MCF10A	12	0.01	Down
0.130598088781977	0.760509007538193	0.887914039321695
YAUCH_HEDGEHOG_SIGNALING_PARACRINE_DN	109	0.01	Up
0.482787658460002	0.760702328237564	0.887914039321695
PID_ILK_PATHWAY 43	0.01	Down	0.218265904024557
0.761084127193999	0.888129063275448
TURASHVILI_BREAST_DUCTAL_CARCINOMA_VS_LOBULAR_NORMAL_DN 51	0.01
Down	0.504733393929056	0.761578103733227
0.888474844609617
TSAI_RESPONSE_TO_RADIATION_THERAPY	29	0.01	Down
0.684352396947892	0.761940015879605	0.888666417067676
ROVERSI_GLIOMA_COPY_NUMBER_DN	38	0.01	Up
0.443528000810583	0.762284907222614	0.888838043570856
MYLLYKANGAS_AMPLIFICATION_HOT_SPOT_22	11	0.01	Down
0.224976304678698	0.762660824255619	0.88904574819217
REACTOME_G2_M_DNA_DAMAGE_CHECKPOINT	68	0.01	Up
0.15375201625397	0.762932180458737	0.889131488504543
WANG_RESPONSE_TO_GSK3_INHIBITOR_SB216763_UP	314	0.01	Up
0.312619059571804	0.763135338298603
0.8891377256745
REACTOME_RNA_POLYMERASE_III_CHAIN_ELONGATION	18	0.01	Up
0.138979188391647	0.763590384682254
0.889437361789772
SEKI_INFLAMMATORY_RESPONSE_LPS_DN	19	0.01	Down
0.550482240660522	0.763932216956207	0.889587853821263
REACTOME_TRNA_PROCESSING	108	0.01	Down
0.148393841798564	0.764186330597495	0.889587853821263
REACTOME_TRANSLESION_SYNTHESIS_BY_Y_FAMILY_DNA_POLYMERASES_BYPASSES_
LESIONS_ON_DNA_TEMPLATE 38	0.01	Up	0.135298778759182
0.764397211083619	0.889587853821263
REACTOME_DNA_DAMAGE_TELOMERE_STRESS_INDUCED_SENESCENCE	44	0.01
Down	0.159096424666249	0.764531550698642
0.889587853821263
HAN_JNK_SINGALING_DN	32	0.01	Up	0.675126556972146
0.764709113941126	0.889587853821263
MATSUDA_NATURAL_KILLER_DIFFERENTIATION	380	0.01	Up
0.350046531779736	0.765010391790287	0.889708075316258
BENPORATH_PROLIFERATION 129	0.01	Up	0.226986178367855
0.765288214095479	0.889800962845106
LIN_TUMOR_ESCAPE_FROM_IMMUNE_ATTACK	13	0.01	Up
0.548943855217865	0.765603913308622	0.889937830442787
NUMATA_CSF3_SIGNALING_VIA_STAT3 18	0.01	Up
0.263018833175353	0.7661781166515 0.890368094424343
REACTOME_PLATELET_CALCIUM_HOMEOSTASIS	16	0.01	Up
0.438199114321372	0.766477799697274	0.890368094424343
FUKUSHIMA_TNFSF11_TARGETS	11	0.01	Down
0.367036917817662	0.766568303764673	0.890368094424343
CHYLA_CBFA2T3_TARGETS_DN	175	0.01	Down
0.386655413522899	0.76698452480494	0.890528919562632

BAE_BRCA1_TARGETS_UP	72	0.01	Down	0.314433717007341
	0.767102998119358	0.890528919562632
FIGUEROA_AML_METHYLATION_CLUSTER_6_DN	19	0.01	Down
0.39545712926741	0.768180722720406	0.891549793087587
REACTOME_EPHRIN_SIGNALING	18	0.01	Down
0.234542693584855	0.768645644883325	0.891859105253109
NAGY_TFTC_COMPONENTS_HUMAN	19	0.01	Down
0.18998992244342	0.768891836289406	0.891864927982999
GARGALOVIC_RESPONSE_TO_OXIDIZED_PHOSPHOLIPIDS_MAGENTA_DN	12
	0.01	Down	0.140317554406017	0.769047488512148
	0.891864927982999
PID_MYC_ACTIV_PATHWAY	72	0.01	Down	0.248547771687617
	0.769456233385097	0.892074456276091
BENPORATH_NOS_TARGETS	158	0.01	Down	0.34261318365722
	0.770136080350272	0.892074456276091
MCCLUNG_DELTA_FOSB_TARGETS_8WK	35	0.01	Down
0.248931499382831	0.77026648800703	0.892074456276091
REACTOME_JOSEPHIN_DOMAIN_DUBS	10	0.01	Down
0.108304972990577	0.770321910799409	0.892074456276091
BONOME_OVARIAN_CANCER_POOR_SURVIVAL_DN	20	0.01	Down
0.134683680982498	0.770336924079924	0.892074456276091
MCCABE_BOUND_BY_HOXC6	292	0.01	Down	0.309257685039634
	0.770717023846028	0.892074456276091
WILLIAMS_ESR2_TARGETS_UP	27	0.01	Down
0.360568271004752	0.7709233955021 0.892074456276091
WEST_ADRENOCORTICAL_CARCINOMA_VS_ADENOMA_UP	15	0.01	Up
	0.187284776653692	0.770966277023776
0.892074456276091
ACEVEDO_METHYLATED_IN_LIVER_CANCER_DN	446	0.01	Down
0.357979481853433	0.771014296469992	0.892074456276091
BOYLAN_MULTIPLE_MYELOMA_PCA3_DN 51	0.01	Down
0.345869675460071	0.771506058478944	0.892413724359972
YANG_BREAST_CANCER_ESR1_BULK_UP 24	0.01	Down
0.229838593274454	0.772520555788458	0.892909712839039
KAYO_CALORIE_RESTRICTION_MUSCLE_DN	70	0.01	Up
0.241737872933774	0.772530564922821	0.892909712839039
MOOTHA_TCA	15	0.01	Down	0.0791906922419629
0.772530783811128	0.892909712839039
KIM_ALL_DISORDERS_CALB1_CORR_UP 478	0.01	Down
0.217772426647125	0.773039766904041	0.893268316769579
KIM_RESPONSE_TO_TSA_AND_DECITABINE_UP	102	0.01	Down
0.515188693484964	0.773305871986408	0.893346156406812
YAO_TEMPORAL_RESPONSE_TO_PROGESTERONE_CLUSTER_9 66	0.01	Down
	0.312420420058996	0.773553126542865
0.893402184946088
KOBAYASHI_EGFR_SIGNALING_24HR_DN	213	0.01	Up
0.279618365782693	0.774141972664	0.893416316168119
HOWLIN_PUBERTAL_MAMMARY_GLAND	45	0.01	Up
0.775227327594623	0.774331744975134	0.893416316168119
MARSON_FOXP3_CORE_DIRECT_TARGETS	13	0.01	Down
0.6312157782925 0.774353545700976	0.893416316168119
REACTOME_NEGATIVE_REGULATION_OF_TCF_DEPENDENT_SIGNALING_BY_WNT_LIGAN
D_ANTAGONISTS	10	0.01	Up	0.684296121335843
0.77436039327942	0.893416316168119

MATZUK_SPERMATOZOA	69	0.01	Down	0.337922087860643
	0.776479681369064	0.895166162847768
ODONNELL_METASTASIS_UP	47	0.01	Down	0.348897431561819
	0.777520162996537	0.895166162847768
FLORIO_NEOCORTEX_BASAL_RADIAL_GLIA_UP	183	0.01	Down
0.559730845446369	0.777693963629402	0.895166162847768
REACTOME_SYNTHESIS_OF_IP2_IP_AND_INS_IN_THE_CYTOSOL	12	0.01
	Up	0.182435916556172	0.777724748639684
0.895166162847768
REACTOME_REGULATION_OF_GENE_EXPRESSION_IN_LATE_STAGE_BRANCHING_MORPH
OGENESIS_PANCREATIC_BUD_PRECURSOR_CELLS 13	0.01	Down
0.285814961444115	0.777896600766022	0.895166162847768
WANG_CLIM2_TARGETS_UP	219	0.01	Down	0.260862676751029
	0.777935498374842	0.895166162847768
MORI_PLASMA_CELL_DN	30	0.01	Down	0.388271540453069
	0.77797122545239	0.895166162847768
REN_ALVEOLAR_RHABDOMYOSARCOMA_UP	67	0.01	Down
0.422178408844584	0.778107608359458	0.895166162847768
ALCALA_APOPTOSIS	80	0.01	Up	0.184822294504567
	0.778112346683671	0.895166162847768
INAMURA_LUNG_CANCER_SCC_DN	10	0.01	Down
0.648207331646516	0.778165143902912	0.895166162847768
BIOCARTA_SKP2E2F_PATHWAY	10	0.01	Down
0.104658525554935	0.778191595768289	0.895166162847768
VANTVEER_BREAST_CANCER_BRCA1_DN 47	0.01	Down
0.486015230627448	0.77876323861043	0.895166162847768
GAUSSMANN_MLL_AF4_FUSION_TARGETS_F_DN	28	0.01	Down
0.479642065942103	0.77887853046599	0.895166162847768
EPPERT_HSC_R	103	0.01	Down	0.375137386633094
0.778936595736359	0.895166162847768
BIOCARTA_BARR_MAPK_PATHWAY	11	0.01	Down
0.223293674134324	0.779054705726795	0.895166162847768
PID_TXA2PATHWAY 44	0.01	Down	0.3979604192567
0.779063410247045	0.895166162847768
BIDUS_METASTASIS_DN	135	0.01	Up	0.391273490219845
	0.780871987450273	0.896772928425463
KEGG_ANTIGEN_PROCESSING_AND_PRESENTATION	54	0.01	Up
	0.53556561131732	0.7810351933206 0.896772928425463
BIOCARTA_IL12_PATHWAY	12	0.01	Down	0.589158581833219
	0.781240247152663	0.896772928425463
SCHLESINGER_METHYLATED_DE_NOVO_IN_CANCER	58	0.01	Up
	0.621936046097356	0.781302582845575
0.896772928425463
CHANGOLKAR_H2AFY_TARGETS_DN	34	0.01	Down
0.330835379328913	0.781517358757915	0.896772928425463
REACTOME_MEIOSIS	64	0.01	Up	0.154180511079974
	0.781848995548689	0.896772928425463
MARTINEZ_RB1_TARGETS_DN 441	0.01	Up	0.366817467364108
	0.781858310678396	0.896772928425463
LIU_SMARCA4_TARGETS	43	0.01	Up	0.609834932439865
	0.782123937079163	0.896848749278275
REACTOME_TP53_REGULATES_TRANSCRIPTION_OF_CELL_DEATH_GENES	40
	0.01	Up	0.261305705217197	0.782564217191309
	0.897007508836056

BECKER_TAMOXIFEN_RESISTANCE_UP 48 0.01 Down
0.359170611018849 0.782699397789521 0.897007508836056
CHUANG_OXIDATIVE_STRESS_RESPONSE_UP 28 0.01 Down
0.287947903000686 0.782861058323437 0.897007508836056
CHANG_POU5F1_TARGETS_UP 15 0.01 Up 0.723761729019152
0.783928330695089 0.897747982968347
REACTOME_DNA_METHYLATION 24 0.01 Up
0.135443334889761 0.78410440303436 0.897747982968347
HOSHIDA_LIVER_CANCER_SURVIVAL_UP 61 0.01 Down
0.629792031645188 0.784106469662676 0.897747982968347
SYED_ESTRADIOL_RESPONSE 17 0.01 Up 0.273707793155696
0.784926684078663 0.898458223818078
REACTOME_XENOBIOTICS 11 0.01 Up 0.661689511844672
0.787097237244157 0.900374585703935
PID_WNT_CANONICAL_PATHWAY 18 0.01 Down
0.162778194507596 0.787103167084501 0.900374585703935
REACTOME_TRANSPORT_OF_BILE_SALTS_AND_ORGANIC_ACIDS_METAL_IONS_AND_AMINE_COMPOUNDS 50 0.01 Up 0.524884864454952
0.787288780167726 0.900374585703935
GAZDA_DIAMOND_BLACKFAN_ANEMIA_PROGENITOR_UP 38 0.01 Up
0.363596232302964 0.787402112658992
0.900374585703935
CHANDRAN_METASTASIS_TOP50_DN 44 0.01 Down
0.299593033396921 0.787727125761574 0.900444182440607
LAMB_CCND1_TARGETS 18 0.01 Down 0.208511371780685
0.787863619474729 0.900444182440607
LIU_CMYB_TARGETS_UP 154 0.01 Down 0.30685201227537
0.788541735664106 0.900990112305581
BOYLAN_MULTIPLE_MYELOMA_C_CLUSTER_DN 22 0.01 Up
0.377017914565692 0.788778171838778 0.901031228059799
KYNG_ENVIRONMENTAL_STRESS_RESPONSE_UP 51 0.01 Up
0.426726250621591 0.789751306778801 0.901913649382802
PARK_HSC_MARKERS 36 0.01 Down 0.267394964408263
0.790210384744518 0.902154335908058
POMEROY_MEDULLOBLASTOMA_DESMOPLASIC_VS_CLASSIC_UP 50 0.01
Down 0.229719522585152 0.790363464917894
0.902154335908058
BARRIER_CANCER_RELAPSE_NORMAL_SAMPLE_DN 30 0.01 Down
0.150973078020873 0.790708319855906 0.902181051788253
RAMASWAMY_METASTASIS_UP 60 0.01 Up 0.238094241155104
0.790788285660893 0.902181051788253
REACTOME_DISEASES_ASSOCIATED_WITH_GLYCOSYLATION_PRECURSOR_BIOSYNTHESIS 18 0.01 Up 0.241609083809998
0.791755953914713 0.903055826654818
REACTOME_THE_NLRP3_INFLAMMASOME 13 0.01 Down
0.381818799538011 0.791985706200949 0.903088723838982
WESTON_VEGFA_TARGETS_3HR 52 0.01 Up
0.679317533039962 0.792325379389954 0.90324691360838
ZHANG_ADIPOGENESIS_BY_BMP7 14 0.01 Up
0.150812518873731 0.792549603986457 0.903273445719859
MATZUK_SPERMATOCYTE 51 0.01 Up 0.195619247625564
0.793164167218414 0.903634152909705
JIANG_AGING_CEREBRAL_CORTEX_DN 41 0.01 Down
0.20816240741464 5 0.793268157370789 0.903634152909705

CONRAD_STEM_CELL	27	0.01	Down	0.474383299831751
	0.794023186182305	0.904176804717671
STEIN_ESRRA_TARGETS_RESPONSIVE_TO_ESTROGEN_UP	27	0.01	Down
	0.208596520203446	0.794146835378279
0.904176804717671
WELCSH_BRCA1_TARGETS_DN 138	0.01	Up	0.177881649631319
	0.795144008050559	0.905082885841292
GENTILE_UV_RESPONSE_CLUSTER_D8	36	0.01	Up
0.307871723207586	0.795625586759735	0.905389937065732
PID_UPA_UPAR_PATHWAY	31	0.01	Down	0.617434726740508
	0.795914240752289	0.905389937065732
REACTOME_COOPERATION_OF_PDCL_PHLP1_AND_TRIC_CCT_IN_G_PROTEIN_BETA_FO
LDING	26	0.01	Up	0.181132215757407
0.79601802698193	0.905389937065732
LE_EGR2_TARGETS_DN	82	0.01	Up	0.310523703872099
	0.796407539373064	0.90560381722285
CHAUHAN_RESPONSE_TO_METHOXYESTRADIOL_DN 98	0.01	Up
0.221070263238581	0.796921868125434	0.90595948336465
KYNG_DNA_DAMAGE_UP	172	0.01	Down	0.33456046370787
	0.797986858644487	0.906859319175917
KASLER_HDAC7_TARGETS_1_UP	166	0.01	Up
0.250613767962808	0.798323834461295	0.906859319175917
REACTOME_MYOGENESIS	21	0.01	Down	0.262399726198368
	0.798641069048567	0.906859319175917
MAGRANGEAS_MULTIPLE_MYELOMA_IGLL_VS_IGLK_DN	20	0.01	Up
	0.295131700236254	0.798934098798496
0.906859319175917
REACTOME_G_ALPHA_Q_SIGNALLING_EVENTS	103	0.01	Down
0.4205555010503 0.799197482904234	0.906859319175917
REACTOME_ACTIVATED_PKN1_STIMULATES_TRANSCRIPTION_OF_AR_ANDROGEN_RECE
PTOR_REGULATED_GENES_KLK2_AND_KLK3	25	0.01	Up
0.158818672695776	0.799198744987769	0.906859319175917
REACTOME_ASSEMBLY_OF_THE_HIV_VIRION	16	0.01	Down
0.186208304899267	0.799203102492034	0.906859319175917
GAVIN_FOXP3_TARGETS_CLUSTER_P7	69	0.01	Down
0.393251991596135	0.79932739100667	0.906859319175917
WATANABE_RECTAL_CANCER_RADIOTHERAPY_RESPONSIVE_UP	105	0.01
	Down	0.185412045608484	0.799812609375521
0.907037790406114
PEART_HDAC_PROLIFERATION_CLUSTER_DN	68	0.01	Down
0.2624541353003 0.80026727990505	0.907037790406114
KYNG_WERNER_SYNDROM_DN	16	0.01	Up	0.345456773807121
	0.800281238973058	0.907037790406114
MENSE_HYPOXIA_UP	95	0.01	Down	0.29224523532928
	0.800430208710725	0.907037790406114
WANG_CISPLATIN_RESPONSE_AND_XPC_UP	144	0.01	Up
0.344737633553768	0.800493640609801	0.907037790406114
DANG_REGULATED_BY_MYC_UP	64	0.01	Up
0.140876301586646	0.800822953344622	0.907182251835704
REACTOME_CA2PLUS_PATHWAY	47	0.01	Down
0.327877896398163	0.801213420037501	0.907217125265524
COLLIS_PRKDC_REGULATORS 19	0.01	Up	0.19142429939232
	0.801257394283455	0.907217125265524
IIZUKA_LIVER_CANCER_PROGRESSION_L1_G1_DN	10	0.01	Up

0.170403142629372	0.801801339098732
0.90760438661516
MARTORIATI_MDM4_TARGETS_NEUROEPITHELIUM_UP	147	0.01	Up
0.302731913122925	0.802341509964719
0.90785164263865
CROONQUIST_IL6_DEPRIVATION_UP	12	0.01	Up
0.57760627666959	0.80242370994513	0.90785164263865
GARGALOVIC_RESPONSE_TO_OXIDIZED_PHOSPHOLIPIDS_GREY_DN	67	0.01
Up	0.339335452547848	0.802716949070773
0.907954878226755
WAKABAYASHI_ADIPOGENESIS_PPARG_RXRA_BOUND_8D	740	0.01	Up
0.241657037224428	0.80324890264822
0.908023594776798
PYEON_HPV_POSITIVE_TUMORS_UP	83	0.01	Down
0.297904854124368	0.803355459181705	0.908023594776798
GAUSSMANN_MLL_AF4_FUSION_TARGETS_C_DN	12	0.01	Up
0.273941656971008	0.80338372334312	0.908023594776798
REACTOME_MITOCHONDRIAL_CALCIUM_ION_TRANSPORT	22	0.01	Down
0.107485483878552	0.804457968949896
0.90900919316988
BROWN_MYELOID_CELL_DEVELOPMENT_DN	103	0.01	Down
0.280142739010913	0.805948943632539	0.910465067008863
PID_INTEGRIN5_PATHWAY	14	0.01	Down	0.636375565594513
0.806316043466612	0.910650908387542
GOBERT_OLIGODENDROCYTE_DIFFERENTIATION_DN	937	0.01	Up
0.295078844553555	0.807232654218127
0.911457116480905
MIKKELSEN_ES_ICP_WITH_H3K27ME3	14	0.01	Up
0.705282584103429	0.807484990409939	0.911513066773651
JAZAG_TGFB1_SIGNALING_VIA_SMAD4_UP	83	0.01	Up
0.31751023419982	0.807691896779044	0.911517719312529
GAZDA_DIAMOND_BLACKFAN_ANEMIA_MYELOID_UP	29	0.01	Down
0.297794493446557	0.808871100952461
0.912157587379443
BACOLOD_RESISTANCE_TO_ALKYLATING_AGENTS_UP	19	0.01	Down
0.447000365405513	0.809048363879825
0.912157587379443
PID_ARF_3PATHWAY	19	0.01	Down	0.186465061769046
0.809407979020103	0.912157587379443
REACTOME_MRNA_SPLICING_MINOR_PATHWAY	52	0.01	Up
0.0913182684887638	0.809446335248962	0.912157587379443
HEIDENBLAD_AMPLICON_8Q24_DN	37	0.01	Up
0.363651970877987	0.809595489808975	0.912157587379443
JISON_SICKLE_CELL_DISEASE_DN	170	0.01	Up
0.202291832157967	0.810180171347954	0.912157587379443
REACTOME_TBC_RABGAPS	42	0.01	Down	0.203313727869097
0.810209044773503	0.912157587379443
REACTOME_LDL_CLEARANCE	16	0.01	Up	0.231849769805227
0.810222012717709	0.912157587379443
REACTOME_SYNTHESIS_OF_PA	29	0.01	Down
0.253792657359846	0.810281664937039	0.912157587379443
JEON_SMAD6_TARGETS_DN	15	0.01	Up	0.213206385333241
0.810288152704364	0.912157587379443
BYSTRYKH_HEMATOPOIESIS_STEM_CELL_QTL_CIS	113	0.01	Up

0.216004706541146	0.810760142822275
0.912460401098179
ONO_FOXP3_TARGETS_DN	26	0.01	Down	0.411997512586164
0.811433935449382	0.912990122614511
ZHAN_V1_LATE_DIFFERENTIATION_GENES_UP	31	0.01	Up
0.418559890255095	0.811694585255607	0.913054845026014
BIOCARTA_PS1_PATHWAY	12	0.01	Down	0.253362010288358
0.813370754240459	0.914073003044752
REACTOME_RNA_POLYMERASE_II_TRANSCRIPTION_ELONGATION	57	0.01
Down	0.119255631989334	0.813396478652625
0.914073003044752
FOURNIER_ACINAR_DEVELOPMENT_EARLY_UP	20	0.01	Down
0.277397089044861	0.813445559993312	0.914073003044752
NEBEN_AML_WITH_FLT3_OR_NRAS_DN	12	0.01	Up
0.32952029485305	0.813719553750971	0.914073003044752
MATZUK_CENTRAL_FOR_FEMALE_FERTILITY	16	0.01	Up
0.720377560414759	0.813982007441398	0.914073003044752
REACTOME_TP53_REGULATES_TRANSCRIPTION_OF_GENES_INVOLVED_IN_G1_CELL_C
YCLE_ARREST	13	0.01	Up	0.289479900100281
0.813992000313688	0.914073003044752
PID_THROMBIN_PAR1_PATHWAY	34	0.01	Down
0.277742860675855	0.814023188250977	0.914073003044752
FIGUEROA_AML_METHYLATION_CLUSTER_2_UP	39	0.01	Down
0.237501956957239	0.814817005531117	0.914735874091501
PARK_TRETINOIN_RESPONSE_AND_PML_RARA_FUSION	28	0.01	Down
0.618300005863364	0.815659835986045
0.915453423909431
FARDIN_HYPOXIA_11	31	0.01	Up	0.292142750497273
0.816228918861646	0.915863452392186
LUI_THYROID_CANCER_PAX8_PPARG_UP	44	0.01	Down
0.401943135918121	0.817975338617821	0.917593997276543
MULLIGHAN_NPM1_MUTATED_SIGNATURE_1_UP	201	0.01	Up
0.28473465865337	0.819556002887633	0.919034458843793
REACTOME_REGULATION_OF_INNATE_IMMUNE_RESPONSES_TO_CYTOSOLIC_DNA 13
0.01	Down	0.155423856078094	0.81985120420146
0.919034458843793
MA_MYELOID_DIFFERENTIATION_UP	31	0.01	Down
0.287881555836138	0.819990672612856	0.919034458843793
HASINA_NOL7_TARGETS_UP	12	0.01	Down	0.438680959138307
0.820077244587865	0.919034458843793
VANTVEER_BREAST_CANCER_BRCA1_UP 35	0.01	Down
0.274282555023401	0.822631143946865	0.921257058164409
TOMLINS_PROSTATE_CANCER_DN	37	0.01	Up
0.435094362623012	0.822728623166908	0.921257058164409
KEGG_NON_HOMOLOGOUS_END_JOINING 12	0.01	Up
0.160269973741265	0.822851059554366	0.921257058164409
BALDWIN_PRKCI_TARGETS_UP	36	0.01	Down
0.383168043156884	0.822966425503932	0.921257058164409
DELYS_THYROID_CANCER_UP 395	0.01	Up	0.546480325829263
0.823085282667023	0.921257058164409
REACTOME_DAP12_INTERACTIONS	29	0.01	Down
0.428577770203465	0.823722830842151	0.921411892217
REACTOME_PROCESSING_OF_DNA_DOUBLE_STRAND_BREAK_ENDS	73	0.01
Up	0.146049594742582	0.823793850152643

0.921411892217
REACTOME_THROMBOXANE_SIGNALLING_THROUGH_TP_RECEPTOR 14 0.01 Down 0.254987212242391 0.824125719296014 0.921411892217
REACTOME_CONDENSATION_OF_PROPHASE_CHROMOSOMES 32 0.01 Up 0.155321651071566 0.824471071937866 0.921411892217
HU_ANGIOGENESIS_DN 35 0.01 Up 0.126352753943349 0.824560846214758 0.921411892217
BOYLAN_MULTIPLE_MYELOMA_C_UP 40 0.01 Up 0.216673853493577 0.824571968205928 0.921411892217
BENPORATH_ES_1 321 0.01 Up 0.28727756791103 0.82501280888289 0.921411892217
REACTOME_RHO_GTPASES_ACTIVATE_PAKS 20 0.01 Down 0.236163241133741 0.825026194276341 0.921411892217
FOURNIER_ACINAR_DEVELOPMENT_LATE_UP 10 0.01 Up 0.487055672486954 0.82523201008721 0.921411892217
JIANG_HYPOXIA_CANCER 55 0.01 Up 0.218016892437375 0.825342174299004 0.921411892217
PID_S1P_S1P2_PATHWAY 21 0.01 Down 0.238957144321887 0.825531170185839 0.921411892217
REACTOME_AXON_GUIDANCE 461 0.01 Down 0.302294978882354 0.825726413375445 0.921411892217
HOLLERN_SOLID_NODULAR_BREAST_TUMOR_DN 26 0.01 Down 0.804627652857986 0.826060386250793 0.921411892217
BERENJENO_TRANSFORMED_BY_RHOA_UP 507 0.01 Down 0.268147343791907 0.826446046449161 0.921411892217
LIN_MELANOMA_COPY_NUMBER_UP 63 0.01 Up 0.20520891638016 0.826453681404809 0.921411892217
REACTOME_SLBP_DEPENDENT_PROCESSING_OF_REPLICATION_DEPENDENT_HISTONE_PRE_MRNAS 11 0.01 Down 0.0522912617630307 0.826533001388295 0.921411892217
BOGNI_TREATMENT_RELATED_MYELOID_LEUKEMIA_DN 31 0.01 Down 0.160001285917706 0.826708378489691 0.921411892217
HELLEBREKERS_SILENCED_DURING_TUMOR_ANGIOGENESIS 68 0.01 Up 0.717744482337324 0.827055191826163 0.921525345601318
GENTILE_UV_RESPONSE_CLUSTER_D7 37 0.01 Down 0.214338039146347 0.827413063107672 0.921525345601318
DUNNE_TARGETS_OF_AML1_MTG8_FUSION_UP 38 0.01 Down 0.637377214324314 0.827454531431193 0.921525345601318
BIOCARTA_VEGF_PATHWAY 23 0.01 Down 0.189441643994507 0.82796610466298 0.921525345601318
FRASOR_TAMOXIFEN_RESPONSE_UP 48 0.01 Down 0.397996311209224 0.827984436863237 0.921525345601318
REACTOME_REGULATION_OF_PYRUVATE_DEHYDROGENASE_PDH_COMPLEX 15 0.01 Up 0.161530598166249 0.828040238238871 0.921525345601318
REACTOME_ADP_SIGNALLING_THROUGH_P2Y_PURINOCEPTOR_1 17 0.01 Down 0.234890402397542 0.82868774342559 0.922017674925254
GARGALOVIC_RESPONSE_TO_OXIDIZED_PHOSPHOLIPIDS_TURQUOISE_DN 49 0.01 Up 0.250325970273244 0.828930567907135

0.922059614635627
ICHIBA_GRAFT_VERSUS_HOST_DISEASE_D7_UP 90 0.01 Up
0.668656459306858 0.829293378726094 0.922234967187974
LEE_EARLY_T_LYMPHOCYTE_DN 44 0.01 Down
0.308323844655927 0.8296802345943 0.922436966238283
OLSSON_E2F3_TARGETS_DN 32 0.01 Down 0.436430496852591
0.830096668159349 0.922625414504484
MCMURRAY_TP53_HRAS_COOPERATION_RESPONSE_UP 21 0.01 Down
0.572856893573044 0.830260245088017
0.922625414504484
FU_INTERACT_WITH_ALKBH8 13 0.01 Down 0.0898993762339667
0.83185301712298 0.924166908543696
KESHELAVA_MULTIPLE_DRUG_RESISTANCE 77 0.01 Up
0.234634406620478 0.832264648358242 0.924298135980822
GALLUZZI_PREVENT_MITOCHONDIAL_PERMEABILIZATION 21 0.01 Down
0.416665398304398 0.832533664515918
0.924298135980822
LEI_MYB_TARGETS 306 0.01 Down 0.45985045581492
0.832588020597631 0.924298135980822
ONO_AML1_TARGETS_UP 14 0.01 Down 0.538840126641175
0.833647743269752 0.924995809708066
REACTOME_POST_CHAPERONIN_TUBULIN_FOLDING_PATHWAY 17 0.01
Up 0.173571759454233 0.833694591535457
0.924995809708066
REACTOME_MITOTIC_G1_G1_S_PHASES 141 0.01 Down
0.189431088183852 0.83383382000825 0.924995809708066
KEGG_VASCULAR_SMOOTH_MUSCLE_CONTRACTION 82 0.01 Up
0.381681070482399 0.834144194877423 0.925111807543552
VISALA_AGING_LYMPHOCYTE_DN 16 0.01 Up
0.402466426268383 0.834617600292774 0.92540851339813
KEGG_STEROID_BIOSYNTHESIS 16 0.01 Down
0.259356754513468 0.83495861024376 0.92553780442585
CHEOK_RESPONSE_TO_MERCAPTOPURINE_AND_LD_MTX_DN 17 0.01 Down
0.367007408572543 0.835302229283079
0.92553780442585
DOANE_RESPONSE_TO_ANDROGEN_DN 225 0.01 Up
0.372317868480249 0.835351918254877 0.92553780442585
PID_RAS_PATHWAY 25 0.01 Up 0.360283895341119
0.835590746814391 0.925574274748814
LE_SKI_TARGETS_UP 16 0.01 Down 0.550415209984727
0.837289733651819 0.927227729185741
REACTOME_SIGNAL_TRANSDUCTION_BY_L1 18 0.01 Down
0.24577564129815 0.837679048724722 0.927430375373799
BREDEMEYER_RAG_SIGNALING_VIA_ATM_NOT_VIA_NFKB_UP 44 0.01
Up 0.392647423912481 0.83831276996381
0.927697984122441
REACTOME_CARBOXYTERMINAL_POST_TRANSLATIONAL_MODIFICATIONS_OF_TUBULIN
26 0.01 Down 0.22751058839157
0.838578833949479 0.927697984122441
KAMIKUBO_MYELOID_CEBPA_NETWORK 22 0.01 Up
0.835759929878921 0.838855827082659 0.927697984122441
REACTOME_PROTEIN_PROTEIN_INTERACTIONS_AT_SYNAPSES 46 0.01
Down 0.330766425686699 0.838913729174662
0.927697984122441

GERHOLD_ADIPOGENESIS_DN 60 0.01 Up 0.407744876496306
0.838952681970573 0.927697984122441
LINDGREN_BLADDER_CANCER_CLUSTER_3_DN 218 0.01 Up
0.249335110744516 0.839872642396646 0.928314550492637
QI_HYPOXIA 117 0.01 Down 0.323623840759309
0.83992330964484 0.928314550492637
KAUFFMANN_DNA_REPLICATION_GENES 130 0.01 Up
0.178547917763987 0.840845055214714 0.929089880513596
WINTER_HYPOXIA_METAGENE 216 0.01 Down 0.435487160196757
0.841137644018114 0.929089880513596
WIKMAN_ASBESTOS_LUNG_CANCER_DN 24 0.01 Up
0.280277798822809 0.841321263020612 0.929089880513596
RUTELLA_RESPONSE_TO_HGF_DN 216 0.01 Down
0.406958325646229 0.841451591450689 0.929089880513596
MULLIGHAN_NPM1_SIGNATURE_3_UP 267 0.01 Down
0.296199912022997 0.842335877628762 0.929494855299624
YAMAZAKI_TCEB3_TARGETS_DN 170 0.01 Down
0.262197580115878 0.842388527678207 0.929494855299624
GHANDHI_DIRECT_IRRADIATION_DN 29 0.01 Down
0.490247207795681 0.842619968127544 0.929494855299624
BOYLAN_MULTIPLE_MYELOMA_C_CLUSTER_UP 30 0.01 Up
0.218903695991606 0.842645502857834 0.929494855299624
WIERENGA_STAT5A_TARGETS_GROUP1 102 0.01 Up
0.344393366499389 0.842858917519966 0.929502167382789
GREGORY_SYNTHETIC_LETHAL_WITH_IMATINIB 121 0.01 Down
0.324488825976402 0.84326411805199 0.929720924857418
BIOCARTA_SM_PATHWAY 14 0.01 Up 0.0567340036556175
0.844159904372096 0.930333736764162
PID_HNF3B_PATHWAY 24 0.01 Up 0.437819457409913
0.844414266948717 0.930333736764162
TURASHVILI_BREAST_DUCTAL_CARCINOMA_VS_LOBULAR_NORMAL_UP 63 0.01
Up 0.401027446420323 0.844544232742431
0.930333736764162
ICHIBA_GRAFT_VERSUS_HOST_DISEASE_D7_DN 23 0.01 Down
0.339483995266452 0.844932232546866 0.930333736764162
REACTOME_POSITIVE_EPIGENETIC_REGULATION_OF_RRNA_EXPRESSION 66
0.01 Down 0.135487116401581 0.84507174759663
0.930333736764162
FIGUEROA_AML_METHYLATION_CLUSTER_5_DN 23 0.01 Down
0.505764095528075 0.845454929219331 0.930333736764162
ODONNELL_METASTASIS_DN 18 0.01 Down 0.310943155229508
0.845468546320714 0.930333736764162
WENG_POR_TARGETS_LIVER_DN 18 0.01 Down
0.54208385517836 0.84552965003927 0.930333736764162
REACTOME_FORMATION_OF_HIV_ELONGATION_COMPLEX_IN_THE_ABSENCE_OF_HIV_T
AT 44 0.01 Up 0.10961098108471
0.845952767285233 0.930333736764162
MARTIN_VIRAL_GPCR_SIGNALING_UP 61 0.01 Up
0.52529161239234 0.846034174502249 0.930333736764162
MARTORIATI_MDM4_TARGETS_FETAL_LIVER_UP 203 0.01 Up
0.270116934389841 0.846096622000422 0.930333736764162
KYNG_DNA_DAMAGE_BY_GAMMA_RADIATION 61 0.01 Down
0.332130625486294 0.846896751930281 0.930666979568638
DACOSTA_UV_RESPONSE_VIA_ERCC3_XPCS_UP 24 0.01 Down

0.204500887629305	0.847212720403975	0.930666979568638
KEGG_FC_GAMMA_R_MEDIATED_PHAGOCYTOSIS	90	0.01	Up
0.363539726643675	0.847291406651932	0.930666979568638
MOREAUX_B_LYMPHOCYTE_MATURATION_BY_TACI_UP	70	0.01	Down
	0.409249830050989	0.847414664107835
0.930666979568638
BOYLAN_MULTIPLE_MYELOMA_D_UP	78	0.01	Down
0.227141319182686	0.847762190202431	0.930666979568638
HONMA_DOCETAXEL_RESISTANCE	35	0.01	Down
0.149279005452818	0.847766500188846	0.930666979568638
KEGG_LEISHMANIA_INFECTION	59	0.01	Down
0.641472893212414	0.848201833788882	0.930666979568638
PURBEY_TARGETS_OF_CTBP1_AND_SATB1_UP	65	0.01	Up
0.224055256574894	0.848262653414663	0.930666979568638
BIOCARTA_PML_PATHWAY	14	0.01	Down	0.273734076758175
	0.848275312912429	0.930666979568638
BLALOCK_ALZHEIMERS_DISEASE_INCIPIENT_UP	336	0.01	Down
0.302272605081601	0.848470140661241	0.930666979568638
REACTOME_AUTOPHAGY	104	0.01	Down	0.145981808135963
	0.848708222775205	0.930701015217015
REACTOME_METABOLISM_OF_STEROID_HORMONES	17	0.01	Down
0.47605286481142	0.849078660869738	0.930880141612066
BREDEMEYER_RAG_SIGNALING_VIA_ATM_NOT_VIA_NFKB_DN	37	0.01
	Down	0.252498760008111	0.849358866618146
0.930960279309575
NAKAMURA_TUMOR_ZONE_PERIPHERAL_VS_CENTRAL_UP	258	0.01	Down
	0.312065673627592	0.84959326121466
0.930990177757167
REACTOME_RNA_POLYMERASE_III_TRANSCRIPTION_INITIATION_FROM_TYPE_3_PRO
MOTER	28	0.01	Up	0.158230876185711
0.849889458627948	0.931087769079363
REACTOME_NOTCH2_ACTIVATION_AND_TRANSMISSION_OF_SIGNAL_TO_THE_NUCLEUS
	21	0.01	Up	0.280201523899654
0.854279855771049	0.935669578872043
SARRIO_EPITHELIAL_MESENCHYMAL_TRANSITION_UP	160	0.01	Up
	0.304227205370047	0.855093565687109
0.936010156797919
REACTOME_HDR_THROUGH_SINGLE_STRAND_ANNEALING_SSA	36	0.01
	Up	0.14420602246563	0.855209653825942
0.936010156797919
REACTOME_TRYPTOPHAN_CATABOLISM	11	0.01	Up
0.74452709042348	0.855215509225596	0.936010156797919
KYNG_RESPONSE_TO_H2O2_VIA_ERCC6_DN	43	0.01	Up
0.195349488340045	0.855869084005545	0.936446331418328
BARIS_THYROID_CANCER_DN	55	0.01	Down	0.385615187340962
	0.856030695394418	0.936446331418328
WANG_BARRETTS_ESOPHAGUS_UP	40	0.01	Down
0.442900126133793	0.85673539584045	0.936732339918085
PID_PTP1B_PATHWAY	46	0.01	Down	0.518139112927766
	0.856924417710622	0.936732339918085
BOYAULT_LIVER_CANCER_SUBCLASS_G123_UP	44	0.01	Down
0.201872084968512	0.857032618817793	0.936732339918085
SPIELMAN_LYMPHOBLAST_EUROPEAN_VS_ASIAN_2FC_DN	15	0.01	Up
	0.428593068456829	0.857125720596904

0.936732339918085
ENGELMANN_CANCER_PROGENITORS_DN 57 0.01 Down
0.354823792897991 0.85760608956435 0.937029502331491
FERREIRA_EWINGS_SARCOMA_UNSTABLE_VS_STABLE_UP 136 0.01 Up
0.242271417456755 0.858130701244631
0.937374848625667
HARRIS_HYPOXIA 72 0.01 Down 0.515725866609259
0.858741432062025 0.937591720312684
SWEET_KRAS_ONCOGENIC_SIGNATURE 85 0.01 Down
0.358436155698172 0.859449312679758 0.937591720312684
ASTIER_INTEGRIN_SIGNALING 45 0.01 Up
0.268511328596658 0.859483540242283 0.937591720312684
SWEET_LUNG_CANCER_KRAS_UP 407 0.01 Up
0.38715156653492 0.859685568252618 0.937591720312684
MURAKAMI_UV_RESPONSE_6HR_UP 31 0.01 Down
0.310775939662015 0.859753708520867 0.937591720312684
LEE_METASTASIS_AND_ALTERNATIVE_SPLICING_DN 37 0.01 Up
0.240761002802328 0.859781910698866
0.937591720312684
MEISSNER_NPC_ICP_WITH_H3K4ME3 16 0.01 Up
0.122276579286369 0.859789337292299 0.937591720312684
FLOTHO_PEDIATRIC_ALL_THERAPY_RESPONSE_DN 27 0.01 Up
0.324146185230713 0.860206744966873
0.937819383610501
RIZ_ERYTHROID_DIFFERENTIATION_APOBEC2 12 0.01 Down
0.444043212645007 0.862077726026999 0.939496592594526
DASU_IL6_SIGNALING_SCAR_UP 29 0.01 Up
0.315292994804108 0.862439730044303 0.939496592594526
REACTOME_TP53_REGULATES_TRANSCRIPTION_OF_DEATH_RECEPTORS_AND_LIGANDS
11 0.01 Up 0.311409176728301
0.86270363358971 0.939496592594526
YANG_BCL3_TARGETS_UP 254 0.01 Up 0.407751154050417
0.86272526204835 0.939496592594526
REACTOME_THROMBIN_SIGNALLING_THROUGH_PROTEINASE_ACTIVATED_RECEPTORS_
PARS 21 0.01 Down 0.248737125098816
0.862790196714172 0.939496592594526
ELVIDGE_HYPOXIA_BY_DMOG_DN 55 0.01 Down
0.271544923140152 0.864063058055399 0.940654745933402
WAKABAYASHI_ADIPOGENESIS_PPARG_RXRA_BOUND_WITH_H4K20ME1_MARK 140
0.01 Up 0.229728634542722 0.865388903481979
0.941870005121427
CHIANG_LIVER_CANCER_SUBCLASS_POLYSOMY7_UP 49 0.01 Down
0.282628819293227 0.866361682229839
0.942700499061516
JOHNSTONE_PARVB_TARGETS_3_UP 421 0.01 Down
0.320261919589843 0.866814148658476 0.942964568785055
MUELLER_PLURINET 271 0.01 Up 0.207463039947634
0.867946499450992 0.943967944600099
FONTAINE_THYROID_TUMOR_UNCERTAIN_MALIGNANCY_UP 29 0.01 Up
0.245390584791546 0.868635739171909
0.944489029409223
ZWANG_DOWN_BY_2ND_EGF_PULSE 205 0.01 Up
0.208965689078005 0.868944347188334 0.944596092046327
REACTOME_NEF_MEDIATES_DOWN_MODULATION_OF_CELL_SURFACE_RECEPTORS_BY_R

ECRUITING_THEM_TO_CLATHRIN_ADAPTERS 19 0.01 Down
0.277789063110619 0.86991974272452 0.944936810848133
ZHAN_V1_LATE_DIFFERENTIATION_GENES_DN 12 0.01 Down
0.249355362598582 0.870098434875359 0.944936810848133
REACTOME_PROTEIN_FOLDING 78 0.01 Up
0.170154403516531 0.870198983302517 0.944936810848133
BREDEMEYER_RAG_SIGNALING_NOT_VIA_ATM_DN 44 0.01 Up
0.386917321304389 0.87023196802990 7 0.944936810848133
KUMAR_TARGETS_OF_MLL_AF9_FUSION 304 0.01 Down
0.38341544598413 0.870549886035778 0.944936810848133
FEVR_CTNNB1_TARGETS_DN 506 0.01 Down 0.221638493657456
0.870566246175173 0.944936810848133
KUROKAWA_LIVER_CANCER_EARLY_RECURRENCE_UP 10 0.01 Down
0.206218922215967 0.870895583804009
0.944936810848133
BROWNE_HCMV_INFECTION_14HR_DN 244 0.01 Up
0.306921873655823 0.870939534448012 0.944936810848133
MAGRANGEAS_MULTIPLE_MYELOMA_IGG_VS_IGA_UP 18 0.01 Up
0.342156547149961 0.871306159265448
0.945106463778521
REACTOME_AURKA_ACTIVATION_BY_TPX2 67 0.01 Up
0.120066964222396 0.872519129169424 0.945831850143564
BAKKER_FOXO3_TARGETS_UP 49 0.01 Down 0.304000362828546
0.872778434835769 0.945831850143564
STAMBOLSKY_RESPONSE_TO_VITAMIN_D3_DN 21 0.01 Down
0.372279841714833 0.872934403845167 0.945831850143564
LIM_MAMMARY_LUMINAL_PROGENITOR_DN 13 0.01 Down
0.270694742852407 0.872994833569467 0.945831850143564
PETRETTO_LEFT_VENTRICLE_MASS_QTL_CIS_DN 10 0.01 Down
0.227760402356427 0.873278815718456 0.945831850143564
RICKMAN_HEAD_AND_NECK_CANCER_D 23 0.01 Down
1.24311953914953 0.873370526555961 0.945831850143564
LIANG_HEMATOPOIESIS_STEM_CELL_NUMBER_LARGE_VS_TINY_DN 35 0.01
Up 0.265632463848639 0.873507513124911
0.945831850143564
LUI_THYROID_CANCER_CLUSTER_2 39 0.01 Up
0.403605598810285 0.873658251790006 0.945831850143564
REACTOME_STIMULI_SENSING_CHANNELS 66 0.01 Down
0.334959599575304 0.874472007039021 0.946484871572454
SENGUPTA_EBNA1_ANTICORRELATED 119 0.01 Up
0.28567033180448 0.875206147100336 0.946984966547184
SHEDDEN_LUNG_CANCER_GOOD_SURVIVAL_A12 188 0.01 Down
0.378664132404215 0.875355402892892 0.946984966547184
WU_CELL_MIGRATION 179 0.01 Down 0.53264461697998
0.875814869612964 0.947254051710846
PID_FANCONI_PATHWAY 45 0.01 Up 0.14053445255767
0.877626615946868 0.948839071269428
MEISSNER_NPC_HCP_WITH_H3K4ME2 307 0.01 Down
0.425091043571857 0.877702526882821 0.948839071269428
HWANG_PROSTATE_CANCER_MARKERS 27 0.01 Up
0.494078215084256 0.877946882523076 0.948875026915418
LI_PROSTATE_CANCER_EPIGENETIC 26 0.01 Down
0.451892841833327 0.878608253172145 0.94933365402441
EBAUER_TARGETS_OF_PAX3_FOXO1_FUSION_DN 37 0.01 Down

0.242181715179379	0.878974108541503	0.94933365402441
JACKSON_DNMT1_TARGETS_DN	15	0.01	Down
0.591244862926893	0.879019096438837	0.94933365402441
FRASOR_TAMOXIFEN_RESPONSE_DN	11	0.01	Up
0.269499922252599	0.879216018176556	0.94933365402441
PEREZ_TP53_TARGETS	961	0.01	Down	0.343459848851295
	0.879582226028779	0.949500986070932
FULCHER_INFLAMMATORY_RESPONSE_LECTIN_VS_LPS_UP	536	0.01	Down
	0.355109426655553	0.88037065357592
0.949944764541944
JAIN_NFKB_SIGNALING	73	0.01	Up	0.183990012297789
	0.880486769921701	0.949944764541944
BIOCARTA_CHEMICAL_PATHWAY	17	0.01	Down
0.247541567237993	0.880627326773366	0.949944764541944
REACTOME_DNA_REPLICATION	117	0.01	Down
0.12512169794012	0.881421675310743	0.950522678859915
BENPORATH_NANOG_TARGETS	915	0.01	Down	0.235769749252475
	0.881585995142822	0.950522678859915
PASQUALUCCI_LYMPHOMA_BY_GC_STAGE_DN	134	0.01	Down
0.299027004446268	0.88252032620855	0.950869124895056
DAVIES_MULTIPLE_MYELOMA_VS_MGUS_DN	21	0.01	Down
0.493700781709089	0.882541080252367	0.950869124895056
REACTOME_G1_PHASE	43	0.01	Down	0.243751476868514
	0.88254280057468	0.950869124895056
MCCABE_HOXC6_TARGETS_CANCER_UP	22	0.01	Up
0.278379739047436	0.882753472344175	0.950869124895056
BARRIER_CANCER_RELAPSE_NORMAL_SAMPLE_UP	25	0.01	Up
0.394741375570356	0.883465850011457	0.951408480067441
MEISSNER_NPC_HCP_WITH_H3K4ME2_AND_H3K27ME3	138	0.01	Down
	0.64343288725878	0.883853495832394
0.951530256533549
KEGG_AMINOACYL_TRNA_BIOSYNTHESIS	41	0.01	Up
0.128931650849883	0.884002302844071	0.951530256533549
OUILLETTE_CLL_13Q14_DELETION_DN	46	0.01	Down
0.385854884867479	0.885061816307535	0.952442629710886
SHIPP_DLBCL_CURED_VS_FATAL_UP	19	0.01	Down
0.726398469996107	0.885590880140153	0.952783869370509
KEGG_CITRATE_CYCLE_TCA_CYCLE	27	0.01	Down
0.110009990796266	0.885989676439266	0.952789820619572
BENPORATH_ES_2	21	0.01	Up	0.219107854652101
0.886038884386048	0.952789820619572
LUI_TARGETS_OF_PAX8_PPARG_FUSION	34	0.01	Up
0.136246020327274	0.886493341049548	0.952789820619572
FISCHER_G1_S_CELL_CYCLE	169	0.01	Up	0.24773185285717
	0.886707736187613	0.952789820619572
TONKS_TARGETS_OF_RUNX1_RUNX1T1_FUSION_SUSTAINDED_IN_ERYTHROCYTE_UP
	37	0.01	Down	0.368543134670361
0.886801947327874	0.952789820619572
TSUNODA_CISPLATIN_RESISTANCE_DN	43	0.01	Down
0.576031793519645	0.886912053689255	0.952789820619572
MASSARWEH_RESPONSE_TO_ESTRADIOL	56	0.01	Up
0.373389986449702	0.887080177818222	0.952789820619572
REACTOME_TRIGLYCERIDE_CATABOLISM	14	0.01	Down
0.341526133062665	0.887552075300269	0.953068938957169

CAIRO_HEPATOBLASTOMA_UP 201 0.01 Down 0.296137652175722
0.888130680553165 0.953462481272147
PID_DNA_PK_PATHWAY 15 0.01 Down 0.14892764499273
0.8884259567133 0.953551737207804
BRACHAT_RESPONSE_TO_CAMPTOTHECIN_UP 25 0.01 Down
0.173433164443358 0.889374787401299 0.954287589419668
PELLICCIOTTA_HDAC_IN_ANTIGEN_PRESENTATION_UP 60 0.01 Up
0.118779367444851 0.88953615120543
0.954287589419668
YAGI_AML_WITH_INV_16_TRANSLOCATION 330 0.01 Down
0.281402884672092 0.890838070672848 0.955431002319326
STEIN_ESR1_TARGETS 70 0.01 Down 0.244275358638956
0.891027088258646 0.955431002319326
MIKKELSEN_MCV6_ICP_WITH_H3K27ME3 26 0.01 Down
0.483224529733164 0.892709843369632 0.956763797837137
PID_HIF1_TFPATHWAY 56 0.01 Down 0.374558578264037
0.892840852108552 0.956763797837137
GAZDA_DIAMOND_BLACKFAN_ANEMIA_PROGENITOR_DN 62 0.01 Down
0.175415656500306 0.892920959653618
0.956763797837137
GNATENKO_PLATELET_SIGNATURE 36 0.01 Up
0.281111929148192 0.893498878141132 0.956763797837137
REACTOME_CENTROSOME_MATURATION 77 0.01 Up
0.124281988564398 0.893712869641473 0.956763797837137
LEE_CALORIE_RESTRICTION_NEOCORTEX_DN 51 0.01 Down
0.234068143580721 0.894045109161952 0.956763797837137
JIANG_VHL_TARGETS 85 0.01 Up 0.189824248241389
0.89407846110715 0.956763797837137
PEDRIOLI_MIR31_TARGETS_UP 145 0.01 Up
0.41223602447349 0.894327580426988 0.956763797837137
ZHAN_MULTIPLE_MYELOMA_MS_DN 38 0.01 Up
0.208408661442932 0.894432024443943 0.956763797837137
SCHLOSSER_SERUM_RESPONSE_AUGMENTED_BY_MYC 98 0.01 Up
0.196784992852859 0.89445460370951
0.956763797837137
GENTILE_UV_RESPONSE_CLUSTER_D9 24 0.01 Down
0.257564848651631 0.894901115463841 0.956763797837137
CLAUS_PGR_POSITIVE_MENINGIOMA_DN 12 0.01 Down
0.500699134769162 0.894952540553801 0.956763797837137
SUNG_METASTASIS_STROMA_DN 47 0.01 Up
0.360586823185292 0.895037101202483 0.956763797837137
REACTOME_SEMA3A_PAK_DEPENDENT_AXON_REPULSION 15 0.01 Up
0.305887757362593 0.89532034385749
0.956839026542895
ROSS_AML_WITH_CBFB_MYH11_FUSION 39 0.01 Down
0.606356734183755 0.895852184324764 0.957179835640555
REACTOME_LYSOSOME_VESICLE_BIOGENESIS 32 0.01 Down
0.153524923926548 0.896368928052391 0.957392044634593
BIOCARTA_FAS_PATHWAY 29 0.01 Down 0.204118709477424
0.896476777723471 0.957392044634593
MIKKELSEN_NPC_HCP_WITH_H3K27ME3 131 0.01 Down
0.623254160898227 0.897156436401265 0.957890304423679
SWEET_KRAS_TARGETS_UP 78 0.01 Up 0.419712097860072
0.898598056992002 0.958922929368249

REACTOME_GLYCOLYSIS 63 0.01 Down 0.210753085298953
0.898611580137262 0.958922929368249
PUJANA_XPRSS_INT_NETWORK 154 0.01 Up
0.158271428839684 0.898763582075291 0.958922929368249
SHETH_LIVER_CANCER_VS_TXNIP_LOSS_PAM2 139 0.01 Down
0.349765849257913 0.899054513331058 0.959005704182038
REACTOME_DAP12_SIGNALING 23 0.01 Down
0.387136038041163 0.900012727884338 0.959420695912443
PARENT_MTOR_SIGNALING_UP 511 0.01 Up
0.228215521292807 0.90006058631226 0.959420695912443
WHITEHURST_PACLITAXEL_SENSITIVITY 26 0.01 Down
0.176632207705724 0.900099002847457 0.959420695912443
DOANE_BREAST_CANCER_CLASSES_DN 32 0.01 Down
0.364381554859477 0.900297329334524 0.959420695912443
FERNANDEZ_BOUND_BY_MYC 151 0.01 Up 0.272811992636423
0.900907111144026 0.959729484769009
KOYAMA_SEMA3B_TARGETS_DN 293 0.01 Up
0.325485761267642 0.901014110283697 0.959729484769009
LINDGREN_BLADDER_CANCER_CLUSTER_1_DN 325 0.01 Up
0.406354932551762 0.901800799639862 0.960134400758908
PLASARI_TGFB1_SIGNALING_VIA_NFIC_1HR_UP 24 0.01 Up
0.425195493043587 0.901821454950859 0.960134400758908
REACTOME_OLFACTORY_SIGNALING_PATHWAY 11 0.01 Up
0.21692114945462 0.902143121416604 0.960249427129442
VERRECCHIA_RESPONSE_TO_TGFB1_C5 20 0.01 Up
0.37127801729276 0.902382554511956 0.960276889803798
HOFMANN_MYELODYSPLASTIC_SYNDROM_HIGH_RISK_UP 10 0.01 Down
0.220306119685514 0.902645406086924
0.960329254523249
BIOCARTA_CSK_PATHWAY 14 0.01 Up 0.688440497299376
0.903046838697598 0.960400613384699
WORSCHECH_TUMOR_REJECTION_DN 10 0.01 Up
0.078049331524848 0.903152499457608 0.960400613384699
SERVITJA_LIVER_HNF1A_TARGETS_DN 84 0.01 Down
0.467328213326284 0.903353457928922 0.960400613384699
WANG_LSD1_TARGETS_DN 22 0.01 Down 0.522190644492284
0.904469050284282 0.961204982145612
KORKOLA_CORRELATED_WITH_POU5F1 18 0.01 Down
0.443109897221224 0.90453772513369 0.961204982145612
PARK_APL_PATHOGENESIS_DN 43 0.01 Up
0.620289199256538 0.904868753081371 0.961329483597438
BOYLAN_MULTIPLE_MYELOMA_D_DN 67 0.01 Down
0.354730517865192 0.905114108468089 0.96136292948111
PID_S1P_S1P4_PATHWAY 11 0.01 Up 0.144499960256927
0.905492995500262 0.961385053441538
PUJANA_BRCA2_PCC_NETWORK 371 0.01 Up
0.185674799147553 0.905613941068197 0.961385053441538
KEGG_O_GLYCAN_BIOSYNTHESIS 22 0.01 Down
0.361133586294256 0.905803634825104 0.961385053441538
GARGALOVIC_RESPONSE_TO_OXIDIZED_PHOSPHOLIPIDS_PURPLE_DN 11 0.01
Up 0.354660849667302 0.906105971993813
0.961385053441538
REACTOME_SPHINGOLIPID_DE_NOVO_BIOSYNTHESIS 41 0.01 Down
0.25376268059116 0.906204331797953

0.961385053441538
BIOCARTA_DREAM_PATHWAY 10 0.01 Down 0.417074536892929
0.906453264987972 0.961422233629291
REACTOME_SIGNALING_BY_RETINOIC_ACID 33 0.01 Up
0.398017619271825 0.908308711379654 0.963162929382294
HUI_MAPK14_TARGETS_UP 14 0.01 Down 0.306386540441428
0.908813657600895 0.963264230104811
REACTOME_SYNTHESIS_OF_PE 12 0.01 Up
0.278241959523366 0.90891736773412 0.963264230104811
REACTOME_TRANSCRIPTION_OF_E2F_TARGETS_UNDER_NEGATIVE_CONTROL_BY_P107_RBL1_AND_P130_RBL2_IN_COMPLEX_WITH_HDAC1 14 0.01 Down
0.129536432785895 0.909047133282449
0.963264230104811
FOSTER_KDM1A_TARGETS_UP 135 0.01 Up 0.51153110167079
0.909429704982665 0.963442499150855
JEON_SMAD6_TARGETS_UP 23 0.01 Up 0.663213757479765
0.909933388541202 0.963748958881409
REACTOME_ACTIVATED_NOTCH1_TRANSMITS_SIGNAL_TO_THE_NUCLEUS 28
0.01 Up 0.271169477473652 0.910460840549285
0.964080442466204
ZHAN_MULTIPLE_MYELOMA_UP 58 0.01 Down
0.211692680874492 0.910874230149167 0.964291018492819
BRACHAT_RESPONSE_TO_CISPLATIN 20 0.01 Down
0.164566094785558 0.911814701255316 0.96491511155042
RIZKI_TUMOR_INVASIVENESS_3D_DN 212 0.01 Down
0.334260280775358 0.911893079836748 0.96491511155042
YAO_HOXA10_TARGETS_VIA_PROGESTERONE_DN 14 0.01 Down
0.654022044066772 0.913056580652296 0.965704280265589
GAUSSMANN_MLL_AF4_FUSION_TARGETS_E_DN 18 0.01 Down
0.587194503147403 0.913068563098721 0.965704280265589
LEE_AGING_CEREBELLUM_DN 52 0.01 Up 0.289184664676107
0.913304711688905 0.965726812289256
LU_AGING_BRAIN_UP 230 0.01 Down 0.408833794411303
0.9137125011986 0.965930783840007
BENPORATH_MYC_MAX_TARGETS 724 0.01 Down
0.163148101792535 0.915122123006876 0.967193497041126
GAUSSMANN_MLL_AF4_FUSION_TARGETS_G_DN 28 0.01 Down
0.291951158945033 0.915577113713915 0.967304893470148
ACEVEDO_LIVER_TUMOR_VS_NORMAL_ADJACENT_TISSUE_DN 209 0.01
Down 0.379923750835171 0.915657913618572
0.967304893470148
LIAN_NEUTROPHIL_GRANULE_CONSTITUENTS 18 0.01 Up
0.934657158484027 0.916181035530515 0.967630111538925
REACTOME_KERATAN_SULFATE_BIOSYNTHESIS 23 0.01 Down
0.811737990136957 0.916798645745563 0.967761958505518
LIU_VMYB_TARGETS_UP 118 0.01 Down 0.281601890473975
0.916894864484072 0.967761958505518
DAZARD_UV_RESPONSE_CLUSTER_G1 52 0.01 Down
0.210174128201489 0.917116073814755 0.967761958505518
ZHAN_V2_LATE_DIFFERENTIATION_GENES 42 0.01 Down
0.251377511776064 0.917174792914578 0.967761958505518
HUANG_GATA2_TARGETS_UP 124 0.01 Down 0.382630372865883
0.917382359330815 0.967761958505518
CHYLA_CBFA2T3_TARGETS_UP 269 0.01 Up

0.421970043812338	0.918097874146771	0.968271708576107
REACTOME_HIV_ELONGATION_ARREST_AND_RECOVERY	32	0.01	Down
	0.0941070165193802	0.918296394585082
0.968271708576107
ZHANG_TARGETS_OF_EWSR1_FLI1_FUSION	75	0.01	Up
0.313092850401753	0.918879170821413	0.968658975807282
REACTOME_ACTIVATION_OF_SMO	16	0.01	Up
0.318449181318838	0.919334531276796	0.968911774464056
REACTOME_INHIBITION_OF_REPLICATION_INITIATION_OF_DAMAGED_DNA_BY_RB1_
E2F1	13	0.01	Down	0.11089002139109
0.919825831129997	0.969202323237069
REACTOME_EPH_EPHRIN_MEDIATED_REPULSION_OF_CELLS 44	0.01	Up
	0.334136539069291	0.920417904352503
0.969598893851535
VANTVEER_BREAST_CANCER_ESR1_UP	190	0.01	Down
0.262677522133087	0.9207488116173 0.969720222169579
REACTOME_G0_AND_EARLY_G1	24	0.01	Down
0.158944828185656	0.921289511560134	0.970062392706208
IWANAGA_CARCINOGENESIS_BY_KRAS_PTEN_DN	263	0.01	Down
0.378229066607147	0.921564264305318	0.970124442166839
ONO_FOXP3_TARGETS_UP	15	0.01	Up	0.499077049767363
	0.922322155884686	0.970694940459299
PETRETTO_HEART_MASS_QTL_CIS_UP	29	0.01	Down
0.290778957136938	0.923020170033881	0.970761685651855
GRADE_COLON_VS_RECTAL_CANCER_DN 47	0.01	Up
0.289052731300569	0.92319100919995	0.970761685651855
LEE_NEURAL_CREST_STEM_CELL_DN	89	0.01	Down
0.395830570505883	0.923689831561484	0.970761685651855
REACTOME_NONHOMOLOGOUS_END_JOINING_NHEJ 45	0.01	Up
0.146384452646487	0.923832934441674	0.970761685651855
REACTOME_NRIF_SIGNALS_CELL_DEATH_FROM_THE_NUCLEUS	16	0.01
	Up	0.159462751117884	0.924054290998505
0.970761685651855
HOLLERN_PAPILLARY_BREAST_TUMOR	14	0.01	Down
0.523191954457227	0.924061711160816	0.970761685651855
LEIN_OLIGODENDROCYTE_MARKERS	53	0.01	Up
0.343420724485201	0.924091179480174	0.970761685651855
KOYAMA_SEMA3B_TARGETS_UP	222	0.01	Up
0.342794421936202	0.924237786197733	0.970761685651855
REACTOME_CONVERSION_FROM_APC_C:CDC20_TO_APC_C:CDH1_IN_LATE_ANAPHASE
	18	0.01	Down	0.10373109207833
0.92432925797329	0.970761685651855
GARGALOVIC_RESPONSE_TO_OXIDIZED_PHOSPHOLIPIDS_BROWN_UP	14	0.01
	Up	0.255615864701468	0.924651173052216
0.970872932228384
TSUNODA_CISPLATIN_RESISTANCE_UP 14	0.01	Up
0.529044818363192	0.925074827301533	0.971090926838018
DORMOY_ELAVL1_TARGETS	16	0.01	Down	0.528570543444617
	0.926246553969952	0.972093920171593
KEGG_PRION_DISEASES	25	0.01	Up	0.606294966623078
	0.926574780398256	0.972211400067731
NOUZOVA_METHYLATED_IN_APL	40	0.01	Up
0.306643691634062	0.92701083267224	0.972441935323622
HOFMANN_CELL_LYMPHOMA_DN	34	0.01	Down

0.310877917116841	0.927280065488049	0.972497408858791
BIOCARTA_INTEGRIN_PATHWAY	32	0.01	Down
0.225032926462211	0.927896858898753	0.972613433483601
CLIMENT_BREAST_CANCER_COPY_NUMBER_UP	19	0.01	Down
0.401108249083784	0.927925293378139	0.972613433483601
FLOTHO_PEDIATRIC_ALL_THERAPY_RESPONSE_UP	48	0.01	Down
	0.179767774988399	0.92803982563096
0.972613433483601
BROWNE_HCMV_INFECTION_10HR_DN	49	0.01	Up
0.239495325575307	0.928314333337812	0.972674342273535
ZHANG_BREAST_CANCER_PROGENITORS_DN	125	0.01	Up
0.355690392643563	0.929408069694737	0.973130555594109
FULCHER_INFLAMMATORY_RESPONSE_LECTIN_VS_LPS_DN	404	0.01	Down
	0.457479172585486	0.929562495379493
0.973130555594109
BONOME_OVARIAN_CANCER_POOR_SURVIVAL_UP	26	0.01	Up
0.345161950679781	0.929631432607037	0.973130555594109
REACTOME_RNA_POLYMERASE_III_TRANSCRIPTION_INITIATION_FROM_TYPE_1_PRO
MOTER	28	0.01	Up	0.129892007602315
0.929771774620698	0.973130555594109
HEIDENBLAD_AMPLICON_8Q24_UP	33	0.01	Up
0.109205523240411	0.929832199394149	0.973130555594109
CAFFAREL_RESPONSE_TO_THC_24HR_5_DN	52	0.01	Up
0.206257441370064	0.930643584767827	0.973741595276692
LIN_MELANOMA_COPY_NUMBER_DN	39	0.01	Up
0.203669525757196	0.930849306986417	0.973741595276692
REACTOME_METABOLISM_OF_COFACTORS	18	0.01	Up
0.233620897050728	0.931363254432413	0.97405254273469
RICKMAN_TUMOR_DIFFERENTIATED_MODERATELY_VS_POORLY_UP	117	0.01
	Up	0.327619725353436	0.932140717644816
0.974068670910598
HAN_SATB1_TARGETS_DN	348	0.01	Up	0.396782972271654
	0.932140781094352	0.974068670910598
MORI_MATURE_B_LYMPHOCYTE_UP	74	0.01	Up
0.376483730674645	0.932229838895824	0.974068670910598
CERIBELLI_GENES_INACTIVE_AND_BOUND_BY_NFY	19	0.01	Down
	0.186288112278983	0.932569120433052
0.974068670910598
ZHAN_MULTIPLE_MYELOMA_MS_UP	39	0.01	Up
0.43116798116485	0.932852705369724	0.974068670910598
KUUSELO_PANCREATIC_CANCER_19Q13_AMPLIFICATION	19	0.01	Up
	0.1494990914486 0.933160829450697	0.974068670910598
KEGG_OLFACTORY_TRANSDUCTION	19	0.01	Down
0.422928649997411	0.933163464589367	0.974068670910598
REACTOME_RMTS_METHYLATE_HISTONE_ARGININES	43	0.01	Down
	0.183455113033201	0.93347082905719
0.974068670910598
SHETH_LIVER_CANCER_VS_TXNIP_LOSS_PAM1	213	0.01	Up
0.309195277854298	0.933545652729477	0.974068670910598
PID_S1P_S1P1_PATHWAY	18	0.01	Up	0.47251513377686
	0.933545680596853	0.974068670910598
NIKOLSKY_OVERCONNECTED_IN_BREAST_CANCER 17	0.01	Down
0.550612890502163	0.934580779150995	0.974922395517225
PID_TAP63_PATHWAY	53	0.01	Up	0.347883161593704

0.935693578087436	0.975856759513463
REACTOME_REPRODUCTION	75	0.01	Up	0.178427380615813
0.936223411373239	0.976012264650511
HORIUCHI_WTAP_TARGETS_DN	266	0.01	Up
0.264392185961334	0.936383268865633	0.976012264650511
REACTOME_TRANSPORT_OF_SMALL_MOLECULES	498	0.01	Down
0.355598375144185	0.936494081743638	0.976012264650511
DELACROIX_RAR_BOUND_ES	363	0.01	Down	0.326107016525136
0.93708027615181	0.976016294854406
DAVICIONI_RHABDOMYOSARCOMA_PAX_FOXO1_FUSION_UP	52	0.01	Up
0.317632794572709	0.937266700342585
0.976016294854406
BILANGES_SERUM_SENSITIVE_VIA_TSC2	30	0.01	Up
0.492988886279657	0.937338713490738	0.976016294854406
KENNY_CTNNB1_TARGETS_UP	47	0.01	Down	0.467864234607229
0.937404158974904	0.976016294854406
REACTOME_REGULATION_OF_INSULIN_SECRETION	45	0.01	Up
0.32801835160087	0.937658827692179
0.976016294854406
BRUINS_UVC_RESPONSE_VIA_TP53_GROUP_B	440	0.01	Up
0.376979024280753	0.937825192031489	0.976016294854406
SCHLOSSER_MYC_TARGETS_AND_SERUM_RESPONSE_DN	48	0.01	Down
0.177697068461586	0.938017885154846
0.976016294854406
KAAB_FAILED_HEART_VENTRICLE_DN	40	0.01	Down
0.339555652190639	0.938664944271757	0.97603720776077
JIANG_HYPOXIA_NORMAL	188	0.01	Up	0.264853854152924
0.938694487111716	0.97603720776077
MEISSNER_BRAIN_HCP_WITH_H3K4ME2_AND_H3K27ME3	28	0.01	Up
0.708526386304706	0.938858091381404
0.97603720776077
MORI_EMU_MYC_LYMPHOMA_BY_ONSET_TIME_DN	17	0.01	Down
0.315596335469423	0.938954229427128	0.97603720776077
PID_HNF3A_PATHWAY	32	0.01	Down	0.661875427471705
0.939123675987838	0.97603720776077
WHITFIELD_CELL_CYCLE_S	135	0.01	Down	0.210238875578888
0.939527701440334	0.976037583733817
REACTOME_TRNA_AMINOACYLATION	42	0.01	Up
0.114436432822359	0.939558314753332	0.976037583733817
WAKABAYASHI_ADIPOGENESIS_PPARG_BOUND_8D	564	0.01	Down
0.257066542414841	0.940264291905074	0.976130051891204
REACTOME_NEUTROPHIL_DEGRANULATION	404	0.01	Down
0.418176388906026	0.94029090116875	0.976130051891204
BHATTACHARYA_EMBRYONIC_STEM_CELL	68	0.01	Up
0.331120591699305	0.940304451977068	0.976130051891204
HU_GENOTOXIN_ACTION_DIRECT_VS_INDIRECT_4HR	30	0.01	Up
0.284266213968831	0.940687871126469
0.976130051891204
REACTOME_SYNTHESIS_OF_IP3_AND_IP4_IN_THE_CYTOSOL	22	0.01
Up	0.313060822078962	0.940733122312057
0.976130051891204
WANG_METASTASIS_OF_BREAST_CANCER_ESR1_UP	18	0.01	Down
0.156180843220279	0.941514225651227
0.976602998944306

LEONARD_HYPOXIA 40 0.01 Down 0.324781813234979
0.941785083212846 0.976602998944306
JAATINEN_HEMATOPOIETIC_STEM_CELL_DN 141 0.01 Down
0.742849854077563 0.941840711996344 0.976602998944306
ZHAN_MULTIPLE_MYELOMA_CD2_DN 44 0.01 Down
0.290570152343158 0.943185390302446 0.977771754937614
INGA_TP53_TARGETS 16 0.01 Up 0.339345943870422
0.943977206974454 0.978062172761845
TRAYNOR_RETT_SYNDROM_UP 33 0.01 Up 0.602105892973187
0.943985606960349 0.978062172761845
REACTOME_RECRUITMENT_OF_NUMA_TO_MITOTIC_CENTROSOMES 84 0.01
Up 0.133708508538086 0.944140326663573
0.978062172761845
KEGG_RNA_POLYMERASE 29 0.01 Up 0.120816299179839
0.944335890942471 0.978062172761845
PHESSE_TARGETS_OF_APC_AND_MBD2_DN 10 0.01 Up
0.346241703309201 0.945019409847274 0.978544631942754
MATTIOLI_MGUS_VS_PCL 94 0.01 Up 0.287843246510069
0.94694597717046 0.980059836033327
AIYAR_COBRA1_TARGETS_UP 36 0.01 Down 0.417302633910939
0.946962411938416 0.980059836033327
MIKKELSEN_NPC_HCP_WITH_H3K4ME3_AND_H3K27ME3 115 0.01 Up
0.546706388318356 0.947322905463708
0.980059836033327
REACTOME_CASPASE_MEDIATED_CLEAVAGE_OF_CYTOSKELETAL_PROTEINS 10
0.01 Up 0.229272789808802 0.947354835943227
0.980059836033327
PLASARI_TGFB1_SIGNALING_VIA_NFIC_10HR_DN 21 0.01 Up
0.514237622060968 0.948181805208846
0.980631691763723
KIM_PTEN_TARGETS_UP 16 0.01 Down 0.295162197372875
0.948431135354947 0.980631691763723
PASINI_SUZ12_TARGETS_DN 292 0.01 Up 0.397406794612488
0.948562090275565 0.980631691763723
FIGUEROA_AML_METHYLATION_CLUSTER_1_UP 89 0.01 Up
0.328214351621925 0.948875346845114 0.980729980241156
DAIRKEE_CANCER_PRONE_RESPONSE_E2 21 0.01 Up
0.280512898941927 0.949335942465007 0.980782295357571
WEI_MIR34A_TARGETS 136 0.01 Up 0.264871966314262
0.949362350856684 0.980782295357571
REACTOME_SEMA3A_PLEXIN_REPULSION_SIGNALING_BY_INHIBITING_INTEGRIN_AD
HESION 13 0.01 Up 0.354259114996154
0.949784761209477 0.980842294675346
BIOCARTA_MTA3_PATHWAY 12 0.01 Up 0.273824561862962
0.949948452662506 0.980842294675346
REACTOME_NEGATIVE_EPIGENETIC_REGULATION_OF_RRNA_EXPRESSION 69
0.01 Down 0.147070352592949 0.950075050281748
0.980842294675346
PID_P73PATHWAY 70 0.01 Up 0.269861497699291
0.952358548360527 0.982959545915103
GOBERT_CORE_OLIGODENDROCYTE_DIFFERENTIATION 34 0.01 Down
0.307606102538756 0.953219893583905
0.982959545915103
SHEPARD_BMYB_MORPHOLINO_UP 182 0.01 Up

0.285681105560765	0.953322280203263	0.982959545915103
REACTOME_TRIGLYCERIDE_METABOLISM	21	0.01	Up
0.294159669727118	0.953945176445051	0.982959545915103
GAZDA_DIAMOND_BLACKFAN_ANEMIA_ERYTHROID_DN	396	0.01	Up
	0.247554383576716	0.954289631243258
0.982959545915103
CROONQUIST_NRAS_SIGNALING_UP	25	0.01	Down
0.566994863635362	0.954303763567634	0.982959545915103
LIAO_METASTASIS 490	0.01	Up	0.283160473245704
0.954311357582048	0.982959545915103
CHESLER_BRAIN_QTL_CIS	60	0.01	Down	0.266708963053869
	0.954542332361828	0.982959545915103
BRACHAT_RESPONSE_TO_METHOTREXATE_DN	23	0.01	Down
0.24342935467738	0.954665187182435	0.982959545915103
CAFFAREL_RESPONSE_TO_THC_8HR_5_UP	15	0.01	Down
0.194227148964688	0.954940842638559	0.982959545915103
AIYAR_COBRA1_TARGETS_DN 28	0.01	Up	0.174566422794175
	0.95528143759604	0.982959545915103
TARTE_PLASMA_CELL_VS_PLASMABLAST_DN	289	0.01	Down
0.229176518451907	0.955418175991503	0.982959545915103
REACTOME_RNA_POLYMERASE_II_TRANSCRIPTION_PRE_INITIATION_AND_PROMOTER
_OPENING	46	0.01	Down	0.129985757684994
0.955763302741919	0.982959545915103
WHITE_NEUROBLASTOMA_WITH_1P36.3_DELETION	16	0.01	Up
	0.183448074044116	0.955862309188583
0.982959545915103
LANDIS_ERBB2_BREAST_TUMORS_65_UP	20	0.01	Down
0.208937285911506	0.955922622409296	0.982959545915103
MAINA_VHL_TARGETS_DN	14	0.01	Down	0.552076377862025
	0.956045254603492	0.982959545915103
MULLIGHAN_MLL_SIGNATURE_1_DN	188	0.01	Down
0.363520064641879	0.956170358600006	0.982959545915103
MUNSHI_MULTIPLE_MYELOMA_UP	69	0.01	Up
0.255457626753794	0.956259137106842	0.982959545915103
GRAESSMANN_APOPTOSIS_BY_SERUM_DEPRIVATION_DN	197	0.01	Down
	0.324546180708112	0.956280777371912
0.982959545915103
MIKKELSEN_ES_LCP_WITH_H3K4ME3	85	0.01	Up
0.434304586547276	0.957167783300731	0.983646361668219
TIEN_INTESTINE_PROBIOTICS_6HR_DN	168	0.01	Down
0.236336629465673	0.957713059043583	0.983981760091635
LI_ADIPOGENESIS_BY_ACTIVATED_PPARG	11	0.01	Up
0.491224798122501	0.95815245948213	0.984092389897936
TONKS_TARGETS_OF_RUNX1_RUNX1T1_FUSION_GRANULOCYTE_DN	14	0.01
	Up	0.648362751343878	0.958258596347779
0.984092389897936
LOPEZ_MESOTHELIOMA_SURVIVAL_UP	10	0.01	Up
0.200600501230137	0.959208564201668	0.98450006308263
MIKKELSEN_IPS_LCP_WITH_H3K4ME3	109	0.01	Up
0.533167048003858	0.95923907467939	0.98450006308263
LEE_LIVER_CANCER_E2F1_UP	58	0.01	Up
0.673275676029291	0.959312630990416	0.98450006308263
NGUYEN_NOTCH1_TARGETS_UP	24	0.01	Up
0.42623103296761	0.959779755719252	0.984754622679305

BRACHAT_RESPONSE_TO_METHOTREXATE_UP 20 0.01 Down
0.186218790359312 0.960311482090674 0.985075333637056
REACTOME_SIRT1_NEGATIVELY_REGULATES_RRNA_EXPRESSION 28 0.01
Up 0.158097571053692 0.960795618739159
0.985347092455515
PID_MYC_PATHWAY 25 0.01 Up 0.195661215504867
0.96130848360286 0.985648182891162
FIGUEROA_AML_METHYLATION_CLUSTER_4_DN 10 0.01 Down
0.336361023453757 0.961899616190178 0.98598828918776
MOHANKUMAR_HOXA1_TARGETS_UP 378 0.01 Down
0.231726512963399 0.962435764203191 0.98598828918776
MEISSNER_BRAIN_HCP_WITH_H3K4ME3_AND_H3K27ME3 698 0.01 Up
0.485917951744358 0.962502266999104
0.98598828918776
REACTOME_RHO_GTPASE_CYCLE 128 0.01 Up
0.334852379723312 0.9626723073656 0.98598828918776
FERRANDO_T_ALL_WITH_MLL_ENL_FUSION_DN 79 0.01 Down
0.301842715661232 0.962902049389807 0.98598828918776
REACTOME_TRANSCRIPTION_OF_E2F_TARGETS_UNDER_NEGATIVE_CONTROL_BY_DREA
M_COMPLEX 19 0.01 Up 0.129788293271945
0.963127093909451 0.98598828918776
STEGER_ADIPOGENESIS_UP 12 0.01 Down 0.649634094480509
0.963175656912893 0.98598828918776
REACTOME_TERMINATION_OF_O_GLYCAN_BIOSYNTHESIS 12 0.01 Down
0.600911474507044 0.963628313208342
0.986003460035981
DAIRKEE_CANCER_PRONE_RESPONSE_BPA 44 0.01 Down
0.196546508168543 0.963629187972874 0.986003460035981
KONDO_EZH2_TARGETS 180 0.01 Down 0.500555030069162
0.963923192897316 0.986079825232917
BANDRES_RESPONSE_TO_CARMUSTIN_MGMT_24HR_DN 24 0.01 Up
0.508821526627747 0.964972515874944
0.986834311983544
KASLER_HDAC7_TARGETS_2_DN 28 0.01 Up
0.32678695372849 0.965324279819966 0.986834311983544
DANG_BOUND_BY_MYC 951 0.01 Down 0.19462101841753
0.965532118739901 0.986834311983544
ELLWOOD_MYC_TARGETS_DN 40 0.01 Down 0.252447288348904
0.965697464577293 0.986834311983544
HERNANDEZ_ABERRANT_MITOSIS_BY_DOCETACEL_2NM_DN 19 0.01 Up
0.567683914853951 0.965758429013484
0.986834311983544
MCCLUNG_CREB1_TARGETS_DN 42 0.01 Up
0.328652543812157 0.966450536277537 0.987087891905801
REACTOME_METABOLISM_OF_NITRIC_OXIDE:_ENOS_ACTIVATION_AND_REGULATION
14 0.01 Up 0.342384681107989
0.966491541268347 0.987087891905801
CHENG_IMPRINTED_BY_ESTRADIOL 92 0.01 Up
0.337396841585693 0.966665383797405 0.987087891905801
PID_E2F_PATHWAY 65 0.01 Down 0.231209564137768
0.967709622251465 0.987649181622168
PASQUALUCCI_LYMPHOMA_BY_GC_STAGE_UP 258 0.01 Up
0.299675041857223 0.967721284824109 0.987649181622168
KLEIN_TARGETS_OF_BCR_ABL1_FUSION 28 0.01 Up

0.35079789527629	0.967874225816608	0.987649181622168
SANSOM_APC_TARGETS	163	0.01	Down	0.30724639542458
	0.968188404241786	0.987745546315667
ZHAN_MULTIPLE_MYELOMA_CD1_UP	37	0.01	Up
0.412086677334218	0.96844658402392	0.987784750439646
LOPEZ_MESOTHELIOMA_SURVIVAL_DN	11	0.01	Up
0.501599760528612	0.968972594058745	0.987896606244356
HOFMANN_MYELODYSPLASTIC_SYNDROM_RISK_UP 19	0.01	Down
0.220423188878678	0.969002046190147	0.987896606244356
REACTOME_NICOTINAMIDE_SALVAGING 15	0.01	Down
0.434261700480651	0.969215580319824	0.987896606244356
CROONQUIST_NRAS_VS_STROMAL_STIMULATION_UP	31	0.01	Down
	0.492886221135325	0.96946169573284
0.987923446456385
KIM_WT1_TARGETS_12HR_UP 157	0.01	Down	0.355979951431891
	0.96986673399891	0.988112187063713
BENPORATH_OCT4_TARGETS	257	0.01	Up	0.297914762772388
	0.97012673862813	0.988126949901715
GUO_HEX_TARGETS_DN	43	0.01	Down	0.305139467607153
	0.970320880281684	0.988126949901715
BIOCARTA_BAD_PATHWAY	20	0.01	Down	0.275419006860051
	0.970554915524203	0.988141414559749
REACTOME_DISASSEMBLY_OF_THE_DESTRUCTION_COMPLEX_AND_RECRUITMENT_OF_A
XIN_TO_THE_MEMBRANE	27	0.01	Down	0.236163149201829
	0.971472811863296	0.988661462248776
WU_ALZHEIMER_DISEASE_DN 16	0.01	Up	0.703845278671004
	0.971518039048889	0.988661462248776
REACTOME_SUMOYLATION_OF_TRANSCRIPTION_COFACTORS 43	0.01	Down
	0.183381019026071	0.971725548434948
0.988661462248776
CAFFAREL_RESPONSE_TO_THC_DN	29	0.01	Down
0.226283002701317	0.972716449182815	0.989292371798525
YAO_TEMPORAL_RESPONSE_TO_PROGESTERONE_CLUSTER_10	67	0.01
	Up	0.141230175366521	0.97281245280907
0.989292371798525
REACTOME_RUNX2_REGULATES_OSTEOBLAST_DIFFERENTIATION	19	0.01
	Down	0.222770914319426	0.973123436213413
0.989292371798525
FAELT_B_CLL_WITH_VH_REARRANGEMENTS_DN	48	0.01	Up
0.166137230846364	0.973461734091309	0.989292371798525
SHETH_LIVER_CANCER_VS_TXNIP_LOSS_PAM3	60	0.01	Down
0.551251981800756	0.973875287504688	0.989292371798525
REACTOME_SIGNALING_BY_MET	71	0.01	Down
0.428226155790202	0.973950062878853	0.989292371798525
BOGNI_TREATMENT_RELATED_MYELOID_LEUKEMIA_UP	21	0.01	Down
	0.147510701626194	0.974005889437492
0.989292371798525
ACEVEDO_NORMAL_TISSUE_ADJACENT_TO_LIVER_TUMOR_UP	146	0.01
	Down	0.266883378413555	0.974242907596879
0.989292371798525
NAKAJIMA_EOSINOPHIL	11	0.01	Up	0.51658111601354
	0.974326436029382	0.989292371798525
MIKKELSEN_MCV6_LCP_WITH_H3K4ME3 113	0.01	Down
0.450955892238752	0.975339392687049	0.989727894352855

PUJANA_BREAST_CANCER_WITH_BRCA1_MUTATED_UP 49 0.01 Down
0.156684834910758 0.975506124142026
0.989727894352855
BANDRES_RESPONSE_TO_CARMUSTIN_MGMT_48HR_DN 101 0.01 Down
0.395897739874856 0.9756143351189 0.989727894352855
KEGG_NITROGEN_METABOLISM 14 0.01 Down
0.365601327629239 0.97582853286862 0.989727894352855
HASLINGER_B_CLL_WITH_17P13_DELETION 23 0.01 Up
0.111042362863977 0.975856290939233 0.989727894352855
FINAK_BREAST_CANCER_SDPP_SIGNATURE 14 0.01 Up
0.843820762861371 0.97628013453117 0.989934402146991
CHOI_ATL_STAGE_PREDICTOR 43 0.01 Down
0.243126028871374 0.976687367687366 0.990026645803292
REACTOME_ASSOCIATION_OF_TRIC_CCT_WITH_TARGET_PROTEINS_DURING_BIOSYNT
HESIS 37 0.01 Down 0.153584964221044
0.976811607149633 0.990026645803292
HOLLEMAN_PREDNISOLONE_RESISTANCE_B_ALL_UP 21 0.01 Down
0.100374458542046 0.977616688638176
0.990619255056042
REACTOME_TRANSFERRIN_ENDOCYTOSIS_AND_RECYCLING 26 0.01 Up
0.210163192041071 0.97788125402021
0.99066401551067
YAGI_AML_RELAPSE_PROGNOSIS 32 0.01 Down
0.260188285879366 0.978320322230955 0.990726736932676
SMIRNOV_RESPONSE_TO_IR_6HR_DN 88 0.01 Up
0.456763060859934 0.978383978919721 0.990726736932676
REACTOME_AQUAPORIN_MEDIATED_TRANSPORT 31 0.01 Up
0.522813193162049 0.979218650159802 0.991348610916286
REACTOME_RRNA_PROCESSING_IN_THE_MITOCHONDRION 10 0.01 Down
0.0933693602671044 0.979981062268027
0.991897067078312
FIGUEROA_AML_METHYLATION_CLUSTER_6_UP 111 0.01 Down
0.265946695896503 0.980854221419929 0.99249017556906
TORCHIA_TARGETS_OF_EWSR1_FLI1_FUSION_DN 270 0.01 Down
0.348961925534425 0.981008642948461 0.99249017556906
GOZGIT_ESR1_TARGETS_DN 638 0.01 Down 0.433556718850025
0.981899610787343 0.993082138145628
REACTOME_CRMPS_IN_SEMA3A_SIGNALING 14 0.01 Up
0.41808001339609 0.982035618255242 0.993082138145628
ZHAN_MULTIPLE_MYELOMA_CD1_AND_CD2_UP 83 0.01 Down
0.221744443083738 0.982553976807573 0.993227508200738
BREDEMEYER_RAG_SIGNALING_NOT_VIA_ATM_UP 49 0.01 Down
0.304665813312937 0.9829546260193 0.993227508200738
BONOME_OVARIAN_CANCER_SURVIVAL_SUBOPTIMAL_DEBULKING 439 0.01
Down 0.3092543151991 0.983070959044644
0.993227508200738
BROWNE_HCMV_INFECTION_2HR_DN 42 0.01 Up
0.424745344033705 0.983389869246277 0.993227508200738
REACTOME_SEMAPHORIN_INTERACTIONS 59 0.01 Down
0.37370931722172 0.98341207477347 0.993227508200738
MARIADASON_REGULATED_BY_HISTONE_ACETYLATION_UP 55 0.01 Down
0.227512428553818 0.983985805943761
0.993227508200738
PID_SYNDECAN_4_PATHWAY 28 0.01 Down 0.547425163899305

0.984112973726971	0.993227508200738
LY_AGING_PREMATURE_DN	26	0.01	Down	0.474307904036249
0.984147445580043	0.993227508200738
HASLINGER_B_CLL_WITH_13Q14_DELETION	21	0.01	Down
0.300443062763235	0.984331444732267	0.993227508200738
JOHANSSON_GLIOMAGENESIS_BY_PDGFB_DN	14	0.01	Down
0.287206277303979	0.984388998672811	0.993227508200738
GAVIN_FOXP3_TARGETS_CLUSTER_P3	131	0.01	Down
0.403809727087387	0.985047700573716	0.993669078563477
HOLLEMAN_PREDNISOLONE_RESISTANCE_ALL_UP	19	0.01	Down
0.0762408988314271	0.985644074129724	0.99404759102829
REACTOME_ERCC6_CSB_AND_EHMT2_G9A_POSITIVELY_REGULATE_RRNA_EXPRESSION
36	0.01	Down	0.124763695146381
0.98671598357474	0.994492393239895
REACTOME_TRANSCRIPTIONAL_REGULATION_BY_E2F6	34	0.01	Up
0.150552473185401	0.986780717935882
0.994492393239895
VANOEVELEN_MYOGENESIS_SIN3A_TARGETS	205	0.01	Up
0.195831110607741	0.986827518673625	0.994492393239895
REACTOME_G_BETA:GAMMA_SIGNALLING_THROUGH_PI3KGAMMA	17	0.01
Up	0.314097551153743	0.986978294568573
0.994492393239895
HALMOS_CEBPA_TARGETS_DN	37	0.01	Down	0.320721604465695
0.9871913367378	0.994492393239895
MYLLYKANGAS_AMPLIFICATION_HOT_SPOT_17	15	0.01	Down
0.375622776618651	0.987499354908159	0.994579789449288
RADMACHER_AML_PROGNOSIS	69	0.01	Down	0.532456117880819
0.98898745953994	0.995855428008968
WINTER_HYPOXIA_UP	93	0.01	Up	0.206716494983636
0.989242241854521	0.995888886256679
BURTON_ADIPOGENESIS_5	112	0.01	Down	0.216093593228285
0.989519935122421	0.99590489968808
ROPERO_HDAC2_TARGETS	78	0.01	Up	0.290105031826282
0.989882491761475	0.99590489968808
REACTOME_REGULATION_OF_PLK1_ACTIVITY_AT_G2_M_TRANSITION	83	0.01
Down	0.129412194396483	0.990149668637636
0.99590489968808
CHIARADONNA_NEOPLASTIC_TRANSFORMATION_CDC25_UP	104	0.01	Down
0.496721686765184	0.990299182922619
0.99590489968808
PID_DELTA_NP63_PATHWAY	44	0.01	Down	0.309357701847276
0.990365940290482	0.99590489968808
BRACHAT_RESPONSE_TO_CAMPTOTHECIN_DN	38	0.01	Down
0.20018092652672	0.990867308354657	0.99618621137423
REACTOME_DNA_DAMAGE_BYPASS	46	0.01	Up
0.125654300251501	0.991111465189447	0.99620886315442
BERENJENO_ROCK_SIGNALING_NOT_VIA_RHOA_DN	46	0.01	Down
0.44690981181822	0.991717234592948
0.996372030889743
HOEGERKORP_CD44_TARGETS_DIRECT_DN	10	0.01	Up
0.544650427771172	0.992017916106684	0.996372030889743
ABBUD_LIF_SIGNALING_1_DN	20	0.01	Down
0.453222822660692	0.992060027941572	0.996372030889743
REACTOME_DEPURINATION	24	0.01	Down	0.121818293330095

0.992446377632016	0.996372030889743
DELACROIX_RARG_BOUND_MEF	317	0.01	Up
0.33045883815411	0.992783270439275	0.996372030889743
MADAN_DPPA4_TARGETS	26	0.01	Down	0.437330488448702
0.992790676727687	0.996372030889743
KEGG_PYRIMIDINE_METABOLISM	90	0.01	Up
0.190671940239587	0.99291132598027	0.996372030889743
AKL_HTLV1_INFECTION_DN	56	0.01	Up	0.476622093016004
0.993047096415139	0.996372030889743
HASLINGER_B_CLL_WITH_CHROMOSOME_12_TRISOMY	21	0.01	Up
0.209356785860131	0.993436863792679
0.996540661180114
LI_CYTIDINE_ANALOG_PATHWAY	13	0.01	Up
0.314199669125474	0.994013393030776	0.996679654543786
BERENJENO_TRANSFORMED_BY_RHOA_REVERSIBLY_DN	29	0.01	Up
0.426280213674015	0.994018885721868
0.996679654543786
BIOCARTA_NOS1_PATHWAY	13	0.01	Up	0.38579380270684
0.99589385235767	0.998336946107878
SMID_BREAST_CANCER_RELAPSE_IN_LUNG_UP	17	0.01	Up
0.601231026252234	0.996676242600045	0.99888457622702
WANG_TUMOR_INVASIVENESS_UP	344	0.01	Up
0.190870392917402	0.996884584861938	0.99888457622702
REACTOME_CARGO_TRAFFICKING_TO_THE_PERICILIARY_MEMBRANE	44	0.01
Up	0.161673761369707	0.997307799246433
0.999014692344183
SHEPARD_CRUSH_AND_BURN_MUTANT_UP	184	0.01	Down
0.310075388620665	0.997458940876684	0.999014692344183
WHITFIELD_CELL_CYCLE_G1_S	117	0.01	Up
0.224419674737384	0.998430118086964	0.999759314737124
RAMPON_ENRICHED_LEARNING_ENVIRONMENT_LATE_UP	18	0.01	Up
0.145059853146253	0.998824413772615
0.999759314737124
REACTOME_SIGNAL_AMPLIFICATION	21	0.01	Up
0.22786035271964	0.998927565958096	0.999759314737124
CHEN_PDGF_TARGETS	17	0.01	Down	0.252831531023561
0.999092067141081	0.999759314737124
VALK_AML_CLUSTER_4	21	0.01	Down	0.239478583912202
0.999660962439922	0.999942502583607
REACTOME_CITRIC_ACID_CYCLE_TCA_CYCLE	22	0.01	Down
0.0894014485755986	0.999790461935736	0.999942502583607
KYNG_DNA_DAMAGE_BY_4NQO_OR_GAMMA_RADIATION	13	0.01	Down
0.354615370253616	0.999942502583607
0.999942502583607

**Supplement table 5:**
edgeR_DE_GSEA_results_c5.CATEGORY.v7.0.entrez.hsa_IPF_effect

| | NGenes | Correlation | Direction | absLog2FC | PValue | FDR |
|---|---|---|---|---|---|---|
| GO_URONIC_ACID_METABOLIC_PROCESS | 11 | 0.01 | Up | 1.23369490479328 | 1.24548037539281e-31 | 7.56131135900977e-28 |
| GO_FLAVONOID_METABOLIC_PROCESS | 14 | 0.01 | Up | 1.02749153476383 | 1.17188749275902e-22 | 3.55726448427001e-19 |
| GO_GLUCURONOSYLTRANSFERASE_ACTIVITY | 20 | 0.01 | Up | 0.786493694659113 | 1.26422956338304e-18 | 2.55837922643281e-15 |
| GO_COMPLEX_OF_COLLAGEN_TRIMERS | 17 | 0.01 | Up | 1.49034253987792 | 2.35172042032805e-15 | 3.56932366795289e-12 |
| GO_EXTRACELLULAR_MATRIX_COMPONENT | 42 | 0.01 | Up | 1.1028164296858 | 5.94056693137404e-14 | 7.21303636807436e-11 |
| GO_EXTRACELLULAR_MATRIX_STRUCTURAL_CONSTITUENT | 110 | 0.01 | Up | 1.07104375595921 | 1.54406349413378e-12 | 1.5623349121477e-09 |
| GO_COLLAGEN_TRIMER | 53 | 0.01 | Up | 1.39671843438991 | 3.02724214294784e-12 | 2.62548386426233e-09 |
| GO_EXTRACELLULAR_MATRIX_STRUCTURAL_CONSTITUENT_CONFERRING_TENSILE_STRENGTH | 29 | 0.01 | Up | 1.27477868588319 | 8.09130363912078e-12 | 6.14028804913778e-09 |
| GO_CORNIFICATION | 63 | 0.01 | Down | 0.903956567833552 | 1.01962191053091e-11 | 6.87791624314797e-09 |
| GO_KERATINIZATION | 70 | 0.01 | Down | 0.864284169314166 | 2.47343706872461e-11 | 1.50162364442271e-08 |
| GO_COLLAGEN_FIBRIL_ORGANIZATION | 41 | 0.01 | Up | 1.22769677972206 | 4.46048697490521e-11 | 2.46178331133178e-08 |
| GO_CORNIFIED_ENVELOPE | 31 | 0.01 | Down | 1.11266412933511 | 1.20500354422069e-10 | 6.09631376413649e-08 |
| GO_RETINOIC_ACID_METABOLIC_PROCESS | 11 | 0.01 | Up | 0.904048292249386 | 1.13073565422501e-09 | 5.2805355052308e-07 |
| GO_LYSOSOMAL_LUMEN | 77 | 0.01 | Up | 0.596943104997967 | 1.44826965286243e-08 | 6.28031790180556e-06 |
| GO_XENOBIOTIC_METABOLIC_PROCESS | 74 | 0.01 | Up | 0.595179459932947 | 2.06111709344614e-08 | 8.34202791620769e-06 |
| GO_RETINOIC_ACID_BINDING | 14 | 0.01 | Up | 0.7219981020834 | 5.77992298636091e-08 | 2.19311952813732e-05 |
| GO_ANTIMICROBIAL_HUMORAL_RESPONSE | 53 | 0.01 | Down | 1.13845692581827 | 7.53479936808951e-08 | 2.69080982139244e-05 |
| GO_COLLAGEN_CONTAINING_EXTRACELLULAR_MATRIX | 271 | 0.01 | Up | 0.852901555752442 | 1.16678175582627e-07 | 3.93529557756739e-05 |
| GO_PLATELET_DERIVED_GROWTH_FACTOR_BINDING | 10 | 0.01 | Up | 1.42384583573847 | 1.30011139015031e-07 | 4.1541980261066e-05 |
| GO_ANTIMICROBIAL_HUMORAL_IMMUNE_RESPONSE_MEDIATED_BY_ANTIMICROBIAL_PEPTIDE | 34 | 0.01 | Down | 1.13512332577899 | 2.23610215157121e-07 | 6.78768808109442e-05 |
| GO_ORGANIC_ACID_BINDING | 123 | 0.01 | Up | 0.465294402695841 | 3.61869807567337e-07 | 0.000104614838178157 |
| GO_EPIDERMAL_CELL_DIFFERENTIATION | 170 | 0.01 | Down | 0.597840957360114 | 4.51849765510383e-07 | 0.000124689996655161 |
| GO_CHEMOKINE_RECEPTOR_BINDING | 31 | 0.01 | Down | 1.20731598738185 | 5.42937330727142e-07 | 0.000143311849341064 |
| GO_ENDOPLASMIC_RETICULUM_LUMEN | 222 | 0.01 | Up | | | |

0.553097109563055	5.97374259032785e-07	0.000151110796941168
GO_EXTRACELLULAR_MATRIX 335	0.01	Up	0.843218264388712
	8.27672876327443e-07	0.000200992081287356
GO_AMINOGLYCAN_CATABOLIC_PROCESS	46	0.01	Up
0.808659500677889	1.33345462588681e-06	0.00031136165514457
GO_EPIDERMIS_DEVELOPMENT	260	0.01	Down
0.551003150507011	2.62067372767502e-06	0.000576367782191635
GO_CHEMOKINE_ACTIVITY	21	0.01	Down	1.47817912865817
	2.65826023741818e-06	0.000576367782191635
GO_NUCLEOTIDE_BINDING_OLIGOMERIZATION_DOMAIN_CONTAINING_2_SIGNALING_
PATHWAY 11	0.01	Down	0.501858242721136
3.61606623440443e-06	0.0007570047623817
GO_DOPAMINE_TRANSPORT	18	0.01	Up	0.817489509652778
	3.97824299320183e-06	0.000805063773724276
GO_MONOSACCHARIDE_BINDING	57	0.01	Up
0.455001201656648	5.73113026900598e-06	0.00112237715687533
GO_MICROVILLUS_ORGANIZATION	18	0.01	Down
0.505077964268478	6.41156825232655e-06	0.00121639471437108
GO_NUCLEOTIDE_BINDING_DOMAIN_LEUCINE_RICH_REPEAT_CONTAINING_RECEPTOR
_SIGNALING_PATHWAY	33	0.01	Down	0.353589487810461
	1.05457112820827e-05	0.00194009130889467
GO_COLLAGEN_METABOLIC_PROCESS	14	0.01	Up
0.619813069827338	1.36240894818728e-05	0.00243270138954264
GO_CYTOPLASMIC_PATTERN_RECOGNITION_RECEPTOR_SIGNALING_PATHWAY	57
	0.01	Down	0.288872541476333	1.69163100840091e-05
	0.0029342548148577
GO_DOPAMINE_SECRETION	13	0.01	Up	0.928204363166641
	1.75114016486213e-05	0.00295310331691056
GO_EYELID_DEVELOPMENT_IN_CAMERA_TYPE_EYE	11	0.01	Down
	0.566598111135408	2.03634305094794e-05
0.00334125369251485
GO_RESPONSE_TO_MURAMYL_DIPEPTIDE	19	0.01	Down
0.42100376132142	2.40013504442352e-05	0.00374137804381372
GO_PIGMENT_METABOLIC_PROCESS	59	0.01	Up
0.380122057605655	2.44596931562276e-05	0.00374137804381372
GO_T_HELPER_17_TYPE_IMMUNE_RESPONSE	19	0.01	Down
0.535646168747897	2.46508189346976e-05	0.00374137804381372
GO_MHC_CLASS_II_PROTEIN_COMPLEX 14	0.01	Up
1.114040502524	2.81612110719983e-05	0.00416991981507565
GO_CHRONIC_INFLAMMATORY_RESPONSE	11	0.01	Down
1.23829649297428	3.49472685747123e-05	0.00505154446469235
GO_GLUTATHIONE_PEROXIDASE_ACTIVITY	16	0.01	Up
0.422978611196357	3.63378822459698e-05	0.00513040193291355
GO_COLLAGEN_ACTIVATED_SIGNALING_PATHWAY 13	0.01	Up
0.885215608537847	4.0784984162001e-05	0.00550196374932374
GO_MONOCARBOXYLIC_ACID_BINDING	45	0.01	Up
0.605118993522346	4.14007819135e-05	0.00550196374932374
GO_MONOAMINE_TRANSPORT	32	0.01	Up	0.72993944441267
	4.16884092355282e-05	0.00550196374932374
GO_MICROVILLUS_ASSEMBLY 15	0.01	Down	0.556813713265613
	4.57829007058473e-05	0.00584907041642131
GO_COLLAGEN_ACTIVATED_TYROSINE_KINASE_RECEPTOR_SIGNALING_PATHWAY
	10	0.01	Up	0.763650359533683
4.62453269623164e-05	0.00584907041642131

GO_XENOBIOTIC_CATABOLIC_PROCESS 10 0.01 Up
0.88492769723411 5.05332618256138e-05 0.0061457032662187
GO_BASEMENT_MEMBRANE 77 0.01 Up 0.763554994111497
5.06152467980457e-05 0.0061457032662187
GO_NEGATIVE_REGULATION_OF_VIRAL_PROCESS 78 0.01 Down
0.365975056228946 5.4496911316522e-05 0.00648726958044324
GO_INTERLEUKIN_1_MEDIATED_SIGNALING_PATHWAY 96 0.01 Down
0.248312843590588 5.62231336590688e-05
0.00656405085469628
GO_RESPONSE_TO_CHEMOKINE 48 0.01 Down
0.958290409619948 6.22292543379842e-05 0.00712818496388494
GO_KERATINOCYTE_MIGRATION 17 0.01 Down
0.64635469186779 7.39587496271328e-05 0.00831488090715413
GO_SKIN_DEVELOPMENT 225 0.01 Down 0.610559577225381
9.8007231721395e-05 0.0108182164323744
GO_L_ASCORBIC_ACID_BINDING 18 0.01 Up
0.324576747924279 0.000107848395118131 0.0116919215493245
GO_OXIDOREDUCTASE_ACTIVITY_ACTING_ON_PEROXIDE_AS_ACCEPTOR 33
0.01 Up 0.407061139069561 0.000113572686228795
0.0120964873350002
GO_ANTIBIOTIC_METABOLIC_PROCESS 59 0.01 Up
0.466755643310264 0.000130304661103284 0.0136230354238158
GO_EXTRACELLULAR_STRUCTURE_ORGANIZATION 306 0.01 Up
0.749940576468806 0.000132393195520529 0.0136230354238158
GO_SULFUR_COMPOUND_CATABOLIC_PROCESS 36 0.01 Up
0.687334881817367 0.000151667161564264 0.0153461889642774
GO_REGULATION_OF_T_HELPER_17_TYPE_IMMUNE_RESPONSE 14 0.01
Down 0.500468128714156 0.000177935710663999
0.0177089786793629
GO_GLUTATHIONE_DERIVATIVE_BIOSYNTHETIC_PROCESS 15 0.01 Up
0.572477641745679 0.000191535315471481
0.0185449432460994
GO_DESMOSOME 22 0.01 Down 0.372734072110955
0.000192444642481347 0.0185449432460994
GO_KERATIN_FILAMENT 23 0.01 Down 0.70957959179153
0.00020309681130417 0.0189712795171701
GO_NIK_NF_KAPPAB_SIGNALING 147 0.01 Down
0.36772269897652 0.000203118624380836 0.0189712795171701
GO_NEGATIVE_REGULATION_OF_VIRAL_GENOME_REPLICATION 49 0.01
Down 0.381887113013992 0.000221267955165037
0.0203532993304081
GO_CHONDROITIN_SULFATE_CATABOLIC_PROCESS 10 0.01 Up
0.961002239811551 0.000247051754884688
0.0223858388642528
GO_T_CELL_CHEMOTAXIS 17 0.01 Down 0.789870087449829
0.000295272702731325 0.0257356570701507
GO_MAP_KINASE_PHOSPHATASE_ACTIVITY 12 0.01 Down
0.458596002921998 0.000296632943008567 0.0257356570701507
GO_LYMPHOCYTE_CHEMOTAXIS 32 0.01 Down
0.777702113783454 0.000302775407454212 0.0257356570701507
GO_CD4_POSITIVE_ALPHA_BETA_T_CELL_ACTIVATION 64 0.01 Down
0.450238346059721 0.000306078558249833
0.0257356570701507
GO_RETINA_VASCULATURE_DEVELOPMENT_IN_CAMERA_TYPE_EYE 15 0.01

Up 0.74185706053668 0.00030805003316621
0.0257356570701507
GO_ALPHA_BETA_T_CELL_DIFFERENTIATION 72 0.01 Down
0.433625139390905 0.000309455273615714 0.0257356570701507
GO_VACUOLAR_LUMEN 136 0.01 Up 0.494571188563793
0.00032023201666486 0.0262720077455725
GO_CELL_MIGRATION_INVOLVED_IN_HEART_DEVELOPMENT 11 0.01 Up
1.029243809815 0.000326544394712014 0.0263138338951276
GO_DNA_BINDING_TRANSCRIPTION_ACTIVATOR_ACTIVITY 258 0.01 Down
0.400458785726472 0.000332791920365728
0.0263138338951276
GO_RAP_PROTEIN_SIGNAL_TRANSDUCTION 13 0.01 Down
0.345853318105013 0.000333744887156123 0.0263138338951276
GO_NEUTROPHIL_MIGRATION 68 0.01 Down 1.01069254708337
0.000405411687594383 0.0315545430177628
GO_GLUTATHIONE_TRANSFERASE_ACTIVITY 17 0.01 Up
0.587953146649051 0.000432702208459774 0.0332523431336619
GO_PROTEIN_SELF_ASSOCIATION 44 0.01 Down
0.390733995491129 0.000450481080723504 0.0338107178242634
GO_DEFENSE_RESPONSE_TO_VIRUS 189 0.01 Down
0.343407475485357 0.000451106595909296 0.0338107178242634
GO_EXTRACELLULAR_MATRIX_ASSEMBLY 16 0.01 Up
0.782776895188944 0.000464297142502412 0.0340666138037366
GO_POSITIVE_REGULATION_OF_CYTOKINE_PRODUCTION_INVOLVED_IN_INFLAMMATO
RY_RESPONSE 13 0.01 Down 0.580639508218623
0.000465743525895263 0.0340666138037366
GO_REGULATION_OF_KERATINOCYTE_MIGRATION 13 0.01 Down
0.686914252554419 0.000495968172945039 0.0357315642354
GO_CELLULAR_RESPONSE_TO_VIRUS 44 0.01 Down
0.297114847946606 0.000500277212981222 0.0357315642354
GO_RESPONSE_TO_INTERLEUKIN_1 159 0.01 Down
0.376518382732205 0.000521896284463334 0.0368422365462431
GO_PULMONARY_VALVE_MORPHOGENESIS 12 0.01 Down
0.596854145936601 0.000535104081541621 0.0373404238970021
GO_NITRIC_OXIDE_MEDIATED_SIGNAL_TRANSDUCTION 19 0.01 Up
0.839254005286768 0.000573725249256096
0.0395805225935654
GO_CELLULAR_RESPONSE_TO_XENOBIOTIC_STIMULUS 115 0.01 Up
0.555621174701821 0.000594783331699354
0.0405722427724357
GO_REGULATION_OF_WNT_SIGNALING_PATHWAY_PLANAR_CELL_POLARITY_PATHWAY
13 0.01 Up 0.710397844477578
0.000602819438506697 0.0406012357874665
GO_NLS_BEARING_PROTEIN_IMPORT_INTO_NUCLEUS 23 0.01 Down
0.243389478661929 0.000608583834073374
0.0406012357874665
GO_NUCLEAR_TRANSCRIBED_MRNA_POLY_A_TAIL_SHORTENING 32 0.01
Down 0.186222776096577 0.00061782053234081
0.0407694396939245
GO_ALPHA_BETA_T_CELL_ACTIVATION 97 0.01 Down
0.434901854787521 0.000645336472118861 0.0421272873358452
GO_ISOPRENOID_BINDING 19 0.01 Up 0.741109479361153
0.00065999095184053 0.042625585836424
GO_WNT_PROTEIN_BINDING 29 0.01 Up 0.509753244349225

0.000675621715823578	0.0431757835448941
GO_OXIDOREDUCTASE_ACTIVITY_ACTING_ON_NAD_P_H_QUINONE_OR_SIMILAR_COMPOUND_AS_ACCEPTOR	51	0.01	Up	0.208078978613079	0.000762536910645862	0.0474685836761545
GO_OLIGOPEPTIDE_BINDING	11	0.01	Up	0.694095079722688	0.000766014100920117	0.0474685836761545
GO_CYTOKINE_RECEPTOR_BINDING	166	0.01	Down	0.627311239212589	0.000766252874363885	0.0474685836761545
GO_BONE_TRABECULA_MORPHOGENESIS	11	0.01	Up	1.17801206616595	0.000788580690889305	0.0483583169130199
GO_CELLULAR_DETOXIFICATION	76	0.01	Up	0.456619033891799	0.000811995219153277	0.0492962297547955
GO_HIPPO_SIGNALING	37	0.01	Down	0.26795410707109	0.000835021932983081	0.050192258961785
GO_GRANULOCYTE_MIGRATION	88	0.01	Down	0.906105716472212	0.000853082835144154	0.0507751558054917
GO_HEXOSAMINIDASE_ACTIVITY	12	0.01	Up	0.433991141956386	0.000863602245675537	0.0509022255679241
GO_RESPONSE_TO_VIRUS	257	0.01	Down	0.371172931892365	0.000914373328067025	0.0533765430259126
GO_ANTIOXIDANT_ACTIVITY	56	0.01	Up	0.40558898661536	0.000967110847211307	0.0555474504647842
GO_ANTIBIOTIC_CATABOLIC_PROCESS	27	0.01	Up	0.426430099292682	0.000969861595991949	0.0555474504647842
GO_CATECHOLAMINE_SECRETION	20	0.01	Up	0.683765193191499	0.00107267738042467	0.0604542421331949
GO_GLUCOCORTICOID_RECEPTOR_BINDING	13	0.01	Down	0.412080588422161	0.00107545019772444	0.0604542421331949
GO_SECONDARY_METABOLIC_PROCESS	45	0.01	Up	0.422223250790462	0.00108678904805759	0.0605311588142903
GO_CHONDROCYTE_DEVELOPMENT_INVOLVED_IN_ENDOCHONDRAL_BONE_MORPHOGENESIS	15	0.01	Up	0.676541461510267	0.00114318996372865	0.0625728866426813
GO_HISTONE_DEMETHYLASE_ACTIVITY	25	0.01	Down	0.256340425893747	0.00114406035535128	0.0625728866426813
GO_UDP_GLYCOSYLTRANSFERASE_ACTIVITY	113	0.01	Up	0.442770448981707	0.00115917835820699	0.0628336768988809
GO_NEGATIVE_REGULATION_OF_CD4_POSITIVE_ALPHA_BETA_T_CELL_ACTIVATION	21	0.01	Down	0.543949989177559	0.00119273794651375	0.064080637816681
GO_FATTY_ACID_METABOLIC_PROCESS	201	0.01	Up	0.346710115408329	0.00124896611569156	0.0665129235821357
GO_MONOCARBOXYLIC_ACID_METABOLIC_PROCESS	346	0.01	Up	0.367952865953768	0.00135445342028875	0.0715033627354173
GO_INTERLEUKIN_1_RECEPTOR_BINDING	11	0.01	Down	0.832731779586931	0.00136849560996302	0.0716218693800476
GO_DEFENSE_RESPONSE_TO_OTHER_ORGANISM	306	0.01	Down	0.475221720074507	0.00145594559418572	0.0755053164862691
GO_T_CELL_DIFFERENTIATION_INVOLVED_IN_IMMUNE_RESPONSE	46	0.01	Down	0.483877484275291	0.00146757162664796	0.0755053164862691
GO_MAP_KINASE_KINASE_KINASE_ACTIVITY	25	0.01	Down	0.265315016350492	0.00155481180738194	0.0793215334673595

GO_LONG_CHAIN_FATTY_ACID_METABOLIC_PROCESS 60 0.01 Up
0.381273583150163 0.00158080504890675
0.0799755620992739
GO_BENZENE_CONTAINING_COMPOUND_METABOLIC_PROCESS 11 0.01
Up 0.763418144751712 0.00159870277997081
0.0802125998115931
GO_DNA_DAMAGE_RESPONSE_SIGNAL_TRANSDUCTION_RESULTING_IN_TRANSCRIPTIO
N 16 0.01 Up 0.257013329544033
0.00161292471063997 0.0802628353958628
GO_REGULATION_OF_VIRAL_GENOME_REPLICATION 82 0.01 Down
0.33052158269571 0.00170059128399688
0.0839373145133746
GO_APICAL_JUNCTION_ASSEMBLY 53 0.01 Down
0.280966998729296 0.00177522503846077 0.086914445229801
GO_REGULATORY_RNA_BINDING 32 0.01 Down
0.186259800633449 0.00179179123054084 0.0870237164849075
GO_3_UTR_MEDIATED_MRNA_DESTABILIZATION 14 0.01 Down
0.310242298819153 0.00181823962895211 0.0876074030743513
GO_REGULATION_OF_CD4_POSITIVE_ALPHA_BETA_T_CELL_ACTIVATION 41
0.01 Down 0.485490843564086 0.00184689640462052
0.0882874651374109
GO_OXIDOREDUCTASE_ACTIVITY_ACTING_ON_THE_CH_CH_GROUP_OF_DONORS 53
0.01 Up 0.269620189045467 0.00187551304986029
0.0889549978570455
GO_REGULATION_OF_CELL_FATE_SPECIFICATION 10 0.01 Up
0.961705377860407 0.00194930540294968
0.091738241095407
GO_GLUTATHIONE_METABOLIC_PROCESS 30 0.01 Up
0.458937243058243 0.00202590455768379 0.0943339794736864
GO_REGULATION_OF_GASTRULATION 27 0.01 Up
0.655328944082905 0.00203553801862179 0.0943339794736864
GO_NEGATIVE_REGULATION_OF_VIRAL_TRANSCRIPTION 22 0.01 Down
0.387430170217177 0.00208148752424539
0.0951994124845014
GO_EXTRACELLULAR_MATRIX_STRUCTURAL_CONSTITUENT_CONFERRING_COMPRESSIO
N_RESISTANCE 10 0.01 Up 1.80983901480472
0.00208729877947553 0.0951994124845014
GO_SCAVENGER_RECEPTOR_ACTIVITY 31 0.01 Up
0.940963667882593 0.00212303158404458 0.0951994124845014
GO_CCR4_NOT_COMPLEX 16 0.01 Down 0.173726255747064
0.00212934007212079 0.0951994124845014
GO_COLLAGEN_BIOSYNTHETIC_PROCESS 37 0.01 Up
0.7543823810267 0.00215624256870607 0.0951994124845014
GO_REGULATION_OF_ALPHA_BETA_T_CELL_ACTIVATION 67 0.01 Down
0.460583292532862 0.00216666150318384
0.0951994124845014
GO_CADHERIN_BINDING 298 0.01 Down 0.190904010679852
0.00216960473748273 0.0951994124845014
GO_COFACTOR_TRANSMEMBRANE_TRANSPORTER_ACTIVITY 19 0.01 Up
0.350120022438833 0.00217966040773278
0.0951994124845014
GO_REGULATION_OF_CELL_CELL_ADHESION 293 0.01 Down
0.501836659405883 0.00222669065173228 0.0965588496190476
GO_TIGHT_JUNCTION 107 0.01 Down 0.321098982564431

0.00226728127727815 0.0976217349954302
GO_POSITIVE_REGULATION_OF_MRNA_CATABOLIC_PROCESS 44 0.01
Down 0.229462518265061 0.00231223998277664
0.0988564009537815
GO_MEMBRANE_BIOGENESIS 28 0.01 Up 0.245150116861477
0.00234120571133566 0.0993061732981178
GO_POSITIVE_REGULATION_OF_DEFENSE_RESPONSE_TO_VIRUS_BY_HOST 21
0.01 Down 0.38845282637405 0.00235547503787333
0.0993061732981178
GO_LEUKOCYTE_CELL_CELL_ADHESION 243 0.01 Down
0.5344385117409 0.0023824074095275 0.0997489336775271
GO_MIRNA_BINDING 26 0.01 Down 0.203140210452335
0.00240420953378129 0.0998744225089579
GO_APICAL_JUNCTION_COMPLEX 121 0.01 Down
0.30808790569218 0.00241830672192667 0.0998744225089579
GO_MODIFIED_AMINO_ACID_BINDING 65 0.01 Up
0.590007958798666 0.00245812392869784 0.100832907913004
GO_ERBB2_SIGNALING_PATHWAY 27 0.01 Down
0.257384924424499 0.00248741871090615 0.101349791905445
GO_DETOXIFICATION 84 0.01 Up 0.452404055404345
0.00262767495105674 0.104730804390943
GO_DERMATAN_SULFATE_BIOSYNTHETIC_PROCESS 10 0.01 Up
0.94530447473033 0.00263617299164231
0.104730804390943
GO_NEGATIVE_REGULATION_OF_ORGAN_GROWTH 24 0.01 Down
0.396530476397089 0.00266417797978955 0.104730804390943
GO_ESTABLISHMENT_OF_ENDOTHELIAL_INTESTINAL_BARRIER 10 0.01
Down 0.358849714010289 0.00266627221405385
0.104730804390943
GO_HYDROLASE_ACTIVITY_ACTING_ON_GLYCOSYL_BONDS 85 0.01 Up
0.28945091115347 0.00267054837253475
0.104730804390943
GO_ENDOCRINE_HORMONE_SECRETION 25 0.01 Down
0.484364426678694 0.00267725048281795 0.104730804390943
GO_REGULATION_OF_VIRAL_LIFE_CYCLE 123 0.01 Down
0.31304634300342 0.00269115557321482 0.104730804390943
GO_CELLULAR_RESPONSE_TO_BIOTIC_STIMULUS 172 0.01 Down
0.58830169372612 0.00275886087906596 0.106681811444646
GO_TIGHT_JUNCTION_ORGANIZATION 48 0.01 Down
0.290712173578114 0.002795849402561 0.107427859005999
GO_CELLULAR_HORMONE_METABOLIC_PROCESS 63 0.01 Up
0.513610965850777 0.00281420474982113 0.107453063120529
GO_STRUCTURAL_CONSTITUENT_OF_RIBOSOME 154 0.01 Up
0.121561991430064 0.00286576940395981 0.108149868453852
GO_INTRACILIARY_TRANSPORT_PARTICLE 27 0.01 Up
0.219173572125282 0.00289333887120783 0.108149868453852
GO_PROTEIN_LOCALIZATION_TO_CILIUM 42 0.01 Up
0.287236003863323 0.00291402955428941 0.108149868453852
GO_EMBRYONIC_CAMERA_TYPE_EYE_MORPHOGENESIS 17 0.01 Down
0.370476741547947 0.00293835959542727
0.108149868453852
GO_CYTOKINE_ACTIVITY 102 0.01 Down 0.907165243187567
0.00294344642294403 0.108149868453852
GO_PULMONARY_VALVE_DEVELOPMENT 15 0.01 Down

0.498280950924749	0.00295924604750098	0.108149868453852
GO_NEGATIVE_REGULATION_OF_CD4_POSITIVE_ALPHA_BETA_T_CELL_DIFFERENTIATION	16	0.01	Down	0.529486986321836
0.00296598931382347	0.108149868453852
GO_NEGATIVE_REGULATION_OF_EMBRYONIC_DEVELOPMENT	22	0.01	Up
0.726454444806772	0.00297496755588756
0.108149868453852
GO_MITOTIC_NUCLEAR_ENVELOPE_DISASSEMBLY	10	0.01	Up
0.405385033524066	0.003039219249160	0.109827976557475
GO_PEPTIDYL_PROLINE_HYDROXYLATION	12	0.01	Up
0.334554473502215	0.00306023326586751	0.109932995012318
GO_PROTEIN_OXIDATION	12	0.01	Up	0.444999254472622
0.0031481037151259	0.111498809955271
GO_POSITIVE_REGULATION_OF_T_CELL_APOPTOTIC_PROCESS	10	0.01
Down	0.884727781541055	0.00315085995266224
0.111498809955271
GO_RESPONSE_TO_OSMOTIC_STRESS	61	0.01	Down
0.431093650186934	0.00317437377241451	0.111498809955271
GO_VIRAL_LIFE_CYCLE	282	0.01	Down	0.285071836560374
0.00318916556563038	0.111498809955271
GO_BONE_GROWTH	31	0.01	Up	0.628908744235517
0.00319565029356236	0.111498809955271
GO_PLATELET_ALPHA_GRANULE_MEMBRANE	14	0.01	Up
0.822182186809472	0.00330988614060396	0.113824161181816
GO_NEGATIVE_REGULATION_OF_CELL_CELL_ADHESION	126	0.01	Down
0.527198766164836	0.00331921025989886
0.113824161181816
GO_REGULATION_OF_DEFENSE_RESPONSE_TO_VIRUS_BY_HOST	31	0.01
Down	0.368821671247618	0.00332642550067675
0.113824161181816
GO_FC_EPSILON_RECEPTOR_SIGNALING_PATHWAY	103	0.01	Down
0.223812534026057	0.00334357507368832
0.113824161181816
GO_CHROMATIN_DNA_BINDING	81	0.01	Down
0.249781716345141	0.00335604099020672	0.113824161181816
GO_REGULATION_OF_ALPHA_BETA_T_CELL_DIFFERENTIATION	45	0.01
Down	0.455789624866224	0.00342690728143214
0.115581967253192
GO_POSITIVE_REGULATION_OF_RECEPTOR_SIGNALING_PATHWAY_VIA_STAT	51
0.01	Down	0.656196709100881	0.00350256571596602
0.117481085423368
GO_COFACTOR_METABOLIC_PROCESS	352	0.01	Up
0.312147315493028	0.0035630733400686	0.118853946415145
GO_POSITIVE_REGULATION_OF_TYROSINE_PHOSPHORYLATION_OF_STAT_PROTEIN
38	0.01	Down	0.691142128433184
0.00362132230091194	0.118935258656912
GO_P_BODY	77	0.01	Down	0.179409568399684
0.00362627005226483	0.118935258656912
GO_GROWTH_PLATE_CARTILAGE_CHONDROCYTE_DIFFERENTIATION	15	0.01
Up	0.70917582490572	0.00363182290510657
0.118935258656912
GO_REGULATION_OF_GENE_EXPRESSION_EPIGENETIC	297	0.01	Down
0.204076402322797	0.00364387384453724
0.118935258656912

GO_REGULATION_OF_MRNA_CATABOLIC_PROCESS 179 0.01 Down
0.181807450966601 0.00370546087735104 0.120298679071648
GO_CARTILAGE_DEVELOPMENT_INVOLVED_IN_ENDOCHONDRAL_BONE_MORPHOGENESIS
30 0.01 Up 0.568444236186959
0.0037506553792647 0.120746377790634
GO_CARBOHYDRATE_BINDING 147 0.01 Up 0.445290682910234
0.00375902905656891 0.120746377790634
GO_REGULATION_OF_DEFENSE_RESPONSE_TO_VIRUS 62 0.01 Down
0.303588684291529 0.00395159125827032
0.126263739626101
GO_GLYCOSAMINOGLYCAN_BINDING 126 0.01 Up
0.75668359545638 0.00400096404707327 0.127172003820847
GO_VIRAL_GENOME_REPLICATION 106 0.01 Down
0.292046273884909 0.00405194035465512 0.12768801230447
GO_REGULATION_OF_SYMBIOSIS_ENCOMPASSING_MUTUALISM_THROUGH_PARASITISM
191 0.01 Down 0.325564182441281
0.0040592631155926 0.12768801230447
GO_RESPONSE_TO_ANTINEOPLASTIC_AGENT 75 0.01 Down
0.361256263793826 0.00410208840897936 0.128369993458318
GO_CARTILAGE_MORPHOGENESIS 15 0.01 Up
0.793296895788173 0.00414438228047085 0.129028434998659
GO_MULTI_ORGANISM_CELLULAR_PROCESS 41 0.01 Down
0.443078072631969 0.00427590733709416 0.131892339246777
GO_RECEPTOR_SIGNALING_PATHWAY_VIA_STAT 95 0.01 Down
0.46095864229493 0.00427982059489623 0.131892339246777
GO_HISTONE_METHYLTRANSFERASE_COMPLEX 84 0.01 Down
0.174811894601887 0.0043465143922198 0.133271155935184
GO_ORGAN_OR_TISSUE_SPECIFIC_IMMUNE_RESPONSE 17 0.01 Down
0.847074945595076 0.00437822399303911
0.133568833476083
GO_ESTABLISHMENT_OF_ENDOTHELIAL_BARRIER 37 0.01 Down
0.397538610549526 0.0044276399786224 0.134401011551083
GO_TOLERANCE_INDUCTION 17 0.01 Down 0.723411275471347
0.00457153336045005 0.137549637390708
GO_PROTEIN_PHOSPHATASE_2A_BINDING 27 0.01 Down
0.240425593651888 0.00457668040733373 0.137549637390708
GO_HISTONE_H4_DEACETYLATION 11 0.01 Down
0.253251838279688 0.00460794695209693 0.137692082821004
GO_LYSINE_ACETYLATED_HISTONE_BINDING 18 0.01 Down
0.198678516604376 0.00464502984209703 0.137692082821004
GO_RNA_DESTABILIZATION 29 0.01 Down 0.214552140347163
0.00466328581445153 0.137692082821004
GO_REGULATION_OF_PROTEIN_DEUBIQUITINATION 12 0.01 Down
0.316547330541561 0.00467442982786999
0.137692082821004
GO_STEROID_CATABOLIC_PROCESS 20 0.01 Up
0.724343640502455 0.00469482146993046 0.137692082821004
GO_DOUBLE_STRANDED_RNA_BINDING 68 0.01 Down
0.231904337963438 0.0047225673515446 0.137839934573208
GO_COFACTOR_CATABOLIC_PROCESS 40 0.01 Up
0.342716256918428 0.00482987897694586 0.140297585019322
GO_CELL_CELL_ADHERENS_JUNCTION 85 0.01 Down
0.245310493759641 0.0048691829976551 0.14052710283877
GO_CELLULAR_RESPONSE_TO_ARSENIC_CONTAINING_SUBSTANCE 18 0.01

Down 0.305913829881871 0.00490954422760271
0.14052710283877
GO_REGULATION_OF_GLIAL_CELL_DIFFERENTIATION 44 0.01 Down
0.479509028317661 0.00491397643561121
0.14052710283877
GO_MHC_PROTEIN_COMPLEX 22 0.01 Up 0.82887001381454
0.00493036944566924 0.14052710283877
GO_PROTEIN_DEMETHYLATION 29 0.01 Down
0.239903106293948 0.0049740721547966 0.141110243232571
GO_CYTOKINE_MEDIATED_SIGNALING_PATHWAY 582 0.01 Down
0.437544517415816 0.00512269690861973 0.144650664801072
GO_RESPONSE_TO_DEXAMETHASONE 30 0.01 Down
0.485689081175613 0.00515321981782522 0.144838877379708
GO_GENE_SILENCING_BY_RNA 164 0.01 Down
0.225158553786089 0.00518107316617297 0.144950669086802
GO_REGULATION_OF_T_CELL_ACTIVATION 229 0.01 Down
0.493964626107401 0.00525651989556758 0.145991840160776
GO_VITAMIN_TRANSPORT 22 0.01 Up 0.513618521160212
0.00526638329685554 0.145991840160776
GO_POSITIVE_REGULATION_OF_LEUKOCYTE_CELL_CELL_ADHESION 160 0.01
Down 0.518649040913531 0.00536905680383395
0.148161562982163
GO_TRABECULA_FORMATION 18 0.01 Up 0.835233531702446
0.00545191826213684 0.149496682619893
GO_GTP_RHO_BINDING 17 0.01 Down 0.298721571205721
0.00547728814047936 0.149496682619893
GO_COPPER_ION_BINDING 42 0.01 Up 0.403306958790863
0.00549131283548609 0.149496682619893
GO_PATTERN_RECOGNITION_RECEPTOR_SIGNALING_PATHWAY 153 0.01
Down 0.455586657602494 0.00559894074951657
0.151477180128481
GO_REGULATION_OF_NUCLEAR_TRANSCRIBED_MRNA_POLY_A_TAIL_SHORTENING
14 0.01 Down 0.219155782191488
0.00562622769142287 0.151477180128481
GO_RESPONSE_TO_BIOTIC_STIMULUS 657 0.01 Down
0.490149597036708 0.00565557372738007 0.151477180128481
GO_REGULATION_OF_NIK_NF_KAPPAB_SIGNALING 91 0.01 Down
0.472773362390845 0.00567022378756737
0.151477180128481
GO_CYTOPLASMIC_PATTERN_RECOGNITION_RECEPTOR_SIGNALING_PATHWAY_IN_RES
PONSE_TO_VIRUS 25 0.01 Down 0.259776286011808
0.00568881519836827 0.151477180128481
GO_TRANSFORMING_GROWTH_FACTOR_BETA_BINDING 17 0.01 Up
0.562195032010194 0.00572811037254917
0.151857458828585
GO_POSITIVE_REGULATION_OF_NEURON_MIGRATION 14 0.01 Down
0.344835426126006 0.00577997409554433
0.152076868076642
GO_NEGATIVE_REGULATION_OF_RNA_CATABOLIC_PROCESS 46 0.01 Down
0.203079585822714 0.00578648600324565
0.152076868076642
GO_POSITIVE_REGULATION_OF_INFLAMMATORY_RESPONSE 91 0.01 Down
0.608986354762371 0.00587626888765857
0.153496075679533

GO_PTERIDINE_CONTAINING_COMPOUND_METABOLIC_PROCESS 23 0.01 Up 0.316430252065684 0.00589105347279379 0.153496075679533
GO_REGULATION_OF_SYNAPSE_ASSEMBLY 62 0.01 Up 0.460909683464898 0.00592535742813084 0.153730106607617
GO_UNSATURATED_FATTY_ACID_METABOLIC_PROCESS 54 0.01 Up 0.390956767165621 0.00602077120297127 0.15554085946059
GO_REGULATION_OF_EPIDERMIS_DEVELOPMENT 65 0.01 Down 0.460146382978612 0.00611095618156125 0.157201758382451
GO_PHOSPHATASE_BINDING 161 0.01 Down 0.264657473416198 0.00622181624262556 0.158456059835308
GO_NEGATIVE_REGULATION_OF_MUSCLE_ADAPTATION 10 0.01 Down 0.707757389616239 0.00626521623235306 0.158456059835308
GO_REGULATION_OF_GENE_SILENCING 111 0.01 Down 0.18589766550592 0.00628846473564293 0.158456059835308
GO_PROTEIN_HYDROXYLATION 25 0.01 Up 0.299078279288846 0.00629882474309065 0.158456059835308
GO_GROWTH_PLATE_CARTILAGE_DEVELOPMENT 24 0.01 Up 0.590289186903964 0.0063032768724455 0.158456059835308
GO_T_CELL_DIFFERENTIATION 172 0.01 Down 0.433719776467254 0.0063163179838815 0.158456059835308
GO_CELLULAR_AMINO_ACID_CATABOLIC_PROCESS 58 0.01 Up 0.306079705302109 0.00639058750320705 0.159659492724156
GO_TOLL_LIKE_RECEPTOR_2_SIGNALING_PATHWAY 13 0.01 Down 0.443672091566954 0.00645677780249213 0.15972717965122
GO_REGULATION_OF_PHOSPHATASE_ACTIVITY 145 0.01 Down 0.348103047600532 0.0064692889070991 0.15972717965122
GO_PROTEIN_MODIFICATION_BY_SMALL_PROTEIN_REMOVAL 265 0.01 Down 0.19064092906809 0.00650813282822636 0.15972717965122
GO_RNA_CAP_BINDING_COMPLEX 12 0.01 Down 0.20310364615474 0.00657236594650368 0.15972717965122
GO_NOSE_DEVELOPMENT 11 0.01 Down 0.394159781871334 0.00661381877366896 0.15972717965122
GO_INTERCELLULAR_BRIDGE 48 0.01 Up 0.33079707569305 0.0066354467605245 0.15972717965122
GO_OXIDOREDUCTASE_ACTIVITY_ACTING_ON_NAD_P_H 88 0.01 Up 0.269030034342796 0.00664452855653653 0.15972717965122
GO_REGULATION_OF_MULTI_ORGANISM_PROCESS 317 0.01 Down 0.340289452439783 0.00665465995033045 0.15972717965122
GO_INTERLEUKIN_6_MEDIATED_SIGNALING_PATHWAY 17 0.01 Down 0.396676464296188 0.00665951247299563 0.15972717965122
GO_POSITIVE_REGULATION_OF_TRANSCRIPTION_BY_RNA_POLYMERASE_II 859 0.01 Down 0.35170031920241 0.00669109512853736 0.15972717965122
GO_EPITHELIAL_CELL_FATE_COMMITMENT 12 0.01 Up 0.320187598888918 0.00669592727333035 0.15972717965122
GO_MODULATION_OF_TRANSCRIPTION_IN_OTHER_ORGANISM_INVOLVED_IN_SYMBIOT

IC_INTERACTION	23	0.01	Down	0.289820726068604
0.00670901512288933	0.15972717965122
GO_RESPONSE_TO_INTERLEUKIN_2	11	0.01	Down
0.398295448737654	0.00678548312196341	0.160916672005624
GO_REGULATION_OF_EPITHELIAL_CELL_DIFFERENTIATION_INVOLVED_IN_KIDNEY_
DEVELOPMENT	12	0.01	Down	0.654372033878043
0.00684139375157322	0.160965487708676
GO_ESTROGEN_METABOLIC_PROCESS	11	0.01	Up
0.335160008561956	0.00685946596487185	0.160965487708676
GO_CALCIUM_DEPENDENT_PHOSPHOLIPID_BINDING	33	0.01	Up
0.606101453117891	0.00686708306976563
0.160965487708676
GO_UBIQUITIN_LIKE_PROTEIN_SPECIFIC_PROTEASE_ACTIVITY	96	0.01
Down	0.181647699952073	0.00707138136406336
0.165116754850879
GO_RIG_I_SIGNALING_PATHWAY	18	0.01	Down
0.250404326197098	0.00711244375633044	0.165279494397334
GO_FATTY_ACID_BETA_OXIDATION	65	0.01	Up
0.221599909330195	0.00713279979115494	0.165279494397334
GO_LIPID_OXIDATION	90	0.01	Up	0.297405339008324
0.00719876603847115	0.165484132537242
GO_TYROSINE_PHOSPHORYLATION_OF_STAT_PROTEIN	48	0.01	Down
0.593609352901365	0.00723123284145352
0.165484132537242
GO_LIPID_CATABOLIC_PROCESS	232	0.01	Up
0.348035424625905	0.007232454790733	0.165484132537242
GO_ORGANIC_ACID_CATABOLIC_PROCESS	189	0.01	Up
0.287402739195717	0.00726128220829561	0.165484132537242
GO_SIGNAL_TRANSDUCTION_IN_ABSENCE_OF_LIGAND	60	0.01	Down
0.373408839319672	0.00729708929721158
0.165484132537242
GO_POSITIVE_REGULATION_OF_SYNAPSE_ASSEMBLY	37	0.01	Up
0.520129705331754	0.00731561705228281
0.165484132537242
GO_CHONDROCYTE_DIFFERENTIATION_INVOLVED_IN_ENDOCHONDRAL_BONE_MORPHOG
ENESIS	19	0.01	Up	0.645129175777481
0.00733243809133883	0.165484132537242
GO_REGULATION_OF_MITOTIC_SPINDLE_ASSEMBLY	20	0.01	Down
0.184616545570835	0.00736482915468047
0.165599547400241
GO_POSITIVE_REGULATION_OF_CELL_CELL_ADHESION	189	0.01	Down
0.501274372312715	0.00739346254458116
0.165629930288385
GO_EPIDERMAL_GROWTH_FACTOR_RECEPTOR_BINDING	29	0.01	Down
0.343561999999275	0.00743279502996589
0.165898892010746
GO_NEGATIVE_REGULATION_OF_MULTI_ORGANISM_PROCESS	131	0.01
Down	0.395378576997767	0.00749115485766453
0.166254203769016
GO_ICOSANOID_METABOLIC_PROCESS	54	0.01	Up
0.471437008888667	0.00750348407720482	0.166254203769016
GO_CELL_CELL_ADHESION_MEDIATED_BY_INTEGRIN	13	0.01	Down
0.678524906388126	0.00753438539616304
0.166331831782203

GO_RESPONSE_TO_BACTERIUM 386 0.01 Down
0.618087268514794 0.00773781893900412 0.170113919303581
GO_REGULATION_OF_CD4_POSITIVE_ALPHA_BETA_T_CELL_DIFFERENTIATION 33
0.01 Down 0.485886983487752 0.00776174528860021
0.170113919303581
GO_OXIDOREDUCTASE_ACTIVITY_ACTING_ON_THE_CH_CH_GROUP_OF_DONORS_NAD_OR_NADP_AS_ACCEPTOR 23 0.01 Up 0.394005318382468
0.00782354210937546 0.170622010115884
GO_REGULATION_OF_NON_CANONICAL_WNT_SIGNALING_PATHWAY 19 0.01
Up 0.550299011383383 0.00784113668626779
0.170622010115884
GO_REGULATION_OF_RECEPTOR_SIGNALING_PATHWAY_VIA_STAT 78 0.01
Down 0.497269317432339 0.00791573810661264
0.171630164447305
GO_REGULATION_OF_HEART_MORPHOGENESIS 29 0.01 Down
0.393263170367921 0.00798073588160463 0.172423656716092
GO_CIRCADIAN_REGULATION_OF_GENE_EXPRESSION 52 0.01 Down
0.278575708646822 0.00805032130694062
0.17331028600864
GO_ACTIVATION_OF_INNATE_IMMUNE_RESPONSE 247 0.01 Down
0.374944386168414 0.00822472726953478 0.176439290647864
GO_PEPTIDYL_PROLINE_DIOXYGENASE_ACTIVITY 11 0.01 Up
0.332911209755805 0.0083628719346696
0.178165411670568
GO_LEUKOCYTE_CHEMOTAXIS 142 0.01 Down 0.754688438706594
0.00839273875160945 0.178165411670568
GO_REGULATION_OF_INFLAMMATORY_RESPONSE 220 0.01 Down
0.553717012389582 0.00842234201792413 0.178165411670568
GO_FIBRONECTIN_BINDING 23 0.01 Up 0.871561174463315
0.00842257834779327 0.178165411670568
GO_MITOCHONDRIAL_MATRIX 424 0.01 Up 0.162263298861426
0.0086404061515474 0.182138561618209
GO_INNATE_IMMUNE_RESPONSE 634 0.01 Down
0.462791462023 0.00869871549003239 0.182234276753264
GO_MIRNA_METABOLIC_PROCESS 20 0.01 Down
0.199377083995215 0.00872122966233701 0.182234276753264
GO_CARBOXYPEPTIDASE_ACTIVITY 28 0.01 Up
0.550868819247448 0.00873499827626418 0.182234276753264
GO_COLLAGEN_BINDING 52 0.01 Up 1.00024120071458
0.00880002586002341 0.182715788555315
GO_BETA_CATENIN_BINDING 66 0.01 Down 0.282534178490422
0.0088182714621491 0.182715788555315
GO_SULFUR_COMPOUND_METABOLIC_PROCESS 153 0.01 Up
0.360987155137896 0.00897623728674511 0.185124166462136
GO_REGULATION_OF_LEUKOCYTE_ADHESION_TO_VASCULAR_ENDOTHELIAL_CELL
14 0.01 Down 0.685061697364494
0.00899733402812689 0.185124166462136
GO_INNATE_IMMUNE_RESPONSE_ACTIVATING_CELL_SURFACE_RECEPTOR_SIGNALING_PATHWAY 90 0.01 Down 0.248598926174975
0.00902598472620527 0.185124166462136
GO_CELLULAR_RESPONSE_TO_STEROL 18 0.01 Down
0.372432693260499 0.00909401628836341 0.185633322294842
GO_3_UTR_MEDIATED_MRNA_STABILIZATION 16 0.01 Down
0.186220358983155 0.00911196343993789 0.185633322294842

GO_COFACTOR_TRANSPORT	37	0.01	Up	0.426689741166481
	0.00930691675413976	0.188398180489068
GO_ORGANIC_ACID_METABOLIC_PROCESS	683	0.01	Up
0.310099444569459	0.00930974372372267	0.188398180489068
GO_OXIDOREDUCTASE_ACTIVITY_ACTING_ON_PAIRED_DONORS_WITH_INCORPORATIO
N_OR_REDUCTION_OF_MOLECULAR_OXYGEN	108	0.01	Up
0.424117775293884	0.00941457397345723	0.189886639843385
GO_RESPONSE_TO_MOLECULE_OF_BACTERIAL_ORIGIN	248	0.01	Down
	0.646939774352917	0.00959988378384582
0.19298309421102
GO_REGULATION_OF_ASTROCYTE_DIFFERENTIATION	23	0.01	Down
	0.593353928425412	0.00965374893545969
0.193425444842164
GO_POSITIVE_REGULATION_OF_INTERFERON_ALPHA_PRODUCTION	20	0.01
	Down	0.310347440654978	0.00975043963449434
0.194604324996764
GO_PALMITOYL_COA_HYDROLASE_ACTIVITY	10	0.01	Up
0.525410235545316	0.00977669562246961	0.194604324996764
GO_COMPLEMENT_ACTIVATION	29	0.01	Up
0.903627156443031	0.0100156825737749	0.197278316954867
GO_GOLGI_ORGANIZATION	122	0.01	Down	0.200178468644568
	0.0100478300212168	0.197278316954867
GO_HYALURONAN_CATABOLIC_PROCESS	12	0.01	Up
0.503554994759894	0.0100505970936386	0.197278316954867
GO_PROSTAGLANDIN_TRANSPORT	11	0.01	Up
0.596038259906902	0.0100627655786451	0.197278316954867
GO_SULFUR_COMPOUND_BINDING	155	0.01	Up
0.561276815477548	0.0100895678691754	0.197278316954867
GO_REGULATION_OF_STEM_CELL_DIFFERENTIATION	93	0.01	Down
	0.177314725866078	0.0101398247429079
0.197278316954867
GO_CILIARY_TRANSITION_ZONE	54	0.01	Up
0.246833812292859	0.0101445435693504	0.197278316954867
GO_CYTOKINE_METABOLIC_PROCESS	78	0.01	Down
0.434453930670823	0.0101709954219854	0.197278316954867
GO_TRANSCRIPTIONAL_REPRESSOR_COMPLEX	72	0.01	Down
0.213957635651552	0.0102795072952069	0.198144808843036
GO_COTRANSLATIONAL_PROTEIN_TARGETING_TO_MEMBRANE	102	0.01
	Up	0.132130861320685	0.0103254909524938
0.198144808843036
GO_NUCLEAR_LOCALIZATION_SEQUENCE_BINDING	20	0.01	Down
	0.163968733284675	0.010363874963839
0.198144808843036
GO_PROTEIN_DEPHOSPHORYLATION	266	0.01	Down
0.288024187456016	0.0104380675182131	0.198144808843036
GO_NEGATIVE_REGULATION_OF_PHOSPHATIDYLINOSITOL_3_KINASE_SIGNALING
	11	0.01	Down	0.437706720370605
0.0104678020909089	0.198144808843036
GO_PERK_MEDIATED_UNFOLDED_PROTEIN_RESPONSE	20	0.01	Down
	0.398409834485392	0.010472809607885
0.198144808843036
GO_NEGATIVE_REGULATION_OF_I_KAPPAB_KINASE_NF_KAPPAB_SIGNALING	35
	0.01	Down	0.298353745255176	0.0104763623269183
	0.198144808843036

GO_RESPONSE_TO_TYPE_I_INTERFERON	78	0.01	Down
0.304422514308146	0.0104767721361579	0.198144808843036
GO_MONOCARBOXYLIC_ACID_CATABOLIC_PROCESS	98	0.01	Up
0.231954846983056	0.0105386926347755
0.198696903682367
GO_SPECIFICATION_OF_SYMMETRY	84	0.01	Up
0.447202159365316	0.01063092233291	0.199497840024289
GO_G_PROTEIN_COUPLED_RECEPTOR_BINDING	146	0.01	Down
0.514192670815873	0.0106468951025975	0.199497840024289
GO_REGULATION_OF_AXON_GUIDANCE	34	0.01	Down
0.519752353775466	0.0107003926031761	0.199848234443883
GO_POSITIVE_REGULATION_OF_DEFENSE_RESPONSE	366	0.01	Down
0.414646674986059	0.0107552790925968
0.199848234443883
GO_CYTOKINE_PRODUCTION_INVOLVED_IN_INFLAMMATORY_RESPONSE	33
0.01	Down	0.674376324108259	0.0107737028456404
0.199848234443883
GO_CELLULAR_RESPONSE_TO_HEAT	119	0.01	Down
0.223304776997944	0.0107972691315424	0.199848234443883
GO_VITAMIN_BINDING	93	0.01	Up	0.340958776025978
0.0108309541439448	0.199862378747383
GO_NEGATIVE_REGULATION_OF_ERK1_AND_ERK2_CASCADE	59	0.01	Down
0.438382360775486	0.0109232874384358
0.200955387996194
GO_GOLGI_LUMEN	56	0.01	Up	0.781505147522087
0.010981650986187	0.201387008176444
GO_STRESS_ACTIVATED_PROTEIN_KINASE_SIGNALING_CASCADE	263	0.01
Down	0.33229248633916	0.0110130928536616
0.201387008176444
GO_INTRACILIARY_TRANSPORT_PARTICLE_B	19	0.01	Up
0.220722587120554	0.0111022134784913	0.202407021104868
GO_LUNG_CELL_DIFFERENTIATION	19	0.01	Down
0.437846644574006	0.0112439159892057	0.203955044943564
GO_NUCLEAR_PORE	79	0.01	Down	0.178165113142893
0.0113197507106317	0.203955044943564
GO_TYPE_I_INTERFERON_PRODUCTION	115	0.01	Down
0.276832666128501	0.0113287151629771	0.203955044943564
GO_POSITIVE_REGULATION_OF_NF_KAPPAB_TRANSCRIPTION_FACTOR_ACTIVITY
129	0.01	Down	0.422886161229241
0.0114183359774198	0.203955044943564
GO_REGULATION_OF_RESPONSE_TO_BIOTIC_STIMULUS	108	0.01	Down
0.353658711786542	0.011418754398577
0.203955044943564
GO_MICRO_RIBONUCLEOPROTEIN_COMPLEX	33	0.01	Down
0.375223019182218	0.0114312130566313	0.203955044943564
GO_LIPOSACCHARIDE_METABOLIC_PROCESS	43	0.01	Up
0.282009239060386	0.0114367049461115	0.203955044943564
GO_REGULATORY_REGION_NUCLEIC_ACID_BINDING	652	0.01	Down
0.323401412691844	0.0114795366318703
0.203955044943564
GO_INTEGRIN_BINDING	102	0.01	Up	0.574274968996476
0.0115112923651627	0.203955044943564
GO_P38MAPK_CASCADE	37	0.01	Down	0.302563691543221
0.0115230736971903	0.203955044943564

GO_REGULATION_OF_RNA_METABOLIC_PROCESS 195 0.01 Down
0.154614345592468 0.0116155916267792 0.20449213578156
GO_THIOESTERASE_BINDING 10 0.01 Down 0.411942201047779
0.0116207851827768 0.20449213578156
GO_DENDRITE_MEMBRANE 16 0.01 Up 0.733222125460952
0.0116924743984431 0.205158994430485
GO_POSITIVE_REGULATION_OF_ERBB_SIGNALING_PATHWAY 25 0.01
Down 0.564645894681667 0.0117456329875153
0.205497803651889
GO_REGULATION_OF_HAIR_CYCLE 21 0.01 Down
0.603122510452097 0.0118149845904251 0.206117159334686
GO_FATTY_ACID_CATABOLIC_PROCESS 90 0.01 Up
0.227280184692404 0.0120221748940343 0.208743304789668
GO_NADH_DEHYDROGENASE_COMPLEX_ASSEMBLY 58 0.01 Up
0.128001063410658 0.0120342870493138 0.208743304789668
GO_RESPONSE_TO_NICOTINE 19 0.01 Down 0.368934637741506
0.0122048465367722 0.211098641950837
GO_REGULATION_OF_DNA_TEMPLATED_TRANSCRIPTION_INITIATION 29 0.01
Down 0.246349474890796 0.0123406575569543
0.2118740725919
GO_REGULATION_OF_ENDOCRINE_PROCESS 21 0.01 Down
0.539537117040086 0.0123595500236876 0.2118740725919
GO_DEMETHYLATION 57 0.01 Down 0.30809560313608
0.012376538260046 0.2118740725919
GO_HYPEROSMOTIC_RESPONSE 17 0.01 Down
0.524218252451031 0.0124240486477499 0.2118740725919
GO_CELLULAR_PROTEIN_CONTAINING_COMPLEX_LOCALIZATION 20 0.01
Down 0.275210211431922 0.0124683939561473
0.2118740725919
GO_NITRIC_OXIDE_SYNTHASE_BIOSYNTHETIC_PROCESS 13 0.01 Down
0.318145543559949 0.0124796446132834
0.2118740725919
GO_POSITIVE_REGULATION_OF_LEUKOCYTE_APOPTOTIC_PROCESS 23 0.01
Down 0.817823026156648 0.0124939743020755
0.2118740725919
GO_RESPONSE_TO_TUMOR_NECROSIS_FACTOR 230 0.01 Down
0.347054790251663 0.0125855311962058 0.212832200256728
GO_MITOCHONDRIAL_RESPIRATORY_CHAIN_COMPLEX_ASSEMBLY 90 0.01
Up 0.127051373568068 0.0126308174617964
0.212852762641196
GO_ACTIVATION_OF_PROTEIN_KINASE_ACTIVITY 263 0.01 Down
0.367879059294139 0.0126664589314557
0.212852762641196
GO_ALPHA_BETA_T_CELL_PROLIFERATION 26 0.01 Down
0.493034625238719 0.012691928854573 0.212852762641196
GO_NEGATIVE_REGULATION_OF_EXTRACELLULAR_MATRIX_ORGANIZATION 11
0.01 Up 0.592016268779257 0.0127834499145341
0.213797037000375
GO_REGULATION_OF_NOTCH_SIGNALING_PATHWAY 78 0.01 Down
0.32690788322116 0.0129518059749196
0.216017621081695
GO_REGULATION_OF_CELLULAR_RESPONSE_TO_HEAT 75 0.01 Down
0.163899259473274 0.0130880633791854
0.217330342718334

GO_HYDROLASE_ACTIVITY_HYDROLYZING_O_GLYCOSYL_COMPOUNDS 65 0.01 Up 0.312941657164922 0.0131021092793461 0.217330342718334
GO_DNA_BINDING_TRANSCRIPTION_FACTOR_ACTIVITY 1239 0.01 Down 0.284380743204955 0.0132337131770413 0.217654114825348
GO_NEGATIVE_REGULATION_OF_NOTCH_SIGNALING_PATHWAY 31 0.01 Down 0.335158105817422 0.0132569298801159 0.217654114825348
GO_EPITHELIAL_CELL_DIFFERENTIATION 461 0.01 Down 0.511056599057473 0.0132604326183781 0.217654114825348
GO_ENGULFMENT_OF_APOPTOTIC_CELL 11 0.01 Up 0.426454152027798 0.0132774729652353 0.217654114825348
GO_REGULATION_OF_T_CELL_DIFFERENTIATION 101 0.01 Down 0.437607492461407 0.0133011961271883 0.217654114825348
GO_ERBB_SIGNALING_PATHWAY 124 0.01 Down 0.325192177873415 0.0133445392385686 0.217654114825348
GO_EXTRACELLULAR_MATRIX_BINDING 47 0.01 Up 0.795525378769766 0.0133725885076355 0.217654114825348
GO_REGULATION_OF_LYMPHOCYTE_CHEMOTAXIS 16 0.01 Down 0.65456254553631 0.0134249980376237 0.217922895952977
GO_RETINOIC_ACID_RECEPTOR_BINDING 22 0.01 Down 0.256265643516369 0.013542747040198 0.219063283273056
GO_I_KAPPAB_KINASE_NF_KAPPAB_SIGNALING 228 0.01 Down 0.329521712719845 0.0135674179724377 0.219063283273056
GO_NEUROTRANSMITTER_UPTAKE 21 0.01 Up 0.660036034454076 0.0137698956764371 0.220805672083338
GO_RESPONSE_TO_INTERFERON_GAMMA 147 0.01 Down 0.511338825960839 0.013795185307312 0.220805672083338
GO_HEPARIN_BINDING 91 0.01 Up 0.731159877045076 0.0137982932285323 0.220805672083338
GO_SULFUR_COMPOUND_BIOSYNTHETIC_PROCESS 143 0.01 Up 0.423835650919483 0.0138208129454239 0.220805672083338
GO_REGULATION_OF_SMAD_PROTEIN_SIGNAL_TRANSDUCTION 19 0.01 Up 0.268909016502441 0.013922118961011 0.221781313715865
GO_IMPORT_INTO_NUCLEUS 137 0.01 Down 0.263373287751309 0.0139549434754506 0.221781313715865
GO_MRNA_3_UTR_AU_RICH_REGION_BINDING 14 0.01 Down 0.283538292177436 0.0139916127582882 0.221783501450568
GO_PROXIMAL_DISTAL_PATTERN_FORMATION 16 0.01 Up 0.737191750028118 0.0140556687867137 0.222218659385779
GO_ENDOCRINE_PROCESS 45 0.01 Down 0.468283377014825 0.0141075602799524 0.222459736258678
GO_CYSTEINE_TYPE_ENDOPEPTIDASE_INHIBITOR_ACTIVITY 29 0.01 Down 0.653475687913337 0.0142364924457056 0.22391125812922
GO_PITUITARY_GLAND_DEVELOPMENT 18 0.01 Down 0.467295260349482 0.0144637317057884 0.226897455260572
GO_ENTRY_OF_BACTERIUM_INTO_HOST_CELL 15 0.01 Down 0.423040493399934 0.0146074301328257 0.228561103959754
GO_POSITIVE_REGULATION_OF_ENDOTHELIAL_CELL_APOPTOTIC_PROCESS 15 0.01 Up 0.65582048082424 0.0146603881404859 0.228800042161671

GO_NEUROPILIN_BINDING	17	0.01	Down	0.36351169984098
	0.0147267133086559	0.22894428881138
GO_COA_HYDROLASE_ACTIVITY	15	0.01	Up
0.40029585575446	0.0147450530267253	0.22894428881138
GO_COFACTOR_BINDING	388	0.01	Up	0.316160908801171
	0.0148198065805213	0.22951797385292
GO_CELLULAR_LIPID_CATABOLIC_PROCESS	159	0.01	Up
0.279521250741776	0.0149477834998754	0.230223749319854
GO_RESPONSE_TO_CYTOKINE 902	0.01	Down	0.417543227010937
	0.0150122159487602	0.230223749319854
GO_EPIDERMAL_GROWTH_FACTOR_RECEPTOR_SIGNALING_PATHWAY	107	0.01
	Down	0.339934719264029	0.015058799266789
0.230223749319854
GO_CORE_PROMOTER_BINDING	49	0.01	Down
0.222427374655589	0.0150867961373664	0.230223749319854
GO_MESSENGER_RIBONUCLEOPROTEIN_COMPLEX	10	0.01	Down
0.274578464522172	0.0151322674529058	0.230223749319854
GO_POSITIVE_REGULATION_OF_INTERLEUKIN_10_PRODUCTION	21	0.01
	Down	0.584758350769866	0.0151463092215982
0.230223749319854
GO_NEGATIVE_REGULATION_OF_ALPHA_BETA_T_CELL_DIFFERENTIATION	18
	0.01	Down	0.540751643336816	0.0152144392021645
	0.230223749319854
GO_REGULATION_OF_ERBB_SIGNALING_PATHWAY 81	0.01	Down
0.338569395216477	0.0152410354065222	0.230223749319854
GO_PROTEIN_TYROSINE_SERINE_THREONINE_PHOSPHATASE_ACTIVITY	34
	0.01	Down	0.276471724673214	0.015244897443171
	0.230223749319854
GO_REGULATION_OF_RESPONSE_TO_CYTOKINE_STIMULUS	144	0.01	Down
	0.36203956577859	0.0152758666345482
0.230223749319854
GO_MITOCHONDRIAL_RESPIRATORY_CHAIN_COMPLEX_I	45	0.01	Up
	0.134880210953307	0.0153684399755768
0.230223749319854
GO_LYMPHOCYTE_MIGRATION 64	0.01	Down	0.628729135040232
	0.0154175888294965	0.230223749319854
GO_REGULATION_OF_DEPHOSPHORYLATION	173	0.01	Down
0.325095435307821	0.0154720504085757	0.230223749319854
GO_POSITIVE_REGULATION_OF_ANIMAL_ORGAN_MORPHOGENESIS	56	0.01
	Down	0.574734046144181	0.0155320781581395
0.230223749319854
GO_LOW_DENSITY_LIPOPROTEIN_PARTICLE_BINDING	11	0.01	Up
	0.955904022077097	0.0155473565057954
0.230223749319854
GO_CD8_POSITIVE_ALPHA_BETA_T_CELL_ACTIVATION	15	0.01	Down
	0.598704146227541	0.0155977803617914
0.230223749319854
GO_PHOSPHATASE_ACTIVATOR_ACTIVITY	16	0.01	Down
0.297723636625236	0.0156039600016171	0.230223749319854
GO_SEMAPHORIN_RECEPTOR_BINDING	19	0.01	Down
0.493321835774267	0.0156547270366884	0.230223749319854
GO_PEPTIDYL_THREONINE_DEPHOSPHORYLATION 15	0.01	Down
0.276540273247524	0.0157183683804345	0.230223749319854
GO_CYTOPLASMIC_MRNA_PROCESSING_BODY_ASSEMBLY	20	0.01	Down

0.147551354425902 0.0157540463953015
0.230223749319854
GO_POSITIVE_REGULATION_OF_TRANSLATIONAL_INITIATION 24 0.01
Down 0.299204265922567 0.0157613511787734
0.230223749319854
GO_CARGO_RECEPTOR_ACTIVITY 59 0.01 Up
0.677394616679585 0.0157636362898604 0.230223749319854
GO_REGULATION_OF_GLIOGENESIS 73 0.01 Down
0.402459465251153 0.0157907485659833 0.230223749319854
GO_PRODUCTION_OF_SMALL_RNA_INVOLVED_IN_GENE_SILENCING_BY_RNA 46
0.01 Down 0.19354744459133 0.0158073465546563
0.230223749319854
GO_SYNAPTIC_TRANSMISSION_CHOLINERGIC 12 0.01 Up
0.523621537000178 0.0158134250479953 0.230223749319854
GO_TRANS_GOLGI_NETWORK_TRANSPORT_VESICLE 27 0.01 Down
0.244681936271884 0.0159635813916022
0.231242405223361
GO_PHOSPHATASE_REGULATOR_ACTIVITY 80 0.01 Down
0.320764920986139 0.0159778929174022 0.231242405223361
GO_PROMOTER_SPECIFIC_CHROMATIN_BINDING 42 0.01 Down
0.269943747273055 0.0160168617003757 0.231242405223361
GO_GOLGI_RIBBON_FORMATION 12 0.01 Down
0.161037393688923 0.0160357523635373 0.231242405223361
GO_PHOSPHATIDATE_PHOSPHATASE_ACTIVITY 10 0.01 Up
0.537556472088085 0.0161820230880967 0.23275239957238
GO_DIENCEPHALON_DEVELOPMENT 36 0.01 Down
0.559729076755618 0.016217141330772 0.23275239957238
GO_PURINE_CONTAINING_COMPOUND_BIOSYNTHETIC_PROCESS 160 0.01
Up 0.250332553543356 0.0163667110790811
0.233872512926912
GO_POSITIVE_REGULATION_OF_INTERLEUKIN_6_PRODUCTION 66 0.01
Down 0.664837325781459 0.0163814967423529
0.233872512926912
GO_MYELOID_DENDRITIC_CELL_DIFFERENTIATION 13 0.01 Down
0.616328475550879 0.0164107544896828
0.233872512926912
GO_NEGATIVE_REGULATION_OF_DEPHOSPHORYLATION 88 0.01 Down
0.27101564290888 0.0165000982271965
0.234595073389485
GO_CALCIUM_INDEPENDENT_CELL_CELL_ADHESION_VIA_PLASMA_MEMBRANE_CELL_A
DHESION_MOLECULES 15 0.01 Down 0.60010041475678
0.0165561260450288 0.234776911659154
GO_CARBOHYDRATE_DERIVATIVE_CATABOLIC_PROCESS 140 0.01 Up
0.466146030275755 0.0166000576938192
0.234776911659154
GO_INSULIN_LIKE_GROWTH_FACTOR_BINDING 22 0.01 Up
0.772168537431198 0.0166289033130352 0.234776911659154
GO_OXIDOREDUCTASE_ACTIVITY 562 0.01 Up
0.310883516711886 0.016829953494692 0.237064147717576
GO_RIBOSOMAL_SUBUNIT 182 0.01 Up 0.113550843197316
0.0169023893794236 0.237179060952473
GO_HUMORAL_IMMUNE_RESPONSE 114 0.01 Down
0.894741114621355 0.0169162466467502 0.237179060952473
GO_CCR_CHEMOKINE_RECEPTOR_BINDING 13 0.01 Down

1.05359879651459	0.0169789972876289	0.237510351458974
GO_CELLULAR_RESPONSE_TO_CHOLESTEROL	15	0.01	Down
0.373243690282026	0.0171978860219515	0.240019232274178
GO_POSITIVE_REGULATION_OF_GLIOGENESIS	45	0.01	Down
0.450812273846894	0.0172479117927492	0.240165303884818
GO_CELLULAR_MODIFIED_AMINO_ACID_BIOSYNTHETIC_PROCESS	38	0.01
Up	0.267502315432486	0.0173144170784301
0.240297382736824
GO_MUSCLE_HYPERTROPHY_IN_RESPONSE_TO_STRESS	15	0.01	Down
0.471141991956576	0.0173512220156891
0.240297382736824
GO_PROXIMAL_PROMOTER_SEQUENCE_SPECIFIC_DNA_BINDING	378	0.01
Down	0.368968555958022	0.0174100391487573
0.240297382736824
GO_POSITIVE_REGULATION_OF_STEROID_BIOSYNTHETIC_PROCESS	12	0.01
Down	0.44968969716166	0.0174157220234232
0.240297382736824
GO_MEMBRANE_DISASSEMBLY	14	0.01	Up	0.308106714681165
0.017469673298131	0.240495207693771
GO_LIPOPROTEIN_METABOLIC_PROCESS	17	0.01	Up
0.685444005360297	0.0175289028356897	0.240764636007856
GO_POSITIVE_REGULATION_OF_FATTY_ACID_TRANSPORT	10	0.01	Up
0.570244791671147	0.0176230301391995
0.240954219564787
GO_PEPTIDE_CROSS_LINKING	28	0.01	Down
1.3888152364248	0.0176604232159291	0.240954219564787
GO_T_CELL_ACTIVATION	334	0.01	Down	0.462545067625135
0.0176617736297694	0.240954219564787
GO_CLATHRIN_COATED_ENDOCYTIC_VESICLE	47	0.01	Up
0.673354528476211	0.0177060155256365	0.241016188915111
GO_NUCLEAR_TRANSPORT	308	0.01	Down	0.207793285214425
0.0178073846866858	0.241853763831923
GO_REGULATION_OF_DEFENSE_RESPONSE	534	0.01	Down
0.437133198577215	0.018180069724319	0.246364293072188
GO_RESPONSE_TO_AXON_INJURY	54	0.01	Up
0.469977069672975	0.0182560654567375	0.246843147857134
GO_CYTOSOLIC_RIBOSOME	104	0.01	Up	0.148924501504796
0.0183567663263529	0.24765317414953
GO_REGULATION_OF_ICOSANOID_SECRETION	11	0.01	Up
0.484531766117747	0.0184075971331799	0.247653552787768
GO_CD40_RECEPTOR_COMPLEX	11	0.01	Down
0.279530959310969	0.0184383801449632	0.247653552787768
GO_MOVEMENT_IN_ENVIRONMENT_OF_OTHER_ORGANISM_INVOLVED_IN_SYMBIOTIC_I
NTERACTION	34	0.01	Down	0.19095468769203?
0.0185525122697554	0.247678703161647
GO_PHOSPHATIDIC_ACID_BINDING	18	0.01	Down
0.277141448780672	0.0185600960222154	0.247678703161647
GO_REGULATION_OF_INNATE_IMMUNE_RESPONSE	338	0.01	Down
0.399575260941481	0.018562643705905	0.247678703161647
GO_APOPTOTIC_CELL_CLEARANCE	37	0.01	Up
0.50589837306613	0.0186499257310988	0.248283779869847
GO_POSITIVE_REGULATION_OF_LEUKOCYTE_ADHESION_TO_VASCULAR_ENDOTHELIAL
_CELL	11	0.01	Down	0.526052350606799
0.0186897854390579	0.248283779869847

GO_PDZ_DOMAIN_BINDING	61	0.01	Down	0.30074274419129
	0.0189519889608547	0.249710859904772
GO_GROWTH_PLATE_CARTILAGE_MORPHOGENESIS	11	0.01	Up
0.873876200944287	0.0189586575540615	0.249710859904772
GO_STRESS_GRANULE_ASSEMBLY	17	0.01	Down
0.15613457421559	0.0189689093062606	0.249710859904772
GO_T_HELPER_1_CELL_DIFFERENTIATION	13	0.01	Down
0.532629315200425	0.0190001634435195	0.249710859904772
GO_FC_RECEPTOR_SIGNALING_PATHWAY	161	0.01	Down
0.303940943559299	0.0190028689303252	0.249710859904772
GO_NEGATIVE_REGULATION_OF_SIGNAL_TRANSDUCTION_IN_ABSENCE_OF_LIGAND
	27	0.01	Down	0.434236023053541
0.0190909384070817	0.250067569909735
GO_POSITIVE_REGULATION_OF_CELL_GROWTH	123	0.01	Down
0.333809317217486	0.019175777342351	0.250067569909735
GO_CYTOSOLIC_TRANSPORT	147	0.01	Down	0.200127440685449
	0.0191825641698043	0.250067569909735
GO_REGULATION_OF_CD8_POSITIVE_ALPHA_BETA_T_CELL_ACTIVATION	13
	0.01	Down	0.637707660121032	0.0191947764088184
	0.250067569909735
GO_RESPONSE_TO_XENOBIOTIC_STIMULUS	180	0.01	Up
0.480338650601783	0.0192821858074665	0.250668415497065
GO_HISTONE_H3_DEACETYLATION	19	0.01	Down
0.21314777496195	0.0194056156724049	0.251691720546823
GO_REGULATION_OF_HEMATOPOIETIC_PROGENITOR_CELL_DIFFERENTIATION	72
	0.01	Down	0.134631899558028	0.0194482637194714
	0.251691720546823
GO_MIRNA_MEDIATED_INHIBITION_OF_TRANSLATION	16	0.01	Down
	0.315876315998021	0.0194852756806138
0.251691720546823
GO_UBIQUITIN_LIKE_PROTEIN_CONJUGATING_ENZYME_ACTIVITY	35	0.01
	Down	0.133203021401224	0.0195782026239507
0.252355134033981
GO_RNA_STABILIZATION	37	0.01	Down	0.208749293664128
	0.0197026310622865	0.253176069000616
GO_POSITIVE_REGULATION_OF_OXIDOREDUCTASE_ACTIVITY	41	0.01
	Down	0.391956971616338	0.0197252974200777
0.253176069000616
GO_VASODILATION	19	0.01	Up	0.708631432519769
0.0197844616239673	0.253191783818848
GO_ACTIVATION_OF_MAPKK_ACTIVITY	47	0.01	Down
0.274390953885054	0.0198099320233821	0.253191783818848
GO_NEGATIVE_REGULATION_BY_HOST_OF_VIRAL_TRANSCRIPTION	10	0.01
	Down	0.468055546728662	0.0200258594221766
0.254790352790322
GO_INTRACELLULAR_RECEPTOR_SIGNALING_PATHWAY	231	0.01	Down
	0.276693765833627	0.020039433394987
0.254790352790322
GO_POSITIVE_REGULATION_OF_PROTEIN_MODIFICATION_BY_SMALL_PROTEIN_CONJ
UGATION_OR_REMOVAL	114	0.01	Down	0.211788929878709
	0.02006091066278 6	0.254790352790322
GO_NEGATIVE_REGULATION_OF_LYMPHOCYTE_DIFFERENTIATION	36	0.01
	Down	0.55185297904006	0.0201658671153945
0.255349121163557

GO_ARACHIDONIC_ACID_METABOLIC_PROCESS	32	0.01	Up
0.484360814599062	0.0202609622828301	0.255349121163557
GO_VESICLE_MEDIATED_TRANSPORT_BETWEEN_ENDOSOMAL_COMPARTMENTS	40
	0.01	Down	0.228195092163217	0.0203187325661221
	0.255349121163557
GO_PROTEIN_SERINE_THREONINE_PHOSPHATASE_COMPLEX	45	0.01	Down
	0.241153819020801	0.0203228355510384
0.255349121163557
GO_REGULATION_OF_SPINDLE_ASSEMBLY	26	0.01	Down
0.174268451757579	0.020350803790919	0.255349121163557
GO_HEMATOPOIETIC_STEM_CELL_DIFFERENTIATION	68	0.01	Down
	0.17131186007619	0.0203572681013279
0.255349121163557
GO_METHYLTRANSFERASE_COMPLEX	111	0.01	Down
0.15305892802534	0.0204578480320546	0.256081640005368
GO_UBIQUITIN_LIKE_PROTEIN_TRANSFERASE_ACTIVITY	363	0.01	Down
	0.195072228002432	0.0210351800140073
0.262750726961162
GO_REGULATION_OF_CHROMATIN_SILENCING	25	0.01	Down
0.172950326584904	0.0210944047294178	0.262750726961162
GO_DETERMINATION_OF_HEART_LEFT_RIGHT_ASYMMETRY	42	0.01	Up
	0.403674423443582	0.0211204669341207
0.262750726961162
GO_INTERFERON_GAMMA_SECRETION	13	0.01	Down
0.718850729714149	0.0211951952521437	0.263141166412606
GO_POSITIVE_REGULATION_OF_TYPE_I_INTERFERON_PRODUCTION	70	0.01
	Down	0.25565195367895	0.0212897710696756
0.263775918702042
GO_MITOCHONDRIAL_PART	924	0.01	Up	0.180588388918817
	0.0213831283077318	0.263989088233409
GO_CARBOXYLIC_ESTER_HYDROLASE_ACTIVITY	90	0.01	Up
0.293630590653741	0.0214105648992972	0.263989088233409
GO_HEART_VALVE_DEVELOPMENT	45	0.01	Down
0.505218707521981	0.0214633452552206	0.263989088233409
GO_MAINTENANCE_OF_GASTROINTESTINAL_EPITHELIUM	13	0.01	Down
	0.340755723504615	0.0215072743538925
0.263989088233409
GO_PROTEIN_IMPORT	163	0.01	Down	0.254847339307584
	0.0215243944449905	0.263989088233409
GO_SINGLE_STRANDED_RNA_BINDING	74	0.01	Down
0.19931797849999	0.0216271936532032	0.264715106186686
GO_PEPTIDYL_TYROSINE_DEPHOSPHORYLATION	84	0.01	Down
0.301109660391148	0.0217986676895251	0.265131282944797
GO_MKS_COMPLEX	13	0.01	Up	0.240217296114541
0.0218089102290752	0.265131282944797
GO_POSITIVE_REGULATION_OF_ALPHA_BETA_T_CELL_ACTIVATION	43	0.01
	Down	0.428642692157985	0.0218307856082206
0.265131282944797
GO_SEQUENCE_SPECIFIC_DOUBLE_STRANDED_DNA_BINDING	593	0.01
	Down	0.331209066070946	0.021835882304793
0.265131282944797
GO_PROTEIN_LOCALIZATION_TO_CILIARY_MEMBRANE	10	0.01	Up
	0.277940768316772	0.0219163197953269
0.265285400735703

GO_NUCLEAR_TRANSCRIBED_MRNA_CATABOLIC_PROCESS_DEADENYLATION_DEPENDEN
T_DECAY 71 0.01 Down 0.180334118674001
0.0219359695551512 0.265285400735703
GO_LIPOPOLYSACCHARIDE_MEDIATED_SIGNALING_PATHWAY 43 0.01
Down 0.566442424000427 0.0220172692163217
0.265499278120918
GO_CYTOPLASMIC_MICROTUBULE_ORGANIZATION 45 0.01 Down
0.211551999349348 0.022041194486811 0.265499278120918
GO_CYTOPLASMIC_STRESS_GRANULE 54 0.01 Down
0.167971269837776 0.0220967527617153 0.265642348547274
GO_MYOFILAMENT 21 0.01 Up 0.661536112866363
0.0221809566305318 0.265752872764567
GO_NEGATIVE_REGULATION_OF_INTERFERON_GAMMA_PRODUCTION 25 0.01
Down 0.898233157469992 0.0221934947276619
0.265752872764567
GO_REGULATION_OF_PHOSPHOPROTEIN_PHOSPHATASE_ACTIVITY 96 0.01
Down 0.340307297873609 0.0222822499746752
0.266290432276089
GO_PROTEIN_LIPID_COMPLEX_BINDING 17 0.01 Up
0.764100129012506 0.0224411438550659 0.2676624446839
GO_TRANSFORMING_GROWTH_FACTOR_BETA_RECEPTOR_BINDING 31 0.01
Down 0.44502673250782 0.0226258310233738
0.269336117927259
GO_ENDONUCLEASE_ACTIVITY_ACTIVE_WITH_EITHER_RIBO_OR_DEOXYRIBONUCLEIC
_ACIDS_AND_PRODUCING_3_PHOSPHOMONOESTERS 16 0.01 Up
0.295301087667167 0.0229267279758276
0.271480610606655
GO_PROTEIN_TYROSINE_PHOSPHATASE_ACTIVITY 83 0.01 Down
0.302894755860022 0.0229581432919847
0.271480610606655
GO_RIBONUCLEOPROTEIN_GRANULE 189 0.01 Down
0.197094432144449 0.0230042711844674 0.271480610606655
GO_CORONARY_VASCULATURE_DEVELOPMENT 38 0.01 Up
0.483192805491052 0.0230283237888984 0.271480610606655
GO_COENZYME_BINDING 239 0.01 Up 0.269287853583785
0.0230295691751651 0.271480610606655
GO_DEFENSE_RESPONSE_TO_BACTERIUM 113 0.01 Down
0.705785431892885 0.0231065998039644 0.271860789554007
GO_AXON_INITIAL_SEGMENT 12 0.01 Up 0.384457399424061
0.0231570635231926 0.271927529302325
GO_POSITIVE_REGULATION_OF_GLIAL_CELL_DIFFERENTIATION 28 0.01
Down 0.493632924417157 0.0232099848165927
0.272007708739311
GO_POSITIVE_REGULATION_OF_NON_CANONICAL_WNT_SIGNALING_PATHWAY 11
0.01 Up 0.382821826055263 0.0232649727016079
0.272007708739311
GO_SEQUENCE_SPECIFIC_DNA_BINDING 754 0.01 Down
0.32023646220453 0.0232983048170716 0.272007708739311
GO_CELLULAR_RESPONSE_TO_DEXAMETHASONE_STIMULUS 24 0.01 Down
0.487466558307871 0.02341105258145
0.27235709113612
GO_RESPONSE_TO_FUNGICIDE 10 0.01 Down
0.270795307812303 0.023417954467642 0.27235709113612
GO_REGULATION_OF_HISTONE_METHYLATION 54 0.01 Down

0.143165357059957	0.0236469738081005	0.273864155427977
GO_TRANSFERASE_ACTIVITY_TRANSFERRING_HEXOSYL_GROUPS	168	0.01	Up	0.343514170029459	0.0236849340601092	0.273864155427977
GO_ELECTRON_TRANSPORT_CHAIN	155	0.01	Up	0.227681767388815	0.0237273620813624	0.273864155427977
GO_RNAI_EFFECTOR_COMPLEX	11	0.01	Down	0.202524139785328	0.023727976569777	0.273864155427977
GO_POSITIVE_REGULATION_OF_REACTIVE_OXYGEN_SPECIES_BIOSYNTHETIC_PROCESS	41	0.01	Down	0.589250548234956	0.0238286851757238	0.274504644595482
GO_RAN_GTPASE_BINDING	37	0.01	Down	0.237657174156406	0.0239392817200268	0.275256400231596
GO_REGULATION_OF_TRANSLATIONAL_INITIATION	70	0.01	Down	0.183192691688356	0.0240711965730741	0.276249970501196
GO_NEGATIVE_REGULATION_OF_CYTOKINE_BIOSYNTHETIC_PROCESS	21	0.01	Down	0.586034994031901	0.0241201501608772	0.276289493635256
GO_CYTOKINE_PRODUCTION	542	0.01	Down	0.474219170960782	0.0243348574049096	0.278223953493797
GO_REGULATION_OF_PSEUDOPODIUM_ASSEMBLY	11	0.01	Down	0.466583476497084	0.0246217331501228	0.280811982046643
GO_POSITIVE_REGULATION_OF_RESPONSE_TO_EXTERNAL_STIMULUS	209	0.01	Down	0.558755510223997	0.0246537286165147	0.280811982046643
GO_ESTABLISHMENT_OF_PROTEIN_LOCALIZATION_TO_VACUOLE	40	0.01	Down	0.217003673220289	0.0247089650948808	0.28091409567607
GO_PROTEIN_LOCALIZATION_TO_GOLGI_APPARATUS	25	0.01	Down	0.247172931427695	0.0248389134149807	0.281389114058617
GO_REGULATION_OF_NUCLEOBASE_CONTAINING_COMPOUND_TRANSPORT	15	0.01	Down	0.125647195888881	0.0248434467361915	0.281389114058617
GO_PML_BODY	92	0.01	Down	0.208131812515849	0.0249658983110192	0.282249476063683
GO_PROTEOGLYCAN_BINDING	25	0.01	Up	0.703629031253903	0.0250774184603546	0.282913639326378
GO_REGULATION_OF_EPITHELIAL_CELL_DIFFERENTIATION	102	0.01	Down	0.44332049432505	0.0251178474051915	0.282913639326378
GO_CELLULAR_RESPONSE_TO_AMMONIUM_ION	36	0.01	Down	0.28235729726747	0.025267336617219	0.28407037148729
GO_POSITIVE_REGULATION_OF_MEMBRANE_PROTEIN_ECTODOMAIN_PROTEOLYSIS	13	0.01	Down	0.463133743014064	0.0255607067752731	0.286837432223073
GO_RETINOID_X_RECEPTOR_BINDING	14	0.01	Down	0.305396964253778	0.0256532686355741	0.286899138227094
GO_RESPONSE_TO_HYPEROXIA	16	0.01	Up	0.603224796497068	0.025668535357016	0.286899138227094
GO_RNA_POLYMERASE_II_CORE_PROMOTER_SEQUENCE_SPECIFIC_DNA_BINDING	30	0.01	Down	0.239531205866709	0.0257079774659099	0.286899138227094

GO_POSITIVE_REGULATION_OF_DNA_BINDING_TRANSCRIPTION_FACTOR_ACTIVITY 210 0.01 Down 0.397959375007885 0.0260403380126665 0.290075031330088
GO_REGULATION_OF_PERK_MEDIATED_UNFOLDED_PROTEIN_RESPONSE 10 0.01 Down 0.412920077501455 0.0261238290537528 0.290472099240537
GO_PROTEIN_LOCALIZATION_TO_VACUOLE 55 0.01 Down 0.200695353053739 0.0261822471643179 0.290589437905985
GO_TRANSCRIPTION_COFACTOR_BINDING 37 0.01 Down 0.292895670542547 0.0263694857968877 0.290605276692449
GO_NEGATIVE_REGULATION_OF_SPROUTING_ANGIOGENESIS 22 0.01 Down 0.344705802735103 0.0264161497940641 0.290605276692449
GO_PHOSPHOPROTEIN_PHOSPHATASE_ACTIVITY 153 0.01 Down 0.266110424433236 0.0264210315256024 0.290605276692449
GO_GROWTH_HORMONE_SECRETION 10 0.01 Down 0.349078740730694 0.0264532916454874 0.290605276692449
GO_INTERACTION_WITH_HOST 178 0.01 Down 0.315945422824022 0.0265055141659815 0.290605276692449
GO_POSITIVE_REGULATION_OF_VASCULAR_SMOOTH_MUSCLE_CELL_PROLIFERATION 28 0.01 Down 0.613258024042013 0.0265680466377319 0.290605276692449
GO_DOSAGE_COMPENSATION 18 0.01 Down 0.151511345385888 0.026575139971488 0.290605276692449
GO_HEAT_GENERATION 12 0.01 Down 0.488086386793593 0.0265881888898818 0.290605276692449
GO_RESPONSE_TO_LEAD_ION 19 0.01 Up 0.435567304057042 0.026741305843101 0.290605276692449
GO_DEFENSE_RESPONSE 1074 0.01 Down 0.47289510462755 0.026783590837539 0.290605276692449
GO_RESPONSE_TO_DSRNA 36 0.01 Down 0.360345474939929 0.0268228693529038 0.290605276692449
GO_SYNAPSE_ASSEMBLY 107 0.01 Up 0.443935166012045 0.0268258951172272 0.290605276692449
GO_POSITIVE_REGULATION_OF_SMAD_PROTEIN_SIGNAL_TRANSDUCTION 11 0.01 Up 0.283819563539894 0.0268300397940509 0.290605276692449
GO_LYS63_SPECIFIC_DEUBIQUITINASE_ACTIVITY 12 0.01 Down 0.241051996377418 0.0268538231303679 0.290605276692449
GO_RIBOSOME 219 0.01 Up 0.139293015194504 0.0270800279560779 0.292046021790659
GO_AMINE_TRANSPORT 36 0.01 Up 0.770920661545745 0.0270831675618746 0.292046021790659
GO_LAMELLAR_BODY 14 0.01 Up 0.964709991268648 0.0271895896676242 0.292673756865508
GO_SNARE_COMPLEX 39 0.01 Down 0.145706475594553 0.0274167204253802 0.294101795436325
GO_PROTEIN_TARGETING_TO_VACUOLE 34 0.01 Down 0.204490986621861 0.0274191428458179 0.294101795436325
GO_DRUG_METABOLIC_PROCESS 230 0.01 Up 0.363079108027855 0.0275608006334617 0.29462985251538
GO_HYDROGEN_PEROXIDE_CATABOLIC_PROCESS 14 0.01 Up 0.282182910919344 0.027612313356025 0.29462985251538

GO_NADH_DEHYDROGENASE_ACTIVITY 39 0.01 Up
0.142593912850498 0.0276139657521415 0.29462985251538
GO_POSITIVE_REGULATION_OF_GENE_EXPRESSION 1472 0.01 Down
0.31729825166361 0.0277640358697921
0.295711336430716
GO_POSITIVE_REGULATION_OF_ASTROCYTE_DIFFERENTIATION 13 0.01
Down 0.638266589328787 0.0278403143660446
0.295722283773609
GO_LYTIC_VACUOLE_ORGANIZATION 55 0.01 Up
0.288040699931507 0.0278624849808112 0.295722283773609
GO_ORGANELLE_ENVELOPE_LUMEN 82 0.01 Up
0.2175693768369 0.0279562132625492 0.296199250814898
GO_PROTEIN_MODIFICATION_BY_SMALL_PROTEIN_CONJUGATION_OR_REMOVAL 960
0.01 Down 0.190731403353127 0.028161684051587
0.297607739292443
GO_POSITIVE_REGULATION_OF_INTERLEUKIN_6_BIOSYNTHETIC_PROCESS 12
0.01 Down 0.598765164529372 0.0281871932289829
0.297607739292443
GO_REGULATION_OF_POSTTRANSCRIPTIONAL_GENE_SILENCING 85 0.01
Down 0.188720140702879 0.0283349880686654
0.298064966857662
GO_POSTSYNAPSE_ASSEMBLY 17 0.01 Up 0.254245853080008
0.0283464393432584 0.298064966857662
GO_CELL_CELL_JUNCTION 342 0.01 Down 0.340328724680512
0.0284164788684333 0.298064966857662
GO_PROTEIN_K48_LINKED_DEUBIQUITINATION 31 0.01 Down
0.189581294753341 0.0284268845018261 0.298064966857662
GO_MRNA_3_UTR_BINDING 67 0.01 Down 0.183910912422944
0.0286154818114134 0.299525155305329
GO_QUINONE_METABOLIC_PROCESS 11 0.01 Up
0.462566323043262 0.0287166159115776 0.300066394490856
GO_PROTEIN_POLYUBIQUITINATION 282 0.01 Down
0.175061276194307 0.0288158203538206 0.300083050509897
GO_MITOCHONDRIAL_TRANSLATION 135 0.01 Up
0.10913368663114 0.0288170677725696 0.300083050509897
GO_MAP_KINASE_ACTIVITY 13 0.01 Down 0.275358756837636
0.0292446100415669 0.302637504335605
GO_NEGATIVE_REGULATION_OF_LEUKOCYTE_CELL_CELL_ADHESION 89 0.01
Down 0.540961674782231 0.0292847159244227
0.302637504335605
GO_REGULATION_OF_TELOMERE_MAINTENANCE 75 0.01 Down
0.147238292519566 0.0292937730963295 0.302637504335605
GO_IMMUNOLOGICAL_SYNAPSE 27 0.01 Down
0.582681709738425 0.0293047818052235 0.302637504335605
GO_ACYL_COA_DEHYDROGENASE_ACTIVITY 11 0.01 Up
0.219637279445436 0.0294281928608948 0.302637504335605
GO_SEMAPHORIN_RECEPTOR_COMPLEX 10 0.01 Up
0.375041897273928 0.029524127044941 0.302637504335605
GO_SEMAPHORIN_RECEPTOR_ACTIVITY 10 0.01 Up
0.375041897273928 0.029524127044941 0.302637504335605
GO_POSTTRANSCRIPTIONAL_REGULATION_OF_GENE_EXPRESSION 545 0.01
Down 0.199587747399095 0.0295573485724808
0.302637504335605
GO_CHEMICAL_HOMEOSTASIS_WITHIN_A_TISSUE 11 0.01 Up

0.928182076141385	0.0295731952561694	0.302637504335605
GO_REGULATION_OF_HIPPO_SIGNALING	12	0.01	Down
0.240231244550625	0.0297040868909923	0.302637504335605
GO_RESPONSE_TO_EPIDERMAL_GROWTH_FACTOR	41	0.01	Down
0.221019714513525	0.0297280117096049	0.302637504335605
GO_PROGRAMMED_NECROTIC_CELL_DEATH	37	0.01	Down
0.340694148388057	0.029747035137879	0.302637504335605
GO_PROTEIN_PHOSPHATASE_BINDING	119	0.01	Down
0.276961868998722	0.0297720033093336	0.302637504335605
GO_REGULATION_OF_ODONTOGENESIS	16	0.01	Down
0.862162050868188	0.0297884982126571	0.302637504335605
GO_POSITIVE_REGULATION_OF_RNA_BIOSYNTHETIC_PROCESS	1213	0.01
Down	0.320038301206967	0.029810118200081
0.302637504335605
GO_ESTABLISHMENT_OR_MAINTENANCE_OF_BIPOLAR_CELL_POLARITY	36
0.01	Down	0.225200554821577	0.029928180181695
0.302926999754942
GO_SERINE_PHOSPHORYLATION_OF_STAT_PROTEIN	10	0.01	Down
0.554604904936626	0.0299384285707404
0.302926999754942
GO_POSITIVE_REGULATION_OF_AUTOPHAGY	105	0.01	Down
0.24122580900291	0.030065676327185	0.303609871357751
GO_VITAMIN_METABOLIC_PROCESS	98	0.01	Up
0.371626002110092	0.0301059368402843	0.303609871357751
GO_PHOTORECEPTOR_CONNECTING_CILIUM	26	0.01	Up
0.276627190248307	0.0302479325949015	0.304535984715832
GO_POSITIVE_REGULATION_OF_HISTONE_METHYLATION	31	0.01	Down
0.151021919112183	0.0303438820568603
0.304996205243707
GO_ENDOSOME_TO_LYSOSOME_TRANSPORT	50	0.01	Down
0.184555912299026	0.0304331666305637	0.305388024155623
GO_NEGATIVE_REGULATION_OF_TYPE_I_INTERFERON_PRODUCTION	42	0.01
Down	0.337968240411634	0.0305509635728611
0.305813306458751
GO_POSITIVE_REGULATION_OF_ACUTE_INFLAMMATORY_RESPONSE	18	0.01
Down	0.594580911694908	0.0305762933652548
0.305813306458751
GO_POSITIVE_REGULATION_OF_DNA_TEMPLATED_TRANSCRIPTION_INITIATION
21	0.01	Down	0.259172566580577
0.0306337363735835	0.305883903822411
GO_INTERMEDIATE_FILAMENT_CYTOSKELETON	100	0.01	Down
0.428493510716034	0.0307752028831447	0.306615314131454
GO_POSITIVE_REGULATION_OF_SMOOTH_MUSCLE_CELL_PROLIFERATION	70
0.01	Down	0.525539538256521	0.0308338197080876
0.306615314131454
GO_ADHERENS_JUNCTION_ORGANIZATION	115	0.01	Down
0.337565596518668	0.0308585005656924	0.306615314131454
GO_CHONDROITIN_SULFATE_PROTEOGLYCAN_METABOLIC_PROCESS	31	0.01
Up	0.615914442316764	0.0309226627924948
0.306750793812477
GO_REGULATION_OF_CELL_FATE_COMMITMENT	17	0.01	Up
0.813029020538244	0.0311518349237881	0.308520048649784
GO_REGULATION_OF_PROTEIN_LOCALIZATION_TO_ENDOSOME	12	0.01
Down	0.316440112020194	0.0312809399345671

0.309294114564751
GO_IMMUNOGLOBULIN_SECRETION 14 0.01 Down
0.593202923160483 0.0313995329375577 0.309961893437257
GO_PROTEOGLYCAN_METABOLIC_PROCESS 56 0.01 Up
0.563446091238486 0.0315259776213613 0.310340405999728
GO_MICROBODY_LUMEN 39 0.01 Up 0.275565762341655
0.0315688496028217 0.310340405999728
GO_PRE_MIRNA_PROCESSING 13 0.01 Down 0.198230562833796
0.031615927591147 0.310340405999728
GO_RHYTHMIC_PROCESS 199 0.01 Down 0.306670232436654
0.0316423507352712 0.310340405999728
GO_UBIQUITIN_LIKE_PROTEIN_BINDING 94 0.01 Down
0.145596251345218 0.0318542761443149 0.311515247441192
GO_PSEUDOPODIUM_ORGANIZATION 14 0.01 Down
0.403816703790568 0.0319146112580688 0.311515247441192
GO_MONOOXYGENASE_ACTIVITY 50 0.01 Up
0.564122621073375 0.0319160737783597 0.311515247441192
GO_NEGATIVE_REGULATION_OF_T_CELL_DIFFERENTIATION 30 0.01
Down 0.480876634018712 0.0321343968526077
0.311814039527574
GO_MEMBRANE_DEPOLARIZATION 53 0.01 Up
0.399262402768015 0.0322263528990623 0.311814039527574
GO_AU_RICH_ELEMENT_BINDING 24 0.01 Down
0.195633658978128 0.0322320331874495 0.311814039527574
GO_POSITIVE_REGULATION_OF_ALCOHOL_BIOSYNTHETIC_PROCESS 14 0.01
Down 0.457242997846491 0.0322586542193202
0.311814039527574
GO_ENDOCARDIAL_CUSHION_DEVELOPMENT 26 0.01 Down
0.501513345643352 0.0323222418522497 0.311814039527574
GO_HISTONE_H3_K4_TRIMETHYLATION 14 0.01 Down
0.24851256255402 0.0323496017244444 0.311814039527574
GO_POSITIVE_REGULATION_OF_VASCULATURE_DEVELOPMENT 134 0.01
Down 0.589257016140403 0.0323837245964827
0.311814039527574
GO_MITOCHONDRIAL_ELECTRON_TRANSPORT_NADH_TO_UBIQUINONE 46 0.01
Up 0.134872209246495 0.032391655018611
0.311814039527574
GO_DENDRITIC_SPINE_DEVELOPMENT 75 0.01 Down
0.344784940493304 0.0324089373977762 0.311814039527574
GO_ORGANIC_ACID_TRANSMEMBRANE_TRANSPORTER_ACTIVITY 86 0.01
Up 0.447512684904474 0.0325609618308233
0.312380287222702
GO_REGULATION_OF_VIRAL_TRANSCRIPTION 60 0.01 Down
0.209887531866994 0.0325974093517106 0.312380287222702
GO_NEGATIVE_REGULATION_OF_GENE_SILENCING 19 0.01 Down
0.285376971243116 0.0326221548507977
0.312380287222702
GO_REGULATION_OF_TOLERANCE_INDUCTION 12 0.01 Down
0.838637286274581 0.0328211470108371 0.313340272943337
GO_RESPONSE_TO_IRON_ION 26 0.01 Up 0.48260088400644
0.0328484279301852 0.313340272943337
GO_MITOCHONDRIAL_PROTEIN_COMPLEX 248 0.01 Up
0.123993534123998 0.032882039002908 0.313340272943337
GO_PEROXISOME_ORGANIZATION 71 0.01 Up

0.231044998073712	0.0329288575420604	0.313340272943337
GO_REGULATION_OF_CENTROSOME_DUPLICATION 41	0.01	Down
0.152029789946355	0.0330942691376933	0.31405282184767
GO_CLATHRIN_COATED_ENDOCYTIC_VESICLE_MEMBRANE	33	0.01	Up
0.586601554427321	0.0331071991405878
0.31405282184767
GO_BONE_MORPHOGENESIS	74	0.01	Up	0.524011159970868
0.033209583762291	0.314342998857036
GO_HISTONE_H4_K20_METHYLATION	10	0.01	Down
0.195081376766594	0.0332608905091672	0.314342998857036
GO_ORGANELLE_INNER_MEMBRANE	461	0.01	Up
0.162966018122108	0.0333113377001161	0.314342998857036
GO_PROTEIN_AUTOUBIQUITINATION	58	0.01	Down
0.164618880023422	0.0333449005540984	0.314342998857036
GO_GERMINAL_CENTER_FORMATION	12	0.01	Down
0.563447472295551	0.0334906216990341	0.315227231526877
GO_POSITIVE_REGULATION_OF_EPITHELIAL_CELL_MIGRATION	120	0.01
Down	0.425851358650909	0.0336103657526473
0.315518597998997
GO_CELL_DIFFERENTIATION_INVOLVED_IN_EMBRYONIC_PLACENTA_DEVELOPMENT
20	0.01	Down	0.368862719412001
0.033673108247923	0.315518597998997
GO_ANTIGEN_PROCESSING_AND_PRESENTATION_OF_ENDOGENOUS_PEPTIDE_ANTIGEN
13	0.01	Down	0.378624853427542
0.0336952952345653	0.315518597998997
GO_WATER_SOLUBLE_VITAMIN_METABOLIC_PROCESS	70	0.01	Up
0.310329737980318	0.0337294630376131
0.315518597998997
GO_REGULATION_OF_T_HELPER_CELL_DIFFERENTIATION	23	0.01	Down
0.499931823119292	0.0338064554363077
0.315752293775114
GO_POSITIVE_REGULATION_OF_GLIAL_CELL_PROLIFERATION	13	0.01
Down	0.402562734829134	0.0339613322842567
0.315951698906988
GO_LIMBIC_SYSTEM_DEVELOPMENT	70	0.01	Down
0.366985240990292	0.0339667617248086	0.315951698906988
GO_RESPONSE_TO_CAFFEINE 13	0.01	Up	0.412721391292915
0.033983933353033	0.315951698906988
GO_REGULATION_OF_REACTIVE_OXYGEN_SPECIES_BIOSYNTHETIC_PROCESS	65
0.01	Down	0.492663171693961	0.0342167713642193
0.317465340569473
GO_ANTIGEN_PROCESSING_AND_PRESENTATION_OF_PEPTIDE_ANTIGEN_VIA_MHC_CL
ASS_I	88	0.01	Down	0.265397732820738
0.0342513256585414	0.317465340569473
GO_POSITIVE_REGULATION_OF_SMOOTHENED_SIGNALING_PATHWAY	27	0.01
Up	0.458838172528231	0.0344122893160863
0.317853855617857
GO_PROTEASOME_BINDING	16	0.01	Down	0.211485377916038
0.0344706383843216	0.317853855617857
GO_OXIDOREDUCTASE_COMPLEX	94	0.01	Up
0.178425266610239	0.0344961222732642	0.317853855617857
GO_NUCLEAR_EXPORT	186	0.01	Down	0.155682957139485
0.0345026669168453	0.317853855617857
GO_POSITIVE_REGULATION_OF_TELOMERASE_ACTIVITY	33	0.01	Down

0.356356392780339	0.0346393595727176
0.318629624190862
GO_HORMONE_RECEPTOR_BINDING	147	0.01	Down
0.233804100743861	0.0348394626111047	0.319621784972337
GO_CELLULAR_RESPONSE_TO_LIPID	439	0.01	Down
0.428423719798378	0.0348525155084314	0.319621784972337
GO_MAGNESIUM_ION_TRANSPORT	17	0.01	Down
0.289911134011985	0.0349629303276841	0.31996300982076
GO_OLFACTORY_LOBE_DEVELOPMENT	17	0.01	Down
0.49265465288435	0.0350316050429084	0.31996300982076
GO_EPHRIN_RECEPTOR_SIGNALING_PATHWAY	75	0.01	Down
0.346888114365952	0.0351846768738855	0.31996300982076
GO_PHOSPHATIDYLINOSITOL_3_KINASE_COMPLEX	22	0.01	Down
0.295079241137167	0.0353182219996059
0.31996300982076
GO_TRANSFERASE_ACTIVITY_TRANSFERRING_GLYCOSYL_GROUPS	221	0.01
Up	0.333166569329115	0.0353262435775047
0.31996300982076
GO_NEGATIVE_REGULATION_OF_ALPHA_BETA_T_CELL_ACTIVATION	29	0.01
Down	0.539764012916196	0.03534699026591
0.31996300982076
GO_CELLULAR_RESPONSE_TO_GLUCOSE_STARVATION	40	0.01	Down
0.223845603025112	0.035404565951085
0.31996300982076
GO_LARGE_RIBOSOMAL_SUBUNIT	115	0.01	Up
0.101631401190603	0.0354215907004315	0.31996300982076
GO_MICROTUBULE_ORGANIZING_CENTER_LOCALIZATION	27	0.01	Down
0.17168986262801З	0.0354240601049134
0.31996300982076
GO_NEGATIVE_REGULATION_OF_STEM_CELL_DIFFERENTIATION	16	0.01
Down	0.177345794439622	0.0354513026748986
0.31996300982076
GO_CELL_CHEMOTAXIS	193	0.01	Down	0.692058883010497
0.0356062329172703	0.31996300982076
GO_REGULATION_OF_NUCLEOBASE_CONTAINING_COMPOUND_METABOLIC_PROCESS
444	0.01	Down	0.186890467392363
0.035613718225669	0.31996300982076
GO_NUCLEAR_HORMONE_RECEPTOR_BINDING	135	0.01	Down
0.226240529677275	0.0356585165365997	0.31996300982076
GO_SMOOTH_ENDOPLASMIC_RETICULUM	24	0.01	Down
0.185761172737956	0.0357097941572849	0.31996300982076
GO_REGULATION_OF_MESODERM_DEVELOPMENT	10	0.01	Up
0.680657482268068	0.0357654315087272	0.31996300982076
GO_REGULATION_OF_RNA_SPLICING	115	0.01	Down
0.13316642137084	0.0357898147944311	0.31996300982076
GO_CELLULAR_RESPONSE_TO_ETHANOL	10	0.01	Up
0.633787864478556	0.0358154337406953	0.31996300982076
GO_ENDOPLASMIC_RETICULUM_PART	1077	0.01	Up
0.337277903827473	0.0358383868684099	0.31996300982076
GO_RESPIRATORY_CHAIN_COMPLEX	69	0.01	Up
0.142737966889493	0.0359233071540369	0.319985317309404
GO_LIVER_MORPHOGENESIS	12	0.01	Down	0.334772236584969
0.0359805958491613	0.319985317309404
GO_MAP_KINASE_KINASE_ACTIVITY	14	0.01	Down

0.224850958545186	0.0359990070371146	0.319985317309404
GO_REGULATION_OF_HEMOPOIESIS	351	0.01	Down
0.357416811335809	0.0360594466029852	0.320053947846087
GO_ORGANELLE_INHERITANCE	13	0.01	Down
0.19200423978168	0.0362047485983793	0.320782237142989
GO_ESCRT_COMPLEX_DISASSEMBLY	10	0.01	Down
0.134607603434157	0.0362471775127805	0.320782237142989
GO_POSITIVE_REGULATION_OF_MEMBRANE_DEPOLARIZATION	10	0.01
Up	0.614729220865436	0.0363362339941683
0.320882015287133
GO_OXIDATION_REDUCTION_PROCESS	757	0.01	Up
0.295340558531676	0.0364135274800349	0.320882015287133
GO_POSITIVE_REGULATION_OF_ADAPTIVE_IMMUNE_RESPONSE	76	0.01
Down	0.456441393473677	0.0364170167242356
0.320882015287133
GO_ANTIBACTERIAL_HUMORAL_RESPONSE	16	0.01	Down
0.667730183552601	0.0365198180682648	0.321321471728167
GO_MYD88_DEPENDENT_TOLL_LIKE_RECEPTOR_SIGNALING_PATHWAY	30	0.01
Down	0.491359412720387	0.0369023919570208
0.324217686788818
GO_REGULATION_OF_LYMPHOCYTE_ACTIVATION	305	0.01	Down
0.477095159394004	0.0369628764834556	0.324279802212513
GO_CILIARY_TIP	42	0.01	Up	0.177096410557114
0.0370756954634169	0.324407475203448
GO_REGULATION_OF_MYELOID_LEUKOCYTE_MEDIATED_IMMUNITY	42	0.01
Down	0.435076426943132	0.0370857965952141
0.324407475203448
GO_THIOLESTER_HYDROLASE_ACTIVITY	29	0.01	Up
0.28332783871715	0.037191736299981	0.324407475203448
GO_NEGATIVE_REGULATION_OF_LIPID_STORAGE	17	0.01	Down
0.406520982015299	0.0372613712237198	0.324407475203448
GO_GROWTH_FACTOR_RECEPTOR_BINDING	97	0.01	Down
0.542776591422047	0.037293320405068	0.324407475203448
GO_EMBRYONIC_HEART_TUBE_DEVELOPMENT	52	0.01	Up
0.379926387309469	0.0373957538684401	0.324407475203448
GO_TUMOR_NECROSIS_FACTOR_SUPERFAMILY_CYTOKINE_PRODUCTION	95
0.01	Down	0.532675098347892	0.0374154877973018
0.324407475203448
GO_RESPIRATORY_ELECTRON_TRANSPORT_CHAIN	97	0.01	Up
0.136028470676742	0.0374600671854065	0.324407475203448
GO_ESTABLISHMENT_OF_PROTEIN_LOCALIZATION_TO_ENDOPLASMIC_RETICULUM
114	0.01	Up	0.13357729464683
0.0375348816216721	0.324407475203448
GO_POSITIVE_REGULATION_OF_IMMUNE_EFFECTOR_PROCESS	153	0.01
Down	0.396982876469848	0.0375477394802908
0.324407475203448
GO_EMBRYONIC_CAMERA_TYPE_EYE_DEVELOPMENT	26	0.01	Down
0.329019848364945	0.0375652207326675
0.324407475203448
GO_KINASE_ACTIVATOR_ACTIVITY	69	0.01	Down
0.376666249968231	0.037635343166729	0.324551375518767
GO_NEGATIVE_REGULATION_OF_CELLULAR_CATABOLIC_PROCESS	202	0.01
Down	0.226253772634307	0.0377442664102164
0.324573381672618

GO_CILIARY_BASAL_BODY 104 0.01 Up 0.223498570319928
0.0377448208632627 0.324573381672618
GO_MITOTIC_SPINDLE_ASSEMBLY 48 0.01 Down
0.175937555566134 0.0378798551549907 0.324786782118297
GO_T_HELPER_1_TYPE_IMMUNE_RESPONSE 30 0.01 Down
0.645906210434689 0.0378868203154825 0.324786782118297
GO_VACUOLAR_TRANSPORT 127 0.01 Down 0.189178986452861
0.0379301315305341 0.324786782118297
GO_MEMBRANE_LIPID_CATABOLIC_PROCESS 27 0.01 Up
0.199089912321927 0.0380400649058166 0.324992294191268
GO_REGULATION_OF_NUCLEOTIDE_BIOSYNTHETIC_PROCESS 23 0.01
Up 0.306754996769994 0.0380611960418368
0.324992294191268
GO_ENTRY_INTO_OTHER_ORGANISM_INVOLVED_IN_SYMBIOTIC_INTERACTION 111
0.01 Down 0.3702777120179 0.0381810997281589
0.325145869943836
GO_LYMPHOCYTE_DIFFERENTIATION 244 0.01 Down
0.446644966344467 0.0383158495550564 0.325145869943836
GO_VENTRICULAR_CARDIAC_MUSCLE_CELL_DIFFERENTIATION 12 0.01
Up 0.306466163929093 0.0383366022159387
0.325145869943836
GO_RESPONSE_TO_COPPER_ION 31 0.01 Up
0.495193457781185 0.0383391068136172 0.325145869943836
GO_CORECEPTOR_ACTIVITY 29 0.01 Up 0.545855984013451
0.0383877771266955 0.325145869943836
GO_FATTY_ACID_BETA_OXIDATION_USING_ACYL_COA_OXIDASE 14 0.01
Up 0.25833086531996 0.038400525242914
0.325145869943836
GO_NEGATIVE_REGULATION_OF_GLUCOSE_TRANSMEMBRANE_TRANSPORT 13
0.01 Down 0.332301855336647 0.0385384642601866
0.325859354489684
GO_INTRINSIC_APOPTOTIC_SIGNALING_PATHWAY_IN_RESPONSE_TO_ENDOPLASMIC_
RETICULUM_STRESS 54 0.01 Down 0.238230740001419
0.038747058180123 0.327167441184321
GO_ATP_SYNTHESIS_COUPLED_ELECTRON_TRANSPORT 80 0.01 Up
0.121509810118721 0.0388565235670944
0.327296601363323
GO_REGULATION_OF_CALCINEURIN_MEDIATED_SIGNALING 29 0.01 Down
0.303691930011834 0.0388701778262157
0.327296601363323
GO_SODIUM_CHANNEL_COMPLEX 11 0.01 Up
0.515495186344257 0.0390173319769991 0.328080640488036
GO_ENHANCER_BINDING 93 0.01 Down 0.256218639515364
0.0391867016810377 0.328626454827486
GO_POSITIVE_REGULATION_OF_CELL_ADHESION_MEDIATED_BY_INTEGRIN 15
0.01 Down 0.943150884512644 0.0392196566551002
0.328626454827486
GO_4_IRON_4_SULFUR_CLUSTER_BINDING 41 0.01 Up
0.185500670278861 0.0392446351095252 0.328626454827486
GO_OXIDOREDUCTASE_ACTIVITY_ACTING_ON_PAIRED_DONORS_WITH_INCORPORATIO
N_OR_REDUCTION_OF_MOLECULAR_OXYGEN_NAD_P_H_AS_ONE_DONOR_AND_INCORPOR
ATION_OF_ONE_ATOM_OF_OXYGEN 26 0.01 Up
0.680756610060338 0.0394470261793203 0.329029521966757
GO_POSITIVE_REGULATION_OF_LYMPHOCYTE_CHEMOTAXIS 12 0.01 Down

0.65550857611039	0.0395068941385239
0.329029521966757
GO_GLYCOLIPID_CATABOLIC_PROCESS 12	0.01	Up
0.229050788426608	0.0395509206185846	0.329029521966757
GO_STRUCTURAL_CONSTITUENT_OF_CYTOSKELETON	69	0.01	Down
0.291133395652307	0.0395582250342379
0.329029521966757
GO_SMOOTHENED_SIGNALING_PATHWAY 110	0.01	Up
0.315062166811781	0.0395637540826441	0.329029521966757
GO_GTPASE_BINDING	469	0.01	Down	0.242573804182057
0.039639814084707	0.3292020344940 3
GO_POSITIVE_REGULATION_OF_FAT_CELL_DIFFERENTIATION	50	0.01
Down	0.460579978675729	0.0397097351471777
0.32920203449403
GO_EXTRACELLULAR_VESICLE_BIOGENESIS	18	0.01	Down
0.170826087519174	0.0397471736590552	0.32920203449403
GO_RESPONSE_TO_PEPTIDOGLYCAN	10	0.01	Down
0.618890885146468	0.0398207649189161	0.329362212292561
GO_EOSINOPHIL_MIGRATION 11	0.01	Down	0.590581125804283
0.0400031253330064	0.330299563240256
GO_VESICLE_DOCKING	56	0.01	Down	0.2104229604399
0.0401306276273824	0.330299563240256
GO_SMALL_GTPASE_BINDING 389	0.01	Down	0.25626104833491
0.0401549874918867	0.330299563240256
GO_HOPS_COMPLEX 13	0.01	Down	0.1602694114412
0.0401767729736651	0.330299563240256
GO_REGULATION_OF_TRANSLATIONAL_INITIATION_BY_EIF2_ALPHA_PHOSPHORYLAT
ION	10	0.01	Down	0.225621290032534
0.0402525956225385	0.330299563240256
GO_MULTIVESICULAR_BODY_ORGANIZATION	31	0.01	Down
0.149264351185501	0.0402605298629204	0.330299563240256
GO_RESPIRASOME	81	0.01	Up	0.14480234370301
0.0404365152643856	0.331295660148563
GO_CELL_CELL_JUNCTION_ASSEMBLY	92	0.01	Down
0.345382556153352	0.0404936191526594	0.331316390668187
GO_NEURAL_RETINA_DEVELOPMENT	33	0.01	Down
0.453771831809754	0.0409037393695312	0.334221536625065
GO_REGULATION_OF_ADAPTIVE_IMMUNE_RESPONSE	120	0.01	Down
0.443881699609317	0.0412296874468473
0.336324473955468
GO_LEUKOCYTE_ADHESION_TO_VASCULAR_ENDOTHELIAL_CELL	22	0.01
Down	0.576726953162112	0.0412719046445105
0.336324473955468
GO_MLL3_4_COMPLEX	11	0.01	Down	0.239825143032703
0.0413300440628254	0.336346779497873
GO_NEGATIVE_REGULATION_OF_RESPONSE_TO_EXTRACELLULAR_STIMULUS	10
0.01	Up	0.301525183737235	0.0417046426271605
0.338940944296508
GO_INTERLEUKIN_6_BIOSYNTHETIC_PROCESS	17	0.01	Down
0.562536420186777	0.0418620505038289	0.339696749372305
GO_CELLULAR_RESPIRATION 167	0.01	Up	0.162134888256311
0.0419588549438224	0.339696749372305
GO_MAGNESIUM_ION_TRANSMEMBRANE_TRANSPORTER_ACTIVITY	16	0.01
Down	0.288153564747389	0.0420096631396397

0.339696749372305
GO_MODIFICATION_DEPENDENT_MACROMOLECULE_CATABOLIC_PROCESS 556
0.01 Down 0.175423129439705 0.0420918780836784
0.339696749372305
GO_POSITIVE_REGULATION_OF_EPITHELIAL_TO_MESENCHYMAL_TRANSITION 42
0.01 Down 0.544729090964174 0.0421454541366507
0.339696749372305
GO_ANCHORED_COMPONENT_OF_EXTERNAL_SIDE_OF_PLASMA_MEMBRANE 10
0.01 Up 0.862066164370382 0.0422909140203765
0.339696749372305
GO_RESPONSE_TO_FUNGUS 23 0.01 Down 0.619427065196014
0.0423012117634234 0.339696749372305
GO_REGULATION_OF_PROTEIN_MODIFICATION_BY_SMALL_PROTEIN_CONJUGATION_OR_REMOVAL 200 0.01 Down 0.216658610325164
0.0424896570873789 0.339696749372305
GO_ACTIVATION_OF_MAPK_ACTIVITY 120 0.01 Down
0.370424358339307 0.0425303233525743 0.339696749372305
GO_PLASMA_MEMBRANE_RECEPTOR_COMPLEX 105 0.01 Down
0.491043084012281 0.0425320750953441 0.339696749372305
GO_VESICLE_DOCKING_INVOLVED_IN_EXOCYTOSIS 36 0.01 Down
0.22948944437476 0.0425799199760982
0.339696749372305
GO_REFLEX 10 0.01 Down 0.423000283843481
0.0425826841371092 0.339696749372305
GO_MITOCHONDRIAL_ENVELOPE 662 0.01 Up
0.187367022374244 0.0425831675022651 0.339696749372305
GO_METALLOENDOPEPTIDASE_INHIBITOR_ACTIVITY 12 0.01 Up
0.596967336761961 0.042656770590552
0.339696749372305
GO_LYMPHOCYTE_COSTIMULATION 36 0.01 Down
0.51533859719607 0.0426648282500992 0.339696749372305
GO_OVULATION_CYCLE_PROCESS 30 0.01 Down
0.512324616178073 0.0426929039319829 0.339696749372305
GO_INTERLEUKIN_6_PRODUCTION 98 0.01 Down
0.600497803561855 0.0429272179596332 0.340902097202371
GO_ENDOSOME_ORGANIZATION 74 0.01 Down
0.176590827695762 0.0429566964848977 0.340902097202371
GO_RESPONSE_TO_INTERLEUKIN_18 10 0.01 Down
0.481049083372834 0.0432225146707197 0.342563820582166
GO_OXIDATIVE_PHOSPHORYLATION 116 0.01 Up
0.125842952333102 0.0433117964885667 0.342823880680689
GO_DERMATAN_SULFATE_PROTEOGLYCAN_BIOSYNTHETIC_PROCESS 13 0.01
Up 0.83434003676435 0.0434690082395315
0.343620246122651
GO_POSITIVE_REGULATION_OF_SMOOTH_MUSCLE_CELL_DIFFERENTIATION 15
0.01 Up 0.622802992179502 0.0436416810653379
0.344536600452102
GO_EXOCYST 14 0.01 Down 0.239204759608964
0.0439145222311456 0.346055525124696
GO_INOSITOL_PHOSPHATE_MEDIATED_SIGNALING 45 0.01 Down
0.306915176326221 0.0439914899162328
0.346055525124696
GO_MACROPHAGE_ACTIVATION_INVOLVED_IN_IMMUNE_RESPONSE 10 0.01
Up 0.856041926192908 0.0440050840712016

0.346055525124696
GO_NOTCH_SIGNALING_PATHWAY 145 0.01 Down
0.329226007796349 0.0444795536260394 0.349334243290666
GO_DOUBLE_STRANDED_DNA_BINDING 668 0.01 Down
0.3138355060888 0.044664777737918 0.350204348714065
GO_METAL_CLUSTER_BINDING 61 0.01 Up
0.16501325065429 0.0447057107977929 0.350204348714065
GO_INSULIN_RECEPTOR_SUBSTRATE_BINDING 10 0.01 Down
0.240973020208032 0.0449861412534508 0.35143893023462
GO_NEGATIVE_REGULATION_OF_CATABOLIC_PROCESS 243 0.01 Down
0.256683018109409 0.0450166738039787
0.35143893023462
GO_FIBROBLAST_GROWTH_FACTOR_RECEPTOR_BINDING 11 0.01 Up
0.679575713334017 0.0452093674658372
0.35143893023462
GO_NEGATIVE_REGULATION_OF_GENE_EXPRESSION_EPIGENETIC 92 0.01
Down 0.17783872981826 0.045228221763214
0.35143893023462
GO_DENDRITIC_CELL_CYTOKINE_PRODUCTION 12 0.01 Down
0.390999915541082 0.0452441103496015 0.35143893023462
GO_ARF_GUANYL_NUCLEOTIDE_EXCHANGE_FACTOR_ACTIVITY 16 0.01
Down 0.205161600626019 0.0452459489669413
0.35143893023462
GO_NEGATIVE_REGULATION_OF_BLOOD_CIRCULATION 22 0.01 Down
0.396258012043738 0.0452685296398407
0.35143893023462
GO_ORGANELLAR_RIBOSOME 87 0.01 Up 0.107086214999393
0.0454069838353201 0.352063600082029
GO_REGULATION_OF_CELLULAR_RESPONSE_TO_VASCULAR_ENDOTHELIAL_GROWTH_FA
CTOR_STIMULUS 15 0.01 Up 0.889433018302085
0.0455746920915134 0.352145882658129
GO_PROTEIN_MODIFICATION_BY_SMALL_PROTEIN_CONJUGATION 789 0.01
Down 0.190219543095302 0.0456207278075115
0.352145882658129
GO_PHAGOLYSOSOME_ASSEMBLY 12 0.01 Up
0.603818455679283 0.0456412331350398 0.352145882658129
GO_RIBOSE_PHOSPHATE_BIOSYNTHETIC_PROCESS 157 0.01 Up
0.230409069414373 0.0456496145036975
0.352145882658129
GO_TAU_PROTEIN_KINASE_ACTIVITY 20 0.01 Down
0.249395891277223 0.0457941633724271 0.352252657716116
GO_PROTEIN_LOCALIZATION_TO_NUCLEUS 220 0.01 Down
0.264286869904357 0.0458128284170657 0.352252657716116
GO_FATTY_ACID_BETA_OXIDATION_USING_ACYL_COA_DEHYDROGENASE 11
0.01 Up 0.251341192623583 0.0460059553848606
0.352252657716116
GO_MICROBODY_PART 83 0.01 Up 0.224480154078146
0.0460177328587229 0.352252657716116
GO_PRI_MIRNA_TRANSCRIPTION_BY_RNA_POLYMERASE_II 39 0.01 Down
0.472118356356427 0.0460497760356021
0.352252657716116
GO_RNA_STEM_LOOP_BINDING 16 0.01 Down
0.291978789162366 0.0460564782777581 0.352252657716116
GO_POSITIVE_REGULATION_OF_STRESS_ACTIVATED_PROTEIN_KINASE_SIGNALING_

CASCADE 143 0.01 Down 0.376337433449106
0.0461523726480192 0.352252657716116
GO_MEMBRANE_DEPOLARIZATION_DURING_CARDIAC_MUSCLE_CELL_ACTION_POTENTIAL 15 0.01 Up 0.532119453461524
0.0461625593615769 0.352252657716116
GO_POSITIVE_REGULATION_OF_CYTOKINE_BIOSYNTHETIC_PROCESS 44 0.01 Down 0.485829253998351 0.046208718006252
0.352252657716116
GO_POSITIVE_REGULATION_OF_NIK_NF_KAPPAB_SIGNALING 65 0.01 Down 0.504566753702589 0.0463002636929984
0.352252657716116
GO_I_KAPPAB_PHOSPHORYLATION 15 0.01 Down
0.246222376253986 0.0463017000259365 0.352252657716116
GO_T_CELL_ACTIVATION_INVOLVED_IN_IMMUNE_RESPONSE 62 0.01 Down 0.520026053567858 0.0464384156910562
0.35257442195112
GO_POSITIVE_REGULATION_OF_LYMPHOCYTE_ACTIVATION 197 0.01 Down 0.470789768553476 0.0464721329526761
0.35257442195112
GO_MRNA_TRANSPORT 138 0.01 Down 0.13860328536964 0.0465910139954841 0.35257442195112
GO_VITAMIN_TRANSMEMBRANE_TRANSPORT 10 0.01 Up
0.570673199221029 0.0465935135237907 0.35257442195112
GO_REGULATION_OF_SPROUTING_ANGIOGENESIS 56 0.01 Down
0.448569363286379 0.0466343700917063 0.35257442195112
GO_RESPONSE_TO_STEROL 25 0.01 Down 0.330773347115261 0.0468447166047814 0.353465878056397
GO_IMMUNE_RESPONSE_TO_TUMOR_CELL 10 0.01 Down
0.51434674308647 0.0469618058589777 0.353465878056397
GO_HISTONE_METHYLATION 114 0.01 Down 0.174247802750753 0.0470825352603041 0.353465878056397
GO_SPLEEN_DEVELOPMENT 23 0.01 Down 0.297108208600317 0.0470876623069215 0.353465878056397
GO_T_CELL_MIGRATION 43 0.01 Down 0.649300535451855 0.047121374312674 0.353465878056397
GO_TRANSLATION_INITIATION_FACTOR_ACTIVITY 49 0.01 Down 0.122807098911045 0.0471532516629782
0.353465878056397
GO_FILOPODIUM_TIP 13 0.01 Down 0.220823515209918 0.0471598354843817 0.353465878056397
GO_RESPONSE_TO_EPINEPHRINE 10 0.01 Down
0.418470078861139 0.0472816824315259 0.353688584418752
GO_INTERLEUKIN_7_MEDIATED_SIGNALING_PATHWAY 17 0.01 Down 0.351079869240167 0.0473060666361434
0.353688584418752
GO_RENAL_VESICLE_DEVELOPMENT 12 0.01 Down
1.13231573679942 0.0473851747319156 0.353844275273628
GO_SARCOLEMMA 95 0.01 Up 0.510740083251923
0.0474453132910323 0.353858104410144
GO_PROTEIN_SERINE_THREONINE_KINASE_ACTIVITY 379 0.01 Down 0.265321739435931 0.0476745808621286
0.355131755109181
GO_CAVEOLA 59 0.01 Up 0.413983235481484
0.0477404177858147 0.355186368109904

GO_NEGATIVE_REGULATION_OF_T_HELPER_CELL_DIFFERENTIATION 12 0.01 Down 0.530870010929941 0.048166720868393 0.357919415412502
GO_RESPONSE_TO_VITAMIN_A 10 0.01 Up 0.422164514101059 0.048431836144098 0.35944948316726
GO_REGULATION_OF_LIPID_STORAGE 36 0.01 Down 0.557758230346088 0.0485237884227016 0.359565783923277
GO_PROTEIN_SERINE_THREONINE_KINASE_ACTIVATOR_ACTIVITY 25 0.01 Down 0.294917146997483 0.0485659599435163 0.359565783923277
GO_REGULATION_OF_TRANSCRIPTION_INITIATION_FROM_RNA_POLYMERASE_II_PROMOTER 22 0.01 Down 0.254832548182559 0.0488928200594395 0.36120014833583
GO_TRANSCYTOSIS 17 0.01 Down 0.388020578259002 0.0489057028384208 0.36120014833583
GO_TRNA_TRANSPORT 35 0.01 Down 0.169095477895065 0.0492323904600063 0.363171133028795
GO_ANION_ANION_ANTIPORTER_ACTIVITY 17 0.01 Up 0.223900287091952 0.0494692726975863 0.363699591609855
GO_REGULATION_OF_RESPONSE_TO_INTERFERON_GAMMA 18 0.01 Down 0.287774979154904 0.0494829618318811 0.363699591609855
GO_TRANSEPITHELIAL_TRANSPORT 14 0.01 Down 0.851223141099825 0.0494837527046187 0.363699591609855
GO_POSITIVE_REGULATION_OF_BIOSYNTHETIC_PROCESS 1491 0.01 Down 0.313953784421981 0.0495638008989689 0.363847442875018
GO_INFLAMMATORY_RESPONSE_TO_WOUNDING 10 0.01 Up 0.616147161097336 0.0496327076265192 0.363913246377534
GO_REGULATION_OF_RECEPTOR_LOCALIZATION_TO_SYNAPSE 14 0.01 Up 0.381409111044055 0.0498862593669374 0.36469106209153
GO_INTRINSIC_COMPONENT_OF_MITOCHONDRIAL_OUTER_MEMBRANE 20 0.01 Up 0.172555176924641 0.0499420199418584 0.36469106209153
GO_NEGATIVE_REGULATION_OF_VASCULAR_SMOOTH_MUSCLE_CELL_PROLIFERATION 15 0.01 Up 0.497954789264005 0.049972933132989 0.36469106209153
GO_REGULATION_OF_TOLL_LIKE_RECEPTOR_4_SIGNALING_PATHWAY 15 0.01 Down 0.479415199612881 0.049979074890488 0.36469106209153
GO_VIRAL_BUDDING_VIA_HOST_ESCRT_COMPLEX 21 0.01 Down 0.177544483889903 0.0501219085628371 0.365027879149316
GO_REGULATION_OF_GLIAL_CELL_PROLIFERATION 22 0.01 Down 0.343846617274066 0.0501454869396359 0.365027879149316
GO_REGULATION_OF_NUCLEAR_TRANSCRIBED_MRNA_CATABOLIC_PROCESS_DEADENYLATION_DEPENDENT_DECAY 19 0.01 Down 0.257820976908128 0.0502274880141509 0.365186921837018
GO_IMMUNE_RESPONSE_REGULATING_SIGNALING_PATHWAY 427 0.01 Down 0.422793202270941 0.0505420023138498 0.365754889540696
GO_POSITIVE_REGULATION_OF_RESPONSE_TO_BIOTIC_STIMULUS 29 0.01 Down 0.347301232785671 0.0505940247547297

0.365754889540696
GO_SMALL_MOLECULE_CATABOLIC_PROCESS 305 0.01 Up
0.303722553711282 0.050651375607875 0.365754889540696
GO_POSITIVE_REGULATION_OF_IMMUNE_SYSTEM_PROCESS 755 0.01 Down
0.445319878372812 0.0508208222972753
0.365754889540696
GO_TOLL_LIKE_RECEPTOR_SIGNALING_PATHWAY 112 0.01 Down
0.487146264823773 0.0508428425108838 0.365754889540696
GO_PHOSPHOPROTEIN_BINDING 73 0.01 Down
0.298221396962238 0.0508498035195043 0.365754889540696
GO_ESTABLISHMENT_OR_MAINTENANCE_OF_CELL_POLARITY 175 0.01
Down 0.284256499868368 0.0508642745084301
0.365754889540696
GO_DEOXYRIBONUCLEOSIDE_MONOPHOSPHATE_BIOSYNTHETIC_PROCESS 10
0.01 Up 0.207692869946472 0.0508821192948544
0.365754889540696
GO_ENDOCYTIC_ADAPTOR_ACTIVITY 10 0.01 Up
0.209891102196174 0.0509195896420928 0.365754889540696
GO_PROTEIN_FOLDING_IN_ENDOPLASMIC_RETICULUM 15 0.01 Down
0.19698206137086 0.0509307088136553
0.365754889540696
GO_THYROID_HORMONE_METABOLIC_PROCESS 13 0.01 Down
0.620452267113129 0.051093435836157 0.365754889540696
GO_RELAXATION_OF_MUSCLE 18 0.01 Up 0.562868316536571
0.0511249984705541 0.365754889540696
GO_ASTROCYTE_DIFFERENTIATION 53 0.01 Down
0.531038584263081 0.0511345540774938 0.365754889540696
GO_SIGNAL_TRANSDUCTION_BY_PROTEIN_PHOSPHORYLATION 697 0.01
Down 0.383494403114787 0.0512774699566085
0.365754889540696
GO_REGENERATION 137 0.01 Up 0.475861637100979
0.0513364189725503 0.365754889540696
GO_NEGATIVE_REGULATION_OF_STEROID_METABOLIC_PROCESS 19 0.01
Up 0.620872641110577 0.0513608278710072
0.365754889540696
GO_INTERFERON_BETA_PRODUCTION 47 0.01 Down
0.321122058398242 0.0514535397832175 0.365754889540696
GO_NUCLEAR_RECEPTOR_BINDING 96 0.01 Down
0.232971187344055 0.0515254448382726 0.365754889540696
GO_TETHERING_COMPLEX 61 0.01 Down 0.168269831175756
0.0515302335141109 0.365754889540696
GO_PROTEIN_KINASE_A_CATALYTIC_SUBUNIT_BINDING 11 0.01 Down
0.23572938706619 0.0516059034395807
0.365754889540696
GO_REGULATION_OF_TELOMERE_MAINTENANCE_VIA_TELOMERE_LENGTHENING 57
0.01 Down 0.149714912194299 0.0516259016649091
0.365754889540696
GO_NEGATIVE_REGULATION_OF_ENDOTHELIAL_CELL_PROLIFERATION 31
0.01 Up 0.612970415910598 0.0516881052205092
0.365754889540696
GO_EPHRIN_RECEPTOR_BINDING 24 0.01 Down
0.370311557327906 0.051691269185623 0.365754889540696
GO_EXTRINSIC_COMPONENT_OF_MEMBRANE 207 0.01 Down
0.31343132245943 0.0521770707843669 0.368476020271342

GO_CELL_MIGRATION_INVOLVED_IN_SPROUTING_ANGIOGENESIS 49 0.01 Down 0.541683509282331 0.0522518006464492 0.368476020271342
GO_VIRAL_RNA_GENOME_REPLICATION 29 0.01 Down 0.250326047638728 0.0522579234810782 0.368476020271342
GO_IMMUNE_SYSTEM_DEVELOPMENT 723 0.01 Down 0.377728436854031 0.0524958388390406 0.36876314363041
GO_PHOSPHATIDYLETHANOLAMINE_BINDING 11 0.01 Up 0.349520226269893 0.0525245209039477 0.36876314363041
GO_CELL_JUNCTION_ORGANIZATION 241 0.01 Down 0.343662656849645 0.052548044937883 0.36876314363041
GO_REGULATION_OF_T_CELL_APOPTOTIC_PROCESS 26 0.01 Down 0.55309111746231 0.0525714945892121 0.36876314363041
GO_ENDOPEPTIDASE_COMPLEX 63 0.01 Down 0.12032530085949 0.052681619890623 0.36876314363041
GO_ACROSOMAL_MEMBRANE 12 0.01 Up 0.302264346926894 0.0527792952916545 0.36876314363041
GO_ALDEHYDE_DEHYDROGENASE_NAD_ACTIVITY 13 0.01 Up 0.399228006048425 0.0527832735719065 0.36876314363041
GO_ANTIGEN_PROCESSING_AND_PRESENTATION_OF_ENDOGENOUS_ANTIGEN 15 0.01 Down 0.374129613636059 0.052788880369633 0.36876314363041
GO_REGULATION_OF_RESPONSE_TO_STRESS 1147 0.01 Down 0.369049500530345 0.0528453195451255 0.36876314363041
GO_HORMONE_CATABOLIC_PROCESS 11 0.01 Up 0.68255990035342 0.0530723993391564 0.369649787366376
GO_T_CELL_APOPTOTIC_PROCESS 36 0.01 Down 0.443939111375452 0.053199141553587 0.369649787366376
GO_PROTEIN_K63_LINKED_UBIQUITINATION 43 0.01 Down 0.189571820829046 0.0532642275314657 0.369649787366376
GO_VESICLE_ORGANIZATION 278 0.01 Down 0.240686599635842 0.0533659160072475 0.369649787366376
GO_REGULATION_OF_LEUKOCYTE_MEDIATED_IMMUNITY 145 0.01 Down 0.423327032086253 0.053484295531704 0.369649787366376
GO_MICROVILLUS_MEMBRANE 19 0.01 Down 0.584735863316388 0.0535135106255982 0.369649787366376
GO_POSITIVE_REGULATION_OF_CYTOKINE_PRODUCTION 324 0.01 Down 0.500828949774238 0.0535202289283936 0.369649787366376
GO_NEGATIVE_REGULATION_OF_PROTEIN_DEPHOSPHORYLATION 50 0.01 Down 0.2943549445978 0.0535583321021436 0.369649787366376
GO_MITOCHONDRIAL_GENE_EXPRESSION 160 0.01 Up 0.110725155228844 0.0535646219416394 0.369649787366376
GO_ALDO_KETO_REDUCTASE_NADP_ACTIVITY 16 0.01 Up 0.492534770642702 0.0535812572693808 0.369649787366376
GO_T_CELL_PROLIFERATION 138 0.01 Down 0.5324972932963 0.0536531368339481 0.369725532030532
GO_ZINC_ION_TRANSMEMBRANE_TRANSPORT 15 0.01 Up 0.225738049275977 0.053794721091516 0.370280897671875
GO_POLY_A_SPECIFIC_RIBONUCLEASE_ACTIVITY 12 0.01 Down 0.17184047407060 3 0.053918231952553

0.37071074313018
GO_RIBOSOMAL_SMALL_SUBUNIT_ASSEMBLY 16 0.01 Up
0.163595001657331 0.0540171692988934 0.370970853861518
GO_TRANSCRIPTION_COACTIVATOR_BINDING 19 0.01 Down
0.299176100828038 0.0541951829712944 0.370978225920549
GO_NEURON_PROJECTION_EXTENSION_INVOLVED_IN_NEURON_PROJECTION_GUIDANC
E 31 0.01 Down 0.556084382364898
0.0542120300201861 0.370978225920549
GO_LATERAL_SPROUTING_FROM_AN_EPITHELIUM 12 0.01 Up
0.578956341858873 0.0543070908194314 0.370978225920549
GO_FOLIC_ACID_METABOLIC_PROCESS 18 0.01 Up
0.301483012397274 0.0543392033362881 0.370978225920549
GO_PEPTIDYL_LYSINE_TRIMETHYLATION 40 0.01 Down
0.177901113101988 0.0543618742118954 0.370978225920549
GO_REGULATION_OF_LYMPHOCYTE_DIFFERENTIATION 124 0.01 Down
0.459962363139665 0.0543848824031113
0.370978225920549
GO_REGULATION_OF_EXTRINSIC_APOPTOTIC_SIGNALING_PATHWAY_IN_ABSENCE_OF
_LIGAND 38 0.01 Down 0.420309623519888
0.054567331088211 0.37137793898901
GO_FIBROBLAST_GROWTH_FACTOR_BINDING 19 0.01 Up
0.297290961004277 0.0545856974352028 0.37137793898901
GO_OLIGOSACCHARIDE_METABOLIC_PROCESS 14 0.01 Up
0.334422573550619 0.0546269971202745 0.37137793898901
GO_RESPONSE_TO_INTERLEUKIN_7 26 0.01 Down
0.27926076498957 0.0547474168731348 0.371382740637596
GO_TUMOR_NECROSIS_FACTOR_MEDIATED_SIGNALING_PATHWAY 137 0.01
Down 0.260273040274079 0.0547500498880989
0.371382740637596
GO_NADP_METABOLIC_PROCESS 10 0.01 Up
0.292403340181858 0.0548120550246269 0.37138837729298
GO_AMINOGLYCAN_BIOSYNTHETIC_PROCESS 92 0.01 Up
0.60715448730547 0.0551028761860686 0.37229116331962
GO_MITOCHONDRIAL_TRANSLATIONAL_TERMINATION 89 0.01 Up
0.109726266764193 0.0551214888094419
0.37229116331962
GO_ALPHA_AMINO_ACID_METABOLIC_PROCESS 131 0.01 Up
0.353697901667408 0.0551292630249281 0.37229116331962
GO_POSITIVE_REGULATION_OF_GROWTH 180 0.01 Down
0.360983831775083 0.0552890582435682 0.372825173941839
GO_RESPONSE_TO_HYDROPEROXIDE 16 0.01 Up
0.474086995985713 0.0553311615420189 0.372825173941839
GO_ADAPTIVE_IMMUNE_RESPONSE 255 0.01 Down
0.485301035038107 0.0553981135136046 0.372862469114294
GO_HISTONE_H3_K4_METHYLATION 47 0.01 Down
0.184827241191975 0.0555412012834313 0.373000102843344
GO_MOLTING_CYCLE 81 0.01 Down 0.432006355810297
0.0555529741039881 0.373000102843344
GO_SINGLE_FERTILIZATION 65 0.01 Up 0.334493199467946
0.0556028814154549 0.373000102843344
GO_POSITIVE_REGULATION_OF_SODIUM_ION_TRANSMEMBRANE_TRANSPORTER_ACTIV
ITY 13 0.01 Down 0.210919275506996
0.0557594019519122 0.373490983754285
GO_POSITIVE_REGULATION_OF_I_KAPPAB_KINASE_NF_KAPPAB_SIGNALING 164

0.01 Down 0.332820506318615 0.0557990977211557
0.373490983754285
GO_PROTEIN_PHOSPHATASE_ACTIVATOR_ACTIVITY 13 0.01 Down
0.228738265252892 0.0559936461608807
0.373736670442
GO_HORMONE_TRANSPORT 196 0.01 Down 0.38360555818621
0.0560383228679759 0.373736670442
GO_VACUOLAR_PART 486 0.01 Up 0.344809020765701
0.0560727223365263 0.373736670442
GO_GROWTH_FACTOR_BINDING 108 0.01 Up
0.626322129269959 0.0560964692931627 0.373736670442
GO_THYROID_HORMONE_RECEPTOR_BINDING 27 0.01 Down
0.180453421752775 0.056164045095428 0.373736670442
GO_NEGATIVE_REGULATION_OF_CELL_ADHESION 206 0.01 Down
0.503830538912291 0.0562051688541502 0.373736670442
GO_PLASMA_MEMBRANE_PHOSPHOLIPID_SCRAMBLING 11 0.01 Up
0.526076974369525 0.0562756268789742
0.373795766720189
GO_NUCLEOPLASM_PART 1007 0.01 Down 0.183385958717864
0.0569603047497107 0.377623827596583
GO_POSITIVE_REGULATION_OF_BMP_SIGNALING_PATHWAY 24 0.01 Up
0.53539687583529 0.0570377980112179
0.377623827596583
GO_NEGATIVE_REGULATION_OF_PROTEIN_COMPLEX_DISASSEMBLY 54 0.01
Down 0.300332350910973 0.0571024053900977
0.377623827596583
GO_GLUTAMINE_FAMILY_AMINO_ACID_METABOLIC_PROCESS 56 0.01
Up 0.378401057471167 0.057123007471355
0.377623827596583
GO_CELL_JUNCTION_ASSEMBLY 198 0.01 Down
0.359631294913916 0.0572097864293589 0.377623827596583
GO_LYASE_ACTIVITY 135 0.01 Up 0.255866310272045
0.0572251558868154 0.377623827596583
GO_MITOCHONDRION_DISTRIBUTION 12 0.01 Down
0.209075602708503 0.0574513407566709 0.378704766269
GO_MODIFICATION_DEPENDENT_PROTEIN_BINDING 133 0.01 Down
0.191701631167686 0.0578216603447049
0.380499447855642
GO_NUCLEOTIDE_EXCISION_REPAIR_DNA_DUPLEX_UNWINDING 22 0.01
Up 0.157453861281498 0.05784895245771
0.380499447855642
GO_PHOSPHATIDYLINOSITOL_PHOSPHATE_KINASE_ACTIVITY 14 0.01
Down 0.209428337385792 0.0580642327290308
0.38150211785492
GO_PODOSOME 28 0.01 Down 0.305567742244055
0.0582118114563207 0.381622396880655
GO_REGULATION_OF_IMMUNE_EFFECTOR_PROCESS 282 0.01 Down
0.441429230165844 0.0582603649637304
0.381622396880655
GO_PROTEIN_SERINE_THREONINE_PHOSPHATASE_ACTIVITY 63 0.01
Down 0.235616903038437 0.0582711187462308
0.381622396880655
GO_REGULATION_OF_STRESS_ACTIVATED_PROTEIN_KINASE_SIGNALING_CASCADE
197 0.01 Down 0.364759940816115

0.0586346637950278	0.383192297672099
GO_MALE_MEIOSIS_I	13	0.01	Up	0.403408929289705
	0.0586447842799483	0.383192297672099
GO_PROTEIN_K48_LINKED_UBIQUITINATION	53	0.01	Down
0.174263299650764	0.0587001872566385	0.383192297672099
GO_INTERLEUKIN_10_PRODUCTION	36	0.01	Down
0.677371123902875	0.0589077465931961	0.383980792601557
GO_POSITIVE_REGULATION_OF_T_CELL_PROLIFERATION	66	0.01	Down
	0.592194103404815	0.0590378837459734
0.383980792601557
GO_OXIDOREDUCTASE_ACTIVITY_ACTING_ON_THE_ALDEHYDE_OR_OXO_GROUP_OF_DONORS	37	0.01	Up	0.340121375676412
0.059045571292709	0.383980792601557
GO_CELLULAR_RESPONSE_TO_NUTRIENT	56	0.01	Up
0.442638765026371	0.0590739680925473	0.383980792601557
GO_INTRACILIARY_TRANSPORT	48	0.01	Up
0.160064409283139	0.0591563531341291	0.38410504799711
GO_REGULATION_OF_RNA_POLYMERASE_II_REGULATORY_REGION_SEQUENCE_SPECIFIC_DNA_BINDING	11	0.01	Down	0.330070725635973
0.0592197365283711	0.384105791093741
GO_AMP_BINDING	12	0.01	Up	0.284555658574897
0.0594637346909305	0.384659311582434
GO_ADENYLATE_CYCLASE_ACTIVATING_G_PROTEIN_COUPLED_RECEPTOR_SIGNALING_PATHWAY	57	0.01	Down	0.538712808969132
0.0594850820779015	0.384659311582434
GO_NCRNA_EXPORT_FROM_NUCLEUS	39	0.01	Down
0.158645922283245	0.0595069033245844	0.384659311582434
GO_NEUROTRANSMITTER_CATABOLIC_PROCESS	12	0.01	Up
0.407692345813653	0.0596207150965165	0.384659311582434
GO_PHOSPHATIDYLINOSITOL_MONOPHOSPHATE_PHOSPHATASE_ACTIVITY	12
	0.01	Down	0.156957660545045	0.0597112019300912
	0.384659311582434
GO_PROSTANOID_METABOLIC_PROCESS	12	0.01	Up
0.402469274645989	0.0597159355752304	0.384659311582434
GO_NEGATIVE_REGULATION_OF_BMP_SIGNALING_PATHWAY	30	0.01	Up
	0.733216620531314	0.0597485967422559
0.384659311582434
GO_PHOSPHATIDYLINOSITOL_3_PHOSPHATE_BIOSYNTHETIC_PROCESS	13
	0.01	Down	0.250810045075139	0.0599103931732546
	0.385292369655539
GO_REGULATION_OF_BMP_SIGNALING_PATHWAY	56	0.01	Up
0.643600409224438	0.0601838747341941	0.385837194244738
GO_REGULATION_OF_TRANSCRIPTION_REGULATORY_REGION_DNA_BINDING	36
	0.01	Down	0.333931355296576	0.0602041809199308
	0.385837194244738
GO_MISMATCHED_DNA_BINDING	13	0.01	Up
0.169954265987237	0.0602591514967279	0.385837194244738
GO_EPITHELIAL_CELL_DIFFERENTIATION_INVOLVED_IN_KIDNEY_DEVELOPMENT
	29	0.01	Down	0.928895784533849
0.0602667477452879	0.385837194244738
GO_SUMO_BINDING	16	0.01	Down	0.170823769291613
0.0603128804708048	0.385837194244738
GO_POSITIVE_REGULATION_OF_TRANSFERASE_ACTIVITY	513	0.01	Down
	0.331995107372037	0.0604035008176255

0.386010161540847
GO_MICROFILAMENT_MOTOR_ACTIVITY 15 0.01 Down
0.281136013999965 0.0606280814040802 0.387037941329307
GO_MULTIVESICULAR_BODY_SORTING_PATHWAY 32 0.01 Down
0.190200573801112 0.0607079002043016 0.387140401407894
GO_CARD_DOMAIN_BINDING 13 0.01 Down 0.361753816607793
0.0608204315572255 0.387451038807887
GO_AMINOGLYCAN_METABOLIC_PROCESS 103 0.01 Up
0.595964794677061 0.061024508655026 0.388343597531093
GO_COVALENT_CHROMATIN_MODIFICATION 405 0.01 Down
0.202669222934148 0.0611641176101648 0.388692978491343
GO_LEUKOCYTE_AGGREGATION 11 0.01 Down
0.829798100577727 0.0612472999992134 0.388692978491343
GO_MYOSIN_BINDING 47 0.01 Down 0.303628253292692
0.0612714841733182 0.388692978491343
GO_SNAP_RECEPTOR_ACTIVITY 29 0.01 Down
0.124227340147205 0.061754080116596 0.39134553276394
GO_REGULATION_OF_EPIDERMAL_GROWTH_FACTOR_ACTIVATED_RECEPTOR_ACTIVITY
23 0.01 Down 0.273306165805674
0.0618932336928019 0.391818375129302
GO_NEGATIVE_REGULATION_OF_INTERLEUKIN_1_SECRETION 10 0.01
Down 0.778119851236217 0.0622853215605963
0.393556045650195
GO_NEGATIVE_REGULATION_OF_VIRAL_RELEASE_FROM_HOST_CELL 14 0.01
Down 0.152712528310973 0.0623387025522915
0.393556045650195
GO_MYELOID_LEUKOCYTE_MIGRATION 131 0.01 Down
0.82939714326066 0.0623621999531358 0.393556045650195
GO_AEROBIC_RESPIRATION 75 0.01 Up 0.137998807169876
0.0624991266204682 0.393816313965179
GO_MITOGEN_ACTIVATED_PROTEIN_KINASE_BINDING 26 0.01 Down
0.251249727142784 0.0625575143352129
0.393816313965179
GO_FERTILIZATION 84 0.01 Up 0.33173561382466
0.0625980469406025 0.393816313965179
GO_RESPONSE_TO_GROWTH_HORMONE 26 0.01 Down
0.375939838688741 0.0627218423967046 0.393890577570357
GO_CYTOSOLIC_LARGE_RIBOSOMAL_SUBUNIT 57 0.01 Up
0.102339080578069 0.0627396126685117 0.393890577570357
GO_REGULATION_OF_NUCLEOCYTOPLASMIC_TRANSPORT 98 0.01 Down
0.28809221865597 0.0628377272108178
0.393896401106656
GO_POSITIVE_REGULATION_OF_CELLULAR_AMIDE_METABOLIC_PROCESS 125
0.01 Down 0.219763185817648 0.0628941252080596
0.393896401106656
GO_POSITIVE_REGULATION_OF_AMYLOID_PRECURSOR_PROTEIN_CATABOLIC_PROCES
S 13 0.01 Down 0.435628511051096
0.0631013892563008 0.393896401106656
GO_NATURAL_KILLER_CELL_ACTIVATION_INVOLVED_IN_IMMUNE_RESPONSE 10
0.01 Down 0.514353878888698 0.0631201282799033
0.393896401106656
GO_PEROXISOMAL_TRANSPORT 61 0.01 Up
0.238554314702019 0.0631325061534378 0.393896401106656
GO_CIRCADIAN_RHYTHM 143 0.01 Down 0.258908053405626

0.0632217521854881	0.393896401106656
GO_INTRINSIC_COMPONENT_OF_MITOCHONDRIAL_MEMBRANE	71	0.01	Up	0.149499463496226	0.0632712447173103	0.393896401106656
GO_RNA_EXPORT_FROM_NUCLEUS	131	0.01	Down	0.132635657540782	0.0633170992872221	0.393896401106656
GO_GUANYL_NUCLEOTIDE_EXCHANGE_FACTOR_ACTIVITY	182	0.01	Down	0.287041114697591	0.0633292218941848	0.393896401106656
GO_ANIMAL_ORGAN_REGENERATION	58	0.01	Up	0.446759184192531	0.06338935659384	0.393896401106656
GO_TRANSLATIONAL_TERMINATION	104	0.01	Up	0.111512529036796	0.0636259480794374	0.394962301421538
GO_CHROMATIN_ORGANIZATION_INVOLVED_IN_REGULATION_OF_TRANSCRIPTION	85	0.01	Down	0.164665034469953	0.0637613241956778	0.395226909060179
GO_SENSORY_PERCEPTION_OF_LIGHT_STIMULUS	87	0.01	Up	0.35878179501209	0.063815864858102	0.395226909060179
GO_ADENYLYLTRANSFERASE_ACTIVITY	26	0.01	Down	0.333470390161386	0.0638638770858237	0.395226909060179
GO_DEPHOSPHORYLATION	395	0.01	Down	0.31132308757032	0.0639760669347317	0.395332634183185
GO_SPINDLE_LOCALIZATION	33	0.01	Down	0.185780721734497	0.0640266206881743	0.395332634183185
GO_STRUCTURAL_CONSTITUENT_OF_SYNAPSE	11	0.01	Down	0.379417848216539	0.0640763156047198	0.395332634183185
GO_POSITIVE_REGULATION_OF_PEPTIDYL_TYROSINE_PHOSPHORYLATION	120	0.01	Down	0.5471949618063	0.0646950534780274	0.39834138331222
GO_GLUTAMATE_METABOLIC_PROCESS	19	0.01	Up	0.308986442324243	0.0647668612767908	0.39834138331222
GO_REGULATION_OF_CELL_ADHESION_MEDIATED_BY_INTEGRIN	37	0.01	Down	0.589994546222824	0.0648497163835897	0.39834138331222
GO_MITOCHONDRIAL_MEMBRANE_PART	207	0.01	Up	0.144363968103374	0.0649735487620548	0.39834138331222
GO_UNSATURATED_FATTY_ACID_BIOSYNTHETIC_PROCESS	31	0.01	Up	0.433739717001023	0.0649740020933443	0.39834138331222
GO_AXON_ENSHEATHMENT_IN_CENTRAL_NERVOUS_SYSTEM	11	0.01	Down	0.26552739773205	0.0650484984707912	0.39834138331222
GO_STRIATED_MUSCLE_ADAPTATION	31	0.01	Down	0.454856331736942	0.0650870822513708	0.39834138331222
GO_REGULATION_OF_PRESYNAPSE_ORGANIZATION	15	0.01	Up	0.260952988358263	0.0651272176685535	0.39834138331222
GO_REGULATION_OF_RESPONSE_TO_EXTERNAL_STIMULUS	528	0.01	Down	0.496181058533388	0.0651545039744744	0.39834138331222
GO_MUSCLE_CELL_PROLIFERATION	140	0.01	Down	0.470072115746367	0.0653061334060415	0.398494623376476
GO_CELLULAR_RESPONSE_TO_GROWTH_HORMONE_STIMULUS	16	0.01	Down	0.314634038836882	0.0653353202820559

0.398494623376476
GO_POSITIVE_REGULATION_OF_TYPE_I_INTERFERON_MEDIATED_SIGNALING_PATHWAY 12 0.01 Down 0.284199218196443 0.0654388028424296 0.398494623376476
GO_CHROMATIN_BINDING 431 0.01 Down 0.228521148195524 0.0654491775320638 0.398494623376476
GO_POSITIVE_REGULATION_OF_LEUKOCYTE_MEDIATED_IMMUNITY 94 0.01 Down 0.398950942242832 0.0655077638164591 0.398494623376476
GO_REGULATION_OF_NUCLEASE_ACTIVITY 21 0.01 Down 0.278947517507178 0.0656070051757051 0.398698827248955
GO_REGULATION_OF_CELLULAR_RESPIRATION 25 0.01 Up 0.255091900268454 0.0657517087277367 0.399178623686089
GO_ZINC_ION_TRANSMEMBRANE_TRANSPORTER_ACTIVITY 14 0.01 Up 0.192857957893661 0.0659373990353426 0.39951555360431
GO_ALPHA_ACTININ_BINDING 15 0.01 Up 0.329307718075014 0.0659501955861896 0.39951555360431
GO_REGULATION_OF_T_CELL_CHEMOTAXIS 11 0.01 Down 0.77947217718565 0.0660081682943538 0.39951555360431
GO_REGULATION_OF_TRANSLATIONAL_INITIATION_IN_RESPONSE_TO_STRESS 14 0.01 Down 0.176757573826843 0.0661126446818501 0.39951555360431
GO_POLY_PURINE_TRACT_BINDING 25 0.01 Down 0.181420192951118 0.0661630911003192 0.39951555360431
GO_ELECTRON_TRANSFER_ACTIVITY 95 0.01 Up 0.269057245629935 0.0662020502266407 0.39951555360431
GO_ESTABLISHMENT_OF_PLANAR_POLARITY_INVOLVED_IN_NEURAL_TUBE_CLOSURE 15 0.01 Up 0.633460639527457 0.0664786228150077 0.400528614731898
GO_RESPONSE_TO_SALT 13 0.01 Down 0.359169233067159 0.0665018684977357 0.400528614731898
GO_REGULATION_OF_CHROMATIN_ORGANIZATION 152 0.01 Down 0.209216291246181 0.0668716734426713 0.401812845677046
GO_INDOLE_CONTAINING_COMPOUND_METABOLIC_PROCESS 10 0.01 Down 0.670563159122368 0.066904236033734 0.401812845677046
GO_FAT_CELL_DIFFERENTIATION 170 0.01 Down 0.376894450559759 0.0669136529368298 0.401812845677046
GO_POLYUBIQUITIN_MODIFICATION_DEPENDENT_PROTEIN_BINDING 47 0.01 Down 0.196787878526001 0.0670292122262165 0.402008829698821
GO_DEFENSE_RESPONSE_TO_FUNGUS 14 0.01 Down 0.772522073341761 0.067142037785672 0.402008829698821
GO_LYMPHOCYTE_ACTIVATION 476 0.01 Down 0.445962767420302 0.0671449437184326 0.402008829698821
GO_PURINE_NUCLEOBASE_BIOSYNTHETIC_PROCESS 11 0.01 Up 0.291847152533582 0.0674548402052336 0.403190250712371
GO_EPITHELIUM_DEVELOPMENT 870 0.01 Down 0.441633636310909 0.0674750938434803 0.403190250712371
GO_RESPONSE_TO_NERVE_GROWTH_FACTOR 41 0.01 Down 0.261056223506027 0.0676859763221527 0.403343005595056
GO_EPITHELIAL_STRUCTURE_MAINTENANCE 19 0.01 Down

0.291420367601691	0.0676870992101873	0.403343005595056
GO_SOLUTE_PROTON_ANTIPORTER_ACTIVITY	16	0.01	Down
0.552162818603423	0.06769997079581	0.403343005595056
GO_GLUCOSE_BINDING	10	0.01	Up	0.414369076641781
	0.0678727993496227	0.403976240050548
GO_ACTIN_FILAMENT_BINDING	147	0.01	Down
0.312985955796488	0.0682335695150297	0.405663869656829
GO_VESICLE_CARGO_LOADING	24	0.01	Down
0.162589784130753	0.0683265398751212	0.405663869656829
GO_SMALL_RIBOSOMAL_SUBUNIT	69	0.01	Up
0.133515846411372	0.0683791372603549	0.405663869656829
GO_REGULATION_OF_VIRAL_INDUCED_CYTOPLASMIC_PATTERN_RECOGNITION_RECEP
TOR_SIGNALING_PATHWAY	19	0.01	Down	0.23878973238586
	0.0685095614482204	0.405663869656829
GO_CELLULAR_RESPONSE_TO_VITAMIN	23	0.01	Up
0.510473783723649	0.0685542145087943	0.405663869656829
GO_CELL_CELL_ADHESION	591	0.01	Down	0.439158115273328
	0.0686218807381688	0.405663869656829
GO_ENDOPLASMIC_RETICULUM	1480	0.01	Up
0.331455513382762	0.0686450898039065	0.405663869656829
GO_REGULATION_OF_FATTY_ACID_METABOLIC_PROCESS	40	0.01	Up
	0.318222911371005	0.0687493741460041
0.405663869656829
GO_NUCLEOSIDE_TRIPHOSPHATE_BIOSYNTHETIC_PROCESS	63	0.01	Up
	0.17324239734011	0.0688065799449957
0.405663869656829
GO_NEURON_PROJECTION_ARBORIZATION	14	0.01	Down
0.386831563104516	0.068897024236712	0.405663869656829
GO_NEGATIVE_REGULATION_OF_HEART_CONTRACTION	15	0.01	Down
	0.450892281549455	0.0689533361347339
0.405663869656829
GO_LYSOSOMAL_TRANSPORT	94	0.01	Down	0.204705254098607
	0.0689581804457005	0.405663869656829
GO_SIN3_COMPLEX	12	0.01	Down	0.120300851811502
0.0691168847447406	0.405671845148242
GO_CHONDROCYTE_DEVELOPMENT	32	0.01	Up
0.727636825793027	0.0692235319972746	0.405671845148242
GO_G_PROTEIN_COUPLED_ACETYLCHOLINE_RECEPTOR_SIGNALING_PATHWAY	10
	0.01	Down	0.287597504584846	0.0693014112613138
	0.405671845148242
GO_MRNA_EXPORT_FROM_NUCLEUS	105	0.01	Down
0.133699890194303	0.069428877528783	0.405671845148242
GO_NEGATIVE_REGULATION_OF_HOMOTYPIC_CELL_CELL_ADHESION	10	0.01
	Down	0.599047895973006	0.0694728940673023
0.405671845148242
GO_PEPTIDASE_COMPLEX	84	0.01	Down	0.134666371753057
	0.0695680458740463	0.405671845148242
GO_NEGATIVE_REGULATION_OF_CHROMATIN_ORGANIZATION	44	0.01
	Down	0.196368414553413	0.069630652816154
0.405671845148242
GO_FERROUS_IRON_BINDING	19	0.01	Up	0.456244002306078
	0.0696516783267753	0.405671845148242
GO_POLYNUCLEOTIDE_ADENYLYLTRANSFERASE_ACTIVITY	11	0.01	Down
	0.20150130023700 7	0.069670211692967 4

0.405671845148242
GO_ANTIGEN_PROCESSING_AND_PRESENTATION_OF_EXOGENOUS_PEPTIDE_ANTIGEN_VIA_MHC_CLASS_I 72 0.01 Down 0.25637937686344 0.0699069831898975 0.405671845148242
GO_POSITIVE_REGULATION_OF_LAMELLIPODIUM_ORGANIZATION 21 0.01 Down 0.126073455680788 0.0699432821271951 0.405671845148242
GO_REGULATION_OF_GOLGI_ORGANIZATION 15 0.01 Down 0.220916169415158 0.0699996069166607 0.405671845148242
GO_HIPPOCAMPUS_DEVELOPMENT 55 0.01 Down 0.328160409165899 0.07007731317922 0.405671845148242
GO_RNA_POLYMERASE_II_DISTAL_ENHANCER_SEQUENCE_SPECIFIC_DNA_BINDING 70 0.01 Down 0.25897994076279 0.0700973091396996 0.405671845148242
GO_DYNEIN_INTERMEDIATE_CHAIN_BINDING 23 0.01 Down 0.299731856782023 0.0701378048760344 0.405671845148242
GO_POSITIVE_REGULATION_OF_LYASE_ACTIVITY 10 0.01 Up 0.524924513196156 0.0701860328552728 0.405671845148242
GO_POSITIVE_REGULATION_OF_TELOMERE_MAINTENANCE 47 0.01 Down 0.135552067582706 0.070200708197186 0.405671845148242
GO_CORE_PROMOTER_SEQUENCE_SPECIFIC_DNA_BINDING 40 0.01 Down 0.21409452449742 0.0702882018076607 0.405671845148242
GO_INTERMEDIATE_FILAMENT_BASED_PROCESS 35 0.01 Down 0.391092555120101 0.0702930663164557 0.405671845148242
GO_COLLAGEN_CATABOLIC_PROCESS 38 0.01 Up 1.01582435628989 0.0703433389539859 0.405671845148242
GO_INTERFERON_GAMMA_PRODUCTION 73 0.01 Down 0.707571974330944 0.0703627825631854 0.405671845148242
GO_ASPARTATE_FAMILY_AMINO_ACID_CATABOLIC_PROCESS 17 0.01 Up 0.337038599699726 0.0705694352667866 0.406200327544657
GO_CHONDROITIN_SULFATE_BIOSYNTHETIC_PROCESS 23 0.01 Up 0.604435835244938 0.0706213486469045 0.406200327544657
GO_POSITIVE_REGULATION_OF_CELL_ACTIVATION 235 0.01 Down 0.506588719537492 0.0706551714523402 0.406200327544657
GO_SELECTIVE_AUTOPHAGY 45 0.01 Down 0.141428050262097 0.0709316976325884 0.407404291700515
GO_POSITIVE_REGULATION_OF_NUCLEOCYTOPLASMIC_TRANSPORT 57 0.01 Down 0.251791335873092 0.0712971404670691 0.40844505625668
GO_ESTABLISHMENT_OF_RNA_LOCALIZATION 181 0.01 Down 0.140840947043202 0.0712989430403463 0.40844505625668
GO_NEGATIVE_REGULATION_OF_NIK_NF_KAPPAB_SIGNALING 19 0.01 Down 0.386264929995827 0.0713147355677924 0.40844505625668
GO_PEPTIDYL_SERINE_MODIFICATION 247 0.01 Down 0.352969092550436 0.0713980981193784 0.408537091124172
GO_PROTEIN_CONTAINING_COMPLEX_SCAFFOLD_ACTIVITY 82 0.01 Down 0.2504269936724 0.0716427953744861 0.409551234198216

GO_PRESYNAPSE_ORGANIZATION	25	0.01	Up
0.32465533460283	0.0720506188343518	0.411172174769581
GO_ER_NUCLEUS_SIGNALING_PATHWAY 45	0.01	Down
0.282525023317524	0.0721264252612574	0.411172174769581
GO_ALPHA_AMINO_ACID_CATABOLIC_PROCESS	55	0.01	Up
0.425773786993244	0.0721295282704008	0.411172174769581
GO_INTERLEUKIN_2_PRODUCTION	48	0.01	Down
0.447707667103013	0.0723198731129333	0.411870496874876
GO_REGULATION_OF_CARDIAC_MUSCLE_CELL_ACTION_POTENTIAL	22	0.01
	Down	0.301815090781579	0.0724414318257851
0.411886492596383
GO_WNT_ACTIVATED_RECEPTOR_ACTIVITY	16	0.01	Up
0.37451852240207	0.0725128717570348	0.411886492596383
GO_REGULATION_OF_VENTRICULAR_CARDIAC_MUSCLE_CELL_ACTION_POTENTIAL
	12	0.01	Down	0.350692303436231
0.0725262165352551	0.411886492596383
GO_MODIFIED_AMINO_ACID_TRANSMEMBRANE_TRANSPORTER_ACTIVITY	12
	0.01	Up	0.333089899840916	0.0725991864325264
	0.411915570870904
GO_TETRAPYRROLE_METABOLIC_PROCESS	23	0.01	Up
0.293742713547803	0.0728391331418362	0.412891108593919
GO_POSITIVE_REGULATION_OF_LEUKOCYTE_PROLIFERATION	92	0.01
	Down	0.551028449182226	0.0729102691031731
0.412908809445302
GO_POSITIVE_REGULATION_OF_ALPHA_BETA_T_CELL_DIFFERENTIATION	34
	0.01	Down	0.427984851033586	0.0733713622802949
	0.415132842873878
GO_POSITIVE_REGULATION_OF_LOCOMOTION	432	0.01	Down
0.464744272208663	0.0737329658183537	0.416790349611941
GO_REGULATION_OF_ALCOHOL_BIOSYNTHETIC_PROCESS	61	0.01	Down
	0.371535025873555	0.074057692254531
0.418236511327682
GO_DRUG_TRANSMEMBRANE_TRANSPORTER_ACTIVITY	62	0.01	Up
	0.346873078845062	0.0742832953191906
0.419044186926862
GO_RESPONSE_TO_ACETYLCHOLINE	13	0.01	Down
0.282217485376448	0.0743387562708336	0.419044186926862
GO_AMINO_ACID_TRANSMEMBRANE_TRANSPORTER_ACTIVITY	44	0.01
	Up	0.413095280183311	0.0744176193481195
0.419099598388157
GO_DETECTION_OF_EXTERNAL_BIOTIC_STIMULUS	10	0.01	Down
	0.902669995937882	0.074519343825169
0.419195980288118
GO_GLYCOLYTIC_PROCESS_THROUGH_FRUCTOSE_6_PHOSPHATE	21	0.01
	Up	0.253965151963036	0.074641113150789
0.419195980288118
GO_CYTOKINE_PRODUCTION_INVOLVED_IN_IMMUNE_RESPONSE	76	0.01
	Down	0.476912937615259	0.0746418801995479
0.419195980288118
GO_REPRODUCTIVE_SYSTEM_DEVELOPMENT	310	0.01	Down
0.377792359642818	0.0748234140227527	0.419827122488107
GO_RIBONUCLEOPROTEIN_COMPLEX_BINDING	124	0.01	Down
0.150678791517097	0.074998691922996	0.420200482427847
GO_RNA_DEPENDENT_DNA_BIOSYNTHETIC_PROCESS	66	0.01	Down

0.138103086342453 0.0750877796288397
0.420200482427847
GO_NEGATIVE_REGULATION_OF_EXOCYTOSIS 24 0.01 Down
0.306416340464036 0.0750975989843871 0.420200482427847
GO_FLAVIN_ADENINE_DINUCLEOTIDE_BINDING 68 0.01 Up
0.299397046497484 0.0752282578437734 0.420543971795164
GO_VITAMIN_D_RECEPTOR_BINDING 15 0.01 Down
0.155414175046477 0.0755325982583306 0.421538047018475
GO_NUCLEAR_CYCLIN_DEPENDENT_PROTEIN_KINASE_HOLOENZYME_COMPLEX 12
0.01 Down 0.147113083147103 0.0756386020855988
0.421538047018475
GO_VIRAL_BUDDING 25 0.01 Down 0.178218760345733
0.0757492362253603 0.421538047018475
GO_POSITIVE_REGULATION_OF_NITRIC_OXIDE_BIOSYNTHETIC_PROCESS 32
0.01 Down 0.603556479709843 0.0757498943400027
0.421538047018475
GO_ENDOCHONDRAL_BONE_MORPHOGENESIS 50 0.01 Up
0.527277427581284 0.0757532547022164 0.421538047018475
GO_MICROTUBULE_ANCHORING 24 0.01 Down
0.210051217771306 0.0758978736827875 0.421956035831687
GO_CELLULAR_RESPONSE_TO_AMINO_ACID_STIMULUS 60 0.01 Up
0.364428794516468 0.0761012219031215
0.42252568451689
GO_POSITIVE_REGULATION_OF_EPITHELIAL_CELL_DIFFERENTIATION 39
0.01 Down 0.4339563411443 0.0761756766816367
0.42252568451689
GO_PROTEIN_LOCALIZATION_TO_ENDOPLASMIC_RETICULUM 134 0.01
Up 0.13278301242484 0.0762217799016844
0.42252568451689
GO_LIPOXYGENASE_PATHWAY 10 0.01 Up 0.513482891967176
0.076278726771621 0.42252568451689
GO_MEMBRANE_DEPOLARIZATION_DURING_ACTION_POTENTIAL 18 0.01
Up 0.512218110789171 0.0764831545839761
0.423271860965651
GO_NEGATIVE_REGULATION_OF_TRANSMEMBRANE_RECEPTOR_PROTEIN_SERINE_THRE
ONINE_KINASE_SIGNALING_PATHWAY 81 0.01 Up
0.481547808511553 0.0767261586203904 0.424229971752632
GO_PROTEIN_METHYLATION 149 0.01 Down 0.162858666506992
0.0770365255340538 0.424725339535809
GO_ENDOCRINE_SYSTEM_DEVELOPMENT 75 0.01 Down
0.443135254427367 0.0771014378739311 0.424725339535809
GO_REGULATION_OF_PROTEIN_LOCALIZATION_TO_SYNAPSE 20 0.01
Up 0.315419341314535 0.0771242715515724
0.424725339535809
GO_BROWN_FAT_CELL_DIFFERENTIATION 29 0.01 Down
0.314308597881971 0.0772318960897754 0.424725339535809
GO_POSITIVE_REGULATION_OF_DEVELOPMENTAL_PROCESS 962 0.01 Down
0.416816007431659 0.0772996573539886
0.424725339535809
GO_REGULATION_OF_SIGNALING_RECEPTOR_ACTIVITY 266 0.01 Down
0.6201676549574 0.0773390077870556 0.424725339535809
GO_NEGATIVE_REGULATION_OF_AXON_GUIDANCE 23 0.01 Down
0.498688724435514 0.0773596031499126 0.424725339535809
GO_NEGATIVE_REGULATION_OF_CELLULAR_AMIDE_METABOLIC_PROCESS 134

0.01 Down 0.261312311532164 0.0773754283522656 0.424725339535809
GO_ADRENERGIC_RECEPTOR_SIGNALING_PATHWAY 12 0.01 Down 0.550793071842853 0.0775311306728174 0.424905004089151
GO_RESPONSE_TO_MAGNESIUM_ION 10 0.01 Up 0.485742259516159 0.0776484837264269 0.424905004089151
GO_CELL_CELL_JUNCTION_ORGANIZATION 120 0.01 Down 0.338432598885892 0.07769101988183 0.424905004089151
GO_ISOPRENOID_METABOLIC_PROCESS 62 0.01 Up 0.491059157392212 0.0777925012313451 0.424905004089151
GO_RRNA_BINDING 62 0.01 Up 0.1318641228387 0.0777983029203484 0.424905004089151
GO_INTERSPECIES_INTERACTION_BETWEEN_ORGANISMS 807 0.01 Down 0.255771774533365 0.0778280949674082 0.424905004089151
GO_PLACENTA_DEVELOPMENT 117 0.01 Down 0.391824611622 0.0779452720108598 0.4250729989853
GO_NEUROMUSCULAR_JUNCTION_DEVELOPMENT 36 0.01 Up 0.268276199279686 0.0779988998302791 0.4250729989853
GO_POSITIVE_REGULATION_OF_NUCLEOTIDE_BIOSYNTHETIC_PROCESS 11 0.01 Up 0.413242803028331 0.0781072128322919 0.425281514892237
GO_NEGATIVE_REGULATION_OF_ERBB_SIGNALING_PATHWAY 48 0.01 Down 0.241355220595534 0.0781999516847483 0.425404934299379
GO_RIBONUCLEOSIDE_TRIPHOSPHATE_BIOSYNTHETIC_PROCESS 54 0.01 Up 0.166615837275091 0.0783604452388863 0.425896385895505
GO_RESPONSE_TO_HEAT 148 0.01 Down 0.282269810879685 0.0784856487006348 0.426041020076591
GO_VESICLE_BUDDING_FROM_MEMBRANE 92 0.01 Down 0.222525091820708 0.078550561914794 0.426041020076591
GO_ESTABLISHMENT_OF_CELL_POLARITY 115 0.01 Down 0.236338626096428 0.0785975856507629 0.426041020076591
GO_POSITIVE_REGULATION_OF_CHROMATIN_SILENCING 14 0.01 Down 0.13656596442841 0.0788454620298628 0.427003389815608
GO_RESPONSE_TO_ARSENIC_CONTAINING_SUBSTANCE 29 0.01 Down 0.264219618598151 0.079338296479309 0.428898207199632
GO_NEGATIVE_REGULATION_OF_TRANSCRIPTION_BY_RNA_POLYMERASE_II 654 0.01 Down 0.305884399092165 0.0793793824922248 0.428898207199632
GO_RESPIRATORY_SYSTEM_PROCESS 17 0.01 Up 0.289577048931811 0.0794072780254302 0.428898207199632
GO_STEROID_HORMONE_RECEPTOR_BINDING 79 0.01 Down 0.243859325702704 0.0795179743241567 0.429114330775071
GO_ORGANELLE_MEMBRANE_FUSION 83 0.01 Down 0.234773737827007 0.0796387653262869 0.42938449759848
GO_INTERFERON_GAMMA_MEDIATED_SIGNALING_PATHWAY 73 0.01 Down 0.478739204424588 0.0797251048878482 0.429468599622117
GO_EXORIBONUCLEASE_ACTIVITY 35 0.01 Down

0.215895497957893	0.0799095763280561	0.430010603417875
GO_NEGATIVE_REGULATION_OF_INTERLEUKIN_2_PRODUCTION	16	0.01	Down	0.556011174118265	0.0800733620860707	0.430010603417875
GO_BRAIN_MORPHOGENESIS	23	0.01	Down	0.322989253480454	0.0800959895295667	0.430010603417875
GO_CELLULAR_RESPONSE_TO_COPPER_ION	19	0.01	Up	0.545795307006792	0.0801090417502251	0.430010603417875
GO_NUCLEAR_BODY	687	0.01	Down	0.194858642149169	0.08059459812469	0.432234810260594
GO_UBIQUITIN_BINDING	74	0.01	Down	0.140092139586194	0.0807524258331924	0.432699009032048
GO_REGULATION_OF_PROTEIN_COMPLEX_DISASSEMBLY	91	0.01	Down	0.286420776033168	0.0808312872160434	0.432739633764197
GO_POSITIVE_REGULATION_OF_EPIDERMIS_DEVELOPMENT	29	0.01	Down	0.485094878556595	0.0811445938695714	0.432940237610557
GO_TRANSFERASE_ACTIVITY_TRANSFERRING_ALKYL_OR_ARYL_OTHER_THAN_METHYL_GROUPS	46	0.01	Up	0.308051853983037	0.0811465506030417	0.432940237610557
GO_INSULIN_SECRETION_INVOLVED_IN_CELLULAR_RESPONSE_TO_GLUCOSE_STIMULUS	52	0.01	Up	0.303287493786264	0.0811743061971938	0.432940237610557
GO_REGULATION_OF_CIRCADIAN_RHYTHM	76	0.01	Down	0.23157294050664	0.0812901824313343	0.432940237610557
GO_POST_GOLGI_VESICLE_MEDIATED_TRANSPORT	101	0.01	Down	0.187790646217013	0.0813688505085245	0.432940237610557
GO_SINGLE_STRANDED_VIRAL_RNA_REPLICATION_VIA_DOUBLE_STRANDED_DNA_INTERMEDIATE	16	0.01	Down	0.337869874660523	0.0813817354127181	0.432940237610557
GO_REGULATION_OF_CELL_ADHESION	516	0.01	Down	0.508383094955039	0.0814040388260267	0.432940237610557
GO_PROTEIN_MONOUBIQUITINATION	64	0.01	Down	0.148722255113848	0.081439260640958	0.432940237610557
GO_POSITIVE_REGULATION_OF_CELL_CYCLE_G2_M_PHASE_TRANSITION	26	0.01	Up	0.254612288632757	0.0817275609863764	0.43362145647709
GO_GOLGI_VESICLE_BUDDING	74	0.01	Down	0.214952604068034	0.08179999381217	0.43362145647709
GO_DENDRITE_MORPHOGENESIS	112	0.01	Down	0.343603786435628	0.0818415774281939	0.43362145647709
GO_REGULATION_OF_LAMELLIPODIUM_ASSEMBLY	27	0.01	Down	0.244121206185606	0.0818531031333791	0.43362145647709
GO_RETINA_LAYER_FORMATION	15	0.01	Down	0.421624748159975	0.0819360508765498	0.433682445398024
GO_STRUCTURAL_CONSTITUENT_OF_NUCLEAR_PORE	23	0.01	Down	0.126097128917729	0.082122693979094	0.434040206591713
GO_RAB_GTPASE_BINDING	153	0.01	Down	0.188329422507613	0.0822098355362651	0.434040206591713
GO_CAMERA_TYPE_EYE_MORPHOGENESIS	73	0.01	Down	0.412312999764572	0.0822181251162032	0.434040206591713

GO_REGULATION_OF_WATER_LOSS_VIA_SKIN 17 0.01 Down
0.331851474219722 0.08229258152872 0.434055831851311
GO_CLATHRIN_BINDING 43 0.01 Up 0.507906173559457
0.0823806480039014 0.434143154541394
GO_POSITIVE_REGULATION_OF_IMMUNE_RESPONSE 568 0.01 Down
0.436576758230885 0.0825992319900292
0.434645461281547
GO_REGULATION_OF_CELLULAR_AMIDE_METABOLIC_PROCESS 347 0.01
Down 0.217432223685685 0.0826825783441688
0.434645461281547
GO_CILIUM_ORGANIZATION 311 0.01 Up 0.215920770477921
0.082733294724407 0.434645461281547
GO_REGULATION_OF_FC_RECEPTOR_MEDIATED_STIMULATORY_SIGNALING_PATHWAY
11 0.01 Down 0.233899082282373
0.082762337875386 0.434645461281547
GO_2_OXOGLUTARATE_DEPENDENT_DIOXYGENASE_ACTIVITY 44 0.01
Up 0.278345857187962 0.0828707107327698
0.434838448451725
GO_GUANOSINE_CONTAINING_COMPOUND_BIOSYNTHETIC_PROCESS 12 0.01
Up 0.279152163514932 0.0832998541811032
0.435453321702411
GO_INTERFERON_ALPHA_PRODUCTION 26 0.01 Down
0.389519007037428 0.0833295160295356 0.435453321702411
GO_REGULATION_OF_EPIDERMAL_CELL_DIFFERENTIATION 47 0.01 Down
0.464075503457311 0.0833755544715933
0.435453321702411
GO_REGULATION_OF_LYSOSOMAL_LUMEN_PH 11 0.01 Up
0.171464516825162 0.0833991363160153 0.435453321702411
GO_ESCRT_III_COMPLEX 11 0.01 Down 0.133870616477969
0.0834051523067079 0.435453321702411
GO_MYELOID_DENDRITIC_CELL_ACTIVATION 19 0.01 Down
0.715346105473664 0.0834182528644216 0.435453321702411
GO_REGULATION_OF_PROTEIN_DEPHOSPHORYLATION 114 0.01 Down
0.326187873924752 0.0838676944477071
0.437423344494871
GO_NEURON_PROJECTION_REGENERATION 43 0.01 Up
0.544303468307314 0.0840510834787104 0.438003543175323
GO_PEPTIDYL_LYSINE_METHYLATION 108 0.01 Down
0.182589665498083 0.0841255372624519 0.438015554648667
GO_POSITIVE_REGULATION_OF_KINASE_ACTIVITY 449 0.01 Down
0.354198149319846 0.0842339514012763
0.438125905877632
GO_REGULATION_OF_SPINDLE_ORGANIZATION 38 0.01 Down
0.190732153995228 0.0842910654035702 0.438125905877632
GO_DENDRITE_DEVELOPMENT 177 0.01 Down 0.343852783207036
0.0846249679884855 0.439485184480834
GO_CUL3_RING_UBIQUITIN_LIGASE_COMPLEX 35 0.01 Down
0.165143833314573 0.0850679255498525 0.441387971709596
GO_HEART_TRABECULA_MORPHOGENESIS 24 0.01 Down
0.293808028318748 0.0851367673977824 0.441387971709596
GO_MUSCLE_HYPERTROPHY 62 0.01 Down 0.40981913721157
0.0852737860807868 0.441721122266601
GO_KERATINOCYTE_PROLIFERATION 34 0.01 Down
0.393324435783559 0.0854111763408399 0.442016767526779

GO_CONVERGENT_EXTENSION 12 0.01 Up 0.533129184335205
0.0855180721590178 0.442016767526779
GO_MAINTENANCE_OF_SYNAPSE_STRUCTURE 11 0.01 Down
0.229560192133446 0.0855492837825671 0.442016767526779
GO_T_TUBULE 33 0.01 Up 0.40508632370171
0.0856520911823375 0.442171637387731
GO_NEGATIVE_REGULATION_OF_PATHWAY_RESTRICTED_SMAD_PROTEIN_PHOSPHORYLATION 11 0.01 Up 0.899040685343313
0.0858334079270093 0.442412575009104
GO_PROTEIN_REFOLDING 33 0.01 Down 0.214873581555764
0.0858525562811045 0.442412575009104
GO_REGULATION_OF_EPITHELIAL_CELL_MIGRATION 185 0.01 Down
0.425243443289693 0.0860291830536337
0.442412575009104
GO_REGULATION_OF_EXOSOMAL_SECRETION 13 0.01 Down
0.160186976803642 0.0860474505066984 0.442412575009104
GO_VESICLE_TARGETING 86 0.01 Down 0.215173105028812
0.0860631281643472 0.442412575009104
GO_POSITIVE_REGULATION_OF_LYMPHOCYTE_APOPTOTIC_PROCESS 13 0.01
Down 0.849699930614398 0.0861916419093191
0.442698357048626
GO_CELLULAR_RESPONSE_TO_PEPTIDE_HORMONE_STIMULUS 232 0.01
Down 0.276868324889625 0.0864561682111057
0.442824602013512
GO_POSITIVE_REGULATION_OF_POTASSIUM_ION_TRANSPORT 21 0.01
Down 0.306067038482853 0.0864664190053087
0.442824602013512
GO_POSITIVE_REGULATION_OF_PROTEIN_SERINE_THREONINE_KINASE_ACTIVITY
273 0.01 Down 0.357248252026787
0.0866501517157183 0.442824602013512
GO_REGULATION_OF_PROTEIN_POLYUBIQUITINATION 18 0.01 Down
0.250419438770069 0.0866956309094116
0.442824602013512
GO_RUFFLE 153 0.01 Down 0.301674077387926
0.0866976669894498 0.442824602013512
GO_REGULATION_OF_LEUKOCYTE_PROLIFERATION 153 0.01 Down
0.591015983646843 0.0868324923657652
0.442824602013512
GO_TETRAHYDROFOLATE_METABOLIC_PROCESS 11 0.01 Up
0.220151210865953 0.0868504932134912 0.442824602013512
GO_SIGNALING_ADAPTOR_ACTIVITY 66 0.01 Down
0.28328517621516 0.0869670670222784 0.442824602013512
GO_TOLL_LIKE_RECEPTOR_4_SIGNALING_PATHWAY 25 0.01 Down
0.588780714534337 0.0869799649670927
0.442824602013512
GO_POSITIVE_REGULATION_OF_DENDRITE_DEVELOPMENT 58 0.01 Down
0.268651999822121 0.0870532816481906
0.442824602013512
GO_HISTONE_DEMETHYLASE_ACTIVITY_H3_K9_SPECIFIC_ 11 0.01 Down
0.243132545044758 0.0870695536646112
0.442824602013512
GO_ACTIVATION_OF_NF_KAPPAB_INDUCING_KINASE_ACTIVITY 17 0.01
Down 0.265767692003348 0.0871078900072259
0.442824602013512

GO_MATING	20	0.01	Down	0.447463294677999
0.0872913207719996	0.442824602013512
GO_IRON_ION_BINDING	96	0.01	Up	0.435733750539527
	0.0873152637594938	0.442824602013512
GO_PEPTIDYL_TYROSINE_AUTOPHOSPHORYLATION	30	0.01	Down
	0.758237057994501	0.0873396474579301
0.442824602013512
GO_ENDOPLASMIC_RETICULUM_TO_GOLGI_VESICLE_MEDIATED_TRANSPORT	186
	0.01	Down	0.198793180343644	0.0873832767603669
	0.442824602013512
GO_PROTEIN_SUMOYLATION	73	0.01	Down	0.181824535163132
	0.0876093810157994	0.443020303332874
GO_POSITIVE_REGULATION_OF_CARDIAC_MUSCLE_CELL_PROLIFERATION	14
	0.01	Up	0.445298129874578	0.0877528077998836
	0.443020303332874
GO_DEMETHYLASE_ACTIVITY	35	0.01	Down	0.271207079905892
	0.0877672256109752	0.443020303332874
GO_POSITIVE_REGULATION_OF_PRI_MIRNA_TRANSCRIPTION_BY_RNA_POLYMERASE_
II	24	0.01	Down	0.482192364389109
0.087815261825627	0.443020303332874
GO_NEGATIVE_REGULATION_OF_MUSCLE_CELL_APOPTOTIC_PROCESS	28	0.01
	Down	0.457471664854634	0.0878389665949832
0.443020303332874
GO_REGULATION_OF_TRANSLATION_IN_RESPONSE_TO_STRESS	18	0.01
	Down	0.170275727020219	0.0878597340162709
0.443020303332874
GO_POSITIVE_REGULATION_OF_INTERLEUKIN_2_PRODUCTION	24	0.01
	Down	0.310450642532997	0.088302283165813
0.444617148777331
GO_REGULATION_OF_CELLULAR_CATABOLIC_PROCESS	712	0.01	Down
	0.214773936415994	0.0883228926742646
0.444617148777331
GO_NUCLEOSIDE_PHOSPHATE_BIOSYNTHETIC_PROCESS	226	0.01	Up
	0.275329900472444	0.0885603446606131
0.445443125463614
GO_ADAPTIVE_IMMUNE_RESPONSE_BASED_ON_SOMATIC_RECOMBINATION_OF_IMMUNE
_RECEPTORS_BUILT_FROM_IMMUNOGLOBULIN_SUPERFAMILY_DOMAINS	183
	0.01	Down	0.495412614568602	0.0886796696567343
	0.445493426353053
GO_CELLULAR_RESPONSE_TO_PEPTIDE	276	0.01	Down
0.2929899743406	0.0887171063187022	0.445493426353053
GO_INTRACILIARY_TRANSPORT_INVOLVED_IN_CILIUM_ASSEMBLY	38	0.01
	Up	0.169693530920233	0.0888705148717581
0.445894955195408
GO_INFLAMMATORY_RESPONSE	460	0.01	Down
0.580941453418219	0.0889942506245832	0.446147064857015
GO_EMBRYONIC_AXIS_SPECIFICATION	21	0.01	Down
0.309186008233133	0.0891418909575718	0.446253186713641
GO_INFLAMMATORY_CELL_APOPTOTIC_PROCESS	18	0.01	Down
0.565391143257697	0.0891624304865172	0.446253186713641
GO_PEPTIDYL_THREONINE_MODIFICATION	103	0.01	Down
0.237211765695204	0.08923881994377	0.446267607807766
GO_CONDENSED_NUCLEAR_CHROMOSOME_KINETOCHORE	10	0.01	Up
	0.299350147452123	0.0893319289403774

0.446365547816487
GO_ACROSOME_REACTION 17 0.01 Up 0.497405838792259
0.0895360938937756 0.446601816721984
GO_NON_MOTILE_CILIUM_ASSEMBLY 50 0.01 Up
0.206112556901562 0.0896397655840891 0.446601816721984
GO_POSITIVE_REGULATION_OF_TRANSFORMING_GROWTH_FACTOR_BETA_PRODUCTION
13 0.01 Down 0.628168130721732
0.0896687314418119 0.446601816721984
GO_POSITIVE_REGULATION_OF_INTERFERON_BETA_PRODUCTION 27 0.01
Down 0.285105312362929 0.0897446011604242
0.446601816721984
GO_CALCINEURIN_MEDIATED_SIGNALING 37 0.01 Down
0.307095912394016 0.0897470295504564 0.446601816721984
GO_GOLGI_TO_ENDOSOME_TRANSPORT 17 0.01 Down
0.149465432433556 0.0899550804354708 0.447270510502656
GO_RECEPTOR_REGULATOR_ACTIVITY 226 0.01 Down
0.703257053961358 0.090151696640727 0.447881301395952
GO_RHO_GUANYL_NUCLEOTIDE_EXCHANGE_FACTOR_ACTIVITY 69 0.01
Down 0.363290226167087 0.0904573672849176
0.448602541779356
GO_GDP_BINDING 58 0.01 Down 0.234494894110779
0.090517552683382 0.448602541779356
GO_GLYCOPROTEIN_METABOLIC_PROCESS 108 0.01 Up
0.413950504429309 0.0906392206074836 0.448602541779356
GO_CELLULAR_RESPONSE_TO_GLUCAGON_STIMULUS 15 0.01 Down
0.48388898275232 0.090702503715738
0.448602541779356
GO_NCRNA_CATABOLIC_PROCESS 28 0.01 Down
0.222360109352959 0.0907485217139293 0.448602541779356
GO_NUCLEAR_PORE_ORGANIZATION 14 0.01 Down
0.126175328137725 0.0908330372860181 0.448602541779356
GO_NUCLEOTIDE_EXCISION_REPAIR_PREINCISION_COMPLEX_ASSEMBLY 29
0.01 Up 0.140346744949657 0.0908656771244576
0.448602541779356
GO_CENTROSOME_DUPLICATION 63 0.01 Down
0.160485393839606 0.0909072072373537 0.448602541779356
GO_SYMPATHETIC_NERVOUS_SYSTEM_DEVELOPMENT 11 0.01 Down
0.412985339450668 0.0909619056054006
0.448602541779356
GO_REGULATION_OF_GLOMERULUS_DEVELOPMENT 10 0.01 Down
0.596945820391014 0.091091066101134 0.448743279917472
GO_REGULATION_OF_CELLULAR_RESPONSE_TO_GROWTH_FACTOR_STIMULUS 193
0.01 Up 0.497487293839536 0.0911859371771244
0.448743279917472
GO_PROTEIN_TYROSINE_KINASE_ACTIVATOR_ACTIVITY 13 0.01 Down
0.767971797205392 0.0913119708345027
0.448743279917472
GO_POSITIVE_REGULATION_OF_CATABOLIC_PROCESS 360 0.01 Down
0.236555154382183 0.0913491251310729
0.448743279917472
GO_POSITIVE_REGULATION_OF_MYOBLAST_DIFFERENTIATION 15 0.01
Down 0.637053341367898 0.0913600220685218
0.448743279917472
GO_METAPHASE_ANAPHASE_TRANSITION_OF_CELL_CYCLE 49 0.01 Up

0.226830482489178	0.0917785762645 0.449790614170231
GO_HISTONE_BINDING	174	0.01	Down	0.186520765335265
0.0918809948723574	0.449790614170231
GO_PRODUCTION_OF_MOLECULAR_MEDIATOR_INVOLVED_IN_INFLAMMATORY_RESPONSE	50	0.01	Down	0.629238807991244
0.0921163782527091	0.449790614170231
GO_POSITIVE_REGULATION_OF_CYTOKINE_PRODUCTION_INVOLVED_IN_IMMUNE_RESPONSE	48	0.01	Down	0.477543905467755
0.0921281723526231	0.449790614170231
GO_POSITIVE_REGULATION_OF_MULTI_ORGANISM_PROCESS	152	0.01	Down	0.251950361737639	0.0921344403130771
0.449790614170231
GO_SODIUM_ION_TRANSMEMBRANE_TRANSPORT	84	0.01	Down
0.508000546943202	0.0921397380408042	0.449790614170231
GO_POSITIVE_REGULATION_OF_PHOSPHATASE_ACTIVITY	28	0.01	Down	0.603223282869435	0.0921654185399539
0.449790614170231
GO_BRANCHED_CHAIN_AMINO_ACID_CATABOLIC_PROCESS	19	0.01	Up	0.184627950250186	0.0921659568485862
0.449790614170231
GO_INTERACTION_WITH_SYMBIONT	57	0.01	Down
0.392838681040126	0.0922458364816773	0.449818854040372
GO_ACTIN_CYTOSKELETON_REORGANIZATION	85	0.01	Up
0.414022386132487	0.0924238237896564	0.449848218237841
GO_CELLULAR_LIPID_METABOLIC_PROCESS	663	0.01	Up
0.339011153335751	0.0924940368840337	0.449848218237841
GO_NEGATIVE_REGULATION_OF_RNA_BIOSYNTHETIC_PROCESS	959	0.01	Down	0.296993393573259	0.0925478823889221
0.449848218237841
GO_ENDOPLASMIC_RETICULUM_TUBULAR_NETWORK_ORGANIZATION	16	0.01	Down	0.159917217199663	0.0925482498071262
0.449848218237841
GO_REGULATION_OF_CHROMATIN_BINDING	20	0.01	Down
0.211708940866426	0.092742251343583	0.450133394431186
GO_NON_MEMBRANE_SPANNING_PROTEIN_TYROSINE_KINASE_ACTIVITY	37	0.01	Down	0.478720142425245	0.0928067404498659	0.450133394431186
GO_RESPONSE_TO_TEMPERATURE_STIMULUS	187	0.01	Down
0.304676707850871	0.092829354279006	0.450133394431186
GO_POSITIVE_REGULATION_OF_CELL_DIFFERENTIATION	704	0.01	Down	0.397521726170467	0.0931323019916596
0.450831077601442
GO_METHYLATION_DEPENDENT_CHROMATIN_SILENCING	17	0.01	Down	0.151993101829764	0.0932039159725204
0.450831077601442
GO_PYRIDINE_CONTAINING_COMPOUND_METABOLIC_PROCESS	32	0.01	Up	0.268756423010311	0.0932132390948136
0.450831077601442
GO_WATER_HOMEOSTASIS	48	0.01	Down	0.514164476673433	0.0934500056548399	0.450831077601442
GO_AUTOPHAGOSOME_ORGANIZATION	89	0.01	Down
0.143906709467432	0.0934926173176803	0.450831077601442
GO_HEPATOCYTE_APOPTOTIC_PROCESS 12	0.01	Down
0.187547685084315	0.0935250967209787	0.450831077601442

GO_MAINTENANCE_OF_CELL_NUMBER 128 0.01 Down
0.319138239472288 0.0935902002356999 0.450831077601442
GO_NEGATIVE_REGULATION_OF_LIPID_METABOLIC_PROCESS 55 0.01
Up 0.463164087411864 0.0936675325259935
0.450831077601442
GO_REGULATION_OF_DNA_BINDING_TRANSCRIPTION_FACTOR_ACTIVITY 336
0.01 Down 0.393712320788193 0.0936693400001944
0.450831077601442
GO_POSITIVE_REGULATION_OF_GTPASE_ACTIVITY 329 0.01 Down
0.308824882066041 0.0937861787814819
0.450831077601442
GO_REGULATION_OF_ENDOPLASMIC_RETICULUM_UNFOLDED_PROTEIN_RESPONSE
25 0.01 Down 0.28463857116765
0.093818426207284 0.450831077601442
GO_S100_PROTEIN_BINDING 12 0.01 Down 0.213900279383507
0.0940061866401053 0.450831077601442
GO_REGULATION_OF_NEURON_MIGRATION 25 0.01 Down
0.418605189497682 0.094060128483301 0.450831077601442
GO_POSITIVE_REGULATION_OF_MULTICELLULAR_ORGANISMAL_PROCESS 1209
0.01 Down 0.430645921170004 0.0940791629757943
0.450831077601442
GO_NEGATIVE_REGULATION_OF_EPIDERMAL_GROWTH_FACTOR_ACTIVATED_RECEPTOR
_ACTIVITY 14 0.01 Down 0.220317459026216
0.0940871314974514 0.450831077601442
GO_REGULATION_OF_TRANSCRIPTION_FROM_RNA_POLYMERASE_II_PROMOTER_INVOL
VED_IN_HEART_DEVELOPMENT 10 0.01 Up
1.01989429793412 0.0942137063053181 0.451081554400304
GO_SPROUTING_ANGIOGENESIS 91 0.01 Down
0.539105611313093 0.0943660635793825 0.451300913280599
GO_1_PHOSPHATIDYLINOSITOL_BINDING 12 0.01 Down
0.40977487982613 0.0944800250273851 0.451300913280599
GO_REGULATION_OF_GENE_EXPRESSION_BY_GENETIC_IMPRINTING 15 0.01
Down 0.251766215467251 0.0944825334837162
0.451300913280599
GO_REGULATION_OF_CAMP_DEPENDENT_PROTEIN_KINASE_ACTIVITY 13 0.01
Down 0.250740860128828 0.0946264622387822
0.451633059946263
GO_GOLGI_VESICLE_TRANSPORT 328 0.01 Down
0.20471646865389 0.0948967701809814 0.45178595084338
GO_CALCIUM_ION_REGULATED_EXOCYTOSIS_OF_NEUROTRANSMITTER 10 0.01
Up 0.652603891091965 0.0949119848773596
0.45178595084338
GO_REGULATION_OF_CYTOKINE_PRODUCTION_INVOLVED_IN_IMMUNE_RESPONSE
65 0.01 Down 0.457109023842852
0.0949212014963206 0.45178595084338
GO_RESPONSE_TO_HYDROGEN_PEROXIDE 113 0.01 Down
0.384682205576606 0.094978352214529 0.45178595084338
GO_FATTY_ACID_DERIVATIVE_TRANSPORT 24 0.01 Up
0.482167506699996 0.0950305813254812 0.45178595084338
GO_CILIARY_MEMBRANE 53 0.01 Up 0.266484084088454
0.0951075501852911 0.451798072906809
GO_POSITIVE_REGULATION_OF_ORGANELLE_ASSEMBLY 64 0.01 Down
0.24232532857377 0.0952072793392451
0.451918211781515

GO_ALPHA_BETA_T_CELL_LINEAGE_COMMITMENT 11 0.01 Down
0.592124915378253 0.095396417056489 0.45222292309249
GO_PEPTIDYL_GLUTAMIC_ACID_MODIFICATION 19 0.01 Up
0.21883379299356 0.0954291874788388 0.45222292309249
GO_CHONDROITIN_SULFATE_PROTEOGLYCAN_BIOSYNTHETIC_PROCESS 25
0.01 Up 0.579384991642269 0.095520965521666
0.45222292309249
GO_RESPONSE_TO_METAL_ION 255 0.01 Up
0.425469063168886 0.0955694301313892 0.45222292309249
GO_MODIFICATION_OF_MORPHOLOGY_OR_PHYSIOLOGY_OF_OTHER_ORGANISM 102
0.01 Down 0.442649006692203 0.0956557012073156
0.452278630864185
GO_NEGATIVE_REGULATION_OF_RESPONSE_TO_ENDOPLASMIC_RETICULUM_STRESS
39 0.01 Down 0.193625198431723
0.0961484596202989 0.454111468398691
GO_GLIAL_CELL_PROLIFERATION 32 0.01 Down
0.413502239552269 0.0962257611570923 0.454111468398691
GO_REGULATION_OF_CELL_ACTIVATION 390 0.01 Down
0.503155117892848 0.0962677416947975 0.454111468398691
GO_REGULATION_OF_ORGANELLE_ASSEMBLY 164 0.01 Down
0.215562776752707 0.0964992630856073 0.454606919059211
GO_RESPONSE_TO_CATECHOLAMINE 27 0.01 Down
0.273950433974532 0.0965225364301306 0.454606919059211
GO_THYMUS_DEVELOPMENT 38 0.01 Down 0.248429111394038
0.096830251001502 0.455702677387689
GO_TRANSFERASE_COMPLEX 716 0.01 Down 0.164911427909942
0.0969130456080234 0.455739039416197
GO_MHC_PROTEIN_COMPLEX_BINDING 18 0.01 Up
0.666929118215727 0.0974159571611923 0.45765255703331
GO_OXIDOREDUCTASE_ACTIVITY_ACTING_ON_THE_ALDEHYDE_OR_OXO_GROUP_OF_DO
NORS_NAD_OR_NADP_AS_ACCEPTOR 31 0.01 Up
0.349605842824285 0.097470722491199 0.45765255703331
GO_DORSAL_VENTRAL_PATTERN_FORMATION 51 0.01 Up
0.495757847399875 0.0975778729050371 0.457742110761895
GO_PYRIMIDINE_NUCLEOSIDE_CATABOLIC_PROCESS 14 0.01 Down
0.472673638846214 0.0976405919019362
0.457742110761895
GO_LUMENAL_SIDE_OF_MEMBRANE 29 0.01 Up
0.566998734156587 0.0979968149499354 0.458500759778821
GO_TRANSCRIPTION_FACTOR_TFIIH_HOLO_COMPLEX 13 0.01 Up
0.214031552736898 0.0980081312701638
0.458500759778821
GO_EXPLORATION_BEHAVIOR 11 0.01 Up 0.445305268347005
0.0980289880073974 0.458500759778821
GO_CYTOSOLIC_SMALL_RIBOSOMAL_SUBUNIT 41 0.01 Up
0.146326421278448 0.0981866778134703 0.458539908313532
GO_POSITIVE_REGULATION_OF_ACTIN_CYTOSKELETON_REORGANIZATION 13
0.01 Up 0.398170945461744 0.0981884171977585
0.458539908313532
GO_MODIFICATION_OF_MORPHOLOGY_OR_PHYSIOLOGY_OF_OTHER_ORGANISM_INVOLV
ED_IN_SYMBIOTIC_INTERACTION 86 0.01 Down
0.363068112892084 0.0983380702109672 0.45888579880921
GO_XENOPHAGY 10 0.01 Down 0.163790077303556
0.0985279065420567 0.459418525819375

GO_PLACENTA_BLOOD_VESSEL_DEVELOPMENT	27	0.01	Down
0.340028244417242	0.0987151500034642	0.459938354313915
GO_PHOSPHATIDYLINOSITOL_ACYL_CHAIN_REMODELING	10	0.01	Up
	0.616741696136949	0.0990450813615981
0.460883713499451
GO_T_CELL_RECEPTOR_SIGNALING_PATHWAY	159	0.01	Down
0.334013358240984	0.0992138917872129	0.460883713499451
GO_DENDRITIC_SPINE_MORPHOGENESIS	44	0.01	Down
0.358611555649956	0.099347138308487	0.460883713499451
GO_ENDOSOMAL_TRANSPORT	205	0.01	Down	0.191701164875225
	0.0993528358354261	0.460883713499451
GO_MRNA_BINDING 207	0.01	Down	0.178394234587334
0.0993769733951142	0.460883713499451
GO_ORGANELLAR_LARGE_RIBOSOMAL_SUBUNIT	57	0.01	Up
0.101806236227098	0.0994640408621099	0.460883713499451
GO_RNA_LOCALIZATION	213	0.01	Down	0.136605561028266
	0.099547303122715	0.460883713499451
GO_EXOPEPTIDASE_ACTIVITY	81	0.01	Up
0.382747645279451	0.0995623358971889	0.460883713499451
GO_COENZYME_METABOLIC_PROCESS	209	0.01	Up
0.280973129992993	0.0996012900858638	0.460883713499451
GO_DNA_METHYLATION_OR_DEMETHYLATION	68	0.01	Down
0.29319482924048	0.0998944332503094	0.461717751294992
GO_SMALL_MOLECULE_METABOLIC_PROCESS	1299	0.01	Up
0.30167012118039	0.0999943793825976	0.461717751294992
GO_HISTONE_DEACETYLASE_ACTIVITY_H3_K14_SPECIFIC_	10	0.01
	Down	0.249494573108799	0.100123656028792
0.461717751294992
GO_POSITIVE_REGULATION_OF_MYELOID_LEUKOCYTE_MEDIATED_IMMUNITY	24
	0.01	Down	0.412747755069047	0.100156793242859
	0.461717751294992
GO_POSITIVE_REGULATION_OF_SMALL_MOLECULE_METABOLIC_PROCESS	65
	0.01	Down	0.343997172499742	0.100161798460798
	0.461717751294992
GO_RESPONSE_TO_ETHANOL	71	0.01	Up	0.461434970592236
	0.100291537823852	0.461965042586196
GO_REGULATION_OF_RAS_PROTEIN_SIGNAL_TRANSDUCTION	192	0.01
	Down	0.325737222127252	0.10038353578586
0.462038245455613
GO_REGULATION_OF_DENDRITE_EXTENSION	10	0.01	Down
0.270122269906007	0.100638505161487	0.462500597668968
GO_TRANSPORT_OF_VIRUS	56	0.01	Down	0.186095410872249
	0.100646644342749	0.462500597668968
GO_UDP_XYLOSYLTRANSFERASE_ACTIVITY	11	0.01	Up
0.30162771771346	0.100712533374794	0.462500597668968
GO_REGULATION_OF_MEMBRANE_LIPID_DISTRIBUTION	37	0.01	Up
	0.420608822632635	0.100819270409769
0.462640809265084
GO_REGULATION_OF_IMMUNE_SYSTEM_PROCESS	1078	0.01	Down
0.436195025774892	0.100926208821186	0.46278173244216
GO_ASYMMETRIC_CELL_DIVISION	11	0.01	Down
0.278019648370405	0.101041999573641	0.462804534564626
GO_EMBRYONIC_PLACENTA_DEVELOPMENT	70	0.01	Down
0.330655918388082	0.101087990616345	0.462804534564626

GO_HEPATOCYTE_GROWTH_FACTOR_RECEPTOR_SIGNALING_PATHWAY 14 0.01 Up 0.576192088746344 0.10123050959244 0.462804534564626
GO_SEMI_LUNAR_VALVE_DEVELOPMENT 28 0.01 Down 0.532239960538133 0.10123610968569 0.462804534564626
GO_POSITIVE_REGULATION_OF_MAP_KINASE_ACTIVITY 209 0.01 Down 0.370989762241905 0.101492818621204 0.463161460639123
GO_POSITIVE_REGULATION_OF_HEART_GROWTH 23 0.01 Up 0.381856601037975 0.101519559702139 0.463161460639123
GO_RNA_POLYMERASE_BINDING 58 0.01 Down 0.169882937278196 0.101543057834076 0.463161460639123
GO_NEGATIVE_REGULATION_OF_CYTOKINE_PRODUCTION 195 0.01 Down 0.520396438957467 0.101666625922161 0.4633769414215
GO_PEPTIDYL_SERINE_DEPHOSPHORYLATION 17 0.01 Down 0.444088797469652 0.101891335580983 0.463681019881939
GO_RHO_GTPASE_BINDING 155 0.01 Down 0.302280690085923 0.101973778196668 0.463681019881939
GO_REGULATION_OF_RNA_POLYMERASE_II_TRANSCRIPTIONAL_PREINITIATION_COMPLEX_ASSEMBLY 12 0.01 Down 0.205057190832167 0.102083291394255 0.463681019881939
GO_MANNOSE_BINDING 11 0.01 Up 0.43270925840639 0.102096872427378 0.463681019881939
GO_ESTABLISHMENT_OF_BLOOD_BRAIN_BARRIER 10 0.01 Up 0.691136628005713 0.102115223782269 0.463681019881939
GO_POSITIVE_REGULATION_OF_PROTEIN_COMPLEX_DISASSEMBLY 30 0.01 Down 0.298128560105683 0.102384637238366 0.464556900354349
GO_SUGAR_TRANSMEMBRANE_TRANSPORTER_ACTIVITY 14 0.01 Up 0.470991545239848 0.102525088309927 0.464562985446648
GO_PROTEIN_ACTIVATION_CASCADE 12 0.01 Up 0.556554880282833 0.102589843840117 0.464562985446648
GO_REGULATION_OF_CELLULAR_AMINO_ACID_METABOLIC_PROCESS 54 0.01 Down 0.154788645295009 0.102615543318062 0.464562985446648
GO_GOLGI_LOCALIZATION 14 0.01 Down 0.194907066667354 0.102790886243298 0.465010037543267
GO_RNA_PHOSPHODIESTER_BOND_HYDROLYSIS_EXONUCLEOLYTIC 39 0.01 Down 0.20409929458388 0.103059047569104 0.465876007291164
GO_REGULATION_OF_SYNCYTIUM_FORMATION_BY_PLASMA_MEMBRANE_FUSION 20 0.01 Down 0.794972443770153 0.103273760047085 0.466499253903167
GO_FLEMMING_BODY 25 0.01 Down 0.191137955710593 0.10372359099742 0.467971500188995
GO_INTERLEUKIN_27_MEDIATED_SIGNALING_PATHWAY 10 0.01 Down 0.601314467947909 0.103753852619731 0.467971500188995
GO_SRP_DEPENDENT_COTRANSLATIONAL_PROTEIN_TARGETING_TO_MEMBRANE_TRANSLOCATION 11 0.01 Up 0.230610664522402 0.104030506119697 0.468850093731211
GO_STEROL_METABOLIC_PROCESS 66 0.01 Up

0.370717040097158	0.104146571954699	0.468850093731211
GO_NEGATIVE_REGULATION_OF_CELL_CYCLE_ARREST	17	0.01	Up
	0.303722986412233	0.104180328849185
0.468850093731211
GO_REGULATION_OF_CATABOLIC_PROCESS	841	0.01	Down
0.225551958916032	0.104394822398751	0.4691794001658
GO_REGULATION_OF_DENDRITE_DEVELOPMENT	114	0.01	Down
0.323342610046823	0.104428079784088	0.4691794001658
GO_REGULATION_OF_PHOSPHATIDYLINOSITOL_3_KINASE_SIGNALING	90
	0.01	Down	0.489287128659168	0.10452248439114
	0.4691794001658
GO_CADHERIN_BINDING_INVOLVED_IN_CELL_CELL_ADHESION	18	0.01
	Down	0.181423115029121	0.104562630279085
0.4691794001658
GO_RESPONSE_TO_PROTOZOAN	12	0.01	Down
0.775015351065089	0.10486767390881	0.470200626514319
GO_PROTEIN_PHOSPHORYLATED_AMINO_ACID_BINDING	44	0.01	Down
	0.345619227280567	0.105012242793364
0.470501347600379
GO_REGULATION_OF_ANIMAL_ORGAN_MORPHOGENESIS	200	0.01	Down
	0.42715717443886	0.105187631365847
0.470939609160809
GO_MALE_MEIOTIC_NUCLEAR_DIVISION	24	0.01	Up
0.304717886957777	0.105432585377814	0.471688449394776
GO_POSTSYNAPTIC_SPECIALIZATION_INTRACELLULAR_COMPONENT	16	0.01
	Down	0.369782887840807	0.105750663714705
0.472664627742638
GO_ESTABLISHMENT_OF_PLANAR_POLARITY_OF_EMBRYONIC_EPITHELIUM	16
	0.01	Up	0.60138853295713	0.105806494663522
	0.472664627742638
GO_REGULATION_OF_ATP_BIOSYNTHETIC_PROCESS	12	0.01	Up
	0.186157251931024	0.106164961554682
0.473917265881229
GO_POSITIVE_REGULATION_OF_CELL_ADHESION	304	0.01	Down
0.50670553397053	0.106724018375126	0.476062832884196
GO_RESPONSE_TO_INTERLEUKIN_6	28	0.01	Down
0.466217942350454	0.106897798284842	0.476487909975975
GO_PROTEIN_PHOSPHORYLATION	1481	0.01	Down
0.346332245208291	0.107010241277341	0.476639159790711
GO_POSITIVE_REGULATION_OF_PROTEIN_MODIFICATION_PROCESS	903	0.01
	Down	0.36252552999048	0.107115883236765
0.476759917251029
GO_SPERM_PART	101	0.01	Up	0.33869787054471
0.107496692877551	0.478056827234334
GO_CALCIUM_ION_BINDING	436	0.01	Up	0.448388823155318
	0.107564754735974	0.478056827234334
GO_DRUG_TRANSMEMBRANE_TRANSPORT	51	0.01	Up
0.495277629406728	0.107798544668601	0.478208218552774
GO_LONG_CHAIN_FATTY_ACID_BIOSYNTHETIC_PROCESS	17	0.01	Up
	0.551165260350537	0.107846906475949
0.478208218552774
GO_REGULATION_OF_DENDRITIC_SPINE_DEVELOPMENT	57	0.01	Down
	0.384421679448865	0.107858929762654
0.478208218552774

GO_RIBOSOME_BINDING	56	0.01	Down	0.168996359920583
	0.107934049728623	0.478208218552774
GO_POSITIVE_REGULATION_OF_P38MAPK_CASCADE	22	0.01	Down
	0.363160260961333	0.108064215079899
0.478208218552774
GO_REGULATION_OF_CHROMOSOME_SEPARATION	53	0.01	Up
0.22502487120834	0.108146994102203	0.478208218552774
GO_RESPONSE_TO_GLUCAGON 22	0.01	Down	0.408124842065869
	0.108235764038319	0.478208218552774
GO_REGULATION_OF_MRNA_SPLICING_VIA_SPLICEOSOME	82	0.01	Down
	0.123097611882883	0.108289241623904
0.478208218552774
GO_NUCLEOTIDE_EXCISION_REPAIR_DNA_DAMAGE_RECOGNITION	23	0.01
	Up	0.121219962393798	0.108307741806962
0.478208218552774
GO_REGULATION_OF_HAIR_FOLLICLE_DEVELOPMENT	15	0.01	Down
	0.590885618481742	0.108503111586074
0.478722667470244
GO_REGULATION_OF_PLASMA_LIPOPROTEIN_PARTICLE_LEVELS	55	0.01
	Up	0.392190372493964	0.108786134558328
0.479281277501596
GO_NEGATIVE_REGULATION_OF_NF_KAPPAB_TRANSCRIPTION_FACTOR_ACTIVITY
	66	0.01	Down	0.417635886404354
0.108823847550834	0.479281277501596
GO_RAS_GUANYL_NUCLEOTIDE_EXCHANGE_FACTOR_ACTIVITY	120	0.01
	Down	0.345047166914423	0.109010274674486
0.479281277501596
GO_QUINONE_BINDING	15	0.01	Up	0.32047660354313
	0.109062326960242	0.479281277501596
GO_CYSTEINE_TYPE_PEPTIDASE_ACTIVITY	148	0.01	Down
0.242345419964702	0.109077769691074	0.479281277501596
GO_HELICASE_ACTIVITY	136	0.01	Down	0.183164276909773
	0.109257502845446	0.479281277501596
GO_ANTIGEN_PROCESSING_AND_PRESENTATION_VIA_MHC_CLASS_IB 10	0.01
	Down	0.286125965197165	0.109279558788814
0.479281277501596
GO_PEPTIDE_ANTIGEN_BINDING	20	0.01	Up
0.682985420528124	0.109289029458048	0.479281277501596
GO_NUCLEOSIDE_BISPHOSPHATE_BIOSYNTHETIC_PROCESS 60	0.01	Up
	0.234778604730181	0.109410431877222
0.479281277501596
GO_REGULATION_OF_BICELLULAR_TIGHT_JUNCTION_ASSEMBLY	17	0.01
	Down	0.310437464127819	0.109452687146412
0.479281277501596
GO_RESPONSE_TO_TUMOR_CELL	13	0.01	Down
0.456294418895678	0.10949812747401	0.479281277501596
GO_SYNAPTIC_TRANSMISSION_DOPAMINERGIC	13	0.01	Up
0.27620420845628	0.109604234129588	0.479400075937126
GO_SPONGIOTROPHOBLAST_LAYER_DEVELOPMENT 11	0.01	Down
0.409176127028763	0.109750299567528	0.479693353977296
GO_ACROSOMAL_VESICLE	58	0.01	Up	0.323716844601903
	0.109899442259809	0.479796191898868
GO_CHONDROCYTE_DIFFERENTIATION	84	0.01	Up
0.678903801706929	0.110101144439492	0.479796191898868

GO_RNA_7_METHYLGUANOSINE_CAP_BINDING	10	0.01	Down
0.182222861018406	0.110112875214625	0.479796191898868
GO_VESICLE_TARGETING_TO_FROM_OR_WITHIN_GOLGI	68	0.01	Down
	0.225191633471669	0.110128484818399
0.479796191898868
GO_PHAGOCYTIC_VESICLE	117	0.01	Down	0.299879530596263
	0.110210056029325	0.479796191898868
GO_ECTODERM_DEVELOPMENT	19	0.01	Down	0.445616463317528
	0.110248013127808	0.479796191898868
GO_PRESYNAPTIC_ACTIVE_ZONE_CYTOPLASMIC_COMPONENT	11	0.01
	Down	0.19166834976713	0.110790018615025
0.481351565335505
GO_REGULATION_OF_KIDNEY_DEVELOPMENT	43	0.01	Down
0.76540261732761	0.110893282946304	0.481351565335505
GO_NEGATIVE_REGULATION_OF_LYMPHOCYTE_ACTIVATION	105	0.01	Down
	0.554603565470843	0.110914802765469
0.481351565335505
GO_COFACTOR_BIOSYNTHETIC_PROCESS	188	0.01	Up
0.284928147872594	0.110997613558953	0.481351565335505
GO_PLASMA_LIPOPROTEIN_PARTICLE_CLEARANCE	37	0.01	Up
	0.463122288575332	0.111001843431017
0.481351565335505
GO_DRUG_TRANSPORT	111	0.01	Up	0.528388363766559
	0.111311743683503	0.481774897869199
GO_REGULATION_OF_MITOCHONDRIAL_MEMBRANE_POTENTIAL	60	0.01
	Up	0.253505263849174	0.111342267588622
0.481774897869199
GO_CELLULAR_RESPONSE_TO_CALCIUM_ION	56	0.01	Up
0.508708872246778	0.111457108154452	0.481774897869199
GO_ANGIOGENESIS_INVOLVED_IN_WOUND_HEALING	17	0.01	Down
	0.545381571627218	0.111489127119704
0.481774897869199
GO_COPII_COATED_VESICLE_CARGO_LOADING	14	0.01	Down
0.158886489773254	0.111496249630411	0.481774897869199
GO_PLATELET_ALPHA_GRANULE	67	0.01	Up
0.582616979861369	0.111752414345982	0.48245443776847
GO_REGULATION_OF_MRNA_PROCESSING	120	0.01	Down
0.1197221386201 0.111859823960573	0.48245443776847
GO_POSITIVE_REGULATION_OF_VIRAL_RELEASE_FROM_HOST_CELL	15	0.01
	Down	0.137267952091563	0.111903144473749
0.48245443776847
GO_CYTOPLASMIC_SIDE_OF_ENDOPLASMIC_RETICULUM_MEMBRANE	12	0.01
	Up	0.222867042010703	0.111971389032412
0.48245443776847
GO_CHROMATIN_REMODELING_AT_CENTROMERE	30	0.01	Up
0.223729586797735	0.112152130393343	0.482890484835451
GO_TELOMERE_MAINTENANCE_VIA_TELOMERE_LENGTHENING	75	0.01
	Down	0.140075949548997	0.112516936994042
0.483826216647685
GO_ANDROGEN_RECEPTOR_BINDING	41	0.01	Down
0.183917969432449	0.112528844985428	0.483826216647685
GO_CARTILAGE_DEVELOPMENT	140	0.01	Up
0.636778115072423	0.11263943857735	0.48395897494911
GO_SUBPALLIUM_DEVELOPMENT	16	0.01	Down

0.425189862819432	0.112741449045962	0.484015286740067
GO_UBIQUITIN_LIKE_PROTEIN_LIGASE_ACTIVITY	195	0.01	Down
	0.209889683383777	0.112837692434807
0.484015286740067
GO_CELLULAR_RESPONSE_TO_ALKALOID	21	0.01	Up
0.361037395021919	0.112891722290222	0.484015286740067
GO_INNER_MITOCHONDRIAL_MEMBRANE_PROTEIN_COMPLEX 121	0.01	Up
	0.135464049210191	0.113259607953999
0.484979453378415
GO_REGULATION_OF_RENAL_SYSTEM_PROCESS	15	0.01	Up
0.436686379234295	0.113276373726008	0.484979453378415
GO_REGULATION_OF_OXIDOREDUCTASE_ACTIVITY	69	0.01	Down
	0.394532883972583	0.113395257031494
0.485146304043832
GO_MITOCHONDRIAL_DEPOLARIZATION 19	0.01	Up
0.410696480797282	0.113590461613922	0.485639220040929
GO_NOTCH_RECEPTOR_PROCESSING	10	0.01	Up
0.204330211463703	0.11437990613197	0.488670239357628
GO_MITOCHONDRIAL_FUSION 21	0.01	Up	0.172733492808843
	0.114543590935144	0.488953403189748
GO_REGULATION_OF_LIPOPROTEIN_LIPASE_ACTIVITY	11	0.01	Down
	0.569662490290402	0.114625521408621
0.488953403189748
GO_POSITIVE_REGULATION_OF_TUMOR_NECROSIS_FACTOR_SUPERFAMILY_CYTOKINE
_PRODUCTION	57	0.01	Down	0.551696949722025
0.114687802032977	0.488953403189748
GO_RNA_POLYADENYLATION	45	0.01	Down	0.124723994589253
	0.115062753755079	0.490207703892693
GO_NEGATIVE_REGULATION_OF_ACTIN_FILAMENT_DEPOLYMERIZATION	29
	0.01	Down	0.331525529638539	0.115223749653539
	0.490227072460784
GO_MEMBRANE_FISSION	10	0.01	Down	0.138702375741176
	0.115229367018395	0.490227072460784
GO_REGULATION_OF_CENTROSOME_CYCLE	54	0.01	Down
0.145306103106971	0.115376248107719	0.490227072460784
GO_LYMPH_VESSEL_DEVELOPMENT	20	0.01	Down
0.384247940572758	0.115405238938081	0.490227072460784
GO_REGULATION_OF_FATTY_ACID_TRANSPORT	17	0.01	Up
0.395112625595293	0.11547104490511	0.490227072460784
GO_MIDBODY_ABSCISSION	17	0.01	Down	0.223322202112635
	0.115627057086941	0.490496371236597
GO_PHOSPHATIDYLINOSITOL_PHOSPHATE_BINDING	124	0.01	Down
	0.253409064728351	0.115696063846287
0.490496371236597
GO_DITERPENOID_METABOLIC_PROCESS	50	0.01	Up
0.529080360005023	0.115969850659397	0.491314000944313
GO_PROTEASOME_ACCESSORY_COMPLEX 24	0.01	Down
0.0918493016154927	0.116145573649007	0.491715326097018
GO_AMYLOID_BETA_FORMATION	25	0.01	Down
0.35942928801932	0.116328735504008	0.491822421059554
GO_VASCULAR_ENDOTHELIAL_GROWTH_FACTOR_SIGNALING_PATHWAY 29	0.01
	Up	0.552777507746068	0.116332893533441
0.491822421059554
GO_POSITIVE_REGULATION_OF_CELL_AGING	14	0.01	Down

0.377393940400675	0.11662079797734	0.492696495838854
GO_CHEMOREPELLENT_ACTIVITY	21	0.01	Down
0.543986462134389	0.11686717511975	0.493394033485398
GO_TUMOR_NECROSIS_FACTOR_SECRETION	14	0.01	Down
0.68497971254029	0.116955964865141	0.493425755869541
GO_ANTIGEN_PROCESSING_AND_PRESENTATION_OF_PEPTIDE_OR_POLYSACCHARIDE_ANTIGEN_VIA_MHC_CLASS_II	89	0.01	Up
0.37779092858873	0.11707945381798	0.493603725089554
GO_ATP_BIOSYNTHETIC_PROCESS	39	0.01	Up
0.131580918826546	0.11729202552514 8	0.494156757087559
GO_PEPTIDE_N_ACETYLTRANSFERASE_ACTIVITY	67	0.01	Down
0.137424049507377	0.117836452882804	0.495685269066201
GO_PROTEIN_HOMOTETRAMERIZATION	64	0.01	Up
0.283659212176618	0.1178763898285	0.495685269066201
GO_SULFURIC_ESTER_HYDROLASE_ACTIVITY	14	0.01	Up
0.414776726271097	0.117966386238939	0.495685269066201
GO_GLIAL_CELL_DIFFERENTIATION	145	0.01	Down
0.393800714250139	0.117992229966139	0.495685269066201
GO_PROTEIN_TARGETING_TO_MEMBRANE	181	0.01	Up
0.17526695227825	0.1180689347602	0.495685269066201
GO_FATTY_ACID_BIOSYNTHETIC_PROCESS	104	0.01	Up
0.352670410012081	0.118144718224147	0.495685269066201
GO_ERROR_PRONE_TRANSLESION_SYNTHESIS	19	0.01	Up
0.135383190710331	0.118356830164186	0.495696155588055
GO_RESPONSE_TO_LAMINAR_FLUID_SHEAR_STRESS	14	0.01	Down
0.263851116263568	0.118367514892427
0.495696155588055
GO_PROTEIN_KINASE_ACTIVITY	488	0.01	Down
0.298046740295476	0.118392262494265	0.495696155588055
GO_REGULATION_OF_GLUCOSE_TRANSMEMBRANE_TRANSPORT	54	0.01
Down	0.324042084827616	0.11862373823889
0.496323028841006
GO_PHOSPHATIDYLINOSITOL_KINASE_ACTIVITY	13	0.01	Down
0.214804565635304	0.118796084303079	0.496701809782364
GO_DYNACTIN_BINDING	11	0.01	Down	0.302142061319171
0.119187663449437	0.497994106938595
GO_INTRA_GOLGI_VESICLE_MEDIATED_TRANSPORT	30	0.01	Down
0.165029231607382	0.11936094548525
0.497994106938595
GO_TRANSCRIPTION_COREGULATOR_ACTIVITY	482	0.01	Down
0.237295992366863	0.119364083902657	0.497994106938595
GO_REGULATION_OF_ACUTE_INFLAMMATORY_RESPONSE	27	0.01	Down
0.508699445329102	0.119438599251485
0.497994106938595
GO_MUSCLE_CELL_CELLULAR_HOMEOSTASIS	17	0.01	Up
0.29451595645152	0.119572332145994	0.497994106938595
GO_DECIDUALIZATION	18	0.01	Down	0.634679945185599
0.119597332880328	0.497994106938595
GO_HYDROLASE_ACTIVITY_HYDROLYZING_N_GLYCOSYL_COMPOUNDS	18	0.01
Up	0.175777359013938	0.119724851737929
0.498183396093877
GO_APICAL_DENDRITE	12	0.01	Down	0.332119243325198
0.119940629458259	0.498688077600502
GO_ERYTHROCYTE_DEVELOPMENT	22	0.01	Down

0.325859308942076	0.120062957676398	0.498688077600502
GO_REGULATION_OF_NEUROTRANSMITTER_RECEPTOR_ACTIVITY	31	0.01	Down	0.391621127966379	0.120107105181892	0.498688077600502
GO_REGULATION_OF_BLOOD_PRESSURE	94	0.01	Up	0.503271906057913	0.120174708866667	0.498688077600502
GO_POSITIVE_REGULATION_OF_VIRAL_LIFE_CYCLE	55	0.01	Down	0.219886796947584	0.120537777996416	0.499590391059131
GO_PHOSPHATIDYLINOSITOL_PHOSPHORYLATION	43	0.01	Down	0.215606817645628	0.120655878788082	0.499590391059131
GO_MAMMARY_GLAND_EPITHELIAL_CELL_DIFFERENTIATION	13	0.01	Down	0.40070790487175	0.120733334054484	0.499590391059131
GO_POSITIVE_REGULATION_OF_SEQUESTERING_OF_CALCIUM_ION	11	0.01	Up	0.392950295371936	0.12075168656002	0.499590391059131
GO_PLASMA_MEMBRANE_ORGANIZATION	74	0.01	Up	0.361502335638401	0.120883759711104	0.499590391059131
GO_LYMPH_VESSEL_MORPHOGENESIS	15	0.01	Down	0.403850036992456	0.120885897622445	0.499590391059131
GO_REGULATION_OF_PHOSPHOLIPID_BIOSYNTHETIC_PROCESS	11	0.01	Up	0.43377497721991	0.121189455676696	0.50029181624656
GO_ORGANIC_CATION_TRANSMEMBRANE_TRANSPORTER_ACTIVITY	16	0.01	Up	0.314074355870935	0.121259573431708	0.50029181624656
GO_EXTRINSIC_COMPONENT_OF_PLASMA_MEMBRANE	112	0.01	Down	0.342757693524241	0.121336141813433	0.50029181624656
GO_CELL_TRAILING_EDGE	11	0.01	Up	0.453544367030469	0.121385248777991	0.50029181624656
GO_POSITIVE_REGULATION_OF_STEM_CELL_DIFFERENTIATION	14	0.01	Down	0.360240145322792	0.121492583012193	0.500309529391098
GO_MONOVALENT_INORGANIC_ANION_HOMEOSTASIS	11	0.01	Down	0.468600782525696	0.121583497606108	0.500309529391098
GO_CARBOHYDRATE_KINASE_ACTIVITY	18	0.01	Up	0.385437229964986	0.12178703570961	0.500309529391098
GO_POSITIVE_REGULATION_OF_TELOMERE_CAPPING	14	0.01	Down	0.166987233349397	0.121888227612364	0.500309529391098
GO_METANEPHRIC_RENAL_VESICLE_MORPHOGENESIS	10	0.01	Down	1.25925081270205	0.121956398217885	0.500309529391098
GO_REGULATION_OF_UBIQUITIN_DEPENDENT_PROTEIN_CATABOLIC_PROCESS	136	0.01	Down	0.200998320631694	0.122010462304103	0.500309529391098
GO_REGULATION_OF_CELL_KILLING	55	0.01	Down	0.548458893782642	0.122020671903455	0.500309529391098
GO_AXONAL_GROWTH_CONE	24	0.01	Down	0.350269233570987	0.122167501787457	0.500309529391098
GO_MAINTENANCE_OF_CELL_POLARITY	12	0.01	Down

0.264341782805527	0.12218228710449	0.500309529391098
GO_MONOCYTE_CHEMOTAXIS	34	0.01	Down	1.03598001586629
	0.122213643895075	0.500309529391098
GO_NAD_DEPENDENT_PROTEIN_DEACETYLASE_ACTIVITY	16	0.01	Down
	0.218849832634511	0.122431077293967
0.500574164657318
GO_RESPONSE_TO_INCREASED_OXYGEN_LEVELS	23	0.01	Up
0.462015194875593	0.122443194616392	0.500574164657318
GO_SCF_DEPENDENT_PROTEASOMAL_UBIQUITIN_DEPENDENT_PROTEIN_CATABOLIC_P
ROCESS	83	0.01	Down	0.13517181230488
0.122722014485836	0.501376413151757
GO_POSITIVE_REGULATION_OF_TELOMERE_MAINTENANCE_VIA_TELOMERE_LENGTHEN
ING	34	0.01	Down	0.139450438683952
0.122812110959851	0.501407078437967
GO_HISTONE_H3_K9_DEMETHYLATION	12	0.01	Down
0.237124007741979	0.122902018657201	0.501436932303675
GO_PHOSPHATIDYLINOSITOL_3_KINASE_REGULATORY_SUBUNIT_BINDING	10
	0.01	Down	0.456609184953851	0.123075430444584
	0.501517109486454
GO_NEGATIVE_REGULATION_OF_BIOSYNTHETIC_PROCESS	1206	0.01	Down
	0.296130958304538	0.12308688735543
0.501517109486454
GO_POSITIVE_REGULATION_OF_CHROMOSOME_ORGANIZATION	153	0.01
	Down	0.173473719179426	0.123197122863688
0.501569413279182
GO_CORTICOSTEROID_RECEPTOR_SIGNALING_PATHWAY	15	0.01	Down
	0.20071417001011	0.123328911378994
0.501569413279182
GO_POSITIVE_REGULATION_OF_MACROAUTOPHAGY	62	0.01	Down
	0.223178790370141	0.123404526565153
0.501569413279182
GO_HYPOTHALAMUS_DEVELOPMENT	11	0.01	Down
0.478561611989	0.123430193285966	0.501569413279182
GO_POSITIVE_REGULATION_OF_FIBROBLAST_MIGRATION	12	0.01	Up
	0.612854063186812	0.123579374336036
0.50157531297749
GO_POSITIVE_REGULATION_OF_HEART_CONTRACTION	23	0.01	Down
	0.464416589573294	0.123657825655583
0.50157531297749
GO_CELLULAR_MODIFIED_AMINO_ACID_CATABOLIC_PROCESS	19	0.01
	Up	0.391277103877666	0.12367949983978
0.50157531297749
GO_REGULATION_OF_DNA_BIOSYNTHETIC_PROCESS	89	0.01	Down
	0.310918150474579	0.123829423720142
0.501727793453873
GO_AXON_EXTENSION	96	0.01	Down	0.349722675258759
	0.123925602024287	0.501727793453873
GO_REGULATION_OF_MEMBRANE_PROTEIN_ECTODOMAIN_PROTEOLYSIS	20
	0.01	Down	0.448859563019633	0.123965028855347
	0.501727793453873
GO_ESC_E_Z_COMPLEX	15	0.01	Down	0.192946640777635
	0.1242541115166	0.502544483662236
GO_POSITIVE_REGULATION_OF_GLUCONEOGENESIS	12	0.01	Down
	0.217195881139441	0.124332369372538

0.502544483662236
GO_REGULATION_OF_TRANSCRIPTION_INVOLVED_IN_G1_S_TRANSITION_OF_MITOTIC_CELL_CYCLE 26 0.01 Up 0.241242925692669
0.124647761732563 0.503484072839913
GO_ARF_PROTEIN_SIGNAL_TRANSDUCTION 16 0.01 Down
0.199604984169504 0.124895481346997 0.503658840673341
GO_REGULATION_OF_PINOCYTOSIS 10 0.01 Up
0.267624166726263 0.124942200590129 0.503658840673341
GO_PRODUCTION_OF_MOLECULAR_MEDIATOR_OF_IMMUNE_RESPONSE 140 0.01
Down 0.387854440382413 0.124954708551681
0.503658840673341
GO_REGULATION_OF_AMYLOID_BETA_FORMATION 19 0.01 Down
0.449336584166417 0.125022874797352 0.503658840673341
GO_N_TERMINAL_PROTEIN_AMINO_ACID_ACETYLATION 15 0.01 Down
0.176419411185101 0.125170674511701
0.503919870663487
GO_POSITIVE_REGULATION_OF_LAMELLIPODIUM_ASSEMBLY 15 0.01
Down 0.129832995381708 0.125378115408292
0.504000774246501
GO_PROTEIN_PHOSPHATASE_TYPE_1_COMPLEX 10 0.01 Down
0.200467442618948 0.12538858331713 0.504000774246501
GO_POSITIVE_REGULATION_OF_PRODUCTION_OF_MOLECULAR_MEDIATOR_OF_IMMUNE_RESPONSE 77 0.01 Down 0.402547478116027
0.125439823733563 0.504000774246501
GO_NEGATIVE_REGULATION_OF_FATTY_ACID_METABOLIC_PROCESS 10 0.01
Up 0.421388287882621 0.125577775003361
0.504047330287709
GO_DNA_N_GLYCOSYLASE_ACTIVITY 13 0.01 Up
0.147072730227426 0.125726590447212 0.504047330287709
GO_ADENYLATE_CYCLASE_BINDING 11 0.01 Down
0.455684960218024 0.125748115050706 0.504047330287709
GO_MITOCHONDRION 1379 0.01 Up 0.199923924958272
0.125783512664451 0.504047330287709
GO_PINOCYTOSIS 19 0.01 Up 0.251802676993463
0.126120382010274 0.504855355711119
GO_REGULATION_OF_PROTEIN_CATABOLIC_PROCESS 328 0.01 Down
0.241196853783629 0.12616133683858
0.504855355711119
GO_CILIARY_PART 303 0.01 Up 0.261428461366232
0.126234628556989 0.504855355711119
GO_REGULATION_OF_POTASSIUM_ION_TRANSMEMBRANE_TRANSPORTER_ACTIVITY
30 0.01 Down 0.419900483967379
0.126500886678949 0.504953041050808
GO_TRANSCRIPTION_FACTOR_BINDING 540 0.01 Down
0.246605872880276 0.12658352909222 0.504953041050808
GO_LYMPHOCYTE_ACTIVATION_INVOLVED_IN_IMMUNE_RESPONSE 119 0.01
Down 0.433504501143261 0.126690422338185
0.504953041050808
GO_ENDOTHELIAL_CELL_MORPHOGENESIS 11 0.01 Up
0.712531885297785 0.126700471378423 0.504953041050808
GO_CILIARY_BASE 29 0.01 Up 0.185148046479335
0.126757834642045 0.504953041050808
GO_REGULATION_OF_PROTEIN_MODIFICATION_PROCESS 1404 0.01 Down
0.332642146671931 0.126758101558463

0.504953041050808
GO_CHEMICAL_SYNAPTIC_TRANSMISSION_POSTSYNAPTIC 53 0.01 Up
0.389056815462079 0.126917325663139
0.505255792853062
GO_LYS48_SPECIFIC_DEUBIQUITINASE_ACTIVITY 14 0.01 Down
0.138934232758881 0.127311276216459
0.50600742164049
GO_EUKARYOTIC_TRANSLATION_INITIATION_FACTOR_4F_COMPLEX 12 0.01
Down 0.130338560498974 0.127317422904483
0.50600742164049
GO_FUCOSYLTRANSFERASE_ACTIVITY 12 0.01 Down
0.296964485336359 0.127374981322581 0.50600742164049
GO_STEROID_BINDING 64 0.01 Up 0.399846006920168
0.127439523585622 0.50600742164049
GO_NEGATIVE_REGULATION_OF_PHOSPHORUS_METABOLIC_PROCESS 449 0.01
Down 0.334977597915213 0.127854600619371
0.506875301739318
GO_RESPONSE_TO_THYROID_HORMONE 20 0.01 Up
0.301292362006763 0.127893643714192 0.506875301739318
GO_MOTILE_CILIUM 102 0.01 Up 0.318614766205845
0.127987843308871 0.506875301739318
GO_POSITIVE_REGULATION_OF_INTERLEUKIN_12_PRODUCTION 24 0.01
Down 0.620036307912561 0.127992066803883
0.506875301739318
GO_TRANSLATION_FACTOR_ACTIVITY_RNA_BINDING 82 0.01 Down
0.136407660243418 0.128644630124431
0.509003651138593
GO_NEGATIVE_REGULATION_OF_CELLULAR_SENESCENCE 15 0.01 Down
0.234751100590035 0.128697184071445
0.509003651138593
GO_TRANSLATIONAL_ELONGATION 129 0.01 Up
0.129978342147445 0.128804068432159 0.509094726205493
GO_SENSORY_PERCEPTION 244 0.01 Up 0.438975411468499
0.129139939392288 0.510090157482487
GO_CELL_DIFFERENTIATION_INVOLVED_IN_KIDNEY_DEVELOPMENT 37 0.01
Down 0.81483236915781 0.129339069331882
0.510286660820404
GO_REGULATION_OF_PROGRAMMED_NECROTIC_CELL_DEATH 17 0.01 Down
0.334834070205765 0.129357794597694
0.510286660820404
GO_DNA_DEMETHYLATION 20 0.01 Down 0.308302135775983
0.129745391437601 0.51148329312836
GO_PHOSPHATIDYLCHOLINE_ACYL_CHAIN_REMODELING 20 0.01 Up
0.492777199059236 0.130092146468018
0.512517469959336
GO_POSITIVE_REGULATION_OF_DNA_BIOSYNTHETIC_PROCESS 59 0.01
Down 0.366582331544753 0.130374751648337
0.512974486553389
GO_MYD88_INDEPENDENT_TOLL_LIKE_RECEPTOR_SIGNALING_PATHWAY 30
0.01 Down 0.344811001479576 0.130377142604493
0.512974486553389
GO_NEGATIVE_REGULATION_OF_CELLULAR_PROTEIN_CATABOLIC_PROCESS 73
0.01 Down 0.224255051634452 0.130653100812233
0.513172237016194

GO_REGULATION_OF_IMMUNE_RESPONSE	704	0.01	Down
0.439339532151373	0.130717296824918	0.513172237016194
GO_REGION_OF_CYTOSOL	18	0.01	Down	0.183030810994426
	0.130838639210395	0.513172237016194
GO_MATERNAL_PLACENTA_DEVELOPMENT	29	0.01	Down
0.517624019656691	0.130923810996333	0.513172237016194
GO_REGULATION_OF_NEUROTRANSMITTER_UPTAKE	11	0.01	Up
	0.747271882694125	0.131089989551817
0.513172237016194
GO_REGULATION_OF_RESPONSE_TO_ENDOPLASMIC_RETICULUM_STRESS	76
	0.01	Down	0.220837163183691	0.131181232026154
	0.513172237016194
GO_DNA_DEALKYLATION	26	0.01	Down	0.268178760481567
	0.131199831759606	0.513172237016194
GO_CELLULAR_RESPONSE_TO_INSULIN_STIMULUS	168	0.01	Down
	0.220451313053465	0.131326383970941
0.513172237016194
GO_ENTRAINMENT_OF_CIRCADIAN_CLOCK	24	0.01	Down
0.202224881557284	0.131409776724234	0.513172237016194
GO_REGULATION_OF_ENDOTHELIAL_CELL_DIFFERENTIATION	26	0.01
	Down	0.359516509002706	0.131467352418478
0.513172237016194
GO_REGULATION_OF_AMYLOID_PRECURSOR_PROTEIN_CATABOLIC_PROCESS	24
	0.01	Down	0.411678891050857	0.131497096905557
	0.513172237016194
GO_LEFT_RIGHT_AXIS_SPECIFICATION	10	0.01	Up
0.257428949186199	0.131634993416794	0.513172237016194
GO_TELOMERASE_RNA_BINDING	21	0.01	Down
0.127914954978601	0.131646423145452	0.513172237016194
GO_PTERIDINE_CONTAINING_COMPOUND_BIOSYNTHETIC_PROCESS	14	0.01
	Up	0.223739922453558	0.131674703989986
0.513172237016194
GO_REGULATION_OF_DNA_METABOLIC_PROCESS	245	0.01	Down
0.205149849103342	0.131811209168336	0.513172237016194
GO_GLUTAMINE_FAMILY_AMINO_ACID_CATABOLIC_PROCESS	20	0.01
	Up	0.406461310641471	0.131907717754829
0.513172237016194
GO_NUCLEOTIDE_EXCISION_REPAIR_DNA_INCISION	39	0.01	Up
	0.120698453619501	0.131911914725248
0.513172237016194
GO_NECROTIC_CELL_DEATH	44	0.01	Down	0.321166086653869
	0.131948914838129	0.513172237016194
GO_REGULATION_OF_INTRACELLULAR_SIGNAL_TRANSDUCTION	1438	0.01
	Down	0.351145813680291	0.13219675940672
0.513806995107682
GO_TRANSCRIPTION_COACTIVATOR_ACTIVITY	276	0.01	Down
0.242606568370836	0.13261042637764	0.514961872133528
GO_REGULATION_OF_FAT_CELL_DIFFERENTIATION	96	0.01	Down
	0.404239884261956	0.132726157557297
0.514961872133528
GO_NEGATIVE_REGULATION_OF_CALCINEURIN_MEDIATED_SIGNALING	12
	0.01	Down	0.197325630515152	0.132907596055106
	0.514961872133528
GO_CYCLIC_NUCLEOTIDE_MEDIATED_SIGNALING	101	0.01	Down

0.520358252696093	0.133026524729463	0.514961872133528
GO_POSITIVE_REGULATION_OF_SMALL_GTPASE_MEDIATED_SIGNAL_TRANSDUCTION	43	0.01	Down	0.379348915616435
0.133053728864831	0.514961872133528
GO_REGULATION_OF_PROTEIN_TYROSINE_KINASE_ACTIVITY	71	0.01	Down	0.44991889831782	0.133061506228367
0.514961872133528
GO_TRANS_GOLGI_NETWORK_TRANSPORT_VESICLE_MEMBRANE	13	0.01	Down	0.287563540124494	0.13308765893222
0.514961872133528
GO_CELLULAR_RESPONSE_TO_PROSTAGLANDIN_STIMULUS	17	0.01	Up	0.518003395447188	0.133258853652435
0.515295860206326
GO_INTRINSIC_COMPONENT_OF_MITOCHONDRIAL_INNER_MEMBRANE	43	0.01	Up	0.134807718543601	0.134010354919596
0.517871969902526
GO_POSITIVE_REGULATION_OF_INTERLEUKIN_8_PRODUCTION	34	0.01	Down	0.440044801363805	0.134143025723193
0.51788312645733
GO_VIRION_ASSEMBLY	38	0.01	Down	0.2043130258406
0.134466702319085	0.51788312645733
GO_REGULATION_OF_INTRINSIC_APOPTOTIC_SIGNALING_PATHWAY_BY_P53_CLASS_MEDIATOR	28	0.01	Up	0.226176121087439
0.134610085979693	0.51788312645733
GO_MYELOID_CELL_DIFFERENTIATION	306	0.01	Down
0.376088968317031	0.134650613208912	0.51788312645733
GO_PATTERN_SPECIFICATION_PROCESS	261	0.01	Up
0.443097093195345	0.134685065460449	0.51788312645733
GO_HISTONE_MRNA_CATABOLIC_PROCESS	12	0.01	Down
0.193758720089167	0.134721926442653	0.51788312645733
GO_REGULATION_OF_DENDRITIC_SPINE_MORPHOGENESIS	33	0.01	Down	0.385319342355487	0.134755798126117
0.51788312645733
GO_IMMUNE_RESPONSE_REGULATING_CELL_SURFACE_RECEPTOR_SIGNALING_PATHWAY	294	0.01	Down	0.394579620048648
0.134897296721614	0.51788312645733
GO_CELL_CELL_SIGNALING_INVOLVED_IN_CARDIAC_CONDUCTION	19	0.01	Up	0.427104684562746	0.13489884390449
0.51788312645733
GO_PLATELET_DEGRANULATION	96	0.01	Up
0.475040837767985	0.134947016979864	0.51788312645733
GO_PRIMARY_MIRNA_PROCESSING	10	0.01	Down
0.102600831292818	0.135029618093428	0.51788312645733
GO_DYNEIN_LIGHT_INTERMEDIATE_CHAIN_BINDING	25	0.01	Down	0.306481158426604	0.135268300311477
0.51788312645733
GO_SARCOPLASM	53	0.01	Up	0.336004747320771
0.135295934974531	0.51788312645733
GO_ATRIOVENTRICULAR_VALVE_DEVELOPMENT	18	0.01	Down
0.358198317832684	0.135308655195106	0.51788312645733
GO_COPII_COATED_VESICLE_BUDDING	66	0.01	Down
0.222130440764416	0.135336397359507	0.51788312645733
GO_PROTEIN_KINASE_A_BINDING	39	0.01	Down
0.31855704676935	0.135652359838117	0.51788312645733

GO_CILIUM 398 0.01 Up 0.258116265460791
0.135692410767526 0.51788312645733
GO_POSITIVE_REGULATION_OF_T_HELPER_1_TYPE_IMMUNE_RESPONSE 12
0.01 Down 0.659303495988875 0.135759081051796
0.51788312645733
GO_CARDIAC_RIGHT_VENTRICLE_MORPHOGENESIS 10 0.01 Down
0.345432333563758 0.135780725533245
0.51788312645733
GO_ACTIVATION_OF_ADENYLATE_CYCLASE_ACTIVITY 19 0.01 Down
0.345004760542713 0.135800835615593
0.51788312645733
GO_CATION_CHANNEL_ACTIVITY 124 0.01 Up
0.47810593641871 0.135804634709285 0.51788312645733
GO_NEGATIVE_REGULATION_OF_CALCIUM_ION_TRANSPORT 38 0.01 Up
0.439508696387763 0.136006156985584
0.518326038329869
GO_REGULATION_OF_MACROPHAGE_MIGRATION 23 0.01 Up
0.719809709267754 0.136169815750791 0.518493914872729
GO_VACUOLAR_ACIDIFICATION 22 0.01 Up
0.300714857931782 0.136221017002471 0.518493914872729
GO_NUCLEOSIDE_MONOPHOSPHATE_BIOSYNTHETIC_PROCESS 40 0.01
Up 0.237544678181425 0.136324296000243
0.51856190539942
GO_MODULATION_BY_SYMBIONT_OF_HOST_CELLULAR_PROCESS 25 0.01
Down 0.280929584858036 0.136644150155675
0.519453121850408
GO_INTESTINAL_EPITHELIAL_CELL_DIFFERENTIATION 13 0.01 Down
0.537821778526084 0.136804656240466
0.519737839822196
GO_NEGATIVE_REGULATION_OF_INTRINSIC_APOPTOTIC_SIGNALING_PATHWAY_IN_RESPONSE_TO_DNA_DAMAGE 30 0.01 Up 0.424535686753211
0.137278501726931 0.520889942787278
GO_PROTEOGLYCAN_BIOSYNTHETIC_PROCESS 50 0.01 Up
0.454581927781244 0.137279510535273 0.520889942787278
GO_TRANSFERASE_ACTIVITY_TRANSFERRING_SULFUR_CONTAINING_GROUPS 45
0.01 Up 0.354039002265184 0.137560994701661
0.521168784853729
GO_NEGATIVE_REGULATION_OF_CARDIAC_MUSCLE_TISSUE_GROWTH 17 0.01
Down 0.41849307010757 0.137570665006511
0.521168784853729
GO_REGULATION_OF_EMBRYONIC_DEVELOPMENT 103 0.01 Up
0.476841816813311 0.137610535681194 0.521168784853729
GO_MODIFIED_AMINO_ACID_TRANSPORT 18 0.01 Up
0.498633948209413 0.137854232798561 0.521766238977597
GO_SPECTRIN_BINDING 19 0.01 Down 0.349158517959533
0.137975806433988 0.521901009882081
GO_POSITIVE_REGULATION_OF_T_CELL_MEDIATED_IMMUNITY 34 0.01
Down 0.431893361760481 0.138204526836434
0.522440649080943
GO_REGULATION_OF_DNA_METHYLATION 17 0.01 Down
0.29180152791879 0.138506226195241 0.523255319994591
GO_REGULATION_OF_LYMPHOCYTE_MIGRATION 43 0.01 Down
0.582616237752245 0.138688027460512 0.523438163575889
GO_POSITIVE_REGULATION_OF_PROTEIN_POLYMERIZATION 108 0.01

Down 0.318667578859553 0.138727063942284
0.523438163575889
GO_SULFURTRANSFERASE_ACTIVITY 11 0.01 Up
0.135073396898243 0.139224411156467 0.524734248698461
GO_POLYSACCHARIDE_BIOSYNTHETIC_PROCESS 58 0.01 Down
0.302539186532744 0.139243431832189 0.524734248698461
GO_LABYRINTHINE_LAYER_MORPHOGENESIS 16 0.01 Down
0.211655887627478 0.139786935423938 0.526370463737044
GO_CELLULAR_RESPONSE_TO_FLUID_SHEAR_STRESS 18 0.01 Down
0.474088372252688 0.139851022567592
0.526370463737044
GO_T_CELL_HOMEOSTASIS 28 0.01 Down 0.441381185368644
0.140062422795148 0.526839509782742
GO_ANTIGEN_BINDING 33 0.01 Up 0.596897946167008
0.140398411785366 0.527550872289745
GO_AXON_REGENERATION 37 0.01 Up 0.438470706023313
0.140425335137577 0.527550872289745
GO_POSITIVE_REGULATION_OF_TRANSCRIPTION_OF_NOTCH_RECEPTOR_TARGET
16 0.01 Down 0.315605396659258
0.140679506834274 0.527930554823401
GO_AXONAL_FASCICULATION 17 0.01 Down 0.444689087049931
0.14070031917382 0.527930554823401
GO_ENDOPLASMIC_RETICULUM_UNFOLDED_PROTEIN_RESPONSE 109 0.01
Down 0.221545902308361 0.140852870883301
0.528176515832315
GO_CHROMOSOME_LOCALIZATION 58 0.01 Down
0.173629947904219 0.141844615967245 0.531310081434144
GO_EXOCYTIC_PROCESS 63 0.01 Down 0.228987338811947
0.141945016716674 0.531310081434144
GO_MANNOSYLATION 32 0.01 Up 0.149744624252832
0.141951071007442 0.531310081434144
GO_SH3_SH2_ADAPTOR_ACTIVITY 41 0.01 Down
0.34019140341863 0.14220995829255 0.531951113243419
GO_AXIS_ELONGATION 26 0.01 Up 0.536799181224703
0.142387964868761 0.5321510920349
GO_GAP_JUNCTION 16 0.01 Down 0.566274361484826
0.142481262409162 0.5321510920349
GO_MOLTING_CYCLE_PROCESS 69 0.01 Down
0.433209335366302 0.142601408391229 0.5321510920349
GO_GLOMERULAR_EPITHELIAL_CELL_DIFFERENTIATION 11 0.01 Down
0.496284072144019 0.142690379671532
0.5321510920349
GO_MACROAUTOPHAGY 276 0.01 Down 0.160895383809351
0.142701692939024 0.5321510920349
GO_NON_MOTILE_CILIUM 75 0.01 Up 0.253027643756615
0.142858419439724 0.532408510999731
GO_REGULATION_OF_CELL_PROJECTION_ASSEMBLY 144 0.01 Down
0.284791881436883 0.143461767958416
0.534329075629168
GO_NEGATIVE_REGULATION_OF_INTERLEUKIN_1_BETA_PRODUCTION 13 0.01
Down 0.716200091662163 0.14368406297289
0.534828906381617
GO_OXIDOREDUCTASE_ACTIVITY_ACTING_ON_CH_OH_GROUP_OF_DONORS 94
0.01 Up 0.35872944922534 0.143806587656399

0.534956981410539
GO_T_CELL_LINEAGE_COMMITMENT 15 0.01 Down
0.542139823117218 0.144119790359857 0.535793782776907
GO_REGULATION_OF_PHOSPHORUS_METABOLIC_PROCESS 1263 0.01 Down
0.357850840424421 0.14435988197361
0.53601544701662
GO_BILE_ACID_METABOLIC_PROCESS 30 0.01 Up
0.414402776489979 0.144573886866948 0.53601544701662
GO_VOLTAGE_GATED_CALCIUM_CHANNEL_ACTIVITY 20 0.01 Up
0.286269099444582 0.144619087558425
0.53601544701662
GO_REGULATION_OF_DNA_TEMPLATED_TRANSCRIPTION_IN_RESPONSE_TO_STRESS
113 0.01 Down 0.195677196723143
0.144770228728644 0.53601544701662
GO_CELLULAR_RESPONSE_TO_HYDROGEN_PEROXIDE 77 0.01 Down
0.359179407420434 0.144813068418736
0.53601544701662
GO_NEGATIVE_REGULATION_OF_NERVOUS_SYSTEM_DEVELOPMENT 209 0.01
Down 0.367347295039098 0.144816138047631
0.53601544701662
GO_RESPONSE_TO_MUSCLE_STRETCH 13 0.01 Down
0.445911975348457 0.14497673490073 0.53601544701662
GO_BASAL_PLASMA_MEMBRANE 24 0.01 Down
0.448166250599046 0.145006203716525 0.53601544701662
GO_POSITIVE_REGULATION_OF_MACROPHAGE_DERIVED_FOAM_CELL_DIFFERENTIATI
ON 10 0.01 Up 0.964575565397481
0.145062583749869 0.53601544701662
GO_POSITIVE_REGULATION_OF_INTERLEUKIN_6_SECRETION 21 0.01
Down 0.464681285561887 0.145136461753204
0.53601544701662
GO_ASSOCIATIVE_LEARNING 44 0.01 Up 0.313835061137757
0.145214299560554 0.53601544701662
GO_CYTOPLASMIC_TRANSLATIONAL_INITIATION 29 0.01 Down
0.134311054529809 0.145238907979302 0.53601544701662
GO_REGULATION_OF_HOMOTYPIC_CELL_CELL_ADHESION 19 0.01 Down
0.547076205736133 0.145577102787405
0.536937175590726
GO_CD4_POSITIVE_ALPHA_BETA_T_CELL_CYTOKINE_PRODUCTION 11 0.01
Down 0.508960718107202 0.145862492224478
0.537663139219674
GO_POSITIVE_REGULATION_OF_STEROID_METABOLIC_PROCESS 15 0.01
Down 0.682522925334821 0.146006551861508
0.537867582737387
GO_ASPARTATE_FAMILY_AMINO_ACID_BIOSYNTHETIC_PROCESS 19 0.01
Up 0.189531208440131 0.146108816025981
0.537917903028338
GO_ESCRT_COMPLEX 27 0.01 Down 0.165776402250053
0.146224413025074 0.538017219075895
GO_POSTSYNAPTIC_DENSITY_INTRACELLULAR_COMPONENT 12 0.01 Down
0.329913883081485 0.146475003084785
0.538070690115292
GO_EMBRYONIC_HINDLIMB_MORPHOGENESIS 17 0.01 Up
0.527616365606342 0.146546050495576 0.538070690115292
GO_EAR_MORPHOGENESIS 70 0.01 Down 0.423751176431893

0.146567698134358	0.538070690115292
GO_DIVALENT_INORGANIC_CATION_TRANSMEMBRANE_TRANSPORTER_ACTIVITY	19	0.01	Up	0.23218841480656	0.146593464248179	0.538070690115292
GO_REGULATION_OF_RAC_PROTEIN_SIGNAL_TRANSDUCTION	12	0.01	Down	0.219190604020961	0.146881505549799	0.538479966393432
GO_POSITIVE_REGULATION_OF_CARBOHYDRATE_METABOLIC_PROCESS	46	0.01	Down	0.324955778100974	0.146882362765199	0.538479966393432
GO_POSITIVE_REGULATION_OF_EXOCYTOSIS	66	0.01	Down	0.299715898414706	0.147327731840277	0.539214876329243
GO_ACYLGLYCEROL_METABOLIC_PROCESS	40	0.01	Up	0.364923163833871	0.147347473379953	0.539214876329243
GO_POTASSIUM_ION_IMPORT_ACROSS_PLASMA_MEMBRANE	21	0.01	Down	0.424557593417993	0.147349280156517	0.539214876329243
GO_POSITIVE_REGULATION_BY_HOST_OF_VIRAL_TRANSCRIPTION	15	0.01	Down	0.172000154571255	0.147476053622408	0.539353687675688
GO_IMMUNE_EFFECTOR_PROCESS	892	0.01	Down	0.413774451451643	0.147791246967611	0.540181011643807
GO_ORGANIC_ACID_TRANSPORT	201	0.01	Up	0.42255501633386	0.148212519869892	0.540423683391823
GO_NEUTROPHIL_HOMEOSTASIS	12	0.01	Down	0.398573411224235	0.148218026725564	0.540423683391823
GO_REGULATION_OF_METHYLATION_DEPENDENT_CHROMATIN_SILENCING	10	0.01	Down	0.140302367769141	0.148296050531732	0.540423683391823
GO_POSITIVE_REGULATION_OF_METHYLATION_DEPENDENT_CHROMATIN_SILENCING	10	0.01	Down	0.140302367769141	0.148296050531732	0.540423683391823
GO_ACUTE_INFLAMMATORY_RESPONSE	66	0.01	Down	0.608732848581635	0.148302727150515	0.540423683391823
GO_MICROBODY	112	0.01	Up	0.225254580398572	0.148704833496665	0.541563913712209
GO_POSITIVE_REGULATION_OF_CALCIUM_ION_TRANSPORT	71	0.01	Up	0.692882201162001	0.148938554174965	0.542041486554951
GO_DORSAL_VENTRAL_NEURAL_TUBE_PATTERNING	21	0.01	Up	0.302060806258753	0.149014534847672	0.542041486554951
GO_REGULATION_OF_PROTEIN_POLYMERIZATION	171	0.01	Down	0.303982424787907	0.149266130424856	0.542631543598385
GO_REGULATION_OF_AUTOPHAGY	284	0.01	Down	0.19883889237469	0.149547770737254	0.543272572741774
GO_POSITIVE_REGULATION_OF_WOUND_HEALING	36	0.01	Down	0.473456447749442	0.149634979896053	0.543272572741774
GO_NITRIC_OXIDE_BIOSYNTHETIC_PROCESS	53	0.01	Down	0.587109175595285	0.149766628988956	0.543272572741774
GO_REGULATION_OF_ACTION_POTENTIAL	34	0.01	Down	0.358290038386176	0.149800409614517	0.543272572741774
GO_VESICLE_COAT	55	0.01	Down	0.162344138191043	0.150088324417236	0.543680160403132

GO_EARLY_ENDOSOME_MEMBRANE 121 0.01 Down
0.274985444918103 0.15009190394262 0.543680160403132
GO_T_CELL_MEDIATED_IMMUNITY 78 0.01 Down
0.42582208003302 0.150215662916512 0.543803989007838
GO_NEGATIVE_REGULATION_OF_DEVELOPMENTAL_GROWTH 82 0.01 Down
0.440452027018466 0.150572215776722
0.544769917747603
GO_ACTIN_FILAMENT 79 0.01 Down 0.328931501491342
0.150670496842019 0.544800825686658
GO_REGULATION_OF_COFACTOR_METABOLIC_PROCESS 30 0.01 Up
0.337863182103507 0.150929559847917
0.545408546416288
GO_FATTY_ACID_TRANSPORT 63 0.01 Up 0.411094857294838
0.151018245186259 0.545408546416288
GO_PROTEIN_LOCALIZATION_TO_ENDOSOME 23 0.01 Down
0.248541238576278 0.151115139978795 0.545434015940108
GO_PHOSPHOLIPASE_ACTIVITY 67 0.01 Up
0.35824326543023 0.151209816358885 0.545451452831128
GO_ACTIVATION_OF_IMMUNE_RESPONSE 443 0.01 Down
0.435272029489521 0.151571079023633 0.546429941064417
GO_CAJAL_BODY 53 0.01 Down 0.115618277161241
0.151792092528501 0.546901954742155
GO_SUBSTRATE_DEPENDENT_CELL_MIGRATION 26 0.01 Down
0.448560916549259 0.15202922525046 0.547431451064972
GO_LIPASE_ACTIVITY 80 0.01 Up 0.352538389408759
0.152177880029939 0.547534859157623
GO_HISTONE_DEACETYLASE_COMPLEX 65 0.01 Down
0.157523271416985 0.152301716272483 0.547534859157623
GO_REGULATION_OF_CELL_PROLIFERATION_INVOLVED_IN_KIDNEY_DEVELOPMENT
11 0.01 Down 0.513997147161731
0.152469832348699 0.547534859157623
GO_NEGATIVE_REGULATION_OF_RECEPTOR_SIGNALING_PATHWAY_VIA_STAT 17
0.01 Down 0.199921255848462 0.152533574712268
0.547534859157623
GO_REGULATION_OF_MACROAUTOPHAGY 158 0.01 Down
0.182795231662899 0.152560601287518 0.547534859157623
GO_CHROMATIN_DISASSEMBLY 18 0.01 Down
0.232984068178408 0.152625149458924 0.547534859157623
GO_REGULATION_OF_GTPASE_ACTIVITY 392 0.01 Down
0.311495577652617 0.152740761182835 0.547534859157623
GO_MYELIN_MAINTENANCE 13 0.01 Down 0.360392618205612
0.152779451723442 0.547534859157623
GO_POSITIVE_REGULATION_OF_PROTEIN_METABOLIC_PROCESS 1243 0.01
Down 0.342105859787377 0.152986337159092
0.547952833565101
GO_REGULATION_OF_PEPTIDYL_SERINE_PHOSPHORYLATION 98 0.01
Down 0.347530348565122 0.153614274065274
0.549628089556818
GO_PERINUCLEAR_REGION_OF_CYTOPLASM 568 0.01 Down
0.304099015820232 0.153635128970173 0.549628089556818
GO_CARBOHYDRATE_TRANSMEMBRANE_TRANSPORTER_ACTIVITY 23 0.01
Up 0.622961103713534 0.153807411261313
0.549638776362476
GO_INTRINSIC_COMPONENT_OF_POSTSYNAPTIC_MEMBRANE 36 0.01 Up

0.420334794273877 0.153819186466784
0.549638776362476
GO_PIGMENT_BIOSYNTHETIC_PROCESS 46 0.01 Up
0.257767144282068 0.153936930588385 0.549735944471814
GO_MICROTUBULE_ANCHORING_AT_CENTROSOME 10 0.01 Down
0.302740602410648 0.154448389265485 0.551103171315384
GO_PROTEIN_HETEROTRIMERIZATION 12 0.01 Up
0.774971487057066 0.15450133381301 0.551103171315384
GO_REGULATION_OF_HORMONE_SECRETION 159 0.01 Down
0.375294432455661 0.154693391971342 0.5514642293940
GO_PHOSPHOTYROSINE_RESIDUE_BINDING 35 0.01 Down
0.394244646107328 0.155603429707822 0.553331164600576
GO_PHOSPHATIDYLINOSITOL_BINDING 204 0.01 Down
0.26728151296281 0.155608320560735 0.553331164600576
GO_POSITIVE_REGULATION_OF_CELL_POPULATION_PROLIFERATION 636 0.01 Down 0.429540671421227 0.155733972201114
0.553331164600576
GO_TRANSMEMBRANE_RECEPTOR_PROTEIN_SERINE_THREONINE_KINASE_BINDING 10 0.01 Down 0.71976424570161
0.155887193470484 0.553331164600576
GO_EPITHELIAL_CILIUM_MOVEMENT 19 0.01 Up
0.379667602368188 0.155959168840639 0.553331164600576
GO_MYOSIN_V_BINDING 14 0.01 Down 0.245198669406287
0.156005475888826 0.553331164600576
GO_POSITIVE_REGULATION_OF_CALCIUM_ION_DEPENDENT_EXOCYTOSIS 14
0.01 Down 0.233908720146204 0.156035673156912
0.553331164600576
GO_REGULATION_OF_TELOMERASE_ACTIVITY 43 0.01 Down
0.310741354235035 0.156142973228577 0.553331164600576
GO_PHAGOPHORE_ASSEMBLY_SITE 31 0.01 Down
0.0852566723867348 0.156185324914176 0.553331164600576
GO_AMINO_ACID_BINDING 35 0.01 Up 0.337220001613551
0.156186080180463 0.553331164600576
GO_POSITIVE_REGULATION_OF_PHOSPHORUS_METABOLIC_PROCESS 778 0.01 Down 0.40220134713499 0.156311685589961
0.553331164600576
GO_CARBON_NITROGEN_LYASE_ACTIVITY 11 0.01 Up
0.281595256006936 0.156335958860868 0.553331164600576
GO_STRIATED_MUSCLE_CELL_PROLIFERATION 42 0.01 Down
0.325971015574716 0.156415235870267 0.553331164600576
GO_T_CELL_CYTOKINE_PRODUCTION 27 0.01 Down
0.494999354625154 0.156559799708908 0.553331164600576
GO_REGULATION_OF_POLYSACCHARIDE_METABOLIC_PROCESS 33 0.01 Down 0.306212997376037 0.156584243252148
0.553331164600576
GO_CYTOKINE_SECRETION 143 0.01 Down 0.599819842930179
0.15687004739658 0.553686399272287
GO_MICROTUBULE_NUCLEATION 26 0.01 Down
0.152120033420674 0.157147250197304 0.553686399272287
GO_CONNECTIVE_TISSUE_DEVELOPMENT 190 0.01 Up
0.601967056136615 0.15717581620415 0.553686399272287
GO_STORE_OPERATED_CALCIUM_ENTRY 16 0.01 Up
0.384348713640858 0.157185981811654 0.553686399272287
GO_OXIDOREDUCTASE_ACTIVITY_ACTING_ON_THE_CH_NH2_GROUP_OF_DONORS_OXYG

EN_AS_ACCEPTOR 11 0.01 Up 0.810772183326664
0.15747215552497 0.553686399272287
GO_CHLORIDE_CHANNEL_REGULATOR_ACTIVITY 13 0.01 Down
0.247387033441208 0.157497385893705 0.553686399272287
GO_REGULATION_OF_PRODUCTION_OF_MOLECULAR_MEDIATOR_OF_IMMUNE_RESPONSE
106 0.01 Down 0.380764117305332
0.157576042943242 0.553686399272287
GO_POSITIVE_REGULATION_OF_RESPONSE_TO_CYTOKINE_STIMULUS 46 0.01
Down 0.326403440344399 0.157592562607108
0.553686399272287
GO_SEGMENT_SPECIFICATION 10 0.01 Up
0.367622811986129 0.157834078834407 0.553686399272287
GO_TOXIN_METABOLIC_PROCESS 16 0.01 Up
0.328098206577711 0.157861335851367 0.553686399272287
GO_G_PROTEIN_COUPLED_RECEPTOR_ACTIVITY 133 0.01 Up
0.503690827607553 0.158012992049857 0.553686399272287
GO_MRNA_CLEAVAGE 21 0.01 Down 0.162668912862612
0.15802142770667 0.553686399272287
GO_REGULATION_OF_LYASE_ACTIVITY 20 0.01 Up
0.421595387069058 0.158024969524943 0.553686399272287
GO_MECHANORECEPTOR_DIFFERENTIATION 41 0.01 Up
0.392801137532582 0.1580464338802 0.553686399272287
GO_REGULATION_OF_CHEMOTAXIS 152 0.01 Down
0.584622998127257 0.158052796893242 0.553686399272287
GO_SMOOTH_MUSCLE_CELL_PROLIFERATION 107 0.01 Down
0.527676093639515 0.158550791935604 0.555110644660353
GO_REGULATION_OF_TRANSCRIPTION_FROM_RNA_POLYMERASE_II_PROMOTER_IN_RE
SPONSE_TO_HYPOXIA 72 0.01 Down 0.169977210472362
0.159047365878042 0.556520214899104
GO_RIBOSOMAL_LARGE_SUBUNIT_BINDING 13 0.01 Down
0.159993945799746 0.159179926684786 0.556520214899104
GO_ADHERENS_JUNCTION_ASSEMBLY 78 0.01 Down
0.377805386924403 0.159258713722452 0.556520214899104
GO_PROTEIN_LOCALIZATION_TO_EARLY_ENDOSOME 10 0.01 Down
0.438419824567551 0.159391089429346
0.556520214899104
GO_TYPE_2_IMMUNE_RESPONSE 25 0.01 Down
0.633723058450532 0.159552074663136 0.556520214899104
GO_POSITIVE_REGULATION_OF_MACROPHAGE_DIFFERENTIATION 12 0.01
Up 0.324285669885885 0.159877022567285
0.556520214899104
GO_NEGATIVE_REGULATION_OF_SECRETION 138 0.01 Down
0.459088382178376 0.159881304754864 0.556520214899104
GO_POSITIVE_REGULATION_OF_MOLECULAR_FUNCTION 1363 0.01 Down
0.334516808917225 0.159901534421107
0.556520214899104
GO_ROUGH_ENDOPLASMIC_RETICULUM 67 0.01 Up
0.309474257209963 0.159931639510243 0.556520214899104
GO_TROPHECTODERMAL_CELL_DIFFERENTIATION 11 0.01 Down
0.376488872828654 0.159935728929884 0.556520214899104
GO_POSITIVE_REGULATION_OF_TRANSCRIPTION_INITIATION_FROM_RNA_POLYMERA
SE_II_PROMOTER 15 0.01 Down 0.262020801549915
0.159961748476188 0.556520214899104
GO_REGULATION_OF_TRANSMEMBRANE_RECEPTOR_PROTEIN_SERINE_THREONINE_KIN

ASE_SIGNALING_PATHWAY 160 0.01 Up 0.481243403433357
0.160232360234554 0.55714241637112
GO_POSITIVE_REGULATION_OF_MUSCLE_HYPERTROPHY 23 0.01 Down
0.245380176797644 0.160539334773015
0.557769759608874
GO_SWI_SNF_SUPERFAMILY_TYPE_COMPLEX 68 0.01 Down
0.139567743625915 0.160683507381417 0.557769759608874
GO_ALTERNATIVE_MRNA_SPLICING_VIA_SPLICEOSOME 65 0.01 Down
0.12319029996176 0.160688405461361
0.557769759608874
GO_CATALYTIC_COMPLEX 1229 0.01 Down 0.167477569894006
0.160900509741281 0.558031674683025
GO_REGULATION_OF_SUPEROXIDE_ANION_GENERATION 16 0.01 Up
0.389231955777035 0.160947695992419
0.558031674683025
GO_CATALYTIC_STEP_2_SPLICEOSOME 86 0.01 Down
0.0975099603135228 0.161238356610209 0.558720355582522
GO_NEGATIVE_REGULATION_OF_TELOMERE_MAINTENANCE_VIA_TELOMERE_LENGTHEN
ING 25 0.01 Down 0.158453937530393
0.161356904205403 0.55881218792413
GO_SCHWANN_CELL_DIFFERENTIATION 29 0.01 Down
0.291316762699043 0.16172205446145 0.559217843234199
GO_PROTEIN_DNA_COMPLEX_DISASSEMBLY 18 0.01 Down
0.224796141192551 0.161759832318784 0.559217843234199
GO_REGULATION_OF_SMOOTH_MUSCLE_CELL_DIFFERENTIATION 30 0.01
Up 0.479621205350071 0.16178606292967
0.559217843234199
GO_GLYCOPROTEIN_COMPLEX 12 0.01 Up 0.335576417407144
0.16184248897422 0.559217843234199
GO_POSITIVE_REGULATION_OF_INSULIN_SECRETION_INVOLVED_IN_CELLULAR_RES
PONSE_TO_GLUCOSE_STIMULUS 25 0.01 Up
0.277307099309938 0.16213517508787 0.55991049371926
GO_MALE_SEX_DIFFERENTIATION 107 0.01 Down
0.381466616091353 0.162363241262402 0.560159686793454
GO_ENZYME_ACTIVATOR_ACTIVITY 427 0.01 Down
0.319650665362917 0.162469585484445 0.560159686793454
GO_CRANIOFACIAL_SUTURE_MORPHOGENESIS 11 0.01 Up
0.358402967308522 0.16248510007945 0.560159686793454
GO_CELL_ADHESION_MOLECULE_BINDING 422 0.01 Down
0.301941201355004 0.162576407203108 0.560159686793454
GO_CHROMATIN_ORGANIZATION 630 0.01 Down
0.204177085217458 0.162880420961057 0.560576196130266
GO_ENDOPLASMIC_RETICULUM_GOLGI_INTERMEDIATE_COMPARTMENT 109 0.01
Down 0.277020428167649 0.162944748267759
0.560576196130266
GO_N_ACETYLTRANSFERASE_ACTIVITY 75 0.01 Down
0.137197486096088 0.162974301790466 0.560576196130266
GO_THIOESTER_BIOSYNTHETIC_PROCESS 46 0.01 Up
0.22047578932513 0.163257093695575 0.561230926288694
GO_POSITIVE_REGULATION_OF_EXCITATORY_POSTSYNAPTIC_POTENTIAL 18
0.01 Up 0.53794678848221 0.163556173923518
0.561623816471604
GO_ORGANIC_ANION_TRANSMEMBRANE_TRANSPORTER_ACTIVITY 125 0.01
Up 0.39565110216708 0.16363777715208

0.561623816471604
GO_MRNA_CLEAVAGE_FACTOR_COMPLEX 19 0.01 Down
0.123965064391583 0.163661257219697 0.561623816471604
GO_POSITIVE_REGULATION_OF_EPIDERMAL_CELL_DIFFERENTIATION 19
0.01 Down 0.412591860633952 0.163741419066832
0.561623816471604
GO_POSITIVE_REGULATION_OF_CD4_POSITIVE_ALPHA_BETA_T_CELL_ACTIVATION
23 0.01 Down 0.461237949200545
0.164391933425087 0.562863723419587
GO_INTERLEUKIN_1_SECRETION 38 0.01 Down
0.692245586056784 0.164416520729697 0.562863723419587
GO_AMIDE_BINDING 240 0.01 Up 0.42564063240031
0.164472777740298 0.562863723419587
GO_MUSCLE_FILAMENT_SLIDING 18 0.01 Up
0.568388141584345 0.164473767970079 0.562863723419587
GO_APICAL_PLASMA_MEMBRANE 204 0.01 Down
0.465655425001174 0.164616579946771 0.563035074285547
GO_POSITIVE_REGULATION_OF_ORGANELLE_ORGANIZATION 533 0.01
Down 0.25752707234916 0.164897750194217
0.563679189993858
GO_ASPARTATE_FAMILY_AMINO_ACID_METABOLIC_PROCESS 40 0.01
Up 0.277745342936252 0.1654293679365
0.565021178719094
GO_SKELETAL_SYSTEM_MORPHOGENESIS 156 0.01 Up
0.536009437198738 0.165476471052965 0.565021178719094
GO_REACTIVE_OXYGEN_SPECIES_BIOSYNTHETIC_PROCESS 82 0.01 Down
0.526592935394631 0.165650980922251
0.565115482581568
GO_POSITIVE_REGULATION_OF_CELL_DEVELOPMENT 383 0.01 Down
0.338260417189327 0.16569025844098
0.565115482581568
GO_MOLECULAR_FUNCTION_REGULATOR 1210 0.01 Down
0.396002097631514 0.166067427697021 0.565761488899867
GO_LYMPHOCYTE_APOPTOTIC_PROCESS 52 0.01 Down
0.434711294633165 0.166134543172998 0.565761488899867
GO_OUTFLOW_TRACT_MORPHOGENESIS 56 0.01 Up
0.550507178137007 0.166210206686841 0.565761488899867
GO_REGULATION_OF_CARBOHYDRATE_BIOSYNTHETIC_PROCESS 72 0.01
Down 0.312293319695193 0.166483004616652
0.565761488899867
GO_REGULATION_OF_DEVELOPMENTAL_GROWTH 229 0.01 Down
0.389953329520307 0.166616210121636 0.565761488899867
GO_REGULATION_OF_PROTEIN_EXPORT_FROM_NUCLEUS 39 0.01 Down
0.187443827558714 0.166643619055603
0.565761488899867
GO_NUCLEAR_INCLUSION_BODY 12 0.01 Down
0.13355628754919 0.166724564309756 0.565761488899867
GO_ENDOPLASMIC_RETICULUM_GOLGI_INTERMEDIATE_COMPARTMENT_MEMBRANE
59 0.01 Down 0.275025990815488
0.166751814476823 0.565761488899867
GO_PCG_PROTEIN_COMPLEX 44 0.01 Down 0.177773266177595
0.16695070174428 0.565761488899867
GO_MRNA_METABOLIC_PROCESS 484 0.01 Down
0.129264813443601 0.166978123600775 0.565761488899867

GO_PLATELET_ALPHA_GRANULE_LUMEN 48 0.01 Up
0.525063558696129 0.167036644700261 0.565761488899867
GO_GUANOSINE_CONTAINING_COMPOUND_METABOLIC_PROCESS 21 0.01
Down 0.285401731258027 0.167038534299664
0.565761488899867
GO_SNRNA_TRANSCRIPTION 71 0.01 Down 0.131649773741848
0.167091146367561 0.565761488899867
GO_COPII_VESICLE_COAT 15 0.01 Down 0.149065156611414
0.167394358421284 0.566008810908047
GO_SIALYLATION 15 0.01 Up 0.452724286243688
0.16748285888734 0.566008810908047
GO_SIALYLTRANSFERASE_ACTIVITY 15 0.01 Up
0.452724286243688 0.16748285888734 0.566008810908047
GO_PROTEIN_K63_LINKED_DEUBIQUITINATION 31 0.01 Down
0.206455046164072 0.167537116323795 0.566008810908047
GO_CELLULAR_RESPONSE_TO_TOPOLOGICALLY_INCORRECT_PROTEIN 145 0.01
Down 0.219467290232612 0.167954442654683
0.566701574729266
GO_INTRA_S_DNA_DAMAGE_CHECKPOINT 13 0.01 Up
0.156441839944101 0.168028178005462 0.566701574729266
GO_INTERLEUKIN_8_PRODUCTION 50 0.01 Down
0.439970735757279 0.168247072485097 0.566701574729266
GO_REGULATION_OF_MAPK_CASCADE 541 0.01 Down
0.412121177336247 0.168252452078865 0.566701574729266
GO_GLIOGENESIS 195 0.01 Down 0.368380519823966
0.168422167715769 0.566701574729266
GO_RNA_CATABOLIC_PROCESS 365 0.01 Down
0.162324888071542 0.168532022275334 0.566701574729266
GO_POSITIVE_REGULATION_OF_BLOOD_VESSEL_ENDOTHELIAL_CELL_MIGRATION
48 0.01 Down 0.417236907476156
0.168587848746909 0.566701574729266
GO_POSITIVE_REGULATION_OF_POSTTRANSCRIPTIONAL_GENE_SILENCING 22
0.01 Down 0.205953034333452 0.168612311191925
0.566701574729266
GO_NEPHRIC_DUCT_DEVELOPMENT 10 0.01 Up
0.622787226827221 0.168654734313679 0.566701574729266
GO_REGULATION_OF_POSITIVE_CHEMOTAXIS 18 0.01 Down
0.677096698441823 0.168675629309139 0.566701574729266
GO_LEUKOCYTE_DIFFERENTIATION 362 0.01 Down
0.448363221385729 0.169199051808768 0.567359974633553
GO_REGULATION_OF_TRANSFERASE_ACTIVITY 778 0.01 Down
0.323771003357045 0.169243754134947 0.567359974633553
GO_POSITIVE_REGULATION_OF_PROTEIN_CATABOLIC_PROCESS 189 0.01
Down 0.233419069127695 0.169307276729873
0.567359974633553
GO_ICOSANOID_BIOSYNTHETIC_PROCESS 26 0.01 Up
0.483606157549795 0.169370600807336 0.567359974633553
GO_CYTOCHROME_COMPLEX 23 0.01 Up 0.220855076781816
0.16937446985322 0.567359974633553
GO_NEGATIVE_REGULATION_OF_CHEMOKINE_PRODUCTION 15 0.01 Down
0.467640865838179 0.169432323177505
0.567359974633553
GO_CD4_POSITIVE_OR_CD8_POSITIVE_ALPHA_BETA_T_CELL_LINEAGE_COMMITMENT
13 0.01 Down 0.609941172508605

0.169551887285938	0.567376573523256
GO_POSITIVE_REGULATION_OF_CELL_PROJECTION_ORGANIZATION	299	0.01	Down	0.327861437753158	0.169624193863401	0.567376573523256
GO_POSITIVE_REGULATION_OF_UBIQUITIN_DEPENDENT_PROTEIN_CATABOLIC_PROCESS	87	0.01	Down	0.198179840136757	0.169789366873651	0.567616325049524
GO_POSITIVE_REGULATION_OF_HEART_RATE	15	0.01	Down	0.527457679221881	0.169952001596638	0.567847331696858
GO_VESICLE_MEDIATED_TRANSPORT_TO_THE_PLASMA_MEMBRANE	83	0.01	Down	0.192830086009434	0.170120538966145	0.567990373162603
GO_C4_DICARBOXYLATE_TRANSPORT	16	0.01	Down	0.342603836017193	0.170331591538483	0.567990373162603
GO_NUCLEOTIDE_EXCISION_REPAIR_PREINCISION_COMPLEX_STABILIZATION	21	0.01	Up	0.127559604731945	0.170417783703518	0.567990373162603
GO_REGULATION_OF_TELOMERE_CAPPING	23	0.01	Down	0.154854218373233	0.170560320870319	0.567990373162603
GO_ALCOHOL_DEHYDROGENASE_NADPPLUS_ACTIVITY	14	0.01	Up	0.468990701036541	0.170564876637342	0.567990373162603
GO_B_CELL_ACTIVATION	161	0.01	Down	0.391934954074986	0.17063981718176	0.567990373162603
GO_CELLULAR_PROTEIN_CATABOLIC_PROCESS	671	0.01	Down	0.204430912111864	0.170719991397137	0.567990373162603
GO_HOMEOSTASIS_OF_NUMBER_OF_CELLS	196	0.01	Down	0.357273971278195	0.170743276399564	0.567990373162603
GO_REGULATION_OF_KINASE_ACTIVITY	691	0.01	Down	0.339476242266599	0.171007004333332	0.568458019882614
GO_RAS_PROTEIN_SIGNAL_TRANSDUCTION	375	0.01	Down	0.330443415936579	0.17107112540364	6	0.568458019882614
GO_ACID_AMINO_ACID_LIGASE_ACTIVITY	12	0.01	Up	0.207342941742637	0.171313027051162	0.568650696822654
GO_REGULATION_OF_ESTABLISHMENT_OR_MAINTENANCE_OF_CELL_POLARITY	22	0.01	Down	0.235653669649548	0.171422510997318	0.568650696822654
GO_NUCLEAR_PORE_OUTER_RING	10	0.01	Down	0.107674106671369	0.17153396573914	0.568650696822654
GO_SUBSTRATE_ADHESION_DEPENDENT_CELL_SPREADING	76	0.01	Up	0.299360719143989	0.171546934999084	0.568650696822654
GO_REGULATION_OF_LAMELLIPODIUM_ORGANIZATION	37	0.01	Down	0.219246195331289	0.171597443020771	0.568650696822654
GO_CELLULAR_ALDEHYDE_METABOLIC_PROCESS	29	0.01	Up	0.451213062097602	0.17195737685154	0.569400354984376
GO_SULFUR_AMINO_ACID_METABOLIC_PROCESS	12	0.01	Up	0.123866562800497	0.172078867851839	0.569400354984376
GO_REGULATION_OF_TRANSPOSITION	15	0.01	Down	0.504195632995103	0.172105032349914	0.569400354984376
GO_INTRAMOLECULAR_OXIDOREDUCTASE_ACTIVITY	36	0.01	Up	0.243181910937619	0.172591085285629	0.570595353837513

GO_NEPHRON_TUBULE_EPITHELIAL_CELL_DIFFERENTIATION 12 0.01 Down 0.483884231729626 0.172654202767174 0.570595353837513
GO_MEMBRANE_FUSION 118 0.01 Down 0.230147932726608 0.172851001776715 0.570934946564983
GO_MITOTIC_METAPHASE_PLATE_CONGRESSION 38 0.01 Down 0.152394802655384 0.174045854670508 0.574121884365754
GO_NEGATIVE_REGULATION_OF_LIPID_LOCALIZATION 32 0.01 Down 0.422561854177175 0.174091977197219 0.574121884365754
GO_INTRACILIARY_RETROGRADE_TRANSPORT 12 0.01 Up 0.160808896361614 0.174125508737597 0.574121884365754
GO_ENDOLYSOSOME 18 0.01 Up 0.357953580530183 0.174194121397088 0.574121884365754
GO_ORGAN_INDUCTION 13 0.01 Down 0.524536459352448 0.174382149001411 0.574429748555381
GO_POSITIVE_REGULATION_OF_BLOOD_VESSEL_DIAMETER 35 0.01 Up 0.53989763532175 0.174698315195319 0.57487180115448
GO_REGULATION_OF_TUMOR_NECROSIS_FACTOR_MEDIATED_SIGNALING_PATHWAY 51 0.01 Down 0.314135494508428 0.17474049534149 0.57487180115448
GO_PROTEIN_POLYMERIZATION 221 0.01 Down 0.28195206458032 0.174814454478343 0.57487180115448
GO_NEURON_PROJECTION_MEMBRANE 26 0.01 Up 0.603825665887942 0.174986945080513 0.57487180115448
GO_CELL_CORTEX 248 0.01 Down 0.283770295167305 0.175066750180403 0.57487180115448
GO_NEGATIVE_REGULATION_OF_PROTEIN_CATABOLIC_PROCESS 107 0.01 Down 0.259209369352445 0.175084493548778 0.57487180115448
GO_CELLULAR_RESPONSE_TO_INORGANIC_SUBSTANCE 155 0.01 Up 0.444821804055496 0.175428637174361 0.575335709274511
GO_REGULATION_OF_CARDIAC_MUSCLE_CELL_DIFFERENTIATION 31 0.01 Up 0.628496945993979 0.175551748301089 0.575335709274511
GO_INOSITOL_LIPID_MEDIATED_SIGNALING 138 0.01 Down 0.43899335089634 0.175695756819713 0.575335709274511
GO_AMMONIUM_ION_METABOLIC_PROCESS 39 0.01 Up 0.361210584711697 0.175696506003048 0.575335709274511
GO_PHOSPHATIDYLSERINE_BINDING 41 0.01 Up 0.622413344073978 0.175699621972483 0.575335709274511
GO_INTERMEDIATE_FILAMENT_BINDING 14 0.01 Down 0.394480420186024 0.175805548259356 0.575372228292479
GO_NEGATIVE_REGULATION_OF_INTERLEUKIN_1_PRODUCTION 16 0.01 Down 0.686704528250285 0.176555855425814 0.577516486147693
GO_GOLGI_TO_PLASMA_MEMBRANE_TRANSPORT 58 0.01 Down 0.198349146293112 0.176966234419322 0.57819660312765
GO_NUCLEOSIDE_MONOPHOSPHATE_KINASE_ACTIVITY 17 0.01 Down 0.2912813361124 0.177305970357713 0.57819660312765
GO_POSITIVE_REGULATION_OF_HEMOPOIESIS 138 0.01 Down 0.40138646906559 0.177362918938514 0.57819660312765

GO_REGULATION_OF_MITOCHONDRIAL_TRANSLATION 25 0.01 Up
0.114964544014709 0.177402101324639
0.57819660312765
GO_NEGATIVE_REGULATION_OF_CELL_DEVELOPMENT 228 0.01 Down
0.395252560640863 0.177555746391148
0.57819660312765
GO_POTASSIUM_ION_TRANSPORT 106 0.01 Down
0.518820783245963 0.177670039642877 0.57819660312765
GO_NEGATIVE_REGULATION_OF_EXTRINSIC_APOPTOTIC_SIGNALING_PATHWAY 81
0.01 Down 0.350605300791814 0.177855490398017
0.57819660312765
GO_HISTONE_H4_K16_ACETYLATION 20 0.01 Down
0.112011367892935 0.177962125413117 0.57819660312765
GO_REGULATION_OF_BONE_DEVELOPMENT 16 0.01 Up
1.0410280965716 0.177974873131623 0.57819660312765
GO_NUCLEAR_RECEPTOR_TRANSCRIPTION_COACTIVATOR_ACTIVITY 59 0.01
Down 0.239973235382785 0.178005023303113
0.57819660312765
GO_FATTY_ACID_LIGASE_ACTIVITY 10 0.01 Up
0.351009595431047 0.178019184898685 0.57819660312765
GO_KINASE_REGULATOR_ACTIVITY 172 0.01 Down
0.297073055715143 0.178160803456096 0.57819660312765
GO_ANION_CHANNEL_ACTIVITY 45 0.01 Down
0.352943951071373 0.178224960778531 0.57819660312765
GO_PROTEIN_AUTOPHOSPHORYLATION 191 0.01 Down
0.373514737867455 0.178327003203098 0.57819660312765
GO_REGULATION_OF_CELLULAR_RESPONSE_TO_STRESS 628 0.01 Down
0.291697093468805 0.178358975924006
0.57819660312765
GO_RESPONSE_TO_LIPID 649 0.01 Down 0.438193581365736
0.178374482229554 0.57819660312765
GO_PHOSPHATIDYLINOSITOL_3_KINASE_SIGNALING 107 0.01 Down
0.463785706239839 0.17838284263846
0.57819660312765
GO_CILIUM_MOVEMENT 37 0.01 Up 0.322292336140827
0.179055844340251 0.580067664116191
GO_MISFOLDED_PROTEIN_BINDING 21 0.01 Down
0.245167933930637 0.179222356823309 0.580067664116191
GO_REGULATION_OF_CELLULAR_PROTEIN_CATABOLIC_PROCESS 223 0.01
Down 0.223512100110708 0.179246736597262
0.580067664116191
GO_INTERLEUKIN_17_PRODUCTION 18 0.01 Down
0.604416943526996 0.179663639600642 0.581107062341766
GO_NEGATIVE_REGULATION_OF_PROTEOLYSIS_INVOLVED_IN_CELLULAR_PROTEIN_C
ATABOLIC_PROCESS 62 0.01 Down 0.203498839818434
0.180119611829535 0.582271652511773
GO_MODULATION_OF_EXCITATORY_POSTSYNAPTIC_POTENTIAL 26 0.01
Up 0.427999299625363 0.180348060933032
0.582434678213422
GO_POSITIVE_REGULATION_OF_INTRACELLULAR_SIGNAL_TRANSDUCTION 746
0.01 Down 0.397066816534251 0.180477649775416
0.582434678213422
GO_EMBRYONIC_EYE_MORPHOGENESIS 24 0.01 Down
0.468956825637788 0.180513052837916 0.582434678213422

GO_REGULATION_OF_EPITHELIAL_TUBE_FORMATION 17 0.01 Up
0.549253232957646 0.180553790874265
0.582434678213422
GO_REGULATION_OF_LONG_TERM_SYNAPTIC_POTENTIATION 26 0.01
Up 0.375938969038067 0.181345687286393
0.58467852762384
GO_REGULATION_OF_CELLULAR_COMPONENT_SIZE 298 0.01 Down
0.341424790974453 0.18211310452833
0.585065210586775
GO_SOLUTE_SODIUM_SYMPORTER_ACTIVITY 29 0.01 Down
0.622727110229428 0.182186644631829 0.585065210586775
GO_GTPASE_REGULATOR_ACTIVITY 261 0.01 Down
0.282751698902494 0.182207889869559 0.585065210586775
GO_NEGATIVE_REGULATION_OF_CELL_DIFFERENTIATION 477 0.01 Down
0.427809738680233 0.182220857409382
0.585065210586775
GO_RESPONSE_TO_COLD 36 0.01 Down 0.382544661116772
0.182261966700253 0.585065210586775
GO_CENTRIOLE_ASSEMBLY 36 0.01 Down 0.129193536278425
0.182297160363024 0.585065210586775
GO_GLYCOSYL_COMPOUND_METABOLIC_PROCESS 51 0.01 Up
0.340882090848491 0.182366602983915 0.585065210586775
GO_RESPONSE_TO_PEPTIDE 362 0.01 Down 0.31022852020253
0.182397679349264 0.585065210586775
GO_PROTEASOMAL_PROTEIN_CATABOLIC_PROCESS 429 0.01 Down
0.186209117954135 0.182414013854524
0.585065210586775
GO_BONE_DEVELOPMENT 152 0.01 Up 0.545661886430201
0.182429326905084 0.585065210586775
GO_ANCHORED_COMPONENT_OF_SYNAPTIC_VESICLE_MEMBRANE 10 0.01
Down 0.171126274755894 0.182971468732821
0.586355642407348
GO_POSITIVE_REGULATION_OF_CALCINEURIN_MEDIATED_SIGNALING 14
0.01 Down 0.379304519094607 0.183148181499233
0.586355642407348
GO_MITOTIC_CYTOKINETIC_PROCESS 23 0.01 Down
0.202413906876764 0.183292578947526 0.586355642407348
GO_POSITIVE_REGULATION_OF_DNA_BINDING 46 0.01 Down
0.337424560351976 0.183333171411415 0.586355642407348
GO_PHOSPHATIDYLINOSITOL_BISPHOSPHATE_BINDING 78 0.01 Down
0.282268854656061 0.183350176207283
0.586355642407348
GO_RENAL_FILTRATION 14 0.01 Up 0.601350927599197
0.183411195014257 0.586355642407348
GO_PHOSPHOLIPASE_A2_ACTIVITY 20 0.01 Up
0.406135238767104 0.183751283943792 0.587046765266549
GO_NEGATIVE_REGULATION_OF_CYTOKINE_SECRETION 40 0.01 Down
0.74987769009314 0.183820771005058
0.587046765266549
GO_MOTOR_NEURON_AXON_GUIDANCE 16 0.01 Down
0.471286310811053 0.183993512658387 0.587289492822853
GO_NEGATIVE_REGULATION_OF_LEUKOCYTE_CHEMOTAXIS 11 0.01 Up
1.11844129525343 0.184453169747582
0.5884472903508

GO_RESPONSE_TO_INTERLEUKIN_15 14 0.01 Down
0.406744486743012 0.184770863105956 0.588708533519889
GO_HISTONE_MRNA_METABOLIC_PROCESS 25 0.01 Down
0.124476153293163 0.184808652604663 0.588708533519889
GO_GLOBAL_GENOME_NUCLEOTIDE_EXCISION_REPAIR 26 0.01 Up
0.15231515798032 0.184825970167832
0.588708533519889
GO_HISTONE_DEACETYLASE_BINDING 96 0.01 Down
0.215607642051815 0.185069337186614 0.588995338861454
GO_LAMININ_BINDING 26 0.01 Up 0.520970721117644
0.185158864308807 0.588995338861454
GO_CELLULAR_RESPONSE_TO_THYROID_HORMONE_STIMULUS 14 0.01
Up 0.375265044254813 0.185386030612262
0.588995338861454
GO_POSITIVE_REGULATION_OF_PROTEIN_LOCALIZATION_TO_NUCLEUS 64
0.01 Down 0.256282238520013 0.185425651413451
0.588995338861454
GO_DOPAMINE_METABOLIC_PROCESS 12 0.01 Up
0.370824066513634 0.185456958803945 0.588995338861454
GO_RESPONSE_TO_INTERFERON_BETA 23 0.01 Down
0.25837364729515 0.185498120227821 0.588995338861454
GO_ENERGY_DERIVATION_BY_OXIDATION_OF_ORGANIC_COMPOUNDS 229 0.01
Up 0.19309798578629 0.185674009626145
0.58920139487092
GO_MYELOID_CELL_HOMEOSTASIS 120 0.01 Down
0.321579158304027 0.185899017553642 0.58920139487092
GO_REGULATION_OF_COAGULATION 56 0.01 Up
0.599256495534612 0.185931835419691 0.58920139487092
GO_POLYSACCHARIDE_METABOLIC_PROCESS 77 0.01 Down
0.293148696027492 0.186028591912637 0.58920139487092
GO_REGULATION_OF_CARBOHYDRATE_METABOLIC_PROCESS 159 0.01 Down
0.312410659472786 0.186090592314985
0.58920139487092
GO_CYTOKINE_SECRETION_INVOLVED_IN_IMMUNE_RESPONSE 14 0.01
Down 0.395386438713126 0.186203198105143
0.58920139487092
GO_REGULATION_OF_ERYTHROCYTE_DIFFERENTIATION 39 0.01 Down
0.358968770261382 0.186242377986706
0.58920139487092
GO_CRISTAE_FORMATION 31 0.01 Up 0.122018153054316
0.186507579402852 0.589733080497248
GO_MITOTIC_SPINDLE_POLE 26 0.01 Down 0.194070231733336
0.186705411820363 0.59005130409236
GO_NEGATIVE_REGULATION_OF_INTRACELLULAR_SIGNAL_TRANSDUCTION 431
0.01 Down 0.335609841208267 0.186870869984738
0.590235669291956
GO_POSITIVE_REGULATION_OF_SPROUTING_ANGIOGENESIS 31 0.01
Down 0.580512503089256 0.186960656872968
0.590235669291956
GO_REGULATION_OF_MYELOID_CELL_DIFFERENTIATION 177 0.01 Down
0.365183749760805 0.187055415535781
0.590235669291956
GO_REGULATION_OF_PROTEIN_DEPOLYMERIZATION 64 0.01 Down
0.312964398923714 0.187265390475874

0.590371348911152
GO_RECEPTOR_SIGNALING_COMPLEX_SCAFFOLD_ACTIVITY 19 0.01 Down
0.293841439083902 0.187308976258689
0.590371348911152
GO_PROCESS_UTILIZING_AUTOPHAGIC_MECHANISM 442 0.01 Down
0.20056444908432 0.187455555918125
0.590371348911152
GO_DNA_REPLICATION_DEPENDENT_NUCLEOSOME_ORGANIZATION 14 0.01
Up 0.158426758165386 0.187572107933428
0.590371348911152
GO_CYTOSOLIC_PART 212 0.01 Up 0.163908575246354
0.187584637135499 0.590371348911152
GO_PROTEASOMAL_UBIQUITIN_INDEPENDENT_PROTEIN_CATABOLIC_PROCESS 21
0.01 Down 0.150019968928619 0.1878857267628
0.591012563304124
GO_LEUKOCYTE_APOPTOTIC_PROCESS 82 0.01 Down
0.491973071553195 0.188143140102796 0.591515796770624
GO_OVULATION 15 0.01 Down 0.631019301717978
0.188313586090289 0.591745228340655
GO_NEGATIVE_REGULATION_OF_METAPHASE_ANAPHASE_TRANSITION_OF_CELL_CYCL
E 32 0.01 Up 0.240329899403831
0.188467732505593 0.591923230233552
GO_TRNA_SPECIFIC_RIBONUCLEASE_ACTIVITY 16 0.01 Up
0.142241025498264 0.188808837878537 0.592075456470727
GO_PROTEIN_TRANSPORT_WITHIN_LIPID_BILAYER 35 0.01 Down
0.239268630361769 0.188809454407386
0.592075456470727
GO_REGULATION_OF_SMOOTHENED_SIGNALING_PATHWAY 65 0.01 Up
0.354613472855773 0.188857715885549
0.592075456470727
GO_OTHER_ORGANISM 65 0.01 Down 0.230762905452254
0.18890630195747 0.592075456470727
GO_SEQUESTERING_OF_METAL_ION 10 0.01 Down
1.27716027467895 0.189113682424237 0.592419590298009
GO_NEUROMUSCULAR_PROCESS_CONTROLLING_BALANCE 26 0.01 Up
0.320928441422065 0.189476107283374
0.592938144089388
GO_OUTER_MEMBRANE 180 0.01 Up 0.235939659429757
0.189566104640349 0.592938144089388
GO_SEX_CHROMOSOME 24 0.01 Down 0.139675288493395
0.189572218362296 0.592938144089388
GO_MEIOSIS_I_CELL_CYCLE_PROCESS 68 0.01 Up
0.24014470173448 0.189926609409091 0.593327326543498
GO_SKELETAL_SYSTEM_DEVELOPMENT 353 0.01 Up
0.535124968147056 0.189985195078413 0.593327326543498
GO_ENDODERM_FORMATION 40 0.01 Up 0.656435063900459
0.189989840685317 0.593327326543498
GO_EUKARYOTIC_TRANSLATION_INITIATION_FACTOR_3_COMPLEX 16 0.01
Down 0.122777575950399 0.190091400280716
0.593339275631994
GO_HISTONE_LYSINE_N_METHYLTRANSFERASE_ACTIVITY 40 0.01 Down
0.208793480867771 0.19026579751803
0.593578446419301
GO_PURINE_NUCLEOSIDE_MONOPHOSPHATE_BIOSYNTHETIC_PROCESS 19 0.01

Up	0.215651436123501	0.190381629348926
0.593634756947782
GO_TRANSMEMBRANE_SIGNALING_RECEPTOR_ACTIVITY	368	0.01	Up
	0.508990158388726	0.190832414972946
0.594489311995676
GO_COATED_MEMBRANE	92	0.01	Down	0.16866775401103
	0.190885417260076	0.594489311995676
GO_INTERLEUKIN_1_BETA_SECRETION 32	0.01	Down
0.72988879020172	0.19105769964231	0.594489311995676
GO_UTERUS_DEVELOPMENT	14	0.01	Down	0.3844715161108
0.191127191835188	0.594489311995676
GO_RAB_GUANYL_NUCLEOTIDE_EXCHANGE_FACTOR_ACTIVITY	35	0.01
	Down	0.202450338222986	0.191145303412215
0.594489311995676
GO_REGULATION_OF_DNA_BINDING	97	0.01	Down
0.314704095288925	0.19175250611328	0.596072434518037
GO_REGULATION_OF_CELL_MIGRATION_INVOLVED_IN_SPROUTING_ANGIOGENESIS
	35	0.01	Down	0.449976745395786
0.192291343976294	0.597174273949696
GO_FATTY_ACYL_COA_BINDING	18	0.01	Up
0.217746955590625	0.192303690589961	0.597174273949696
GO_CELLULAR_MACROMOLECULE_CATABOLIC_PROCESS	1019	0.01	Down
	0.196262737220205	0.192423667617805
0.597241352815795
GO_BLEB_ASSEMBLY	11	0.01	Up	0.691111438813541
	0.192538022590347	0.597290922404701
GO_CALCIUM_MEDIATED_SIGNALING	131	0.01	Down
0.434882294458444	0.193030486554419	0.598512810966231
GO_MEGAKARYOCYTE_DIFFERENTIATION	64	0.01	Down
0.324121343629525	0.193481004731222	0.599482692031068
GO_CARBON_OXYGEN_LYASE_ACTIVITY 56	0.01	Up
0.227168562880679	0.193785526398694	0.599482692031068
GO_REGULATION_OF_PROTEIN_COMPLEX_ASSEMBLY	352	0.01	Down
	0.271659803431857	0.193802577448665
0.599482692031068
GO_MICROTUBULE_END	24	0.01	Down	0.180001311094665
	0.193817517557465	0.599482692031068
GO_REGULATION_OF_PROTEASOMAL_UBIQUITIN_DEPENDENT_PROTEIN_CATABOLIC_P
ROCESS	114	0.01	Down	0.20046583598207
0.193837016052872	0.599482692031068
GO_CARDIAC_EPITHELIAL_TO_MESENCHYMAL_TRANSITION 23	0.01	Down
	0.496714896089314	0.19413198045163
0.600089232852262
GO_REGULATION_OF_CELLULAR_AMINE_METABOLIC_PROCESS	64	0.01
	Down	0.187583513893096	0.194295593567662
0.600289337684111
GO_PROTEIN_BINDING_INVOLVED_IN_HETEROTYPIC_CELL_CELL_ADHESION	10
	0.01	Down	0.211027881830433	0.194457318340664
	0.600327041711437
GO_RESPONSE_TO_AMMONIUM_ION	71	0.01	Down
0.305616609740473	0.194505565977005	0.600327041711437
GO_ENERGY_COUPLED_PROTON_TRANSPORT_DOWN_ELECTROCHEMICAL_GRADIENT
	22	0.01	Up	0.10212837552163
0.19468364048592	0.600571332007123

GO_NEGATIVE_REGULATION_OF_HORMONE_SECRETION 38 0.01 Down
0.435747587047909 0.195393860078235
0.6017706851565
GO_REGULATION_OF_INTRINSIC_APOPTOTIC_SIGNALING_PATHWAY_IN_RESPONSE_T
O_DNA_DAMAGE 38 0.01 Up 0.355970695098949
0.195396240383144 0.6017706851565
GO_OVULATION_CYCLE 44 0.01 Down 0.500862160428795
0.195467239817734 0.6017706851565
GO_REGULATION_OF_ORGAN_GROWTH 62 0.01 Down
0.357504410835799 0.195560089792863 0.6017706851565
GO_CERAMIDE_CATABOLIC_PROCESS 16 0.01 Up
0.141796395253736 0.195656684434427 0.6017706851565
GO_REPLICATIVE_SENESCENCE 12 0.01 Up
0.347620696204361 0.195690850988988 0.6017706851565
GO_HAIR_FOLLICLE_MATURATION 14 0.01 Down
0.443883995989081 0.195839112255319 0.6017706851565
GO_POSITIVE_REGULATION_OF_RNA_SPLICING 28 0.01 Down
0.130009034872582 0.195963153850764 0.6017706851565
GO_MICROTUBULE_ANCHORING_AT_MICROTUBULE_ORGANIZING_CENTER 12
0.01 Down 0.253867571835822 0.19596452718735
0.6017706851565
GO_GLUCOSE_IMPORT 44 0.01 Down 0.249416093130943
0.196146636206441 0.602008496688282
GO_ORGANIC_ACID_TRANSMEMBRANE_TRANSPORT 91 0.01 Up
0.406927134663267 0.196270070723908 0.602008496688282
GO_NEGATIVE_REGULATION_OF_MACROAUTOPHAGY 30 0.01 Down
0.165051876825372 0.196431785267899
0.602008496688282
GO_NEGATIVE_REGULATION_OF_AXON_EXTENSION 34 0.01 Down
0.400216696497599 0.196544769274692
0.602008496688282
GO_T_CELL_DIFFERENTIATION_IN_THYMUS 57 0.01 Down
0.369055460439534 0.196636999173152 0.602008496688282
GO_MEMBRANE_LIPID_METABOLIC_PROCESS 174 0.01 Up
0.304504345595646 0.196817068031727 0.602008496688282
GO_CENTRAL_NERVOUS_SYSTEM_NEURON_AXONOGENESIS 28 0.01 Down
0.358330892938225 0.196827917918289
0.602008496688282
GO_ZINC_ION_TRANSPORT 18 0.01 Up 0.212733454581825
0.196862398777046 0.602008496688282
GO_TRANSLATION_PREINITIATION_COMPLEX 18 0.01 Down
0.114139787067868 0.197127600560395 0.602008496688282
GO_CHLORIDE_CHANNEL_COMPLEX 20 0.01 Down
0.420704603106179 0.197239001844322 0.602008496688282
GO_NEGATIVE_REGULATION_OF_AUTOPHAGY 66 0.01 Down
0.181386532032105 0.197249755433166 0.602008496688282
GO_PROTEIN_N_TERMINUS_BINDING 98 0.01 Down
0.220105641969054 0.19725759307678 0.602008496688282
GO_CARBOHYDRATE_DERIVATIVE_METABOLIC_PROCESS 752 0.01 Up
0.316291157632917 0.197380495876725
0.602008496688282
GO_NEGATIVE_REGULATION_OF_INTERLEUKIN_10_PRODUCTION 14 0.01
Down 1.03528192064136 0.197488288623514
0.602008496688282

GO_PROTEIN_CATABOLIC_PROCESS 789 0.01 Down
0.221982486037891 0.197656673552719 0.602008496688282
GO_AMYLOID_BETA_BINDING 55 0.01 Up 0.545126307144948
0.197801017684748 0.602008496688282
GO_POSITIVE_REGULATION_OF_ORGAN_GROWTH 28 0.01 Up
0.364865241919309 0.19786070563584 2 0.602008496688282
GO_REGULATION_OF_EARLY_ENDOSOME_TO_LATE_ENDOSOME_TRANSPORT 17
0.01 Down 0.168805438401126 0.197917644262244
0.602008496688282
GO_NEGATIVE_REGULATION_OF_AUTOPHAGOSOME_ASSEMBLY 11 0.01
Down 0.141899068022451 0.19803545366328
0.602008496688282
GO_REGULATION_OF_MYELINATION 27 0.01 Down
0.202166687051519 0.198061049845035 0.602008496688282
GO_EXPORT_ACROSS_PLASMA_MEMBRANE 14 0.01 Down
0.789438135840525 0.198221195070452 0.602008496688282
GO_CHROMATIN 440 0.01 Down 0.196344883282401
0.198223519169803 0.602008496688282
GO_LIPID_MODIFICATION 194 0.01 Up 0.29407122120856
0.198604260039109 0.602580645993017
GO_NEGATIVE_REGULATION_OF_FIBROBLAST_PROLIFERATION 25 0.01
Up 0.302188471861326 0.198987957961309
0.602580645993017
GO_CELLULAR_RESPONSE_TO_CORTICOSTEROID_STIMULUS 45 0.01 Down
0.447041995159436 0.199001378169503
0.602580645993017
GO_BRANCHING_INVOLVED_IN_SALIVARY_GLAND_MORPHOGENESIS 20 0.01
Up 0.611403556891571 0.199019286368692
0.602580645993017
GO_NEGATIVE_REGULATION_OF_CALCIUM_ION_TRANSPORT_INTO_CYTOSOL 13
0.01 Up 0.54082730520023 0.199020298564361
0.602580645993017
GO_STRUCTURAL_MOLECULE_ACTIVITY 579 0.01 Up
0.438451875253998 0.199075613938429 0.602580645993017
GO_INNER_MITOCHONDRIAL_MEMBRANE_ORGANIZATION 44 0.01 Up
0.110188404776601 0.199106700026683
0.602580645993017
GO_UBIQUITIN_LIKE_MODIFIER_ACTIVATING_ENZYME_ACTIVITY 10 0.01
Down 0.116568953216308 0.199331988792034
0.602839616810907
GO_HEMIDESMOSOME_ASSEMBLY 12 0.01 Down
0.364434679720374 0.199390866505732 0.602839616810907
GO_TELOMERE_ORGANIZATION 144 0.01 Down
0.160805402987904 0.200278544191315 0.605222021794662
GO_HETEROCHROMATIN 68 0.01 Down 0.189474650634741
0.200627683068332 0.605975454680518
GO_POSITIVE_REGULATION_OF_CELL_MATRIX_ADHESION 42 0.01 Down
0.495811158084959 0.201020552499786
0.606860156253706
GO_REGULATION_OF_NECROTIC_CELL_DEATH 23 0.01 Down
0.332011644664185 0.201249321182916 0.607248821521613
GO_POTASSIUM_ION_HOMEOSTASIS 13 0.01 Down
0.52417819645474 0.201869950416784 0.608266509824021
GO_MUSCLE_CELL_APOPTOTIC_PROCESS 50 0.01 Down

0.391890395550707	0.201936922549761	0.608266509824021
GO_MANNOSYLTRANSFERASE_ACTIVITY 23	0.01	Up
0.137148688388298	0.202005269191253	0.608266509824021
GO_NEGATIVE_REGULATION_OF_PROTEIN_METABOLIC_PROCESS	843	0.01
	Down	0.353614527556053	0.202072435138629
0.608266509824021
GO_INSULIN_RECEPTOR_SIGNALING_PATHWAY	109	0.01	Down
0.21851089268841	0.202087555644054	0.608266509824021
GO_ESTABLISHMENT_OF_MITOCHONDRION_LOCALIZATION	21	0.01	Down
	0.238513170884908	0.202542489534307
0.609047855008979
GO_REGULATION_OF_INTRINSIC_APOPTOTIC_SIGNALING_PATHWAY_IN_RESPONSE_T
O_DNA_DAMAGE_BY_P53_CLASS_MEDIATOR	18	0.01	Up
0.260468027338573	0.202618302902592	0.609047855008979
GO_NUCLEOSIDE_CATABOLIC_PROCESS 23	0.01	Down
0.404950453398453	0.202648108568298	0.609047855008979
GO_TRANSCRIPTION_COREPRESSOR_ACTIVITY	204	0.01	Down
0.229649925509066	0.203083158514362	0.609601831896922
GO_ADENYL_NUCLEOTIDE_BINDING	1256	0.01	Down
0.259054764703865	0.203085517026997	0.609601831896922
GO_ACTIVATED_T_CELL_PROLIFERATION	30	0.01	Down
0.463880041303948	0.203139949217491	0.609601831896922
GO_INTERLEUKIN_35_MEDIATED_SIGNALING_PATHWAY	10	0.01	Down
	0.552051698825765	0.203234081330814
0.609601831896922
GO_DEVELOPMENTAL_PROCESS_INVOLVED_IN_REPRODUCTION	443	0.01
	Down	0.371769804013797	0.203964216467044
0.610700474176781
GO_REGULATION_OF_CELL_DIFFERENTIATION	1309	0.01	Down
0.387956601650475	0.204088323389069	0.610700474176781
GO_REGULATION_OF_PEPTIDYL_TYROSINE_PHOSPHORYLATION	175	0.01
	Down	0.49399430352591	0.204311259447824
0.610700474176781
GO_PHOSPHATIDYLCHOLINE_METABOLIC_PROCESS	28	0.01	Up
	0.406314017006248	0.20436925335815
0.610700474176781
GO_POSITIVE_REGULATION_OF_EPITHELIAL_CELL_APOPTOTIC_PROCESS	26
	0.01	Up	0.485339812842932	0.20443997770483
	0.610700474176781
GO_STEM_CELL_DIFFERENTIATION	186	0.01	Down
0.356528745679487	0.204491942724766	0.610700474176781
GO_UBIQUITIN_CONJUGATING_ENZYME_BINDING 28	0.01	Down
0.18362276914756	0.204509014009756	0.610700474176781
GO_NUCLEAR_CHROMATIN	290	0.01	Down	0.195811043816564
	0.204550217271414	0.610700474176781
GO_POSITIVE_REGULATION_OF_ALPHA_BETA_T_CELL_PROLIFERATION	13
	0.01	Down	0.555523967649806	0.204660206743183
	0.610700474176781
GO_HYDRO_LYASE_ACTIVITY 41	0.01	Up	0.250330395788132
	0.204661446699054	0.610700474176781
GO_LATERAL_PLASMA_MEMBRANE	45	0.01	Down
0.343906068323336	0.204706879418506	0.610700474176781
GO_POSITIVE_REGULATION_OF_HORMONE_SECRETION	78	0.01	Down
	0.37326481288775	0.204954444500967

0.610996528392216
GO_NEGATIVE_REGULATION_OF_CELL_AGING 20 0.01 Down
0.236275829522922 0.205186767440538 0.610996528392216
GO_L_ALPHA_AMINO_ACID_TRANSMEMBRANE_TRANSPORT 31 0.01 Up
0.393513824176872 0.205202106728968
0.610996528392216
GO_POSITIVE_REGULATION_OF_SMOOTH_MUSCLE_CONTRACTION 20 0.01
Down 0.464787661256076 0.205263618196449
0.610996528392216
GO_HORMONE_MEDIATED_SIGNALING_PATHWAY 166 0.01 Down
0.260120960448512 0.205309325962794 0.610996528392216
GO_HISTONE_METHYLTRANSFERASE_ACTIVITY_H3_K4_SPECIFIC_ 14 0.01
Down 0.194901298977045 0.205431980569834
0.611062005898806
GO_REGULATION_OF_MAP_KINASE_ACTIVITY 277 0.01 Down
0.379932018362588 0.205666221015402 0.611333653705141
GO_REGULATION_OF_CELL_MATRIX_ADHESION 93 0.01 Down
0.470010393316656 0.205816913261318 0.611333653705141
GO_CELLULAR_RESPONSE_TO_OSMOTIC_STRESS 31 0.01 Down
0.422890381182691 0.205844122833955 0.611333653705141
GO_NEUROTRANSMITTER_BINDING 14 0.01 Up
0.295358327204715 0.205958505403956 0.611333653705141
GO_DENDRITE_SELF_AVOIDANCE 10 0.01 Up
0.583955015138617 0.206026792205686 0.611333653705141
GO_GOLGI_TO_VACUOLE_TRANSPORT 14 0.01 Down
0.1751147654855 0.206342659960367 0.611971806848749
GO_POSITIVE_REGULATION_OF_LYMPHOCYTE_MEDIATED_IMMUNITY 72 0.01
Down 0.408877678901208 0.206449682190833
0.611990244424096
GO_PROTEIN_TRIMERIZATION 43 0.01 Up
0.48834985550186 0.206563032989134 0.61202741497171
GO_REGULATION_OF_POSTSYNAPTIC_MEMBRANE_POTENTIAL 56 0.01
Up 0.416069290974898 0.206879579924537
0.612564615771687
GO_ATP_DEPENDENT_MICROTUBULE_MOTOR_ACTIVITY_MINUS_END_DIRECTED 13
0.01 Down 0.335758457346374 0.20694897984374
0.612564615771687
GO_PHAGOPHORE_ASSEMBLY_SITE_MEMBRANE 16 0.01 Down
0.0916039967063452 0.207095958987131 0.612564615771687
GO_RAC_GUANYL_NUCLEOTIDE_EXCHANGE_FACTOR_ACTIVITY 15 0.01
Down 0.340581182818243 0.207147942048966
0.612564615771687
GO_L_AMINO_ACID_TRANSPORT 38 0.01 Up
0.406296736645709 0.207388749442915 0.612621096243705
GO_ESTABLISHMENT_OF_SPINDLE_ORIENTATION 23 0.01 Down
0.160467695482744 0.207436808829682 0.612621096243705
GO_JNK_CASCADE 185 0.01 Down 0.359604237158112
0.207469770034106 0.612621096243705
GO_NEGATIVE_REGULATION_OF_ANOIKIS 16 0.01 Down
0.368312305654565 0.207644780836182 0.612839797985638
GO_EXOSOME_RNASE_COMPLEX_ 25 0.01 Down
0.147973481169512 0.207935172304238 0.61322344751158
GO_REGULATION_OF_CHROMOSOME_ORGANIZATION 292 0.01 Down
0.1954285232167 0.207976787749357 0.61322344751158

GO_TOOTH_MINERALIZATION 15 0.01 Down 0.68141119476548
0.208447961107658 0.614094517572626
GO_POSITIVE_REGULATION_OF_VIRAL_PROCESS 97 0.01 Down
0.190579667004565 0.208600351927929 0.614094517572626
GO_POSITIVE_REGULATION_OF_INTERLEUKIN_13_PRODUCTION 12 0.01
Up 0.526026037912793 0.208672854288336
0.614094517572626
GO_POSITIVE_REGULATION_OF_PODOSOME_ASSEMBLY 10 0.01 Down
0.35131039501978 0.208683438502733
0.614094517572626
GO_NUCLEAR_SPECK 362 0.01 Down 0.203271717643394
0.208777974677961 0.614094517572626
GO_SIGNAL_PEPTIDE_PROCESSING 11 0.01 Down
0.220600141168391 0.209207884638058 0.614938365243229
GO_POSITIVE_REGULATION_OF_BLOOD_PRESSURE 16 0.01 Up
0.331325969511647 0.209270743099363
0.614938365243229
GO_CHROMATIN_REMODELING 136 0.01 Down 0.213281694570694
0.209368736774461 0.614938365243229
GO_DEFENSE_RESPONSE_TO_GRAM_NEGATIVE_BACTERIUM 33 0.01 Down
0.765818480811085 0.209489703769482
0.614996127458667
GO_ALCOHOL_CATABOLIC_PROCESS 40 0.01 Up
0.398337438830008 0.21007082497585 0.615644533703607
GO_POSITIVE_REGULATION_OF_NUCLEOBASE_CONTAINING_COMPOUND_METABOLIC_P
ROCESS 203 0.01 Down 0.195742567850592
0.210153668421728 0.615644533703607
GO_VENTRICULAR_CARDIAC_MUSCLE_CELL_ACTION_POTENTIAL 22 0.01
Down 0.494900064023335 0.21018833506701
0.615644533703607
GO_MUCOPOLYSACCHARIDE_METABOLIC_PROCESS 12 0.01 Up
0.992019679673218 0.210237178885845 0.615644533703607
GO_CHANNEL_INHIBITOR_ACTIVITY 31 0.01 Down
0.289088668722556 0.21031416368563 0.615644533703607
GO_PHOSPHATIDYLINOSITOL_BISPHOSPHATE_PHOSPHATASE_ACTIVITY 22
0.01 Down 0.159014538886663 0.210319018761535
0.615644533703607
GO_COATED_VESICLE 238 0.01 Down 0.331360140046948
0.210536451584956 0.615983998830008
GO_POSITIVE_REGULATION_OF_CATALYTIC_ACTIVITY 1110 0.01 Down
0.331668096893661 0.210771043335901
0.616373316036731
GO_POSITIVE_REGULATION_OF_BLOOD_CIRCULATION 37 0.01 Down
0.433061426718085 0.211094908369906
0.617023201113961
GO_5S_RRNA_BINDING 11 0.01 Up 0.138846844727089
0.211293431585617 0.617306267158941
GO_SERINE_FAMILY_AMINO_ACID_METABOLIC_PROCESS 30 0.01 Up
0.374722998157512 0.211399505777435
0.617319095514578
GO_WNT_SIGNALOSOME 10 0.01 Up 0.477330477142641
0.21175512737641 0.618060278029896
GO_DNA_METHYLATION 51 0.01 Down 0.282309178777179
0.212404082325099 0.619627402037547

GO_POSITIVE_REGULATION_OF_DENDRITIC_SPINE_DEVELOPMENT 33 0.01 Down 0.340120905957208 0.212496170489569 0.619627402037547
GO_GTP_DEPENDENT_PROTEIN_BINDING 20 0.01 Down 0.253334391402496 0.212612430238394 0.619668777713533
GO_MHC_CLASS_II_PROTEIN_COMPLEX_BINDING 15 0.01 Up 0.625056399980128 0.212901465310191 0.619790329768716
GO_CELLULAR_RESPONSE_TO_ORGANIC_CYCLIC_COMPOUND 414 0.01 Down 0.363126458504583 0.212921726201864 0.619790329768716
GO_POSITIVE_REGULATION_OF_SODIUM_ION_TRANSMEMBRANE_TRANSPORT 16 0.01 Down 0.362582305771632 0.213059222077597 0.619790329768716
GO_REGULATION_OF_MUSCLE_HYPERTROPHY 47 0.01 Down 0.419458562932904 0.21309318932089 0.619790329768716
GO_REGULATION_OF_HISTONE_MODIFICATION 121 0.01 Down 0.217952521087983 0.213439461444122 0.619790329768716
GO_POSITIVE_REGULATION_OF_CHONDROCYTE_DIFFERENTIATION 12 0.01 Up 0.495909184724357 0.2134521739306 0.619790329768716
GO_TRANSCRIPTION_PREINITIATION_COMPLEX_ASSEMBLY 38 0.01 Down 0.160747582394704 0.213638052937945 0.619790329768716
GO_COENZYME_BIOSYNTHETIC_PROCESS 129 0.01 Up 0.275795829227831 0.213754656301675 0.619790329768716
GO_LONG_CHAIN_FATTY_ACID_TRANSPORT 47 0.01 Up 0.392635801870416 0.213852347243765 0.619790329768716
GO_PROTEIN_HEXAMERIZATION 12 0.01 Down 0.159849994282348 0.213910421834997 0.619790329768716
GO_REGULATION_OF_AUTOPHAGOSOME_MATURATION 10 0.01 Down 0.135015741890682 0.213939323337654 0.619790329768716
GO_RIBONUCLEOSIDE_CATABOLIC_PROCESS 17 0.01 Down 0.424972766250229 0.213949741610755 0.619790329768716
GO_REGULATION_OF_CLATHRIN_DEPENDENT_ENDOCYTOSIS 13 0.01 Down 0.33466544273032 0.214063874551496 0.619790329768716
GO_PHOTOPERIODISM 22 0.01 Down 0.197770164914055 0.214083400020589 0.619790329768716
GO_ERYTHROCYTE_HOMEOSTASIS 99 0.01 Down 0.316736665104771 0.214314133177176 0.620132675368062
GO_POSITIVE_REGULATION_OF_EXTRINSIC_APOPTOTIC_SIGNALING_PATHWAY_VIA_DEATH_DOMAIN_RECEPTORS 15 0.01 Down 0.302051184317062 0.214478636355667 0.620132675368062
GO_REGULATION_OF_MICROTUBULE_BASED_PROCESS 178 0.01 Down 0.208974639622441 0.214508090639587 0.620132675368062
GO_MRNA_3_END_PROCESSING 96 0.01 Down 0.116801407465958 0.214662940757922 0.620266358033168
GO_NEGATIVE_REGULATION_OF_DNA_DAMAGE_RESPONSE_SIGNAL_TRANSDUCTION_BY_P53_CLASS_MEDIATOR 13 0.01 Up 0.257176322755142 0.214758669837872 0.620266358033168
GO_HORMONE_METABOLIC_PROCESS 143 0.01 Up 0.529062510292028 0.215069133374765 0.620512464764931

GO_PODOSOME_ASSEMBLY	18	0.01	Down	0.388944302562799
	0.215131576112293	0.620512464764931
GO_TRANSPOSITION	18	0.01	Down	0.459975469031495
	0.215199405914828	0.620512464764931
GO_CORTICAL_CYTOSKELETON_ORGANIZATION	43	0.01	Down
0.360412402182248	0.215252717969848	0.620512464764931
GO_AMINOACYL_TRNA_EDITING_ACTIVITY	11	0.01	Up
0.125350043450929	0.215464576418208	0.620533891572552
GO_AMINOACYL_TRNA_METABOLISM_INVOLVED_IN_TRANSLATIONAL_FIDELITY 11
	0.01	Up	0.125350043450929	0.215464576418208
	0.620533891572552
GO_AMYLOID_PRECURSOR_PROTEIN_CATABOLIC_PROCESS	33	0.01	Down
	0.332874417260384	0.215627021108021
0.620707276029776
GO_POSTSYNAPTIC_NEUROTRANSMITTER_RECEPTOR_INTERNALIZATION	12
	0.01	Down	0.236016165066989	0.215797873985078
	0.62090468860825
GO_TRNA_WOBBLE_BASE_MODIFICATION	21	0.01	Up
0.147325119957332	0.216294124419211	0.622037721150653
GO_LUNG_EPITHELIUM_DEVELOPMENT	30	0.01	Down
0.548487227485306	0.216533933897947	0.622432534419715
GO_NUCLEOBASE_CONTAINING_COMPOUND_TRANSPORT	216	0.01	Down
	0.166411557042048	0.216881897292323
0.623137718155085
GO_LONG_TERM_SYNAPTIC_DEPRESSION	15	0.01	Down
0.324574086178203	0.217200172298182	0.623483240122664
GO_MACROMOLECULE_METHYLATION	256	0.01	Down
0.175008333849913	0.217241739133503	0.623483240122664
GO_NEGATIVE_REGULATION_OF_NEURON_DIFFERENTIATION	161	0.01
	Down	0.376046867377641	0.217310251375318
0.623483240122664
GO_SCHWANN_CELL_DEVELOPMENT	22	0.01	Down
0.201649151790634	0.217437991271547	0.623555051964838
GO_ISWI_TYPE_COMPLEX	10	0.01	Down	0.159268465324959
	0.21761567172771	0.62376994478703
GO_CELLULAR_RESPONSE_TO_STARVATION	131	0.01	Down
0.216241254610932	0.217766129769963	0.623906641733574
GO_GENERATION_OF_PRECURSOR_METABOLITES_AND_ENERGY	408	0.01
	Up	0.230525741846278	0.218043595728131
0.624403239987636
GO_ATRIAL_SEPTUM_MORPHOGENESIS	10	0.01	Down
0.313963004536212	0.218246180126362	0.624403239987636
GO_MICROTUBULE_POLYMERIZATION	63	0.01	Down
0.232103596549695	0.218248011077872	0.624403239987636
GO_HEAT_SHOCK_PROTEIN_BINDING	106	0.01	Down
0.233833425104987	0.218542867443844	0.624952307231078
GO_SECRETORY_GRANULE_LOCALIZATION	11	0.01	Down
0.265563078624831	0.218751892042877	0.625017852021383
GO_2_IRON_2_SULFUR_CLUSTER_BINDING	20	0.01	Up
0.111174226708251	0.218771690915078	0.625017852021383
GO_HOMOPHILIC_CELL_ADHESION_VIA_PLASMA_MEMBRANE_ADHESION_MOLECULES
	119	0.01	Up	0.371147493032626
0.218986342095112	0.625088197652441
GO_MESODERM_MORPHOGENESIS	49	0.01	Down

0.456000821357976	0.219246701752276	0.625088197652441
GO_TRANSFERASE_ACTIVITY_TRANSFERRING_PHOSPHORUS_CONTAINING_GROUPS
	767	0.01	Down	0.262948511092572
0.219278419384408	0.625088197652441
GO_DNA_TEMPLATED_TRANSCRIPTION_INITIATION	206	0.01	Down
	0.218181081801962	0.219322075965822
0.625088197652441
GO_INCLUSION_BODY_ASSEMBLY	20	0.01	Down
0.224806949784885	0.219623267036195	0.625088197652441
GO_REGULATION_OF_CALCIUM_MEDIATED_SIGNALING	66	0.01	Down
	0.409009733775652	0.219629965977277
0.625088197652441
GO_REGULATION_OF_MYOTUBE_DIFFERENTIATION	41	0.01	Down
	0.530190857882005	0.219659664162875
0.625088197652441
GO_NUCLEOBASE_CONTAINING_SMALL_MOLECULE_CATABOLIC_PROCESS	39
	0.01	Down	0.295291592877716	0.21974515613439
	0.625088197652441
GO_AROMATASE_ACTIVITY	10	0.01	Up	0.658370426367769
	0.219796178100516	0.625088197652441
GO_PROTEIN_DOMAIN_SPECIFIC_BINDING	577	0.01	Down
0.268762638664737	0.219825943335194	0.625088197652441
GO_DEOXYRIBONUCLEOTIDE_BIOSYNTHETIC_PROCESS	14	0.01	Up
	0.188983729134177	0.220033989274784
0.625377682094732
GO_ACTIVIN_BINDING	13	0.01	Down	0.343263679722794
	0.22013376818258	0.625377682094732
GO_SMALL_GTPASE_MEDIATED_SIGNAL_TRANSDUCTION	478	0.01	Down
	0.33079309482594	0.220340457777096
0.625672085671071
GO_INORGANIC_ANION_TRANSMEMBRANE_TRANSPORT	56	0.01	Down
	0.373382847294554	0.220734917613765
0.626499151394656
GO_GATED_CHANNEL_ACTIVITY	136	0.01	Up
0.447490328797811	0.221086532314986	0.627203896114149
GO_SODIUM_CHANNEL_ACTIVITY	21	0.01	Up
0.483592909565312	0.221278271516394	0.627454640997678
GO_ZINC_ION_BINDING	626	0.01	Down	0.318734943213108
	0.221494676382061	0.627717596889991
GO_SEX_DIFFERENTIATION	175	0.01	Down	0.372925856393884
	0.221577797749177	0.627717596889991
GO_CARDIAC_MUSCLE_CELL_PROLIFERATION	33	0.01	Down
0.342882230287381	0.221741707720193	0.627888949425975
GO_TRANSCRIPTION_INITIATION_FROM_RNA_POLYMERASE_II_PROMOTER	160
	0.01	Down	0.227617868530427	0.222273139365415
	0.628652338907648
GO_SYNAPTIC_MEMBRANE_ADHESION	14	0.01	Up
0.501262214372084	0.222282529069145	0.628652338907648
GO_RNA_POLYMERASE_II_PREINITIATION_COMPLEX_ASSEMBLY	23	0.01
	Down	0.172391931171847	0.222321952171754
0.628652338907648
GO_NADP_BINDING	42	0.01	Up	0.313045126892929
0.222859980872495	0.62977799529562
GO_REGULATION_OF_LYMPHOCYTE_MEDIATED_IMMUNITY	103	0.01	Down

0.432046678324085 0.222967468736166
0.62977799529562
GO_CYTOKINETIC_PROCESS 38 0.01 Down 0.190308334226058
0.223095416172033 0.62977799529562
GO_NEGATIVE_REGULATION_OF_INTRINSIC_APOPTOTIC_SIGNALING_PATHWAY_BY_P
53_CLASS_MEDIATOR 22 0.01 Up 0.231747874273439
0.223368731589646 0.62977799529562
GO_BLOOD_VESSEL_ENDOTHELIAL_CELL_MIGRATION 97 0.01 Down
0.475785220398197 0.223538239669792
0.62977799529562
GO_SYNAPTIC_VESICLE_MEMBRANE 63 0.01 Up
0.292259837593008 0.223555531767077 0.62977799529562
GO_GAMMA_TUBULIN_BINDING 24 0.01 Up
0.223306850514115 0.223627832504432 0.62977799529562
GO_REGULATION_OF_EXTRACELLULAR_MATRIX_ORGANIZATION 33 0.01
Up 0.556438184071538 0.22367352679747
0.62977799529562
GO_NEGATIVE_REGULATION_OF_SINGLE_STRANDED_VIRAL_RNA_REPLICATION_VIA_
DOUBLE_STRANDED_DNA_INTERMEDIATE 12 0.01 Down
0.21246926572788 0.223757123705809 0.62977799529562
GO_O_ACYLTRANSFERASE_ACTIVITY 36 0.01 Up
0.239936651677457 0.223853709860204 0.62977799529562
GO_POSITIVE_REGULATION_OF_TRANSCRIPTION_FROM_RNA_POLYMERASE_II_PROMO
TER_INVOLVED_IN_CELLULAR_RESPONSE_TO_CHEMICAL_STIMULUS 19 0.01
Down 0.25842001455325 0.223861128948764
0.62977799529562
GO_AMINO_ACID_TRANSMEMBRANE_TRANSPORT 53 0.01 Up
0.451316117984305 0.224238136250331 0.630341559974858
GO_NEGATIVE_REGULATION_OF_SIGNAL_TRANSDUCTION_BY_P53_CLASS_MEDIATOR
30 0.01 Up 0.23202600312721
0.224400080385571 0.630341559974858
GO_CELLULAR_RESPONSE_TO_ACID_CHEMICAL 154 0.01 Up
0.397691712664709 0.224426066420055 0.630341559974858
GO_RESPONSE_TO_PAIN 11 0.01 Up 0.225871045313419
0.224572735833825 0.630341559974858
GO_REGULATION_OF_CELLULAR_RESPONSE_TO_TRANSFORMING_GROWTH_FACTOR_BET
A_STIMULUS 91 0.01 Up 0.417426674447888
0.224580595326242 0.630341559974858
GO_CATENIN_COMPLEX 16 0.01 Down 0.290046921623746
0.22475595219064 0.630542230013573
GO_BLOC_COMPLEX 19 0.01 Up 0.151165201605375
0.22497025702501 0.630851930900156
GO_NEGATIVE_REGULATION_OF_SMALL_GTPASE_MEDIATED_SIGNAL_TRANSDUCTION
52 0.01 Down 0.288367223594724
0.225328435055508 0.631346410059632
GO_FIBRILLAR_CENTER 129 0.01 Down 0.19466743359073
0.225354582539816 0.631346410059632
GO_POSITIVE_REGULATION_OF_RESPONSE_TO_WOUNDING 45 0.01 Down
0.420603449360172 0.225632293157506
0.631400821737208
GO_LIPID_PHOSPHORYLATION 54 0.01 Down
0.21223080293308 0.225649699059798 0.631400821737208
GO_REGULATION_OF_PHOSPHOLIPASE_C_ACTIVITY 24 0.01 Up
0.700567918843573 0.225794398961157

0.631400821737208
GO_REGULATION_OF_MITOCHONDRIAL_FISSION 20 0.01 Up
0.318677000933236 0.226012061748229 0.631400821737208
GO_RESPONSE_TO_CADMIUM_ION 49 0.01 Up
0.492281169855204 0.226182021603807 0.631400821737208
GO_PRIMARY_ALCOHOL_CATABOLIC_PROCESS 10 0.01 Up
0.570550387827598 0.226274535162031 0.631400821737208
GO_NUCLEOSIDE_TRIPHOSPHATASE_REGULATOR_ACTIVITY 295 0.01 Down
0.271835338452662 0.226288570449645
0.631400821737208
GO_NUCLEAR_PERIPHERY 123 0.01 Down 0.184703324330041
0.226492401846125 0.631400821737208
GO_B_CELL_APOPTOTIC_PROCESS 19 0.01 Down
0.264729804234798 0.226502531876525 0.631400821737208
GO_REGULATION_OF_EXTRINSIC_APOPTOTIC_SIGNALING_PATHWAY 128 0.01
Down 0.35960088518816 0.226561185271029
0.631400821737208
GO_POSITIVE_REGULATION_OF_JNK_CASCADE 121 0.01 Down
0.379556344875744 0.226568550871706 0.631400821737208
GO_UBIQUITIN_LIGASE_COMPLEX 262 0.01 Down
0.15459286220182 0.22665662272964 0.631400821737208
GO_REGULATION_OF_ACTIN_CYTOSKELETON_REORGANIZATION 28 0.01
Up 0.432190827623482 0.22692795601334
0.631400821737208
GO_ANTIGEN_RECEPTOR_MEDIATED_SIGNALING_PATHWAY 184 0.01 Down
0.347971239276643 0.227050779571342
0.631400821737208
GO_SPERMATID_NUCLEUS_DIFFERENTIATION 11 0.01 Down
0.285361012152295 0.227082722178006 0.631400821737208
GO_POSITIVE_REGULATION_OF_CALCIUM_ION_TRANSPORT_INTO_CYTOSOL 33
0.01 Up 0.765213047900183 0.227185388825086
0.631400821737208
GO_REGULATION_OF_SMALL_MOLECULE_METABOLIC_PROCESS 270 0.01
Down 0.307336204538792 0.227417767973453
0.631400821737208
GO_ORGANIC_ANION_TRANSPORT 312 0.01 Up
0.401816299987006 0.227443571059533 0.631400821737208
GO_REGULATION_OF_HYDROLASE_ACTIVITY 971 0.01 Down
0.359364931733754 0.22752818341403 0.631400821737208
GO_CELLULAR_RESPONSE_TO_DRUG 269 0.01 Down
0.372278155791506 0.227557626052765 0.631400821737208
GO_REGULATION_OF_METALLOPEPTIDASE_ACTIVITY 15 0.01 Up
0.342351413231289 0.227558062586232
0.631400821737208
GO_PROTON_TRANSPORTING_ATP_SYNTHASE_COMPLEX_COUPLING_FACTOR_F_O_
10 0.01 Up 0.0967806983165701
0.227772884817945 0.631708169817152
GO_RESPONSE_TO_INSULIN 202 0.01 Down 0.216894816288597
0.227974197899522 0.631748178959495
GO_TRANSCRIPTION_ELONGATION_FACTOR_COMPLEX 51 0.01 Down
0.150993550740068 0.228111973138494
0.631748178959495
GO_RESPONSE_TO_DOPAMINE 11 0.01 Down 0.0966836863061876
0.228170960463742 0.631748178959495

GO_MAINTENANCE_OF_SISTER_CHROMATID_COHESION 11 0.01 Down 0.120305375378174 0.228272368220753 0.631748178959495
GO_NEGATIVE_REGULATION_OF_SIGNALING 1031 0.01 Down 0.352217676989375 0.228399673876632 0.631748178959495
GO_REGULATION_OF_MONOOXYGENASE_ACTIVITY 42 0.01 Down 0.397908620291892 0.22841167069405 0.631748178959495
GO_NEGATIVE_REGULATION_OF_POSTTRANSCRIPTIONAL_GENE_SILENCING 10 0.01 Down 0.31625512846625 0.228564147401341 0.631882030452432
GO_RETROGRADE_VESICLE_MEDIATED_TRANSPORT_GOLGI_TO_ENDOPLASMIC_RETICULUM 77 0.01 Down 0.13691124064662 0.228796455658539 0.632072001654067
GO_ENDOCARDIAL_CUSHION_MORPHOGENESIS 19 0.01 Down 0.532312829754585 0.228929882960528 0.632072001654067
GO_NEGATIVE_REGULATION_OF_HISTONE_MODIFICATION 35 0.01 Down 0.182302830611242 0.228945203695815 0.632072001654067
GO_CDC42_PROTEIN_SIGNAL_TRANSDUCTION 10 0.01 Up 0.471996003455914 0.229094695779681 0.632197226399292
GO_NEGATIVE_REGULATION_OF_OSTEOCLAST_DIFFERENTIATION 19 0.01 Up 0.491788006290402 0.229499817988212 0.633027439803014
GO_STRIATUM_DEVELOPMENT 14 0.01 Down 0.408650086426064 0.229850177447403 0.633286739860694
GO_ACUTE_INFLAMMATORY_RESPONSE_TO_ANTIGENIC_STIMULUS 13 0.01 Down 0.535878002738788 0.229919598602448 0.633286739860694
GO_HOMOTYPIC_CELL_CELL_ADHESION 58 0.01 Down 0.369003391037588 0.230003850209645 0.633286739860694
GO_RESPONSE_TO_NUTRIENT 159 0.01 Up 0.452322195679298 0.230042903480693 0.633286739860694
GO_KINASE_ACTIVITY 629 0.01 Down 0.283885424624122 0.230202861263668 0.633286739860694
GO_SODIUM_ION_TRANSPORT 130 0.01 Down 0.451098357281387 0.230240036119646 0.633286739860694
GO_POSITIVE_REGULATION_OF_INTERLEUKIN_4_PRODUCTION 17 0.01 Up 0.581633534569355 0.230411021846094 0.633286739860694
GO_REGULATION_OF_VASOCONSTRICTION 27 0.01 Down 0.408706907457388 0.230428332787395 0.633286739860694
GO_REGULATION_OF_KERATINOCYTE_DIFFERENTIATION 33 0.01 Down 0.471778119798266 0.230599387637703 0.63347008251063
GO_LABYRINTHINE_LAYER_DEVELOPMENT 35 0.01 Down 0.242869080024318 0.230846364827461 0.633625722120706
GO_REGULATION_OF_LEUKOCYTE_DIFFERENTIATION 197 0.01 Down 0.446201725038221 0.230864782956844 0.633625722120706
GO_NEGATIVE_REGULATION_OF_INTERLEUKIN_6_PRODUCTION 27 0.01 Down 0.671567712028364 0.231169666868008 0.633773611275777
GO_REGULATION_OF_T_CELL_MEDIATED_IMMUNITY 49 0.01 Down 0.405754987777463 0.231237739680076

0.633773611275777
GO_NEGATIVE_REGULATION_OF_MAPK_CASCADE 144 0.01 Down
0.410598021040823 0.231277469819877 0.633773611275777
GO_REGULATION_OF_T_HELPER_2_CELL_DIFFERENTIATION 10 0.01
Up 0.467832541719517 0.231336241572578
0.633773611275777
GO_MEMBRANE_RAFT_ORGANIZATION 19 0.01 Up
0.411172620544183 0.231525387283501 0.634005695172818
GO_LIPID_METABOLIC_PROCESS 916 0.01 Up
0.331365089166795 0.231740658106687 0.634309078163074
GO_BRANCHING_INVOLVED_IN_BLOOD_VESSEL_MORPHOGENESIS 24 0.01
Up 0.660491399629808 0.232944828389258
0.637008462827818
GO_DNA_REPLICATION_INDEPENDENT_NUCLEOSOME_ORGANIZATION 39 0.01
Up 0.177623514937853 0.233025730739937
0.637008462827818
GO_REGULATION_OF_MEMBRANE_DEPOLARIZATION 34 0.01 Up
0.405075709662797 0.233041639917737
0.637008462827818
GO_SERTOLI_CELL_DEVELOPMENT 10 0.01 Down
0.15978967426954 0.233537589439207 0.637948585232264
GO_POSITIVE_REGULATION_OF_PROTEIN_COMPLEX_ASSEMBLY 206 0.01
Down 0.304279156571944 0.233618107272032
0.637948585232264
GO_RNA_POLYMERASE_CORE_ENZYME_BINDING 44 0.01 Down
0.16972300480071 0.233700815937499 0.637948585232264
GO_GENETIC_IMPRINTING 22 0.01 Down 0.228368913581938
0.233933868478452 0.63829775978997
GO_REGULATION_OF_TRANSCRIPTION_FROM_RNA_POLYMERASE_II_PROMOTER_IN_RE
SPONSE_TO_OXIDATIVE_STRESS 10 0.01 Up
0.212155294628045 0.234112596522414 0.638498460686243
GO_PROTEASOME_CORE_COMPLEX 18 0.01 Down
0.137916186997782 0.234428708863968 0.639073503149146
GO_RNA_BINDING 1500 0.01 Down 0.159024996047704
0.234583434118687 0.639208271335076
GO_BETA_CATENIN_TCF_COMPLEX 10 0.01 Down
0.119696613437323 0.234886899385794 0.639350140508319
GO_PHAGOCYTIC_VESICLE_MEMBRANE 69 0.01 Down
0.293508718857761 0.234988451206667 0.639350140508319
GO_NEGATIVE_REGULATION_OF_SYSTEMIC_ARTERIAL_BLOOD_PRESSURE 10
0.01 Down 0.358695954443261 0.23510633759718
0.639350140508319
GO_METANEPHRIC_NEPHRON_MORPHOGENESIS 17 0.01 Down
1.03386229381622 0.235122331717234 0.639350140508319
GO_RIBONUCLEOTIDE_BINDING 1529 0.01 Down
0.262230736674368 0.235382384519564 0.639350140508319
GO_REGULATION_OF_RYANODINE_SENSITIVE_CALCIUM_RELEASE_CHANNEL_ACTIVIT
Y 19 0.01 Up 0.424755866801797
0.235443424491188 0.639350140508319
GO_POSITIVE_REGULATION_OF_RNA_METABOLIC_PROCESS 60 0.01 Down
0.146010818941935 0.23589726531006
0.639350140508319
GO_REGULATION_OF_ANTIGEN_PROCESSING_AND_PRESENTATION 14 0.01
Up 0.715521904968916 0.236023626331969

0.639350140508319
GO_LYSINE_CATABOLIC_PROCESS 11 0.01 Up
0.349387559888984 0.236034603126615 0.639350140508319
GO_MEDIATOR_COMPLEX 36 0.01 Down 0.204678927614175
0.23605802691608 0.639350140508319
GO_GLOMERULAR_EPITHELIUM_DEVELOPMENT 13 0.01 Down
0.640629440084324 0.236201670246426 0.639350140508319
GO_CGMP_MEDIATED_SIGNALING 17 0.01 Up
0.734949181244418 0.236286489428206 0.639350140508319
GO_CARBON_SULFUR_LYASE_ACTIVITY 10 0.01 Up
0.230264626717515 0.236327093368704 0.639350140508319
GO_SEMAPHORIN_PLEXIN_SIGNALING_PATHWAY 33 0.01 Down
0.416743795003271 0.236343835125039 0.639350140508319
GO_DYNEIN_HEAVY_CHAIN_BINDING 11 0.01 Up
0.218701933000022 0.236359274833953 0.639350140508319
GO_IRON_SULFUR_CLUSTER_ASSEMBLY 21 0.01 Up
0.116998242831163 0.236401562539652 0.639350140508319
GO_TRNA_WOBBLE_URIDINE_MODIFICATION 17 0.01 Up
0.150628055304081 0.23643148887344 0.639350140508319
GO_MICROTUBULE_CYTOSKELETON_ORGANIZATION_INVOLVED_IN_MITOSIS 111
0.01 Down 0.197298196747453 0.236628099815705
0.639350140508319
GO_ORGANELLAR_SMALL_RIBOSOMAL_SUBUNIT 28 0.01 Up
0.114757504641725 0.23670475259068 0.639350140508319
GO_REGULATION_OF_DNA_DAMAGE_RESPONSE_SIGNAL_TRANSDUCTION_BY_P53_CLAS
S_MEDIATOR 28 0.01 Up 0.241235009629262
0.236741742029764 0.639350140508319
GO_HISTONE_METHYLTRANSFERASE_ACTIVITY 50 0.01 Down
0.184238053665522 0.237101075381234 0.639941214211641
GO_EXTRINSIC_APOPTOTIC_SIGNALING_PATHWAY 190 0.01 Down
0.334935979818739 0.237171426779146
0.639941214211641
GO_PRE_MRNA_BINDING 32 0.01 Down 0.124237704874405
0.237341255751531 0.639961295521925
GO_NEGATIVE_REGULATION_OF_COAGULATION 34 0.01 Up
0.570302887302936 0.237389694863346 0.639961295521925
GO_CELL_FATE_COMMITMENT 141 0.01 Down 0.503424588942844
0.238033033331458 0.641410805750237
GO_REGULATION_OF_AUTOPHAGOSOME_ASSEMBLY 33 0.01 Down
0.152837633873222 0.238221269448056 0.641618288903185
GO_ENDOTHELIUM_DEVELOPMENT 92 0.01 Down
0.467203083106597 0.238433153233947 0.641618288903185
GO_N_TERMINAL_PROTEIN_AMINO_ACID_MODIFICATION 25 0.01 Down
0.160339213180754 0.238455892339699
0.641618288903185
GO_ADRENERGIC_RECEPTOR_BINDING 16 0.01 Down
0.332991618970689 0.238538702266324 0.641618288903185
GO_FATTY_ACID_DERIVATIVE_BIOSYNTHETIC_PROCESS 40 0.01 Up
0.316361883872806 0.238642766586635
0.641618288903185
GO_INTERLEUKIN_1_PRODUCTION 63 0.01 Down
0.595310888595303 0.238744146702734 0.641618288903185
GO_NEGATIVE_REGULATION_OF_CELL_CYCLE_G2_M_PHASE_TRANSITION 94
0.01 Down 0.141836269502204 0.238983623956604

0.641977690725903
GO_HISTONE_H3_K36_METHYLATION 12 0.01 Down
0.196316271505712 0.239134107129859 0.642097817065623
GO_REGULATION_OF_DIGESTIVE_SYSTEM_PROCESS 15 0.01 Up
0.4885269013088 0.239261658563084 0.642156290511265
GO_REGULATION_OF_VASCULATURE_DEVELOPMENT 237 0.01 Down
0.568580589460093 0.239371417720773
0.642166980549189
GO_POSITIVE_REGULATION_OF_MAPK_CASCADE 382 0.01 Down
0.428808960073899 0.239915201694459 0.643341514791104
GO_AXONEME_ASSEMBLY 37 0.01 Up 0.383917356365241
0.240344462584641 0.643729449178349
GO_PYRIMIDINE_CONTAINING_COMPOUND_CATABOLIC_PROCESS 28 0.01
Down 0.322044649824724 0.240552983716535
0.643729449178349
GO_VACUOLE 651 0.01 Up 0.345998779351282
0.240605655924314 0.643729449178349
GO_CYTOPLASMIC_SIDE_OF_MEMBRANE 132 0.01 Down
0.333328269041609 0.240614756913953 0.643729449178349
GO_RNA_POLYMERASE_II_SPECIFIC_DNA_BINDING_TRANSCRIPTION_FACTOR_BINDI
NG 232 0.01 Down 0.246863453311755
0.240648368507512 0.643729449178349
GO_ANKYRIN_BINDING 12 0.01 Up 0.721826564530039
0.240696071427253 0.643729449178349
GO_ANTIBIOTIC_BIOSYNTHETIC_PROCESS 15 0.01 Up
0.562779230213336 0.241041013077493 0.644363573239107
GO_NEUROTRANSMITTER_BIOSYNTHETIC_PROCESS 59 0.01 Down
0.565420951299809 0.241340661393759
0.644363573239107
GO_POSITIVE_REGULATION_OF_GLYCOLYTIC_PROCESS 11 0.01 Up
0.485385189652367 0.241356773599429
0.644363573239107
GO_NEGATIVE_REGULATION_OF_CATION_CHANNEL_ACTIVITY 27 0.01
Up 0.599700145971766 0.241357727811848
0.644363573239107
GO_CATECHOLAMINE_METABOLIC_PROCESS 18 0.01 Up
0.283691604955973 0.241905397239811 0.645541831491382
GO_POSITIVE_REGULATION_OF_NITRIC_OXIDE_SYNTHASE_ACTIVITY 16
0.01 Down 0.515030847188212 0.242072697017188
0.645704456762455
GO_RETROGRADE_TRANSPORT_ENDOSOME_TO_GOLGI 77 0.01 Down
0.181268958959808 0.242241344351247
0.645870532084505
GO_UBIQUITIN_LIKE_PROTEIN_LIGASE_BINDING 278 0.01 Down
0.201208554693062 0.242349495758112
0.645875236500219
GO_REGULATION_OF_COLLATERAL_SPROUTING 16 0.01 Up
0.434424735897474 0.242531149600321 0.645958538958789
GO_NLRP3_INFLAMMASOME_COMPLEX_ASSEMBLY 11 0.01 Down
0.658237978226077 0.242593554410482 0.645958538958789
GO_INNER_EAR_MORPHOGENESIS 58 0.01 Down
0.402406865115181 0.242758803730589 0.646033283597531
GO_EPITHELIAL_CELL_DEVELOPMENT 161 0.01 Down
0.432216386091723 0.242924279904498 0.646033283597531

GO_HISTONE_DEACETYLATION	68	0.01	Down
0.176162046498597	0.242957967578221	0.646033283597531
GO_NEGATIVE_REGULATION_OF_TELOMERE_MAINTENANCE_VIA_TELOMERASE	19
	0.01	Down	0.146659382446195	0.243047277176208
	0.646033283597531
GO_MYOBLAST_MIGRATION	10	0.01	Down	0.212322195501466
	0.243244533900203	0.646274645649073
GO_INTERMEDIATE_FILAMENT_ORGANIZATION	12	0.01	Down
0.388157182134092	0.243429406324797	0.646482907173159
GO_ENDOMEMBRANE_SYSTEM_ORGANIZATION	363	0.01	Down
0.236551662029284	0.243666605102257	0.646570948906923
GO_POSITIVE_REGULATION_OF_AXON_EXTENSION	35	0.01	Down
	0.375823063444452	0.243878170427556
0.646570948906923
GO_NEURAL_TUBE_PATTERNING	30	0.01	Up
0.267998775128763	0.244066443187048	0.646570948906923
GO_ATPASE_COMPLEX	82	0.01	Down	0.170104905992446
	0.244089091270135	0.646570948906923
GO_POSITIVE_REGULATION_OF_SUPEROXIDE_ANION_GENERATION	13	0.01
	Up	0.342501422900107	0.244452602884827
0.646570948906923
GO_PHOSPHATIDYLINOSITOL_4_PHOSPHATE_BINDING	26	0.01	Down
	0.176594533785959	0.244486309323492
0.646570948906923
GO_LOW_DENSITY_LIPOPROTEIN_RECEPTOR_PARTICLE_METABOLIC_PROCESS	20
	0.01	Up	0.275835838757112	0.244565950060161
	0.646570948906923
GO_TAXIS	432	0.01	Down	0.523160717962992
0.24463321676689	0.646570948906923
GO_BLOC_1_COMPLEX	14	0.01	Up	0.153438119499934
	0.244667469402662	0.646570948906923
GO_CHROMOSOME_SEPARATION	73	0.01	Up
0.201191293339988	0.244711738275672	0.646570948906923
GO_AORTA_MORPHOGENESIS	26	0.01	Down	0.672458500700444
	0.244748598876235	0.646570948906923
GO_ACTIN_FILAMENT_DEPOLYMERIZATION	41	0.01	Down
0.332586732636967	0.244763908409482	0.646570948906923
GO_REGULATION_OF_CELL_CELL_ADHESION_MEDIATED_BY_CADHERIN	13
	0.01	Up	0.509774780332658	0.244864282430376
	0.646570948906923
GO_PROTEIN_COFACTOR_LINKAGE	10	0.01	Up
0.182750187653957	0.244953579720956	0.646570948906923
GO_CARBON_CARBON_LYASE_ACTIVITY	37	0.01	Up
0.201976213005763	0.245489013781308	0.647702652180061
GO_SENSORY_ORGAN_MORPHOGENESIS	159	0.01	Down
0.443712914782492	0.245674131694125	0.647819356886635
GO_NEGATIVE_REGULATION_OF_NUCLEOBASE_CONTAINING_COMPOUND_METABOLIC_P
ROCESS	132	0.01	Down	0.155711640216639
0.245746660996528	0.647819356886635
GO_DICARBOXYLIC_ACID_METABOLIC_PROCESS	51	0.01	Up
0.208652979311052	0.246259882291216	0.648631433702536
GO_NEGATIVE_REGULATION_OF_MEIOTIC_CELL_CYCLE	11	0.01	Down
	0.450330895230825	0.246268399717402
0.648631433702536

GO_RECEPTOR_CLUSTERING 40 0.01 Up 0.380495681841418
0.246569867256597 0.649143826589245
GO_DRUG_CATABOLIC_PROCESS 64 0.01 Up
0.43338694213537 0.246792756060532 0.64944899091612
GO_POSITIVE_REGULATION_OF_ERYTHROCYTE_DIFFERENTIATION 24 0.01
Down 0.387008479921506 0.247365994728507
0.650675456671043
GO_NEGATIVE_REGULATION_OF_TUMOR_NECROSIS_FACTOR_MEDIATED_SIGNALING_P
ATHWAY 15 0.01 Up 0.240461876217876
0.247623985087504 0.651071985043844
GO_MONOCARBOXYLIC_ACID_TRANSPORT 101 0.01 Up
0.468671472859248 0.248101035917232 0.65204389136516
GO_SECONDARY_METABOLITE_BIOSYNTHETIC_PROCESS 20 0.01 Up
0.341001302665645 0.248607198515862
0.653091433227952
GO_POSITIVE_REGULATION_OF_CREB_TRANSCRIPTION_FACTOR_ACTIVITY 12
0.01 Down 0.271254600476666 0.248829492142026
0.653288363021347
GO_TUMOR_NECROSIS_FACTOR_RECEPTOR_SUPERFAMILY_BINDING 39 0.01
Down 0.395712896430488 0.248897378301495
0.653288363021347
GO_RIBOSOMAL_SMALL_SUBUNIT_BIOGENESIS 62 0.01 Up
0.118777744536774 0.249213228670113 0.653369786083633
GO_CEREBELLAR_PURKINJE_CELL_LAYER_DEVELOPMENT 17 0.01 Up
0.231368573879254 0.249268453704159
0.653369786083633
GO_CELL_ACTIVATION 1020 0.01 Down 0.440369275118395
0.249476626033668 0.653369786083633
GO_REGULATION_OF_CENTRIOLE_REPLICATION 21 0.01 Down
0.124489829382469 0.24954877016266 0.653369786083633
GO_O_GLYCAN_PROCESSING 42 0.01 Down 0.449767952969531
0.249680492624174 0.653369786083633
GO_CELLULAR_COMPONENT_MAINTENANCE 45 0.01 Down
0.276629563455327 0.249698137894899 0.653369786083633
GO_CAMP_RESPONSE_ELEMENT_BINDING 13 0.01 Down
0.256420909164598 0.249754430634678 0.653369786083633
GO_POSITIVE_REGULATION_OF_SIGNALING 1342 0.01 Down
0.382009143126603 0.250024461632129 0.653369786083633
GO_THIOESTER_METABOLIC_PROCESS 35 0.01 Up
0.269633256017314 0.250133162105896 0.653369786083633
GO_NEGATIVE_REGULATION_OF_HEMOPOIESIS 101 0.01 Down
0.430651796573458 0.250373048161208 0.653369786083633
GO_EUKARYOTIC_48S_PREINITIATION_COMPLEX 15 0.01 Down
0.113976513972887 0.250680035340059 0.653369786083633
GO_REGULATION_OF_SUPEROXIDE_METABOLIC_PROCESS 26 0.01 Up
0.498407065103043 0.250687403307758
0.653369786083633
GO_NEGATIVE_REGULATION_OF_UBIQUITIN_DEPENDENT_PROTEIN_CATABOLIC_PROC
ESS 43 0.01 Down 0.213691530563873
0.250753027137754 0.653369786083633
GO_INTRACELLULAR_LIGAND_GATED_ION_CHANNEL_ACTIVITY 20 0.01
Up 0.56241810883846 0.250815044822871
0.653369786083633
GO_NEGATIVE_REGULATION_OF_NEUROTRANSMITTER_TRANSPORT 17 0.01

Up 0.670113088933811 0.250885627664202
0.653369786083633
GO_METALLOCARBOXYPEPTIDASE_ACTIVITY 14 0.01 Up
0.679327427046665 0.250978840974894 0.653369786083633
GO_POSITIVE_REGULATION_OF_TRANSCRIPTION_BY_RNA_POLYMERASE_I 21
0.01 Down 0.113625518040845 0.250980349121522
0.653369786083633
GO_ACTININ_BINDING 21 0.01 Up 0.377264364214149
0.25109874542879 0.653369786083633
GO_RECYCLING_ENDOSOME 156 0.01 Down 0.284939359406812
0.251120578387022 0.653369786083633
GO_NUCLEAR_MATRIX 98 0.01 Down 0.205343651168473
0.251205224367388 0.653369786083633
GO_HISTONE_ACETYLTRANSFERASE_BINDING 26 0.01 Down
0.271277612914463 0.251245288893833 0.653369786083633
GO_POSITIVE_REGULATION_OF_NEURON_PROJECTION_DEVELOPMENT 216 0.01
Down 0.336013103995459 0.251296071570628
0.653369786083633
GO_ISOPRENOID_BIOSYNTHETIC_PROCESS 18 0.01 Down
0.398117949890667 0.251669367354313 0.653688316137692
GO_POSITIVE_REGULATION_OF_ESTABLISHMENT_OF_PROTEIN_LOCALIZATION_TO_M
ITOCHONDRION 56 0.01 Down 0.183811174147353
0.251672640236501 0.653688316137692
GO_POSITIVE_REGULATION_OF_KERATINOCYTE_DIFFERENTIATION 14 0.01
Down 0.346468330754449 0.251793684040569
0.653688316137692
GO_EXCRETION 32 0.01 Up 0.446194491102857
0.252012648402533 0.653688316137692
GO_MICROTUBULE_PLUS_END_BINDING 14 0.01 Down
0.120506043778049 0.252014986554738 0.653688316137692
GO_MULTI_ORGANISM_LOCALIZATION 69 0.01 Down
0.192226453866157 0.252158709163697 0.653688316137692
GO_POSITIVE_REGULATION_OF_PROTEOLYSIS_INVOLVED_IN_CELLULAR_PROTEIN_C
ATABOLIC_PROCESS 110 0.01 Down 0.195420482478881
0.252328107354313 0.653688316137692
GO_MRNA_CLEAVAGE_INVOLVED_IN_MRNA_PROCESSING 12 0.01 Down
0.13491983797729 0.252330691581651
0.653688316137692
GO_GABA_ERGIC_SYNAPSE 33 0.01 Up 0.325959994653168
0.252387648332524 0.653688316137692
GO_NUCLEOTIDYLTRANSFERASE_ACTIVITY 121 0.01 Down
0.175232388764473 0.252656363514396 0.653758512154356
GO_HEPATICOBILIARY_SYSTEM_DEVELOPMENT 111 0.01 Down
0.335277981214926 0.252673264461459 0.653758512154356
GO_REGULATION_OF_MRNA_3_END_PROCESSING 27 0.01 Down
0.123955099387958 0.252771283247146 0.653758512154356
GO_SEX_DETERMINATION 11 0.01 Down 0.440607229397503
0.253030674787697 0.653758512154356
GO_K63_LINKED_POLYUBIQUITIN_MODIFICATION_DEPENDENT_PROTEIN_BINDING
21 0.01 Down 0.250768545219959
0.253064525092732 0.653758512154356
GO_EMBRYONIC_DIGESTIVE_TRACT_DEVELOPMENT 22 0.01 Down
0.804321003322057 0.253259066574959
0.653758512154356

GO_FORMATION_OF_CYTOPLASMIC_TRANSLATION_INITIATION_COMPLEX 14 0.01 Down 0.118964482329355 0.253261913040727 0.653758512154356
GO_BARBED_END_ACTIN_FILAMENT_CAPPING 14 0.01 Down 0.127327881340906 0.25327623465443 0.653758512154356
GO_PROTON_TRANSPORTING_ATP_SYNTHASE_ACTIVITY_ROTATIONAL_MECHANISM 16 0.01 Up 0.131450947291574 0.253613581379759 0.65435106355993
GO_LEUKOCYTE_MIGRATION 291 0.01 Down 0.647109411790366 0.253779863693417 0.654501933934892
GO_REGULATION_OF_PROTON_TRANSPORT 11 0.01 Down 0.260598452726318 0.25389687273934 0.654525653673262
GO_MONOCARBOXYLIC_ACID_BIOSYNTHETIC_PROCESS 135 0.01 Up 0.394556724211023 0.254026991349526 0.654583134330633
GO_NEGATIVE_REGULATION_OF_RNA_METABOLIC_PROCESS 44 0.01 Down 0.140438812923862 0.254145686143807 0.654611141526964
GO_ENVELOPE 1041 0.01 Up 0.202279624447296 0.254739121425538 0.655671495538757
GO_REGULATION_OF_CARBOHYDRATE_CATABOLIC_PROCESS 68 0.01 Down 0.253251239332243 0.254773358256618 0.655671495538757
GO_EMBRYONIC_PATTERN_SPECIFICATION 38 0.01 Down 0.355658533883319 0.255134040322537 0.656210851622296
GO_POSITIVE_REGULATION_OF_PROTEIN_IMPORT 35 0.01 Down 0.284199300804942 0.255327030504325 0.656210851622296
GO_PROTEIN_TRANSPORT_ALONG_MICROTUBULE 61 0.01 Up 0.190680647782474 0.255391657545397 0.656210851622296
GO_ACTIVATION_OF_CYSTEINE_TYPE_ENDOPEPTIDASE_ACTIVITY 13 0.01 Up 0.218091312731467 0.255499029613651 0.656210851622296
GO_BILE_ACID_BIOSYNTHETIC_PROCESS 23 0.01 Up 0.429056416400853 0.255542246336889 0.656210851622296
GO_TRANSMEMBRANE_RECEPTOR_PROTEIN_TYROSINE_KINASE_ACTIVITY 41 0.01 Up 0.449218585522335 0.255631471600516 0.656210851622296
GO_POSITIVE_REGULATION_OF_NEUTROPHIL_MIGRATION 18 0.01 Down 0.724345465032127 0.256022342760451 0.656936450929288
GO_REGULATION_OF_HISTONE_H3_K9_METHYLATION 18 0.01 Down 0.148101908409153 0.256165430003617 0.656956137690681
GO_NEGATIVE_REGULATION_OF_TRANSCRIPTION_FROM_RNA_POLYMERASE_II_PROMOTER_IN_RESPONSE_TO_STRESS 12 0.01 Down 0.16301019564906 0.256246439474804 0.656956137690681
GO_POSITIVE_REGULATION_OF_CARDIAC_MUSCLE_CELL_DIFFERENTIATION 16 0.01 Up 0.703015491409322 0.256633982781738 0.657671975292499
GO_POSITIVE_REGULATION_OF_CHROMATIN_BINDING 11 0.01 Down 0.251107060597571 0.257207800733852 0.658684472060871
GO_RESPONSE_TO_MECHANICAL_STIMULUS 148 0.01 Down

0.460132212489394	0.257246068729423	0.658684472060871
GO_POSITIVE_REGULATION_OF_BINDING	153	0.01	Down
0.335456824229187	0.257528018930272	0.658815067681885
GO_G_PROTEIN_COUPLED_PURINERGIC_RECEPTOR_SIGNALING_PATHWAY	11
	0.01	Down	0.385774039115586	0.257583387099958
	0.658815067681885
GO_NEGATIVE_REGULATION_OF_NATURAL_KILLER_CELL_MEDIATED_IMMUNITY	13
	0.01	Down	0.777783141252824	0.257648576063951
	0.658815067681885
GO_POSITIVE_REGULATION_OF_DEVELOPMENTAL_GROWTH	112	0.01	Down
	0.378871783742156	0.257731145732907
0.658815067681885
GO_PROTEIN_MANNOSYLATION	22	0.01	Up
0.157170875224782	0.257959062883364	0.659120147628326
GO_RECYCLING_ENDOSOME_MEMBRANE	72	0.01	Down
0.276041861431401	0.258173631242459	0.659265818124259
GO_BASIC_AMINO_ACID_TRANSPORT	10	0.01	Up
0.5386682678596	0.258233259018199	0.659265818124259
GO_MISMATCH_REPAIR	29	0.01	Up	0.158817214458849
	0.258343405675924	0.659269783883368
GO_RNA_SPLICING	395	0.01	Down	0.112686756580331
0.258677315035756	0.659418842579262
GO_POSITIVE_REGULATION_OF_NEUROBLAST_PROLIFERATION	14	0.01
	Up	0.413147523921704	0.258686700705004
0.659418842579262
GO_RESPONSE_TO_FLUID_SHEAR_STRESS	32	0.01	Down
0.523438476599532	0.258727669745314	0.659418842579262
GO_PROTEIN_EXIT_FROM_ENDOPLASMIC_RETICULUM	40	0.01	Down
	0.17713684430983	0.258846365525832
0.659444517460062
GO_VITAMIN_B6_BINDING	38	0.01	Up	0.288747211053816
	0.259357131562824	0.66038143618946
GO_INTERLEUKIN_6_SECRETION	26	0.01	Down
0.515931275016285	0.259562784058363	0.66038143618946
GO_ENERGY_HOMEOSTASIS	29	0.01	Down	0.40300489300535
	0.259590499808669	0.66038143618946
GO_REGULATION_OF_PHAGOCYTOSIS	68	0.01	Up
0.563971217376102	0.259649232117318	0.66038143618946
GO_ESTABLISHMENT_OF_EPITHELIAL_CELL_POLARITY	26	0.01	Down
	0.273054927180259	0.260249580674228
0.661194959752537
GO_G_PROTEIN_ALPHA_SUBUNIT_BINDING	16	0.01	Up
0.496709529005861	0.260332449779178	0.661194959752537
GO_SOLUTE_CATION_SYMPORTER_ACTIVITY	53	0.01	Down
0.479974438936845	0.260453695740863	0.661194959752537
GO_FATTY_ACID_BINDING	22	0.01	Up	0.693888242780563
	0.26067076498028	0.661194959752537
GO_DICARBOXYLIC_ACID_TRANSMEMBRANE_TRANSPORTER_ACTIVITY	20	0.01
	Up	0.315281633943985	0.260690446777645
0.661194959752537
GO_ATP_DEPENDENT_DNA_HELICASE_ACTIVITY	36	0.01	Down
0.149604697403748	0.260713463762497	0.661194959752537
GO_SENSORY_PERCEPTION_OF_PAIN	50	0.01	Up
0.444991516828922	0.260736762226274	0.661194959752537

GO_POSITIVE_REGULATION_OF_CHROMATIN_ORGANIZATION 88 0.01 Down 0.208911120811474 0.260903117464183 0.661194959752537
GO_NEGATIVE_REGULATION_OF_EPITHELIAL_CELL_APOPTOTIC_PROCESS 26 0.01 Down 0.341395334689769 0.260994300065393 0.661194959752537
GO_POSITIVE_REGULATION_OF_PROTEIN_TARGETING_TO_MEMBRANE 27 0.01 Up 0.313750929942762 0.261078748088061 0.661194959752537
GO_CELLULAR_SENESCENCE 58 0.01 Down 0.274328997120522 0.261246478092522 0.661194959752537
GO_MISMATCH_REPAIR_COMPLEX_BINDING 11 0.01 Up 0.191001660539136 0.261463901554535 0.661194959752537
GO_LYMPHOCYTE_MEDIATED_IMMUNITY 167 0.01 Down 0.485343397783906 0.26146920793868 0.661194959752537
GO_CELL_CELL_ADHESION_MEDIATOR_ACTIVITY 40 0.01 Down 0.319898502708985 0.26157396968982 0.661194959752537
GO_SERINE_TYPE_ENDOPEPTIDASE_INHIBITOR_ACTIVITY 47 0.01 Down 0.787158404829732 0.261655608294797 0.661194959752537
GO_PROTEIN_DEACETYLASE_ACTIVITY 29 0.01 Down 0.184850070204543 0.261712744818407 0.661194959752537
GO_CORTICAL_CYTOSKELETON 85 0.01 Down 0.275758120956177 0.262129871915578 0.661194959752537
GO_GLYCOPROTEIN_CATABOLIC_PROCESS 17 0.01 Up 0.536715137781624 0.26216598219273 0.661194959752537
GO_SET1C_COMPASS_COMPLEX 12 0.01 Down 0.127950898158635 0.262236792244667 0.661194959752537
GO_MANGANESE_ION_BINDING 54 0.01 Down 0.271598784515947 0.262244518600517 0.661194959752537
GO_SNARE_BINDING 78 0.01 Down 0.279972480660772 0.262311642131333 0.661194959752537
GO_ADAPTATION_OF_SIGNALING_PATHWAY 12 0.01 Down 0.294989714914257 0.262365122392334 0.661194959752537
GO_POSITIVE_REGULATION_OF_MICROTUBULE_POLYMERIZATION_OR_DEPOLYMERIZATION 27 0.01 Down 0.214214528908486 0.262611962481394 0.661542416690681
GO_NEURON_MATURATION 32 0.01 Down 0.35400172033431 0.262840137657139 0.661637471658868
GO_NEURAL_NUCLEUS_DEVELOPMENT 44 0.01 Down 0.335006715588329 0.262943169796441 0.661637471658868
GO_MAST_CELL_ACTIVATION 43 0.01 Down 0.434780223934423 0.263060991671152 0.661637471658868
GO_MONOCARBOXYLIC_ACID_TRANSMEMBRANE_TRANSPORTER_ACTIVITY 26 0.01 Up 0.613420791409525 0.263085629481882 0.661637471658868
GO_LEFT_RIGHT_PATTERN_FORMATION 16 0.01 Up 0.35412457920034 0.26368300269865 0.662647319091895
GO_CARBOHYDRATE_DERIVATIVE_TRANSMEMBRANE_TRANSPORTER_ACTIVITY 36 0.01 Up 0.254643715289044 0.263705472397631 0.662647319091895
GO_POSITIVE_REGULATION_OF_NEURON_DIFFERENTIATION 267 0.01 Down 0.332786199118345 0.263896048238225 0.66285184478869

GO_HYDROLASE_ACTIVITY_ACTING_ON_CARBON_NITROGEN_BUT_NOT_PEPTIDE_BONDS_IN_CYCLIC_AMIDINES 25 0.01 Down 0.49774085681145 0.264243676501876 0.66345052110955
GO_CLATHRIN_COAT 42 0.01 Down 0.183995075945463 0.264601232954053 0.663461315382825
GO_NEGATIVE_REGULATION_OF_PHOSPHORYLATION 365 0.01 Down 0.348290288097962 0.264623044784391 0.663461315382825
GO_PLASMA_MEMBRANE_RAFT 78 0.01 Up 0.397132389327716 0.264794342675275 0.663461315382825
GO_THREONINE_TYPE_PEPTIDASE_ACTIVITY 18 0.01 Down 0.134705309901596 0.264809930504245 0.663461315382825
GO_CELLULAR_IRON_ION_HOMEOSTASIS 48 0.01 Up 0.401430410373512 0.264970063249155 0.663461315382825
GO_COHESIN_COMPLEX 10 0.01 Down 0.220618776489807 0.264980389319902 0.663461315382825
GO_REGULATION_OF_MAST_CELL_ACTIVATION 31 0.01 Down 0.466103697464382 0.265012961588429 0.663461315382825
GO_GLUCAN_METABOLIC_PROCESS 52 0.01 Down 0.244629656828759 0.265342737582697 0.664013091452825
GO_HEART_GROWTH 57 0.01 Down 0.359025768778169 0.265587373354475 0.664225426164858
GO_LYSOPHOSPHOLIPASE_ACTIVITY 15 0.01 Up 0.322569055136031 0.265646406642773 0.664225426164858
GO_SOLUTE_CATION_ANTIPORTER_ACTIVITY 22 0.01 Down 0.511715687879296 0.265837488673583 0.664429556911207
GO_NEGATIVE_REGULATION_OF_RESPONSE_TO_STIMULUS 1226 0.01 Down 0.36351931302448 0.266396160474274 0.665551888987373
GO_TRIGLYCERIDE_METABOLIC_PROCESS 25 0.01 Up 0.471783058708041 0.266516873464928 0.665579571701184
GO_NUCLEAR_OUTER_MEMBRANE 20 0.01 Up 0.201823765958468 0.266827020997206 0.665701103596287
GO_CENTROMERE_COMPLEX_ASSEMBLY 38 0.01 Up 0.216858950965218 0.266833168655721 0.665701103596287
GO_APICAL_PART_OF_CELL 254 0.01 Down 0.457471320377391 0.2668944961544 0.665701103596287
GO_NEGATIVE_REGULATION_OF_CELLULAR_RESPONSE_TO_GROWTH_FACTOR_STIMULUS 106 0.01 Up 0.538174577709194 0.267368858244181 0.665867682427842
GO_NUCLEUS_ORGANIZATION 108 0.01 Down 0.193933379854709 0.267388442401178 0.665867682427842
GO_CELL_SUBSTRATE_JUNCTION_ASSEMBLY 86 0.01 Down 0.391280209379061 0.267430805570775 0.665867682427842
GO_ENZYME_REGULATOR_ACTIVITY 794 0.01 Down 0.339594427444618 0.267485328719155 0.665867682427842
GO_HEMATOPOIETIC_PROGENITOR_CELL_DIFFERENTIATION 138 0.01 Down 0.303973797407877 0.267510170036022 0.665867682427842
GO_WASH_COMPLEX 11 0.01 Down 0.154112682262736 0.267619361740065 0.665867682427842
GO_POSITIVE_REGULATION_OF_LEUKOCYTE_CHEMOTAXIS 63 0.01 Down 0.585056175512725 0.268031917816119 0.666620963974459

GO_LUNG_MORPHOGENESIS 43 0.01 Down 0.4284643794434
0.268593894130552 0.667713171918741
GO_ENDOSOME_LUMEN 20 0.01 Up 0.63270069520749
0.268705764741497 0.667713171918741
GO_ACYLGLYCEROL_HOMEOSTASIS 14 0.01 Up
0.398979292609411 0.269098397698079 0.667713171918741
GO_NEGATIVE_REGULATION_OF_HYDROLASE_ACTIVITY 311 0.01 Down
0.423164935535239 0.269100065361639
0.667713171918741
GO_ENSHEATHMENT_OF_NEURONS 91 0.01 Down
0.307954044328098 0.269226717848817 0.667713171918741
GO_REGULATION_OF_GRANULOCYTE_CHEMOTAXIS 31 0.01 Down
0.745506198611303 0.269514021411837 0.667713171918741
GO_INTERLEUKIN_1_BETA_PRODUCTION 53 0.01 Down
0.610453450770917 0.269532770929072 0.667713171918741
GO_POSITIVE_REGULATION_OF_CHEMOTAXIS 97 0.01 Down
0.579258617455619 0.269566297237018 0.667713171918741
GO_NEGATIVE_CHEMOTAXIS 33 0.01 Down 0.46433156259182
0.269579718260537 0.667713171918741
GO_CELL_DIFFERENTIATION_INVOLVED_IN_METANEPHROS_DEVELOPMENT 15
0.01 Down 0.999349998686686 0.269597285835271
0.667713171918741
GO_DEOXYRIBONUCLEASE_ACTIVITY 49 0.01 Up
0.184958767854675 0.269785793347168 0.667713171918741
GO_POSITIVE_REGULATION_OF_T_HELPER_CELL_DIFFERENTIATION 15 0.01
Down 0.495550287704882 0.269790876415199
0.667713171918741
GO_REGULATION_OF_EXTRINSIC_APOPTOTIC_SIGNALING_PATHWAY_VIA_DEATH_DOM
AIN_RECEPTORS 45 0.01 Down 0.332607363169868
0.270103279543414 0.667876535635443
GO_RNA_SPLICING_VIA_TRANSESTERIFICATION_REACTIONS 313 0.01
Down 0.103772207841337 0.2701229478092
0.667876535635443
GO_REGULATION_OF_CELL_SHAPE 128 0.01 Down
0.394754280019427 0.2701869167387 0.667876535635443
GO_REGULATION_OF_PROTEASOMAL_PROTEIN_CATABOLIC_PROCESS 169 0.01
Down 0.208829863313367 0.270867590930639
0.669023271272768
GO_CHROMOSOME_ORGANIZATION 971 0.01 Down
0.192673259284049 0.270871223980969 0.669023271272768
GO_TRANSLATION_REGULATOR_ACTIVITY_NUCLEIC_ACID_BINDING 97 0.01
Down 0.153301051440465 0.271472222116577
0.669631751959171
GO_POSITIVE_REGULATION_OF_MITOCHONDRIAL_FISSION 14 0.01 Up
0.348601150810326 0.271536475093053
0.669631751959171
GO_PROTEIN_ACETYLATION 169 0.01 Down 0.213725731569983
0.271757861074066 0.669631751959171
GO_SIN3_TYPE_COMPLEX 15 0.01 Down 0.131089895215477
0.271778582431736 0.669631751959171
GO_REGULATION_OF_UBIQUITIN_PROTEIN_LIGASE_ACTIVITY 19 0.01
Up 0.258263388182594 0.271812057753172
0.669631751959171
GO_NEGATIVE_REGULATION_OF_SMOOTH_MUSCLE_CELL_PROLIFERATION 35

0.01 Up 0.649306676285142 0.271999342510337 0.669631751959171
GO_NEGATIVE_REGULATION_OF_PROGRAMMED_NECROTIC_CELL_DEATH 12 0.01 Down 0.370956638065423 0.272103326692501 0.669631751959171
GO_PEPTIDE_RECEPTOR_ACTIVITY 42 0.01 Up 0.496273867226936 0.272149243040859 0.669631751959171
GO_LAMELLIPODIUM_MEMBRANE 20 0.01 Up 0.383576119526521 0.272186815810205 0.669631751959171
GO_COSTAMERE 16 0.01 Down 0.285118349030852 0.272220583731714 0.669631751959171
GO_REGULATION_OF_NEUROTRANSMITTER_TRANSPORT 83 0.01 Up 0.374275557933602 0.272406694475058 0.669818162072936
GO_REGULATION_OF_SISTER_CHROMATID_COHESION 21 0.01 Down 0.158298716540533 0.272686145665398 0.670233842240742
GO_REGULATION_OF_GRANULOCYTE_DIFFERENTIATION 11 0.01 Up 0.485020319385577 0.272801445473458 0.670245882423861
GO_P53_BINDING 59 0.01 Down 0.20346886156561 0.272991204960465 0.670315142415957
GO_DNA_HELICASE_ACTIVITY 45 0.01 Down 0.151276467982251 0.273054156326808 0.670315142415957
GO_CALCIUM_ION_TRANSMEMBRANE_TRANSPORTER_ACTIVITY 68 0.01 Up 0.38228624823864 0.273171259975908 0.670315142415957
GO_NUCLEOTIDE_EXCISION_REPAIR 105 0.01 Up 0.135207809787765 0.273400374760741 0.670315142415957
GO_PROTEIN_KINASE_COMPLEX 94 0.01 Down 0.232369108419833 0.273586251121451 0.670315142415957
GO_NEGATIVE_REGULATION_OF_KERATINOCYTE_PROLIFERATION 12 0.01 Down 0.347112188288495 0.273651744402204 0.670315142415957
GO_NEURONAL_ACTION_POTENTIAL 12 0.01 Up 0.380623951988231 0.273808059042267 0.670315142415957
GO_REGULATION_OF_SMALL_GTPASE_MEDIATED_SIGNAL_TRANSDUCTION 282 0.01 Down 0.326991118058457 0.273926479709619 0.670315142415957
GO_REGULATION_OF_MRNA_BINDING 10 0.01 Down 0.148789831161733 0.273929051242474 0.670315142415957
GO_HOST_CELL_PART 10 0.01 Up 0.519501343103448 0.274027367100241 0.670315142415957
GO_ANATOMICAL_STRUCTURE_MATURATION 109 0.01 Down 0.42489807758842 0.27411838584084 0.670315142415957
GO_CELL_PROLIFERATION_INVOLVED_IN_KIDNEY_DEVELOPMENT 17 0.01 Down 0.783667535149466 0.274154587155135 0.670315142415957
GO_POSITIVE_REGULATION_OF_INTRINSIC_APOPTOTIC_SIGNALING_PATHWAY 51 0.01 Down 0.33657698151339 0.27441032785883 0.670670330286214
GO_POSITIVE_REGULATION_OF_GRANULOCYTE_CHEMOTAXIS 17 0.01 Down 0.870930759137731 0.274612860047333 0.670895240783645

GO_HSP70_PROTEIN_BINDING 38 0.01 Down
0.270183158647645 0.274998236508319 0.671566489880129
GO_REGULATION_OF_VESICLE_FUSION 21 0.01 Down
0.214095319495563 0.275164939485787 0.671703396710178
GO_PEPTIDE_ALPHA_N_ACETYLTRANSFERASE_ACTIVITY 11 0.01 Down
0.169155645773305 0.275429516465072
0.67196277553166
GO_DEFINITIVE_HEMOPOIESIS 12 0.01 Down
0.399338215872533 0.275492562724148 0.67196277553166
GO_THYROID_GLAND_DEVELOPMENT 18 0.01 Down
0.545844910896564 0.275834019401967 0.67212336148549
GO_C21_STEROID_HORMONE_METABOLIC_PROCESS 24 0.01 Up
0.557327390595469 0.275867685716247
0.67212336148549
GO_LEUKOCYTE_HOMEOSTASIS 64 0.01 Down
0.480734940012487 0.275902066886472 0.67212336148549
GO_REGULATION_OF_MULTICELLULAR_ORGANISM_GROWTH 46 0.01 Down
0.316807097475655 0.276003139830255
0.67212336148549
GO_REGULATION_OF_PROTEIN_IMPORT 51 0.01 Down
0.380742021079462 0.276161862819706 0.67212336148549
GO_RENAL_SYSTEM_PROCESS 70 0.01 Up 0.447844154183513
0.276313722435257 0.67212336148549
GO_REGULATION_OF_HUMORAL_IMMUNE_RESPONSE 37 0.01 Up
0.838655503339342 0.276333373458703
0.67212336148549
GO_SIGNAL_SEQUENCE_BINDING 38 0.01 Down
0.182790181895323 0.276480103697334 0.672210936942938
GO_ACTIVIN_RECEPTOR_SIGNALING_PATHWAY 26 0.01 Down
0.343919970171061 0.276647852041255 0.672349523515797
GO_POSITIVE_REGULATION_OF_CELL_CYCLE_PHASE_TRANSITION 76 0.01
Up 0.232472436440696 0.277376029174091
0.673849489041978
GO_WNT_SIGNALING_PATHWAY_CALCIUM_MODULATING_PATHWAY 31 0.01
Down 0.278737630645451 0.277505542445762
0.673884033216054
GO_NEPHRON_TUBULE_FORMATION 12 0.01 Up
0.933796399401128 0.277670762784175 0.673884033216054
GO_9PLUS0_NON_MOTILE_CILIUM 57 0.01 Up
0.256683170169516 0.277723250058733 0.673884033216054
GO_NEGATIVE_REGULATION_OF_EPITHELIAL_CELL_DIFFERENTIATION 28
0.01 Down 0.553257780019634 0.277867903836646
0.673965658886248
GO_CELL_PROLIFERATION_INVOLVED_IN_HEART_MORPHOGENESIS 13 0.01
Down 0.2595866345532 0.278331373025005
0.674062756103843
GO_INTRINSIC_APOPTOTIC_SIGNALING_PATHWAY_BY_P53_CLASS_MEDIATOR 71
0.01 Up 0.233161082451925 0.278536513509288
0.674062756103843
GO_EAR_DEVELOPMENT 139 0.01 Down 0.478815744803319
0.278589751205931 0.674062756103843
GO_REGULATION_OF_PROTEIN_LOCALIZATION_TO_NUCLEUS 96 0.01
Down 0.323380591498949 0.278660111813345
0.674062756103843

GO_CELLULAR_RESPONSE_TO_STEROID_HORMONE_STIMULUS 196 0.01 Down 0.301242792169006 0.278795247080758 0.674062756103843
GO_MITOTIC_DNA_REPLICATION 10 0.01 Up 0.229738699032074 0.27880655053985 0.674062756103843
GO_HISTONE_H3_K27_METHYLATION 16 0.01 Down 0.146143111307428 0.278819924626072 0.674062756103843
GO_NEGATIVE_REGULATION_OF_GLIAL_CELL_DIFFERENTIATION 13 0.01 Down 0.312106658338043 0.278858064333619 0.674062756103843
GO_ORGANIC_ACID_BIOSYNTHETIC_PROCESS 208 0.01 Up 0.369353590778276 0.278966341229353 0.674062756103843
GO_REGULATION_OF_CELL_MATURATION 13 0.01 Down 0.207101807692008 0.279018235231256 0.674062756103843
GO_IMMUNOGLOBULIN_PRODUCTION 71 0.01 Down 0.323547592530067 0.279321818071005 0.674261157216977
GO_ENERGY_COUPLED_PROTON_TRANSMEMBRANE_TRANSPORT_AGAINST_ELECTROCHEMICAL_GRADIENT 25 0.01 Up 0.247051118793341 0.279364763182957 0.674261157216977
GO_NEUROPEPTIDE_SIGNALING_PATHWAY 21 0.01 Up 0.58098089850254 0.279530298689062 0.674261157216977
GO_POSITIVE_REGULATION_OF_LYMPHOCYTE_DIFFERENTIATION 71 0.01 Down 0.415941749973449 0.279638786345345 0.674261157216977
GO_TRANSFERASE_COMPLEX_TRANSFERRING_PHOSPHORUS_CONTAINING_GROUPS 227 0.01 Down 0.194733761763528 0.279663957302266 0.674261157216977
GO_POSITIVE_REGULATION_OF_APOPTOTIC_SIGNALING_PATHWAY 159 0.01 Down 0.304143490595391 0.27992680229569 0.674261157216977
GO_UBIQUITIN_SPECIFIC_PROTEASE_BINDING 13 0.01 Down 0.0950991693428254 0.280106442544071 0.674261157216977
GO_NEGATIVE_REGULATION_OF_CELL_DEATH 737 0.01 Down 0.355258382631574 0.280421673199038 0.674261157216977
GO_PEPTIDE_METABOLIC_PROCESS 88 0.01 Up 0.425876371612584 0.280658398963509 0.674261157216977
GO_DEOXYRIBOSE_PHOSPHATE_BIOSYNTHETIC_PROCESS 11 0.01 Up 0.189150405312275 0.280692761105872 0.674261157216977
GO_DNA_SECONDARY_STRUCTURE_BINDING 22 0.01 Up 0.204523510417565 0.280763260333936 0.674261157216977
GO_EXTRINSIC_COMPONENT_OF_SYNAPTIC_MEMBRANE 10 0.01 Down 0.247754366429415 0.280838604517941 0.674261157216977
GO_POSTSYNAPTIC_MEMBRANE_ORGANIZATION 26 0.01 Up 0.236408014460107 0.280890888624199 0.674261157216977
GO_DETECTION_OF_BIOTIC_STIMULUS 13 0.01 Down 0.779592130795725 0.280987574734915 0.674261157216977
GO_LIGAND_GATED_CALCIUM_CHANNEL_ACTIVITY 17 0.01 Up 0.383385444133576 0.28100288662416 0.674261157216977
GO_PANCREATIC_JUICE_SECRETION 10 0.01 Up 0.588909740694726 0.281054304532545 0.674261157216977
GO_GROWTH_FACTOR_ACTIVITY 81 0.01 Down

0.554657799506004	0.28105477474464	0.674261157216977
GO_LEUKOCYTE_PROLIFERATION	197	0.01	Down
0.565975448601872	0.281298923586859	0.674261157216977
GO_REGULATION_OF_CELL_PROJECTION_ORGANIZATION	520	0.01	Down
	0.349814010714693	0.281320310689616
0.674261157216977
GO_INFLAMMATORY_RESPONSE_TO_ANTIGENIC_STIMULUS	27	0.01	Down
	0.422257228570782	0.281531322696769
0.674261157216977
GO_TRNA_PROCESSING	128	0.01	Up	0.128571804315454
	0.28161382319966	0.674261157216977
GO_RESPONSE_TO_CORTICOSTERONE	10	0.01	Down
0.305952264996698	0.281634787997521	0.674261157216977
GO_MEMBRANOUS_SEPTUM_MORPHOGENESIS	10	0.01	Up
0.536721931827131	0.281664151376194	0.674261157216977
GO_GPI_ANCHOR_BIOSYNTHETIC_PROCESS	30	0.01	Up
0.150442270070906	0.281765863261319	0.674261157216977
GO_POSITIVE_REGULATION_OF_REPRODUCTIVE_PROCESS	46	0.01	Down
	0.478073277758342	0.281948443857302
0.674432231149597
GO_VENTRICULAR_SYSTEM_DEVELOPMENT	24	0.01	Up
0.369842947189365	0.282233700704255	0.674848679391703
GO_PLATELET_DERIVED_GROWTH_FACTOR_RECEPTOR_BETA_SIGNALING_PATHWAY
	13	0.01	Up	0.503108474745901
0.283198884185691	0.676398754229129
GO_GLAND_DEVELOPMENT	323	0.01	Down	0.383465512749279
	0.283283328236221	0.676398754229129
GO_SMOOTH_MUSCLE_CELL_DIFFERENTIATION	48	0.01	Up
0.444405665565194	0.283389757074056	0.676398754229129
GO_MORPHOGENESIS_OF_AN_EPITHELIAL_FOLD	19	0.01	Up
0.528693799140199	0.283407710784677	0.676398754229129
GO_SPLICEOSOMAL_COMPLEX	174	0.01	Down	0.0974950681232847
	0.283439043116275	0.676398754229129
GO_REGULATION_OF_INTERLEUKIN_4_PRODUCTION	21	0.01	Up
	0.502647795909812	0.284210731137229
0.677755373507335
GO_ENDOSOME_TRANSPORT_VIA_MULTIVESICULAR_BODY_SORTING_PATHWAY	24
	0.01	Down	0.170144392958458	0.284297980708083
	0.677755373507335
GO_REGULATION_OF_OLIGODENDROCYTE_DIFFERENTIATION	22	0.01
	Down	0.354003623529238	0.284342437213504
0.677755373507335
GO_NEGATIVE_REGULATION_OF_T_CELL_PROLIFERATION	47	0.01	Down
	0.599101415291051	0.284456990343253
0.677762318828058
GO_METAPHASE_PLATE_CONGRESSION	49	0.01	Down
0.179406744660643	0.285263956453366	0.679418391380301
GO_REGULATION_OF_CELLULAR_RESPONSE_TO_DRUG	23	0.01	Down
	0.230765502973726	0.285651641953724
0.680074948353355
GO_RETINA_DEVELOPMENT_IN_CAMERA_TYPE_EYE	88	0.01	Up
	0.406219202787023	0.286132134882895
0.680951858437498
GO_BETA_CATENIN_TCF_COMPLEX_ASSEMBLY	28	0.01	Down

0.24176530083559	0.286353781117809	0.681212306099616
GO_RESPONSE_TO_VITAMIN	68	0.01	Up	0.528918588722218
	0.286570600746684	0.681353140808964
GO_POSITIVE_REGULATION_OF_PLASMA_MEMBRANE_BOUNDED_CELL_PROJECTION_ASSEMBLY	81	0.01	Down	0.2847970602108	0.286637443852099
	0.681353140808964
GO_ESTABLISHMENT_OF_MITOTIC_SPINDLE_LOCALIZATION	23	0.01
	Down	0.171730136381373	0.286824610929029
0.681531198806315
GO_NUCLEOSIDE_DIPHOSPHATE_KINASE_ACTIVITY	14	0.01	Up
	0.248781024016308	0.287293390390247
0.681716492231999
GO_NEGATIVE_REGULATION_OF_LEUKOCYTE_MIGRATION	28	0.01	Up
	0.924033429294131	0.28740129927713
0.681716492231999
GO_NEURON_PROJECTION_ORGANIZATION	67	0.01	Down
0.352641528582121	0.287421694951196	0.681716492231999
GO_SMOOTHENED_SIGNALING_PATHWAY_INVOLVED_IN_DORSAL_VENTRAL_NEURAL_TUBE_PATTERNING	14	0.01	Up	0.197354441335599
0.287448750130397	0.681716492231999
GO_REGULATION_OF_GLUCOSE_METABOLIC_PROCESS	57	0.01	Down
	0.368554084322936	0.287509618911655
0.681716492231999
GO_CELLULAR_RESPONSE_TO_NITROGEN_COMPOUND	466	0.01	Down
	0.338573236663382	0.287726305939643
0.681716492231999
GO_CELL_SURFACE	487	0.01	Up	0.566754348651968
0.287823964111494	0.681716492231999
GO_RESPONSE_TO_BRONCHODILATOR	31	0.01	Down
0.565832361921703	0.287848929306846	0.681716492231999
GO_INTERLEUKIN_4_PRODUCTION	25	0.01	Up
0.445763611079017	0.287935297708071	0.681716492231999
GO_REGULATION_OF_TOLL_LIKE_RECEPTOR_SIGNALING_PATHWAY	53	0.01
	Down	0.437764328106811	0.288025498694627
0.681716492231999
GO_TUBE_FORMATION	126	0.01	Up	0.429329426766206
	0.288206361356296	0.681821963069128
GO_POSITIVE_REGULATION_OF_MYOTUBE_DIFFERENTIATION	23	0.01
	Down	0.556573201213579	0.288294676198065
0.681821963069128
GO_N_ACYLTRANSFERASE_ACTIVITY	84	0.01	Down
0.163434479970996	0.288591951130266	0.681866298254546
GO_NONRIBOSOMAL_PEPTIDE_BIOSYNTHETIC_PROCESS	18	0.01	Up
	0.25469161222942	0.28864766706804
0.681866298254546
GO_REGULATION_OF_LYMPHOCYTE_APOPTOTIC_PROCESS	40	0.01	Down
	0.531337933841645	0.288650368393046
0.681866298254546
GO_VACUOLAR_MEMBRANE	367	0.01	Up	0.284901991853074
	0.288863011987967	0.682103207226351
GO_APICOLATERAL_PLASMA_MEMBRANE	15	0.01	Down
0.293874332295497	0.289036206365054	0.682246815257481
GO_NEURON_TO_NEURON_SYNAPSE	225	0.01	Down
0.27109769496706	0.289261568088771	0.682513400647853

GO_SULFATION	10	0.01	Up	0.371414888898835
0.289592658678932	0.68281388855923
GO_MAMMARY_GLAND_MORPHOGENESIS	37	0.01	Up
0.464443056544124	0.289698281635298	0.68281388855923
GO_GROWTH_CONE_PART	15	0.01	Up	0.392394709107471
	0.289726334529497	0.68281388855923
GO_CELL_CORTEX_PART	139	0.01	Down	0.258977171784372
	0.289847417399528	0.682834175798422
GO_CELL_VOLUME_HOMEOSTASIS	24	0.01	Down
0.522395520463043	0.290081914823905	0.682863109170243
GO_TRNA_METABOLIC_PROCESS	175	0.01	Up
0.128258523588217	0.290106568982265	0.682863109170243
GO_TISSUE_MIGRATION	245	0.01	Down	0.474631774582144
	0.290381043392999	0.682863109170243
GO_ACTIN_FILAMENT_POLYMERIZATION	143	0.01	Down
0.316860891497656	0.290554232400146	0.682863109170243
GO_CELLULAR_AMINO_ACID_BIOSYNTHETIC_PROCESS	48	0.01	Up
	0.248487091506503	0.290556305280495
0.682863109170243
GO_POSITIVE_REGULATION_OF_MORPHOGENESIS_OF_AN_EPITHELIUM	24
	0.01	Down	0.60498912983185	0.29060998781214
	0.682863109170243
GO_CARBOHYDRATE_PHOSPHORYLATION	22	0.01	Up
0.339960277119092	0.290647055525598	0.682863109170243
GO_PHOTORECEPTOR_CELL_MAINTENANCE	19	0.01	Up
0.246617085495165	0.290841720470973	0.683056125717322
GO_REGULATION_OF_GROWTH	498	0.01	Down	0.370669022413189
	0.291101030460719	0.68340075635229
GO_DRUG_BINDING	1340	0.01	Down	0.268262524710783
0.291760199053865	0.684279306070975
GO_NEUROTRANSMITTER_RECEPTOR_INTERNALIZATION	13	0.01	Down
	0.230120723502301	0.291885162831236
0.684279306070975
GO_REGULATION_OF_CELL_POPULATION_PROLIFERATION	1154	0.01	Down
	0.416330432131968	0.291929855895561
0.684279306070975
GO_NEGATIVE_REGULATION_OF_ORGANELLE_ASSEMBLY	33	0.01	Down
	0.123099068587469	0.292120143861974
0.684279306070975
GO_PROTON_TRANSPORTING_TWO_SECTOR_ATPASE_COMPLEX_PROTON_TRANSPORTING
_DOMAIN	20	0.01	Up	0.149625313521418
0.292340067214702	0.684279306070975
GO_POSITIVE_CHEMOTAXIS	46	0.01	Down	0.670295821346699
	0.292376922913526	0.684279306070975
GO_HISTONE_H2B_UBIQUITINATION	12	0.01	Down
0.118673720052837	0.292387429421399	0.684279306070975
GO_RNA_INTERFERENCE	14	0.01	Down	0.170768997557957
	0.292479072662843	0.684279306070975
GO_NEGATIVE_REGULATION_OF_MITOTIC_CELL_CYCLE	280	0.01	Down
	0.231487826143266	0.29266451826752
0.684279306070975
GO_POSITIVE_REGULATION_OF_DNA_METABOLIC_PROCESS	140	0.01	Down
	0.21163440223643	0.292786168452954
0.684279306070975

GO_REGULATION_OF_DENDRITE_MORPHOGENESIS 70 0.01 Down
0.326705633780278 0.292926934061901 0.684279306070975
GO_RNA_3_END_PROCESSING 132 0.01 Down 0.121667529743013
0.292961913816453 0.684279306070975
GO_NEGATIVE_REGULATION_OF_PROTEIN_MODIFICATION_PROCESS 500 0.01
Down 0.314573957131444 0.293129731977851
0.684279306070975
GO_NEGATIVE_REGULATION_OF_CALCIUM_ION_TRANSMEMBRANE_TRANSPORT 25
0.01 Up 0.330279159286964 0.293130949381598
0.684279306070975
GO_ANAPHASE_PROMOTING_COMPLEX 20 0.01 Up
0.117674341570708 0.29334787882943 0.684279306070975
GO_MITOCHONDRIAL_DNA_METABOLIC_PROCESS 11 0.01 Up
0.180285413023278 0.293354329607903 0.684279306070975
GO_CYCLOSPORIN_A_BINDING 15 0.01 Down
0.154155908685461 0.293391374353936 0.684279306070975
GO_REGULATION_OF_MUSCLE_ADAPTATION 66 0.01 Down
0.435580494716325 0.293767378988756 0.684893148172326
GO_CYCLIN_BINDING 27 0.01 Down 0.168650872218606
0.294043512795488 0.68490079871729
GO_POSITIVE_REGULATION_OF_NERVOUS_SYSTEM_DEVELOPMENT 372 0.01
Down 0.359431114885122 0.294064036620843
0.68490079871729
GO_REGULATION_OF_MORPHOGENESIS_OF_A_BRANCHING_STRUCTURE 41 0.01
Up 0.678258136962109 0.294160095988706
0.68490079871729
GO_POSITIVE_REGULATION_OF_HISTONE_H3_K4_METHYLATION 14 0.01
Down 0.147303432776179 0.294221921109322
0.68490079871729
GO_BASOLATERAL_PLASMA_MEMBRANE 142 0.01 Down
0.430651576810571 0.294949104654295 0.685959582953976
GO_PROTEIN_DENEDDYLATION 10 0.01 Up
0.0914998912018144 0.294953161141895 0.685959582953976
GO_ACTIVATION_OF_JUN_KINASE_ACTIVITY 35 0.01 Down
0.273896465328063 0.295015725760638 0.685959582953976
GO_RESPONSE_TO_UV_B 14 0.01 Up 0.261403201199705
0.295154943518311 0.68602054444857
GO_DEVELOPMENT_OF_PRIMARY_SEXUAL_CHARACTERISTICS 146 0.01
Down 0.362958749081751 0.295729876682668
0.687093793088587
GO_PHOSPHATIDYLINOSITOL_DEPHOSPHORYLATION 23 0.01 Down
0.206098969466771 0.296209322144478
0.687655949756304
GO_REGULATION_OF_T_CELL_DIFFERENTIATION_IN_THYMUS 20 0.01
Down 0.50559326579277 0.296424608035203
0.687655949756304
GO_LIPOPROTEIN_LOCALIZATION 14 0.01 Up
0.579323423814353 0.296437903122509 0.687655949756304
GO_RESPONSE_TO_PROSTAGLANDIN 20 0.01 Up
0.478108759107922 0.296599855461103 0.687655949756304
GO_CARBOHYDRATE_DERIVATIVE_BIOSYNTHETIC_PROCESS 543 0.01 Up
0.316232614825746 0.296631408721735
0.687655949756304
GO_TETRAPYRROLE_BIOSYNTHETIC_PROCESS 26 0.01 Up

0.180452892106935 0.296651446616992 0.687655949756304
GO_PLATELET_DERIVED_GROWTH_FACTOR_RECEPTOR_BINDING 14 0.01
Down 0.589538328932899 0.29677115205383
0.687670864167483
GO_HUMORAL_IMMUNE_RESPONSE_MEDIATED_BY_CIRCULATING_IMMUNOGLOBULIN
25 0.01 Up 0.808754878656441
0.296903699167226 0.687715512264109
GO_POSITIVE_REGULATION_OF_VIRAL_GENOME_REPLICATION 32 0.01
Down 0.206846319831109 0.297244845285858
0.688243118127552
GO_PEPTIDYL_LYSINE_MODIFICATION 346 0.01 Down
0.204051192075923 0.298115409404113 0.68999140489268
GO_REGULATION_OF_LEUKOCYTE_DEGRANULATION 36 0.01 Down
0.393728657136615 0.298227218981781
0.68999140489268
GO_REGULATION_OF_CELL_SIZE 142 0.01 Down
0.386643519380522 0.298438327113614 0.690216793869238
GO_ENAMEL_MINERALIZATION 11 0.01 Down
0.579678597359913 0.29863523775947 0.690319152725778
GO_AMINOPEPTIDASE_ACTIVITY 36 0.01 Up
0.313405169233545 0.298710000693563 0.690319152725778
GO_SERINE_THREONINE_PROTEIN_KINASE_COMPLEX 77 0.01 Down
0.234902158712642 0.29892453676025
0.69036454269758
GO_POSITIVE_REGULATION_OF_PEPTIDYL_SERINE_PHOSPHORYLATION 71
0.01 Down 0.387183358548446 0.298957071775974
0.69036454269758
GO_CELL_GROWTH 364 0.01 Down 0.373642323926169
0.299108549851595 0.69045171336465
GO_REGULATION_OF_ESTABLISHMENT_OF_PROTEIN_LOCALIZATION_TO_MITOCHONDR
ION 66 0.01 Down 0.174265270088847
0.29929482302628 0.690619107028713
GO_MODIFICATION_BY_SYMBIONT_OF_HOST_MORPHOLOGY_OR_PHYSIOLOGY 38
0.01 Down 0.313117464142905 0.299427131780927
0.690661898572191
GO_PHOSPHOLIPASE_A2_ACTIVITY_CONSUMING_1_2_DIPALMITOYLPHOSPHATIDYLCH
OLINE_ 14 0.01 Up 0.446937944065501
0.299632017507479 0.690843312738168
GO_SYNTAXIN_1_BINDING 13 0.01 Down 0.183638977658758
0.299733369420579 0.690843312738168
GO_NEGATIVE_REGULATION_OF_ERAD_PATHWAY 12 0.01 Down
0.160346786263135 0.299872389271527 0.690901432739067
GO_PURINE_NUCLEOSIDE_MONOPHOSPHATE_METABOLIC_PROCESS 19 0.01
Down 0.322632353848529 0.300139318332162
0.691254097721759
GO_POSITIVE_REGULATION_OF_TRANSCRIPTION_REGULATORY_REGION_DNA_BINDIN
G 16 0.01 Down 0.220183530895378
0.300265581215557 0.691282648297173
GO_MOLECULAR_CARRIER_ACTIVITY 29 0.01 Down
0.251600694770317 0.300449107029587 0.691352525442987
GO_NEGATIVE_REGULATION_OF_MUSCLE_HYPERTROPHY 24 0.01 Down
0.556615075291561 0.300523688789992
0.691352525442987
GO_NEGATIVE_REGULATION_OF_ENDOPLASMIC_RETICULUM_UNFOLDED_PROTEIN_RES

PONSE	11	0.01	Down	0.238126437958994
0.300944282433371	0.691921780667831
GO_TRANSLATION_REGULATOR_ACTIVITY	122	0.01	Down
0.161019106677659	0.301198123127724	0.691921780667831
GO_REGULATION_OF_PROTEIN_SERINE_THREONINE_KINASE_ACTIVITY	424
	0.01	Down	0.351227978463945	0.301222914899078
	0.691921780667831
GO_TELOMERE_CAPPING	40	0.01	Down	0.157639169368902
	0.30122702459316	0.691921780667831
GO_POLY_PYRIMIDINE_TRACT_BINDING	26	0.01	Down
0.164018534983957	0.30176811039638	0.692526645750316
GO_PRIMARY_ALCOHOL_METABOLIC_PROCESS	45	0.01	Up
0.462665104461493	0.30182722773644	0.692526645750316
GO_ATPASE_ACTIVITY_COUPLED_TO_TRANSMEMBRANE_MOVEMENT_OF_IONS_ROTATIO
NAL_MECHANISM	33	0.01	Up	0.145708387782639
0.301832565418438	0.692526645750316
GO_NEGATIVE_REGULATION_OF_APOPTOTIC_SIGNALING_PATHWAY	191	0.01
	Down	0.324854346883759	0.302348574762949
0.6927207983487
GO_RACEMASE_AND_EPIMERASE_ACTIVITY_ACTING_ON_CARBOHYDRATES_AND_DERIV
ATIVES	12	0.01	Down	0.318906575742417
0.30240849744105	0.6927207983487
GO_PROTEIN_DEPOLYMERIZATION	82	0.01	Down
0.295504201497912	0.302431774290698	0.6927207983487
GO_MATURATION_OF_5_8S_RRNA_FROM_TRICISTRONIC_RRNA_TRANSCRIPT_SSU_RRN
A_5_8S_RRNA_LSU_RRNA_	21	0.01	Up	0.118424521642439
	0.302533851589602	0.6927207983487
GO_MICROBODY_MEMBRANE	53	0.01	Up	0.180101734593873
	0.30259372836682	0.6927207983487
GO_NEGATIVE_REGULATION_OF_UBIQUITIN_PROTEIN_TRANSFERASE_ACTIVITY
	16	0.01	Up	0.236363998789416
0.302601804846113	0.6927207983487
GO_RECEPTOR_SERINE_THREONINE_KINASE_BINDING	12	0.01	Down
	0.614273080979834	0.303281525201404
0.693277635459435
GO_SYNAPTIC_VESICLE_CYTOSKELETAL_TRANSPORT	17	0.01	Up
	0.202807646438382	0.303302346769032
0.693277635459435
GO_POSITIVE_REGULATION_OF_SMOOTH_MUSCLE_CELL_MIGRATION	31	0.01
	Down	0.622723092510718	0.303397558529585
0.693277635459435
GO_RESPONSE_TO_IMMOBILIZATION_STRESS	17	0.01	Up
0.505216351546575	0.30359515523602	0.693277635459435
GO_POSITIVE_REGULATION_OF_RESPONSE_TO_ENDOPLASMIC_RETICULUM_STRESS
	36	0.01	Down	0.257527276596866
0.303632701161947	0.693277635459435
GO_DEAMINASE_ACTIVITY	23	0.01	Down	0.517725774049087
	0.303645259855687	0.693277635459435
GO_COPI_COATED_VESICLE_MEMBRANE	18	0.01	Down
0.128368419909122	0.303905855971182	0.693277635459435
GO_CLATHRIN_COATED_VESICLE	147	0.01	Down
0.411404963889278	0.303907475917629	0.693277635459435
GO_CYTOCHROME_COMPLEX_ASSEMBLY	34	0.01	Up
0.107967437672794	0.303918932873195	0.693277635459435

GO_NEURON_PROJECTION_EXTENSION 124 0.01 Down
0.353911429689697 0.30398699812107 0.693277635459435
GO_FAT_SOLUBLE_VITAMIN_METABOLIC_PROCESS 24 0.01 Up
0.511561550163078 0.304282995011758
0.693692100156358
GO_REGULATION_OF_POTASSIUM_ION_TRANSPORT 50 0.01 Down
0.496974845078471 0.304715134992568
0.693745911114228
GO_REGULATION_OF_TUBE_SIZE 73 0.01 Up
0.480398631089743 0.3047587501449 0.693745911114228
GO_NUCLEOCYTOPLASMIC_CARRIER_ACTIVITY 18 0.01 Down
0.134106795860506 0.304775247304676 0.693745911114228
GO_CUL2_RING_UBIQUITIN_LIGASE_COMPLEX 13 0.01 Down
0.164780071995167 0.304849608392406 0.693745911114228
GO_RESPONSE_TO_FOLLICLE_STIMULATING_HORMONE 12 0.01 Down
0.370283515565756 0.304877959290522
0.693745911114228
GO_INTERLEUKIN_13_PRODUCTION 16 0.01 Up
0.476713199775592 0.305460261554684 0.694810508766761
GO_RESPONSE_TO_COBALT_ION 10 0.01 Up
0.201225413838312 0.30559937366195 0.694866590824607
GO_CELLULAR_RESPONSE_TO_LITHIUM_ION 10 0.01 Up
0.329381709255428 0.305764416986088 0.694981570768454
GO_INTRINSIC_APOPTOTIC_SIGNALING_PATHWAY_IN_RESPONSE_TO_DNA_DAMAGE_BY_P53_CLASS_MEDIATOR 43 0.01 Up 0.265973426594064
0.305950245659416 0.695143690643081
GO_NEGATIVE_REGULATION_OF_DEVELOPMENTAL_PROCESS 643 0.01 Down
0.434010205572055 0.306277328565022
0.695626510182658
GO_PHOSPHATIDYLINOSITOL_3_KINASE_REGULATOR_ACTIVITY 13 0.01
Down 0.362499310238374 0.306633182286495
0.696174289327341
GO_EXOCYTIC_VESICLE 141 0.01 Up 0.29620962311266
0.306774179199388 0.696234034362423
GO_PROTEASOME_REGULATORY_PARTICLE_BASE_SUBCOMPLEX 12 0.01
Down 0.0918283806295652 0.307082215020736
0.696672693344876
GO_INTRINSIC_COMPONENT_OF_PLASMA_MEMBRANE 906 0.01 Up
0.490835114771875 0.307409392278569
0.697154434263427
GO_NEGATIVE_REGULATION_OF_TUMOR_NECROSIS_FACTOR_SUPERFAMILY_CYTOKINE_PRODUCTION 33 0.01 Down 0.585645994343851
0.307822679387196 0.697396437354242
GO_POSITIVE_REGULATION_OF_URINE_VOLUME 10 0.01 Up
0.41014690907369 0.307937448988053 0.697396437354242
GO_DEVELOPMENTAL_INDUCTION 20 0.01 Down
0.459571838595012 0.308009015947618 0.697396437354242
GO_EXOGENOUS_PROTEIN_BINDING 58 0.01 Down
0.426313189789537 0.308070304245195 0.697396437354242
GO_OMEGA_PEPTIDASE_ACTIVITY 10 0.01 Up
0.411600520307598 0.308090470265867 0.697396437354242
GO_CELLULAR_DEFENSE_RESPONSE 28 0.01 Down
0.597232954243492 0.308552327833524 0.698037244379918
GO_SMAD_BINDING 71 0.01 Down 0.367045597735413

0.308603519010987	0.698037244379918
GO_TRANSFERASE_ACTIVITY_TRANSFERRING_NITROGENOUS_GROUPS 18	0.01
	Up	0.28208201940954	0.308874940934287
0.698285388558668
GO_CYTOKINE_RECEPTOR_ACTIVITY	56	0.01	Down
0.589560128830732	0.30902743810118	0.698285388558668
GO_POSITIVE_REGULATION_OF_SUPRAMOLECULAR_FIBER_ORGANIZATION	166
	0.01	Down	0.299495798537341	0.30911149583336
	0.698285388558668
GO_ACID_THIOL_LIGASE_ACTIVITY	19	0.01	Up
0.306141768856314	0.309173303318349	0.698285388558668
GO_REGULATION_OF_HISTONE_H3_K4_METHYLATION	24	0.01	Down
	0.140142399508119	0.309391282164775
0.698471241579732
GO_REGULATION_OF_INFLAMMATORY_RESPONSE_TO_ANTIGENIC_STIMULUS	13
	0.01	Down	0.54732276227203	0.309485692612334
	0.698471241579732
GO_POLY_A_BINDING	19	0.01	Down	0.179940655785764
	0.309632941921778	0.69854388346604
GO_ATRIAL_SEPTUM_DEVELOPMENT	12	0.01	Down
0.272387292321921	0.309947594165322	0.698773102250376
GO_RESPONSE_TO_INTERLEUKIN_12	42	0.01	Down
0.255928225214403	0.309964744582484	0.698773102250376
GO_ENDOPLASMIC_RETICULUM_ORGANIZATION	49	0.01	Down
0.179558336522606	0.31025411654649	0.699165828342145
GO_VESICLE_CYTOSKELETAL_TRAFFICKING	50	0.01	Down
0.213427735262419	0.310650083047757	0.699767571674123
GO_POSITIVE_REGULATION_OF_ACTIN_FILAMENT_POLYMERIZATION 80	0.01
	Down	0.325993042087275	0.310751667473799
0.699767571674123
GO_CELL_ADHESION_MEDIATED_BY_INTEGRIN	57	0.01	Down
0.630915241249274	0.310935089298214	0.699920996340176
GO_BASAL_TRANSCRIPTION_MACHINERY_BINDING	55	0.01	Down
	0.174868056275806	0.311666751288305
0.701307949248072
GO_POSITIVE_REGULATION_OF_T_CELL_CYTOKINE_PRODUCTION	17	0.01
	Down	0.48822667749492	0.312004605702359
0.70157967410586
GO_NEGATIVE_REGULATION_OF_DENDRITE_MORPHOGENESIS	11	0.01
	Down	0.432371007591817	0.312018632858808
0.70157967410586
GO_AUTOPHAGOSOME	80	0.01	Down	0.196553159614692
	0.312371143685227	0.702112259649393
GO_LOW_DENSITY_LIPOPROTEIN_PARTICLE_RECEPTOR_CATABOLIC_PROCESS	13
	0.01	Up	0.263622616567032	0.312602882827516
	0.702373094613564
GO_EXCITATORY_SYNAPSE_ASSEMBLY	14	0.01	Up
0.248745483298503	0.313438947223285	0.703448828983758
GO_SALIVARY_GLAND_DEVELOPMENT	33	0.01	Up
0.522758666069164	0.313450920810047	0.703448828983758
GO_CELL_CELL_ADHESION_MEDIATED_BY_CADHERIN	26	0.01	Up
	0.47795259055519	0.31357813756601
0.703448828983758
GO_TRANSCRIPTION_FACTOR_COMPLEX 273	0.01	Down

0.260589799128026	0.31370751586178	0.703448828983758
GO_NUCLEAR_DNA_REPLICATION	50	0.01	Up
0.162764994261989	0.313710468729293	0.703448828983758
GO_SISTER_CHROMATID_COHESION	47	0.01	Down
0.198419024250794	0.313776878420033	0.703448828983758
GO_NEGATIVE_REGULATION_OF_CELL_MIGRATION_INVOLVED_IN_SPROUTING_ANGIOGENESIS 14	0.01	Down	0.259805862180441
0.314140292113005	0.704003585610209
GO_PROTON_TRANSPORTING_TWO_SECTOR_ATPASE_COMPLEX_CATALYTIC_DOMAIN	15	0.01	Up	0.129321504254202
0.314594397842559	0.704464151002176
GO_NEGATIVE_REGULATION_OF_ENDOCYTOSIS	32	0.01	Up
0.366828584645391	0.314724099137108	0.704464151002176
GO_PHOSPHATIDYLINOSITOL_3_4_5_TRISPHOSPHATE_BINDING	34	0.01	Down	0.207908077203833	0.314803984672769
0.704464151002176
GO_REGULATION_OF_RNA_BINDING	12	0.01	Down
0.140109789319647	0.31480995580117	0.704464151002176
GO_REGULATION_OF_EXIT_FROM_MITOSIS	16	0.01	Up
0.254201197430059	0.315083287301167	0.704795008885022
GO_MESODERM_DEVELOPMENT 82	0.01	Down	0.424859168420184	0.315191868624709	0.704795008885022
GO_RESPONSE_TO_ABIOTIC_STIMULUS 927	0.01	Down
0.300883037484856	0.315366906582834	0.704795008885022
GO_INTERLEUKIN_12_PRODUCTION	39	0.01	Down
0.594469661128465	0.315422177423918	0.704795008885022
GO_PROTEIN_TRANSPORT_WITHIN_PLASMA_MEMBRANE	27	0.01	Down	0.222901503755332	0.316846099944881
0.707716215145465
GO_REGULATION_OF_INTRACELLULAR_TRANSPORT	347	0.01	Down	0.255090395137182	0.317236719555428
0.707931620209062
GO_ESTABLISHMENT_OF_ORGANELLE_LOCALIZATION	400	0.01	Down	0.242515944038308	0.317506621426846
0.707931620209062
GO_CELLULAR_RESPONSE_TO_CARBOHYDRATE_STIMULUS	106	0.01	Up	0.284304327051259	0.317678472239459
0.707931620209062
GO_SULFUR_COMPOUND_TRANSMEMBRANE_TRANSPORTER_ACTIVITY	21	0.01	Up	0.232931646619046	0.317740398556806
0.707931620209062
GO_PEPTIDYL_PROLINE_MODIFICATION	46	0.01	Up
0.234975849410866	0.317767770985981	0.707931620209062
GO_SITE_OF_POLARIZED_GROWTH	140	0.01	Down
0.331014302931675	0.317926544870462	0.707931620209062
GO_POSITIVE_REGULATION_OF_ENDOPLASMIC_RETICULUM_UNFOLDED_PROTEIN_RESPONSE	13	0.01	Down	0.339208818470316
0.317959155596437	0.707931620209062
GO_AMYLOID_PRECURSOR_PROTEIN_METABOLIC_PROCESS	46	0.01	Down	0.361216251302629	0.317984270002737
0.707931620209062
GO_INTRINSIC_COMPONENT_OF_SYNAPTIC_MEMBRANE	56	0.01	Up	0.374182531580878	0.318028752510428
0.707931620209062

GO_REGULATION_OF_BEHAVIOR	29	0.01	Down	0.495535008223331	0.31820116792329	0.707931620209062
GO_FIBROBLAST_APOPTOTIC_PROCESS	18	0.01	Down	0.252767772090418	0.318225233330675	0.707931620209062
GO_PROTEIN_ACYLATION	207	0.01	Down	0.207105823342332	0.318756364107835	0.708853438277901
GO_MAGNESIUM_DEPENDENT_PROTEIN_SERINE_THREONINE_PHOSPHATASE_ACTIVITY	14	0.01	Down	0.179959427496223	0.319040745106337	0.709125158265886
GO_PURINE_NTP_DEPENDENT_HELICASE_ACTIVITY	72	0.01	Down	0.187686570550678	0.319112161486147	0.709125158265886
GO_REGULATION_OF_PROTEIN_KINASE_C_SIGNALING	10	0.01	Down	0.449740600478119	0.319436848619903	0.709306213144494
GO_PROTEOLYSIS	1352	0.01	Down	0.314924279209261	0.31961996760395	0.709306213144494
GO_TRANSITION_METAL_ION_TRANSMEMBRANE_TRANSPORTER_ACTIVITY	30	0.01	Up	0.331006542667421	0.319711880209283	0.709306213144494
GO_CATALYTIC_ACTIVITY_ACTING_ON_A_GLYCOPROTEIN	20	0.01	Down	0.452410528958586	0.319814264552142	0.709306213144494
GO_REGULATION_OF_STEROID_BIOSYNTHETIC_PROCESS	70	0.01	Down	0.333198479133188	0.31984760893524	0.709306213144494
GO_MITOCHONDRIAL_RESPIRATORY_CHAIN_COMPLEX_IV_ASSEMBLY	20	0.01	Up	0.105477208149471	0.319964935722149	0.709306213144494
GO_REGULATION_OF_PROTEIN_STABILITY	235	0.01	Down	0.199852724563104	0.320051397485506	0.709306213144494
GO_REGULATION_OF_PROTEOLYSIS	545	0.01	Down	0.3571694675052	0.320358653224851	0.709306213144494
GO_NEGATIVE_REGULATION_OF_CARDIAC_MUSCLE_TISSUE_DEVELOPMENT	24	0.01	Down	0.464049425481773	0.32061221688622	0.709306213144494
GO_EXCITATORY_SYNAPSE	15	0.01	Up	0.304906961934776	0.320710986017477	0.709306213144494
GO_PEPTIDYL_LYSINE_DIMETHYLATION	14	0.01	Up	0.162769608899578	0.320719836878501	0.709306213144494
GO_RAB_PROTEIN_SIGNAL_TRANSDUCTION	64	0.01	Down	0.259735463886676	0.320880178134607	0.709306213144494
GO_PORE_COMPLEX_ASSEMBLY	15	0.01	Down	0.229859229946486	0.32089673286415	0.709306213144494
GO_POSITIVE_REGULATION_OF_REGULATED_SECRETORY_PATHWAY	37	0.01	Down	0.333016867542831	0.320994545257513	0.709306213144494
GO_REGULATION_OF_COENZYME_METABOLIC_PROCESS	12	0.01	Up	0.191693423971912	0.32107989425983	0.709306213144494
GO_KETONE_BIOSYNTHETIC_PROCESS	30	0.01	Up	0.316936809612296	0.321163539381722	0.709306213144494
GO_EMBRYONIC_PLACENTA_MORPHOGENESIS	21	0.01	Down	0.270031937694122	0.321179835271654	0.709306213144494

GO_PYRIMIDINE_NUCLEOSIDE_TRIPHOSPHATE_BIOSYNTHETIC_PROCESS 17 0.01 Up 0.270233703530899 0.321476715452377 0.709350042464741
GO_NEGATIVE_REGULATION_OF_CELL_MOTILITY 216 0.01 Up 0.528405883572335 0.321571380151623 0.709350042464741
GO_VOLTAGE_GATED_POTASSIUM_CHANNEL_ACTIVITY 27 0.01 Down 0.624812150798026 0.321647353451378 0.709350042464741
GO_REGULATION_OF_MYOBLAST_FUSION 16 0.01 Down 0.766236764859893 0.321667051046851 0.709350042464741
GO_CELLULAR_RESPONSE_TO_NITRIC_OXIDE 13 0.01 Down 0.689722830379924 0.321916584787166 0.709374476497643
GO_PROTEIN_SERINE_THREONINE_TYROSINE_KINASE_ACTIVITY 39 0.01 Down 0.224325209800247 0.321919698175245 0.709374476497643
GO_INTRINSIC_COMPONENT_OF_ENDOPLASMIC_RETICULUM_MEMBRANE 142 0.01 Up 0.263044499442182 0.32202867027302 0.709374476497643
GO_STEROID_HORMONE_MEDIATED_SIGNALING_PATHWAY 149 0.01 Down 0.257969497941436 0.322566230018748 0.710188949484465
GO_REGULATION_OF_VITAMIN_METABOLIC_PROCESS 10 0.01 Down 0.687820074187981 0.3226323707261 0.710188949484465
GO_HOMOLOGOUS_RECOMBINATION 37 0.01 Up 0.192926285169053 0.322841902974338 0.710386901142596
GO_MAMMARY_GLAND_ALVEOLUS_DEVELOPMENT 11 0.01 Up 0.374695802791009 0.323012185338413 0.710386901142596
GO_REGULATION_OF_SISTER_CHROMATID_SEGREGATION 71 0.01 Up 0.207151927947762 0.323073337844623 0.710386901142596
GO_PROTEIN_CONTAINING_COMPLEX_LOCALIZATION 240 0.01 Down 0.178023660641053 0.3233169072326 0.710665077410975
GO_NEGATIVE_REGULATION_OF_MITOCHONDRION_ORGANIZATION 41 0.01 Up 0.21707700523506 0.323606513888489 0.711044207678979
GO_RIBOSOME_ASSEMBLY 57 0.01 Up 0.120914986287661 0.323726980517622 0.711051555254155
GO_REGIONALIZATION 201 0.01 Up 0.46753144073582 0.324122801296482 0.711653540169721
GO_POSITIVE_REGULATION_OF_ADHERENS_JUNCTION_ORGANIZATION 28 0.01 Down 0.352165290153507 0.324235495323579 0.711653540169721
GO_CENTRAL_NERVOUS_SYSTEM_PROJECTION_NEURON_AXONOGENESIS 19 0.01 Down 0.429043885288077 0.324508760532907 0.711748619760716
GO_PROTEIN_TETRAMERIZATION 110 0.01 Up 0.266754877956855 0.32451328932592 0.711748619760716
GO_POSITIVE_REGULATION_OF_CELLULAR_PROTEIN_CATABOLIC_PROCESS 129 0.01 Down 0.21783878351093 0.32493173628923 0.712409018061362
GO_ENDOTHELIAL_CELL_DEVELOPMENT 52 0.01 Down 0.480609313591793 0.325244356769875 0.712601983233942
GO_ACTIVATION_OF_PROTEIN_KINASE_A_ACTIVITY 13 0.01 Down 0.317352622811503 0.325444648423894

0.712601983233942
GO_TETRAPYRROLE_BINDING 81 0.01 Up 0.50520102500848 0.325474362846496 0.712601983233942
GO_MULTICELLULAR_ORGANISM_GROWTH 110 0.01 Down 0.288345301690797 0.325489260337295 0.712601983233942
GO_POSITIVE_REGULATION_OF_FATTY_ACID_METABOLIC_PROCESS 11 0.01 Up 0.368849873318034 0.325688386958426 0.712709101571784
GO_NEGATIVE_REGULATION_OF_MORPHOGENESIS_OF_AN_EPITHELIUM 15 0.01 Down 0.491174137094176 0.325772979222813 0.712709101571784
GO_NATURAL_KILLER_CELL_ACTIVATION 52 0.01 Down 0.49230213066231 0.326042176737413 0.712926867893479
GO_CELLULAR_AMINO_ACID_METABOLIC_PROCESS 245 0.01 Up 0.26848792396782 0.326107381344126 0.712926867893479
GO_REGULATION_OF_PRODUCTION_OF_SMALL_RNA_INVOLVED_IN_GENE_SILENCING_BY_RNA 17 0.01 Down 0.255931083762865 0.326767086315769 0.714111944212755
GO_REGULATION_OF_ORGANIC_ACID_TRANSPORT 33 0.01 Up 0.420840177275123 0.326982275389271 0.71432507876512
GO_PHAGOCYTIC_CUP 22 0.01 Down 0.43758852298879 0.327415963544791 0.714769110370849
GO_SYNAPTIC_VESICLE_RECYCLING 49 0.01 Down 0.273287160202806 0.32745607259706 0.714769110370849
GO_DOPAMINERGIC_NEURON_DIFFERENTIATION 20 0.01 Up 0.680189219330039 0.327538735801631 0.714769110370849
GO_CELLULAR_RESPONSE_TO_REACTIVE_NITROGEN_SPECIES 16 0.01 Down 0.612782072766094 0.328726563991032 0.717007463269546
GO_RESPONSE_TO_PROGESTERONE 31 0.01 Down 0.354444529313248 0.329214500108956 0.717007463269546
GO_B_CELL_DIFFERENTIATION 85 0.01 Down 0.406588753061068 0.329388674904557 0.717007463269546
GO_NUCLEAR_OUTER_MEMBRANE_ENDOPLASMIC_RETICULUM_MEMBRANE_NETWORK 890 0.01 Up 0.28173583233305 0.329404144642816 0.717007463269546
GO_NEGATIVE_REGULATION_OF_REGULATED_SECRETORY_PATHWAY 18 0.01 Down 0.300127215123467 0.329477613474364 0.717007463269546
GO_POSITIVE_REGULATION_OF_SECRETION 271 0.01 Down 0.449404524342378 0.329522283941971 0.717007463269546
GO_RENAL_SYSTEM_VASCULATURE_DEVELOPMENT 18 0.01 Up 0.942412632711218 0.329598425060056 0.717007463269546
GO_POSITIVE_REGULATION_OF_UBIQUITIN_PROTEIN_TRANSFERASE_ACTIVITY 28 0.01 Down 0.175255396199986 0.329648008471795 0.717007463269546
GO_POSITIVE_REGULATION_OF_GENE_EXPRESSION_EPIGENETIC 50 0.01 Down 0.174216631292331 0.329771359779179 0.717007463269546
GO_MESODERMAL_CELL_DIFFERENTIATION 22 0.01 Down 0.72244618783112 0.329959516101214 0.717007463269546
GO_REGULATION_OF_LEUKOCYTE_APOPTOTIC_PROCESS 65 0.01 Down 0.56381461433257 0.330098016815554

0.717007463269546
GO_U2_TYPE_CATALYTIC_STEP_2_SPLICEOSOME 30 0.01 Down
0.0832283358066084 0.330219015616516 0.717007463269546
GO_CORTICAL_ACTIN_CYTOSKELETON 65 0.01 Down
0.275310008245827 0.330271964408143 0.717007463269546
GO_PALLIUM_DEVELOPMENT 122 0.01 Down 0.325687246606321
0.330297159186429 0.717007463269546
GO_RESPONSE_TO_ALKALOID 64 0.01 Up 0.328010825521554
0.330336003090911 0.717007463269546
GO_RESPONSE_TO_ANTIMETABOLITE 11 0.01 Up
0.24893673350999 0.330610204515863 0.717346158547464
GO_PIGMENT_GRANULE_MATURATION 10 0.01 Down
0.145245860718002 0.330857185288628 0.717470327088945
GO_NUCLEOBASE_BIOSYNTHETIC_PROCESS 18 0.01 Up
0.265964919058236 0.330903791113333 0.717470327088945
GO_CELL_PROLIFERATION_IN_HINDBRAIN 10 0.01 Up
0.423936519560292 0.331059396247898 0.717551443991785
GO_PROTEIN_LOCALIZATION_TO_PLASMA_MEMBRANE 210 0.01 Down
0.26805703628523 0.331216835544877
0.7176364770139
GO_REGULATION_OF_SPINDLE_CHECKPOINT 13 0.01 Up
0.239564367677139 0.33164388439324 0.718127769512004
GO_ENDODEOXYRIBONUCLEASE_ACTIVITY 33 0.01 Up
0.179349912574353 0.331725340424598 0.718127769512004
GO_SPHINGOLIPID_METABOLIC_PROCESS 69 0.01 Up
0.335239733282558 0.331798450581646 0.718127769512004
GO_MACROPHAGE_MIGRATION 32 0.01 Up 0.629235641221284
0.332624627553881 0.719659342081116
GO_REGULATION_OF_SECRETION 503 0.01 Down
0.431137530521815 0.333204455259004 0.720657017412688
GO_BETA_CATENIN_DESTRUCTION_COMPLEX 11 0.01 Down
0.226273797700431 0.333440786017936 0.72091132902952
GO_SYNAPTONEMAL_STRUCTURE 19 0.01 Up
0.242053367739372 0.333826917678885 0.721489219376473
GO_AMYLOID_BETA_METABOLIC_PROCESS 34 0.01 Down
0.413245872760629 0.33410082157369 0.721693472021007
GO_GOLGI_ASSOCIATED_VESICLE_MEMBRANE 98 0.01 Down
0.318026874984619 0.334272287377632 0.721693472021007
GO_PATTERN_RECOGNITION_RECEPTOR_ACTIVITY 13 0.01 Up
1.19496889337285 0.334366756234032
0.721693472021007
GO_CELLULAR_RESPONSE_TO_OXYGEN_CONTAINING_COMPOUND 816 0.01
Down 0.393288189877824 0.334396925843369
0.721693472021007
GO_NEGATIVE_REGULATION_OF_CELL_CYCLE 541 0.01 Down
0.249494248095867 0.334985842992275 0.722670024559682
GO_NEGATIVE_REGULATION_OF_TELOMERE_MAINTENANCE 34 0.01 Down
0.14318682728475 0.335144611485792
0.722670024559682
GO_NUCLEAR_ENVELOPE_REASSEMBLY 18 0.01 Down
0.132533081936763 0.335206521027848 0.722670024559682
GO_NEUTRAL_LIPID_METABOLIC_PROCESS 74 0.01 Up
0.33932655057617 0.33539217388536 0.722813591642889
GO_POSITIVE_REGULATION_OF_T_CELL_MEDIATED_CYTOTOXICITY 15 0.01

Down 0.403398662870254 0.335619459087367
0.722974512751964
GO_NUCLEOTIDE_ACTIVATED_PROTEIN_KINASE_COMPLEX 12 0.01 Down
0.127136500329054 0.335796863769745
0.722974512751964
GO_INTRACELLULAR_ESTROGEN_RECEPTOR_SIGNALING_PATHWAY 48 0.01
Down 0.193070035906372 0.335824102447791
0.722974512751964
GO_RELAXATION_OF_CARDIAC_MUSCLE 11 0.01 Up
0.522873437295073 0.335988153168494 0.723071278938648
GO_OLIGOSACCHARIDE_CATABOLIC_PROCESS 10 0.01 Up
0.307125909837532 0.336309595601856 0.723506575088189
GO_RESPONSE_TO_ATP 19 0.01 Up 0.37072025676660 5
0.336705593645966 0.724101898343841
GO_NEGATIVE_REGULATION_OF_PEPTIDYL_THREONINE_PHOSPHORYLATION 16
0.01 Down 0.148133861382123 0.336992843167415
0.724452164613426
GO_RESPONSE_TO_KETONE 137 0.01 Down 0.381441831619212
0.33711626788076 0.724452164613426
GO_SYMPORTER_ACTIVITY 81 0.01 Down 0.425802746408232
0.337226456464757 0.724452164613426
GO_POSITIVE_REGULATION_OF_RUFFLE_ASSEMBLY 11 0.01 Down
0.15539850699648 0.337562282231372
0.724917090706283
GO_ONE_CARBON_METABOLIC_PROCESS 21 0.01 Up
0.189369844920688 0.337819152998445 0.724965614892305
GO_POSITIVE_REGULATION_OF_PROTEIN_TYROSINE_KINASE_ACTIVITY 38
0.01 Down 0.56119906028565 0.337910866785291
0.724965614892305
GO_PHAGOSOME_MATURATION 39 0.01 Up 0.366045877860217
0.337958797857206 0.724965614892305
GO_NUCLEOTIDE_SALVAGE 13 0.01 Down 0.247410813089405
0.33814970727119 0.724965614892305
GO_CHROMOSOME_ORGANIZATION_INVOLVED_IN_MEIOTIC_CELL_CYCLE 33
0.01 Up 0.198822379828548 0.3381819504818
0.724965614892305
GO_CELL_REDOX_HOMEOSTASIS 64 0.01 Up
0.265842808449645 0.338558759682859 0.725122402453741
GO_ANCHORED_COMPONENT_OF_MEMBRANE 89 0.01 Down
0.646145309770713 0.338620306162633 0.725122402453741
GO_NEURON_PROJECTION_TERMINUS 69 0.01 Up
0.285149740811375 0.33875870568345 0.725122402453741
GO_POSITIVE_REGULATION_OF_CYTOSKELETON_ORGANIZATION 183 0.01
Down 0.288093109678513 0.338802909053668
0.725122402453741
GO_POSITIVE_REGULATION_OF_EXTRACELLULAR_MATRIX_ORGANIZATION 18
0.01 Up 0.537274284804887 0.338852290522362
0.725122402453741
GO_POSITIVE_REGULATION_OF_CANONICAL_WNT_SIGNALING_PATHWAY 122
0.01 Down 0.294664770397001 0.338983160299998
0.725146852072336
GO_GLUCOSYLTRANSFERASE_ACTIVITY 16 0.01 Up
0.207557215636971 0.339493985624553 0.725827367950053
GO_POSITIVE_REGULATION_OF_ENDOTHELIAL_CELL_MIGRATION 85 0.01

Down 0.423616484306561 0.339598746376894
0.725827367950053
GO_LIPID_DROPLET_ORGANIZATION 15 0.01 Up
0.180830517149574 0.339659949324016 0.725827367950053
GO_RESPONSE_TO_ENDOPLASMIC_RETICULUM_STRESS 251 0.01 Down
0.223212989548933 0.339816698605103
0.72590681816734
GO_ATP_HYDROLYSIS_COUPLED_TRANSMEMBRANE_TRANSPORT 25 0.01
Up 0.244133381915776 0.340305892430539
0.726455857899943
GO_SENSORY_PERCEPTION_OF_MECHANICAL_STIMULUS 95 0.01 Up
0.476567516754375 0.340389903002968
0.726455857899943
GO_NEGATIVE_REGULATION_OF_SMOOTH_MUSCLE_CELL_MIGRATION 21 0.01
Up 0.540419949708042 0.340461306464933
0.726455857899943
GO_NEUROTRANSMITTER_TRANSPORT 157 0.01 Up
0.401451255172723 0.340552359015522 0.726455857899943
GO_NF_KAPPAB_BINDING 28 0.01 Down 0.246608810952334
0.34073952263286 0.726599803970527
GO_TRANSLATION_INITIATION_FACTOR_BINDING 28 0.01 Down
0.222300002784262 0.340864882049872
0.726611902712351
GO_SYNAPSE_ORGANIZATION 273 0.01 Up 0.400032005766324
0.341263329542979 0.727078582514463
GO_PYRIMIDINE_NUCLEOTIDE_CATABOLIC_PROCESS 12 0.01 Up
0.139366389116862 0.341323333909771
0.727078582514463
GO_FAT_SOLUBLE_VITAMIN_BIOSYNTHETIC_PROCESS 11 0.01 Down
0.592949650218 0.341575964489904 0.727361515404493
GO_COATED_VESICLE_MEMBRANE 146 0.01 Down
0.338366504059915 0.341834870857372 0.727657609037554
GO_MICROTUBULE_BUNDLE_FORMATION 62 0.01 Up
0.366046160004247 0.342157338349287 0.727711826178793
GO_PHOSPHATIDYLINOSITOL_PHOSPHATE_PHOSPHATASE_ACTIVITY 25 0.01
Down 0.194988139410998 0.3422187017223
0.727711826178793
GO_PHOSPHATASE_INHIBITOR_ACTIVITY 32 0.01 Down
0.250952283848043 0.342219941317815 0.727711826178793
GO_REGULATION_OF_ER_TO_GOLGI_VESICLE_MEDIATED_TRANSPORT 14 0.01
Down 0.206127666202173 0.342427345335744
0.727897903898215
GO_REGULATION_OF_SYNAPSE_STRUCTURE_OR_ACTIVITY 147 0.01 Up
0.385899597319216 0.342861984775271
0.728228237713078
GO_PEPTIDYL_AMINO_ACID_MODIFICATION 986 0.01 Down
0.300364225648182 0.343077757671704 0.728228237713078
GO_AMINO_ACID_TRANSPORT 86 0.01 Up 0.419600794667453
0.343085079614009 0.728228237713078
GO_OXIDOREDUCTASE_ACTIVITY_ACTING_ON_THE_CH_NH_GROUP_OF_DONORS 22
0.01 Up 0.264131331063838 0.343152045155301
0.728228237713078
GO_INTRACELLULAR_STEROID_HORMONE_RECEPTOR_SIGNALING_PATHWAY 112
0.01 Down 0.222170076612843 0.343182505039881

0.728228237713078
GO_VITAMIN_TRANSMEMBRANE_TRANSPORTER_ACTIVITY 16 0.01 Up
0.358211220087471 0.343425136586432
0.728488471074852
GO_DNA_BINDING_TRANSCRIPTION_REPRESSOR_ACTIVITY_RNA_POLYMERASE_II_SPECIFIC 170 0.01 Down 0.380057089393064
0.344330836234131 0.730154560523022
GO_RUFFLE_MEMBRANE 82 0.01 Down 0.361759883835841
0.344491357466643 0.730239885188545
GO_REGULATION_OF_ANION_TRANSPORT 61 0.01 Up
0.426075846117227 0.344770534475963 0.730568851852927
GO_ANION_TRANSPORT 386 0.01 Up 0.395457948918194
0.344921638388508 0.730568851852927
GO_POST_EMBRYONIC_DEVELOPMENT 75 0.01 Down
0.357826044498058 0.345114326651041 0.730568851852927
GO_REGULATION_OF_CALCIUM_ION_DEPENDENT_EXOCYTOSIS 66 0.01
Up 0.339233222410435 0.345249316385046
0.730568851852927
GO_POSITIVE_REGULATION_OF_CALCIUM_MEDIATED_SIGNALING 30 0.01
Down 0.436958679529505 0.345502181419864
0.730568851852927
GO_PROTEIN_PHOSPHATASE_TYPE_2A_COMPLEX 19 0.01 Down
0.350786185176463 0.345509500303648 0.730568851852927
GO_AZUROPHIL_GRANULE_LUMEN 73 0.01 Up
0.342738318213798 0.345518954847575 0.730568851852927
GO_TRANSMEMBRANE_TRANSPORTER_ACTIVITY 592 0.01 Up
0.369212942741407 0.345875873881502 0.730568851852927
GO_SNARE_COMPLEX_ASSEMBLY 17 0.01 Down
0.133396696072622 0.345888751514164 0.730568851852927
GO_ENDOPLASMIC_RETICULUM_MANNOSE_TRIMMING 16 0.01 Up
0.161606794850824 0.345966937963579
0.730568851852927
GO_RESPIRATORY_CHAIN_COMPLEX_IV_ASSEMBLY 24 0.01 Up
0.0950626323309415 0.345970260101658
0.730568851852927
GO_VOLTAGE_GATED_ANION_CHANNEL_ACTIVITY 13 0.01 Down
0.190747880435827 0.346133304407579 0.730659002454247
GO_ENDOCARDIUM_DEVELOPMENT 10 0.01 Down
0.172305599080536 0.346417683768185 0.730796965065864
GO_PROTEASOME_CORE_COMPLEX_BETA_SUBUNIT_COMPLEX 10 0.01 Down
0.202099568816893 0.346500118410676
0.730796965065864
GO_FATTY_ACYL_COA_BIOSYNTHETIC_PROCESS 25 0.01 Up
0.21402904327989 0.346867177566318 0.730796965065864
GO_BLASTOCYST_FORMATION 30 0.01 Down 0.298084692839432
0.346895388495002 0.730796965065864
GO_LYMPHANGIOGENESIS 12 0.01 Down 0.421919293254172
0.346942508616289 0.730796965065864
GO_INTRINSIC_APOPTOTIC_SIGNALING_PATHWAY 261 0.01 Down
0.296284745695604 0.347065463813969
0.730796965065864
GO_PROTEIN_CONTAINING_COMPLEX_REMODELING 10 0.01 Up
0.790606062985207 0.347298728795237
0.730796965065864

GO_CELLULAR_RESPONSE_TO_X_RAY	11	0.01	Up
0.528823907972148	0.347325368577293	0.730796965065864
GO_POSITIVE_REGULATION_OF_HYDROLASE_ACTIVITY	602	0.01	Down
	0.335325717446774	0.347371736278122
0.730796965065864
GO_ANCHORING_JUNCTION	500	0.01	Down	0.311867249904542
	0.34749492683568	0.730796965065864
GO_RACEMASE_AND_EPIMERASE_ACTIVITY	18	0.01	Down
0.40516303221296	0.347522786714734	0.730796965065864
GO_REGULATION_OF_CELL_DEVELOPMENT	662	0.01	Down
0.370132897583499	0.347787205255114	0.731099765617658
GO_NUCLEOBASE_CONTAINING_COMPOUND_TRANSMEMBRANE_TRANSPORTER_ACTIVITY
	38	0.01	Up	0.253281525745057
0.348140066447402	0.731475350275945
GO_SMOOTH_MUSCLE_CONTRACTION	59	0.01	Up
0.532134857702307	0.348206846038129	0.731475350275945
GO_PEPTIDYL_METHIONINE_MODIFICATION	11	0.01	Down
0.159767674508513	0.348372113983297	0.731476495028929
GO_REGULATION_OF_PEPTIDE_SECRETION	316	0.01	Down
0.452907111199464	0.34854488215682	0.731476495028929
GO_RESPONSE_TO_TOPOLOGICALLY_INCORRECT_PROTEIN	178	0.01	Down
	0.213407182601261	0.348748975751023
0.731476495028929
GO_CELL_JUNCTION	956	0.01	Down	0.330045627559795
	0.348815450340361	0.731476495028929
GO_EARLY_ENDOSOME	296	0.01	Down	0.282645368635976
	0.348900242033894	0.731476495028929
GO_POSITIVE_REGULATION_OF_PROTEIN_EXPORT_FROM_NUCLEUS	20	0.01
	Down	0.180645135214667	0.348930312898003
0.731476495028929
GO_RESPONSE_TO_PEPTIDE_HORMONE	305	0.01	Down
0.293934539136764	0.349207176014664	0.731618471608637
GO_POSITIVE_REGULATION_OF_MUSCLE_CONTRACTION	30	0.01	Down
	0.408492095029628	0.349239059581919
0.731618471608637
GO_REGULATION_OF_CELL_DEATH	1264	0.01	Down
0.345657779574041	0.349535056531802	0.731912503121979
GO_REGULATION_OF_PODOSOME_ASSEMBLY	14	0.01	Down
0.415587056752508	0.349620533528865	0.731912503121979
GO_AXIS_SPECIFICATION	53	0.01	Up	0.422806477740962
	0.349845107113161	0.732130177622888
GO_POSITIVE_REGULATION_OF_ENDOCYTOSIS	74	0.01	Up
0.457683742758084	0.350051381530415	0.73230942014857
GO_VENTRICULAR_TRABECULA_MYOCARDIUM_MORPHOGENESIS	12	0.01
	Down	0.310430957878222	0.350368903585143
0.732721189688393
GO_IN_UTERO_EMBRYONIC_DEVELOPMENT	278	0.01	Down
0.26189763071969	0.350748976198985	0.733263441633622
GO_NEURON_REMODELING	10	0.01	Down	0.552756530108522
	0.351118824213682	0.733783952427286
GO_CARBOHYDRATE_DERIVATIVE_TRANSPORT	59	0.01	Up
0.307799656140645	0.351458059497601	0.73406037907341
GO_NUCLEAR_TRANSCRIPTIONAL_REPRESSOR_COMPLEX	26	0.01	Down
	0.216056292937066	0.351492920765344

0.73406037907341
GO_NUCLEAR_MIGRATION 18 0.01 Down 0.262497255515615 0.351645891501488 0.734127306501214
GO_VITAMIN_BIOSYNTHETIC_PROCESS 18 0.01 Down 0.434127736556509 0.351848537592903 0.734297378136061
GO_REGULATION_OF_APOPTOTIC_SIGNALING_PATHWAY 347 0.01 Down 0.310431884980707 0.351969258833131 0.734297378136061
GO_REGULATION_OF_MACROPHAGE_CHEMOTAXIS 20 0.01 Up 0.6786181953617 0.352375906265303 0.734441889911469
GO_PIGMENTATION 77 0.01 Up 0.340834571633603 0.352442932825219 0.734441889911469
GO_REGULATION_OF_PLATELET_AGGREGATION 14 0.01 Down 0.489254239247292 0.352551619412964 0.734441889911469
GO_NEGATIVE_REGULATION_OF_PROTEASOMAL_PROTEIN_CATABOLIC_PROCESS 48 0.01 Down 0.201896469100298 0.352600118574307 0.734441889911469
GO_NEGATIVE_REGULATION_OF_GROWTH 193 0.01 Down 0.465564798711256 0.352643404561346 0.734441889911469
GO_REGULATION_OF_CALCIUM_ION_TRANSPORT 151 0.01 Up 0.533428310278131 0.353031011255995 0.734997005944837
GO_REGULATION_OF_ANATOMICAL_STRUCTURE_MORPHOGENESIS 813 0.01 Down 0.431937973544899 0.353392896023589 0.735329389221304
GO_RESPONSE_TO_NITRIC_OXIDE 14 0.01 Down 0.641886780847064 0.353494621643895 0.735329389221304
GO_NEGATIVE_REGULATION_OF_TRANSPORT 321 0.01 Down 0.417255568180528 0.353554025224343 0.735329389221304
GO_NEGATIVE_REGULATION_OF_DENDRITE_DEVELOPMENT 20 0.01 Down 0.397653131279707 0.353814207242939 0.735618511017769
GO_SH3_DOMAIN_BINDING 107 0.01 Down 0.276050695579941 0.354019240360632 0.73577437335667
GO_HEPARAN_SULFATE_PROTEOGLYCAN_METABOLIC_PROCESS 29 0.01 Up 0.42332298615342 0.354247690486489 0.73577437335667
GO_NEGATIVE_REGULATION_OF_B_CELL_ACTIVATION 22 0.01 Down 0.528955301294886 0.354369424292397 0.73577437335667
GO_REGULATION_OF_CELL_MORPHOGENESIS 381 0.01 Down 0.366447727734241 0.354373952840537 0.73577437335667
GO_NCRNA_TRANSCRIPTION 101 0.01 Down 0.1329905685762 0.354988664551093 0.736666873919859
GO_SENSORY_PERCEPTION_OF_CHEMICAL_STIMULUS 30 0.01 Up 0.576452682401279 0.355046495320294 0.736666873919859
GO_INNER_EAR_AUDITORY_RECEPTOR_CELL_DIFFERENTIATION 23 0.01 Up 0.3353045020142 0.355281888940219 0.736903432783079
GO_NEGATIVE_REGULATION_OF_MUSCLE_CONTRACTION 20 0.01 Down 0.557687021496658 0.355719776978742 0.736959813439427
GO_DNA_REPAIR_COMPLEX 36 0.01 Up 0.171265522529602 0.355723285509685 0.736959813439427

GO_PEPTIDYL_HISTIDINE_MODIFICATION	10	0.01	Up
0.117892715573689	0.355910386968866	0.736959813439427
GO_APOPTOTIC_PROCESS	1490	0.01	Down	0.332812480606685
	0.356101659343869	0.736959813439427
GO_RNA_HELICASE_ACTIVITY	47	0.01	Down
0.211256967842076	0.356168714850209	0.736959813439427
GO_MORPHOGENESIS_OF_EMBRYONIC_EPITHELIUM	126	0.01	Up
	0.440523682126073	0.356193696179478
0.736959813439427
GO_GLYCOLIPID_BIOSYNTHETIC_PROCESS	52	0.01	Up
0.264389593513118	0.356284329148275	0.736959813439427
GO_NATURAL_KILLER_CELL_MEDIATED_IMMUNITY	42	0.01	Down
	0.502317450684829	0.356319929679366
0.736959813439427
GO_KILLING_OF_CELLS_OF_OTHER_ORGANISM	19	0.01	Down
0.734523570249957	0.356569511791607	0.736959813439427
GO_POSITIVE_REGULATION_OF_MONOOXYGENASE_ACTIVITY	20	0.01
	Down	0.447849579619654	0.356692243814548
0.736959813439427
GO_MATING_BEHAVIOR	13	0.01	Down	0.320126136251838
	0.356753512901795	0.736959813439427
GO_CYTOPLASMIC_REGION	381	0.01	Down	0.276920905668909
	0.356765753862374	0.736959813439427
GO_HEXOSE_TRANSMEMBRANE_TRANSPORTER_ACTIVITY	11	0.01	Up
	0.535246303161143	0.35709731039386
0.737393799796301
GO_MIDBRAIN_DOPAMINERGIC_NEURON_DIFFERENTIATION	12	0.01	Up
	0.585238227485267	0.357669335864053
0.738323882363368
GO_ATP_DEPENDENT_MICROTUBULE_MOTOR_ACTIVITY	31	0.01	Down
	0.254209235429115	0.357995730924966
0.738746459022932
GO_PROTEIN_LOCALIZATION_TO_PHAGOPHORE_ASSEMBLY_SITE	13	0.01
	Down	0.123525127116463	0.358127564303898
0.738767394797474
GO_NUCLEOTIDE_PHOSPHORYLATION	16	0.01	Up
0.232037900672838	0.358823441328804	0.73995146477825
GO_PEROXISOME_PROLIFERATOR_ACTIVATED_RECEPTOR_BINDING	10	0.01
	Down	0.285659780003114	0.359662807913387
0.740670598232914
GO_REGULATION_OF_HETEROTYPIC_CELL_CELL_ADHESION	17	0.01	Down
	0.655001472479836	0.359669268964217
0.740670598232914
GO_CELL_KILLING	84	0.01	Down	0.504332907694067
0.359732095969131	0.740670598232914
GO_CHROMOSOME	861	0.01	Down	0.185770722010293
0.359765063909021	0.740670598232914
GO_POSITIVE_REGULATION_OF_CELL_MORPHOGENESIS_INVOLVED_IN_DIFFERENTIA
TION	125	0.01	Down	0.326444016987001
0.359782176608279	0.740670598232914
GO_NEGATIVE_REGULATION_OF_REACTIVE_OXYGEN_SPECIES_METABOLIC_PROCESS
	23	0.01	Up	0.380887107055454
0.359914473127473	0.740689566581029
GO_REGULATION_OF_INCLUSION_BODY_ASSEMBLY	15	0.01	Down

0.203354083974795 0.360035399601485
0.740689566581029
GO_PEPTIDE_SECRETION 403 0.01 Down 0.433890426730982
0.360327974994038 0.741040357787535
GO_DEVELOPMENTAL_MATURATION 171 0.01 Down
0.375200035467883 0.360763544008638 0.741684888478307
GO_REGULATION_OF_TRANSLATIONAL_FIDELITY 17 0.01 Up
0.113564063204389 0.361368414421895 0.742135021558987
GO_POSITIVE_REGULATION_OF_MITOTIC_SISTER_CHROMATID_SEGREGATION 16
0.01 Down 0.109226963408207 0.361393116371899
0.742135021558987
GO_TRANSFERASE_ACTIVITY_TRANSFERRING_AMINO_ACYL_GROUPS 11 0.01
Down 0.568319245418096 0.36143790201074
0.742135021558987
GO_REGULATION_OF_INTRACELLULAR_PROTEIN_TRANSPORT 209 0.01
Down 0.255435267916751 0.36147146413275
0.742135021558987
GO_NEGATIVE_REGULATION_OF_DNA_REPAIR 28 0.01 Down
0.166289539702508 0.361888121993407 0.742548513612395
GO_MITOCHONDRIAL_FISSION 30 0.01 Up
0.272223833518027 0.361917485056676 0.742548513612395
GO_MYELOID_CELL_APOPTOTIC_PROCESS 27 0.01 Down
0.519743666704768 0.362208300015449 0.742618046106542
GO_RESPONSE_TO_ORGANIC_CYCLIC_COMPOUND 643 0.01 Down
0.378597768188022 0.362289030962125 0.742618046106542
GO_KINASE_BINDING 609 0.01 Down 0.291381559799802
0.362318341717605 0.742618046106542
GO_MUSCLE_CONTRACTION 207 0.01 Up 0.463513758008063
0.362474184311835 0.742686727288947
GO_INTRACELLULAR_TRANSPORT 1606 0.01 Down
0.212800920847828 0.363344518759203 0.743061127226645
GO_CELLULAR_RESPONSE_TO_REACTIVE_OXYGEN_SPECIES 133 0.01 Down
0.41085940118526 0.363350336076811
0.743061127226645
GO_REGULATION_OF_EXOCYTOSIS 147 0.01 Down
0.338501081796491 0.36335604284401 0.743061127226645
GO_REGULATION_OF_IMMUNOGLOBULIN_PRODUCTION 47 0.01 Down
0.328198270793367 0.363509371393163
0.743061127226645
GO_CALMODULIN_DEPENDENT_PROTEIN_KINASE_ACTIVITY 20 0.01 Down
0.243956902244496 0.363556595989254
0.743061127226645
GO_DNA_MODIFICATION 89 0.01 Down 0.249742757838821
0.363561453203845 0.743061127226645
GO_SECONDARY_ALCOHOL_METABOLIC_PROCESS 61 0.01 Up
0.383029362369224 0.363806549802964 0.743061127226645
GO_POSITIVE_REGULATION_OF_LONG_TERM_SYNAPTIC_POTENTIATION 14
0.01 Up 0.228375160598894 0.363943039402555
0.743061127226645
GO_POSITIVE_REGULATION_OF_INTRACELLULAR_ESTROGEN_RECEPTOR_SIGNALING_
PATHWAY 10 0.01 Down 0.252818796087267
0.364035597799228 0.743061127226645
GO_RESPONSE_TO_PLATELET_DERIVED_GROWTH_FACTOR 18 0.01 Down
0.496247881320655 0.364216574812948

0.743061127226645
GO_CELLULAR_RESPONSE_TO_KETONE 73 0.01 Down
0.435554012365304 0.364388301254082 0.743061127226645
GO_SMALL_NUCLEOLAR_RIBONUCLEOPROTEIN_COMPLEX_ASSEMBLY 11 0.01
Up 0.179025787072623 0.364442655589951
0.743061127226645
GO_OUTFLOW_TRACT_SEPTUM_MORPHOGENESIS 20 0.01 Up
0.495647449988469 0.364470033113041 0.743061127226645
GO_QUINONE_BIOSYNTHETIC_PROCESS 16 0.01 Up
0.146678279402966 0.364480800691471 0.743061127226645
GO_ACTIN_POLYMERIZATION_OR_DEPOLYMERIZATION 168 0.01 Down
0.320605918353274 0.364492840863276
0.743061127226645
GO_HINDLIMB_MORPHOGENESIS 20 0.01 Up
0.488240162294307 0.364937937810577 0.743259766264672
GO_PHAGOCYTOSIS 207 0.01 Up 0.480709813191829
0.365022184141794 0.743259766264672
GO_PROTEIN_TARGETING_TO_LYSOSOME 20 0.01 Down
0.200575952688336 0.365114276559053 0.743259766264672
GO_PROTEIN_KINASE_B_SIGNALING 182 0.01 Down
0.385426591921417 0.365126070003209 0.743259766264672
GO_NEGATIVE_REGULATION_OF_CELL_ACTIVATION 135 0.01 Down
0.57422771572248 0.365288032572107
0.743259766264672
GO_CELLULAR_RESPONSE_TO_MECHANICAL_STIMULUS 63 0.01 Down
0.383249757631852 0.365504402646566
0.743259766264672
GO_ENDOCYTIC_VESICLE_MEMBRANE 135 0.01 Up
0.431659782406928 0.365524276651089 0.743259766264672
GO_POSITIVE_REGULATION_OF_BEHAVIOR 10 0.01 Down
0.523596944984446 0.365569702201666 0.743259766264672
GO_RETINA_MORPHOGENESIS_IN_CAMERA_TYPE_EYE 28 0.01 Down
0.477563790263998 0.366026659284348
0.743711845065326
GO_DETECTION_OF_STIMULUS_INVOLVED_IN_SENSORY_PERCEPTION 28 0.01
Up 0.575643331799333 0.3660370602957
0.743711845065326
GO_PROTEIN_LOCALIZATION_TO_CHROMOSOME_TELOMERIC_REGION 28 0.01
Down 0.0890632505708235 0.36646373173717
0.743951870828128
GO_DELAYED_RECTIFIER_POTASSIUM_CHANNEL_ACTIVITY 10 0.01 Down
0.875779115585911 0.366471640750609
0.743951870828128
GO_INCLUSION_BODY 66 0.01 Down 0.22632842605048
0.366522820893911 0.743951870828128
GO_REGULATION_OF_PROTEIN_ACETYLATION 63 0.01 Down
0.281407575479024 0.366882410270046 0.744093300837683
GO_NEGATIVE_REGULATION_OF_ANIMAL_ORGAN_MORPHOGENESIS 25 0.01
Down 0.760654788741569 0.366918189203416
0.744093300837683
GO_FIBROBLAST_PROLIFERATION 70 0.01 Up
0.379822591770535 0.36696019481272 0.744093300837683
GO_METALLOPEPTIDASE_ACTIVITY 124 0.01 Up
0.542260709026432 0.367272606939301 0.744338376251551

GO_MYOSIN_II_COMPLEX	10	0.01	Up	0.249930939250369
	0.367346174856448	0.744338376251551
GO_NEGATIVE_REGULATION_OF_CARDIAC_MUSCLE_CELL_DIFFERENTIATION	15
	0.01	Up	0.549010497550947	0.367482866549454
	0.744338376251551
GO_LONG_TERM_SYNAPTIC_POTENTIATION	47	0.01	Up
0.40192530054644	0.367571479493024	0.744338376251551
GO_REGULATION_OF_MONOCYTE_CHEMOTAXIS	22	0.01	Down
1.13625736871022	0.367807135422115	0.744399511403795
GO_PURINE_NUCLEOSIDE_BIOSYNTHETIC_PROCESS	21	0.01	Up
	0.277820482210873	0.367846900710161
0.744399511403795
GO_NEGATIVE_REGULATION_OF_CHROMOSOME_SEGREGATION	38	0.01
	Up	0.241611951003452	0.36847470741217
0.745272116665553
GO_SECOND_MESSENGER_MEDIATED_SIGNALING	238	0.01	Down
0.466789813366883	0.368523619540436	0.745272116665553
GO_NUCLEAR_TRANSCRIBED_MRNA_CATABOLIC_PROCESS_NONSENSE_MEDIATED_DECA
Y	119	0.01	Up	0.13053555840799
0.368732866006891	0.745446962879732
GO_RESPIRATORY_GASEOUS_EXCHANGE	40	0.01	Up
0.64147790240706	0.369019088525714	0.745777259134357
GO_FEMALE_MEIOTIC_NUCLEAR_DIVISION	22	0.01	Up
0.198497495319603	0.369182505700606	0.745859231982821
GO_CERAMIDE_METABOLIC_PROCESS	22	0.01	Down
0.239977882769995	0.369435595332219	0.746122255243479
GO_CAMP_BINDING	12	0.01	Down	0.404155500256664
0.36985850262174	0.746727957903753
GO_HIPPOCAMPAL_MOSSY_FIBER_TO_CA3_SYNAPSE	18	0.01	Up
	0.240228166617359	0.370177693475598
0.747123928553975
GO_RESPONSE_TO_STEROL_DEPLETION	10	0.01	Down
0.186139247030854	0.370619132725799	0.747766286067906
GO_RESPIRATORY_BURST_INVOLVED_IN_DEFENSE_RESPONSE	10	0.01
	Up	0.511639276336415	0.370768171228714
0.747818460973264
GO_REGULATION_OF_ACTIN_FILAMENT_ORGANIZATION	210	0.01	Down
	0.312086817192489	0.370961552286813
0.747845436897767
GO_REGULATION_OF_CELLULAR_COMPONENT_MOVEMENT	753	0.01	Down
	0.464814782992344	0.371027912359755
0.747845436897767
GO_MEMBRANE_INVAGINATION	56	0.01	Up
0.528912199807217	0.37119248001409	0.747928823818633
GO_CELL_COMMUNICATION_BY_ELECTRICAL_COUPLING	18	0.01	Up
	0.398563236040022	0.371425614812146
0.747948403645669
GO_L_AMINO_ACID_TRANSMEMBRANE_TRANSPORTER_ACTIVITY	30	0.01
	Up	0.280209809358126	0.371448597758473
0.747948403645669
GO_NUCLEOID	41	0.01	Up	0.108095763864783
0.371685855580844	0.7479548725937
GO_SODIUM_ION_IMPORT_ACROSS_PLASMA_MEMBRANE	10	0.01	Down
	0.650040758478868	0.371840481942271

0.7479548725937
GO_DNA_PACKAGING_COMPLEX 45 0.01 Down
0.149169371766399 0.371855622440527 0.7479548725937
GO_PENTOSE_PHOSPHATE_SHUNT 15 0.01 Up
0.144212343463576 0.372062000783333 0.7479548725937
GO_POSITIVE_REGULATION_OF_FOCAL_ADHESION_ASSEMBLY 22 0.01
Down 0.424551541360079 0.372096242761394
0.7479548725937
GO_CHLORIDE_TRANSMEMBRANE_TRANSPORT 39 0.01 Down
0.431396798394843 0.372191017971598 0.7479548725937
GO_MACROMOLECULE_CATABOLIC_PROCESS 1196 0.01 Down
0.228874185109145 0.37247541173769 0.748278697769528
GO_CELL_LEADING_EDGE 332 0.01 Down 0.329272939537106
0.372798195201336 0.748567548377932
GO_REGULATION_OF_TRANSCRIPTION_BY_RNA_POLYMERASE_I 30 0.01
Down 0.129965587013369 0.372873642555682
0.748567548377932
GO_REGULATION_OF_T_CELL_MEDIATED_CYTOTOXICITY 22 0.01 Down
0.418069714960301 0.372989101275448
0.748567548377932
GO_RIBONUCLEOPROTEIN_COMPLEX_SUBUNIT_ORGANIZATION 236 0.01
Down 0.126707279318044 0.373328069978215
0.748823791725747
GO_NURD_COMPLEX 14 0.01 Down 0.113043192907122
0.373366611818496 0.748823791725747
GO_PEPTIDE_HORMONE_SECRETION 156 0.01 Down
0.343562061303299 0.373486812937829 0.748823791725747
GO_CYSTEINE_TYPE_ENDOPEPTIDASE_ACTIVITY 84 0.01 Down
0.275475263337972 0.373787143018192 0.749178522701699
GO_POSITIVE_REGULATION_OF_SODIUM_ION_TRANSPORT 27 0.01 Down
0.37698410538764 0.374309007538465
0.749976892662053
GO_NEGATIVE_REGULATION_OF_LYASE_ACTIVITY 11 0.01 Up
0.37297158351254 0.374651012769687
0.750088525338242
GO_ORGANOPHOSPHATE_BIOSYNTHETIC_PROCESS 464 0.01 Up
0.268347244630621 0.374787626528588 0.750088525338242
GO_NEGATIVE_REGULATION_OF_MICROTUBULE_DEPOLYMERIZATION 19 0.01
Down 0.235580530254754 0.374910692739543
0.750088525338242
GO_HISTONE_H3_K9_TRIMETHYLATION 15 0.01 Down
0.155596185483273 0.375136925142989 0.750088525338242
GO_NEGATIVE_REGULATION_OF_ENDOTHELIAL_CELL_APOPTOTIC_PROCESS 18
0.01 Down 0.334732718915448 0.37515507718345
0.750088525338242
GO_POSITIVE_REGULATION_OF_PROTEIN_LOCALIZATION_TO_CELL_PERIPHERY
50 0.01 Down 0.320981797769055
0.375163656493526 0.750088525338242
GO_CELLULAR_EXTRAVASATION 40 0.01 Down
0.65572836910781 0.375229591739786 0.750088525338242
GO_MEMBRANE_LIPID_BIOSYNTHETIC_PROCESS 119 0.01 Up
0.266415250895667 0.375638129926881 0.750574984965787
GO_CALCIUM_RELEASE_CHANNEL_ACTIVITY 16 0.01 Up
0.386029341916237 0.375774883820804 0.750574984965787

GO_TUBE_CLOSURE 82	0.01	Up	0.369277243895114
0.375847532897056	0.750574984965787
GO_REGULATION_OF_PROTEIN_LOCALIZATION	771	0.01	Down
0.322987541170507	0.376040048260573	0.750574984965787
GO_CYCLIC_NUCLEOTIDE_BIOSYNTHETIC_PROCESS	13	0.01	Up
	0.536895437931701	0.376091105957161
0.750574984965787
GO_MITOCHONDRION_LOCALIZATION	35	0.01	Down
0.204955786539635	0.376642838216439	0.751429073549787
GO_ORGANELLE_FUSION	104	0.01	Down	0.223462706574587
	0.376787628652383	0.75147098999626
GO_POSITIVE_REGULATION_OF_CARDIAC_MUSCLE_TISSUE_DEVELOPMENT	30
	0.01	Up	0.56158500618917	0.377173350701614
	0.751993238788013
GO_POSTSYNAPTIC_CYTOSOL 15	0.01	Down	0.173698170493957
	0.377352998893188	0.752104417688951
GO_LIPOPOLYSACCHARIDE_BINDING	16	0.01	Down
0.834285185684748	0.37749626866427	0.752141042979798
GO_PROTEIN_POLYGLUTAMYLATION	10	0.01	Up
0.216274453533965	0.377619156482033	0.752141042979798
GO_PASSIVE_TRANSMEMBRANE_TRANSPORTER_ACTIVITY	190	0.01	Up
	0.467315402774881	0.377899718526612
0.752452998089557
GO_UBIQUITIN_DEPENDENT_PROTEIN_CATABOLIC_PROCESS_VIA_THE_MULTIVESICU
LAR_BODY_SORTING_PATHWAY	22	0.01	Down
0.178159055342654	0.378473316800414	0.753348034850923
GO_TROPOMYOSIN_BINDING	11	0.01	Up	0.58426528102692
	0.378660549272504	0.753388154343377
GO_REGULATION_OF_HORMONE_METABOLIC_PROCESS	30	0.01	Down
	0.51726030094521	0.378946246920645
0.753388154343377
GO_PROTEIN_TARGETING	393	0.01	Up	0.178170213490899
	0.379019053511384	0.753388154343377
GO_NUCLEOSIDE_BISPHOSPHATE_METABOLIC_PROCESS	50	0.01	Up
	0.290013955361034	0.379286158272768
0.753388154343377
GO_CIRCULATORY_SYSTEM_PROCESS	302	0.01	Up
0.460018895042694	0.379359247652879	0.753388154343377
GO_METANEPHROS_MORPHOGENESIS	21	0.01	Down
0.88919929259393	0.379755802346576	0.753388154343377
GO_REGULATION_OF_CELLULAR_COMPONENT_BIOGENESIS	729	0.01	Down
	0.291706868079885	0.37978883878564
0.753388154343377
GO_REGULATION_OF_CELL_CYCLE	1020	0.01	Down
0.239383359450999	0.379804777052914	0.753388154343377
GO_ION_TRANSMEMBRANE_TRANSPORTER_ACTIVITY	483	0.01	Up
	0.378585823833353	0.379811649973473
0.753388154343377
GO_PROTEIN_TAG	10	0.01	Up	0.0943761749043402
0.379831781840925	0.753388154343377
GO_PHOSPHATIDYLETHANOLAMINE_ACYL_CHAIN_REMODELING	18	0.01
	Up	0.511268911356153	0.38000532576563
0.753388154343377
GO_GALACTOSYLTRANSFERASE_ACTIVITY	27	0.01	Down

0.400242788911039	0.380476564141552	0.753388154343377
GO_POSITIVE_REGULATION_OF_CELLULAR_COMPONENT_ORGANIZATION	963
	0.01	Down	0.312454429716652	0.380480978783595
	0.753388154343377
GO_ORGANOPHOSPHATE_METABOLIC_PROCESS	415	0.01	Up
0.298698022925256	0.380495869624465	0.753388154343377
GO_RESPONSE_TO_ACID_CHEMICAL	243	0.01	Up
0.390844619321363	0.38049624691406	0.753388154343377
GO_REGULATION_OF_RHO_PROTEIN_SIGNAL_TRANSDUCTION	109	0.01
	Down	0.381590760636076	0.380521258624433
0.753388154343377
GO_MICROTUBULE_MINUS_END_BINDING	10	0.01	Down
0.161437525532141	0.380704201741592	0.753388154343377
GO_COPI_VESICLE_COAT	15	0.01	Down	0.127910453464683
	0.380888056913914	0.753388154343377
GO_LYMPH_NODE_DEVELOPMENT	13	0.01	Down
0.584594071067687	0.380936746577896	0.753388154343377
GO_PROTEIN_O_LINKED_MANNOSYLATION	18	0.01	Up
0.14075896728667	0.380975396777165	0.753388154343377
GO_POSITIVE_REGULATION_OF_TRANSPORTER_ACTIVITY	63	0.01	Down
	0.356320767465551	0.381124858307365
0.753438298529473
GO_NUCLEOSOMAL_DNA_BINDING	35	0.01	Down
0.133184976153249	0.381257823971052	0.753455810328209
GO_MICROTUBULE_ORGANIZING_CENTER_ORGANIZATION	122	0.01	Down
	0.174679617415429	0.381724626968218
0.754132837723413
GO_MITOCHONDRIA_ASSOCIATED_ENDOPLASMIC_RETICULUM_MEMBRANE	12
	0.01	Down	0.129554481638129	0.381956555148125
	0.754345558329299
GO_REGULATION_OF_MEGAKARYOCYTE_DIFFERENTIATION	50	0.01	Down
	0.288442277877449	0.382691824098083
0.755152908522714
GO_PROTEIN_SERINE_THREONINE_KINASE_INHIBITOR_ACTIVITY	31	0.01
	Up	0.224356106538514	0.382797651822443
0.755152908522714
GO_POSITIVE_REGULATION_OF_PHOSPHOPROTEIN_PHOSPHATASE_ACTIVITY	17
	0.01	Down	0.477234538479011	0.3828396388037
0.755152908522714
GO_NAD_BINDING	51	0.01	Up	0.238916828875705
0.382862897781735	0.755152908522714
GO_REPRODUCTIVE_BEHAVIOR	18	0.01	Down
0.290197145337056	0.383072548851087	0.755321027630707
GO_MOLECULAR_TRANSDUCER_ACTIVITY	560	0.01	Up
0.47793142434758	0.383206604819438	0.75534003177234
GO_REGULATION_OF_TRANSMEMBRANE_TRANSPORT	313	0.01	Down
	0.443085512519003	0.38334856772987
0.755374603923415
GO_POSITIVE_REGULATION_OF_AXONOGENESIS	71	0.01	Down
0.392612265352307	0.383487951684929	0.755404073549384
GO_AXO_DENDRITIC_PROTEIN_TRANSPORT	10	0.01	Down
0.292241942206946	0.384159958782459	0.756482358017616
GO_9PLUS2_MOTILE_CILIUM	53	0.01	Up	0.351388775035801
	0.38430303607855	0.75651871985503

GO_TUMOR_NECROSIS_FACTOR_RECEPTOR_BINDING 25 0.01 Down
0.456202292547433 0.384487377771497
0.75663626270689
GO_BASAL_PART_OF_CELL 38 0.01 Down 0.473243281011833
0.384668806461005 0.756747998711848
GO_SECRETION_BY_TISSUE 21 0.01 Up 0.55963040135182
0.385045021421645 0.756867098237551
GO_MATURE_B_CELL_DIFFERENTIATION_INVOLVED_IN_IMMUNE_RESPONSE 15
0.01 Up 0.521135221946823 0.385076177186315
0.756867098237551
GO_CATALYTIC_ACTIVITY_ACTING_ON_A_TRNA 120 0.01 Up
0.138927636939279 0.385300341368259 0.756867098237551
GO_AMINE_BIOSYNTHETIC_PROCESS 12 0.01 Down
0.304847694000881 0.38535269323872 0.756867098237551
GO_POLYAMINE_BIOSYNTHETIC_PROCESS 12 0.01 Down
0.304847694000881 0.38535269323872 0.756867098237551
GO_REGULATION_OF_NATURAL_KILLER_CELL_DIFFERENTIATION 10 0.01
Down 0.613754678270455 0.385774154018136
0.757449834749064
GO_ENERGY_RESERVE_METABOLIC_PROCESS 59 0.01 Down
0.280566361107333 0.386031549861835 0.757710164633431
GO_REGULATION_OF_CALCIUM_ION_IMPORT 21 0.01 Up
0.718310866178119 0.386637722738874 0.75865469125653
GO_NEGATIVE_REGULATION_OF_OXIDATIVE_STRESS_INDUCED_NEURON_DEATH 16
0.01 Up 0.260522146837212 0.387258590920085
0.759516797934396
GO_SULFOTRANSFERASE_ACTIVITY 29 0.01 Up
0.447549112326215 0.387327294746317 0.759516797934396
GO_POSITIVE_REGULATION_OF_JUN_KINASE_ACTIVITY 68 0.01 Down
0.336853302679562 0.388239174662807
0.75994524276831
GO_MYOBLAST_DIFFERENTIATION 61 0.01 Down
0.538354349194395 0.388530840333166 0.75994524276831
GO_MITOCHONDRIAL_RNA_METABOLIC_PROCESS 23 0.01 Up
0.0939925437731104 0.388555417591466 0.75994524276831
GO_CELLULAR_RESPONSE_TO_HORMONE_STIMULUS 493 0.01 Down
0.306817457445096 0.388562069108847
0.75994524276831
GO_RNA_CAP_BINDING 17 0.01 Down 0.159883326373558
0.388696653048897 0.75994524276831
GO_NUCLEOBASE_CONTAINING_SMALL_MOLECULE_INTERCONVERSION 25 0.01
Up 0.200277402999684 0.388714130589181
0.75994524276831
GO_REGULATION_OF_INTEGRIN_MEDIATED_SIGNALING_PATHWAY 12 0.01
Up 0.411677401197307 0.388718634397162
0.75994524276831
GO_POSITIVE_REGULATION_OF_LIPID_STORAGE 15 0.01 Down
0.832938649002802 0.388758335930327 0.75994524276831
GO_REGULATION_OF_SYNAPTIC_TRANSMISSION_GABAERGIC 17 0.01
Down 0.385045801273932 0.38884434115685
0.75994524276831
GO_REGULATION_OF_CYTOKINESIS 70 0.01 Down
0.231242253336913 0.388909670474584 0.75994524276831
GO_POSITIVE_REGULATION_OF_CELL_MIGRATION_INVOLVED_IN_SPROUTING_ANGIO

GENESIS 20	0.01	Down	0.597760142023236
0.388922725956373	0.75994524276831
GO_TRNA_MODIFICATION	85	0.01	Up	0.133167670087119
	0.389203679356964	0.760249529400298
GO_REGULATION_OF_ESTABLISHMENT_OF_PROTEIN_LOCALIZATION	559	0.01
	Down	0.344649847863164	0.389563110015177
0.76065481421716
GO_REGULATION_OF_ENDOPLASMIC_RETICULUM_STRESS_INDUCED_INTRINSIC_APOPTOTIC_SIGNALING_PATHWAY 27	0.01	Down	0.198013495347196
	0.389976785451825	0.76065481421716
GO_INTRINSIC_COMPONENT_OF_POSTSYNAPTIC_DENSITY_MEMBRANE 17	0.01
	Up	0.327254039531201	0.39005996000795
0.76065481421716
GO_PROTON_TRANSPORTING_TWO_SECTOR_ATPASE_COMPLEX_ASSEMBLY	10
	0.01	Up	0.153071029642934	0.390092899456975
	0.76065481421716
GO_PEPTIDYL_LYSINE_ACETYLATION	147	0.01	Down
0.202205536982337	0.39021690528609	0.76065481421716
GO_POSITIVE_REGULATION_OF_MITOCHONDRIAL_TRANSLATION	17	0.01
	Up	0.113305594060901	0.390278375634618
0.76065481421716
GO_POSITIVE_REGULATION_OF_LEUKOCYTE_MIGRATION	91	0.01	Down
	0.582013827258668	0.390288213850511
0.76065481421716
GO_BASE_CONVERSION_OR_SUBSTITUTION_EDITING	13	0.01	Down
	0.400796315789999	0.390632185937317
0.761080873178899
GO_NUCLEOLUS	863	0.01	Down	0.178639649192385
0.391102699565605	0.761519889448449
GO_REGULATION_OF_BINDING	307	0.01	Down
0.306807269942545	0.391108386641454	0.761519889448449
GO_FACE_DEVELOPMENT	44	0.01	Down	0.347293727521416
	0.391356060784792	0.761757821424966
GO_DNA_ENDOREDUPLICATION	10	0.01	Down
0.263370041353965	0.391712897425888	0.762010266943446
GO_MICROTUBULE_BASED_MOVEMENT	218	0.01	Up
0.231382980600883	0.391806205096373	0.762010266943446
GO_LAYER_FORMATION_IN_CEREBRAL_CORTEX	10	0.01	Down
0.230719817649746	0.391917360888192	0.762010266943446
GO_CENTRIOLE_CENTRIOLE_COHESION 12	0.01	Up
0.274824273564035	0.391987821390938	0.762010266943446
GO_NEURON_DEATH_IN_RESPONSE_TO_OXIDATIVE_STRESS 23	0.01	Up
	0.266438239375991	0.392377742088618
0.762303984700679
GO_REGULATION_OF_OSSIFICATION	134	0.01	Down
0.525009141935701	0.392405896656187	0.762303984700679
GO_STRUCTURAL_CONSTITUENT_OF_MUSCLE	27	0.01	Up
0.422780994546118	0.392515608001041	0.762303984700679
GO_C_ACYLTRANSFERASE_ACTIVITY	13	0.01	Up
0.26112694756458	0.393036180218784	0.763070882669728
GO_LOCOMOTORY_BEHAVIOR	108	0.01	Up	0.422092288409641
	0.393212389641436	0.763168931430038
GO_REGULATION_OF_PROTEIN_EXIT_FROM_ENDOPLASMIC_RETICULUM	22
	0.01	Down	0.222217668543794	0.393622211765402

0.763393916306019
GO_NEGATIVE_REGULATION_OF_MULTICELLULAR_ORGANISMAL_PROCESS 815
0.01 Down 0.470415535827275 0.393641431854877
0.763393916306019
GO_EMBRYO_IMPLANTATION 40 0.01 Up 0.624820873453755
0.393986804267416 0.763393916306019
GO_POSITIVE_REGULATION_OF_ORGANIC_ACID_TRANSPORT 19 0.01
Up 0.4123913002577 0.394105563866716
0.763393916306019
GO_TRANSCRIPTION_COUPLED_NUCLEOTIDE_EXCISION_REPAIR 73 0.01
Up 0.106022519459199 0.394214104069038
0.763393916306019
GO_REGULATION_OF_TRANSPORTER_ACTIVITY 155 0.01 Down
0.423939508976421 0.394269918381656 0.763393916306019
GO_SYNAPTIC_VESICLE_LOCALIZATION 109 0.01 Up
0.272795413163844 0.394331867442822 0.763393916306019
GO_POSITIVE_REGULATION_OF_BONE_RESORPTION 13 0.01 Up
0.411289309054981 0.394334264789273
0.763393916306019
GO_PHOSPHATASE_ACTIVITY 224 0.01 Down 0.299216548799811
0.394743384461019 0.763942329315539
GO_ZINC_ION_HOMEOSTASIS 24 0.01 Down 0.545932183296633
0.394895370324339 0.763992923275673
GO_CELLULAR_RESPONSE_TO_LEUCINE_STARVATION 10 0.01 Up
0.186581260063659 0.395258941865438
0.764446066287025
GO_PROTEIN_INSERTION_INTO_MITOCHONDRIAL_MEMBRANE_INVOLVED_IN_APOPTOT
IC_SIGNALING_PATHWAY 29 0.01 Down 0.271007588244608
0.3954050007736 0.764446066287025
GO_GOLGI_MEMBRANE 635 0.01 Down 0.282811919562396
0.395538343850824 0.764446066287025
GO_NUCLEOTIDE_SUGAR_METABOLIC_PROCESS 17 0.01 Down
0.410987272449441 0.395633263098968 0.764446066287025
GO_CELLULAR_RESPONSE_TO_TOXIC_SUBSTANCE 176 0.01 Up
0.431775461005039 0.396362394773555 0.76513174669962
GO_CELL_MIGRATION_IN_HINDBRAIN 10 0.01 Down
0.115955262955425 0.396528237562602 0.76513174669962
GO_LENS_MORPHOGENESIS_IN_CAMERA_TYPE_EYE 16 0.01 Down
0.287610143896242 0.396596719319737
0.76513174669962
GO_PROTEIN_INSERTION_INTO_MITOCHONDRIAL_INNER_MEMBRANE 11 0.01
Up 0.0757987328288385 0.396751202580258
0.76513174669962
GO_DNA_BINDING_TRANSCRIPTION_FACTOR_BINDING 294 0.01 Down
0.247010399600432 0.396838612234307
0.76513174669962
GO_DEVELOPMENTAL_CELL_GROWTH 165 0.01 Down
0.378498812256104 0.396959369209593 0.76513174669962
GO_MESENCHYMAL_CELL_DIFFERENTIATION 159 0.01 Down
0.511799983606388 0.396961431032839 0.76513174669962
GO_CLATHRIN_VESICLE_COAT 26 0.01 Down
0.193568445494493 0.396996376561325 0.76513174669962
GO_REGULATION_OF_INTRACELLULAR_ESTROGEN_RECEPTOR_SIGNALING_PATHWAY
34 0.01 Down 0.206230612470049

0.39736964645052	0.765410147251357
GO_POSITIVE_REGULATION_OF_GLUCOSE_TRANSMEMBRANE_TRANSPORT	29
	0.01	Down	0.338246620109266	0.397392980421063
	0.765410147251357
GO_NEGATIVE_REGULATION_OF_CELL_KILLING	17	0.01	Down
0.66333238342247	0.397585820101685	0.765538697696584
GO_PEPTIDASE_ACTIVATOR_ACTIVITY 35	0.01	Up
0.405819016697254	0.397733174410485	0.765579613774907
GO_REGULATION_OF_IRE1_MEDIATED_UNFOLDED_PROTEIN_RESPONSE	12
	0.01	Down	0.312788316390717	0.398005870028485
	0.765672781087056
GO_REGULATION_OF_RETINOIC_ACID_RECEPTOR_SIGNALING_PATHWAY	12
	0.01	Down	0.337161692457192	0.398099020333188
	0.765672781087056
GO_RESPONSE_TO_PH	22	0.01	Up	0.207620140328943
	0.39831783550309	0.765672781087056
GO_ESTABLISHMENT_OR_MAINTENANCE_OF_CYTOSKELETON_POLARITY	15
	0.01	Down	0.472347288608197	0.398341932746529
	0.765672781087056
GO_POSITIVE_REGULATION_OF_CELLULAR_PROTEIN_LOCALIZATION 281	0.01
	Down	0.259832500710865	0.398412175169496
0.765672781087056
GO_TRANSCRIPTION_ELONGATION_FROM_RNA_POLYMERASE_I_PROMOTER	29
	0.01	Up	0.150318819010568	0.398755980459315
	0.766090999167247
GO_HEART_MORPHOGENESIS	176	0.01	Up	0.39890978150834
	0.398983059852093	0.766156348241905
GO_FORMATION_OF_EXTRACHROMOSOMAL_CIRCULAR_DNA	13	0.01	Down
	0.131414177542162	0.399130890175189
0.766156348241905
GO_INTRINSIC_COMPONENT_OF_ORGANELLE_MEMBRANE	328	0.01	Up
	0.221730141361283	0.399168593228323
0.766156348241905
GO_EXOGENOUS_DRUG_CATABOLIC_PROCESS	12	0.01	Up
0.597302528377577	0.399365883936247	0.766292756440252
GO_POSITIVE_REGULATION_OF_MYELOID_LEUKOCYTE_CYTOKINE_PRODUCTION_INVO
LVED_IN_IMMUNE_RESPONSE 19	0.01	Down	0.605028951715758
	0.399625188885889	0.766338565092051
GO_CHAPERONE_MEDIATED_PROTEIN_TRANSPORT 11	0.01	Up
0.227982089959843	0.399642216616939	0.766338565092051
GO_DICARBOXYLIC_ACID_CATABOLIC_PROCESS	11	0.01	Up
0.186258821257	0.400000665740854	0.766596063748327
GO_POLYMERIC_CYTOSKELETAL_FIBER 475	0.01	Down
0.298553190614889	0.400059426620556	0.766596063748327
GO_IRON_ION_HOMEOSTASIS 68	0.01	Up	0.354020606379388
	0.400155316425374	0.766596063748327
GO_REGULATION_OF_GTP_BINDING	11	0.01	Down
0.225881533455056	0.400415691348937	0.766852890277413
GO_POSITIVE_T_CELL_SELECTION	22	0.01	Down
0.549782622545582	0.400747463185107	0.766853546873067
GO_INNER_EAR_RECEPTOR_CELL_DEVELOPMENT	29	0.01	Up
0.349813105488714	0.400986538802998	0.766853546873067
GO_POSITIVE_REGULATION_OF_PROTEOLYSIS	300	0.01	Down
0.317985925390357	0.400989377008193	0.766853546873067

GO_LIPOPROTEIN_PARTICLE_RECEPTOR_BINDING 18 0.01 Up
0.313761576781511 0.40102059619725
0.766853546873067
GO_RESPONSE_TO_AUDITORY_STIMULUS 14 0.01 Down
0.615420198402046 0.401047605225167 0.766853546873067
GO_SODIUM_ION_HOMEOSTASIS 19 0.01 Down
0.535508418890598 0.401516845040729 0.767159511400205
GO_AMEBOIDAL_TYPE_CELL_MIGRATION 321 0.01 Down
0.470210530080877 0.401562919821854 0.767159511400205
GO_STEROID_DEHYDROGENASE_ACTIVITY 24 0.01 Up
0.719173069512958 0.401586711782219 0.767159511400205
GO_NOTCH_BINDING 18 0.01 Up 0.348249687790578
0.401993898034689 0.767605373969198
GO_AMMONIUM_TRANSPORT 51 0.01 Up 0.700144112999061
0.402148185286954 0.767605373969198
GO_NEURON_FATE_COMMITMENT 27 0.01 Up
0.564104085394834 0.402225328815154 0.767605373969198
GO_RRNA_CATABOLIC_PROCESS 18 0.01 Down
0.202527127107938 0.402417901819521 0.767605373969198
GO_CHEMOKINE_BIOSYNTHETIC_PROCESS 12 0.01 Down
0.445936538401275 0.402452298689501 0.767605373969198
GO_NEGATIVE_REGULATION_OF_SODIUM_ION_TRANSPORT 13 0.01 Down
0.410196052578379 0.402644549420667
0.767606585266539
GO_MAST_CELL_MEDIATED_IMMUNITY 37 0.01 Down
0.431191868989846 0.402789567647625 0.767606585266539
GO_NEUTRAL_AMINO_ACID_TRANSMEMBRANE_TRANSPORTER_ACTIVITY 19
0.01 Up 0.252529972561825 0.402835735236417
0.767606585266539
GO_MEMBRANE_REPOLARIZATION_DURING_ACTION_POTENTIAL 16 0.01
Down 0.574345877269155 0.402958686747564
0.767606585266539
GO_AMINE_METABOLIC_PROCESS 97 0.01 Down
0.289709231921196 0.403379600344307 0.768167363139989
GO_PROTEIN_KINASE_A_REGULATORY_SUBUNIT_BINDING 17 0.01 Down
0.290204657460441 0.403725979210558
0.768176168468153
GO_PYRIMIDINE_CONTAINING_COMPOUND_SALVAGE 11 0.01 Down
0.226145441810443 0.403938818958513
0.768176168468153
GO_REGULATION_OF_NEURON_DIFFERENTIATION 479 0.01 Down
0.359574282279772 0.4039662651675 0.768176168468153
GO_RESPONSE_TO_AMINO_ACID 90 0.01 Up
0.320473012239661 0.404198666024482 0.768176168468153
GO_REGULATION_OF_NITRIC_OXIDE_SYNTHASE_ACTIVITY 34 0.01 Down
0.412005936951158 0.404505700490203
0.768176168468153
GO_NATURAL_KILLER_CELL_DIFFERENTIATION 17 0.01 Down
0.490343955089693 0.404512546312926 0.768176168468153
GO_REGULATION_OF_ACTIN_NUCLEATION 24 0.01 Down
0.263332442450254 0.404516960682978 0.768176168468153
GO_METALLOEXOPEPTIDASE_ACTIVITY 38 0.01 Up
0.441302400378084 0.404574588789988 0.768176168468153
GO_POSITIVE_REGULATION_OF_DNA_TEMPLATED_TRANSCRIPTION_ELONGATION

29	0.01	Down	0.1135146447952 0.404714354622484
0.768176168468153
GO_HISTONE_H2A_UBIQUITINATION	25	0.01	Down
0.141210996776519	0.404746874669405	0.768176168468153
GO_ACTIN_FILAMENT_BUNDLE_ORGANIZATION	128	0.01	Down
0.284641639945726	0.404804976957359	0.768176168468153
GO_BASE_EXCISION_REPAIR_AP_SITE_FORMATION	10	0.01	Up
0.114849952517445	0.404946457851433
0.768176168468153
GO_NUCLEOTIDE_DIPHOSPHATASE_ACTIVITY	14	0.01	Up
0.177107471874174	0.405029141042095	0.768176168468153
GO_CELLULAR_RESPONSE_TO_ANTIBIOTIC	117	0.01	Down
0.391869263597895	0.405361050013815	0.768565563595838
GO_SYNAPTIC_SIGNALING	384	0.01	Up	0.369136613918718
0.406090749436952	0.769708691798857
GO_POSITIVE_REGULATION_OF_ANION_TRANSPORT	31	0.01	Up
0.424921789826124	0.406420243656947
0.769883592773869
GO_INVADOPODIUM 15	0.01	Down	0.335904068236783
0.406722207184617	0.769883592773869
GO_TERMINATION_OF_RNA_POLYMERASE_I_TRANSCRIPTION	31	0.01
Up	0.157700276471849	0.40679387649841
0.769883592773869
GO_REGULATION_OF_RECEPTOR_MEDIATED_ENDOCYTOSIS	67	0.01	Down
0.394876332373527	0.40687084547358
0.769883592773869
GO_MITOTIC_SISTER_CHROMATID_COHESION	20	0.01	Down
0.1612106352164 0.407000155907044	0.769883592773869
GO_CLEAVAGE_INVOLVED_IN_RRNA_PROCESSING 21	0.01	Up
0.128887508885497	0.407119149547674	0.769883592773869
GO_REGULATION_OF_REACTIVE_OXYGEN_SPECIES_METABOLIC_PROCESS	85
0.01	Up	0.392430565874758	0.407164587740606
0.769883592773869
GO_PEPTIDE_BINDING	191	0.01	Up	0.448501719847016
0.40743403643234	0.769883592773869
GO_VENTRICULAR_CARDIAC_MUSCLE_CELL_MEMBRANE_REPOLARIZATION	17
0.01	Down	0.648307893130963	0.407441517867599
0.769883592773869
GO_ANION_TRANSMEMBRANE_TRANSPORTER_ACTIVITY	188	0.01	Up
0.380770071145992	0.407640747911079
0.769883592773869
GO_POSITIVE_REGULATION_OF_CYCLASE_ACTIVITY	11	0.01	Up
0.495011336044831	0.407702244840657
0.769883592773869
GO_ACTIN_CYTOSKELETON	387	0.01	Down	0.312477492297887
0.407704785170152	0.769883592773869
GO_POSITIVE_REGULATION_OF_GLYCOPROTEIN_BIOSYNTHETIC_PROCESS	15
0.01	Down	0.196210127388608	0.407886247987047
0.76992289935611
GO_POSITIVE_REGULATION_OF_ENDOTHELIAL_CELL_CHEMOTAXIS	12	0.01
Up	0.461213308699132	0.407979240195784
0.76992289935611
GO_MITOTIC_SPINDLE_ORGANIZATION 93	0.01	Down
0.199302929674527	0.408701009590362	0.770935649992076

GO_MHC_CLASS_I_PROTEIN_BINDING 16 0.01 Down
0.368479633914726 0.408840479927301 0.770935649992076
GO_STEROL_BIOSYNTHETIC_PROCESS 65 0.01 Down
0.257865423922994 0.408896852738344 0.770935649992076
GO_PROTEIN_LOCALIZATION_TO_CYTOSKELETON 53 0.01 Down
0.180214386395612 0.409258298787988 0.770943552864694
GO_NEGATIVE_REGULATION_OF_ESTABLISHMENT_OF_PROTEIN_LOCALIZATION 129
0.01 Down 0.474580485541363 0.409391579055983
0.770943552864694
GO_NEGATIVE_REGULATION_OF_LEUKOCYTE_DIFFERENTIATION 71 0.01
Down 0.522714441991722 0.409440675948081
0.770943552864694
GO_NEGATIVE_REGULATION_OF_PROTEOLYSIS 230 0.01 Down
0.449349103218315 0.409526582067716 0.770943552864694
GO_RETINOIC_ACID_RECEPTOR_SIGNALING_PATHWAY 22 0.01 Down
0.270363390935419 0.409559848470612
0.770943552864694
GO_CELLULAR_CARBOHYDRATE_BIOSYNTHETIC_PROCESS 63 0.01 Down
0.302937140464196 0.409662971757784
0.770943552864694
GO_TERMINATION_OF_RNA_POLYMERASE_II_TRANSCRIPTION 34 0.01
Down 0.0800706475233375 0.411577550940203
0.773866576678963
GO_REGULATION_OF_MICROTUBULE_POLYMERIZATION 39 0.01 Down
0.23358994134172 0.41163515083826
0.773866576678963
GO_MACROMOLECULE_DEACYLATION 91 0.01 Down
0.172661178075105 0.411808561059292 0.773866576678963
GO_HISTONE_UBIQUITINATION 45 0.01 Down
0.126559516978518 0.411901797612253 0.773866576678963
GO_MATURATION_OF_SSU_RRNA_FROM_TRICISTRONIC_RRNA_TRANSCRIPT_SSU_RRNA
_5_8S_RRNA_LSU_RRNA_ 34 0.01 Up 0.118485973733063
0.411968204998324 0.773866576678963
GO_SNORNA_BINDING 25 0.01 Up 0.135755168373603
0.412044055996611 0.773866576678963
GO_NEURONAL_STEM_CELL_POPULATION_MAINTENANCE 19 0.01 Down
0.50116490694831 0.412148733950294
0.773866576678963
GO_OXIDOREDUCTASE_ACTIVITY_ACTING_ON_PAIRED_DONORS_WITH_INCORPORATIO
N_OR_REDUCTION_OF_MOLECULAR_OXYGEN_REDUCED_FLAVIN_OR_FLAVOPROTEIN_AS
_ONE_DONOR_AND_INCORPORATION_OF_ONE_ATOM_OF_OXYGEN 15 0.01
Up 0.52770615146847 0.412235959311442
0.773866576678963
GO_SUPEROXIDE_ANION_GENERATION 23 0.01 Up
0.458536727717722 0.412631662055047 0.774329464218954
GO_CELLULAR_RESPONSE_TO_ATP 14 0.01 Up
0.369293712028743 0.412737629091177 0.774329464218954
GO_RETROGRADE_TRANSPORT_ENDOSOME_TO_PLASMA_MEMBRANE 24 0.01
Down 0.165118039648084 0.412901359020658
0.774397327962438
GO_RESPIRATORY_CHAIN_COMPLEX_III 11 0.01 Up
0.0929885298199586 0.41331684919722 0.774937180814182
GO_ENDONUCLEASE_COMPLEX 29 0.01 Down 0.18843690215675
0.413572013046303 0.775176193641279

GO_PHOSPHATIDYLINOSITOL_BIOSYNTHETIC_PROCESS 105 0.01 Down
0.219275118623626 0.413713433966206
0.775201931360752
GO_REGULATION_OF_B_CELL_PROLIFERATION 39 0.01 Down
0.520013110740049 0.413952491052715 0.775226767085345
GO_DENDRITIC_CELL_DIFFERENTIATION 30 0.01 Down
0.577992890897971 0.413982075257897 0.775226767085345
GO_RESPONSE_TO_NITROGEN_COMPOUND 682 0.01 Down
0.345330907539449 0.414430215946494 0.775691609020459
GO_REGULATION_OF_GLUCONEOGENESIS 35 0.01 Down
0.234993795807934 0.414646364274948 0.775691609020459
GO_ACETYLTRANSFERASE_ACTIVITY 91 0.01 Down
0.155997216179894 0.414683642899647 0.775691609020459
GO_TELOMERIC_DNA_BINDING 35 0.01 Down
0.145276160515341 0.414755739466347 0.775691609020459
GO_SPINDLE_ASSEMBLY 91 0.01 Down 0.200792592851947
0.414898400177697 0.775691609020459
GO_HISTONE_H3_K9_MODIFICATION 41 0.01 Down
0.186542909302812 0.414996927375795 0.775691609020459
GO_PHOSPHATIDYLINOSITOL_3_5_BISPHOSPHATE_BINDING 22 0.01
Down 0.18766889118061 0.415168449867155
0.775773363848415
GO_POSITIVE_REGULATION_OF_MRNA_3_END_PROCESSING 11 0.01 Down
0.144296316634779 0.415297304539972
0.775775364880669
GO_REGULATION_OF_PROTEIN_KINASE_A_SIGNALING 14 0.01 Up
0.473219881156318 0.415828165188315
0.776528080854587
GO_MEMORY 68 0.01 Up 0.383680339912533
0.416103750455742 0.776716333970497
GO_REGULATION_OF_BROWN_FAT_CELL_DIFFERENTIATION 12 0.01 Down
0.238223393607663 0.416184851656404
0.776716333970497
GO_MUSCLE_ADAPTATION 82 0.01 Down 0.41286651186549
0.416703896601901 0.776935426622959
GO_PHOTORECEPTOR_CELL_DEVELOPMENT 22 0.01 Up
0.485293069713422 0.416789046825474 0.776935426622959
GO_REGULATION_OF_INTRACELLULAR_STEROID_HORMONE_RECEPTOR_SIGNALING_PA
THWAY 66 0.01 Down 0.207520057466397
0.416963554000107 0.776935426622959
GO_SYNTAXIN_BINDING 49 0.01 Down 0.220086867278055
0.417010413161142 0.776935426622959
GO_REGULATION_OF_CELL_AGING 43 0.01 Down
0.310514236273511 0.417066578167147 0.776935426622959
GO_CYTOKINESIS 139 0.01 Down 0.213011377616205
0.417070096419737 0.776935426622959
GO_REGULATION_OF_BLOOD_CIRCULATION 154 0.01 Down
0.431799770902003 0.41751652582017 0.777353753788519
GO_POSITIVE_REGULATION_OF_PROTEIN_BINDING 82 0.01 Down
0.421647190820878 0.417584925079789
0.777353753788519
GO_MAST_CELL_GRANULE 16 0.01 Down 0.454701008695183
0.417678791773703 0.777353753788519
GO_SPERM_PRINCIPAL_PIECE 15 0.01 Up

0.195462381458267	0.418106178658889	0.777748459079133
GO_MULTIVESICULAR_BODY	31	0.01	Up	0.536903784331246
	0.418147087865968	0.777748459079133
GO_DENDRITE_TERMINUS	12	0.01	Down	0.157622336665886
	0.418336127776465	0.777861755507172
GO_RESPONSE_TO_WOUNDING	473	0.01	Up	0.455809366644444
	0.418603234645212	0.777932512235041
GO_REGULATION_OF_T_HELPER_1_TYPE_IMMUNE_RESPONSE	18	0.01
	Down	0.786948850076808	0.418780217427867
0.777932512235041
GO_POSITIVE_REGULATION_OF_MRNA_PROCESSING	25	0.01	Down
	0.116325435426251	0.418821629917328
0.777932512235041
GO_ALCOHOL_METABOLIC_PROCESS	169	0.01	Up
0.372265563183748	0.41888673735733	0.777932512235041
GO_SOMATIC_STEM_CELL_POPULATION_MAINTENANCE	54	0.01	Down
	0.366099733418297	0.419988758758511
0.779738651327239
GO_RESPONSE_TO_MUSCLE_ACTIVITY	17	0.01	Up
0.323982817760576	0.42029497945483	0.779738651327239
GO_SIGNAL_RELEASE	286	0.01	Down	0.354039615244362
	0.420346449457991	0.779738651327239
GO_PROTEIN_IMPORT_INTO_MITOCHONDRIAL_MATRIX	18	0.01	Down
	0.102522299572626	0.420373020226331
0.779738651327239
GO_POSITIVE_REGULATION_OF_KIDNEY_DEVELOPMENT	31	0.01	Down
	0.734686341877222	0.420715609185191
0.780135755456106
GO_NEGATIVE_REGULATION_OF_IMMUNE_RESPONSE	101	0.01	Down
	0.520282040690996	0.42110009515752
0.780526810460396
GO_RIBOSOMAL_LARGE_SUBUNIT_BIOGENESIS	67	0.01	Up
0.0959237310133311	0.421271713902494	0.780526810460396
GO_NUCLEAR_EUCHROMATIN	21	0.01	Down	0.200706301455196
	0.421312198629339	0.780526810460396
GO_HISTONE_MONOUBIQUITINATION	29	0.01	Down
0.127901802410471	0.421667904127811	0.780620358331231
GO_HETEROTYPIC_CELL_CELL_ADHESION	42	0.01	Down
0.550916892685448	0.421700080757152	0.780620358331231
GO_REGULATION_OF_MACROPHAGE_DERIVED_FOAM_CELL_DIFFERENTIATION	20
	0.01	Up	0.640567300775398	0.421748439355368
	0.780620358331231
GO_TAU_PROTEIN_BINDING	43	0.01	Down	0.323944695831154
	0.422222804712975	0.781163550893598
GO_MATURATION_OF_SSU_RRNA	44	0.01	Up
0.12481835820147	0.422422697279737	0.781163550893598
GO_POSITIVE_REGULATION_OF_CELL_KILLING	38	0.01	Down
0.450603660340164	0.422495747537718	0.781163550893598
GO_ATP_DEPENDENT_3_5_DNA_HELICASE_ACTIVITY	10	0.01	Down
	0.155915044134341	0.422685764027885
0.781163550893598
GO_CELL_SUBSTRATE_ADHESION	264	0.01	Up
0.520239788388794	0.42273053430946	0.781163550893598
GO_V_D_J_RECOMBINATION	11	0.01	Down	0.246800376561426

0.422900394284276	0.781163550893598
GO_REGULATION_OF_TRANSPORT	1230	0.01	Down
0.393524993096379	0.422985560367776	0.781163550893598
GO_GUANYL_NUCLEOTIDE_EXCHANGE_FACTOR_COMPLEX	10	0.01	Down
0.23761423361217	0.423071282381511
0.781163550893598
GO_ALKALI_METAL_ION_BINDING	12	0.01	Up
0.368490448819809	0.423443749344447	0.781613561042912
GO_NEGATIVE_REGULATION_OF_DNA_METABOLIC_PROCESS 82	0.01	Down
0.164182885754091	0.423942917328547
0.782145036820693
GO_FC_RECEPTOR_MEDIATED_STIMULATORY_SIGNALING_PATHWAY	77	0.01
Down	0.402111925286366	0.424069310425054
0.782145036820693
GO_PROTEIN_ACETYLTRANSFERASE_COMPLEX	88	0.01	Down
0.144489932983008	0.424118178424266	0.782145036820693
GO_NEGATIVE_REGULATION_OF_INTRACELLULAR_PROTEIN_TRANSPORT	33
0.01	Down	0.364115368470989	0.424449486412983
0.782189827572616
GO_PIGMENT_ACCUMULATION 12	0.01	Down	0.152806591813843
0.424787449742487	0.782189827572616
GO_OXIDOREDUCTASE_ACTIVITY_ACTING_ON_A_HEME_GROUP_OF_DONORS	20
0.01	Up	0.209539225681482	0.424916279336636
0.782189827572616
GO_NEGATIVE_REGULATION_OF_CELL_MORPHOGENESIS_INVOLVED_IN_DIFFERENTIA
TION	77	0.01	Down	0.43531102029695
0.424934078863036	0.782189827572616
GO_NUCLEAR_TRANSCRIPTION_FACTOR_COMPLEX 167	0.01	Down
0.240676419904322	0.425091397930027	0.782189827572616
GO_NEGATIVE_REGULATION_OF_STRESS_ACTIVATED_PROTEIN_KINASE_SIGNALING_
CASCADE 40	0.01	Down	0.401361079257674
0.425095790488458	0.782189827572616
GO_FANCONI_ANAEMIA_NUCLEAR_COMPLEX	12	0.01	Up
0.219948916978917	0.425172273557232	0.782189827572616
GO_RESPONSE_TO_REACTIVE_OXYGEN_SPECIES	181	0.01	Down
0.445078580878281	0.425173189093993	0.782189827572616
GO_ENDOPLASMIC_RETICULUM_EXIT_SITE	22	0.01	Down
0.144678806550811	0.425420181225714	0.782311209487824
GO_SOMATIC_DIVERSIFICATION_OF_IMMUNE_RECEPTORS_VIA_GERMLINE_RECOMBIN
ATION_WITHIN_A_SINGLE_LOCUS	48	0.01	Down
0.269316372269441	0.425816115205014	0.782311209487824
GO_NEGATIVE_REGULATION_OF_NITRIC_OXIDE_BIOSYNTHETIC_PROCESS	10
0.01	Down	0.500767282401451	0.425922113562403
0.782311209487824
GO_CEREBRAL_CORTEX_CELL_MIGRATION	34	0.01	Down
0.322256128357967	0.425930221520314	0.782311209487824
GO_PROTECTION_FROM_NON_HOMOLOGOUS_END_JOINING_AT_TELOMERE	10
0.01	Down	0.128144652802037	0.426067043362812
0.782311209487824
GO_REGULATION_OF_ORGANELLE_ORGANIZATION 1063	0.01	Down
0.261633224214851	0.426075907172449	0.782311209487824
GO_DEFENSE_RESPONSE_TO_GRAM_POSITIVE_BACTERIUM	41	0.01	Down
0.77181163205615	0.426141190870736
0.782311209487824

GO_RESPONSE_TO_ANTIBIOTIC 232 0.01 Down
0.419762784027013 0.426402219298757 0.782553770665886
GO_NEGATIVE_REGULATION_OF_PROTEASOMAL_UBIQUITIN_DEPENDENT_PROTEIN_CA
TABOLIC_PROCESS 31 0.01 Down 0.224624886422438
0.426714519815763 0.782580950974289
GO_ENDONUCLEOLYTIC_CLEAVAGE_INVOLVED_IN_RRNA_PROCESSING 11 0.01
Up 0.136633187223049 0.426795217159112
0.782580950974289
GO_REGULATION_OF_CHONDROCYTE_DIFFERENTIATION 35 0.01 Up
0.733145107660764 0.426803743810883
0.782580950974289
GO_CYTOSKELETON_ORGANIZATION 1054 0.01 Down
0.315748475465625 0.427132523689274 0.78268553314526
GO_REGULATION_OF_CARDIAC_MUSCLE_CONTRACTION_BY_REGULATION_OF_THE_REL
EASE_OF_SEQUESTERED_CALCIUM_ION 13 0.01 Up
0.331058047158071 0.427234386764504 0.78268553314526
GO_REGULATION_OF_KERATINOCYTE_PROLIFERATION 26 0.01 Down
0.365302664501894 0.427331268180366
0.78268553314526
GO_ACTIN_MEDIATED_CELL_CONTRACTION 68 0.01 Up
0.47542553982034 0.427595109140812 0.78268553314526
GO_PYRIMIDINE_NUCLEOTIDE_BIOSYNTHETIC_PROCESS 41 0.01 Up
0.253188093027713 0.427681568117006
0.78268553314526
GO_PRENYLTRANSFERASE_ACTIVITY 15 0.01 Down
0.138163094173562 0.427762909924737 0.78268553314526
GO_POSITIVE_REGULATION_OF_OLIGODENDROCYTE_DIFFERENTIATION 11
0.01 Down 0.412761452608799 0.427865125519847
0.78268553314526
GO_GENERAL_TRANSCRIPTION_INITIATION_FACTOR_BINDING 41 0.01
Down 0.190782559332715 0.427970136604239
0.78268553314526
GO_HISTONE_METHYLTRANSFERASE_ACTIVITY_H3_K9_SPECIFIC_ 10 0.01
Up 0.202524646477138 0.428021078906648
0.78268553314526
GO_STEROID_METABOLIC_PROCESS 215 0.01 Up
0.405164682754559 0.428726293866004 0.782861918381811
GO_ANDROGEN_RECEPTOR_SIGNALING_PATHWAY 52 0.01 Down
0.227793209953861 0.428749433661897 0.782861918381811
GO_14_3_3_PROTEIN_BINDING 26 0.01 Down
0.232041100484942 0.428818930384275 0.782861918381811
GO_PHOTORECEPTOR_INNER_SEGMENT 24 0.01 Up
0.373175502521396 0.428884908965803 0.782861918381811
GO_NEGATIVE_REGULATION_OF_CYSTEINE_TYPE_ENDOPEPTIDASE_ACTIVITY 74
0.01 Down 0.445624625556886 0.428926780603383
0.782861918381811
GO_EXTRINSIC_COMPONENT_OF_ORGANELLE_MEMBRANE 38 0.01 Down
0.239201831065205 0.429051752619029
0.782861918381811
GO_KIDNEY_MORPHOGENESIS 69 0.01 Down 0.582867437310649
0.429122216444423 0.782861918381811
GO_INACTIVATION_OF_MAPK_ACTIVITY 23 0.01 Down
0.448012901293432 0.429216197464821 0.782861918381811
GO_GOLGI_STACK 122 0.01 Down 0.279412939516834

0.429386978740434	0.782861918381811
GO_1_PHOSPHATIDYLINOSITOL_3_KINASE_REGULATOR_ACTIVITY	10	0.01	Down	0.361288245349145	0.429486905121422	0.782861918381811
GO_PROTEIN_COMPLEX_INVOLVED_IN_CELL_ADHESION	26	0.01	Up	0.699906197166365	0.429535999033077	0.782861918381811
GO_HEART_DEVELOPMENT	402	0.01	Up	0.426908544941941	0.43021973527795	0.782901614922612
GO_ORGANIC_HYDROXY_COMPOUND_TRANSMEMBRANE_TRANSPORTER_ACTIVITY	20	0.01	Up	0.894481743823096	0.430424333778974	0.782901614922612
GO_NADPLUS_BINDING	14	0.01	Down	0.267587243445422	0.430468068582926	0.782901614922612
GO_REGULATION_OF_POSTSYNAPTIC_MEMBRANE_NEUROTRANSMITTER_RECEPTOR_LEVELS	44	0.01	Down	0.253888850019492	0.430516037080883	0.782901614922612
GO_REGULATION_OF_GLYCOGEN_METABOLIC_PROCESS	26	0.01	Down	0.277933620803738	0.430539974760717	0.782901614922612
GO_REGULATION_OF_T_CELL_CYTOKINE_PRODUCTION	22	0.01	Down	0.427866636492955	0.430632840571428	0.782901614922612
GO_REGULATION_OF_CELLULAR_PROTEIN_LOCALIZATION	449	0.01	Down	0.266855682430616	0.430730299504703	0.782901614922612
GO_ION_CHANNEL_REGULATOR_ACTIVITY	69	0.01	Down	0.391528924226777	0.430758635180312	0.782901614922612
GO_OXIDOREDUCTASE_ACTIVITY_ACTING_ON_THE_CH_NH_GROUP_OF_DONORS_NAD_OR_NADP_AS_ACCEPTOR	17	0.01	Up	0.219966775382713	0.43092055362378	0.782901614922612
GO_BONE_REMODELING	60	0.01	Up	0.528319656531628	0.430961367411442	0.782901614922612
GO_NEGATIVE_REGULATION_OF_DNA_BIOSYNTHETIC_PROCESS	28	0.01	Down	0.205525193433652	0.431084475946938	0.782901614922612
GO_LIGASE_ACTIVITY_FORMING_CARBON_NITROGEN_BONDS	38	0.01	Up	0.195136689057889	0.431105270743912	0.782901614922612
GO_CATION_CATION_ANTIPORTER_ACTIVITY	17	0.01	Down	0.583304094228208	0.43227368283853	0.784788734603084
GO_APOPTOTIC_SIGNALING_PATHWAY	511	0.01	Down	0.303049165463478	0.432436692975242	0.784849974006785
GO_DEOXYRIBOSE_PHOSPHATE_CATABOLIC_PROCESS	19	0.01	Up	0.172301766884708	0.43289155181902	0.785440708635167
GO_ENDODERM_DEVELOPMENT	57	0.01	Up	0.537647420686095	0.43373968990375	0.785832116294373
GO_LATE_ENDOSOME_TO_VACUOLE_TRANSPORT	16	0.01	Down	0.0975114943241398	0.433772911996291	0.785832116294373
GO_CILIUM_MOVEMENT_INVOLVED_IN_CELL_MOTILITY	14	0.01	Up	0.275026668391716	0.433960777766468	0.785832116294373
GO_INTEGRIN_MEDIATED_SIGNALING_PATHWAY	84	0.01	Up

0.573267002302966	0.434083164767929	0.785832116294373
GO_CARDIAC_MUSCLE_CELL_DIFFERENTIATION	78	0.01	Up
0.4793961593009 0.434093926347236	0.785832116294373
GO_NEGATIVE_REGULATION_OF_CYTOKINE_PRODUCTION_INVOLVED_IN_IMMUNE_RES
PONSE	15	0.01	Down	0.557543739774112
0.43410696580181	0.785832116294373
GO_REGULATION_OF_RESPONSE_TO_DRUG	47	0.01	Up
0.496729321908109	0.434169757337493	0.785832116294373
GO_POSITIVE_REGULATION_OF_MULTICELLULAR_ORGANISM_GROWTH 19	0.01
Down	0.381591364053841	0.434178007839518
0.785832116294373
GO_COPPER_ION_TRANSPORT 18	0.01	Down	0.379549264403492
0.434361474028938	0.785832116294373
GO_NEGATIVE_REGULATION_OF_EPITHELIAL_CELL_PROLIFERATION 104	0.01
Up	0.438478375221525	0.434608256164812
0.785832116294373
GO_ATP_METABOLIC_PROCESS	33	0.01	Down
0.170156804205573	0.434618751236741	0.785832116294373
GO_TRANSFORMING_GROWTH_FACTOR_BETA1_PRODUCTION	10	0.01	Up
0.938494645227394	0.434679954824844
0.785832116294373
GO_REGULATION_OF_PHOSPHOLIPASE_ACTIVITY 42	0.01	Up
0.530760597537985	0.434827726132771	0.785832116294373
GO_REGULATION_OF_SODIUM_ION_TRANSMEMBRANE_TRANSPORT	46	0.01
Down	0.500519805647897	0.434919438436682
0.785832116294373
GO_MICROTUBULE_POLYMERIZATION_OR_DEPOLYMERIZATION	89	0.01
Down	0.241069145095167	0.435184369891918
0.785948242397939
GO_ALPHA_AMINO_ACID_BIOSYNTHETIC_PROCESS	52	0.01	Up
0.258391795845862	0.435242627399419
0.785948242397939
GO_HYDROLASE_ACTIVITY_ACTING_ON_CARBON_NITROGEN_BUT_NOT_PEPTIDE_BOND
S_IN_LINEAR_AMIDES	59	0.01	Down	0.249653698781032
0.435464440176714	0.786114961734414
GO_CILIARY_PLASM	71	0.01	Up	0.270231499056981
0.435638750408115	0.78619079739247
GO_ION_ANTIPORTER_ACTIVITY	34	0.01	Up
0.40360219066008	0.435821739430679	0.78619079739247
GO_RESPONSE_TO_STARVATION	158	0.01	Down
0.230285419416085	0.435894947129477	0.78619079739247
GO_RETINAL_GANGLION_CELL_AXON_GUIDANCE	12	0.01	Down
0.375029817803287	0.436214691572384	0.786533826117002
GO_REGULATION_OF_ARP2_3_COMPLEX_MEDIATED_ACTIN_NUCLEATION	14
0.01	Down	0.184624065305593	0.43687985200229
0.787372691273695
GO_RESPONSE_TO_OXYGEN_RADICAL	24	0.01	Up
0.411721519979896	0.43705021711666	0.787372691273695
GO_LOCOMOTION	1291	0.01	Down	0.454846800039707
0.437069011627797	0.787372691273695
GO_CLATHRIN_COAT_ASSEMBLY	15	0.01	Down
0.215811050082166	0.437367155708764	0.787389942746482
GO_REGULATION_OF_PROTEIN_LOCALIZATION_TO_PLASMA_MEMBRANE	77
0.01	Down	0.307322360719318	0.437434849417336

0.787389942746482
GO_NEGATIVE_REGULATION_OF_SUPRAMOLECULAR_FIBER_ORGANIZATION 99 0.01 Down 0.308486949526002 0.437467678617013 0.787389942746482
GO_UBIQUITIN_LIKE_PROTEIN_CONJUGATING_ENZYME_BINDING 34 0.01 Down 0.183751319464485 0.437772686049455 0.787705387375887
GO_REGULATION_OF_EPITHELIAL_TO_MESENCHYMAL_TRANSITION 73 0.01 Down 0.570319453001553 0.438063629696252 0.787995346929169
GO_NEGATIVE_REGULATION_OF_CELLULAR_RESPONSE_TO_TRANSFORMING_GROWTH_FACTOR_BETA_STIMULUS 57 0.01 Up 0.366344647126564 0.438541716952829 0.788396880936472
GO_POSITIVE_REGULATION_OF_EXECUTION_PHASE_OF_APOPTOSIS 12 0.01 Up 0.550058347394037 0.438548952699351 0.788396880936472
GO_MYELOID_LEUKOCYTE_ACTIVATION 522 0.01 Down 0.446976546725186 0.438676439433932 0.788396880936472
GO_ATPASE_ACTIVITY 363 0.01 Down 0.222847073305651 0.439014307647548 0.788581294736991
GO_POSITIVE_REGULATION_OF_RESPONSE_TO_DRUG 12 0.01 Down 0.282331731542125 0.439055914132262 0.788581294736991
GO_TRANSMEMBRANE_RECEPTOR_PROTEIN_PHOSPHATASE_ACTIVITY 15 0.01 Down 0.310947679037646 0.439236467844493 0.788581294736991
GO_ACETYL_COA_BIOSYNTHETIC_PROCESS 20 0.01 Up 0.187876455718499 0.439321971605267 0.788581294736991
GO_POSITIVE_REGULATION_OF_ACTIN_NUCLEATION 16 0.01 Down 0.288207565992843 0.439630378834459 0.788581294736991
GO_PROTEIN_STABILIZATION 144 0.01 Down 0.201587307717658 0.439641218790832 0.788581294736991
GO_INTEGRATOR_COMPLEX 16 0.01 Down 0.106742754271006 0.439957978158161 0.788581294736991
GO_LYSINE_N_METHYLTRANSFERASE_ACTIVITY 54 0.01 Down 0.188119797217968 0.439971722274887 0.788581294736991
GO_ERK1_AND_ERK2_CASCADE 201 0.01 Down 0.491610466687416 0.440032218135841 0.788581294736991
GO_LYSOSOMAL_PROTEIN_CATABOLIC_PROCESS 10 0.01 Down 0.373642754634575 0.440077981645351 0.788581294736991
GO_CELL_PROJECTION_ASSEMBLY 477 0.01 Up 0.271973199015221 0.44066996934515 0.789409083474301
GO_OSTEOBLAST_DEVELOPMENT 14 0.01 Up 0.448769790338254 0.440897327418789 0.789583384884798
GO_DNA_DAMAGE_RESPONSE_DETECTION_OF_DNA_DAMAGE 38 0.01 Up 0.122727943964815 0.441123946867321 0.789756261112211
GO_CHLORIDE_CHANNEL_ACTIVITY 34 0.01 Down 0.380636276163494 0.441551174138116 0.790288083193544
GO_REGULATION_OF_MUSCLE_ORGAN_DEVELOPMENT 93 0.01 Down 0.447288899339242 0.441817784535432 0.790532204513588
GO_PROTEIN_LOCALIZATION_TO_LYSOSOME 33 0.01 Down

0.172824951811553	0.442324435613163	0.791205553508401
GO_CHLORIDE_TRANSPORT	51	0.01	Down	0.413051968393373
	0.442489107335104	0.791227265116925
GO_REGULATION_OF_B_CELL_ACTIVATION	86	0.01	Down
0.436663349688937	0.442597231483623	0.791227265116925
GO_AMELOGENESIS 16	0.01	Down	0.480686639030158
0.442787893655693	0.791335090486815
GO_POSITIVE_REGULATION_OF_MONONUCLEAR_CELL_MIGRATION	20	0.01
	Down	0.855974614071262	0.442925624055524
0.791348282413504
GO_LOW_DENSITY_LIPOPROTEIN_PARTICLE_RECEPTOR_BINDING	17	0.01
	Up	0.325053185785909	0.44320778231091
0.791619431129607
GO_CLATHRIN_ADAPTOR_COMPLEX	26	0.01	Down
0.179600191211363	0.443475389221787	0.791864437636902
GO_ACTIN_FILAMENT_ORGANIZATION	319	0.01	Down
0.320857129326556	0.443617925028122	0.79188604023691
GO_PH_REDUCTION 42	0.01	Up	0.279466610444432
0.444220818536374	0.792536735836788
GO_POSITIVE_REGULATION_OF_TORC1_SIGNALING	10	0.01	Up
	0.113606033785773	0.444243536823026
0.792536735836788
GO_HEME_BIOSYNTHETIC_PROCESS	20	0.01	Up
0.190157171715873	0.444385312599461	0.792556766389933
GO_CARDIAC_MUSCLE_CELL_MEMBRANE_REPOLARIZATION	23	0.01	Down
	0.568879876075453	0.444637174923349
0.79277306577376
GO_DETECTION_OF_MECHANICAL_STIMULUS_INVOLVED_IN_SENSORY_PERCEPTION
	12	0.01	Up	0.573046351887776
0.44530628730598	0.793661950062233
GO_FATTY_ACID_ELONGATION	11	0.01	Up
0.19968443523356	0.445427439603435	0.793661950062233
GO_TERMINAL_BOUTON	24	0.01	Up	0.204730161817775
	0.445527907397808	0.793661950062233
GO_NEGATIVE_REGULATION_OF_VIRAL_ENTRY_INTO_HOST_CELL	14	0.01
	Down	0.363209098331474	0.445804781407191
0.793922214116473
GO_DNA_BIOSYNTHETIC_PROCESS	162	0.01	Down
0.247649462268358	0.446057762545103	0.793995051364093
GO_NUCLEOLAR_PART	174	0.01	Down	0.182511689657675
	0.446107250898192	0.793995051364093
GO_FILOPODIUM_MEMBRANE	14	0.01	Up	0.223352359189037
	0.446534120707323	0.794521803787864
GO_POSITIVE_REGULATION_OF_B_CELL_MEDIATED_IMMUNITY	28	0.01
	Down	0.415263612262148	0.446866886025003
0.794521803787864
GO_LEUKOCYTE_MEDIATED_IMMUNITY	603	0.01	Down
0.428678410846444	0.446909951478788	0.794521803787864
GO_PHENOL_CONTAINING_COMPOUND_METABOLIC_PROCESS 34	0.01	Down
	0.420594317676944	0.44692669410897
0.794521803787864
GO_SIGNAL_TRANSDUCTION_INVOLVED_IN_G2_DNA_DAMAGE_CHECKPOINT	13
	0.01	Up	0.171234860351977	0.447675919328692
	0.795516855762328

GO_PHOSPHOLIPASE_C_ACTIVITY 20 0.01 Up
0.474033993196565 0.447748492198958 0.795516855762328
GO_MUSCLE_CELL_DEVELOPMENT 108 0.01 Up
0.409380406490509 0.448127588167364 0.795614130104878
GO_SKELETAL_MUSCLE_ADAPTATION 17 0.01 Down
0.415562144876995 0.448239499902295 0.795614130104878
GO_RESPONSE_TO_INTERLEUKIN_4 28 0.01 Down
0.423880465519516 0.448323923529814 0.795614130104878
GO_REGULATION_OF_T_CELL_MIGRATION 32 0.01 Down
0.695418169771025 0.448327448375686 0.795614130104878
GO_CHROMATIN_SILENCING_AT_RDNA 19 0.01 Down
0.183980772283713 0.448702363858071 0.796046771181283
GO_REGULATION_OF_DEFENSE_RESPONSE_TO_BACTERIUM 15 0.01 Down
0.595546285530629 0.448845062718805
0.796067302297944
GO_PHOSPHOLIPID_METABOLIC_PROCESS 204 0.01 Up
0.320172894951395 0.449357083040414 0.796742655122182
GO_MULTICELLULAR_ORGANISMAL_HOMEOSTASIS 331 0.01 Down
0.426364989847168 0.449537734880097 0.796830244804984
GO_RESPONSE_TO_TESTOSTERONE 32 0.01 Down
0.311576015370157 0.449857698880589 0.797164649709299
GO_NEGATIVE_REGULATION_OF_RESPONSE_TO_CYTOKINE_STIMULUS 50 0.01
Down 0.447522267279626 0.450316912596168
0.797561056701384
GO_NEGATIVE_REGULATION_OF_CHROMOSOME_ORGANIZATION 118 0.01
Down 0.195115905647941 0.450547052248929
0.797561056701384
GO_NEGATIVE_REGULATION_OF_DEFENSE_RESPONSE_TO_VIRUS 21 0.01
Up 0.167174040680508 0.450647615802635
0.797561056701384
GO_TRANSFORMING_GROWTH_FACTOR_BETA_PRODUCTION 29 0.01 Down
0.637531043686855 0.450823274687836
0.797561056701384
GO_RIBONUCLEOPROTEIN_COMPLEX_DISASSEMBLY 15 0.01 Down
0.14937176749836 0.450946772863334
0.797561056701384
GO_NEGATIVE_REGULATION_OF_NUCLEAR_DIVISION 49 0.01 Up
0.284505380528594 0.451000603706351
0.797561056701384
GO_EMBRYONIC_MORPHOGENESIS 401 0.01 Up
0.434700761944778 0.45100100603786 0.797561056701384
GO_OXIDIZED_DNA_BINDING 11 0.01 Up 0.17626339989973
0.451534327053558 0.798271665562653
GO_MAINTENANCE_OF_LOCATION_IN_CELL 78 0.01 Up
0.284739739765441 0.452214651945142 0.79924167451498
GO_MICROTUBULE_BINDING 187 0.01 Down 0.271115392332993
0.452550799202656 0.799533941960661
GO_NEGATIVE_REGULATION_OF_IMMUNE_EFFECTOR_PROCESS 89 0.01
Down 0.473191413940141 0.452643412702815
0.799533941960661
GO_AMINOACYL_TRNA_SYNTHETASE_MULTIENZYME_COMPLEX 12 0.01
Down 0.0842669247663698 0.452787498244666
0.799555817871834
GO_POSITIVE_REGULATION_OF_FILOPODIUM_ASSEMBLY 22 0.01 Up

0.239500510607586 0.453222133718285
0.800090600117391
GO_REGULATION_OF_CARDIOCYTE_DIFFERENTIATION 40 0.01 Up
0.597312924749995 0.45363571265801
0.800587910333365
GO_POLYSOMAL_RIBOSOME 29 0.01 Up 0.124753797410614
0.453908828986538 0.800837111530738
GO_REGULATION_OF_GLYCOPROTEIN_BIOSYNTHETIC_PROCESS 28 0.01
Down 0.246664146149254 0.45418733265518
0.801095670119
GO_GLYCOLIPID_BINDING 15 0.01 Up 0.593800442201541
0.454487550740488 0.801215676580393
GO_ER_TO_GOLGI_TRANSPORT_VESICLE_MEMBRANE 55 0.01 Down
0.398509629705815 0.454612334964123
0.801215676580393
GO_POSITIVE_REGULATION_OF_MONOCYTE_CHEMOTAXIS 15 0.01 Down
0.950093412757142 0.454834502168475
0.801215676580393
GO_RIBONUCLEOSIDE_MONOPHOSPHATE_BIOSYNTHETIC_PROCESS 30 0.01
Up 0.247495280926409 0.454836472010538
0.801215676580393
GO_CHOLESTEROL_TRANSPORTER_ACTIVITY 11 0.01 Up
0.432334264886707 0.454915242492606 0.801215676580393
GO_CIS_GOLGI_NETWORK 48 0.01 Up 0.216063354779863
0.45522453024539 0.801527877934965
GO_POSITIVE_REGULATION_BY_HOST_OF_VIRAL_PROCESS 13 0.01 Up
0.210255378409378 0.455414044955968
0.801629071304053
GO_RETINOL_DEHYDROGENASE_ACTIVITY 11 0.01 Down
0.877378412430769 0.455965726688736 0.802108760820697
GO_PROTEIN_METHYLTRANSFERASE_ACTIVITY 77 0.01 Down
0.157765752553033 0.456152101422941 0.802108760820697
GO_POSITIVE_REGULATION_OF_LYMPHOCYTE_MIGRATION 27 0.01 Down
0.711655646026882 0.456284781281108
0.802108760820697
GO_POSITIVE_REGULATION_OF_SIGNAL_TRANSDUCTION_BY_P53_CLASS_MEDIATOR
18 0.01 Up 0.200652448739109
0.456336758905882 0.802108760820697
GO_REGULATION_OF_MICROTUBULE_POLYMERIZATION_OR_DEPOLYMERIZATION 62
0.01 Down 0.239153523391541 0.456347168485371
0.802108760820697
GO_RRNA_METABOLIC_PROCESS 201 0.01 Up
0.132647383976659 0.456541812519096 0.802218623387389
GO_TRANSITION_METAL_ION_TRANSPORT 93 0.01 Up
0.331851796009293 0.456881442160815 0.802583112082843
GO_METHYLOSOME 12 0.01 Up 0.0892807612769026
0.457175412959074 0.802867206269753
GO_PHOSPHATIDYLINOSITOL_4_5_BISPHOSPHATE_BINDING 51 0.01
Down 0.27323979941692 0.457414343605625
0.803054505503108
GO_REGULATION_OF_COLLAGEN_BIOSYNTHETIC_PROCESS 30 0.01 Up
0.697523702494308 0.457709916923131
0.803341111778065
GO_REGULATION_OF_FATTY_ACID_OXIDATION 26 0.01 Up

0.315046933073203	0.457906114754522	0.803453185744133
GO_PHOSPHOLIPID_BINDING 322	0.01	Down	0.33544686988661
	0.458084438489045	0.803510409084432
GO_PHOSPHATIDYLCHOLINE_BIOSYNTHETIC_PROCESS	34	0.01	Up
	0.332584269870001	0.458444681633561
0.803510409084432
GO_REGULATION_OF_MITOTIC_CELL_CYCLE	561	0.01	Down
0.226663022461297	0.458464523828948	0.803510409084432
GO_POSITIVE_REGULATION_OF_TRANSPORT	684	0.01	Down
0.410059228348679	0.458542543322037	0.803510409084432
GO_REGULATION_OF_CYTOPLASMIC_TRANSPORT	22	0.01	Down
0.182965875191308	0.458858705751611	0.803510409084432
GO_REGULATION_OF_MEMBRANE_POTENTIAL	220	0.01	Up
0.373538893013939	0.458912655417446	0.803510409084432
GO_ENDOPLASMIC_RETICULUM_TO_CYTOSOL_TRANSPORT	27	0.01	Down
	0.182313479510128	0.45900492590173
0.803510409084432
GO_GLYCOGEN_BIOSYNTHETIC_PROCESS	35	0.01	Down
0.249603066759668	0.459113141957175	0.803510409084432
GO_ATPASE_ACTIVITY_COUPLED	294	0.01	Down
0.229010956586308	0.459196415315028	0.803510409084432
GO_PHYSIOLOGICAL_CARDIAC_MUSCLE_HYPERTROPHY	18	0.01	Down
	0.423140967410917	0.459262249962606
0.803510409084432
GO_SYNAPTIC_VESICLE_CYCLE	123	0.01	Down
0.272881083079642	0.459884773430392	0.804158041991558
GO_POSITIVE_REGULATION_OF_RESPONSE_TO_OXIDATIVE_STRESS	16	0.01
	Down	0.461280415478411	0.459914352199054
0.804158041991558
GO_CELL_MOTILITY	1137	0.01	Down	0.474168482977141
	0.460029794076212	0.804158041991558
GO_LEUKOCYTE_MEDIATED_CYTOTOXICITY	69	0.01	Down
0.478513462713062	0.460329664629839	0.804275995761335
GO_INTRACELLULAR_MRNA_LOCALIZATION	12	0.01	Down
0.1441447389915 0.460362227848894	0.804275995761335
GO_REGULATION_OF_MRNA_POLYADENYLATION	16	0.01	Down
0.0966282333372887	0.461082233743788	0.804933991236779
GO_ENTRAINMENT_OF_CIRCADIAN_CLOCK_BY_PHOTOPERIOD	20	0.01
	Down	0.177462580004874	0.461106165557963
0.804933991236779
GO_REGULATION_OF_MACROPHAGE_DIFFERENTIATION	16	0.01	Up
	0.486676834213941	0.461136620247326
0.804933991236779
GO_GLUTAMATE_RECEPTOR_SIGNALING_PATHWAY 38	0.01	Down
0.336794251963015	0.461615948192809	0.805476162489726
GO_MEMBRANE_PROTEIN_ECTODOMAIN_PROTEOLYSIS	38	0.01	Down
	0.350200002674203	0.461769348125324
0.805476162489726
GO_REGULATION_OF_TRANS_SYNAPTIC_SIGNALING	260	0.01	Up
	0.365288213727467	0.461845251462154
0.805476162489726
GO_REGULATION_OF_CYTOPLASMIC_TRANSLATION	25	0.01	Down
	0.203628619445502	0.462059298444493
0.805498424453196

GO_RNA_POLYMERASE_II_CTD_HEPTAPEPTIDE_REPEAT_KINASE_ACTIVITY 16 0.01 Down 0.160708089199294 0.462165547078357 0.805498424453196
GO_CHROMOSOME_CENTROMERIC_REGION 159 0.01 Down 0.160189188535093 0.46243177578747 0.805498424453196
GO_GABA_RECEPTOR_BINDING 11 0.01 Up 0.287481845746562 0.46246201400386 0.805498424453196
GO_POSITIVE_REGULATION_OF_EXTRINSIC_APOPTOTIC_SIGNALING_PATHWAY 46 0.01 Down 0.308548315482387 0.462531642678572 0.805498424453196
GO_SINGLE_STRANDED_DNA_ENDODEOXYRIBONUCLEASE_ACTIVITY 10 0.01 Up 0.175615797933878 0.462770961439844 0.805498424453196
GO_CYCLASE_ACTIVITY 14 0.01 Up 0.509921538045711 0.462786773924024 0.805498424453196
GO_WHITE_FAT_CELL_DIFFERENTIATION 10 0.01 Down 0.37679859313843 0.463099236086464 0.805811253161629
GO_CELL_FATE_SPECIFICATION 41 0.01 Up 0.677966076675667 0.463339115253076 0.805995227927058
GO_NUCLEAR_HETEROCHROMATIN 29 0.01 Down 0.125004756797333 0.463664040054416 0.805995227927058
GO_EXON_EXON_JUNCTION_COMPLEX 20 0.01 Down 0.111421407524405 0.463773842089723 0.805995227927058
GO_REGULATION_OF_MITOCHONDRIAL_MEMBRANE_PERMEABILITY_INVOLVED_IN_APOPTOTIC_PROCESS 57 0.01 Up 0.243264899885618 0.463837792065087 0.805995227927058
GO_PURINE_NUCLEOSIDE_METABOLIC_PROCESS 29 0.01 Down 0.321296185717646 0.463868773904981 0.805995227927058
GO_POSITIVE_REGULATION_OF_NOTCH_SIGNALING_PATHWAY 40 0.01 Down 0.334138621005157 0.464479471722667 0.806811205113425
GO_LOW_DENSITY_LIPOPROTEIN_PARTICLE_CLEARANCE 26 0.01 Up 0.276013392097063 0.464604179389974 0.806811205113425
GO_REGULATION_OF_ANATOMICAL_STRUCTURE_SIZE 363 0.01 Down 0.371958128057573 0.46492126794202 0.807130974456965
GO_SYNCYTIUM_FORMATION 33 0.01 Down 0.633670558495485 0.465480454463852 0.807852863225457
GO_NEGATIVE_REGULATION_OF_PROTEIN_COMPLEX_ASSEMBLY 102 0.01 Down 0.260183705045622 0.465603223262374 0.807852863225457
GO_OUTER_MITOCHONDRIAL_MEMBRANE_PROTEIN_COMPLEX 16 0.01 Up 0.0754672756566801 0.46628746451881 0.808808913455342
GO_ANAPHASE_PROMOTING_COMPLEX_DEPENDENT_CATABOLIC_PROCESS 75 0.01 Down 0.136334261927323 0.466574366977341 0.809075401862164
GO_CONDENSED_NUCLEAR_CHROMOSOME_CENTROMERIC_REGION 17 0.01 Up 0.263866053902761 0.466708058587601 0.80907613469027
GO_ASTROCYTE_ACTIVATION 16 0.01 Down 0.561504961814097 0.467077849963387 0.809143935898995
GO_INSULIN_LIKE_GROWTH_FACTOR_I_BINDING 11 0.01 Up

0.672283393937413	0.467116259415343	0.809143935898995
GO_NUCLEOSIDE_PHOSPHATE_METABOLIC_PROCESS	166	0.01	Up
0.258458096450343	0.46714700960731
0.809143935898995
GO_NEGATIVE_REGULATION_OF_GLIOGENESIS	22	0.01	Down
0.275660610460878	0.467679110261117	0.809639801737841
GO_ARP2_3_COMPLEX_MEDIATED_ACTIN_NUCLEATION	30	0.01	Down
0.169580913805162	0.46776759564044
0.809639801737841
GO_REGULATION_OF_CHOLESTEROL_BIOSYNTHETIC_PROCESS	41	0.01
Down	0.246880623254906	0.467947452288221
0.809639801737841
GO_NEGATIVE_REGULATION_OF_AXONOGENESIS	54	0.01	Down
0.440383739701627	0.467966737654107	0.809639801737841
GO_ORGANIC_HYDROXY_COMPOUND_TRANSPORT	132	0.01	Up
0.564110865690407	0.468147971493624	0.80972260254638
GO_POSITIVE_REGULATION_OF_B_CELL_PROLIFERATION	26	0.01	Down
0.483169177294284	0.468527989384169
0.80972419019184
GO_TRANSMEMBRANE_RECEPTOR_PROTEIN_TYROSINE_KINASE_ADAPTOR_ACTIVITY
10	0.01	Up	0.267137930324428
0.468628640126773	0.80972419019184
GO_FOUR_WAY_JUNCTION_DNA_BINDING	15	0.01	Up
0.202668161713671	0.468692333455029	0.80972419019184
GO_MITOCHONDRIAL_ELECTRON_TRANSPORT_UBIQUINOL_TO_CYTOCHROME_C	13
0.01	Up	0.096188003043785	0.468881108724598
0.80972419019184
GO_SOLUTE_PROTON_SYMPORTER_ACTIVITY	19	0.01	Down
0.335919582592938	0.469050811657609	0.80972419019184
GO_DE_NOVO_PROTEIN_FOLDING	38	0.01	Down
0.26827319313212	0.46915157177623	0.80972419019184
GO_EXTRINSIC_COMPONENT_OF_MITOCHONDRIAL_INNER_MEMBRANE	13	0.01
Up	0.127241434225637	0.469231496439733
0.80972419019184
GO_HORMONE_ACTIVITY	21	0.01	Down	0.688546826811118
0.469299949495911	0.80972419019184
GO_POSITIVE_REGULATION_OF_INTRACELLULAR_TRANSPORT	183	0.01
Down	0.233633084145162	0.469349271171979
0.80972419019184
GO_PROTEIN_C_TERMINUS_BINDING	164	0.01	Down
0.237289172645056	0.469555081869568	0.809849119894927
GO_CARDIAC_CHAMBER_DEVELOPMENT	121	0.01	Up
0.402022525430438	0.469957891176629	0.810313648774018
GO_NEGATIVE_REGULATION_OF_MYOBLAST_DIFFERENTIATION	17	0.01
Up	0.511517836222137	0.470423640080971
0.810570210156239
GO_POSITIVE_REGULATION_OF_AUTOPHAGY_OF_MITOCHONDRION	19	0.01
Up	0.19220106531704	0.47062190699636
0.810570210156239
GO_POSITIVE_REGULATION_OF_MESENCHYMAL_CELL_PROLIFERATION	20
0.01	Up	0.483643601686272	0.470642574071861
0.810570210156239
GO_REGULATION_OF_PHOSPHOLIPID_METABOLIC_PROCESS	48	0.01	Down
0.433295060897127	0.47074200830303

0.810570210156239
GO_PROTEIN_MATURATION_BY_IRON_SULFUR_CLUSTER_TRANSFER 14 0.01 Up 0.077883381643215 0.470774264702833 0.810570210156239
GO_PATHWAY_RESTRICTED_SMAD_PROTEIN_PHOSPHORYLATION 45 0.01 Up 0.540387005889114 0.470910244785824 0.81057445310313
GO_PIGMENT_GRANULE_ORGANIZATION 24 0.01 Up 0.255731852604319 0.471494830405211 0.811274250158202
GO_MATURE_B_CELL_DIFFERENTIATION 20 0.01 Up 0.549800615014964 0.471652607140382 0.811274250158202
GO_TRANSMEMBRANE_RECEPTOR_PROTEIN_SERINE_THREONINE_KINASE_ACTIVITY 13 0.01 Down 0.193912781351379 0.471878547449781 0.811274250158202
GO_NEGATIVE_REGULATION_OF_TISSUE_REMODELING 13 0.01 Up 1.00934056539722 0.471920282081456 0.811274250158202
GO_GLUCOSE_METABOLIC_PROCESS 91 0.01 Down 0.369311414739025 0.471984953312267 0.811274250158202
GO_RIBOSOMAL_SMALL_SUBUNIT_BINDING 16 0.01 Down 0.136873363942343 0.472182610513232 0.811384270712095
GO_MONOSACCHARIDE_CATABOLIC_PROCESS 33 0.01 Up 0.239857753181724 0.472815497185787 0.812210248292969
GO_INORGANIC_ION_IMPORT_ACROSS_PLASMA_MEMBRANE 47 0.01 Down 0.421000589527434 0.4730092620246 0.812210248292969
GO_MORPHOGENESIS_OF_AN_ENDOTHELIUM 14 0.01 Down 0.651007080786952 0.473083712792621 0.812210248292969
GO_RNA_METABOLIC_PROCESS 1008 0.01 Down 0.149319046384041 0.473333232987387 0.812210248292969
GO_NEGATIVE_REGULATION_OF_PRI_MIRNA_TRANSCRIPTION_BY_RNA_POLYMERASE_II 12 0.01 Down 0.424564514214066 0.473363410963393 0.812210248292969
GO_CELL_CYCLE 1513 0.01 Down 0.226878698225168 0.47346599715184 0.812210248292969
GO_CILIARY_BASAL_BODY_PLASMA_MEMBRANE_DOCKING 91 0.01 Up 0.153898533871578 0.473732029240861 0.812437047887364
GO_MYELOID_CELL_DEVELOPMENT 51 0.01 Down 0.427315351691127 0.473902499153753 0.812499879232543
GO_PROTEIN_DISULFIDE_OXIDOREDUCTASE_ACTIVITY 20 0.01 Down 0.182560787437693 0.474064603222332 0.812548336014336
GO_REGULATION_OF_URINE_VOLUME 14 0.01 Up 0.334517047055802 0.474636389509562 0.813242766156185
GO_3_5_DNA_HELICASE_ACTIVITY 13 0.01 Down 0.139594861852673 0.47483686093733 0.813242766156185
GO_ORGANIC_CYCLIC_COMPOUND_CATABOLIC_PROCESS 522 0.01 Down 0.230904604761062 0.474874990132908 0.813242766156185
GO_EPITHELIAL_CELL_APOPTOTIC_PROCESS 75 0.01 Down 0.378356820484105 0.47500557548836 0.813242766156185
GO_FEEDING_BEHAVIOR 36 0.01 Down 0.35448707166745 0.475503552204095 0.813865820533143
GO_FOREBRAIN_DEVELOPMENT 246 0.01 Down

0.370528429686592	0.475762704896049	0.814079870750821
GO_REGULATION_OF_B_CELL_APOPTOTIC_PROCESS	13	0.01	Down
	0.329152520656486	0.476096124265724
0.814368063560114
GO_ALCOHOL_BIOSYNTHETIC_PROCESS	134	0.01	Down
0.324882460192583	0.47619941124006	0.814368063560114
GO_NEGATIVE_REGULATION_OF_NEURON_PROJECTION_REGENERATION	13
	0.01	Up	0.569904243928811	0.477225450906016
	0.815719152537757
GO_NEGATIVE_REGULATION_OF_INTRACELLULAR_STEROID_HORMONE_RECEPTOR_SIGNALING_PATHWAY	29	0.01	Down	0.236761869529043
0.477491567578582	0.815719152537757
GO_REGULATION_OF_NEUROTRANSMITTER_SECRETION	66	0.01	Up
	0.289955438644951	0.47752199958164
0.815719152537757
GO_REGULATION_OF_LEUKOCYTE_MIGRATION	130	0.01	Down
0.633444167343333	0.477570047086745	0.815719152537757
GO_MIDBODY	155	0.01	Down	0.185407030414764
0.477661272816953	0.815719152537757
GO_NEGATIVE_REGULATION_OF_RELEASE_OF_CYTOCHROME_C_FROM_MITOCHONDRIA
	15	0.01	Down	0.264832981859789
0.477797228570679	0.815721871387118
GO_RESPONSE_TO_ISCHEMIA	36	0.01	Up	0.484671091816477
	0.478507026888264	0.816671940100269
GO_PHARYNGEAL_SYSTEM_DEVELOPMENT	20	0.01	Down
0.40604113715391	0.478746633918197	0.816671940100269
GO_REGULATION_OF_NATURAL_KILLER_CELL_MEDIATED_IMMUNITY	29	0.01
	Down	0.553768967495114	0.478757278012989
0.816671940100269
GO_PEPTIDYL_TYROSINE_MODIFICATION	256	0.01	Down
0.473365014410506	0.478964988746225	0.816744280328853
GO_TFIID_CLASS_TRANSCRIPTION_FACTOR_COMPLEX_BINDING	11	0.01
	Down	0.187399349938343	0.479130653915611
0.816744280328853
GO_CONTRACTILE_RING	10	0.01	Up	0.154179217096911
	0.479203282248621	0.816744280328853
GO_SECRETORY_GRANULE_ORGANIZATION	25	0.01	Down
0.326310134549426	0.479402982129465	0.816855319817003
GO_PROTEIN_DNA_COMPLEX	118	0.01	Down	0.187025049610034
	0.479780724800967	0.817269579199403
GO_PROTEIN_LIPID_COMPLEX_ASSEMBLY	19	0.01	Up
0.387420590781764	0.480105888489665	0.817370204334949
GO_BLOOD_VESSEL_ENDOTHELIAL_CELL_PROLIFERATION_INVOLVED_IN_SPROUTING_ANGIOGENESIS	15	0.01	Down	0.556629584493555
0.480109067477916	0.817370204334949
GO_CARBOHYDRATE_TRANSPORT	103	0.01	Down
0.364046796451783	0.480576506119107	0.817838460633834
GO_SECONDARY_LYSOSOME	12	0.01	Up	0.453759161538258
	0.480653537727149	0.817838460633834
GO_EPITHELIAL_TO_MESENCHYMAL_TRANSITION_INVOLVED_IN_ENDOCARDIAL_CUSHION_FORMATION	11	0.01	Up	0.547731881835562
0.480895437161258	0.818020789858783
GO_POSITIVE_REGULATION_OF_TRANSMEMBRANE_TRANSPORT	121	0.01
	Down	0.457737366871106	0.481151139310873

0.818122676932724
GO_NEGATIVE_REGULATION_OF_PROTEIN_ACETYLATION 18 0.01 Down 0.239093566596363 0.481224852466934 0.818122676932724
GO_SPECIFIC_GRANULE_MEMBRANE 73 0.01 Up 0.428635191522606 0.481395374478435 0.818183459814832
GO_REGULATION_OF_PROTEIN_TARGETING_TO_MEMBRANE 31 0.01 Up 0.297124735136239 0.482128877461426 0.818912347512056
GO_LOCOMOTOR_RHYTHM 10 0.01 Up 0.385745724027407 0.482256457098991 0.818912347512056
GO_PHOSPHATIDYLINOSITOL_3_KINASE_BINDING 28 0.01 Down 0.525526688122956 0.48240677556801 0.818912347512056
GO_OXIDATIVE_DEMETHYLATION 13 0.01 Down 0.370683370533294 0.482483865080631 0.818912347512056
GO_B_CELL_HOMEOSTASIS 24 0.01 Down 0.5374221370089 0.482498676832585 0.818912347512056
GO_ORGANIC_HYDROXY_COMPOUND_CATABOLIC_PROCESS 46 0.01 Up 0.445726652544662 0.482784361237896 0.818987817992388
GO_SIGNAL_RELEASE_FROM_SYNAPSE 106 0.01 Up 0.306888237862331 0.482812946894211 0.818987817992388
GO_NUCLEOSOME_BINDING 62 0.01 Down 0.12260558004789 0.483308914656002 0.819195317666881
GO_NEGATIVE_REGULATION_OF_MYELOID_CELL_APOPTOTIC_PROCESS 11 0.01 Down 0.483351556427173 0.483323455905849 0.819195317666881
GO_LIPID_STORAGE 52 0.01 Down 0.514700722684047 0.483340080362835 0.819195317666881
GO_SIGNALING_RECEPTOR_BINDING 1015 0.01 Down 0.447381764023272 0.483585970128257 0.819383316954688
GO_MICROTUBULE_CYTOSKELETON_ORGANIZATION 439 0.01 Down 0.241652185020993 0.483916700281654 0.819532313817709
GO_CELL_CELL_SIGNALING_BY_WNT 421 0.01 Down 0.338344776788246 0.483943888162821 0.819532313817709
GO_BHLH_TRANSCRIPTION_FACTOR_BINDING 22 0.01 Down 0.166257585143138 0.484311731983957 0.819875910196308
GO_REGULATION_OF_SUPRAMOLECULAR_FIBER_ORGANIZATION 280 0.01 Down 0.324925845988414 0.484416881876817 0.819875910196308
GO_NEUROTRANSMITTER_RECEPTOR_LOCALIZATION_TO_POSTSYNAPTIC_SPECIALIZATION_MEMBRANE 12 0.01 Up 0.273881364488416 0.48464603562062 0.819999780524496
GO_POSITIVE_REGULATION_OF_B_CELL_DIFFERENTIATION 10 0.01 Up 0.36161958522182 0.484760206276135 0.819999780524496
GO_SUPEROXIDE_METABOLIC_PROCESS 48 0.01 Up 0.448548578282237 0.485276402540849 0.820644300787045
GO_COCHLEA_DEVELOPMENT 30 0.01 Up 0.329519330620886 0.486087631731135 0.821513225591953
GO_MYELIN_ASSEMBLY 17 0.01 Up 0.441877288358427 0.486146475938607 0.821513225591953

GO_RESPONSE_TO_WATER	10	0.01	Up	0.38144967465815
	0.48628562352473	0.821513225591953
GO_GOLGI_TO_PLASMA_MEMBRANE_PROTEIN_TRANSPORT	36	0.01	Down
	0.200517089837688	0.486331499386835
0.821513225591953
GO_REGULATION_OF_EXTENT_OF_CELL_GROWTH	86	0.01	Down
0.378154040293646	0.486642696628033	0.821574754657616
GO_RESPONSE_TO_GONADOTROPIN	20	0.01	Down
0.416856005777728	0.486829384342861	0.821574754657616
GO_SERTOLI_CELL_DIFFERENTIATION	14	0.01	Down
0.149745319636779	0.486847418596387	0.821574754657616
GO_POSITIVE_REGULATION_OF_DOUBLE_STRAND_BREAK_REPAIR_VIA_NONHOMOLOGO
US_END_JOINING	10	0.01	Down	0.226788275607481
0.487125355029391	0.821574754657616
GO_PROTEIN_LIPID_COMPLEX	22	0.01	Up
0.727546106211681	0.487176055880607	0.821574754657616
GO_CYTOPLASMIC_UBIQUITIN_LIGASE_COMPLEX	11	0.01	Down
0.0926968766339103	0.487280618811758	0.821574754657616
GO_CELLULAR_RESPONSE_TO_ABIOTIC_STIMULUS	250	0.01	Down
	0.268843809125211	0.487315218501413
0.821574754657616
GO_RECEPTOR_COMPLEX	223	0.01	Down	0.487180769079557
	0.487879277840253	0.821942158401849
GO_NUCLEAR_CHROMOSOME	470	0.01	Down	0.190042636108584
	0.48797265093399	0.821942158401849
GO_PHOSPHOLIPID_TRANSPORT	68	0.01	Up
0.401902540207905	0.48812775239802	0.821942158401849
GO_NEGATIVE_REGULATION_OF_INTERLEUKIN_8_PRODUCTION	13	0.01
	Down	0.440551590295931	0.488134408770624
0.821942158401849
GO_TEMPERATURE_HOMEOSTASIS	127	0.01	Down
0.398852121898007	0.488210084532543	0.821942158401849
GO_REGULATION_OF_CYTOSKELETON_ORGANIZATION	436	0.01	Down
	0.306990049175285	0.488585720554296
0.822151661352118
GO_OLIGOSACCHARYLTRANSFERASE_COMPLEX	13	0.01	Up
0.106608452498913	0.488605368828602	0.822151661352118
GO_MITOCHONDRION_ORGANIZATION	479	0.01	Up
0.178258843589171	0.488923175623963	0.822154727708473
GO_REGULATION_OF_MESENCHYMAL_CELL_PROLIFERATION	26	0.01	Up
	0.417662850007339	0.488952025214768
0.822154727708473
GO_INTEGRAL_COMPONENT_OF_SYNAPTIC_VESICLE_MEMBRANE	18	0.01
	Up	0.256344288733966	0.489013461004002
0.822154727708473
GO_VASCULAR_ENDOTHELIAL_CELL_PROLIFERATION	16	0.01	Up
	0.489581977760022	0.48927348312895
0.822273465689144
GO_REGULATION_OF_DEVELOPMENTAL_PIGMENTATION	13	0.01	Down
	0.608369981785356	0.489473745790209
0.822273465689144
GO_HISTONE_DEUBIQUITINATION	23	0.01	Down
0.17947661731721	0.489502698910336	0.822273465689144
GO_POSITIVE_REGULATION_OF_MAST_CELL_ACTIVATION	14	0.01	Down

0.550153643788242 0.48964824209902
0.822273465689144
GO_NUCLEAR_UBIQUITIN_LIGASE_COMPLEX 42 0.01 Up
0.152119807131001 0.489761299939375 0.822273465689144
GO_NUCLEOTIDE_EXCISION_REPAIR_DNA_GAP_FILLING 23 0.01 Up
0.125292167976222 0.490230603123989
0.822721521006164
GO_NEGATIVE_REGULATION_OF_DENDRITIC_SPINE_DEVELOPMENT 10 0.01
Down 0.497321775655945 0.490299203261456
0.822721521006164
GO_RESPONSE_TO_FATTY_ACID 59 0.01 Up
0.414718691771311 0.490686391705573 0.82314370932427
GO_HYDROLASE_ACTIVITY_ACTING_ON_ACID_ANHYDRIDES 741 0.01 Down
0.243153825371699 0.490886995533043
0.82325274858594
GO_POTASSIUM_CHANNEL_COMPLEX 39 0.01 Down
0.555046821752485 0.491367511265527 0.823622288852547
GO_POSITIVE_REGULATION_OF_NEUROINFLAMMATORY_RESPONSE 11 0.01
Up 0.883195715852551 0.491378674060933
0.823622288852547
GO_RESPONSE_TO_AMPHETAMINE 17 0.01 Down
0.503199311041994 0.491952023548938 0.824240562224385
GO_POSTSYNAPTIC_SPECIALIZATION_MEMBRANE 31 0.01 Down
0.308675360122498 0.492122123263383 0.824240562224385
GO_MONOSACCHARIDE_BIOSYNTHETIC_PROCESS 72 0.01 Up
0.226635946083642 0.492258828657986 0.824240562224385
GO_ADENYLATE_CYCLASE_INHIBITING_G_PROTEIN_COUPLED_RECEPTOR_SIGNALING
_PATHWAY 33 0.01 Up 0.522206948033753
0.492346795106873 0.824240562224385
GO_REGULATION_OF_PROTEIN_LOCALIZATION_TO_CENTROSOME 10 0.01
Down 0.130803874147914 0.49242637443384
0.824240562224385
GO_POSTSYNAPTIC_DENSITY_MEMBRANE 25 0.01 Down
0.308484679786369 0.493269352061998 0.825423990178718
GO_FEAR_RESPONSE 21 0.01 Up 0.334212373758573
0.493610904730979 0.825480314879099
GO_POSITIVE_REGULATION_OF_MITOTIC_NUCLEAR_DIVISION 42 0.01
Down 0.336544267547711 0.493644001893762
0.825480314879099
GO_POSITIVE_REGULATION_OF_CENTROSOME_CYCLE 11 0.01 Down
0.108045228353576 0.493994532016073
0.825480314879099
GO_POSITIVE_REGULATION_OF_CELL_DEATH 562 0.01 Down
0.368683715874707 0.494365613750889 0.825480314879099
GO_SUBSTANTIA_NIGRA_DEVELOPMENT 35 0.01 Down
0.258616906314497 0.494574474116513 0.825480314879099
GO_REGULATION_OF_NERVOUS_SYSTEM_DEVELOPMENT 642 0.01 Down
0.369446001547888 0.494624340898369
0.825480314879099
GO_BETA_CATENIN_DESTRUCTION_COMPLEX_DISASSEMBLY 17 0.01 Up
0.293803293974968 0.494664890248689
0.825480314879099
GO_LONG_CHAIN_FATTY_ACYL_COA_BIOSYNTHETIC_PROCESS 14 0.01
Up 0.253499216859207 0.494737368984719

0.825480314879099
GO_RHO_PROTEIN_SIGNAL_TRANSDUCTION 165 0.01 Down
0.390085316174972 0.494820333473382 0.825480314879099
GO_MONONUCLEAR_CELL_MIGRATION 54 0.01 Down
0.88792375318389 0.494864087802634 0.825480314879099
GO_CYSTEINE_TYPE_ENDOPEPTIDASE_ACTIVATOR_ACTIVITY_INVOLVED_IN_APOPTO
TIC_PROCESS 15 0.01 Up 0.288084606041036
0.49491068608673 0.825480314879099
GO_SCAFFOLD_PROTEIN_BINDING 45 0.01 Down
0.325130416541239 0.494939902186492 0.825480314879099
GO_RNA_POLYMERASE_II_GENERAL_TRANSCRIPTION_INITIATION_FACTOR_BINDING
19 0.01 Down 0.169426747727334
0.495070635228924 0.825480314879099
GO_REGULATION_OF_NEURON_PROJECTION_DEVELOPMENT 379 0.01 Down
0.370662857828625 0.495443212345647
0.825793084638862
GO_ACIDIC_AMINO_ACID_TRANSPORT 40 0.01 Down
0.317264209349395 0.495530259815414 0.825793084638862
GO_PROTEIN_LOCALIZATION_TO_MICROTUBULE 16 0.01 Down
0.187783025794034 0.496122732074284 0.826553541828479
GO_NUCLEOBASE_CONTAINING_SMALL_MOLECULE_METABOLIC_PROCESS 282
0.01 Up 0.277973558610537 0.496424848390847
0.826829973821901
GO_GLUTAMATERGIC_SYNAPSE 219 0.01 Down
0.278943468254609 0.496778862874228 0.827192670463368
GO_CARBOHYDRATE_TRANSMEMBRANE_TRANSPORT 78 0.01 Down
0.338658856877364 0.49696872319161 0.827281908005557
GO_MITOGEN_ACTIVATED_PROTEIN_KINASE_KINASE_BINDING 14 0.01
Down 0.230060087736079 0.497441541597288
0.82773330973028
GO_ORGANELLE_LOCALIZATION 573 0.01 Down
0.227809778849021 0.497512575721593 0.82773330973028
GO_GTP_METABOLIC_PROCESS 12 0.01 Down
0.174727429271435 0.498089941565313 0.828106525786004
GO_NEUROMUSCULAR_PROCESS 53 0.01 Up
0.311254258912047 0.498282514262379 0.828106525786004
GO_REGULATION_OF_NON_MOTILE_CILIUM_ASSEMBLY 11 0.01 Down
0.192241684763419 0.49848707984465
0.828106525786004
GO_NUCLEOSIDE_DIPHOSPHATASE_ACTIVITY 11 0.01 Down
0.247650906344814 0.498852858710606 0.828106525786004
GO_REGULATION_OF_CELLULAR_CARBOHYDRATE_METABOLIC_PROCESS 89
0.01 Down 0.370536772405928 0.498965809955435
0.828106525786004
GO_OXIDOREDUCTASE_ACTIVITY_OXIDIZING_METAL_IONS 15 0.01 Up
0.490858867976242 0.499110868760107
0.828106525786004
GO_INTRACELLULAR_PROTEIN_TRANSPORT 1049 0.01 Down
0.201658595804148 0.499313619414074 0.828106525786004
GO_MITOCHONDRIAL_TRANSPORT 234 0.01 Up
0.160156761431643 0.499344073282893 0.828106525786004
GO_PYRIMIDINE_RIBONUCLEOSIDE_TRIPHOSPHATE_BIOSYNTHETIC_PROCESS 13
0.01 Up 0.254789801844556 0.499550893306664
0.828106525786004

GO_ORGANIC_CATION_TRANSPORT 23 0.01 Up
0.30671969111741 0.499610834263074 0.828106525786004
GO_HISTONE_ACETYLTRANSFERASE_ACTIVITY_H4_K5_SPECIFIC_ 10 0.01
Down 0.0856929067867065 0.499744304052433
0.828106525786004
GO_NEGATIVE_REGULATION_OF_INCLUSION_BODY_ASSEMBLY 10 0.01
Down 0.159453587393979 0.499902489685904
0.828106525786004
GO_DEVELOPMENTAL_GROWTH 466 0.01 Down 0.392891253240234
0.500005422586564 0.828106525786004
GO_REGULATION_OF_HEART_RATE_BY_CARDIAC_CONDUCTION 26 0.01
Up 0.539053969863728 0.500040551296683
0.828106525786004
GO_REGULATION_OF_NEUTROPHIL_MIGRATION 22 0.01 Down
0.780571364872254 0.500077286405464 0.828106525786004
GO_RECEPTOR_BIOSYNTHETIC_PROCESS 20 0.01 Down
0.37395247436456 0.500166465850519 0.828106525786004
GO_RESPONSE_TO_SALT_STRESS 15 0.01 Down
0.494965571846092 0.500169468960225 0.828106525786004
GO_REGULATION_OF_NEUROTRANSMITTER_LEVELS 204 0.01 Up
0.405000267701684 0.500192164397509
0.828106525786004
GO_EXTRACELLULAR_MATRIX_DISASSEMBLY 64 0.01 Up
0.639287989023525 0.500481550788995 0.828359731417663
GO_VASCULAR_PROCESS_IN_CIRCULATORY_SYSTEM 87 0.01 Up
0.434944158904068 0.500817773655977
0.828552879545262
GO_PEPTIDYL_ASPARAGINE_MODIFICATION 31 0.01 Down
0.122507097150923 0.500871202097037 0.828552879545262
GO_PYRIMIDINE_NUCLEOSIDE_MONOPHOSPHATE_BIOSYNTHETIC_PROCESS 15
0.01 Up 0.229612101573583 0.501367533975045
0.829048425113787
GO_COBALAMIN_METABOLIC_PROCESS 16 0.01 Up
0.254350542873095 0.501577576399671 0.829048425113787
GO_ATTACHMENT_OF_MITOTIC_SPINDLE_MICROTUBULES_TO_KINETOCHORE 10
0.01 Down 0.114407702375913 0.501691150709329
0.829048425113787
GO_POSITIVE_REGULATION_OF_CELLULAR_EXTRAVASATION 10 0.01
Up 0.88439214757196 0.501754473239073
0.829048425113787
GO_POST_TRANSLATIONAL_PROTEIN_MODIFICATION 297 0.01 Up
0.271959850767303 0.501853559923105
0.829048425113787
GO_REGULATION_OF_PEPTIDYL_LYSINE_ACETYLATION 53 0.01 Down
0.277886892864042 0.502053491882523
0.829130554698264
GO_EXOCRINE_SYSTEM_DEVELOPMENT 39 0.01 Up
0.528355078356289 0.502176420626835 0.829130554698264
GO_SPINAL_CORD_DEVELOPMENT 53 0.01 Up
0.363495404364899 0.502329965559927 0.829158570123523
GO_NERVOUS_SYSTEM_PROCESS 494 0.01 Up
0.396623235323811 0.502780901595886 0.829617492996198
GO_POSITIVE_REGULATION_OF_DENDRITE_MORPHOGENESIS 33 0.01
Down 0.27035600369642 0.502881300317247

0.829617492996198
GO_NEGATIVE_REGULATION_OF_CARTILAGE_DEVELOPMENT 20 0.01 Up
0.849551852616086 0.503160572879693
0.829852713380227
GO_REGULATION_OF_AXONOGENESIS 145 0.01 Down
0.394763975350798 0.504076793415488 0.831138026296967
GO_NEGATIVE_REGULATION_OF_T_CELL_APOPTOTIC_PROCESS 16 0.01
Down 0.591989855849735 0.504567292314567
0.831429565023169
GO_POSITIVE_REGULATION_OF_UBIQUITIN_PROTEIN_LIGASE_ACTIVITY 10
0.01 Down 0.199434768107713 0.504583861163036
0.831429565023169
GO_CYCLIN_DEPENDENT_PROTEIN_SERINE_THREONINE_KINASE_INHIBITOR_ACTIVI
TY 12 0.01 Up 0.339749554247668
0.504736567938984 0.831429565023169
GO_CELLULAR_PROTEIN_COMPLEX_DISASSEMBLY 190 0.01 Up
0.191513499419594 0.504843174983434 0.831429565023169
GO_GROWTH 725 0.01 Down 0.37858413692503
0.504943480010912 0.831429565023169
GO_AMMONIUM_ION_BINDING 25 0.01 Up 0.306211463256216
0.505075314743114 0.831429565023169
GO_PSEUDOURIDINE_SYNTHASE_ACTIVITY 13 0.01 Up
0.124583987686506 0.505757136569839 0.832326260806586
GO_CALMODULIN_BINDING 131 0.01 Down 0.354768975263612
0.506088261854973 0.832443005888575
GO_ORGANOPHOSPHATE_ESTER_TRANSPORT 89 0.01 Up
0.358719607336414 0.506102311766551 0.832443005888575
GO_REGULATION_OF_SYNAPTIC_TRANSMISSION_GLUTAMATERGIC 31 0.01
Down 0.361407210171406 0.506251630687934
0.83246306877206
GO_CYTOSKELETAL_PROTEIN_BINDING 735 0.01 Down
0.30726564320026 0.507429533507151 0.833381568251521
GO_MRNA_METHYLATION 14 0.01 Down 0.123736321958166
0.507528774451207 0.833381568251521
GO_REGULATION_OF_FIBROBLAST_MIGRATION 30 0.01 Up
0.462260850580709 0.507531734299555 0.833381568251521
GO_AMINO_SUGAR_METABOLIC_PROCESS 21 0.01 Down
0.522648661302406 0.507553498219184 0.833381568251521
GO_UDP_N_ACETYLGLUCOSAMINE_BIOSYNTHETIC_PROCESS 11 0.01 Up
0.421441696927658 0.507810665648965
0.833381568251521
GO_NEGATIVE_REGULATION_OF_VASCULATURE_DEVELOPMENT 87 0.01
Down 0.573720550469837 0.508117921851857
0.833381568251521
GO_REGULATION_OF_DOUBLE_STRAND_BREAK_REPAIR_VIA_NONHOMOLOGOUS_END_JO
INING 20 0.01 Down 0.169705304683997
0.508249398927426 0.833381568251521
GO_SYNAPTIC_VESICLE_EXOCYTOSIS 77 0.01 Up
0.291474329635209 0.508515893100212 0.833381568251521
GO_MODULATION_BY_HOST_OF_VIRAL_GENOME_REPLICATION 12 0.01
Down 0.331199040853408 0.508603172747705
0.833381568251521
GO_MULTI_ORGANISM_BEHAVIOR 36 0.01 Down
0.202699929191676 0.508666900027994 0.833381568251521

GO_INTRACELLULAR_STEROL_TRANSPORT 20 0.01 Down
0.254074582891504 0.508791100514594 0.833381568251521
GO_POSITIVE_REGULATION_OF_SYNAPTIC_TRANSMISSION 100 0.01 Up
0.436164942560725 0.508849823110121
0.833381568251521
GO_POSITIVE_REGULATION_OF_SYNAPTIC_TRANSMISSION_GLUTAMATERGIC 16
0.01 Down 0.302252630951732 0.50906377699531
0.833381568251521
GO_MESONEPHRIC_TUBULE_MORPHOGENESIS 46 0.01 Down
0.651996718119361 0.509363803184246 0.833381568251521
GO_ADA2_GCN5_ADA3_TRANSCRIPTION_ACTIVATOR_COMPLEX 15 0.01
Up 0.126267991533685 0.509464858450788
0.833381568251521
GO_REGULATION_OF_LIPID_LOCALIZATION 86 0.01 Down
0.483046912235725 0.509468231278972 0.833381568251521
GO_GANGLION_DEVELOPMENT 10 0.01 Down 0.662197328925474
0.509908013869792 0.833381568251521
GO_POSITIVE_REGULATION_OF_PHAGOCYTOSIS 51 0.01 Up
0.582480390484092 0.509913187549635 0.833381568251521
GO_MEIOTIC_CELL_CYCLE_PROCESS 115 0.01 Up
0.256266385603236 0.510394778119851 0.833381568251521
GO_PHOTOTRANSDUCTION 22 0.01 Down 0.164470015203785
0.51046771003437 0.833381568251521
GO_TELOMERE_MAINTENANCE_IN_RESPONSE_TO_DNA_DAMAGE 13 0.01
Down 0.153500192838042 0.510684791072945
0.833381568251521
GO_G_PROTEIN_COUPLED_CHEMOATTRACTANT_RECEPTOR_ACTIVITY 10 0.01
Up 0.729156152198493 0.510798620984729
0.833381568251521
GO_REGULATION_OF_ANION_TRANSMEMBRANE_TRANSPORT 20 0.01 Down
0.398146269808254 0.511198146181049
0.833381568251521
GO_METHYLATED_HISTONE_BINDING 60 0.01 Down
0.19166316713088 0.511560373282782 0.833381568251521
GO_EPITHELIAL_TO_MESENCHYMAL_TRANSITION 107 0.01 Down
0.500097395037597 0.511602744710222 0.833381568251521
GO_POSITIVE_REGULATION_OF_LEUKOCYTE_DIFFERENTIATION 109 0.01
Down 0.412301551652563 0.511615801334373
0.833381568251521
GO_REGULATION_OF_BONE_REMODELING 31 0.01 Up
0.684634197375566 0.511681338418799 0.833381568251521
GO_REGULATION_OF_DENDRITIC_CELL_DIFFERENTIATION 11 0.01 Up
0.635562879319016 0.511769644144172
0.833381568251521
GO_SWI_SNF_COMPLEX 17 0.01 Down 0.155376157374093
0.511874227370043 0.833381568251521
GO_RESPONSE_TO_UV_C 15 0.01 Down 0.160802655777884
0.511875848428766 0.833381568251521
GO_FIBROBLAST_MIGRATION 39 0.01 Up 0.437570191221083
0.512373933159118 0.833381568251521
GO_MAMMARY_GLAND_DUCT_MORPHOGENESIS 28 0.01 Up
0.437542844073752 0.512392007729396 0.833381568251521
GO_DENDRITIC_TREE 393 0.01 Down 0.303414039668657
0.512393315998334 0.833381568251521

GO_FEMALE_SEX_DIFFERENTIATION 77 0.01 Down
0.366912020148912 0.512401971758986 0.833381568251521
GO_REGULATION_OF_VASCULAR_SMOOTH_MUSCLE_CELL_PROLIFERATION 43
0.01 Down 0.573035965398522 0.512531181412082
0.833381568251521
GO_NEUROMUSCULAR_PROCESS_CONTROLLING_POSTURE 10 0.01 Up
0.282049666023688 0.512670086161734
0.833381568251521
GO_REGULATION_OF_SKELETAL_MUSCLE_CELL_DIFFERENTIATION 15 0.01
Up 0.365429256566193 0.512742113912723
0.833381568251521
GO_RESPONSE_TO_STEROID_HORMONE 280 0.01 Down
0.352624747477102 0.513053597875356 0.833381568251521
GO_FATTY_ACID_DERIVATIVE_BINDING 23 0.01 Up
0.468544291466238 0.513083047115743 0.833381568251521
GO_NEGATIVE_REGULATION_OF_CHONDROCYTE_DIFFERENTIATION 16 0.01
Up 0.960413453109309 0.513130358195157
0.833381568251521
GO_CLATHRIN_COATED_PIT 57 0.01 Down 0.21755463230824
0.513258628379795 0.833381568251521
GO_ENDOSOME 755 0.01 Down 0.28988241729262
0.513344912666626 0.833381568251521
GO_NEUTRAL_AMINO_ACID_TRANSPORT 23 0.01 Up
0.430758319064254 0.513531424224034 0.833381568251521
GO_POSITIVE_REGULATION_OF_VASCULAR_ASSOCIATED_SMOOTH_MUSCLE_CELL_MIG
RATION 14 0.01 Down 0.439824728728675
0.513557772602151 0.833381568251521
GO_RELEASE_OF_SEQUESTERED_CALCIUM_ION_INTO_CYTOSOL_BY_ENDOPLASMIC_RE
TICULUM 20 0.01 Up 0.346915324746502
0.513639727477997 0.833381568251521
GO_CEREBELLAR_CORTEX_MORPHOGENESIS 20 0.01 Down
0.30784910757669 0.51366323833872 0.833381568251521
GO_PROTEIN_KINASE_C_ACTIVITY 13 0.01 Up
0.239606471818545 0.51372921424657 0.833381568251521
GO_MESENCHYMAL_TO_EPITHELIAL_TRANSITION 13 0.01 Down
1.05160065402839 0.513742182913455 0.833381568251521
GO_NEGATIVE_REGULATION_OF_LIPID_BIOSYNTHETIC_PROCESS 37 0.01
Up 0.46527816770854 0.513863493272931
0.833381568251521
GO_PHOSPHOLIPID_TRANSPORTER_ACTIVITY 52 0.01 Down
0.290571549412645 0.513981985502431 0.833381568251521
GO_CELL_COMMUNICATION_INVOLVED_IN_CARDIAC_CONDUCTION 35 0.01
Up 0.394711153979478 0.514312621959798
0.833381568251521
GO_ACTIVATION_OF_GTPASE_ACTIVITY 78 0.01 Down
0.21213541096046 0.514337170485412 0.833381568251521
GO_POSITIVE_REGULATION_OF_TRANSCRIPTION_FROM_RNA_POLYMERASE_II_PROMO
TER_IN_RESPONSE_TO_ENDOPLASMIC_RETICULUM_STRESS 10 0.01 Down
0.227689615978276 0.51461700845162
0.833381568251521
GO_MESENCHYME_MORPHOGENESIS 29 0.01 Down
0.511066855817375 0.514808046518276 0.833381568251521
GO_ENDOCYTOSIS 409 0.01 Up 0.391589396172867
0.514811011889308 0.833381568251521

GO_IRON_ION_TRANSMEMBRANE_TRANSPORT	14	0.01	Down
0.407061631891607	0.514947377588445	0.833381568251521
GO_SOMATIC_RECOMBINATION_OF_IMMUNOGLOBULIN_GENE_SEGMENTS	41
	0.01	Down	0.273659248385464	0.515003004991491
	0.833381568251521
GO_TRANSFORMING_GROWTH_FACTOR_BETA_RECEPTOR_CYTOPLASMIC_MEDIATOR_ACTIVITY	10	0.01	Down	0.308386820782057
0.51504700232069	0.833381568251521
GO_MORPHOGENESIS_OF_AN_EPITHELIAL_BUD	13	0.01	Up
0.643591074107611	0.515155965824219	0.833381568251521
GO_REGULATION_OF_DOUBLE_STRAND_BREAK_REPAIR	66	0.01	Down
	0.208894373563845	0.515259741288478
0.833381568251521
GO_EXONUCLEASE_ACTIVITY_ACTIVE_WITH_EITHER_RIBO_OR_DEOXYRIBONUCLEIC_ACIDS_AND_PRODUCING_5_PHOSPHOMONOESTERS	51	0.01	Down
0.171530293040614	0.515492671726582	0.833381568251521
GO_TISSUE_REMODELING	111	0.01	Up	0.515798532578878
	0.515522866083821	0.833381568251521
GO_BRUSH_BORDER_MEMBRANE	29	0.01	Up
0.558185252019773	0.515637034944279	0.833381568251521
GO_NEGATIVE_REGULATION_OF_HISTONE_ACETYLATION	14	0.01	Down
	0.268018953162525	0.516277577498321
0.833381568251521
GO_MEGAKARYOCYTE_DEVELOPMENT	13	0.01	Down
0.446726769070673	0.516409908075809	0.833381568251521
GO_NEGATIVE_REGULATION_OF_NEUROLOGICAL_SYSTEM_PROCESS	11	0.01
	Up	0.229190050788963	0.516438392153433
0.833381568251521
GO_IMPORT_INTO_CELL	492	0.01	Up	0.404461740273398
	0.516568815520324	0.833381568251521
GO_POSITIVE_REGULATION_OF_EXTRINSIC_APOPTOTIC_SIGNALING_PATHWAY_IN_ABSENCE_OF_LIGAND	11	0.01	Down	0.32873493195739
	0.51685591276985	0.833381568251521
GO_NEUROMUSCULAR_JUNCTION	54	0.01	Down
0.380795862040362	0.516911087662454	0.833381568251521
GO_PHOSPHORYLATION_OF_RNA_POLYMERASE_II_C_TERMINAL_DOMAIN	10
	0.01	Down	0.160974326395721	0.516917111249653
	0.833381568251521
GO_RIBONUCLEOPROTEIN_COMPLEX	808	0.01	Down
0.152917729379687	0.517303453326683	0.833381568251521
GO_METALLOENDOPEPTIDASE_ACTIVITY	73	0.01	Up
0.665871695693866	0.517481848516545	0.833381568251521
GO_DAMAGED_DNA_BINDING	63	0.01	Up	0.159854129559784
	0.517614348969768	0.833381568251521
GO_POSITIVE_REGULATION_OF_ERK1_AND_ERK2_CASCADE	122	0.01	Up
	0.540168875044836	0.517648472087086
0.833381568251521
GO_RESPONSE_TO_ELECTRICAL_STIMULUS	24	0.01	Down
0.382000667330811	0.5177120580229	0.833381568251521
GO_MITOCHONDRIAL_CALCIUM_ION_TRANSMEMBRANE_TRANSPORT	23	0.01
	Down	0.105010475420285	0.517797836408829
0.833381568251521
GO_NEGATIVE_REGULATION_OF_GTPASE_ACTIVITY	40	0.01	Down
	0.196641342074105	0.517871177296246

0.833381568251521
GO_STEROID_DEHYDROGENASE_ACTIVITY_ACTING_ON_THE_CH_OH_GROUP_OF_DONORS_NAD_OR_NADP_AS_ACCEPTOR	20	0.01	Up
0.708450082366975	0.518250699742139	0.833381568251521
GO_RESPONSE_TO_NITROSATIVE_STRESS	10	0.01	Up
0.191245480464553	0.518342488144774	0.833381568251521
GO_SAGA_COMPLEX 13	0.01	Up	0.13718129902567
0.518559292390759	0.833381568251521
GO_TRANSMEMBRANE_RECEPTOR_PROTEIN_KINASE_ACTIVITY	54	0.01
Up	0.387756077110809	0.51858356216237
0.833381568251521
GO_MACROPHAGE_DIFFERENTIATION	30	0.01	Up
0.605940788587875	0.518586221502499	0.833381568251521
GO_FICOLIN_1_RICH_GRANULE	117	0.01	Down
0.291138784204735	0.518653414226534	0.833381568251521
GO_PROTEIN_DESTABILIZATION	40	0.01	Down
0.225970317226229	0.518836346917797	0.833381568251521
GO_SODIUM_ION_TRANSMEMBRANE_TRANSPORTER_ACTIVITY	69	0.01
Down	0.563094199201576	0.518875002942598
0.833381568251521
GO_NEGATIVE_REGULATION_OF_FAT_CELL_DIFFERENTIATION	35	0.01
Down	0.347859739692352	0.518888472829001
0.833381568251521
GO_ANCHORED_COMPONENT_OF_PLASMA_MEMBRANE	33	0.01	Up
0.699394149019619	0.519098446387386
0.833381568251521
GO_PROTEIN_FOLDING_CHAPERONE	19	0.01	Down
0.250143968241236	0.519134757714279	0.833381568251521
GO_NEURAL_TUBE_DEVELOPMENT	135	0.01	Up
0.339807501516989	0.519155276887871	0.833381568251521
GO_REGULATION_OF_ADHERENS_JUNCTION_ORGANIZATION 59	0.01	Down
0.402486179753198	0.519164732519725
0.833381568251521
GO_EARLY_PHAGOSOME	11	0.01	Down	0.234625396356988
0.519973121965	0.834458584046925
GO_MITOCHONDRIAL_MEMBRANE_ORGANIZATION	121	0.01	Up
0.179917975234601	0.520182183447984	0.834573476668264
GO_CARDIAC_MYOFIBRIL_ASSEMBLY	14	0.01	Up
0.54993958467706	0.520437820197986	0.834763013585727
GO_NEGATIVE_REGULATION_OF_BLOOD_PRESSURE	26	0.01	Down
0.516428980983612	0.52067868765638
0.834928767237686
GO_NEGATIVE_REGULATION_OF_PEPTIDYL_SERINE_PHOSPHORYLATION	23
0.01	Down	0.236953871122637	0.520960984917694
0.835160850128154
GO_ANOIKIS	32	0.01	Down	0.25593147887256
0.521379047265907	0.83541881197328
GO_NEPHRON_MORPHOGENESIS	56	0.01	Down
0.659795214826472	0.521514739525011	0.83541881197328
GO_RESPONSE_TO_LITHIUM_ION	17	0.01	Up
0.369520953415962	0.521628934874356	0.83541881197328
GO_NEGATIVE_REGULATION_OF_STRIATED_MUSCLE_CELL_APOPTOTIC_PROCESS
20	0.01	Down	0.447187260718368
0.521719086847158	0.83541881197328

GO_REGULATION_OF_TYPE_2_IMMUNE_RESPONSE 22 0.01 Down
0.632143263728336 0.521809938231375 0.83541881197328
GO_MRNA_CIS_SPLICING_VIA_SPLICEOSOME 31 0.01 Down
0.0825317683418823 0.522129593205525 0.83568140120462
GO_REGULATION_OF_CELL_CYCLE_CHECKPOINT 31 0.01 Up
0.191406935641781 0.522249256493218 0.83568140120462
GO_POSITIVE_REGULATION_OF_FIBROBLAST_PROLIFERATION 42 0.01
Up 0.435781977720628 0.522577830726188
0.835986827493725
GO_NEGATIVE_REGULATION_OF_CYTOSKELETON_ORGANIZATION 107 0.01
Down 0.261550877463831 0.522951944700196
0.836259052381309
GO_INTRACELLULAR_LIPID_TRANSPORT 36 0.01 Down
0.258760256082361 0.523023492322818 0.836259052381309
GO_ARACHIDONIC_ACID_SECRETION 16 0.01 Up
0.361276188519679 0.523688949138443 0.836843322287327
GO_PHOSPHOLIPID_CATABOLIC_PROCESS 28 0.01 Up
0.424066258152328 0.523879760263157 0.836843322287327
GO_ORGANONITROGEN_COMPOUND_BIOSYNTHETIC_PROCESS 1478 0.01 Up
0.25713795044619 0.523933336755491
0.836843322287327
GO_MAMMARY_GLAND_EPITHELIAL_CELL_PROLIFERATION 23 0.01 Down
0.235819307908895 0.523940284634184
0.836843322287327
GO_NEGATIVE_REGULATION_OF_TYROSINE_PHOSPHORYLATION_OF_STAT_PROTEIN
10 0.01 Down 0.212890481291089
0.524280331284346 0.836961339898874
GO_CARBOXY_LYASE_ACTIVITY 26 0.01 Up
0.198591297237552 0.524289898803396 0.836961339898874
GO_GLIAL_CELL_PROJECTION 16 0.01 Down
0.292081385534495 0.525364532789281 0.838191196395425
GO_GRANULOCYTE_DIFFERENTIATION 23 0.01 Up
0.449446244232276 0.525389432691201 0.838191196395425
GO_NEGATIVE_REGULATION_OF_RESPONSE_TO_DNA_DAMAGE_STIMULUS 74
0.01 Up 0.267557477391015 0.525474500655739
0.838191196395425
GO_MELANOCYTE_DIFFERENTIATION 16 0.01 Up
0.586580019866385 0.525757782817919 0.838235952900721
GO_PEPTIDASE_ACTIVATOR_ACTIVITY_INVOLVED_IN_APOPTOTIC_PROCESS 17
0.01 Up 0.277839357383836 0.525778703450164
0.838235952900721
GO_CHROMOSOME_SEGREGATION 240 0.01 Down
0.175745040230623 0.526130722813848 0.838576954109444
GO_PLATELET_DERIVED_GROWTH_FACTOR_RECEPTOR_SIGNALING_PATHWAY 50
0.01 Down 0.39703598283127 0.526590975550374
0.8386957988062
GO_TRANSCRIPTION_COREPRESSOR_BINDING 14 0.01 Down
0.28464228554141 0.526672675725295 0.8386957988062
GO_POSITIVE_REGULATION_OF_OXIDATIVE_STRESS_INDUCED_CELL_DEATH 13
0.01 Down 0.399684308222127 0.526963855556243
0.8386957988062
GO_WW_DOMAIN_BINDING 24 0.01 Up 0.236240932796888
0.527299577232308 0.8386957988062
GO_NEGATIVE_REGULATION_OF_DNA_BINDING_TRANSCRIPTION_FACTOR_ACTIVITY

125 0.01 Down 0.373502945157351
0.527323752582738 0.8386957988062
GO_UV_DAMAGE_EXCISION_REPAIR 12 0.01 Up
0.15590173471137 0.527335652765631 0.8386957988062
GO_REGULATION_OF_INTRINSIC_APOPTOTIC_SIGNALING_PATHWAY 146 0.01
Down 0.349116787902129 0.527346480301653
0.8386957988062
GO_NEURAL_TUBE_FORMATION 95 0.01 Up
0.366081979104401 0.527379178792248 0.8386957988062
GO_CELLULAR_MACROMOLECULE_LOCALIZATION 1656 0.01 Down
0.225938852529005 0.527495874586006 0.8386957988062
GO_TELOMERIC_LOOP_DISASSEMBLY 10 0.01 Down
0.184404876474161 0.527586765877265 0.8386957988062
GO_CARDIAC_MUSCLE_CONTRACTION 84 0.01 Up
0.501738811937901 0.527947869932669 0.839050135696657
GO_INTEGRAL_COMPONENT_OF_PRESYNAPTIC_ACTIVE_ZONE_MEMBRANE 10
0.01 Up 0.289217215426281 0.528128600281516
0.839117700159405
GO_POSITIVE_REGULATION_OF_PEPTIDE_SECRETION 182 0.01 Down
0.47707317952929 0.528624878385259
0.839686456482707
GO_CYTOPLASMIC_SEQUESTERING_OF_TRANSCRIPTION_FACTOR 16 0.01
Up 0.222666462694332 0.529120045211539
0.84016054789196
GO_CELLULAR_RESPONSE_TO_DSRNA 21 0.01 Down
0.403574015231685 0.529200121090241 0.84016054789196
GO_PROTEIN_AUTOPROCESSING 13 0.01 Up
0.410078558478077 0.529574888142926 0.840524231081894
GO_PROTON_TRANSPORTING_TWO_SECTOR_ATPASE_COMPLEX 43 0.01
Up 0.149623576279636 0.529839046973062
0.840524231081894
GO_REGULATION_OF_TYPE_I_INTERFERON_MEDIATED_SIGNALING_PATHWAY 29
0.01 Down 0.204689028941795 0.529908184029245
0.840524231081894
GO_PROTEIN_TRANSPORTER_ACTIVITY 90 0.01 Down
0.141803430530106 0.529982994001234 0.840524231081894
GO_PROTEIN_LIPID_COMPLEX_SUBUNIT_ORGANIZATION 25 0.01 Up
0.404318535602152 0.530231558683784
0.840698822870006
GO_NEGATIVE_REGULATION_OF_LIPID_TRANSPORT 19 0.01 Up
0.393106254377177 0.530779295726937
0.841347546829826
GO_CELLULAR_GLUCOSE_HOMEOSTASIS 109 0.01 Up
0.293268360123497 0.531067229654849 0.841584221152334
GO_TRNA_5_LEADER_REMOVAL 12 0.01 Up
0.115348460574062 0.531414912009146 0.841731062212918
GO_NEGATIVE_REGULATION_OF_ORGANIC_ACID_TRANSPORT 11 0.01
Up 0.437566362835288 0.531437186865774
0.841731062212918
GO_T_CELL_RECEPTOR_COMPLEX 10 0.01 Down
0.80796456292933 0.531904906964416 0.841945562205803
GO_HYDROLASE_ACTIVITY_ACTING_ON_CARBON_NITROGEN_BUT_NOT_PEPTIDE_BOND
S 97 0.01 Down 0.32018117293251
0.531989340949958 0.841945562205803

GO_NEGATIVE_REGULATION_OF_NEURON_PROJECTION_DEVELOPMENT 120 0.01
Down 0.395092769890994 0.53222445362041
0.841945562205803
GO_CHEMOKINE_C_X_C_MOTIF_LIGAND_2_PRODUCTION 12 0.01 Down
0.472291226900006 0.532401234149803
0.841945562205803
GO_MULTI_ORGANISM_MEMBRANE_ORGANIZATION 10 0.01 Up
0.346154136247813 0.532447234658596 0.841945562205803
GO_REGULATION_OF_SODIUM_ION_TRANSPORT 63 0.01 Down
0.495565194521146 0.532478672237999 0.841945562205803
GO_DENDRITE_EXTENSION 19 0.01 Down 0.23093056700398
0.532650159732549 0.841945562205803
GO_NEGATIVE_REGULATION_OF_PROTEIN_TYROSINE_KINASE_ACTIVITY 29
0.01 Down 0.340316885299692 0.532682079465078
0.841945562205803
GO_ANDROGEN_METABOLIC_PROCESS 10 0.01 Up
0.625499816585344 0.532845732988117 0.841985019513497
GO_MUSCLE_TISSUE_DEVELOPMENT 256 0.01 Down
0.440873679277523 0.533131659212856 0.84202899806689
GO_MITOTIC_CELL_CYCLE 856 0.01 Down 0.218052797040686
0.533150958420215 0.84202899806689
GO_DIACYLGLYCEROL_METABOLIC_PROCESS 11 0.01 Up
0.185449885672668 0.533473609828165 0.842309702736367
GO_BRUSH_BORDER 62 0.01 Down 0.400271669159876
0.533704309302533 0.842309702736367
GO_CALCIUM_IMPORT_INTO_THE_MITOCHONDRION 11 0.01 Down
0.0898081490278933 0.53374492281779
0.842309702736367
GO_EYE_MORPHOGENESIS 98 0.01 Down 0.451560718439729
0.53401821614941 0.842507645457467
GO_POSITIVE_REGULATION_OF_TRANSCRIPTION_OF_NUCLEOLAR_LARGE_RRNA_BY_R
NA_POLYMERASE_I 10 0.01 Down 0.126476538471167
0.534147904359379 0.842507645457467
GO_IRE1_MEDIATED_UNFOLDED_PROTEIN_RESPONSE 62 0.01 Down
0.209825509907716 0.534831984972179
0.842956760850296
GO_ENDOPEPTIDASE_ACTIVITY 273 0.01 Down
0.442490378312046 0.53486662575562 0.842956760850296
GO_ARTERY_MORPHOGENESIS 50 0.01 Down 0.703299450477711
0.534957537419835 0.842956760850296
GO_SENSORY_PERCEPTION_OF_SMELL 14 0.01 Up
0.38087486069525 0.535278550662271 0.842956760850296
GO_POSITIVE_REGULATION_OF_HORMONE_METABOLIC_PROCESS 12 0.01
Down 0.454015942397524 0.535337344931414
0.842956760850296
GO_STAGA_COMPLEX 12 0.01 Up 0.126781708497504
0.535457904625415 0.842956760850296
GO_NEGATIVE_REGULATION_OF_MOLECULAR_FUNCTION 817 0.01 Down
0.357382733048029 0.535642696920156
0.842956760850296
GO_OSTEOCLAST_DIFFERENTIATION 71 0.01 Up
0.453276985138238 0.535727397176947 0.842956760850296
GO_GENERAL_TRANSCRIPTION_INITIATION_FACTOR_ACTIVITY 20 0.01
Down 0.148160314725109 0.535760327519862

0.842956760850296
GO_TRANSCRIPTION_INITIATION_FROM_RNA_POLYMERASE_I_PROMOTER 35 0.01 Up 0.160356659550743 0.535821139865144 0.842956760850296
GO_NEGATIVE_REGULATION_OF_INTRACELLULAR_TRANSPORT 49 0.01 Down 0.339678920536873 0.536244878403782 0.84340483336512
GO_RESPONSE_TO_MORPHINE 19 0.01 Up 0.455492535677681 0.53642310284847 0.843410444459197
GO_NEGATIVE_REGULATION_OF_BLOOD_VESSEL_ENDOTHELIAL_CELL_MIGRATION 27 0.01 Up 0.514628805511973 0.53665512467199 0.843410444459197
GO_EPITHELIAL_CELL_DIFFERENTIATION_INVOLVED_IN_PROSTATE_GLAND_DEVELOPMENT 10 0.01 Up 0.36165285704258 0.536762986266605 0.843410444459197
GO_O_METHYLTRANSFERASE_ACTIVITY 23 0.01 Up 0.125539602167231 0.536804143862681 0.843410444459197
GO_ACID_SECRETION 64 0.01 Up 0.448671412390262 0.537077771346243 0.843622031007255
GO_HEAD_MORPHOGENESIS 33 0.01 Down 0.334476713365514 0.537540326332355 0.844130191713328
GO_CELL_COMMUNICATION_BY_ELECTRICAL_COUPLING_INVOLVED_IN_CARDIAC_CONDUCTION 16 0.01 Up 0.426188423409755 0.537772723322975 0.844276752855904
GO_MAINTENANCE_OF_PROTEIN_LOCATION_IN_NUCLEUS 21 0.01 Down 0.213273543578437 0.538186057298 0.844707226953505
GO_SMOOTH_MUSCLE_CELL_APOPTOTIC_PROCESS 19 0.01 Down 0.342995280724255 0.538446055304947 0.844708233709037
GO_COLLATERAL_SPROUTING 23 0.01 Up 0.454685936359627 0.538639209978818 0.844708233709037
GO_CARDIOBLAST_DIFFERENTIATION 11 0.01 Down 0.78661643944038 0.538773496432801 0.844708233709037
GO_ESCRT_I_COMPLEX 10 0.01 Down 0.238298863620868 0.538856272354762 0.844708233709037
GO_INTRINSIC_COMPONENT_OF_POSTSYNAPTIC_SPECIALIZATION_MEMBRANE 21 0.01 Up 0.352084038414111 0.538882389911892 0.844708233709037
GO_POSITIVE_REGULATION_OF_VIRAL_TRANSCRIPTION 39 0.01 Down 0.118583708149133 0.53947810385858 0.8452147093731
GO_SOMATIC_DIVERSIFICATION_OF_IMMUNE_RECEPTORS 53 0.01 Down 0.249012475295501 0.539483939848585 0.8452147093731
GO_HISTONE_H3_K9_METHYLATION 32 0.01 Down 0.192472992513414 0.540000139289567 0.845805171730382
GO_RIBOSOME_BIOGENESIS 273 0.01 Up 0.122544518489396 0.540318825342414 0.846010609363291
GO_SERINE_HYDROLASE_ACTIVITY 102 0.01 Down 0.592446748321865 0.540410005453936 0.846010609363291
GO_SCHAFFER_COLLATERAL__CA1_SYNAPSE 50 0.01 Down 0.338605660848531 0.540625868037884 0.846043745195967
GO_POSITIVE_REGULATION_OF_SISTER_CHROMATID_COHESION 10 0.01 Down 0.119388103716914 0.540709888216167 0.846043745195967

GO_UV_PROTECTION	12	0.01	Up	0.161535839720147
	0.540922161541721	0.846111209721142
GO_NUCLEOSIDE_DIPHOSPHATE_METABOLIC_PROCESS	22	0.01	Down
	0.335281888871131	0.5410317437222 0.846111209721142
GO_POSITIVE_REGULATION_OF_AUTOPHAGOSOME_ASSEMBLY	11	0.01
	Down	0.146397840159783	0.541446800549902
0.84628126222692
GO_SOMITOGENESIS	44	0.01	Down	0.405628345528358
	0.541457959696363	0.84628126222692
GO_AXON_CYTOPLASM	50	0.01	Down	0.239199508799448
	0.541927424958802	0.84628126222692
GO_EATING_BEHAVIOR	15	0.01	Up	0.24322267723466
	0.542316770597035	0.84628126222692
GO_IONOTROPIC_GLUTAMATE_RECEPTOR_BINDING	22	0.01	Down
	0.222369708102368	0.542506198461996
0.84628126222692
GO_HEXOSE_CATABOLIC_PROCESS	29	0.01	Up
0.261541981857702	0.542533808881378	0.84628126222692
GO_PROGRAMMED_CELL_DEATH_IN_RESPONSE_TO_REACTIVE_OXYGEN_SPECIES 11
	0.01	Down	0.171341060823056	0.542722848642845
	0.84628126222692
GO_REGULATION_OF_VASCULAR_SMOOTH_MUSCLE_CELL_DIFFERENTIATION	14
	0.01	Up	0.596135331132857	0.542746664839504
	0.84628126222692
GO_RESPONSE_TO_INORGANIC_SUBSTANCE	405	0.01	Up
0.418341689222776	0.542777280946993	0.84628126222692
GO_PARTURITION	13	0.01	Up	0.598197516526532
0.543148893352827	0.84628126222692
GO_RESPONSE_TO_DIETARY_EXCESS	13	0.01	Down
0.412438955340232	0.543318889815188	0.84628126222692
GO_REGULATION_OF_ENDOTHELIAL_CELL_DEVELOPMENT	11	0.01	Down
	0.430722913235956	0.543375426814554
0.84628126222692
GO_CRANIAL_NERVE_DEVELOPMENT	27	0.01	Down
0.540276606225724	0.543424264034515	0.84628126222692
GO_SECRETORY_VESICLE	716	0.01	Up	0.406047787942465
	0.54350962524603	0.84628126222692
GO_NEGATIVE_REGULATION_OF_IMMUNE_SYSTEM_PROCESS 316	0.01	Down
	0.495976653679677	0.543603626631608
0.84628126222692
GO_ENDOCYTIC_RECYCLING	43	0.01	Down	0.232408316914316
	0.54365837600725	0.84628126222692
GO_EXONUCLEASE_ACTIVITY 71	0.01	Down	0.178426405489297
	0.543769287375091	0.84628126222692
GO_NEGATIVE_REGULATION_OF_CELL_MATRIX_ADHESION	30	0.01	Down
	0.373896698313323	0.543942292077698
0.84628126222692
GO_OSTEOCLAST_DEVELOPMENT	14	0.01	Up
0.524150283780092	0.544165363967344	0.84628126222692
GO_REGULATION_OF_CELLULAR_RESPONSE_TO_INSULIN_STIMULUS	49	0.01
	Down	0.209072403139631	0.544295050276443
0.84628126222692
GO_RESPONSE_TO_STIMULUS_INVOLVED_IN_REGULATION_OF_MUSCLE_ADAPTATION
	10	0.01	Up	0.436547860460448

0.54434114355804	0.84628126222692
GO_SMOOTH_MUSCLE_TISSUE_DEVELOPMENT	17	0.01	Up
0.803935470261243	0.544345598876061	0.84628126222692
GO_METALLOAMINOPEPTIDASE_ACTIVITY	17	0.01	Down
0.283544565991334	0.544355028715886	0.84628126222692
GO_NEGATIVE_REGULATION_OF_JUN_KINASE_ACTIVITY	12	0.01	Up
0.789493890705043	0.544486017173176
0.84628126222692
GO_RESPONSE_TO_OXYGEN_CONTAINING_COMPOUND	1138	0.01	Down
0.395698824010163	0.544896534539939
0.846374519503552
GO_POSITIVE_REGULATION_OF_LEUKOCYTE_DEGRANULATION	19	0.01
Down	0.442843347439136	0.54498434569315
0.846374519503552
GO_NEGATIVE_REGULATION_OF_ORGANELLE_ORGANIZATION	303	0.01
Down	0.230187846692296	0.545035069468738
0.846374519503552
GO_B_CELL_PROLIFERATION	47	0.01	Down	0.473278127968799
0.545122002870093	0.846374519503552
GO_PROTEIN_LOCALIZATION_TO_CELL_JUNCTION	12	0.01	Down
0.10935940972032	0.545345549168998
0.846374519503552
GO_REGULATION_OF_CALCIUM_ION_TRANSPORT_INTO_CYTOSOL	64	0.01
Up	0.601552192417862	0.545382493872162
0.846374519503552
GO_HETEROCHROMATIN_ORGANIZATION	17	0.01	Up
0.420367255099083	0.547532472171257	0.849160571474753
GO_PYRIMIDINE_CONTAINING_COMPOUND_BIOSYNTHETIC_PROCESS	50	0.01
Up	0.241153284855305	0.547645290157046
0.849160571474753
GO_NEGATIVE_REGULATION_OF_PROTEIN_KINASE_B_SIGNALING	40	0.01
Down	0.260847545452577	0.547656007087644
0.849160571474753
GO_POSITIVE_REGULATION_OF_CARDIOCYTE_DIFFERENTIATION	23	0.01
Up	0.659710284666609	0.547938988870206
0.849160571474753
GO_DETECTION_OF_STIMULUS	117	0.01	Up
0.35254250047665	0.548050438751047	0.849160571474753
GO_REGULATION_OF_HORMONE_BIOSYNTHETIC_PROCESS	22	0.01	Down
0.544554345810928	0.548132176824101
0.849160571474753
GO_CELL_CYCLE_DNA_REPLICATION	60	0.01	Up
0.179532502110652	0.548156857125606	0.849160571474753
GO_TISSUE_REGENERATION	44	0.01	Up	0.442209628094819
0.548687946985638	0.849579270396708
GO_90S_PRERIBOSOME	29	0.01	Up	0.128567676394402
0.548724387675502	0.849579270396708
GO_TRANSMEMBRANE_RECEPTOR_PROTEIN_TYROSINE_KINASE_SIGNALING_PATHWAY
553	0.01	Down	0.389061783443027
0.548846960714197	0.849579270396708
GO_NEGATIVE_REGULATION_OF_PEPTIDE_SECRETION	87	0.01	Down
0.538480695671452	0.54906943962888
0.849707001781017
GO_NEGATIVE_REGULATION_OF_CELL_DIVISION	13	0.01	Down

0.392981368039633	0.549425775413534	0.849950150334669
GO_NEUTRAL_LIPID_CATABOLIC_PROCESS	18	0.01	Up
0.3705354234375 0.549623344585272	0.849950150334669
GO_NEGATIVE_REGULATION_OF_LYMPHOCYTE_APOPTOTIC_PROCESS	24	0.01
Down	0.50007579256829	0.549646564027988
0.849950150334669
GO_REGULATION_OF_SODIUM_ION_TRANSMEMBRANE_TRANSPORTER_ACTIVITY	37
0.01	Down	0.362267740829171	0.550197087998404
0.850386973271639
GO_REGULATION_OF_ANOIKIS	22	0.01	Down
0.303461718209014	0.550209196345083	0.850386973271639
GO_SARCOPLASMIC_RETICULUM_MEMBRANE	22	0.01	Up
0.273902832693534	0.55041813271996	0.850493378402361
GO_TELOMERE_MAINTENANCE_VIA_SEMI_CONSERVATIVE_REPLICATION	27
0.01	Up	0.115326809010273	0.55067259086804
0.850546773471905
GO_COMMISSURAL_NEURON_AXON_GUIDANCE	10	0.01	Up
0.313982497098924	0.550899109050542	0.850546773471905
GO_NERVE_DEVELOPMENT	47	0.01	Down	0.545008593014282
0.551001542417206	0.850546773471905
GO_NEGATIVE_REGULATION_OF_CELL_CYCLE_PROCESS	296	0.01	Down
0.215144155139533	0.55101308846401
0.850546773471905
GO_EXODEOXYRIBONUCLEASE_ACTIVITY	22	0.01	Up
0.173648968268748	0.551276152489765	0.850736533239797
GO_RAC_PROTEIN_SIGNAL_TRANSDUCTION	31	0.01	Down
0.228127475006864	0.551788076928283	0.851310143591259
GO_VESICLE_MEDIATED_TRANSPORT_IN_SYNAPSE	140	0.01	Down
0.260493683352017	0.552046571836058
0.851324494945215
GO_MORPHOGENESIS_OF_A_BRANCHING_STRUCTURE	149	0.01	Down
0.53144286668889	0.552077835051772
0.851324494945215
GO_KINESIN_COMPLEX	39	0.01	Down	0.23847908400211
0.552710080379795	0.852060679394361
GO_BUNDLE_OF_HIS_CELL_TO_PURKINJE_MYOCYTE_COMMUNICATION	10	0.01
Down	0.329657795343914	0.552835944018183
0.852060679394361
GO_NEGATIVE_REGULATION_OF_PEPTIDYL_LYSINE_ACETYLATION	16	0.01
Down	0.246601369401559	0.553008542260134
0.852110370573928
GO_H4_HISTONE_ACETYLTRANSFERASE_ACTIVITY	21	0.01	Down
0.0956052423522918	0.55317000200195
0.852142877988795
GO_CLATHRIN_DEPENDENT_ENDOCYTOSIS	36	0.01	Down
0.292015396756373	0.553794348351961	0.852733799244558
GO_TRANSCRIPTION_ELONGATION_FROM_RNA_POLYMERASE_II_PROMOTER	80
0.01	Down	0.141227635548353	0.554045283533411
0.852733799244558
GO_CELL_ACTIVATION_INVOLVED_IN_IMMUNE_RESPONSE	554	0.01	Down
0.415112395808508	0.554202135435672
0.852733799244558
GO_ENTEROENDOCRINE_CELL_DIFFERENTIATION 17	0.01	Up
0.326492178394516	0.55424985860443	0.852733799244558

GO_CHEMOKINE_PRODUCTION 62 0.01 Down 0.608786011384551
0.554413642325131 0.852733799244558
GO_NEURON_DIFFERENTIATION 956 0.01 Down
0.356419239196635 0.554453685874858 0.852733799244558
GO_RECEPTOR_TYROSINE_KINASE_BINDING 56 0.01 Up
0.2913848903371 0.554668185469099 0.852733799244558
GO_MULTIMERIC_RIBONUCLEASE_P_COMPLEX 10 0.01 Up
0.121133019488521 0.554677281043775 0.852733799244558
GO_REGULATION_OF_RUFFLE_ASSEMBLY 23 0.01 Down
0.150146232948702 0.555285211081575 0.85320187480318
GO_TYPE_B_PANCREATIC_CELL_PROLIFERATION 15 0.01 Up
0.628326504218446 0.555435476429444 0.85320187480318
GO_HIGH_DENSITY_LIPOPROTEIN_PARTICLE 11 0.01 Up
0.844879128895225 0.555462110759792 0.85320187480318
GO_CARNITINE_SHUTTLE 10 0.01 Up 0.2777098541337
0.555543899044139 0.85320187480318
GO_CULLIN_FAMILY_PROTEIN_BINDING 17 0.01 Down
0.117032448581692 0.555852356199181 0.853253672341864
GO_NEGATIVE_REGULATION_OF_STEM_CELL_PROLIFERATION 10 0.01
Down 0.603363604089139 0.555925932577443
0.853253672341864
GO_REGULATION_OF_GENERATION_OF_PRECURSOR_METABOLITES_AND_ENERGY 122
0.01 Down 0.242845409000508 0.556387320074174
0.853253672341864
GO_SYNAPTIC_TRANSMISSION_GABAERGIC 20 0.01 Down
0.389515482712786 0.556444753418936 0.853253672341864
GO_MODIFICATION_OF_POSTSYNAPTIC_STRUCTURE 13 0.01 Up
0.181820877403174 0.556461368054466
0.853253672341864
GO_CELL_CELL_RECOGNITION 39 0.01 Down
0.324889039213354 0.556575635821708 0.853253672341864
GO_POSITIVE_REGULATION_OF_ESTABLISHMENT_OF_PROTEIN_LOCALIZATION 345
0.01 Down 0.339269867858998 0.556727050730441
0.853253672341864
GO_NEGATIVE_REGULATION_OF_SMALL_MOLECULE_METABOLIC_PROCESS 54
0.01 Up 0.431230733981236 0.557006189646224
0.853253672341864
GO_ATP_HYDROLYSIS_COUPLED_CATION_TRANSMEMBRANE_TRANSPORT 46
0.01 Up 0.225585852797111 0.557162863743565
0.853253672341864
GO_HSP90_PROTEIN_BINDING 35 0.01 Down
0.230749860007026 0.557182476957633 0.853253672341864
GO_DNA_REPLICATION_INITIATION 29 0.01 Up
0.144041248625981 0.557233794838999 0.853253672341864
GO_DEVELOPMENT_OF_PRIMARY_FEMALE_SEXUAL_CHARACTERISTICS 67 0.01
Down 0.363497179203377 0.557296538024806
0.853253672341864
GO_CELL_AGING 94 0.01 Down 0.280688349540346
0.55770034483095 0.853253672341864
GO_POSITIVE_REGULATION_OF_MUSCLE_TISSUE_DEVELOPMENT 48 0.01
Up 0.51492904288296 0.557901848863559
0.853253672341864
GO_2_OXOGLUTARATE_METABOLIC_PROCESS 16 0.01 Up
0.18712199596857 0.557933272953925 0.853253672341864

GO_POTASSIUM_ION_TRANSMEMBRANE_TRANSPORTER_ACTIVITY 58 0.01 Down 0.621903755084605 0.558013656784102 0.853253672341864
GO_TOLL_LIKE_RECEPTOR_9_SIGNALING_PATHWAY 19 0.01 Down 0.521382390321704 0.558062713020838 0.853253672341864
GO_CARDIAC_CHAMBER_MORPHOGENESIS 88 0.01 Up 0.411027041068462 0.558133102445086 0.853253672341864
GO_MITOTIC_SPINDLE 89 0.01 Down 0.200305756594741 0.558247996465472 0.853253672341864
GO_REGULATION_OF_ENDOTHELIAL_CELL_MIGRATION 132 0.01 Down 0.450184595836327 0.55883639662756 0.853351921498398
GO_PROTEIN_ADP_RIBOSYLATION 28 0.01 Down 0.152472844681436 0.558952968980022 0.853351921498398
GO_POSITIVE_REGULATION_OF_DEPHOSPHORYLATION 47 0.01 Down 0.480527213999813 0.558982394963904 0.853351921498398
GO_DNA_POLYMERASE_COMPLEX 13 0.01 Up 0.184245381558142 0.558998906714792 0.853351921498398
GO_PHOSPHATIDYLINOSITOL_PHOSPHATE_5_PHOSPHATASE_ACTIVITY 11 0.01 Down 0.131703205885562 0.559015086773041 0.853351921498398
GO_NEGATIVE_REGULATION_OF_PRODUCTION_OF_MOLECULAR_MEDIATOR_OF_IMMUNE_RESPONSE 23 0.01 Down 0.485203042586319 0.559178157100072 0.853386272437037
GO_POSITIVE_REGULATION_OF_INSULIN_SECRETION 48 0.01 Up 0.278968094812346 0.55943147780405 0.85339565597597
GO_FACE_MORPHOGENESIS 28 0.01 Down 0.350247474566767 0.559465444042886 0.85339565597597
GO_PEPTIDE_CATABOLIC_PROCESS 21 0.01 Up 0.482933558287678 0.559651975269025 0.853465747766453
GO_POSITIVE_REGULATION_OF_INTRACELLULAR_STEROID_HORMONE_RECEPTOR_SIGNALING_PATHWAY 12 0.01 Down 0.237037926341459 0.559837641164835 0.853534485060702
GO_DIVALENT_INORGANIC_CATION_TRANSPORT 289 0.01 Up 0.427163332910316 0.560248760341915 0.853868966480196
GO_CONDENSED_NUCLEAR_CHROMOSOME 62 0.01 Up 0.185667832835524 0.560338323580481 0.853868966480196
GO_POSITIVE_REGULATION_OF_TRANSCRIPTION_ELONGATION_FROM_RNA_POLYMERASE_II_PROMOTER 17 0.01 Down 0.1174600400306 0.561678010266088 0.855695658801862
GO_PROTEIN_PALMITOYLATION 29 0.01 Up 0.184720191170981 0.561925754834333 0.855858318514611
GO_PROTON_TRANSPORTING_ATP_SYNTHASE_COMPLEX 20 0.01 Up 0.121160341869474 0.562074165054727 0.855869640342926
GO_PROTEIN_K11_LINKED_UBIQUITINATION 27 0.01 Down 0.0884481539735955 0.562540150699999 0.856364406945761
GO_COMPLEMENT_BINDING 11 0.01 Up 0.767837563900298 0.562760484884584 0.85648505984816
GO_MYELIN_SHEATH 40 0.01 Down 0.322687783759283 0.563050827552287 0.856712173952364

GO_ANION_TRANSMEMBRANE_TRANSPORT 170 0.01 Up
0.361772900010358 0.564357835396271 0.858485697492048
GO_VENTRICULAR_CARDIAC_MUSCLE_TISSUE_DEVELOPMENT 34 0.01
Down 0.306346989650879 0.564781808697661
0.85891542099286
GO_MOLECULAR_ADAPTOR_ACTIVITY 176 0.01 Down
0.282971096234985 0.565196070724788 0.859004032751925
GO_REGULATION_OF_LIPOPROTEIN_PARTICLE_CLEARANCE 12 0.01 Up
0.424479787860513 0.565241018597205
0.859004032751925
GO_OLIGOSACCHARIDE_LIPID_INTERMEDIATE_BIOSYNTHETIC_PROCESS 20
0.01 Up 0.171747101789851 0.565264554578149
0.859004032751925
GO_METHYL_CPG_BINDING 16 0.01 Down 0.15027945699412
0.565749212600977 0.859037508485779
GO_NUCLEAR_INNER_MEMBRANE 41 0.01 Up
0.232577578141783 0.565840403996231 0.859037508485779
GO_MELANIN_BIOSYNTHETIC_PROCESS 13 0.01 Up
0.358993542372166 0.566053198637663 0.859037508485779
GO_CELLULAR_COMPONENT_DISASSEMBLY_INVOLVED_IN_EXECUTION_PHASE_OF_APO
PTOSIS 31 0.01 Down 0.256496135883128
0.566512606642339 0.859037508485779
GO_METANEPHROS_DEVELOPMENT 63 0.01 Up
0.615716594552167 0.566542357708791 0.859037508485779
GO_CELLULAR_PIGMENTATION 47 0.01 Up
0.237228607521961 0.566725143229107 0.859037508485779
GO_PROTEIN_HOMOTRIMERIZATION 22 0.01 Up
0.435324876818062 0.566767133529576 0.859037508485779
GO_SECRETORY_GRANULE_MEMBRANE 228 0.01 Up
0.432224165475626 0.566959130409754 0.859037508485779
GO_I_BAND 83 0.01 Up 0.376356041757144
0.566993307403669 0.859037508485779
GO_GOLGI_CISTERNA 92 0.01 Down 0.274644125297294
0.567070547081784 0.859037508485779
GO_NEGATIVE_REGULATION_OF_LYMPHOCYTE_MEDIATED_IMMUNITY 27 0.01
Down 0.522748246078019 0.567164606799522
0.859037508485779
GO_OOCYTE_MATURATION 19 0.01 Down 0.254065610908691
0.567210625837089 0.859037508485779
GO_NEGATIVE_REGULATION_OF_NECROTIC_CELL_DEATH 14 0.01 Down
0.35313881116553 0.567334014987342
0.859037508485779
GO_REGULATION_OF_PROTEIN_GLYCOSYLATION 11 0.01 Down
0.159841112674443 0.567365253910557 0.859037508485779
GO_NEGATIVE_REGULATION_OF_INTRINSIC_APOPTOTIC_SIGNALING_PATHWAY 89
0.01 Down 0.373058251069809 0.567420205084296
0.859037508485779
GO_NEPHRON_EPITHELIUM_DEVELOPMENT 74 0.01 Down
0.6115760366541 0.567555014403969 0.859037508485779
GO_VASCULAR_SMOOTH_MUSCLE_CELL_DIFFERENTIATION 22 0.01 Up
0.487368805822202 0.567798791667978
0.859037508485779
GO_LATE_ENDOSOME_MEMBRANE 119 0.01 Up
0.286453353932905 0.567833556506906 0.859037508485779

GO_AMIDE_TRANSMEMBRANE_TRANSPORTER_ACTIVITY	38	0.01	Up
	0.357983481827322	0.56852423691084
0.859812606282431
GO_NEGATIVE_REGULATION_OF_HISTONE_H3_K9_METHYLATION	10	0.01
	Up	0.0931316934171064	0.568629157342112
0.859812606282431
GO_NCRNA_METABOLIC_PROCESS	424	0.01	Up
0.140987890383041	0.568927951441062	0.860050197509633
GO_TRABECULA_MORPHOGENESIS	37	0.01	Up
0.562274472582416	0.569425952714485	0.860551569556216
GO_CLUSTER_OF_ACTIN_BASED_CELL_PROJECTIONS	92	0.01	Down
	0.388474126821117	0.569543107639084
0.860551569556216
GO_POSTSYNAPSE	379	0.01	Down	0.302117748766777
0.570525867356009	0.861821980770921
GO_NUCLEOSIDE_TRIPHOSPHATE_METABOLIC_PROCESS	45	0.01	Down
	0.164693634704572	0.571207429255399
0.86254254707464
GO_M_BAND	15	0.01	Up	0.238031397843285
0.571287033732025	0.86254254707464
GO_AUTONOMIC_NERVOUS_SYSTEM_DEVELOPMENT	25	0.01	Down
0.425981794276599	0.571631049890166	0.862847365460766
GO_PROTEIN_PHOSPHATASE_1_BINDING	18	0.01	Down
0.400323104567727	0.571944515923345	0.863105929945471
GO_NEGATIVE_REGULATION_OF_HISTONE_METHYLATION	19	0.01	Down
	0.115562605487047	0.57212605683137
0.863165330771185
GO_MORPHOGENESIS_OF_AN_EPITHELIUM	426	0.01	Down
0.405011025249862	0.57228457326854	0.863189973742436
GO_PHOSPHORUS_OXYGEN_LYASE_ACTIVITY	13	0.01	Up
0.495539594304987	0.572563672046246	0.863396436411516
GO_POSITIVE_REGULATION_OF_CHROMOSOME_SEGREGATION	26	0.01
	Down	0.122645255649821	0.573578760124996
0.864712354784914
GO_RESPONSE_TO_ANGIOTENSIN	19	0.01	Down
0.36537524235105	0.573997243961698	0.864954225766842
GO_METENCEPHALON_DEVELOPMENT	74	0.01	Down
0.320952568110339	0.574126363106573	0.864954225766842
GO_PROTEIN_N_LINKED_GLYCOSYLATION	70	0.01	Down
0.242022015116846	0.574166616676062	0.864954225766842
GO_NEGATIVE_REGULATION_OF_T_CELL_MEDIATED_IMMUNITY	12	0.01
	Up	0.41120897589098	0.574623788486218
0.865389604343774
GO_ACUTE_PHASE_RESPONSE	27	0.01	Down	0.649935806826749
	0.574790181712414	0.865389604343774
GO_FILOPODIUM	82	0.01	Down	0.335284602883131
0.574883260470835	0.865389604343774
GO_GTPASE_ACTIVITY	250	0.01	Down	0.283292432080692
	0.575085563778233	0.865479538348451
GO_RNA_POLYMERASE_COMPLEX	104	0.01	Down
0.139937340007655	0.575535395493486	0.865941855276568
GO_REGULATION_OF_JUN_KINASE_ACTIVITY	80	0.01	Down
0.405150130048638	0.575706799974479	0.865951806161471
GO_POSITIVE_REGULATION_OF_CELLULAR_COMPONENT_BIOGENESIS	413	0.01

Down 0.310320285403971 0.575827284051039
0.865951806161471
GO_MUSCLE_FIBER_DEVELOPMENT 38 0.01 Up
0.365697639641102 0.576377534439697 0.866294869814944
GO_AMINE_CATABOLIC_PROCESS 17 0.01 Down
0.58455264201896 0.576397169313483 0.866294869814944
GO_NEGATIVE_REGULATION_OF_MYELOID_CELL_DIFFERENTIATION 59 0.01
Down 0.368968161579505 0.576483491031523
0.866294869814944
GO_EMBRYO_DEVELOPMENT_ENDING_IN_BIRTH_OR_EGG_HATCHING 464 0.01
Down 0.3216644018517 0.576834230130092
0.866437435111767
GO_REGULATION_OF_ION_TRANSMEMBRANE_TRANSPORT 252 0.01 Down
0.470275926348129 0.576863797186915
0.866437435111767
GO_ATPASE_ACTIVITY_COUPLED_TO_TRANSMEMBRANE_MOVEMENT_OF_IONS_PHOSPHORYLATIVE_MECHANISM 19 0.01 Down 0.358300604951356
0.577140799820554 0.866439132419991
GO_POSITIVE_REGULATION_OF_PHOSPHATIDYLINOSITOL_3_KINASE_SIGNALING
58 0.01 Down 0.585939515352473
0.577150362626658 0.866439132419991
GO_THYMIC_T_CELL_SELECTION 14 0.01 Up
0.437248979399168 0.577576954347506 0.866710457987492
GO_PHOSPHORIC_ESTER_HYDROLASE_ACTIVITY 288 0.01 Down
0.303498967672576 0.577730089895965 0.866710457987492
GO_NADPLUS_KINASE_ACTIVITY 13 0.01 Up
0.187817205456051 0.577841533316609 0.866710457987492
GO_ENDOLYSOSOME_MEMBRANE 13 0.01 Up
0.275776460626298 0.577902146917043 0.866710457987492
GO_CORPUS_CALLOSUM_DEVELOPMENT 14 0.01 Up
0.343975080979062 0.57850469301163 0.867040474757821
GO_LYMPHOCYTE_HOMEOSTASIS 46 0.01 Down
0.464351714445896 0.578750124498775 0.867040474757821
GO_INNERVATION 17 0.01 Down 0.680407295988562
0.579140522145662 0.867040474757821
GO_PHOSPHATIDYLGLYCEROL_METABOLIC_PROCESS 17 0.01 Up
0.222044143354524 0.579149185580471
0.867040474757821
GO_REPRESSING_TRANSCRIPTION_FACTOR_BINDING 65 0.01 Down
0.223323606556511 0.579176165624397
0.867040474757821
GO_CELLULAR_CARBOHYDRATE_METABOLIC_PROCESS 199 0.01 Down
0.320071054554018 0.579228832297578
0.867040474757821
GO_POSITIVE_REGULATION_OF_EPITHELIAL_CELL_PROLIFERATION 135 0.01
Down 0.428050220395279 0.579315400639913
0.867040474757821
GO_ESTABLISHMENT_OR_MAINTENANCE_OF_ACTIN_CYTOSKELETON_POLARITY 12
0.01 Down 0.562491682946456 0.579349375323105
0.867040474757821
GO_ATP_DEPENDENT_CHROMATIN_REMODELING 58 0.01 Down
0.185865157587633 0.579407545065472 0.867040474757821
GO_POSITIVE_REGULATION_OF_NEURON_APOPTOTIC_PROCESS 42 0.01
Down 0.359312240391556 0.580028009267217

0.867596048326014
GO_REGULATION_OF_VASCULAR_PERMEABILITY 19 0.01 Down
0.200165097798081 0.580064628587595 0.867596048326014
GO_ADP_BINDING 36 0.01 Up 0.194411821742074
0.580212950948068 0.867604144139341
GO_POSITIVE_REGULATION_OF_NEUROTRANSMITTER_TRANSPORT 22 0.01
Down 0.273472161695286 0.580658854320821
0.868057105289757
GO_SEMAPHORIN_PLEXIN_SIGNALING_PATHWAY_INVOLVED_IN_NEURON_PROJECTION
_GUIDANCE 12 0.01 Up 0.443071044288959
0.581065377453342 0.868139257411263
GO_ENDOCRINE_PANCREAS_DEVELOPMENT 28 0.01 Down
0.429448053658216 0.581094730901849 0.868139257411263
GO_CLEAVAGE_FURROW 48 0.01 Up 0.222731441113453
0.581142800546758 0.868139257411263
GO_NEGATIVE_REGULATION_OF_MYOTUBE_DIFFERENTIATION 12 0.01
Down 0.64011045504215 0.581592440693782
0.868445221119809
GO_MYOSIN_COMPLEX 38 0.01 Down 0.292119389477298
0.581633712579994 0.868445221119809
GO_AP_TYPE_MEMBRANE_COAT_ADAPTOR_COMPLEX 43 0.01 Down
0.179574405361074 0.582256055973494
0.869160687439164
GO_SPLICEOSOMAL_TRI_SNRNP_COMPLEX 31 0.01 Up
0.0823435652799279 0.582711472254781 0.869564446654335
GO_REGULATION_OF_CELL_JUNCTION_ASSEMBLY 76 0.01 Down
0.413304424487981 0.582813001719073 0.869564446654335
GO_ESTROGEN_RECEPTOR_BINDING 35 0.01 Down
0.268192181725563 0.583149725691545 0.869815794771032
GO_GLYCOSYL_COMPOUND_CATABOLIC_PROCESS 32 0.01 Down
0.345092048702102 0.583959395136318 0.869815794771032
GO_POSITIVE_REGULATION_OF_PROTEIN_KINASE_B_SIGNALING 116 0.01
Down 0.431451996794802 0.583994655639593
0.869815794771032
GO_SIGNAL_RECOGNITION_PARTICLE 11 0.01 Down
0.134683067144715 0.584006850241886 0.869815794771032
GO_NEGATIVE_REGULATION_OF_CHEMOTAXIS 43 0.01 Down
0.648981974521628 0.584080178990955 0.869815794771032
GO_NUCLEAR_CHROMOSOME_TELOMERIC_REGION 103 0.01 Up
0.181462746893175 0.584117463799871 0.869815794771032
GO_REGULATION_OF_DNA_TEMPLATED_TRANSCRIPTION_ELONGATION 45 0.01
Down 0.137112485661689 0.584170348586358
0.869815794771032
GO_NON_CANONICAL_WNT_SIGNALING_PATHWAY 131 0.01 Down
0.275245799151363 0.584292087623933 0.869815794771032
GO_REGULATION_OF_LEUKOCYTE_CHEMOTAXIS 80 0.01 Down
0.655123803301397 0.584383031393058 0.869815794771032
GO_REGULATION_OF_CELLULAR_RESPONSE_TO_HYPOXIA 12 0.01 Up
0.152260492375254 0.584414203075447
0.869815794771032
GO_MATURATION_OF_LSU_RRNA 20 0.01 Up
0.106896582178404 0.585014779816357 0.869955833998901
GO_MEMBRANE_PROTEIN_COMPLEX 822 0.01 Down
0.28693610127624 0.585059242574701 0.869955833998901

GO_CALCIUM_ION_IMPORT 45 0.01 Up 0.471706301262678
0.58506472643848 0.869955833998901
GO_REGULATION_OF_HORMONE_LEVELS 324 0.01 Down
0.435255403532019 0.58513244632794 0.869955833998901
GO_AZUROPHIL_GRANULE_MEMBRANE 54 0.01 Down
0.394282220546449 0.585363938683076 0.869955833998901
GO_NEGATIVE_REGULATION_OF_CELL_CYCLE_PHASE_TRANSITION 218 0.01
Down 0.213875590706329 0.585368074762891
0.869955833998901
GO_PROSTATE_GLAND_MORPHOGENESIS 24 0.01 Up
0.448144046469069 0.585802105610772 0.870034155120515
GO_PARAXIAL_MESODERM_DEVELOPMENT 12 0.01 Up
0.273007867529765 0.585835524059322 0.870034155120515
GO_PROTON_TRANSPORTING_ATPASE_ACTIVITY_ROTATIONAL_MECHANISM 22
0.01 Up 0.162997226535652 0.585943039974
0.870034155120515
GO_CELLULAR_RESPONSE_TO_GAMMA_RADIATION 29 0.01 Up
0.148217075374372 0.585994014213109 0.870034155120515
GO_STRIATED_MUSCLE_CELL_APOPTOTIC_PROCESS 28 0.01 Down
0.371722078463482 0.586382390430555
0.870397919878705
GO_LEUKOCYTE_TETHERING_OR_ROLLING 13 0.01 Down
0.774089503815903 0.58685257128823 0.870882904006562
GO_NEUROGENESIS 1131 0.01 Down 0.358594338920442
0.587198588405839 0.871012363063028
GO_RESPONSE_TO_X_RAY 29 0.01 Up 0.329670511588441
0.587226750455769 0.871012363063028
GO_SMALL_MOLECULE_BIOSYNTHETIC_PROCESS 501 0.01 Up
0.319248521907654 0.587638457719258 0.871342644288574
GO_SPLICEOSOMAL_TRI_SNRNP_COMPLEX_ASSEMBLY 12 0.01 Down
0.0845913325576247 0.587736473128267
0.871342644288574
GO_RESPONSE_TO_LEPTIN 15 0.01 Down 0.310791678881641
0.587926707875232 0.871371652476007
GO_ENDOTHELIAL_CELL_PROLIFERATION 107 0.01 Up
0.494661085434928 0.588168529074629 0.871371652476007
GO_POSITIVE_REGULATION_OF_MACROPHAGE_ACTIVATION 18 0.01 Up
0.599280297959912 0.58826242936153
0.871371652476007
GO_HOMEOSTATIC_PROCESS 1318 0.01 Down 0.351214704761107
0.588330160352356 0.871371652476007
GO_NEGATIVE_REGULATION_OF_CARDIOCYTE_DIFFERENTIATION 17 0.01
Up 0.512892967215753 0.588986142991703
0.872130457098202
GO_RNA_POLYMERASE_III_TRANSCRIPTION_FACTOR_COMPLEX 11 0.01
Down 0.10094070382972 0.589229674422476
0.872173521791122
GO_REGULATION_OF_CELL_GROWTH_INVOLVED_IN_CARDIAC_MUSCLE_CELL_DEVELOP
MENT 14 0.01 Up 0.402344182390243
0.589302550879128 0.872173521791122
GO_LIPOPROTEIN_CATABOLIC_PROCESS 11 0.01 Up
0.314238203538441 0.589483671925489 0.8722289476626
GO_POSITIVE_REGULATION_OF_LIPID_CATABOLIC_PROCESS 11 0.01
Up 0.27496281665003 0.589756415190829

0.872397518549661
GO_NEGATIVE_REGULATION_OF_REACTIVE_OXYGEN_SPECIES_BIOSYNTHETIC_PROCE
SS	20	0.01	Down	0.342256224140278
0.590287703328375	0.872397518549661
GO_NAD_BIOSYNTHETIC_PROCESS	23	0.01	Down
0.654211129579258	0.590596262417921	0.872397518549661
GO_ENDOTHELIAL_CELL_CHEMOTAXIS	22	0.01	Up
0.498432736555493	0.590674671779949	0.872397518549661
GO_EPITHELIAL_CELL_MORPHOGENESIS	30	0.01	Up
0.471111963454897	0.590848621293184	0.872397518549661
GO_RECEPTOR_MEDIATED_ENDOCYTOSIS	198	0.01	Up
0.487473684693721	0.591195581391206	0.872397518549661
GO_GLYCEROLIPID_METABOLIC_PROCESS	181	0.01	Up
0.290435354387609	0.59122418635116	0.872397518549661
GO_DNA_REPLICATION_ORIGIN_BINDING	15	0.01	Up
0.129882343170756	0.591241973400058	0.872397518549661
GO_ANTERIOR_POSTERIOR_PATTERN_SPECIFICATION	115	0.01	Up
0.464250208801519	0.591328261914912
0.872397518549661
GO_VASCULAR_ASSOCIATED_SMOOTH_MUSCLE_CELL_MIGRATION	19	0.01
Up	0.51604919683047	0.591385039373533
0.872397518549661
GO_CELL_JUNCTION_MAINTENANCE	13	0.01	Down
0.304061577305828	0.591471194445457	0.872397518549661
GO_POSITIVE_REGULATION_OF_INTERLEUKIN_1_SECRETION	23	0.01
Up	0.659178479752737	0.591578781393261
0.872397518549661
GO_CALCIUM_MEDIATED_SIGNALING_USING_INTRACELLULAR_CALCIUM_SOURCE
16	0.01	Down	0.42481438978665
0.591666351607653	0.872397518549661
GO_CARDIAC_ATRIUM_DEVELOPMENT	20	0.01	Down
0.318296528179614	0.591718073269376	0.872397518549661
GO_NEGATIVE_REGULATION_OF_TRANSCRIPTION_ELONGATION_FROM_RNA_POLYMERA
SE_II_PROMOTER	11	0.01	Up	0.220891005777827
0.591753085387499	0.872397518549661
GO_REGULATION_OF_NATURAL_KILLER_CELL_ACTIVATION	24	0.01	Down
0.555957895585093	0.5920019273822	0.872552488744194
GO_NEGATIVE_REGULATION_OF_DNA_REPLICATION	36	0.01	Down
0.18268335992042	0.592395687777347
0.872920927304921
GO_NUCLEOTIDE_TRANSPORT	17	0.01	Up	0.27617324231938
0.592880297713247	0.8729797864303
GO_PORE_COMPLEX	10	0.01	Down	0.213115228170977
0.592881286957916	0.8729797864303
GO_DNA_UNWINDING_INVOLVED_IN_DNA_REPLICATION	10	0.01	Down
0.193673759938787	0.592883136669842
0.8729797864303
GO_CHAPERONE_BINDING	86	0.01	Down	0.193709627145794
0.593010811931899	0.8729797864303
GO_REGULATION_OF_CILIUM_ASSEMBLY	45	0.01	Up
0.227073952826468	0.593190598864848	0.873032757747513
GO_MACROMOLECULE_TRANSMEMBRANE_TRANSPORTER_ACTIVITY	21	0.01
Up	0.171850009276319	0.593362477400354
0.873074066964991

GO_REGULATION_OF_PROTEIN_MATURATION 49 0.01 Up
0.418117233040363 0.594175803745808 0.873499126475626
GO_AEROBIC_ELECTRON_TRANSPORT_CHAIN 15 0.01 Up
0.095743431492516 0.594294707968396 0.873499126475626
GO_NEGATIVE_REGULATION_OF_CATALYTIC_ACTIVITY 582 0.01 Down
0.366721357906559 0.594340520649262
0.873499126475626
GO_NEGATIVE_REGULATION_OF_TOLL_LIKE_RECEPTOR_SIGNALING_PATHWAY 32
0.01 Down 0.447731942448201 0.594464075237448
0.873499126475626
GO_MAGNESIUM_ION_BINDING 182 0.01 Up
0.275612079805566 0.594499996186722 0.873499126475626
GO_NEURON_CELLULAR_HOMEOSTASIS 21 0.01 Down
0.314587716643851 0.594551040851271 0.873499126475626
GO_POSITIVE_REGULATION_OF_CHEMOKINE_SECRETION 12 0.01 Up
0.809992870883047 0.594879799109287
0.873499126475626
GO_VENTRICULAR_SEPTUM_DEVELOPMENT 57 0.01 Down
0.354364542893527 0.595073886767766 0.873499126475626
GO_CHANNEL_REGULATOR_ACTIVITY 87 0.01 Down
0.422186725229386 0.595181998419583 0.873499126475626
GO_RESPONSE_TO_INTERFERON_ALPHA 18 0.01 Down
0.240917324299946 0.595228677578313 0.873499126475626
GO_MITOCHONDRIAL_DNA_REPLICATION 12 0.01 Up
0.104572974995942 0.595234044840992 0.873499126475626
GO_TRANSFERASE_ACTIVITY_TRANSFERRING_ONE_CARBON_GROUPS 196 0.01
Up 0.190982779460957 0.595829778798865
0.873698139042141
GO_APPENDAGE_DEVELOPMENT 126 0.01 Up
0.404723817032288 0.595929382047793 0.873698139042141
GO_REGULATION_OF_HEART_RATE 60 0.01 Down
0.469023124632713 0.595943024721767 0.873698139042141
GO_INTRINSIC_COMPONENT_OF_PEROXISOMAL_MEMBRANE 15 0.01 Up
0.138311952946592 0.595945312761243
0.873698139042141
GO_CELL_MATURATION 105 0.01 Down 0.353619496105947
0.596292306127002 0.873995796836559
GO_DNA_STRAND_ELONGATION 19 0.01 Up
0.130528922838626 0.596736326314139 0.874435490478672
GO_SPERM_CAPACITATION 13 0.01 Down 0.416058504283485
0.597024464416773 0.874646603154978
GO_REGULATION_OF_PEPTIDYL_THREONINE_PHOSPHORYLATION 38 0.01
Down 0.289448544806886 0.597335428805232
0.87469382548432
GO_NEGATIVE_REGULATION_OF_LEUKOCYTE_PROLIFERATION 59 0.01
Down 0.736771014015259 0.597344852653268
0.87469382548432
GO_POSITIVE_REGULATION_OF_MRNA_SPLICING_VIA_SPLICEOSOME 15 0.01
Down 0.0959645572377207 0.597588444567426
0.87480254457967
GO_T_CELL_SELECTION 31 0.01 Down 0.479849036913067
0.597809663773071 0.87480254457967
GO_REGULATION_OF_HISTONE_H3_K9_TRIMETHYLATION 10 0.01 Down
0.123649383991113 0.597851384856045

0.87480254457967
GO_REGULATION_OF_ANIMAL_ORGAN_FORMATION 24 0.01 Down
0.536190913050143 0.598922798181606 0.875437644784918
GO_IMMUNOGLOBULIN_PRODUCTION_INVOLVED_IN_IMMUNOGLOBULIN_MEDIATED_IMMUNE_RESPONSE 42 0.01 Down 0.32889972668575
0.598932844322326 0.875437644784918
GO_SECRETION 1173 0.01 Down 0.411258169276951
0.598995545352415 0.875437644784918
GO_POSTSYNAPTIC_MODULATION_OF_CHEMICAL_SYNAPTIC_TRANSMISSION 11
0.01 Down 0.141798048173462 0.599158599085383
0.875437644784918
GO_ACTIVATION_OF_JNKK_ACTIVITY 11 0.01 Down
0.407314003885456 0.599231100808344 0.875437644784918
GO_OXIDOREDUCTASE_ACTIVITY_ACTING_ON_THE_CH_NH2_GROUP_OF_DONORS 14
0.01 Up 0.696955225013693 0.599293383640486
0.875437644784918
GO_SPINDLE_POLE 146 0.01 Down 0.220725324648367
0.599294819918649 0.875437644784918
GO_NEUROTROPHIN_SIGNALING_PATHWAY 33 0.01 Down
0.301629839301791 0.599477067003005 0.875483177865378
GO_INNER_DYNEIN_ARM_ASSEMBLY 10 0.01 Up
0.602532650810165 0.599614405133296 0.875483177865378
GO_CELL_DEATH_IN_RESPONSE_TO_HYDROGEN_PEROXIDE 23 0.01 Down
0.212062892628864 0.600028245125024
0.875876767529218
GO_METHYLATION 303 0.01 Down 0.197129053116136
0.600228903503821 0.875959056050889
GO_CELL_CYCLE_G2_M_PHASE_TRANSITION 244 0.01 Down
0.170526354675439 0.600585107907763 0.876268250446535
GO_EPITHELIAL_CELL_PROLIFERATION_INVOLVED_IN_PROSTATE_GLAND_DEVELOPMENT 11 0.01 Up 0.319213682584179
0.600884473751854 0.87639510493387
GO_INSULIN_SECRETION 136 0.01 Down 0.33182959520455
0.600960767886625 0.87639510493387
GO_REGULATION_OF_HELICASE_ACTIVITY 13 0.01 Up
0.173899430081991 0.601601315173676 0.877118536123773
GO_COMPACT_MYELIN 13 0.01 Up 0.306103631015717
0.602099534608962 0.877392492308112
GO_POSITIVE_REGULATION_OF_DNA_DAMAGE_RESPONSE_SIGNAL_TRANSDUCTION_BY_P53_CLASS_MEDIATOR 12 0.01 Up 0.243155553277986
0.602130172983606 0.877392492308112
GO_I_SMAD_BINDING 11 0.01 Down 0.163941632136002
0.6022227829761 0.877392492308112
GO_EMBRYONIC_DIGIT_MORPHOGENESIS 47 0.01 Up
0.364989847646461 0.602744135729191 0.87794137428309
GO_REGULATION_OF_NEUROBLAST_PROLIFERATION 18 0.01 Up
0.3737744079174 0.603125277606295 0.878239284244309
GO_FICOLIN_1_RICH_GRANULE_MEMBRANE 46 0.01 Up
0.43058402079705 0.603292004969313 0.878239284244309
GO_CELLULAR_RESPONSE_TO_RETINOIC_ACID 44 0.01 Down
0.395863017190291 0.60338264776528 0.878239284244309
GO_REGULATION_OF_RESPONSE_TO_WOUNDING 117 0.01 Down
0.487472556120887 0.603659943727176 0.878410603725151
GO_POSITIVE_REGULATION_OF_RECEPTOR_INTERNALIZATION 17 0.01

Up 0.59347712175752 0.603842232791799
0.878410603725151
GO_STEROL_TRANSPORTER_ACTIVITY 25 0.01 Down
0.282291962897381 0.603934419362342 0.878410603725151
GO_CANONICAL_WNT_SIGNALING_PATHWAY 262 0.01 Down
0.347560839512132 0.604089495097165 0.878425706523327
GO_CDP_DIACYLGLYCEROL_BIOSYNTHETIC_PROCESS 12 0.01 Down
0.233467955776276 0.604431163354521
0.878712067223491
GO_G_PROTEIN_COUPLED_RECEPTOR_SIGNALING_PATHWAY_COUPLED_TO_CYCLIC_NU
CLEOTIDE_SECOND_MESSENGER 97 0.01 Down
0.517028625915137 0.604660902549355 0.878824768481391
GO_LEUKOCYTE_ACTIVATION_INVOLVED_IN_INFLAMMATORY_RESPONSE 25
0.01 Up 0.781180804937143 0.604798201732046
0.878824768481391
GO_UNFOLDED_PROTEIN_BINDING 107 0.01 Down
0.195950050971568 0.605371613340694 0.879296804156187
GO_LRR_DOMAIN_BINDING 16 0.01 Down 0.147010321070643
0.605479305390864 0.879296804156187
GO_RNA_POLYMERASE_I_COMPLEX 12 0.01 Up
0.122041887983731 0.605557558586233 0.879296804156187
GO_MONOVALENT_INORGANIC_CATION_TRANSPORT 300 0.01 Down
0.394574073244158 0.605941934635306
0.879413472919623
GO_PHOSPHATIDYLGLYCEROL_BINDING 11 0.01 Up
0.262351468704141 0.606063741192793 0.879413472919623
GO_MANNOSIDASE_ACTIVITY 15 0.01 Up 0.225905267637867
0.606178901957298 0.879413472919623
GO_GLYCOSPHINGOLIPID_BIOSYNTHETIC_PROCESS 15 0.01 Down
0.43528263389177 0.606291262577389
0.879413472919623
GO_MITOTIC_NUCLEAR_ENVELOPE_REASSEMBLY 10 0.01 Down
0.171238580568883 0.606383859983601 0.879413472919623
GO_POTASSIUM_CHANNEL_ACTIVITY 39 0.01 Up
0.646789090478018 0.606507035268401 0.879413472919623
GO_REACTIVE_OXYGEN_SPECIES_METABOLIC_PROCESS 128 0.01 Up
0.400221242026453 0.606792734713026
0.879617643849757
GO_NEGATIVE_REGULATION_OF_ANTIGEN_RECEPTOR_MEDIATED_SIGNALING_PATHWA
Y 22 0.01 Up 0.439895073426344
0.607438424945821 0.879895167413238
GO_PROTEIN_CONTAINING_COMPLEX_BINDING 879 0.01 Down
0.342288634181722 0.607462847926877 0.879895167413238
GO_NCRNA_PROCESSING 360 0.01 Up 0.131796007219782
0.607560488043276 0.879895167413238
GO_REGULATION_OF_THE_FORCE_OF_HEART_CONTRACTION 14 0.01 Up
0.56236583008474 0.607563917278256
0.879895167413238
GO_MOTOR_ACTIVITY 93 0.01 Down 0.272285094199842
0.607741334656208 0.879942199546349
GO_NUCLEUS_LOCALIZATION 20 0.01 Down 0.262834435633948
0.607973331496399 0.880014377114577
GO_AROMATIC_AMINO_ACID_FAMILY_CATABOLIC_PROCESS 10 0.01 Down
0.723434413561649 0.608151175604532

0.880014377114577
GO_RNA_PHOSPHODIESTER_BOND_HYDROLYSIS 129 0.01 Down
0.202869776721115 0.608273067615771 0.880014377114577
GO_MUSCLE_CELL_MIGRATION 70 0.01 Down
0.472257207137197 0.608442547345338 0.880014377114577
GO_POSITIVE_REGULATION_OF_CALCIUM_ION_TRANSMEMBRANE_TRANSPORT 44
0.01 Up 0.606420924132667 0.608738087994242
0.880014377114577
GO_POSITIVE_REGULATION_OF_ERAD_PATHWAY 15 0.01 Up
0.155656745883545 0.608994797220566 0.880014377114577
GO_POSITIVE_REGULATION_OF_PROTEIN_AUTOPHOSPHORYLATION 25 0.01
Down 0.604389162174054 0.609114475614984
0.880014377114577
GO_TUMOR_NECROSIS_FACTOR_BIOSYNTHETIC_PROCESS 15 0.01 Down
0.583057095345112 0.609211171019688
0.880014377114577
GO_CELL_CYCLE_ARREST 207 0.01 Down 0.271230491072735
0.609361168596966 0.880014377114577
GO_LIPID_TRANSPORTER_ACTIVITY 101 0.01 Down
0.409799874937356 0.609384250376198 0.880014377114577
GO_ODONTOGENESIS_OF_DENTIN_CONTAINING_TOOTH 61 0.01 Up
0.452839383431722 0.609385676394281
0.880014377114577
GO_RNA_PHOSPHODIESTER_BOND_HYDROLYSIS_ENDONUCLEOLYTIC 65 0.01
Up 0.218376562863567 0.609787703091294
0.880225373634737
GO_REGULATION_OF_PEPTIDE_HORMONE_SECRETION 130 0.01 Down
0.339816036706235 0.609821762725696
0.880225373634737
GO_LIPOPROTEIN_BIOSYNTHETIC_PROCESS 82 0.01 Up
0.17899267197629 0.610331620687699 0.880751906155223
GO_G_PROTEIN_COUPLED_RECEPTOR_SIGNALING_PATHWAY 421 0.01 Down
0.489805727858341 0.610843914244732
0.881274417114025
GO_CYCLIN_DEPENDENT_PROTEIN_KINASE_ACTIVITY 27 0.01 Up
0.262899418578002 0.611056602152731
0.881274417114025
GO_REGULATION_OF_FEEDING_BEHAVIOR 10 0.01 Down
0.333511497426434 0.611177081902272 0.881274417114025
GO_RESPONSE_TO_ALCOHOL 151 0.01 Up 0.41062779478742
0.611274348619199 0.881274417114025
GO_PROTEIN_CONTAINING_COMPLEX_ASSEMBLY 1489 0.01 Down
0.247378353215634 0.611515582816495 0.881412892516368
GO_BINDING_OF_SPERM_TO_ZONA_PELLUCIDA 19 0.01 Down
0.179511472675776 0.611797085341362 0.881609329481939
GO_POSITIVE_REGULATION_OF_OSSIFICATION 65 0.01 Down
0.543531868996579 0.612529526915055 0.882455329354841
GO_PRESYNAPTIC_ENDOCYTOSIS 40 0.01 Down
0.312913801722341 0.61366176430306 0.883660473147539
GO_POSTSYNAPTIC_ENDOSOME 10 0.01 Down
0.149463257366004 0.613890726917554 0.883660473147539
GO_G1_DNA_DAMAGE_CHECKPOINT 60 0.01 Down
0.229029295659445 0.613936102121188 0.883660473147539
GO_PROTEIN_NITROSYLATION 12 0.01 Down

0.911153691912381	0.614105764433517	0.883660473147539
GO_PROTEIN_HOMODIMERIZATION_ACTIVITY	633	0.01	Up
0.36181041789034	0.61446156229845	0.883660473147539
GO_REGULATION_OF_FATTY_ACID_BIOSYNTHETIC_PROCESS	31	0.01
Down	0.309017068402865	0.61463422449778
0.883660473147539
GO_POLYOL_CATABOLIC_PROCESS	20	0.01	Down
0.189931894570984	0.615394073206528	0.883660473147539
GO_RETINOL_METABOLIC_PROCESS	21	0.01	Down
0.539800505679025	0.615490073417131	0.883660473147539
GO_PHOTOTRANSDUCTION_VISIBLE_LIGHT	15	0.01	Down
0.148255281565665	0.615653774015637	0.883660473147539
GO_CELLULAR_RESPONSE_TO_ENDOGENOUS_STIMULUS	986	0.01	Down
0.350402498779997	0.615655098723356
0.883660473147539
GO_VESICLE_LOCALIZATION	243	0.01	Down	0.250198991512532
0.615664669123222	0.883660473147539
GO_COAGULATION	228	0.01	Up	0.413099216953534
0.615790250477537	0.883660473147539
GO_U4_SNRNP	10	0.01	Up	0.0716514590869248
0.615815883084925	0.883660473147539
GO_GLIAL_CELL_DEVELOPMENT	79	0.01	Down
0.401741788384531	0.616055814662075	0.883660473147539
GO_INTRASPECIES_INTERACTION_BETWEEN_ORGANISMS	29	0.01	Down
0.209651890989184	0.616459730166991
0.883660473147539
GO_TRANSPORTER_ACTIVITY	762	0.01	Up	0.343908297087996
0.616643318480296	0.883660473147539
GO_POSITIVE_REGULATION_OF_AMINO_ACID_TRANSPORT	10	0.01	Down
0.388031612613801	0.616713798460953
0.883660473147539
GO_ROOF_OF_MOUTH_DEVELOPMENT	64	0.01	Up
0.431158810042444	0.616719949147826	0.883660473147539
GO_NEGATIVE_REGULATION_OF_SMOOTH_MUSCLE_CONTRACTION	12	0.01
Up	0.525530291610456	0.616733441817249
0.883660473147539
GO_LENS_DEVELOPMENT_IN_CAMERA_TYPE_EYE	52	0.01	Down
0.283059897334434	0.616797351940438	0.883660473147539
GO_POSITIVE_REGULATION_OF_DNA_REPLICATION	32	0.01	Up
0.23763702411983	0.616902851949013
0.883660473147539
GO_POSITIVE_REGULATION_OF_PROTEIN_TARGETING_TO_MITOCHONDRION	31
0.01	Down	0.122230025222928	0.616952620633965
0.883660473147539
GO_UBIQUITINATION_LIKE_MODIFICATION_DEPENDENT_PROTEIN_BINDING	10
0.01	Up	0.118451147535649	0.617043238030633
0.883660473147539
GO_EPHRIN_RECEPTOR_ACTIVITY	11	0.01	Up
0.266241512962709	0.617197036877773	0.883660473147539
GO_RESPONSE_TO_MANGANESE_ION	13	0.01	Down
0.24214191874379	0.617209239567121	0.883660473147539
GO_CELL_CYCLE_PROCESS	1117	0.01	Down	0.221728866611202
0.617489193129087	0.883660473147539
GO_ORGANELLE_SUBCOMPARTMENT	334	0.01	Down

0.265330759814723	0.61749303866912	0.883660473147539
GO_MAINTENANCE_OF_PROTEIN_LOCATION	91	0.01	Up
0.294079838086389	0.617503782263284	0.883660473147539
GO_BRANCH_ELONGATION_OF_AN_EPITHELIUM	17	0.01	Up
0.470663135693219	0.617972943533651	0.883660473147539
GO_GOLGI_TRANSPORT_COMPLEX	12	0.01	Down
0.137892157567984	0.618257381174246	0.883660473147539
GO_PROSTANOID_BIOSYNTHETIC_PROCESS	18	0.01	Up
0.551116354816884	0.618418123234812	0.883660473147539
GO_PRIMARY_ACTIVE_TRANSMEMBRANE_TRANSPORTER_ACTIVITY	90	0.01
Up	0.23329974249956	0.618670894852619
0.883660473147539
GO_PROTEASOME_ASSEMBLY	13	0.01	Down	0.0951687498422824
0.618724223158122	0.883660473147539
GO_POSITIVE_REGULATION_OF_MYELOID_LEUKOCYTE_DIFFERENTIATION	39
0.01	Up	0.420866433745984	0.61874928589561
0.883660473147539
GO_RNA_POLYMERASE_I_ACTIVITY	11	0.01	Up
0.130680231888352	0.618813696370771	0.883660473147539
GO_CARDIAC_SEPTUM_DEVELOPMENT	87	0.01	Up
0.37543474346062	0.619258680592933	0.883660473147539
GO_VENTRICULAR_SEPTUM_MORPHOGENESIS	36	0.01	Down
0.370275330382414	0.619564673342734	0.883660473147539
GO_NEGATIVE_REGULATION_OF_MRNA_PROCESSING	28	0.01	Down
0.104081399998906	0.619619665397471
0.883660473147539
GO_CATION_CHANNEL_COMPLEX	91	0.01	Up
0.459058136622932	0.619686520951364	0.883660473147539
GO_NUCLEAR_TRANSCRIBED_MRNA_CATABOLIC_PROCESS	201	0.01	Down
0.154651810162376	0.619701894489145
0.883660473147539
GO_TUBULIN_DEACETYLATION	10	0.01	Up
0.188042978934324	0.619722517541269	0.883660473147539
GO_MYELOID_LEUKOCYTE_MEDIATED_IMMUNITY	458	0.01	Down
0.409145492136876	0.619860568682953	0.883660473147539
GO_CENTRAL_NERVOUS_SYSTEM_DEVELOPMENT	655	0.01	Down
0.36953552408295	0.620022668536311	0.883660473147539
GO_STRIATED_MUSCLE_CELL_DIFFERENTIATION	184	0.01	Up
0.465513912805147	0.620054645599408	0.883660473147539
GO_CELLULAR_RESPONSE_TO_NITROGEN_STARVATION	10	0.01	Up
0.145867438472856	0.620061364847084
0.883660473147539
GO_NEGATIVE_REGULATION_OF_MUSCLE_TISSUE_DEVELOPMENT	38	0.01
Down	0.471307178826965	0.6201459619879
0.883660473147539
GO_TELENCEPHALON_GLIAL_CELL_MIGRATION	18	0.01	Down
0.206209800908024	0.62020709538489	0.883660473147539
GO_PRERIBOSOME_LARGE_SUBUNIT_PRECURSOR	22	0.01	Up
0.109027152827958	0.620658046038955	0.883786252061832
GO_CELLULAR_RESPONSE_TO_UV	73	0.01	Up
0.235371461159284	0.620726321720004	0.883786252061832
GO_TELOMERASE_HOLOENZYME_COMPLEX	19	0.01	Down
0.0744749363567318	0.62073209994921	0.883786252061832
GO_PYRUVATE_METABOLIC_PROCESS	19	0.01	Up

0.167160644824596	0.620902839532579	0.883801385733586
GO_POSITIVE_REGULATION_OF_CARTILAGE_DEVELOPMENT 21	0.01	Up
	0.454763736453004	0.621309045051176
0.883801385733586
GO_PHOTORECEPTOR_OUTER_SEGMENT_MEMBRANE 10	0.01	Down
0.171777727569865	0.621376787121037	0.883801385733586
GO_CENTRAL_NERVOUS_SYSTEM_NEURON_DIFFERENTIATION	101	0.01
	Down	0.424098143614115	0.621427451199963
0.883801385733586
GO_NEGATIVE_REGULATION_OF_EPIDERMIS_DEVELOPMENT 12	0.01	Up
	0.682672855650529	0.621721917667718
0.883801385733586
GO_GUANYL_NUCLEOTIDE_BINDING	308	0.01	Down
0.271501650778351	0.621793819673234	0.883801385733586
GO_GTPASE_INHIBITOR_ACTIVITY	12	0.01	Down
0.271421083662852	0.62179735082092	0.883801385733586
GO_AXON 410	0.01	Down	0.317837323606059
0.621907349671204	0.883801385733586
GO_ION_CHANNEL_BINDING	88	0.01	Up	0.358338936964736
	0.622057517573275	0.883807907602938
GO_UBIQUITIN_UBIQUITIN_LIGASE_ACTIVITY	13	0.01	Down
0.137339451478434	0.622314422176584	0.883966040485269
GO_REGULATION_OF_SPHINGOLIPID_BIOSYNTHETIC_PROCESS	13	0.01
	Down	0.259912545551549	0.622539232481642
0.88407852173007
GO_REGULATION_OF_SIGNAL_TRANSDUCTION_BY_P53_CLASS_MEDIATOR	166
	0.01	Down	0.182062204261017	0.623010921779819
	0.884209384637875
GO_PYRIMIDINE_CONTAINING_COMPOUND_METABOLIC_PROCESS	20	0.01
	Up	0.224653100448931	0.623019248875789
0.884209384637875
GO_CHEMOKINE_BINDING	16	0.01	Up	0.583681076724585
	0.623273724847531	0.884209384637875
GO_RNA_POLYMERASE_III_REGULATORY_REGION_DNA_BINDING	10	0.01
	Up	0.0797086427905494	0.623338464751741
0.884209384637875
GO_POSITIVE_REGULATION_OF_PROTEIN_LOCALIZATION_TO_MEMBRANE	101
	0.01	Down	0.30169489523221	0.623359605707479
	0.884209384637875
GO_PEPTIDYL_L_CYSTEINE_S_PALMITOYLATION 18	0.01	Up
0.203719779112697	0.623909866162611	0.884783180909416
GO_ANATOMICAL_STRUCTURE_HOMEOSTASIS	317	0.01	Down
0.294372007328674	0.624311316090739	0.88508225456735
GO_REGULATION_OF_DEFENSE_RESPONSE_TO_VIRUS_BY_VIRUS	25	0.01
	Up	0.344426725186264	0.624412336733974
0.88508225456735
GO_NEGATIVE_REGULATION_OF_INFLAMMATORY_RESPONSE 83	0.01	Down
	0.439924056989814	0.625399044485165
0.886273949362613
GO_MEMBRANE_PROTEIN_PROTEOLYSIS 53	0.01	Down
0.34797449560155	0.625610261929642	0.886286599804351
GO_DNA_REPLICATION_FACTOR_A_COMPLEX	15	0.01	Down
0.131256126882598	0.625828322526159	0.886286599804351
GO_REGULATION_OF_B_CELL_MEDIATED_IMMUNITY	38	0.01	Down

0.382271861164865	0.625845932031173
0.886286599804351
GO_SMN_COMPLEX	12	0.01	Down	0.0855362071248993
0.626292266080883	0.886325510188221
GO_POSITIVE_REGULATION_OF_CAMP_MEDIATED_SIGNALING	11	0.01
Down	0.46746151303776	0.626341120041673
0.886325510188221
GO_DNA_APURINIC_OR_APYRIMIDINIC_SITE_ENDONUCLEASE_ACTIVITY	12
0.01	Down	0.25146604524423	0.626374013244512
0.886325510188221
GO_RESPONSE_TO_MINERALOCORTICOID	19	0.01	Down
0.342857836737394	0.626558787618179	0.886325510188221
GO_ESTABLISHMENT_OF_PROTEIN_LOCALIZATION_TO_PLASMA_MEMBRANE	51
0.01	Down	0.198585743184079	0.626687657676102
0.886325510188221
GO_GLIAL_CELL_APOPTOTIC_PROCESS	14	0.01	Up
0.240943851144964	0.626886602178728	0.886325510188221
GO_REGULATION_OF_ACTIN_FILAMENT_BUNDLE_ASSEMBLY	77	0.01	Down
0.251993551309878	0.626927511921754
0.886325510188221
GO_AXONAL_TRANSPORT	48	0.01	Down	0.246458233975257
0.627245539453336	0.886325510188221
GO_CALCIUM_ION_TRANSMEMBRANE_TRANSPORT	188	0.01	Up
0.42880528108105	0.627260460095306	0.886325510188221
GO_ANTIPORTER_ACTIVITY	55	0.01	Up	0.317988314411082
0.627333341670036	0.886325510188221
GO_DETECTION_OF_MECHANICAL_STIMULUS	17	0.01	Up
0.493474921199175	0.627681837943539	0.886537124395797
GO_KINASE_INHIBITOR_ACTIVITY	58	0.01	Down
0.209995651253047	0.627860417784771	0.886537124395797
GO_SPINAL_CORD_MOTOR_NEURON_DIFFERENTIATION	14	0.01	Up
0.25760535215144	0.627921204892427
0.886537124395797
GO_PROTEIN_CATABOLIC_PROCESS_IN_THE_VACUOLE	13	0.01	Down
0.303481208441659	0.628185634342049
0.886651229529168
GO_POSITIVE_REGULATION_OF_SKELETAL_MUSCLE_TISSUE_DEVELOPMENT	17
0.01	Up	0.495406374426577	0.628412080938969
0.886651229529168
GO_MULTICELLULAR_ORGANISMAL_RESPONSE_TO_STRESS	38	0.01	Up
0.429971438714977	0.628440164826883
0.886651229529168
GO_REGULATION_OF_CELL_CYCLE_PROCESS	654	0.01	Down
0.223134386330308	0.628649495787237	0.886671594919646
GO_MESODERMAL_CELL_FATE_COMMITMENT	11	0.01	Down
0.880864135149843	0.628746700070676	0.886671594919646
GO_TROPHOBLAST_CELL_MIGRATION	10	0.01	Up
0.614997204840215	0.629159895593916	0.887048241093977
GO_REGULATION_OF_TORC1_SIGNALING	30	0.01	Up
0.164271547682559	0.62939713934889	0.887144263285137
GO_KINETOCHORE	111	0.01	Down	0.16476462471306
0.629520257985895	0.887144263285137
GO_REGULATION_OF_MEMBRANE_INVAGINATION	12	0.01	Up
0.729699196376326	0.629765402761552	0.887255383490925

GO_REGULATION_OF_DELAYED_RECTIFIER_POTASSIUM_CHANNEL_ACTIVITY 11 0.01 Down 0.62800352188754 0.629891402214773 0.887255383490925
GO_PALMITOYLTRANSFERASE_COMPLEX 11 0.01 Down 0.239497285594862 0.630094683686941 0.887335844273583
GO_RIBONUCLEASE_P_ACTIVITY 11 0.01 Up 0.114891494435433 0.63041919886099 0.887418160438173
GO_REGULATION_OF_PROTEIN_LOCALIZATION_TO_CHROMOSOME_TELOMERIC_REGION 14 0.01 Down 0.0556952195743495 0.630445482782052 0.887418160438173
GO_RESPONSE_TO_FIBROBLAST_GROWTH_FACTOR 109 0.01 Down 0.379791935849645 0.630914002921118 0.887730060795593
GO_VASCULAR_ENDOTHELIAL_GROWTH_FACTOR_RECEPTOR_SIGNALING_PATHWAY 78 0.01 Down 0.390791439841806 0.631000632900157 0.887730060795593
GO_ENDODERMAL_CELL_FATE_COMMITMENT 10 0.01 Up 0.282879217006886 0.631105739152328 0.887730060795593
GO_GERM_CELL_NUCLEUS 12 0.01 Up 0.134405263432851 0.631555403395682 0.888078632912782
GO_RESPONSE_TO_LIGHT_STIMULUS 226 0.01 Down 0.229114269250953 0.631807926569675 0.888078632912782
GO_DISULFIDE_OXIDOREDUCTASE_ACTIVITY 36 0.01 Down 0.157409210322205 0.631900106875877 0.888078632912782
GO_POSITIVE_REGULATION_OF_MEIOTIC_CELL_CYCLE 13 0.01 Up 0.548115102272325 0.631938674713098 0.888078632912782
GO_POSITIVE_REGULATION_OF_CYTOPLASMIC_TRANSPORT 12 0.01 Down 0.218901830030879 0.632117158310468 0.88808718953843
GO_RESPONSE_TO_REDOX_STATE 12 0.01 Up 0.276610143155578 0.632316782144295 0.88808718953843
GO_CARBOHYDRATE_CATION_SYMPORTER_ACTIVITY 12 0.01 Up 0.322202295367417 0.632476528821235 0.88808718953843
GO_POSITIVE_REGULATION_OF_MYELOID_CELL_DIFFERENTIATION 65 0.01 Down 0.439562985161868 0.632809046871126 0.88808718953843
GO_PROTEIN_LOCALIZATION_TO_CELL_PERIPHERY 248 0.01 Down 0.268688166814132 0.632814291141364 0.88808718953843
GO_RESPONSE_TO_CARBOHYDRATE 165 0.01 Up 0.309519621160028 0.632977219627673 0.88808718953843
GO_MITOTIC_SISTER_CHROMATID_SEGREGATION 128 0.01 Up 0.173466984420661 0.633209063219324 0.88808718953843
GO_SENSORY_PERCEPTION_OF_TASTE 14 0.01 Up 0.73386381162849 0.633250858287621 0.88808718953843
GO_PHAGOSOME_ACIDIFICATION 23 0.01 Up 0.244564729562228 0.633261315024191 0.88808718953843
GO_ALPHA_TUBULIN_BINDING 29 0.01 Up 0.333420745621252 0.633482407297171 0.888192077298182
GO_NEGATIVE_REGULATION_OF_VASCULAR_ENDOTHELIAL_GROWTH_FACTOR_RECEPTOR_SIGNALING_PATHWAY 11 0.01 Up 0.514721131932914 0.634007610371998 0.888412345648575
GO_NEUROTRANSMITTER_RECEPTOR_COMPLEX 17 0.01 Down

0.276632271417329 0.634069459498211 0.888412345648575
GO_MITOTIC_G2_DNA_DAMAGE_CHECKPOINT 20 0.01 Down
0.164371850392156 0.634179407474074 0.888412345648575
GO_NUCLEOSIDE_MONOPHOSPHATE_METABOLIC_PROCESS 26 0.01 Down
0.298750551433517 0.63429436885934
0.888412345648575
GO_REGULATION_OF_SYSTEMIC_ARTERIAL_BLOOD_PRESSURE_BY_RENIN_ANGIOTENS
IN 15 0.01 Up 0.393967159028029
0.634470824422645 0.888412345648575
GO_NUCLEAR_EXOSOME_RNASE_COMPLEX_ 17 0.01 Down
0.138817192997934 0.634517531005143 0.888412345648575
GO_RAL_GTPASE_BINDING 15 0.01 Up 0.369693692558606
0.634677419077539 0.888421785550247
GO_NUCLEAR_TRANSCRIBED_MRNA_CATABOLIC_PROCESS_EXONUCLEOLYTIC 35
0.01 Down 0.132271542828348 0.634865253716531
0.888421785550247
GO_ENDORIBONUCLEASE_ACTIVITY 52 0.01 Up
0.237723920193308 0.634963288997285 0.888421785550247
GO_BRANCHING_INVOLVED_IN_MAMMARY_GLAND_DUCT_MORPHOGENESIS 22
0.01 Up 0.445005392285396 0.635332635794267
0.888733740070736
GO_GLUTAMATE_SECRETION 25 0.01 Up 0.380543368095554
0.635490206085692 0.88874937598393
GO_REGULATION_OF_SYSTEMIC_ARTERIAL_BLOOD_PRESSURE_BY_CIRCULATORY_REN
IN_ANGIOTENSIN 12 0.01 Up 0.476031491052603
0.635999162620445 0.889092001250761
GO_NEGATIVE_REGULATION_OF_PEPTIDASE_ACTIVITY 153 0.01 Down
0.564601481580967 0.6360280944543 0.889092001250761
GO_ER_ASSOCIATED_MISFOLDED_PROTEIN_CATABOLIC_PROCESS 12 0.01
Up 0.157458884401505 0.636866135285804
0.890058542200763
GO_POSITIVE_REGULATION_OF_MITOCHONDRIAL_OUTER_MEMBRANE_PERMEABILIZAT
ION_INVOLVED_IN_APOPTOTIC_SIGNALING_PATHWAY 34 0.01 Down
0.242334675492916 0.637201153377889
0.890321795663329
GO_SKELETAL_MUSCLE_CONTRACTION 25 0.01 Down
0.582915856740894 0.637417915770706 0.890419734616649
GO_NEGATIVE_REGULATION_OF_MICROTUBULE_POLYMERIZATION_OR_DEPOLYMERIZA
TION 29 0.01 Down 0.256154613471936
0.637702449609781 0.890463276431624
GO_PEPTIDYL_ARGININE_METHYLATION 12 0.01 Down
0.121043075773382 0.637742435500692 0.890463276431624
GO_VASCULAR_ENDOTHELIAL_GROWTH_FACTOR_RECEPTOR_BINDING 12 0.01
Up 0.739724949279242 0.637965019486502
0.890529112225895
GO_RIBOSOMAL_LARGE_SUBUNIT_ASSEMBLY 28 0.01 Up
0.104074589322341 0.638161344165102 0.890529112225895
GO_POSITIVE_REGULATION_OF_STEROL_TRANSPORT 17 0.01 Down
0.32640753600405 0.638299863655596
0.890529112225895
GO_ATRIAL_CARDIAC_MUSCLE_CELL_TO_AV_NODE_CELL_SIGNALING 11 0.01
Up 0.461717547391056 0.638476889951457
0.890529112225895
GO_REGULATION_OF_TRANSCRIPTION_ELONGATION_FROM_RNA_POLYMERASE_II_PRO

MOTER	28	0.01	Down	0.154493497634252
0.638523015239552	0.890529112225895
GO_NUCLEAR_ENVELOPE	400	0.01	Down	0.230737042348139
	0.638740442964452	0.890551069540937
GO_MULTICELLULAR_ORGANISMAL_MOVEMENT	34	0.01	Down
0.504874352175991	0.638832137679259	0.890551069540937
GO_POSITIVE_REGULATION_OF_ION_TRANSMEMBRANE_TRANSPORT	88	0.01
	Down	0.504115135958522	0.639110113412659
0.890594220271144
GO_REGULATION_OF_MAST_CELL_ACTIVATION_INVOLVED_IN_IMMUNE_RESPONSE
	23	0.01	Down	0.429795577943612
0.639285184348276	0.890594220271144
GO_INSULIN_LIKE_GROWTH_FACTOR_RECEPTOR_SIGNALING_PATHWAY	26
	0.01	Up	0.455098643813484	0.639303181014931
	0.890594220271144
GO_POSITIVE_REGULATION_OF_WNT_SIGNALING_PATHWAY	149	0.01	Down
	0.340552478830056	0.639591410953584
0.890679642810588
GO_SPINDLE_ORGANIZATION	147	0.01	Down	0.209521131432453
	0.639657921702218	0.890679642810588
GO_VENOUS_BLOOD_VESSEL_DEVELOPMENT	11	0.01	Down
0.486282930715816	0.640236129128852	0.8909733221024
GO_ION_TRANSPORT	1002	0.01	Up	0.397638246735871
	0.64030907675549	0.8909733221024
GO_DESMOSOME_ORGANIZATION	10	0.01	Down
0.268437081440317	0.640309109591957	0.8909733221024
GO_COPPER_ION_HOMEOSTASIS	16	0.01	Down
0.149525252117579	0.641068885088301	0.891652765483364
GO_CELLULAR_RESPONSE_TO_EXTERNAL_STIMULUS	278	0.01	Down
	0.323936947464936	0.641102686337811
0.891652765483364
GO_MEMBRANE_REGION	244	0.01	Down	0.435241962194861
	0.641319292568295	0.891652765483364
GO_INTRINSIC_COMPONENT_OF_PRESYNAPTIC_ACTIVE_ZONE_MEMBRANE	12
	0.01	Up	0.371373786684162	0.641467810039144
	0.891652765483364
GO_NEGATIVE_REGULATION_OF_KIDNEY_DEVELOPMENT	11	0.01	Down
	0.812953889245687	0.64153175418075
0.891652765483364
GO_MITOGEN_ACTIVATED_PROTEIN_KINASE_KINASE_KINASE_BINDING	17
	0.01	Down	0.152025935368935	0.641781575782459
	0.891795822058894
GO_REGULATION_OF_CELL_MORPHOGENESIS_INVOLVED_IN_DIFFERENTIATION	235
	0.01	Down	0.359792552757749	0.642102734071199
	0.891847977671202
GO_CYSTEINE_TYPE_ENDOPEPTIDASE_REGULATOR_ACTIVITY_INVOLVED_IN_APOPTO
TIC_PROCESS	32	0.01	Up	0.442742326824118
0.642345324143328	0.891847977671202
GO_RNA_POLYMERASE_II_TRANSCRIPTION_FACTOR_BINDING	40	0.01
	Down	0.327757885973756	0.642382162170554
0.891847977671202
GO_PRESYNAPTIC_MEMBRANE	82	0.01	Down	0.283663551633315
	0.642618657063015	0.891847977671202
GO_SYNAPTIC_TRANSMISSION_GLUTAMATERGIC	39	0.01	Down

0.350595573413131	0.642654103915771	0.891847977671202
GO_NEURON_PROJECTION_CYTOPLASM	71	0.01	Down
0.240019068268509	0.642700527476776	0.891847977671202
GO_OOCYTE_DIFFERENTIATION	35	0.01	Down
0.329339869449585	0.643167304819379	0.892291752184289
GO_CELLULAR_RESPONSE_TO_OXYGEN_RADICAL	21	0.01	Up
0.434968708782322	0.643561544036738	0.892634711868183
GO_OLIGOSACCHARIDE_BIOSYNTHETIC_PROCESS 14	0.01	Down
0.532177640146441	0.643740832216297	0.892679440928538
GO_STEROL_BINDING	44	0.01	Up	0.358231592353355
	0.643956934337054	0.892775188024721
GO_AMINO_ACID_IMPORT	15	0.01	Up	0.548086554592248
	0.64417386996808	0.892872046706898
GO_REGULATION_OF_CELLULAR_LOCALIZATION	704	0.01	Down
0.295207313104542	0.644519879227801	0.893010114905874
GO_U6_SNRNA_BINDING	11	0.01	Down	0.0963002044807211
	0.644636295355052	0.893010114905874
GO_NEUROEPITHELIAL_CELL_DIFFERENTIATION 37	0.01	Up
0.356111154943445	0.644971997579739	0.893010114905874
GO_PLATELET_AGGREGATION 44	0.01	Down	0.340062493429437
	0.645221811371729	0.893010114905874
GO_REGULATION_OF_WOUND_HEALING	93	0.01	Down
0.523889973030634	0.645236924330733	0.893010114905874
GO_REGULATION_OF_CYSTEINE_TYPE_ENDOPEPTIDASE_ACTIVITY_INVOLVED_IN_AP
OPTOTIC_SIGNALING_PATHWAY	15	0.01	Down
0.478261349891457	0.64538099338606	0.893010114905874
GO_BONE_RESORPTION	41	0.01	Up	0.485012077738566
	0.645443413300175	0.893010114905874
GO_NEGATIVE_REGULATION_OF_CELL_POPULATION_PROLIFERATION 520	0.01
	Down	0.43578983896058	0.645450236239001
0.893010114905874
GO_VESICLE_LUMEN	260	0.01	Up	0.407622186642253
	0.645821057391729	0.893198316511623
GO_PRC1_COMPLEX 15	0.01	Up	0.201238499204446
0.646067269303786	0.893198316511623
GO_REGULATION_OF_VIRAL_ENTRY_INTO_HOST_CELL	24	0.01	Down
	0.352788174980168	0.646127306958396
0.893198316511623
GO_CHROMATIN_ASSEMBLY	97	0.01	Up	0.187571333186962
	0.646174766285463	0.893198316511623
GO_BLOOD_VESSEL_REMODELING	27	0.01	Down
0.464878037556001	0.647020622876704	0.89399118319751
GO_ACTOMYOSIN_STRUCTURE_ORGANIZATION	148	0.01	Down
0.329756588265035	0.647112116633465	0.89399118319751
GO_ESTABLISHMENT_OF_PROTEIN_LOCALIZATION_TO_MEMBRANE	297	0.01
	Up	0.189952393903852	0.647190125210519
0.89399118319751
GO_SMAD_PROTEIN_COMPLEX_ASSEMBLY	12	0.01	Down
0.176189217296331	0.647579111756174	0.894325019898028
GO_INORGANIC_ANION_TRANSMEMBRANE_TRANSPORTER_ACTIVITY	75	0.01
	Down	0.344258154155547	0.648196955905223
0.894974691676281
GO_MAMMARY_GLAND_DEVELOPMENT	105	0.01	Down
0.356686399653022	0.64847766120298	0.895158681483241

GO_NEGATIVE_REGULATION_OF_MULTICELLULAR_ORGANISM_GROWTH 10 0.01 Down 0.319871172245412 0.648732770346309 0.895262090660731
GO_APOPTOTIC_PROCESS_INVOLVED_IN_MORPHOGENESIS 19 0.01 Up 0.404541398863337 0.648847504349731 0.895262090660731
GO_GLUTAMINE_FAMILY_AMINO_ACID_BIOSYNTHETIC_PROCESS 15 0.01 Up 0.299014410254568 0.649132108001838 0.895352023662341
GO_REGULATION_OF_PEPTIDASE_ACTIVITY 306 0.01 Down 0.453160044923819 0.649207644236802 0.895352023662341
GO_LIVER_REGENERATION 25 0.01 Down 0.235472935809368 0.649378908004457 0.895384817282548
GO_REGULATION_OF_CYCLASE_ACTIVITY 19 0.01 Up 0.42520362207575 0.650014814366453 0.896058114899803
GO_REGULATION_OF_SKELETAL_MUSCLE_TISSUE_DEVELOPMENT 34 0.01 Down 0.358489602656838 0.65037109456346 0.896219507371731
GO_CILIUM_OR_FLAGELLUM_DEPENDENT_CELL_MOTILITY 20 0.01 Up 0.254750199542201 0.650427137124007 0.896219507371731
GO_RESPONSE_TO_CORTICOSTEROID 106 0.01 Down 0.486707506847924 0.650944151839008 0.89652092348334
GO_DYNEIN_LIGHT_CHAIN_BINDING 19 0.01 Down 0.303381679985913 0.651025797667063 0.89652092348334
GO_RESPONSE_TO_LEUKEMIA_INHIBITORY_FACTOR 80 0.01 Down 0.322137713446643 0.651223503832561 0.89652092348334
GO_POSITIVE_REGULATION_OF_IMMUNOGLOBULIN_PRODUCTION 31 0.01 Down 0.343224240437179 0.651237745430122 0.89652092348334
GO_ERROR_FREE_TRANSLESION_SYNTHESIS 19 0.01 Up 0.112426591036128 0.651384251932962 0.89652092348334
GO_CELLULAR_RESPONSE_TO_INTERFERON_BETA 15 0.01 Down 0.19463869007577 0.651586085036753 0.89659544928788
GO_COP9_SIGNALOSOME 33 0.01 Up 0.198056176470966 0.651838202945966 0.896739118532735
GO_REGULATION_OF_RESPONSE_TO_OXIDATIVE_STRESS 74 0.01 Up 0.361753756111769 0.652208958453895 0.89701307832617
GO_PEPTIDYL_ARGININE_N_METHYLATION 10 0.01 Down 0.10599984574956 0.652514814803323 0.89701307832617
GO_PROTEIN_ARGININE_N_METHYLTRANSFERASE_ACTIVITY 10 0.01 Down 0.10599984574956 0.652514814803323 0.89701307832617
GO_NEGATIVE_REGULATION_OF_ANION_TRANSPORT 17 0.01 Up 0.387891374452162 0.652829398664385 0.89701307832617
GO_POSITIVE_REGULATION_OF_BONE_MINERALIZATION 27 0.01 Up 0.652031907216516 0.652848359922987 0.89701307832617
GO_AGING 240 0.01 Down 0.358092087257504 0.652923866434417 0.89701307832617
GO_ESTABLISHMENT_OF_PROTEIN_LOCALIZATION_TO_CHROMOSOME 25 0.01

Up 0.108118391094394 0.653074539033103
0.897017087436644
GO_MUSCLE_ORGAN_MORPHOGENESIS 50 0.01 Up
0.473957682192096 0.653253015429775 0.897059275429578
GO_BICARBONATE_TRANSPORT 20 0.01 Up
0.463653669496394 0.653973267820333 0.897845253038725
GO_NEGATIVE_REGULATION_OF_CELL_GROWTH 141 0.01 Down
0.447395052573176 0.654229179318957 0.897993521963686
GO_GASTRULATION 132 0.01 Up 0.456038661651106
0.654567489939453 0.89823136538151
GO_POSITIVE_REGULATION_OF_G1_S_TRANSITION_OF_MITOTIC_CELL_CYCLE 31
0.01 Up 0.277937003966959 0.654698367948144
0.89823136538151
GO_RNA_POLYMERASE_II_TRANSCRIPTION_FACTOR_COMPLEX 136 0.01
Down 0.205176063556851 0.654875237827029
0.898271027755963
GO_REGULATION_OF_MICROTUBULE_BINDING 10 0.01 Down
0.12135415417311 0.6555280657962 0.898351786556688
GO_BLASTOCYST_DEVELOPMENT 77 0.01 Down
0.247601997362374 0.655610377152566 0.898351786556688
GO_HEART_FORMATION 20 0.01 Up 0.489855059890928
0.655860092400884 0.898351786556688
GO_MITOTIC_G2_M_TRANSITION_CHECKPOINT 28 0.01 Down
0.154350812857914 0.655883641751523 0.898351786556688
GO_REGULATION_OF_DNA_REPAIR 108 0.01 Down
0.21114359837252 0.656012059170409 0.898351786556688
GO_POSTSYNAPTIC_MEMBRANE 148 0.01 Down
0.311817708608245 0.656101790911759 0.898351786556688
GO_REGULATION_OF_ACTIVIN_RECEPTOR_SIGNALING_PATHWAY 18 0.01
Down 0.346198510266445 0.656218059156965
0.898351786556688
GO_TORC2_COMPLEX 12 0.01 Up 0.180369928485722
0.656230424679282 0.898351786556688
GO_INORGANIC_ANION_TRANSPORT 94 0.01 Down
0.364983869443112 0.656265882618829 0.898351786556688
GO_RESPONSE_TO_CALCIUM_ION 102 0.01 Up
0.433252127857213 0.656506078691393 0.898477999038649
GO_PROTEIN_LOCALIZATION_TO_MICROTUBULE_ORGANIZING_CENTER 31
0.01 Down 0.176310420634675 0.657010899298722
0.89878514432749
GO_NEGATIVE_REGULATION_OF_ADAPTIVE_IMMUNE_RESPONSE 32 0.01
Down 0.496261292214807 0.657026597022797
0.89878514432749
GO_TRANSITION_METAL_ION_BINDING 808 0.01 Down
0.335456176286994 0.657297014079727 0.898952505626947
GO_G_RICH_STRAND_TELOMERIC_DNA_BINDING 10 0.01 Down
0.0779839702840842 0.657709402175153 0.899014078319263
GO_TIM23_MITOCHONDRIAL_IMPORT_INNER_MEMBRANE_TRANSLOCASE_COMPLEX
14 0.01 Down 0.126080713421953
0.657892272726674 0.899014078319263
GO_PROTEIN_KINASE_C_ACTIVATING_G_PROTEIN_COUPLED_RECEPTOR_SIGNALING_
PATHWAY 18 0.01 Up 0.331144469972721
0.658035476688616 0.899014078319263
GO_REGULATION_OF_LIPID_CATABOLIC_PROCESS 35 0.01 Up

0.441490102468932 0.658039559193884
0.899014078319263
GO_NEGATIVE_REGULATION_OF_PROTEIN_BINDING 81 0.01 Up
0.272951130834766 0.65808245166378
0.899014078319263
GO_REGULATION_OF_PH 74 0.01 Down 0.36560756296762
0.658863742071735 0.89987891521203
GO_REPLICATION_FORK_PROCESSING 31 0.01 Up
0.195118798946953 0.65909414232609 0.899991124170421
GO_HYDROLASE_ACTIVITY_ACTING_ON_ESTER_BONDS 566 0.01 Up
0.275374112403762 0.659952276122101
0.900637718696128
GO_SMALL_NUCLEOLAR_RIBONUCLEOPROTEIN_COMPLEX 21 0.01 Up
0.114895230824395 0.660001043930656
0.900637718696128
GO_POSITIVE_REGULATION_OF_B_CELL_ACTIVATION 55 0.01 Down
0.3979673626614 0.660012717917818 0.900637718696128
GO_MCM_COMPLEX 11 0.01 Up 0.126497418633457
0.661015143716418 0.901488746559377
GO_RECEPTOR_LOCALIZATION_TO_SYNAPSE 32 0.01 Up
0.300626807486492 0.661166121916956 0.901488746559377
GO_NEGATIVE_REGULATION_OF_ACTIN_FILAMENT_POLYMERIZATION 42 0.01
Down 0.310152776918292 0.661247520068355
0.901488746559377
GO_ACTIN_BINDING 324 0.01 Down 0.329292777492347
0.661484779329381 0.901488746559377
GO_NEGATIVE_REGULATION_OF_EXTRINSIC_APOPTOTIC_SIGNALING_PATHWAY_VIA_
DEATH_DOMAIN_RECEPTORS 26 0.01 Down 0.36760268313989
0.661538073541808 0.901488746559377
GO_CELLULAR_COPPER_ION_HOMEOSTASIS 13 0.01 Down
0.12244123645473 0.661539713593382 0.901488746559377
GO_REGULATION_OF_MICROTUBULE_DEPOLYMERIZATION 24 0.01 Down
0.266170985198778 0.661675812002732
0.901488746559377
GO_REGULATION_OF_CELL_CYCLE_ARREST 90 0.01 Up
0.2700355507911 0.662415967114398 0.902108042055415
GO_NEGATIVE_REGULATION_OF_BIOMINERAL_TISSUE_DEVELOPMENT 15 0.01
Up 0.867921360958535 0.662427549247742
0.902108042055415
GO_REGULATION_OF_UBIQUITIN_PROTEIN_TRANSFERASE_ACTIVITY 46 0.01
Down 0.202752916488172 0.662803457762142
0.902167167549885
GO_LOW_DENSITY_LIPOPROTEIN_PARTICLE_RECEPTOR_ACTIVITY 11 0.01
Down 0.443935524684382 0.662850226260993
0.902167167549885
GO_IRES_DEPENDENT_VIRAL_TRANSLATIONAL_INITIATION 10 0.01
Down 0.113241683906597 0.662916773915341
0.902167167549885
GO_VASCULAR_ENDOTHELIAL_GROWTH_FACTOR_PRODUCTION 26 0.01
Down 0.686229784978767 0.663106904465983
0.902223670330117
GO_NEGATIVE_REGULATION_OF_INTERFERON_BETA_PRODUCTION 14 0.01
Down 0.426292893245649 0.663500444480619
0.90255684482228

GO_RUFFLE_ASSEMBLY	36	0.01	Down	0.241130396991123
	0.663709876480062	0.902639484791768
GO_PROTEIN_KINASE_A_SIGNALING	20	0.01	Up
0.405243389865372	0.664085526157192	0.902818766843374
GO_POSITIVE_REGULATION_OF_ENDOTHELIAL_CELL_DIFFERENTIATION	11
	0.01	Up	0.387747507592912	0.664139122504119
	0.902818766843374
GO_CERAMIDE_BINDING	11	0.01	Up	0.349643379154805
	0.664428021655431	0.902836661662597
GO_REGULATION_OF_CATION_TRANSMEMBRANE_TRANSPORT	192	0.01	Down
	0.491412944415745	0.664449712454041
0.902836661662597
GO_REGULATION_OF_STORE_OPERATED_CALCIUM_ENTRY	11	0.01	Up
	0.363713090423784	0.665028142942458
0.903063481813179
GO_SARCOPLASMIC_RETICULUM_CALCIUM_ION_TRANSPORT	22	0.01	Up
	0.323371759729696	0.665108789400155
0.903063481813179
GO_INTERCALATED_DISC	35	0.01	Up	0.378089451955523
	0.665170043346771	0.903063481813179
GO_VCP_NPL4_UFD1_AAA_ATPASE_COMPLEX	10	0.01	Up
0.146461189088357	0.665211643990864	0.903063481813179
GO_CELL_MORPHOGENESIS_INVOLVED_IN_NEURON_DIFFERENTIATION	436
	0.01	Down	0.351213035705437	0.665828190835295
	0.903698400751415
GO_CONTRACTILE_ACTIN_FILAMENT_BUNDLE_ASSEMBLY	81	0.01	Down
	0.265426299211109	0.666689064707151
0.904288539319122
GO_SOMITE_DEVELOPMENT	59	0.01	Down	0.409582962692567
	0.666917637613471	0.904288539319122
GO_ENDOPEPTIDASE_REGULATOR_ACTIVITY	97	0.01	Down
0.665871464534868	0.666928551466326	0.904288539319122
GO_MALE_GAMETE_GENERATION	307	0.01	Up
0.275437766830474	0.666950741587812	0.904288539319122
GO_REGULATION_OF_CYSTEINE_TYPE_ENDOPEPTIDASE_ACTIVITY	189	0.01
	Down	0.392047542100572	0.667009111615351
0.904288539319122
GO_HOMOLOGOUS_CHROMOSOME_SEGREGATION	25	0.01	Up
0.252015426238401	0.667175358775333	0.904288539319122
GO_SUPRAMOLECULAR_COMPLEX	631	0.01	Up
0.376945749510186	0.667305659059408	0.904288539319122
GO_TRANSCRIPTION_FACTOR_TFIID_COMPLEX	36	0.01	Down
0.139049447203081	0.667606494244663	0.904292984177178
GO_NEGATIVE_REGULATION_OF_LIPID_CATABOLIC_PROCESS	15	0.01
	Down	0.646416913618041	0.667719023032641
0.904292984177178
GO_LINOLEIC_ACID_METABOLIC_PROCESS	12	0.01	Up
0.352537161169924	0.667755797737817	0.904292984177178
GO_DNA_TOPOLOGICAL_CHANGE	10	0.01	Down
0.206980162465732	0.66810881497341	0.904468094170856
GO_GOLGI_ASSOCIATED_VESICLE	154	0.01	Down
0.299173509522652	0.668317011715427	0.904468094170856
GO_MAST_CELL_MIGRATION	10	0.01	Down	0.356268611501577
	0.668332049160016	0.904468094170856

GO_REGULATION_OF_INSULIN_LIKE_GROWTH_FACTOR_RECEPTOR_SIGNALING_PATHWAY 13 0.01 Up 0.687343810544439 0.668500783725978 0.904494820147184
GO_MICROTUBULE_BASED_PROCESS 606 0.01 Down 0.23427360391212 0.669007901960069 0.9047695830818
GO_REGULATION_OF_SYNAPTIC_PLASTICITY 103 0.01 Up 0.341628287295087 0.669083457319387 0.9047695830818
GO_HETEROCHROMATIN_ASSEMBLY 10 0.01 Down 0.396095416999981 0.669150951743911 0.9047695830818
GO_POSITIVE_REGULATION_OF_CYCLIN_DEPENDENT_PROTEIN_KINASE_ACTIVITY 30 0.01 Down 0.277225264329967 0.66982293180018 0.905121045182862
GO_CARDIAC_MUSCLE_CELL_ACTION_POTENTIAL_INVOLVED_IN_CONTRACTION 33 0.01 Up 0.520149746467621 0.669834992644025 0.905121045182862
GO_PRESYNAPSE 296 0.01 Up 0.289513670300871 0.669858154506111 0.905121045182862
GO_NEGATIVE_REGULATION_OF_CELLULAR_PROTEIN_LOCALIZATION 89 0.01 Down 0.309769042652068 0.67021387349515 0.905400183798188
GO_REGULATION_OF_ACTOMYOSIN_STRUCTURE_ORGANIZATION 74 0.01 Down 0.244145247332548 0.670524165013762 0.905505750699365
GO_POSITIVE_REGULATION_OF_AMINE_TRANSPORT 16 0.01 Down 0.432652326390057 0.670693525930381 0.905505750699365
GO_NEGATIVE_REGULATION_OF_INTERLEUKIN_12_PRODUCTION 11 0.01 Up 0.692539971365521 0.670739476345749 0.905505750699365
GO_PROTEIN_DISULFIDE_ISOMERASE_ACTIVITY 14 0.01 Down 0.172811187367255 0.671140956776062 0.905846320272893
GO_APOPTOTIC_PROCESS_INVOLVED_IN_DEVELOPMENT 27 0.01 Up 0.448974672499271 0.671469974488546 0.906088956461428
GO_MEMBRANE_ORGANIZATION 673 0.01 Down 0.269646678116185 0.67209189756364 0.906439881182634
GO_AMYLOID_FIBRIL_FORMATION 12 0.01 Down 0.534880200771364 0.672147205270339 0.906439881182634
GO_MEMBRANE_DOCKING 163 0.01 Down 0.178206056496158 0.672177951751642 0.906439881182634
GO_SMALL_RIBOSOMAL_SUBUNIT_RRNA_BINDING 11 0.01 Up 0.12403337540468 0.672374762616276 0.906503927124897
GO_REGULATION_OF_LIPID_KINASE_ACTIVITY 45 0.01 Down 0.454312800530923 0.6728056164854 0.90688341422799
GO_ACETYL_COA_BIOSYNTHETIC_PROCESS_FROM_PYRUVATE 15 0.01 Up 0.158550923918722 0.673389716220869 0.90746924909587
GO_PRONUCLEUS 14 0.01 Down 0.217267591077881 0.673652961500516 0.907622532017229
GO_MITOPHAGY 20 0.01 Down 0.117224221473298 0.674038598010757 0.907940609834325
GO_RESPONSE_TO_ESTROGEN 50 0.01 Up 0.364842246489722 0.674766497126221 0.908643696630142
GO_REGULATION_OF_CHOLESTEROL_METABOLIC_PROCESS 11 0.01 Up

0.40349866393324 0.674859895915881
0.908643696630142
GO_RENAL_SYSTEM_DEVELOPMENT 225 0.01 Up
0.54401050369593 0.675232979014428 0.908834010284245
GO_NEGATIVE_REGULATION_OF_NEURON_APOPTOTIC_PROCESS 108 0.01
Up 0.39490305592323 0.675300645757244
0.908834010284245
GO_OLIGODENDROCYTE_DEVELOPMENT 31 0.01 Up
0.310505706761531 0.676094128576002 0.909700233728924
GO_RESPONSE_TO_RADIATION 346 0.01 Down
0.251214086510929 0.676491699333065 0.909832274594559
GO_PROTEIN_TRANSMEMBRANE_IMPORT_INTO_INTRACELLULAR_ORGANELLE 32
0.01 Down 0.109414011593772 0.676565768847723
0.909832274594559
GO_RESPONSE_TO_COCAINE 25 0.01 Up 0.350172921076832
0.676641857979646 0.909832274594559
GO_S_ADENOSYLMETHIONINE_METABOLIC_PROCESS 11 0.01 Up
0.206747740305728 0.67738911798565
0.910498966521624
GO_RENAL_WATER_HOMEOSTASIS 22 0.01 Down
0.474736728441117 0.677447303830429 0.910498966521624
GO_DNA_METABOLIC_PROCESS 716 0.01 Down
0.181108161032062 0.677621457955595 0.910498966521624
GO_NEGATIVE_REGULATION_OF_DNA_TEMPLATED_TRANSCRIPTION_ELONGATION
12 0.01 Up 0.20793554618942
0.677737576957868 0.910498966521624
GO_FOREBRAIN_CELL_MIGRATION 41 0.01 Down
0.330347411627594 0.678383750126562 0.910916363545746
GO_PANCREAS_DEVELOPMENT 48 0.01 Down 0.477132204424914
0.678589438090159 0.910916363545746
GO_ALPHA_LINOLENIC_ACID_METABOLIC_PROCESS 11 0.01 Up
0.124626662857752 0.678593023602938
0.910916363545746
GO_PROTEIN_IMPORT_INTO_PEROXISOME_MATRIX 12 0.01 Up
0.125064265961534 0.678735708751228
0.910916363545746
GO_LYSOSOME_LOCALIZATION 56 0.01 Down
0.322832851913158 0.678826604086138 0.910916363545746
GO_POSITIVE_REGULATION_OF_DOUBLE_STRAND_BREAK_REPAIR 27 0.01
Up 0.235212903491804 0.679151459468731
0.910916363545746
GO_PROFILIN_BINDING 10 0.01 Up 0.235539122198575
0.679251197417391 0.910916363545746
GO_HETEROPHILIC_CELL_CELL_ADHESION_VIA_PLASMA_MEMBRANE_CELL_ADHESION
_MOLECULES 30 0.01 Up 0.480250535055454
0.67992766179806 0.910916363545746
GO_LIPID_BIOSYNTHETIC_PROCESS 519 0.01 Up
0.312873718952432 0.680139907306407 0.910916363545746
GO_AMYLOID_BETA_CLEARANCE 29 0.01 Down
0.714329182675347 0.680354123240523 0.910916363545746
GO_ASTROCYTE_DEVELOPMENT 28 0.01 Down
0.620953102208512 0.680783196159162 0.910916363545746
GO_CYTOSKELETAL_PART 1254 0.01 Down 0.265645877788205
0.680851611331333 0.910916363545746

GO_ATPASE_ACTIVITY_COUPLED_TO_MOVEMENT_OF_SUBSTANCES 98 0.01
Up 0.281420888991043 0.680956007573919
0.910916363545746
GO_PEPTIDASE_ACTIVITY 423 0.01 Down 0.398424902144314
0.681106292236202 0.910916363545746
GO_CELLULAR_RESPONSE_TO_DNA_DAMAGE_STIMULUS 757 0.01 Down
0.196727601880965 0.681115145196044
0.910916363545746
GO_EPITHELIAL_CELL_MATURATION 11 0.01 Down
0.606012528712223 0.681458806980914 0.910916363545746
GO_SCHMIDT_LANTERMAN_INCISURE 10 0.01 Down
0.164189041100904 0.681545573932136 0.910916363545746
GO_SOMATODENDRITIC_COMPARTMENT 531 0.01 Down
0.311909280694426 0.681585522355418 0.910916363545746
GO_LENS_FIBER_CELL_DEVELOPMENT 11 0.01 Up
0.177156486968156 0.681703561148327 0.910916363545746
GO_CIRCULATORY_SYSTEM_DEVELOPMENT 781 0.01 Up
0.468716748248806 0.681710777766485 0.910916363545746
GO_CARDIAC_MUSCLE_TISSUE_DEVELOPMENT 137 0.01 Down
0.426397695660825 0.681968732278711 0.910916363545746
GO_CALCIUM_ACTIVATED_CATION_CHANNEL_ACTIVITY 13 0.01 Up
0.320147276397946 0.681990667786874
0.910916363545746
GO_REGULATION_OF_LONG_TERM_NEURONAL_SYNAPTIC_PLASTICITY 17 0.01
Down 0.195304148123293 0.682021709067124
0.910916363545746
GO_REGULATION_OF_CARDIAC_MUSCLE_CONTRACTION_BY_CALCIUM_ION_SIGNALING
17 0.01 Down 0.334042256665346
0.682227267716616 0.910916363545746
GO_LIGAND_GATED_CATION_CHANNEL_ACTIVITY 39 0.01 Up
0.471920639861346 0.68222994697167 0.910916363545746
GO_RNA_MODIFICATION 150 0.01 Up 0.149233796256285
0.682290328306028 0.910916363545746
GO_NEGATIVE_REGULATION_OF_PHAGOCYTOSIS 16 0.01 Up
0.3802613351695 0.682317569888875 0.910916363545746
GO_ORGANOPHOSPHATE_ESTER_TRANSMEMBRANE_TRANSPORTER_ACTIVITY 22
0.01 Down 0.207671302532796 0.682326626373829
0.910916363545746
GO_NEGATIVE_REGULATION_OF_KINASE_ACTIVITY 210 0.01 Down
0.313631507877297 0.682399542316925
0.910916363545746
GO_RETROGRADE_AXONAL_TRANSPORT 14 0.01 Down
0.369164660651645 0.68292963587072 0.91123059633182
GO_PROTEIN_TARGETING_TO_MITOCHONDRION 96 0.01 Up
0.117976216123051 0.682935136437124 0.91123059633182
GO_DNA_LIGATION 17 0.01 Up 0.258228658447328
0.683473248753249 0.911748207686436
GO_NEGATIVE_REGULATION_OF_LIPASE_ACTIVITY 12 0.01 Down
0.456966253913983 0.683871650149518
0.911967182130232
GO_MESENCHYME_DEVELOPMENT 187 0.01 Down
0.502169233080987 0.683937832356934 0.911967182130232
GO_REGULATION_OF_VASCULAR_ENDOTHELIAL_GROWTH_FACTOR_RECEPTOR_SIGNALI
NG_PATHWAY 22 0.01 Up 0.37337826969186

0.684146590289153	0.912045223901065
GO_CELLULAR_COMPONENT_ASSEMBLY_INVOLVED_IN_MORPHOGENESIS	72
	0.01	Up	0.423397820015237	0.68430211433386
	0.912051685912532
GO_PSEUDOURIDINE_SYNTHESIS	17	0.01	Up
0.124821277931341	0.684451899360483	0.912051685912532
GO_VISUAL_BEHAVIOR	35	0.01	Up	0.271090457192735
	0.684985540782287	0.912206762597037
GO_NEGATIVE_REGULATION_OF_TRANSMEMBRANE_TRANSPORT	78	0.01
	Down	0.418359477622857	0.685110256978754
0.912206762597037
GO_OXYGEN_BINDING	11	0.01	Down	0.658044287130844
	0.685195827717436	0.912206762597037
GO_SKELETAL_MUSCLE_CELL_PROLIFERATION	10	0.01	Down
0.400144339172215	0.685220224367547	0.912206762597037
GO_CHLORIDE_TRANSMEMBRANE_TRANSPORTER_ACTIVITY	45	0.01	Down
	0.418693403592787	0.68535098180245
0.912206762597037
GO_REGULATION_OF_ENDOTHELIAL_CELL_CHEMOTAXIS	15	0.01	Up
	0.443897564311482	0.685586430134802
0.912206762597037
GO_BILE_ACID_AND_BILE_SALT_TRANSPORT	13	0.01	Up
0.819011492438038	0.685834835625441	0.912206762597037
GO_REGULATION_OF_T_CELL_RECEPTOR_SIGNALING_PATHWAY	31	0.01
	Down	0.453314187208543	0.686085380454559
0.912206762597037
GO_CELLULAR_RESPONSE_TO_VITAMIN_D	16	0.01	Up
0.351578373086526	0.686118629118337	0.912206762597037
GO_NEURON_PART	1117	0.01	Down	0.294534940460631
0.686156843975103	0.912206762597037
GO_GLANDULAR_EPITHELIAL_CELL_DIFFERENTIATION	32	0.01	Down
	0.418368126926397	0.686321859038883
0.912206762597037
GO_NEGATIVE_REGULATION_OF_MICROTUBULE_POLYMERIZATION	10	0.01
	Up	0.29524537158458	0.68637135423213
0.912206762597037
GO_REGULATION_OF_ASPARTIC_TYPE_PEPTIDASE_ACTIVITY	11	0.01
	Down	0.397342717337346	0.68707659191606
0.912705781580434
GO_POSITIVE_REGULATION_OF_TRANSCRIPTION_BY_RNA_POLYMERASE_III	10
	0.01	Down	0.175828530811113	0.687172268120007
	0.912705781580434
GO_MRNA_TRANSCRIPTION	20	0.01	Down	0.276033713734886
	0.687247502292029	0.912705781580434
GO_CELLULAR_TRANSITION_METAL_ION_HOMEOSTASIS	81	0.01	Up
	0.40639056680006	0.687399477917998
0.912705781580434
GO_NEUROTRANSMITTER_METABOLIC_PROCESS	79	0.01	Down
0.495761921400084	0.68797247149794	0.912705781580434
GO_NEURON_DEVELOPMENT	805	0.01	Down	0.360377592852965
	0.688036573732888	0.912705781580434
GO_EMBRYONIC_DIGESTIVE_TRACT_MORPHOGENESIS	11	0.01	Up
	0.709838098112591	0.688080463213882
0.912705781580434

GO_POSITIVE_REGULATION_OF_DNA_DEPENDENT_DNA_REPLICATION 11 0.01 Up 0.305918923602002 0.688130102724475 0.912705781580434
GO_CELLULAR_MODIFIED_AMINO_ACID_METABOLIC_PROCESS 29 0.01 Down 0.375445230842535 0.688167943735348 0.912705781580434
GO_ENDONUCLEASE_ACTIVITY 99 0.01 Up 0.21422341587715 0.688426077649756 0.912705781580434
GO_ARTERY_DEVELOPMENT 72 0.01 Up 0.609038279508829 0.688536223898069 0.912705781580434
GO_ORGANELLE_FISSION 356 0.01 Up 0.233016919689082 0.688550894356512 0.912705781580434
GO_LYSOPHOSPHOLIPID_ACYLTRANSFERASE_ACTIVITY 16 0.01 Down 0.265622456998103 0.688945166101594 0.913029055534332
GO_RESPONSE_TO_ANESTHETIC 39 0.01 Up 0.352723959631205 0.689446095978875 0.913493506915702
GO_HISTONE_EXCHANGE 36 0.01 Up 0.20530131630494 0.690445863142846 0.914519855383477
GO_RRNA_CONTAINING_RIBONUCLEOPROTEIN_COMPLEX_EXPORT_FROM_NUCLEUS 16 0.01 Down 0.113166789001726 0.690663668038867 0.914519855383477
GO_METANEPHRIC_MESENCHYME_DEVELOPMENT 11 0.01 Up 0.625678272864777 0.690672630033477 0.914519855383477
GO_INTRINSIC_APOPTOTIC_SIGNALING_PATHWAY_IN_RESPONSE_TO_DNA_DAMAGE 98 0.01 Up 0.286643993692205 0.69086120102102 0.914550324299775
GO_INOSITOL_PHOSPHATE_METABOLIC_PROCESS 40 0.01 Up 0.274723856379649 0.691311866514666 0.914550324299775
GO_ANIMAL_ORGAN_FORMATION 44 0.01 Up 0.489033493123346 0.691350643489982 0.914550324299775
GO_POSITIVE_REGULATION_OF_VASCULAR_ENDOTHELIAL_GROWTH_FACTOR_PRODUCTION 22 0.01 Up 0.679499574040781 0.691404276686742 0.914550324299775
GO_STEROID_BIOSYNTHETIC_PROCESS 139 0.01 Down 0.37908708371897 0.691448853324982 0.914550324299775
GO_DETECTION_OF_LIGHT_STIMULUS 28 0.01 Down 0.187723111260794 0.691619103328124 0.914576252734707
GO_CELL_PART_MORPHOGENESIS 513 0.01 Down 0.345092047455784 0.692151993711968 0.914591268918618
GO_NEGATIVE_REGULATION_OF_CELLULAR_RESPONSE_TO_DRUG 14 0.01 Down 0.248766035205878 0.692191898642494 0.914591268918618
GO_POSITIVE_REGULATION_OF_INTRACELLULAR_PROTEIN_TRANSPORT 138 0.01 Down 0.224137644178638 0.692337166642593 0.914591268918618
GO_MYOBLAST_FUSION 24 0.01 Down 0.665964340050585 0.692383463286042 0.914591268918618
GO_CYTOKINE_BINDING 83 0.01 Up 0.673525014259904 0.692619644271692 0.914591268918618
GO_PROLINE_RICH_REGION_BINDING 18 0.01 Down 0.182816298912239 0.692661060465738 0.914591268918618
GO_MULTICELLULAR_ORGANISM_AGING 23 0.01 Up 0.268613911480626 0.692685003209983 0.914591268918618

GO_REGULATION_OF_CHROMOSOME_SEGREGATION 88 0.01 Up
0.201979711595773 0.693118035019306 0.914660017352129
GO_INTERLEUKIN_2_BIOSYNTHETIC_PROCESS 14 0.01 Down
0.396943922782653 0.693318378831349 0.914660017352129
GO_RESPONSE_TO_ESTRADIOL 85 0.01 Up
0.419473980459907 0.69340904038228 0.914660017352129
GO_TRANSCRIPTION_BY_RNA_POLYMERASE_I 60 0.01 Down
0.144639076821902 0.693444533140169 0.914660017352129
GO_POST_EMBRYONIC_ANIMAL_ORGAN_DEVELOPMENT 15 0.01 Up
0.412756821145353 0.693490373887638
0.914660017352129
GO_3_5_EXODEOXYRIBONUCLEASE_ACTIVITY 11 0.01 Up
0.194562591807873 0.694181359849756 0.915372509914828
GO_NUCLEAR_ENVELOPE_ORGANIZATION 48 0.01 Up
0.190532931598101 0.694521268689632 0.915621850643812
GO_POSITIVE_REGULATION_OF_REGULATORY_T_CELL_DIFFERENTIATION 10
0.01 Up 0.430207223097536 0.694819852846981
0.915816614553631
GO_RESPONSE_TO_RETINOIC_ACID 71 0.01 Down
0.425289938044286 0.694971136480721 0.915817184626537
GO_FOREBRAIN_REGIONALIZATION 11 0.01 Up
0.489473232120498 0.695290391989419 0.916037538125293
GO_NEGATIVE_REGULATION_OF_LOCOMOTION 255 0.01 Up
0.497389510130083 0.695440127362786 0.916037538125293
GO_DNA_HELICASE_COMPLEX 15 0.01 Up 0.0918927576167503
0.69606248843676 0.916226224982215
GO_DEATH_RECEPTOR_BINDING 16 0.01 Down
0.278113800361684 0.696367482077768 0.916226224982215
GO_SECRETORY_GRANULE 616 0.01 Up 0.421429610085657
0.696378752887752 0.916226224982215
GO_REGULATION_OF_HISTONE_UBIQUITINATION 10 0.01 Down
0.109785204357193 0.696466306211104 0.916226224982215
GO_SEQUESTERING_OF_CALCIUM_ION 92 0.01 Up
0.556985866111642 0.696473114992161 0.916226224982215
GO_NEGATIVE_REGULATION_OF_COLD_INDUCED_THERMOGENESIS 39 0.01
Down 0.337977378766044 0.696493564167992
0.916226224982215
GO_RIBONUCLEASE_ACTIVITY 93 0.01 Down
0.23548721015769 0.696639804730341 0.916226224982215
GO_AXON_PART 266 0.01 Down 0.314905002481493
0.696891520455366 0.916311709163368
GO_LAMIN_BINDING 15 0.01 Down 0.245455734925115
0.697019835943081 0.916311709163368
GO_PROTEIN_PEPTIDYL_PROLYL_ISOMERIZATION 33 0.01 Down
0.205164555036184 0.697157599180629
0.916311709163368
GO_ORGANIC_ACID_SODIUM_SYMPORTER_ACTIVITY 13 0.01 Down
0.642830495074113 0.697767732372682
0.916727027094769
GO_TELENCEPHALON_DEVELOPMENT 172 0.01 Down
0.365940465890371 0.697922250576052 0.916727027094769
GO_NUCLEOSIDE_PHOSPHATE_CATABOLIC_PROCESS 56 0.01 Down
0.320609375287896 0.698224824836923
0.916727027094769

GO_RRNA_MODIFICATION	32	0.01	Up	0.109575786579305
	0.698239371853551	0.916727027094769
GO_PREASSEMBLY_OF_GPI_ANCHOR_IN_ER_MEMBRANE	17	0.01	Up
	0.135740447102998	0.698344834817502
0.916727027094769
GO_MLL1_2_COMPLEX	29	0.01	Down	0.120235229589035
	0.698810982588413	0.916727027094769
GO_NEGATIVE_REGULATION_OF_LEUKOCYTE_MEDIATED_IMMUNITY	34	0.01
	Down	0.499014778600382	0.698846301374292
0.916727027094769
GO_ENDOTHELIAL_CELL_APOPTOTIC_PROCESS	39	0.01	Up
0.493791037942612	0.698880540505293	0.916727027094769
GO_ESTABLISHMENT_OF_PIGMENT_GRANULE_LOCALIZATION	23	0.01
	Up	0.199831010533402	0.699157464841123
0.916727027094769
GO_MAINTENANCE_OF_PROTEIN_LOCALIZATION_IN_ORGANELLE	36	0.01
	Down	0.175922917929348	0.699249856886989
0.916727027094769
GO_RNA_POLYMERASE_II_ACTIVATING_TRANSCRIPTION_FACTOR_BINDING	28
	0.01	Down	0.283622182027116	0.699330131592699
	0.916727027094769
GO_NOTOCHORD_DEVELOPMENT	13	0.01	Up
0.361736134018402	0.699410816079314	0.916727027094769
GO_EMBRYONIC_ORGAN_DEVELOPMENT	285	0.01	Down
0.407494759598826	0.699822354815886	0.916727027094769
GO_RESPONSE_TO_FOOD	17	0.01	Up	0.25514620891501
	0.699843237001483	0.916727027094769
GO_VOLTAGE_GATED_ION_CHANNEL_ACTIVITY	78	0.01	Up
0.456506641763301	0.699885063809527	0.916727027094769
GO_CELLULAR_RESPONSE_TO_ALCOHOL	69	0.01	Down
0.423814801264834	0.699889601479864	0.916727027094769
GO_POLYSACCHARIDE_CATABOLIC_PROCESS	20	0.01	Up
0.235262913803767	0.700120018380337	0.91683102493249
GO_ENDOPLASMIC_RETICULUM_TUBULAR_NETWORK	19	0.01	Down
	0.0952802382463054	0.700500192125859
0.917131047314231
GO_CELLULAR_BIOGENIC_AMINE_METABOLIC_PROCESS	35	0.01	Down
	0.417084588999882	0.701123881579048
0.917741641069738
GO_NEGATIVE_REGULATION_OF_CELL_JUNCTION_ASSEMBLY	20	0.01
	Down	0.415689594776096	0.701601502303255
0.917741641069738
GO_RESPONSE_TO_GROWTH_FACTOR	521	0.01	Down
0.412187915708773	0.701652306895973	0.917741641069738
GO_DETERMINATION_OF_ADULT_LIFESPAN	12	0.01	Up
0.327721131804311	0.701692071067898	0.917741641069738
GO_POSITIVE_REGULATION_OF_PEPTIDYL_THREONINE_PHOSPHORYLATION	25
	0.01	Down	0.369665583259348	0.701722401226441
	0.917741641069738
GO_OXIDOREDUCTASE_ACTIVITY_ACTING_ON_A_SULFUR_GROUP_OF_DONORS_NAD_P_
AS_ACCEPTOR	11	0.01	Up	0.141154858607157
0.702384105864262	0.918016687049634
GO_U2_TYPE_SPLICEOSOMAL_COMPLEX	81	0.01	Down
0.0801018677997368	0.702477145226201	0.918016687049634

GO_POSITIVE_REGULATION_OF_PROTEIN_HOMODIMERIZATION_ACTIVITY 10 0.01 Down 0.269119516017916 0.702519325144039 0.918016687049634
GO_CELLULAR_RESPONSE_TO_CADMIUM_ION 28 0.01 Up 0.555489105278484 0.702687553273942 0.918016687049634
GO_ADRENAL_GLAND_DEVELOPMENT 15 0.01 Down 0.407742738408783 0.702986848584389 0.918016687049634
GO_AXON_DEVELOPMENT 378 0.01 Down 0.360213763322733 0.70306913494441 0.918016687049634
GO_NEGATIVE_REGULATION_OF_PROTEIN_LOCALIZATION_TO_CELL_PERIPHERY 22 0.01 Up 0.344737050557285 0.703127468067258 0.918016687049634
GO_SPINDLE_POLE_CENTROSOME 12 0.01 Up 0.118175326258601 0.703418433786883 0.918016687049634
GO_NUCLEOTIDE_EXCISION_REPAIR_COMPLEX 12 0.01 Up 0.145750176348783 0.7036297942413 0.918016687049634
GO_NEURAL_CREST_CELL_MIGRATION 38 0.01 Up 0.535970647972872 0.703697448858753 0.918016687049634
GO_CALCIUM_ION_REGULATED_EXOCYTOSIS 98 0.01 Up 0.302448737624639 0.703814642062271 0.918016687049634
GO_NUCLEOSIDE_SALVAGE 15 0.01 Down 0.243537842325394 0.70381848105793 0.918016687049634
GO_RETINAL_METABOLIC_PROCESS 10 0.01 Down 0.551849188149973 0.703948655469078 0.918016687049634
GO_HAIR_CELL_DIFFERENTIATION 28 0.01 Up 0.322123044282332 0.704128326281015 0.918016687049634
GO_CELLULAR_RESPONSE_TO_FATTY_ACID 41 0.01 Up 0.441322556705623 0.704200907855402 0.918016687049634
GO_MESONEPHROS_DEVELOPMENT 74 0.01 Down 0.565175236459849 0.704473885003028 0.918175387688575
GO_EXECUTION_PHASE_OF_APOPTOSIS 64 0.01 Down 0.299500308811143 0.704939807210808 0.918585440990946
GO_POSITIVE_REGULATION_OF_PROTEIN_MATURATION 17 0.01 Up 0.424645226084468 0.705175658045628 0.918598716063724
GO_ALCOHOL_BINDING 58 0.01 Up 0.422061872104643 0.705252613337674 0.918598716063724
GO_CELLULAR_RESPONSE_TO_OXYGEN_LEVELS 193 0.01 Down 0.271731118338469 0.705491064653481 0.918708980321909
GO_MITOCHONDRIAL_TRANSMEMBRANE_TRANSPORT 84 0.01 Up 0.135467410331845 0.705667776957061 0.918708980321909
GO_OLIGODENDROCYTE_DIFFERENTIATION 64 0.01 Down 0.301410022622068 0.705791250901232 0.918708980321909
GO_INTRACELLULAR_CALCIUM_ACTIVATED_CHLORIDE_CHANNEL_ACTIVITY 10 0.01 Up 0.347657188229113 0.706589509055547 0.918964966440129
GO_ANION_HOMEOSTASIS 30 0.01 Down 0.403756936391044 0.706659445525353 0.918964966440129
GO_ACTIN_CORTICAL_PATCH 11 0.01 Down 0.261860541461519 0.706833284136761 0.918964966440129
GO_POSITIVE_REGULATION_OF_TRANSMEMBRANE_RECEPTOR_PROTEIN_SERINE_THREONINE_KINASE_SIGNALING_PATHWAY 75 0.01 Up 0.39554782843596 0.707203523390406 0.918964966440129

GO_HEMATOPOIETIC_STEM_CELL_PROLIFERATION	17	0.01	Up
	0.437320706584558	0.70737341757235
0.918964966440129
GO_NUCLEOTIDE_SUGAR_BIOSYNTHETIC_PROCESS	23	0.01	Up
	0.275223084351368	0.707416971933515
0.918964966440129
GO_TRNA_METHYLATION	39	0.01	Up	0.140005461858129
	0.707484443676497	0.918964966440129
GO_PRESYNAPTIC_ACTIVE_ZONE	48	0.01	Up
0.274902145241987	0.707632749136539	0.918964966440129
GO_PROTEIN_HETERODIMERIZATION_ACTIVITY	342	0.01	Down
0.324195441187925	0.70775290540297	0.918964966440129
GO_ESTABLISHMENT_OF_PROTEIN_LOCALIZATION_TO_POSTSYNAPTIC_MEMBRANE
	12	0.01	Down	0.234957640242721
0.707766524369001	0.918964966440129
GO_POSITIVE_REGULATION_OF_ACTIN_FILAMENT_BUNDLE_ASSEMBLY	52
	0.01	Down	0.245725048690124	0.707830649541662
	0.918964966440129
GO_CELL_CELL_SIGNALING	1042	0.01	Down	0.395262843920028
	0.707976790149526	0.918964966440129
GO_MITOCHONDRIAL_GENOME_MAINTENANCE	20	0.01	Up
0.172545711642263	0.708103493179117	0.918964966440129
GO_SYNAPSIS	20	0.01	Up	0.229936814315923
0.708162455386274	0.918964966440129
GO_RNA_SPLICING_VIA_ENDONUCLEOLYTIC_CLEAVAGE_AND_LIGATION	16
	0.01	Up	0.147096618645522	0.708299784860156
	0.918964966440129
GO_REGULATION_OF_CHEMOKINE_C_X_C_MOTIF_LIGAND_2_PRODUCTION	10
	0.01	Down	0.536521685037577	0.708409824236502
	0.918964966440129
GO_NEGATIVE_REGULATION_OF_PROTEIN_MODIFICATION_BY_SMALL_PROTEIN_CONJ
UGATION_OR_REMOVAL	76	0.01	Down	0.236636051100633
	0.709338423559597	0.919972990692226
GO_ORGANONITROGEN_COMPOUND_CATABOLIC_PROCESS	1046	0.01	Down
	0.275117797226888	0.709616406479663
0.920002617671709
GO_NEGATIVE_REGULATION_OF_CELLULAR_COMPONENT_ORGANIZATION	547
	0.01	Down	0.297034676679028	0.709664348304499
	0.920002617671709
GO_REGULATION_OF_VACUOLE_ORGANIZATION	39	0.01	Down
0.159337995482465	0.710112820723795	0.920387475365961
GO_MITOCHONDRIAL_PROTEIN_PROCESSING	12	0.01	Up
0.154348068252689	0.710447643628309	0.920624897431689
GO_3_PHOSPHOADENOSINE_5_PHOSPHOSULFATE_METABOLIC_PROCESS	10
	0.01	Down	0.388958783528302	0.71073970407029
	0.920806816775658
GO_CELL_CYCLE_G1_S_PHASE_TRANSITION	225	0.01	Down
0.239504357338214	0.711125882788236	0.921110568467544
GO_REGULATION_OF_MUSCLE_SYSTEM_PROCESS	155	0.01	Down
0.459857369586909	0.711504579741624	0.921329403266787
GO_MACROMOLECULE_DEPALMITOYLATION	10	0.01	Up
0.14584184353308	0.711770829791137	0.921329403266787
GO_PALMITOYL_PROTEIN_HYDROLASE_ACTIVITY	10	0.01	Up
0.14584184353308	0.711770829791137	0.921329403266787

GO_ACTIN_FILAMENT_BUNDLE 56 0.01 Up
0.240229815057211 0.71192886783423 0.921329403266787
GO_INTERLEUKIN_8_SECRETION 19 0.01 Up
0.62975716243583 0.71219686955948 0.921329403266787
GO_CYTOSKELETAL_ADAPTOR_ACTIVITY 13 0.01 Down
0.3115362393094 0.712300111962365 0.921329403266787
GO_PHOSPHORIC_DIESTER_HYDROLASE_ACTIVITY 61 0.01 Up
0.329323376862935 0.712662302504712
0.921329403266787
GO_INNER_EAR_RECEPTOR_CELL_STEREOCILIUM_ORGANIZATION 20 0.01
Down 0.282864681720573 0.712715240655462
0.921329403266787
GO_NEGATIVE_REGULATION_OF_ATPASE_ACTIVITY 12 0.01 Down
0.291275662199992 0.712983587867207
0.921329403266787
GO_DICARBOXYLIC_ACID_TRANSPORT 55 0.01 Up
0.348064526888071 0.713053356200975 0.921329403266787
GO_KINESIN_BINDING 35 0.01 Down 0.278664860642663
0.713097710948826 0.921329403266787
GO_REGULATION_OF_CARTILAGE_DEVELOPMENT 49 0.01 Up
0.619844875194474 0.713135812747174 0.921329403266787
GO_RRNA_METHYLATION 23 0.01 Up 0.106077954565295
0.713267698131098 0.921329403266787
GO_METANEPHRIC_NEPHRON_EPITHELIUM_DEVELOPMENT 14 0.01 Up
0.413848533467457 0.713460027667131
0.921381797057467
GO_POSITIVE_REGULATION_OF_INTERLEUKIN_5_PRODUCTION 11 0.01
Up 0.532982731156302 0.714332237832532
0.921635329882941
GO_NEGATIVE_REGULATION_OF_CAMP_MEDIATED_SIGNALING 15 0.01
Down 0.400558458062907 0.714413874708641
0.921635329882941
GO_REGULATION_OF_AMINO_ACID_TRANSPORT 17 0.01 Up
0.524596503112486 0.714518544050739 0.921635329882941
GO_RETINA_HOMEOSTASIS 38 0.01 Down 0.391373553270866
0.714591106585853 0.921635329882941
GO_NEGATIVE_REGULATION_OF_DEFENSE_RESPONSE 139 0.01 Down
0.424711061479482 0.714804011561913
0.921635329882941
GO_EYE_PHOTORECEPTOR_CELL_DIFFERENTIATION 23 0.01 Down
0.598658763424068 0.714811363883416
0.921635329882941
GO_IRON_ION_TRANSPORT 60 0.01 Up 0.370193191635278
0.715218574173573 0.921635329882941
GO_RESPONSE_TO_TRANSFORMING_GROWTH_FACTOR_BETA 198 0.01 Up
0.40933526627364 9 0.715290129281718
0.921635329882941
GO_NEGATIVE_REGULATION_OF_EPITHELIAL_CELL_MIGRATION 57 0.01
Down 0.458195705046104 0.715335427849946
0.921635329882941
GO_CYCLIC_NUCLEOTIDE_BINDING 19 0.01 Up
0.467399705397376 0.71543106350721 0.921635329882941
GO_TOR_COMPLEX 15 0.01 Up 0.156932737622145
0.715645498716105 0.921635329882941

GO_REGULATION_OF_STRIATED_MUSCLE_CONTRACTION 57 0.01 Down
0.423231390176545 0.715889332228515
0.921635329882941
GO_SEH1_ASSOCIATED_COMPLEX 17 0.01 Up
0.136101700968432 0.715969697486113 0.921635329882941
GO_REGULATION_OF_REGULATED_SECRETORY_PATHWAY 104 0.01 Up
0.359122794963296 0.716130137928514
0.921635329882941
GO_ANATOMICAL_STRUCTURE_FORMATION_INVOLVED_IN_MORPHOGENESIS 780
0.01 Down 0.479311995977188 0.716142914631319
0.921635329882941
GO_PHOSPHATIDYLGLYCEROL_ACYL_CHAIN_REMODELING 11 0.01 Up
0.300963475865701 0.716479360583497
0.921635329882941
GO_MICROTUBULE_PLUS_END 17 0.01 Down 0.124994195426908
0.716781401810595 0.921635329882941
GO_LAMELLIPODIUM_ASSEMBLY 55 0.01 Down
0.272587551558893 0.717056738868129 0.921635329882941
GO_ENZYME_LINKED_RECEPTOR_PROTEIN_SIGNALING_PATHWAY 771 0.01
Down 0.395353350313556 0.717152662646249
0.921635329882941
GO_ANTIGEN_PROCESSING_AND_PRESENTATION 192 0.01 Down
0.332218981033668 0.717347311688922 0.921635329882941
GO_RESPONSE_TO_LEUCINE 13 0.01 Down 0.197813920918004
0.717633515505263 0.921635329882941
GO_3_5_EXONUCLEASE_ACTIVITY 50 0.01 Down
0.153400769909629 0.717635183305554 0.921635329882941
GO_CALCIUM_DEPENDENT_CYSTEINE_TYPE_ENDOPEPTIDASE_ACTIVITY 15
0.01 Down 0.49075099400627 0.717911259108289
0.921635329882941
GO_DNA_BINDING_BENDING 16 0.01 Up 0.276274649772021
0.718110257741562 0.921635329882941
GO_REGULATION_OF_ADENYLATE_CYCLASE_ACTIVITY 14 0.01 Up
0.33751386352678 0.71820643865109
0.921635329882941
GO_BETA_TUBULIN_BINDING 35 0.01 Up 0.234710251583054
0.718227173604996 0.921635329882941
GO_HYALURONAN_BIOSYNTHETIC_PROCESS 11 0.01 Down
0.514751264241783 0.718328927585707 0.921635329882941
GO_NEGATIVE_REGULATION_OF_POTASSIUM_ION_TRANSMEMBRANE_TRANSPORT 13
0.01 Up 0.684540749853063 0.718364344995473
0.921635329882941
GO_ENDOSOMAL_PART 451 0.01 Down 0.289875005058071
0.718401195643907 0.921635329882941
GO_GOLGI_APPARATUS_PART 797 0.01 Down 0.312968584049279
0.718420495903029 0.921635329882941
GO_POSITIVE_REGULATION_OF_T_CELL_MIGRATION 25 0.01 Down
0.725909672666866 0.718697727539628
0.921635329882941
GO_PROTEIN_MATURATION 224 0.01 Up 0.345103340676193
0.718805625904646 0.921635329882941
GO_CONTRACTILE_FIBER 149 0.01 Up 0.416938155344632
0.718819604008599 0.921635329882941
GO_NUCLEOSIDE_METABOLIC_PROCESS 42 0.01 Up

0.301897949119356	0.718891633629627	0.921635329882941
GO_ESTABLISHMENT_OF_EPITHELIAL_CELL_APICAL_BASAL_POLARITY	12
	0.01	Up	0.282296768857389	0.718969679183925
	0.921635329882941
GO_DEVELOPMENTAL_PIGMENTATION	32	0.01	Down
0.41129085314038	0.719408671453144	0.921713531757235
GO_PHOSPHATIDIC_ACID_BIOSYNTHETIC_PROCESS	31	0.01	Down
	0.237245708999158	0.719438842672514
0.921713531757235
GO_PYRIMIDINE_NUCLEOSIDE_BIOSYNTHETIC_PROCESS	14	0.01	Down
	0.227911448181427	0.719486151704421
0.921713531757235
GO_AZOLE_TRANSPORT	16	0.01	Down	0.581265780805749
	0.719854898025295	0.921991368335774
GO_INTRACELLULAR_PROTEIN_TRANSMEMBRANE_TRANSPORT	50	0.01
	Up	0.136966683329147	0.72008294761565
0.922088921108334
GO_RESPONSE_TO_PLATELET_AGGREGATION_INHIBITOR	10	0.01	Down
	0.418475530002556	0.720238831826139
0.922094042179774
GO_POSITIVE_REGULATION_OF_STRESS_FIBER_ASSEMBLY	44	0.01	Down
	0.207923531039233	0.720771370961789
0.922581276219485
GO_POSITIVE_REGULATION_OF_ACTIN_FILAMENT_DEPOLYMERIZATION	10
	0.01	Down	0.349435378652788	0.721094228652458
	0.922693854101084
GO_RESPIRATORY_SYSTEM_DEVELOPMENT	157	0.01	Up
0.484327212998143	0.721163290678578	0.922693854101084
GO_POSITIVE_REGULATION_OF_CATION_CHANNEL_ACTIVITY	33	0.01
	Down	0.369566369180857	0.721460560719966
0.922879701671074
GO_PROTEIN_PROCESSING	161	0.01	Up	0.400990205202974
	0.72210050782754	0.923252165688203
GO_DIOXYGENASE_ACTIVITY	75	0.01	Up	0.325573308405633
	0.722106883699822	0.923252165688203
GO_REGULATION_OF_METANEPHROS_DEVELOPMENT	16	0.01	Up
	0.679009631882602	0.722207961596653
0.923252165688203
GO_CENTRAL_NERVOUS_SYSTEM_NEURON_DEVELOPMENT	54	0.01	Down
	0.386709648261575	0.722369524191747
0.923264290814336
GO_POSITIVE_REGULATION_OF_NEURON_DEATH	74	0.01	Down
0.369765287241198	0.723059985334469	0.923334944955135
GO_NEGATIVE_REGULATION_OF_TRANSPORTER_ACTIVITY	50	0.01	Down
	0.463931111887255	0.723170953930695
0.923334944955135
GO_NEGATIVE_REGULATION_OF_RNA_SPLICING	24	0.01	Down
0.0986854136792974	0.723183038454769	0.923334944955135
GO_NEGATIVE_REGULATION_OF_ANION_TRANSMEMBRANE_TRANSPORT	12	0.01
	Up	0.412021063911509	0.723455604696922
0.923334944955135
GO_KIDNEY_EPITHELIUM_DEVELOPMENT	101	0.01	Down
0.566621743786893	0.723472214824097	0.923334944955135
GO_REGULATION_OF_ENDOCYTOSIS	152	0.01	Up

0.359528686711367	0.723497157500085	0.923334944955135
GO_DISTAL_AXON	193	0.01	Down	0.323227651470387
0.723748479926772	0.923334944955135
GO_AORTA_DEVELOPMENT	46	0.01	Up	0.573542326298537
	0.72386120546098	0.923334944955135
GO_INTERSTRAND_CROSS_LINK_REPAIR	50	0.01	Up
0.165558287803592	0.723939398624969	0.923334944955135
GO_MALE_GENITALIA_DEVELOPMENT	11	0.01	Up
0.326277204814056	0.724081868196968	0.923334944955135
GO_ACTIVATION_OF_PHOSPHOLIPASE_C_ACTIVITY	13	0.01	Up
	0.552848117666777	0.7240977883267	0.923334944955135
GO_TRANSPORT_VESICLE	290	0.01	Up	0.324644810626476
	0.724553121840981	0.923683560942243
GO_NEGATIVE_REGULATION_OF_OXIDOREDUCTASE_ACTIVITY	14	0.01
	Down	0.471929308530062	0.724675473689327
0.923683560942243
GO_RESPONSE_TO_UV	127	0.01	Down	0.217205943255242
	0.725131770520057	0.924008930473073
GO_DOPAMINE_RECEPTOR_BINDING	13	0.01	Down
0.218347976709126	0.725263753165107	0.924008930473073
GO_PURINE_CONTAINING_COMPOUND_CATABOLIC_PROCESS	41	0.01	Down
	0.377884453523309	0.72550353495105
0.924008930473073
GO_ACETYLGLUCOSAMINYLTRANSFERASE_ACTIVITY	46	0.01	Up
	0.353813229242935	0.725539543990305
0.924008930473073
GO_MYOBLAST_PROLIFERATION	11	0.01	Up
0.440862623255578	0.725700100768405	0.924019570420509
GO_KINETOCHORE_ORGANIZATION	18	0.01	Up
0.209856448148733	0.726675726816322	0.924891095698107
GO_POSITIVE_REGULATION_OF_PHOSPHOLIPASE_ACTIVITY	32	0.01
	Up	0.630583121462466	0.726689264780098
0.924891095698107
GO_PROTON_TRANSPORTING_V_TYPE_ATPASE_V0_DOMAIN	10	0.01	Up
	0.202469928726265	0.726958826346187
0.925040250418717
GO_GLIAL_CELL_MIGRATION	38	0.01	Up	0.234571286991322
	0.728001118489854	0.926172420442562
GO_CHROMOSOMAL_REGION	285	0.01	Down	0.169639753350692
	0.72822029050330l	0.926249567721141
GO_TRANSCRIPTION_INITIATION_FROM_RNA_POLYMERASE_III_PROMOTER	12
	0.01	Down	0.141939502494933	0.728400784812997
	0.926249567721141
GO_REGULATION_OF_CARDIAC_MUSCLE_CELL_MEMBRANE_REPOLARIZATION	17
	0.01	Down	0.597463980669084	0.728519467281906
	0.926249567721141
GO_RNA_METHYLTRANSFERASE_ACTIVITY	64	0.01	Up
0.129112337420736	0.728800872782038	0.926283978541154
GO_RECOMBINATIONAL_REPAIR	108	0.01	Up
0.175007435406205	0.728851682670251	0.926283978541154
GO_NUCLEAR_MEMBRANE_PART	10	0.01	Up
0.319751559096855	0.72928295354926	0.926455925359551
GO_CELL_DIVISION	497	0.01	Down	0.235546888397895
	0.729619326876616	0.926455925359551

GO_POSITIVE_REGULATION_OF_PHAGOCYTOSIS_ENGULFMENT 10 0.01 Down 0.677486184950393 0.729693792911487 0.926455925359551
GO_ADAPTIVE_THERMOGENESIS 112 0.01 Down 0.376295013508675 0.729697222323212 0.926455925359551
GO_POSITIVE_REGULATION_OF_CYTOKINE_SECRETION 87 0.01 Down 0.5920747709616 0.729920837886571 0.926455925359551
GO_TOXIN_TRANSPORT 35 0.01 Up 0.302040905652763 0.729945598086878 0.926455925359551
GO_GLYCOSYL_COMPOUND_BIOSYNTHETIC_PROCESS 39 0.01 Up 0.240871504113147 0.730055204565984 0.926455925359551
GO_RESPONSE_TO_BMP 100 0.01 Up 0.5135751154447 0.730280565350685 0.926548236623617
GO_EMBRYONIC_ORGAN_MORPHOGENESIS 176 0.01 Up 0.436559788286374 0.730552384523091 0.926699441378957
GO_NEGATIVE_REGULATION_OF_PROTEIN_MATURATION 24 0.01 Down 0.478933938698562 0.730990885046304 0.92696579235595
GO_TRNA_THREONYLCARBAMOYLADENOSINE_METABOLIC_PROCESS 13 0.01 Up 0.098226080681151 0.731104861223113 0.92696579235595
GO_ERAD_PATHWAY 91 0.01 Up 0.189378150824304 0.731322114148133 0.92696579235595
GO_PROTEIN_LOCALIZATION_TO_CELL_SURFACE 49 0.01 Up 0.377988447567171 0.731373109106408 0.92696579235595
GO_MEIOTIC_CELL_CYCLE 153 0.01 Up 0.242830775634579 0.732013969467633 0.92746800332161
GO_POSITIVE_REGULATION_OF_LIPASE_ACTIVITY 38 0.01 Up 0.564790503357176 0.732088415954432 0.92746800332161
GO_WALKING_BEHAVIOR 18 0.01 Down 0.437758726004529 0.732227662645442 0.92746800332161
GO_PALMITOYLTRANSFERASE_ACTIVITY 35 0.01 Up 0.211959505303859 0.733017639718643 0.928274945918206
GO_NEGATIVE_REGULATION_OF_CILIUM_ASSEMBLY 10 0.01 Up 0.102767459108686 0.733382947669098 0.92854387388928
GO_REGULATION_OF_NEURONAL_SYNAPTIC_PLASTICITY 27 0.01 Up 0.25370527437658 0.733736940282323 0.928626791334369
GO_REGULATION_OF_PROTEIN_TARGETING_TO_MITOCHONDRION 40 0.01 Down 0.12125665564931 0.733754359748142 0.928626791334369
GO_LONG_TERM_MEMORY 21 0.01 Up 0.341128951302947 0.733956471067824 0.928688982045178
GO_FOREBRAIN_GENERATION_OF_NEURONS 35 0.01 Down 0.432844460613043 0.734431552176238 0.928946540318071
GO_EXTRINSIC_COMPONENT_OF_CYTOPLASMIC_SIDE_OF_PLASMA_MEMBRANE 63 0.01 Down 0.379887743175167 0.73466050655039 0.928946540318071
GO_NEGATIVE_REGULATION_OF_MYELOID_LEUKOCYTE_DIFFERENTIATION 34 0.01 Up 0.463266328520535 0.73540646168843 0.929632787835835

GO_REGULATION_OF_DNA_DEPENDENT_DNA_REPLICATION 46 0.01 Down
0.236219731282795 0.73560821670024
0.929632787835835
GO_REGULATION_OF_PLATELET_DERIVED_GROWTH_FACTOR_RECEPTOR_SIGNALING_PATHWAY 20 0.01 Down 0.378197265279007
0.735626194299208 0.929632787835835
GO_REGULATION_OF_ACTIN_FILAMENT_BASED_PROCESS 302 0.01 Down
0.342213988263186 0.735671563688564
0.929632787835835
GO_TRANS_GOLGI_NETWORK 216 0.01 Down 0.268571871726598
0.735916201725778 0.929632787835835
GO_4_HYDROXYPROLINE_METABOLIC_PROCESS 10 0.01 Up
0.277028400219179 0.735927388953883 0.929632787835835
GO_ZYMOGEN_ACTIVATION 37 0.01 Up 0.296555021715658
0.736599092116013 0.929697242575756
GO_VENTRAL_SPINAL_CORD_DEVELOPMENT 20 0.01 Down
0.387426592202179 0.736627550454864 0.929697242575756
GO_FIBRINOLYSIS 15 0.01 Up 0.486732948802019
0.736643218434212 0.929697242575756
GO_POSITIVE_REGULATION_OF_G_PROTEIN_COUPLED_RECEPTOR_SIGNALING_PATHWAY 18 0.01 Up 0.603136408430368
0.736799506764839 0.929697242575756
GO_DIHYDROLIPOYL_DEHYDROGENASE_COMPLEX 10 0.01 Up
0.146706197688874 0.737083659442862 0.929697242575756
GO_CELL_RECOGNITION 87 0.01 Up 0.48779023761163
0.737299213842984 0.929697242575756
GO_CATION_TRANSMEMBRANE_TRANSPORTER_ACTIVITY 340 0.01 Up
0.38401675686182 0.737603559362936
0.929697242575756
GO_POSITIVE_REGULATION_OF_RHO_PROTEIN_SIGNAL_TRANSDUCTION 14
0.01 Down 0.590957954618417 0.737670564290479
0.929697242575756
GO_COCHLEA_MORPHOGENESIS 16 0.01 Up
0.437463812731334 0.737679947514043 0.929697242575756
GO_POSITIVE_REGULATION_OF_TYPE_2_IMMUNE_RESPONSE 14 0.01
Up 0.789824817611524 0.737828043483859
0.929697242575756
GO_NEGATIVE_REGULATION_OF_ANDROGEN_RECEPTOR_SIGNALING_PATHWAY 12
0.01 Down 0.32597701842478 0.737852627997663
0.929697242575756
GO_CELLULAR_RESPONSE_TO_OXIDATIVE_STRESS 251 0.01 Down
0.375831352375862 0.738003941534158
0.929697242575756
GO_FILOPODIUM_ASSEMBLY 49 0.01 Down 0.308051142472065
0.738067453196883 0.929697242575756
GO_POSITIVE_REGULATION_OF_LIPID_LOCALIZATION 50 0.01 Down
0.577355157270147 0.738159343520866
0.929697242575756
GO_NEGATIVE_REGULATION_OF_RESPONSE_TO_BIOTIC_STIMULUS 36 0.01
Up 0.245796121082719 0.73827547462654
0.929697242575756
GO_EPITHELIAL_CELL_CELL_ADHESION 13 0.01 Down
0.47565987005658 0.739008857983947 0.930000423879142
GO_REGULATION_OF_TISSUE_REMODELING 47 0.01 Up

0.543794585044465	0.73921898608609	0.930000423879142
GO_NEGATIVE_REGULATION_OF_PEPTIDE_HORMONE_SECRETION	28	0.01
	Up	0.376478938577451	0.739228702910892
0.930000423879142
GO_RESPONSE_TO_HORMONE	680	0.01	Down	0.332244699839929
	0.739328696389311	0.930000423879142
GO_REGULATION_OF_MORPHOGENESIS_OF_AN_EPITHELIUM	155	0.01	Down
	0.424746496511119	0.739529566508773
0.930000423879142
GO_DNA_CATABOLIC_PROCESS	32	0.01	Up
0.229889154483962	0.73953426662555	0.930000423879142
GO_STRIATED_MUSCLE_CONTRACTION	102	0.01	Down
0.477094997929199	0.739601780938838	0.930000423879142
GO_NEGATIVE_REGULATION_OF_SYNAPTIC_TRANSMISSION	35	0.01	Down
	0.307444277780433	0.739856216871678
0.930000423879142
GO_CELLULAR_AMIDE_METABOLIC_PROCESS	837	0.01	Up
0.212957322488299	0.740127681654738	0.930000423879142
GO_NADPLUS_ADP_RIBOSYLTRANSFERASE_ACTIVITY	18	0.01	Down
	0.155226177066777	0.740463319723144
0.930000423879142
GO_PURINERGIC_RECEPTOR_SIGNALING_PATHWAY	14	0.01	Down
	0.46003750196215	0.740505851168621
0.930000423879142
GO_EMBRYONIC_SKELETAL_SYSTEM_DEVELOPMENT	76	0.01	Up
	0.489819884243949	0.740610631365356
0.930000423879142
GO_LIPID_DROPLET	68	0.01	Up	0.268495220101977
	0.740711423407059	0.930000423879142
GO_LABYRINTHINE_LAYER_BLOOD_VESSEL_DEVELOPMENT	17	0.01	Down
	0.271379724244512	0.740952168752707
0.930000423879142
GO_HINDBRAIN_DEVELOPMENT	102	0.01	Up
0.374306648853439	0.741169961995329	0.930000423879142
GO_PERICENTRIOLAR_MATERIAL	21	0.01	Down
0.265741725448398	0.74131459107344	0.930000423879142
GO_REGULATION_OF_OSTEOCLAST_DIFFERENTIATION	44	0.01	Up
	0.482797417016469	0.741326744995086
0.930000423879142
GO_PROTEIN_LOCALIZATION_TO_ORGANELLE	823	0.01	Down
0.21016866083329	0.74151383354943	0.930000423879142
GO_RRNA_TRANSCRIPTION	30	0.01	Up	0.136163783017498
	0.74155625436558	0.930000423879142
GO_NUCLEAR_MEMBRANE	263	0.01	Down	0.239002281759514
	0.741652584597506	0.930000423879142
GO_MRNA_MODIFICATION	19	0.01	Down	0.117168712054586
	0.741733166269611	0.930000423879142
GO_MACROPHAGE_CHEMOTAXIS	27	0.01	Up
0.602854947350573	0.74204188660823l	0.930053379780359
GO_PROTEIN_HETEROOLIGOMERIZATION	88	0.01	Down
0.305017557858577	0.74208179404646	0.930053379780359
GO_NEURON_PROJECTION	838	0.01	Down	0.30619027431161
	0.742781106924588	0.930737688367219
GO_SOMATIC_STEM_CELL_DIVISION	17	0.01	Down

0.419251709883184	0.745009825440583	0.933337732201771
GO_DISORDERED_DOMAIN_SPECIFIC_BINDING	29	0.01	Down
0.216914803281289	0.745247141729938	0.933442417462854
GO_NEGATIVE_REGULATION_OF_ALCOHOL_BIOSYNTHETIC_PROCESS	12	0.01
	Up	0.741161497521955	0.745731624072801
0.933480095514496
GO_POSITIVE_REGULATION_OF_CELL_CYCLE_G1_S_PHASE_TRANSITION	39
	0.01	Up	0.244123585308109	0.74574051066303
	0.933480095514496
GO_DNA_TEMPLATED_TRANSCRIPTION_ELONGATION	106	0.01	Down
	0.137859810965154	0.745986009537515
0.933480095514496
GO_METAL_ION_TRANSPORT	517	0.01	Down	0.421012438076113
	0.746082673394746	0.933480095514496
GO_NEGATIVE_REGULATION_OF_RESPONSE_TO_OXIDATIVE_STRESS	40	0.01
	Up	0.309040210276598	0.746315858367138
0.933480095514496
GO_SPINAL_CORD_PATTERNING	11	0.01	Up
0.590833337565799	0.746541703758902	0.933480095514496
GO_MODULATION_BY_HOST_OF_VIRAL_PROCESS	20	0.01	Up
0.383545153552759	0.746650466434203	0.933480095514496
GO_MORPHOGENESIS_OF_A_POLARIZED_EPITHELIUM	128	0.01	Down
	0.279161572282587	0.746739153868373
0.933480095514496
GO_CARDIOCYTE_DIFFERENTIATION	99	0.01	Up
0.459523351547042	0.74691441806911	0.933480095514496
GO_ORGANIC_HYDROXY_COMPOUND_METABOLIC_PROCESS	338	0.01	Up
	0.358643233672897	0.746941758397863
0.933480095514496
GO_PRERIBOSOME	72	0.01	Up	0.113258416835194
0.747066388001496	0.933480095514496
GO_N_METHYLTRANSFERASE_ACTIVITY	84	0.01	Down
0.20531384881546	0.74734161469105	0.933480095514496
GO_RESPONSE_TO_CAMP	66	0.01	Down	0.524550261998071
	0.74734257645543	0.933480095514496
GO_BASEMENT_MEMBRANE_ORGANIZATION	17	0.01	Up
0.670661375055086	0.747434994704353	0.933480095514496
GO_FRIZZLED_BINDING	23	0.01	Up	0.348257577706558
	0.747583631097262	0.933480095514496
GO_SPHINGOLIPID_BIOSYNTHETIC_PROCESS	82	0.01	Up
0.29859018905218	0.748329190502052	0.934218900994851
GO_RENAL_TUBULE_DEVELOPMENT	67	0.01	Down
0.600168284394142	0.749178554631909	0.935011923010935
GO_POSITIVE_REGULATION_OF_TRIGLYCERIDE_BIOSYNTHETIC_PROCESS	10
	0.01	Up	0.179793161908535	0.74927244365808
	0.935011923010935
GO_VIRAL_LATENCY	11	0.01	Down	0.234265093482002
	0.749761504750647	0.935311224365066
GO_RESPONSE_TO_AMINE	24	0.01	Down	0.473175031679554
	0.749930808975578	0.935311224365066
GO_INTRAMOLECULAR_TRANSFERASE_ACTIVITY	25	0.01	Up
0.147342711163264	0.750043578947177	0.935311224365066
GO_CELLULAR_CARBOHYDRATE_CATABOLIC_PROCESS	35	0.01	Up
	0.251123129594163	0.750716614565653

0.935311224365066
GO_PHOSPHATE_ION_TRANSPORT 18 0.01 Up
0.300274846288681 0.75078292478767 0.935311224365066
GO_MITOTIC_RECOMBINATION 23 0.01 Up
0.157526420912998 0.750847047216354 0.935311224365066
GO_AMMONIUM_TRANSMEMBRANE_TRANSPORTER_ACTIVITY 16 0.01 Down
0.766840991814783 0.751075739024993
0.935311224365066
GO_ENDODEOXYRIBONUCLEASE_ACTIVITY_PRODUCING_5_PHOSPHOMONOESTERS 12
0.01 Down 0.154558323957492 0.751091627782422
0.935311224365066
GO_PURINERGIC_RECEPTOR_ACTIVITY 10 0.01 Up
0.482085187710121 0.751233721352182 0.935311224365066
GO_IDENTICAL_PROTEIN_BINDING 1319 0.01 Down
0.331903842667149 0.751327884004763 0.935311224365066
GO_REGULATION_OF_CELLULAR_KETONE_METABOLIC_PROCESS 111 0.01
Down 0.26698015891482 0.751351765694741
0.935311224365066
GO_EXTRINSIC_APOPTOTIC_SIGNALING_PATHWAY_VIA_DEATH_DOMAIN_RECEPTORS
71 0.01 Down 0.30992527220208
0.751361034628303 0.935311224365066
GO_NEGATIVE_REGULATION_OF_POTASSIUM_ION_TRANSPORT 16 0.01
Up 0.773220462278131 0.751616458445721
0.935437375814672
GO_SIDE_OF_MEMBRANE 318 0.01 Down 0.517684598728751
0.752342090049923 0.935795680108133
GO_POSITIVE_REGULATION_OF_SUBSTRATE_ADHESION_DEPENDENT_CELL_SPREADIN
G 28 0.01 Down 0.211434603926315
0.752394556546142 0.935795680108133
GO_FEMALE_GAMETE_GENERATION 86 0.01 Down
0.340021049850358 0.752543197569155 0.935795680108133
GO_REGULATION_OF_ANTIGEN_RECEPTOR_MEDIATED_SIGNALING_PATHWAY 45
0.01 Down 0.413117268211713 0.752781443389708
0.935795680108133
GO_PROTEIN_INSERTION_INTO_MITOCHONDRIAL_MEMBRANE 45 0.01
Down 0.20383709616482 0.753004078628088
0.935795680108133
GO_NEUROBLAST_PROLIFERATION 37 0.01 Down
0.351592255378538 0.753072200534876 0.935795680108133
GO_PROTEIN_LOCALIZATION_TO_SYNAPSE 46 0.01 Up
0.290236560979362 0.753105621178577 0.935795680108133
GO_DENDRITE_CYTOPLASM 26 0.01 Down 0.254303957788003
0.753137488553507 0.935795680108133
GO_CHOLESTEROL_STORAGE 15 0.01 Up 0.630524695981297
0.754041489604826 0.936694759614848
GO_ADIPOSE_TISSUE_DEVELOPMENT 34 0.01 Up
0.338491667145301 0.754403169195591 0.936694759614848
GO_EMBRYONIC_FORELIMB_MORPHOGENESIS 18 0.01 Up
0.480810460391509 0.754429283053617 0.936694759614848
GO_PEPTIDE_DISULFIDE_OXIDOREDUCTASE_ACTIVITY 14 0.01 Down
0.0892656246683148 0.754611376552384
0.936694759614848
GO_REGULATION_OF_NMDA_RECEPTOR_ACTIVITY 13 0.01 Down
0.42715873924083 0.754632526647376 0.936694759614848

GO_ORGANELLE_MEMBRANE_CONTACT_SITE 19 0.01 Up
0.213318022080412 0.754969357297613 0.93679854191909
GO_U5_SNRNP 16 0.01 Down 0.0796680019210685
0.755243352456471 0.93679854191909
GO_WIDE_PORE_CHANNEL_ACTIVITY 13 0.01 Down
0.611484183619072 0.755247070929253 0.93679854191909
GO_LAMELLIPODIUM 171 0.01 Down 0.294577186691052
0.755333365622459 0.93679854191909
GO_REGULATION_OF_SYSTEMIC_ARTERIAL_BLOOD_PRESSURE_MEDIATED_BY_A_CHEMICAL_SIGNAL 25 0.01 Down 0.488863064016877
0.755568048725435 0.936898207477965
GO_NEGATIVE_REGULATION_OF_CELLULAR_RESPONSE_TO_INSULIN_STIMULUS 25
0.01 Down 0.208318009354153 0.756447013698842
0.937796573446124
GO_POSITIVE_REGULATION_OF_ESTABLISHMENT_OF_PROTEIN_LOCALIZATION_TO_TELOMERE 10 0.01 Down 0.0564332223818512
0.756836849871232 0.93808830452598
GO_ENDOTHELIAL_CELL_MIGRATION 173 0.01 Down
0.496373493449661 0.757387311115677 0.938346689103883
GO_RESOLUTION_OF_MEIOTIC_RECOMBINATION_INTERMEDIATES 12 0.01
Down 0.173167948995536 0.757719665829923
0.938346689103883
GO_REGULATION_OF_EPITHELIAL_CELL_APOPTOTIC_PROCESS 56 0.01
Down 0.423478114418525 0.757789277174475
0.938346689103883
GO_POSITIVE_REGULATION_OF_DNA_REPAIR 59 0.01 Up
0.242173157831724 0.757795123781443 0.938346689103883
GO_NEGATIVE_REGULATION_OF_PROTEIN_IMPORT_INTO_NUCLEUS 10 0.01
Up 0.777790142141555 0.757904603617958
0.938346689103883
GO_COPII_COATED_ER_TO_GOLGI_TRANSPORT_VESICLE 84 0.01 Down
0.348216735297409 0.757972683802577
0.938346689103883
GO_INOSITOL_PHOSPHATE_PHOSPHATASE_ACTIVITY 14 0.01 Up
0.202797295375447 0.7582584129943 0.93839453102875
GO_TRANSITION_METAL_ION_HOMEOSTASIS 105 0.01 Up
0.368917294963396 0.758698037678904 0.93839453102875
GO_RESPONSE_TO_AMYLOID_BETA 28 0.01 Up
0.438801922711636 0.758777688579343 0.93839453102875
GO_ACTIVATING_TRANSCRIPTION_FACTOR_BINDING 54 0.01 Down
0.308172976304896 0.75898814914152
0.93839453102875
GO_AUDITORY_RECEPTOR_CELL_DEVELOPMENT 14 0.01 Up
0.275466796647792 0.759024237960291 0.93839453102875
GO_POST_ANAL_TAIL_MORPHOGENESIS 11 0.01 Down
0.533656869782261 0.759151313247197 0.93839453102875
GO_ACTOMYOSIN 62 0.01 Up 0.232868960616641
0.7592370902083 0.93839453102875
GO_NEUROTRANSMITTER_RECEPTOR_ACTIVITY 16 0.01 Down
0.455817677799446 0.759247889377898 0.93839453102875
GO_DNA_DEPENDENT_ATPASE_ACTIVITY 78 0.01 Down
0.158279733902424 0.75951160600939 0.938529403639936
GO_NEGATIVE_REGULATION_OF_TORC1_SIGNALING 15 0.01 Up
0.161350013794285 0.759826591382222

0.938727561310841
GO_POSITIVE_REGULATION_OF_STEM_CELL_PROLIFERATION 26 0.01 Up 0.334012849409563 0.760088520998617 0.938860103963907
GO_NEGATIVE_REGULATION_OF_RESPONSE_TO_WOUNDING 61 0.01 Up 0.477508707380564 0.760323163801899 0.938958894922971
GO_ADHESION_OF_SYMBIONT_TO_HOST 11 0.01 Down 0.58033449159866 0.760974334180994 0.939357385534621
GO_NEURON_SPINE 107 0.01 Down 0.307291136241203 0.761019814096124 0.939357385534621
GO_AXO_DENDRITIC_TRANSPORT 58 0.01 Down 0.243849905591028 0.761110027910526 0.939357385534621
GO_OXIDOREDUCTASE_ACTIVITY_ACTING_ON_NAD_P_H_OXYGEN_AS_ACCEPTOR 12 0.01 Up 0.424248816248363 0.761517366651954 0.93945679023269
GO_DOUBLE_STRAND_BREAK_REPAIR 198 0.01 Down 0.182563164168563 0.761548742382132 0.93945679023269
GO_ORGANELLE_ASSEMBLY 688 0.01 Up 0.221762650421809 0.761779402451727 0.93945679023269
GO_TISSUE_MORPHOGENESIS 504 0.01 Down 0.415513331421424 0.762137174460161 0.93945679023269
GO_NUCLEAR_TRANSCRIBED_MRNA_CATABOLIC_PROCESS_EXONUCLEOLYTIC_3_5_ 10 0.01 Up 0.12199154805576 0.762211426141462 0.93945679023269
GO_PROTEIN_DEMANNOSYLATION 20 0.01 Up 0.167175208532114 0.762308303455103 0.93945679023269
GO_INNER_CELL_MASS_CELL_PROLIFERATION 13 0.01 Up 0.150560190530591 0.762368120845486 0.93945679023269
GO_CYSTEINE_TYPE_ENDOPEPTIDASE_INHIBITOR_ACTIVITY_INVOLVED_IN_APOPTOTIC_PROCESS 16 0.01 Down 0.545289877447373 0.76242852997471 0.93945679023269
GO_AXONEMAL_DYNEIN_COMPLEX_ASSEMBLY 17 0.01 Up 0.448283128970197 0.762636472236413 0.939522326085078
GO_NBAF_COMPLEX 12 0.01 Down 0.194540700088486 0.762890707318994 0.939644853749972
GO_FORMATION_OF_PRIMARY_GERM_LAYER 87 0.01 Up 0.55455719176583 0.763111185114946 0.939725761629378
GO_REGULATION_OF_AUTOPHAGY_OF_MITOCHONDRION_IN_RESPONSE_TO_MITOCHONDRIAL_DEPOLARIZATION 13 0.01 Up 0.141847287807075 0.763303534070543 0.93977200473378
GO_PROTEIN_QUALITY_CONTROL_FOR_MISFOLDED_OR_INCOMPLETELY_SYNTHESIZED_PROTEINS 27 0.01 Down 0.130617899942691 0.763975548860747 0.940229046272812
GO_NEGATIVE_REGULATION_OF_DNA_DEPENDENT_DNA_REPLICATION 25 0.01 Down 0.206266820801447 0.764026077550678 0.940229046272812
GO_MYELOID_LEUKOCYTE_CYTOKINE_PRODUCTION 29 0.01 Down 0.540973779137184 0.76413936984188 0.940229046272812
GO_LIGASE_ACTIVITY 127 0.01 Up 0.17586841876793 0.764804866377423 0.94048306686771
GO_MEMBRANE_PROTEIN_INTRACELLULAR_DOMAIN_PROTEOLYSIS 18 0.01 Down 0.203488993185308 0.764967753497778

0.94048306686771
GO_PLATELET_ACTIVATION 110 0.01 Down 0.430892409309098
0.765077376410689 0.94048306686771
GO_REGULATION_OF_DNA_REPLICATION 90 0.01 Down
0.221017417175057 0.765093138624723 0.94048306686771
GO_NEGATIVE_REGULATION_OF_CYCLIN_DEPENDENT_PROTEIN_KINASE_ACTIVITY
30 0.01 Up 0.334321052303798
0.765666818755275 0.94048306686771
GO_RIBONUCLEOPROTEIN_COMPLEX_BIOGENESIS 430 0.01 Down
0.122808028032823 0.765715241566018 0.94048306686771
GO_CYTOLYSIS 18 0.01 Up 0.822743935994345
0.765752739011565 0.94048306686771
GO_POSITIVE_REGULATION_OF_CHOLESTEROL_EFFLUX 13 0.01 Down
0.361278573026065 0.765962010936739
0.94048306686771
GO_POSITIVE_REGULATION_OF_PROTEIN_DEPOLYMERIZATION 14 0.01
Down 0.303654288105719 0.766013719243822
0.94048306686771
GO_CELLULAR_COMPONENT_MORPHOGENESIS 844 0.01 Down
0.368534566284703 0.766076441953753 0.94048306686771
GO_CELL_CYCLE_PHASE_TRANSITION 522 0.01 Down
0.202367044299228 0.766290173587528 0.94048306686771
GO_REGULATION_OF_SMOOTH_MUSCLE_CONTRACTION 37 0.01 Up
0.49503815521613 0.766610459989143
0.94048306686771
GO_PLATELET_DENSE_GRANULE 15 0.01 Up
0.247557822308844 0.766763931507293 0.94048306686771
GO_CYTOPLASMIC_EXOSOME_RNASE_COMPLEX_ 14 0.01 Down
0.128230961424784 0.766797978614643 0.94048306686771
GO_MACROPHAGE_CYTOKINE_PRODUCTION 16 0.01 Up
0.572119669438539 0.766845390705898 0.94048306686771
GO_ADULT_LOCOMOTORY_BEHAVIOR 43 0.01 Up
0.413080412363484 0.766963727820415 0.94048306686771
GO_MIDDLE_EAR_MORPHOGENESIS 14 0.01 Down
0.672141808575789 0.767067363269097 0.94048306686771
GO_OXIDOREDUCTASE_ACTIVITY_ACTING_ON_A_SULFUR_GROUP_OF_DONORS 54
0.01 Up 0.185121614619847 0.767219869660068
0.94048306686771
GO_INTESTINAL_ABSORPTION 18 0.01 Up
0.559352765838338 0.7672891830334 0.94048306686771
GO_NEURAL_CREST_CELL_DIFFERENTIATION 65 0.01 Down
0.445433874856405 0.767533755851246 0.940499787170767
GO_CELLULAR_RESPONSE_TO_PH 12 0.01 Up
0.161342401122644 0.767612657788033 0.940499787170767
GO_CALCIUM_ION_TRANSMEMBRANE_TRANSPORT_VIA_HIGH_VOLTAGE_GATED_CALCIU
M_CHANNEL 10 0.01 Up 0.392718275947338
0.767803796453051 0.940544158245858
GO_HEART_PROCESS 159 0.01 Down 0.441148012276088
0.768170587732639 0.94080363891968
GO_NAD_BIOSYNTHESIS_VIA_NICOTINAMIDE_RIBOSIDE_SALVAGE_PATHWAY 12
0.01 Down 0.558294906493509 0.768359624531568
0.940820111667358
GO_PLATELET_MORPHOGENESIS 16 0.01 Down
0.251249178553846 0.768493976899757 0.940820111667358

GO_LATERAL_VENTRICLE_DEVELOPMENT	11	0.01	Up
0.522370768090477	0.769164529622463	0.941241075109464
GO_POSITIVE_REGULATION_OF_NEUROTRANSMITTER_SECRETION	15	0.01
	Down	0.22647110794665	0.76948606885541
0.941241075109464
GO_NEGATIVE_REGULATION_OF_PROTEIN_POLYMERIZATION	53	0.01
	Down	0.304301243657238	0.769545684672046
0.941241075109464
GO_GLYCOSYLATION	215	0.01	Down	0.288145494957828
	0.769701223005916	0.941241075109464
GO_REGULATION_OF_RESPIRATORY_BURST	10	0.01	Up
0.367175808738784	0.769754524009666	0.941241075109464
GO_CARDIAC_MUSCLE_CELL_CONTRACTION	43	0.01	Up
0.479428031633832	0.769768067520753	0.941241075109464
GO_REGULATION_OF_TRANSLATIONAL_ELONGATION	17	0.01	Down
	0.152012379497075	0.770254675655803
0.941646422856702
GO_RNA_POLYMERASE_II_GENERAL_TRANSCRIPTION_INITIATION_FACTOR_ACTIVIT
Y	13	0.01	Down	0.157543461011144
0.770545892150558	0.941722637038855
GO_REGULATION_OF_HISTONE_H4_ACETYLATION	13	0.01	Up
0.299175018527434	0.770736774850821	0.941722637038855
GO_CELLULAR_RESPONSE_TO_LIGHT_STIMULUS	91	0.01	Up
0.213357781253052	0.77086414412313	0.941722637038855
GO_CARDIAC_CELL_DEVELOPMENT	53	0.01	Up
0.422377750547326	0.77093749070715	0.941722637038855
GO_CELLULAR_PROTEIN_CONTAINING_COMPLEX_ASSEMBLY	890	0.01	Down
	0.209823693410943	0.771218681310093
0.941876607168291
GO_CHEMOKINE_SECRETION	15	0.01	Up	0.705738493012726
	0.771996393672771	0.942636787205831
GO_SUSCEPTIBILITY_TO_NATURAL_KILLER_CELL_MEDIATED_CYTOTOXICITY	11
	0.01	Up	0.398436395593896	0.772404112997522
	0.942944976876725
GO_APOLIPOPROTEIN_BINDING	11	0.01	Up
0.237801580261279	0.772612894458252	0.942986195463661
GO_NEGATIVE_REGULATION_OF_OXIDATIVE_STRESS_INDUCED_INTRINSIC_APOPTOT
IC_SIGNALING_PATHWAY	17	0.01	Up	0.351748943094741
	0.772748529473187	0.942986195463661
GO_STEREOCILIUM_BUNDLE	34	0.01	Down	0.359037553262692
	0.773222407415012	0.943374846345768
GO_NEGATIVE_REGULATION_OF_OSSIFICATION	48	0.01	Up
0.543508180505107	0.77375575424682	0.943539019470076
GO_POSITIVE_REGULATION_OF_TOLL_LIKE_RECEPTOR_SIGNALING_PATHWAY	18
	0.01	Down	0.218281605501423	0.773783371622274
	0.943539019470076
GO_LEUCINE_ZIPPER_DOMAIN_BINDING	10	0.01	Up
0.133290380664056	0.774059689921813	0.943539019470076
GO_CALCIUM_CHANNEL_REGULATOR_ACTIVITY	24	0.01	Down
0.272438642442638	0.774062698160002	0.943539019470076
GO_PHOSPHATIDYLSERINE_ACYL_CHAIN_REMODELING	13	0.01	Up
	0.445061660521868	0.774134056330168
0.943539019470076
GO_SNORNA_3_END_PROCESSING	10	0.01	Up

0.152524526762368	0.774471934219692	0.943574501674902
GO_KINETOCHORE_ASSEMBLY 15	0.01	Up	0.238044454909078
	0.774521724905636	0.943574501674902
GO_RNA_POLYMERASE_II_HOLOENZYME 78	0.01	Down
0.141601279831331	0.774629437711697	0.943574501674902
GO_POSITIVE_REGULATION_OF_MEMBRANE_POTENTIAL	12	0.01	Up
	0.1550150185435 0.775481678912634	0.944127817142785
GO_NEGATIVE_REGULATION_OF_CARBOHYDRATE_METABOLIC_PROCESS	33
	0.01	Up	0.368191785387535	0.775617251883836
	0.944127817142785
GO_POST_MRNA_RELEASE_SPLICEOSOMAL_COMPLEX	10	0.01	Down
	0.128611439075143	0.775651856073607
0.944127817142785
GO_CELLULAR_CHEMICAL_HOMEOSTASIS	504	0.01	Up
0.399473309558527	0.775712101747326	0.944127817142785
GO_TRNA_BINDING 57	0.01	Up	0.153917850830786
0.776356965241443	0.944127817142785
GO_NEGATIVE_REGULATION_OF_EPITHELIAL_TO_MESENCHYMAL_TRANSITION	23
	0.01	Up	0.53528307779309	0.776374319286922
	0.944127817142785
GO_CELL_AGGREGATION	10	0.01	Down	0.20510449332759
	0.776451025054154	0.944127817142785
GO_FLUID_TRANSPORT	15	0.01	Down	0.858431395657363
	0.776536539466246	0.944127817142785
GO_ACTIVATION_OF_PROTEIN_KINASE_B_ACTIVITY	25	0.01	Down
	0.351456365452696	0.776794664848302
0.944127817142785
GO_NEGATIVE_REGULATION_OF_SIGNALING_RECEPTOR_ACTIVITY	30	0.01
	Up	0.289417044696534	0.776810451744536
0.944127817142785
GO_MUSCLE_CELL_DIFFERENTIATION	245	0.01	Up
0.427202553207745	0.776941430237784	0.944127817142785
GO_POSITIVE_REGULATION_OF_CELL_CYCLE	315	0.01	Down
0.269556647783044	0.777272794217188	0.944127817142785
GO_POSITIVE_REGULATION_OF_MACROPHAGE_MIGRATION	15	0.01	Up
	0.626859206451742	0.777587334760717
0.944127817142785
GO_POSITIVE_REGULATION_OF_MITOCHONDRION_ORGANIZATION	111	0.01
	Down	0.264516904087619	0.777644833606578
0.944127817142785
GO_ORGANELLE_TRANSPORT_ALONG_MICROTUBULE	74	0.01	Down
	0.195447593941478	0.777798303087595
0.944127817142785
GO_PHOSPHATIDYLINOSITOL_METABOLIC_PROCESS	85	0.01	Down
	0.320381208968289	0.777818247557424
0.944127817142785
GO_REGULATION_OF_PROTEIN_TARGETING	72	0.01	Up
0.198367828577298	0.778071234025362	0.944127817142785
GO_COENZYME_CATABOLIC_PROCESS	11	0.01	Down
0.221704245641774	0.778337335070024	0.944127817142785
GO_METAL_ION_HOMEOSTASIS	389	0.01	Up
0.426258418651072	0.778341658913323	0.944127817142785
GO_NEUROTRANSMITTER_TRANSPORTER_ACTIVITY	23	0.01	Up
	0.553324205149499	0.778382544570927

0.944127817142785
GO_PROTEIN_LOCALIZATION_TO_NUCLEOLUS 11 0.01 Down
0.127095100035856 0.778466896096046 0.944127817142785
GO_COPI_COATED_VESICLE 25 0.01 Down 0.204335069632461
0.778504999607442 0.944127817142785
GO_ACETYLGALACTOSAMINYLTRANSFERASE_ACTIVITY 36 0.01 Up
0.403095546035109 0.779076831918475
0.944632603670274
GO_NEGATIVE_REGULATION_OF_PROTEIN_LOCALIZATION_TO_MEMBRANE 25
0.01 Up 0.307045269102442 0.77936341354354
0.94464745712682
GO_PLASMA_MEMBRANE_TUBULATION 19 0.01 Down
0.209550987372059 0.779400282119624 0.94464745712682
GO_CELLULAR_ION_HOMEOSTASIS 400 0.01 Up
0.431763881420147 0.779803788452967 0.944753522143186
GO_POSITIVE_REGULATION_OF_CYSTEINE_TYPE_ENDOPEPTIDASE_ACTIVITY 118
0.01 Up 0.356452290090576 0.779830166936559
0.944753522143186
GO_AP_3_ADAPTOR_COMPLEX 10 0.01 Down 0.228213887119067
0.779972570500796 0.944753522143186
GO_QUATERNARY_AMMONIUM_GROUP_BINDING 16 0.01 Up
0.335934456429088 0.780110262972129 0.944753522143186
GO_PROTEIN_KINASE_B_BINDING 12 0.01 Up
0.3182901438279 0.780533222840805 0.945077222949048
GO_SIGNAL_TRANSDUCTION_BY_P53_CLASS_MEDIATOR 246 0.01 Down
0.197045704052969 0.780744248352792
0.945080975104513
GO_CELLULAR_HOMEOSTASIS 643 0.01 Up 0.366260892405337
0.781066752216287 0.945080975104513
GO_NEGATIVE_REGULATION_OF_ADHERENS_JUNCTION_ORGANIZATION 16
0.01 Up 0.337619714760199 0.78119633125514
0.945080975104513
GO_POSITIVE_REGULATION_OF_MUSCLE_CELL_DIFFERENTIATION 71 0.01
Up 0.508497765169291 0.781321826582038
0.945080975104513
GO_PHOSPHATIDYLETHANOLAMINE_BIOSYNTHETIC_PROCESS 13 0.01
Up 0.27013328231388 0.781845269920452
0.945080975104513
GO_REGULATION_OF_REPRODUCTIVE_PROCESS 93 0.01 Down
0.454874632248549 0.781860651214381 0.945080975104513
GO_BLASTODERM_SEGMENTATION 11 0.01 Down
0.424722353176306 0.782179282004858 0.945080975104513
GO_POSITIVE_REGULATION_OF_RECEPTOR_RECYCLING 13 0.01 Down
0.283784534297552 0.782376293543976
0.945080975104513
GO_SPINDLE_MIDZONE 30 0.01 Down 0.228581251846142
0.782378486733131 0.945080975104513
GO_RNA_POLYMERASE_III_COMPLEX 17 0.01 Up
0.145565016824515 0.782461711897688 0.945080975104513
GO_AUTOPHAGY_OF_MITOCHONDRION 71 0.01 Up
0.161035626807209 0.782476953212494 0.945080975104513
GO_PROTEIN_NEDDYLATION 14 0.01 Down 0.0989665210223452
0.782554499591743 0.945080975104513
GO_FOAM_CELL_DIFFERENTIATION 25 0.01 Up

0.562059547406008	0.782629076989837	0.945080975104513
GO_REGULATION_OF_CARDIAC_MUSCLE_CONTRACTION	53	0.01	Down
	0.43844278255651	0.782749505270689
0.945080975104513
GO_REGULATION_OF_SYSTEM_PROCESS 330	0.01	Down
0.427538845809268	0.782994066202077	0.945080975104513
GO_REGULATION_OF_SYSTEMIC_ARTERIAL_BLOOD_PRESSURE_BY_HORMONE	19
	0.01	Down	0.450967431574382	0.783027063873447
	0.945080975104513
GO_POSITIVE_REGULATION_OF_OSTEOBLAST_DIFFERENTIATION	50	0.01
	Down	0.520446659316994	0.783322450441678
0.945249571980009
GO_APOPTOTIC_DNA_FRAGMENTATION	20	0.01	Down
0.246851154239373	0.78353380811467	0.945316722786996
GO_PROTEIN_LOCALIZATION_TO_CHROMOSOME	75	0.01	Down
0.134108546035621	0.784041994365752	0.945415027405324
GO_POSITIVE_REGULATION_OF_MUSCLE_CELL_APOPTOTIC_PROCESS 19	0.01
	Up	0.331019410297591	0.784186942204138
0.945415027405324
GO_PROTON_TRANSMEMBRANE_TRANSPORT	113	0.01	Down
0.271979588364284	0.784513043793587	0.945415027405324
GO_CELLULAR_RESPONSE_TO_ESTROGEN_STIMULUS	11	0.01	Up
	0.228837385334241	0.784513337573622
0.945415027405324
GO_GAMMA_TUBULIN_COMPLEX	20	0.01	Down
0.136600181371817	0.784738022379673	0.945415027405324
GO_GLUTAMATE_RECEPTOR_BINDING	34	0.01	Down
0.232604745004902	0.784788420520781	0.945415027405324
GO_TRANSMEMBRANE_RECEPTOR_PROTEIN_SERINE_THREONINE_KINASE_SIGNALING_
PATHWAY 240	0.01	Up	0.440084810589592
0.784820256063829	0.945415027405324
GO_SEMINIFEROUS_TUBULE_DEVELOPMENT	10	0.01	Down
0.246839591475399	0.784966318601928	0.945415027405324
GO_POSITIVE_REGULATION_OF_PROTEIN_ACETYLATION	37	0.01	Down
	0.324194472055371	0.785052485672103
0.945415027405324
GO_NUCLEAR_NUCLEOSOME	10	0.01	Down	0.207164251825835
	0.785236282411795	0.945415027405324
GO_MITOCHONDRIAL_OUTER_MEMBRANE_PERMEABILIZATION	52	0.01
	Up	0.240189611139453	0.785396227603245
0.945415027405324
GO_BLOOD_MICROPARTICLE	61	0.01	Up	0.466110728324959
	0.785484005638685	0.945415027405324
GO_NEGATIVE_REGULATION_OF_ACTIN_FILAMENT_BUNDLE_ASSEMBLY	17
	0.01	Down	0.267901913942288	0.785931896426615
	0.94544055716453
GO_MATURATION_OF_5_8S_RRNA	27	0.01	Up
0.122558581223345	0.786077190840745	0.94544055716453
GO_TORC1_SIGNALING	38	0.01	Up	0.160269412973108
	0.786422263841263	0.94544055716453
GO_NPBAF_COMPLEX	12	0.01	Down	0.189484976555563
	0.786438078857755	0.94544055716453
GO_CELL_SURFACE_RECEPTOR_SIGNALING_PATHWAY_INVOLVED_IN_CELL_CELL_SIG
NALING	470	0.01	Down	0.350121939792331

0.786522049002885	0.94544055716453
GO_REGULATION_OF_RESPONSE_TO_DNA_DAMAGE_STIMULUS	192	0.01
	Down	0.23726131481325	0.786603835945868
0.94544055716453
GO_DNA_DEPENDENT_DNA_REPLICATION	131	0.01	Up
0.18609853237551	0.786719744756554	0.94544055716453
GO_NEGATIVE_REGULATION_OF_MACROPHAGE_DERIVED_FOAM_CELL_DIFFERENTIATI
ON	10	0.01	Down	0.316559036153315
0.786751061570615	0.94544055716453
GO_PRERIBOSOME_SMALL_SUBUNIT_PRECURSOR	14	0.01	Down
0.10323797371149	0.786982365369045	0.945455746919374
GO_INSULIN_RECEPTOR_BINDING	17	0.01	Down
0.272767489748282	0.787296793442977	0.945455746919374
GO_REPLICATION_FORK	61	0.01	Up	0.152448630943487
	0.787566234008691	0.945455746919374
GO_ACROSOME_ASSEMBLY	10	0.01	Down	0.259828879507214
	0.787734734108699	0.945455746919374
GO_MULTICELLULAR_ORGANISM_REPRODUCTION	500	0.01	Down
0.313860178475699	0.787761611833305	0.945455746919374
GO_SKELETAL_MUSCLE_ORGAN_DEVELOPMENT	113	0.01	Down
0.403675521422883	0.787938483353238	0.945455746919374
GO_NEUROTRANSMITTER_SODIUM_SYMPORTER_ACTIVITY	11	0.01	Down
	0.663923612014953	0.788091169279561
0.945455746919374
GO_RESPONSE_TO_OXYGEN_LEVELS	313	0.01	Down
0.320116920284943	0.788274851623457	0.945455746919374
GO_PHOSPHORYLATED_CARBOHYDRATE_DEPHOSPHORYLATION	13	0.01
	Down	0.143741900754887	0.788383808362967
0.945455746919374
GO_NEURON_DEATH 262	0.01	Down	0.342684888710163
0.788439794640486	0.945455746919374
GO_ACTIN_FILAMENT_BASED_MOVEMENT	89	0.01	Up
0.423282121626004	0.788580301226064	0.945455746919374
GO_NEURON_APOPTOTIC_PROCESS	180	0.01	Up
0.362387088585929	0.788632499159893	0.945455746919374
GO_MUSCLE_STRUCTURE_DEVELOPMENT 438	0.01	Down
0.425330217843435	0.788952987248452	0.945531326498951
GO_PLASMA_MEMBRANE_REGION	741	0.01	Down
0.381467303905527	0.78900703344485	0.945531326498951
GO_RETROMER_COMPLEX	20	0.01	Down	0.135184992760982
	0.789176652993745	0.945547949541154
GO_SNORNA_PROCESSING	11	0.01	Up	0.139375572040051
	0.789527582824473	0.945781759141155
GO_PERINUCLEAR_ENDOPLASMIC_RETICULUM	19	0.01	Up
0.265790375632977	0.789770117101296	0.945885654157027
GO_METANEPHRIC_NEPHRON_DEVELOPMENT	25	0.01	Down
0.875575581193251	0.790217635765027	0.946092151928606
GO_TRNA_5_END_PROCESSING	16	0.01	Up
0.123315889049025	0.790254208932624	0.946092151928606
GO_MICROTUBULE_ASSOCIATED_COMPLEX	114	0.01	Down
0.212860388526199	0.790722814719205	0.94646652369091
GO_PRECATALYTIC_SPLICEOSOME	44	0.01	Down
0.0801733158846421	0.791010433917605	0.946624156182492
GO_U2_SNRNP	19	0.01	Up	0.0833901916448301

0.791475972164263	0.946994605244233
GO_MAIN_AXON	39	0.01	Up	0.467089345763199
0.791839437619552	0.947165495540593
GO_CELLULAR_RESPONSE_TO_ESTRADIOL_STIMULUS	21	0.01	Up
	0.412454603826458	0.791930827765451
0.947165495540593
GO_POSITIVE_REGULATION_OF_LIPID_BIOSYNTHETIC_PROCESS	51	0.01
	Down	0.337371219069393	0.792482512330428
0.9476386315458
GO_ER_OVERLOAD_RESPONSE	12	0.01	Down	0.250101496194104
	0.792954168946637	0.947938810574767
GO_REGULATION_OF_NEUROINFLAMMATORY_RESPONSE	20	0.01	Up
	1.0069171713753	0.793045827525818	0.947938810574767
GO_MESENCHYMAL_CELL_PROLIFERATION	32	0.01	Up
0.582809138669495	0.793502765267111	0.947940538348673
GO_ANIMAL_ORGAN_MATURATION	16	0.01	Up
0.772555522082861	0.793537080970571	0.947940538348673
GO_RNA_SURVEILLANCE	15	0.01	Up	0.147719095032135
	0.793620205302621	0.947940538348673
GO_REGULATION_OF_MEMBRANE_REPOLARIZATION	23	0.01	Up
	0.513797565717497	0.793671842600281
0.947940538348673
GO_NON_RECOMBINATIONAL_REPAIR	79	0.01	Down
0.16880714119459	0.793999382924149	0.948087328971557
GO_PROTEIN_KINASE_C_SIGNALING	16	0.01	Down
0.498246924268	0.794476514769311	0.948087328971557
GO_CRANIAL_SKELETAL_SYSTEM_DEVELOPMENT	54	0.01	Down
0.473331332663811	0.794729103741284	0.948087328971557
GO_OTIC_VESICLE_DEVELOPMENT	10	0.01	Down
0.233604230462253	0.794763336651848	0.948087328971557
GO_TOR_SIGNALING	106	0.01	Down	0.186836404016724
	0.794910744345294	0.948087328971557
GO_REGULATION_OF_ANDROGEN_RECEPTOR_SIGNALING_PATHWAY	23	0.01
	Down	0.227768855974827	0.795057349708477
0.948087328971557
GO_TRAPP_COMPLEX	11	0.01	Down	0.138539931217008
	0.795198655441249	0.948087328971557
GO_MAINTENANCE_OF_PROTEIN_LOCATION_IN_CELL	64	0.01	Up
	0.250589162301646	0.795333029976959
0.948087328971557
GO_WHOLE_MEMBRANE	1343	0.01	Up	0.319848778391164
	0.795421794318888	0.948087328971557
GO_PHOSPHATIDYLINOSITOL_3_PHOSPHATE_BINDING	34	0.01	Down
	0.186667036605621	0.795489616359322
0.948087328971557
GO_POLYOL_BIOSYNTHETIC_PROCESS	59	0.01	Up
0.352613941199997	0.795512576804663	0.948087328971557
GO_L_GLUTAMATE_TRANSMEMBRANE_TRANSPORT	12	0.01	Down
0.285365364604774	0.796133398604047	0.948616572968267
GO_TRNA_METHYLTRANSFERASE_ACTIVITY	34	0.01	Up
0.147080024396889	0.796351695949628	0.948616572968267
GO_CHAPERONE_MEDIATED_PROTEIN_COMPLEX_ASSEMBLY	18	0.01	Up
	0.214071283576273	0.796598943377625
0.948616572968267

GO_REGULATION_OF_CYTOSOLIC_CALCIUM_ION_CONCENTRATION	186	0.01	Up	0.509085940328134	0.796737146817179	0.948616572968267
GO_MITOTIC_SPINDLE_MIDZONE	10	0.01	Down	0.207265338027643	0.796787567512127	0.948616572968267
GO_EXOCYTOSIS	709	0.01	Down	0.386473981356611	0.796982242213072	0.948616572968267
GO_OLIGOSACCHARYL_TRANSFERASE_ACTIVITY	10	0.01	Up	0.0806257057326079	0.79705042640605	0.948616572968267
GO_NEGATIVE_REGULATION_OF_INTRACELLULAR_ESTROGEN_RECEPTOR_SIGNALING_PATHWAY	10	0.01	Down	0.182483786909018	0.797351665062581	0.948620384209779
GO_TRANSLATIONAL_INITIATION	183	0.01	Down	0.146861375575555	0.79767308783067	0.948620384209779
GO_NEGATIVE_REGULATION_OF_CELL_PROJECTION_ORGANIZATION	142	0.01	Down	0.364688205144338	0.798055137494	0.948620384209779
GO_MULTI_MULTICELLULAR_ORGANISM_PROCESS	156	0.01	Down	0.516673632653562	0.798170179206442	0.948620384209779
GO_NUCLEOSOME_ORGANIZATION	107	0.01	Down	0.174744649853381	0.798254818396743	0.948620384209779
GO_CELL_PROJECTION_PART	993	0.01	Up	0.309928789073247	0.798261321221478	0.948620384209779
GO_REGULATION_OF_TRANSCRIPTION_BY_RNA_POLYMERASE_III	22	0.01	Down	0.1380341617869	0.798469438259321	0.948620384209779
GO_DEVELOPMENTAL_GROWTH_INVOLVED_IN_MORPHOGENESIS	176	0.01	Down	0.389118812549466	0.798487429970773	0.948620384209779
GO_DETECTION_OF_ABIOTIC_STIMULUS	52	0.01	Up	0.330951267406033	0.79875828513928	0.948620384209779
GO_RECEPTOR_CATABOLIC_PROCESS	32	0.01	Up	0.24236675155793	0.799105815533452	0.948620384209779
GO_COGNITION	174	0.01	Up	0.373610049017341	0.799481026717159	0.948620384209779
GO_POSITIVE_REGULATION_OF_COLLAGEN_BIOSYNTHETIC_PROCESS	20	0.01	Up	0.613858729540392	0.79949127414403	0.948620384209779
GO_CYCLIN_DEPENDENT_PROTEIN_SERINE_THREONINE_KINASE_REGULATOR_ACTIVITY	42	0.01	Up	0.298280754102784	0.799512051646823	0.948620384209779
GO_REGULATION_OF_NEURAL_PRECURSOR_CELL_PROLIFERATION	54	0.01	Down	0.292348448242142	0.79971351664529	0.948620384209779
GO_RESPONSE_TO_INSECTICIDE	11	0.01	Up	0.480163362550993	0.799725207960339	0.948620384209779
GO_MACROPHAGE_ACTIVATION	62	0.01	Down	0.652265076906324	0.79982200476639	0.948620384209779
GO_ORGANELLE_DISASSEMBLY	92	0.01	Down	0.164446012456079	0.799874098969052	0.948620384209779
GO_RDNA_BINDING	10	0.01	Up	0.118190309214799	0.800303174812827	0.948620384209779
GO_LYMPHOID_PROGENITOR_CELL_DIFFERENTIATION	18	0.01	Up	0.465621771444355	0.800314752111206

0.948620384209779
GO_CATALYTIC_ACTIVITY_ACTING_ON_A_RRNA 20 0.01 Up
0.10502608894828 0.800489759922542 0.948620384209779
GO_POLY_A_PLUS_MRNA_EXPORT_FROM_NUCLEUS 12 0.01 Up
0.170122628884202 0.80054525574055 0.948620384209779
GO_TRANSPORTER_COMPLEX 144 0.01 Down 0.420850700027166
0.800637982653371 0.948620384209779
GO_INTEGRIN_ACTIVATION 15 0.01 Up 0.419285877395913
0.800647479606474 0.948620384209779
GO_LIPOPROTEIN_PARTICLE_RECEPTOR_ACTIVITY 14 0.01 Down
0.442785368894638 0.800998238461148
0.948643262444312
GO_CHONDROCYTE_PROLIFERATION 13 0.01 Down
0.572712642678964 0.801240736233886 0.948643262444312
GO_RECEPTOR_RECYCLING 30 0.01 Up 0.257205576887213
0.801245435883785 0.948643262444312
GO_AZUROPHIL_GRANULE 133 0.01 Up 0.357254911971364
0.801291821745089 0.948643262444312
GO_ION_TRANSMEMBRANE_TRANSPORT 671 0.01 Up
0.390685186131209 0.801537883769387 0.948749559829196
GO_BODY_FLUID_SECRETION 60 0.01 Up 0.397034750364434
0.801992218979604 0.949102292675473
GO_CALCIUM_DEPENDENT_CELL_CELL_ADHESION_VIA_PLASMA_MEMBRANE_CELL_ADH
ESION_MOLECULES 26 0.01 Up 0.393557537037455
0.802176534692462 0.949135400919497
GO_CORONARY_VASCULATURE_MORPHOGENESIS 11 0.01 Down
0.517741539538721 0.802805398269574 0.949328360801003
GO_GENITALIA_DEVELOPMENT 26 0.01 Down
0.325481626672913 0.80304444370964 0.949328360801003
GO_LEARNING 77 0.01 Up 0.304996067754226
0.803044453847232 0.949328360801003
GO_POSITIVE_REGULATION_OF_LIPID_TRANSPORT 37 0.01 Up
0.450953498645801 0.803067457628719
0.949328360801003
GO_REGULATION_OF_DNA_CATABOLIC_PROCESS 10 0.01 Up
0.167392105738502 0.803203646276941 0.949328360801003
GO_CELL_CELL_ADHESION_VIA_PLASMA_MEMBRANE_ADHESION_MOLECULES 185
0.01 Up 0.399287535758056 0.803277843754695
0.949328360801003
GO_MORPHOGENESIS_OF_AN_EPITHELIAL_SHEET 47 0.01 Down
0.401223396370476 0.803536208981089 0.949448875968119
GO_PROTEASE_BINDING 105 0.01 Down 0.515506421371952
0.804030721257697 0.949848318496883
GO_NEGATIVE_REGULATION_OF_PROTEIN_SERINE_THREONINE_KINASE_ACTIVITY
121 0.01 Up 0.364530936602648
0.804241455194288 0.949878328370964
GO_REGULATION_OF_MICROTUBULE_BASED_MOVEMENT 17 0.01 Up
0.286135111584018 0.804369047299477
0.949878328370964
GO_AMINO_ACID_ACTIVATION 49 0.01 Up
0.119493974770562 0.805237829497914 0.950719343228673
GO_ION_HOMEOSTASIS 500 0.01 Up 0.409432586904079
0.805407670706802 0.950734973529262
GO_NEURON_RECOGNITION 32 0.01 Down 0.505494795908402

0.805875746075364	0.951102576676425
GO_MITOCHONDRIAL_TRNA_PROCESSING	11	0.01	Up
0.0760101710340331	0.806361949456001	0.951491427628257
GO_STEM_CELL_DIVISION	30	0.01	Up	0.468206578669723
0.807098009808906	0.95194150036961
GO_CELL_PROJECTION_MEMBRANE	231	0.01	Up
0.408461779417205	0.807113576562887	0.95194150036961
GO_VINCULIN_BINDING	10	0.01	Down	0.144014335798887
0.807250321550757	0.95194150036961
GO_PROTON_TRANSPORTING_V_TYPE_ATPASE_COMPLEX	21	0.01	Up
0.179448214776747	0.807489353972846
0.95194150036961
GO_REGULATION_OF_MEIOTIC_NUCLEAR_DIVISION	17	0.01	Up
0.459582054139364	0.807595887467795
0.95194150036961
GO_LIPID_LOCALIZATION	267	0.01	Down	0.413793373137599
0.807798208267733	0.95194150036961
GO_REGULATION_OF_FIBROBLAST_GROWTH_FACTOR_RECEPTOR_SIGNALING_PATHWAY
24	0.01	Up	0.452703442062768
0.807840983347756	0.95194150036961
GO_FOREBRAIN_NEURON_DIFFERENTIATION	28	0.01	Down
0.49157569566866	0.80803597114981	0.951986489588686
GO_NEGATIVE_REGULATION_OF_ION_TRANSMEMBRANE_TRANSPORT	61	0.01
Up	0.449793909661526	0.808594550662185
0.952238508199081
GO_SPERM_MIDPIECE	12	0.01	Down	0.470893062918943
0.808645796300502	0.952238508199081
GO_REGULATION_OF_MAMMARY_GLAND_EPITHELIAL_CELL_PROLIFERATION	15
0.01	Down	0.242451072977366	0.808889543807877
0.952238508199081
GO_REGULATION_OF_LIPID_METABOLIC_PROCESS	294	0.01	Up
0.349214446752831	0.808971131263933
0.952238508199081
GO_TRANSLATION_ELONGATION_FACTOR_ACTIVITY	18	0.01	Up
0.130336378206872	0.809087953431681
0.952238508199081
GO_MICROTUBULE_MOTOR_ACTIVITY	61	0.01	Down
0.264352240878286	0.809190983989303	0.952238508199081
GO_POSITIVE_REGULATION_OF_TUMOR_NECROSIS_FACTOR_BIOSYNTHETIC_PROCESS
10	0.01	Up	0.60337423537099
0.809632028118695	0.952315674004745
GO_ANIMAL_ORGAN_MORPHOGENESIS	720	0.01	Up
0.451151449626947	0.809661965234911	0.952315674004745
GO_ORGANOPHOSPHATE_CATABOLIC_PROCESS	100	0.01	Down
0.324810211828081	0.809909192839118	0.952315674004745
GO_GLYCEROPHOSPHOLIPID_METABOLIC_PROCESS	143	0.01	Up
0.272369428263845	0.809925969178817
0.952315674004745
GO_POLYSOME	65	0.01	Up	0.150895876917859
0.810190248618203	0.952315674004745
GO_REGULATION_OF_LIPID_BIOSYNTHETIC_PROCESS	140	0.01	Down
0.327667575764494	0.810197736161178
0.952315674004745
GO_REGULATION_OF_FILOPODIUM_ASSEMBLY	34	0.01	Down

0.291550424074963	0.810729538125088	0.952645698104968
GO_TUBULIN_BINDING	257	0.01	Down	0.272960933898381
	0.810792344277445	0.952645698104968
GO_CHOLESTEROL_EFFLUX	26	0.01	Up	0.447027434947432
	0.81142684218985	0.953206725800035
GO_ACTIN_FILAMENT_BASED_PROCESS	584	0.01	Down
0.362434114177834	0.812101194213911	0.953606619839411
GO_REGULATION_OF_NEUROLOGICAL_SYSTEM_PROCESS	70	0.01	Down
	0.366066239645408	0.812287795910376
0.953606619839411
GO_NEGATIVE_REGULATION_OF_NEUROTRANSMITTER_SECRETION	10	0.01
	Down	0.321271844399319	0.812343578179072
0.953606619839411
GO_COLUMNAR_CUBOIDAL_EPITHELIAL_CELL_DIFFERENTIATION	76	0.01
	Down	0.391676850807299	0.812473160693859
0.953606619839411
GO_HETEROTRIMERIC_G_PROTEIN_COMPLEX	17	0.01	Down
0.259872549995733	0.812575656918371	0.953606619839411
GO_SPINDLE	300	0.01	Down	0.201775468936034
0.812777596249122	0.953606619839411
GO_ACTION_POTENTIAL	70	0.01	Up	0.421188304283121
	0.81300749788499	0.953606619839411
GO_NEGATIVE_REGULATION_OF_OSTEOBLAST_DIFFERENTIATION	30	0.01
	Up	0.552884808684364	0.813023861684861
0.953606619839411
GO_VOLTAGE_GATED_CATION_CHANNEL_ACTIVITY	48	0.01	Down
	0.482964455776515	0.813597680313428
0.953773302096432
GO_REGULATION_OF_MUSCLE_CONTRACTION	102	0.01	Up
0.488014974005094	0.813822316033175	0.953773302096432
GO_SULFUR_COMPOUND_TRANSPORT	29	0.01	Up
0.345000837305921	0.81389822396451	0.953773302096432
GO_REGULATION_OF_CALCIUM_ION_TRANSMEMBRANE_TRANSPORT	95	0.01
	Up	0.481720279014425	0.813956514073697
0.953773302096432
GO_NUCLEOBASE_CONTAINING_SMALL_MOLECULE_BIOSYNTHETIC_PROCESS	87
	0.01	Up	0.284249076177378	0.814059950212803
	0.953773302096432
GO_DIOL_BIOSYNTHETIC_PROCESS	18	0.01	Up
0.164533374492622	0.814108590259218	0.953773302096432
GO_POST_EMBRYONIC_ANIMAL_MORPHOGENESIS	11	0.01	Up
0.577515863853354	0.814347003535286	0.953868543018082
GO_CUL4_RING_E3_UBIQUITIN_LIGASE_COMPLEX	30	0.01	Up
	0.102428268076437	0.814751542960854
0.954158298093238
GO_STEROL_TRANSPORT	72	0.01	Down	0.429419464728526
	0.815098740067504	0.954164031126602
GO_CYSTEINE_TYPE_ENDOPEPTIDASE_ACTIVITY_INVOLVED_IN_APOPTOTIC_PROCES
S	11	0.01	Down	0.248104908427636
0.815131065305059	0.954164031126602
GO_CERAMIDE_BIOSYNTHETIC_PROCESS	45	0.01	Up
0.321896341724732	0.815227940941144	0.954164031126602
GO_REGULATION_OF_PROTEIN_LOCALIZATION_TO_MEMBRANE	150	0.01
	Down	0.291423944277497	0.815642363934342

0.954222059892636
GO_PROTEIN_TRANSMEMBRANE_TRANSPORT 59 0.01 Up
0.168772883979982 0.815724350420632 0.954222059892636
GO_REGULATION_OF_CELL_DIVISION 127 0.01 Down
0.349695279783702 0.815749051365966 0.954222059892636
GO_OXIDOREDUCTASE_ACTIVITY_ACTING_ON_SINGLE_DONORS_WITH_INCORPORATIO
N_OF_MOLECULAR_OXYGEN 14 0.01 Up 0.504483624030443
0.816559242680341 0.95498577582592
GO_ISOMERASE_ACTIVITY 128 0.01 Up 0.231345601980965
0.817119963429252 0.955457491906585
GO_POSITIVE_REGULATION_OF_PROTEIN_LOCALIZATION_TO_CELL_SURFACE 15
0.01 Up 0.508907896947491 0.817733881726246
0.95570800110712
GO_ORGANIC_HYDROXY_COMPOUND_BIOSYNTHETIC_PROCESS 184 0.01
Down 0.337184896899286 0.81778756698799
0.95570800110712
GO_ENDOPLASMIC_RETICULUM_SUBCOMPARTMENT 22 0.01 Down
0.120973887384141 0.817806467756793 0.95570800110712
GO_WOUND_HEALING 390 0.01 Up 0.457362345457339
0.818153753371019 0.95579882379518
GO_REGULATION_OF_ACTIN_FILAMENT_BASED_MOVEMENT 29 0.01 Down
0.399846506943618 0.818324924215026
0.95579882379518
GO_CARBOHYDRATE_BIOSYNTHETIC_PROCESS 157 0.01 Down
0.304224178550681 0.818626010120261 0.95579882379518
GO_REGULATION_OF_OSTEOBLAST_DIFFERENTIATION 88 0.01 Down
0.516990299380161 0.818702681360917
0.95579882379518
GO_DNA_DEPENDENT_DNA_REPLICATION_MAINTENANCE_OF_FIDELITY 41
0.01 Up 0.19367507063924 0.819089856124685
0.95579882379518
GO_REGULATION_OF_PROTEIN_HOMOOLIGOMERIZATION 17 0.01 Up
0.284773926024422 0.81913753313554
0.95579882379518
GO_CALCIUM_ION_TRANSMEMBRANE_IMPORT_INTO_CYTOSOL 91 0.01
Up 0.554266039670258 0.819221377020123
0.95579882379518
GO_ARP2_3_PROTEIN_COMPLEX 11 0.01 Down
0.141410004823639 0.81947008563687 0.95579882379518
GO_EXTRACELLULAR_LIGAND_GATED_ION_CHANNEL_ACTIVITY 12 0.01
Down 0.57796094093984 0.819591867090312
0.95579882379518
GO_ACTIVE_ION_TRANSMEMBRANE_TRANSPORTER_ACTIVITY 51 0.01
Up 0.228451253615949 0.81971698454083
0.95579882379518
GO_POSITIVE_REGULATION_OF_PATHWAY_RESTRICTED_SMAD_PROTEIN_PHOSPHORYL
ATION 29 0.01 Down 0.448682165111329
0.819752566172126 0.95579882379518
GO_REGULATION_OF_SKELETAL_MUSCLE_ADAPTATION 10 0.01 Down
0.478568930400436 0.819784458579412
0.95579882379518
GO_MATERNAL_PROCESS_INVOLVED_IN_FEMALE_PREGNANCY 51 0.01
Down 0.520155628734527 0.820028470679818
0.95579882379518

GO_SNRNA_BINDING	37	0.01	Down	0.136747450410523	0.820088300634013	0.95579882379518
GO_SYNAPTONEMAL_COMPLEX_ORGANIZATION	12	0.01	Up	0.242864103186338	0.82045052107731	0.956037449800451
GO_REGULATION_OF_RECEPTOR_BIOSYNTHETIC_PROCESS	17	0.01	Down	0.300466099701234	0.821266096652714	0.956804159044066
GO_SMALL_SUBUNIT_PROCESSOME	36	0.01	Up	0.109491596051297	0.821578663849461	0.956984663896792
GO_VIRAL_GENE_EXPRESSION	187	0.01	Down	0.15779312778302	0.821777358005672	0.95703248426097
GO_REGULATION_OF_CELL_CYCLE_G1_S_PHASE_TRANSITION	145	0.01	Down	0.264732021776696	0.822191908852629	0.957238072278429
GO_MEIOTIC_SPINDLE_ORGANIZATION	10	0.01	Down	0.163377632539103	0.822269238499754	0.957238072278429
GO_CELL_CYCLE_CHECKPOINT	200	0.01	Down	0.207073812541292	0.822524910062697	0.957352133625505
GO_GLANDULAR_EPITHELIAL_CELL_DEVELOPMENT	18	0.01	Up	0.303320313126297	0.822756292458329	0.957437886048402
GO_TELOMERASE_RNA_LOCALIZATION	19	0.01	Down	0.0950507243333605	0.822939128364265	0.957467123093035
GO_CYCLIC_NUCLEOTIDE_PHOSPHODIESTERASE_ACTIVITY	15	0.01	Down	0.327887178824754	0.82335954160783	0.95772734095552
GO_RESPONSE_TO_PURINE_CONTAINING_COMPOUND	101	0.01	Up	0.467474084087366	0.823478293491651	0.95772734095552
GO_POSITIVE_REGULATION_OF_PROTEIN_OLIGOMERIZATION	19	0.01	Down	0.42661427612938	0.823695682798766	0.957796684595156
GO_REGULATION_OF_NUCLEAR_DIVISION	169	0.01	Up	0.283953310595447	0.824198342175649	0.958011798095961
GO_CELL_FATE_COMMITMENT_INVOLVED_IN_FORMATION_OF_PRIMARY_GERM_LAYER	23	0.01	Down	0.621306392817622	0.824250549950212	0.958011798095961
GO_POSITIVE_REGULATION_OF_RELEASE_OF_SEQUESTERED_CALCIUM_ION_INTO_CYTOSOL	28	0.01	Up	0.761289861213247	0.82435408223576	0.958011798095961
GO_CALCIUM_CHANNEL_COMPLEX	33	0.01	Down	0.290850314857503	0.824893542817458	0.958333798428529
GO_ATP_DEPENDENT_MICROTUBULE_MOTOR_ACTIVITY_PLUS_END_DIRECTED	18	0.01	Down	0.195312575155538	0.82494686716974	0.958333798428529
GO_ATTACHMENT_OF_SPINDLE_MICROTUBULES_TO_KINETOCHORE	25	0.01	Down	0.141256719469491	0.825311252651907	0.958443168925451
GO_POSITIVE_REGULATION_OF_STRIATED_MUSCLE_CELL_DIFFERENTIATION	42	0.01	Up	0.598305533428197	0.825512023009121	0.958443168925451
GO_POSITIVE_REGULATION_OF_COLD_INDUCED_THERMOGENESIS	69	0.01	Down	0.416259320163484	0.825684433112867	0.958443168925451

GO_RESPONSE_TO_TOXIC_SUBSTANCE 342 0.01 Up
0.432117239367084 0.82582574773547 0.958443168925451
GO_POSTSYNAPTIC_SPECIALIZATION_ORGANIZATION 20 0.01 Up
0.358169309557925 0.825830376651134
0.958443168925451
GO_C21_STEROID_HORMONE_BIOSYNTHETIC_PROCESS 17 0.01 Up
0.379506512708092 0.826452500552996
0.95866234375332
GO_REGULATION_OF_PROTEIN_AUTOPHOSPHORYLATION 41 0.01 Down
0.488872156631508 0.826656713005426
0.95866234375332
GO_MUSCLE_SYSTEM_PROCESS 276 0.01 Up
0.444545514894215 0.826938261032068 0.95866234375332
GO_CELLULAR_RESPONSE_TO_VASCULAR_ENDOTHELIAL_GROWTH_FACTOR_STIMULUS
44 0.01 Down 0.617250783726976
0.826945625831658 0.95866234375332
GO_RNA_POLYMERASE_II_ACTIVITY 11 0.01 Up
0.13127936279362 0.827098158450579 0.95866234375332
GO_REGULATION_OF_PLATELET_ACTIVATION 23 0.01 Up
0.623929062621353 0.827464637358476 0.95866234375332
GO_HISTONE_H3_K9_ACETYLATION 11 0.01 Down
0.149720310910865 0.827545264450057 0.95866234375332
GO_DNA_POLYMERASE_BINDING 17 0.01 Up
0.130147058856234 0.827623265430537 0.95866234375332
GO_PROTEIN_LOCALIZATION_TO_POSTSYNAPTIC_MEMBRANE 24 0.01
Up 0.254419502365568 0.827623964521227
0.95866234375332
GO_NEGATIVE_REGULATION_OF_CYTOKINE_PRODUCTION_INVOLVED_IN_INFLAMMATO
RY_RESPONSE 14 0.01 Up 0.718115448005359
0.827747934422605 0.95866234375332
GO_SECONDARY_ACTIVE_TRANSMEMBRANE_TRANSPORTER_ACTIVITY 138 0.01
Down 0.424030833531135 0.82775621906686
0.95866234375332
GO_SPECIFIC_GRANULE 124 0.01 Up 0.438195883499661
0.828102426542372 0.958880379847176
GO_PLASMA_MEMBRANE_PROTEIN_COMPLEX 324 0.01 Down
0.451756747357615 0.828794065341068 0.959073552485312
GO_REGULATION_OF_MESONEPHROS_DEVELOPMENT 18 0.01 Down
0.840265922704257 0.828803775409118
0.959073552485312
GO_RESPONSE_TO_ENDOGENOUS_STIMULUS 1157 0.01 Down
0.354389985049662 0.828881084269082 0.959073552485312
GO_REGULATION_OF_TRIGLYCERIDE_BIOSYNTHETIC_PROCESS 14 0.01
Up 0.256439516011461 0.829049083322186
0.959073552485312
GO_BONE_MATURATION 14 0.01 Up 0.865513521891247
0.829184560028766 0.959073552485312
GO_REGULATION_OF_WNT_SIGNALING_PATHWAY 285 0.01 Down
0.361181074033097 0.829217110359974 0.959073552485312
GO_TUBE_MORPHOGENESIS 627 0.01 Down 0.505713753000014
0.82945544592955 0.95916647852158
GO_ACTIN_NUCLEATION 42 0.01 Down 0.270443384329701
0.830324402766274 0.959742867067599
GO_EPITHELIAL_TUBE_BRANCHING_INVOLVED_IN_LUNG_MORPHOGENESIS 24

0.01 Down 0.463502903362144 0.830332925051837
0.959742867067599
GO_PROTEIN_LOCALIZATION_TO_CELL_CORTEX 10 0.01 Down
0.281914657040789 0.830428147044325 0.959742867067599
GO_MULTICELLULAR_ORGANISMAL_SIGNALING 99 0.01 Down
0.431211186422389 0.830690856412193 0.959863758903392
GO_NEGATIVE_REGULATION_OF_NEURON_DEATH 148 0.01 Down
0.348714552789502 0.830913114343151 0.959937871965227
GO_LACTATION 35 0.01 Down 0.309678260760056
0.831543009007288 0.960482802070632
GO_MITOTIC_CELL_CYCLE_CHECKPOINT 152 0.01 Down
0.20956062264236 0.832351402889113 0.96069487970819
GO_CELLULAR_RESPONSE_TO_LOW_DENSITY_LIPOPROTEIN_PARTICLE_STIMULUS
15 0.01 Up 0.552380664069065
0.832384791822498 0.96069487970819
GO_NEGATIVE_REGULATION_OF_T_CELL_RECEPTOR_SIGNALING_PATHWAY 17
0.01 Up 0.438647927190677 0.83240414206304
0.96069487970819
GO_NEGATIVE_REGULATION_OF_BONE_MINERALIZATION 10 0.01 Up
1.17404017887585 0.83240993451418
0.96069487970819
GO_EMBRYO_DEVELOPMENT 726 0.01 Down 0.36970849006951
0.832562846552239 0.96069487970819
GO_RESPONSE_TO_VITAMIN_D 29 0.01 Up
0.53453223058822 0.832676075938807 0.96069487970819
GO_MITOCHONDRIAL_RNA_PROCESSING 15 0.01 Up
0.0736287988666586 0.833183557987499 0.961057810850034
GO_SIGNAL_TRANSDUCTION_INVOLVED_IN_CELL_CYCLE_CHECKPOINT 69
0.01 Down 0.234878779227829 0.833464673913755
0.961057810850034
GO_REGULATION_OF_KETONE_BIOSYNTHETIC_PROCESS 12 0.01 Up
0.565506830210473 0.833569966553671
0.961057810850034
GO_REGULATION_OF_SYSTEMIC_ARTERIAL_BLOOD_PRESSURE 48 0.01
Down 0.435136649043558 0.834367515006334
0.961057810850034
GO_ACTIN_BASED_CELL_PROJECTION 151 0.01 Down
0.346139610222273 0.834461764949296 0.961057810850034
GO_REGULATION_OF_MYOBLAST_DIFFERENTIATION 37 0.01 Down
0.520850154517965 0.834580151095208
0.961057810850034
GO_DENDRITIC_SHAFT 23 0.01 Up 0.349309321714655
0.834946715812942 0.961057810850034
GO_NEGATIVE_REGULATION_OF_FIBROBLAST_GROWTH_FACTOR_RECEPTOR_SIGNALIN
G_PATHWAY 14 0.01 Down 0.552129834871022
0.83497605717151 0.961057810850034
GO_NUCLEOTIDASE_ACTIVITY 11 0.01 Down
0.259246726355255 0.835002820881361 0.961057810850034
GO_HISTONE_H4_ACETYLATION 61 0.01 Down
0.199623511519724 0.83519510526233 0.961057810850034
GO_MRNA_METHYLTRANSFERASE_ACTIVITY 11 0.01 Down
0.119652930833264 0.835287934751075 0.961057810850034
GO_REGULATION_OF_APOPTOTIC_PROCESS_INVOLVED_IN_MORPHOGENESIS 10
0.01 Up 0.391155392419997 0.83572463303434

0.961057810850034
GO_REGULATION_OF_APOPTOTIC_PROCESS_INVOLVED_IN_DEVELOPMENT 10
0.01 Up 0.391155392419997 0.83572463303434
0.961057810850034
GO_METHIONINE_BIOSYNTHETIC_PROCESS 10 0.01 Up
0.115021523645746 0.835740553067963 0.961057810850034
GO_CENTRIOLAR_SATELLITE 26 0.01 Up 0.20185789149778
0.835741127912813 0.961057810850034
GO_CHROMATIN_ASSEMBLY_OR_DISASSEMBLY 120 0.01 Up
0.186845741036981 0.83586594297128 0.961057810850034
GO_ENDOCARDIAL_CUSHION_FORMATION 15 0.01 Down
0.489263385638395 0.835985021737529 0.961057810850034
GO_DNA_REPLICATION_CHECKPOINT 13 0.01 Up
0.168113392174853 0.836261262998119 0.961057810850034
GO_CELLULAR_COMPONENT_DISASSEMBLY 474 0.01 Up
0.275324145350077 0.836307470664913 0.961057810850034
GO_CYTOPLASMIC_TRANSLATIONAL_ELONGATION 10 0.01 Up
0.160710588472929 0.836346500310208 0.961057810850034
GO_PROTEIN_DNA_COMPLEX_SUBUNIT_ORGANIZATION 198 0.01 Up
0.183921079273693 0.836773615107922
0.961057810850034
GO_MONOVALENT_INORGANIC_CATION_HOMEOSTASIS 98 0.01 Down
0.375113832504876 0.837059927871015
0.961057810850034
GO_N_GLYCAN_PROCESSING 14 0.01 Down 0.219472859674963
0.837328315625121 0.961057810850034
GO_INTRINSIC_COMPONENT_OF_SYNAPTIC_VESICLE_MEMBRANE 28 0.01
Down 0.225909283741797 0.837451434684985
0.961057810850034
GO_UROGENITAL_SYSTEM_DEVELOPMENT 253 0.01 Up
0.514003845034637 0.837591035944767 0.961057810850034
GO_MYELOID_LEUKOCYTE_DIFFERENTIATION 149 0.01 Down
0.461551898557418 0.83768858251992 0.961057810850034
GO_NUCLEOSIDE_MONOPHOSPHATE_PHOSPHORYLATION 11 0.01 Up
0.13931059499098 0.837699682240988
0.961057810850034
GO_DNA_DEAMINATION 10 0.01 Down 0.37177363642274
0.837751421026184 0.961057810850034
GO_ESTABLISHMENT_OF_PROTEIN_LOCALIZATION_TO_ORGANELLE 501 0.01
Down 0.200250155973143 0.837805405977283
0.961057810850034
GO_REGULATION_OF_BIOMINERAL_TISSUE_DEVELOPMENT 54 0.01 Down
0.689317865783441 0.838045703683328
0.961057810850034
GO_PROTON_TRANSMEMBRANE_TRANSPORTER_ACTIVITY 100 0.01 Down
0.296038826580594 0.838178153504849
0.961057810850034
GO_TRANS_GOLGI_NETWORK_MEMBRANE 83 0.01 Up
0.312242446754173 0.838187532172524 0.961057810850034
GO_REGULATION_OF_PROTEIN_OLIGOMERIZATION 32 0.01 Up
0.355132936412295 0.838711085255993
0.961057810850034
GO_MICROTUBULE_DEPOLYMERIZATION 32 0.01 Up
0.27216730544538 0.8387180417918 0.961057810850034

GO_REGULATION_OF_RESPIRATORY_GASEOUS_EXCHANGE 11 0.01 Up
0.318927083988362 0.83880144288993
0.961057810850034
GO_NUCLEAR_CHROMOSOME_SEGREGATION 194 0.01 Down
0.181661765825283 0.83895459511055 0.961057810850034
GO_CELLULAR_KETONE_METABOLIC_PROCESS 153 0.01 Up
0.272147254759913 0.839002213589132 0.961057810850034
GO_PEPTIDE_BIOSYNTHETIC_PROCESS 601 0.01 Down
0.169282094797641 0.839147957431664 0.961057810850034
GO_LATE_ENDOSOME 217 0.01 Up 0.305254736540176
0.839210708004176 0.961057810850034
GO_REGULATION_OF_FERTILIZATION 13 0.01 Up
0.268504525482422 0.839488330667448 0.961057810850034
GO_POSITIVE_REGULATION_OF_MAST_CELL_ACTIVATION_INVOLVED_IN_IMMUNE_RESPONSE 10 0.01 Up 0.510388656669689
0.839617553100482 0.961057810850034
GO_SLEEP 12 0.01 Down 0.481207811565655
0.839861899400068 0.961057810850034
GO_BASE_EXCISION_REPAIR 39 0.01 Up 0.171470252559154
0.840106733827387 0.961057810850034
GO_C_TERMINAL_PROTEIN_AMINO_ACID_MODIFICATION 11 0.01 Down
0.0817062362061099 0.840122400155113
0.961057810850034
GO_CHAPERONE_COMPLEX 20 0.01 Down 0.136968508179084
0.84026559808679 0.961057810850034
GO_FUCOSYLATION 19 0.01 Down 0.28285124825264
0.840272584416403 0.961057810850034
GO_RESPONSE_TO_ACIDIC_PH 12 0.01 Down
0.226868592637732 0.840780914776591 0.961349658915019
GO_PLASMA_MEMBRANE_BOUNDED_CELL_PROJECTION_CYTOPLASM 143 0.01
Up 0.25423407217066 0.841124214501645
0.961349658915019
GO_NITRIC_OXIDE_SYNTHASE_BINDING 13 0.01 Up
0.368338459854343 0.841229719911046 0.961349658915019
GO_REGULATION_OF_OXIDATIVE_STRESS_INDUCED_INTRINSIC_APOPTOTIC_SIGNALING_PATHWAY 24 0.01 Up 0.292416560511919
0.841305258426822 0.961349658915019
GO_PRESYNAPTIC_ACTIVE_ZONE_MEMBRANE 21 0.01 Up
0.30759718831285 0.841319508781996 0.961349658915019
GO_SNRNA_PROCESSING 25 0.01 Up 0.120520297055584
0.841748467510961 0.961423080595095
GO_MYOFIBRIL_ASSEMBLY 40 0.01 Up 0.484288008225583
0.841805581875172 0.961423080595095
GO_CRANIAL_NERVE_MORPHOGENESIS 16 0.01 Up
0.667140226548002 0.84185885298032 0.961423080595095
GO_REGULATION_OF_RELEASE_OF_SEQUESTERED_CALCIUM_ION_INTO_CYTOSOL
54 0.01 Up 0.600048414289643
0.842432061581357 0.961896754910742
GO_PYRIDINE_CONTAINING_COMPOUND_BIOSYNTHETIC_PROCESS 29 0.01
Down 0.551918601279142 0.84275001512655
0.961908132341829
GO_ANNEALING_ACTIVITY 12 0.01 Up 0.217679410910222
0.842758912193409 0.961908132341829
GO_MEIOTIC_CHROMOSOME_SEPARATION 15 0.01 Down

0.193984953667971	0.843245472754032	0.962282568625889
GO_REGULATION_OF_CATION_CHANNEL_ACTIVITY	86	0.01	Down
	0.461867794079308	0.843878011751094
0.962313285049697
GO_ORGANISM_EMERGENCE_FROM_PROTECTIVE_STRUCTURE 20	0.01	Up
	0.267384593192503	0.844166835168161
0.962313285049697
GO_NEGATIVE_REGULATION_OF_CALCIUM_MEDIATED_SIGNALING	23	0.01
	Down	0.333206622753652	0.844221672950006
0.962313285049697
GO_KERATINOCYTE_DEVELOPMENT	10	0.01	Down
0.395267487045625	0.844483045139511	0.962313285049697
GO_POSITIVE_REGULATION_OF_LIPID_KINASE_ACTIVITY 23	0.01	Up
	0.640607069793554	0.844649249799219
0.962313285049697
GO_EUCHROMATIN	28	0.01	Down	0.220168751294866
0.844690790744283	0.962313285049697
GO_5_3_EXONUCLEASE_ACTIVITY	16	0.01	Up
0.160670606393474	0.844751543754336	0.962313285049697
GO_ASPARTIC_TYPE_PEPTIDASE_ACTIVITY	14	0.01	Up
0.44335722786216	0.844846511107289	0.962313285049697
GO_PLASMINOGEN_ACTIVATION	15	0.01	Down
0.264036560320166	0.844923900238326	0.962313285049697
GO_UBIQUITIN_DEPENDENT_ERAD_PATHWAY	70	0.01	Down
0.171252552605394	0.845401108900895	0.962313285049697
GO_GLIAL_CELL_ACTIVATION	33	0.01	Up
0.650082168045872	0.845445702235262	0.962313285049697
GO_REGULATION_OF_HISTONE_PHOSPHORYLATION	12	0.01	Down
	0.174789411144407	0.845644628402202
0.962313285049697
GO_PEPTIDYL_CYSTEINE_MODIFICATION	40	0.01	Up
0.415248607319287	0.845665701925045	0.962313285049697
GO_CENTROSOME	458	0.01	Up	0.207486412947267
0.845740178697477	0.962313285049697
GO_POSITIVE_REGULATION_OF_ANTIGEN_RECEPTOR_MEDIATED_SIGNALING_PATHWA
Y	17	0.01	Up	0.43486873028099
0.845759875018718	0.962313285049697
GO_POSITIVE_REGULATION_OF_GLUCOSE_METABOLIC_PROCESS	18	0.01
	Down	0.32067814472947	0.845855904199706
0.962313285049697
GO_REGULATION_OF_CELLULAR_EXTRAVASATION 17	0.01	Down
0.799779526440375	0.846241713019981	0.962313285049697
GO_REGULATION_OF_PROTEIN_SUMOYLATION	19	0.01	Down
0.176258003295965	0.846252381336084	0.962313285049697
GO_TRICARBOXYLIC_ACID_CYCLE	31	0.01	Up
0.116855008195979	0.84628407657393	0.962313285049697
GO_MYOTUBE_DIFFERENTIATION	70	0.01	Up
0.47103851518187	0.846676820411466	0.962369500709466
GO_SECONDARY_PALATE_DEVELOPMENT 20	0.01	Down
0.569868031531543	0.846738198884988	0.962369500709466
GO_POSITIVE_REGULATION_OF_TOR_SIGNALING 31	0.01	Up
0.155689927745422	0.846809071452803	0.962369500709466
GO_PHENOL_CONTAINING_COMPOUND_BIOSYNTHETIC_PROCESS	21	0.01
	Down	0.348780890060281	0.847342317237288

0.962795285036042
GO_REGULATION_OF_RESPONSE_TO_REACTIVE_OXYGEN_SPECIES 30 0.01 Down 0.389895898303853 0.847964582957715 0.963160570607384
GO_POSITIVE_REGULATION_OF_NATURAL_KILLER_CELL_MEDIATED_IMMUNITY 20 0.01 Down 0.353018285330534 0.848324867366848 0.963160570607384
GO_TRANSLESION_SYNTHESIS 39 0.01 Up 0.132723591619751 0.848577435426356 0.963160570607384
GO_PYRIMIDINE_RIBONUCLEOTIDE_BIOSYNTHETIC_PROCESS 18 0.01 Up 0.264058398614252 0.848684480100828 0.963160570607384
GO_PRP19_COMPLEX 13 0.01 Down 0.0804311735842122 0.848693356983789 0.963160570607384
GO_POSITIVE_REGULATION_OF_PROTEIN_DEPHOSPHORYLATION 36 0.01 Up 0.383542063660839 0.848774667067912 0.963160570607384
GO_ACYLGLYCEROL_O_ACYLTRANSFERASE_ACTIVITY 17 0.01 Down 0.23551782436377 0.849142718079144 0.963160570607384
GO_CELLULAR_RESPONSE_TO_ZINC_ION 13 0.01 Down 0.393341641607378 0.849175979443253 0.963160570607384
GO_TRANSFERASE_ACTIVITY_TRANSFERRING_ACYL_GROUPS 198 0.01 Down 0.210428548465232 0.849232205395544 0.963160570607384
GO_PROTEIN_COMPLEX_OLIGOMERIZATION 401 0.01 Down 0.31507966960937 0.849317587980793 0.963160570607384
GO_CELLULAR_RESPONSE_TO_CAMP 34 0.01 Down 0.609435953363415 0.849408943342436 0.963160570607384
GO_GLUCOSAMINE_CONTAINING_COMPOUND_METABOLIC_PROCESS 13 0.01 Down 0.48436053740796 0.849919103028404 0.963517809587066
GO_PROTEIN_LOCALIZATION_TO_CHROMOSOME_CENTROMERIC_REGION 18 0.01 Up 0.154839275954428 0.85005092112403 0.963517809587066
GO_WOUND_HEALING_SPREADING_OF_EPIDERMAL_CELLS 14 0.01 Down 0.487266042244357 0.850296563976089 0.963517809587066
GO_SUMO_TRANSFERASE_ACTIVITY 19 0.01 Down 0.201616619003033 0.850511019011336 0.963517809587066
GO_REGULATION_OF_NUCLEAR_CELL_CYCLE_DNA_REPLICATION 13 0.01 Up 0.205900652901928 0.850517532791482 0.963517809587066
GO_NEGATIVE_REGULATION_OF_PROTEIN_OLIGOMERIZATION 12 0.01 Down 0.185547057320043 0.850785265189605 0.963641295702629
GO_HISTONE_H2A_MONOUBIQUITINATION 17 0.01 Up 0.139387952994866 0.851310596350302 0.964056450371701
GO_POSITIVE_REGULATION_OF_CELL_SUBSTRATE_ADHESION 90 0.01 Up 0.553170513770603 0.851532130594301 0.964127483185006
GO_METAL_ION_TRANSMEMBRANE_TRANSPORTER_ACTIVITY 211 0.01 Up 0.463049310027898 0.851781295877256 0.964160087809976

GO_CULLIN_RING_UBIQUITIN_LIGASE_COMPLEX 143 0.01 Down
0.158745960202957 0.851926095590951 0.964160087809976
GO_DICARBOXYLIC_ACID_BIOSYNTHETIC_PROCESS 11 0.01 Down
0.522006680124147 0.852037369642649
0.964160087809976
GO_GLYCOPROTEIN_BIOSYNTHETIC_PROCESS 270 0.01 Down
0.327584146570785 0.852429704315237 0.96421501376657
GO_CYCLIN_DEPENDENT_PROTEIN_KINASE_HOLOENZYME_COMPLEX 36 0.01
Down 0.230076461129971 0.852713831755788
0.96421501376657
GO_POSITIVE_REGULATION_OF_HETEROTYPIC_CELL_CELL_ADHESION 10
0.01 Down 0.846343311248219 0.852853624815827
0.96421501376657
GO_NAD_METABOLIC_PROCESS 18 0.01 Up
0.275782767696817 0.852957691252441 0.96421501376657
GO_IMPORT_ACROSS_PLASMA_MEMBRANE 69 0.01 Up
0.46819795822912 0.852985393913404 0.96421501376657
GO_HISTONE_H4_K5_ACETYLATION 16 0.01 Up
0.16980608516539 0.853038846802874 0.96421501376657
GO_LIGAND_GATED_ION_CHANNEL_ACTIVITY 47 0.01 Up
0.517182350609241 0.853889124915891 0.964223215325546
GO_REGULATION_OF_DOUBLE_STRAND_BREAK_REPAIR_VIA_HOMOLOGOUS_RECOMBINATION 39 0.01 Up 0.224076272525623
0.854032844433863 0.964223215325546
GO_MONOVALENT_INORGANIC_CATION_TRANSMEMBRANE_TRANSPORTER_ACTIVITY
197 0.01 Up 0.432881582564605
0.854062266972373 0.964223215325546
GO_CARDIAC_MUSCLE_TISSUE_MORPHOGENESIS 38 0.01 Down
0.336810383781147 0.854202729353929 0.964223215325546
GO_NEGATIVE_REGULATION_OF_GLYCOPROTEIN_BIOSYNTHETIC_PROCESS 10
0.01 Down 0.28944661729465 0.854387214423235
0.964223215325546
GO_MRNA_SPLICE_SITE_SELECTION 28 0.01 Up
0.117541535154665 0.85475651428066 0.964223215325546
GO_DIGESTIVE_SYSTEM_DEVELOPMENT 97 0.01 Down
0.582122918705418 0.854912148885847 0.964223215325546
GO_TERTIARY_GRANULE 128 0.01 Down 0.461413057035004
0.854925146580094 0.964223215325546
GO_REGULATION_OF_HEART_GROWTH 42 0.01 Up
0.387115757322737 0.85500562786624 0.964223215325546
GO_TELOMERE_CAP_COMPLEX 11 0.01 Down 0.120987397526499
0.855160670411705 0.964223215325546
GO_POSITIVE_REGULATION_OF_ENDOTHELIAL_CELL_PROLIFERATION 69
0.01 Down 0.428194474384435 0.855247648941249
0.964223215325546
GO_APOPTOTIC_MITOCHONDRIAL_CHANGES 107 0.01 Up
0.271479044773831 0.855378128405292 0.964223215325546
GO_NUCLEOSIDE_TRIPHOSPHATE_CATABOLIC_PROCESS 10 0.01 Up
0.130728672718184 0.855458170331544
0.964223215325546
GO_CATALYTIC_ACTIVITY_ACTING_ON_DNA 157 0.01 Down
0.164157728219336 0.855555036680975 0.964223215325546
GO_STEROID_HORMONE_RECEPTOR_ACTIVITY 38 0.01 Down
0.405093969467563 0.855597422691268 0.964223215325546

GO_REGULATION_OF_MEIOTIC_CELL_CYCLE 29 0.01 Down
0.411859931993985 0.855702477476634 0.964223215325546
GO_REGULATION_OF_CYCLIN_DEPENDENT_PROTEIN_KINASE_ACTIVITY 90
0.01 Down 0.275009302489843 0.855746118295839
0.964223215325546
GO_REGULATION_OF_MONOCYTE_DIFFERENTIATION 15 0.01 Down
0.377288448094773 0.856302589535873
0.96455373311408
GO_REGULATION_OF_RIBOSOME_BIOGENESIS 14 0.01 Down
0.0973664625834686 0.856447691673278 0.96455373311408
GO_PHOTORECEPTOR_OUTER_SEGMENT 39 0.01 Down
0.278916949970287 0.856622489848529 0.96455373311408
GO_CHROMOSOME_TELOMERIC_REGION 133 0.01 Up
0.173837420819315 0.856674967707317 0.96455373311408
GO_ANATOMICAL_STRUCTURE_ARRANGEMENT 12 0.01 Up
0.61338305928726 0.857376760159592 0.96464178223542
GO_CUL4A_RING_E3_UBIQUITIN_LIGASE_COMPLEX 11 0.01 Down
0.0873851596587533 0.857384186265857
0.96464178223542
GO_STEM_CELL_PROLIFERATION 80 0.01 Down
0.427022776784035 0.857471441915034 0.96464178223542
GO_ENDORIBONUCLEASE_ACTIVITY_PRODUCING_5_PHOSPHOMONOESTERS 27
0.01 Down 0.147144639572936 0.857631179302366
0.96464178223542
GO_ACTIVE_TRANSMEMBRANE_TRANSPORTER_ACTIVITY 229 0.01 Up
0.351249115719127 0.857678215644783
0.96464178223542
GO_DIGESTION 64 0.01 Up 0.537824799006062
0.857768571954327 0.96464178223542
GO_RECEPTOR_INTERNALIZATION 74 0.01 Down
0.421021825962419 0.858013245675003 0.96464178223542
GO_POSITIVE_REGULATION_OF_NATURAL_KILLER_CELL_ACTIVATION 17
0.01 Up 0.407234385458066 0.858288535364359
0.96464178223542
GO_CYTOSKELETON_DEPENDENT_INTRACELLULAR_TRANSPORT 161 0.01
Up 0.207998931256016 0.858301093513048
0.96464178223542
GO_POSITIVE_REGULATION_OF_COAGULATION 19 0.01 Down
0.416081023966887 0.858479185512432 0.96464178223542
GO_GLYCEROLIPID_BIOSYNTHETIC_PROCESS 189 0.01 Down
0.236063803880972 0.858500996445062 0.96464178223542
GO_DETECTION_OF_VISIBLE_LIGHT 22 0.01 Down
0.176382025668597 0.859052386489902 0.964900550190941
GO_REGULATION_OF_SYNAPTIC_VESICLE_RECYCLING 20 0.01 Down
0.200608189682346 0.85919359090161
0.964900550190941
GO_AORTIC_VALVE_DEVELOPMENT 22 0.01 Down
0.506378005478332 0.859208099873534 0.964900550190941
GO_PHOSPHATIDYLSERINE_METABOLIC_PROCESS 17 0.01 Down
0.377821083414997 0.859670831061715 0.965241652556995
GO_NEGATIVE_REGULATION_OF_TRANSLATIONAL_INITIATION 17 0.01
Down 0.151136124935229 0.860033238881483
0.965462699061556
GO_A_BAND 21 0.01 Up 0.218874671794391

0.860567628705628	0.965462699061556
GO_PERIKARYON	66	0.01	Up	0.301448187417909
0.860921158520289	0.965462699061556
GO_EPITHELIAL_CELL_PROLIFERATION	292	0.01	Down
0.447227979398862	0.860923413341696	0.965462699061556
GO_BIOLOGICAL_ADHESION	1008	0.01	Down	0.500887151709762
0.860955083958359	0.965462699061556
GO_FAD_BINDING	26	0.01	Up	0.286314608835505
0.861252019859518	0.965462699061556
GO_SMOOTH_MUSCLE_CELL_MIGRATION	60	0.01	Down
0.515579709076485	0.861311110244966	0.965462699061556
GO_CELLULAR_RESPONSE_TO_EXTRACELLULAR_STIMULUS	223	0.01	Down
0.307513903185059	0.861478796485814
0.965462699061556
GO_POSITIVE_REGULATION_OF_VACUOLE_ORGANIZATION	13	0.01	Down
0.149796621260639	0.861516899537338
0.965462699061556
GO_DNA_TEMPLATED_TRANSCRIPTION_TERMINATION	74	0.01	Down
0.115487893909292	0.861574486822212
0.965462699061556
GO_REGULATION_OF_MICROGLIAL_CELL_ACTIVATION	14	0.01	Up
0.809246869434535	0.861617015897795
0.965462699061556
GO_EMBRYONIC_HEMOPOIESIS	16	0.01	Up
0.345281154059894	0.862196074766693	0.96566975685429
GO_CENTRIOLE	120	0.01	Up	0.210790745949371
0.862262625774415	0.96566975685429
GO_RECEPTOR_ANTAGONIST_ACTIVITY	12	0.01	Down
0.76628688026822	0.86241545074283	0.96566975685429
GO_EPITHELIAL_TUBE_MORPHOGENESIS	248	0.01	Up
0.460758706232096	0.862438053313121	0.96566975685429
GO_TRNA_GUANINE_METHYLTRANSFERASE_ACTIVITY	13	0.01	Down
0.133905787240917	0.862641774814869
0.965719751964055
GO_REGULATION_OF_MITOCHONDRIAL_OUTER_MEMBRANE_PERMEABILIZATION_INVOLVED_IN_APOPTOTIC_SIGNALING_PATHWAY	42	0.01	Up
0.24028584308095	0.86322500130775	0.966076733275373
GO_LUNG_ALVEOLUS_DEVELOPMENT	34	0.01	Down
0.56658321765773	0.863278912538116	0.966076733275373
GO_DYNEIN_COMPLEX_BINDING	18	0.01	Down
0.248270999440676	0.863511641713462	0.966159081614897
GO_APPENDAGE_MORPHOGENESIS	104	0.01	Up
0.444483703755537	0.863803059726867	0.966163995496091
GO_REGULATION_OF_B_CELL_RECEPTOR_SIGNALING_PATHWAY	17	0.01
Up	0.342969045881239	0.863862517122841
0.966163995496091
GO_TRANSFORMING_GROWTH_FACTOR_BETA_RECEPTOR_SIGNALING_PATHWAY	153
0.01	Up	0.40437034108074	0.86405404988082
0.966163995496091
GO_DNA_STRAND_ELONGATION_INVOLVED_IN_DNA_REPLICATION	13	0.01
Up	0.15445499023479	0.864254866050552
0.966163995496091
GO_GATOR2_COMPLEX	12	0.01	Down	0.136786292086037
0.864516000129613	0.966163995496091

GO_ESTABLISHMENT_OF_PROTEIN_LOCALIZATION_TO_MITOCHONDRIAL_MEMBRANE 18 0.01 Up 0.0993716775465664 0.864848499189129 0.966163995496091
GO_REGULATION_OF_EXTRACELLULAR_MATRIX_DISASSEMBLY 14 0.01 Up 0.530601561945432 0.865015901664796 0.966163995496091
GO_CELLULAR_RESPONSE_TO_GONADOTROPIN_STIMULUS 12 0.01 Down 0.388131374464477 0.865116462311656 0.966163995496091
GO_POSITIVE_REGULATION_OF_TISSUE_REMODELING 22 0.01 Up 0.353132045852124 0.865148298173996 0.966163995496091
GO_SPLICEOSOMAL_SNRNP_ASSEMBLY 38 0.01 Up 0.0822383780623685 0.865185602479896 0.966163995496091
GO_RNA_5_END_PROCESSING 20 0.01 Up 0.114305501321073 0.865266618928059 0.966163995496091
GO_REGULATION_OF_SYNAPTIC_VESICLE_CYCLE 73 0.01 Down 0.228136686176607 0.865947991180839 0.966567420808047
GO_HISTONE_H2A_ACETYLATION 15 0.01 Up 0.103477220469234 0.86631046835282 0.966567420808047
GO_NEGATIVE_REGULATION_OF_G_PROTEIN_COUPLED_RECEPTOR_SIGNALING_PATHWAY 35 0.01 Down 0.563302203051803 0.866558257622983 0.966567420808047
GO_ODONTOGENESIS 88 0.01 Up 0.539137900143056 0.866641287944175 0.966567420808047
GO_CATION_TRANSMEMBRANE_TRANSPORT 490 0.01 Down 0.412087150923929 0.866660586850998 0.966567420808047
GO_TISSUE_HOMEOSTASIS 147 0.01 Down 0.419126595558297 0.866734524757592 0.966567420808047
GO_NEURON_MIGRATION 101 0.01 Up 0.408894951053385 0.867047092669233 0.966567420808047
GO_NEGATIVE_REGULATION_OF_TRANSFERASE_ACTIVITY 236 0.01 Down 0.30583271938432 0.867194266392834 0.966567420808047
GO_POSITIVE_REGULATION_OF_NEUROLOGICAL_SYSTEM_PROCESS 38 0.01 Up 0.410049191238549 0.867699919229139 0.966567420808047
GO_DNA_GEOMETRIC_CHANGE 88 0.01 Up 0.171061283900323 0.867715041182519 0.966567420808047
GO_DNA_PACKAGING 126 0.01 Up 0.194075407757428 0.867742267077015 0.966567420808047
GO_H4_HISTONE_ACETYLTRANSFERASE_COMPLEX 41 0.01 Up 0.109537295403737 0.8679992137634 0.966567420808047
GO_POSITIVE_REGULATION_OF_CILIUM_ASSEMBLY 22 0.01 Up 0.3008633830759 0.868133640924412 0.966567420808047
GO_ACTIN_MONOMER_BINDING 19 0.01 Up 0.326612794261509 0.868155731949016 0.966567420808047
GO_POSITIVE_REGULATION_OF_LIPID_METABOLIC_PROCESS 88 0.01 Down 0.437927848337806 0.868155820555731 0.966567420808047
GO_MICROTUBULE_CYTOSKELETON 1004 0.01 Down 0.23332009494478 0.868566843897149 0.966567420808047
GO_LOCALIZATION_WITHIN_MEMBRANE 110 0.01 Up 0.324101581316275 0.868592881447096 0.966567420808047

GO_NUCLEOTIDE_SUGAR_TRANSMEMBRANE_TRANSPORT 14 0.01 Down
0.160821709632905 0.868716181376837
0.966567420808047
GO_REGULATORY_T_CELL_DIFFERENTIATION 21 0.01 Down
0.413331613368635 0.868880814810301 0.966567420808047
GO_PEPTIDASE_REGULATOR_ACTIVITY 125 0.01 Up
0.603217623445423 0.869005078530894 0.966567420808047
GO_REGULATION_OF_PROTEIN_LOCALIZATION_TO_CELL_PERIPHERY 93 0.01
Down 0.308189657951415 0.869036996978638
0.966567420808047
GO_PSEUDOPODIUM 13 0.01 Down 0.133125995332488
0.869139319905401 0.966567420808047
GO_PIGMENT_GRANULE_LOCALIZATION 25 0.01 Up
0.193950511110211 0.869297328761842 0.966567420808047
GO_DNA_INTEGRATION 11 0.01 Up 0.194508875473784
0.869549242859622 0.966567420808047
GO_HINDBRAIN_MORPHOGENESIS 28 0.01 Down
0.292155575129767 0.869758235171009 0.966567420808047
GO_REGULATION_OF_THYMOCYTE_APOPTOTIC_PROCESS 12 0.01 Down
0.428679604807204 0.869780964920828
0.966567420808047
GO_INDUCTION_OF_POSITIVE_CHEMOTAXIS 11 0.01 Up
0.75364359331641 0.869926599785071 0.966567420808047
GO_OXIDOREDUCTASE_ACTIVITY_ACTING_ON_NAD_P_H_HEME_PROTEIN_AS_ACCEPTO
R 11 0.01 Up 0.369673867541682
0.870154922003553 0.966632604376358
GO_CHAPERONE_MEDIATED_AUTOPHAGY 14 0.01 Down
0.241225588190818 0.870303708700572 0.966632604376358
GO_POSITIVE_REGULATION_OF_NEURAL_PRECURSOR_CELL_PROLIFERATION 31
0.01 Up 0.316666016965981 0.870747098590896
0.966781009944514
GO_CATALYTIC_ACTIVITY_ACTING_ON_RNA 329 0.01 Down
0.170319586483016 0.870879380951693 0.966781009944514
GO_NEGATIVE_REGULATION_OF_MRNA_SPLICING_VIA_SPLICEOSOME 19 0.01
Down 0.0906117851163014 0.87091506232689
0.966781009944514
GO_MITOTIC_CELL_CYCLE_ARREST 14 0.01 Down
0.362672824882571 0.871172496382695 0.966889986387449
GO_REGULATION_OF_REMOVAL_OF_SUPEROXIDE_RADICALS 10 0.01 Up
0.673087240024655 0.872012176731107
0.96707912569799
GO_PROTEIN_LOCALIZATION_TO_MITOCHONDRION 134 0.01 Down
0.152265277005587 0.872168771317453
0.96707912569799
GO_REGULATION_OF_PROTEIN_BINDING 181 0.01 Down
0.342862958722764 0.872237734355825 0.96707912569799
GO_NUCLEOLAR_LARGE_RRNA_TRANSCRIPTION_BY_RNA_POLYMERASE_I 14
0.01 Down 0.125569228598635 0.872311555105333
0.96707912569799
GO_POSITIVE_REGULATION_OF_PEPTIDASE_ACTIVITY 157 0.01 Up
0.39172274873786 0.872910109946471
0.96707912569799
GO_SEXUAL_REPRODUCTION 476 0.01 Up 0.303392133689473
0.87294244201824 0.96707912569799

GO_REGULATION_OF_AUTOPHAGY_OF_MITOCHONDRION 39 0.01 Up
0.152164610010331 0.873042755942376
0.96707912569799
GO_CELLULAR_RESPONSE_TO_EXOGENOUS_DSRNA 16 0.01 Down
0.357307479560173 0.873370540150604 0.96707912569799
GO_PROTEIN_LOCALIZATION_TO_CHROMATIN 25 0.01 Down
0.173586769288484 0.873452130527303 0.96707912569799
GO_CYTOSOLIC_CALCIUM_ION_TRANSPORT 112 0.01 Up
0.493932034457269 0.873511598347853 0.96707912569799
GO_U1_SNRNP 19 0.01 Up 0.0796594894954556
0.873719623507122 0.96707912569799
GO_SPHINGOLIPID_BINDING 14 0.01 Up 0.403640199621351
0.873857987767745 0.96707912569799
GO_POSITIVE_REGULATION_OF_ACTIVATED_T_CELL_PROLIFERATION 16
0.01 Down 0.375994737681446 0.873866297885611
0.96707912569799
GO_RESPONSE_TO_DRUG 670 0.01 Down 0.398600529007759
0.874048262300312 0.96707912569799
GO_POSITIVE_REGULATION_OF_ION_TRANSPORT 154 0.01 Up
0.506769002325492 0.874294990751975 0.96707912569799
GO_CELL_BODY 353 0.01 Up 0.327933671641159
0.874344298460874 0.96707912569799
GO_NEUROINFLAMMATORY_RESPONSE 38 0.01 Up
0.759470285323305 0.87435754272932 0.96707912569799
GO_POSITIVE_REGULATION_OF_OSTEOCLAST_DIFFERENTIATION 16 0.01
Down 0.575854743516801 0.874433310227815
0.96707912569799
GO_TRANSPORT_VESICLE_MEMBRANE 144 0.01 Down
0.33795616951354 0.874576454640679 0.96707912569799
GO_ADULT_BEHAVIOR 70 0.01 Down 0.353511473189678
0.8747437215773 0.96707912569799
GO_KIDNEY_MESENCHYME_DEVELOPMENT 15 0.01 Down
0.721045973401572 0.874769936400567 0.96707912569799
GO_HEAD_DEVELOPMENT 537 0.01 Down 0.352569875355546
0.874953204966238 0.96707912569799
GO_RELEASE_OF_CYTOCHROME_C_FROM_MITOCHONDRIA 53 0.01 Down
0.333735741451189 0.875006693701048
0.96707912569799
GO_LENS_FIBER_CELL_DIFFERENTIATION 23 0.01 Up
0.278127265894787 0.875401569414695 0.96733944811005
GO_RNA_POLYMERASE_III_ACTIVITY 15 0.01 Up
0.142844380830288 0.875628082203001 0.967413664614089
GO_RNA_CAPPING 34 0.01 Up 0.114149802306729
0.875891621502824 0.967419534064763
GO_PROTEIN_TYROSINE_KINASE_BINDING 69 0.01 Up
0.313408712013898 0.875952096648658 0.967419534064763
GO_RESPONSE_TO_MITOCHONDRIAL_DEPOLARISATION 19 0.01 Down
0.125891613109675 0.876204850201825
0.967522671075897
GO_ANTERIOR_POSTERIOR_AXIS_SPECIFICATION 31 0.01 Up
0.513982897577091 0.876752742211604
0.967792831307962
GO_PHOSPHATIDYLINOSITOL_3_4_BISPHOSPHATE_BINDING 24 0.01
Down 0.27898431874541 0.876909868715228

0.967792831307962
GO_ATPASE_DEPENDENT_TRANSMEMBRANE_TRANSPORT_COMPLEX 10 0.01
Up 0.395076562913551 0.87699502871628
0.967792831307962
GO_POSITIVE_REGULATION_OF_TRANSCRIPTION_FROM_RNA_POLYMERASE_II_PROMOTER_IN_RESPONSE_TO_STRESS 25 0.01 Up
0.236942558555321 0.877500691978139 0.967792831307962
GO_PROTEIN_FOLDING 190 0.01 Down 0.17448036524235
0.877592780964821 0.967792831307962
GO_NUCLEASE_ACTIVITY 164 0.01 Up 0.207503564244549
0.877695402330752 0.967792831307962
GO_NEGATIVE_REGULATION_OF_SMOOTHENED_SIGNALING_PATHWAY 25 0.01
Up 0.321836866026343 0.877725117775859
0.967792831307962
GO_REGULATION_OF_CELL_CYCLE_G2_M_PHASE_TRANSITION 193 0.01
Down 0.155157769927287 0.87777802297566
0.967792831307962
GO_EYE_PHOTORECEPTOR_CELL_DEVELOPMENT 15 0.01 Down
0.623112752392183 0.878040467872789 0.967792831307962
GO_MICROTUBULE_ORGANIZING_CENTER 640 0.01 Up
0.229054010854133 0.878180426043667 0.967792831307962
GO_G0_TO_G1_TRANSITION 44 0.01 Down 0.187539593577042
0.878404741393188 0.967792831307962
GO_PROTEIN_LOCALIZATION_TO_MEMBRANE 521 0.01 Up
0.239468190008266 0.878526807014965 0.967792831307962
GO_REGULATION_OF_RECEPTOR_INTERNALIZATION 36 0.01 Down
0.382663535465672 0.878561447537803
0.967792831307962
GO_PIGMENT_GRANULE 94 0.01 Down 0.21389258040043
0.878681285812796 0.967792831307962
GO_INTRINSIC_COMPONENT_OF_EXTERNAL_SIDE_OF_PLASMA_MEMBRANE 14
0.01 Up 0.798918340965582 0.879011889113474
0.967981349321223
GO_NEGATIVE_REGULATION_OF_INSULIN_SECRETION 26 0.01 Up
0.386617110174113 0.87945927452792
0.968255950193711
GO_MULTI_ORGANISM_REPRODUCTIVE_PROCESS 601 0.01 Up
0.348415484958775 0.879580228186183 0.968255950193711
GO_TRIGLYCERIDE_CATABOLIC_PROCESS 12 0.01 Down
0.435427042583368 0.879793433597939 0.968267535712995
GO_TRANSCRIPTION_EXPORT_COMPLEX 13 0.01 Up
0.142611883056815 0.879909733903573 0.968267535712995
GO_NEGATIVE_REGULATION_OF_JNK_CASCADE 32 0.01 Up
0.414785267902553 0.880424843216546 0.968658793615015
GO_DNA_POLYMERASE_ACTIVITY 27 0.01 Down
0.136631920658924 0.880725052645367 0.968813515964853
GO_ENDOPLASMIC_RETICULUM_CHAPERONE_COMPLEX 10 0.01 Up
0.269852541028687 0.880991441226706
0.968930985450604
GO_NEGATIVE_REGULATION_OF_RECEPTOR_MEDIATED_ENDOCYTOSIS 16 0.01
Down 0.427196410461104 0.881168750902894
0.968950459469565
GO_SPERMATID_DIFFERENTIATION 83 0.01 Up
0.309203408628516 0.881609994472646 0.969219974371921

GO_GASTRULATION_WITH_MOUTH_FORMING_SECOND 21 0.01 Down 0.333164649240548 0.881733144202952 0.969219974371921
GO_CELL_CORTEX_REGION 33 0.01 Up 0.249347598252103 0.881944643448081 0.969276960603421
GO_CYTOPLASMIC_DYNEIN_COMPLEX 19 0.01 Down 0.131034532786131 0.882324521102101 0.969347776976864
GO_TAT_PROTEIN_BINDING 10 0.01 Up 0.238069732187346 0.88236598806373 0.969347776976864
GO_NEGATIVE_REGULATION_OF_WNT_SIGNALING_PATHWAY 163 0.01 Down 0.35651631121341 0.882488084887354 0.969347776976864
GO_PROTEIN_LOCALIZATION_TO_KINETOCHORE 14 0.01 Down 0.137922976375254 0.882899358709907 0.969568662310082
GO_VOLTAGE_GATED_CALCIUM_CHANNEL_COMPLEX 18 0.01 Up 0.417708813745728 0.883131966604189 0.969568662310082
GO_MODIFICATION_OF_SYNAPTIC_STRUCTURE 18 0.01 Down 0.180288096884686 0.883168292303534 0.969568662310082
GO_GLYCEROLIPID_CATABOLIC_PROCESS 35 0.01 Up 0.461743012779135 0.88348244323482 0.969738187105152
GO_NEGATIVE_REGULATION_OF_REPRODUCTIVE_PROCESS 32 0.01 Down 0.484102290803439 0.884252814709253 0.970408322143867
GO_TRANSCRIPTION_FACTOR_TFTC_COMPLEX 14 0.01 Down 0.146745033193316 0.884743052662948 0.970770842710421
GO_CELL_ADHESION_MEDIATOR_ACTIVITY 48 0.01 Down 0.452446956677349 0.884905316499751 0.97077343268341
GO_GAMMA_DELTA_T_CELL_ACTIVATION 13 0.01 Up 0.524305000921672 0.885110571858005 0.970823176467922
GO_REPRODUCTION 904 0.01 Down 0.348052257256584 0.885502958976328 0.971044374236594
GO_NEGATIVE_REGULATION_OF_PLATELET_ACTIVATION 13 0.01 Up 0.649977939668727 0.885632136410479 0.971044374236594
GO_REGULATION_OF_RECEPTOR_RECYCLING 21 0.01 Up 0.271294472828419 0.8859410891031 0.971137990997813
GO_PHOSPHOTRANSFERASE_ACTIVITY_PHOSPHATE_GROUP_AS_ACCEPTOR 31 0.01 Up 0.259462562797828 0.886180878315671 0.971137990997813
GO_HYDROGEN_PEROXIDE_METABOLIC_PROCESS 11 0.01 Up 0.586796294197781 0.886197409014641 0.971137990997813
GO_SKELETAL_MUSCLE_TISSUE_REGENERATION 23 0.01 Down 0.342046383433921 0.886475750517745 0.971267692003831
GO_GLUTAMINE_METABOLIC_PROCESS 18 0.01 Down 0.282317181467649 0.886698559901686 0.971336513382017
GO_REGULATION_OF_ESTABLISHMENT_OF_PLANAR_POLARITY 103 0.01 Down 0.284118729177485 0.887133565010582 0.971499917070482
GO_NEGATIVE_REGULATION_OF_TRANSCRIPTION_REGULATORY_REGION_DNA_BINDING 14 0.01 Down 0.398997815087543 0.887167771411423 0.971499917070482
GO_DIVALENT_INORGANIC_CATION_HOMEOSTASIS 284 0.01 Up 0.466480063080375 0.887350176603428

0.971524422391238
GO_DICHOTOMOUS_SUBDIVISION_OF_AN_EPITHELIAL_TERMINAL_UNIT 10
0.01 Up 0.534436030485358 0.888041439661235
0.971740424213915
GO_HISTONE_PHOSPHORYLATION 32 0.01 Up
0.20877672792407 0.88821024517853 0.971740424213915
GO_NEGATIVE_REGULATION_OF_BLOOD_VESSEL_DIAMETER 38 0.01 Down
0.465313649596315 0.888347430208108
0.971740424213915
GO_EPIDERMIS_MORPHOGENESIS 20 0.01 Down
0.549573081242352 0.888515206772824 0.971740424213915
GO_TRANSFERASE_ACTIVITY_TRANSFERRING_PENTOSYL_GROUPS 42 0.01
Up 0.233505492295875 0.888801975615004
0.971740424213915
GO_VESICLE_FUSION_TO_PLASMA_MEMBRANE 15 0.01 Down
0.137261898896149 0.888933964654261 0.971740424213915
GO_POSTREPLICATION_REPAIR 49 0.01 Down
0.12944960264014 0.889251838335801 0.971740424213915
GO_GERM_CELL_DEVELOPMENT 145 0.01 Down
0.318697728646239 0.889287511772467 0.971740424213915
GO_CELLULAR_RESPONSE_TO_LEPTIN_STIMULUS 12 0.01 Down
0.30421447215977 0.889331660017006 0.971740424213915
GO_PROTEIN_ADP_RIBOSYLASE_ACTIVITY 15 0.01 Down
0.146995398227927 0.889477438118039 0.971740424213915
GO_GOLGI_CISTERNA_MEMBRANE 70 0.01 Down
0.302427062230171 0.889491428449885 0.971740424213915
GO_LAMELLIPODIUM_ORGANIZATION 71 0.01 Down
0.251057350346479 0.889870923020498 0.971740424213915
GO_RECEPTOR_METABOLIC_PROCESS 143 0.01 Down
0.349195836509314 0.889889129568995 0.971740424213915
GO_TRICARBOXYLIC_ACID_CYCLE_ENZYME_COMPLEX 12 0.01 Up
0.135208821464402 0.889967336462789
0.971740424213915
GO_T_HELPER_2_CELL_DIFFERENTIATION 13 0.01 Up
0.490953082718719 0.890070151246475 0.971740424213915
GO_STEROL_HOMEOSTASIS 50 0.01 Down 0.377190698123885
0.890108466324095 0.971740424213915
GO_TRANSLATION_REPRESSOR_ACTIVITY 22 0.01 Down
0.301374926855927 0.890601176647935 0.971759318161707
GO_S_ADENOSYLMETHIONINE_DEPENDENT_METHYLTRANSFERASE_ACTIVITY 143
0.01 Up 0.170976011910081 0.890742399436938
0.971759318161707
GO_REGULATION_OF_SYNAPTIC_VESICLE_ENDOCYTOSIS 14 0.01 Down
0.224776807118248 0.890848896411198
0.971759318161707
GO_DNA_CONFORMATION_CHANGE 215 0.01 Up
0.183397259039536 0.891033196842846 0.971759318161707
GO_NEUTRAL_LIPID_BIOSYNTHETIC_PROCESS 28 0.01 Down
0.241015727405259 0.891273103007513 0.971759318161707
GO_POTASSIUM_CHANNEL_REGULATOR_ACTIVITY 31 0.01 Down
0.444826137289736 0.891306179263966 0.971759318161707
GO_POSITIVE_REGULATION_OF_CYTOSOLIC_CALCIUM_ION_CONCENTRATION_INVOLV
ED_IN_PHOSPHOLIPASE_C_ACTIVATING_G_PROTEIN_COUPLED_SIGNALING_PATHWAY
13 0.01 Down 0.535427066207501

0.891590516644348	0.971759318161707
GO_PROTEIN_KINASE_C_BINDING	44	0.01	Up
0.361593607474324	0.891704506343265	0.971759318161707
GO_POSITIVE_REGULATION_OF_SMOOTH_MUSCLE_CELL_APOPTOTIC_PROCESS	12
	0.01	Up	0.416541479070015	0.891705933228602
	0.971759318161707
GO_NEGATIVE_REGULATION_OF_RESPONSE_TO_DRUG	22	0.01	Down
	0.377303194700163	0.891726430815165
0.971759318161707
GO_REGULATION_OF_MYELOID_LEUKOCYTE_DIFFERENTIATION	83	0.01
	Up	0.425199069534824	0.892080219721535
0.971970390152447
GO_PHOSPHATE_ION_TRANSMEMBRANE_TRANSPORT	11	0.01	Down
	0.200975128809151	0.892347135496926
0.972086750332288
GO_LONG_CHAIN_FATTY_ACID_IMPORT_INTO_CELL	13	0.01	Up
	0.394339224249574	0.892862523080216
0.972473695303191
GO_REGULATION_OF_CYTOLYSIS	10	0.01	Down
0.885508710939703	0.893120941215223	0.972475637633971
GO_FILAMENTOUS_ACTIN	18	0.01	Up	0.265856937928011
	0.893184673932963	0.972475637633971
GO_U12_TYPE_SPLICEOSOMAL_COMPLEX	26	0.01	Up
0.103746721651838	0.893598632874784	0.972636185668145
GO_NEGATIVE_REGULATION_OF_DNA_RECOMBINATION	31	0.01	Up
	0.184274274553202	0.893684280325745
0.972636185668145
GO_INOSITOL_PHOSPHATE_BIOSYNTHETIC_PROCESS	30	0.01	Down
	0.435335896789854	0.893812762286704
0.972636185668145
GO_RHYTHMIC_BEHAVIOR	22	0.01	Down	0.354111670015698
	0.894043103553056	0.972712487754589
GO_RESPIRATORY_BURST	26	0.01	Up	0.5819189022295
0.894551985056537	0.973091757978541
GO_NEGATIVE_REGULATION_OF_SYNAPSE_ORGANIZATION	13	0.01	Up
	0.573010998140473	0.894746516883305
0.973129004657568
GO_GOLGI_CIS_CISTERNA	10	0.01	Down	0.357931785245504
	0.895246657299506	0.973360144041426
GO_POSITIVE_REGULATION_OF_DNA_RECOMBINATION	34	0.01	Up
	0.276854750494343	0.895279697632569
0.973360144041426
GO_REGULATION_OF_G_PROTEIN_COUPLED_RECEPTOR_SIGNALING_PATHWAY	89
	0.01	Up	0.423779547518812	0.895507292726722
	0.973433263051733
GO_MONOSACCHARIDE_METABOLIC_PROCESS	119	0.01	Down
0.345885866326077	0.895995387536775	0.973637346424934
GO_GDP_DISSOCIATION_INHIBITOR_ACTIVITY	13	0.01	Up
0.25789117855167	0.896145135500591	0.973637346424934
GO_MACROPHAGE_APOPTOTIC_PROCESS	10	0.01	Up
0.661518728566372	0.896176164029407	0.973637346424934
GO_CIS_TRANS_ISOMERASE_ACTIVITY	34	0.01	Down
0.191470210721699	0.896953603555742	0.974307626979228
GO_SMAD_PROTEIN_SIGNAL_TRANSDUCTION	47	0.01	Up

0.355416720540941	0.897300059291768	0.974497399772821
GO_MAINTENANCE_OF_PROTEIN_LOCALIZATION_IN_ENDOPLASMIC_RETICULUM 10
	0.01	Up	0.113187947988794	0.897656254011121
	0.974497399772821
GO_R_SMAD_BINDING	21	0.01	Up	0.390459515933505
	0.897763457354122	0.974497399772821
GO_NEGATIVE_REGULATION_OF_PLATELET_DERIVED_GROWTH_FACTOR_RECEPTOR_SI
GNALING_PATHWAY 11	0.01	Up	0.426689993461807
0.897839347548639	0.974497399772821
GO_NEGATIVE_REGULATION_OF_DNA_BINDING	41	0.01	Up
0.283621468970478	0.89793089348199	0.974497399772821
GO_3_5_RNA_HELICASE_ACTIVITY	19	0.01	Down
0.196958192682353	0.898128936332299	0.974538118404538
GO_B_CELL_RECEPTOR_SIGNALING_PATHWAY	36	0.01	Up
0.42151133086004	0.89854115094072	0.974628302043847
GO_POSITIVE_REGULATION_OF_PEPTIDE_HORMONE_SECRETION	57	0.01
	Down	0.302518073460989	0.898671493304632
0.974628302043847
GO_INTRONIC_TRANSCRIPTION_REGULATORY_REGION_DNA_BINDING 11	0.01
	Down	0.227118681340389	0.898808874065803
0.974628302043847
GO_CLATHRIN_COAT_OF_COATED_PIT	17	0.01	Down
0.151028083325718	0.898871515063259	0.974628302043847
GO_RESPONSE_TO_IONIZING_RADIATION	138	0.01	Down
0.226888704375215	0.899014740808029	0.974628302043847
GO_TYPE_I_TRANSFORMING_GROWTH_FACTOR_BETA_RECEPTOR_BINDING	11
	0.01	Down	0.361279514119295	0.899378574415743
	0.974848656539542
GO_CHAPERONE_MEDIATED_PROTEIN_FOLDING	53	0.01	Down
0.249105346154043	0.900105188546585	0.975462084910089
GO_CARDIAC_MUSCLE_CELL_ACTION_POTENTIAL 45	0.01	Up
0.459303705044514	0.900714358056177	0.975533284773038
GO_AMINO_SUGAR_BIOSYNTHETIC_PROCESS	13	0.01	Up
0.439762624416836	0.900810525414529	0.975533284773038
GO_REGULATION_OF_B_CELL_DIFFERENTIATION 23	0.01	Up
0.490034118784764	0.900828025662351	0.975533284773038
GO_CELLULAR_RESPONSE_TO_LIPOPROTEIN_PARTICLE_STIMULUS	26	0.01
	Down	0.472508185492861	0.900885249076698
0.975533284773038
GO_LIGASE_ACTIVITY_FORMING_CARBON_OXYGEN_BONDS	41	0.01	Up
	0.123200995608062	0.900974325106642
0.975533284773038
GO_REGULATION_OF_AMYLOID_BETA_CLEARANCE 12	0.01	Down
0.499916919686801	0.901519906539741	0.975601792895882
GO_AGGRESOME	36	0.01	Up	0.249926884653058
0.901544623829824	0.975601792895882
GO_CYTOPLASMIC_MICROTUBULE	55	0.01	Up
0.206527583006278	0.90159286464781	0.975601792895882
GO_DNA_DIRECTED_DNA_POLYMERASE_ACTIVITY 22	0.01	Up
0.148321341278294	0.901680392017591	0.975601792895882
GO_DNA_REPLICATION	240	0.01	Up	0.181755395419846
	0.902287014460582	0.975934585326592
GO_DIGESTIVE_TRACT_MORPHOGENESIS	34	0.01	Up
0.630662421673467	0.902384074266136	0.975934585326592

GO_PERICARDIUM_DEVELOPMENT 14 0.01 Down
0.442920941486071 0.902489647834122 0.975934585326592
GO_POSITIVE_REGULATION_OF_NUCLEAR_DIVISION 49 0.01 Down
0.371508724256028 0.902765583205355
0.975934585326592
GO_CYTOPLASMIC_SEQUESTERING_OF_PROTEIN 23 0.01 Up
0.25206864028783 0.902791736319246 0.975934585326592
GO_RIBONUCLEASE_P_COMPLEX 13 0.01 Up
0.117918042881847 0.903093663278287 0.975967567582307
GO_CELL_DIFFERENTIATION_IN_SPINAL_CORD 18 0.01 Up
0.470222053970873 0.903197119870336 0.975967567582307
GO_CELLULAR_PROCESS_INVOLVED_IN_REPRODUCTION_IN_MULTICELLULAR_ORGANI
SM 205 0.01 Up 0.323957969119107
0.903304523512598 0.975967567582307
GO_POSITIVE_REGULATION_OF_CELLULAR_CARBOHYDRATE_METABOLIC_PROCESS
30 0.01 Down 0.333278859776895
0.903483028532423 0.975986737761626
GO_SERINE_FAMILY_AMINO_ACID_BIOSYNTHETIC_PROCESS 16 0.01
Down 0.305672013702018 0.903696573730066
0.976043746506891
GO_REGULATION_OF_ATPASE_ACTIVITY 59 0.01 Up
0.195350680062362 0.904147988287158 0.976122294638995
GO_REGULATION_OF_STEROL_TRANSPORT 31 0.01 Up
0.421380539162174 0.904227204876334 0.976122294638995
GO_RESPONSE_TO_PROSTAGLANDIN_E 15 0.01 Up
0.423908978744346 0.904301544668593 0.976122294638995
GO_POSITIVE_REGULATION_OF_VASOCONSTRICTION 14 0.01 Down
0.376920934074764 0.904458422029082
0.976122294638995
GO_ER_MEMBRANE_PROTEIN_COMPLEX 10 0.01 Up
0.083452705659693 0.90457322181502 0.976122294638995
GO_INTRINSIC_COMPONENT_OF_GOLGI_MEMBRANE 59 0.01 Down
0.203785200662628 0.904840485836513
0.976237176028696
GO_CARBOHYDRATE_METABOLIC_PROCESS 431 0.01 Up
0.299823204491966 0.905020948140393 0.976258382402332
GO_AXONEME_PART 20 0.01 Up 0.289671813866045
0.905578259098464 0.97668602078287
GO_NEGATIVE_REGULATION_OF_FATTY_ACID_BIOSYNTHETIC_PROCESS 13
0.01 Down 0.402828411711826 0.905786941362206
0.976737570339245
GO_SNRNA_3_END_PROCESSING 17 0.01 Down
0.143715059742584 0.906121822736893 0.976888208625073
GO_REGULATION_OF_RESPONSE_TO_EXTRACELLULAR_STIMULUS 15 0.01
Up 0.322413781552776 0.906248458404944
0.976888208625073
GO_TUBE_DEVELOPMENT 765 0.01 Down 0.49488490747899
0.906900747321332 0.977114973013425
GO_SARCOMERE_ORGANIZATION 27 0.01 Down
0.414468310309134 0.906950451297988 0.977114973013425
GO_LIPID_BINDING 509 0.01 Up 0.377704798503106
0.907100079435452 0.977114973013425
GO_MITOCHONDRIAL_CALCIUM_ION_HOMEOSTASIS 20 0.01 Up
0.217083358760983 0.907102617015922

0.977114973013425
GO_METANEPHRIC_EPITHELIUM_DEVELOPMENT 15 0.01 Up
0.428181268918549 0.908001686731927 0.977791842855923
GO_CHAPERONE_COFACTOR_DEPENDENT_PROTEIN_REFOLDING 29 0.01
Down 0.298579826277819 0.908053106575802
0.977791842855923
GO_REGULATION_OF_DNA_RECOMBINATION 80 0.01 Down
0.234060386200986 0.908442423978696 0.978027322075677
GO_RIBONUCLEOTIDE_CATABOLIC_PROCESS 25 0.01 Up
0.29998591067252 0.908593987235516 0.978027322075677
GO_RESPONSE_TO_ACTIVITY 42 0.01 Down 0.337023198174816
0.908787168626987 0.978061850865881
GO_AUTOPHAGOSOME_MEMBRANE 27 0.01 Down
0.173172532532392 0.909545524850923 0.978704516371846
GO_NEGATIVE_REGULATION_OF_DOUBLE_STRAND_BREAK_REPAIR_VIA_HOMOLOGOUS_
RECOMBINATION 14 0.01 Down 0.191685783095152
0.91022057256506 0.979094391091949
GO_EPIBOLY 27 0.01 Up 0.34101732831284
0.910323448979869 0.979094391091949
GO_PROTEIN_HETEROTETRAMERIZATION 29 0.01 Up
0.212434113096141 0.910617258028406 0.979094391091949
GO_CELLULAR_METABOLIC_COMPOUND_SALVAGE 30 0.01 Down
0.229266447483347 0.910768972975786 0.979094391091949
GO_NEGATIVE_REGULATION_OF_ION_TRANSPORT 88 0.01 Up
0.472818182927815 0.911146836613923 0.979094391091949
GO_CARBOXY_TERMINAL_DOMAIN_PROTEIN_KINASE_COMPLEX 21 0.01
Up 0.192958147371017 0.911344638087843
0.979094391091949
GO_UBIQUITIN_PROTEIN_TRANSFERASE_REGULATOR_ACTIVITY 16 0.01
Up 0.148634144399172 0.911416165595143
0.979094391091949
GO_FIBROBLAST_GROWTH_FACTOR_RECEPTOR_SIGNALING_PATHWAY 84 0.01
Up 0.336230398663421 0.911560907175969
0.979094391091949
GO_POLYPEPTIDE_N_ACETYLGALACTOSAMINYLTRANSFERASE_ACTIVITY 13
0.01 Up 0.328503688501522 0.911646488956182
0.979094391091949
GO_NEURON_PROJECTION_GUIDANCE 196 0.01 Down
0.367220018719648 0.911710528400032 0.979094391091949
GO_REGULATION_OF_CELL_SUBSTRATE_ADHESION 161 0.01 Up
0.510088489669879 0.911733011236359
0.979094391091949
GO_SPERM_AXONEME_ASSEMBLY 10 0.01 Down
0.246758408426398 0.911843137412927 0.979094391091949
GO_NEURAL_PRECURSOR_CELL_PROLIFERATION 96 0.01 Down
0.311572063842446 0.912440359959245 0.979562409427511
GO_CHEMOATTRACTANT_ACTIVITY 24 0.01 Up
0.776576271262578 0.912724604296412 0.979611346564907
GO_G2_DNA_DAMAGE_CHECKPOINT 30 0.01 Up
0.167804018939644 0.913051017202361 0.979611346564907
GO_GLAND_MORPHOGENESIS 97 0.01 Down 0.47781663172643
0.91312721105588 0.979611346564907
GO_SYNAPTIC_VESICLE_PRIMING 13 0.01 Down
0.187089798965549 0.913131380367453 0.979611346564907

GO_NEGATIVE_REGULATION_OF_RHO_PROTEIN_SIGNAL_TRANSDUCTION 18 0.01 Down 0.286211641569489 0.913316211712241 0.979636523198766
GO_FORELIMB_MORPHOGENESIS 21 0.01 Down 0.447311621002063 0.914102758162212 0.98030698548009
GO_CELL_DIVISION_SITE 61 0.01 Up 0.228588468892537 0.914266537776222 0.980309457937026
GO_TRANSCRIPTION_BY_RNA_POLYMERASE_III 53 0.01 Up 0.145101230397364 0.915116128902881 0.980930375913282
GO_REGULATION_OF_CALCIUM_ION_TRANSMEMBRANE_TRANSPORTER_ACTIVITY 55 0.01 Up 0.393447157443282 0.915171146083121 0.980930375913282
GO_NEGATIVE_REGULATION_OF_CELL_SUBSTRATE_ADHESION 50 0.01 Down 0.431442830605552 0.915330354068315 0.980930375913282
GO_NEGATIVE_REGULATION_OF_NEURAL_PRECURSOR_CELL_PROLIFERATION 15 0.01 Down 0.199106016000975 0.91611219872326 0.981322964119636
GO_RESPONSE_TO_ORGANOPHOSPHORUS 90 0.01 Up 0.487150547027354 0.916292299704968 0.981322964119636
GO_POSTTRANSLATIONAL_PROTEIN_TARGETING_TO_ENDOPLASMIC_RETICULUM_MEMBRANE 11 0.01 Down 0.101300715228639 0.916333474410219 0.981322964119636
GO_NEGATIVE_REGULATION_OF_BINDING 132 0.01 Up 0.267310248580192 0.91657088216844 0.981322964119636
GO_KERATAN_SULFATE_BIOSYNTHETIC_PROCESS 23 0.01 Down 0.811737990136957 0.916798645745563 0.981322964119636
GO_REGULATION_OF_LIPID_TRANSPORT 64 0.01 Up 0.406683688059053 0.917014037115397 0.981322964119636
GO_RAC_GTPASE_BINDING 62 0.01 Up 0.272806050696317 0.917179153114998 0.981322964119636
GO_REGULATION_OF_OXIDATIVE_PHOSPHORYLATION 19 0.01 Up 0.1876979357589 0.917246984065419 0.981322964119636
GO_B_CELL_ACTIVATION_INVOLVED_IN_IMMUNE_RESPONSE 56 0.01 Down 0.339036383349755 0.917443810865155 0.981322964119636
GO_TRICARBOXYLIC_ACID_METABOLIC_PROCESS 10 0.01 Down 0.162415433146785 0.917562252593982 0.981322964119636
GO_CAMERA_TYPE_EYE_DEVELOPMENT 206 0.01 Up 0.382109013848038 0.917604250429571 0.981322964119636
GO_SINGLE_STRANDED_DNA_BINDING 95 0.01 Down 0.159032226887507 0.917680855517009 0.981322964119636
GO_PURINE_CONTAINING_COMPOUND_SALVAGE 13 0.01 Down 0.283843949410331 0.917900049746267 0.981322964119636
GO_ENDONUCLEASE_ACTIVITY_ACTIVE_WITH_EITHER_RIBO_OR_DEOXYRIBONUCLEIC_ACIDS_AND_PRODUCING_5_PHOSPHOMONOESTERS 36 0.01 Down 0.139582240519578 0.918113433208266 0.981322964119636
GO_BLASTOCYST_GROWTH 17 0.01 Up 0.128915011058827 0.918121303936671 0.981322964119636
GO_MUSCLE_ORGAN_DEVELOPMENT 265 0.01 Up 0.455966207305647 0.918391184510122 0.981352858196362
GO_VACUOLE_ORGANIZATION 149 0.01 Down 0.196446802891694 0.91847256469638 0.981352858196362

GO_SEPTIN_CYTOSKELETON 12 0.01 Up 0.218612130962991
0.919382238179145 0.981969188579348
GO_REGULATION_OF_VOLTAGE_GATED_CALCIUM_CHANNEL_ACTIVITY 22 0.01
Up 0.388433834340103 0.919547278118949
0.981969188579348
GO_PROTEIN_DIMERIZATION_ACTIVITY 934 0.01 Up
0.334373441362303 0.920022330052295 0.981969188579348
GO_INTERFERON_GAMMA_BIOSYNTHETIC_PROCESS 10 0.01 Down
0.702546517397314 0.920401784244796
0.981969188579348
GO_MOTILE_CILIUM_ASSEMBLY 18 0.01 Up
0.369415959955519 0.92043344401189 0.981969188579348
GO_BLOOD_VESSEL_MORPHOGENESIS 440 0.01 Down
0.543482625841653 0.920480135289448 0.981969188579348
GO_GLYCOLYTIC_PROCESS 86 0.01 Up 0.271491063857336
0.9206427771037 0.981969188579348
GO_REGULATION_OF_MONONUCLEAR_CELL_MIGRATION 39 0.01 Down
0.892827159806344 0.92082712835526
0.981969188579348
GO_RESPONSE_TO_ZINC_ION 30 0.01 Down 0.448414207153774
0.921008466770515 0.981969188579348
GO_DNA_SYNTHESIS_INVOLVED_IN_DNA_REPAIR 47 0.01 Down
0.136843007313876 0.921053074402608 0.981969188579348
GO_DNA_DOUBLE_STRAND_BREAK_PROCESSING 22 0.01 Down
0.198314175779125 0.921140512597549 0.981969188579348
GO_NEGATIVE_REGULATION_OF_PROTEIN_LOCALIZATION_TO_NUCLEUS 22
0.01 Up 0.555580839979331 0.921309785832399
0.981969188579348
GO_ESTABLISHMENT_OF_PROTEIN_LOCALIZATION_TO_TELOMERE 17 0.01
Down 0.0737840482158014 0.921338480441867
0.981969188579348
GO_MITOTIC_CHROMOSOME_CONDENSATION 14 0.01 Up
0.135240662651391 0.921595478121428 0.981969188579348
GO_MICROTUBULE_ORGANIZING_CENTER_PART 161 0.01 Up
0.2091476689026 0.921761324854359 0.981969188579348
GO_COENZYME_A_BIOSYNTHETIC_PROCESS 10 0.01 Up
0.238952535118314 0.922355996409671 0.981969188579348
GO_BRANCHING_MORPHOGENESIS_OF_AN_EPITHELIAL_TUBE 116 0.01
Down 0.546447052629441 0.922742039913693
0.981969188579348
GO_ORGAN_GROWTH 117 0.01 Down 0.466079466336679
0.922995046690202 0.981969188579348
GO_POSITIVE_REGULATION_OF_ISOTYPE_SWITCHING 22 0.01 Down
0.308998322865537 0.923283297032289
0.981969188579348
GO_NEGATIVE_REGULATION_OF_SUBSTRATE_ADHESION_DEPENDENT_CELL_SPREADIN
G 13 0.01 Up 0.389674404024612
0.923493964840072 0.981969188579348
GO_H4_H2A_HISTONE_ACETYLTRANSFERASE_COMPLEX 18 0.01 Down
0.117074402437206 0.923582421819757
0.981969188579348
GO_RNA_POLYMERASE_II_CORE_COMPLEX 15 0.01 Up
0.107255706810944 0.923589821981773 0.981969188579348
GO_CEREBRAL_CORTEX_RADIALLY_ORIENTED_CELL_MIGRATION 23 0.01

Up 0.272030345168097 0.923721879026862
0.981969188579348
GO_REGULATION_OF_METAL_ION_TRANSPORT 231 0.01 Down
0.506802539922221 0.92382996336234 0.981969188579348
GO_MICROTUBULE 332 0.01 Down 0.24429815957445
0.924061431807379 0.981969188579348
GO_RNA_METHYLATION 77 0.01 Up 0.123635668634634
0.924310514898157 0.981969188579348
GO_REGULATION_OF_CARDIAC_MUSCLE_TISSUE_DEVELOPMENT 56 0.01
Up 0.505269745687861 0.924484459194974
0.981969188579348
GO_LIGASE_ACTIVITY_FORMING_CARBON_SULFUR_BONDS 29 0.01 Up
0.240771832428726 0.924608357861996
0.981969188579348
GO_NADPH_BINDING 13 0.01 Down 0.229603127574588
0.924757717654062 0.981969188579348
GO_REPLISOME 27 0.01 Down 0.125905692010368
0.924784249534086 0.981969188579348
GO_POSITIVE_REGULATION_OF_CELL_JUNCTION_ASSEMBLY 29 0.01
Down 0.419951720205335 0.924799115211405
0.981969188579348
GO_SPECIFIC_GRANULE_LUMEN 44 0.01 Up
0.500211589813354 0.924911693364641 0.981969188579348
GO_POSITIVE_REGULATION_OF_INTERFERON_GAMMA_PRODUCTION 41 0.01
Up 0.72794280705167 0.924916086180299
0.981969188579348
GO_VASCULOGENESIS 54 0.01 Down 0.368148123819418
0.924976451054366 0.981969188579348
GO_BODY_MORPHOGENESIS 44 0.01 Down 0.336627074085048
0.925213553164319 0.981969188579348
GO_B_CELL_MEDIATED_IMMUNITY 77 0.01 Up
0.505624858837583 0.925267375495058 0.981969188579348
GO_UDP_GALACTOSYLTRANSFERASE_ACTIVITY 17 0.01 Down
0.429477979036984 0.92530325884719 0.981969188579348
GO_POSITIVE_REGULATION_OF_COLLATERAL_SPROUTING 10 0.01 Up
0.300526248133573 0.92537624844648
0.981969188579348
GO_FILAMIN_BINDING 10 0.01 Up 0.363639881384489
0.925517472673099 0.981969188579348
GO_REPLACEMENT_OSSIFICATION 20 0.01 Down
0.555793488125922 0.925519304406363 0.981969188579348
GO_CELL_SURFACE_RECEPTOR_SIGNALING_PATHWAY_INVOLVED_IN_HEART_DEVELOP
MENT 19 0.01 Down 0.437451760070956
0.926349982114686 0.982414647379104
GO_FOREBRAIN_NEURON_DEVELOPMENT 19 0.01 Down
0.489099739019637 0.926395423398751 0.982414647379104
GO_POSTSYNAPSE_ORGANIZATION 116 0.01 Down
0.335419421253106 0.926424618060512 0.982414647379104
GO_CALCIUM_DEPENDENT_PROTEIN_BINDING 43 0.01 Down
0.355933921051131 0.926749032931517 0.982511020618091
GO_MRNA_TRANSCRIPTION_BY_RNA_POLYMERASE_II 14 0.01 Down
0.293468678687381 0.92683917230766
0.982511020618091
GO_PEROXISOMAL_MEMBRANE_TRANSPORT 17 0.01 Down

0.134075773932832	0.927250729046985	0.982775694141803
GO_OSTEOBLAST_PROLIFERATION	19	0.01	Down
0.808977141400929	0.9280033249147	0.983088455894509
GO_CONDENSED_CHROMOSOME	168	0.01	Down	0.173967246701769
	0.928121238784556	0.983088455894509
GO_REGULATION_OF_ESTABLISHMENT_OF_PROTEIN_LOCALIZATION_TO_CHROMOSOME
	12	0.01	Up	0.0779961785377606
0.928278518110065	0.983088455894509
GO_T_CELL_MEDIATED_CYTOTOXICITY	36	0.01	Down
0.410445952403699	0.928471647794371	0.983088455894509
GO_POSITIVE_REGULATION_OF_INTERLEUKIN_8_SECRETION	11	0.01
	Down	0.649262525775747	0.928577523328366
0.983088455894509
GO_PERICENTRIC_HETEROCHROMATIN	16	0.01	Up
0.0956608710929128	0.929392109788754	0.983088455894509
GO_LEADING_EDGE_MEMBRANE	124	0.01	Down
0.422171777185682	0.929606271382261	0.983088455894509
GO_MICROVILLUS	60	0.01	Down	0.421075281998775
0.929658201094787	0.983088455894509
GO_LIPID_HOMEOSTASIS	86	0.01	Down	0.325571847781087
	0.929680917023969	0.983088455894509
GO_THYMOCYTE_APOPTOTIC_PROCESS	17	0.01	Down
0.355087357806444	0.929683320368671	0.983088455894509
GO_MRNA_5_UTR_BINDING	24	0.01	Up	0.176330127878837
	0.929710485957709	0.983088455894509
GO_CHEMICAL_HOMEOSTASIS	751	0.01	Down	0.393089037974893
	0.929816513030181	0.983088455894509
GO_RESPONSE_TO_HEPATOCYTE_GROWTH_FACTOR	14	0.01	Up
0.529582929833029	0.929881003422474	0.983088455894509
GO_POSITIVE_REGULATION_OF_CELL_CYCLE_PROCESS	229	0.01	Up
	0.269967350854613	0.931007853222162
0.983088455894509
GO_REGULATION_OF_PROTEIN_DEACETYLATION	34	0.01	Up
0.168248745183307	0.931057754060647	0.983088455894509
GO_PROTON_EXPORTING_ATPASE_ACTIVITY	27	0.01	Down
0.209209523783469	0.931233523872365	0.983088455894509
GO_POSITIVE_REGULATION_OF_DENDRITIC_SPINE_MORPHOGENESIS	16	0.01
	Down	0.384698115839256	0.931371492385117
0.983088455894509
GO_PIGMENT_CELL_DIFFERENTIATION	26	0.01	Down
0.416836112501623	0.931546778760878	0.983088455894509
GO_ENDOPLASMIC_RETICULUM_QUALITY_CONTROL_COMPARTMENT	22	0.01
	Down	0.164677650339521	0.931619807841104
0.983088455894509
GO_INOSITOL_TRISPHOSPHATE_BIOSYNTHETIC_PROCESS	19	0.01	Down
	0.401558529248387	0.931696093810162
0.983088455894509
GO_SPERM_MOTILITY	37	0.01	Up	0.286134566328118
	0.932027906310035	0.983088455894509
GO_POSITIVE_REGULATION_OF_SIGNALING_RECEPTOR_ACTIVITY	24	0.01
	Down	0.526525266439395	0.932146764070715
0.983088455894509
GO_VESICLE_MEMBRANE	600	0.01	Up	0.373574843066764
	0.932288577892527	0.983088455894509

GO_REGULATION_OF_RHODOPSIN_MEDIATED_SIGNALING_PATHWAY 11 0.01 Up 0.158000775541708 0.932383906900689 0.983088455894509
GO_POSITIVE_REGULATION_OF_MEMBRANE_PERMEABILITY 58 0.01 Down 0.239862942253548 0.93241233839123 0.983088455894509
GO_CEREBELLAR_CORTEX_FORMATION 13 0.01 Down 0.201151532669729 0.932462415996271 0.983088455894509
GO_VIRAL_TRANSLATION 14 0.01 Down 0.114067763021112 0.932536324164782 0.983088455894509
GO_RESPONSE_TO_GAMMA_RADIATION 52 0.01 Down 0.217455640520308 0.932562540015087 0.983088455894509
GO_GOLGI_APPARATUS 1269 0.01 Down 0.311000305708348 0.932733172246308 0.983088455894509
GO_SM_LIKE_PROTEIN_FAMILY_COMPLEX 74 0.01 Up 0.084742559400037 0.932742514246637 0.983088455894509
GO_EXIT_FROM_MITOSIS 28 0.01 Up 0.190968225418163 0.932774207386815 0.983088455894509
GO_REGULATION_OF_BODY_FLUID_LEVELS 335 0.01 Up 0.410142570859249 0.932817139812787 0.983088455894509
GO_RECEPTOR_INHIBITOR_ACTIVITY 17 0.01 Down 0.646067465325435 0.932889572460594 0.983088455894509
GO_TOLL_LIKE_RECEPTOR_3_SIGNALING_PATHWAY 18 0.01 Up 0.438683410212245 0.933195447852884 0.983240118694005
GO_DEACETYLASE_ACTIVITY 40 0.01 Up 0.215263302319141 0.933876020169754 0.983403579227047
GO_REGULATION_OF_DNA_DAMAGE_CHECKPOINT 17 0.01 Up 0.161947466684756 0.934014286948017 0.983403579227047
GO_REGULATION_OF_SENSORY_PERCEPTION 17 0.01 Down 0.50077423758417 0.934081707475969 0.983403579227047
GO_NEURAL_CREST_FORMATION 10 0.01 Down 0.216596918124966 0.934363469060363 0.983403579227047
GO_POSITIVE_REGULATION_OF_CELL_DIVISION 59 0.01 Down 0.391075395125603 0.934396172097785 0.983403579227047
GO_SEQUENCE_SPECIFIC_MRNA_BINDING 10 0.01 Up 0.275279015979913 0.93460571772623 0.983403579227047
GO_REGULATION_OF_ION_TRANSPORT 364 0.01 Up 0.486348710960066 0.934799524466348 0.983403579227047
GO_DNA_INTEGRITY_CHECKPOINT 148 0.01 Up 0.200260226438834 0.934977165214629 0.983403579227047
GO_CHROMOSOME_CONDENSATION 28 0.01 Down 0.217101825604594 0.935169969480915 0.983403579227047
GO_TERTIARY_GRANULE_LUMEN 42 0.01 Down 0.522542466373573 0.935454545554446 0.983403579227047
GO_SPLICEOSOMAL_COMPLEX_ASSEMBLY 54 0.01 Down 0.110365860115111 0.935676722413189 0.983403579227047
GO_OVARIAN_FOLLICLE_DEVELOPMENT 39 0.01 Down 0.352109257616211 0.935953070301875 0.983403579227047
GO_HORMONE_BINDING 51 0.01 Down 0.421409842657827 0.935992683910652 0.983403579227047
GO_REGULATION_OF_POSTSYNAPSE_ORGANIZATION 66 0.01 Up 0.365223608213792 0.936001113299726 0.983403579227047

GO_NEUROTROPHIN_TRK_RECEPTOR_SIGNALING_PATHWAY	27	0.01	Down
	0.311137754243829	0.936107646117416
0.983403579227047
GO_DNA_DAMAGE_RESPONSE_SIGNAL_TRANSDUCTION_BY_P53_CLASS_MEDIATOR
	98	0.01	Down	0.241157939708994
0.936122582825233	0.983403579227047
GO_POSITIVE_REGULATION_OF_EMBRYONIC_DEVELOPMENT	28	0.01	Up
	0.361158387936925	0.936248123157711
0.983403579227047
GO_DENDRITIC_CELL_MIGRATION	19	0.01	Up
0.443210323571164	0.936349449665419	0.983403579227047
GO_REGULATION_OF_CARDIAC_CONDUCTION	40	0.01	Down
0.339423023400221	0.936857837530369	0.983403579227047
GO_DIGESTIVE_SYSTEM_PROCESS	50	0.01	Up
0.528521375982432	0.93697679023166	0.983403579227047
GO_REGULATION_OF_CELL_CYCLE_PHASE_TRANSITION	394	0.01	Down
	0.202951636563387	0.937134632199651
0.983403579227047
GO_CELL_BODY_MEMBRANE	10	0.01	Down	0.701409048330109
	0.937135255673846	0.983403579227047
GO_INORGANIC_ION_TRANSMEMBRANE_TRANSPORT	482	0.01	Down
	0.405073236816366	0.937198911966296
0.983403579227047
GO_MITOCHONDRIAL_TRANSCRIPTION	15	0.01	Up
0.162062579511781	0.937238199540058	0.983403579227047
GO_POSITIVE_REGULATION_OF_CHEMOKINE_PRODUCTION	40	0.01	Down
	0.512916798941844	0.937476626758822
0.983433885234343
GO_REGULATION_OF_INSULIN_SECRETION	116	0.01	Down
0.334608589557549	0.937591060407903	0.983433885234343
GO_SPECIFICATION_OF_ANIMAL_ORGAN_IDENTITY	20	0.01	Down
	0.529063717390028	0.937829991471493
0.983514575612961
GO_CELLULAR_RESPONSE_TO_PROSTAGLANDIN_E_STIMULUS	14	0.01
	Up	0.45118864876176	0.938140007054798
0.983669772509444
GO_REGULATION_OF_VESICLE_MEDIATED_TRANSPORT	385	0.01	Down
	0.359972031104908	0.93831564555196
0.983684041468822
GO_REGULATION_OF_STEROID_METABOLIC_PROCESS	89	0.01	Up
	0.384743397442476	0.938885032853703
0.984111021142063
GO_NEGATIVE_REGULATION_OF_TYPE_I_INTERFERON_MEDIATED_SIGNALING_PATHW
AY	10	0.01	Down	0.238942846608471
0.939564437059053	0.984502339950139
GO_POSITIVE_REGULATION_OF_PEPTIDYL_LYSINE_ACETYLATION	29	0.01
	Up	0.322502959343932	0.939824342048925
0.984502339950139
GO_REGULATION_OF_HIGH_VOLTAGE_GATED_CALCIUM_CHANNEL_ACTIVITY	10
	0.01	Down	0.356459588724531	0.940040124426573
	0.984502339950139
GO_OUTER_DYNEIN_ARM_ASSEMBLY	10	0.01	Down
0.508421184373425	0.940060720449032	0.984502339950139
GO_REGULATION_OF_NEURON_PROJECTION_REGENERATION	25	0.01	Up

0.522054814330983	0.940184675282859
0.984502339950139
GO_TONGUE_DEVELOPMENT	13	0.01	Down	0.335458848202971
0.940231356783216	0.984502339950139
GO_PHOSPHATIDYLINOSITOL_5_PHOSPHATE_BINDING	13	0.01	Up
0.15445469031982	0.941074946385293
0.985101679826076
GO_PHOTORECEPTOR_CELL_DIFFERENTIATION	33	0.01	Up
0.505439367479837	0.941311550735772	0.985101679826076
GO_PHOSPHATIDYLINOSITOL_4_5_BISPHOSPHATE_PHOSPHATASE_ACTIVITY	12
0.01	Down	0.150282115835814	0.941373366464868
0.985101679826076
GO_AUDITORY_RECEPTOR_CELL_MORPHOGENESIS	10	0.01	Up
0.272768293551451	0.941837861108827	0.985101679826076
GO_PROTEIN_O_LINKED_GLYCOSYLATION	85	0.01	Up
0.335786108736726	0.94189068395737	0.985101679826076
GO_SYNAPSE_MATURATION	12	0.01	Up	0.206349012170215
0.941910426891912	0.985101679826076
GO_POSITIVE_REGULATION_OF_MITOTIC_CELL_CYCLE	127	0.01	Up
0.271964584611116	0.94193959008242
0.985101679826076
GO_SCF_UBIQUITIN_LIGASE_COMPLEX	58	0.01	Up
0.171239023747452	0.942526272542103	0.985438827899798
GO_PEPTIDYL_ARGININE_MODIFICATION	17	0.01	Up
0.153223866641068	0.942586604120264	0.985438827899798
GO_SINGLE_STRANDED_DNA_DEPENDENT_ATPASE_ACTIVITY	11	0.01
Down	0.128129890760703	0.942935575479901
0.985595707686634
GO_CEREBELLAR_CORTEX_DEVELOPMENT	33	0.01	Down
0.267270783902306	0.943296957203657	0.985595707686634
GO_REGULATION_OF_STRIATED_MUSCLE_CELL_DIFFERENTIATION	76	0.01
Down	0.561694997872972	0.943572677931445
0.985595707686634
GO_REGULATION_OF_NEUTROPHIL_ACTIVATION	10	0.01	Up
0.423550584789863	0.943624491256537	0.985595707686634
GO_ANTIGEN_PROCESSING_AND_PRESENTATION_OF_PEPTIDE_ANTIGEN	167
0.01	Down	0.330832984840319	0.943737754771559
0.985595707686634
GO_NEGATIVE_REGULATION_OF_ENDOPLASMIC_RETICULUM_STRESS_INDUCED_INTRINSIC_APOPTOTIC_SIGNALING_PATHWAY	16	0.01	Up
0.181721430636416	0.943771088595158	0.985595707686634
GO_TRACHEA_DEVELOPMENT	15	0.01	Up	0.312848169454444
0.943873076015498	0.985595707686634
GO_DOPAMINE_RECEPTOR_SIGNALING_PATHWAY	22	0.01	Up
0.32523941384762	0.944143900216637	0.985616936377353
GO_MIDBRAIN_DEVELOPMENT	65	0.01	Down	0.354739244977468
0.944276791148165	0.985616936377353
GO_TRANSCRIPTIONALLY_ACTIVE_CHROMATIN	21	0.01	Down
0.159546587266112	0.944623312732607	0.985616936377353
GO_NEGATIVE_REGULATION_OF_VIRAL_INDUCED_CYTOPLASMIC_PATTERN_RECOGNITION_RECEPTOR_SIGNALING_PATHWAY	11	0.01	Up
0.101243112293706	0.944779581662655	0.985616936377353
GO_CELL_MORPHOGENESIS_INVOLVED_IN_DIFFERENTIATION	553	0.01
Down	0.355807236211244	0.945070665734445

0.985616936377353
GO_SISTER_CHROMATID_SEGREGATION 152 0.01 Up
0.180851675579286 0.945686311801079 0.985616936377353
GO_REGULATION_OF_PROTEIN_FOLDING 10 0.01 Down
0.102908716470153 0.945761440422422 0.985616936377353
GO_PROTEIN_TYROSINE_KINASE_ACTIVITY 98 0.01 Up
0.408187777858095 0.945851954387674 0.985616936377353
GO_TELOMERE_MAINTENANCE_VIA_RECOMBINATION 14 0.01 Up
0.158992662382979 0.9459647363074 0.985616936377353
GO_HISTONE_H3_ACETYLATION 54 0.01 Up
0.213789972967123 0.946026610300482 0.985616936377353
GO_RESPONSE_TO_AMINO_ACID_STARVATION 47 0.01 Up
0.172603561132843 0.946280719496469 0.985616936377353
GO_POSITIVE_REGULATION_OF_CELLULAR_RESPONSE_TO_TRANSFORMING_GROWTH_FACTOR_BETA_STIMULUS 26 0.01 Down 0.278483460883377
0.946976180837048 0.985616936377353
GO_REGULATION_OF_RETROGRADE_PROTEIN_TRANSPORT_ER_TO_CYTOSOL 12
0.01 Up 0.241908566952732 0.946981429885576
0.985616936377353
GO_POSITIVE_REGULATION_OF_CYTOKINESIS 25 0.01 Down
0.251384297248371 0.947162578867264 0.985616936377353
GO_TERTIARY_GRANULE_MEMBRANE 57 0.01 Up
0.52098496678822 0.947277536923652 0.985616936377353
GO_REGULATION_OF_MITOPHAGY 15 0.01 Up
0.125258156004035 0.947340123695443 0.985616936377353
GO_PHOSPHOLIPID_TRANSLOCATING_ATPASE_ACTIVITY 19 0.01 Up
0.386332148913076 0.947605504002421
0.985616936377353
GO_STEREOCILIUM_TIP 10 0.01 Down 0.303610909902107
0.947772152108541 0.985616936377353
GO_EXTERNAL_SIDE_OF_PLASMA_MEMBRANE 173 0.01 Down
0.647754472254136 0.947888776224524 0.985616936377353
GO_CARDIAC_SEPTUM_MORPHOGENESIS 58 0.01 Up
0.388295940111014 0.948140729273307 0.985616936377353
GO_RNA_POLYMERASE_II_REPRESSING_TRANSCRIPTION_FACTOR_BINDING 33
0.01 Down 0.181872072629729 0.9484163873147
0.985616936377353
GO_NEGATIVE_REGULATION_OF_MAP_KINASE_ACTIVITY 65 0.01 Up
0.458541877143595 0.948487439869329
0.985616936377353
GO_SIGNAL_TRANSDUCTION_IN_RESPONSE_TO_DNA_DAMAGE 122 0.01
Up 0.232252757106386 0.948869683034541
0.985616936377353
GO_FATTY_ACID_TRANSMEMBRANE_TRANSPORT 16 0.01 Down
0.218805681099946 0.948901920346225 0.985616936377353
GO_POSITIVE_REGULATION_OF_CELL_CYCLE_ARREST 71 0.01 Down
0.269012442444331 0.949063472872017
0.985616936377353
GO_MEMBRANE_REPOLARIZATION 28 0.01 Up
0.523773616152059 0.949180201007319 0.985616936377353
GO_POSITIVE_REGULATION_OF_BIOMINERAL_TISSUE_DEVELOPMENT 32 0.01
Up 0.62117270226819 0.949452238509067
0.985616936377353
GO_ENDOCYTIC_VESICLE 243 0.01 Down 0.404395882480083

0.949460262226901 0.985616936377353
GO_N_ACETYLGLUCOSAMINE_METABOLIC_PROCESS 11 0.01 Down
0.54940925465526 0.949554753248644
0.985616936377353
GO_SEQUENCE_SPECIFIC_SINGLE_STRANDED_DNA_BINDING 14 0.01
Down 0.106965068066262 0.949577778649595
0.985616936377353
GO_CELLULAR_MONOVALENT_INORGANIC_CATION_HOMEOSTASIS 79 0.01
Down 0.35266037892435 0.949675782454448
0.985616936377353
GO_REGULATION_OF_MACROPHAGE_ACTIVATION 41 0.01 Down
0.637208110105143 0.949702866634455 0.985616936377353
GO_TRANSLATION_REPRESSOR_ACTIVITY_MRNA_REGULATORY_ELEMENT_BINDING
12 0.01 Up 0.32735048274278
0.949788962486548 0.985616936377353
GO_NEGATIVE_REGULATION_OF_TOR_SIGNALING 44 0.01 Down
0.186392783412937 0.949814518283969 0.985616936377353
GO_NEGATIVE_REGULATION_OF_AMYLOID_PRECURSOR_PROTEIN_CATABOLIC_PROCES
S 12 0.01 Down 0.602145071472034
0.949898752272612 0.985616936377353
GO_POSITIVE_REGULATION_OF_CELLULAR_RESPONSE_TO_INSULIN_STIMULUS 17
0.01 Down 0.266916154086063 0.950201161508947
0.985616936377353
GO_REGULATION_OF_SYNAPTIC_VESICLE_TRANSPORT 49 0.01 Up
0.200098340120444 0.950209877330894
0.985616936377353
GO_CARDIAC_CONDUCTION 79 0.01 Up 0.420409769170508
0.950228386447321 0.985616936377353
GO_PROTEIN_INSERTION_INTO_MEMBRANE 55 0.01 Up
0.208606560828505 0.950503599100261 0.985616936377353
GO_GLOMERULAR_MESANGIUM_DEVELOPMENT 11 0.01 Down
0.96021021441804 0.950588015141579 0.985616936377353
GO_REGULATION_OF_EXECUTION_PHASE_OF_APOPTOSIS 23 0.01 Down
0.3528971954894 0.950964391667379 0.985616936377353
GO_NEGATIVE_REGULATION_OF_NUCLEOCYTOPLASMIC_TRANSPORT 18 0.01
Down 0.508673060867742 0.95102724447943
0.985616936377353
GO_CLATHRIN_COATED_VESICLE_MEMBRANE 86 0.01 Up
0.440506795998778 0.951188001359538 0.985616936377353
GO_REGULATION_OF_MEMBRANE_PERMEABILITY 73 0.01 Down
0.233602655007001 0.951193717055787 0.985616936377353
GO_RESPONSE_TO_OXIDATIVE_STRESS 363 0.01 Up
0.365960267470473 0.951199082562166 0.985616936377353
GO_REGULATION_OF_ATTACHMENT_OF_SPINDLE_MICROTUBULES_TO_KINETOCHORE
11 0.01 Down 0.130284325873046
0.951543091987442 0.985696070961821
GO_AUTOPHAGOSOME_MATURATION 33 0.01 Up
0.141441008919462 0.951674554402007 0.985696070961821
GO_MHC_PROTEIN_BINDING 22 0.01 Up 0.463963255119972
0.951839504111097 0.985696070961821
GO_GLUCAN_CATABOLIC_PROCESS 19 0.01 Down
0.219027340361523 0.951924899365699 0.985696070961821
GO_REGULATION_OF_OXIDATIVE_STRESS_INDUCED_CELL_DEATH 55 0.01
Up 0.325113482074127 0.952457547739905

0.985914136217622
GO_HOMEOSTASIS_OF_NUMBER_OF_CELLS_WITHIN_A_TISSUE 22 0.01 Down 0.429454882806633 0.952588415573609 0.985914136217622
GO_NUCLEAR_REPLICATION_FORK 38 0.01 Up 0.154644804345621 0.952622685398216 0.985914136217622
GO_CELL_MATRIX_ADHESION 173 0.01 Up 0.555660315775118 0.952929483386918 0.986063557805007
GO_MITOTIC_NUCLEAR_DIVISION 240 0.01 Up 0.218721225641092 0.953391546476402 0.986373564870183
GO_NCRNA_3_END_PROCESSING 34 0.01 Down 0.136687772480717 0.953844946628876 0.986401379529394
GO_MAINTENANCE_OF_LOCATION 249 0.01 Down 0.470289823645105 0.953904796524636 0.986401379529394
GO_GEMINI_OF_COILED_BODIES 12 0.01 Up 0.092138451162696 0.95393685635962 0.986401379529394
GO_POSITIVE_REGULATION_OF_GLYCOGEN_METABOLIC_PROCESS 13 0.01 Up 0.383443448709648 0.954551047641039 0.986401379529394
GO_RESPONSE_TO_EXTRACELLULAR_STIMULUS 382 0.01 Up 0.335950033615404 0.954785734218084 0.986401379529394
GO_MEIOTIC_CHROMOSOME_SEGREGATION 47 0.01 Up 0.222028304548442 0.954821929676351 0.986401379529394
GO_REGULATION_OF_STEM_CELL_POPULATION_MAINTENANCE 23 0.01 Down 0.381098078040457 0.954839622390972 0.986401379529394
GO_REGULATION_OF_RELEASE_OF_CYTOCHROME_C_FROM_MITOCHONDRIA 43 0.01 Down 0.340114554888135 0.954865245521675 0.986401379529394
GO_CELL_CELL_CONTACT_ZONE 52 0.01 Up 0.376849715250584 0.954945893383675 0.986401379529394
GO_OSTEOBLAST_DIFFERENTIATION 162 0.01 Up 0.471658367393506 0.955043206864401 0.986401379529394
GO_NUCLEOBASE_METABOLIC_PROCESS 12 0.01 Up 0.702846266257559 0.955561966245482 0.986445747875696
GO_LAMELLIPODIUM_MORPHOGENESIS 15 0.01 Down 0.17675439958744 0.955678845931297 0.986445747875696
GO_PHOSPHOLIPID_BIOSYNTHETIC_PROCESS 206 0.01 Down 0.244437518736331 0.956238737600888 0.986445747875696
GO_PURINE_NUCLEOTIDE_TRANSPORT 12 0.01 Down 0.185589910978779 0.95632808824259 0.986445747875696
GO_PURINE_NUCLEOTIDE_TRANSMEMBRANE_TRANSPORTER_ACTIVITY 12 0.01 Down 0.185589910978779 0.95632808824259 0.986445747875696
GO_CYTOPLASMIC_TRANSLATION 89 0.01 Up 0.137713034944209 0.956420629634414 0.986445747875696
GO_REGULATION_OF_CAMP_MEDIATED_SIGNALING 28 0.01 Down 0.440372792528967 0.956781381438357 0.986445747875696
GO_NEGATIVE_REGULATION_OF_STRIATED_MUSCLE_CELL_DIFFERENTIATION 28 0.01 Up 0.581239970038756 0.956958342317097 0.986445747875696
GO_PHOSPHOLIPASE_BINDING 10 0.01 Down 0.267758849778404 0.956984461577993 0.986445747875696

GO_E_BOX_BINDING 33 0.01 Down 0.240300716661176
0.957041079678477 0.986445747875696
GO_POSITIVE_REGULATION_OF_RELEASE_OF_CYTOCHROME_C_FROM_MITOCHONDRIA
29 0.01 Up 0.392118417101445
0.957075739319242 0.986445747875696
GO_POLARIZED_EPITHELIAL_CELL_DIFFERENTIATION 16 0.01 Up
0.248447804614268 0.957414403519048
0.986445747875696
GO_TRIGLYCERIDE_LIPASE_ACTIVITY 10 0.01 Down
0.384841707162616 0.957774813628344 0.986445747875696
GO_ATPASE_BINDING 64 0.01 Up 0.276364780397841
0.957921968337871 0.986445747875696
GO_CELLULAR_RESPONSE_TO_RADIATION 151 0.01 Down
0.230366053332942 0.957984097707037 0.986445747875696
GO_PEPTIDE_HORMONE_BINDING 21 0.01 Down
0.511193216131736 0.958235513694058 0.986445747875696
GO_NUCLEOTIDE_TRANSMEMBRANE_TRANSPORTER_ACTIVITY 16 0.01
Up 0.184225986342268 0.958261464335387
0.986445747875696
GO_TBP_CLASS_PROTEIN_BINDING 21 0.01 Up
0.21044582584481 0.958461361417077 0.986445747875696
GO_FATTY_ACID_HOMEOSTASIS 11 0.01 Down
0.295802386899194 0.95856304852924 0.986445747875696
GO_BONE_MINERALIZATION 72 0.01 Up 0.697154418551535
0.958775807008697 0.986445747875696
GO_ATF6_MEDIATED_UNFOLDED_PROTEIN_RESPONSE 10 0.01 Up
0.213834398187487 0.958829395689408
0.986445747875696
GO_INTRINSIC_COMPONENT_OF_PRESYNAPTIC_MEMBRANE 32 0.01 Down
0.363837967900116 0.958893687500534
0.986445747875696
GO_POSITIVE_REGULATION_OF_PROTEIN_EXIT_FROM_ENDOPLASMIC_RETICULUM
11 0.01 Up 0.172571072885424
0.959643216473539 0.986445747875696
GO_PURINE_CONTAINING_COMPOUND_METABOLIC_PROCESS 121 0.01 Down
0.303074866324013 0.95967553875333
0.986445747875696
GO_HORMONE_BIOSYNTHETIC_PROCESS 53 0.01 Up
0.517809021306106 0.959690392456002 0.986445747875696
GO_DERLIN_1_RETROTRANSLOCATION_COMPLEX 10 0.01 Up
0.156738433667052 0.959721212671238 0.986445747875696
GO_PEROXISOME_PROLIFERATOR_ACTIVATED_RECEPTOR_SIGNALING_PATHWAY 16
0.01 Up 0.320398153611089 0.959732119049276
0.986445747875696
GO_MITOTIC_CYTOKINESIS 66 0.01 Down 0.211950570998919
0.959773996576684 0.986445747875696
GO_DNA_CATABOLIC_PROCESS_ENDONUCLEOLYTIC 26 0.01 Up
0.221254235978442 0.960032526823536
0.986445747875696
GO_CELL_CELL_JUNCTION_MAINTENANCE 10 0.01 Down
0.348807555072744 0.960040957684603 0.986445747875696
GO_INTERLEUKIN_5_PRODUCTION 13 0.01 Up
0.478838857854043 0.960161144321162 0.986445747875696
GO_OSSIFICATION 279 0.01 Down 0.4976922668039

0.960477274113165	0.986445747875696
GO_CELL_PROJECTION_ORGANIZATION 1184	0.01	Up
0.324350389103621	0.960619578576	0.986445747875696
GO_CENTROSOME_SEPARATION	13	0.01	Up
0.0873791982010246	0.961222639664764	0.986445747875696
GO_SAGA_TYPE_COMPLEX	26	0.01	Down	0.130813505765571
	0.961625552007955	0.986445747875696
GO_CARBOHYDRATE_CATABOLIC_PROCESS	148	0.01	Down
0.253322497570537	0.961795237985073	0.986445747875696
GO_SWR1_COMPLEX 11	0.01	Down	0.15093858651023
0.96192257382987	0.986445747875696
GO_REGULATION_OF_STEM_CELL_PROLIFERATION	43	0.01	Down
	0.389668382186837	0.962141115899288
0.986445747875696
GO_GLYCEROPHOSPHOLIPID_CATABOLIC_PROCESS	18	0.01	Down
	0.553675746673335	0.962192335048931
0.986445747875696
GO_RUFFLE_ORGANIZATION	45	0.01	Up	0.269927814119611
	0.962212981230857	0.986445747875696
GO_MAMMARY_GLAND_EPITHELIUM_DEVELOPMENT 57	0.01	Up
0.356570830325062	0.962697922364281	0.986445747875696
GO_INOSITOL_PHOSPHATE_CATABOLIC_PROCESS 14	0.01	Down
0.144013176019102	0.962777122770515	0.986445747875696
GO_BEHAVIOR	339	0.01	Up	0.35514863346674
0.962822016457619	0.986445747875696
GO_BIOMINERAL_TISSUE_DEVELOPMENT	99	0.01	Down
0.646776689783481	0.962859565892174	0.986445747875696
GO_CELL_FATE_DETERMINATION	19	0.01	Up
0.565448155694527	0.962868306104876	0.986445747875696
GO_NUCLEAR_RECEPTOR_ACTIVITY	34	0.01	Up
0.413298043447207	0.963001914330556	0.986445747875696
GO_MITOCHONDRION_MORPHOGENESIS	20	0.01	Up
0.162589958593512	0.963028922245053	0.986445747875696
GO_POSITIVE_REGULATION_OF_MACROPHAGE_CHEMOTAXIS 13	0.01	Down
	0.692149195255416	0.963349779763332
0.986445747875696
GO_POSITIVE_REGULATION_OF_RECEPTOR_MEDIATED_ENDOCYTOSIS 36	0.01
	Down	0.460079324727315	0.963372254394654
0.986445747875696
GO_RENAL_ABSORPTION	12	0.01	Up	0.465101544316808
	0.963381365426485	0.986445747875696
GO_PROTEIN_AUTO_ADP_RIBOSYLATION	11	0.01	Up
0.12011753731331	0.963469649546531	0.986445747875696
GO_G_PROTEIN_BETA_GAMMA_SUBUNIT_COMPLEX_BINDING 12	0.01	Down
	0.267306642495754	0.963535378834274
0.986445747875696
GO_OOGENESIS	52	0.01	Down	0.309505154346939
0.963805375194454	0.986555797134636
GO_REGULATION_OF_RECEPTOR_BINDING	17	0.01	Down
0.620086539547657	0.964103797774762	0.986694901599896
GO_SPINDLE_MICROTUBULE	45	0.01	Down	0.211846370113516
	0.965125011194864	0.987425768468479
GO_NEGATIVE_REGULATION_OF_SMOOTH_MUSCLE_CELL_DIFFERENTIATION	13
	0.01	Up	0.363637196090538	0.965143223536807

0.987425768468479
GO_SUPRAMOLECULAR_FIBER_ORGANIZATION 517 0.01 Up
0.393493153790671 0.965884620137981 0.987893831404117
GO_PROTEIN_DEGLYCOSYLATION 29 0.01 Up
0.154117090558917 0.965926170847445 0.987893831404117
GO_NEGATIVE_REGULATION_OF_PLASMA_MEMBRANE_BOUNDED_CELL_PROJECTION_ASSEMBLY 20 0.01 Down 0.191127136475344
0.966194563139801 0.988001885265578
GO_NUCLEAR_BODY_ORGANIZATION 13 0.01 Down
0.256045780882858 0.966835989203089 0.988330668438594
GO_EMBRYONIC_CRANIAL_SKELETON_MORPHOGENESIS 39 0.01 Up
0.48550195016028 0.966841680095011
0.988330668438594
GO_ENZYME_INHIBITOR_ACTIVITY 244 0.01 Up
0.411525614912846 0.967135672561435 0.98846475894284
GO_PERIPHERAL_NERVOUS_SYSTEM_DEVELOPMENT 49 0.01 Down
0.500708453344003 0.967708628446279
0.988766748522586
GO_DENDRITIC_SPINE_MAINTENANCE 15 0.01 Up
0.34449628242107 0.967756880204449 0.988766748522586
GO_ROUGH_ENDOPLASMIC_RETICULUM_MEMBRANE 24 0.01 Down
0.26649318039531 0.968195702406986 0.989015030437659
GO_AMIDE_BIOSYNTHETIC_PROCESS 698 0.01 Up
0.182988862458417 0.968325702671956 0.989015030437659
GO_POSITIVE_REGULATION_OF_PROTEIN_DEACETYLATION 19 0.01 Up
0.167481216492988 0.9686483520016 0.989140268378724
GO_POSITIVE_REGULATION_OF_REACTIVE_OXYGEN_SPECIES_METABOLIC_PROCESS
37 0.01 Up 0.405508556584186
0.968890646582396 0.989140268378724
GO_SITE_OF_DOUBLE_STRAND_BREAK 53 0.01 Down
0.169819154498953 0.968941964374892 0.989140268378724
GO_HISTONE_KINASE_ACTIVITY 14 0.01 Up
0.242322513242398 0.96916738532662 0.989140268378724
GO_ATPASE_REGULATOR_ACTIVITY 34 0.01 Down
0.188036218528956 0.969262964352665 0.989140268378724
GO_MATURATION_OF_LSU_RRNA_FROM_TRICISTRONIC_RRNA_TRANSCRIPT_SSU_RRNA_5_8S_RRNA_LSU_RRNA_ 13 0.01 Up 0.110498591612127
0.969715176990711 0.989329889207255
GO_SEGMENTATION 65 0.01 Down 0.405051955209313
0.970030948195334 0.989329889207255
GO_C4_DICARBOXYLATE_TRANSMEMBRANE_TRANSPORTER_ACTIVITY 10 0.01
Down 0.208894653642287 0.970209121478207
0.989329889207255
GO_PROTEIN_HOMOOLIGOMERIZATION 240 0.01 Down
0.318923338008805 0.970363500429639 0.989329889207255
GO_CLATHRIN_COAT_OF_ENDOCYTIC_VESICLE 13 0.01 Down
0.15832085425258 0.970413265037227 0.989329889207255
GO_CELLULAR_RESPONSE_TO_LEUCINE 10 0.01 Up
0.153058163113347 0.970687811131295 0.989329889207255
GO_CELLULAR_RESPONSE_TO_INCREASED_OXYGEN_LEVELS 13 0.01 Down
0.278459751282693 0.97085086665213
0.989329889207255
GO_POSITIVE_REGULATION_OF_INTERLEUKIN_1_PRODUCTION 35 0.01
Up 0.598549895771482 0.970871356841351

0.989329889207255
GO_RNA_POLYMERASE_ACTIVITY	43	0.01	Down
0.120599684133423	0.971131266696793	0.989329889207255
GO_TRANSMEMBRANE_TRANSPORT	988	0.01	Up
0.35780330996372	0.971175748527691	0.989329889207255
GO_SMN_SM_PROTEIN_COMPLEX	17	0.01	Up
0.0514620989159234	0.971388099417245	0.989329889207255
GO_NEGATIVE_REGULATION_OF_INNATE_IMMUNE_RESPONSE	44	0.01
Down	0.542586317254827	0.971511147505604
0.989329889207255
GO_SODIUM_CHANNEL_REGULATOR_ACTIVITY	26	0.01	Down
0.502260645014641	0.971567254069124	0.989329889207255
GO_PHOSPHOTRANSFERASE_ACTIVITY_FOR_OTHER_SUBSTITUTED_PHOSPHATE_GROUP
S	19	0.01	Up	0.178780743834265
0.972315036687287	0.989893534355072
GO_CEREBRAL_CORTEX_DEVELOPMENT	87	0.01	Down
0.305888045855077	0.972446885009661	0.989893534355072
GO_CARDIAC_VENTRICLE_DEVELOPMENT	89	0.01	Up
0.420327401429889	0.972833728078922	0.990121301452998
GO_DNA_RECOMBINATION	222	0.01	Up	0.189620990951311
0.973043212111882	0.990168511688106
GO_ECTODERMAL_PLACODE_DEVELOPMENT	10	0.01	Up
0.323370391652086	0.9739325186057 0.990872560708486
GO_REGULATION_OF_MUSCLE_CELL_DIFFERENTIATION	122	0.01	Up
0.481364714369097	0.974210614679027
0.990872560708486
GO_RESPONSE_TO_MISFOLDED_PROTEIN	25	0.01	Down
0.175553556275846	0.97439314381251	0.990872560708486
GO_PHOSPHOLIPID_DEPHOSPHORYLATION	32	0.01	Up
0.304468506187826	0.974558927609399	0.990872560708486
GO_REGULATION_OF_BONE_MINERALIZATION	45	0.01	Up
0.753298885760887	0.974635458106081	0.990872560708486
GO_PRESPLICEOSOME	15	0.01	Up	0.0941277801141299
0.974714368728558	0.990872560708486
GO_REGULATION_OF_CHOLESTEROL_EFFLUX	19	0.01	Down
0.359176336782023	0.975229084912464	0.991164072294552
GO_POSITIVE_REGULATION_OF_T_CELL_RECEPTOR_SIGNALING_PATHWAY	10
0.01	Down	0.461465468516584	0.975327650780375
0.991164072294552
GO_GLOMERULUS_DEVELOPMENT	42	0.01	Up
0.7726444226549 0.975761588993148	0.991169978392403
GO_NUCLEOBASE_CONTAINING_COMPOUND_KINASE_ACTIVITY	37	0.01
Down	0.243335030180647	0.975775342826416
0.991169978392403
GO_CONDENSED_CHROMOSOME_CENTROMERIC_REGION	95	0.01	Down
0.180628192038249	0.975823251663876
0.991169978392403
GO_POSITIVE_REGULATION_OF_PROTEIN_SUMOYLATION	11	0.01	Down
0.162496278621062	0.976552265262732
0.991744530346277
GO_REGULATION_OF_ERAD_PATHWAY	28	0.01	Down
0.176558641550128	0.976827426154613	0.991858053886043
GO_PARASYMPATHETIC_NERVOUS_SYSTEM_DEVELOPMENT	13	0.01	Down
0.562670200365448	0.977440308144414

0.992092051676208
GO_POSITIVE_REGULATION_OF_TELOMERASE_RNA_LOCALIZATION_TO_CAJAL_BODY
15 0.01 Up 0.0816439386272547
0.977842119044233 0.992092051676208
GO_NEUROMUSCULAR_SYNAPTIC_TRANSMISSION 11 0.01 Down
0.316077121199294 0.977917977391491 0.992092051676208
GO_SENSORY_ORGAN_DEVELOPMENT 349 0.01 Down
0.415047646631004 0.977941816822323 0.992092051676208
GO_MYOTUBE_CELL_DEVELOPMENT 20 0.01 Up
0.324311617521013 0.977979205683525 0.992092051676208
GO_NEGATIVE_REGULATION_OF_GLUCONEOGENESIS 12 0.01 Up
0.161744756066446 0.978147318515702
0.992092051676208
GO_ACTIVATION_OF_CYSTEINE_TYPE_ENDOPEPTIDASE_ACTIVITY_INVOLVED_IN_APOPTOTIC_PROCESS 75 0.01 Up 0.372487783513178
0.978210803894922 0.992092051676208
GO_CELL_PROLIFERATION_IN_FOREBRAIN 14 0.01 Down
0.433467807800164 0.978437982588323 0.992092051676208
GO_SYNAPSE_PART 589 0.01 Up 0.293163191009695
0.978739799748688 0.992092051676208
GO_INO80_TYPE_COMPLEX 24 0.01 Up 0.114945064935435
0.978821951240984 0.992092051676208
GO_CATION_TRANSPORT 673 0.01 Down 0.41080883906017
0.979092948288485 0.992092051676208
GO_REGULATION_OF_PLASMINOGEN_ACTIVATION 11 0.01 Up
0.266334019795016 0.979104664942262 0.992092051676208
GO_REGULATION_OF_PROTEIN_HOMODIMERIZATION_ACTIVITY 20 0.01
Up 0.292801359006725 0.979182272054659
0.992092051676208
GO_SITE_OF_DNA_DAMAGE 66 0.01 Down 0.161747262522928
0.979762922555905 0.992334650897831
GO_STEROID_HYDROXYLASE_ACTIVITY 13 0.01 Up
0.581944213478277 0.979798162636167 0.992334650897831
GO_C2H2_ZINC_FINGER_DOMAIN_BINDING 10 0.01 Down
0.156270697309382 0.980105676021058 0.992334650897831
GO_DYNEIN_COMPLEX 36 0.01 Down 0.214962634163105
0.980148282203909 0.992334650897831
GO_NEGATIVE_REGULATION_OF_MUSCLE_CELL_DIFFERENTIATION 42 0.01
Down 0.502875024521108 0.980246779256946
0.992334650897831
GO_SPERM_EGG_RECOGNITION 27 0.01 Down
0.21912750829823 0.980543483026383 0.992334650897831
GO_RENAL_SYSTEM_PROCESS_INVOLVED_IN_REGULATION_OF_SYSTEMIC_ARTERIAL_BLOOD_PRESSURE 10 0.01 Up 0.390857359231635
0.98056589865526 0.992334650897831
GO_CARDIAC_VENTRICLE_MORPHOGENESIS 43 0.01 Up
0.446956893258459 0.981106656949568 0.992708334190216
GO_POSITIVE_REGULATION_OF_RESPONSE_TO_DNA_DAMAGE_STIMULUS 86
0.01 Up 0.223607947504105 0.981262183079474
0.992708334190216
GO_TRANSMISSION_OF_NERVE_IMPULSE 24 0.01 Up
0.475191031096303 0.981694208067825 0.992979929553443
GO_SENSORY_SYSTEM_DEVELOPMENT 246 0.01 Down
0.399154443457936 0.982129434874329 0.993254672517417

GO_SYNAPSE 768 0.01 Down 0.294804322440311
0.982319594033959 0.993281521548995
GO_CELL_SUBSTRATE_JUNCTION 388 0.01 Up
0.313876621686707 0.982688796007933 0.993352762074771
GO_REGULATION_OF_HISTONE_DEACETYLATION 22 0.01 Up
0.163451028318149 0.982717293530073 0.993352762074771
GO_RESPIRATORY_CHAIN_COMPLEX_IV 11 0.01 Up
0.249850769031133 0.982989881153336 0.993363272425391
GO_SYNAPTIC_MEMBRANE 214 0.01 Up 0.298343938638041
0.983266771217746 0.993363272425391
GO_CARBOHYDRATE_HOMEOSTASIS 168 0.01 Up
0.301635818622849 0.983394475723746 0.993363272425391
GO_NUCLEIC_ACID_PHOSPHODIESTER_BOND_HYDROLYSIS 249 0.01 Down
0.190252987839164 0.983427096267606
0.993363272425391
GO_REGULATION_OF_GLUTAMATE_RECEPTOR_SIGNALING_PATHWAY 24 0.01
Up 0.365308988454281 0.983951580318235
0.993363272425391
GO_POSITIVE_REGULATION_OF_HISTONE_DEACETYLATION 16 0.01 Down
0.174039440230722 0.984158528962647
0.993363272425391
GO_ATPASE_ACTIVATOR_ACTIVITY 20 0.01 Up
0.144482442369513 0.98431612933526 0.993363272425391
GO_NUCLEOTIDE_SUGAR_TRANSMEMBRANE_TRANSPORTER_ACTIVITY 14 0.01
Up 0.177197311134505 0.984569552210089
0.993363272425391
GO_GAMETE_GENERATION 405 0.01 Up 0.29010276591591
0.984667620531112 0.993363272425391
GO_PROSTATE_GLAND_DEVELOPMENT 40 0.01 Down
0.39143127824032 0.984702176037316 0.993363272425391
GO_SPHINGOID_BIOSYNTHETIC_PROCESS 11 0.01 Up
0.0997196634788184 0.984771936151289 0.993363272425391
GO_COLUMNAR_CUBOIDAL_EPITHELIAL_CELL_DEVELOPMENT 42 0.01
Down 0.37433251162796 0.98493189553495
0.993363272425391
GO_S_ACYLTRANSFERASE_ACTIVITY 28 0.01 Down
0.165459037357136 0.985085839452344 0.993363272425391
GO_CARDIOVASCULAR_SYSTEM_DEVELOPMENT 538 0.01 Up
0.515415060785639 0.985130627044967 0.993363272425391
GO_CYTOSKELETON_DEPENDENT_CYTOKINESIS 85 0.01 Down
0.20633227629466 0.985182056213684 0.993363272425391
GO_TRANSFERRIN_TRANSPORT 30 0.01 Down
0.194434194535511 0.985377948936917 0.993395803386918
GO_INTRINSIC_APOPTOTIC_SIGNALING_PATHWAY_IN_RESPONSE_TO_OXIDATIVE_ST
RESS 40 0.01 Up 0.298847044671351
0.985823461664672 0.993522455569464
GO_NEPHRON_DEVELOPMENT 101 0.01 Down 0.65869709850805
0.985830879978661 0.993522455569464
GO_POSITIVE_REGULATION_OF_VASCULAR_ENDOTHELIAL_CELL_PROLIFERATION
12 0.01 Down 0.457923507175754
0.986269178957495 0.993724880389007
GO_CYTOPLASMIC_VESICLE_PART 1175 0.01 Down
0.351551052585726 0.986359105456128 0.993724880389007
GO_DNA_DEALKYLATION_INVOLVED_IN_DNA_REPAIR 10 0.01 Down

0.121669088099848 0.98677224730135
0.993976159509954
GO_PROTEIN_CONTAINING_COMPLEX_DISASSEMBLY 288 0.01 Down
0.198244204738066 0.98722420992332
0.994113460998398
GO_FEMALE_GENITALIA_DEVELOPMENT 11 0.01 Up
0.381835440800295 0.987321949932181 0.994113460998398
GO_NEGATIVE_REGULATION_OF_CELL_CYCLE_G1_S_PHASE_TRANSITION 97
0.01 Up 0.276683358586364 0.987399797367871
0.994113460998398
GO_EMBRYONIC_SKELETAL_SYSTEM_MORPHOGENESIS 57 0.01 Up
0.502375081833928 0.988103007434166
0.994391262725475
GO_POSITIVE_REGULATION_OF_MESONEPHROS_DEVELOPMENT 14 0.01
Up 0.731287258834146 0.988154969070873
0.994391262725475
GO_VACUOLAR_PROTON_TRANSPORTING_V_TYPE_ATPASE_COMPLEX 12 0.01
Down 0.157183488637376 0.988326743349554
0.994391262725475
GO_NEGATIVE_REGULATION_OF_B_CELL_PROLIFERATION 11 0.01 Down
0.574668143172663 0.988330897592738
0.994391262725475
GO_NEGATIVE_REGULATION_OF_PEPTIDYL_TYROSINE_PHOSPHORYLATION 49
0.01 Up 0.419062378772487 0.988565025297643
0.994462016335044
GO_CEREBRAL_CORTEX_NEURON_DIFFERENTIATION 12 0.01 Up
0.442870526486884 0.988748199485847
0.994481497527929
GO_REGULATION_OF_LIPOPOLYSACCHARIDE_MEDIATED_SIGNALING_PATHWAY 16
0.01 Up 0.453137948198706 0.989090455740717
0.994628370735848
GO_POSITIVE_REGULATION_OF_ATPASE_ACTIVITY 41 0.01 Up
0.147025145559418 0.989479263981674
0.994628370735848
GO_ENDOPLASMIC_RETICULUM_CALCIUM_ION_HOMEOSTASIS 21 0.01
Up 0.2031732482243 0.98948780343298
0.994628370735848
GO_POSITIVE_REGULATION_OF_CALCIUM_ION_TRANSMEMBRANE_TRANSPORTER_ACTI
VITY 26 0.01 Up 0.409126574747099
0.989549556785459 0.994628370735848
GO_SKELETAL_MUSCLE_CELL_DIFFERENTIATION 45 0.01 Up
0.37611046096048 0.991036083901134 0.995875569880282
GO_PROTEIN_LOCALIZATION_TO_NUCLEOPLASM 12 0.01 Down
0.0630883171412429 0.991273657124057 0.995875569880282
GO_BONE_CELL_DEVELOPMENT 27 0.01 Down
0.486872295216298 0.991282501859091 0.995875569880282
GO_REGULATION_OF_MITOCHONDRION_ORGANIZATION 162 0.01 Up
0.235645231285672 0.991578490002427
0.995937054211885
GO_NEGATIVE_REGULATION_OF_RESPONSE_TO_EXTERNAL_STIMULUS 221 0.01
Up 0.44016723397865 0.991671799161727
0.995937054211885
GO_CELLULAR_RESPONSE_TO_IONIZING_RADIATION 69 0.01 Up
0.232991991645566 0.992324158465785

0.996373394467326
GO_PHOSPHOLIPASE_C_ACTIVATING_G_PROTEIN_COUPLED_RECEPTOR_SIGNALING_PATHWAY 33 0.01 Down 0.524944200335873
0.992434511010364 0.996373394467326
GO_NEGATIVE_REGULATION_OF_CENTROSOME_CYCLE 12 0.01 Up
0.110726369382486 0.992919582144221
0.996695566004888
GO_NERVE_GROWTH_FACTOR_SIGNALING_PATHWAY 11 0.01 Down
0.434022348656115 0.993440717432987
0.996994505094985
GO_PEPTIDE_HORMONE_PROCESSING 21 0.01 Down
0.476116313356236 0.993545833606434 0.996994505094985
GO_TYPE_B_PANCREATIC_CELL_DEVELOPMENT 14 0.01 Up
0.344750307998607 0.993929394812137 0.99700647502621
GO_DNA_REPAIR 476 0.01 Up 0.165811725393473
0.99395505045055 0.99700647502621
GO_REGULATION_OF_FATTY_ACID_BETA_OXIDATION 14 0.01 Up
0.221206260141335 0.994050435403336
0.99700647502621
GO_POLYOL_METABOLIC_PROCESS 56 0.01 Down
0.286508637537073 0.994725335563327 0.997518584771219
GO_ACTIVATION_OF_MAPKKK_ACTIVITY 10 0.01 Down
0.290230916433763 0.995033490260265 0.997662810796048
GO_REGULATION_OF_LIPASE_ACTIVITY 56 0.01 Up
0.515897191688714 0.995553267924778 0.998019136322874
GO_INSULIN_LIKE_GROWTH_FACTOR_RECEPTOR_BINDING 11 0.01 Up
0.315394485626044 0.995732420427687
0.998033931718093
GO_REGULATION_OF_ISOTYPE_SWITCHING 29 0.01 Up
0.283852865131145 0.996581325479144 0.998469235631909
GO_MITOTIC_DNA_INTEGRITY_CHECKPOINT 100 0.01 Down
0.20985832392394 0.996617446881459 0.998469235631909
GO_SH2_DOMAIN_BINDING 29 0.01 Up 0.408165408917595
0.996660116608363 0.998469235631909
GO_REGULATION_OF_SUBSTRATE_ADHESION_DEPENDENT_CELL_SPREADING 43
0.01 Down 0.252802080171774 0.997033449810832
0.998499334301346
GO_VASCULAR_SMOOTH_MUSCLE_CONTRACTION 14 0.01 Down
0.504949115805927 0.997472613655322 0.998499334301346
GO_NEGATIVE_REGULATION_OF_LEUKOCYTE_APOPTOTIC_PROCESS 37 0.01
Down 0.538574565991019 0.997565091162676
0.998499334301346
GO_MONOCYTE_CHEMOTACTIC_PROTEIN_1_PRODUCTION 11 0.01 Up
0.431176818401309 0.99760627221649
0.998499334301346
GO_ESTABLISHMENT_OR_MAINTENANCE_OF_MONOPOLAR_CELL_POLARITY 18
0.01 Up 0.270359856730932 0.997785238436456
0.998499334301346
GO_MONONUCLEAR_CELL_DIFFERENTIATION 29 0.01 Down
0.51980992834618 0.997821449428132 0.998499334301346
GO_GTPASE_ACTIVATING_PROTEIN_BINDING 13 0.01 Up
0.219645849729568 0.997841453007126 0.998499334301346
GO_REGULATION_OF_SECONDARY_METABOLIC_PROCESS 11 0.01 Up
0.367744630597106 0.998236334110967

0.998684895753245
GO_CELL_DEATH_IN_RESPONSE_TO_OXIDATIVE_STRESS 72 0.01 Down 0.317989176194477 0.998355893975695 0.998684895753245
GO_REGULATION_OF_PROTEIN_LOCALIZATION_TO_CELL_SURFACE 32 0.01 Down 0.375221062494744 0.998750594220022 0.998915133032909
GO_PEPTIDE_TRANSMEMBRANE_TRANSPORTER_ACTIVITY 28 0.01 Down 0.263052606515644 0.999063625525033 0.999063625525033

**Supplement table 6:** edgeR_DE_GSEA_results_regulons_IPF_effect

| | NGenes | Correlation | Direction | absLog2FC | PValue | FDR |
|---|---|---|---|---|---|---|
| GRHL3_E | 318 | 0.01 | Down | 0.624206487356476 | 1.08947856949779e-15 | 1.45227493314055e-12 |
| ZNF654_E | 397 | 0.01 | Down | 0.228528706875837 | 3.41228861679034e-15 | 2.27429036309076e-12 |
| OSR1_E | 189 | 0.01 | Up | 0.877881568975381 | 1.76930782393035e-14 | 6.7693727813836e-12 |
| MIER3_E | 451 | 0.01 | Down | 0.209452094263078 | 2.03131966433116e-14 | 6.7693727813836e-12 |
| BARX2_E | 415 | 0.01 | Down | 0.590540793171815 | 7.67007918793639e-14 | 2.04484311150384e-11 |
| ZBTB43_E | 330 | 0.01 | Down | 0.24515367139328 | 1.03729048880489e-13 | 2.3045137026282e-11 |
| HAND2_E | 232 | 0.01 | Up | 0.609103973321204 | 2.50402184562524e-13 | 4.7683730288835e-11 |
| PURB_E | 405 | 0.01 | Down | 0.194805299238808 | 3.31633078641157e-12 | 5.52583617285827e-10 |
| ZFHX4_E | 260 | 0.01 | Up | 0.699361869552749 | 1.37706203263899e-11 | 2.01243885957124e-09 |
| RC3H2_E | 806 | 0.01 | Down | 0.193460729959787 | 1.50970657132126e-11 | 2.01243885957124e-09 |
| IKZF2_E | 172 | 0.01 | Down | 0.325809965758677 | 1.88152863093197e-11 | 2.28007060457484e-09 |
| ZNF805_E | 113 | 0.01 | Down | 0.225000517489119 | 2.87402524875953e-10 | 3.19256304716371e-08 |
| ZNF430_E | 268 | 0.01 | Down | 0.224503704371386 | 4.02172609364891e-10 | 4.12381606371846e-08 |
| ZBED2_E | 175 | 0.01 | Down | 0.655427765425168 | 4.61567955977779e-10 | 4.39478632370271e-08 |
| ADNP2_E | 256 | 0.01 | Down | 0.192653363732834 | 5.66128661358236e-10 | 5.03099670393686e-08 |
| ARID4B_E | 492 | 0.01 | Down | 0.191812697510855 | 6.85305694780273e-10 | 5.70945306963815e-08 |
| ZNF562_E | 418 | 0.01 | Down | 0.181856931411713 | 7.78624733939033e-10 | 6.10533394318077e-08 |
| ZNF148_E | 626 | 0.01 | Down | 0.190622404814515 | 8.31061414357151e-10 | 6.15447147410046e-08 |
| NFKBIZ_E | 233 | 0.01 | Down | 0.440345407993603 | 1.23540794470319e-09 | 8.66736205415449e-08 |
| HIVEP1_E | 210 | 0.01 | Down | 0.223518785288327 | 2.25954306163971e-09 | 1.50598545058287e-07 |
| KAT6A_E | 465 | 0.01 | Down | 0.213908945938011 | 3.38683642055288e-09 | 2.14983473742714e-07 |
| ZNF267_E | 617 | 0.01 | Down | 0.296512757002855 | 4.36352284643138e-09 | 2.55420718386152e-07 |
| FOXN1_E | 201 | 0.01 | Down | 0.67038622381182 | 4.42460611183047e-09 | 2.55420718386152e-07 |
| ZNF800_E | 726 | 0.01 | Down | 0.18886260583763 | 4.59872261160363e-09 | 2.55420718386152e-07 |
| TSC22D2_E | 273 | 0.01 | Down | 0.258780641461329 | 5.01590151152061e-09 | 2.67447868594279e-07 |
| ZNF292_E | 355 | 0.01 | Down | 0.2087794942054 | 9.32522522974395e-09 | 4.78097124278796e-07 |

HEYL_E 321 0.01 Up 0.69588490279495
1.00519521315854e-08 4.94109094248793e-07
BNC2_E 494 0.01 Up 0.648939664324279
1.03788857006498e-08 4.94109094248793e-07
TSHZ3_E 772 0.01 Up 0.558509027509548
1.36675114838875e-08 6.28234234759381e-07
GRHL1_E 492 0.01 Down 0.366938069842394
1.41459918303752e-08 6.28553570329672e-07
CNOT4_E 310 0.01 Down 0.172740597102938
1.46850437182416e-08 6.3145687988439e-07
ZNF791_E 348 0.01 Down 0.196842512315897
1.56635319232681e-08 6.43301090168415e-07
ZNF131_E 634 0.01 Down 0.166177715205853
1.6199942631217e-08 6.43301090168415e-07
ZBTB5_E 275 0.01 Down 0.21702055422929
1.64082798692619e-08 6.43301090168415e-07
ZFPM2_E 452 0.01 Up 0.633610250194522
1.97120710801748e-08 7.5074830713923e-07
HMGXB4_E 551 0.01 Down 0.162372500377065
2.77730808095152e-08 1.02837546441899e-06
ZNF507_E 439 0.01 Down 0.168816291692483
3.48637760574139e-08 1.23134468654763e-06
PBRM1_E 930 0.01 Down 0.167370512824574
3.51020990913802e-08 1.23134468654763e-06
MXD1_E 376 0.01 Down 0.401878186260545
4.48991589368071e-08 1.53463022725036e-06
ELMSAN1_E 165 0.01 Down 0.255936894245488
5.40219248987678e-08 1.80028064725144e-06
PRDM8_E 244 0.01 Up 0.587235185748204
5.66902359225233e-08 1.84312401182253e-06
IRF6_E 476 0.01 Down 0.351343280247077
5.92176654692065e-08 1.87945590643934e-06
RBAK_E 266 0.01 Down 0.171791598560976
6.66164269455788e-08 2.0261182278946e-06
ZNF521_E 444 0.01 Up 0.593439198773284
6.68786211758159e-08 2.0261182278946e-06
ZNF423_E 756 0.01 Up 0.591790845556011
6.91288250200287e-08 2.04774941670441e-06
ZBTB34_E 228 0.01 Down 0.212375451767585
7.1117708736932e-08 2.06086751622457e-06
OVOL2_E 393 0.01 Down 0.529078080472775
7.60399328880811e-08 2.15662192637898e-06
PRDM2_E 296 0.01 Down 0.182414616019159
1.00814742259816e-07 2.79970940484029e-06
REL_A 13 0.01 Down 0.715523169274427
1.12290810917198e-07 3.05476838678827e-06
SALL2_E 517 0.01 Up 0.353399692910622
1.56211386508526e-07 4.1645955643173e-06
ZFP91_E 987 0.01 Down 0.148745906524536
1.59679184559298e-07 4.17357554936361e-06
ZNF853_E 405 0.01 Up 0.365923518638418
1.75215616701524e-07 4.49158494352175e-06
ZBTB44_E 425 0.01 Down 0.196813329253129
1.90709278790257e-07 4.79651827598892e-06

ZBTB41_E	500	0.01	Down	0.189609605461517
1.97530704993328e-07	4.87608203252048e-06
SP6_E	126	0.01	Down	0.340506293079276
2.05319749688805e-07	4.97620411518505e-06
FOXS1_E 264	0.01	Up	0.688009243257746
2.88291408317829e-07	6.8623651301369e-06
CSRNP1_E	132	0.01	Down	0.408726810748586
3.19873275816713e-07	7.48054520462594e-06
ZNF721_E	283	0.01	Down	0.183153423426574
3.48757724113665e-07	8.01541459040544e-06
NPAS4_E 208	0.01	Up	0.539087930110716
3.82939030154772e-07	8.54568498258147e-06
EBF2_E	282	0.01	Up	0.800395410059057
3.84651987212969e-07	8.54568498258147e-06
IRX4_E	148	0.01	Down	0.584488014098806
4.75546960179301e-07	1.03918704576887e-05
ZNF81_E 320	0.01	Down	0.187653292008689
5.42660582652929e-07	1.1667202527038e-05
PURA_E	400	0.01	Down	0.182593641558719
5.79916030232054e-07	1.2270286798402e-05
IKZF5_E 419	0.01	Down	0.177929345308849
6.095573987175e-07	1.26959376951629e-05
OSR2_E	316	0.01	Up	0.688541121062023
7.77432895802758e-07	1.59433546170012e-05
ZBTB21_E	354	0.01	Down	0.270970748178862
9.63813017228849e-07	1.9466102302516e-05
KMT2A_C 47	0.01	Down	0.252177914949224
1.07866712819874e-06	2.13045010389261e-05
BNC1_E	204	0.01	Down	0.570751333193245
1.08680125329856e-06	2.13045010389261e-05
RFX7_E	385	0.01	Down	0.195897106089084
1.1830101041633e-06	2.28543836065172e-05
TBX18_E 219	0.01	Up	0.76943889833723
1.57196035097136e-06	2.99346163977832e-05
RBPJL_E 51	0.01	Down	1.23919215483945
1.82817027113068e-06	3.4323253118552e-05
RCOR2_E 252	0.01	Up	0.359492232639281
2.19835293427581e-06	4.07000619637451e-05
NPAS3_E 168	0.01	Up	0.544507388223891
2.35447362083224e-06	4.29933333776626e-05
PURG_E	149	0.01	Up	0.455696251358978
3.40068635358456e-06	6.12583095855165e-05
ZNF438_E	433	0.01	Up	0.338521868658458
3.45550723728026e-06	6.14158819639279e-05
RFX5_A	14	0.01	Up	1.19149676676848
4.04425669621398e-06	7.09341338954372e-05
ZNF460_E	165	0.01	Down	0.250520626819944
4.12258839888204e-06	7.11674902231399e-05
ZNF431_E	289	0.01	Down	0.221010120960237
4.1643392628694e-06	7.11674902231399e-05
UBP1_E	807	0.01	Down	0.155082271029378
4.94838720549616e-06	8.34962043661567e-05
ZNF451_E	528	0.01	Down	0.17688332178382
5.16647052100104e-06	8.60863150561798e-05

TAF1_C	49	0.01	Down	0.203188630895447
5.97832366064721e-06	9.78381314220546e-05
PRDM16_E	294	0.01	Up	0.537310197166
6.12341979511803e-06	9.78381314220546e-05
ZNF655_E	591	0.01	Down	0.159108103849693
6.14199876637662e-06	9.78381314220546e-05
ZBTB2_E	311	0.01	Down	0.153151895813632
6.16534361549332e-06	9.78381314220546e-05
KDM2A_E	531	0.01	Down	0.171908527674276
6.49434274901549e-06	0.000101846575111031
ZNF614_E	169	0.01	Down	0.182849492869074
6.5731708911407e-06	0.000101884148812681
ZNF469_E	407	0.01	Up	0.623861259752398
6.84410577910672e-06	0.000104864287397118
ZNF468_E	379	0.01	Down	0.166760927671296
7.10682111359358e-06	0.000107652188004776
ETV7_E	332	0.01	Down	0.456367572568172
8.08161361207837e-06	0.000121042594886522
ZNF699_E	241	0.01	Down	0.189606398636447
8.19971793450628e-06	0.000121446933407743
RELA_A	105	0.01	Down	0.710046175176087
8.77564840050701e-06	0.000128548783712921
ZBTB47_E	632	0.01	Up	0.369980994599525
9.02364692651908e-06	0.00013074479731576
PRDM10_E	203	0.01	Down	0.1928137387288
9.54839950399767e-06	0.000136860392890633
AEBP2_E	665	0.01	Down	0.213393331013143
1.04394925609543e-05	0.000148040889188853
ZNF484_E	162	0.01	Down	0.185771571567805
1.07709774848887e-05	0.000151133820919544
MESP1_E	420	0.01	Up	0.216373237977376
1.09436877466855e-05	0.000151957664232623
DMRT1_E	18	0.01	Down	1.72816260953292
1.13310666011039e-05	0.000155714554425479
TOX3_E	94	0.01	Down	0.530638617446739
1.14786194783238e-05	0.00015613265065924
ZBED6_E	251	0.01	Down	0.203966886384051
1.22425939994943e-05	0.000164842200013392
ZNF573_E	104	0.01	Down	0.210863815756784
1.31119487598514e-05	0.000174560507401041
IRX5_E	88	0.01	Down	0.416936153919489
1.33103701110468e-05	0.000174560507401041
NFKB1_A	118	0.01	Down	0.712864335656626
1.33572181207098e-05	0.000174560507401041
OVOL1_E	630	0.01	Down	0.477061192171396
1.46690894435716e-05	0.000189843652701757
ARNTL2_E	633	0.01	Down	0.345720901906271
1.52304381547625e-05	0.000194089233962001
FOXE1_E	84	0.01	Down	0.876156554163703
1.53418572246151e-05	0.000194089233962001
THAP7_E	694	0.01	Up	0.153487190335128
1.54339525881261e-05	0.000194089233962001
HOPX_E	263	0.01	Down	0.585481602283068
1.73182647021848e-05	0.000215749970542172

TCF21_E 353 0.01 Up 0.668419557505567
1.98639904654692e-05 0.00024517314157843
ELF3_C 49 0.01 Down 0.448012792952796
2.47522803009761e-05 0.000302704492121111
ETV3_E 696 0.01 Down 0.17748249575800З
2.51296483372643e-05 0.000304525647577939
FOXL2_A 13 0.01 Down 0.939223325238589
3.32659199333722e-05 0.000399490732172839
THAP8_E 666 0.01 Up 0.205689586286784
3.46069761215012e-05 0.00041188481401751
ZNF837_E 316 0.01 Up 0.204002843630143
3.56914460847902e-05 0.000421032722398454
DMRTB1_E 20 0.01 Down 1.32796952446775
3.62957516385084e-05 0.000424405587141506
ZNF362_E 565 0.01 Up 0.273722806007149
4.06597855049468e-05 0.000471299948505166
ZNF32_E 771 0.01 Up 0.158390155040701
4.11206813226608e-05 0.000472533346578508
ZNF771_E 355 0.01 Up 0.215085066913848
4.38782918621489e-05 0.000499912504720038
STAT2_B 33 0.01 Down 0.338695548296308
4.67273097267316e-05 0.000527860202251977
ZNF567_E 123 0.01 Down 0.171555436499007
4.73026788269678e-05 0.000529869503162589
RFXANK_A 12 0.01 Up 1.1629924820314
5.14465907696613e-05 0.0005667628553385
RFXAP_A 12 0.01 Up 1.1629924820314 5.14465907696613e-05
0.0005667628553385
ZNF254_E 173 0.01 Down 0.186061860087706
5.26069932647881e-05 0.000574796082147234
POU2F3_E 450 0.01 Down 0.411310111924752
5.69373996498029e-05 0.000617053282383636
ZNF770_E 1221 0.01 Down 0.195064744405636
7.02049273603125e-05 0.000751466207366673
ZNF414_E 690 0.01 Up 0.163539166373683
7.04675738340841e-05 0.000751466207366673
ZNF280C_E 334 0.01 Down 0.168391686312293
7.1045601876274e-05 0.000751617359532327
ZFP92_E 130 0.01 Up 0.458185943521967
7.77944499197048e-05 0.000816535446795012
NFXL1_E 429 0.01 Down 0.150466925504102
8.50523690937208e-05 0.000885740687515076
KDM5C_E 305 0.01 Down 0.138919346070276
8.5793924814184e-05 0.000886537223079901
MKX_E 132 0.01 Up 0.622991456621875
8.74667562723468e-05 0.000896870662392602
ZNF518B_E 226 0.01 Down 0.174127292209919
8.930917054431e-05 0.000908771941492864
VDR_A 13 0.01 Down 0.501368274128157
0.000100686740566544 0.00101678352405457
ZBTB11_C 22 0.01 Down 0.202385550457981
0.000104033570820941 0.00103976951028448
HOXB6_E 32 0.01 Up 0.813528591823031
0.000104522966525221 0.00103976951028448

FOXN2_E 345	0.01	Down	0.237674894268432
0.00011544664524873	0.00113992872678931
MEIS3_E 779	0.01	Up	0.49123209352165
0.000116379947892624	0.00114069463632991
ETV6_D	31	0.01	Down	0.488523837056695
0.000139665424694168	0.0013589343877177
ZNF611_E	198	0.01	Down	0.164286761195967
0.000141144903569515	0.00136337794534901
HIVEP2_E	377	0.01	Down	0.22041873571965
0.00014508147927513	0.001391320948732
HMG20A_E	489	0.01	Down	0.154499772201564
0.000147577332801458	0.00140514703303102
JDP2_E	365	0.01	Up	0.43312759076698
0.00015228691792503	0.00143970540137635
ZNF426_E	96	0.01	Down	0.185820790866235
0.000158432441826159	0.00148725665460754
FOXJ3_E 1065	0.01	Down	0.227595096283217
0.000163181071200865	0.00150963570574057
ZNF510_E	256	0.01	Down	0.167578019799654
0.000164105715080137	0.00150963570574057
LRRFIP2_E	249	0.01	Down	0.184315113829961
0.000164213936483408	0.00150963570574057
EHF_C	20	0.01	Down	0.412961454571448
0.000165513641319329	0.00151116221834703
ZNF845_E	224	0.01	Down	0.144571718582747
0.000177324786700549	0.00159740575915898
ZNF575_E	436	0.01	Up	0.274298146021448
0.000177356378361237	0.00159740575915898
IRX2_E	79	0.01	Down	0.336846365408976
0.000192524641200491	0.00172238487731714
HMG20B_E	741	0.01	Up	0.173942767877312
0.000216032640058564	0.00191981006132044
ZNF678_E	323	0.01	Down	0.186743066295938
0.000222013727979526	0.00195989602249476
ZNF398_E	347	0.01	Down	0.138085494280529
0.000239507409034146	0.00209562478900529
NFATC4_E	893	0.01	Up	0.381021381456598
0.000240533077807808	0.00209562478900529
HOXC4_E 48	0.01	Up	0.597632095248883
0.000272847466377194	0.00236172514727792
ZNF580_E	428	0.01	Up	0.195227695978752
0.000279372243918739	0.00240260129770115
ZFP30_E 157	0.01	Down	0.168012516745191
0.000285242790363395	0.00243254773231731
RCOR1_C 17	0.01	Down	0.268366214345715
0.000287321355083514	0.00243254773231731
PRDM4_E 709	0.01	Down	0.121096894792609
0.000288328988526733	0.00243254773231731
ZNF440_E	382	0.01	Down	0.214040045042915
0.000307118895831574	0.00257477665499049
ZNF417_E	104	0.01	Down	0.200801802412848
0.000311642823777496	0.00259637427559626
NFAT5_E 733	0.01	Down	0.237823334029991
0.000313836732048614	0.00259841219764473

RLF_E 456 0.01 Down 0.208948781080614
0.000321927251169636 0.00264012675211967
ZNF511_E 660 0.01 Up 0.170253336400579
0.000322836204497754 0.00264012675211967
GZF1_E 199 0.01 Down 0.184907230031879
0.000364596692456209 0.00296345970148858
ZNF28_E 388 0.01 Down 0.14274010820621
0.000371302454843255 0.00298758329792473
ZSCAN29_E 403 0.01 Down 0.143020973264359
0.000372047132374722 0.00298758329792473
TOX_E 214 0.01 Up 0.467586291787338
0.000381281960069151 0.00304340630402502
KLF10_E 108 0.01 Down 0.293089315304635
0.000421650053800785 0.00334190553446835
TSHZ2_E 403 0.01 Up 0.584099597340069
0.000423692449606265 0.00334190553446835
SOX12_E 333 0.01 Up 0.221608706523244
0.000432874803349789 0.00339424772273687
THAP5_E 290 0.01 Down 0.157199435187835
0.000439441040630826 0.00342558425240287
ARID4A_E 474 0.01 Down 0.178469298295418
0.000442206216351347 0.00342709817672294
ZNF841_E 364 0.01 Down 0.178608031540635
0.000448773599406011 0.00345789137577002
ZNF91_E 389 0.01 Down 0.187558015179771
0.000454144191195621 0.00347916210841243
ZNF146_E 997 0.01 Down 0.179333472261307
0.000472098149317451 0.00359603904594379
ZNF304_E 68 0.01 Down 0.183142729918685
0.00048475732040401 0.00367148584146901
ZNF548_E 187 0.01 Down 0.197805851449802
0.000626822588967535 0.00472064695533177
ZNF664_E 656 0.01 Down 0.160625700368669
0.000639940282424861 0.0047923617779345
GATAD2B_E 360 0.01 Down 0.160438145072736
0.000661270590848505 0.00492443406481038
TFAM_E 1279 0.01 Down 0.13942011864277
0.000672498207092173 0.00498022283363259
VEZF1_E 786 0.01 Down 0.21975082003491
0.00071024413632067 0.00523069300395278
SFPQ_E 497 0.01 Down 0.142125201455793
0.000718062237422151 0.00525921407958092
ZNF597_E 75 0.01 Down 0.143954478453219
0.000757521136760738 0.0055178998650386
SIX6_E 15 0.01 Down 1.66390451050734
0.000773941738333565 0.00560687139781871
ATOH8_E 203 0.01 Up 0.490520596105642
0.000803093878279714 0.00578661697160464
GLIS2_E 863 0.01 Up 0.420555972423722
0.000865206806410063 0.00620064877927212
ZNF397_E 378 0.01 Down 0.155343479177036
0.000902533753120082 0.00643356948079716
ZNF34_E 203 0.01 Up 0.2079727212044
0.000933175447388896 0.00661661101792233

| | | | | | | |
|---|---|---|---|---|---|---|
| ZBTB1_E | 227 | 0.01 | Down | 0.174289603545194 | 0.000978843349834944 | 0.00688935315919999 |
| ZNF581_E | 477 | 0.01 | Up | 0.147341365312596 | 0.00098197831976594 | 0.00688935315919999 |
| ARID1A_C | 21 | 0.01 | Down | 0.188519489093544 | 0.000997684915170861 | 0.00693371483861126 |
| ZNF570_E | 131 | 0.01 | Down | 0.155451877301232 | 0.000998704612913249 | 0.00693371483861126 |
| MGA_E | 691 | 0.01 | Down | 0.212108189072602 | 0.00102987380098806 | 0.00711306620060666 |
| ZNF205_E | 493 | 0.01 | Up | 0.161268707903161 | 0.00104028362327764 | 0.00714792819499531 |
| ZNF536_E | 133 | 0.01 | Up | 0.574238316096756 | 0.00104934004203042 | 0.00717318090270024 |
| ZNF524_E | 785 | 0.01 | Up | 0.159330144779105 | 0.00106319434212417 | 0.00723080641863021 |
| GABPA_C | 51 | 0.01 | Down | 0.187466877808793 | 0.00110614177380309 | 0.00748470550497221 |
| KLF5_C | 80 | 0.01 | Down | 0.287509235990421 | 0.00112198203684764 | 0.00755354573291868 |
| ZNF219_E | 413 | 0.01 | Up | 0.177652830053565 | 0.00114515269410677 | 0.00767079668966999 |
| HOXC6_E | 26 | 0.01 | Up | 0.599763982054285 | 0.00120055331687759 | 0.00799990524434111 |
| MEOX2_E | 630 | 0.01 | Up | 0.534447634574171 | 0.00120628728740627 | 0.00799990524434111 |
| ZNF649_E | 304 | 0.01 | Down | 0.122476908719182 | 0.00125974617793835 | 0.00828021088278191 |
| NFX1_E | 594 | 0.01 | Down | 0.147988693298304 | 0.00126097735124136 | 0.00828021088278191 |
| ZNF517_E | 560 | 0.01 | Up | 0.185973015108606 | 0.00128955730537649 | 0.00842637200032772 |
| ZNF283_E | 201 | 0.01 | Down | 0.164039249798609 | 0.0013262046451052 | 0.00862356483865967 |
| CDC5L_E | 1215 | 0.01 | Down | 0.132193035553153 | 0.00137456551731288 | 0.00889463997367997 |
| THAP4_E | 1072 | 0.01 | Up | 0.125246084648954 | 0.00146729307733846 | 0.00944880034827135 |
| NCOR1_C | 15 | 0.01 | Down | 0.268317418365436 | 0.00148202535640614 | 0.00949778750042973 |
| FEZF2_E | 23 | 0.01 | Down | 1.34308413527714 | 0.00162333878423887 | 0.0103536392315331 |
| ZNF135_E | 202 | 0.01 | Up | 0.239267271819386 | 0.001655635092179 | 0.0105093408470219 |
| ZNF208_E | 21 | 0.01 | Up | 0.590542922598041 | 0.00167400092396876 | 0.0105261196555633 |
| THAP3_E | 416 | 0.01 | Up | 0.159918932739611 | 0.00167407154312034 | 0.0105261196555633 |
| MTF1_E | 1175 | 0.01 | Down | 0.22249100372862 | 0.00176908967864043 | 0.0110546623237227 |
| ZBTB45_E | 628 | 0.01 | Up | 0.13889701605234 | 0.00177471698220305 | 0.0110546623237227 |
| ZNF781_E | 186 | 0.01 | Up | 0.291067785092604 | 0.00182495833764888 | 0.011314741693423 |

JUND_A	12	0.01	Down	0.63695058126955
0.00189027250015718	0.0116654316792108
ZNF37A_E	201	0.01	Down	0.163622684517876
0.00192759393987972	0.0118409342021183
MYCL_E	349	0.01	Down	0.35461129809276
0.00196704163724277	0.0120278279928652
ZNF714_E	149	0.01	Down	0.194375261931674
0.00201094542019182	0.0122401381055511
NFIX_E	729	0.01	Up	0.393142439886742
0.00211540174500415	0.0127686127669107
ARNT2_E	336	0.01	Up	0.400881461396662
0.00211692679781491	0.0127686127669107
DNAJC2_E	1146	0.01	Down	0.142970986862401
0.00216840144552718	0.013020176247242
SMARCA5_E	1582	0.01	Down	0.176212035660832
0.0021929387815552	0.0131084636583546
MIER1_C	18	0.01	Down	0.249924148980568
0.00231337186937944	0.0137666281334053
ZSCAN12_E	196	0.01	Down	0.14541738710033
0.0023512392107071	0.0139297860794336
ZNF775_E	810	0.01	Up	0.225782817438422
0.00244467847243801	0.0144164556527848
ZNF541_E	57	0.01	Up	0.380938072729524
0.00246188398548777	0.0144164556527848
PLAGL1_E	385	0.01	Up	0.460899785332474
0.00246583037421976	0.0144164556527848
ZFX_C	55	0.01	Down	0.206918163670883
0.00249983486544123	0.0145225079258003
ZNF835_E	156	0.01	Up	0.260502616448298
0.00250575905696479	0.0145225079258003
ZNF621_E	279	0.01	Down	0.176218192621374
0.00253956868067586	0.0146547404819953
ZBTB12_E	315	0.01	Up	0.206295011241733
0.00255876459275376	0.0147018672506067
PEG3_E	191	0.01	Up	0.36436717776159
0.00258675949166261	0.0147989287656063
BBX_E	682	0.01	Down	0.196120245170605
0.00261488357452933	0.0148958966019128
CEBPB_A	23	0.01	Down	0.829884766982715
0.00265572560212703	0.0150548057667002
ZNF808_E	160	0.01	Down	0.200025111105685
0.00268020275955104	0.0150548057667002
ARID2_C	29	0.01	Down	0.192378804614678
0.00268191470931719	0.0150548057667002
ZNF587_E	256	0.01	Down	0.181699535269345
0.00268795481806049	0.0150548057667002
SOHLH1_E	14	0.01	Down	1.14550689896414
0.00270490513215946	0.0150863537287387
ZNF525_E	151	0.01	Down	0.163966485632446
0.00287260390967779	0.0159549208816687
ZNF281_E	648	0.01	Down	0.241152093018754
0.00306230139742815	0.0168902523918772
NFYC_D	18	0.01	Up	0.857747691667467
0.00306634739597471	0.0168902523918772

ZNF550_E 94 0.01 Down 0.178784175502285
0.00315170605403854 0.0172889883540468
ZNFX1_E 584 0.01 Down 0.20361510863615
0.00338786759996828 0.0185083094703185
HLX_E 252 0.01 Up 0.453389232539148
0.00354431556349191 0.0192174603407334
ZNF487_E 215 0.01 Up 0.270082064881357
0.00354650805988029 0.0192174603407334
ZNF444_E 523 0.01 Up 0.174639849661511
0.00363156863841417 0.019598708481806
FOXK2_C 50 0.01 Down 0.207183503136469
0.00369133845451443 0.019798611849452
HOXA3_E 59 0.01 Up 0.377192759458406
0.00370448632158027 0.019798611849452
IRF1_A 21 0.01 Down 0.501898294199187
0.00373219665789061 0.019798611849452
ZNF470_E 57 0.01 Down 0.196344712349949
0.00373483548241233 0.019798611849452
ZNF618_E 432 0.01 Up 0.325396467899519
0.0037428733578859 0.019798611849452
ZNF141_E 265 0.01 Down 0.170874992657442
0.00383575531762284 0.0201838332057072
ZNF823_E 157 0.01 Down 0.187816527748816
0.00384598172111751 0.0201838332057072
ZNF358_E 600 0.01 Up 0.242606925386807
0.00392587912929706 0.020522340703345
KLF6_C 47 0.01 Down 0.307904803693626
0.00396774234695898 0.0206601583925637
SMAD9_E 243 0.01 Up 0.396416436379261
0.0040330452772963 0.0208898780015651
SMARCC2_C 16 0.01 Down 0.190250859580541
0.0040432021938513 0.0208898780015651
PLAGL2_E 210 0.01 Down 0.215202121032606
0.00406832593553877 0.0209385269192015
EMX2_E 295 0.01 Up 0.508139940503156
0.00412552171379313 0.0211497484253221
EBF4_E 499 0.01 Up 0.26764063332911
0.00414109852888902 0.0211497484253221
ZNF677_E 108 0.01 Down 0.238506586939063
0.00420754073827803 0.0214070679546741
PKNOX2_E 409 0.01 Up 0.457601124256413
0.00423318294041885 0.0214556382493473
ZKSCAN8_E 346 0.01 Down 0.166095940427086
0.00430896935591647 0.021757030876654
ZNF264_E 502 0.01 Down 0.267084353138826
0.0044130031729383 0.0221982386019878
BCL11B_E 286 0.01 Down 0.474422422011978
0.00443076455713973 0.0222037938145386
ZNF623_E 362 0.01 Down 0.143170765839877
0.00448214163425203 0.0223771340766216
ZBTB24_E 615 0.01 Down 0.138484583470456
0.0045336659391019 0.0225499130478464
CSRNP3_E 295 0.01 Up 0.392149797864091
0.0045649799177386 0.0226212573617568

| | | | | | | |
|---|---|---|---|---|---|---|
| SP5_E | 42 | 0.01 | Up | 0.460675766322195 | 0.00459131576268076 | 0.0226278204503147 |
| ZNF576_E | 845 | 0.01 | Up | 0.125847358548129 | 0.00462267516457171 | 0.0226278204503147 |
| ZNF432_E | 250 | 0.01 | Down | 0.157826546306441 | 0.00462820751801039 | 0.0226278204503147 |
| ZNF334_E | 173 | 0.01 | Up | 0.255478411227949 | 0.00464721135811662 | 0.0226278204503147 |
| ZNF615_E | 177 | 0.01 | Down | 0.177909697313514 | 0.00465117989751404 | 0.0226278204503147 |
| TCF15_E | 91 | 0.01 | Up | 0.548788193434139 | 0.00484219966596109 | 0.0234714623808223 |
| SALL3_E | 11 | 0.01 | Down | 1.14517603703946 | 0.00486716242444809 | 0.023506983738367 |
| ZNF599_E | 258 | 0.01 | Up | 0.190993604549504 | 0.00512247128870488 | 0.024650737284634 |
| ZBTB33_B | 12 | 0.01 | Down | 0.200840091254651 | 0.00532549424188942 | 0.0255355533253187 |
| PATZ1_E | 567 | 0.01 | Up | 0.271285172345493 | 0.00547064077446161 | 0.0261375059224277 |
| NCOA1_C | 20 | 0.01 | Down | 0.164327012866843 | 0.00556304754223831 | 0.0264840799064417 |
| ZBTB37_E | 366 | 0.01 | Down | 0.201694727819994 | 0.00585298720546414 | 0.0277652382380203 |
| KDM5B_C | 17 | 0.01 | Down | 0.184910412279485 | 0.0058877028902375 | 0.027830879264846 |
| PRRX1_E | 707 | 0.01 | Up | 0.550634424536055 | 0.00593922061956659 | 0.0279290502843227 |
| ZBTB22_E | 411 | 0.01 | Up | 0.160401886996403 | 0.00595037530438684 | 0.0279290502843227 |
| IRX6_E | 47 | 0.01 | Up | 0.46798874113676 | 0.00605843054290077 | 0.0283364488199534 |
| ZNF420_E | 176 | 0.01 | Down | 0.155215817379915 | 0.00609395483620664 | 0.0284029433449771 |
| SNAI1_D | 13 | 0.01 | Down | 0.679137733563081 | 0.00628623213907155 | 0.0291970294124822 |
| ZNF260_E | 552 | 0.01 | Down | 0.210576177387261 | 0.00641274740161013 | 0.0295825538093832 |
| ZNF669_E | 110 | 0.01 | Down | 0.159437376742704 | 0.00641362194366972 | 0.0295825538093832 |
| ZNF829_E | 71 | 0.01 | Down | 0.158622841176478 | 0.00655392480792871 | 0.030125454375755 |
| ZNF674_E | 133 | 0.01 | Down | 0.166840783573762 | 0.00658396479947315 | 0.0301595363494767 |
| SOX7_E | 180 | 0.01 | Down | 0.482393475299536 | 0.00687402202485382 | 0.0313803813668841 |
| ZNF776_E | 195 | 0.01 | Down | 0.172634765335112 | 0.0069920032453387 | 0.0318100352424454 |
| PHTF2_E | 417 | 0.01 | Down | 0.200194000510866 | 0.00702139120901779 | 0.0318350832708187 |
| ZNF670_E | 54 | 0.01 | Down | 0.142980012328747 | 0.00706301701344593 | 0.0318751952961903 |
| ZBTB38_E | 497 | 0.01 | Down | 0.194269486225842 | 0.00707806287147212 | 0.0318751952961903 |

LRRFIP1_E 576 0.01 Down 0.23307566854687
0.00719611549022481 0.0322977169982144
ZBED3_E 415 0.01 Up 0.263319825164785
0.00736308071768291 0.0329361966331252
ZNF579_E 327 0.01 Up 0.153604057929755
0.00741733753982774 0.0330679295671919
JUNB_C 50 0.01 Down 0.265615883180537
0.00747369619911841 0.0332081234447495
ZNF33A_E 480 0.01 Down 0.162619421860822
0.00752127153476229 0.0333084882253759
ZNF383_E 205 0.01 Down 0.144393920808866
0.00771438321559206 0.0340505722727954
ZBTB26_E 172 0.01 Down 0.165754461536893
0.0078620983743594 0.0345880433433039
ZNF160_E 183 0.01 Down 0.170032650342744
0.00801335863156071 0.0351375232101001
ZNF675_E 342 0.01 Down 0.219899097968306
0.00812019310336847 0.035489237399312
ZNF813_E 136 0.01 Down 0.142246533852753
0.00820798949658926 0.0357085742113158
NFKBIL1_E 566 0.01 Up 0.151413273139135
0.00822395520095571 0.0357085742113158
MAFF_C 37 0.01 Down 0.319673025801051
0.00827240424745267 0.0357813456465851
ZXDB_E 226 0.01 Down 0.171926206580981
0.00829440045370952 0.0357813456465851
CREBZF_E 441 0.01 Down 0.16316550807129
0.00837379321000006 0.0360073108030003
ZNF701_E 208 0.01 Down 0.155727182364966
0.00842664857370041 0.0361180789348638
FIZ1_E 698 0.01 Up 0.132401986324523
0.0085338154539023 0.0364601794873454
FOXM1_A 21 0.01 Up 0.588504643506747
0.00862728034373135 0.0367417402498207
ZNF749_E 191 0.01 Down 0.140096568283213
0.00868235119469917 0.0368585163774968
TET1_E 175 0.01 Up 0.376103754073497
0.0087571106701485 0.0370578683279618
TOX4_E 1088 0.01 Down 0.121732849190755
0.00889202850312457 0.0375097278312185
ZFHX2_E 318 0.01 Up 0.246291861338569
0.00906222823450892 0.0379874518014373
TFEC_E 718 0.01 Up 0.531736355126212
0.0090622728228485 0.0379874518014373
ZNF765_E 95 0.01 Down 0.142557457340442
0.0092969144912958 0.0388488621219351
ZNF100_E 87 0.01 Down 0.164015855981596
0.00945921650219621 0.0394035487419611
NFKB2_B 10 0.01 Down 0.68968089732703
0.0101303576462433 0.0418894791980947
ZNF639_C 26 0.01 Down 0.158954500677176
0.0101724938014437 0.0418894791980947
ZNF681_E 170 0.01 Down 0.202840924639772
0.010180675192613 0.0418894791980947

TRERF1_E 551 0.01 Up 0.408112811102436
0.0101816888673538 0.0418894791980947
ZNF467_E 431 0.01 Up 0.340647399965828
0.010332475082788 0.0423790439549428
CASZ1_E 217 0.01 Down 0.2729211192333 0.0104087212325146
0.0425608141194537
HES6_E 164 0.01 Up 0.174412590397192
0.0105521302415908 0.0430152587524174
ZBTB39_E 275 0.01 Down 0.147603032271778
0.0106737126679703 0.0433782286170864
EBF3_E 205 0.01 Up 0.593587389854726
0.0107185859543868 0.0434281917240049
ZSCAN1_E 74 0.01 Up 0.303719552372619
0.0108547580717771 0.043846643968724
HES2_E 622 0.01 Down 0.361569376603874
0.0111110816912678 0.0447464407687614
ZNF69_E 236 0.01 Up 0.242587933077262
0.0111737657232617 0.0448205700989807
ZNF227_E 207 0.01 Down 0.150019213718172
0.011196736566362 0.0448205700989807
ZNF8_E 408 0.01 Down 0.257243830149258
0.0113956581808674 0.0454802765122642
PPP1R13L_E 559 0.01 Down 0.259182053034323
0.0115241767386587 0.0458559032615883
RELB_C 62 0.01 Down 0.417770256182729
0.0116062280514058 0.0459129441808366
ZNF230_E 82 0.01 Down 0.151203490700339
0.0116073984913293 0.0459129441808366
ZNF234_E 187 0.01 Down 0.139978626597333
0.0117006563054891 0.0460568023707596
ZNF189_E 78 0.01 Down 0.175719420653722
0.0117128702203207 0.0460568023707596
ZNF280B_E 70 0.01 Up 0.280560989070228
0.0120922105709129 0.0474085785030203
ZNF461_E 135 0.01 Down 0.169782894160943
0.0121691165398696 0.0475701828376721
SMAD7_D 16 0.01 Up 0.896467375011969
0.0122598798661457 0.04778485339641
SOX21_E 394 0.01 Down 0.464867303772146
0.012406837973179 0.0482166618607801
NR4A1_C 35 0.01 Down 0.252579229914732
0.0124669201080503 0.0483093154186949
NFE2L3_E 216 0.01 Down 0.217047734698544
0.0126469994241393 0.0488650731373266
NRL_E 600 0.01 Up 0.225237654983493
0.0127666039944698 0.0491575171335977
ZNF500_E 270 0.01 Up 0.153514866446094
0.012796442944755 0.0491575171335977
ZNF83_D 111 0.01 Down 0.290508134354737
0.012866079089488 0.0492829983514009
ZNF17_E 76 0.01 Down 0.161951367614107
0.0129788209369062 0.0495281775023338
ZNF491_E 140 0.01 Up 0.188813664719535
0.0130043976937861 0.0495281775023338

ZFPM1_E 240 0.01 Up 0.204715748583922
0.013141201368381 0.0498334325411696
RXRG_D 36 0.01 Up 0.458912534823288
0.0131634341652615 0.0498334325411696
ZNF846_E 364 0.01 Up 0.242647057562083
0.0131967004403847 0.0498334325411696
ZNF45_E 280 0.01 Down 0.14260582645795
0.0133712908643404 0.050350086785779
ATF2_A 14 0.01 Down 0.520995481232631
0.0136516845279556 0.0512611140162389
ZNF121_E 674 0.01 Down 0.223398935992885
0.0142267191419629 0.053270271393923
ZNF555_E 47 0.01 Down 0.153358201356355
0.0150658917593014 0.0562544361768871
MTA2_C 33 0.01 Down 0.182347892228681
0.0151953776556734 0.0565794369134432
ELF1_C 52 0.01 Down 0.171927744103574
0.0153954035991911 0.057164548740172
ATF3_C 45 0.01 Down 0.202432598246022
0.0155445430542844 0.0575579885871143
ZSCAN2_E 110 0.01 Up 0.159884269255725
0.0158573782175987 0.0584892387867011
FOXF1_E 132 0.01 Up 0.60697103406021
0.0158837992804094 0.0584892387867011
ZNF891_E 203 0.01 Down 0.186515697406976
0.0160342802966612 0.0588807042298882
ZBED4_E 756 0.01 Down 0.157787865821424
0.0161491600909742 0.0591396439595291
ZNF529_E 400 0.01 Down 0.16928524345417
0.0162320122820838 0.0592801982795004
VENTX_E 658 0.01 Up 0.528426056984207
0.0167120495274539 0.0608665628964372
KLF3_C 11 0.01 Down 0.276485385881904
0.0168298522252046 0.0611285913247896
ZNF407_E 398 0.01 Down 0.142642664913552
0.0172852364952251 0.0626120115438454
ZNF445_E 388 0.01 Down 0.186306368814918
0.0178307050795728 0.0642926169675298
ZBTB6_E 676 0.01 Down 0.20958558941768
0.0178456626241456 0.0642926169675298
ZNF668_E 476 0.01 Up 0.14907147385989
0.018015983819071 0.0646316412852456
ZNF101_E 309 0.01 Down 0.259191488305029
0.0180367371028592 0.0646316412852456
ZBTB46_E 468 0.01 Up 0.312132172062665
0.0182318692579095 0.0650086777908444
ZNF213_E 433 0.01 Up 0.156714324827468
0.0182394939938303 0.0650086777908444
ZNF773_E 59 0.01 Down 0.140740505631504
0.0183288492153673 0.0651529493442256
HIF3A_E 324 0.01 Up 0.421485158331884
0.0185483474849722 0.0657578382911383
FOS_A 63 0.01 Down 0.614931747358598
0.0192100537952119 0.0679230814562798

| | | | | | | |
|---|---|---|---|---|---|---|
| JUN_A | 84 | 0.01 | Down | 0.705089037029953 | 0.0193840643675554 | 0.0683570312220934 |
| NHLH2_E | 81 | 0.01 | Down | 0.609777837142864 | 0.0194374868037303 | 0.0683645644046767 |
| ZBTB4_E | 597 | 0.01 | Up | 0.255626787622915 | 0.0198408800285877 | 0.0695402030192262 |
| ZNF497_E | 302 | 0.01 | Up | 0.218761942925293 | 0.0198760820332522 | 0.0695402030192262 |
| ZNF184_E | 264 | 0.01 | Down | 0.153566609730237 | 0.0199309881510838 | 0.0695497570821852 |
| GRHL2_C | 71 | 0.01 | Down | 0.298961958180461 | 0.0202961019231545 | 0.0706389134818929 |
| TBX2_E | 635 | 0.01 | Up | 0.450147662724632 | 0.0204634413559438 | 0.071026493257306 |
| ZNF551_E | 58 | 0.01 | Down | 0.16101658299952 | 0.0205140284351559 | 0.071026493257306 |
| NEUROD1_C | 23 | 0.01 | Down | 0.183530622900036 | 0.0207151954989482 | 0.071537190673829 |
| RORC_E | 732 | 0.01 | Down | 0.331190935702782 | 0.0212666477239828 | 0.0730891060938258 |
| ELK1_A | 27 | 0.01 | Down | 0.528414313260469 | 0.0213166258684387 | 0.0730891060938258 |
| RXRB_D | 39 | 0.01 | Up | 0.44005892237404 | 0.0213729478645244 | 0.0730891060938258 |
| NR4A3_E | 118 | 0.01 | Down | 0.389497513683586 | 0.0214231369548741 | 0.0730891060938258 |
| FOXC2_E | 467 | 0.01 | Up | 0.470917424060879 | 0.0214387400470262 | 0.0730891060938258 |
| ZNF253_E | 151 | 0.01 | Down | 0.167647240244782 | 0.0216521479048495 | 0.0734913594411096 |
| ZNF217_C | 10 | 0.01 | Down | 0.329999252452088 | 0.0216669949440031 | 0.0734913594411096 |
| ZNF768_E | 744 | 0.01 | Up | 0.241422055729642 | 0.0217867505352618 | 0.0737099961002638 |
| ZNF556_E | 151 | 0.01 | Down | 0.23132635064092 | 0.0221864478693058 | 0.0748722405311002 |
| ZNF506_E | 284 | 0.01 | Down | 0.163179005342641 | 0.0222429016568384 | 0.0748732017893071 |
| BACH2_C | 42 | 0.01 | Down | 0.321175084438138 | 0.0225216555117408 | 0.075620571277457 |
| TWIST1_A | 12 | 0.01 | Up | 0.747189590654753 | 0.022618374329937 | 0.0757107232896518 |
| ZNF391_E | 218 | 0.01 | Up | 0.219110834833936 | 0.0226620994692956 | 0.0757107232896518 |
| TSC22D4_D | 49 | 0.01 | Up | 0.560474989628269 | 0.0227466401841591 | 0.0758031784137103 |
| RARA_A | 22 | 0.01 | Down | 0.603420463557216 | 0.0228206593535976 | 0.0758601968038544 |
| TEAD2_C | 29 | 0.01 | Up | 0.370819274144254 | 0.0232142027089406 | 0.0769764482861141 |
| MYCN_A | 13 | 0.01 | Up | 0.201742734893466 | 0.0233643997211945 | 0.0772822452316435 |
| ZNF780A_E | 195 | 0.01 | Down | 0.160815121451211 | 0.0236723037829962 | 0.0779474705855889 |

SP110_E 368 0.01 Down 0.332103287228046
0.0236824648065743 0.0779474705855889
ZNF274_C 18 0.01 Down 0.26236764540205
0.0239780279028653 0.0786136713288736
ZFP2_E 168 0.01 Up 0.23797249427221
0.0240028238791084 0.0786136713288736
TLX1_E 33 0.01 Down 1.08101540382078
0.0242985903873926 0.0792567668591739
ZNF410_E 401 0.01 Down 0.272881396330777
0.0243180927572409 0.0792567668591739
ZNF155_E 73 0.01 Down 0.133818522124874
0.0255225740129519 0.0829794906323535
GLI4_E 892 0.01 Up 0.171007807715315
0.0260139717060897 0.0843713486234006
ZNF229_E 40 0.01 Down 0.205731268602201
0.0264832575202996 0.0856849084333965
ZNF57_E 200 0.01 Down 0.234223875087648
0.0270549306872297 0.0869961582245548
SP100_E 517 0.01 Down 0.290294179509313
0.0270739518148092 0.0869961582245548
ATF5_E 308 0.01 Up 0.158210968999762
0.0270843253287249 0.0869961582245548
SP2_C 22 0.01 Down 0.330260010517666
0.0271902866052944 0.0870742029748116
HEY2_E 319 0.01 Up 0.376189677211712
0.0272392667970716 0.0870742029748116
ZNF691_E 434 0.01 Up 0.151332553423563
0.0274465286914554 0.0875268486739473
ELF2_E 557 0.01 Down 0.162019083344414
0.0290432208196287 0.0923976452328523
ZNF543_E 38 0.01 Down 0.240608528197599
0.0291252009504524 0.0924378401594119
ZNF311_E 182 0.01 Up 0.196855012925649
0.0295324941808365 0.0935078734989432
IRF4_C 59 0.01 Down 0.359074456951935
0.0299211411484069 0.0945139363763657
ZNF74_E 408 0.01 Up 0.170928104103818
0.0303301816350258 0.0955795085567126
ZNF628_E 256 0.01 Up 0.131822056716099
0.0304809625965483 0.0958281206160351
PHOX2B_E 179 0.01 Down 0.309261507431598
0.0315628752603091 0.098996029934099
ZNF425_E 180 0.01 Up 0.16164036555095
0.0317007346063244 0.0991925268116209
BCL11A_C 30 0.01 Down 0.476985676866917
0.0317743502989963 0.0991925268116209
KLF2_D 26 0.01 Up 0.646495458394244
0.0327466147503888 0.101988872575393
ZNF571_E 125 0.01 Down 0.168338826543992
0.0331833830675543 0.10310827419359
GMEB1_E 724 0.01 Down 0.180694370971417
0.0332616678244245 0.10311170255716
NCOA3_C 21 0.01 Down 0.26242659093325
0.0338190125712303 0.104595693172738

| | | | | | | |
|---|---|---|---|---|---|---|
| HOXB2_E | 458 | 0.01 | Up | 0.396309248391654 | 0.0339107281406466 | 0.104636575489541 |
| IRF9_C | 36 | 0.01 | Down | 0.203095694644786 | 0.0348503110508928 | 0.107287447184388 |
| ZNF710_E | 337 | 0.01 | Down | 0.2579402623544 | 0.0353324254082898 | 0.10852102089689 |
| DMRTC2_E | 25 | 0.01 | Down | 1.48553888531024 | 0.0358426730054138 | 0.109835133600498 |
| PRDM5_E | 394 | 0.01 | Up | 0.356255988017584 | 0.0366739558861171 | 0.11211275522222 |
| ZKSCAN7_E | 157 | 0.01 | Up | 0.190589999347426 | 0.0367541440600975 | 0.11211275522222 |
| GTF3A_E | 1113 | 0.01 | Up | 0.132220571428317 | 0.0374732835590022 | 0.114045404073402 |
| ZNF799_E | 108 | 0.01 | Down | 0.140214427646571 | 0.0378781462949379 | 0.115014963578934 |
| ZNF354B_E | 358 | 0.01 | Down | 0.146423000299658 | 0.0382736081433322 | 0.115829964361471 |
| ZFP41_E | 388 | 0.01 | Up | 0.146670313055046 | 0.0383203407977559 | 0.115829964361471 |
| FOXL1_E | 402 | 0.01 | Up | 0.468758808393239 | 0.0385022521020311 | 0.116090853742441 |
| ZNF585A_E | 219 | 0.01 | Down | 0.171678037770571 | 0.038580831363767 | 0.116090853742441 |
| MXD4_E | 545 | 0.01 | Up | 0.197687182195486 | 0.0394147852653286 | 0.118227089035094 |
| TOX2_E | 69 | 0.01 | Up | 0.513974426746004 | 0.0394681580049638 | 0.118227089035094 |
| TONSL_E | 975 | 0.01 | Up | 0.197598433933625 | 0.0396617130485412 | 0.118540501106963 |
| ZNF41_E | 662 | 0.01 | Down | 0.231086296881821 | 0.0400803687348866 | 0.119519481458102 |
| ZNF251_E | 354 | 0.01 | Up | 0.207785652217274 | 0.0401685879169016 | 0.119519481458102 |
| SALL1_E | 48 | 0.01 | Down | 0.402449547957428 | 0.0405053188483856 | 0.120252984465252 |
| ZNF787_E | 614 | 0.01 | Up | 0.127757448512032 | 0.0408000851611847 | 0.120672287853117 |
| SMARCA1_E | 956 | 0.01 | Up | 0.348524429370366 | 0.0408276082683838 | 0.120672287853117 |
| ZNF19_E | 151 | 0.01 | Up | 0.176220755885421 | 0.0411951852456434 | 0.121489340558501 |
| ARID3A_C | 21 | 0.01 | Down | 0.33526791675204 | 0.0416541530732474 | 0.122510713428982 |
| ZNF568_E | 156 | 0.01 | Down | 0.163729345449474 | 0.0417253292548822 | 0.122510713428982 |
| SPDEF_E | 474 | 0.01 | Down | 0.351373632065168 | 0.041876153209119 | 0.122683323577485 |
| ZNF256_E | 41 | 0.01 | Down | 0.146623211812811 | 0.0421153932601741 | 0.122956960787174 |
| MEF2A_C | 96 | 0.01 | Down | 0.330417146833494 | 0.0421540368190087 | 0.122956960787174 |
| MITF_A | 27 | 0.01 | Up | 0.550395098599483 | 0.0423768116748223 | 0.123336877647463 |

| | | | | | | |
|---|---|---|---|---|---|---|
| HMBOX1_C | 24 | 0.01 | Down | 0.200630752662578 | 0.0434366421307826 | 0.126146065273057 |
| ZNF112_E | 69 | 0.01 | Down | 0.147350399726685 | 0.0436254176355787 | 0.126418873278753 |
| ZNF317_E | 602 | 0.01 | Down | 0.218463457891947 | 0.0447696102500367 | 0.128877365288902 |
| FOXO3_A | 31 | 0.01 | Down | 0.324019289701847 | 0.0448302533211443 | 0.128877365288902 |
| ZNF48_E | 455 | 0.01 | Up | 0.145186826613764 | 0.0448415070559947 | 0.128877365288902 |
| TFAP2C_A | 10 | 0.01 | Down | 0.491304127160828 | 0.0448605382551015 | 0.128877365288902 |
| PRDM6_E | 414 | 0.01 | Up | 0.418649373249737 | 0.0451948970390007 | 0.129558704845135 |
| DMAP1_E | 828 | 0.01 | Up | 0.159454943942222 | 0.0453453142256584 | 0.129708108357109 |
| ZNF35_E | 558 | 0.01 | Down | 0.123802268505564 | 0.0454416253584169 | 0.129708108357109 |
| ZNF174_E | 333 | 0.01 | Up | 0.163975357308543 | 0.0458721069275886 | 0.130657090885632 |
| BACH1_B | 27 | 0.01 | Down | 0.278409526504361 | 0.0462585336518645 | 0.131476813129926 |
| ZNF883_E | 35 | 0.01 | Up | 0.217672465766726 | 0.0470765304820783 | 0.133517053473639 |
| NFKBIE_E | 567 | 0.01 | Down | 0.377214976310639 | 0.0472620621920376 | 0.133758660089142 |
| RREB1_E | 525 | 0.01 | Down | 0.270854185695058 | 0.0477827941566877 | 0.13494589959929 |
| ZNF26_E | 474 | 0.01 | Down | 0.14728050528265 | 0.0490042846494419 | 0.137892459945884 |
| RCOR3_E | 519 | 0.01 | Down | 0.170063517913242 | 0.049054732011351 | 0.137892459945884 |
| ZNF566_E | 129 | 0.01 | Down | 0.155331365413715 | 0.0492240884228467 | 0.137892459945884 |
| ZNF660_E | 70 | 0.01 | Up | 0.283115639026849 | 0.0492399181802258 | 0.137892459945884 |
| ISX_E | 320 | 0.01 | Down | 0.446213528919685 | 0.0497083898975951 | 0.138912544514663 |
| DMTF1_E | 633 | 0.01 | Down | 0.18257731866614 | 0.0500217695836422 | 0.139495855345178 |
| ZNF697_E | 102 | 0.01 | Down | 0.241873175116666 | 0.0513977024672908 | 0.143033689747179 |
| ZBTB20_E | 128 | 0.01 | Down | 0.212214860817561 | 0.0520711378070242 | 0.144574094061899 |
| ZNF394_E | 587 | 0.01 | Down | 0.24632979102078 | 0.0521681464694475 | 0.144574094061899 |
| ARID3B_E | 583 | 0.01 | Down | 0.226473460444052 | 0.0524062082764066 | 0.144932522059025 |
| TCF12_C | 71 | 0.01 | Down | 0.25103727917134 | 0.0530380390416963 | 0.146376202986711 |
| SNAI3_E | 223 | 0.01 | Up | 0.575855010221686 | 0.0539330537617628 | 0.148538761703367 |
| PPARD_D | 28 | 0.01 | Up | 0.44734613602022 | 0.0541563086191419 | 0.148846101833642 |

HBP1_C 38 0.01 Down 0.228308755051936
0.0563681987318854 0.154606602694657
ZNF527_E 164 0.01 Down 0.168995808999279
0.0567592394277889 0.155251749736265
RFX8_E 142 0.01 Up 0.545339316043502
0.0568363494908459 0.155251749736265
DLX5_E 19 0.01 Down 0.667192425094349
0.0572087512317565 0.15594941797941
ZNF14_E 113 0.01 Down 0.135531207274661
0.057372678924327 0.156077104094139
ZSCAN32_E 39 0.01 Down 0.118896956005747
0.0582386788835659 0.158110303364141
ZEB2_C 47 0.01 Up 0.347880290267083
0.058866592154473 0.159490177524212
ZNF140_E 587 0.01 Down 0.230030114226423
0.0602202901091915 0.162826869605583
ZNF561_E 404 0.01 Down 0.128042361840735
0.0605130891960408 0.162962885815495
ZNF778_E 402 0.01 Down 0.134148256006026
0.060515100134036 0.162962885815495
ZBED5_E 349 0.01 Down 0.140918488984441
0.0611704255479934 0.164395518660232
SMAD6_E 91 0.01 Up 0.495229902622797
0.0621130595315105 0.166592974558357
WT1_A 18 0.01 Down 0.512972691118077
0.0623903807811807 0.166804951062918
NR6A1_E 445 0.01 Down 0.2970682690371 0.0624423635261785
0.166804951062918
HNF1B_C 27 0.01 Down 0.451651460211734
0.0630979339275576 0.168219091850868
STAT1_A 149 0.01 Down 0.463357084793489
0.0635267413048607 0.168809744013038
ZNF415_E 92 0.01 Up 0.347133198536169
0.0635727618113616 0.168809744013038
THAP9_E 189 0.01 Down 0.14911705115152
0.0638259234146751 0.169145041574079
TCFL5_E 358 0.01 Up 0.207048253185339
0.0639923608417005 0.169249636908704
TSHZ1_E 253 0.01 Up 0.277582941976785
0.0644086138163041 0.170013232113135
MAFB_C 45 0.01 Down 0.489587865944055
0.0662466052026763 0.174519218844205
KAT7_E 572 0.01 Down 0.132539159616764
0.0674630119690314 0.177373165591161
LYL1_C 75 0.01 Up 0.431404381664971
0.0676564185432998 0.177531507713029
ZNF92_D 194 0.01 Up 0.440084465803256
0.0686246665321291 0.179718429248189
SMAD4_A 16 0.01 Down 0.306251824149729
0.0689962534814408 0.180337266452472
ZBTB40_E 322 0.01 Down 0.132886689370033
0.0703153717691225 0.183425421855656
HNF4A_A 38 0.01 Up 0.566788203047612
0.0707294730709452 0.183699587023937

NR1D2_E 94 0.01 Down 0.215918504968062
0.0707847258605784 0.183699587023937
ZNF239_E 597 0.01 Up 0.131392209016462
0.0708338992725458 0.183699587023937
ZSCAN25_E 344 0.01 Down 0.129343776498941
0.071879107543442 0.186048253117297
ZFP64_E 716 0.01 Up 0.176607902021727
0.0720705759393881 0.186182321176753
ARID1B_C 17 0.01 Down 0.208369971605855
0.0725970705472291 0.187179680927382
KLF12_E 824 0.01 Up 0.299480636562948
0.0728535155338782 0.187478255225212
SMAD5_C 11 0.01 Down 0.168404964318848
0.0732722786631296 0.187729310647764
DDIT3_D 17 0.01 Down 0.42025712279211
0.073396050868541 0.187729310647764
ZNF790_E 88 0.01 Down 0.180681984671453
0.0735470096978952 0.187729310647764
ZNF107_E 117 0.01 Down 0.237093628855349
0.0736258153213342 0.187729310647764
ZNF696_E 247 0.01 Up 0.130883266553628
0.0736552359105631 0.187729310647764
ZNF784_E 346 0.01 Up 0.173503389914851
0.0739554749252282 0.188092002902396
NR2C2_C 84 0.01 Down 0.134537820679693
0.0743274832535353 0.188092002902396
ZNF225_E 168 0.01 Down 0.144927050171123
0.07434966947662 0.188092002902396
ARID5B_E 533 0.01 Down 0.32190899476217
0.074361954635831 0.188092002902396
MYRF_E 225 0.01 Down 0.415579435813945
0.075089030677458 0.189571359645931
CUX2_E 288 0.01 Down 0.332339539934966
0.0755267069116104 0.190058487219124
GATAD2A_E 881 0.01 Down 0.175107124569296
0.0755671404547153 0.190058487219124
ZNF347_E 134 0.01 Down 0.175158646022563
0.0772783697467004 0.193996359458289
YBX2_E 96 0.01 Up 0.286830598940566
0.0775674814966406 0.194356114351545
THAP6_E 439 0.01 Down 0.153254881672514
0.0777750013169919 0.194510462956004
SIX5_C 27 0.01 Up 0.18978381886705
0.0790531817470203 0.197336875035165
ZNF700_E 453 0.01 Down 0.165021030910294
0.0793948644101993 0.197819353754758
ZNF396_E 301 0.01 Up 0.202031113183044
0.0796082953641025 0.197981077836471
HOXA4_E 40 0.01 Up 0.426680460981789
0.0809112783954675 0.20084680465765
ZNF557_E 179 0.01 Down 0.123494718721912
0.0811593624933566 0.201088160229822
ZNF827_E 197 0.01 Up 0.238705119649379
0.0823601653140222 0.203684787316496

ZNF708_E 453 0.01 Down 0.274888075709385
0.0826627575679142 0.204054547848203
ZNF766_D 201 0.01 Down 0.283320824923571
0.0829597307758057 0.204105026710007
ZNF18_E 712 0.01 Up 0.221883951374106
0.0830309940607273 0.204105026710007
SIM1_E 15 0.01 Down 0.916422550708289
0.0832623312276343 0.204105026710007
NCOA2_C 16 0.01 Down 0.21272016596213
0.0833494051137013 0.204105026710007
ZNF133_E 430 0.01 Up 0.150872283727728
0.0834487918656818 0.204105026710007
ID1_E 78 0.01 Down 0.300999638105291
0.0836770522527988 0.204288481049415
HOXB13_C 32 0.01 Down 0.289116201015284
0.0846158977772893 0.206202909939902
ZNF496_E 381 0.01 Up 0.198840621746906
0.085223081749496 0.207303591189924
LMX1B_E 154 0.01 Up 0.399849094226325
0.0867116998461239 0.210540429681026
GMEB2_E 528 0.01 Down 0.162927064103854
0.0872076540770357 0.211108878617577
SP140L_E 513 0.01 Down 0.324966521813624
0.0872663736399705 0.211108878617577
ELF4_E 874 0.01 Down 0.308936114100635
0.0874209309804222 0.211108878617577
NR3C2_D 13 0.01 Down 0.474880798714013
0.0879027643860263 0.211888580337384
FOSL1_A 10 0.01 Down 1.14625198522022
0.0890415410530081 0.214246162858592
MZF1_E 820 0.01 Up 0.209482871159195
0.0896154524342372 0.215238555125835
NEUROD6_E 20 0.01 Down 1.0709779701644
0.0907248263932839 0.217511139536416
ZNF610_E 121 0.01 Up 0.197134850731383
0.0912183242912756 0.218301662980737
NFATC1_C 29 0.01 Down 0.392317779002022
0.0923898629921088 0.220447669137962
ZNF446_E 365 0.01 Up 0.148270332517446
0.0924457967352744 0.220447669137962
ZNF595_E 80 0.01 Down 0.144041213611025
0.0927546123465193 0.220696363276907
ZNF200_E 235 0.01 Down 0.145927970675014
0.0928876940923961 0.220696363276907
ZNF493_E 150 0.01 Down 0.159947121287234
0.0930467788159202 0.220696363276907
ZNF286A_E 253 0.01 Down 0.165562364806325
0.0932285591424653 0.220734759035357
UBTF_C 20 0.01 Down 0.16771629924601
0.0935221093283099 0.221037183926661
IRX3_E 89 0.01 Down 0.322404089338345
0.0954100231809154 0.224910754484948
HIC1_E 982 0.01 Up 0.441052560617721
0.0954984899013356 0.224910754484948

| | | | | | | |
|---|---|---|---|---|---|---|
| FOXA3_E | 381 | 0.01 | Down | 0.383412083326088 | 0.0963268092990603 | 0.226461440556697 |
| ZNF181_E | 150 | 0.01 | Down | 0.149870805802186 | 0.0972123081142783 | 0.228140856894952 |
| TWIST2_D | 21 | 0.01 | Up | 0.412864211380531 | 0.0979406503802624 | 0.22944619851826 |
| ZNF653_E | 438 | 0.01 | Up | 0.12511213413683 2 | 0.0991108567637087 | 0.231780301870217 |
| NFKBIA_D | 200 | 0.01 | Down | 0.39606558250679 | 0.100947895852713 | 0.235662951263865 |
| TEAD1_C | 119 | 0.01 | Down | 0.307766777637331 | 0.101490753659556 | 0.236516039559769 |
| FOXP2_C | 30 | 0.01 | Down | 0.262824827862877 | 0.102810734100175 | 0.238969556949706 |
| ZIC2_E | 12 | 0.01 | Up | 0.866156061152241 | 0.102916965041608 | 0.238969556949706 |
| TSC22D3_E | 161 | 0.01 | Up | 0.451116562706394 | 0.103081391782506 | 0.238969556949706 |
| NCOR2_C | 22 | 0.01 | Down | 0.153362141854396 | 0.104594874232046 | 0.242057234984926 |
| ZGLP1_E | 457 | 0.01 | Up | 0.178846495873157 | 0.104885658509552 | 0.242309502241305 |
| ZNF44_E | 147 | 0.01 | Down | 0.158865943989827 | 0.105355586770227 | 0.242974043537566 |
| DLX3_E | 598 | 0.01 | Down | 0.31776541516267 | 0.106349068037332 | 0.244841636776793 |
| ZNF763_E | 110 | 0.01 | Up | 0.19267174937514 | 0.10782426896233 | 0.247809914701356 |
| PBX1_E | 935 | 0.01 | Up | 0.329812444467753 | 0.108398038452052 | 0.248699802507032 |
| ZNF782_E | 108 | 0.01 | Down | 0.157470240389795 | 0.109350109155913 | 0.250453085059849 |
| DMBX1_E | 268 | 0.01 | Down | 0.278605333564155 | 0.110594346936042 | 0.252868378157365 |
| ZNF740_C | 35 | 0.01 | Down | 0.217299566991845 | 0.110841944879627 | 0.253000535144766 |
| POU6F1_E | 785 | 0.01 | Up | 0.245556241307283 | 0.112225136075366 | 0.255344988481231 |
| ZNF577_E | 179 | 0.01 | Up | 0.188846446671206 | 0.112252185483872 | 0.255344988481231 |
| CSRNP2_E | 201 | 0.01 | Down | 0.162547865437499 | 0.113137600386036 | 0.256920649598955 |
| ZNF236_E | 325 | 0.01 | Down | 0.191707495978612 | 0.114130163173058 | 0.258733856308991 |
| IRF5_E | 975 | 0.01 | Up | 0.401192878839899 | 0.114872701772419 | 0.259193424867875 |
| HIVEP3_E | 163 | 0.01 | Down | 0.25106180725810 1 | 0.114999108619487 | 0.259193424867875 |
| PROX1_C | 20 | 0.01 | Down | 0.295056358857634 | 0.115093381174168 | 0.259193424867875 |
| ZNF777_E | 369 | 0.01 | Up | 0.121736702120006 | 0.115110658305913 | 0.259193424867875 |
| NFIB_E | 229 | 0.01 | Up | 0.446376694871832 | 0.115545599507804 | 0.259416773892973 |

| | | | | | | |
|---|---|---|---|---|---|---|
| PHOX2A_E | 225 | 0.01 | Down | 0.320608853117756 | 0.115680810774192 | 0.259416773892973 |
| ZNF518A_E | 375 | 0.01 | Down | 0.180746668512732 | 0.115793683770682 | 0.259416773892973 |
| ZFP42_E | 300 | 0.01 | Down | 0.159451280877515 | 0.116356917326153 | 0.260241226167386 |
| TBP_C | 37 | 0.01 | Down | 0.19517990736539 | 0.116832021507249 | 0.260334382951027 |
| ZNF408_E | 430 | 0.01 | Up | 0.114752920171707 | 0.116971785636887 | 0.260334382951027 |
| NR5A2_C | 25 | 0.01 | Down | 0.304816859396825 | 0.116984467657663 | 0.260334382951027 |
| ZNF84_E | 316 | 0.01 | Down | 0.175115155136453 | 0.117767838651167 | 0.261640881536675 |
| KAT6B_E | 387 | 0.01 | Down | 0.200770715761957 | 0.118645364981601 | 0.263151866090639 |
| ELK4_C | 12 | 0.01 | Down | 0.277401514917877 | 0.12046271990086 | 0.266673471859533 |
| SREBF1_A | 20 | 0.01 | Down | 0.247596544898107 | 0.120633235957463 | 0.266673471859533 |
| ZNF134_E | 458 | 0.01 | Down | 0.236651145488813 | 0.121552613355851 | 0.268260982787001 |
| LHX3_E | 240 | 0.01 | Down | 0.400745138459161 | 0.122945510227244 | 0.270794398739934 |
| ZBTB7C_E | 612 | 0.01 | Down | 0.302883163526969 | 0.123106830935034 | 0.270794398739934 |
| KLF15_E | 523 | 0.01 | Up | 0.372549403817159 | 0.123803698485009 | 0.271878632752087 |
| GATA3_A | 44 | 0.01 | Down | 0.412395206747925 | 0.124176062202711 | 0.272247846901668 |
| ZNF143_C | 43 | 0.01 | Down | 0.189821549404328 | 0.124482029161281 | 0.272470517031178 |
| ZNF589_D | 142 | 0.01 | Down | 0.37035331899466 | 0.12509349318975 | 0.27336004331465 |
| ZNF71_E | 251 | 0.01 | Up | 0.157202125930048 | 0.125974365879797 | 0.274834418523355 |
| MNT_C | 55 | 0.01 | Down | 0.287991675107611 | 0.128147909733663 | 0.279119548488518 |
| ZFP69_E | 153 | 0.01 | Up | 0.17094814820898 | 0.128673260982837 | 0.279806618091552 |
| TBPL1_E | 473 | 0.01 | Down | 0.135302607861965 | 0.129564647081969 | 0.281286114918997 |
| XBP1_C | 10 | 0.01 | Down | 0.151048777235636 | 0.130018528867644 | 0.281812518667593 |
| ZNF519_E | 313 | 0.01 | Up | 0.233981856018959 | 0.130530161994834 | 0.282462184966093 |
| ZNF516_E | 224 | 0.01 | Down | 0.248940423527019 | 0.131518633933303 | 0.284139933603068 |
| SATB2_E | 134 | 0.01 | Up | 0.317207306107251 | 0.131744819838826 | 0.284168033730025 |
| ZFP90_E | 462 | 0.01 | Up | 0.206708193947934 | 0.133381902815273 | 0.28688840353006 |
| ZNF124_E | 211 | 0.01 | Down | 0.194914375414129 | 0.133564294374833 | 0.28688840353006 |

ZNF24_C 18 0.01 Down 0.431888468052864
0.133651686865842 0.28688840353006
ZNF772_E 111 0.01 Down 0.154066489217145
0.134546032676246 0.288343828870476
ZSCAN26_E 331 0.01 Up 0.199714835916245
0.135705519986335 0.290361891078307
ZNF287_E 475 0.01 Up 0.134057087816228
0.137281763596089 0.293263767425619
CXXC1_E 829 0.01 Up 0.141908110234556
0.137515919864399 0.29329395388679
ZNF322_E 658 0.01 Down 0.236013155748635
0.138706324528092 0.295360272517487
ZNF273_E 179 0.01 Up 0.292481414152066
0.146009910131813 0.310416603198895
HSF2_E 597 0.01 Down 0.257578890675491
0.147401235618949 0.312395534622028
NR2F6_C 28 0.01 Down 0.20778711202392 8
0.147409445819396 0.312395534622028
PRRX2_E 981 0.01 Up 0.435987683000718
0.147935674721668 0.312671303953479
HOXD3_E 267 0.01 Down 0.296477666257742
0.148008696770176 0.312671303953479
ZNF142_E 492 0.01 Down 0.12746607908589
0.149953374064993 0.316278239918728
ERF_E 573 0.01 Down 0.17087311742364
0.150422373941418 0.316766231380584
NFKBID_E 418 0.01 Down 0.367415947224925
0.150869682864021 0.317207077693597
MAX_C 52 0.01 Down 0.222427697297541
0.151137766689895 0.317270303933275
ZNF480_E 210 0.01 Down 0.110127381137288
0.151795284510141 0.31814955071072
THAP1_C 51 0.01 Down 0.210057145794556
0.152043853765958 0.318170262276329
ZNF221_E 29 0.01 Down 0.170601215094292
0.152609912619594 0.318533060950495
ZNF585B_E 150 0.01 Down 0.170839586907784
0.152695143246336 0.318533060950495
NFE2L1_C 28 0.01 Up 0.267673508173233
0.154506933051817 0.321808971496987
ZNF442_E 28 0.01 Down 0.215360461031334
0.155084243790867 0.322507483577575
ZNF384_C 61 0.01 Down 0.179848821476893
0.156569479905467 0.324714294476063
WIZ_E 849 0.01 Up 0.136237911815218
0.15669529005719 0.324714294476063
POU3F4_E 256 0.01 Down 0.325693347206083
0.15687622328776 0.324714294476063
ZNF586_E 367 0.01 Down 0.273719415076347
0.158810702250529 0.328208784651094
SCX_E 255 0.01 Up 0.192307509002589
0.159410841149009 0.328463562599826
ATF4_A 14 0.01 Down 0.584160912091628
0.159426800451679 0.328463562599826

ZNF25_E 683 0.01 Up 0.2658479207936 0.159690834886269
0.328499819295364
BCL6_C 43 0.01 Down 0.390560105834358
0.160982268011504 0.330646168350284
EOMES_C 15 0.01 Up 0.550718180231862
0.162059074026591 0.332345762580686
HOXC13_E 306 0.01 Down 0.461969798267918
0.164556188739549 0.336948386466696
HOXA7_E 30 0.01 Up 0.253151058445965
0.165061257055921 0.337464195790711
SREBF2_A 13 0.01 Down 0.207979128283966
0.166436466202354 0.339754685218588
ESRRA_C 60 0.01 Down 0.245507011855096
0.168301859428664 0.343037276174937
YY1_A 57 0.01 Down 0.153865607848475
0.168606512341513 0.343133558704177
ZNF30_E 191 0.01 Up 0.196211311904421
0.16910399647935 0.343424565686115
FOXQ1_E 458 0.01 Down 0.350974797158616
0.169409957760205 0.343424565686115
ZNF554_E 493 0.01 Up 0.232732224892471
0.16952240376704 0.343424565686115
ZNF354C_E 317 0.01 Down 0.186974312936385
0.171882019251895 0.347190425670179
ZNF761_E 215 0.01 Down 0.135074394991897
0.171902236265805 0.347190425670179
ZSCAN5A_E 448 0.01 Up 0.123764259564529
0.175382670576955 0.353683963508443
ZFP1_E 364 0.01 Down 0.158888432029118
0.175694912178722 0.353778425882533
SMAD1_C 15 0.01 Up 0.639824498429491
0.17729168299019 0.35645522387017
ZNF737_E 123 0.01 Down 0.177597300170972
0.1782850821235 0.35791267239552
ZNF233_E 33 0.01 Up 0.229801784339171
0.179091862577213 0.358991658369061
FOXA1_A 17 0.01 Down 0.771094875282048
0.180652055204273 0.361575359740685
TFAP2B_E 542 0.01 Down 0.331195722211384
0.182624433199726 0.364975066649527
ZBTB7A_C 82 0.01 Down 0.273773060698663
0.183373519889292 0.365923506006626
HES1_D 129 0.01 Down 0.30048000309995
0.184299255833608 0.366814646534441
ZNF879_E 156 0.01 Down 0.136859176480975
0.184370452496681 0.366814646534441
ZNF680_E 637 0.01 Down 0.23945781598481
0.18466428341371 0.366851698644525
NR2F1_C 54 0.01 Up 0.425266794235027
0.185760495196227 0.368480268000848
PRDM15_E 354 0.01 Down 0.130858128579571
0.186054190255293 0.368514465988567
ST18_E 78 0.01 Up 0.76808865828001
0.188283728767937 0.37237716683629

MYT1_E 61 0.01 Up 0.550059711345557
0.189151529074798 0.373510229864882
ZNF616_E 247 0.01 Down 0.150403374585015
0.189417040801695 0.373510229864882
ZNF350_E 366 0.01 Down 0.239334234216157
0.190268711553894 0.37463543944068
LHX8_E 121 0.01 Down 0.316288266046277
0.191550089634616 0.375494513945505
PAX4_E 121 0.01 Down 0.316288266046277
0.191550089634616 0.375494513945505
VSX2_E 121 0.01 Down 0.316288266046277
0.191550089634616 0.375494513945505
TFE3_E 1223 0.01 Up 0.23586197153932
0.193147777279854 0.378070465659391
ZNF626_E 99 0.01 Down 0.164064325252204
0.193976397154472 0.378566726096526
ZFP3_E 288 0.01 Down 0.147694761480617
0.194095767511286 0.378566726096526
SP140_E 362 0.01 Down 0.612388986039472
0.194475590158534 0.378566726096526
TAL1_A 13 0.01 Down 0.394567236269573
0.194537289854554 0.378566726096526
POU4F1_E 243 0.01 Down 0.281809185759735
0.194845932237272 0.378614617598082
SPI1_A 72 0.01 Up 0.729860554578591
0.196003090372346 0.380308761959734
PRDM14_B 147 0.01 Down 0.425682375506298
0.1967057141158 0.381117321099362
ZNF319_E 336 0.01 Up 0.162008296578951
0.197248069893526 0.381613464685153
ZNF165_D 84 0.01 Up 0.268070230640151
0.198649004486862 0.382176904984594
ZNF592_C 12 0.01 Down 0.199468067917289
0.198711349784098 0.382176904984594
RFX4_E 167 0.01 Up 0.292446404195406
0.198725848615637 0.382176904984594
TGIF1_E 771 0.01 Down 0.321203141359944
0.198866302661673 0.382176904984594
ZNF136_E 471 0.01 Down 0.269208568644172
0.199494641756833 0.382176904984594
IRF2_C 70 0.01 Down 0.246049714056561
0.199543716498972 0.382176904984594
E2F5_C 11 0.01 Up 0.271666172374201
0.199546230959698 0.382176904984594
ZSCAN20_E 102 0.01 Down 0.132487243298982
0.201241934836314 0.384871591301014
ZNF282_E 673 0.01 Up 0.193037793856168
0.20267878863356 0.386936849840798
ZNF180_E 298 0.01 Down 0.13075513136788
0.202989314248042 0.386936849840798
YY2_E 361 0.01 Down 0.198495429210165
0.203367655039556 0.386936849840798
ZNF736_E 307 0.01 Down 0.15940618118268
0.203482919533683 0.386936849840798

TADA2B_E 431 0.01 Down 0.122007927667267
0.205284923868019 0.389807412416053
SOX5_E 562 0.01 Up 0.422153374895223
0.2068275702696 0.392178024423011
ETV4_A 12 0.01 Down 0.661420402701771
0.207332206889496 0.392576465601843
ZNF513_E 736 0.01 Up 0.138318112609915
0.208915126244843 0.395012572034575
MTA1_C 53 0.01 Up 0.169758538585531
0.210079737315309 0.39665196861375
KLF11_E 202 0.01 Down 0.227027995926463
0.210957452940033 0.397745805896837
ZNF85_E 425 0.01 Up 0.253419482102941
0.211821387866858 0.398798707498874
ZNF692_E 668 0.01 Up 0.159864518248281
0.212114241272844 0.398798707498874
EGR3_E 351 0.01 Down 0.33219275492663
0.212477197245041 0.398918456236111
ZNF16_E 287 0.01 Up 0.110193970000686
0.213705695506879 0.400364609019468
MLXIPL_E 291 0.01 Up 0.320922226047696
0.21384816325721 0.400364609019468
ZNF549_E 396 0.01 Down 0.270817176782872
0.214739982431254 0.401470401936692
ZNF596_E 172 0.01 Up 0.149906108215587
0.215335189005839 0.402019337457679
POU2F1_A 15 0.01 Up 0.684320497158578
0.215776537975044 0.402073879977959
HIC2_E 413 0.01 Down 0.299368729020254
0.21596766546453 0.402073879977959
HMGA2_E 67 0.01 Down 0.358677108476722
0.216810551585422 0.402552921026546
BARHL2_E 296 0.01 Down 0.294391804980524
0.216828955211598 0.402552921026546
HOXD10_E 10 0.01 Down 0.770017839642564
0.217574138087521 0.403374584242928
PDX1_C 41 0.01 Down 0.316369844760507
0.220691606473882 0.408585988096784
ZNF337_E 140 0.01 Down 0.157520500252585
0.224350097393585 0.414783189772052
FOXD1_E 285 0.01 Down 0.30275109751037
0.225456214675944 0.416250878342151
ID4_E 438 0.01 Up 0.444476830472503
0.226247397189405 0.416334738628483
ALX1_E 470 0.01 Down 0.337404120241782
0.226285146401356 0.416334738628483
ZNF565_E 45 0.01 Down 0.108542014116541
0.226438623785184 0.416334738628483
NONO_C 26 0.01 Down 0.145729313102235
0.227543046348082 0.417789091986216
SOX11_C 19 0.01 Down 0.245447841468375
0.229318853771184 0.420470470532309
FOXJ2_C 28 0.01 Down 0.264621511014699
0.229681446533511 0.420556824490618

ETS1_A	108	0.01	Down	0.577084727999628
0.231071792644813	0.422522221667402
FOXN4_E 24	0.01	Down	0.395721145211784
0.233869000137248	0.427051201620481
ASCL1_C 38	0.01	Down	0.256498187592355
0.235269033506584	0.429020002276713
NKX6-1_E	542	0.01	Down	0.346080561472896
0.237679844184698	0.432243790989253
PROP1_E 209	0.01	Down	0.348179200922556
0.237685445457706	0.432243790989253
E4F1_E	762	0.01	Up	0.123820737905417
0.24053589229377	0.436831531917705
ZNF263_B	259	0.01	Up	0.301837693881905
0.241453235861641	0.4374548058382
GLI1_D	63	0.01	Down	0.485194201150375
0.241719536480815	0.4374548058382
ZNF512B_E	481	0.01	Up	0.152951306122449
0.241863609829523	0.4374548058382
MEIS1_C 45	0.01	Down	0.385631239296135
0.242651560166835	0.438285270599446
THAP10_E	427	0.01	Up	0.142894087987759
0.243349088226395	0.438751429478255
ONECUT3_E	277	0.01	Down	0.341842560060861
0.243567935344268	0.438751429478255
ZFP14_E 366	0.01	Down	0.179360117396814
0.245516106241713	0.441663926612961
ZNF718_E	35	0.01	Down	0.141847454368135
0.247845863279429	0.444907193824333
ZKSCAN1_C	41	0.01	Down	0.309778240039968
0.247986530391207	0.444907193824333
E2F2_A	12	0.01	Up	0.158662148099317
0.248661087310835	0.445517781431914
ATRX_D	12	0.01	Down	0.210520563872519
0.252869235795165	0.451833250259503
GATA5_E 308	0.01	Up	0.351552341539463
0.253084260603268	0.451833250259503
ZNF619_E	264	0.01	Down	0.128343004275013
0.253202879177681	0.451833250259503
KAT8_E	535	0.01	Up	0.130268843731673
0.257846532112437	0.459504581959731
ZNF223_E	13	0.01	Down	0.123700194449048
0.260272179018615	0.463208030216039
ZNF454_E	33	0.01	Up	0.242998647903638
0.26125478830656	0.464336843750192
SIX2_C	34	0.01	Up	0.395079552212187
0.261818564023162	0.464586341361964
NR1H2_C 13	0.01	Down	0.223464791809878
0.26209219583043	0.464586341361964
ZKSCAN5_E	371	0.01	Down	0.122999392593648
0.263884230561001	0.467141672427375
SPIB_C	49	0.01	Down	0.431208078184729
0.264529213618625	0.46766238959367
HIF1A_A 56	0.01	Up	0.369194604258174
0.264973138747371	0.467734186936047

SOX18_E 134 0.01 Up 0.531138385101766
0.265581631836783 0.467734186936047
ZNF70_E 275 0.01 Up 0.185222495475389
0.265622490255504 0.467734186936047
SMARCE1_E 968 0.01 Down 0.139137851627027
0.266125701337033 0.46800205789217
ZNF76_E 1113 0.01 Up 0.171423155057631
0.270099504478822 0.474062293098845
ZNF689_E 419 0.01 Up 0.143838220309827
0.270283077835801 0.474062293098845
SNAI2_C 64 0.01 Down 0.283918307525465
0.270994259193831 0.474685082135843
RFX1_C 36 0.01 Up 0.314668674245543
0.274219385299868 0.479703990294913
ZNF224_E 417 0.01 Down 0.148161284931304
0.274656665269754 0.479839233033529
ZNF512_E 397 0.01 Up 0.179990627905117
0.276501984882281 0.48243081917288
ZNF75A_E 186 0.01 Down 0.141876994961983
0.278246019365707 0.484839142241161
ZNF366_E 136 0.01 Down 0.463766777171551
0.279680048754229 0.486701703641498
ZNF583_E 80 0.01 Down 0.158032694552233
0.281821348860784 0.489170060073543
POU5F1B_E 243 0.01 Down 0.285127947370072
0.281832412705537 0.489170060073543
GLIS1_E 349 0.01 Up 0.358447150704765
0.282467321622501 0.48963451199323
IKZF3_E 239 0.01 Down 0.588588426715778
0.284746853152623 0.492944876951228
NR2E1_E 295 0.01 Down 0.385598460671106
0.28657151031025 0.495070405998783
NEUROG2_E 207 0.01 Down 0.344521736479101
0.286717444434404 0.495070405998783
ZNF600_E 77 0.01 Down 0.150976719483869
0.287287796213096 0.495413495927629
ZNF490_E 254 0.01 Down 0.370997089349741
0.288339044058181 0.496583909211312
EGR2_D 12 0.01 Down 0.864015155500318
0.289882199325244 0.49859738283942
MXI1_C 75 0.01 Down 0.21818022034473
0.290912529033128 0.499128796680913
MYF6_E 410 0.01 Up 0.378479356166861
0.290940041276121 0.499128796680913
TFEB_E 688 0.01 Up 0.208976503609331
0.291320608338427 0.499139294235377
IKZF4_E 123 0.01 Down 0.184117447423197
0.293217627053766 0.501321466578235
ZNF786_E 249 0.01 Up 0.145036581958471
0.293817868809758 0.501321466578235
MIER2_E 546 0.01 Up 0.160921019967375
0.294033921151451 0.501321466578235
NKX6-2_E 282 0.01 Down 0.317313177398873
0.294098564789332 0.501321466578235

ZNF117_E	154	0.01	Down	0.234265823205543
0.294982811885703	0.502186575023809
ZNF23_E 130	0.01	Down	0.135467299604841
0.295850119761195	0.503020675563358
ZBTB8B_E	202	0.01	Down	0.249980362605626
0.297591513040577	0.504900733491104
TBX21_C 63	0.01	Down	0.442014382106095
0.297713410745692	0.504900733491104
SIX4_E	58	0.01	Down	0.294004766075401
0.298735594251053	0.505551291802627
ZEB1_B	15	0.01	Down	0.39855391530825
0.299129648386978	0.505551291802627
ZHX3_E	431	0.01	Up	0.316483076219622
0.29923478562061	0.505551291802627
ZFAT_E	232	0.01	Up	0.133099827421331
0.300269003414073	0.506656432342987
HSF1_B	11	0.01	Down	0.305832774718626
0.301339011036348	0.507561438473001
RFX3_E	569	0.01	Down	0.245554758130695
0.301566886174506	0.507561438473001
THRA_D	10	0.01	Down	0.324268169874854
0.302685345199887	0.508801469295649
YBX1_D	22	0.01	Up	0.292505616486562
0.305078607100991	0.512178568344612
KMT2B_D 83	0.01	Down	0.189746949884915
0.307494099775722	0.515584446542186
NR1H3_C 12	0.01	Up	0.363488678410973
0.308290877359062	0.516060281942955
E2F7_C	29	0.01	Down	0.275188806434491
0.308886027127762	0.516060281942955
ZC3H4_E 401	0.01	Up	0.134471425013249
0.308939313571251	0.516060281942955
LEF1_A	10	0.01	Up	0.45699189338727
0.310225766368879	0.51756063400465
ZNF684_E	216	0.01	Up	0.165247959790423
0.311203425729148	0.518542708121193
ISL2_E	262	0.01	Down	0.333217018884902
0.31220106418145	0.519160285261804
TBX15_E 387	0.01	Up	0.410192677843794
0.312352999834934	0.519160285261804
ZNF268_E	289	0.01	Down	0.152549161821875
0.313690733413277	0.520533605852987
ELF5_C	19	0.01	Down	0.191254711514188
0.313960254392949	0.520533605852987
ZNF789_E	491	0.01	Up	0.17781783663299
0.314356177734713	0.520542589963196
MAFG_C	34	0.01	Down	0.285314852373287
0.315861811105734	0.521161682007535
MAZ_C	67	0.01	Up	0.21990533611795
0.315892510470441	0.521161682007535
E2F3_A	11	0.01	Up	0.335183106115859
0.315902955035325	0.521161682007535
NKX3-1_E	706	0.01	Down	0.266526696795106
0.31781780599595	0.523672602463042

KAT5_E 694 0.01 Up 0.114719840266274
0.318426658449329 0.524028068781426
LHX6_E 319 0.01 Up 0.488025248391376
0.319039471703971 0.524389168657699
ZNF532_E 302 0.01 Up 0.236827478550821
0.319558229407487 0.524594975123374
FOXO6_D 30 0.01 Up 0.268803282707527
0.321130360742226 0.52652739344328
T_C 22 0.01 Down 0.426938022667801
0.325682295512525 0.533334766484271
TCF3_B 13 0.01 Down 0.209873914097729
0.328034503600772 0.536527599140895
STAT3_A 50 0.01 Down 0.623100872257836
0.328735748693526 0.536551638868173
ARNT_C 55 0.01 Down 0.253191460834917
0.329162489699197 0.536551638868173
ETV2_E 840 0.01 Up 0.229277550820259
0.329256744631782 0.536551638868173
SOX8_E 517 0.01 Up 0.409001580660415
0.329966068525234 0.536992146975146
ZBTB48_E 641 0.01 Up 0.14277906623115
0.330679568764841 0.536992146975146
ZXDA_E 145 0.01 Down 0.200309642075023
0.330735598399546 0.536992146975146
NFATC3_D 11 0.01 Up 0.594515473447331
0.331965917630663 0.538334024576245
MTA3_C 19 0.01 Up 0.277679673592084
0.334605733611175 0.541955580684929
TCF7L2_A 12 0.01 Up 0.385391916907615
0.338646380172482 0.547834496079998
OTX1_E 337 0.01 Down 0.315175492868414
0.341694046083048 0.552094743549943
ZNF793_E 246 0.01 Down 0.210259996434588
0.343069810341774 0.553115109464413
TBX1_E 358 0.01 Up 0.364815937870673
0.343155435504178 0.553115109464413
ZNF540_E 200 0.01 Up 0.195479369511332
0.344717431037077 0.55496175793771
USF1_A 41 0.01 Up 0.482852597832805
0.347728971304302 0.559134763267352
SMARCC1_D 213 0.01 Down 0.401695485975982
0.348244780234104 0.559289508496459
TFCP2L1_E 380 0.01 Down 0.493067263662444
0.350286859641699 0.561892158727298
ZNF671_E 66 0.01 Down 0.236939539718975
0.351434517637941 0.563051927818398
POU4F2_C 19 0.01 Down 0.293649344081822
0.351854655568436 0.563051927818398
ZC3H8_D 199 0.01 Down 0.29855927267448
0.352304341723253 0.56309554858165
PBX2_C 39 0.01 Down 0.311559739862962
0.353050986513543 0.563613131763537
SOX15_E 544 0.01 Up 0.34401250984229
0.353767141438266 0.564080860690441

BATF2_E 216	0.01	Down	0.375459297344691
0.356176997428723	0.567244847756856
ONECUT2_E	291	0.01	Down	0.335756701459108
0.356842251674367	0.567626159286313
ISL1_E	289	0.01	Down	0.344150260778671
0.357592399983912	0.56814144121401
HDX_E	235	0.01	Up	0.33862098623582 8
0.358740641494342	0.569287232276141
L3MBTL1_E	576	0.01	Up	0.177364491322466
0.361983211274487	0.573749846169907
ZNF404_E	140	0.01	Up	0.190723033808642
0.363110402807253	0.574612272680144
RORB_E	298	0.01	Down	0.371741161491048
0.363389456766213	0.574612272680144
TEAD4_C 61	0.01	Down	0.272507303598865
0.36523292144108	0.576772104251287
MAF_C	33	0.01	Down	0.306767860616398
0.365620726250816	0.576772104251287
ZNF343_E	337	0.01	Down	0.125299058061962
0.36731685140841	0.578059630561756
ZNF528_E	374	0.01	Down	0.301349052381947
0.367542073143281	0.578059630561756
GLI2_A	13	0.01	Up	0.996531493047751
0.368012670909434	0.578059630561756
ZNF354A_E	571	0.01	Down	0.193791295195459
0.368341288382912	0.578059630561756
ZNF836_E	107	0.01	Down	0.169066665860708
0.368605165774563	0.578059630561756
MECOM_E 626	0.01	Down	0.36936249891104
0.370566584644346	0.580229448279023
IKZF1_C 36	0.01	Down	0.395220956136845
0.370897127956701	0.580229448279023
MXD3_E	422	0.01	Up	0.163998848061915
0.37148993460232	0.580229448279023
ZNF79_E 159	0.01	Down	0.100867171025775
0.371729894096239	0.580229448279023
TP73_C	38	0.01	Up	0.359103059710989
0.374356429520756	0.583645755030605
RUNX2_C 58	0.01	Down	0.327112669241872
0.375136720576809	0.584179028655241
RARG_D	27	0.01	Down	0.756260377917581
0.378868852821683	0.589302428017857
MEF2B_C 31	0.01	Up	0.486874696079743
0.379338816328301	0.589345736789774
SIM2_E	14	0.01	Down	0.192220551930432
0.379974429114588	0.589646000011346
HMX1_E	269	0.01	Down	0.288633981256273
0.380662825733812	0.590027379887409
ZBED1_C 22	0.01	Down	0.141533310880128
0.381326370221911	0.590369397800009
HLF_E	710	0.01	Up	0.355389680476533
0.3835953818671 0.593193322539263
ATF1_A	12	0.01	Down	0.440735220104659
0.386361388673977	0.59677836744196

| | | | | | | |
|---|---|---|---|---|---|---|
| RARB_D | 34 | 0.01 | Down | 0.637098971668096 | 0.387097284139451 | 0.597223008979038 |
| ZSCAN4_E | 267 | 0.01 | Up | 0.379882980833499 | 0.388815269619638 | 0.599180062893616 |
| VSX1_E | 131 | 0.01 | Down | 0.326378945920905 | 0.391191047452038 | 0.602145111147306 |
| POU3F3_E | 275 | 0.01 | Down | 0.368320595114784 | 0.392763606224658 | 0.603868381888661 |
| NOTO_E | 209 | 0.01 | Down | 0.298715429520365 | 0.394579705972984 | 0.605961691315654 |
| ZNF75D_E | 256 | 0.01 | Down | 0.155084595552631 | 0.399070394849292 | 0.611123958718513 |
| ZNF324B_E | 304 | 0.01 | Up | 0.108369857575389 | 0.399497870027176 | 0.611123958718513 |
| POU6F2_E | 273 | 0.01 | Down | 0.285905491873896 | 0.399687807753515 | 0.611123958718513 |
| ZNF250_E | 474 | 0.01 | Down | 0.251084519736708 | 0.399775012755096 | 0.611123958718513 |
| PAX3_E | 120 | 0.01 | Down | 0.34452936425878 | 0.401157673789932 | 0.612535142224489 |
| LBX2_E | 245 | 0.01 | Down | 0.321441422378271 | 0.40177779878293 | 0.612634856909611 |
| ZFP62_E | 381 | 0.01 | Down | 0.149869991822953 | 0.402584515822391 | 0.612634856909611 |
| DBP_E | 786 | 0.01 | Up | 0.296672741839989 | 0.402601751427472 | 0.612634856909611 |
| HSF4_E | 751 | 0.01 | Up | 0.250625688902595 | 0.40542845728034 | 0.616232763460312 |
| SATB1_D | 195 | 0.01 | Down | 0.382299954340511 | 0.406501177588634 | 0.617159532717141 |
| ZNF419_E | 76 | 0.01 | Down | 0.170273479580616 | 0.407042636806393 | 0.617278537955542 |
| ZNF341_E | 724 | 0.01 | Up | 0.218026835455345 | 0.409499244588342 | 0.620298287541204 |
| ZSCAN16_E | 166 | 0.01 | Up | 0.116354213958502 | 0.410276851706604 | 0.620770764273443 |
| DMRT3_E | 265 | 0.01 | Down | 0.402939212930062 | 0.415172726343948 | 0.627466263283993 |
| DLX2_E | 274 | 0.01 | Down | 0.300791538622014 | 0.415773802568518 | 0.627663056425634 |
| ATF6_D | 13 | 0.01 | Down | 0.19655239502839 | 0.42128715541136 | 0.635266717379346 |
| NKX2-1_E | 284 | 0.01 | Down | 0.373655522586877 | 0.425307969848675 | 0.640322418242356 |
| BHLHE41_E | 298 | 0.01 | Up | 0.285435030491651 | 0.425600647083816 | 0.640322418242356 |
| ZBTB14_E | 768 | 0.01 | Up | 0.206490953093548 | 0.432165864079203 | 0.649466850978104 |
| DLX4_E | 277 | 0.01 | Down | 0.287020918955345 | 0.434793517658894 | 0.652679908828048 |
| ZNF175_D | 173 | 0.01 | Down | 0.336640475111144 | 0.436278415428777 | 0.65414753429233 |
| ZNF646_E | 401 | 0.01 | Down | 0.134419303882566 | 0.436752667306957 | 0.65414753429233 |

PRDM1_C 36 0.01 Down 0.41406369691446
0.437557815725553 0.654617921843055
ZNF620_E 25 0.01 Down 0.118929498153567
0.438082743375822 0.654668494304899
NR1I2_D 102 0.01 Down 0.291445471569676
0.442351104806004 0.659659087684065
NKX3-2_E 528 0.01 Down 0.296490156579035
0.44241202129749 0.659659087684065
NOBOX_E 367 0.01 Down 0.280448505023558
0.443921001504035 0.660769523926223
PSPC1_E 776 0.01 Down 0.10697732194451
0.444300099680885 0.660769523926223
ZNF3_E 424 0.01 Down 0.114974588305667
0.444643858185913 0.660769523926223
ZNF277_E 736 0.01 Down 0.140153816074133
0.445240415247438 0.660919235551041
ZNF22_E 412 0.01 Down 0.196324548395142
0.447604830608929 0.663689921247722
ATOH1_E 506 0.01 Down 0.421458736472014
0.451003356868602 0.667986083006495
ZNF672_E 504 0.01 Up 0.12528906440886
0.452641032870137 0.669667588030958
ZNF214_E 449 0.01 Up 0.256424576072441
0.453787516435488 0.670219628644686
YBX3_E 600 0.01 Down 0.26332474090893
0.454019748436723 0.670219628644686
STAT5A_A 12 0.01 Down 0.39673529391226
0.455678635440292 0.671924359559635
CEBPD_B 12 0.01 Down 0.685820558769749
0.457376670644545 0.673682985601302
ZNF485_E 369 0.01 Up 0.137005201316974
0.458551370040166 0.674667744220244
GBX2_E 256 0.01 Down 0.311172481858873
0.460256432901439 0.675796006460011
ZHX1_D 272 0.01 Down 0.261082229438683
0.460741517266435 0.675796006460011
BHLHE22_C 17 0.01 Down 0.534796962032233
0.460839137188409 0.675796006460011
GLI3_D 18 0.01 Up 0.614926224185211
0.462672665924407 0.677739190854104
CLOCK_C 42 0.01 Down 0.270856880954883
0.463207182018623 0.67777735854097
ZNF280D_E 574 0.01 Down 0.183852792588999
0.463879132462766 0.678016319707092
MSX2_E 292 0.01 Down 0.296072835243392
0.466279881904069 0.680489142580001
FOXD3_E 338 0.01 Up 0.437516355276961
0.466591955227397 0.680489142580001
DNMT3B_C 10 0.01 Down 0.246956226733516
0.468997312128556 0.683249636139197
ZFHX3_E 349 0.01 Up 0.332364497788763
0.470542744889559 0.683501267840061
ZNF195_E 635 0.01 Down 0.144447070434923
0.470880358434608 0.683501267840061

ZNF792_E 114 0.01 Down 0.382774420192847
0.471237249846102 0.683501267840061
ZSCAN22_E 515 0.01 Down 0.199769006388597
0.472257276638559 0.683501267840061
HOXA9_C 18 0.01 Down 0.356275026379012
0.472498765459697 0.683501267840061
ZBTB25_E 330 0.01 Down 0.176713919618507
0.472661693147184 0.683501267840061
ZKSCAN4_E 224 0.01 Up 0.177670927460419
0.472759316540537 0.683501267840061
HOXB3_E 411 0.01 Up 0.337778907315649
0.474060051624968 0.684002857444297
RAX_E 270 0.01 Down 0.343376308757052
0.474132513337232 0.684002857444297
THAP11_C 112 0.01 Up 0.154748478606739
0.475285782246309 0.684925348901978
HES4_E 271 0.01 Up 0.295415848710004
0.47985726988595 0.690766458701912
ZNF687_E 431 0.01 Down 0.127373062504898
0.481801002118462 0.692618379064982
LYAR_E 963 0.01 Down 0.133785402973232
0.482477912336243 0.692618379064982
BCL3_C 35 0.01 Down 0.285491634435609
0.482702531246338 0.692618379064982
ZNF296_E 567 0.01 Down 0.257258758893772
0.484346623239946 0.693325495643636
FOXO1_A 30 0.01 Up 0.39936318507709
0.484502338678986 0.693325495643636
LBX1_E 267 0.01 Down 0.290126705848296
0.485213189628382 0.693325495643636
LHX4_E 267 0.01 Down 0.217119498751224
0.485679785838622 0.693325495643636
ZNF138_E 376 0.01 Down 0.16791397610813
0.486877542496401 0.693325495643636
TBR1_E 275 0.01 Down 0.346522852195585
0.487966870973896 0.693325495643636
SNAPC4_C 28 0.01 Up 0.126169843035281
0.488251126913791 0.693325495643636
HOXB7_D 52 0.01 Up 0.37402329031103
0.488478510174561 0.693325495643636
POU2F2_C 72 0.01 Down 0.483990625542029
0.488575468977341 0.693325495643636
ZNF439_E 104 0.01 Down 0.186221171501992
0.489417968010998 0.693325495643636
KDM5D_E 40 0.01 Down 0.432000687915743
0.490323933210341 0.693325495643636
BSX_E 263 0.01 Down 0.300698662655726
0.491006973314001 0.693325495643636
LHX9_E 263 0.01 Down 0.300698662655726
0.491006973314001 0.693325495643636
RAX2_E 263 0.01 Down 0.300698662655726
0.491006973314001 0.693325495643636
CREB3L2_D 12 0.01 Up 0.483213986037758
0.492836531693487 0.693325495643636

CREB3L3_D	12	0.01	Up	0.483213986037758
0.492836531693487	0.693325495643636
CREB3L4_D	12	0.01	Up	0.483213986037758
0.492836531693487	0.693325495643636
TP53_A	205	0.01	Up	0.34158193943924
0.493021637103821	0.693325495643636
ZNF547_E	341	0.01	Up	0.293237320340883
0.493077696826832	0.693325495643636
ZNF33B_E	445	0.01	Up	0.175840141831342
0.493914606511559	0.693770464151642
SCRT1_E	299	0.01	Up	0.328855117542118
0.494708977827825	0.694154807836306
ZNF443_E	128	0.01	Down	0.151064646606856
0.496145550935227	0.695438506200482
ZBTB18_E	485	0.01	Up	0.321229477709394
0.496746563062757	0.695549546809511
TFDP2_E	585	0.01	Up	0.211034505082537
0.498596804985981	0.697407703091619
ZNF436_E	320	0.01	Down	0.360863921852594
0.499158893480047	0.69746205975776
ZNF683_E	128	0.01	Down	0.622180545043061
0.502419431369469	0.700785132955333
HMGXB3_E	783	0.01	Down	0.133393428818637
0.502588587475842	0.700785132955333
HESX1_E	226	0.01	Down	0.342035441093184
0.503412484667925	0.701200461925124
SOX1_E	216	0.01	Up	0.34000268319744
0.50486140624746	0.701591067268322
ZBTB42_E	569	0.01	Up	0.178301928186808
0.505608494424494	0.701591067268322
ZNF514_E	136	0.01	Up	0.143268808998387
0.505937646360291	0.701591067268322
NFE2L2_B	10	0.01	Up	0.137585513952238
0.506490603022818	0.701591067268322
UNCX_E	300	0.01	Down	0.368519992463799
0.507568310858085	0.701591067268322
ZNF471_E	285	0.01	Up	0.206658754002313
0.508016213387815	0.701591067268322
FOXP1_B	152	0.01	Down	0.247894458091119
0.508065590141024	0.701591067268322
SHOX_E	265	0.01	Down	0.300313639863191
0.508197002701267	0.701591067268322
ASCL2_E	373	0.01	Up	0.370913630969634
0.508969670492286	0.701591067268322
EN1_E	272	0.01	Down	0.315475434009224
0.509303393234195	0.701591067268322
CEBPA_A	94	0.01	Up	0.549363125670105
0.509482485458166	0.701591067268322
SOX2_B	201	0.01	Down	0.386836194828252
0.513165281753031	0.704825607540203
ZBTB32_E	52	0.01	Down	0.622757501343191
0.513745676421129	0.704825607540203
AR_A	109	0.01	Down	0.273335482691588
0.514133063967933	0.704825607540203

BHLHA15_E	654	0.01	Down	0.30606205445434
0.514180268353991	0.704825607540203
ZBTB49_E	81	0.01	Down	0.118701446982523
0.514475105878933	0.704825607540203
POU3F1_E	552	0.01	Down	0.374013513661568
0.515676393390605	0.705746029147511
ZNF552_E	250	0.01	Down	0.216361637789065
0.516714884597641	0.70644199094221
NR4A2_E 560	0.01	Down	0.349665844445588
0.51862548244905	0.70707161878131
FIGLA_E 277	0.01	Up	0.291275539187095
0.518668253966627	0.70707161878131
CTCF_A	82	0.01	Down	0.373406996949641
0.518766724057105	0.70707161878131
HMX3_E	277	0.01	Down	0.278380701481486
0.519609225397731	0.707496524469024
CREM_C	47	0.01	Down	0.315141474292864
0.520277212955748	0.707683188642869
HOXD13_E	281	0.01	Down	0.440818067714328
0.5223007103549 0.709711362796209
THAP2_E 284	0.01	Down	0.188824488095168
0.524223931895019	0.711599288407394
ZNF785_E	415	0.01	Up	0.156772366117069
0.527198969836756	0.714351615434806
ETV1_C	27	0.01	Down	0.332693476147886
0.527323323021642	0.714351615434806
TBX20_E 265	0.01	Down	0.321897602207342
0.528193736487811	0.714804315470306
ZNF788_E	104	0.01	Up	0.163340347833581
0.531893121562214	0.717261268481796
ZNF232_E	50	0.01	Up	0.104867277075708
0.532182522935534	0.717261268481796
FOXB1_E 261	0.01	Down	0.304093272858484
0.532503323504394	0.717261268481796
PAX9_E	337	0.01	Down	0.369825211132604
0.533378691899256	0.717261268481796
ZNF558_E	297	0.01	Down	0.142343919271825
0.533519666329977	0.717261268481796
FOXO4_A 11	0.01	Down	0.277578839649286
0.53354582390628	0.717261268481796
HOXB8_E 225	0.01	Down	0.302722448369077
0.533775827707383	0.717261268481796
KLF13_C 28	0.01	Down	0.277239702997073
0.537199920299262	0.72113544185188
MYBL1_E 322	0.01	Down	0.311553721057106
0.539626303630833	0.723663845814788
ADNP_D	187	0.01	Down	0.408840667769915
0.540183582279161	0.723683130832283
TSC22D1_E	195	0.01	Up	0.361860936453536
0.540902628012115	0.723918878654768
NPAS2_E 398	0.01	Down	0.327959474104221
0.54150190072303	0.723994015710931
SMAD2_C 27	0.01	Down	0.245993789195709
0.542455613160947	0.724542417177898

ETS2_A 21 0.01 Down 1.12088425406178
0.545143270170219 0.727403382519422
KLF1_C 45 0.01 Down 0.240157485462828
0.546777154258461 0.728853946626528
MEOX1_E 353 0.01 Up 0.469721444604966
0.551312723812385 0.734165695146763
KLF16_E 698 0.01 Down 0.201585187861041
0.55336920744595 0.736168815893664
KLF14_E 286 0.01 Down 0.237292617613377
0.557453583324051 0.740863037458585
PROX2_E 107 0.01 Down 0.170539852221067
0.560890873033946 0.744688778639692
FOXG1_E 279 0.01 Down 0.37911544920541
0.561661318544737 0.744969689174264
HES5_E 202 0.01 Up 0.238896019756464
0.565356481379624 0.749125437056699
ZNF222_E 30 0.01 Down 0.114102889743178
0.567809525245883 0.75162869627881
HOXB4_E 363 0.01 Down 0.326773095122774
0.569655352183098 0.753323992519911
CIC_E 395 0.01 Down 0.156887507938746
0.570828618305358 0.75411886817688
ALX3_E 237 0.01 Down 0.298591562110073
0.571387889616391 0.75411886817688
ARX_E 203 0.01 Up 0.336204354625611
0.574162017568457 0.757030632461675
GSC2_E 292 0.01 Down 0.296792277650788
0.574819848077076 0.757149068662789
ZNF90_E 220 0.01 Up 0.186394762330146
0.57559708330856 0.757424394916397
TEAD3_E 741 0.01 Up 0.300628180985175
0.578324259863938 0.760172067571943
MYBL2_C 22 0.01 Up 0.336891705236401
0.578952699955181 0.760172067571943
ZNF333_E 389 0.01 Down 0.169959898422341
0.579452209287239 0.760172067571943
CREBL2_E 981 0.01 Up 0.191183282040301
0.579966236099525 0.760172067571943
ZNF569_E 277 0.01 Down 0.156286875062053
0.581168285567159 0.760605481034577
BHLHE40_C 54 0.01 Down 0.27861525508727
0.581972980880627 0.760605481034577
ZNF10_E 178 0.01 Up 0.209540980588927
0.582008695165243 0.760605481034577
HOXB5_E 283 0.01 Down 0.358019612809191
0.585649528596364 0.764613929107692
ZNF584_D 122 0.01 Down 0.388602019337407
0.587652292461688 0.766477990069893
ZNF324_E 542 0.01 Down 0.231477141907458
0.588906830899583 0.767175981709888
BARHL1_E 285 0.01 Down 0.338540930232895
0.590053986019849 0.767175981709888
PHF5A_E 1597 0.01 Up 0.111337324075163
0.59017122698376 0.767175981709888

| | | | | | | |
|---|---|---|---|---|---|---|
| NFE2_C | 41 | 0.01 | Up | 0.333923071738047 | 0.591194970688806 | 0.767175981709888 |
| ZNF816_E | 434 | 0.01 | Down | 0.256004466764086 | 0.591331515181239 | 0.767175981709888 |
| TFAP2E_E | 241 | 0.01 | Up | 0.201388778825678 | 0.592133059562003 | 0.767175981709888 |
| SP8_E | 298 | 0.01 | Down | 0.240548746430418 | 0.592429017858869 | 0.767175981709888 |
| EGR1_A | 88 | 0.01 | Down | 0.397508555755417 | 0.592791643781834 | 0.767175981709888 |
| ZNF544_E | 139 | 0.01 | Down | 0.176997105112156 | 0.595670706830515 | 0.770154269840036 |
| ZNF665_E | 80 | 0.01 | Down | 0.187820388377776 | 0.596552561436656 | 0.770529332163728 |
| DLX6_E | 287 | 0.01 | Down | 0.311778005632189 | 0.597116879313677 | 0.770529332163728 |
| SOX6_C | 27 | 0.01 | Up | 0.388247434938974 | 0.600346637930094 | 0.773947841741601 |
| ZNF624_E | 256 | 0.01 | Down | 0.199638991368475 | 0.601158463296165 | 0.77424563437081 |
| FOXD2_E | 340 | 0.01 | Up | 0.338686244088122 | 0.603120092431783 | 0.776022281092246 |
| NHLH1_E | 441 | 0.01 | Up | 0.316023513638117 | 0.604736489364661 | 0.776171357309013 |
| TBX5_E | 273 | 0.01 | Up | 0.330621673087346 | 0.60489790374831 | 0.776171357309013 |
| TBX6_E | 313 | 0.01 | Up | 0.181317522786455 | 0.604982775877018 | 0.776171357309013 |
| ZNF682_E | 95 | 0.01 | Up | 0.160559747674933 | 0.607772640822465 | 0.779000894438794 |
| CREB1_A | 44 | 0.01 | Down | 0.601142612906095 | 0.609303136747793 | 0.779184698761559 |
| HES7_E | 159 | 0.01 | Up | 0.217582533720524 | 0.610601880085117 | 0.779184698761559 |
| ARID5A_E | 666 | 0.01 | Down | 0.32780795899495 | 0.610844209513061 | 0.779184698761559 |
| MAFA_E | 10 | 0.01 | Up | 0.180925948986914 | 0.611224939516696 | 0.779184698761559 |
| DUX4_C | 29 | 0.01 | Up | 0.325986036806265 | 0.611361966812697 | 0.779184698761559 |
| PLAG1_E | 418 | 0.01 | Down | 0.292199423382341 | 0.611423251991441 | 0.779184698761559 |
| CRX_E | 363 | 0.01 | Down | 0.308120814215433 | 0.612704229517376 | 0.779420691791081 |
| POU4F3_E | 228 | 0.01 | Down | 0.294680972727794 | 0.612777858212343 | 0.779420691791081 |
| CUX1_C | 36 | 0.01 | Up | 0.227672580838317 | 0.61923317425197 | 0.786880668520378 |
| GATA4_D | 17 | 0.01 | Down | 0.324464645003702 | 0.621108031931638 | 0.788511434823689 |
| HINFP_C | 14 | 0.01 | Up | 0.251182145065234 | 0.622252779790859 | 0.78887252531772 |
| ZNF416_E | 132 | 0.01 | Down | 0.105516493804207 | 0.62257606649230 4 | 0.78887252531772 |

ZNF286B_E 12 0.01 Down 0.143016582871302
0.629124434004101 0.795724261760283
ZNF429_E 144 0.01 Down 0.159117393904298
0.629177323252317 0.795724261760283
ZNF285_E 74 0.01 Up 0.211468277728434
0.631114472593976 0.796643219262892
ELK3_E 1102 0.01 Down 0.306014780183422
0.631354539524305 0.796643219262892
NR1I3_E 308 0.01 Up 0.329521135835659
0.631719986247568 0.796643219262892
DNMT1_D 31 0.01 Up 0.440559629139907
0.63229446810213 0.796643219262892
SHOX2_E 317 0.01 Up 0.354972898183741
0.634849131623696 0.799106602884218
AHRR_E 15 0.01 Up 0.556207320567752
0.636398820999869 0.800301536219646
ZSCAN31_E 357 0.01 Up 0.291767257819705
0.639400250620617 0.8033181282538
HKR1_E 395 0.01 Up 0.139059030416156
0.640959721103659 0.803722466812377
ZNF418_E 369 0.01 Up 0.291442608067567
0.642237906209578 0.803722466812377
OLIG1_E 191 0.01 Up 0.353472775644191
0.642710808551763 0.803722466812377
GATA6_C 47 0.01 Up 0.420322872547748
0.64314796766401 0.803722466812377
HOXA1_E 106 0.01 Up 0.376646748862888
0.643153983745726 0.803722466812377
EMX1_E 269 0.01 Down 0.302092667608481
0.643508388276417 0.803722466812377
ZKSCAN3_E 146 0.01 Up 0.175214574120204
0.643942681587111 0.803722466812377
PAX7_E 112 0.01 Down 0.298885159292061
0.644772411761304 0.804005261812739
ZNF502_E 323 0.01 Down 0.237465940877428
0.647102117438821 0.806156189295279
ZNF546_E 268 0.01 Up 0.18073029444324
0.652958635259778 0.812592958371747
FOXN3_E 645 0.01 Down 0.217439651477841
0.653488110558524 0.812592958371747
DMRTA1_E 154 0.01 Up 0.271642874950695
0.654399050749392 0.812967320269282
OLIG2_E 492 0.01 Up 0.359831053066034
0.65543777216918 0.813499581286328
FOXF2_E 339 0.01 Up 0.421854973227843
0.658368629942506 0.815816900805384
IRF3_C 61 0.01 Up 0.216827475502686
0.659297563559431 0.815816900805384
ESRRG_E 513 0.01 Up 0.353570546440989
0.659315276306161 0.815816900805384
FEV_E 657 0.01 Down 0.171199111168472
0.659848066422651 0.815816900805384
ZNF275_E 261 0.01 Down 0.213472987744956
0.660380125412305 0.815816900805384

| | | | | | | |
|---|---|---|---|---|---|---|
| ZNF860_E | 128 | 0.01 | Down | 0.279519297451817 | 0.660976933885832 | 0.815816900805384 |
| ALX4_E | 270 | 0.01 | Down | 0.342779781130099 | 0.662971796013532 | 0.816773122161297 |
| ZNF627_E | 182 | 0.01 | Down | 0.13344951962752 | 0.663278071045369 | 0.816773122161297 |
| TCF7L1_D | 10 | 0.01 | Up | 0.432537032795194 | 0.663589865941999 | 0.816773122161297 |
| MYB_A | 16 | 0.01 | Up | 0.631167453532019 | 0.665928843162458 | 0.818679986914536 |
| TBX19_E | 615 | 0.01 | Up | 0.247221913380737 | 0.666367431209506 | 0.818679986914536 |
| ZGPAT_E | 517 | 0.01 | Up | 0.12819138998064 | 0.668660481903055 | 0.820595436198263 |
| PTF1A_E | 272 | 0.01 | Down | 0.289150677722402 | 0.669253881161015 | 0.820595436198263 |
| ZBTB17_E | 1055 | 0.01 | Up | 0.16954086683472 | 0.669773319267599 | 0.820595436198263 |
| BARX1_E | 266 | 0.01 | Down | 0.297939136765032 | 0.679129535636287 | 0.831294463731103 |
| PBX4_E | 348 | 0.01 | Down | 0.378419059239296 | 0.681458198765611 | 0.833124434162347 |
| ZNF717_E | 202 | 0.01 | Up | 0.16127926969314 | 0.681874536887562 | 0.833124434162347 |
| ZNF530_E | 170 | 0.01 | Up | 0.122477319726563 | 0.68326731189415 | 0.833808305289665 |
| HOXD12_E | 282 | 0.01 | Down | 0.336835356386183 | 0.683685279581098 | 0.833808305289665 |
| MSX1_E | 452 | 0.01 | Up | 0.408136330945864 | 0.686068249503728 | 0.835949704377029 |
| MYC_A | 321 | 0.01 | Up | 0.23967881235324 | 0.686942275747617 | 0.836250277234313 |
| ZIK1_E | 96 | 0.01 | Down | 0.162013111009326 | 0.689203843014585 | 0.838237885710258 |
| ZIC3_E | 502 | 0.01 | Up | 0.319910819248732 | 0.691543730386491 | 0.839637446969161 |
| ZC3H6_E | 438 | 0.01 | Down | 0.172716386527482 | 0.691614341164395 | 0.839637446969161 |
| POU5F1_C | 59 | 0.01 | Down | 0.361180751461636 | 0.692552800097074 | 0.840011722046769 |
| ZNF248_E | 349 | 0.01 | Up | 0.188191351801196 | 0.695173332111685 | 0.84105287876417 |
| SOX13_C | 38 | 0.01 | Up | 0.41429111862182 | 0.696240624958224 | 0.84105287876417 |
| ZNF667_E | 371 | 0.01 | Up | 0.296064769542256 | 0.696495250953845 | 0.84105287876417 |
| EBF1_C | 67 | 0.01 | Down | 0.310375671208103 | 0.696536428999253 | 0.84105287876417 |
| ZNF182_E | 354 | 0.01 | Down | 0.152233905716123 | 0.69699621854711 | 0.84105287876417 |
| PPARA_A | 17 | 0.01 | Down | 0.682860848122453 | 0.698361671097444 | 0.84105287876417 |
| OTX2_C | 15 | 0.01 | Up | 0.452976827556967 | 0.698714735533698 | 0.84105287876417 |

SRY_E 441 0.01 Down 0.282361732037231
0.699190534745132 0.84105287876417
GSX1_E 261 0.01 Down 0.305649940939848
0.69924031275998 0.84105287876417
NRF1_C 75 0.01 Down 0.139598157910776
0.70018055381738 0.84105287876417
ZXDC_E 331 0.01 Down 0.165792573761613
0.70035160947354 0.84105287876417
FOXJ1_E 81 0.01 Down 0.390337725905907
0.701567441574912 0.841053310606292
BHLHE23_E 172 0.01 Down 0.309982947069489
0.701613864511776 0.841053310606292
DEAF1_D 113 0.01 Up 0.191884337036452
0.709715011450681 0.849534262881795
ZNF215_E 249 0.01 Up 0.239126615007628
0.70996336747961 0.849534262881795
TCF7_C 47 0.01 Down 0.310356019014281
0.715895651190302 0.854580316753775
NR0B1_E 16 0.01 Down 1.22412511455381
0.716254086353683 0.854580316753775
ZNF441_E 166 0.01 Down 0.157733212600224
0.716719029912709 0.854580316753775
ONECUT1_C 40 0.01 Down 0.403463190749291
0.717291352119561 0.854580316753775
CEBPG_C 39 0.01 Down 0.218995391831901
0.717385877304932 0.854580316753775
ERG_A 11 0.01 Up 1.19496299056791
0.718447475771392 0.854824194848436
ZNF257_E 305 0.01 Down 0.27660939715593
0.719134016073554 0.854824194848436
ETV5_E 750 0.01 Up 0.303025292838941
0.719514438574603 0.854824194848436
ESX1_E 245 0.01 Down 0.331385402922397
0.720946648501466 0.855596069129071
ZNF707_E 350 0.01 Up 0.135459388197124
0.721447848237866 0.855596069129071
ZFP28_E 481 0.01 Down 0.265668401312163
0.728297272185537 0.862041718739656
ZNF572_E 97 0.01 Up 0.177351814821348
0.728518589824123 0.862041718739656
ZFP37_E 85 0.01 Down 0.206610579395974
0.728822968506821 0.862041718739656
ZNF169_E 242 0.01 Down 0.145186323224382
0.732456763386867 0.865571689356998
KLF8_E 549 0.01 Down 0.293747997833001
0.733308873527685 0.865811096910898
ZNF43_E 192 0.01 Down 0.169242766371566
0.73577447400647 0.867953428186393
HOMEZ_E 430 0.01 Up 0.121826490030109
0.738386016026912 0.870263978217395
TCF23_E 30 0.01 Up 0.671390542220608
0.740221284294124 0.870908896332066
ZNF202_E 567 0.01 Down 0.120947863089596
0.740239894631831 0.870908896332066

BATF_C 41 0.01 Down 0.424724047690293
0.741089064917164 0.87113908600933
ZNF320_E 501 0.01 Down 0.237142004732908
0.742022361414127 0.871467672039675
ZNF449_E 449 0.01 Down 0.294091533537959
0.745098384203722 0.872959173441877
GLIS3_E 672 0.01 Up 0.390568196857617
0.745213807833539 0.872959173441877
SOX4_E 703 0.01 Down 0.346508558104276
0.745256968774836 0.872959173441877
ZNF2_E 197 0.01 Down 0.12832496315911
0.746739804651673 0.873928147147217
HNF4G_C 48 0.01 Down 0.486332930555433
0.748449575876533 0.875160776002998
POGZ_E 665 0.01 Up 0.186332648862858
0.750882909710213 0.877236563228496
CREB5_D 21 0.01 Down 0.455542490574223
0.751998095204773 0.877240019343401
EVX2_E 438 0.01 Down 0.291619131503061
0.752420264990178 0.877240019343401
HOXC12_E 279 0.01 Down 0.317673873356188
0.753339901933655 0.877240019343401
ZNF814_E 244 0.01 Down 0.169153940172465
0.753518246172688 0.877240019343401
GBX1_E 247 0.01 Down 0.338739115977831
0.754835613223477 0.878006869482456
ZIC1_E 572 0.01 Up 0.287765651666915
0.756090674844958 0.878034284424492
ZNF726_E 133 0.01 Up 0.22452533945299
0.756493199072823 0.878034284424492
PBX3_C 48 0.01 Up 0.374052908668978
0.758322971244457 0.878034284424492
E2F6_C 66 0.01 Down 0.197540362883797
0.758373954947108 0.878034284424492
FOXC1_E 760 0.01 Up 0.417334638209923
0.75843775156825 0.878034284424492
NFIC_A 17 0.01 Down 0.530617943157541
0.758889595395494 0.878034284424492
MIXL1_E 238 0.01 Down 0.348885004692239
0.759470014959819 0.878034284424492
TBX3_E 353 0.01 Up 0.345808565819391
0.761148309314476 0.879212041868454
FOSB_E 305 0.01 Down 0.382985305564463
0.764900908005977 0.88278174058179
ZNF276_E 715 0.01 Down 0.15994761266071
0.76782586158895 0.884408876215001
TCF4_C 57 0.01 Up 0.357005916899059
0.768160439288243 0.884408876215001
EVX1_E 186 0.01 Down 0.322455313192114
0.768707942350971 0.884408876215001
HOXD8_E 267 0.01 Down 0.408370272612979
0.768964656814093 0.884408876215001
ZNF746_E 453 0.01 Up 0.159397070305117
0.771872294938298 0.88679500902011

| | | | | | | |
|---|---|---|---|---|---|---|
| MBD1_C | 41 | 0.01 | Down | 0.134179372960268 | 0.772369846565902 | 0.88679500902011 |
| SP1_A | 227 | 0.01 | Up | 0.505109805983359 | 0.774353799633131 | 0.888307758098936 |
| KLF7_E | 313 | 0.01 | Down | 0.310453437563019 | 0.778079896850793 | 0.891102873737402 |
| HMGA1_D | 11 | 0.01 | Down | 0.430797561799266 | 0.778127340607904 | 0.891102873737402 |
| PAX2_E | 322 | 0.01 | Up | 0.337083067123824 | 0.780699839512023 | 0.89328144726998 |
| ZNF662_E | 291 | 0.01 | Up | 0.180827545169457 | 0.781915711246702 | 0.893318137862343 |
| MLX_E | 1276 | 0.01 | Up | 0.15859716330364 | 0.78207221821857 | 0.893318137862343 |
| ZNF613_E | 101 | 0.01 | Down | 0.114440884867987 | 0.784035820742729 | 0.894334193850559 |
| ZNF658_E | 122 | 0.01 | Up | 0.150480077451946 | 0.78440203640933 | 0.894334193850559 |
| FOXK1_D | 63 | 0.01 | Up | 0.189065992769449 | 0.785692586752681 | 0.894334193850559 |
| TBX4_E | 260 | 0.01 | Up | 0.311538074549695 | 0.785698567932584 | 0.894334193850559 |
| ZNF212_E | 285 | 0.01 | Up | 0.137651825594442 | 0.786440277539942 | 0.894334193850559 |
| ZFP69B_E | 360 | 0.01 | Down | 0.17411854968202 | 0.787330483844093 | 0.894334193850559 |
| TFAP4_C | 37 | 0.01 | Down | 0.182658773826501 | 0.787658172228474 | 0.894334193850559 |
| ZNF582_E | 317 | 0.01 | Up | 0.253308342244407 | 0.788973971168121 | 0.895065790269877 |
| ZNF300_E | 265 | 0.01 | Down | 0.176600681584358 | 0.792175175316631 | 0.896686086500637 |
| ZNF606_E | 238 | 0.01 | Down | 0.146375205015803 | 0.792474405975772 | 0.896686086500637 |
| RUNX1_A | 16 | 0.01 | Up | 0.591941942150976 | 0.792778274067447 | 0.896686086500637 |
| FOXI1_E | 481 | 0.01 | Down | 0.296085829634443 | 0.793092945224494 | 0.896686086500637 |
| ZNF433_E | 37 | 0.01 | Down | 0.222518522541499 | 0.795278521247346 | 0.898395143070095 |
| USF2_A | 28 | 0.01 | Up | 0.483203971268212 | 0.796191706236214 | 0.898665151916065 |
| GATAD1_C | 13 | 0.01 | Up | 0.302250109959974 | 0.798176543072256 | 0.900143258811605 |
| ZSCAN21_E | 51 | 0.01 | Down | 0.120749310249943 | 0.800154865003445 | 0.901611525823832 |
| TRPS1_E | 239 | 0.01 | Down | 0.395583136852361 | 0.801331778948682 | 0.902175051806244 |
| NKX2-5_E | 477 | 0.01 | Down | 0.325166159512848 | 0.803014053229965 | 0.903306103759952 |
| BATF3_E | 949 | 0.01 | Up | 0.363888401994521 | 0.804819598903952 | 0.90428550823423 |
| ZNF197_D | 268 | 0.01 | Down | 0.221620445075253 | 0.805241484076542 | 0.90428550823423 |

ZNF711_E 353 0.01 Up 0.337763773808037
0.806526496709385 0.904966178546809
TADA2A_E 296 0.01 Down 0.11351364247779
0.807925736056793 0.905243147235523
HOXD9_E 289 0.01 Down 0.351703751229322
0.808131541793152 0.905243147235523
SP4_C 53 0.01 Up 0.249212089453 0.812503056292337
0.907574178034047
ZNF880_E 42 0.01 Up 0.161138813460969
0.813170381086932 0.907574178034047
ZNF331_E 389 0.01 Down 0.332874041180278
0.813547322018659 0.907574178034047
ESR1_A 111 0.01 Down 0.431829949372563
0.81373071811093 0.907574178034047
MEIS2_C 51 0.01 Up 0.318077106450029
0.814297397408141 0.907574178034047
OLIG3_E 210 0.01 Up 0.344604961126765
0.814632986425052 0.907574178034047
SSRP1_C 24 0.01 Up 0.143024618605485
0.814978462945802 0.907574178034047
NEUROG1_E 210 0.01 Down 0.290468755902947
0.817347288606559 0.90945236703885
SMAD3_A 25 0.01 Up 0.949421465826656
0.820800740293722 0.912006815563551
ZNF329_E 365 0.01 Up 0.265261150934602
0.823206663624603 0.912006815563551
NR1D1_E 306 0.01 Up 0.317191253961251
0.824327422578111 0.912006815563551
ZNF266_E 100 0.01 Up 0.112467629560381
0.825503930874941 0.912006815563551
ZBTB16_D 13 0.01 Down 0.304083358900332
0.826003337039237 0.912006815563551
MEF2D_E 795 0.01 Up 0.285956247042269
0.82674386899067 0.912006815563551
PITX3_E 278 0.01 Down 0.325628109845028
0.826973715144551 0.912006815563551
HOXC9_E 516 0.01 Down 0.319406317721987
0.82705026337371 0.912006815563551
INSM1_E 436 0.01 Up 0.399005926953881
0.82955503139475 0.912006815563551
ZNF526_E 267 0.01 Down 0.118384244206968
0.829660166373895 0.912006815563551
AHR_C 38 0.01 Up 0.439042346521528
0.829763982425313 0.912006815563551
ZNF652_E 133 0.01 Down 0.198209851169062
0.829939057777276 0.912006815563551
ZIC4_E 276 0.01 Up 0.306584913208048
0.830212998844802 0.912006815563551
ID2_E 342 0.01 Up 0.2919247999614 0.830312833125928
0.912006815563551
PKNOX1_C 28 0.01 Up 0.230600017708738
0.830535923161606 0.912006815563551
RERE_E 347 0.01 Down 0.189860381732753
0.830589853033872 0.912006815563551

TFAP2A_A	45	0.01	Down	0.404367727985793
0.835703783776893	0.915380889404467
ZNF607_E	158	0.01	Up	0.186301077863515
0.835923451599942	0.915380889404467
HOXA13_E	409	0.01	Down	0.355689448867878
0.835993326290754	0.915380889404467
JARID2_D	74	0.01	Up	0.779569731448712
0.836771225410365	0.915380889404467
ZHX2_D	331	0.01	Up	0.225739473783225
0.837238368158296	0.915380889404467
GCM1_E	312	0.01	Down	0.310411549039878
0.837843128047131	0.915380889404467
ZNF844_E	77	0.01	Down	0.181416318932204
0.838469666888863	0.915380889404467
MNX1_E	332	0.01	Down	0.338946818465999
0.839843741769686	0.916130693763496
ZNF132_E	127	0.01	Down	0.163115564546061
0.841750819536205	0.917460214588521
HOXC10_E	288	0.01	Up	0.402033254029495
0.845455677540954	0.920556399507146
ZNF462_E	120	0.01	Down	0.326277758913404
0.846467834568415	0.920556399507146
ZNF235_E	40	0.01	Up	0.162662861866368
0.846663275165612	0.920556399507146
FEZF1_E	298	0.01	Up	0.308921367079856
0.852884241596908	0.926564542826959
E2F4_A	204	0.01	Up	0.216712476747168
0.858561352089751	0.931246863109358
ZNF226_E	277	0.01	Up	0.149377874673689
0.858591443932033	0.931246863109358
ID3_D	184	0.01	Up	0.384666927181274
0.859777549972778	0.931775182206271
ZNF77_E	48	0.01	Up	0.12949877444313
0.865236165891747	0.936929170701624
ZNF563_E	420	0.01	Up	0.266160874397341
0.866287706991708	0.93724402380695
NEUROD2_E	473	0.01	Down	0.360464760460144
0.867564722853252	0.93724402380695
EN2_E	258	0.01	Down	0.320519248183729
0.867636253096606	0.93724402380695
NFATC2_D	13	0.01	Up	0.655084746821278
0.868420089685862	0.93733115753138
ATF7_C	27	0.01	Down	0.188775119037254
0.869954726022109	0.937865507213557
PPARG_A	27	0.01	Down	0.606852506803238
0.870322304893601	0.937865507213557
SALL4_E	249	0.01	Down	0.298983840197576
0.871243086040727	0.938099381011542
ZNF382_E	448	0.01	Up	0.330554922162259
0.87547745930098	0.941897863800005
ZNF594_E	179	0.01	Down	0.172916680408806
0.876993053318355	0.942767532317232
ZNF345_E	271	0.01	Up	0.171991614509006
0.878611834244298	0.943664300846107

CREB3_C 22 0.01 Up 0.263912979117359
0.88015987699701 0.943664300846107
ZNF284_E 73 0.01 Up 0.115939435740754
0.880195116949797 0.943664300846107
ZFY_E 37 0.01 Down 0.541497843728453
0.880658957428775 0.943664300846107
RFX2_C 18 0.01 Up 0.174949517398656
0.882834852804614 0.945236031155462
ZNF574_E 268 0.01 Down 0.11797460815141
0.885541296025074 0.94737283114079
ZNF211_E 167 0.01 Up 0.160765536654808
0.887305172808724 0.948498633002429
ZSCAN9_E 185 0.01 Up 0.150124982036977
0.888038910174388 0.948522329537226
RHOXF1_E 291 0.01 Down 0.256741464505682
0.889134865015389 0.948932566105295
MAFK_C 45 0.01 Up 0.367806196677525
0.891993958093174 0.951018595059157
ZNF644_D 167 0.01 Down 0.345918848441445
0.892516325895728 0.951018595059157
NPAS1_E 178 0.01 Up 0.319566875599839
0.893328188764178 0.951123383085183
E2F1_A 257 0.01 Down 0.29666074374424
0.89416757309498 0.951257282470557
TEF_E 338 0.01 Up 0.306110593233359
0.896674682929618 0.952316879255661
PAX1_E 285 0.01 Down 0.291804803124291
0.897612238355458 0.952316879255661
LMX1A_E 82 0.01 Up 0.305854387357516
0.897647560413207 0.952316879255661
HOXD11_E 274 0.01 Down 0.309627579472355
0.899023808182725 0.952316879255661
AIRE_E 275 0.01 Down 0.346633114250014
0.899384116053532 0.952316879255661
HOXA5_E 277 0.01 Down 0.319050139501904
0.900219441502128 0.952316879255661
GFI1B_C 25 0.01 Down 0.233124946055214
0.900225753776385 0.952316879255661
MLXIP_E 552 0.01 Down 0.253699389057541
0.900878908283112 0.952316879255661
ZNF605_E 173 0.01 Down 0.173017523929935
0.902206535690574 0.952964589600266
ESRRB_E 370 0.01 Up 0.366232367413438
0.904096971180855 0.95377476087323
ZNF316_E 453 0.01 Up 0.130084082728161
0.904404574451435 0.95377476087323
SIX1_E 506 0.01 Down 0.365677777538435
0.90553824621734 0.95389073577487
KDM5A_D 177 0.01 Down 0.132695628492942
0.906093153569197 0.95389073577487
ZBTB7B_E 925 0.01 Down 0.226144843977611
0.906661337004321 0.95389073577487
ZNF483_E 221 0.01 Up 0.22086428405451
0.908069344135889 0.953909280733442

SCRT2_E 279 0.01 Down 0.327831655880366
0.908110185484424 0.953909280733442
BCL6B_E 466 0.01 Up 0.404941602285825
0.909808085346304 0.954940297454034
ZSCAN30_E 270 0.01 Up 0.192628975068517
0.913022405128208 0.957242485370168
ZNF154_E 44 0.01 Up 0.361498041507887
0.913437690465757 0.957242485370168
NKX2-3_E 296 0.01 Down 0.368217795192932
0.917056766470661 0.960051180815784
NR2C1_E 701 0.01 Up 0.238652742748964
0.917789787157228 0.960051180815784
HMX2_E 284 0.01 Down 0.288040666414777
0.918278511282914 0.960051180815784
GSX2_E 283 0.01 Down 0.306270471690517
0.92033452146797 0.960348443585193
MEF2C_C 61 0.01 Down 0.324746432252696
0.920743336392155 0.960348443585193
NKX2-2_E 24 0.01 Up 1.0831830243363
0.921374010580817 0.960348443585193
CTCFL_C 44 0.01 Up 0.322234410359408
0.921444605660511 0.960348443585193
ZNF629_E 526 0.01 Up 0.146042140331763
0.922575061008682 0.96066688899547
ZNF676_E 10 0.01 Up 0.497344641821892
0.923191511480268 0.96066688899547
FOXP4_E 277 0.01 Up 0.193099324940685
0.924082123191423 0.960843580510271
KLF9_C 64 0.01 Down 0.226593060760767
0.926748112493123 0.961902914786225
DUXA_E 251 0.01 Down 0.310133944626335
0.927620969638182 0.961902914786225
DLX1_E 291 0.01 Up 0.38724686733119
0.927934022951679 0.961902914786225
NKX2-8_E 548 0.01 Down 0.317079082341222
0.927987358150852 0.961902914786225
GSC_E 290 0.01 Down 0.3239580813981 0.933201372952882
0.965763731021601
MYOG_E 482 0.01 Up 0.350400733477454
0.934305712839232 0.965763731021601
ZNF559_E 109 0.01 Down 0.228626609233159
0.934821889833953 0.965763731021601
SPIC_E 289 0.01 Down 0.423829118595553
0.935155754340075 0.965763731021601
GFI1_E 659 0.01 Up 0.413335558258998
0.935334566203216 0.965763731021601
ZNF774_E 233 0.01 Up 0.228408514067461
0.938119472789631 0.967889517978776
MYNN_D 305 0.01 Up 0.221422262618149
0.939010601590115 0.968059653456785
ZNF302_E 183 0.01 Down 0.154096172412458
0.940215448484662 0.968552699250428
TCF25_C 41 0.01 Up 0.127396425357742
0.941029182118671 0.968595892189644

| | | | | | | |
|---|---|---|---|---|---|---|
| SIX3_E | 261 | 0.01 | Up | 0.331241082264607 | 0.942609807373754 | 0.968595892189644 |
| FOXH1_E | 320 | 0.01 | Down | 0.322894660453079 | 0.942745572537036 | 0.968595892189644 |
| HOXA11_E | 277 | 0.01 | Down | 0.313696626479299 | 0.943163892019623 | 0.968595892189644 |
| GATA2_A | 26 | 0.01 | Down | 0.585525279495226 | 0.94454560882881 | 0.96926812668884 |
| ZNF335_E | 798 | 0.01 | Down | 0.170335537139788 | 0.946406693138669 | 0.970430863041419 |
| NFKBIB_E | 698 | 0.01 | Up | 0.139903895481428 | 0.947919834149936 | 0.971235310470304 |
| ZKSCAN2_E | 187 | 0.01 | Down | 0.15543189199268 | 0.954632867909255 | 0.977362221907095 |
| ZNF93_E | 50 | 0.01 | Down | 0.235987782057055 | 0.956319186900347 | 0.978337280228827 |
| DNAJC1_E | 977 | 0.01 | Down | 0.147601391562975 | 0.959904077976896 | 0.980421118071592 |
| ZNF713_E | 75 | 0.01 | Down | 0.200137399293003 | 0.960866676802313 | 0.980421118071592 |
| TP63_B | 14 | 0.01 | Up | 0.281688853079013 | 0.961248964555316 | 0.980421118071592 |
| NFYB_C | 69 | 0.01 | Down | 0.275823274029432 | 0.961298125521058 | 0.980421118071592 |
| MYOD1_C | 40 | 0.01 | Up | 0.268107946779304 | 0.962454277596474 | 0.980849810425152 |
| E2F8_E | 522 | 0.01 | Up | 0.307293015691818 | 0.966250378778485 | 0.983966199321406 |
| SOX17_E | 691 | 0.01 | Up | 0.379155170323667 | 0.967286585019196 | 0.984269479259991 |
| GCM2_E | 318 | 0.01 | Up | 0.295543377504709 | 0.970739343198962 | 0.986485419531957 |
| ZNF7_E | 362 | 0.01 | Down | 0.114801379352504 | 0.970944388916674 | 0.986485419531957 |
| ZNF764_E | 138 | 0.01 | Down | 0.104751692062465 | 0.972072441668823 | 0.986732899797633 |
| SP3_A | 44 | 0.01 | Up | 0.622897029535204 | 0.974598694848057 | 0.986732899797633 |
| PHTF1_E | 521 | 0.01 | Down | 0.23860663242121 | 0.974897120189985 | 0.986732899797633 |
| VAX1_E | 253 | 0.01 | Down | 0.296252140307395 | 0.975611320734552 | 0.986732899797633 |
| VAX2_E | 253 | 0.01 | Down | 0.296252140307395 | 0.975611320734552 | 0.986732899797633 |
| HOXA2_E | 232 | 0.01 | Down | 0.317947705013854 | 0.975629378794659 | 0.986732899797633 |
| ZFP82_E | 374 | 0.01 | Up | 0.376759571449914 | 0.977998197649632 | 0.988378769876391 |
| ZNF780B_E | 313 | 0.01 | Down | 0.193049082974895 | 0.979328716341001 | 0.988973620365572 |
| EGR4_E | 299 | 0.01 | Up | 0.289712785591485 | 0.981300331573622 | 0.990214490528113 |
| NANOG_C | 36 | 0.01 | Up | 0.403577755177413 | 0.984495271802692 | 0.99268698737745 |

ZNF501_E 141 0.01 Up 0.17704276079828
0.985325917313667 0.992773581087769
ZNF641_E 276 0.01 Up 0.144857255738949
0.986544404233497 0.992807172970669
NR2E3_E 520 0.01 Up 0.299625748381455
0.986848840349689 0.992807172970669
MYT1L_E 68 0.01 Down 0.579024558911545
0.988949693783111 0.99414650284765
IRX1_E 14 0.01 Down 0.681130897744409
0.989831990521872 0.99414650284765
SOX3_E 559 0.01 Up 0.312006266246933
0.990417521216564 0.99414650284765
ZIM3_E 256 0.01 Down 0.342537151831222
0.99146376968575 0.994447859286008
TGIF2_C 16 0.01 Down 0.178092654424847
0.992797307718289 0.995036700141714
CREB3L1_C 18 0.01 Down 0.284871501627734
0.995471959713659 0.99696778534809
PAX5_C 46 0.01 Up 0.362281624522003
0.996537460615697 0.997285611862405
RUNX3_B 11 0.01 Down 0.652526828567196
0.997996414064255 0.997996414064255

**Supplement table 7:** edgeR_DE_results_IPF_effect

| ENTREZID | SYMBOL | GENENAME | LENGTH | logFC | logCPM | LR | PValue | FDR |
|---|---|---|---|---|---|---|---|---|
| 90019 | SYT8 | synaptotagmin 8 | 4704 | 3.83031642933146 | 4.76137271588789 | 82.0438399390938 | 1.331013652902e-19 | 1.86435082361983e-15 |
| 140828 | LINC00261 | long intergenic non-protein coding RNA 261 | 4912 | 3.9058709305555 | 0.804366847797816 | 72.7850866223606 | 1.44565140074608e-17 | 1.01246195851252e-13 |
| 91156 | IGFN1 | immunoglobulin like and fibronectin type III domain containing 1 | 12116 | -2.92914830968497 | 1.04954759081783 | 71.9160930378613 | 2.24545176950911e-17 | 1.0484014311838e-13 |
| 8000 | PSCA | prostate stem cell antigen | 1763 | -2.2484598041392 | 3.32175816521053 | 66.5175758607993 | 3.46787691751244e-16 | 1.21436379958992e-12 |
| 6558 | SLC12A2 | solute carrier family 12 member 2 | 10664 | -1.49376842598675 | 6.41760032583251 | 51.7045264957122 | 6.45138430180897e-13 | 1.80729079830876e-09 |
| 80114 | BICC1 | BicC family RNA binding protein 1 | 6569 | 2.99426345972451 | 4.45253233674204 | 47.2613803950031 | 6.21242423257672e-12 | 1.45029043709504e-08 |
| 9615 | GDA | guanine deaminase | 7904 | -4.35355698811671 | 2.85280000317873 | 44.4740537311561 | 2.57750617206039e-11 | 4.52508998913839e-08 |
| 23221 | RHOBTB2 | Rho related BTB domain containing 2 | 6580 | 1.01014813394304 | 5.84854357215035 | 44.4687688042876 | 2.58447347134341e-11 | 4.52508998913839e-08 |
| 56967 | C14orf132 | chromosome 14 open reading frame 132 | 7966 | 2.06736118021292 | 4.36308984147712 | 43.7974900776276 | 3.6417175825803e-11 | 5.6677264643558e-08 |
| 25984 | KRT23 | keratin 23 | 2902 | -3.03947980427182 | 6.07061853379088 | 42.4944105826774 | 7.08844995120787e-11 | 9.92879184665687e-08 |
| 9047 | SH2D2A | SH2 domain containing 2A | 2512 | -2.2973199482849 | 0.198527581398444 | 41.5542948632042 | 1.14642349849457e-10 | 1.45981399485576e-07 |
| 5521 | PPP2R2B | protein phosphatase 2 regulatory subunit Bbeta | 12857 | 2.0269903689329 | 2.6151052574474 | 40.7196338955174 | 1.75712806519346e-10 | 2.05100773409706e-07 |
| 10783 | NEK6 | NIMA related kinase 6 | 4419 | 1.40771914374428 | 5.14700873602658 | 38.990328876342 | 4.25910240187443e-10 | 4.58901902638885e-07 |
| 10371 | SEMA3A | semaphorin 3A | 8331 | -1.3999781542996 | 4.74982592223833 | 38.5548614396701 | 5.32360008155255e-10 | 5.32626188159333e-07 |
| 54578 | UGT1A6 | UDP glucuronosyltransferase family 1 member A6 | 5180 | 1.45361427700307 | 7.45786146944106 | 38.0190853628091 | 7.00559809470401e-10 | 6.5418275008346e-07 |
| 91461 | PKDCC | protein kinase domain containing, cytoplasmic | 3925 | 2.15185096964272 | 0.223978834195268 | 36.7687455253893 | 1.33005541532257e-09 | 1.16438038765145e-06 |
| 115749 | C12orf56 | chromosome 12 open reading frame 56 | 5504 | -2.31694772947584 | -0.699228974691679 | 36.0651625947336 | 1.90828115415248e-09 | 1.57231141918905e-06 |

7136	TNNI2	troponin I2, fast skeletal type 1082
3.658363799431	1.78855287393166	34.5749501749979
4.10150541282659e-09	3.19165479541456e-06
23316	CUX2	cut like homeobox 2	7648	2.94262619253436
	4.62730191689765	34.1907556113165
4.99657488611308e-09	3.49999210649128e-06
85301	COL27A1 collagen type XXVII alpha 1 chain	13988
0.79282709074905	6.35041010625252	34.1903992441519
	4.99748997856969e-09	3.49999210649128e-06
6581	SLC22A3 solute carrier family 22 member 3	3348
1.83654100027912	4.43042872620132	33.8394450859256
	5.98529327142384e-09	3.9921906120397e-06
10736	SIX2	SIX homeobox 2	2205	-2.36953389284089
2.22482826530701	33.0780050258322	8.85346047477273e-09
	5.63683731227917e-06
23767	FLRT3	fibronectin leucine rich transmembrane protein 3
	5190	1.75903585800718	5.2460093749324
32.9778710742611	9.32138229635185e-09	5.67672181847828e-06
131873	COL6A6	collagen type VI alpha 6 chain
9720	-2.20778745337283	-0.237409200597925
32.7146209655865	1.06731357106268e-08	6.22910882911458e-06
54657	UGT1A4	UDP glucuronosyltransferase family 1 member A4	2522
	1.36371386844351	7.17249606972986
32.4382447175982	1.23040990179757e-08	6.25366848499782e-06
54575	UGT1A10 UDP glucuronosyltransferase family 1 member A10 3157
	1.36021750222018	7.19447409710169
32.3810000727146	1.26719429904992e-08	6.25366848499782e-06
54579	UGT1A5	UDP glucuronosyltransferase family 1 member A5	2345
	1.36186998998636	7.15752630797695
32.3801703453482	1.26773548734907e-08	6.25366848499782e-06
54600	UGT1A9	UDP glucuronosyltransferase family 1 member A9	2416
	1.36137415469824	7.15676168365737
32.3498558944997	1.28766758477079e-08	6.25366848499782e-06
54576	UGT1A8	UDP glucuronosyltransferase family 1 member A8	2407
	1.36118188431188	7.15718132336632
32.3246640246167	1.30447013250967e-08	6.25366848499782e-06
54659	UGT1A3	UDP glucuronosyltransferase family 1 member A3	2364
	1.3606762805606 7.15961310964801	32.2733150908981
	1.33940211715524e-08	6.25366848499782e-06
54658	UGT1A1	UDP glucuronosyltransferase family 1 member A1	4467
	1.35527899670806	7.220941223639	32.1438883074102
	1.4316635080958e-08	6.46880992190256e-06
54577	UGT1A7	UDP glucuronosyltransferase family 1 member A7	4167
	1.35532591840753	7.19434418388932
32.0130847238087	1.53137618670368e-08	6.70312070223702e-06
9315	NREP	neuronal regeneration related protein	9337
2.14886405711659	4.38297822862843	31.6383180701197
	1.85728473336838e-08	7.88332947281542e-06
8527	DGKD	diacylglycerol kinase delta	9771
0.830322812417789	5.68015891530781	30.8471392421572
	2.79173363264963e-08	1.15011214683892e-05
100507533	SOX21-AS1	SOX21 antisense divergent transcript
1	3287	-2.00163248647716	0.370157742538742
30.3325817634858	3.63960623737388e-08	1.45657041619703e-05

654463	FER1L6	fer-1 like family member 6
6058	-2.63214636512789	2.49615163758075
29.9631014894254	4.40346484101748e-08	1.71331477855922e-05
58524	DMRT3	doublesex and mab-3 related transcription factor 3
	2183	-3.6148021188473	0.878532737500385
28.4354907927969	9.68730447373691e-08	3.667299290909e-05
80149	ZC3H12A	zinc finger CCCH-type containing 12A
3117	-1.6282446803702	6.25991061510752
27.9973076887204	1.21484353447718e-07	4.47797720721627e-05
7025	NR2F1	nuclear receptor subfamily 2 group F member 1	4201
	1.65940106335351	2.87108495273772
27.8361200644816	1.32037904634389e-07	4.74219212875356e-05
57139	RGL3	ral guanine nucleotide dissociation stimulator like
3	4358	3.22856809159606	2.37979897799412
26.8643630059516	2.18246317115384e-07	7.64244040958798e-05
4837	NNMT	nicotinamide N-methyltransferase	2815
2.94675035577442	4.43924672170923	26.684814561207
2.39496347709548e-07	8.18201303016497e-05
107075116	MIR3142HG	MIR3142 host gene
2301	-1.82488469353181	0.451758539904696
26.2524146017597	2.99580019742131e-07	9.81435417956977e-05
128553	TSHZ2	teashirt zinc finger homeobox 2
12581	-2.83449059787295	2.03968046619322
26.2414213883102	3.01290233255872e-07	9.81435417956977e-05
1289	COL5A1	collagen type V alpha 1 chain	11189
1.58060995457755	7.30501378194219	26.06405988972
3.30274955579502e-07	0.000105140029609138
55075	UACA	uveal autoantigen with coiled-coil domains and
ankyrin repeats	11031	-0.826293220284411	4.90656339951855
	25.9400830428882	3.52179982356837e-07
0.000109621889174938
22837	COBLL1	cordon-bleu WH2 repeat protein like 1	17178
0.827608771747435	6.68262007772591	25.4120511499997
	4.63016617835918e-07	0.000140988560131037
55612	FERMT1	fermitin family member 1
5359	-1.67501140856345	8.34448781104499
25.0285759300853	5.64869082343316e-07	0.000168343005029422
2706	GJB2	gap junction protein beta 2
2250	-2.04953302531315	5.2391996563143	24.8631219449432
	6.15486530124801e-07	0.000179606663072044
5654	HTRA1	HtrA serine peptidase 1	2120	0.951537823581377
	4.24871591436869	24.3596876773575
7.99237950778116e-07	0.00022846787707243
10544	PROCR	protein C receptor	1696	0.928692581928044
	4.22469943355717	24.2721079050365
8.36415121450179e-07	0.000234313332123053
3170	FOXA2	forkhead box A2	2603	2.90243001863281
2.705364499604	24.1217828961054	9.04315515718091e-07
0.000248367596640457
9697	TRAM2	translocation associated membrane protein 2	6908
	0.805525272264177	5.64326956911266
24.0654838666226	9.3114482025505e-07	0.000250818182640625
100126311	MIR147B	microRNA 147b	80	-1.0397833441396
	5.82788766800658	23.8698574312558

1.03073112163003e-06	0.000272404732465507
8560	DEGS1	delta 4-desaturase, sphingolipid 1	2293
0.824117329054976	6.04761763156333	23.6421732611306
	1.16016810910578e-06	0.00030093471674527
3696	ITGB8	integrin subunit beta 8 14113	-1.38809817653538
	9.95722482428052	23.4504641459398
1.28171796544306e-06	0.000326418609853836
7128	TNFAIP3 TNF alpha induced protein 3
5051	-1.23768786745253	8.51486238394146
23.1166866342701	1.5246182865072e-06	0.000381345148912613
91683	SYT12	synaptotagmin 12	10830	1.63666052440125
	4.00388921783767	22.9542790361429
1.65900506965564e-06	0.000402362394098938
1847	DUSP5	dual specificity phosphatase 5
2615	-1.28901995910715	5.54682871701079
22.946079161543 1.6660968699749e-06	0.000402362394098938
4318	MMP9	matrix metallopeptidase 9
2336	-3.88646646358111	6.41736269852419
22.8702287096639	1.73315620113508e-06	0.000411463032361
55351	STK32B	serine/threonine kinase 32B	5324
2.44662356566195	0.639554063087618	22.8128415297943
	1.78568557969265e-06	0.000416868298579249
3854	KRT6B	keratin 6B	2282	-1.64254790893414
6.7040339088129 22.739797476075 1.85486304041514e-06
0.000417315496194087
84419	C15orf48	chromosome 15 open reading frame 48
1405	-1.02540330735382	8.30821918805363
22.7259079036816	1.86831844270053e-06	0.000417315496194087
441094	NR2F1-AS1	NR2F1 antisense RNA 1	7720
1.29892808941862	2.72402182801731	22.7170185740828
	1.87698124225226e-06	0.000417315496194087
10620	ARID3B	AT-rich interaction domain 3B
6212	-0.967671059134849	3.82330701524784
22.5625101320139	2.0341557569473e-06	0.000445194057618138
7447	VSNL1	visinin like 1	3833	1.06796667725149
3.86662266879682	22.3281712112933	2.29810450273921e-06
	0.000495223842613356
9672	SDC3	syndecan 3	7067	0.815701452086522
5.97395642701785	22.2063436435037	2.44863425222345e-06
	0.000519666969255968
4940	OAS3	2'-5'-oligoadenylate synthetase 3
8251	-1.1554826864508	5.64716494857164
22.1516619852463	2.51937771190441e-06	0.000526700352397687
389376	SFTA2	surfactant associated 2 1204	2.61318833391824
	1.63846857957059	21.8633778509669
2.92768338725149e-06	0.000603059723606347
6372	CXCL6	C-X-C motif chemokine ligand 6
2183	-2.73079954216321	6.7981495884013 21.5800992108452
	3.3935479802519e-06	0.000688890239991137
51297	BPIFA1	BPI fold containing family A member 1
1081	-4.0732416064031	4.43859718272277
21.5013308155515	3.53583196798933e-06	0.000703661535897401
347	APOD	apolipoprotein D	2197	2.37630488137662
	2.62489611586971	21.4846163888623

3.56678582485296e-06	0.000703661535897401
2192	FBLN1	fibulin 1	7285	1.65826005308169
7.27342314470593	21.45555976572	3.62124513110919e-06
0.000704483063214534
26037	SIPA1L1 signal induced proliferation associated 1 like 1
	11897	-0.646755780748389	7.09755929882032
21.2996094049378	3.92811528747793e-06	0.000753713847009636
11166	SOX21	SRY-box transcription factor 21
2778	-2.03031492446528	0.388426189508704
21.0689855200237	4.4304121654448e-06	0.0008386051783971
9407	TMPRSS11D	transmembrane serine protease 11D	3294
	-3.89015113249265	0.891692428757327
20.94698514043	4.72170324988659e-06	0.000874108565055315
147920	IGFL2	IGF like family member 2	2426
1.40346051160291	1.98316015749713	20.9384491021899
	4.74278938703534e-06	0.000874108565055315
80731	THSD7B	thrombospondin type 1 domain containing 7B	6761
	2.08913435722768	0.750987915244901
20.7651908294294	5.19181037521975e-06	0.000937733602364328
83987	CCDC8	coiled-coil domain containing 8 3213
1.75181513687072	5.6624101713292 20.7541223473142
5.22190483218516e-06	0.000937733602364328
5999	RGS4	regulator of G protein signaling 4	4836
3.04498082460595	0.618167615084196	20.7209898474172
	5.31304039361018e-06	0.000942022237889845
3656	IRAK2	interleukin 1 receptor associated kinase 2
3483	-1.17564654853379	4.48822673554006
20.61053232684	5.62856088904462e-06	0.000985490654660599
81848	SPRY4	sprouty RTK signaling antagonist 4
5516	-0.707577687212106	4.40490055575631
20.5364358431192	5.85069019728204e-06	0.00101173601967073
53827	FXYD5	FXYD domain containing ion transport regulator 5
	2993	-1.11479773646359	6.37829723646139
20.3667235390078	6.39319030556803e-06	0.00108819805961159
8644	AKR1C3	aldo-keto reductase family 1 member C3	4532
1.33579913946811	7.05301404178933	20.3503183896377
	6.44823580693665e-06	0.00108819805961159
6706	SPRR2G	small proline rich protein 2G
568	-3.06066439574784	1.88449024176279
20.3069630737529	6.59601027507379e-06	0.00109988471336855
4488	MSX2	msh homeobox 2	2270	1.99577830619329
0.100822861839864	20.2095396880411	6.94059530174156e-06
	0.00114372845166464
57476	GRAMD1B GRAM domain containing 1B
10218	-1.43943147060233	0.815984897094341
20.1534759777938	7.14702495585111e-06	0.00115747935123885
4060	LUM	lumican 3008	3.96284714765671
3.21376244956127	20.1421919286834	7.18931274061399e-06
	0.00115747935123885
219790	RTKN2	rhotekin 2	9548	-1.63002641893973
4.42412444318217	19.789413857701 8.64602145119417e-06
0.00136148647947641
79960	JADE1	jade family PHD finger 1	9129
1.09300778467647	3.81129277810418	19.7883492386338

8.65083862878565e-06    0.00136148647947641
9592    IER2    immediate early response 2
2103    -0.679423991011737      5.966688437308  19.7593864034157
        8.78292824955853e-06    0.00136691639990629
3918    LAMC2   laminin subunit gamma 2 6543    -1.19298872453531
        12.7761040638834        19.6894328173551
9.11037559106812e-06    0.0014022970429021
27122   DKK3    dickkopf WNT signaling pathway inhibitor 3      5059
        1.10873389230933        8.65790102020753
19.5752979919717        9.67115665636125e-06    0.001472433600931
59      ACTA2   actin alpha 2, smooth muscle    3027
2.33308674896258        5.25161833226589        19.4798663883416
        1.01665648649018e-05    0.00152689292616324
2615    LRRC32  leucine rich repeat containing 32       4459
2.70583013264087        0.390411351237698       19.4511968168321
        1.03203150977811e-05    0.00152689292616324
148641  SLC35F3 solute carrier family 35 member F3      3195
2.61047711750322        0.992669509748176       19.4374015319499
        1.03951268161504e-05    0.00152689292616324
4151    MB      myoglobin       2168    -1.81250180615869
1.2004438816388 19.4246266408291        1.04648904770237e-05
0.00152689292616324
3576    CXCL8   C-X-C motif chemokine ligand 8
2274    -2.30844412996995       10.0558188491315
19.3872646247291        1.06716342453013e-05    0.00154100598839109
4017    LOXL2   lysyl oxidase like 2    5659    1.80494194076071
        6.00927317294208        19.2826484011773
1.12726143203958e-05    0.00161117866107943
2348    FOLR1   folate receptor alpha   1577    1.86966872304686
        4.02443337881273        19.2066553184431
1.17303714275992e-05    0.00165966982410487
23138   N4BP3   NEDD4 binding protein 3 6080    -1.41671719785138
        3.0150973480594 19.167019934719 1.19764821216638e-05
0.00167754585078145
51332   SPTBN5  spectrin beta, non-erythrocytic 5
12338   -1.50940104040111       3.24658993965546
18.913706756259 1.367664685125e-05      0.00187154674788888
11145   PLAAT3  phospholipase A and acyltransferase 3   3104
0.963496626677586       5.29265007554904        18.9045872250906
        1.37421825493969e-05    0.00187154674788888
3783    KCNN4   potassium calcium-activated channel subfamily N
member 4        3289    1.3170591772566 5.4150640043312
18.9017888601641        1.37623556102345e-05    0.00187154674788888
340719  NANOS1  nanos C2HC-type zinc finger 1   4017
1.35832898844589        1.37224009770492        18.7126935422007
        1.51967483505257e-05    0.00204673898217128
5099    PCDH7   protocadherin 7 12058   0.990391015026088
6.30916659325239        18.601121450501 1.61125736988511e-05
0.00213958348844788
5270    SERPINE2        serpin family E member 2
8891    -1.09802706205912       6.16989558160617
18.5917934559295        1.61916077515153e-05    0.00213958348844788
8710    SERPINB7        serpin family B member 7
2637    -1.39990411889805       6.26190547220277

18.5311800760294	1.67147443502779e-05	0.00218502668607449
57381	RHOJ	ras homolog family member J	3793
2.2988894685168 0.693610218036585	18.5161018333488
1.68474964015168e-05	0.00218502668607449
56999	ADAMTS9 ADAM metallopeptidase with thrombospondin type 1
motif 9 10680	-1.40713152162016	7.33576267390319
18.4731275467242	1.72316906659048e-05	0.00219571489797209
650	BMP2	bone morphogenetic protein 2
3601	-1.67501114211606	5.20511478991167
18.4627355084292	1.73259103936337e-05	0.00219571489797209
55076	TMEM45A transmembrane protein 45A	6005
1.17344057324701	5.87740320564795	18.4545835619061
1.7400182314193e-05	0.00219571489797209
5924	RASGRF2 Ras protein specific guanine nucleotide releasing
factor 2	10702	0.996528365856032	5.35290421792029
18.3842716885564	1.8054212468315e-05
0.00225790494681865
5265	SERPINA1	serpin family A member 1	4902
3.09102000470936	5.04888779628524	18.0667257273293
2.13296583038695e-05	0.00263347544590247
1278	COL1A2	collagen type I alpha 2 chain	11156
2.15119986529311	7.97524204364668	18.0574971175886
2.14332976963576e-05	0.00263347544590247
162394	SLFN5	schlafen family member 5
10709	-0.857391432784056	7.90139129804924
18.0020701779037	2.20664869016602e-05	0.00268769810462221
3159	HMGA1	high mobility group AT-hook 1
2327	-0.696020441538297	7.60967146257742
17.9527992877704	2.26451137328012e-05	0.00272134246253944
29126	CD274	CD274 molecule	4108	-1.85725855089006
5.35192521702518	17.9309729036603	2.29062872660318e-05
0.00272134246253944
284418	FAM71E2 family with sequence similarity 71 member E2
4686	-1.55796119567281	0.920301406937178
17.9293716491961	2.29255665438462e-05	0.00272134246253944
79625	NDNF	neuron derived neurotrophic factor	3556
1.80705132072122	2.97237624340148	17.8040740377023
2.44858391206724e-05	0.00288212729885091
85360	SYDE1	synapse defective Rho GTPase homolog 1	3965
1.42855701598516	2.54374801694254	17.706813483731
2.57701892555033e-05	0.00300802534084862
101928969	RHOXF1-AS1	RHOXF1 antisense RNA 1	2495
2.91327054343034	0.160168731167929	17.6676088092703
2.63068464342708e-05	0.00304528923970935
6678	SPARC	secreted protein acidic and cysteine rich	4831
3.35140061193802	6.7633603188507 17.517487107936
2.84677254250548e-05	0.00326842155761264
338382	RAB7B	RAB7B, member RAS oncogene family	3686
1.50546452876009	3.23012927256677	17.4925040611062
2.88442697832313e-05	0.00328472916141236
6700	SPRR2A	small proline rich protein 2A
709	-4.67050883384125	2.96468366451314
17.4262833571606	2.98667441113821e-05	0.00337373778038814
169834	ZNF883	zinc finger protein 883 2319

2.24612840829452	-0.154921870361588	17.3122338298269
	3.17138531633982e-05	0.00355372753007775
1306	COL15A1 collagen type XV alpha 1 chain	6653
3.33283721530466	2.94700877740005	17.296806556001
3.19723718072076e-05	0.00355426199923458
10205	MPZL2	myelin protein zero like 2
4753	-0.72611050444764	7.44414135860264
17.2260100129366	3.31861584418209e-05	0.00366014583696524
7185	TRAF1	TNF receptor associated factor 1
6449	-2.07190096642509	4.32899400862876
17.136575452832 3.47859300065321e-05	0.00377487598077459
92949	ADAMTSL1	ADAMTS like 1	13172	2.31326694890692
	-0.800727837853213	17.1325704277135
3.48593566220645e-05	0.00377487598077459
6913	TBX15	T-box transcription factor 15	3806
2.29474743162809	-0.526636222289884	17.1230296210353
	3.50349023702932e-05	0.00377487598077459
51719	CAB39	calcium binding protein 39
5360	-0.649611187852555	7.6390843996938 17.1073698630309
	3.53249617904134e-05	0.00377707434960551
4811	NID1	nidogen 1	5864	2.47358726029305
3.2723992321818 17.0826773219642	3.5787243702061e-05
0.00379751456465734
6518	SLC2A5	solute carrier family 2 member 5	6693
1.38924615854264	4.23948482744286	17.0388189189047
	3.66233842283612e-05	0.00385702062320794
4792	NFKBIA	NFKB inhibitor alpha	2373	-0.752584138807818
	7.15191630617609	16.9510552158474
3.8355994311881e-05	0.00400934636064565
64220	STRA6	stimulated by retinoic acid 6
7690	-1.41772365074492	6.23989338092109
16.9090430206227	3.9214319560718e-05	0.00406870351175539
3119	HLA-DQB1	major histocompatibility complex, class II,
DQ beta 1	4904	1.09971392022316	5.74212540674199
	16.8796981281339	3.98252756819399e-05
0.0040721225369148
387700	SLC16A12	solute carrier family 16 member 12	4835
	1.67277302458063	2.47028600972657
16.8795342978385	3.98287133260032e-05	0.0040721225369148
338707	B4GALNT4	beta-1,4-N-acetyl-galactosaminyltransferase
4	3761	1.20954820323529	2.11878293962191
16.7853147047793	4.18559701437961e-05	0.00424838096959531
182	JAG1	jagged canonical Notch ligand 1
8534	-0.621368052689773	8.07101045363817
16.733547875703 4.30136312041674e-05	0.00433447433292642
84675	TRIM55	tripartite motif containing 55	3081
1.94202357701708	1.61419385260643	16.6559673520452
	4.48091210030118e-05	0.00448315255635133
639	PRDM1	PR/SET domain 1 6369	-1.44224906716919
3.27168533262699	16.6421018967283	4.51378656755422e-05
	0.00448401478381078
114897	C1QTNF1 C1q and TNF related 1	4854	-1.29206996404363
	2.91571239795905	16.5057762437845
4.85020886900102e-05	0.0047720557399885

94122	SYTL5	synaptotagmin like 5	4791	-1.80300910139448
	1.70629660397611	16.497324188933 4.87187813820486e-05
	0.0047720557399885
200150	PLD5	phospholipase D family member 5 9545
2.34694768629476	1.74873763979676	16.4564563223657
	4.97803588838601e-05	0.00484217699226548
285704	RGMB	repulsive guidance molecule BMP co-receptor b
6671	-0.87279873021083	5.60801824647833
16.3870094102008	5.1637924864995e-05	0.00495405851212566
150759	LINC00342	long intergenic non-protein coding RNA 342
	2618	-1.30436162511439	3.30953821544929
16.3870090716646	5.16379340879808e-05	0.00495405851212566
4249	MGAT5	alpha-1,6-mannosylglycoprotein 6-beta-N-
acetylglucosaminyltransferase	8255	-0.569294464068935
6.55688179941132	16.3262713720827	5.33196048581571e-05
	0.00501073778449277
2308	FOXO1	forkhead box O1 5811	-0.86347893957303
4.88981638279068	16.3200659513082	5.3494488112415e-05
	0.00501073778449277
3340	NDST1	N-deacetylase and N-sulfotransferase 1	9006
0.607660484057146	9.61982672299425	16.3146158789013
	5.36485595863503e-05	0.00501073778449277
374407	DNAJB13 DnaJ heat shock protein family (Hsp40) member B13
	3475	1.57877686215584	0.934190536879062
16.3142243035485	5.3659646439203e-05	0.00501073778449277
83716	CRISPLD2	cysteine rich secretory protein LCCL domain
containing 2	7965	1.26926747500117	2.87279363775079
	16.285650652289 5.44749147726665e-05	0.00505317967695854
83849	SYT15	synaptotagmin 15	12684	2.69269046446334
	1.00089179988913	16.1313864805853
5.90964956070808e-05	0.00544581982873935
9721	GPRIN2	G protein regulated inducer of neurite outgrowth 2
	1966	1.16037891057039	4.32024843286344
15.9935762217678	6.35577747375794e-05	0.00579102359869603
1846	DUSP4	dual specificity phosphatase 4
7257	-1.14932434338029	7.56011709352863
15.9902534573466	6.36694248732198e-05	0.00579102359869603
139818	DOCK11	dedicator of cytokinesis 11	6721
1.04050539833333	3.62976962206458	15.9610058950309
	6.46607388775631e-05	0.00584324496424534
4669	NAGLU	N-acetyl-alpha-glucosaminidase	3219
0.704326827334655	4.89163967040876	15.9474470077613
	6.51255547288485e-05	0.00584752336594219
6549	SLC9A2	solute carrier family 9 member A2
5984	-1.56874707831276	1.78527421518258
15.9340588638099	6.55878148483668e-05	0.00585151925210875
65062	TMEM237 transmembrane protein 237	7647
0.675291192887756	5.1240382555418 15.9108831722021
6.63958302679323e-05	0.00586290316069255
2947	GSTM3	glutathione S-transferase mu 3	4750
1.50968815908945	3.59681466600706	15.9064208906972
	6.6552552477341e-05	0.00586290316069255
8942	KYNU	kynureninase	17600	-2.09512760452669
4.87867432193224	15.8554462228175	6.83694855316889e-05

0.00598532114901479
729582 DIRC3 disrupted in renal carcinoma 3
4665 -2.73115882950034 1.15184522594568
15.8432666962021 6.88109455691952e-05 0.00598655226451998
1594 CYP27B1 cytochrome P450 family 27 subfamily B member 1
3440 -1.15513040685168 2.46234287674638
15.7948179703497 7.05955954332391e-05 0.00608567212626361
6695 SPOCK1 SPARC (osteonectin), cwcv and kazal like domains proteoglycan 1 6222 -1.08206489409377 5.21795777076359
15.7888346647061 7.08192015835631e-05
0.00608567212626361
4293 MAP3K9 mitogen-activated protein kinase kinase kinase 9
12386 -0.519198634222875 7.36538583264577
15.6681862396876 7.54832204186897e-05 0.00641359311868827
2919 CXCL1 C-X-C motif chemokine ligand 1
1296 -2.18095600409499 8.84642677639806
15.6664887428308 7.55510005414125e-05 0.00641359311868827
6703 SPRR2D small proline rich protein 2D
1057 -2.75708688628554 6.46804862372935
15.6542037075889 7.60433637531271e-05 0.00641650238608465
27295 PDLIM3 PDZ and LIM domain 3 9117 2.11920856866262
0.674789725471059 15.4814530506886
8.33187957612104e-05 0.00698830163010344
3624 INHBA inhibin subunit beta A 7532 -1.36957712510235
7.78017112113819 15.4595753431013
8.42888198140656e-05 0.00702758035199772
4319 MMP10 matrix metallopeptidase 10
1758 -2.30440147905255 7.45162868326031
15.3707660721789 8.83445884532886e-05 0.00732214586074091
5796 PTPRK protein tyrosine phosphatase receptor type K
9000 -0.621329714834622 9.44269309313022
15.3552303359364 8.9073999758977e-05 0.00733917361543524
864 RUNX3 RUNX family transcription factor 3
4523 -1.56351430865102 3.15770199980948
15.3256004446933 9.04819821600577e-05 0.00741158552114578
1826 DSCAM DS cell adhesion molecule
8577 -2.12623666443806 0.875847812776289
15.3101759820221 9.12237785741694e-05 0.00742890387493251
1295 COL8A1 collagen type VIII alpha 1 chain 9408
3.03431222475315 3.19236526518064 15.2905724296921
9.21753946270611e-05 0.00746301013029621
162514 TRPV3 transient receptor potential cation channel subfamily V member 3 9780 -2.29522490893332
3.86825105267539 15.1878099550647 9.7329698248631e-05
0.0078350407090148
387104 SOGA3 SOGA family member 3 12233 1.96643030802339
0.713599200928293 15.1287541380873
0.000100422034395435 0.00803777963301064
5209 PFKFB3 6-phosphofructo-2-kinase/fructose-2,6-biphosphatase 3 5481 0.69219063707667 6.7675277311556
15.1045367451024 0.0001017185415886 0.00809529325017905
4715 NDUFB9 NADH:ubiquinone oxidoreductase subunit B9 2369
0.616960002650837 6.54882650811263
15.0647595442912 0.000103884724708389 0.00821877928154697

346389	MACC1	MET transcriptional regulator MACC1
9898	-0.729582043677746	8.59563174814159
15.0546309162012	0.000104443686165157	0.00821877928154697
10893	MMP24	matrix metallopeptidase 24	4414
1.72997489447734	5.28197671914709	15.0106618929286
	0.000106905478968114	0.0083116556768207
501	ALDH7A1 aldehyde dehydrogenase 7 family member A1	6911
	0.826270072564206	5.80297942424795
15.0070725806248	0.000107109001882059	0.0083116556768207
2152	F3	coagulation factor III, tissue factor
2875	-0.647499366568253	10.803409528844 15.0018798664236
	0.000107404132041439	0.0083116556768207
8910	SGCE	sarcoglycan epsilon	2547	1.14422894776407
	3.87438268345642	14.9434595480282
0.000110781420495617	0.00852590855429729
284297	SSC5D	scavenger receptor cysteine rich family member with
5 domains	7314	2.64887810032483	0.251719316812683
	14.8189080993643	0.000118343002360106
0.00905808980359566
80323	CCDC68	coiled-coil domain containing 68
5180	-0.951595229552941	4.07237095148537
14.765356227107 0.000121751896808876	0.00926836314457569
23024	PDZRN3	PDZ domain containing ring finger 3	6136
1.67231649991402	0.0974434218941073	14.6980315205631
	0.000126178138097065	0.0095055548197288
1475	CSTA	cystatin A	1373	-1.68183728758523
3.40849421510933	14.6973319286441	0.000126224972975623
	0.0095055548197288
6415	SELENOW selenoprotein W 3151	0.73864704740918
6.64132593610726	14.6095797171999	0.000132240596936037
	0.00990531572878645
3169	FOXA1	forkhead box A1 3677	-0.636078269723743
6.43600501029452	14.5976859641604	0.000133077857598663
	0.00991500825204507
80820	EEPD1	endonuclease/exonuclease/phosphatase family domain
containing 1	4847	0.977440287969563	2.2268483288884
14.5593150511633	0.000135815535034921	0.0100544468570871
1284	COL4A2	collagen type IV alpha 2 chain	7885
1.01355040372591	9.19685303329237	14.5514311944365
	0.000136385014838763	0.0100544468570871
100506895	LUARIS	lncRNA upregulator of antiviral response
interferon signaling	2916	1.13179949967267
0.511382887452289	14.3995649452093	0.000147836461302772
	0.0108415984998321
25797	QPCT	glutaminyl-peptide cyclotransferase
2599	-0.587650726675206	5.50584277498371
14.3600313686094	0.000150973171809415	0.0110139646746587
389337	ARHGEF37	Rho guanine nucleotide exchange factor 37
	5324	-0.788493729508391	7.09548968362664
14.3186611185339	0.000154327403036849	0.0112003312659956
25891	PAMR1	peptidase domain containing associated with muscle
regeneration 1	3863	1.21294488904272	3.10785676124057
	14.2882964151286	0.000156836989347382
0.0112661597600517

3212	HOXB2	homeobox B2	1598	2.27889143072263
0.0890855393218446	14.2795954055064	0.00015756365823699
	0.0112661597600517
56650	CLDND1	claudin domain containing 1
5499	-0.511299161120423	6.56919699090374
14.2785951393163	0.000157647412934256	0.0112661597600517
10455	ECI2	enoyl-CoA delta isomerase 2	4579
0.869504052833646	3.96221082900996	14.2464817667131
	0.00016036030230715	0.0114018616975444
3995	FADS3	fatty acid desaturase 3 6283	0.646354426204267
	6.32116742078207	14.2199834084954
0.000162634214776298	0.0115051386180384
9843	HEPH	hephaestin	7113	2.60457114595512
0.495002150132266	14.205571551521 0.00016388454923616
0.0115353310610598
5990	RFX2	regulatory factor X2	6154	1.04675247932486
	2.22924055025838	14.1875400558593
0.00016546255685892	0.0115881701696145
196996	GRAMD2A GRAM domain containing 2A	4593
1.41712548250657	1.46242504077415	14.1680924138504
	0.000167181654900412	0.0116503156228362
405753	DUOXA2	dual oxidase maturation factor 2
2554	-2.60411477700238	1.88993301053788
14.1338108625143	0.000170255937250732	0.0118058164013416
4139	MARK1	microtubule affinity regulating kinase 1
5417	-0.563351182752376	4.87760986309442
14.1186497089544	0.000171633649255123	0.0118427217986035
9729	KIAA0408	KIAA0408	7229	1.95594062747519
	0.207319878336522	14.0583768892662
0.000177222676357094	0.0121232886586912
6035	RNASE1	ribonuclease A family member 1, pancreatic	1506
	1.48408430536927	6.06434753743362
14.0494669698838	0.000178064294552485	0.0121232886586912
59342	SCPEP1	serine carboxypeptidase 1	3853
0.934392761400686	6.3366653304706 14.0454802593149
0.000178442176031286	0.0121232886586912
10331	B3GNT3	UDP-GlcNAc:betaGal beta-1,3-N-
acetylglucosaminyltransferase 3 2301	-0.947205233650489
4.90359012399401	14.0379105081177	0.000179161901359968
	0.0121232886586912
223117	SEMA3D	semaphorin 3D	8062	-1.45740359893703
1.01597107905656	13.9791664209045	0.000184847555183491
	0.0123754378813922
22936	ELL2	elongation factor for RNA polymerase II 2
7328	-0.585951351150878	6.60819326917732
13.9785012344339	0.000184912966487486	0.0123754378813922
6704	SPRR2E	small proline rich protein 2E
734	-3.31939981302453	3.75620972306957
13.9721489203752	0.000185538798821472	0.0123754378813922
390	RND3	Rho family GTPase 3	3594	-0.517083161834302
	9.20002564800277	13.936190480565 0.00018912187658199
	0.0125126091923682
103611157	TGFB2-OT1	TGFB2 overlapping transcript 1
1371	-0.78110685880599	5.99078075474087

13.930359145321 0.000189709473446024    0.0125126091923682
3213    HOXB3   homeobox B3     6681
2.23457867994972        -0.470296818546235      13.9247612900716
        0.000190275273647065    0.0125126091923682
285672  SREK1IP1        SREK1 interacting protein 1
7537    -0.710576012747388      6.35850507304885
13.90343784226  0.00019244614452882     0.0125962296561457
7127    TNFAIP2 TNF alpha induced protein 2
4897    -1.57088331977725       7.29940382029439
13.8768795690019        0.000195184881708271    0.012716068084129
56300   IL36G   interleukin 36 gamma    1380    -2.5318037019756
        5.63615142896339        13.8534234991373
0.000197636351172526    0.0128161683836739
9734    HDAC9   histone deacetylase 9   13807   -0.764840079916164
        6.35071120686774        13.8168116885206
0.000201524915317896    0.0130081082435842
7058    THBS2   thrombospondin 2        6397    3.91015919386122
        3.42944982479328        13.77879974323  0.000205643777810131
        0.0132130843843417
7739    ZNF185  zinc finger protein 185 with LIM domain
4812    -1.01812454172483       7.5353935094117 13.7660309305781
        0.000207046315012571    0.0132424554081328
64175   P3H1    prolyl 3-hydroxylase 1  6766    0.626022590122604
        5.35826301666322        13.7451970267552
0.000209355450353109    0.0133292808777091
6423    SFRP2   secreted frizzled related protein 2     2032
3.89604091240734        1.74983694574799        13.7241098417472
        0.00021171908343242     0.0133706696424434
55841   WWC3    WWC family member 3     7114    -0.649872225846398
        6.1553111714929 13.7223756294487        0.000211914661285245
        0.0133706696424434
9509    ADAMTS2 ADAM metallopeptidase with thrombospondin type 1
motif 2 7687    3.60114617163784        3.31536894458115
13.6145411862646        0.000224440329749459    0.0140974695013483
123872  DNAAF1  dynein axonemal assembly factor 1
5912    -1.40360041507465       -0.571900115993694
13.6029350157494        0.000225832174711829    0.014121568174949
54869   EPS8L1  EPS8 like 1     5341    -0.78645961377242
7.72182736566456        13.5893522218343        0.000227472111853694
        0.0141608972032653
1528    CYB5A   cytochrome b5 type A    9313    0.711337245975705
        6.12042010118618        13.5267075431441
0.000235192299183448    0.0145767191799228
170689  ADAMTS15        ADAM metallopeptidase with thrombospondin
type 1 motif 15 5673    1.64767768353749        4.5623087139744
13.5011839664255        0.000238412981661335    0.0147112362737018
9118    INA     internexin neuronal intermediate filament protein
alpha   3231    1.60461277013143        4.59788510932343
13.4905624802865        0.000239766311998163    0.0147298540884135
8140    SLC7A5  solute carrier family 7 member 5        4745
1.11254233340395        8.87995841137554        13.4809398830112
        0.000240999052404427    0.0147409333058027
91624   NEXN    nexilin F-actin binding protein 4436
0.690790339963014       6.18871329265694        13.458982815332

0.000243835918453485	0.0148496074338172
27111	SDCBP2	syndecan binding protein 2
2653	-0.72795386428946	5.57728747411777
13.4379793627479	0.000246581067317211	0.0149517792636891
101	ADAM8	ADAM metallopeptidase domain 8
4780	-1.05834586194776	6.71009729645405
13.3957208463134	0.000252198999124815	0.0152265145721607
3976	LIF	LIF interleukin 6 family cytokine
3969	-1.30910351798507	8.46306543759471
13.385491185292 0.000253578235154088	0.0152440787116022
201799	TMEM154 transmembrane protein 154
11587	-0.808518417068866	5.30399084347524
13.3271338312241	0.000261593058129989	0.0156586921590887
10516	FBLN5	fibulin 5	4973	1.45235636933214
2.89837749432711	13.1948128334892	0.00028072503567256
0.0166946008613366
2200	FBN1	fibrillin 1	16057	2.32306134555673
4.1448782067761 13.191094118098 0.000281282630347357
0.0166946008613366
9806	SPOCK2	SPARC (osteonectin), cwcv and kazal like domains proteoglycan 2	6569	1.13043333909307	5.15439934464626
13.090605913534 0.00029678034018784	0.0175400937764181
6699	SPRR1B	small proline rich protein 1B
620	-1.63178980856303	7.5025828773493 13.0442957035164
0.000304209536722868	0.0179036259700723
1292	COL6A2	collagen type VI alpha 2 chain	5386
2.02800501081777	5.13340120192225	12.9794810074845
0.000314923249435892	0.0184566106897429
4495	MT1G	metallothionein 1G	750	-2.00764207279082
-0.106947889603115	12.9623006576323
0.000317826373362486	0.0185491417153681
54952	TRNAU1AP	tRNA selenocysteine 1 associated protein 1
4311	0.562390555509352	4.81006643259613
12.9404737268349	0.000321553628526179	0.018569792213071
79628	SH3TC2	SH3 domain and tetratricopeptide repeats 2
15047	-1.01025154123422	4.32093582932372
12.9338078248425	0.000322700687239542	0.018569792213071
146223	CMTM4	CKLF like MARVEL transmembrane domain containing 4
3796	-0.514653443683289	6.23321035189136
12.9316357561449	0.000323075344217213	0.018569792213071
1544	CYP1A2	cytochrome P450 family 1 subfamily A member 2	2728
1.17242872031298	1.4093680231849 12.9282034021081
0.00032366828037083	0.018569792213071
857	CAV1	caveolin 1	4082	0.652387315023269
8.49432612526194	12.9216181017243	0.000324808959249117
0.018569792213071
3720	JARID2	jumonji and AT-rich interaction domain containing 2
6015	-0.549578578861835	5.77618749760062
12.8946806690513	0.000329517313183381	0.0187623943323562
64283	ARHGEF28	Rho guanine nucleotide exchange factor 28
15617	-0.840389924802509	6.87653048262287
12.8852842913024	0.000331175833161065	0.0187804854052107
1739	DLG1	discs large MAGUK scaffold protein 1
11001	-0.58493133463056	8.58760946824704

12.8748754693356	0.000333022885210207	0.0188090788433039
5549	PRELP	proline and arginine rich end leucine rich repeat protein 5747	2.68693145581563	-0.298821159967023
12.8598016327785	0.000335716178477762	0.0188758725739563
79627	OGFRL1	opioid growth factor receptor like 1
9051	-0.631714515886181	6.33440209297239
12.853210652871 0.000336900702754985	0.0188758725739563
57624	NYAP2	neuronal tyrosine-phosphorylated phosphoinositide-3-kinase adaptor 2	4828
1.69217746908895	-0.121437443446036	12.8277280732667
0.000341520191994791	0.0190042472698295
25960	ADGRA2	adhesion G protein-coupled receptor A2	7061
2.00942251519326	1.24821421843757	12.8256183304288
0.000341905498107877	0.0190042472698295
8611	PLPP1	phospholipid phosphatase 1	2016
0.894831826155897	5.4867011982029 12.7354350713265
0.000358792351404823	0.0198129448135985
100132994	CXorf49B	chromosome X open reading frame 49B
3512	-1.44197227864826	3.46908216995201
12.7328751486593	0.000359283785439709	0.0198129448135985
1282	COL4A1	collagen type IV alpha 1 chain	8188
0.964642120697475	8.64718320468568	12.6723044096439
0.000371111858467879	0.0203369242942817
2791	GNG11	G protein subunit gamma 11	3055
1.94906841210423	0.466991577706719	12.6693972224533
0.000371689342424226	0.0203369242942817
399668	SMIM10L2A	small integral membrane protein 10 like 2A
5230	1.39094887448329	2.15361092099983
12.6494588829881	0.000375674396801222	0.0204749855097071
3091	HIF1A	hypoxia inducible factor 1 subunit alpha	5151
0.706522895205367	9.55065544072964
12.6250689192917	0.000380607848386126	0.0206634656292421
6617	SNAPC1	small nuclear RNA activating complex polypeptide 1
2811	0.577083225109219	6.006599928904
12.5809507170915	0.00038969844307033	0.0210753130968576
5371	PML	promyelocytic leukemia	12657	-0.58036132791081
6.91983433721897	12.5427832187192
0.000397739435960142	0.0214274472288219
10974	ADIRF	adipogenesis regulatory factor
998	-1.03841672725584	6.147565997622	12.4969073417149
0.000407626249164371	0.0218611064129856
2065	ERBB3	erb-b2 receptor tyrosine kinase 3
8658	-0.496380092895878	7.25540067745159
12.4877184206362	0.000409636185660195	0.0218611064129856
2944	GSTM1	glutathione S-transferase mu 1	2512
1.86117668160982	1.25343152132024	12.4835864324863
0.000410543251116108	0.0218611064129856
7070	THY1	Thy-1 cell surface antigen	5078
3.31536934036245	2.26855777399413	12.476824645587
0.000412031990649546	0.0218611064129856
112399	EGLN3	egl-9 family hypoxia inducible factor 3 3444
0.797627772532704	5.69801752672977	12.4639445230169
0.000414882886929877	0.0219293003668935
134266	GRPEL2	GrpE like 2, mitochondrial

4419 -0.555756299852636 4.87264836164799
12.4542871315865 0.000417033509304659 0.0219601066347006
199713 NLRP7 NLR family pyrin domain containing 7 3914
1.81113753910431 -0.683853300974768 12.4437788643753
0.000419386394764875 0.0220012930017663
3676 ITGA4 integrin subunit alpha 4 7901
1.48800612372511 3.41846804802105 12.4038805785372
0.00042844256781458 0.0223925188335031
5317 PKP1 plakophilin 1 6211 -0.689056270716404
8.97073823186769 12.374244400825 0.00043529692905876
0.0226175196399988
10411 RAPGEF3 Rap guanine nucleotide exchange factor 3
12078 -1.0389653601153 3.55415152700176
12.3638416142888 0.000437729069890862 0.0226175196399988
100131096 TNRC6C-AS1 TNRC6C antisense RNA 1 3192
0.578491351967511 4.95708521060552 12.3601418026885
0.000438597376551773 0.0226175196399988
27018 BEX3 brain expressed X-linked 3 1080
1.08994891123249 4.9787739320313 12.3533914866869
0.000440186085006212 0.0226175196399988
85455 DISP2 dispatched RND transporter family member 2
12754 2.00356893459288 0.838025909669669
12.3506997063198 0.000440821222368792 0.0226175196399988
1953 MEGF6 multiple EGF like domains 6 7039
0.871262939980956 5.15509329588433 12.3385898494997
0.00044369005363434 0.0226816298585993
23764 MAFF MAF bZIP transcription factor F
2977 -0.758098097426438 5.7911702035747 12.3128530620169
0.000449849835924422 0.0228643697426903
80321 CEP70 centrosomal protein 70 4420 0.609741109770646
3.7850244379502 12.3077042470298 0.000451092462127123
0.0228643697426903
871 SERPINH1 serpin family H member 1 4914
0.662427094049081 7.09641342713568 12.2854993764902
0.000456491237242022 0.0228643697426903
54873 PALMD palmdelphin 6125 1.2473206696931
4.7797691974485 12.2844827989524 0.000456739957544968
0.0228643697426903
3620 IDO1 indoleamine 2,3-dioxygenase 1
3261 -2.82591618163088 3.24053948523789
12.2841333397193 0.000456825489348962 0.0228643697426903
100289019 SLC25A25-AS1 SLC25A25 antisense RNA 1 6391
-0.778984365155897 3.22131572847886
12.2831801655509 0.000457058865421096 0.0228643697426903
23508 TTC9 tetratricopeptide repeat domain 9
5217 -0.772662714477311 7.87433505293452
12.2664544379207 0.000461173642901788 0.0229881110894141
56938 ARNTL2 aryl hydrocarbon receptor nuclear translocator like
2 6983 -0.834263285815043 7.0024573407484
12.2584427943775 0.000463157847917154 0.023005148850268
4128 MAOA monoamine oxidase A 5528 -1.0057396871454
5.92312521848801 12.2403174161294
0.000467678725569447 0.0231297595778681
2135 EXTL2 exostosin like glycosyltransferase 2 4834

0.828672221027974	3.22538869295021	12.2351759602214
	0.00046896920968905	0.0231297595778681
23286	WWC1	WW and C2 domain containing 1
8086	-0.54960692104609	7.63323767473795
12.2210859980136	0.000472524180277562	0.0232233199759572
23048	FNBP1	formin binding protein 1
6600	-0.527831016297701	6.04034006917219
12.200391991058 0.000477794751584453	0.0234002485505015
389015	SLC9A4	solute carrier family 9 member A4
4478	-1.53410060602383	0.657877717328077
12.1645001729204	0.000487077158931287	0.0237717413419879
80206	FHOD3	formin homology 2 domain containing 3	7965
0.742743633416462	6.17691017099586	12.1150632943621
	0.000500161403760939	0.0243255582724982
1326	MAP3K8	mitogen-activated protein kinase kinase kinase 8
	3601	-1.34208948928842	4.2973322348173
12.1003352976284	0.000504127567127734	0.0244329956621546
8829	NRP1	neuropilin 1	11522	0.932548441293193
5.30475994931278	12.0881517082008	0.000507432519120029
	0.0244329956621546
5271	SERPINB8	serpin family B member 8
5310	-0.45517606868419	6.20628517995098
12.0845959971434	0.000508401166814069	0.0244329956621546
1839	HBEGF	heparin binding EGF like growth factor
3018	-1.15937325056222	6.82093382766339
12.0811276716812	0.000509347807049986	0.0244329956621546
5631	PRPS1	phosphoribosyl pyrophosphate synthetase 1	2280
	0.851683086551074	5.87350395524737
12.0596600071566	0.000515246881798415	0.0246115899495921
25802	LMOD1	leiomodin 1	3971	2.14863824163524
2.32683512979393	12.0548245620179	0.000516585096393238
	0.0246115899495921
7433	VIPR1	vasoactive intestinal peptide receptor 1	7384
	1.34664876778622	6.07661248070547
12.0273986990059	0.000524241890108519	0.0248216069577513
79888	LPCAT1	lysophosphatidylcholine acyltransferase 1	6494
	0.808450624456518	6.44743312621576
12.026348257319 0.000524537421253258	0.0248216069577513
79180	EFHD2	EF-hand domain family member D2
2531	-0.79518814792136	5.81889779776341
11.984168520419 0.000536544268825627	0.0253042948600692
23621	BACE1	beta-secretase 1	7190	0.68308481114776
	4.52409663030853	11.9580947032152
0.000544104803904195	0.0255747516385438
2512	FTL	ferritin light chain	878	1.13589449837113
	9.72297319355758	11.9198069851359
0.000555402349789258	0.0260184639247429
1290	COL5A2	collagen type V alpha 2 chain	7156
1.17157365962155	7.16985607979789	11.8882885590697
	0.000564880017359271	0.0263742480105043
4938	OAS1	2'-5'-oligoadenylate synthetase 1
4947	-1.17392664935179	4.05018901319785
11.8236618562266	0.000584827293682888	0.0272148701083595
64127	NOD2	nucleotide binding oligomerization domain containing

2	5747	-1.15716139585293	1.45203294463814
11.801327981144 0.000591884866016256	0.0274520904579129
79670	TUT7	terminal uridylyl transferase 7
6707	-0.454243852065114	5.9424798123673 11.7428084691095
	0.000610787864559297	0.0282353320755184
330	BIRC3	baculoviral IAP repeat containing 3
7518	-1.59049369271943	5.90484323033748
11.7362607137539	0.000612940496375828	0.0282416366208428
54507	ADAMTSL4	ADAMTS like 4	6432	1.18406459972354
	5.16087355766444	11.7094367825186
0.000621839380092103	0.0284008659577846
23129	PLXND1	plexin D1	9738	1.22042688709757
3.75484439389318	11.7085001925466	0.00062215244187484
	0.0284008659577846
84301	DDI2	DNA damage inducible 1 homolog 2
12136	-0.766152658163851	6.18071034395521
11.6967418544301	0.000626096318481281	0.0284008659577846
10317	B3GALT5 beta-1,3-galactosyltransferase 5
13688	-1.71682956800569	1.19922159557213
11.6916053178293	0.00062782708425274	0.0284008659577846
7368	UGT8	UDP glycosyltransferase 8	4596
1.44482744032455	1.06501033240702	11.6896934374059
	0.000628472529246735	0.0284008659577846
1277	COL1A1	collagen type I alpha 1 chain	9819
1.24898381224278	10.177735236014 11.6869053572645
0.000629414977972366	0.0284008659577846
9575	CLOCK	clock circadian regulator
11762	-0.621245268904661	6.73492614377094
11.683436160998 0.000630589656091311	0.0284008659577846
57188	ADAMTSL3	ADAMTS like 3	9754	-1.92610871381302
	2.46762911373062	11.6628075834523
0.000637620410492679	0.02862547785183
5420	PODXL	podocalyxin like	6874	-1.60835788615003
	6.45103952512784	11.6528760768615
0.000641033506190523	0.0286867614096187
10186	LHFPL6	LHFPL tetraspan subfamily member 6	2327
0.768060876811376	3.85430120385149	11.6253455683437
	0.0006505915207807	0.0289513243049353
280636	SELENOH selenoprotein H 2265	0.604674817819135
4.98680694641875	11.6206103602186	0.000652249934780702
	0.0289513243049353
129642	MBOAT2	membrane bound O-acyltransferase domain containing 2
	7751	-0.700587454619048	7.96615069709473
11.6180564162299	0.000653146175509357	0.0289513243049353
196883	ADCY4	adenylate cyclase 4	8194	-0.736792792353395
	3.20355498526176	11.608296204674 0.00065658274482115
	0.0290118438697472
2304	FOXE1	forkhead box E1 3462	-1.46525828593222
4.06367937784756	11.586843570017 0.000664200541692975
0.029244375347469
51458	RHCG	Rh family C glycoprotein
2607	-3.05634583266561	2.81731507818718
11.5797461507683	0.000666740427681292	0.029244375347469
4791	NFKB2	nuclear factor kappa B subunit 2

4484	-0.709206471646936	6.56633399352817
11.5759336732762	0.000668108810679667	0.029244375347469
3557	IL1RN	interleukin 1 receptor antagonist
2769	-1.61907182880962	8.71492634149089
11.5669708050195	0.000671336967706924	0.0292941336656414
6320	CLEC11A C-type lectin domain containing 11A	1469
1.52712592804547	0.319263908428758	11.5508141545355
0.000677195962137254	0.0294580243529706
84327	ZBED3	zinc finger BED-type containing 3	6514
0.940319171034198	2.50712914002093	11.544094496588
0.000679647939765091	0.0294731538460979
286887	KRT6C	keratin 6C	2896	-1.25783126211471
6.62538646424777	11.531107609595 0.000684412250126242
0.0295881555170317
84919	PPP1R15B	protein phosphatase 1 regulatory subunit 15B
5227	-0.448920172857605	6.48137079663382
11.5176906958376	0.000689369753775069	0.0297107758188535
8463	TEAD2	TEA domain transcription factor 2	2980
0.436845320217484	6.61535289691236	11.5032407722165
0.000694749532623376	0.0298447752707684
4481	MSR1	macrophage scavenger receptor 1 6488
3.62074908344951	3.5848959550244 11.4979222400902
0.000696740309383969	0.0298447752707684
9414	TJP2	tight junction protein 2
7306	-0.555025559519694	8.41946641949662
11.4726664909865	0.000706272719651395	0.030160859707796
9351	SLC9A3R2	SLC9A3 regulator 2
3550	-0.648802254594396	5.70364935691964
11.4626030554568	0.000710107625494147	0.0302061463976108
55103	RALGPS2 Ral GEF with PH domain and SH3 binding motif 2
9178	-0.594702415801154	5.31652380338023
11.4575017584033	0.000712059620747568	0.0302061463976108
729238	SFTPA2	surfactant protein A2	2685	3.6898922297117
2.459491795949	11.4529583565196	0.00071380270276356
0.0302061463976108
222865	TMEM130 transmembrane protein 130	4663
1.18243563254931	-0.422365450752564	11.4179989403803
0.000727359935394121	0.0306871404068236
266977	ADGRF1	adhesion G protein-coupled receptor F1
12401	-1.03535394663885	6.95247897448055
11.4064166892045	0.000731908658449755	0.0307451650134129
23704	KCNE4	potassium voltage-gated channel subfamily E
regulatory subunit 4	4124	2.45734601010474
0.444239385652105	11.4033312985407	0.000733125231275785
0.0307451650134129
3752	KCND3	potassium voltage-gated channel subfamily D member 3
7863	-1.09190746072814	1.4002708091209
11.3618734914717	0.000749671658206489	0.0312997711672267
1397	CRIP2	cysteine rich protein 2 4837	0.582777325923447
5.41332370166553	11.3590328994097
0.000750819098464208	0.0312997711672267
23529	CLCF1	cardiotrophin like cytokine factor 1
1975	-0.83910743524623	5.88085850918375
11.3366526971459	0.000759921738758668	0.0314102457706776

22808	MRAS	muscle RAS oncogene homolog	6365
0.83042772146852	5.51711552189062	11.3326333551276
	0.000761568288037103	0.0314102457706776
10769	PLK2	polo like kinase 2	4768	-0.580129693221558
	8.92839372597866	11.3261686453877
0.000764224163335193	0.0314102457706776
23057	NMNAT2	nicotinamide nucleotide adenylyltransferase 2	7165
	1.24461951191212	1.88590945360616
11.3190866493362	0.000767144379843246	0.0314102457706776
7286	TUFT1	tuftelin 1	3878	-0.673027760186275
7.40851188805948	11.3188807910728	0.000767229432633919
	0.0314102457706776
55741	EDEM2	ER degradation enhancing alpha-mannosidase like
protein 2	1919	0.448858089966991	5.52367543391907
	11.318507540323 0.00076738366992306	0.0314102457706776
3486	IGFBP3	insulin like growth factor binding protein 3
4571	-1.32673512832855	9.47299188194116
11.3104100398513	0.000770737499591253	0.0314102457706776
7039	TGFA	transforming growth factor alpha
4963	-0.854607110368319	8.5577075080285 11.3016189417739
	0.000774395375675439	0.0314102457706776
81607	NECTIN4 nectin cell adhesion molecule 4
3502	-0.728021290595388	6.81728337355215
11.2981772355834	0.000775832204284663	0.0314102457706776
5922	RASA2	RAS p21 protein activator 2
5993	-0.894898229360405	5.33881437009484
11.2980297372557	0.000775893841411755	0.0314102457706776
54502	RBM47	RNA binding motif protein 47
8544	-0.605101581463538	6.38524064949593
11.2906323163531	0.000778991457677396	0.0314447646907415
23135	KDM6B	lysine demethylase 6B	7258	-0.697749788437174
	6.24242228247585	11.2755844213272
0.000785331276070648	0.0316095838618436
79646	PANK3	pantothenate kinase 3	10580	-0.601655073585136
	7.80829951692761	11.2642762506892
0.000790129806133267	0.031689116333151
4645	MYO5B	myosin VB	10873	-0.652036213177184
8.21211247041771	11.2597609693125	0.000792054096504752
	0.031689116333151
6318	SERPINB4	serpin family B member 4
2013	-3.04156486600095	1.68556311783855
11.254080838686 0.000794481541743274	0.031689116333151
30846	EHD2	EH domain containing 2	3923	0.574538745236027
	7.88365638081821	11.2497049241386
0.000796356746574509	0.031689116333151
23062	GGA2	golgi associated, gamma adaptin ear containing, ARF
binding protein 2	8231	0.484272241666511
6.60148299904925	11.2361345940732	0.000802200513907316
	0.0317643918650124
3714	JAG2	jagged canonical Notch ligand 2 6463
0.597429007427184	5.88939004086769	11.2347852454123
	0.000802783945185578	0.0317643918650124
58504	ARHGAP22	Rho GTPase activating protein 22	6783
	0.907115093305458	3.25060059249942

11.2237757353179	0.000807560287862248	0.0317795052815179
8876	VNN1	vanin 1 3106	-2.38897777826404
1.93106398772229	11.2224071907798	0.000808156017784659
	0.0317795052815179
22854	NTNG1	netrin G1	9138	-1.15464514646594
2.11840331596057	11.2182407826291	0.000809972398479466
	0.0317795052815179
1849	DUSP7	dual specificity phosphatase 7
5648	-0.655000382508113	6.54288701331964
11.2072697755301	0.000814775062130179	0.0318530126840423
10162	LPCAT3	lysophosphatidylcholine acyltransferase 3	4725
	0.54810368745365	5.24995070542317
11.1997130644836	0.000818099814073987	0.0318530126840423
84256	FLYWCH1 FLYWCH-type zinc finger 1	8419
0.57369283178107	5.67594233295108	11.1984244548528
	0.000818668134950755	0.0318530126840423
124976	SPNS2	sphingolipid transporter 2
4111	-1.57672228600441	5.32685933406938
11.1848777151527	0.000824666910524576	0.0319975330075284
10608	MXD4	MAX dimerization protein 4	7058
0.596619262540425	6.1481905997533 11.1550321856955
0.000838040364632951	0.0324266060425794
219833	C11orf45	chromosome 11 open reading frame 45	3750
	0.878906943077509	2.8614699727345 11.1147064226112
	0.000856458803052105	0.0330479847227296
25791	NGEF	neuronal guanine nucleotide exchange factor
4678	-1.92325023872125	1.03239539985656
11.0949776858583	0.000865618254360078	0.0333096562879714
2117	ETV3	ETS variant transcription factor 3
6707	-0.461305658959461	5.6834411437269 11.0830260371839
	0.000871215137966281	0.0334207234778779
5794	PTPRH	protein tyrosine phosphatase receptor type H
4344	-1.89468368625117	2.52695645523824
11.0786435975042	0.000873276561212487	0.0334207234778779
55501	CHST12	carbohydrate sulfotransferase 12	16143
0.624114961719445	4.72034061214169	11.0621547930611
	0.000881076881455701	0.0335548489350634
79974	CPED1	cadherin like and PC-esterase domain containing 1
	7705	1.84350613168236	0.525321925469294
11.0581337927339	0.000882989745254945	0.0335548489350634
2591	GALNT3	polypeptide N-acetylgalactosaminyltransferase 3
4686	-0.728388300066327	6.74987787197027
11.0560808986572	0.00088396796294984	0.0335548489350634
3310	HSPA6	heat shock protein family A (Hsp70) member 6
2371	-1.90441892865542	1.00313809763033
11.0247161902128	0.000899050424548387	0.0340351332341872
4477	MSMB	microseminoprotein beta 492	-3.13252841095032
	2.51504413373716	10.9911096429441
0.000915499811396275	0.0345326601880362
6744	ITPRID2 ITPR interacting domain containing 2
9582	-1.21861215360406	8.29246064443967
10.9835784187799	0.000919227658972317	0.0345326601880362
116496	NIBAN1	niban apoptosis regulator 1
8372	-1.22816914275613	5.07371884004926

10.9828501835019	0.000919588937683838	0.0345326601880362
1830	DSG3	desmoglein 3	5525	-1.1409126988199
8.09954909373812	10.95782019731	0.000932094007762003
0.0349086651516641
100126784	LOC100126784	uncharacterized LOC100126784	1686
	1.42505180903816	2.4710807671172	10.9465085731437
	0.000937801607066737	0.0349768006609683
152007	GLIPR2	GLI pathogenesis related 2
3055	-0.930418077473776	4.49455030312586
10.9443249500907	0.000938907478298284	0.0349768006609683
83464	APH1B	aph-1 homolog B, gamma-secretase subunit	5165
	0.553505247405538	4.26287230659381
10.9261080857906	0.000948184737980484	0.0352287098803518
220	ALDH1A3	aldehyde dehydrogenase 1 family member A3
4771	-0.510537309320366	12.527218901267	10.9107871732858
	0.000956058907412299	0.0354272939580002
64332	NFKBIZ	NFKB inhibitor zeta	5902	-0.831093262365942
	6.34033005197601	10.9017825749007
0.000960717629123258	0.0355059942773865
5010	CLDN11	claudin 11	4150	1.27640430028654
0.467445839291642	10.8727563823451	0.000975891791631135
	0.0359718850667824
1672	DEFB1	defensin beta 1	478	-2.1301115188962
0.935596397322972	10.8671314851207	0.000978860262860158
	0.0359866028920793
84436	ZNF528	zinc finger protein 528	7868	-0.68720594858607
	4.31248084203655	10.8606340616786
0.000982300566399152	0.0360185445904527
7124	TNF	tumor necrosis factor	1676	-1.45008543232244
	3.97403219551427	10.8338304044284
0.000996622512447066	0.0364482807620002
64225	ATL2	atlastin GTPase 2	4884	-0.498123645568091
	5.96340985253112	10.8079940995952
0.00101062744247476	0.0368642150696456
163732	CITED4	Cbp/p300 interacting transactivator with Glu/Asp rich carboxy-terminal domain 4	1316	-0.839889900958189
3.59224382593048	10.7878426680515	0.00102168883970267
	0.0371708976044554
10006	ABI1	abl interactor 1	4218	-0.463912595935671
	7.48039222941307	10.7764217214465
0.00102801223450559	0.0373040605407249
4973	OLR1	oxidized low density lipoprotein receptor 1	3057
	2.90664433012198	2.26312769578352
10.722686569504	0.00105829901633378	0.0382217655009495
50854	SNHG32	small nucleolar RNA host gene 32	1801
0.498130661993691	5.64198205044597	10.7218815448123
	0.00105875954982283	0.0382217655009495
2982	GUCY1A1	guanylate cyclase 1 soluble subunit alpha 1	5766
	1.39121293380286	2.52827774162079
10.667711604661	0.00109021911343913	0.0392321191863512
57198	ATP8B2	ATPase phospholipid transporting 8B2	7572
0.828847822653942	4.93757819326369	10.6641022403827
	0.0010923485744754	0.0392321191863512
57475	PLEKHH1	pleckstrin homology, MyTH4 and FERM domain

containing H1	10817	-0.752466334664307	4.00546082684182
	10.6460859070373	0.00110304097781736
0.0393341130514471
131177	FAM3D	family with sequence similarity 3 member D
1640	-2.10914182536439	0.243018623614396
10.6444752755668	0.00110400200217355	0.0393341130514471
10395	DLC1	DLC1 Rho GTPase activating protein	10868
1.22820817900671	3.83726063745361	10.6372822761652
	0.0011083042374697	0.0393341130514471
3913	LAMB2	laminin subunit beta 2	6865	0.458006612247556
	8.50701451573907	10.6357226972248
0.00110923927889077	0.0393341130514471
1577	CYP3A5	cytochrome P450 family 3 subfamily A member 5
6731	-1.3130505128792	0.0223418056054831
10.6356731053532	0.0011092690246538	0.0393341130514471
51313	GASK1B	golgi associated kinase 1B	7612
1.6397263229392 5.33517907785886	10.6280406937564
0.00111385666343609	0.0393341130514471
26585	GREM1	gremlin 1, DAN family BMP antagonist	14606
5.96580187423699	4.81526002209773	10.6263995095
0.00111484564013882	0.0393341130514471
27071	DAPP1	dual adaptor of phosphotyrosine and 3-
phosphoinositides 1	4548	-0.853734294951017
7.7084704687473 10.6102323205373	0.00112463558376976
0.0395798256830729
51450	PRRX2	paired related homeobox 2
1311	-1.64452986067489	1.06117944092622
10.5880439586561	0.00113821337521086	0.0399572800666129
5013	OTX1	orthodenticle homeobox 1
4833	-1.35674874633771	3.02384212880592
10.5825314536522	0.00114161230251701	0.0399764088033894
2303	FOXC2	forkhead box C2 2478	-1.31960958955639
0.939344570700911	10.5690827251012	0.00114994773615293
	0.040167875162828
4299	AFF1	AF4/FMR2 family member 1
10733	-0.733886359923579	7.84808807253042
10.5441237718352	0.00116558067008792	0.0406126578256754
6234	RPS28	ribosomal protein S28	2119	0.598145202283922
	4.37911341983778	10.5250999031719
0.00117764045974683	0.0409310419843023
22891	ZNF365	zinc finger protein 365 9073	-1.50966091183822
	2.89359722117474	10.5178575799104
0.00118226469361133	0.0409900533747868
4208	MEF2C	myocyte enhancer factor 2C	10414
1.37947588496615	0.646092806306348	10.5101980363544
	0.00118717531655422	0.0410186775542717
53373	TPCN1	two pore segment channel 1
8728	-0.741781922128396	6.01008703658857
10.4989845031481	0.0011944016939865	0.0410186775542717
57545	CC2D2A	coiled-coil and C2 domain containing 2A 7606
0.882538014987551	3.1194771521841 10.4988320678512
0.00119450023444272	0.0410186775542717
8406	SRPX	sushi repeat containing protein X-linked	1987
	0.995284766962354	2.29336749372106

10.4983621536328	0.001194804058124	0.0410186775542717
114569	MAL2	mal, T cell differentiation protein 2 (gene/
pseudogene)	3430	-0.615068385117732	8.94438938154493
10.4760315721262	0.00120933244083593
0.0414159400948382
2012	EMP1	epithelial membrane protein 1
7838	-1.47949820844909	8.31853787178005
10.4453261596939	0.00122960192560565	0.0420074004194105
57136	APMAP	adipocyte plasma membrane associated protein	2454
0.472963904883873	6.55213627116524
10.427474385241	0.00124154407022791	0.0423121844079862
7042	TGFB2	transforming growth factor beta 2
5151	-0.642915499637555	8.32791007917383
10.4153620226256	0.00124971354465148	0.0424872272328477
57106	NAT14	N-acetyltransferase 14 (putative)	2163
0.686744132260504	3.64464321050992	10.3769697149923
0.00127596945518068	0.043195110681344
9314	KLF4	Kruppel like factor 4	3659	-0.759564441595063
5.70166447421072	10.3750181602932
0.00127731890737825	0.043195110681344
6707	SPRR3	small proline rich protein 3
1441	-2.32034998355085	4.38570421098906
10.3714548135679	0.00127978660189603	0.043195110681344
51274	KLF3	Kruppel like factor 3	6297	-0.58597051028513
6.6577964742707	10.3523324906938	0.00129311188821608
0.0435071054611737
9244	CRLF1	cytokine receptor like factor 1	3018
2.26904370499876	2.62670767386411	10.3492933519376
0.00129524259136928	0.0435071054611737
10308	ZNF267	zinc finger protein 267	3711	-0.611997441745022
3.97739729565963	10.3435194331643
0.00129930040925128	0.043538997206657
6604	SMARCD3	SWI/SNF related, matrix associated, actin dependent
regulator of chromatin, subfamily d, member 3	5198
0.943726312544256	2.7318073101381	10.3102056161274
0.00132296541932049	0.0442261972038714
8870	IER3	immediate early response 3
1244	-0.615605942802225	9.38826823730264
10.2753355033412	0.00134820355486842	0.044962588554862
89872	AQP10	aquaporin 10	2292	1.89400178888244
0.103447333920343	10.2694795938265	0.00135248946968608
0.0449983848025959
51561	IL23A	interleukin 23 subunit alpha
1187	-0.883420037572228	3.66344655201285
10.2582852839806	0.00136072096045077	0.0450979571969298
3490	IGFBP7	insulin like growth factor binding protein 7	1930
0.635640195485382	8.73002622942928
10.2469579783163	0.00136910187535824	0.0450979571969298
8851	CDK5R1	cyclin dependent kinase 5 regulatory subunit 1
4325	-0.869367435273539	4.16899130056648
10.2460451800396	0.0013697795130102	0.0450979571969298
80271	ITPKC	inositol-trisphosphate 3-kinase C
3385	-0.445526787280877	6.82303477565191
10.2421736272397	0.00137265742917352	0.0450979571969298

2185	PTK2B	protein tyrosine kinase 2 beta
7693	-0.835776071123158	5.1856668781902 10.2407049221297
	0.00137375079099441	0.0450979571969298
157693	FAM87A	family with sequence similarity 87 member A	4013
	1.28868725273556	0.820601936316443
10.2392962444828	0.00137480029435918	0.0450979571969298
2524	FUT2	fucosyltransferase 2	3380	-0.843837318963793
	6.26129104113164	10.2240545725698
0.00138620779492335	0.0453659172511479
6578	SLCO2A1 solute carrier organic anion transporter family
member 2A1	5283	1.78251834181632	7.15100570601482
	10.2139716196069	0.00139380691650892
0.0455082831690919
113189	CHST14	carbohydrate sulfotransferase 14	3574
0.58289691453236	4.51684044575545	10.2059890059438
	0.00139985299836325	0.0455827356100392
8204	NRIP1	nuclear receptor interacting protein 1
13782	-0.518359448794115	6.75362915362938
10.2023792305117	0.00140259577696344	0.0455827356100392
1291	COL6A1	collagen type VI alpha 1 chain	5343
1.23834737640651	5.58502685773375	10.179382862563
0.00142019707421672	0.0460335265098688
83742	MARVELD1	MARVEL domain containing 1	3379
0.51100763922759	6.38700238863836	10.1746744320962
	0.00142382837446708	0.0460335265098688
8809	IL18R1	interleukin 18 receptor 1
4353	-1.33267360769375	2.29074855035916
10.1714427979791	0.00142632615872657	0.0460335265098688
6439	SFTPB	surfactant protein B	4110	4.82929852674052
	4.51701040608097	10.1469337208171
0.00144541463011812	0.0465423510898034
254427	PROSER2 proline and serine rich 2
3663	-0.706755821061936	5.69012899008361
10.1370902572614	0.00145315366065244	0.0466521104854334
1796	DOK1	docking protein 1	3049	1.0539804153554
2.59071564450913	10.1317203449041	0.00145739321433874
	0.0466521104854334
29094	LGALSL	galectin like	4003	-1.10608330980991
6.77963043454657	10.1299227785267	0.00145881519187691
	0.0466521104854334
4664	NAB1	NGFI-A binding protein 1
4509	-0.496099540341522	4.77195105851072
10.1191702854013	0.00146735041238022	0.0468181713581086
3426	CFI	complement factor I	3875	1.33628111084851
	3.0842527253144 10.1068962474731	0.00147715524928006
	0.0470238944924223
10695	CNPY3	canopy FGF signaling regulator 3	1712
0.451444402032387	5.06962003986479	10.0929990888963
	0.00148833679437735	0.0472724115166521
55914	ERBIN	erbb2 interacting protein
10216	-0.561377256886442	7.5763168242563 10.0796107483284
	0.00149919001125774	0.0474966227184336
28514	DLL1	delta like canonical Notch ligand 1	3226
0.85031208138354	3.43012326028844	10.0759423418014

0.00150217775856829	0.0474966227184336
744	MPPED2	metallophosphoesterase domain containing 2	8197
	1.58219411386688	0.618078302146243
10.0695112176036	0.00150743017119023	0.047555347765454
202374	STK32A	serine/threonine kinase 32A	7776
0.687687322600699	4.27447516601558	10.0596698259118
	0.00151550384135332	0.0477026119232268
5820	PVT1	Pvt1 oncogene	5493	0.679509576632876
3.24874493008188	10.0395906744436	0.00153211244571164
	0.0481172623925627
83604	TMEM47	transmembrane protein 47	4054
0.997433371827085	5.1426333420894 10.0318896040868
0.00153853124064088	0.048210754111089
157506	RDH10	retinol dehydrogenase 10
5321	-0.739029976672008	6.11719355206897
10.0186134545204	0.00154966084777099	0.0484511149435898
7086	TKT	transketolase	9507	0.538016503876059
7.65151896034212	10.012113683586 0.00155513940782373
0.0485141151122204
326624	RAB37	RAB37, member RAS oncogene family
5074	-1.37435176413653	1.69642111564759
10.0063422682121	0.00156002048926039	0.0485582377623784
57205	ATP10D	ATPase phospholipid transporting 10D (putative) 7965
	0.720063048491205	5.97501584955302
9.97684695786599	0.00158520904531915	0.0492328671791248
10170	DHRS9	dehydrogenase/reductase 9
3750	-2.72105593752196	2.85135511734391
9.95761773815754	0.00160185196399025	0.0496396912823263
342897	NCCRP1	NCCRP1, F-box associated domain containing
1978	-1.47245016000142	5.63575338891731
9.95295131768468	0.00160591737678173	0.0496558161072442
196374	KRT78	keratin 78	3260	-2.30796773263641
1.71407468285667	9.93750874612054	0.00161944571145249
	0.0499638239654517
9902	MRC2	mannose receptor C type 2	6860
0.80795486108936	6.62907529269615	9.90632213695983
	0.00164711954475412	0.0506736295144399
284194	LGALS9B galectin 9B	2670	-1.83300997178464
1.42131853943175	9.90345499973764	0.00164968766035444
	0.0506736295144399
65251	ZNF649	zinc finger protein 649 4365	-0.807958472420374
	2.84578303097856	9.87346815983986
0.00167679140317017	0.0513934730507758
148170	CDC42EP5	CDC42 effector protein 5
903	-0.796439187749375	3.03494047222317
9.85875292678456	0.00169025641523633	0.0516589406308891
6666	SOX12	SRY-box transcription factor 12 4824
1.18874492838503	2.49370580843106	9.85595515526303
	0.00169282885340031	0.0516589406308891
222962	SLC29A4 solute carrier family 29 member 4	5811
0.922815647422936	1.12233302806094	9.85136846650727
	0.00169705471385807	0.0516753160369781
51430	SUCO	SUN domain containing ossification factor
7894	-0.409325988296496	6.46684189739757

9.84549994331219	0.00170247715909126	0.0517279773696123
2946	GSTM2	glutathione S-transferase mu 2	5864
1.25945821843127	2.51590996273337	9.82832406519478
	0.00171844855841871	0.0521002358393309
9945	GFPT2	glutamine-fructose-6-phosphate transaminase 2	4780
	2.01361039329306	0.980397335138861
9.7964645920289	0.00174847697793881	0.0528137610576008
5592	PRKG1	protein kinase cGMP-dependent 1	7507
1.8783240662491	0.787498018892988	9.7953635677865
0.00174952417582115	0.0528137610576008
1514	CTSL	cathepsin L	2366	0.779250424382446
5.65895389778096	9.77153892322654	0.00177234042134884
	0.0533874672727596
114786	XKR4	XK related 4	19885	1.69443426854791
0.322083825861523	9.76060953599387	0.00178290787770804
	0.0535903832919795
6820	SULT2B1	sulfotransferase family 2B member 1
2822	-1.59545608310104	3.37422244015195
9.75335102035372	0.00178996128750143	0.0535903832919795
10085	EDIL3	EGF like repeats and discoidin domains 3	5738
	0.868530281141582	6.71565130889636
9.75274169664002	0.00179055467842125	0.0535903832919795
3635	INPP5D	inositol polyphosphate-5-phosphatase D	8229
1.09317508214907	6.31289850121459	9.740441168495
0.00180257629445184	0.0537359646151239
25805	BAMBI	BMP and activin membrane bound inhibitor
2314	-0.994475457339587	3.94783754229483
9.73860164187608	0.00180438112798707	0.0537359646151239
9001	HAP1	huntingtin associated protein 1
6223	-1.08692104990511	1.48343134319394
9.73600906731918	0.00180692791702173	0.0537359646151239
3925	STMN1	stathmin 1	6261	0.668717276761601
7.60388651316655	9.72852065205154	0.00181430456191732
	0.0538410254211354
51170	HSD17B11	hydroxysteroid 17-beta dehydrogenase 11	2435
	1.12964041206194	4.71026927062468
9.69702530222077	0.00184566546669318	0.0546558904692841
688	KLF5	Kruppel like factor 5	3893	-0.615086017215768
	7.19291084864153	9.68794612641697
0.00185480750080771	0.0547109582584507
1040	CDS1	CDP-diacylglycerol synthase 1
4461	-0.43237283240464	6.75815770759523
9.68742167152725	0.00185533698670408	0.0547109582584507
10105	PPIF	peptidylprolyl isomerase F
2674	-0.719002963648964	5.3497558668267	9.6737809351161
0.00186916252006524	0.0550028559213315
4130	MAP1A	microtubule associated protein 1A	10629
1.19037214154251	2.07539676995582	9.66546777631088
	0.00187763946727571	0.055096019275981
51309	ARMCX1	armadillo repeat containing X-linked 1	2141
0.660622783443562	4.06000840755684	9.66296867426065
	0.00188019541757114	0.055096019275981
642475	MROH6	maestro heat like repeat family member 6
4408	-0.846485306751941	3.88274456792722

9.64408349324373	0.0018996245619568	0.0555491466374298
1509	CTSD	cathepsin D	3313	0.691716034129754
9.33475159282403	9.61670730774409	0.001928151250157
	0.0562311351161321
122402	TDRD9	tudor domain containing 9	5480
2.01058790838007	0.262935404450088	9.6135483313565
0.0019314708099303	0.0562311351161321
5151	PDE8A	phosphodiesterase 8A	7886	-0.509379046810123
	5.34439842034485	9.60784851237683
0.00193747503813966	0.0562311351161321
55315	SLC29A3 solute carrier family 29 member 3	2684
0.652907901559763	3.7909715063917 9.60334622332737
0.00194223115369403	0.0562311351161321
26525	IL36RN	interleukin 36 receptor antagonist
2831	-2.59503664201342	1.73039628270915
9.59892586499259	0.00194691223450404	0.0562311351161321
5774	PTPN3	protein tyrosine phosphatase non-receptor type 3
	9846	-0.421560600016351	6.81900346304755
9.59539594916231	0.00195065856498283	0.0562311351161321
80342	TRAF3IP3	TRAF3 interacting protein 3
7750	-1.15862529507833	0.751981342482155
9.59502924851332	0.00195104816637683	0.0562311351161321
28983	TMPRSS11E	transmembrane serine protease 11E	2138
	-1.62876971065345	2.22928339368839
9.59063256864507	0.00195572557013055	0.056250201356917
8061	FOSL1	FOS like 1, AP-1 transcription factor subunit
1887	-0.51171349304285	4.73692830851848
9.58432347621294	0.00196245736057892	0.0563281562492395
1029	CDKN2A	cyclin dependent kinase inhibitor 2A	3493
1.83258728012977	3.00771973558642	9.57960949310223
	0.00196750250152988	0.0563574796297118
224	ALDH3A2 aldehyde dehydrogenase 3 family member A2	4987
	0.39147631732237	5.80727680991058
9.5689975903191 0.00197890806555795	0.0565685005597351
2535	FZD2	frizzled class receptor 2	2112
1.57154092212257	2.8449805543591 9.55633130157197
0.00199260945758148	0.0568441561554862
79875	THSD4	thrombospondin type 1 domain containing 4
15837	0.678730443180327	5.88974500610062
9.53650540889155	0.0020142489160324	0.0573446840789955
79444	BIRC7	baculoviral IAP repeat containing 7	1562
1.50342860697452	1.5706101000187 9.52306063938539
0.00202905897700491	0.0575028472965649
3454	IFNAR1	interferon alpha and beta receptor subunit 1
6301	-0.342639067920532	6.98028396359822
9.52038750089276	0.00203201670109655	0.0575028472965649
3853	KRT6A	keratin 6A	2928	-1.16079535535733
11.220041174738 9.5198435238708 0.00203261912569465
0.0575028472965649
54587	MXRA8	matrix remodeling associated 8	3622
1.09492638806919	4.21570465194172	9.51410604558995
	0.00203898410549113	0.0575028472965649
570	BAAT	bile acid-CoA:amino acid N-acyltransferase	3389
	2.64868403089703	2.36164986349488

9.51271247559843	0.00204053313681343	0.0575028472965649
115362	GBP5	guanylate binding protein 5
8143	-2.09822211125793	1.37326773724413
9.50920586980757	0.00204443620715994	0.0575028472965649
285386	TPRG1	tumor protein p63 regulated 1
8605	-1.3457058208285	1.92749490107637
9.5040753099545 0.00205016048440612	0.0575482923949428
9507	ADAMTS4 ADAM metallopeptidase with thrombospondin type 1
motif 4 10334	2.45550680082011	0.590258089357929
9.49631064037826	0.0020588546213697	0.0576738844551777
6788	STK3	serine/threonine kinase 3
11906	-0.457167275180506	5.14391047055753
9.49273600479274	0.00206286971600229	0.0576738844551777
51703	ACSL5	acyl-CoA synthetase long chain family member 5	4236
0.987817880567008	3.52042257334738
9.48077619295408	0.00207636100476582	0.0579354354457268
80111	C3orf36 chromosome 3 open reading frame 36	1668
2.20002335922399	2.77254280557027	9.4739120795343
0.00208414445810016	0.0580370008441529
5724	PTAFR	platelet activating factor receptor
4706	-0.98321932226258	5.83066942955982
9.46772812965719	0.00209118198378667	0.0580938030638041
11009	IL24	interleukin 24	1979	-2.43254371947462
1.02253219683606	9.46483806201077	0.00209447922804463
0.0580938030638041
284069	FAM171A2	family with sequence similarity 171 member
A2	3592	1.03184994577651	1.68160827229064
9.45017476539202	0.00211128982556643	0.0583211276371498
4681	NBL1	NBL1, DAN family BMP antagonist
3465	-0.421787983310511	6.58975514568512
9.44752667628978	0.00211434025751806	0.0583211276371498
6928	HNF1B	HNF1 homeobox B 5962	1.57442409603394
3.63377114354546	9.44681034734656	0.00211516619116671
0.0583211276371498
4051	CYP4F3	cytochrome P450 family 4 subfamily F member 3
5860	-1.24307074955372	3.16041449946215
9.43114827821298	0.00213330671201805	0.0587057507175576
165	AEBP1	AE binding protein 1	4955	2.85850294389137
3.11623583605417	9.42664186214642
0.00213855542126734	0.058734795658219
8581	LY6D	lymphocyte antigen 6 family member D
1049	-2.19332940606294	5.03808247512611
9.40695839921085	0.00216163510820367	0.0592524911166513
3934	LCN2	lipocalin 2	1598	-1.89692039835745
8.9496235690684 9.39162400723058	0.0021797902752204
0.0596054068137915
55715	DOK4	docking protein 4	4788	-0.450765940061415
5.71692975840972	9.38890891031723
0.00218302089637146	0.0596054068137915
10802	SEC24A	SEC24 homolog A, COPII coat complex component
7201	-0.51230367385289	5.99808888387244
9.37528790358105	0.00219930161403942	0.0599331083810314
3814	KISS1	KiSS-1 metastasis suppressor
709	-1.83413453243361	1.4755063319498 9.37049381947159

0.00220506107281886	0.059973379508687
221416	C6orf223	chromosome 6 open reading frame 223
3917	-1.21648611946017	0.949259415511466
9.36222238511257	0.00221503410179554	0.0601278733795546
55930	MYO5C	myosin VC	10045	-0.63979306969542
6.56480354902482	9.35730218985407	0.00222098816683603
0.0601728844349559
873	CBR1	carbonyl reductase 1	2450	0.47038884295613
5.08004253945996	9.35036274084938
0.0022294133940263	0.0602845432627923
619208	FAM229B	family with sequence similarity 229 member B	2543
0.94802529333243	1.90984131475165
9.33426542401471	0.00224908229357693	0.0606992209752062
3910	LAMA4	laminin subunit alpha 4	11575	2.13926924548977
3.81393948587245	9.31945983442871
0.00226732825423023	0.0609899604269992
2214	FCGR3A	Fc fragment of IgG receptor IIIa	2883
2.87025117923858	2.13055479160865	9.31846184790797
0.0022685635312677	0.0609899604269992
51099	ABHD5	abhydrolase domain containing 5
3172	-0.431718484780974	5.34908508376761
9.30484088591177	0.00228549155031181	0.0613273566000337
64398	MPP5	membrane palmitoylated protein 5
8743	-0.432115970760446	6.53193687988265
9.30063155376166	0.00229074875038949	0.0613508943531655
2246	FGF1	fibroblast growth factor 1
5275	-0.849230295117164	4.44996827833533
9.29570030159318	0.00229692319205297	0.0613988609753549
100505812	CARD8-AS1	CARD8 antisense RNA 1	1969
0.948707593407539	2.05597290370466	9.28829081654125
0.00230623239947552	0.0614375523464687
7059	THBS3	thrombospondin 3	4623	0.537420672180959
4.92118089248334	9.28756764115105
0.00230714303806972	0.0614375523464687
6776	STAT5A	signal transducer and activator of transcription 5A
5754	-0.591830082022776	4.29712310952198
9.27928819955362	0.00231759472894182	0.0615987654047211
102524628	RASGRF2-AS1	RASGRF2 antisense RNA 1	1166
1.43888277528263	0.14718764977654	9.26989465978484
0.00232951100374543	0.061711542223774
374462	PTPRQ	protein tyrosine phosphatase receptor type Q
11619	1.98095818476237	-0.00970692517099546
9.26899983753548	0.00233064937790936	0.061711542223774
3786	KCNQ3	potassium voltage-gated channel subfamily Q member 3
12478	1.53398869542755	1.49075226875293
9.23138594516284	0.002379015035051	0.0628733275395459
6248	RSC1A1	regulator of solute carriers 1
1854	-0.698951610656553	4.01555877918845
9.19330056909192	0.00242902500064802	0.0640665948756753
84900	RNFT2	ring finger protein, transmembrane 2	6487
1.03712752698075	-0.10688156569114	9.19007142470344
0.00243331394830151	0.0640665948756753
90007	MIDN	midnolin	4492	-0.452461162537181
6.2663637989601	9.18155450301914	0.00244466302133322

0.064244643414286
2035	EPB41	erythrocyte membrane protein band 4.1
8287	-0.463483266495121	5.59725192881548
9.16616202859463	0.00246531044293883	0.0644216634486987
25803	SPDEF	SAM pointed domain containing ETS transcription
factor	1914	-1.10375922512819	3.93876988964559
9.16509088456642	0.00246675384183796	0.0644216634486987
9368	SLC9A3R1	SLC9A3 regulator 1
2315	-0.858952969628487	5.87318821802195
9.16445155482884	0.00246761576663505	0.0644216634486987
7465	WEE1	WEE1 G2 checkpoint kinase
9239	-0.71023946216932	5.35059173009937
9.16283534836651	0.00246979604997153	0.0644216634486987
7483	WNT9A	Wnt family member 9A	3971	-0.579057748986558
5.58002684430684	9.15855911351352
0.00247557418018581	0.064452356025767
121457	IKBIP	IKBKB interacting protein	4163
0.654498046532299	4.66651156536413	9.15149074754384
0.00248515518826711	0.0645369884272875
255239	ANKK1	ankyrin repeat and kinase domain containing 1
2598	-0.968943775351667	0.875233043106523
9.14936803379875	0.00248803981942852	0.0645369884272875
5277	PIGA	phosphatidylinositol glycan anchor biosynthesis
class A	4889	-0.602535635865522	4.32103453989671
9.13021951674551	0.00251421550670804	0.0650954096163761
64106	NPFFR1	neuropeptide FF receptor 1	8921
1.22396491339935	1.04842622393293	9.11079300404068
0.00254105688495058	0.0656689737776805
55601	DDX60	DExD/H-box helicase 60	6750	-0.730613771720084
4.33286499244856	9.0995133120743	0.00255677518279824
0.0659534990524032
157285	PRAG1	PEAK1 related, kinase-activating pseudokinase 1
4904	-0.429016424121417	6.50237259021139
9.09216284147013	0.00256707117050137	0.0660973637595821
9537	TP53I11	tumor protein p53 inducible protein 11	7345
0.761764724103422	5.29250089979723	9.08315614297092
0.00257974447714998	0.0663017998008069
9182	RASSF9	Ras association domain family member 9
5682	-0.807221317514315	4.22847325222361
9.07333073712186	0.00259364223323161	0.0665368988294417
11117	EMILIN1	elastin microfibril interfacer 1	3958
2.57831084746863	2.06002519136904	9.03070189646575
0.00265482525089144	0.067981969450158
51338	MS4A4A	membrane spanning 4-domains A4A	2508
2.23106650959318	0.453254948202919	9.01985367409213
0.00267062771658176	0.068191951486997
713	C1QB	complement C1q B chain	1601	3.13661232172674
3.55354445850488	9.01839331204724
0.00267276228788187	0.068191951486997
50618	ITSN2	intersectin 2	12022	-0.594738506129036
7.1071317664097	9.0083447292356	0.0026874970682312
0.068279718385265
57121	LPAR5	lysophosphatidic acid receptor 5
3258	-0.899807911575579	2.69325093048678

9.00505788245735	0.00269233462826694	0.068279718385265
9495	AKAP5	A-kinase anchoring protein 5	6487
0.920024809218337	0.438447302510327	9.00453428364337
	0.002693106073599	0.068279718385265
5325	PLAGL1	PLAG1 like zinc finger 1	8719
0.669196873218883	3.5158657474633 9.00277414313558
0.00269570102570511	0.068279718385265
10595	ERN2	endoplasmic reticulum to nucleus signaling 2
5015	-1.29922081840385	3.16624955095067
8.99296085677495	0.00271021522853056	0.0685234380975226
80201	HKDC1	hexokinase domain containing 1	4259
1.13665341153581	4.43215111258199	8.97884173301341
	0.00273123724884365	0.0689305227829784
54886	PLPPR1	phospholipid phosphatase related 1	3669
1.62357041570952	-0.251969415690049	8.97526800997575
	0.00273658438685237	0.0689308351625274
6279	S100A8	S100 calcium binding protein A8
1006	-2.95870121334242	3.13153506763764
8.9722609029063 0.00274109196726835	0.0689308351625274
5653	KLK6	kallikrein related peptidase 6
1976	-1.62729962580114	5.73846412295162
8.96023916672893	0.00275918770125244	0.0692469849559235
253039	CUTALP	cutA divalent cation tolerance homolog-like, pseudogene	7181	0.668431630717526	3.58648894748969
	8.95735215764007	0.00276355140932114
0.0692469849559235
3672	ITGA1	integrin subunit alpha 1	17668
1.7178718504074 5.45714605961249	8.9424299615515
0.00278621825321891	0.0696119634180776
81849	ST6GALNAC5	ST6 N-acetylgalactosaminide alpha-2,6-sialyltransferase 5	4263	-1.39698294622601
1.92048769284755	8.94122501190882	0.00278805679142868
	0.0696119634180776
3855	KRT7	keratin 7	3748	0.63097557503351
11.9884571523384	8.93457076915274	0.0027982321892534
	0.0696216869497032
5155	PDGFB	platelet derived growth factor subunit B
3761	-0.851008777851744	7.32074699424309
8.93204109780504	0.00280211034605445	0.0696216869497032
90362	FAM110B family with sequence similarity 110 member B	4943
	1.05724385581641	2.11250099699737
8.92935463423449	0.00280623485227433	0.0696216869497032
6352	CCL5	C-C motif chemokine ligand 5
1512	-1.71688648259334	1.15661783166856
8.92799268143781	0.00280832820208341	0.0696216869497032
57214	CEMIP	cell migration inducing hyaluronidase 1 7621
2.02399346051073	3.36845267888133	8.91845468890129
	0.00282303280393345	0.0698625803616534
55283	MCOLN3	mucolipin 3	6623	1.23123650631377
4.09952027231579	8.90993919601279	0.0028362270897227
	0.069965603150125
57333	RCN3	reticulocalbin 3	2093	1.65032599457836
	1.53831340315333	8.90932197021474
0.00283718587772335	0.069965603150125

79784	MYH14	myosin heavy chain 14	8753	-0.619692810557454
	8.59951180299773	8.90391522800746
0.00284559870191151	0.0700384953015394
3073	HEXA	hexosaminidase subunit alpha	12611
0.562350308021962	6.10140889616111	8.90092614707679
	0.00285026054491869	0.0700384953015394
7298	TYMS	thymidylate synthetase	2470	0.763813731503244
	3.54104750502793	8.89519132696666
0.00285922643715171	0.0700384953015394
9712	USP6NL	USP6 N-terminal like	14202	-0.591025120042811
	6.25202514235236	8.89460626667081
0.00286014273666599	0.0700384953015394
131368	ZPLD1	zona pellucida like domain containing 1 4607
1.38371577096475	3.56425211306155	8.8744377379729
0.00289191295021443	0.0706928877725193
11035	RIPK3	receptor interacting serine/threonine kinase 3
3256	-0.453418653409605	4.90740508638702
8.86815482906622	0.00290188306138562	0.0708130244613735
3993	LLGL2	LLGL scribble cell polarity complex component 2
7909	-0.395882573733514	7.60968319483056
8.86131448026152	0.00291277745441539	0.0709552587895588
23213	SULF1	sulfatase 1	9333	2.66892939291234
3.98102290420786	8.83410017906676	0.00295653378346251
	0.0718961262238878
5625	PRODH	proline dehydrogenase 1 11181	1.23357715563373
	3.3784057827067 8.82873770760595	0.00296523423361977
	0.0719827312137126
117145	THEM4	thioesterase superfamily member 4	4390
0.864382472260633	2.49894107706908	8.81336665357487
	0.00299031771013048	0.072466055650169
358	AQP1	aquaporin 1 (Colton blood group)	3895
3.37746413169056	2.87775069141285	8.80183511885486
	0.00300927703608317	0.0727995568988204
7538	ZFP36	ZFP36 ring finger protein
2010	-0.53034070112422	4.91974440758061
8.78136950181068	0.00304322635812903	0.0734127274559917
1948	EFNB2	ephrin B2	4461	-0.685264401217142
6.98319399026502	8.78024340387206	0.00304510563660535
	0.0734127274559917
348	APOE	apolipoprotein E	2191	1.6340047294748
5.00026052987832	8.77206193460817	0.0030587946557074
	0.0735799058558992
84815	MGC12916	uncharacterized protein MGC12916	1652
	-1.30343237678169	0.229870636259061
8.76982623465297	0.00306254623502458	0.0735799058558992
2057	EPOR	erythropoietin receptor 3079	1.4328996532291
1.3556503865746 8.74987689175419	0.00309622956123567
0.0742617936031303
4842	NOS1	nitric oxide synthase 1 13113	2.60932189723218
	1.06995447884978	8.74439009054799
0.00310555954059355	0.0743582435642631
6801	STRN	striatin	8345	-0.516918266134884
7.00277265060511	8.74015456843068	0.00311278132702769
	0.0744039727776055

221079	ARL5B	ADP ribosylation factor like GTPase 5B
7196	-0.622089997590336	5.99708510542922
8.71236160349197	0.00316059504158372	0.0753815071319953
149345	SHISA4	shisa family member 4	1746	0.61462981492191
	4.28863590842861	8.71014459017104
0.0031644410790043	0.0753815071319953
4052	LTBP1	latent transforming growth factor beta binding
protein 1	7329	0.524497072392696	6.8233633003145
8.70048699838225	0.00318125046594481	0.0756532687206944
26032	SUSD5	sushi domain containing 5	5008
1.03281456043653	0.87115568071033	8.69494767689498
	0.0031909327728802	0.0757549073724289
5352	PLOD2	procollagen-lysine,2-oxoglutarate 5-dioxygenase 2
	7646	0.883948375910905	7.30720142493341
8.67077733697488	0.00323353232978289	0.076636357602824
84680	ACCS	1-aminocyclopropane-1-carboxylate synthase homolog
(inactive)	5826	0.866021661976088	3.69608271199495
	8.65988569853693	0.00325291700555609
0.076965554893284
8537	BCAS1	breast carcinoma amplified sequence 1
8186	-1.28112040601279	4.38582023204521
8.65527643809284	0.00326115598293873	0.0770303741197687
368	ABCC6	ATP binding cassette subfamily C member 6	7559
	1.51928966623164	0.388495060915008
8.63903841437303	0.00329035052678011	0.0775891242905875
5027	P2RX7	purinergic receptor P2X 7	5546
1.74732933795318	0.244030414182512	8.62806614371485
	0.00331022796681489	0.0779266607246658
9818	NUP58	nucleoporin 58	10422	-0.444578166453407
6.48435485337656	8.6139178509173	0.00333603939346144
0.0783276332337259
124359	CDYL2	chromodomain Y like 2	8557	0.822372706574381
	3.55420489012765	8.61171632222754
0.00334007410382633	0.0783276332337259
84458	LCOR	ligand dependent nuclear receptor corepressor
11116	-0.593819027144121	6.22230831240716
8.60781997953347	0.00334722703708768	0.0783276332337259
23208	SYT11	synaptotagmin 11	5240	1.10494069542154
	4.61525389084364	8.60651352381288
0.00334962892175354	0.0783276332337259
79413	ZBED2	zinc finger BED-type containing 2
2311	-1.03267394435993	5.62738756435431
8.58856071181657	0.00338281270602334	0.0789717626221148
3955	LFNG	LFNG O-fucosylpeptide 3-beta-N-
acetylglucosaminyltransferase	3418	1.19594112155653
4.98186397162511	8.58107210695877	0.00339675314985357
	0.0791130753412322
2525	FUT3	fucosyltransferase 3 (Lewis blood group)
3109	-0.550609184443217	6.4337796370654 8.57737918278093
	0.00340364921430503	0.0791130753412322
1612	DAPK1	death associated protein kinase 1	8907
0.79482499072877	7.28932941668477	8.57622347775134
	0.00340581026849168	0.0791130753412322
654346	LGALS9C galectin 9C	4754	-1.58683480022551

1.23162732308888	8.55250552653942	0.00345046986417439
	0.0799338159046019
85463	ZC3H12C	zinc finger CCCH-type containing 12C
9719	-0.719528141950465	4.90347373635034
8.54935881875381	0.00345643950879619	0.0799338159046019
914	CD2	CD2 molecule	1679	-2.20553855219953
0.966678564898894	8.5483986274545	0.00345826318542077
0.0799338159046019
56261	GPCPD1	glycerophosphocholine phosphodiesterase 1
7527	-0.800454050564274	5.27215966368451
8.52544570786129	0.00350214971961033	0.0807607572005724
5293	PIK3CD	phosphatidylinositol-4,5-bisphosphate 3-kinase
catalytic subunit delta	6332	-0.422872202313071
5.73969980333515	8.52366838869929	0.0035055715269471
	0.0807607572005724
51752	ERAP1	endoplasmic reticulum aminopeptidase 1
7945	-0.517687396223382	6.26650395679502
8.50001782641797	0.00355143001317588	0.0815217437267183
10211	FLOT1	flotillin 1	3186	0.405148220846201
6.03391958450445	8.49972820543915	0.00355199535340507
	0.0815217437267183
9245	GCNT3	glucosaminyl (N-acetyl) transferase 3, mucin type
	6190	-1.49738006242675	4.92175662085322
8.49764535025137	0.00355606378360997	0.0815217437267183
10577	NPC2	NPC intracellular cholesterol transporter 2	2749
	0.635298528700607	7.97815647476185
8.48603200035809	0.00357883510062675	0.081909711200129
6327	SCN2B	sodium voltage-gated channel beta subunit 2	5826
	1.18573448407853	1.02283806807298
8.47776823418066	0.00359512883737876	0.0821484006935796
2534	FYN	FYN proto-oncogene, Src family tyrosine kinase	7411
	0.765409573684148	5.02526585867708
8.46173775443446	0.00362695169710295	0.0827405739760928
629	CFB	complement factor B	6817	-1.54191753503146
	7.14792175973368	8.43498162170789
0.00368070601486323	0.083830323821446
6483	ST3GAL2	ST3 beta-galactoside alpha-2,3-sialyltransferase 2
	9744	0.779469488411911	3.55235087741516
8.42586952433576	0.00369919701900365	0.0841146958525717
440335	SMIM22	small integral membrane protein 22	1673
1.21509650826596	2.95854234084236	8.42249887899658
	0.00370606091834449	0.0841341900863068
1604	CD55	CD55 molecule (Cromer blood group)
5568	-0.768625564474393	6.38336998679588
8.41546751304202	0.00372042114385106	0.0843235258283525
6575	SLC20A2	solute carrier family 20 member 2
5554	-0.607931339893337	7.93079865111372
8.40151757187365	0.00374907898962588	0.0848357825649269
2882	GPX7	glutathione peroxidase 7	1691
1.01609333811943	2.58660219449709	8.39681300938715
	0.0037587942803727	0.0849019703337691
3488	IGFBP5	insulin like growth factor binding protein 5	6442
	2.17464895990052	5.8024568361796	8.39423604571712
	0.00376412676356612	0.0849019703337691

26468	LHX6	LIM homeobox 6	5748	1.55328566618145
1.50601601168657	8.38811621425256	0.0037768213214899
	0.084977765462069
57826	RAP2C	RAP2C, member of RAS oncogene family
4118	-0.484005389081866	5.4952176167823 8.38676947170681
	0.00377962075268573	0.084977765462069
249	ALPL	alkaline phosphatase, biomineralization associated
	3007	-1.85449868467101	1.78551668432146
8.37795549848092	0.00379799426302267	0.0852414177066108
4790	NFKB1	nuclear factor kappa B subunit 1
6257	-0.518868920497936	6.17025996371334
8.37531385764639	0.00380351867399384	0.0852414177066108
27161	AGO2	argonaute RISC catalytic component 2
15833	-0.662121576909754	6.82134122066861
8.36425719271123	0.00382673010546559	0.0856246143566399
55312	RFK	riboflavin kinase	3877	-0.417305554155798
	5.26854495338408	8.36096409650698
0.00383367116516869	0.0856431132544144
5937	RBMS1	RNA binding motif single stranded interacting
protein 1	11765	-0.576057290836134	6.82219942967083
	8.35563883874762	0.00384492264758501
0.0857576935107057
4126	MANBA	mannosidase beta	8437	0.543738809206389
	5.23843846064014	8.35191654955881
0.00385280723590587	0.0857969331531534
643365	LINC00452	long intergenic non-protein coding RNA 452
	3578	-1.08205421301718	1.42939502162565
8.34538983718804	0.00386667190695443	0.085859319411786
201134	CEP112	centrosomal protein 112 7849	0.73134978064315
	2.09809312720421	8.34482778289707
0.00386786824793581	0.085859319411786
7051	TGM1	transglutaminase 1	3735	-1.15320211329742
	4.54033332380276	8.34049317634833
0.00387710718199731	0.0859282283199942
341	APOC1	apolipoprotein C1	988	3.05967757619643
	2.76382329209799	8.33468873004165
0.00388951415666734	0.0860670217890038
9425	CDYL	chromodomain Y like	4531	-0.412556772688943
	6.18582696336543	8.32583257379892
0.00390852202341642	0.0863512113280658
51361	HOOK1	hook microtubule tethering protein 1
6758	-0.586033480544742	5.04974151938063
8.32102889351978	0.00391887161209389	0.0863722216644824
8740	TNFSF14 TNF superfamily member 14
4778	-1.41200321004481	1.1055291587317 8.31966938988622
	0.00392180573846011	0.0863722216644824
10517	FBXW10	F-box and WD repeat domain containing 10
4390	-1.26549918633135	2.97941845238045
8.31408655078765	0.00393387826277972	0.0865020923496948
23105	FSTL4	follistatin like 4	6734	1.10106648825818
	0.323031382778336	8.30609977230163
0.00395121491860369	0.0867471275311629
10435	CDC42EP2	CDC42 effector protein 2
2224	-0.612367757706336	4.62555568905573

8.2953212396928 0.00397473486689851     0.0870671852408482
56905   C15orf39        chromosome 15 open reading frame 39
5093    -0.425299084734817      5.87384428210827
8.29372730908634        0.00397822506990382     0.0870671852408482
5275    SERPINB13       serpin family B member 13
4535    -1.11240921302944       5.29488907320011
8.28378935481282        0.00400005648526972     0.0874084105915336
9053    MAP7    microtubule associated protein 7
5447    -0.336774061429964      6.06650974572087
8.26573447647211        0.00404003135479622     0.0881277677468359
2309    FOXO3   forkhead box O3 7721    -0.496438854862369
6.55731467241932        8.26325188797199        0.00404555969595313
        0.0881277677468359
118738  ZNF488  zinc finger protein 488 3496    1.69260403110437
        2.09981935167505        8.25931274924828
0.00405434735174494     0.0881820549004524
30850   CDR2L   cerebellar degeneration related protein 2 like  3536
        0.377133544833268       6.72786835423129
8.22439370674753        0.00413310112371983     0.0897555774262692
4248    MGAT3   beta-1,4-mannosyl-glycoprotein 4-beta-N-
acetylglucosaminyltransferase   5457    -1.9585294175721
2.01872955377527        8.21408061749938        0.00415665698743772
        0.0901273907477402
80210   ARMC9   armadillo repeat containing 9   9661
0.770339315897936       4.83122240856611        8.20788217532766
        0.00417088038514433     0.0902960147677227
64780   MICAL1  microtubule associated monooxygenase, calponin and
LIM domain containing 1 7056    0.504329807719211
6.17960532810022        8.20275625335497        0.00418268011435369
        0.0903040387981138
116984  ARAP2   ArfGAP with RhoGAP domain, ankyrin repeat and PH
domain 2        10451   -0.485534541946544      5.94664282365807
        8.20212085238981        0.00418414515456385
0.0903040387981138
345079  SOWAHB  sosondowah ankyrin repeat domain family member B
        3220    -0.481030968979149      6.67021508633207
8.19919729695653        0.00419089271180973     0.0903105141758751
7764    ZNF217  zinc finger protein 217 5796    -0.462229240651078
        7.08022909118418        8.18313803904742
0.00422815544228937     0.0909735380647423
22801   ITGA11  integrin subunit alpha 11       10864
2.8444851903503 2.87440755297   8.17856044993132
0.00423883858325498     0.0910489649418861
51661   FKBP7   FKBP prolyl isomerase 7 3080    0.969498632429656
        1.51576902610925        8.17402959643954
0.00424943970989251     0.0910489649418861
6545    SLC7A4  solute carrier family 7 member 4        2447
0.770010519434001       3.22323952383487        8.17329468055613
        0.00425116178139455     0.0910489649418861
4908    NTF3    neurotrophin 3  2251    1.73558551766303
0.26115330934947        8.1618901098524 0.00427797645931473
0.0913651685075708
283422  LINC01559       long intergenic non-protein coding RNA 1559
        3953    -2.78759478046931       3.79609065516992

8.16146840840945	0.00427897126729253	0.0913651685075708
33	ACADL	acyl-CoA dehydrogenase long chain	4383
0.90763415820697	1.49703162073729	8.15558698199841
	0.004292870321857	0.0914898562070453
143903	LAYN	layilin 3144	0.441796410441417
7.30049638086794	8.15134162163553	0.00430293155858304
	0.0914898562070453
3606	IL18	interleukin 18	2628	0.438717836354382
6.51942698913824	8.14952882119144	0.00430723509183544
	0.0914898562070453
10965	ACOT2	acyl-CoA thioesterase 2 2393	0.696302283412706
	2.71107554873858	8.14797060462548
0.00431093775231312	0.0914898562070453
80781	COL18A1 collagen type XVIII alpha 1 chain	8525
0.809587566995018	6.09719101969433	8.13726330318134
	0.00433646840729757	0.0918189854503831
54328	GPR173	G protein-coupled receptor 173	4608
0.934961206767578	0.231830342076216	8.13597252655862
	0.0043395565337441	0.0918189854503831
9394	HS6ST1	heparan sulfate 6-O-sulfotransferase 1	4683
0.412487411373727	6.46642046004259	8.12078982018904
	0.00437604898441959	0.092450415497775
148252	DIRAS1	DIRAS family GTPase 1	3455	1.18211339969606
	0.97261472317708	8.11807798595901
0.00438259983511977	0.092450415497775
128414	NKAIN4	sodium/potassium transporting ATPase interacting 4
	2668	1.32937945859083	5.50772704671224
8.11258754450948	0.00439589342330625	0.0925549942138934
81035	COLEC12 collectin subfamily member 12	7340
1.51618906624114	0.915364716547775	8.11057648713954
	0.00440077290972035	0.0925549942138934
3887	KRT81	keratin 81	1929	-1.72259560781059
3.58788973474271	8.10358312975036	0.00441778406614159
	0.0927734653889734
85012	TCEAL3	transcription elongation factor A like 3	2305
	0.518845319535924	3.70489070001637
8.09611097854732	0.00443603386972695	0.0929638597351246
5357	PLS1	plastin 1	5294	-0.429299104289811
5.21453031826156	8.09426936785158	0.00444054355439153
	0.0929638597351246
1593	CYP27A1 cytochrome P450 family 27 subfamily A member 1	2983
	1.88123787858174	3.37912751853241
8.09173334578105	0.00444676133522763	0.0929638597351246
100132215	LOC100132215	uncharacterized LOC100132215
2324	-1.3134145284637	0.541753582212178
8.08792670260824	0.00445611104976912	0.0930204880389211
25890	ABI3BP	ABI family member 3 binding protein	10848
2.22202846337114	0.90375138128144	8.07753216681758
	0.00448174375127819	0.093363302739433
1281	COL3A1	collagen type III alpha 1 chain 6811
2.86181505904068	7.98003152315587	8.07586681428313
	0.004485864406628	0.093363302739433
3614	IMPDH1	inosine monophosphate dehydrogenase 1	3662
0.485576009086357	5.13898616148867	8.06289569711143

0.00451809174505292	0.0938614830467885
2825	GPR1	G protein-coupled receptor 1	2834
1.9964534545746 1.5965088159485 8.06084723410876
0.00452320275980454	0.0938614830467885
151354	LRATD1	LRAT domain containing 1
6598	-0.558905168964479	6.42699661109268
8.05764584020672	0.00453120218660785	0.0938883861358226
5641	LGMN	legumain	3899	0.69989380743757
5.95757264940439	8.05004054202981	0.00455026360603337
0.0941324887958089
79718	TBL1XR1 transducin beta like 1 X-linked receptor 1
4492	-0.508433103686421	7.06574386517626
8.04758964869013	0.00455642374552427	0.0941324887958089
64866	CDCP1	CUB domain containing protein 1
6507	-0.575432715488647	8.37243775793357
8.03879100467694	0.00457860854489512	0.094451502044692
100616211	MIR4709 microRNA 4709	72	0.671194295760632
5.26393371426352	8.03166887250924
0.00459664674604854	0.094495070265464
7145	TNS1	tensin 1	15290	1.03921228879903
6.44281413931354	8.03092609130076	0.00459853214834333
0.094495070265464
23348	DOCK9	dedicator of cytokinesis 9
13945	-0.548756285668407	7.62314467579656
8.02997030840319	0.00460095937181741	0.094495070265464
9962	SLC23A2 solute carrier family 23 member 2	7515
0.419704882123329	4.83811139568169	8.01313186179313
0.00464393546070425	0.0951023740565981
80139	ZNF703	zinc finger protein 703 3349	-0.744547986958441
5.26537969116759	8.01306449329993
0.00464410822122604	0.0951023740565981
100133941	CD24	CD24 molecule	3097	-0.419698635163598
11.865056679619 8.00491379809773	0.00466505833375327
0.0952800112773765
26499	PLEK2	pleckstrin 2	1603	-0.718973681914483
5.57126066382472	8.00439799563506	0.00466638735891199
0.0952800112773765
26471	NUPR1	nuclear protein 1, transcriptional regulator	5674
0.804195430609441	7.09349180729525
7.99851558198952	0.00468157138816706	0.0954509031063406
563	AZGP1	alpha-2-glycoprotein 1, zinc-binding	2699
1.97713484588731	1.09441277752472	7.98994043808765
0.00470379625117528	0.0957646425729828
23002	DAAM1	dishevelled associated activator of morphogenesis 1
10712	-0.440096222521823	6.10489142271352
7.97154791622531	0.00475182873653522	0.0966021264334527
151987	PPP4R2	protein phosphatase 4 regulatory subunit 2
5964	-0.34392603278752	6.62721038450198
7.96337883884943	0.00477332241459789	0.0968984450163371
9936	CD302	CD302 molecule	4075	1.11587068667137
1.67708108969724	7.95521098473784	0.00479491189555996
0.0971243243889604
272	AMPD3	adenosine monophosphate deaminase 3
5969	-0.495426719522152	5.53178868741126

7.95392599915263	0.00479831744678808	0.0971243243889604
102724344	LINC02188	long intergenic non-protein coding
RNA 2188	3891	-1.48251010043494	2.50136933723756
	7.95082933128592	0.00480653456211434
0.097150259179705
79192	IRX1	iroquois homeobox 1	1858	1.16113349679992
	0.873179619585119	7.94716823215152
0.00481626791041077	0.097206721356086
1019	CDK4	cyclin dependent kinase 4	3168
0.678108133258834	5.1372280653309 7.94414069476046
0.00482433204659826	0.0972071654177044
80320	SP6	Sp6 transcription factor
4075	-0.879322856327839	3.93293513515494
7.94195226228869	0.00483016971019649	0.0972071654177044
2070	EYA4	EYA transcriptional coactivator and phosphatase 4
	8968	-2.21904177109648	3.976018375969
7.93666643354264	0.00484429940679407	0.097351652497797
23198	PSME4	proteasome activator subunit 4
10241	-0.395891489770839	7.50039401110496
7.9048095045471 0.00493035333975061	0.0987397654828061
2268	FGR	FGR proto-oncogene, Src family tyrosine kinase	3474
	2.32630629488904	1.68416423581323
7.90157676149509	0.00493917242281725	0.0987397654828061
323	APBB2	amyloid beta precursor protein binding family B
member 2	12956	0.584212384961627	5.08032672114959
	7.90129851024459	0.00493993225686571
0.0987397654828061
85441	HELZ2	helicase with zinc finger 2
12066	-0.640219109286718	5.68020210136617
7.90069879635949	0.00494157032936725	0.0987397654828061
5581	PRKCE	protein kinase C epsilon	9431
0.99721775695047	3.52946797300427	7.89376285853943
	0.00496055557579794	0.0989779230059854
57224	NHSL1	NHS like 1	8561	-0.561691340399511
4.62175548910374	7.87216388356125	0.00502015436738111
	0.0997409368378061
84766	CRACR2A calcium release activated channel regulator 2A
12466	1.1441469392244 1.45433896873214	7.87014130467416
	0.00502577256905154	0.0997409368378061
8510	MMP23B	matrix metallopeptidase 23B	2612
1.14137290539057	0.573559168235404	7.86846428709024
	0.00503043574905658	0.0997409368378061
1634	DCN	decorin 12676	2.93654095403227
1.85439850744899	7.86813566536259	0.00503135004386914
	0.0997409368378061
342132	ZNF774	zinc finger protein 774 8511	0.93402532186528
	2.30066317773103	7.86703983081219
0.00503440011025408	0.0997409368378061
8999	CDKL2	cyclin dependent kinase like 2	3781
0.961217590771841	1.74382943147693	7.85613498814294
	0.00506485471950649	0.100202570700745
8639	AOC3	amine oxidase copper containing 3	4524
2.27565151296566	1.31419726348955	7.85281182264498
	0.00507417279737263	0.100245329157685

103695436	LINC01269	long intergenic non-protein coding
RNA 1269	602	-1.11934940294725	1.62781168822607
	7.84246511783646	0.00510329674359423
0.100678700686654
79971	WLS	Wnt ligand secretion mediator	4469
0.550863928733234	6.49334034980512	7.83746908249168
	0.00511742053744936	0.100735617719992
203447	NRK	Nik related kinase	9573	1.12781826592058
	3.76354010027528	7.83635855095525
0.00512056541847895	0.100735617719992
128434	VSTM2L	V-set and transmembrane domain containing 2 like
	1959	1.15449377222763	3.81596445557659
7.83037080173941	0.00513755591929556	0.100928114672613
801	CALM1	calmodulin 1	7705	-0.436002349176469
8.8181745768538 7.8191747327304 0.00516947958009333
0.101413025880066
57491	AHRR	aryl-hydrocarbon receptor repressor	7678
0.859713184388522	2.46564073871565	7.79752397995757
	0.00523178770727288	0.102491818763316
123036	TC2N	tandem C2 domains, nuclear
6147	-0.608878717630935	5.4311224608895 7.77513197592507
	0.00529703462621868	0.103572278522127
1364	CLDN4	claudin 4	1831	-0.537241681889797
9.49929516549872	7.77353164133478	0.005301729399612
	0.103572278522127
79905	TMC7	transmembrane channel like 7	5638
0.6389388513888 4.86703893208845	7.76762566680445
0.00531909202703837	0.103601100779711
6362	CCL18	C-C motif chemokine ligand 18	1397
3.58629458310753	2.92991389416033	7.76657140283548
	0.00532219748755637	0.103601100779711
1896	EDA	ectodysplasin A 6404	0.715759910598272
2.7224376081319 7.76548691163464	0.00532539391457073
0.103601100779711
84514	GHDC	GH3 domain containing	4019	0.457661032359808
	4.279147566004	7.75823197567905	0.00534682749427377
	0.103873804039241
154064	RAET1L	retinoic acid early transcript 1L
1346	-0.946232508731514	5.41890064670172
7.74852671664415	0.00537563776271122	0.104154475400693
79689	STEAP4	STEAP4 metalloreductase 11245	-1.11177180706179
	5.85759119240938	7.74727596732789
0.00537936213347381	0.104154475400693
55227	LRRC1	leucine rich repeat containing 1
4897	-0.416683416549968	5.21149985122439
7.74585971245591	0.00538358250803896	0.104154475400693
8522	GAS7	growth arrest specific 7
10100	-0.971375754503239	4.90257442890652
7.73885762564667	0.00540449806240694	0.104293695604702
2318	FLNC	filamin C	9280	1.19794211707196
4.76669282884433	7.73846602426245	0.00540567023695392
	0.104293695604702
374286	CDRT1	CMT1A duplicated region transcript 1
8649	-0.807519913296049	7.76567381162234

7.73409840864588	0.00541876133568514	0.104402462213125
8976	WASL	WASP like actin nucleation promoting factor
4435	-0.410401229044374	6.95172213997514
7.72750709913155	0.00543857876163509	0.104510738801355
3667	IRS1	insulin receptor substrate 1	9953
0.55919314980509	6.09095453472691	7.72485080786004
	0.00544658604095613	0.104510738801355
337876	CHSY3	chondroitin sulfate synthase 3	3938
1.06255922276659	0.42723477252416	7.72479143983922
	0.00544676514064317	0.104510738801355
51673	TPPP3	tubulin polymerization promoting protein family
member 3	2618	-1.48302476350674	0.280283687617101
	7.72027044270138	0.00546042160587583
0.104564152245521
22807	IKZF2	IKAROS family zinc finger 2
12240	-0.851423032167493	4.98580762041506
7.71479122130539	0.00547701936199876	0.104564152245521
3164	NR4A1	nuclear receptor subfamily 4 group A member 1
7295	-1.000916046791 2.97232309156292	7.71277716680676
	0.00548313329408904	0.104564152245521
6274	S100A3	S100 calcium binding protein A3 955
1.21893657155322	4.0252257122937 7.71267393366636
0.00548344685953063	0.104564152245521
10551	AGR2	anterior gradient 2, protein disulphide isomerase
family member	2845	-1.82828087201831	4.10263245838934
	7.71016863258567	0.00549106219357761
0.104564152245521
5174	PDZK1	PDZ domain containing 1 2480	-1.25605481410139
	0.621295877016187	7.70909148810529
0.00549433969106186	0.104564152245521
84166	NLRC5	NLR family CARD domain containing 5
11353	-0.941549197972325	4.65528448894754
7.70463401078537	0.00550792396812735	0.1046804491473
10418	SPON1	spondin 1	5102	2.30537420836313
0.862092212162385	7.69422531152784	0.00553977826203574
	0.105143189859532
55286	C4orf19 chromosome 4 open reading frame 19
2770	-0.979929997509719	5.47857360696869
7.68917836789237	0.00555529125581019	0.105211231032515
5326	PLAGL2	PLAG1 like zinc finger 2
5705	-0.493742957145043	5.37794554990474
7.68817329446579	0.00555838587592355	0.105211231032515
127933	UHMK1	U2AF homology motif kinase 1
9246	-0.429333735130238	7.82007534211743
7.68306351332023	0.00557414609608056	0.105229316725746
151636	DTX3L	deltex E3 ubiquitin ligase 3L
5868	-0.384163533444811	6.42222598282872
7.68299212503125	0.00557436660316294	0.105229316725746
4059	BCAM	basal cell adhesion molecule (Lutheran blood group)
	5024	0.850699623092869	6.55811927124489
7.67871986225884	0.00558757915281556	0.10533677145826
79921	TCEAL4	transcription elongation factor A like 4	2518
	0.425691386035019	5.40174108821037
7.67527031000824	0.00559827065997157	0.105396474642771

57447	NDRG2	NDRG family member 2	7550	-1.51892019188653
	6.15681842966694	7.66915920181759
0.00561726261259665	0.105498316251937
284217	LAMA1	laminin subunit alpha 1	15872	0.91059646306649
	2.10753531693563	7.66868349202376
0.00561874376554187	0.105498316251937
4033	LRMP	lymphoid restricted membrane protein	4531
1.26755676479797	2.56884495208838	7.66169010546042
	0.00564056414781001	0.105766240988454
7163	TPD52	tumor protein D52	8142	-0.632647461055903
	6.79933370716742	7.65808360527043
0.00565185070868824	0.105836193685289
1041	CDSN	corneodesmosin	2552	-1.94569271820149
1.46276329104307	7.65427861651629	0.00566378341153233
	0.105918043051179
338442	HCAR2	hydroxycarboxylic acid receptor 2
2051	-1.33975774571338	3.46857952284696
7.65153480480507	0.00567240413330593	0.105937819593621
6442	SGCA	sarcoglycan alpha	1727	1.04697029195096
	2.87385158899918	7.63906815528467
0.00571174168336338	0.10642862216013
127544	RNF19B	ring finger protein 19B	2610	-0.435731031909579
	5.88862936975038	7.63839285179077
0.00571388047864763	0.10642862216013
2528	FUT6	fucosyltransferase 6	3782	-0.83828457320276
	2.57288290580165	7.62094946694292
0.00576941048011798	0.107320229209844
55353	LAPTM4B	lysosomal protein transmembrane 4 beta	3197
0.40161690648212	6.28445825170059	7.61331832687914
	0.00579387656241429	0.107505157486112
148808	MFSD4A	major facilitator superfamily domain containing 4A
	5790	-1.776347446697	3.19657388765928
7.61306137434932	0.00579470221332298	0.107505157486112
5311	PKD2	polycystin 2, transient receptor potential cation
channel	5498	0.680395670319272	5.54777122674851
7.60683640305811	0.00581474127464292	0.107601845751355
338773	TMEM119	transmembrane protein 119	3189
2.09604128155335	-0.0724689929581767	7.60667001266051
	0.00581527787775941	0.107601845751355
441478	NRARP	NOTCH regulated ankyrin repeat protein
1770	-1.2788789193372	0.510771802981054
7.60161877585566	0.00583159200307672	0.107607441323355
144501	KRT80	keratin 80	4350	-0.816740556162166
8.34532441784439	7.60062971838605	0.00583479185237232
	0.107607441323355
50506	DUOX2	dual oxidase 2	9834	-0.978201790781689
6.4072472562815	7.59944486742114	0.00583862750094593
0.107607441323355
83605	CCM2	CCM2 scaffold protein	8196	0.347614253185478
	6.05868500115607	7.59330974868803
0.00585852954229828	0.107832356503248
8434	RECK	reversion inducing cysteine rich protein with kazal
motifs	6045	0.706026008887692	2.93351591306426
7.57484397952889	0.00591885029728782	0.108694671009896

827	CAPN6	calpain 6	3568	1.65593303489069
2.21940811379694	7.57422015699603	0.0059208991204791
	0.108694671009896
8321	FZD1	frizzled class receptor 1	6963
0.658993743371512	4.8627934480692 7.56877272193992
0.00593882093098048	0.108875306294204
6665	SOX15	SRY-box transcription factor 15
1396	-1.31504298367623	5.00689920785518
7.56650902281709	0.00594628466588607	0.108875306294204
3643	INSR	insulin receptor	10451	-0.678652140209163
	5.53418386562662	7.56049465895814
0.00596616139709391	0.109096635364353
6330	SCN4B	sodium voltage-gated channel beta subunit 4	5649
	1.11103959974673	2.01121505629761
7.54846315463317	0.00600612773464858	0.109684264901203
80315	CPEB4	cytoplasmic polyadenylation element binding protein
4	10272	-0.632938569635671	6.10602375490993
7.54180571301739	0.00602835976263777	0.109924362040595
644	BLVRA	biliverdin reductase A	1821	0.492125927618393
	4.88464477046888	7.53983088532438
0.00603497068674359	0.109924362040595
100192386	FLJ16779	uncharacterized LOC100192386	7638
	1.42617585600949	6.15705271218254
7.53404311683533	0.00605438843646039	0.110134829648702
4354	MPP1	membrane palmitoylated protein 1	5396
1.73784186042676	1.95301656131057	7.52925846084367
	0.00607048890980732	0.110279376739006
55236	UBA6	ubiquitin like modifier activating enzyme 6
11432	-0.427217407324562	6.7522862382867 7.5251257643305
0.00608443068841388	0.110279376739006
29995	LMCD1	LIM and cysteine rich domains 1 9130
0.830621334210693	4.64151573465403	7.52467479043575
	0.00608595403864149	0.110279376739006
54809	SAMD9	sterile alpha motif domain containing 9
6855	-0.672378745564334	7.01112243056297
7.51810986629243	0.00610817389881254	0.110539007494402
2762	GMDS	GDP-mannose 4,6-dehydratase
4304	-0.575305221335873	3.60915281825047
7.51458769534954	0.0061201292601791	0.110612452319134
93517	SDR42E1 short chain dehydrogenase/reductase family 42E,
member 1	11911	-0.512211055404326	5.31500519788093
	7.50443497271132	0.00615472460768353
0.111061126364827
2113	ETS1	ETS proto-oncogene 1, transcription factor
7149	-0.725885880388191	7.60046679379997
7.50100164693868	0.00616646874846428	0.111061126364827
9839	ZEB2	zinc finger E-box binding homeobox 2	18847
2.29435238488738	2.532210299651	7.50033811553582
0.00616874108030524	0.111061126364827
9641	IKBKE	inhibitor of nuclear factor kappa B kinase subunit
epsilon 3791	-0.646780857108966	5.13560258544339
7.49635788448047	0.00618238974197329	0.111163970623646
5754	PTK7	protein tyrosine kinase 7 (inactive)	9024
0.426241103531364	7.79444753938051	7.47346276714967

0.00626150042688634	0.1123909661524
5329	PLAUR	plasminogen activator, urokinase receptor
4457	-0.812595849136235	7.85673375009366
7.47197494563363	0.00626667698757936	0.1123909661524
9685	CLINT1	clathrin interactor 1	6368	-0.371536871307243
6.59546399511435	7.46774223934116
0.00628142769785103	0.11251145494092
5520	PPP2R2A protein phosphatase 2 regulatory subunit Balpha
7881	-0.407388207530608	6.79832002844929
7.45610030328174	0.00632218215741906	0.113087778251294
54439	RBM27	RNA binding motif protein 27
6900	-0.374156484091147	5.58945126492245
7.45181404312637	0.00633725478684951	0.113087778251294
5241	PGR	progesterone receptor	15404	1.96507308406828
0.629046365532926	7.45165227321753
0.00633782436833483	0.113087778251294
401093	MBNL1-AS1	MBNL1 antisense RNA 1	6585
0.881333686407145	2.42502721393589	7.4465510443991
0.00635581229890594	0.113264456578595
6469	SHH	sonic hedgehog signaling molecule	5038
1.05052622327676	0.518435201548207	7.44198826369905
0.0063719457070924	0.113402341233017
7292	TNFSF4	TNF superfamily member 4	3942
1.72710519228655	1.37732249536745	7.43948795873233
0.00638080418947439	0.113402341233017
10404	CPQ	carboxypeptidase Q	3457	0.713894771027456
3.19003870991067	7.43750518213789
0.0063878380261905	0.113402341233017
64410	KLHL25	kelch like family member 25
3758	-0.557215566549906	4.40113029382365
7.42953952569325	0.0064161758060147	0.113427129400396
55790	CSGALNACT1	chondroitin sulfate N-
acetylgalactosaminyltransferase 1	6580	-0.818572923676403
3.06550553386716	7.42906087410283
0.00641788268833134	0.113427129400396
91860	CALML4	calmodulin like 4	5255	-0.878901283578602
2.29837553627132	7.42856727408457
0.00641964336257079	0.113427129400396
27189	IL17C	interleukin 17C 1379	-1.85011809781885
0.450578935945912	7.42801164808176	0.00642162587381409
0.113427129400396
5017	OVOL1	ovo like transcriptional repressor 1
3849	-0.945076748078974	5.68013448770068
7.40909205532908	0.00648950658115669	0.114428458228977
379	ARL4D	ADP ribosylation factor like GTPase 4D	1612
0.61006224274888	4.09152584001372	7.40766551369929
0.00649465440794152	0.114428458228977
23120	ATP10B	ATPase phospholipid transporting 10B (putative)
11310	-1.41748703041048	2.62914910070836
7.402596902333	0.00651297877671411	0.114472474473406
2719	GPC3	glypican 3	2711	1.45230402333762
1.09779373036913	7.40091949907275	0.00651905465054963
0.114472474473406
345275	HSD17B13	hydroxysteroid 17-beta dehydrogenase 13 2584

1.14273003056761	1.60601098321309
7.40019789718051	0.00652167020987918	0.114472474473406
100170841	EPOP	elongin BC and polycomb repressive complex 2 associated protein	3719	0.653989174023272
2.52514767790251	7.38505557136652	0.00657680378298825
0.115260026541941
645638	WFDC21P WAP four-disulfide core domain 21, pseudogene
1099	-1.66112526780572	4.81514569159027
7.38240443713414	0.00658650547142098	0.115260026541941
51199	NIN	ninein	13683	-0.469042237242716
6.55205898423566	7.38111633778998	0.00659122447776786
0.115260026541941
26063	DECR2	2,4-dienoyl-CoA reductase 2	4046
0.746330429676491	2.95970875735776	7.37792866778649
0.00660291748991197	0.115294438327512
113612	CYP2U1	cytochrome P450 family 2 subfamily U member 1	6096
0.610217811468095	4.4337706583843 7.3760946197862
0.00660965474241395	0.115294438327512
56929	FEM1C	fem-1 homolog C 5816	-0.403953568238778
5.79315643507282	7.37190218446957	0.00662508174721504
0.11541980103637
1645	AKR1C1	aldo-keto reductase family 1 member C1	8834
0.900935290529902	7.33885034998284	7.36767141059724
0.00664068710230811	0.115547955580161
254827	NAALADL2	N-acetylated alpha-linked acidic dipeptidase like 2	11741	-0.805426697747427	2.83380824326325
7.3645527883746 0.00665221427534826	0.115604919795041
8566	PDXK	pyridoxal kinase	11697	0.360510767756088
8.5709130721765 7.35706819428935	0.0066799625603512
0.115942761673062
415116	PIM3	Pim-3 proto-oncogene, serine/threonine kinase
2461	-0.439356543008599	4.89972267368983
7.35405686439954	0.00669116004305754	0.115942761673062
100499467	LINC00673	long intergenic non-protein coding RNA 673 4699	0.489543698125602	6.22968700718236
7.35151957146341	0.00670060972108743	0.115942761673062
6317	SERPINB3	serpin family B member 3
2182	-3.47933233001731	3.82614152572569
7.35040513415427	0.00670476454309849	0.115942761673062
83468	GLT8D2	glycosyltransferase 8 domain containing 2	3271
1.24428421330674	-0.08130132886257
7.34141600411543	0.00673837397591627	0.116380276548285
57556	SEMA6A	semaphorin 6A	10956	-1.11139160525166
1.41141337574273	7.33173614110316	0.00677475843103369
0.116864582935331
79633	FAT4	FAT atypical cadherin 4 16439	0.811087709318274
4.01239906942759	7.32082362705247
0.00681601683097406	0.117431670050988
1601	DAB2	DAB adaptor protein 2	7332	0.888845213238091
4.37375402549852	7.30457689415165
0.00687791903643309	0.118352594524961
199675	MCEMP1	mast cell expressed membrane protein 1	1906
2.70011921216226	1.39097574212992	7.28994245390664
0.00693416971493271	0.119085324432261

1048	CEACAM5 CEA cell adhesion molecule 5
3523	-1.30680147569838	7.32518621944136
7.28811416732105	0.00694123007993233	0.119085324432261
6705	SPRR2F	small proline rich protein 2F
681	-2.32676302134145	1.28062032393248
7.28492243082658	0.00695357333722317	0.119085324432261
10318	TNIP1	TNFAIP3 interacting protein 1
5793	-0.648916778622304	8.68391998332475
7.28468094645791	0.00695450813061964	0.119085324432261
8459	TPST2	tyrosylprotein sulfotransferase 2	6957
0.562105028122201	4.88705190712163	7.28185168947702
	0.00696546983861347	0.119127394419364
2319	FLOT2	flotillin 2	3336	0.409668139481193
6.44120524637112	7.27206504409565	0.00700352364618483
	0.11963214111233
29114	TAGLN3	transgelin 3	1791	-0.992916191976405
0.676001668707901	7.26894386362167	0.00701570447250703
	0.119694241834843
135228	CD109	CD109 molecule	9563	0.580271544163387
8.0806552644939 7.26106065479847	0.00704656621181437
0.120074516945114
5236	PGM1	phosphoglucomutase 1	4090	0.52963546171069
	5.61334909127364	7.25675458617062
0.00706348244752054	0.12021652325932
283383	ADGRD1	adhesion G protein-coupled receptor D1	14718
1.87394269671196	2.06224518533861	7.25267214234525
	0.00707955849439683	0.120343902707544
647024	C6orf132	chromosome 6 open reading frame 132
6852	-0.513554802194214	7.63240550149336
7.23915500721603	0.00713305422305848	0.121106291518036
79616	CCNJL	cyclin J like	4473	0.599187563236813
5.46758469760208	7.22866021636972	0.00717487334983694
	0.121572476763465
84552	PARD6G	par-6 family cell polarity regulator gamma	4224
	0.399136465535876	4.84795246117955
7.22651279888386	0.00718346110117319	0.121572476763465
5083	PAX9	paired box 9	6035	-1.05022822873377
4.3983476830987 7.22574095546807	0.00718655035054963
0.121572476763465
23187	PHLDB1	pleckstrin homology like domain family B member 1
	21722	0.662095389295915	6.71271634130105
7.21717974241265	0.00722090718225065	0.12200632919395
8796	SCEL	sciellin	4205	-0.668573886815321
7.92519479086247	7.21336931447581	0.00723625266373805
	0.122118302483107
51368	TEX264	testis expressed 264, ER-phagy receptor 5688
0.381829307419707	5.02899118869624	7.20585586014894
	0.00726660880825607	0.122421237338657
8605	PLA2G4C phospholipase A2 group IVC	6613
1.07568990534807	2.07090138276671	7.20460297538104
	0.00727168340585153	0.122421237338657
10148	EBI3	Epstein-Barr virus induced 3
1284	-1.42544494789642	0.406257563322216
7.20126266204249	0.00728523048507542	0.122502068912907

5175	PECAM1	platelet and endothelial cell adhesion molecule 1
	7759	2.59553474581942	3.05308264140127
7.19215496665056	0.00732229912363472	0.12297775038939
23333	DPY19L1	dpy-19 like C-mannosyltransferase 1	5849
0.580426602487582	5.88189931420872	7.18923764914914
	0.00733421342215252	0.123030332220467
23586	DDX58	DExD/H-box helicase 58	4353	-0.34616679910372
	6.43814053411881	7.16258803547817
0.00744397163484319	0.124722141972785
1741	DLG3	discs large MAGUK scaffold protein 3
7844	-0.336168732210921	5.95408221329086
7.15925780246007	0.00745780492865809	0.124792715688621
2878	GPX3	glutathione peroxidase 3	2729
0.646258252299921	5.69791333611396	7.15377120288719
	0.0074806527795816	0.124792715688621
1238	ACKR2	atypical chemokine receptor 2
8856	-1.14117404467467	3.2291020328564 7.15208697857978
	0.00748768073887432	0.124792715688621
79812	MMRN2	multimerin 2	5334	-1.02490541147553
2.17348561913809	7.15159987159409	0.00748971460497936
	0.124792715688621
8754	ADAM9	ADAM metallopeptidase domain 9
6913	-0.367318202702089	10.1369170998301
7.15087785735082	0.00749273034155284	0.124792715688621
29933	GPR132	G protein-coupled receptor 132
4200	-1.54833068456602	0.797225191989866
7.14337527473647	0.00752414097118449	0.125167033947009
151556	GPR155	G protein-coupled receptor 155	7769
0.775630539119682	3.56361939240104	7.12697336193388
	0.00759327955565138	0.126167338951375
11000	SLC27A3 solute carrier family 27 member 3	5470
0.685198106584677	3.13870960013623	7.11185393757766
	0.00765758734249034	0.127085101784671
387914	SHISA2	shisa family member 2	2860	-1.1803882878316
	0.657181367420224	7.0989026675525 0.00771311591952745
	0.12785516530748
2817	GPC1	glypican 1	6516	0.50261734683612
8.18255310636968	7.09526056040102	0.0077288054979379
	0.127963804503092
6051	RNPEP	arginyl aminopeptidase	3479	0.384796417924471
	6.72263556466055	7.09029718070224
0.00775023944446153	0.128167182879071
5450	POU2AF1 POU class 2 homeobox associating factor 1	4583
	1.38514312635078	0.742166188971	7.08789685234618
	0.00776062685839878	0.12818761840282
2683	B4GALT1 beta-1,4-galactosyltransferase 1
4901	-0.598415386003047	8.61040504605769
7.0786593926415 0.00780073492176433	0.128698343991935
2941	GSTA4	glutathione S-transferase alpha 4	2175
1.04436773692123	1.46987244393115	7.07256628306594
	0.0078273065056827	0.128785026615016
3087	HHEX	hematopoietically expressed homeobox	2525
0.986955135947621	0.0763002475907495	7.07195145388103
	0.00782999286416985	0.128785026615016

81792	ADAMTS12	ADAM metallopeptidase with thrombospondin type 1 motif 12 9778	3.91380478551184	2.8457325582808	7.06927143283846	0.00784171363193034	0.128785026615016
80024	SLC8B1	solute carrier family 8 member B1	6405	-0.390546394252944	5.99176714111127	7.06793848747246	0.00784754979617752	0.128785026615016
80824	DUSP16	dual specificity phosphatase 16	5724	-0.323811724100729	6.04809171356284	7.0669317485147 0.0078519606431944	0.128785026615016
2114	ETS2	ETS proto-oncogene 2, transcription factor	4595	-0.530800161744329	6.62919509025505	7.0634264812205 0.00786733813631209	0.128886321959443
79070	POGLUT2 protein O-glucosyltransferase 2 2367	0.695089740265654	1.72057669050438	7.06042042928872	0.00788055011616449	0.128951945650836
23351	KHNYN	KH and NYN domain containing	8015	-0.341225011760969	7.45600082710502	7.05220501673779	0.00791677375743584	0.129393523944462
159195	USP54	ubiquitin specific peptidase 54	9498	-0.48878501735328	7.15419515821775	7.04967793609649	0.00792795044633965	0.129425177041818
6280	S100A9	S100 calcium binding protein A9	577	-2.44190399345351	7.03420224667314	7.04267017487695	0.00795902863658893	0.129705642354656
7020	TFAP2A	transcription factor AP-2 alpha	8245	-0.613363675236436	7.08088780863806	7.04052464873905	0.00796856855933585	0.129705642354656
28231	SLCO4A1 solute carrier organic anion transporter family member 4A1	4051	-1.63451269771023	3.82591837512049	7.03954900602393	0.00797291054953656	0.129705642354656
56944	OLFML3	olfactomedin like 3	2821	0.975870142571973	0.66422629078221	7.03535576637781	0.00799159966522276	0.129858859061224
2022	ENG	endoglin	3935	1.6629415041469	3.92260862782678	7.03230734902031	0.00800521447099684	0.129872464878076
23471	TRAM1	translocation associated membrane protein 1	3878	0.335057264459647	10.5014626986446	7.02982395369054	0.00801632331529464	0.129872464878076
80023	NRSN2	neurensin 2	2933	0.51409427652977	4.83326466568325	7.02806407382583	0.00802420523970658	0.129872464878076
100528031	FKBP1A-SDCBP2	FKBP1A-SDCBP2 readthrough (NMD candidate)	1375	-0.601232485008661	3.53578177898403	7.02687730754833	0.00802952485074707	0.129872464878076
8843	HCAR3	hydroxycarboxylic acid receptor 3	2137	-1.21384637910343	3.24001736371371	7.00962652848678	0.00810725904010211	0.130978520616736
6573	SLC19A1 solute carrier family 19 member 1	8369	0.643254211287274	2.44475579520017	6.98840108622363	0.00820396084866165	0.13212160439058
4638	MYLK	myosin light chain kinase	22167

1.05311813115082	4.68611336112379	6.98776499276127
	0.00820687699302716	0.13212160439058
8447	DOC2B	double C2 domain beta	6448	-0.979402193822605
	4.50353720512378	6.98586877054461
0.00821557643967445	0.13212160439058
6655	SOS2	SOS Ras/Rho guanine nucleotide exchange factor 2
	6938	-0.342597296713185	6.33732089104002
6.98583240412518	0.00821574337289889	0.13212160439058
5396	PRRX1	paired related homeobox 1	8246
2.70231051556143	2.55310383483938	6.95089783435876
	0.00837771170694225	0.134571894404977
100506237	NKX2-1-AS1	NKX2-1 antisense RNA 1	1775
0.79567942286313	0.440704207239121	6.94795557389228
	0.00839150782266753	0.13463900351902
57514	ARHGAP31	Rho GTPase activating protein 31	9384
	-0.50102565051003	4.91447788758471
6.93598283091589	0.00844786299738433	0.135265070347016
11254	SLC6A14 solute carrier family 6 member 14
4520	-1.85309749578729	7.06584121556062
6.93556388065679	0.00844984197570062	0.135265070347016
9914	ATP2C2	ATPase secretory pathway Ca2+ transporting 2
6693	-1.50754648819461	4.0304669680967 6.92303948644828
	0.00850922253393836	0.135926839978782
57007	ACKR3	atypical chemokine receptor 3	2258
0.863373319870409	5.99710107086397	6.92275205066466
	0.00851059032350907	0.135926839978782
602	BCL3	BCL3 transcription coactivator
3242	-0.529414643146733	3.95357188667573
6.91803418604447	0.00853307289700262	0.136130697116533
9124	PDLIM1	PDZ and LIM domain 1	2239	0.462256220224911
	8.43701438236056	6.91189527660356
0.00856241836404207	0.136416960973137
4097	MAFG	MAF bZIP transcription factor G
5220	-0.394834617925928	6.57047893369489
6.91020943456259	0.00857049515644752	0.136416960973137
140609	NEK7	NIMA related kinase 7	5485	-0.588759230933633
	6.96022684027985	6.90562040881349
0.00859252049198062	0.13661229799225
2331	FMOD	fibromodulin	3310	2.89136713901478
2.43317430269193	6.89134165302288	0.00866142360122038
	0.137514635712808
30001	ERO1A	endoplasmic reticulum oxidoreductase 1 alpha
7548	-0.653084373644944	7.65838897993739
6.88811840926303	0.00867705566471667	0.137514635712808
6414	SELENOP selenoprotein P 5688	1.11958260485199
3.06732938507283	6.88745539268457	0.00868027473666477
	0.137514635712808
134147	CMBL	carboxymethylenebutenolidase homolog	3140
0.915154398238151	2.17303420399263	6.88461075614791
	0.00869409986794114	0.137514635712808
84957	RELT	RELT TNF receptor	6322	-0.491350339061058
	4.05072603383347	6.88373456331896
0.00869836276444264	0.137514635712808
3572	IL6ST	interleukin 6 signal transducer

9292	-0.43732892489032	7.86643204921474
6.87970500377228	0.00871799513699835	0.137573931229812
9919	SEC16A	SEC16 homolog A, endoplasmic reticulum export factor
	9975	-0.407327935969222	7.7990020523082
6.87893392969175	0.00872175704519691	0.137573931229812
1891	ECH1	enoyl-CoA hydratase 1	2467	0.333219230785832
	5.92784229573759	6.87437726723209
0.00874402199009207	0.137629405244489
5671	PSG3	pregnancy specific beta-1-glycoprotein 3	3486
	1.33880321562266	1.24422106074959
6.87419262085155	0.00874492544210717	0.137629405244489
4071	TM4SF1	transmembrane 4 L six family member 1
3668	-0.48832097830552	8.16427785698655
6.86181752652655	0.00880569357125755	0.138348005412341
55605	KIF21A	kinesin family member 21A
10114	-0.40109639237682	5.65898086567989
6.86087507460653	0.0088103391752558	0.138348005412341
58475	MS4A7	membrane spanning 4-domains A7	4348
2.42883814687255	2.44694138809743	6.85815461807321
	0.00882376312999228	0.138403639598882
51560	RAB6B	RAB6B, member RAS oncogene family	6005
0.946574927652276	4.0512699094866 6.84690302177022
0.00887950632043777	0.139108015238745
202052	DNAJC18 DnaJ heat shock protein family (Hsp40) member C18
	5878	0.505452893534672	4.41334484710373
6.84508793464479	0.00888853242226576	0.139108015238745
9590	AKAP12	A-kinase anchoring protein 12
8741	-1.12824349168411	8.24484966792944
6.83630933934489	0.00893231955448405	0.139637276785333
114294	LACTB	lactamase beta	3972	0.365647057435423
5.18880477746982	6.83006431846957	0.00896360374235614
	0.139970119976792
646	BNC1	basonuclin 1	4797	-0.393063641714166
6.44531787473659	6.82164044767951	0.00900598059910071
	0.140475245269046
4815	NINJ2	ninjurin 2	2048	1.19534549169794
1.24054160299153	6.81669788253826	0.009030939940177
	0.140509146221118
150946	GAREM2	GRB2 associated regulator of MAPK1 subtype 2	5939
	1.69037952448253	1.07962607788081
6.81148524652105	0.00905733985148753	0.140509146221118
22846	VASH1	vasohibin 1	6502	1.12468517514395
2.7031034862001 6.80965613579174	0.00906662228177303
0.140509146221118
5909	RAP1GAP RAP1 GTPase activating protein
5755	-0.769825948600025	2.65904303228178
6.80937906799105	0.00906802920367604	0.140509146221118
2687	GGT5	gamma-glutamyltransferase 5	2590
1.48329020167523	0.975958140343283	6.80897091868328
	0.00907010215125811	0.140509146221118
51393	TRPV2	transient receptor potential cation channel
subfamily V member 2	3742	2.31540887628186
1.8217953309396 6.80776271490162	0.00907624133628589
0.140509146221118

6383	SDC2	syndecan 2	4254	1.17648966970016
3.77401623241332	6.80734330061991	0.00907837347969670
	0.140509146221118
2550	GABBR1	gamma-aminobutyric acid type B receptor subunit 1
	8470	-0.890538742783012	7.1646801132592
6.80145440091201	0.00910836462969838	0.140817729986959
64135	IFIH1	interferon induced with helicase C domain 1
4528	-0.62149800157067	4.92433038338605
6.797745628381	0.00912730481755722	0.140954970870478
115701	ALPK2	alpha kinase 2	8561	1.4535058811458
1.31821885768142	6.79362083049826	0.00914841696528938
	0.141125414573577
8214	DGCR6	DiGeorge syndrome critical region gene 6	6475
	0.614672023055149	3.68647676947042
6.78635984837694	0.0091857027691795	0.141310347475805
10536	P3H3	prolyl 3-hydroxylase 3	4601	0.907750353990895
	2.943830372529	6.78615819707391	0.00918674048505188
	0.141310347475805
79096	C11orf49	chromosome 11 open reading frame 49	5671
	0.458846473670582	4.73962606547777
6.78539465169817	0.00919067084675223	0.141310347475805
57561	ARRDC3	arrestin domain containing 3
4934	-0.591377466091986	4.60939434683373
6.77871897440478	0.00922510742864998	0.141684297974891
55500	ETNK1	ethanolamine kinase 1	8191	-0.393993964494882
	5.46310940522624	6.7725115097619	0.00925724729258194
	0.142011040907302
894	CCND2	cyclin D2	7157	0.813771366308355
7.37522201041519	6.77069793533906	0.00926665891263469
	0.142011040907302
4008	LMO7	LIM domain 7	16059	-0.507809142524716
9.29830883247726	6.76619973992092	0.00929004480483463
	0.14218745967066
64759	TNS3	tensin 3	11797	0.716557477195324
5.98579289634052	6.76445039849771	0.00929915585960366
	0.14218745967066
7466	WFS1	wolframin ER transmembrane glycoprotein	4513
0.543575220332006	5.67848507469391	6.75858558290777
	0.00932976827221232	0.14218745967066
50509	COL5A3	collagen type V alpha 3 chain	6750
2.82240295503363	3.41623433472429	6.7583941073838
0.00933076944970333	0.14218745967066
976	ADGRE5	adhesion G protein-coupled receptor E5	4624
1.02419663983355	4.13197249408698	6.75746315493129
	0.00933563873492344	0.14218745967066
366	AQP9	aquaporin 9	3235	2.53716919768516
1.41399178953055	6.7568059419897	0.00933907781088079
0.14218745967066
57169	ZNFX1	zinc finger NFX1-type containing 1
9172	-0.433417900403993	7.16497793545839
6.7532017914878	0.00935796075946093	0.14232025663167
220002	CYB561A3	cytochrome b561 family member A3
10597	0.456772122673597	7.37514226317016
6.74891841765476	0.00938045317486164	0.142435677127429

8470	SORBS2	sorbin and SH3 domain containing 2
16730	-0.903566426028098	5.06233659477674
6.74535979748883	0.00939918194699251	0.142435677127429
9071	CLDN10	claudin 10	3380	-1.60058286293834
1.06941350688871	6.74504495101552	0.0094008408054148
	0.142435677127429
4534	MTM1	myotubularin 1	4016	-0.485373049136518
5.01993793908119	6.744023333421	0.00940622555457068
0.142435677127429
619279	ZNF704	zinc finger protein 704	15008	0.638182185563904
	4.62138174583246	6.73175905492743
0.00947111536681854	0.143263404906077
56983	POGLUT1	protein O-glucosyltransferase 1	7097
0.528325982623417	4.26644605595371	6.72144056383912
	0.00952606521933626	0.14383949122951
8482	SEMA7A	semaphorin 7A (John Milton Hagen blood group)
4309	-0.625227546764891	7.71016435596047
6.7207325726086	0.00952984749168916	0.14383949122951
51111	KMT5B	lysine methyltransferase 5B
8632	-0.342079845415052	6.01672599995745
6.71883214734093	0.00954000766418324	0.14383949122951
3726	JUNB	JunB proto-oncogene, AP-1 transcription factor subunit	1820	-0.587545880481235	6.79533830561108
6.71410745307142	0.009565315237014	0.143924457795889
10892	MALT1	MALT1 paracaspase	10850	-0.487702028411706
	7.06148306495001	6.71195636785589
0.00957686018541342	0.143924457795889
60485	SAV1	salvador family WW domain containing protein 1
4760	-0.446062402398907	5.75135114231188
6.71146567228125	0.00957949576450719	0.143924457795889
5154	PDGFA	platelet derived growth factor subunit A	4151
	0.525847789292142	5.35476371682412
6.71011695342067	0.00958674370840041	0.143924457795889
157773	C8orf48	chromosome 8 open reading frame 48	1445
1.05247221302755	1.16638150443294	6.70714449920888
	0.00960273736243921	0.1440102165264З
6781	STC1	stanniocalcin 1	4149	1.22740168078202
0.313662893590309	6.70261881190928	0.00962714089090412
	0.144221778030903
4929	NR4A2	nuclear receptor subfamily 4 group A member 2
3967	-0.828198040278608	2.93655102538109
6.68948717928367	0.00969830991015353	0.144978352417645
56477	CCL28	C-C motif chemokine ligand 28
4524	-1.74541679293938	4.75202565625945
6.6894807579875	0.00969834484295949	0.144978352417645
55917	CTTNBP2NL	CTTNBP2 N-terminal like
5964	-0.371256951816642	7.15500486216512
6.6789596460512	0.00975575477623495	0.145599882838479
2247	FGF2	fibroblast growth factor 2	6775
1.62577433706765	3.48460473926151	6.677128753854
0.00976578081415402	0.145599882838479
10611	PDLIM5	PDZ and LIM domain 5	11090	-0.611899747986229
	8.04041939278429	6.67615697899458
0.00977110658015065	0.145599882838479

5698	PSMB9	proteasome 20S subunit beta 9
3072	-1.14232766963044	2.47815207095375
6.66759555157468	0.00981815588203339	0.145937437559585
1312	COMT	catechol-O-methyltransferase	5195
0.318763443999377	6.90679249333216	6.66609862370717
	0.00982640605882683	0.145937437559585
11259	FILIP1L filamin A interacting protein 1 like	5903
0.752711810577051	7.55180645723316	6.66321550733294
	0.00984231610692229	0.145937437559585
3664	IRF6	interferon regulatory factor 6
4915	-0.509686841956816	9.34879074146548
6.66309539543204	0.00984297949904619	0.145937437559585
85465	SELENOI selenoprotein I 8552	-0.52571111332428
5.97278137962055	6.66257502266777	0.00984585410821787
	0.145937437559585
9526	MPDU1	mannose-P-dolichol utilization defect 1 2876
0.401340239881342	6.14214297137824	6.65006607930337
	0.00991521471820372	0.14677200547573
100528030	POC1B-GALNT4	POC1B-GALNT4 readthrough	5500
	-0.479485969703365	4.2866760663151 6.64850766806473
	0.00992389093345964	0.14677200547573
5194	PEX13	peroxisomal biogenesis factor 13
5910	-0.308462485246592	6.08627702032854
6.64665999036151	0.00993418767724063	0.14677200547573
653499	LGALS7B galectin 7B	644	-0.737985465422177
4.51364057745097	6.64488795062516	0.00994407319172324
	0.14677200547573
25959	KANK2	KN motif and ankyrin repeat domains 2	6684
0.429772102286557	5.07248402160682	6.63838805522832
	0.00998041995369067	0.147153412938258
958	CD40	CD40 molecule	2431	-0.708728094656938
3.65537634935541	6.63502039679128	0.00999930508156269
	0.147276830996266
113251	LARP4	La ribonucleoprotein 4	8112	-0.376674986318504
	6.37897788275027	6.6257305191195 0.0100515908864376
	0.147891421792365
30061	SLC40A1 solute carrier family 40 member 1	4330
0.726095644594034	3.47663882323647	6.62308873694235
	0.0100665106263176	0.147955523969392
1452	CSNK1A1 casein kinase 1 alpha 1 7714	-0.356978107220627
	8.67102253691873	6.62041414442156
0.010081638790983	0.148022551934276
65056	GPBP1	GC-rich promoter binding protein 1
6472	-0.296057418461701	6.31242344359036
6.61435746548887	0.0101159830813933	0.148371282744582
55281	TMEM140 transmembrane protein 140	2313
1.11882961787328	2.89625925262146	6.60765657519015
	0.0101541201325076	0.148774854284554
101927668	LOC101927668	uncharacterized LOC101927668
2564	-0.617386548380535	3.47998346774477
6.59245594291814	0.0102411794638354	0.149893626697955
50486	G0S2	G0/G1 switch 2	866	-1.04547588834936
2.22465061767031	6.58409741495468	0.0102893773821457
	0.150441867423502

134285	TMEM171 transmembrane protein 171
1535	-1.14513521880185	3.53251538493534
6.58064705993037	0.0103093409763952	0.150575333941662
29113	C6orf15 chromosome 6 open reading frame 15
1140	-1.40772278233422	2.38117681041816
6.5787853004226 0.01032012952128	0.150575333941662
26035	GLCE	glucuronic acid epimerase
6630	-0.475166654318131	5.73304747066486
6.57695349955576	0.010330755759116	0.150575333941662
3775	KCNK1	potassium two pore domain channel subfamily K member
1	3627	-0.680503510676712	4.78131860246397
6.56719926207969	0.0103875290968427	0.151229118668939
8291	DYSF	dysferlin	8386	1.0029823900761
5.58811792398529	6.56554240861658	0.0103972043462689
0.151229118668939
3991	LIPE	lipase E, hormone sensitive type
5206	-0.803277784337655	2.15133973029105
6.55794326040724	0.0104416983395125	0.151680368675802
80336	PABPC1L poly(A) binding protein cytoplasmic 1 like	4984
0.523322613153445	4.43123914011938
6.55391035615369	0.0104653907698257	0.151680368675802
55655	NLRP2	NLR family pyrin domain containing 2	5770
3.20989342593198	4.6349258767497 6.5533462592677
0.0104687091073763	0.151680368675802
26586	CKAP2	cytoskeleton associated protein 2	4547
0.666098377403331	5.1592079112324 6.55286449386346
0.0104715439786893	0.151680368675802
10626	TRIM16	tripartite motif containing 16
5639	-0.720473059038319	8.36403127466108
6.54897494439558	0.0104944602617453	0.151695062943392
144108	SPTY2D1 SPT2 chromatin protein domain containing 1
6005	-0.373781705991466	5.770842673796	6.54747890193422
0.0105032882699832	0.151695062943392
22998	LIMCH1	LIM and calponin homology domains 1	10862
0.54336410027489	4.86619212445783	6.54718079764682
0.0105050482655165	0.151695062943392
84220	RGPD5	RANBP2 like and GRIP domain containing 5
13276	-0.545859647183986	5.42730167466104
6.5345299992901 0.0105800174899772	0.152478190559096
5756	TWF1	twinfilin actin binding protein 1
3932	-0.353530542241319	6.6496024768963 6.53435600268624
0.0105810524183224	0.152478190559096
1889	ECE1	endothelin converting enzyme 1
7265	-0.587270316442504	9.28964818704427
6.53217451782939	0.0105940366610802	0.152508398264903
101927181	NA	NA	2067	0.716901807553366
1.95937070926427	6.52506299327383	0.0106364781772965
0.152962166149273
51542	VPS54	VPS54 subunit of GARP complex
5091	-0.328053986224132	5.45484248903885
6.51923850570806	0.0106713685447247	0.153306522262522
54520	CCDC93	coiled-coil domain containing 93
8065	-0.525158910445038	6.72762020608352
6.5161140776686 0.0106901331241826	0.153342313766754

394	ARHGAP5 Rho GTPase activating protein 5
10277	-0.463471729398006	7.82258976789831
6.51517908958135	0.0106957550189276	0.153342313766754
7076	TIMP1	TIMP metallopeptidase inhibitor 1	1541
1.44603669857147	6.16553742335461	6.50987110145731
0.0107277284043514	0.153619680565998
25758	KIAA1549L	KIAA1549 like	12318	-1.18338460647361
3.44147983776736	6.50722068647834
0.0107437302183394	0.153619680565998
5184	PEPD	peptidase D	4157	0.442190378166385
4.96028464789992	6.50651355582128	0.010748003637801
0.153619680565998
2948	GSTM4	glutathione S-transferase mu 4	5460
1.42148341487485	3.29964316229368	6.50358275089602
0.0107657340370661	0.153716245318232
4651	MYO10	myosin X	13963	-0.455036303998049
7.55862478096786	6.49998565141351	0.0107875363123803
0.153870693612536
219285	SAMD9L	sterile alpha motif domain containing 9 like
7135	-0.638088533000977	5.36336517150137
6.49550247680749	0.0108147725646651	0.154102257694063
27346	TMEM97	transmembrane protein 97	3034
0.937598962690647	2.8042190997202 6.48717600527331
0.0108655449936377	0.154559484081826
250	ALPP	alkaline phosphatase, placental 3059
1.86473779410579	3.73545433840858	6.48597832931448
0.0108728681663555	0.154559484081826
100533464	PRH1-PRR4	PRH1-PRR4 readthrough	1633
0.680714024931248	1.30941916255733	6.48481867470117
0.0108799636827787	0.154559484081826
25849	PARM1	prostate androgen-regulated mucin-like protein 1
5034	-0.75424816236152	4.06212764901012
6.47640229987772	0.0109316030521544	0.155135728421
401647	GOLGA7B golgin A7 family member B
6456	-0.548803646378136	2.99564373790613
6.47416073609908	0.0109453987080804	0.155137530627845
11320	MGAT4A	alpha-1,3-mannosyl-glycoprotein 4-beta-N-acetylglucosaminyltransferase A 9630	-0.674157073602385
4.80860739059602	6.47234483289859	0.0109565877400555
0.155137530627845
2876	GPX1	glutathione peroxidase 1	1425
0.504858943578962	7.00095585245363	6.47098777231871
0.0109649571872326	0.155137530627845
7518	XRCC4	X-ray repair cross complementing 4	2676
0.720013646175009	3.63128984156066	6.46867977376384
0.0109792064539971	0.155182386277636
116068	LYSMD3	LysM domain containing 3
4459	-0.412483872308959	5.55601760902208
6.46301521013011	0.0110142592814035	0.155520896930059
10565	ARFGEF1 ADP ribosylation factor guanine nucleotide exchange factor 1	10964	-0.33110691248216	7.51270790118042
6.45948890053723	0.0110361383678156
0.155530747124242
92181	UBTD2	ubiquitin domain containing 2

3301	-0.361058602693471	6.02403620679973
6.45932369812277	0.0110371644635894	0.155530747124242
157769	FAM91A1	family with sequence similarity 91 member A1
7246	-0.345165429656642	6.8207077615219	6.45683114221688
	0.0110526579692594	0.155592542889865
30818	KCNIP3	potassium voltage-gated channel interacting protein
3	7870	1.10026798047276	1.86033659460371
6.44453311676155	0.0111294287136404	0.156515971879479
221710	SMIM13	small integral membrane protein 13
4688	-0.448174839632572	5.13340777813872
6.43254792307076	0.0112047731165065	0.157417509571622
79873	NUDT18	nudix hydrolase 18	1756	0.525757335263305
	3.36120584315538	6.42866040267075
0.0112293241060404	0.157604351456221
1363	CPE	carboxypeptidase E	2958	1.23747724764679
	3.479069918814	6.41470540811322	0.0113179102510854
	0.158688657544497
59084	ENPP5	ectonucleotide pyrophosphatase/phosphodiesterase family member 5	4264	0.684881506444398	4.17518379306481
	6.41283232339656	0.0113298550444777
0.158697279607999
105370333	PCCA-DT	PCCA divergent transcript	913
0.746522290095315	1.09715086827507	6.40335810107972
	0.0113904713588569	0.159386945378131
9486	CHST10	carbohydrate sulfotransferase 10	4487
0.747164623854442	2.34496343239678	6.3988438886287
0.0114194703859724	0.15951158538394
3290	HSD11B1	hydroxysteroid 11-beta dehydrogenase 1
1513	-2.08013391342551	1.70788439262805
6.39842663094392	0.0114221546469688	0.15951158538394
219988	PATL1	PAT1 homolog 1, processing body mRNA decay factor
	4278	-0.289985314497758	6.14479357594865
6.38260544573541	0.0115244132274721	0.160642197614935
251	ALPG	alkaline phosphatase, germ cell	2493
1.50794435542328	0.924706021605823	6.3823530931493
0.0115260518742779	0.160642197614935
479	ATP12A	ATPase H+/K+ transporting non-gastric alpha2 subunit
	4442	-1.84849526005878	2.27281723134516
6.38031956257052	0.0115392652992984	0.160666490106634
51132	RLIM	ring finger protein, LIM domain interacting
8379	-0.334784243388019	6.27028362030521
6.37736832986788	0.0115584694109814	0.160774062601406
282809	POC1B	POC1 centriolar protein B
8746	-0.351000646632512	5.31023404330256
6.37400620994102	0.0115803872097018	0.160919130601481
1955	MEGF9	multiple EGF like domains 9	6298
0.385473379303161	6.20164595864638	6.35347686039231
	0.0117151477425821	0.162570642151974
2920	CXCL2	C-X-C motif chemokine ligand 2
1500	-1.31574229855784	4.97705522611972
6.35076393732865	0.0117330761953075	0.162570642151974
10938	EHD1	EH domain containing 1	7101	-0.342334808414152
	7.22126433644895	6.34919916657076
0.0117434298480981	0.162570642151974

54434	SSH1	slingshot protein phosphatase 1 14537
0.360005982055746	7.797388296853	6.34750624789898
0.0117546419846933	0.162570642151974
80004	ESRP2	epithelial splicing regulatory protein 2
5563	-0.440285458639941	6.24652254825775
6.34710990377981	0.0117572685442957	0.162570642151974
9048	ARTN	artemin 2658	-0.937579123573047
4.02639966365192	6.3446985512517 0.0117732614898255
0.162631236378684
10912	GADD45G growth arrest and DNA damage inducible gamma	1322
0.73532409517075	-0.062332759515471
6.34059845506207	0.01180050607954	0.162846983897652
1435	CSF1	colony stimulating factor 1	5418
0.534605646845408	5.04579308451579	6.33685081601465
0.0118254652645827	0.163030799174222
8511	MMP23A	matrix metallopeptidase 23A (pseudogene)	1275
1.02697458031342	0.269535368915177
6.33332710974475	0.0118489825069311	0.163194393288677
400619	LINC00511	long intergenic non-protein coding RNA 511
2326	0.746694525460756	1.71388233940443
6.33074840122749	0.0118662232542272	0.163271305620787
2140	EYA3	EYA transcriptional coactivator and phosphatase 3
6483	-0.368407248552913	5.35035813202669
6.31806692913586	0.0119513847233665	0.164281693641015
2977	GUCY1A2 guanylate cyclase 1 soluble subunit alpha 2
16361	1.73959290665322	0.484042006151185
6.30969225848785	0.0120079680611858	0.164897655522577
6732	SRPK1	SRSF protein kinase 1	6692	-0.377108387518016
7.31316915958678	6.30473748110948
0.012041574403459	0.165197191644711
64746	ACBD3	acyl-CoA binding domain containing 3
3578	-0.293484874941499	6.55733333229751
6.29776472136202	0.0120890316338683	0.165619038932048
22853	LMTK2	lemur tyrosine kinase 2 9139	-0.499743577744915
6.71229843344661	6.29674738055473
0.012095971787498	0.165619038932048
55647	RAB20	RAB20, member RAS oncogene family	1518
0.809110051108496	2.94344112021993	6.29095719628236
0.012135549594836	0.165998674975457
64101	LRRC4	leucine rich repeat containing 4
4036	-0.498523930475247	4.43251276905729
6.28879932195436	0.0121503333722015	0.166026537874779
55568	GALNT10 polypeptide N-acetylgalactosaminyltransferase 10
10496	1.02014272421966	6.48385683034235
6.28720140243053	0.0121612927721513	0.166026537874779
331	XIAP	X-linked inhibitor of apoptosis
9025	-0.418708268549545	6.54264182371829
6.27749088770466	0.012228111445542	0.166631355983005
481	ATP1B1	ATPase Na+/K+ transporting subunit beta 1
2962	-0.635189363409782	8.53995268283602
6.27730594065565	0.0122293877311722	0.166631355983005
6670	SP3	Sp3 transcription factor
10704	-0.345463055115726	6.60106023547557
6.2713670269313 0.0122704439118202	0.166919141770802

83595	SOX7	SRY-box transcription factor 7
3701	-0.549049020157942	4.48581411010647
6.27080413817759	0.0122743425447224	0.166919141770802
29085	PHPT1	phosphohistidine phosphatase 1	1612
0.49913831059322	5.82045119881714	6.26617107414038
	0.0123064800625016	0.16710148245362
57128	LYRM4	LYR motif containing 4	9443	0.418316676387832
	3.99917197204712	6.26543258515661
0.0123116106155591	0.16710148245362
1286	COL4A4	collagen type IV alpha 4 chain	9895
1.22766533925041	5.42103494686093	6.24783699923441
	0.0124345056594989	0.168606118850533
123606	NIPA1	NIPA magnesium transporter 1
8700	-0.352325585850501	6.51494992253722
6.23695296482585	0.0125111546167672	0.169213057273875
2619	GAS1	growth arrest specific 1
2827	-1.08702352064772	0.0721031995230924
6.23655469341437	0.0125139685644089	0.169213057273875
195827	PRXL2C	peroxiredoxin like 2C	3084	0.522367490908022
	3.54900321199753	6.23439001892456
0.0125292742362368	0.169213057273875
200844	C3orf67	chromosome 3 open reading frame 67
9455	-0.726947997857518	2.82740044784541
6.23390684757033	0.0125326932003671	0.169213057273875
23705	CADM1	cell adhesion molecule 1	11017
0.610754935706112	4.43113984068693	6.232921342193
0.0125396696973143	0.169213057273875
5324	PLAG1	PLAG1 zinc finger	7833	-0.734216340564505
	3.06865730226441	6.23010275269438
0.0125596448256409	0.169312068673409
948	CD36	CD36 molecule	9761	2.24058638735978
1.47603266307704	6.22847684620401	0.0125711823674124
	0.169312068673409
5744	PTHLH	parathyroid hormone like hormone
3377	-0.819172813354136	5.16582388952278
6.21872561946481	0.0126406066190422	0.170083551309244
51293	CD320	CD320 molecule	2277	0.543036844575665
3.30998322903626	6.20508395138948	0.0127383907413638
	0.171230983114734
1462	VCAN	versican	14831	0.718373325261193
9.59566164438992	6.2034252786259 0.01275033307551
0.171230983114734
730755	KRTAP2-3	keratin associated protein 2-3
875	-1.44065682066485	4.45397518842029
6.19612856836962	0.0128030057074129	0.17177365990779
85403	EAF1	ELL associated factor 1 4690	-0.328939284172089
	6.06913820154982	6.17826899916948
0.0129328739272023	0.173350014448156
7187	TRAF3	TNF receptor associated factor 3
8263	-0.378802236705923	6.57170997142746
6.17507488214921	0.0129562427274137	0.173497219773311
93129	ORAI3	ORAI calcium release-activated calcium modulator 3
	2876	0.596105805122825	4.16049045714125
6.16701876848662	0.0130153758550264	0.174122607069107

7453	WARS1	tryptophanyl-tRNA synthetase 1
6358	-0.689841509624572	7.31909484236512
6.16429571928336	0.0130354261355597	0.174190474589756
83988	NCALD	neurocalcin delta	8042	-0.984208249433683
	0.828369026703607	6.16295350678214
0.0130453207570967	0.174190474589756
54503	ZDHHC13 zinc finger DHHC-type containing 13
7763	-0.338005101995228	6.12836812212144
6.16020026094201	0.013065641507872	0.17429565771 5013
100528016	TMX2-CTNND1	TMX2-CTNND1 readthrough (NMD
candidate)	5044	-0.366551349264864	8.85843523724512
	6.1540547288384 0.013111170072054	0.174359013909453
11074	TRIM31	tripartite motif containing 31
3535	-1.91357571111143	-0.145831484576894
6.15252303841835	0.0131224764680937	0.174359013909453
8324	FZD7	frizzled class receptor 7	3859
0.805384900134702	3.49671232136902	6.15201869656088
	0.013126219027585	0.174359013909453
127703	C1orf216	chromosome 1 open reading frame 216	3629
	0.55042192424207	4.30133683670219
6.1515527000743 0.0131296780137036	0.174359013909453
9069	CLDN12	claudin 12	6018	-0.352287190710472
6.95563367300156	6.15115499838151	0.0131326308042031
	0.174359013909453
200197	TMEM51-AS1	TMEM51 antisense RNA 1
7100	-0.662175679011924	5.29354663715998
6.14558587011242	0.0131740513478525	0.174743311770237
5652	PRSS8	serine protease 8	3010	-0.323365407726587
	7.40843196223145	6.14391134568077
0.0131865319145499	0.174743379874267
84612	PARD6B	par-6 family cell polarity regulator beta
4726	-0.972483605187933	3.52972619720317
6.14010433973933	0.0132149515480051	0.174954467233371
57469	PNMA8B	PNMA family member 8B	5308	0.75594110179949
	0.550563557895938	6.13521038128794
0.0132515779126095	0.175273703325704
84941	HSH2D	hematopoietic SH2 domain containing
3398	-0.670485420322639	2.80996404427476
6.12926324297732	0.0132962267922817	0.175698347810839
51166	AADAT	aminoadipate aminotransferase	2981
0.683818509953583	2.95876704092586	6.1124273242896
0.0134234652284444	0.177212514095024
3912	LAMB1	laminin subunit beta 1	8140	0.543602864239576
	8.15426076423635	6.10773268560786
0.0134591679987182	0.177516540638462
54814	QPCTL	glutaminyl-peptide cyclotransferase like	2396
	0.572852375628222	3.17590885140346
6.09813568845269	0.0135324574305554	0.177636414836351
467	ATF3	activating transcription factor 3
3348	-0.884035646210795	5.48449170998989
6.09741225091483	0.0135379987176864	0.177636414836351
400986	ANKRD36C	ankyrin repeat domain 36C
8574	-0.993557963975121	3.65140397825108
6.09631783843543	0.0135463859832885	0.177636414836351

3096	HIVEP1	HIVEP zinc finger 1	10979	-0.675665551356933
	6.81169024754652	6.09309139804195
0.0135711436182371	0.177636414836351
3489	IGFBP6	insulin like growth factor binding protein 6	1279
	0.85365075900293	3.29075544120292
6.09284451467325	0.0135730399585298	0.177636414836351
441631	TSPAN11	tetraspanin 11	7113	1.79682100392339
0.757955048741258	6.0921715348726	0.013578210579877
0.177636414836351
84525	HOPX	HOP homeobox	2981	1.43674948713799
4.73056867560939	6.09096515590656	0.0135874844713999
	0.177636414836351
84640	USP38	ubiquitin specific peptidase 38
7787	-0.304219265441266	6.01229046438425
6.09013094283947	0.0135939011925535	0.177636414836351
51278	IER5	immediate early response 5
4188	-0.34144661474535	6.34437685890133
6.09001728914185	0.0135947756517613	0.177636414836351
1385	CREB1	cAMP responsive element binding protein 1
9575	-0.401292625046643	5.97330124999114
6.08997818798529	0.0135950765120703	0.177636414836351
63917	GALNT11	polypeptide N-acetylgalactosaminyltransferase 11
	8295	0.361147822143379	5.48098449028497
6.08562827057527	0.0136285893510383	0.177908342068959
3815	KIT	KIT proto-oncogene, receptor tyrosine kinase	5563
	1.08310297739105	1.83783213187366
6.08337112387233	0.013646012438891	0.177969922003302
26273	FBXO3	F-box protein 3	12446	-0.44827114420787
6.21319244954199	6.07258721022494	0.013729571047391
	0.178893117824005
3914	LAMB3	laminin subunit beta 3	4688	-0.504247111133116
	11.5977361806112	6.06688835272462
0.01377394072707	0.179222394366583
10762	NUP50	nucleoporin 50	8318	-0.618722215916875
6.7464755177648	6.06605612451034	0.0137804325503541
0.179222394366583
9849	ZNF518A	zinc finger protein 518A
10891	-0.430428938819968	5.02278163248311
6.06427568397903	0.0137943315061573	0.179236735998836
8840	CCN4	cellular communication network factor 4	3844
2.98660228240249	0.785154652143733	6.06030922796797
	0.0138253473988896	0.179331308189888
117177	RAB3IP	RAB3A interacting protein
12612	-0.440398499864116	5.84789456663832
6.05914904471472	0.0138344330613381	0.179331308189888
1808	DPYSL2	dihydropyrimidinase like 2	7052
0.461670662008785	5.41950534887533	6.05843616173642
	0.0138400188586613	0.179331308189888
3074	HEXB	hexosaminidase subunit beta	4977
0.497709346638613	6.6747041342955	6.05281088710333
0.0138841771869441	0.179737217982926
57532	NUFIP2	nuclear FMR1 interacting protein 2
10850	-0.407328932068992	7.59187215166075
6.05084421454671	0.013899649712927	0.179771369832843

168667	BMPER	BMP binding endothelial regulator	5888
0.822600459207777	1.52687450544555	6.04902787014684
	0.0139139553095454	0.179790380092991
9440	MED17	mediator complex subunit 17
10773	-0.358152279603521	5.67100405576207
6.04014900767302	0.0139841038805818	0.180446691537934
55765	INAVA	innate immunity activator
5413	-0.411604160661806	7.92227392347562
6.03839555081339	0.0139980002564707	0.180446691537934
54811	ZNF562	zinc finger protein 562 13007	-0.415272307562125
	5.88851215134882	6.03771532047608
0.0140033949954833	0.180446691537934
23515	MORC3	MORC family CW-type zinc finger 3
6408	-0.335441183726428	6.58875496185802
6.02974673799069	0.0140667514003871	0.180970568131606
1285	COL4A3	collagen type IV alpha 3 chain	9124
1.20326974227074	3.68415735708396	6.0293529494032
0.0140698899618276	0.180970568131606
55228	PNMA8A	PNMA family member 8A	3696	1.29975000021032
	4.96043485956854	6.02674424863819
0.014090699952273	0.181071958010539
83872	HMCN1	hemicentin 1	18579	1.21995558176449
0.824352031826935	6.02082540960999	0.0141380329312059
	0.181453888857189
51367	POP5	POP5 homolog, ribonuclease P/MRP subunit	2072
	0.556211359424132	3.19468578822944
6.01813758590853	0.0141595814563152	0.181453888857189
152078	SLC66A1L	solute carrier family 66 member 1 like	3180
	0.8370900018186 0.267470118991405	6.01674825766461
	0.0141707330727963	0.181453888857189
27198	HCAR1	hydroxycarboxylic acid receptor 1	4768
0.889548818929196	3.67528037563355	6.01656070306802
	0.0141722391953855	0.181453888857189
8794	TNFRSF10C	TNF receptor superfamily member 10c	1868
	0.500739017449879	4.56475659998577
6.01388684818249	0.0141937290167057	0.181563070627395
9143	SYNGR3	synaptogyrin 3	3420	-0.81609781044466
1.9150970856165 6.00986192020403	0.0142261406859881
0.181811635573572
4628	MYH10	myosin heavy chain 10	9011	0.336653803564323
	5.90578805779475	5.99106028254925
0.0143785572352987	0.183592024790181
29965	CDIP1	cell death inducing p53 target 1	4268
0.602263487100888	5.81391300158691	5.98551295059634
	0.0144238473282352	0.184002576982323
4360	MRC1	mannose receptor C-type 1	5198
2.55048649365983	3.48382249844171	5.98228733173003
	0.0144502497405306	0.184171654336317
727851	RGPD8	RANBP2 like and GRIP domain containing 8
25009	-0.473286107097103	6.58861796628487
5.97936182470522	0.014474238687149	0.184309692082633
84914	ZNF587	zinc finger protein 587 7760	-0.312251509635026
	6.1356682494578 5.97548184154848	0.0145061175184055
	0.184393117143807

3294	HSD17B2 hydroxysteroid 17-beta dehydrogenase 2
9550	-1.15015393680387	1.35190107838287
5.9753600995547 0.0145071189471319	0.184393117143807
7068	THRB	thyroid hormone receptor beta
8709	-0.633706061201088	4.34763934098748
5.96893621915778	0.0145600617579706	0.184898263865724
124411	ZNF720	zinc finger protein 720 9816	-0.373647302054156
	4.91062890237371	5.9673281370874 0.0145733459651288
	0.184899326932572
79622	SNRNP25 small nuclear ribonucleoprotein U11/U12 subunit 25
	3060	0.600386587254153	3.00781317303872
5.9631478233287 0.0146079375293696	0.185068786433753
5213	PFKM	phosphofructokinase, muscle	7284
0.449335174950552	4.96654245666662	5.96252149838743
	0.0146131275644843	0.185068786433753
5290	PIK3CA	phosphatidylinositol-4,5-bisphosphate 3-kinase
catalytic subunit alpha 9411	-0.33609942239607
6.26427855159591	5.94603092328774	0.0147504616928456
	0.186639310688066
51429	SNX9	sorting nexin 9 4646	-0.327084270817407
6.81143650010796	5.94290727650643	0.0147766248678426
	0.186801610581111
84141	EVA1A	eva-1 homolog A, regulator of programmed cell death
	2744	0.480510592902381	5.12513026821961
5.9394932551627 0.0148052748285784	0.18699502662209
493	ATP2B4	ATPase plasma membrane Ca2+ transporting 4
9721	-0.412093454523783	8.33632369638394
5.93111829502654	0.0148757987484628	0.187594245889485
5806	PTX3	pentraxin 3	1940	1.5494485861339
0.468073039116896	5.92980673994331	0.0148868743865465
	0.187594245889485
26108	PYGO1	pygopus family PHD finger 1	8594
1.0956578937027 1.64490924715865	5.92778644695716
0.0149039516921531	0.187594245889485
1466	CSRP2	cysteine and glycine rich protein 2	5873
0.739159709548163	5.1823465939452 5.92700033785803
0.0149106020328684	0.187594245889485
7275	TUB	TUB bipartite transcription factor	6722
1.07657644361204	-0.0764440948548938	5.92592758054311
	0.0149196822960582	0.187594245889485
79001	VKORC1	vitamin K epoxide reductase complex subunit 1	2514
	0.473759703740779	4.36839238832608
5.91463386006654	0.0150156232754745	0.188631242349391
222584	FAM83B	family with sequence similarity 83 member B
3167	-0.476731540871677	4.47392296646593
5.90255006871529	0.0151189797725186	0.189759453112606
10529	NEBL	nebulette	13654	-0.70467121915048
5.99198153423588	5.89923124562958	0.015147494769517
	0.189822226130236
79048	SECISBP2	SECIS binding protein 2
9813	-0.343541888860266	5.64451508948741
5.89815731790396	0.0151567337062989	0.189822226130236
161582	DNAAF4	dynein axonemal assembly factor 4	2934
0.740868173074564	1.53126817134279	5.89723917452924

0.0151646370414603	0.189822226130236
7871	SLMAP	sarcolemma associated protein
14111	-0.306510447451363	7.25020713039222
5.89423981047607	0.0151904850208709	0.189923190685451
142891	SAMD8	sterile alpha motif domain containing 8
7316	-0.392823239648004	6.17588629286706
5.89315773509597	0.0151998212863847	0.189923190685451
3845	KRAS	KRAS proto-oncogene, GTPase
7302	-0.395260098786534	6.42048541012228
5.88827836338446	0.0152419945133952	0.190022033682514
127845	GOLT1A	golgi transport 1A	1136	-1.10501788964145
1.76943472223689	5.88803330067361
0.0152441158073713	0.190022033682514
4690	NCK1	NCK adaptor protein 1	3727	-0.597304445934424
5.54325830352283	5.88753495498651
0.0152484304889802	0.190022033682514
23580	CDC42EP4	CDC42 effector protein 4
4488	-0.718713169952025	6.04143818610992
5.88523686660972	0.0152683436406175	0.190074559872169
57630	SH3RF1	SH3 domain containing ring finger 1
5624	-0.348790560959837	6.5318118284336 5.88391781749305
0.0152797854227217	0.190074559872169
84886	C1orf198	chromosome 1 open reading frame 198	5161
0.42211511572761	5.735958577037	5.87085582327742
0.0153935665376413	0.191283029935399
54469	ZFAND6	zinc finger AN1-type containing 6
5153	-0.295274740439785	6.3952306613073 5.86963506768586
0.015404244860936	0.191283029935399
8693	GALNT4	polypeptide N-acetylgalactosaminyltransferase 4
5399	-0.467062018179848	4.25434616486652
5.86747705459156	0.0154231403347073	0.191297776158914
5921	RASA1	RAS p21 protein activator 1
5232	-0.36533458470037	6.64870514277641
5.86638094267227	0.0154327469879041	0.191297776158914
55342	STRBP	spermatid perinuclear RNA binding protein
8083	-0.324614672427947	5.20697436593646
5.86401662638031	0.0154534895584186	0.191385524531184
112724	RDH13	retinol dehydrogenase 13
6713	-0.558296216617756	5.3391738377991 5.85737660099697
0.0155118974286523	0.191823261596211
4921	DDR2	discoidin domain receptor tyrosine kinase 2
10730	1.70991221357591	2.06977624373769
5.85655514956039	0.0155191389803694	0.191823261596211
8034	SLC25A16	solute carrier family 25 member 16	7813
0.384790513259753	4.98732564215151
5.8549431521727 0.0155333597601936	0.191823261596211
64089	SNX16	sorting nexin 16	4610	-0.592789690879403
4.71443099513639	5.85378172039608
0.0155436140438137	0.191823261596211
54549	SDK2	sidekick cell adhesion molecule 2	11607
0.769286084941297	4.98733865276962	5.85172173272059
0.0155618188380699	0.191878870127505
55222	LRRC20	leucine rich repeat containing 20	3625
0.837944772907747	3.56926949903213	5.84836544654554

0.0155915265187198	0.192052635462707
55785	FGD6	FYVE, RhoGEF and PH domain containing 6
10044	-0.613061120510914	6.70964589696889
5.84557903774831	0.0156162344020703	0.192052635462707
728340	GTF2H2C GTF2H2 family member C	6964	0.613262527808416
3.53182688689696	5.84548768036054
0.0156170451768418	0.192052635462707
3749	KCNC4	potassium voltage-gated channel subfamily C member 4
24423	-0.664335076407891	2.89532840093106
5.84113091571351	0.0156557608093601	0.192257155476241
23225	NUP210	nucleoporin 210 8182	-0.975673046053237
2.68397708977431	5.83962866202386	0.0156691332431975
0.192257155476241
8630	HSD17B6 hydroxysteroid 17-beta dehydrogenase 6	2272
1.22539325644531	2.52999510457362	5.83898646548689
0.0156748533985769	0.192257155476241
3932	LCK	LCK proto-oncogene, Src family tyrosine kinase
3339	-1.36862148762227	0.845671741596883
5.83522638017625	0.0157083883356117	0.192499908501236
339324	ZNF260	zinc finger protein 260 5952	-0.539813054393529
5.56620757582612	5.83188910733604
0.0157382142751005	0.192696824607808
22955	SCMH1	Scm polycomb group protein homolog 1	4447
0.426875281015306	4.17048688876411	5.83023713577702
0.0157529998940515	0.192709405690811
23258	DENND5A DENN domain containing 5A	8648
0.381368540906491	5.59384137549706	5.82490609623975
0.015800811930927	0.193120589663856
6553	SLC9A5	solute carrier family 9 member A5	5419
0.758251984239192	1.09112698149163	5.82341775974358
0.0158141869311375	0.193120589663856
27244	SESN1	sestrin 1	5121	0.520252417180786
3.88575885638991	5.82070309305023	0.0158386124566086
0.19325038735167
3569	IL6	interleukin 6	2069	-1.14989917505024
5.17780344228865	5.81735120512235	0.0158688251014737
0.193450507568618
8685	MARCO	macrophage receptor with collagenous structure	2079
2.85585452690851	2.12431860588999
5.81503067344853	0.0158897762800375	0.193537475090857
101927482	HECW2-AS1	HECW2 antisense RNA 1
1607	-0.929354945337812	1.08271667136448
5.80991754821492	0.0159360413723563	0.193932347091741
4810	NHS	NHS actin remodeling regulator
10168	-0.469119597612201	4.63138081395863
5.80445527058126	0.0159856191025569	0.194336320870656
1437	CSF2	colony stimulating factor 2
787	-0.887447394178708	5.61223691063116
5.8032054102186 0.0159969856474524	0.194336320870656
7431	VIM	vimentin	4965	0.785482789558287
8.76651048849632	5.7951908882269 0.0160700699750363
0.195055000121606
10781	ZNF266	zinc finger protein 266 5267	-0.452299908793979
4.89122682538224	5.79068878443023

0.01611127548462	0.195361063878855
83959	SLC4A11 solute carrier family 4 member 11
4081	-0.454018926722502	6.36383783894494
5.7879830248818 0.016136092332586	0.195361063878855
5554	PRH1	proline rich protein HaeIII subfamily 1 3144
0.678071132554588	1.01786115359173	5.78787020790301
	0.016137127929452	0.195361063878855
4939	OAS2	2'-5'-oligoadenylate synthetase 2
6264	-0.725498592269284	3.65924821012404
5.78162267292686	0.0161945838943603	0.195887337312871
8751	ADAM15	ADAM metallopeptidase domain 15
6584	-0.477289065397932	7.13780365643838
5.7628686089341 0.0163683272190824	0.197818084001456
91768	CABLES1 Cdk5 and Abl enzyme substrate 1 6654
0.608064447135211	2.51091550170551	5.75911530014966
	0.0164033291747385	0.198003733175852
84547	PGBD1	piggyBac transposable element derived 1 3100
0.723132510083577	2.72716267923387	5.75819099352827
	0.0164119607494227	0.198003733175852
2568	GABRP	gamma-aminobutyric acid type A receptor pi subunit
	4621	-1.58984640025376	6.81504238572886
5.74613287863974	0.0165249946679612	0.199195869461388
84084	RAB6C	RAB6C, member RAS oncogene family
3075	-0.373646482209983	4.263265027808	5.73884176776287
	0.0165937313894024	0.199727572832685
5602	MAPK10	mitogen-activated protein kinase 10
9816	-0.782167589659482	1.14070165909374
5.73842998490774	0.0165976222443954	0.199727572832685
214	ALCAM	activated leukocyte cell adhesion molecule
7179	-0.647708424519842	8.71002659808144
5.73610620902718	0.0166195967863145	0.199820336640264
25870	SUMF2	sulfatase modifying factor 2	4101
0.30584730393203	6.74902252546166	5.72826272501085
	0.0166939896645776	0.200317329973334
23075	SWAP70	switching B cell complex subunit SWAP70
7190	-0.298510639400447	7.10527705781856
5.72757715699605	0.0167005083516755	0.200317329973334
57088	PLSCR4	phospholipid scramblase 4	4227
0.863766399623151	2.74541552474363	5.7272272165573
0.0167038367536842	0.200317329973334
84230	LRRC8C	leucine rich repeat containing 8 VRAC subunit C
10038	-0.556019415768153	4.97844751499971
5.72501390267713	0.0167249041847075	0.200398402835926
401089	FOXL2NB FOXL2 neighbor	3546	1.34195831534589
0.709780666770963	5.72327705504157	0.0167414555597011
	0.200425271816011
168850	ZNF800	zinc finger protein 800 5793	-0.411632813189938
	4.85073419227794	5.71287061348057
0.0168409787262637	0.201444567906725
80256	FAM214B family with sequence similarity 214 member B
6409	-0.307808012433426	7.03591933379678
5.71012730390217	0.0168673161713567	0.201587455300506
23165	NUP205	nucleoporin 205 7619	-0.283357645505624
6.47159181035547	5.70483085442507	0.016918285578706

0.201989726275911
57124 CD248 CD248 molecule 2558 2.49334857815597
0.80221893387567 5.70363491792979 0.0169298164237823
0.201989726275911
153478 PLEKHG4B pleckstrin homology and RhoGEF domain
containing G4B 12203 1.15552113608561 2.5996799633656
5.70203539636017 0.016945251182838 6 0.20200181558980 5
1293 COL6A3 collagen type VI alpha 3 chain 19633
3.35991930837622 6.02056900716859 5.68924387236671
0.0170692078070737 0.203306457273539
4860 PNP purine nucleoside phosphorylase
4139 -0.543874799117622 6.09226675124335
5.68708959750635 0.0170901757264428 0.203383255225391
102800447 LINC01828 long intergenic non-protein coding
RNA 1828 5087 -1.46110946043214 0.528973684720988
5.67536107687379 0.0172047984665905
0.204464235753323
6331 SCN5A sodium voltage-gated channel alpha subunit 5
10017 1.08163359975794 -0.275036667935293
5.67480991232507 0.0172102044658505 0.204464235753323
9610 RIN1 Ras and Rab interactor 1 5469
0.623130278833003 2.3848468594404 5.67202298865239
0.01723756635823 0.204615755915023
7225 TRPC6 transient receptor potential cation channel
subfamily C member 6 5537 1.33576019604015
4.46397653350509 5.66923252565909 0.0172650079716818
0.204767964995213
28984 RGCC regulator of cell cycle 1032 0.993916366528298
0.646145008118316 5.66294164045061
0.0173270384722455 0.20532980362161
101928075 LOC101928075 uncharacterized LOC101928075
2941 -0.803911344824749 0.823958174023673
5.6602194924695 0.0173539511182868 0.205332824695747
120224 TMEM45B transmembrane protein 45B
3027 -1.1551152311268 3.85883961898964
5.65854479860508 0.0173705294866346 0.205332824695747
4919 ROR1 receptor tyrosine kinase like orphan receptor 1 6715
0.941847575749145 3.28242812655154
5.65846990034992 0.0173712713118055 0.205332824695747
11107 PRDM5 PR/SET domain 5 9517 0.720349559583817
1.62122050137688 5.6551444739544 0.0174042407628945
0.205435948279471
256987 SERINC5 serine incorporator 5 9583 -0.461345736357989
7.11787194029573 5.65455335961539
0.0174101080348551 0.205435948279471
51339 DACT1 dishevelled binding antagonist of beta catenin 1
4490 0.743300787003626 2.75086714246028
5.65315503632981 0.017423995613337 0.205435948279471
1746 DLX2 distal-less homeobox 2 2970 -0.870725019509457
0.629780584088792 5.65132525144314
0.0174421855618113 0.205477454301338
80028 FBXL18 F-box and leucine rich repeat protein 18
3605 -0.479369230249501 5.11658425027067
5.64369769403457 0.0175182227128848 0.206199786167544

2060 EPS15 epidermal growth factor receptor pathway substrate 15 7440 -0.288530282474328 5.86950090176662 5.63683523388121 0.0175869251429039 0.206707289468335
157 GRK3 G protein-coupled receptor kinase 3 9303 -0.462311974899442 4.44532903805549 5.63621424628703 0.0175931557661031 0.206707289468335
154 ADRB2 adrenoceptor beta 2 3443 -0.313335560655784 5.77510830364135 5.63380978941312 0.0176173021643977 0.206707289468335
84868 HAVCR2 hepatitis A virus cellular receptor 2 3804 1.68628931198937 1.28776988502914 5.63350467786996 0.0176203686460479 0.206707289468335
2678 GGT1 gamma-glutamyltransferase 1 4434 -1.04463339536034 2.82049494770372 5.62315107711135 0.0177247532911317 0.207757840459316
57820 CCNB1IP1 cyclin B1 interacting protein 1 3680 0.382210560694364 4.28613809090324 5.6199897161736 0.0177567530632807 0.207958896452653
3002 GZMB granzyme B 1411 -2.2067310785948 0.961778055380548 5.60891385215902 0.0178693358757286 0.209061407086935
282974 STK32C serine/threonine kinase 32C 3543 0.657551519011687 1.58154960821808 5.60654797619165 0.0178934796519037 0.209061407086935
9113 LATS1 large tumor suppressor kinase 1 8643 -0.451521775393679 6.01967223195046 5.60633366249878 0.0178956683870376 0.209061407086935
7110 TMF1 TATA element modulatory factor 1 6999 -0.349594595867329 6.40771079613973 5.6043278210412 0.0179161669760785 0.209126459028276
57604 TRMT9B tRNA methyltransferase 9B (putative) 14668 1.06863959682695 1.10190579805218 5.59794122089375 0.0179815961069777 0.209669232808305
7379 UPK2 uroplakin 2 1106 -1.05976617602698 0.201939845003323 5.59605910789213 0.0180009249109143 0.209669232808305
130026 ICA1L islet cell autoantigen 1 like 10598 0.574513095489478 2.91565528219825 5.59541215875218 0.0180075738608118 0.209669232808305
200315 APOBEC3A apolipoprotein B mRNA editing enzyme catalytic subunit 3A 1544 -1.79192237349411 2.91391800917426 5.58653847390778 0.0180990285403734 0.210559047147019
153572 IRX2 iroquois homeobox 2 3384 0.530995504428221 4.7850714815188 5.5840996550712 0.018124247608057 0.21058022348155
1646 AKR1C2 aldo-keto reductase family 1 member C2 6001 0.822815230102926 7.23169537878237 5.58345530880501 0.018130916650157 0.21058022348155
147166 TRIM16L tripartite motif containing 16 like 3910 -0.439826256393527 5.95838626581203 5.5735331718936 0.0182339322039886 0.211601233124497
83699 SH3BGRL2 SH3 domain binding glutamate rich protein like 2 4653 -0.3403118195516 5.90179203001083

5.5692753043392 0.0182783242272803 0.211894338643353
6565 SLC15A2 solute carrier family 15 member 2
6210 -1.18052234240969 5.04596520248635
5.5682103311549 0.0182894449503687 0.211894338643353
5376 PMP22 peripheral myelin protein 22 2925
1.59503421267281 1.69178079986562 5.56189908031615
0.0183554922058336 0.212440307329539
84804 MFSD9 major facilitator superfamily domain containing 9
4049 -0.402430661804764 5.39911648089274
5.56081106737574 0.0183669031324389 0.212440307329539
53358 SHC3 SHC adaptor protein 3 9929 1.21933066130813
3.64238172308025 5.55708543705457
0.0184060325447809 0.212504755309723
8445 DYRK2 dual specificity tyrosine phosphorylation regulated kinase 2 9494 -0.416397653326064 5.40734216799855
5.5543306974798 0.0184350202794287 0.212504755309723
84287 ZDHHC16 zinc finger DHHC-type containing 16 3078
0.277342903620336 6.01696705847167 5.55385307561527
0.018440051020993 0.212504755309723
5095 PCCA propionyl-CoA carboxylase subunit alpha 4489
0.500161882939758 3.15820355607553 5.55314829344646
0.0184474770084396 0.212504755309723
221955 DAGLB diacylglycerol lipase beta 5756
0.332990852980979 6.03650891674942 5.55306719331328
0.0184483317238969 0.212504755309723
93082 NEURL3 neuralized E3 ubiquitin protein ligase 3
2497 -1.45400152850931 4.38267666296013
5.54943177812211 0.0184866874570044 0.212524698786356
148304 C1orf74 chromosome 1 open reading frame 74
4684 -0.582512459733488 7.14044919843129
5.54903137290032 0.0184909169961521 0.212524698786356
9181 ARHGEF2 Rho/Rac guanine nucleotide exchange factor 2
7879 -0.441618057665623 6.53094628451283
5.54858991498622 0.0184955813393709 0.212524698786356
25994 HIGD1A HIG1 hypoxia inducible domain family member 1A
1991 -0.402345692354584 7.55155711302152
5.54494700448411 0.0185341179193207 0.212635892206017
90523 MLIP muscular LMNA interacting protein
6904 -1.6222334122785 0.722638230771695
5.54480520281499 0.0185356196461446 0.212635892206017
51582 AZIN1 antizyme inhibitor 1 6426 -0.379452322683826
7.7629544998681 5.54299262624932 0.0185548265093581
0.21268204166595
1021 CDK6 cyclin dependent kinase 6
12260 -0.489134364136755 7.70582935475287
5.54117593498464 0.01857409760212 0.212728851277918
3344 FOXN2 forkhead box N2 5481 -0.5427266799486
4.36451186263875 5.53501573597665 0.0186395977973618
0.213304613029123
116071 BATF2 basic leucine zipper ATF-like transcription factor 2
2343 -0.945372366500983 0.823927632059657
5.53255587454025 0.0186658196430734 0.213430314890228
80352 RNF39 ring finger protein 39 2206 -0.881397373263468
3.01443340242342 5.52716100802346

0.0187234618302365      0.213914787810866
80310   PDGFD   platelet derived growth factor D        4266
0.97906426858133        1.77648030328508        5.51807805459837
        0.0188209256755371      0.214853061073552
10400   PEMT    phosphatidylethanolamine N-methyltransferase    1720
        0.570227873967654       2.80569448018832
5.51105456930148        0.018896649846641       0.215541835832166
10753   CAPN9   calpain 9       2593    -1.30125237241583
2.03635503546499        5.49616463175787        0.0190582296067142
        0.217207991945685
25843   MOB4    MOB family member 4, phocein
4296    -0.30078681330258       5.95441443820905
5.49341561233142        0.0190882166644404      0.217237291363782
9424    KCNK6   potassium two pore domain channel subfamily K member
6       6169    -0.378569812415277      6.40823845123394
5.49308569384183        0.0190918187812391      0.217237291363782
2013    EMP2    epithelial membrane protein 2   8718
0.626183067578966       7.17202459658319        5.48859871129571
        0.0191408783654347      0.217606955269807
54536   EXOC6   exocyst complex component 6
6193    -0.414801818930715      4.36383875454352
5.48712165231781        0.0191570566585934      0.217606955269807
5296    PIK3R2  phosphoinositide-3-kinase regulatory subunit 2  5719
        0.68443160969171        6.65256085523482
5.48500633306733        0.0191802504593931      0.217606955269807
23132   RAD54L2 RAD54 like 2    9776    -0.362100225486336
5.63115738916227        5.48444147087966        0.0191864489011359
        0.217606955269807
1824    DSC2    desmocollin 2   12371   -0.453732995646735
8.58243694456841        5.47980270635717        0.0192374301532543
        0.21800864413967
85456   TNKS1BP1        tankyrase 1 binding protein 1
7149    -0.301158163154632      8.4802082894967 5.47799035724336
        0.0192573863663415      0.218058375774734
1500    CTNND1  catenin delta 1 9089    -0.343581771879692
9.0861531820475 5.47590567611715        0.0192803677550775
0.218142254560074
3487    IGFBP4  insulin like growth factor binding protein 4    2200
        0.393642067121879       7.39746918780998
5.47256000117156        0.0193173095245715      0.21838382123541
403     ARL3    ADP ribosylation factor like GTPase 3   3844
0.374908964017495       4.28964750351847        5.46900989847428
        0.0193565885213019      0.218521323457876
124583  CANT1   calcium activated nucleotidase 1
4421    -0.3200988741958        6.79202872368551
5.46864106262134        0.0193606741208841      0.218521323457876
5066    PAM     peptidylglycine alpha-amidating monooxygenase   8846
        0.325570533648056       8.07585148249686
5.46689481468675        0.0193800294353818      0.218563665299028
402     ARL2    ADP ribosylation factor like GTPase 2   1879
0.363902832147789       5.01965047524817        5.45732827023189
        0.0194864199803487      0.219520230420729
1488    CTBP2   C-terminal binding protein 2
9735    -0.288255944239836      6.9889148261371 5.45645225360452

0.0194961923783384	0.219520230420729
6452	SH3BP2	SH3 domain binding protein 2
13295	-0.317832419201072	6.34715381593158
5.44960579008769	0.0195727427958705	0.220205147262456
2252	FGF7	fibroblast growth factor 7	6657
1.12920797641702	0.205930969032816	5.44776549968769
0.0195933720496254	0.220260322872475
7150	TOP1	DNA topoisomerase I	3738	-0.264695549453678
7.00913193793821	5.44370523932388
0.0196389662267207	0.220595829942002
119385	AGAP11	ArfGAP with GTPase domain, ankyrin repeat and PH domain 11	5672	-1.22227889379397	3.2649153101763
5.43878605967561	0.0196943525151896	0.221040701666876
881	CCIN	calicin
1941	-1.28835504502718	-0.0166732140488373
5.43484508117599	0.0197388414936178	0.221362652362774
10123	ARL4C	ADP ribosylation factor like GTPase 4C	4013
0.97789779849482	7.19437312665971	5.43280188023673
0.0197619476992193	0.221443670139724
2770	GNAI1	G protein subunit alpha i1	4280
0.4060219711926	5.22021198401888	5.43141171477995
0.0197776848250728	0.221443670139724
5742	PTGS1	prostaglandin-endoperoxide synthase 1	5790
1.45613887309912	1.84005993394105	5.42806592459689
0.0198156134038175	0.221576497437176
9175	MAP3K13	mitogen-activated protein kinase kinase kinase 13
12623	-0.438754561382252	5.17688731382365
5.42629952586229	0.0198356679969131	0.221576497437176
80736	SLC44A4	solute carrier family 44 member 4
3369	-0.622691381938511	7.41350424943759
5.42618183679997	0.0198370049108459	0.221576497437176
22873	DZIP1	DAZ interacting zinc finger protein 1	8470
0.950847905624662	2.12580471065672	5.41530136614618
0.0199610073135105	0.22278392784091
22904	SBNO2	strawberry notch homolog 2
6626	-0.34332283666354	6.80845727569043
5.41157663268348	0.0200036412459212	0.223082008703518
10505	SEMA4F	ssemaphorin 4F	6227	0.348924796972422
4.71192766947563	5.40885007377377	0.02003490959143
0.223252966306413
54726	OTUD4	OTU deubiquitinase 4	9085	-0.35875957966999
6.59418302860785	5.4039154865331	0.0200916283294954
0.223707025446138
54836	BSPRY	B-box and SPRY domain containing
2339	-0.900095673188182	4.31931112357495
5.40171593602894	0.0201169637101768	0.223729127986217
374354	NHLRC2	NHL repeat containing 2	11223	-0.47045435068485
5.54491417010571	5.40097038918198
0.0201255587393899	0.223729127986217
3799	KIF5B	kinesin family member 5B
5877	-0.315935017509921	8.79983531391019
5.39244317221409	0.0202241352554861	0.224646679241549
476	ATP1A1	ATPase Na+/K+ transporting subunit alpha 1	6971
0.34799412278032	9.06050699978957

5.37864139952766	0.0203847464597945	0.226186123854112
471	ATIC	5-aminoimidazole-4-carboxamide ribonucleotide formyltransferase/IMP cyclohydrolase	3928	0.384172524730256	5.44069275350058	5.37776221777347	0.0203950220909362	0.226186123854112
9953	HS3ST3B1	heparan sulfate-glucosamine 3-sulfotransferase 3B1	5367	-1.22594214582974	3.3433102234119 5.37506655250218	0.0204265616894593	0.226356684797671
2191	FAP	fibroblast activation protein alpha	6595	2.12376493637654	3.45821354571271	5.37111444332223	0.0204728930708654	0.226690761457401
84079	ANKRD27 ankyrin repeat domain 27	7855	-0.319324688333466	5.43883627173864	5.3696503095202 0.0204900849966198	0.226701911964971
57154	SMURF1	SMAD specific E3 ubiquitin protein ligase 1	6276	-0.322814637152408	7.77197133723277	5.36490962151078	0.0205458529144934	0.227139512054703
100532736	MICOS10-NBL1	MICOS10-NBL1 readthrough	3188	-0.311224197157007	6.54271576692723	5.36042699122036	0.0205987295759678	0.227461573877108
55830	GLT8D1	glycosyltransferase 8 domain containing 1	3916	0.278125155130856	5.45214431087194	5.35968774085592	0.0206074632148247	0.227461573877108
26046	LTN1	listerin E3 ubiquitin protein ligase 1	8876	-0.34910835397173	5.90918674192033	5.35801498738052	0.0206272395758321	0.22750058640841
633	BGN	biglycan	2453	3.48085447110071	4.53122623003522	5.35368687732069	0.0206785004482971	0.227886511234695
64063	PRSS22	serine protease 22	2387	-0.369316898388542	6.99178935045	5.35035949441772	0.0207179986077945	0.228142300707058
81493	SYNC	syncoilin, intermediate filament protein	3487	0.433058187208115	5.03025108533411	5.34527329001386	0.0207785260371817	0.228629076357269
5627	PROS1	protein S	4429	0.883797775709856	5.02537218161885	5.34304812960168	0.0208050636537918	0.228634715395735
8798	DYRK4	dual specificity tyrosine phosphorylation regulated kinase 4	6701	0.638120920987345	3.03143048358112	5.34249345002739	0.0208116843099566	0.228634715395735
54854	FAM83E	family with sequence similarity 83 member E	2034	-0.697726812858507	4.26834607095379	5.3382818725442 0.0208620248814391	0.229008136766707
51806	CALML5	calmodulin like 5	859	-1.31081853716123	3.35239527488443	5.33184616752787	0.0209391937660546	0.229595610143786
84910	TMEM87B transmembrane protein 87B	5256	-0.340711205666475	7.2349165828284 5.33043449531422	0.0209561602353007	0.229595610143786
5966	REL	REL proto-oncogene, NF-kB subunit	11255	-0.680574891052906	5.4223435251294 5.32972302337719

0.0209647165969803      0.229595610143786
345274  SLC10A6 solute carrier family 10 member 6
1502    -1.61465482550138       1.6601633270025 5.32657625240855
0.0210026039327743      0.22975503741986
729540  RGPD6   RANBP2 like and GRIP domain containing 6
8902    -0.574173199339829      4.63481825420393
5.32579013614551        0.0210120798839752      0.22975503741986
54442   KCTD5   potassium channel tetramerization domain containing
5       3420    -0.320937997079052      6.09343970576665
5.32326779804934        0.0210425143745088      0.229908345431938
116349  EXOC3-AS1       EXOC3 antisense RNA 1   1663
0.748722326132429       -0.0374889493240903     5.32006375825445
0.0210812400674249      0.230132664667581
4520    MTF1    metal regulatory transcription factor 1
8024    -0.374031876903535      5.69610031902085
5.31881325371477        0.0210963743227887      0.230132664667581
64651   CSRNP1  cysteine and serine rich nuclear protein 1
3214    -0.564297906828092      4.97561857553731
5.31749548288629        0.0211123348395689      0.230132664667581
375791  CYSRT1  cysteine rich tail 1    1297    -0.950392843795534
2.12357680441229        5.31442593756219
0.0211495609016256      0.230193044331244
55179   FAIM    Fas apoptotic inhibitory molecule       4072
0.42840520857405        4.32325063970379        5.31103139379152
0.0211908075217085      0.230193044331244
114801  TMEM200A        transmembrane protein 200A
6137    -1.71327923354094       3.47895130916572
5.31008136710615        0.0212023660675125      0.230193044331244
5875    RABGGTA Rab geranylgeranyltransferase subunit alpha
3545    -0.296333626551233      5.99241626297357
5.30903306663048        0.0212151278401536      0.230193044331244
10427   SEC24B  SEC24 homolog B, COPII coat complex component
5219    -0.279768317716268      6.26601031365538
5.30887503658511        0.0212170523517079      0.230193044331244
169044  COL22A1 collagen type XXII alpha 1 chain
7830    -1.58351482342124       5.58965533143511
5.30809462908856        0.0212265589113442      0.230193044331244
55213   RCBTB1  RCC1 and BTB domain containing protein 1        4534
0.489147003329192       6.48045403864029
5.30757320910169        0.0212329130631804      0.230193044331244
55328   RNLS    renalase, FAD dependent amine oxidase
3933    -1.28355919925438       1.64767065549632
5.30554597997508        0.0212576360896001      0.230282837360424
10913   EDAR    ectodysplasin A receptor        4426
0.60610673966947        2.73351604588694        5.29621251660622
0.0213718471645143      0.231341161695016
5033    P4HA1   prolyl 4-hydroxylase subunit alpha 1    3093
0.572146044120983       6.10013930350593        5.2943773330104
0.0213943784436152      0.231406223057697
9693    RAPGEF2 Rap guanine nucleotide exchange factor 2
11811   -0.415995620614798      6.3753496761543 5.28964767637049
0.0214525597520803      0.231856484913109
166968  MIER3   MIER family member 3    5763    -0.433680382143518
5.21965954590617        5.28301753517758

0.0215343955565354	0.232391112966293
6890	TAP1	transporter 1, ATP binding cassette subfamily B member	4578	-0.396967552437408	7.19668341514999	5.28295180816848	0.0215352084407974	0.232391112966293
3065	HDAC1	histone deacetylase 1	3963	-0.359307682345313	6.91654077730691	5.28106539715309	0.0215585523328698	0.232463928041961
10537	UBD	ubiquitin D	1087	-1.13759103640243	6.79776851788013	5.27690072761749	0.0216101819598375	0.232841399008803
26576	SRPK3	SRSF protein kinase 3	4744	0.479202831409759	3.9712737136252	5.27492710668481	0.0216346937389143	0.23290504859774
79983	POF1B	POF1B actin binding protein	4029	-0.516100847186802	4.618823621647	5.27338265489642	0.0216538954449866	0.23290504859774
51524	TMEM138	transmembrane protein 138	6348	0.375420299308755	5.20296906362917	5.2724119781541	0.0216659726081856	0.23290504859774
9043	SPAG9	sperm associated antigen 9	11624	-0.33011444549267	7.73290821466924	5.26744986629013	0.0217278202807604	0.233390781190653
92270	ATP6AP1L	ATPase H+ transporting accessory protein 1 like	5735	0.68661681321545	1.78861307492604	5.26166986462986	0.0218000926962569	0.233987661606491
114771	PGLYRP3	peptidoglycan recognition protein 3	1173	-1.32397225303802	0.894946516127974	5.25602602129111	0.0218709028605954	0.234567945151883
5170	PDPK1	3-phosphoinositide dependent protein kinase 1	10926	-0.292478018219279	6.64988457047586	5.24895765717744	0.0219599218196617	0.235342482729917
8935	SKAP2	src kinase associated phosphoprotein 2	5520	-0.586272378174975	4.3385864542051	5.24518897843656	0.022007537853593	0.2356724638496
643699	GOLGA8N	golgin A8 family member N	6532	1.34423830031578	1.0063119201012	5.24292119233992	0.022036242121393	0.235729007860745
51379	CRLF3	cytokine receptor like factor 3	3535	-0.349255594194045	4.37995230951314	5.24211334089364	0.0220464767828639	0.235729007860745
3908	LAMA2	laminin subunit alpha 2	10094	1.34646164196884	1.78987512994624	5.23807069393685	0.0220977670559768	0.236036963191844
155066	ATP6V0E2	ATPase H+ transporting V0 subunit e2	3335	0.651463318709052	3.9610750966563	5.23718812372359	0.0221089809172342	0.236036963191844
831	CAST	calpastatin	11891	-0.294163754809087	8.79218312301424	5.23166892252189	0.0221792412413995	0.23644475531402
84725	PLEKHA8	pleckstrin homology domain containing A8	10572	-0.394033813047019	5.48864978564243	5.2315358045941	0.0221809387079762	0.23644475531402
9306	SOCS6	suppressor of cytokine signaling 6	5984	-0.390139095056751	5.72958672790016

5.22845409440658	0.022220273088582	0.236683927872067
5129	CDK18	cyclin dependent kinase 18	6831
0.473296417236709	5.61874884464497	5.22432123638504
	0.022273137596662	0.237066746441068
55616	ASAP3	ArfGAP with SH3 domain, ankyrin repeat and PH domain
3	4988	0.408551896870367	4.47545850615635
5.22042731747154	0.022323064991956	0.237337818457984
10335	MRVI1	murine retrovirus integration site 1 homolog	7700
	0.958379202663848	1.42410776216599
5.21969293970677	0.022332494090642	0.237337818457984
5118	PCOLCE	procollagen C-endopeptidase enhancer	3532
0.931124205612363	2.07305958709358	5.21385914624704
	0.022407544268231	0.23795486926847
29109	FHOD1	formin homology 2 domain containing 1	5383
0.331675204589774	4.32241259116627	5.20792328091231
	0.0224841758324307	0.238489062129182
2310	FOXO3B	forkhead box O3B	1833	-0.463792773689672
	3.52092121630384	5.20603973749122
0.0225085489475678	0.238489062129182
863	CBFA2T3	CBFA2/RUNX1 partner transcriptional co-repressor 3
	5972	0.774017171698503	2.31557027584716
5.20473911094701	0.0225253950767311	0.238489062129182
5284	PIGR	polymeric immunoglobulin receptor
4376	-1.58136494249678	5.17249094059131
5.20469602257998	0.0225259533945104	0.238489062129182
8120	AP3B2	adaptor related protein complex 3 subunit beta 2
	8866	1.44192756845203	1.32016144848484
5.20190253289684	0.0225621806726472	0.238692193868406
2026	ENO2	enolase 2	4281	0.490128904882667
4.68244178434481	5.19108779539504	0.0227030015314956
	0.239778821867178
84775	ZNF607	zinc finger protein 607	4931	-0.513864448303833
	3.39183177186046	5.19057071393656
0.0227097573256759	0.239778821867178
3956	LGALS1	galectin 1	1099	0.675548161809781
7.44992014939184	5.18860940066088	0.0227354013022104
	0.239778821867178
11118	BTN3A2	butyrophilin subfamily 3 member A2
4837	-0.693913209449539	3.73072268379193
5.18750936863921	0.022749797246291	0.239778821867178
9747	TCAF1	TRPM8 channel associated factor 1	6310
0.378008182176675	5.95392549593802	5.18745677436479
	0.0227504857757893	0.239778821867178
84069	PLEKHN1	pleckstrin homology domain containing N1
2833	-0.43010373256435	5.081912215055	5.1847306368624
0.0227862041558524	0.239974707978214
221037	JMJD1C	jumonji domain containing 1C
9305	-0.494541813907959	6.62852547983471
5.17663485044268	0.022892619704709	0.240914293165934
25996	REXO2	RNA exonuclease 2	3699	0.320696532920556
	6.86397410137165	5.16631192516377
0.0230290576741369	0.242168176307534
647979	NORAD	non-coding RNA activated by DNA damage
5339	-0.395915775455696	8.71166054489811

5.160079874084	0.0231118344083046	0.242856312495966
1545	CYP1B1	cytochrome P450 family 1 subfamily B member 1	5909
	0.83149064026756	8.55397952430169
5.15542662002727	0.0231738419922324	0.243223835354895
57534	MIB1	mindbomb E3 ubiquitin protein ligase 1
10033	-0.334640516512298	7.00379188095508
5.15484989358436	0.0231815392445766	0.243223835354895
200931	SLC51A	solute carrier family 51 alpha subunit
4191	-1.28543499398304	0.583064576103064
5.14054359179271	0.0233733284750167	0.24505255385446
154664	ABCA13	ATP binding cassette subfamily A member 13
19074	-1.10973291966869	4.25137884874451
5.1392320361884	0.0233909932255047	0.245054332168769
9338	TCEAL1	transcription elongation factor A like 1	1602
	0.50775016454517	3.71149218504169
5.13736743829944	0.0234161304861976	0.245134334618961
104	ADARB1	adenosine deaminase RNA specific B1	10049
0.490077196617349	3.35672232031952	5.13605963190386
	0.0234337781690043	0.245135870659628
51053	GMNN	geminin DNA replication inhibitor	2476
0.955083045644577	3.765884274116	5.13459617314215
0.0234535425944704	0.245159530687124
196446	MYRFL	myelin regulatory factor like
3747	-0.693541719153726	2.52096500382338
5.13154949145472	0.0234947443664961	0.245235142533787
923	CD6	CD6 molecule	5958	-1.6773804958195
0.407458293000252	5.13039529613206	0.0235103727135597
	0.245235142533787
113201	CASC4	cancer susceptibility 4	6065	-0.292116448977075
	6.51718109545814	5.13017917914284
0.0235133002372296	0.245235142533787
9784	SNX17	sorting nexin 17	3765	0.264866469401619
	6.47252171574905	5.1261626673051	0.0235677768145641
	0.245505624130669
55964	SEPTIN3	septin 3	5427	0.97398941724225
0.996208502348823	5.1256831820179	0.0235742888881095
0.245505624130669
140707	BRI3BP	BRI3 binding protein	6648	-0.387682678448877
	5.04385661153195	5.12126884798981
0.0236343293959091	0.245801339267315
339105	PRSS53	serine protease 53	5908	0.472861084145688
	3.72041774712311	5.12101540175506
0.0236377813945223	0.245801339267315
374378	GALNT18	polypeptide N-acetylgalactosaminyltransferase 18
	3627	0.379160017698884	5.87190584690116
5.11835635938137	0.0236740297285203	0.245852781402082
347902	AMIGO2	adhesion molecule with Ig like domain 2	3956
0.647629015746439	9.24559904451859	5.11807763508364
	0.0236778326630548	0.245852781402082
56948	SDR39U1	short chain dehydrogenase/reductase family 39U
member 1	3099	0.381436289773666	4.19816926077822
	5.11465775014091	0.0237245454349216
0.245994516687464
83660	TLN2	talin 2	15723	0.565989529442106

2.81754965868482	5.1145069413193 0.023726607556562
0.245994516687464
3660	IRF2	interferon regulatory factor 2
3116	-0.421585226260323	5.03974745627488
5.1108814944233 0.0237762369938134	0.24616931040828
66035	SLC2A11 solute carrier family 2 member 11	9247
0.444017905138115	6.58890956247964	5.11016481385644
0.0237860605111281	0.24616931040828
320	APBA1	amyloid beta precursor protein binding family A
member 1	6534	0.720480204422731	1.81745971704285
5.10942606490566	0.0237961909254523
0.24616931040828
56996	SLC12A9 solute carrier family 12 member 9	8894
0.3291499517957 5.47926485808934	5.10785461900434
0.023817754944047	0.246210548709422
2517	FUCA1	alpha-L-fucosidase 1	2047	0.922396779863616
5.68443858589252	5.10255040985234
0.0238906912511488	0.246782383742508
3678	ITGA5	integrin subunit alpha 5
7577	-0.413723758995633	7.68906540843495
5.10062477253675	0.0239172272989375	0.2468744309331
221692	PHACTR1 phosphatase and actin regulator 1	8623
1.5063687833229 0.877696428281412	5.09530822825879
0.0239906502515466	0.247121239646433
283234	CCDC88B coiled-coil domain containing 88B
8331	-0.54384952287655	3.81738020484873
5.09504675726691	0.0239942672655256	0.247121239646433
56829	ZC3HAV1 zinc finger CCCH-type containing, antiviral 1
8235	-0.344802745287362	6.19167938976735
5.09490755762704	0.0239961930905452	0.247121239646433
11112	HIBADH	3-hydroxyisobutyrate dehydrogenase	2323
0.395686629175583	4.05447412047684	5.09378648636959
0.0240117089425855	0.247121239646433
9353	SLIT2	slit guidance ligand 2	9712	-0.812797800447412
2.33999103418962	5.08798302072493
0.0240921966398426	0.247591998902411
83931	STK40	serine/threonine kinase 40
5590	-0.420459889085477	5.2467911234277 5.08681265824836
0.0241084621694814	0.247591998902411
57326	PBXIP1	PBX homeobox interacting protein 1	3851
0.438611937459745	7.51833618382045	5.08666756052526
0.0241104795104511	0.247591998902411
55681	SCYL2	SCY1 like pseudokinase 2
7481	-0.287153992671273	7.38000054326714
5.08148725522827	0.0241826178506002	0.247718996042699
152831	KLB	klotho beta	6082	0.712518507395719
0.934127476035139	5.08115424831713	0.0241872627918654
0.247718996042699
57162	PELI1	pellino E3 ubiquitin protein ligase 1
4125	-0.37966278839979	5.09823595558455
5.0808079770705 0.0241920937322358	0.247718996042699
1316	KLF6	Kruppel like factor 6	6139	-0.381955512044116
9.19278085532861	5.08070088846034
0.0241935879621912	0.247718996042699

1573 CYP2J2 cytochrome P450 family 2 subfamily J member 2
2990 -0.563838793944185 2.75095480712683
5.07562379637705 0.0242645397021434 0.248263993869922
22937 SCAP SREBF chaperone 6301 0.283453737595394
5.99775245618399 5.0653228683469 0.0244091581173993
0.249543901208787
114789 SLC25A25 solute carrier family 25 member 25
4936 -0.422291370078314 4.75595144775344
5.06417951491348 0.0244252651215283 0.249543901208787
51110 LACTB2 lactamase beta 2 1931 0.328622252502874
5.21367902755345 5.05821974540482
0.0245094022146438 0.250220988936236
84733 CBX2 chromobox 2 5687 0.93998644147768
1.4771552317338 5.05688624284231 0.0245282690922845
0.250231220084216
400680 LINC00664 long intergenic non-protein coding RNA 664
6085 1.204052037238 0.209628348866843
5.05396549367468 0.0245696455905873 0.250383187209743
307 ANXA4 annexin A4 11048 0.309157521304807
7.13580327647837 5.05331176927513 0.0245789164284569
0.250383187209743
135114 HINT3 histidine triad nucleotide binding protein 3
3314 -0.297955069316247 6.07460977018592
5.05140854137821 0.0246059278640607 0.250476185749926
84290 CAPNS2 calpain small subunit 2 1016 -1.18874898718384
2.47306129843497 5.04823420772628
0.0246510479503058 0.250729287952062
8542 APOL1 apolipoprotein L1 3519 0.968085152966527
7.52927670414993 5.04714201020295
0.0246665923322583 0.250729287952062
5266 PI3 peptidase inhibitor 3 577 -2.56192024804111
2.73247686646985 5.04140283816101
0.0247484407765281 0.251378832456004
54 ACP5 acid phosphatase 5, tartrate resistant 2792
1.55239880227613 5.12150453356569 5.03950854657995
0.0247755178031551 0.251452441852048
1123 CHN1 chimerin 1 4971 0.942177000378726
1.21522767260737 5.03800894892191 0.0247969748768239
0.251452441852048
164 AP1G1 adaptor related protein complex 1 subunit gamma 1
10980 -0.268457172113557 7.43992800123968
5.03713118090756 0.0248095434168295 0.251452441852048
10052 GJC1 gap junction protein gamma 1 8660
1.10578631323174 0.524581852154116 5.03564620484074
0.0248308214756959 0.251486129002221
253461 ZBTB38 zinc finger and BTB domain containing 38
13706 -0.540121372452763 8.02263178906839
5.03409402986358 0.0248530826704837 0.251526760997204
1063 CENPF centromere protein F 11690 0.741502764492073
1.43972040099614 5.03286334720661
0.0248707477676253 0.251526760997204
84978 FRMD5 FERM domain containing 5 6325
0.597654007337672 4.28642154899593 5.02836110812576
0.0249354836052454 0.251967427087451

3988	LIPA	lipase A, lysosomal acid type	3341
0.396253749928202	6.76990690877329	5.02619873656712
	0.024966637612732	0.251967427087451
387856	CCDC184	coiled-coil domain containing 184	2343
0.944272068066599	0.769357208026488	5.0260843678858
0.0249682864851418	0.251967427087451
4166	CHST6	carbohydrate sulfotransferase 6
8469	-1.11733096119832	2.70726812104793
5.02412082880353	0.024996612795649	0.252071674174253
2921	CXCL3	C-X-C motif chemokine ligand 3
2219	-1.54839079257341	4.34765365628062
5.02205871444876	0.0250263970879206	0.252090894221274
5519	PPP2R1B	protein phosphatase 2 scaffold subunit Abeta
6720	-0.306990800808114	6.21656945319591
5.02122081560455	0.0250385098633256	0.252090894221274
84179	SLC49A3	solute carrier family 49 member 3	2644
0.672618592442537	3.09289365592284	5.02025279826708
	0.0250525112269589	0.252090894221274
728689	EIF3CL	eukaryotic translation initiation factor 3 subunit C
like	4067	-0.397019347423428	7.05256674217447
5.01855293839263	0.0250771176052661	0.2521573483826
7096	TLR1	toll like receptor 1	4267	-0.767860443186326
	2.77040497338476	5.01639091034879
0.0251084503730758	0.252178100428509
55466	DNAJA4	DnaJ heat shock protein family (Hsp40) member A4
	6591	-0.603615263181581	5.87930341050199
5.01592630763841	0.0251151888411344	0.252178100428509
126969	SLC44A3	solute carrier family 44 member 3	2869
0.369288956877121	5.71346892868531	5.01274352133075
	0.0251614016063174	0.25242815746783
23170	TTLL12	tubulin tyrosine ligase like 12
4622	-0.442218022645623	5.47866621383701
5.01173000831611	0.0251761359307888	0.25242815746783
29087	THYN1	thymocyte nuclear protein 1	2119
0.427165986338448	4.2314878759457 5.01004436239146
0.0252006614833397	0.252493322887796
4216	MAP3K4	mitogen-activated protein kinase kinase kinase 4
	7522	-0.366287653771807	5.40554978351337
5.00496280233594	0.0252747466196961	0.253054593210925
714	C1QC	complement C1q C chain	1335	2.08599487780809
	3.02983838929627	5.00140401700639
0.0253267655310634	0.25327855308301
9022	CLIC3	chloride intracellular channel 3
1177	-0.816340657305749	3.51887534548583
5.00095887602705	0.0253332799935245	0.25327855308301
54812	AFTPH	aftiphilin	5767	-0.289756017763544
6.10703948900554	4.99611962077153	0.0254042129721896
	0.25380656997251
8773	SNAP23	synaptosome associated protein 23
5900	-0.256809558049257	6.66884915763473
4.99099543454842	0.0254795472939342	0.25437777544272
64078	SLC28A3	solute carrier family 28 member 3
5063	-0.867862799927865	3.37694113266778
4.98582989232427	0.0255557244810867	0.254956576073064

51105	PHF20L1 PHD finger protein 20 like 1
20571	-0.297605936578199	6.38110793432995
4.97575777976457	0.0257049408312367	0.25626270905561
585	BBS4	Bardet-Biedl syndrome 4 4149	0.413994077318887
	3.94734353710464	4.97165065006673
0.0257660466369482	0.256633096093619
91298	C12orf29	chromosome 12 open reading frame 29
6454	-0.462060593386628	4.98158193410483
4.97079897053734	0.0257787367890142	0.256633096093619
38	ACAT1	acetyl-CoA acetyltransferase 1	5584
0.478983787708151	3.85121928949877	4.96717536207293
	0.0258328017172411	0.256988674469741
6048	RNF5	ring finger protein 5	1178	0.394311132121871
	5.24007502912157	4.96520207372153
0.0258622930810884	0.257099460033219
892	CCNC	cyclin C	5765	-0.31728755026818
6.43392456784869	4.96389148144371	0.0258818995887533
	0.257111891872105
5365	PLXNB3	plexin B3	7007	0.425849921678789
5.60288328488052	4.956100160077	0.0259987772572845
0.257953211458122
23648	SSBP3	single stranded DNA binding protein 3
4884	-0.419076888607176	5.12160347771433
4.95551010530167	0.0260076509804503	0.257953211458122
6556	SLC11A1 solute carrier family 11 member 1	8676
2.2491159854768 2.65638132876935	4.95454015292505
0.0260222447511258	0.257953211458122
668	FOXL2	forkhead box L2 2917	1.12184732859828
1.3499528420468 4.95334395957191	0.0260402542301553
0.257953211458122
114757	CYGB	cytoglobin	2949	1.48592139075401
3.68643979760776	4.95167222013618	0.0260654450677626
	0.258020274956997
5097	PCDH1	protocadherin 1 5903	-0.290704782323788
7.84003434013836	4.94931836211464	0.0261009574370503
	0.258102259497418
977	CD151	CD151 molecule (Raph blood group)	3762
0.317902729873597	9.70730300104266	4.94868108887067
	0.0261105805460014	0.258102259497418
10112	KIF20A	kinesin family member 20A	3958
0.826815652386277	-0.1450212142853	4.94708949082878
	0.02613463048776	0.258157760827847
8777	MPDZ	multiple PDZ domain crumbs cell polarity complex
component	11273	0.398882290780174	5.48044066214877
	4.94587110640216	0.0261530565156504
0.258157760827847
374897	SBSN	suprabasin	1986	-2.07545520822087
1.18505336719231	4.94165028346044	0.0262169938621384
	0.258341227863664
8436	CAVIN2	caveolae associated protein 2	3229
0.62338651875753	3.45058651505329	4.9409308729347
0.0262279077388735	0.258341227863664
2776	GNAQ	G protein subunit alpha q
6629	-0.255420730601811	6.35741737402869

4.94007551740592	0.0262408901245769	0.258341227863664
85460	ZNF518B zinc finger protein 518B
10905	-0.362181542860684	5.25223280596286
4.93888650035249	0.0262589478409033	0.258341227863664
23710	GABARAPL1	GABA type A receptor associated protein like
1	4086	0.354451166001858	6.96143614972667
4.93823397529138	0.0262688632892651	0.258341227863664
283464	GXYLT1	glucoside xylosyltransferase 1
7497	-0.47653786251098	4.26777105674421
4.93734978247177	0.0262823052549241	0.258341227863664
29904	EEF2K	eukaryotic elongation factor 2 kinase	8020
0.338905092522511	4.49632604335004	4.93586348821986
0.0263049167931379	0.258382166564854
80176	SPSB1	splA/ryanodine receptor domain and SOCS box
containing 1	3403	0.325654730806869	6.91153488560195
4.9287266297228 0.0264137743833397	0.259166894139574
4582	MUC1	mucin 1, cell surface associated	4717
0.709750007616453	7.56824794481396	4.92720699427025
0.0264370135855478	0.259166894139574
3554	IL1R1	interleukin 1 receptor type 1
5516	-0.538256712325143	7.81927998165766
4.92699122623594	0.0264403149657636	0.259166894139574
728492	SERF1B	small EDRK-rich factor 1B	5941
0.43284457620997	5.15644444225761	4.92373984450191
0.0264901149643351	0.25947345475905
162333	MARCHF10	membrane associated ring-CH-type finger 10
4920	1.09912088216699	0.502968980421938
4.92228479232861	0.026512432931951	0.259510585658866
440193	CCDC88C coiled-coil domain containing 88C
12426	-0.427458588452909	5.13622753944539
4.91940386872166	0.026556678959237	0.259762152361755
6374	CXCL5	C-X-C motif chemokine ligand 5
2538	-1.63909977171895	6.21409232516006
4.90547609880537	0.0267716706008602	0.26168233782711
253982	ASPHD1	aspartate beta-hydroxylase domain containing 1	2312
0.934803031103223	0.450146772324087
4.89952270974204	0.0268641194656544	0.26240287402749
11149	BVES	blood vessel epicardial substance	5808
0.761296406567444	2.70659608002547	4.89682082847042
0.0269061858210482	0.262606678722139
56971	CEACAM19	CEA cell adhesion molecule 19
5203	-0.39824930363387	4.74560760833148
4.89577538816914	0.0269224809484537	0.262606678722139
2081	ERN1	endoplasmic reticulum to nucleus signaling 1
8965	-0.605022185649209	5.3113637890419 4.89235287385378
0.0269758990032626	0.262826710464186
253143	PRR14L	proline rich 14 like	11809	-0.41474025927858
6.48487372403574	4.891187976357	0.026994105664122
0.262826710464186
84333	PCGF5	polycomb group ring finger 5
8306	-0.406740981555904	5.41542971103318
4.89072594352299	0.0270013305031744	0.262826710464186
23400	ATP13A2 ATPase cation transporting 13A2
5253	-0.390266627032844	7.13034276186835

4.88671676679374	0.0270641067550489	0.26325482174859
51599	LSR	lipolysis stimulated lipoprotein receptor
2916	-0.357991506607571	8.24630812069116
4.87940362514115	0.0271790081589118	0.264189012686938
7082	TJP1	tight junction protein 1
11265	-0.385333862846915	8.05418383455914
4.87113067813118	0.0273096014190647	0.265108589713321
6314	ATXN7	ataxin 7	9172	-0.411235805942644
5.22513737584563	4.87101289896669	0.0273114653356409
	0.265108589713321
1510	CTSE	cathepsin E	2671	1.31699768483516
4.69548717534841	4.86926192031016	0.0273391910937811
	0.265193940201241
11177	BAZ1A	bromodomain adjacent to zinc finger domain 1A
7698	-0.328605430691929	7.09695997715694
4.8677633614678	0.0273629431735737	0.265240654001555
55284	UBE2W	ubiquitin conjugating enzyme E2 W
9392	-0.362430792025233	5.87952239685586
4.86436756481982	0.0274168458287028	0.265565946338854
441869	ANKRD65	ankyrin repeat domain 65	2644
0.33665799776686	6.00273110022524	4.86263621841181
	0.027444370585812	0.265565946338854
718	C3	complement C3	8328	-1.20913702329624
8.7448410995615	4.86206992081827	0.0274533797600244
0.265565946338854
4067	LYN	LYN proto-oncogene, Src family tyrosine kinase
6185	-0.557295365198762	6.71973754766171
4.85657384404476	0.0275409762881242	0.266099275440525
80223	RAB11FIP1	RAB11 family interacting protein 1
9602	-0.539259834590728	9.98456745656843
4.85622732505294	0.0275465088447749	0.266099275440525
23443	SLC35A3	solute carrier family 35 member A3
6241	-0.419141809062713	5.49539204892857
4.85499306010905	0.0275662246358303	0.266106208459046
78999	LRFN4	leucine rich repeat and fibronectin type III domain containing 4	2431	0.595793471966599	4.22712569374673
	4.85132092154467	0.0276249691612586
0.266329028311659
83938	LRMDA	leucine rich melanocyte differentiation associated
	6560	1.02591503774333	1.45521525183469
4.85117321673425	0.0276273347709602	0.266329028311659
50853	VILL	villin like	6476	-0.430656737459414
5.87175224217259	4.84575238632027	0.0277142996179823
	0.2669039949357
144406	WDR66	WD repeat domain 66	17370	0.446037117828157
	5.268301776609	4.84508111961162	0.027725088358067
	0.2669039949357
9766	SUSD6	sushi domain containing 6
5830	-0.429087227500998	7.29601608835802
4.84137988749325	0.0277846538555002	0.267293713292577
7378	UPP1	uridine phosphorylase 1	4577	-0.613296024725572
	4.68224952406206	4.83681743916908
0.0278582625317469	0.267692531848019
7508	XPC	XPC complex subunit, DNA damage recognition and

repair factor	4798	0.330922949424763	5.73656020318167
	4.83644174043548	0.02786433293599
0.267692531848019
55924	INKA2	inka box actin regulator 2	6414
0.553344132962441	4.4131318036787	4.83198621921828
0.0279364286683084	0.268198566789047
100506714	NUP50-DT	NUP50 divergent transcript	2263
	-0.563355555928452	3.35733673910285
4.83082185528661	0.027955301457272	0.268198566789047
92305	TMEM129	transmembrane protein 129	3442
0.395316930783736	5.81511785331982	4.82906692966269
	0.0279837715340528	0.268287945159123
25998	IBTK	inhibitor of Bruton tyrosine kinase
8722	-0.396773581121263	6.67993129417771
4.82574129496084	0.0280378059391384	0.268443783421131
10260	DENND4A	DENN domain containing 4A
10231	-0.374923734869021	4.57198238504195
4.82570746323179	0.0280383561894135	0.268443783421131
5618	PRLR	prolactin receptor	13108	-1.34264764132986
	-0.235285597879059	4.822568127189	0.0280894643059582
	0.268701723031894
5359	PLSCR1	phospholipid scramblase 1
3856	-0.497987442652356	6.03298344700009
4.8216969491344	0.0281036641851734	0.268701723031894
1182	CLCN3	chloride voltage-gated channel 3
8123	-0.298933034203855	6.71491438020253
4.81393211758057	0.0282305585551538	0.269730855171923
148022	TICAM1	toll like receptor adaptor molecule 1
2676	-0.419569301941112	6.06363362589905
4.81270200079279	0.0282507160086986	0.269739454078965
401541	CENPP	centromere protein P	9794	0.765081690420163
	0.985533242035724	4.80945291510615
0.0283040295994661	0.26990904918863
1512	CTSH	cathepsin H	5892	0.627309560923399
7.15029323720832	4.80927101602151	0.0283070174382878
	0.26990904918863
388536	ZNF790	zinc finger protein 790	3851	-0.467883679253378
	3.2594923449303	4.79891203819972	0.0284777148212767
	0.271036334945342
10314	LANCL1	LanC like 1	5254	0.354856545057711
5.33290397058093	4.7989023278284	0.0284778753324515
0.271036334945342
55317	AP5S1	adaptor related protein complex 5 subunit sigma 1
	5490	0.33582636818526	4.12714813107581
4.79833857686544	0.0284871956771169	0.271036334945342
79912	PYROXD1	pyridine nucleotide-disulphide oxidoreductase domain
1	5725	-0.521710206638729	4.69443914568271
4.79740464799707	0.0285026430623609	0.271036334945342
6789	STK4	serine/threonine kinase 4
9075	-0.302749493131337	5.60135361222247
4.79617596652135	0.0285229790017873	0.271045703445071
100507431	LINC02551	long intergenic non-protein coding
RNA 2551	2255	1.43463942337273	0.915878582849881
	4.79314649766781	0.0285731843750554

0.271338707485696
406937  MIR145  microRNA 145    2549
1.61870926134289        -0.504053217761644      4.78436096440772
        0.0287193020731349      0.272323854180562
220001  VWCE    von Willebrand factor C and EGF domains 4874
0.781879355738113       1.31309368073248        4.78368800220846
        0.0287305265555866      0.272323854180562
128     ADH5    alcohol dehydrogenase 5 (class III), chi polypeptide
        4866    0.333429101933001       7.22797657146401
4.78286750296471        0.0287442180195915      0.272323854180562
2353    FOS     Fos proto-oncogene, AP-1 transcription factor
subunit 2938    1.22929059933547        2.81495081120583
4.78224005449197        0.0287546926774507      0.272323854180562
54939   COMMD4  COMM domain containing 4        4943
0.434842439923133       4.98115218379641        4.7787431320416
0.0288131433232048      0.272693039546033
6648    SOD2    superoxide dismutase 2  14697   -0.789410732769273
        7.60589935199221        4.77662445761595
0.0288486168824643      0.272844413688506
10644   IGF2BP2 insulin like growth factor 2 mRNA binding protein 2
        4735    -0.300044835505133      6.45567052423831
4.77228786487176        0.0289213675360078      0.273107514943962
30845   EHD3    EH domain containing 3  4636    0.446129730081518
        5.38483026757256        4.77079836292008
0.028946399464504       0.273107514943962
23242   COBL    cordon-bleu WH2 repeat protein
14349   -0.390459161619752      6.34381773865374
4.77036874076389        0.0289536237005585      0.273107514943962
156     GRK2    G protein-coupled receptor kinase 2
6665    -0.271967545158577      6.46992999418952
4.77008068434851        0.0289584685140662      0.273107514943962
9915    ARNT2   aryl hydrocarbon receptor nuclear translocator 2
        7741    1.02620472234979        4.20460490377474
4.76916202981447        0.0289739249808473      0.273107514943962
399948  COLCA1  colorectal cancer associated 1  8733
1.72317976412345        3.27617389110563        4.76771703904576
        0.0289982544999391      0.273144571537395
63898   SH2D4A  SH2 domain containing 4A
4148    -0.398258049426541      5.60174781727315
4.76661301428383        0.0290168574603873      0.273144571537395
140606  SELENOM selenoprotein M 3187    -0.648664627785459
3.7075529781831 4.76524191284089        0.0290399779993405
0.273178624470627
558     AXL     AXL receptor tyrosine kinase    5154
0.523271357998914       6.66334738795827        4.7594393829566
0.0291380371951872      0.27354240475033
11076   TPPP    tubulin polymerization promoting protein
6022    -0.974811740263507      2.84105468184685
4.75779646926733        0.0291658640204913      0.27354240475033
4947    OAZ2    ornithine decarboxylase antizyme 2      7671
0.296723574660383       6.3172726233664 4.75777043974685
0.0291663051177385      0.27354240475033
23059   CLUAP1  clusterin associated protein 1  7909
0.638090198696766       3.80490202387655        4.75744943024242

0.0291717455282443	0.27354240475033
94031	HTRA3	HtrA serine peptidase 3 3321	2.34826646139182
0.822734238294454	4.7571810860049
0.0291762941884053	0.27354240475033
118932	ANKRD22 ankyrin repeat domain 22
1733	-0.896739076316699	2.92524068802812
4.75372174828353	0.0292349990519824	0.27354368156393
81788	NUAK2	NUAK family kinase 2	3441	-0.445488290099321
4.61642520641229	4.75288702735189
0.0292491826569971	0.27354368156393
571	BACH1	BTB domain and CNC homolog 1
7723	-0.469858312303595	5.89100108650866
4.75250734802924	0.0292556365514584	0.27354368156393
55048	VPS37C	VPS37C subunit of ESCRT-I
3960	-0.311537930353646	5.04421419643405
4.75026455120068	0.0292937905035716	0.27354368156393
7305	TYROBP	TYRO protein tyrosine kinase binding protein	907
1.99092387722408	2.16130694453242
4.75019839927938	0.0292949166516053	0.27354368156393
6507	SLC1A3	solute carrier family 1 member 3	6265
1.3526998520141 1.2462769008965 4.74954315671136
0.0293060737160321	0.27354368156393
340277	FAM221A family with sequence similarity 221 member A
7458	-0.516345766972484	2.9993951807716 4.74912865593959
0.029313133863601	0.27354368156393
54542	RC3H2	ring finger and CCCH-type domains 2
11545	-0.349885384235007	7.53924664803877
4.74564412985124	0.0293725555159731	0.273741762424003
10116	FEM1B	fem-1 homolog B 7569	-0.502130208757955
7.00140778611101	4.7455919219491 0.0293734467711342
0.273741762424003
100534611	TM4SF19-TCTEX1D2	TM4SF19-TCTEX1D2 readthrough (NMD candidate) 1736	0.631631394909527	0.426929143242902
4.74229344767378	0.0294298130625748
0.274084701840083
23635	SSBP2	single stranded DNA binding protein 2	11284
0.562339748459653	2.63197485963936	4.73998790854569
0.02946927837655	0.27424665467838
6482	ST3GAL1 ST3 beta-galactoside alpha-2,3-sialyltransferase 1
8371	0.454472977137171	5.89330760775393
4.73891728342947	0.0294876236569687	0.27424665467838
57631	LRCH2	leucine rich repeats and calponin homology domain containing 2	4929	1.34758539374385	0.375005870655672
4.73651447565072	0.0295288393586338
0.27424665467838
1836	SLC26A2 solute carrier family 26 member 2	8606
0.684253132198639	6.49801123226729	4.73588369803505
0.0295396691178945	0.27424665467838
8869	ST3GAL5 ST3 beta-galactoside alpha-2,3-sialyltransferase 5
5445	0.50238339757212	5.74518671306364
4.73515051770064	0.029552262219223	0.27424665467838
64062	RBM26	RNA binding motif protein 26
10122	-0.331817794110962	5.58562599054194
4.73442796380358	0.0295646782726033	0.27424665467838

10526	IPO8	importin 8	6925	-0.29246756051687
5.99780451258765	4.7315896154028	0.0296135038480003
0.274517768629345
8500	PPFIA1	PTPRF interacting protein alpha 1
10467	-0.291659507329994	7.45908676787389
4.73010811608705	0.02963902211757	0.274572607672489
54492	NEURL1B	neuralized E3 ubiquitin protein ligase 1B
6424	-0.479692559743361	7.29866976468933
4.72213076075159	0.0297768234156314	0.275591766230036
343990	KIAA1211L	KIAA1211 like	4705	0.660855935985562
4.61265519484856	4.72073072636137
0.0298010764628483	0.275591766230036
26298	EHF	ETS homologous factor	4386	-0.482763033394065
8.06969938580029	4.72032769280016
0.0298080621002717	0.275591766230036
1952	CELSR2	cadherin EGF LAG seven-pass G-type receptor 2
12349	-0.498346799143602	6.90727569906417
4.7141420424792	0.0299154900815022	0.276402552487864
125115	KRT40	keratin 40	2225	-1.83655298806785
2.16625278594997	4.71144178823172	0.0299625125994839
0.27643904814002
220963	SLC16A9	solute carrier family 16 member 9
4429	-1.03963919806374	1.43007242899821
4.71125759097279	0.0299657230351604	0.27643904814002
11182	SLC2A6	solute carrier family 2 member 6
3540	-0.393958169679219	6.00401200507758
4.71033931762759	0.0299817332773272	0.27643904814002
140901	STK35	serine/threonine kinase 35
7055	-0.301216502494745	5.78340767208277
4.70938489828941	0.029998383177899	0.27643904814002
6387	CXCL12	C-X-C motif chemokine ligand 12	6077
1.29930667189368	0.157591115921102	4.7073099107567
0.0300346147092087	0.276444822410772
283149	BCL9L	BCL9 like	10470	-0.314764284461447
7.72723818912258	4.70708857473312	0.0300384821667163
0.276444822410772
4043	LRPAP1	LDL receptor related protein associated protein 1
8724	0.328409622923421	6.96430366693632
4.70333613780667	0.030104128439128	0.27686705649827
5337	PLD1	phospholipase D1	9560	-0.332423936210087
4.53143427062312	4.69894880466991
0.0301810714170205	0.277208266050423
83641	FAM107B	family with sequence similarity 107 member B
7019	-0.49887875783479	6.64750585656877
4.69823181753657	0.0301936650699333	0.277208266050423
730394	GTF2H2C_2	GTF2H2 family member C, copy 2	7489
0.521395616127537	2.82471420646762	4.69783708846026
0.0302006006991465	0.277208266050423
10491	CRTAP	cartilage associated protein	6695
0.352630374712731	6.29065460531536	4.69183556171996
0.0303062559494034	0.277851601034083
2030	SLC29A1	solute carrier family 29 member 1 (Augustine blood
group)	3333	0.663027637939015	4.11211914933314
4.69160274281307	0.0303103624173684	0.277851601034083

4987	OPRL1	opioid related nociceptin receptor 1	3549
0.847335547484179	0.200145097739922	4.68973708060097
	0.030343289984211	0.277971525708858
9076	CLDN1	claudin 1	3663	-0.388834543972406
10.6317510413361	4.68529057980012	0.0304219178711143
	0.278355066916762
29903	CCDC106	coiled-coil domain containing 106	2413
0.619728212448094	3.31517281347846	4.68512203907829
	0.0304249023666426	0.278355066916762
140876	RIPOR3	RIPOR family member 3	6009	-1.40992807030632
	3.5719720692498	4.68020990502909	0.0305120199582334
	0.278890938383081
9912	ARHGAP44	Rho GTPase activating protein 44	4331
	0.453583275713543	4.23960721470921
4.67957516726221	0.030523296104895	0.278890938383081
54875	CNTLN	centlein	11064	0.770156446861884
1.45849039242297	4.67491360603771	0.0306062421850714
	0.279310675261289
1543	CYP1A1	cytochrome P450 family 1 subfamily A member 1	3435
	0.783119419319606	9.43080330794662
4.67397679448595	0.030622939798208З	0.279310675261289
53405	CLIC5	chloride intracellular channel 5	8997
0.455762289322085	7.68800988009864	4.6736337419064
0.0306290567003169	0.279310675261289
4642	MYO1D	myosin ID	7731	-0.306928415067639
7.78303018957323	4.67045598105364	0.0306857793088202
	0.279645875587928
201292	TRIM65	tripartite motif containing 65	3906
0.30070437419723	5.09411321305595	4.66650786755333
	0.0307564052088612	0.279985859370734
170959	ZNF431	zinc finger protein 431	14132	-0.532671877774781
	6.46422907382803	4.66482446276829
0.0307865703441646	0.279985859370734
55916	NXT2	nuclear transport factor 2 like export factor 2
3064	-0.399555731380019	3.74245949154652
4.66306672042816	0.0308181004658543	0.279985859370734
55266	TMEM19	transmembrane protein 19	6683
0.259481814259904	5.64049764509171	4.66301763148775
	0.0308189814991247	0.279985859370734
108	ADCY2	adenylate cyclase 2	15784	-1.27925677399202
	0.0531043684212749	4.66176259206142
0.030841515483695	0.279985859370734
867	CBL	Cbl proto-oncogene	11231	-0.467820402964722
	6.6414433565758	4.66167883198783	0.0308430199906506
	0.279985859370734
3589	IL11	interleukin 11	2671	-0.844380599175299
5.209795288684	4.65831709634932	0.0309034670447033
0.280352890476139
284207	METRNL	meteorin like, glial cell differentiation regulator
	2619	-0.418153476939939	4.74790162855802
4.65370337807347	0.0309866270577273	0.280570036777184
5366	PMAIP1	phorbol-12-myristate-13-acetate-induced protein 1
	2235	-0.413862049747696	5.68604301596939
4.65309791732789	0.0309975574997529	0.280570036777184

9689	BZW1	basic leucine zipper and W2 domains 1
7962	-0.401504671536903	8.11215163735143
4.65281036493025	0.031002750119486	0.280570036777184
85294	KRTAP2-4	keratin associated protein 2-4
764	-1.30971062473589	2.11278859782942
4.65254593370809	0.0310075260177826	0.280570036777184
8293	SERF1A	small EDRK-rich factor 1A	4846
0.425308313772888	5.13184312703491	4.65074503927283
	0.0310400724252518	0.280683211401228
79572	ATP13A3 ATPase 13A3	11864	-0.39477805042488
7.93933943797102	4.64813928075546	0.0310872277042051
	0.280908077908867
201626	PDE12	phosphodiesterase 12	8773	-0.269007160226659
	5.55330504375139	4.64715553552148
0.0311050495349934	0.280908077908867
57822	GRHL3	grainyhead like transcription factor 3
3940	-0.680413190405134	6.0544928440956 4.64401861936983
	0.0311619500670177	0.281097240278532
7444	VRK2	VRK serine/threonine kinase 2
3509	-0.334106578848729	5.17153875325799
4.64350240244417	0.0311713241246253	0.281097240278532
54813	KLHL28	kelch like family member 28
7820	-0.486697192660836	4.84017099960373
4.64257994555202	0.0311880824764544	0.281097240278532
196051	PLPP4	phospholipid phosphatase 4	1785
1.55303514714536	1.5841415993223 4.64092196265517
0.0312182268179425	0.281097240278532
222	ALDH3B2 aldehyde dehydrogenase 3 family member B2
3235	-0.968107720038521	1.21156263668714
4.64047616203092	0.0312263372509028	0.281097240278532
687	KLF9	Kruppel like factor 9	5175	0.47571108305644
	3.81351592801836	4.63764446087937
0.0312779056184559	0.281177017699035
114818	KLHL29	kelch like family member 29	11071
0.486046815359619	4.6736003199452 4.63689232670656
0.0312916177268399	0.281177017699035
644943	RASSF10 Ras association domain family member 10
2530	-0.507122814081677	4.05547163667858
4.63583332178307	0.0313109350057447	0.281177017699035
3352	HTR1D	5-hydroxytryptamine receptor 1D 2835
1.1048262051168 1.61531036394794	4.6336815095007
0.0313502244776595	0.281177017699035
3852	KRT5	keratin 5	4292	-0.492798038193311
11.4671693815201	4.63313373372921	0.0313602344056107
	0.281177017699035
4327	MMP19	matrix metallopeptidase 19	4672
1.09060119165957	0.124410664089197	4.63294769771377
	0.0313636347431351	0.281177017699035
83737	ITCH	itchy E3 ubiquitin protein ligase
6666	-0.310965599005342	7.07749022709943
4.63228679341925	0.031375717759948	0.281177017699035
60559	SPCS3	signal peptidase complex subunit 3
6831	-0.500345980844822	7.19907518685984
4.62937050668489	0.0314290929369436	0.281475258802922

57463 AMIGO1 adhesion molecule with Ig like domain 1 5144
0.867148431697069 2.16328943079655 4.6256425061729
0.0314974624266777 0.281768319772469
4088 SMAD3 SMAD family member 3 8831 -0.319040594906385
8.1771579138834 4.62539276016615 0.0315020481733196
0.281768319772469
7106 TSPAN4 tetraspanin 4 5841 0.487119165372638
4.17648437481922 4.62224110954638 0.0315599775423301
0.28182477576146
285 ANGPT2 angiopoietin 2
6775 -1.82496287795172 -0.304808618951622
4.62147737435587 0.0315740322084166 0.28182477576146
55031 USP47 ubiquitin specific peptidase 47
11158 -0.254219824719559 6.67602929239845
4.62089131365886 0.0315848216405754 0.28182477576146
51351 ZNF117 zinc finger protein 117 10691 -0.514554073105399
4.82968097225122 4.62067303538779
0.0315888411469617 0.28182477576146
84765 ZNF577 zinc finger protein 577 6234 -0.540583924268661
2.32371166514449 4.61710590637551
0.0316546040661926 0.282231724478141
100506305 LINC00958 long intergenic non-protein coding
RNA 958 2870 -1.69281048877961 4.79079536358154
4.61397635545987 0.0317124175076495 0.282567323174075
56950 SMYD2 SET and MYND domain containing 2 4889
0.28713591843411 6.15577982682841 4.61085469843356
0.0317701948820986 0.282704237355333
3108 HLA-DMA major histocompatibility complex, class II, DM alpha
2534 1.19753846004333 2.81405339350308
4.61000353731368 0.031785967697154 0.282704237355333
51088 KLHL5 kelch like family member 5
5144 -0.3996618862622 7.68970670482374
4.60769788682197 0.0318287346237698 0.282704237355333
55080 TAPBPL TAP binding protein like
3188 -0.423936412874657 3.73860495316654
4.60665829984044 0.0318480372941269 0.282704237355333
599 BCL2L2 BCL2 like 2 3658 -0.306019705609451
6.17025635928804 4.6059188593511 0.0318617743881178
0.282704237355333
91351 DDX60L DExD/H-box 60 like 11099 -0.486777559005606
5.48221770568467 4.60479581096007
0.0318826498570152 0.282704237355333
81539 SLC38A1 solute carrier family 38 member 1
9977 -0.322561440142891 9.34711962793976
4.60248814182052 0.031925590122173 0.282704237355333
124401 ANKS3 ankyrin repeat and sterile alpha motif domain
containing 3 7868 0.409637236603476 3.73943827402964
4.60227050714235 0.0319296429017747
0.282704237355333
3560 IL2RB interleukin 2 receptor subunit beta
4700 -1.49375976495644 1.7315556159866 4.60117848256251
0.0319499866233523 0.282704237355333
2707 GJB3 gap junction protein beta 3
2387 -0.409829073015835 6.93159077617757

4.60099959945422	0.0319533203907037	0.282704237355333
79815	NIPAL2	NIPA like domain containing 2
4581	-0.280775604930359	6.15434144047523
4.59969985966598	0.0319775539840626	0.282704237355333
57664	PLEKHA4	pleckstrin homology domain containing A4	3851
	0.592139466986116	3.4516552203246	4.59934782939971
	0.0319841208737507	0.282704237355333
2122	MECOM	MDS1 and EVI1 complex locus	9103
0.361267471291445	5.8859875379379	4.59902398493296
0.0319901632189764	0.282704237355333
283463	MUC19	mucin 19, oligomeric	29490	1.20734481219564
	0.637272256600585	4.59418561935887
0.0320805802575499	0.283027525609752
5590	PRKCZ	protein kinase C zeta	7636	0.293896079324604
	5.41338112783776	4.5940457586219	0.0320831978701237
	0.283027525609752
8702	B4GALT4	beta-1,4-galactosyltransferase 4
7145	-0.63875144737035	6.29682185292113
4.59309386661162	0.0321010192604541	0.283027525609752
1727	CYB5R3	cytochrome b5 reductase 3	5973
0.300576031962064	7.73606841014472	4.5921131104278
0.0321193918543584	0.283027525609752
3675	ITGA3	integrin subunit alpha 3	7697
0.374342117155932	9.93470497326673	4.59166571913943
	0.032127776520276	0.283027525609752
4585	MUC4	mucin 4, cell surface associated
21116	-1.3157022682338	9.96289891919966
4.5870560460362	0.0322143006396179	0.283500715421274
7782	SLC30A4	solute carrier family 30 member 4	7157
0.741554579103025	2.866404894777	4.58664805598288
0.0322219703684349	0.283500715421274
4615	MYD88	MYD88 innate immune signal transduction adaptor
3989	-0.302215126154274	7.37136297655812
4.58142793490859	0.0323202708236644	0.284187089408077
1272	CNTN1	contactin 1	7492	-0.845951426070697
4.59332453809951	4.57964920317835	0.0323538377021423
	0.284303767060168
29978	UBQLN2	ubiquilin 2	4230	-0.26304613974092
6.13762885781744	4.57805691468458	0.0323839170335664
	0.284389671403865
55810	FOXJ2	forkhead box J2	6296	-0.454135412788351
5.9363357733259	4.57476427058482	0.0324462097843329
0.284758183238816
5567	PRKACB	protein kinase cAMP-activated catalytic subunit beta
	6730	0.507665221510537	4.27572032693159
4.57254415537842	0.0324882822103684	0.284850779969112
6638	SNRPN	small nuclear ribonucleoprotein polypeptide N	3208
	0.334799304159355	6.92536141329319
4.57206166688772	0.0324974331684615	0.284850779969112
4856	CCN3	cellular communication network factor 3	2673
1.0864019424716	1.20378094988777	4.56726481630121
0.0325885575425991	0.285470872732449
26148	NA	NA	4929	-0.470177269579726
4.05289393281009	4.56553452720371	0.0326214927732062

0.285580780796437
79901	CYBRD1	cytochrome b reductase 1	5176
1.07510485741371	5.24973932578332	4.55975262381437
0.0327318008721603	0.286367479585477
342926	ZNF677	zinc finger protein 677 5547	-0.738761367411142
2.18894317142541	4.55745822293672
0.0327756816999715	0.286487484639479
60677	CELF6	CUGBP Elav-like family member 6 5134
0.347790910828442	4.77536350272783	4.55654357367904
0.0327931916508961	0.286487484639479
929	CD14	CD14 molecule	1886	1.69184188966301
3.09796805552826	4.55556892718575	0.032811860932824
0.286487484639479
85359	DGCR6L	DiGeorge syndrome critical region gene 6 like	1154
0.411497449839724	3.92717435452154
4.55476178360578	0.0328273301096854	0.286487484639479
643338	C15orf62	chromosome 15 open reading frame 62
2370	-0.835951081178968	2.6966410092599 4.55234322895445
0.0328737281326549	0.286713767094705
23683	PRKD3	protein kinase D3	10071	-0.288118533465599
6.15029840526066	4.55023586183077
0.0329142121029799	0.286872255769611
347734	SLC35B2 solute carrier family 35 member B2	2363
0.273498841144423	7.18416569080284	4.54926600382309
0.032932861232065	0.286872255769611
6373	CXCL11	C-X-C motif chemokine ligand 11
1908	-2.09562086508819	1.02948750018576
4.54305905912799	0.0330524744262555	0.287469291215919
100132111	C2CD4D-AS1	C2CD4D and THEM5 antisense RNA 1
506	0.550205965186428	1.27176644262747
4.54263335390123	0.0330606947422513	0.287469291215919
55704	CCDC88A coiled-coil domain containing 88A	14537
0.810821925598926	4.38476402511794	4.54251551753615
0.0330629705253691	0.287469291215919
140733	MACROD2 mono-ADP ribosylhydrolase 2	11344
0.736316713606406	0.65016153851727	4.53711827105127
0.033167383542563	0.288198226600918
694	BTG1	BTG anti-proliferation factor 1
4854	-0.408599756286756	6.96590970528317
4.53354744203611	0.0332366526533591	0.288621074839182
10650	PRELID3A	PRELI domain containing 3A	2643
0.652919201735523	0.359536869816451	4.53230200049066
0.0332608480126592	0.288652229314323
204851	HIPK1	homeodomain interacting protein kinase 1
9280	-0.38052080165356	6.71945767048422
4.53069018881994	0.0332921882081529	0.288745312836903
4211	MEIS1	Meis homeobox 1 10934	-0.584871335574119
3.0609132969663 4.52480626603551	0.0334068577584711
0.289375518769554
7216	TRO	trophinin	6716	1.48496389609512
2.76307860596748	4.52432669112071	0.0334162221790711
0.289375518769554
729830	FAM160A1	family with sequence similarity 160 member
A1	4932	-0.509961958415225	4.67521719827581

4.52378367558414	0.0334268286834538	0.289375518769554
9413	FAM189A2	family with sequence similarity 189 member
A2	4015	0.651029229586624	4.13280217858256
4.51881135848748	0.033524114658519	0.289781040573397
6564	SLC15A1 solute carrier family 15 member 1
4192	-1.02862958334375	1.25042100173896
4.51723367529051	0.0335550446331818	0.289781040573397
5721	PSME2	proteasome activator subunit 2
3417	-0.30107929679088	5.54898473771641
4.51719866720781	0.0335557312934622	0.289781040573397
5493	PPL	periplakin	7686	-0.561670207869871
8.78837880199749	4.51716329032615	0.0335564252024024
	0.289781040573397
56895	AGPAT4	1-acylglycerol-3-phosphate O-acyltransferase 4	8769
	0.457389897120299	4.0612677681633 4.51293690937641
	0.033639432614781	0.290318874082094
64432	MRPS25	mitochondrial ribosomal protein S25	5383
0.315336275463606	4.70284365462026	4.50704363679661
	0.0337555365921272	0.291079520538002
4255	MGMT	O-6-methylguanine-DNA methyltransferase
3557	-0.525953376864575	3.3515087836455 4.5063549788186
0.0337691312111268	0.291079520538002
6676	SPAG4	sperm associated antigen 4	2240
0.613280037340981	0.889847832210887	4.50459303753171
	0.0338039392877381	0.291176452323607
643988	FNDC10	fibronectin type III domain containing 10	2085
	0.660376352822015	1.58032497635596
4.50204480142999	0.0338543473666159	0.291176452323607
23046	KIF21B	kinesin family member 21B
12150	-1.07385499478086	1.22554141891543
4.50109956745619	0.0338730655331097	0.291176452323607
23552	CDK20	cyclin dependent kinase 20	5727
0.614177576780106	2.48061598380145	4.49929244448565
	0.033908881520978	0.291176452323607
100506930	LINC00665	long intergenic non-protein coding
RNA 665 5707	-1.08719112589292	3.72351054749832
4.49895268642839	0.0339156197205125	0.291176452323607
51161	C3orf18 chromosome 3 open reading frame 18	3248
0.747192259834416	2.25897140631912	4.49874510138642
	0.0339197373076095	0.291176452323607
5332	PLCB4	phospholipase C beta 4	6900	-0.496205078615684
	5.36413145923111	4.49792007198942
0.0339361074781184	0.291176452323607
3101	HK3	hexokinase 3	3290	1.69111568130734
0.602031522498445	4.49738745587571	0.0339466799917505
	0.291176452323607
23514	SPIDR	scaffold protein involved in DNA repair 7477
0.313359813065618	5.45548451253802	4.49347567171051
	0.0340244350774217	0.291652748888061
4314	MMP3	matrix metallopeptidase 3	2048
2.72464114307675	3.25365498598516	4.49169526643993
	0.0340598860575287	0.291652748888061
85480	TSLP	thymic stromal lymphopoietin
4282	-1.11414632316097	0.728207806965222

4.4902188386072 0.0340893135949793 0.291652748888061
27065 NSG1 neuronal vesicle trafficking associated 1 4200
0.555939167318453 5.26734846370961
4.48784165906415 0.0341367503848426 0.291652748888061
79033 ERI3 ERI1 exoribonuclease family member 3 2642
0.331007897871765 4.90973140660526 4.48628317186438
0.0341678875481306 0.291652748888061
6239 RREB1 ras responsive element binding protein 1
17254 -0.422249984054635 6.19047157120973
4.48622408506412 0.034169068632791 0.291652748888061
285349 ZNF660 zinc finger protein 660 6513
0.702164534836852 -0.0561376733838586 4.48563841686332
0.0341807778486342 0.291652748888061
55731 FAM222B family with sequence similarity 222 member B
5514 -0.330072689390217 5.55034217360657
4.48526441701615 0.0341882573907749 0.291652748888061
54498 SMOX spermine oxidase 2946 -0.370338626721669
4.96245726218429 4.48519698287367
0.0341896061736415 0.291652748888061
91947 ARRDC4 arrestin domain containing 4
4072 -0.576690205442943 5.31888112791907
4.475824939842 0.0343776030392587 0.293077958472852
3755 KCNG1 potassium voltage-gated channel modifier subfamily G
member 1 2865 -1.20725869675016 1.62837848460071
4.47192268550595 0.0344561979685661
0.293363396155494
54739 XAF1 XIAP associated factor 1
6615 -0.983123322010919 2.76502727405629
4.47010147716603 0.0344929430200987 0.293363396155494
22795 NID2 nidogen 2 6392 2.47935288414997
1.11532370898657 4.46941357343597 0.0345068329558581
0.293363396155494
6309 SC5D sterol-C5-desaturase 5566 0.565108513313805
8.05945403702104 4.46814053673572
0.0345325531385703 0.293363396155494
27297 CRCP CGRP receptor component 3540 -0.28786370659667
5.73508865854444 4.46651767722502
0.034565370136214 0.293363396155494
123 PLIN2 perilipin 2 3596 0.855179740864725
4.87701110602697 4.46615669305976 0.0345726742859578
0.293363396155494
54790 TET2 tet methylcytosine dioxygenase 2
16474 -0.481493152416085 6.00380808728395
4.46537674781507 0.0345884612000691 0.293363396155494
158056 MAMDC4 MAM domain containing 4 5469 0.483728942817122
3.57206376313803 4.46504551023249
0.0345951680779206 0.293363396155494
285172 FAM126B family with sequence similarity 126 member B
11071 -0.494733527802271 4.928909348669 4.46424671834328
0.0346113475598403 0.293363396155494
5980 REV3L REV3 like, DNA directed polymerase zeta catalytic
subunit 12425 0.429780471689151 4.94895830288813
4.46379379486991 0.034620525012139 0.293363396155494
6697 SPR sepiapterin reductase 1480 0.347743197389019

4.60725837288128	4.45953700791833
0.0347069034370991	0.293917531102447
84083	ZRANB3	zinc finger RANBP2-type containing 3	9115
0.573500305938002	1.35489043832974	4.45810797098808
0.0347359518520619	0.293985787064551
101928123	LOC101928123	uncharacterized LOC101928123
3050	-0.920271126667456	1.31287436320776
4.45502306232248	0.0347987463209191	0.294277579702497
131034	CPNE4	copine 4	6427	0.904124086892439
1.2707623565629 4.45435074603706	0.0348124473168442
0.294277579702497
493869	GPX8	glutathione peroxidase 8 (putative)	3868
0.358098333156367	5.40130264354573	4.45004388145946
0.034900349943363	0.294766983628944
55802	DCP1A	decapping mRNA 1A	6127	-0.351057429923808
5.22649252076526	4.4486851143063 0.0349281303693904
0.294766983628944
55206	SBNO1	strawberry notch homolog 1
11124	-0.280772885810552	6.5191200995978 4.44822392002684
0.0349375648886845	0.294766983628944
55057	CRYBG2	crystallin beta-gamma domain containing 2
5871	-0.627452641595923	5.38457011552432
4.44672872312078	0.0349681700333734	0.294766983628944
29886	SNX8	sorting nexin 8 5773	-0.37000700230402
4.58059873072633	4.44636769392275	0.0349755641315988
0.294766983628944
54972	TMEM132A	transmembrane protein 132A
6882	-0.427126794638644	8.85643855313776
4.44424778855159	0.0350190140881497	0.294955700741258
8416	ANXA9	annexin A9	1826	-1.35847652212048
1.12621496174431	4.44312094179554	0.0350421331141044
0.294973052000758
9911	TMCC2	transmembrane and coiled-coil domain family 2	4922
0.650173463288676	4.53329448801685
4.43691094400547	0.0351698276501384	0.295870135672966
5660	PSAP	prosaposin	3239	0.385167700999102
10.6214370793004	4.43465474511254	0.0352163417199011
0.296083612527403
55144	LRRC8D	leucine rich repeat containing 8 VRAC subunit D
5177	-0.310555124494322	5.03208977372986
4.43302751653973	0.0352499287972358	0.296163796714154
317	APAF1	apoptotic peptidase activating factor 1
7908	-0.397471320164722	5.39973653629422
4.43214459814677	0.0352681668393809	0.296163796714154
9501	RPH3AL	rabphilin 3A like (without C2 domains)	6066
0.507747867976793	3.86156365971309	4.43061629502034
0.0352997596328365	0.296251487823332
4282	MIF	macrophage migration inhibitory factor	1109
0.417201324041463	6.53331354654687	4.42938507780804
0.0353252326525596	0.296287744769103
712	C1QA	complement C1q A chain	1272	1.87588612241235
2.43358197912868	4.42732204034598
0.0353679585614315	0.296468579036488
653238	GTF2H2B general transcription factor IIH subunit 2B

(pseudogene)	4463	0.517767143393148	2.52801660629799
	4.42438716716702	0.0354288335342605
0.296801238824394
100505783	OSER1-DT	OSER1 divergent transcript	2186
	0.644883035950188	1.35612309326159
4.42209019257361	0.0354765537098386	0.297023363905385
23219	FBXO28	F-box protein 28	6044	-0.239870032220645
	6.37627132044612	4.41946533527346
0.0355311680609535	0.29730290981468
81553	FAM49A	family with sequence similarity 49 member A	5448
	0.613003521906776	3.74189948915172
4.41776225849831	0.0355666503391348	0.297422132119559
51390	AIG1	androgen induced 1	6016	0.344895453429301
	4.37100899056115	4.41495854304525
0.0356251443401932	0.297733530294204
144811	LACC1	laccase domain containing 1
5696	-0.351410458935321	5.36634253603853
4.40634246344698	0.0358055326534431	0.2990626689784
51092	SIDT2	SID1 transmembrane family member 2	9061
0.274009396202525	5.17422096006051	4.40284860564487
	0.035878953027913	0.299497315293193
51341	ZBTB7A	zinc finger and BTB domain containing 7A
5725	-0.31803506430154	6.18179303235745
4.39977398166299	0.035943693892941	0.299636086369813
79714	CCDC51	coiled-coil domain containing 51	2410
0.454185299511098	4.14033982303184	4.39912553609479
	0.0359573635002054	0.299636086369813
317649	EIF4E3	eukaryotic translation initiation factor 4E family
member 3	10779	-0.332852837889349	4.77248648548902
	4.39800003615132	0.0359811026074055
0.299636086369813
54680	ZNHIT6	zinc finger HIT-type containing 6
6197	-0.494812321024899	5.53761537599975
4.39799802946925	0.0359811449470997	0.299636086369813
25776	CBY1	chibby family member 1, beta catenin antagonist 2348
	0.349051233090378	4.16581123656543
4.39668119867743	0.0360089404721929	0.299689381576949
115361	GBP4	guanylate binding protein 4
6127	-1.4205829354798	3.35922856855897
4.39465480709253	0.0360517572368975	0.299867555592176
25907	TMEM158 transmembrane protein 158 (gene/pseudogene)	1813
	0.860720905603902	0.685031885995328
4.39104277098092	0.0361282101649287	0.300237658390642
26225	ARL5A	ADP ribosylation factor like GTPase 5A
6327	-0.268406097428934	5.75574369451635
4.39052785697578	0.0361391227276806	0.300237658390642
57562	CEP126	centrosomal protein 126 8191	0.694915713165591
	2.1276166558301 4.38936358916535	0.0361638097545058
	0.300264661073718
5753	PTK6	protein tyrosine kinase 6
2947	-0.713843811545306	4.39560578296506
4.38753776255323	0.0362025599640995	0.300277625986791
27333	GOLIM4	golgi integral membrane protein 4	4647
0.355404270327762	5.00483447792871	4.38703729965037

0.036213189064926 0.300277625986791
89910 UBE3B ubiquitin protein ligase E3B
12453 -0.23212765279628 6.22861134890479
4.38626093901879 0.0362296842948295 0.300277625986791
29966 STRN3 striatin 3 4965 -0.290911248158035
5.93653408823366 4.38457214193216 0.036265593106261
0.300397494168775
8301 PICALM phosphatidylinositol binding clathrin assembly
protein 5833 -0.447327451684239 8.46702874706888
4.37999506489587 0.0363631027135164 0.30062067074403
100505989 SMIM31 small integral membrane protein 31
3190 -1.0906662570324 -0.0616212736971192
4.37849766926171 0.0363950626339545 0.30062067074403
64838 FNDC4 fibronectin type III domain containing 4 1982
0.520525348647441 4.22615073032249
4.3779824051255 0.0364060670614881 0.30062067074403
728 C5AR1 complement C5a receptor 1 2311
1.66170229946775 2.41053915046753 4.37782521059166
0.036409424937575 0.30062067074403
374655 ZNF710 zinc finger protein 710 4930 -0.311677960849261
5.65521033064223 4.37696786519876
0.036427744637843 0.30062067074403
166929 SGMS2 sphingomyelin synthase 2
7176 -0.369773128985321 6.42836948737345
4.37640730659695 0.0364397278343724 0.30062067074403
9371 KIF3B kinesin family member 3B
6103 -0.236547728428804 6.68195779823568
4.3762649634775 0.036442771394543 0.30062067074403
160760 PPTC7 PTC7 protein phosphatase homolog
5687 -0.28880605060252 6.02690667052627
4.37464901624897 0.0364773420119745 0.300728740177591
11236 RNF139 ring finger protein 139 2700 0.276523476801271
5.97286841881653 4.37039505184862
0.0365685130606292 0.301303036729549
84659 RNASE7 ribonuclease A family member 7
1499 -0.7697830777429 3.59453227724168
4.36808098260223 0.0366182083957017 0.301332692149367
2877 GPX2 glutathione peroxidase 2 1499
0.658045695088668 5.21760525744376 4.36807546062359
0.0366183270663871 0.301332692149367
65258 MPPE1 metallophosphoesterase 1 7630
0.326127654939843 4.60872875586583 4.36544890755201
0.0366748189643769 0.301332692149367
84133 ZNRF3 zinc and ring finger 3 6968 0.570627324909213
3.78290268222169 4.36343367548246
0.0367182244329113 0.301332692149367
112812 FDX2 ferredoxin 2 2489 0.382375599473668
4.18492249456815 4.36340406173666 0.0367188626752183
0.301332692149367
9891 NUAK1 NUAK family kinase 1 7008 -0.39285298991826
7.53690584832956 4.3627501493807 0.0367329589063793
0.301332692149367
9949 AMMECR1 AMMECR nuclear protein 1
6516 -0.36863397693339 4.83811610199425

4.36226746432935	0.0367433676666732	0.301332692149367
9771	RAPGEF5 Rap guanine nucleotide exchange factor 5
10245	-0.704390118547803	2.31933972806982
4.3622281154215 0.0367442163340557	0.301332692149367
3687	ITGAX	integrin subunit alpha X	7074
1.50340377401933	1.78321047149763	4.35714257732196
	0.0368540728825061	0.301740673091576
4891	SLC11A2 solute carrier family 11 member 2
9823	-0.375646797474491	6.6886710271231 3
4.35689067991223	0.0368595232403838	0.301740673091576
152519	NIPAL1	NIPA like domain containing 1
5302	-0.642260292711514	4.3925548551867 4.35680356214108
	0.0368614084225794	0.301740673091576
9121	SLC16A5 solute carrier family 16 member 5	3089
0.474978507957577	4.13230920444074	4.35555178465895
	0.0368885073721584	0.301740673091576
54332	GDAP1	ganglioside induced differentiation associated
protein 1	4052	0.464973003572762	3.35401777037931
	4.35494384493855	0.0369016758053737
0.301740673091576
11128	POLR3A	RNA polymerase III subunit A
8917	-0.256325770922542	5.48366500594053
4.35228247423377	0.0369593810764271	0.302036202297267
5802	PTPRS	protein tyrosine phosphatase receptor type S	9425
	0.624509487067378	5.0255343494453 4.34493485124288
	0.0371191873369512	0.303064161795642
26090	ABHD12	abhydrolase domain containing 12	3429
0.474795614725312	5.29643261046241	4.34389893571998
	0.0371417759999244	0.303064161795642
9792	SERTAD2 SERTA domain containing 2
6274	-0.373864143755541	5.34557934233568
4.34157116260768	0.0371925868138628	0.303064161795642
4828	NMB	neuromedin B	1022	-0.898691988695464
3.68496087288188	4.34119860470282	0.0372007257933739
	0.303064161795642
8821	INPP4B	inositol polyphosphate-4-phosphatase type II B
13779	-0.348385275182833	8.21656374555434
4.34043200288301	0.0372174790096678	0.303064161795642
26286	ARFGAP3 ADP ribosylation factor GTPase activating protein 3
	3588	-0.413751698608536	6.80741565338984
4.33990182923232	0.0372290699780316	0.303064161795642
56975	FAM20C	FAM20C golgi associated secretory pathway kinase
	4629	0.505656979873835	5.21929323188798
4.33955630132201	0.0372366261476619	0.303064161795642
115273	RAB42	RAB42, member RAS oncogene family
2180	-0.940552159212 3.54654673501572	4.33491382517
0.0373383059745018	0.303715244938935
84707	BEX2	brain expressed X-linked 2	1176
0.752607461555992	2.08407377772648	4.33283266878259
	0.0373839819775193	0.303910293417942
3115	HLA-DPB1	major histocompatibility complex, class II,
DP beta 1	2659	1.67796951565822	3.24686012255744
	4.33106956960303	0.0374227232449565
0.304048772907254

55665   URGCP   upregulator of cell proliferation
7692    -0.338447941797962      6.33688317275494
4.32857794747508        0.0374775443642013      0.304255816798123
374618  TEX9    testis expressed 9      8290    0.56087935538439
        2.73752964895933        4.32793746903984
0.0374916498746028      0.304255816798123
8073    PTP4A2  protein tyrosine phosphatase 4A2
6431    -0.345973362318725      7.30863868174158
4.3248598039996 0.0375595081304723      0.304482235221897
51768   TM7SF3  transmembrane 7 superfamily member 3    3888
0.349063235778445       7.73533565573489        4.32470042107724
        0.037563025805914       0.304482235221897
84285   EIF1AD  eukaryotic translation initiation factor 1A domain
containing      3327    -0.301134268158896      5.18089489595042
        4.31882179259635        0.0376930120679861
0.305257394062167
25978   CHMP2B  charged multivesicular body protein 2B
3106    -0.308032660168821      6.55270750193145
4.31840521547809        0.0377022411456807      0.305257394062167
9991    PTBP3   polypyrimidine tract binding protein 3
8239    -0.294806628632806      8.1450695263641 4.31622565464858
        0.0377505669476245      0.305472091990397
5341    PLEK    pleckstrin      3314    1.70932035793564
0.952800540085197       4.3151670714505 0.0377740615485046
0.305485727546134
29842   TFCP2L1 transcription factor CP2 like 1
9372    -1.0347836766164        3.12749465468312
4.30907628846703        0.0379095407521515      0.30622554316292
26579   MYEOV   myeloma overexpressed   2667    -0.914894499044295
        5.65762809291868        4.30877268254534
0.0379163072632653      0.30622554316292
51364   ZMYND10 zinc finger MYND-type containing 10
4009    -0.948218533198454      -0.404428214856889
4.30745921673322        0.0379455952600187      0.30622554316292
4864    NPC1    NPC intracellular cholesterol transporter 1
10102   -0.274575832626166      7.33562052860795
4.30712771768932        0.0379529908567737      0.30622554316292
375449  MAST4   microtubule associated serine/threonine kinase
family member 4 14675   -0.650136776548656      6.58099915627578
        4.30348431544712        0.0380343731547467
0.306705506493113
64854   USP46   ubiquitin specific peptidase 46
8917    -0.24542301864636       5.68022042377525
4.29849236148907        0.0381461749910611      0.307322651695534
117289  TAGAP   T cell activation RhoGTPase activating protein  4089
        0.956473782655312       0.612640545837773
4.29810850392183        0.0381547862710454      0.307322651695534
9510    ADAMTS1 ADAM metallopeptidase with thrombospondin type 1
motif 1 7063    -0.557034162071078      8.81675039301256
4.29551927614131        0.0382129250385028      0.307469382153464
51326   ARL17A  ADP ribosylation factor like GTPase 17A
9153    -0.555098891350306      3.53974425463163
4.2953421593289 0.0382169054279418      0.307469382153464
11030   RBPMS   RNA binding protein, mRNA processing factor     8931

0.274557452408463 5.68643379385093
4.2918319129133 0.0382958817758735 0.307927908171447
8553 BHLHE40 basic helix-loop-helix family member e40
3837 -0.251848268952067 8.7024388033611 4.28808959683833
0.0383802678597248 0.308429381475137
4240 MFGE8 milk fat globule-EGF factor 8 protein 4521
0.563799667924139 8.00126215922893 4.2867757383454
0.0384099404760044 0.30849084647213
23760 PITPNB phosphatidylinositol transfer protein beta
5129 -0.251895314745536 6.63563180624795
4.28113361772799 0.0385376376524078 0.309339077706176
400966 RGPD1 RANBP2 like and GRIP domain containing 1
13692 -0.450371906184555 3.91738467964036
4.27943568233714 0.0385761538068646 0.309470897120706
4084 MXD1 MAX dimerization protein 1
6351 -0.609749749957952 7.84663453245618
4.27772559403479 0.0386149864348362 0.309605103029623
79794 C12orf49 chromosome 12 open reading frame 49 8664
0.381483227685895 7.34509512911346
4.27513153595777 0.0386739704775813 0.30982214980349
11176 BAZ2A bromodomain adjacent to zinc finger domain 2A
10918 -0.317186855861419 7.64610710592353
4.2745900213344 0.0386862954241667 0.30982214980349
80162 PGGHG protein-glucosylgalactosylhydroxylysine glucosidase
5260 -0.982213579657084 6.30152591000217
4.27146302199652 0.0387575469572745 0.310215405846025
643314 KIAA0754 KIAA0754 7410 -0.566829445700838
6.04113932039237 4.27045447649194
0.0387805569231305 0.310222307722609
6278 S100A7 S100 calcium binding protein A7
536 -2.92549155444195 1.47498249431095
4.26215172110337 0.0389705295456885 0.311359691218842
51564 HDAC7 histone deacetylase 7 9928 -0.279995627797863
7.49792234074124 4.26178107731073
0.0389790328464955 0.311359691218842
7580 ZNF32 zinc finger protein 32 1436 0.343898446311995
4.04028021628997 4.260607092523 0.0390059792214738
0.311359691218842
92359 CRB3 crumbs cell polarity complex component 3
1592 -0.336308010835104 5.54870963627628
4.26035990156346 0.0390116554643441 0.311359691218842
11309 SLCO2B1 solute carrier organic anion transporter family
member 2B1 9379 2.03470423959131 3.11162898727791
4.24993431474487 0.039251848624138
0.313098316445501
8663 EIF3C eukaryotic translation initiation factor 3 subunit C
6749 -0.359393865536454 7.15261731167721
4.24208008733496 0.0394338249458698 0.314370851460898
55754 TMEM30A transmembrane protein 30A
5123 -0.265342680852219 8.55629168388069
4.2377972261844 0.0395334281774015 0.314968191987814
8289 ARID1A AT-rich interaction domain 1A
9246 -0.370882457032777 7.48223251924395
4.23692579478931 0.0395537266871254 0.314968191987814

3097	HIVEP2	HIVEP zinc finger 2	9774	-0.538429315860135
	6.2308215778361 4.23448725870223	0.0396105863630458
	0.315188279567867
7138	TNNT1	troponin T1, slow skeletal type
2171	-1.04517323537189	1.62192322639895
4.23381101146862	0.039626369695082	0.315188279567867
10773	ZBTB6	zinc finger and BTB domain containing 6
4106	-0.363072527760415	4.59288819528268
4.22799873099529	0.0397622983219713	0.31608996174566
54863	TOR4A	torsin family 4 member A
4148	-0.359393178035863	5.51092780267305
4.22636722636362	0.0398005413042473	0.316214510521039
925	CD8A	CD8a molecule	3634	-1.41204369968316
0.14341547947258	4.22477394333028	0.0398379256118805
	0.316332099799099
5307	PITX1	paired like homeodomain 1
3511	-1.26559742374496	3.92144632737684
4.22200199874068	0.039903053520562	0.316669728420687
79828	METTL8	methyltransferase like 8	8132
0.360769730756258	4.45300244672997	4.22025839502525
	0.0399440773997315	0.316815793962649
261729	STEAP2	STEAP2 metalloreductase 8715	-0.589430106697824
	2.45483938965735	4.21756066183331
0.0400076374811984	0.316828437466297
127733	UBXN10	UBX domain protein 10	5571	1.14684060136978
	0.886949382780333	4.21662647453921
0.0400296721829578	0.316828437466297
728239	MAGED4	MAGE family member D4	5599	0.889920189411653
	4.99459822087431	4.21614326342565
0.040041074694424	0.316828437466297
65975	STK33	serine/threonine kinase 33	7599
0.847789517711478	1.49682059225006	4.2150614788914
0.0400666143242853	0.316828437466297
4015	LOX	lysyl oxidase	5468	1.09174400411978
4.48556372442233	4.21468295302713	0.0400755549012827
	0.316828437466297
22874	PLEKHA6	pleckstrin homology domain containing A6
7679	-0.526819779033427	6.0280586455357 4.21443607105794
	0.0400813872485385	0.316828437466297
2627	GATA6	GATA binding protein 6	3945	0.488563149560097
	5.37591625622819	4.21148596036123
0.0401511497206317	0.317200876557749
57472	CNOT6	CCR4-NOT transcription complex subunit 6
6586	-0.252642052865761	6.35995557019236
4.20987543034066	0.0401892883480932	0.317323202870204
100913187	APOBEC3A_B	APOBEC3A and APOBEC3B deletion
hybrid	1103	-1.18062926141187	2.7532755576736
4.20753624952586	0.0402447497384261	0.317582089907681
79768	KATNBL1	katanin regulatory subunit B1 like 1
5877	-0.368297396069515	5.29286892912632
4.2045546495032 0.0403155592126589	0.317744223887009
100093630	SNHG8	small nucleolar RNA host gene 8 923
0.409804527897806	3.84059776152431	4.20436363674141
	0.0403200999949404	0.317744223887009

51310	SLC22A17	solute carrier family 22 member 17	4278
	0.913363068908265	2.34735016038211
4.20380640059941	0.0403333497587707	0.317744223887009
258010	SVIP	small VCP interacting protein
5263	-0.44971802693777	4.18001600761489
4.19948432926467	0.0404362737787076	0.318319630835893
94134	ARHGAP12	Rho GTPase activating protein 12	5938
	-0.271323051786234	5.61699650096095
4.19747078081553	0.0404843176997684	0.318319630835893
4317	MMP8	matrix metallopeptidase 8	3438
1.96623720285831	1.59380321013967	4.19699251680438
	0.0404957380396135	0.318319630835893
79042	TSEN34	tRNA splicing endonuclease subunit 34	3542
0.328012086196329	5.21993441860971	4.19604403965228
	0.0405183964803893	0.318319630835893
445	ASS1	argininosuccinate synthase 1
3050	-0.969178715849218	6.02705851563295
4.19597615572135	0.0405200186892552	0.318319630835893
100506343	INKA2-AS1	INKA2 antisense RNA 1	2131
0.899038988683059	0.0421511956375143	4.19427609769683
	0.0405606668834538	0.318460348114651
6364	CCL20	C-C motif chemokine ligand 20
1343	-1.63474585650424	5.06083109434899
4.1927196331377 0.0405979193141575	0.318574260971095
55117	SLC6A15 solute carrier family 6 member 15
8781	-0.923241879805852	1.74232106975402
4.19043721330757	0.0406526117352224	0.318824822270582
6443	SGCB	sarcoglycan beta	4541	0.386111414435202
	6.20157525735361	4.18513484524743
0.040779968413244	0.319563651683998
5330	PLCB2	phospholipase C beta 2	7350	1.45209803406521
	1.25909741397132	4.18148266289541
0.0408679331698217	0.319563651683998
83637	ZMIZ2	zinc finger MIZ-type containing 2
7339	-0.267466842910499	7.07399053169755
4.17981533339846	0.040908157912217	0.319563651683998
79924	ADM2	adrenomedullin 2	4276	-0.498824794842528
	4.28550709200667	4.17841540350896
0.0409419636808243	0.319563651683998
79755	ZNF750	zinc finger protein 750 3713	-0.712259884624645
	3.93584567546621	4.1783785606402 0.0409428537654079
	0.319563651683998
1054	CEBPG	CCAAT enhancer binding protein gamma
4411	-0.31043815572923	6.00275995296357
4.17835785026898	0.0409433541149051	0.319563651683998
27151	CPAMD8	C3 and PZP like alpha-2-macroglobulin domain
containing 8	10199	-1.05923869556556	2.67500873872245
	4.17782746655279	0.0409561700388965
0.319563651683998
4599	MX1	MX dynamin like GTPase 1
8616	-0.50345535238344	5.00869852969143
4.17351162952247	0.0410606124909058	0.319563651683998
163259	DENND2C DENN domain containing 2C
7413	-0.522782553402743	5.40656759797455

4.17331338323411	0.0410654167283462	0.319563651683998
79720	VPS37B	VPS37B subunit of ESCRT-I
5953	-0.278490696060046	6.96094394608814
4.17284595420943	0.0410767465916284	0.319563651683998
100506115	SMIM25	small integral membrane protein 25	2451
	1.22663898894414	0.649697269903589
4.17207186880216	0.041095516622018	0.319563651683998
152273	FGD5	FYVE, RhoGEF and PH domain containing 5 8046
1.64591643635152	0.720589676641044	4.17203493966815
	0.0410964123051058	0.319563651683998
5055	SERPINB2	serpin family B member 2
3041	-1.06363875341646	10.0777512038105
4.17172955174192	0.0411038200017544	0.319563651683998
24141	LAMP5	lysosomal associated membrane protein family member
5	2042	1.58334123834563	-0.0907474226000534
4.17129698931748	0.0411143149293301	0.319563651683998
58526	MID1IP1 MID1 interacting protein 1
3790	-0.378316516044335	4.58113520053728
4.17125548191929	0.0411153221392064	0.319563651683998
22878	TRAPPC8 trafficking protein particle complex 8
8845	-0.270350590582142	6.14212657583864
4.17082949601105	0.0411256605223829	0.319563651683998
10207	PATJ	PATJ crumbs cell polarity complex component
11867	-0.40040843621493	6.95123699564781
4.17014889562969	0.0411421838888432	0.319563651683998
43849	KLK12	kallikrein related peptidase 12
1769	-1.39192217141417	0.694988367871432
4.16950607057109	0.041157796562791	0.319563651683998
26094	DCAF4	DDB1 and CUL4 associated factor 4	5581
0.380718294145678	3.94211127478681	4.16775921673199
	0.0412002548790514	0.319563651683998
11151	CORO1A	coronin 1A	4801	-0.76379092700574
4.4013760925401 4.16764183697553	0.0412031095124794
0.319563651683998
81557	MAGED4B MAGE family member D4B	9565	0.881543429868172
	5.05212186350926	4.16633885305973
0.0412348115685444	0.319632543243277
10042	HMGXB4	HMG-box containing 4	6006	-0.266783015279214
	5.43907973033654	4.16480533916673
0.0412721553233524	0.319745066158295
29887	SNX10	sorting nexin 10	3518	1.16552629806034
	2.03801790919743	4.16371912408502
0.0412986280584558	0.319773290887115
56288	PARD3	par-3 family cell polarity regulator
6513	-0.269912690444305	6.45128051262327
4.16194051609957	0.0413420139962142	0.319932370190592
5971	RELB	RELB proto-oncogene, NF-kB subunit
3035	-0.465707340210365	4.78321705856836
4.15432713490051	0.0415282705469151	0.321196292407863
5144	PDE4D	phosphodiesterase 4D	15979	-0.630367882934144
	2.31942057245044	4.15103503376506
0.0416090825339902	0.321643719124503
5903	RANBP2	RAN binding protein 2	12302	-0.324401574872321
	7.67886734190744	4.14931302426339

0.041651418905882	0.321793394713011
581	BAX	BCL2 associated X, apoptosis regulator	2587
0.302275303963326	6.35103982310302	4.1468763757144
0.0417114023436954	0.321878723269437
57614	RELCH	RAB11 binding and LisH domain, coiled-coil and HEAT repeat containing	8299	-0.284658571358157
5.75738181194171	4.14571036393687	0.0417401383780425
	0.321878723269437
2256	FGF11	fibroblast growth factor 11
7215	-0.424613519447385	5.30554942739894
4.14528391563816	0.0417506532715146	0.321878723269437
29957	SLC25A24	solute carrier family 25 member 24
4594	-0.270828407833574	7.03031689346254
4.14416157608239	0.0417783399905642	0.321878723269437
9788	MTSS1	MTSS I-BAR domain containing 1
7727	-0.632451340687299	5.7484153409202 4.14413226256531
	0.0417790633769869	0.321878723269437
84146	ZNF644	zinc finger protein 644 7567	-0.342626969459608
	6.27470771538095	4.1432702144688 0.0418003425163923
	0.321878723269437
100422894	MIR4260 microRNA 4260	67	-0.514830159962319
	3.38992846397433	4.14219108427575
0.0418269962491634	0.321906998056061
8444	DYRK3	dual specificity tyrosine phosphorylation regulated kinase 3	4218	0.58383079130989	2.73090035241341
	4.14038195684408	0.0418717204581217
0.322074238581499
247	ALOX15B arachidonate 15-lipoxygenase type B	3028
1.13862774980198	5.1094407991796 4.13347747206592
0.0420428715820112	0.322884044095605
11135	CDC42EP1	CDC42 effector protein 1
2573	-0.340210938804292	7.08286892953465
4.1334378909981 0.0420438548499837	0.322884044095605
3394	IRF8	interferon regulatory factor 8	5162
1.83592483846145	1.14953767357958	4.13277925510652
	0.0420602201485259	0.322884044095605
8242	KDM5C	lysine demethylase 5C	8727	-0.296802920776914
	7.44830380845039	4.1324176915523 0.0420692068590333
	0.322884044095605
6548	SLC9A1	solute carrier family 9 member A1
5974	-0.228430334503964	6.93692977644149
4.13052693570055	0.0421162347289793	0.323067962677334
79991	STN1	STN1 subunit of CST complex	1617
0.437950007403562	5.0481760503758 4.12736403915368
0.0421950275106392	0.323492460876706
6696	SPP1	secreted phosphoprotein 1	2321
1.68068360229918	3.3750384279111 4.1264525112958
0.042217763866825	0.323492460876706
55088	CCDC186 coiled-coil domain containing 186
7366	-0.309427717578485	6.73802984435545
4.12207717829666	0.0423270777486153	0.323919362508581
55624	POMGNT1 protein O-linked mannose N-acetylglucosaminyltransferase 1 (beta 1,2-)	6198
0.27424889189367	6.65141520750055	4.12142180385956

        0.0423434773102879      0.323919362508581
80020   FOXRED2 FAD dependent oxidoreductase domain containing 2
        5577    0.626482253207061       2.17980188071535
4.12062684347509        0.0423633787230267      0.323919362508581
5111    PCNA    proliferating cell nuclear antigen      1471
0.338243100360196       5.36889563833383        4.12052299245866
        0.0423659793043279      0.323919362508581
359845  RFLNB   refilin B       3621    -0.737368763374484
4.48455463412321        4.11714592505075        0.0424506375322768
        0.324342101362994
255488  RNF144B ring finger protein 144B
5175    -0.310912293249352      7.195492859566  4.11647087084671
        0.0424675814878083      0.324342101362994
23092   ARHGAP26        Rho GTPase activating protein 26
13347   -0.402169838320426      4.78646876130048
4.1151590471357 0.0425005287833769      0.324403911362683
401145  CCSER1  coiled-coil serine rich protein 1
10428   -0.751500649561645      2.60570412787364
4.11430493477377        0.0425219948070169      0.324403911362683
23331   TTC28   tetratricopeptide repeat domain 28      12645
0.597507167279972       2.43779026161765        4.11208366419836
        0.0425778743905081      0.324653394985218
221749  PXDC1   PX domain containing 1  2474    0.322575156813656
        7.19545549558182        4.10757339471017
0.042691575036838       0.325066597296624
1999    ELF3    E74 like ETS transcription factor 3
7191    -0.468591098696108      8.39724020061297
4.10669360295196        0.0427137911379665      0.325066597296624
84752   B3GNT9  UDP-GlcNAc:betaGal beta-1,3-N-acetylglucosaminyltransferase 9 2917    0.35095709090238
3.74641206300933        4.10630581320363        0.0427235872876184
        0.325066597296624
7802    DNALI1  dynein axonemal light intermediate chain 1      4456
        0.614949488689631       3.07217878356067
4.10625404942128        0.0427248950969576      0.325066597296624
2548    GAA     glucosidase alpha, acid 5156    0.601851018291926
        6.11449154119641        4.10201865304668
0.0428320449268528      0.325704914924227
23507   LRRC8B  leucine rich repeat containing 8 VRAC subunit B
7691    -0.676163123301229      4.9818766719753 4.10005864913565
        0.04288172598914        0.325905770987457
7357    UGCG    UDP-glucose ceramide glucosyltransferase
4615    -0.338640662280468      6.97549048584234
4.09192189395002        0.0430886206911944      0.32715827060421
220441  RNF152  ring finger protein 152 9924    -0.533208840572783
        4.7327986668685 4.09174070456827        0.0430932397561768
        0.32715827060421
55288   RHOT1   ras homolog family member T1
6372    -0.279595746824545      5.20342603799813
4.08985693608613        0.043141293572139 4      0.327264810010132
10449   ACAA2   acetyl-CoA acyltransferase 2    5072
0.488244659126966       6.19439948220139        4.08935912449353
        0.0431540018625483      0.327264810010132
51246   SHISA5  shisa family member 5   4664    0.291223748070357

6.5693113681105 4.0882429248013 0.0431825108717375
0.327303803993737
79949 PLEKHS1 pleckstrin homology domain containing S1
5402 -1.54092668789322 2.54951010500861
4.08480948080672 0.0432703291844399 0.327596370884305
644815 FAM83G family with sequence similarity 83 member G
7876 -0.36707016299915 6.53981427272414
4.08412145684835 0.0432879495781958 0.327596370884305
92737 DNER delta/notch like EGF repeat containing 3935
0.764602389442525 1.87230789290831 4.08348264494407
0.0433043164009849 0.327596370884305
84433 CARD11 caspase recruitment domain family member 11
4707 -0.263769272808373 6.42562826321495
4.0830789548774 0.0433146625885438 0.327596370884305
7182 NR2C2 nuclear receptor subfamily 2 group C member 2
11821 -0.319007830185118 6.24550081409169
4.07866176977169 0.043428040818651 0.328133149834651
283078 MKX mohawk homeobox
4106 -0.626547009350378 -0.289352372920291
4.07803475542466 0.0434441600125621 0.328133149834651
6688 SPI1 Spi-1 proto-oncogene 2069 1.89156216698337
1.99495368906649 4.07757768045031
0.0434559143958933 0.328133149834651
11326 VSIG4 V-set and immunoglobulin domain containing 4 2192
2.27084420771433 2.79595179234742
4.07541070472062 0.0435116870443118 0.328312341257771
60489 APOBEC3G apolipoprotein B mRNA editing enzyme
catalytic subunit 3G 5454 -0.609059664768037
2.41644967140973 4.07392068663277 0.0435500801476636
0.328312341257771
5045 FURIN furin, paired basic amino acid cleaving enzyme
4728 -0.251698689707408 8.46560447441905
4.07315043610549 0.0435699410653068 0.328312341257771
196463 PLBD2 phospholipase B domain containing 2 4804
0.260679159509584 6.18333379397435 4.07178976270121
0.0436050493311209 0.328312341257771
5523 PPP2R3A protein phosphatase 2 regulatory subunit B''alpha
7837 -0.357399284043072 5.40330067184803
4.0716057673966 0.0436097990849767 0.328312341257771
163351 GBP6 guanylate binding protein family member 6
2987 -0.50985031617563 5.74168912404845
4.07119981958962 0.043620280365582 4 0.328312341257771
1109 AKR1C4 aldo-keto reductase family 1 member C4 1414
0.798573987767884 1.27800838988075 4.06984554418128
0.0436552659564407 0.328335178685236
85464 SSH2 slingshot protein phosphatase 2
12608 -0.418386573737913 5.69677203830506
4.06926794999827 0.0436701961798097 0.328335178685236
5586 PKN2 protein kinase N2 6897 -0.302697441209374
6.79344631758759 4.06501307655782
0.0437803460323518 0.328817800841142
6897 TARS1 threonyl-tRNA synthetase 1
5718 -0.262733764758934 7.98630586708999
4.06497481768486 0.0437813378002948 0.328817800841142

100507127	LINC00707	long intergenic non-protein coding
RNA 707 3070	-1.35870641462869	0.75123057571364
4.06393138720943	0.043808395305799	0.328844690808321
9334	B4GALT5 beta-1,4-galactosyltransferase 5
4722	-0.487934312068244	7.78994745766155
4.05996447472478	0.0439114232045206	0.329441513029309
64699	TMPRSS3 transmembrane serine protease 3 5660
1.24539352751559	1.79244654028181	4.05457249244478
	0.0440518717389423	0.330318290924714
6596	HLTF	helicase like transcription factor	6024
0.416043995321419	3.89056138913194	4.05040122146804
	0.0441608478872415	0.330958264503259
64792	IFT22	intraflagellar transport 22	4132
0.378958507549477	4.14715619500304	4.04491513527589
	0.0443046062772064	0.331858085628251
10457	GPNMB	glycoprotein nmb	5956	1.19877378414258
	6.39125772774766	4.04200842326827
0.0443809738335928	0.332252432114983
10966	RAB40B	RAB40B, member RAS oncogene family	5108
0.559816255803771	2.97608869528982	4.0397576939406
0.0444402020460949	0.33237935703052
388284	C16orf86	chromosome 16 open reading frame 86	1789
	0.786683860115574	0.246546229784671
4.03890655466112	0.0444626215550854	0.33237935703052
26256	CABYR	calcium binding tyrosine phosphorylation regulated
	3322	0.546923702263958	2.09794660865948
4.03866006280555	0.0444691165185403	0.33237935703052
100271835	LIMS3-LOC440895 LIMS3-LOC440895 readthrough
9684	-0.621063320769548	4.5931294205527 4.03643205518156
	0.0445278689770388	0.332640992406071
6942	TCF20	transcription factor 20 7548	-0.256270804410101
	6.3478679713969 4.03384481835901	0.044596196850164
	0.33296700305905
10678	B3GNT2	UDP-GlcNAc:betaGal beta-1,3-N-
acetylglucosaminyltransferase 2 2861	-0.286261614843419
5.84355512430978	4.0329803296613 0.0446190522411535
0.33296700305905
8303	SNN	stannin 3312	0.295886897714576
6.81379525532523	4.02992436188488	0.0446999449517372
	0.333393039903611
9969	MED13	mediator complex subunit 13
11027	-0.445652843761658	7.58135994674054
4.02896813276794	0.0447252884299214	0.333404531685955
63928	CHP2	calcineurin like EF-hand protein 2
2382	-1.4419420140475	0.162047972676874
4.02599904958165	0.0448040761727847	0.33379047958021
84969	TOX2	TOX high mobility group box family member 2	3223
	0.975409976629861	0.419767131265029
4.0252218995405 0.0448247227879185	0.33379047958021
101929378	LINC01876	long intergenic non-protein coding
RNA 1876	2006	-1.1756719489862	-0.589539353244063
	4.02356675064861	0.0448687286735907
0.3338180045753
5908	RAP1B	RAP1B, member of RAS oncogene family

16942	-0.2971887407418	6.3134397862376 4.02287005409648
	0.0448872655327577	0.3338180045753
3909	LAMA3	laminin subunit alpha 3 13522	-0.555307854463609
	9.54775695286228	4.02239477538706
0.0448999158006615	0.3338180045753
115548	FCHO2	FCH and mu domain containing endocytic adaptor 2
	6127	-0.315842216683328	6.12633941343031
4.02007265081558	0.0449617766897204	0.333926689440507
25782	RAB3GAP2	RAB3 GTPase activating non-catalytic protein
subunit 2	10344	-0.263555703096904	6.39447325444411
	4.01910101100448	0.0449876875884216
0.333926689440507
220323	OAF	out at first homolog	2525	-0.577379473355575
	3.9415674363344 4.0187458840942 0.0449971617468875
0.333926689440507
5597	MAPK6	mitogen-activated protein kinase 6
6388	-0.294834064921397	6.92031745566668
4.018268743813	0.045009894314534	0.333926689440507
27345	KCNMB4	potassium calcium-activated channel subfamily M
regulatory beta subunit 4	4874	0.722998951001646
3.2017170130558 4.01718718246495	0.0450387700082892
0.333964029383857
3949	LDLR	low density lipoprotein receptor
7610	-0.468642652275667	8.65654601346791
4.01407183247744	0.0451220531877682	0.334374192150182
1105	CHD1	chromodomain helicase DNA binding protein 1
10437	-0.350704239580331	5.71835506853521
4.01299438973219	0.0451508943800056	0.334374192150182
6874	TAF4	TATA-box binding protein associated factor 4
7570	-0.364284471603789	4.13517531165446
4.01244153519883	0.0451657008315945	0.334374192150182
81618	ITM2C	integral membrane protein 2C	2611
0.569763752936811	5.98676556492969	4.01128719498891
	0.0451966326535096	0.334426430838727
4069	LYZ	lysozyme	1764	1.99201014355059
5.46910766804487	4.00900356941798	0.045257890671151
	0.334702890512572
48	ACO1	aconitase 1	7555	0.519010635322457
4.56247897649659	4.0017404494186 0.0454533051675131
0.335673320677978
55272	IMP3	IMP U3 small nucleolar ribonucleoprotein 3	1237
	0.319479320331757	5.01353192324727
4.00131084908849	0.0454648913834085	0.335673320677978
54456	MOV10L1 Mov10 like RISC complex RNA helicase 1	6872
1.07706516307509	1.51786201320415	4.00080717158629
	0.0454784794046481	0.335673320677978
345651	ACTBL2	actin beta like 2	2794	-0.723234479367301
	2.75490356749994	4.00011393874663
0.0454971881743463	0.335673320677978
29844	TFPT	TCF3 fusion partner	1205	0.317072809781741
	4.1823959359	3.99967887034974	0.0455089338164833
	0.335673320677978
11279	KLF8	Kruppel like factor 8	9022	0.395608090581014
	3.94264346354994	3.99798262731186

0.0455547581957038	0.335834472656433
79004	CUEDC2	CUE domain containing 2 1412	0.336550232129154
	4.96044591587994	3.99214715298676
0.0457127764469843	0.336822125035723
128077	LIX1L	limb and CNS expressed 1 like	3975
0.551065354350797	2.91703831411315	3.98197737337843
	0.0459895447297801	0.338593310790046
25932	CLIC4	chloride intracellular channel 4
4495	-0.325988822875047	8.87978985101047
3.98153937749046	0.0460015042788219	0.338593310790046
5881	RAC3	Rac family small GTPase 3	1527
0.525232468489568	3.0604535068265 3.98023912155942
0.046037027288768	0.33867680737068
219855	SLC37A2 solute carrier family 37 member 2	6371
0.370997171418611	4.42975094914437	3.96975790427837
	0.0463244322007521	0.340612242433561
80313	LRRC27	leucine rich repeat containing 27	9910
0.484343860780671	3.19980085842753	3.96814273732392
	0.0463688894986942	0.34076024932225
8853	ASAP2	ArfGAP with SH3 domain, ankyrin repeat and PH domain
2	7248	-0.334451170229583	7.17510929799777
3.95493454979466	0.0467341355218911	0.343264308471488
28981	IFT81	intraflagellar transport 81	7005
0.33552832500529	4.04467300578866	3.9534551590822
0.0467752335925406	0.343386109502472
4862	NPAS2	neuronal PAS domain protein 2	12353
0.252030626310976	6.21007841298676	3.95049861672749
	0.0468574817471268	0.343809715469882
83543	AIF1L	allograft inflammatory factor 1 like
5299	-0.584868131658163	2.90335357586819
3.94843405915736	0.0469150061566519	0.344051566092264
3106	HLA-B	major histocompatibility complex, class I, B
3012	-0.730998644138555	9.23153492627706
3.94556305059268	0.0469951243758023	0.344458768776485
3953	LEPR	leptin receptor 11370	-0.501249267331922
4.72761766756313	3.94377163951907	0.0470451884883435
	0.344522089197348
84803	GPAT3	glycerol-3-phosphate acyltransferase 3
3468	-0.820556292207767	0.575096517607518
3.94189445116519	0.0470977101470057	0.344522089197348
100616358	MIR4800 microRNA 4800	80	0.728908302150104
	0.0706223420575232	3.94116687277805
0.0471180836119616	0.344522089197348
10103	TSPAN1	tetraspanin 1	2366	-0.377206340177869
8.94945639584047	3.94040108191689	0.0471395371323483
	0.344522089197348
23517	MTREX	Mtr4 exosome RNA helicase
6578	-0.253204173570613	5.88809717016195
3.93982234713766	0.0471557571377658	0.344522089197348
160518	DENND5B DENN domain containing 5B	11310
0.859187705838958	2.47173940930275	3.9394653348044
0.0471657659378555	0.344522089197348
308	ANXA5	annexin A5	3034	0.38231206545776
8.26947769639924	3.93789576083265	0.0472097953492788

0.344522089197348
23325 WASHC4 WASH complex subunit 4 7698 -0.303582509704166 6.72645579638558 3.93705359018023 0.0472334376384885 0.344522089197348
80853 KDM7A lysine demethylase 7A 9430 -0.514666500443181 5.81309275860094 3.93552047287112 0.0472765089625472 0.344522089197348
554202 MIR31HG MIR31 host gene 745 0.593107891130106 2.68116675783754 3.9353795099674 0.0472804712465102 0.344522089197348
382 ARF6 ADP ribosylation factor 6 3865 -0.253958542758422 6.95161568842177 3.93497035682311 0.0472919739932824 0.344522089197348
8837 CFLAR CASP8 and FADD like apoptosis regulator 22529 -0.359404587132053 8.94788940364632 3.93472332500849 0.0472989203631399 0.344522089197348
55633 TBC1D22B TBC1 domain family member 22B 3462 -0.281785668799732 5.24894966166335 3.93247554314989 0.047362175956802 0.344803533589878
56928 SPPL2B signal peptide peptidase like 2B 6523 0.239224400758465 6.63630716206116 3.93141560100177 0.0473920351009369 0.344841680861727
84002 B3GNT5 UDP-GlcNAc:betaGal beta-1,3-N-acetylglucosaminyltransferase 5 5822 -0.447983444630312 6.22421041646923 3.9303224992211 0.0474228491677477 0.344853608005452
8609 KLF7 Kruppel like factor 7 9262 -0.508036498374343 6.69012072124894 3.92961110394504 0.047442914444671 0.344853608005452
29780 PARVB parvin beta 7403 0.857240653258303 2.25562255896763 3.92722491225329 0.0475102836555892 0.345069374306961
27075 TSPAN13 tetraspanin 13 2224 -0.368325670145529 6.38165602624029 3.92681491347953 0.0475218692823679 0.345069374306961
1073 CFL2 cofilin 2 7179 -0.393894918296925 4.99186510916951 3.92498516223534 0.0475736102042802 0.345266092296038
79690 GAL3ST4 galactose-3-O-sulfotransferase 4 3397 0.575945264148029 2.66128312647043 3.92194971397994 0.0476595764212049 0.345537945898154
341277 OVCH2 ovochymase 2 (gene/pseudogene) 2088 0.648647810692999 1.33308681030746 3.92192043821237 0.0476604063307798 0.345537945898154
23308 ICOSLG inducible T cell costimulator ligand 11467 -0.655706411379268 2.9468922435963 3.92075648844075 0.0476934143393147 0.345598372814683
4301 AFDN afadin, adherens junction formation factor 13720 -0.395625172554783 8.05696986201595 3.91962144790978 0.0477256257486051 0.345652967870068
55668 GPATCH2L G-patch domain containing 2 like 14327 -0.38635816872003 6.22133656961164 3.91776477004805 0.0477783660525306 0.345728948670766
79594 MUL1 mitochondrial E3 ubiquitin protein ligase 1 2438

0.31529173993397	6.23466795437119
3.91751442908198	0.0477854818752482	0.345728948670766
79172	CENPO	centromere protein O	4405	0.291166623454825
4.25056482695017	3.91344097485342
0.0479014251271745	0.346239493969819
25833	POU2F3	POU class 2 homeobox 3	3347	-1.15135887173797
2.50144838943883	3.91300945167504
0.0479137250026703	0.346239493969819
1028	CDKN1C	cyclin dependent kinase inhibitor 1C	2586
0.485426077537763	3.33269544828412	3.912431464785
0.0479302048124137	0.346239493969819
27324	TOX3	TOX high mobility group box family member 3
5031	-1.16097838449279	2.23438892285659
3.90944139163369	0.0480155546138355	0.346518260368248
158586	ZXDB	zinc finger X-linked duplicated B	5894
0.473558245653482	5.48378081207735	3.90801989265092
0.0480561866595466	0.346518260368248
10866	HCP5	HLA complex P5	3138	-0.996018458285312
4.39011737001533	3.90798560632493	0.0480571671459394
0.346518260368248
9663	LPIN2	lipin 2	6664	0.306346884064937
5.07569034936518	3.90739719470737	0.0480739972488528
0.346518260368248
81624	DIAPH3	diaphanous related formin 3	7259
0.398570442694132	4.0434222185151	3.90675092755717
0.0480924893378935	0.346518260368248
57563	KLHL8	kelch like family member 8
5855	-0.32208217598519	4.49921221701524
3.90501816598691	0.0481421070965942	0.34669742627352
400221	FLJ22447	uncharacterized LOC400221	2653
1.31387862648671	3.4723504656431	3.90390737466191
0.0481739431037531	0.34674841780795
6487	ST3GAL3	ST3 beta-galactoside alpha-2,3-sialyltransferase 3
4443	-0.879655407189359	3.59841906281713
3.90000759166141	0.0482858894844334	0.347375682592942
55540	IL17RB	interleukin 17 receptor B	3038
0.636582062127133	1.40892970515524	3.89817578043827
0.0483385678245108	0.347576139382917
56952	PRTFDC1	phosphoribosyl transferase domain containing 1	2329
0.614567940771154	2.52084575118565
3.89375902049247	0.0484658324733547	0.348312424553248
161	AP2A2	adaptor related protein complex 2 subunit alpha 2
9638	0.254620826173568	6.0061747470955
3.88938896399237	0.0485920995321017	0.349040788792897
387923	SERP2	stress associated endoplasmic reticulum protein family member 2	1927	0.712763163808923	0.744006341681071
3.88569040458777	0.0486992358795661
0.349386900320084
26297	SERGEF	secretion regulating guanine nucleotide exchange factor	3787	0.428023103791074	3.05504299546696
3.88391566250647	0.048750733430385	0.349386900320084
9651	PLCH2	phospholipase C eta 2	10781	0.363531062750184
7.28755474417955	3.88260118069077
0.0487889127041592	0.349386900320084

1960	EGR3	early growth response 3 4855	-1.03553537475683
	0.905600722077315	3.88213486630463
0.0488024644430773	0.349386900320084
397	ARHGDIB Rho GDP dissociation inhibitor beta	1499
1.08326701474412	4.34695882662101	3.88212168461487
	0.0488028475789064	0.349386900320084
5781	PTPN11	protein tyrosine phosphatase non-receptor type 11
	6617	-0.234825141607448	8.42937922789467
3.8818975325524 0.0488093632150928	0.349386900320084
388931	MFSD2B	major facilitator superfamily domain containing 2B
	3883	-0.55928157093054	2.02659860764182
3.88170742463324	0.0488148899783254	0.349386900320084
79148	MMP28	matrix metallopeptidase 28	2858
0.780278399303661	2.44765567223282	3.87723399470197
	0.0489451311398536	0.35014016949741
374875	HSD11B1L	hydroxysteroid 11-beta dehydrogenase 1 like
	3288	0.598942759922681	2.30300666397093
3.87398633670184	0.0490399145519819	0.350482798998196
80017	DGLUCY	D-glutamate cyclase	7212	0.495209744451064
	3.97105717446391	3.87387832137911
0.0490430703245852	0.350482798998196
9882	TBC1D4	TBC1 domain family member 4	6658
0.410899879145026	3.84592585720893	3.87248523356044
	0.0490837899616967	0.35051663102812
79820	CATSPERB	cation channel sperm associated auxiliary
subunit beta	5635	-0.735323189213299	0.692590374677731
	3.87200438909611	0.0490978532217585
0.35051663102812
57573	ZNF471	zinc finger protein 471 6566	-0.449093607115722
	3.18875738920501	3.86980777657821
0.0491621516955352	0.350796871522854
80209	PROSER1 proline and serine rich 1
7315	-0.270724415138627	5.64259572237769
3.86873337887179	0.0491936334673251	0.350842782065592
23592	LEMD3	LEM domain containing 3 5562	-0.285378005807675
	6.12311830600609	3.86779282688468
0.0492212107972754	0.350860814064853
2803	GOLGA4	golgin A4	10737	-0.265474157113986
8.0083802111201 3.86468760447251	0.0493123730482362
0.351181552993737
128486	FITM2	fat storage inducing transmembrane protein 2	4597
	0.321252577226672	4.14752786272392
3.86455227653568	0.0493163500206097	0.351181552993737
254102	EHBP1L1 EH domain binding protein 1 like 1
6006	-0.296400011323261	6.80970966993908
3.86348751453234	0.0493476527122518	0.351225900173024
84557	MAP1LC3A	microtubule associated protein 1 light chain
3 alpha 1762	0.311547019986919	4.04434234965409
3.86211088862312	0.0493881549262681	0.351322056690367
10254	STAM2	signal transducing adaptor molecule 2
7005	-0.250166803449886	5.99301899478341
3.86132384493552	0.049411326599559	0.351322056690367
9172	MYOM2	myomesin 2	7062	0.610361190372302
0.675444809267807	3.85955814112831	0.0494633531909979

0.351375332137117
643749 TRAF3IP2-AS1 TRAF3IP2 antisense RNA 1 4819
0.597517145641143 -0.0300906065631167 3.85936692498514
0.049468990859884 0.351375332137117
3659 IRF1 interferon regulatory factor 1
7799 -0.55605067584057 5.62943638163962
3.85604320183702 0.049567093449604 0.351893703978005
55806 HR HR lysine demethylase and nuclear receptor
corepressor 8662 -0.511879451365882 5.13618173736159
3.85405261974432 0.0496259455926644
0.352133039471353
23405 DICER1 dicer 1, ribonuclease III
12294 -0.375746541932021 6.67293474986396
3.85258025868147 0.0496695238677095 0.352263808007598
1102 RCBTB2 RCC1 and BTB domain containing protein 2 3545
0.461315055081379 3.48407304668382
3.85061619944784 0.0497277181423653 0.352415049621696
92999 ZBTB47 zinc finger and BTB domain containing 47 3781
0.360951848344242 4.6738572902038 3.84875851610479
0.0497828268290114 0.352415049621696
85446 ZFHX2 zinc finger homeobox 2 10128 -0.472988782273256
4.2362171197948 3.84848003947759 0.0497910934739313
0.352415049621696
4313 MMP2 matrix metallopeptidase 2 4933
0.459920541395731 7.83579775161841 3.84809207745805
0.0498026126364267 0.352415049621696
115817 DHRS1 dehydrogenase/reductase 1
4872 -0.293869153639402 5.24284745540595
3.84761947528023 0.0498166486935788 0.352415049621696
8548 BLZF1 basic leucine zipper nuclear factor 1
2320 -0.266406795750417 5.46715473936685
3.84646400082191 0.0498509833319671 0.352479921015074
285148 IAH1 isoamyl acetate hydrolyzing esterase 1 (putative)
5906 0.327263359915946 4.58815559867437
3.84305724732993 0.0499523597755556 0.353018518353283
34 ACADM acyl-CoA dehydrogenase medium chain 4396
0.332703666037863 4.45384283201804 3.83903098857786
0.0500724520565653 0.353688772544786
133022 TRAM1L1 translocation associated membrane protein 1 like 1
2023 0.556360620137983 1.8970193366896
3.83785336499684 0.0501076349814059 0.353758892734149
5912 RAP2B RAP2B, member of RAS oncogene family
4904 -0.293860241814014 6.42796160787944
3.83177248155729 0.0502897248977008 0.354766914530705
113763 ZBED6CL ZBED6 C-terminal like 2933 0.84106260834775
4.56839726948919 3.83139437144507
0.0503010703403998 0.354766914530705
100529211 TMEM256-PLSCR3 TMEM256-PLSCR3 readthrough (NMD
candidate) 2743 0.255373165501086 6.57612903316701
3.82689097300356 0.0504364059416876
0.355164492716518
653602 NA NA 2340 -0.464862074740312
4.63467664134728 3.8268377660678 0.050438007208636
0.355164492716518

115207	KCTD12	potassium channel tetramerization domain containing
12	6235	-0.615671014310808	6.57231646414945
3.82657616598986	0.0504458808652846	0.355164492716518
4953	ODC1	ornithine decarboxylase 1
2968	-0.708574130548687	4.22054431428127
3.82526238447264	0.0504854427990151	0.355164492716518
64421	DCLRE1C	DNA cross-link repair 1C
6326	-0.390913015495217	3.75514998621755
3.82450037989929	0.0505084040858296	0.355164492716518
84866	TMEM25	transmembrane protein 25	5529
0.314569353282263	5.7226531309801	3.82446140690302
0.0505095787457203	0.355164492716518
23215	PRRC2C	proline rich coiled-coil 2C
13921	-0.386842988734805	8.4020612610639	3.82221217320907
	0.0505774203911855	0.355463084505437
256356	GK5	glycerol kinase 5	12158	-0.297029466867369
	5.08809259026671	3.82136743751126
0.0506029242801151	0.355463972112122
84336	TMEM101	transmembrane protein 101	2654
0.306550897378593	4.65830301096576	3.81872023130493
	0.0506829356564011	0.355847558766521
116987	AGAP1	ArfGAP with GTPase domain, ankyrin repeat and PH
domain 1	13910	-0.330700587958272	5.17719482115371
	3.81700200779407	0.0507349402461463
0.355936438890199
64760	FAM160B2	family with sequence similarity 160 member
B2	8278	-0.26481069279585	6.66149374797676
3.8166230553566	0.0507464173958541	0.355936438890199
59339	PLEKHA2	pleckstrin homology domain containing A2
6298	-0.272100331869063	6.25149705576453
3.80821254119011	0.0510018503440308	0.357547450879469
23067	SETD1B	SET domain containing 1B, histone lysine
methyltransferase	8314	-0.337430326678043
5.46124292923239	3.80711334780967	0.0510353338887373
	0.357547450879469
7511	XPNPEP1	X-prolyl aminopeptidase 1
6691	-0.298693120810662	6.13149021546867
3.80654418131291	0.0510526809280316	0.357547450879469
93	ACVR2B	activin A receptor type 2B
15744	-0.489139115346303	3.30080789693426
3.80408476870328	0.0511277105386938	0.357893973770857
6253	RTN2	reticulon 2	3238	0.382149253632256
3.57712352073334	3.79991969879716	0.0512550409136113
	0.358606072965511
100287314	LINC00941	long intergenic non-protein coding
RNA 941	2218	-0.699477128834874	2.19175844340178
3.79702802401398	0.0513436395196996	0.359046609462023
7410	VAV2	vav guanine nucleotide exchange factor 2	5682
	0.257515044315578	6.8117425428303	3.7954032566002
0.0513934921523127	0.359215890507706
56904	SH3GLB2	SH3 domain containing GRB2 like, endophilin B2
6814	-0.330366290767396	6.68329659035424
3.79444272397331	0.0514229881882556	0.35924279079945
2203	FBP1	fructose-bisphosphatase 1	1668

1.26152613381922	3.35443790807622	3.79227186961044
	0.0514897167624319	0.359529642418437
9320	TRIP12	thyroid hormone receptor interactor 12
12594	-0.241162949173647	8.57952624963968
3.7903208046267 0.0515497675109897	0.35976960315218
442213	PTCHD4	patched domain containing 4	5177
0.670093234754395	2.5751348498494 3.78848739664487
0.0516062644741707	0.359984535104437
9728	SECISBP2L	SECIS binding protein 2 like
8699	-0.293573688931674	7.10448658725777
3.78582679745524	0.0516883680743288	0.360025048662188
29841	GRHL1	grainyhead like transcription factor 1
7952	-0.693599061143791	5.63071672541422
3.78582354137414	0.051688468638554	0.360025048662188
51068	NMD3	NMD3 ribosome export adaptor
4869	-0.278644503294606	5.98301352411398
3.78580044290634	0.0516891820418119	0.360025048662188
841	CASP8	caspase 8	7243	0.333447968140644
3.95324706192081	3.78152790690221	0.0518213199455436
	0.360477431287892
54933	RHBDL2	rhomboid like 2 2795	-0.567943685769283
4.89782051443409	3.78095465951284	0.0518390760972347
	0.360477431287892
26580	BSCL2	BSCL2 lipid droplet biogenesis associated, seipin
	3965	0.37729221155782	5.58535667734998
3.77900473912165	0.0518995224470785	0.360477431287892
9034	CCRL2	C-C motif chemokine receptor like 2	1959
0.902453729272789	0.61373084839963	3.77794121696221
	0.051932522411788	0.360477431287892
51755	CDK12	cyclin dependent kinase 12
9285	-0.263826653390271	6.77383577081268
3.77792757701826	0.0519329457889374	0.360477431287892
10057	ABCC5	ATP binding cassette subfamily C member 5
11441	0.31327857168566	6.74244683382688
3.77641191358158	0.0519800139919415	0.360477431287892
7326	UBE2G1	ubiquitin conjugating enzyme E2 G1
4539	-0.234588625481158	6.25515833117351
3.77640456989444	0.0519802421564731	0.360477431287892
4363	ABCC1	ATP binding cassette subfamily C member 1	7575
	0.234874517499971	7.4103399308792 3.77612945415335
	0.0519887906226183	0.360477431287892
100506311	HOTAIRM1	HOXA transcript antisense RNA,
myeloid-specific 1	1199	-0.812935634909191
0.21685337631445	3.77535537046369	0.0520128511288563
	0.360477431287892
200933	FBXO45	F-box protein 45	5976	-0.282799983560692
	4.9608020285363 3.77459577619739	0.0520364727198211
	0.360477431287892
5274	SERPINI1	serpin family I member 1	2418
0.531259909422084	2.15477893824801	3.77457169344559
	0.0520372218222402	0.360477431287892
9823	ARMCX2	armadillo repeat containing X-linked 2	3589
0.442341534426451	4.37484569174191	3.77182699821255
	0.0521226713270827	0.360890883479213

92689	FAM114A1	family with sequence similarity 114 member A1	4871	-0.310509692917083	6.82887677972062	3.7681965940012	0.0522359233941779	0.361496333489254
55190	NUDT11	nudix hydrolase 11	2385	0.557875998290145	2.06842709769176	3.76121482147231	0.0524544554601828	0.362367050255895
85477	SCIN	scinderin	11623	-1.26786094465084	1.96644730655697	3.76076308221824	0.052468628352742	0.362367050255895
51762	RAB8B	RAB8B, member RAS oncogene family	5200	-0.454228800884633	5.77068525275499	3.75937536055208	0.0525121921644822	0.362367050255895
2621	GAS6	growth arrest specific 6	4059	0.490602827473736	9.56288383754371	3.75903209405169	0.0525229740034287	0.362367050255895
23433	RHOQ	ras homolog family member Q	4619	0.321250523230804	6.27509775212276	3.75793897992762	0.0525573238162532	0.362367050255895
2239	GPC4	glypican 4	4960	0.494696897889971	5.24902305838064	3.75742374990496	0.0525735225517198	0.362367050255895
5205	ATP8B1	ATPase phospholipid transporting 8B1	6180	-0.355270422758213	6.87829841571137	3.75674173983526	0.0525949729482241	0.362367050255895
23413	NCS1	neuronal calcium sensor 1	5289	0.300871395975818	7.46726341220174	3.75566857081272	0.0526287447292652	0.362367050255895
3728	JUP	junction plakoglobin	4942	-0.300362660006478	10.3248151083698	3.75565712471769	0.0526291050525179	0.362367050255895
951	CD37	CD37 molecule	3854	1.28139425307773	0.332151850515962	3.75488356992929	0.0526534626220191	0.362367050255895
100	ADA	adenosine deaminase	2231	-0.656592413816271	2.75255168314143	3.75339322868026	0.0527004238959955	0.362367050255895
891	CCNB1	cyclin B1	2558	0.474172411518169	3.73229496461107	3.75236454155185	0.0527328641174234	0.362367050255895
6905	TBCE	tubulin folding cofactor E	2568	0.407101110916389	3.9013574631558	3.75181739352455	0.0527501273461614	0.362367050255895
25816	TNFAIP8	TNF alpha induced protein 8	9591	-0.31542850049071	6.24221653611385	3.75181543193405	0.052750189247643	0.362367050255895
2146	EZH2	enhancer of zeste 2 polycomb repressive complex 2 subunit	4522	-0.358236237315152	3.6995016397926	3.75136899335411	0.0527642794124991	0.362367050255895
65981	CAPRIN2	caprin family member 2	8360	-0.398920281938072	5.07851899512246	3.75100824296426	0.0527756680604002	0.362367050255895
124739	USP43	ubiquitin specific peptidase 43	5068	-0.362263025417841	4.7575916947815	3.74590329876835	0.05293710698165	0.36329743140224

266727	MDGA1	MAM domain containing glycosylphosphatidylinositol anchor 1	14187	0.837006749786872	1.6811405146402	3.74491462069217	0.0529684333559689	0.363334400596012
7994	KAT6A	lysine acetyltransferase 6A	11852	-0.363366912232948	6.43410568534721	3.73790351732274	0.0531911448190922	0.364683487753805
23466	CBX6	chromobox 6	8030	-0.385922059242052	6.5557695976258	3.73489090124374	0.0532871465498524	0.365150355158467
5783	PTPN13	protein tyrosine phosphatase non-receptor type 13	9496	-0.388509328965487	7.06596512019102	3.73356076931142	0.0533295916607493	0.365150355158467
6038	RNASE4	ribonuclease A family member 4	2717	0.403823941303215	3.32804080567627	3.73331470943862	0.0533374474658545	0.365150355158467
64754	SMYD3	SET and MYND domain containing 3	5866	0.519389884916952	3.27904332863138	3.72952778080065	0.053458505141914	0.365760155558364
1476	CSTB	cystatin B	2490	-0.663554267538542	7.81937770683432	3.72889548465812	0.0534787462399893	0.365760155558364
57669	EPB41L5	erythrocyte membrane protein band 4.1 like 5	8932	-0.306774284745656	5.86354394576	3.71905929104418	0.0537946704944023	0.367612861051605
4907	NT5E	5'-nucleotidase ecto	4533	-0.569714727322722	7.16321214730908	3.71882793647096	0.0538021250200464	0.367612861051605
440515	ZNF506	zinc finger protein 506	6021	-0.411874131846783	3.61838670375388	3.71333925927733	0.0539792981902727	0.368100764327648
11041	B4GAT1	beta-1,4-glucuronyltransferase 1	2072	0.533440241567614	2.87582870191098	3.71279074757015	0.0539970379556834	0.368100764327648
284837	AATBC	apoptosis associated transcript in bladder cancer	4874	0.440830572388412	3.96344751200887	3.71157842349917	0.0540362684141547	0.368100764327648
5813	PURA	purine rich element binding protein A	11497	-0.413689351882016	5.0116989060158	3.71147464691461	0.0540396279974059	0.368100764327648
26049	FAM169A	family with sequence similarity 169 member A	6860	0.403493958460107	3.5665633885985	3.71085880054828	0.0540595694873878	0.368100764327648
84264	HAGHL	hydroxyacylglutathione hydrolase like	3107	0.835319903167833	1.63741100885402	3.71047759878299	0.0540719169450949	0.368100764327648
1968	EIF2S3	eukaryotic translation initiation factor 2 subunit gamma	3472	-0.334822030668318	6.71580940495139	3.71035293431652	0.054075955584251	0.368100764327648
7456	WIPF1	WAS/WASL interacting protein family member 1	9035	0.625812540555107	3.96462058020503	3.70991184057525	0.0540902478554387	0.368100764327648
51290	ERGIC2	ERGIC and golgi 2	5826	-0.256471361617374	5.69656475169094	3.70930089547735	0.0541100502427806	0.368100764327648

23526 ARHGAP45 Rho GTPase activating protein 45 5928
0.726543363027827 3.80288361532856
3.70810510802679 0.05414883119034 0.368185766253928
92521 SPECC1 sperm antigen with calponin homology and coiled-coil domains 1 13676 0.229840254236292 6.63638863056998
3.7059972903138 0.0542172621954351 0.368413105169387
374969 SVBP small vasohibin binding protein 1208
0.40670075633076 3.84689135028253 3.70509823086137
0.0542464783331951 0.368413105169387
51147 ING4 inhibitor of growth family member 4 2736
0.307079243766148 4.42744664331899 3.70464626147498
0.0542611719828975 0.368413105169387
54504 CPVL carboxypeptidase vitellogenic like 3340
1.54479092226616 0.963419044581516 3.70171836611494
0.0543564608835384 0.368865757079153
22887 FOXJ3 forkhead box J3 6893 -0.218068821867811
5.87841795146571 3.70077553121021 0.0543871833536867
0.368865757079153
6916 TBXAS1 thromboxane A synthase 1 6271
1.21781175762999 1.5380376376981 3.69912021297267
0.0544411667634841 0.368865757079153
284119 CAVIN1 caveolae associated protein 1 3828
0.30481605144266 9.29249349271348 3.69905315942549
0.0544433547170493 0.368865757079153
309 ANXA6 annexin A6 5427 1.15194254498087
3.96720303314872 3.69855807817653 0.0544595120753686
0.368865757079153
153339 TMEM167A transmembrane protein 167A
4908 -0.252311016046039 6.92069463252914
3.69757436335938 0.0544916314490399 0.368904921076221
402778 IFITM10 interferon induced transmembrane protein 10 4784
0.713485263798109 2.41270427916689
3.69639519709872 0.0545301589859317 0.368969045427827
51596 CUTA cutA divalent cation tolerance homolog 1876
0.336467099188729 5.73647673229474 3.69480128572732
0.0545822836077925 0.368969045427827
100128927 ZBTB42 zinc finger and BTB domain containing 42
3800 -0.379759626479796 4.04277887931418
3.69423842696059 0.0546007030174594 0.368969045427827
1513 CTSK cathepsin K 1887 0.86163428769526
2.51610324813079 3.6940622285339 0.0546064704199247
0.368969045427827
57530 CGN cingulin 6616 -0.38736241810136
7.61274245757435 3.69322580077001 0.0546338575412842
0.36897610539092
29767 TMOD2 tropomodulin 2 11636 0.355123529586107
5.08870383640115 3.69026502090263 0.0547309192768266
0.369453487378559
55252 ASXL2 ASXL transcriptional regulator 2
13023 -0.395311995757534 6.31188783358028
3.68775167326532 0.0548134565041736 0.369832411008651
57045 TWSG1 twisted gastrulation BMP signaling modulator 1 4435
0.318814600041727 6.01603712561287
3.68694087240139 0.054840110985905 0.369834104275194

143662	MUC15	mucin 15, cell surface associated
3356	-0.911570467896445	1.51611577819542
3.68369069076139	0.0549470963269169	0.370177102709992
91039	DPP9	dipeptidyl peptidase 9	8324	0.248554623481181
	7.06704264885089	3.6832152049581
0.054962766305905	0.370177102709992
27	ABL2	ABL proto-oncogene 2, non-receptor tyrosine kinase
	13257	-0.458090503701365	7.81518332631284
3.68298799503657	0.0549702558461329	0.370177102709992
23363	OBSL1	obscurin like cytoskeletal adaptor 1	8939
0.431626010740082	5.60193389484686	3.68178556740035
	0.055009909589871	0.37026612379881
54977	SLC25A38	solute carrier family 25 member 38	2151
	0.326191276468787	5.18660299766856
3.67855406933069	0.0551166285041156	0.370708219156754
10020	GNE	glucosamine (UDP-N-acetyl)-2-epimerase/N-acetylmannosamine kinase	5477	-0.922476710880016
6.45908594720413	3.67814655994891	0.0551301019137161
	0.370708219156754
92747	BPIFB1	BPI fold containing family B member 1
3101	-1.95926684816242	4.01172040411163
3.6773941210708	0.0551549888429124	0.370708219156754
9990	SLC12A6	solute carrier family 12 member 6
10056	-0.302446327479998	6.84750884812484
3.67392572531934	0.0552698601955367	0.371302125543829
51495	HACD3	3-hydroxyacyl-CoA dehydratase 3	4472
0.234502325154911	6.78350312137892	3.66848371766274
	0.0554506080593428	0.372337807807869
7799	PRDM2	PR/SET domain 2	10923	-0.281334004044911
6.23102364797808	3.66616020968935	0.0555279706332027
	0.37267862226127
5159	PDGFRB	platelet derived growth factor receptor beta	7137
	2.71258396779495	3.25437242906702
3.66339715958468	0.0556201170455321	0.373118285180445
123016	TTC8	tetratricopeptide repeat domain 8	6257
0.364216372048604	4.06699407070251	3.65911334504643
	0.0557633010132578	0.373899740207133
10810	WASF3	WASP family member 3	5162	0.441370714197344
	2.91708118897022	3.65770647517959
0.0558104100483348	0.374036561505754
27344	PCSK1N	proprotein convertase subtilisin/kexin type 1 inhibitor	1363	1.173582200652	1.57374355745004
3.65648592195921	0.0558513144700339	0.374131689039582
84918	LRP11	LDL receptor related protein 11
4381	-0.392617287877748	6.21499748237139
3.65477627650615	0.0559086633256439	0.374230308054833
7389	UROD	uroporphyrinogen decarboxylase	2908
0.259441075248798	5.63681861538023	3.65392178415481
	0.0559373500809848	0.374230308054833
161424	NOP9	NOP9 nucleolar protein	6033	-0.245999908618088
	6.21682354593899	3.65261045353446
0.055981404042617	0.374230308054833
10095	ARPC1B	actin related protein 2/3 complex subunit 1B	3168
	0.279936911445541	7.78930276090646

3.65221400333638	0.0559947299740389	0.374230308054833
6590	SLPI	secretory leukocyte peptidase inhibitor
596	-1.0364191592093	7.09934723738597
3.65206844613924	0.0559996234513408	0.374230308054833
4241	MELTF	melanotransferrin	8393	0.466370004255757
	6.90702952143344	3.6508019667433	0.0560422202859353
	0.374336375557986
3122	HLA-DRA	major histocompatibility complex, class II, DR alpha
	1280	1.728480168702	5.50691370674979
3.64966413797714	0.0560805193933581	0.374398979792414
9476	NAPSA	napsin A aspartic peptidase	3581
1.11611327577144	0.993305174029908	3.64890325445014
	0.0561061460855649	0.374398979792414
10564	ARFGEF2	ADP ribosylation factor guanine nucleotide exchange factor 2	8945	-0.349022241831866	6.85149875280802
	3.64814249327449	0.056131781078323
0.374398979792414
57572	DOCK6	dedicator of cytokinesis 6
10161	-0.260539658648347	5.8438099266207	3.6460594290841
0.056202036678727	0.374689161236996
10276	NET1	neuroepithelial cell transforming 1
4175	-0.387145132507715	8.13813190601728
3.64514908729116	0.0562327691076609	0.374715697854903
149483	CCDC17	coiled-coil domain containing 17
3323	-0.651299561525869	0.831084216204195
3.64264004159123	0.0563175648674519	0.37477131141099
55016	MARCHF1	membrane associated ring-CH-type finger 1
10421	1.43249103094704	0.289839051619867
3.64260150795821	0.0563188682082548	0.37477131141099
6385	SDC4	syndecan 4	2613	-0.440933572386787
8.78775325323189	3.64204963282384	0.0563375380469314
	0.37477131141099
56901	NDUFA4L2	NDUFA4 mitochondrial complex associated like 2	2806	0.98707507761001	1.10585822118056
3.6408368442101	0.0563785895500748	0.37477131141099
131	ADH7	alcohol dehydrogenase 7 (class IV), mu or sigma polypeptide	3108	0.945712274645123	1.67392953898211
	3.64057039653925	0.0563876127501128
0.37477131141099
11190	CEP250	centrosomal protein 250	16295	0.413361163921753
	5.84613701597626	3.64015595083583
0.0564016509212798	0.37477131141099
58985	IL22RA1	interleukin 22 receptor subunit alpha 1
2798	-0.609489975747911	2.06458704087761
3.63777657074611	0.0564823174626211	0.375129360217607
79930	DOK3	docking protein 3	4821	0.586826344372955
	3.37591156078543	3.6359802847676	0.0565432968509089
	0.375133876298678
10691	GMEB1	glucocorticoid modulatory element binding protein 1
	6459	-0.30341318101919	4.53938499805369
3.63515589332794	0.056571306252058	0.375133876298678
114796	PSMG3-AS1	PSMG3 antisense RNA 1 (head to head)	6355
	0.40814324150472	3.78681373930968
3.6346720823294	0.0565877509988879	0.375133876298678

5874	RAB27B	RAB27B, member RAS oncogene family
7432	-0.518923946872706	6.60606858082794
3.63271006706836	0.0566544919644547	0.375133876298678
329	BIRC2	baculoviral IAP repeat containing 2
4532	-0.306261222547626	6.76456369935379
3.63202860433515	0.0566776925101848	0.375133876298678
376267	RAB15	RAB15, member RAS oncogene family	4379
0.279923130582938	5.83653634775264	3.631879870486
0.0566827575266042	0.375133876298678
7703	PCGF2	polycomb group ring finger 2	3511
0.350303284086768	5.25037832024236	3.63170449384933
	0.0566887304597851	0.375133876298678
147040	KCTD11	potassium channel tetramerization domain containing
11	3056	-0.307402757780419	6.74234547434957
3.629721747139	0.0567563048338121	0.375133876298678
7188	TRAF5	TNF receptor associated factor 5	9022
0.496856456178185	4.36321506529972	3.62802583496854
	0.0568141713827126	0.375133876298678
2313	FLI1	Fli-1 proto-oncogene, ETS transcription factor	8172
	1.30826589990642	0.289852199201844
3.62802231139407	0.0568142916764509	0.375133876298678
1820	ARID3A	AT-rich interaction domain 3A
7210	-0.427562658326222	3.74211023889231
3.62697612479706	0.0568500201215373	0.375133876298678
154881	KCTD7	potassium channel tetramerization domain containing
7	4873	0.419517486299183	3.89657531122283
3.62694279270181	0.0568511588419083	0.375133876298678
199777	ZNF626	zinc finger protein 626	6872	-0.429969339271006
	3.3241235824254	3.62616177046566	0.0568778477419985
	0.375133876298678
51633	OTUD6B	OTU deubiquitinase 6B	3977	-0.403332558808993
	3.9389398535515	3.6259990009343	0.0568834115369238
0.375133876298678
245812	CNPY4	canopy FGF signaling regulator 4	2165
0.512684798340205	2.88561725850543	3.62580338754122
	0.0568900987664455	0.375133876298678
9759	HDAC4	histone deacetylase 4	13368	-0.453671061787704
	3.65498965800949	3.6251773046994	0.0569115076129571
	0.375133876298678
137492	VPS37A	VPS37A subunit of ESCRT-I
6050	-0.231636947291748	5.85707654068369
3.6226840777416	0.0569968481629393	0.375519685897597
1263	PLK3	polo like kinase 3	3479	-0.284949380625487
	5.10665980008971	3.62182012878605
0.0570264519157274	0.375538087439395
23186	RCOR1	REST corepressor 1	6202	-0.310895614484344
	6.65557206943532	3.61891065399141
0.057126266999341	0.376018619295005
113220	KIF12	kinesin family member 12	2975
0.685442573470227	1.94343629321865	3.61060356204705
	0.0574122798501747	0.377723721870078
10082	GPC6	glypican 6	6468	1.35426076908234
3.52879848393355	3.60816711143032	0.0574964548326804
	0.378086882335236

25814	ATXN10	ataxin 10	5098	0.266121497955986
6.69582550697129	3.60744394917712	0.0575214640005989
	0.378086882335236
5031	P2RY6	pyrimidinergic receptor P2Y6
3976	-0.770823839534645	2.73372026124172
3.60513024419443	0.0576015566479324	0.378435742949151
53917	RAB24	RAB24, member RAS oncogene family
3516	-0.46403221132596	4.46990955714151
3.60324183287969	0.0576670148442686	0.378688174835288
440895	LOC440895	two pore channel 3 pseudogene
2431	-0.764625725527633	2.5175275429499 3.60210236128583
	0.0577065506783475	0.378750210989757
26039	SS18L1	SS18L1 subunit of BAF chromatin remodeling complex
	5094	-0.342271479666161	4.94801665777983
3.60141130863455	0.0577305419049855	0.378750210989757
151393	RMDN2	regulator of microtubule dynamics 2	4969
0.470711242441769	2.69409747596173	3.59871412563081
	0.0578242812294206	0.379187596994614
3485	IGFBP2	insulin like growth factor binding protein 2	4767
	0.865403059406051	6.88178612938184
3.59791287962359	0.0578521592927889	0.379192884985538
23035	PHLPP2	PH domain and leucine rich repeat protein
phosphatase 2	12493	-0.402085216128219	4.29261397643502
	3.59517163916744	0.0579476443575208
0.379641091915713
653269	POTEI	POTE ankyrin domain family member I
7023	-0.414132728134616	5.91215000372552
3.5938229236918 0.0579946853173233	0.379771649013439
57655	GRAMD1A GRAM domain containing 1A	5942
0.257746307758628	6.30422858399893	3.59044254904609
	0.0581127655208289	0.380225858929655
93589	CACNA2D4	calcium voltage-gated channel auxiliary
subunit alpha2delta 4	7470
0.744983776080886	-0.0731009825169372	3.59028319430284
	0.0581183382571851	0.380225858929655
166614	DCLK2	doublecortin like kinase 2	4606
1.15853297999839	2.75453570019011	3.58266337017749
	0.0583854720963425	0.381795194982946
9525	VPS4B	vacuolar protein sorting 4 homolog B
4315	-0.236225445851575	7.55864282371774
3.58087573702366	0.0584483312161973	0.381924303092402
152137	CCDC50	coiled-coil domain containing 50
9202	-0.326088342455691	7.13213152014353
3.58055124584115	0.0584597491133084	0.381924303092402
81571	MIR600HG	MIR600 host gene
5984	-0.573017500408102	1.7936176031576 3.57924928326061
	0.0585055852127495	0.382016872929452
170954	PPP1R18 protein phosphatase 1 regulatory subunit 18
4599	-0.337861724669485	7.42726931834212
3.57859979419737	0.0585284650036842	0.382016872929452
388722	C1orf53 chromosome 1 open reading frame 53	611
0.692476561163076	0.191541311887663	3.57745845995212
	0.0585686942444062	0.382101397429621
1184	CLCN5	chloride voltage-gated channel 5

10619 -0.409130465465561 4.70241600339984
3.57578060518288 0.0586278878807331 0.382298100565135
142 PARP1 poly(ADP-ribose) polymerase 1 7169
0.250567944522492 5.85067130248941 3.57505711620027
0.0586534317208877 0.382298100565135
9648 GCC2 GRIP and coiled-coil domain containing 2
13282 -0.335081783815937 6.01967552897195
3.57360760539405 0.0587046444197474 0.38245393227321
5292 PIM1 Pim-1 proto-oncogene, serine/threonine kinase
2977 -0.411975822979016 7.2242606113458 3.57261435523088
0.0587397644009796 0.382459925878804
2553 GABPB1 GA binding protein transcription factor subunit beta
1 5677 -0.344056370468095 4.75548944740633
3.57203742191019 0.0587601742336821 0.382459925878804
9321 TRIP11 thyroid hormone receptor interactor 11
10249 -0.319513644192071 6.26763793312215
3.5708598385651 0.058801856306847 0.382553460887137
101927811 LOC101927811 uncharacterized LOC101927811
896 -0.554769764277625 1.58999169084466
3.56994522829877 0.0588342517924513 0.382586520360662
54460 MRPS21 mitochondrial ribosomal protein S21 1641
0.327684481303675 5.37799098965144 3.56840725202176
0.0588887696715258 0.382763339577291
9742 IFT140 intraflagellar transport 140 8168
0.331904684519125 4.03743174392769 3.5660698939764
0.0589717267144614 0.382984872020289
1770 DNAH9 dynein axonemal heavy chain 9
16980 -1.33280355761197 -0.566005569358384
3.56590630067133 0.0589775375846193 0.382984872020289
54793 KCTD9 potassium channel tetramerization domain containing
9 4045 -0.275795133291093 5.98635281535302
3.5625705833716 0.0590961558243284 0.383186176495876
10640 EXOC5 exocyst complex component 5
12575 -0.242898213339568 6.67000195059562
3.56247722848378 0.0590994791736788 0.383186176495876
90427 BMF Bcl2 modifying factor 5447 0.614895270424757
5.06617519303252 3.5621746630362 0.05911025159473
0.383186176495876
51097 SCCPDH saccharopine dehydrogenase (putative) 2503
0.494414087251441 5.49328303732234 3.56195806903935
0.0591179644040543 0.383186176495876
51209 RAB9B RAB9B, member RAS oncogene family 3748
0.859636096040494 0.893148456443722 3.55826092490097
0.0592497830745093 0.383687075927607
6775 STAT4 signal transducer and activator of transcription 4
4250 0.723135309010558 2.21330786818519
3.55825405749862 0.0592500282167069 0.383687075927607
54438 GFOD1 glucose-fructose oxidoreductase domain containing 1
10686 -0.516526942402432 5.3268215892589 3
3.555948983261 0.0593323722461291 0.383866395718196
1436 CSF1R colony stimulating factor 1 receptor 5151
1.23964080085619 3.13151898783865 3.55584913468322
0.0593359418791246 0.383866395718196
57048 PLSCR3 phospholipid scramblase 3 2648

0.246703192021033	6.56447845462464	3.55514775625377
0.0593610229197061	0.383866395718196
55165	CEP55	centrosomal protein 55	3271	0.769022792096236
0.176728303533914	3.55441213503981
0.059387340581233	0.383866395718196
23530	NNT	nicotinamide nucleotide transhydrogenase	8245
0.277729826714	6.09809367519816	3.55310838527937
0.0594340141514206	0.383876772002754
55161	TMEM33	transmembrane protein 33
8240	-0.273705656423047	6.70499452495882
3.55281185922544	0.0594446350746684	0.383876772002754
51385	ZNF589	zinc finger protein 589 5521	0.468477618796232
3.09929798751479	3.55207144074514
0.0594711640783876	0.383876772002754
127534	GJB4	gap junction protein beta 4
2840	-0.731303070920347	5.09701476259108
3.55056564639571	0.0595251551452715	0.384048294850216
92579	G6PC3	glucose-6-phosphatase catalytic subunit 3	2048
0.369511878929169	4.6637378876496 3.54849845714572
0.0595993602351971	0.384191835736565
5947	RBP1	retinol binding protein 1
3905	-0.695060066010072	6.56552680579687
3.54788369037806	0.0596214472514244	0.384191835736565
4794	NFKBIE	NFKB inhibitor epsilon	2581	-0.420195779417287
4.58763026098452	3.54765435862537
0.059629688790697	0.384191835736565
2593	GAMT	guanidinoacetate N-methyltransferase	1743
0.745888345040544	2.48521599453739	3.54273821686366
0.0598066527902417	0.385012767098795
8915	BCL10	BCL10 immune signaling adaptor
2819	-0.355457233897593	5.85840617341874
3.54194226311185	0.0598353568522282	0.385012767098795
65083	NOL6	nucleolar protein 6	5906	0.248526036833115
6.0148697083193 3.54182559286564	0.059839565501112
0.385012767098795
23253	ANKRD12 ankyrin repeat domain 12
12314	-0.370560801161073	6.0478934060238 3.54035405096785
0.0598926755081581	0.385177550891998
135250	RAET1E	retinoic acid early transcript 1E	2637
1.01654022597495	1.44579487936492	3.53931677783233
0.0599301422714208	0.385200445223827
126299	ZNF428	zinc finger protein 428 1803	0.331392402255394
3.84772223662627	3.53873308244683
0.0599512365665698	0.385200445223827
284371	ZNF841	zinc finger protein 841 4277	-0.56956857317466
3.7138701007633 3.53418929776215	0.0601157156735819
0.385813252920832
123920	CMTM3	CKLF like MARVEL transmembrane domain containing 3
3163	0.31325087969894	6.55270678595109
3.53325041409369	0.0601497618450715	0.385813252920832
2009	EML1	EMAP like 1	8899	-0.696514536616154
1.96200345191684	3.53251182307952	0.0601765593395784
0.385813252920832
10735	STAG2	stromal antigen 2	8187	-0.248692366771932

6.77448781390126	3.53233461170177
0.0601829907962376	0.385813252920832
157378	TMEM65	transmembrane protein 65
9029	-0.354575823869413	5.08956791565109
3.53229762093746	0.0601843333784549	0.385813252920832
158219	TTC39B	tetratricopeptide repeat domain 39B
12208	-0.58448795151023	4.2539592084208 3.53058380887892
0.0602465712477669	0.385890203837904
55783	CMTR2	cap methyltransferase 2 7582	-0.308605998831909
5.45541062871359	3.53044990701336
0.0602514368382591	0.385890203837904
54884	RETSAT	retinol saturase	4005	0.282800213371878
7.0722837838062 3.52788938092313	0.0603445591194629
0.386309981529395
22822	PHLDA1	pleckstrin homology like domain family A member 1
8069	-0.431113486221758	7.51563587097948
3.52433193618788	0.0604741919971383	0.386963000138838
1002	CDH4	cadherin 4	7351	0.670910424077802
3.57586108232786	3.52237184294948	0.0605457441451866
0.387199880638926
30817	ADGRE2	adhesion G protein-coupled receptor E2
8622	-0.774932514506719	2.36561222440087
3.5218037728509 0.0605664980709564	0.387199880638926
253943	YTHDF3	YTH N6-methyladenosine RNA binding protein 3
6163	-0.240528901087805	7.10115103931839
3.51844184165774	0.0606894782486355	0.387809088425473
9764	KIAA0513	KIAA0513	8837	-0.283845463270852
5.11148017768826	3.51506690617502
0.0608132014379454	0.388422486338943
386593	CHKB-CPT1B	CHKB-CPT1B readthrough (NMD candidate)	5022
0.328027324101435	5.77118329707736
3.51424314354838	0.0608434408374284	0.388438503103856
2125	EVPL	envoplakin	7307	-0.452946950823505
7.43349470326095	3.51091822432852	0.0609656576340408
0.388846354501803
80380	PDCD1LG2	programmed cell death 1 ligand 2	2365
-0.687219114041744	4.10391392769894
3.51082858445529	0.0609689562157125	0.388846354501803
85458	DIXDC1	DIX domain containing 1 8509	0.450644848102017
4.56996522311982	3.50967229506897
0.0610115225613103	0.388846354501803
389161	ANKUB1	ankyrin repeat and ubiquitin domain containing 1
5844	0.704520005132745	3.17573385064038
3.50910454485872	0.0610324346609737	0.388846354501803
1982	EIF4G2	eukaryotic translation initiation factor 4 gamma 2
8655	-0.261569655329853	10.7595624716972
3.50871274894138	0.0610468702367566	0.388846354501803
3055	HCK	HCK proto-oncogene, Src family tyrosine kinase	3092
1.64206424948139	1.86901992236641
3.50782727624497	0.0610795085410488	0.388846354501803
85415	RHPN2	rhophilin Rho GTPase binding protein 2
4804	-0.387322511735109	5.18992051919539
3.50722683972404	0.0611016510500798	0.388846354501803
5610	EIF2AK2 eukaryotic translation initiation factor 2 alpha

kinase 2 10753 -0.246220774489451 6.35252466359518
3.50620331026224 0.0611394158057324
0.388909989641641
1719 DHFR dihydrofolate reductase 4409 0.44171812762739
2.97752672109483 3.50007840944356
0.0613659236578207 0.389642482597285
710 SERPING1 serpin family G member 1 3603
0.943085636961967 5.25360112770455 3.49986514080143
0.0613738267233798 0.389642482597285
64900 LPIN3 lipin 3 4568 0.282855548089657
6.05931503045747 3.4995538815292 0.0613853629585556
0.389642482597285
254251 LCORL ligand dependent nuclear receptor corepressor like
10032 -0.509372948736247 3.24789296076707
3.49943802410418 0.0613896575840621 0.389642482597285
57720 GPR107 G protein-coupled receptor 107
8888 -0.217558826879862 7.9557725061481 3.49933012781863
0.0613936573921759 0.389642482597285
3111 HLA-DOA major histocompatibility complex, class II, DO alpha
4012 1.46955814371162 0.928779444200838
3.497151651398 0.0614744747699649 0.389978699321965
200424 TET3 tet methylcytosine dioxygenase 3
12271 -0.490429670332053 6.04654226046273
3.49379178266484 0.0615993417285102 0.390519333056117
254225 RNF169 ring finger protein 169 7823 -0.303642958803655
5.69565873294245 3.49335864991132
0.0616154584175069 0.390519333056117
100008587 RNA5-8SN5 RNA, 5.8S ribosomal N5 1224
0.723189344283361 3.85777195281587 3.49195511369424
0.0616677142702156 0.390673755668435
55000 TUG1 taurine up-regulated 1 6793 -0.274602780459835
7.94806256277651 3.4905942475952 0.0617184265112187
0.39077588509934
57701 NCKAP5L NCK associated protein 5 like 8090
0.414084220782485 5.93351466765297 3.49002553412797
0.0617396325926207 0.39077588509934
83548 COG3 component of oligomeric golgi complex 3
5704 -0.309919582089653 5.44713790700651
3.48632231671266 0.061877907373634 0.391474186351622
130507 UBR3 ubiquitin protein ligase E3 component n-recognin 3
10063 -0.266447866970229 5.85483962766365
3.48523346969965 0.0619186266481738 0.391554945129106
79739 TTLL7 tubulin tyrosine ligase like 7 10543
0.54496162768739 2.98727051862251 3.482864678341
0.0620073102138965 0.39193880603161
84695 LOXL3 lysyl oxidase like 3 4363 0.784197078745345
0.353081291830107 3.48009849533226
0.0621110427189651 0.392417399803584
140775 SMCR8 SMCR8-C9orf72 complex subunit
8279 -0.338209656540106 6.60760752682635
3.47749953513632 0.0622086727601401 0.392857024053779
414919 C8orf82 chromosome 8 open reading frame 82 3001
0.390622557440757 3.41792682811746 3.4753722077247
0.0622887075612971 0.392894460690245

55092	TMEM51	transmembrane protein 51
2240	-0.343440662250531	6.03515762325437
3.47439345436278	0.0623255672818639	0.392894460690245
79084	WDR77	WD repeat domain 77	2805	0.344513174239089
	5.57700633365078	3.47373457202282
0.0623503937884591	0.392894460690245
64420	SUSD1	sushi domain containing 1	4183
0.287670498945151	6.02399724313697	3.47361707291125
	0.0623548222296838	0.392894460690245
140612	ZFP28	ZFP28 zinc finger protein
6910	-0.482665454245849	3.16478281717602
3.47361633201422	0.0623548501545237	0.392894460690245
54532	USP53	ubiquitin specific peptidase 53
11950	-0.413196892186028	6.68445709860182
3.47229252044406	0.062404766667484	0.39303217927673
25909	AHCTF1	AT-hook containing transcription factor 1
9609	-0.247327905455531	5.69531188731139
3.46968569313815	0.062503185917249	0.393475112423778
100128252	ZNF667-AS1	ZNF667 antisense RNA 1 (head to
head)	2765	-0.937619743529445	2.60484721932427
3.46843055164117	0.062550632023709	0.393596901507678
162	AP1B1	adaptor related protein complex 1 subunit beta 1
	6721	-0.267046053552742	7.08398786619208
3.46712718714062	0.0625999416437675	0.393699378407845
57608	JCAD	junctional cadherin 5 associated	9515
0.782168157482003	1.68137822983036	3.4665145821174
0.0626231323690068	0.393699378407845
1017	CDK2	cyclin dependent kinase 2
3894	-0.269670746789837	4.75317868025292
3.46529502441747	0.0626693271008357	0.393779116997636
100505738	MIR4458HG	MIR4458 host gene	4339
0.756309327013993	0.920260045415098	3.46469569670545
	0.0626920419008159	0.393779116997636
7552	ZNF711	zinc finger protein 711	5004	0.61554161182012
	2.25932560750073	3.46338105061836
0.0627418983346381	0.39391562975046
100506746	AFF1-AS1	AFF1 antisense RNA 1
4180	-0.510722236303946	3.86820278945909
3.45959320338122	0.0628857844542691	0.394642107012073
55349	CHDH	choline dehydrogenase	7758	0.487156672136935
	4.12124592394514	3.45827398138539
0.0629359792817335	0.394780233676328
51374	ATRAID	all-trans retinoic acid induced differentiation
factor	2019	0.276066734630985	5.34138759174342
3.45683248823413	0.0629908752036798	0.394947712165596
28986	MAGEH1	MAGE family member H1	1462	0.43185928347873
	2.76286669163013	3.45077622891457
0.0632220720354547	0.396219938702735
64324	NSD1	nuclear receptor binding SET domain protein 1
15242	-0.341049207871282	7.21978783922424
3.4471635540724	0.0633604159471117	0.396909367697314
101927189	LOC101927189	uncharacterized LOC101927189
3449	-1.40962819620714	2.22932927176272
3.44634434984375	0.0633918314491791	0.396928646897028

283373	ANKRD52	ankyrin repeat domain 52
8820	-0.257609910360676	7.58026529674122
3.44495045318985	0.0634453238899252	0.397057493435089
3658	IREB2	iron responsive element binding protein 2
10806	-0.253127974763191	6.51954963587771
3.44433121965572	0.0634691031485088	0.397057493435089
169200	TMEM64	transmembrane protein 64	5523
0.318618433756491	4.92403000663253	3.44204236554536
0.0635570802029528	0.397430367144089
55327	LIN7C	lin-7 homolog C, crumbs cell polarity complex
component	4688	-0.361318420616206	4.98273718717874
3.43604212115304	0.0637883303483928
0.398698412846916
118788	PIK3AP1	phosphoinositide-3-kinase adaptor protein 1	5691
1.42195370011764	2.39972823743124
3.4339759090089	0.0638681699461415	0.399019382888316
6338	SCNN1B	sodium channel epithelial 1 beta subunit
3399	-0.852930920197935	2.2541251582576	3.4329207522126
0.0639089828960013	0.399096354625185
5892	RAD51D	RAD51 paralog D	11389	0.372292229316408
3.36076236864873	3.42825576153186	0.0640897560578075
0.400046886408961
333	APLP1	amyloid beta precursor like protein 1	3656
0.725611524750586	1.48759106985421	3.42612830411781
0.0641723782026655	0.400384187743757
100125288	ZGLP1	zinc finger GATA like protein 1	2431
0.576396199712085	1.0039447191714	3.42347067581407
0.064275749753024	0.400525183011524
9580	SOX13	SRY-box transcription factor 13
7420	-0.304012113831767	4.18317242328977
3.42319697278948	0.0642864058347529	0.400525183011524
3732	CD82	CD82 molecule	4054	-0.603390259383492
7.54732638594463	3.4229450801833	0.0642962144365091
0.400525183011524
114880	OSBPL6	oxysterol binding protein like 6	11486
0.505722712942515	1.96229469022653	3.42260768238273
0.0643093550790974	0.400525183011524
147463	ANKRD29	ankyrin repeat domain 29	6143
0.810309721418603	1.37946688381693	3.41861997700787
0.0644648815734323	0.401315376088474
102606465	LOC102606465	uncharacterized LOC102606465	2840
0.600283691397926	0.931509504248193
3.4169184424089	0.0645313660652967	0.401550797190853
6535	SLC6A8	solute carrier family 6 member 8	5498
0.457476438271246	5.34774750403784	3.41466462921034
0.0646195425212354	0.401920928994203
10466	COG5	component of oligomeric golgi complex 5
8467	-0.2565143545189	5.93557984615649
3.41245262001506	0.064706208473745	0.402281341363402
9532	BAG2	BCL2 associated athanogene 2	6651
0.661147734557696	1.93274460944647	3.41012784800702
0.0647974261240156	0.402669719484954
440107	PLEKHG7	pleckstrin homology and RhoGEF domain containing G7
3998	-1.1986359688736	1.31422824573161

3.40721451522626	0.0649119310407682	0.403202402699796
23043	TNIK	TRAF2 and NCK interacting kinase	8445
1.25816452220743	1.9988553535907 3.40358260870195
0.0650549810737112	0.403862112478177
55814	BDP1	B double prime 1, subunit of RNA polymerase III
transcription initiation factor IIIB	12436	-0.40064769448677
	6.26147632215356	3.40305464254136
0.0650758040881877	0.403862112478177
2207	FCER1G	Fc fragment of IgE receptor Ig	964
1.67809115971904	2.38009188773267	3.40098624183828
	0.0651574504742446	0.404189729314767
780	DDR1	discoidin domain receptor tyrosine kinase 1
9001	-0.300232996421073	9.71927546674972
3.39956355349182	0.0652136719873489	0.404315964529877
8914	TIMELESS	timeless circadian regulator	5818
0.399713723354878	4.50973229129754	3.39893553029157
	0.0652385065532764	0.404315964529877
27293	SMPDL3B sphingomyelin phosphodiesterase acid like 3B	2670
	0.267154712762443	4.98543400352315
3.39828110072799	0.0652643960735384	0.404315964529877
4752	NEK3	NIMA related kinase 3	4272	0.458415100112632
	2.38507841012928	3.39727285151251
0.0653043043191849	0.404384345976491
11285	B4GALT7 beta-1,4-galactosyltransferase 7	4927
0.292267096986686	4.30037466928491	3.39485629126818
	0.065400062022179	0.404689672253226
10554	AGPAT1	1-acylglycerol-3-phosphate O-acyltransferase 1	3631
	0.241236586822498	5.93792910861423
3.39342240248867	0.0654569516012711	0.404689672253226
3068	HDGF	heparin binding growth factor
3745	-0.273734076507266	8.255113507237	3.39318903067634
	0.0654662156322085	0.404689672253226
22850	ADNP2	ADNP homeobox 2 5893	-0.350156864817988
4.74291515234613	3.39311437468699	0.0654691795049483
	0.404689672253226
10225	CD96	CD96 molecule	5299	-1.60323495347825
1.11905366444239	3.3913781727518 0.0655381478366786
0.404937290140431
266655	BRD3OS	BRD3 opposite strand	2785	0.307865793767549
	4.40330884398557	3.38872655197116
0.0656436297061545	0.405245882886336
50651	SLC45A1 solute carrier family 45 member 1	2888
0.580798389945616	0.417837911763633	3.38866811986773
	0.0656459561839863	0.405245882886336
2171	FABP5	fatty acid binding protein 5
2077	-0.854423379978865	-0.273496208856152
3.38518129854737	0.0657849436808313	0.40583944402143
94025	MUC16	mucin 16, cell surface associated
43830	-1.25197119834148	8.2425700310069 3.38480265798262
	0.0658000554988697	0.40583944402143
8826	IQGAP1	IQ motif containing GTPase activating protein 1
11305	-0.259233229659116	9.34090401536319
3.38241233944629	0.0658955404369641	0.406045144783359
55432	YOD1	YOD1 deubiquitinase	6521	-0.459416087292465

5.32184286414894 3.382065061029 0.0659094253105831
0.406045144783359
132720 FAM241A family with sequence similarity 241 member A
8901 -0.623498967357184 2.16690420839334
3.38179131000802 0.0659203726163603 0.406045144783359
51547 SIRT7 sirtuin 7 6152 -0.306576234636049
5.34300073090936 3.37857880167839 0.0660489860453913
0.40665852639024
153571 C5orf38 chromosome 5 open reading frame 38 1925
0.554992358193424 2.39966475663322 3.37780991030595
0.0660798084955362 0.406669542002186
6787 NEK4 NIMA related kinase 4 6391 -0.290815995411987
4.56045856431594 3.37674909959874
0.066122358297311 0.406711474200557
4285 MIPEP mitochondrial intermediate peptidase 3407
0.320859565797779 3.88515808677565 3.37567134746231
0.0661656175978233 0.406711474200557
388650 DIPK1A divergent protein kinase domain 1A 3091
0.439291136917042 4.5293538630188 3.37546929659536
0.0661737309704483 0.406711474200557
51321 AMZ2 archaelysin family metallopeptidase 2 4214
0.25386045822271 5.2750368181623 3.37372237193287
0.0662439232278667 0.406964312566986
10172 ZNF256 zinc finger protein 256 2176 -0.538437442497182
1.6041504450009 3.37231243172208 0.0663006332825049
0.407127655079525
7067 THRA thyroid hormone receptor alpha 6959
0.455778107487982 3.67583061721607 3.37112888407263
0.0663482774994297 0.407127655079525
3595 IL12RB2 interleukin 12 receptor subunit beta 2 4233
1.1329196209189 0.848019253012593 3.37089468864288
0.0663577094700189 0.407127655079525
3177 SLC29A2 solute carrier family 29 member 2 2720
0.538046290408186 3.24824272456778 3.36915745686243
0.0664277193547722 0.407267040088938
149371 EXOC8 exocyst complex component 8
5100 -0.279525117691389 5.21488774310566
3.36713095527146 0.0665094864348371 0.407267040088938
3892 KRT86 keratin 86 2394 -0.686499139432939
1.13398291150998 3.36606364837541 0.0665525942974322
0.407267040088938
84913 ATOH8 atonal bHLH transcription factor 8 6076
0.532411785020886 3.03256981599394 3.36584244588016
0.0665615322623107 0.407267040088938
3669 ISG20 interferon stimulated exonuclease gene 20
7363 -0.482407353772652 3.88769524240449
3.36555912125808 0.0665729822229487 0.407267040088938
55293 UEVLD UEV and lactate/malate dehyrogenase domains
5515 -0.221011049847271 5.80970753794272
3.36539777513271 0.066579503622965 0.407267040088938
9885 OSBPL2 oxysterol binding protein like 2
8845 -0.208942336869498 6.66050393270227
3.36528753995296 0.066583959577616 0.407267040088938
83734 ATG10 autophagy related 10 5449 0.468234739728433

3.0767884400135 3.36347221608182 0.0666573849048423
0.407538188722011
167153 TENT2 terminal nucleotidyltransferase 2
5508 -0.225011052188604 5.668935291851 3.3621067411008
0.0667126719693561 0.407698253173984
57223 PPP4R3B protein phosphatase 4 regulatory subunit 3B
8217 -0.268886313803337 6.49390654994359
3.36029079155392 0.0667862743286426 0.407956150636777
730098 LOC730098 uncharacterized LOC730098 3889
0.504446587086526 0.235284684302157 3.35908858669672
0.0668350486939356 0.407956150636777
56548 CHST7 carbohydrate sulfotransferase 7 2289
0.49680314750423 1.87209275398997 3.35891121904815
0.0668422478554583 0.407956150636777
5529 PPP2R5E protein phosphatase 2 regulatory subunit B'epsilon
9188 -0.199762339791948 6.11885455822234
3.35567296946872 0.0669738306763743 0.408581204827515
11123 RCAN3 RCAN family member 3 7077 -0.268297608550185
6.03961922223037 3.35431598841783
0.0670290524263194 0.408610838802748
2119 ETV5 ETS variant transcription factor 5 5933
0.479144893765433 4.82219643444931 3.35412000208443
0.0670370320246101 0.408610838802748
57415 C3orf14 chromosome 3 open reading frame 14 4430
0.490388090544476 2.88546888580807 3.35300372831341
0.067082500556215 0.408710128443194
148229 ATP8B3 ATPase phospholipid transporting 8B3 7014
0.723045268913346 -0.051785654976565 3.35144114009395
0.0671462039200115 0.40892038187287
2023 ENO1 enolase 1 3958 0.336718982927444
9.62144610458577 3.35029121707583 0.0671931251034023
0.409028293491246
79153 GDPD3 glycerophosphodiester phosphodiesterase domain
containing 3 1891 -0.809283802496624 1.14942421761713
3.34873725416565 0.0672565883150375
0.409118093251091
60386 SLC25A19 solute carrier family 25 member 19 3316
0.474115972525169 2.28029133789324
3.34797631220324 0.0672876882373132 0.409118093251091
5504 PPP1R2 protein phosphatase 1 regulatory inhibitor subunit 2
5126 -0.257107928187878 5.57335238547461
3.34732112497581 0.0673144782469119 0.409118093251091
10008 KCNE3 potassium voltage-gated channel subfamily E
regulatory subunit 3 3677 -1.06812632941382
1.3368412569112 3.34707098066767 0.067324709426984
0.409118093251091
4322 MMP13 matrix metallopeptidase 13
2834 -0.863990809221028 5.2378384018048 3.34528460923811
0.0673978223377894 0.409384777747362
653583 PHLDB3 pleckstrin homology like domain family B member 3
4553 -0.487724934735386 3.24008651564512
3.34398017275089 0.0674512641438969 0.409531797513465
5814 PURB purine rich element binding protein B
9069 -0.290835473963305 6.91038095375804

3.3425894101622 0.0675082925623081      0.409645273142968
6546    SLC8A1  solute carrier family 8 member A1       21922
1.20689837746637        0.973968818448184       3.34145907193465
        0.0675546802101112      0.409645273142968
90701   SEC11C  SEC11 homolog C, signal peptidase complex subunit
        3903    -0.389185083229236      4.70610542716057
3.34088706778162        0.0675781675185361      0.409645273142968
7263    TST     thiosulfate sulfurtransferase   1937
0.347551581227222       4.76145256854367        3.34032743223467
        0.0676011554028875      0.409645273142968
57703   CWC22   CWC22 spliceosome associated protein homolog
3324    -0.282712192468197      4.61490891793619
3.33996170483497        0.0676161827305306      0.409645273142968
153769  SH3RF2  SH3 domain containing ring finger 2
7232    -0.413467768392753      6.79062476430767
3.33737619124273        0.0677225204930764      0.409832672545318
257397  TAB3    TGF-beta activated kinase 1 (MAP3K7) binding protein
3       12711   -0.272002230874756      6.6048089094957
3.33719873155572        0.0677298256610154      0.409832672545318
254170  FBXO33  F-box protein 33        3853    -0.259131013453645
        5.0248314664057 3.33707566215168        0.0677348923354331
        0.409832672545318
28996   HIPK2   homeodomain interacting protein kinase 2
15402   0.324370391591332       6.73135294705904
3.3355878874805 0.0677961749646532      0.410026348328971
5347    PLK1    polo like kinase 1      4760    0.60852353982902
        0.82626176491286        3.33374369759907
0.0678722209157325      0.410228478446983
79029   SPATA5L1        spermatogenesis associated 5 like 1
3406    -0.441621868106476      3.42833148873587
3.33301617759441        0.0679022455927872      0.410228478446983
147948  ZNF582  zinc finger protein 582 3350    -0.527359028116417
        1.83951022200832        3.33264768841027
0.0679174585220642      0.410228478446983
55596   ZCCHC8  zinc finger CCHC-type containing 8
9124    -0.329020054930668      4.91222294891389
3.32861725080142        0.0680840916699831      0.410662439799625
64598   MOSPD3  motile sperm domain containing 3        1726
0.404678897576792       3.62087933307177        3.32861541180598
        0.0680841678005381      0.410662439799625
55196   RESF1   retroelement silencing factor 1
6646    -0.42804735368813       5.35735300568202
3.32825489645859        0.0680990941382719      0.410662439799625
1906    EDN1    endothelin 1    2033    -0.626204929105707
7.37587149647937        3.32785697699285        0.0681155731733815
        0.410662439799625
960     CD44    CD44 molecule (Indian blood group)      9116
0.313095293041416       9.5520686585347 3.3273663576471
0.0681358970582087      0.410662439799625
440556  PRDM16-DT       PRDM16 divergent transcript     4881
0.858419323456177       0.0335319887067746      3.32580714971022
        0.0682005302437008      0.410816594822061
306     ANXA3   annexin A3      5123    -0.281700323170761
8.84449260783735        3.32533457184609        0.0682201327590572

0.410816594822061
80222	TARS2	threonyl-tRNA synthetetase 2, mitochondrial	4077
0.269205181115668	4.13188922166149
3.32425993643844	0.0682647310367529	0.410908503494542
25864	ABHD14A abhydrolase domain containing 14A	2378
0.57869007049506	1.74828009740385	3.32203766695017
0.0683570560425172	0.411287493121795
441150	C6orf226	chromosome 6 open reading frame 226	552
0.557896344578574	0.527255842522984
3.31757059544207	0.0685430464738402	0.412066792510294
652	BMP4	bone morphogenetic protein 4	2652
0.792404172341162	4.15088038188149	3.3175137947365
0.0685454149031902	0.412066792510294
114793	FMNL2	formin like 2	6519	0.313556456344754
6.43835666029663	3.31623419645244	0.068598793752234
0.412210769664325
9060	PAPSS2	3'-phosphoadenosine 5'-phosphosulfate synthase 2
4083	0.853948655226583	2.96734788414415
3.31429647379331	0.0686797111389794	0.412270904842631
6198	RPS6KB1 ribosomal protein S6 kinase B1
7687	-0.253923494107667	5.43096274235926
3.31414780462539	0.0686859236328936	0.412270904842631
11056	DDX52	DExD-box helicase 52	9546	-0.212802987595942
6.43208694195645	3.31320428115109
0.0687253650171624	0.412270904842631
10826	FAXDC2	fatty acid hydroxylase domain containing 2	7222
0.829904465128397	3.55851815940428
3.31272104866893	0.0687455745937242	0.412270904842631
141	ADPRH	ADP-ribosylarginine hydrolase	4615
0.529578617029672	3.46718930975019	3.31201640221995
0.0687750554619956	0.412270904842631
11248	NXPH3	neurexophilin 3 2906	0.976946783548316
0.636539593671566	3.31159892794229	0.068792528054109
0.412270904842631
79691	QTRT2	queuine tRNA-ribosyltransferase accessory subunit 2
8054	-0.316049839491837	4.62545240681823
3.31094419036468	0.0688199404158438	0.412270904842631
6297	SALL2	spalt like transcription factor 2	5382
0.849195906539446	2.57115560479637	3.31003649294966
0.0688579629593065	0.412270904842631
8692	HYAL2	hyaluronidase 2 4870	0.289973959202234
6.22999750464799	3.30966096314299	0.0688737001022172
0.412270904842631
151742	PPM1L	protein phosphatase, Mg2+/Mn2+ dependent 1L
12447	-0.438010232090378	4.12710335168588
3.30586953872966	0.069032801191123	0.413046751936805
5887	RAD23B	RAD23 homolog B, nucleotide excision repair protein
4615	-0.286579215581427	8.16702738705466
3.30378667070067	0.0691203727567356	0.413318549591193
149837	LINC00654	long intergenic non-protein coding RNA 654
6294	0.905865185069717	1.89779286504974
3.30338574224996	0.0691372429279764	0.413318549591193
56649	TMPRSS4 transmembrane serine protease 4
9948	-0.60574677615999	7.27972203638682

3.29933129871227	0.0693080926290281	0.413919010101534
3475	IFRD1	interferon related developmental regulator 1
7489	-0.393716741250284	6.40406477891633
3.29846808936378	0.0693445256081732	0.413919010101534
7850	IL1R2	interleukin 1 receptor type 2
3073	-1.05761988686507	2.21554159809402
3.29765467273336	0.0693788757727653	0.413919010101534
7706	TRIM25	tripartite motif containing 25
6666	-0.245216768184338	6.87234269612895
3.29757343694412	0.06938230731828	0.413919010101534
54985	HCFC1R1 host cell factor C1 regulator 1 1433
0.654773905896202	4.26861397400392	3.29749931777335
	0.0693854384035412	0.413919010101534
340061	STING1	stimulator of interferon response cGAMP interactor 1
	3850	-0.638598413894224	5.95205213355598
3.29501412297833	0.0694905104007406	0.414369339797009
51742	ARID4B	AT-rich interaction domain 4B
11073	-0.234152748504418	6.97737302134267
3.29322847158541	0.0695661114155863	0.414565347561628
27350	APOBEC3C	apolipoprotein B mRNA editing enzyme
catalytic subunit 3C	2818	0.415790643365556
5.1958441587284 3.29269232359401	0.0695888280627612
0.414565347561628
6018	RLF	rearranged L-myc fusion 6246	-0.365065958765963
	5.48583268154648	3.29214155855262
0.0696121723042015	0.414565347561628
440695	ETV3L	ETS variant transcription factor 3 like
1976	-0.578622212820286	0.358342900709325
3.29125058550121	0.0696499540530142	0.414614069876995
54468	MIOS	meiosis regulator for oocyte development
6714	-0.36777817259397	5.22438252928401
3.28967643121944	0.0697167597916732	0.414664060261729
10746	MAP3K2	mitogen-activated protein kinase kinase kinase 2
	11508	-0.335602332884028	6.39094931379972
3.28913689385702	0.0697396730698592	0.414664060261729
388389	CCDC103 coiled-coil domain containing 103	3895
0.589808399779543	2.22824657591526	3.28881060526876
	0.0697535339297213	0.414664060261729
57448	BIRC6	baculoviral IAP repeat containing 6
17723	-0.456885387652233	7.23275598418769
3.28826382318528	0.0697767680471831	0.414664060261729
219333	USP12	ubiquitin specific peptidase 12
5195	-0.264769714565369	5.88624599358767
3.28163945033995	0.0700589131613234	0.41616420553463
84168	ANTXR1	ANTXR cell adhesion molecule 1	8185
0.429486531730949	6.29059002828785	3.27641282794808
	0.0702823874201179	0.416792613816083
1308	COL17A1 collagen type XVII alpha 1 chain	6426
0.503970583162843	9.46045982530177	3.27572694780403
	0.0703117701527958	0.416792613816083
3801	KIFC3	kinesin family member C3
7188	-0.287430284295595	6.6118090773079 3.27544354705003
	0.0703239147276394	0.416792613816083
8907	AP1M1	adaptor related protein complex 1 subunit mu 1

14371 0.239260390925159 5.96238961994156
3.27397477820163 0.0703868918931712 0.416792613816083
80018 NAA25 N(alpha)-acetyltransferase 25, NatB auxiliary
subunit 10128 -0.255393983628044 5.56574312687565
3.2731545893684 0.0704220858357536 0.416792613816083
10009 ZBTB33 zinc finger and BTB domain containing 33
5313 -0.283290331860586 5.6156068928033 3.27207064748936
0.0704686262206591 0.416792613816083
8722 CTSF cathepsin F 2896 0.680881184996753
3.96970133503055 3.27173543340892 0.0704830257198854
0.416792613816083
1964 EIF1AX eukaryotic translation initiation factor 1A X-linked
4427 -0.33332208230378 5.7689002693618
3.27067833196751 0.0705284553484044 0.416792613816083
2199 FBLN2 fibulin 2 4812 0.747207022968173
3.49939938089714 3.27053580322767 0.0705345830135544
0.416792613816083
8556 CDC14A cell division cycle 14A 5579 -0.447760476066321
2.94299034977775 3.27048282107176
0.0705368609936418 0.416792613816083
84140 FAM161A FAM161 centrosomal protein A 5252
0.518490426015009 1.52892154463421 3.26989223238562
0.0705622588266934 0.416792613816083
7430 EZR ezrin 3342 -0.319957045293124
10.2219105868384 3.26976278776735 0.0705678268061104
0.416792613816083
7534 YWHAZ tyrosine 3-monooxygenase/tryptophan 5-monooxygenase
activation protein zeta 8555 -0.313943413238384
10.3367346290964 3.26969883321112 0.0705705779450988
0.416792613816083
114915 EPB41L4A-AS1 EPB41L4A antisense RNA 1 3344
0.479899064694221 2.66181873157129 3.26944992147769
0.0705812864975904 0.416792613816083
90592 ZNF700 zinc finger protein 700 3319 -0.408181856197857
3.6183494964757 3.26606493839151 0.0707270864061196
0.417407964237885
836 CASP3 caspase 3 2899 -0.211034673648577
6.42424502951769 3.26564742200791 0.0707450922467865
0.417407964237885
6611 SMS spermine synthase 2160 -0.322019999144993
6.02642048054122 3.26302574678132
0.0708582670807335 0.417774730296799
23095 KIF1B kinesin family member 1B
17181 -0.277210127383171 7.19187891650727
3.26270018442217 0.0708723347822726 0.417774730296799
26167 PCDHB5 protocadherin beta 5 3408 1.71381283039094
1.37714268132988 3.26213572040478
0.0708967326276498 0.417774730296799
1652 DDT D-dopachrome tautomerase 1753
0.475109210142056 3.76720443428181 3.25743930633656
0.0711000751117626 0.418796783890017
865 CBFB core-binding factor subunit beta
4129 -0.261603534566645 6.03167672729068
3.25630291051454 0.071149371924848 0.418910993085896

10512 SEMA3C semaphorin 3C 6268 -0.381678155320199
8.77597277750004 3.25502351593718 0.0712049158497612
0.419061872398153
253260 RICTOR RPTOR independent companion of MTOR complex 2
13550 -0.45730871071911 6.1122177811063 3.25380102548923
0.0712580327378118 0.419145840963653
253430 IPMK inositol polyphosphate multikinase
6133 -0.472324047126882 3.5238724294383 3.25331800791294
0.0712790314253888 0.419145840963653
5593 PRKG2 protein kinase cGMP-dependent 2 5116
0.539719899390184 1.44990469892105 3.25228016865005
0.0713241728441516 0.419235287044915
7450 VWF von Willebrand factor 10391 -0.899688171347952
2.51933587000559 3.2484571259976 0.071490722768898
0.420037983986558
9826 ARHGEF11 Rho guanine nucleotide exchange factor 11
8950 -0.257717375743512 6.17820457872148
3.24694467648967 0.0715567272874183 0.420215360348619
9824 ARHGAP11A Rho GTPase activating protein 11A 6428
0.548273229721581 2.38906556143412
3.24597500271749 0.0715990789933458 0.420215360348619
9024 BRSK2 BR serine/threonine kinase 2 6903
0.744847127226195 0.810490957389588 3.24570415318427
0.0716109134826982 0.420215360348619
120376 COLCA2 colorectal cancer associated 2 2068
0.772291895705296 2.2286423196218 3.24501435299417
0.0716410630568539 0.420216235442777
10538 BATF basic leucine zipper ATF-like transcription factor
980 -1.11944937118289 1.44181153082461
3.24343267011184 0.0717102461433691 0.420274760150847
155382 VPS37D VPS37D subunit of ESCRT-I 1626
0.598462870868888 0.640026672087536 3.24341430269485
0.0717110499579157 0.420274760150847
7186 TRAF2 TNF receptor associated factor 2
3663 -0.336993795730808 4.49402310055576
3.24185112410734 0.0717794948522333 0.420499951650034
11260 XPOT exportin for tRNA 7364 -0.238712979502916
6.64097253545244 3.23599733193803
0.0720364301705093 0.421828711286925
1368 CPM carboxypeptidase M 7524 0.941915237087147
3.00464349411336 3.23517863835158
0.0720724428842519 0.42186322920172
64847 SPATA20 spermatogenesis associated 20 5337
0.352996893463152 5.50152942426749 3.23392586550445
0.0721275872604557 0.422009655287052
7940 LST1 leukocyte specific transcript 1 2281
1.60833421565741 0.406901593863078 3.23284420518605
0.0721752360225354 0.422112121489625
7566 ZNF18 zinc finger protein 18 3012 0.404825195641094
2.76121245440204 3.23187237534518
0.0722180753725421 0.422186386370283
56915 EXOSC5 exosome component 5 1488 0.502606188743154
2.0016754852809 3.22930569022461 0.0723313488838706
0.42267217514241

10553	HTATIP2 HIV-1 Tat interactive protein 2 2797
0.229829400360671	6.43848158135507	3.22673634878524
	0.0724449304647497	0.423159358223415
25894	PLEKHG4 pleckstrin homology and RhoGEF domain containing G4
	7328	0.499184192472102	1.77383588808777
3.22350932801604	0.0725878568005473	0.423817469864638
23462	HEY1	hes related family bHLH transcription factor with
YRPW motif 1	3232	-0.648533365107592	0.158353150609293
	3.2227504116185 0.0726215134861899	0.423837308083776
55036	CCDC40	coiled-coil domain containing 40	9961
0.523694620269501	2.4734106926032 3.22125637630987
0.0726878203981262	0.424014087017458
3134	HLA-F	major histocompatibility complex, class I, F
4342	-0.575934082844261	4.84151347728178
3.22016512310814	0.0727362924375183	0.424014087017458
129303	TMEM150A	transmembrane protein 150A	3094
0.352884227081574	4.02289703571571	3.22002277172877
	0.0727426180554689	0.424014087017458
4833	NME4	NME/NM23 nucleoside diphosphate kinase 4	3670
	0.614688848284955	4.65539648836796
3.2181115183641 0.072827604917409	0.424211517728146
55628	ZNF407	zinc finger protein 407 10320	-0.308374149044151
	4.78652862765091	3.21730785914257
0.0728633726815917	0.424211517728146
79865	TREML2	triggering receptor expressed on myeloid cells like
2	3302	-1.2151507762212	0.42220014637706
3.21664661594203	0.0728928161903711	0.424211517728146
92565	FANK1	fibronectin type III and ankyrin repeat domains 1
	3384	0.415925689475594	3.40681102389942
3.21653850283797	0.0728976314108408	0.424211517728146
284900	TTC28-AS1	TTC28 antisense RNA 1	10818
0.419912033361763	2.66844511455064	3.21347361005203
	0.0730342799022622	0.424830215361706
10293	TRAIP	TRAF interacting protein	3263
0.51364967307936	0.471302637036339	3.21174802765179
	0.0731113360262755	0.424978175400602
205717	USF3	upstream transcription factor family member 3
13738	-0.505213241547776	4.94127713554524
3.2115452427977 0.0731203971382488	0.424978175400602
1270	CNTF	ciliary neurotrophic factor
1894	-0.526362766360933	0.380911008139587
3.20422542065884	0.0734482796033236	0.426548683796724
9877	ZC3H11A zinc finger CCCH-type containing 11A
9700	-0.251344887933061	7.66384372798557
3.2041532797781 0.0734515189060968	0.426548683796724
58533	SNX6	sorting nexin 6 4105	-0.216539867086828
6.88857588809785	3.19982615911201	0.0736460979363371
	0.427501406462608
100506084	ARL17B	ADP ribosylation factor like GTPase 17B
2875	-0.505222425636851	2.03651949197571
3.1949295218709 0.0738669540615737	0.428588050749378
3673	ITGA2	integrin subunit alpha 2
7910	-0.431283635946464	10.3298004076586
3.19432008672514	0.0738944915085135	0.428588050749378

100381270	ZBED6	zinc finger BED-type containing 6
4250	-0.424453217797712	5.6279659621593 3.19059505906485
	0.0740630473342185	0.428810416283181
29902	FAM216A family with sequence similarity 216 member A	2753
	0.604657249682219	0.886839052798941
3.19039232318492	0.0740722328772305	0.428810416283181
8013	NR4A3	nuclear receptor subfamily 4 group A member 3
6314	-1.10569734277248	1.62813724562789
3.18974207345388	0.0741017025995991	0.428810416283181
6335	SCN9A	sodium voltage-gated channel alpha subunit 9
10364	0.657391365747671	4.04002946616509
3.18921443449092	0.074125624776669	0.428810416283181
9618	TRAF4	TNF receptor associated factor 4
4596	-0.252107420599538	7.37529352133084
3.1885997838742 0.074153502357008	0.428810416283181
5214	PFKP	phosphofructokinase, platelet	4836
0.397815417286367	7.6196368207434 3.18833974268529
0.0741652999564306	0.428810416283181
439994	LINC00863	long intergenic non-protein coding RNA 863
	3287	0.463463667612864	2.30365084968362
3.18770237010524	0.0741942249238428	0.428810416283181
9332	CD163	CD163 molecule	5813	1.79658969948889
2.18404833679303	3.18747627828064	0.0742044882354801
	0.428810416283181
117584	RFFL	ring finger and FYVE like domain containing E3
ubiquitin protein ligase	8512	-0.382118961439683
4.47540140672531	3.18680390512689	0.0742350192461046
	0.428810416283181
9957	HS3ST1	heparan sulfate-glucosamine 3-sulfotransferase 1
	8166	-0.612510768264475	4.78567298215985
3.18462782856909	0.0743339226747229	0.428810416283181
51157	ZNF580	zinc finger protein 580 2692	0.380686753488862
	3.55628408952863	3.18399241828794
0.0743628289723853	0.428810416283181
121551	BTBD11	BTB domain containing 11
7924	-0.370935593432272	3.33939089143061
3.18395444629509	0.0743645567891856	0.428810416283181
79137	RETREG2 reticulophagy regulator family member 2 4618
0.232521273115059	6.66640461400546	3.18360475532434
	0.0743804705941647	0.428810416283181
80194	TMEM134 transmembrane protein 134	5892
0.267769463171756	4.87509706515066	3.18328718666839
	0.0743949257389711	0.428810416283181
9889	ZBED4	zinc finger BED-type containing 4
6893	-0.239040340989712	6.18752980556383
3.18263316172152	0.0744247052636471	0.428810416283181
2006	ELN	elastin 6542	1.82146717958085
0.320147963894873	3.18172490162858	0.0744660820354144
	0.428810416283181
79982	DNAJB14 DnaJ heat shock protein family (Hsp40) member B14
	13078	-0.27314627999248	6.06592267037942
3.18098479087924	0.0744998168477697	0.428810416283181
57786	RBAK	RB associated KRAB zinc finger
6960	-0.344738924106531	4.92896354466011

3.1802226846316 0.0745345713829203 0.428810416283181
1591 CYP24A1 cytochrome P450 family 24 subfamily A member 1 3972
0.64973478958404 7.21725955198424
3.17978672501192 0.0745544603946273 0.428810416283181
64600 PLA2G2F phospholipase A2 group IIF
6440 -1.18017315702663 0.551724180892548
3.17971355839687 0.0745577989037049 0.428810416283181
9917 FAM20B FAM20B glycosaminoglycan xylosylkinase
6039 -0.241401932656935 6.35213632779397
3.17932076043164 0.074575724570988 0.428810416283181
9659 PDE4DIP phosphodiesterase 4D interacting protein
27018 0.266111283112178 5.92340429237503
3.17793866853428 0.0746388343066872 0.42899719004258
8508 NIPSNAP1 nipsnap homolog 1 3067
0.283589046250437 5.80685036949621 3.176714587817
0.0746947767581024 0.429068245029587
55346 TCP11L1 t-complex 11 like 1 5370 0.294444605959819
4.504672536253 3.1763278293739 0.0747124615997117
0.429068245029587
7980 TFPI2 tissue factor pathway inhibitor 2
2663 -0.957260404616499 8.45940204426095
3.17380159977404 0.0748280861635631 0.429556148726651
51552 RAB14 RAB14, member RAS oncogene family
4304 -0.226006052079389 7.40419662640201
3.17256173888325 0.0748849044232602 0.429652220868988
9046 DOK2 docking protein 2 2211 1.3714044922276
0.450683800067556 3.17209795122176 0.0749061700122844
0.429652220868988
9063 PIAS2 protein inhibitor of activated STAT 2 17123
0.336309656399664 3.63689776527951 3.16907335935934
0.075045012776864 0.430272408500014
7139 TNNT2 troponin T2, cardiac type 7259
0.761253993782481 1.6064520063646 3.16840364228225
0.0750757932591129 0.430272764394597
94115 CGB8 chorionic gonadotropin subunit beta 8 882
0.873734444456338 0.153481207974467 3.167687806479
0.0751087083741858 0.430285348955918
9576 SPAG6 sperm associated antigen 6 3887
0.868270590809994 0.735934855098697 3.16442445202272
0.0752589584669772 0.430851430601353
728730 MAP4K3-DT MAP4K3 divergent transcript 6741
0.476827866843424 1.81712259753446 3.16420572551168
0.0752690405284152 0.430851430601353
7701 ZNF142 zinc finger protein 142 6994 -0.303348574362506
5.42475177040606 3.16070081112626
0.0754307954587469 0.431600960780501
4355 MPP2 membrane palmitoylated protein 2 6114
1.19433818979861 1.10805467001419 3.1558133610982
0.0756569795485996 0.432718379966205
64319 FBRS fibrosin 7218 -0.270339706698936
6.77899909668849 3.15335095732573 0.0757712120411277
0.43319484369799
25831 HECTD1 HECT domain E3 ubiquitin protein ligase 1
12557 -0.29574756079317 8.03373078763108

3.15150118141139	0.0758571462607873	0.433435108843751
79602	ADIPOR2 adiponectin receptor 2	5427	-0.245914782314841
	6.9324561945365 3.15111444829463	0.0758751257860269
	0.433435108843751
89891	WDR34	WD repeat domain 34	2604	0.320407815360848
	4.76844522234402	3.15003900185942
0.0759251482162252	0.433544048538388
829	CAPZA1	capping actin protein of muscle Z-line subunit alpha
1	4130	-0.273811615805243	7.83768112501262
3.14859568631095	0.0759923371252998	0.433750352997538
79801	SHCBP1	SHC binding and spindle associated 1	4459
0.92218987934554	-0.358487771119387	3.14793293026026
	0.0760232110094207	0.433750352997538
10725	NFAT5	nuclear factor of activated T cells 5
16401	-0.472221624585924	7.45306822064961
3.14511954040356	0.0761544203894383	0.434322054721035
55686	MREG	melanoregulin	3056	-0.396582280415295
4.79563211934834	3.14243643871176	0.0762797802819261
	0.434859944000382
23179	RGL1	ral guanine nucleotide dissociation stimulator like
1	5305	0.49016275145226	3.7504476748416
3.14130325452288	0.0763327915410374	0.434985114367498
56262	LRRC8A	leucine rich repeat containing 8 VRAC subunit A
5230	-0.255649407649844	9.09567111179638
3.14004782330798	0.076391567867403	0.43501135139317
54149	C21orf91	chromosome 21 open reading frame 91
6235	-0.366312608555561	5.17528721538658
3.13987828172154	0.0763995091330904	0.43501135139317
1654	DDX3X	DEAD-box helicase 3 X-linked
7082	-0.27595149534495	9.04880014075747
3.13898009527634	0.0764415946666273	0.435074122915664
10658	CELF1	CUGBP Elav-like family member 1
10441	-0.261078301690358	6.99487819404852
3.13830609082083	0.0764731922728603	0.435077174722158
1365	CLDN3	claudin 3	1274	0.722695832959829
1.53381401939077	3.13675106465913	0.0765461461315921
	0.435315415698421
55789	DEPDC1B DEP domain containing 1B	3806
0.878812111082931	1.70649658899117	3.13543980964523
	0.0766077216583732	0.435316319138655
92	ACVR2A	activin A receptor type 2A
6770	-0.349578968986593	4.10895233513427
3.13542405176309	0.0766084619602187	0.435316319138655
94039	ZNF101	zinc finger protein 101 6557	-0.355074439333247
	3.44552087275903	3.1343414193417 0.0766593422197639
	0.435428794189875
84282	RNF135	ring finger protein 135 3029	0.276981467980986
	4.26453135394763	3.12792484845508
0.0769616475024945	0.436968705540105
54165	DCUN1D1 defective in cullin neddylation 1 domain containing
1	8412	-0.239299787442984	5.72452377487057
3.12719830385434	0.0769959580816193	0.436986379598558
1832	DSP	desmoplakin	10093	-0.300662492973209
10.5832012488002	3.12559570419126	0.0770716979574969

0.437239073831778
10643 IGF2BP3 insulin like growth factor 2 mRNA binding protein 3
8511 0.564313700563321 2.13217744415052
3.12411264767676 0.077141859551257 0.43740984994546
642361 LOC642361 uncharacterized LOC642361 1701
0.553184534721387 0.0888463535300997 3.12363953501122
0.0771642563871801 0.43740984994546
2643 GCH1 GTP cyclohydrolase 1 2951 -0.361247060206073
4.77465163099577 3.12267815546181
0.0772097890019119 0.437490903944086
2005 ELK4 ETS transcription factor ELK4
11873 -0.324460737485463 5.85892938339102
3.12144505783834 0.0772682329762915 0.437645021956698
4283 CXCL9 C-X-C motif chemokine ligand 9
2740 -2.90362515186167 4.50993647718721
3.1195921991591 0.0773561406913898 0.437732098265056
11215 AKAP11 A-kinase anchoring protein 11
9920 -0.288090967209279 6.73097430391781
3.11733079295649 0.0774635775217149 0.437732098265056
222171 PRR15 proline rich 15 1982 -0.666610760391226
3.70775690226432 3.11712772459877 0.077473232920549
0.437732098265056
55764 IFT122 intraflagellar transport 122 9636
0.461085226610036 2.56565895853441 3.11659871698434
0.0774983920071811 0.437732098265056
92597 MOB1B MOB kinase activator 1B 8801 -0.290771548453321
6.0151882008101 3.11652400454557 0.0775019459663752
0.437732098265056
126661 CCDC163 coiled-coil domain containing 163 2757
0.651300867312972 0.964682442375 3.11640170106671
0.0775077641388202 0.437732098265056
3146 HMGB1 high mobility group box 1
7723 -0.235283466548295 6.85338974216184
3.11606735497717 0.0775236719166661 0.437732098265056
2039 DMTN dematin actin binding protein
4473 -0.48028332539448 5.7511747582114 3.11465275948589
0.0775910155258872 0.437732098265056
65059 RAPH1 Ras association (RalGDS/AF-6) and pleckstrin
homology domains 1 11697 -0.597230539109269
6.39243307367181 3.11433964209512 0.0776059303829738
0.437732098265056
51499 TRIAP1 TP53 regulated inhibitor of apoptosis 1 1442
0.27556505332044 4.89330781171889 3.11219339414012
0.0777082466332537 0.437732098265056
57484 RNF150 ring finger protein 150 11007 1.15631618011611
0.953529872973284 3.11137325123293
0.077747382933546 0.437732098265056
719 C3AR1 complement C3a receptor 1 2145
1.62224763340673 1.62657754033217 3.11129885806649
0.0777509339412078 0.437732098265056
55274 PHF10 PHD finger protein 10 5534 -0.290312201639175
4.85954684208466 3.11108784391149
0.0777610072257575 0.437732098265056
54665 RSBN1 round spermatid basic protein 1

6819    -0.369601684286896      4.35192503692245
3.11058005654607        0.0777852534732822      0.437732098265056
9026    HIP1R   huntingtin interacting protein 1 related        7102
        0.191886351367  6.72370197713218        3.11032787863171
        0.0777972976946352      0.437732098265056
57194   ATP10A  ATPase phospholipid transporting 10A (putative) 8128
        0.904270209596581       5.22832792990612
3.10986612145064        0.0778193567939525      0.437732098265056
2799    GNS     glucosamine (N-acetyl)-6-sulfatase      6284
0.247944406352431       7.42220609309318        3.10947729306686
        0.077837937153966       0.437732098265056
645369  TMEM200C        transmembrane protein 200C      1920
0.773163851109096       0.483803860625868       3.1087042472973
0.0778748917190621      0.437732098265056
8202    NCOA3   nuclear receptor coactivator 3
8402    -0.278372546279573      6.93753191324754
3.10865231825986        0.0778773748037781      0.437732098265056
2444    FRK     fyn related Src family tyrosine kinase
13230   -0.465874517849689      5.99888478329344
3.10776522968754        0.0779198057783349      0.437794913572859
1207    CLNS1A  chloride nucleotide-sensitive channel 1A        4296
        0.262519890846701       5.55033563758332
3.10503834102641        0.0780503934405186      0.438352791067099
6772    STAT1   signal transducer and activator of transcription 1
        6053    -0.386227843245258      7.34870295435491
3.10185991919373        0.0782029019086636      0.439033285384629
64781   CERK    ceramide kinase 4938    -0.247423207024938
5.80060439083709        3.10019621072259        0.0782828585939357
        0.439306089873901
100507266       STX18-AS1       STX18 antisense RNA 1 (head to head)
        6107    0.567902288569805       0.960674358691788
3.0991214707547 0.0783345566439729      0.439420158154637
254065  BRWD3   bromodomain and WD repeat domain containing 3
12694   -0.37508556388759       5.69041312875587
3.0966237582296 0.0784548457518539      0.439918744774307
4358    MPV17   mitochondrial inner membrane protein MPV17      4654
        0.309428889655012       3.93223995872933
3.09590670573283        0.0784894155220293      0.439936471875576
8544    PIR     pirin   2512    0.501344258681774
2.04779822850154        3.09449207896833        0.0785576641036708
        0.440142880440047
100507373       LOC100507373    uncharacterized LOC100507373
3186    -0.645545377311179      0.431207253784436
3.09116270939697        0.0787185416348745      0.440683549424852
651     BMP3    bone morphogenetic protein 3
5734    -1.49680107100487       -0.203078394639706
3.09082523943168        0.0787348682178965      0.440683549424852
64282   TENT4B  terminal nucleotidyltransferase 4B
8144    -0.335394631279389      4.47031962729049
3.09001925546151        0.0787738759679188      0.440683549424852
113540  CMTM1   CKLF like MARVEL transmembrane domain containing 1
        1465    0.55018368003797        0.672616945320597
3.08989254094136        0.0787800105489991      0.440683549424852
22861   NLRP1   NLR family pyrin domain containing 1

11180	-0.342208371601955	6.38442826975794
3.08897742596641	0.078824328933668	0.440755439271013
56100	PCDHGB6	protocadherin gamma subfamily B, 6	6782
0.258494261878439	6.53291179637894	3.08668569004689
	0.0789354339787932	0.440972303192361
57688	ZSWIM6	zinc finger SWIM-type containing 6
5501	-0.325767314990968	5.51912479758181
3.08660182926747	0.078939502808651	0.440972303192361
100128893	GATA6-AS1	GATA6 antisense RNA 1 (head to head)
	1788	0.774405795214458	0.397212515197769
3.08552780367067	0.0789916332889136	0.440972303192361
6419	SETMAR	SET domain and mariner transposase fusion gene	3567
	0.355567801036662	3.30252490240959
3.08533464779827	0.0790010125185577	0.440972303192361
3998	LMAN1	lectin, mannose binding 1
6767	-0.265607025067258	6.95469333651192
3.08493290546016	0.0790205240960109	0.440972303192361
57094	CPA6	carboxypeptidase A6	2395	-0.81932734149666
	3.19997160848983	3.08235763793039
0.0791457214664945	0.441495069924806
594	BCKDHB	branched chain keto acid dehydrogenase E1 subunit
beta	4071	0.384516198238998	3.31307359542473
3.08026184051883	0.0792477669570573	0.441757740599745
259236	TMIE	transmembrane inner ear	1824	-0.566741004026238
	0.491125236604798	3.07819142814939
0.0793487156179651	0.441757740599745
2669	GEM	GTP binding protein overexpressed in skeletal muscle
	2648	-0.867129487482029	1.63884617819586
3.07780582255954	0.0793675321796341	0.441757740599745
101928100	LOC101928100	uncharacterized LOC101928100	2830
	0.942304488769417	4.336789227049	3.07763035352866
	0.0793760962087247	0.441757740599745
146198	ZFP90	ZFP90 zinc finger protein	6087
0.278552820831777	5.63651406694996	3.07744658790223
	0.0793850662328771	0.441757740599745
54470	ARMCX6	armadillo repeat containing X-linked 6	2489
0.279159326231276	5.2474989046373	3.07672033492437
0.0794205270039489	0.441757740599745
401474	SAMD12	sterile alpha motif domain containing 12
10756	-0.261724903067669	5.85305941832524
3.07640793072348	0.0794357860200925	0.441757740599745
7874	USP7	ubiquitin specific peptidase 7
5923	-0.218976181591874	6.95074885122792
3.07621693320455	0.079445116625313	0.441757740599745
84261	FBXW9	F-box and WD repeat domain containing 9	2114
0.422316970250363	2.9230555083427	3.07360940127584
0.0795726178695165	0.442268134981334
10766	TOB2	transducer of ERBB2, 2	4565	-0.328900222325436
	6.21062599060345	3.07304887531743
0.0796000548502851	0.442268134981334
375035	SFT2D2	SFT2 domain containing 2
11221	-0.260590042459329	7.43251165250455
3.06788071056666	0.0798535103003112	0.443360608094609
6821	SUOX	sulfite oxidase	5008	0.355749452854821

3.78113795862516	3.06774891148651	0.0798599853089669
	0.443360608094609
10863	ADAM28	ADAM metallopeptidase domain 28
9351	-0.377419994580124	4.56127609843123
3.06444075345999	0.0800226935632614	0.444026857042603
65983	GRAMD2B GRAM domain containing 2B
4825	-0.275983543965264	7.3845174288185 3.06402040370168
	0.0800433935912452	0.444026857042603
254295	PHYHD1	phytanoyl-CoA dioxygenase domain containing 1	2915
	0.552152227422653	3.01221412351576
3.06236996424911	0.0801247249389312	0.444302067387019
669	BPGM	bisphosphoglycerate mutase
2539	-0.460356350767247	8.47031926548757
3.05981468568825	0.0802508212578021	0.444825189298787
10459	MAD2L2	mitotic arrest deficient 2 like 2	3324
0.372883678830256	3.46843533746035	3.05898975902693
	0.0802915748946817	0.444875035423183
54464	XRN1	5'-3' exoribonuclease 1 11982	-0.355441783957558
	5.90793882571013	3.05779610988418
0.0803505839335158	0.444927590739658
113246	C12orf57	chromosome 12 open reading frame 57	1806
	0.292867130159991	4.85095898274754
3.05751294145275	0.0803645894603652	0.444927590739658
84312	BRMS1L	BRMS1 like transcriptional repressor	3897
0.318498781040384	3.93432596031137	3.05616119654067
	0.0804314831182228	0.445122000804799
2280	FKBP1A	FKBP prolyl isomerase 1A
2577	-0.307029130313575	7.746978035641	3.05385766482499
	0.0805456160998027	0.445577584798553
128346	C1orf162	chromosome 1 open reading frame 162	2768
	1.21537952772838	0.498863847479237
3.05235219554223	0.080620301860296	0.445737184054869
64798	DEPTOR	DEP domain containing MTOR interacting protein	2622
	0.658515935460969	0.30011055585803
3.05195466634272	0.0806400355922155	0.445737184054869
100506119	LINC01503	long intergenic non-protein coding
RNA 1503	1594	0.54885969916381	3.08120729663647
	3.05099722874575	0.0806875850964418
0.445737184054869
4041	LRP5	LDL receptor related protein 5	6164
0.220969681691918	5.99929681227751	3.05066058946288
	0.0807043108946582	0.445737184054869
197358	NLRC3	NLR family CARD domain containing 3
6686	-1.25658729222868	0.206933656283457
3.05007170009969	0.0807335786354825	0.445737184054869
5469	MED1	mediator complex subunit 1
7330	-0.256577140246902	6.32718592708471
3.04728751773081	0.080872107270683	0.446326086107351
9829	DNAJC6	DnaJ heat shock protein family (Hsp40) member C6
	6954	-0.448633038119182	5.01979502299053
3.04512524110991	0.0809798694555362	0.446568903875678
847	CAT	catalase	3347	0.454980311104293
5.42460523768507	3.04510690700164	0.0809807838411351
	0.446568903875678

414	ARSD	arylsulfatase D 7030	0.249484554423705
5.89886472138446	3.04448614078817	0.0810117501783464
	0.446568903875678
9052	GPRC5A	G protein-coupled receptor class C group 5 member A
	8440	-0.246697516829564	11.0893806245058
3.04379532106395	0.0810462260833442	0.446583197777105
80204	FBXO11	F-box protein 11	7074	-0.223614228853358
	6.58061149332799	3.0429232293938 0.0810897711067889
	0.44664743369752
57402	S100A14 S100 calcium binding protein A14
1277	-0.743962802236716	6.78208487385416
3.04017292190815	0.081227264005756	0.447228886371315
51012	PRELID3B	PRELI domain containing 3B
2645	-0.41518606430633	6.26705817475429
3.0374004978766 0.0813661170898474	0.447817368203337
28951	TRIB2	tribbles pseudokinase 2 4588	-0.568457064643899
	5.38106418416695	3.03540251728856
0.0814663419726455	0.448141373861672
322	APBB1	amyloid beta precursor protein binding family B
member 1	5244	0.827295196603659	3.97674752594469
	3.0349516863615 0.0814889754569628	0.448141373861672
57018	CCNL1	cyclin L1	8729	-0.323900599884666
6.46231473235504	3.03361365385132	0.0815561899022946
	0.448191270834162
84303	CHCHD6	coiled-coil-helix-coiled-coil-helix domain
containing 6	2763	0.428642370275374	2.96784839856624
	3.03236994841288	0.0816187195804749
0.448191270834162
7172	TPMT	thiopurine S-methyltransferase
3181	-0.296365733028393	4.92940511643646
3.03225533127205	0.0816244847797715	0.448191270834162
10114	HIPK3	homeodomain interacting protein kinase 3
7958	-0.317783511652849	7.20852797291695
3.03216235285991	0.081629161881888	0.448191270834162
81786	TRIM7	tripartite motif containing 7
7338	-0.56827358212661	3.01569002968237
3.03078728530248	0.0816983658094652	0.448191270834162
27106	ARRDC2	arrestin domain containing 2
4637	-0.523473463797332	5.2985893305392 3.0303418382555
0.0817207976881969	0.448191270834162
27440	HDHD5	haloacid dehalogenase like hydrolase domain
containing 5	4936	0.285008374188192	3.93916149511987
	3.03031732646552	0.0817220322489077
0.448191270834162
60	ACTB	actin beta	3256	-0.391099052529207
12.3540836567093	3.02946155033169	0.0817651468845636
	0.448252216208252
5624	PROC	protein C, inactivator of coagulation factors Va and
VIIIa	2495	1.20049206327972	1.76380129269219
3.02825489031984	0.0818259810173589	0.44841021757048
63976	PRDM16	PR/SET domain 16	9767	0.561885655483037
	3.35698300116535	3.0238815950223 0.0820468716567378
	0.44944486949391
701	BUB1B	BUB1 mitotic checkpoint serine/threonine kinase B

4476	0.512904388391028	0.0924040542288981
3.02246715120068	0.0821184514128025	0.449596611875151
2932	GSK3B	glycogen synthase kinase 3 beta
8006	-0.236141355590102	7.5841762767615 3.02206592092488
0.0821387684578075	0.449596611875151
80746	TSEN2	tRNA splicing endonuclease subunit 2	4954
0.440413407957458	2.12763119198079	3.02006462553302
0.0822401888426306	0.449902167897882
28970	C11orf54	chromosome 11 open reading frame 54
6269	-0.589396630320595	4.30223589013756
3.01917864776576	0.0822851310216477	0.449902167897882
54778	RNF111	ring finger protein 111 7181	-0.256452578562756
5.93650970397851	3.01816960165852
0.0823363482665139	0.449902167897882
2633	GBP1	guanylate binding protein 1
4992	-0.550578666585244	5.7477660526109 3.01797935546526
0.0823460086533885	0.449902167897882
6872	TAF1	TATA-box binding protein associated factor 1
10860	-0.294018532875518	5.14296092247014
3.01779854845413	0.0823551908681494	0.449902167897882
2118	ETV4	ETS variant transcription factor 4
3607	-0.404689438370872	4.18184542173352
3.01700273279231	0.0823956192017195	0.449947539243074
4867	NPHP1	nephrocystin 1	11763	0.474809838403434
2.11248409251228	3.01573681360026	0.082459973476472
0.450123479534273
8774	NAPG	NSF attachment protein gamma
5586	-0.237562742444375	5.67289673385391
3.01466813681601	0.0825143429731029	0.450145689060805
207107	SFTA1P	surfactant associated 1, pseudogene	824
0.914223220623273	3.44163064307821	3.01439359813281
0.0825283165208929	0.450145689060805
55837	EAPP	E2F associated phosphoprotein	2030
0.242239409542339	4.97267222208763	3.0136871274851
0.0825642864110876	0.450166586126938
1147	CHUK	component of inhibitor of nuclear factor kappa B
kinase complex	6928	-0.235294816301284	5.69390343674112
3.01283820979266	0.082607531369807
0.450201058579243
3840	KPNA4	karyopherin subunit alpha 4
9413	-0.233991693158308	7.38410266074509
3.01131668936088	0.0826851007819771	0.450201058579243
26355	FAM162A family with sequence similarity 162 member A	3793
0.327900777242609	4.20499190332953
3.00913855382985	0.0827962823301795	0.450201058579243
5793	PTPRG	protein tyrosine phosphatase receptor type G
14633	0.308244876315762	4.58529094768387
3.00892501742462	0.0828071908457452	0.450201058579243
84879	MFSD2A	major facilitator superfamily domain containing 2A
4007	-0.654529674373376	4.01074460936393
3.0086705729944 0.082820191175672	0.450201058579243
55258	THNSL2	threonine synthase like 2	6431
1.13223327304627	4.46365412283804	3.00858492510673
0.0828245676791629	0.450201058579243

91775	NXPE3	neurexophilin and PC-esterase domain family member 3
	9474	0.374476285327345	4.7245137826123
3.00765545827027	0.0828720783737657	0.450201058579243
1004	CDH6	cadherin 6	12379	-0.53708658111232
8.98104579448833	3.0072615285585	0.0828922233922517
0.450201058579243
148327	CREB3L4	cAMP responsive element binding protein 3 like 4
	2904	0.40260085835213	2.73207925009904
3.0064066095977	0.0829359609539681	0.450201058579243
2352	FOLR3	folate receptor gamma	1396	0.710728626297497
	2.99170554095264	3.00627740155106
0.0829425733911989	0.450201058579243
10743	RAI1	retinoic acid induced 1	9693	-0.275163159538308
	6.3755649662014	3.00601459588083	0.082956024666263
	0.450201058579243
11043	MID2	midline 2	3097	-0.34991218224353
3.38241184899699	3.00587022836558	0.0829634148820315
	0.450201058579243
51026	GOLT1B	golgi transport 1B	3789	-0.312739110666323
	6.10903825356769	3.00538139146112
0.0829884438674666	0.450201058579243
84246	MED10	mediator complex subunit 10
1880	-0.226385874924742	5.46018367101658
3.00292880197982	0.0831141422865786	0.450385362428495
9213	XPR1	xenotropic and polytropic retrovirus receptor 1	8903
	0.26722993389996	6.56432749815178
3.00281701525646	0.0831198763996091	0.450385362428495
196951	FAM227B	family with sequence similarity 227 member B	6020
	0.508093879489388	0.592743189587304
3.00230476468754	0.0831461578175622	0.450385362428495
2115	ETV1	ETS variant transcription factor 1	9616
0.779894281407679	0.101525165644261	3.00197056321472
	0.0831633091243032	0.450385362428495
4292	MLH1	mutL homolog 1	3532	0.261223333990468
4.71657870257441	3.00158328144366	0.083183189305527
	0.450385362428495
2063	NR2F6	nuclear receptor subfamily 2 group F member 6
2676	-0.264532125205069	5.49388996765789
2.99963339248107	0.083283360188932	0.450751715773687
10867	TSPAN9	tetraspanin 9	5714	0.370407695463517
4.68274654609803	2.99901388327542	0.0833152132603752
	0.450751715773687
5973	RENBP	renin binding protein	2057	0.90539143862404
	1.18495103075472	2.99680133561003
0.0834290824572202	0.45116403429224
83882	TSPAN10	tetraspanin 10	4745	0.717461207074443
0.520351397073606	2.99591497755214	0.0834747461996473
	0.45116403429224
79618	HMBOX1	homeobox containing 1	5861	-0.3716729118934
	3.99604096019482	2.99565675823296
0.0834880543218023	0.45116403429224
81890	QTRT1	queuine tRNA-ribosyltransferase catalytic subunit 1
	3430	0.417589911679433	3.15326716327104
2.99455944495978	0.0835446332866763	0.451295672366554

7786	MAP3K12 mitogen-activated protein kinase kinase kinase 12
	7971	0.420926241808102	3.93937090384539
2.99205229289414	0.0836740609585822	0.451820575114441
127294	MYOM3	myomesin 3	9254	-0.815377781924007
1.51931327575088	2.99121981677412	0.0837170840845356
	0.451878688544158
55276	PGM2	phosphoglucomutase 2	3704	-0.34593341619557
	5.56810678771116	2.98945168796	0.0838085419078536
	0.452115625142755
115286	SLC25A26	solute carrier family 25 member 26	5850
	0.310044438239396	3.79054291233836
2.98912338142874	0.083825535696133	0.452115625142755
152579	SCFD2	sec1 family domain containing 2 3185
0.436118594746885	3.29861721011036	2.98746297746487
	0.0839115385041802	0.452405280919189
2241	FER	FER tyrosine kinase	13614	-0.46655718158452
	5.49753507046241	2.98158194710274
0.0842169219964293	0.453806841167001
23336	SYNM	synemin 7819	-0.805624144014474
2.16788397898747	2.98120963827727	0.08423629521198
	0.453806841167001
1777	DNASE2	deoxyribonuclease 2, lysosomal	2389
0.287784841407966	5.91365569684466	2.97960694667594
	0.0843197468241528	0.454081773843102
80034	CSRNP3	cysteine and serine rich nuclear protein 3
12565	0.714490343080971	1.48460886812348
2.97869249470015	0.0843674020650141	0.454163797357668
4147	MATN2	matrilin 2	6614	0.521042712437588
2.99454552455387	2.97730085541841	0.0844399810510627
	0.4543586630468
254128	NIFK-AS1	NIFK antisense RNA 1	2551
0.538868991289701	1.09463395980451	2.97675482199366
	0.0844684770881606	0.4543586630468
202781	PAXIP1-AS1	PAXIP1 antisense RNA 1 (head to head)	2256
	0.50943753864864	0.245961908890806
2.97578376114342	0.08451917986476	0.454456872309287
340526	RTL5	retrotransposon Gag like 5	4339
0.60628955896739	2.5204789912683 2.97508892443389
0.0845554801218685	0.454477594039529
79694	MANEA	mannosidase endo-alpha	5187	0.408873316404835
	3.19707013880714	2.97358005486101
0.0846343658349665	0.454553895731135
83481	EPPK1	epiplakin 1	16002	-0.506942955529881
7.22332208990775	2.97357596325537	0.0846345798576996
	0.454553895731135
7576	ZNF28	zinc finger protein 28	6892	-0.277267236243275
	5.12676699903031	2.96917934550413
0.0848648955689593	0.455616171803148
100506098	LOC100506098	uncharacterized LOC100506098
901	-0.473466576632223	1.6986859422105 2.96689579380243
	0.0849847861585481	0.455947762262128
10636	RGS14	regulator of G protein signaling 14	3797
0.719544931398369	2.79372688146341	2.96676303662487
	0.0849917617809964	0.455947762262128

58191	CXCL16	C-X-C motif chemokine ligand 16
2794	-0.373735741035208	7.33935974633637
2.96529826880987	0.085068767979382	0.456186153555591
196872	LINC00638	long intergenic non-protein coding RNA 638
	2518	0.561952914511905	0.189724302198297
2.96456175529648	0.0851075166846302	0.456219282893844
10001	MED6	mediator complex subunit 6
4513	-0.291600680947716	4.89441873449294
2.96251663405678	0.0852152126579959	0.456621837681924
8563	THOC5	THO complex 5	7878	0.232323460063075
6.03528845578919	2.96151855012474	0.0852678252107699
	0.456729035459753
54749	EPDR1	ependymin related 1	2864	0.511691965332202
	4.61544197926132	2.96032544065523
0.085330764356853	0.456891443557508
3075	CFH	complement factor H	8145	0.492039661505968
	5.52169950983288	2.95951038493752
0.0853737892230698	0.456947140102231
55769	ZNF83	zinc finger protein 83	8028	-0.362525089872798
	5.64055760058275	2.95850176526858
0.0854270643599904	0.457057635787007
1687	GSDME	gasdermin E	8897	0.293543096547034
6.12813789172902	2.95285218085579	0.085726140161707
	0.458482644232543
28988	DBNL	drebrin like	12587	-0.197946712674262
6.89551462442335	2.94919721016655	0.0859202291043074
	0.459115052506501
10100	TSPAN2	tetraspanin 2	3410	-0.5041610622338
7.03897286242587	2.94901995929337	0.0859296536901671
	0.459115052506501
79649	MAP7D3	MAP7 domain containing 3	7067
0.366843958269882	3.10091299003583	2.94877426753319
	0.0859427191884091	0.459115052506501
374986	MIGA1	mitoguardin 1	9594	-0.253985982002868
5.70110331180428	2.94708895875982	0.0860323991998383
	0.459418915589834
54891	INO80D	INO80 complex subunit D	14373	-0.461168653085936
	5.33100994941964	2.94643737994407
0.0860670986758932	0.45942905912852
4775	NFATC3	nuclear factor of activated T cells 3	9070
0.268981233443702	5.32769858530862	2.94456373366672
	0.0861669631061894	0.459777510552534
136895	C7orf31	chromosome 7 open reading frame 31	3837
0.539103374446805	2.39413822013272	2.94330339977184
	0.0862342088253093	0.459777510552534
50863	NTM	neurotrimin	6588	1.13285228781459
3.06666290387467	2.94220749769987	0.0862927273693381
	0.459777510552534
4659	PPP1R12A	protein phosphatase 1 regulatory subunit 12A
	11815	-0.227498018164604	6.74490523628905
2.94131934607663	0.0863401840659691	0.459777510552534
9748	SLK	STE20 like kinase	7974	-0.253152134192373
	7.62742500670832	2.94110489703547
0.0863516469765382	0.459777510552534

131544	CRYBG3	crystallin beta-gamma domain containing 3
10929	-0.342647773927161	5.78223331453813
2.94063579276539	0.0863767276753929	0.459777510552534
285527	FRYL	FRY like transcription coactivator
18152	-0.391695722260269	7.08304311513034
2.94050832255601	0.0863835442429455	0.459777510552534
10602	CDC42EP3	CDC42 effector protein 3	6777
0.379518591960473	8.28024333970257	2.93973176132528
0.0864250840322887	0.459777510552534
255374	MBLAC1	metallo-beta-lactamase domain containing 1	1571
0.506093263377377	0.569808740995561
2.9395067505089 0.0864371243434039	0.459777510552534
89846	FGD3	FYVE, RhoGEF and PH domain containing 3 5890
0.631049100587541	3.83850463589611	2.93864509187165
0.086483248420432	0.459777510552534
84101	USP44	ubiquitin specific peptidase 44 4845
0.6851140412372 0.992882070343327	2.93806161415999
0.0865144967643635	0.459777510552534
57551	TAOK1	TAO kinase 1	16980	-0.394308871747508
7.88432930770628	2.93784176834917	0.0865262738499665
0.459777510552534
29901	SAC3D1	SAC3 domain containing 1	1772
0.438085128033749	2.70335254024454	2.935035728482
0.0866767452388257	0.460402415836265
100132707	PAXIP1-AS2	PAXIP1 antisense RNA 2	2974
0.677454775772804	1.58531768432436	2.93260466049893
0.0868073381238622	0.460921298370333
56888	KCMF1	potassium channel modulatory factor 1
7968	-0.199420020017367	6.58580387055616
2.93164745568276	0.0868588159912675	0.461019869492112
29922	NME7	NME/NM23 family member 7	5737
0.387674111123689	3.79109563253673	2.93034651011712
0.0869288330096476	0.461052814427406
55640	FLVCR2	feline leukemia virus subgroup C cellular receptor family member 2 4867	0.737731291775133	1.76212296404611
2.93030896163132	0.0869308547799514
0.461052814427406
55183	RIF1	replication timing regulatory factor 1
16623	-0.344321883448341	6.14843516779765
2.92861312364764	0.0870222190111993	0.46122191128523
399474	TMEM200B	transmembrane protein 200B	2639
0.774348532357413	1.35120896353523	2.92849487372251
0.0870285936693697	0.46122191128523
2966	GTF2H2	general transcription factor IIH subunit 2	1817
0.545567449755857	2.57430570190516
2.92689086311982	0.087115131966757	0.461505820932616
1643	DDB2	damage specific DNA binding protein 2	4897
0.321896072323963	5.66089804169801	2.92431494850884
0.0872542516914957	0.462068167653225
284340	CXCL17	C-X-C motif chemokine ligand 17
1189	-0.892427738571638	4.956035682647	2.91968711433766
0.0875048299618291	0.463119146827663
9473	THEMIS2	thymocyte selection associated family member 2	4621
0.32049721203492	4.81929561616155

2.91942880558376	0.0875188392698525	0.463119146827663
84144	SYDE2	synapse defective Rho GTPase homolog 2	6451
0.372382987898571	3.2817116243726	2.91800446137777
0.087596131850389	0.46335310378716
987	LRBA	LPS responsive beige-like anchor protein
14929	-0.285200273268314	7.11167969874257
2.91654195687181	0.0876755721581609	0.46355189755767
57335	ZNF286A	zinc finger protein 286A
9184	-0.305382893136357	5.15684306382285
2.91609432742022	0.0876999020866585	0.46355189755767
90843	TCEAL8	transcription elongation factor A like 8	1222
	0.295566688264322	4.92924702428202
2.91480276065715	0.0877701434033254	0.463748169992599
84321	THOC3	THO complex 3	3453	0.493114941358245
4.28268595661951	2.91317990805769	0.0878584879729183
	0.463922167762158
85352	SHISAL1	shisa like 1	6751	0.540141264476429
5.97529066856656	2.91298112094009	0.0878693161328625
	0.463922167762158
116362	RBP7	retinol binding protein 7
861	-0.739967036097152	0.930096330407385
2.90857816860008	0.0881095209959971	0.464912571716509
92170	MTG1	mitochondrial ribosome associated GTPase 1	4477
	0.280379100299082	5.22719847381285
2.90832623611203	0.088123286778563	0.464912571716509
9481	SLC25A27	solute carrier family 25 member 27	3288
	0.699442612977167	1.10417545956741
2.90744146078969	0.0881716500412704	0.464980762613219
11188	NISCH	nischarin	9161	0.30589229864996
6.99108734681634	2.90687543792717	0.0882026048592363
	0.464980762613219
80155	NAA15	N(alpha)-acetyltransferase 15, NatA auxiliary subunit	9712	-0.296153907516511	5.90986615794591
2.90564052365487	0.0882701810810033	0.465161936193233
3398	ID2	inhibitor of DNA binding 2	2205
0.525093924788787	3.76140427650881	2.904722903887
0.0883204308407725	0.465251701687364
154810	AMOTL1	angiomotin like 1	10035	-0.293524362480052
	7.50994521161847	2.90406233465886
0.0883566234507232	0.465267377697098
1909	EDNRA	endothelin receptor type A	4238
0.818466207061089	1.62393807230728	2.90192495627601
	0.0884738406708333	0.465709540126404
26227	PHGDH	phosphoglycerate dehydrogenase
5622	-0.372378144086853	4.7105538756612	2.89891931935527
	0.0886389598024246	0.466403422221097
79605	PGBD5	piggyBac transposable element derived 5
10662	-0.291999049967672	6.7718709973623	2.8974239928035
0.0887212321898634	0.466578278124931
6450	SH3BGR	SH3 domain binding glutamate rich protein	1495
	0.503769554281406	0.792845694631659
2.89567849409826	0.0888173736926329	0.466578278124931
1503	CTPS1	CTP synthase 1	5462	-0.369624671507484
5.98146751094165	2.89484114075749	0.0888635349231236

0.466578278124931
79767 ELMO3 engulfment and cell motility 3 2575
0.236458314741526 5.14323872462839 2.8940961268565
0.0889046275727785 0.466578278124931
115811 IQCD IQ motif containing D 2460 -0.740281161826887
1.78537718459467 2.89319147541191
0.0889545530301142 0.466578278124931
154313 CFAP206 cilia and flagella associated protein 206 3976
0.516807263067273 0.723502440744631
2.892723790923 0.0889803752889054 0.466578278124931
5868 RAB5A RAB5A, member RAS oncogene family
3785 -0.200437086893991 6.97747133553392
2.89215712826546 0.0890116733175366 0.466578278124931
81688 C6orf62 chromosome 6 open reading frame 62
4134 -0.210304450666084 7.69486175372015
2.89201274818706 0.0890196496556006 0.466578278124931
125965 COX6B2 cytochrome c oxidase subunit 6B2
3187 -0.952089825607609 1.99427747858468
2.89185608231289 0.0890283056042884 0.466578278124931
79754 ASB13 ankyrin repeat and SOCS box containing 13 3003
0.619073932286988 3.12956401409312
2.89172978628482 0.0890352842524812 0.466578278124931
29954 POMT2 protein O-mannosyltransferase 2 10008
0.235134032938723 5.89000875799637 2.89166969647916
0.0890386047995959 0.466578278124931
220213 OTUD1 OTU deubiquitinase 1 2933 0.625821274782338
5.75714476014821 2.89026645251201
0.0891161858899334 0.466810177920829
84898 PLXDC2 plexin domain containing 2 12567
0.487110495308288 5.5718693432649 2.885554325855
0.0893772418408248 0.467833046254992
23136 EPB41L3 erythrocyte membrane protein band 4.1 like 3
11000 1.17999647632377 2.55532541323293
2.88541539915233 0.0893849510764083 0.467833046254992
26130 GAPVD1 GTPase activating protein and VPS9 domains 1
13625 -0.229524940452783 6.68849623546324
2.88490233234418 0.089413428115716 0.467833046254992
6591 SNAI2 snail family transcriptional repressor 2 2282
0.528931089075424 5.74963027140076
2.88393064312857 0.0894673873018435 0.467833046254992
338651 KRTAP5-AS1 KRTAP5-1/KRTAP5-2 antisense RNA 1 2931
0.317338256274969 3.98141769363567
2.8837314010975 0.08947845583759 0.467833046254992
81025 GJA9 gap junction protein alpha 9
1799 -0.542359499692979 1.04383316247932
2.88252440363568 0.089545540187 0.467983333351696
22941 SHANK2 SH3 and multiple ankyrin repeat domains 2
16873 0.367362898884822 5.27919723916033
2.88182121322271 0.0895846483241844 0.467983333351696
27289 RND1 Rho family GTPase 1 2058 -0.803274413730611
7.03767493914369 2.88104726650414
0.0896277130100061 0.467983333351696
10915 TCERG1 transcription elongation regulator 1
8714 -0.209183793333633 5.9098908523857 2.88081138395347

0.0896408426774185	0.467983333351696
171024	SYNPO2	synaptopodin 2	14034	0.629918806654048
1.41349393509286	2.87769610066043	0.0898144410805729
0.468302965261472
7482	WNT2B	Wnt family member 2B	12561	-0.537394532831597
2.32946106263375	2.87756689879058
0.0898216486948168	0.468302965261472
25836	NIPBL	NIPBL cohesin loading factor
12529	-0.330364585124374	6.8339886102862 2.87740802717788
0.089830512315998	0.468302965261472
55819	RNF130	ring finger protein 130 12822	0.340108083543733
4.61319650186668	2.87731323696295
0.0898358012177893	0.468302965261472
4214	MAP3K1	mitogen-activated protein kinase kinase kinase 1
7716	-0.35385343031373	4.87142638886329
2.87544680253782	0.0899400093753547	0.468540086511999
3983	ABLIM1	actin binding LIM protein 1	8312
0.302812824148236	7.82625661970203	2.87530038767255
0.0899481896645082	0.468540086511999
54505	DHX29	DExH-box helicase 29	4734	-0.201501539412817
5.85895929617054	2.87171313734411
0.0901488639348894	0.469254718246299
6098	ROS1	ROS proto-oncogene 1, receptor tyrosine kinase	7856
1.19283078795919	2.85944660090098
2.87165028481364	0.090152384293624	0.469254718246299
644596	SMIM10L2B	small integral membrane protein 10 like 2B
2696	0.649503532392354	0.752333765370786
2.86850617903161	0.0903286755896203	0.469943392203314
8440	NCK2	NCK adaptor protein 2	2990	-0.229235734440094
6.43601008116956	2.86752861362378
0.0903835642895883	0.469943392203314
3945	LDHB	lactate dehydrogenase B 2983	0.358933596110655
6.9427569051931 2.86742976460505	0.0903891165150914
0.469943392203314
11060	WWP2	WW domain containing E3 ubiquitin protein ligase 2
9015	-0.195896272645786	6.00913386955574
2.86663424803959	0.0904338131614599	0.469943392203314
10970	CKAP4	cytoskeleton associated protein 4	3638
0.245496086699581	8.29320629895446	2.86630277712001
0.0904524441622142	0.469943392203314
27063	ANKRD1	ankyrin repeat domain 1 1979	0.704204642844467
8.71178369787845	2.86402742508415
0.0905804474693104	0.470046427337167
3916	LAMP1	lysosomal associated membrane protein 1 5739
0.274994573259198	8.84596072632425	2.86396627705813
0.0905838901538182	0.470046427337167
84976	DISP1	dispatched RND transporter family member 1	6338
0.347970260734043	3.27528623392753
2.86339969813676	0.0906157957711915	0.470046427337167
79709	COLGALT1	collagen beta(1-O)galactosyltransferase 1
4900	0.298941082450836	6.38648523699443
2.86337194718524	0.0906173588168993	0.470046427337167
84300	UQCC2	ubiquinol-cytochrome c reductase complex assembly
factor 2	1414	0.367301735415089	4.13080120814108

2.8625715037404 0.0906624556228664 0.470046427337167
3671 ISLR immunoglobulin superfamily containing leucine rich repeat 2671 1.4437811079733 1.34911128566704
2.8623733439232 0.0906736236642412 0.470046427337167
100862671 TMEM265 transmembrane protein 265
995 -0.292852808111314 5.764842425636 2.86001915270107
0.0908064172686417 0.470312846351624
55291 PPP6R3 protein phosphatase 6 regulatory subunit 3
8305 -0.217751540680213 6.99675018662038
2.85873964720878 0.0908786792717388 0.470312846351624
285636 C5orf51 chromosome 5 open reading frame 51
5572 -0.270336191413491 5.47099678243319
2.858230756978 0.0909074369640636 0.470312846351624
285315 C3orf33 chromosome 3 open reading frame 33 2134
0.43807547954576 1.91863879158921 2.85818581213185
0.0909099772990182 0.470312846351624
23063 WAPL WAPL cohesin release factor
7280 -0.235129400027454 6.67410355014028
2.85790585689555 0.0909258024294946 0.470312846351624
54881 TEX10 testis expressed 10 7261 0.264085188237731
4.44743494637441 2.85629279900453
0.0910170426105713 0.470312846351624
3985 LIMK2 LIM domain kinase 2 6170 0.218071064345736
8.48840369529302 2.85621518496166
0.0910214352371606 0.470312846351624
6667 SP1 Sp1 transcription factor
8083 -0.258286196866885 7.12894109209904
2.85602123632709 0.0910324129158264 0.470312846351624
9044 BTAF1 B-TFIID TATA-box binding protein associated factor 1
7582 -0.30945059821806 5.94401713519069
2.855704852123 0.0910503236500944 0.470312846351624
7422 VEGFA vascular endothelial growth factor A
14431 -0.539036537210504 7.68149422347074
2.85552005882067 0.0910607866999075 0.470312846351624
101410538 MMP24OS MMP24 opposite strand 4253
0.318407541888207 5.78986162264522 2.85291848378245
0.0912082272624492 0.470758833776316
22806 IKZF3 IKAROS family zinc finger 3
10008 -1.13889110039712 0.612658581785146
2.8528104537541 0.0912143553129807 0.470758833776316
100289341 MAN1B1-DT MAN1B1 divergent transcript 1872
0.540987969510358 0.170109321141687
2.85197049845547 0.0912620173978491 0.470831336166362
7957 EPM2A EPM2A glucan phosphatase, laforin 5662
0.394564186249724 2.69125439208496 2.85027660250273
0.0913582174243641 0.471041528497395
55626 AMBRA1 autophagy and beclin 1 regulator 1
7430 -0.233532418728903 5.42645191164631
2.85006896217426 0.0913700173432871 0.471041528497395
401138 AMTN amelotin 1037 -0.979971143861751
5.76015921351075 2.84763794478363 0.0915082919604788
0.471580811438715
729082 OIP5-AS1 OIP5 antisense RNA 1
12673 -0.248558948061739 7.5308393748238 2.8464248717812

0.0915773757055109	0.471763259105219
113828	FAM83F	family with sequence similarity 83 member F
16389	-0.282369062781972	5.49619182081813
2.8447891281129 0.0916706199790383	0.471919357492392
6692	SPINT1	serine peptidase inhibitor, Kunitz type 1
3542	-0.24938197292951	9.42169210821407
2.84471127593079	0.0916750604509743	0.471919357492392
11178	LZTS1	leucine zipper tumor suppressor 1	5706
0.746077126099582	2.51151334428143	2.84380911269942
	0.0917265343754533	0.47201086223254
64582	GPR135	G protein-coupled receptor 135	4578
0.521594781075268	-0.0630311934649817	2.84216346605369
	0.0918205094477973	0.472320923920417
196441	ZFC3H1	zinc finger C3H1-type containing
9670	-0.321493026625534	6.50337979822965
2.83985663121845	0.091952417934597	0.472825814247393
1949	EFNB3	ephrin B3	3222	0.887428131095218
2.71132297138972	2.83862376112099	0.0920229998040177
	0.473015103946744
5782	PTPN12	protein tyrosine phosphatase non-receptor type 12
	5547	-0.276772642373411	7.3468165169252
2.83747369173601	0.0920888943919417	0.473180170120296
9654	TTLL4	tubulin tyrosine ligase like 4
7685	-0.2703698589568	4.67038853788108
2.83635950691429	0.0921527818220399	0.473198697573769
5331	PLCB3	phospholipase C beta 3	4964	-0.195089763611908
	6.3578809467186 2.83596457338203	0.0921754388945693
	0.473198697573769
23646	PLD3	phospholipase D family member 3 5184
0.281526655122112	7.59512608043654	2.83564374263491
	0.0921938491953178	0.473198697573769
57132	CHMP1B	charged multivesicular body protein 1B
3054	-0.212932811947543	7.17828995825716
2.83465233588092	0.092250764557452	0.473317384306311
4644	MYO5A	myosin VA	12988	-0.264821410096362
7.19757186920831	2.83366408550857	0.0923075367125694
	0.473435249627594
5646	PRSS3	serine protease 3	1363	-1.31371343107303
	0.888037387996879	2.83150952284191
0.0924314418223194	0.473897220206892
100316904	SAP25	Sin3A associated protein 25
1388	-0.58773178527132	0.109722176352604
2.82903555098504	0.0925739385453903	0.474140032690135
133	ADM	adrenomedullin	2718	-0.382702019023722
4.40385348739384	2.82864746510314	0.0925963133099787
	0.474140032690135
29985	SLC39A3 solute carrier family 39 member 3	4659
0.2898283175967 4.11911100301026	2.82783837107661
0.0926429798461201	0.474140032690135
51172	NAGPA	N-acetylglucosamine-1-phosphodiester alpha-N-
acetylglucosaminidase	5255	0.358232223437347
4.29808800791935	2.8273988787186 0.0926683393875373
0.474140032690135
10487	CAP1	cyclase associated actin cytoskeleton regulatory

protein 1	3814	-0.371679730835676	9.70301158663452
	2.82733780057788	0.0926718643089971
0.474140032690135
55603	TENT5A	terminal nucleotidyltransferase 5A
6053	-0.524151475505987	3.14100277410983
2.82695820766471	0.092693774517057	0.474140032690135
200958	MUC20	mucin 20, cell surface associated
5429	-0.43729600291464	5.84075583416046
2.82657753323406	0.0927157528049032	0.474140032690135
5998	RGS3	regulator of G protein signaling 3	12982
0.453321681928943	4.84760763366039	2.82513046109793
	0.0927993514055977	0.474299131453083
5303	PIN4	peptidylprolyl cis/trans isomerase, NIMA-interacting
4	4011	-0.316621636416914	3.72682519012448
2.82486689016837	0.0928145869431642	0.474299131453083
51302	CYP39A1 cytochrome P450 family 39 subfamily A member 1	3111
	0.485977223838188	1.13754001642371
2.82260522810179	0.0929454325915687	0.474794556641175
1116	CHI3L1	chitinase 3 like 1	3408	0.753738693853778
	2.10020623237422	2.82121556757622
0.0930259291176403	0.474817028713112
9791	PTDSS1	phosphatidylserine synthase 1	6295
0.201113700412577	6.36014427224954	2.8209396973132
0.0930419180038725	0.474817028713112
9540	TP53I3	tumor protein p53 inducible protein 3	2107
0.386214400697751	7.2427360192023 2.82072275398832
0.0930544937015724	0.474817028713112
79026	AHNAK	AHNAK nucleoprotein	19465	-0.428366838977191
	10.9350854097688	2.81986940806178
0.0931039781037362	0.474817028713112
93145	OLFM2	olfactomedin 2	2500	0.509995692227381
3.42456134160182	2.81959000464665	0.0931201865546788
	0.474817028713112
6197	RPS6KA3 ribosomal protein S6 kinase A3
8837	-0.250153413354384	6.68963121775832
2.81902068129406	0.0931532230244615	0.474817028713112
4234	METTL1	methyltransferase like 1	2621
0.416481314030995	2.53374835880479	2.81606532028422
	0.0933249204258932	0.475519156204251
55023	PHIP	pleckstrin homology domain interacting protein
13252	-0.34008994508202	6.10805884820933
2.81367143142988	0.0934642502713144	0.475960554148166
8497	PPFIA4	PTPRF interacting protein alpha 4	9707
0.850303177466074	1.87362227106358	2.8131989214556
0.0934917781601435	0.475960554148166
5600	MAPK11	mitogen-activated protein kinase 11
2688	-0.613604316148644	1.79934837261453
2.81254012183559	0.0935301737868307	0.475960554148166
51228	GLTP	glycolipid transfer protein
2696	-0.348744091724087	5.7867666741554 2.81176100354827
	0.0935756038613635	0.475960554148166
6594	SMARCA1 SWI/SNF related, matrix associated, actin dependent
regulator of chromatin, subfamily a, member 1	4363
0.292996003378	6.71681828626083	2.81066085565735

0.0936397939143784	0.475960554148166
8729	GBF1	golgi brefeldin A resistant guanine nucleotide exchange factor 1	6641	-0.238814165000108
7.08623000932156	2.80970824482355	0.0936954143451622
	0.475960554148166
123099	DEGS2	delta 4-desaturase, sphingolipid 2
4466	-0.855624834655838	0.980443672412573
2.8093012950153 0.0937191860221006	0.475960554148166
79866	BORA	BORA aurora kinase A activator
3189	-0.448518778224914	1.91539472270651
2.80872710724286	0.093752737940566	0.475960554148166
139716	GAB3	GRB2 associated binding protein 3	5272
0.694461341422716	0.286095867440349	2.80706777352794
	0.0938497724080374	0.475960554148166
2359	FPR3	formyl peptide receptor 3	2772
1.66144802150924	2.49724090293176	2.80628132305232
	0.0938958005865225	0.475960554148166
196528	ARID2	AT-rich interaction domain 2
12127	-0.371374599186609	5.66193077531501
2.80583869999568	0.0939217165544507	0.475960554148166
84858	ZNF503	zinc finger protein 503 3769	0.416111634422816
	4.62901796261523	2.8049409752102 0.093974303024197
	0.475960554148166
1636	ACE	angiotensin I converting enzyme 11230
1.06594285568191	0.905777256239044	2.80481089827408
	0.0939819252615017	0.475960554148166
92370	PXYLP1	2-phosphoxylose phosphatase 1
7052	-0.365114513288303	3.04637865021486
2.8046191042975 0.0939931652155837	0.475960554148166
4601	MXI1	MAX interactor 1, dimerization protein	4607
0.283871472530725	4.94960812951768	2.80343355221772
	0.0940626761443886	0.475960554148166
26036	ZNF451	zinc finger protein 451 16416	-0.250394214060194
	5.58597630034288	2.80338084653823
0.0940657676657513	0.475960554148166
5063	PAK3	p21 (RAC1) activated kinase 3	10312
0.6913276121903 1.28085903194197	2.80335175805038
0.0940674739368264	0.475960554148166
91833	WDR20	WD repeat domain 20	7328	-0.237010626791316
	4.80382979312562	2.80321429589393
0.094075537640688	0.475960554148166
89853	MVB12B	multivesicular body subunit 12B 7884
0.397976164720757	4.47228754358823	2.80311427084181
	0.0940814057093486	0.475960554148166
22797	TFEC	transcription factor EC 8163	1.33925219885016
	0.70949484225164	2.80267092063611
0.0941074200821705	0.475960554148166
6520	SLC3A2	solute carrier family 3 member 2	5178
0.372169813705198	8.29164707301574	2.80236912792459
	0.0941251327900636	0.475960554148166
3508	IGHMBP2 immunoglobulin mu DNA binding protein 2
6465	-0.25683402275536	5.01899552794495
2.79919119020477	0.0943118714976549	0.476732726116078
10437	IFI30	IFI30 lysosomal thiol reductase 2481

1.18503668075402	5.22504754674116	2.79489620711958
	0.0945648896913816	0.477839253213269
79644	SRD5A3	steroid 5 alpha-reductase 3
4420	-0.855874741303417	6.31875772877078
2.79333323341677	0.0946571479686897	0.478132950449851
25898	RCHY1	ring finger and CHY zinc finger domain containing 1
	6565	-0.290473421052524	4.65394327205993
2.79072196992481	0.094811502677756	0.478739984862051
196394	AMN1	antagonist of mitotic exit network 1 homolog	3195
	0.489097855946057	2.36360130237076
2.7888741957559 0.0949208921803668	0.479119616854197
152926	PPM1K	protein phosphatase, Mg2+/Mn2+ dependent 1K
9516	-0.334918919612097	3.75344696008363
2.78580422918288	0.0951029401628936	0.479865591808952
2000	ELF4	E74 like ETS transcription factor 4
4316	-0.231265722352251	5.94885509433644
2.782763603554	0.0952836229478109	0.480604143546988
54619	CCNJ	cyclin J	4148	-0.303212386799579
4.18138331257467	2.78129576405813	0.0953709799467351
	0.480871604072685
1522	CTSZ	cathepsin Z	2038	0.319099023783904
7.92648407614694	2.78054985025483	0.0954154056726863
	0.480922485519006
102724105	ZNF528-AS1	ZNF528 antisense RNA 1
1713	-0.696709523650132	0.956777411381545
2.77900931834046	0.0955072292356908	0.480939485505468
23118	TAB2	TGF-beta activated kinase 1 (MAP3K7) binding protein
2	6763	-0.184663984039913	6.96939961824337
2.77891704088422	0.0955127324965156	0.480939485505468
5451	POU2F1	POU class 2 homeobox 1	15753	-0.413204896646981
	4.76818169215544	2.77836443094833
0.0955456963814423	0.480939485505468
100132741	LOC100132741	uncharacterized LOC100132741	4113
	0.714878401581269	-0.0197716600440061
2.7779634074012 0.0955696257025835	0.480939485505468
57414	RHBDD2	rhomboid domain containing 2	2441
0.256613688483604	6.52073575681038	2.77761439586593
	0.0955904567464284	0.480939485505468
56052	ALG1	ALG1 chitobiosyldiphosphodolichol beta-
mannosyltransferase	3540	0.241462926223341
4.28184207873498	2.77545015458388	0.095719741734952
	0.481251624704694
55679	LIMS2	LIM zinc finger domain containing 2	8489
0.58113186934897	4.75686163851087	2.77498691454896
	0.0957474389760894	0.481251624704694
8479	HIRIP3	HIRA interacting protein 3	3931
0.38022117897513	2.59229632537579	2.77485093898627
	0.0957555706469609	0.481251624704694
9958	USP15	ubiquitin specific peptidase 15
19418	-0.24939580390235	6.29019631463316
2.77350018507735	0.0958363898895754	0.481280712999935
6047	RNF4	ring finger protein 4	7270	-0.213598980817412
	6.511888300589	2.77312559791404	0.0958588156036051
	0.481280712999935

283651	HMGN2P46	high mobility group nucleosomal binding domain 2 pseudogene 46	702	-0.606454949704498
1.95635009042275	2.77303169197154	0.0958644384429084
	0.481280712999935
100529251	CKLF-CMTM1	CKLF-CMTM1 readthrough	840
0.356188631792547	3.0429722069278 2.77207900235356
0.0959215032890546	0.48133194686683
56107	PCDHGA9 protocadherin gamma subfamily A, 9	4749
0.236597321141033	6.48641268396074	2.77171326567595
	0.0959434202588148	0.48133194686683
6091	ROBO1	roundabout guidance receptor 1
10875	-0.278926550088766	6.37166632350988
2.77080655436745	0.0959977789551873	0.481403845526947
29956	CERS2	ceramide synthase 2	4644	0.220608254931639
	7.83637472813527	2.77032870973473
0.0960264399516163	0.481403845526947
11169	WDHD1	WD repeat and HMG-box DNA binding protein 1	6385
	0.36501152758917	3.09359329435646
2.76908261276002	0.0961012243786044	0.481559679563929
2288	FKBP4	FKBP prolyl isomerase 4 5180	-0.23754883890691
	6.2392941065091 2.76809465558754	0.0961605616531489
	0.481559679563929
9679	FAM53B	family with sequence similarity 53 member B	8766
	0.303220869877767	5.61873593229013
2.76809294842627	0.0961606642207689	0.481559679563929
10261	IGSF6	immunoglobulin superfamily member 6	3532
1.53139621615235	1.14343294989165	2.76699095188447
	0.0962268979341079	0.481719142016815
3157	HMGCS1	3-hydroxy-3-methylglutaryl-CoA synthase 1
4702	-0.619882418177381	7.56307045335349
2.76630256727856	0.0962682973919036	0.481731759073502
23659	PLA2G15 phospholipase A2 group XV	4464
0.238157146467322	5.20292841399805	2.76580550622146
	0.0962982027133438	0.481731759073502
23054	NCOA6	nuclear receptor coactivator 6
10000	-0.242806986658371	6.59801959636043
2.76431509005093	0.096387933220607	0.482008490046784
23382	AHCYL2	adenosylhomocysteinase like 2
6858	-0.343141879750434	5.6819334666559 2.76328412525159
	0.0964500557568997	0.482147013200176
440275	EIF2AK4 eukaryotic translation initiation factor 2 alpha kinase 4	10713	0.229778985718538	6.55174396013952
	2.7623563519475 0.0965059976088503	0.482198337464483
283299	LOC283299	uncharacterized LOC283299	2869
0.677088175751081	0.593734945620891	2.76197215473522
	0.096529173859528	0.482198337464483
56886	UGGT1	UDP-glucose glycoprotein glucosyltransferase 1
11349	-0.308125843789077	6.82368305728946
2.76118566077766	0.0965766371117516	0.482263442432907
161725	OTUD7A	OTU deubiquitinase 7A	14235	0.71775054872774
	0.565824198130123	2.75899722745687
0.0967088383287404	0.482620264413503
54915	YTHDF1	YTH N6-methyladenosine RNA binding protein 1	3301
	0.284299413478344	6.72181798029535

2.75886215295423	0.096717004512651	0.482620264413503
5351	PLOD1	procollagen-lysine,2-oxoglutarate 5-dioxygenase 1
	5320	0.337612780355426	6.72529070117085
2.75815995005777	0.0967594696231974	0.482660217596911
196074	METTL15 methyltransferase like 15
7778	-0.313662466406881	3.44242524393467
2.75673998207219	0.0968454030568809	0.482916895912328
79071	ELOVL6	ELOVL fatty acid elongase 6
7297	-0.468681181987337	4.81152997019796
2.75445675923774	0.096983753282589	0.483434673391183
6603	SMARCD2 SWI/SNF related, matrix associated, actin dependent regulator of chromatin, subfamily d, member 2
3688	-0.20467866528952	5.90507954481422
2.75359520909285	0.097036014253282	0.483523106241807
10058	ABCB6	ATP binding cassette subfamily B member 6 (Langereis blood group)	7613	0.193183136508525	6.24401876045858
	2.75288507029232	0.0970791138063481
0.483565841780056
3113	HLA-DPA1	major histocompatibility complex, class II, DP alpha 1	5748	1.48972145806556	3.98006960157442
	2.75063780708182	0.0972156416597727
0.483907789879459
84904	ARHGEF39	Rho guanine nucleotide exchange factor 39
	6496	0.675172912460692	0.104611525797725
2.75061781235813	0.0972168573371028	0.483907789879459
84129	ACAD11	acyl-CoA dehydrogenase family member 11 5674
0.303725185038372	3.85996383098356	2.74822641251936
	0.0973623738168184	0.484337064724145
9559	VPS26A	VPS26, retromer complex component A
3565	-0.185874279160655	6.79445250120302
2.748064169462	0.0973722548913539	0.484337064724145
463	ZFHX3	zinc finger homeobox 3	17503	-0.478060968390571
	6.19359582978538	2.74635105590262
0.0974766552408533	0.484684242086842
10015	PDCD6IP programmed cell death 6 interacting protein
9038	-0.221790045482436	8.01003105626673
2.74488868439767	0.0975658714607788	0.484882883672394
55107	ANO1	anoctamin 1	6021	-0.725472179729465
4.7088961063165 2.74456158720647	0.097585839157027
0.484882883672394
158158	RASEF	RAS and EF-hand domain containing
6649	-0.402206642484622	5.75987617730158
2.7367321147951 0.0980651229168122	0.487091552019783
6726	SRP9	signal recognition particle 9
1728	-0.257681339236442	6.88866329184822
2.73605892999686	0.0981064518501847	0.48712409467052
132671	SPATA18 spermatogenesis associated 18	4941
0.299902663409674	4.39652198680342	2.73506524069125
	0.098167492288566	0.487177325647863
3903	LAIR1	leukocyte associated immunoglobulin like receptor 1
	7847	1.49919309893217	1.45186043461476
2.73448076061266	0.0982034151339333	0.487177325647863
23133	PHF8	PHD finger protein 8	9016	-0.267412263936274
	5.24103172218025	2.73418634823757

0.098221515501504	0.487177325647863
55362	TMEM63B transmembrane protein 63B	4425
0.21229986942333	7.98122333292233	2.73333104717057
	0.0982741197522424	0.487265697475985
55697	VAC14	VAC14 component of PIKFYVE complex	5317
0.336196456780438	5.37201104907213	2.7323129622461
0.0983367758943925	0.487336846479567
8934	RAB29	RAB29, member RAS oncogene family	3882
0.402988674908519	5.55454842297547	2.73156423260198
	0.0983828828772295	0.487336846479567
55660	PRPF40A pre-mRNA processing factor 40 homolog A
9536	-0.221365626782663	7.37773765630465
2.73140248292442	0.0983928465655897	0.487336846479567
653489	RGPD3	RANBP2 like and GRIP domain containing 3
5818	-0.417677232899379	3.2526152653616 2.72916131070824
	0.0985310148669936	0.48779489167343
55100	WDR70	WD repeat domain 70	7039	0.265902400569102
	4.35668025344156	2.72877300198853
0.0985549756147502	0.48779489167343
6944	VPS72	vacuolar protein sorting 72 homolog
4545	-0.21938838627237	5.53587281453402
2.7279232682965 0.0986074309598701	0.487882121319286
389136	VGLL3	vestigial like family member 3
11574	-0.441302817101763	6.84864264255794
2.7251272161825 0.0987802506443907	0.488194035832708
6662	SOX9	SRY-box transcription factor 9
3935	-0.366441149338713	5.94723155761134
2.72498405042674	0.0987891083902877	0.488194035832708
678655	CD27-AS1	CD27 antisense RNA 1	3411
0.2986844900304 3.67419769431149	2.72421700721942
0.0988365805504293	0.488194035832708
347240	KIF24	kinesin family member 24	6226
0.590431804039876	1.39049267315958	2.72403633735699
	0.0988477657964242	0.488194035832708
1375	CPT1B	carnitine palmitoyltransferase 1B	4102
0.499204792980632	4.16879277877352	2.72379329304271
	0.0988628148152171	0.488194035832708
10474	TADA3	transcriptional adaptor 3	4234
0.224593994625198	6.2258008877456 2.72345948545828
0.0988834878647137	0.488194035832708
121274	ZNF641	zinc finger protein 641 8474	0.291914386683097
	4.51335869470334	2.72200886889923
0.0989733808660894	0.488194035832708
2769	GNA15	G protein subunit alpha 15
2613	-0.30617293208817	6.79947332917328
2.72125712214816	0.0990200008108438	0.488194035832708
10606	PAICS	phosphoribosylaminoimidazole carboxylase and phosphoribosylaminoimidazolesuccinocarboxamide synthase 7517
0.302611675893866	5.27576825715972	2.72120450332031
	0.0990232648907525	0.488194035832708
10990	LILRB5	leukocyte immunoglobulin like receptor B5	7017
	1.62110801914483	0.624244320279641
2.72114771394876	0.0990267878118773	0.488194035832708
50515	CHST11	carbohydrate sulfotransferase 11

6098    -0.462535122414902      4.60373019168834
2.72071134698696        0.0990538623428682      0.488194035832708
6675    UAP1    UDP-N-acetylglucosamine pyrophosphorylase 1
3437    -0.385397601741888      6.01163999304286
2.71926133261655        0.0991438870053789      0.488294063885929
83990   BRIP1   BRCA1 interacting protein C-terminal helicase 1
7991    -0.590744209990714      2.076719343226  2.71913877903585
        0.0991514998804098      0.488294063885929
5468    PPARG   peroxisome proliferator activated receptor gamma
        4046    0.764509835533423       2.95553985845639
2.71870034560877        0.0991787400410843      0.488294063885929
55728   N4BP2   NEDD4 binding protein 2 9835    -0.385809221831354
        4.7959732226261 2.71627612260648        0.0993295061692894
        0.488864509105143
80233   FAAP100 FA core complex associated protein 100  5354
0.236396561386215       5.27337948455324        2.71517584237545
        0.0993980168069244      0.489029863510569
9232    PTTG1   PTTG1 regulator of sister chromatid separation,
securin 2013    0.429351477453613       2.9077313334237
2.71435410652989        0.0994492171017453      0.489090360352613
2329    FMO4    flavin containing dimethylaniline monoxygenase 4
        2517    0.49745611949288        1.37762667998533
2.71385790269663        0.099480148257628       0.489090360352613
7644    ZNF91   zinc finger protein 91  5990    -0.415306384643323
        4.82884265015523        2.71303963860854
0.0995311781716156      0.489119723948126
55357   TBC1D2  TBC1 domain family member 2     3681
0.307021131933784       6.75607276328407        2.71264242605321
        0.099555960089677       0.489119723948126
9819    TSC22D2 TSC22 domain family member 2
11676   -0.266901005956043      5.26963363961739
2.71012248940762        0.0997133348707117      0.489549393129574
1120    CHKB    choline kinase beta     3021    0.254588228371442
        5.38535126400182        2.71009127713397
0.0997152858381881      0.489549393129574
26051   PPP1R16B        protein phosphatase 1 regulatory subunit 16B
        6400    -1.35948841606444       0.925607099579353
2.70956374263384        0.0997482664376243      0.489549393129574
5441    POLR2L  RNA polymerase II subunit L     1142
0.347964534913377       5.63249479789821        2.70797788684087
        0.0998474833125235      0.489672539303419
221927  BRAT1   BRCA1 associated ATM activator 1        6639
0.33001278798367        4.55946181059476        2.70728727310775
        0.0998907242868206      0.489672539303419
118812  MORN4   MORN repeat containing 4        2512
0.347415388359607       3.67694799601528        2.70697239934099
        0.099910446072223       0.489672539303419
6195    RPS6KA1 ribosomal protein S6 kinase A1  5884
0.221696142752801       6.30599066426082        2.7063834096757
0.0999473482346196      0.489672539303419
25981   DNAH1   dynein axonemal heavy chain 1
13872   -0.490739243217231      2.55338176621046
2.70629142464306        0.0999531127467644      0.489672539303419
9107    MTMR6   myotubularin related protein 6

5779	-0.222246692847938	6.2339848979915 2.7058128019883
0.0999831129012479	0.489672539303419
199870	FAM76A	family with sequence similarity 76 member A	4034
	0.327231711681538	3.25979675175209
2.70472565497422	0.100051292016221	0.489680844127025
401934	RNF223	ring finger protein 223 1902	-0.569249232106858
	3.51973846115835	2.70467088716151
0.100054728056796	0.489680844127025
196403	DTX3	deltex E3 ubiquitin ligase 3	4087
0.63348050117671	3.81374288472332	2.70051375972655
	0.100315915825883	0.49061772100408
55653	BCAS4	breast carcinoma amplified sequence 4	7583
0.348311161065597	4.36625925244376	2.70050908483225
	0.100316209963282	0.49061772100408
550112	UBA6-AS1	UBA6 antisense RNA 1 (head to head)	4986
	0.374003404685013	3.63303250407655
2.69984303585532	0.100358126405435	0.490651405431391
85865	GTPBP10 GTP binding protein 10	7997	-0.290076392222262
	4.0476794862414 2.69849298825725	0.100443147622627
	0.490731722396677
80852	GRIP2	glutamate receptor interacting protein 2
11704	-1.77546160649956	2.41958754136427
2.69806881911593	0.100469876536042	0.490731722396677
51065	RPS27L	ribosomal protein S27 like	9127
0.206795188318654	6.37170860493955	2.69791361335903
	0.100479658730183	0.490731722396677
5576	PRKAR2A protein kinase cAMP-dependent type II regulatory subunit alpha	6472	-0.274106825900069	6.20601263497685
	2.69319908089572	0.100777299269036
0.492013813475564
9214	FCMR	Fc fragment of IgM receptor
2348	-0.543700678916563	0.169544377874486
2.69214914474452	0.100843715513438	0.492113984420252
399473	SPRED3	sprouty related EVH1 domain containing 3
5864	-0.380109055424954	3.36180726930182
2.69176411059221	0.100868083763157	0.492113984420252
10507	SEMA4D	semaphorin 4D	14160	-0.46260843906403
3.61206441328945	2.69045905066599	0.100950726982997
	0.492345693889569
55636	CHD7	chromodomain helicase DNA binding protein 7
13860	-0.438330607502907	4.92440455746988
2.6871874795897 0.101158225428875	0.493185960174818
83591	THAP2	THAP domain containing 2
6527	-0.423020883163019	3.94643478658547
2.68639803763267	0.101208365339702	0.493187006860166
339263	LINC02693	long intergenic non-protein coding RNA 2693
	8796	0.530737724559435	2.48757339240202
2.68607547371119	0.101228860192973	0.493187006860166
10567	RABAC1	Rab acceptor 1	1916	0.246821104946836
5.73327472993923	2.68515618669672	0.101287294135315
	0.493300114378775
915	CD3D	CD3d molecule	929	-1.53516937035609
0.0636803374792238	2.6838622425061 0.101369605425393
0.493529392837497

730091	LINC00886	long intergenic non-protein coding RNA 886
	3822	0.483354614416108	1.81220014521356
2.68305900005451	0.10142073863355	0.493606770688023
92714	ARRDC1	arrestin domain containing 1
3821	-0.214646439558262	5.84757644803784
2.68072563844651	0.101569436925898	0.494158771455732
8996	NOL3	nucleolar protein 3	2402	0.27748780312159
	3.85395120567258	2.67910813457255
0.101672655582026	0.494489196783834
57591	MRTFA	myocardin related transcription factor A	5617
	0.207668202652051	6.79578282660785
2.67844912777062	0.101714742019523	0.494522176837019
10069	RWDD2B	RWD domain containing 2B	3395
0.273069348661783	4.2786921074554 2.67714079780184
0.101798352924469	0.494756949831034
79637	ARMC7	armadillo repeat containing 7	2823
0.31344543567444	4.22347435633562	2.6764102145249
0.101845074707407	0.494812334868766
152816	ODAPH	odontogenesis associated phosphoprotein
2054	-0.85391473024482	2.03493177947532
2.67576795805082	0.101886167240685	0.494840341380121
23467	NPTXR	neuronal pentraxin receptor	5784
0.66783656531073	3.69531480183008	2.67417919025121
	0.101987896911804	0.495162728611313
6586	SLIT3	slit guidance ligand 3	14260	0.810676205714836
	1.73890538105983	2.67312460994433
0.102055483600433	0.495319181840355
51084	CRYL1	crystallin lambda 1	2196	-0.371540722030382
	3.31476078154989	2.6702423212357 0.102240455788153
	0.496045051688484
7159	TP53BP2 tumor protein p53 binding protein 2
5991	-0.273000399534451	6.59547077202745
2.6691157072046 0.102312856406947	0.496068687359165
5522	PPP2R2C protein phosphatase 2 regulatory subunit Bgamma
9198	-1.28150438506976	2.23901561220579
2.66906433837939	0.102316158904878	0.496068687359165
728769	SCAMP1-AS1	SCAMP1 antisense RNA 1	1792
0.488243874072238	1.5640191834416 2.66752971913534
0.102414873258186	0.496215320757278
7157	TP53	tumor protein p53	4576	0.258602057080817
	6.19197331516722	2.66694577160197
0.102452463032294	0.496215320757278
23143	LRCH1	leucine rich repeats and calponin homology domain
containing 1	7307	-0.260130395221765	4.92706472980202
	2.66694238082051	0.102452681347187
0.496215320757278
253970	SFTA3	surfactant associated 3 3243	0.445577145481477
	3.57838680240223	2.66232019879522
0.102750753514776	0.497486970093837
5287	PIK3C2B phosphatidylinositol-4-phosphate 3-kinase catalytic
subunit type 2 beta	9414	-0.400058383179344
4.34736582436492	2.66087083962369	0.102844414119539
	0.497768385823215
814	CAMK4	calcium/calmodulin dependent protein kinase IV

13910	-0.563399534144587	0.963086647006916
2.65725509569279	0.103078478518405	0.498728928707186
6092	ROBO2	roundabout guidance receptor 2
16663	-0.67912724977094	1.248359805683	2.65646131364898
	0.103129941829558	0.498805626797866
4212	MEIS2	Meis homeobox 2 10798	-0.371117464023127
3.10492224563886	2.65519348283932	0.103212197503998
	0.499031153068174
7026	NR2F2	nuclear receptor subfamily 2 group F member 2	6459
	0.409222631052543	5.77733110428874
2.65174949845004	0.103436002682373	0.499940679631471
51780	KDM3B	lysine demethylase 3B	7961	-0.223359553362676
	6.84471474244594	2.65035566289659
0.103526731175695	0.500206596611922
55907	CMAS	cytidine monophosphate N-acetylneuraminic acid
synthetase	2018	-0.22648885410429	5.57804379956671
	2.64746768407803	0.103714994859237
0.500943425170114
3371	TNC	tenascin C	9636	0.330802270366521
10.9054440844654	2.64657848450997	0.10377303610066
	0.500998466859957
90355	C5orf30 chromosome 5 open reading frame 30
3568	-0.423839701235463	3.42172755033963
2.64619733327269	0.103797926096066	0.500998466859957
137075	CLDN23	claudin 23	2169	-0.321803855227495
4.03222975782017	2.64239642836679	0.104046491301082
	0.501816803131872
64132	XYLT2	xylosyltransferase 2	3667	0.287715248674912
	5.14050659546351	2.64059617011281
0.104164448994607	0.501816803131872
23527	ACAP2	ArfGAP with coiled-coil, ankyrin repeat and PH
domains 2	11139	-0.269897604352805	5.73151671570634
	2.64059098890989	0.104164788692049
0.501816803131872
163702	IFNLR1	interferon lambda receptor 1
4923	-0.319531131544075	5.52456172498156
2.64039329184655	0.104177751296288	0.501816803131872
9821	RB1CC1	RB1 inducible coiled-coil 1
7615	-0.273717315250953	7.11729138396969
2.64013051236998	0.104194983960188	0.501816803131872
5547	PRCP	prolylcarboxypeptidase	6482	0.239955237885066
	4.87827530481492	2.63990905935884
0.104209508923789	0.501816803131872
284656	EPHA10	EPH receptor A10	8022	0.832501977086034
	0.630505371481584	2.63888088049586
0.104276975585251	0.501816803131872
144348	ZNF664	zinc finger protein 664 5781	-0.23090800551531
	8.1503686184446 2.63881225560937	0.104281480290544
	0.501816803131872
55199	FAM86C1 family with sequence similarity 86 member C1	2972
	0.462068112436107	1.99848351022643
2.63868390605671	0.104289906041042	0.501816803131872
11012	KLK11	kallikrein related peptidase 11
3113	-0.505073681124368	5.05979772801343

2.63755207752361	0.104364239367059	0.502002026378569
64750	SMURF2	SMAD specific E3 ubiquitin protein ligase 2
6961	-0.465382648128476	7.45708411802569
2.63666576170428	0.104422489056548	0.502109785175103
6674	SPAG1	sperm associated antigen 1
5055	-0.305097111869081	5.72971641507071
2.63546410257814	0.10450152035011	0.502187985823632
7087	ICAM5	intercellular adhesion molecule 5	3111
0.665241066779874	2.69068726898053	2.63532827667973
0.104510457533797	0.502187985823632
55159	RFWD3	ring finger and WD repeat domain 3
6465	-0.270263754285975	5.56939904565123
2.63357895331755	0.104625635811541	0.502433039863742
84059	ADGRV1	adhesion G protein-coupled receptor V1
23499	-0.882118568551707	0.740118910479857
2.63346420226571	0.104633196065006	0.502433039863742
3713	IVL	involucrin	2153	-0.651789501244614
8.40254385370957	2.62982877901095	0.104873022266155
0.503412070898571
9200	HACD1	3-hydroxyacyl-CoA dehydratase 1	3857
0.493674709570628	4.37461563194063	2.62905015656018
0.104924465691644	0.503486464865658
23389	MED13L	mediator complex subunit 13L
15909	-0.328860328529725	6.70058174086023
2.62752195036791	0.105025514376113	0.503583273768395
6137	RPL13	ribosomal protein L13	5444	0.284061188778552
7.19504341996431	2.62746780470437
0.105029096573756	0.503583273768395
101928649	CTC-338M12.4	uncharacterized LOC101928649	1844
0.444839359558761	1.52652548257135
2.62711415165979	0.105052497033715	0.503583273768395
64110	MAGEF1	MAGE family member F1	1636	0.294998729621019
4.27819601680396	2.62545965950624
0.105162047130406	0.503722926071585
130574	LYPD6	LY6/PLAUR domain containing 6
5033	-0.513210004624174	0.269758568944149
2.62506796119131	0.105188001261954	0.503722926071585
200407	CREG2	cellular repressor of E1A stimulated genes 2
7150	-0.573792701218603	0.842675570300518
2.62490901032956	0.105198535430051	0.503722926071585
2644	GCHFR	GTP cyclohydrolase I feedback regulator	1531
0.681430324018386	1.94287185436113	2.62450253898741
0.105225478809556	0.503722926071585
30008	EFEMP2	EGF containing fibulin extracellular matrix protein 2	5397	0.428191199442655	5.23116740193328
2.62313669248309	0.105316070800085	0.503984353842428
587	BCAT2	branched chain amino acid transaminase 2	4553
0.333080712751366	3.41508326412525
2.62205201963039	0.105388074391457	0.5041566796452
11031	RAB31	RAB31, member RAS oncogene family
5586	-0.376111928221658	6.70142306557069
2.62126061108836	0.10544064433265	0.504235952600693
151195	CCNYL1	cyclin Y like 1	4707	-0.285673862704858
5.35255011217924	2.61832045922374	0.105636198170692

0.504998712551836
51148 CERCAM cerebral endothelial cell adhesion molecule 6059
0.241470730698701 7.17017486353531
2.61728873660637 0.105704913774779 0.505154802880699
57409 MIF4GD MIF4G domain containing 2283 0.241623396669122
4.11093207166515 2.6164284562677 0.10576224831293
0.505195365078792
79651 RHBDF2 rhomboid 5 homolog 2 6924 -0.332850922663523
5.96801259051916 2.61542975213778
0.10582885106106 0.505195365078792
57695 USP37 ubiquitin specific peptidase 37
9290 -0.367786828710137 4.31731573827536
2.61529800106831 0.105837640863315 0.505195365078792
293 SLC25A6 solute carrier family 25 member 6 4328
0.240942704849882 7.2231157667772 2.61499781378942
0.105857670914989 0.505195365078792
286676 ILDR1 immunoglobulin like domain containing receptor 1
2953 -0.401518045882861 4.07047745448228
2.61315616396518 0.10598064634864 5 0.505609984129929
79783 SUGCT succinyl-CoA:glutarate-CoA transferase 2857
0.792516561840529 3.24431807558885 2.61244657254108
0.106028070833629 0.505664006866408
8650 NUMB NUMB endocytic adaptor protein
9546 -0.190489231449936 7.32818901716468
2.61076528177365 0.10614053031538 0.506005280805167
29994 BAZ2B bromodomain adjacent to zinc finger domain 2B
23094 -0.35878917089255 6.9668350115942 2.60985434113314
0.106201516652576 0.506005280805167
9037 SEMA5A semaphorin 5A 12097 -0.521051141554576
3.16468012292852 2.60932363467354 0.106237064502511
0.506005280805167
55209 SETD5 SET domain containing 5 10808 -0.260440263980994
7.02166632296611 2.60897070495432
0.106260711712087 0.506005280805167
4291 MLF1 myeloid leukemia factor 1 3569
0.432105088665306 2.4697258577955 2.60816110482584
0.106314978840969 0.506005280805167
112487 DTD2 D-aminoacyl-tRNA deacylase 2 3302
0.359205166792335 3.75970929091535 2.60814020003857
0.106316380481874 0.506005280805167
26517 TIMM13 translocase of inner mitochondrial membrane 13 1973
0.249294037531308 5.35521910278243
2.60749788443044 0.106359456852934 0.506038353308101
3694 ITGB6 integrin subunit beta 6 5281 -0.222620392280804
10.1715939489955 2.60446468032046
0.106563135031282 0.506791257914047
53826 FXYD6 FXYD domain containing ion transport regulator 6
4339 1.22990208063115 0.263220096902482
2.60337547963722 0.106636378701144 0.506791257914047
1318 SLC31A2 solute carrier family 31 member 2
2504 -0.239962933064731 6.34117260783069
2.60314567098446 0.10665183931634 0.506791257914047
27131 SNX5 sorting nexin 5 5281 0.249309254419455
5.36100330136735 2.60224152834317 0.106712690323357

0.506791257914047
2870 GRK6 G protein-coupled receptor kinase 6
4355 -0.264828252376434 5.07698942771202
2.60214431213024 0.106719235479421 0.506791257914047
9635 CLCA2 chloride channel accessory 2 4496
0.714094115391179 3.53849206013291 2.60179140400743
0.106742998990941 0.506791257914047
90865 IL33 interleukin 33 3253 1.39070901036241
0.797734131193128 2.60137609460357 0.106770971807264
0.506791257914047
23392 ECPAS Ecm29 proteasome adaptor and scaffold
8449 -0.238663887892454 7.20176953215331
2.60014787738197 0.106853744421057 0.507012329981621
8804 CREG1 cellular repressor of E1A stimulated genes 1 2688
0.367784615906205 5.13197704922979
2.59849540859607 0.106965219508906 0.507369397108448
374654 KIF7 kinesin family member 7 4654 0.52769689097599
1.48191091217521 2.59784384430626
0.107009208928053 0.507406225272594
441518 RTL8B retrotransposon Gag like 8B 2026
0.369883447085624 3.67346729846441 2.59652910598845
0.107098032034951 0.50765554474232
8398 PLA2G6 phospholipase A2 group VI 6758
0.46559604140325 2.65758687194778 2.5955917216983
0.107161410686052 0.507741900698727
58486 ZBED5 zinc finger BED-type containing 5
8096 -0.215601415659414 5.74091747123567
2.59518757703775 0.107188748509041 0.507741900698727
56898 BDH2 3-hydroxybutyrate dehydrogenase 2 3564
0.293053338795539 5.2201578963628 2.59405200103104
0.107265603998908 0.507934183641886
154043 CNKSR3 CNKSR family member 3 22895 -0.450185995455717
2.74458735555795 2.59216097914861
0.107393722119795 0.508164249259602
79729 SH3D21 SH3 domain containing 21
4474 -0.44791860796847 3.82054745375417
2.59201091574101 0.107403896216459 0.508164249259602
727800 RNF208 ring finger protein 208 1077 -0.473507409650305
1.6641836334154 2.59145954133775 0.107441287743303
0.508164249259602
132946 ARL9 ADP ribosylation factor like GTPase 9 1258
0.792175035326419 0.344538618928205 2.5907453348863
0.107489742989316 0.508164249259602
1138 CHRNA5 cholinergic receptor nicotinic alpha 5 subunit 3623
0.703951311455899 1.72606203150622
2.59065923834343 0.107495585818248 0.508164249259602
9943 OXSR1 oxidative stress responsive kinase 1
6075 -0.267881024410703 7.19118060628725
2.58952708324655 0.10757245048144 0.50835604382373
79600 TCTN1 tectonic family member 1 9605
0.324342128577598 3.61421644721919 2.58497260525355
0.107882274820656 0.509648237238761
25999 CLIP3 CAP-Gly domain containing linker protein 3 3772
0.8929412926218 1.87664681343195 2.58367398536408

0.107970794542072	0.509694119565606
9401	RECQL4	RecQ like helicase 4	4236	0.49912844240252
	2.10950533010169	2.583614562868	0.107974846952009
	0.509694119565606
222663	SCUBE3	signal peptide, CUB domain and EGF like domain containing 3	7356	0.543880042505006	1.37671245756499
	2.58288240558474	0.108024791269824
0.509694119565606
64928	MRPL14	mitochondrial ribosomal protein L14	959
0.272544043054458	5.8823284609816	2.58257844590938
0.108045533417086	0.509694119565606
2697	GJA1	gap junction protein alpha 1	3037
0.663564004082652	7.87098330183267	2.58188940472098
	0.108092569645445	0.509694119565606
26958	COPG2	coatomer protein complex subunit gamma 2	4139
	0.245507148995892	5.18360538695914
2.58162948657661	0.108110318357208	0.509694119565606
23099	ZBTB43	zinc finger and BTB domain containing 43
5974	-0.249731292016739	5.82497692513163
2.5807800700703	0.108168343743867	0.509796093815727
100507602	TRIM52-AS1	TRIM52 antisense RNA 1 (head to head)	1316	0.50613284721868	0.862030106410653
2.57940702241416	0.108262211789274	0.51006686866208
6232	RPS27	ribosomal protein S27	946	0.339506450139327
	4.4327820158318	2.57859448748265	0.108317802660475
	0.510157182873328
10096	ACTR3	actin related protein 3	12049	-0.323427917676863
	8.56676786431911	2.57742025457119
0.108398195073268	0.510364207862609
4924	NUCB1	nucleobindin 1	3937	0.221350642369586
6.91251595815038	2.5761150407173	0.108487631850828
0.510521483703991
23272	TASOR	transcription activation suppressor
9824	-0.211356524118622	6.38230792058542
2.57556066594578	0.108525643628774	0.510521483703991
5578	PRKCA	protein kinase C alpha	11152	0.412070128770514
	2.67592953350137	2.57525805987983
0.108546398570754	0.510521483703991
7018	TF	transferrin
8565	-0.979097641700454	-0.187546785907805
2.57480632645279	0.10857738987322	0.510521483703991
64768	IPPK	inositol-pentakisphosphate 2-kinase
5166	-0.249871234691505	5.10264003143713
2.57403728298519	0.108630172684163	0.510598264693648
11120	BTN2A1	butyrophilin subfamily 2 member A1
3787	-0.215893945451371	6.33063971901594
2.57269495115783	0.108722370329616	0.510860194970457
55089	SLC38A4	solute carrier family 38 member 4	4145
0.520013598697897	2.92979812135898	2.5712178598736
	0.108823923241456	0.51116589297219
282991	BLOC1S2	biogenesis of lysosomal organelles complex 1 subunit 2	2943	0.228098085221928	6.07877759807255
2.57038983381672	0.108880897233514	0.511262060861494
79088	ZNF426	zinc finger protein 426	7527	-0.280273306989745

5.20971039195094	2.56981139277748
0.108920717471122	0.511277643973864
147339	C18orf25	chromosome 18 open reading frame 25
9576	-0.366366146170985	7.56353939546925
2.56913082383346	0.10896758874603	0.511326303372074
349136	WDR86	WD repeat domain 86	4044
0.801799739850347	-0.0065376492466463	2.56819703151438
0.109031935766736	0.511456906994195
9805	SCRN1	secernin 1	6046	-0.30192199875649
7.46041023505154	2.56680270403035	0.109128095698407
0.511736604100295
211	ALAS1	5'-aminolevulinate synthase 1
2536	-0.25128223748748	5.52362668641707
2.56522156859839	0.109237251633236	0.511867677207616
8742	TNFSF12 TNF superfamily member 12	1879
0.381151435543579	3.03425391576278	2.56459366321797
0.109280633256513	0.511867677207616
2043	EPHA4	EPH receptor A4 7554	-0.485278215869619
6.29828602850995	2.56438753186336	0.109294878881592
0.511867677207616
146439	BICDL2	BICD family like cargo adaptor 2
4690	-0.345204230029506	5.2693840303116 2.56376334194126
0.109338028754049	0.511867677207616
57610	RANBP10 RAN binding protein 10	6460	-0.201937274349411
5.31572464028633	2.56333559747591
0.10936760928199	0.511867677207616
84987	COX14	cytochrome c oxidase assembly factor COX14	2085
0.324366507182288	4.40381941146397
2.56300170734187	0.109390705456959	0.511867677207616
5272	SERPINB9	serpin family B member 9
4118	-0.78729088125967	4.04835930598225
2.56269605496554	0.109411853042022	0.511867677207616
7134	TNNC1	troponin C1, slow skeletal and cardiac type	1168
0.721882939270138	1.83667670623586
2.56206377483061	0.109455613727638	0.51190142954358
55839	CENPN	centromere protein N	3568	0.378356617513243
3.08295773414357	2.56139641500126
0.109501823180298	0.511946607906019
10964	IFI44L	interferon induced protein 44 like
10240	-1.21830881266978	0.616754699355139
2.55910980885698	0.10966031546802	0.512181848029569
56106	PCDHGA10	protocadherin gamma subfamily A, 10	6643
0.221494519591435	6.51314844284219
2.55869656904584	0.109688985391039	0.512181848029569
5452	POU2F2	POU class 2 homeobox 2	8903	-0.568610460445901
1.75328220866205	2.55855414238416
0.109698868636343	0.512181848029569
55738	ARFGAP1 ADP ribosylation factor GTPase activating protein 1
6613	0.196152963108248	6.764113125873
2.55820792513416	0.109722897363898	0.512181848029569
22876	INPP5F	inositol polyphosphate-5-phosphatase F
6332	-0.268440249824351	5.62488018290556
2.5580340071722 0.109734970081869	0.512181848029569
339745	SPOPL	speckle type BTB/POZ protein like

6468	-0.297088873980524	5.58398126289025
2.55743470903499	0.109776582247609	0.512205392252584
51700	CYB5R2	cytochrome b5 reductase 2
8596	-0.390209422352985	3.22812914365353
2.55657836313354	0.109836072589286	0.512243816780464
126298	IRGQ	immunity related GTPase Q
9869	-0.17733479164339	6.40335071823239
2.55626344928701	0.109857958564183	0.512243816780464
10983	CCNI	cyclin I	3741	-0.188169393370278
7.95852253926155	2.55564617146634	0.109900872230673
	0.512273383472557
5770	PTPN1	protein tyrosine phosphatase non-receptor type 1
	4041	0.216438379106609	6.5064633861005
2.55481260086397	0.109958852002899	0.512373133734067
79369	B3GNT4	UDP-GlcNAc:betaGal beta-1,3-N-
acetylglucosaminyltransferase 4 3057	0.515967160941148
1.44152274981353	2.55358772667626	0.110044110285679
	0.512430912665385
2653	GCSH	glycine cleavage system protein H	1880
0.47630508898182	0.013262667374013	2.55358328481254
	0.110044419597164	0.512430912665385
3481	IGF2	insulin like growth factor 2	6259
1.33221674533609	4.73066386363937	2.55181111127433
	0.110167902133345	0.51266833659169
51530	ZC3HC1	zinc finger C3HC-type containing 1	3011
0.282628644784969	3.6744181439701 2.55180098632295
0.110168608063182	0.51266833659169
79977	GRHL2	grainyhead like transcription factor 2
6060	-0.2808008821033	6.15955734791202
2.55088473419936	0.110232511408408	0.512795346163258
118672	PSTK	phosphoseryl-tRNA kinase	6118
0.402660697776831	1.82740188934241	2.54583070785772
	0.110585734252347	0.514267722334868
55132	LARP1B	La ribonucleoprotein 1B 8792	-0.386002949531173
	4.21908883821604	2.54519128610301
0.110630511696339	0.514305203229543
202	CRYBG1	crystallin beta-gamma domain containing 1
10374	-0.250712153537649	7.40126753906457
2.54447535803291	0.110680670394444	0.514367667622753
401264	TRAM2-AS1	TRAM2 antisense RNA 1 (head to head)	2546
	0.454043193708913	1.38066144526543
2.54231230308642	0.110832368412026	0.514901819020645
126133	ALDH16A1	aldehyde dehydrogenase 16 family member A1
	4643	0.321921314182497	4.55530154752454
2.54041613954625	0.110965537127139	0.515129723110132
58515	SELENOK selenoprotein K 2165	-0.292202165366794
5.39106681476365	2.54038350364485	0.110967830706802
	0.515129723110132
6795	AURKC	aurora kinase C 1948	0.548644890492835
0.019085288623853	2.54004313137702	0.110991754433239
	0.515129723110132
81846	SBF2	SET binding factor 2	15946	-0.234098833270247
	5.27835341418733	2.5380485390972 0.111132062348535
	0.515610068670396

27165	GLS2	glutaminase 2	4141	0.565523103457851
1.21589582022803	2.53283084389475	0.111500020707502
	0.517145956970193
11275	KLHL2	kelch like family member 2
3872	-0.253856485500343	5.01622636528866
2.53132827308411	0.111606232386235	0.517467228412443
66000	TMEM108 transmembrane protein 108	8425
0.703456391864093	0.218617604099376	2.52988565038595
	0.111708311332729	0.517678446864121
26019	UPF2	UPF2 regulator of nonsense mediated mRNA decay
5906	-0.238892033189759	5.77171845694527
2.52963998555838	0.111725704638412	0.517678446864121
93380	MMGT1	membrane magnesium transporter 1
3738	-0.211329251380878	5.59780849387073
2.52892057451023	0.111776656782122	0.517703581487733
83692	CD99L2	CD99 molecule like 2	5762	0.305247247642622
	6.43950376461984	2.52822391105233
0.111826022213075	0.517703581487733
284338	PRR19	proline rich 19 2575	0.520616127052822
1.90876431586622	2.52795375292389	0.111845172021152
	0.517703581487733
4048	LTA4H	leukotriene A4 hydrolase	7350
0.326961838904945	6.70368556954501	2.5274770598331
0.111878970597799	0.517703581487733
5706	PSMC6	proteasome 26S subunit, ATPase 6
7482	-0.217115946076174	6.25320275671481
2.52497780807185	0.112056357110436	0.518353168443156
9136	RRP9	ribosomal RNA processing 9, U3 small nucleolar RNA
binding protein 1578	0.280615268409471	3.78241770895053
	2.52410200281284	0.112118591384384
0.518469828168063
54989	ZNF770	zinc finger protein 770 5439	-0.240945279084018
	6.17565359578477	2.52265504844074
0.112221494621637	0.518618800538093
8178	ELL	elongation factor for RNA polymerase II
4929	-0.276788208587394	4.33561632568301
2.52259059977325	0.112226080444358	0.518618800538093
58508	KMT2C	lysine methyltransferase 2C
20114	-0.456231517139007	6.76145878623845
2.52182964486887	0.112280241533719	0.518618800538093
81559	TRIM11	tripartite motif containing 11
8493	-0.220077590535458	5.53906785208625
2.52042011121577	0.112380641379356	0.518618800538093
65249	ZSWIM4	zinc finger SWIM-type containing 4
4713	-0.340020057197644	6.15819189838171
2.51959928706417	0.112439153513639	0.518618800538093
2564	GABRE	gamma-aminobutyric acid type A receptor epsilon
subunit 9612	-0.582431954854701	3.40807014210758
2.5195875455426 0.112439990746865	0.518618800538093
130367	SGPP2	sphingosine-1-phosphate phosphatase 2
3336	-0.343067812949875	7.46839496402491
2.51949292009692	0.112446738297389	0.518618800538093
26872	STEAP1	STEAP family member 1	1625	-0.486327226744358
	2.30414553781465	2.51948908176246

0.112447012010722	0.518618800538093
58517	RBM25	RNA binding motif protein 25
11859	-0.203445226910978	6.89664699581225
2.51822566336713	0.112537146823489	0.518671657708095
9422	ZNF264	zinc finger protein 264 12532	-0.371458316353846
	5.1939331090317 2.5180707406294 0.112548204799707
0.518671657708095
23271	CAMSAP2 calmodulin regulated spectrin associated protein family member 2 7777	-0.236846692524287	7.49644369618622
	2.51753402350766	0.112586523507763
0.518671657708095
4793	NFKBIB	NFKB inhibitor beta	2324	-0.251947773434381
	4.95805679554073	2.51725303427776
0.112606590354131	0.518671657708095
8775	NAPA	NSF attachment protein alpha	5839
0.204154686268274	6.2309280300791 2.51655210306557
0.112656664531084	0.518695793741578
1191	CLU	clusterin	6616	-0.622601284899297
4.49543652902525	2.51548455423414	0.112732976821491
	0.518695793741578
8916	HERC3	HECT and RLD domain containing E3 ubiquitin protein ligase 3	7104	-0.271794458860478	5.27255408045662
	2.51515029726396	0.112756882435359
0.518695793741578
55032	SLC35A5 solute carrier family 35 member A5	4853
0.206790011404049	5.49287083369305	2.5151073389578
0.112759955161213	0.518695793741578
9201	DCLK1	doublecortin like kinase 1
14375	-0.658027666024077	4.63788137256327
2.51410486268495	0.112831686579706	0.518855362416923
23140	ZZEF1	zinc finger ZZ-type and EF-hand domain containing 1
	15878	-0.311887728864709	6.43145770194174
2.5131976133428 0.112896647444075	0.518983702247836
109	ADCY3	adenylate cyclase 3	6058	0.397775293607701
	3.92448867517466	2.51112013840234
0.113045554056708	0.519497728238946
4883	NPR3	natriuretic peptide receptor 3	9866
0.593166210909923	2.19400715794631	2.50902380556563
	0.113196031726607	0.519670038597326
9767	JADE3	jade family PHD finger 3
5137	-0.309583481905217	4.12425608653092
2.50836423688262	0.113243422114616	0.519670038597326
3257	HPS1	HPS1 biogenesis of lysosomal organelles complex 3 subunit 1	4193	0.251350823033679	5.74751726688207
	2.50815619338995	0.113258374694192
0.519670038597326
3071	NCKAP1L NCK associated protein 1 like	10596
1.44187288376406	2.36954291647124	2.50701888165898
	0.113340154454116	0.519670038597326
7080	NKX2-1	NK2 homeobox 1	3165	0.286795654916028
5.19804083779522	2.50687253043335	0.113350682738867
	0.519670038597326
56997	COQ8A	coenzyme Q8A	7499	0.453727147901163
2.87312202814812	2.50613704395121	0.1134036088395

0.519670038597326
90649	ZNF486	zinc finger protein 486	4261	0.635090307437778
	4.03410428141021	2.506081366986	0.113407616500015
	0.519670038597326
84288	EFCAB2	EF-hand calcium binding domain 2	8607
0.560899145003711	2.59188550269036	2.50547947634373
	0.113450950906226	0.519670038597326
54497	HEATR5B	HEAT repeat containing 5B
8206	-0.247530060398715	5.99288959639263
2.5051923604963	0.113471628855522	0.519670038597326
10847	SRCAP	Snf2 related CREBBP activator protein
15230	-0.267114031353861	7.94784068395863
2.50471850637157	0.113505764693664	0.519670038597326
32	ACACB	acetyl-CoA carboxylase beta	14436
0.794998271086351	2.00500115203578	2.50457309044654
	0.11351624253826	0.519670038597326
4779	NFE2L1	nuclear factor, erythroid 2 like 1
5825	-0.211857980731402	8.86072694760705
2.5042051093512	0.113542761925761	0.519670038597326
92822	ZNF276	zinc finger protein 276	7427	-0.283751362424751
	4.66810756925337	2.50389162414282
0.113565359330793	0.519670038597326
221035	REEP3	receptor accessory protein 3
5248	-0.254005453069749	7.28980550579887
2.4995878931793	0.113876093076352	0.520921762155603
5423	POLB	DNA polymerase beta	3651	-0.563326937021927
	5.9321398819023	2.4984744724304	0.113956635709704
0.521120011879147
84532	ACSS1	acyl-CoA synthetase short chain family member 1	8691
	0.610435895236078	1.28364548111784
2.49777651663113	0.114007156461716	0.5211321348733
222658	KCTD20	potassium channel tetramerization domain containing
20	6239	-0.200508236527135	6.48213741227295
2.49736631685402	0.114036859835996	0.5211321348733
93010	B3GNT7	UDP-GlcNAc:betaGal beta-1,3-N-
acetylglucosaminyltransferase 7	3806	-0.496013760889923
5.00250618917836	2.49681914161452	0.114076495156462
	0.5211321348733
133619	PRRC1	proline rich coiled-coil 1
9064	-0.223855277634618	6.59986842957835
2.49616967456564	0.114123559826727	0.5211321348733
4725	NDUFS5	NADH:ubiquinone oxidoreductase subunit S5	548
	0.497156710996051	6.60936343985476
2.49535963737861	0.114182290476438	0.5211321348733
826	CAPNS1	calpain small subunit 1	3478	0.210735259292955
	8.27492077263157	2.49414504930383
0.114270415000805	0.5211321348733
729515	TMEM242	transmembrane protein 242	4644
0.225977100982002	4.68821990446733	2.49406531213635
	0.114276202959579	0.5211321348733
24138	IFIT5	interferon induced protein with tetratricopeptide
repeats 5	4015	0.317715389111813	4.50744865124063
	2.49385436464934	0.114291516767539
0.5211321348733

93622	LOC93622	Morf4 family associated protein 1 like 1
pseudogene	2311	0.350763182361842	3.12339623882763
	2.493578508925	0.11431154602685	0.5211321348733
56833	SLAMF8	SLAM family member 8	2996	1.54347201044827
	1.91873987017722	2.49330597449501
0.114331337935722	0.5211321348733
10804	GJB6	gap junction protein beta 6
2244	-0.853537736154891	0.580924245780964
2.4907650940867	0.114516042914094	0.521804233278374
9197	SLC33A1	solute carrier family 33 member 1
9627	-0.23908038306494	5.98478171011138
2.48887387652228	0.114653735397566	0.522261746898766
55567	DNAH3	dynein axonemal heavy chain 3
14960	-0.607166367880987	0.125793016197652
2.48700830724862	0.114789739463388	0.522711274598075
643008	SMIM5	small integral membrane protein 5
5117	-0.671507467764643	2.94232088644907
2.48591249704894	0.114869709377029	0.522905433618472
79098	C1orf116	chromosome 1 open reading frame 116
5502	-0.284743363823813	9.25024244050178
2.48506210809614	0.114931811321485	0.523018155029252
144455	E2F7	E2F transcription factor 7	6303
0.911943783742166	4.7238620770192	2.48451235092145
0.114971978532742	0.523031017638232
55279	ZNF654	zinc finger protein 654	4966	-0.269955659854346
	4.58356608962796	2.48339587417442
0.115053599961868	0.523232394372041
11080	DNAJB4	DnaJ heat shock protein family (Hsp40) member B4
	3783	0.387484149602456	4.28893104794632
2.48224152194893	0.115138057552992	0.523298510306199
6470	SHMT1	serine hydroxymethyltransferase 1	4382
0.35528013911621	3.3618690079463	2.48217594146773
0.115142857768523	0.523298510306199
3425	IDUA	alpha-L-iduronidase	4000	0.404201086133803
	3.97422456840827	2.48109450508208
0.115222046218353	0.523465061415974
6873	TAF2	TATA-box binding protein associated factor 2
6547	-0.187459705617058	5.50061145723675
2.47990103027485	0.115309508473263	0.523465061415974
10256	CNKSR1	connector enhancer of kinase suppressor of Ras 1
	4043	-0.247717913108283	5.44798464866812
2.47969317152021	0.11532474862232	0.523465061415974
116447	TOP1MT	DNA topoisomerase I mitochondrial	6237
0.340188898940592	2.95144711132244	2.4796353130562
0.1153289911851	0.523465061415974
11057	ABHD2	abhydrolase domain containing 2
10345	-0.343092186574349	7.78452084555729
2.47787998811606	0.11545778505793	0.523792275682774
517	ATP5MC2	ATP synthase membrane subunit c locus 2	3991
0.255610637443759	5.37296829190757	2.47763214606394
	0.115475982816335	0.523792275682774
7693	ZNF134	zinc finger protein 134	5138	-0.259711988821417
	4.80879520796668	2.47534600154939
0.115643991922946	0.524385042041019

64399 HHIP hedgehog interacting protein 12074
0.918818351224969 0.0574227284080412 2.47394863114697
0.11574681773146 0.524681448532222
56270 WDR45B WD repeat domain 45B 3542 -0.178255669474276
6.78886702635354 2.47227849555185
0.115869847444341 0.52506921810187
3123 HLA-DRB1 major histocompatibility complex, class II,
DR beta 1 1304 1.23550937641904 4.9147538112689
2.47128359410259 0.115943204975498 0.52523171801158
9021 SOCS3 suppressor of cytokine signaling 3
2734 -0.450199629430872 4.49338574432689
2.47059130109058 0.115994280389285 0.52529320575904
57522 SRGAP1 SLIT-ROBO Rho GTPase activating protein 1
26641 -0.30180432580227 4.35538660299617
2.46955110647722 0.116071069704437 0.525471064431174
6286 S100P S100 calcium binding protein P
1280 -1.2018953701857 4.89797007614644
2.46850120937683 0.11614863219572 0.525652307323246
374899 ZNF829 zinc finger protein 829 7742 -0.420511265166238
1.7412026873447 2.46605053279192 0.116329901858103
0.526295209284851
155435 RBM33 RNA binding motif protein 33
13132 -0.258142250508145 5.86961388218624
2.46549342571606 0.116371153069333 0.526295209284851
3552 IL1A interleukin 1 alpha 2124 -0.984801422041204
5.43462754860179 2.46505794568974
0.116403409606944 0.526295209284851
85450 ITPRIP inositol 1,4,5-trisphosphate receptor interacting
protein 7026 -0.353303935376161 6.48617529346861
2.46173087504408 0.116650175951919 0.527240727511626
8864 PER2 period circadian regulator 2
6877 -0.405761091409297 3.17561164393763
2.46050428255752 0.116741297126336 0.527310400155685
2011 MARK2 microtubule affinity regulating kinase 2
6154 -0.188926294521514 6.56663014073828
2.46045523532526 0.11674494238442 0.527310400155685
3613 IMPA2 inositol monophosphatase 2 3825
0.332835751294703 3.8341316918198 2.45945808641213
0.116819079138377 0.527310400155685
90060 CCDC120 coiled-coil domain containing 120
4537 -0.230080965437786 5.82685823957734
2.4594231929552 0.116821674364128 0.527310400155685
83891 SNX25 sorting nexin 25 4567 0.299017788659968
4.98298245441142 2.45899103352957
0.116853821809327 0.527310400155685
8558 CDK10 cyclin dependent kinase 10 6791
0.265473319747173 5.50635727455602 2.45790165698719
0.116934901650535 0.527506334112412
10568 SLC34A2 solute carrier family 34 member 2
4699 -0.872678553063031 7.010874246298 2.45567968755816
0.1171004705658 0.527525822554465
2582 GALE UDP-galactose-4-epimerase
3978 -0.234970071696705 5.1733802437921 2.45541186782956
0.117120444500425 0.527525822554465

64785	GINS3	GINS complex subunit 3	2511	-0.355542090031416	2.94213226855419	2.45522295057695	0.117134536170436	0.527525822554465
123169	LEO1	LEO1 homolog, Paf1/RNA polymerase II complex component	2679	-0.212233792936068	5.05352171360186	2.45521928393596	0.117134809690164	0.527525822554465
2896	GRN	granulin precursor	3680	0.290174722073213	8.21947479090242	2.4550435075821	0.117147922873678	0.527525822554465
51192	CKLF	chemokine like factor	1489	0.300570408129454	3.96352247100857	2.45481206576806	0.117165191259152	0.527525822554465
92304	SCGB3A1	secretoglobin family 3A member 1	846	1.18876942313793	0.661918744251203	2.45323068533778	0.117283256840655	0.527747454027708
115557	ARHGEF25	Rho guanine nucleotide exchange factor 25	3503	0.933565218797274	0.766515945525961	2.45314348344108	0.117289771142161	0.527747454027708
54897	CASZ1	castor zinc finger 1	11336	-0.380131302707412	6.1882767962685	2.45178110068469	0.117391598068295	0.5280360032571
54555	DDX49	DEAD-box helicase 49	2780	0.261585620201583	4.47018236773963	2.450803682082	0.11746471226926	0.528195256743348
729857	RGPD2	RANBP2 like and GRIP domain containing 2	6104	-0.444892658396239	1.76570309064004	2.44954188352509	0.117559173476916	0.528450366781505
4259	MGST3	microsomal glutathione S-transferase 3	4445	0.381380102799547	6.17501455761873	2.4486397624287	0.117626759851797	0.528584544512068
340481	ZDHHC21	zinc finger DHHC-type containing 21	9426	-0.295224085765701	6.43599080112162	2.44747091337175	0.11771439320568	0.528729081876993
342908	ZNF404	zinc finger protein 404	1857	0.663180404044279	0.556195108878141	2.44703642666024	0.11774698683256	0.528729081876993
90693	CCDC126	coiled-coil domain containing 126	3550	-0.299267730456324	4.0402998424095	2.44670086282258	0.117772166449362	0.528729081876993
27327	TNRC6A	trinucleotide repeat containing adaptor 6A	12522	-0.316870231141313	6.08613230455769	2.44582272013648	0.117838087587626	0.528855524780478
3077	HFE	homeostatic iron regulator	5675	0.433584293444031	2.76959609175762	2.44523255629039	0.117882413410147	0.528884998281847
1759	DNM1	dynamin 1	8851	0.586903103776551	0.960010492478585	2.44338775062383	0.118021091344387	0.529225810176432
10492	SYNCRIP	synaptotagmin binding cytoplasmic RNA interacting protein	8205	-0.225780860064596	7.66716215420684	2.44277024875063	0.118067550544649	0.529225810176432
100507582	BHLHE40-AS1	BHLHE40 antisense RNA 1	2717	0.602236295528509	0.396345646385329	2.44271477359878

0.118071725337428 0.529225810176432
4316 MMP7 matrix metallopeptidase 7 2323
1.08153616801744 7.64845094178091 2.44152620716324
0.11816121034356 0.529382982609489
8576 STK16 serine/threonine kinase 16 4368
0.251235425989815 4.53754691176142 2.44124518057995
0.118182379283206 0.529382982609489
63932 CXorf56 chromosome X open reading frame 56 2580
0.296428574491242 4.14586150636068 2.43978601331558
0.118292361670929 0.529679514051937
9666 DZIP3 DAZ interacting zinc finger protein 3 6972
0.347238821548986 3.31815129751538 2.43901186744926
0.118350757618014 0.529679514051937
25946 ZNF385A zinc finger protein 385A 3179
0.363948356036988 5.68094712981738 2.43886255106709
0.118362024629297 0.529679514051937
26057 ANKRD17 ankyrin repeat domain 17
12041 -0.250891812816916 7.44362813820511
2.4380657168151 0.11842217165021 0.529779418174541
25901 CCDC28A coiled-coil domain containing 28A 1507
0.305109367521939 3.62296271406348 2.43720981169266
0.118486815140829 0.52989936771315
65985 AACS acetoacetyl-CoA synthetase
16094 -0.225768093524146 5.86126356034544
2.43659253549233 0.118533460052778 0.52993877272878
89845 ABCC10 ATP binding cassette subfamily C member 10 8083
0.182251533842953 5.94964935866811
2.43436699155178 0.118701803539315 0.530522068339244
5962 RDX radixin 6592 -0.205249591440452
6.98546467610466 2.43314612574597 0.118794263972185
0.530765950704434
55906 ZC4H2 zinc finger C4H2-type containing 5173
0.420778707138765 3.58555920836414 2.43210131200253
0.118873454631121 0.530902904067415
440145 MZT1 mitotic spindle organizing protein 1
2299 -0.326196255334129 3.87543042820973
2.43163592798314 0.118908746765054 0.530902904067415
80273 GRPEL1 GrpE like 1, mitochondrial
4177 -0.263159902655483 5.31518694586164
2.43124206139827 0.118938624470875 0.530902904067415
54509 RHOF ras homolog family member F, filopodia associated
4442 -0.327712754392757 6.46882797255134
2.43032836611998 0.119007967029085 0.531043196615609
83695 RHNO1 RAD9-HUS1-RAD1 interacting nuclear orphan 1 2643
0.298469667176839 3.8105900149779 2.42848085236645
0.119148316037402 0.531500147368116
3078 CFHR1 complement factor H related 1 2197
0.528976595497867 0.819206033490567 2.42658744153111
0.119292341721133 0.531973202956993
22976 PAXIP1 PAX interacting protein 1
9643 -0.262693314057753 4.07169065670313
2.42499359190985 0.119413730088804 0.532300851487066
57526 PCDH19 protocadherin 19 9756 -0.962211782477481
2.73885459460139 2.42400726642435

0.119488917584441	0.532300851487066
10097	ACTR2	actin related protein 2	4347	-0.242222588154788
	8.63280857973879	2.42392108454496
0.119495489709141	0.532300851487066
388325	SCIMP	SLP adaptor and CSK interacting membrane protein
	4208	1.52512015690765	0.0168295061125851
2.42362823266185	0.119517825225018	0.532300851487066
90411	MCFD2	multiple coagulation factor deficiency 2
5994	-0.192944223991113	7.46326777713648
2.4225319321122	0.119601479987797	0.532408552944082
5552	SRGN	serglycin	1220	1.20230928302459
2.34117996685064	2.4218740351812	0.11965171285511
0.532408552944082
4357	MPST	mercaptopyruvate sulfurtransferase	6760
0.248317836411082	5.19143027491883	2.42181740292955
	0.119656038028698	0.532408552944082
3480	IGF1R	insulin like growth factor 1 receptor
13509	-0.288671146438191	7.50931950718221
2.41935067020118	0.11984459778089	0.533078209309917
11006	LILRB4	leukocyte immunoglobulin like receptor B4	4735
	1.33344055621579	0.612515592506216
2.41674032456361	0.120044493738627	0.533797848824428
23521	RPL13A	ribosomal protein L13a	2369	0.258081498844112
	7.8861884209528	2.41624262044058	0.120082648981069
	0.533798052769862
1213	CLTC	clathrin heavy chain	11836	-0.186731542523417
	9.43822810067711	2.41565748426191
0.120127524156228	0.533828118926489
94121	SYTL4	synaptotagmin like 4	7428	-0.467673722701416
	3.98284391157059	2.41108924478007
0.120478509938896	0.535218042725695
3275	PRMT2	protein arginine methyltransferase 2	8031
0.176373740561807	6.49266112692552	2.40548971282462
	0.120910281830546	0.536610930722537
8241	RBM10	RNA binding motif protein 10
3889	-0.214332860789342	5.43149350201711
2.40408520227255	0.12101885031681	0.536610930722537
57541	ZNF398	zinc finger protein 398	6041	-0.215565170208773
	4.72980518384302	2.40339187695442
0.121072484086824	0.536610930722537
51135	IRAK4	interleukin 1 receptor associated kinase 4
5629	-0.210913212222729	4.72442726929802
2.40316156242059	0.12109030642222	0.536610930722537
7529	YWHAB	tyrosine 3-monooxygenase/tryptophan 5-monooxygenase activation protein beta	7933	-0.223863468930719
8.61415067991341	2.40247174034357	0.121143704034481
	0.536610930722537
2960	GTF2E1	general transcription factor IIE subunit 1
3370	-0.235319321202102	3.96245662249112
2.40203062083448	0.12117786380226	0.536610930722537
116254	GINM1	glycoprotein integral membrane 1
2248	-0.197761846880252	5.81910501355304
2.40195298161477	0.121183877195085	0.536610930722537
2124	EVI2B	ecotropic viral integration site 2B	2691

1.23425781543039        1.62429466705364        2.40174081909382
        0.121200311511537       0.536610930722537
51421   AMOTL2  angiomotin like 2       5984    -0.264369251556732
        8.26861449288973        2.40170846761794
0.121202817705486       0.536610930722537
10285   SMNDC1  survival motor neuron domain containing 1
5417    -0.217634278731634      5.21268856199303
2.40068872166255        0.121281844502157       0.536610930722537
3176    HNMT    histamine N-methyltransferase   4756
0.458975908450149       3.55788484324634        2.39901318976277
        0.121411816455924       0.536610930722537
55890   GPRC5C  G protein-coupled receptor class C group 5 member C
        5593    0.30350181562993        6.04146499617416
2.39896240940942        0.121415757925541       0.536610930722537
10188   TNK2    tyrosine kinase non receptor 2
11297   -0.234100724896361      5.67370299937325
2.39895336653763        0.121416459830062       0.536610930722537
9533    POLR1C  RNA polymerase I and III subunit C      2867
0.254638326174753       4.26658318336497        2.39700766802103
        0.121567588888154       0.536610930722537
25900   IFFO1   intermediate filament family orphan 1   5203
0.926915158347244       0.839681576667964       2.39700555566708
        0.121567753075155       0.536610930722537
2957    GTF2A1  general transcription factor IIA subunit 1
6819    -0.283478721402423      5.45984832456117
2.39682351807431        0.121581903239922       0.536610930722537
384     ARG2    arginase 2      2418    -0.514285795718332
4.66090187856421        2.39649790816347        0.121607218135632
        0.536610930722537
91875   TTC5    tetratricopeptide repeat domain 5       8764
0.245228417680432       3.94169218889631        2.3964660518523
0.121609695152801       0.536610930722537
7003    TEAD1   TEA domain transcription factor 1
10379   -0.304891428717514      8.0500018295092 2.3961969821253
0.121630619151171       0.536610930722537
54622   ARL15   ADP ribosylation factor like GTPase 15
3797    -0.243107696939314      4.59887968920317
2.39590722102072        0.121653156660706       0.536610930722537
124540  MSI2    musashi RNA binding protein 2   14660
0.262216385901863       5.11928155870175        2.3956739911095
0.121671300557401       0.536610930722537
54621   VSIG10  V-set and immunoglobulin domain containing 10
7339    -0.366864901855283      5.16897639559571
2.39500368679107        0.121723462936336       0.536610930722537
2690    GHR     growth hormone receptor 6610    0.652315853917225
        0.0546581174369599      2.39457086697931
0.121757157689461       0.536610930722537
267004  PGBD3   piggyBac transposable element derived 3 2212
0.411185550239435       3.08677909748659        2.39454112841108
        0.121759473198038       0.536610930722537
9583    ENTPD4  ectonucleoside triphosphate diphosphohydrolase 4
        7064    -0.213163052011965      7.20757894248805
2.39447477909715        0.121764639473573       0.536610930722537
27125   AFF4    AF4/FMR2 family member 4

11297	-0.268233765903605	8.68657193777799
2.39387031078388	0.12181171746482	0.536610930722537
6715	SRD5A1	steroid 5 alpha-reductase 1
7715	-0.269732018997532	5.52810748584825
2.39340071696719	0.121848304909633	0.536610930722537
55061	SUSD4	sushi domain containing 4	4710
0.378120078464317	2.29088315595345	2.39318986514453
	0.121864736962118	0.536610930722537
8481	OFD1	OFD1 centriole and centriolar satellite protein
5172	-0.207545985347084	5.04168765179215
2.39189251947039	0.121965895457528	0.53688758569252
8085	KMT2D	lysine methyltransferase 2D
20476	-0.451932117501718	7.81750350589164
2.39092052849821	0.122041745927077	0.536989906483227
11119	BTN3A1	butyrophilin subfamily 3 member A1
5547	-0.364507059142997	3.87491598901202
2.38991632207898	0.122120165295491	0.536989906483227
26994	RNF11	ring finger protein 11	3122	-0.201889321164333
	7.40612951597877	2.38990125683638
0.122121342178878	0.536989906483227
3604	TNFRSF9	TNF receptor superfamily member 9
6331	-0.607784294370761	2.7655514568108	2.3896305858047
0.122142488902375	0.536989906483227
6988	TCTA	T cell leukemia translocation altered	2747
0.280027802835577	4.81017340588594	2.38835691469887
	0.122242051557882	0.537258994719564
9331	B4GALT6	beta-1,4-galactosyltransferase 6
5306	-0.396220204555635	3.12289149548731
2.38699802640428	0.122348374855413	0.537557618130417
54545	MTMR12	myotubularin related protein 12
5545	-0.243486505483085	5.49073519225924
2.38617184279432	0.122413067941195	0.537673202462314
51585	PCF11	PCF11 cleavage and polyadenylation factor subunit
	8703	-0.272673673645548	5.64219259363171
2.38507832812992	0.122498752327076	0.537873844318297
3636	INPPL1	inositol polyphosphate phosphatase like 1	6398
	0.38516558352324	8.60265761835841
2.38460898262451	0.122535549169678	0.537873844318297
171177	RHOV	ras homolog family member V
1705	-0.653776535377955	5.27083916268602
2.3824786252742	0.122702724058802	0.538136713897969
55454	CSGALNACT2	chondroitin sulfate N-
acetylgalactosaminyltransferase 2	3730	-0.357316739822003
	6.10513336366945	2.3824333581028	0.122706279039398
	0.538136713897969
28959	TMEM176B	transmembrane protein 176B	1979
1.29728981577435	1.81844493602412	2.38237716668485
	0.122710692096103	0.538136713897969
55776	SAYSD1	SAYSVFN motif domain containing 1	2925
0.277712473575907	3.47734279824332	2.38074309526909
	0.122839102774475	0.538531240238518
54625	PARP14	poly(ADP-ribose) polymerase family member 14
10495	-0.324040346229189	7.3154591736846	2.37946954289671
	0.122939286042238	0.538801808383489

221002	RASGEF1A	RasGEF domain family member 1A
6073	-1.12154903970571	2.12812075752347
2.37839560889512	0.123023836888534	0.538900719746511
26233	FBXL6	F-box and leucine rich repeat protein 6
3375	-0.28030726806875	4.43865761733278
2.3782056108629 0.123038802152448	0.538900719746511
3964	LGALS8	galectin 8	9710	-0.185415004481242
6.86115480473822	2.37622713347357	0.123194757713899
	0.539209039231889
79781	IQCA1	IQ motif containing with AAA domain 1	5975
0.346481547592735	4.44190051882128	2.37612039304554
	0.123203177873648	0.539209039231889
84276	NICN1	nicolin 1	2929	0.318810659714067
3.97484450564918	2.37584781441705	0.123224682985741
	0.539209039231889
157638	LRATD2	LRAT domain containing 2	5404
0.314220518925926	7.65081214165917	2.37298381293397
	0.123450890569912	0.539796682959656
2101	ESRRA	estrogen related receptor alpha
3030	-0.194662358679952	5.53312854254098
2.37275529261524	0.123468959643189	0.539796682959656
51398	WDR83OS WD repeat domain 83 opposite strand	2656
0.190173362782031	5.94285866758265	2.37268410180521
	0.123474589291264	0.539796682959656
3842	TNPO1	transportin 1	11977	-0.229785330468928
8.08074503498491	2.3718809556903 0.123538120457038
0.539905913647964
221	ALDH3B1 aldehyde dehydrogenase 3 family member B1	6626
	0.455258197292006	1.68806654152794
2.36998179304187	0.123688494046502	0.540394490364741
3628	INPP1	inositol polyphosphate-1-phosphatase	3676
0.261971896240187	4.72454157578503	2.36443008953278
	0.1241292380397 0.542150993832892
22903	BTBD3	BTB domain containing 3 6228	-0.253806886284366
	4.25883679303791	2.36286614402915
0.124253712608273	0.542440041376405
79848	CSPP1	centrosome and spindle pole associated protein 1
	7096	0.296064594950404	3.35949821361074
2.36262559538418	0.124272870191824	0.542440041376405
171568	POLR3H	RNA polymerase III subunit H	6594
0.245585393904117	5.42496069256088	2.35992512533923
	0.124488164109896	0.543210503017854
201176	ARHGAP27	Rho GTPase activating protein 27	7727
	-0.199999652458493	6.12376622241299
2.35477459790063	0.124899937253582	0.54467835691212
339803	LOC339803	uncharacterized LOC339803	3109
0.425814788548339	2.42714937557788	2.35402841004291
	0.124959718669305	0.54467835691212
1201	CLN3	CLN3 lysosomal/endosomal transmembrane protein,
battenin	5799	0.334229913561632	6.04499690420588
	2.35344709040537	0.125006313567672
0.54467835691212
4678	NASP	nuclear autoantigenic sperm protein	8304
0.229811969728524	5.15027783988542	2.35324025862996

0.125022896547136	0.54467835691212
3627	CXCL10	C-X-C motif chemokine ligand 10
1176	-1.77505166177163	3.06013615240928
2.35309158847327	0.125034817860444	0.54467835691212
5569	PKIA	cAMP-dependent protein kinase inhibitor alpha	4216
0.422525190245381	2.91708971088778
2.35265653329157	0.12506971059141	0.54467835691212
60481	ELOVL5	ELOVL fatty acid elongase 5	3813
0.233103556778014	6.86528644600306	2.35231940264075
0.12509675692056	0.54467835691212
219347	TMEM254-AS1	TMEM254 antisense RNA 1	3138
0.634974159610827	0.316603206841733	2.35137823650721
0.125172296434122	0.544837898120804
7572	ZNF24	zinc finger protein 24	8056	-0.208576110076404
6.91937834740381	2.35070744545223
0.12522616613891	0.544840161117882
1106	CHD2	chromodomain helicase DNA binding protein 2
18730	-0.267178076909418	7.10792248461125
2.34963786731318	0.125312114817381	0.544840161117882
27301	APEX2	apurinic/apyrimidinic endodeoxyribonuclease 2
3245	-0.207321481172255	5.12360999799243
2.34912184521175	0.125353604527224	0.544840161117882
6310	ATXN1	ataxin 1	16487	-0.330690580628043
6.56773495343494	2.34843307263656	0.125409007674649
0.544840161117882
221336	BEND6	BEN domain containing 6	5588	0.566564440703838
0.636419520930584	2.34790712714053
0.125451331739761	0.544840161117882
726	CAPN5	calpain 5	6166	-0.575018533214214
3.08843560325614	2.34781236139394	0.125458959448722
0.544840161117882
5538	PPT1	palmitoyl-protein thioesterase 1	3306
0.296959735433756	5.89474217328644	2.34772858200169
0.125465703293616	0.544840161117882
80759	KHDC1	KH domain containing 1	4267	0.560598519989163
1.41687553132781	2.34750133061108
0.125483997984314	0.544840161117882
26060	APPL1	adaptor protein, phosphotyrosine interacting with PH domain and leucine zipper 1	7033	-0.215184648320589
5.65608349188848	2.34609612748235	0.125597188595271
0.545142898707925
83594	NUDT12	nudix hydrolase 12	4304	-0.344236094723872
3.63733850552688	2.34558966188661
0.12563801275398	0.545142898707925
51311	TLR8	toll like receptor 8	4353	1.50608441343797
0.551208136956801	2.34436563013507
0.125736737945801	0.545142898707925
375775	PNPLA7	patatin like phospholipase domain containing 7	9976
0.784338304507975	1.90002215685047
2.34397119293681	0.125768569938403	0.545142898707925
257396	LOC257396	uncharacterized LOC257396	3194
0.44644401938971	0.381397078072512	2.34366916925953
0.125792950001627	0.545142898707925
10367	MICU1	mitochondrial calcium uptake 1	3846

0.21236809947759	5.72808527121973	2.34352146559954
	0.125804874901882	0.545142898707925
22834	ZNF652	zinc finger protein 652 6196	-0.364688652086941
	3.68289689059317	2.34220666548901
0.125911081085452	0.545142898707925
282890	ZNF311	zinc finger protein 311 3176	0.847110792384996
	0.13805922366087	2.3419146816811З
0.125934680310973	0.545142898707925
11097	NUP42	nucleoporin 42	4159	0.282244411943821
4.30132029944362	2.34167980151108	0.125953667775003
	0.545142898707925
26985	AP3M1	adaptor related protein complex 3 subunit mu 1
5912	-0.203402483877632	6.1960670409079 2.34121371317396
	0.125991355286104	0.545142898707925
55848	PLGRKT	plasminogen receptor with a C-terminal lysine
1493	-0.354467660737622	3.54494796353928
2.34077043547295	0.126027209979329	0.545142898707925
400745	SH2D5	SH2 domain containing 5 4423	-0.485199420842562
	2.24517437870196	2.34013216897445
0.126078856345785	0.545142898707925
9388	LIPG	lipase G, endothelial type
11970	-0.552770831110007	8.39691496227555
2.33960835770915	0.12612125896093	0.545142898707925
57167	SALL4	spalt like transcription factor 4	5772
0.741349872271864	1.13907716421078	2.33946988219919
	0.126132471227604	0.545142898707925
2086	ERV3-1	endogenous retrovirus group 3 member 1, envelope
	4574	-0.634538169802558	4.27799932549254
2.33940762767384	0.126137512294737	0.545142898707925
54558	SPATA6	spermatogenesis associated 6	5939
0.421625972858066	1.78693130572649	2.33855295355833
	0.126206742278076	0.545145564231875
388242	LOC388242	SAGA complex associated factor 29 pseudogene
	3982	-0.530923784850959	1.55501116385046
2.3384390964822 0.12621596807339	0.545145564231875
3145	HMBS	hydroxymethylbilane synthase	4247
0.397555754561765	3.61348775222727	2.33780159223664
	0.126267638653736	0.545200621030483
151242	PPP1R1C protein phosphatase 1 regulatory inhibitor subunit
1C	4887	0.605011744775487	1.49241230714918
2.33686454671724	0.126343630156805	0.545360624840173
22882	ZHX2	zinc fingers and homeoboxes 2
4716	-0.216779245057191	5.01839669765389
2.33474953782503	0.126515337858719	0.545769635880052
23639	LRRC6	leucine rich repeat containing 6	2001
0.43660718591827	1.88562819770474	2.33473752552839
	0.126516313821841	0.545769635880052
54210	TREM1	triggering receptor expressed on myeloid cells 1
	2799	1.01909233954924	1.49278986609751
2.33342652961211	0.126622878651162	0.545770018145882
100529241	HSPE1-MOB4	HSPE1-MOB4 readthrough
890	-0.213718401024356	5.19115398053967
2.33336049874503	0.126628248636615	0.545770018145882
151011	SEPTIN10	septin 10	3071	0.231875080776274

6.0861228092729 2.33329845365618 0.12663329470794 0.545770018145882
27250 PDCD4 programmed cell death 4 4547 0.357554094247229 6.28759500894745 2.33254986292015 0.126694194549917 0.545811380048792
27248 ERLEC1 endoplasmic reticulum lectin 1 2592 -0.189508572607417 6.12358490956646 2.33222263221174 0.126720825866972 0.545811380048792
255967 PAN3 poly(A) specific ribonuclease subunit PAN3 6352 -0.243698563910667 5.25371505531523 2.331197754045 0.126804274763424 0.546002913191295
92283 ZNF461 zinc finger protein 461 3219 -0.415152592559646 2.35232669256719 2.32996015617847 0.126905125467463 0.5462692355325
23200 ATP11B ATPase phospholipid transporting 11B (putative) 9196 -0.286716421126338 6.78501775657959 2.3292932562488 0.12695950751591 0.546335429116853
23082 PPRC1 PPARG related coactivator 1 6095 -0.300821275053944 5.42980272361188 2.32855420772466 0.127019803172649 0.54634825868997
126626 GABPB2 GA binding protein transcription factor subunit beta 2 9189 -0.332046653581861 3.36793037266619 2.32830061940841 0.127040499646836 0.54634825868997
344595 DUBR DPPA2 upstream binding RNA 4206 0.556103215089205 3.00043974239532 2.32752889046571 0.127103506974151 0.546377072825397
3730 ANOS1 anosmin 1 7132 0.413046551872516 7.42735411281357 2.32706798359112 0.127141154009394 0.546377072825397
284184 NDUFAF8 NADH:ubiquinone oxidoreductase complex assembly factor 8 1376 0.373824275821305 3.2656304059337 2.3267856408636 0.127164221989776 0.546377072825397
57549 IGSF9 immunoglobulin superfamily member 9 6628 0.271611211664889 5.27721458165665 2.32583391796061 0.1272420140141 0.546543664610704
4209 MEF2D myocyte enhancer factor 2D 6400 -0.211615749600856 5.6491871315798 2.32340356209705 0.127440907054644 0.546984494075883
474343 SPIN2B spindlin family member 2B 1478 0.47019473213175 1.13486045974818 2.32333568668375 0.127446466738486 0.546984494075883
905 CCNT2 cyclin T2 9081 -0.267946785735509 5.66408415288972 2.32295409082721 0.12747772832875 0.546984494075883
57466 SCAF4 SR-related CTD associated factor 4 5901 -0.228409821390934 5.42298060012392 2.32267195077381 0.127500847658868 0.546984494075883
91289 LMF2 lipase maturation factor 2 3054 0.226552484918994 5.66342153979843 2.32186721269348 0.127566815743766 0.547099935126435
4082 MARCKS myristoylated alanine rich protein kinase C substrate 4275 -0.205797120257431 7.96872336375984 2.31859334918569 0.127835581295278 0.548042689130465

100506451	RASSF8-AS1	RASSF8 antisense RNA 1	3981
0.550677418461912	0.456844656445133	2.31823680522539
	0.127864889560817	0.548042689130465
1478	CSTF2	cleavage stimulation factor subunit 2
2187	-0.222545946674927	4.48136043313422
2.31581559612461	0.128064113309922	0.548728673946798
54477	PLEKHA5	pleckstrin homology domain containing A5
12194	-0.239419938722527	5.93527764344291
2.31382817473687	0.12822790206907	0.549242826541791
9144	SYNGR2	synaptogyrin 2	3146	0.352470337157098
8.00555826106351	2.31337587885125	0.128265209563291
	0.549242826541791
10920	COPS8	COP9 signalosome subunit 8	4743
0.200380187248085	5.6847865938968	2.31293309682533
0.128301744016902	0.549242826541791
1723	DHODH	dihydroorotate dehydrogenase (quinone)	6921
0.456708307776313	1.15625433946645	2.31209167989942
	0.128371202228799	0.549372266916829
9774	BCLAF1	BCL2 associated transcription factor 1
9297	-0.179482586856434	7.14428655306643
2.30906607113169	0.128621309725429	0.5502619826802
101927043	EPCAM-DT	EPCAM divergent transcript	3552
	0.615286506333857	1.28420882313933
2.30822363563818	0.128691044966264	0.5502619826802
9545	RAB3D	RAB3D, member RAS oncogene family
4417	-0.378285476259186	5.63383488097446
2.30815225726965	0.128696955469432	0.5502619826802
1656	DDX6	DEAD-box helicase 6	11444	-0.195843680313621
	7.86752127740875	2.30482824052198
0.128972536668424	0.551176256824537
84330	ZNF414	zinc finger protein 414	2758	0.366204236909592
	2.23899671641496	2.30462402474641
0.128989488817793	0.551176256824537
90557	CCDC74A	coiled-coil domain containing 74A	3077
0.484035629282703	1.78297733215403	2.30286687223439
	0.129135454359858	0.551441751041405
8899	PRPF4B	pre-mRNA processing factor 4B
7732	-0.202726027066806	6.58913574331627
2.30234567352962	0.129178785384188	0.551441751041405
283335	LOC283335	uncharacterized LOC283335
5408	-0.490541888128608	0.241874843043521
2.30214028422588	0.129195865334364	0.551441751041405
9972	NUP153	nucleoporin 153	6030	-0.210423928722615
6.15979549223535	2.30198118749578	0.12920909737402
	0.551441751041405
27107	ZBTB11	zinc finger and BTB domain containing 11
7416	-0.245246590987166	5.50097171443548
2.30101405479087	0.129289566080887	0.551617103897345
1911	PHC1	polyhomeotic homolog 1	6090	0.311339587894617
	3.83560590180889	2.29958743667552
0.129408367479169	0.551898397829176
23005	MAPKBP1	mitogen-activated protein kinase binding protein 1
	10357	-0.212497624253601	5.97703299720037
2.29927622466266	0.129434299769318	0.551898397829176

26353	HSPB8	heat shock protein family B (small) member 8
2860	-0.389441049217135	6.72068916926679
2.29663032777472	0.129655007856952	0.552610415399802
57507	ZNF608	zinc finger protein 608 6869	0.56523499117145
	3.87628021965633	2.29632874575041
0.129680191005865	0.552610415399802
55119	PRPF38B pre-mRNA processing factor 38B
5817	-0.194520312888973	6.03373671313659
2.29573139312267	0.129730088124535	0.552654910085268
285590	SH3PXD2B	SH3 and PX domains 2B
7977	-0.259989166343603	8.10570933460601
2.29415522663214	0.129861848733844	0.552910379380537
253959	RALGAPA1	Ral GTPase activating protein catalytic alpha subunit 1 13265	-0.332074009352263	4.6170151308093
2.29362310835359	0.1299063651637 0.552910379380537
55095	SAMD4B	sterile alpha motif domain containing 4B
5467	-0.188968288707302	6.4991277530205 2.29359785189925
	0.129908478513697	0.552910379380537
728130	NUTM2D	NUT family member 2D	6985	0.507410710194543
	1.8068663645623 2.29233252316583	0.130014404778674
	0.553113612087476
80124	VCPIP1	valosin containing protein interacting protein 1
	9942	-0.266490054458758	5.5600639726502
2.2919468003649 0.130046714487023	0.553113612087476
55905	RNF114	ring finger protein 114 4344	-0.248501098669154
	6.1403360175126 2.29161285956296	0.130074693954176
	0.553113612087476
26031	OSBPL3	oxysterol binding protein like 3
8100	-0.292998700899393	7.72005923544843
2.2909577572669 0.130129601663257	0.553140999425007
8880	FUBP1	far upstream element binding protein 1
3896	-0.235531704225596	6.2092174035325 2.29059383391478
	0.130160115235584	0.553140999425007
114787	GPRIN1	G protein regulated inducer of neurite outgrowth 1
	4211	-0.271789389291798	3.74468128207277
2.28996027431119	0.130213255803858	0.553198991217662
10782	ZNF274	zinc finger protein 274 6913	-0.305899719868293
	4.84995285101338	2.28884840012109
0.130306574066074	0.553352881648146
5019	OXCT1	3-oxoacid CoA-transferase 1	4194
0.372278048347465	4.28996698746298	2.28858740597715
	0.130328489794905	0.553352881648146
27230	SERP1	stress associated endoplasmic reticulum protein 1
	4244	-0.316546781265926	6.77445538012327
2.28714377854664	0.130449785733645	0.553700045082171
3710	ITPR3	inositol 1,4,5-trisphosphate receptor type 3
9870	-0.352001668700595	9.06405521387181
2.28623252445186	0.130526415601058	0.553857468441086
2014	EMP3	epithelial membrane protein 3	1667
0.48621362595346	4.28589363641528	2.28557552292739
	0.130581695813072	0.553924231754605
651746	ANKRD33B	ankyrin repeat domain 33B	9783
0.520920129415018	2.01035717274848	2.28490861616683
	0.130637836157812	0.553994602198752

26287	ANKRD2	ankyrin repeat domain 2	1451	0.722982524125798
	1.18234308507084	2.28407074167831
0.130708406757829	0.554044615712842
5655	KLK10	kallikrein related peptidase 10
3853	-0.524147615818021	7.65575478015046
2.28382942696572	0.130728739553862	0.554044615712842
145407	ARMH4	armadillo like helical domain containing 4	7450
	0.416897633851942	2.25502357756509
2.28306564642662	0.13079311775084	0.554149788365399
169714	QSOX2	quiescin sulfhydryl oxidase 2	4530
0.260232229579361	3.92419573901807	2.28156959573201
	0.130919320665396	0.554516759770244
101669762	BLACAT1	bladder cancer associated transcript 1
972	-0.675788356204443	0.000467179826138593
2.28052800313624	0.131007266894012	0.554532476931648
101928979	LINC01719	long intergenic non-protein coding
RNA 1719	2458	0.539860116606733	0.456439830428014
	2.28016266092729	0.131038129948807
0.554532476931648
643155	SMIM15	small integral membrane protein 15
2889	-0.216607160288085	5.67866945193533
2.28009181699318	0.131044115578645	0.554532476931648
134957	STXBP5	syntaxin binding protein 5
10504	-0.267132999303219	5.51618386536338
2.27918875114906	0.131120442673386	0.554532476931648
57693	ZNF317	zinc finger protein 317	5065	-0.202379073903074
	5.7588294152553	2.2791823985902	0.131120979767089
0.554532476931648
2526	FUT4	fucosyltransferase 4	6059	-0.406771774477329
	3.16059057601061	2.27666801865388
0.131333757478836	0.55520201116852
56995	TULP4	TUB like protein 4	11544	-0.306949408746424
	6.12331797006342	2.27636901592873
0.131359086004143	0.55520201116852
9923	ZBTB40	zinc finger and BTB domain containing 40
8992	-0.263712949980962	6.0650592358891	2.27584619177571
	0.131403387544856	0.55520201116852
84959	UBASH3B	ubiquitin associated and SH3 domain containing B
	9011	0.187140432974929	6.29931927793226
2.27507013480489	0.131469177529834	0.55520201116852
57633	LRRN1	leucine rich repeat neuronal 1	4264
0.540175735721275	8.78840696011183	2.27497222256443
	0.131477480620117	0.55520201116852
54930	HAUS4	HAUS augmin like complex subunit 4	2589
0.339962431644701	3.42246392786814	2.27069919514443
	0.131840409830176	0.556416268417126
9716	AQR	aquarius intron-binding spliceosomal factor
13528	-0.182992985781845	5.81573240356261
2.26979924335645	0.131916989624789	0.556416268417126
7748	ZNF195	zinc finger protein 195	5709	-0.226660880030751
	5.30170218332347	2.26936111586434
0.131954289272881	0.556416268417126
80830	APOL6	apolipoprotein L6	10121	-0.480464303322311
	6.59539930367693	2.2693059651383	0.131958985321851

0.556416268417126
2212 FCGR2A Fc fragment of IgG receptor IIa 5232
1.56485158021403 2.11837667209821 2.26891012927546
0.131992695967191 0.556416268417126
147664 ERVV-1 endogenous retrovirus group V member 1, envelope
2202 0.815573720928677 0.148049747771002
2.26850853731824 0.132026906644542 0.556416268417126
201627 DENND6A DENN domain containing 6A
5124 -0.24710187953516 5.08779166968368
2.26831850877358 0.132043098180804 0.556416268417126
9887 SMG7 SMG7 nonsense mediated mRNA decay factor
8317 -0.193842626919518 6.96425441404468
2.26680790911806 0.132171888932236 0.556791473165061
7405 UVRAG UV radiation resistance associated
6352 -0.214773655167921 4.72200401079168
2.26619566393393 0.132224127685717 0.556844063888705
55508 SLC35E3 solute carrier family 35 member E3 18389
0.291228944154856 3.89535675462295 2.26539149728917
0.132292776830758 0.556965712374037
7023 TFAP4 transcription factor AP-4 7318
0.47131308848793 0.524776050463892 2.26415087087066
0.132398763227782 0.557176531575368
56181 MTFR1L mitochondrial fission regulator 1 like 5940
0.206810311623822 5.80541632293172 2.26373990063319
0.132433893213391 0.557176531575368
84836 ABHD14B abhydrolase domain containing 14B 3402
0.316909950302094 3.68217586109524 2.26287963803379
0.132507462686423 0.557176531575368
51279 C1RL complement C1r subcomponent like 4243
0.383775487236734 4.91378877446074 2.26257217651225
0.132533767799709 0.557176531575368
26524 LATS2 large tumor suppressor kinase 2
5578 -0.264747749127479 5.95864035780746
2.26247896608764 0.132541743643116 0.557176531575368
6324 SCN1B sodium voltage-gated channel beta subunit 1 4576
0.532503286790993 1.30202934663543
2.25937020400547 0.132808061989912 0.558128570144825
10483 SEC23B SEC23 homolog B, coat complex II component
4039 -0.254687906754271 6.38998923097421
2.25792934044829 0.132931699124936 0.558480596773537
5932 RBBP8 RB binding protein 8, endonuclease 5091
0.202246216629587 6.06230288804056 2.25580301089313
0.133114389440145 0.558908020832006
80725 SRCIN1 SRC kinase signaling inhibitor 1 11397
0.439336121997185 0.404653800082258 2.25574474458306
0.133119399520903 0.558908020832006
26091 HERC4 HECT and RLD domain containing E3 ubiquitin protein
ligase 4 15847 -0.224461689071134 6.3637319908914
2.25458679104634 0.133219010900428 0.558908020832006
2317 FLNB filamin B 13952 -0.322939749417842
10.6109385811341 2.25452784247045 0.133224084098775
0.558908020832006
55040 EPN3 epsin 3 4582 -0.552593976216034
2.98812543038682 2.25401773345748 0.133267993823576

0.558908020832006
9867 PJA2 praja ring finger ubiquitin ligase 2
5217 -0.198838717943765 7.1261353341386 2.25396134660354
0.133272848545648 0.558908020832006
9778 KIAA0232 KIAA0232 8499 -0.174969130390339
6.89617038137487 2.25247637738838
0.133400770649321 0.559277041150866
28978 TMEM14A transmembrane protein 14A 992
0.293417450965312 3.88977760831301 2.25054294951075
0.133567530717883 0.55980861842172
7586 ZKSCAN1 zinc finger with KRAB and SCAN domains 1
10319 -0.265536946866758 7.54362777623582
2.24962484330564 0.133646799855265 0.559973295115971
93099 DMKN dermokine 7444 0.368705105952928
5.61394646154269 2.2485842168183 0.133736710951006
0.560182449249624
5926 ARID4A AT-rich interaction domain 4A
7568 -0.292283691026508 4.78092180972206
2.24755365007029 0.13382581951088 0.56022886427513
51322 WAC WW domain containing adaptor with coiled-coil
12188 -0.172965985443385 7.54634841953425
2.24753093107765 0.133827784669421 0.56022886427513
580 BARD1 BRCA1 associated RING domain 1
6067 -0.517736481445235 2.63899196505973
2.24521727305047 0.134028081609087 0.56075466429738
9330 GTF3C3 general transcription factor IIIC subunit 3
7105 -0.199022354587467 5.53585251616687
2.24515524771524 0.134033455848335 0.56075466429738
54882 ANKHD1 ankyrin repeat and KH domain containing 1
11124 -0.269781069079719 7.169220678966 2.24242642931203
0.134270135330185 0.561215103208574
28227 PPP2R3B protein phosphatase 2 regulatory subunit B''beta
18030 0.361429415203158 3.6735359889691
2.24204858659501 0.134302943698212 0.561215103208574
10717 AP4B1 adaptor related protein complex 4 subunit beta 1
5369 -0.286434531848913 3.69520417520016
2.24192935751402 0.134313298309662 0.561215103208574
10406 WFDC2 WAP four-disulfide core domain 2
1382 -0.334517603937101 6.81888608873486
2.24165710606624 0.13433694570286 0.561215103208574
83666 PARP9 poly(ADP-ribose) polymerase family member 9
8938 -0.272244193751362 5.75797707889201
2.24157768143999 0.134343845295806 0.561215103208574
128272 ARHGEF19 Rho guanine nucleotide exchange factor 19
3745 -0.445738232619133 3.75946772762614
2.23976334605595 0.134501564019471 0.561628599648954
9 NAT1 N-acetyltransferase 1 3168 -0.302413069058417
4.03702929490076 2.23937527888041
0.134535325262339 0.561628599648954
57103 TIGAR TP53 induced glycolysis regulatory phosphatase 4017
0.233894679394519 5.71945997124441
2.23905590915099 0.134563117043042 0.561628599648954
10018 BCL2L11 BCL2 like 11 6658 -0.327494612374851
4.40757761334986 2.23802225738041 0.134653110182866

0.561743582658304
9472 AKAP6 A-kinase anchoring protein 6 19748
0.345495278573916 2.53099754006413 2.2378183002247
0.134670875317097 0.561743582658304
7045 TGFBI transforming growth factor beta induced 7810
0.234917273516764 11.1096464592376 2.23720107585143
0.134724652942285 0.56180059951253
27342 RABGEF1 RAB guanine nucleotide exchange factor 1
6150 -0.220914460865959 6.38978452708025
2.23346845568289 0.135050381591377 0.562991278259052
6929 TCF3 transcription factor 3 6080 0.178960980782863
7.11320666835462 2.23137616663874
0.135233351332501 0.56342863336808
84320 ACBD6 acyl-CoA binding domain containing 6 2356
0.235767255922019 4.54518746757568 2.23134882774894
0.135235743941135 0.56342863336808
9813 EFCAB14 EF-hand calcium binding domain 14
6285 -0.181649940416041 7.60908060087011
2.23062578470164 0.135299039456204 0.563524723658357
10730 YME1L1 YME1 like 1 ATPase 7335 -0.19712905017361
7.90718761743847 2.22875943261299
0.135462574031381 0.564038131527215
2696 GIPR gastric inhibitory polypeptide receptor
4736 -0.455084827762286 3.22320486777727
2.22735303235646 0.135585952588742 0.564384082588563
55214 P3H2 prolyl 3-hydroxylase 2 5605 0.414132685584158
3.77223585165298 2.22644667095756
0.135665531130079 0.564547562251641
23369 PUM2 pumilio RNA binding family member 2
6978 -0.239695365603963 7.16266421382351
2.22542483348964 0.13575531116728 0.564647246783624
80267 EDEM3 ER degradation enhancing alpha-mannosidase like
protein 3 7804 -0.221438218663693 5.89910366441301
2.2247190669125 0.135817359611236 0.564647246783624
115992 RNF166 ring finger protein 166 4394 0.473726000735937
3.86851650969058 2.22463464363569
0.135824783925749 0.564647246783624
5611 DNAJC3 DnaJ heat shock protein family (Hsp40) member C3
5591 -0.237699910487292 7.03792023214734
2.22400189242318 0.135880443517577 0.564647246783624
106707243 PRH1-TAS2R14 PRH1-TAS2R14 readthrough 1662
0.44130998679268 0.860288050559939
2.22359965137288 0.135915839691248 0.564647246783624
3429 IFI27 interferon alpha inducible protein 27 4147
1.03293786761066 6.373364069092 2.22241132103795
0.136020469996003 0.564647246783624
444882 IGFL4 IGF like family member 4 1190
1.11486568393034 0.568491573740752 2.2223252959015
0.136028047850671 0.564647246783624
29078 NDUFAF4 NADH:ubiquinone oxidoreductase complex assembly
factor 4 2540 0.356544533542673 3.14320834110393
2.22215900350513 0.136042697695208
0.564647246783624
988 CDC5L cell division cycle 5 like

6423	-0.166968374485972	6.30764310238109
2.22205010497774	0.136052292274915	0.564647246783624
7022	TFAP2C	transcription factor AP-2 gamma
3005	-0.379557456758621	4.31487817230993
2.22113143254913	0.136133262700284	0.564815939171469
8926	SNURF	SNRPN upstream reading frame	1734
0.241191359710818	4.83841755265832	2.2191724547522
0.136306104350897	0.565196294003188
84706	GPT2	glutamic--pyruvic transaminase 2
6354	-0.227011566004854	5.42288847366453
2.21876451840272	0.136342127694539	0.565196294003188
25975	EGFL6	EGF like domain multiple 6	2607
0.688358073708076	-0.167405059932788	2.21849971097925
0.136365517560466	0.565196294003188
404734	ANKHD1-EIF4EBP3 ANKHD1-EIF4EBP3 readthrough
11385	-0.258635524023783	7.21370894980259
2.21826400580354	0.136386340667579	0.565196294003188
55729	ATF7IP	activating transcription factor 7 interacting
protein 13189	-0.252645276265962	6.63685161013153
2.2158877713327 0.136596465427407	0.565770643496053
23353	SUN1	Sad1 and UNC84 domain containing 1
9969	-0.241835156896177	8.08200057070889
2.21565820107182	0.136616784976762	0.565770643496053
8562	DENR	density regulated re-initiation and release factor
3576	-0.217432390715877	6.71141679520468
2.215214885648	0.13665603295763	0.565770643496053
10409	BASP1	brain abundant membrane attached signal protein 1
2217	0.266893333158396	6.82553267988033
2.21456346522416	0.136713727982863	0.565770643496053
541468	LURAP1	leucine rich adaptor protein 1	1854
0.690994500603721	0.568246819420463	2.21402045931751
0.136761841797516	0.565770643496053
5498	PPOX	protoporphyrinogen oxidase	4607
0.315329199084416	3.77019355484387	2.21395901103887
0.13676728770455	0.565770643496053
8295	TRRAP	transformation/transcription domain associated
protein 13218	-0.312601044419973	6.70560889053501
2.21305571884707	0.136847370795738	0.565934786754029
9020	MAP3K14 mitogen-activated protein kinase kinase kinase 14
5852	-0.429265680750581	2.3361718744713
2.21250245013072	0.136896447826783	0.565970644837589
284004	HEXD	hexosaminidase D	4153	0.351026314102845
3.32496953857849	2.21050881032941
0.137073454824471	0.566535226239705
100507178	SLFNL1-AS1	SLFNL1 antisense RNA 1
4800	-0.610075471138645	0.699312840774823
2.20944903725502	0.137167652040603	0.566586171180416
54839	LRRC49	leucine rich repeat containing 49
9991	-0.388194520885204	2.81575529105366
2.20868215691869	0.137235860912851	0.566586171180416
2178	FANCE	FA complementation group E
2554	-0.395081350471045	1.84596809103115
2.20859722267503	0.13724341758253	0.566586171180416
9475	ROCK2	Rho associated coiled-coil containing protein kinase

2	8444	-0.221134825589717	6.65082808704374
2.20850034136485	0.137252037761779	0.566586171180416
2201	FBN2	fibrillin 2	13196	0.661427729266497
1.50847878240754	2.20809587885936	0.137288032054425
	0.566586171180416
93624	TADA2B	transcriptional adaptor 2B
4687	-0.207955410038179	5.25454548679181
2.20737416616902	0.137352285640491	0.566637155652638
26277	TINF2	TERF1 interacting nuclear factor 2
2712	-0.222723314810231	6.14380683282685
2.20686093500242	0.13739799876134	0.566637155652638
286527	TMSB15B thymosin beta 15B	6690	0.368347585187675
	3.74219971428398	2.20642538814308
0.137436805971792	0.566637155652638
1729	DIAPH1	diaphanous related formin 1
7082	-0.194809473013026	8.48278696264715
2.20614044107202	0.137462201392708	0.566637155652638
8767	RIPK2	receptor interacting serine/threonine kinase 2
3117	-0.417941491249989	4.94504799396403
2.2052390117487 0.137542574379536	0.566801659115671
84304	NUDT22	nudix hydrolase 22	2575	0.222939074415159
	4.69814377157247	2.20471696491891
0.137589145052747	0.566815176743641
339123	JMJD8	jumonji domain containing 8	2627
0.28248832105975	5.35985175349225	2.20429514333598
	0.137626787756488	0.566815176743641
23587	ELP5	elongator acetyltransferase complex subunit 5	2768
	0.331067116232731	3.73973173358054
2.20304848344202	0.13773810523465	0.567106890071061
7074	TIAM1	TIAM Rac1 associated GEF 1
8751	-0.285408053915791	4.22479585584682
2.19906495101602	0.138094482174097	0.568407114843545
60681	FKBP10	FKBP prolyl isomerase 10	4273
0.446439443682377	4.73130167426954	2.1976603372104
0.138220388911557	0.568758221940124
165679	SPTSSB	serine palmitoyltransferase small subunit B
3745	-1.05716975576199	1.14770062548362
2.19685160591646	0.138292940332592	0.568889637368171
140700	SAMD10	sterile alpha motif domain containing 10
2453	-0.334431884495211	3.06289781123044
2.19573262445946	0.138393394774584	0.569135725369231
1108	CHD4	chromodomain helicase DNA binding protein 4
7472	-0.204882044232833	8.13934080096074
2.19346403279243	0.138597304510982	0.569725717326886
1780	DYNC1I1 dynein cytoplasmic 1 intermediate chain 1	4568
	0.557076545633154	2.13250687456205
2.19323167908267	0.138618208370817	0.569725717326886
55	ACPP	acid phosphatase, prostate
4193	-0.740699282376977	0.726404208352309
2.19211290494375	0.138718909165127	0.569848943947707
5562	PRKAA1	protein kinase AMP-activated catalytic subunit alpha
1	6563	-0.190157022752669	6.41905035095498
2.19199466677478	0.138729556568978	0.569848943947707
55365	TMEM176A	transmembrane protein 176A	3479

1.27741116869755	0.982256935878963	2.19091915236284
	0.13882644926139	0.570079763941451
6641	SNTB1	syntrophin beta 1	6499	-0.602178451926883
	4.31521872904332	2.18972011032272
0.138934560011364	0.570356501195538
23344	ESYT1	extended synaptotagmin 1	5493
0.207881104846908	8.40413019522979	2.18811951073086
	0.139079024123788	0.570782270993816
5051	PAFAH2	platelet activating factor acetylhydrolase 2
3835	-0.248524637777483	5.03660119022994
2.18641604837618	0.1392329573949 0.570871099401043
2037	EPB41L2 erythrocyte membrane protein band 4.1 like 2
10247	0.383467949216277	3.68875933825523
2.18624532204368	0.139248395606032	0.570871099401043
84376	HOOK3	hook microtubule tethering protein 3
15250	-0.339208251042792	7.34949719942017
2.18613530168256	0.139258345396955	0.570871099401043
79608	RIC3	RIC3 acetylcholine receptor chaperone	7657
0.558982586564522	3.36759471488204	2.1854672513166
0.139318778267431	0.570871099401043
4850	CNOT4	CCR4-NOT transcription complex subunit 4
6999	-0.219119901853592	5.00318507633432
2.18517643285361	0.139345095349194	0.570871099401043
4233	MET	MET proto-oncogene, receptor tyrosine kinase
7039	-0.254453829688872	9.13147459665035
2.18515423771652	0.139347104086396	0.570871099401043
23403	FBXO46	F-box protein 46	3535	-0.31160476651037
	3.7313793250573 2.18406635176545	0.139445601394549
	0.570871099401043
4085	MAD2L1	mitotic arrest deficient 2 like 1	5748
0.452176282838496	1.32871222021208	2.18291822835491
	0.139549637358524	0.570871099401043
115290	FBXO17	F-box protein 17	4960	0.645874838041378
	1.5124764386442 2.18279154817	0.139561121681703
0.570871099401043
56102	PCDHGB3 protocadherin gamma subfamily B, 3	4757
0.211275483878105	6.45280519338835	2.18272525500934
	0.139567131979521	0.570871099401043
112616	CMTM7	CKLF like MARVEL transmembrane domain containing 7
	3751	0.224167115025098	5.97875940512234
2.18223272442225	0.139611795088206	0.570871099401043
55667	DENND4C DENN domain containing 4C
8745	-0.227859733830278	6.52467675280473
2.1821704204477 0.139617446010502	0.570871099401043
2263	FGFR2	fibroblast growth factor receptor 2	7486
0.408327598377885	5.2115194008192 2.18202652634078
0.139630498075817	0.570871099401043
100130476	WAKMAR2 wound and keratinocyte migration associated
lncRNA 2	2981	-0.497758795218085	0.759918578384043
	2.18151733292962	0.13967669599495
0.570893341348487
64207	IRF2BPL interferon regulatory factor 2 binding protein like
	4157	0.257327685605396	5.05595816811988
2.18031483094705	0.139785864282063	0.57104007273019

112574	SNX18	sorting nexin 18	5559	-0.258245549183656
5.42875318787682	2.18022380202947
0.139794132176185	0.571040072730191
55630	SLC39A4 solute carrier family 39 member 4	5033
0.455835029150546	3.22944261505946	2.17862979988137
0.139938999800158	0.571341279520426
5596	MAPK4	mitogen-activated protein kinase 4
7601	-0.590459172249529	2.39144354351387
2.17851489891496	0.139949448849474	0.571341279520426
10142	AKAP9	A-kinase anchoring protein 9
18210	-0.343097468791547	6.95074433587754
2.17725064458827	0.14006447750618	0.571561063427989
1459	CSNK2A2 casein kinase 2 alpha 2 6905	-0.160008986529623
6.4661454971215 2.17682416496618	0.140103304855343
0.571561063427989
26112	CCDC69	coiled-coil domain containing 69
4993	-0.275177062665683	5.66673199837974
2.17646771032832	0.140135766295376	0.571561063427989
283106	CSNK2A3 casein kinase 2 alpha 3 1416	-0.188727229282693
5.36296072524754	2.17587644243096
0.140189630219636	0.571561063427989
7718	ZNF165	zinc finger protein 165 2107	-0.344902663747328
3.99389849416712	2.17562911251242
0.140212168627566	0.571561063427989
54877	ZCCHC2	zinc finger CCHC-type containing 2
9103	-0.314129433175506	4.2117307336112 2.17504529975328
0.140265385811017	0.571561063427989
8412	BCAR3	BCAR3 adaptor protein, NSP family member
5042	-0.334008761300221	7.19568916994025
2.17478717372683	0.140288922400616	0.571561063427989
114991	ZNF618	zinc finger protein 618 10384	0.362647624493005
5.46523367666729	2.17403729602999
0.140357323321727	0.571594974583864
767	CA8	carbonic anhydrase 8	3812	-1.21746945845756
0.512602700624838	2.17380125879651
0.140378861467016	0.571594974583864
5927	KDM5A	lysine demethylase 5A	12106	-0.243494497887366
6.76479883921591	2.17009627420993
0.140717424206196	0.572807021463584
3716	JAK1	Janus kinase 1	5047	-0.202401809201563
7.81225250230931	2.1662989181269 0.141065379942959
0.573870670001679
10200	MPHOSPH6	M-phase phosphoprotein 6
2190	-0.28078323657396	4.55150910880053
2.16562627266779	0.14112711587483	0.573870670001679
140469	MYO3B	myosin IIIB	8236	0.689465760140256
2.10608367889968	2.16491628230942	0.14119231228471
0.573870670001679
647135	SRGAP2B SLIT-ROBO Rho GTPase activating protein 2B	7123
0.313625137979918	4.23268925391819
2.16427334354133	0.14125138073627	0.573870670001679
11343	MGLL	monoglyceride lipase	10102	0.239303291061201
8.38684051805642	2.16408897954857
0.141268323856667	0.573870670001679

9149	DYRK1B	dual specificity tyrosine phosphorylation regulated kinase 1B	3192	0.330453950097285	5.30365124341158	2.16397480370223	0.141278817805021	0.573870670001679
653361	NCF1	neutrophil cytosolic factor 1	3462	1.13631053498518	0.752364789345683	2.16320373021373	0.141349710419779	0.573870670001679
595135	PGM5P2	phosphoglucomutase 5 pseudogene 2	1980	-0.655647525043603	0.531736881473244	2.16195314886408	0.141464774072234	0.573870670001679
3930	LBR	lamin B receptor	5257	-0.26251013475436	4.85034525925157	2.16189217906585	0.141470386479846	0.573870670001679
9668	ZNF432	zinc finger protein 432	2701	-0.416672441508815	3.38076450106764	2.16113854516731	0.141539780849769	0.573870670001679
56894	AGPAT3	1-acylglycerol-3-phosphate O-acyltransferase 3	10961	-0.206791253836164	7.16664778551287	2.16086940244118	0.141564572696613	0.573870670001679
9262	STK17B	serine/threonine kinase 17b	6227	-0.469910553395369	5.20843823934256	2.16071230962069	0.14157904542008	0.573870670001679
9466	IL27RA	interleukin 27 receptor subunit alpha	2962	0.630698333077836	1.78394773494271	2.16055942869106	0.141593131705127	0.573870670001679
1843	DUSP1	dual specificity phosphatase 1	2019	-0.358703944154063	7.03125224766077	2.1603676336864	0.141610805719062	0.573870670001679
10385	BTN2A2	butyrophilin subfamily 2 member A2	6398	-0.21856828823647	5.16056396901645	2.16032559377709	0.141614680050863	0.573870670001679
284111	SLC13A5	solute carrier family 13 member 5	5623	-1.00609168016056	1.17666896136608	2.16011328835173	0.14163424760447	0.573870670001679
301	ANXA1	annexin A1	4561	-0.314330973378139	9.12035575765904	2.15891685521922	0.141744576089427	0.573946195246532
10288	LILRB2	leukocyte immunoglobulin like receptor B2	4514	1.24176763848353	0.216615338882382	2.15876193458251	0.14175886908153	0.573946195246532
6654	SOS1	SOS Ras/Rac guanine nucleotide exchange factor 1	9263	-0.241681965077288	7.16667176235674	2.15857828581527	0.141775814632184	0.573946195246532
348938	NIPAL4	NIPA like domain containing 4	4208	-1.28219036476934	2.2037764990749	2.15754092138571	0.141871576591713	0.574071491552054
535	ATP6V0A1	ATPase H+ transporting V0 subunit a1	8182	-0.193200316871062	6.70289901765102	2.15735513767505	0.141888734472279	0.574071491552054
2332	FMR1	FMRP translational regulator 1	8827	-0.239483899851421	5.85880004022788	2.15667946091853	0.141951155638901	0.574158197237679
7805	LAPTM5	lysosomal protein transmembrane 5	3274	1.31868783059228	4.59511022335672	2.15536473222585

0.142072702954997	0.574388004010888
103908605	LOC103908605	uncharacterized LOC103908605	4853
0.609793145178387	1.38234570443858
2.15517789114118	0.142089985999695	0.574388004010888
337867	UBAC2	UBA domain containing 2	4603	0.176389442305021
6.42988608799651	2.15464502309294
0.142139289977197	0.574421533384478
400242	DICER1-AS1	DICER1 antisense RNA 1	1826
0.436500523599823	0.388192044047514	2.15232569544234
0.142354111538243	0.574913023338754
51094	ADIPOR1	adiponectin receptor 1	3193	-0.200939288800089
7.57189014146648	2.15229087954776
0.142357339048114	0.574913023338754
83850	ESYT3	extended synaptotagmin 3	4407
0.501035790654899	1.86991013009687	2.15174956905815
0.142407530306441	0.574913023338754
666	BOK	BCL2 family apoptosis regulator BOK	2657
0.199365876405615	6.54694423588261	2.15156027356698
0.142425086812699	0.574913023338754
57560	IFT80	intraflagellar transport 80	8582
0.231785538784846	4.35708525341497	2.15032502233444
0.14253971190615	0.575209952368034
5143	PDE4C	phosphodiesterase 4C	7900	-0.608151105144058
0.920892726721896	2.14869455154948
0.142691170278458	0.575271384560355
100130449	LOC100130449	uncharacterized LOC100130449
2032	-0.872244978943763	3.18978864402679
2.14816393363115	0.142740499694086	0.575271384560355
150771	ITPRIPL1	ITPRIP like 1	4652	-0.599646495914017
2.23450790681525	2.14815191025861
0.142741617680951	0.575271384560355
123745	PLA2G4E	phospholipase A2 group IVE
5823	-1.36672063221902	0.926489008197002
2.14782452754449	0.142772062973325	0.575271384560355
6451	SH3BGRL	SH3 domain binding glutamate rich protein like	2304
0.307511566269826	5.52545818628688
2.14782101739824	0.142772389442564	0.575271384560355
441250	TYW1B	tRNA-yW synthesizing protein 1 homolog B	3258
0.615271809389232	2.85809070232805
2.14750960128025	0.142801356758503	0.575271384560355
140467	ZNF358	zinc finger protein 358	2285	0.322434999184271
5.15722963939033	2.14543914075362
0.142994115121716	0.575843938259595
3248	HPGD	15-hydroxyprostaglandin dehydrogenase	5529
0.727788306558133	1.70178159298983	2.1447786210293
0.143055670600893	0.575843938259595
348825	TPRXL	tetrapeptide repeat homeobox like
3636	-0.603103119423681	0.0129538494544028
2.14431172201015	0.143099200069318	0.575843938259595
7976	FZD3	frizzled class receptor 3
13787	-0.300808423104196	5.02612972585267
2.14421808646372	0.143107931600085	0.575843938259595
100506881	MKLN1-AS	MKLN1 antisense RNA	2715
0.480240625056838	1.15232310708362	2.14333492576939

0.143190315984338 0.575843938259595
57115 PGLYRP4 peptidoglycan recognition protein 4
2272 -1.50551562880494 3.32752308148564
2.14292349557022 0.143228713883236 0.575843938259595
4162 MCAM melanoma cell adhesion molecule 7422
0.350954115668776 7.45298718704047 2.14285637578226
0.143234979128431 0.575843938259595
55620 STAP2 signal transducing adaptor family member 2
2766 -0.200947396874732 6.01153981861753
2.14245584552196 0.143272372730398 0.575843938259595
100616124 MIR4647 microRNA 4647 80 0.348408367636259
2.68746704432708 2.14142723537956
0.143368454363887 0.576064813618749
84316 NAA38 N(alpha)-acetyltransferase 38, NatC auxiliary subunit 1466 0.293797130062401 4.39676870970551
2.14006707269539 0.143495617408536 0.576378892810294
80217 CFAP43 cilia and flagella associated protein 43
7184 -0.569582991388798 0.397622388337522
2.13971108860934 0.143528919691747 0.576378892810294
10795 ZNF268 zinc finger protein 268 14628 -0.224005677784939
4.66413398742985 2.13878183975545
0.143615891846757 0.576562853854263
2622 GAS8 growth arrest specific 8 5615
0.237679429349505 4.77144515321004 2.13803924769095
0.143685436682081 0.576676765503126
285195 SLC9A9 solute carrier family 9 member A9 4087
0.446107298838137 0.643461976426089 2.13685036263881
0.143796856450779 0.576958627415084
23199 GSE1 Gse1 coiled-coil protein
10173 -0.26879233985988 5.81472895248839
2.1363111357549 0.143847423700704 0.576996238194662
53635 PTOV1 PTOV1 extended AT-hook containing adaptor protein
4931 0.284671665632743 6.67050013010266
2.13521805904899 0.143949990933311 0.577242348411936
27291 R3HCC1L R3H domain and coiled-coil containing 1 like
3571 -0.268815500395101 4.79025912588626
2.13471718457392 0.143997017253221 0.577265661323945
55316 RSAD1 radical S-adenosyl methionine domain containing 1
4198 0.225503228703537 4.11061709422473
2.13273583761047 0.144183212437434 0.57761650476427
56913 C1GALT1 core 1 synthase, glycoprotein-N-acetylgalactosamine 3-beta-galactosyltransferase 1 7805 -0.162615588043607
5.95329811875185 2.13250436168427 0.144204982845887
0.57761650476427
116238 TLCD1 TLC domain containing 1 1311 -0.632921977523729
2.49701481716534 2.13204113299597
0.144248560815742 0.57761650476427
401115 C4orf48 chromosome 4 open reading frame 48 540
0.473007459684598 1.27892582717606 2.13171097476164
0.14427962931114 0.57761650476427
11214 AKAP13 A-kinase anchoring protein 13
19875 -0.322132429879701 7.39859951637327
2.13159310913414 0.144290722508045 0.57761650476427
56479 KCNQ5 potassium voltage-gated channel subfamily Q member 5

	7570	-0.609375929792385	0.843625202492483
2.12922984121407	0.14451334978328	0.578054331652536
10422	UBAC1	UBA domain containing 1 2979	0.230880521624711
	4.91754574030628	2.12918740495969
0.144517350946094	0.578054331652536
26240	FAM50B	family with sequence similarity 50 member B	1628
	0.385073694428642	2.87203095707476
2.12911794705209	0.144523900153293	0.578054331652536
5495	PPM1B	protein phosphatase, Mg2+/Mn2+ dependent 1B
7103	-0.169301335726792	5.87936744280397
2.12738681661465	0.144687237011127	0.57838140189942
8165	AKAP1	A-kinase anchoring protein 1	5399
0.200980732169611	5.54900522259423	2.12737600070605
	0.144688258174882	0.57838140189942
1907	EDN2	endothelin 2	1639	-0.734536289145704
3.76151318621045	2.12666998800707	0.144754932605759
	0.578482836236483
5316	PKNOX1	PBX/knotted 1 homeobox 1
7621	-0.219072618734917	4.9930612341348 2.12494004162026
	0.144918451649126	0.578566285200713
348995	NUP43	nucleoporin 43	4432	-0.201053496914525
5.23243676585533	2.12490058818938	0.144922183316567
	0.578566285200713
90594	ZNF439	zinc finger protein 439 3043	-0.52767283988355
	2.86354972962419	2.12429162812301
0.144979794992663	0.578566285200713
57403	RAB22A	RAB22A, member RAS oncogene family
9285	-0.192508103256296	6.74924415947314
2.12416227373386	0.144992036101422	0.578566285200713
3399	ID3	inhibitor of DNA binding 3, HLH protein 1797
0.355682855258655	5.7397110156543 2.12406509345166
0.145001233265144	0.578566285200713
23593	HEBP2	heme binding protein 2	10356	-0.191523562239615
	6.88922699536441	2.12352834910673
0.145052042712981	0.578566285200713
23294	ANKS1A	ankyrin repeat and sterile alpha motif domain
containing 1A	6777	-0.174671719842615	6.08252782560151
	2.12339200257519	0.14506495278253
0.578566285200713
10849	CD3EAP	CD3e molecule associated protein	3661
0.325138719499769	3.78712450165725	2.1227257394049
0.145128057014202	0.578612113175863
100506658	OCLN	occludin	6761	-0.280902556895044
	7.06716396966199	2.12228212277793
0.145170090727657	0.578612113175863
143187	VTI1A	vesicle transport through interaction with t-SNAREs
1A	8408	-0.177453016140646	5.66537249366519
2.12196265378634	0.145200369658968	0.578612113175863
135112	NCOA7	nuclear receptor coactivator 7
7820	-0.609749529303282	7.61979512690783
2.12128849796193	0.145264288788348	0.578702188014332
3895	KTN1	kinectin 1	17575	-0.177472252329135
8.38660965110078	2.12071296382814	0.145318882332579
	0.578755071035665

7675	ZNF121	zinc finger protein 121 7251	-0.384552673321376
	4.53174994293793	2.12023272857451
0.145364453790295	0.578772002342427
2123	EVI2A	ecotropic viral integration site 2A	2071
1.06634888298811	0.22676825569794	2.1177055815906
0.145604530494392	0.579436193252173
26025	PCDHGA12	protocadherin gamma subfamily A, 12	4815
	0.206158230497586	6.44391276358205
2.11732817729853	0.145640421903889	0.579436193252173
123688	HYKK	hydroxylysine kinase	5707	-0.425907790185531
	0.69626748741982	2.1170752811639 0.145664478081634
	0.579436193252173
7035	TFPI	tissue factor pathway inhibitor 11714
0.41046131148403	4.36063255578282	2.11652409680659
	0.14571692377839	0.579436193252173
5164	PDK2	pyruvate dehydrogenase kinase 2 5481
0.496142703079673	3.93844228229707	2.11630149953099
	0.145738110146884	0.579436193252173
57016	AKR1B10 aldo-keto reductase family 1 member B10
1744	-0.573342287956748	2.04949516127936
2.1144626810764 0.145913258139231	0.57996793608292
2033	EP300	E1A binding protein p300
10611	-0.331146653864576	7.5535037795416 2.11308529639825
	0.146044609891752	0.580325347731567
150372	NFAM1	NFAT activating protein with ITAM motif 1	5817
	1.02156191644434	0.0913883612928138
2.11131299814926	0.14621381803613	0.580411559254203
10146	G3BP1	G3BP stress granule assembly factor 1
12406	-0.17884522608285	7.74955513399002
2.11126152270606	0.146218735897626	0.580411559254203
219749	ZNF25	zinc finger protein 25	4244	0.330576238651552
	2.98027792630154	2.1112528683735 0.146219562733667
	0.580411559254203
644873	LINC01184	long intergenic non-protein coding RNA 1184
	5825	-0.319537961604536	3.43603037778579
2.11112212228635	0.146232054873141	0.580411559254203
10045	SH2D3A	SH2 domain containing 3A
5716	-0.240374397265572	6.31440166159469
2.11011399804372	0.146328416566421	0.580615153325109
1288	COL4A6	collagen type IV alpha 6 chain	9282
0.63312111590281	4.56431698244681	2.10971836212573
	0.146366253044261	0.580615153325109
375061	FAM89A	family with sequence similarity 89 member A	2394
	0.458252492389003	1.8521032742219 2.10918568394438
	0.146417212932237	0.580652860006183
285362	SUMF1	sulfatase modifying factor 1	4279
0.180192119572937	5.88890773744205	2.10854646070145
	0.146478392116171	0.580731060959867
23533	PIK3R5	phosphoinositide-3-kinase regulatory subunit 5	7506
	1.16860874005342	1.39545224256747
2.10762722303507	0.146566421649036	0.580915638946815
1305	COL13A1 collagen type XIII alpha 1 chain	5761
0.976513294530292	0.555184741078271	2.10654181056599
	0.146670441521987	0.581163472248505

1080	CFTR	CF transmembrane conductance regulator
7023	-0.843864713804433	5.44541093500798
2.10496723195666	0.1468214884118 0.581597451409525
79961	DENND2D DENN domain containing 2D	5515
0.254483582939381	4.61306455306729	2.10370895000329
	0.146942319610468	0.581846997306627
10120	ACTR1B	actin related protein 1B	2892
0.18258612349582	6.26243137552595	2.10333782496602
	0.146977979693439	0.581846997306627
2289	FKBP5	FKBP prolyl isomerase 5 10628	0.767797439311962
	1.82611460235345	2.10301398534173
0.14700910426702	0.581846997306627
3120	HLA-DQB2	major histocompatibility complex, class II,
DQ beta 2	2028	1.06681216587042	0.956549794398562
	2.09860245127791	0.147433843151699
0.583232987448549
5253	PHF2	PHD finger protein 2	5569	-0.234039896614431
	5.99305465307315	2.0981628024571 0.147476248012501
	0.583232987448549
2590	GALNT2	polypeptide N-acetylgalactosaminyltransferase 2 7743
	0.237543955948242	7.9980047646111 2.09808033444642
	0.147484203722622	0.583232987448549
4750	NEK1	NIMA related kinase 1	7755	0.226081179161831
	4.34314566109189	2.09709254963917
0.14757953323629	0.583445250364299
56113	PCDHGA2 protocadherin gamma subfamily A, 2	6724
0.203466125322351	6.45015754597168	2.09558771994051
	0.147724895639923	0.583695828433685
80012	PHC3	polyhomeotic homolog 3	16919	-0.319688404156069
	5.8150832427791 2.09522648888041	0.147759813617007
	0.583695828433685
5463	POU6F1	POU class 6 homeobox 1	6021	0.512088720007717
	2.0428626480441 2.09514252868892	0.147767930864985
	0.583695828433685
9133	CCNB2	cyclin B2	2801	0.392701131552578
0.542574088150982	2.09440578951819	0.147839180197519
	0.583812629553607
1958	EGR1	early growth response 1 3138	-0.548010085971844
	5.21760555450538	2.09297860926463
0.147977311789249	0.584041213733647
1997	ELF1	E74 like ETS transcription factor 1
4443	-0.211534859124455	7.43609984617507
2.0929461256828 0.147980457452753	0.584041213733647
2260	FGFR1	fibroblast growth factor receptor 1	12155
0.781186114130627	2.97957536784641	2.09199872719175
	0.148072235387183	0.584238817202332
78988	MRPL57	mitochondrial ribosomal protein L57	2246
0.245893017484496	3.86241479687339	2.09122244924344
	0.148147484162853	0.584246215228748
8634	RTCA	RNA 3'-terminal phosphate cyclase
3037	-0.287166635279252	5.87017090053561
2.09111882365931	0.148157532411831	0.584246215228748
29937	NENF	neudesin neurotrophic factor	1297
0.242823143371766	4.11485300486491	2.09036321116339

0.148230825064584 0.584370719583347
11108 PRDM4 PR/SET domain 4 4777 -0.211311753446903
6.01781006355553 2.08834291506333 0.148426990266683
0.584956992292945
84283 TMEM79 transmembrane protein 79
3067 -0.354603205873148 5.03803681930901
2.08797174359921 0.148463061869167 0.584956992292945
670 BPHL biphenyl hydrolase like 5780 0.261968409591383
4.11533782806639 2.08729587306034
0.148528770513171 0.584997441815311
53981 CPSF2 cleavage and polyadenylation specific factor 2
13035 -0.183862316828919 6.62906441695481
2.08700707881749 0.148556857323985 0.584997441815311
81577 GFOD2 glucose-fructose oxidoreductase domain containing 2
7243 -0.192846643140394 5.49714729244761
2.08606874568627 0.148648156747729 0.585192448444473
54676 GTPBP2 GTP binding protein 2 4624 -0.201727676462204
6.89917661249098 2.08336203658457
0.148911873765093 0.586065921839746
56114 PCDHGA1 protocadherin gamma subfamily A, 1 6252
0.202751014122466 6.44453397349633 2.0821657765802
0.149028594821207 0.586360541477709
3717 JAK2 Janus kinase 2 5285 -0.267552114026204
4.13423370068897 2.08142984521308 0.149100452215002
0.586478526867603
23511 NUP188 nucleoporin 188 8135 -0.163048591425491
6.50950976173297 2.08085432008034 0.149156674596357
0.586500054011513
80328 ULBP2 UL16 binding protein 2 1349 -0.439602092734148
4.97527866212822 2.08051668513692
0.149189668911474 0.586500054011513
5743 PTGS2 prostaglandin-endoperoxide synthase 2
5269 -0.596244163295802 9.84052002499755
2.07956549663606 0.149282665180858 0.586639044506246
56926 NCLN nicalin 4047 0.223105858376781
6.5193450693071 2.07912523990223 0.149325730597291
0.586639044506246
10776 ARPP19 cAMP regulated phosphoprotein 19
6917 -0.176667733301559 8.46351783133027
2.07887035203186 0.14935066985859 0.586639044506246
164832 LONRF2 LON peptidase N-terminal domain and ring finger 2
13933 -0.696532432815443 1.03927277049338
2.07827120997553 0.149409310884091 0.58670485493509
79143 MBOAT7 membrane bound O-acyltransferase domain containing 7
5292 0.243491315867029 7.53610006616787
2.07681864860155 0.149551588382588 0.586992487414887
171586 ABHD3 abhydrolase domain containing 3
4366 -0.270407489434822 4.60261370132208
2.07659997082862 0.149573020997302 0.586992487414887
51375 SNX7 sorting nexin 7 2067 0.283520399029194
5.32832786549312 2.07602694844467 0.149629199409219
0.586992487414887
56922 MCCC1 methylcrotonoyl-CoA carboxylase 1 4321
0.321772460398511 4.41687768788735 2.07539797367808

0.149690890786059	0.586992487414887
200634	KRTCAP3	keratinocyte associated protein 3	1270
0.380375389390919	2.94129930026951	2.0750196394657
0.14972801257659	0.586992487414887
1043	CD52	CD52 molecule	935	1.53738485203551
1.6649365316108	2.07469814318581	0.149759565663403
0.586992487414887
29919	RMC1	regulator of MON1-CCZ1	3090	-0.206513384118579
5.2190095364236	2.0745316554	0.14977590847582
0.586992487414887
7109	TRAPPC10	trafficking protein particle complex 10
11941	-0.191376462810954	6.88637379889785
2.07357600753241	0.149869756015741	0.587151335921208
54899	PXK	PX domain containing serine/threonine kinase like
5354	0.399763397426468	3.26596335552375
2.07279884382974	0.149946124886578	0.587151335921208
25966	C2CD2	C2 calcium dependent domain containing 2
8602	-0.356890907683303	5.31439451401607
2.0722473025133	0.150000349412861	0.587151335921208
7205	TRIP6	thyroid hormone receptor interactor 6	2841
0.25594337022089	6.00342950160819	2.07194961255575
0.150029625875016	0.587151335921208
920	CD4	CD4 molecule	4583	1.1794893842192
2.13204645676427	2.07187152233757	0.150037306764853
0.587151335921208
7171	TPM4	tropomyosin 4	5817	-0.344453184704984
10.3551426652043	2.07156001805165	0.150067950496032
0.587151335921208
163049	ZNF791	zinc finger protein 791	6642	-0.228041199246233
5.14873715681574	2.06996702706505
0.15022476909745	0.587600765358274
63875	MRPL17	mitochondrial ribosomal protein L17	2276
0.256059246482011	4.69183330295141	2.06936812036898
0.15028377511958	0.587667458989381
219902	TLCD5	TLC domain containing 5	4468	0.43535709139685
1.72250395125057	2.06875330483309
0.150344375794545	0.587740349359251
606500	SNORD68	small nucleolar RNA, C/D box 68	2367
0.272974863907058	6.86829574558961	2.06818022394947
0.150400887689094	0.587758905914211
10859	LILRB1	leukocyte immunoglobulin like receptor B1	5008
1.29365451324356	0.476830239728246
2.06760030246083	0.150458098599925	0.587758905914211
8732	RNGTT	RNA guanylyltransferase and 5'-phosphatase
4916	-0.226920292095194	4.99477250258841
2.06742894749125	0.150475007968042	0.587758905914211
147179	WIPF2	WAS/WASL interacting protein family member 2
8123	-0.160039578228607	6.80784162873503
2.06617506444133	0.150598807022028	0.588038879164703
2232	FDXR	ferredoxin reductase	3599	0.324940024029573
3.34491859461493	2.06585274703091
0.150630648850072	0.588038879164703
5720	PSME1	proteasome activator subunit 1
1781	-0.243141760013084	6.34896643589465

2.06488398431238	0.150726399050525	0.588248724296656
125058	TBC1D16 TBC1 domain family member 16	11555
0.228737113474377	6.69585308606599	2.06393714293913
	0.150820049142636	0.58845025859078
7323	UBE2D3	ubiquitin conjugating enzyme E2 D3
10069	-0.191788740601952	8.00186910501774
2.06343461383835	0.150869779982556	0.588480369873477
622	BDH1	3-hydroxybutyrate dehydrogenase 1
9114	-0.401341305309362	3.6402018839103 2.06281414787845
	0.150931207584016	0.588556075899028
64800	EFCAB6	EF-hand calcium binding domain 6	8557
0.448378908160743	0.330542748762283	2.06186096653169
	0.151025629938536	0.588760366977196
55116	TMEM39B transmembrane protein 39B	2820
0.304670449923826	2.87421359976286	2.05972748263153
	0.151237215679426	0.589353887139929
9901	SRGAP3	SLIT-ROBO Rho GTPase activating protein 3
18924	0.525052165707409	2.92348117948352
2.05940351023123	0.151269374588797	0.589353887139929
6804	STX1A	syntaxin 1A	3693	-0.269357765103042
4.41304830332375	2.0590537345735 0.151304103530748
0.589353887139929
1802	DPH2	diphthamide biosynthesis 2	2930
0.222729920707304	4.8235365042344 2.05791381955586
0.151417347415665	0.5895643285543
23770	FKBP8	FKBP prolyl isomerase 8 2902	0.204171445281883
	7.2164717365067 2.05709600971952	0.151498651141418
	0.5895643285543
728673	LOC728673	uncharacterized LOC728673	2541
0.528331225310201	0.899695170207716	2.05669570270276
	0.151538466245806	0.5895643285543
6647	SOD1	superoxide dismutase 1	2019	0.211408544643348
	6.26399362738201	2.05558286319388
0.151649213095018	0.5895643285543
147	ADRA1B	adrenoceptor alpha 1B	1738	0.450631396538046
	3.40094719996809	2.05487553910493
0.15171965174752	0.5895643285543
2580	GAK	cyclin G associated kinase
15155	-0.197268003156647	6.93395835940052
2.0548702634479 0.151720177261357	0.5895643285543
54414	SIAE	sialic acid acetylesterase	8139
0.344291712959505	4.23955794495468	2.05477716026712
	0.15172945170786	0.5895643285543
4909	NTF4	neurotrophin 4	2078	0.447136879754801
2.14367446726695	2.05465918245474	0.151741204955647
	0.5895643285543
56097	PCDHGC5 protocadherin gamma subfamily C, 5	4990
0.202475634482136	6.44139844214239	2.05413389596848
	0.151793547836242	0.5895643285543
170463	SSBP4	single stranded DNA binding protein 4	2974
0.344221090287275	3.34349016496071	2.05291049837833
	0.151915534210878	0.5895643285543
276	AMY1A	amylase alpha 1A	6471	-0.677153082689079
	1.62105225171889	2.05274727642991

0.151931817649433	0.5895643285543
79872	CBLL1	Cbl proto-oncogene like 1
5481	-0.221574860712719	4.88328360900997
2.05263619987008	0.151942900060938	0.5895643285543
56098	PCDHGC4	protocadherin gamma subfamily C, 4	4974
0.202817595194102	6.4393574308627	2.05260302035221
0.151946210649535	0.5895643285543
11345	GABARAPL2	GABA type A receptor associated protein like
2	1363	-0.186388885710375	6.29469967408901
2.05259110468663	0.151947399591706	0.5895643285543
216	ALDH1A1	aldehyde dehydrogenase 1 family member A1	2669
	0.861993827919333	3.33152909258787
2.05063505826784	0.152142716310242	0.590146950071431
128344	PIFO	primary cilia formation	4261	0.516308629879062
	2.28583795073706	2.05024376451287
0.152181822207326	0.590146950071431
53834	FGFRL1	fibroblast growth factor receptor like 1
4102	-0.253699907329436	5.50386432874909
2.04967583406246	0.152238601429357	0.590203733800442
100533195	FAM24B-CUZD1	FAM24B-CUZD1 readthrough	2903
	0.546586346124401	0.913175129813981
2.04852965875668	0.152353264153033	0.590452683725845
286006	LSMEM1	leucine rich single-pass membrane protein 1
2592	-0.672711287498486	0.849941258495899
2.0475819730599	0.152448143970126	0.590452683725845
56998	CTNNBIP1	catenin beta interacting protein 1
3339	-0.240380554473358	5.33431051440073
2.0474026705635	0.152466102796552	0.590452683725845
254896	LOC254896	uncharacterized LOC254896	1491
0.381902322931778	2.87664419073388	2.04721340365429
	0.152485062252418	0.590452683725845
126014	OSCAR	osteoclast associated Ig-like receptor	2084
0.595530489358416	0.323710536734399	2.04597347887019
	0.152609335462698	0.590452683725845
202243	CCDC125	coiled-coil domain containing 125
7932	-0.296853193435604	4.04970517574405
2.04535808094605	0.152671057190822	0.590452683725845
5269	SERPINB6	serpin family B member 6	4729
0.304301849249945	7.37319705186821	2.04522546793362
	0.152684361401241	0.590452683725845
4680	CEACAM6	CEA cell adhesion molecule 6
3052	-0.678188158792639	10.2100497067814
2.04457445933519	0.152749691960647	0.590452683725845
27112	FAM155B	family with sequence similarity 155 member B	4013
	0.442791237587362	2.2720774062967	2.0444544379739
0.152761739902165	0.590452683725845
56104	PCDHGB1	protocadherin gamma subfamily B, 1	4773
0.204207057817878	6.45515248167916	2.04389121562165
	0.152818291470924	0.590452683725845
26095	PTPN20	protein tyrosine phosphatase non-receptor type 20
	4506	0.89478331870618	0.793659407715628
2.04383528661103	0.152823908439823	0.590452683725845
57496	MRTFB	myocardin related transcription factor B
16578	-0.23560050240802	6.49500353164658

2.04356763051167	0.152850792468549	0.590452683725845
80221	ACSF2	acyl-CoA synthetase family member 2	5797
0.371558059876457	5.50581447817003	2.04356735967099
	0.152850819675156	0.590452683725845
1382	CRABP2	cellular retinoic acid binding protein 2
1100	-0.491039863820543	7.44578893432486
2.04067778749982	0.153141396870925	0.591255057743824
1783	DYNC1LI2	dynein cytoplasmic 1 light intermediate
chain 2 8278	-0.182329410604611	8.49334368156498
2.04066232485582	0.153142953487156	0.591255057743824
56655	POLE4	DNA polymerase epsilon 4, accessory subunit	2151
	0.326855901417279	3.32859136274732
2.03965586309561	0.153244312044588	0.591483350457026
8629	JRK	Jrk helix-turn-helix protein
9526	-0.330993410489418	4.7008619674564 2.03894522859758
	0.153315924283835	0.591596735934898
163859	SDE2	SDE2 telomere maintenance homolog
3975	-0.224498178554272	5.12915838224303
2.03834896263562	0.153376040615235	0.591665712172293
8645	KCNK5	potassium two pore domain channel subfamily K member
5	3756	-0.787088945938386	1.80321478152988
2.03720005931976	0.153491949943579	0.591796941096332
79669	C3orf52 chromosome 3 open reading frame 52
6021	-0.270945401290877	6.36109550698507
2.03592035902503	0.153621171895931	0.591796941096332
54874	FNBP1L	formin binding protein 1 like
5798	-0.221052129598262	7.86337793241171
2.03546542974487	0.153667139591491	0.591796941096332
9500	MAGED1	MAGE family member D1	6016	0.271319694048958
	7.25628734911634	2.03513994362751
0.153700037453229	0.591796941096332
100500810	MIR3620 microRNA 3620	79	-0.384575686190288
	1.51834739386205	2.03484810063669
0.15372954169295	0.591796941096332
89839	ARHGAP11B	Rho GTPase activating protein 11B	2497
	0.501561152186647	-0.0517319083064456
2.03444694306687	0.153770107710288	0.591796941096332
23435	TARDBP	TAR DNA binding protein 6421	-0.163806451424733
	7.13171630940582	2.03435058175548
0.153779853805458	0.591796941096332
116843	SLC18B1 solute carrier family 18 member B1
2444	-0.242791702053288	4.06208835049256
2.03400298479669	0.153815015987063	0.591796941096332
374879	ZNF699	zinc finger protein 699 3623	-0.374776868986493
	2.66663509627348	2.03366676557386
0.153849035894715	0.591796941096332
55757	UGGT2	UDP-glucose glycoprotein glucosyltransferase 2	8439
	0.223876160718994	4.66217764909734
2.03353224165591	0.153862649913281	0.591796941096332
80008	TMEM156 transmembrane protein 156
2562	-0.713630066210978	1.74552288544035
2.0334120988164 0.153874809700353	0.591796941096332
43	ACHE	acetylcholinesterase (Cartwright blood group)	3305
	0.577342813657106	0.41432002538541

2.03292481472396	0.153924139426015	0.591824161663516
9770	RASSF2	Ras association domain family member 2	5759
0.750421823217964	2.79287493167819	2.03162488733799
	0.154055824054129	0.591933503960855
60678	EEFSEC	eukaryotic elongation factor, selenocysteine-tRNA
specific	2421	0.30996323419911	2.92174736978608
	2.03138643118229	0.154079993892017
0.591933503960855
124220	ZG16B	zymogen granule protein 16B
2665	-0.524427511196956	1.88486044406895
2.03082113298354	0.15413730951689	0.591933503960855
55332	DRAM1	DNA damage regulated autophagy modulator 1	4180
	0.348394398406201	7.93737492811548
2.03060619471751	0.154159108467762	0.591933503960855
58476	TP53INP2	tumor protein p53 inducible nuclear protein
2	4228	-0.218603902405722	7.19163566601329
2.02975087800974	0.154245888987458	0.591933503960855
27242	TNFRSF21	TNF receptor superfamily member 21
3595	-0.270735637104662	8.17388376610706
2.02949236130866	0.154272129021664	0.591933503960855
79832	QSER1	glutamine and serine rich 1
10537	-0.229180898486839	5.87259014025335
2.02878158220257	0.154344300823501	0.591933503960855
84081	NSRP1	nuclear speckle splicing regulatory protein 1
4806	-0.199308042228658	4.72244856637186
2.02820545853103	0.154402827950519	0.591933503960855
56108	PCDHGA7	protocadherin gamma subfamily A, 7	5443
0.201999942565519	6.44786486147133	2.02767970639701
	0.154456259915952	0.591933503960855
90627	STARD13	StAR related lipid transfer domain containing 13
	9313	-0.434648309024368	4.64119058057829
2.02712102008385	0.154513061957678	0.591933503960855
10670	RRAGA	Ras related GTP binding A	1589
0.174161311841782	6.29129689226001	2.02694420683277
	0.154531043625409	0.591933503960855
121512	FGD4	FYVE, RhoGEF and PH domain containing 4
10629	-0.269812284417561	5.91450460106504
2.02647147915647	0.154579131048798	0.591933503960855
9696	CROCC	ciliary rootlet coiled-coil, rootletin	8871
0.420421146634808	3.23271318913354	2.02637403027511
	0.154589045981873	0.591933503960855
728118	NUTM2A	NUT family member 2A	5082	0.560059188541019
	1.5140727674132	2.02605417984965	0.154621594223411
	0.591933503960855
10569	SLU7	SLU7 homolog, splicing factor
5082	-0.165237909725884	6.28770391753402
2.02496760715517	0.154732222838	0.591933503960855
440533	PSG8	pregnancy specific beta-1-glycoprotein 8	4090
	0.730584603560446	0.54300187480283
2.02417625080768	0.15481285073823	0.591933503960855
84928	TMEM209	transmembrane protein 209
4759	-0.309366849564141	3.93457687823501
2.02410985179496	0.15481961801608	0.591933503960855
5261	PHKG2	phosphorylase kinase catalytic subunit gamma 2

7782	-0.194078052986756	5.85691729830562
2.02377269846118	0.154853985307708	0.591933503960855
89958	SAPCD2	suppressor APC domain containing 2
3848	-0.555466186266077	1.58614202853218
2.02329650915032	0.154902539802218	0.591933503960855
79656	BEND5	BEN domain containing 5	2138	0.558650901913985
	0.420463192079974	2.02296030158318
0.154936831506986	0.591933503960855
7038	TG	thyroglobulin	11132	0.544021471797602
1.57073946437686	2.02284092661222	0.154949009305514
	0.591933503960855
399959	MIR100HG	mir-100-let-7a-2-mir-125b-1 cluster host
gene	5789	1.05118230105411	0.787573239028875
2.02280236511856	0.154952943311153	0.591933503960855
154860	FEZF1-AS1	FEZF1 antisense RNA 1	3313
0.921076348637373	-0.388127987160256	2.02270650892623
	0.15496272295667	0.591933503960855
729678	LINC00847	long intergenic non-protein coding RNA 847
	2453	0.326378345869669	2.76147980652742
2.02266641545484	0.154966813666342	0.591933503960855
576	ADGRB2	adhesion G protein-coupled receptor B2	7756
0.727415875255059	1.9156867370474	2.02219004503763
0.15501542679616	0.591957765303658
5718	PSMD12	proteasome 26S subunit, non-ATPase 12
5364	-0.179411445904632	6.52600070365048
2.02177265226964	0.155058035541311	0.591959090713311
5891	MOK	MOK protein kinase	11009	0.39408723163063
	2.62274475756546	2.02078581095901
0.155158828230855	0.592022629043861
729683	LOC729683	uncharacterized LOC729683	1824
0.437098983196776	0.837534609907956	2.02078206281222
	0.155159211195832	0.592022629043861
23049	SMG1	SMG1 nonsense mediated mRNA decay associated PI3K
related kinase	25988	-0.447047704684398	7.85628902174807
	2.02031959367824	0.155206471976746
0.592041681094302
54014	BRWD1	bromodomain and WD repeat domain containing 1
19794	-0.412484815839318	6.8764442421427	2.01912011971476
	0.155329125138213	0.592068432048935
717	C2	complement C2	5536	0.936195455359214
1.72591723405286	2.01903182950611	0.155338157670056
	0.592068432048935
2908	NR3C1	nuclear receptor subfamily 3 group C member 1
9172	-0.222863215249261	5.74129410035103
2.01830374477234	0.155412667132573	0.592068432048935
56103	PCDHGB2	protocadherin gamma subfamily B, 2	4769
0.200301220816709	6.45741835545933	2.01821680782055
	0.155421566645495	0.592068432048935
60682	SMAP1	small ArfGAP 1	4161	-0.180140587248165
5.71032452405072	2.0181849088999	0.155424832201323
0.592068432048935
71	ACTG1	actin gamma 1	2976	-0.304332976101829
12.0567937820929	2.01715218153731	0.155530596646005
	0.592101091172254

28982	FLVCR1	feline leukemia virus subgroup C cellular receptor 1
	6442	0.325558491272697	2.66282742348699
2.01693387839505	0.155552964130397	0.592101091172254
64975	MRPL41	mitochondrial ribosomal protein L41	1262
0.278075837865044	3.77070210753136	2.01686306016502
	0.155560220997636	0.592101091172254
6389	SDHA	succinate dehydrogenase complex flavoprotein subunit
A	8383	0.162695994518585	6.43833658966808
2.01495846370824	0.155755532567622	0.592683440552753
22908	SACM1L	SAC1 like phosphatidylinositide phosphatase
7352	-0.181981885565172	5.92798215915842
2.01414696955003	0.155838833831233	0.592839365962543
91147	TMEM67	transmembrane protein 67	8281
0.337668983078692	3.06256584462782	2.01194642069764
	0.156064978669693	0.593538462184736
55610	VPS50	VPS50 subunit of EARP/GARPII complex
9454	-0.186853904451467	5.34158663440468
2.01091903481133	0.156170688128867	0.593779269441106
79858	NEK11	NIMA related kinase 11	3515	0.277301915291427
	3.56027707597075	2.01025036351264
0.15623953253327	0.593879818777072
955	ENTPD6	ectonucleoside triphosphate diphosphohydrolase 6
	2777	-0.306177298416477	5.59748850235361
2.00981375045502	0.156284503516272	0.593889593258931
23080	AVL9	AVL9 cell migration associated
14751	-0.189645622589744	6.16258820248782
2.00841782942754	0.156428381553739	0.59399481634408
9354	UBE4A	ubiquitination factor E4A
6576	-0.186224557950331	6.45496485587122
2.00838804191778	0.156431453401465	0.59399481634408
25911	DPCD	deleted in primary ciliary dyskinesia homolog
(mouse)	1718	0.32281997475808	3.47452973488672
2.008310847725	0.156439414399465	0.59399481634408
56109	PCDHGA6	protocadherin gamma subfamily A, 6	7910
0.199498576127623	6.45657317042039	2.00620699261616
	0.156656561239078	0.594556789239396
79778	MICALL2	MICAL like 2	8843	0.301436429522765
5.93119554141172	2.00605449412778	0.156672314491512
	0.594556789239396
53838	C11orf24	chromosome 11 open reading frame 24	3295
	0.304753736307304	4.84808262060868
2.00557704150208	0.156721647485617	0.594582913415775
100526664	LY75-CD302	LY75-CD302 readthrough	5650
0.420198974397519	4.98870472854005	2.00394084416099
	0.156890842214444	0.594783247078383
153396	TMEM161B	transmembrane protein 161B
9598	-0.234795615167793	4.32238354041636
2.00365684467436	0.156920230968227	0.594783247078383
54482	TRMT13	tRNA methyltransferase 13 homolog
4133	-0.362999360937172	3.10450335869034
2.00361946213237	0.156924099843709	0.594783247078383
890	CCNA2	cyclin A2	2798	0.411524704716235
0.745783055539746	2.00342424853262	0.156944305076155
	0.594783247078383

100534612	C1QTNF3-AMACR	C1QTNF3-AMACR readthrough (NMD candidate)	3424	0.232613259357093	4.8239749711201
2.00207232117305	0.15708431505728	0.595152826888645
64426	SUDS3	SDS3 homolog, SIN3A corepressor complex component
	5731	-0.162760027807646	6.15482411518812
2.00155704231759	0.157137716368458	0.595194157158732
57616	TSHZ3	teashirt zinc finger homeobox 3
5817	-0.374496003262625	3.98938248849045
2.00094885137471	0.15720077324734	0.59519658364800 2
904	CCNT1	cyclin T1	7371	-0.209431089886816
5.87919279627688	2.00073123493365	0.15722334257854
	0.595196583648002
79587	CARS2	cysteinyl-tRNA synthetase 2, mitochondrial
10460	0.246648573262778	5.0304360710859 1.99961390303344
	0.157339280788648	0.595381505570187
51473	DCDC2	doublecortin domain containing 2	4996
0.799227762100934	2.96541043189392	1.9994412707947
0.157357202371731	0.595381505570187
121227	LRIG3	leucine rich repeats and immunoglobulin like domains 3	5893	0.279239366205096	4.10786465589244
1.99875613330398	0.157428351882518	0.595489852773004
91664	ZNF845	zinc finger protein 845 4311	-0.252604334187369
	3.83730755791649	1.99826250413555
0.157479636486895	0.595523020591774
23550	PSD4	pleckstrin and Sec7 domain containing 4
13442	-0.374359247623511	6.86177645959398
1.99784660874857	0.157522859841103	0.595525694411426
81614	NIPA2	NIPA magnesium transporter 2
3505	-0.17599455076628	6.65639137742831
1.99659689777722	0.157652821525274	0.595856198355239
9832	JAKMIP2 janus kinase and microtubule interacting protein 2
	9275	1.1602186922454 0.101654869175666
1.99610667953068	0.157703834273557	0.595888213290995
2131	EXT1	exostosin glycosyltransferase 1 8325
0.189431933351052	6.992354690915	1.99466320392513
0.157854153267559	0.596126598379265
7262	PHLDA2	pleckstrin homology like domain family A member 2
	955	-0.358469829136163	5.22734898634396
1.99464857000655	0.157855678031757	0.596126598379265
57210	SLC45A4 solute carrier family 45 member 4
8208	-0.226039487012369	4.39346839498793
1.9942750587983 0.15789460126987	0.596126598379265
7167	TPI1	triosephosphate isomerase 1	2700
0.303766052393844	8.04405566728271	1.99336793999942
	0.157989176687651	0.596314821429193
400931	MIRLET7BHG	MIRLET7B host gene	5728
0.318624034752793	3.58359555517849	1.99259323540417
	0.158069997727757	0.596314821429193
10362	HMG20B	high mobility group 20B 3857	0.202350359615226
	5.29063441030974	1.992572387453	0.158072173339516
	0.596314821429193
54918	CMTM6	CKLF like MARVEL transmembrane domain containing 6
	4413	-0.192252557833008	7.79331928395861
1.98786224229556	0.158564580302506	0.597953877104439

1240	CMKLR1	chemerin chemokine-like receptor 1	5690
1.35289628322916	0.767984296345239	1.98760009979353
	0.1585920363706 0.597953877104439
654434	SNHG20	small nucleolar RNA host gene 20	3500
0.403307551800538	1.53224520487333	1.98637319914038
	0.158720610425015	0.598194850183817
201501	ZBTB7C	zinc finger and BTB domain containing 7C
7648	-0.574434567837361	3.62123626424271
1.98617528561723	0.158741362042782	0.598194850183817
55367	PIDD1	p53-induced death domain protein 1	5320
0.41754922367394	4.26443376202054	1.9855658120365
0.158805285930066	0.59827478214697
79647	AKIRIN1 akirin 1	2741	-0.174894569955049
7.21660436263407	1.98507077669874	0.158857228664485
	0.598309546088583
10423	CDIPT	CDP-diacylglycerol--inositol 3-
phosphatidyltransferase 3325	0.197924131388485
6.30084452403606	1.98368982890885	0.159002230061854
	0.598561512877621
57050	UTP3	UTP3 small subunit processome component
2073	-0.207241928440227	5.42470957718259
1.98335447207274	0.159037465716217	0.598561512877621
114327	EFHC1	EF-hand domain containing 1	7162
0.298334673469885	3.59698599925322	1.98321305001826
	0.159052327474156	0.598561512877621
8139	GAN	gigaxonin	15406	-0.354530258389345
5.64900935919845	1.98264550430036	0.159111985611155
	0.59862519002295
359948	IRF2BP2 interferon regulatory factor 2 binding protein 2
	4724	-0.191280786201132	7.30463643172468
1.98209149363895	0.159170245566582	0.598637301759824
646851	FAM227A family with sequence similarity 227 member A
10781	0.442808727023241	2.22212664591489
1.98051563701463	0.1593360960285 0.598637301759824
6505	SLC1A1	solute carrier family 1 member 1
3998	-0.511463499734606	4.94151656013725
1.98029959832091	0.159358848273981	0.598637301759824
81031	SLC2A10 solute carrier family 2 member 10
5000	-0.196120674948951	5.9438676901928 1.98027108345756
	0.159361851610357	0.598637301759824
8564	KMO	kynurenine 3-monooxygenase
5946	-0.502868745164484	0.764627086937688
1.98013801735332	0.159375867689031	0.598637301759824
170960	ZNF721	zinc finger protein 721 4860	-0.288516065708718
	4.59867536402027	1.97877836584532
0.159519162560139	0.598637301759824
2746	GLUD1	glutamate dehydrogenase 1
3519	-0.181812340925824	6.66673915119064
1.97753228848266	0.1596506165963 0.598637301759824
79613	TANGO6	transport and golgi organization 6 homolog	6839
	0.246428722987335	3.79804491092773
1.97743923261762	0.159660438406948	0.598637301759824
23122	CLASP2	cytoplasmic linker associated protein 2
15211	-0.195458151942125	6.15620333027896

1.97737745976444	0.159666958755003	0.598637301759824
26986	PABPC1	poly(A) binding protein cytoplasmic 1
4705	-0.196578896800221	8.89416203368718
1.97644317385362	0.159765612998721	0.598637301759824
23141	ANKLE2	ankyrin repeat and LEM domain containing 2
17517	-0.206402349666687	8.80111982413437
1.9762739847335 0.159783485643087	0.598637301759824
83706	FERMT3	fermitin family member 3	3066
0.982861161885343	1.63849617839461	1.97620362392093
	0.159790919024151	0.598637301759824
89953	KLC4	kinesin light chain 4	6421	0.283220237994174
	5.11413730088639	1.97544749494813
0.159870826347606	0.598637301759824
79092	CARD14	caspase recruitment domain family member 14
7950	-0.636501069974971	1.90646210330154
1.97497981389191	0.159920273423301	0.598637301759824
339500	ZNF678	zinc finger protein 678 8915	-0.409051761803857
	3.10373978805023	1.9743902243306 0.159982634490185
	0.598637301759824
26260	FBXO25	F-box protein 25	11438	0.193569057631269
	5.12563509540704	1.97367297268299
0.160058535763489	0.598637301759824
144363	ETFRF1	electron transfer flavoprotein regulatory factor 1
	2466	-0.36535883081238	2.44809005272503
1.97259108602288	0.160173101195477	0.598637301759824
51655	RASD1	ras related dexamethasone induced 1	1748
0.741782682961694	3.43778135709498	1.97249188752174
	0.160183610407682	0.598637301759824
9985	REC8	REC8 meiotic recombination protein	3844
0.508567504981806	4.50385761560411	1.97215230762147
	0.160219591871013	0.598637301759824
27249	MMADHC	metabolism of cobalamin associated D
2191	-0.182748934174922	6.95669770935793
1.97095817467549	0.160346193854065	0.598637301759824
93986	FOXP2	forkhead box P2 16141	-0.70284547025908
1.97702465728378	1.97016809974138	0.160430020257228
	0.598637301759824
115908	CTHRC1	collagen triple helix repeat containing 1	1776
	0.699034759569475	2.42252471191	1.96981712512629
	0.160467274437738	0.598637301759824
7030	TFE3	transcription factor binding to IGHM enhancer 3
4283	-0.188930824432375	6.63455401617355
1.96976579429028	0.160472723772617	0.598637301759824
23012	STK38L	serine/threonine kinase 38 like
6921	-0.369969496522081	7.3155963230728 1.96951871140175
	0.160498957299356	0.598637301759824
2533	FYB1	FYN binding protein 1	6100	0.652186120147229
	5.28169392078421	1.96934175238539
0.160517748568829	0.598637301759824
9187	SLC24A1 solute carrier family 24 member 1	9084
0.239204681546761	3.86825348344203	1.96884967998146
	0.160570014893651	0.598637301759824
253827	MSRB3	methionine sulfoxide reductase B3	5791
0.43353860432791	5.28225487968597	1.96874691299328

0.160580932915628       0.598637301759824
161882  ZFPM1   zinc finger protein, FOG family member 1        6110
        0.353829791832523       2.41593072175263
1.96847495024501        0.160609830471324       0.598637301759824
4501    MT1X    metallothionein 1X      1053    0.289672913161969
        3.86362490857164        1.9676097435711 0.160701802920325
        0.598637301759824
56112   PCDHGA3 protocadherin gamma subfamily A, 3      4876
0.198890758694265       6.44022423756325        1.9674242981226
0.160721523791824       0.598637301759824
29789   OLA1    Obg like ATPase 1       6100    -0.254941102686988
        6.0363433997003 1.96738178296345        0.160726045380617
        0.598637301759824
9444    QKI     QKI, KH domain containing RNA binding
17368   -0.431994745919708      7.26258641157186
1.96675285518854        0.160792950290919       0.598637301759824
261734  NPHP4   nephrocystin 4  13167   0.302027673894797
3.56531653474645        1.96642921887811        0.160827390871428
        0.598637301759824
91452   ACBD5   acyl-CoA binding domain containing 5
4371    -0.174095026678848      5.90021862689046
1.96613176091499        0.160859053032094       0.598637301759824
64859   NABP1   nucleic acid binding protein 1
13806   -0.315736004453211      7.5684477520424 1.96567602553486
        0.160907576414465       0.598637301759824
2872    MKNK2   MAPK interacting serine/threonine kinase 2
5221    -0.265341706877702      7.16103541247756
1.96487569796999        0.160992829843868       0.598637301759824
23196   FAM120A family with sequence similarity 120A
7204    -0.189811592642889      7.98922391850809
1.96450735383888        0.161032084329585       0.598637301759824
51727   CMPK1   cytidine/uridine monophosphate kinase 1
3844    -0.219146215855348      7.69612036451731
1.96392196742994        0.161094491492116       0.598637301759824
8554    PIAS1   protein inhibitor of activated STAT 1
9805    -0.188883602722972      5.5638279676272 1.96384921714754
        0.161102249216027       0.598637301759824
201161  CENPV   centromere protein V    2302    0.662405332975384
        0.180089859825836       1.96342356188877
0.161147647486849       0.598637301759824
3140    MR1     major histocompatibility complex, class I-related
        9348    0.190811962283387       5.81597125212539
1.96303696605013        0.161188892513383       0.598637301759824
1024    CDK8    cyclin dependent kinase 8
3538    -0.227295333582637      4.79717635493758
1.96203529566842        0.161295814439664       0.598637301759824
91179   SCARF2  scavenger receptor class F member 2     3782
0.401840830783933       1.94893890021056        1.96103393324022
        0.161402784330628       0.598637301759824
51804   SIX4    SIX homeobox 4  6658    -0.408825568081058
4.57160399133022        1.96072344773962        0.161435968169802
        0.598637301759824
10787   NCKAP1  NCK associated protein 1
22520   -0.178807081984799      8.90817492270198

1.9607177481902 0.161436577394732 0.598637301759824
4783 NFIL3 nuclear factor, interleukin 3 regulated 2085
0.235185942482542 4.52773189331098 1.9604110136112
0.161469368126211 0.598637301759824
401321 LINC00997 long intergenic non-protein coding RNA 997
3118 0.400231120154892 1.05179122412318
1.9603672432576 0.161474047911067 0.598637301759824
10561 IFI44 interferon induced protein 44
1759 -0.594951872120114 1.73263934825845
1.95939241875022 0.161578313091012 0.598637301759824
4311 MME membrane metalloendopeptidase 9620
0.808239645797669 2.22169983790874 1.95910566596323
0.16160899817872 0.598637301759824
124565 SLC38A10 solute carrier family 38 member 10 6815
0.169513630838539 6.43714696585171
1.95910105589191 0.161609491551454 0.598637301759824
25878 MXRA5 matrix remodeling associated 5 9793
0.766205505253852 7.11253529171814 1.95902861508739
0.16161724443753 0.598637301759824
978 CDA cytidine deaminase 963 -0.492419124816703
5.83090964704149 1.9583585277577 0.161688979804416
0.598637301759824
115024 NT5C3B 5'-nucleotidase, cytosolic IIIB 2730
0.197759193799095 4.60956154828587 1.95809028261232
0.161717706632909 0.598637301759824
83593 RASSF5 Ras association domain family member 5
6654 -0.511134467045001 5.06007409042335
1.95784384259495 0.161744103443789 0.598637301759824
8575 PRKRA protein activator of interferon induced protein
kinase EIF2AK2 3941 0.20959486534062 5.09691445865126
1.95784142966937 0.161744361922617
0.598637301759824
9765 ZFYVE16 zinc finger FYVE-type containing 16
11461 -0.198233283747035 6.18115921292395
1.95775884431591 0.161753208963261 0.598637301759824
2820 GPD2 glycerol-3-phosphate dehydrogenase 2
7155 -0.321914483341105 6.40498111598602
1.9576285322662 0.161767169897732 0.598637301759824
9975 NR1D2 nuclear receptor subfamily 1 group D member 2
5820 -0.275571316400473 6.39434919806351
1.95736791990767 0.161795094631787 0.598637301759824
100379345 MIR181A2HG MIR181A2 host gene 662
0.415228407566866 0.585850982147282 1.9572123309742
0.161811768678617 0.598637301759824
6782 HSPA13 heat shock protein family A (Hsp70) member 13
4194 -0.200651819861654 5.76693974923365
1.95710338336928 0.16182344546665 0.598637301759824
64284 RAB17 RAB17, member RAS oncogene family 4675
0.534472792025231 3.91337448614299 1.9567241277091
0.161864100832445 0.598637301759824
163486 DENND1B DENN domain containing 1B
13203 -0.287060644910283 4.75743065082005
1.95575737864586 0.161967786900203 0.598637301759824
79752 ZFAND1 zinc finger AN1-type containing 1

4214	-0.302477456086628	4.74340889450359
1.955227704370509	0.162024698727122	0.598637301759824
57528	KCTD16	potassium channel tetramerization domain containing
16	14294	-0.565158296975161	1.57178882056146
1.95379455390137	0.162178537556593	0.598637301759824
64981	MRPL34	mitochondrial ribosomal protein L34	1348
0.261287996797569	4.39628597449212	1.95353716848409
0.162206196507836	0.598637301759824
57186	RALGAPA2	Ral GTPase activating protein catalytic
alpha subunit 2 11366	-0.254187601177119	6.46841749989825
1.9533942998378 0.162221551669829	0.598637301759824
56105	PCDHGA11	protocadherin gamma subfamily A, 11	5814
0.195088482120417	6.46130247370847
1.95296872109728	0.162267301629418	0.598637301759824
84717	HDGFL2	HDGF like 2	4299	0.244828664341877
5.91432860328223	1.95281095949288	0.162284264832705
0.598637301759824
57705	WDFY4	WDFY family member 4	13688	0.858603892474266
0.962150780196291	1.9524485059649 0.162323245053143
0.598637301759824
2879	GPX4	glutathione peroxidase 4	2019
0.266983508039597	7.1627660235699 1.95197729652341
0.162373937449026	0.598637301759824
79363	CPLANE2 ciliogenesis and planar polarity effector 2	1545
0.378409949565423	1.44555496017465
1.9519562101603 0.162376206328068	0.598637301759824
7691	ZNF132	zinc finger protein 132 3031	0.35392450075689
1.93604603976388	1.95131288190447
0.162445445424576	0.598637301759824
7169	TPM2	tropomyosin 2	4047	0.471970696648891
6.47412996812487	1.95130279720735	0.162446531072277
0.598637301759824
10451	VAV3	vav guanine nucleotide exchange factor 3	7271
0.704776847578774	3.3127331903639 1.95121575333553
0.16245590194865	0.598637301759824
83694	RPS6KL1 ribosomal protein S6 kinase like 1	7617
0.601346239656371	0.862747857068239	1.95118542652377
0.162459166985127	0.598637301759824
80829	ZFP91	ZFP91 zinc finger protein
5220	-0.172505202925722	7.35231812842113
1.95086733803171	0.162493417448973	0.598637301759824
25809	TTLL1	tubulin tyrosine ligase like 1	1749
0.338668922837961	2.57651203153891	1.95048817642773
0.162534254780943	0.598637301759824
5889	RAD51C	RAD51 paralog C 6068	0.333178552346801
3.11802916774595	1.95035634513255	0.162548456321217
0.598637301759824
204962	SLC44A5 solute carrier family 44 member 5	6299
1.31408968718306	3.47383024564101	1.94983850286892
0.162604254642366	0.598637301759824
23461	ABCA5	ATP binding cassette subfamily A member 5
13158	-0.279931593872727	4.45345022867283
1.94965729517792	0.162623785219878	0.598637301759824
8985	PLOD3	procollagen-lysine,2-oxoglutarate 5-dioxygenase 3

5496	0.199106955458036	6.23737098697294
1.94957598472864	0.162632549735934	0.598637301759824
6001	RGS10	regulator of G protein signaling 10
1252	-0.375945950166845	3.88181712863352
1.94909654340386	0.162684240048814	0.598637301759824
54518	APBB1IP amyloid beta precursor protein binding family B member 1 interacting protein	3981	1.1269443485217
0.207859955771853	1.94881442279348	0.162714665261027
0.598637301759824
6541	SLC7A1	solute carrier family 7 member 1
7692	-0.334764093158808	8.28202970127283
1.94873937475424	0.162722759887775	0.598637301759824
100289635	ZNF605	zinc finger protein 605
10824	-0.252452576550954	4.97563065914379
1.94779552123746	0.16282460248994	0.598637301759824
26154	ABCA12	ATP binding cassette subfamily A member 12
9416	-0.440940544411281	6.17111297660913
1.9476591633009 0.162839321641858	0.598637301759824
64581	CLEC7A	C-type lectin domain containing 7A	4190
0.798193230169251	3.35618211941522	1.94744184292389
0.162862783420535	0.598637301759824
1329	COX5B	cytochrome c oxidase subunit 5B 1441
0.207750483767245	5.07993669377789	1.94731770131225
0.162876187406774	0.598637301759824
400569	MED11	mediator complex subunit 11	1477
0.309089226066633	3.22642050670091	1.94652210030302
0.162962121019887	0.598743302860159
7102	TSPAN7	tetraspanin 7	3388	0.39073087157141
0.838001557408591	1.9462592785939 0.162990520011836
0.598743302860159
4729	NDUFV2	NADH:ubiquinone oxidoreductase core subunit V2
2854	-0.238806129877272	5.60863981538208
1.94584962816823	0.1630347957315 0.598748920768515
367	AR	androgen receptor	12775	0.627212821767589
2.4691004675007 1.94536709554767	0.163086966297351
0.598783522130275
84064	HDHD2	haloacid dehalogenase like hydrolase domain containing 2	2727	0.270160931992367	4.37814285864591
1.94459126664481	0.163170887497677
0.598875448209175
5702	PSMC3	proteasome 26S subunit, ATPase 3
2276	-0.190177664148061	6.66748805276944
1.94428873897989	0.163203625193322	0.598875448209175
23654	PLXNB2	plexin B2	7257	-0.257459791944641
9.23247172217555	1.94395018845728	0.16324026995521
0.598875448209175
7454	WAS	WASP actin nucleation promoting factor	2381
0.77884570885409	-0.2372871640685	1.943379690545
0.16330204199608	0.598945195663548
3954	LETM1	leucine zipper and EF-hand containing transmembrane protein 1	8184	-0.176042757564937	6.17889724647373
1.94237605735612	0.163410777600298
0.599125587570923
9493	KIF23	kinesin family member 23	6377

0.281283365242117	3.83418461676695	1.94213623804371
	0.163436772335868	0.599125587570923
64098	PARVG	parvin gamma	8656	1.28208531301251
1.35119722340148	1.93960678212073	0.16371123544311
	0.599974692534918
23368	PPP1R13B	protein phosphatase 1 regulatory subunit 13B
	8943	-0.230738969574176	4.87978016558391
1.93874665116165	0.163804685397302	0.600160143437093
10040	TOM1L1	target of myb1 like 1 membrane trafficking protein
	2507	-0.315482156248126	5.54583661298164
1.93773273144399	0.163914922173769	0.600406986110875
3693	ITGB5	integrin subunit beta 5	6259	0.264648659708167
	7.79328414128369	1.93570027065982
0.164136153499766	0.601027779491116
11322	TMC6	transmembrane channel like 6	8445
0.212857417241943	6.85222685213221	1.93520001997783
	0.164190657591251	0.601027779491116
29108	PYCARD	PYD and CARD domain containing	1661
0.320946973445178	3.37123158271444	1.93499381831822
	0.164213130014457	0.601027779491116
10914	PAPOLA	poly(A) polymerase alpha
12766	-0.169786949398686	7.68212563493928
1.9345144391764	0.164265387654169	0.601061986643665
9169	SCAF11	SR-related CTD associated factor 11
14035	-0.216636970341447	7.24930240988663
1.93374883125585	0.164348886818428	0.601210461652054
6815	STYX	serine/threonine/tyrosine interacting protein
4859	-0.283370897034877	4.50093102643334
1.93275569274986	0.164457273330551	0.601391530585804
342184	FMN1	formin 1	20084	-0.401266060265222
4.49721894881191	1.93250858175816	0.164484254563733
	0.601391530585804
58477	SRPRB	SRP receptor subunit beta	4180
0.261953410730419	5.82164121921037	1.93209195242083
	0.164529756401465	0.601400912817152
167227	DCP2	decapping mRNA 2	9951	-0.305585912350017
	5.24240442108701	1.93020943657606
0.164735533303254	0.601995986167148
56160	NSMCE3	NSE3 homolog, SMC5-SMC6 complex component	1681
	0.221894155422673	4.5622417463456	1.9281089068913
0.1649654885502	0.602518067448242
207	AKT1	AKT serine/threonine kinase 1
11128	-0.173356155801776	7.5589945244714	1.92750121082307
	0.165032084434428	0.602518067448242
10555	AGPAT2	1-acylglycerol-3-phosphate O-acyltransferase 2	1798
	0.210969228766317	5.77898843647314
1.92703093105176	0.16508364231061	0.602518067448242
2356	FPGS	folylpolyglutamate synthase	3931
0.20968693491207	5.90857656319307	1.92696920530811
	0.16509041081607	0.602518067448242
26140	TTLL3	tubulin tyrosine ligase like 3
10065	-0.218793998734743	5.24356484113213
1.9265932773196	0.16513163986279	0.602518067448242
51566	ARMCX3	armadillo repeat containing X-linked 3	4239

0.206606402110737	6.49205134599646	1.92589957115279
	0.1652077514201 0.602518067448242
113835	ZNF257	zinc finger protein 257 5029	0.765049845755783
	0.567711706500261	1.92489484982906
0.165318057852176	0.602518067448242
23102	TBC1D2B TBC1 domain family member 2B
9244	-0.214612465978479	6.91704970018265
1.92426874605119	0.165386839183976	0.602518067448242
51019	WASHC3	WASH complex subunit 3	2593	0.269618514500997
	4.18693726688185	1.92390639633813
0.165426660468469	0.602518067448242
388591	RNF207	ring finger protein 207 6585	-0.318821774835861
	4.93104464429425	1.92378877899244
0.165439588664035	0.602518067448242
149041	RC3H1	ring finger and CCCH-type domains 1
11508	-0.26132328215774	5.9212627498457 1.92329421994573
	0.165493961966297	0.602518067448242
353116	RILPL1	Rab interacting lysosomal protein like 1	5259
	0.285524815317119	3.50553926381882
1.92314075631898	0.165510838371825	0.602518067448242
51343	FZR1	fizzy and cell division cycle 20 related 1
5802	-0.16112155505098	6.96943809954118
1.92247897657148	0.165583636885999	0.602518067448242
50512	PODXL2	podocalyxin like 2	2186	0.418770080916257
	3.07222024420989	1.92244234423987
0.1655876676639 0.602518067448242
4967	OGDH	oxoglutarate dehydrogenase
5210	-0.170857377924294	7.18399550507885
1.92220561839235	0.16561371810647	0.602518067448242
116154	PHACTR3 phosphatase and actin regulator 3	6095
0.620616796768508	2.44076694290481	1.92184094260043
	0.165653857945634	0.602518067448242
121506	ERP27	endoplasmic reticulum protein 27	2426
0.674144935431284	0.129713250580615	1.92165277266743
	0.165674574149625	0.602518067448242
170384	FUT11	fucosyltransferase 11	4910	0.213799834516746
	4.79639580288704	1.92137094247346
0.165705607241963	0.602518067448242
5328	PLAU	plasminogen activator, urokinase
2950	-0.319205795307338	10.7271000623806
1.92125126060159	0.165718787751154	0.602518067448242
203859	ANO5	anoctamin 5	6900	0.723482956067752
0.903747540954079	1.92107038105843	0.165738710207616
	0.602518067448242
286451	YIPF6	Yip1 domain family member 6
8850	-0.160463993529982	6.48447589119096
1.92012354975829	0.165843040934344	0.602562949339896
54935	DUSP23	dual specificity phosphatase 23 825
0.3042717323165 3.30699821116275	1.92004798980799
0.165851370072919	0.602562949339896
58490	RPRD1B	regulation of nuclear pre-mRNA domain containing 1B
	5511	-0.188414349285603	4.91388050498498
1.9195647087479 0.165904654522072	0.602562949339896
163227	ZNF100	zinc finger protein 100 6612	0.391511135274555

3.30481986379152 1.91939717184284
0.165923130977653 0.602562949339896
23582 CCNDBP1 cyclin D1 binding protein 1 6654
0.179696345360686 6.58973456676245 1.91782598820956
0.166096520281324 0.603036277755446
345462 ZNF879 zinc finger protein 879 3883 -0.326979973522662
2.13824094929587 1.91672380759925
0.166218276093154 0.603115984938125
162073 ITPRIPL2 ITPRIP like 2 7812 -0.196780840891744
7.074586002577 1.91651684471103 0.166241150280944
0.603115984938125
203260 CCDC107 coiled-coil domain containing 107 1459
0.227979049171436 4.18999422350386 1.91635171204641
0.166259403844986 0.603115984938125
8225 GTPBP6 GTP binding protein 6 (putative) 5928
0.236129961489 4.41953362811249 1.91606857269596
0.166290707062971 0.603115984938125
54491 OTULINL OTU deubiquitinase with linear linkage specificity like 3597 0.508623545124526 3.51712781055868
1.91535618083358 0.166369497264746 0.60311995110925
80219 COQ10B coenzyme Q10B 2722 -0.193203695371838
5.97307288008067 1.91526122679771 0.166380002362613
0.60311995110925
4026 LPP LIM domain containing preferred translocation partner in lipoma 25125 -0.399864171730714
7.52683494263349 1.91481864392301 0.166428976874691
0.60311995110925
27166 PRELID1 PRELI domain containing 1
1875 -0.339261442377029 5.78500813321653
1.91450191970998 0.166464034481928 0.60311995110925
2330 FMO5 flavin containing dimethylaniline monoxygenase 5
5986 0.465540844638744 0.828293041283769
1.91337015061545 0.166589376959169 0.603309430423665
54901 CDKAL1 CDK5 regulatory subunit associated protein 1 like 1
4602 -0.252851597865934 4.04494620992077
1.91299645555031 0.166630787095826 0.603309430423665
94234 FOXQ1 forkhead box Q1 1715 0.453731268896443
1.79858290728795 1.91286327820529 0.166645547676816
0.603309430423665
51011 FAHD2A fumarylacetoacetate hydrolase domain containing 2A
5665 0.284721716239377 4.00917152964153
1.91027678909395 0.166932515723407 0.603315700612795
285958 SNHG15 small nucleolar RNA host gene 15 3647
0.36618088340983 3.10812171294517 1.90984237780175
0.166980768637678 0.603315700612795
55231 CCDC87 coiled-coil domain containing 87 2908
0.47737048955419 0.24424619470958 1.90958938448224
0.167008877625389 0.603315700612795
8621 CDK13 cyclin dependent kinase 13
8921 -0.200825817896104 6.27272474901384
1.90923568451004 0.167048184774374 0.603315700612795
4723 NDUFV1 NADH:ubiquinone oxidoreductase core subunit V1 4154
0.195891959639841 5.85799808642045
1.90922912000173 0.167048914397047 0.603315700612795

80896	NPL	N-acetylneuraminate pyruvate lyase	4975
0.669195870213736	1.07344670612714	1.90907448544782
	0.167066102554939	0.603315700612795
100131089	SRP14-AS1	SRP14 antisense RNA1 (head to head)
	2992	0.394484239272114	0.30944046053996
1.90880778788808	0.167095751656605	0.603315700612795
23038	WDTC1	WD and tetratricopeptide repeats 1
6024	-0.193635486185875	6.60298387064621
1.90867499985961	0.16711051611484	0.603315700612795
10985	GCN1	GCN1 activator of EIF2AK4
9654	-0.179779783408392	7.73345530745261
1.90863644301243	0.167114803460083	0.603315700612795
55322	C5orf22 chromosome 5 open reading frame 22
4172	-0.199762020221577	4.94425559558228
1.90863422185334	0.167115050446661	0.603315700612795
51510	CHMP5	charged multivesicular body protein 5
2390	-0.169161825531946	6.71112797468468
1.9085800306075 0.16712107648873	0.603315700612795
723961	INS-IGF2	INS-IGF2 readthrough	2413
1.08792818896453	1.93675654839044	1.90799355620376
	0.167186308098819	0.603318184403317
27175	TUBG2	tubulin gamma 2 2772	0.239282052572499
3.92347575490852	1.90779940114028	0.16720790974896
	0.603318184403317
7464	CORO2A	coronin 2A	5717	-0.232394463157406
6.49318532578423	1.90459101210336	0.167565337696254
	0.604397139312851
348110	ARPIN	actin related protein 2/3 complex inhibitor	8594
	0.263112651892011	4.41517646844493
1.90417288760783	0.167611982939788	0.604397139312851
403341	ZBTB34	zinc finger and BTB domain containing 34
6564	-0.31986152616969	4.56319652209759
1.9039541758857 0.167636387965334	0.604397139312851
11101	ATE1	arginyltransferase 1	5720	-0.202247644749157
	5.18115606038088	1.90316884658034
0.167724052774161	0.604465255950516
3064	HTT	huntingtin	18128	-0.219618080480718
7.64317816234338	1.90246895068762	0.167802225078682
	0.604465255950516
199	AIF1	allograft inflammatory factor 1 1276
1.53084421626823	0.490458608566755	1.90226353821029
	0.167825175794732	0.604465255950516
7915	ALDH5A1 aldehyde dehydrogenase 5 family member A1	5525
	0.320661028592293	3.85776406932695
1.90201641807716	0.167852791269305	0.604465255950516
30833	NT5C	5', 3'-nucleotidase, cytosolic	1571
0.288518858150161	3.17939304070227	1.90137443793877
	0.167924556364743	0.604465255950516
64417	TMEM267 transmembrane protein 267
3158	-0.32694365897477	4.05243046032662
1.90113970240071	0.167950805546101	0.604465255950516
64219	PJA1	praja ring finger ubiquitin ligase 1
3035	-0.200144894055726	5.29360158512491
1.90107889974805	0.167957605538229	0.604465255950516

252839	TMEM9	transmembrane protein 9 3915	0.169071809783611
	6.01923295614846	1.90045613376274
0.168027272044598	0.604465255950516
4300	MLLT3	MLLT3 super elongation complex subunit	8129
0.252608899298256	4.37136492653854	1.90030957805043
	0.168043671497916	0.604465255950516
100507034	PITRM1-AS1	PITRM1 antisense RNA 1	4490
0.42651383898512	3.76332358915281	1.89859480235332
	0.168235689717349	0.604726484149437
56110	PCDHGA5	protocadherin gamma subfamily A, 5	5733
0.19549829288216	6.44663149986715	1.89851549176474
	0.168244576890025	0.604726484149437
150726	FBXO41	F-box protein 41	7466	0.344792296019134
	3.9614088192772	1.89850445704902	0.168245813431167
	0.604726484149437
9536	PTGES	prostaglandin E synthase
1896	-0.514505591823699	5.32462402072082
1.89797027699882	0.168305685665838	0.604786490282555
56099	PCDHGB7	protocadherin gamma subfamily B, 7	4966
0.194164447540349	6.52440397187632	1.89707653730102
	0.168405912862065	0.604991439204654
6291	SAA4	serum amyloid A4, constitutive
727	-0.742713910525756	2.62240642239754
1.89610338929106	0.168515123004958	0.605138897711163
4038	LRP4	LDL receptor related protein 4	8597
0.397495078616053	3.99589613913104	1.89594091155538
	0.168533364744146	0.605138897711163
64105	CENPK	centromere protein K	4312	0.477979642019446
	1.15796774335919	1.89546829846815
0.168586438943963	0.60517433374887
23180	RFTN1	raftlin, lipid raft linker 1
4776	-0.246076940765601	5.53214673138632
1.89281050568937	0.168885264792239	0.606091699704046
10659	CELF2	CUGBP Elav-like family member 2 10427
0.376123405567623	5.47669744865087	1.89166948797924
	0.169013740099382	0.606269826781525
240	ALOX5	arachidonate 5-lipoxygenase	3730
0.72065272158518	3.20012589828823	1.89160090467915
	0.169021465951443	0.606269826781525
2040	STOM	stomatin	3198	0.292738000975737
8.46954880929958	1.89076739222844	0.169115392818576
	0.606314982476098
51569	UFM1	ubiquitin fold modifier 1
4382	-0.201187752007159	6.88132676507069
1.89072095045943	0.16912062800986	0.606314982476098
26005	C2CD3	C2 domain containing 3 centriole elongation
regulator	12571	-0.27605724014586	5.64008052170775
	1.88972671297982	0.169232748942693
0.606447128719032
23108	RAP1GAP2	RAP1 GTPase activating protein 2	7039
	-0.325830501700684	5.54591713621816
1.88947403616795	0.169261257088031	0.606447128719032
7371	UCK2	uridine-cytidine kinase 2
6756	-0.327165534065348	5.26015899495046

1.8892425802208 0.169287375832899 0.606447128719032
150356 CHADL chondroadherin like 2599 0.406893898486663
0.675943985476595 1.88807818118642
0.169418843087707 0.60659376852294
8153 RND2 Rho family GTPase 2 4353 0.531823939293683
0.382354794115282 1.88763696943005
0.169468688990602 0.60659376852294
23780 APOL2 apolipoprotein L2 3430 0.328624151027336
6.47899311781053 1.88761537902813
0.169471128598225 0.60659376852294
3837 KPNB1 karyopherin subunit beta 1
7678 -0.189932884596596 8.35524139269019
1.88699202840377 0.16954158148317 0.60659376852294
10935 PRDX3 peroxiredoxin 3 3417 0.189939676563803
6.77647557495646 1.88673351485258 0.169570809286574
0.60659376852294
5289 PIK3C3 phosphatidylinositol 3-kinase catalytic subunit type
3 15182 -0.196064956679619 5.22880853297412
1.88658017650163 0.169588148606828 0.60659376852294
9403 SELENOF selenoprotein F 2103 -0.185041403304474
7.19589324698464 1.8856504006101 0.169693330143022
0.606650743307442
150709 ANKAR ankyrin and armadillo repeat containing 5758
0.417305136457987 0.623924537773484 1.88555336649383
0.16970431150665 0.606650743307442
10600 USP16 ubiquitin specific peptidase 16
5694 -0.173969494449819 5.67845862020354
1.88525348789333 0.16973825396111 0.606650743307442
54566 EPB41L4B erythrocyte membrane protein band 4.1 like
4B 7879 -0.240601440416564 6.24192765178495
1.88459983557006 0.169812266112391 0.606650743307442
79809 TTC21B tetratricopeptide repeat domain 21B 7222
0.258435097186007 3.45326990625277 1.8838773299802
0.169894117515472 0.606650743307442
51444 RNF138 ring finger protein 138 3795 -0.318009010512463
3.40565603080739 1.88352810843015
0.169933696447607 0.606650743307442
203069 R3HCC1 R3H domain and coiled-coil containing 1 2630
0.225354326290539 4.38351028786101 1.88338026651335
0.169950455260027 0.606650743307442
23327 NEDD4L NEDD4 like E3 ubiquitin protein ligase
16120 -0.189895726818706 7.56161426802664
1.88285092422041 0.170010475097879 0.606650743307442
2968 GTF2H4 general transcription factor IIH subunit 4 2353
0.215974814350383 4.53692884696916
1.88254288797631 0.170045413194641 0.606650743307442
6597 SMARCA4 SWI/SNF related, matrix associated, actin dependent
regulator of chromatin, subfamily a, member 4
10984 -0.184338222152092 7.46372910187083
1.88225299113039 0.170078301410848 0.606650743307442
10295 BCKDK branched chain keto acid dehydrogenase kinase 3873
0.208432618788081 4.68227269876361
1.88223367319698 0.170080493251112 0.606650743307442
26030 PLEKHG3 pleckstrin homology and RhoGEF domain containing G3

11910 -0.204043915862803 7.06396249821544
1.88160848090382 0.170151445965404 0.606749313553313
7423 VEGFB vascular endothelial growth factor B 1545
0.39284596303744 4.86456296721475 1.88112420371201
0.170206429656215 0.606778100020876
1519 CTSO cathepsin O 2954 -0.308900711064992
4.35253714764446 1.88077440330526 0.170246157855504
0.606778100020876
9802 DAZAP2 DAZ associated protein 2
5731 -0.171540129517622 7.67579896631322
1.88019565554612 0.170311911876615 0.60680478628351
64093 SMOC1 SPARC related modular calcium binding 1
4369 -0.806699525146345 4.26875329403257
1.87994597283841 0.170340288403424 0.60680478628351
107133515 SMURF2P1-LRRC37BP1 SMURF2P1-LRRC37BP1
readthrough transcribed pseudogene 2971 0.261641256315565
3.31023397453756 1.87765307995235
0.170601130419059 0.607579464474893
23210 JMJD6 jumonji domain containing 6, arginine demethylase
and lysine hydroxylase 5969 -0.19971502339283
4.75717483819615 1.87493989751726 0.170910377651138
0.608526095515885
3705 ITPK1 inositol-tetrakisphosphate 1-kinase
6783 -0.225320636884085 5.5001872970473 1.87161727433156
0.171289965972615 0.609714847524026
22978 NT5C2 5'-nucleotidase, cytosolic II
5156 -0.210427491992978 7.77522738097271
1.8712559035155 0.171331308621016 0.609714847524026
29984 RHOD ras homolog family member D 1265
0.211875639683577 7.02380479784346 1.86942328775172
0.171541145689995 0.610306534843729
55379 LRRC59 leucine rich repeat containing 59
3331 -0.264900071962686 8.04948248995759
1.86887529551863 0.171603948903475 0.610374939637119
6307 MSMO1 methylsterol monooxygenase 1
2892 -0.418583308675053 8.16354420240438
1.86761689714291 0.171748268966178 0.610733181875922
9718 ECE2 endothelin converting enzyme 2 6552
0.347065753242525 2.43484671196273 1.86648394044141
0.171878322011267 0.611040521931933
51696 HECA hdc homolog, cell cycle regulator
5614 -0.217690815427812 5.86175699315376
1.8655655104279 0.171983832425239 0.611260477234288
5805 PTS 6-pyruvoyltetrahydropterin synthase
3130 -0.224094199126601 4.74926605067414
1.86511412659119 0.172035715259969 0.611289767540939
2077 ERF ETS2 repressor factor 3295 0.289199141742051
5.42901356523586 1.86392851550059
0.172172077146249 0.611526576789535
79968 WDR76 WD repeat domain 76 4078 0.389106119858129
0.507046204279911 1.86340803898287
0.172231978478124 0.611526576789535
1387 CREBBP CREB binding protein 15160 -0.304812446978782
6.94245378663008 1.8633962410134 0.172233336577048

0.611526576789535
168002 DACT2 dishevelled binding antagonist of beta catenin 2
4223 1.69317843810486 1.48576063405001
1.86278630000865 0.172303565451337 0.611620892366163
7403 KDM6A lysine demethylase 6A 7139 -0.329196794553287
6.16517355045665 1.86166859771589
0.172432343639288 0.611913935129476
26084 ARHGEF26 Rho guanine nucleotide exchange factor 26
5489 0.44155329398848 2.87567738034369
1.86131165344447 0.172473492960032 0.611913935129476
51592 TRIM33 tripartite motif containing 33
9574 -0.230501878644849 5.99861821715238
1.85967689020872 0.172662096394564 0.612427952443316
5021 OXTR oxytocin receptor 4514 0.710862279798906
1.1678488695591 1.85923620195888 0.172712979389292
0.612448730193421
84950 PRPF38A pre-mRNA processing factor 38A
5658 -0.215341695089758 4.67697004318781
1.85886889103511 0.172755403226544 0.612448730193421
6722 SRF serum response factor 4202 -0.255442321354982
6.02809446218882 1.85766522822308
0.172894508405356 0.612648244982323
3491 CCN1 cellular communication network factor 1
2455 -0.238779388030262 9.35080886797675
1.85762501110148 0.172899158450426 0.612648244982323
664 BNIP3 BCL2 interacting protein 3 5325
0.459583900401816 4.37450879040996 1.85724027507506
0.172943650254133 0.612650912774315
55243 KIRREL1 kirre like nephrin family adhesion molecule 1 7872
0.210157900775409 5.99292186428336
1.85625810983936 0.173057289946798 0.6126603598372
4320 MMP11 matrix metallopeptidase 11 5176
0.48290508383072 2.14653211584954 1.85592879623076
0.173095411821842 0.6126603598372
11257 TP53TG1 TP53 target 1 1023 0.398796038907297
4.24679834498038 1.85591524612651 0.17309698061069
0.6126603598372
8792 TNFRSF11A TNF receptor superfamily member 11a 8993
0.730355835441525 2.12567745791911
1.85535696099222 0.173161631345731 0.6126603598372
5128 CDK17 cyclin dependent kinase 17
6547 -0.187889891278139 5.43663017805554
1.85532774869236 0.173165014963624 0.6126603598372
596 BCL2 BCL2 apoptosis regulator
7485 -0.650517647826663 0.785638329475136
1.85449396034004 0.173261623522436 0.612847363807769
8473 OGT O-linked N-acetylglucosamine (GlcNAc) transferase
9812 -0.299363187286178 7.86158303280441
1.85411304520082 0.173305779652372 0.612848789596258
137392 FAM92A family with sequence similarity 92 member A 7423
0.39634599713444 2.37464949676963
1.85366939747041 0.173357224139849 0.612875981455544
55773 TBC1D23 TBC1 domain family member 23
5168 -0.195853275739749 6.20553011402087

1.85312318152287	0.173420586363539	0.612876496956947
728431	LINC01137	long intergenic non-protein coding RNA 1137
	1395	-0.463219768242448	1.88520566436203
1.85291381853384	0.173444879984104	0.612876496956947
5476	CTSA	cathepsin A	5856	0.272753765849166
7.28459388360196	1.85107671739445	0.17365821766926
	0.613475575004621
140885	SIRPA	signal regulatory protein alpha
5068	-0.24581565705413	7.46385394147352
1.85009877395015	0.173771906802175	0.613558076606356
11059	WWP1	WW domain containing E3 ubiquitin protein ligase 1
	6336	-0.171905910402571	6.62184461505366
1.8494844678148 0.1738433657098 0.613558076606356
56911	MAP3K7CL	MAP3K7 C-terminal like	4497
0.545643374467027	1.8204746792053 1.84939607887659
0.173853650299607	0.613558076606356
126792	B3GALT6 beta-1,3-galactosyltransferase 6	2777
0.244912455743736	4.48540140468777	1.84936912810629
	0.173856786324739	0.613558076606356
51024	FIS1	fission, mitochondrial 1	1795
0.198249837436323	5.90149287704908	1.84878166037385
	0.173925160981289	0.61364476822794
64288	ZSCAN31 zinc finger and SCAN domain containing 31
4913	-0.328323594380975	4.56347287874736
1.847831697721	0.174035791486916	0.613880466219398
1499	CTNNB1	catenin beta 1	6614	-0.199928230203983
9.60989496820445	1.84638795643247	0.174204081486117
	0.61431937798994
84197	POMK	protein O-mannose kinase
1877	-0.356237652642855	3.01356746197573
1.8456268569644 0.174292874614464	0.614400365330765
219654	ZCCHC24 zinc finger CCHC-type containing 24	5051
0.331845086187161	3.0709476339758 1.8445218510542
0.17442188210922	0.614400365330765
2186	BPTF	bromodomain PHD finger transcription factor
15098	-0.254505710907671	7.17816987755544
1.84448163431395	0.174426579416085	0.614400365330765
6608	SMO	smoothened, frizzled class receptor	4311
0.413812697347489	3.87996054308438	1.84423514670907
	0.174455372299432	0.614400365330765
4218	RAB8A	RAB8A, member RAS oncogene family
5191	-0.169635272106499	5.80134205172332
1.84419638611627	0.174459900527013	0.614400365330765
10007	GNPDA1	glucosamine-6-phosphate deaminase 1	3961
0.349363393862225	5.23857229428921	1.84393680160028
	0.174490230119639	0.614400365330765
9578	CDC42BPB	CDC42 binding protein kinase beta	8009
	-0.171406667296968	7.85590369551985
1.84315408620705	0.174581718558824	0.614568015042334
1121	CHM	CHM Rab escort protein	8645	-0.194762322901638
	5.59974866430385	1.84170635471085
0.174751083670768	0.61495204579308
55510	DDX43	DEAD-box helicase 43	2807	1.31165779558368
	1.48802527453464	1.84086221209296

0.174849924339992	0.61495204579308
8514	KCNAB2	potassium voltage-gated channel subfamily A regulatory beta subunit 2	9534	0.966943318382639
2.40034475028108	1.84066189565157	0.17487338884705
	0.61495204579308
221294	NT5DC1	5'-nucleotidase domain containing 1	7309
0.203307911449682	4.82295592950939	1.84038471467003
	0.174905863033292	0.61495204579308
3866	KRT15	keratin 15	7069	-0.4230485996258
9.0292097625716 1.84034661735973	0.174910327010754
0.61495204579308
57095	PITHD1	PITH domain containing 1	2186
0.199216863782176	4.76176108081357	1.83973776253796
	0.174981686189838	0.615048551684082
55901	THSD1	thrombospondin type 1 domain containing 1	3348
	0.562570423825873	2.78795389564357
1.83802464794298	0.17518264715287	0.615520093853056
10163	WASF2	WASP family member 2	5676	-0.180798255673779
	7.73808162793568	1.83784507545788
0.175203727721292	0.615520093853056
92922	CCDC102A	coiled-coil domain containing 102A	2582
	0.466787950296069	2.53741555433989
1.83741835444984	0.175253833543953	0.615541736823007
51442	VGLL1	vestigial like family member 1
1541	-0.511489761020261	3.57190138716202
1.83661095187042	0.175348684395307	0.615720486920297
92667	MGME1	mitochondrial genome maintenance exonuclease 1	2336
	0.239564794165786	4.17331379821579
1.83576612780661	0.175447994728646	0.615914802547404
6734	SRPRA	SRP receptor subunit alpha
4167	-0.181785153015789	7.29555944728585
1.83529900319968	0.175502933752017	0.615953293175773
9960	USP3	ubiquitin specific peptidase 3
7764	-0.19906536687183	5.25809197653879
1.83344714913282	0.17572092747356	0.616370826008256
64855	NIBAN2	niban apoptosis regulator 2
4629	-0.301074251961842	9.69943015833051
1.83262630005788	0.175817654745143	0.616370826008256
1212	CLTB	clathrin light chain B	2039	0.232695891116648
	8.25811861360899	1.83224942354067
0.175862085704574	0.616370826008256
3109	HLA-DMB major histocompatibility complex, class II, DM beta
	4829	1.12806150088693	2.55819490055598
1.83215044167347	0.175873757082082	0.616370826008256
84444	DOT1L	DOT1 like histone lysine methyltransferase
11272	-0.251801298561074	5.07033228575166
1.8320597922212 0.175884446732065	0.616370826008256
389119	INKA1	inka box actin regulator 1	1040
0.381691758685021	2.30346091822568	1.83204723257608
	0.175885927861426	0.616370826008256
23190	UBXN4	UBX domain protein 4	5112	-0.166850256860636
	7.91600183799956	1.83103130451351
0.176005781499288	0.616499461803239
1959	EGR2	early growth response 2 3148	0.842279486657681

0.874657190271442	1.83098994921494
0.176010662365328	0.616499461803239
10106	CTDSP2	CTD small phosphatase 2 5442	-0.209909376137367
7.46872593658691	1.83019266088847
0.176104791037208	0.616672553535986
7040	TGFB1	transforming growth factor beta 1	3238
0.226368710289768	5.75020537876326	1.82964162447045
0.176169880876723	0.616672553535986
2861	GPR37	G protein-coupled receptor 37	3021
0.254575733001154	3.85593348829062	1.82924496819047
0.176216752107935	0.616672553535986
342371	ATXN1L	ataxin 1 like	8542	-0.209076240522283
6.79704780349563	1.82908055643685	0.176236184179664
0.616672553535986
246175	CNOT6L	CCR4-NOT transcription complex subunit 6 like
9367	-0.239049816279439	4.51592498012295
1.82760076981925	0.176411193630196	0.616967934599266
29092	LINC00339	long intergenic non-protein coding RNA 339
2406	0.37477451888166	2.11468842808494
1.82737601923295	0.176437791663365	0.616967934599266
51014	TMED7	transmembrane p24 trafficking protein 7
3757	-0.258486133490936	6.9703224744119 1.82724971436097
0.176452741201161	0.616967934599266
5509	PPP1R3D protein phosphatase 1 regulatory subunit 3D
3630	-0.25268803716195	4.19407434940385
1.8262801049753 0.176567553549236	0.617215303859283
9324	HMGN3	high mobility group nucleosomal binding domain 3
1014	0.265564781956841	4.54503990266447
1.82564243769777	0.176643107289631	0.617234801118694
23411	SIRT1	sirtuin 1	4551	-0.264026376982028
4.92102529958023	1.8254705762433 0.176663476601157
0.617234801118694
29800	ZDHHC1	zinc finger DHHC-type containing 1	3206
0.371440603162416	3.46247241392298	1.82511752229188
0.176705329655598	0.617234801118694
285190	RGPD4	RANBP2 like and GRIP domain containing 4
5464	-0.356289955157085	2.93611828620132
1.82279488511496	0.176980953795757	0.617906989285218
152006	RNF38	ring finger protein 38	6138	-0.24449285261538
5.73207783219099	1.82265507799582
0.176997560332821	0.617906989285218
5727	PTCH1	patched 1	16591	-0.391726278991234
3.0902956350397 1.82238109440972	0.177030109802354
0.617906989285218
57508	INTS2	integrator complex subunit 2
7138	-0.275554720033504	4.28115764937075
1.82145046965845	0.177140720349946	0.617986557848183
26996	GPR160	G protein-coupled receptor 160
4153	-0.499309505768272	2.4403446686113 1.82144689109366
0.177141145838542	0.617986557848183
347733	TUBB2B	tubulin beta 2B class IIb
2194	-0.445157139047934	5.0991895539422 1.82056850909149
0.177245620462343	0.618054777901935
84293	PRXL2A	peroxiredoxin like 2A	6520	0.2630538115546

6.9358290296276 1.82054052534902 0.177248950012999
0.618054777901935
53833 IL20RB interleukin 20 receptor subunit beta
2473 -0.481432367218914 3.86527704870545
1.81961453181956 0.17735916693354 0.618285179501117
575 ADGRB1 adhesion G protein-coupled receptor B1
7356 -0.835543130822128 0.318149916981293
1.81921665950364 0.177406548209818 0.618296471951959
63935 PCIF1 PDX1 C-terminal inhibiting factor 1 3219
0.234316003061485 4.5560741520236 1.81818647127301
0.177529297762142 0.618570366605555
84343 HPS3 HPS3 biogenesis of lysosomal organelles complex 2
subunit 1 6256 0.207514851060455 4.76537096408967
1.81737667912684 0.177625855377775
0.61864098009344
9147 NEMF nuclear export mediator factor
9426 -0.160095337273304 6.08342157655809
1.8172757270428 0.177637896904106 0.61864098009344
93210 PGAP3 post-GPI attachment to proteins phospholipase 3 3669
0.221338564574425 4.48561319927582
1.81670253270026 0.177706285175197 0.618725313559281
284252 KCTD1 potassium channel tetramerization domain containing
1 7410 0.185090711737749 5.5232880240577
1.81548990899382 0.177851064460745 0.618860400456241
3419 IDH3A isocitrate dehydrogenase (NAD(+)) 3 catalytic
subunit alpha 6832 -0.203438509868066 5.22720283903042
1.81534359789612 0.17786854224709
0.618860400456241
91614 DEPDC7 DEP domain containing 7 3326 0.458777238978253
0.251240855766435 1.81526752323584
0.177877630630172 0.618860400456241
7113 TMPRSS2 transmembrane serine protease 2
5933 -0.304464041601796 7.04501841779982
1.8145317512618 0.177965558501384 0.619012559704217
328 APEX1 apurinic/apyrimidinic endodeoxyribonuclease 1 1886
0.267166142537039 6.15391334505936
1.81397513884415 0.178032109389086 0.619090306904898
57643 ZSWIM5 zinc finger SWIM-type containing 5
6046 -0.439616670981211 0.0278989754509169
1.81322132463757 0.178122284357222 0.619171217530152
23041 MON2 MON2 homolog, regulator of endosome-to-Golgi
trafficking 16014 -0.208128752109506 5.69712415744687
1.81304164983913 0.178143785724746
0.619171217530152
221662 RBM24 RNA binding motif protein 24
3444 -0.540942886386551 0.405869023446871
1.81158777697741 0.178317878431143 0.619518915157523
117247 SLC16A10 solute carrier family 16 member 10
11237 1.23225735782236 0.8514731726479 1.81145965661191
0.178333229507991 0.619518915157523
25807 RHBDD3 rhomboid domain containing 3 3014
0.291839625490495 4.20843472844607 1.811098499896
0.178376510661119 0.619518915157523
9061 PAPSS1 3'-phosphoadenosine 5'-phosphosulfate synthase 1

3478	0.189915343986852	7.10807234463145
1.81036427485341	0.178464537868958	0.619670992050197
6892	TAPBP	TAP binding protein	5056	-0.209788479838724
8.06577115274124	1.80869186650028
0.17866523249417	0.619905083507378
339230	CCDC137	coiled-coil domain containing 137
2762	-0.198045762991661	5.0434933394175	1.80858674281423
0.178677856396604	0.619905083507378
114781	BTBD9	BTB domain containing 9	9537	-0.207628772513711
5.17784322872559	1.80846327527957
0.178692684457714	0.619905083507378
4170	MCL1	MCL1 apoptosis regulator, BCL2 family member
4371	-0.191273724222861	8.84112392498672
1.80830336244395	0.178711891597457	0.619905083507378
55220	KLHDC8A	kelch domain containing 8A	3651
0.439434490944879	-0.116538211625417	1.80795917575715
0.178753239971893	0.619905083507378
29899	GPSM2	G protein signaling modulator 2	3551
0.267917025355477	3.8323562948918	1.80750867609924
0.178807376804827	0.619939338342874
6642	SNX1	sorting nexin 1	9780	0.160742936290883
6.80049354880512	1.80693419451185	0.178876440108804
0.62002531467558
494513	PJVK	pejvakin	2456	0.393117653144683
0.40563335030355	1.80651958672727	0.178926302776111
0.620044711277828
1376	CPT2	carnitine palmitoyltransferase 2	3401
0.235399285267165	4.46516935818576	1.80537506428322
0.17906403177122	0.620184933164052
6948	TCN2	transcobalamin 2	3122	0.349935388357414
2.9802490000952	1.80533098430456	0.179069338696151
0.620184933164052
104355295	SGMS1-AS1	SGMS1 antisense RNA 1	12420
0.390500196660727	1.68108893694043	1.80501671607612
0.179107179687613	0.620184933164052
221491	SMIM29	small integral membrane protein 29	2229
0.286169719855187	3.98792116774037	1.8047120463469
0.179143873747537	0.620184933164052
54908	SPDL1	spindle apparatus coiled-coil protein 1
4759	-0.293074233755237	3.42573491076331
1.80428126761269	0.179195771053391	0.620211308412366
1806	DPYD	dihydropyrimidine dehydrogenase	6443
0.638180589370151	2.12838349116313	1.80344686494872
0.179296343669832	0.620406098266634
2595	GANC	glucosidase alpha, neutral C	10954
0.193568879236332	5.64307962363581	1.80286330890799
0.179366719891471	0.620496331321272
389432	SAMD5	sterile alpha motif domain containing 5	6320
0.562852180055052	0.638398209262522	1.80234417074021
0.179429354225925	0.620559744356179
140735	DYNLL2	dynein light chain LC8-type 2
6805	-0.156625332755047	6.59707428082513
1.80174947776021	0.179501135365856	0.620654752670832
56606	SLC2A9	solute carrier family 2 member 9	6397

0.352072497316013	4.88807355935008	1.80078588722682
	0.179617513976242	0.620903879137518
58529	MYOZ1	myozenin 1	1541	0.467455738025023
0.695968816431561	1.79985906015277	0.179729534753039
	0.6211378221776
2149	F2R	coagulation factor II thrombin receptor 4066
0.735479204728935	5.69706864885245	1.79748523389338
	0.18001681540264	0.621778153070869
254394	MCM9	minichromosome maintenance 9 homologous
recombination repair factor	12496	-0.257040465878933
3.75288867377105	1.79730277997386	0.180038917957112
	0.621778153070869
57606	SLAIN2	SLAIN motif family member 2
6524	-0.186265012994728	6.59805530550747
1.79722790302152	0.180047989494927	0.621778153070869
5978	REST	RE1 silencing transcription factor
7955	-0.213141478959848	6.62743505954343
1.79615566258614	0.180177952219766	0.621814908601833
400706	NA	NA	2719	1.0102801063928 1.68739651414022
	1.79549635986431	0.18025791799429
0.621814908601833
54858	PGPEP1	pyroglutamyl-peptidase I
7520	-0.284872516358068	6.38203704172675
1.79531894433595	0.18027944344934	0.621814908601833
79575	ABHD8	abhydrolase domain containing 8 2210
0.295999771467235	3.77753092356108	1.79496883064649
	0.180321930734832	0.621814908601833
4773	NFATC2	nuclear factor of activated T cells 2
8286	-0.721298399702782	2.49171275042678
1.79493158326323	0.180326451491647	0.621814908601833
10808	HSPH1	heat shock protein family H (Hsp110) member 1
8390	-0.325886708626704	7.83656000017165
1.7948132053386 0.18034082002401	0.621814908601833
4674	NAP1L2	nucleosome assembly protein 1 like 2	2550
0.377055992242224	0.345312911113157	1.79457790087949
	0.180369384853948	0.621814908601833
79892	MCMBP	minichromosome maintenance complex binding protein
	6000	-0.167605468053225	6.12121223077454
1.79412280060554	0.180424646690714	0.621852368650793
5358	PLS3	plastin 3	4099	-0.249014693276369
9.18869674314545	1.79364078478199	0.180483198183447
	0.62190114562252
10198	MPHOSPH9	M-phase phosphoprotein 9	10106
0.40778275019485	2.32765737140714	1.79258110172908
	0.180611997451688	0.622191895795816
10855	HPSE	heparanase	4721	-0.510374342275011
1.65926651734353	1.79183993527373	0.180702145794679
	0.622349386807493
627	BDNF	brain derived neurotrophic factor	7740
0.806766468232951	2.74439262532687	1.79030960600027
	0.180888445076655	0.622594104163361
11082	ESM1	endothelial cell specific molecule 1
2398	-0.848546554400432	0.217881954481957
1.78923456431201	0.181019451458457	0.622594104163361

54946	SLC41A3 solute carrier family 41 member 3	6592
0.215985008974583	5.30497162977616	1.78921386242899
	0.181021975302015	0.622594104163361
100506603	VASH1-AS1	VASH1 antisense RNA 1	1843
0.381589330611281	1.48585307773392	1.78891597981327
	0.181058295789099	0.622594104163361
8065	CUL5	cullin 5	6742	-0.187185713229965
5.87211632028911	1.78885518400971	0.181065709588288
	0.622594104163361
27286	SRPX2	sushi repeat containing protein X-linked 2	2368
	0.196035335722642	5.64991738619718
1.78882782937957	0.181069045487982	0.622594104163361
285966	TCAF2	TRPM8 channel associated factor 2	6466
0.43251655877459	0.678340230520789	1.7886347776339
0.181092590188916	0.622594104163361
286343	LURAP1L leucine rich adaptor protein 1 like	2132
0.212336641845501	6.63573858721821	1.78833801103273
	0.181128790923517	0.622594104163361
5869	RAB5B	RAB5B, member RAS oncogene family
6588	-0.170040138429972	8.36500185780246
1.78766121720767	0.181211380181907	0.622627258657489
8887	TAX1BP1 Tax1 binding protein 1	8722	-0.153437392679053
	7.59787409494783	1.78753048295162
0.181227338726821	0.622627258657489
57157	PHTF2	putative homeodomain transcription factor 2	8646
	0.311285375278024	5.37683269291475
1.78685176340666	0.181310215208731	0.622759240909439
7189	TRAF6	TNF receptor associated factor 6
4871	-0.209024677390517	5.04665595839438
1.7846836692568 0.181575249001886	0.623398280597526
55827	DCAF6	DDB1 and CUL4 associated factor 6	6306
0.16838519624232	6.11132532334554	1.78442634651181
	0.181606734609321	0.623398280597526
4054	LTBP3	latent transforming growth factor beta binding
protein 3	10543	-0.260237921109993	6.00869882604422
	1.78407541892421	0.18164968374832
0.623398280597526
143872	ARHGAP42	Rho GTPase activating protein 42	6944
	-0.311939740487357	4.95207799974049
1.78351587905424	0.181718188745694	0.623398280597526
64895	PAPOLG	poly(A) polymerase gamma
8501	-0.277007630445487	4.41769731911253
1.7834915594595 0.181721166895076	0.623398280597526
3872	KRT17	keratin 17	2537	-0.234042763125395
12.7137633634677	1.78232122684403	0.181864551299879
	0.623398280597526
84187	TMEM164 transmembrane protein 164
6583	-0.214184210848803	4.73666420131522
1.78216972755196	0.181883121954838	0.623398280597526
11164	NUDT5	nudix hydrolase 5	4545	0.179217956239482
	5.3867336060932 1.78208145144379	0.181893943778081
	0.623398280597526
2801	GOLGA2	golgin A2	6832	-0.156766105923301
6.99181261871863	1.78195823811681	0.181909049823627

0.623398280597526
2055 CLN8 CLN8 transmembrane ER and ERGIC protein
7838 -0.190075036205553 6.06641162824877
1.78151459359041 0.181963453008036 0.623398280597526
2631 NIPSNAP2 nipsnap homolog 2 5651
0.203299219055353 5.33784835273003 1.78133213107838
0.181985833466358 0.623398280597526
8131 NPRL3 NPR3 like, GATOR1 complex subunit 6609
0.278795942080933 4.41439736279962 1.78021234151435
0.182123254285534 0.623716484786667
255743 NPNT nephronectin 6117 -0.713174172521143
3.01568691527211 1.7788564679636 0.182289808297978
0.62399195792535
101927365 LOC101927365 uncharacterized LOC101927365 1638
0.537012398435051 1.01847081857492
1.77856507243009 0.182325626035872 0.62399195792535
6416 MAP2K4 mitogen-activated protein kinase kinase 4
5714 -0.146120256039065 6.0123127007212 1.7772495202889
0.1824874325253 0.62399195792535
51282 SCAND1 SCAN domain containing 1 1510
0.271956571488409 4.27815873187793 1.77712029399094
0.182503335701164 0.62399195792535
1230 CCR1 C-C motif chemokine receptor 1 2731
1.20247316277781 1.35103863612217 1.77683277411413
0.182538724966149 0.62399195792535
170691 ADAMTS17 ADAM metallopeptidase with thrombospondin type 1 motif 17 8242 -0.468359727358114 0.458442083031529
1.7764393219204 0.182587165752911 0.62399195792535
9400 RECQL5 RecQ like helicase 5 8334 -0.226459035159383
5.1474999668837 1.77634958737128 0.182598215717643
0.62399195792535
60468 BACH2 BTB domain and CNC homolog 2
9786 -0.684277180124536 3.96546258517633
1.77615759995247 0.182621859763021 0.62399195792535
4734 NEDD4 NEDD4 E3 ubiquitin protein ligase
9097 -0.253628385097187 5.96193092643464
1.77569394864021 0.182678974972628 0.62399195792535
26043 UBXN7 UBX domain protein 7 11291 -0.230161419306437
6.48673917217315 1.77561635628626
0.182688535262559 0.62399195792535
283871 PGP phosphoglycolate phosphatase 2727
0.212353024695106 4.67655045891359 1.77557423003211
0.182693725955013 0.62399195792535
440519 ZNF724 zinc finger protein 724 3322 -0.43744922446981
0.366876346228224 1.77375732084431
0.182917763739435 0.624604855362816
8991 SELENBP1 selenium binding protein 1
3504 -0.47842167032046 1.41433587912749
1.77258485566868 0.183062506051939 0.624775568503794
100132077 LOC100132077 uncharacterized LOC100132077 2162
0.468342750619535 0.184298356569571
1.77239362230569 0.183086126643901 0.624775568503794
150465 TTL tubulin tyrosine ligase 15732 0.173775477785028
6.2486203765915 1.77201303812977 0.183133145813756

0.624775568503794
246330 PELI3 pellino E3 ubiquitin protein ligase family member 3
3926 0.3444014029216 2.67324943365614
1.77190759050209 0.183146175789004 0.624775568503794
203228 C9orf72 C9orf72-SMCR8 complex subunit
6062 -0.295395595776537 3.31693849611574
1.77045915795365 0.183325264835507 0.624934747887237
54516 MTRF1L mitochondrial translational release factor 1 like
4273 -0.203693711511753 4.56533885966926
1.769879822421 0.183396952658208 0.624934747887237
25903 OLFML2B olfactomedin like 2B 3415 1.25882144788534
0.534837972703009 1.76985409549293
0.183400136899429 0.624934747887237
100272228 LINC00894 long intergenic non-protein coding RNA 894 6323 -0.444612256716244 1.71860502637067
1.76977579431662 0.183409828689333 0.624934747887237
27148 STK36 serine/threonine kinase 36 6458
0.25912678410045 5.09303768011362 1.76972660904369
0.18341591693899 0.624934747887237
60509 AGBL5 ATP/GTP binding protein like 5 5283
0.296669967007918 4.39625872757648 1.76910654625301
0.183492689630759 0.625044285909055
285598 ARL10 ADP ribosylation factor like GTPase 10 11459
0.32301475459339 5.37069187549244 1.76802664972698
0.183626485331902 0.625347964999746
125988 MICOS13 mitochondrial contact site and cristae organizing system subunit 13 1854 0.25053065616633
4.3414988944428 1.76691434885927 0.183764414123034
0.625665568454384
284836 LINC00319 long intergenic non-protein coding RNA 319
8195 0.560755550803369 2.30005898185032
1.76538098522632 0.183954753068356 0.626079509468128
2747 GLUD2 glutamate dehydrogenase 2
2493 -0.204779706934077 4.5273621714033 1.76489304250819
0.18401537015601 0.626079509468128
5825 ABCD3 ATP binding cassette subfamily D member 3
7519 -0.141086989869389 5.98770966297223
1.76485509208191 0.184020085705739 0.626079509468128
4804 NGFR nerve growth factor receptor
4150 -0.776112439951971 3.28172230517547
1.76402726710063 0.184122982412887 0.62618664274325
10712 FAM189B family with sequence similarity 189 member B 4468
0.235174934617831 5.62100970045515
1.76371583847756 0.184161709539345 0.62618664274325
10048 RANBP9 RAN binding protein 9 3347 -0.174414927094255
6.43153739334273 1.76352303019652
0.184185690590575 0.62618664274325
8270 LAGE3 L antigen family member 3 949
0.371861569672972 1.67651300709251 1.762861057422
0.184268052821909 0.626234830079737
2549 GAB1 GRB2 associated binding protein 1
9075 -0.284189421627011 5.13584957263582
1.76249958791075 0.184313044665848 0.626234830079737
1056 CEL carboxyl ester lipase 2681 0.399411530520888

1.39786888879705	1.76216741598737
0.184354401099671	0.626234830079737
131096	KCNH8	potassium voltage-gated channel subfamily H member 8
6148	0.884495299130754	1.5688650739433
1.76197229595719	0.18437869916819	0.626234830079737
286410	ATP11C	ATPase phospholipid transporting 11C
7407	-0.23908722799413	5.02641115046371
1.76089606262735	0.184512788040348	0.626496758392294
56111	PCDHGA4	protocadherin gamma subfamily A, 4	4887
0.185080087336686	6.48335531058977	1.75996793378224
0.18462851532678	0.626496758392294
375444	C5orf34	chromosome 5 open reading frame 34
3011	-0.393424863256114	0.495072739946076
1.75986944538382	0.184640800671633	0.626496758392294
57221	ARFGEF3	ARFGEF family member 3	14877	-0.354801700896157
5.69251352947145	1.75976504333387
0.184653824715443	0.626496758392294
10681	GNB5	G protein subunit beta 5	13302
0.205481143828085	5.3560701312106	1.75939694682454
0.184699752868077	0.626496758392294
8898	MTMR2	myotubularin related protein 2
5776	-0.193407283525783	6.33693433195431
1.75920120291632	0.184724181634909	0.626496758392294
64285	RHBDF1	rhomboid 5 homolog 1	4914	-0.222560855430838
6.28267861015608	1.75586618018097
0.185140968092875	0.627491926024572
4502	MT2A	metallothionein 2A	1299	-0.325931101632446
7.80961743705743	1.75563396861989
0.185170028806346	0.627491926024572
57506	MAVS	mitochondrial antiviral signaling protein
11771	-0.176330722740006	7.82334548763902
1.75541028548501	0.185198027216221	0.627491926024572
80764	THAP7	THAP domain containing 7	2879
0.274925786271942	3.46717547755249	1.75456963831743
0.185303294952383	0.627491926024572
81839	VANGL1	VANGL planar cell polarity protein 1
9062	-0.308412459708475	7.17318828523248
1.75447101579573	0.185315649240306	0.627491926024572
201232	SLC16A13	solute carrier family 16 member 13
1748	-0.394033109432147	3.69082304614763
1.75443378800761	0.185320312955338	0.627491926024572
96626	LIMS3	LIM zinc finger domain containing 3	4534
0.278960032197791	4.04451472159055	1.75434689879329
0.185331198540991	0.627491926024572
64403	CDH24	cadherin 24	3748	-0.393333307557362
3.45664895694844	1.75349828234882	0.185437553282628
0.627700292128994
100616439	MIR4658	microRNA 4658	65	-0.390448045119857
0.451235378549749	1.75313697316868
0.18548285667033	0.627701950563256
84981	MIR22HG	MIR22 host gene	2986	-0.208569087586679
5.81399701454761	1.75271739120566	0.185535482825241
0.627728383558733
126353	MISP	mitotic spindle positioning

2966	-0.199775295239481	7.90674436572621
1.7502657463239 0.185843327841191	0.628550823226801
27341	RRP7A	ribosomal RNA processing 7 homolog A
4082	-0.222855916638879	5.72617807591525
1.75006694600788	0.185868316542115	0.628550823226801
1992	SERPINB1	serpin family B member 1
3094	-0.458809042952317	7.66582203726479
1.74909072363187	0.185991081907924	0.628814164683634
114902	C1QTNF5 C1q and TNF related 5	4194	0.906063438092969
	0.477616156054458	1.74772431295522
0.186163073864753	0.629229303677457
100500827	MIR3614 microRNA 3614	86	-0.333201368331057
	1.6052930271555 1.74720614806548	0.186228344302722
	0.629229303677457
5860	QDPR	quinoid dihydropteridine reductase	3059
0.518838461402615	3.92708281476146	1.74704507307091
	0.186248639469318	0.629229303677457
9158	FIBP	FGF1 intracellular binding protein	3353
0.219345295944111	5.27476189586754	1.74616376070473
	0.186359728795771	0.629452790268232
9550	ATP6V1G1	ATPase H+ transporting V1 subunit G1
1305	-0.185585608832257	7.79904990855454
1.7445426864179 0.186564266189135	0.629953403451562
11158	RABL2B	RAB, member of RAS oncogene family like 2B	5870
	0.224233349061734	4.26045577934695
1.74427638192115	0.186597891834121	0.629953403451562
54737	MPHOSPH8	M-phase phosphoprotein 8
4905	-0.169514231924922	5.48233870877962
1.74390778755642	0.186644445005189	0.62995873281631
79147	FKRP	fukutin related protein 5081	0.180017186948933
	4.88939979005647	1.7430327177643 0.186755019698316
	0.630072023247626
401109	LINC00885	long intergenic non-protein coding RNA 885
	1828	0.439373401541094	-0.0479877063625354
1.74249381438472	0.186823153958337	0.630072023247626
55120	FANCL	FA complementation group L	2412
0.331346873782265	2.43608992061128	1.74222901102295
	0.186856643982851	0.630072023247626
1031	CDKN2C	cyclin dependent kinase inhibitor 2C
3318	-0.718581142263416	0.831973666250014
1.7422187538929 0.186857941355796	0.630072023247626
122509	IFI27L1 interferon alpha inducible protein 27 like 1	2380
	0.420628364203538	3.29281546908266
1.74156600742415	0.186940525527762	0.630198782447019
23545	ATP6V0A2	ATPase H+ transporting V0 subunit a2
13729	-0.233292981019821	5.2771443973361 1.74080473018031
	0.187036894409327	0.630289174067745
5140	PDE3B	phosphodiesterase 3B	10919	0.764669034876404
	0.589340732653764	1.74064331301156
0.187057335375142	0.630289174067745
23167	EFR3A	EFR3 homolog A	6106	-0.16866519395495
7.33491209051443	1.73862116574064	0.187313629021962
	0.630823242581514
23366	KIAA0895	KIAA0895	6728	-0.264814635621445

3.38618240561872	1.73849380953561
0.187329784234871	0.630823242581514
8945	BTRC	beta-transducin repeat containing E3 ubiquitin
protein ligase	6613	-0.167619799385968	5.61526903481401
1.73832701102044	0.187350945180203
0.630823242581514
3117	HLA-DQA1	major histocompatibility complex, class II,
DQ alpha 1	6518	1.13629181396966	3.30181311007287
1.73786848589179	0.187409130440447
0.630867505426423
55876	GSDMB	gasdermin B	5905	-0.347087849892208
3.33965223775054	1.73689303284384	0.187532982007877
0.631132743629104
154141	MBOAT1	membrane bound O-acyltransferase domain containing 1
1794	-0.446689599470865	2.89389912458216
1.73591622945346	0.187657100444079	0.631398752322897
164284	APCDD1L	APC down-regulated 1 like	3942
0.44397033200031	1.53776798023587	1.73436034436754
0.187854997840261	0.631912813340186
7694	ZNF135	zinc finger protein 135	6418	-0.708920561007141
2.52006073532523	1.73365416547827
0.187944898826714	0.632063432860931
728780	ANKDD1B	ankyrin repeat and death domain containing 1B
3287	-0.426676870233483	1.66812911728382
1.73295930902367	0.18803340726556	0.632209298024172
84329	HVCN1	hydrogen voltage gated channel 1	3688
0.4316034160305	1.4167679395174	1.7322302051955	0.188126330171485
0.632369932016316
147694	ZNF548	zinc finger protein 548	7071	-0.241226456349004
4.02531894384136	1.73035970050149
0.188364966997687	0.633020175800529
100008589	RNA28SN5	RNA, 28S ribosomal N5
593	-0.454753546864125	2.64513775724944
1.72874539869931	0.188571201056722	0.633561240873473
113130	CDCA5	cell division cycle associated 5	3925
0.425188293979205	1.1834862329474	1.726414974612
0.188869386697283	0.634410911143607
22856	CHSY1	chondroitin sulfate synthase 1	4990
0.208742432879561	5.62169016271866	1.72549010864756
0.188987878584963	0.634600428733105
79789	CLMN	calmin	15701	0.42883045173164
4.68877997977385	1.72509854321583	0.189038071210355
0.634600428733105
2551	GABPA	GA binding protein transcription factor subunit
alpha	5429	-0.229915826109913	4.79956787578595
1.72491405204919	0.189061725501767	0.634600428733105
80131	LRRC8E	leucine rich repeat containing 8 VRAC subunit E
4329	-0.303300079609706	4.39576972925328
1.72376531795151	0.189209086508654	0.634746991039831
80218	NAA50	N(alpha)-acetyltransferase 50, NatE catalytic
subunit	6436	-0.16078028032717	7.28560731073942
1.72374753043097	0.189211369364982	0.634746991039831
6924	ELOA	elongin A	5391	-0.160151301004459
6.61296838965224	1.72325157069834	0.189275033922812

0.634746991039831
10794	ZNF460	zinc finger protein 460 3505	-0.50236894302217
3.0553932781969 1.72316105836427	0.189286655356134
0.634746991039831
60312	AFAP1	actin filament associated protein 1
10036	-0.218310114591199	7.22774291288193
1.72153047842573	0.189496157945965	0.635297435219995
65986	ZBTB10	zinc finger and BTB domain containing 10
10216	-0.318346076770229	4.89736835105525
1.72113209429868	0.189547384780896	0.635317113813354
375	ARF1	ADP ribosylation factor 1
3281	-0.195841522468026	8.43232413840148
1.72045279548137	0.189634770678982	0.635337964156403
10298	PAK4	p21 (RAC1) activated kinase 4
7115	-0.210425061434082	5.05485889775601
1.72037856441932	0.189644322705642	0.635337964156403
55238	SLC38A7 solute carrier family 38 member 7	6991
0.19462648425704	6.44743586105375	1.71973266073806
0.189727461028004	0.635464501821916
125150	ZSWIM7	zinc finger SWIM-type containing 7	2559
0.253351037328413	3.41585105864325	1.71891093344316
0.189833292105884	0.635547133838076
9015	TAF1A	TATA-box binding protein associated factor, RNA polymerase I subunit A	2895	-0.364467554634674
1.78364327618472	1.71842856494199	0.189895448870295
0.635547133838076
9080	CLDN9	claudin 9	2050	-0.469057373681681
2.63226317773046	1.71794735009863	0.189957480611855
0.635547133838076
1665	DHX15	DEAH-box helicase 15	8162	-0.164480910846706
7.83363771999673	1.71744719850812
0.190021978454218	0.635547133838076
2250	FGF5	fibroblast growth factor 5
5998	-0.691751163759765	4.73488747124623
1.71726797612551	0.190045096572663	0.635547133838076
25840	METTL7A methyltransferase like 7A	4083
0.72293892516395	4.58911692877105	1.71723912757651
0.190048818089238	0.635547133838076
10522	DEAF1	DEAF1 transcription factor	3938
0.197146154590988	5.26476077996949	1.71677280051581
0.190108986934586	0.635547133838076
55014	STX17	syntaxin 17	4740	-0.192098423833338
4.50165540937447	1.7166430026322 0.190125738333845
0.635547133838076
10101	NUBP2	nucleotide binding protein 2	3269
0.23670001863714	4.83334442672499	1.71637374241354
0.190160493889868	0.635547133838076
3148	HMGB2	high mobility group box 2	2775
0.439897173563879	1.99229712153676	1.71595945265692
0.190213984025562	0.635574254352589
339400	FLG-AS1 FLG antisense RNA 1	4897	0.713407116328359
1.65893282315192	1.71489626528441
0.190351335471558	0.6357697441866
26999	CYFIP2	cytoplasmic FMR1 interacting protein 2	10582

0.345506256588339	5.3869086302004 1.71480393540425
0.190363268874034	0.6357697441866
60672	MIIP	migration and invasion inhibitory protein	2821
	0.242997423440775	3.73255181172235
1.71211506590149	0.190711181324293	0.636754434854125
4702	NDUFA8	NADH:ubiquinone oxidoreductase subunit A8	844
	0.204106483233717	5.13152476622318
1.71182292442931	0.190749026104656	0.636754434854125
57455	REXO1	RNA exonuclease 1 homolog
5199	-0.196904888044129	5.23797129199638
1.70851634665708	0.19117798013463	0.638034302536516
271	AMPD2	adenosine monophosphate deaminase 2	6823
0.314942181143682	5.71640142839848	1.7074790279281
0.191312780688342	0.638332091258124
64223	MLST8	MTOR associated protein, LST8 homolog	4135
0.205455499352934	4.58302950152467	1.7069276359468
0.191384479737805	0.638419244507606
115416	MALSU1	mitochondrial assembly of ribosomal large subunit 1
	3851	0.232594259643959	3.80092250769175
1.70600742132906	0.19150420766834	0.638666532573915
100507053	LOC100507053	uncharacterized LOC100507053	6244
	0.381940821944754	1.95992740021198
1.70496271273822	0.191640239300927	0.638968062815539
80110	ZNF614	zinc finger protein 614 5813	-0.315361318981381
	3.20157981202652	1.70454342864019
0.191694866042153	0.638998093444177
23624	CBLC	Cbl proto-oncogene C	1579	-0.339623029580338
	4.33435379832737	1.70413450382587
0.191748160621975	0.639023670195575
55114	ARHGAP17	Rho GTPase activating protein 17	9135
	-0.168419746103779	5.89455024856637
1.70162773067695	0.192075242957407	0.639817343395489
54853	WDR55	WD repeat domain 55	7675	0.169135295957814
	5.96411555514881	1.70155808954304
0.19208433898691	0.639817343395489
8812	CCNK	cyclin K	8220	-0.18679680566075
5.80235407422813	1.70120894048083	0.192129949935143
	0.639817343395489
3156	HMGCR	3-hydroxy-3-methylglutaryl-CoA reductase
6215	-0.281295316907083	7.75923047346925
1.70090988350853	0.192169027176756	0.639817343395489
10501	SEMA6B	semaphorin 6B	4412	1.56022655804339
2.06801940689673	1.70038779901557	0.192237269269437
	0.639892450251188
9477	MED20	mediator complex subunit 20	3278
0.200728451312479	4.57466951012383	1.69953375500376
	0.192348963074274	0.639987875036482
6651	SON	SON DNA binding protein 13172	-0.209768426599261
	8.72654123820606	1.69946989366895
0.192357318048375	0.639987875036482
7772	ZNF229	zinc finger protein 229 5591	-0.543732599396548
	3.25345875046595	1.69819955592151
0.192523604342364	0.640236780814154
51528	JKAMP	JNK1/MAPK8 associated membrane protein	4350

0.15623923418044	6.21995698439789	1.69813960165215
	0.192531456460295	0.640236780814154
25820	ARIH1	ariadne RBR E3 ubiquitin protein ligase 1
22880	-0.160329489014429	6.9045864314052 1.69785103211589
	0.192569255198831	0.640236780814154
51657	STYXL1	serine/threonine/tyrosine interacting like 1
2028	-0.194395264632647	4.90808104600769
1.69731639698151	0.192639308135454	0.640287692191387
80228	ORAI2	ORAI calcium release-activated calcium modulator 2
	11103	0.300603617194006	5.58395268535379
1.69703651951586	0.19267599218867	0.640287692191387
65982	ZSCAN18 zinc finger and SCAN domain containing 18
8307	-0.245701372483623	4.60371371944578
1.69633199433225	0.192768371724256	0.640442737841948
338322	NLRP10	NLR family pyrin domain containing 10
2350	-0.593414881856075	0.0924375393368708
1.69530535863124	0.19290308003502	0.640659574351663
79673	ZNF329	zinc finger protein 329 4309	-0.209281541862576
	4.31616632900345	1.69505261394532
0.19293626037869	0.640659574351663
80854	SETD7	SET domain containing 7, histone lysine
methyltransferase	8861	-0.20381517628317
8.22998640711933	1.69419610001029	0.19304875319351
	0.640659574351663
26512	INTS6	integrator complex subunit 6
25509	-0.188246530147739	5.63713406922524
1.69396232805948	0.193079469640824	0.640659574351663
63934	ZNF667	zinc finger protein 667 8703	-0.44869425439849
	2.66320423788148	1.69373933710289
0.193108774841607	0.640659574351663
55233	MOB1A	MOB kinase activator 1A 6440	-0.16844179853375
	7.45357809193496	1.69359794708287
0.193127358838577	0.640659574351663
23131	GPATCH8 G-patch domain containing 8
9734	-0.182389332601936	6.62594185506261
1.69319604306936	0.193180195641796	0.640659574351663
2886	GRB7	growth factor receptor bound protein 7
3597	-0.349212976070329	5.07716846381685
1.6930488857099 0.193199546088486	0.640659574351663
10745	PHTF1	putative homeodomain transcription factor 1	7170
	0.180601195735111	5.36298326457561
1.69205271438096	0.193330597144767	0.640815354564521
4646	MYO6	myosin VI	8785	-0.181940059921354
8.3305942293039 1.69199629099512	0.193338023016325
0.640815354564521
1163	CKS1B	CDC28 protein kinase regulatory subunit 1B	2338
	0.334764952220105	2.53659780437758
1.6912291319801 0.193439021758289	0.640998433349504
8243	SMC1A	structural maintenance of chromosomes 1A
10484	-0.160763949940532	6.2455195494274 1.68964505511662
	0.193647765322949	0.6413184576603
4351	MPI	mannose phosphate isomerase	8025
0.309310636257923	3.96159714010355	1.6894290649752
0.19367624806672	0.6413184576603

4065    LY75    lymphocyte antigen 75   8798    0.369087452721576
        5.27031012743183        1.68908559534449
0.193721551689881       0.6413184576603
259173  ALS2CL  ALS2 C-terminal like    5717    -0.26540748633259
        7.52411513594302        1.68846980050876
0.19380280600866        0.6413184576603
55200   PLEKHG6 pleckstrin homology and RhoGEF domain containing G6
        4260    -0.328559557385162      4.21908832417745
1.68782524620332        0.193887897828247       0.6413184576603
7155    TOP2B   DNA topoisomerase II beta
6044    -0.178495592231005      6.92537616818724
1.68773038339515        0.193900424973405       0.6413184576603
252969  NEIL2   nei like DNA glycosylase 2      3367
0.324494404577479       2.77985652815403        1.68745278856061
        0.193937088304013       0.6413184576603
54970   TTC12   tetratricopeptide repeat domain 12
5302    -0.354019881698977      3.34901653412803
1.68726895352614        0.193961372772506       0.6413184576603
63925   ZNF335  zinc finger protein 335 7521    -0.185296887093229
        5.1401925326999 1.68718046764445        0.193973062957777
        0.6413184576603
115825  WDFY2   WD repeat and FYVE domain containing 2  16197
0.27412439433149        4.82389269139836        1.68653306441426
        0.194058618773736       0.6413184576603
152815  THAP6   THAP domain containing 6
7127    -0.218105606555643      3.93923242337787
1.6863382974781 0.194084366302536       0.6413184576603
27235   COQ2    coenzyme Q2, polyprenyltransferase      2774
0.259201951081891       3.72039377175956        1.6863333168664
0.194085024774899       0.6413184576603
92291   CAPN13  calpain 13      4952    -0.670523166248723
3.32439525555742        1.68450942071081        0.194326332582864
        0.641964372756644
222234  FAM185A family with sequence similarity 185 member A    2729
        0.328222470016285       2.19598732466553
1.68384977701767        0.19441369212163        0.64209685225052
5300    PIN1    peptidylprolyl cis/trans isomerase, NIMA-interacting
1       4570    0.199811433355004       5.26400423478423
1.68326186876046        0.19449159013254        0.64209685225052
51635   DHRS7   dehydrogenase/reductase 7
2977    -0.191931526733837      6.74818725444914
1.68316854953378        0.194503958313626       0.64209685225052
253635  GPATCH11        G-patch domain containing 11
4040    -0.196114550859479      4.01758422757492
1.68266207193767        0.194571101007454       0.642167156411735
4522    MTHFD1  methylenetetrahydrofolate dehydrogenase,
cyclohydrolase and formyltetrahydrofolate synthetase 1  9377
0.243087676692883       4.92656850994425        1.68221604951695
        0.194630251734587       0.642211056783594
101927528       LINC01204       long intergenic non-protein coding
RNA 1204        5260    -0.708195310878101      0.499201112432202
        1.68126634715767        0.194756269725969
0.642255840723733
6769    STAC    SH3 and cysteine rich domain    4396

0.691145412265661	1.63318155791389	1.68124056788984
	0.194759691761382	0.642255840723733
114825	PWWP2A	PWWP domain containing 2A
6510	-0.197817305577529	4.3215835387001	1.68107718420843
	0.194781381551682	0.642255840723733
80264	ZNF430	zinc finger protein 430	6845	-0.26004271161878
	4.35801852440966	1.68034445589375
0.194878688719544	0.64242546314301
84154	RPF2	ribosome production factor 2 homolog
3922	-0.216942075220478	4.21887793211645
1.67874719939639	0.195091003434562	0.642470914830985
7494	XBP1	X-box binding protein 1	3028	-0.381119343405797
	6.49932899201622	1.67828010533076
0.195153142879731	0.642470914830985
54753	ZNF853	zinc finger protein 853	3857	0.627281877098464
	1.56309744519041	1.67793286913006
0.195199352150127	0.642470914830985
5663	PSEN1	presenilin 1	9947	-0.158979816186795
7.61633591600704	1.67751159335363	0.195255431603644
	0.642470914830985
6738	RO60	Ro60, Y RNA binding protein
9102	-0.237383616340473	5.63704758941395
1.67735571296456	0.195276186892492	0.642470914830985
84911	ZNF382	zinc finger protein 382	11318	-0.476513777321234
	0.925421151822939	1.67731274511147
0.195281908465125	0.642470914830985
729288	ZNF286B	zinc finger protein 286B (pseudogene)
5884	-0.268171301599275	4.05060110297649
1.67722963140973	0.195292976390879	0.642470914830985
399823	FOXI2	forkhead box I2	3217	-0.876915423593749
2.11745392380201	1.676958991663	0.195329021512874
0.642470914830985
124790	HEXIM2	HEXIM P-TEFb complex subunit 2	2038
0.407646578819739	2.24107969377576	1.67612062732223
	0.195440728398711	0.642470914830985
321	APBA2	amyloid beta precursor protein binding family A
member 2	6869	0.483891659117181	1.6506369108048
1.67571708706208	0.195494524312405	0.642470914830985
79830	ZMYM1	zinc finger MYM-type containing 1
4828	-0.295299807177891	3.87604994597519
1.67570637126206	0.195495953070923	0.642470914830985
10999	SLC27A4	solute carrier family 27 member 4	3110
0.174393798602218	6.32210533140058	1.67569105668427
	0.195497995014668	0.642470914830985
23557	SNAPIN	SNAP associated protein	1544	0.197798105736923
	4.42732494406603	1.67546946762717
0.195527543017174	0.642470914830985
8209	GATD3A	glutamine amidotransferase like class 1 domain
containing 3A	6440	0.181295586938488	5.16356054584252
	1.67488374286917	0.195605672216735
0.642470914830985
55852	TEX2	testis expressed 2	9094	0.163652731088851
	5.76801252406026	1.67479665902824
0.195617291357809	0.642470914830985

55269	PSPC1	paraspeckle component 1 4817	0.171823692403413
	5.51796860728612	1.67472868045691
0.195626361944324	0.642470914830985
9352	TXNL1	thioredoxin like 1	10905	-0.164785370704577
	6.19571472332108	1.67425394625853
0.195689720893983	0.642528345185657
23588	KLHDC2	kelch domain containing 2	6841
0.186889141282368	5.33158100194589	1.67054967129816
	0.196184926704478	0.643893427530523
5606	MAP2K3	mitogen-activated protein kinase kinase 3
3783	-0.176809673747029	6.02895734105506
1.6701158533526 0.196243017581244	0.643893427530523
83607	AMMECR1L	AMMECR1 like	4761	-0.152229474058732
	6.4302429758833 1.67011313746644	0.196243381318469
	0.643893427530523
5899	RALB	RAS like proto-oncogene B
2915	-0.227180890496734	6.42709064980944
1.66851826549419	0.196457118162005	0.644294717623191
100129250	SMIM27	small integral membrane protein 27	1885
	0.411684017790473	0.763937250606451
1.66844918407582	0.19646638227166	0.644294717623191
150684	COMMD1	copper metabolism domain containing 1	1323
0.247719839246566	3.8885288578334 1.66817110416366
0.196503679138022	0.644294717623191
3663	IRF5	interferon regulatory factor 5	4375
0.227833588958997	4.5893477859649 1.66773139769044
0.196562670746653	0.644337310823394
9817	KEAP1	kelch like ECH associated protein 1	3761
0.166113572292033	6.12321734741587	1.66618696717344
	0.196770038109921	0.644708382764748
79733	E2F8	E2F transcription factor 8
4140	-0.434875664481438	0.0265468233505709
1.66608242821586	0.196784083591164	0.644708382764748
57602	USP36	ubiquitin specific peptidase 36
7414	-0.221976171801683	5.42293381137448
1.66564727129035	0.196842562347795	0.644708382764748
389602	LOC389602	uncharacterized LOC389602	2536
0.484912751164142	0.983519970567536	1.66551768388313
	0.196859980944158	0.644708382764748
64756	ATPAF1	ATP synthase mitochondrial F1 complex assembly
factor 1	5677	0.186038534433559	4.62906211154589
	1.66448272128517	0.19699916112342
0.644998958559253
5670	PSG2	pregnancy specific beta-1-glycoprotein 2
2798	-0.619835091366737	2.79002074275159
1.66396489260266	0.197068841212777	0.644998958559253
399512	SLC25A35	solute carrier family 25 member 35	3897
	0.403007749232539	1.72733991032432
1.66382013599314	0.197088325125074	0.644998958559253
84888	SPPL2A	signal peptide peptidase like 2A
8144	-0.183876219791881	6.71374384894695
1.66329352752533	0.197159224484426	0.644998958559253
79890	RIN3	Ras and Rab interactor 3
8683	-0.238822334821656	5.47930591303472

1.66314706117349	0.197178949136198	0.644998958559253
9880	ZBTB39	zinc finger and BTB domain containing 39
6170	-0.298999076655876	3.82765631956209
1.66248511258347	0.197268122736161	0.645061563512837
84302	PGAP4	post-GPI attachment to proteins GalNAc transferase 4
	5370	0.205697747510388	4.97070984327713
1.66232133344803	0.197290193338259	0.645061563512837
55915	LANCL2	LanC like 2	5235	0.181579866359405
4.80838373547749	1.66072237539614	0.197505818410899
	0.645347157233988
100507421	TMEM178B	transmembrane protein 178B
10656	-1.08628214945027	2.8879324118168 1.66070332578066
	0.197508388983019	0.645347157233988
80174	DBF4B	DBF4 zinc finger B	7146	0.362097423566921
	1.89574077910842	1.66058470171265
0.197524397103396	0.645347157233988
114990	VASN	vasorin 2806	-0.299430454246958
5.02228545152907	1.66030732846719	0.197561834098618
	0.645347157233988
2078	ERG	ETS transcription factor ERG	7588
0.524961436988652	-0.222946399555612	1.65882228291101
	0.197762411891634	0.645851737786458
10361	NPM2	nucleophosmin/nucleoplasmin 2	2063
0.474334161067943	3.4256356780027 1.65780674249747
0.197899713435393	0.646149483936956
149175	MANEAL	mannosidase endo-alpha like
3018	-0.444100408997226	2.01421067825374
1.65594597890589	0.198151579778264	0.646666672353438
6840	SVIL	supervillin	9937	-0.21322900689758
8.04401511404237	1.65584943757693	0.198164657526359
	0.646666672353438
29099	COMMD9	COMM domain containing 9	4048
0.211733563328836	5.5640495169203 1.65524763097129
0.198246202700893	0.646666672353438
63939	FAM217B family with sequence similarity 217 member B
5323	-0.223651488243753	4.17451828834134
1.6549561865593 0.198285707751013	0.646666672353438
10454	TAB1	TGF-beta activated kinase 1 (MAP3K7) binding protein
1	3986	0.203632974613282	4.61565482386937
1.65493225173985	0.198288952506462	0.646666672353438
79269	DCAF10	DDB1 and CUL4 associated factor 10
8492	-0.193762157226291	5.61851459390363
1.65428434327204	0.198376810747141	0.646741252948544
5050	PAFAH1B3	platelet activating factor acetylhydrolase
1b catalytic subunit 3	1419	0.190036784019626
5.48320156273054	1.65408267569103	0.198404166768037
	0.646741252948544
6050	RNH1	ribonuclease/angiogenin inhibitor 1	6152
0.168219647578294	7.06513957088656	1.65293882717696
	0.198559412542641	0.64699556787813
10201	NME6	NME/NM23 nucleoside diphosphate kinase 6
4716	-0.205672353926552	4.43500662065871
1.65235775590451	0.198638331481936	0.64699556787813
728978	UNC5B-AS1	UNC5B antisense RNA 1	707

0.532728466117872	0.18513866741589	1.65202656034853
	0.198683329700549	0.64699556787813
83986	FAM234A family with sequence similarity 234 member A	5570
	0.17716698178096	6.30114175456103
1.65178845738251	0.198715687192406	0.64699556787813
55573	CDV3	CDV3 homolog	4367	-0.221795236223781
7.93152396754751	1.65155837821985	0.198746960148817
	0.64699556787813
122830	NAA30	N(alpha)-acetyltransferase 30, NatC catalytic
subunit 7653	-0.210309884118531	5.51802281293382
1.65146738689526	0.198759329519497	0.64699556787813
26276	VPS33B	VPS33B late endosome and lysosome associated
4012	-0.214424047278183	4.19140989952925
1.64832568851405	0.199186967101714	0.648168117404795
101927832	PAQR9-AS1	PAQR9 antisense RNA 1
4767	-0.31079630180184	2.93874748698545
1.64767941364275	0.199275069797057	0.648168117404795
7409	VAV1	vav guanine nucleotide exchange factor 1
4305	-0.546459005878137	3.68524968364803
1.64739012395708	0.199314521712443	0.648168117404795
22885	ABLIM3	actin binding LIM protein family member 3
7225	-0.505850502971333	6.57541645186813
1.64689222306583	0.199382444484228	0.648168117404795
10765	KDM5B	lysine demethylase 5B	12092	-0.18892856086016
	7.8909278716537 1.64680588322723	0.199394225579635
	0.648168117404795
283554	GPR137C G protein-coupled receptor 137C 6193
0.622149949673688	0.421693566611262	1.64678417062882
	0.199397188398462	0.648168117404795
22992	KDM2A	lysine demethylase 2A	10380	-0.217253875496545
	8.0554256447923 1.64561230194636	0.199557173876567
	0.648358174840891
85451	UNK	unk zinc finger 5575	0.209873600296246
4.89331881482095	1.64538072417425	0.199588807098417
	0.648358174840891
9202	ZMYM4	zinc finger MYM-type containing 4
7647	-0.165185783687356	6.32915385068469
1.64533890192693	0.199594520590699	0.648358174840891
8216	LZTR1	leucine zipper like transcription regulator 1	7556
	0.162195993217149	6.51861147392602
1.64437407493786	0.199726382533117	0.648636086283646
1979	EIF4EBP2	eukaryotic translation initiation factor 4E
binding protein 2	7257	-0.166996115245245
7.65332098832506	1.64391620679282	0.199788994715403
	0.648689023870804
57520	HECW2	HECT, C2 and WW domain containing E3 ubiquitin
protein ligase 2	12692	0.533220506255402
0.243425193136314	1.64228871968266	0.200011735719688
	0.649261734003632
23163	GGA3	golgi associated, gamma adaptin ear containing, ARF
binding protein 3	6640	-0.193976865819682
5.93770367156694	1.64114731405849	0.200168124745265
	0.649618842286127
200205	IBA57	iron-sulfur cluster assembly factor IBA57

10219	-0.263190531572009	3.41739726528016
1.64057160156171	0.200247060166531	0.649623867897802
1428	CRYM	crystallin mu	3782	-0.452287085738502
0.0973739708990399	1.6401336046005	0.200309915335655
0.649623867897802
119710	C11orf74	chromosome 11 open reading frame 74	1474
0.292863231383771	2.74823203609439
1.63982811533965	0.200349053019474	0.649623867897802
28232	SLCO3A1	solute carrier organic anion transporter family member 3A1	6688	0.190953849988638	5.46385455599642
1.63936456110522	0.200412675167924
0.649623867897802
5494	PPM1A	protein phosphatase, Mg2+/Mn2+ dependent 1A
11860	-0.147171402261654	6.16091734247227
1.63895748693407	0.200468565091059	0.649623867897802
22879	MON1B	MON1 homolog B, secretory trafficking associated
6942	-0.146094388914604	6.36547012704025
1.63851386232809	0.200529494127619	0.649623867897802
115399	LRRC56	leucine rich repeat containing 56	2769
0.413585125692404	1.61881055659436	1.63845641769153
0.200537385377049	0.649623867897802
3054	HCFC1	host cell factor C1	9004	-0.18135548591884
6.71663675682221	1.63843227922928
0.200540701420011	0.649623867897802
389558	FAM180A	family with sequence similarity 180 member A	2526
1.20251985035157	-0.2333469263581
1.63581678974757	0.200900388810244	0.650382778327687
79919	MAB21L4	mab-21 like 4	2691	-0.312699818193509
7.10328206496836	1.63542637635742	0.200954144277621
0.650382778327687
284309	ZNF776	zinc finger protein 776	5684	-0.225510039423195
4.53062351678227	1.63523810567085
0.200980073046048	0.650382778327687
6693	SPN	sialophorin	7382	1.09181004224535
2.44231810725459	1.63437089359062	0.20109955689313
0.650382778327687
8607	RUVBL1	RuvB like AAA ATPase 1	3993	0.219894102849305
4.66142425133671	1.63436141816861
0.201100862871084	0.650382778327687
7078	TIMP3	TIMP metallopeptidase inhibitor 3	4603
0.288414712520398	8.10766892376457	1.63403942581743
0.201145248343918	0.650382778327687
6649	SOD3	superoxide dismutase 3	1624	0.672400175878232
0.694721219826408	1.63366200685471
0.201197288839425	0.650382778327687
788	SLC25A20	solute carrier family 25 member 20	1982
0.231326143163983	3.75436116232976
1.63342468843719	0.201230019649028	0.650382778327687
353189	SLCO4C1	solute carrier organic anion transporter family member 4C1	5334	-0.630722276309663	0.726485814466855
1.63317923688349	0.201263878763481
0.650382778327687
23013	SPEN	spen family transcriptional repressor
13294	-0.337898376372422	7.59112941077622

1.63308972369703	0.201276228436605	0.650382778327687
55553	SOX6	SRY-box transcription factor 6	19503
0.420158285261829	1.46807301408844	1.63302079451203
	0.20128573884847	0.650382778327687
7741	ZSCAN26 zinc finger and SCAN domain containing 26	3198
	0.271396766427303	3.44856446771661
1.63186887161122	0.201444752111637	0.65074645821672
54834	GDAP2	ganglioside induced differentiation associated
protein 2	10496	-0.190959797345072	4.85202048492644
	1.63040843609867	0.201646565412139
0.651248199614441
92154	MTSS2	MTSS I-BAR domain containing 2	4999
0.3360581722658 5.88361757020293	1.62961157706145
0.201756781316083	0.651371236537578
6768	ST14	suppression of tumorigenicity 14
4350	-0.230358930222838	8.99338147938803
1.62946062460458	0.201777667975766	0.651371236537578
4297	KMT2A	lysine methyltransferase 2A
21961	-0.376300995410925	6.95608704375121
1.62909043778262	0.201828899947243	0.651386498055538
129450	TYW5	tRNA-yW synthesizing protein 5
10496	-0.199090105186299	4.24792904467429
1.6276285397707 0.202031368817192	0.651721355515788
10430	TMEM147 transmembrane protein 147	1932
0.177555368919525	5.45808976524581	1.62753209622324
	0.202044734386194	0.651721355515788
8540	AGPS	alkylglycerone phosphate synthase
8197	-0.147696495832729	6.33708940898332
1.6273070947825 0.202075920116593	0.651721355515788
6652	SORD	sorbitol dehydrogenase	6159	-0.248101751612334
	4.2087282270577 1.62699802662232	0.202118766928006
	0.651721355515788
84930	MASTL	microtubule associated serine/threonine kinase like
	3631	-0.257654578652141	3.77982125702154
1.62571621857641	0.202296581014996	0.652123443379088
8128	ST8SIA2 ST8 alpha-N-acetyl-neuraminide alpha-2,8-
sialyltransferase 2	5903	0.730145697198255
0.148990183661988	1.62542805611573	0.202336580632935
	0.652123443379088
55146	ZDHHC4	zinc finger DHHC-type containing 4	3557
0.179055982893996	5.10727764129166	1.62474507314692
	0.202431422180793	0.652152815593602
221656	KDM1B	lysine demethylase 1B	4595	-0.228847837566981
	5.05382551132121	1.62467741195515
0.202440820713399	0.652152815593602
9227	LRAT	lecithin retinol acyltransferase
5799	-0.602991244926285	1.09872663767689
1.62435673750727	0.202485371244133	0.652152815593602
374819	LRRC37A3	leucine rich repeat containing 37 member A3
	7641	0.315753037285805	3.28958063305916
1.62310290902181	0.202659673451524	0.652564148513907
9948	WDR1	WD repeat domain 1	7827	-0.252550015696043
	9.24771101818886	1.62269625069086
0.20271624339596	0.652596281601288

164312	LRRN4	leucine rich repeat neuronal 4	2692
0.582767271696285	2.10263709669221	1.62224714789188
	0.202778739365877	0.652647472954468
92421	CHMP4C	charged multivesicular body protein 4C
1866	-0.18159417058114	5.80548138364718
1.62160163300267	0.202868607272913	0.652786717682447
253512	SLC25A30	solute carrier family 25 member 30
6794	-0.192373362374376	4.87556744971024
1.62040554941681	0.203035248775241	0.653151082167838
51090	PLLP	plasmolipin	8640	0.558227726191512
3.62717189750215	1.61942351284566	0.203172189215638
	0.653151082167838
1362	CPD	carboxypeptidase D	12532	-0.230149712470151
	7.71292288142532	1.61888866566639
0.203246816930538	0.653151082167838
100302156	MIR1229 microRNA 1229	69	0.337491986686083
	1.35730617015269	1.61859750322495
0.20328745666502	0.653151082167838
51704	GPRC5B	G protein-coupled receptor class C group 5 member B
	6667	-0.661164040452552	5.19253504664424
1.61786705049871	0.203389453591353	0.653151082167838
6560	SLC12A4 solute carrier family 12 member 4	7882
0.185155007858055	7.4158503277781 1.61755542247156
0.203432986185193	0.653151082167838
56126	PCDHB10 protocadherin beta 10	3290	0.5272991134811
0.108399969299167	1.61705133578291	0.203503427359182
	0.653151082167838
11273	ATXN2L	ataxin 2 like	6508	-0.171852794745042
7.34410377747227	1.61689473961232	0.203525315989217
	0.653151082167838
78992	YIPF2	Yip1 domain family member 2	2087
0.178256941440094	5.61734894574709	1.61669261691595
	0.203553572306729	0.653151082167838
85313	PPIL4	peptidylprolyl isomerase like 4
2646	-0.196293990848685	4.98929371106746
1.61636179720752	0.203599830163801	0.653151082167838
100130705	ATP6V1FNB	ATP6V1F neighbor	3631
0.595957092544072	1.10538966173224	1.61593612085349
	0.203659369859942	0.653151082167838
55656	INTS8	integrator complex subunit 8
7866	-0.166352503954456	5.27344991153335
1.61591203874153	0.203662738857569	0.653151082167838
101927027	CHROMR	cholesterol induced regulator of metabolism
RNA	4001	0.292879070251049	2.5318728861161
1.61584346471496	0.203672332467824	0.653151082167838
205564	SENP5	SUMO specific peptidase 5
7035	-0.15490463384363	5.74093613040933
1.61545605527268	0.203726541639555	0.653151082167838
140460	ASB7	ankyrin repeat and SOCS box containing 7
5397	-0.182322426206372	5.07955977440908
1.6153746921452 0.203737928725607	0.653151082167838
4763	NF1	neurofibromin 1 21093	-0.260225245187355
7.19262554947995	1.61476290492965	0.203823574752675
	0.653151082167838

372	ARCN1	archain 1	4177	-0.172972195919395
8.01937271123235	1.61473189620121	0.203827916892292
	0.653151082167838
134492	NUDCD2	NudC domain containing 2	8847
0.256138014854165	3.58951873836419	1.61454714665534
	0.203853789555752	0.653151082167838
23236	PLCB1	phospholipase C beta 1	15755	0.565880713599141
	1.36545099526106	1.61444698266231
0.20386781832211	0.653151082167838
25874	MPC2	mitochondrial pyruvate carrier 2	2552
0.211256820890009	4.26070526981242	1.61316498063191
	0.204047473453926	0.653577169144557
55625	ZDHHC7	zinc finger DHHC-type containing 7	3947
0.152469459294489	7.70362534860299	1.61179988143501
	0.204238978710323	0.65400440301809
2635	GBP3	guanylate binding protein 3
4297	-0.773056592714049	6.83220253911794
1.6114601931082	0.20428666538049	0.65400440301809
26065	LSM14A	LSM14A mRNA processing body assembly factor
4748	-0.158042641064152	6.73900820418289
1.61121617175601	0.20432093007833	0.65400440301809
79896	THNSL1	threonine synthase like 1	3788
0.390113353449121	1.474834095665	1.61057278114586
0.204411305420242	0.654119990563912
163071	ZNF114	zinc finger protein 114	3087	1.0264775510944
6.00468632856236	1.61029427957361	0.204450440400429
	0.654119990563912
153527	ZMAT2	zinc finger matrin-type 2	1812
0.167123450916598	6.14606186208422	1.60929051620985
	0.204591562330631	0.654143357621295
9708	PCDHGA8	protocadherin gamma subfamily A, 8	8661
0.175090168525922	6.46631026216211	1.60875105644687
	0.204667453959782	0.654143357621295
55384	MEG3	maternally expressed 3	12831	0.970106746818137
	3.26152279760232	1.60873245395591
0.204670071565117	0.654143357621295
6199	RPS6KB2	ribosomal protein S6 kinase B2
5822	-0.16082227528313	5.42747426735106
1.6079700316093	0.204777387992764	0.654143357621295
80095	ZNF606	zinc finger protein 606	7635	-0.244434739851332
	3.54992614846487	1.60791079972545
0.204785728084174	0.654143357621295
56892	TCIM	transcriptional and immune response regulator
1854	-0.707096068658072	5.27039847883482
1.60769548280966	0.204816048961571	0.654143357621295
4650	MYO9B	myosin IXB	9800	-0.172965100369853
7.08534688924205	1.60736533765844	0.204862550208511
	0.654143357621295
79887	PLBD1	phospholipase B domain containing 1	2841
0.732625738646997	2.28693535499478	1.60734904642328
	0.20486484516629	0.654143357621295
10594	PRPF8	pre-mRNA processing factor 8
11355	-0.163905817922156	8.20962138377275
1.60711739729598	0.204897481023549	0.654143357621295

101243544	NA	NA	3809	0.250863913579337
4.12544189786349	1.60692383510413	0.204924755710876
	0.654143357621295
54843	SYTL2	synaptotagmin like 2	11258	-0.332888359688725
	7.0818161640784	1.60578654653994	0.205085096630366
	0.654506026088297
373156	GSTK1	glutathione S-transferase kappa 1	4228
0.198128458455904	6.34283201889484	1.60529476647198
	0.205154476202088	0.654578302542745
201229	LYRM9	LYR motif containing 9	3110	0.427909899984485
	1.03686169786274	1.60443415392522
0.20527595673295	0.654790922832443
79155	TNIP2	TNFAIP3 interacting protein 2
2686	-0.167717476958194	5.12881631584507
1.6041604506918	0.205314609343906	0.654790922832443
7468	NSD2	nuclear receptor binding SET domain protein 2
19776	-0.189526057854821	5.94611072723337
1.60291783274479	0.205490200981202	0.654900696582775
5480	PPIC	peptidylprolyl isomerase C	2098
0.173762861489853	5.62774260097417	1.60288609256171
	0.205494688437936	0.654900696582775
130576	LYPD6B	LY6/PLAUR domain containing 6B
3771	-0.453461228422612	1.03154324190366
1.60271321995399	0.205519131359044	0.654900696582775
10559	SLC35A1	solute carrier family 35 member A1	2730
0.249382914174035	3.57600286032422	1.60254166389412
	0.205543391522058	0.654900696582775
5516	PPP2CB	protein phosphatase 2 catalytic subunit beta
4516	-0.176728774908647	7.53520609402159
1.60183813147461	0.205642915122309	0.654900696582775
54187	NANS	N-acetylneuraminate synthase	4922
0.221217341225795	5.22449715590145	1.60166503611495
	0.205667410377799	0.654900696582775
23387	SIK3	SIK family kinase 3	10920	-0.19365495403772
	5.71904742890134	1.60143735534923
0.205699635425845	0.654900696582775
51586	MED15	mediator complex subunit 15
10374	-0.171716579446566	7.58607396176868
1.60127179682852	0.205723071675891	0.654900696582775
776	CACNA1D	calcium voltage-gated channel subunit alpha1 D
10657	0.518637152703113	0.553848788864832
1.60079131296586	0.205791106199012	0.654968421842662
55038	CDCA4	cell division cycle associated 4	2513
0.206561507682869	4.94074344607069	1.59991078253997
	0.205915854603904	0.655075559857675
23229	ARHGEF9	Cdc42 guanine nucleotide exchange factor 9	7391
	0.251137168856487	3.42769871362507
1.59989349691845	0.205918304423027	0.655075559857675
91748	ELMSAN1	ELM2 and Myb/SANT domain containing 1
13484	-0.249862209541478	4.93872720071564
1.59866668102283	0.206092263846452	0.655082376226141
51296	SLC15A3	solute carrier family 15 member 3	6119
0.466610981205977	2.78684981414108	1.59828644571883
	0.206146215479372	0.655082376226141

4855	NOTCH4	notch receptor 4	9748	-0.735295999074732
	4.09370388227934	1.59804046019277
0.206181127279182	0.655082376226141
1844	DUSP2	dual specificity phosphatase 2
1932	-0.680279366316078	0.62028879792297
1.59772114178189	0.206226457353397	0.655082376226141
23019	CNOT1	CCR4-NOT transcription complex subunit 1
10852	-0.187970243512341	7.99985433040357
1.59764334230282	0.206237503460761	0.655082376226141
64773	PCED1A	PC-esterase domain containing 1A	2341
0.28867061550351	4.00586378537067	1.59699700967773
	0.20632929792533	0.655082376226141
648987	LOC648987	uncharacterized LOC648987
2784	-0.393257313395768	1.40548935113623
1.59695898992184	0.206334699129033	0.655082376226141
51608	GET4	guided entry of tail-anchored proteins factor 4
6525	-0.168422445503938	7.18185805865353
1.59693535237193	0.206338057236403	0.655082376226141
84294	UTP23	UTP23 small subunit processome component
5252	-0.23471887281517	3.60335030047241
1.59691209759611	0.206341361027325	0.655082376226141
22927	HABP4	hyaluronan binding protein 4	4056
0.302583638539635	3.6676379686726 1.59603259877165
0.206466356727929	0.655129793264172
885	CCK	cholecystokinin 2085	0.587562480077519
0.739039813246788	1.59536073672028	0.206561903030745
	0.655129793264172
30844	EHD4	EH domain containing 4	7879	-0.260072585284843
	6.86160090342039	1.595176060151	0.20658817525558
	0.655129793264172
10846	PDE10A	phosphodiesterase 10A	10693	0.616635144456958
	1.99674017088539	1.59454391237368
0.206678134947721	0.655129793264172
23410	SIRT3	sirtuin 3	5202	0.261905230588143
3.2222870992566 1.59452090038483	0.206681410610857
0.655129793264172
60592	SCOC	short coiled-coil protein
6294	-0.185474114676954	6.45868485441698
1.59451308237711	0.206682523486454	0.655129793264172
11227	GALNT5	polypeptide N-acetylgalactosaminyltransferase 5
6641	-0.224746704007205	9.78687824104754
1.59450483214164	0.206683697896365	0.655129793264172
85021	REPS1	RALBP1 associated Eps domain containing 1
9602	-0.162735024146502	5.3717225447685 1.5935641965691
0.206817647804832	0.655406061719972
10527	IPO7	importin 7	6688	-0.199977942424614
7.35071706166622	1.59311825259957	0.206881187674119
	0.65545912593336
133957	CCDC127 coiled-coil domain containing 127	9701
0.214878965096396	4.98780925724239	1.59273938988002
	0.206935187661311	0.655481947890543
23600	AMACR	alpha-methylacyl-CoA racemase	4838
0.209123079327893	5.01170991105988	1.59233569562128
	0.206992745260184	0.655516025968662

160335	TMTC2	transmembrane O-mannosyltransferase targeting cadherins 2	8323	-0.218969618551505	4.64036285987159	1.59133509767807	0.207135489286978	0.65581980073298
8658	TNKS	tankyrase	11700	-0.221387407539154	6.03543609056563	1.59054951209964	0.207247641443907	0.656026601967189
23175	LPIN1	lipin 1	11861	-0.214859917220206	5.65098921487901	1.5894172248296	0.20740941588874	0.65621188689188
387103	CENPW	centromere protein W	795	0.440655031081906	1.1287674281325	1.58934963320931	0.207419077695706	0.65621188689188
80775	TMEM177	transmembrane protein 177	2588	0.335026277929162	2.28550461507921	1.5891562609731	0.207446722007371	0.65621188689188
1716	DGUOK	deoxyguanosine kinase	1455	0.222731825943136	5.08311566836571	1.58821658371531	0.207581119396021	0.656488764818258
100126791	EGOT	eosinophil granule ontogeny transcript	1464	-0.568279753234623	1.25191265015498	1.58766015199287	0.207660751592599	0.65650202650619З
84795	PYROXD2	pyridine nucleotide-disulphide oxidoreductase domain 2	3867	0.567622181476814	1.13225510740349	1.5874192741406	0.207695235364941	0.656502026506193
79177	ZNF576	zinc finger protein 576	5952	0.237455715166484	3.59920290775859	1.58704462292918	0.207748883418859	0.656502026506193
5141	PDE4A	phosphodiesterase 4A	6646	0.299255440470209	3.85978385043298	1.58675071027986	0.207790981609834	0.656502026506193
84085	FBXO30	F-box protein 30	9009	-0.220533816203114	5.03602971206619	1.58608241197382	0.20788674198445	0.656502026506193
41	ASIC1	acid sensing ion channel subunit 1	5745	0.435534153898293	0.557943725086473	1.58535464761749	0.207991082601618	0.656502026506193
5734	PTGER4	prostaglandin E receptor 4	3961	-0.449745845530874	3.65450168462406	1.58521771395892	0.208010721897838	0.656502026506193
7351	UCP2	uncoupling protein 2	3237	0.544304843578487	5.90807095408103	1.5849629160644	0.20804727135297	0.656502026506193
65095	KRI1	KRI1 homolog	4267	0.204379934829275	4.24346342867354	1.58445599868037	0.208120008647533	0.656502026506193
114034	TOE1	target of EGR1, exonuclease	2742	0.282727472524281	2.45121600419311	1.584103050229	0.208170670785834	0.656502026506193
339122	RAB43	RAB43, member RAS oncogene family	4991	-0.166117110904597	6.98483805470406	1.58391143350909	0.208198181513924	0.656502026506193
404672	GTF2H5	general transcription factor IIH subunit 5	7481	0.233397372978789	4.20568396336257

1.58388799627428	0.208201546731462	0.656502026506193
65998	C11orf95	chromosome 11 open reading frame 95	5948
	0.250876074789776	5.00379106148817
1.58367696861275	0.208231849875423	0.656502026506193
101927021	LOC101927021	uncharacterized LOC101927021	2002
	0.449079696785025	0.706422007387671
1.58360986668902	0.208241486668595	0.656502026506193
9456	HOMER1	homer scaffold protein 1
6154	-0.238784997044975	4.11996924578842
1.58219475543049	0.208444839749812	0.656670447777756
100532735	INO80B-WBP1	INO80B-WBP1 readthrough (NMD
candidate)	2040	0.210116400602473	5.6680730982319
1.58170204501722	0.208515697918418	0.656670447777756
79893	GGNBP2	gametogenetin binding protein 2
9729	-0.149101041779537	6.39715337003249
1.58149546808755	0.208545414847034	0.656670447777756
29110	TBK1	TANK binding kinase 1	4762	-0.154784617128156
	5.57592537131482	1.58148802255162
0.208546486011102	0.656670447777756
3936	LCP1	lymphocyte cytosolic protein 1	5061
0.777927098472537	4.3349962434563 1.58139655492914
0.208559645679712	0.656670447777756
102724826	ZNF337-AS1	ZNF337 antisense RNA 1	5308
0.383264094113262	0.411158342436913	1.58109898371658
	0.208602464769242	0.656670447777756
9873	FCHSD2	FCH and double SH3 domains 2
5188	-0.169689197032314	5.19401287977745
1.58095573848393	0.208623080788964	0.656670447777756
54742	LY6K	lymphocyte antigen 6 family member K	4049
0.437916925072585	1.13100429559978	1.57975251562542
	0.208796345219922	0.657068166141417
9744	ACAP1	ArfGAP with coiled-coil, ankyrin repeat and PH
domains 1	3763	-0.655859322700228	0.610904516503512
	1.57914210393756	0.20888430977605
0.657197333116156
1536	CYBB	cytochrome b-245 beta chain	4423
1.31731836318793	3.71827474926285	1.57849077678221
	0.208978218903152	0.657344001168907
638	BIK	BCL2 interacting killer 953	-0.425873590410327
	3.85008944132446	1.57768249577399
0.209094827312313	0.657344001168907
53354	PANK1	pantothenate kinase 1	7638	0.338049724057892
	2.74519997626759	1.57760744558645
0.209105658496882	0.657344001168907
4326	MMP17	matrix metallopeptidase 17
4003	-0.637581322623264	2.98549508373658
1.57750507204352	0.209120434037579	0.657344001168907
29035	C16orf72	chromosome 16 open reading frame 72
15875	-0.169394922735478	7.3548286868282 1.57719236014863
	0.209165575298766	0.657344001168907
283120	H19	H19 imprinted maternally expressed transcript
2825	-0.997698663144544	2.52149690731237
1.57648616957792	0.209267559284117	0.657516981357699
10628	TXNIP	thioredoxin interacting protein

3625 -0.235026246396644 7.95950786442172
1.57578149062348 0.209369383650645 0.657537342359584
26022 TMEM98 transmembrane protein 98 5187
0.33824723147217 7.03285031705454 1.5756435760679
0.209389318825374 0.657537342359584
143279 HECTD2 HECT domain E3 ubiquitin protein ligase 2
11550 -0.328981897105426 3.68222695485962
1.5750706635846 0.209472155831461 0.657537342359584
8969 H2AC11 H2A clustered histone 11
2250 -0.380635264564788 0.395528055014753
1.57502569284618 0.209478659760541 0.657537342359584
2175 FANCA FA complementation group A 9736
0.596456324941294 1.54056899483441 1.57481761032244
0.209508756975143 0.657537342359584
641638 SNHG6 small nucleolar RNA host gene 6 1247
0.264832467542666 3.53311979763011 1.57266655723485
0.209820187183848 0.658367240565449
1969 EPHA2 EPH receptor A2 4159 -0.271560837502193
8.9423323751837 1.57195553360447 0.209923249967946
0.658543104658683
23576 DDAH1 dimethylarginine dimethylaminohydrolase 1 7653
0.274286851424782 6.19596713245924
1.57112195878352 0.21004415289502 0.658698358207083
4135 MAP6 microtubule associated protein 6 6433
0.600223856290224 1.83713946590625 1.57096592965477
0.210066792754411 0.658698358207083
54756 IL17RD interleukin 17 receptor D
9354 -0.295115304500727 4.89810331448554
1.56922892380785 0.210319027998194 0.659341679760676
26528 DAZAP1 DAZ associated protein 1
6773 -0.144845667351105 6.48473444496463
1.56854291327923 0.210418744244875 0.659433259077203
6891 TAP2 transporter 2, ATP binding cassette subfamily B
member 7031 -0.208913728162488 6.74378664526876
1.56838024115252 0.210442397949247 0.659433259077203
55041 PLEKHB2 pleckstrin homology domain containing B2
6728 -0.28095065966126 7.98734294257928
1.56735796101754 0.210591117016817 0.65960172465087
7090 TLE3 TLE family member 3, transcriptional corepressor
12634 -0.198856899419275 4.94362408778286
1.5672992489462 0.210599662101207 0.65960172465087
6398 SECTM1 secreted and transmembrane 1
2961 -0.417555129662185 3.30519344272631
1.56661608268824 0.21069912184034 0.65960172465087
7041 TGFB1I1 transforming growth factor beta 1 induced transcript
1 5403 0.282845941713096 5.74533364917694
1.56654499950256 0.210709473785027 0.65960172465087
51043 ZBTB7B zinc finger and BTB domain containing 7B
4303 -0.18765317082825 6.19917578443813
1.5663929840248 0.210731614036742 0.65960172465087
9208 LRRFIP1 LRR binding FLII interacting protein 1
11629 -0.212679449254125 8.10887788479745
1.56383691713648 0.211104305006932 0.660286096527558
112770 GLMP glycosylated lysosomal membrane protein 3323

0.311779756750829	5.75225380546621	1.56327803402722
	0.211185897825944	0.660286096527558
6839	SUV39H1 suppressor of variegation 3-9 homolog 1 3368
0.293888191216315	2.06682781347218	1.56310980060838
	0.211210465987686	0.660286096527558
9406	ZRANB2	zinc finger RANBP2-type containing 2
4993	-0.26839036390482	5.75887719265007
1.56308881258623	0.21121353123551	0.660286096527558
6723	SRM	spermidine synthase	1898	0.307492803860738
	4.60259050847923	1.56297135716741
0.211230686277473	0.660286096527558
9117	SEC22C	SEC22 homolog C, vesicle trafficking protein	8159
	0.144032931172842	5.97868484360229
1.56295484824983	0.211233097632611	0.660286096527558
3880	KRT19	keratin 19	2336	-0.192578760970303
10.8835258132534	1.56232061505125	0.211325760859098
	0.660337226602967
1893	ECM1	extracellular matrix protein 1
3360	-0.585202277342469	6.492996815368	1.56219758496786
	0.211343741476483	0.660337226602967
1164	CKS2	CDC28 protein kinase regulatory subunit 2	613
	0.328267176082074	4.01316388239494
1.56162431846929	0.211427547231832	0.660451751578116
79796	ALG9	ALG9 alpha-1,2-mannosyltransferase	8998
0.173763029419575	4.89656186922866	1.56123257761485
	0.211484838444014	0.660483418525151
196	AHR	aryl hydrocarbon receptor
6661	-0.25787203391413	7.05249456101766
1.56065483533872	0.211569365546021	0.660600112171894
79176	FBXL15	F-box and leucine rich repeat protein 15	2593
	0.346951302805618	2.84168012701872
1.56025576409302	0.211627775420882	0.660635224051772
2217	FCGRT	Fc fragment of IgG receptor and transporter	4173
	0.332065818282772	5.54931135681021
1.55836509751518	0.211904762059475	0.661352496026529
24145	PANX1	pannexin 1	2769	0.18044804857821
6.24324694628384	1.55683691084147	0.212128958979156
	0.661904728986642
51477	ISYNA1	inositol-3-phosphate synthase 1 3284
0.406692529524448	5.23648875917157	1.55592666856229
	0.212262632381154	0.662174318878136
55361	PI4K2A	phosphatidylinositol 4-kinase type 2 alpha	4864
	0.179764248440728	6.78057830038455
1.55504326125539	0.212392460536975	0.662431795756269
9827	RGP1	RGP1 homolog, RAB6A GEF complex partner 1
7116	-0.141005029815325	6.71006192218209
1.55449643822571	0.212472870493832	0.662535061666764
23137	SMC5	structural maintenance of chromosomes 5
6637	-0.208735926463419	5.75723732914204
1.55368883604433	0.212591693949841	0.662695743945262
56889	TM9SF3	transmembrane 9 superfamily member 3
7095	-0.167335168559584	8.2691380687992 1.55350315688139
	0.212619024294282	0.662695743945262
11340	EXOSC8	exosome component 8	3630	-0.229438421158346

3.62448952844259	1.55278991798532
0.212724045594626	0.662865118560709
9735	KNTC1	kinetochore associated 1
11325	-0.275214690553989	3.39091363268595
1.55249143578499	0.212768014067891	0.662865118560709
1673	DEFB4A	defensin beta 4A	380	-1.33831535049593
1.50723371881784	1.5515405934072
0.212906166207999	0.663112940752603
221527	ZBTB12	zinc finger and BTB domain containing 12	1875
0.448154920288135	0.854230786381296
1.55130938877622	0.212942243699951	0.663112940752603
3112	HLA-DOB major histocompatibility complex, class II, DO beta
1858	0.495417730452164	1.10480249244908
1.55090513281159	0.213001868498867	0.663151182943682
54931	TRMT10C tRNA methyltransferase 10C, mitochondrial RNase P
subunit 1888	-0.194487186188169	4.66999032480344
1.54986354132923	0.213155586998062	0.663236774593411
84807	NFKBID	NFKB inhibitor delta	2456	-0.356855302694189
2.16657263000148	1.54984815050875
0.2131578593692 0.663236774593411
64231	MS4A6A	membrane spanning 4-domains A6A 5791
1.14941680696267	-0.000511300547285725	1.54973206790416
0.213174999262788	0.663236774593411
51773	RSF1	remodeling and spacing factor 1
15740	-0.202278316543846	6.51302500737584
1.5494357298259 0.213218761761557	0.663236774593411
404217	CTXN1	cortexin 1	1274	0.336698006324633
3.43949128102001	1.54900772382333	0.213281987500738
0.663286145409157
23066	CAND2	cullin associated and neddylation dissociated 2
(putative)	5478	0.659443930250202	0.0176270551221487
1.54818349127611	0.213403807210806
0.663491622057672
57148	RALGAPB Ral GTPase activating protein non-catalytic beta
subunit 12633	-0.171100984381416	7.11239903665881
1.54791980844114	0.21344279638694	0.663491622057672
89795	NAV3	neuron navigator 3	11153	0.787641286784036
0.996703505091412	1.54577081177748
0.213760871175707	0.664201284834995
5888	RAD51	RAD51 recombinase	3046	0.326216436714034
1.16299101986562	1.54496861369724
0.213879749514308	0.664201284834995
79611	ACSS3	acyl-CoA synthetase short chain family member 3
10572	-0.579801963304969	0.868092933495821
1.54490233151714	0.213889575433825	0.664201284834995
3267	AGFG1	ArfGAP with FG repeats 1
9321	-0.206685517298817	7.01951054677061
1.54483756339589	0.213899177421767	0.664201284834995
84236	RHBDD1	rhomboid domain containing 1
6926	-0.209006040739658	4.8833989537845 1.54477678447666
0.213908188469384	0.664201284834995
10133	OPTN	optineurin	5138	-0.146589077411591
8.23421070749254	1.54445009946737	0.213956630330656
0.66420358224397

290	ANPEP	alanyl aminopeptidase, membrane	4871
1.28066360106713	2.58069994743546	1.5441322997491
0.214003767164064	0.66420358224397
10232	MSLN	mesothelin	2642	1.03026125221373
4.93721377173507	1.54376127578468	0.214058813964427
	0.664227250155012
1657	DMXL1	Dmx like 1	13680	-0.250293271946196
5.98013122547873	1.54248034916366	0.21424898744387
	0.664661846634085
64777	RMND5B	required for meiotic nuclear division 5 homolog B
	7701	-0.205420603933736	4.61989256886792
1.54199491947797	0.214321109500982	0.664661846634085
84818	IL17RC	interleukin 17 receptor C
4540	-0.320386445202118	3.89923457153981
1.54185955245653	0.214341226618559	0.664661846634085
114907	FBXO32	F-box protein 32	7395	-0.302185851493898
	7.18158191965715	1.54111977893121
0.214451205253003	0.664855695435772
2180	ACSL1	acyl-CoA synthetase long chain family member 1
6284	-0.276830118637137	6.63550780736317
1.54051004576195	0.214541901841039	0.664989692207885
130916	MTERF4	mitochondrial transcription termination factor 4
	9938	0.183312616941794	4.86786378845844
1.53928740706478	0.214723904355303	0.665281926802857
100419583	LOC100419583	ring finger protein 4 pseudogene
	3425	-0.407388275632367	3.86388053273618
1.53923857968516	0.214731176631378	0.665281926802857
92715	DPH7	diphthamide biosynthesis 7	4475
0.193836504429878	4.67091233028006	1.53825510120038
	0.214877716813469	0.665378324832564
151188	ARL6IP6	ADP ribosylation factor like GTPase 6 interacting
protein 6	3349	-0.26015249676294	4.27140264422382
	1.53809992325318	0.214900849487434
0.665378324832564
10724	OGA	O-GlcNAcase	7247	-0.208113636321212
7.67481956351954	1.53770485237598	0.214959756831056
	0.665378324832564
259282	BOD1L1	biorientation of chromosomes in cell division 1 like
1	15860	-0.242426365688476	6.79691114519331
1.53693139721337	0.21507513906816	0.665378324832564
79663	HSPBAP1	HSPB1 associated protein 1	5434
0.39927720996985	1.60955270127328	1.53679249813795
	0.21509586751015	0.665378324832564
51714	SELENOT	selenoprotein T	4135	-0.207076986874882
7.22552557188914	1.53677381914551	0.215098655229137
	0.665378324832564
125061	AFMID	arylformamidase	4048	0.304169560310032
3.19460765991762	1.53674921002022	0.215102328047765
	0.665378324832564
57458	TMCC3	transmembrane and coiled-coil domain family 3
6515	-0.448259510585594	4.98803555265489
1.53648130338114	0.215142316924872	0.665378324832564
55229	PANK4	pantothenate kinase 4 (inactive)
5107	-0.186839157574177	4.49369423611797

1.53559471776571	0.215274715515427	0.665640825656643
6778	STAT6	signal transducer and activator of transcription 6
	7298	-0.165750344122062	7.61290971613907
1.53512598964005	0.215344752373912	0.665696161652961
101059938	NPIPA7	nuclear pore complex interacting protein
family member A7	3007	0.257589296140547
3.7319200063332 1.53460196633622	0.215423083477686
0.665696161652961
8036	SHOC2	SHOC2 leucine rich repeat scaffold protein
4429	-0.138805598451493	6.53834108063669
1.53360575219069	0.215572091271406	0.665696161652961
101929147	WARS2-AS1	WARS2 antisense RNA 1	5462
0.295873322388918	2.5714447369339 1.53353841171634
0.215582168084281	0.665696161652961
3551	IKBKB	inhibitor of nuclear factor kappa B kinase subunit
beta	10239	-0.223154209336992	6.18063407500911
1.53324291677375	0.215626392505729	0.665696161652961
492311	IGIP	IgA inducing protein	3458	-0.389603812506103
	2.25858169311744	1.53315838990923
0.215639044967559	0.665696161652961
83452	RAB33B	RAB33B, member RAS oncogene family
4562	-0.249023609049566	3.74667783446398
1.53304183336076	0.215656493263852	0.665696161652961
4436	MSH2	mutS homolog 2	4462	0.21511748593651
4.45188797549601	1.53293277860827	0.215672820076875
	0.665696161652961
283450	HECTD4	HECT domain E3 ubiquitin protein ligase 4
21572	-0.338209048620766	7.25318383844156
1.5324370022405 0.215747062332468	0.665696161652961
22823	MTF2	metal response element binding transcription factor
2	5909	-0.190743057207643	4.81251308115443
1.53229807985758	0.215767871343217	0.665696161652961
9546	APBA3	amyloid beta precursor protein binding family A
member 3	5556	-0.191414515016272	4.24902386708077
	1.53047206822406	0.216041609545893
0.66639392752903
22821	RASA3	RAS p21 protein activator 3
4763	-0.35689349773987	2.94051840828353
1.52983258991192	0.216137571747266	0.666543145632752
93974	ATP5IF1 ATP synthase inhibitory factor subunit 1	2225
	0.241387316286531	5.8933527267542 1.52748721645461
	0.2164899608288 0.667380230882124
686	BTD	biotinidase	3704	0.21670336100758
4.27907430371045	1.52739185789951	0.216504302786348
	0.667380230882124
8532	CPZ	carboxypeptidase Z	8396	-0.670332287888163
	1.84125667891062	1.52663232948643
0.216618576515145	0.667475016798827
4703	NEB	nebulin 30567	-0.612767939713502
0.822678947373788	1.5265540492693 0.216630358132896
0.667475016798827
57026	PDXP	pyridoxal phosphatase	2203	0.367382050047845
	2.25257249160375	1.52501792045866
0.216861708776089	0.66785125400011

55034	MOCOS	molybdenum cofactor sulfurase	6281
0.270375431522684	4.38955828662117	1.52418116779557
	0.216987852806546	0.66785125400011
80208	SPG11	SPG11 vesicle trafficking associated, spatacsin
11853	-0.163498299882479	6.28816643255484
1.52393507258516	0.217024969343401	0.66785125400011
55193	PBRM1	polybromo 1	9654	-0.189773858254573
6.99514867898576	1.52380870731061	0.21704403092822
	0.66785125400011
113174	SAAL1	serum amyloid A like 1	3070	0.271893243079053
	3.16837627805546	1.52369942656949
0.217060517003069	0.66785125400011
388335	TMEM220	transmembrane protein 220	3508
0.827508621947815	-0.152503334891452	1.52336767604835
	0.217110573973654	0.66785125400011
114783	LMTK3	lemur tyrosine kinase 3	5163	-0.320391139937447
	4.00858789564732	1.52305410136125
0.21715790108583	0.66785125400011
5025	P2RX4	purinergic receptor P2X 4	8159
0.341083079610999	4.52975142982495	1.52266268515588
	0.217216993891662	0.66785125400011
645426	TMEM191C	transmembrane protein 191C	4299
0.402882696444553	0.478254615936625	1.52247820667203
	0.217244851582721	0.66785125400011
728190	NUTM2A-AS1	NUTM2A antisense RNA 1	6851
0.420349224146461	4.88228580021265	1.52232240346396
	0.217268382399647	0.66785125400011
8720	MBTPS1	membrane bound transcription factor peptidase, site
1	10238	0.154290480922485	7.38435098944334
1.52226571536919	0.217276944704684	0.66785125400011
3778	KCNMA1	potassium calcium-activated channel subfamily M
alpha 1	18328	-0.500029764342767	1.81539503680516
1.5211718580358	0.217442242298178	0.668212700278758
10067	SCAMP3	secretory carrier membrane protein 3	2442
0.157752001466287	6.01695753984359	1.52033841112515
	0.21756828874014	0.668355856488222
10901	DHRS4	dehydrogenase/reductase 4	2305
0.327075222571751	2.52675693522451	1.52023285814767
	0.217584258269886	0.668355856488222
3251	HPRT1	hypoxanthine phosphoribosyltransferase 1	1624
	0.409842623794026	2.57482121800004
1.519228951947	0.217736213096138	0.668579449523683
11234	HPS5	HPS5 biogenesis of lysosomal organelles complex 2
subunit 2	6433	-0.200255484096361	6.01371101017769
	1.51885104962015	0.217793446514277
0.668579449523683
10137	RBM12	RNA binding motif protein 12
6702	-0.181767519517604	6.47272727366922
1.51880616993382	0.217800244747381	0.668579449523683
26175	TMEM251	transmembrane protein 251	1604
0.260814075328868	3.26987111538322	1.51826162624687
	0.217882750666437	0.668686171907271
9785	DHX38	DEAH-box helicase 38	6361	-0.153329284162824
	6.43679246508988	1.51682452864347

0.218100669897854	0.66920834244452
85462	FHDC1	FH2 domain containing 1 6593	-0.28081558842552
	5.32565644583737	1.51610278569993
0.218210211911073	0.669397818273849
5046	PCSK6	proprotein convertase subtilisin/kexin type 6
13239	-0.555288060329945	2.75570552616761
1.51554394079139	0.218295075268597	0.669511521630663
57111	RAB25	RAB25, member RAS oncogene family
1699	-0.229045710131697	6.44184105849302
1.51416003674373	0.218505397434884	0.670009873439235
5412	UBL3	ubiquitin like 3	4384	-0.275109953067249
	6.6662233728069 1.51363795553539	0.218584804782767
	0.670106666796284
11072	DUSP14	dual specificity phosphatase 14
2499	-0.30728334382357	7.25717872691861
1.51312352348726	0.218663082388468	0.670199955145573
79003	MIS12	MIS12 kinetochore complex component
3157	-0.190936923340089	4.42253181289346
1.51178680955421	0.218866637423207	0.670381958825976
8309	ACOX2	acyl-CoA oxidase 2	3859	0.574001017033687
	0.54567630853689	1.511530812573	0.21890564648282
	0.670381958825976
57634	EP400	E1A binding protein p400
18705	-0.230269179097131	6.12553321340723
1.51151637255485	0.218907847113493	0.670381958825976
81928	CABLES2 Cdk5 and Abl enzyme substrate 2
3785	-0.273387487911845	3.4527598167465 1.51147661725246
	0.218913905880632	0.670381958825976
55207	ARL8B	ADP ribosylation factor like GTPase 8B
3576	-0.173199602776469	7.52915251439682
1.51107633606557	0.218974920480684	0.670422231950369
9825	SPATA2	spermatogenesis associated 2
4270	-0.220065241772223	3.73837603603804
1.50986085947255	0.219160319209049	0.670800493084923
785	CACNB4	calcium voltage-gated channel auxiliary subunit beta
4	13587	-0.43682937673028	1.01234411493919
1.50963854295807	0.219194249792939	0.670800493084923
51	ACOX1	acyl-CoA oxidase 1	8472	-0.184276615154495
	6.503003571576	1.50916148327464	0.219267081176825
	0.670876803417166
5777	PTPN6	protein tyrosine phosphatase non-receptor type 6
	5009	0.204813843203921	4.89438900475671
1.50855405432901	0.219359857488288	0.671014091251027
10994	ILVBL	ilvB acetolactate synthase like 5615
0.186120300370746	5.65225671124531	1.50756913438597
	0.219510389913562	0.671261210511369
9700	ESPL1	extra spindle pole bodies like 1, separase
8890	-0.399454732579749	0.0202562841061219
1.50739845063444	0.219536489280544	0.671261210511369
10521	DDX17	DEAD-box helicase 17	9995	-0.192993772572456
	8.75544363508592	1.50664981991389
0.21965100668151	0.671464786247906
23594	ORC6	origin recognition complex subunit 6	4309
0.329055861975616	2.20135660371186	1.50612221962828

0.219731756096775	0.671565068218968
26098	EDRF1	erythroid differentiation regulatory factor 1
7741	-0.220855161755977	4.66704950287002
1.50379536806471	0.220088304815624	0.672412735602718
7280	TUBB2A	tubulin beta 2A class IIa
1727	-0.379496686971592	5.73093798628549
1.50368575343033	0.220105118350715	0.672412735602718
7589	ZSCAN21	zinc finger and SCAN domain containing 21	2140
0.307450914211138	2.8278360457168	1.50256598868727
0.220276964405125	0.67279098133942
57680	CHD8	chromodomain helicase DNA binding protein 8
9613	-0.193963685770311	7.36069915380095
1.50187946073753	0.220382402515187	0.672966276876002
114803	MYSM1	Myb like, SWIRM and MPN domains 1
8938	-0.265936712580931	5.63515137839324
1.49924545830511	0.220787496986445	0.674056336157178
144165	PRICKLE1	prickle planar cell polarity protein 1
6785	-0.331228432021294	4.17162220488492
1.49826476573647	0.220938549198645	0.674370507436354
4199	ME1	malic enzyme 1	3371	0.235404218780576
5.76055653608231	1.49768456924825	0.221027972740378
0.674496473676357
134549	SHROOM1	shroom family member 1	4245	0.316250171780975
2.98030155992299	1.49720603522906
0.221101759984009	0.674574679175783
1859	DYRK1A	dual specificity tyrosine phosphorylation regulated kinase 1A	10213	-0.198977814195547	6.26526619065953
1.49656770603056	0.221200232591851
0.674728148500448
6376	CX3CL1	C-X3-C motif chemokine ligand 1
5983	-0.668555189825377	6.68246229945893
1.49558288624206	0.221352259862894	0.674939453634605
11267	SNF8	SNF8 subunit of ESCRT-II	5270
0.172498902873631	5.4238406979236	1.49549471003096
0.221365877775211	0.674939453634605
60684	TRAPPC11	trafficking protein particle complex 11
5841	-0.150770598870915	5.5927658450655	1.49474508039223
0.221481690866513	0.675128188323294
11261	CHP1	calcineurin like EF-hand protein 1
3950	-0.163708470888976	7.39192197860409
1.49419556578876	0.221566633467314	0.675128188323294
51114	ZDHHC9	zinc finger DHHC-type containing 9	5243
0.158810098678739	7.06047401771199	1.49415841990299
0.22157237679176	0.675128188323294
5125	PCSK5	proprotein convertase subtilisin/kexin type 5
12705	-0.364868472953384	2.3174981719758	1.49312819982944
0.221731735616628	0.675265409558922
8704	B4GALT2	beta-1,4-galactosyltransferase 2	3772
0.220192308326662	5.59526457074409	1.4928043143244
0.221781863928582	0.675265409558922
89765	RSPH1	radial spoke head component 1
2988	-0.465562830695043	-0.0430948601478595
1.49273476323993	0.221792630236663	0.675265409558922
83658	DYNLRB1	dynein light chain roadblock-type 1	2460

0.198449715631501	6.09853542493451	1.49262095739586
	0.221810248402984	0.675265409558922
5595	MAPK3	mitogen-activated protein kinase 3
3669	-0.242320598560808	5.57900817532916
1.49201203608989	0.221904543337296	0.675394544364851
255104	TMCO4	transmembrane and coiled-coil domains 4 4054
0.234537629199244	3.97659127038964	1.49154412899286
	0.221977034004351	0.675394544364851
4713	NDUFB7	NADH:ubiquinone oxidoreductase subunit B7	531
	0.201110684911029	5.08698332426224
1.49141321158937	0.221997321500377	0.675394544364851
101928841	LOC101928841	collagen alpha-1(II) chain-like 9759
	0.368039682740669	0.300954663133888
1.49099431336889	0.22206225058887	0.675445373289535
81563	C1orf21 chromosome 1 open reading frame 21
10971	-0.274334184195593	2.74041330366493
1.49022633374909	0.222181346208787	0.67566090237657
102465801	MIR7703 microRNA 7703	643	-0.24635029711839
	3.8475178249787 1.48930565154392	0.222324223177009
	0.675948642075181
27032	ATP2C1	ATPase secretory pathway Ca2+ transporting 1
8533	-0.173113270159338	8.3943514366512 1.48865369923887
	0.222425463537969	0.67610969352785
23262	PPIP5K2 diphosphoinositol pentakisphosphate kinase 2
17270	-0.19328767322384	5.55820552642213
1.48793083543747	0.222537780120225	0.67620611085294
57505	AARS2	alanyl-tRNA synthetase 2, mitochondrial 4298
0.258255201355294	3.68137319599207	1.48782818382426
	0.222553735348901	0.67620611085294
125144	SNHG29	small nucleolar RNA host gene 29	3759
0.26063383256313	7.46852207259224	1.4866873197557
0.222731153051776	0.676511455050746
91749	MFSD4B	major facilitator superfamily domain containing 4B
	5958	0.244270340882448	3.46436161995754
1.48654738511199	0.222752926181355	0.676511455050746
673	BRAF	B-Raf proto-oncogene, serine/threonine kinase
10380	-0.26379205891897	4.70805416844577
1.48625052300056	0.222799124876782	0.676511455050746
3881	KRT31	keratin 31	1596	-0.661646749562687
0.77813906425784	1.48574406709532	0.222877967758122
	0.676604181705249
9337	CNOT8	CCR4-NOT transcription complex subunit 8
3930	-0.156066557749655	6.617294305259	1.48512913969105
	0.222973741928626	0.676629827220686
373	TRIM23	tripartite motif containing 23
4807	-0.252795754025907	4.7408318201906 1.48506953014342
	0.222983028660719	0.676629827220686
8654	PDE5A	phosphodiesterase 5A	9775	0.315576193582009
	4.26767142183538	1.48423635581904
0.223112879629272	0.676877215717394
10499	NCOA2	nuclear receptor coactivator 2
8447	-0.223984719911648	5.90963875410975
1.48364774063909	0.223204670425624	0.677009055576377
23551	RASD2	RASD family member 2	3412	0.62759763764652

3.23356320920159	1.48237039619217
0.223404019804419	0.677467007014612
1595	CYP51A1	cytochrome P450 family 51 subfamily A member 1
4741	-0.267308867749644	7.53620392884436
1.48183799331957	0.223487172485316	0.677572472944118
7336	UBE2V2	ubiquitin conjugating enzyme E2 V2
6606	-0.179491429284264	5.65949533242198
1.4813466387944	0.223563946983739	0.677658559922362
138050	HGSNAT	heparan-alpha-glucosaminide N-acetyltransferase	8943
0.14483044413906	7.31009029490629
1.48020306383462	0.223742753786106	0.677910939217464
4190	MDH1	malate dehydrogenase 1	4990	0.171222104999673
6.82389928871565	1.47995816377464
0.223781068082232	0.677910939217464
9448	MAP4K4	mitogen-activated protein kinase kinase kinase
kinase 4	11961	-0.152460064096033	8.48021676834365
1.47988572474953	0.223792402580249
0.677910939217464
9465	AKAP7	A-kinase anchoring protein 7	4394
0.367895263486155	0.427918060602798	1.47850639317329
0.224008357231661	0.678418391295973
3105	HLA-A	major histocompatibility complex, class I, A
2710	-0.382045573976167	9.19911422630389
1.47804786659233	0.224080201620906	0.678480915601433
340120	ANKRD34B	ankyrin repeat domain 34B	4024
0.536367616353587	3.10456121762496	1.47775643095294
0.22412587966644	0.678480915601433
7978	MTERF1	mitochondrial transcription termination factor 1
4890	-0.228592935862949	3.43328157290528
1.47738679733592	0.224183830096232	0.67850970357777
64852	TUT1	terminal uridylyl transferase 1, U6 snRNA-specific
4432	0.227171731536271	3.5678824352821
1.47645802767255	0.224329520059694	0.67869400222479
140893	RBBP8NL	RBBP8 N-terminal like	2793	-0.568165497778056
0.887916350836217	1.47638085108255
0.224341631348667	0.67869400222479
4647	MYO7A	myosin VIIA	10120	0.602504260540483
2.12489842650376	1.47532808134177	0.224506920361518
0.679047383611268
92856	IMP4	IMP U3 small nucleolar ribonucleoprotein 4	3861
0.172474018249325	4.90711815822879
1.47477429144039	0.224593926172998	0.679163886853449
254531	LPCAT4	lysophosphatidylcholine acyltransferase 4
4068	-0.32772349042957	6.09077674868678
1.4730035637212	0.224872396299718	0.679709343884199
91272	BOD1	biorientation of chromosomes in cell division 1	2018
0.190179170480719	4.8257810684847	1.47285871243429
0.224895194379532	0.679709343884199
1477	CSTF1	cleavage stimulation factor subunit 1	4800
0.188945579526275	4.87549626142344	1.47252250342688
0.224948120852243	0.679709343884199
9994	CASP8AP2	caspase 8 associated protein 2
8171	-0.23940040681973	4.67401448160301
1.47198982651609	0.225032006157272	0.679709343884199

375287	RBM43	RNA binding motif protein 43	4717
0.337630359183309	2.84448817699628	1.47183557290596
	0.225056304832443	0.679709343884199
1838	DTNB	dystrobrevin beta	7146	0.316736604586894
	3.51923899635135	1.47177744142225
0.225065462763969	0.679709343884199
50640	PNPLA8	patatin like phospholipase domain containing 8
5668	-0.150329656665585	6.25451396625214
1.47101785019198	0.225185168524459	0.679924262884694
256281	NUDT14	nudix hydrolase 14	1916	0.226673407188169
	4.04070255875404	1.47003768493406
0.225339747962696	0.680060759207524
53335	BCL11A	BAF chromatin remodeling complex subunit BCL11A
8375	-0.288160473188877	2.99818929200915
1.46988884155664	0.225363232817245	0.680060759207524
51690	LSM7	LSM7 homolog, U6 small nuclear RNA and mRNA
degradation associated	1698	0.195687860030178
4.10844778901959	1.4698077464299 0.225376029431093
0.680060759207524
22859	ADGRL1	adhesion G protein-coupled receptor L1	8487
0.400553800098727	4.22423780089679	1.46942746016138
	0.225436049312346	0.680095357035974
3653	IPW	imprinted in Prader-Willi syndrome	9063
0.182547654350389	5.40237095142369	1.46862366192589
	0.225562974437851	0.680331736208221
83982	IFI27L2	interferon alpha inducible protein 27 like 2	1662
	0.291281700901284	4.03197685113995
1.4679733989985 0.225665718069134	0.680495094293725
3032	HADHB	hydroxyacyl-CoA dehydrogenase trifunctional
multienzyme complex subunit beta	4280	0.181892195161382
	6.6472494845572 1.46732974523934	0.225767472753448
	0.680509212178381
4946	OAZ1	ornithine decarboxylase antizyme 1	2533
0.176416424229014	8.92967837582982	1.46723293364209
	0.225782782382206	0.680509212178381
971	CD72	CD72 molecule	3386
0.583854567155763	-0.0263004252025736	1.46702195595678
	0.225816150368039	0.680509212178381
84727	SPSB2	splA/ryanodine receptor domain and SOCS box
containing 2	2248	0.365488701406331	3.34565021118946
	1.46632948821447	0.225925711871475
0.680539073143152
4905	NSF	N-ethylmaleimide sensitive factor, vesicle fusing
ATPase	4503	-0.170995277981271	5.53248374984044
1.46587191040437	0.22599814435939	0.680539073143152
114882	OSBPL8	oxysterol binding protein like 8
9198	-0.214212797447878	7.22010284602524
1.4658224365395 0.226005977517416	0.680539073143152
4893	NRAS	NRAS proto-oncogene, GTPase
4449	-0.17636677888558	6.87627965372095
1.46573133870913	0.226020401817801	0.680539073143152
11070	TMEM115 transmembrane protein 115	2197
0.156535058505406	6.06433339237148	1.46522004390107
	0.226101380021811	0.680636584991511

158228	FAM201A family with sequence similarity 201 member A	3847
	0.382988372534074	1.51928294049986
1.46328105031675	0.22640879203716	0.681415545780943
2773	GNAI3	G protein subunit alpha i3
27174	-0.201650573533293	7.0313801624615 1.46249559399251
	0.226533462728468	0.681644299127314
8237	USP11	ubiquitin specific peptidase 11 5269
0.19689617151633	5.46897916927766	1.46191622333231
	0.226625475544823	0.681774707035297
79874	RABEP2	rabaptin, RAB GTPase binding effector protein 2 4566
	0.19592051581142	5.02842882220271
1.46120844109116	0.226737942991928	0.681904750813897
51720	UIMC1	ubiquitin interaction motif containing 1	4739
	0.17927070842351	4.97669700179422
1.46088633891649	0.226789147613816	0.681904750813897
11124	FAF1	Fas associated factor 1 7545	0.17257687328455
	5.01986088383175	1.46064436592994
0.226827623227539	0.681904750813897
80199	FUZ	fuzzy planar cell polarity protein	2950
0.360855903448524	2.83709565395454	1.46025653083798
	0.226889308426803	0.681904750813897
254042	METAP1D methionyl aminopeptidase type 1D, mitochondrial 3523
	0.36863965191584	0.901690991073898
1.45972845548006	0.22697333125961	0.681904750813897
25855	BRMS1	BRMS1 transcriptional repressor and anoikis
regulator	2514	0.163058474720132	5.5645237333921
1.45954070542791	0.227003213450718	0.681904750813897
102465538	MIR6892 microRNA 6892	115	-0.36134870532173
	0.362918766371745	1.45950130650522
0.227009484760848	0.681904750813897
51447	IP6K2	inositol hexakisphosphate kinase 2	5624
0.186406545664673	6.94360756056613	1.4590044084519
0.227088596213167	0.681996133610168
9231	DLG5	discs large MAGUK scaffold protein 5
10271	-0.173469759991601	7.50346034373769
1.45835617430749	0.227191851790098	0.682037602545928
8428	STK24	serine/threonine kinase 24
12077	-0.194864364210061	8.03383204890699
1.45810536770549	0.227231817277465	0.682037602545928
79734	KCTD17	potassium channel tetramerization domain containing
17	2786	0.262250126282516	3.59025049277712
1.45779424282919	0.227281406093648	0.682037602545928
54454	ATAD2B	ATPase family AAA domain containing 2B
10492	-0.259339227056323	4.57706617193475
1.45769532502101	0.227297174890012	0.682037602545928
54947	LPCAT2	lysophosphatidylcholine acyltransferase 2	7439
	0.32034847450821	7.426395592016	1.45660256135153
	0.227471463292253	0.682357021222823
90480	GADD45GIP1	GADD45G interacting protein 1	1782
0.228903662814533	4.60788112642369	1.45637650586222
	0.227507537662341	0.682357021222823
80741	LY6G5C	lymphocyte antigen 6 family member G5C	2306
0.341571433824172	0.613940940847581	1.45611190905663
	0.227549771266638	0.682357021222823

29072 SETD2 SET domain containing 2, histone lysine methyltransferase 10019 -0.191379905364538 6.99773277901071 1.45528377264257 0.227682015142828 0.682607445656164
101929243 LOC101929243 uncharacterized LOC101929243 771 0.394695806025475 -0.108763031669615 1.45437766295603 0.22782681633766 0.682895402619645
158358 KIAA2026 KIAA2026 7233 -0.239646289948635 5.37572283088446 1.45350151891969 0.22796693417844 0.683166880181658
6191 RPS4X ribosomal protein S4 X-linked 2982 0.227708260513127 7.45071654723048 1.45300541624488 0.228046319639933 0.683166880181658
9411 ARHGAP29 Rho GTPase activating protein 29 14016 -0.245221243149514 5.65234376754161 1.45269486055471 0.228096031126437 0.683166880181658
23071 ERP44 endoplasmic reticulum protein 44 4854 -0.204823341817843 5.63657289588098 1.45173153744103 0.228250315799031 0.683166880181658
10102 TSFM Ts translation elongation factor, mitochondrial 3195 0.204182437006379 4.07933717001837 1.45155622317036 0.22827840742184 0.683166880181658
4087 SMAD2 SMAD family member 2 36366 -0.157398654304136 6.816458094719 1.45144553265447 0.228296146147422 0.683166880181658
283358 B4GALNT3 beta-1,4-N-acetyl-galactosaminyltransferase 3 6357 0.254652959620028 5.87280361278652 1.45119591046572 0.22833615547619 0.683166880181658
91404 SESTD1 SEC14 and spectrin domain containing 1 11349 -0.16843722344903 6.22449303956904 1.45101145395617 0.228365725499691 0.683166880181658
51149 MRNIP MRN complex interacting protein 8496 0.230214128167318 4.32088125943117 1.45087243594037 0.22838801437187 0.683166880181658
9196 KCNAB3 potassium voltage-gated channel subfamily A regulatory beta subunit 3 4676 0.251186560092162 3.24767365019397 1.45076576452764 0.228405118861334 0.683166880181658
9811 CTIF cap binding complex dependent translation initiation factor 8100 -0.17975295522588 6.52127596479419 1.44988907116188 0.228545752810781 0.683342664189469
6303 SAT1 spermidine/spermine N1-acetyltransferase 1 2498 -0.285757346695917 10.2854265639454 1.44979118980814 0.228561460821565 0.683342664189469
54361 WNT4 Wnt family member 4 4527 -0.628839565233825 4.30002916083443 1.4491554745158 0.228663512107548 0.683501880941193
219899 TBCEL tubulin folding cofactor E like 7020 0.239866912490827 4.80932312871831 1.44607563324918 0.229158695669386 0.684835897213802
4694 NDUFA1 NADH:ubiquinone oxidoreductase subunit A1 766 0.2184695325175 5.3758576801844 1.44481439692333 0.229361852476739 0.685096375853986
254228 CALHM5 calcium homeostasis modulator family member 5 9784

0.499802902975504	3.27719251140843
1.44472218762765	0.229376713825334	0.685096375853986
55959	SULF2	sulfatase 2	6561	-0.408136451541512
7.38251637697928	1.44354757636211	0.229566127123889
0.685096375853986
6727	SRP14	signal recognition particle 14	2005
0.188274761541047	7.40903242910419	1.4427586103935
0.229693458499945	0.685096375853986
10326	SIRPB1	signal regulatory protein beta 1
6559	-0.652793878972211	2.88864866068288
1.44256705282607	0.22972438684663	0.685096375853986
85439	STON2	stonin 2	6675	-0.190608417060814
5.21056302978153	1.44246711176988	0.229740525039964
0.685096375853986
3433	IFIT2	interferon induced protein with tetratricopeptide repeats 2	3489	0.630136684704093	4.09691337982312
1.44240493890643	0.229750565223511
0.685096375853986
25819	NOCT	nocturnin	2504	-0.209598397218125
4.40792111097512	1.44231621090926	0.229764894663024
0.685096375853986
441389	LINC01239	long intergenic non-protein coding RNA 1239
2568	0.425466547182988	1.72881626254242
1.44209782878261	0.229800167637148	0.685096375853986
122809	SOCS4	suppressor of cytokine signaling 4
7043	-0.171721652378936	5.98892962901159
1.4420726024969	0.229804242594688	0.685096375853986
1264	CNN1	calponin 1	2887	0.781149797167062
4.13113341683402	1.44203100319084	0.22981096257732
0.685096375853986
4482	MSRA	methionine sulfoxide reductase A	3566
0.25219141792051	3.62969663566548	1.44189590163296
0.229832788615541	0.685096375853986
10274	STAG1	stromal antigen 1	7846	-0.194454549079961
5.96472887429391	1.44138884559012
0.229914727226941	0.685108777709335
57674	RNF213	ring finger protein 213	28570	-0.389036264019034
8.29723541331805	1.44120689602289
0.229944138265787	0.685108777709335
388963	C2orf81	chromosome 2 open reading frame 81	2653
0.377143095835425	1.84467556359508	1.44096228812117
0.229983684785414	0.685108777709335
253980	KCTD13	potassium channel tetramerization domain containing
13	4861	-0.194302107611689	3.98200573294289
1.44025343355867	0.230098333793365	0.6852291518197
4800	NFYA	nuclear transcription factor Y subunit alpha
3811	-0.13727974031348	6.01132048338649
1.440107570189	0.230121934044397	0.6852291518197
79648	MCPH1	microcephalin 1	9775	-0.169016878492514
4.72475616250302	1.4395179651084	0.230217360085945
0.685349663358304
5338	PLD2	phospholipase D2	4888	-0.198735760631333
5.51350687397653	1.43906176478902
0.230291227636437	0.685349663358304

90639	COX19	cytochrome c oxidase assembly factor COX19
4356	-0.19148601343189	4.49997027994081
1.43895083914809	0.230309192934071	0.685349663358304
11153	FICD	FIC domain containing	4053	-0.280310233384346
	3.18132604661219	1.4373499073129 0.230568664884509
	0.685520305659535
4795	NFKBIL1 NFKB inhibitor like 1	1779	0.205169735634718
	3.79064364569565	1.43688593475478
0.230643929348017	0.685520305659535
83439	TCF7L1	transcription factor 7 like 1	3950
0.254209408648072	4.56517340787807	1.4368858312247
0.230643946145711	0.685520305659535
199990	FAAP20	FA core complex associated protein 20	10916
0.247894535167804	4.31849587503605	1.43683458690181
	0.230652260687794	0.685520305659535
10825	NEU3	neuraminidase 3 7556	0.25013074287407
3.40171788408391	1.4366375168112 0.23068423925625
0.685520305659535
89874	SLC25A21	solute carrier family 25 member 21	5022
	0.515042142117354	0.403830191191827
1.43646610308383	0.230712058912295	0.685520305659535
55632	G2E3	G2/M-phase specific E3 ubiquitin protein ligase
9237	-0.226945743904343	4.21429145738319
1.43630936778373	0.230737499867204	0.685520305659535
9690	UBE3C	ubiquitin protein ligase E3C
9220	-0.165217776377259	7.83677016797151
1.43618267554362	0.230758066765525	0.685520305659535
23345	SYNE1	spectrin repeat containing nuclear envelope protein
1	46064	0.379391624256367	4.4042440137994
1.4358747482472 0.230808064119662	0.68552344235032
89857	KLHL6	kelch like family member 6	10546
0.842002197853375	0.768156565830193	1.43444906871558
	0.231039718287087	0.68594657143768
547	KIF1A	kinesin family member 1A
12357	-0.772495135499742	3.47182128463314
1.43439523936496	0.231048470339329	0.68594657143768
390916	NUDT19	nudix hydrolase 19	2967	-0.175919552646609
	4.8956080665037 1.43374526181232	0.231154181007995
	0.68611498482284
4074	M6PR	mannose-6-phosphate receptor, cation dependent	3777
	0.152845765684234	6.58420922949183
1.43337271691082	0.231214797043547	0.68614950470105
64084	CLSTN2	calsyntenin 2	14636	0.751186455771649
1.87440609132851	1.43227270670754	0.231393889409286
	0.686493489006451
79745	CLIP4	CAP-Gly domain containing linker protein family
member 4	7775	-0.296250078763548	8.01281826753462
	1.43205881538424	0.23142873242582
0.686493489006451
26229	B3GAT3	beta-1,3-glucuronyltransferase 3	2428
0.217486067134851	4.68425313111883	1.43158809702044
	0.23150543500395	0.686575614672945
5067	CNTN3	contactin 3	5109	-0.225517967307288
4.87227619892856	1.43085695514258	0.231624633871879

0.686728956578249
57662 CAMSAP3 calmodulin regulated spectrin associated protein family member 3 4845 -0.215581854941811 5.12329056038065 1.4306695709214 0.231655195247535 0.686728956578249
51027 BOLA1 bolA family member 1 865 0.33493255727224 2.23387320222074 1.42973313627527 0.231807995715447 0.687036520521852
51073 MRPL4 mitochondrial ribosomal protein L4 4070 0.180427059309028 4.84744945937407 1.4289102355676 0.231942370958089 0.687126663623411
57730 ANKRD36B ankyrin repeat domain 36B 6771 -0.405596583960114 2.71522485143241 1.42832044375208 0.232038738610553 0.687126663623411
80331 DNAJC5 DnaJ heat shock protein family (Hsp40) member C5 5287 -0.155621982939395 7.80544407746154 1.42819630178519 0.23205902865198 0.687126663623411
7204 TRIO trio Rho guanine nucleotide exchange factor 17500 -0.238901211119449 9.37689674377626 1.42795186474773 0.232098986248988 0.687126663623411
123283 TARS3 threonyl-tRNA synthetase 3 4584 -0.348731455868027 4.56300333566445 1.42753898168267 0.232166498217469 0.687126663623411
114898 C1QTNF2 C1q and TNF related 2 2414 0.362202037151687 -0.047789027595923 1.42739683750895 0.232189746198065 0.687126663623411
730094 MOSMO modulator of smoothened 6045 -0.215979565384255 3.80321474165845 1.42694895441 0.232263016805582 0.687126663623411
153642 ARSK arylsulfatase family member K 4314 0.320544947438273 3.11335286399129 1.42644724001106 0.232345126992161 0.687126663623411
123263 MTFMT mitochondrial methionyl-tRNA formyltransferase 3432 0.236516367553383 3.65773619301993 1.42614690319451 0.232394296648326 0.687126663623411
4090 SMAD5 SMAD family member 5 8779 -0.200274993060341 6.19599623769024 1.42577367833546 0.232455416680229 0.687126663623411
94241 TP53INP1 tumor protein p53 inducible nuclear protein 1 5676 0.227960311823148 6.8090369110844 1.42576987381629 0.232456039815635 0.687126663623411
100507347 VIM-AS1 VIM antisense RNA 1 1992 0.372201969804874 1.66519251623584 1.42572383174107 0.232463581124872 0.687126663623411
159090 FAM122B family with sequence similarity 122B 4404 -0.172985443058014 4.55535445049598 1.42564717469159 0.232476137567741 0.687126663623411
151313 FAHD2B fumarylacetoacetate hydrolase domain containing 2B 2662 0.327204018441551 2.13499966783521 1.42493885889236 0.232592198613338 0.687308835487116
8925 HERC1 HECT and RLD domain containing E3 ubiquitin protein ligase family member 1 17053 -0.280686508734695 6.30252318101573 1.42460739523201 0.232646534602622 0.687308835487116
1767 DNAH5 dynein axonemal heavy chain 5 16475

0.308789918007898	4.79409296740385	1.42437293217172
	0.232684978787742	0.687308835487116
3787	KCNS1	potassium voltage-gated channel modifier subfamily S member 1	4538	-0.686937795043113	0.412735864892572
	1.42292231702693	0.232923002323621
0.687547721904829
348013	TMEM255B	transmembrane protein 255B	7287
0.52736915832757	0.519744605982091	1.42285632430026
	0.23293383770217	0.687547721904829
27314	RAB30	RAB30, member RAS oncogene family
11832	-0.229212682726263	4.30218605108623
1.42281340350473	0.23294088521564	0.687547721904829
55255	WDR41	WD repeat domain 41	7008	0.231020707012975
	5.58627727421369	1.42268362219913
0.232962196627423	0.687547721904829
100506428	CBR3-AS1	CBR3 antisense RNA 1
7912	-0.381038153034634	2.72013852062519
1.42113120470291	0.233217302059292	0.688155624593322
9879	DDX46	DEAD-box helicase 46	9222	-0.133424471753148
	6.32508413658178	1.42038716383511
0.23333968832769	0.688162142877736
90233	ZNF551	zinc finger protein 551 4930	-0.337657578405693
	3.03519396505053	1.42022989189586
0.23336556770859	0.688162142877736
678	ZFP36L2 ZFP36 ring finger protein like 2
3696	-0.211193753957771	5.99153001222439
1.42015391772532	0.233378070637536	0.688162142877736
3338	DNAJC4	DnaJ heat shock protein family (Hsp40) member C4
	3153	0.188835031308764	4.15150616728279
1.41992328368437	0.23341603061413	0.688162142877736
27236	ARFIP1	ADP ribosylation factor interacting protein 1
4152	-0.148301472543054	5.72545879364449
1.41908556541238	0.233553973155157	0.688423927185243
3090	HIC1	HIC ZBTB transcriptional repressor 1
7693	-0.384687567645937	1.34232831496482
1.41853400973796	0.233644848633416	0.688546895604516
51465	UBE2J1	ubiquitin conjugating enzyme E2 J1
4290	-0.240289349560582	7.43520334121003
1.4180864669448 0.233718618137401	0.688619411916402
51144	HSD17B12	hydroxysteroid 17-beta dehydrogenase 12
8075	-0.255980908399417	7.24339516363939
1.41771909307566	0.233779194253801	0.688653033420189
283987	HID1	HID1 domain containing	6196	-0.176957509757857
	5.65473606098866	1.41721505935639
0.23386233503611	0.688753096478299
100505881	MAGI2-AS3	MAGI2 antisense RNA 3	11625
0.362800764504538	3.79555084498844	1.41674881686222
	0.233939273949322	0.688783090322805
8086	AAAS	aladin WD repeat nucleoporin	4138
0.189639313117666	4.23597510400212	1.41655741547526
	0.233970867691576	0.688783090322805
10087	CERT1	ceramide transporter 1	6253	-0.155532563400014
	7.15882183241264	1.41592714982113
0.234074939195774	0.688944667643456

80762	NDFIP1	Nedd4 family interacting protein 1
6024	-0.159357808343049	7.07419079779925
1.41516638530828	0.234200633655581	0.689169805801203
10157	AASS	aminoadipate-semialdehyde synthase	4591
0.28845781384733	4.54450256585128	1.41483681270789
	0.23425511138621	0.689185327701457
342979	PALM3	paralemmin 3	2533	0.919081705558384
1.95700378642484	1.41416717631493	0.23436584822834
	0.689366324261729
4695	NDUFA2	NADH:ubiquinone oxidoreductase subunit A2	1488
	0.224560122199202	4.9564134953577 1.41375667807044
	0.234433763053011	0.689421314105295
858	CAV2	caveolin 2	6555	0.211297242788354
8.00650060130729	1.41323495675647	0.234520113515769
	0.689530484889876
4335	MNT	MAX network transcriptional repressor
6328	-0.23424397255019	4.9500635990345 1.41293036992383
	0.234570543682791	0.689534020013611
8933	RTL8C	retrotransposon Gag like 8C	1181
0.203755498746796	6.09824779574589	1.41199826641991
	0.234724952732241	0.689750193676894
27247	NFU1	NFU1 iron-sulfur cluster scaffold	2197
0.194154184514782	4.30539512686553	1.41189196745027
	0.234742569662151	0.689750193676894
10383	TUBB4B	tubulin beta 4B class IVb
2111	-0.207776488311289	7.92780419296512
1.41112883662711	0.234869090319748	0.689904294197285
11142	PKIG	cAMP-dependent protein kinase inhibitor gamma	2496
	0.230301414199497	4.52017197431911
1.41079478438045	0.234924499327232	0.689904294197285
4830	NME1	NME/NM23 nucleoside diphosphate kinase 1	3438
	0.301148815667375	5.51582370883088
1.40997774345462	0.235060087987017	0.689904294197285
5192	PEX10	peroxisomal biogenesis factor 10	3789
0.209045565713426	4.29677582185163	1.40970165728368
	0.235105926133869	0.689904294197285
10062	NR1H3	nuclear receptor subfamily 1 group H member 3	4533
	0.374424900754042	1.38734367707933
1.40966102612696	0.235112672973114	0.689904294197285
135932	TMEM139 transmembrane protein 139	2564
0.807194902947983	5.12198904662661	1.40946925019328
	0.235144520698263	0.689904294197285
84340	GFM2	G elongation factor mitochondrial 2
4197	-0.153945559301866	5.72276367828648
1.40879449486595	0.235256617050079	0.689904294197285
113802	HENMT1	HEN methyltransferase 1 2426	0.371573634851876
	1.95599879409012	1.40869096406686
0.235273822228921	0.689904294197285
3135	HLA-G	major histocompatibility complex, class I, G	1763
	0.34670334521383	6.04591014831548
1.40863351015648	0.235283370815805	0.689904294197285
954	ENTPD2	ectonucleoside triphosphate diphosphohydrolase 2
	2633	-0.41079119971735	0.489340430897647
1.40855289110578	0.235296770137839	0.689904294197285

102723553 SMIM11B small integral membrane protein 11B 8524
0.281925199902477 3.02622824410106
1.40831200972635 0.235336811428188 0.689904294197285
4524 MTHFR methylenetetrahydrofolate reductase
10055 -0.178173642811214 5.42953292365487
1.40747709114141 0.235475662336622 0.690166897331882
6289 SAA2 serum amyloid A2 2227 -0.878972179727541
7.05618869301443 1.40658843405285
0.23562355917906 0.690455898205249
64061 TSPYL2 TSPY like 2 4948 0.294748998454087
3.33682874095984 1.40607876846947 0.235708432215518
0.690560135963765
390980 ZNF805 zinc finger protein 805 2523 -0.323605168288666
2.36660384160334 1.4047686293038 0.235926775586646
0.690756429347267
139285 AMER1 APC membrane recruitment protein 1 8443
0.298632513890199 2.26980541040038 1.40465807553802
0.23594521130745 0.690756429347267
54878 DPP8 dipeptidyl peptidase 8 10211 -0.15311592231827
6.12838102504988 1.40458314881543
0.235957706926976 0.690756429347267
6733 SRPK2 SRSF protein kinase 2 5856 0.131859032715686
6.09625126639249 1.40418955982507
0.236023359399661 0.690756429347267
84848 MIR503HG MIR503 host gene 1678
0.353424920153709 2.42936364381132 1.40398155820463
0.236058063971713 0.690756429347267
23499 MACF1 microtubule actin crosslinking factor 1
35647 -0.296590379973875 9.38391333023222
1.40390209750774 0.236071323430097 0.690756429347267
1486 CTBS chitobiase 6939 -0.211200943652395
3.76226071732158 1.40347980568486 0.236141805617438
0.690818352398381
9572 NR1D1 nuclear receptor subfamily 1 group D member 1
2772 -0.300444633791283 4.19991352835776
1.40242182714663 0.236318498404701 0.690826068540342
11073 TOPBP1 DNA topoisomerase II binding protein 1
6515 -0.173703975736424 5.23730285607397
1.4024098144578 0.236320505560875 0.690826068540342
55861 DBNDD2 dysbindin domain containing 2
2764 -0.164113956367888 5.53087871367359
1.40239110144298 0.236323632291122 0.690826068540342
1737 DLAT dihydrolipoamide S-acetyltransferase
4831 -0.264267117337153 6.0371049483002 1.40228283440491
0.236341723455795 0.690826068540342
55323 LARP6 La ribonucleoprotein 6, translational regulator
5554 -0.326025756421793 4.85372806754862
1.40188040762898 0.236408982723443 0.690852564330919
5708 PSMD2 proteasome 26S subunit, non-ATPase 2
5453 -0.187629391210113 9.19997851264351
1.40139772366501 0.236489686301145 0.690852564330919
7643 ZNF90 zinc finger protein 90 4619 0.267593914399135
4.4496152186284 1.40096744710739 0.236561655630637
0.690852564330919

126003	TRAPPC5 trafficking protein particle complex 5	5833
0.247145145946804	3.88363337197217	1.40067303003688
	0.236610915992836	0.690852564330919
3691	ITGB4	integrin subunit beta 4	6553	0.233258515177593
	10.2901272237833	1.40055091400927
0.23663135147043	0.690852564330919
7941	PLA2G7	phospholipase A2 group VII	2026
1.05546502245185	1.76978648249972	1.40045908634028
	0.236646719758674	0.690852564330919
324	APC	APC regulator of WNT signaling pathway
12440	-0.204706552905037	5.94774273806238
1.39981501626563	0.236754545393189	0.690941150384413
58485	TRAPPC1 trafficking protein particle complex 1	1171
0.184991712742836	5.80350763643699	1.39968857165286
	0.236775720842806	0.690941150384413
7414	VCL	vinculin	9992	-0.283283390584048
9.04108720221829	1.39914803353065	0.236866269664135
	0.691061412036147
6814	STXBP3	syntaxin binding protein 3
5895	-0.152891529129491	5.7620152815453 1.39782956382622
	0.237087310631143	0.691491759336557
1810	DR1	down-regulator of transcription 1
10949	-0.183173891242681	6.11280279132296
1.39755817992652	0.237132839068809	0.691491759336557
23232	TBC1D12 TBC1 domain family member 12
6921	-0.216483302992738	3.82208458032049
1.39721448577099	0.237190513773917	0.691491759336557
2328	FMO3	flavin containing dimethylaniline monoxygenase 3
	3463	-0.821354594216661	0.442816188161291
1.39683711366015	0.237253859505798	0.691491759336557
128866	CHMP4B	charged multivesicular body protein 4B
1646	-0.156540816215396	7.53102673265875
1.39647594955952	0.237314503768829	0.691491759336557
49854	ZBTB21	zinc finger and BTB domain containing 21
8062	-0.238746611507076	4.83800295553738
1.39537992307413	0.237498656318851	0.691491759336557
84740	AFAP1-AS1	AFAP1 antisense RNA 1	6795
0.586980156658216	0.22501086300958	1.39490031530488
	0.237579293722916	0.691491759336557
4037	LRP3	LDL receptor related protein 3	5095
0.34978366729619	4.0752212331927 1.39463073393149
0.237624633550173	0.691491759336557
360	AQP3	aquaporin 3 (Gill blood group)
2928	-0.335626350336526	7.9110915524083 1.39440382665944
	0.237662804322856	0.691491759336557
257364	SNX33	sorting nexin 33	8008	0.196451334234975
	7.72057453005993	1.39434667622157
0.237672419450552	0.691491759336557
10975	UQCR11	ubiquinol-cytochrome c reductase, complex III
subunit XI	2151	0.179751883368882	6.04817154541977
	1.39424356396002	0.237689768500056
0.691491759336557
5029	P2RY2	purinergic receptor P2Y2
2861	-0.306355074012463	4.59029062301281

1.39421406451626	0.237694732181772	0.691491759336557
84926	SPRYD3	SPRY domain containing 3
3479	-0.241848263541223	5.48499173749726
1.39368835279076	0.237783211402522	0.691491759336557
79089	TMUB2	transmembrane and ubiquitin like domain containing 2
	3511	0.149975573289397	6.15945262712549
1.39336654134406	0.237837393167928	0.691491759336557
84068	SLC10A7	solute carrier family 10 member 7
6687	-0.291084581997563	2.48120923344921
1.39315545046424	0.237872941605375	0.691491759336557
23172	ABRAXAS2	abraxas 2, BRISC complex subunit	2992
	-0.161462822988783	5.25824189320801
1.392680043784	0.237953025349313	0.691491759336557
6494	SIPA1	signal-induced proliferation-associated 1	4692
	0.242837196488001	4.09371826414529
1.39232138787583	0.23801346370713	0.691491759336557
63941	NECAB3	N-terminal EF-hand calcium binding protein 3	4342
	0.332824838529712	2.84889687297074
1.39202163059482	0.23806399114017	0.691491759336557
89970	RSPRY1	ring finger and SPRY domain containing 1
6064	-0.1275262805009	5.90827371535564
1.39151738752699	0.238149016296478	0.691491759336557
8502	PKP4	plakophilin 4	9525	-0.152686192780103
6.62142835795817	1.39149622693514	0.238152585187976
	0.691491759336557
10625	IVNS1ABP	influenza virus NS1A binding protein	7929
	0.21814323801938	7.55324196397529
1.39131539598276	0.238183086322699	0.691491759336557
100037417	DDTL	D-dopachrome tautomerase like	1545
0.334245687693567	2.07392399259002	1.3907884036692
0.238272002258328	0.691491759336557
54929	TMEM161A	transmembrane protein 161A	3281
0.245090798960892	3.51216552455921	1.39075143623397
	0.238278241041485	0.691491759336557
375033	PEAR1	platelet endothelial aggregation receptor 1
5530	-0.517200837555236	1.59004766560046
1.39033601313434	0.23834836326081	0.691491759336557
4697	NDUFA4	NDUFA4 mitochondrial complex associated 3205
0.210618982221471	6.04866800755975	1.38978679919818
	0.238441107432777	0.691491759336557
1565	CYP2D6	cytochrome P450 family 2 subfamily D member 6
2443	-0.41733055098074	0.590331843288032
1.38965664592447	0.238463092459693	0.691491759336557
79786	KLHL36	kelch like family member 36
9995	-0.153736061399259	6.51923177229354
1.38948593718559	0.238491931701055	0.691491759336557
593	BCKDHA	branched chain keto acid dehydrogenase E1 subunit
alpha	3579	0.215501571372784	4.28478085699294
1.38919633218656	0.238540866755999	0.691491759336557
27316	RBMX	RNA binding motif protein X-linked	5882
0.140078958019432	6.72021133809109	1.38909713180189
	0.238557631617938	0.691491759336557
56916	SMARCAD1	SWI/SNF-related, matrix-associated actin-
dependent regulator of chromatin, subfamily a, containing DEAD/H box

1	6313	-0.142971506369406	5.99826686469642
1.38862440997117	0.238637541242212	0.691491759336557
2677	GGCX	gamma-glutamyl carboxylase	9098
0.142431924941239	5.94567755334181	1.38828039454669
0.238695714565794	0.691491759336557
54708	MARCHF5 membrane associated ring-CH-type finger 5
4632	-0.142780876286388	6.46584498159509
1.3882141833789 0.238706912910405	0.691491759336557
9087	TMSB4Y	thymosin beta 4 Y-linked	1669
0.338694541098256	0.0859984895817925	1.38818115641402
0.238712499024232	0.691491759336557
124989	EFCAB13 EF-hand calcium binding domain 13
6672	-0.41637672476071	1.47466614150886
1.38793543433053	0.238754064941984	0.691491759336557
55671	PPP4R3A protein phosphatase 4 regulatory subunit 3A
5912	-0.14429567759099	6.77083848599675
1.38778035641823	0.238780302172987	0.691491759336557
472	ATM	ATM serine/threonine kinase	22317
0.279765690641814	5.27304745624686	1.38759882668014
0.238811019167204	0.691491759336557
6341	SCO1	SCO cytochrome c oxidase assembly protein 1
10230	0.147177454763467	5.34392758288273
1.38599135461398	0.239083232048111	0.691491759336557
115106	HAUS1	HAUS augmin like complex subunit 1	2165
0.199468426551319	3.93853089403477	1.38562160287464
0.239145899919994	0.691491759336557
440026	TMEM41B transmembrane protein 41B
4422	-0.16380671025872	6.0291183596634 1.38438441709653
0.239355731123521	0.691491759336557
7428	VHL	von Hippel-Lindau tumor suppressor
4213	-0.149628669604286	5.46620594320138
1.38427499380129	0.239374300473865	0.691491759336557
768	CA9	carbonic anhydrase 9	1670	1.07446124233083
0.294435408975165	1.38416114503717
0.23939362269263	0.691491759336557
6899	TBX1	T-box transcription factor 1
3496	-0.391931156009076	2.98562366525405
1.38411136110438	0.239402072532467	0.691491759336557
5993	RFX5	regulatory factor X5	4586	0.201559657848055
6.22760341527726	1.38403537230862
0.23941497082906	0.691491759336557
116224	FAM122A family with sequence similarity 122A
4554	-0.174888139376942	4.15490629433183
1.38367760833811	0.239475708834157	0.691491759336557
55250	ELP2	elongator acetyltransferase complex subunit 2
11068	0.154046522175516	5.7060374594401 1.38363824785923
0.239482392251328	0.691491759336557
267002	PGBD2	piggyBac transposable element derived 2
3651	-0.274164491058435	2.83092434423777
1.38362421426331	0.239484775213514	0.691491759336557
83985	SPNS1	sphingolipid transporter 1 (putative)	6987
0.185669222114374	6.15701955467801	1.38362024153952
0.23948544980355	0.691491759336557
10166	SLC25A15	solute carrier family 25 member 15	1837

0.34442658037332 1.9621612550672 1.38348672530481 0.239508122926588 0.691491759336557
53615 MBD3 methyl-CpG binding domain protein 3 8080 0.16605991657374 6.81685512826878 1.38346232010094 0.239512267589808 0.691491759336557
10959 TMED2 transmembrane p24 trafficking protein 2 2556 -0.204302096724289 7.71521168554491 1.38317457356089 0.239561141300005 0.691491759336557
784 CACNB3 calcium voltage-gated channel auxiliary subunit beta 3 5737 0.139442891051398 6.49849503187156 1.38300972592839 0.239589146106628 0.691491759336557
4542 MYO1F myosin IF 7685 0.974503220803674 2.02038673863268 1.38283956842991 0.239618057140905 0.691491759336557
56605 ERO1B endoplasmic reticulum oxidoreductase 1 beta 5361 -0.285065504121688 2.95472262685762 1.38276777715755 0.239630256287545 0.691491759336557
23315 SLC9A8 solute carrier family 9 member A8 6946 -0.234814298416616 5.37364949196368 1.38229062154325 0.239711356230874 0.691583309315313
54926 UBE2R2 ubiquitin conjugating enzyme E2 R2 4075 -0.13696992865815 6.55227831594231 1.38173225240798 0.239806302015188 0.691714759540102
6792 CDKL5 cyclin dependent kinase like 5 13702 -0.272009758009358 5.05921483929913 1.3791239248187 0.240250431166757 0.692560023231557
152503 SH3D19 SH3 domain containing 19 7407 -0.164111444759223 6.89077520423877 1.37897072429485 0.24027654823416 0.692560023231557
55677 IWS1 interacts with SUPT6H, CTD assembly factor 1 3990 -0.136723541744905 6.62830456174314 1.37872998814583 0.240317595029643 0.692560023231557
23279 NUP160 nucleoporin 160 8576 -0.165650688385413 5.54331535050109 1.37871304854738 0.240320483642411 0.692560023231557
411 ARSB arylsulfatase B 7189 0.281871909901633 4.25841585828097 1.37837246123583 0.240378571004793 0.692560023231557
54453 RIN2 Ras and Rab interactor 2 5053 0.148986255861322 5.7642918934907 1.37827026271115 0.240396004351526 0.692560023231557
65980 BRD9 bromodomain containing 9 9572 -0.160195570878763 5.31772853339781 1.37711180059983 0.240593725990849 0.692689008974736
23376 UFL1 UFM1 specific ligase 1 4539 -0.198234358750397 5.31981145382322 1.37708001988235 0.240599152981728 0.692689008974736
80351 TNKS2 tankyrase 2 6157 -0.200379153041874 7.37998407355388 1.37685092728944 0.240638278070469 0.692689008974736
260425 MAGI3 membrane associated guanylate kinase, WW and PDZ domain containing 3 8180 -0.182466485121593 4.6084817523286 1.37679476226497 0.240647871275016 0.692689008974736

22894	DIS3	DIS3 homolog, exosome endoribonuclease and 3'-5' exoribonuclease	10672	-0.147766643436739	6.1268244719829	1.37655960459198	0.240688042170346	0.692689008974736
767557	SYS1-DBNDD2	SYS1-DBNDD2 readthrough (NMD candidate)	1402	-0.162274773454641	5.59855903229431	1.37555025295521	0.240860557685652	0.692734420411395
84000	TMPRSS13	transmembrane serine protease 13	5367	-0.379553049251533	4.4094363834338	1.37540605906021	0.240885215172152	0.692734420411395
90379	DCAF15	DDB1 and CUL4 associated factor 15	2961	-0.180115193190053	4.48028609771565	1.37534553630918	0.240895565616096	0.692734420411395
201514	ZNF584	zinc finger protein 584	2680	0.275375722560051	2.85098247431877	1.37530998097203	0.240901646449911	0.692734420411395
4689	NCF4	neutrophil cytosolic factor 4	1759	0.727250669615763	1.5816222120361	1.37397071713113	0.241130829417818	0.693251134576228
55922	NKRF	NFKB repressing factor	4004	-0.171722700033429	4.23054476325131	1.37351292951334	0.241209229641983	0.693334225240151
56132	PCDHB3	protocadherin beta 3	3355	0.993522252600533	0.549362974151045	1.37248904832936	0.241384690741053	0.693599588705324
64745	METTL17	methyltransferase like 17	4449	0.184210421256234	5.23844164499725	1.37229654518863	0.241417697080497	0.693599588705324
79441	HAUS3	HAUS augmin like complex subunit 3	7067	-0.221297742208332	3.95967620851622	1.37200562722892	0.241467587919914	0.693599588705324
132989	C4orf36	chromosome 4 open reading frame 36	2538	-0.310371587140996	3.33823703044763	1.37165715399263	0.241527365692303	0.693599588705324
386724	AMIGO3	adhesion molecule with Ig like domain 3	2862	0.329980061530447	2.28468568184422	1.37153025148991	0.241549139266408	0.693599588705324
54925	ZSCAN32	zinc finger and SCAN domain containing 32	4013	0.170708274798013	4.29411638806234	1.37082038238938	0.241670980676689	0.69380721999147
6650	CAPN15	calpain 15	7686	-0.193157733671561	5.12703062013875	1.37031895655119	0.24175709012345	0.69391220519655
51133	KCTD3	potassium channel tetramerization domain containing 3	4929	0.156891198729954	5.45909285681859	1.36984662248197	0.24183823782794	0.693995908587859
5226	PGD	phosphogluconate dehydrogenase	2966	0.211952884443796	7.29085904636346	1.36949563341012	0.241898559754817	0.693995908587859
391059	FRRS1	ferric chelate reductase 1	3868	0.345748104413459	2.72207807299233	1.36928428648347	0.241934891242558	0.693995908587859
389541	LAMTOR4	late endosomal/lysosomal adaptor, MAPK and MTOR activator 4	1936	0.218685815469784	4.78060402719471	1.36887364273005	0.242005501748137

0.69404505292918
7084	TK2	thymidine kinase 2	6860	0.222682793084953
	5.46562092613785	1.36840136034131
0.242086742090678	0.69404505292918
554251	FBXO48	F-box protein 48	5666	-0.343354888181226
	2.10005186856783	1.36805388149238
0.242146535375504	0.69404505292918
57547	ZNF624	zinc finger protein 624 4812	-0.284718168892386
	2.9479220192844 1.36803245484614	0.242150223007418
	0.69404505292918
57406	ABHD6	abhydrolase domain containing 6 3241
0.25577266856631	3.98450849869952	1.36747861022138
	0.242245566144248	0.694176277615074
54910	SEMA4C	semaphorin 4C	6157	-0.171416886728624
5.53172927370178	1.36708847786421	0.242312754018037
	0.694226783704366
100505806	SNHG18	small nucleolar RNA host gene 18	1951
	0.378678952516222	2.14679328560387
1.36522779533898	0.242633509769123	0.695003593320267
1398	CRK	CRK proto-oncogene, adaptor protein
4090	-0.160692004860034	7.73247018999998
1.36493329876911	0.242684324273589	0.695007019034996
55339	WDR33	WD repeat domain 33	14580	-0.138982351506451
	6.1848788552697 1.36464509441105	0.242734065628912
	0.695007370659069
6668	SP2	Sp2 transcription factor
3184	-0.177533528013426	4.69569255852011
1.36419468179609	0.242811827478225	0.695087935313201
85440	DOCK7	dedicator of cytokinesis 7
12723	-0.160536551921286	6.74388723899149
1.36214002212638	0.243166941174377	0.69586138941821
23479	ISCU	iron-sulfur cluster assembly enzyme	4439
0.214089833747088	5.54298871045563	1.36205659410797
	0.243181373684739	0.69586138941821
29127	RACGAP1 Rac GTPase activating protein 1 4487
0.219650306754986	4.18930045940124	1.36079990219401
	0.243398899672345	0.696185877862687
85395	FAM207A family with sequence similarity 207 member A	1402
	0.345204274272704	2.21128437578802
1.36066031524687	0.243423075986434	0.696185877862687
3727	JUND	JunD proto-oncogene, AP-1 transcription factor
subunit 1863	0.191376868645027	5.85017689779878
1.36028389934266	0.243488285422987	0.696185877862687
1848	DUSP6	dual specificity phosphatase 6
4632	-0.321645055167089	6.96241460129839
1.36025332551957	0.243493582897787	0.696185877862687
55805	LRP2BP	LRP2 binding protein	7360
0.32246484404277	-0.0606917965096499	1.35894704054126
	0.243720052018748	0.696556999464569
55194	EVA1B	eva-1 homolog B 1211	0.272998671945965
3.01765712580448	1.35893095435541	0.243722842462758
	0.696556999464569
5264	PHYH	phytanoyl-CoA 2-hydroxylase	3381
0.259777288538676	4.287318216874	1.35860334221898

0.243779681274915	0.696577314487502
50937	CDON	cell adhesion associated, oncogene regulated
10600	-0.338940719360677	3.68900214953716
1.35806156062732	0.243873712725035	0.696703873983187
149076	ZNF362	zinc finger protein 362 3331	0.293804145099038
	4.18196722566745	1.35731272396261
0.244003753427521	0.696804522297506
2802	GOLGA3	golgin A3	10550	-0.183903664238062
7.72280610151654	1.35708031932183	0.244044129308712
	0.696804522297506
79158	GNPTAB	N-acetylglucosamine-1-phosphate transferase subunits alpha and beta	9620	0.16801253275643	7.38884341936254
	1.35699942987593	0.244058184221572
0.696804522297506
2571	GAD1	glutamate decarboxylase 1	8242
0.472577860179223	3.20029527040663	1.35614140967099
	0.244207329977301	0.69695475210744
1774	DNASE1L1	deoxyribonuclease 1 like 1	4130
0.219924922899226	4.90993131270001	1.35612422677822
	0.244210317936983	0.69695475210744
112849	L3HYPDH trans-L-3-hydroxyproline dehydratase	3504
0.286547714329197	2.46756841267457	1.35540846708098
	0.244334822127663	0.696970941814064
65009	NDRG4	NDRG family member 4	8936	-0.316124904063601
	5.52551292173321	1.35516325197914
0.244377494348064	0.696970941814064
6192	RPS4Y1	ribosomal protein S4 Y-linked 1 2275
0.258161343483727	6.07347259118862	1.35493733902115
	0.244416815666224	0.696970941814064
57542	KLHL42	kelch like family member 42
7622	-0.185656272560182	5.40820650374509
1.35481976166457	0.244437283667498	0.696970941814064
339451	KLHL17	kelch like family member 17
3395	-0.251326957268647	3.98278655387084
1.35466180322935	0.244464784545763	0.696970941814064
4809	SNU13	small nuclear ribonucleoprotein 13	2780
0.168980001344782	5.74513393418656	1.3536702876177
0.244637495649739	0.697258035704301
6491	STIL	STIL centriolar assembly protein	7291
0.28681102330433	3.51590639051224	1.3535122251187
0.244665042156539	0.697258035704301
7784	ZP3	zona pellucida glycoprotein 3
2716	-0.312142988601604	2.55774436155023
1.3523750091615 0.24486334329715	0.697619381338259
27352	SGSM3	small G protein signaling modulator 3
4932	-0.150961391542246	6.25823397359443
1.35221393123523	0.244891446993662	0.697619381338259
9188	DDX21	DExD-box helicase 21	5020	-0.239012250339824
	7.32728108865879	1.35130404685381
0.24505027091828	0.697929878965503
5037	PEBP1	phosphatidylethanolamine binding protein 1	1780
	0.180394822701382	7.11787070188509
1.35089980069445	0.245120874032903	0.697989038946712
5394	EXOSC10 exosome component 10	6003	-0.150911694825068

6.31064738527382	1.35058791444206
0.245175363115633	0.698002299016397
6300	MAPK12	mitogen-activated protein kinase 12	4838
0.400721916339502	1.84789581034771	1.34998650618392
0.245280475827426	0.698088631289294
400720	ZNF772	zinc finger protein 772 6572	-0.422199700155931
3.22442774393503	1.34984414924917
0.245305364689506	0.698088631289294
349565	NMNAT3	nicotinamide nucleotide adenylyltransferase 3	3451
0.491226573535595	0.10345259482129
1.34947639784277	0.245369674491671	0.698129805119813
9265	CYTH3	cytohesin 3	5808	0.164413432545792
6.61869593386767	1.34891415642335	0.245468035159754
0.698249594051443
51126	NAA20	N(alpha)-acetyltransferase 20, NatB catalytic subunit 2376	-0.178843080508901	6.11410962977084
1.34866592834268	0.245511476454869	0.698249594051443
3320	HSP90AA1	heat shock protein 90 alpha family class A member 1	5248	-0.176842495331032	10.1398724102309
1.34829211716396	0.245576913210146
0.698293914603026
116138	KLHDC3	kelch domain containing 3	2019
0.148835257370053	5.66895708732558	1.34754288192906
0.245708133214415	0.698525232785532
79831	KDM8	lysine demethylase 8	4039	0.334322311249057
2.15559759282242	1.34612121121982
0.245957358174843	0.698955279950642
23426	GRIP1	glutamate receptor interacting protein 1
6378	-0.233924913565296	4.22838644134587
1.34593347896481	0.245990291523936	0.698955279950642
80173	IFT74	intraflagellar transport 74	3755
0.244051846029619	3.11462943371319	1.34582625044054
0.246009104744532	0.698955279950642
54206	ERRFI1	ERBB receptor feedback inhibitor 1
4186	-0.471502887244946	8.15913925223239
1.3447327195322 0.246201064930862	0.699228758977611
9820	CUL7	cullin 7	5628	0.163651086114118
5.41650108563966	1.34470917376673	0.246205200205438
0.699228758977611
92745	SLC38A5 solute carrier family 38 member 5	4014
0.287652367386765	7.40391310529286	1.34338148772156
0.246438515278042	0.699632463320428
79101	TAF1D	TATA-box binding protein associated factor, RNA polymerase I subunit D	4606	-0.220624537695392
4.86674603409793	1.34300892720058	0.246504034118163
0.699632463320428
9757	KMT2B	lysine methyltransferase 2B
10279	-0.223434058192059	6.09552744846133
1.34296613218712	0.246511561452503	0.699632463320428
57619	SHROOM3 shroom family member 3	12608	0.210195873076443
7.56839242537191	1.34276385988086
0.246547143496083	0.699632463320428
2639	GCDH	glutaryl-CoA dehydrogenase	3529
0.219319227956292	3.36663139283843	1.34215637972894

0.246654044050602	0.699794043957218
4148	MATN3	matrilin 3	3118	0.465173079094644
3.61508775159831	1.34040050492768	0.246963350720165
0.700083808828176
91869	RFT1	RFT1 homolog	6409	0.236215739230367
3.67904513577878	1.34036890876079	0.246968920895835
0.700083808828176
643650	LINC00842	long intergenic non-protein coding RNA 842
4440	0.211975994907465	4.22094698620041
1.33988767773255	0.247053777440246	0.700083808828176
11196	SEC23IP SEC23 interacting protein
7867	-0.135182083430639	6.05292936238255
1.33976581522714	0.247075271386859	0.700083808828176
388695	LYSMD1	LysM domain containing 1
2508	-0.199918263465156	3.76010176348193
1.33959675273158	0.247105094192771	0.700083808828176
5286	PIK3C2A phosphatidylinositol-4-phosphate 3-kinase catalytic subunit type 2 alpha	8909	-0.201365944861152
6.89099641626416	1.33950937564396	0.247120509330806
0.700083808828176
8939	FUBP3	far upstream element binding protein 3
5507	-0.128990986298927	6.20284406136661
1.33932744051141	0.24715261025193	0.700083808828176
10518	CIB2	calcium and integrin binding family member 2
2146	-0.319010291542765	2.69760109749139
1.33883706617694	0.247239158076904	0.700083808828176
54541	DDIT4	DNA damage inducible transcript 4
2058	-0.198368669718875	6.03676464774483
1.33875724755524	0.247253249043671	0.700083808828176
284759	SIRPB2	signal regulatory protein beta 2	5034
0.452306390330386	1.89135685753101	1.33845925188281
0.247305865087176	0.700083808828176
56125	PCDHB11 protocadherin beta 11	4198	-0.67083656545438
0.384266082296969	1.33841505209089
0.24731367045483	0.700083808828176
23064	SETX	senataxin	12140	-0.224924537090766
7.27319059008952	1.33750515952039	0.247474417971648
0.700083808828176
84679	SLC9A7	solute carrier family 9 member A7
10033	-0.223396865826245	6.33011033911729
1.33663043152183	0.247629073628909	0.700083808828176
127602	DNAH14	dynein axonemal heavy chain 14	19693
0.407726421264922	2.04188098660028	1.33650690284283
0.24765092355775	0.700083808828176
100820829	MYZAP	myocardial zonula adherens protein
2561	-0.499351015235231	4.396879420705	1.33646770922229
0.247657856674272	0.700083808828176
2296	FOXC1	forkhead box C1 3926	-0.27352318169258
5.43173605100629	1.33634084556728	0.24768029972316
0.700083808828176
7272	TTK	TTK protein kinase	4579	0.360715184967838
-0.118426034086916	1.33626627154261
0.247693493540119	0.700083808828176
85028	SNHG12	small nucleolar RNA host gene 12

3377	-0.234056427107469	4.48250606336139
1.33607625231165	0.247727116090748	0.700083808828176
583	BBS2	Bardet-Biedl syndrome 2 8337	0.229845707399071
	5.07505915107974	1.33563211057088
0.247805725599638	0.700083808828176
20	ABCA2	ATP binding cassette subfamily A member 2
11514	0.22803026239672	5.47699442588039
1.33535842704775	0.247854180580826	0.700083808828176
9527	GOSR1	golgi SNAP receptor complex member 1
8115	-0.137197805306516	5.92793268208507
1.33528661894798	0.247866895945244	0.700083808828176
10460	TACC3	transforming acidic coiled-coil containing protein 3
	4698	0.345236973794797	3.31423766463489
1.3349908311705 0.247919280759083	0.700083808828176
285676	ZNF454	zinc finger protein 454 2662	-0.477953384176539
	0.497846374421405	1.33449039421476
0.248007940358574	0.700083808828176
4140	MARK3	microtubule affinity regulating kinase 3
8891	-0.14211104639453	6.75560440565758
1.33432762879256	0.248036784957003	0.700083808828176
56271	BEX4	brain expressed X-linked 4	1328
0.199550474888043	4.16308209262581	1.33427495775931
	0.248046119973848	0.700083808828176
35	ACADS	acyl-CoA dehydrogenase short chain	2648
0.270949210245612	3.02299484665973	1.33422100619436
	0.248055682388394	0.700083808828176
1829	DSG2	desmoglein 2	6190	0.206134222714755
10.1991235550656	1.33388715174306	0.248114865031778
	0.700109773267549
1627	DBN1	drebrin 1	5119	0.152467137441877
8.47828362310885	1.33349815960502	0.24818384378372
	0.700163363520357
23304	UBR2	ubiquitin protein ligase E3 component n-recognin 2
	8840	-0.17166910764842	6.30105402468152
1.33313736446485	0.248247843440018	0.700198398410649
961	CD47	CD47 molecule	7895	-0.194398019318412
7.1960341034652 1.33286460216836	0.248296240801434
0.700198398410649
27086	FOXP1	forkhead box P1 13628	-0.226830587125549
5.27130793347766	1.3324240307656 0.248374437642875
0.700277928354217
7923	HSD17B8 hydroxysteroid 17-beta dehydrogenase 8	1171
0.316006687584021	1.64901272167314	1.33165117074863
	0.248511685124833	0.700523882781956
9101	USP8	ubiquitin specific peptidase 8
20639	-0.143552929130883	6.36747037418113
1.33108347678409	0.248612557596186	0.700667222183053
23598	PATZ1	POZ/BTB and AT hook containing zinc finger 1	5693
	0.189046813335982	4.69333584046456
1.33009254638715	0.248788754355522	0.700823196856762
170622	COMMD6	COMM domain containing 6	2897
0.253162927073497	4.83731214090095	1.33005380321038
	0.248795646364055	0.700823196856762
112937	GLB1L3	galactosidase beta 1 like 3	8945

0.599647971943162	2.38511517819631	1.32992813927288
	0.248818002282336	0.700823196856762
4853	NOTCH2	notch receptor 2	16804	-0.264712430706632
	8.15472562399222	1.32894593344082
0.248992823894741	0.701174604803707
84058	WDR54	WD repeat domain 54	2860	0.242311134361458
	3.23494827322711	1.32851842348843
0.249068962873254	0.701227406372897
3638	INSIG1	insulin induced gene 1	3650	-0.312755200362098
	6.14734999094958	1.3282785336496 0.249111699443959
	0.701227406372897
412	STS	steroid sulfatase	6521	-0.439910846245032
	6.16236048362344	1.32778869842859
0.249198991899413	0.701332183953199
5696	PSMB8	proteasome 20S subunit beta 8
2006	-0.283869773746241	5.41977368418854
1.32731122724846	0.249284117067802	0.701430821166875
24142	NAA80	N(alpha)-acetyltransferase 80, NatH catalytic
subunit 2188	0.367416339162727	2.18348611527871
1.3267345598191 0.249386974795858	0.701579304271054
9582	APOBEC3B	apolipoprotein B mRNA editing enzyme
catalytic subunit 3B	1896	-0.42531782061795
3.52128480963228	1.32640480698536	0.249445814790374
	0.701603921238709
11212	PLPBP	pyridoxal phosphate binding protein
3428	-0.165841232255869	4.89271320750928
1.32611629745568	0.249497309425396	0.701607872539955
2224	FDPS	farnesyl diphosphate synthase
3718	-0.353940656321026	7.79027285716913
1.32522386141432	0.249656678428055	0.701800914879208
11230	PRAF2	PRA1 domain family member 2	2411
0.20009601034655	5.38651431825278	1.32517076616387
	0.249666163978232	0.701800914879208
201164	PLD6	phospholipase D family member 6 2560
0.471595372086089	1.34447134948944	1.32469126924117
	0.24975184687602	0.701870033464085
79682	CENPU	centromere protein U	2953	-0.42099652321745
	1.71977890259968	1.32426837193323
0.249827445724103	0.701870033464085
6672	SP100	SP100 nuclear antigen	12618	0.145951231796661
	6.30124315143317	1.32419212724116
0.249841078521591	0.701870033464085
4998	ORC1	origin recognition complex subunit 1	3157
0.390008950669088	0.635602648415182	1.32305334981065
	0.250044804215849	0.702037839439779
972	CD74	CD74 molecule	3184	0.573845085088788
6.79813128940777	1.32288868337105	0.250074279669004
	0.702037839439779
1069	CETN2	centrin 2	1928	0.195911132362865
5.01069492525913	1.32288679971717	0.250074616869536
	0.702037839439779
100506144	TMEM35B transmembrane protein 35B	922
0.230531894860081	3.78812863662644	1.32272705444821
	0.250103215548095	0.702037839439779

541565	C8orf58 chromosome 8 open reading frame 58	3685
0.198664896411914	4.1718636639583 1.3224578800012
0.250151414053254	0.702037839439779
9114	ATP6V0D1	ATPase H+ transporting V0 subunit d1	8212
	0.168731703436421	7.16909959273777
1.32137054293084	0.250346229156714	0.702443836498016
134429	STARD4	StAR related lipid transfer domain containing 4
5684	-0.213992399850039	6.44480129813526
1.32040975240352	0.250518526187991	0.702786500363547
54868	TMEM104 transmembrane protein 104	4745
0.154358387795705	5.50240280186686	1.31875673988015
	0.250815298926663	0.703478152195789
8615	USO1	USO1 vesicle transport factor
4588	-0.1754650913949	7.16765817464916
1.31629099219884	0.251258787973443	0.704580949578382
55748	CNDP2	carnosine dipeptidase 2 9779	0.184304681147295
	7.33485461857445	1.3154863801264 0.251403713687444
	0.704846240516418
8353	H3C6	H3 clustered histone 6	462	-0.466609977370215
	0.892923934082121	1.31478588980805
0.251529968712932	0.705059089806292
605	BCL7A	BAF chromatin remodeling complex subunit BCL7A	6565
	0.187120505695563	4.81580061956397
1.31384618607822	0.251699461424789	0.705393028446781
10231	RCAN2	regulator of calcineurin 2	3712
0.388375903700753	3.13392973362238	1.31339103815822
	0.251781606083263	0.705482087699193
23264	ZC3H7B	zinc finger CCCH-type containing 7B
10525	-0.163450432091613	7.53265793869711
1.31305473466528	0.251842322970432	0.705511083569369
80345	ZSCAN16 zinc finger and SCAN domain containing 16
1328	-0.255930116155205	2.81565440525918
1.31208212804309	0.252018020524443	0.705783089520589
11037	STON1	stonin 1	5629	0.63075664047858
2.04029170714007	1.31170047845471	0.252087005158286
	0.705783089520589
5069	PAPPA	pappalysin 1	11573	0.592924004130335
3.1016985745103 1.31168068764335	0.252090583056436
0.705783089520589
837	CASP4	caspase 4	7256	-0.201290361429657
5.52304838111402	1.31136993419795	0.252146771057149
	0.705799324979514
29970	SCHIP1	schwannomin interacting protein 1	8987
0.22554897023313	5.08849906553266	1.31097105171081
	0.252218916416254	0.705860212236258
8714	ABCC3	ATP binding cassette subfamily C member 3	8634
	0.390746975830169	4.33575986393667
1.31001281274752	0.252392335555483	0.706048911598801
79161	TMEM243 transmembrane protein 243	3127
0.279046673557682	3.03761982948187	1.30964685649823
	0.252458603860475	0.706048911598801
9322	TRIP10	thyroid hormone receptor interactor 10
3982	-0.177682874861187	6.94198585153124
1.30959045735553	0.252468818665081	0.706048911598801

51761	ATP8A2	ATPase phospholipid transporting 8A2
10415	-0.518029327058322	-0.181147219192106
1.30948472343306	0.252487970171942	0.706048911598801
1831	TSC22D3 TSC22 domain family member 3	4766
0.258175812972705	5.6794155526288 1.30799353833916
0.252758257713003	0.706663655845514
10224	ZNF443	zinc finger protein 443 2753	-0.280442132684948
	2.42379590295509	1.30636736742613
0.253053417986786	0.707061752456804
114044	MCM3AP-AS1	MCM3AP antisense RNA 1
7605	-0.339008826162259	0.913729080166642
1.30581223220316	0.253154275566084	0.707061752456804
84662	GLIS2	GLIS family zinc finger 2	4469
0.305260729305394	5.54958887262184	1.30558582328038
	0.253195423982244	0.707061752456804
138716	RPP25L	ribonuclease P/MRP subunit p25 like	1092
0.281680093275243	2.5327482601005 1.30530706682241
0.253246097538161	0.707061752456804
1523	CUX1	cut like homeobox 1	17788	-0.179347565439812
	7.24320566757685	1.30524072497911
0.253258159283323	0.707061752456804
26048	ZNF500	zinc finger protein 500 6710	0.246823729664206
	3.81415597485318	1.30505967283961
0.25329108033618	0.707061752456804
8832	CD84	CD84 molecule	9255	0.868430450432286
1.2740919601012 1.30498951170881	0.253303839282449
0.707061752456804
79818	ZNF552	zinc finger protein 552 2989	0.184674947649426
	4.61796116204193	1.30498597971298
0.253304481604072	0.707061752456804
23324	MAN2B2	mannosidase alpha class 2B member 2	8898
0.132209501003539	6.29106123841775	1.30443808026214
	0.253404145758361	0.707199017660364
6249	CLIP1	CAP-Gly domain containing linker protein 1
16293	-0.145562543158589	8.2621182221429 1.30415190938291
	0.253456220069291	0.707203441137562
29943	PADI1	peptidyl arginine deiminase 1	4903
0.58323971425042	2.47212276978186	1.30369032472815
	0.253540242047046	0.707296986726343
5119	CHMP1A	charged multivesicular body protein 1A
3208	-0.146989312326595	6.68702156898921
1.30328362121108	0.25361430244643	0.707362710945271
57535	KIAA1324	KIAA1324	9300	0.360204535661194
	0.237260583793848	1.30266919789604
0.25372623895369	0.707534029270225
3006	H1-2	H1.2 linker histone, cluster member	642
0.358370887508153	5.05184736497772	1.30194535739496
	0.25385818725077	0.70770917883726
3174	HNF4G	hepatocyte nuclear factor 4 gamma
4479	-0.392504499133665	1.98557863073398
1.30164106179049	0.253913682261792	0.70770917883726
80025	PANK2	pantothenate kinase 2	6037	0.163272214080164
	5.6015780648834 1.301265406337	0.253982211910266
0.70770917883726

9632	SEC24C	SEC24 homolog C, COPII coat complex component
5341	-0.13162107538478	7.71936128675483
1.30112771392325	0.25400733640769	0.70770917883726
23268	DNMBP	dynamin binding protein 7072	-0.267286835069648
	5.20886387381094	1.29952631208673
0.25429976627186	0.70770917883726
1296	COL8A2	collagen type VIII alpha 2 chain
4670	-0.471670351263372	3.13170728387852
1.29946166713448	0.254311579701875	0.70770917883726
51171	HSD17B14	hydroxysteroid 17-beta dehydrogenase 14
1281	-0.697734045057908	2.1452038104405 1.29921002228202
	0.254357572540511	0.70770917883726
1374	CPT1A	carnitine palmitoyltransferase 1A	6565
0.142625644568735	5.9532194154587 1.29913089326892
0.254372036975254	0.70770917883726
132864	CPEB2	cytoplasmic polyadenylation element binding protein
2	7177	-0.348857628657289	5.2972583729848
1.29904388773227	0.254387942372848	0.70770917883726
5870	RAB6A	RAB6A, member RAS oncogene family
4896	-0.16308247416719	7.97823519163415
1.29886077570103	0.254421420906271	0.70770917883726
6238	RRBP1	ribosome binding protein 1
6980	-0.181947300213619	8.28333944755643
1.29870418527884	0.25445005477425	0.70770917883726
11044	TENT4A	terminal nucleotidyltransferase 4A
6785	-0.180260961070554	5.38403754214933
1.298511220217	0.254485345507544	0.70770917883726
114791	TUBGCP5 tubulin gamma complex associated protein 5	6478
	0.198149909583456	4.12297041677505
1.29846169913454	0.254494403224185	0.70770917883726
56256	SERTAD4 SERTA domain containing 4
5467	-0.241233979245048	5.36687217667654
1.29825024645928	0.254533083712088	0.70770917883726
64919	BCL11B	BAF chromatin remodeling complex subunit BCL11B 7855
	0.338609489664073	2.33222061232003
1.2980617840657 0.254567564749763	0.70770917883726
55739	NAXD	NAD(P)HX dehydratase	2847	0.158543814246097
	5.65164158674645	1.29789843890269
0.254597454998283	0.70770917883726
6478	SIAH2	siah E3 ubiquitin protein ligase 2
2571	-0.223276816818425	5.57427368275886
1.29682838773687	0.254793368751121	0.70793811624712
7390	UROS	uroporphyrinogen III synthase	6643
0.181474687851826	5.38671116003112	1.2962721378303
0.254895284906404	0.70793811624712
11253	MAN1B1	mannosidase alpha class 1B member 1	8331
0.163976427921038	6.54091910700362	1.29617757844252
	0.254912615079208	0.70793811624712
2702	GJA5	gap junction protein alpha 5
3292	-0.567854463641639	0.811325569023258
1.29592183512697	0.254959493169512	0.70793811624712
6878	TAF6	TATA-box binding protein associated factor 6	4086
	0.171187990109658	5.32381808867328
1.29587402671681	0.254968257692998	0.70793811624712

23032 USP33 ubiquitin specific peptidase 33
6918 -0.141123324674858 6.26320367202925
1.29566705648777 0.255006204994486 0.70793811624712
8814 CDKL1 cyclin dependent kinase like 1 6149
0.352419774088786 0.181415098459369 1.29551762494486
0.255033607095236 0.70793811624712
9903 KLHL21 kelch like family member 21
6670 -0.216639033848601 7.0662670970718 1.29513193294821
0.255104350369579 0.707994181816265
145389 SLC38A6 solute carrier family 38 member 6 5147
0.312954618559705 2.50398620010313 1.29413230050544
0.255287814618586 0.708033508765128
90317 ZNF616 zinc finger protein 616 4697 -0.217275069478153
3.82090799223223 1.29392114796754
0.255326588601695 0.708033508765128
440 ASNS asparagine synthetase (glutamine-hydrolyzing)
4455 -0.24309189957662 5.48941133941564
1.29368879244044 0.255369264481893 0.708033508765128
3688 ITGB1 integrin subunit beta 1 5473 -0.237398678888696
10.7387692062003 1.29362488392644
0.255381003863708 0.708033508765128
124491 TMEM170A transmembrane protein 170A
5761 -0.193856453137386 4.70980416041572
1.29345604484336 0.255412021185737 0.708033508765128
3029 HAGH hydroxyacylglutathione hydrolase 5975
0.17394540209368 5.68717941646664 1.29340275531148
0.255421811936189 0.708033508765128
196264 MPZL3 myelin protein zero like 3
3968 -0.359719926943609 6.37650946106498
1.29253549287841 0.255581216907034 0.708335200873926
55175 KLHL11 kelch like family member 11
2383 -0.330923206302156 2.64854699408592
1.29198514976398 0.255682434927809 0.70847335962728
165545 DQX1 DEAQ-box RNA dependent ATPase 1 2940
0.335865246751132 1.49336743941419 1.29171451757507
0.255732227191799 0.70847335962728
386607 ZFP91-CNTF ZFP91-CNTF readthrough (NMD candidate)
2417 -0.157455330887597 5.57368576360207
1.29137329661607 0.255795023770617 0.708507197539062
23001 WDFY3 WD repeat and FYVE domain containing 3
16856 -0.269868212397946 7.12432817543127
1.29097531106969 0.255868291038006 0.708520190043164
55366 LGR4 leucine rich repeat containing G protein-coupled
receptor 4 6340 -0.273645676847514 5.90822140194361
1.2907983280576 0.255900881090053 0.708520190043164
163786 SASS6 SAS-6 centriolar assembly protein
3972 -0.279170475334252 2.54078055703617
1.29039839833983 0.25597454390601 0.708584078358001
7756 ZNF207 zinc finger protein 207 15929 -0.13286087238538
7.85140159728544 1.28856728136945
0.256312150402475 0.709378441155398
118426 BORCS5 BLOC-1 related complex subunit 5
4980 -0.268633795308624 2.49177722160328
1.28809813751162 0.256398735708263 0.709477892348012

55230	USP40	ubiquitin specific peptidase 40 9352
0.16522957284192	6.11389914075874	1.28780035477404
	0.25645371328404	0.709489860155946
6515	SLC2A3	solute carrier family 2 member 3	6159
0.466546562392166	2.8639465171436 1.28713719101954
0.25657620090857	0.709688555712152
84536	LINC01547	long intergenic non-protein coding RNA 1547
	2656	0.468585383240695	-0.0578961385795121
1.28637635971241	0.256716816842855	0.709937305729097
340252	ZNF680	zinc finger protein 680 5586	-0.248748653929981
	3.19983932226563	1.28488825304547
0.256992121954564	0.710511562910794
79695	GALNT12 polypeptide N-acetylgalactosaminyltransferase 12
	3137	-0.185258376873657	4.18017075091223
1.28470568935032	0.257025921986792	0.710511562910794
23212	RRS1	ribosome biogenesis regulator 1 homolog 1706
0.290548405779286	3.73583483154361	1.28407573548876
	0.257142594442354	0.710693828009875
152518	NFXL1	nuclear transcription factor, X-box binding like 1
	4484	-0.22166147950882	4.30080808221367
1.28350356549546	0.25724862153057	0.710846605203925
57181	SLC39A10	solute carrier family 39 member 10	6917
	0.228441364005233	5.41671267449174
1.28306734272471	0.257329492913542	0.710929823913211
116841	SNAP47	synaptosome associated protein 47	4537
0.184725465204954	4.73434723331462	1.28276830500592
	0.257384949681272	0.710942810133224
1731	SEPTIN1 septin 1	3371	-0.455301902104712
0.185264219562548	1.28223618902942	0.257483667483544
	0.711075262311121
2316	FLNA	filamin A	10585	-0.227947198955112
11.1714226970604	1.28142115756111	0.257634962250005
	0.711352831901404
57519	STARD9	StAR related lipid transfer domain containing 9
20706	0.44862985248065	2.47498543378615
1.2810255678693 0.257708435371699	0.711415462012493
7544	ZFY	zinc finger protein Y-linked
5527	-0.221252947978089	4.96074210058292
1.28028665010402	0.257845744364457	0.711480748340612
158471	PRUNE2	prune homolog 2 with BCH domain 14656
0.617723653277424	1.04256441847598	1.28026056959147
	0.257850592412436	0.711480748340612
6940	ZNF354A zinc finger protein 354A	2947
0.180578913124162	5.01599066731128	1.27998701901087
	0.257901448901298	0.711480748340612
5210	PFKFB4	6-phosphofructo-2-kinase/fructose-2,6-biphosphatase
4	6107	-0.264382116451005	6.88678348457758
1.27975176494304	0.25794519549824	0.711480748340612
7730	ZNF177	zinc finger protein 177 2952	0.314573354512055
	1.15013331797345	1.27918766187592
0.258050130460347	0.711480748340612
7443	VRK1	VRK serine/threonine kinase 1	2744
0.252484144180237	2.78715537658613	1.27898444162508
	0.258087946597215	0.711480748340612

79850	TLCD3A	TLC domain containing 3A	2740
0.197320233098464	5.26221314947795	1.27885989436139
	0.258111126295505	0.711480748340612
112464	CAVIN3	caveolae associated protein 3	1672
0.30123311976522	3.85521730469531	1.27871313929356
	0.258138442426429	0.711480748340612
106865373	WRB-SH3BGR	WRB-SH3BGR readthrough	1314
0.285976724697058	2.57971025079357	1.27792621679739
	0.258284976529617	0.711586559499685
54059	YBEY	ybeY metalloendoribonuclease	1807
0.290995436194674	2.30720812082969	1.2776234549363
0.258341381683472	0.711586559499685
90231	KIAA2013	KIAA2013	3281	0.135584410176906
	5.7923478751323	1.27759892233023	0.258345952824847
	0.711586559499685
90634	N4BP2L1	NEDD4 binding protein 2 like 1	6053
0.460179337854166	1.90107384498754	1.27741599049427
	0.258380041523195	0.711586559499685
29855	UBN1	ubinuclein 1	8551	-0.154424927545376
6.67027232495004	1.27692252838404	0.258472024163792
	0.711699949373351
3005	H1-0	H1.0 linker histone	2344	-0.185634298824454
	7.78475062518022	1.27657833857819
0.258536206013691	0.711722246268431
100128398	LOC100128398	uncharacterized LOC100128398
8787	-0.309143391961147	2.26776433572942
1.27615203888388	0.258615726391738	0.711722246268431
55325	UFSP2	UFM1 specific peptidase 2	2781
0.172457182002666	4.12977934245431	1.27606182977472
	0.258632557543108	0.711722246268431
7750	ZMYM2	zinc finger MYM-type containing 2
13618	-0.161570404624482	7.32439573993702
1.27566731178497	0.258706182341777	0.711785011993963
56852	RAD18	RAD18 E3 ubiquitin protein ligase
6983	-0.187657822895582	3.74756316353097
1.27515600201209	0.258801641335361	0.71188116362721
50804	MYEF2	myelin expression factor 2	14067
0.587570356014838	3.57488379395852	1.27482187445263
	0.258864044818048	0.71188116362721
9241	NOG	noggin	1892	-0.447821693437616
1.46141643954028	1.27466366454939	0.258893599451489
	0.71188116362721
55065	SLC52A1	solute carrier family 52 member 1	3934
0.502028996097982	1.62174254838457	1.27331627791036
	0.259145469152544	0.7122405096221
79863	RBFA	ribosome binding factor A	4250
0.333343207480714	3.09419046968911	1.27323266152209
	0.259161109716804	0.7122405096221
1495	CTNNA1	catenin alpha 1	8830	-0.149877294351794
10.1484406159837	1.2729417974972	0.259215525341185
0.7122405096221
8382	NME5	NME/NM23 family member 5	2392
0.308569828640813	1.59482535794216	1.27287683673115
	0.259227680306523	0.7122405096221

6093 ROCK1 Rho associated coiled-coil containing protein kinase
1 11113 -0.156808727868125 7.15089108005655
1.2724991125049 0.259298371165088 0.712295015671579
285489 DOK7 docking protein 7 4031 0.558669512484383
2.2261110221453 1.27210126946589 0.259372853033118
0.712335045983492
23321 TRIM2 tripartite motif containing 2
9032 -0.274435052502676 7.333860488386 1.2716412633518
0.259459005706756 0.712335045983492
29988 SLC2A8 solute carrier family 2 member 8 3923
0.19063874009501 4.25463295200387 1.27160653950825
0.259465510431054 0.712335045983492
201158 TVP23C trans-golgi network vesicle protein 23 homolog C
4823 -0.184994483365664 5.12433335865617
1.27095768485129 0.259587095747382 0.712365650217102
29088 MRPL15 mitochondrial ribosomal protein L15 1191
0.23542075968757 5.23160005875792 1.27077318390804
0.259621681238791 0.712365650217102
339229 OXLD1 oxidoreductase like domain containing 1 1600
0.266863002242868 3.24336572056176 1.27073291055162
0.259629231409888 0.712365650217102
64224 HERPUD2 HERPUD family member 2 3246 -0.133113254270375
5.68719538786478 1.26933071080896
0.259892275643208 0.712871488479837
9261 MAPKAPK2 MAPK activated protein kinase 2
4195 -0.140405983972316 7.65331243635346
1.26920764690765 0.259915377430322 0.712871488479837
10154 PLXNC1 plexin C1 9887 0.689713807784043
2.37771447853751 1.26841643942867 0.260063965103428
0.713139381206679
10210 TOPORS TOP1 binding arginine/serine rich protein
4098 -0.149337803515205 5.48255086699537
1.26808833933049 0.260125612689627 0.713168811302332
9927 MFN2 mitofusin 2 4973 0.132597878544615
7.29542317947068 1.2675656587444 0.260223857882109
0.71320100049023
3978 LIG1 DNA ligase 1 6033 0.2542750703913
3.27600544925165 1.26697286070889 0.260335338240238
0.71320100049023
22983 MAST1 microtubule associated serine/threonine kinase 1
6533 0.43570891142608 0.0121679330479152
1.26674072277696 0.260379009725145 0.71320100049023
27134 TJP3 tight junction protein 3
4217 -0.366968645603056 4.65479760992666
1.26654871960943 0.260415137619041 0.71320100049023
23269 MGA MAX dimerization protein MGA
17651 -0.303399402250977 5.77367942444088
1.26633106445247 0.260456099775196 0.71320100049023
963 CD53 CD53 molecule 2080 1.08110303936256
2.80223123498866 1.26618650988562 0.260483308992235
0.71320100049023
10005 ACOT8 acyl-CoA thioesterase 8 4213 0.193802708019712
4.16371458214187 1.26613090746169
0.260493775862641 0.71320100049023

113622	ADPRHL1 ADP-ribosylhydrolase like 1	2313
0.372812964549035	0.843729243654778	1.26554829622159
	0.260603480760003	0.713361922025672
170371	TMEM273 transmembrane protein 273	4123
0.986530136754827	0.244291799603972	1.26325431022451
	0.261035991815143	0.714406240202171
84838	ZNF496	zinc finger protein 496 10531	0.161743461410191
	4.75709778537731	1.26239093164279
0.261199004622665	0.714712728609038
63967	CLSPN	claspin 5411	-0.341747962029722
0.435775591233376	1.26207950359871	0.261257835692277
	0.714734082918306
408029	C2orf27B	chromosome 2 open reading frame 27B	1325
	0.442972212733567	1.03705107631871
1.26170975928625	0.26132770446052	0.71478561928891
27163	NAAA	N-acylethanolamine acid amidase 4459
0.281494818982699	3.59310449886984	1.2610411978209
0.261454098041651	0.714977862173323
90326	THAP3	THAP domain containing 3	4072
0.295257068400789	3.45620645161201	1.26037619662473
	0.261579893717155	0.714977862173323
112942	CFAP36	cilia and flagella associated protein 36	3505
	0.190154781642314	4.75382800625772
1.26021379192883	0.261610626576092	0.714977862173323
56672	AKIP1	A-kinase interacting protein 1	2300
0.174903432344278	4.24807538283811	1.26001053861474
	0.261649095781871	0.714977862173323
79963	ABCA11P ATP binding cassette subfamily A member 11,
pseudogene	1909	0.240690874301476	3.1935219266422
1.25975120144352	0.26169819001051	0.714977862173323
6664	SOX11	SRY-box transcription factor 11
8719	-0.893234693892217	1.53151835584653
1.25971916786354	0.261704254969846	0.714977862173323
57728	WDR19	WD repeat domain 19	8873	0.223188586969636
	4.41739764115653	1.25899367998163
0.261841659210184	0.715140345787131
10273	STUB1	STIP1 homology and U-box containing protein 1	2252
	0.153042164800867	6.13067195435809
1.2587909019127 0.261880080485378	0.715140345787131
5954	RCN1	reticulocalbin 1	5038	0.289720626813728
	6.47107738615918	1.25859662888411
0.261916896829298	0.715140345787131
51251	NT5C3A	5'-nucleotidase, cytosolic IIIA
3493	-0.178149422304507	4.35710246182815
1.25688179097949	0.262242151319406	0.715394381646906
200350	FOXD4L1 forkhead box D4 like 1	2491
0.520640541718971	-0.115017947915115	1.25665079103218
	0.262286003516094	0.715394381646906
10580	SORBS1	sorbin and SH3 domain containing 1	10819
0.399795220077357	1.39412405538367	1.25658869804211
	0.262297792576404	0.715394381646906
7702	ZNF143	zinc finger protein 143 4233	-0.183343648237146
	4.80440470047219	1.25657188193316
0.262300985419037	0.715394381646906

158 ADSL adenylosuccinate lyase 5866 0.189590934703757
5.21204610583116 1.2564317110681 0.262327601270521
0.715394381646906
91975 ZNF300 zinc finger protein 300 4411 -0.269148144899562
3.59311719867491 1.25634095294873
0.262344836343511 0.715394381646906
51562 MBIP MAP3K12 binding inhibitory protein 1 3468
0.173909654601751 4.13376872849949 1.256084954928
0.26239345823659 0.715394381646906
10771 ZMYND11 zinc finger MYND-type containing 11
5300 -0.141435599224072 6.59563360522984
1.25566877698589 0.262472527082715 0.715394381646906
57180 ACTR3B actin related protein 3B 6024
0.286750008679301 2.1068804996925 1.25530114719182
0.262542396949682 0.715394381646906
6431 SRSF6 serine and arginine rich splicing factor 6 4593
0.133097189909806 6.80058405234816
1.25520007250928 0.262561610739023 0.715394381646906
27000 DNAJC2 DnaJ heat shock protein family (Hsp40) member C2
6564 -0.160328004494305 4.65239982369623
1.25514673315806 0.262571750984989 0.715394381646906
1855 DVL1 dishevelled segment polarity protein 1
3814 -0.153547162193902 7.10155571953471
1.2543881941491 0.262716008046105 0.715502719741946
6400 SEL1L SEL1L adaptor subunit of ERAD E3 ubiquitin ligase
9559 -0.151856501036634 7.46077435865431
1.25436854264401 0.262719746627189 0.715502719741946
9698 PUM1 pumilio RNA binding family member 1
6621 -0.167400026442961 7.36284421431687
1.25383871415266 0.262820568213253 0.715502719741946
54993 ZSCAN2 zinc finger and SCAN domain containing 2 4777
0.201669799244984 4.01663783823891
1.25364794092001 0.262856882396742 0.715502719741946
9039 UBA3 ubiquitin like modifier activating enzyme 3
4001 -0.141850675342816 6.66170340576349
1.25345091886784 0.262894392596157 0.715502719741946
1022 CDK7 cyclin dependent kinase 7
2419 -0.212250783573011 5.01105224073798
1.25332691185157 0.262918005176826 0.715502719741946
10664 CTCF CCCTC-binding factor 4301 -0.148064490539898
6.10425388137683 1.25248676650659
0.263078049344561 0.715799191369321
26470 SEZ6L2 seizure related 6 homolog like 2 4110
0.560671021167321 3.67688817680944 1.25201939949765
0.263167133151116 0.715902511953326
2909 ARHGAP35 Rho GTPase activating protein 35
11245 0.140847737995937 6.80740280132142
1.25158729762278 0.263249528466894 0.715987601016658
55001 TTC22 tetratricopeptide repeat domain 22
6055 -0.211465308860597 3.92753659549708
1.25127278795339 0.263309520861952 0.716011737276911
84247 RTL6 retrotransposon Gag like 6 5495
0.145296834839303 6.43880024165194 1.25069667695345
0.263419457474813 0.716033964318332

23644	EDC4	enhancer of mRNA decapping 4
6173	-0.14334595993868	6.47489123932908
1.25056969965862	0.263443695633006	0.716033964318332
55105	GPATCH2 G-patch domain containing 2
8833	-0.186294981848339	4.04318353647702
1.25042639432244	0.263471053908523	0.716033964318332
285282	RABL3	RAB, member of RAS oncogene family like 3
4684	-0.166086975473825	4.83137775626705
1.25010292777318	0.263532819548248	0.716057169015994
151887	CCDC80	coiled-coil domain containing 80	12988
0.354019264060828	7.26245589848734	1.24984630070747
	0.26358183504294	0.716057169015994
55529	PIP4P2	phosphatidylinositol-4,5-bisphosphate 4-phosphatase
2	3066	0.376664985402552	2.33190719160798
1.24943989534257	0.263659481197707	0.716129213328735
2774	GNAL	G protein subunit alpha L
8445	-0.354004413413296	4.23469856263022
1.24873304839267	0.26379459618692	0.716184164059375
3601	IL15RA	interleukin 15 receptor subunit alpha	3740
0.343722188929261	2.63933388997499	1.2481573505086
0.263904705368202	0.716184164059375
55246	CCDC25	coiled-coil domain containing 25	4415
0.187251011472328	5.10900248960254	1.24796108128133
	0.263942257281383	0.716184164059375
94015	TTYH2	tweety family member 2	5375	0.532176291403805
	0.652253628793495	1.24707880462452
0.264111144043097	0.716184164059375
727764	MAFIP	MAFF interacting protein (pseudogene)
2216	-0.617359387307259	2.28974259790875
1.24699004918527	0.264128141178608	0.716184164059375
9203	ZMYM3	zinc finger MYM-type containing 3	6916
0.18295141703797	5.15139373701953	1.24695666563116
	0.26413453465735	0.716184164059375
3455	IFNAR2	interferon alpha and beta receptor subunit 2	4614
	0.361713915267674	5.62524772826369
1.24652625753083	0.264216981843086	0.716184164059375
22890	ZBTB1	zinc finger and BTB domain containing 1
5142	-0.1672494939671	6.5667228231443 1.24652259947527
	0.26421768270204	0.716184164059375
90956	ADCK2	aarF domain containing kinase 2
2913	-0.173127205670368	4.7310626599243 1.2462213706252
0.264275404061949	0.716184164059375
400818	NBPF9	NBPF member 9	9046	0.256926582244319
4.47351755746724	1.24621239785254	0.264277123661633
	0.716184164059375
11051	NUDT21	nudix hydrolase 21	5474	-0.140500054071756
	5.80639780343252	1.24615079139028
0.264288930694569	0.716184164059375
3631	INPP4A	inositol polyphosphate-4-phosphatase type I A
13187	0.173398034934358	6.2911899728263 1.24612810932635
	0.264293277934098	0.716184164059375
949	SCARB1	scavenger receptor class B member 1	7741
0.436080304213542	3.51293939995495	1.24502667923089
	0.264504484772847	0.716589657887226

170958	ZNF525	zinc finger protein 525 7193	-0.314575231403265
	3.44061998040567	1.24481428990097
0.264545236020193	0.716589657887226
64844	MARCHF7 membrane associated ring-CH-type finger 7
6288	-0.15561297097011	7.17033627219401
1.24449010288152	0.264607452995542	0.716619604429341
100507334	LOC100507334	two pore channel 3 pseudogene
1715	-0.464521322713691	-0.0415779018985033
1.24384176241629	0.264731935079569	0.716757837304063
23033	DOP1A	DOP1 leucine zipper like protein A
10045	-0.203977635120562	5.34759382063668
1.2434780135202 0.264801807152133	0.716757837304063
10468	FST	follistatin	2969	-0.592072990415532
7.15475127961065	1.24275862539048	0.264940061054123
	0.716757837304063
143379	C10orf82	chromosome 10 open reading frame 82	2411
	0.641347318426014	0.291425058161561
1.24261426772125	0.264967814902643	0.716757837304063
59349	KLHL12	kelch like family member 12	3552
0.142228832920483	5.21964958560981	1.242528727989
0.264984262265972	0.716757837304063
9441	MED26	mediator complex subunit 26
5518	-0.171285537286706	4.60193188040517
1.2423690620667 0.265014965822432	0.716757837304063
140691	TRIM69	tripartite motif containing 69
6627	-0.388096408763337	1.45193185059196
1.24236007338134	0.265016694466891	0.716757837304063
125919	ZNF543	zinc finger protein 543 3659	-0.251944509148803
	2.63730917849388	1.24106343270543
0.265266202805694	0.717035200308876
140576	S100A16 S100 calcium binding protein A16
1701	-0.273543137088185	9.17178169008488
1.2407157506475 0.265333155803284	0.717035200308876
7314	UBB	ubiquitin B	1621	0.185056049326689
8.40546334633681	1.24056691932424	0.265361822618244
	0.717035200308876
9369	NRXN3	neurexin 3	15535	0.384275175089973
3.23682436788806	1.23979062510822	0.265511409284434
	0.717035200308876
152185	SPICE1	spindle and centriole associated protein 1
7941	-0.216037043106938	4.10008302724239
1.23966923649784	0.265534809536124	0.717035200308876
7584	ZNF35	zinc finger protein 35	2663	-0.172825515043384
	4.02726487917026	1.23944360451983
0.26557831174519	0.717035200308876
64840	PORCN	porcupine O-acyltransferase	3012
0.232605515221431	7.08434688221265	1.23889041278859
	0.265685005533076	0.717035200308876
114609	TIRAP	TIR domain containing adaptor protein
3591	-0.228003955800471	3.15681510482625
1.23848456871604	0.2657633143801 0.717035200308876
8780	RIOK3	RIO kinase 3	5320	-0.158563106936907
7.04029759120622	1.23834373884933	0.265790494643704
	0.717035200308876

2539	G6PD	glucose-6-phosphate dehydrogenase	3709
0.181445197015176	5.42723209024971	1.23828856299835
	0.265801144565082	0.717035200308876
3052	HCCS	holocytochrome c synthase
2335	-0.134437566648278	5.6345922479756 1.23809622074905
	0.265838274195798	0.717035200308876
113177	IZUMO4	IZUMO family member 4	2770	-0.530571498385132
	1.25179090073955	1.23784530317346
0.265886720875787	0.717035200308876
11342	RNF13	ring finger protein 13	3635	-0.131919867655533
	6.19220658562264	1.23745983656683
0.265961167432347	0.717035200308876
957	ENTPD5	ectonucleoside triphosphate diphosphohydrolase 5 (inactive)	5842	-0.217314528979228	4.86241392585021
	1.23741037517702	0.265970721967129
0.717035200308876
6950	TCP1	t-complex 1	4929	-0.153103269928607
7.38958181273672	1.23693715674551	0.26606215592118
	0.717035200308876
494115	RBMXL1	RBMX like 1	5087	-0.209107802215248
4.60642652874221	1.23630797292575	0.266183785683537
	0.717035200308876
6305	SBF1	SET binding factor 1	9441	-0.152817310491442
	7.46819944676424	1.23629445271655
0.266186400082922	0.717035200308876
4943	TBC1D25 TBC1 domain family member 25	4223
0.171314908640697	4.91540399554779	1.23623529725143
	0.26619783933598	0.717035200308876
55770	EXOC2	exocyst complex component 2
4983	-0.15712543522753	5.67652301576247
1.23618148291195	0.266208246274654	0.717035200308876
146722	CD300LF CD300 molecule like family member f	2161
0.976948151851538	0.207673715153524	1.23595943023245
	0.266251193507074	0.717035200308876
4089	SMAD4	SMAD family member 4	13384	-0.156964548867303
	6.11772563728564	1.23549250464816
0.26634152978037	0.717035200308876
9283	GPR37L1 G protein-coupled receptor 37 like 1
6588	-0.405380187120142	1.21945276496019
1.23514970305146	0.266407876033993	0.717035200308876
23371	TNS2	tensin 2	8093	0.304875660951207
2.39591181088619	1.2350248336538 0.266432048528367
0.717035200308876
79998	ANKRD53 ankyrin repeat domain 53	2526
0.491925929666008	-0.0686213571684952	1.23464742999365
	0.266505123768833	0.717035200308876
338809	C12orf74	chromosome 12 open reading frame 74
2369	-0.734259724729954	-0.0620151843791402
1.23454324780093	0.266525300568029	0.717035200308876
4154	MBNL1	muscleblind like splicing regulator 1
9989	-0.191644837627155	7.22783890788139
1.23441127529986	0.266550862203351	0.717035200308876
121536	AEBP2	AE binding protein 2	6572	-0.160828013891456
	4.69997341196694	1.23402646636708

0.266625412933312	0.717035200308876
10193	RNF41	ring finger protein 41	6537	0.132153212969353
	6.19820033099252	1.23395660721581
0.26663894983961	0.717035200308876
54958	TMEM160 transmembrane protein 160	663
0.346328647639339	1.14007530344412	1.23375333642069
	0.266678343359904	0.717035200308876
83878	USHBP1	USH1 protein network component harmonin binding protein 1	2829	-0.280015533381623	2.82852225408557
	1.2336239560075 0.266703420830525	0.717035200308876
5378	PMS1	PMS1 homolog 1, mismatch repair system component
	3880	0.240005825638704	3.23100366508411
1.23360974739851	0.266706175027432	0.717035200308876
55186	SLC25A36	solute carrier family 25 member 36
9277	-0.212424261516519	5.73417007512697
1.23326531676211	0.266772950305581	0.717077089796638
5715	PSMD9	proteasome 26S subunit, non-ATPase 9	8959
0.146907477677666	5.63945696875768	1.23267673401639
	0.266887107926109	0.717246300982543
23559	WBP1	WW domain binding protein 1	2438
0.196141270862204	5.46366913341476	1.23203814591577
	0.267011033098682	0.717353708700336
10497	UNC13B	unc-13 homolog B	6940	-0.160563170487967
	5.79881735406852	1.23194300605968
0.267029502189159	0.717353708700336
84269	CHCHD5	coiled-coil-helix-coiled-coil-helix domain containing 5	3759	0.248011639639218	2.79270817347218
	1.22995242575294	0.26741629032821
0.718200163668681
79086	SMIM7	small integral membrane protein 7	6821
0.142911458468815	5.90177773671009	1.229541550602
0.2674962140863 0.718200163668681
195814	SDR16C5 short chain dehydrogenase/reductase family 16C member 5	3279	-0.225608563695771	4.36417042850343
	1.22953025475404	0.267498411784073
0.718200163668681
6405	SEMA3F	semaphorin 3F	4826	-0.166455404428628
7.38893268784229	1.22903704437999	0.267594391766988
	0.718221675544432
9318	COPS2	COP9 signalosome subunit 2
7510	-0.14145310803968	6.44988058572111
1.22896212077585	0.267608975845391	0.718221675544432
25945	NECTIN3 nectin cell adhesion molecule 3 5816
0.245542339389272	3.38460310029151	1.22740915937431
	0.267911487176797	0.718745224841925
5108	PCM1	pericentriolar material 1
14445	-0.155620455974382	7.11229673617561
1.22735989008903	0.26792109163439	0.718745224841925
790	CAD	carbamoyl-phosphate synthetase 2, aspartate transcarbamylase, and dihydroorotase	8422	0.168385340021142
	5.85135285449361	1.2271706286964 0.267957989871102
	0.718745224841925
9833	MELK	maternal embryonic leucine zipper kinase	3087
	0.253143224607629	3.28802030874413

1.22573472306175	0.268238139335444	0.719358915885807
90522	YIF1B	Yip1 interacting factor homolog B, membrane trafficking protein	3193	0.224129238131124
3.91295839735202	1.22525272837831	0.268332259827972
	0.719473576456816
55352	COPRS	coordinator of PRMT5 and differentiation stimulator
	1432	0.181206698338076	4.48555873028688
1.22428075600691	0.268522185028877	0.719845023100379
25824	PRDX5	peroxiredoxin 5 893	0.169910681612083
7.65856787244722	1.22354833491505	0.268665412295776
	0.719862383224916
56942	CMC2	C-X9-C motif containing 2	12781
0.210809106631234	4.45354287628433	1.22347743197883
	0.268679282650441	0.719862383224916
27300	ZNF544	zinc finger protein 544 7130	-0.285132038052035
	5.00716558030362	1.22345924827681
0.268682839972861	0.719862383224916
406	ARNTL	aryl hydrocarbon receptor nuclear translocator like
	6726	0.268538176765771	4.3783052629339
1.22316293731284	0.26874081630061	0.719880017961876
23117	NPIPB3	nuclear pore complex interacting protein family member B3	4305	-0.305295538528931	3.47343969534391
	1.22238112456943	0.268893861074027
0.719968320940275
23030	KDM4B	lysine demethylase 4B	9744	0.154553014545408
	6.16708835961198	1.2223470658307 0.268900530759343
	0.719968320940275
55024	BANK1	B cell scaffold protein with ankyrin repeats 1	5742
	0.311733418516021	2.87122287750272
1.22220689309565	0.268927982805705	0.719968320940275
102724550	LINC02015	long intergenic non-protein coding RNA 2015	3005	-0.413063105844682	1.46113957117576
	1.22106977521639	0.269150810343476
0.720427173797261
9627	SNCAIP	synuclein alpha interacting protein	7325
0.584409494596467	0.742896142763296	1.2204707534012
0.269268286261192	0.720493930830862
84189	SLITRK6 SLIT and NTRK like family member 6	4318
0.487514025662185	4.58399185201641	1.22001843031352
	0.269357035090658	0.720493930830862
81562	LMAN2L	lectin, mannose binding 2 like	2628
0.18419626937888	5.01288677094348	1.21943694480946
	0.269471180124795	0.720493930830862
81617	CAB39L	calcium binding protein 39 like
5006	-0.198945576004065	4.55836140246251
1.21943244556927	0.269472063556723	0.720493930830862
55006	TRMT61B tRNA methyltransferase 61B	2056
0.202513450670057	3.64788971173115	1.2193265647824
0.269492854437373	0.720493930830862
7485	GET1	guided entry of tail-anchored proteins factor 1 4522
	0.187329722913671	4.51306670288141
1.21907526820965	0.269542207360128	0.720493930830862
84529	C15orf41	chromosome 15 open reading frame 41
8968	-0.315894147378186	2.4879671539446 1.21905142702193

0.269546890192087	0.720493930830862
84720	PIGO	phosphatidylinositol glycan anchor biosynthesis class O 7209	0.143719679066194	5.70044810225495
1.21884593966942	0.269587255763872	0.720493930830862
4221	MEN1	menin 1 4246	0.18432226990289
4.91258130973196	1.218250001758	0.26970436346149
0.720590228048762
23011	RAB21	RAB21, member RAS oncogene family
16036	-0.159461115044615	6.44351792717009
1.21786824927284	0.269779414996385	0.720590228048762
401409	RAB19	RAB19, member RAS oncogene family
997	-0.350996933772031	0.460135267151226
1.2168896569902 0.269971922870082	0.720590228048762
64924	SLC30A5 solute carrier family 30 member 5
5784	-0.139809524041607	5.76788372336193
1.21671212523679	0.270006865162439	0.720590228048762
79696	ZC2HC1C zinc finger C2HC-type containing 1C	4812
0.394974660327941	1.02864248361976	1.21669092842519
0.270011037555357	0.720590228048762
84671	ZNF347	zinc finger protein 347 9131	-0.204514725481828
4.36883731092721	1.21660101352455
0.270028737353266	0.720590228048762
23020	SNRNP200	small nuclear ribonucleoprotein U5 subunit 200	10926	-0.161992924412701	7.8745139635263
1.21659062692295	0.270030782054946	0.720590228048762
9179	AP4M1	adaptor related protein complex 4 subunit mu 1	4722
0.16197916344218	4.63647291224316
1.21656997640492	0.270034847364029	0.720590228048762
23528	ZNF281	zinc finger protein 281 4895	-0.156106769842446
5.54885594770838	1.21600608893826
0.270145885127855	0.720620623091409
9135	RABEP1	rabaptin, RAB GTPase binding effector protein 1 6531
0.133095902986795	6.36737206189707
1.21584581850671	0.270177455154061	0.720620623091409
3191	HNRNPL	heterogeneous nuclear ribonucleoprotein L	7223
0.128623833881718	7.37358053740712
1.21572844046986	0.270200579172991	0.720620623091409
26190	FBXW2	F-box and WD repeat domain containing 2 11263
0.125645959351481	6.64265239090485	1.21523875809834
0.270297075565808	0.720633606609157
254863	TMEM256 transmembrane protein 256	707
0.298295706862378	2.86804463311911	1.21476225892521
0.270391015442101	0.720633606609157
11066	SNRNP35 small nuclear ribonucleoprotein U11/U12 subunit 35
5140	-0.183878197901466	4.26961474408389
1.21472106043304	0.270399139474086	0.720633606609157
91754	NEK9	NIMA related kinase 9	10787	0.14207470051745
5.93529014089067	1.21465970010974
0.270411239832778	0.720633606609157
56912	IFT46	intraflagellar transport 46	3814
0.17595731821224	4.13719079224711	1.21398739625429
0.270543863630716	0.720849894973453
23148	NACAD	NAC alpha domain containing	4917
0.324378662636652	3.86489641512299	1.21311365645261

0.270716345766659	0.720914344713615
23774	BRD1	bromodomain containing 1
5850	-0.182208970880653	5.62247168715734
1.21297564851866	0.270743602036948	0.720914344713615
201475	RAB12	RAB12, member RAS oncogene family
2467	-0.128733578513962	5.78969032327201
1.21286210253838	0.270766029697045	0.720914344713615
148362	BROX	BRO1 domain and CAAX motif containing
4980	-0.151744283321111	6.28981335348435
1.21259574167625	0.270818650548637	0.720914344713615
10494	STK25	serine/threonine kinase 25	7584
0.13049406713228	7.0853423923993 1.21230087377137
0.270876918017266	0.720914344713615
203547	VMA21	vacuolar ATPase assembly factor VMA21
5109	-0.175969913152825	4.94675169071665
1.21207776287223	0.270921016338926	0.720914344713615
10936	GPR75	G protein-coupled receptor 75
2115	-0.365892899934719	1.05586778215204
1.21204076734557	0.270928329447595	0.720914344713615
154214	RNF217	ring finger protein 217 12779	-0.262774145590338
5.38070940444308	1.21138974443835
0.271057061126233	0.720937615201858
6526	SLC5A3	solute carrier family 5 member 3
11574	-0.864159055969745	7.76364555144201
1.21132559813032	0.271069749400516	0.720937615201858
27348	TOR1B	torsin family 1 member B	2911
0.164591283287373	6.07568455158875	1.21120822649048
0.271092967680468	0.720937615201858
770	CA11	carbonic anhydrase 11	1929	0.449609102687488
3.4897016919571 1.21095558127815	0.271142954014663
0.720937615201858
10890	RAB10	RAB10, member RAS oncogene family
3631	-0.162772839791853	7.70436752731232
1.21055950021546	0.27122134261265	0.721009175550464
9980	DOP1B	DOP1 leucine zipper like protein B
9030	-0.261249091329864	6.01284650978021
1.20952642551591	0.271425932389268	0.721270257518226
4179	CD46	CD46 molecule	8708	-0.159403170787601
8.55792633780126	1.20923543591141	0.271483594727772
0.721270257518226
80213	TM2D3	TM2 domain containing 3 5580	0.155987431744108
5.06221266413721	1.20915903660735
0.271498736511955	0.721270257518226
64129	TINAGL1 tubulointerstitial nephritis antigen like 1	5281
0.184084732299793	9.39246837401845
1.20902387169241	0.271525527799929	0.721270257518226
7105	TSPAN6	tetraspanin 6	4535	0.195372665036844
6.84635918477403	1.20716411849358	0.271894489097749
0.721829487794605
55659	ZNF416	zinc finger protein 416 2561	-0.28434885424849
2.63995916293684	1.20685143744777
0.27195658431925	0.721829487794605
728752	LOC728752	uncharacterized LOC728752
1392	-0.308523590564258	0.801792743048472

1.20655231054855	0.272016004444187	0.721829487794605
25844	YIPF3	Yip1 domain family member 3	3053
0.130244810958057	7.42268946744897	1.20648585770035
	0.272029207190468	0.721829487794605
54942	ABITRAM actin binding transcription modulator
2758	-0.167391644646302	4.02609258456764
1.20633404909342	0.272059371293298	0.721829487794605
55347	ABHD10	abhydrolase domain containing 10	3021
0.240173323436212	3.11982019045705	1.20622827820964
	0.272080390256972	0.721829487794605
3187	HNRNPH1 heterogeneous nuclear ribonucleoprotein H1
9307	-0.193855442826206	8.44712226854666
1.20614577418705	0.272096787003321	0.721829487794605
26047	CNTNAP2 contactin associated protein like 2
10864	-0.63431063303076	5.14574855409896
1.20557182135895	0.272210887920118	0.721848235536496
57146	TMEM159 transmembrane protein 159
2558	-0.164353501382542	6.36684938648805
1.20545246734413	0.272234622834393	0.721848235536496
145783	LOC145783	uncharacterized LOC145783	5229
0.285878839727484	1.91860997522755	1.20476424404979
	0.272371534495847	0.721848235536496
10014	HDAC5	histone deacetylase 5	7618	0.257853683802548
	4.59265794878687	1.2046363688152 0.272396982847156
	0.721848235536496
5090	PBX3	PBX homeobox 3	3922	0.195247536938087
3.76564078427651	1.20455976378466	0.272412229381227
	0.721848235536496
284697	BTBD8	BTB domain containing 8 1955	-0.411572081338949
	0.282377042293048	1.20455557562819
0.272413062971794	0.721848235536496
80279	CDK5RAP3	CDK5 regulatory subunit associated protein 3
	6216	-0.140696169175801	6.38198523741081
1.20421335841976	0.272481187040093	0.721892185903269
81875	ISG20L2 interferon stimulated exonuclease gene 20 like 2
	5262	-0.148660565520182	5.43397400202733
1.20339605804481	0.272643970729926	0.722083031917982
169792	GLIS3	GLIS family zinc finger 3
11558	-0.215527025275249	5.32128246318808
1.20332289506398	0.272658548717505	0.722083031917982
54872	PIGG	phosphatidylinositol glycan anchor biosynthesis
class G 10895	0.121407508848973	6.63233688760215
1.20307536403885	0.272707877407448	0.722083031917982
25798	BRI3	brain protein I3	2212	0.221935673919565
	5.96570228380306	1.20221040454337
0.27288033682911	0.722225908497796
79658	ARHGAP10	Rho GTPase activating protein 10
10759	0.23635953312673	3.72084330891771
1.20203917205359	0.272914494117038	0.722225908497796
57648	KIAA1522	KIAA1522	5965	-0.154439249093868
	8.22446022253519	1.2020056666418 0.272921178370196
	0.722225908497796
10073	SNUPN	snurportin 1	4040	0.216730737876208
3.77184029781464	1.20165372348945	0.27299140262196

0.722225908497796
9557 CHD1L chromodomain helicase DNA binding protein 1 like
3912 0.195004761321755 4.96046418480084
1.20151220661043 0.273019646283703 0.722225908497796
83903 HASPIN histone H3 associated protein kinase 2797
0.324096121862917 0.618340466692639 1.20068268837448
0.2731852735751 0.72226106906466
100128731 OST4 oligosaccharyltransferase complex subunit 4, non-catalytic 719 0.203092268678306 6.29680959156164
1.20057698402652 0.27320638827414
0.72226106906466
4673 NAP1L1 nucleosome assembly protein 1 like 1 17355
0.193901974468022 7.15895598609592 1.20011372506897
0.273298949501783 0.72226106906466
55300 PI4K2B phosphatidylinositol 4-kinase type 2 beta
3631 -0.19112236501156 5.18255008705286
1.20007928244718 0.273305832862653 0.72226106906466
441459 ANKRD18B ankyrin repeat domain 18B
5857 -0.369224308609291 1.55334061302432
1.19984040517672 0.273353578486171 0.72226106906466
100130086 HSFX2 heat shock transcription factor family, X-linked 2 3328 0.417971376097626 0.425695050950226
1.19979834145999 0.273361987059353
0.72226106906466
29015 SLC43A3 solute carrier family 43 member 3 3664
0.159028560574968 5.75994545230139 1.19963870112606
0.273393902250166 0.72226106906466
23277 CLUH clustered mitochondria homolog
6900 -0.171368064318569 6.08433175156287
1.19873939828986 0.27357377758928 0.722600019365086
11165 NUDT3 nudix hydrolase 3 9905 0.147539754453894
5.71748426766903 1.19818260747887
0.273685219276365 0.722758081901215
5997 RGS2 regulator of G protein signaling 2 2096
0.568009892366727 0.834334448791087 1.19739832975087
0.273842289057716 0.723020225412574
26523 AGO1 argonaute RISC component 1
13886 -0.172799099283672 5.86734220010057
1.19717158221356 0.27388772157058 0.723020225412574
6272 SORT1 sortilin 1 9160 -0.153384155350903
7.53617514817185 1.19636481847068 0.274049446331874
0.723177897107803
80178 TEDC2 tubulin epsilon and delta complex 2 2854
0.315946446595083 0.812805594422145 1.19635852311111
0.274050708777627 0.723177897107803
91 ACVR1B activin A receptor type 1B
5690 -0.164488010895338 5.11288903886858
1.19585226135276 0.274152256335761 0.723230021015089
222484 LNX2 ligand of numb-protein X 2
4627 -0.161324120951779 6.13834465106162
1.19574524401703 0.274173728249456 0.723230021015089
10290 SPEG striated muscle enriched protein kinase
17066 -0.531639173061964 3.59400607688719
1.19528671574022 0.274265751062326 0.723265010459849

8723	SNX4	sorting nexin 4 2709	-0.152387934677595
5.47049420813427	1.19516460328561	0.274290264550776
	0.723265010459849
152002	XXYLT1	xyloside xylosyltransferase 1	5409
0.273109599366987	4.63816956900823	1.19434941551766
	0.274453980033428	0.723560492815401
3301	DNAJA1	DnaJ heat shock protein family (Hsp40) member A1
	2730	-0.146946629550532	7.98368385818708
1.1940843134856	0.274507247326428	0.723564737166217
4976	OPA1	OPA1 mitochondrial dynamin like GTPase
8448	-0.146347738416712	6.58163328962252
1.19306461910266	0.274712256810786	0.723811788336473
645212	EIF3J-DT	EIF3J divergent transcript	3330
0.262624839720532	2.31152760107175	1.19277810238243
	0.274769895560143	0.723811788336473
6447	SCG5	secretogranin V 2196
0.672227829022927	-0.109811580105425	1.19275425427769
	0.274774693782761	0.723811788336473
7559	ZNF12	zinc finger protein 12	5297	-0.191430567795858
	4.84811787987071	1.19239535705979
0.274846916365751	0.723811788336473
3021	H3-3B	H3.3 histone B	3160	-0.169993732411784
9.05555475648834	1.19233358699557	0.27485934905131
	0.723811788336473
23008	KLHDC10	kelch domain containing 10
6622	-0.128564728430452	6.26339814949459
1.19131167250574	0.275065136003991	0.724002458351669
80157	CWH43	cell wall biogenesis 43 C-terminal homolog
3138	-0.807117000718017	0.455376197321005
1.19115986265905	0.275095723053448	0.724002458351669
100506710	EBLN3P	endogenous Bornavirus like nucleoprotein 3,
pseudogene	4614	-0.145885935164693	5.65420876411075
	1.19070223015898	0.27518795390756
0.724002458351669
6237	RRAS	RAS related	1160	0.205698789299025
5.33391514044213	1.19057246044143	0.275214114649746
	0.724002458351669
221264	AK9	adenylate kinase 9	10098	-0.259155160719866
	2.96475112650996	1.19053782506808
0.275221097446755	0.724002458351669
440093	H3-5	H3.5 histone	1057	-0.150461469262552
5.85826944775604	1.19043472058677	0.27524188553742
	0.724002458351669
89932	PAPLN	papilin, proteoglycan like sulfated glycoprotein
	7699	-0.405940141287348	3.51332469800536
1.1894973887871	0.275430962363718	0.724363779539728
7857	SCG2	secretogranin II	2719	0.466057757250829
	0.283810086723239	1.18751504037458
0.275831375728592	0.725082131808761
140825	NEURL2	neuralized E3 ubiquitin protein ligase 2	1301
	0.338927619076449	0.247847987621003
1.18707314317059	0.275920733953746	0.725082131808761
79962	DNAJC22	DnaJ heat shock protein family (Hsp40) member C22
	4412	0.537859250190419	0.824807326264463

1.18699203255187	0.275937139685563	0.725082131808761
23309	SIN3B	SIN3 transcription regulator family member B
8777	-0.128441576627179	6.6418907349011 1.18651284367763
	0.276034087207581	0.725082131808761
83943	IMMP2L	inner mitochondrial membrane peptidase subunit 2
	3411	0.302045745521622	1.92423774355492
1.18623550556378	0.276090216672775	0.725082131808761
55083	KIF26B	kinesin family member 26B	14403
0.359374240561551	4.3844587608872 1.18596708413017
0.276144555185236	0.725082131808761
79665	DHX40	DEAH-box helicase 40	5939	0.14981522913082
	5.68175085641851	1.18550811114641
0.27623749958988	0.725082131808761
84061	MAGT1	magnesium transporter 1 4862	-0.228478529702527
	6.92478768944954	1.1854213906618 0.276255065353721
	0.725082131808761
6645	SNTB2	syntrophin beta 2	9875	-0.171280889539824
	5.69642281537058	1.18528637886728
0.276282415612301	0.725082131808761
6275	S100A4	S100 calcium binding protein A4
1025	-0.40394492625733	7.15634529382157
1.18514958054789	0.276310131251461	0.725082131808761
5162	PDHB	pyruvate dehydrogenase E1 beta subunit	4717
0.140702922788159	6.2009245918143 1.1849951470373
0.276341424019507	0.725082131808761
4820	NKTR	natural killer cell triggering receptor
17356	-0.222491507158908	6.3094624347193 1.18496351472676
	0.276347834205067	0.725082131808761
6448	SGSH	N-sulfoglucosamine sulfohydrolase	6315
0.203131685881688	4.87768484463807	1.18481929817552
	0.276377061592559	0.725082131808761
6760	SS18	SS18 subunit of BAF chromatin remodeling complex
	6039	-0.132279451001865	6.84290196995384
1.18384366294442	0.276574889069112	0.72546525677735
997	CDC34	cell division cycle 34	1942	-0.144230862590131
	6.09302994127638	1.18239239882161
0.276869488374314	0.725994822879736
23086	EXPH5	exophilin 5	10843	-0.353036131881316
3.89296773014565	1.18233847874436	0.276880441480942
	0.725994822879736
130535	KCTD18	potassium channel tetramerization domain containing
18	2940	0.168101925687147	4.12026245472406
1.18099040791061	0.277154460407562	0.726119320687037
284434	NWD1	NACHT and WD repeat domain containing 1
8495	-0.549673124254426	0.604592599117686
1.18041762674642	0.277270991457823	0.726119320687037
51143	DYNC1LI1	dynein cytoplasmic 1 light intermediate
chain 1 6132	0.127470830807868	6.22037584447744
1.18031874660054	0.277291114642902	0.726119320687037
1797	DXO	decapping exoribonuclease	2483
0.252346274246294	3.71468819346291	1.17903370182307
	0.277552802464797	0.726119320687037
200576	PIKFYVE	phosphoinositide kinase, FYVE-type zinc finger
containing	10832	-0.226566841264828	6.13329641162141

1.17902948719627	0.277553661246952
0.726119320687037
9119	KRT75	keratin 75	2324	-0.425644114135821
5.51101026247715	1.17898669546325	0.277562380779321
0.726119320687037
112869	SGF29	SAGA complex associated factor 29	2124
0.258700768016377	2.35740499188155	1.17898381276848
0.277562968189063	0.726119320687037
103	ADAR	adenosine deaminase RNA specific
8356	-0.149339531369001	8.24183513397091
1.17895347390242	0.277569150466563	0.726119320687037
146330	FBXL16	F-box and leucine rich repeat protein 16	3989
0.585526885439429	-0.16879362384136
1.17893279715263	0.277573363953581	0.726119320687037
819	CAMLG	calcium modulating ligand	2221
0.159412874093259	4.95615661477321	1.17878262293734
0.27760396871824	0.726119320687037
4507	MTAP	methylthioadenosine phosphorylase
13999	-0.194808254037797	5.96554989035835
1.17866202547432	0.277628548961858	0.726119320687037
201931	TMEM192	transmembrane protein 192	10241
0.179841408710619	5.28186856171072	1.17858662508986
0.277643918503143	0.726119320687037
100506394	GAS6-DT	GAS6 divergent transcript	1952
0.359023041081222	2.47455281966057	1.17844187250447
0.277673427731906	0.726119320687037
5792	PTPRF	protein tyrosine phosphatase receptor type F
10712	-0.160779825077555	9.41417773678512
1.17834563826662	0.277693048206503	0.726119320687037
1060	CENPC	centromere protein C	8613	-0.206057514210179
3.86146005591416	1.17815268152153
0.277732393963702	0.726119320687037
65108	MARCKSL1	MARCKS like 1	1699	0.140174525751884
7.84865306692848	1.17786570314365
0.277790924628708	0.726119320687037
27241	BBS9	Bardet-Biedl syndrome 9	6195	0.179940309583779
3.96307772100268	1.17772839779399
0.277818934229147	0.726119320687037
53407	STX18	syntaxin 18	5332	-0.138106713944768
5.47670081168315	1.17742296895595	0.277881252928789
0.726119320687037
2184	FAH	fumarylacetoacetate hydrolase	7063
0.281383960302426	3.2302286975157	1.17674806283046
0.278019020996026	0.726119320687037
5087	PBX1	PBX homeobox 1	19029	0.321919875050101
4.41550811827081	1.17649248196351	0.278071214973895
0.726119320687037
59271	EVA1C	eva-1 homolog C	4353	-0.395029770585989
2.20763095153846	1.1764602641577	0.278077795275836
0.726119320687037
10282	BET1	Bet1 golgi vesicular membrane trafficking protein
5247	-0.214494522644832	4.16502290388495
1.17640530595574	0.278089020625777	0.726119320687037
140890	SREK1	splicing regulatory glutamic acid and lysine rich

protein 1	18240	-0.195512875960487	6.05308177216705
	1.17551389046209	0.27827117419219
0.726119320687037
5930	RBBP6	RB binding protein 6, ubiquitin ligase
8744	-0.165719701061478	6.0512332697685	1.1754819874755
0.278277696095217	0.726119320687037
22974	TPX2	TPX2 microtubule nucleation factor	3605
0.316155704780569	2.50454238602501	1.17504895224739
	0.278366240200916	0.726119320687037
59269	HIVEP3	HIVEP zinc finger 3	14224	-0.28412778853086
	2.99293068636785	1.17504349972673
0.278367355321581	0.726119320687037
55589	BMP2K	BMP2 inducible kinase	9938	-0.216614669184588
	5.4812976844023	1.17463054718775	0.278451826544153
	0.726119320687037
29097	CNIH4	cornichon family AMPA receptor auxiliary protein 4
	5098	0.140111120805826	6.39348160221412
1.17443824033974	0.278491174759898	0.726119320687037
79635	CCDC121	coiled-coil domain containing 121	2844
0.33696834870865	0.618210768406733	1.17434124763987
	0.278511023250995	0.726119320687037
2138	EYA1	EYA transcriptional coactivator and phosphatase 1
	5170	-0.568717714309102	1.61103389098087
1.17425991696585	0.278527668054317	0.726119320687037
7358	UGDH	UDP-glucose 6-dehydrogenase	4045
0.190819829167528	6.433352008339	1.17422431466936
0.278534954669198	0.726119320687037
23160	WDR43	WD repeat domain 43	4130	-0.187768755220546
	5.41206984403715	1.17395892751357
0.278589278228445	0.726125794593568
283232	TMEM80	transmembrane protein 80	4071
0.233678354721657	4.17699787895246	1.1727509335891
0.278836717917338	0.726515696487735
5291	PIK3CB	phosphatidylinositol-4,5-bisphosphate 3-kinase
catalytic subunit beta	7412	-0.131791711547148
5.9356904412948	1.17272220386148	0.278842606148216
0.726515696487735
79149	ZSCAN5A	zinc finger and SCAN domain containing 5A	3562
	0.288969514465318	1.02932141593298
1.1717908133085	0.279033581940637	0.726650351985008
6256	RXRA	retinoid X receptor alpha	10814
0.133991937712724	6.24024196376975	1.17169502108349
	0.279053232884586	0.726650351985008
9611	NCOR1	nuclear receptor corepressor 1
15191	-0.194924376803259	7.3432822255329	1.1714762679097
	0.279098114741661	0.726650351985008
5445	PON2	paraoxonase 2	3843	0.207613553651607
7.70736272101876	1.17121456556404	0.279151820489202
	0.726650351985008
8690	JRKL	JRK like	3511	-0.241830409185606
3.91685237977177	1.17115344476707	0.27916436538715
	0.726650351985008
821	CANX	calnexin	6235	-0.205811152108322
9.47122611465712	1.17095279161893	0.279205553964683

0.726650351985008
3198 HOXA1 homeobox A1
2539 -0.879849210364444 -0.124997019750434
1.16989441557268 0.279422936325379 0.727081008565779
7629 ZNF76 zinc finger protein 76 6789 0.197958755741133
4.49133482455822 1.16945692084234
0.27951285674488 0.727175305269262
55829 SELENOS selenoprotein S 3766 -0.187965566116493
6.44927066725409 1.1690139045225 0.279603949234365
0.727175305269262
124637 CYB5D1 cytochrome b5 domain containing 1
3980 -0.194106826939035 3.90540380895314
1.1689605585503 0.279614920695384 0.727175305269262
81790 RNF170 ring finger protein 170 6055 -0.138363846336246
5.6439231540964 1.16843449877661 0.27972314247196
0.727299601055915
54887 UHRF1BP1 UHRF1 binding protein 1
9908 -0.237490831000128 5.71630746495693
1.16800279673194 0.27981199231239 0.727299601055915
134430 WDR36 WD repeat domain 36 7656 -0.164442792082686
5.1107033763013 1.1679712463102 0.279818487191428
0.727299601055915
3188 HNRNPH2 heterogeneous nuclear ribonucleoprotein H2
2223 -0.122948991361003 6.66057872515908
1.16767687600126 0.279879094582431 0.727322166570708
79680 RTL10 retrotransposon Gag like 10
6803 -0.180162733962574 5.27947541605596
1.1672552820139 0.279965924683162 0.727412856063263
29959 NRBP1 nuclear receptor binding protein 1
2533 -0.139831760238666 7.38654930329785
1.16632095299755 0.280158477156873 0.727735925779126
1267 CNP 2',3'-cyclic nucleotide 3' phosphodiesterase 6187
0.142382070133686 6.88651281278089
1.16575467126653 0.280275261443771 0.727735925779126
1677 DFFB DNA fragmentation factor subunit beta 3870
0.2881970503667 2.36524727186674 1.16535499296416
0.280357724090689 0.727735925779126
9657 IQCB1 IQ motif containing B1 3049 -0.31259543574043
4.12970355620611 1.16498034373686
0.280435050483837 0.727735925779126
64789 EXO5 exonuclease 5 3092 0.217336514055752
3.43919650685702 1.1649478337448 0.280441761710743
0.727735925779126
144717 PHETA1 PH domain containing endocytic trafficking adaptor 1
3361 0.200415457964246 4.9383921957246
1.16494147219779 0.280443074985642 0.727735925779126
80742 PRR3 proline rich 3 4281 0.286007460002974
2.70976529190482 1.16442058124426 0.280550633826926
0.727735925779126
8943 AP3D1 adaptor related protein complex 3 subunit delta 1
8601 -0.137380081836879 7.95469591044246
1.16419017532265 0.280598226988029 0.727735925779126
439 GET3 guided entry of tail-anchored proteins factor 3,
ATPase 1652 0.141755273452239 6.1283617787415

1.16384199503732	0.280670167226514	0.727735925779126
292	SLC25A5 solute carrier family 25 member 5	1492
0.167876208050717	7.1757831255292 1.16373269383091
0.280692755601854	0.727735925779126
116150	NUS1	NUS1 dehydrodolichyl diphosphate synthase subunit
	4676	-0.178480573970685	5.73356685612525
1.16330452808703	0.280781263181473	0.727735925779126
84698	CAPS2	calcyphosine 2	6689	0.363596839613257
1.14861373255264	1.16326429430349	0.280789581853201
	0.727735925779126
84912	SLC35B4 solute carrier family 35 member B4	7011
0.164421188520929	5.12580622787712	1.16320567097334
	0.280801703274801	0.727735925779126
6576	SLC25A1 solute carrier family 25 member 1	2096
0.220918591006631	5.72362084905284	1.16306132100769
	0.280831553027698	0.727735925779126
285513	GPRIN3	GPRIN family member 3	14244	0.775561759612345
	1.84699433895462	1.16266721098979
0.280913070417321	0.727735925779126
9360	PPIG	peptidylprolyl isomerase G
7368	-0.116514950910356	5.95184935513587
1.16262622331973	0.280921549988415	0.727735925779126
26007	TKFC	triokinase and FMN cyclase	8215
0.18000071197769	3.95315070735145	1.16166459131326
	0.281120586197939	0.727941759938002
200728	TMEM17	transmembrane protein 17	2982
0.27754674037663	2.45785757808662	1.16166453910371
	0.281120597008978	0.727941759938002
10458	BAIAP2	BAR/IMD domain containing adaptor protein 2
10288	-0.156492275908169	6.71219991507157
1.16144641406612	0.281165768771956	0.727941759938002
4485	MST1	macrophage stimulating 1	5154
0.36933357031386	2.80472159760898	1.16101872213656
	0.281254366599544	0.727941759938002
130162	CLHC1	clathrin heavy chain linker domain containing 1
4173	-0.390067225864689	1.84195226615759
1.16098740407148	0.281260855628219	0.727941759938002
440854	CAPN14	calpain 14	2197	0.494250485197472
3.10955445876244	1.16070754471488	0.281318850219527
	0.727957349902996
81502	HM13	histocompatibility minor 13
9914	-0.186224221099981	7.24096874935889
1.16027507400893	0.281408499814748	0.727973628929969
130589	GALM	galactose mutarotase	2790	0.570551596936929
	2.39481378896868	1.16004620293704
0.281455958557118	0.727973628929969
126364	LRRC25	leucine rich repeat containing 25	2415
0.872401200553811	-0.316993573892756	1.15968876529164
	0.281530097117459	0.727973628929969
2519	FUCA2	alpha-L-fucosidase 2	2793	0.160952299094506
	6.0066084355762 1.15963743195012	0.281540746539535
	0.727973628929969
6236	RRAD	RRAD, Ras related glycolysis inhibitor and calcium
channel regulator	1709	-0.32623612187181

4.06720271020085	1.15942413442428	0.281585001894951
	0.727973628929969
2305	FOXM1	forkhead box M1 4899	0.39952460937644
0.918666902131327	1.15904204584132	0.281664300304707
	0.727990754606314
54916	TMEM260 transmembrane protein 260	8077
0.165536654667815	4.61699957853998	1.15866238120033
	0.281743123608359	0.727990754606314
92126	DSEL	dermatan sulfate epimerase like
9531	-0.294449562792237	4.34903595476103
1.15864108270353	0.281747546278348	0.727990754606314
5310	PKD1	polycystin 1, transient receptor potential channel interacting	18532	-0.270267494145026	6.03922302243999
	1.15686066413744	0.282117563795727
0.728643201360185
54058	C21orf58	chromosome 21 open reading frame 58	5315
	0.363118185421295	0.981344219936567
1.15664668445751	0.282162075711626	0.728643201360185
2963	GTF2F2	general transcription factor IIF subunit 2	3619
	0.145627836872917	6.05723964247639
1.15655834400263	0.282180454826249	0.728643201360185
139735	ZFP92	ZFP92 zinc finger protein	6105
0.343278960321196	1.30817137936549	1.15642530409763
	0.282208136458842	0.728643201360185
54845	ESRP1	epithelial splicing regulatory protein 1
4604	-0.153408531665107	7.13254344216058
1.15577531730665	0.282343428688658	0.728664843493617
113178	SCAMP4	secretory carrier membrane protein 4	4820
0.150396077264993	6.42755172176214	1.15573524741307
	0.282351771765442	0.728664843493617
3695	ITGB7	integrin subunit beta 7 4481	-0.614288615698283
	1.7488647292286 1.15563530219126	0.282372583028725
	0.728664843493617
219972	MPEG1	macrophage expressed 1	4442	1.10077336988714
	2.30343012313829	1.15536403139612
0.282429078629462	0.728676386878408
131408	FAM131A family with sequence similarity 131 member A
6597	-0.172213560290792	3.84892106196947
1.15494664211226	0.282516033192248	0.728704030315474
8266	UBL4A	ubiquitin like 4A	2630	-0.145921775177715
	5.30989114066427	1.15464128356196
0.282579669881605	0.728704030315474
84953	MICALCL MICAL C-terminal like	3199	-0.392530417744087
	3.5457627405913 1.1542785583224 0.282655285346786
0.728704030315474
4070	TACSTD2 tumor associated calcium signal transducer 2
2351	-0.150402823782596	10.9682081889195
1.15423061540242	0.282665281675844	0.728704030315474
9609	RAB36	RAB36, member RAS oncogene family	4107
0.304498462216355	2.25685441550424	1.15403311592651
	0.282706465981445	0.728704030315474
388507	ZNF788P zinc finger family member 788, pseudogene	7434
	0.417477305564222	0.135429558475877
1.15366369896937	0.282783520395095	0.728704030315474

655	BMP7	bone morphogenetic protein 7
5418	-0.353513593203161	1.55194058059364
1.15356571401906	0.282803962932457	0.728704030315474
440119	FZD10-AS1	FZD10 antisense divergent transcript
11498	-0.984182879327315	1.81202751047718
1.15231242412604	0.283065600993964	0.729189881669083
901	CCNG2	cyclin G2	7161	-0.191538404179896
6.91388065835108	1.15189543376515	0.283152720231819
	0.729189881669083
51701	NLK	nemo like kinase	5290	-0.168112381420125
	4.65065637088562	1.15175154124682
0.283182790688392	0.729189881669083
221395	ADGRF5	adhesion G protein-coupled receptor F5	6650
0.941121663363204	5.06547233747397	1.15148718077216
	0.283238046904902	0.729189881669083
148867	SLC30A7	solute carrier family 30 member 7
8376	-0.206353541529441	5.56310659186327
1.15134262823442	0.283268266823741	0.729189881669083
84326	METTL26	methyltransferase like 26	1830
0.189529666680603	3.61740801259118	1.15100463014523
	0.283338944086241	0.729189881669083
6094	ROM1	retinal outer segment membrane protein 1	2248
	0.291040534584907	0.248279670992791
1.15091863106861	0.283356930529365	0.729189881669083
28973	MRPS18B	mitochondrial ribosomal protein S18B	1888
0.161749194863874	4.44271898401815	1.150608034783
0.283421902828524	0.729223106708143
647087	STMP1	short transmembrane mitochondrial protein 1	4930
	0.16583407454731	5.39611319277264
1.15014914976722	0.283517929530713	0.72929423873219
2805	GOT1	glutamic-oxaloacetic transaminase 1	2424
0.141237029018855	5.43451120320565	1.14997834876427
	0.283553682025809	0.72929423873219
51693	TRAPPC2L	trafficking protein particle complex 2 like
	4463	0.212164164711381	4.6149017951463
1.14961030400235	0.283630741486238	0.729354057184536
149603	RNF187	ring finger protein 187	3285	0.152906678842079
	6.68757639271959	1.14936994444452
0.283681081140955	0.729354057184536
3796	KIF2A	kinesin family member 2A
4346	-0.147585998081588	5.63878346551822
1.14883966437264	0.283792180261374	0.729505793525612
393	ARHGAP4	Rho GTPase activating protein 4	5580
0.190160779018605	3.82542761857018	1.14832017483654
	0.283901072093727	0.729609299784346
23303	KIF13B	kinesin family member 13B
11112	-0.213135920881852	6.58284982407099
1.14815062198428	0.283936624054007	0.729609299784346
55824	PAG1	phosphoprotein membrane anchor with
glycosphingolipid microdomains 1	11023	-0.501740433518403
	2.69549632144315	1.14762446808702
0.284046984279336	0.729759007483614
5000	ORC4	origin recognition complex subunit 4
8915	-0.147197312974164	5.27002068339282

1.1465118288843 0.28428053819768 0.730225105177866
28977 MRPL42 mitochondrial ribosomal protein L42
16111 -0.154809155493815 5.48042671228814
1.14618748570054 0.284348666822955 0.730266185586565
9858 PPP1R26 protein phosphatase 1 regulatory subunit 26
5219 -0.172912383246996 5.54682213443481
1.14491306973013 0.28461655968876 0.730632336431239
4901 NRL neural retina leucine zipper 3113
0.346389934131631 0.315045194781217 1.14485057229811
0.284629705396848 0.730632336431239
90161 HS6ST2 heparan sulfate 6-O-sulfotransferase 2 4561
0.464437320506112 5.62860139597656 1.14476491750132
0.284647723274453 0.730632336431239
6508 SLC4A3 solute carrier family 4 member 3 5091
0.239964798917628 5.09926850103104 1.14352133443816
0.284909479607967 0.731151556887014
3037 HAS2 hyaluronan synthase 2 4190 -0.941834307417842
3.49488743649881 1.14307633518835
0.285003219679626 0.731151556887014
26123 TCTN3 tectonic family member 3 3223
0.160122723879725 5.03219042571186 1.14306027329508
0.285006603883994 0.731151556887014
50636 ANO7 anoctamin 7 5695 0.348440168720273
1.2844534408111 1.14226467218281 0.285174299023952
0.731328277623853
493812 HCG11 HLA complex group 11 3472 -0.307860323615971
2.94252600895622 1.14222378671331
0.28518292015776 0.731328277623853
8881 CDC16 cell division cycle 16 3901 0.16049298273193
6.16662816506907 1.14188759363333
0.285253822560892 0.731328277623853
130271 PLEKHH2 pleckstrin homology, MyTH4 and FERM domain
containing H2 9835 0.326511760040532 3.12646063535072
1.14153123635163 0.28532900196266
0.731328277623853
339665 SLC35E4 solute carrier family 35 member E4
5531 -0.33507583305302 1.98644502684332
1.14149547084429 0.285336548669548 0.731328277623853
2475 MTOR mechanistic target of rapamycin kinase
12119 -0.136969213221263 6.85320808687651
1.1411280405266 0.285414093029038 0.731393194485498
102466730 MIR6764 microRNA 6764 2742 -0.13503750585122
5.78645094197427 1.14043314731856
0.285560819922579 0.731567599274028
9091 PIGQ phosphatidylinositol glycan anchor biosynthesis
class Q 6886 0.225893816853588 5.76943950365476
1.14014576216728 0.285621529349013 0.731567599274028
23125 CAMTA2 calmodulin binding transcription activator 2
5999 -0.14124533382574 6.10830584197918
1.13900158692183 0.285863396094014 0.731567599274028
63910 SLC17A9 solute carrier family 17 member 9 5521
0.458839727130252 1.76001994476409 1.13897362742166
0.285869309694125 0.731567599274028
100505385 IQCJ-SCHIP1 IQCJ-SCHIP1 readthrough 3761

0.211691120245401	4.96269998985692	1.13870274968461
	0.285926609981314	0.731567599274028
26093	CCDC9	coiled-coil domain containing 9
3028	-0.202979102315536	3.84641758881483
1.13826502601848	0.286019234938173	0.731567599274028
10540	DCTN2	dynactin subunit 2	4217	0.129081029061442
	7.4191667850735 1.13824561686108	0.286023342914199
	0.731567599274028
1193	CLIC2	chloride intracellular channel 2	2840
0.431482894947717	0.0596099605984356	1.13824008376064
	0.286024514016569	0.731567599274028
7077	TIMP2	TIMP metallopeptidase inhibitor 2	5633
0.221170587903975	8.47660512334241	1.13799323953862
	0.286076765755657	0.731567599274028
64428	CIAO3	cytosolic iron-sulfur assembly component 3	8101
	0.172926736544338	4.84489425594809
1.13775943146402	0.286126269190403	0.731567599274028
85474	LBX2	ladybird homeobox 2	2331	0.357830476601131
	0.0460239970046664	1.13749450531273
0.286182374298172	0.731567599274028
440465	BAIAP2-DT	BAIAP2 divergent transcript	4773
0.174035084581219	4.56231976053193	1.13707815654606
	0.286270575368159	0.731567599274028
90990	KIFC2	kinesin family member C2	4447
0.24430027939286	5.19073636270642	1.13701424562327
	0.286284117581553	0.731567599274028
161835	FSIP1	fibrous sheath interacting protein 1	3084
0.36915024434172	0.886873217323361	1.13675378221353
	0.286339316107753	0.731567599274028
10342	TFG	trafficking from ER to golgi regulator
4994	-0.159478065619654	7.29280942107074
1.13637264915462	0.286420111822097	0.731567599274028
51533	PHF7	PHD finger protein 7	2335	0.311997908452335
	1.26732819385999	1.13620040094119
0.286456635910097	0.731567599274028
80205	CHD9	chromodomain helicase DNA binding protein 9
16304	-0.2693601804244	7.01232185758218
1.13608542307269	0.286481019494022	0.731567599274028
140688	NOL4L	nucleolar protein 4 like
9448	-0.210914607478528	4.74889881455138
1.13578193636502	0.286545393181848	0.731567599274028
162993	ZNF846	zinc finger protein 846 3426	0.239716857743191
	3.10672795668875	1.1357759064146 0.286546672403041
	0.731567599274028
403313	PLPP6	phospholipid phosphatase 6
2944	-0.167645544423932	4.37935668036822
1.13559738756109	0.286584547493072	0.731567599274028
402055	SRRD	SRR1 domain containing	4319	0.157524082386243
	4.87713205818607	1.13552961198388
0.286598928625903	0.731567599274028
29057	FAM156A family with sequence similarity 156 member A	5432
	0.206008747911197	3.48450308053736
1.13526752474255	0.286654548898026	0.731567599274028
10158	PDZK1IP1	PDZK1 interacting protein 1

1438    -0.54008988965605       4.99754819343713
1.13454197214856        0.286808597526358       0.731567599274028
649446  DLGAP1-AS1      DLGAP1 antisense RNA 1  2105
0.298718992326396       1.55567549732448        1.1340667717248
0.286909548618685       0.731567599274028
5867    RAB4A   RAB4A, member RAS oncogene family       3310
0.140474851909614       5.28117355649923        1.13400539373338
        0.286922590988036       0.731567599274028
153443  SRFBP1  serum response factor binding protein 1
3111    -0.199944078986215      3.62920004460138
1.13386531636186        0.286952359214994       0.731567599274028
84153   RNASEH2C        ribonuclease H2 subunit C       3697
0.222188621945316       3.76650054100889        1.13372711724816
        0.286981732133614       0.731567599274028
284131  ENDOV   endonuclease V  7253    -0.237945707351494
3.35135144409303        1.1335538241902 0.287018569337738
0.731567599274028
6167    RPL37   ribosomal protein L37   8153    0.200332588165911
        7.22194130977935        1.13338711143287
0.287054013426184       0.731567599274028
1794    DOCK2   dedicator of cytokinesis 2      12890
0.888671432810203       1.72896997193357        1.13307602178277
        0.287120167743294       0.731567599274028
26152   ZNF337  zinc finger protein 337 3823    0.164453611987757
        4.24033906227162        1.13278395012388
0.287182295450812       0.731567599274028
1534    CYB561  cytochrome b561 5320    -0.170589974484631
6.43381201231506        1.13275605666863        0.287188229670636
        0.731567599274028
54948   MRPL16  mitochondrial ribosomal protein L16     2271
0.269215614599539       3.54570542847312        1.13267394650943
        0.287205699179545       0.731567599274028
5130    PCYT1A  phosphate cytidylyltransferase 1, choline, alpha
        7978    -0.144099155901664      7.12168698830274
1.13217998019813        0.287310822461452       0.731643215560381
406988  MIR205  microRNA 205    110     0.349156693990004
1.56530256401126        1.1317234423221 0.287408023911434
0.731643215560381
83442   SH3BGRL3        SH3 domain binding glutamate rich protein
like 3  1786    -0.221499725839219      8.29573047323226
1.13159864980094        0.287434600753014       0.731643215560381
2300    FOXL1   forkhead box L1 3655    0.424703585906435
2.43748877000979        1.13155300801154        0.287444321784021
        0.731643215560381
2734    GLG1    golgi glycoprotein 1    14111   0.134031413132987
        7.50809465523841        1.13085595967272
0.287592834787779       0.731702019877167
3162    HMOX1   heme oxygenase 1        2405    0.265629191837862
        4.89517476638642        1.13077909102084
0.287609218381113       0.731702019877167
11174   ADAMTS6 ADAM metallopeptidase with thrombospondin type 1
motif 6 8253    -0.481140731429536      2.9078681439364
1.13058952691961        0.287649626673875       0.731702019877167
57143   ADCK1   aarF domain containing kinase 1 5317

0.250152860280225	2.98941961183275	1.13046405031363
	0.287676377772796	0.731702019877167
8648	NCOA1	nuclear receptor coactivator 1
8592	-0.175725074129529	5.43091110678355
1.12915864231701	0.287954873150524	0.732035147248916
89927	BMERB1	bMERB domain containing 1	3577
0.206068161552615	4.60133613406463	1.12912857198459
	0.287961292385688	0.732035147248916
3725	JUN	Jun proto-oncogene, AP-1 transcription factor
subunit 3540	-0.176447095430938	7.27495971732148
1.12896687100996	0.287995814465877	0.732035147248916
23119	HIC2	HIC ZBTB transcriptional repressor 2
7219	-0.247444700050067	2.45169602211162
1.1288704664904 0.288016398692709	0.732035147248916
828	CAPS	calcyphosine	4669	-0.242389389183712
4.06200478487629	1.12855594819773	0.288083567434844
	0.73207302776848
1386	ATF2	activating transcription factor 2	5751
0.159291714974332	5.57358149463067	1.12737832828365
	0.288335237790931	0.732579661842476
84132	USP42	ubiquitin specific peptidase 42
7072	-0.188630926957838	4.35579835155032
1.1270020591934 0.288415709601153	0.732651223138075
84295	PHF6	PHD finger protein 6	8016	-0.146524238114389
	5.37936484864487	1.12655885804202
0.288510532674943	0.732663659425483
6525	SMTN	smoothelin	7943	0.191413158929049
6.65378902641336	1.12606955624341	0.28861526508077
	0.732663659425483
9830	TRIM14	tripartite motif containing 14
5469	-0.338512758995193	4.53074404353399
1.12588236098976	0.288655346008741	0.732663659425483
30849	PIK3R4	phosphoinositide-3-kinase regulatory subunit 4
5376	-0.155503921477248	5.19089309569639
1.12578333784349	0.288676551011433	0.732663659425483
6217	RPS16	ribosomal protein S16	2528	0.204043400364465
	6.85271217766476	1.12575723945656
0.288682140099182	0.732663659425483
7265	TTC1	tetratricopeptide repeat domain 1	1938
0.13189631956126	6.086012429266	1.12325773713101
0.289218058763199	0.733478413962509
84286	TMEM175 transmembrane protein 175	3623
0.236646381737899	3.41594585133836	1.12273614408747
	0.28933005331317	0.733478413962509
402415	XKRX	XK related X-linked	3372	-0.385005801179862
	0.970427763673618	1.12236301082636
0.289410205010556	0.733478413962509
4124	MAN2A1	mannosidase alpha class 2A member 1
11815	-0.216874623874881	7.22585492012739
1.12227707607558	0.289428668416883	0.733478413962509
1540	CYLD	CYLD lysine 63 deubiquitinase
12954	-0.170077324735323	7.25443627693365
1.12208322914586	0.2894703226725 0.733478413962509
7392	USF2	upstream transcription factor 2, c-fos interacting

2805 0.175378740756656 5.69639447037397
1.12205314943033 0.289476786952415 0.733478413962509
57732 ZFYVE28 zinc finger FYVE-type containing 28 8785
0.384740709733067 1.98505451725498 1.1219802071186
0.289492463380223 0.733478413962509
4982 TNFRSF11B TNF receptor superfamily member 11b 2827
0.636004934725375 1.65437295139741
1.12190836281114 0.289507904887036 0.733478413962509
113655 MFSD3 major facilitator superfamily domain containing 3
1823 0.230689052818301 3.58577558156423
1.12183559379667 0.289523546210742 0.733478413962509
6277 S100A6 S100 calcium binding protein A6 971
0.261095007617526 9.59116493926348 1.12180764377547
0.289529554209519 0.733478413962509
65110 UPF3A UPF3A regulator of nonsense mediated mRNA decay 5772
0.173456317094074 5.11231520384899
1.12140233513376 0.289616695209238 0.733478413962509
89890 KBTBD6 kelch repeat and BTB domain containing 6
5178 -0.241379514691662 3.03916077288623
1.12087832408202 0.28972940665017 0.733478413962509
10509 SEMA4B semaphorin 4B 6565 0.196815823763791
7.25378260634661 1.12075790956638 0.289755314941771
0.733478413962509
25801 GCA grancalcin 3664 0.183332716349969
3.65983706358595 1.12066637992557 0.289775010361243
0.733478413962509
9730 DCAF1 DDB1 and CUL4 associated factor 1
6849 -0.160395716175505 5.24749188324112
1.12060302345899 0.289788644454096 0.733478413962509
6223 RPS19 ribosomal protein S19 2440 0.208318040519167
8.43093533511613 1.12023441627349
0.289867983660855 0.733488068515682
79730 NSUN7 NOP2/Sun RNA methyltransferase family member 7
4728 -0.324144717529634 2.85686209073753
1.11995463165142 0.289928223129948 0.733488068515682
152330 CNTN4 contactin 4 9120 -0.938638454258887
2.42625896654567 1.1198555665598 0.289949556319792
0.733488068515682
112840 WDR89 WD repeat domain 89 3208 -0.216907779515225
3.26063753381836 1.11919462201727
0.290091938748006 0.733715743236426
26271 FBXO5 F-box protein 5 3833 0.307907018962285
1.81507074674357 1.11826691013854 0.290291939262746
0.734089040125164
3384 ICAM2 intercellular adhesion molecule 2 3243
0.418725832477307 0.793385444498652 1.11775067824101
0.290403307096697 0.734238108755133
58472 SQOR sulfide quinone oxidoreductase
3267 -0.264207573896417 6.90210560511765
1.11708359183194 0.290547299790869 0.734398985264306
9126 SMC3 structural maintenance of chromosomes 3
4275 -0.13420253017582 5.87055068655272
1.11687424479984 0.290592506770992 0.734398985264306
9931 HELZ helicase with zinc finger

15677	-0.220674044942683	6.96723542517565
1.11608974300407	0.290761994076787	0.734398985264306
3298	HSF2	heat shock transcription factor 2	2702
0.273394381151014	3.95213539017832	1.11590795740506
	0.290801285847169	0.734398985264306
2170	FABP3	fatty acid binding protein 3	993
0.647689508187691	0.0740623464069446	1.11584009003601
	0.29081595667195	0.734398985264306
400073	C12orf76	chromosome 12 open reading frame 76	3043
	0.292594428795101	1.92499751223915
1.11577534672572	0.290829953051745	0.734398985264306
8437	RASAL1	RAS protein activator like 1
4813	-0.553131078379129	1.20779760393921
1.11553697778983	0.290881491683308	0.734398985264306
3033	HADH	hydroxyacyl-CoA dehydrogenase	3055
0.205407132678061	5.96212133565422	1.11528170288799
	0.290936698535618	0.734398985264306
5228	PGF	placental growth factor 5962	0.387814951776237
	2.53999449530416	1.11523082252773
0.290947703739376	0.734398985264306
9755	TBKBP1	TBK1 binding protein 1	4280	0.26486650193652
	3.4279989165893 1.11499388018847	0.290998960346527
	0.734398985264306
23358	USP24	ubiquitin specific peptidase 24
11772	-0.199590840424157	6.82131784292361
1.11478721882768	0.291043675819388	0.734398985264306
93474	ZNF670	zinc finger protein 670 1994	-0.262207641938069
	2.26697020566869	1.11439061289545
0.291129514297197	0.734483268508796
284029	LINC00324	long intergenic non-protein coding RNA 324
	2082	0.366435452920018	0.411161354637562
1.11367232041708	0.291285058492475	0.734619100329206
928	CD9	CD9 molecule	3299	0.183110700188012
9.63064017031732	1.1136575987548 0.291288247535405
0.734619100329206
51303	FKBP11	FKBP prolyl isomerase 11
3599	-0.154632701813923	4.86648269381557
1.11238628686105	0.291563810410188	0.735181690803871
25825	BACE2	beta-secretase 2	9619	-0.209654975859132
	7.41821319025148	1.11187134002461
0.29167552225317	0.735331000755968
10301	DLEU1	deleted in lymphocytic leukemia 1	10603
0.288643296616905	1.65050688342035	1.11047860621102
	0.29197793383943	0.735960935628739
112939	NACC1	nucleus accumbens associated 1
4474	-0.124219578940209	6.88389152835464
1.11016972241979	0.292045057628285	0.735997683015364
23625	FAM89B	family with sequence similarity 89 member B	1617
	0.127519437560222	6.15716934656446
1.10963518794132	0.29216126431906	0.736030481886901
8706	B3GALNT1	beta-1,3-N-acetylgalactosaminyltransferase 1 (globoside blood group) 4183	0.212958275856257
3.87162092695456	1.10962643842349	0.292163166937329
	0.736030481886901

101927752	LOC101927752	uncharacterized LOC101927752	1322
	0.335813422638562	1.65169132017688
1.10924762188193	0.292245557313406	0.73610565029471
4688	NCF2	neutrophil cytosolic factor 2
2928	-0.224896480200864	7.97626915866075
1.10751263010243	0.292623287338416	0.736924556948793
90324	CCDC97	coiled-coil domain containing 97
3821	-0.153963247116892	5.17142107025361
1.10672968906278	0.29279394793434	0.737221791967696
23177	CEP68	centrosomal protein 68	8623	0.289219016763579
	3.78332421631363	1.10640125799125
0.292865575156442	0.737238934706722
113791	PIK3IP1	phosphoinositide-3-kinase interacting protein 1	3805
	0.205997363601655	6.52166407333514
1.10621583609267	0.292906023534155	0.737238934706722
145567	TTC7B	tetratricopeptide repeat domain 7B	21268
0.337075720932901	2.49972921357412	1.10486691766593
	0.293200494876451	0.737747054818384
11235	PDCD10	programmed cell death 10
3275	-0.150734783341249	5.95174128154574
1.10480857324445	0.293213240106657	0.737747054818384
4437	MSH3	mutS homolog 3	4709	-0.161997321307102
4.22245772724874	1.10408103996111	0.293372227904354
	0.737830296841093
9796	PHYHIP	phytanoyl-CoA 2-hydroxylase interacting protein	4170
	0.304028081256292	1.35640266345023
1.1037726834854	0.29343964628529	0.737830296841093
23531	MMD	monocyte to macrophage differentiation associated
	3177	-0.30088979228608	4.54125736194485
1.10369524929359	0.293456579438835	0.737830296841093
57649	PHF12	PHD finger protein 12	12227	-0.17606277593074
	5.46978450947513	1.10369320057733
0.293457027464962	0.737830296841093
60676	PAPPA2	pappalysin 2
10930	-0.408398363144786	-0.0586790011300345
1.10309007512099	0.293588960769327	0.738029535803296
1003	CDH5	cadherin 5	5972	-0.544929136268144
3.4578159849496	1.10284446171832	0.293642710280334
0.738032198617735
55796	MBNL3	muscleblind like splicing regulator 3
12199	-0.319557439501612	2.75081147619314
1.1020882250132	0.293808282563476	0.738216161780448
1266	CNN3	calponin 3	2838	-0.164586742556433
8.77554619375725	1.1020287423317	0.293821310910687
0.738216161780448
100528017	SAA2-SAA4	SAA2-SAA4 readthrough
743	-0.751968483521264	6.08768612321464
1.10149468100863	0.293938318180599	0.738377694181429
4267	CD99	CD99 molecule (Xg blood group)	3120
0.187673265775507	6.849307966617	1.10046932659336
0.294163129814635	0.738571022091011
25841	ABTB2	ankyrin repeat and BTB domain containing 2
4973	-0.362201684161231	5.08797323778957
1.09971776957809	0.294328050421716	0.738571022091011

25942 SIN3A SIN3 transcription regulator family member A
12453 -0.143141177048198 6.0438953507389 1.09927680546896
0.294424870054209 0.738571022091011
9277 WDR46 WD repeat domain 46 3115 0.141063437074052
5.4739294400373 1.09922834196656 0.294435513357885
0.738571022091011
8776 MTMR1 myotubularin related protein 1
6323 -0.146421014602847 5.98533915104615
1.0991914861569 0.294443607769846 0.738571022091011
8452 CUL3 cullin 3 11505 -0.116720017105907
6.67069180744641 1.09898804727908 0.294488292910145
0.738571022091011
348235 SKA2 spindle and kinetochore associated complex subunit 2
3148 -0.199161639224742 5.00667942486007
1.09872727312776 0.294545584381909 0.738571022091011
165324 UBXN2A UBX domain protein 2A 6884 -0.144852095308284
5.53654937856616 1.09852866370396
0.29458922798129 0.738571022091011
26027 ACOT11 acyl-CoA thioesterase 11
9768 -0.205849614165854 3.88663187267391
1.09837309360917 0.294623419649051 0.738571022091011
11173 ADAMTS7 ADAM metallopeptidase with thrombospondin type 1
motif 7 7730 0.321621195382469 4.53141876010868
1.09834841622742 0.294628843785854 0.738571022091011
10425 ARIH2 ariadne RBR E3 ubiquitin protein ligase 2 8017
0.116549416126498 6.52898973227887
1.09827615868283 0.294644726870903 0.738571022091011
84253 GARNL3 GTPase activating Rap/RanGAP domain like 3 9695
0.405628531628443 0.869881832816912
1.09826115978359 0.294648023948352 0.738571022091011
64863 METTL4 methyltransferase like 4
5597 -0.176561685039712 3.82600223626361
1.09711385812004 0.294900365370103 0.738894893290697
57176 VARS2 valyl-tRNA synthetase 2, mitochondrial 8448
0.185395841340197 5.48063834813241 1.09680627859351
0.294968062495938 0.738894893290697
9255 AIMP1 aminoacyl tRNA synthetase complex interacting
multifunctional protein 1 3328 -0.141912039546068
5.38389803062216 1.0967506276346 0.294980313194333
0.738894893290697
7184 HSP90B1 heat shock protein 90 beta family member 1
4688 -0.328150872185204 9.07364731939171
1.09671463145395 0.294988237544198 0.738894893290697
113878 DTX2 deltex E3 ubiquitin ligase 2
5042 -0.219340133487272 5.8926951846093 1.09613261734101
0.295116402423379 0.738921890286631
23589 CARHSP1 calcium regulated heat stable protein 1
5577 -0.137869467694101 5.62234804752044
1.09604698638371 0.295135265169537 0.738921890286631
23383 MAU2 MAU2 sister chromatid cohesion factor
11145 -0.130797949681674 6.24712808754041
1.09562353659681 0.29522856519231 0.738921890286631
152405 TEX55 testis expressed 55 1997 -0.781356672505059
1.1196694401344 1.09553122000202 0.29524891061465

0.738921890286631
51373 MRPS17 mitochondrial ribosomal protein S17 2159
0.22572007916924 3.43206440256868 1.095468273267
0.295262784317432 0.738921890286631
51382 ATP6V1D ATPase H+ transporting V1 subunit D
3931 -0.149853811442616 7.14770388456858
1.09412866548459 0.295558237187538 0.739463604936608
134637 ADAT2 adenosine deaminase tRNA specific 2 7175
0.324883088062534 1.6024214739911б 1.09391434122509
0.295605541933951 0.739463604936608
5695 PSMB7 proteasome 20S subunit beta 7
1787 -0.161307559900263 6.800373673177 1.09376901537617
0.295637623163062 0.739463604936608
100534599 ISY1-RAB43 ISY1-RAB43 readthrough
4868 -0.137170081516419 7.15861016939212
1.09339626112167 0.295719930475729 0.73948760272446
388327 C17orf100 chromosome 17 open reading frame 100 1756
0.350815203916119 0.59729892030353
1.09324741228195 0.295752805773001 0.73948760272446
55975 KLHL7 kelch like family member 7
7066 -0.120957538139877 6.01269860957037
1.09271709704554 0.295869971216585 0.739566236944602
54904 NSD3 nuclear receptor binding SET domain protein 3
15252 -0.176101182187727 6.87359121186405
1.09262712792474 0.295889854489723 0.739566236944602
10493 VAT1 vesicle amine transport 1 3501
0.166376514765937 8.99866777691821 1.09231869171245
0.295958032231609 0.739604666809661
401588 ZNF674-AS1 ZNF674 antisense RNA 1 (head to head) 2078
0.338103116084465 0.460684825586143
1.092019248722 0.296024241352882 0.73963816422223
255928 SYT14 synaptotagmin 14 12436 0.696635499894473
3.28933941485777 1.0910071519817 0.296248164268701
0.739995365712878
95681 CEP41 centrosomal protein 41 11491 0.175480911123206
4.1684837268072 1.09054471162395 0.296350549906995
0.739995365712878
6426 SRSF1 serine and arginine rich splicing factor 1
3720 -0.139769008141925 7.13191303368492
1.09045716848712 0.296369937320692 0.739995365712878
25788 RAD54B RAD54 homolog B 5771 0.257525287402101
2.3476675625724 1.08990126572197 0.296493086224172
0.739995365712878
401397 SMIM30 small integral membrane protein 30 1896
0.153590870861215 4.63621067594209 1.08960036507309
0.296559771999989 0.739995365712878
100130776 AGAP2-AS1 AGAP2 antisense RNA 1
1500 -0.305058383115961 1.76948979254649
1.08946659616041 0.296589424122366 0.739995365712878
2907 GRINA glutamate ionotropic receptor NMDA type subunit
associated protein 1 2527 0.164744334831204
7.04631663725172 1.08941725241596 0.296600362950997
0.739995365712878
10046 MAMLD1 mastermind like domain containing 1

5958	-0.45974435696439	2.45129875394488
1.08920404656384	0.296647633702994	0.739995365712878
11159	RABL2A	RAB, member of RAS oncogene family like 2A	6008
	0.193869979609743	4.05212067529648
1.08904691809911	0.296682477492408	0.739995365712878
51366	UBR5	ubiquitin protein ligase E3 component n-recognin 5
	13085	-0.190115855844824	7.2836426077109
1.08886513191223	0.2967227957708 0.739995365712878
54629	MINDY2	MINDY lysine 48 deubiquitinase 2
9454	-0.329697638343695	5.59889750214916
1.08836301893353	0.296834195722767	0.739995365712878
5411	PNN	pinin, desmosome associated protein
4201	-0.123948502469987	6.20197232640523
1.08812838372616	0.29688627083501	0.739995365712878
11337	GABARAP GABA type A receptor-associated protein 2351
0.155938765348699	7.45156038789337	1.08808823021104
	0.296895183710846	0.739995365712878
101928527	PIGBOS1 PIGB opposite strand 1	1632
0.246903558777033	2.36399427522056	1.088035768712
0.29690682910733	0.739995365712878
85378	TUBGCP6 tubulin gamma complex associated protein 6
7689	-0.183063927245807	5.92403555958199
1.08775009257842	0.296970253745968	0.740021765561249
84620	ST6GAL2 ST6 beta-galactoside alpha-2,6-sialyltransferase 2
	7708	0.689340054716546	1.91221575128818
1.08645554974441	0.297257880872086	0.740354559016291
723790	H2AC19	H2A clustered histone 19
1113	-0.247833711395135	4.23090755134726
1.08637341239944	0.297276142565462	0.740354559016291
6845	VAMP7	vesicle associated membrane protein 7
5334	-0.130656320187396	5.89373240588114
1.08597133183801	0.297365558400702	0.740354559016291
55506	MACROH2A2	macroH2A.2 histone	1980
0.184588409794556	4.97409416002363	1.08571960799916
	0.29742155505169	0.740354559016291
6242	RTKN	rhotekin	5005	-0.160865337734841
6.4544437910399 1.08557714327297	0.297453252716688
0.740354559016291
100129550	LINC02035	long intergenic non-protein coding
RNA 2035	5476	0.260617412163193	3.36413199187049
	1.08516299030541	0.297545424249981
0.740354559016291
9794	MAML1	mastermind like transcriptional coactivator 1
10690	-0.181207678012937	5.94501657456856
1.08486804175541	0.297611088688237	0.740354559016291
84798	C19orf48	chromosome 19 open reading frame 48	3256
	0.235921940583191	3.61148360650869
1.08484813194952	0.29761552188172	0.740354559016291
6288	SAA1	serum amyloid A1	765	-0.718604048044202
	6.88810673358722	1.08461818040433
0.297666729918927	0.740354559016291
5229	PGGT1B	protein geranylgeranyltransferase type I subunit
beta	10064	-0.174643651197045	4.69451174618001
1.08444983873787	0.297704225193045	0.740354559016291

6567	SLC16A2 solute carrier family 16 member 2
4161	-0.173605783687889	5.65344578032542
1.08431818091991	0.297733553984734	0.740354559016291
5481	PPID	peptidylprolyl isomerase D
2509	-0.152697221420858	4.83299487490975
1.08419027552991	0.297762050420024	0.740354559016291
54553	MIR4453HG	MIR4453 host gene
3000	-0.299240438163099	0.240071294270851
1.08406065716045	0.297790932069521	0.740354559016291
5500	PPP1CB	protein phosphatase 1 catalytic subunit beta
5760	-0.172289468954193	8.68189789669609
1.08371246796062	0.297868533633148	0.740416069316681
6498	SKIL	SKI like proto-oncogene 7782	-0.227631975303087
8.19158766245675	1.0826735111246 0.298100242315625
0.740837433187235
80346	REEP4	receptor accessory protein 4	2266
0.146491088595868	5.18774698814507	1.08247818968216
0.298143828862458	0.740837433187235
133522	PPARGC1B	PPARG coactivator 1 beta	12889
0.50417058809286	1.04948312844542	1.08084156854894
0.298509367650199	0.74149028280169
56980	PRDM10	PR/SET domain 10	6626	-0.225776715236738
4.51982272526797	1.08082783721286
0.298512436975707	0.74149028280169
7461	CLIP2	CAP-Gly domain containing linker protein 2
7315	-0.276031448831452	6.00405158159782
1.08019047321656	0.29865494975034	0.741712744885286
6907	TBL1X	transducin beta like 1 X-linked
7009	-0.243321088942674	4.16493186226279
1.07945468423943	0.298819578871841	0.741967403263249
57531	HACE1	HECT domain and ankyrin repeat containing E3
ubiquitin protein ligase 1	4775	0.186755479025238
3.82340395623223	1.079258777056	0.29886343179919
0.741967403263249
80705	TSGA10	testis specific 10	7166	0.318838291692054
2.2601091280288 1.07860053914863	0.299010835980048
0.742081157775223
9130	FAM50A	family with sequence similarity 50 member A	2567
0.182205492082504	6.13298880955894
1.07858101065003	0.299015210572097	0.742081157775223
5970	RELA	RELA proto-oncogene, NF-kB subunit
5464	-0.132383018558077	6.94450842645602
1.07794780339646	0.299157100373572	0.742207374791501
92345	NAF1	nuclear assembly factor 1 ribonucleoprotein
4808	-0.242755528366236	2.61600229071751
1.07779975184105	0.299190288426233	0.742207374791501
55304	SPTLC3	serine palmitoyltransferase long chain base subunit
3	8520	0.316038200571892	1.90095907873192
1.07764477667947	0.29922503358607	0.742207374791501
81627	TRMT1L	tRNA methyltransferase 1 like
4945	-0.167872260498074	4.92146573454012
1.07711224825408	0.299344465101531	0.742372153448503
79650	USB1	U6 snRNA biogenesis phosphodiesterase 1
7298	-0.14044110160817	6.14458963992793

1.07644732663475	0.299493674892956	0.742553340436043
26119	LDLRAP1 low density lipoprotein receptor adaptor protein 1
	5397	-0.14788520841117	5.44284489506337
1.07631424306884	0.299523550614953	0.742553340436043
5902	RANBP1	RAN binding protein 1	4622	0.185546108544548
	4.77054932950639	1.07547864028626
0.299711221447712	0.742887113576023
54978	SLC35F6 solute carrier family 35 member F6
3902	-0.130877517165356	6.25020335048826
1.07445637932123	0.299941020361358	0.74332517201018
4192	MDK	midkine 2196	0.143178432845357
6.86994556081288	1.07396502626191	0.300051554672331
	0.743467561700927
100533105	C8orf44-SGK3	C8orf44-SGK3 readthrough	3398
	-0.205125351176156	3.31560867675003
1.07307697409481	0.300251463129852	0.743732124331314
4603	MYBL1	MYB proto-oncogene like 1	5720
0.437357748790866	0.114223733155906	1.07298087921982
	0.300273105225215	0.743732124331314
682	BSG	basigin (Ok blood group)	3146
0.12550888184884	8.65241178086918	1.07278325606386
	0.300317619420141	0.743732124331314
5305	PIP4K2A phosphatidylinositol-5-phosphate 4-kinase type 2
alpha	4530	0.185913685290379	4.26268915213848
1.07164786801768	0.300573527920078	0.744234294781075
3339	HSPG2	heparan sulfate proteoglycan 2
17684	-0.201093936216413	9.50783784266708
1.06940111632285	0.301080758613175	0.74528061107353
6811	STX5	syntaxin 5	2257	-0.14512907437636
5.11168247126837	1.06930484072983	0.301102518602492
	0.74528061107353
85019	TMEM241 transmembrane protein 241	4548
0.242561843179546	2.88367275202659	1.06899733319658
	0.301172034330943	0.745320969058926
10742	RAI2	retinoic acid induced 2 2622	0.604248212021649
	2.37056590187398	1.06850918504675
0.301282428509392	0.74546245824608
975	CD81	CD81 molecule	7394	0.17089775878968
8.15391589532499	1.06808576517882	0.301378226710151
	0.745567789037281
339929	LPP-AS2 LPP antisense RNA 2	2852	0.268356274945662
	1.35912000147752	1.06758932734059
0.301490595156785	0.745714067872344
51320	MEX3C	mex-3 RNA binding family member C
3824	-0.160500415968869	6.63390796492155
1.06731294916658	0.301553176623732	0.745737172487397
100887750	MRPS31P5	mitochondrial ribosomal protein S31
pseudogene 5	3796	0.247140706326408	2.32954896367161
	1.06677471713382	0.30167509884314
0.745860579902869
6919	TCEA2	transcription elongation factor A2	4802
0.240456908023285	3.39664248050636	1.06657708139081
	0.301719883947209	0.745860579902869
147372	CCBE1	collagen and calcium binding EGF domains 1	6577

0.766869584878012	2.18983133235109
1.06638761372937	0.30176282618045	0.745860579902869
1066	CES1	carboxylesterase 1	2937	1.03756514577417
2.87966415475371	1.06503258072712
0.302070169966455	0.746488509301365
29980	DONSON	downstream neighbor of SON
3050	-0.218371518223229	3.00710618940318
1.0647717486295	0.302129377324126	0.746503120158587
7798	LUZP1	leucine zipper protein 1
9658	-0.158525432457808	7.88768699489797
1.06425686874283	0.302246296003837	0.746660294202072
144132	DNHD1	dynein heavy chain domain 1
19629	-0.425394119372166	3.10377737104415
1.06393894838045	0.30231851836678	0.74670701582851
64788	LMF1	lipase maturation factor 1	11055
0.202188824116079	3.9677577913078	1.06327018769341
0.302470514282409	0.746886037711652
991	CDC20	cell division cycle 20	2038	0.36914960129348
0.208854677754057	1.06298306841673
0.30253580101095	0.746886037711652
81689	ISCA1	iron-sulfur cluster assembly 1	3050
0.151340897110461	4.88808429583158	1.06265253200173
0.302610982689591	0.746886037711652
56603	CYP26B1	cytochrome P450 family 26 subfamily B member 1	5015
0.331206656524541	2.17916713144602
1.06246517194595	0.30265360910901	0.746886037711652
387921	NHLRC3	NHL repeat containing 3	6409	0.205653157102031
4.24066965399458	1.06243111119463
0.302661359127455	0.746886037711652
57478	USP31	ubiquitin specific peptidase 31
12017	-0.167565629729537	6.20084049589444
1.06204732507358	0.302748701674693	0.746886037711652
7804	LRP8	LDL receptor related protein 8
10258	-0.275046841296637	3.3779314897386	1.06168762978665
0.302830591145317	0.746886037711652
5673	PSG5	pregnancy specific beta-1-glycoprotein 5	4678
0.606813536771549	4.35988161119529
1.06150768249634	0.302871569300279	0.746886037711652
10587	TXNRD2	thioredoxin reductase 2	8373	0.181446461171109
4.30555736793347	1.06122239234251
0.302936551135544	0.746886037711652
85407	NKD1	NKD inhibitor of WNT signaling pathway 1	8915
0.509414579054725	1.52837786314284
1.06089553867341	0.303011022214576	0.746886037711652
59307	SIGIRR	single Ig and TIR domain containing
3496	-0.198827836088529	4.77635541007171
1.06064494539698	0.303068133967028	0.746886037711652
23194	FBXL7	F-box and leucine rich repeat protein 7	4854
0.462809851815799	3.4106969804095	1.06057124595649
0.303084933169566	0.746886037711652
10671	DCTN6	dynactin subunit 6	1984	-0.151644297451653
5.0729453112349	1.06054187817994	0.30309162765628
0.746886037711652
3292	HSD17B1	hydroxysteroid 17-beta dehydrogenase 1	5658

0.281523961964862	3.31851446389801	1.0603406144722
0.303137511557846	0.746886037711652
101929331	LOC101929331	uncharacterized LOC101929331	583
	0.37145205284746	0.437025867206409
1.05994632333824	0.303227427672201	0.746905324161262
221468	TMEM217	transmembrane protein 217
2737	-0.193730133501853	4.77067405905574
1.05969941257128	0.303283751977141	0.746905324161262
23037	PDZD2	PDZ domain containing 2	15642	-0.348951912332305
	4.57381494939654	1.05960492152687
0.303305310475424	0.746905324161262
29785	CYP2S1	cytochrome P450 family 2 subfamily S member 1
2874	-0.47105515137115	5.23992295382411
1.05918254415273	0.303401701720567	0.747011361574966
5238	PGM3	phosphoglucomutase 3	7224	-0.214152903348893
	6.02788985328786	1.05798570518678
0.303675048846421	0.747552971738457
84932	RAB2B	RAB2B, member RAS oncogene family
3610	-0.140969037982685	5.09387705685578
1.05766433746977	0.303748500506126	0.747602397924671
116461	TSEN15	tRNA splicing endonuclease subunit 15	2443
0.17120389430787	4.28387688786251	1.05668997786252
	0.303971340107587	0.748019423908462
5563	PRKAA2	protein kinase AMP-activated catalytic subunit alpha
2	9354	-0.202798150871878	3.4315083701464
1.05639862342227	0.304038014976951	0.748052077249631
7621	ZNF70	zinc finger protein 70	6924	0.26220884842384
	2.32200239735917	1.0557399710322 0.304188813680002
	0.748291660206495
64778	FNDC3B	fibronectin type III domain containing 3B
13095	-0.152467794061302	8.6023752407416 1.05525267983404
	0.304300441427399	0.748434817045405
2673	GFPT1	glutamine--fructose-6-phosphate transaminase 1
8866	-0.181801735439036	7.28303174163828
1.05488373107218	0.304384994755947	0.748511345074885
81544	GDPD5	glycerophosphodiester phosphodiesterase domain
containing 5	9867	-0.265951792017858	3.36334314145881
	1.05457641541597	0.304455446599866
0.748512387339153
64748	PLPPR2	phospholipid phosphatase related 2
3273	-0.244220984817278	3.89260638445813
1.05441571468673	0.304492295499286	0.748512387339153
56169	GSDMC	gasdermin C	2934	-0.61379899946315
2.28477228410043	1.05401836334863	0.304583433482238
	0.748605045233499
9772	TMEM94	transmembrane protein 94	6409
0.181770694681283	5.7992593441857 1.0535780091864
0.304684475982545	0.748665119236586
8797	TNFRSF10A	TNF receptor superfamily member 10a
2951	-0.18951982695054	5.90660290153693
1.05314043065239	0.304784924557721	0.748665119236586
58528	RRAGD	Ras related GTP binding D	5072
0.326715575546417	3.63744936924521	1.05275122654746
	0.304874304523907	0.748665119236586

55049	REX1BD	required for excision 1-B domain containing	2968
	0.237737132193751	3.12535651335015
1.05267804975241	0.304891113220596	0.748665119236586
11247	NXPH4	neurexophilin 4 2150	0.533223842391283
0.908721977376593	1.0524617202167 0.304940811089786
0.748665119236586
8227	AKAP17A A-kinase anchoring protein 17A
8150	-0.165649111449451	5.80316555761319
1.05235487589328	0.304965360543318	0.748665119236586
493856	CISD2	CDGSH iron sulfur domain 2	2607
0.170448586861607	4.45846368417358	1.05228237177497
	0.304982021158275	0.748665119236586
203111	ERICH5	glutamate rich 5	1761	0.23655051067283
	3.07382991910232	1.05188527080193
0.305073291305471	0.748727541547266
100506649	PXN-AS1 PXN antisense RNA 1	2332
0.317038570070667	0.566592789230992	1.05168011205079
	0.305120459080918	0.748727541547266
79763	ISOC2	isochorismatase domain containing 2	2969
0.179229442086975	4.03198902804447	1.05121113074012
	0.305228317408843	0.748727541547266
79660	PPP1R3B protein phosphatase 1 regulatory subunit 3B	5776
	0.284944111419336	4.98611083634637
1.05086320749488	0.305308366177467	0.748727541547266
51706	CYB5R1	cytochrome b5 reductase 1
4504	-0.178825307946491	7.64093667463591
1.0507864858759 0.305326021633594	0.748727541547266
342892	ZNF850	zinc finger protein 850 7850	-0.271321499641837
	1.52280138093243	1.05077713811321
0.305328172864852	0.748727541547266
80862	ZNRD1ASP	zinc ribbon domain containing 1 antisense,
pseudogene	19531	0.339866346248001	1.9227678688325
1.04931066128658	0.305665901170627	0.749153995488833
84270	CARD19	caspase recruitment domain family member 19	3081
	0.182225669740942	5.01836922425421
1.04924136052859	0.305681873041411	0.749153995488833
101929125	LINC01730	long intergenic non-protein coding
RNA 1730	1193	0.29841148119835	0.715136518189631
	1.0491403428392 0.305705156706853	0.749153995488833
23322	RPGRIP1L	RPGRIP1 like	12857	0.277035613244157
	2.73554027096262	1.04871657358924
0.305802856727461	0.749153995488833
64949	MRPS26	mitochondrial ribosomal protein S26	1127
0.153942964456873	4.21244518424595	1.04840389527432
	0.3058749706597 0.749153995488833
339942	H1FX-AS1	H1FX antisense RNA 1	3420
0.358484335982161	0.849047248153658	1.04826198709752
	0.305907706633513	0.749153995488833
10010	TANK	TRAF family member associated NFKB activator
5003	-0.118452521127957	6.11998572566924
1.04820098209239	0.305921780919099	0.749153995488833
6683	SPAST	spastin 5213	-0.170808053625146
4.99906150370319	1.04816556108655	0.305929953180276
	0.749153995488833

25948	KBTBD2	kelch repeat and BTB domain containing 2
4987	-0.14787280099704	5.8588203177074 1.04770225678297
	0.306036871838979	0.74928482150823
100190939	TPT1-AS1	TPT1 antisense RNA 1
10402	-0.240891595356791	3.35763820860707
1.0469190343567 0.306217729462397	0.749596598493498
57189	DENND11 DENN domain containing 11
7356	-0.185890082798206	6.59862969867247
1.04626141527707	0.306369690400205	0.749740513064936
7284	TUFM	Tu translation elongation factor, mitochondrial 2271
	0.134877837908656	7.17217667071299
1.04620136181798	0.306383572269843	0.749740513064936
64172	OSGEPL1 O-sialoglycoprotein endopeptidase like 1	3583
	0.280844146057537	2.08355936141148
1.04553370383995	0.306537962043428	0.749843034356641
55030	FBXO34	F-box protein 34	3809	-0.132812831045103
	5.68652874382683	1.04535613538564
0.30657904009377	0.749843034356641
834	CASP1	caspase 1	3698	0.426562217837535
2.89299973505008	1.04516872072846	0.30662240367056
	0.749843034356641
10079	ATP9A	ATPase phospholipid transporting 9A (putative)
8317	-0.173949821846669	7.44327800853383
1.04509440150775	0.306639601684504	0.749843034356641
112970	KTI12	KTI12 chromatin associated homolog	1714
0.211314483256726	3.41258212377338	1.04425772189098
	0.30683330182089	0.750185731996021
3981	LIG4	DNA ligase 4	4387	-0.131031868575317
5.62316787858734	1.04384585151112	0.306928712424872
	0.750288041000904
100529207	RAD51L3-RFFL	RAD51L3-RFFL readthrough	2197
	-0.240059966104495	3.02281489194451
1.04355952193953	0.306995063919757	0.750319291628693
57609	DIP2B	disco interacting protein 2 homolog B
10361	-0.195162260915324	7.68082091062612
1.04329334508977	0.30705676208651	0.750339160248735
9445	ITM2B	integral membrane protein 2B	10208
0.124535667232461	8.52423519299088	1.0430459997652
0.30711410963806	0.750348392412403
8089	YEATS4	YEATS domain containing 4	1714
0.22072860537066	3.39350637037028	1.04262289215881
	0.307212240270229	0.750350123858351
171546	SPTSSA	serine palmitoyltransferase small subunit A
2777	-0.175360037430469	4.14930729259583
1.04258100364095	0.307221957616024	0.750350123858351
80003	PCNX2	pecanex 2	13815	0.215402987428872
4.59494906726459	1.04212114608962	0.30732866211339
	0.750378177356265
392	ARHGAP1 Rho GTPase activating protein 1 6138
0.140435010329566	6.47336465885084	1.04206976661078
	0.3073405870988 0.750378177356265
126295	ZNF57	zinc finger protein 57	2327	0.242391082123377
	2.60622179490028	1.04177152600906
0.307409819488013	0.750379876192923

148103	ZNF599	zinc finger protein 599 6796	0.297278394663544
	1.55103018760536	1.04127580311548
0.307524939410105	0.750379876192923
8239	USP9X	ubiquitin specific peptidase 9 X-linked
13962	-0.146626352488653	7.93521110896755
1.04110819970157	0.307563873994669	0.750379876192923
5613	PRKX	protein kinase X-linked 6452	-0.171314387853345
	5.44242630708123	1.04076122836607
0.307644496412711	0.750379876192923
378708	CENPS	centromere protein S	1739	0.279515742653181
	2.84810708330667	1.04073548174071
0.307650480003421	0.750379876192923
152485	ZNF827	zinc finger protein 827 8433	-0.203072797741238
	5.20929953077826	1.04068284429106
0.307662713570069	0.750379876192923
84062	DTNBP1	dystrobrevin binding protein 1	3052
0.20058159766316	3.4594154606964 1.03983997157292
0.307858693204352	0.750591756395622
656	BMP8B	bone morphogenetic protein 8b
5143	-0.410492666004465	1.15168952677752
1.03971648869798	0.307887418298736	0.750591756395622
90141	EFCAB11 EF-hand calcium binding domain 11	7968
0.249267797248109	2.68826700799196	1.03921675680882
	0.308003703527739	0.750591756395622
124935	SLC43A2 solute carrier family 43 member 2	10606
0.433217171189941	3.93580819898588	1.03916298072208
	0.308016220364738	0.750591756395622
9096	TBX18	T-box transcription factor 18	7631
0.688257282745199	0.576687177401493	1.03901072043235
	0.308051663811669	0.750591756395622
7125	TNNC2	troponin C2, fast skeletal type
876	-0.54240277172248	-0.0558809597958761
1.03892720115754	0.308071107840254	0.750591756395622
11162	NUDT6	nudix hydrolase 6	6216	0.356867432376475
	0.788114823332245	1.03830560940908
0.30821586999557	0.750708668164817
84939	PWWP3A	PWWP domain containing 3A, DNA repair factor	9906
	0.155813936394833	5.36622206228704
1.03766316409359	0.308365581573237	0.750708668164817
9736	USP34	ubiquitin specific peptidase 34
18125	-0.208242261962703	7.69285449730111
1.03747488510976	0.308409474866075	0.750708668164817
55629	PNRC2	proline rich nuclear receptor coactivator 2
2564	-0.140438812488492	5.98861589920516
1.03714434322139	0.308486553400832	0.750708668164817
51400	PPME1	protein phosphatase methylesterase 1	7917
0.136871044165868	6.22015245918636	1.03700525397941
	0.308518994879056	0.750708668164817
30815	ST6GALNAC6	ST6 N-acetylgalactosaminide alpha-2,6-
sialyltransferase 6	3115	0.183673345038685
5.62681864589021	1.0369511343863 0.308531619048724
0.750708668164817
8661	EIF3A	eukaryotic translation initiation factor 3 subunit A
	7101	-0.130972362123589	8.45002002844244

1.03613640176549	0.308721748216145	0.750708668164817
100049716	NINJ2-AS1	NINJ2 antisense RNA 1	3530
0.321886274038584	1.63858843625113	1.03591312581734
	0.308773879332144	0.750708668164817
139231	FAM199X	family with sequence similarity 199, X-linked
7457	-0.154911914941735	5.96922206984923
1.03589629840459	0.308777808708783	0.750708668164817
4201	MEA1	male-enhanced antigen 1	1089	0.163244940481499
	5.62492296821925	1.03580886451226
0.308798226481828	0.750708668164817
7260	EIPR1	EARP complex and GARP complex interacting protein 1
	5775	-0.16935408012703	4.81360935864709
1.03580164090819	0.308799913433701	0.750708668164817
57140	RNPEPL1	arginyl aminopeptidase like 1
7767	-0.138585194217113	6.34026860651938
1.03578450457221	0.308803915386161	0.750708668164817
5784	PTPN14	protein tyrosine phosphatase non-receptor type 14
	18756	-0.21563196551858	7.1834465121057
1.03527617930101	0.308922658327523	0.750708668164817
23451	SF3B1	splicing factor 3b subunit 1
9164	-0.129548243244756	8.4481626981612	1.03504195875846
	0.308977391367257	0.750708668164817
7570	ZNF22	zinc finger protein 22	2551	0.207076164060146
	3.90822879788637	1.03495092280676
0.308998668198701	0.750708668164817
53339	BTBD1	BTB domain containing 1	4169	-0.121809125005227
	6.71907014880326	1.03476948211421
0.309041080015916	0.750708668164817
2729	GCLC	glutamate-cysteine ligase catalytic subunit	7934
	0.119079306016303	5.84292227278771
1.03443742085362	0.309118719034068	0.750708668164817
83715	ESPN	espin	4615	0.292502330609286
3.06214121385805	1.03436097523248	0.309136596323042
	0.750708668164817
4928	NUP98	nucleoporin 98	9905	-0.12286194029749
7.06268875646195	1.03435752769674	0.309137402582613
	0.750708668164817
6653	SORL1	sortilin related receptor 1	18789
0.265845618973374	5.69204783611233	1.03313612057613
	0.309423219668683	0.751272497469101
90993	CREB3L1	cAMP responsive element binding protein 3 like 1
	4037	-0.396064089425093	3.21379776638842
1.03257029386423	0.309555743198426	0.751455400156121
51479	ANKFY1	ankyrin repeat and FYVE domain containing 1
9729	-0.127186209219789	6.75744515191415
1.03232165056534	0.30961400184151	0.751455400156121
285343	TCAIM	T cell activation inhibitor, mitochondrial
8743	-0.139357845097957	4.97901119126425
1.03200304866684	0.309688673055973	0.751455400156121
4835	NQO2	N-ribosyldihydronicotinamide:quinone reductase 2
	11558	0.213183429449465	3.38122574628062
1.03168876848856	0.309762354332916	0.751455400156121
2072	ERCC4	ERCC excision repair 4, endonuclease catalytic
subunit	8460	-0.175672896240878	4.31914162506776

1.03166983556571	0.309766793781783	0.751455400156121
51098	IFT52	intraflagellar transport 52	2542
0.168776107149188	4.29924932740984	1.03106148503185
0.309909485716927	0.751671370811601
57187	THOC2	THO complex 2	10277	-0.132106935946863
6.22129932886827	1.03030356158715	0.310087380453955
0.751853023465469
23424	TDRD7	tudor domain containing 7
4320	-0.140081863172585	5.30389805325835
1.03018263539891	0.310115775732051	0.751853023465469
84334	COA8	cytochrome c oxidase assembly factor 8	3932
0.15343568449322	5.28061721792888	1.03005644456188
0.310145410836259	0.751853023465469
23505	TMEM131 transmembrane protein 131	12985
0.120265436984994	6.74650928604513	1.02957799840776
0.310257804310565	0.751995339155231
2950	GSTP1	glutathione S-transferase pi 1	1915
0.186717900601468	9.29709712987078	1.0292261275902
0.310340497355558	0.752065630875312
2806	GOT2	glutamic-oxaloacetic transaminase 2	3766
0.153351106993167	7.23599455887986	1.02820924751681
0.310579635314839	0.752315598400489
387119	CEP85L	centrosomal protein 85 like
9103	-0.303949599347021	2.29622251596428
1.0281486361713 0.310593896756117	0.752315598400489
84687	PPP1R9B protein phosphatase 1 regulatory subunit 9B	4298
0.147350751834232	6.57005648199559
1.0277719300835 0.310682552280169	0.752315598400489
4238	MFAP3	microfibril associated protein 3
3945	-0.138576679679514	5.64734733631667
1.02767144327708	0.310706206815487	0.752315598400489
389834	LOC389834	ankyrin repeat domain 57 pseudogene
3027	-0.655437922747907	0.29759932148405
1.02764599997252	0.31071219652651	0.752315598400489
5587	PRKD1	protein kinase D1	5187	0.326111762430673
2.30072713563364	1.02724988975869
0.310805465825056	0.752353558252029
102465978	MIR6850 microRNA 6850	61	0.302724053906834
1.68127783490477	1.0270079245007 0.310862457628028
0.752353558252029
7321	UBE2D1	ubiquitin conjugating enzyme E2 D1
2974	-0.177600890409648	5.2548110340364 1.02689520335097
0.310889012291193	0.752353558252029
10750	GRAP	GRB2 related adaptor protein
3107	-0.408769652439845	-0.0233463186263973
1.02650229990777	0.310981594936712	0.752447607579638
23174	ZCCHC14 zinc finger CCHC-type containing 14
7390	-0.259783052053341	5.30741936635573
1.02608235962406	0.311080588172073	0.752557132733373
56101	PCDHGB5 protocadherin gamma subfamily B, 5	4851
0.169677407810636	6.73978355508964	1.02563874710238
0.311185206292511	0.752593651109967
4150	MAZ	MYC associated zinc finger protein	3763
0.131471122620279	6.99381470769073	1.02513473302585

0.311304124673652	0.752593651109967
84277	DNAJC30 DnaJ heat shock protein family (Hsp40) member C30
	2485	0.214675772302467	3.27814283021817
1.02479684270013	0.311383880542163	0.752593651109967
6622	SNCA	synuclein alpha 4014	0.353989377896232
3.04219462899737	1.02465384687678	0.311417641394216
	0.752593651109967
7296	TXNRD1	thioredoxin reductase 1 6708	-0.149810649337606
	7.37517680146034	1.0245929112281l
0.311432029553408	0.752593651109967
100505636	MALINC1 mitosis associated long intergenic non-coding RNA 1	3439	-0.274614755716938	0.815341296401011
	1.02435489113992	0.311488239289786
0.752593651109967
129138	ANKRD54 ankyrin repeat domain 54	3729
0.17207904060354	4.38830020772502	1.02434787576812
	0.311489896208498	0.752593651109967
2971	GTF3A	general transcription factor IIIA	3001
0.153779489096008	5.31726315291643	1.02392875151427
	0.311588907505074	0.752593651109967
9103	FCGR2C	Fc fragment of IgG receptor IIc (gene/pseudogene)
	3521	1.05875769961654	0.901582238989554
1.0238686806523 0.31160310163085	0.752593651109967
9056	SLC7A7	solute carrier family 7 member 7	5131
0.54415376212854	3.16377542283462	1.02374223653896
	0.311632981826073	0.752593651109967
124512	METTL23 methyltransferase like 23	2155
0.172747608084587	3.89872029859109	1.02172918854549
	0.312109191652744	0.753496318463423
133746	JMY	junction mediating and regulatory protein, p53 cofactor	9042	-0.276415550095639	5.27482134342323
	1.02152106245784	0.31215848039403
0.753496318463423
1465	CSRP1	cysteine and glycine rich protein 1
10318	-0.247773757811289	8.429701997542	1.02111744920933
	0.312254093640583	0.753496318463423
51292	GMPR2	guanosine monophosphate reductase 2	3640
0.143314322277766	5.6170709034711 1.02105706783532
0.312268400863633	0.753496318463423
2987	GUK1	guanylate kinase 1	4725	-0.143381215876686
	7.23222404791251	1.02086033798539
0.312315021465325	0.753496318463423
7581	ZNF33A	zinc finger protein 33A 7523	-0.175004083571922
	5.70382384256601	1.02052750321812
0.312393916581777	0.753496318463423
84056	KATNAL1 katanin catalytic subunit A1 like 1
8105	-0.167164080696939	5.2801707889824 1.02038008137797
	0.312428869745943	0.753496318463423
51639	SF3B6	splicing factor 3b subunit 6	1197
0.154013876193189	5.48642169547523	1.01992591418703
	0.312536583153445	0.753496318463423
728294	D2HGDH	D-2-hydroxyglutarate dehydrogenase	5279
0.300285044678778	3.82029411907385	1.01987894300205
	0.312547725926134	0.753496318463423

57460	PPM1H	protein phosphatase, Mg2+/Mn2+ dependent 1H
7063	-0.303556003135467	4.90809080363956
1.01961261361375	0.312610915906303	0.753496318463423
114823	LENG8	leukocyte receptor cluster member 8
6469	-0.167799827733881	7.52063471812748
1.01952341918716	0.312632082126043	0.753496318463423
51776	MAP3K20 mitogen-activated protein kinase kinase kinase 20
	13799	-0.377338960017676	5.2120155300684
1.01925929478391	0.312694770947086	0.753496318463423
11238	CA5B	carbonic anhydrase 5B	8520	0.263536811184646
	3.00205630462844	1.01898027281639
0.312761013460018	0.753496318463423
9581	PREPL	prolyl endopeptidase like
8153	-0.141491149370534	5.45450504353765
1.01883882200003	0.312794602246356	0.753496318463423
6609	SMPD1	sphingomyelin phosphodiesterase 1	2750
0.186171733323915	5.02695986546801	1.01875852685744
	0.312813671154766	0.753496318463423
8148	TAF15	TATA-box binding protein associated factor 15	3905
	0.131832390395459	6.63209356128562
1.01833342085227	0.312914652823775	0.753602059026272
7936	NELFE	negative elongation factor complex member E	2719
	0.149844623191064	5.59281910208388
1.01798186187798	0.312998195930738	0.753602059026272
113263	GLCCI1	glucocorticoid induced 1
6885	-0.201994619164032	4.61317831422133
1.01767072440516	0.313072157715773	0.753602059026272
51060	TXNDC12 thioredoxin domain containing 12	3630
0.13948967050155	5.34668567494435	1.01758608205452
	0.31309228234889	0.753602059026272
157697	ERICH1	glutamate rich 1	9370	-0.168363945130101
	4.21886504538001	1.01744185242535
0.313126578391726	0.753602059026272
5255	PHKA1	phosphorylase kinase regulatory subunit alpha 1
6350	-0.221808762404318	3.92494264867571
1.01711464708171	0.313204402008713	0.75365986238379
164091	PAQR7	progestin and adipoQ receptor family member 7	3297
	0.157983980029789	6.2420351578116 1.01617396403569
	0.313428277928895	0.754069029362768
55294	FBXW7	F-box and WD repeat domain containing 7
10287	-0.156701177861637	4.56308978607873
1.01528876177561	0.3136391406985 0.754446753179442
23518	R3HDM1	R3H domain containing 1 7898	0.156430152879845
	4.75034754225565	1.01409125004841
0.313924693254888	0.755003979811334
122011	CSNK1A1L	casein kinase 1 alpha 1 like
2406	-0.282735767263868	2.13411830700039
1.01342736814553	0.314083145516791	0.755195904306181
5424	POLD1	DNA polymerase delta 1, catalytic subunit	4858
	0.241945668768697	3.55966242790691
1.01330515925815	0.314112325157979	0.755195904306181
54938	SARS2	seryl-tRNA synthetase 2, mitochondrial	3706
0.178889412548537	4.15714148365713	1.0124127485253
0.31432512226973	0.755490510831083

65018	PINK1	PTEN induced kinase 1	4836	0.178648662982089
	7.19047398549623	1.01227753955959
0.314357828651156	0.755490510831083
9173	IL1RL1	interleukin 1 receptor like 1
5746	-0.860227201895078	2.02773917315397
1.01211504570492	0.314396672209208	0.755490510831083
641339	ZNF674	zinc finger protein 674 6203	-0.238385162502134
	2.64013432892165	1.01157535037574
0.314525729428241	0.755623098756889
445582	POTEE	POTE ankyrin domain family member E
4192	-0.324813765589709	2.01672284984824
1.01123880550155	0.31460624238917	0.755623098756889
10054	UBA2	ubiquitin like modifier activating enzyme 2
4815	-0.128524372222016	6.35133936455443
1.01119133144451	0.314617601964209	0.755623098756889
23339	VPS39	VPS39 subunit of HOPS complex
7710	-0.118678194365606	6.63466212173268
1.01098226799131	0.314667632972723	0.755623098756889
255738	PCSK9	proprotein convertase subtilisin/kexin type 9	4981
	0.511895716807845	3.49073280594394
1.00936904927531	0.315054042726647	0.75630472257898
23126	POGZ	pogo transposable element derived with ZNF domain
	10191	-0.141241994220625	6.04925456069554
1.00934639157486	0.315059474280599	0.75630472257898
342510	CD300E	CD300e molecule 3752	0.832316572399652
0.586242439059246	1.00804662894156	0.315371261447087
	0.756469769233422
130502	TTC32	tetratricopeptide repeat domain 32	1986
0.362423472736626	1.18962837137468	1.0077809853064
0.315435033747425	0.756469769233422
25818	KLK5	kallikrein related peptidase 5
1653	-0.615741594503859	5.6565443130244 1.00771428442994
	0.315451049084894	0.756469769233422
80349	WDR61	WD repeat domain 61	4670	0.162422229268393
	4.6923717909793 1.00742579654801	0.315520329258304
	0.756469769233422
90874	ZNF697	zinc finger protein 697 5041	-0.295409672467067
	3.87507344850344	1.00730544573884
0.315549237312007	0.756469769233422
145694	LOC145694	uncharacterized LOC145694	2265
0.383132044472746	1.86737983177678	1.00719236351137
	0.31557640262406	0.756469769233422
254048	UBN2	ubinuclein 2	16386	-0.312737420979911
5.24339278525371	1.00693800052039	0.315637518462531
	0.756469769233422
222255	ATXN7L1 ataxin 7 like 1 8295	-0.275363255993706
1.95685515083348	1.00693704348098	0.315637748439803
	0.756469769233422
7132	TNFRSF1A	TNF receptor superfamily member 1A	4005
	0.1144661220517 7.28853744951909	1.00688774776544
	0.315649594533096	0.756469769233422
8669	EIF3J	eukaryotic translation initiation factor 3 subunit J
	3195	-0.173718516461376	5.9326706161511
1.00680993559379	0.315682945077 0.756469769233422

128387	TATDN3	TatD DNase domain containing 3	3302
0.233428946808493	2.64335277450876	1.00571545002655
	0.315931477174871	0.756774028652708
84265	POLR3GL	RNA polymerase III subunit G like	2512
0.242185731484562	2.57434859414944	1.00569109348824
	0.315937337274607	0.756774028652708
64114	TMBIM1	transmembrane BAX inhibitor motif containing 1
2407	-0.142209593281398	9.95284225637872
1.00554700478	0.31597200743626	0.756774028652708
26953	RANBP6	RAN binding protein 6	4583	-0.123598307936017
	5.8893310385695	1.00530767756007	0.316029604592937
	0.756774028652708
80235	PIGZ	phosphatidylinositol glycan anchor biosynthesis
class Z	3211	0.273558505486074	2.78006122551959
1.00514940680638	0.316067702027068	0.756774028652708
57713	SFMBT2	Scm like with four mbt domains 2	9953
0.561580085787544	0.98196286293188	1.00456684065522
	0.31620798364919	0.756774028652708
23431	AP4E1	adaptor related protein complex 4 subunit epsilon 1
	6872	-0.181501311844591	5.87354123419592
1.00408719865801	0.316323542405727	0.756774028652708
4012	LNPEP	leucyl and cystinyl aminopeptidase
13604	-0.172395054036755	6.78517120133682
1.00396865667945	0.316352110906307	0.756774028652708
29123	ANKRD11	ankyrin repeat domain 11
15355	-0.168469143908436	7.66712184553593
1.00374509829765	0.316405997455616	0.756774028652708
8805	TRIM24	tripartite motif containing 24
9223	-0.170591990546986	4.82934933750226
1.00367269823782	0.316423451355718	0.756774028652708
80755	AARSD1	alanyl-tRNA synthetase domain containing 1	3704
	0.208900618526603	3.56261905228099
1.00361589612036	0.316437145852894	0.756774028652708
57418	WDR18	WD repeat domain 18	2357	0.184447331037025
	4.02686374939149	1.00284032352071
0.316624207380767	0.756774028652708
254428	SLC41A1	solute carrier family 41 member 1	6357
0.144397414542325	4.91873730404012	1.00238858736
0.316733229089391	0.756774028652708
25953	PNKD	PNKD metallo-beta-lactamase domain containing	4990
	0.28579457549078	4.63568299496824
1.00217123422465	0.316785702482289	0.756774028652708
80224	NUBPL	nucleotide binding protein like	7319
0.244953640759732	2.9368508687261	1.00200820238516
0.316825069106512	0.756774028652708
118429	ANTXR2	ANTXR cell adhesion molecule 2
9864	-0.164067377241518	6.57198998258995
1.00193846993702	0.316841909070201	0.756774028652708
5934	RBL2	RB transcriptional corepressor like 2	7320
0.14190855651852	6.16434694846849	1.00190716087073
	0.316849470401619	0.756774028652708
152217	NCBP2AS2	NCBP2 antisense 2 (head to head)	918
	0.283705575484651	2.73932605426776
1.00180612349356	0.316873873160756	0.756774028652708

567	B2M	beta-2-microglobulin	3151	-0.287083475327973
	10.0593077180488	1.00179860814205
0.316875688382637	0.756774028652708
283267	LINC00294	long intergenic non-protein coding RNA 294
	3306	0.204767683534133	3.94692387467361
1.00168195127462	0.316903866869598	0.756774028652708
751071	CSKMT	citrate synthase lysine methyltransferase	2874
	0.294847422609307	0.321929158068955
1.00157438232596	0.316929853078945	0.756774028652708
23650	TRIM29	tripartite motif containing 29
9746	-0.195819101887509	8.42585255963381
1.00083096617669	0.317109521881368	0.756982773143295
9567	GTPBP1	GTP binding protein 1	7946	-0.154781216744894
	5.44210952433971	1.00076546062231
0.317125359663372	0.756982773143295
8718	TNFRSF25	TNF receptor superfamily member 25	3918
	-0.254419468291921	5.03434904253455
1.00042793855982	0.317206981411024	0.757048592370799
23381	SMG5	SMG5 nonsense mediated mRNA decay factor
5304	-0.130174654402162	7.82031849566684
0.999591593387459	0.317409350490841	0.757367762684432
91137	SLC25A46	solute carrier family 25 member 46
8283	-0.138858266847106	5.96473017480456
0.999171307184916	0.31751111038272	0.757367762684432
55691	FRMD4A	FERM domain containing 4A	13328
0.323650906366045	2.66029739574528	0.999081389925159
	0.31753288674802	0.757367762684432
7855	FZD5	frizzled class receptor 5	6708
0.202440715976674	4.96823698908839	0.998569115444051
	0.317656987922124	0.757367762684432
3718	JAK3	Janus kinase 3	6813	-0.464601133515268
1.18573545382675	0.998529398002841	0.317666612337645
	0.757367762684432
84126	ATRIP	ATR interacting protein 5431	0.21613378479724
	3.68788114321698	0.998490197593947
0.317676111839896	0.757367762684432
79867	TCTN2	tectonic family member 2	5106
0.268694439824037	4.19384502393006	0.998094642536948
	0.317771988231315	0.757367762684432
4548	MTR	5-methyltetrahydrofolate-homocysteine
methyltransferase	11374	0.142499706102133
5.51124637701259	0.997912297708033	0.317816198564997
	0.757367762684432
7220	TRPC1	transient receptor potential cation channel
subfamily C member 1	4768	0.230761467584542
2.99612247725102	0.997422777083052	0.317934924990664
	0.757367762684432
2972	BRF1	BRF1 RNA polymerase III transcription initiation
factor subunit	13167	0.136309319599087	5.02358089476609
	0.997397050008873	0.317941166344631
0.757367762684432
7402	UTRN	utrophin	14401	0.20636461599389
6.61606147450348	0.997244157177988	0.317978261259211
	0.757367762684432

192286	HIGD2A	HIG1 hypoxia inducible domain family member 2A	668
	0.172655039575422	4.71164791683776
0.997048483970149	0.318025743850713	0.757367762684432
57691	KIAA1586	KIAA1586	3109	0.236766755138712
	2.68889896453589	0.996808692672833
0.318083944947274	0.757367762684432
148223	C19orf25	chromosome 19 open reading frame 25	4531
	0.144755042856403	4.80899225012659
0.996652897337253	0.318121766402704	0.757367762684432
24139	EML2	EMAP like 2	7405	-0.203875363528861
4.15617140629926	0.996529301325856	0.318151775229185
	0.757367762684432
84231	TRAF7	TNF receptor associated factor 7
4489	-0.124331040411559	6.61878928156272
0.994995455388334	0.31852449888779	0.758034472731687
23647	ARFIP2	ADP ribosylation factor interacting protein 2	4788
	0.118739639804509	6.05273214299918
0.994931385004543	0.318540080423981	0.758034472731687
57673	BEND3	BEN domain containing 3	6839	-0.221268945178381
	3.13494270785672	0.993848139313515
0.318803670923202	0.758333644540394
94027	CGB7	chorionic gonadotropin subunit beta 7	3386
0.351101301913654	1.95113242406155	0.993618089268025
	0.318859686795802	0.758333644540394
4286	MITF	melanocyte inducing transcription factor	7193
	0.219815932318423	5.74959906235422
0.993511517547329	0.318885640783178	0.758333644540394
9074	CLDN6	claudin 6	1739	-0.430888498276029
2.41893686486886	0.99328727295156	0.318940261345787
	0.758333644540394
2	A2M	alpha-2-macroglobulin	6384	0.883023137378225
	1.42442308449087	0.993116977219323
0.318981749480508	0.758333644540394
9683	N4BP1	NEDD4 binding protein 1	7839	-0.13259548300378
	7.08303371012303	0.992962807872029
0.319019314955076	0.758333644540394
79693	YRDC	yrdC N6-threonylcarbamoyltransferase domain
containing	1826	-0.215434460075667	5.01549244528424
	0.99273639407474	0.319074494316655
0.758333644540394
84541	KBTBD8	kelch repeat and BTB domain containing 8
4736	-0.262681691426611	2.25281597312083
0.99243433289731	0.319148129245712	0.758333644540394
79711	IPO4	importin 4	5888	0.131475781533376
5.73385048960374	0.992414132002134	0.319153054513145
	0.758333644540394
3858	KRT10	keratin 10	2124	0.15406277665718
4.34658736992704	0.99182377455633	0.319297036232
0.758486243555316
79758	DHRS12	dehydrogenase/reductase 12	3525
0.216287692266256	2.90950082104307	0.991706786326085
	0.319325578514007	0.758486243555316
54986	ULK4	unc-51 like kinase 4	4813	0.258510527892108
	2.02986726211369	0.991157859493079

0.319459548096669	0.758675803694479
90390	MED30	mediator complex subunit 30
1440	-0.291007466752449	1.68138247089851
0.990855229699013	0.319533438689753	0.758722643791722
84826	SFT2D3	SFT2 domain containing 3	3735
0.195420715572641	3.20135700176476	0.990451517456057
	0.319632044721604	0.758828144138222
51187	RSL24D1	ribosomal L24 domain containing 1
3009	-0.152360021107678	5.93812698667406
0.990142717715514	0.319707495539322	0.75885322796003
130074	FAM168B	family with sequence similarity 168 member B
944	-0.155393955684956	4.37517940239616
0.989444073108235	0.31987828584259	0.75885322796003
10239	AP3S2	adaptor related protein complex 3 subunit sigma 2
	8142	0.134548542554856	5.68438765695616
0.989363537030393	0.319897981362376	0.75885322796003
55904	KMT2E	lysine methyltransferase 2E
9233	-0.202422070592206	6.61200032033269
0.989344584670064	0.319902616493192	0.75885322796003
51388	NIP7	nucleolar pre-rRNA processing protein NIP7
3274	-0.129071951780543	5.25150659284572
0.989300109392529	0.319913494046118	0.75885322796003
7249	TSC2	TSC complex subunit 2	11121	-0.133023108265517
	6.6720675880601	0.989061677849023	0.319971816799175
	0.758863060939054
26272	FBXO4	F-box protein 4	3014	0.323072031627625
1.35252130840706	0.987582365428281	0.320333984231052
	0.759478732883218
65993	MRPS34	mitochondrial ribosomal protein S34	1097
0.1663335217619	4.94316613734864	0.987558402330746
0.320339855349043	0.759478732883218
28985	MCTS1	MCTS1 re-initiation and release factor
2205	-0.189968917319043	4.55173924535309
0.987124553068966	0.320446175789017	0.759487235402618
154091	SLC2A12	solute carrier family 2 member 12
4468	-0.365559014594032	5.56104211650328
0.987101259184492	0.320451885573604	0.759487235402618
8367	H4C5	H4 clustered histone 5	1487	0.442735082918092
	0.019642201493687	0.986860457616254
0.320510918572096	0.759496513676313
220869	CBWD5	COBW domain containing 5	14553
0.161859506437941	4.19137603971946	0.986642981234297
	0.320564245652485	0.759496513676313
10233	LRRC23	leucine rich repeat containing 23	3054
0.233534429846533	2.15014720142834	0.986304790488762
	0.320647196133547	0.759564565574597
102724814	LOC102724814	uncharacterized LOC102724814
3243	-0.279610264121555	1.98797593275241
0.985854329390283	0.320757727853726	0.759697919182811
122953	JDP2	Jun dimerization protein 2	6235
0.16123381269386	4.13208856767773	0.985569008646429
	0.320827764267111	0.75973533289762
57192	MCOLN1	mucolipin 1	3956	0.238055372668474
5.74688292008978	0.984116410918162	0.321184639304007

0.760408542033607
644961	ACTG1P20	actin gamma 1 pseudogene 20
191	-0.274132491500867	2.73630513405237
0.983788576181325	0.321265254219504	0.760408542033607
23031	MAST3	microtubule associated serine/threonine kinase 3
6772	-0.220888312193382	3.91570910777063
0.983499541204438	0.321336350337709	0.760408542033607
122622	ADSS1	adenylosuccinate synthase 1	5775
0.346664110804014	2.33683046081984	0.983335361531022
0.321376744082039	0.760408542033607
2194	FASN	fatty acid synthase	9067	-0.249573675881711
8.14105181415883	0.983307938431039
0.321383491742626	0.760408542033607
56252	YLPM1	YLP motif containing 1	14639	-0.138798138872031
6.6967433553043 0.982895075813559	0.321485102262029
0.760431870239309
217	ALDH2	aldehyde dehydrogenase 2 family member	9824
0.285746588152066	7.0002007613086 0.982826717552342
0.321501930146155	0.760431870239309
55004	LAMTOR1 late endosomal/lysosomal adaptor, MAPK and MTOR activator 1	2778	0.14920149551992	6.42106251207009
0.982477978704033	0.321587797920009
0.760506548280527
79170	PRR15L	proline rich 15 like	1519	0.368047755496389
4.73451252312657	0.982135529745065
0.321672146379039	0.760577608766239
100288413	ERVMER34-1	endogenous retrovirus group MER34 member 1, envelope	4077	0.32302564832252
1.5957822355621 0.981529925256852	0.321821383939482
0.760802046386553
5018	OXA1L	OXA1L mitochondrial inner membrane protein	4497
0.147911088574523	5.72316178797161
0.981110179151419	0.321924874435277	0.760887621461896
1294	COL7A1	collagen type VII alpha 1 chain
11071	-0.226192993757868	9.71571117978719
0.980942510065688	0.321966226344304	0.760887621461896
1861	TOR1A	torsin family 1 member A	3766
0.135323383966682	6.16560689860412	0.980393588521963
0.322101654815552	0.760985419348837
6689	SPIB	Spi-B transcription factor	4707
0.28565178810085	0.309692548688999	0.980334380449642
0.322116266960752	0.760985419348837
100288637	LOC100288637	OTU deubiquitinase 7A pseudogene
6722	0.342563417655204	-0.293911983609282
0.979916126735491	0.322219513996723	0.761100966703557
4306	NR3C2	nuclear receptor subfamily 3 group C member 2
6607	-0.192403277160711	7.35102983812715
0.979200049013969	0.322396380996488	0.76139034035033
6812	STXBP1	syntaxin binding protein 1
6009	-0.166978677387551	5.18681639705608
0.978794998540565	0.32249648277799	0.761498353720719
1903	S1PR3	sphingosine-1-phosphate receptor 3	4206
0.301323255612105	5.54432945538458	0.977990607699681
0.322695396744796	0.761698145942117

25780	RASGRP3 RAS guanyl releasing protein 3	11118
0.290733190380696	3.18017924085726	0.977740040525076
	0.322757391359235	0.761698145942117
8263	F8A1	coagulation factor VIII associated 1	3472
0.201128875979764	4.46731818227104	0.977716531185269
	0.322763208779102	0.761698145942117
55285	RBM41	RNA binding motif protein 41
6101	-0.158999946318349	4.67211869163761
0.977040276428488	0.322930608225925	0.761698145942117
54928	IMPAD1	inositol monophosphatase domain containing 1
7594	-0.138177175888827	6.93499081247475
0.976957937926898	0.322950998040739	0.761698145942117
27130	INVS	inversin	5943	-0.168274013816711
4.10093248455457	0.976873410125624	0.322971931767743
	0.761698145942117
7707	ZNF148	zinc finger protein 148 10574	-0.15603568481764
	6.02472729984352	0.976540903372429
0.323054296082547	0.761698145942117
163590	TOR1AIP2	torsin 1A interacting protein 2
18876	-0.146098633172418	6.79818876819045
0.976523107833103	0.323058704947019	0.761698145942117
23248	RPRD2	regulation of nuclear pre-mRNA domain containing 2
	8738	-0.1487586884277	5.86051982908439
0.976446907733528	0.323077584503588	0.761698145942117
83443	SF3B5	splicing factor 3b subunit 5	693
0.170062047117569	5.45330173384273	0.976255988916165
	0.323124893495256	0.761698145942117
401561	LINC01451	long intergenic non-protein coding RNA 1451
	3406	0.359841910022136	-0.047270598265787
0.975808905905685	0.323235714825061	0.761813277144046
23261	CAMTA1	calmodulin binding transcription activator 1
10033	0.154204164833639	4.71372341728498
0.975551914528902	0.323299439630319	0.761813277144046
139341	FUNDC1	FUN14 domain containing 1	1399
0.208215995143793	4.09092221545332	0.975400881779107
	0.323336898166728	0.761813277144046
28972	SPCS1	signal peptidase complex subunit 1	5324
0.133431900767724	5.81029423726014	0.974905546003704
	0.323459789571996	0.761942254083722
29923	HILPDA	hypoxia inducible lipid droplet associated	1867
	0.218842844807185	2.98918334194019
0.974578610575689	0.323540935109939	0.761942254083722
7477	WNT7B	Wnt family member 7B	5051	0.177893252613146
	6.26326081203792	0.974522632157807
0.323554831676303	0.761942254083722
10190	TXNDC9	thioredoxin domain containing 9
2546	-0.158139926993384	5.93295753152538
0.973922409387669	0.323703885708133	0.762165124745977
4738	NEDD8	NEDD8 ubiquitin like modifier	3879
0.158972571195602	6.4322173780254 0.973492366459936
0.32381073453925	0.762222961633369
100526739	CENPS-CORT	CENPS-CORT readthrough	2244
0.268936804057143	2.53430835202197	0.973071147503878
	0.323915436119986	0.762222961633369

23762	OSBP2	oxysterol binding protein 2
6700	-0.325949139268197	2.32140180654068
0.972992863783769	0.323934899880097	0.762222961633369
5534	PPP3R1	protein phosphatase 3 regulatory subunit B, alpha
	3363	-0.126491599098187	6.17615001795581
0.972821298467959	0.323977561740487	0.762222961633369
29063	ZCCHC4	zinc finger CCHC-type containing 4
4400	-0.2198860285784	3.34076668823602
0.972610576027911	0.324029970665992	0.762222961633369
4172	MCM3	minichromosome maintenance complex component 3	3650
	0.13593726910554	4.95664533663539
0.97251014168174	0.324054953703628	0.762222961633369
7317	UBA1	ubiquitin like modifier activating enzyme 1
6437	-0.123299736125293	8.85186126227541
0.971831382599447	0.324223861717828	0.762412155386024
23564	DDAH2	dimethylarginine dimethylaminohydrolase 2	2538
	0.136152459042143	6.04930412545694
0.97174948365301	0.324244249991757	0.762412155386024
5819	NECTIN2 nectin cell adhesion molecule 2
3747	-0.11526478485631	7.27726649639425
0.970502773756888	0.324554820589163	0.763014328968179
399909	PCNX3	pecanex 3	7972	-0.137266681093939
6.57298050073568	0.970206244334896	0.324628747548
0.763026269299208
10461	MERTK	MER proto-oncogene, tyrosine kinase	4700
0.297344383755803	2.91709623637216	0.969936943677519
	0.324695905465009	0.763026269299208
8986	RPS6KA4 ribosomal protein S6 kinase A4
3807	-0.131291757633279	5.63342034671136
0.96979598433364	0.324731065082198	0.763026269299208
8405	SPOP	speckle type BTB/POZ protein	5271
0.174033830289184	5.5619370676373 0.96932893867907
0.324847596679122	0.763026269299208
142678	MIB2	mindbomb E3 ubiquitin protein ligase 2	8767
0.250198423572634	4.7834614337682 0.969233886989642
0.324871319601048	0.763026269299208
5426	POLE	DNA polymerase epsilon, catalytic subunit
16609	-0.229179870865891	4.4390928109434 0.968972150691179
	0.324936655369585	0.763026269299208
84034	EMILIN2 elastin microfibril interfacer 2	6435
0.324095453567207	3.90251739190594	0.96895385933994
	0.324941221986848	0.763026269299208
22925	PLA2R1	phospholipase A2 receptor 1	9805
0.195978177451824	4.38318144793424	0.968270245929911
	0.325111953733489	0.763154531815778
50649	ARHGEF4 Rho guanine nucleotide exchange factor 4
9926	-0.343921729329698	4.74534983703834
0.967957796969813	0.325190027046413	0.763154531815778
11213	IRAK3	interleukin 1 receptor associated kinase 3
10004	-0.463616477148726	4.18251363479314
0.967734967186345	0.325245721881005	0.763154531815778
9049	AIP	aryl hydrocarbon receptor interacting protein	1640
	0.163443037370381	4.87284623282631
0.967692565349101	0.325256321369471	0.763154531815778

79630	C1orf54 chromosome 1 open reading frame 54	1351
0.28543768608932	1.22443551610686	0.96764479804111
	0.325268262650117	0.763154531815778
22847	ZNF507	zinc finger protein 507 11239	-0.192504944106446
	3.97453927192845	0.96719516977436
0.325380693002081	0.763290465061154
7812	CSDE1	cold shock domain containing E1
5778	-0.119868827298077	9.57261468783797
0.966834050808103	0.32547102867741	0.763374530925063
4131	MAP1B	microtubule associated protein 1B	12630
0.425186310162486	6.1851815825765 0.965958590530136
0.32569016713712	0.763690830110834
283726	SAXO2	stabilizer of axonemal microtubules 2	4160
0.33991753801891	0.177762573945735	0.965812545933757
	0.325726742895052	0.763690830110834
23042	PDXDC1	pyridoxal dependent decarboxylase domain containing
1	9410	-0.129748029499209	7.08288675728786
0.96530376564594	0.32585420551531	0.763690830110834
27339	PRPF19	pre-mRNA processing factor 19	2739
0.130513589619146	5.74460919060918	0.964868926195194
	0.325963196377158	0.763690830110834
29979	UBQLN1	ubiquilin 1	7117	-0.12988590053509
7.68162621639902	0.964802130947611	0.32597994262881
	0.763690830110834
57118	CAMK1D	calcium/calmodulin dependent protein kinase ID
8762	-0.20641892075857	5.37396594494486
0.964702735813653	0.326004864113919	0.763690830110834
100506190	LINC00963	long intergenic non-protein coding
RNA 963 12761	0.169393200892989	5.0376633095611
0.964511614303817	0.326052791374219	0.763690830110834
1183	CLCN4	chloride voltage-gated channel 4	6789
0.266733302333441	4.3235959229076 0.963932436594156
0.326198087902776	0.763690830110834
23612	PHLDA3	pleckstrin homology like domain family A member 3
	3138	0.137120318528491	7.9976867796259
0.963748581371533	0.326244229023931	0.763690830110834
113451	AZIN2	antizyme inhibitor 2	6559	0.488893639797899
	1.5814208852149 0.963420842296555	0.326326501312916
	0.763690830110834
79598	CEP97	centrosomal protein 97	9550	-0.209128430606397
	5.20382782801013	0.963113908788102
0.326403575702433	0.763690830110834
133584	EGFLAM	EGF like, fibronectin type III and laminin G domains
	5724	0.958049852699936	0.494912877657608
0.963051320562062	0.32641929525586	0.763690830110834
10950	BTG3	BTG anti-proliferation factor 3
2306	-0.180391713977043	5.94412998627302
0.96300701919202	0.326430422519422	0.763690830110834
9702	CEP57	centrosomal protein 57	7101	-0.152365308805882
	4.73810242944102	0.962882159909884
0.326461786378255	0.763690830110834
26001	RNF167	ring finger protein 167 2744	0.133960078710881
	6.39194607239242	0.962824809775015
0.326476193705757	0.763690830110834

84986 ARHGAP19 Rho GTPase activating protein 19 5687
-0.19802556520346 3.04939843192544
0.962816669345881 0.326478238788011 0.763690830110834
4735 SEPTIN2 septin 2 6079 0.156089115937047
9.01854721291204 0.96218312948406 0.326637452137441
0.763935680762922
11116 FGFR1OP FGFR1 oncogene partner 15159 -0.169634092293918
4.38751361751158 0.961690408528128
0.326761347750308 0.7640978276103
400916 CHCHD10 coiled-coil-helix-coiled-coil-helix domain
containing 10 1360 0.309362476538366 1.57669282361224
0.960811111229216 0.32698260348126
0.764269441499069
55187 VPS13D vacuolar protein sorting 13 homolog D
20023 -0.20840704489744 6.99360291925323
0.960591355082613 0.327037931268577 0.764269441499069
55176 SEC61A2 SEC61 translocon alpha 2 subunit 3916
0.228418705393969 3.61148166510706 0.960549352455082
0.327048507640961 0.764269441499069
91252 SLC39A13 solute carrier family 39 member 13 4584
0.135479715483631 6.95532955337956
0.960492930492407 0.32706271555351 0.764269441499069
285464 NA NA 4438 0.275140944426192
3.52598784293391 0.960288030673913 0.327114319340548
0.764269441499069
140686 WFDC3 WAP four-disulfide core domain 3 2405
0.330812884399352 1.79883126336484 0.960098338948795
0.327162102607869 0.764269441499069
5676 PSG7 pregnancy specific beta-1-glycoprotein 7 (gene/
pseudogene) 5961 -0.511090029413713 2.35070519270863
0.959813514032524 0.327233867265711
0.764309617940774
7337 UBE3A ubiquitin protein ligase E3A
11766 -0.111438554941129 6.78553962305981
0.958865439429573 0.327472895127173 0.76456262621077
23633 KPNA6 karyopherin subunit alpha 6
7480 -0.113158953651104 7.07409010428647
0.958496845041465 0.327565887371888 0.76456262621077
51422 PRKAG2 protein kinase AMP-activated non-catalytic subunit
gamma 2 6740 -0.124915948210493 5.41345594310363
0.958471421132856 0.327572302830914 0.76456262621077
253832 ZDHHC20 zinc finger DHHC-type containing 20
5574 -0.138263589227941 7.23373163158937
0.958418844984692 0.327585570403952 0.76456262621077
5573 PRKAR1A protein kinase cAMP-dependent type I regulatory
subunit alpha 7389 -0.124339724140635 8.57371508955656
0.957520990991284 0.327812253610426
0.76456262621077
158135 TTLL11 tubulin tyrosine ligase like 11 3535
0.37900913817064 3.54634770344212 0.957396805585617
0.327843623356168 0.76456262621077
51093 RRNAD1 ribosomal RNA adenine dimethylase domain containing
1 2742 0.19027537883017 3.96312244922743
0.957305348570808 0.327866728321844 0.76456262621077

7690	ZNF131	zinc finger protein 131 6232	-0.14165036915184
	4.75069937245427	0.957301453807966
0.327867712311403	0.764562626210077
23028	KDM1A	lysine demethylase 1A	3404	0.121291852067608
	6.11562287366885	0.957252546132757
0.327880068889368	0.764562626210077
8882	ZPR1	ZPR1 zinc finger	7069	-0.188846682280454
	5.51669069328228	0.957221020339716
0.327888034243159	0.764562626210077
6095	RORA	RAR related orphan receptor A
17040	-0.29944164420046	4.41845109512185
0.95667609542393	0.328025756364931	0.764578035819504
25962	VIRMA	vir like m6A methyltransferase associated
10771	-0.121686285746201	6.16266517413902
0.956343295701779	0.328109904577319	0.764578035819504
1307	COL16A1 collagen type XVI alpha 1 chain 9269
0.236573387697127	5.75827886713873	0.956202935228248
	0.328145403228303	0.764578035819504
197407	ZNF48	zinc finger protein 48	3640	0.213567641345014
	4.5728346481189 0.956139817674586	0.32816136798772
	0.764578035819504
414328	IDNK	IDNK gluconokinase	1814	0.293197063852878
	0.0140705703871507	0.955866988770145
0.328230388348733	0.764578035819504
22926	ATF6	activating transcription factor 6
7890	-0.136437509000384	6.60248725694671
0.955700930968199	0.328272407147523	0.764578035819504
84883	AIFM2	apoptosis inducing factor mitochondria associated 2
	5135	0.195405591506911	5.05076881796176
0.955367459743606	0.328356809334986	0.764578035819504
29766	TMOD3	tropomodulin 3	9334	-0.127127335798059
8.61452767810325	0.955301390095521	0.32837353510463
	0.764578035819504
7462	LAT2	linker for activation of T cells family member 2
	4545	0.41364899750102	0.951220407732526
0.955252503781778	0.32838591157922	0.764578035819504
11277	TREX1	three prime repair exonuclease 1	2197
0.304670482372909	3.01528588379992	0.954463416528773
	0.328585769310711	0.764916215844296
10208	USPL1	ubiquitin specific peptidase like 1
3814	-0.159259983045343	4.06649044853047
0.953902551255876	0.328727921839378	0.765118733608951
3516	RBPJ	recombination signal binding protein for
immunoglobulin kappa J region	9253	-0.153068328094512
6.70123864402234	0.953679080928822	0.328784583633207
	0.765118733608951
8815	BANF1	barrier to autointegration factor 1	1645
0.198485213164076	4.89613015444644	0.953473830475062
	0.328836637133282	0.765118733608951
1130	LYST	lysosomal trafficking regulator
19614	-0.275416470934832	6.09243600929695
0.952489772699161	0.329086355806468	0.765495064034134
54554	WDR5B	WD repeat domain 5B	3720	-0.227475619237661
	2.63154057006736	0.952262184434904

0.329144145434411	0.765495064034134
4277	MICB	MHC class I polypeptide-related sequence B	2798
	0.274418495718452	4.30436230219366
0.952190576558058	0.329162331025743	0.765495064034134
80135	RPF1	ribosome production factor 1 homolog
1728	-0.157268159305817	4.88972558349377
0.951892269318407	0.329238103727225	0.765504368659374
54539	NDUFB11 NADH:ubiquinone oxidoreductase subunit B11	1418
	0.14555965423123	5.15359511370625
0.951159241704594	0.32942439776255	0.765504368659374
334	APLP2	amyloid beta precursor like protein 2	7451
0.14511053455909	9.98098958805549	0.950903011630771
	0.329489549952557	0.765504368659374
23429	RYBP	RING1 and YY1 binding protein
7512	-0.185238737813665	6.05986283871089
0.950788881934265	0.329518575476963	0.765504368659374
90378	SAMD1	sterile alpha motif domain containing 1
2263	-0.143235187191878	4.70488845529455
0.9507483986932 0.32952887201265	0.765504368659374
139886	SPIN4	spindlin family member 4
4117	-0.510530489082966	2.64685216499696
0.950684821183739	0.329545043223792	0.765504368659374
317762	CCDC85C coiled-coil domain containing 85C
17449	-0.148723236273911	5.16808767138702
0.950457860917254	0.329602780141822	0.765504368659374
25906	ANAPC15 anaphase promoting complex subunit 15	2883
0.225097028644134	4.351437529185	0.950454856746575
0.329603544469528	0.765504368659374
10463	SLC30A9 solute carrier family 30 member 9
7225	-0.131956430939713	6.38936379254474
0.950185321117658	0.329672129906304	0.765536724734349
26018	LRIG1	leucine rich repeats and immunoglobulin like domains
1	7032	-0.294439957775069	3.91410994812188
0.949524218463598	0.329840432974258	0.765646954510494
64374	SIL1	SIL1 nucleotide exchange factor 3722
0.16194038398376	4.82554401428269	0.949519374586977
	0.329841666546621	0.765646954510494
9656	MDC1	mediator of DNA damage checkpoint 1	8064
0.173983785705833	4.50281995614017	0.949354788734514
	0.329883584669867	0.765646954510494
23077	MYCBP2	MYC binding protein 2	15945	-0.221650077972135
	6.39996864538838	0.948328297170804
0.330145180195978	0.765834247180864
8411	EEA1	early endosome antigen 1
10150	-0.199495064590792	6.02576714798509
0.94823814998616	0.330168166868816	0.765834247180864
27173	SLC39A1 solute carrier family 39 member 1	3388
0.120601323565824	7.5944849319573 0.948005180590961
0.33022758170453	0.765834247180864
2181	ACSL3	acyl-CoA synthetase long chain family member 3	7240
	0.16376613286251	7.30895426835324
0.94759609131313	0.330331947305103	0.765834247180864
51735	RAPGEF6 Rap guanine nucleotide exchange factor 6
11406	-0.197970530151915	4.84810350018139

0.947094497257986	0.330459972334966	0.765834247180864
727957	MROH1	maestro heat like repeat family member 1
10903	-0.186807578903457	6.49943810762928
0.94703605260316	0.330474893828531	0.765834247180864
10075	HUWE1	HECT, UBA and WWE domain containing E3 ubiquitin protein ligase 1	17380	-0.22214165306773
8.66869210827755	0.946638375454995	0.330576448512278
	0.765834247180864
100129781	MIR193BHG	MIR193B host gene
1392	-0.338328862295364	0.812473688386562
0.94659086562859	0.330588583857972	0.765834247180864
64744	SMAP2	small ArfGAP2	3631	-0.18064586328311
4.49282509160474	0.946518982211103	0.330606946031299
	0.765834247180864
84102	SLC41A2	solute carrier family 41 member 2
4538	-0.254099362821646	4.96922628908224
0.946516555793597	0.330607565868287	0.765834247180864
210	ALAD	aminolevulinate dehydratase	4983
0.201972331615461	5.25134130106874	0.946407188383647
	0.3306355057683	0.765834247180864
29982	NRBF2	nuclear receptor binding factor 2
1901	-0.139087886389814	5.02370720386574
0.946061806263527	0.330723760565508	0.765834247180864
8675	STX16	syntaxin 16	5088	-0.124690393206694
6.6499316921204	0.946053435857261	0.330725899827656
0.765834247180864
6713	SQLE	squalene epoxidase	4305	-0.206478459964202
	7.32689810880355	0.946038440273128
0.330729732362179	0.765834247180864
6183	MRPS12	mitochondrial ribosomal protein S12	2144
0.176739563923894	4.43846064431788	0.945677231548782
	0.330822067063472	0.7658493636999
105377348	UBE2D3-AS1	UBE2D3 antisense RNA 1
883	-0.327713013336254	0.602084820567395
0.945585142928135	0.330845612889848	0.7658493636999
57190	SELENON	selenoprotein N	4737	0.1605041914108
6.33277196444103	0.943512227719637	0.331376220427017
	0.76695087896914
58487	CREBZF	CREB/ATF bZIP transcription factor
5587	-0.225936410217697	5.60801054155641
0.943245476423602	0.33144458347585	0.766982369196469
5725	PTBP1	polypyrimidine tract binding protein 1
6883	-0.118748319624359	7.92520638275992
0.942849910435454	0.331545993755377	0.766997593169982
8325	FZD8	frizzled class receptor 8	4030
0.229972451545134	3.0072543378726	0.942733271685214
0.331575904035534	0.766997593169982
54906	TASOR2	transcription activation suppressor family member 2
	8626	-0.131255423227649	6.72910577814656
0.942579130127175	0.331615436869952	0.766997593169982
124045	SPATA33	spermatogenesis associated 33	5746
0.256892675666024	2.4929992944157	0.941988507091049
0.331766972674753	0.767029787987608
3689	ITGB2	integrin subunit beta 2	6797	0.893045747420908

3.65691546479731 0.941164875318133
0.331978445490311 0.767029787987608
5536 PPP5C protein phosphatase 5 catalytic subunit 7059
0.118738287723239 5.93962708446533 0.94111397820695
0.331991519547659 0.767029787987608
8123 PWAR5 Prader Willi/Angelman region RNA 5
3369 -0.383391465080071 2.81780522473972
0.941055289782156 0.332006595829445 0.767029787987608
65094 JMJD4 jumonji domain containing 4 4315
0.157953124234688 4.20841242987989 0.940865264661792
0.33205541703621 0.767029787987608
10370 CITED2 Cbp/p300 interacting transactivator with Glu/Asp rich carboxy-terminal domain 2 2535 0.295163258746683
4.94395145416763 0.940588125318659 0.332126636764256
0.767029787987608
84067 FAM160A2 family with sequence similarity 160 member A2 4736 -0.148940761895988 4.83180342731054
0.940418034428804 0.33217035708707 0.767029787987608
122616 CLBA1 clathrin binding box of aftiphilin containing 1 5225
0.240269026974299 2.90040656221939
0.940402595490367 0.33217432590617 0.767029787987608
55008 HERC6 HECT and RLD domain containing E3 ubiquitin protein ligase family member 6 5793 -0.269623676781922
4.84970301400538 0.939698836342103 0.33235530527623
0.767029787987608
23244 PDS5A PDS5 cohesin associated factor A
12204 -0.12546583742864 7.7021011273909 0.939542587828356
0.332395504125915 0.767029787987608
388630 TRABD2B TraB domain containing 2B
7034 -0.266769883214689 0.100460365012246
0.939503863554614 0.332405467919245 0.767029787987608
220988 HNRNPA3 heterogeneous nuclear ribonucleoprotein A3
6033 -0.140869799978518 6.75952108654017
0.938998193712727 0.332535613827139 0.767029787987608
4705 NDUFA10 NADH:ubiquinone oxidoreductase subunit A10 5021
0.131246030982594 5.40738132322869
0.938787324661234 0.332589905965175 0.767029787987608
552891 DNAJC25-GNG10 DNAJC25-GNG10 readthrough
1448 -0.1563603152001 5.65898498253222
0.938569401917867 0.332646026629622 0.767029787987608
6542 SLC7A2 solute carrier family 7 member 2
7905 -0.845489053243485 5.17509512868442
0.938327305658163 0.332708387402371 0.767029787987608
51504 TRMT112 tRNA methyltransferase subunit 11-2 1625
0.156712437920549 5.44604501424418 0.938318843740319
0.332710567361771 0.767029787987608
9670 IPO13 importin 13 5083 0.131586075084888
5.98249890194213 0.938239810684402 0.332730928778841
0.767029787987608
9055 PRC1 protein regulator of cytokinesis 1 5525
0.287295903164509 1.88478915180425 0.938230775746629
0.332733256570692 0.767029787987608
9569 GTF2IRD1 GTF2I repeat domain containing 1 6498
-0.18666131209628 5.69938466539557

0.937834795042477	0.332835299720079	0.767029787987608
9881	TRANK1	tetratricopeptide repeat and ankyrin repeat containing 1	11177	0.265535964770842	4.3813120460727
0.937648421404194	0.332883342148755	0.767029787987608
4814	NINJ1	ninjurin 1	2648	0.207118463958571
5.98590980685145	0.937603699806335	0.332894871626385
	0.767029787987608
8099	CDK2AP1	cyclin dependent kinase 2 associated protein 1	3117
	0.138106610671533	7.41725859999524
0.937254262367048	0.332984976897728	0.767029787987608
9828	ARHGEF17	Rho guanine nucleotide exchange factor 17
	9539	-0.148086185993181	6.41514059839123
0.937197694050042	0.332999566556334	0.767029787987608
55527	FEM1A	fem-1 homolog A	9593	-0.135528875467685
5.70964469655208	0.936998372658343	0.333050980771671
	0.767029787987608
170506	DHX36	DEAH-box helicase 36	10562	-0.105492509440037
	6.09931757087175	0.93679098299273
0.333104487413922	0.767029787987608
6786	STIM1	stromal interaction molecule 1
6735	-0.132904674855473	6.92503916222357
0.936766726079441	0.333110746459562	0.767029787987608
85016	CFAP300	cilia and flagella associated protein 300	2647
	0.308631932844416	0.074104214530111
0.93674330750392	0.333116789336631	0.767029787987608
10329	RXYLT1	ribitol xylosyltransferase 1	4534
0.190664401107272	3.44448421286252	0.936441659039176
	0.333194639114941	0.767029787987608
8568	RRP1	ribosomal RNA processing 1
4251	-0.14354133775705	4.56826193269302
0.936157059813932	0.333268111069928	0.767029787987608
4176	MCM7	minichromosome maintenance complex component 7	4386
	0.169272171591839	6.20782410881485
0.935960819305393	0.333318784991379	0.767029787987608
56260	C8orf44	chromosome 8 open reading frame 44	2323
0.272458465706562	0.616171278773179	0.935929277517821
	0.333326930783221	0.767029787987608
222194	RSBN1L	round spermatid basic protein 1 like
7206	-0.171788510736774	4.32755881991102
0.935465230516066	0.333446803483559	0.767047828539391
64326	COP1	COP1 E3 ubiquitin ligase
4299	-0.129022720902223	5.66363365994217
0.935456635350842	0.333449024330903	0.767047828539391
7993	UBXN8	UBX domain protein 8	2739	0.244415844713667
	2.59986181593847	0.935210082409675
0.333512737907499	0.767047828539391
26053	AUTS2	activator of transcription and developmental regulator AUTS2	10420	-0.17507879399487	5.23050659344388
	0.935051149192841	0.333553817636427
0.767047828539391
9031	BAZ1B	bromodomain adjacent to zinc finger domain 1B
7053	-0.140500992723178	7.20027694369769
0.933590394402074	0.333931697951488	0.76767905483266
221061	FAM171A1	family with sequence similarity 171 member

A1	3955	-0.144332861335751	5.75857330742339
0.933437742139589	0.333971220311877	0.76767905483266
144402	CPNE8	copine 8	5178	0.199155050022396
3.82950826511495	0.933354675133156	0.333992729360336
	0.76767905483266
389641	LOC389641	uncharacterized LOC389641
2024	-0.265987934007628	1.66022859995185
0.932772173122914	0.334143612173747	0.76789162815205
2935	GSPT1	G1 to S phase transition 1
8018	-0.124502732310467	7.82001318785098
0.932574415102067	0.334194857229592	0.76789162815205
5178	PEG3	paternally expressed 3	9278	-0.752545263777892
	2.3103063201167	0.932168040569799	0.334300194068707
	0.767968524914866
8506	CNTNAP1	contactin associated protein 1
6045	-0.270179172691684	2.2240579866334	0.931848923738948
	0.334382943835085	0.767968524914866
284129	SLC26A11	solute carrier family 26 member 11	4903
	0.193930512638167	4.47757234899381
0.931810898300807	0.334392805986704	0.767968524914866
203054	ADCK5	aarF domain containing kinase 5	2742
0.229463604060353	3.26646526214591	0.930536480044161
	0.334723559870975	0.76842745048253
28966	SNX24	sorting nexin 24	3550	-0.150678182372582
	5.14975363890791	0.930424891979778
0.334752541501489	0.76842745048253
92335	STRADA	STE20 related adaptor alpha	5259
0.13653441760364	5.10106787674716	0.930193631221238
	0.334812615183528	0.76842745048253
8365	H4C8	H4 clustered histone 8	397	0.341738625252376
	2.23032509427027	0.929899743612983
0.334888977992549	0.76842745048253
57412	AS3MT	arsenite methyltransferase	2404
0.360976643253527	1.48658761300526	0.929769977929146
	0.334922703301687	0.76842745048253
23524	SRRM2	serine/arginine repetitive matrix 2
12857	-0.178345515362256	9.50893838848733
0.929266030015015	0.335053719366957	0.76842745048253
23398	PPWD1	peptidylprolyl isomerase domain and WD repeat
containing 1	3777	-0.153098690556	4.02914859391834
0.929092198691319	0.335098927825913	0.76842745048253
23039	XPO7	exportin 7	6909	-0.106405253504349
6.58984954530806	0.928923864251814	0.335142714481867
	0.76842745048253
10610	ST6GALNAC2	ST6 N-acetylgalactosaminide alpha-2,6-
sialyltransferase 2	7558	0.255254486932732
6.21254472388966	0.928810834986081	0.335172119614433
	0.76842745048253
23554	TSPAN12	tetraspanin 12	3368	-0.252962763401852
4.53907457523855	0.928785314206582	0.335178759451287
	0.76842745048253
160777	CCDC60	coiled-coil domain containing 60	3440
0.379396226939595	0.246777731621037	0.928718681305363
	0.335196096412384	0.76842745048253

2357	FPR1	formyl peptide receptor 1	2594
0.77020059211309	-0.168772581915607	0.927819190677623
	0.335430248905218	0.76881151612525
5826	ABCD4	ATP binding cassette subfamily D member 4	8769
	0.18341893828213	5.66955341202634
0.927548766111267	0.335500687740385	0.76881151612525
25972	UNC50	unc-50 inner nuclear membrane RNA binding protein
	2230	0.131385526750581	5.05102994188504
0.927442806910828	0.335528292858832	0.76881151612525
79573	TTC13	tetratricopeptide repeat domain 13
4212	-0.143154920128868	4.25523286700728
0.926556159171625	0.335759406763173	0.768987985367518
55022	PID1	phosphotyrosine interaction domain containing 1 3364
	0.171152048940395	4.56997919418364
0.92652638609575	0.335767171120325	0.768987985367518
51186	TCEAL9	transcription elongation factor A like 9	1096
	0.150121854992846	5.86784556278515
0.926515503569867	0.335770009174537	0.768987985367518
84222	TMEM191A	transmembrane protein 191A (pseudogene) 2707
	0.264093913255528	1.71206809831848
0.925916273125563	0.335926332076451	0.769014873196694
375616	KCP	kielin cysteine rich BMP regulator	8259
0.346409010626255	1.4412678936939 0.925616167134446
0.336004658173891	0.769014873196694
5195	PEX14	peroxisomal biogenesis factor 14	2995
0.143106453676773	4.76022064993668	0.92536701932159
	0.336069703030659	0.769014873196694
205251	MTLN	mitoregulin	2051	0.639401190738882
2.54649436472186	0.925261271809504	0.336097315561266
	0.769014873196694
115939	TSR3	TSR3 ribosome maturation factor 1243
0.164438822179431	4.654560840712	0.925192001765115
0.33611540483192	0.769014873196694
9373	PLAA	phospholipase A2 activating protein
4515	-0.140557785059029	5.74380752650966
0.925129115359749	0.336131828199729	0.769014873196694
6160	RPL31	ribosomal protein L31	5791	0.196926842922933
	7.20746871168833	0.924968249159448
0.33617384479012	0.769014873196694
79877	DCAKD	dephospho-CoA kinase domain containing	2756
0.292001070388153	3.1595701676908 0.924092517031674
0.336402700396461	0.769014873196694
56658	TRIM39	tripartite motif containing 39
4144	-0.151447634809465	5.01779813380171
0.923974813469567	0.336433475846643	0.769014873196694
162466	PHOSPHO1	phosphoethanolamine/phosphocholine
phosphatase 1	2375	1.07162064733398	0.516878017529534
	0.923929320535635	0.336445371701299
0.769014873196694
283635	FAM177A1	family with sequence similarity 177 member
A1	4699	-0.148700990227415	6.53214190255103
0.923869356761628	0.336461052367293	0.769014873196694
9589	WTAP	WT1 associated protein	5830	-0.117421236256409
	6.20965585093735	0.92373319937835

0.33649666147446	0.769014873196694
6894	TARBP1	TAR (HIV-1) RNA binding protein 1
7743	-0.191071265152318	4.2096523474993 0.923374315306777
	0.336590544269998	0.769014873196694
55791	LRIF1	ligand dependent nuclear receptor interacting factor
1	3621	-0.229342572274461	4.10345141962475
0.923342547361923	0.336598856341887	0.769014873196694
147650	SPACA6	sperm acrosome associated 6
3648	-0.394637384422937	2.55540695371828
0.923231674995323	0.336627868213618	0.769014873196694
9684	LRRC14	leucine rich repeat containing 14	6136
0.154750679606368	4.93802872578878	0.922656041529553
	0.336778547569873	0.769014873196694
285267	ZNF619	zinc finger protein 619 4514	-0.264986979798645
	1.62785893199313	0.922254457641832
0.336883720769105	0.769014873196694
23365	ARHGEF12	Rho guanine nucleotide exchange factor 12
	24404	-0.164681101764442	8.06095575887693
0.922178650690171	0.336903579242621	0.769014873196694
11276	SYNRG	synergin gamma	14277	-0.142943385963998
6.17776633217405	0.921794868779504	0.337004139261844
	0.769014873196694
23052	ENDOD1	endonuclease domain containing 1	4687
0.189686274991389	6.44064406631942	0.921729261893709
	0.337021333858171	0.769014873196694
18	ABAT	4-aminobutyrate aminotransferase
9744	-0.191239120556353	5.13151917647495
0.921684650783689	0.337033026443294	0.769014873196694
23476	BRD4	bromodomain containing 4
11637	-0.145007726693654	6.61141708604941
0.921649646989394	0.337042201328862	0.769014873196694
255252	LRRC57	leucine rich repeat containing 57
8114	-0.118704117101754	5.479963222084	0.921640878703768
	0.337044499646927	0.769014873196694
2790	GNG10	G protein subunit gamma 10
1491	-0.155238301928436	5.66251985947296
0.9212615245129 0.337143955008351	0.769116511042666
400	ARL1	ADP ribosylation factor like GTPase 1
4979	-0.117046427840345	6.42447656681988
0.920683724571951	0.337295512547177	0.769162421076457
102723354	LINC02298	long intergenic non-protein coding
RNA 2298	1177	0.313470825531501	0.467110390764081
	0.920301313531281	0.337395869611841
0.769162421076457
6257	RXRB	retinoid X receptor beta	3920
0.140849035433901	5.59299698380436	0.920147056399557
	0.337436363022139	0.769162421076457
5321	PLA2G4A phospholipase A2 group IVA	2875
0.264572291185664	3.5959639493191 0.92006003167416
0.337459210403449	0.769162421076457
10269	ZMPSTE24	zinc metallopeptidase STE24
3572	-0.11819922856877	7.15151087602155
0.920053499543769	0.337460925425906	0.769162421076457
115509	ZNF689	zinc finger protein 689 4159	0.188447260885995

3.28515907651375 0.919500645501828
0.33760612061304 0.769162421076457
55262 MAP11 microtubule associated protein 11
3255 -0.145236200932198 5.28245916242954
0.919450833877733 0.337619206676152 0.769162421076457
29068 ZBTB44 zinc finger and BTB domain containing 44
10803 0.159483161731475 5.74592604791941
0.919394157274425 0.337634097072561 0.769162421076457
548593 SLX1A SLX1 homolog A, structure-specific endonuclease subunit 4926 -0.181150092108783 4.95375474616209
0.919302063964828 0.337658294224255 0.769162421076457
3430 IFI35 interferon induced protein 35
1973 -0.32748290304172 2.43004605653915
0.918801864868618 0.337789760192375 0.769336775774731
65244 SPATS2 spermatogenesis associated serine rich 2 6840
0.140612675029033 5.21025112002336
0.917911796138501 0.338023864417224 0.769513780917081
57486 NLN neurolysin 10948 0.219134721863965
5.18848605175166 0.917765423399242 0.338062383947611
0.769513780917081
396 ARHGDIA Rho GDP dissociation inhibitor alpha
2903 -0.166867965107859 7.89968627154196
0.917576429120345 0.338112128500143 0.769513780917081
7533 YWHAH tyrosine 3-monooxygenase/tryptophan 5-monooxygenase activation protein eta 2453 0.159182226518206
7.30381768036198 0.917303465113861 0.338183991786797
0.769513780917081
84074 QRICH2 glutamine rich 2 5457 -0.265541825555236
2.47468785883344 0.917177425730682
0.338217181118313 0.769513780917081
55026 TMEM255A transmembrane protein 255A
3858 -0.580918722351132 1.03629397339779
0.916994968851865 0.338265234338044 0.769513780917081
1845 DUSP3 dual specificity phosphatase 3 5012
0.117002020542769 7.68310369979354 0.916720395769744
0.338337565250987 0.769513780917081
10333 TLR6 toll like receptor 6 6033 -0.412801169896151
2.05038485548582 0.916695766951463
0.338344054246586 0.769513780917081
51227 PIGP phosphatidylinositol glycan anchor biosynthesis class P 7923 -0.217685971182839 3.34582029436261
0.916516951966201 0.338391171937942 0.769513780917081
4493 MT1E metallothionein 1E 1052 0.311710929411255
3.50253309121203 0.916419496329752
0.338416855175927 0.769513780917081
57037 ANKMY2 ankyrin repeat and MYND domain containing 2 2854
0.190076972808144 3.90186279477288
0.91578036623509 0.338585354959789 0.769771963467256
7004 TEAD4 TEA domain transcription factor 4 2272
0.238958643750568 2.5236741346325 0.915409411095505
0.338683204970823 0.769869466411281
344787 ZNF860 zinc finger protein 860 3263 -0.278631200333197
1.72634714830242 0.915043268059185
0.338779822898667 0.769955583207064

51439	FAM8A1	family with sequence similarity 8 member A1	4677
	0.129584127396081	4.76846542662154
0.914849276664907	0.338831028406393	0.769955583207064
100506469	TMEM147-AS1	TMEM147 antisense RNA 1 3579
0.306791488231121	1.77346814627129	0.914040137478281
	0.339044719021935	0.770159500360855
8218	CLTCL1	clathrin heavy chain like 1	10049
0.199368147454131	4.59783305494954	0.914010844683461
	0.33905245853447	0.770159500360855
79068	FTO	FTO alpha-ketoglutarate dependent dioxygenase
12723	0.138004947235429	5.04889995417223
0.9138849768405 0.339085717050431	0.770159500360855
7803	PTP4A1	protein tyrosine phosphatase 4A1
5796	-0.13239547436429	7.82005466667329
0.912870792430674	0.339353858710675	0.770596999010715
200942	KLHDC8B kelch domain containing 8B	2158
0.193317998706398	4.37659447787602	0.912454391774423
	0.339464033996929	0.770596999010715
9780	PIEZO1	piezo type mechanosensitive ion channel component 1
	10855	-0.164455955478353	9.48105287950216
0.91235664113929	0.339489904767366	0.770596999010715
1287	COL4A5	collagen type IV alpha 5 chain	8738
0.241091185249345	6.33749415695718	0.912288947810012
	0.339507822097646	0.770596999010715
4716	NDUFB10 NADH:ubiquinone oxidoreductase subunit B10	1070
	0.169809593129394	5.18195752577939
0.911925591147586	0.339604018462762	0.770596999010715
65267	WNK3	WNK lysine deficient protein kinase 3
11472	-0.500567785509931	2.05234142085453
0.911908932145451	0.339608429706086	0.770596999010715
4055	LTBR	lymphotoxin beta receptor	6280
0.15191579755703	7.2329271328924 0.911047919549077
0.339836527650545	0.770905172794004
138474	TAF1L	TATA-box binding protein associated factor 1 like
	6216	-0.272174473441013	1.42358271845796
0.910980795473574	0.339854318698006	0.770905172794004
23034	SAMD4A	sterile alpha motif domain containing 4A
12170	-0.160769548702767	6.88574648813756
0.910548330945128	0.339968972243435	0.770989847953808
23317	DNAJC13 DnaJ heat shock protein family (Hsp40) member C13
	9562	-0.154529611255701	7.02613505581401
0.910351794369255	0.340021094555029	0.770989847953808
175	AGA	aspartylglucosaminidase 2675	-0.181044811838789
	4.1882546213076 0.910217269236336	0.340056777372644
	0.770989847953808
51501	HIKESHI heat shock protein nuclear import factor hikeshi
	3351	0.160032108338198	4.26982825297673
0.90960356534525	0.340219626373784	0.771123655027261
7170	TPM3	tropomyosin 3	8389	-0.172335069065136
8.68013462735295	0.909142526041023	0.340342034186314
	0.771123655027261
286257	PAXX	PAXX non-homologous end joining factor
1367	-0.227213109326716	2.43807646657011
0.908630700284836	0.340477995432596	0.771123655027261

283971  CLEC18C C-type lectin domain family 18 member C 4894
0.278561530953067       -0.11564220610455       0.90846753291495
        0.34052135453076        0.771123655027261
57234   LINC00869       long intergenic non-protein coding RNA 869
        8501    -0.193902307979387      3.6105473208043
0.908439108539321       0.340528908610399       0.771123655027261
607     BCL9    BCL9 transcription coactivator
6420    -0.177227221101595      4.4630864221901 0.908056484607414
        0.340630616939828       0.771123655027261
25977   NECAP1  NECAP endocytosis associated 1
4155    -0.132474143078138      5.19220987317016
0.907722224646932       0.340719502721582       0.771123655027261
246721  POLR2J2 RNA polymerase II subunit J2
3233    -0.373408071407498      5.2221694468821 0.907674912628607
        0.340732086367114       0.771123655027261
1738    DLD     dihydrolipoamide dehydrogenase
5299    -0.114236368355077      6.43910377994447
0.907582778819034       0.340756593124443       0.771123655027261
9874    TLK1    tousled like kinase 1   9405    -0.116129242949321
        6.09181378740169        0.907550987823626
0.340765049791398       0.771123655027261
103091866       LINC01145       long intergenic non-protein coding
RNA 1145        2290    -0.243929389996406      1.44367986080369
        0.907480614532432       0.340783770672211
0.771123655027261
5193    PEX12   peroxisomal biogenesis factor 12        2861
0.22033257536104        2.4225995655844 0.907411142559515
0.340802253140228       0.771123655027261
26995   TRUB2   TruB pseudouridine synthase family member 2     6125
        0.139475216144026       5.35817853389956
0.907301292753589       0.340831480564987       0.771123655027261
10993   SDS     serine dehydratase      3006    1.04189907336908
        0.18286892894001        0.906809905851217
0.340962263799679       0.771254263320144
51005   AMDHD2  amidohydrolase domain containing 2      4533
0.249982773655987       4.02752267303233        0.906520756268364
        0.34103925290249        0.771254263320144
55334   SLC39A9 solute carrier family 39 member 9
7136    -0.104821470864039      6.89402504189388
0.906262172893684       0.341108123263188       0.771254263320144
10656   KHDRBS3 KH RNA binding domain containing, signal
transduction associated 3       8258    -0.300100752679002
1.84549850175741        0.906257166672307       0.341109456790768
        0.771254263320144
51517   NCKIPSD NCK interacting protein with SH3 domain 3431
0.14506653033536        4.36392692780875        0.905909449298067
        0.341202096867411       0.771339214141676
54842   MFSD6   major facilitator superfamily domain containing 6
        6031    -0.12316877700543       7.56472440819542
0.905518561798437       0.341306278944932       0.771446623840454
254359  ZDHHC24 zinc finger DHHC-type containing 24     6266
0.16631443378434        4.17500257035993        0.90525089801792
        0.34137764329204        0.771446623840454
51308   REEP2   receptor accessory protein 2    3284

0.46477148535091	1.88584717627503	0.904803521822224
	0.341496967336228	0.771446623840454
79929	MAP6D1	MAP6 domain containing 1	2408
0.312031516644791	1.23092414525762	0.904518234756197
	0.34157308836206	0.771446623840454
6367	CCL22	C-C motif chemokine ligand 22
2929	-0.806660901098338	0.67229390230535
0.904301930869789	0.341630818359471	0.771446623840454
9267	CYTH1	cytohesin 1	5508	-0.125149242126473
6.88866840831709	0.904185331991954	0.341661943240344
	0.771446623840454
81889	FAHD1	fumarylacetoacetate hydrolase domain containing 1
	3039	0.161070941862313	4.68968545561902
0.903895958417434	0.341739205076896	0.771446623840454
81932	HDHD3	haloacid dehalogenase like hydrolase domain
containing 3	2105	0.204871915994257	4.30506200663912
	0.903873394953512	0.341745230440545
0.771446623840454
29992	PILRA	paired immunoglobin like type 2 receptor alpha	2006
	0.429672923788101	2.38452671233993
0.903873165531532	0.341745291706291	0.771446623840454
7515	XRCC1	X-ray repair cross complementing 1	3628
0.15865091180204	4.60589236654059	0.903487270011649
	0.341848363718215	0.771541435131631
10806	SDCCAG8 serologically defined colon cancer antigen 8	5322
	0.163705308629053	4.56914439471943
0.903107753235267	0.341949772853484	0.771541435131631
202020	TAPT1-AS1	TAPT1 antisense RNA 1 (head to head)	2763
	0.308352642025466	0.305265639913426
0.902692017516535	0.342060906455989	0.771541435131631
54517	PUS7	pseudouridine synthase 7	4421
0.242526246774176	3.7498329420064 0.902660833360182
0.34206924450262	0.771541435131631
8871	SYNJ2	synaptojanin 2	9921	-0.293535497815427
6.58601425794752	0.902502114572762	0.342111687116161
	0.771541435131631
202018	TAPT1	transmembrane anterior posterior transformation 1
	7828	0.136752195447439	4.85573389955488
0.902479302611757	0.342117787792001	0.771541435131631
79174	CRELD2	cysteine rich with EGF like domains 2
5165	-0.223408335675846	4.05650649419339
0.902007220312342	0.342244071210951	0.771635051170415
55608	ANKRD10 ankyrin repeat domain 10
4563	-0.196887093224404	6.08766991881272
0.901585203967301	0.34235701505906	0.771635051170415
7917	BAG6	BCL2 associated athanogene 6
5310	-0.121786352215128	8.08731959586032
0.901397388985785	0.342407295975179	0.771635051170415
128240	NAXE	NAD(P)HX epimerase	1933	0.148221315794738
	4.91159618866877	0.901375597920321
0.342413130418404	0.771635051170415
3911	LAMA5	laminin subunit alpha 5 13596	0.166415011631641
	8.41771734044731	0.90093386075273
0.342531432140825	0.771635051170415

5898	RALA	RAS like proto-oncogene A
3665	-0.261735571393107	7.70667663460549
0.900482112809328	0.342652471884467	0.771635051170415
29080	CCDC59	coiled-coil domain containing 59
3784	-0.171216092873976	3.90543683311456
0.90044842550709	0.342661500252756	0.771635051170415
143458	LDLRAD3	low density lipoprotein receptor class A domain
containing 3	4575	0.205505624317293	3.41510987571491
	0.900064001759617	0.34276455050561
0.771635051170415
5048	PAFAH1B1	platelet activating factor acetylhydrolase
1b regulatory subunit 1	7622	-0.120306934402233
7.96801004834444	0.900018313104669	0.342776800779533
	0.771635051170415
253769	WDR27	WD repeat domain 27	10244	-0.255645988438939
	3.76553169214682	0.899669413380785
0.342870369028805	0.771635051170415
9897	WASHC5	WASH complex subunit 5	4784	-0.116062652212253
	6.22765934575202	0.899482962031854
0.342920385855436	0.771635051170415
79759	ZNF668	zinc finger protein 668	3935	0.164395742112688
	4.03985916328076	0.898922904351977
0.343070684318359	0.771635051170415
55552	ZNF823	zinc finger protein 823	2459	-0.189313307813743
	3.23469994838363	0.89866637473643
0.343139556932145	0.771635051170415
23678	SGK3	serum/glucocorticoid regulated kinase family member
3	6007	-0.166911353905288	3.8781234107748
0.898376300901246	0.343217457926011	0.771635051170415
55225	RAVER2	ribonucleoprotein, PTB binding 2	6070
0.169920653328592	4.29488062878417	0.898143969675971
	0.343279869031775	0.771635051170415
319089	TTC6	tetratricopeptide repeat domain 6
8241	-0.323672205160403	1.66385640146462
0.898130405854057	0.34328351315281	0.771635051170415
119548	PNLIPRP3	pancreatic lipase related protein 3	2354
	0.823365849246904	3.104976798427	0.898011253274039
	0.343315527497675	0.771635051170415
2710	GK	glycerol kinase	5359	-0.43164096095991
2.00895469770055	0.897818854677111	0.343367230357365
	0.771635051170415
8841	HDAC3	histone deacetylase 3	4631	0.118352819870561
	5.82333379941474	0.897808508579651
0.343370010939688	0.771635051170415
54585	LZTFL1	leucine zipper transcription factor like 1	7201
	0.187894087362203	4.2734061650646	0.897772167821685
	0.343379778000427	0.771635051170415
163782	KANK4	KN motif and ankyrin repeat domains 4	5980
0.425907964890923	2.5198119154271	0.897754208265833
0.343384605007797	0.771635051170415
1000	CDH2	cadherin 2	4950	0.406375308252166
6.75345008206868	0.897616173620406	0.343421707772966
	0.771635051170415
100526830	SLX1A-SULT1A3	SLX1A-SULT1A3 readthrough (NMD

candidate)	8665	-0.210053737375187	5.64013710981472
	0.896843197989043	0.343629578396799
0.771635051170415
677825	SNORA44 small nucleolar RNA, H/ACA box 44
132	-0.296113881142922	0.301197002104124
0.896773189501948	0.343648413653773	0.771635051170415
100507567	RAB11B-AS1	RAB11B antisense RNA 1	1156
0.289676855319567	1.72645340737095	0.896764752654764
	0.343650683618216	0.771635051170415
11226	GALNT6	polypeptide N-acetylgalactosaminyltransferase 6
7347	-0.260614671822426	4.5664591247761 0.89661414449453
	0.343691208700047	0.771635051170415
50862	RNF141	ring finger protein 141 5146	-0.188758584250775
	5.95619856729491	0.896599778045214
0.343695074707882	0.771635051170415
26064	RAI14	retinoic acid induced 14	7687
0.324790838251153	8.52262024070306	0.896525015058614
	0.343715194356765	0.771635051170415
57599	WDR48	WD repeat domain 48	7488	-0.109084766434285
	5.93927859967867	0.896370106567582
0.343756887220918	0.771635051170415
89790	SIGLEC10	sialic acid binding Ig like lectin 10	3536
	1.32594889347187	0.195134593423487
0.895415506206744	0.344013964096458	0.771857691005854
8641	PCDHGB4 protocadherin gamma subfamily B, 4	6906
0.130187369581436	6.50809890089349	0.89530510666725
	0.3440437117962 0.771857691005854
55705	IPO9	importin 9	11915	0.116845043243412
6.83001221414639	0.895195761761268	0.344073178746444
	0.771857691005854
53836	GPR87	G protein-coupled receptor 87	1608
0.127194697912986	6.89875670864037	0.89507354303251
	0.344106119052007	0.771857691005854
200232	FAM209A family with sequence similarity 209 member A	1137
	0.154679918592487	4.09326908063132
0.894979022498732	0.344131597082284	0.771857691005854
4141	MARS1	methionyl-tRNA synthetase 1
6623	-0.116095879582291	7.66392727919225
0.89425358497477	0.344327223828802	0.77210002398743
79746	ECHDC3	enoyl-CoA hydratase domain containing 3 2185
0.445052625176763	2.19951197804074	0.894169584336201
	0.344349885760654	0.77210002398743
11017	SNRNP27 small nuclear ribonucleoprotein U4/U6.U5 subunit 27
	2275	-0.176702251034898	5.03994694650631
0.892821338627584	0.34471389543012	0.772541186491908
653160	LOC653160	uncharacterized LOC653160	3346
0.317413074371797	0.65599137142337	0.892806729839733
	0.344717842471139	0.772541186491908
6923	ELOB	elongin B	1128	0.17280247178594
6.27772150321434	0.892740587822928	0.3447357136614
0.772541186491908
8565	YARS1	tyrosyl-tRNA synthetase 1
8212	-0.121487199973844	6.45072392894824
0.892623859113546	0.344767256140567	0.772541186491908

222183	SRRM3	serine/arginine repetitive matrix 3
6873	-0.364920039207469	1.58080404238264
0.892289754675172	0.344857559556318	0.772619935493497
23379	ICE1	interactor of little elongation complex ELL subunit
1	8678	-0.136926191439974	6.05133234266455
0.891675335589442	0.345023711355696	0.772785946010873
79184	BRCC3	BRCA1/BRCA2-containing complex subunit 3
3182	-0.123541938904535	5.14039297460727
0.891607726508758	0.345042000881845	0.772785946010873
6015	RING1	ring finger protein 1	1818	0.134026287996994
	5.42149727098735	0.891351606550522
0.345111298039131	0.772817578198899
3336	HSPE1	heat shock protein family E (Hsp10) member 1	1977
	0.213521272317816	4.77612125999035
0.890801467961097	0.345260210098327	0.772962821635709
100526794	NT5C1B-RDH14	NT5C1B-RDH14 readthrough	2324
	0.228492448811613	1.96620379960804
0.89061657090452	0.34531027773977	0.772962821635709
256691	MAMDC2	MAM domain containing 2	3854	0.424467174158601
	2.85672723794412	0.890500510291874
0.345341710415954	0.772962821635709
219931	TPCN2	two pore segment channel 2	6130
0.16545631418579	4.7077429560352	0.889550679368227
0.345599098566351	0.773097743542276
26127	FGFR1OP2	FGFR1 oncogene partner 2
5971	-0.147043834902856	5.1646224909105
0.889524688960435	0.345606145177881	0.773097743542276
81704	DOCK8	dedicator of cytokinesis 8	13784
0.254551714310552	3.92828037286874	0.88951184101921
	0.345609628629114	0.773097743542276
57184	FAM219B	family with sequence similarity 219 member B	3771
	0.148018593910167	6.21592080766213
0.889463391804895	0.345622765050456	0.773097743542276
7873	MANF	mesencephalic astrocyte derived neurotrophic factor
	2111	-0.224782303222504	5.64206596651936
0.889151157986781	0.345707439711166	0.773163676837667
1464	CSPG4	chondroitin sulfate proteoglycan 4	8290
0.423889722654996	3.5259822288379	0.888429471824839
0.345903261240931	0.773478125830414
152110	NEK10	NIMA related kinase 10	9582	0.335543595917775
	0.714133983317117	0.888070391296986
0.346000749723736	0.773488271168445
899	CCNF	cyclin F	4932	-0.231310476715407
2.63662810211947	0.888005978528768	0.346018241389411
	0.773488271168445
613037	NPIPB13	nuclear pore complex interacting protein family,
member B13	2295	-0.312330561117056	0.457176715546254
	0.887648952401733	0.346115215654289
0.773581590181845
9900	SV2A	synaptic vesicle glycoprotein 2A	4824
0.537044033618102	1.1078883122067	0.887349635900406
0.346196543393103	0.773639914375749
10021	HCN4	hyperpolarization activated cyclic nucleotide gated
potassium channel 4	7228	0.487972848083407

0.197511656452232	0.886712703575892	0.346369691424096
	0.773851455497715
1366	CLDN7	claudin 7	2712	-0.150388591439729
8.81025837184581	0.886512906592827	0.346424029776883
	0.773851455497715
100271836	SMG1P3	SMG1 pseudogene 3
937	-0.286943798361532	2.31224652416716
0.886391888273961	0.34645694848477	0.773851455497715
54432	YIPF1	Yip1 domain family member 1
2075	-0.112568637633236	6.01195676948603
0.886009662304012	0.346560947241098	0.773886630567698
5981	RFC1	replication factor C subunit 1	6713
0.138492954803518	6.12034470493349	0.885867643995439
	0.346599599372128	0.773886630567698
80212	CCDC92	coiled-coil domain containing 92	8223
0.147111002087213	5.60597603735433	0.885426230281382
	0.346719773171696	0.773886630567698
257000	TINCR	TINCR ubiquitin domain containing
3925	-0.485681998387158	3.48674551175202
0.885093228093648	0.346810469581123	0.773886630567698
114932	MRFAP1L1	Morf4 family associated protein 1 like 1
	1593	0.119205120591243	5.8628132285483
0.884945054765467	0.346850836379523	0.773886630567698
8727	CTNNAL1	catenin alpha like 1	3131	0.183053050323699
	4.82566079630579	0.884895438808137
0.346864354657897	0.773886630567698
659	BMPR2	bone morphogenetic protein receptor type 2
11893	-0.22066336470608	8.17139411574298
0.884859766312957	0.346874074365997	0.773886630567698
93487	MAPK1IP1L	mitogen-activated protein kinase 1
interacting protein 1 like	6988	-0.10688685629969
6.49203687859109	0.883998812105631	0.347108771185402
	0.773886630567698
9711	RUBCN	rubicon autophagy regulator
8613	-0.124123997765695	5.87004404090992
0.883929440832164	0.347127691225024	0.773886630567698
11180	WDR6	WD repeat domain 6	5304	0.130318241984663
	7.71886390349722	0.883782496173332
0.347167772934605	0.773886630567698
114625	ERMAP	erythroblast membrane associated protein (Scianna
blood group)	4631	0.161530888634491	4.30759831987669
	0.883553683476535	0.347230198072938
0.773886630567698
6522	SLC4A2	solute carrier family 4 member 2	7590
0.139207423027112	7.7554455132916	0.883429793662202
0.347264004294119	0.773886630567698
60487	TRMT11	tRNA methyltransferase 11 homolog
2690	-0.177048265996782	3.42156391873895
0.883377124034935	0.347278377781083	0.773886630567698
80169	CTC1	CST telomere replication complex component 1
8367	-0.144051030971187	4.7653781300626	0.883364191363603
	0.347281907217161	0.773886630567698
91419	ATP23	ATP23 metallopeptidase and ATP synthase assembly
factor homolog	1373	0.276534315443327	0.722621451408749

0.88329259849108	0.347301446401695
0.773886630567698
114984	FLYWCH2	FLYWCH family member 2	1365	0.183142555787203
3.53758604163668	0.882259928824325
0.347583449426675	0.774391820601151
9263	STK17A	serine/threonine kinase 17a	4943
0.147669206261692	6.90821336345488	0.881693686020185
0.347738211792234	0.774613411668865
23373	CRTC1	CREB regulated transcription coactivator 1	6965
0.202540092286622	3.19347760813952
0.881356996589393	0.347830278233537	0.774695294516958
79813	EHMT1	euchromatic histone lysine methyltransferase 1
16291	-0.115861645776467	6.28431796266271
0.881109062934769	0.347898095900535	0.774723152508553
3759	KCNJ2	potassium inwardly rectifying channel subfamily J
member 2	5577	-0.318032242245524	1.37997742613772
0.880879295790276	0.347960960480711
0.774739973526199
84816	RTN4IP1	reticulon 4 interacting protein 1
1974	-0.303950439914857	2.23681852845139
0.880344480315229	0.348107346407423	0.774942721094847
7091	TLE4	TLE family member 4, transcriptional corepressor
8187	-0.16059951153705	5.0618275077727
0.879252438755696	0.348406512152179	0.775458073532018
132160	PPM1M	protein phosphatase, Mg2+/Mn2+ dependent 1M	3571
0.257284145629155	3.06659821013802
0.878940814725784	0.348491945867781	0.775458073532018
51167	CYB5R4	cytochrome b5 reductase 4
9635	-0.108473489805998	5.93913383007011
0.87889345871236	0.348504931311776	0.775458073532018
1977	EIF4E	eukaryotic translation initiation factor 4E
14067	-0.166741038545608	4.70403292866602
0.878113007250489	0.348719032830886	0.775811228218268
100130581	LINC00910	long intergenic non-protein coding
RNA 910	4168	0.357546645293514	0.278924575866844
0.877303952785505	0.348941169739714	0.775906004346052
340371	NRBP2	nuclear receptor binding protein 2
4924	-0.125648692332079	5.47744874824027
0.877161497960168	0.348980302560381	0.775906004346052
4253	MIA2	MIA SH3 domain ER export factor 2
9550	-0.124939947626162	6.01783335212092
0.876720524326169	0.349101477324858	0.775906004346052
10840	ALDH1L1	aldehyde dehydrogenase 1 family member L1
13712	-0.334611578854153	0.111938448234216
0.876670620298536	0.349115194011981	0.775906004346052
65992	DDRGK1	DDRGK domain containing 1	1832
0.12405522823933	6.37122247549179	0.87659169098373
0.349136890123833	0.775906004346052
782	CACNB1	calcium voltage-gated channel auxiliary subunit beta
1	7094	-0.192381993658768	3.54486950629337
0.876216651538116	0.349240006122615	0.775906004346052
57555	NLGN2	neuroligin 2	5550	0.313989478695726
5.22882244937561	0.876198507854554	0.349244995725331
0.775906004346052

79074	C2orf49 chromosome 2 open reading frame 49
4768	-0.144633486644168	4.8638302051994 0.875783041976841
	0.349359277423762	0.775906004346052
2237	FEN1	flap structure-specific endonuclease 1	2814
0.18895224797085	3.78613093067707	0.875759759231412
	0.349365683285087	0.775906004346052
10891	PPARGC1A	PPARG coactivator 1 alpha	10612
0.591010890013301	2.04693827403992	0.875304231959424
	0.349491046201655	0.775906004346052
54776	PPP1R12C	protein phosphatase 1 regulatory subunit 12C
	4062	-0.130691412054151	6.24757046959637
0.875222944153396	0.34951342336149	0.775906004346052
5911	RAP2A	RAP2A, member of RAS oncogene family	5570
0.147406311016211	5.65639401302651	0.875165927780554
	0.349529120168569	0.775906004346052
66008	TRAK2	trafficking kinesin protein 2
7627	-0.121263248162887	6.48821354927543
0.875130419508916	0.349538896204581	0.775906004346052
7569	ZNF182	zinc finger protein 182 3587	0.259191880840361
	3.33624657148591	0.87510880490313
0.349544847258254	0.775906004346052
26020	LRP10	LDL receptor related protein 10
7112	-0.116026163325303	9.21817095836523
0.874935577702104	0.349592546114652	0.775906004346052
128308	MRPL55	mitochondrial ribosomal protein L55	1886
0.173639658229867	3.68360602549106	0.874537076321629
	0.349702308804127	0.775978662914756
60674	GAS5	growth arrest specific 5	3631
0.173226238638967	4.83131602386943	0.874358388945979
	0.349751541440985	0.775978662914756
54902	TTC19	tetratricopeptide repeat domain 19
5966	-0.128315894183403	6.29160132494009
0.874213454618058	0.349791481233938	0.775978662914756
221472	FGD2	FYVE, RhoGEF and PH domain containing 2 10167
0.390871214543976	1.76389313124287	0.87376744264941
	0.349914428429705	0.776128487571635
3839	KPNA3	karyopherin subunit alpha 3
4474	-0.109966294796968	6.53245442125841
0.87305944833841	0.350109714197548	0.776369584900674
100859930	HEIH	hepatocellular carcinoma up-regulated EZH2-associated long non-coding RNA	1652	0.220693702141732
3.79742381491649	0.872867999088381	0.350162547039662
	0.776369584900674
54890	ALKBH5	alkB homolog 5, RNA demethylase 3190
0.107956592898889	6.12592574942373	0.872770678788751
	0.35018940796762	0.776369584900674
79797	ZNF408	zinc finger protein 408 3078	0.169818447689558
	4.37025377280285	0.872318000411241
0.350314386515539	0.776473129097356
4076	CAPRIN1 cell cycle associated protein 1
7689	-0.109345949155838	8.51611383117526
0.871892086860683	0.350432031096224	0.776473129097356
283537	SLC46A3 solute carrier family 46 member 3	3776
0.236991689006124	4.05865678812699	0.871833602107561

	0.350448189781816	0.776473129097356
5603	MAPK13	mitogen-activated protein kinase 13
7197	-0.121642417256719	7.31101567583472
0.871798635218596	0.350457851228206	0.776473129097356
160418	TMTC3	transmembrane O-mannosyltransferase targeting
cadherins 3	7327	-0.113083163391233	6.47494694324132
	0.871409626664175	0.350565359814313
0.776588485674376
9439	MED23	mediator complex subunit 23
7412	-0.130632114000787	5.94210960938512
0.870383656809132	0.35084911812507	0.777094180515157
64595	TTTY15	testis-specific transcript, Y-linked 15
5284	-0.213889540893523	4.42852017607696
0.86991086145947	0.350979987134324	0.777157378997268
6429	SRSF4	serine and arginine rich splicing factor 4
8893	-0.105493087772312	6.44389521201232
0.869879686992301	0.350988618514794	0.777157378997268
84446	BRSK1	BR serine/threonine kinase 1	3841
0.38752631194772	0.792686535103122	0.868196245921277
	0.351455148439195	0.777865705192992
7126	TNFAIP1	TNF alpha induced protein 1
3889	-0.13412608504554	8.02620472669395
0.867994218994738	0.351511192701097	0.777865705192992
55638	SYBU	syntabulin	7024	-0.266514857800998
2.53395978958114	0.867791028312807	0.351567572090457
	0.777865705192992
84457	PHYHIPL	phytanoyl-CoA 2-hydroxylase interacting protein like
	4093	0.482488232964645	0.965001641764563
0.867787850195585	0.351568454021673	0.777865705192992
160	AP2A1	adaptor related protein complex 2 subunit alpha 1
	7386	0.122405094034189	6.94774700319906
0.86766525256283	0.351602477309639	0.777865705192992
9743	ARHGAP32	Rho GTPase activating protein 32
12841	-0.286605533975112	6.50997052608834
0.867146742321077	0.35174642390069	0.777865705192992
283578	TMED8	transmembrane p24 trafficking protein family member
8	7784	-0.143255739552549	5.16290378656692
0.867041810940677	0.351775564277463	0.777865705192992
54991	C1orf159	chromosome 1 open reading frame 159
6322	-0.17087729133417	4.29426453287095
0.866985169916941	0.351791295361292	0.777865705192992
80335	WDR82	WD repeat domain 82	5726	-0.104176980678767
	7.09123276110479	0.866694307342947
0.351872092624018	0.777865705192992
898	CCNE1	cyclin E1	2043	0.23086267986794
2.63679348311674	0.866317448045708	0.351976816133386
	0.777865705192992
28960	DCPS	decapping enzyme, scavenger	3352
0.202241800890948	2.93525477827215	0.866316515840166
	0.35197707523194	0.777865705192992
9677	PPIP5K1	diphosphoinositol pentakisphosphate kinase 1	8075
	0.143623745827662	4.79621282187726
0.866266707799054	0.351990919329138	0.777865705192992
843	CASP10	caspase 10	7598	0.282023047493361

4.25931206648423	0.865971670980854	0.352072939762916
	0.777865705192992
5431	POLR2B	RNA polymerase II subunit B
5709	-0.111311610700692	7.36603495273223
0.865729120266224	0.3521403885543 0.777865705192992
8269	TMEM187 transmembrane protein 187	1924
0.275698132976883	2.61894699782245	0.865725009437307
	0.352141531850415	0.777865705192992
100134938	UPK3BL1 uroplakin 3B like 1
1342	-0.384741976523119	4.97006866215351
0.864847219219186	0.352385777292844	0.778181444181605
8526	DGKE	diacylglycerol kinase epsilon
10933	-0.27410850174653	3.45516780346982
0.864812003820626	0.352395580812731	0.778181444181605
51449	PCYOX1	prenylcysteine oxidase 1	5870
0.134065751732194	7.46060073935661	0.864424378372515
	0.352503515419376	0.77823051427553
8292	COLQ	collagen like tail subunit of asymmetric
acetylcholinesterase	4851	0.342860548587059
0.133291005759185	0.864333158625064	0.352528922187352
	0.77823051427553
79596	OBI1	ORC ubiquitin ligase 1	3529	-0.155474184027725
	4.11560786834449	0.864111405942686
0.352590695762288	0.778244228733433
5915	RARB	retinoic acid receptor beta	4281
0.322811418875364	5.43505077594888	0.863911566732781
	0.352646377566979	0.778244495128514
100885850	PTGES3L-AARSD1	PTGES3L-AARSD1 readthrough	2383
	0.203490762835954	3.45408062532528
0.863573354840683	0.352740641925532	0.778329894683511
7579	ZSCAN20 zinc finger and SCAN domain containing 20
5308	-0.267367976666961	3.06097739272211
0.86261761088049	0.353007207114461	0.778633545946054
2274	FHL2	four and a half LIM domains 2	6626
0.223321935652746	8.31339688686885	0.862509787490367
	0.353037297249227	0.778633545946054
9289	ADGRG1	adhesion G protein-coupled receptor G1
8659	-0.13648931984475	9.00746459749352
0.862326018210069	0.353088589579681	0.778633545946054
7844	RNF103	ring finger protein 103 7896	-0.107609078515648
	6.77199234326193	0.862251341155005
0.353109435798825	0.778633545946054
3712	IVD	isovaleryl-CoA dehydrogenase	6533
0.182249966082693	5.45671969033023	0.861808248439331
	0.353233160425709	0.778633545946054
10904	BLCAP	BLCAP apoptosis inducing factor 3252
0.10684183214013	6.93452961237402	0.861774918030794
	0.353242469659004	0.778633545946054
25875	LETMD1	LETM1 domain containing 1	4400
0.144633311979086	4.37324930053403	0.86168573777153
	0.353267379487911	0.778633545946054
150864	FAM117B family with sequence similarity 117 member B	5986
	0.196199993337373	3.42846267269272
0.861392823150279	0.353349213284455	0.778691383020038

9446	GSTO1	glutathione S-transferase omega 1
1691	-0.206646169795542	6.37900830853427
0.861057051210933	0.353443052323432	0.778737162848962
80142	PTGES2	prostaglandin E synthase 2
3039	-0.148931421539672	5.34457361790978
0.860130203276984	0.353702257688295	0.778737162848962
201595	STT3B	STT3 oligosaccharyltransferase complex catalytic
subunit B	5012	0.12203781473114	7.61171975721171
	0.860111618410343	0.353707457850364
0.778737162848962
51109	RDH11	retinol dehydrogenase 11
5976	-0.147782974058023	7.4737195833767 0.860078803554543
	0.353716639907677	0.778737162848962
79078	C1orf50	chromosome 1 open reading frame 50	2105
0.203013936723	3.01629139125124	0.859921345905907
0.35376070330113	0.778737162848962
440345	NPIPB4	nuclear pore complex interacting protein family
member B4	7747	-0.233254481396372	3.87973413827142
	0.859751830336333	0.353808149405132
0.778737162848962
1154	CISH	cytokine inducible SH2 containing protein
5112	-0.297623353663589	2.2331672720059 0.859678731713765
	0.353828611831754	0.778737162848962
30837	SOCS7	suppressor of cytokine signaling 7
8621	-0.143567002779698	5.37759105721471
0.859663868138277	0.353832772775976	0.778737162848962
4591	TRIM37	tripartite motif containing 37
4841	-0.125020200190797	5.81912167136751
0.859529634435282	0.353870353504222	0.778737162848962
84617	TUBB6	tubulin beta 6 class V	4117	0.18768635651417
	8.52609720769605	0.859160804245903
0.353973641120793	0.778828964140499
157680	VPS13B	vacuolar protein sorting 13 homolog B
16716	-0.218285212282971	5.98337136495464
0.858814141279794	0.3540707585871 0.778828964140499
3836	KPNA1	karyopherin subunit alpha 1
8451	-0.103193522734831	6.75136489221742
0.858785164870987	0.354078877964353	0.778828964140499
10412	NSA2	NSA2 ribosome biogenesis factor 4962
0.173874412771287	5.41113340573263	0.858069976837559
	0.354279359008102	0.778931964221713
54521	WDR44	WD repeat domain 44	4202	-0.110090951002739
	5.89851476818073	0.858031550480952
0.35429013505388	0.778931964221713
57661	PHRF1	PHD and ring finger domains 1
5769	-0.13297244079234	6.43732917441333
0.858022991111771	0.354292535450599	0.778931964221713
23352	UBR4	ubiquitin protein ligase E3 component n-recognin 4
	24675	-0.250554567792667	8.76245426417269
0.857427811889465	0.354459502587049	0.779176750272567
102724571	LNCTAM34A	long non coding transcriptional
activator of miR34a	1662	0.279085870684403
1.18818826678277	0.857071706401783	0.35455945323883
	0.779274166250791

8761	PABPC4	poly(A) binding protein cytoplasmic 4
6782	-0.149664312326909	6.66013011165513
0.856637008553932	0.354681515243359	0.77932125325694
120227	CYP2R1	cytochrome P450 family 2 subfamily R member 1	4516
	0.147621110208216	3.92567953687385
0.856599133525363	0.354692153174341	0.77932125325694
148413	MRPL20-AS1	MRPL20 antisense RNA 1	2525
0.15792166416135	3.84444030051366	0.856127539462892
	0.354824646021643	0.779398176815671
3632	INPP5A	inositol polyphosphate-5-phosphatase A
3706	-0.130697388377812	5.1021811287250 7
0.856078418429071	0.354838450314381	0.779398176815671
55556	ENOSF1	enolase superfamily member 1	10616
0.2139572009568	4.07478198396107	0.855657538805154
0.354956758614214	0.779535797728018
79623	GALNT14	polypeptide N-acetylgalactosaminyltransferase 14
	4567	0.211535393424056	3.81501948805495
0.855155708354232	0.355097892595343	0.779621880251674
116228	COX20	cytochrome c oxidase assembly factor COX20	4864
	0.153862594067449	4.77031559913449
0.855122358243207	0.355107274648796	0.779621880251674
1	A1BG	alpha-1-B glycoprotein	6535	0.345142539519813
	0.564021292035503	0.854556810094859
0.355266426352496	0.779846983394893
29796	UQCR10	ubiquinol-cytochrome c reductase, complex III
subunit X	975	0.147424687911053	5.61022951719178
	0.854362403530404	0.355321157137589
0.779846983394893
27152	INTU	inturned planar cell polarity protein
16066	-0.210523882991106	3.78740038026594
0.854024329320023	0.355416361814175	0.779933727076791
11147	HHLA3	HERV-H LTR-associating 3	1222
0.295895480139526	1.67608625517474	0.853317790031551
	0.355615442351703	0.780248355422979
10327	AKR1A1	aldo-keto reductase family 1 member A1	3121
0.139020937497797	6.02388592604967	0.852815365024455
	0.355757102862852	0.780436920877052
50999	TMED5	transmembrane p24 trafficking protein 5
9423	-0.154969796133436	6.62443187985211
0.852352675973847	0.35588762808264	0.780601003218531
55072	RNF31	ring finger protein 31	8046	-0.125544936980273
	6.37223933155956	0.852147112736512
0.355945638811845	0.780606006393849
641649	TMEM91	transmembrane protein 91	2796
0.326089498820238	0.743795165140196	0.851611061630334
	0.356096975456303	0.780815644210462
7760	ZNF213	zinc finger protein 213	6443	-0.16345635718686
	4.06316709020278	0.850889161053374
0.356300920083463	0.781023792800477
8460	TPST1	tyrosylprotein sulfotransferase 1
2871	-0.170808420477849	5.09427063449272
0.850751381952245	0.356339862349975	0.781023792800477
145376	PPP1R36	protein phosphatase 1 regulatory subunit 36
3093	-0.296418656352093	0.194304268367796

0.850683036644567	0.356359181822506	0.781023792800477
1662	DDX10	DEAD-box helicase 10	8461	0.16303078176987
	3.86757785282345	0.850170991877	0.3565039695007
0.781218883103302
64979	MRPL36	mitochondrial ribosomal protein L36	1394
0.153532135606513	4.14054089708503	0.849929641743353
	0.356572242536384	0.781221779211748
79837	PIP4K2C phosphatidylinositol-5-phosphate 4-kinase type 2
gamma	3299	-0.120914213114148	7.84616085509609
0.849641747944311	0.356653705283664	0.781221779211748
5042	PABPC3	poly(A) binding protein cytoplasmic 3
3387	-0.151786822699813	3.93074849886936
0.84957494302459	0.356672612126731	0.781221779211748
2686	GGT7	gamma-glutamyltransferase 7	2966
0.225900324042586	3.5813791329543 0.849043358898312
0.356823107808517	0.781243602630984
23107	MRPS27	mitochondrial ribosomal protein S27	6511
0.126856051334213	5.23832954394799	0.848977267979279
	0.356841824760532	0.781243602630984
55449	DHRS4-AS1	DHRS4 antisense RNA 1	3593
0.215606657022596	3.2515844144649 0.848948750064533
0.356849901451634	0.781243602630984
1520	CTSS	cathepsin S	4228	-0.398595049640562
6.22500088287597	0.848634225070484	0.356938996177236
	0.781316536873659
64431	ACTR6	actin related protein 6 3813	-0.143558121907281
	4.49467232383598	0.84831009820347
0.357030842741048	0.781395470980291
6881	TAF10	TATA-box binding protein associated factor 10	6103
	0.144303238540901	4.65316402181084
0.847928856349899	0.357138915271461	0.781447874153873
349075	ZNF713	zinc finger protein 713 6825	0.167747035377516
	3.59255156598724	0.847720955559721
0.357197868865016	0.781447874153873
23144	ZC3H3	zinc finger CCCH-type containing 3
3731	-0.156714194832309	4.49197865785883
0.847635320510592	0.357222155936835	0.781447874153873
11314	CD300A	CD300a molecule 2686	0.876587811590446
0.561891020033673	0.847372512065208	0.357296705549511
	0.781458827516516
66036	MTMR9	myotubularin related protein 9
8804	-0.143561713067315	5.35255736022164
0.846775093759579	0.3574662518193 0.781458827516516
22883	CLSTN1	calsyntenin 1	5559	-0.119745202882676
9.77991140613567	0.846698856304158	0.35748789582293
	0.781458827516516
100129973	CERNA1	competing endogenous lncRNA 1 for
miR-4707-5p and miR-4767	2096	0.391760557555599
0.453381587027175	0.846512752311466	0.357540738769007
	0.781458827516516
59272	ACE2	angiotensin I converting enzyme 2
3926	-0.343697546390678	1.52544960537176
0.84644346243023	0.35756041589345	0.781458827516516
55122	AKIRIN2 akirin 2	1999	-0.127272641060152

5.28850258565528	0.846438213404767	0.357561906586232
	0.781458827516516
51118	UTP11	UTP11 small subunit processome component
3385	-0.140091376869913	4.95669352287456
0.845751179183269	0.357757094012144	0.781629810003515
387590	TPTEP1	TPTE pseudogene 1	10678	-0.442562272904597
	4.69842522189051	0.845612938411875
0.357796386102192	0.781629810003515
2888	GRB14	growth factor receptor bound protein 14 2958
0.322213460004409	-0.069640944841153	0.844864881426985
	0.358009108749733	0.781629810003515
79634	SCRN3	secernin 3	4144	-0.204535012389665
4.59961097862966	0.84483116527714l	0.358018700598611
	0.781629810003515
23139	MAST2	microtubule associated serine/threonine kinase 2
	7877	0.181955613441572	6.40668643710214
0.844565008863846	0.358094431371518	0.781629810003515
54943	DNAJC28 DnaJ heat shock protein family (Hsp40) member C28
	2484	0.24503312131209	1.39812995424585
0.844536267507338	0.358102610605636	0.781629810003515
283870	BRICD5	BRICHOS domain containing 5	2083
0.324237467707474	1.8066838521403 0.844519335558526
0.358107429230922	0.781629810003515
92249	LINC01278	long intergenic non-protein coding RNA 1278
	3748	0.235346231128693	2.89650179805423
0.844475109435356	0.358120015864301	0.781629810003515
80303	EFHD1	EF-hand domain family member D1 3052
0.537005650027961	2.64312945980688	0.844396582183542
	0.358142366002896	0.781629810003515
9110	MTMR4	myotubularin related protein 4
7475	-0.143992316852617	5.50665821195907
0.84415203582148	0.358211980173072	0.781638052195959
57153	SLC44A2 solute carrier family 44 member 2
5216	-0.16512583907825	10.0295485879845
0.843868737061641	0.358292649107377	0.781638052195959
9294	S1PR2	sphingosine-1-phosphate receptor 2	3582
0.195945910930607	3.25256748677573	0.843795340555733
	0.358313552734365	0.781638052195959
9266	CYTH2	cytohesin 2	9630	-0.126854241906898
7.4710282366818 0.843303826899536	0.358453581058258
0.781821754886798
220382	FAM181B family with sequence similarity 181 member B
3924	-0.428159053784952	0.143437962227134
0.842841227399504	0.358585440699669	0.781907287039432
84080	ENKD1	enkurin domain containing 1	2679
0.304375028882427	2.54451337718626	0.842774586738344
	0.358604441489349	0.781907287039432
55421	NCBP3	nuclear cap binding subunit 3
14718	-0.118988336136738	6.18034020019271
0.842410776872246	0.358708196487097	0.782011783376617
51741	WWOX	WW domain containing oxidoreductase	4886
0.235717104149341	2.68075622727746	0.841834549707714
	0.35887261530242	0.782248478453314
55163	PNPO	pyridoxamine 5'-phosphate oxidase	2861

0.173807681527163	3.77369106921602	0.841605516472771
	0.358937995700457	0.782269255605462
4776	NFATC4	nuclear factor of activated T cells 4	8437
0.261384211669027	4.6254303954707 0.84129553201057
0.359026510734829	0.782340438062032
100505894	TMEM161B-AS1	TMEM161B antisense RNA 1	4439
	0.261148557764698	1.2582653715859 0.840670987625919
	0.359204938533354	0.782583124221118
284439	SLC25A42	solute carrier family 25 member 42	3120
	0.308484563341953	2.12034495069858
0.840514642190313	0.35924962438365	0.782583124221118
5832	ALDH18A1	aldehyde dehydrogenase 18 family member A1
	3766	0.176700382031088	5.68430952728522
0.840021809323076	0.359390533418456	0.782670951017613
81542	TMX1	thioredoxin related transmembrane protein 1	4499
	0.147494837148974	6.15077644653578
0.839867415245408	0.359434692875464	0.782670951017613
6230	RPS25	ribosomal protein S25	1477	0.188440181949816
	6.42101834011414	0.839606142056029
0.359509438676922	0.782670951017613
339231	ARL16	ADP ribosylation factor like GTPase 16	2597
0.149751654578052	4.91293516854903	0.839592121458974
	0.359513450335355	0.782670951017613
83732	RIOK1	RIO kinase 1	3237	-0.153421156757522
3.69812439276565	0.839333361652631	0.359587499327985
	0.782699097241301
218	ALDH3A1 aldehyde dehydrogenase 3 family member A1	3918
	0.296392235024449	3.43952085584167
0.839153600979018	0.359638953590629	0.782699097241301
677838	SNORA61 small nucleolar RNA, H/ACA box 61
130	-0.317089133394834	0.121652953428616
0.838854179555873	0.359724681764342	0.782699097241301
22902	RUFY3	RUN and FYVE domain containing 3
7925	-0.152772504320145	4.83837809189843
0.838766134198903	0.359749895626436	0.782699097241301
24140	FTSJ1	FtsJ RNA 2'-O-methyltransferase 1	2872
0.112182916463167	6.97135886932253	0.83855369743771
	0.359810741917281	0.782709902474818
3931	LCAT	lecithin-cholesterol acyltransferase	2625
0.230086837778451	3.95496655392586	0.837163443290704
	0.36020928960791	0.78324934948941
27076	LYPD3	LY6/PLAUR domain containing 3
2519	-0.427747966004626	4.59407076994844
0.837117613139885	0.360222438204845	0.78324934948941
2058	EPRS1	glutamyl-prolyl-tRNA synthetase 1
5450	-0.152399921757712	7.4369014054341 0.837103524697849
	0.360226480289197	0.78324934948941
79934	COQ8B	coenzyme Q8B	5488	0.165693568276318
3.99680388037827	0.836445456918257	0.360415354742426
	0.783536723347563
84248	FYTTD1	forty-two-three domain containing 1
8025	-0.125814497806276	6.53763120885616
0.836100857434445	0.360514313949156	0.783536723347563
5327	PLAT	plasminogen activator, tissue type

6618	-0.274045709985062	4.59551493040091
0.83605855687339	0.360526464051906	0.783536723347563
83648	FAM167A family with sequence similarity 167 member A	5033
	0.359704351735181	2.72287014782679
0.834316798583437	0.361027244447274	0.784183522447215
100128338	FAM83H-AS1	FAM83H antisense RNA 1 (head to head)	3229	-0.180090074908271	4.65754726311527
0.834170428696467	0.361069371556593	0.784183522447215
23162	MAPK8IP3	mitogen-activated protein kinase 8 interacting protein 3	10285	0.199933982319345
6.30879518856506	0.834042403929516	0.361106224265977
	0.784183522447215
151473	SLC16A14	solute carrier family 16 member 14
4946	-0.525959262439138	1.12945246092096
0.833666019669469	0.361214598819025	0.784183522447215
147007	TMEM199 transmembrane protein 199
5115	-0.125489617127556	5.15939488929612
0.833394527939031	0.361292798896949	0.784183522447215
6745	SSR1	signal sequence receptor subunit 1
11349	-0.159978326556051	7.31116965410809
0.833123719777053	0.361370825348969	0.784183522447215
7328	UBE2H	ubiquitin conjugating enzyme E2 H
6392	-0.122637351457031	9.2599963179951 0.832958754320344
	0.361418367324492	0.784183522447215
54764	ZRANB1	zinc finger RANBP2-type containing 1
5993	-0.121888732943741	6.42883032260514
0.832590114627763	0.36152463808852	0.784183522447215
9870	AREL1	apoptosis resistant E3 ubiquitin protein ligase 1
	7308	-0.101923905465215	6.88370272294877
0.832453229929818	0.361564109932028	0.784183522447215
60313	GPBP1L1 GC-rich promoter binding protein 1 like 1
7433	-0.104572119766351	7.3420910704049 0.832215167503122
	0.361632771383455	0.784183522447215
54822	TRPM7	transient receptor potential cation channel subfamily M member 7	12333	-0.137217947631042
6.5582262924501 0.832021355694911	0.361688683455794
0.784183522447215
2157	F8	coagulation factor VIII 9609	0.255309728667185
	2.1561535503495 0.83187406824893	0.361731181860887
	0.784183522447215
4670	HNRNPM	heterogeneous nuclear ribonucleoprotein M
4283	-0.106754902086384	6.90770262234728
0.831652666363638	0.361795078257522	0.784183522447215
4430	MYO1B	myosin IB	9533	0.117391888332602
8.72562566504602	0.8315844277829 0.361814774994939
0.784183522447215
140465	MYL6B	myosin light chain 6B	3259	0.132030478238573
	5.85587926508988	0.831580167505592
0.361816004752539	0.784183522447215
64963	MRPS11	mitochondrial ribosomal protein S11	4033
0.165544211768409	4.34811696430259	0.831564051798104
	0.361820656711718	0.784183522447215
22919	MAPRE1	microtubule associated protein RP/EB family member 1
	2623	-0.133981052802085	7.21230386531746

0.831498536291363	0.361839569264152	0.784183522447215
3841	KPNA5	karyopherin subunit alpha 5
5556	-0.199432901239281	3.77511160157873
0.831492651761377	0.361841268035377	0.784183522447215
10360	NPM3	nucleophosmin/nucleoplasmin 3	928
0.290271195120568	2.66178536098925	0.831331511265901
	0.361887791040108	0.784183522447215
56675	NRIP3	nuclear receptor interacting protein 3
4123	-0.350358989297537	2.11592796934642
0.830907823733345	0.362010153622595	0.784207345640807
23300	ATMIN	ATM interactor	6262	-0.1030118060425
6.71437485311948	0.830722635419733	0.362063654657703
	0.784207345640807
79883	PODNL1	podocan like 1	3883	0.277950056494965
1.25456724426664	0.830444662602632	0.362143981697108
	0.784207345640807
7699	ZNF140	zinc finger protein 140	5374	-0.142717887822265
	4.57616046501702	0.830405152044907
0.362155401235929	0.784207345640807
84517	ACTRT3	actin related protein T3	2005
0.229949685929391	3.22651202242273	0.829955365679858
	0.362285435799746	0.784207345640807
100288332	NPIPA5	nuclear pore complex interacting protein
family member A5	1520	0.164243383442332
4.42179441969965	0.829846658296999	0.362316873100096
	0.784207345640807
253650	ANKRD18A	ankyrin repeat domain 18A
5303	-0.335075163955528	1.07183462891799
0.829838942745112	0.362319104518816	0.784207345640807
219771	CCNY	cyclin Y	5743	-0.122139767097728
6.47110885362243	0.829743602149818	0.36234667958787
	0.784207345640807
56648	EIF5A2	eukaryotic translation initiation factor 5A2	5593
	0.234758774763292	3.26439336189228
0.829284298741278	0.362479563140014	0.784373743380531
101928464	ZNF433-AS1	ZNF433 and ZNF878 antisense RNA 1
	2891	-0.293846144650952	1.18222107661521
0.829043742827304	0.362549186535923	0.78440095261485
137964	GPAT4	glycerol-3-phosphate acyltransferase 4
7939	-0.102535369564778	7.21261724103854
0.828853923213259	0.36260413851511	0.78440095261485
54819	ZCCHC10	zinc finger CCHC-type containing 10
2559	-0.141462331969366	4.11292219191223
0.828581752486901	0.362682950877434	0.784441357419078
51337	THEM6	thioesterase superfamily member 6	2259
0.208839478612641	4.67668067736923	0.828402660277564
	0.362734823445661	0.784441357419078
4040	LRP6	LDL receptor related protein 6
10337	-0.200514999780711	6.2345392095402	0.828150654627009
	0.362807832150604	0.784478126726383
26973	CHORDC1	cysteine and histidine rich domain containing 1
10655	-0.151061282830969	5.17619991212722
0.827654382449651	0.36295166684431	0.784521199153186
6737	TRIM21	tripartite motif containing 21

1929	-0.15843328834008	4.71795265047275
0.827424658358243	0.363018274525506	0.784521199153186
140685	ZBTB46	zinc finger and BTB domain containing 46
6084	-0.422579549501499	2.92906611739401
0.827387239586741	0.363029125565206	0.784521199153186
79943	ZNF696	zinc finger protein 696 4012	0.254080046576099
	1.79567750975101	0.826998478644194
0.363141888575751	0.784521199153186
4303	FOXO4	forkhead box O4 3447	0.209483249537035
4.15852974851769	0.826989307334385	0.363144549367203
	0.784521199153186
1571	CYP2E1	cytochrome P450 family 2 subfamily E member 1
9621	-0.389002515037113	2.29252323918554
0.826750146980221	0.363213944375143	0.784521199153186
554249	LOC554249	uncharacterized LOC554249	2541
0.307219279305578	0.64219679788626	0.826729910737472
	0.363219816984966	0.784521199153186
5283	PIGH	phosphatidylinositol glycan anchor biosynthesis
class H 2915	0.18485428082039	3.29614284948082
0.826311733527824	0.363341202486421	0.784662384093016
116372	LYPD1	LY6/PLAUR domain containing 1	2292
0.48613434266224	3.13694469724935	0.82573612832601
	0.363508376841036	0.784702297038737
285550	FAM200B family with sequence similarity 200 member B
4817	-0.16870434928193	3.63617852585479
0.825720768070722	0.363512839412887	0.784702297038737
51182	HSPA14	heat shock protein family A (Hsp70) member 14
4818	-0.138669776589547	4.6190017772075 0.825669444683804
	0.363527750802054	0.784702297038737
3099	HK2	hexokinase 2	6064	-0.204468218267978
5.81012637843644	0.824700747133104	0.363809352939024
	0.785189153561927
100302736	TMED7-TICAM2	TMED7-TICAM2 readthrough	3865
	-0.137467058773 5.23081376439708	0.823913397575886
	0.364038459304593	0.785483905609308
513	ATP5F1D ATP synthase F1 subunit delta	1505
0.184402168716279	4.18862955598206	0.82359931895158
	0.364129906973434	0.785483905609308
115704	EVI5L	ecotropic viral integration site 5 like 6252
0.152441838674874	4.75142419390131	0.823556017234175
	0.364142517269642	0.785483905609308
124923	RSKR	ribosomal protein S6 kinase related
4979	-0.300122089075977	1.83372636514837
0.823196640957143	0.364247197914377	0.785483905609308
2281	FKBP1B	FKBP prolyl isomerase 1B	1887
0.153536743689951	4.36503385860716	0.823181890750465
	0.364251495307101	0.785483905609308
128439	SNHG11	small nucleolar RNA host gene 11	2813
0.207324280163943	3.30020160996303	0.822956946168435
	0.364317040390261	0.785483905609308
60598	KCNK15	potassium two pore domain channel subfamily K member
15	2599	0.490536786275029	2.48455614212774
0.822883417474333	0.364338468961496	0.785483905609308
10763	NES	nestin	5558	0.496341454189422

1.48063227765141	0.821878152924883	0.364631609764695
	0.785994915046797
55182	RNF220	ring finger protein 220 9687	0.131374389955969
	5.38744604235012	0.8212834916161 0.364805170149925
	0.786248040973996
92815	H2AW	H2A.W histone	895	-0.278994236276624
1.41554825254538	0.820661905301205	0.364986711353382
	0.786518287065665
5707	PSMD1	proteasome 26S subunit, non-ATPase 1
5187	-0.107474321854092	7.32560081938029
0.820164112218229	0.36513218765176	0.786710744875896
51540	SCLY	selenocysteine lyase	11733	0.179606178622477
	3.18235508169698	0.819792034570287
0.365240977045366	0.78682411034673
6386	SDCBP	syndecan binding protein
4778	-0.141773894414068	7.64785131268952
0.819526746421531	0.365318570382949	0.786870246863597
5914	RARA	retinoic acid receptor alpha	4731
0.196391071732452	4.7467421252561 0.819259507782933
0.365396757320662	0.786917647569267
26268	FBXO9	F-box protein 9 8164	0.121146553571605
5.97311946815836	0.81883926790357	0.365519755299641
	0.786922752674718
83869	TTTY14	testis-specific transcript, Y-linked 14 3140
0.246297890233152	1.64698743338163	0.81875228501892
	0.365545221070777	0.786922752674718
729359	PLIN4	perilipin 4	6502	0.370848509761773
1.30237426875647	0.818675614160887	0.365567669854672
	0.786922752674718
91283	MSANTD3 Myb/SANT DNA binding domain containing 3	3253
	0.131190162841038	6.45080375593359
0.818286958164698	0.365681495497986	0.787046820442578
10061	ABCF2	ATP binding cassette subfamily F member 2	4448
	0.106318519458488	6.38966992312744
0.817644817643902	0.365869666858341	0.787330837868303
100526783	ARPIN-AP3S2	ARPIN-AP3S2 readthrough 2673
0.135900697321521	5.2765130544125 0.817145422157905
0.366016101413398	0.787369363313472
53346	TM6SF1	transmembrane 6 superfamily member 1	2661
0.495838716702034	0.855414209674147	0.817035329291413
	0.366048394181779	0.787369363313472
56907	SPIRE1	spire type actin nucleation factor 1	6544
0.108403520280827	5.70995634552261	0.816778031424231
	0.366123880978916	0.787369363313472
83851	SYT16	synaptotagmin 16	14067	0.376678779784071
	1.44648347950435	0.816677114710563
0.366153494116034	0.787369363313472
1616	DAXX	death domain associated protein 3394
0.109496637700041	6.06712381792837	0.816625530638227
	0.366168632299847	0.787369363313472
79987	SVEP1	sushi, von Willebrand factor type A, EGF and
pentraxin domain containing 1	14190	0.491345695464776
3.91483911257977	0.81604558666232	0.366338886074036
	0.787614547542444

83448	PUS7L	pseudouridine synthase 7 like
14037	-0.208472899576148	4.63970415617854
0.814969231785195	0.366655162229554	0.788126360047156
154791	FMC1	formation of mitochondrial complex V assembly factor
1 homolog	1524	0.294225977547448	1.64130455065116
	0.814852532723923	0.366689475864019
0.788126360047156
79041	TMEM38A transmembrane protein 38A	3005
0.23555389087039	3.84624855925208	0.814149464213054
	0.366896297169778	0.788306166056854
483	ATP1B3	ATPase Na+/K+ transporting subunit beta 3	2584
	0.254202891054957	7.91143384142334
0.81383980553008	0.366987440705329	0.788306166056854
23061	TBC1D9B TBC1 domain family member 9B	11473
0.111618251924488	7.89422326320093	0.813805861029437
	0.366997433688865	0.788306166056854
51222	ZNF219	zinc finger protein 219 4643	0.186243974909452
	5.04866825292834	0.813659784325225
0.367040441790564	0.788306166056854
56681	SAR1A	secretion associated Ras related GTPase 1A	6914
	0.134508006038402	6.99150589974439
0.813611935767213	0.367054530950439	0.788306166056854
4774	NFIA	nuclear factor I A	10872	0.293685994866698
	3.29553173954322	0.812479251442994
0.367388273012333	0.788768184821132
3416	IDE	insulin degrading enzyme	6482
0.136289409722795	5.75565044864115	0.812375384767558
	0.367418898124459	0.788768184821132
9474	ATG5	autophagy related 5	3953	-0.115976024894049
	5.57158535875017	0.812308586736023
0.367438595413571	0.788768184821132
375190	FAM228B family with sequence similarity 228 member B	6569
	0.321321522378091	1.29126540871756
0.81187833621574	0.367565502132303	0.788822119870396
1730	DIAPH2	diaphanous related formin 2	9550
0.189045310209575	3.96069083171706	0.811809370664836
	0.367585849881193	0.788822119870396
942	CD86	CD86 molecule	3433	0.713885379926286
1.04912162110422	0.811606436424823	0.367645733133291
	0.788822119870396
91949	COG7	component of oligomeric golgi complex 7
3869	-0.137564505538265	4.64951873798352
0.811300599860626	0.367736007148397	0.788822119870396
2098	ESD	esterase D	3401	0.168839951114569
5.54657616334724	0.811269116616657	0.36774530183149
	0.788822119870396
143689	PIWIL4	piwi like RNA-mediated gene silencing 4
4558	-0.306122153984302	1.08004788285615
0.810843140800031	0.367871093234426	0.788971122788946
340075	ARSI	arylsulfatase family member I	3269
0.456574891602806	2.93178430806756	0.809951120068522
	0.368134702453459	0.789405092257203
100534593	STX16-NPEPL1	STX16-NPEPL1 readthrough (NMD
candidate)	4070	-0.130574677782393	5.93635586436109

	0.809609444056946	0.368235743865341
0.789405092257203
9459	ARHGEF6 Rac/Cdc42 guanine nucleotide exchange factor 6	6434
	0.345154980034267	2.71212569936543
0.809586560227276	0.368242512515782	0.789405092257203
9138	ARHGEF1 Rho guanine nucleotide exchange factor 1
8469	-0.150176299766268	6.40644755366356
0.809156902560559	0.368369630144802	0.789556757373871
54893	MTMR10	myotubularin related protein 10 8817
0.158843101427032	5.07068325985805	0.808939439447656
	0.36843399160665	0.789573886235364
54020	SLC37A1 solute carrier family 37 member 1
5423	-0.181695869512743	6.01836778561096
0.808241850910292	0.368640559031459	0.789895718273465
7743	ZNF189	zinc finger protein 189 3433	0.14782045221075
	5.01221444470564	0.807965213146886
0.368722520659887	0.789950496617167
84240	ZCCHC9	zinc finger CCHC-type containing 9
5127	-0.162075364775465	3.96004605749962
0.807205620563172	0.368947701597777	0.790223527022201
51193	ZNF639	zinc finger protein 639 6859	-0.137098566506744
	4.53081091386138	0.807025609988411
0.369001093732082	0.790223527022201
54961	SSH3	slingshot protein phosphatase 3 3639
0.128058659990068	5.7372809878639 0.806964535776177
0.369019211126738	0.790223527022201
6159	RPL29	ribosomal protein L29	1912	0.157095071537653
	6.40225879030791	0.806625841353686
0.369119705816983	0.790304754175618
84331	MCRIP2	MAPK regulated corepressor interacting protein 2
	3404	0.209642895558792	2.71071559659319
0.806456428624629	0.369169986904481	0.790304754175618
842	CASP9	caspase 9	5269	-0.155783476142024
3.83676056638972	0.805933973108872	0.369325109915685
	0.790516016899297
7348	UPK1B	uroplakin 1B	2266	-0.802016167424537
6.56185303462403	0.804880514815636	0.369638170020548
	0.791065217338093
641	BLM	BLM RecQ like helicase	6551	-0.275585049883665
	0.662972096037102	0.804190428991001
0.369843445905644	0.791250421995565
4152	MBD1	methyl-CpG binding domain protein 1
6734	-0.154461983429817	7.05959923876973
0.804082428875915	0.369875586463406	0.791250421995565
10302	SNAPC5	small nuclear RNA activating complex polypeptide 5
	4094	0.185241591049615	3.30695666593679
0.803994590038274	0.369901729948265	0.791250421995565
84279	PRADC1	protease associated domain containing 1 1407
0.196854117065807	3.20373928808669	0.803321043362033
	0.370102283388302	0.791250421995565
57587	CFAP97	cilia and flagella associated protein 97
6886	-0.146015974411744	5.28553667961498
0.803277139146896	0.370115361452489	0.791250421995565
126820	WDR63	WD repeat domain 63	3923	0.346763907608806

0.304723625297159	0.803254083562861
0.37012222944314	0.791250421995565
54664	TMEM106B	transmembrane protein 106B
13867	-0.219957563275111	6.6587095863836 0.80320654501989
0.370136391183921	0.791250421995565
79735	TBC1D17 TBC1 domain family member 17	4745
0.21867444837929	5.80930248555937	0.802642188223899
0.370304570935617	0.791250421995565
9317	PTER	phosphotriesterase related
4003	-0.175827588564229	4.08922469997838
0.802630763386358	0.370307976666728	0.791250421995565
29799	YPEL1	yippee like 1	4713	0.27673486323777
2.18395579075969	0.80236165731437	0.370388209519171
0.791250421995565
115560	ZNF501	zinc finger protein 501 3910	-0.306419174620363
1.02301102872004	0.802327922267815
0.370398269193101	0.791250421995565
2323	FLT3LG	fms related tyrosine kinase 3 ligand	3804
0.242576084845608	2.09679435195554	0.802313447493333
0.370402585637532	0.791250421995565
7517	XRCC3	X-ray repair cross complementing 3
7507	-0.253924795805693	1.41309493350833
0.801936034791137	0.370515156619176	0.791280178027692
30011	SH3KBP1 SH3 domain containing kinase binding protein 1
5780	-0.208076935850681	6.55095468906834
0.801887962245473	0.370529498656645	0.791280178027692
5315	PKM	pyruvate kinase M1/2	8571	0.145463465158337
11.589924485674 0.800627975501357	0.370905681480706
0.791962786661622
79823	CAMKMT	calmodulin-lysine N-methyltransferase	3991
0.236602659463748	1.84846208588835	0.800433003529652
0.370963940118412	0.791966454692668
344558	SH3RF3	SH3 domain containing ring finger 3	5803
0.282618789258764	0.903083102040455	0.79990264336412
0.371122479148468	0.79210860100242
2176	FANCC	FA complementation group C
7912	-0.255506421590693	2.50009922489594
0.799710426597581	0.371179961308305	0.79210860100242
28964	GIT1	GIT ArfGAP 1	5635	-0.144698917269213
5.83106184742823	0.79926693611823	0.371312633973327
0.79210860100242
818	CAMK2G	calcium/calmodulin dependent protein kinase II gamma
6780	-0.189861729723328	6.48943480254646
0.799251687214984	0.37131719694600 2	0.79210860100242
7014	TERF2	telomeric repeat binding factor 2
5621	-0.129127102457987	4.6517603973088 0.799225639410976
0.37132499148561	0.79210860100242
51069	MRPL2	mitochondrial ribosomal protein L2	2539
0.154540419208974	3.88941192748955	0.799075819037169
0.37136982814185	0.79210860100242
8626	TP63	tumor protein p63	7454	0.195675094245599
7.75956914657338	0.798558830216166
0.371524605173139	0.792318079272253
1981	EIF4G1	eukaryotic translation initiation factor 4 gamma 1

8257	-0.114258074889302	9.32853326929795
0.798227934584915	0.371623716597762	0.792408798658068
65109	UPF3B	UPF3B regulator of nonsense mediated mRNA decay
2531	-0.197642625704411	4.12240168441047
0.797826552141082	0.371743990159362	0.792455086739067
10726	NUDC	nuclear distribution C, dynein complex regulator
2856	0.127385379823014	6.22826232428353
0.79777788772158	0.371758576066424	0.792455086739067
23154	NCDN	neurochondrin	4917	0.150700183970148
5.6478460329367 0.797472098474678	0.37185024682711
0.792529885469771
55973	BCAP29	B cell receptor associated protein 29
8409	-0.147051786163086	5.76908499801745
0.797111508368346	0.371958386614719	0.792639756779609
92659	MAFG-DT MAFG divergent transcript
1895	-0.30040320549242	0.252366801265116
0.796703767673531	0.372080719675258	0.792779835791198
55885	LMO3	LIM domain only 3	10268	0.283785103597455
6.13709616111214	0.796360628889644
0.372183714034666	0.792872387453228
10535	RNASEH2A	ribonuclease H2 subunit A	2039
0.201013094597517	2.32255756085751	0.796181924984385
0.372237368450948	0.792872387453228
90	ACVR1	activin A receptor type 1	3990
0.110609816615242	6.34242186849591	0.795773731516231
0.372359965847674	0.792900886760295
1027	CDKN1B	cyclin dependent kinase inhibitor 1B
2811	-0.172680232292389	5.17839865417777
0.795459120017426	0.372454495242847	0.792900886760295
79364	ZXDC	ZXD family zinc finger C
7357	-0.154795405660255	5.01472392693322
0.794737835923353	0.372671341803629	0.792900886760295
55311	ZNF444	zinc finger protein 444 7239	0.168605460879315
4.22833812789088	0.794654615028116
0.372696372664226	0.792900886760295
9731	CEP104	centrosomal protein 104 7876	-0.112450626749816
5.8717248129087 0.794634520713068	0.372702416907529
0.792900886760295
4838	NODAL	nodal growth differentiation factor	1693
0.403462550023152	-0.0908896510560139	0.794269795570759
0.37281214777296	0.792900886760295
91966	CXorf40A	chromosome X open reading frame 40A
4104	-0.149401495448678	4.18129705896775
0.794226285752476	0.37282524111426	0.792900886760295
23390	ZDHHC17 zinc finger DHHC-type containing 17	8468
0.19130239997808	6.31230040477031	0.794011575274993
0.37288986302514	0.792900886760295
8417	STX7	syntaxin 7	16387	-0.112988515171261
6.49255665023939	0.79397108395532	0.372902051548406
0.792900886760295
23350	U2SURP	U2 snRNP associated SURP domain containing
10739	-0.126587285058716	6.33671525347822
0.793817936462752	0.372948156392253	0.792900886760295
221302	ZUP1	zinc finger containing ubiquitin peptidase 1

2656	-0.132798349667879	4.27444842045344
0.793723872237477	0.372976478254059	0.792900886760295
545	ATR	ATR serine/threonine kinase
12496	-0.155315921884252	5.20004281945026
0.793558401748911	0.37302630718246	0.792900886760295
3459	IFNGR1	interferon gamma receptor 1
2571	-0.202558761182147	7.45263994888177
0.793369548844993	0.373083188757633	0.792900886760295
29	ABR	ABR activator of RhoGEF and GTPase
10028	-0.131785688020653	7.75801263842129
0.793333458322813	0.373094060429025	0.792900886760295
55718	POLR3E	RNA polymerase III subunit E
9488	-0.1251349049234	5.16235257651214
0.793238199295664	0.373122757761609	0.792900886760295
6880	TAF9	TATA-box binding protein associated factor 9	2577
0.133428247767649	5.6436953279945	0.793115650671649
0.373159680787875	0.792900886760295
26267	FBXO10	F-box protein 10	6337	0.136083235702744
4.62207179581743	0.79293846255486
0.373213075348799	0.792900886760295
23239	PHLPP1	PH domain and leucine rich repeat protein
phosphatase 1	7070	-0.189270485532589	3.75481336468576
0.792615483862104	0.373310430535976
0.792924272284376
11321	GPN1	GPN-loop GTPase 1	3746	-0.135556986821234
6.0981504916766	0.792512564432723	0.373341460921208
0.792924272284376
130733	TMEM178A	transmembrane protein 178A	2470
0.396802639669599	-0.165518902535657	0.792206556051751
0.373433744312729	0.792924272284376
10026	PIGK	phosphatidylinositol glycan anchor biosynthesis
class K	5513	0.136902459634188	4.18660234415519
0.792150942076102	0.373450519330337	0.792924272284376
435	ASL	argininosuccinate lyase	3554	-0.170573198767338
4.89142796222839	0.79165872726184
0.373599033642575	0.793119379240914
400657	LINC00909	long intergenic non-protein coding RNA 909
3742	0.217773720629286	2.9924140302472
0.791258435821677	0.373719873229382	0.793121365774334
169270	ZNF596	zinc finger protein 596	5385	0.246734415215189
1.28533060200019	0.791100901426565
0.373767444572693	0.793121365774334
7374	UNG	uracil DNA glycosylase	2567	0.151134515947012
4.4552898591182	0.790797877027333	0.373858974020659
0.793121365774334
125950	RAVER1	ribonucleoprotein, PTB binding 1	5138
0.142129582718768	5.26648540397267	0.790771061808428
0.373867075152145	0.793121365774334
514	ATP5F1E	ATP synthase F1 subunit epsilon	5126
0.174559280111688	7.11534169170093	0.790408732829263
0.373976562288291	0.793121365774334
9786	KIAA0586	KIAA0586	10184	-0.153096829236222
3.80967971329653	0.790277846172344
0.374016124147386	0.793121365774334

64359	NXN	nucleoredoxin	5431	0.189580754835249
6.91243678221434	0.790245334976674	0.374025951905223
	0.793121365774334
100379661	GRIK1-AS2	GRIK1 antisense RNA 2
1402	-0.285952144165691	2.34581699762504
0.790156011847529	0.374052955115674	0.793121365774334
162681	C18orf54	chromosome 18 open reading frame 54	6062
	0.265632440947278	1.68117024585196
0.789886463462208	0.37413445865864	0.79317411267316
4125	MAN2B1	mannosidase alpha class 2B member 1
5658	-0.124401669716846	6.66524084300899
0.789633362136115	0.374211011754735	0.793216350128415
338799	LINC01089	long intergenic non-protein coding RNA 1089
	4510	-0.143047175123917	4.64593202341428
0.789120313799678	0.37436625587746	0.79342535119921
90271	OLMALINC	oligodendrocyte maturation-associated long
intergenic non-coding RNA	2551	-0.234652654522048
1.88384319637756	0.788366149933271	0.374594623519913
	0.793789242306116
163081	ZNF567	zinc finger protein 567	4724	-0.224440373236741
	2.8711614826216	0.787660331595287	0.374808528730503
	0.794122381172009
5890	RAD51B	RAD51 paralog B	4167	0.242121871049563
1.46373684213716	0.786615640684325	0.375125447270028
	0.794673644874664
10947	AP3M2	adaptor related protein complex 3 subunit mu 2	6276
	0.164530844591462	4.81474606075064
0.786335129773402	0.375210607396929	0.794733854197608
9349	RPL23	ribosomal protein L23	5656	0.159728507546355
	7.50516183817982	0.78557348885343
0.375441970391757	0.794991089178679
50488	MINK1	misshapen like kinase 1	7701	-0.109653623378835
	7.49364182321082	0.785561654596783
0.375445566853499	0.794991089178679
3709	ITPR2	inositol 1,4,5-trisphosphate receptor type 2
12750	-0.226990538373504	4.68943062537492
0.785035367444301	0.375605555827271	0.795118662809328
4189	DNAJB9	DnaJ heat shock protein family (Hsp40) member B9
	2837	-0.200958461854034	5.87156779950664
0.784963600384133	0.375627380114295	0.795118662809328
51291	GMIP	GEM interacting protein	3740	-0.130880451196279
	4.96116414221107	0.784629881390918
0.3757288871049	0.795118662809328
92255	LMBRD2	LMBR1 domain containing 2
8421	-0.183176644291088	5.30305023067406
0.784616761689943	0.37573287849896	0.795118662809328
55253	TYW1	tRNA-yW synthesizing protein 1 homolog	3890
0.144385811812357	4.48088301520107	0.783638638905622
	0.376030619684306	0.795605601493268
388610	TRNP1	TMF1 regulated nuclear protein 1	1976
0.206685199803464	4.77963883636541	0.783340864767943
	0.376121328134498	0.795605601493268
55331	ACER3	alkaline ceramidase 3	8442	-0.152243981501108
	5.59590972966032	0.782976662067995

0.376232313971397	0.795605601493268
3860	KRT13	keratin 13	2913	0.281930601589743
8.48787296183793	0.782971659598843	0.376233838726485
	0.795605601493268
168374	ZNF92	zinc finger protein 92	3512	-0.223993754304117
	3.29930100991708	0.782928533452913
0.376246983957407	0.795605601493268
166824	RASSF6	Ras association domain family member 6
6632	-0.2811842447949	5.81184105938731
0.781329964795681	0.376734697690893	0.796463327919787
5787	PTPRB	protein tyrosine phosphatase receptor type B
14542	-0.281399485305504	3.04499815820582
0.781226378563785	0.376766331890948	0.796463327919787
51316	PLAC8	placenta associated 8	3853	-0.511456845077305
	0.365874017104359	0.780944847623109
0.376852327503665	0.79652490589163
63977	PRDM15	PR/SET domain 15	13004	-0.162487461617228
	4.00855909581149	0.780566014636037
0.376968088289153	0.796649368235692
9937	DCLRE1A DNA cross-link repair 1A
4600	-0.169040827111138	3.92773937291504
0.780373868226945	0.377026821988346	0.796653295457952
54799	MBTD1	mbt domain containing 1 7201	-0.162378174084557
	4.45965622848847	0.779610122065364
0.3772604048347 0.79702661998788
3303	HSPA1A	heat shock protein family A (Hsp70) member 1A
2483	-0.209218328560638	6.479672615815	0.779336129593091
	0.377344251865493	0.797083537306585
64776	C11orf1 chromosome 11 open reading frame 1	3831
0.226592607038406	2.51915783587406	0.778266791114262
	0.377671741333366	0.797437573670819
153562	MARVELD2	MARVEL domain containing 2
4898	-0.10202501880086	6.16728270575611
0.778165461480685	0.377702794717861	0.797437573670819
3607	FOXK2	forkhead box K2 7088	-0.0940128125972133
6.58104215998367	0.77811767021629	0.377717442031638
	0.797437573670819
25828	TXN2	thioredoxin 2	2050	0.155690649795725
5.00053376251909	0.778045442403013	0.377739580303126
	0.797437573670819
51063	CALHM2	calcium homeostasis modulator family member 2	3108
	0.345917967231422	1.89534456828742
0.777187449451892	0.378002700284601	0.797646950548545
23708	GSPT2	G1 to S phase transition 2	2802
0.234743388177844	2.98217538252364	0.776952726127067
	0.378074727660816	0.797646950548545
55064	SPATA6L spermatogenesis associated 6 like
7361	-0.284716408931228	1.01017874708691
0.776870572400793	0.378099941982829	0.797646950548545
10634	GAS2L1	growth arrest specific 2 like 1 4481
0.14528440769753	4.46974022596518	0.776841177773317
	0.378108964250123	0.797646950548545
2651	GCNT2	glucosaminyl (N-acetyl) transferase 2 (I blood
group)	9052	-0.1935537236834	5.74600228370738

0.776759780299685	0.378133949642186	0.797646950548545
4072	EPCAM	epithelial cell adhesion molecule	2519
0.159850250725178	7.94403141929572	0.776608349753076
	0.378180438251795	0.797646950548545
5604	MAP2K1	mitogen-activated protein kinase kinase 1
3759	-0.131336897948959	6.73380619307286
0.776381855258464	0.378249986242649	0.797673525639986
10263	CDK2AP2	cyclin dependent kinase 2 associated protein 2	2019
	0.134606235262095	4.74445115223637
0.775965648616621	0.378377834735949	0.797811649053307
2120	ETV6	ETS variant transcription factor 6
7075	-0.142146293648735	6.12878760271827
0.775797837850847	0.378429399322494	0.797811649053307
115426	UHRF2	ubiquitin like with PHD and ring finger domains 2
	9919	-0.124014179148271	5.68920125665561
0.774073231032659	0.378959907861737	0.798190803698166
8795	TNFRSF10B	TNF receptor superfamily member 10b	5158
	0.107455166582682	9.11473349525982
0.77383843503992	0.379032214852205	0.798190803698166
57718	PPP4R4	protein phosphatase 4 regulatory subunit 4
4974	-0.306478408126153	3.26790603751189
0.773736786467779	0.379063524243044	0.798190803698166
388403	YPEL2	yippee like 2	8667	0.175048416427437
5.00384267315202	0.773729300769538	0.379065830102399
	0.798190803698166
57396	CLK4	CDC like kinase 4	7924	0.202769025116223
	4.57829882188078	0.773709281492614
0.379071996843716	0.798190803698166
645027	EVPLL	envoplakin like	2156	0.310672157822655
1.20411199940756	0.773619904688779	0.379099530215141
	0.798190803698166
101927178	LOC101927178	uncharacterized LOC101927178
4165	-0.157067993759778	4.78942350154711
0.773586494249386	0.379109823341972	0.798190803698166
9797	TATDN2	TatD DNase domain containing 2
5621	-0.114898766453836	6.36622983312568
0.773461599547787	0.379148304537739	0.798190803698166
8938	BAIAP3	BAI1 associated protein 3
6146	-0.332618744334183	0.256502930354215
0.773432267250842	0.379157342884728	0.798190803698166
3321	IGSF3	immunoglobulin superfamily member 3
7400	-0.153114930303069	7.99931760296999
0.773255276736734	0.379211886547055	0.798190803698166
2273	FHL1	four and a half LIM domains 1	5997
0.552956307771883	3.43140864804073	0.77317677607355
	0.379236081859877	0.798190803698166
144404	TMEM120B	transmembrane protein 120B	7510
0.200048061961169	4.22810197142444	0.772865489865077
	0.379332047300044	0.798272834514981
100533184	ARHGAP19-SLIT1	ARHGAP19-SLIT1 readthrough (NMD
candidate)	2356	-0.256916824251341	1.04882018172418
	0.772550723178928	0.379429120571033
0.798317263816687
23195	MDN1	midasin AAA ATPase 1	18602	-0.2261737733173

5.63320917089127 0.77242744752407
0.379467148032519 0.798317263816687
124222 PAQR4 progestin and adipoQ receptor family member 4
2972 -0.17278739332693 4.86349161994279
0.772206354526816 0.379535363176694 0.798340866799212
10631 POSTN periostin 5464 1.06612632377318
4.58925056894776 0.771750204184286 0.379676156670229
0.798488110009131
389677 RBM12B RNA binding motif protein 12B
7377 -0.186665425505658 4.40836022597948
0.771610227503539 0.379719376081304 0.798488110009131
407004 MIR22 microRNA 22 111 0.259125295521748
0.864343790280257 0.771252497757836 0.379829860818519
0.79860054945737
2004 ELK3 ETS transcription factor ELK3
4648 -0.150245485240865 7.64982796429144
0.770701463023869 0.380000136576818 0.798779763341689
3770 KCNJ14 potassium inwardly rectifying channel subfamily J
member 14 4340 -0.263790543738356 0.791470833572527
0.770607597565684 0.380029152774257
0.798779763341689
11316 COPE coatomer protein complex subunit epsilon
2885 -0.102506729867474 6.68749582963622
0.769233440023926 0.38045429828946 0.799201162984425
5599 MAPK8 mitogen-activated protein kinase 8
9629 -0.138887156276614 5.53465461476423
0.769218316124153 0.380458981154442 0.799201162984425
147837 ZNF563 zinc finger protein 563 2847 0.276977074697449
1.00961788072712 0.768852656092545
0.380572226520367 0.799201162984425
5645 PRSS2 serine protease 2 1716 -0.572193362254236
4.2397837556968 0.768760655829215 0.380600726616513
0.799201162984425
388558 ZNF808 zinc finger protein 808 3600 -0.240718300594488
3.29311791088761 0.768732638961486
0.380609406358483 0.799201162984425
3192 HNRNPU heterogeneous nuclear ribonucleoprotein U
8838 -0.106127807250873 8.37436272356829
0.7687014472748 0.380619070001664 0.799201162984425
153129 SLC38A9 solute carrier family 38 member 9
8823 -0.141364222374782 4.94058544194552
0.768536937913559 0.380670043158822 0.799201162984425
161145 TMEM229B transmembrane protein 229B
6271 -0.364140443088884 3.01202283184782
0.768485132923701 0.380686096910979 0.799201162984425
7006 TEC tec protein tyrosine kinase
4030 -0.269565749761025 1.94207677162441
0.768004275720564 0.38083515453539 0.799367593061904
84317 CCDC115 coiled-coil domain containing 115 1646
0.159567451855207 4.85303643150517 0.766967123190028
0.381156935383012 0.799367593061904
81706 PPP1R14C protein phosphatase 1 regulatory inhibitor
subunit 14C 2152 0.121921827516295 6.26971716471449
0.766956456704989 0.381160246701042

0.799367593061904
27037 TRMT2A tRNA methyltransferase 2 homolog A 3963
0.137007803666223 4.81820061584883 0.766935512611998
0.381166748731952 0.799367593061904
80198 MUS81 MUS81 structure-specific endonuclease subunit
5476 -0.138813449512881 5.26663414235528
0.766930791144823 0.381168214518956 0.799367593061904
4649 MYO9A myosin IXA 20253 -0.214050504402511
5.31512127449396 0.766783395320374 0.381213977797021
0.799367593061904
339761 CYP27C1 cytochrome P450 family 27 subfamily C member 1
4491 -0.239397383248703 5.41489265553639
0.766735351722901 0.381228895994649 0.799367593061904
64218 SEMA4A semaphorin 4A 6822 -0.308181943810397
3.29941178979958 0.766575231080459 0.381278621618534
0.799367593061904
5865 RAB3B RAB3B, member RAS oncogene family
12844 -0.311978171166604 5.75415980168174
0.766503240270154 0.381300981425665 0.799367593061904
10398 MYL9 myosin light chain 9 2821 0.257053968886776
8.4166686642936 0.766332469221531 0.3813540290553
0.799367593061904
7376 NR1H2 nuclear receptor subfamily 1 group H member 2 3667
0.104514377475505 5.89423514883071
0.766100842876135 0.381425997214723 0.799367593061904
80329 ULBP1 UL16 binding protein 1 3142 0.349927300207804
3.27633859201151 0.766022951839908
0.38145020290039 0.799367593061904
9564 BCAR1 BCAR1 scaffold protein, Cas family member 9178
0.128778648807558 6.78456967047518
0.765472893820288 0.381621202705827 0.799606310590952
200734 SPRED2 sprouty related EVH1 domain containing 2 5154
0.127512568800116 5.74852769566791
0.764084108013097 0.382053425866535 0.800392212999187
10539 GLRX3 glutaredoxin 3 4725 -0.115545987916503
6.33755278370157 0.763572695864617 0.382212764095253
0.800562178228277
3996 LLGL1 LLGL scribble cell polarity complex component 1
6638 -0.119437996781355 6.18569473983602
0.763375249579241 0.382274306668346 0.800562178228277
613 BCR BCR activator of RhoGEF and GTPase
12389 -0.106776108373437 7.31207521599872
0.762602969180001 0.38251515572654 8 0.800562178228277
79692 ZNF322 zinc finger protein 322 5059 -0.161390863623309
4.25998470341613 0.762080826914801
0.382678116692849 0.800562178228277
2648 KAT2A lysine acetyltransferase 2A 4010
0.113325061692794 5.99661711940541 0.762049087801977
0.38268802566368 7 0.800562178228277
51081 MRPS7 mitochondrial ribosomal protein S7 3324
0.128752618356799 5.54455010676924 0.762008474283137
0.382700705759951 0.800562178228277
50859 SPOCK3 SPARC (osteonectin), cwcv and kazal like domains
proteoglycan 3 4482 -0.280561669567654 4.04069310530054

0.761999822371594	0.382703407082042
0.800562178228277
9801	MRPL19	mitochondrial ribosomal protein L19
4601	-0.125373150894877	5.36710715738549
0.761965478981617	0.382714130130175	0.800562178228277
23660	ZKSCAN5	zinc finger with KRAB and SCAN domains 5	6056
0.158942863829972	4.76955821207904
0.761534339782999	0.382848781100266	0.800562178228277
7248	TSC1	TSC complex subunit 1	9911	-0.160465424302804
6.09552815505025	0.761270209469927
0.382931305966033	0.800562178228277
79006	METRN	meteorin, glial cell differentiation regulator	3692
0.318166475177702	3.2203313136048	0.761191199226345
0.382955996823415	0.800562178228277
25902	MTHFD1L	methylenetetrahydrofolate dehydrogenase (NADP+ dependent) 1 like	5021	0.130532820308298
5.51116068057696	0.761002472851061	0.383014983323682
0.800562178228277
10979	FERMT2	fermitin family member 2
5185	-0.224297075621367	5.90416874335533
0.760911200769568	0.383043515068543	0.800562178228277
9703	KIAA0100	KIAA0100	9235	-0.108183321711751
8.29985603760017	0.760830715598786
0.383068677314479	0.800562178228277
84142	ABRAXAS1	abraxas 1, BRCA1 A complex subunit	7777
0.216727719283633	2.54332187348045
0.760101315954532	0.383296817941935	0.800562178228277
8936	WASF1	WASP family member 1	3260	0.157590508683505
4.62733063283962	0.760043218480291
0.383314997874959	0.800562178228277
27040	LAT	linker for activation of T cells	4430
0.339603751589969	2.87358707610641	0.759981975348902
0.383334163474984	0.800562178228277
7639	ZNF85	zinc finger protein 85	4668	0.279942121500155
1.92138407452436	0.759942302035602
0.383346579680085	0.800562178228277
5876	RABGGTB	Rab geranylgeranyltransferase subunit beta
6302	-0.139002751242062	5.6068259624559	0.759766979001439
0.383401455806298	0.800562178228277
999	CDH1	cadherin 1	6177	-0.144180240325959
10.3636867872	0.759543815093139	0.383471322250235
0.800562178228277
219	ALDH1B1	aldehyde dehydrogenase 1 family member B1	3155
0.291695513359057	6.33560415190414
0.759259338311495	0.383560410246425	0.800562178228277
4708	NDUFB2	NADH:ubiquinone oxidoreductase subunit B2	4103
0.164080266523384	5.45889604062761
0.759016532667193	0.383636471554673	0.800562178228277
7341	SUMO1	small ubiquitin like modifier 1	2164
0.135575642800764	6.16702801904062	0.758850783977518
0.38368840628893	0.800562178228277
23522	KAT6B	lysine acetyltransferase 6B
13400	-0.179045382608537	4.9969323073797	0.758838671127834
0.383692202050235	0.800562178228277

8660	IRS2	insulin receptor substrate 2
8138	-0.274482426494554	5.89628647782924
0.758641530283029	0.383753986878274	0.800562178228277
56834	GPR137	G protein-coupled receptor 137
4031	-0.164021702259714	5.0865434829106	0.758399001197056
	0.38383001595578	0.800562178228277
11331	PHB2	prohibitin 2	2499	0.125291306344922
6.19299555722556	0.758389373712518	0.383833034462815
	0.800562178228277
55743	CHFR	checkpoint with forkhead and ring finger domains
	16314	-0.124865955974486	5.6529586312234
0.758244281691032	0.383878529271656	0.800562178228277
6046	BRD2	bromodomain containing 2
7337	-0.117781415349763	8.12037462165732
0.75819655566179	0.383893495837447	0.800562178228277
63035	BCORL1	BCL6 corepressor like 1 7909	-0.213138783864668
	4.86704446593531	0.758078341684737
0.38393057052203	0.800562178228277
6227	RPS21	ribosomal protein S21	1023	0.212153763762127
	6.41721855378494	0.758043152945135
0.383941607520261	0.800562178228277
7692	ZNF133	zinc finger protein 133 5426	0.151016165334957
	4.28164755408155	0.757961956734437
0.383967076545853	0.800562178228277
6938	TCF12	transcription factor 12 7163	-0.115907013948047
	6.82418181741279	0.757772758755742
0.384026432071151	0.800562178228277
550631	CCDC157 coiled-coil domain containing 157	6586
0.287356495595647	2.41660127566844	0.757518446502656
	0.384106235885247	0.800562178228277
51119	SBDS	SBDS ribosome maturation factor
1732	-0.139411369219597	7.47995573185387
0.757426925760171	0.384134961081763	0.800562178228277
57325	KAT14	lysine acetyltransferase 14	3752
0.15967140307506	4.87021253001324	0.757083219083682
	0.384242865963653	0.800583741580189
10865	ARID5A	AT-rich interaction domain 5A
3565	-0.165292814775934	4.42124247778288
0.757029865782613	0.384259619807497	0.800583741580189
84750	FUT10	fucosyltransferase 10	5606	0.212310353041603
	3.41246734503953	0.756167997229591
0.384530404991512	0.801028760070808
54883	CWC25	CWC25 spliceosome associated protein homolog
3848	-0.126863952684701	4.47181681831229
0.755978541707269	0.38458996519455	0.801031283225542
100507642	LOC100507642	uncharacterized LOC100507642	5008
	0.41948471007794	0.241544182108143
0.755800363405346	0.384645992073606	0.801031283225542
10552	ARPC1A	actin related protein 2/3 complex subunit 1A
2557	-0.138982289252542	7.75931645917194
0.755500943877252	0.384740168499675	0.801108300903069
29098	RANGRF	RAN guanine nucleotide release factor	1577
0.156192441565865	3.76478860544726	0.755290831124237
	0.384806274828554	0.801126856647378

57585	CRAMP1	cramped chromatin regulator homolog 1
9543	-0.131265078170417	6.08828844349739
0.754496295416772	0.385056400112919	0.801528458371476
79932	KIAA0319L	KIAA0319 like	9629	0.0985700232789782
	6.81150061847843	0.753961603034737
0.385224855048412	0.801759962059898
23542	MAPK8IP2	mitogen-activated protein kinase 8
interacting protein 2	5901	0.296452806875903
0.94992415143192	0.753771392253715	0.385284806257236
	0.801765604107133
92070	CTBP1-DT	CTBP1 divergent transcript	739
0.238995741078906	0.687189761202972	0.753444937686542
	0.385387730122506	0.801860655945624
8940	TOP3B	DNA topoisomerase III beta
6641	-0.170927912933983	5.3222128722458
0.752839021090384	0.385578865816671	0.801878985799118
22839	DLGAP4	DLG associated protein 4
8789	-0.134821724122723	7.29841989445095
0.752763660067991	0.385602647791865	0.801878985799118
79631	EFL1	elongation factor like GTPase 1
4254	-0.114085479587614	5.36796807175052
0.752652213791522	0.385637821157746	0.801878985799118
3652	IPP	intracisternal A particle-promoted polypeptide
3580	-0.16210946079803	3.52352538886662
0.752154628826657	0.385794918781914	0.801878985799118
56851	EMC7	ER membrane protein complex subunit 7	1538
0.126229975993233	5.97885155723892	0.75203510815132
	0.385832667435547	0.801878985799118
64318	NOC3L	NOC3 like DNA replication regulator
4464	-0.149939894446554	3.97966155070015
0.752011399798491	0.385840155956304	0.801878985799118
8550	MAPKAPK5	MAPK activated protein kinase 5
13813	-0.115813498039581	5.35151448500068
0.751908537255712	0.385872648516277	0.801878985799118
8323	FZD6	frizzled class receptor 6
4065	-0.112713016605033	7.21620688899473
0.75187317381884	0.385883820135016	0.801878985799118
57583	TMEM181	transmembrane protein 181
5390	-0.162944750281874	6.94614093572758
0.751784687432803	0.385911775774388	0.801878985799118
5763	PTMS	parathymosin	1475	0.123349785165323
7.08652010984105	0.751179707810337	0.386102985154272
	0.802157299474324
1946	EFNA5	ephrin A5	5703	-0.214597281667488
5.04870533195021	0.750598545043893	0.386286794018796
	0.802241844077349
2733	GLE1	GLE1 RNA export mediator	3527
0.137940519295532	4.77483060114211	0.750221432828391
	0.38640613288573	0.802241844077349
112950	MED8	mediator complex subunit 8
2704	-0.150685706104202	4.83152707155584
0.750216902778078	0.386407566759733	0.802241844077349
53343	NUDT9	nudix hydrolase 9	3184	-0.124259741503672
	5.3229319600208	0.750163014293316

0.386424624391514	0.802241844077349
64115	VSIR	V-set immunoregulatory receptor
5128	-0.161490384707918	5.83205890834114
0.750024452191298	0.38646848917363	0.802241844077349
401237	CASC15	cancer susceptibility 15	6785
0.272315119339161	3.85581210910804	0.749763971499051
0.386550969077504	0.802241844077349
387521	TMEM189	transmembrane protein 189
7931	-0.105537295140726	7.00444934711028
0.749706253497479	0.386569248586043	0.802241844077349
9741	LAPTM4A	lysosomal protein transmembrane 4 alpha	1946
0.123489870586149	8.52544813474486	0.749603247963591
0.386601873886064	0.802241844077349
84958	SYTL1	synaptotagmin like 1	5584	0.150266127461793
5.14478544868756	0.749321688534579
0.386691073195706	0.80227008531405
6503	SLA	Src like adaptor	7637	0.805637141047138
1.84154347916133	0.74908099606731
0.386767348911021	0.80227008531405
643853	TMPPE	transmembrane protein with metallophosphoesterase domain	3966	-0.251510296970942	1.84036251708932
0.749018010977749	0.386787312495594	0.80227008531405
29968	PSAT1	phosphoserine aminotransferase 1
2188	-0.199376923012629	5.8917858072489	0.748625292908514
0.386911820480298	0.802380017467207
25862	USP49	ubiquitin specific peptidase 49	10921
0.208606535283341	3.34133231413639	0.748489515097151
0.386954880987434	0.802380017467207
55706	NDC1	NDC1 transmembrane nucleoporin	5135
0.123306546521826	4.65866921772591	0.748266696592494
0.387025560337218	0.802407789171612
93058	COQ10A	coenzyme Q10A	3243	0.270884444275358
2.27742828361485	0.747944502369617	0.38712779479536
0.802410733425045
26088	GGA1	golgi associated, gamma adaptin ear containing, ARF binding protein 1	9369	-0.102841483006572
6.01361927471719	0.747898909744322	0.387142264759757
0.802410733425045
1241	LTB4R	leukotriene B4 receptor	4465	-0.2112284338986
3.0868820347698	0.74762135025819	0.387230371866885
0.802410733425045
140838	NANP	N-acetylneuraminic acid phosphatase
3964	-0.158578740334617	3.5348867839112	0.747271012904926
0.387341622034912	0.802410733425045
5910	RAP1GDS1	Rap1 GTPase-GDP dissociation stimulator 1
5424	-0.13504146029842	5.93551177658993
0.747182021917098	0.38736988851815	0.802410733425045
25822	DNAJB5	DnaJ heat shock protein family (Hsp40) member B5
5891	0.170663175920898	3.87272556831929
0.747058543341932	0.387409114272526	0.802410733425045
3958	LGALS3	galectin 3	2397	-0.150637555828342
7.86620271949124	0.746858564458307	0.387472654090374
0.802410733425045
4665	NAB2	NGFI-A binding protein 2

3010	-0.195683651620183	4.69670024684224
0.74637992823121	0.387624792804685	0.802410733425045
79777	ACBD4	acyl-CoA binding domain containing 4	3776
0.230791428651205	2.76053880836491	0.746026149308914
	0.387737299297841	0.802410733425045
80019	UBTD1	ubiquitin domain containing 1	1706
0.184205122548719	5.27161555580245	0.746002898455211
	0.387744695015661	0.802410733425045
131616	TMEM42	transmembrane protein 42	3106
0.20916353413461	2.61494048176073	0.745369862441516
	0.387946130814215	0.802410733425045
6774	STAT3	signal transducer and activator of transcription 3
	6484	-0.108996933882262	7.90333038285954
0.745356575294238	0.387950360462635	0.802410733425045
55815	TSNAXIP1	translin associated factor X interacting
protein 1	3852	0.330258148778746	0.911977222269896
	0.745198681546611	0.388000627232283
0.802410733425045
5127	CDK16	cyclin dependent kinase 16	6524
0.103974694819999	6.86593283794178	0.745197268763514
	0.38800107704532	0.802410733425045
10655	DMRT2	doublesex and mab-3 related transcription factor 2
	3560	-0.25271024708237	0.0660757914013285
0.745066849494018	0.388042604167385	0.802410733425045
6729	SRP54	signal recognition particle 54
3903	-0.0983272966556276	6.21726770847127
0.744975835544789	0.388071587901672	0.802410733425045
9529	BAG5	BCL2 associated athanogene 5
5271	-0.101210772739239	6.55972903345785
0.744936412669105	0.388084143223577	0.802410733425045
6539	SLC6A12	solute carrier family 6 member 12
5280	-1.09311843634058	3.24365225200307
0.744910125181001	0.388092515534575	0.802410733425045
902	CCNH	cyclin H	4660	0.147690838953921
4.41179519947123	0.744661137724133	0.38817182841965
	0.802410733425045
10573	MRPL28	mitochondrial ribosomal protein L28	2404
0.148217490897706	5.20593966057456	0.744353128671261
	0.388269974158688	0.802410733425045
283212	KLHL35	kelch like family member 35	3101
0.277011527416691	-0.0607845239162423	0.744226244503808
	0.388310415546288	0.802410733425045
4899	NRF1	nuclear respiratory factor 1
4125	-0.14785871668958	3.94345600891022
0.744161985245913	0.388330898989271	0.802410733425045
113278	SLC52A3	solute carrier family 52 member 3
3603	-0.349976062134788	2.09206749643062
0.744116642805285	0.38834535338815	0.802410733425045
25974	MMACHC	metabolism of cobalamin associated C	5296
0.215544325440364	2.72577059607925	0.7438218364935
0.388439351361363	0.802410733425045
648791	PPP1R3G	protein phosphatase 1 regulatory subunit 3G	4907
	0.311648725604745	1.7333594800652	0.743759784167204
	0.388459140669325	0.802410733425045

55333	SYNJ2BP synaptojanin 2 binding protein
7189	-0.0998531648045666	6.15682587242094
0.743327936722819	0.388596902413786	0.802576940741654
79676	OGFOD2	2-oxoglutarate and iron dependent oxygenase domain
containing 2	7126	0.100807587640871	6.21520881639722
	0.742821314429008	0.388758606707901
0.802792540786904
11232	POLG2	DNA polymerase gamma 2, accessory subunit	3388
	0.231129552098388	2.3144282692081 0.742328920272144
	0.388915861769206	0.8029305632110 2
91408	BTF3L4	basic transcription factor 3 like 4
6022	-0.134423411325021	5.11296585352184
0.742045283682309	0.389006487545874	0.8029305632110 2
5919	RARRES2 retinoic acid receptor responder 2	1921
0.487538002952291	1.38934527186707	0.742028140264448
	0.389011966069715	0.8029305632110 2
1259	CNGA1	cyclic nucleotide gated channel subunit alpha 1 3581
	0.362591012531014	0.935068834120853
0.74189430616056	0.389054739238466	0.8029305632110 2
5447	POR	cytochrome p450 oxidoreductase
6112	-0.161477591130992	6.41863263037405
0.74093721293076	0.389360820719963	0.802980755611273
9851	KIAA0753	KIAA0753	6299	-0.123653625936648
	5.02778458420783	0.740721899517621
0.389429726064512	0.802980755611273
28958	COA3	cytochrome c oxidase assembly factor 3	1922
0.144715266010362	4.54190978258195	0.740498875279695
	0.389501117436828	0.802980755611273
399665	FAM102A family with sequence similarity 102 member A
6100	-0.213170300432117	6.67387195513934
0.740368167980137	0.389542966310727	0.802980755611273
2810	SFN	stratifin	1320	-0.151192578282793
10.1150740065176	0.740295363274072	0.389566279158426
	0.802980755611273
10125	RASGRP1 RAS guanyl releasing protein 1	6097
0.21284248179916	4.61896746097568	0.740197948241864
	0.389597475617221	0.802980755611273
8635	RNASET2 ribonuclease T2 7809	0.175854548559077
3.51405850823537	0.74009672540728	0.389629895281904
	0.802980755611273
22826	DNAJC8	DnaJ heat shock protein family (Hsp40) member C8
	2888	0.107652322561437	6.25152658588724
0.740057135771906	0.389642576105784	0.802980755611273
58495	OVOL2	ovo like zinc finger 2	2175	0.224573125715934
	2.46368727073998	0.740030707169968
0.389651041691511	0.802980755611273
90589	ZNF625	zinc finger protein 625 1904	-0.244260785387863
	0.999064896779416	0.740026683153334
0.389652330683931	0.802980755611273
8681	JMJD7-PLA2G4B	JMJD7-PLA2G4B readthrough
7862	-0.159636879080747	4.36832792999943
0.739685657307167	0.389761591889903	0.802999137377734
54956	PARP16	poly(ADP-ribose) polymerase family member 16	3715
	0.210230518801104	2.61862559975981

0.73964098588027	0.389775907405669	0.802999137377734
2302	FOXJ1	forkhead box J1
2641	-0.648579764842759	-0.240121235158005
0.739186051156988	0.389921739751473	0.803181442455718
4728	NDUFS8	NADH:ubiquinone oxidoreductase core subunit S8	4523
	0.150918197596781	5.204869469284	0.738826158193282
	0.39003716113542	0.803301061023943
84299	MIEN1	migration and invasion enhancer 1	1200
0.142339328206078	5.05603770701852	0.738119007724261
	0.390264094027805	0.803455548595669
23288	IQCE	IQ motif containing E	9240	0.127576425710366
	5.29983011317929	0.738101383119954
0.390269752383527	0.803455548595669
10799	RPP40	ribonuclease P/MRP subunit p40	1962
0.241403368211755	2.1191407748397	0.737836971133065
0.390354655596637	0.803455548595669
1939	EIF2D	eukaryotic translation initiation factor 2D	8064
	0.128163796674087	4.65226817575093
0.737522928475011	0.390455529670159	0.803455548595669
160428	ALDH1L2	aldehyde dehydrogenase 1 family member L2
8507	-0.451195097766885	3.3543988963713	0.737342492078596
	0.390513504773963	0.803455548595669
84306	PDCD2L	programmed cell death 2 like	1973
0.169165162547839	4.20538951090839	0.737250016914196
	0.390543222277084	0.803455548595669
340267	COL28A1	collagen type XXVIII alpha 1 chain	6619
0.33344656288866	0.765859096891668	0.737227716535291
	0.390550389135267	0.803455548595669
6404	SELPLG	selectin P ligand	2476	0.495951691014974
	2.06615797901685	0.737163401965638
0.390571059497959	0.803455548595669
23636	NUP62	nucleoporin 62	6016	-0.105660467850879
5.8639674638799	0.736871956213765	0.390664748257848
0.803530268553257
84449	ZNF333	zinc finger protein 333	11247	-0.232693656139041
	2.49241253130063	0.73649091968413
0.390787285606633	0.803664294448995
3313	HSPA9	heat shock protein family A (Hsp70) member 9	4751
	0.109664204044892	7.66062633980561
0.736198512469059	0.390881358032424	0.803739750728151
4194	MDM4	MDM4 regulator of p53	11008	-0.241796879805872
	3.69795549779775	0.735400099641176
0.391138386438129	0.804055695562809
84318	CCDC77	coiled-coil domain containing 77	3614
0.19444082250076	3.02556949504365	0.735364605037687
	0.39114981863104	0.804055695562809
22864	R3HDM2	R3H domain containing 2	6133	-0.118211086823171
	5.50241839603657	0.735029093103616
0.391257904833188	0.804159863976297
50484	RRM2B	ribonucleotide reductase regulatory TP53 inducible
subunit M2B	5548	0.198970865048128	7.03314174953554
	0.734825756091936	0.391323431309561
0.804161705862501
1340	COX6B1	cytochrome c oxidase subunit 6B1	936

0.153689012758916	6.34488581880209	0.734604390933541
	0.391394785339827	0.804161705862501
285600	KIAA0825	KIAA0825	10415	0.295387538167678
	1.18738599867411	0.734465043586997
0.391439711615645	0.804161705862501
9652	TTC37	tetratricopeptide repeat domain 37
8092	-0.11261137884941	6.75016632518839
0.734313908565419	0.391488446653559	0.804161705862501
5268	SERPINB5	serpin family B member 5
3939	-0.157929993996259	8.86422395374082
0.733991052006957	0.391592584205121	0.804193595461913
6176	RPLP1	ribosomal protein lateral stalk subunit P1	3271
	0.144376931241628	7.82983387399282
0.733901941564316	0.391621333818544	0.804193595461913
5165	PDK3	pyruvate dehydrogenase kinase 3	2695
0.257001995957168	2.06184495846299	0.733731869244843
	0.391676212482414	0.804193595461913
653519	GPR89A	G protein-coupled receptor 89A	5557
0.127768202605808	4.31507277327606	0.733404889940946
	0.391781752626841	0.804292394700888
79085	SLC25A23	solute carrier family 25 member 23
4859	-0.170290853456976	7.4369796841016	0.732851373619297
	0.391960505823404	0.804341309154685
84232	MAF1	MAF1 homolog, negative regulator of RNA polymerase
III	2377	0.109914198061346	6.16807073406872
0.732637081348986	0.392029741028576	0.804341309154685
23516	SLC39A14	solute carrier family 39 member 14	6942
	0.168114729764644	6.87340066805647
0.732562445600905	0.392053859043068	0.804341309154685
381	ARF5	ADP ribosylation factor 5	1871
0.105723163082385	6.77315648064634	0.732479846891877
	0.392080552713651	0.804341309154685
100506516	LINC01176	long intergenic non-protein coding
RNA 1176	818	0.276874740077575	0.0523855998671393
	0.732296957251747	0.392139666982529
0.804341309154685
741	ZNHIT2	zinc finger HIT-type containing 2	1306
0.23494109080575	2.434228910087	0.732259246436682
0.392151857602478	0.804341309154685
728392	LOC728392	uncharacterized LOC728392
2865	-0.240382064197176	2.77766426938844
0.732086989862612	0.392207549191583	0.804341309154685
10652	YKT6	YKT6 v-SNARE homolog	3707	-0.117613677342463
	7.37119882541861	0.731786550929755
0.392304710073325	0.804371872911745
51411	BIN2	bridging integrator 2	4010	0.695507209483309
	0.293042906582467	0.731493681064975
0.392399456397227	0.804371872911745
221143	EEF1AKMT1	EEF1A lysine methyltransferase 1	1339
	0.260915068954761	1.57519261576967
0.731374039750293	0.392438171005356	0.804371872911745
79065	ATG9A	autophagy related 9A	4724	0.0973177670359054
	6.93193911229974	0.73126789870274
0.392472521665848	0.804371872911745

1890	TYMP	thymidine phosphorylase 3320	-0.333826848507236
	5.41813438570826	0.731128763806648
0.392517556735487	0.804371872911745
221914	GPC2	glypican 2	3828	0.261929983059673
1.58422758816737	0.730864662779818	0.392603061147918
	0.804371872911745
79007	DBNDD1	dysbindin domain containing 1	3522
0.175531947391033	5.8468383250953 0.730798649869257
0.392624437431113	0.804371872911745
2934	GSN	gelsolin	11091	0.122731132061582
9.57175683905938	0.730362186399475	0.392765814977077
	0.804543838897911
4854	NOTCH3	notch receptor 3	9394	-0.213276316456243
	8.07245709315565	0.730108902619918
0.392847891193639	0.804594299159132
401494	HACD4	3-hydroxyacyl-CoA dehydratase 4
8267	-0.158800651426292	4.1981524498679 0.729904077962577
	0.392914282321823	0.804612624631839
3268	AGFG2	ArfGAP with FG repeats 2	5478
0.280796969310756	4.1350831029463 0.729444360553375
0.393063352160251	0.80480023004073
10241	CALCOCO2	calcium binding and coiled-coil domain 2
	7451	-0.0960848915672623	6.11201915317675
0.728873876817238	0.393248452536189	0.80506154262999
10352	WARS2	tryptophanyl tRNA synthetase 2, mitochondrial	3847
	0.211537638988963	3.59364510875342
0.728528291722475	0.393360642739745	0.805061901232658
5032	P2RY11	purinergic receptor P2Y11	2295
0.24236728170303	2.90769775018764	0.728436931236142
	0.393390309552112	0.805061901232658
839	CASP6	caspase 6	1943	0.138973674241231
5.52963857340789	0.728233461928873	0.393456392186349
	0.805061901232658
390940	PINLYP	phospholipase A2 inhibitor and LY6/PLAUR domain
containing	1514	0.281693229174249	0.927298652346297
	0.728042403798588	0.393518458452489
0.805061901232658
8287	USP9Y	ubiquitin specific peptidase 9 Y-linked
11237	-0.150317283771673	6.80902812585379
0.727988394793201	0.393536006121226	0.805061901232658
825	CAPN3	calpain 3	6489	0.213087389299063
4.51759039261324	0.727372603853567	0.393736157780864
	0.805227175208027
51101	ZC2HC1A	zinc finger C2HC-type containing 1A	4181
0.165420683747691	3.91790844180166	0.727344565139823
	0.393745274738954	0.805227175208027
5791	PTPRE	protein tyrosine phosphatase receptor type E
11235	0.164263373303887	6.53575894569026
0.727209307167868	0.39378925895445	0.805227175208027
83451	ABHD11	abhydrolase domain containing 11
2179	-0.12833733929083	4.14133902586288
0.726863719269534	0.393901671947908	0.805270441403564
5931	RBBP7	RB binding protein 7, chromatin remodeling factor
	7392	-0.107482343627407	5.9411983084188

0.726790793877246	0.393925399050276	0.805270441403564
10053	AP1M2	adaptor related protein complex 1 subunit mu 2
2214	-0.122378450307781	6.51579821079619
0.726539510134914	0.394007172808933	0.805320074352068
1122	CHML	CHM like Rab escort protein
7078	-0.314284957863847	4.42755106159577
0.725832066883669	0.394237523002741	0.805673327210299
140739	UBE2F	ubiquitin conjugating enzyme E2 F (putative)
4238	-0.130610905401035	5.75708577305242
0.725223160535961	0.394435943845741	0.805961234930312
493900	TMEM9B-AS1	TMEM9B antisense RNA 1	1162
0.277531193375686	0.725901298858127	0.724629864266402
0.394629416199315	0.806127667137069
3295	HSD17B4	hydroxysteroid 17-beta dehydrogenase 4	8231
0.117400566190042	6.58157912273543	0.724519794494874
0.394665324684505	0.806127667137069
5959	RDH5	retinol dehydrogenase 5	2210	0.336508009940976
0.0183744253148317	0.724262149562805
0.394749395576896	0.806127667137069
56970	ATXN7L3	ataxin 7 like 3	5301	-0.113527380606173
6.93634992819247	0.724116620722327	0.394796893796542
0.806127667137069
6834	SURF1	SURF1 cytochrome c oxidase assembly factor	1761
0.139265086594712	4.70150794260457
0.723586961069381	0.394969835496497	0.806127667137069
140680	C20orf96	chromosome 20 open reading frame 96	1730
0.182480722016018	3.08199627620585
0.723584791843972	0.394970544004942	0.806127667137069
445329	SULT1A4	sulfotransferase family 1A member 4
4540	-0.207693714179307	4.52056999726171
0.723434444302576	0.395019654705074	0.806127667137069
26168	SENP3	SUMO specific peptidase 3	3659
0.113420919588705	6.08324636649741	0.723402887028396
0.395029963937393	0.806127667137069
339290	LINC00667	long intergenic non-protein coding RNA 667
5584	0.178651135834455	4.53754905688092
0.723270125919207	0.395073339010516	0.806127667137069
260293	CYP4X1	cytochrome P450 family 4 subfamily X member 1	2548
0.357044093799411	1.59958432947967
0.723114291533148	0.395124261227025	0.806127667137069
113000	RPUSD1	RNA pseudouridine synthase domain containing 1	2769
0.165902161127116	4.57313465720123
0.723034113029854	0.395150464950604	0.806127667137069
29801	ZDHHC8	zinc finger DHHC-type containing 8	5542
0.143671071770365	5.69706779700857	0.722767833544804
0.395237507663965	0.806143798564291
1942	EFNA1	ephrin A1	2539	-0.215255000908021
6.86396187474919	0.722657817938483	0.395273478156604
0.806143798564291
55623	THUMPD1	THUMP domain containing 1
5092	-0.108157382325717	5.82064965843882
0.722251681656111	0.395406308556787	0.80625753561697
100131434	LINC00893	long intergenic non-protein coding
RNA 893	4153	-0.156867331505069	4.97387560155537

0.722135347490418	0.395444368507788	0.80625753561697
205	AK4	adenylate kinase 4	7578	0.226966099804483
	5.47701964020086	0.721655549647622
0.395601395246173	0.806297066466787
51616	TAF9B	TATA-box binding protein associated factor 9b
2791	-0.156837144228256	3.82039653697224
0.721581977283336	0.395625481726711	0.806297066466787
2581	GALC	galactosylceramidase	6825	0.143581125463196
	5.35271214578972	0.72143401689029
0.395673928125261	0.806297066466787
29089	UBE2T	ubiquitin conjugating enzyme E2 T	1065
0.221356313119388	2.56220583423074	0.721372684362894
	0.395694012628878	0.806297066466787
39	ACAT2	acetyl-CoA acetyltransferase 2
4299	-0.241386544202329	6.2929946186942 0.721063043294578
	0.39579543293575	0.806386418782698
129531	MITD1	microtubule interacting and trafficking domain
containing 1	4305	-0.167591825921898	3.87986260014224
	0.720598241715369	0.395947745126507
0.806452585706503
27077	B9D1	B9 domain containing 1	2554	0.260749368444173
	1.6778601879762 0.72047580819183	0.395987879790997
	0.806452585706503
55588	MED29	mediator complex subunit 29	3835
0.111911200467643	5.42160605240355	0.720270726404704
	0.396055120347559	0.806452585706503
11096	ADAMTS5 ADAM metallopeptidase with thrombospondin type 1
motif 5 9056	0.53151656311401	4.5321862692384
0.720172517064881	0.396087326262552	0.806452585706503
55152	DALRD3	DALR anticodon binding domain containing 3	3682
	0.135791303191104	3.92220405059067
0.720085746384476	0.396115784226511	0.806452585706503
10528	NOP56	NOP56 ribonucleoprotein 5364	0.127985089897297
	6.12076901844495	0.719790579432046
0.396212611480921	0.806532487867063
8021	NUP214	nucleoporin 214 13062	-0.140003575493152
5.48095296182616	0.718893379744234	0.396507140699702
	0.806978658473838
3588	IL10RB	interleukin 10 receptor subunit beta	2977
0.144451693501391	5.08691480156059	0.718723229466477
	0.396563032609095	0.806978658473838
114769	CARD16	caspase recruitment domain family member 16	1688
	0.412553428464902	1.99694713589511
0.718596611014981	0.396604632321975	0.806978658473838
5747	PTK2	protein tyrosine kinase 2
13742	-0.123487918775326	7.75806531906154
0.718181768447195	0.396740970429811	0.807138819580299
114928	GPRASP2 G protein-coupled receptor associated sorting
protein 2	3975	0.197209078056889	3.09297331045958
	0.717148834938438	0.397080738422015
0.807502592736767
84985	FAM83A	family with sequence similarity 83 member A
6175	-0.345646999809378	6.76528441316839
0.717115965790811	0.39709155713286	0.807502592736767

84441	MAML2	mastermind like transcriptional coactivator 2
7109	-0.218385077859476	4.84988046939425
0.71711240473428	0.397092729261858	0.807502592736767
57147	SCYL3	SCY1 like pseudokinase 3
6883	-0.143035252109945	4.17018421991154
0.716904618484238	0.397161131182669	0.807524454126236
10955	SERINC3 serine incorporator 3	4683	-0.102647401395104
	8.67675609661543	0.716707576893796
0.397226011753145	0.807539150453746
26257	NKX2-8	NK2 homeobox 8	1261	0.293446440014518
0.21468195014309	0.716505333463475	0.397292621075884
	0.807544305876271
8890	EIF2B4	eukaryotic translation initiation factor 2B subunit
delta	5341	0.120760838915511	5.25300194303503
0.716349811643198	0.397343853508907	0.807544305876271
1778	DYNC1H1 dynein cytoplasmic 1 heavy chain 1
15826	-0.148272114158349	9.97261123985268
0.715506333884676	0.397621880673286	0.807992119917412
94097	SFXN5	sideroflexin 5	11087	0.136898387102169
4.16442488337702	0.714285541753527	0.398024776489121
	0.808693508019019
60686	C14orf93	chromosome 14 open reading frame 93	4243
	0.211136615602843	1.74367855437417
0.713720794543032	0.398211358722324	0.808927403914989
11007	CCDC85B coiled-coil domain containing 85B	1524
0.283251417689522	0.869677685199982	0.713584488167354
	0.3982564108282 0.808927403914989
27436	EML4	EMAP like 4	8357	-0.143849324865964
5.37136574722146	0.713325086619761	0.398342168832262
	0.808927403914989
738	VPS51	VPS51 subunit of GARP complex	4878
0.128846071808968	5.45578622666916	0.713238190652916
	0.398370902563403	0.808927403914989
4034	LRCH4	leucine rich repeats and calponin homology domain
containing 4	8954	-0.171519937934602	6.19913134838769
	0.712575651393081	0.398590081853961
0.80907399840517
654364	NME1-NME2	NME1-NME2 readthrough	1241
0.186565273299645	7.51882657672548	0.712548466882026
	0.398599078680757	0.80907399840517
79780	CCDC82	coiled-coil domain containing 82
6850	-0.133361193030005	5.33189300133673
0.712441880955915	0.398634356570998	0.80907399840517
9846	GAB2	GRB2 associated binding protein 2
6771	-0.114852618190881	5.40801665521504
0.712321686205982	0.398674144141678	0.80907399840517
120379	PIH1D2	PIH1 domain containing 2	2474
0.23652640576531	0.27296843360768	0.712049927917037
	0.398764124357026	0.809139372717495
7789	ZXDA	zinc finger X-linked duplicated A	4113
0.21766285086764	3.49451876884418	0.711737627693267
	0.398867564472642	0.809232035858676
9416	DDX23	DEAD-box helicase 23	5419	-0.110685482888112
	7.1460040491224 0.710886595764109	0.399149640635848

0.809467393212604
22852 ANKRD26 ankyrin repeat domain 26 7941
0.153848733117013 4.07338022125307 0.710391694197684
0.399313809582671 0.809467393212604
100506100 LOC100506100 uncharacterized LOC100506100 544
0.286193676403989 0.723454121808415
0.710194777598661 0.399379158039746 0.809467393212604
10434 LYPLA1 lysophospholipase 1 4694 -0.178873397164591
6.59726920480238 0.710034243168025
0.399432444228499 0.809467393212604
27079 RPUSD2 RNA pseudouridine synthase domain containing 2 2358
0.196792856277965 3.0330696893086 0.709890882826683
0.39948003852996 0.809467393212604
1457 CSNK2A1 casein kinase 2 alpha 1 6047 -0.095839574321053
7.04152460051793 0.709664944493142
0.399555064640579 0.809467393212604
10196 PRMT3 protein arginine methyltransferase 3
5467 -0.156605714021255 3.38535841515344
0.709408487810352 0.399640249559053 0.809467393212604
355 FAS Fas cell surface death receptor 3688
0.120122115362886 5.9840128438302 0.709316935122381
0.399670666170673 0.809467393212604
57658 CALCOCO1 calcium binding and coiled-coil domain 1
8898 0.180003292525924 7.10178909623355
0.709100516681801 0.399742580358157 0.809467393212604
23347 SMCHD1 structural maintenance of chromosomes flexible hinge
domain containing 1 11520 -0.137699047572918
5.87742985941736 0.709065034356236 0.399754372652366
0.809467393212604
410 ARSA arylsulfatase A 2633 0.155428697859139
5.5646353413654 0.70898920576545 0.399779575426347
0.809467393212604
805 CALM2 calmodulin 2 6466 -0.137495733421061
9.68382475891714 0.708772584649381 0.399851585406364
0.809467393212604
257019 FRMD3 FERM domain containing 3 6486
0.387900383316321 0.789010447673007 0.708433788916562
0.399964246827467 0.809467393212604
54841 BIVM basic, immunoglobulin-like variable motif containing
5461 0.152465840003167 4.01869546414992
0.708405463530826 0.399973668092888 0.809467393212604
152100 CMC1 C-X9-C motif containing 1 10536
0.191485051106075 3.07657935009245 0.708352420288677
0.399991311594174 0.809467393212604
221322 TBC1D32 TBC1 domain family member 32 6912
0.209800766544429 2.14534631319233 0.708274663733793
0.400017177397588 0.809467393212604
11167 FSTL1 follistatin like 1 8727 -0.134004095038559
9.10191471605312 0.708191451345776
0.400044860776698 0.809467393212604
56943 ENY2 ENY2 transcription and export complex 2 subunit 6783
0.120456152479158 5.51040196662965
0.707883351138551 0.400147384788652 0.809467393212604
286749 STON1-GTF2A1L STON1-GTF2A1L readthrough 4294

0.482963836477498	1.00649167205156	0.707766367323007
	0.400186322527476	0.809467393212604
9577	BABAM2	BRISC and BRCA1 A complex member 2	3595
0.171306192965321	4.1116888623735 0.707764679870214
0.400186884231669	0.809467393212604
23361	ZNF629	zinc finger protein 629 6087	0.175902186716124
	5.80237713305273	0.7074344676629 0.400296824329341
	0.809467393212604
81488	POLR2M	RNA polymerase II subunit M	6356
0.127422856013403	5.70598113726432	0.707098419451761
	0.400408752440585	0.809467393212604
604	BCL6	BCL6 transcription repressor
5938	-0.340657973623451	5.40308187960244
0.706958709079174	0.400455299359141	0.809467393212604
10078	TSSC4	tumor suppressing subtransferable candidate 4	2659
	0.119806014207655	5.27757402864069
0.706776522023944	0.400516009912086	0.809467393212604
55174	INTS10	integrator complex subunit 10	6478
0.111181158659931	5.22026657475636	0.706712585072587
	0.400537318914832	0.809467393212604
57410	SCYL1	SCY1 like pseudokinase 1	4109
0.163586267388299	6.17030669146183	0.706676592033119
	0.400549315452598	0.809467393212604
51329	ARL6IP4 ADP ribosylation factor like GTPase 6 interacting protein 4	2781	0.103675485047792	5.95611690607522
	0.70653042891476	0.400598037212659
0.809467393212604
9516	LITAF	lipopolysaccharide induced TNF factor
5060	-0.137866554056459	8.42950673488933
0.706519450784498	0.400601696990774	0.809467393212604
159686	CFAP58	cilia and flagella associated protein 58
3548	-0.284903681416575	0.0650087650340173
0.70618823651242	0.400712136678097	0.809573762938137
23313	KIAA0930	KIAA0930	10019	0.160947963515672
	7.82550966450881	0.705727374074741
0.400865879633814	0.809767576583621
2275	FHL3	four and a half LIM domains 3	2004
0.145757103481674	4.38418589972399	0.705168125790369
	0.401052558984403	0.810027857778591
80737	VWA7	von Willebrand factor A domain containing 7
3638	-0.253253083247649	1.10420694501599
0.704591189165392	0.401245275098001	0.810300254944882
157657	C8orf37 chromosome 8 open reading frame 37	3334
0.225196708503031	1.26787143860593	0.703926200788681
	0.401467570356102	0.810404408416283
84908	FAM136A family with sequence similarity 136 member A
3086	-0.143746133858709	5.17462248469712
0.70368048498616	0.401549754595391	0.810404408416283
7474	WNT5A	Wnt family member 5A	7038	0.266722932880815
	3.78904489643112	0.703678344629218
0.401550470585141	0.810404408416283
101928725	LOC101928725	uncharacterized LOC101928725	1396
	0.239397903067525	0.841409136609685
0.703604677854496	0.40157511464091	0.810404408416283

9920 KBTBD11 kelch repeat and BTB domain containing 11 6710
0.484690535615799 3.05316701380185
0.703571736328374 0.401586135419249 0.810404408416283
57546 PDP2 pyruvate dehyrogenase phosphatase catalytic subunit
2 8364 -0.148941827877189 4.30280908896392
0.703395381618809 0.401645143384506 0.810406730536844
23536 ADAT1 adenosine deaminase tRNA specific 1 4486
0.129914500476193 5.07425604870548 0.702727280059101
0.401868803290129 0.810741225361492
3556 IL1RAP interleukin 1 receptor accessory protein
11243 0.184619811364208 4.77388383781191
0.702301129279661 0.402011560083706 0.810742436962126
10000 AKT3 AKT serine/threonine kinase 3
9171 -0.321230249476762 4.32314315998934
0.702242329015833 0.402031263447655 0.810742436962126
81619 TSPAN14 tetraspanin 14 17888 -0.115180452471929
8.59926791932803 0.702136562530235 0.402066708244432
0.810742436962126
55128 TRIM68 tripartite motif containing 68
4282 -0.164937765374686 4.26985505865123
0.702034462043282 0.402100928791025 0.810742436962126
84162 KIAA1109 KIAA1109 20470 -0.217875948644451
7.30471585619014 0.701674426574364
0.402221634017654 0.810797852976599
4904 YBX1 Y-box binding protein 1 1930 0.145873215799701
7.76230133621008 0.701607183724207
0.402244183646347 0.810797852976599
54802 TRIT1 tRNA isopentenyltransferase 1 2663
0.134712207730619 4.01538727064514 0.701282327418895
0.402353148851687 0.810846071639184
54537 SHLD2 shieldin complex subunit 2
4242 -0.132619508789843 4.64629475740331
0.701190724419334 0.402383882627541 0.810846071639184
55437 STRADB STE20 related adaptor beta
3242 -0.197742970671765 2.86414457173307
0.70100654398855 0.402445687449142 0.81085396203972
197335 WDR90 WD repeat domain 90 10534 -0.237324190930607
4.18255454887741 0.700257692698365
0.402697119316518 0.810918274424231
79699 ZYG11B zyg-11 family member B, cell cycle regulator
8198 -0.110176686856477 5.91221187370496
0.700219953605131 0.402709796521297 0.810918274424231
57343 ZNF304 zinc finger protein 304 4609 -0.141853741057894
4.19967178714288 0.700191836491721
0.40271924191661 0.810918274424231
11200 CHEK2 checkpoint kinase 2 4142 0.228884002106021
2.08653117677604 0.700070549666266
0.402759989553272 0.810918274424231
283899 INO80E INO80 complex subunit E 6076 0.120803659132721
5.14841978654439 0.70002944107587
0.402773801795865 0.810918274424231
7056 THBD thrombomodulin 4109 0.294750903555832
4.58887706044457 0.699877170231616 0.402824969903891
0.810918274424231

11133 KPTN kaptin, actin binding protein 2124
0.186693700365881 3.13483838104415 0.699505001560725
0.402950070887465 0.810933904844789
22868 FASTKD2 FAST kinase domains 2 6079 -0.122051924039663
5.03692606266155 0.69945627501475
0.40296645404297 0.810933904844789
84307 ZNF397 zinc finger protein 397 8710 -0.144672040176729
4.20105786097534 0.699228144652617
0.403043170414891 0.810933904844789
100134229 KDM7A-DT KDM7A divergent transcript 2380
-0.250659163523007 1.75729198786691
0.699165281536091 0.403064313952268 0.810933904844789
840 CASP7 caspase 7 3528 -0.170265923905841
6.02207730153125 0.698576205972024 0.40326252360715
0.811097519783352
101928143 LOC101928143 uncharacterized LOC101928143
1431 -0.213028249219525 2.44155085517977
0.698556401609261 0.403269189761675 0.811097519783352
5553 PRG2 proteoglycan 2, pro eosinophil major basic protein
3289 0.140460060276574 4.91596951474241
0.698008283177484 0.403453750310017 0.811097519783352
4671 NAIP NLR family apoptosis inhibitory protein 9513
0.281882061309021 2.45792197280936 0.697998694928724
0.403456979927074 0.811097519783352
51534 VTA1 vesicle trafficking 1 7262 -0.099624323754193
6.10906934385774 0.697656519301567
0.403572259863981 0.811097519783352
597 BCL2A1 BCL2 related protein A1 1089 -0.659331462429449
3.37090989657684 0.697543184646477
0.403610453211161 0.811097519783352
128710 SLX4IP SLX4 interacting protein
6492 -0.184130716026722 3.29552806525822
0.697434322943881 0.403647144150085 0.811097519783352
1104 RCC1 regulator of chromosome condensation 1
3888 -0.121181863762178 6.35287239114737
0.697388548440593 0.403662573520489 0.811097519783352
92285 ZNF585B zinc finger protein 585B
7571 -0.163397793799153 4.22683324854428
0.697376022249134 0.403666795917023 0.811097519783352
9956 HS3ST2 heparan sulfate-glucosamine 3-sulfotransferase 2
2605 -0.26623724607568 4.89857928072767
0.696926274127888 0.403818442109655 0.811285846045602
7625 ZNF74 zinc finger protein 74 6251 -0.159735576835424
3.79685006833075 0.696669058038876
0.403905207627859 0.811334449625611
55840 EAF2 ELL associated factor 2 1770 -0.274333775667619
0.198329154752845 0.696511167452195
0.403958481594132 0.811334449625611
283219 KCTD21 potassium channel tetramerization domain containing
21 4164 -0.124235225032088 4.54001718832431
0.696320877823248 0.404022700956865 0.81134709280327
23204 ARL6IP1 ADP ribosylation factor like GTPase 6 interacting
protein 1 3884 -0.108489591978113 8.12189431789519
0.695898278271812 0.404165374033543

0.811464416034125
60492 CCDC90B coiled-coil domain containing 90B 6561
0.124992567197049 5.66910622538824 0.69577444780224
0.404207194134774 0.811464416034125
10807 ENTR1 endosome associated trafficking regulator 1
3133 -0.13412495684649 5.08932541122548
0.695633146553537 0.404254922187915 0.811464416034125
1854 DUT deoxyuridine triphosphatase 3674
0.137947335166361 4.09348740388457 0.695296769046129
0.404368575238235 0.811576247795094
139728 PNCK pregnancy up-regulated nonubiquitous CaM kinase 4584
0.361019861453479 -0.310630628572211
0.694721746856717 0.40456296798088 0.811850070559913
374907 B3GNT8 UDP-GlcNAc:betaGal beta-1,3-N-acetylglucosaminyltransferase 8 1874 0.210587605712767
2.26725664003005 0.694488352839654 0.404641908367477
0.811892165950902
10452 TOMM40 translocase of outer mitochondrial membrane 40 3610
0.126214677858399 5.00585581117357
0.694252181920815 0.404721810862681 0.811910335487354
80014 WWC2 WW and C2 domain containing 2
9966 -0.158961796096007 6.36917589144266
0.693772960692591 0.404884014108489 0.811910335487354
55794 DDX28 DEAD-box helicase 28 2169 0.172605673705102
3.21025892266821 0.693404624401579
0.405008750350739 0.811910335487354
399687 MYO18A myosin XVIIIA 10562 -0.156926383982816
6.53839708256237 0.693258094131945 0.405058388090976
0.811910335487354
8808 IL1RL2 interleukin 1 receptor like 2
2644 -0.307991160428201 0.812045153844905
0.693209953407777 0.405074697900702 0.811910335487354
3304 HSPA1B heat shock protein family A (Hsp70) member 1B
2521 -0.20310722676537 8.0571750781324 0.693068306191833
0.405122692752135 0.811910335487354
26873 OPLAH 5-oxoprolinase, ATP-hydrolysing
4319 -0.295289231413827 2.36578058253945
0.693000402839698 0.405145703641298 0.811910335487354
267 AMFR autocrine motility factor receptor 4969
0.106503713268303 7.46974843166939 0.692925678079166
0.405171028355317 0.811910335487354
28968 SLC6A16 solute carrier family 6 member 16 3599
0.324030828089347 1.1415768194074 0.692920906755077
0.405172645467024 0.811910335487354
637 BID BH3 interacting domain death agonist
4446 -0.256093773174791 5.93404479644201
0.69254606604926 0.405299717085981 0.81196575775473
92675 DTD1 D-aminoacyl-tRNA deacylase 1 3329
0.147765199770397 4.6225003114095 0.692497337781941
0.4053162403242 0.81196575775473
352909 DNAAF3 dynein axonemal assembly factor 3 3302
0.336513108837203 1.08614108741045 0.691766564062478
0.405564156017215 0.81234622241286
1871 E2F3 E2F transcription factor 3

5164 -0.122438280669973 5.44683605842181
0.691496259230957 0.40565591332661 0.812386035429629
9340 GLP2R glucagon like peptide 2 receptor 6052
0.337894181207514 1.47705990454629 0.691233393655295
0.405745174415313 0.812386035429629
55798 METTL2B methyltransferase like 2B
6933 -0.143785387671415 5.08282910743623
0.691195546817454 0.405758028404775 0.812386035429629
5098 PCDHGC3 protocadherin gamma subfamily C, 3 5432
0.109056692807223 6.96239580577191 0.690768189680639
0.405903213790694 0.81256057103991
10121 ACTR1A actin related protein 1A 4327
0.106267333578767 8.02694833038335 0.689972572611339
0.406173709940717 0.812918868557342
22938 SNW1 SNW domain containing 1 2432 -0.101686135052466
5.99138651692118 0.689900373302265
0.406198269510447 0.812918868557342
57567 ZNF319 zinc finger protein 319 4274 -0.166541352442495
4.5850282009269 0.689184368962231 0.406441945670296
0.813209771123711
57182 ANKRD50 ankyrin repeat domain 50
8794 -0.153903040013932 6.49834565581014
0.689132101528671 0.406459742103027 0.813209771123711
93323 HAUS8 HAUS augmin like complex subunit 8
3755 -0.210645777310695 1.74452347784738
0.688445481248545 0.406693633955189 0.813540093841015
51335 NGRN neugrin, neurite outgrowth associated 8130
0.106947460092825 7.34941562830643 0.688306483465157
0.406741006437398 0.813540093841015
1432 MAPK14 mitogen-activated protein kinase 14
6860 -0.0970963540394387 6.82350656039366
0.68655739168014 0.407337813572388 0.814617469261628
63905 MANBAL mannosidase beta like 2045 0.115088492812936
5.67128230267934 0.685996752501049
0.40752938053224 0.814884230280526
7486 WRN WRN RecQ like helicase 5634 -0.142895049219437
4.15064970049229 0.685565728148926
0.40767674866392 0.815062549034473
100144748 KLLN killin, p53 regulated DNA replication
inhibitor 4277 0.327289160851368 -0.120217622616252
0.685125520039517 0.407827337362541
0.815247254807637
388115 CCDC9B coiled-coil domain containing 9B 7510
0.135359773974875 8.13843907556846 0.684699553157088
0.407973131977842 0.81542232585811
835 CASP2 caspase 2 8020 -0.129421089287898
5.78858111941065 0.684237985762302 0.408131197763735
0.815516127745101
6208 RPS14 ribosomal protein S14 4544 0.145529167510601
7.51047564921588 0.683740655932155
0.408301611081676 0.815516127745101
79939 SLC35E1 solute carrier family 35 member E1
7257 -0.109731349931689 7.29308168833332
0.68373890884538 0.40830220991631 0.815516127745101

5704	PSMC4	proteasome 26S subunit, ATPase 4
2387	-0.106270351402115	6.46518633067881
0.683692565164215	0.408318095230888	0.815516127745101
25839	COG4	component of oligomeric golgi complex 4
4786	-0.0981815071328207	6.01418957256219
0.683619934392297	0.408342992846825	0.815516127745101
406991	MIR21	microRNA 21	4325	-0.118574749707455
7.36086874838852	0.683528825018648	0.408374228022159
0.815516127745101
8505	PARG	poly(ADP-ribose) glycohydrolase
5129	-0.107865030121366	5.37806585214804
0.683205827642015	0.408484989981891	0.815516127745101
5976	UPF1	UPF1 RNA helicase and ATPase
8600	-0.148435641294269	7.24055214013929
0.682730382425575	0.408648109416623	0.815516127745101
7153	TOP2A	DNA topoisomerase II alpha	6437
0.27785673646647	1.66837602484581	0.682439430568774
0.408747978584973	0.815516127745101
158405	KIAA1958	KIAA1958	13119	-0.257566753386543
3.15684293255775	0.682362692481263
0.408774324888015	0.815516127745101
10379	IRF9	interferon regulatory factor 9
3794	-0.141590867939508	5.59055067669464
0.681991277237639	0.40890187722342	0.815516127745101
10150	MBNL2	muscleblind like splicing regulator 2	4908
0.219863777093941	6.57848913968623	0.681852899709469
0.408949414112395	0.815516127745101
10446	LRRN2	leucine rich repeat neuronal 2
3694	-0.562691539906171	1.11378650824211
0.681732042249354	0.40899093896266	0.815516127745101
4736	RPL10A	ribosomal protein L10a	1501	0.179969100170481
7.05192639489798	0.681716819408507
0.408996169746393	0.815516127745101
5074	PAWR	pro-apoptotic WT1 regulator
11611	-0.158309225608033	7.34714528220501
0.68166976989778	0.409012337245598	0.815516127745101
83596	BCL2L12	BCL2 like 12	2216	0.178255440978003
2.79257126171022	0.68166879322951	0.409012672865479
0.815516127745101
9097	USP14	ubiquitin specific peptidase 14
5987	-0.124231366147572	7.33997513163375
0.681523144546366	0.409062727747086	0.815516127745101
662	BNIP1	BCL2 interacting protein 1
2440	-0.205306453823846	2.55755241167121
0.681353696408772	0.409120973073858	0.815516127745101
128178	EDARADD	EDAR associated death domain	3726
0.128339737941977	5.26434958058266	0.681287756328096
0.409143642306732	0.815516127745101
64221	ROBO3	roundabout guidance receptor 3	7026
0.384618164323217	3.43440501773754	0.680899781332489
0.4092770597837	0.815516127745101
83590	TMUB1	transmembrane and ubiquitin like domain containing 1
2005	0.124769804293845	4.61741456701167
0.680768612630353	0.40932218074460	0.815516127745101

79135	APOO	apolipoprotein O	2353	-0.282191171732662
	2.95409320089449	0.680442977719689
0.409434228081142	0.815516127745101
401612	SLC25A53	solute carrier family 25 member 53
6845	-0.220901090374648	1.11939048638131
0.68024654163122	0.409501841334871	0.815516127745101
157695	TDRP	testis development related protein	3613
0.148648796727252	4.95501964163342	0.680140872341651
	0.409538219464181	0.815516127745101
23178	PASK	PAS domain containing serine/threonine kinase
10686	-0.185675125713561	3.8521701704619 0.679988583949152
	0.409590655223901	0.815516127745101
3400	ID4	inhibitor of DNA binding 4, HLH protein
2344	-0.300903020609839	3.87367630290681
0.679884429691974	0.40962652317557	0.815516127745101
728888	NPIPB11 nuclear pore complex interacting protein family
member B11	4398	-0.244171662168121	2.85208695970586
	0.679855845170897	0.409636367730321
0.815516127745101
284306	ZNF547	zinc finger protein 547 3723	-0.223848841422094
	2.16031732776847	0.679815450815196
0.409650280203793	0.815516127745101
3654	IRAK1	interleukin 1 receptor associated kinase 1
4001	-0.112790308971253	7.53597829075526
0.679548109219393	0.409742374507351	0.815583549769001
57185	NIPAL3	NIPA like domain containing 3
2018	-0.192366995518937	6.61682683905379
0.679277260998475	0.409835707811515	0.815653418487624
23223	RRP12	ribosomal RNA processing 12 homolog	7179
0.151310867665673	5.05987853666231	0.678851924388816
	0.409982340329866	0.815672351056493
25921	ZDHHC5	zinc finger DHHC-type containing 5
5429	-0.107096640342893	7.93856465349389
0.678745658738851	0.41001898688484	0.815672351056493
7596	ZNF45	zinc finger protein 45	4393	0.143402874249054
	4.43624172981135	0.678742952225281
0.410019920310471	0.815672351056493
63027	SLC22A23	solute carrier family 22 member 23	8898
	-0.1358007180638	6.72593211767894
0.678025656449989	0.410267412323188	0.81604879926312
1174	AP1S1	adaptor related protein complex 1 subunit sigma 1
	1539	0.154909285144925	6.41562903646076
0.677617059057923	0.410408490877899	0.816151088873778
705	BYSL	bystin like	2248	0.152147519023479
4.60415714723339	0.677282147850526	0.410524180644083
	0.816151088873778
23214	XPO6	exportin 6	7065	-0.130281077348713
6.4541987497258 0.677259503977858	0.41053200434238
0.816151088873778
64397	ZNF106	zinc finger protein 106 12208	-0.141764843077385
	7.22558997617461	0.677201900989418
0.410551907775015	0.816151088873778
59274	TLNRD1	talin rod domain containing 1
4845	-0.134690836321356	4.56137879736129

0.676873821006195	0.410665295599707	0.816260649278431
5204	PFDN5	prefoldin subunit 5	3380	0.124749801033672
	6.6685655448917	0.676271237158474	0.410873674878815
	0.816558961978938
7543	ZFX	zinc finger protein X-linked
9037	-0.144430461538579	5.43278102317571
0.675730132034474	0.411060926883058	0.816737237513995
94101	ORMDL1	ORMDL sphingolipid biosynthesis regulator 1	3958
	0.111832333314189	5.31661983296845
0.675675043833033	0.411079997463673	0.816737237513995
153020	RASGEF1B	RasGEF domain family member 1B
9765	-0.278858917709201	3.28050970504276
0.674494325819527	0.411489055017455	0.817434008456885
522	ATP5PF	ATP synthase peripheral stalk subunit F6	2043
	0.140531446495579	5.83471477352513
0.674084032539824	0.411631340747814	0.817453894288113
10609	P3H4	prolyl 3-hydroxylase family member 4 (inactive)	3739
	0.130814541455826	4.89627385251639
0.673901703259041	0.411694594041408	0.817453894288113
55145	THAP1	THAP domain containing 1
2462	-0.165846822802078	3.65220304932752
0.673870275925522	0.411705498193573	0.817453894288113
375748	ERCC6L2	ERCC excision repair 6 like 2
10352	-0.15571872949102	4.22796717174928
0.673792438222906	0.411732506903879	0.817453894288113
9655	SOCS5	suppressor of cytokine signaling 5
5034	-0.139320849856725	4.93450283003097
0.673297509237238	0.411904302186483	0.8176790760662
5638	PRRG1	proline rich and Gla domain 1	6180
0.173945238848842	4.68692419343861	0.672716821065951
	0.412106000175605	0.81795391843987
735301	SNHG9	small nucleolar RNA host gene 9
275	-0.243042190767872	0.55643860137014
0.67256273403139	0.412159545680632	0.81795391843987
55140	ELP3	elongator acetyltransferase complex subunit 3	5390
	0.130183371175623	4.81178360792103
0.671935700642276	0.412377546706157	0.818045041081404
54974	THG1L	tRNA-histidine guanylyltransferase 1 like	3220
	0.195513213073839	2.49192822039468
0.671818575398493	0.412418286551476	0.818045041081404
9746	CLSTN3	calsyntenin 3	6865	0.136866609254619
6.26779882935476	0.671710714614853	0.412455809177664
	0.818045041081404
56925	LXN	latexin	1504	0.220464832983625
2.62290068009564	0.67165243758119	0.412476084696894
	0.818045041081404
286076	BREA2	breast cancer estrogen-induced apoptosis 2
1963	-0.269482900164285	0.517185911016595
0.671590962424721	0.412497474488324	0.818045041081404
201562	HACD2	3-hydroxyacyl-CoA dehydratase 2
4986	-0.152946736474566	8.0578619655464	0.671107752436853
	0.412665660561246	0.818123094836052
8766	RAB11A	RAB11A, member RAS oncogene family
5426	-0.122479937688732	7.92108694103812

0.671072658894474	0.412677879160967	0.818123094836052
200916	RPL22L1 ribosomal protein L22 like 1	2326
0.276523219373792	3.41026056488569	0.670963809569329
	0.412715780878275	0.818123094836052
54914	FOCAD	focadhesin	8190	-0.114997678788933
6.09408929925611	0.670806787494449	0.412770465567672
	0.818123094836052
219743	TYSND1	trypsin domain containing 1	4597
0.161046578317284	4.45689737781764	0.670465578584768
	0.412889332251525	0.818242908439037
6598	SMARCB1 SWI/SNF related, matrix associated, actin dependent regulator of chromatin, subfamily b, member 1	2481
0.0914903471663947	6.08980918368653	0.669999195628025
	0.413051887474832	0.818415468281194
7072	TIA1	TIA1 cytotoxic granule associated RNA binding protein	6607	-0.137699830249425	6.48887665549531
0.66972547191493	0.413147336380906	0.818415468281194
23295	MGRN1	mahogunin ring finger 1	8311	-0.129991812277558
	6.19251992369309	0.669556270861563
0.413206353972885	0.818415468281194
200558	APLF	aprataxin and PNKP like factor	4872
0.198058428395757	2.51816581141434	0.669545466666134
	0.413210122916014	0.818415468281194
57001	SDHAF3	succinate dehydrogenase complex assembly factor 3
	2303	0.231795048764625	1.81521851094844
0.669242465027833	0.413315842902169	0.818455625057425
51234	EMC4	ER membrane protein complex subunit 4	3012
0.121829041816448	6.50686172855413	0.669152439001607
	0.413347261487558	0.818455625057425
9050	PSTPIP2 proline-serine-threonine phosphatase interacting protein 2	4120	0.13825384462615	5.33342585390534
	0.66894629356338	0.413419218395144
0.818482401704704
84991	RBM17	RNA binding motif protein 17	6419
0.097309903926344	6.19619671645863	0.668221898211934
	0.413672221897634	0.818867554002283
55627	SMPD4	sphingomyelin phosphodiesterase 4	7375
0.0888950914728297	6.14325594420623	0.667444739373098
	0.41394390816174	0.819202843542847
4213	MEIS3P1 Meis homeobox 3 pseudogene 1
690	-0.204625703128759	2.58391518348371
0.667380814268517	0.41396626742101	0.819202843542847
360023	ZBTB41	zinc finger and BTB domain containing 41
8562	-0.160801697166567	5.22231287013142
0.667235623410422	0.414017057859628	0.819202843542847
7473	WNT3	Wnt family member 3	3846	0.183054521608287
	3.45561055395253	0.666649277614081
0.41422226619858	0.819458052091094
7029	TFDP2	transcription factor Dp-2	11897
0.156071693548683	5.51065866038975	0.666138764183003
	0.414401057375005	0.819458052091094
4701	NDUFA7	NADH:ubiquinone oxidoreductase subunit A7	892
	0.185208315953871	3.87254441448529
0.665927461083726	0.414475093004217	0.819458052091094

10206	TRIM13	tripartite motif containing 13
7812	-0.120001509273139	5.57956572593226
0.66563909333531	0.414576161851696	0.819458052091094
57710	KIAA1614	KIAA1614	10195	-0.322632685228697
	0.955905215989253	0.665345070237136
0.414679250371504	0.819458052091094
1841	DTYMK	deoxythymidylate kinase 2451	0.148259083516293
	3.67728938439755	0.665256106904222
0.414710449608283	0.819458052091094
54732	TMED9	transmembrane p24 trafficking protein 9 2440
0.159682630215349	7.91241524099101	0.66505376856955
	0.414781422135469	0.819458052091094
165631	PARP15	poly(ADP-ribose) polymerase family member 15	5507
	0.492871832569027	0.767050918312518
0.664993447687353	0.414802583865878	0.819458052091094
79827	CLMP	CXADR like membrane protein	5097
0.819063248446417	2.1844668034138 0.664820050876036
0.414863423716245	0.819458052091094
10490	VTI1B	vesicle transport through interaction with t-SNAREs
1B	5789	-0.0966074383820965	6.58008896298244
0.664803294816285	0.414869303625553	0.819458052091094
1474	CST6	cystatin E/M	759	-0.156871562223052
6.97783570117347	0.664758491757851	0.41488502618069
	0.819458052091094
90624	LYRM7	LYR motif containing 7	6351	-0.168171639063729
	3.28759258148005	0.664731993019364
0.41489432569037	0.819458052091094
93436	ARMC6	armadillo repeat containing 6	4052
0.124443777943585	4.68250868543142	0.664697067980526
	0.41490658281074	0.819458052091094
3655	ITGA6	integrin subunit alpha 6
6533	-0.201498642199362	8.61241009416058
0.663912043967912	0.415182232508298	0.819544334739128
54103	GSAP	gamma-secretase activating protein
9001	-0.329252149327635	4.193890981677	0.663749145595286
	0.415239465862458	0.819544334739128
55156	ARMC1	armadillo repeat containing 1
3421	-0.120270937148303	5.38658123482975
0.663564716319335	0.415304278073889	0.819544334739128
90293	KLHL13	kelch like family member 13	6983
0.281543710485156	0.87898085380586	0.66341247034461
	0.415357791707158	0.819544334739128
346653	FAM71F2 family with sequence similarity 71 member F2
5875	-0.265152843454886	1.72697339582523
0.663319024684749	0.415390642407897	0.819544334739128
10287	RGS19	regulator of G protein signaling 19
2028	-0.170586708157869	3.56443605715969
0.663118026900256	0.415461315963404	0.819544334739128
5929	RBBP5	RB binding protein 5, histone lysine
methyltransferase complex subunit	4404	-0.104166790219598
	5.24437811678936	0.662923267796637
0.415529812908676	0.819544334739128
85026	ARRDC1-AS1	ARRDC1 antisense RNA 1	1841
0.270039374827646	1.78051543728267	0.662858501761658

0.415552594886554	0.819544334739128
4831	NME2	NME/NM23 nucleoside diphosphate kinase 2	2632
0.176024681253153	7.37717535215046
0.662738241723559	0.4155949022391 0.819544334739128
10495	ENOX2	ecto-NOX disulfide-thiol exchanger 2
4279	-0.140569397915743	3.88192151278692
0.662652977817281	0.415624901856895	0.819544334739128
8507	ENC1	ectodermal-neural cortex 1
5858	-0.158713696961109	9.14372655437035
0.662461681598245	0.415692220015258	0.819544334739128
51523	CXXC5	CXXC finger protein 5	3852	-0.152848408278679
5.37415277587454	0.662410883159339
0.415710098971663	0.819544334739128
7757	ZNF208	zinc finger protein 208
6867	-0.486253389530678	-0.0973817692847603
0.66196169832401	0.415868243125891	0.819544334739128
140459	ASB6	ankyrin repeat and SOCS box containing 6
4622	-0.12787887925076	5.59409014581794
0.661866263023279	0.415901854443745	0.819544334739128
285346	ZNF852	zinc finger protein 852 1975	-0.249052568537414
1.95888672242661	0.661831695062193
0.415914029912736	0.819544334739128
2931	GSK3A	glycogen synthase kinase 3 alpha
4281	-0.0971493768176501	6.31651599321701
0.66164101370498	0.415981200876753	0.819544334739128
79169	C1orf35 chromosome 1 open reading frame 35	4308
0.165761388071782	3.60513793267486	0.661614793041622
0.415990438838518	0.819544334739128
2314	FLII	FLII actin remodeling protein
6465	-0.106604302907914	8.70928955603257
0.661577885246928	0.416003442564089	0.819544334739128
6714	SRC	SRC proto-oncogene, non-receptor tyrosine kinase
6468	-0.111551106750588	7.29466683632854
0.661324257035545	0.416092819711782	0.819605136507233
5717	PSMD11	proteasome 26S subunit, non-ATPase 11
7200	-0.103345947730343	7.11651434469513
0.661137003834256	0.416158824954134	0.819619890485454
57537	SORCS2	sortilin related VPS10 domain containing receptor 2
6317	-0.270995142885334	6.25199309326834
0.660363899801851	0.41643150236513	0.820041621485784
94056	SYAP1	synapse associated protein 1
6418	-0.0930671435206476	6.20388746230361
0.659791657474667	0.416633505651725	0.820218064260317
84280	BTBD10	BTB domain containing 10
3495	-0.100424775744758	6.75250847552786
0.659778311521811	0.416638218548737	0.820218064260317
9649	RALGPS1 Ral GEF with PH domain and SH3 binding motif 1
10482	-0.160022377587132	4.28011843095495
0.658805208795139	0.416982066619087	0.820573490628502
25936	NSL1	NSL1 component of MIS12 kinetochore complex	4091
0.123642232951838	3.99259221611238
0.658725947114421	0.417010092467362	0.820573490628502
10277	UBE4B	ubiquitination factor E4B
9128	-0.0954858533738286	6.25599293610075

0.658698257297864	0.417019883867695	0.820573490628502
145853	C15orf61	chromosome 15 open reading frame 61	4811
	0.229532773028477	1.28878837129155
0.658311956190171	0.41715651947327	0.820573490628502
7556	ZNF10	zinc finger protein 10	5541	0.265875364074502
	2.74369371842642	0.658302374079261
0.417159909533261	0.820573490628502
3784	KCNQ1	potassium voltage-gated channel subfamily Q member 1
	3971	-0.375343843679952	2.71685687225014
0.658273119895789	0.417170259639149	0.820573490628502
23	ABCF1	ATP binding cassette subfamily F member 1
4523	-0.101648742051422	6.78100432020777
0.657528994555975	0.417433658865874	0.820944185856639
9517	SPTLC2	serine palmitoyltransferase long chain base subunit
2	9143	-0.134239441049124	7.08246527949153
0.657409622936619	0.417475936021763	0.820944185856639
5364	PLXNB1	plexin B1	9733	0.127297459441102
8.39200607680939	0.656499820300326	0.417798364707878
	0.82146289927895
255231	MCOLN2	mucolipin 2	3442	0.563885072170027
0.305275857143585	0.656250581771941	0.41788675795768
	0.821521378065013
4598	MVK	mevalonate kinase	6473	-0.19083970192024
	5.08770451560391	0.655847275612189
0.418029850697384	0.821687358787294
10322	SMYD5	SMYD family member 5	3380	0.173140824835892
	4.15788100934084	0.655425759095134
0.418179482340013	0.821866144119063
7158	TP53BP1	tumor protein p53 binding protein 1	12991
0.112570445525504	6.62157536963234	0.65472766737372
	0.418427469044005	0.822077330475594
257194	NEGR1	neuronal growth regulator 1
13417	-0.459576398739793	3.82413724721021
0.654710179542352	0.418433684144681	0.822077330475594
7541	ZBTB14	zinc finger and BTB domain containing 14
4120	-0.144476890659421	3.93509652956495
0.654627671665777	0.418463008944884	0.822077330475594
55094	GPATCH1	G-patch domain containing 1
4488	-0.167328197152326	3.3303079931659	0.654084582536219
	0.418656109034971	0.822317405852595
493911	PHOSPHO2	phosphatase, orphan 2	1488
0.244708324815728	0.980934641835751	0.653953799190187
	0.418702630009332	0.822317405852595
10902	BRD8	bromodomain containing 8	7046
0.125565527466245	5.84400499332617	0.653705026711449
	0.418791142165924	0.822375932751731
10693	CCT6B	chaperonin containing TCP1 subunit 6B	4334
0.293824654386816	0.906913016869078	0.652933492229662
	0.419065827979861	0.822551073064863
22990	PCNX1	pecanex 1	16715	-0.124437433837555
6.61355202446209	0.652906872431213	0.419075310089355
	0.822551073064863
7955	RNF217-AS1	RNF217 antisense RNA 1 (head to head)	4507
	0.299740293004058	0.91838170868711

0.652808847897635	0.419110229688459	0.822551073064863
8744	TNFSF9	TNF superfamily member 9	1665
0.233468593774162	4.87249062148015	0.652784039088552
	0.41911906809851	0.822551073064863
53371	NUP54	nucleoporin 54	4062	-0.112434560245132
4.51903566687956	0.652629998412487	0.419173952990433
	0.822551073064863
440138	ALG11	ALG11 alpha-1,2-mannosyltransferase
7716	-0.112628525523479	5.07479524287017
0.652108166860875	0.419359961829514	0.822688967371665
1173	AP2M1	adaptor related protein complex 2 subunit mu 1	4756
	0.111740420926376	8.66156832881359
0.651851634679975	0.419451448786421	0.822688967371665
50619	DEF6	DEF6 guanine nucleotide exchange factor
2545	-0.165051806209263	2.9787366434964 0.651648447255635
	0.419523932533496	0.822688967371665
391	RHOG	ras homolog family member G
1782	-0.130879720485601	5.37170354607844
0.651368589035037	0.419623797879702	0.822688967371665
10637	LEFTY1	left-right determination factor 1	2634
0.173990581796367	3.28346873850725	0.651341426777513
	0.419633492417475	0.822688967371665
131583	FAM43A	family with sequence similarity 43 member A	2494
	0.415790277237451	1.50280029415665
0.651268468568783	0.419659533735881	0.822688967371665
100288798	LOC100288798	uncharacterized LOC100288798	2675
	0.312185796942133	0.544865653079906
0.651128677621831	0.419709436708915	0.822688967371665
100616483	MIR4751 microRNA 4751	74	0.271033312539682
	0.359281482877543	0.651115623391824
0.419714097296917	0.822688967371665
219699	UNC5B	unc-5 netrin receptor B 6841	-0.256765379466869
	5.21663483304108	0.650843368622951
0.419811314586527	0.822764388332655
54932	EXD3	exonuclease 3'-5' domain containing 3
7626	-0.170601413091435	3.04506181027036
0.650619509956118	0.419891275647859	0.822783464799084
127428	TCEANC2 transcription elongation factor A N-terminal and central domain containing 2	12165	-0.129831484283991
4.13905639285913	0.650487246710078	0.419938530009899
	0.822783464799084
84993	UBL7	ubiquitin like 7	1885	0.12687756435738
	4.55348795330088	0.650207962255713
0.420038337439676	0.822786235781464
7048	TGFBR2	transforming growth factor beta receptor 2
4704	-0.102383643039476	8.20681389888656
0.649976446724537	0.420121100585861	0.822786235781464
256643	BCLAF3	BCLAF1 and THRAP3 family member 3
8630	-0.125023814006117	4.17317189434802
0.649792355515245	0.420186927645586	0.822786235781464
56474	CTPS2	CTP synthase 2	5510	-0.124936973244558
4.95248709779618	0.649695613497006	0.420221526678705
	0.822786235781464
6490	PMEL	premelanosome protein	3901	0.236218351870817

0.224212886231573 0.649595141817798
0.420257464094912 0.822786235781464
56882 CDC42SE1 CDC42 small effector 1
4184 -0.115397618877937 8.17996939258332
0.649497508287709 0.420292390734374 0.822786235781464
51333 ZNF771 zinc finger protein 771 5637 0.284728668235454
0.715358397255634 0.649265076079254
0.420375556604328 0.822790310867277
6741 SSB small RNA binding exonuclease protection factor La
5113 0.120375088098983 6.08708455437048
0.649067197365039 0.420446378342837 0.822790310867277
54981 NMRK1 nicotinamide riboside kinase 1
3865 -0.174089009355209 3.71725273980394
0.648999263087685 0.420470696450915 0.822790310867277
9986 RCE1 Ras converting CAAX endopeptidase 1 2995
0.117965689068522 4.83398566270011 0.648357355387279
0.4207005807335 0.823011326829416
55700 MAP7D1 MAP7 domain containing 1
5271 -0.113198084244586 8.46355005479507
0.648355748127301 0.420701156571616 0.823011326829416
23228 PLCL2 phospholipase C like 2 5146 0.355032356765955
1.45806677244356 0.647977504244857
0.420836703930039 0.823156325578174
64222 TOR3A torsin family 3 member A 3917
0.151308646827228 4.8616168406166 0.647820991085847
0.42089281100813 0.823156325578174
151903 CCDC12 coiled-coil domain containing 12 3437
0.132560025212807 4.31055710766382 0.647314867579816
0.421074323421587 0.823366210522308
6671 SP4 Sp4 transcription factor
6234 -0.222262713438817 2.47153169615057
0.64696326245113 0.421200489325159 0.823366210522308
6659 SOX4 SRY-box transcription factor 4
5851 -0.136926534989874 9.28200786256375
0.646781569398854 0.421265708081314 0.823366210522308
4832 NME3 NME/NM23 nucleoside diphosphate kinase 3 1445
0.206218390669081 2.79960271371228
0.646529777739925 0.421356113688369 0.823366210522308
23216 TBC1D1 TBC1 domain family member 1
9032 -0.161136952159536 5.90570333908579
0.646497907013668 0.42136755891548 0.823366210522308
51203 NUSAP1 nucleolar and spindle associated protein 1 2875
0.268093721600769 0.663642157857757
0.646392218797605 0.421405516368264 0.823366210522308
5920 PLAAT4 phospholipase A and acyltransferase 4 1299
0.356791001510094 5.30453180145423 0.646375264632795
0.421411605856674 0.823366210522308
51522 TMEM14C transmembrane protein 14C 1321
0.0994231217906455 6.42548294915156 0.646176849018222
0.421482881282145 0.823390616195119
150696 PROM2 prominin 2 5086 0.229062208722705
7.59974579718388 0.645967265068521 0.4215581881951
0.823422889701403
79937 CNTNAP3 contactin associated protein like 3 9521

0.234553644888527	3.35137226402206	0.645422510301643
	0.421754021513414	0.823690543689123
8519	IFITM1	interferon induced transmembrane protein 1
1111	-0.353012789998391	3.6723198885304 0.64476980760476
	0.421988840070837	0.823979058542109
374918	IGFL1	IGF like family member 1
769	-0.52592378727926	7.10030031197952
0.644684897043703	0.422019402154715	0.823979058542109
146712	B3GNTL1 UDP-GlcNAc:betaGal beta-1,3-N-acetylglucosaminyltransferase like 1	6891	0.235418220738187
	0.0912377417121393	0.644066399669683
0.422242119474458	0.824223017930532
692201	SNORD86 small nucleolar RNA, C/D box 86 2980
0.126113381254064	5.72458133480498	0.644011106618809
	0.422262038742735	0.824223017930532
79753	SNIP1	Smad nuclear interacting protein 1
4888	-0.121493031809542	4.61389007879501
0.643454189561442	0.422462745994913	0.82449988618514
5862	RAB2A	RAB2A, member RAS oncogene family
5654	-0.118079600726962	6.83918251444697
0.643091002278844	0.422593711918757	0.824524513517463
84548	TMEM185A	transmembrane protein 185A	5158
0.133166230576942	4.47878289092964	0.642950288058589
	0.422644470028209	0.824524513517463
79868	ALG13	ALG13 UDP-N-acetylglucosaminyltransferase subunit
	7925	-0.155595503245687	4.47388583900584
0.642929525458385	0.422651960238123	0.824524513517463
29070	CCDC113 coiled-coil domain containing 113	5927
0.24569518873594	2.24989275545023	0.642377499622397
	0.422851179169269	0.824617745022076
3141	HLCS	holocarboxylase synthetase
7889	-0.151804543733652	4.96772035951731
0.642271482086219	0.422889455609583	0.824617745022076
79083	MLPH	melanophilin	7514	-0.304889070932548
4.33399131901254	0.642270467333145	0.422889821999846
	0.824617745022076
57650	CIP2A	cell proliferation regulating inhibitor of protein phosphatase 2A	4579	0.215587728408995	2.39280316530967
	0.642144693236264	0.422935238112272
0.824617745022076
196743	PAOX	polyamine oxidase	2277	-0.187525957391931
	2.8809331545879 0.641498389645662	0.423168728891499
	0.824915863801954
9367	RAB9A	RAB9A, member RAS oncogene family
2043	-0.12383631370368	5.4843085164872 0.641333432410917
	0.423228354095113	0.824915863801954
6188	RPS3	ribosomal protein S3	5894	0.158036724884852
	7.99118686213143	0.641232568195505
0.423264818529638	0.824915863801954
7417	VDAC2	voltage dependent anion channel 2
2827	-0.149095567419559	7.28195890699773
0.64089787651983	0.423385850005638	0.825014758218346
340307	CTAGE6	CTAGE family member 6	2608	-0.35111562132974
	2.08389282026529	0.640600091046768

0.423493579033553	0.825014758218346
3987	LIMS1	LIM zinc finger domain containing 1	7771
0.167618966998066	6.88304083622819	0.640403805104569
	0.423564611321983	0.825014758218346
11011	TLK2	tousled like kinase 2	6300	-0.106486518165125
	5.20675412460497	0.64021798473636
0.423631872767114	0.825014758218346
51602	NOP58	NOP58 ribonucleoprotein 3299	-0.137646879913331
	5.15487215616251	0.640113345359225
0.423669756155433	0.825014758218346
388121	TNFAIP8L3	TNF alpha induced protein 8 like 3
2262	-0.418458660894071	1.69722127091748
0.640096398845674	0.423675891908254	0.825014758218346
51667	NUB1	negative regulator of ubiquitin like proteins 1 5411
	0.107959723659389	6.1294045104033 0.639952874401409
	0.423727862541785	0.825014758218346
121260	SLC15A4 solute carrier family 15 member 4	5496
0.137751578108584	5.75210463144798	0.638997772564833
	0.424073951382405	0.825400818240876
22872	SEC31A	SEC31 homolog A, COPII coat complex component
9145	-0.146903947482479	8.01231226407237
0.638970962573097	0.424083672315113	0.825400818240876
9918	NCAPD2	non-SMC condensin I complex subunit D2	6487
0.117989138156006	5.33153471911739	0.638377068281024
	0.424299095931141	0.825400818240876
7001	PRDX2	peroxiredoxin 2 2093	0.122037558314182
7.17350466538497	0.638325417108909	0.424317839150249
	0.825400818240876
8835	SOCS2	suppressor of cytokine signaling 2
6311	-0.266562776253521	5.36526028661427
0.638322769747077	0.424318799860522	0.825400818240876
10180	RBM6	RNA binding motif protein 6
7302	-0.105484832441442	6.62147757811557
0.638115486004736	0.424394031907411	0.825400818240876
25934	NIPSNAP3A	nipsnap homolog 3A	2585
0.15037299716059	4.39092920568319	0.638093759479439
	0.424401918542337	0.825400818240876
80067	DCAF17	DDB1 and CUL4 associated factor 17	7657
0.12903911071058	4.86494746678901	0.637815609633513
	0.424502905172064	0.825400818240876
58492	ZNF77	zinc finger protein 77	2437	-0.244545275192908
	1.98675721817139	0.637782783819794
0.424514825473743	0.825400818240876
10330	CNPY2	canopy FGF signaling regulator 2	3176
0.0910395368049452	6.35677800364617	0.637781145354468
	0.424515420475996	0.825400818240876
25961	NUDT13	nudix hydrolase 13	2521	0.213682106952369
	0.991162850297604	0.637196818840597
0.424727696214939	0.825592155930304
23357	ANGEL1	angel homolog 1 6414	0.142135403116395
4.18052579066059	0.637185771726696	0.424731710975496
	0.825592155930304
80761	NA	NA	2660	0.308809079628261
3.02346556820693	0.63676492861418	0.424884696924915

0.825769004956191
84678 KDM2B lysine demethylase 2B 13623 -0.119762488092333
4.69957953401907 0.636524590328492
0.424972102430777 0.825769004956191
9311 ASIC3 acid sensing ion channel subunit 3
2748 -0.34979195859797 0.581807746935383
0.636449121511333 0.424999554274947 0.825769004956191
100129461 LYRM4-AS1 LYRM4 antisense RNA 1 1489
0.268159440706411 -0.0164878969333237 0.636035011310507
0.425150234619345 0.8258664775746
51202 DDX47 DEAD-box helicase 47 5574 0.113478011002966
6.41854249026532 0.635987183715912
0.425167642592307 0.8258664775746
147645 VSIG10L V-set and immunoglobulin domain containing 10 like
3771 -0.322644228786167 1.85587386880317
0.635591317040088 0.4253117686629 0.826015533185089
114904 C1QTNF6 C1q and TNF related 6 6956 0.182583615236355
5.10397146981407 0.635452512146003
0.425362321757981 0.826015533185089
4726 NDUFS6 NADH:ubiquinone oxidoreductase subunit S6 1451
0.119839938637375 4.67281081763455
0.634956436011878 0.425543067776523 0.826251975373685
6499 SKIV2L Ski2 like RNA helicase 5565 -0.103231919792338
5.88813578290386 0.634723875042407
0.425627841432864 0.826302033950122
6168 RPL37A ribosomal protein L37a 6596 0.181270654889324
8.01403715602545 0.634367143635018
0.425757927332764 0.826330475696632
4005 LMO2 LIM domain only 2 3109 -0.346906862397316
2.0296689837911 0.634347481344525 0.425765099134227
0.826330475696632
79762 C1orf115 chromosome 1 open reading frame 115
3413 -0.26114980483439 2.83506541835817
0.634198422562964 0.425819474089976 0.826330475696632
127700 OSCP1 organic solute carrier partner 1 2788
0.199242790497503 2.38983030985458 0.633878552554972
0.425936194303532 0.826341791067277
64328 XPO4 exportin 4 10500 -0.137618952641905
4.92813179361101 0.633859097760961 0.42594329488868
0.826341791067277
286077 FAM83H family with sequence similarity 83 member H
5654 -0.119303216097365 7.75742888034494
0.633516023818515 0.426068538825028 0.826470298202765
83643 CCDC3 coiled-coil domain containing 3
3683 -0.406990294946361 1.95635578489543
0.633242804844613 0.426168320861056 0.826483638971731
10228 STX6 syntaxin 6 6477 -0.0976401711211621
6.46710541384453 0.633083869965154 0.426226381489518
0.826483638971731
7270 TTF1 transcription termination factor 1
3215 -0.127669829648566 4.09085080817232
0.633005785989354 0.426254910769693 0.826483638971731
4330 MN1 MN1 proto-oncogene, transcriptional regulator 7693
0.295976680220956 4.52533033160673

0.632851099505045	0.426311436536786	0.826483638971731
1434	CSE1L	chromosome segregation 1 like
3800	-0.109604949241723	6.25391761834855
0.631984445862336	0.426628339182227	0.826983552024004
101928378	PTOV1-AS2	PTOV1 antisense RNA 2
1023	-0.231507711264335	1.92142623931313
0.631287491263528	0.426883446634336	0.827132056381921
582	BBS1	Bardet-Biedl syndrome 1 6381	0.136969727201766
	4.83546000419364	0.631240347858698
0.426900710907897	0.827132056381921
623	BDKRB1	bradykinin receptor B1	2704	-0.348341828387527
	1.32532222744001	0.631127344052548
0.426942098043464	0.827132056381921
728378	POTEF	POTE ankyrin domain family member F
4371	-0.238882609098606	2.05032183600712
0.631031768943937	0.42697710671461	0.827132056381921
285193	DUSP28	dual specificity phosphatase 28 3709
0.238942699491763	1.39506190108034	0.63096705447951
	0.427000813745681	0.827132056381921
5822	PWP2	PWP2 small subunit processome component
14587	-0.11228920420174	4.85354929848907
0.630657986266772	0.427114062924335	0.827132056381921
84447	SYVN1	synoviolin 1	5496	-0.0968648523368301
6.23906160253537	0.630439220240078	0.427194250812117
	0.827132056381921
84572	GNPTG	N-acetylglucosamine-1-phosphate transferase subunit
gamma	4102	0.118141888079169	5.41456830526007
0.630309632572311	0.427241761397348	0.827132056381921
31	ACACA	acetyl-CoA carboxylase alpha
13008	-0.138459607521544	6.74094939057643
0.630290180757466	0.427248893682296	0.827132056381921
27340	UTP20	UTP20 small subunit processome component
10024	-0.143879483154012	4.73726926599236
0.629948773518779	0.427374104726994	0.827132056381921
79856	SNX22	sorting nexin 22	5767	0.241943195881406
	-0.0149115461069956	0.629847694319122
0.427411186116705	0.827132056381921
1152	CKB	creatine kinase B	2669	0.136427629192145
	6.8855787756949 0.629841206509957	0.42741356636627
	0.827132056381921
90025	UBE3D	ubiquitin protein ligase E3D	6394
0.251264227268567	1.55572635538611	0.629462328821089
	0.427552603801342	0.827166158098905
3682	ITGAE	integrin subunit alpha E	6657
0.183593951234968	4.29172275605929	0.629302279264984
	0.42761135796469	0.827166158098905
5533	PPP3CC	protein phosphatase 3 catalytic subunit gamma
2899	-0.133693096947268	4.65963421626636
0.629240701224566	0.427633966495152	0.827166158098905
90333	ZNF468	zinc finger protein 468 5281	-0.146820947132908
	5.75848244559311	0.629149639809091
0.42766740322355	0.827166158098905
100528007	BORCS7-ASMT	BORCS7-ASMT readthrough (NMD
candidate)	2670	0.21392396603216	2.9532431930063

0.628902584510692	0.427758139012438	0.82722742691526
88455	ANKRD13A	ankyrin repeat domain 13A	7350
0.112252358127071	7.60827769960875	0.628454462225591
	0.427922794610004	0.827381135265443
168451	THAP5	THAP domain containing 5
4961	-0.153149011269319	4.13301781307477
0.628289345735793	0.427983488211691	0.827381135265443
4215	MAP3K3	mitogen-activated protein kinase kinase kinase 3
	6107	0.133374079894535	5.23715412110349
0.628053066453344	0.428070362459056	0.827381135265443
4686	NCBP1	nuclear cap binding protein subunit 1
5548	-0.118591955892103	5.73377225719301
0.627536611300979	0.428260343344244	0.827381135265443
115294	PCMTD1	protein-L-isoaspartate (D-aspartate) O-methyltransferase domain containing 1	7323	-0.128707865341524
	6.15326940258151	0.627337671607052
0.428333558359555	0.827381135265443
199857	ALG14	ALG14 UDP-N-acetylglucosaminyltransferase subunit
	9633	0.170435382443066	3.48692512348096
0.626920340522944	0.428487208524605	0.827381135265443
6342	SCP2	sterol carrier protein 2	8305
0.104845510651355	6.51662245442347	0.626817578409302
	0.428525055558317	0.827381135265443
357	SHROOM2	shroom family member 2	8191	-0.171024621509652
	5.00358468403929	0.626703207493165
0.4285671840203	0.827381135265443
56478	EIF4ENIF1	eukaryotic translation initiation factor 4E nuclear import factor 1	5585	-0.11795903286316
4.88916918501116	0.626638555385368	0.428591001354125
	0.827381135265443
144097	SPINDOC	spindlin interactor and repressor of chromatin binding	2807	0.148984266593508	4.77861165030436
0.626618887636276	0.42859824719644	0.827381135265443
64779	MTHFSD	methenyltetrahydrofolate synthetase domain containing	5544	-0.140243333985441	3.6089873402196
0.626386990967385	0.428683694750936	0.827381135265443
311	ANXA11	annexin A11	8513	-0.150168764659931
8.70103975609551	0.626269217078459	0.428727101045466
	0.827381135265443
219927	MRPL21	mitochondrial ribosomal protein L21	2825
0.164252889107524	4.00548929639912	0.626239897652061
	0.428737907933116	0.827381135265443
51409	HEMK1	HemK methyltransferase family member 1	19425
0.126207882694201	4.06594950171233	0.625869067114064
	0.428874628388487	0.827381135265443
6097	RORC	RAR related orphan receptor C
6109	-0.492936279896025	-0.230553520524901
0.625785196294101	0.428905559613939	0.827381135265443
4692	NDN	necdin, MAGE family member	1931
0.204777070509628	4.40463737516775	0.625759608898832
	0.428914996814615	0.827381135265443
7320	UBE2B	ubiquitin conjugating enzyme E2 B
3291	-0.113484893959889	6.22760526554468
0.625723035458648	0.428928486457713	0.827381135265443

643836	ZFP62	ZFP62 zinc finger protein
4197	-0.170391368115745	4.08210528295918
0.625514440830859	0.429005436157637	0.827381135265443
5669	PSG1	pregnancy specific beta-1-glycoprotein 1	4961
	0.684017074846809	3.14156442926068
0.625357243476138	0.429063439397203	0.827381135265443
6623	SNCG	synuclein gamma 1247	-0.200930980077702
4.05438114312054	0.624994212914297	0.42919743699782
	0.827381135265443
55920	RCC2	regulator of chromosome condensation 2
4126	-0.100985770603334	7.47865434031304
0.624758414999405	0.429284505852201	0.827381135265443
11245	GPR176	G protein-coupled receptor 176	5128
0.237437812065741	4.75962787748333	0.624611961783899
	0.429338597437392	0.827381135265443
5395	PMS2	PMS1 homolog 2, mismatch repair system component
	2855	0.142509854805302	4.16394081392009
0.624553858447772	0.429360060398121	0.827381135265443
4242	MFNG	MFNG O-fucosylpeptide 3-beta-N-
acetylglucosaminyltransferase	3042	0.257571689310112
2.04605925614002	0.624471960619635	0.429390315629989
	0.827381135265443
23251	MINAR1	membrane integral NOTCH2 associated receptor 1
6800	-0.237583217312601	1.8275052080215 0.624426419190207
	0.429407141237418	0.827381135265443
8613	PLPP3	phospholipid phosphatase 3	5270
0.204683030035513	3.77612126758223	0.624297073305627
	0.429454934437239	0.827381135265443
50808	AK3	adenylate kinase 3	4483	0.111294887930432
	7.00491439344682	0.624253419156517
0.429471066425185	0.827381135265443
23181	DIP2A	disco interacting protein 2 homolog A	13557
0.0938741448739816	6.08159433884046	0.62398689448861
	0.429569578032707	0.827381135265443
6041	RNASEL	ribonuclease L	6125	0.166934101430785
4.13520282903524	0.62395219632962	0.429582405518334
	0.827381135265443
5905	RANGAP1 Ran GTPase activating protein 1
5348	-0.130936716084168	7.9605726324831 0.623878384105353
	0.429609694922936	0.827381135265443
79772	MCTP1	multiple C2 and transmembrane domain containing 1
	12089	0.255229189452194	1.55626318724133
0.622637236463868	0.4300689574558 0.827669588481351
84975	MFSD5	major facilitator superfamily domain containing 5
	2246	0.128310063981222	5.08633022927218
0.622528515520424	0.430109222904164	0.827669588481351
1192	CLIC1	chloride intracellular channel 1
1942	-0.125509085651977	8.2913867552842 0.622273127005315
	0.430203830004716	0.827669588481351
6813	STXBP2	syntaxin binding protein 2
6417	-0.0876673969088953	6.93705330703631
0.622224519665103	0.430221839861775	0.827669588481351
51428	DDX41	DEAD-box helicase 41	4259	0.096784358808652
	5.74065560442516	0.622095089641221

0.430269801481156	0.827669588481351
642587	MIR205HG	MIR205 host gene	3518
0.167522394910315	6.43184497127862	0.622076244426779
	0.430276785443124	0.827669588481351
64393	ZMAT3	zinc finger matrin-type 3	9646
0.129714820447428	7.0731448454173 0.621809966405037
0.430375485370517	0.827669588481351
1012	CDH13	cadherin 13	11217	0.286779527403947
5.68820292769663	0.621799032149426	0.430379539048316
	0.827669588481351
7766	ZNF223	zinc finger protein 223 3329	-0.18225394463973
	2.86172434529797	0.621714475935764
0.430410888689483	0.827669588481351
6142	RPL18A	ribosomal protein L18a	3527	0.166174106637893
	5.3809250937123 0.621682448436488	0.430422763948571
	0.827669588481351
56731	SLC2A4RG	SLC2A4 regulator	2244
0.177716238549186	3.98188659719585	0.621585926870456
	0.430458555525367	0.827669588481351
221806	VWDE	von Willebrand factor D and EGF domains 6490
0.402747440037443	2.2166720445199 0.621558981609763
0.430468548017894	0.827669588481351
79632	FAM184A family with sequence similarity 184 member A	7791
	0.349118010792705	0.246799676165254
0.620906317307973	0.430710692098038	0.827904844263762
57684	ZBTB26	zinc finger and BTB domain containing 26
4645	-0.188354714666402	2.98589848733989
0.620890796663037	0.430716452901989	0.827904844263762
653145	ANXA8	annexin A8	4854	0.130727212766614
9.49085767030331	0.620751331744486	0.430768223387899
	0.827904844263762
9913	SUPT7L	SPT7 like, STAGA complex gamma subunit	5309
0.107415777684781	6.00380174108735	0.620157644974825
	0.43098871014511	0.828214962683847
169611	OLFML2A olfactomedin like 2A	6746	0.279013411805681
	5.35323030249072	0.61976642359913
0.431134097641345	0.828372267534607
5525	PPP2R5A protein phosphatase 2 regulatory subunit B'alpha
	3825	-0.103509421180905	5.18764457509325
0.619542286643883	0.431217425946729	0.828372267534607
91746	YTHDC1	YTH domain containing 1 7032	-0.113691802124083
	6.59855439008444	0.619460095777839
0.43124798849592	0.828372267534607
92312	MEX3A	mex-3 RNA binding family member A	6124
0.148542507682125	6.18361036206292	0.618484752141653
	0.431610919414395	0.828851626663126
4171	MCM2	minichromosome maintenance complex component 2	4902
	0.177015414437042	3.83809552674413
0.618471403998707	0.431615889546715	0.828851626663126
54431	DNAJC10 DnaJ heat shock protein family (Hsp40) member C10
	21826	-0.163067955994014	7.70664021015385
0.617965067642899	0.431804486165136	0.829036827434639
100616361	MIR4775 microRNA 4775	75	0.235276347757212
	0.0384651648014924	0.61789470142887

0.431830705573151	0.829036827434639
102465248	MIR6501 microRNA 6501	67	-0.262165360298258
	1.59653932999091	0.617632487451438
0.43192843134115	0.829100627076285
376497	SLC27A1 solute carrier family 27 member 1	8249
0.172829155800047	3.48249694399632	0.616686743667879
	0.432281184045978	0.829100627076285
84792	FAM220A family with sequence similarity 220 member A	2395
	0.114766969510509	4.31334148110149
0.616671213508866	0.432286980284613	0.829100627076285
129607	CMPK2	cytidine/uridine monophosphate kinase 2
4302	-0.273905194051231	2.13302951854079
0.616643212018712	0.432297431430004	0.829100627076285
113510	HELQ	helicase, POLQ like	3713	-0.162688646749732
	3.47234855130973	0.616620158184491
0.43230603622334	0.829100627076285
64400	AKTIP	AKT interacting protein 3995	-0.117460533802378
	5.68177629871752	0.616463107276758
0.432364662052689	0.829100627076285
9637	FEZ2	fasciculation and elongation protein zeta 2
6828	-0.111739458503596	5.78560397787573
0.616251714288367	0.432443592401909	0.829100627076285
124044	SPATA2L spermatogenesis associated 2 like
2482	-0.192914468637424	3.35901474882341
0.616205352680179	0.432460905920218	0.829100627076285
124446	TMEM219 transmembrane protein 219	2428
0.108685297884344	5.91750777076785	0.616192018854052
	0.432465885567995	0.829100627076285
56850	GRIPAP1 GRIP1 associated protein 1
6841	-0.0889717930927054	6.04973024233688
0.615870462468766	0.43258600032215	0.829100627076285
90459	ERI1	exoribonuclease 1	6234	-0.126625174477439
	4.13543675220295	0.615869178672913
0.432586479975123	0.829100627076285
10111	RAD50	RAD50 double strand break repair protein	9771
	0.12892430967344	6.32629095687132
0.615845429110337	0.432595353453971	0.829100627076285
2787	GNG5	G protein subunit gamma 5
1112	-0.115146014405312	5.76084292857233
0.615624475773927	0.432677920848938	0.829100627076285
55020	TTC38	tetratricopeptide repeat domain 38
3578	-0.171616099862194	3.85548906625181
0.615509219387963	0.432721000166063	0.829100627076285
53947	A4GALT	alpha 1,4-galactosyltransferase (P blood group)
2943	-0.177161376047893	5.57126363017773
0.615062641256692	0.432887978988717	0.829100627076285
65987	KCTD14	potassium channel tetramerization domain containing
14	1917	0.316005459745703	1.12647177835255
0.615039670790283	0.432896570461096	0.829100627076285
91893	FDXACB1 ferredoxin-fold anticodon binding domain containing
1	3765	0.211890350848697	1.88304555896981
0.615030084225507	0.432900156129117	0.829100627076285
83746	L3MBTL2 L3MBTL histone methyl-lysine binding protein 2
6451	-0.202384017947928	4.98499580930644

0.614807838237823	0.432983295569986	0.829100627076285
22979	EFR3B	EFR3 homolog B	10591	-0.235616484240119
1.90654312500309	0.614758484082261	0.433001761634245
	0.829100627076285
1203	CLN5	CLN5 intracellular trafficking protein	4226
0.127184137467192	5.52668043070628	0.614469327823997
	0.433109974715097	0.829100627076285
9665	MARF1	meiosis regulator and mRNA stability factor 1
10919	-0.129472159608222	6.41598821530805
0.613930702800332	0.433311657977509	0.829100627076285
199704	ZNF585A	zinc finger protein 585A
11329	-0.171515512123524	3.68325478938284
0.613766989737094	0.433372987148849	0.829100627076285
57082	KNL1	kinetochore scaffold 1	10250	0.256799306185445
	1.11991388269828	0.613619740678839
0.433428159953047	0.829100627076285
55974	SLC50A1	solute carrier family 50 member 1	2340
0.127769152385342	4.84879213935541	0.613607040926016
	0.433432918928504	0.829100627076285
23334	SZT2	SZT2 subunit of KICSTOR complex
17401	-0.173665071822535	5.64840573448512
0.613447747617904	0.433492617623603	0.829100627076285
23613	ZMYND8	zinc finger MYND-type containing 8
7625	-0.109575990025874	5.62892737493361
0.613363630125766	0.433524147562552	0.829100627076285
123811	FOPNL	FGFR1OP N-terminal like	2391	-0.108458322910335
	4.92468566976234	0.613359957777051
0.433525524156316	0.829100627076285
26155	NOC2L	NOC2 like nucleolar associated transcriptional
repressor	5539	-0.113065333766559	6.09211881597416
	0.6132633407491	0.433561743802284	0.829100627076285
27257	LSM1	LSM1 homolog, mRNA degradation associated	1690
	0.110016185964467	5.42419082498303
0.613213307154517	0.43358050212992	0.829100627076285
57130	ATP13A1	ATPase 13A1	8376	0.0979320636222771
6.41897773469884	0.61291307550475	0.433693089310126
	0.829202716621204
149840	SHLD1	shieldin complex subunit 1	1702
0.212057767061518	1.62230757479038	0.612444121782079
	0.433869036462881	0.829352256741693
55761	TTC17	tetratricopeptide repeat domain 17
11284	-0.122129764415976	6.36463377631598
0.6122324333545	0.433948495612646	0.829352256741693
390010	NKX1-2	NK1 homeobox 2	3364	0.283206874564971
6.6296622609345	0.612231271094991	0.433948931938307
0.829352256741693
85414	SLC45A3	solute carrier family 45 member 3	3702
0.158698264076632	3.46119726921201	0.611704946187324
	0.434146589039868	0.829616817555448
11018	TMED1	transmembrane p24 trafficking protein 1	1513
0.142194763032894	4.70238640076564	0.611348126967965
	0.434280667606253	0.829759829649542
51004	COQ6	coenzyme Q6, monooxygenase	6686
0.132416001112321	4.0766035735204	0.610906422729423

0.434446729967298	0.829963904344238
101926996	RUNDC3A-AS1	RUNDC3A antisense RNA 1 5403
0.222746740743858	1.87681113557599	0.60999304776476
	0.434790427793453	0.830507230615422
51649	MRPS23	mitochondrial ribosomal protein S23	6379
0.130561743018508	5.5222650100688 0.609707730056229
0.434897876159565	0.830599202531638
11237	RNF24	ring finger protein 24	7668	-0.124189397513811
	5.56241647524318	0.608615244231931
0.435309672030532	0.831272334850942
6422	SFRP1	secreted frizzled related protein 1	4761
0.651400469047676	0.812353991978324	0.607370284115447
	0.435779666161574	0.831768752657298
9275	BCL7B	BAF chromatin remodeling complex subunit BCL7B	3542
	0.105173320211372	5.3974115133193 0.607298338033077
	0.43580685072346	0.831768752657298
50617	ATP6V0A4	ATPase H+ transporting V0 subunit a4	4034
	0.292515645574642	0.739397848372309
0.607298173083509	0.435806913052019	0.831768752657298
84626	KRBA1	KRAB-A domain containing 1	5337
0.122441852722775	5.6928825087187 0.607297522262453
0.435807158974221	0.831768752657298
65117	RSRC2	arginine and serine rich coiled-coil 2
7604	-0.0988394936067167	6.37870654610531
0.606955017981651	0.435936608591074	0.831808880346801
6633	SNRPD2	small nuclear ribonucleoprotein D2 polypeptide	1498
	0.142061161697578	5.69794831619009
0.606892739803975	0.435960152957278	0.831808880346801
503538	A1BG-AS1	A1BG antisense RNA 1	2793
0.38957698816012	2.03907846802359	0.606735743655419
	0.436019514221933	0.831808880346801
378805	LINC-PINT	long intergenic non-protein coding RNA, p53 induced transcript	11134	-0.262654825400335
3.11649285247988	0.606613550260739	0.436065724879457
	0.831808880346801
56623	INPP5E	inositol polyphosphate-5-phosphatase E
3394	-0.127148672076997	4.55396882659832
0.606325201967689	0.436174801045823	0.831903654445648
84967	LSM10	LSM10, U7 small nuclear RNA associated	1496
0.145695696953711	3.72935099288469	0.606164383265137
	0.436235653518021	0.831906439595224
54838	WBP1L	WW domain binding protein 1 like	4281
0.0913353402426794	6.93851624599985	0.605959191374414
	0.436313315266738	0.831941275107704
64131	XYLT1	xylosyltransferase 1	11004	-0.282630094834728
	4.12906656503882	0.605774081914014
0.436383394266604	0.83196164468386
11168	PSIP1	PC4 and SFRS1 interacting protein 1	6768
0.188681857954415	2.90422976834955	0.605342450680614
	0.436546868633604	0.832072514512544
55257	MRGBP	MRG domain binding protein	2781
0.12552372105247	4.19625717122021	0.605262094635596
	0.4365773127062 0.832072514512544
55762	ZNF701	zinc finger protein 701 6294	-0.186061928314688

3.78425040903594 0.605150070175691
0.436619760238966 0.832072514512544
26207 PITPNC1 phosphatidylinositol transfer protein cytoplasmic 1
7819 -0.130679390737841 4.49017785622396
0.603976671527853 0.437064755661785 0.832659635343368
26173 INTS1 integrator complex subunit 1 9062
0.116044179012416 6.82448546628756 0.603817168510318
0.437125298533035 0.832659635343368
29920 PYCR2 pyrroline-5-carboxylate reductase 2 3602
0.130081977078926 4.95939345958048 0.603808438321849
0.437128612637868 0.832659635343368
9673 SLC25A44 solute carrier family 25 member 44 7293
-0.099792239456507 5.75382003002517
0.60371093609421 0.437165628493976 0.832659635343368
8608 RDH16 retinol dehydrogenase 16
3464 -0.288424895343849 0.953318343477388
0.60284425442058 0.437494867330393 0.833173433949261
150223 YDJC YdjC chitooligosaccharide deacetylase homolog 1754
0.144110568432945 3.8659741979046 0.602468012580758
0.437637913837007 0.833271519489189
55720 TSR1 TSR1 ribosome maturation factor
5545 -0.108150127611507 6.20614278113532
0.602395869172458 0.437665350816161 0.833271519489189
90075 ZNF30 zinc finger protein 30 2745 0.226424524152392
1.65763960219371 0.6017611400138 0.437906859244252
0.833461284317284
11001 SLC27A2 solute carrier family 27 member 2
2802 -0.499540075095896 2.35207580430886
0.60166933089906 0.437941808630937 0.833461284317284
4851 NOTCH1 notch receptor 1 9880 0.183503534398135
6.88170280535152 0.601504579907157
0.438004535838225 0.833461284317284
64761 PARP12 poly(ADP-ribose) polymerase family member 12
6285 -0.13462056939466 5.58358655303327
0.601326056238662 0.438072522379966 0.833461284317284
7079 TIMP4 TIMP metallopeptidase inhibitor 4 1650
0.368368628874775 0.909653505531824 0.601163418419873
0.438134473238396 0.833461284317284
83935 NA NA 1953 -0.238335201017137
2.64924272386 0.600960764310576 0.438211685600522
0.833461284317284
3434 IFIT1 interferon induced protein with tetratricopeptide
repeats 1 4613 -0.286582561446343 2.16957005449935
0.600947816417042 0.438216619529565
0.833461284317284
10589 DRAP1 DR1 associated protein 1
1459 -0.12066874786577 5.53306942386032
0.600835171632149 0.43825954758324 0.833461284317284
63940 GPSM3 G protein signaling modulator 3
1854 -0.174294602437451 3.36693238906314
0.600727592492611 0.438300551173064 0.833461284317284
3118 HLA-DQA2 major histocompatibility complex, class II,
DQ alpha 2 1524 0.807570889655329 0.134101227060657
0.600483595290811 0.438393572039634

0.833525012021061
51275	MAPKAPK5-AS1	MAPKAPK5 antisense RNA 1	2290
0.165352725591923	3.19238318715222	0.600177757174423
	0.438510211704594	0.833543575143971
3155	HMGCL	3-hydroxy-3-methylglutaryl-CoA lyase	5658
0.127563952501172	4.5721369420102 0.599762650812167
0.438668599877727	0.833543575143971
27284	SULT1B1 sulfotransferase family 1B member 1
7356	-0.448681160976256	1.1474091587843 0.599691192541471
	0.438695874374687	0.833543575143971
10267	RAMP1	receptor activity modifying protein 1	1234
0.262615454027376	1.79296551422798	0.59955686987548
	0.438747150258042	0.833543575143971
5189	PEX1	peroxisomal biogenesis factor 1
7849	-0.120371656575293	4.51632088731968
0.599526274005106	0.438758831107555	0.833543575143971
7384	UQCRC1	ubiquinol-cytochrome c reductase core protein 1 3079
	0.0887838351957113	7.05433508060556
0.599423949388857	0.438797899923187	0.833543575143971
9563	H6PD	hexose-6-phosphate dehydrogenase/glucose 1-
dehydrogenase	9321	0.115679497030065	5.54883924319724
	0.599232544747053	0.438870994925622
0.833543575143971
55526	DHTKD1	dehydrogenase E1 and transketolase domain containing
1	5281	-0.114656944893038	4.67286895813551
0.598801479406708	0.439035681695594	0.833543575143971
113419	TEX261	testis expressed 261	7767	0.0987855970499883
	6.6656453036808 0.598702344787535	0.439073569080194
	0.833543575143971
6418	SET	SET nuclear proto-oncogene
4332	-0.100897345235338	8.05705194794573
0.5984047795151 0.439187323047372	0.833543575143971
11335	CBX3	chromobox 3	4641	0.106918103401426
6.70675174572999	0.598185880406561	0.439271033188643
	0.833543575143971
283687	ST20-AS1	ST20 antisense RNA 1
4223	-0.198855110698018	3.19647688410046
0.598167913239379	0.439277905176671	0.833543575143971
26960	NBEA	neurobeachin	11724	-0.17167854409583
3.81462009215427	0.597935453169313	0.439366830161924
	0.833543575143971
1937	EEF1G	eukaryotic translation elongation factor 1 gamma
	5096	0.123284094243921	8.05336511449707
0.597736090894273	0.439443115963109	0.833543575143971
51559	NT5DC3	5'-nucleotidase domain containing 3	8722
0.265529159890107	3.88605862064158	0.59766101493327
	0.439471848982922	0.833543575143971
8394	PIP5K1A phosphatidylinositol-4-phosphate 5-kinase type 1
alpha	5405	-0.0934823495193484	7.26889689783268
0.597628520847405	0.439484285990997	0.833543575143971
8552	INE1	inactivation escape 1	945	-0.228593391418272
	1.4911129775493 0.597499859632471	0.439533535966371
	0.833543575143971
5530	PPP3CA	protein phosphatase 3 catalytic subunit alpha

5213	-0.0961981999194228	6.42480976144166
0.597424388585508	0.439562429323112	0.833543575143971
51747	LUC7L3	LUC7 like 3 pre-mRNA splicing factor
8151	-0.114256302641831	6.3252430280709 0.597331054808691
	0.439598165284594	0.833543575143971
220929	ZNF438	zinc finger protein 438 4267	0.216821620530303
	2.03845453811361	0.597253423534669
0.439627892414569	0.833543575143971
9682	KDM4A	lysine demethylase 4A	5172	0.0947232932715695
	6.73087391445878	0.597187794790809
0.439653025855905	0.833543575143971
51295	ECSIT	ECSIT signaling integrator	2370
0.150957097445128	3.53217793076458	0.596585736425227
	0.43988369547204	0.83386803660534
57677	ZFP14	ZFP14 zinc finger protein
7817	-0.242902466273993	2.85113495319144
0.596175224433529	0.440041083552076	0.834053512491735
3460	IFNGR2	interferon gamma receptor 2
2960	-0.177409742592844	6.61177513417836
0.595871926225129	0.440157422015113	0.834152376789259
10845	CLPX	caseinolytic mitochondrial matrix peptidase chaperone subunit	5461	0.0901109014740452
5.75863488835377	0.595617619812227	0.440255004736941
	0.834152376789259
22832	CEP162	centrosomal protein 162 5474	0.155530957599989
	3.46305573765262	0.595573595136497
0.440271901306703	0.834152376789259
51386	EIF3L	eukaryotic translation initiation factor 3 subunit L
	4296	0.102167924838635	7.10664090711899
0.595243110217485	0.44039877248574	0.834279903463317
284018	C17orf58	chromosome 17 open reading frame 58
1752	-0.19773966972218	2.23378470820315
0.594568896260355	0.440657773495959	0.834657665092346
10654	PMVK	phosphomevalonate kinase	1280
0.142710484502529	3.99435757278823	0.594080727465527
	0.44084545097897	0.834805070180325
79899	PRR5L	proline rich 5 like	6616	0.182161736621877
	5.80084380770519	0.594038829790946
0.440861564352278	0.834805070180325
728661	SLC35E2B	solute carrier family 35 member E2B
8395	-0.142672176591219	7.18880822747259
0.593762020965208	0.440968044671961	0.834805070180325
84128	WDR75	WD repeat domain 75	5066	-0.0912003688765082
	5.46483665541707	0.593746561634461
0.440973992594005	0.834805070180325
26747	NUFIP1	nuclear FMR1 interacting protein 1	3729
0.165344917730103	3.06145910871251	0.593369446033135
	0.441119124703585	0.834896420081265
10253	SPRY2	sprouty RTK signaling antagonist 2	3038
0.177856397209833	4.56445309335128	0.593311430954396
	0.44114145820103	0.834896420081265
6731	SRP72	signal recognition particle 72
4697	-0.103332809022317	6.72352501453059
0.592748742337132	0.441358161240762	0.83500652117768

81545	FBXO38	F-box protein 38	8722	-0.106613886623202
	5.9074467920064 0.592523974319931	0.441444769854464
	0.83500652117768
7879	RAB7A	RAB7A, member RAS oncogene family
2717	-0.0991812820823952	8.43663156910681
0.592230718376129	0.441557807881002	0.83500652117768
79993	ELOVL7	ELOVL fatty acid elongase 7	4576
0.150494309919084	4.75340209677055	0.592158843337472
	0.441585519532384	0.83500652117768
94104	PAXBP1	PAX3 and PAX7 binding protein 1
7212	-0.122804974534158	4.99499811778353
0.591678159869593	0.441770917444133	0.83500652117768
64969	MRPS5	mitochondrial ribosomal protein S5	2539
0.121073834844189	4.9262564512831 0.591597798723342
0.441801924148721	0.83500652117768
26058	GIGYF2	GRB10 interacting GYF protein 2
13069	-0.107685490915463	6.57040297433234
0.591527225750099	0.441829156929565	0.83500652117768
9123	SLC16A3 solute carrier family 16 member 3	6175
0.155861026690253	7.58375585618564	0.591265540193795
	0.441930159028137	0.83500652117768
81491	GPR63	G protein-coupled receptor 63	5952
0.252035232100644	-0.087863797215765	0.591211900214312
	0.441950866704942	0.83500652117768
9857	CEP350	centrosomal protein 350 14301	-0.193589522382691
	6.4897518473113 0.591171433621594	0.441966489794165
	0.83500652117768
241	ALOX5AP arachidonate 5-lipoxygenase activating protein	1715
	0.658176850498268	3.2352803131612 0.591120529430455
	0.4419861437745 0.83500652117768
25886	POC1A	POC1 centriolar protein A	2247
0.193724101091022	1.88717245423319	0.590918310007723
	0.442064233486764	0.83500652117768
100132247	NPIPB5	nuclear pore complex interacting protein
family member B5	9035	-0.1931415017922
4.55589314081058	0.59087261278065	0.442081883021746
	0.83500652117768
10286	BCAS2	BCAS2 pre-mRNA processing factor	1299
0.111950945316784	5.10382322617577	0.590871402061278
	0.442082350649871	0.83500652117768
123228	SENP8	SUMO peptidase family member, NEDD8 specific	5007
	0.211435748474987	1.72423162188389
0.590841666331242	0.442093836014398	0.83500652117768
51614	ERGIC3	ERGIC and golgi 3	6012	0.095000014716793
	7.1395095467668 0.59038616039955	0.442269831730276
	0.835226308891192
83856	FSD1L	fibronectin type III and SPRY domain containing 1
like	9877	0.266753212821539	2.63018762998356
0.589735415907423	0.442521449982889	0.835588831209265
4643	MYO1E	myosin IE	9106	-0.178945360454042
7.7825096152664 0.58934570879245	0.442672240648214
0.835760894292968
4174	MCM5	minichromosome maintenance complex component 5	6807
	0.165935807092367	3.81486481840887

0.588547330350529	0.44298140745707	0.83611982225915
64860	ARMCX5	armadillo repeat containing X-linked 5
4849	-0.148372102774527	3.97304689925236
0.588455795187332	0.443016875132508	0.83611982225915
6509	SLC1A4	solute carrier family 1 member 4
6939	-0.366997494034399	5.27723491352278
0.588292158673568	0.443080291267838	0.83611982225915
126328	NDUFA11	NADH:ubiquinone oxidoreductase subunit A11	4962
	0.123095788241427	5.924601271356	0.588188172706168
	0.443120597549581	0.83611982225915
378938	MALAT1	metastasis associated lung adenocarcinoma transcript
1	8708	-0.359873146580885	8.75716754351775
0.587908697938474	0.443228953565824	0.83611982225915
7528	YY1	YY1 transcription factor
6988	-0.0879189319499961	6.76663938303717
0.587746008521947	0.443292049210997	0.83611982225915
51575	ESF1	ESF1 nucleolar pre-rRNA processing protein homolog
	3216	-0.126818475450084	4.12696437161015
0.58767846782829	0.443318247511815	0.83611982225915
6133	RPL9	ribosomal protein L9	3259	0.177258195794384
	5.47344412814799	0.587622662074068
0.443339895760599	0.83611982225915
388566	ZNF470	zinc finger protein 470	8036	-0.19521475551211
	3.34244915089311	0.58744589974561
0.443408476431234	0.836129574525463
56006	SMG9	SMG9 nonsense mediated mRNA decay factor	7927
	0.119462154079085	5.47679743431465
0.587301652035201	0.443464454140763	0.836129574525463
9906	SLC35E2A	solute carrier family 35 member E2A
3264	-0.162934698050002	3.3457248980029	0.58656720899191
	0.443749636221776	0.83655466413976
5134	PDCD2	programmed cell death 2	6476	0.107137538267206
	5.52787643586617	0.58638579324996
0.443820123051078	0.836574951362731
8904	CPNE1	copine 1	3897	0.085273619105607
6.96515877369119	0.58545488362957	0.44418208870578
	0.836645870192171
23640	HSPBP1	HSPA (Hsp70) binding protein 1	2159
0.127137026042812	4.42812972377543	0.585399773395739
	0.444203531525514	0.836645870192171
23142	DCUN1D4	defective in cullin neddylation 1 domain containing
4	7625	0.127413836512183	5.13522280939485
0.585383128971046	0.444210008012959	0.836645870192171
8867	SYNJ1	synaptojanin 1	8070	-0.131915754709247
4.38313913575551	0.585036256053126	0.444345012449716
	0.836645870192171
51115	RMDN1	regulator of microtubule dynamics 1	5646
0.130358796124055	4.78683574901893	0.585031092230114
	0.444347022711117	0.836645870192171
79891	ZNF671	zinc finger protein 671	2914	0.271866558766179
	1.25023571994268	0.584797588656726
0.444437939676276	0.836645870192171
147807	ZNF524	zinc finger protein 524	1162	-0.174776056776972
	2.91855027804802	0.584674035808762

0.444486057858497 0.836645870192171
4155 MBP myelin basic protein 18615 0.184194853330972
6.48027270556541 0.584388552734916
0.444597271277812 0.836645870192171
9841 ZBTB24 zinc finger and BTB domain containing 24 5519
0.178742856309665 3.35936474741637
0.584225510102717 0.444660805838229 0.836645870192171
93343 MVB12A multivesicular body subunit 12A 4053
0.126709078085903 4.24541906871995 0.58364303300824
0.444887900664068 0.836645870192171
64689 GORASP1 golgi reassembly stacking protein 1
6256 -0.0993678629356079 6.05865499595605
0.583459425808549 0.444959522218294 0.836645870192171
65003 MRPL11 mitochondrial ribosomal protein L11 2539
0.176984937535072 3.38013842570405 0.583458203652424
0.444959999017122 0.836645870192171
9988 DMTF1 cyclin D binding myb like transcription factor 1
8698 -0.136744123417551 5.68107246740961
0.583294968995894 0.445023688734776 0.836645870192171
79852 EPHX3 epoxide hydrolase 3 2034 -0.315381768487363
3.77015285036622 0.583032290321384
0.445126208427944 0.836645870192171
54707 GPN2 GPN-loop GTPase 2 4979 0.103850718205951
5.3932524773612 0.58301972943793 0.44513111167524
0.836645870192171
6850 SYK spleen associated tyrosine kinase 5210
0.134559572275867 5.4240901905665 0.582985121608523
0.445144621567271 0.836645870192171
84527 ZNF559 zinc finger protein 559 7013 0.168625134567504
3.26870516017495 0.582976069147509
0.445148155490018 0.836645870192171
5471 PPAT phosphoribosyl pyrophosphate amidotransferase 5439
0.175885978927108 2.90637672730776
0.582882308009948 0.445184760756111 0.836645870192171
6709 SPTAN1 spectrin alpha, non-erythrocytic 1
15023 -0.116331077550081 9.50439718377143
0.582845630001771 0.445199081477697 0.836645870192171
9364 RAB28 RAB28, member RAS oncogene family
2221 -0.140629942169888 3.56566186470756
0.582501484637609 0.445333485846774 0.836645870192171
55892 MYNN myoneurin 5730 -0.121955936006765
4.61128178779791 0.582458574150099 0.445350248743264
0.836645870192171
6311 ATXN2 ataxin 2 13933 -0.0953742184778503
6.06524615998977 0.582444284516148 0.445355831177187
0.836645870192171
81566 CSRNP2 cysteine and serine rich nuclear protein 2
5230 -0.0960092238972341 6.82691406440616
0.582288733691939 0.445416606168785 0.836645870192171
255031 LINC00957 long intergenic non-protein coding RNA 957
2703 0.242900209995713 0.388081679048056
0.582230006429418 0.445439554735411 0.836645870192171
6794 STK11 serine/threonine kinase 11 8639
0.116897683765206 5.61000995189383 0.582089290974459

0.445494548880455	0.836645870192171
285175	UNC80	unc-80 homolog, NALCN channel complex subunit
19258	-0.414948151089615	0.822276118205238
0.582040278362825	0.44551370636509	0.836645870192171
55148	UBR7	ubiquitin protein ligase E3 component n-recognin 7
(putative)	3976	0.112912751229929	5.03182970911198
0.581953001393934	0.445547823340145
0.836645870192171
57628	DPP10	dipeptidyl peptidase like 10
8445	-0.668347773502123	0.371514328448876
0.581493409973133	0.445727546567254	0.836645870192171
2720	GLB1	galactosidase beta 1	4251	0.132907685019615
6.77608465602305	0.581269653567745
0.445815087170275	0.836645870192171
375593	TRIM73	tripartite motif containing 73	1799
0.267273155264472	-0.132297489399104	0.581205257042837
0.445840286070775	0.836645870192171
55055	ZWILCH	zwilch kinetochore protein
4849	-0.153180692309075	3.93348232603141
0.581122148394169	0.445872810445151	0.836645870192171
81931	ZNF93	zinc finger protein 93	4888	-0.194707132570981
2.91581662169333	0.58109412019999
0.445883780042284	0.836645870192171
10963	STIP1	stress induced phosphoprotein 1
4657	-0.101491739525181	7.01152295422095
0.581088006077081	0.445886173026376	0.836645870192171
9640	ZNF592	zinc finger protein 592	8134	-0.128736517834098
6.06542443744725	0.581081841500653
0.445888585777437	0.836645870192171
1945	EFNA4	ephrin A4	1263	0.12641338795392
4.38019641574503	0.580371642713779	0.446166685720435
0.83681182827174
23112	TNRC6B	trinucleotide repeat containing adaptor 6B
19998	-0.162540767911109	5.67201162483941
0.580208489735869	0.44623061121808	0.83681182827174
51441	YTHDF2	YTH N6-methyladenosine RNA binding protein 2
3859	-0.0869588431839957	7.08816032163458
0.580152229443076	0.446252658039193	0.83681182827174
29934	SNX12	sorting nexin 12	2595	0.0911607625157236
6.36354101916465	0.580104424583858
0.446271392741986	0.83681182827174
285629	LINC02159	long intergenic non-protein coding RNA 2159
2323	0.249941820228525	0.857315363766147
0.579860604832248	0.446366964571721	0.83681182827174
440307	TTLL13P	tubulin tyrosine ligase like 13, pseudogene	3838
0.101146838692795	6.94127751488209
0.579851729649704	0.446370444041374	0.83681182827174
91316	GUSBP11	GUSB pseudogene 11	6797	0.189132899745952
4.42268804898583	0.579622309028512
0.446460401817274	0.83681182827174
79102	RNF26	ring finger protein 26	2787	0.111036328262711
5.16331684449518	0.57957721770701
0.446478085814584	0.83681182827174
83539	CHST9	carbohydrate sulfotransferase 9

11522 -0.491866384175687 3.40446391093808
0.579410856914944 0.446543338896717 0.83681182827174
64112 MOAP1 modulator of apoptosis 1 2537
0.110259901989732 5.21099552654625 0.579331536039028
0.446574456795264 0.83681182827174
55643 BTBD2 BTB domain containing 2 5505 0.136212884565165
5.36732484370454 0.57868380154494
0.446828691620905 0.837130967120856
7770 ZNF227 zinc finger protein 227 4608 0.169321674456823
3.49643247145526 0.578411937860594
0.446935464645546 0.837130967120856
55343 SLC35C1 solute carrier family 35 member C1 4505
0.110625921531952 6.42520784259449 0.578264739618639
0.446993292505961 0.837130967120856
9295 SRSF11 serine and arginine rich splicing factor 11
8259 -0.135162026113669 7.11364827913274
0.578187358494233 0.447023696881263 0.837130967120856
7499 XG Xg glycoprotein (Xg blood group)
2517 -0.448245018496265 1.31379715025323
0.577570349602894 0.447266245223796 0.837130967120856
6210 RPS15A ribosomal protein S15a 4806 0.186580964676687
6.52129685207056 0.577558275974912
0.447270993447629 0.837130967120856
65979 PHACTR4 phosphatase and actin regulator 4
7322 -0.113093279680077 6.86984341050876
0.577542028936548 0.447277383081862 0.837130967120856
79094 CHAC1 ChaC glutathione specific gamma-glutamylcyclotransferase 1 2058 -0.21121922404541
3.91375151479478 0.577479744311475 0.447301879688994
0.837130967120856
100528018 ARL2-SNX15 ARL2-SNX15 readthrough (NMD candidate) 2392 0.1014489778283 5.2323466903693
0.577375315070003 0.447342956493607 0.837130967120856
113675 SDSL serine dehydratase like 2216 0.272224415195416
1.31982549406692 0.577325779958746
0.447362442971058 0.837130967120856
255043 TMEM86B transmembrane protein 86B
1999 -0.228390593021668 0.894359264269765
0.577224762474975 0.447402186040318 0.837130967120856
10651 MTX2 metaxin 2 1717 0.136980761303789
4.84375594066143 0.576704038737716 0.447607140192612
0.837331565012347
80148 SLC66A2 solute carrier family 66 member 2
5225 -0.141783729527009 5.05555066899985
0.576648638536611 0.447628954009599 0.837331565012347
23060 ZNF609 zinc finger protein 609 9227 -0.116106590377556
7.00116159903362 0.576376156730849
0.447736267711617 0.837420470267943
9296 ATP6V1F ATPase H+ transporting V1 subunit F 789
0.109841899187678 6.53924128760633 0.575618352435427
0.448034930243336 0.837775578723216
55198 APPL2 adaptor protein, phosphotyrosine interacting with PH domain and leucine zipper 2 5124 -0.109176972251998
5.22284840519232 0.575590906689662 0.44804575285326

0.837775578723216
10036 CHAF1A chromatin assembly factor 1 subunit A 4299
0.154137527452167 3.31117462322257 0.575411983412422
0.44811631717182 0.837794092687531
6449 SGTA small glutamine rich tetratricopeptide repeat
containing alpha 3801 -0.0962912361177657
6.09404060938523 0.574792905820255 0.448360604341699
0.837794092687531
23597 ACOT9 acyl-CoA thioesterase 9 5046 0.0945217668295404
6.60257923738161 0.574773357499641
0.448368321456143 0.837794092687531
7266 DNAJC7 DnaJ heat shock protein family (Hsp40) member C7
3969 -0.101226499089789 6.33463418802108
0.574769891900038 0.448369689596704 0.837794092687531
55002 TMCO3 transmembrane and coiled-coil domains 3
10407 -0.0885608622728463 7.21735324050651
0.574724689788141 0.448387534965891 0.837794092687531
160851 DGKH diacylglycerol kinase eta
19637 -0.243555433890063 6.66980540565827
0.574532068105341 0.448463592587139 0.837794092687531
57569 ARHGAP20 Rho GTPase activating protein 20 7125
-0.349092620964403 1.74600491927339
0.574365632983872 0.448529326484067 0.837794092687531
822 CAPG capping actin protein, gelsolin like
2607 -0.132458778846977 7.52687898820433
0.574238564160453 0.44857952267443 0.837794092687531
57636 ARHGAP23 Rho GTPase activating protein 23 8133
-0.114624389981799 6.83123372643397
0.5742020033762 0.448593966956271 0.837794092687531
2495 FTH1 ferritin heavy chain 1 2030 0.214975896410754
10.2319366033801 0.573078463413594
0.449038203694629 0.838511947627073
28962 OSTM1 osteoclastogenesis associated transmembrane protein
1 4932 -0.130591959910312 6.37704919080498
0.572912428099087 0.449103910533123 0.838522857216404
7991 TUSC3 tumor suppressor candidate 3 5190
0.0955966553021541 6.87141211469678 0.572734354939554
0.449174397876245 0.838542688398316
284349 ZNF283 zinc finger protein 283 7229 -0.174497238371873
3.04946449193688 0.572272320667842
0.449357366970448 0.83875058990347
26262 TSPAN17 tetraspanin 17 5215 -0.0893709818612339
6.73004007274846 0.572150762614994 0.44940552418259
0.83875058990347
200081 TXLNA taxilin alpha 5103 0.0985773303803162
6.83135737137374 0.571863696569526 0.449519282025131
0.838825936788238
9726 ZNF646 zinc finger protein 646 7022 -0.134605329274062
5.22584053005092 0.571571606973393
0.449635076662769 0.838825936788238
5817 PVR PVR cell adhesion molecule
3519 -0.163320464502738 7.63312155592653
0.571400746397387 0.449702833390251 0.838825936788238
254013 ETFBKMT electron transfer flavoprotein subunit beta lysine

methyltransferase	6868	-0.222427046622478
0.656645312998246	0.571230329882659	0.44977042987878
	0.838825936788238
100874058	COX10-AS1	COX10 antisense RNA 1	2422
0.216782374019161	0.750239933455004	0.571120700847873
	0.449813923110496	0.838825936788238
23432	GPR161	G protein-coupled receptor 161	11224
0.118394485200841	7.07713627065615	0.570810817769704
	0.449936898798661	0.838825936788238
9220	TIAF1	TGFB1-induced anti-apoptotic factor 1
5337	-0.144959605697117	4.4758997802374 0.570771120275232
	0.449952656357307	0.838825936788238
90850	ZNF598	zinc finger protein 598 4701	-0.135184281283271
	5.48644660699983	0.570596471255348
0.450021991919003	0.838825936788238
51035	UBXN1	UBX domain protein 1	2532	0.0880864780599398
	5.84998629080972	0.570393991791359
0.45010239701522	0.838825936788238
7301	TYRO3	TYRO3 protein tyrosine kinase	10394
0.152878207912726	4.04819916603543	0.570191518650373
	0.450182822013993	0.838825936788238
346171	ZFP57	ZFP57 zinc finger protein
2149	-0.437542122506059	3.1717039287829 0.570069989485432
	0.450231105767714	0.838825936788238
56204	FAM214A family with sequence similarity 214 member A
9832	-0.155218592517854	6.39160318375628
0.569977531931862	0.450267844731056	0.838825936788238
2863	GPR39	G protein-coupled receptor 39	3387
0.139335193501025	3.93353055489365	0.569743455989885
	0.450360878141107	0.838825936788238
23233	EXOC6B	exocyst complex component 6B	5918
0.094490925760046	6.49930686600829	0.569688916736704
	0.450382559059538	0.838825936788238
25987	TSKU	tsukushi, small leucine rich proteoglycan
4131	-0.109848040749014	6.16603037302432
0.569671642455532	0.450389426421779	0.838825936788238
91057	CCDC34	coiled-coil domain containing 34	3733
0.224521337574057	1.44399653489897	0.569321551753055
	0.450528639638671	0.838825936788238
58493	INIP	INTS3 and NABP interacting protein
4423	-0.106427893768206	4.70381925181742
0.569274413337549	0.450547389313385	0.838825936788238
257106	ARHGAP30	Rho GTPase activating protein 30	5064
	-0.365286899750651	2.8944881793005 0.569191249508333
	0.450580471349768	0.838825936788238
114799	ESCO1	establishment of sister chromatid cohesion N-
acetyltransferase 1	5288	-0.123182394446275
4.78779896688194	0.569183050623415	0.450583733018826
	0.838825936788238
23768	FLRT2	fibronectin leucine rich transmembrane protein 2
	34465	0.229700249578585	5.36651239327505
0.568896330308032	0.450697818888702	0.838857431330287
3818	KLKB1	kallikrein B1	2953	0.328673725053315
0.695141926824838	0.568839523822732	0.450720427514224

0.838857431330287
51276 ZNF571 zinc finger protein 571 5652 -0.219916913494064
2.12627527118251 0.568462468889435
0.45087053792207 0.839025325451632
4718 NDUFC2 NADH:ubiquinone oxidoreductase subunit C2 2518
0.100297179804748 6.11615236354309
0.568270811603881 0.450946869082271 0.839055897347951
9524 TECR trans-2,3-enoyl-CoA reductase 2846
0.128079838141859 5.55580320442234 0.568077445613603
0.45102390122672 0.839087765238765
28956 LAMTOR2 late endosomal/lysosomal adaptor, MAPK and MTOR
activator 2 1438 0.118853215071353 5.18822053506873
0.567710975122559 0.451169950265438
0.839150262045668
283455 KSR2 kinase suppressor of ras 2
18294 -0.360126667293651 -0.0529151224977818
0.567392367741961 0.451296984542737 0.839150262045668
375757 SWI5 SWI5 homologous recombination repair protein 1649
0.154763226183101 3.43852180472369
0.567281878314422 0.451341051645474 0.839150262045668
3309 HSPA5 heat shock protein family A (Hsp70) member 5
3908 -0.184730483295635 8.85090234930884
0.567226674348035 0.451363071465166 0.839150262045668
9896 FIG4 FIG4 phosphoinositide 5-phosphatase 3592
0.121526214421261 4.41498041771383 0.567210304945752
0.451369601233418 0.839150262045668
146923 RUNDC1 RUN domain containing 1 3828 -0.103925457443246
5.39788972183134 0.567091616881693
0.451416950418656 0.839150262045668
11315 PARK7 Parkinsonism associated deglycase 2397
0.121920932719491 6.58387647981975 0.56690193379405
0.451492638343571 0.839179589341613
8970 H2BC11 H2B clustered histone 11
381 -0.259557799589275 0.974528939095287
0.566648851774725 0.451593654805817 0.83925598021296
100507062 PSMD6-AS2 PSMD6 antisense RNA 2
2456 -0.300960414716676 -0.0447342164714888
0.566129125059767 0.451801212060126 0.839492227104124
80823 BHLHB9 basic helix-loop-helix family member b9
5518 -0.197380972317782 3.12972409707641
0.56597981203774 0.451860869069061 0.839492227104124
7257 TSNAX translin associated factor X
5908 -0.0996334536580784 5.23729501579013
0.565827364886067 0.451921791013313 0.839492227104124
22906 TRAK1 trafficking kinesin protein 1
10992 -0.0901749987028354 7.33949033367863
0.565730489941398 0.451960511500835 0.839492227104124
3566 IL4R interleukin 4 receptor 5894 -0.124505499467377
7.35142574537105 0.56520681702797
0.452169911159098 0.839619616357999
8349 H2BC21 H2B clustered histone 21
2194 -0.143157584454818 4.72304536067919
0.565002320398631 0.452251723894844 0.839619616357999
546 ATRX ATRX chromatin remodeler

19060	-0.13435739407973	7.38384522834959
0.564826995366559	0.452321884433339	0.839619616357999
51809	GALNT7	polypeptide N-acetylgalactosaminyltransferase 7
6865	-0.122491950930082	6.62168703483074
0.56474703331736	0.452353888833232	0.839619616357999
147657	ZNF480	zinc finger protein 480	4836	-0.137269358495159
	4.41121563140539	0.564718106596645
0.45236546747803	0.839619616357999
55066	PDPR	pyruvate dehydrogenase phosphatase regulatory
subunit	9255	-0.227108823216525	5.82175735904312
0.564636148202162	0.452398275899081	0.839619616357999
10813	UTP14A	UTP14A small subunit processome component
4087	-0.147815209525703	4.33396327321625
0.56451021605568	0.452448694529177	0.839619616357999
79567	RIPOR1	RHO family interacting cell polarization regulator 1
	6729	0.120814644131999	7.1924403390712
0.564298621061084	0.452533429248229	0.839665617098946
4983	OPHN1	oligophrenin 1	8224	0.270813826086565
4.0463592791727	0.563912733996611	0.452688024470836
0.83972282695181
9470	EIF4E2	eukaryotic translation initiation factor 4E family
member 2	7177	-0.103457277583131	6.26618741787995
	0.563706966482424	0.452770493483776
0.83972282695181
50861	STMN3	stathmin 3	3032	-0.343541857075689
1.18082055758611	0.563692382614049	0.452776339406641
	0.83972282695181
1795	DOCK3	dedicator of cytokinesis 3
8755	-0.216964401253407	4.00021764115307
0.56362322409904	0.452804063108947	0.83972282695181
4801	NFYB	nuclear transcription factor Y subunit beta
4956	-0.114434602524181	4.56857989656025
0.5632988842511	0.452934117319207	0.839852817221357
541578	CXorf40B	chromosome X open reading frame 40B
2509	-0.114265006675121	4.32077916664453
0.56245141162464	0.453274214450154	0.840359353870694
10525	HYOU1	hypoxia up-regulated 1	7694	-0.142739179111174
	7.91617368069037	0.56223282372779
0.453362000364113	0.840359353870694
81691	REXO5	RNA exonuclease 5	4114	0.221354687463012
	0.325808937830655	0.562169888353364
0.453387280445551	0.840359353870694
57670	KIAA1549	KIAA1549	12427	0.247739870175991
	4.74321024297554	0.561741462109271
0.453559430826802	0.840465946634389
5691	PSMB3	proteasome 20S subunit beta 3	1179
0.113172573961031	5.92030268205941	0.561541446522696
	0.453639836267885	0.840465946634389
6856	SYPL1	synaptophysin like 1	2804	-0.104753323734268
	7.35794330314936	0.561434956230414
0.453682654048675	0.840465946634389
728806	NSFP1	N-ethylmaleimide-sensitive factor pseudogene 1
1017	-0.255580784655629	1.44162680083078
0.561308705392732	0.453733425395342	0.840465946634389

3181	HNRNPA2B1	heterogeneous nuclear ribonucleoprotein A2/
B1	7541	-0.092489042652462	8.92665688155537
0.561280410424814	0.453744805343703	0.840465946634389
92799	SHKBP1	SH3KBP1 binding protein 1
3936	-0.0858967438077727	6.11860164764569
0.561059471049059	0.453833680317826	0.840502998368124
5918	RARRES1	retinoic acid receptor responder 1
2236	-0.445745635633313	3.65405395263122
0.56089236773979	0.453900917365489	0.840502998368124
9420	CYP7B1	cytochrome P450 family 7 subfamily B member 1
2282	-0.261116632657487	2.35659134646979
0.560783262317582	0.453944826347887	0.840502998368124
6453	ITSN1	intersectin 1	21748	-0.083752209137379
6.38337544150275	0.560449580117591	0.45407915662067
	0.84064059566293
7776	ZNF236	zinc finger protein 236	8764	-0.136206244906839
	4.29811553426494	0.559844619232585
0.454322854607506	0.840980603209639
2794	GNL1	G protein nucleolar 1 (putative)	8646
0.112240475384859	6.23501866303654	0.559627308627483
	0.454410444554546	0.841031593138944
147184	TMEM99	transmembrane protein 99 (putative)
2362	-0.1782982743276	2.80108596303012
0.559378895657803	0.454510603220667	0.841042875453413
140710	SOGA1	suppressor of glucose, autophagy associated 1
15128	-0.171679492437825	6.10321467325183
0.559314359893193	0.454536629341211	0.841042875453413
3735	KARS1	lysyl-tRNA synthetase 1	4065	-0.106568176446971
	6.83975128316431	0.558983843491845
0.454669957409794	0.84113442038909
8603	FAM193A	family with sequence similarity 193 member A
6174	-0.108685359285453	5.21515513712228
0.558894009958678	0.454706206267308	0.84113442038909
23367	LARP1	La ribonucleoprotein 1, translational regulator
8284	-0.093466228372847	7.89299724965122
0.558447253669371	0.454886545013721	0.841177341795308
6619	SNAPC3	small nuclear RNA activating complex polypeptide 3
	6684	-0.0860791048331764	5.96489078187038
0.558419766869671	0.454897644074046	0.841177341795308
9675	TTI1	TELO2 interacting protein 1
4122	-0.110739619664902	4.2762925934718	0.558390230489062
	0.454909571221493	0.841177341795308
7005	TEAD3	TEA domain transcription factor 3
3046	-0.0950363228724829	6.5187246151401	0.557543877712888
	0.455251547711573	0.8413956140206
22849	CPEB3	cytoplasmic polyadenylation element binding protein
3	7928	-0.233659387695159	1.80290244337168
0.557511555511642	0.455264615802848	0.8413956140206
93429	LOC93429	uncharacterized LOC93429
2803	-0.312930393287091	0.837807017497434
0.557468841975024	0.455281886088506	0.8413956140206
9019	MPZL1	myelin protein zero like 1	6714
0.119753108889575	7.7180551090688	0.557387861553764
0.455314631582183	0.8413956140206

57134	MAN1C1	mannosidase alpha class 1C member 1
9026	-0.281992415286253	-0.188664332896398
0.557354899391948	0.455327961324777	0.8413956140206
7705	ZNF146	zinc finger protein 146 4146	-0.0966409916467712
	6.54515229986868	0.557094545664683
0.455433268719912	0.841479197330143
50717	DCAF8	DDB1 and CUL4 associated factor 8	8274
0.0960912108696831	6.64503790557584	0.556680606738496
	0.455600776884612	0.841556561468419
55898	UNC45A	unc-45 myosin chaperone A	7498
0.0998324795898748	6.69884298336606	0.556292768185958
	0.455757811036869	0.841556561468419
81573	ANKRD13C	ankyrin repeat domain 13C
4595	-0.117546057932234	5.1772981267336 0.556061508670858
	0.4558514875641 0.841556561468419
83667	SESN2	sestrin 2	3453	-0.113618420988488
5.85781773160649	0.555963454645806	0.455891215558913
	0.841556561468419
79886	CAAP1	caspase activity and apoptosis inhibitor 1
3014	-0.109797204798204	4.50269231979194
0.555784163392158	0.455963872080435	0.841556561468419
84460	ZMAT1	zinc finger matrin-type 1	7514
0.304817150334896	0.618297402052599	0.555767208874412
	0.455970743723274	0.841556561468419
5515	PPP2CA	protein phosphatase 2 catalytic subunit alpha
5832	-0.0932329498721528	7.92323332025809
0.555766567287947	0.455971003760592	0.841556561468419
196527	ANO6	anoctamin 6	8235	-0.108598179634475
8.46585921185028	0.555714000483903	0.455992310070072
	0.841556561468419
497661	C18orf32	chromosome 18 open reading frame 32
5740	-0.0906123183093344	6.09296075167667
0.555605075621751	0.456036464345585	0.841556561468419
79090	TRAPPC6A	trafficking protein particle complex 6A 933
	0.221798966335008	2.21769731674315
0.555502275343859	0.45607814210801	0.841556561468419
10939	AFG3L2	AFG3 like matrix AAA peptidase subunit 2
6624	-0.0860043381458422	5.73572014486693
0.555359509042135	0.456136033031215	0.841556561468419
27445	PCLO	piccolo presynaptic cytomatrix protein
22873	-0.255607992216884	3.48954352160548
0.555178444782241	0.456209470207648	0.841581199683725
57665	RDH14	retinol dehydrogenase 14	1587
0.112151902898622	4.39860567680353	0.554926209839636
	0.456311804284123	0.841588699370627
55125	CEP192	centrosomal protein 192 9901	-0.150363364765509
	5.18768247153075	0.554872246142079
0.456333702557287	0.841588699370627
57584	ARHGAP21	Rho GTPase activating protein 21
10473	-0.102009452026997	8.12328611706157
0.55421479311201	0.456600627656406	0.841970114742401
4205	MEF2A	myocyte enhancer factor 2A
6925	-0.0958937556369394	6.14100520323971
0.553728549976785	0.456798200097505	0.842059616331275

1649	DDIT3	DNA damage inducible transcript 3	1178
0.173584735351144	4.19377153007893	0.5534170857247
0.456924826448504	0.842059616331275
23250	ATP11A	ATPase phospholipid transporting 11A
11737	-0.156852687709121	6.5299818833978 0.553410914742592
	0.456927335831546	0.842059616331275
55572	FOXRED1 FAD dependent oxidoreductase domain containing 1
	5657	0.134866090940069	3.95208295245063
0.553386029135986	0.456937455595165	0.842059616331275
100288198	HMGN3-AS1	HMGN3 antisense RNA 1	2946
0.247126950806078	0.839035754638692	0.55333834748598
	0.456956846386327	0.842059616331275
200035	NUDT17	nudix hydrolase 17	1657	0.24403163487205
	1.05017624011519	0.553138707438285
0.457038048494618	0.842059616331275
4925	NUCB2	nucleobindin 2	4786	-0.195838387185138
4.39887037177299	0.553060208003746	0.457069983791439
	0.842059616331275
121441	NEDD1	NEDD1 gamma-tubulin ring complex targeting factor
	4463	-0.102390703712114	5.20557771910859
0.552301843072297	0.457378684700356	0.842403608912035
6626	SNRPA	small nuclear ribonucleoprotein polypeptide A	2361
	0.13104393320983	4.31992035883488
0.552054042069967	0.457479626184226	0.842403608912035
642273	FAM110C family with sequence similarity 110 member C
3968	-0.114545260375127	4.55611130265466
0.551921827089134	0.457533498207147	0.842403608912035
55571	CNOT11	CCR4-NOT transcription complex subunit 11
2786	-0.0938246424575064	5.82598385822252
0.551823019355055	0.457573764726037	0.842403608912035
5002	SLC22A18	solute carrier family 22 member 18	5316
	0.278217867189283	2.94468058654549
0.551733870487009	0.457610099826652	0.842403608912035
55701	ARHGEF40	Rho guanine nucleotide exchange factor 40
	8189	-0.229745003753377	6.4546768303504
0.551715586180194	0.457617552667357	0.842403608912035
255394	TCP11L2 t-complex 11 like 2	6558	0.231496208199067
	5.12455721197059	0.551134362316816
0.457854564428463	0.84263755191522
85406	DNAJC14 DnaJ heat shock protein family (Hsp40) member C14
	5751	0.0884180540086053	6.56463684006415
0.551108895378597	0.457864953782162	0.84263755191522
26521	TIMM8B	translocase of inner mitochondrial membrane 8
homolog B	1166	0.143355269005786	4.49256393559924
	0.550766637380558	0.458004615632422
0.842783848024611
9131	AIFM1	apoptosis inducing factor mitochondria associated 1
	3306	0.119326371015511	4.67463233508832
0.550527755916648	0.458102133520125	0.842852565902586
51196	PLCE1	phospholipase C epsilon 1	13329
0.217906685103441	3.73433862915596	0.550342477336741
	0.458177791849604	0.842881052066903
144100	PLEKHA7	pleckstrin homology domain containing A7
9545	-0.209483810110815	6.73190249975816

0.54954633297703	0.458503121484602	0.84318135315585
51377	UCHL5	ubiquitin C-terminal hydrolase L5
6661	-0.117043860115758	4.64979468442612
0.549528713480385	0.458510325494986	0.84318135315585
8799	PEX11B	peroxisomal biogenesis factor 11 beta	1838
0.112095382452022	4.6822929485872 0.549501084149567
0.458521622545021	0.84318135315585
400506	KNOP1	lysine rich nucleolar protein 1
7761	-0.123732055409297	5.10887085475531
0.549273352656492	0.458614753910044	0.843241908377262
222068	TMED4	transmembrane p24 trafficking protein 4
3769	-0.0941097289184133	6.70124107201759
0.548962057714434	0.458742107155934	0.843365362243492
10189	ALYREF	Aly/REF export factor	1574	-0.133979015692906
4.64027661303074	0.548770972630621
0.458820309295942	0.843389701969214
55911	APOBR	apolipoprotein B receptor	3792
0.478106042900328	1.14418164137277	0.548463211810281
0.458946305670288	0.843389701969214
83461	CDCA3	cell division cycle associated 3	5653
0.276189467958572	0.494428002841071	0.548187831631281
0.459059091916946	0.843389701969214
441172	FLJ46906	uncharacterized LOC441172
1220	-0.197729657787584	1.58056621122727
0.547994270690294	0.459138394069868	0.843389701969214
246269	AFG1L	AFG1 like ATPase	8296	-0.18512349504391
2.34469513475643	0.547796952542591
0.459219257873752	0.843389701969214
83479	DDX59	DEAD-box helicase 59	4470	-0.120315029879129
4.25127376336109	0.547657021555545
0.459276617259364	0.843389701969214
22845	DOLK	dolichol kinase 2090	0.130002892546013
4.62175789822476	0.547576511272853	0.459309624527382
0.843389701969214
106821730	BUB1B-PAK6	BUB1B-PAK6 readthrough
3745	-0.165237544558945	3.42908958415714
0.547514552426161	0.459335028712575	0.843389701969214
29940	DSE	dermatan sulfate epimerase
16550	-0.116092801387591	7.2504232080665 0.547501669613411
0.45934031116514	0.843389701969214
1894	ECT2	epithelial cell transforming 2
5499	-0.118338581369654	5.67207922730382
0.547022835227658	0.459536720070946	0.843389701969214
285367	RPUSD3	RNA pseudouridine synthase D3	5348
0.154481592610668	3.87372315418826	0.546898430750829
0.459587770224052	0.843389701969214
8844	KSR1	kinase suppressor of ras 1
11511	-0.16644103691294	3.55005723200119
0.546536819597007	0.459736210629148	0.843389701969214
51728	POLR3K	RNA polymerase III subunit K
1889	-0.156040468377618	3.08888042982581
0.546496249532085	0.459752869269726	0.843389701969214
7727	ZNF174	zinc finger protein 174 2876	0.179620992980902
2.98170059672697	0.54614288362964

0.459898006740473	0.843389701969214
1399	CRKL	CRK like proto-oncogene, adaptor protein
5325	-0.0855830957934069	6.76337999923863
0.546112051508636	0.459910673816724	0.843389701969214
27128	CYTH4	cytohesin 4	6161
0.432925914301301	-0.29447110689962	0.546002502782454
0.459955685314851	0.843389701969214
53340	SPA17	sperm autoantigenic protein 17	5205
0.179528277639019	2.54815989472459	0.545982805744628
0.45996377919532	0.843389701969214
8531	YBX3	Y-box binding protein 3 8610	-0.103781744826663
6.9657739812063 0.545947641677941	0.4599782293278
0.843389701969214
55795	PCID2	PCI domain containing 2 6243	0.107789640031658
4.68178671926292	0.545917215041973
0.459990733266553	0.843389701969214
79871	RPAP2	RNA polymerase II associated protein 2
17142	-0.167650312369218	4.83821064324256
0.54582358976598	0.46002921229978	0.843389701969214
386758	ZNF582-AS1	ZNF582 antisense RNA 1 (head to head)
2072	-0.249134575280307	0.298239769386399
0.545730162265528	0.460067615134926	0.843389701969214
100137047	JMJD7	jumonji domain containing 7
1971	-0.157736331362914	3.22670118749425
0.545700008125178	0.46008001090503	0.843389701969214
51191	HERC5	HECT and RLD domain containing E3 ubiquitin protein ligase 5	4766	0.240871441103282	0.425568430666737
0.545065080873466	0.460341139919006
0.84375796216246
123775	C16orf46	chromosome 16 open reading frame 46	2719
0.210944064796151	1.52262235075074
0.544670800741608	0.46050341535012	0.843944961246779
400793	C1orf226	chromosome 1 open reading frame 226
4647	-0.262417168531897	1.86964412374791
0.544286565688266	0.460661643800097	0.8439641711261
130399	ACVR1C	activin A receptor type 1C
9195	-0.2907857950839	0.348022666473333
0.544162745331144	0.46071265154579	0.8439641711261
54949	SDHAF2	succinate dehydrogenase complex assembly factor 2
3118	0.104381269113916	5.18957895068967
0.544132252263701	0.460725214527675	0.8439641711261
7404	UTY	ubiquitously transcribed tetratricopeptide repeat containing, Y-linked	9191	-0.145462982362276
4.4130252343773 0.544060181536334	0.460754909445369
0.8439641711261
8803	SUCLA2	succinate-CoA ligase ADP-forming beta subunit
3840	-0.107603385824991	5.61576161789292
0.543902781047642	0.460819772889986	0.843972614915014
10560	SLC19A2 solute carrier family 19 member 2
3671	-0.145238954546691	5.06380672867711
0.543286824177898	0.461073743016369	0.844268948752212
353322	ANKRD37 ankyrin repeat domain 37	2580
0.232790910759952	2.11070583091393	0.543218023657989
0.46110212450592	0.844268948752212

11010	GLIPR1	GLI pathogenesis related 1
6273	-0.291281711196953	6.78246994298064
0.542436862993767	0.461424562412861	0.844549572669489
114821	ZBED9	zinc finger BED-type containing 9	5480
0.193530782841275	2.46285350434061	0.542414675204294
	0.461433726045837	0.844549572669489
203102	ADAM32	ADAM metallopeptidase domain 32
10044	-0.335062899972436	1.05581672002338
0.542326376399839	0.461470196618796	0.844549572669489
284161	GDPD1	glycerophosphodiester phosphodiesterase domain
containing 1	2970	-0.168665000084119	2.94345826752821
	0.542035463350488	0.461590386544932
0.844549572669489
5188	GATB	glutamyl-tRNA amidotransferase subunit B
11757	0.11870537758462	4.25863598757702
0.541897768362096	0.461647292193534	0.844549572669489
79906	MORN1	MORN repeat containing 1	9098
0.217557056460122	2.44355897090711	0.541879646343895
	0.46165478236811	0.844549572669489
64793	CEP85	centrosomal protein 85	5422	0.132773215864529
	3.90996305995189	0.541761842145225
0.461703477791056	0.844549572669489
9242	MSC	musculin	2113	0.385494056476307
0.871946740187658	0.541678948837937	0.461737747376522
	0.844549572669489
136	ADORA2B adenosine A2b receptor	1735	0.266048723955291
	3.9691583989109 0.541120129960937	0.46196887869565
	0.844647169058672
57465	TBC1D24 TBC1 domain family member 24
7666	-0.119589545024381	7.40778307031523
0.540970550620017	0.462030776942584	0.844647169058672
3312	HSPA8	heat shock protein family A (Hsp70) member 8
4849	-0.151416804629731	10.1065070759912
0.54086323550564	0.462075193723087	0.844647169058672
11067	DEPP1	DEPP1 autophagy regulator
2979	-0.284558817537827	3.47660551643879
0.540775210178115	0.462111631699808	0.844647169058672
23053	ZSWIM8	zinc finger SWIM-type containing 8
7351	-0.120995413592931	7.3253019459009 0.540769636206505
	0.462113939192372	0.844647169058672
399979	SNX19	sorting nexin 19	8152	-0.0969224964152576
	7.29389499926782	0.540656790855422
0.462160658442658	0.844647169058672
9886	RHOBTB1 Rho related BTB domain containing 1	4825
0.204562692074927	3.83474497131162	0.540482741671944
	0.462232731500572	0.844647169058672
23176	SEPTIN8 septin 8	7711	-0.0855308200998192
6.52444167819233	0.540384260095088	0.46227352024015
	0.844647169058672
4237	MFAP2	microfibril associated protein 2
2915	-0.234663714047944	4.12747627720338
0.540160871072095	0.462366063915493	0.84470607242263
283314	C1RL-AS1	C1RL antisense RNA 1
6094	-0.272413312367299	3.6361317458995 0.539705415054989

	0.462554837637214	0.844940742147164
122786	FRMD6	FERM domain containing 6
9416	-0.161071888241244	8.33737585181714
0.53949279030428	0.462643006618018	0.844979882168646
100131211	NEMP2	nuclear envelope integral membrane protein 2
	6169	0.229620462715686	1.66217418666178
0.539362817450197	0.4626969155589 0.844979882168646
5872	RAB13	RAB13, member RAS oncogene family	2751
0.106032530407152	5.21752197675541	0.539084487990468
	0.462812392122696	0.845080586163813
84078	KBTBD7	kelch repeat and BTB domain containing 7
4734	-0.186724223432446	3.10465234060415
0.538936941590411	0.46287362649865	0.845082232320984
333926	PPM1J	protein phosphatase, Mg2+/Mn2+ dependent 1J	3307
	0.13986565645872	6.8086969836399 0.538403520739507
	0.463095113289569	0.845138531368813
10199	MPHOSPH10	M-phase phosphoprotein 10
4673	-0.111929372427565	4.87344985418763
0.53828589132501	0.463143978042941	0.845138531368813
1438	CSF2RA	colony stimulating factor 2 receptor alpha subunit
	7660	0.67837425063691	0.07061792892898
0.538276712096605	0.463147791556508	0.845138531368813
7862	BRPF1	bromodomain and PHD finger containing 1
5477	-0.0994823889375372	4.97612772095069
0.538159507269461	0.463196488738996	0.845138531368813
79183	TTPAL	alpha tocopherol transfer protein like
7044	-0.115682992834595	6.05818723860238
0.538066145674687	0.463235285184738	0.845138531368813
27074	LAMP3	lysosomal associated membrane protein 3
3816	-0.428758180764929	2.58248594701622
0.537991075989154	0.463266484175751	0.845138531368813
1902	LPAR1	lysophosphatidic acid receptor 1
4159	-0.193385909327857	5.39739753095473
0.537031853771467	0.46366543223851	0.845756180409534
55650	PIGV	phosphatidylinositol glycan anchor biosynthesis
class V 3382	0.126319819547573	4.45619310009605
0.53656061310694	0.463861625591666	0.8460038788623
26504	CNNM4	cyclin and CBS domain divalent metal cation
transport mediator 4	5105	-0.131993305418304
4.86755632178075	0.536333783283695	0.463956109685886
	0.846029966999633
6885	MAP3K7	mitogen-activated protein kinase kinase kinase 7
	7227	-0.0805943718695198	5.84299599604124
0.536183123107968	0.464018882907507	0.846029966999633
8819	SAP30	Sin3A associated protein 30	1108
0.215980591994209	2.22213601920557	0.535993723920276
	0.464097816131864	0.846029966999633
415117	STX19	syntaxin 19	1158	-0.282553511387362
0.786527593219444	0.535759518792662	0.464195452121733
	0.846029966999633
22866	CNKSR2	connector enhancer of kinase suppressor of Ras 2
	7160	-0.387394359915468	0.589399856064321
0.535620447961991	0.464253443809848	0.846029966999633
3329	HSPD1	heat shock protein family D (Hsp60) member 1

4245	-0.141231832148184	6.77989810823476
0.535497981546314	0.464304521131982	0.846029966999633
1051	CEBPB	CCAAT enhancer binding protein beta
1956	-0.103472404662358	5.06967975511949
0.535430997700176	0.464332462017937	0.846029966999633
146691	TOM1L2	target of myb1 like 2 membrane trafficking protein
	7863	0.114308003626016	7.50959073504958
0.535367062474933	0.464359133739786	0.846029966999633
23287	AGTPBP1	ATP/GTP binding protein 1
4939	-0.16845676186407	4.24197831530714
0.534674143294669	0.464648354311879	0.846446806977045
1186	CLCN7	chloride voltage-gated channel 7	8119
0.173719607483296	7.07810773775626	0.5340720425556
0.464899901413281	0.846794917957844
374887	YJEFN3	YjeF N-terminal domain containing 3	3683
0.120304500729126	5.67251792055133	0.533861410643738
	0.464987951084678	0.846845173689128
9230	RAB11B	RAB11B, member RAS oncogene family	2470
0.100052356362257	6.11288158431412	0.533163964089894
	0.465279692340603	0.847084334298734
23151	GRAMD4	GRAM domain containing 4	4885
0.1051629553094 4.9901393142675 0.533158906700252
0.465281808910337	0.847084334298734
4321	MMP12	matrix metallopeptidase 12
1874	-0.518677024211355	3.24229387460864
0.533113775720665	0.465300697372347	0.847084334298734
116832	RPL39L	ribosomal protein L39 like	1060
0.231067247570595	1.09862316447948	0.532846326066629
	0.465412657010634	0.847091879667835
114885	OSBPL11 oxysterol binding protein like 11
4191	-0.0997137440612309	5.30525930183178
0.532814949517018	0.465425794668641	0.847091879667835
133686	NADK2	NAD kinase 2, mitochondrial	4885
0.189594525351359	3.94041703031339	0.532656102737572
	0.465492314417732	0.847100471081496
57538	ALPK3	alpha kinase 3	10940	0.257847426466594
4.9828970703524 0.532514874183697	0.465551469007308
0.847100471081496
79944	L2HGDH	L-2-hydroxyglutarate dehydrogenase
6879	-0.151774153079593	3.0356152297707 0.53174966117466
	0.465872193110967	0.847573945824823
65124	SOWAHC	sosondowah ankyrin repeat domain family member C
	4657	0.12870832179227	5.17742287009488
0.531476659355803	0.465986702057373	0.847659473574987
7551	ZNF3	zinc finger protein 3	5424	-0.101448263536917
	5.33125334415506	0.53105124626429
0.466165228788422	0.847659473574987
23556	PIGN	phosphatidylinositol glycan anchor biosynthesis
class N 6796	-0.0982689804085416	5.65731406502884
0.530982315524967	0.466194166224636	0.847659473574987
7700	ZNF141	zinc finger protein 141 5104	-0.200857392769285
	2.13499332518232	0.530943688489289
0.46621038328621	0.847659473574987
55336	FBXL8	F-box and leucine rich repeat protein 8 2866

0.210341467411118	1.90203530779806	0.530860063353108
	0.466245495312806	0.847659473574987
3575	IL7R	interleukin 7 receptor	6060	0.513108588311302
	3.25745103197668	0.530570872420579
0.466366951731448	0.847659473574987
147991	DPY19L3	dpy-19 like C-mannosyltransferase 3	10781
0.16054482211349	3.54198692582452	0.530462421640436
	0.466412512706814	0.847659473574987
8091	HMGA2	high mobility group AT-hook 2
15403	-0.237457656222229	5.8306186835168	0.53035173289247
	0.466459021219706	0.847659473574987
57707	MEAK7	MTOR associated protein, eak-7 homolog	6155
0.119362978663228	6.09797395523132	0.530333294101001
	0.466466769438641	0.847659473574987
51513	ETV7	ETS variant transcription factor 7
2374	-0.204956246026048	2.36785928318564
0.530196229484115	0.466524372227427	0.847659473574987
54617	INO80	INO80 complex ATPase subunit
7014	-0.100504315492974	5.67626400949398
0.529827315253165	0.466679468775443	0.847831299499044
3963	LGALS7	galectin 7	795	-0.25395881272454
5.36726602660454	0.529447156943078	0.466839378930557
	0.848011824754288
120071	LARGE2	LARGE xylosyl- and glucuronyltransferase 2	3115
	0.189916927905721	4.92213070231093
0.529267567086123	0.466914952320462	0.848014581780217
677844	SNORA78	small nucleolar RNA, H/ACA box 78
471	-0.218555647838371	0.616831905357739
0.529084819376093	0.46699187470039	0.848014581780217
57492	ARID1B	AT-rich interaction domain 1B
16869	-0.111851655406998	6.5509596056505	0.528871708738368
	0.467081603139634	0.848014581780217
29763	PACSIN3	protein kinase C and casein kinase substrate in
neurons 3	2752	-0.108620831153823	6.08925564566147
	0.528868236088954	0.467083065498278
0.848014581780217
10060	ABCC9	ATP binding cassette subfamily C member 9
11421	0.448149794596565	0.416100365439783
0.527457575238817	0.467677713212005	0.84898415357187
1956	EGFR	epidermal growth factor receptor
12961	-0.15279454738229	8.86208446226749
0.52684020334209	0.467938341393673	0.849302441319113
3980	LIG3	DNA ligase 3	12919	-0.122982246497293
4.11571492589825	0.526755030975567	0.467974315849283
	0.849302441319113
102724428	SIK1B	salt inducible kinase 1B (putative)	9417
	0.157551510325425	9.03538104807473
0.526236123851476	0.468193583867957	0.849364922334258
114299	NA	NA	4832	0.226713074411277
1.59703262681543	0.526173174083477	0.468220194976838
	0.849364922334258
55251	PCMTD2	protein-L-isoaspartate (D-aspartate) O-
methyltransferase domain containing 2	5997	0.117010592215502
	4.90891534845376	0.526086124912444

0.468256997764496	0.849364922334258
8533	COPS3	COP9 signalosome subunit 3	2788
0.108564826462621	5.43195720559878	0.525810119944095
	0.468373718316517	0.849364922334258
10844	TUBGCP2 tubulin gamma complex associated protein 2	7697
	0.084485396097508	6.16230124593631
0.525788500896589	0.468382862832979	0.849364922334258
347404	LANCL3	LanC like 3	10298	-0.233410319057576
0.430854937239317	0.525676014607479	0.468430447390434
	0.849364922334258
55359	STYK1	serine/threonine/tyrosine kinase 1
3424	-0.26162303268309	2.70081454464884
0.525669475701676	0.468433213752562	0.849364922334258
347273	CAVIN4	caveolae associated protein 4	2615
0.265451211253878	0.957662083261218	0.525404220297053
	0.468545455352776	0.849458476977263
5830	PEX5	peroxisomal biogenesis factor 5 5488
0.116567254442912	4.86843980506413	0.524983348413087
	0.468723634067699	0.849501397573455
4609	MYC	MYC proto-oncogene, bHLH transcription factor
3001	-0.167286773294568	5.86164058813897
0.524915959135399	0.468752173851036	0.849501397573455
89832	CHRFAM7A	CHRNA7 (exons 5-10) and FAM7A (exons A-E)
fusion	6837	0.33208915517942	0.391311740460188
0.524869485099849	0.468771857525318	0.849501397573455
6721	SREBF2	sterol regulatory element binding transcription
factor 2	7557	-0.104888911720941	8.00792475066616
	0.524775370679436	0.468811722941587
0.849501397573455
26009	ZZZ3	zinc finger ZZ-type containing 3
9008	-0.097481678462091	5.59857906671574
0.524334023455243	0.468998743557898	0.849644958280813
64839	FBXL17	F-box and leucine rich repeat protein 17
8125	-0.116405546998919	4.96828915563544
0.524215846212414	0.469048841457383	0.849644958280813
25800	SLC39A6 solute carrier family 39 member 6	3682
0.0831370759341238	7.83898558975635	0.524016689643318
	0.469133287729656	0.849644958280813
93587	TRMT10A tRNA methyltransferase 10A
4379	-0.141820502918357	3.34553513536809
0.523969639026532	0.469153241680879	0.849644958280813
122553	TRAPPC6B	trafficking protein particle complex 6B
4533	-0.09368923434012	5.62054872350987
0.523872971245247	0.469194242328985	0.849644958280813
10278	EFS	embryonal Fyn-associated substrate
3237	-0.157690474421584	4.91343720065052
0.523548688473518	0.469331825657036	0.849784240172971
4739	NEDD9	neural precursor cell expressed, developmentally
down-regulated 9	10359	0.185610819360327
8.49513167998934	0.523404600468893	0.469392978651863
	0.849785117225882
94239	H2AZ2	H2A.Z variant histone 2 4749	0.0947715430761252
	5.96093328871178	0.522771951282722
0.469661635786919	0.850044452986872

23567	ZNF346	zinc finger protein 346 4686	0.119624573375059
	3.85739792020388	0.522719978567352
0.469683717221496	0.850044452986872
8476	CDC42BPA	CDC42 binding protein kinase alpha
14055	-0.119115690426959	6.9391760748032 0.522638616760787
	0.469718288435667	0.850044452986872
80143	SIKE1	suppressor of IKBKE 1	5894	-0.108476899737173
	5.63950269953028	0.521827736568703
0.470063061304007	0.850284962820331
55249	YY1AP1	YY1 associated protein 1
5001	-0.0856106762224671	5.88163497943638
0.521686806273689	0.47012302412015	0.850284962820331
4660	PPP1R12B	protein phosphatase 1 regulatory subunit 12B
	18917	0.213982604604517	6.271734500335
0.521677770948688	0.470126868877437	0.850284962820331
115004	CGAS	cyclic GMP-AMP synthase 4155	-0.206432935723351
	2.85015949301172	0.521430620964546
0.470232057042846	0.850284962820331
8504	PEX3	peroxisomal biogenesis factor 3 4124
0.128666706440246	3.66535949387153	0.521302268039433
	0.470286699599907	0.850284962820331
79713	IGFLR1	IGF like family receptor 1	2926
0.201935131650689	2.43400450627655	0.521080192451807
	0.470381266053362	0.850284962820331
91151	TIGD7	tigger transposable element derived 7
4197	-0.241431960806886	1.63458551702347
0.520744206159897	0.470524397388242	0.850284962820331
100508120	GMDS-DT GMDS divergent transcript
12882	-0.208215503786006	1.1934713826785 0.520303077977216
	0.47071242619151	0.850284962820331
7525	YES1	YES proto-oncogene 1, Src family tyrosine kinase
	5100	-0.0864038535011529	7.01878682476842
0.520270584140015	0.470726281327713	0.850284962820331
6218	RPS17	ribosomal protein S17	3812	0.14670912263592
	7.6433866049646 0.520233597068454	0.470742053146929
	0.850284962820331
9976	CLEC2B	C-type lectin domain family 2 member B
2638	-0.273290865838946	0.952107990823011
0.520183456134131	0.470763435323369	0.850284962820331
22984	PDCD11	programmed cell death 11
7382	-0.121566365661427	5.90103974411063
0.520046784892513	0.470821725571478	0.850284962820331
389831	LOC389831	uncharacterized LOC389831	2195
0.487114247714497	3.37430692216297	0.520040478162748
	0.470824415671062	0.850284962820331
9982	FGFBP1	fibroblast growth factor binding protein 1
1359	-0.437951701496415	5.54502466664258
0.519895801147243	0.470886133635642	0.850284962820331
140564	APOBEC3D	apolipoprotein B mRNA editing enzyme
catalytic subunit 3D	2545	-0.225383618673039
0.341817192622547	0.519843740837903	0.470908345307117
	0.850284962820331
94005	PIGS	phosphatidylinositol glycan anchor biosynthesis
class S 5990	0.097200531203313	5.80858636420347

0.519810372552584	0.47092258286507	0.850284962820331
4920	ROR2	receptor tyrosine kinase like orphan receptor 2 5818
	0.311312462521245	2.15043830048525
0.519452731334942	0.471075224611153	0.850284962820331
1315	COPB1	coatomer protein complex subunit beta 1
4721	-0.10609236618675	8.0397885669867 0.519398394844529
	0.471098422483937	0.850284962820331
6125	RPL5	ribosomal protein L5	2890	0.139450388310875
	7.78719160604156	0.519356462226483
0.47111632603308	0.850284962820331
23113	CUL9	cullin 9	12251	0.176123278482835
5.70517925736686	0.519354710420487	0.471117074008003
	0.850284962820331
54876	DCAF16	DDB1 and CUL4 associated factor 16
4816	-0.118601158727144	4.50862180652201
0.51924653475734	0.471163265874765	0.850284962820331
693197	MIR612	microRNA 612	100	-0.572249923850938
0.421119027613769	0.518927348919824	0.471299603378231
	0.850284962820331
284613	CYB561D1	cytochrome b561 family member D1	5690
	-0.109535709782823	5.17635751884451
0.518879993463996	0.471319836281963	0.850284962820331
59348	ZNF350	zinc finger protein 350 4539	-0.164423622039003
	2.97871191155356	0.518763916001066
0.471369437009334	0.850284962820331
171017	ZNF384	zinc finger protein 384 5181	-0.095139747636107
	5.8393182687387 0.51857382638093	0.471450681863785
	0.850284962820331
57217	TTC7A	tetratricopeptide repeat domain 7A
6949	-0.0937874999723076	6.5116566023739 0.518524077603111
	0.471471948369015	0.850284962820331
8875	VNN2	vanin 2 2633	-0.570187259899295
0.912072584255535	0.518401733340019	0.47152425443318
	0.850284962820331
378	ARF4	ADP ribosylation factor 4
2013	-0.116113941251844	8.30893336294949
0.518334620543996	0.471552951269017	0.850284962820331
79056	PRRG4	proline rich and Gla domain 4
5530	-0.250099296244427	6.67308897583489
0.518025326661302	0.471685239052374	0.850284962820331
79683	ZDHHC14 zinc finger DHHC-type containing 14	11121
0.177345194899729	2.18201434703578	0.517987234588526
	0.471701535521717	0.850284962820331
51194	IPO11	importin 11	6749	-0.106986495995796
4.73707435960194	0.517913641598238	0.471733022494238
	0.850284962820331
23452	ANGPTL2 angiopoietin like 2	3889	-0.439084418876895
	0.897082846499856	0.517477888824436
0.471919530177149	0.850293807870751
7498	XDH	xanthine dehydrogenase	6037	-0.205473625784212
	3.89622688333038	0.517389012047168
0.471957585189579	0.850293807870751
9166	EBAG9	estrogen receptor binding site associated antigen 9
	3681	0.121684650182957	3.78998598803201

0.517230440527724	0.47202549422371	0.850293807870751
84223	IQCG	IQ motif containing G	7474	0.165943231499781
	2.46720700945853	0.517225415861423
0.472027646319953	0.850293807870751
327	APEH	acylaminoacyl-peptide hydrolase 4669
0.0922160093720957	6.17502979349999	0.517144207999254
	0.472062430358823	0.850293807870751
84549	MAK16	MAK16 homolog	4846	0.144053732397718
4.42993212726259	0.517051467702753	0.472102159192606
	0.850293807870751
11093	ADAMTS13	ADAM metallopeptidase with thrombospondin
type 1 motif 13 5708	0.273869191612932	2.20708277108706
	0.516788441329732	0.472214865955547
0.850387455314906
85358	SHANK3	SH3 and multiple ankyrin repeat domains 3
7455	-0.187390372702988	3.5266626322425 0.516550884955046
	0.472316696223358	0.850461494279544
55850	USE1	unconventional SNARE in the ER 1	2431
0.142331150284385	3.24514709872526	0.516381794553411
	0.472389199689948	0.850482714660296
158399	ZNF483	zinc finger protein 483 5738	0.294962155294079
	0.380982876586945	0.516138359681051
0.472493612585437	0.850561371479786
1787	TRDMT1	tRNA aspartic acid methyltransferase 1
3527	-0.194621979457609	2.0170950473181 0.515882512253569
	0.47260338965672	0.850598349161936
8932	MBD2	methyl-CpG binding domain protein 2
5810	-0.0978815536163898	6.32284248131778
0.515807443761361	0.472635607305444	0.850598349161936
23114	NFASC	neurofascin	21271	0.335323993887625
1.30862180805308	0.515305275695817	0.472851217583682
	0.850648067412838
28974	C19orf53	chromosome 19 open reading frame 53	1174
	0.114836930668966	5.53292217481005
0.515301725099384	0.472852742630226	0.850648067412838
283174	MIR4697HG	MIR4697 host gene
5164	-0.395924763704647	1.13375803702918
0.515025406154376	0.472971451129975	0.850648067412838
91300	R3HDM4	R3H domain containing 4 2957	-0.110880965746027
	5.28525361546593	0.514789994393951
0.473072623691355	0.850648067412838
84820	POLR2J4 RNA polymerase II subunit J4, pseudogene
7103	-0.202367382480818	4.66467048309489
0.514753455607142	0.473088330052798	0.850648067412838
54944	LINC01521	long intergenic non-protein coding RNA 1521
	1943	0.259608308276801	0.274360810957295
0.514639884660355	0.473137154425476	0.850648067412838
83752	LONP2	lon peptidase 2, peroxisomal
9957	-0.0772996492914166	6.82825362541658
0.514566326803596	0.473168781446476	0.850648067412838
54627	MAP10	microtubule associated protein 10	3516
0.209191994559413	1.74326076285242	0.514543679163
0.473178519741523	0.850648067412838
80381	CD276	CD276 molecule	5542	0.0863047774472128

7.22209395187788	0.51438186968106	0.473248105903777
	0.850648067412838
4486	MST1R	macrophage stimulating 1 receptor	5252
0.125756934495441	5.71732986174891	0.514329722334509
	0.473270535400032	0.850648067412838
7514	XPO1	exportin 1	11708	-0.103507736780549
7.58641448265091	0.513740043993934	0.473524286372236
	0.850905430141977
57719	ANO8	anoctamin 8	4979	-0.119347590248515
5.1043253635232 0.513714647190747	0.473535220101143
0.850905430141977
729614	FLJ37453	uncharacterized LOC729614	2473
0.184352684548751	1.0734627084806 0.513524210649917
0.473617219107349	0.850943610574223
26100	WIPI2	WD repeat domain, phosphoinositide interacting 2
	8977	0.0833526581929865	6.55668761600186
0.51318859960849	0.473761784018968	0.851094178370358
7709	ZBTB17	zinc finger and BTB domain containing 17
4240	-0.121276391900404	4.96264109353636
0.512931416832038	0.473872614266723	0.851184112340854
5682	PSMA1	proteasome 20S subunit alpha 1
3708	-0.0945457917357682	7.20389749086519
0.512500367919884	0.474058464624671	0.851282403304746
23275	POFUT2	protein O-fucosyltransferase 2	5862
0.0956852889731812	5.38951851765624	0.512381731059224
	0.474109636648896	0.851282403304746
84817	TXNDC17 thioredoxin domain containing 17	2780
0.153841906023127	6.00275160780089	0.51238167352415
	0.474109661467861	0.851282403304746
8453	CUL2	cullin 2	5147	-0.0841831939074297
5.8033042772907 0.512201209005577	0.474187519006924
0.851313070844653
645840	NA	NA	1134	0.270895700871371
1.08388989496016	0.511670250214296	0.474416710107833
	0.851338724244541
101927550	LOC101927550	uncharacterized LOC101927550
3831	-0.278921516841663	0.397293846084398
0.511629442890483	0.474434332237947	0.851338724244541
5594	MAPK1	mitogen-activated protein kinase 1
11336	-0.101702380941608	7.68470647124232
0.511597135163107	0.474448284677604	0.851338724244541
1234	CCR5	C-C motif chemokine receptor 5 (gene/pseudogene)
	3450	-0.668520707974189	0.925826427897197
0.511412446537902	0.474528057234035	0.851338724244541
6158	RPL28	ribosomal protein L28	6933	0.118418067905126
	7.21510042861393	0.511224800063815
0.474609129673856	0.851338724244541
6996	TDG	thymine DNA glycosylase 4316	-0.0991003700789934
	6.15567428755517	0.511201271643539
0.474619296686762	0.851338724244541
663	BNIP2	BCL2 interacting protein 2
7969	-0.0915846685531521	6.93819942274034
0.51118283254198	0.474627264769445	0.851338724244541
137872	ADHFE1	alcohol dehydrogenase iron containing 1 3750

0.216739488310562	1.74815648036368	0.510524491019513
	0.474911896050154	0.851654212030662
414189	AGAP6	ArfGAP with GTPase domain, ankyrin repeat and PH
domain 6	3160	0.148728234152064	3.84963457464309
	0.510436431733645	0.474949989143395
0.851654212030662
84915	FAM222A family with sequence similarity 222 member A
3722	-0.250945333780424	1.09429832753801
0.510232526025011	0.475038214673009	0.851654212030662
56963	RGMA	repulsive guidance molecule BMP co-receptor a
13141	-0.278318226064026	4.71837198403065
0.510213704729551	0.475046359577037	0.851654212030662
126433	FBXO27	F-box protein 27	2833	0.239249920612803
	3.02709503349883	0.510036124571897
0.475123218451463	0.851682994733765
404550	C16orf74	chromosome 16 open reading frame 74
2181	-0.28967233958617	3.08979691083033
0.509660466325659	0.475285874555759	0.851698628577565
29888	STRN4	striatin 4	6590	-0.0816438319262373
6.56434722135668	0.509274369032847	0.475453144989128
	0.851698628577565
283349	RASSF3	Ras association domain family member 3
3705	-0.146310068039848	6.08419905840024
0.50915026852266	0.475506929860343	0.851698628577565
220074	LRTOMT	leucine rich transmembrane and O-methyltransferase
domain containing	6144	0.197489386719949
1.55406159748065	0.508668964253723	0.475715619624862
	0.851698628577565
9619	ABCG1	ATP binding cassette subfamily G member 1	6121
	0.272944978412881	3.69954779641055
0.508664652544379	0.475717489821401	0.851698628577565
2202	EFEMP1	EGF containing fibulin extracellular matrix protein
1	3962	0.28152420852675	5.86111242752271
0.508451840345394	0.475809811611753	0.851698628577565
262	AMD1	adenosylmethionine decarboxylase 1
7222	-0.104687807128218	6.8125016689200８
0.508328880336464	0.475863167178244	0.851698628577565
767558	LUZP6	leucine zipper protein 6
3775	-0.0896378983307019	8.13663875986047
0.50826659138739	0.475890199698133	0.851698628577565
830	CAPZA2	capping actin protein of muscle Z-line subunit alpha
2	8425	-0.101037278577129	7.23901775435177
0.508220614817908	0.475910154479449	0.851698628577565
8027	STAM	signal transducing adaptor molecule
3246	-0.0955051997420098	5.87522046710956
0.507959420253606	0.476023544159251	0.851698628577565
201191	SAMD14	sterile alpha motif domain containing 14	5400
	0.311974863238908	1.136193606527	0.50779377331846
	0.476095477523219	0.851698628577565
9129	PRPF3	pre-mRNA processing factor 3
5373	-0.097426862491854	5.40597645599809
0.507680432510456	0.476144706772132	0.851698628577565
81606	LBH	LBH regulator of WNT signaling pathway
4535	-0.195082898532212	9.40254452567785

0.507389845756961	0.476270960140004	0.851698628577565
8031	NCOA4	nuclear receptor coactivator 4
4079	-0.0865242342359523	8.20538723608395
0.507288355174129	0.476315068342528	0.851698628577565
30851	TAX1BP3 Tax1 binding protein 3	1383	0.112479380669503
	8.55608920468353	0.507231339710643
0.476339850402606	0.851698628577565
51684	SUFU	SUFU negative regulator of hedgehog signaling	6865
	0.125881965535308	5.09161257115248
0.507215495523631	0.476346737531516	0.851698628577565
79847	MFSD13A major facilitator superfamily domain containing 13A
	3526	0.275185672324189	1.34139014698318
0.507113470660663	0.476391089438977	0.851698628577565
5093	PCBP1	poly(rC) binding protein 1
1750	-0.082772067846604	7.88513457782662
0.506987561801723	0.476445833387663	0.851698628577565
7375	USP4	ubiquitin specific peptidase 4
6659	-0.102414038479333	5.87050686969693
0.506970003003218	0.476453468596585	0.851698628577565
2021	ENDOG	endonuclease G	1172	0.158724605640488
3.39103159311799	0.506916275394486	0.476476832567066
	0.851698628577565
388564	TMEM238 transmembrane protein 238	1000
0.236870371757464	0.540756176261238	0.506894228050309
	0.476486420607913	0.851698628577565
2992	GYG1	glycogenin 1	2983	0.123177366831696
5.24436719693104	0.506885734514569	0.476490114395944
	0.851698628577565
84925	SLC49A4 solute carrier family 49 member 4	3596
0.0975487608714201	5.21530222113471	0.506792970555925
	0.476530459924493	0.851698628577565
26146	TRAF3IP1	TRAF3 interacting protein 1
4617	-0.113504449850159	4.62200812370349
0.506558949673226	0.476632266597866	0.851712857003149
4616	GADD45B growth arrest and DNA damage inducible beta	2133
	0.123763680204641	6.25476527833185
0.506383780565152	0.476708494028088	0.851712857003149
100101467	ZSCAN30 zinc finger and SCAN domain containing 30
	12081	0.11618359828807	4.06918117280164
0.506355415206888	0.476720839502013	0.851712857003149
4636	MYL5	myosin light chain 5	4033	0.238976459341341
	2.82797716158927	0.506080423002748
0.47684055155313	0.851720072894081
23355	VPS8	VPS8 subunit of CORVET complex	10243
0.0898780042872101	5.83911058043769	0.506066780346437
	0.476846491870878	0.851720072894081
653784	MZT2A	mitotic spindle organizing protein 2A
4093	-0.111799375681317	4.381556258102	0.504916711096255
	0.477347691782091	0.852506581511125
10336	PCGF3	polycomb group ring finger 3
7425	-0.0810278174286217	7.33465323409404
0.503479426162035	0.477975269170285	0.853248729638399
220972	MARCHF8 membrane associated ring-CH-type finger 8	7539
	0.113252497950292	4.44904938494144

0.503361672175323	0.478026745056675	0.853248729638399
374928	ZNF773	zinc finger protein 773 8073	-0.154865434285467
	3.0834748638315 0.503348746153403	0.478032396188981
	0.853248729638399
22836	RHOBTB3 Rho related BTB domain containing 3
8833	-0.099346370696282	7.61423872115578
0.503321203482285	0.478044437943422	0.853248729638399
10175	CNIH1	cornichon family AMPA receptor auxiliary protein 1
	5060	-0.115706109868847	6.73311290295391
0.503267713460762	0.47806782538746	0.853248729638399
6904	TBCD	tubulin folding cofactor D
13546	-0.0924038485447473	5.5545618615409 0.503090417202087
	0.478145357998821	0.85327838316849
790955	UQCC3	ubiquinol-cytochrome c reductase complex assembly
factor 3	2398	0.156924330315189	2.64679441743983
	0.502750816110346	0.478293924743809
0.853372534221022
55832	CAND1	cullin associated and neddylation dissociated 1
13655	-0.0843751952636606	7.48736676220791
0.50269130709534	0.47831996617186	0.853372534221022
57509	MTUS1	microtubule associated scaffold protein 1
12010	0.153253783541122	7.58405498938698
0.502274853903728	0.478502272917205	0.853589064792574
55884	WSB2	WD repeat and SOCS box containing 2
3976	-0.0993290895688774	7.54728471899429
0.50173112453475	0.478740467107386	0.8538476906586
255783	INAFM1	InaF motif containing 1 1255	0.21357789513634
	1.79524725077715	0.501665634160896
0.478769169874537	0.8538476906586
57451	TENM2	teneurin transmembrane protein 2	11070
0.21060172872294	6.20212537894379	0.500796566743531
	0.479150326642658	0.854110854164212
284391	ZNF844	zinc finger protein 844 6620	-0.257964040861489
	2.46533433991733	0.500468851262056
0.479294185435032	0.854110854164212
84874	ZNF514	zinc finger protein 514 6813	-0.144004437961382
	4.22154915486745	0.500302399213476
0.479367280733203	0.854110854164212
84033	OBSCN	obscurin, cytoskeletal calmodulin and titin-
interacting RhoGEF	30286	-0.269726743656 5.39121392472779
	0.500265888911841	0.479383316206982
0.854110854164212
374383	NCR3LG1 natural killer cell cytotoxicity receptor 3 ligand 1
	6438	-0.170949647944004	3.33829450991421
0.500060119331978	0.479473707467166	0.854110854164212
202134	FAM153B family with sequence similarity 153 member B
8225	-0.640179778084581	-0.255023392364562
0.500048562625771	0.479478784969332	0.854110854164212
693199	MIR614	microRNA 614	90	-0.172015179179103
2.99880505027462	0.500037625068157	0.47948359052624
	0.854110854164212
9552	SPAG7	sperm associated antigen 7	2831
0.0938473247040082	5.79446754351713	0.499973788597728
	0.479511639475211	0.854110854164212

64858	DCLRE1B DNA cross-link repair 1B	3940
0.150464792291717	3.27158284112929	0.499956994330294
	0.479519019113131	0.854110854164212
80325	ABTB1	ankyrin repeat and BTB domain containing 1	6149
	0.193772515862396	4.74742564542014
0.499858692683407	0.479562217972125	0.854110854164212
8721	EDF1	endothelial differentiation related factor 1
1200	-0.10868965844413	7.11237008178663
0.499135407590245	0.479880263102444	0.854110854164212
4898	NRDC	nardilysin convertase	6773	-0.0877990508232139
	7.42041056173125	0.498920693199342
0.479974744471875	0.854110854164212
55614	KIF16B	kinesin family member 16B	6280
0.0976977862523821	4.96832701063216	0.498827139359371
	0.480015920759736	0.854110854164212
9601	PDIA4	protein disulfide isomerase family A member 4
3277	-0.15800838235709	6.99218755913547
0.498656243974011	0.480091152686474	0.854110854164212
9139	CBFA2T2 CBFA2/RUNX1 partner transcriptional co-repressor 2
	9457	-0.130379846262084	5.57711506428695
0.498614378839818	0.480109585590264	0.854110854164212
10613	ERLIN1	ER lipid raft associated 1	5792
0.0955615256633516	5.1267405289251 0.498600349218296
0.480115762986561	0.854110854164212
55696	RBM22	RNA binding motif protein 22
3615	-0.0867313252378978	5.81617103225822
0.498598708007009	0.480116485638408	0.854110854164212
25941	TPGS2	tubulin polyglutamylase complex subunit 2
10571	-0.0999154105748829	7.2151140838347 0.498439584261888
	0.480186558859459	0.854110854164212
8539	API5	apoptosis inhibitor 5	4651	-0.0848273030177723
	6.92409563805691	0.498181914990191
0.480300064062134	0.854110854164212
134553	C5orf24 chromosome 5 open reading frame 24
5741	-0.138489712713141	6.47435529327755
0.498105473759722	0.480333745442976	0.854110854164212
64146	PDF	peptide deformylase, mitochondrial	1158
0.148804231866748	3.64672890564541	0.497618185136492
	0.480548544622549	0.854110854164212
407977	TNFSF12-TNFSF13 TNFSF12-TNFSF13 readthrough	1816
0.156663851245292	4.73352963382424	0.497615154700009
	0.480549880946352	0.854110854164212
3959	LGALS3BP	galectin 3 binding protein	3584
0.145651906296617	8.19276155694703	0.497584368021649
	0.48056345721438	0.854110854164212
55435	AP1AR	adaptor related protein complex 1 associated
regulatory protein	3444	-0.127359154873464
4.55419277560588	0.497507333171464	0.480597430695191
	0.854110854164212
64342	HS1BP3	HCLS1 binding protein 3 9059	0.123394112664217
	4.92438904714023	0.497484704079808
0.48060741119969	0.854110854164212
5906	RAP1A	RAP1A, member of RAS oncogene family
5532	-0.0839946437604875	6.05875721994699

0.497414700207699	0.480638288385482	0.854110854164212
7259	TSPYL1	TSPY like 1	3326	-0.0856119314768701
6.98169970434843	0.497368060019038	0.480658862164525
	0.854110854164212
374900	ZNF568	zinc finger protein 568 9437	-0.192396007062001
	2.91079373516535	0.497320259658816
0.480679949213761	0.854110854164212
8665	EIF3F	eukaryotic translation initiation factor 3 subunit F
	9423	0.0828210907227147	5.92595350789565
0.497135551406387	0.480761447214105	0.854110854164212
79741	CCDC7	coiled-coil domain containing 7
8250	-0.249387905167188	0.440444175879278
0.49709800367083	0.480778017006589	0.854110854164212
7755	ZNF205	zinc finger protein 205 2330	0.141890589521066
	3.5719132591349 0.49703226174681	0.480807031133348
	0.854110854164212
5528	PPP2R5D protein phosphatase 2 regulatory subunit B'delta
	3314	0.0816654759615221	5.86848547830669
0.496179411013042	0.481183683458679	0.854226575342645
1456	CSNK1G3 casein kinase 1 gamma 3 5000	-0.11525271467026
	5.60932228984618	0.496153969940426
0.481194926672058	0.854226575342645
9419	CRIPT	CXXC repeat containing interactor of PDZ3 domain
	2561	-0.12434699071067	3.81776264212892
0.495678106736438	0.481405305160983	0.854226575342645
6627	SNRPA1	small nuclear ribonucleoprotein polypeptide A'
3248	-0.141067076126784	4.01516247666315
0.495671959986694	0.481408023618869	0.854226575342645
84163	GTF2IRD2	GTF2I repeat domain containing 2	7549
	0.139534603338337	3.64957174903288
0.49563908693442	0.481422562463376	0.854226575342645
4298	MLLT1	MLLT1 super elongation complex subunit	4740
0.117520781572556	6.09758477390558	0.49551746794026
	0.48147635746039	0.854226575342645
79697	RIOX1	ribosomal oxygenase 1	2429	0.117753024741565
	4.03057118604255	0.495508061437533
0.481480518593404	0.854226575342645
81856	ZNF611	zinc finger protein 611 5609	-0.112567166184266
	4.38941072062159	0.495393239267102
0.48153131697493	0.854226575342645
2053	EPHX2	epoxide hydrolase 2	4473	0.263567984344737
	0.745686027875457	0.495385404028481
0.481534783677228	0.854226575342645
4824	NKX3-1	NK3 homeobox 1	3271	-0.15952998484723
4.05476555133374	0.495317326094749	0.481564906493137
	0.854226575342645
79066	METTL16 methyltransferase like 16
6318	-0.0976227652059976	4.91791578261495
0.495301271392073	0.481572010756365	0.854226575342645
1445	CSK	C-terminal Src kinase	4570	-0.0940378296485265
	6.30199135180131	0.49514659231393
0.481640465621688	0.854226575342645
23276	KLHL18	kelch like family member 18
7064	-0.0871809470917183	5.97354065021544

0.494899975883088	0.481749641407077	0.854226575342645
10426	TUBGCP3 tubulin gamma complex associated protein 3
7014	-0.107027013905141	4.324561383766	0.49481947893514
	0.481785285781779	0.854226575342645
9455	HOMER2	homer scaffold protein 2
2970	-0.118342074536482	5.8777025408004 0.494815698796124
	0.48178695974919	0.854226575342645
84260	TCHP	trichoplein keratin filament binding
5307	-0.11714848237519	4.21178077731769
0.494380722423838	0.481979645163139	0.854447493796865
92906	HNRNPLL heterogeneous nuclear ribonucleoprotein L like
5923	-0.0962295722045721	5.32275702417155
0.494259060526473	0.482033561503736	0.854447493796865
284186	TMEM105 TMEM105 long non-coding RNA	3743
0.221545281793744	0.00164819076746435	0.493939107900257
	0.482175400769922	0.854477897336617
2804	GOLGB1	golgin B1	11865	-0.131970547292436
8.03863937809187	0.493922202738872	0.482182896962195
	0.854477897336617
79946	C10orf95	chromosome 10 open reading frame 95
1469	-0.259861421026454	1.2922365323295 0.493807589680355
	0.482233724455341	0.854477897336617
23285	KIAA1107	KIAA1107	4556	-0.198320906536345
	2.35163773881772	0.49329587872317
0.482460760631959	0.854772055928642
129787	TMEM18	transmembrane protein 18	4961
0.103059086415882	4.39931829894336	0.493043893120869
	0.48257262635734	0.854862119310391
116931	MED12L	mediator complex subunit 12L	12619
0.30166045093398	0.351975711473398	0.492591788165217
	0.482773439486833	0.855039418803258
85363	TRIM5	tripartite motif containing 5	4804
0.12620135523514	5.58568310352923	0.492543715974008
	0.482794799979651	0.855039418803258
58473	PLEKHB1 pleckstrin homology domain containing B1	4933
	0.493826320318808	1.74990343396065
0.492282753401433	0.482910783760646	0.855061584649875
9409	PEX16	peroxisomal biogenesis factor 16	3013
0.114686921386721	3.89163933389142	0.492092520991546
	0.482995360714696	0.855061584649875
728568	C12orf73	chromosome 12 open reading frame 73	3096
	0.145125862808522	3.30860639679639
0.492025617784001	0.483025111546733	0.855061584649875
8559	PRPF18	pre-mRNA processing factor 18
2995	-0.0965970729304858	4.83830369733594
0.49196628812841	0.483051497061074	0.855061584649875
317648	NOP14-AS1	NOP14 antisense RNA 1	5648
0.146221542695174	4.7759977729731 0.491693141539509
0.483173003419598	0.855168594755915
60496	AASDHPPT	aminoadipate-semialdehyde dehydrogenase-
phosphopantetheinyl transferase 6183	0.105743700418901
5.25603301626085	0.491209816712544	0.483388128741124
	0.855271527865107
1717	DHCR7	7-dehydrocholesterol reductase	4160

0.144714663280086	6.93696736975429	0.491089688910883
	0.483441621477927	0.855271527865107
84759	PCGF1	polycomb group ring finger 1	2629
0.135288835751366	3.47304582022594	0.491025980053985
	0.483469994898978	0.855271527865107
1200	TPP1	tripeptidyl peptidase 1 4901	0.12170317669787
	6.36261463948517	0.49101383951313
0.483475402130072	0.855271527865107
80864	EGFL8	EGF like domain multiple 8
1393	-0.250778581056162	1.76861093785275
0.490857742897616	0.483544934311463	0.855286512804731
90203	SNX21	sorting nexin family member 21	4066
0.127659037173227	4.17309390991019	0.49037226441718
	0.483761292811211	0.855312426309526
79034	C7orf26 chromosome 7 open reading frame 26	2728
0.117524857268109	4.72727094317621	0.490143326915168
	0.483863376564209	0.855312426309526
57787	MARK4	microtubule affinity regulating kinase 4
6668	-0.104252148961866	5.76876947721237
0.490107317842856	0.483879436324805	0.855312426309526
3665	IRF7	interferon regulatory factor 7
2628	-0.146692160068236	3.53600218672659
0.490073285920477	0.483894615100274	0.855312426309526
5885	RAD21	RAD21 cohesin complex component
6192	-0.0857925469889359	7.16728515467846
0.489896137559484	0.483973638755454	0.855312426309526
11332	ACOT7	acyl-CoA thioesterase 7 4124	-0.149098303896513
	5.99977068242193	0.489544376290219
0.484130618153134	0.855312426309526
90861	JPT2	Jupiter microtubule associated homolog 2	4768
	0.137509984840652	7.52439377445033
0.489507309094762	0.484147164906591	0.855312426309526
2069	EREG	epiregulin	5717	-0.545845859486947
4.73211725188413	0.489417930585958	0.484187067208328
	0.855312426309526
55602	CDKN2AIP	CDKN2A interacting protein
3193	-0.12491418147315	4.67861115751588
0.489369531149933	0.484208677004363	0.855312426309526
100506696	PCAT6	prostate cancer associated transcript 6 890
	0.268547400077088	0.181512399271684
0.489273266264204	0.484251662908941	0.855312426309526
7227	TRPS1	transcriptional repressor GATA binding 1
11072	0.211013078649585	5.04422676201572
0.489233525135347	0.484269410659487	0.855312426309526
79736	TEFM	transcription elongation factor, mitochondrial	2724
	0.139465003803527	3.46159028311588
0.489102384776523	0.484327983456538	0.855312426309526
3420	IDH3B	isocitrate dehydrogenase (NAD(+)) 3 non-catalytic
subunit beta	2454	0.094638025693118	5.7556429650962
0.488915838436062	0.484411323044189	0.855312426309526
3679	ITGA7	integrin subunit alpha 7	6373
0.316159691820231	0.0351591288368593	0.488908795905179
	0.48441446975894	0.855312426309526
256714	MAP7D2	MAP7 domain containing 2	4376

0.267381659401988	1.06166668634118	0.48866405238462
	0.484523846022347	0.855352548966713
10156	RASA4	RAS p21 protein activator 4
7677	-0.269241376185369	4.25193923769318
0.488468662442756	0.484611195368187	0.855352548966713
60488	MRPS35	mitochondrial ribosomal protein S35
1979	-0.118091275790498	5.56958723477314
0.488448094413755	0.484620391846922	0.855352548966713
151295	SLC23A3	solute carrier family 23 member 3
3601	-0.264907620312049	1.75451109685745
0.487980918148063	0.484829355658522	0.855613554832925
29775	CARD10	caspase recruitment domain family member 10
5652	-0.135174888498592	6.18957214568644
0.487814848400134	0.484903673048405	0.855636904558958
129880	BBS5	Bardet-Biedl syndrome 5	5493	0.173423724950396
	2.86302019662482	0.48732643341193
0.485122351341353	0.855675070780398
642819	ZNF487	zinc finger protein 487	2699	0.177238351226853
	1.29146762620342	0.487087523697289
0.485229377879638	0.855675070780398
653464	SRGAP2C	SLIT-ROBO Rho GTPase activating protein 2C	7311
	0.18656051881444	3.43725908390524
0.486978787459844	0.485278102357992	0.855675070780398
1798	DPAGT1	dolichyl-phosphate N-
acetylglucosaminephosphotransferase 1	5280	0.095011735436473
	5.15775532166714	0.486914226325614
0.485307035893152	0.855675070780398
619505	SNORA21	small nucleolar RNA, H/ACA box 21	3081
0.136587975693041	5.91055735700475	0.486784616320534
	0.485365130157576	0.855675070780398
10204	NUTF2	nuclear transport factor 2	2649
0.0981283952200947	6.53534849343216	0.4865472746661З
	0.485471542104605	0.855675070780398
9218	VAPA	VAMP associated protein A
7762	-0.0840412989372121	6.78491465697007
0.486478786832954	0.485502255744753	0.855675070780398
8972	MGAM	maltase-glucoamylase	10248	0.298086165454691
	2.96335471447339	0.486457932460098
0.485511608609166	0.855675070780398
157381	LINC00964	long intergenic non-protein coding RNA 964
	4335	0.287333038554005	0.264603872395286
0.486334386082167	0.48556702336603	0.855675070780398
27183	VPS4A	vacuolar protein sorting 4 homolog A
5763	-0.104101121011904	6.51253356547603
0.486270847822311	0.485595526512711	0.855675070780398
10940	POP1	POP1 homolog, ribonuclease P/MRP subunit
4852	-0.163966164940221	4.36195881705429
0.486065350559649	0.485687731161906	0.855675070780398
100131067	CKMT2-AS1	CKMT2 antisense RNA 1	3884
0.218639870244065	1.19896620930572	0.485850951600866
	0.485783960795945	0.855675070780398
127253	TYW3	tRNA-yW synthesizing protein 3 homolog	4611
0.113357086297413	4.50484173682676	0.485774270406765
	0.485818385615563	0.855675070780398

5335	PLCG1	phospholipase C gamma 1 9153	0.12274955844269
	7.4711418812064 0.485621832486913	0.485886832220159
	0.855675070780398
10363	HMG20A	high mobility group 20A 5483	-0.100777868704529
	4.96957263117601	0.485545749561814
0.485921000411962	0.855675070780398
100529215	ZNF559-ZNF177	ZNF559-ZNF177 readthrough	4117
	0.190130913337941	1.8452751321168 0.48551042507808б
	0.485936865687733	0.855675070780398
114794	ELFN2	extracellular leucine rich repeat and fibronectin
type III domain containing 2	10752	-0.304339689607491
2.2263301856092 0.485412513159254	0.485980845323352
0.855675070780398
8614	STC2	stanniocalcin 2 6128	0.233228905612313
5.03166935729196	0.485167107261073	0.486091104599666
	0.855675070780398
51015	ISOC1	isochorismatase domain containing 1
1940	-0.132246896974752	3.67267727854645
0.4851049269994 0.486119048363658	0.855675070780398
11325	DDX42	DEAD-box helicase 42	7795	-0.0802180523921839
	7.2974981179786 0.484906799305136	0.486208104546233
	0.855675070780398
2041	EPHA1	EPH receptor A1 4362	0.125103418217392
5.53968135622836	0.484906645585829	0.48620817365183
	0.855675070780398
9925	ZBTB5	zinc finger and BTB domain containing 5
4627	-0.131797883371172	5.23954428319835
0.484105419298976	0.486568591728373	0.856201792002427
387893	KMT5A	lysine methyltransferase 5A
4579	-0.0880862874637869	5.44751077293252
0.483806367692566	0.486703228482894	0.856307640794665
63893	UBE2O	ubiquitin conjugating enzyme E2 O
6472	-0.110365123524132	4.97646954223179
0.483601281266015	0.486795596639325	0.856307640794665
55695	NSUN5	NOP2/Sun RNA methyltransferase 5
3414	-0.108029978077419	4.48184293596141
0.483433191369329	0.486871323728641	0.856307640794665
118987	PDZD8	PDZ domain containing 8 9881	-0.111643936846817
	6.2494115881483 0.483420370417008	0.486877100566875
	0.856307640794665
6773	STAT2	signal transducer and activator of transcription 2
	6109	-0.162616534186793	7.31819315342102
0.483293180767419	0.486934415572893	0.856307640794665
56647	BCCIP	BRCA2 and CDKN1A interacting protein
4498	-0.111130959491658	4.91894674805938
0.483146993747241	0.487000305113991	0.856316002225918
112483	SAT2	spermidine/spermine N1-acetyltransferase family
member 2	1551	0.126580770286329	4.56013033006301
	0.482942018319966	0.487092716719263
0.856339025104813
9099	USP2	ubiquitin specific peptidase 2
4538	-0.302546335911094	2.26753717388883
0.48284676334475	0.487135671595284	0.856339025104813
23645	PPP1R15A	protein phosphatase 1 regulatory subunit 15A

2378	-0.13396939815272	7.55870484750564
0.482434058416004	0.487321851953082	0.856362249653828
8731	RNMT	RNA guanine-7 methyltransferase
6982	-0.100756985555324	6.19282263264071
0.482154876337994	0.487447864141749	0.856362249653828
60314	C12orf10	chromosome 12 open reading frame 10	2509
0.117274701882518	3.90088939145364
0.482039628094022	0.487499898591626	0.856362249653828
285237	C3orf38 chromosome 3 open reading frame 38
3102	-0.0976226810771853	5.0790239512681 0.48201476299927
0.487511126362415	0.856362249653828
221786	FAM200A family with sequence similarity 200 member A
2717	-0.160923773862583	3.20742917090266
0.481998339242824	0.48751854270356	0.856362249653828
133383	SETD9	SET domain containing 9 2535	0.187982599107467
2.06112233576723	0.481901822126421
0.48756212993112	0.856362249653828
54785	BORCS6	BLOC-1 related complex subunit 6	2575
0.155041479139543	3.72565108086395	0.48186922958461
0.487576850216983	0.856362249653828
8863	PER3	period circadian regulator 3
7144	-0.250621253729807	3.85450086804579
0.481452587991285	0.487765089675813	0.856364350949284
9180	OSMR	oncostatin M receptor	7174	-0.103222929971581
8.72308837186128	0.481202472349025
0.487878150364866	0.856364350949284
126789	PUSL1	pseudouridine synthase like 1
2489	-0.131492703939461	3.50462389051801
0.481108215187563	0.48792076906615	0.856364350949284
800	CALD1	caldesmon 1	9843	-0.171702072343714
10.142544311693 0.480877757547205	0.488024997331334
0.856364350949284
79642	ARSJ	arylsulfatase family member J
5024	-0.131695768846304	4.24051498089852
0.480842915550289	0.488040758417035	0.856364350949284
9462	RASAL2	RAS protein activator like 2
16044	-0.14530658906308	6.41190326961012
0.480809337761931	0.488055948426958	0.856364350949284
6888	TALDO1	transaldolase 1 3005	-0.118923332201745
7.30075921264294	0.480646418501724	0.488129661431065
0.856364350949284
84324	SARNP	SAP domain containing ribonucleoprotein 3396
0.126421925817828	5.48294477590598	0.480312422556022
0.488280836123598	0.856364350949284
9701	PPP6R2	protein phosphatase 6 regulatory subunit 2
5349	-0.0848172772294973	6.34223740102275
0.480244735173724	0.488311482561879	0.856364350949284
7975	MAFK	MAF bZIP transcription factor K
4458	-0.133635548776367	6.98280606929578
0.480137190875723	0.48836018136883	0.856364350949284
10694	CCT8	chaperonin containing TCP1 subunit 8
3341	-0.103833359236448	7.18493581677581
0.480083338161421	0.488384570287692	0.856364350949284
27246	RNF115	ring finger protein 115 11041	0.0979476473914454

5.46596330979656	0.479905590981719
0.488465083137453	0.856364350949284
124997	WDR81	WD repeat domain 81	7913	0.118252678585726
5.53148328545335	0.479818200490342
0.488504675865636	0.856364350949284
404201	WDFY3-AS2	WDFY3 antisense RNA 2	6203
0.275067970286608	0.349498269006708	0.479782246255144
0.488520966672817	0.856364350949284
57827	C6orf47 chromosome 6 open reading frame 47
2475	-0.100091804834086	5.23737120071491
0.479177033562761	0.488795323238114	0.856364350949284
29102	DROSHA	drosha ribonuclease III 7780	-0.0914789563154919
5.63665552019829	0.479114588838076
0.48882364542489	0.856364350949284
400410	ST20	suppressor of tumorigenicity 20
1614	-0.178424861695646	2.89133149145612
0.479112878257559	0.488824421307561	0.856364350949284
10194	TSHZ1	teashirt zinc finger homeobox 1 5697
0.214293668302877	2.66542874095252	0.479091432510343
0.488834148812835	0.856364350949284
3382	ICA1	islet cell autoantigen 1	4891
0.134177470916508	4.71435373706437	0.478925275174984
0.488909526481435	0.856364350949284
116115	ZNF526	zinc finger protein 526 4271	-0.1527792009525
3.72296870424312	0.478866836565963
0.488936041891557	0.856364350949284
100533970	P2RX5-TAX1BP3	P2RX5-TAX1BP3 readthrough (NMD
candidate)	5905	0.109424660622335	8.36751823544118
0.47883284972113	0.488951463878023
0.856364350949284
55766	H2AJ	H2A.J histone	3356	0.13756169673584
4.31800589491688	0.478716563803111	0.489004236295156
0.856364350949284
689	BTF3	basic transcription factor 3	2467
0.103930797260772	7.69675535678503	0.478584262761316
0.489064288159595	0.856364350949284
81608	FIP1L1	factor interacting with PAPOLA and CPSF1
5636	-0.0931863693640376	5.3117932879697 0.478315654655061
0.489186247997252	0.856364350949284
5106	PCK2	phosphoenolpyruvate carboxykinase 2, mitochondrial
4189	-0.152469699825157	4.81815563727365
0.478312566738133	0.489187650341018	0.856364350949284
26472	PPP1R14B	protein phosphatase 1 regulatory inhibitor
subunit 14B	1655	0.10387982481163	6.4002060891555
0.478282224810606	0.489201430152134	0.856364350949284
1314	COPA	coatomer protein complex subunit alpha
6232	-0.115195688184703	7.84203587715913
0.47821174260195	0.489233442198573	0.856364350949284
3843	IPO5	importin 5	9671	0.127753844481695
6.91667788336002	0.478087413259203	0.489289919372251
0.856364350949284
9361	LONP1	lon peptidase 1, mitochondrial	5085
0.0771101136084345	5.9594775374724 0.477900837629541
0.489374692576632	0.856405712009105

79366	HMGN5	high mobility group nucleosome binding domain 5 2416
	0.201454787250374	1.3185455631835 0.47747108153138
	0.489570051252381	0.856515683406188
8396	PIP4K2B phosphatidylinositol-5-phosphate 4-kinase type 2
beta	6135	0.0850557809904431	6.36619327873509
0.477368575691454	0.489616667579262	0.856515683406188
10741	RBBP9	RB binding protein 9, serine hydrolase	3861
0.0980179951582028	5.19085783350118	0.477210412294852
	0.489688609663678	0.856515683406188
2327	FMO2	flavin containing dimethylaniline monoxygenase 2
	4786	0.529041122425336	5.12901035852707
0.477174381128204	0.489705001236227	0.856515683406188
400713	ZNF880	zinc finger protein 880 3719	-0.218176476324458
	3.02150852598269	0.477040569475214
0.48976588385441	0.856515683406188
288	ANK3	ankyrin 3	25614	0.202656336722428
4.59571984913343	0.476955868880911	0.489804428077644
	0.856515683406188
5893	RAD52	RAD52 homolog, DNA repair protein	7304
0.173301133361439	3.03781026509392	0.476793764080362
	0.489878210294423	0.856537772012731
221184	CPNE2	copine 2	7781	-0.142396083425033
5.20689764983811	0.476215378413285	0.490141614331727
	0.856781120354617
27109	DMAC2L	distal membrane arm assembly complex 2 like
11297	0.128012343285783	3.56981711749667
0.4759202350725 0.490276117292412	0.856781120354617
2764	GMFB	glia maturation factor beta
7966	-0.124781134642382	6.84730674742042
0.475751271284075	0.490353145315226	0.856781120354617
10971	YWHAQ	tyrosine 3-monooxygenase/tryptophan 5-monooxygenase
activation protein theta	2713	-0.10409052075048
8.82325403175916	0.475678799599649	0.490386190240598
	0.856781120354617
6209	RPS15	ribosomal protein S15	2137	0.107819484184051
	6.31543936134567	0.475498380866107
0.490468471936589	0.856781120354617
9722	NOS1AP	nitric oxide synthase 1 adaptor protein
7996	-0.261282899431955	0.383621726093452
0.475379818877486	0.49052255581226	0.856781120354617
22803	XRN2	5'-3' exoribonuclease 2 3437	0.0793662089107259
	6.94943947005835	0.475314476604268
0.490552366949071	0.856781120354617
144983	HNRNPA1L2	heterogeneous nuclear ribonucleoprotein A1
like 2	985	0.199272843529733	1.57354307111286
0.475258760294789	0.490577788815746	0.856781120354617
2649	NR6A1	nuclear receptor subfamily 6 group A member 1
7124	-0.226872154288395	1.87784117460874
0.475175075877964	0.490615975915092	0.856781120354617
3762	KCNJ5	potassium inwardly rectifying channel subfamily J
member 5	6109	0.519023402782027	3.18400975127766
	0.47514638626788	0.490629068777353
0.856781120354617
2744	GLS	glutaminase	9827	0.300527216226414

9.92839668595445	0.474964624532277	0.490712031569423
	0.856819175541375
340393	TMEM249 transmembrane protein 249
2929	-0.18943198611297	1.59356077613372
0.474489700264722	0.490928915168791	0.857067469276355
79829	NAA40	N(alpha)-acetyltransferase 40, NatD catalytic
subunit 5321	0.108189135175872	5.119199252384
0.474385307179592	0.490976609800348	0.857067469276355
10951	CBX1	chromobox 1	3087	0.0873660187871151
6.84837639855098	0.474220985840756	0.491051699853256
	0.857068040360679
100616453	MIR4687 microRNA 4687	80	-0.221249815651586
	0.709214913206143	0.474116812301394
0.491099314052603	0.857068040360679
115353	LRRC42	leucine rich repeat containing 42
2411	-0.0793699129462895	6.01478651781265
0.473524222255658	0.491370313682662	0.857434157686938
59343	SENP2	SUMO specific peptidase 2
7169	-0.0827318491531907	6.06687693765951
0.472840853062298	0.491683138258818	0.857873158643655
9993	DGCR2	DiGeorge syndrome critical region gene 2	5636
	0.0858276119357395	7.71330697343374
0.472698049466711	0.491748551237369	0.857880428095881
255919	CNEP1R1 CTD nuclear envelope phosphatase 1 regulatory
subunit 1	4005	-0.116762076645142	4.59672110395569
	0.472322903838458	0.491920460746415
0.8579670117553
2355	FOSL2	FOS like 2, AP-1 transcription factor subunit
7157	-0.124086152420175	8.2749938228227 0.472322408907075
	0.491920687613823	0.8579670117553
84897	TBRG1	transforming growth factor beta regulator 1	6627
	0.0992428351823925	5.70387522304513
0.471979606945524	0.492077863755948	0.858037653922579
150590	C2orf15 chromosome 2 open reading frame 15	3346
0.119581123398209	3.89493766882039	0.471896388154008
	0.492116032596211	0.858037653922579
56994	CHPT1	choline phosphotransferase 1	5702
0.184078545363559	3.29763955901216	0.471833316628683
	0.492144964061826	0.858037653922579
54859	ELP6	elongator acetyltransferase complex subunit 6	2807
	0.111259046450251	4.65778655699859
0.471369503372926	0.492357806756283	0.85821131372946
146664	MGAT5B	alpha-1,6-mannosylglycoprotein 6-beta-N-
acetylglucosaminyltransferase B 6386	0.472021641479089
1.65258566039171	0.471190558870548	0.492439965114325
	0.85821131372946
54205	CYCS	cytochrome c, somatic	4457	-0.118174527035347
	6.29073712965892	0.471169126804984
0.492449806705963	0.85821131372946
6538	SLC6A11 solute carrier family 6 member 11
6700	-0.321669193483633	-0.0563264692720855
0.471082365366373	0.492489650871521	0.85821131372946
23404	EXOSC2	exosome component 2	4240	0.139021505285761
	3.46850194529786	0.470891285698074

0.492577420988736	0.858257486725864
106660612	LINC00680	long intergenic non-protein coding RNA 680	2969	0.198262643796317	2.73747287906792
0.470216513172261	0.492887579230917	0.858635549165093
4772	NFATC1	nuclear factor of activated T cells 1
9068	-0.232015997231543	3.77024408798798
0.4701452419774	0.492920358038027	0.858635549165093
5861	RAB1A	RAB1A, member RAS oncogene family
2951	-0.104204125590598	7.64865250591868
0.469695865701965	0.493127117813148	0.858635549165093
54840	APTX	aprataxin	3238	0.100925754500979
4.88003324755279	0.469613886661435	0.493164852366957
0.858635549165093
2976	GTF3C2	general transcription factor IIIC subunit 2
5839	-0.0751591536450439	6.32746415527019
0.469583718684476	0.493178739758454	0.858635549165093
84984	CEP19	centrosomal protein 19	2201	0.322667498301732
1.72223814160914	0.469503717070012
0.493215570518204	0.858635549165093
4712	NDUFB6	NADH:ubiquinone oxidoreductase subunit B6	1388
0.118888549636611	4.65326677456036
0.469486484488613	0.493223504575023	0.858635549165093
10450	PPIE	peptidylprolyl isomerase E	8071
0.105957665288769	4.98528289807588	0.469313991833125
0.493302933750996	0.858667104890046
64170	CARD9	caspase recruitment domain family member 9
6699	-0.177252683270961	2.64201135674185
0.468604713689655	0.493629766717334	0.858941290542849
26092	TOR1AIP1	torsin 1A interacting protein 1
9615	-0.0967603672299945	6.84131739131991
0.468583641047218	0.493639482475298	0.858941290542849
84947	SERAC1	serine active site containing 1
8873	-0.128372821207326	4.7617703565153	0.468341355669641
0.493751213690208	0.858941290542849
4133	MAP2	microtubule associated protein 2	11530
0.134836792837467	8.91685432704975	0.468333783549951
0.493754706298059	0.858941290542849
10250	SRRM1	serine and arginine repetitive matrix 1
7825	-0.0821177840090995	6.15768761940565
0.468306991328525	0.493767064428573	0.858941290542849
6118	RPA2	replication protein A2	2019	0.110180594157185
4.74023966499659	0.467668151427258
0.494061888381702	0.859278944706774
3093	UBE2K	ubiquitin conjugating enzyme E2 K
5245	-0.0917767860428273	6.66849132416103
0.46756212171433	0.494110849579649	0.859278944706774
8443	GNPAT	glyceronephosphate O-acyltransferase	3173
0.0894299265841219	5.79821705735156	0.467448850982223
0.494163163454852	0.859278944706774
84224	NBPF3	NBPF member 3	6545	-0.200060005752737
3.80722701296905	0.46735492004934	0.494206552335102
0.859278944706774
55521	TRIM36	tripartite motif containing 36	6113
0.212306040900824	2.05898221621555	0.467175335049665

0.494289524631172	0.859316541083384
51124	IER3IP1	immediate early response 3 interacting protein 1
3685	-0.115281095856543	5.44985677073421
0.466705507536664	0.494506706202337	0.859433023312051
200894	ARL13B	ADP ribosylation factor like GTPase 13B	3977
0.164336110682495	3.4842851430077	0.466470443788589
0.494615426507925	0.859433023312051
4091	SMAD6	SMAD family member 6	4364	0.252382981664693
1.46864733721917	0.466239487308242
0.494722286275739	0.859433023312051
81929	SEH1L	SEH1 like nucleoporin	10481	-0.104249630305529
4.75372793420174	0.466210591927741
0.494735658424685	0.859433023312051
113235	SLC46A1	solute carrier family 46 member 1	7324
0.176903951894413	5.25672218911987	0.46617699216476
0.494751208422811	0.859433023312051
22985	ACIN1	apoptotic chromatin condensation inducer 1
9110	-0.0971353762008858	7.79043328981105
0.466057371614369	0.494806575582802	0.859433023312051
150383	CDPF1	cysteine rich DPF motif domain containing 1	2005
0.180913241839197	1.55287576465221
0.465997412960465	0.49483433176027	0.859433023312051
7009	TMBIM6	transmembrane BAX inhibitor motif containing 6
5952	-0.0878633641917352	9.93773461886896
0.465969215674932	0.494847385807931	0.859433023312051
26275	HIBCH	3-hydroxyisobutyryl-CoA hydrolase	7175
0.144109929160927	3.89527744504652	0.465741732963345
0.494952721034591	0.859501507089609
9282	MED14	mediator complex subunit 14
9656	-0.111764578364475	6.47098784350949
0.465619055351752	0.495009542206888	0.859501507089609
55751	TMEM184C	transmembrane protein 184C
4712	-0.0754649080418905	6.12477006785748
0.465373419726451	0.495123347408888	0.859592554183973
23166	STAB1	stabilin 1	10588	0.784797998934792
2.95679048752816	0.464900932016896	0.495342379031748
0.859866241553811
55180	LINS1	lines homolog 1	7790	-0.137565663236122
3.54754744523043	0.46450291933877	0.495527012746926
0.859932857949927
25956	SEC31B	SEC31 homolog B, COPII coat complex component	8426
0.0928485975241604	5.10788144656572
0.464365999942725	0.495590554934597	0.859932857949927
145741	C2CD4A	C2 calcium dependent domain containing 4A
3445	-0.348811637519296	1.04013138984325
0.464352253283793	0.495596935305523	0.859932857949927
79947	DHDDS	dehydrodolichyl diphosphate synthase subunit	5596
0.110245766039969	5.39739151577462
0.464215106284655	0.495660598253888	0.859932857949927
6945	MLX	MAX dimerization protein MLX
4259	-0.091124462341117	5.90438071342138
0.464156438901753	0.495687835563586	0.859932857949927
79925	SPEF2	sperm flagellar 2
14887	-0.239772434087935	-0.0191586935262182

0.464024470824006	0.495749113153827	0.859932857949927
7289	TULP3	TUB like protein 3	3181	0.0961864597059955
	5.07622011450017	0.46381810838327
0.495844960335181	0.859979102502063
221895	JAZF1	JAZF zinc finger 1	4020	-0.229853470990436
	2.53332085839047	0.463622193460431
0.49593598394421	0.859979102502063
1967	EIF2B1	eukaryotic translation initiation factor 2B subunit
alpha	3627	0.088583854663124	5.25131067503358
0.463570593953962	0.495959962162609	0.859979102502063
64145	RBSN	rabenosyn, RAB effector 7921	-0.095190602741701
	5.35044338655888	0.462945342209041
0.496250671093361	0.860190406842201
64167	ERAP2	endoplasmic reticulum aminopeptidase 2
7422	-0.7074203381515	5.16775486506182
0.462908942093009	0.496267604064176	0.860190406842201
4796	TONSL	tonsoku like, DNA repair protein	7111
0.185269448933232	2.27337544454139	0.462778955456956
	0.496328080500328	0.860190406842201
120	ADD3	adducin 3	6541	-0.133290410495397
6.79714885909295	0.462347888135326	0.496528724034451
	0.860190406842201
51075	TMX2	thioredoxin related transmembrane protein 2	2246
	0.098896616147666	6.32616240434559
0.462312820730077	0.4965450524383 0.860190406842201
9662	CEP135	centrosomal protein 135 8640	-0.170892025360705
	3.83438574742888	0.462290916854595
0.496555251979299	0.860190406842201
7347	UCHL3	ubiquitin C-terminal hydrolase L3
2061	-0.125300732171728	5.4396864502988 0.462270116964731
	0.496564937776403	0.860190406842201
10472	ZBTB18	zinc finger and BTB domain containing 18
3906	-0.13963626634148	3.75629336942964
0.462194152262754	0.496600314646323	0.860190406842201
29062	WDR91	WD repeat domain 91	6032	0.221791459426618
	4.76539025189702	0.462106619996206
0.496641083819078	0.860190406842201
11282	MGAT4B	alpha-1,3-mannosyl-glycoprotein 4-beta-N-
acetylglucosaminyltransferase B 5050	0.0955773384877095
8.06906509603621	0.46198886497686	0.496695938497874
	0.860190406842201
10903	MTMR11	myotubularin related protein 11
4591	-0.128421794652111	5.19083536264272
0.461830949939284	0.496769517256308	0.860211475857227
64425	POLR1E	RNA polymerase I subunit E
2512	-0.180179769170442	3.1487526793148 0.461472044978416
	0.496936813451971	0.86039480173322
8653	DDX3Y	DEAD-box helicase 3 Y-linked
5738	-0.103250181753447	6.19047032344323
0.460981711698171	0.497165526009207	0.860676307455805
55054	ATG16L1 autophagy related 16 like 1
6047	-0.0819949355302487	5.93894385870094
0.460736251706741	0.497280085876416	0.860676307455805
55356	SLC22A15	solute carrier family 22 member 15

5292	-0.164441834452317	4.16992890385807
0.46044593792594	0.497415637233845	0.860676307455805
146542	ZNF688	zinc finger protein 688 3023	0.158044782544673
	2.39810795545276	0.460427755336036
0.497424129003069	0.860676307455805
55857	KIZ	kizuna centrosomal protein	5207
0.178098453935167	2.29449469879832	0.460323041177936
	0.497473038159333	0.860676307455805
10419	PRMT5	protein arginine methyltransferase 5
3467	-0.115809523911079	6.20911820209087
0.460268886285306	0.497498335634795	0.860676307455805
4236	MFAP1	microfibril associated protein 1	2373
0.0812595895786575	5.57327338797905	0.460202124158122
	0.497529525342304	0.860676307455805
641298	SMG1P1	SMG1 pseudogene 1	685	-0.276687434658994
	0.569614922513047	0.459997477116682
0.497625152250669	0.860687193881732
8802	SUCLG1	succinate-CoA ligase alpha subunit	4286
0.10518060911654	5.17211136643965	0.459925672437755
	0.49765871230443	0.860687193881732
9862	MED24	mediator complex subunit 24
6962	-0.0768213780632809	6.08266177668872
0.459685893156379	0.497770808004498	0.860767309708065
23475	QPRT	quinolinate phosphoribosyltransferase	2410
0.439453408007603	1.7838779048306 0.459501014220006
0.497857267189086	0.860767309708065
23007	PLCH1	phospholipase C eta 1	7173	0.29835130373452
	0.210211421130253	0.459318776350113
0.49794251611883	0.860767309708065
1763	DNA2	DNA replication helicase/nuclease 2	4867
0.196747302905605	2.25334856033252	0.459224512358329
	0.497986621492581	0.860767309708065
11021	RAB35	RAB35, member RAS oncogene family
4022	-0.0807264596378645	6.27236738198068
0.459169638851876	0.498012299412734	0.860767309708065
9343	EFTUD2	elongation factor Tu GTP binding domain containing 2
	10375	0.0850746767312905	6.68874149205253
0.458792850969814	0.49818867684857	0.860895541424434
392490	FLJ44635	TPT1-like protein
2269	-0.168065835475892	2.88286343600357
0.45874856634036	0.498209413777858	0.860895541424434
5467	PPARD	peroxisome proliferator activated receptor delta
	4457	-0.15643557935933	6.9140458419002
0.457327468121147	0.498875640251338	0.861566583120394
55177	RMDN3	regulator of microtubule dynamics 3	5114
0.10432324910488	5.75725969659276	0.457261245673365
	0.498906722838974	0.861566583120394
54106	TLR9	toll like receptor 9	4333	-0.116057247462118
	4.39653127062511	0.457181648294764
0.498944087509669	0.861566583120394
84464	SLX4	SLX4 structure-specific endonuclease subunit
8591	-0.145662678328954	3.99722007758717
0.457102564202174	0.498981215927488	0.861566583120394
124995	MRPL10	mitochondrial ribosomal protein L10	2764

0.10198080619988	4.55594497147947	0.45681746046111
	0.499115105354969	0.861566583120394
4046	LSP1	lymphocyte specific protein 1
5399	-0.446080406519654	2.82009242530182
0.456700534851494	0.499170033154136	0.861566583120394
9585	KIF20B	kinesin family member 20B	6851
0.215611888608121	1.81342677428425	0.456695481291316
	0.499172407381119	0.861566583120394
10252	SPRY1	sprouty RTK signaling antagonist 1	3963
0.213175886205518	1.56298741285655	0.456642360429306
	0.499197365396938	0.861566583120394
10392	NOD1	nucleotide binding oligomerization domain containing
1	6820	-0.169144965964823	3.0850275209359
0.456416209111985	0.499303642766554	0.861566583120394
8079	MLF2	myeloid leukemia factor 2	3567
0.096725621501313	7.34380446300801	0.456333194918187
	0.499342664012513	0.861566583120394
138151	NACC2	NACC family member 2	7212	0.0858697499420149
	6.37247868276504	0.45632974491172
0.499344285817551	0.861566583120394
11163	NUDT4	nudix hydrolase 4	10078	0.149056986501121
	5.4030946201832 0.456157976877599	0.499425043119535
	0.861566583120394
5147	PDE6D	phosphodiesterase 6D	2850	0.118481281105182
	4.27109520857121	0.455850695927573
0.499569567508311	0.861566583120394
55011	PIH1D1	PIH1 domain containing 1	3247
0.108800711251919	5.05564083488343	0.455832732345403
	0.499578018568752	0.861566583120394
149420	PDIK1L	PDLIM1 interacting kinase 1 like
4535	-0.174071181478655	2.91425242052074
0.455762294824977	0.499611158604856	0.861566583120394
10905	MAN1A2	mannosidase alpha class 1A member 2
9667	-0.0918868917981612	6.39845690575915
0.455676646184287	0.499651460319486	0.861566583120394
79864	JHY	junctional cadherin complex regulator	4680
0.12904349457799	3.39671692739947	0.45565755409786
	0.499660444795787	0.861566583120394
106614088	MIR34AHG	MIR34A host gene
4211	-0.119851366004288	4.38649642887518
0.45556303629084	0.499704927626907	0.861566583120394
203427	SLC25A43	solute carrier family 25 member 43
3115	-0.118546843765762	4.4658868328606 0.455392892516491
	0.49978501917021	0.861598617048262
90864	SPSB3	splA/ryanodine receptor domain and SOCS box
containing 3	3192	0.120512281657307	5.36077169878195
	0.455095691200036	0.499924972587132
0.861615199743254
54936	ADPRHL2 ADP-ribosylhydrolase like 2	1668
0.091852202480662	5.34990518427202	0.454994555150908
	0.499972613165528	0.861615199743254
23765	IL17RA	interleukin 17 receptor A	9036
0.111155394754556	5.86555076792586	0.454980620378468
	0.499979177804896	0.861615199743254

5537	PPP6C	protein phosphatase 6 catalytic subunit
4443	-0.0787832650997725	6.23373953062973
0.454659088134768	0.500130691546237	0.861752169941432
400960	PCBP1-AS1	PCBP1 antisense RNA 1	18619
0.157683925228531	3.41538442269269	0.454490349764098
	0.500210236316923	0.861752169941432
813	CALU	calumenin	4537	0.149042522275745
8.44953969637431	0.454319561407829	0.500290769342451
	0.861752169941432
552889	ATXN7L3B	ataxin 7 like 3B
7604	-0.0953720709155143	6.68000603095327
0.454289915115914	0.500304750907669	0.861752169941432
10675	CSPG5	chondroitin sulfate proteoglycan 5	4366
0.23833916754408	0.325994381669452	0.45395514728794
	0.500462677698683	0.861883498273296
989	SEPTIN7 septin 7	10906	-0.0901737618986262
7.31542412912316	0.453732386050234	0.500567812278493
	0.861883498273296
3386	ICAM4	intercellular adhesion molecule 4 (Landsteiner-Wiener blood group)	1427	0.219999684350729
2.68226112684944	0.453626782434948	0.500617666158608
	0.861883498273296
147495	APCDD1	APC down-regulated 1	5943	0.393056568384889
	5.12912735343525	0.453493438641488
0.500680627811489	0.861883498273296
222389	BEND7	BEN domain containing 7	9815	-0.121043080779284
	6.06910864558509	0.453455603725075
0.500698494976906	0.861883498273296
29969	MDFIC	MyoD family inhibitor domain containing
5966	-0.18763549562657	4.71764763613046
0.453222631487126	0.500808537694275	0.861883498273296
3265	HRAS	HRas proto-oncogene, GTPase	1854
0.102919300894757	5.38814825870649	0.453149126749146
	0.500843265644651	0.861883498273296
56957	OTUD7B	OTU deubiquitinase 7B	12177	-0.117007954244795
	6.19784875518675	0.453085659664396
0.500873254511646	0.861883498273296
6397	SEC14L1 SEC14 like lipid binding 1
10356	-0.11742308931514	7.46556868975407
0.452640709194327	0.501083584043222	0.862063651509903
80298	MTERF2	mitochondrial transcription termination factor 2
	3626	0.171538174284464	2.01740190019149
0.452603798044241	0.501101038808712	0.862063651509903
134145	ATPSCKMT	ATP synthase c subunit lysine N-methyltransferase	2462	0.140530520498721
3.5560934405793 0.452109216297151	0.501335019372946
0.862297609015297
79228	THOC6	THO complex 6	1860	0.155868663117882
3.48049161828387	0.452056103105837	0.501360157622659
	0.862297609015297
2873	GPS1	G protein pathway suppressor 1	4772
0.0919338540209559	6.42743096417872	0.451689414560175
	0.501533768294969	0.862490299878161
80007	C10orf88	chromosome 10 open reading frame 88

3173	-0.118108654640799	3.99647667310008
0.451297027695303	0.501719659317572	0.862580808815841
10063	COX17	cytochrome c oxidase copper chaperone COX17
1809	-0.156796624300001	4.22096605407128
0.451139478918853	0.501794330146763	0.862580808815841
50855	PARD6A	par-6 family cell polarity regulator alpha
1488	-0.192896349325566	1.11363100711396
0.450837275031178	0.501937613773201	0.862580808815841
51729	WBP11	WW domain binding protein 11
5315	-0.0887652957858174	5.71307081411105
0.450733038254663	0.501987051612636	0.862580808815841
57449	PLEKHG5 pleckstrin homology and RhoGEF domain containing G5
	6920	-0.108798645525877	7.09834406026338
0.450484799223489	0.502104820823224	0.862580808815841
56935	SMCO4	single-pass membrane protein with coiled-coil
domains 4	1223	0.2349239799008 1.09168649768322
0.45046394683142	0.502114715728381	0.862580808815841
7140	TNNT3	troponin T3, fast skeletal type
2567	-0.263571036076256	0.0678934260739751
0.450284513515211	0.502199874397302	0.862580808815841
388569	ZNF324B zinc finger protein 324B	4042
0.168127749062251	2.31948584900955	0.450070470152435
	0.502301491111219	0.862580808815841
460	ASTN1	astrotactin 1	9201	-0.420214391679102
0.154074193392807	0.449949853004	0.502358769329315
0.862580808815841
55007	FAM118A family with sequence similarity 118 member A
6512	-0.277276044724548	4.92091845300389
0.449712240059	0.502471638642983	0.862580808815841
4123	MAN2C1	mannosidase alpha class 2C member 1
8557	-0.0906383691302575	5.99071933273871
0.449710669915035	0.502472384626039	0.862580808815841
5257	PHKB	phosphorylase kinase regulatory subunit beta
11593	0.0894851008316066	6.50131177849995
0.449679002091102	0.502487430566693	0.862580808815841
50650	ARHGEF3 Rho guanine nucleotide exchange factor 3
6594	-0.0892347486192104	5.42063639158891
0.449496763766497	0.50257403013273	0.862580808815841
10981	RAB32	RAB32, member RAS oncogene family	1217
0.108955136205643	6.40755095740601	0.449438630983394
	0.502601660157133	0.862580808815841
5334	PLCL1	phospholipase C like 1 (inactive)	6739
0.426856683535865	0.504154483181096	0.449416472712951
	0.50261219247906	0.862580808815841
51106	TFB1M	transcription factor B1, mitochondrial
2736	-0.163936359991085	3.11313092658788
0.449172737365515	0.502728070188617	0.862580808815841
5339	PLEC	plectin 16991	-0.210824323533534
10.2844356897678	0.449109388553097	0.502758195213898
	0.862580808815841
80311	KLHL15	kelch like family member 15
6272	-0.178137817693062	2.24949398981535
0.44905311826564	0.502784956684899	0.862580808815841
100128788	SRRM2-AS1	SRRM2 antisense RNA 1

5858	-0.21452799538939	-0.00504754412674478
0.448995602818266	0.502812312849979	0.862580808815841
3030	HADHA	hydroxyacyl-CoA dehydrogenase trifunctional
multienzyme complex subunit alpha	3811	0.0980331515431825
	7.2674442990373 0.448983559690475	0.50281804126375
	0.862580808815841
10402	ST3GAL6 ST3 beta-galactoside alpha-2,3-sialyltransferase 6
	5967	-0.303007556638274	2.3212773476368
0.448835873100782	0.502888298671333	0.862595689381504
51606	ATP6V1H ATPase H+ transporting V1 subunit H	5090
0.0975545414121559	6.7259867643915 0.44845178964917
0.503071093142005	0.862803575565087
8717	TRADD	TNFRSF1A associated via death domain
2567	-0.105455060481861	4.42996668222403
0.448305533551897	0.503140729715106	0.862817360568009
7353	UFD1	ubiquitin recognition factor in ER associated
degradation 1	8736	-0.0854654663542505	6.39044568158026
	0.44816365295716	0.503208298712888
0.862827597021842
54987	CZIB	CXXC motif containing zinc binding protein	4487
	0.125801507464487	3.59672874128173
0.447943584081118	0.503313134628864	0.862901722979988
65010	SLC26A6 solute carrier family 26 member 6	6373
0.110369165043615	4.6542274254022 0.447608627685366
0.503472771992677	0.863069773259262
7851	MALL	mal, T cell differentiation protein like
3167	-0.253131470004145	6.96403617984287
0.447432158645483	0.503556910413698	0.863108375448443
1998	ELF2	E74 like ETS transcription factor 2
8560	-0.112780620112864	4.84586272145793
0.446746271905674	0.503884161600074	0.863542417847033
9122	SLC16A4 solute carrier family 16 member 4	3167
0.107346142007529	5.74621759618414	0.446643050700814
	0.503933442099068	0.863542417847033
11046	SLC35D2 solute carrier family 35 member D2
2203	-0.145216968024148	4.57592080611685
0.446414925630247	0.504042384193393	0.863623446531726
4926	NUMA1	nuclear mitotic apparatus protein 1	15224
0.12642449376801	8.2091726011511 0.445889388681167
0.504293509880748	0.863948042184398
64083	GOLPH3	golgi phosphoprotein 3	2785	-0.0799113051023696
	6.96340893768559	0.445664282919719
0.504401141214648	0.864026756144499
22877	MLXIP	MLX interacting protein 13905	-0.0843404839487468
	7.65183322435919	0.445390485714427
0.50453210669491	0.864145416785963
25842	ASF1A	anti-silencing function 1A histone chaperone	2517
	0.097250875444678	4.50386257384609
0.444825027345013	0.504802766789767	0.864503283338338
100128537	MIR29B2CHG	MIR29B2 and MIR29C host gene
15645	-0.272491951173867	2.42613784955523
0.444479667702835	0.504968197577396	0.86457684740835
27089	UQCRQ	ubiquinol-cytochrome c reductase complex III subunit
VII	1472	0.126886736318165	6.00170737125586

0.444477634564748	0.504969171746106	0.86457684740835
7841	MOGS	mannosyl-oligosaccharide glucosidase	3234
0.0789110135320666	6.60457502294957	0.444081438586739
	0.505159068662387	0.864632793606905
23512	SUZ12	SUZ12 polycomb repressive complex 2 subunit
4491	-0.11661907820113	5.75026453664646
0.444038779655422	0.5051795224023 0.864632793606905
55081	IFT57	intraflagellar transport 57	3766
0.132328939740055	4.34486713231748	0.443894466289231
	0.505248727061893	0.864632793606905
128218	TMEM125 transmembrane protein 125
2127	-0.13842922499853	4.48250955496287
0.443894392497356	0.505248762452525	0.864632793606905
56924	PAK6	p21 (RAC1) activated kinase 6
5919	-0.131399830717932	4.06869486385471
0.443750226412504	0.50531791271258	0.864645492715014
677850	SNORD1C small nucleolar RNA, C/D box 1C
2665	-0.198938313239698	0.266446780876212
0.443352411526079	0.505508811525868	0.864802484885335
158427	TSTD2	thiosulfate sulfurtransferase like domain containing
2	5484	0.0870346474074696	5.32964363448603
0.443301723739552	0.505533143873858	0.864802484885335
2868	GRK4	G protein-coupled receptor kinase 4	4292
0.188424210179362	2.5642872601314 0.442901742799679
0.505725222629779	0.865025423540764
100528064	NEDD8-MDP1	NEDD8-MDP1 readthrough	1051
0.121100372957043	5.07403803902431	0.442597317164712
	0.505871497729154	0.865169971757296
53944	CSNK1G1 casein kinase 1 gamma 1 11221	-0.08579658436677
	6.33946106068788	0.442306000381535
0.506011542052243	0.865254250372904
6059	ABCE1	ATP binding cassette subfamily E member 1
6265	-0.134245137326818	5.85038322171159
0.44211053834556	0.506105543545392	0.865254250372904
9013	TAF1C	TATA-box binding protein associated factor, RNA
polymerase I subunit C	5577	0.131747146452991
5.34368359446387	0.442109391767019	0.506106095045706
	0.865254250372904
55771	PRR11	proline rich 11 4080	-0.270027565077755
1.80960712615747	0.441750332765506	0.506278851933687
	0.865402028090709
283951	C16orf91	chromosome 16 open reading frame 91	978
	0.143521012798747	3.06232317310914
0.441672942596429	0.506316100535686	0.865402028090709
285440	CYP4V2	cytochrome P450 family 4 subfamily V member 2	9807
	0.255354895249201	4.17994153165215
0.441524260928184	0.506387675597287	0.865418761846169
10262	SF3B4	splicing factor 3b subunit 4
2245	-0.0821611500229056	6.32135808902077
0.441381561756096	0.506456387036747	0.865430598172956
2821	GPI	glucose-6-phosphate isomerase	6766
0.10851889689928	7.18983069058846	0.44120052059148
	0.5065435836699 0.865474015182275
57161	PELI2	pellino E3 ubiquitin protein ligase family member 2

6278	-0.214343674239056	1.29064636097716
0.441055580867535	0.506613410997475	0.865487748242668
643837	LINC01128	long intergenic non-protein coding RNA 1128
9470	0.121909084664418	3.83706592486042
0.440465742027271	0.506897747155704	0.86586789565975
101930370	LOC101930370	uncharacterized LOC101930370	3367
0.203676696321951	-0.0427649762355958
0.439733022472585	0.507251342186346	0.866303417194927
100616113	MIR4680	microRNA 4680	66	0.217382898634999
0.366805791401838	0.43968111724373
0.507276406641879	0.866303417194927
2859	GPR35	G protein-coupled receptor 35	3892
0.225232237602396	1.95685913845705	0.439169548894211
0.507523551292021	0.86636015612815
90673	PPP1R3E	protein phosphatase 1 regulatory subunit 3E	5548
0.190413342283688	1.29529714850344
0.439137540733423	0.507539021694374	0.86636015612815
8772	FADD	Fas associated via death domain	1841
0.0886271905431017	5.46675289982369	0.438948554851205
0.507630380180171	0.86636015612815
10165	SLC25A13	solute carrier family 25 member 13
3710	-0.0963805162760191	5.17811460635339
0.43892969134528	0.507639500622747	0.86636015612815
5016	OVGP1	oviductal glycoprotein 1	2567
0.278374847162927	1.67391691516519	0.43876829065397
0.50771754885288	0.86636015612815
57653	SUGT1P4-STRA6LP-CCDC180	SUGT1P4-STRA6LP-CCDC180 readthrough
7116	0.326733340785748	-0.056389107418541
0.438763848155022	0.507719697395833	0.86636015612815
54888	NSUN2	NOP2/Sun RNA methyltransferase 2
5506	-0.089788783785902	6.56591420145944
0.438637682835406	0.507780721751511	0.86636015612815
2771	GNAI2	G protein subunit alpha i2	7650
0.113783274541168	8.29884886451638	0.438394631282151
0.507898317900268	0.86636015612815
58489	ABHD17C	abhydrolase domain containing 17C
3627	-0.115560590063255	4.69938858560662
0.438388661712651	0.50790120675968	0.86636015612815
5591	PRKDC	protein kinase, DNA-activated, catalytic subunit
15417	-0.13432767066643	7.55256590716714
0.438253815812203	0.507966470402356	0.86636015612815
8439	NSMAF	neutral sphingomyelinase activation associated
factor	6165	0.100418901545111	5.31564882564362
0.438205202175919	0.507990002304598	0.86636015612815
55015	PRPF39	pre-mRNA processing factor 39
4932	-0.125469291156668	4.78198538603487
0.438003412182269	0.508087700781536	0.866421283765154
101730217	SPECC1L-ADORA2A	SPECC1L-ADORA2A readthrough (NMD
candidate)	6276	-0.104515789573109	6.24801466750898
0.437830581667981	0.508171403993265
0.86645853386898
1874	E2F4	E2F transcription factor 4
3582	-0.0833719236805864	5.36064506215413
0.437432198517218	0.508364434682615	0.866478916120875

402569	KPNA7	karyopherin subunit alpha 7
1725	-0.143670268917207	3.74143857180188
0.437143867072527	0.508504220270059	0.866478916120875
10477	UBE2E3	ubiquitin conjugating enzyme E2 E3	2866
0.104964458769819	6.01464880019482	0.436978158157478
	0.508584587373417	0.866478916120875
28971	AAMDC	adipogenesis associated Mth938 domain containing
	1943	0.172101415114533	2.86334748899403
0.436970254940645	0.508588420898584	0.866478916120875
55714	TENM3	teneurin transmembrane protein 3	11984
0.370509288620991	0.677148935121376	0.436620721927625
	0.508758014815656	0.866478916120875
9057	SLC7A6	solute carrier family 7 member 6	7588
0.0759610554119694	6.6865048264655	0.43635805184308
0.508885526909155	0.866478916120875
151009	LINC01106	long intergenic non-protein coding RNA 1106
	2774	-0.206486628274968	3.61346078866481
0.436350564034321	0.508889162642654	0.866478916120875
55663	ZNF446	zinc finger protein 446	4923	0.150470693283803
	3.69537553184191	0.436176313460434
0.508973783315587	0.866478916120875
150368	PHETA2	PH domain containing endocytic trafficking adaptor 2
	2451	0.12176114270152	3.86677613596658
0.436117467482005	0.509002365957617	0.866478916120875
91612	CHURC1	churchill domain containing 1	3615
0.14693206944712	5.93632804993551	0.436092848823534
	0.509014324542921	0.866478916120875
57711	ZNF529	zinc finger protein 529	7075	-0.150274861543357
	2.99319926242108	0.435917784694317
0.50909937643756	0.866478916120875
79465	ULBP3	UL16 binding protein 3	988	0.191910605596621
	2.40970218611957	0.435902536349341
0.509106785744562	0.866478916120875
23312	DMXL2	Dmx like 2	13807	0.169685017238866
5.23727679753045	0.435879165892239	0.509118142019547
	0.866478916120875
96459	FNIP1	folliculin interacting protein 1
6720	-0.149216640660121	5.94631362274972
0.435789395567554	0.509161767674245	0.866478916120875
51258	MRPL51	mitochondrial ribosomal protein L51	1551
0.0915286919593609	5.66635052026725	0.435686596390113
	0.509211732880788	0.866478916120875
54998	AURKAIP1	aurora kinase A interacting protein 1	1669
	0.10336838520104	5.49375140402648
0.435413056615086	0.509344727231272	0.866478916120875
80005	DOCK5	dedicator of cytokinesis 5
15479	-0.134814343972372	6.97683508788455
0.435402165853077	0.509350023533551	0.866478916120875
389333	PROB1	proline rich basic protein 1
5122	-0.206837792221417	1.96335276669137
0.435176962639494	0.509459563750979	0.866478916120875
96610	BMS1P20	BMS1 pseudogene 20	2042	-0.129415968476637
	3.68921342801662	0.435166709421303
0.509464551946998	0.866478916120875

2965	GTF2H1	general transcription factor IIH subunit 1	6472
	0.113995954420876	5.91269643704229
0.435152106321794	0.509471656508155	0.866478916120875
148534	TLCD4	TLC domain containing 4	7162	0.172994161383307
	3.38962569734566	0.435129969007267
0.509482426870245	0.866478916120875
102466854	MIR7705	microRNA 7705	57	-0.20579210143731
	0.907865806939639	0.434407960603236
0.509833917879229	0.866756757678793
51466	EVL	Enah/Vasp-like	10621	0.100713249701158
4.64163937000065	0.434359695570715	0.509857429393191
	0.866756757678793
3898	LAD1	ladinin 1	3482	0.103207293801191
6.95079837385471	0.434266357544743	0.509902902787739
	0.866756757678793
1468	SLC25A10	solute carrier family 25 member 10	3322
	0.123055006946639	4.99161007228423
0.434246893548917	0.509912386343758	0.866756757678793
11132	CAPN10	calpain 10	6080	-0.0973070217209701
4.71535321120509	0.434007471775768	0.51002906615138
	0.866756757678793
79719	AAGAB	alpha and gamma adaptin binding protein
3833	-0.0809370106337474	5.97392344854584
0.433949483686547	0.510057333009527	0.866756757678793
94103	ORMDL3	ORMDL sphingolipid biosynthesis regulator 3	3752
	0.0838767416994719	6.67366155699882
0.433905126451762	0.510078957203276	0.866756757678793
10667	FARS2	phenylalanyl-tRNA synthetase 2, mitochondrial	2351
	0.161681646233748	3.27095597893479
0.433356153310331	0.510346713390124	0.866868039165439
2889	RAPGEF1	Rap guanine nucleotide exchange factor 1	7002
	0.109345161147745	7.21096522909535
0.433325809032695	0.510361520598552	0.866868039165439
339559	ZFP69	ZFP69 zinc finger protein	3248
0.180688119789519	1.88060340511398	0.433169123625646
	0.510437990778232	0.866868039165439
4232	MEST	mesoderm specific transcript	4631
0.205641475485074	3.29650202615338	0.43287252830925
	0.51058279815741	0.866868039165439
729873	TBC1D3	TBC1 domain family member 3
2116	-0.288936519593716	1.04023441569164
0.432647308772383	0.510692805074623	0.866868039165439
143888	POGLUT3	protein O-glucosyltransferase 3	4993
0.135316340938924	4.31475808909888	0.432545290696504
	0.510742648594268	0.866868039165439
1642	DDB1	damage specific DNA binding protein 1	13950
0.092237624821094	8.29856330318427	0.432504432899471
	0.510762613069273	0.866868039165439
1611	DAP	death associated protein	3361
0.119156273524092	6.95627018238251	0.432451058674189
	0.510788695521395	0.866868039165439
9482	STX8	syntaxin 8	3779	0.140049539460643
4.17463053022009	0.432409237889583	0.510809133749719
	0.866868039165439

5465	PPARA	peroxisome proliferator activated receptor alpha
	11986	-0.148303692349721	5.38635726608795
0.432317317428947	0.510854061157935	0.866868039165439
22931	RAB18	RAB18, member RAS oncogene family
6223	-0.102489043576853	7.37036911187874
0.432217075212954	0.510903063739125	0.866868039165439
643866	CBLN3	cerebellin 3 precursor	2562	-0.322414351922423
	2.24494329410612	0.432185836915854
0.510918335986515	0.866868039165439
3133	HLA-E	major histocompatibility complex, class I, E
2975	-0.113683644555554	8.3497139253094 0.432123137038616
	0.510948992028976	0.866868039165439
9910	RABGAP1L		RAB GTPase activating protein 1 like
16071	0.18099257826697	6.3872459292377 0.43186627149862
	0.511074615343451	0.866948662773973
84520	GON7	GON7 subunit of KEOPS complex	1154
0.15649859070861	2.46715411209185	0.431772883701505
	0.511120301077138	0.866948662773973
55812	SPATA7	spermatogenesis associated 7	6872
0.15358329681818	2.56829475222561	0.431302094227418
	0.511350721193585	0.867234477752578
1718	DHCR24	24-dehydrocholesterol reductase 4250
0.153671309178578	9.54390056260346	0.430800887396202
	0.511596226483602	0.86744529083357
101928269	LOC101928269	uncharacterized LOC101928269	1748
	0.220467282879437	0.843634579328218
0.430795467276178	0.511598882528459	0.86744529083357
548644	POLR2J3 RNA polymerase II subunit J3
8814	-0.219255051053844	5.57413495660739
0.430313973211341	0.511834926590569	0.867684652068925
8895	CPNE3	copine 3	6793	-0.0932291077838015
6.98258357540866	0.430254806286435	0.511863945173522
	0.867684652068925
51622	CCZ1	CCZ1 homolog, vacuolar protein trafficking and biogenesis associated	6431	-0.0952942171548962
5.48338804513559	0.429962864397172	0.512007170677642
	0.867743398036248
105371433	LOC105371433	uncharacterized LOC105371433	484
	0.211052562364772	1.6546995915073 0.429931622493718
	0.512022501946854	0.867743398036248
286097	MICU3	mitochondrial calcium uptake family member 3	4606
	0.189565567028243	1.66624272859484
0.429498070221618	0.512235340343411	0.867999082045748
9793	CKAP5	cytoskeleton associated protein 5
8538	-0.0791730890263741	7.09727209654385
0.429101550648078	0.512430133230344	0.868170539201871
54996	MARC2	mitochondrial amidoxime reducing component 2
2597	-0.189062452690886	2.34112259325356
0.42903978959311	0.512460485337408	0.868170539201871
10919	EHMT2	euchromatic histone lysine methyltransferase 2	7289
	0.120039591254235	5.68150971397066
0.428899510792704	0.512529436128254	0.868182345126189
54968	TMEM70	transmembrane protein 70	2610
0.115453268312297	3.9691714023311 0.428471515723619

0.512739906791397	0.868347193997767
6844	VAMP2	vesicle associated membrane protein 2
3298	-0.0960366087437108	5.03671518032295
0.428449490671978	0.512750741883025	0.868347193997767
8906	AP1G2	adaptor related protein complex 1 subunit gamma 2
	6897	-0.0944041807273731	6.66180070783538
0.427905489117805	0.513018486506902	0.8686185449574
5728	PTEN	phosphatase and tensin homolog
10129	-0.0836217168526443	5.93088064335069
0.42778417316962	0.513078228470763	0.8686185449574
757	TMEM50B transmembrane protein 50B	4495
0.0950445483223978	4.75844175898213	0.427595872389822
	0.513170981033926	0.8686185449574
80700	UBXN6	UBX domain protein 6	3401	0.0992275639570483
	6.42877945319905	0.427540746799252
0.513198140128546	0.8686185449574
51507	RTF2	replication termination factor 2	4063
0.0792631302867588	6.36459754949288	0.427494274450176
	0.513221037914432	0.8686185449574
54964	C1orf56 chromosome 1 open reading frame 56	2331
0.206536160413434	0.39839264479986	0.426937934411554
	0.513495294968646	0.868977720988985
79712	GTDC1	glycosyltransferase like domain containing 1
14162	0.137447031664373	3.91451688793001
0.426699951954292	0.51361269024351	0.869071388287127
113277	TMEM106A	transmembrane protein 106A
3577	-0.220907134635577	2.8190374089438 0.426522411248057
	0.513700300399568	0.86911464037888
55746	NUP133	nucleoporin 133 6080	0.076629479557043
5.73955181337799	0.425985856222242	0.513965229796551
	0.869270833260238
387522	TMEM189-UBE2V1	TMEM189-UBE2V1 readthrough
2757	-0.0949195921687298	7.32250293260589
0.425973649565776	0.513971259722537	0.869270833260238
56704	JPH1	junctophilin 1	4378	0.192639788127288
2.98424575743658	0.42590420573132	0.514005566397662
	0.869270833260238
51201	ZDHHC2	zinc finger DHHC-type palmitoyltransferase 2
6735	-0.101847261458757	6.02241034764971
0.425675446342375	0.514118606429957	0.869270833260238
516	ATP5MC1 ATP synthase membrane subunit c locus 1 2615
0.153430382357121	3.90384772832007	0.425583471627618
	0.514164067370234	0.869270833260238
1235	CCR6	C-C motif chemokine receptor 6
6528	-0.184308307997265	3.43768248078466
0.425581628453614	0.514164978479409	0.869270833260238
30834	ZNRD1	zinc ribbon domain containing 1
3672	-0.137642950326191	3.14559933572139
0.424807620358772	0.514547831349155	0.869643287181958
23051	ZHX3	zinc fingers and homeoboxes 3	11621
0.0934940376601118	6.42747095578648	0.424656116287053
	0.514622828988842	0.869643287181958
202051	SPATA24 spermatogenesis associated 24	1204
0.209074194325634	0.196788398445826	0.424650109668872

0.514625802781496	0.869643287181958
4312	MMP1	matrix metallopeptidase 1
1970	-0.533279597589831	6.90693546591067
0.424551520759451	0.514674617054081	0.869643287181958
23266	ADGRL2	adhesion G protein-coupled receptor L2
7540	-0.142882244226816	6.28875213902147
0.424487612756593	0.514706264101906	0.869643287181958
9595	CYTIP	cytohesin 1 interacting protein 3428
0.488683157529694	0.933400256639663	0.424355322242336
0.514771784744828	0.869643287181958
158293	FAM120AOS	family with sequence similarity 120A
opposite strand 2771	0.126352183562887	5.49747063617994
0.424258222165655	0.514819885579553
0.869643287181958
55184	DZANK1	double zinc ribbon and ankyrin repeat domains 1 6619
0.15342636824857	2.47865492955003
0.423918665360382	0.514988154791863	0.86982263163748
7226	TRPM2	transient receptor potential cation channel
subfamily M member 2	6705	0.686059328628394
1.29006985361818	0.423733235106134	0.515080086272561
0.869825448203306
55847	CISD1	CDGSH iron sulfur domain 1	2857
0.13055061748914	4.0648418664816 0.423664801608112
0.515114021049934	0.869825448203306
54480	CHPF2	chondroitin polymerizing factor 2	4205
0.0792007057329581	6.85876819483809	0.422724905818079
0.515580491434375	0.87039729416099
51076	CUTC	cutC copper transporter 1806	0.134239852173665
3.23609661255174	0.422676907573289
0.515604332765361	0.87039729416099
79643	CHMP6	charged multivesicular body protein 6	2118
0.095461642982133	4.72062009723557	0.422606938061469
0.515639090950803	0.87039729416099
10811	NOXA1	NADPH oxidase activator 1	1678
0.192780782777593	2.43432729807322	0.422138776382827
0.515871761196511	0.870685113758228
65084	TMEM135 transmembrane protein 135	5030
0.133694087378105	3.59085023615065	0.421961265422598
0.5159600298053 0.870729173190703
29948	OSGIN1	oxidative stress induced growth inhibitor 1
3460	-0.207382258886732	4.01863048132075
0.421632172691965	0.516123743411	0.87089765833004
79095	C9orf16 chromosome 9 open reading frame 16
1662	-0.11472474437295	4.70129761088597
0.421510651870621	0.516184219280074	0.87089765833004
9789	SPCS2	signal peptidase complex subunit 2
2982	-0.114130223475188	5.24056112257994
0.421237954275114	0.516319974673002	0.870899715262453
388	RHOB	ras homolog family member B
2372	-0.188940143276616	6.31855380522159
0.421168322342069	0.516354649126069	0.870899715262453
51550	CINP	cyclin dependent kinase 2 interacting protein	4786
0.123560772305103	3.7966466865631 0.421133549235357
0.516371966534924	0.870899715262453

868	CBLB	Cbl proto-oncogene B	7380	-0.192713833534148
	5.88416062601613	0.420888292295047
0.516494136187101	0.870904676057385
7415	VCP	valosin containing protein
5438	-0.0840308644118878	9.29662168450574
0.420878007084639	0.516499260656007	0.870904676057385
8549	LGR5	leucine rich repeat containing G protein-coupled
receptor 5	5251	-0.402130517441521	4.458086415204
0.420599008169539	0.516638302094148	0.870962044965187
90196	SYS1	SYS1 golgi trafficking protein
6117	-0.0919452785001796	5.56145792386112
0.420560205606407	0.516657644864406	0.870962044965187
6536	SLC6A9	solute carrier family 6 member 9
5159	-0.195603870331111	6.32093109614126
0.420354112575239	0.516760401857933	0.871030439088335
253264	ZNF503-AS1	ZNF503 antisense RNA 1
3272	-0.339119607837902	0.590769106780127
0.420169703587545	0.516852377626447	0.871061867627745
5202	PFDN2	prefoldin subunit 2	677	-0.105756770837813
	4.98536440943082	0.419785824319312
0.517043933105272	0.871061867627745
5433	POLR2D	RNA polymerase II subunit D
5226	-0.107213202360373	4.76958229487969
0.419742316066964	0.517065651537099	0.871061867627745
79922	MRM1	mitochondrial rRNA methyltransferase 1	1857
0.16335003706778	1.71098221500015	0.419683438448892
	0.517095044587403	0.871061867627745
874	CBR3	carbonyl reductase 3	1169	-0.209968290399984
	0.36964972900667	0.419591946392692
0.517140725318143	0.871061867627745
113146	AHNAK2	AHNAK nucleoprotein 2	18771	-0.223457612391209
	10.1850276021345	0.419528207231451
0.517172553587093	0.871061867627745
22905	EPN2	epsin 2	7368	-0.0915416566934302
5.81547775474623	0.419444494529412	0.517214360895263
	0.871061867627745
84245	MRI1	methylthioribose-1-phosphate isomerase 1	3456
	0.15389262646165	4.60267270354952
0.419268859677983	0.517302094675024	0.871104885803445
4013	VWA5A	von Willebrand factor A domain containing 5A
6403	-0.115133596995707	5.66591423312717
0.419138108058572	0.517367425151028	0.87111017238736
6461	SHB	SH2 domain containing adaptor protein B	3986
0.0785156062422295	6.23325630757821	0.418968233419378
	0.517452325174587	0.87114840369236
100113407	TMEM170B	transmembrane protein 170B	8362
	-0.168219727815717	2.24057626018121
0.418761640536121	0.517555609176726	0.871217572135369
540	ATP7B	ATPase copper transporting beta	9425
0.115663966976959	4.37658504829302	0.418460736858673
	0.517706107579199	0.871257460384286
56977	STOX2	storkhead box 2	10952	0.166166805415129
3.5353010702754	0.418350373299575	0.517761325675747
0.871257460384286

10788 IQGAP2 IQ motif containing GTPase activating protein 2 7904
0.54141503373924 0.753996186807144
0.418341211770827 0.51776590991924 0.871257460384286
10550 ARL6IP5 ADP ribosylation factor like GTPase 6 interacting protein 5 2522 -0.118008441980597 8.44183488117542
0.417778376823605 0.518047677758836
0.871537243060691
4711 NDUFB5 NADH:ubiquinone oxidoreductase subunit B5
5871 -0.102767461750148 5.38626911770212
0.417760522036829 0.518056620669901 0.871537243060691
399726 MIR1915HG MIR1915 host gene 3801
0.198593677720185 3.03312625361384 0.417225972521631
0.518324485565005 0.871597177447093
23609 MKRN2 makorin ring finger protein 2
2848 -0.0922403915085121 5.25837007066229
0.417169787975034 0.51835265398785 0.871597177447093
57597 BAHCC1 BAH domain and coiled-coil containing 1
12030 -0.200027367838404 3.67039286142071
0.417043371630328 0.518416043328253 0.871597177447093
84937 ZNRF1 zinc and ring finger 1 6939 0.0975497195496117
4.24637068372483 0.417008996265447
0.518433282628573 0.871597177447093
55763 EXOC1 exocyst complex component 1
5347 -0.0757584595345997 6.06816280455176
0.41699303792571 0.518441286103614 0.871597177447093
83941 TM2D1 TM2 domain containing 1 5516 0.0952547314286674
5.11702146388361 0.41689087911643
0.518492526241796 0.871597177447093
5116 PCNT pericentrin 11660 0.12063334379467
5.39012722510655 0.416820508405181 0.518527827491014
0.871597177447093
811 CALR calreticulin 2714 -0.12388578263063
9.11184415072673 0.416114978823593 0.518881987760192
0.872011892733485
9423 NTN1 netrin 1 6383 -0.30708652176697
7.50414532936061 0.416029284126694 0.518925033560276
0.872011892733485
5156 PDGFRA platelet derived growth factor receptor alpha
10749 0.103419634187755 4.46346296284574
0.415849735440805 0.519015244061392 0.872011892733485
376940 ZC3H6 zinc finger CCCH-type containing 6
11803 -0.172962068471705 3.64532212857549
0.41566013006371 0.519110537275911 0.872011892733485
150967 LINC01963 long intergenic non-protein coding RNA 1963
3148 0.160806845305181 3.04539130806396
0.415549200019999 0.519166303551164 0.872011892733485
145788 C15orf65 chromosome 15 open reading frame 65 843
0.209133940884053 0.982497277787097
0.415513226723875 0.519184390154586 0.872011892733485
7164 TPD52L1 TPD52 like 1 5772 -0.144936435577283
4.83000297524084 0.415399070859461 0.519241792619558
0.872011892733485
11137 PWP1 PWP1 homolog, endonuclein 2876
0.126588653372863 5.78726458613897 0.415337829343343

0.519272592081816	0.872011892733485
4957	ODF2	outer dense fiber of sperm tails 2	6393
0.105629678935795	4.49390989138927	0.415103167975253
0.519390637270694	0.87204442424577
26061	HACL1	2-hydroxyacyl-CoA lyase 1	3055
0.129069253951636	3.47418265782775	0.41489741250232
0.51949418035512	0.87204442424577
7283	TUBG1	tubulin gamma 1	2411	0.0981698970399973
5.47197946810347	0.414890222878242	0.519497799072943
0.87204442424577
6189	RPS3A	ribosomal protein S3A	2341	0.12383519378866
5.56139325061655	0.414684426517439
0.519601400347781	0.87204442424577
101927415	TMED2-DT	TMED2 divergent transcript
8279	-0.200486455647006	-0.0667852258289516
0.414588413281531	0.519649747432987	0.87204442424577
114548	NLRP3	NLR family pyrin domain containing 3
4757	-0.389514561245204	0.699796637252877
0.414557112047252	0.519665510757439	0.87204442424577
2937	GSS	glutathione synthetase	1922	0.0978744779675039
5.42521110183707	0.414423928067966
0.519732591719177	0.87205251703528
10428	CFDP1	craniofacial development protein 1	3402
0.109701180816959	4.63896602890268	0.414010236277072
0.519941053713283	0.872202399494061
54757	FAM20A	FAM20A golgi associated secretory pathway
pseudokinase	6203	0.339553542305773	0.229751690057534
0.413999515584239	0.519946457897866
0.872202399494061
100527950	GJA9-MYCBP	GJA9-MYCBP readthrough
5092	-0.127694902363443	4.77393920162303
0.413792817121649	0.520050671691183	0.872272752769537
81537	SGPP1	sphingosine-1-phosphate phosphatase 1	3312
0.157190777148744	4.48092615795221	0.413053572101873
0.520423687643997	0.872727411996475
83719	YPEL3	yippee like 3	3181	-0.130742631889711
5.17904788938868	0.413008683036857	0.520446353423757
0.872727411996475
7316	UBC	ubiquitin C	3599	0.106850168700344
11.2674145219269	0.412651281693343	0.520626877758379
0.872805605182104
5684	PSMA3	proteasome 20S subunit alpha 3
3119	-0.0897769077028145	5.99308885940671
0.41252356585027	0.520691414142435	0.872805605182104
285331	CCDC66	coiled-coil domain containing 66
9267	-0.14135179922476	3.42055202139028
0.412386562965656	0.520760659072291	0.872805605182104
54407	SLC38A2	solute carrier family 38 member 2
6602	-0.192629756262878	10.5384907467699
0.412355135317512	0.520776545738747	0.872805605182104
100653515	CEP295NL	CEP295 N-terminal like	2758
0.147316109363624	3.96558002198076	0.412214348604838
0.520847723874794	0.872805605182104
100133612	LINC01134	long intergenic non-protein coding

RNA 1134	4064	0.23575989954692	1.10841883492885
	0.412001869561693	0.520955180316068
0.872805605182104
84629	TNRC18	trinucleotide repeat containing 18
12572	-0.110893860956702	7.05829354213797
0.41181645999929	0.521048978954069	0.872805605182104
29880	ALG5	ALG5 dolichyl-phosphate beta-glucosyltransferase
	2194	0.0951286633239565	5.0744859897653
0.411769110835517	0.521072937659363	0.872805605182104
55654	TMEM127 transmembrane protein 127
6367	-0.0776106427634735	7.51502550104679
0.411727450306994	0.521094019520146	0.872805605182104
1350	COX7C	cytochrome c oxidase subunit 7C 1993
0.11309959935897	6.51594199072133	0.41158215775852
	0.521167555030458	0.872805605182104
2121	EVC	EvC ciliary complex subunit 1	7148
0.117568708531116	5.51096135754324	0.411362910807348
	0.521278555056967	0.872805605182104
51002	TPRKB	TP53RK binding protein	2047	-0.125300863820656
	3.54225997047737	0.411270514165391
0.521325345998809	0.872805605182104
284325	C19orf54	chromosome 19 open reading frame 54	3375
	0.184775819322392	2.75692590574259
0.411209677946555	0.521356158354419	0.872805605182104
84060	RBM48	RNA binding motif protein 48
4353	-0.110579728456748	3.75422151206444
0.411002003898218	0.52146136543677	0.872805605182104
91603	ZNF830	zinc finger protein 830 2246	-0.117752885782174
	3.63998551594379	0.410926117978481
0.521499818384226	0.872805605182104
54677	CROT	carnitine O-octanoyltransferase 4979
0.246404944485194	7.06523980315006	0.410903928276667
	0.521511063306524	0.872805605182104
8313	AXIN2	axin 2	5526	0.271435078606486
1.28438113451806	0.410820052592879	0.521553572279238
	0.872805605182104
7372	UMPS	uridine monophosphate synthetase
7579	-0.101279143552736	5.09880934319814
0.410699655503372	0.521614601340715	0.872805605182104
84973	SNHG7	small nucleolar RNA host gene 7 4345
0.127901931300154	3.79209478122167	0.410530369811198
	0.521700433293529	0.872844955702636
728855	LINC00623	long intergenic non-protein coding RNA 623
	2075	-0.141444160051415	3.6643034803245
0.409782633816057	0.522079852455481	0.873226640840534
219541	MED19	mediator complex subunit 19	1775
0.121303661439174	3.77044817555315	0.409727013884051
	0.522108094839425	0.873226640840534
23036	ZNF292	zinc finger protein 292 13128	-0.193391168305497
	6.17386125812203	0.409712247237053
0.522115593420395	0.873226640840534
7175	TPR	translocated promoter region, nuclear basket protein
	12350	-0.0910066300515712	7.41665433700323
0.409558780400221	0.522193535966383	0.87325272902114

84105 PCBD2 pterin-4 alpha-carbinolamine dehydratase 2
6587 -0.132723309201823 2.94979299034195
0.409405441078678 0.522271434310956 0.873278737065006
26149 ZNF658 zinc finger protein 658 4843 0.212089737209695
1.54017134128949 0.409279265017545
0.522335548791788 0.873281693951609
84072 HORMAD1 HORMA domain containing 1
2405 -0.959062541956316 1.76135420936598
0.408930548269581 0.522512816462839 0.873433866929082
55364 IMPACT impact RWD domain protein 4918
0.109180353835163 4.98922518041239 0.408752517172687
0.522603358366668 0.873433866929082
54934 KANSL2 KAT8 regulatory NSL complex subunit 2 3386
0.100982582658344 4.88658778019518 0.408529758293948
0.522716686766426 0.873433866929082
79188 TMEM43 transmembrane protein 43 4241
0.0777661177654371 7.47536446212154 0.408482504235373
0.522740732825651 0.873433866929082
51496 CTDSPL2 CTD small phosphatase like 2
9665 -0.126559174437795 4.58883048275463
0.408370403776099 0.522797784953867 0.873433866929082
771 CA12 carbonic anhydrase 12 3655 -0.528464098278727
2.72011140785898 0.408364657652172
0.522800709668981 0.873433866929082
105416157 NKILA NF-kappaB interacting lncRNA
2598 -0.159278993303823 3.28298822720019
0.408236724772311 0.522865833631507 0.873438489168338
90736 FAM104B family with sequence similarity 104 member B 2499
0.188364885768554 1.95877014451267
0.407985679675656 0.522993669385235 0.873545181824379
135293 PM20D2 peptidase M20 domain containing 2
4708 -0.126827057576472 5.32383127197356
0.407866390045305 0.523054432780829 0.873545181824379
8664 EIF3D eukaryotic translation initiation factor 3 subunit D
4070 -0.092899072854888 6.9699874178235
0.407724498899825 0.523126725160728 0.873561759576338
5933 RBL1 RB transcriptional corepressor like 1
5954 -0.164554593899168 2.71917445887993
0.407464441744744 0.523259268138674 0.873606520384501
7094 TLN1 talin 1 10076 0.110495414583039
8.95807642988878 0.407427171601654 0.523278268439063
0.873606520384501
2547 XRCC6 X-ray repair cross complementing 6 3288
0.0903112086610556 7.79106955389526 0.407286483260044
0.523350002310044 0.873622152586913
388677 NOTCH2NLA notch 2 N-terminal like A
6318 -0.157463531699953 2.8379391116783 0.406789644287784
0.523603469087327 0.873911982961534
55277 FGGY FGGY carbohydrate kinase domain containing
12174 0.175798622775077 1.77806847587219
0.406579605624412 0.523710687653551 0.873911982961534
57717 PCDHB16 protocadherin beta 16 5001 0.402556386754107
0.569097027706703 0.406579384268861
0.523710800669602 0.873911982961534

23474	ETHE1	ETHE1 persulfide dioxygenase
1604	-0.0966355725269389	5.6035440238117 0.406142434387249
	0.523933975698976	0.874147911665164
23673	STX12	syntaxin 12	3846	-0.108601777289501
6.53352498178722	0.405978792875715	0.524017600130007
	0.874147911665164
1537	CYC1	cytochrome c1	1975	-0.0925088614915516
5.71419957527472	0.405790782129493	0.524113707035524
	0.874147911665164
5073	PARN	poly(A)-specific ribonuclease	4382
0.084250555510104	5.07775762218358	0.40574647591737
	0.524136359941187	0.874147911665164
51259	TMEM216 transmembrane protein 216	2364
0.161471241756148	2.69489535813212	0.405371880223967
	0.524327952963195	0.874147911665164
3422	IDI1	isopentenyl-diphosphate delta isomerase 1
3589	-0.190234539390292	7.24049784586958
0.405337731085979	0.524345425278996	0.874147911665164
9372	ZFYVE9	zinc finger FYVE-type containing 9	5809
0.0981228915566422	5.06600959218343	0.405203481753716
	0.524414123620868	0.874147911665164
6884	TAF13	TATA-box binding protein associated factor 13
1400	-0.106123401665874	6.42155243006434
0.404996049482534	0.524520302735735	0.874147911665164
9054	NFS1	NFS1 cysteine desulfurase	6024
0.0993108561553419	4.24212406721663	0.404924549513861
	0.524556910540082	0.874147911665164
4848	CNOT2	CCR4-NOT transcription complex subunit 2
13993	-0.0880087687262388	6.09087983130256
0.404922934713873	0.524557737366052	0.874147911665164
64764	CREB3L2 cAMP responsive element binding protein 3 like 2
	8788	-0.132473979306492	6.33382948396647
0.404913870540394	0.524562378537409	0.874147911665164
5991	RFX3	regulatory factor X3	11169	-0.19504931723289
	2.94926934382663	0.404838289923321
0.524601081277745	0.874147911665164
113829	SLC35A4 solute carrier family 35 member A4
4023	-0.0800173050831001	6.38742669311571
0.404617550756242	0.524714144754658	0.87422935191819
57580	PREX1	phosphatidylinositol-3,4,5-trisphosphate dependent
Rac exchange factor 1	7044	0.557698478602609
2.09386696931381	0.404420141758678	0.52481529512611
	0.87422935191819
60490	PPCDC	phosphopantothenoylcysteine decarboxylase
3335	-0.14942432985311	3.47954217723002
0.404377405854961	0.524837197135722	0.87422935191819
3184	HNRNPD	heterogeneous nuclear ribonucleoprotein D
6981	-0.0715845450159195	6.59129285817911
0.404128011119063	0.524965043542264	0.874303004116907
55585	UBE2Q1	ubiquitin conjugating enzyme E2 Q1	4161
0.0797983640586953	6.75777603339155	0.403852799350091
	0.525106188753739	0.874303004116907
64422	ATG3	autophagy related 3	6111	0.0890322271357525
	5.15730524894872	0.403848317841258

0.525108487700659	0.874303004116907
79016	DDA1	DET1 and DDB1 associated 1	4596
0.0831410851592591	6.81839231186513	0.403694961924558
	0.525187167782205	0.874303004116907
51058	ZNF691	zinc finger protein 691 2036	0.194735696664839
	2.08926931907201	0.403682604537016
0.525193508719901	0.874303004116907
374291	NDUFS7	NADH:ubiquinone oxidoreductase core subunit S7	6133
	0.122176452945824	4.42767509661732
0.403396534277762	0.525340337847086	0.874443507097342
1666	DECR1	2,4-dienoyl-CoA reductase 1	3507
0.104099463152755	5.35869023417821	0.402991708886235
	0.52554824443051	0.874685629721739
55010	PARPBP	PARP1 binding protein	4745	0.18214859429431
	1.60370192134782	0.402735183870071
0.525680064088055	0.874752350816453
2054	STX2	syntaxin 2	3723	0.176208764010942
4.23029135138906	0.402514029166426	0.5257937553986
0.874752350816453
1153	CIRBP	cold inducible RNA binding protein	5810
0.0957396915115078	6.45225473523001	0.402321384431886
	0.525892825994996	0.874752350816453
84243	ZDHHC18	zinc finger DHHC-type containing 18
5436	-0.106592611422521	6.29847280451804
0.402109718172589	0.526001717068468	0.874752350816453
285613	RELL2	RELT like 2	2596	0.195612967426033
4.05202078947938	0.402106294734814	0.526003478575981
	0.874752350816453
16	AARS1	alanyl-tRNA synthetase 1
4498	-0.0855073058430202	6.57474443792998
0.402069701302742	0.526022308143933	0.874752350816453
23409	SIRT4	sirtuin 4	1222	0.240016659433222
0.760305359476629	0.402063516172031	0.526025490892195
	0.874752350816453
9658	ZNF516	zinc finger protein 516 8217	-0.0987506675151854
	5.98073900145683	0.401620240745095
0.526253681251163	0.874874079220891
339366	ADAMTSL5	ADAMTS like 5	3625	-0.224616531638921
	3.38422451722317	0.401538904836923
0.526295570703636	0.874874079220891
9804	TOMM20	translocase of outer mitochondrial membrane 20
3542	-0.0867725270194308	7.16410788089525
0.401537903606195	0.526296086392478	0.874874079220891
26220	DGCR5	DiGeorge syndrome critical region gene 5	6905
	0.367608881788333	0.395253986793977
0.401353021099791	0.52639132649156	0.874874079220891
22980	TCF25	transcription factor 25 7799	0.068741121529815
	6.5678570216911 0.401235544650178	0.526451859108197
	0.874874079220891
27020	NPTN	neuroplastin	4032	0.0808167760098064
7.54966033871562	0.401193648805586	0.526473449971649
	0.874874079220891
64215	DNAJC1	DnaJ heat shock protein family (Hsp40) member C1
	2501	0.084620526721087З	5.43422481896443

0.400674879943885	0.526740926476911	0.87510783572108
554203	JPX	JPX transcript, XIST activator	4427
0.100601469378994	4.1890645588775	0.400672424293603
0.526742193183451	0.87510783572108
83444	INO80B	INO80 complex subunit B	1522	0.131868143495397
3.94187619147477	0.400469150934963
0.526847067237095	0.87510783572108
100303453	TSNAX-DISC1	TSNAX-DISC1 readthrough (NMD candidate)	3890	-0.107055583724135	3.73371180614336
0.400226256007175	0.52697243198977
0.87510783572108
148932	MOB3C	MOB kinase activator 3C	5916	-0.128333746631456
4.30149770228055	0.400099665137226
0.527037790135964	0.87510783572108
716	C1S	complement C1s	6462	0.411754005841719
3.56322224708389	0.400048388911236	0.527064267879365
0.87510783572108
9519	TBPL1	TATA-box binding protein like 1
5039	-0.0957275269752785	4.15429989079989
0.399872122506054	0.527155305507053	0.87510783572108
55735	DNAJC11	DnaJ heat shock protein family (Hsp40) member C11
5083	0.0758722176991551	6.01807803013327
0.399763223029389	0.527211563671973	0.87510783572108
79990	PLEKHH3	pleckstrin homology, MyTH4 and FERM domain containing H3	4803	0.138688826355851	4.70313069238861
0.399710334569029	0.527238890057911
0.87510783572108
29998	BICRA	BRD4 interacting chromatin remodeling complex associated protein	6144	-0.11989686699747
3.81046551931082	0.399622534374029	0.527284260211092
0.87510783572108
728113	ANXA8L1	annexin A8 like 1	3124	0.0907627693125236
8.20068056453876	0.399589448627125
0.52730135885528	0.87510783572108
343099	CCDC18	coiled-coil domain containing 18	5633
0.161125348707554	1.43364058002193	0.399286772122836
0.527457827439433	0.875231782812696
64122	FN3K	fructosamine 3 kinase	3834	0.221864973308697
2.18680217503707	0.399084324695897
0.527562528935229	0.875231782812696
348793	WDR53	WD repeat domain 53	1797	0.140792535670419
3.083594763732	0.398820469822477	0.527699044836881
0.875231782812696
57215	THAP11	THAP domain containing 11	2114
0.128364229374936	3.89474444781897	0.398658705889353
0.527782771134553	0.875231782812696
317749	DHRS4L2	dehydrogenase/reductase 4 like 2	2516
0.154920596043551	2.3192137462612	0.398558062128277
0.527834874646805	0.875231782812696
5429	POLH	DNA polymerase eta	3540	0.135487734707905
4.92729715773172	0.398477853342682
0.527876405512554	0.875231782812696
114112	TXNRD3	thioredoxin reductase 3	3055	0.170692973152062
3.39243760571173	0.398430510457794

0.527900921667855	0.875231782812696
54433	GAR1	GAR1 ribonucleoprotein	2046	-0.133962290859724
	4.06059070298698	0.397763723926076
0.528246428570343	0.875231782812696
84448	ABLIM2	actin binding LIM protein family member 2	6049
	0.19368576460238	1.68695778841529
0.397722096209603	0.528268012109271	0.875231782812696
84971	ATG4D	autophagy related 4D cysteine peptidase 2278
0.0960098228001097	4.85782181743375	0.397709136754509
	0.528274731774483	0.875231782812696
1490	CCN2	cellular communication network factor 2 2339
0.189845259596792	8.44764037445144	0.397598425201778
	0.528332143556802	0.875231782812696
10682	EBP	EBP cholestenol delta-isomerase
2409	-0.125395948802758	6.33808297421781
0.39744452466698	0.528411970446553	0.875231782812696
60526	LDAH	lipid droplet associated hydrolase	4433
0.130828932995673	4.14948249317495	0.39722429084101
	0.52852624142261	0.875231782812696
84861	KLHL22	kelch like family member 22
3989	-0.110547427497499	4.81103840528754
0.397211555095767	0.528532850871291	0.875231782812696
79751	SLC25A22	solute carrier family 25 member 22
4647	-0.137697456802546	3.88711729734715
0.397193791532633	0.528542069845579	0.875231782812696
7710	ZNF154	zinc finger protein 154 5713	0.301319214514752
	2.65095974345851	0.397088462229252
0.528596739804443	0.875231782812696
55245	UQCC1	ubiquinol-cytochrome c reductase complex assembly
factor 1	6782	0.11750665474304	4.95921576169722
	0.397064224292368	0.528609321659472
0.875231782812696
92092	ZC3HAV1L	zinc finger CCCH-type containing, antiviral
1 like	1753	-0.213042943526282	1.50827408452428
0.39699487660908	0.528645322839494	0.875231782812696
5526	PPP2R5B protein phosphatase 2 regulatory subunit B'beta
2855	-0.0863568293199072	6.07680111341614
0.396916804357062	0.528685858551875	0.875231782812696
10533	ATG7	autophagy related 7	8833	-0.101763491011651
	4.5997548467153 0.396734376967419	0.528780597966201
	0.875231782812696
721	C4B	complement C4B (Chido blood group)
12576	-0.170485709046197	1.24813068327167
0.396732295345437	0.528781679183623	0.875231782812696
25915	NDUFAF3 NADH:ubiquinone oxidoreductase complex assembly
factor 3	2434	0.116775977181148	3.80963068359189
	0.396563843343703	0.528869188146929
0.875231782812696
80227	PAAF1	proteasomal ATPase associated factor 1	7937
0.15293012776621	3.11415980075867	0.396529214703566
	0.528887180561199	0.875231782812696
55727	BTBD7	BTB domain containing 7 11333	-0.0964527702469072
	6.53305539916451	0.396252803365281
0.529030838230643	0.875231782812696

64430	PCNX4	pecanex 4	22042	0.112043924514163
6.08520325104061	0.396211392504547	0.529052366483697
	0.875231782812696
724102	SNHG4	small nucleolar RNA host gene 4
1926	-0.219878014053785	0.349250552895386
0.39620549972358	0.529055430089822	0.875231782812696
6205	RPS11	ribosomal protein S11	1390	0.116238709275418
	8.20533476190086	0.39612626263004 8
0.529096627862822	0.875231782812696
51362	CDC40	cell division cycle 40	5809	-0.0835943301431239
	5.02426110755178	0.396070483316258
0.529125632673514	0.875231782812696
56937	PMEPA1	prostate transmembrane protein, androgen induced 1
	5619	0.102258235702933	9.2684481064525
0.395840043349651	0.529245489899594	0.875326671038329
9467	SH3BP5	SH3 domain binding protein 5
4159	-0.115825420508997	3.76574603838456
0.395562342912704	0.529389993123189	0.875377446592641
728489	DNLZ	DNL-type zinc finger	3144	-0.171313390884305
	2.39288826106077	0.395540847104087
0.52940118155824	0.875377446592641
28512	NKIRAS1 NFKB inhibitor interacting Ras like 1
3473	-0.106810360449128	4.01199874219638
0.395317928296437	0.529517234435238	0.875442169093413
6468	FBXW4	F-box and WD repeat domain containing 4 5038
0.099868645711287	5.14370238438225	0.395225580962471
	0.52956532438984	0.875442169093413
7716	VEZF1	vascular endothelial zinc finger 1
4913	-0.0827436223112071	6.76000306570186
0.395055499262979	0.529653915100032	0.875485294879178
55787	TXLNG	taxilin gamma	4460	-0.0958113844114234
4.13816250905664	0.394765588163343	0.529804982870287
	0.875611305541362
3098	HK1	hexokinase 1	4940	0.0846136054140592
7.89357585134773	0.394610254561528	0.529885956418477
	0.875611305541362
29952	DPP7	dipeptidyl peptidase 7	2820	0.111872221520522
	4.77418721572367	0.394549397259153
0.529917686661964	0.875611305541362
50814	NSDHL	NAD(P) dependent steroid dehydrogenase-like
1996	-0.146291436629865	5.74866136965825
0.393947746932675	0.530231563457319	0.876026599356767
25914	RTTN	rotatin 9042	0.153866998656324
4.80824982703508	0.39376289909552	0.530328064384646
	0.876064681144596
80216	ALPK1	alpha kinase 1	12325	0.143697249275799
3.56109282286727	0.39366401051344	0.530379702727648
	0.876064681144596
29058	TMEM230 transmembrane protein 230	3051
0.0913141376304768	5.9911063943922 0.393420295034467
0.530507006409575	0.876121028173915
100506686	IQCH-AS1	IQCH antisense RNA 1	6504
0.215472425478524	1.6343503124624 0.393359227307826
0.530538913469776	0.876121028173915

29929	ALG6	ALG6 alpha-1,3-glucosyltransferase
4950	-0.147557095881761	3.24355161267438
0.393176737009839	0.530634282748436	0.876175220848444
54682	MANSC1	MANSC domain containing 1
6015	-0.150495190565927	4.42824646064127
0.393007522139353	0.530722741855138	0.87621799212222
64925	CCDC71	coiled-coil domain containing 71	1781
0.107948868142015	4.25495403190987	0.392486038677691
	0.53099552054582	0.876536215405194
26099	SZRD1	SUZ RNA binding domain containing 1
4747	-0.0779960350824482	7.47571728522743
0.392399817387769	0.531040645672055	0.876536215405194
388969	C2orf68	chromosome 2 open reading frame 68	4413
0.110898476919169	4.25075741498324	0.392117400896105
	0.531188500670816	0.876676956391672
79583	TMEM231	transmembrane protein 231	5403
0.137230208695632	3.51723066334482	0.391897112695716
	0.531303880783531	0.876764073767073
285381	DPH3	diphthamide biosynthesis 3
3562	-0.101841457102639	5.56379474970096
0.391581463664508	0.531469286634356	0.87693371396954
342909	ZNF284	zinc finger protein 284	6209	-0.107047105747796
	3.7764563897167	0.391364558132856	0.531583002877075
	0.876933716537969
51668	HSPB11	heat shock protein family B (small) member 11	3236
	0.103735585687788	3.87546863326553
0.391285283120813	0.531624575028784	0.876933716537969
26284	ERAL1	Era like 12S mitochondrial rRNA chaperone 1	2318
	0.0880054846002061	5.41959038668469
0.391184247286269	0.531677567141015	0.876933716537969
8682	PEA15	proliferation and apoptosis adaptor protein 15
2806	-0.104750175610755	9.14517389419752
0.391103899390956	0.531719715467764	0.876933716537969
105376382	LINC02657	long intergenic non-protein coding
RNA 2657	697	0.242083638964735	1.70067180818801
	0.390882396290369	0.531835941182306
0.877022136583537
7099	TLR4	toll like receptor 4	5256	-0.434799989051066
	2.9176948667147	0.390699271936821	0.531932063633796
	0.877077388501304
57621	ZBTB2	zinc finger and BTB domain containing 2
3090	-0.114068292385727	4.61089266253855
0.390381874226627	0.532098740750583	0.877248947939432
10155	TRIM28	tripartite motif containing 28	4034
0.0827577542365135	7.59286891346594	0.390201049611676
	0.532193740482371	0.877302309395855
55110	MAGOHB	mago homolog B, exon junction complex subunit
4856	-0.178867163222595	2.92724874422019
0.389922766250319	0.532340001866137	0.877412322654624
51665	ASB1	ankyrin repeat and SOCS box containing 1
7410	-0.0944573332918198	5.60669717392372
0.389782287602632	0.532413862802423	0.877412322654624
30832	ZNF354C	zinc finger protein 354C
5411	-0.135666895476037	3.06420933359366

0.38971661168023	0.532448400268745	0.877412322654624
254552	NUDT8	nudix hydrolase 8	1475	0.199721173700453
	0.740043494170715	0.3893992190455	0.532615366820871
	0.877415925673087
22870	PPP6R1	protein phosphatase 6 regulatory subunit 1
6583	-0.0818779018430366	7.1983387350058	0.389034118925691
	0.532807547186525	0.877415925673087
100499177	THAP9-AS1	THAP9 antisense RNA 1
4436	-0.197237007676193	4.57796862268051
0.389008904240773	0.532820824244605	0.877415925673087
57446	NDRG3	NDRG family member 3	3248	0.0835231832931984
	5.88071115077699	0.388900802279885
0.532877753246329	0.877415925673087
51604	PIGT	phosphatidylinositol glycan anchor biosynthesis
class T	2428	0.0723049869288864	6.44632409217863
0.388773241041625	0.532944944091458	0.877415925673087
7388	UQCRH	ubiquinol-cytochrome c reductase hinge protein	1408
	0.10253268407956	5.81779272615222
0.38863365844329	0.533018484538816	0.877415925673087
221937	FOXK1	forkhead box K1	13093	-0.128649563799873
6.10724983926699	0.388570252251895	0.533051896764053
	0.877415925673087
85007	PHYKPL	5-phosphohydroxy-L-lysine phospho-lyase	8402
0.10516910588555	4.23838550810414	0.388568105275242
	0.533053028190921	0.877415925673087
157922	CAMSAP1	calmodulin regulated spectrin associated protein 1
	8642	-0.111326242635318	7.21218111819665
0.388537188020621	0.533069321635872	0.877415925673087
1337	COX6A1	cytochrome c oxidase subunit 6A1	861
0.0845103413472786	6.80023175968455	0.388522620268901
	0.533076999177409	0.877415925673087
196294	IMMP1L	inner mitochondrial membrane peptidase subunit 1
	2854	-0.164644459419798	2.08334585529312
0.388307838138761	0.533190217520884	0.877494949719843
23254	KAZN	kazrin, periplakin interacting protein
12644	-0.118870501432903	4.09442890717882
0.387981427374306	0.533362362048914	0.877494949719843
10513	APPBP2	amyloid beta precursor protein binding protein 2
	6617	-0.0840575701806673	5.81964952294359
0.387860268769581	0.533426285001793	0.877494949719843
25940	FAM98A	family with sequence similarity 98 member A
4921	-0.107583212394617	5.80911425742969
0.387762516332934	0.533477869024063	0.877494949719843
6161	RPL32	ribosomal protein L32	3720	0.133543654805505
	7.08880683387681	0.387747263376014
0.533485918832832	0.877494949719843
54828	BCAS3	BCAS3 microtubule associated cell migration factor
	8006	0.109556675174105	4.35458028773948
0.387718893321479	0.533500891826529	0.877494949719843
10171	RCL1	RNA terminal phosphate cyclase like 1	2785
0.161696378422213	3.09515023161842	0.387315272622951
	0.533713994953904	0.877742389024227
1647	GADD45A	growth arrest and DNA damage inducible alpha
2069	-0.0935661516874755	8.55204943071049

0.387082504739425	0.533836961481266	0.8778415495971
24147	FJX1	four-jointed box kinase 1	2450
0.153253105623751	4.15897007359025	0.386750506674101
	0.534012438057277	0.877973724703536
80774	LIMD2	LIM domain containing 2 4745	0.127694259609196
	4.02762800174029	0.386470721097234
0.534160398997752	0.877973724703536
8310	ACOX3	acyl-CoA oxidase 3, pristanoyl
3846	-0.0966090432749594	4.9016501987779 0.386435424814215
	0.534179070249644	0.877973724703536
84545	MRPL43	mitochondrial ribosomal protein L43	3573
0.0904124030655495	5.11236243216958	0.38637374515407
	0.534211700786067	0.877973724703536
28965	SLC27A6 solute carrier family 27 member 6
3493	-0.450098458027155	2.78401848417541
0.386337748698125	0.534230745744859	0.877973724703536
100302197	MIR1306 microRNA 1306	109	-0.176653869526861
	0.29695865461372	0.386017403107043
0.534400287981445	0.878022700631926
56259	CTNNBL1 catenin beta like 1	4148	0.0865415491227928
	4.96139042492238	0.385896433944267
0.534464336021937	0.878022700631926
90135	BTBD6	BTB domain containing 6 2392	-0.0940500252558402
	5.61977304170186	0.385848460475302
0.534489739791675	0.878022700631926
197370	NSMCE1	NSE1 homolog, SMC5-SMC6 complex component	3770
	0.135616031323152	3.81516083301428
0.385807776998174	0.534511284949556	0.878022700631926
26145	IRF2BP1 interferon regulatory factor 2 binding protein 1
	2564	0.147122303261428	3.99846260193489
0.385597771799805	0.534622524555456	0.878082080784306
100529855	ZNF625-ZNF20	ZNF625-ZNF20 readthrough (NMD
candidate)	3083	-0.139719932226464	3.20433801671821
	0.385468257599019	0.534691149062877
0.878082080784306
55759	WDR12	WD repeat domain 12	9098	0.091066627249922
	4.99158987834285	0.385344060663318
0.534756971163612	0.878082080784306
84298	LLPH	LLP homolog, long-term synaptic facilitation factor
	7993	-0.1102449694255	4.93725100866529
0.385266302877113	0.534798188846357	0.878082080784306
9404	LPXN	leupaxin	2467	-0.294956879478583
3.82453189259184	0.385099337715481	0.534886712858754
	0.8781244944928
150275	CCDC117 coiled-coil domain containing 117
4326	-0.106803269140606	4.96012792009457
0.384979912034984	0.534950048006378	0.878125550501035
100996928	FMC1-LUC7L2	FMC1-LUC7L2 readthrough 1661
0.071776098520214	5.99849374691408	0.384581808644828
	0.535161272913526	0.878332809095103
10629	TAF6L	TATA-box binding protein associated factor 6 like
	3149	0.138600872213991	3.61393754020773
0.384505604537191	0.535201722397851	0.878332809095103
55486	PARL	presenilin associated rhomboid like	2034

0.096715064994893	4.89472037497468	0.384117149257875
	0.535408002499277	0.878470162414236
533	ATP6V0B ATPase H+ transporting V0 subunit b
3535	-0.0861230553816778	7.43104383149375
0.384111788825436	0.535410850041432	0.878470162414236
9733	SART3	spliceosome associated factor 3, U4/U6 recycling
protein 6634	-0.0781877310432423	5.70236014079354
0.383826862519953	0.535562246765219	0.878578324667382
11270	NRM	nurim	2045	0.150595057101297
2.58865340370706	0.383751655393954	0.535602221341813
	0.878578324667382
6154	RPL26	ribosomal protein L26	2472	0.132831480414291
	7.14328864117765	0.383545179760858
0.535711996475934	0.878620745288265
84267	C9orf64 chromosome 9 open reading frame 64	2729
0.110713927008581	3.99724936325118	0.383399332807169
	0.535789562318423	0.878620745288265
401431	ATP6V0E2-AS1	ATP6V0E2 antisense RNA 1	3291
0.202368126085715	0.655919223817982	0.383349135029704
	0.535816263743297	0.878620745288265
528	ATP6V1C1	ATPase H+ transporting V1 subunit C1
5851	-0.0805781782470545	6.71233408273925
0.383040926858815	0.535980260231138	0.878674016635241
65243	ZFP69B	ZFP69 zinc finger protein B
2386	-0.195005199123709	1.43271787415407
0.382998744916176	0.536002712193696	0.878674016635241
81490	PTDSS2	phosphatidylserine synthase 2	6113
0.1252398062087 5.22821918443567	0.382934437937578
0.536036943824383	0.878674016635241
6301	SARS1	seryl-tRNA synthetase 1 3836	-0.0876750937692421
	7.4256430071787 0.382386515295085	0.53632877311472
	0.879049511469446
54471	MIEF1	mitochondrial elongation factor 1
7265	-0.0729838984158108	5.90675725525783
0.381773156415328	0.536655797491744	0.879198785754561
140766	ADAMTS14	ADAM metallopeptidase with thrombospondin
type 1 motif 14 5269	-0.461746310255839	5.77381515728148
	0.381637805448463	0.536728011415056
0.879198785754561
23731	TMEM245 transmembrane protein 245	9484
0.0784394718716904	7.48131830738552	0.381629076859491
	0.536732668994254	0.879198785754561
161291	TMEM30B transmembrane protein 30B
4471	-0.0762655349951186	5.90084004310789
0.381628571088152	0.536732938876335	0.879198785754561
63892	THADA	THADA armadillo repeat containing	14220
0.0927802610418521	4.99290269054905	0.381627160619861
	0.536733691510477	0.879198785754561
51070	NOSIP	nitric oxide synthase interacting protein	4248
	0.0874814478629968	5.02059388350025
0.381494516281407	0.536804479895682	0.879211921176196
51280	GOLM1	golgi membrane protein 1	4233
0.102773273000655	7.15207045571639	0.381198524907774
	0.536962503110105	0.87936791547565

105369635	MIR200CHG	MIR200C and MIR141 host gene	350	-0.187082822609357	1.00591191867793	0.380912680149404	0.53711518972269	0.879476665332896
10641	NPRL2	NPR2 like, GATOR1 complex subunit	3677	0.116304383916456	3.53648348957099	0.38082516489885	0.537161952619118	0.879476665332896
55845	BRK1	BRICK1 subunit of SCAR/WAVE actin nucleating complex	1233	-0.100632067703778	6.96095355471259	0.380658903452797	0.537250813195819	0.879476665332896
25976	TIPARP	TCDD inducible poly(ADP-ribose) polymerase	5048	0.1448241971021 7.0885767566159 0.380604189239762	0.537280061772942	0.879476665332896
2768	GNA12	G protein subunit alpha 12	6120	0.108793078574872	6.58578727177215	0.380402932017109	0.537387672826906	0.879543857190717
207063	DHRSX	dehydrogenase/reductase X-linked	7240	0.102554412248239	5.23653221521947	0.380292576039404	0.537446696201567	0.879543857190717
950	SCARB2	scavenger receptor class B member 2	5685	0.0800123995495044	8.68635976340387	0.379787491122563	0.537716989437664	0.879863836336438
54856	GON4L	gon-4 like	11967	-0.0833147082137927	6.20961668764725	0.37969249830337	0.537767851993735	0.879863836336438
11052	CPSF6	cleavage and polyadenylation specific factor 6	7190	-0.0699150468771686	6.17098704907542	0.379416892758615	0.537915470765182	0.879886700846482
259232	NALCN	sodium leak channel, non-selective	8951	-0.330141055710965	0.865649880769013	0.379351146666365	0.537950696357225	0.879886700846482
6231	RPS26	ribosomal protein S26	1943	0.148317245646792	1.95630388198979	0.379314599359364	0.537970279578016	0.879886700846482
51451	LCMT1	leucine carboxyl methyltransferase 1	2606	-0.102948640390324	4.83245667364133	0.379100474854115	0.538085040517031	0.879971647696679
83786	FRMD8	FERM domain containing 8	4091	0.0792815480733401	6.32644010756318	0.378927644611115	0.538177702223163	0.880020438365613
64097	EPB41L4A	erythrocyte membrane protein band 4.1 like 4A	8582	-0.116218047603804	5.07816886230057	0.378551615683122	0.538379408168204	0.880247504402012
55160	ARHGEF10L	Rho guanine nucleotide exchange factor 10 like	7817	-0.132722108149507	5.38175194985513	0.378160452133386	0.538589379093586	0.880409119130342
1075	CTSC	cathepsin C	11421	-0.280315334726712	6.56424639386626	0.378049884634265	0.538648757255945	0.880409119130342
65065	NBEAL1	neurobeachin like 1	16376	-0.114443661506792	6.77544802143566	0.377969950574661	0.538691691767727	0.880409119130342
10417	SPON2	spondin 2	4348	-0.207911501434776	3.69362685196001	0.377765647501192	0.53880145631211	0.880409119130342

889	KRIT1	KRIT1 ankyrin repeat containing
6204	-0.0983814604936984	4.92553979480244
0.377486544155296	0.538951474409845	0.880409119130342
2852	GPER1	G protein-coupled estrogen receptor 1	3149
0.334216516278905	2.47552809790636	0.377402178096005
	0.538996836218063	0.880409119130342
199953	TMEM201 transmembrane protein 201	6966
0.131933087758045	3.25001167493931	0.377175018430671
	0.539119009790682	0.880409119130342
1660	DHX9	DExH-box helicase 9	5645	-0.0734118003394374
	7.48550659926416	0.376922678955353
0.539254785269404	0.880409119130342
79641	ROGDI	rogdi atypical leucine zipper	3683
0.0946098431857818	5.07125074012444	0.376642488812654
	0.539405619606659	0.880409119130342
8578	SCARF1	scavenger receptor class F member 1	4961
0.292207956421134	0.908500566733407	0.376504835699766
	0.539479750494643	0.880409119130342
8161	COIL	coilin	3030	-0.0894620201689793
4.53151219818926	0.376054430982838	0.539722439208916
	0.880409119130342
203197	TMEM268 transmembrane protein 268	4792
0.109647115024082	3.83261748190994	0.375965132194253
	0.539770579263246	0.880409119130342
7763	ZFAND5	zinc finger AN1-type containing 5
7053	-0.0779843977154157	7.51602228829094
0.37584657894069	0.539834502253104	0.880409119130342
3600	IL15	interleukin 15	7860	-0.153932533742508
3.74347713023172	0.375718435284767	0.539903611914265
	0.880409119130342
7111	TMOD1	tropomodulin 1	3708	0.267615446489192
1.45600976619861	0.37571182938165	0.539907175009397
	0.880409119130342
79885	HDAC11	histone deacetylase 11	5281	0.107684533025786
	4.67175010715693	0.375701697715959
0.539912639915737	0.880409119130342
5801	PTPRR	protein tyrosine phosphatase receptor type R
5142	-0.235337241436525	0.945051701194814
0.375614894072455	0.539959464978365	0.880409119130342
55106	SLFN12	schlafen family member 12
3560	-0.133681560567113	3.97166246251399
0.375534476308637	0.540002851905904	0.880409119130342
5158	PDE6B	phosphodiesterase 6B	5086	0.244727068859443
	1.06340477110829	0.375528495335063
0.540006079011271	0.880409119130342
54851	ANKRD49 ankyrin repeat domain 49
3526	-0.128516020650571	3.73737281137323
0.375211735982396	0.540177040815743	0.880409119130342
2743	GLRB	glycine receptor beta	3459	0.163883403099436
	2.77784622507691	0.375058296096896
0.540259891289988	0.880409119130342
22924	MAPRE3	microtubule associated protein RP/EB family member 3
	3459	0.125100424308069	5.1885787629394
0.375025848315779	0.540277414583026	0.880409119130342

63826 SRR serine racemase 3498 -0.140170550698488
3.3814358766497 0.375005133933092 0.540288601846801
0.880409119130342
80153 EDC3 enhancer of mRNA decapping 3 5103
0.0732285952903003 6.04364287668827 0.374964749238774
0.540310413722121 0.880409119130342
100128750 RBPMS-AS1 RBPMS antisense RNA 1
1171 -0.18008404205936 0.681703190931617
0.374911126643455 0.540339377917212 0.880409119130342
90120 TMEM250 transmembrane protein 250 3396
0.076710772798316 5.73516711773241 0.37487541306859
0.540358670151581 0.880409119130342
4931 NVL nuclear VCP like 4813 0.0990272624288933
4.44376776126953 0.374747370157452
0.540427848462976 0.880409119130342
63891 RNF123 ring finger protein 123 7458 -0.077944147437073
5.76036747164548 0.374651000949974
0.54047992499864 0.880409119130342
22888 UBOX5 U-box domain containing 5 4738
0.133081817447873 2.98073520368505 0.374554234915698
0.540532225238529 0.880409119130342
94107 TMEM203 transmembrane protein 203 1557
0.0926735822416679 4.73881050863695 0.374526854253517
0.540547025662225 0.880409119130342
10749 KIF1C kinesin family member 1C
8580 -0.0871381515907831 9.21119606618115
0.374355334724328 0.540639756256248 0.880409119130342
119392 SFR1 SWI5 dependent homologous recombination repair
protein 1 1812 0.176320192491105 2.09440191702887
0.374288547830067 0.540675871927874
0.880409119130342
9435 CHST2 carbohydrate sulfotransferase 2
4582 -0.187057770729629 2.69341939664151
0.374042264735459 0.540809090223716 0.880409119130342
54976 C20orf27 chromosome 20 open reading frame 27 2415
0.157144572635552 2.70760826947163
0.373933624546451 0.540867874520591 0.880409119130342
63943 FKBPL FKBP prolyl isomerase like 1345
0.173181246240751 3.01933598652813 0.373746912925037
0.540968930030454 0.880409119130342
23394 ADNP activity dependent neuroprotector homeobox
7081 -0.0856676891586963 7.31036849235958
0.373675180243083 0.541007763737021 0.880409119130342
642852 LINC00205 long intergenic non-protein coding RNA 205
1549 0.173701418057094 1.96085603524974
0.373648204784333 0.54102236868489 0.880409119130342
3066 HDAC2 histone deacetylase 2 15461 -0.0725932755176255
7.43253655850571 0.373628616333928
0.541032974641335 0.880409119130342
80727 TTYH3 tweety family member 3 6129 0.10573520457246
7.90240224333647 0.373568331688205
0.541065617512776 0.880409119130342
84233 TMEM126A transmembrane protein 126A
971 -0.125775903781513 3.45867607865252

0.373504107691763	0.541100397440403	0.880409119130342
7745	ZKSCAN8 zinc finger with KRAB and SCAN domains 8	7565
	0.164012710158065	5.01454031530992
0.373380984487381	0.541167085189865	0.880409119130342
122525	C14orf28	chromosome 14 open reading frame 28
3864	-0.172858107206959	2.20340375419896
0.373276334619629	0.541223779019874	0.880409119130342
196383	RILPL2	Rab interacting lysosomal protein like 2	6141
	0.129603697698283	4.04411612920726
0.373169950373594	0.541281423633019	0.880409119130342
23509	POFUT1	protein O-fucosyltransferase 1	6426
0.0933231295485681	5.8076066083485 0.373135253714217
0.541300226558869	0.880409119130342
5532	PPP3CB	protein phosphatase 3 catalytic subunit beta	5094
	0.0842023748878362	5.547639010553	0.373069209613262
	0.541336020709006	0.880409119130342
57679	ALS2	alsin Rho guanine nucleotide exchange factor ALS2
	10384	-0.0808050521167556	5.84004191718304
0.373007291617725	0.541369582570831	0.880409119130342
100534012	TNFAIP8L2-SCNM1 TNFAIP8L2-SCNM1 readthrough	1290
	0.0935999071071552	4.6125982188502 0.372645236042249
	0.541565906789748	0.880626150035291
93185	IGSF8	immunoglobulin superfamily member 8
2592	-0.0837295223482159	5.89344581226585
0.372446949330119	0.541673483019614	0.880651817021556
4722	NDUFS3	NADH:ubiquinone oxidoreductase core subunit S3	3670
	0.0784745096495547	5.66048082522179
0.372384381614586	0.541707435957573	0.880651817021556
11078	TRIOBP	TRIO and F-actin binding protein	13026
0.0801258904188312	6.7972652297456 0.371960467702337
0.541937579846393	0.88080870242868
147687	ZNF417	zinc finger protein 417 4204	-0.13536563160719
	3.85629383193398	0.371805092089431
0.542021978704292	0.88080870242868
51100	SH3GLB1 SH3 domain containing GRB2 like, endophilin B1
6881	-0.0805994253766909	7.51557663198605
0.371336417079327	0.542276706046816	0.88080870242868
25804	LSM4	LSM4 homolog, U6 small nuclear RNA and mRNA
degradation associated	3029	0.102176939776971
5.75604767271787	0.371105127386782	0.542402494463331
	0.88080870242868
162387	MFSD6L	major facilitator superfamily domain containing 6
like	2188	-0.383290918577326	1.64193488323776
0.371042075232706	0.542436795113816	0.88080870242868
85025	TMEM60	transmembrane protein 60	1023
0.118950897299888	3.48586036186838	0.370853220624092
	0.542539556768473	0.88080870242868
3927	LASP1	LIM and SH3 protein 1	7061	-0.0882782060180815
	8.81806528512213	0.370698104696345
0.542623986992485	0.88080870242868
53918	PELO	pelota mRNA surveillance and ribosome rescue factor
	5247	0.0826222338298997	5.58023959774865
0.370610327868846	0.5426717750012 0.88080870242868
8425	LTBP4	latent transforming growth factor beta binding

protein 4	8108	-0.0972755562993935	5.78610597121921
	0.370459214750515	0.542754063122796
0.88080870242868
5356	PLRG1	pleiotropic regulator 1 5049	-0.0679945449030284
	6.26831114324784	0.370443565717633
0.542762586063308	0.88080870242868
3326	HSP90AB1	heat shock protein 90 alpha family class B
member 1	2980	-0.102253996004382	9.70830638080595
	0.370390391559567	0.542791548174794
0.88080870242868
23161	SNX13	sorting nexin 13	16418	-0.0848922645302926
	5.77633901117826	0.37034993250484
0.54281358672104	0.88080870242868
9854	C2CD2L	C2CD2 like	8370	-0.125093721698238
4.84034834760943	0.370246061919886	0.54287017386495
	0.88080870242868
965	CD58	CD58 molecule	1783	-0.0953530008510518
4.54703973468094	0.37020087524138	0.542894794287819
	0.88080870242868
3897	L1CAM	L1 cell adhesion molecule
7423	-0.31300806973061	3.79518941412211
0.370011011355693	0.542998266068037	0.88080870242868
22863	ATG14	autophagy related 14	5052	-0.100753552230019
	5.07855945915784	0.369937836851143
0.54303815432696	0.88080870242868
1349	COX7B	cytochrome c oxidase subunit 7B 3024
0.108626851932346	5.83025337644713	0.369904570194947
	0.543056290148535	0.88080870242868
26292	MYCBP	MYC binding protein	1030	-0.147396580812779
	3.61714814344525	0.369800957364617
0.543112783411939	0.88080870242868
79005	SCNM1	sodium channel modifier 1	3042
0.0916034702359259	4.67374034893023	0.369669400289378
	0.543184528457686	0.88080870242868
5701	PSMC2	proteasome 26S subunit, ATPase 2
3538	-0.0726487385666285	6.67398277652336
0.369569159289789	0.543239206926057	0.88080870242868
377	ARF3	ADP ribosylation factor 3
5408	-0.0866413815372086	7.91884267807075
0.369515360198662	0.543268556909988	0.88080870242868
51030	TVP23B	trans-golgi network vesicle protein 23 homolog B
	5862	-0.0748529844492874	6.00170065754244
0.369168987823663	0.543457589743421	0.88080870242868
387509	GPR153	G protein-coupled receptor 153
4082	-0.138458135295883	4.38221165948041
0.369116825308911	0.543486067971198	0.88080870242868
286144	TRIQK	triple QxxK/R motif containing
6724	-0.117343594692454	4.0230170784161 0.369062820244714
	0.543515555045293	0.88080870242868
196792	FAM24B	family with sequence similarity 24 member B	1177
	0.202254420297419	1.07245353139624
0.369052444940796	0.543521220358835	0.88080870242868
155061	ZNF746	zinc finger protein 746 8260	-0.0891104897138015
	4.90290467045532	0.369043232677342

0.543526250699092	0.88080870242868
55149	MTPAP	mitochondrial poly(A) polymerase
7888	-0.0999477347273881	4.11305935635739
0.368967578749643	0.543567564645012	0.88080870242868
57539	WDR35	WD repeat domain 35	7836	0.125861501017674
	3.35144636337094	0.36882033844229
0.543647987911261	0.88080870242868
51233	DRICH1	aspartate rich 1	2231	0.298503685038901
	0.466161239036985	0.368769455785248
0.54367578533848	0.88080870242868
47	ACLY	ATP citrate lyase	5217	-0.115356536393869
	8.41847595929683	0.368416120660982
0.543868886359413	0.88080870242868
10247	RIDA	reactive intermediate imine deaminase A homolog 1567
	0.172081099451228	2.81154009006824
0.368240142955221	0.543965107131757	0.88080870242868
26574	AATF	apoptosis antagonizing transcription factor
2940	-0.0882344201791829	5.27287827896637
0.368231274502763	0.543969957040752	0.88080870242868
55278	QRSL1	glutaminyl-tRNA amidotransferase subunit QRSL1
5453	-0.0968333833980065	4.42301409934527
0.368206888204014	0.543983293638523	0.88080870242868
29843	SENP1	SUMO specific peptidase 1
6342	-0.106479441192114	4.48659279764846
0.368194497028316	0.543990070467344	0.88080870242868
100526693	ARPC4-TTLL3	ARPC4-TTLL3 readthrough
3158	-0.0892942350089094	6.44059779921405
0.36816745452127	0.544004860763226	0.88080870242868
64386	MMP25	matrix metallopeptidase 25
4126	-0.206100007677826	1.03041938418226
0.367910684359096	0.544145332515286	0.880851605408187
10927	SPIN1	spindlin 1	4914	0.0758246992986168
7.55247794364673	0.367592248986654	0.54431963274518
	0.880851605408187
22997	IGSF9B	immunoglobulin superfamily member 9B
5753	-0.303841275142133	0.787786458674045
0.3675474497308	0.544344162553095	0.880851605408187
55471	NDUFAF7	NADH:ubiquinone oxidoreductase complex assembly factor 7	3651	0.107052761398004	3.53934271803543
	0.367503855864166	0.544368034313505
0.880851605408187
1265	CNN2	calponin 2	3016	-0.121715195775356
8.26673462378935	0.367464245960619	0.544389726156661
	0.880851605408187
6388	SDF2	stromal cell derived factor 2	3332
0.0779113575895818	5.84393563228083	0.367412025592838
	0.544418326398169	0.880851605408187
90338	ZNF160	zinc finger protein 160	4344	-0.103032883234728
	5.11618175999191	0.366941830575826
0.544675969766251	0.880851605408187
1487	CTBP1	C-terminal binding protein 1	8361
0.0779305907618248	6.67330600486811	0.366885238348289
	0.544706994690313	0.880851605408187
64710	NUCKS1	nuclear casein kinase and cyclin dependent kinase

substrate 1	6546	0.0748272043263276	7.40224963898105
	0.366869331660382	0.544715715624801
0.880851605408187
79677	SMC6	structural maintenance of chromosomes 6
9490	-0.0802671179451574	5.18715231972382
0.366687313190376	0.544815526714937	0.880851605408187
203238	CCDC171 coiled-coil domain containing 171	15144
0.18983213028598	1.48855293501748	0.366666707823764
	0.544826827943792	0.880851605408187
2052	EPHX1	epoxide hydrolase 1	2508	0.129457390480933
	6.1091780494494 0.366655695303191	0.544832868053844
	0.880851605408187
112597	CYTOR	cytoskeleton regulator RNA	2862
0.167775520715246	4.22898749306927	0.366626496777933
	0.544848883362357	0.880851605408187
4121	MAN1A1	mannosidase alpha class 1A member 1
5014	-0.162477927280988	4.22247835867081
0.366136568137918	0.545117737917328	0.881115212563882
9333	TGM5	transglutaminase 5	3607	-0.271862674096689
	2.15056727028575	0.366066436510168
0.545156243641787	0.881115212563882
440957	SMIM4	small integral membrane protein 4	2482
0.205975497213923	2.16934404339345	0.365905356731421
	0.545244703469851	0.881115212563882
4134	MAP4	microtubule associated protein 4	14943
0.0831246275937614	8.82762145087336	0.365871029351013
	0.545263558399638	0.881115212563882
84138	SLC7A6OS	solute carrier family 7 member 6 opposite
strand	5234	-0.0887081864907138	4.48551955710026
0.365140158602256	0.545665289238002	0.881368929010319
64108	RTP4	receptor transporter protein 4	1554
0.352914815297751	2.25236107733809	0.365109625486863
	0.545682084028908	0.881368929010319
8546	AP3B1	adaptor related protein complex 3 subunit beta 1
	7310	-0.0672045356717685	6.49786453373327
0.36506731876439	0.545705356496445	0.881368929010319
100533952	RBAK-RBAKDN	RBAK-RBAKDN readthrough
1049	-0.189252461489049	1.43378062110988
0.365024272606751	0.54572903760862	0.881368929010319
2001	ELF5	E74 like ETS transcription factor 5
4494	-0.462982717059879	2.25084981392842
0.36501310074523	0.545735183929928	0.881368929010319
79590	MRPL24	mitochondrial ribosomal protein L24	1695
0.150558054140799	3.83544253428321	0.364686637844557
	0.545914847768762	0.881557444396709
55709	KBTBD4	kelch repeat and BTB domain containing 4	3302
	0.097762871313277	4.5097300619475 0.364399447900091
	0.546072989097376	0.881711165220398
9587	MAD2L1BP	MAD2L1 binding protein
1880	-0.0856377055484086	4.90097130557314
0.363965154664033	0.546312294529072	0.881792157309283
8536	CAMK1	calcium/calmodulin dependent protein kinase I
2720	-0.242453197603301	1.93951640552261
0.36387337967799	0.546362889552882	0.881792157309283

10325 RRAGB Ras related GTP binding B 2725
0.124720466166454 3.51657444813521 0.363731003990829
0.546441397677618 0.881792157309283
93109 TMEM44 transmembrane protein 44
4561 -0.103578037987917 4.49510304018954
0.363635266528018 0.546494200557568 0.881792157309283
100129480 MKRN2OS MKRN2 opposite strand 1462
0.161429580154288 1.72835199759578 0.363606200764927
0.546510233315551 0.881792157309283
56654 NPDC1 neural proliferation, differentiation and control 1
2737 -0.14536431508057 5.5841150868514
0.363534846734382 0.546549596108647 0.881792157309283
80055 PGAP1 post-GPI attachment to proteins inositol deacylase 1
2443 0.184636257695706 1.85465873089473
0.363509054296202 0.546563825926979 0.881792157309283
79621 RNASEH2B ribonuclease H2 subunit B 8519
0.155530245381321 3.35382336072483 0.363194509211159
0.546737417443958 0.881911192337348
284040 CDRT4 CMT1A duplicated region transcript 4 3245
0.123343961958288 3.68355763405775 0.363147205910302
0.546763532109483 0.881911192337348
1608 DGKG diacylglycerol kinase gamma 10814
0.219464388928029 1.45748674918248 0.362766506541284
0.546973788720388 0.882084976713933
7852 CXCR4 C-X-C motif chemokine receptor 4 2224
0.385236828651529 1.46272692592886 0.362621691190412
0.547053808325123 0.882084976713933
138162 C9orf116 chromosome 9 open reading frame 116 1864
0.181316485270506 0.475614777275334
0.36257914654115 0.547077321061927 0.882084976713933
26509 MYOF myoferlin 8473 -0.1057171209274
10.3521174990334 0.362463515406034 0.547141235292768
0.882084976713933
57727 NCOA5 nuclear receptor coactivator 5
3220 -0.0847227700159587 5.64743868466652
0.362382277714453 0.547186147116967 0.882084976713933
255027 MPV17L MPV17 mitochondrial inner membrane protein like 6053
0.173121084547525 0.544546200530666
0.362109852122096 0.547336806230526 0.882175460173386
9555 MACROH2A1 macroH2A.1 histone
17730 -0.0733700815428135 7.71988122511463
0.361936144002684 0.547432912078041 0.882175460173386
51108 METTL9 methyltransferase like 9
3677 -0.0771856288939191 6.65688853790129
0.361788144756261 0.547514819004669 0.882175460173386
51260 PBDC1 polysaccharide biosynthesis domain containing 1
1201 -0.108140079112744 4.70046625266973
0.361720324370371 0.547552360327986 0.882175460173386
1580 CYP4B1 cytochrome P450 family 4 subfamily B member 1 3364
0.485269776394589 3.95386024509936
0.361711614073243 0.547557182176585 0.882175460173386
22982 DIP2C disco interacting protein 2 homolog C 8563
0.156317845642026 5.04817244585121 0.361532937268628
0.547656111577775 0.882233370312811

6181	RPLP2	ribosomal protein lateral stalk subunit P2	1955
	0.110401015939689	7.15466249907833
0.361217724176981	0.547830719288601	0.882276421607479
80776	B9D2	B9 domain containing 2	1316	0.152244127870472
	1.72885265449349	0.361066905014727
0.547914300054629	0.882276421607479
3791	KDR	kinase insert domain receptor
6259	-0.312647757396794	6.2175368599795	0.361016078626676
	0.547942472308462	0.882276421607479
7571	ZNF23	zinc finger protein 23	5529	0.107179689348871
	3.83275551068116	0.361011391277717
0.547945070566942	0.882276421607479
79053	ALG8	ALG8 alpha-1,3-glucosyltransferase	3448
0.102331479267721	4.54618422131175	0.360916315259884
	0.54799777739595	0.882276421607479
51710	ZNF44	zinc finger protein 44	5815	-0.134196972591258
	3.29389451400221	0.360574423141123
0.548187388569879	0.882480261084738
23380	SRGAP2	SLIT-ROBO Rho GTPase activating protein 2
11282	-0.10540938265389	5.63821728842197
0.360258349954179	0.548362789562429	0.882661180579285
222229	LRWD1	leucine rich repeats and WD repeat domain containing
1	5926	0.119306434331536	4.28063718584679
0.3597546606393	0.54864252226698	0.88290887354898
9905	SGSM2	small G protein signaling modulator 2
9444	-0.104863790385307	6.70178334048153
0.359741348230472	0.548649919158149	0.88290887354898
10677	AVIL	advillin	5889	-0.280264670242619
1.60614199842588	0.359535493386014	0.548764323796236
	0.88290887354898
27068	PPA2	inorganic pyrophosphatase 2	4654
0.112317871527759	5.23066395172129	0.359493750270129
	0.548787528126722	0.88290887354898
55167	MSL2	MSL complex subunit 2	5095	-0.109891165883533
	4.17191343428123	0.359414047861005
0.548831838508672	0.88290887354898
10413	YAP1	Yes associated protein 1
6296	-0.0850016123483884	8.02936941198866
0.358980964026102	0.549072727338107	0.882988669932724
8543	LMO4	LIM domain only 4	5833	0.123870973224502
	4.90781060112596	0.358943336424156
0.549093665793329	0.882988669932724
2335	FN1	fibronectin 1	17315	-0.168787171316669
13.5590603482338	0.358861365580196	0.54913928488843
	0.882988669932724
55240	STEAP3	STEAP3 metalloreductase	4562	0.17972844812613
	5.81080475348661	0.35864297899397
0.549260857771974	0.882988669932724
29997	NOP53	NOP53 ribosome biogenesis factor	5341
0.0934847466940103	4.87054402489595	0.358579011976822
	0.549296476874858	0.882988669932724
4682	NUBP1	nucleotide binding protein 1	1955
0.090786360109484	4.15020323261585	0.358460085943957
	0.549362710579655	0.882988669932724

27158	NDOR1	NADPH dependent diflavin oxidoreductase 1
2851	-0.113090702016573	3.90651941030725
0.358428610571083	0.549380242717631	0.882988669932724
23604	DAPK2	death associated protein kinase 2
13612	-0.197436316280041	2.45678348233804
0.358376010832312	0.549409543699931	0.882988669932724
51072	MEMO1	mediator of cell motility 1
5354	-0.101402256854089	4.73240900297186
0.358194168570829	0.549510862469714	0.882988669932724
27067	STAU2	staufen double-stranded RNA binding protein 2
8998	-0.0764089254896227	5.55212620177062
0.358116541896575	0.549554125095495	0.882988669932724
374868	ATP9B	ATPase phospholipid transporting 9B (putative)
14930	0.0971536768890559	4.09846007081445
0.35807932049504	0.549574871455236	0.882988669932724
1195	CLK1	CDC like kinase 1	5170	-0.162086892836301
	5.47727277397755	0.357720163138605
0.549775132840327	0.883140476560998
199745	THAP8	THAP domain containing 8	1825
0.190642938890777	1.65469047479397	0.35764689739727
	0.549816001608849	0.883140476560998
55582	KIF27	kinesin family member 27
5140	-0.172354586303032	2.76723395571422
0.357419916905149	0.54994265093966	0.883140476560998
57122	NUP107	nucleoporin 107	7694	-0.0946501658485038
4.78608772672653	0.357380752477084	0.549964509214418
	0.883140476560998
53938	PPIL3	peptidylprolyl isomerase like 3	2914
0.116971403976393	3.85335482224325	0.357257017812041
	0.550033578133547	0.883140476560998
6610	SMPD2	sphingomyelin phosphodiesterase 2	2014
0.13108226843684	2.97629499788881	0.357231801360591
	0.550047655994727	0.883140476560998
87769	GGACT	gamma-glutamylamine cyclotransferase
4216	-0.213616137603735	1.47518523146448
0.356815950698486	0.550279914756701	0.883348219139121
283209	PGM2L1	phosphoglucomutase 2 like 1
8515	-0.125402387023472	7.81263124865071
0.356774323737	0.55030317414207	0.883348219139121
25926	NOL11	nucleolar protein 11	4713	-0.0844895171078539
	5.17398797182777	0.356533452275762
0.550437799084899	0.883371662024238
143098	MPP7	membrane palmitoylated protein 7
6167	-0.11447590304314	5.12591808577782
0.356522518823116	0.550443911340583	0.883371662024238
3013	H2AC7	H2A clustered histone 7	393	0.191798450600289
	1.02969041072682	0.356324972250334
0.550554370013934	0.883447710022359
27154	BRPF3	bromodomain and PHD finger containing 3
6613	-0.0983138800498086	6.60039024620319
0.35589804521436	0.55079322929056	0.883683071683484
64901	RANBP17	RAN binding protein 17	10993	0.157634122409997
	2.75522192781483	0.355837316237888
0.550827222022453	0.883683071683484

25813	SAMM50	SAMM50 sorting and assembly machinery component 6191
	0.0874138489198598	5.51420524029691
0.355623195135969	0.550947106594141	0.883771910376099
3690	ITGB3	integrin subunit beta 3 4913	0.267009474431874
	5.17351570303799	0.355513062471543
0.551008787985612	0.883771910376099
51271	UBAP1	ubiquitin associated protein 1
2772	-0.0823491304495726	7.56184802685432
0.35515869934336	0.551207342091356	0.883847598707383
7567	ZNF19	zinc finger protein 19	7985	-0.161165303488051
	1.70490473611923	0.355114224031989
0.551232271646502	0.883847598707383
84057	MND1	meiotic nuclear divisions 1	1511
0.253608782538221	0.549932440064449	0.355004577790337
	0.551293740220792	0.883847598707383
387680	WASHC2A WASH complex subunit 2A 5159	0.0881867496580095
	6.23207767928277	0.354839697987515
0.551386197396213	0.883847598707383
51611	DPH5	diphthamide biosynthesis 5	2970
0.144819494587136	2.87266772648358	0.354807143761395
	0.551404455781847	0.883847598707383
23354	HAUS5	HAUS augmin like complex subunit 5	5075
0.152810198174088	2.26021782416788	0.354753437100191
	0.551434580217307	0.883847598707383
7840	ALMS1	ALMS1 centrosome and basal body associated protein
	14054	-0.133891836580401	4.80045646203491
0.354497326983902	0.551578276646628	0.883976764415254
100133991	MAP3K14-AS1	MAP3K14 antisense RNA 1 3775
0.150033606920213	2.2067400442028 0.354285483320688
0.551697189511822	0.884066186190606
2542	SLC37A4 solute carrier family 37 member 4	6186
0.104160235528029	4.11811757123782	0.35399505705834
	0.551860290924813	0.884226389268344
26097	CHTOP	chromatin target of PRMT1
5625	-0.0717410246731153	5.89568870245355
0.353826953944118	0.551954737887589	0.884276565662983
79080	CCDC86	coiled-coil domain containing 86	2044
0.119779264026287	4.49920039657166	0.353593732494613
	0.552085821215505	0.884385418317197
100126338	MIR937	microRNA 937	86	-0.181833007448994
	0.09888001305281	0.353442362806628
0.552170930653158	0.884420608994715
55170	PRMT6	protein arginine methyltransferase 6
2616	-0.146590522557911	4.04007337739648
0.353152664373322	0.552333885901777	0.884580464192338
8975	USP13	ubiquitin specific peptidase 13 8440
0.243113250057924	1.16916063297194	0.352837036120742
	0.552511529641791	0.884763804240605
124751	KRBA2	KRAB-A domain containing 2	3695
0.1223307212605 6.403745628399	0.352290158702289
0.552819581016754	0.884813166120311
8620	NPFF	neuropeptide FF-amide peptide precursor
1406	-0.194960714728261	0.774560555970753
0.35223456165627	0.552850916461538	0.884813166120311

83939	EIF2A	eukaryotic translation initiation factor 2A
4131	-0.0910857916971445	5.90166797182849
0.352214626850326	0.552862152873174	0.884813166120311
153241	CEP120	centrosomal protein 120 6057	-0.10728336217669
	4.59616493113511	0.352198253860939
0.552871381960235	0.884813166120311
54620	FBXL19	F-box and leucine rich repeat protein 19
4340	-0.105184287062429	5.67560492131494
0.351995889863019	0.552985473964091	0.884813166120311
5584	PRKCI	protein kinase C iota	5841	-0.0754678276262749
	6.72844682515142	0.351969069587653
0.553000598454497	0.884813166120311
2213	FCGR2B	Fc fragment of IgG receptor IIb 6651
0.61178598711667	0.848599545107255	0.351908697601186
	0.553034646277513	0.884813166120311
89941	RHOT2	ras homolog family member T2	5174
0.0843259824440452	6.15904017290407	0.35160396200839
	0.553206567457786	0.884813166120311
529	ATP6V1E1	ATPase H+ transporting V1 subunit E1	3212
	0.0984087939457201	7.46947224520968
0.351539239568382	0.553243094562675	0.884813166120311
56946	EMSY	EMSY transcriptional repressor, BRCA2 interacting
	11783	-0.113861381466087	4.78719636248504
0.351126549219984	0.553476109641752	0.884813166120311
171023	ASXL1	ASXL transcriptional regulator 1	9454
0.0780069398573301	6.50655023203427	0.351012172014824
	0.553540722557658	0.884813166120311
7762	ZNF215	zinc finger protein 215 4025	-0.172833484318664
	0.984471972515002	0.350985713254209
0.553555671422211	0.884813166120311
10809	STARD10 StAR related lipid transfer domain containing 10
	3710	-0.0899239903262968	6.4120255184917
0.350820299688699	0.553649145250176	0.884813166120311
8315	BRAP	BRCA1 associated protein
4686	-0.0782961135831653	5.42677588670887
0.350802677876661	0.553659104943815	0.884813166120311
11240	PADI2	peptidyl arginine deiminase 2	5984
0.366276723470664	0.861163045225432	0.350773845978505
	0.553675401208565	0.884813166120311
284390	ZNF763	zinc finger protein 763 3280	-0.162784844438589
	2.02696301138457	0.350736486334857
0.553696518841482	0.884813166120311
5790	PTPRCAP protein tyrosine phosphatase receptor type C
associated protein	1292	-0.266003276384076
0.381141711049292	0.3505263970832 0.553815300620983
0.884813166120311
100534595	HNRNPUL2-BSCL2	HNRNPUL2-BSCL2 readthrough (NMD
candidate)	4188	0.0752813378250045	7.51634539496957
	0.350438064658995	0.553865257001464
0.884813166120311
81858	SHARPIN SHANK associated RH domain interactor
2783	-0.0796731492305796	5.18934191874914
0.350434839795412	0.553867080983126	0.884813166120311
527	ATP6V0C ATPase H+ transporting V0 subunit c

1498	-0.101747192560177	7.29540956030378
0.350421862679013	0.553874420949978	0.884813166120311
29028	ATAD2	ATPase family AAA domain containing 2
7118	-0.122835820896746	3.92837285776646
0.350325519688607	0.5539289191006 0.884813166120311
100507458	ZNF213-AS1	ZNF213 antisense RNA 1 (head to
head)	2741	-0.158735377282658	2.1827981215961
0.350188989655268	0.554006167110294	0.884813166120311
105371743	SNHG30	small nucleolar RNA host gene 30	1298
	0.187397243659537	1.0632351330224 0.350142372935276
	0.554032547273459	0.884813166120311
362	AQP5	aquaporin 5	1885	-0.37442502911476
2.89003012789814	0.350079830681153	0.554067943339387
	0.884813166120311
8801	SUCLG2	succinate-CoA ligase GDP-forming beta subunit
3090	-0.112356952577283	4.8835451294015 0.349782766014552
	0.554236126715933	0.884813166120311
23534	TNPO3	transportin 3	4676	-0.0665771135826069
6.45804237907163	0.349661391302833	0.554304870837898
	0.884813166120311
54148	MRPL39	mitochondrial ribosomal protein L39
1199	-0.119526526773411	4.25973617837606
0.349551340976199	0.55436721498378	0.884813166120311
6135	RPL11	ribosomal protein L11	1987	0.130755531418823
	7.58549433909926	0.34953427625328
0.55437688343398	0.884813166120311
100532746	PPT2-EGFL8	PPT2-EGFL8 readthrough (NMD
candidate)	6487	-0.192160316103374	3.98604242957438
	0.349479584137956	0.554407872781035
0.884813166120311
29941	PKN3	protein kinase N3	3375	0.178113332671486
	1.63089720931213	0.349427415216526
0.554437435499249	0.884813166120311
81894	SLC25A28	solute carrier family 25 member 28	1755
	0.0998753330035704	4.16741581004604
0.349237473593183	0.554545095450406	0.884826314927069
253558	LCLAT1	lysocardiolipin acyltransferase 1
7517	-0.101614784683204	4.08100097245858
0.349123003666303	0.554609996709005	0.884826314927069
55009	FAM174C	family with sequence similarity 174 member C	2193
	0.134180170606324	4.13475906539585
0.348977497796342	0.55469251520554	0.884826314927069
9469	CHST3	carbohydrate sulfotransferase 3
6970	-0.206345398315408	5.52673603332289
0.348967200343374	0.554698355920225	0.884826314927069
57488	ESYT2	extended synaptotagmin 2
7098	-0.0730749523099048	7.20596252033177
0.348640169412626	0.554883908339622	0.884869129436117
92979	MARCHF9	membrane associated ring-CH-type finger 9	4005
	0.123370680902934	4.58868530441989
0.348579187183769	0.554918521719598	0.884869129436117
9694	EMC2	ER membrane protein complex subunit 2	3781
0.0889369182188531	4.68231250813855	0.348476163971904
	0.554977006746582	0.884869129436117

221504	ZBTB9	zinc finger and BTB domain containing 9
3014	-0.116710318457272	3.47545786610806
0.348474608644714	0.554977889776275	0.884869129436117
441951	ZFAS1	ZNFX1 antisense RNA 1	1838	0.111071971064722
	5.88983114288638	0.348139942740545
0.555167956598301	0.884969267648517
548645	DNAJC25 DnaJ heat shock protein family (Hsp40) member C25
	2586	0.125067802078542	3.33845713894948
0.347894800106877	0.555307258629406	0.884969267648517
10432	RBM14	RNA binding motif protein 14
3923	-0.0761278366163203	5.31541679237913
0.347867569921061	0.555322736234478	0.884969267648517
200312	RNF215	ring finger protein 215 3105	0.0828499398453543
	4.64904492598851	0.347611261901337
0.555468461432962	0.884969267648517
122416	ANKRD9	ankyrin repeat domain 9 7215	0.103786681311321
	5.06359825398707	0.347282070534328
0.555655731065919	0.884969267648517
389114	ZNF662	zinc finger protein 662 5070	0.181260185494075
	2.36115850311534	0.347198114203814
0.555703511089073	0.884969267648517
339448	C1orf174	chromosome 1 open reading frame 174
4388	-0.0856053758894696	4.2678847272215 0.347176058354122
	0.555716064488858	0.884969267648517
122961	ISCA2	iron-sulfur cluster assembly 2	3061
0.110482778734365	3.74097603858056	0.347120442516923
	0.555747721416985	0.884969267648517
3633	INPP5B	inositol polyphosphate-5-phosphatase B	8930
0.112491774673856	3.68294835582568	0.347117212414285
	0.555749560119235	0.884969267648517
1915	EEF1A1	eukaryotic translation elongation factor 1 alpha 1
	5948	0.102396510209648	10.1375900337508
0.346941167821115	0.555849789102983	0.884969267648517
65263	PYCR3	pyrroline-5-carboxylate reductase 3	3244
0.178407302083948	2.22528339114542	0.346939080942304
	0.555850977449068	0.884969267648517
9221	NOLC1	nucleolar and coiled-body phosphoprotein 1
3970	-0.0864719539437409	6.06677699431902
0.346923720669963	0.555859724305142	0.884969267648517
9894	TELO2	telomere maintenance 2	4239	0.0950780561629588
	4.11547882630864	0.346899503789031
0.555873515057322	0.884969267648517
91373	UAP1L1	UDP-N-acetylglucosamine pyrophosphorylase 1 like 1
	5283	-0.1550892651415	4.77959761244924
0.346808715147397	0.55592522210604	0.884969267648517
92104	TTC30A	tetratricopeptide repeat domain 30A	5975
0.11796901513068	3.80351167389036	0.346679687533985
	0.555998723162663	0.884985694924934
55422	ZNF331	zinc finger protein 331 7728	0.141073613129718
	3.09934919156058	0.346285779252089
0.556223228480216	0.885242445326938
2760	GM2A	GM2 ganglioside activator	4479
0.146570788130625	6.69945812027665	0.346028140293441
	0.556370161041039	0.885375692535995

752	FMNL1	formin like 1	7373	-0.305280128439512
2.65911430495395	0.345731864050741	0.556539219650378
	0.885544115601822
84418	CYSTM1	cysteine rich transmembrane module containing 1 2185
	0.125612587001502	4.99092577583828
0.345228106378187	0.556826893446625	0.885793330862014
745	MYRF	myelin regulatory factor
10680	-0.226751872131212	2.10032902085399
0.345003283279483	0.556955371138686	0.885793330862014
1001	CDH3	cadherin 3	5438	0.109176773969213
9.42107127611525	0.34478467263898	0.557080352654604
	0.885793330862014
84342	COG8	component of oligomeric golgi complex 8 6546
0.0757269104525441	5.425737261427	0.344657577495996
0.557153038497616	0.885793330862014
100268168	LOC100268168	uncharacterized LOC100268168
2780	-0.205552123310333	0.788795785803517
0.344540392273966	0.557220072816243	0.885793330862014
2395	FXN	frataxin	2939	0.160503611625784
2.27002660233572	0.344417499612455	0.557290388470731
	0.885793330862014
80262	C16orf70	chromosome 16 open reading frame 70	5963
	0.0957967059079971	4.83176948767741
0.34437403706	0.557315260533198	0.885793330862014
100505576	LINC00672	long intergenic non-protein coding
RNA 672 2365	-0.214997949544772	0.153767586299565
0.344352252122079	0.557327728066972	0.885793330862014
85302	FBF1	Fas binding factor 1	7890	0.0819995345820784
	4.73925175415505	0.344292778265384
0.557361767692456	0.885793330862014
143884	CWF19L2	CWF19 like cell cycle control factor 2	3581
0.124193855824087	3.85227581908269	0.344267515648973
	0.55737622784651	0.885793330862014
2737	GLI3	GLI family zinc finger 3
9239	-0.211211575636558	3.54900664660768
0.344240875975743	0.557391476991346	0.885793330862014
387066	SNHG5	small nucleolar RNA host gene 5 2794
0.256625494147177	3.68895196967664	0.344120318050688
	0.55746049696255	0.885802516273901
22889	KHDC4	KH domain containing 4, pre-mRNA splicing factor
	5635	-0.11894020384105	5.97504252271682
0.343725014326662	0.557686924235885	0.885933857140603
6690	SPINK1	serine peptidase inhibitor, Kazal type 1	1480
	0.442141207101999	4.15355865963887
0.343702639244423	0.557699745761236	0.885933857140603
9028	RHBDL1	rhomboid like 1 1884	0.225621259888447
0.0360323829259505	0.343644782966501	0.557732901568204
	0.885933857140603
200186	CRTC2	CREB regulated transcription coactivator 2
3612	-0.0772357844813242	5.81366649051847
0.343355302880031	0.557898850815656	0.886061272911697
10381	TUBB3	tubulin beta 3 class III
5622	-0.151946266457872	8.80604426535845
0.343250237682241	0.557959104422105	0.886061272911697

6643	SNX2	sorting nexin 2 4254	0.0916780808288025
6.02058602950487	0.343173900592006	0.558002890580001
	0.886061272911697
10549	PRDX4	peroxiredoxin 4 1904	0.138186195658779
5.46529551088232	0.342742295365781	0.558250577944559
	0.886332651830491
9871	SEC24D	SEC24 homolog D, COPII coat complex component
9920	-0.152567645084564	6.25323154936648
0.342551656332695	0.558360047562098	0.886332651830491
84960	CCDC183 coiled-coil domain containing 183	5694
0.122063374874814	2.94895257170734	0.342523269093186
	0.558376351710692	0.886332651830491
83478	ARHGAP24	Rho GTPase activating protein 24	7908
	0.193347958959923	2.56775891230605
0.342327242565877	0.55848896387978	0.886332651830491
353091	RAET1G	retinoic acid early transcript 1G
1486	-0.198903524578761	3.69894540724804
0.342325121850056	0.558490182412787	0.886332651830491
65008	MRPL1	mitochondrial ribosomal protein L1	1943
0.126004713926596	3.58327925289027	0.341987578290464
	0.558684194728278	0.886394194051117
2530	FUT8	fucosyltransferase 8	6700	-0.0959117662831169
	5.15367510211933	0.34197522397923
0.558691298140898	0.886394194051117
6926	TBX3	T-box transcription factor 3
5950	-0.320610940872211	2.1966935429372 0.341749571938109
	0.558821072590538	0.886394194051117
55062	WIPI1	WD repeat domain, phosphoinositide interacting 1
	3402	-0.0995002573902035	5.73707729780348
0.341671383107176	0.558866053077345	0.886394194051117
746	TMEM258 transmembrane protein 258	3068
0.106196438079193	4.85773517183989	0.341601740382526
	0.558906122973075	0.886394194051117
92344	GORAB	golgin, RAB6 interacting
3773	-0.124541242359301	3.32624981168567
0.34159734104351	0.558908654377059	0.886394194051117
55646	LYAR	Ly1 antibody reactive	2319	-0.162890399265929
	2.65310599619636	0.341374563574975
0.559036870365278	0.886404967645393
23555	TSPAN15 tetraspanin 15	2646	-0.170850549474613
5.72811186000575	0.341365629189482	0.559042013577455
	0.886404967645393
131118	DNAJC19 DnaJ heat shock protein family (Hsp40) member C19
	3586	0.110144201586662	3.54353361269549
0.340998749071606	0.559253291596313	0.886583168366481
432369	ATP5F1EP2	ATP synthase F1 subunit epsilon pseudogene 2
	385	0.167929215061291	3.04874388031855
0.340950664910027	0.559280993480847	0.886583168366481
79666	PLEKHF2 pleckstrin homology and FYVE domain containing 2
	3137	-0.097124644700694	4.84409953608552
0.340674001253412	0.559440433772752	0.886735561373196
127829	ARL8A	ADP ribosylation factor like GTPase 8A
1975	-0.0762621937134683	6.19288577341972
0.340465660316328	0.559560557223509	0.886806588210014

55809	TRERF1	transcriptional regulating factor 1
7732	-0.145321883652821	3.66986812511685
0.34037669367752	0.559611867865232	0.886806588210014
5900	RALGDS	ral guanine nucleotide dissociation stimulator	7624
	0.0734646072651098	5.97017630755623
0.340137261218011	0.559750002860547	0.886925145935258
100287765	LINC00630	long intergenic non-protein coding
RNA 630	4077	-0.173904612497361	1.71864262496185
0.33993824212534	0.559864871899106	0.886985995729408
57099	AVEN	apoptosis and caspase activation inhibitor	1808
	0.147042869555951	4.1818084924604	0.339797172666962
	0.559946321091374	0.886985995729408
9859	CEP170	centrosomal protein 170	10939	0.0727355021107346
	6.14042047630072	0.339659609994831
0.560025767389887	0.886985995729408
653645	TBC1D3P1-DHX40P1	TBC1D3P1-DHX40P1 readthrough,
transcribed pseudogene	2013	0.0987142828221172
3.674299466375	0.339632020174925	0.560041703878838
0.886985995729408
79836	LONRF3	LON peptidase N-terminal domain and ring finger 3
	4041	0.204978009607826	0.63242073304584
0.339226655473672	0.560275951498117	0.887224637820659
79917	MAGIX	MAGI family member, X-linked	5107
0.16247960471314	1.19615429010136	0.339114632592008
	0.560340719084097	0.887224637820659
200845	KCTD6	potassium channel tetramerization domain containing
6	3068	0.14487275504024	2.36280103833614
0.339042542219794	0.560382406710885	0.887224637820659
51264	MRPL27	mitochondrial ribosomal protein L27	3027
0.0974925844676218	5.03697471639283	0.338755879843688
	0.560548233414203	0.887386879004605
56945	MRPS22	mitochondrial ribosomal protein S22
4894	-0.131817033534608	4.62568062323323
0.338599600817943	0.560638676318719	0.887429759204012
55969	RAB5IF	RAB5 interacting factor	1517	-0.105591116714932
	6.40394474752449	0.338053359318863
0.560955020714389	0.887809037613818
133015	PACRGL	parkin coregulated like	4032	0.11349238957699
	3.25381302623077	0.337881081417718
0.56105486281347	0.887809037613818
83892	KCTD10	potassium channel tetramerization domain containing
10	9865	-0.0910512757636236	7.21622606924218
0.337762506169327	0.561123601813298	0.887809037613818
85377	MICALL1	MICAL like 1	5968	0.0971905277796009
8.58698041705249	0.337748330710383	0.561131820517964
	0.887809037613818
51078	THAP4	THAP domain containing 4	3147
0.079086626009606	5.11678498919349	0.337345352435612
	0.561365557447043	0.888065900163889
9775	EIF4A3	eukaryotic translation initiation factor 4A3	4967
	0.0739324027795022	6.28922803426463
0.337249862260762	0.561420971367976	0.888065900163889
23682	RAB38	RAB38, member RAS oncogene family
1465	-0.180730994737737	3.70536996173303

0.33669313257656	0.561744256309445	0.888428224383829
9810	RNF40	ring finger protein 40	6523	-0.0731912248668878
	6.68085613194979	0.33652919503832
0.561839520577707	0.888428224383829
90102	PHLDB2	pleckstrin homology like domain family B member 2
	12336	-0.114872097468864	8.5529756768443
0.336508608386566	0.561851485693378	0.888428224383829
5797	PTPRM	protein tyrosine phosphatase receptor type M
9900	-0.0763886112303669	5.97739669380553
0.336418717481834	0.561903736689965	0.888428224383829
729920	CRPPA	CDP-L-ribitol pyrophosphorylase A
6309	-0.16884455935184	2.14032691535757
0.336293817607484	0.561976352882158	0.888442751108396
22950	SLC4A1AP	solute carrier family 4 member 1 adaptor
protein 5207	-0.0745703209639468	5.45975170449112
0.336055900408341	0.562114726677848	0.8885612206948
79954	NOL10	nucleolar protein 10	3824	0.0950143349168469
	4.7111361045671 0.335832399024693	0.562244775793421
	0.888586080844088
54529	ASNSD1	asparagine synthetase domain containing 1
2504	-0.0983280623257963	5.24419353303854
0.335810827100275	0.562257330943182	0.888586080844088
103344931	LOC103344931	uncharacterized LOC103344931
4237	-0.131734529588223	3.05445363338321
0.335376677253406	0.562510126569504	0.888798756565328
57594	HOMEZ	homeobox and leucine zipper encoding
5534	-0.102501099441615	4.8941573955167 0.335344124292185
	0.562529090226485	0.888798756565328
5879	RAC1	Rac family small GTPase 1
3114	-0.0777042852527494	8.29044200034939
0.335149676422361	0.562642391016054	0.888798756565328
51601	LIPT1	lipoyltransferase 1	2279	-0.169848659368762
	0.914076834446903	0.335093251097878
0.56267527711114	0.888798756565328
51218	GLRX5	glutaredoxin 5	2845	0.103169663609917
4.19227459518132	0.335012036739016	0.562722617367468
	0.888798756565328
51435	SCARA3	scavenger receptor class A member 3
4031	-0.272452932022457	4.29947869845492
0.334906219010676	0.562784310545453	0.888798756565328
83999	KREMEN1 kringle containing transmembrane protein 1
7977	-0.147042065221967	5.99827403713985
0.334817440148612	0.56283607986967	0.888798756565328
152687	ZNF595	zinc finger protein 595 3933	-0.15457045712646
	2.09949784722102	0.33457006414427
0.56298037976568	0.888832756297465
23259	DDHD2	DDHD domain containing 2
10686	-0.0715287947691001	6.2538609760871 0.334562963096526
	0.562984523015001	0.888832756297465
79607	FAM118B family with sequence similarity 118 member B	3846
	0.0832215237619502	4.70574023973599
0.334290247474531	0.563143688856154	0.888917091246069
7385	UQCRC2	ubiquinol-cytochrome c reductase core protein 2
5217	-0.0860794698549131	6.9411091599374 0.334253974888966

0.563164865261485 0.888917091246069
55166 CENPQ centromere protein Q 1733 -0.147915900492553
2.02430714819342 0.333925941392774
0.563356445121381 0.889100589853914
57690 TNRC6C trinucleotide repeat containing adaptor 6C
11058 -0.190126323445794 3.33748445271224
0.333796921339811 0.563431830491258 0.889100589853914
4836 NMT1 N-myristoyltransferase 1
6669 -0.0647335812206276 7.12416273569712
0.333728963738672 0.563471545379681 0.889100589853914
57715 SEMA4G semaphorin 4G 5253 0.193293312653067
1.72655587369592 0.333321867109837 0.563709568424617
0.889176527280011
7297 TYK2 tyrosine kinase 2 6682 -0.0796389161521332
6.68258855589615 0.333299244552819
0.563722801163899 0.889176527280011
1632 ECI1 enoyl-CoA delta isomerase 1 3234
0.147839056802541 4.15270509754655 0.333281053922708
0.563733441945601 0.889176527280011
3418 IDH2 isocitrate dehydrogenase (NADP(+)) 2
2814 -0.0979411621478952 5.50027135002309
0.332985290868972 0.563906505696193 0.889176527280011
5434 POLR2E RNA polymerase II subunit E 4865
0.0832668130504188 6.61528324893392 0.332938878618465
0.563933672810165 0.889176527280011
2562 GABRB3 gamma-aminobutyric acid type A receptor beta3
subunit 8995 -0.524513236478037 1.4950768240973
0.332908105241344 0.563951687201245 0.889176527280011
84661 DPY30 dpy-30 histone methyltransferase complex regulatory
subunit 631 0.113229072354899 4.07240394781406
0.332871680054247 0.563973011533022 0.889176527280011
54919 DNAAF5 dynein axonemal assembly factor 5
7137 -0.0876214767695197 4.76907902575799
0.332778586796497 0.564027518018341 0.889176527280011
58509 CACTIN cactin, spliceosome C complex subunit 3849
0.0837030930605879 4.66109125111826 0.332581763567625
0.564142792288435 0.889197689716198
55578 SUPT20H SPT20 homolog, SAGA complex component
10933 -0.074740680811092 5.96051078308782
0.332446983205848 0.564221755880483 0.889197689716198
833 CARS1 cysteinyl-tRNA synthetase 1
10037 -0.0726306263411036 7.00810146965431
0.332403336243942 0.564247331840031 0.889197689716198
84271 POLDIP3 DNA polymerase delta interacting protein 3
3650 -0.0629574283172245 6.46985794923542
0.332251042718031 0.564336589324162 0.889197689716198
1062 CENPE centromere protein E 9444 -0.162158305443832
2.3218207133447 0.332116332827929 0.564415563959291
0.889197689716198
5163 PDK1 pyruvate dehydrogenase kinase 1 14883
0.252170235336027 3.0384602646655 0.332105635646954
0.564421836172393 0.889197689716198
26056 RAB11FIP5 RAB11 family interacting protein 5
5932 -0.084365061069923 6.79414366145355

0.331909220509615	0.564537026663432	0.889253810519205
131601	TPRA1	transmembrane protein adipocyte associated 1	3596
	0.087469511402252	6.68069582481306
0.331824202454158	0.564586900806615	0.889253810519205
284370	ZNF615	zinc finger protein 615 4650	-0.135919869854428
	2.96879534232826	0.331720208696268
0.564647918237867	0.889253810519205
27309	ZNF330	zinc finger protein 330 2536	0.088246534840965
	4.2147976062297	0.331481182419509
0.564788213120082	0.889293949093554
79915	ATAD5	ATPase family AAA domain containing 5
7075	-0.169195853033853	0.550847583475871
0.33136761226861	0.564854895967998	0.889293949093554
6228	RPS23	ribosomal protein S23	4204	-0.1099967710999
	6.5087797877165 0.331352325604371	0.564863872712598
	0.889293949093554
8408	ULK1	unc-51 like autophagy activating kinase 1
7608	-0.107748362870711	6.93485133542057
0.33094454152652	0.565103436425264	0.889437756261439
6129	RPL7	ribosomal protein L7	2907	0.116213012046126
	4.12101026433088	0.330736855824668
0.565225522480545	0.889437756261439
9014	TAF1B	TATA-box binding protein associated factor, RNA polymerase I subunit B	4064	0.141760577637838
4.03261007107022	0.330708601582646	0.565242135408343
	0.889437756261439
147968	CAPN12	calpain 12	4570	0.214566890166528
2.3752474503301 0.330682080960055	0.565257729861919
0.889437756261439
162989	DEDD2	death effector domain containing 2
2631	-0.091257836482411	4.7824039935951 0.330602274919712
	0.565304661819236	0.889437756261439
57704	GBA2	glucosylceramidase beta 2
4803	-0.0929633283226196	6.33180082061781
0.330525895908252	0.565349585484977	0.889437756261439
54675	CRLS1	cardiolipin synthase 1	5370	0.0900305899223271
	5.00926150441746	0.330397852244644
0.56542491212487	0.889437756261439
2523	FUT1	fucosyltransferase 1 (H blood group)
4343	-0.133823672094341	4.42471776586662
0.330260374824759	0.565505810157507	0.889437756261439
22993	HMGXB3	HMG-box containing 3	6263	0.0755648013582724
	6.70630974644378	0.33022486032354
0.565526712162803	0.889437756261439
54832	VPS13C	vacuolar protein sorting 13 homolog C
18992	-0.149050816446095	7.3527604990614 0.329760236334891
	0.565800303719478	0.889680132408703
221830	TWISTNB TWIST neighbor	3997	-0.102548420525471
4.27066245095376	0.32974741928545	0.565807854608177
	0.889680132408703
64926	RASAL3	RAS protein activator like 3
4681	-0.436223609937234	0.649420037460255
0.329396004541159	0.566014959097265	0.88982633392722
10635	RAD51AP1	RAD51 associated protein 1	2558

0.195926853582874	0.226322087357313	0.329374074070337
	0.566027888576535	0.88982633392722
100506123	LOC100506123	uncharacterized LOC100506123
6083	-0.185367489839287	3.46858134722231
0.328868385051024	0.566326184909175	0.89002726492441
84681	HINT2	histidine triad nucleotide binding protein 2	2395
	0.120763290631971	3.29487358650666
0.328862722715529	0.566329526738168	0.89002726492441
80833	APOL3	apolipoprotein L3	5583	-0.318392620224679
	1.39782889823158	0.328834256425196
0.566346327712663	0.89002726492441
6584	SLC22A5 solute carrier family 22 member 5
12788	-0.124138077030714	5.01573016855017
0.328515428664927	0.566534567811912	0.890069207426668
80145	THOC7	THO complex 7	1784	0.0921886795372038
5.14775387422115	0.32830593578953	0.566658321234667
	0.890069207426668
2193	FARSA	phenylalanyl-tRNA synthetase subunit alpha	2435
	0.076063206869367	5.6603135672895 0.328132259288111
	0.566760956672099	0.890069207426668
54953	ODR4	odr-4 GPCR localization factor homolog
4073	-0.08155669102833	5.53970180669996
0.328130925969259	0.566761744746449	0.890069207426668
57613	FAM234B family with sequence similarity 234 member B	6807
	0.13897289457407	2.36115874823045
0.328093820024762	0.566783677520079	0.890069207426668
692087	SNORD49B	small nucleolar RNA, C/D box 49B	72
	-0.188262032193033	0.34696066240562
0.328077701137828	0.566793205668338	0.890069207426668
10227	MFSD10	major facilitator superfamily domain containing 10
	3159	0.0861374308142531	7.0010914240445
0.328036048044069	0.566817828960225	0.890069207426668
339287	MSL1	MSL complex subunit 1	7397	-0.0745358116560371
	6.8654209577468 0.327824091523752	0.566943159561567
	0.890166218582992
9878	TOX4	TOX high mobility group box family member 4
7091	-0.0619962935310271	6.58588614600634
0.327668714278175	0.56703506882674	0.890210738517838
80856	LNPK	lunapark, ER junction formation factor
9122	-0.10403661666259	5.44692283908322
0.327523611762853	0.56712092647805	0.890245748871237
10818	FRS2	fibroblast growth factor receptor substrate 2
7242	-0.0875546611196718	5.76693187375956
0.327256068599901	0.567279298891421	0.89039456965174
6599	SMARCC1 SWI/SNF related, matrix associated, actin dependent regulator of chromatin subfamily c member 1
7645	-0.0848149726940628	7.4171536526064 0.327032750162175
	0.56741155824549	0.890502374940569
1508	CTSB	cathepsin B	7428	-0.118696892245582
11.1083054457151	0.326868602545346	0.567508812160828
	0.890555224281505
192670	AGO4	argonaute RISC component 4
7258	-0.100668393976173	5.68693942148613
0.326729604754876	0.567591190686203	0.890584721400431

51160	VPS28	VPS28 subunit of ESCRT-I
3769	-0.0804639411334461	5.79821305432119
0.326170634017277	0.567922705568941	0.891005078058261
124460	SNX20	sorting nexin 20	5754	0.487215181527349
	-0.241305893835945	0.326016833986003
0.568013987519527	0.89104848506955
79846	CFAP69	cilia and flagella associated protein 69	9139
	0.175480026472498	0.65445345142466
0.325778305171047	0.568155613394694	0.891073527784395
6640	SNTA1	syntrophin alpha 1	2342	0.16101850380683
	2.54994050395281	0.325696653497426
0.568204109660262	0.891073527784395
23155	CLCC1	chloride channel CLIC like 1	4964
0.113731728851675	5.19656898162637	0.325668555073463
	0.568220800326281	0.891073527784395
114038	LINC00313	long intergenic non-protein coding RNA 313
	3984	-0.168158126979404	1.54690972228988
0.325425301109008	0.568365334846186	0.89113286145999
90871	DMAC1	distal membrane arm assembly complex 1	3000
0.147392594884353	3.29656182719915	0.325308215889635
	0.56843492904793	0.89113286145999
29107	NXT1	nuclear transport factor 2 like export factor 1	1123
	0.114217382826352	3.27574982444515
0.325283708793648	0.568449497904263	0.89113286145999
57639	CCDC146	coiled-coil domain containing 146
4907	-0.150883009607847	3.444188348579	0.325024780104737
	0.568603468995606	0.891191293870704
387263	C6orf120	chromosome 6 open reading frame 120	2667
	0.140322045922372	3.78797580622665
0.32498317624399	0.568628216171467	0.891191293870704
11329	STK38	serine/threonine kinase 38
3576	-0.0719526240662223	7.66540037504541
0.324891833230716	0.568682556992795	0.891191293870704
57617	VPS18	VPS18 core subunit of CORVET and HOPS complexes
4049	-0.0767581600125575	6.20602554060035
0.324741783309026	0.568771845080463	0.891191293870704
83475	DOHH	deoxyhypusine hydroxylase	2387
0.105042029806419	3.65094725447707	0.324624471522666
	0.568841671184396	0.891191293870704
80167	ABHD18	abhydrolase domain containing 18
6491	-0.107275207869081	3.64951133010238
0.324579371609875	0.568868519918467	0.891191293870704
100128071	FAM229A family with sequence similarity 229 member A
	2845	0.204869990920989	2.37473424249995
0.324437518101295	0.568952983800102	0.891223936936707
9647	PPM1F	protein phosphatase, Mg2+/Mn2+ dependent 1F
9828	-0.105286884763863	4.76743476847955
0.324284070160331	0.569044378898727	0.891267428741414
101927151	LOC101927151	uncharacterized LOC101927151	4071
	0.153980424913503	2.27527127528128
0.324161847645803	0.569117196317357	0.891281816728223
148266	ZNF569	zinc finger protein 569	5016	0.156359202597242
	2.28019502943881	0.323731469859672
0.569373750481071	0.891425166389623

84172 POLR1B RNA polymerase I subunit B
6329 -0.0996791710610759 4.61105561217171
0.323710914917724 0.569386009214211 0.891425166389623
8884 SLC5A6 solute carrier family 5 member 6 5606
0.0902973803413367 4.37872859005107 0.323577838757208
0.569465386786934 0.891425166389623
51103 NDUFAF1 NADH:ubiquinone oxidoreductase complex assembly
factor 1 1790 0.162988382095263 4.11620503803525
0.323406360168267 0.569567702589594
0.891425166389623
162427 RETREG3 reticulophagy regulator family member 3 5145
0.0657451868325458 6.66600870776254 0.323375439489869
0.569586155791802 0.891425166389623
3851 KRT4 keratin 4 3312 -0.507128508295179
4.91612912450153 0.323319384151603 0.569619612125801
0.891425166389623
51634 RBMX2 RNA binding motif protein X-linked 2
2487 -0.104844393268202 3.65144117837126
0.323004374225668 0.569807695560738 0.891425166389623
84311 MRPL45 mitochondrial ribosomal protein L45
1705 -0.136154669130844 4.5949492097829
0.323000751320797 0.569809859399503 0.891425166389623
10885 WDR3 WD repeat domain 3 10275 0.101026907595835
4.71259930715241 0.32292768851346
0.569853500774694 0.891425166389623
57605 PITPNM2 phosphatidylinositol transfer protein membrane
associated 2 9161 0.214938199972087 2.69737899158147
0.322905398153092 0.569866816400181
0.891425166389623
5621 PRNP prion protein 2851 -0.0766595627539444
7.63094261919936 0.322798679361334 0.56993057558726
0.891425166389623
79152 FA2H fatty acid 2-hydroxylase
3010 -0.28068860764688 2.83187857580299
0.322728641405803 0.569972427370991 0.891425166389623
51021 MRPS16 mitochondrial ribosomal protein S16 2794
0.0696512962392805 6.0995658827453 0.322539902509504
0.570085239836652 0.891502060331806
6229 RPS24 ribosomal protein S24 4880 0.13481344111002
7.41473808860852 0.322196952481534
0.570290338704307 0.891604956607262
10605 PAIP1 poly(A) binding protein interacting protein 1
3391 -0.064045137520299 6.43106706178666
0.322168850235741 0.57030715145301 0.891604956607262
9141 PDCD5 programmed cell death 5 2253 -0.122631350465725
4.77978736040618 0.322110604543809
0.570342001228034 0.891604956607262
81622 UNC93B1 unc-93 homolog B1, TLR signaling regulator 3548
0.109921519245086 4.46697583707332
0.321655019279078 0.570614732483045 0.891834903170642
51232 CRIM1 cysteine rich transmembrane BMP regulator 1 6884
0.101673085537506 7.65770080196214
0.321616894622913 0.570637566974498 0.891834903170642
6730 SRP68 signal recognition particle 68

7641 -0.066659740126187 6.52010238500384
0.321545879985059 0.570680105455734 0.891834903170642
166 TLE5 TLE family member 5, transcriptional modulator 5298
0.0753542690343143 8.23009813589433
0.321227241540484 0.5708710495343 0.892033778539373
90987 ZNF251 zinc finger protein 251 4939 0.11517271194224
4.42309459029446 0.3210234838834 0.570993216903147
0.892125152165351
51020 HDDC2 HD domain containing 2 2558 0.105643888239392
3.90948825204386 0.320617401533838
0.571236845304421 0.892406256098486
101752400 CAPN10-DT CAPN10 divergent transcript 4000
0.192098900272973 0.468983983125021
0.320394252654327 0.571370809846387 0.892515995708525
100533179 UBE2F-SCLY UBE2F-SCLY readthrough (NMD
candidate) 3195 -0.0842871851388362 4.69333502512785
0.319738038235962 0.571765116908317
0.893032336366502
7050 TGIF1 TGFB induced factor homeobox 1
8222 -0.0936565837060075 6.96402974624893
0.319292680117108 0.572033028635426 0.893135833864843
84455 EFCAB7 EF-hand calcium binding domain 7 5704
0.147165254474856 2.3053750632757 0.319153037477589
0.572117083482811 0.893135833864843
8555 CDC14B cell division cycle 14B 10462 0.0739465549307157
5.13021726419276 0.319145378135696
0.572121694558661 0.893135833864843
138199 CARNMT1 carnosine N-methyltransferase 1
4610 -0.105355841610926 4.00691808178842
0.319033679288452 0.572188947777007 0.893135833864843
284996 RNF149 ring finger protein 149 4738 -0.0755670224894508
6.17051283023065 0.319011292624893
0.57220242852339 0.893135833864843
79073 TMEM109 transmembrane protein 109 2421
0.0865194066104695 6.15428042350181 0.31899213962371
0.572213962526101 0.893135833864843
79654 HECTD3 HECT domain E3 ubiquitin protein ligase 3
5077 -0.0642195518828489 6.04691018992814
0.318745459304612 0.57236255509267 0.893268223864404
23171 GPD1L glycerol-3-phosphate dehydrogenase 1 like
4802 -0.116548329070543 5.00048451669563
0.318558593917373 0.57247516748533 0.893344437496326
55871 CBWD1 COBW domain containing 1
14170 -0.0922094132915927 5.33083223370983
0.318325085912669 0.572615949773157 0.893464588222415
64794 DDX31 DEAD-box helicase 31 5078 0.105172083853156
3.94381402435723 0.318212314283464
0.572683964316635 0.893471183802975
151827 LRRC34 leucine rich repeat containing 34 4429
0.258979955841305 0.465706995294687 0.317550422260751
0.573083483545809 0.893845951090634
101928492 OR2A1-AS1 OR2A1 antisense RNA 1 5760
0.228729903128393 2.46984350904242 0.317520531984788
0.573101538313743 0.893845951090634

7818	DAP3	death associated protein 3
3978	-0.0789584355271731	6.08923775474973
0.317262870307218	0.57325722138524	0.893845951090634
79047	KCTD15	potassium channel tetramerization domain containing
15	6388	0.115306259041415	4.37010546594881
0.317192997784005	0.573299453770538	0.893845951090634
6446	SGK1	serum/glucocorticoid regulated kinase 1 9877
0.131628104401686	6.34586863382777	0.317146171835724
	0.573327759762579	0.893845951090634
10467	ZNHIT1	zinc finger HIT-type containing 1	5510
0.0742014383341388	5.65709130691051	0.317100164047474
	0.573355573862809	0.893845951090634
652965	SNORA48 small nucleolar RNA, H/ACA box 48
1473	-0.0993384466096535	5.96585267751796
0.31707485315976	0.573370876743724	0.893845951090634
51545	ZNF581	zinc finger protein 581 1736	0.122588975733724
	3.55889925483698	0.316802382740716
0.573535662440096	0.893886050854989
57003	CCDC47	coiled-coil domain containing 47	4537
0.071995248681884	7.20450758216963	0.316674086039573
	0.573613286522387	0.893886050854989
26578	OSTF1	osteoclast stimulating factor 1
1319	-0.0966311428443073	5.44937208126483
0.316635117134684	0.573636868199717	0.893886050854989
116840	CNTROB	centrobin, centriole duplication and spindle
assembly protein	6234	0.0800761111373994
4.92255793386354	0.316255235172022	0.573866850453942
	0.893886050854989
10614	HEXIM1	HEXIM P-TEFb complex subunit 1	3600
0.0741476698102925	6.80782208479529	0.316147270686479
	0.573932245818409	0.893886050854989
154865	IQUB	IQ motif and ubiquitin domain containing
5893	-0.222442465360977	-0.196672611604832
0.316117749382655	0.57395012977958	0.893886050854989
8455	ATRN	attractin	8753	-0.0760955521425065
6.91758764194325	0.316114799955876	0.573951916598244
	0.893886050854989
1978	EIF4EBP1	eukaryotic translation initiation factor 4E
binding protein 1	1073	0.141481525287504
5.03148475149217	0.316113674483148	0.573952598433854
	0.893886050854989
26052	DNM3	dynamin 3	11329	-0.151033801000957
1.36587518230161	0.316073639710938	0.573976853399944
	0.893886050854989
389	RHOC	ras homolog family member C	4505
0.101998220167207	8.41879480450566	0.315908459313192
	0.574076948911335	0.893886050854989
90423	ATP6V1E2	ATPase H+ transporting V1 subunit E2	5283
	0.143785638377483	1.75342595193124
0.315862169874329	0.574105005420841	0.893886050854989
7011	TEP1	telomerase associated protein 1
11328	-0.125414985372411	5.94694735781069
0.315767483109703	0.574162404479356	0.893886050854989
103021294	ABALON	apoptotic BCL2L1-antisense long non-coding

RNA	1903	-0.163598685576703	0.549214287900284
0.315534934350421	0.574303423464303	0.893942362716349
824	CAPN2	calpain 2	8792	0.0797948512106999
10.563717264058 0.315497356956712	0.574326217040367
0.893942362716349
165918	RNF168	ring finger protein 168 5347	0.106539243181347
	4.72113530230143	0.315263848764346
0.574467897735734	0.894063538176048
8402	SLC25A11	solute carrier family 25 member 11	2669
	0.088211659974865	5.18550236868355
0.315119877560686	0.574555286412376	0.894100199619837
192669	AGO3	argonaute RISC catalytic component 3
21157	-0.133220965063008	4.9786831620537 0.314796955850895
	0.574751391276704	0.894212591158287
8525	DGKZ	diacylglycerol kinase zeta
10787	-0.0821907175756904	5.64086661191829
0.31479070132827	0.574755190847295	0.894212591158287
116225	ZMYND19 zinc finger MYND-type containing 19
2372	-0.0832871324486652	4.32161509517562
0.314633356255634	0.574850793005651	0.894262001069542
81533	ITFG1	integrin alpha FG-GAP repeat containing 1	7770
	0.0700781241904509	6.69270624436688
0.313755721430869	0.575384617198309	0.894930189072561
2242	FES	FES proto-oncogene, tyrosine kinase	4237
0.184271031496296	1.74295486935193	0.313706388106546
	0.575414653458643	0.894930189072561
26608	TBL2	transducin beta like 2	5067	-0.0975108457604731
	4.76271792551826	0.313415624610197
0.575591745902249	0.894930189072561
9274	BCL7C	BAF chromatin remodeling complex subunit BCL7C	4398
	0.0928348025791201	4.21558307392978
0.313410173028245	0.575595067273571	0.894930189072561
51816	ADA2	adenosine deaminase 2	5800	0.290292262747617
	5.36049376488442	0.313402443117276
0.575599776779803	0.894930189072561
80306	MED28	mediator complex subunit 28	10982
0.0913229538223947	5.29511425898455	0.313115988963716
	0.575774354935417	0.895102262994494
5955	RCN2	reticulocalbin 2	10019	-0.0655888509177563
	6.92236849564844	0.312735841714861
0.576006195901445	0.895182336105825
23074	UHRF1BP1L	UHRF1 binding protein 1 like
6616	-0.0826309059982657	5.01197682928354
0.312584595048767	0.576098488370135	0.895182336105825
550	AUP1	AUP1 lipid droplet regulating VLDL assembly factor
	2634	-0.074965810032442	7.07413675017464
0.312531038281033	0.57613117635702	0.895182336105825
54965	PIGX	phosphatidylinositol glycan anchor biosynthesis
class X 4446	0.0980387321548668	3.87103338608824
0.312521476134259	0.576137012931092	0.895182336105825
149473	CCDC24	coiled-coil domain containing 24	3554
0.106780065488761	3.71795489540528	0.312507719178816
	0.57614541015164	0.895182336105825
84865	CCDC142 coiled-coil domain containing 142

4897	-0.0953391201838355	4.47175359647843
0.312299114063292	0.576272772072404	0.895280034890059
10946	SF3A3	splicing factor 3a subunit 3	4740
0.0729568491318277	5.70880555147555	0.311904042863702
	0.576514132176598	0.895280034890059
1947	EFNB1	ephrin B1	3314	-0.0784280654425424
6.85991985348076	0.311751353869866	0.576607467907813
	0.895280034890059
79022	TMEM106C	transmembrane protein 106C	3475
0.101388642209653	5.1544887359897 0.31158134007334
0.576711429229873	0.895280034890059
55900	ZNF302	zinc finger protein 302 5032	-0.107044451818247
	3.89819274781023	0.311342075724154
0.576857799437082	0.895280034890059
195828	ZNF367	zinc finger protein 367 3646	0.19090260213091
	0.249863365102988	0.311237186767984
0.576921988615809	0.895280034890059
166815	TIGD2	tigger transposable element derived 2	2083
0.127929754562041	3.14673083117016	0.311150950630003
	0.576974773403248	0.895280034890059
8879	SGPL1	sphingosine-1-phosphate lyase 1
7486	-0.0786234218879299	7.85332188100554
0.311128153647957	0.576988728945524	0.895280034890059
57337	SENP7	SUMO specific peptidase 7
6396	-0.125727429130339	3.98822349549052
0.311094817093167	0.577009137665095	0.895280034890059
222236	NAPEPLD N-acyl phosphatidylethanolamine phospholipase D 7561
	0.160570233335712	3.19962980972947
0.311071535881808	0.577023391329311	0.895280034890059
27335	EIF3K	eukaryotic translation initiation factor 3 subunit K
	2798	0.0871786483652852	5.86888543372529
0.310982546321398	0.577077880649594	0.895280034890059
6187	RPS2	ribosomal protein S2	2413	0.115696802244156
	7.43933251576409	0.310892646324191
0.577132937819774	0.895280034890059
55204	GOLPH3L golgi phosphoprotein 3 like
3329	-0.0801693829763413	5.28432288584433
0.310882672555856	0.577139046666143	0.895280034890059
221078	NSUN6	NOP2/Sun RNA methyltransferase 6	2476
0.133024674862958	2.38059140215711	0.310873390665333
	0.577144731858769	0.895280034890059
1535	CYBA	cytochrome b-245 alpha chain	3357
0.110485658799048	5.76397214952571	0.310750845147465
	0.577219801879308	0.895280034890059
79541	OR2A4	olfactory receptor family 2 subfamily A member 4
	933	-0.344397637249148	1.3685952061183
0.310732640334654	0.577230955600209	0.895280034890059
84284	NTPCR	nucleoside-triphosphatase, cancer-related
15854	0.107993546868883	4.87456150823867
0.310564317435956	0.577334103929139	0.895340876188601
154743	BMT2	base methyltransferase of 25S rRNA 2 homolog
4138	-0.0933308279818383	4.47023402085391
0.310382048646051	0.577445839625439	0.895415019997069
199223	TTC21A	tetratricopeptide repeat domain 21A

7677	-0.171340823123875	2.31972665448632
0.310091540840745	0.577624017662276	0.89550049332384
144715	RAD9B	RAD9 checkpoint clamp component B	3178
0.145693514825275	1.03685170528108	0.310083704626172
0.577628825385942	0.89550049332384
284440	LINC00663	long intergenic non-protein coding RNA 663
1645	0.16868651897161	0.162163524735413
0.309485909897629	0.577995822995784	0.895959229432605
51231	VRK3	VRK serine/threonine kinase 3
6144	-0.0771923157004631	4.86700496361357
0.309300005237173	0.578110048033476	0.895959229432605
285154	CYP1B1-AS1	CYP1B1 antisense RNA 1	7167
0.275874071772988	1.53709675394438e-05	0.30919719217003
0.578173238568549	0.895959229432605
151525	WDSUB1	WD repeat, sterile alpha motif and U-box domain containing 1	2135	0.108321229006038	3.36770534940558
0.309185238328122	0.578180586481139
0.895959229432605
80317	ZKSCAN3	zinc finger with KRAB and SCAN domains 3	4842
0.140550710671048	2.93559478831171
0.308940871762959	0.578330837049815	0.896092924176633
90024	FLJ20021	uncharacterized LOC90024	790
0.149868470702201	1.80065228858246	0.30861782039121
0.578529586972812	0.89626625087977
51652	CHMP3	charged multivesicular body protein 3
7018	-0.0773195888311154	7.57249983987699
0.308551063320181	0.578570674695144	0.89626625087977
199964	TMEM61	transmembrane protein 61	1097
0.174962845494418	0.941181040467316	0.308335564354
0.578703349962135	0.89637264435692
55744	COA1	cytochrome c oxidase assembly factor 1 homolog	5119
0.107564644601621	4.5229336140542	0.307970624754432
0.578928169069734	0.896579270811443
115950	ZNF653	zinc finger protein 653	3557	0.13776772061691
2.57051680706731	0.307743533863986
0.579068155165247	0.896579270811443
23658	LSM5	LSM5 homolog, U6 small nuclear RNA and mRNA degradation associated	4701	0.0903619973982157
4.01436995273377	0.307719374765214	0.579083051582016
0.896579270811443
64418	TMEM168	transmembrane protein 168
7995	-0.0797700481376148	4.91533546363219
0.307571039115324	0.579174531596335	0.896579270811443
54760	PCSK4	proprotein convertase subtilisin/kexin type 4
3792	-0.215852514783959	1.36196101949839
0.307533666522149	0.579197584181101	0.896579270811443
9094	UNC119	unc-119 lipid binding chaperone
5128	-0.0941969165609892	4.35817502133058
0.307496023504038	0.579220805423913	0.896579270811443
54467	ANKIB1	ankyrin repeat and IBR domain containing 1
7130	-0.0702273747017823	7.31337783926902
0.307184171853947	0.579413252121408	0.896716936891193
5982	RFC2	replication factor C subunit 2
2211	-0.101275697218919	3.60633508374781

0.307144439022611	0.579437780809752	0.896716936891193
131566	DCBLD2	discoidin, CUB and LCCL domain containing 2
11274	0.132630310253126	9.48959409668955
0.306892962476311	0.579593075599754	0.89674788130648
4627	MYH9	myosin heavy chain 9	8959	-0.0961937054001961
11.8203019940731	0.306879649490838
0.579601299110316	0.89674788130648
11221	DUSP10	dual specificity phosphatase 10 2914
0.174363503195683	6.82970325197859	0.306801073682863
0.579649840604617	0.89674788130648
3704	ITPA	inosine triphosphatase	2071	0.114217792858243
5.00334960316222	0.306521089208964
0.579822871673061	0.896916506187142
64423	INF2	inverted formin, FH2 and WH2 domain containing
9488	-0.101275717775258	7.35800026336957
0.306370669575358	0.579915874002419	0.896961312627195
440270	GOLGA8B golgin A8 family member B	7086
0.13187648170664	6.78955233641081	0.306118438195554
0.580071892052526	0.897080467070203
79833	GEMIN6	gem nuclear organelle associated protein 6	4361
0.127972695947664	2.90971567743057
0.305906822455068	0.580202852022841	0.897080467070203
60560	NAA35	N(alpha)-acetyltransferase 35, NatC auxiliary subunit 2726	-0.0678876556851234	5.41323153857698
0.305884202169306	0.580216854253698	0.897080467070203
4193	MDM2	MDM2 proto-oncogene	13267	0.0850073241988108
7.4519306902182 0.305832126973201	0.580249092000859
0.897080467070203
79159	NOL12	nucleolar protein 12	3589	-0.098085171831455
3.4893003190559 0.305445303131348	0.580488671989679
0.897351818624814
84650	EBPL	EBP like	1500	0.236503540049305
2.8219051892075 0.305188642416027	0.580647744548043
0.897498671141518
1951	CELSR3	cadherin EGF LAG seven-pass G-type receptor 3
13873	0.187259061212034	3.3836223819939 0.305017733109525
0.580753718735407	0.897563426936649
23211	ZC3H4	zinc finger CCCH-type containing 4
6502	-0.0997510993536197	5.34501191946424
0.304764835079794	0.580910602035897	0.897659144833651
5644	PRSS1	serine protease 1	2081	-0.454506815307118
3.42958344842942	0.304711297784365
0.580943824367605	0.897659144833651
23649	POLA2	DNA polymerase alpha 2, accessory subunit
5726	-0.109502482816425	3.48011529574097
0.304485082164627	0.581084243520311	0.897777079085484
5007	OSBP	oxysterol binding protein
5492	-0.0610011355018801	6.78256748616511
0.304362420266036	0.581160412073588	0.897795730882844
63933	MCUR1	mitochondrial calcium uniporter regulator 1
6057	-0.0807268116733946	5.01405377555334
0.304126811465506	0.581306772698319	0.897868430666311
9375	TM9SF2	transmembrane 9 superfamily member 2
3885	-0.0690888817329698	8.48092953732419

0.304011311709814	0.581378548368977	0.897868430666311
55777	MBD5	methyl-CpG binding domain protein 5
14352	-0.103113713323741	4.1694692124941 0.30387128463002
	0.581465590041986	0.897868430666311
55666	NPLOC4	NPL4 homolog, ubiquitin recognition factor
9574	-0.0660248432774111	7.87919376892138
0.303560595956469	0.581658809399081	0.897868430666311
8711	TNK1	tyrosine kinase non receptor 1
3860	-0.0977905723993094	4.63176033980084
0.303555451843796	0.581662009641611	0.897868430666311
84777	DLGAP1-AS2	DLGAP1 antisense RNA 2	2261
0.183067274341633	1.27752920937567	0.30332826308927
	0.581803383008039	0.897868430666311
11113	CIT	citron rho-interacting serine/threonine kinase
13230	-0.247449765976313	1.41725351842059
0.303265128773422	0.581842682023003	0.897868430666311
79157	MFSD11	major facilitator superfamily domain containing 11
	6548	0.0782310086291994	5.30512220795425
0.303224603669279	0.581867910363919	0.897868430666311
388796	SNHG17	small nucleolar RNA host gene 17
4186	-0.18352074637605	3.27572908641284
0.303185886797372	0.581892015067223	0.897868430666311
8737	RIPK1	receptor interacting serine/threonine kinase 1
4231	-0.063690348655138	6.25376311068772
0.303103911572826	0.581943058570179	0.897868430666311
2874	GPS2	G protein pathway suppressor 2	4280
0.082313818186403	5.51153427214482	0.303056385995923
	0.581972655434367	0.897868430666311
9326	ZNHIT3	zinc finger HIT-type containing 3	6687
0.0861597427662831	4.8050543100323 0.302950922190082
0.582038344512472	0.897868430666311
2634	GBP2	guanylate binding protein 2
6276	-0.15932802823776	4.8847216216289 0.302827019196862
	0.582115537639242	0.897868430666311
2671	GFER	growth factor, augmenter of liver regeneration
2687	-0.103790442768169	4.17665087222972
0.302765776925426	0.582153699944558	0.897868430666311
105378663	LOC105378663	uncharacterized LOC105378663	976
	0.157637753053965	0.33228934470712
0.302635273889721	0.582235037948742	0.897868430666311
8745	ADAM23	ADAM metallopeptidase domain 23 6421
0.475123181087887	0.268738474358209	0.302565410759625
	0.58227859059715	0.897868430666311
117854	TRIM6	tripartite motif containing 6	3947
0.174002269334497	2.80262993083967	0.302535568918476
	0.582297195985776	0.897868430666311
9847	C2CD5	C2 calcium dependent domain containing 5
6759	-0.0937607465523167	4.72533002288098
0.302390692464279	0.582387538593455	0.897908888616238
92105	INTS4	integrator complex subunit 4
6074	-0.0795841635904	4.34547868560212
0.302124797448673	0.582553419775216	0.898065788112641
22796	COG2	component of oligomeric golgi complex 2 10938
0.071101647980531	5.07180804187543	0.301806212897365

0.582752297046799	0.898264359287039
6164	RPL34	ribosomal protein L34	3393	0.120342442801025
6.34580587057136	0.301610695856581
0.582874416763974	0.898264359287039
283	ANG	angiogenin	1430	0.165379069417496
1.54187132429455	0.301610375199079	0.582874617088591
0.898264359287039
7779	SLC30A1 solute carrier family 30 member 1	5474
0.0889224859084945	6.23554698964766	0.301504166777937
0.582940976387515	0.898267794968088
284023	RNF227	ring finger protein 227 2529	0.16620212550958
1.33422430123609	0.301304504301967
0.583065767213161	0.898319742936536
5496	PPM1G	protein phosphatase, Mg2+/Mn2+ dependent 1G	4092
0.0676157953698437	6.68085741321308
0.301245019953484	0.583102955863425	0.898319742936536
64376	IKZF5	IKAROS family zinc finger 5
4934	-0.0998920712491788	4.21472799803056
0.301024986120464	0.583240558984387	0.898348942622601
51651	PTRH2	peptidyl-tRNA hydrolase 2	2612
0.116454745737551	4.43522092940747	0.301009604100003
0.583250180924533	0.898348942622601
51594	NBAS	NBAS subunit of NRZ tethering complex
8948	-0.0879478194515893	5.5517663122	0.300551746926065
0.583536731621322	0.898541213256708
54458	PRR13	proline rich 13 2897	0.0764784993125475
5.57950998629064	0.300516551722101	0.583558770346908
0.898541213256708
339175	METTL2A methyltransferase like 2A
3843	-0.0967961145198774	4.47301107370309
0.300502674790965	0.583567460340992	0.898541213256708
284695	ZNF326	zinc finger protein 326 10984	-0.0765227656262284
5.4807547576306 0.300396612373126	0.583633887233051
0.898544719550818
84501	SPIRE2	spire type actin nucleation factor 2	7677
0.253604407642788	2.05030472875995	0.300170702754313
0.583775425274515	0.898630500358611
112936	VPS26B	VPS26, retromer complex component B	6483
0.0766058906488356	5.92066333285035	0.300035021358735
0.583860466359817	0.898630500358611
6472	SHMT2	serine hydroxymethyltransferase 2	4838
0.070351267425053	6.46376024404952	0.299806165752941
0.584003962942154	0.898630500358611
6780	STAU1	staufen double-stranded RNA binding protein 1
4094	-0.0658285635300781	7.48214708025903
0.299754813100247	0.584036171758169	0.898630500358611
154467	CCDC167 coiled-coil domain containing 167
619	-0.144709866292464	3.35635818824241
0.299725094926345	0.584054812885095	0.898630500358611
51574	LARP7	La ribonucleoprotein 7, transcriptional regulator
4083	0.0822369512826846	5.26249061008125
0.299345025584849	0.584293322497966	0.898630500358611
100507217	LINC01578	long intergenic non-protein coding
RNA 1578	5395	0.0916311507181315	5.52565249010512

0.299221778452726	0.584370707537714
0.898630500358611
80255	SLC35F5	solute carrier family 35 member F5
7445	-0.0727259908046626	6.26789313471317
0.299120789103021	0.584434132683191	0.898630500358611
55017	C14orf119	chromosome 14 open reading frame 119	3676
0.0767459508762332	6.05751356807657
0.298950748575066	0.58454095602356	0.898630500358611
7089	TLE2	TLE family member 2, transcriptional corepressor
4038	0.153086718960061	5.39543085273295
0.298764672809384	0.584657898295511	0.898630500358611
54805	CNNM2	cyclin and CBS domain divalent metal cation transport mediator 2	16324	0.171032215514202
3.8040907860603	0.298702088716567	0.584697240892267
0.898630500358611
5635	PRPSAP1	phosphoribosyl pyrophosphate synthetase associated protein 1	5958	0.0797154433568143	5.76334536689484
0.298534690774687	0.584802499531477
0.898630500358611
100506334	LINC00649	long intergenic non-protein coding RNA 649	11854	0.224431343561248	3.22086483529159
0.298476650928052	0.584839003522317	0.898630500358611
127687	C1orf122	chromosome 1 open reading frame 122	2231
0.102109341867564	4.74807557226932
0.298457381787351	0.584851123810681	0.898630500358611
6777	STAT5B	signal transducer and activator of transcription 5B
6447	-0.075335053865382	5.4172687680436
0.298351637582967	0.584917645954185	0.898630500358611
144568	A2ML1	alpha-2-macroglobulin like 1
7303	-0.391412254219788	3.39297252246429
0.298236875985744	0.584989858130121	0.898630500358611
81029	WNT5B	Wnt family member 5B	4305	-0.150355892378152
5.78916071795317	0.298213316763615
0.585004684682022	0.898630500358611
9631	NUP155	nucleoporin 155	9233	-0.0893468594845734
4.85187504021367	0.298190956284692	0.585018757530787
0.898630500358611
2297	FOXD1	forkhead box D1	2107	0.472658020438787
1.5357004505904	0.298141182894387	0.585050085499783
0.898630500358611
91304	TMEM259	transmembrane protein 259	3936
0.070633504156214	7.24601285786651	0.298077446644569
0.585090206817166	0.898630500358611
150094	SIK1	salt inducible kinase 1	986	0.105130826690565
5.40934814381897	0.298018172800631
0.585127524092939	0.898630500358611
23541	SEC14L2	SEC14 like lipid binding 2
5985	-0.10672427985526	7.4963203900774	0.298018104184223
0.585127567294842	0.898630500358611
10152	ABI2	abl interactor 2	23331	0.0935897501628545
5.00279251639175	0.297851281454442
0.585232620440034	0.898630500358611
79173	C19orf57	chromosome 19 open reading frame 57
3840	-0.179136893186658	0.634982265169143

0.297736734065072	0.58530477638847	0.898630500358611
91582	RPS19BP1	ribosomal protein S19 binding protein 1 3763
	0.0771376976017475	5.58861036511646
0.297669528005006	0.585347119365388	0.898630500358611
51474	LIMA1	LIM domain and actin binding 1	6317
0.0703390061834974	8.79578175783322	0.297352495368202
	0.585546948514819	0.898630500358611
26122	EPC2	enhancer of polycomb homolog 2
5582	-0.0994009354489742	5.04948504929925
0.297225696446368	0.585626909959599	0.898630500358611
9112	MTA1	metastasis associated 1 7212	0.0705834820739457
	5.37621948309432	0.297183152196107
0.585653744007148	0.898630500358611
1954	MEGF8	multiple EGF like domains 8	11372
0.12328945431786	6.26906246116332	0.297177451501177
	0.585657339810839	0.898630500358611
51715	RAB23	RAB23, member RAS oncogene family
4926	-0.0957209624326043	4.59218319382474
0.297063948652738	0.585728942846311	0.898630500358611
5986	RFNG	RFNG O-fucosylpeptide 3-beta-N-
acetylglucosaminyltransferase	3305	0.0842111782101599
5.36354644964199	0.297033271638625	0.585748298422476
	0.898630500358611
60493	FASTKD5 FAST kinase domains 5	2874	-0.0695640154724553
	5.34641227895429	0.296746617212555
0.585929224911978	0.898630500358611
11344	TWF2	twinfilin actin binding protein 2
1824	-0.0750734119511512	5.8282127080423
0.296442156585272	0.586121514205827	0.898630500358611
7746	ZSCAN9	zinc finger and SCAN domain containing 9
3705	-0.0970490682147506	4.15710226978188
0.296360636362445	0.586173021953202	0.898630500358611
3364	HUS1	HUS1 checkpoint clamp component
4177	-0.104808105532821	4.36725600539306
0.296329937298704	0.586192421233714	0.898630500358611
55256	ADI1	acireductone dioxygenase 1	5560
0.089729943889516	6.74887247843325	0.29631671979277
	0.586200774010293	0.898630500358611
6152	RPL24	ribosomal protein L24	2682	-0.0962067619962581
	7.29960645723645	0.296229551442366
0.586255865924654	0.898630500358611
25853	DCAF12	DDB1 and CUL4 associated factor 12
4390	-0.066533169059456	6.68064287864446
0.296085899439163	0.586346679419719	0.898630500358611
8541	PPFIA3	PTPRF interacting protein alpha 3
6451	-0.204910550958366	1.98158131140736
0.295869957429659	0.586483246731117	0.898630500358611
124944	C17orf49	chromosome 17 open reading frame 49
2370	-0.0967607002252775	4.11304995021539
0.295738035029528	0.586566709650808	0.898630500358611
7553	ZNF7	zinc finger protein 7	10296	-0.0868354463115048
	4.09102689219339	0.295601459316247
0.586653141983049	0.898630500358611
91689	SMDT1	single-pass membrane protein with aspartate rich

tail 1  1731    -0.0849034691171542     4.15396974370554
0.295255740120161       0.586872047047742       0.898630500358611
23362   PSD3    pleckstrin and Sec7 domain containing 3
14812   -0.0928069423446807     6.31991085517452
0.295190316169339       0.586913491351879       0.898630500358611
4094    MAF     MAF bZIP transcription factor   6976
0.161238166884074       5.18579198441459        0.295181839207679
        0.586918861713213       0.898630500358611
10961   ERP29   endoplasmic reticulum protein 29        3083
0.0672736533756973      6.09247731718172        0.295041404472606
        0.58700784506858        0.898630500358611
254887  ZDHHC23 zinc finger DHHC-type containing 23     7138
0.122011399235065       3.42871786925875        0.294972681503896
        0.587051399848241       0.898630500358611
8678    BECN1   beclin 1        6414    -0.0616199571484253
6.8119605109625 0.294955450773443       0.58706232125458
0.898630500358611
84451   MAP3K21 mitogen-activated protein kinase kinase kinase 21
        7916    -0.140797560162014      2.87804887899774
0.294910904609374       0.58709055799996        0.898630500358611
10817   FRS3    fibroblast growth factor receptor substrate 3   2858
        0.166229596119301       1.32329277748516
0.294667612501995       0.5872448237787 0.898630500358611
56624   ASAH2   N-acylsphingosine amidohydrolase 2      6985
0.12904389071783        2.20322080989272        0.294649224055352
        0.58725648680945        0.898630500358611
27349   MCAT    malonyl-CoA-acyl carrier protein transacylase   2388
        0.119670937996051       3.36906389188104
0.294493378458913       0.587355352144824       0.898630500358611
7335    UBE2V1  ubiquitin conjugating enzyme E2 V1
4818    -0.0823664535514696     7.08311913548784
0.294480721445879       0.587363382987941       0.898630500358611
23558   WBP2    WW domain binding protein 2
5399    -0.0877383937446845     6.92650564623594
0.294337762330017       0.587454105705172       0.898630500358611
9704    DHX34   DExH-box helicase 34    5696    -0.101158335334103
        4.16835397874856        0.294317213590294
0.587467148403318       0.898630500358611
57470   LRRC47  leucine rich repeat containing 47       4516
0.0674609425370143      5.68190700153642        0.294117355176022
        0.587594033305122       0.898630500358611
54784   ALKBH4  alkB homolog 4, lysine demethylase      2312
0.127293399643659       3.44848148504178        0.294098377881054
        0.58760608439545        0.898630500358611
29916   SNX11   sorting nexin 11        3606    0.0870190149958024
        4.90796993906246        0.294090773881985
0.587610913279617       0.898630500358611
100533975       SLMO2-ATP5E     SLMO2-ATP5E readthrough
612     -0.147179039026194      3.34881169213849
0.294021418688168       0.587654960701369       0.898630500358611
116236  ABHD15  abhydrolase domain containing 15        3592
0.0917541236451559      4.62205376172527        0.293967880424143
        0.587688967403009       0.898630500358611
286826  LIN9    lin-9 DREAM MuvB core complex component

4442	-0.132018645109086	2.71898536210721
0.293683784568693	0.587869487900992	0.898630500358611
115752	DIS3L	DIS3 like exosome 3'-5' exoribonuclease
5436	-0.113823647439261	3.91422870717457
0.293676475775762	0.587874133552324	0.898630500358611
2738	GLI4	GLI family zinc finger 4
2912	-0.151336286788618	3.00075809889534
0.293488040632345	0.587993933463068	0.898630500358611
148479	PHF13	PHD finger protein 13	3681	0.110036150651069
	5.09109205897983	0.293461418522	0.588010862800518
	0.898630500358611
29086	BABAM1	BRISC and BRCA1 A complex member 1	2964
0.0714585966747579	6.02466526026474	0.293398477730747
	0.588050891597934	0.898630500358611
221935	SDK1	sidekick cell adhesion molecule 1
12062	-0.234418278410318	3.21781208376244
0.293388936362124	0.588056960158761	0.898630500358611
9491	PSMF1	proteasome inhibitor subunit 1
5118	-0.0699954534291624	6.34717439542257
0.293370271876318	0.588068831630558	0.898630500358611
100526825	INMT-MINDY4	INMT-MINDY4 readthrough (NMD
candidate)	2880	0.147843047708933	1.37298568792787
	0.293269885120059	0.588132690606099
0.898630500358611
23235	SIK2	salt inducible kinase 2	9796	-0.0866160548371386
	6.14036863668822	0.293067045639567
0.588261765926904	0.898630500358611
1974	EIF4A2	eukaryotic translation initiation factor 4A2	6133
	0.0686544520718856	8.31508455509961
0.293041902486564	0.588277769599411	0.898630500358611
5699	PSMB10	proteasome 20S subunit beta 10
1960	-0.133575769790312	3.95528989660663
0.29282830886431	0.588413758228281	0.898630500358611
84273	NOA1	nitric oxide associated 1	3415
0.0833535179569637	4.56268528040898	0.292808078342091
	0.5884266417166	0.898630500358611
3685	ITGAV	integrin subunit alpha V	8211
0.11194744109542	10.6398607124657	0.292746554075165
	0.588465826009589	0.898630500358611
341640	FREM2	FRAS1 related extracellular matrix 2
16223	-0.225424322801801	0.124157449233525
0.292717820195412	0.588484128201732	0.898630500358611
55260	TMEM143	transmembrane protein 143	3701
0.169877492792336	1.47681520581479	0.292664563494851
	0.588518053401363	0.898630500358611
11284	PNKP	polynucleotide kinase 3'-phosphatase	4025
0.086290858660405	4.93097173652994	0.292544835729518
	0.588594336090317	0.898630500358611
23503	ZFYVE26	zinc finger FYVE-type containing 26
15706	-0.111433941735893	6.43615187807682
0.292459577102221	0.588648669602963	0.898630500358611
55619	DOCK10	dedicator of cytokinesis 10	11065
0.364076872254681	1.69572328127163	0.29242773847443
	0.588668962309311	0.898630500358611

64769	MEAF6	MYST/Esa1 associated factor 6
3487	-0.0713712310955965	5.61603854174628
0.292264953506272	0.588772737450386	0.898630500358611
8741	TNFSF13	TNF superfamily member 13	2276
0.11796761413248	5.05203561352747	0.292206878750017
	0.588809769047612	0.898630500358611
54681	P4HTM	prolyl 4-hydroxylase, transmembrane
6326	-0.0892331549307444	4.6821425936474	0.292138772866393
	0.588853203091436	0.898630500358611
5687	PSMA6	proteasome 20S subunit alpha 6
2292	-0.0987203401858318	6.62674095737363
0.292086999354726	0.588886225658006	0.898630500358611
4666	NACA	nascent polypeptide associated complex subunit alpha
	10008	-0.0805936623021358	7.54567375983595
0.291765156839688	0.589091590507187	0.898716043333805
55212	BBS7	Bardet-Biedl syndrome 7	4829	0.0756677957100393
	4.62387053394411	0.291760598528826
0.589094500175498	0.898716043333805
51573	GDE1	glycerophosphodiester phosphodiesterase 1
4416	-0.10286878195011	7.90361858183524
0.291391728934016	0.589330054988665	0.898716043333805
3612	IMPA1	inositol monophosphatase 1
3720	-0.0871082400536387	4.4376421058755	0.291387355142877
	0.589332849183533	0.898716043333805
84188	FAR1	fatty acyl-CoA reductase 1
7501	-0.0915232219496235	6.59326608187525
0.291387191640013	0.589332953637804	0.898716043333805
100507463	PSMB8-AS1	PSMB8 antisense RNA 1 (head to head)
	1951	-0.270536633743096	0.505519775006954
0.291234826220327	0.589430309173786	0.898716043333805
112609	MRAP2	melanocortin 2 receptor accessory protein 2	2153
	0.152890372857195	0.909896340117536
0.291218289125269	0.589440877707977	0.898716043333805
5690	PSMB2	proteasome 20S subunit beta 2
4843	-0.0837448058292063	6.96857695825862
0.291160544875623	0.589477783967965	0.898716043333805
102800317	TPTEP2-CSNK1E	TPTEP2-CSNK1E readthrough
2321	-0.0652981450663085	6.63615239066321
0.291030709911112	0.589560783048144	0.898716043333805
10618	TGOLN2	trans-golgi network protein 2
7081	-0.0806076268653573	8.1190411006641	0.290994550961102
	0.589583902491207	0.898716043333805
51765	STK26	serine/threonine kinase 26	3354
0.0969811365432247	5.48755325679218	0.290452905513341
	0.589930443891315	0.899146433904859
10284	SAP18	Sin3A associated protein 18	3396
0.072509140586756	6.49966705680709	0.290226476422589
	0.5900754354206	0.899269570660032
28511	NKIRAS2	NFKB inhibitor interacting Ras like 2
5566	-0.0715871140680931	6.11997919559445
0.289807750899051	0.590343754345764	0.899560096410584
8857	FCGBP	Fc fragment of IgG binding protein
12961	-0.351392071494243	3.13610793635005
0.289702548656207	0.590411207141357	0.899560096410584

10630	PDPN	podoplanin	3707	0.138609107377692
6.68863903158809	0.28956551281783	0.590499094458201
	0.899560096410584
7849	PAX8	paired box 8	7400	-0.395055597992895
4.82480451393405	0.289482748349428	0.590552188067284
	0.899560096410584
6156	RPL30	ribosomal protein L30	3374	0.111787679305795
	7.18859976710593	0.289427348775689
0.590587732497764	0.899560096410584
55210	ATAD3A	ATPase family AAA domain containing 3A	3069
0.0854514776148204	4.84510512647641	0.289328128434814
	0.590651403347479	0.899560096410584
55565	ZNF821	zinc finger protein 821	3402	0.113849494389073
	3.59448600436183	0.289219126489936
0.590721367394699	0.899568840301973
26974	ZNF285	zinc finger protein 285	6866	-0.233887038148142
	0.596957485912219	0.288970019188361
0.590881323373315	0.899714609902166
245711	SPDYA	speedy/RINGO cell cycle regulator family member A
	3483	-0.0857481401537301	6.36627002343663
0.288797189310969	0.590992352562686	0.899785856776689
55651	NHP2	NHP2 ribonucleoprotein	1510	0.112331559488421
	4.58282535100218	0.288527839435204
0.591165473420814	0.899951612455748
23130	ATG2A	autophagy related 2A	7870	-0.103277125275933
	5.85859596896124	0.288231861424322
0.591355829252142	0.900036194425162
199692	ZNF627	zinc finger protein 627	3514	-0.0973831962641075
	4.36274543080956	0.288133859532058
0.591418886112349	0.900036194425162
2108	ETFA	electron transfer flavoprotein subunit alpha
5031	-0.0960367476538328	6.46427953758585
0.288091129056099	0.591446384290236	0.900036194425162
65080	MRPL44	mitochondrial ribosomal protein L44
1748	-0.0770811171952146	5.22390616590529
0.28778786213681	0.591641619991597	0.900036194425162
2997	GYS1	glycogen synthase 1	3983	-0.0855258404228645
	6.27447373352102	0.287759505948173
0.591659881772287	0.900036194425162
400746	NCMAP	non-compact myelin associated protein
4122	-0.311022769441059	1.06045465143208
0.287747500613535	0.591667613724244	0.900036194425162
3297	HSF1	heat shock transcription factor 1
5307	-0.0680248651632141	6.04256005660504
0.287735980188206	0.591675033569041	0.900036194425162
255057	CBARP	CACN subunit beta associated regulatory protein	4543
	0.189824185426543	1.65430360564728
0.287511647482689	0.591819555425626	0.900036194425162
8834	TMEM11	transmembrane protein 11
3309	-0.0855932659527286	4.59280805087482
0.287475296079343	0.591842980924023	0.900036194425162
285830	HLA-F-AS1	HLA-F antisense RNA 1	3972
0.150827695178028	2.01246139449013	0.287345993072663
	0.591926321565703	0.900036194425162

3127 HLA-DRB5 major histocompatibility complex, class II, DR beta 5 1260 0.490076806589786 3.1636971909799 0.287343618945023 0.591927852005754 0.900036194425162
83638 C11orf68 chromosome 11 open reading frame 68 1619 0.0815004873109206 4.77094668162533 0.287069024257754 0.592104919627301 0.900168374833441
105376805 LOC105376805 uncharacterized LOC105376805 508 -0.18356133628106 0.705157657939031 0.287009504218192 0.592143314465291 0.900168374833441
8399 PLA2G10 phospholipase A2 group X 1116 0.207086943953831 0.563323212692344 0.286799152208849 0.59227904817107 0.900274869569761
23201 FAM168A family with sequence similarity 168 member A 7887 0.0752573585237584 6.43998258223983 0.286701759167592 0.592341914610903 0.900274869569761
4704 NDUFA9 NADH:ubiquinone oxidoreductase subunit A9 10878 -0.0718340848814217 5.89622249610775 0.28636208238618 0.592561280869994 0.900425874183412
60625 DHX35 DEAH-box helicase 35 4930 -0.114265861317181 3.44014935468287 0.28633065908065 0.592581582773919 0.900425874183412
55661 DDX27 DEAD-box helicase 27 5457 -0.0651059144534643 6.14748525187339 0.286249350523754 0.592634121089232 0.900425874183412
79038 ZFYVE21 zinc finger FYVE-type containing 21 8390 0.0701455779044817 5.67939609311809 0.285832293242855 0.592903757678216 0.900661822111715
136647 MPLKIP M-phase specific PLK1 interacting protein 7668 0.0961026986195376 3.64038674602006 0.285810248953637 0.592918016826738 0.900661822111715
100630923 LOC100630923 LOC100289561-PRKRIP1 readthrough 3455 0.0824819379097307 5.35988366283802 0.28513289302356 0.593356503400087 0.901113056869462
84622 ZNF594 zinc finger protein 594 4651 -0.184791374170559 1.81203766663713 0.285063421652052 0.593401513549664 0.901113056869462
10672 GNA13 G protein subunit alpha 13 5104 -0.0704747470871778 6.55307145272162 0.28496220416204 0.593467104465244 0.901113056869462
55716 LMBR1L limb development membrane protein 1 like 6047 -0.0784124798679972 5.38732413112606 0.284725085529171 0.593620820608991 0.901113056869462
50485 SMARCAL1 SWI/SNF related, matrix associated, actin dependent regulator of chromatin, subfamily a like 1 5092 0.0764056321184294 4.72481967949056 0.284556589423419 0.593730101030838 0.901113056869462
51175 TUBE1 tubulin epsilon 1 3853 -0.124727480681748 2.80248539029111 0.284417373105908 0.593820423010965 0.901113056869462
6428 SRSF3 serine and arginine rich splicing factor 3 6434 -0.0745088289099646 7.57567945518196 0.284373082078645 0.593849164484384 0.901113056869462
401505 TOMM5 translocase of outer mitochondrial membrane 5 3644 0.107074454366902 4.77829500599108

0.284294467106506	0.593900186641062	0.901113056869462
2588	GALNS	galactosamine (N-acetyl)-6-sulfatase	8402
0.0961896953438549	4.68320667400565	0.284133194104264
	0.594004883315911	0.901113056869462
55536	CDCA7L	cell division cycle associated 7 like	4074
0.197551956866309	2.92243890244545	0.28366770073286
	0.594307290687787	0.901113056869462
64976	MRPL40	mitochondrial ribosomal protein L40	1965
0.10460449200378	3.84336243341452	0.283655660130126
	0.594315117079965	0.901113056869462
7123	CLEC3B	C-type lectin domain family 3 member B
1330	-0.13400295499011	2.03508499844851
0.283629519241866	0.594332109392212	0.901113056869462
7088	TLE1	TLE family member 1, transcriptional corepressor
	4906	0.0875748111714823	5.26121728398114
0.283576405660082	0.594366637809661	0.901113056869462
80324	PUS1	pseudouridine synthase 1	5732
0.108267888651283	3.57923297181522	0.283555064545029
	0.594380512547322	0.901113056869462
6224	RPS20	ribosomal protein S20	3002	0.112403613377152
	7.52042819050185	0.283466863565053
0.594437862745753	0.901113056869462
127396	ZNF684	zinc finger protein 684	3039	0.147365184573373
	1.12905475571279	0.283463456073398
0.59444007860079	0.901113056869462
57724	EPG5	ectopic P-granules autophagy protein 5 homolog
13675	-0.0960907821414931	6.35069251653396
0.283385905559015	0.594490513485136	0.901113056869462
2272	FHIT	fragile histidine triad diadenosine triphosphatase
	5439	0.155335885290496	1.64940293233801
0.283321971984541	0.594532099269746	0.901113056869462
389856	USP27X	ubiquitin specific peptidase 27 X-linked
2618	-0.135099354408446	2.17903889073362
0.283224705113719	0.594595378347024	0.901113056869462
89849	ATG16L2	autophagy related 16 like 2
8617	-0.148459924278229	3.36880490986737
0.283058876346332	0.594703294013747	0.901113056869462
54880	BCOR	BCL6 corepressor	7642	-0.127857178372935
	3.89429618389637	0.282869077862668
0.594826858136235	0.901113056869462
83874	TBC1D10A	TBC1 domain family member 10A
7047	-0.0676724008188661	5.4558201125804	0.282719686026294
	0.594924153809516	0.901113056869462
55621	TRMT1	tRNA methyltransferase 1	3364
0.0812820910004413	4.51639276223582	0.282698756003736
	0.594937787714029	0.901113056869462
78994	PRR14	proline rich 14	3860	0.0834627068877983
5.16799992128051	0.282583921453916	0.595012602943637
	0.901113056869462
25930	PTPN23	protein tyrosine phosphatase non-receptor type 23
	5981	-0.0727128786489312	6.37794703293288
0.282539847644856	0.595041322407567	0.901113056869462
22841	RAB11FIP2	RAB11 family interacting protein 2	6169
	0.0980689530723375	4.48445633626501

0.282393880392338	0.595136458399217	0.901113056869462
55856	ACOT13	acyl-CoA thioesterase 13	4394
0.116465004270373	3.97049182352733	0.282367515407495
	0.595153645471318	0.901113056869462
64795	RMND5A	required for meiotic nuclear division 5 homolog A
	6804	-0.104760038027505	5.62398295264068
0.282175564538122	0.595278807364751	0.901113056869462
23607	CD2AP	CD2 associated protein	6751	-0.0903471493674061
	7.46825208887185	0.282152781726722
0.595293666559486	0.901113056869462
286336	FAM78A	family with sequence similarity 78 member A	5499
	0.222594211360708	1.09795325496608
0.282136714119931	0.595304146488048	0.901113056869462
2091	FBL	fibrillarin	2173	0.107026906641885
4.35886933850846	0.282078640803661	0.595342027392263
	0.901113056869462
27005	USP21	ubiquitin specific peptidase 21	3764
0.13046830668511	4.18590593624463	0.282032819925339
	0.595371919627318	0.901113056869462
10585	POMT1	protein O-mannosyltransferase 1	6067
0.105055571240304	5.24488934624645	0.281776478364983
	0.59553920700256	0.901113056869462
57533	TBC1D14	TBC1 domain family member 14
6241	-0.0662168958905188	6.7296360498581	0.281754416384516
	0.595553609111932	0.901113056869462
5250	SLC25A3	solute carrier family 25 member 3
8896	-0.0746387760480587	8.46863716473043
0.28172816827211	0.595570744878909	0.901113056869462
84225	ZMYND15	zinc finger MYND-type containing 15
3303	-0.210674727806384	3.09250376270012
0.281710791656115	0.595582089558193	0.901113056869462
642	BLMH	bleomycin hydrolase	4012	0.128736393722694
	3.30738487247623	0.281690414508533
0.595595393767222	0.901113056869462
4287	ATXN3	ataxin 3	28202	-0.0942860425763151
4.90554629365086	0.281454198441383	0.595749663914194
	0.90124911355936
10129	FRY	FRY microtubule binding protein	18452
0.153010754420674	2.57814908218052	0.281175722321436
	0.595931640226267	0.901355439703449
22893	BAHD1	bromo adjacent homology domain containing 1	5196
	0.0886772540405184	5.80303220561658
0.2809190383106	0.596099478290667	0.901355439703449
54921	CHTF8	chromosome transmission fidelity factor 8
3383	-0.069556970259914	6.4764028223436	0.280902324041376
	0.596110410661463	0.901355439703449
6868	ADAM17	ADAM metallopeptidase domain 17	5491
0.0656369774435429	6.64679998506336	0.280828662562598
	0.596158595692199	0.901355439703449
10195	ALG3	ALG3 alpha-1,3- mannosyltransferase	3464
0.0853727247812374	4.49490036875302	0.280795108048629
	0.596180547775894	0.901355439703449
58525	WIZ	WIZ zinc finger	7965	0.0761670364050735
6.19436103572665	0.28066084711859	0.596268400972162

0.901355439703449
23263 MCF2L MCF.2 cell line derived transforming sequence like
14430 -0.210964655276554 4.1768724206537
0.280580364286635 0.596321077549338 0.901355439703449
144245 ALG10B ALG10 alpha-1,2-glucosyltransferase B
11449 -0.197935089683793 3.05660431061762
0.280469657611436 0.596393551652569 0.901355439703449
30000 TNPO2 transportin 2 7029 -0.0683175624338175
6.88705180029952 0.280461181132111 0.596399101532917
0.901355439703449
10612 TRIM3 tripartite motif containing 3 6710
0.0996446041252455 3.50620541300718 0.280021615823637
0.596687049305652 0.901519352053722
100885848 PTGES3L prostaglandin E synthase 3 like 2856
0.216118400471017 0.396580368052205 0.280003431340872
0.596698967716268 0.901519352053722
706 TSPO translocator protein 1455 0.0815145079554997
6.94299944743962 0.280000874645907
0.596700643456133 0.901519352053722
23761 PISD phosphatidylserine decarboxylase
5002 -0.0700782643555472 5.54537946083555
0.279679844965526 0.596911134458656 0.901692861013915
3797 KIF3C kinesin family member 3C
6403 -0.0771434647901507 8.08806899361268
0.279488677117399 0.597036551807005 0.901692861013915
11103 KRR1 KRR1 small subunit processome component homolog
10749 0.0678056788845863 6.04298760912689
0.27930031726288 0.597160180634769 0.901692861013915
3587 IL10RA interleukin 10 receptor subunit alpha 6548
0.49241471107546 1.82131718332326 0.279281775258163
0.597172353448221 0.901692861013915
162461 TMEM92 transmembrane protein 92 3101
0.107678115753603 4.64162994120888 0.279254610100637
0.597190188289102 0.901692861013915
100130275 TFAP2A-AS1 TFAP2A antisense RNA 1
1692 -0.17397624865793 0.167889766703417
0.278934182051386 0.597400643884601 0.901692861013915
101060321 TBC1D3G TBC1 domain family member 3G
4713 -0.183612136937848 1.33997711964138
0.278809498895673 0.597482576963012 0.901692861013915
80063 ATF7IP2 activating transcription factor 7 interacting
protein 2 4831 0.155808074235585 1.17145815408043
0.278762590659401 0.597513407851085
0.901692861013915
1936 EEF1D eukaryotic translation elongation factor 1 delta
9390 -0.0720409841506129 6.94545185350012
0.278541090810165 0.597659035579309 0.901692861013915
408 ARRB1 arrestin beta 1 5828 0.155361672487997
4.18522585121673 0.278488917044523 0.597693348618536
0.901692861013915
6480 ST6GAL1 ST6 beta-galactoside alpha-2,6-sialyltransferase 1
8741 0.117008221079011 6.79154428880794
0.27841743848677 0.597740364482312 0.901692861013915
28957 MRPS28 mitochondrial ribosomal protein S28 1817

0.104801232616708	3.92516735111504	0.278384507946285
	0.597762027529567	0.901692861013915
84268	RPAIN	RPA interacting protein 7430	0.0878036869475374
	5.38913743821687	0.278276910114371
0.597832821175623	0.901692861013915
51548	SIRT6	sirtuin 6	1722	-0.0768793605984902
4.30915699935948	0.278266050927428	0.5978399669145
0.901692861013915
79577	CDC73	cell division cycle 73	5352	-0.0677658451379074
	6.11975540991306	0.278182466101736
0.597894974727863	0.901692861013915
63926	ANKEF1	ankyrin repeat and EF-hand domain containing 1	5421
	0.156474453370538	3.3132045761667 0.278132705857949
	0.597927727341271	0.901692861013915
55070	DET1	DET1 partner of COP1 E3 ubiquitin ligase	2337
	0.148878294827928	1.53728503632866
0.278095457379042	0.597952247056028	0.901692861013915
7016	TESK1	testis associated actin remodelling kinase 1	3298
	0.0859399496387598	5.49697909931422
0.278006451732466	0.598010845682189	0.901692861013915
2287	FKBP3	FKBP prolyl isomerase 3 2262	0.0918037208324261
	5.80645271372044	0.277942556498971
0.598052919764237	0.901692861013915
51571	FAM49B	family with sequence similarity 49 member B	9131
	0.0934559764540775	6.33017379216861
0.277866552925218	0.598102975061061	0.901692861013915
79701	OGFOD3	2-oxoglutarate and iron dependent oxygenase domain
containing 3	5586	0.0961583941951785	4.46510103588636
	0.277218989424895	0.598529809949189
0.901794876421383
768211	RELL1	RELT like 1	1947	-0.153991738327495
2.77457219129136	0.277112860779731	0.598599824281888
	0.901794876421383
64599	GIGYF1	GRB10 interacting GYF protein 1
6709	-0.12696985201639	5.97522206904732
0.277099349354842	0.598608739158651	0.901794876421383
8793	TNFRSF10D	TNF receptor superfamily member 10d	3538
	0.114227871017383	6.72080856123828
0.276947536496095	0.598708924802054	0.901794876421383
2617	GARS1	glycyl-tRNA synthetase 1
4152	-0.0808431633887378	7.6899711641063 0.276862202976957
	0.598765254224505	0.901794876421383
100505648	RAD51-AS1	RAD51 antisense RNA 1	1696
0.161364887377989	1.37371720557463	0.276808370350285
	0.598800795326993	0.901794876421383
6843	VAMP1	vesicle associated membrane protein 1	3580
0.278862376359273	1.93952172445015	0.276743071981896
	0.598843912205956	0.901794876421383
3937	LCP2	lymphocyte cytosolic protein 2
7244	-0.436909750171047	1.21539734667212
0.276673273759613	0.598890007545543	0.901794876421383
9946	CRYZL1	crystallin zeta like 1	6177	-0.101661466308702
	3.67212628780321	0.27648714361253
0.599012965778876	0.901794876421383

266812	NAP1L5	nucleosome assembly protein 1 like 5	2321
0.163456059293194	0.431119520048488	0.276447667421566
	0.599039050680583	0.901794876421383
10384	BTN3A3	butyrophilin subfamily 3 member A3
5658	-0.158472527839106	3.39730395828042
0.276415781463434	0.599060121877696	0.901794876421383
685	BTC	betacellulin	2816	-0.131399041781966
3.06786564214606	0.276271389847149	0.599155559608949
	0.901794876421383
219738	FAM241B family with sequence similarity 241 member B
1349	-0.115941356729802	3.76418666807198
0.276073878702391	0.599286159024541	0.901794876421383
2068	ERCC2	ERCC excision repair 2, TFIIH core complex helicase
subunit 6450	-0.0956975053853712	4.35028633850681
0.275782230279148	0.599479113462505	0.901794876421383
57019	CIAPIN1 cytokine induced apoptosis inhibitor 1	3509
0.0640148236428612	6.07837561596842	0.275721976261906
	0.599518993710358	0.901794876421383
54487	DGCR8	DGCR8 microprocessor complex subunit
7315	-0.0643018258090464	6.09541961705471
0.275599223396913	0.599600257190852	0.901794876421383
253725	WASHC2C WASH complex subunit 2C 5702	0.102739584370975
	6.20335556580059	0.275585501512197
0.599609342634147	0.901794876421383
10179	RBM7	RNA binding motif protein 7	8183
0.0647092991613874	5.77620223482174	0.275562245950567
	0.599624741111241	0.901794876421383
50813	COPS7A	COP9 signalosome subunit 7A	3113
0.0696626805181975	5.55047188404327	0.27554097111604
	0.599638828792275	0.901794876421383
375056	MIA3	MIA SH3 domain ER export factor 3	10171
0.0702884487716287	6.1642263977631 0.275495652585793
0.599668839940178	0.901794876421383
3597	IL13RA1 interleukin 13 receptor subunit alpha 1
4372	-0.0775799414960811	7.06265806989595
0.275424412347846	0.599716023504498	0.901794876421383
3149	HMGB3	high mobility group box 3	3967
0.0737025263628311	8.40727374836074	0.2753382243869
0.599773117592849	0.901794876421383
51071	DERA	deoxyribose-phosphate aldolase
2844	-0.0809329562557466	5.27990846734805
0.275278010005401	0.59981301258557	0.901794876421383
256364	EML3	EMAP like 3	5490	-0.0771179877197798
5.1830189023354 0.275262637887487	0.599823198262209
0.901794876421383
26269	FBXO8	F-box protein 8 2810	-0.0817246176881797
4.12468187964221	0.275203185972006	0.599862594946152
	0.901794876421383
55218	EXD2	exonuclease 3'-5' domain containing 2	6263
0.068220661056779	6.08343342209212	0.275166960229219
	0.599886603126762	0.901794876421383
8317	CDC7	cell division cycle 7	3557	0.182077872337527
	2.53368413683482	0.275133244398972
0.599908949703323	0.901794876421383

112	ADCY6	adenylate cyclase 6	8410	0.060744666755309
	7.08640962580616	0.27502256941478
0.599982316451954	0.901808381429608
550643	NBDY	negative regulator of P-body association	1529
	0.0761425614887661	5.22870430664031
0.274564239407504	0.600286345179169	0.902119454934129
80011	PSME3IP1	proteasome activator subunit 3 interacting
protein 1	4110	-0.0650321575997423	5.6345538785736
0.274450994672193	0.600361514792818	0.902119454934129
54971	BANP	BTG3 associated nuclear protein
7721	-0.0960161643524187	4.09988953338959
0.274419398048842	0.600382491532516	0.902119454934129
22907	DHX30	DExH-box helicase 30	7744	0.0650651843531399
	6.70914925540815	0.274255323202869
0.600491444232721	0.902186384143272
26790	SNORD58B	small nucleolar RNA, C/D box 58B	66
	-0.150131986158046	1.72996787871368
0.273771035111058	0.600813274566542	0.902573094900639
6204	RPS10	ribosomal protein S10	1642	0.108403190687158
	5.22627597727281	0.273282410212978
0.601138354570872	0.902964604018682
3308	HSPA4	heat shock protein family A (Hsp70) member 4	5156
	0.0671255383580154	7.5525653146454 0.272705926694911
	0.601522362700357	0.903057053944552
124808	CCDC43	coiled-coil domain containing 43
2855	-0.0791015401071048	4.89538478272886
0.272554333891511	0.601623427716372	0.903057053944552
150290	DUSP18	dual specificity phosphatase 18
4450	-0.120087912789496	3.73249727794379
0.272480219792469	0.601672851672508	0.903057053944552
2647	BLOC1S1	biogenesis of lysosomal organelles complex 1 subunit
1	1762	0.0985499826153106	4.11147841085439
0.272452164233639	0.601691563120498	0.903057053944552
9217	VAPB	VAMP associated protein B and C
10585	-0.0730899887958124	6.3053850262632 0.27242289078276
	0.601711088139509	0.903057053944552
5036	PA2G4	proliferation-associated 2G4
3826	-0.0730057826115295	6.49404533645233
0.272395194788146	0.60172956224418	0.903057053944552
51527	GSKIP	GSK3B interacting protein
4567	-0.0691957845073234	5.84554493167696
0.272222016493593	0.601845104750865	0.903057053944552
9941	EXOG	exo/endonuclease G	5223	0.104315935822661
	3.35964532018217	0.272122639291305
0.601911429158546	0.903057053944552
163126	EID2	EP300 interacting inhibitor of differentiation 2
	1981	0.0952535304368573	3.88679834219229
0.271794563767633	0.602130496410695	0.903057053944552
55168	MRPS18A	mitochondrial ribosomal protein S18A
1284	-0.0791419218987501	4.58354435020831
0.271691338551886	0.602199458195275	0.903057053944552
51526	OSER1	oxidative stress responsive serine rich 1
1712	-0.0859549824698703	5.2833743344258 0.271550982765611
	0.602293252572841	0.903057053944552

22900	CARD8	caspase recruitment domain family member 8	9833
	0.126609102668629	3.70973484311013
0.271523313825664	0.602311746298795	0.903057053944552
9088	PKMYT1	protein kinase, membrane associated tyrosine/
threonine 1	3991	0.171644541552869	0.258147363389402
	0.271398973819303	0.602394869082469
0.903057053944552
401074	LINC00960	long intergenic non-protein coding RNA 960
	3181	0.369578530369326	2.35132717414967
0.271334900448238	0.602437712350667	0.903057053944552
118881	COMTD1	catechol-O-methyltransferase domain containing 1
	1580	0.135903204750986	2.6671252278797
0.271240521405494	0.602500831508759	0.903057053944552
8790	FPGT	fucose-1-phosphate guanylyltransferase
4180	-0.130599659735044	3.19829724181462
0.27117430252634	0.602545125944514	0.903057053944552
2995	GYPC	glycophorin C (Gerbich blood group)	3242
0.321297638711504	0.386401867637422	0.271118853562093
	0.602582221570952	0.903057053944552
51454	GULP1	GULP PTB domain containing engulfment adaptor 1
11979	-0.182172305941496	5.28971922884465
0.271116496907027	0.602583798291459	0.903057053944552
1655	DDX5	DEAD-box helicase 5	6994	-0.0654209003312309
	9.89448084649237	0.271045916770845
0.602631023982065	0.903057053944552
100379224	LOC100379224	uncharacterized LOC100379224	3488
	0.163358862741587	1.2056573388478 0.270986528937715
	0.602670766875023	0.903057053944552
634	CEACAM1 CEA cell adhesion molecule 1
5524	-0.128448546057621	4.68858899030367
0.270951563089042	0.602694168936606	0.903057053944552
51287	COA4	cytochrome c oxidase assembly factor 4 homolog	1480
	0.0950620288745577	4.35245526314007
0.270910658051355	0.602721548434633	0.903057053944552
4035	LRP1	LDL receptor related protein 1	20839
0.121804223706744	8.32131474649648	0.270886525936021
	0.602737702327656	0.903057053944552
2051	EPHB6	EPH receptor B6 5325	-0.0947008404060816
5.77574475656149	0.270849989454288	0.602762161369441
	0.903057053944552
100526837	EEF1E1-BLOC1S5	EEF1E1-BLOC1S5 readthrough (NMD
candidate)	768	0.10869381937046	3.32122893773051
	0.270776002056744	0.60281169803538
0.903057053944552
6240	RRM1	ribonucleotide reductase catalytic subunit M1	4177
	0.0779863368928075	5.66446386780927
0.270565021499031	0.602953002760607	0.903172143050777
101927597	NCK1-DT NCK1 divergent transcript
1429	-0.164510764920823	1.11601434237263
0.270302710105996	0.603128784083688	0.903338845023548
57697	FANCM	FA complementation group M
8468	-0.151119692238116	2.87964903234904
0.270056321574145	0.603293993814349	0.903403821797996
715	C1R	complement C1r	6038	0.292682493870143

4.11094938354686	0.270045637277782	0.603301160069855
	0.903403821797996
6261	RYR1	ryanodine receptor 1	15965	0.161922611825474
	2.4130582682775 0.269911962337936	0.60339083480125
	0.903441520369974
286827	TRIM59	tripartite motif containing 59	7865
0.148684543236204	2.83997666399029	0.269507212009009
	0.603662530290443	0.903751716735595
23111	SPART	spartin 6387	0.08113600040137
5.84969133597323	0.269142772554841	0.6039073881091
0.903986162100726
118472	ZNF511	zinc finger protein 511 2728	0.105933249830651
	3.17284632218477	0.269082052544014
0.603948204821775	0.903986162100726
284454	LOC284454	uncharacterized LOC284454	1774
0.102899339357223	4.1614212393766 0.268895695656465
0.604073512640945	0.904026535465507
79876	UBA5	ubiquitin like modifier activating enzyme 5	4595
	0.0700840201924368	5.14451683596993
0.268849980693851	0.604104260152577	0.904026535465507
100529240	ZNF816-ZNF321P	ZNF816-ZNF321P readthrough
1252	-0.134427391705174	1.78810989344672
0.268592737445058	0.604277341740599	0.904114504555772
23753	SDF2L1	stromal cell derived factor 2 like 1
1099	-0.140438009234616	2.75458247879674
0.268330758310296	0.604453717887279	0.904114504555772
284114	TMEM102 transmembrane protein 102	2129
0.0745794542902886	4.93507036714148	0.268221919651147
	0.604527025081736	0.904114504555772
30835	CD209	CD209 molecule	4413	0.476506440555362
0.0667272159743649	0.268152137601925	0.604574036010152
	0.904114504555772
55201	MAP1S	microtubule associated protein 1S
5040	-0.108121243115686	4.88077801438111
0.268085808048504	0.604618728244924	0.904114504555772
100506243	KRBOX1	KRAB box domain containing 1	3845
0.34934461490432	0.754829203702767	0.268014356941764
	0.604666879179256	0.904114504555772
23085	ERC1	ELKS/RAB6-interacting/CAST family member 1
13484	-0.0848757141179996	6.21550375471069
0.267965561484846	0.604699767137613	0.904114504555772
388886	LRRC75B leucine rich repeat containing 75B
3422	-0.115970829976435	2.9172113233431 0.267905672206524
	0.604740137480882	0.904114504555772
55113	XKR8	XK related 8	2743	0.0905929445053993
3.81506887686967	0.267899986546301	0.604743970385023
	0.904114504555772
155060	LOC155060	AI894139 pseudogene	3239
0.144470956020178	2.89389186827744	0.267705274028176
	0.604875264096813	0.904183974781641
135295	SRSF12	serine and arginine rich splicing factor 12	4548
	0.19226408639099	0.0915152733261344
0.267639628496228	0.604919542205951	0.904183974781641
7565	ZNF17	zinc finger protein 17	3328	-0.104334510336481

3.04757020206781	0.267538778602083
0.604987579175933	0.904189182833685
10638	NA	NA	648	0.193751481588345
0.282940693823106	0.267437383608822	0.605056000277574
0.904194963820333
286075	ZNF707	zinc finger protein 707 5203	0.0833512549527391
4.240916492104	0.267322349722334	0.605133644779834
0.904214525542046
27143	PALD1	phosphatase domain containing paladin 1
4555	-0.216288501218358	1.24157345381109
0.266585423177117	0.605631551476013	0.904362411743007
3428	IFI16	interferon gamma inducible protein 16	5294
0.123529207351269	5.97760200419116	0.266582226381135
0.605633713299084	0.904362411743007
84313	VPS25	vacuolar protein sorting 25 homolog	2404
0.0666034622692281	5.51245904081589	0.2665194002543
0.605676202598841	0.904362411743007
9929	JOSD1	Josephin domain containing 1
3958	-0.0671544211024023	6.85987901478341
0.26649128445375	0.605695219364746	0.904362411743007
79230	ZNF557	zinc finger protein 557 6018	-0.123908985333859
3.22354586225902	0.2664392747599 0.605730400661993
0.904362411743007
138639	PTPDC1	protein tyrosine phosphatase domain containing 1
6342	-0.117407662917185	3.34727147237562
0.266404981596802	0.605753600215917	0.904362411743007
51042	ZNF593	zinc finger protein 593 1956	0.140595037395368
3.09136834886781	0.266250635647076
0.605858039696211	0.904362411743007
100271722	LINC00899	long intergenic non-protein coding
RNA 899 2827	0.150636908558255	0.662411952756535
0.266136277895708	0.605935445551431	0.904362411743007
23623	RUSC1	RUN and SH3 domain containing 1 5427
0.0654439368647066	5.87937226841332	0.266113352510335
0.605950965695736	0.904362411743007
55631	LRRC40	leucine rich repeat containing 40
2884	-0.0942078203468581	4.47252268564138
0.265925780753326	0.606077980784105	0.904362411743007
146857	SLFN13	schlafen family member 13	9329
0.158202255596373	3.19728389396895	0.265758205272295
0.60619150327182	0.904362411743007
90121	TSR2	TSR2 ribosome maturation factor 1534
0.071969970203598	4.71822363575193	0.265715772538493
0.60622025612987	0.904362411743007
23384	SPECC1L sperm antigen with calponin homology and coiled-coil
domains 1 like	7267	-0.0752796037477455	6.95104140160152
0.265638281632683	0.606272772266536
0.904362411743007
3094	HINT1	histidine triad nucleotide binding protein 1	1936
0.0931568272349679	7.55128713953344
0.265463167577622	0.606391484021473	0.904362411743007
161394	SAMD15	sterile alpha motif domain containing 15	2616
0.152141910771188	0.786976027780155
0.264970605896782	0.606725662935222	0.904362411743007

56257	MEPCE	methylphosphate capping enzyme	3420
0.0927365931589719	5.05070130884019	0.264722537917105
	0.6068941136699	0.904362411743007
245972	ATP6V0D2	ATPase H+ transporting V0 subunit d2	2446
	0.239411088687828	1.33439866522028
0.264641337527898	0.60694927453555	0.904362411743007
29914	UBIAD1	UbiA prenyltransferase domain containing 1	5082
	0.0834729473047394	3.92846639611159
0.264548566900395	0.607012308369136	0.904362411743007
80207	OPA3	outer mitochondrial membrane lipid metabolism regulator OPA3	9999	-0.0686718071641863	5.51588476929221
	0.26440289383946	0.607111315443845
0.904362411743007
55280	CWF19L1	CWF19 like cell cycle control factor 1
3985	-0.0997716301141032	4.15683903167913
0.264375843791218	0.60712970387567	0.904362411743007
51134	CEP83	centrosomal protein 83	6925	-0.156563003635694
	1.84755363132019	0.264351508387194
0.607146247927915	0.904362411743007
51079	NDUFA13	NADH:ubiquinone oxidoreductase subunit A13	3344
	0.0800349264579968	6.48904400864219
0.264277996156125	0.607196229941695	0.904362411743007
3035	HARS1	histidyl-tRNA synthetase 1
8436	-0.0715111987884573	6.65477443339465
0.264179645513011	0.607263113692395	0.904362411743007
10957	PNRC1	proline rich nuclear receptor coactivator 1	2992
	0.0824429070603399	5.37983688495211
0.264156434607138	0.607278900657836	0.904362411743007
473	RERE	arginine-glutamic acid dipeptide repeats
11503	-0.0950234632794332	7.51357583066782
0.264090934148335	0.607323455715096	0.904362411743007
11016	ATF7	activating transcription factor 7
8305	-0.0742104876965884	6.05864388410128
0.264076510670446	0.607333267865369	0.904362411743007
23469	PHF3	PHD finger protein 3	9789	-0.0954818199093106
	6.14215321427843	0.264056756001366
0.607346707322297	0.904362411743007
23473	CAPN7	calpain 7	5598	-0.0754222932418326
5.48226620914059	0.263858946898704	0.607481315466335
	0.904362411743007
27092	CACNG4	calcium voltage-gated channel auxiliary subunit gamma 4	3380	0.159375424726049	5.80702431552754
0.263770421465267	0.607541577254728	0.904362411743007
51460	SFMBT1	Scm like with four mbt domains 1
6123	-0.127453909713976	3.06755039265619
0.263683757134832	0.607600584518036	0.904362411743007
2990	GUSB	glucuronidase beta	3992	0.0940180693912291
	5.28748021575128	0.263670583967922
0.60760955482336	0.904362411743007
832	CAPZB	capping actin protein of muscle Z-line subunit beta
	3238	-0.0848517919416414	7.6387774768732
0.263654676550686	0.607620387401623	0.904362411743007
136051	ZNF786	zinc finger protein 786	3271	-0.102054675141178
	3.51625080334572	0.263646014137273

0.607626286475772	0.904362411743007
7024	TFCP2	transcription factor CP2
3827	-0.0746418684196858	5.15343829192635
0.263635428075975	0.607633495713878	0.904362411743007
5034	P4HB	prolyl 4-hydroxylase subunit beta
5303	-0.0944960156811658	9.49672787874445
0.263476341247269	0.607741857836704	0.904362411743007
142679	DUSP19	dual specificity phosphatase 19	5376
0.168522464442809	-0.163536163785185	0.263443999603427
	0.60776389242919	0.904362411743007
285971	ZNF775	zinc finger protein 775	2537	0.182527958816445
	0.811880412732611	0.263425264001848
0.607776657909817	0.904362411743007
9907	AP5Z1	adaptor related protein complex 5 subunit zeta 1
	6873	-0.0721132925757178	6.15076093768494
0.26342367194087	0.607777742685175	0.904362411743007
65264	UBE2Z	ubiquitin conjugating enzyme E2 Z
5753	-0.0641622760394435	7.73488571803788
0.263394798781668	0.60779741657722	0.904362411743007
908	CCT6A	chaperonin containing TCP1 subunit 6A
3901	-0.0766761458489784	7.58638164822305
0.263322353137994	0.607846786337873	0.904362411743007
93550	ZFAND4	zinc finger AN1-type containing 4	6232
0.140866545117171	2.43726612361311	0.263205960586285
	0.607926122704277	0.904362411743007
84255	SLC37A3	solute carrier family 37 member 3
8199	-0.0677872577845216	6.20128618170823
0.263179228676265	0.607944347038777	0.904362411743007
494470	RNF165	ring finger protein 165	9867	-0.203332299651195
	1.12493801815005	0.262878917203986
0.60814916314031	0.904571023479486
145438	FRMD6-AS1	FRMD6 antisense RNA 1
2229	-0.220267023422162	0.152371299322891
0.262777569841121	0.60821831662613	0.904577825544935
152225	LINC02085	long intergenic non-protein coding RNA 2085
	2324	-0.22116320361532	0.376871955650026
0.262623208730537	0.608323675942268	0.904588622068704
340348	TSPAN33	tetraspanin 33	2476	0.377801638596381
2.34856394106533	0.262577710388108	0.608354738337059
	0.904588622068704
151963	MB21D2	Mab-21 domain containing 2
3368	-0.17217250346885	4.613505930011	0.262304915304234
	0.608541050889268	0.904727306442615
91056	AP5B1	adaptor related protein complex 5 subunit beta 1
	4811	-0.0775981297839162	6.03078636281731
0.262232022108448	0.60859085591209	0.904727306442615
54963	UCKL1	uridine-cytidine kinase 1 like 1
3184	-0.0672911241619066	5.27482905078303
0.262157504703293	0.608641779725049	0.904727306442615
740	MRPL49	mitochondrial ribosomal protein L49
3080	-0.0583361038427992	5.96283165482696
0.261895814121958	0.608820686615682	0.904808770149478
7264	TSTA3	tissue specific transplantation antigen P35B
3622	-0.0857744769583427	5.77395876109066

0.261888370707812	0.608825777015695	0.904808770149478
5198	PFAS	phosphoribosylformylglycinamidine synthase	6471
	0.104345142369519	3.13233678299794
0.261542797499452	0.609062208087108	0.904872085311346
80184	CEP290	centrosomal protein 290	10388	0.0864636314912704
	4.65847976204817	0.261486871710922
0.609100489397294	0.904872085311346
30811	HUNK	hormonally up-regulated Neu-associated kinase
8294	-0.224104487047402	1.08298784512397
0.261482543319863	0.60910345240463	0.904872085311346
275	AMT	aminomethyltransferase	4561	0.179264496254
2.66995676251245	0.261448458853431	0.609126786064159
	0.904872085311346
353376	TICAM2	toll like receptor adaptor molecule 2
3244	-0.0996020392031635	3.67312002919175
0.260861141048162	0.609529155759967	0.905373794775171
25793	FBXO7	F-box protein 7	8521	0.0648488240990275
6.78683497133935	0.260641324367842	0.609679898403231
	0.905501679242292
100137049	PLA2G4B	phospholipase A2 group IVB
5042	-0.106055507948594	3.55445423938943
0.26039882559059	0.609846288632537	0.9055850693623
4879	NPPB	natriuretic peptide B	694	0.488174071036487
	0.359548861125623	0.260371054253472
0.60986535013169	0.9055850693623
105376575	NA	NA	3256	-0.178920108947325
1.83354432797582	0.260119679686861	0.610037945186855
	0.905745335831278
6717	SRI	sorcin	4237	0.112426989942894
6.55363145038419	0.259922278477688	0.610173555541257
	0.905811317418771
2869	GRK5	G protein-coupled receptor kinase 5	7000
0.123399269207513	3.33531570007483	0.259757240459767
	0.61028698289025	0.905811317418771
60561	RINT1	RAD50 interactor 1	3655	-0.0810291838802168
	4.66261173249894	0.259756660017318
0.610287381896903	0.905811317418771
55129	ANO10	anoctamin 10	4092	-0.0697848399261712
5.95747564038397	0.259631410662735	0.610373493710406
	0.905811317418771
5880	RAC2	Rac family small GTPase 2	2532
0.193097573779991	5.39567749846936	0.259584536503439
	0.610405727501662	0.905811317418771
1020	CDK5	cyclin dependent kinase 5	2938
0.117308350152796	4.48004502014	0.259406740307515
0.610528025329894	0.905896827414813
56954	NIT2	nitrilase family member 2	9264
0.0796725698536572	5.18791594526014	0.259036281448772
	0.610783016609744	0.906179187972957
146456	TMED6	transmembrane p24 trafficking protein 6	1692
0.132212273325618	1.50219378952909	0.258697226774622
	0.611016593293931	0.906393994975851
474170	LRRC37A2	leucine rich repeat containing 37 member A2
	6756	-0.140707497696312	1.57717307523941

0.258542991219125	0.611122910801123	0.906393994975851
10464	PIBF1	progesterone immunomodulatory binding factor 1
4253	-0.094213641837651	3.7435102425701 0.258470305683072
	0.611173028162844	0.906393994975851
25851	TECPR1	tectonin beta-propeller repeat containing 1
9066	-0.0769181399982057	5.01029874017954
0.258395670544502	0.611224499022968	0.906393994975851
55888	ZKSCAN7 zinc finger with KRAB and SCAN domains 7	4208
	0.143765473019501	2.16421012468187
0.258213172510687	0.611350395115542	0.906393994975851
7071	KLF10	Kruppel like factor 10	4003	-0.0797080503795255
	6.61132851302243	0.258004406422337
0.611494480862357	0.906393994975851
84954	MPND	MPN domain containing	2049	0.115991360990991
	3.24875976058937	0.257883554867349
0.61157792345501	0.906393994975851
283518	KCNRG	potassium channel regulator
1873	-0.128501368342468	2.61815863146311
0.257879262759309	0.611580887415826	0.906393994975851
55870	ASH1L	ASH1 like histone lysine methyltransferase
13296	-0.113958011780461	6.79435578108895
0.25770495332273	0.611701284874055	0.906393994975851
65977	PLEKHA3 pleckstrin homology domain containing A3
14193	-0.0801821017348503	4.71976911988292
0.257663637054154	0.611729829985896	0.906393994975851
10049	DNAJB6	DnaJ heat shock protein family (Hsp40) member B6
	14981	-0.0861081821007531	7.26036308673392
0.257566263964776	0.611797115737795	0.906393994975851
58500	ZNF250	zinc finger protein 250 7357	0.106054117086915
	3.85591418018057	0.257379725261975
0.611926060478786	0.906393994975851
55297	CCDC91	coiled-coil domain containing 91
4708	-0.0828859178841066	4.67835209334476
0.257347764445711	0.61194815926486	0.906393994975851
56180	MOSPD1	motile sperm domain containing 1	2675
0.101753329969566	5.26238960421289	0.257304276284211
	0.611978231217383	0.906393994975851
55844	PPP2R2D protein phosphatase 2 regulatory subunit Bdelta 7756
	0.0622391801726278	5.53611747851313
0.257169256007979	0.612071617757994	0.906393994975851
7918	GPANK1	G-patch domain and ankyrin repeats 1
3528	-0.101966533450425	3.91301092523978
0.257160440913112	0.612077715774793	0.906393994975851
101928399	LINC01679	long intergenic non-protein coding
RNA 1679	2798	0.18477326312796	0.34082766175993
	0.256978272584216	0.612203763755127
0.906393994975851
990	CDC6	cell division cycle 6	5120	0.140915577441983
	3.05550109987766	0.256942158118301
0.612228759156625	0.906393994975851
55320	MIS18BP1	MIS18 binding protein 1 6250
0.120714254094933	4.23483360374798	0.256873226407262
	0.612276474029954	0.906393994975851
347051	SLC10A5 solute carrier family 10 member 5

2568	-0.164526242973239	0.652082320446033
0.256856947447496	0.61228774358142	0.906393994975851
9749	PHACTR2 phosphatase and actin regulator 2
11694	-0.0884160243180133	5.47295684706572
0.256818492315862	0.612314366971989	0.906393994975851
80301	PLEKHO2 pleckstrin homology domain containing O2
4386	-0.147926531208121	3.62448420449356
0.256717938039122	0.612383994900423	0.906393994975851
163087	ZNF383	zinc finger protein 383 8144	-0.122846821617852
	3.14026453354064	0.256671534307145
0.612416132537407	0.906393994975851
64856	VWA1	von Willebrand factor A domain containing 1
5065	-0.0971911314301712	4.9029400584374 0.256365090233356
	0.612628456927548	0.906542956168116
665	BNIP3L	BCL2 interacting protein 3 like
4307	-0.0875272286787881	7.23375138582425
0.255929288168261	0.612930683707379	0.906542956168116
55503	TRPV6	transient receptor potential cation channel
subfamily V member 6	11925	-0.233648414781612
4.01013563167365	0.255921468353641	0.612936109664491
	0.906542956168116
7251	TSG101	tumor susceptibility 101	3839
0.0647484550422266	6.33000713231629	0.255734776929067
	0.613065680700479	0.906542956168116
23583	SMUG1	single-strand-selective monofunctional uracil-DNA
glycosylase 1	5353	0.0902179821981145	4.59720454241601
	0.255720354735978	0.613075692728672
0.906542956168116
3185	HNRNPF	heterogeneous nuclear ribonucleoprotein F
3253	-0.0582441770172765	7.55195353183186
0.255661775684452	0.613116362526798	0.906542956168116
4288	MKI67	marker of proliferation Ki-67
12807	-0.26794785271784	1.03892705500642
0.255512554145978	0.61321998930992	0.906542956168116
64978	MRPL38	mitochondrial ribosomal protein L38	4176
0.0748865143260143	5.09074774200926	0.255466038051358
	0.613252300143393	0.906542956168116
112752	IFT43	intraflagellar transport 43	4172
0.0935475111168476	3.91111273807959	0.255435783288849
	0.613273317574192	0.906542956168116
5829	PXN	paxillin	7374	-0.0764196828852342
8.54082585603497	0.255328843000516	0.613347619564378
	0.906542956168116
100527963	PMF1-BGLAP	PMF1-BGLAP readthrough	1815
0.085517609200675	3.79981766128607	0.255240632683516
	0.613408922694266	0.906542956168116
6136	RPL12	ribosomal protein L12	2452	0.116294096330933
	6.55971894485839	0.25520678862614
0.613432446677159	0.906542956168116
84749	USP30	ubiquitin specific peptidase 30 5135
0.091044615619019	3.83983249725902	0.255090541173871
	0.613513261687707	0.906542956168116
84830	ADTRP	androgen dependent TFPI regulating protein	4150
	0.212859566798971	2.68751031797426

0.255088198531229	0.613514890526022	0.906542956168116
51360	MBTPS2	membrane bound transcription factor peptidase, site
2	8927	-0.0960950653836542	4.5082285591569
0.255066161482553	0.613530213335001	0.906542956168116
4613	MYCN	MYCN proto-oncogene, bHLH transcription factor
2602	-0.374780919947242	0.744858413232175
0.255032494036859	0.613553624600219	0.906542956168116
80152	CENPT	centromere protein T	5539	-0.0789440148828892
	6.1272689987548	0.25490737927304	0.61364064240516
	0.906542956168116
353274	ZNF445	zinc finger protein 445	18871	0.101150292763415
	4.86134028501537	0.254759155889353
0.613743767032485	0.906542956168116
23087	TRIM35	tripartite motif containing 35
4269	-0.0723072502854393	5.18817557771285
0.254755266649894	0.613746473430588	0.906542956168116
84254	CAMKK1	calcium/calmodulin dependent protein kinase kinase 1
	7424	0.133362177858003	1.70615604123212
0.25463714219652	0.613828684834857	0.906568788325795
100532724	NPHP3-ACAD11	NPHP3-ACAD11 readthrough (NMD
candidate)	7789	0.0859122352701525	5.10945998270379
	0.254521807843723	0.613908977491593
0.906570523547394
51181	DCXR	dicarbonyl and L-xylulose reductase	1902
0.125930066364578	3.0687647329691	0.254449532751956
0.613959305088212	0.906570523547394
1302	COL11A2	collagen type XI alpha 2 chain	6740
0.199718508101849	0.229184013373863	0.254171200671777
	0.614153200942761	0.906761240181855
728448	PPIEL	peptidylprolyl isomerase E like pseudogene
3059	-0.173129791685251	1.45983103280741
0.254048175826988	0.614238946784101	0.906792256282135
26249	KLHL3	kelch like family member 3
11430	-0.147646602275961	2.06341590480341
0.253575126246973	0.614568895548511	0.907159641626925
6103	RPGR	retinitis pigmentosa GTPase regulator	7143
0.117895317994704	2.51754387232115	0.253428297590908
	0.614671385962964	0.907159641626925
23468	CBX5	chromobox 5	15040	-0.0729024473208199
6.15329697299261	0.253412953283643	0.614682098856368
	0.907159641626925
94120	SYTL3	synaptotagmin like 3	2896	0.163047032590344
	2.15878767403396	0.253096523005361
0.614903110811716	0.907197599872109
93273	LEMD1	LEM domain containing 1	1039	-0.25978983119696
	0.114683723040139	0.25304998121071
0.614935632711753	0.907197599872109
10514	MYBBP1A	MYB binding protein 1a	6203	0.0931872796419965
	5.39038877832194	0.253007074146531
0.614965618099925	0.907197599872109
92482	BBIP1	BBSome interacting protein 1	3828
0.0934143024107442	3.51104578171106	0.252908524525198
	0.61503450107605	0.907197599872109
10621	POLR3F	RNA polymerase III subunit F

2928 -0.0950324168004181 4.19690588735923
0.252811236892976 0.61510251845809 0.907197599872109
6941 TCF19 transcription factor 19 3538 -0.166045250673049
2.62530228609539 0.25275579241368
0.615141289079249 0.907197599872109
409 ARRB2 arrestin beta 2 4372 0.198450613636849
2.7046955483037 0.252727334219516 0.615161191089119
0.907197599872109
80000 GREB1L GREB1 like retinoic acid receptor coactivator
10572 0.176647597679309 2.07171398184578
0.252563538052115 0.615275767923795 0.907267967190728
9646 CTR9 CTR9 homolog, Paf1/RNA polymerase II complex
component 5562 -0.0576523506546513 6.05817896127202
0.25240421650834 0.615387259376804
0.907267967190728
84790 TUBA1C tubulin alpha 1c 4141 -0.127539471392508
8.46767958032124 0.252381408411111
0.615403223836589 0.907267967190728
5001 ORC5 origin recognition complex subunit 5 3724
0.0933648358166934 3.74817037998215 0.251786437088553
0.615819991760508 0.907721429464448
617 BCS1L BCS1 homolog, ubiquinol-cytochrome c reductase
complex chaperone 3379 0.0844929326516885
3.83991792454769 0.251757298561204 0.615840418662144
0.907721429464448
1725 DHPS deoxyhypusine synthase 3006 -0.0727741412633229
5.13344929469957 0.251534647839764
0.615996551755338 0.907856029086387
9762 LZTS3 leucine zipper tumor suppressor family member 3
5395 -0.0772241235404991 6.09660980855849
0.251332790480824 0.616138178334332 0.907969222927827
29100 TMEM208 transmembrane protein 208 1787
0.0789714722591346 5.11671918203413 0.251104925879297
0.616298137577932 0.908039050597314
441263 DTX2P1-UPK3BP1-PMS2P11 DTX2P1-UPK3BP1-PMS2P11 readthrough,
transcribed pseudogene 2043 -0.120598577118027
3.38608689995083 0.251080602631282 0.616315217678922
0.908039050597314
51177 PLEKHO1 pleckstrin homology domain containing O1 6174
0.0955036189961274 5.01120269072206
0.250889541526181 0.616449419280517 0.908080310239354
100272147 CMC4 C-X9-C motif containing 4 1573
0.130972742726053 2.03341806304751 0.25074373027903
0.616551880326739 0.908080310239354
221393 ADGRF4 adhesion G protein-coupled receptor F4
3391 -0.105494881075476 7.53320765011211
0.250607635787674 0.616647547068064 0.908080310239354
29081 METTL5 methyltransferase like 5 2865
0.0981007471755834 4.3709977762448 0.250403189668233
0.616791322218913 0.908080310239354
22954 TRIM32 tripartite motif containing 32
3780 -0.0775457766651193 5.33253086253239
0.250342589137325 0.616833953174039 0.908080310239354
8273 SLC10A3 solute carrier family 10 member 3 2247

0.0808391996133257	4.56194965419584	0.250069918529366
	0.617025849974393	0.908080310239354
143384	CACUL1	CDK2 associated cullin domain 1
12406	-0.0666476235502114	6.61189795930832
0.250068387022393	0.617026928169003	0.908080310239354
64968	MRPS6	mitochondrial ribosomal protein S6
4646	-0.304324526105297	5.91639911394184
0.249993464053233	0.617079679650042	0.908080310239354
5580	PRKCD	protein kinase C delta	3396	0.082321489202412
	5.6363103111237 0.249991522816373	0.617081046560073
	0.908080310239354
391356	PTRHD1	peptidyl-tRNA hydrolase domain containing 1	1917
	0.0953689731196519	3.54959250491604
0.249991028227761	0.617081394822672	0.908080310239354
6399	TRAPPC2 trafficking protein particle complex 2
6067	-0.0942019268199958	3.67486688128635
0.249989416998203	0.617082529366525	0.908080310239354
10291	SF3A1	splicing factor 3a subunit 1
6329	-0.0639879470769485	6.6408823837714 0.249868295367193
	0.617167829992377	0.908080310239354
56171	DNAH7	dynein axonemal heavy chain 7	21324
0.189218972110283	0.493806630266368	0.24979310847148
	0.61722079380858	0.908080310239354
440823	MIAT	myocardial infarction associated transcript
11505	0.336739343126901	1.97717443699442
0.249750449541651	0.617250848417855	0.908080310239354
6259	RYK	receptor like tyrosine kinase	8047
0.0630300309182671	6.27738395684972	0.249550184910134
	0.617391984284163	0.908146834419079
8538	BARX2	BARX homeobox 2 1813	-0.207621667774952
1.39092789057693	0.249502305782083	0.617425737429349
	0.908146834419079
59277	NTN4	netrin 4	4166	0.0837743989814488
9.38756708600402	0.249207361755225	0.617633752156651
	0.908329266641216
197259	MLKL	mixed lineage kinase domain like pseudokinase
5692	-0.127054117868056	2.95422524389649
0.249142574882715	0.617679464893095	0.908329266641216
158234	TRMT10B tRNA methyltransferase 10B	2866
0.134968235577466	2.1032572788641 0.248824599758925
0.617903931518849	0.908420590657955
220136	CFAP53	cilia and flagella associated protein 53
1851	-0.166865771999874	-0.135515752313519
0.248764622789235	0.617946290844535	0.908420590657955
84196	USP48	ubiquitin specific peptidase 48 9497
0.0587081759489446	6.08880261270156	0.248613797070208
	0.618052841181877	0.908420590657955
10127	ZNF263	zinc finger protein 263 4649	0.0724533233551942
	5.05620776550806	0.248545938740245
0.618100792640817	0.908420590657955
63874	ABHD4	abhydrolase domain containing 4 4070
0.131449520052983	5.91516855659788	0.248529159732385
	0.61811265063122	0.908420590657955
64065	PERP	p53 apoptosis effector related to PMP22

4290	-0.104731186668333	10.9869482914899
0.248340277031275	0.618246171516239	0.908420590657955
26135	SERBP1	SERPINE1 mRNA binding protein 1
7676	-0.0678288485521305	7.4369513895514 0.248305293109654
0.618270908552439	0.908420590657955
6829	SUPT5H	SPT5 homolog, DSIF elongation factor subunit
8191	-0.0654254907169424	7.64423696315684
0.248160112853789	0.618373588368989	0.908420590657955
101927391	LOC101927391	uncharacterized LOC101927391
3385	-0.15798638228376	1.4059196353382 0.248133869688227
0.618392153043739	0.908420590657955
8655	DYNLL1	dynein light chain LC8-type 1
2708	-0.0870948519017637	7.70399190002664
0.248088065361401	0.618424558413252	0.908420590657955
28976	ACAD9	acyl-CoA dehydrogenase family member 9	4327
0.071581251644288	5.18813374994594	0.24804508524411
0.6184549691236 0.908420590657955
8900	CCNA1	cyclin A1	2400	-0.237662358719235
1.81291000185128	0.247719916792548	0.618685149640288
0.908636411567987
5478	PPIA	peptidylprolyl isomerase A	4432
0.0916232327368571	8.27974539022235	0.247639620927998
0.618742018562922	0.908636411567987
54440	SASH3	SAM and SH3 domain containing 3 2711
0.426027570625351	1.23904496079181	0.247523349959401
0.618824386965046	0.908636411567987
116113	FOXP4	forkhead box P4 6170	-0.0957335504017672
5.6065339774975 0.247426168297087	0.618893250684421
0.908636411567987
11005	SPINK5	serine peptidase inhibitor, Kazal type 5	4381
0.352624847719249	1.71023829000067
0.247348094347345	0.618948586744335	0.908636411567987
1762	DMWD	DM1 locus, WD repeat containing
4809	-0.079471129357511	5.4633283166707 0.247288092240012
0.618991121523647	0.908636411567987
6687	SPG7	SPG7 matrix AAA peptidase subunit, paraplegin
10724	0.0645734332618371	6.469267427025	0.247002866148364
0.619193402967059	0.908695867281367
83855	KLF16	Kruppel like factor 16	2955	-0.0848647993422354
4.49721353937175	0.246948216933967
0.619232176641473	0.908695867281367
55577	NAGK	N-acetylglucosamine kinase
8536	-0.0623818241509115	6.61679264721963
0.246735754170359	0.619382970041048	0.908695867281367
51426	POLK	DNA polymerase kappa	9105	-0.0919636806675324
5.21737016355235	0.246718199297746
0.61939543306264	0.908695867281367
11224	RPL35	ribosomal protein L35	1535	0.0956069141656576
6.847539604379	0.246635956511795	0.61945382840872
0.908695867281367
25827	FBXL2	F-box and leucine rich repeat protein 2 7664
0.0708923263877938	4.85848944843002	0.246592223802349
0.619484885153905	0.908695867281367
29896	TRA2A	transformer 2 alpha homolog

3516	-0.0703272171871048	6.09176269189916
0.24657343351322	0.619498230115632	0.908695867281367
101867536	PRNCR1	prostate cancer associated non-coding RNA 1
	12727	-0.130064486414305	2.2599177834449
0.246440871248585	0.619592394544106	0.908695867281367
10475	TRIM38	tripartite motif containing 38
9413	-0.0693933265020677	5.45489889434587
0.246408358817117	0.61961549428174	0.908695867281367
101927755	RNFT1-DT	RNFT1 divergent transcript	2286
	-0.144478291925271	1.37768549475523
0.24610888064198	0.619828359726927	0.908912880516653
55780	ERMARD	ER membrane associated RNA degradation
4251	-0.0955259685088355	3.7765132330212 0.245977914381655
	0.619921499675627	0.908925826591156
439921	MXRA7	matrix remodeling associated 7
9257	-0.0923973782516553	7.68330828520596
0.24591399342544	0.619966969890191	0.908925826591156
24149	ZNF318	zinc finger protein 318 8697	-0.112796620711888
	5.03465955138655	0.245745189196811
0.620087084296226	0.908960931554552
444	ASPH	aspartate beta-hydroxylase
20047	-0.112289963832114	8.18872424344999
0.245697958183882	0.620120701216199	0.908960931554552
84619	ZGPAT	zinc finger CCCH-type and G-patch domain containing
	3335	-0.06797030067737	5.23230033510561
0.245499568823757	0.620261949854955	0.908997863072501
132660	LIN54	lin-54 DREAM MuvB core complex component
6262	-0.105733525414232	3.80555368669517
0.245480277028826	0.620275688958875	0.908997863072501
80128	TRIM46	tripartite motif containing 46	4533
0.182531634543237	3.20434903285513	0.245205662426524
	0.620471335130751	0.909189454040844
27141	CIDEB	cell death inducing DFFA like effector b
3190	-0.194414611154521	1.42300448917205
0.245060439441367	0.620574852810769	0.909207767748809
9861	PSMD6	proteasome 26S subunit, non-ATPase 6
9123	-0.0627168609049476	6.24368851782564
0.245006018201622	0.620613655132888	0.909207767748809
126917	IFFO2	intermediate filament family orphan 2	6188
0.12043079365739	8.50899105328556	0.244554114185291
	0.620936069942162	0.909486657283703
117178	SSX2IP	SSX family member 2 interacting protein 8639
0.0849476623271071	4.14299048072344	0.244407990920564
	0.621040402162284	0.909486657283703
79744	ZNF419	zinc finger protein 419 8218	-0.107108821291924
	3.59047396161377	0.244386282439251
0.621055905360057	0.909486657283703
2664	GDI1	GDP dissociation inhibitor 1
5256	-0.0749139416094095	7.07567710699609
0.24435068653227	0.621081328172266	0.909486657283703
445571	CBWD3	COBW domain containing 3	11089
0.147725173555306	2.04367227341988	0.244223042891859
	0.621172510983049	0.909486657283703
134728	IRAK1BP1	interleukin 1 receptor associated kinase 1

binding protein 1 7542 0.172392252210769
1.92613581559142 0.244193517524214 0.621193606784692
0.909486657283703
22930 RAB3GAP1 RAB3 GTPase activating protein catalytic
subunit 1 6629 -0.0573111547635253 7.05914999157125
0.243736734207801 0.621520179672513
0.909553545831845
100506211 MIR210HG MIR210 host gene 2321
0.168329590443491 2.08964017957851 0.243730118134602
0.621524912566532 0.909553545831845
23189 KANK1 KN motif and ankyrin repeat domains 1 11932
0.105971969364281 5.61454211207423 0.243705889422813
0.6215422455727 0.909553545831845
59344 ALOXE3 arachidonate lipoxygenase 3
3760 -0.166679538624361 1.69303172285927
0.243698547683014 0.621547497999983 0.909553545831845
101929145 PPP3CB-AS1 PPP3CB antisense RNA 1 (head to
head) 5443 0.146512058072899 0.57941592572515
0.243397834760631 0.621762718635502 0.909553545831845
5994 RFXAP regulatory factor X associated protein
2306 -0.131137620009042 0.764072914160287
0.243377227166353 0.621777473569393 0.909553545831845
8483 CILP cartilage intermediate layer protein 5695
0.312351443559444 2.38924072748311 0.24312398958131
0.621958853854605 0.909553545831845
79706 PRKRIP1 PRKR interacting protein 1 2400
0.0770842233564818 5.15137867757659 0.242940393577077
0.62209042711253 0.909553545831845
7922 SLC39A7 solute carrier family 39 member 7 2845
0.0640602608756604 7.83166843517342 0.242893457908679
0.622124073267819 0.909553545831845
100130733 LRRC70 leucine rich repeat containing 70
2421 -0.161790257403618 0.172984766942586
0.242883006848958 0.622131565731538 0.909553545831845
148156 ZNF558 zinc finger protein 558 8184 0.0808046526567975
4.53205493067883 0.242865187955393
0.622144340727151 0.909553545831845
80230 RUFY1 RUN and FYVE domain containing 1 5911
0.0663375719306457 5.72349801150724 0.242735335404426
0.622237454210558 0.909553545831845
57380 MRS2 magnesium transporter MRS2
5133 -0.0860896873972336 5.49324159147542
0.242664125600895 0.622288529828638 0.909553545831845
7554 ZNF8 zinc finger protein 8 11466 0.0610264169894419
5.6410721263943 0.242563862494702 0.622360459875046
0.909553545831845
138429 PIP5KL1 phosphatidylinositol-4-phosphate 5-kinase like 1
3341 -0.134146778358087 2.42712240820179
0.242407740348694 0.622472500701868 0.909553545831845
8666 EIF3G eukaryotic translation initiation factor 3 subunit G
3486 -0.0591731323329914 6.24926542827325
0.242355953569014 0.62250967530815 0.909553545831845
2162 F13A1 coagulation factor XIII A chain 4690
0.284435686108506 -0.598532277921035 0.242276233621444

0.622566911096791	0.909553545831845
145781	GCOM1	GRINL1A complex locus 1 4648	-0.140184640940143
5.34787264575395	0.24227516684239
0.62256767708153	0.909553545831845
54495	TMX3	thioredoxin related transmembrane protein 3	7637
0.0678441923187963	6.62460597100447
0.242142744795768	0.622662777025524	0.909553545831845
6155	RPL27	ribosomal protein L27	2026	0.11043499657405
7.20766946848391	0.241940911033566
0.622807787697602	0.909553545831845
26071	RTL8A	retrotransposon Gag like 8A	1265
0.0868194693654253	4.53654695325055	0.241891825322414
0.622843065461965	0.909553545831845
221756	SERPINB9P1	serpin family B member 9 pseudogene 1
1460	-0.487704074858623	2.65016623806826
0.24184619286507	0.622875865369712	0.909553545831845
163050	ZNF564	zinc finger protein 564 7094	0.0856843240047754
3.99591419983471	0.241819309027491
0.622895190857126	0.909553545831845
5064	PALM	paralemmin	4908	0.291378874910558
5.0216757295869 0.241735251703879	0.622955624199213
0.909553545831845
10666	CD226	CD226 molecule	13638	-0.263155995548737
1.16824642044677	0.2416770842022 0.62299745159095
0.909553545831845
374	AREG	amphiregulin	1648	-0.185155362960297
5.7952471695183 0.241660991151861	0.62300902496966
0.909553545831845
7083	TK1	thymidine kinase 1	1861	-0.205695554086429
1.58462025931049	0.241570617922736
0.623074026112681	0.909553545831845
80314	EPC1	enhancer of polycomb homolog 1
11286	-0.089882772069918	5.3534254285422 0.241548379035262
0.623090023795617	0.909553545831845
81831	NETO2	neuropilin and tolloid like 2	7481
0.132237235184491	4.40830097726497	0.241466733375532
0.623148763953452	0.909553545831845
9562	MINPP1	multiple inositol-polyphosphate phosphatase 1
3016	-0.089583453630931	4.78082458326236
0.241360746326158	0.623225034972885	0.909553545831845
9710	KIAA0355	KIAA0355	7459	-0.094734623639312
5.1127834183291 0.241322868844827	0.623252297629332
0.909553545831845
84952	CGNL1	cingulin like 1 7768	0.247571891695518
4.78929107427762	0.241127746325319	0.623392780905599
0.909655697815537
5435	POLR2F	RNA polymerase II subunit F	5469
0.0799082086307936	4.55176776458349	0.241045272779196
0.62345218098302	0.909655697815537
5425	POLD2	DNA polymerase delta 2, accessory subunit
3695	-0.115043144384883	4.19031609889917
0.240939610036676	0.623528301090574	0.909672004309517
56172	ANKH	ANKH inorganic pyrophosphate transport regulator
9374	-0.133936274810309	4.47580132415021

0.240723957294344	0.623683722987563	0.909720354147131
83707	TRPT1	tRNA phosphotransferase 1	2210
0.0801464374722571	4.20420391258117	0.240538149465305
	0.623817704809448	0.909720354147131
55248	PACC1	proton activated chloride channel 1
2914	-0.0702608128215543	4.67072322543531
0.240425763143726	0.623898775203661	0.909720354147131
6434	TRA2B	transformer 2 beta homolog
11769	-0.0734083248677039	6.22720487358431
0.240394979111435	0.623920985508727	0.909720354147131
7439	BEST1	bestrophin 1	8930	0.163536990643967
2.64483092462239	0.240221822210516	0.624045948984365
	0.909720354147131
55197	RPRD1A	regulation of nuclear pre-mRNA domain containing 1A
	7964	-0.0612702022984974	6.48034913741769
0.240085000486175	0.624144729684437	0.909720354147131
8824	CES2	carboxylesterase 2	5513	0.0842396553788579
	5.96330407468238	0.23997799765327
0.624222006469934	0.909720354147131
6827	SUPT4H1	SPT4 homolog, DSIF elongation factor subunit	2079
	0.0672358588899777	6.00161709408756
0.239974702959273	0.624224386216594	0.909720354147131
10591	DNPH1	2'-deoxynucleoside 5'-phosphate N-hydrolase 1
1250	-0.131350433625856	2.92279746400676
0.239931054340426	0.62425591538271	0.909720354147131
10424	PGRMC2	progesterone receptor membrane component 2	4626
	0.0875621321430656	6.57768993866231
0.239919923300249	0.624263956353123	0.909720354147131
100616329	MIR4700	microRNA 4700	74	-0.152399335076075
	0.114402876832158	0.239866494938663
0.624302555774774	0.909720354147131
55308	DDX19A	DEAD-box helicase 19A	8113	0.0650383740696915
	5.56263922242532	0.239813547839255
0.624340812766214	0.909720354147131
64763	ZNF574	zinc finger protein 574	4682	-0.093938769746924
	4.20474319551467	0.239600210205861
0.624495013207088	0.909796879476734
6898	TAT	tyrosine aminotransferase	5407
0.258826660099642	0.00705112671195341	0.239484435431566
	0.624578730837973	0.909796879476734
124936	CYB5D2	cytochrome b5 domain containing 2	2182
0.100790782881309	4.09635095794067	0.239458252706198
	0.624597667245827	0.909796879476734
285368	PRRT3	proline rich transmembrane protein 3
3939	-0.181307367399092	1.23619819521751
0.239381556912242	0.624653144137057	0.909796879476734
51104	ABHD17B	abhydrolase domain containing 17B
2959	-0.0753626487251149	4.4218751114099	0.239237608111228
	0.624757297347996	0.909822893779741
727936	GXYLT2	glucoside xylosyltransferase 2
3667	-0.237278064834515	1.25613283891618
0.239167925164308	0.624807729936117	0.909822893779741
55726	INTS13	integrator complex subunit 13
3345	-0.0716192520133115	4.48144705235419

0.239070648892552	0.624878148244619	0.909822893779741
10391	CORO2B	coronin 2B	3998	-0.190388088846018
2.87874293746926	0.238997897402655	0.624930824663018
	0.909822893779741
100134934	TEN1	TEN1 subunit of CST complex	1303
0.0829385482686676	4.6826062205585 0.238634359368512
0.625194196824021	0.91011173507733
11092	SPACA9	sperm acrosome associated 9	2823
0.154298408060857	1.50666202408591	0.23837300063156
	0.625383696795721	0.910255036114107
84561	SLC12A8 solute carrier family 12 member 8	5586
0.12229278830638	3.20959329428443	0.238228276405004
	0.625488685452594	0.910255036114107
116540	MRPL53	mitochondrial ribosomal protein L53
1344	-0.0897654778586074	3.91096917591427
0.238107937362324	0.625576014208373	0.910255036114107
100130958	SYCE1L	synaptonemal complex central element protein
1 like	1244	0.152572187790756	1.37424963811196
0.237982983771237	0.625666720611383	0.910255036114107
11243	PMF1	polyamine modulated factor 1	1712
0.0678828095703675	4.69788591199174	0.237932794028225
	0.625703162685392	0.910255036114107
8260	NAA10	N(alpha)-acetyltransferase 10, NatA catalytic
subunit 4742	0.0710169455289128	5.17395092625023
0.237919206855729	0.625713028960665	0.910255036114107
8028	MLLT10	MLLT10 histone lysine methyltransferase DOT1L
cofactor	6945	-0.0788117590593997	5.19650517393124
	0.237871689264021	0.625747536427696
0.910255036114107
57158	JPH2	junctophilin 2	5943	0.270236624596845
2.52647737773507	0.237663131256296	0.625899042572291
	0.910343043926215
283385	MORN3	MORN repeat containing 3
3856	-0.175237245798808	0.556889955862517
0.237592570635807	0.625950319684163	0.910343043926215
286053	NSMCE2	NSE2 (MMS21) homolog, SMC5-SMC6 complex SUMO ligase
	2991	0.0834900736051439	4.47402390967677
0.237445045492658	0.626057558142092	0.910343043926215
80086	TUBA4B	tubulin alpha 4b	1678	-0.122232684224205
	2.30088804665523	0.237360780427942
0.626118830305847	0.910343043926215
22809	ATF5	activating transcription factor 5	2954
0.118545459065387	4.95795656703477	0.23725446898972
	0.626196152384448	0.910343043926215
79639	TMEM53	transmembrane protein 53	3635
0.099218389708591	3.2887404717297 0.237251944516595
0.62619798873628	0.910343043926215
10645	CAMKK2	calcium/calmodulin dependent protein kinase kinase 2
	6931	-0.0599260080975028	6.12736067562757
0.236904972366922	0.626450498244655	0.910465745430812
6138	RPL15	ribosomal protein L15	6528	0.0800161137198419
	8.16697235700266	0.236892012404951
0.62645993430676	0.910465745430812
79000	AUNIP	aurora kinase A and ninein interacting protein

2408	-0.199857987098321	0.0198154195471368
0.236739227102817	0.626571200343628	0.910465745430812
8886	DDX18	DEAD-box helicase 18	7851	0.0610090273314234
	6.15230010720052	0.236671877228282
0.626620262053297	0.910465745430812
100506233	RAB30-DT	RAB30 divergent transcript	2123
	-0.182900945557504	1.3109803873655 0.236660108011595
	0.62662883634647	0.910465745430812
26085	KLK13	kallikrein related peptidase 13
2007	-0.263377558768278	1.88979061481155
0.236583363679867	0.626684753789135	0.910465745430812
4289	MKLN1	muskelin 1	12835	-0.0788246256484049
6.01524998906995	0.236432109010053	0.626794993784451
	0.910465745430812
92002	CCNQ	cyclin Q	1730	0.0885217776561174
3.64216411909063	0.236421952755158	0.626802397600437
	0.910465745430812
10248	POP7	POP7 homolog, ribonuclease P/MRP subunit
988	-0.0866735564822824	4.04625213352022
0.236299883070083	0.626891400660595	0.910500606496574
25852	ARMC8	armadillo repeat containing 8
9848	-0.0605136024561204	5.3551340180436 0.236060060446829
	0.627066342169867	0.910660264880593
339487	ZBTB8OS	zinc finger and BTB domain containing 8 opposite
strand	3531	0.0871687725230595	4.33227135250416
0.235896806459358	0.627185492609343	0.910738875697601
151230	KLHL23	kelch like family member 23	5080
0.118528737448156	3.14444892194476	0.235668569588642
	0.627352156059304	0.9108864569216
55374	TMCO6	transmembrane and coiled-coil domains 6
3502	-0.128118519861878	1.9481854501856 0.235327130544252
	0.627601668439934	0.91109468830862
65990	ANTKMT	adenine nucleotide translocase lysine
methyltransferase	1923	0.114979268308973
2.29288195378212	0.235238732561285	0.627666303204742
	0.91109468830862
100532732	MSH5-SAPCD1	MSH5-SAPCD1 readthrough (NMD
candidate)	4140	0.166757182193093	0.971481361469867
	0.235205361050904	0.627690707658898
0.91109468830862
8671	SLC4A4	solute carrier family 4 member 4	8361
0.306862550441684	3.00845354306415	0.235003105747211
	0.627838661940021	0.911167703424384
65265	C8orf33	chromosome 8 open reading frame 33	3378
0.0689635259742784	5.3661660353402 0.234958759738587
0.627871112547451	0.911167703424384
79035	NABP2	nucleic acid binding protein 2	2218
0.0770035464898442	4.48863350795692	0.234758231218733
	0.628017898404905	0.911199451411657
11274	USP18	ubiquitin specific peptidase 18
2129	-0.163387727868038	2.1465360365915 0.234660249956153
	0.628089648344848	0.911199451411657
2531	KDSR	3-ketodihydrosphingosine reductase	9287
0.0818986913221085	6.95864014167101	0.234645802253496

0.628100229705672	0.911199451411657
285512	FAM13A-AS1	FAM13A antisense RNA 1
4510	-0.175294165985561	1.62324161818892
0.234539962883134	0.628177757727635	0.911199451411657
390595	UBAP1L	ubiquitin associated protein 1 like
8536	-0.21061342220245	1.31616706295073
0.234484688272668	0.62821825532108	0.911199451411657
29128	UHRF1	ubiquitin like with PHD and ring finger domains 1
5145	0.119886133542253	2.53630606360388
0.234337837570882	0.628325875845025	0.911250468255531
2592	GALT	galactose-1-phosphate uridylyltransferase	4609
0.095367007687425	4.24874856312508
0.234230101937063	0.628404857100959	0.911250468255531
9935	MAFB	MAF bZIP transcription factor B
3393	-0.116942023949876	4.15596326968005
0.234170448538006	0.628448598796918	0.911250468255531
55223	TRIM62	tripartite motif containing 62	3640
0.111746074562894	3.86755868235479	0.234042782759209
0.628542234638985	0.911291903590546
5984	RFC4	replication factor C subunit 4	2997
0.115510423433618	2.7163517147949	0.233839729733321
0.628691227835054	0.91136626966393
823	CAPN1	calpain 1	7553	-0.0635410453450257
9.35187460734142	0.233733271379663	0.628769375140169
0.91136626966393
2672	GFI1	growth factor independent 1 transcriptional repressor	3301	0.255949260113675	1.29271622018958
0.233706920101307	0.628788722069838
0.91136626966393
9328	GTF3C5	general transcription factor IIIC subunit 5	3592
0.0589234703595929	5.84786844281278
0.233555154413181	0.628900173607384	0.911382535478648
57461	ISY1	ISY1 splicing factor homolog	4102
0.0759182033760086	4.9069393354029	0.233404745953118
0.629010672522537	0.911382535478648
1892	ECHS1	enoyl-CoA hydratase, short chain 1	1617
0.0847240060457996	6.39393343159626	0.233248115813339
0.629125788935603	0.911382535478648
100532737	ATP6V1G2-DDX39B	ATP6V1G2-DDX39B readthrough (NMD candidate)	2065	-0.054326964012808	6.94464833500942
0.23321627607100З	0.629149195605597
0.911382535478648
25829	TMEM184B	transmembrane protein 184B
4708	-0.0621540593297413	7.40456556938466
0.233107955411082	0.629228841219216	0.911382535478648
6657	SOX2	SRY-box transcription factor 2
2513	-0.149704963229932	3.2249441722155	0.232965851746573
0.629333361285873	0.911382535478648
286262	TPRN	taperin	5338	-0.114557254294636
3.70188563947847	0.232886223796235	0.629391946407867
0.911382535478648
64327	LMBR1	limb development membrane protein 1
11044	-0.0602509981888731	6.19058397414232
0.232801020746436	0.629454647006139	0.911382535478648

1317	SLC31A1	solute carrier family 31 member 1	5127
0.0725300823185339	6.06006478085786	0.23272221239905
	0.629512654361104	0.911382535478648
63979	FIGNL1	fidgetin like 1	5034	-0.109694456915658
4.20331976383047	0.23266057021534	0.629558034898094
	0.911382535478648
3028	HSD17B10	hydroxysteroid 17-beta dehydrogenase 10
1115	-0.074640932486076	4.97848254964039
0.232565179821375	0.629628275236035	0.911382535478648
1793	DOCK1	dedicator of cytokinesis 1
8142	-0.0752553051447346	6.97278940894884
0.232548350836595	0.629640669034635	0.911382535478648
25871	NEPRO	nucleolus and neural progenitor protein
6494	-0.0619518836726221	5.19769286585565
0.232541376655603	0.629645805370663	0.911382535478648
347862	GATD1	glutamine amidotransferase like class 1 domain
containing 1	5510	0.0645935443231445	6.08217287532947
	0.232293323860681	0.629828552746576
0.911552855788519
83852	SETDB2	SET domain bifurcated histone lysine
methyltransferase 2	6156	0.10291122292842
3.46585646654516	0.232094810915459	0.629974889376619
	0.9115890824463
64708	COPS7B	COP9 signalosome subunit 7B	6066
0.0683198242536226	5.30074479629236	0.231846635280231
	0.630157944056188	0.9115890824463
90853	SPOCD1	SPOC domain containing 1
7085	-0.188936124353022	5.94717134169253
0.23182690615592	0.630172501462396	0.9115890824463
51631	LUC7L2	LUC7 like 2, pre-mRNA splicing factor	6316
0.0542377452626517	6.42593896434779	0.231788635935464
	0.630200741847636	0.9115890824463
57222	ERGIC1	endoplasmic reticulum-golgi intermediate compartment
1	14139	0.0743951387565116	7.06836970037881
0.231718287043591	0.630252661238514	0.9115890824463
55860	ACTR10	actin related protein 10
4039	-0.0606114949329714	6.33295635503997
0.231595439261463	0.630343349516647	0.9115890824463
102724827	MIATNB	MIAT neighbor	2847	0.151695683749148
	1.08205753153392	0.231423044275179
0.630470664332568	0.9115890824463
92609	TIMM50	translocase of inner mitochondrial membrane 50	5463
	0.0773863204875432	5.09677339282454
0.231197613336473	0.630637234678511	0.9115890824463
126075	CCDC159	coiled-coil domain containing 159	2968
0.125851484028812	2.10725014579039	0.231080045236672
	0.630724145119828	0.9115890824463
6241	RRM2	ribonucleotide reductase regulatory subunit M2	4955
	0.288160547502516	0.820342583538147
0.231038441738471	0.630754906397155	0.9115890824463
10144	FAM13A	family with sequence similarity 13 member A
13215	0.127032088781713	2.94008845025643
0.231034996235024	0.630757454126352	0.9115890824463
25943	C20orf194	chromosome 20 open reading frame 194

8541 -0.0711272181482628 5.55365645786171
0.231005165117899 0.630779513305344 0.9115890824463
81794 ADAMTS10 ADAM metallopeptidase with thrombospondin type 1 motif 10 5703 0.35153856327994 1.69733013493566
0.230979227693059 0.630798694645598
0.9115890824463
51529 ANAPC11 anaphase promoting complex subunit 11 2528
0.0747485985568299 4.92305832555407 0.230954987272975
0.630816622209118 0.9115890824463
2107 ETF1 eukaryotic translation termination factor 1 4601
0.06701695288803 7.74951388372903
0.230858314544435 0.63088813028327 0.9115890824463
23116 TOGARAM1 TOG array regulator of axonemal microtubules 1 10274 -0.108549447898948 4.77544948970521
0.230817729509422 0.630918156214082 0.9115890824463
29890 RBM15B RNA binding motif protein 15B
6641 -0.0581431309535386 6.94718927489111
0.230761232579958 0.630959959614255 0.9115890824463
64782 AEN apoptosis enhancing nuclease
6018 -0.0852040216628499 5.34185550943613
0.230640136649602 0.631049582548703 0.91162453617571
339044 PKD1P1 polycystin 1, transient receptor potential channel interacting pseudogene 1 7190 0.105368298814797
5.29173306502094 0.230244193735757 0.631342821618823
0.91175358944019
6016 RIT1 Ras like without CAAX 1 3919 -0.0527445946431817
6.11192495925806 0.230176784805643
0.631392776206069 0.91175358944019
25970 SH2B1 SH2B adaptor protein 1 7145 0.0845525133451664
5.63946094209175 0.230121674481332
0.631433623378668 0.91175358944019
84922 FIZ1 FLT3 interacting zinc finger 1 6790
0.113289769994813 3.51745287221533 0.230073056520567
0.631469663477574 0.91175358944019
23090 ZNF423 zinc finger protein 423 8997
0.257509412263897 -0.062982063204784 0.229874519294168
0.631616886146056 0.91175358944019
83759 RBM4B RNA binding motif protein 4B 4241
0.0641912099988654 5.04939193262564 0.229816672086933
0.631659796691362 0.91175358944019
5996 RGS1 regulator of G protein signaling 1 4117
0.423293200843133 0.433359733724429 0.22975686985127
0.631704164442114 0.91175358944019
6464 SHC1 SHC adaptor protein 1 4342 -0.0791601323881632
7.56998976506391 0.229684047677296
0.631758201376909 0.91175358944019
2959 GTF2B general transcription factor IIB
2225 -0.0684301251321507 5.06771513923558
0.229662702767049 0.631774042141239 0.91175358944019
445815 PALM2AKAP2 PALM2 and AKAP2 fusion 7727
0.109369398628876 7.44463929133127 0.229641412053979
0.631789843585813 0.91175358944019
6875 TAF4B TATA-box binding protein associated factor 4b
5395 -0.15949089431545 1.81335560572385

0.229273609927907	0.632062959582138	0.911930271920244
2195	FAT1	FAT atypical cadherin 1 16177	-0.119776791568861
	9.24325293782539	0.229256862961734
0.632075401656473	0.911930271920244
7052	TGM2	transglutaminase 2	8036	0.292590057728655
	6.03559523440425	0.229213541404036
0.632107589781798	0.911930271920244
348327	ZNF530	zinc finger protein 530 2962	-0.125880552815586
	1.49548723237886	0.22841718822358
0.632699951195182	0.912273841519091
9852	EPM2AIP1	EPM2A interacting protein 1	8126
0.0868863743783153	4.463071428153	0.228322407797059
0.632770537348686	0.912273841519091
9715	FAM131B	family with sequence similarity 131 member B	6430
	0.163066932337524	2.10375135497868
0.228313392487493	0.632777252288539	0.912273841519091
64411	ARAP3	ArfGAP with RhoGAP domain, ankyrin repeat and PH
domain 3	6186	0.0777213167949959	5.60800736484338
	0.228214967768011	0.63285057329782
0.912273841519091
64682	ANAPC1	anaphase promoting complex subunit 1	13327
0.0677133475094939	5.16541119512296	0.22817512100378
	0.632880262470641	0.912273841519091
79792	GSDMD	gasdermin D	5116	0.113654117955995
4.57829759785988	0.228157274446289	0.6328935606813
0.912273841519091
441294	CTAGE15	CTAGE family member 15	2587	-0.217707999109478
	1.89353399862867	0.22801210655282
0.633001755038414	0.912273841519091
962	CD48	CD48 molecule
3274	-0.389458977626659	-0.058878929445983
0.227977224565693	0.633027759062452	0.912273841519091
57536	KIAA1328	KIAA1328	6002	-0.121663455744192
	2.49709280688541	0.227885508777046
0.633096143528231	0.912273841519091
3190	HNRNPK	heterogeneous nuclear ribonucleoprotein K
5114	-0.0593590604568756	9.00734593760344
0.227767066074328	0.633184480928238	0.912273841519091
9442	MED27	mediator complex subunit 27	2107
0.0836530536789478	3.9762635266063 0.227720443845854
0.633219260635987	0.912273841519091
10728	PTGES3	prostaglandin E synthase 3
3399	-0.0736262846285019	7.34280341790126
0.227498611915053	0.633384804901079	0.912273841519091
6185	RPN2	ribophorin II	2819	-0.0861229757745892
7.89762199500901	0.227476637351728	0.633401209028028
	0.912273841519091
5245	PHB	prohibitin	3838	0.0728865326811322
6.42990284868764	0.227462193554683	0.633411991928125
	0.912273841519091
132884	EVC2	EvC ciliary complex subunit 2	5509
0.125696719110986	3.13089308793038	0.227326442252209
	0.633513356507762	0.912273841519091
9325	TRIP4	thyroid hormone receptor interactor 4

2045	-0.0650480490073916	5.36619478941484
0.227285633994289	0.633543835019489	0.912273841519091
64787	EPS8L2	EPS8 like 2	5693	0.0876484459260893
8.98069562791548	0.227248061975928	0.633571899446444
	0.912273841519091
23360	FNBP4	formin binding protein 4
7596	-0.0724341680355256	6.45195321192856
0.227233768997142	0.633582576337518	0.912273841519091
345778	MTX3	metaxin 3	7916	-0.0971993719758464
4.80080858296312	0.227203435777042	0.633605236694527
	0.912273841519091
11266	DUSP12	dual specificity phosphatase 12 1516
0.0742552320442212	4.25134223080634	0.227145753428932
	0.633648333271881	0.912273841519091
81926	ABHD17A abhydrolase domain containing 17A
6139	-0.0775580454902284	5.10012169011818
0.226785609087592	0.633917562144591	0.912435140463598
127262	TPRG1L	tumor protein p63 regulated 1 like	2414
0.069687133710112	6.98144869818381	0.226709048772875
	0.633974829236659	0.912435140463598
84962	AJUBA	ajuba LIM protein	5979	0.0903280192976013
	7.70925640360475	0.226607954011087
0.634050466263216	0.912435140463598
254122	SNX32	sorting nexin 32	3966	0.154957000790697
	0.232453829298473	0.226540813245878
0.634100711043098	0.912435140463598
84705	GTPBP3	GTP binding protein 3, mitochondrial	4526
0.0860556502383892	3.98620545150956	0.226438169625174
	0.634177542034244	0.912435140463598
114826	SMYD4	SET and MYND domain containing 4
6777	-0.0764776909024622	4.04805919996017
0.22637655740407	0.634223670384665	0.912435140463598
57678	GPAM	glycerol-3-phosphate acyltransferase, mitochondrial
	9808	-0.0787166984134361	4.03765965311969
0.226357502005862	0.634237938497418	0.912435140463598
84899	TMTC4	transmembrane O-mannosyltransferase targeting
cadherins 4	6272	-0.10732561314319	4.23979550849166
	0.22629933143007	0.634281499442711
0.912435140463598
27185	DISC1	DISC1 scaffold protein	12906	0.167479046001494
	2.20351835729615	0.22596794856116
0.634529786036549	0.912651258271052
57552	NCEH1	neutral cholesterol ester hydrolase 1
4808	-0.0918147291870852	6.35107373440987
0.225841783515108	0.634624373092912	0.912651258271052
79582	SPAG16	sperm associated antigen 16
8257	-0.124580075238751	2.74437478665998
0.225838007302835	0.634627204651963	0.912651258271052
85457	CIPC	CLOCK interacting pacemaker
5300	-0.0808660630369556	4.96067667254946
0.22555895377398	0.634836530803467	0.912797639357813
25873	RPL36	ribosomal protein L36	2117	0.0849586045331974
	6.74997902155856	0.225528575951373
0.634859327666439	0.912797639357813

91694	LONRF1	LON peptidase N-terminal domain and ring finger 1
	5685	-0.108830202469559	3.4245417312109
0.225366421960258	0.634981047038511	0.912878941380317
351	APP	amyloid beta precursor protein	6316
0.102562440292476	11.4925555724377	0.225208929450673
	0.635099318690379	0.912904093515797
2218	FKTN	fukutin 8032	-0.0856355220471049
4.59804624440531	0.225159405305568	0.635136520152334
	0.912904093515797
11127	KIF3A	kinesin family member 3A	6860
0.062798156200268	5.74625443968231	0.224923357296021
	0.635313903202469	0.912904093515797
23243	ANKRD28 ankyrin repeat domain 28
11279	-0.057009770758086	5.94143311026028
0.224895949533181	0.635334506706875	0.912904093515797
101927726	BEAN1-AS1	BEAN1 antisense RNA 1	1311
0.136858182246664	1.59459096535876	0.224725642732841
	0.635462567574135	0.912904093515797
51754	TMEM8B	transmembrane protein 8B
5444	-0.118858946291773	3.391524011279	0.224544464835224
	0.635598868135917	0.912904093515797
100289137	FAM95C	family with sequence similarity 95 member C
	2774	-0.167172516015945	1.10314975074409
0.224429602033579	0.635685314574246	0.912904093515797
389084	SNORC	secondary ossification center associated regulator of chondrocyte maturation	3508	-0.144070559027771
0.521169510139345	0.224203699485329	0.635855409247779
	0.912904093515797
7903	ST8SIA4 ST8 alpha-N-acetyl-neuraminide alpha-2,8-sialyltransferase 4	7794	-0.282302281575419
0.90524895706537	0.224118925456104	0.635919267430088
	0.912904093515797
153830	RNF145	ring finger protein 145 4614	0.0634535799471764
	7.41224973533639	0.224108632415851
0.635927021929529	0.912904093515797
23016	EXOSC7	exosome component 7	3899	0.0813866765574319
	4.10764003380375	0.22408636959501
0.635943794885095	0.912904093515797
8760	CDS2	CDP-diacylglycerol synthase 2	11893
0.0560466971689338	6.67917441410829	0.224065156016557
	0.635959778283505	0.912904093515797
652995	UCA1	urothelial cancer associated 1
2299	-0.352812350672635	6.82647293203052
0.224056572304335	0.635966245955893	0.912904093515797
8495	PPFIBP2 PPFIA binding protein 2 9598	0.13532064963187
	2.76863573635084	0.22371113100528
0.636226669194996	0.912904093515797
151126	ZNF385B zinc finger protein 385B
5310	-0.318660318250881	2.03631140361004
0.223641084642654	0.63627948989988	0.912904093515797
729991	BORCS8	BLOC-1 related complex subunit 8	4039
0.0756432771977111	4.2750382883741 0.223622564522691
0.636293460852639	0.912904093515797
10209	EIF1	eukaryotic translation initiation factor 1

3623	-0.0630208553663433	8.83890689544557
0.223606424526565	0.636305636897685	0.912904093515797
55101	DMAC2	distal membrane arm assembly complex 2	3024
0.063140912498214	5.50989898139262	0.223382373979311
	0.636474716639689	0.912904093515797
116151	FAM210B family with sequence similarity 210 member B
3577	-0.113773206870208	7.965927945848	0.22337237502478
	0.636482264768235	0.912904093515797
79844	ZDHHC11 zinc finger DHHC-type containing 11
6953	-0.158534839188588	1.20929006242805
0.223257126547648	0.636569279819034	0.912904093515797
79064	TMEM223 transmembrane protein 223	2280
0.101711426670465	2.60743790025619	0.22324888140394
	0.636575506130841	0.912904093515797
8743	TNFSF10 TNF superfamily member 10
3473	-0.161466479176543	7.08088461432504
0.223207345213496	0.636606874281498	0.912904093515797
85365	ALG2	ALG2 alpha-1,3/1,6-mannosyltransferase
2983	-0.0770741041565436	5.3575212663245 0.223092140583915
	0.636693895559185	0.912904093515797
80097	MZT2B	mitotic spindle organizing protein 2B	1815
0.0822728922513111	5.17669386637549	0.223060952058617
	0.636717458934126	0.912904093515797
55755	CDK5RAP2	CDK5 regulatory subunit associated protein 2
	10080	-0.0709738862236433	5.308752156508
0.222930362944849	0.636816142753817	0.912904093515797
23481	PES1	pescadillo ribosomal biogenesis factor 1	4557
	0.0727887874091483	5.41102866281506
0.222728493784953	0.636968761185289	0.912904093515797
51122	COMMD2	COMM domain containing 2	4901
0.0609460836668759	5.6741104712308 0.222698372540599
0.636991540893744	0.912904093515797
340542	BEX5	brain expressed X-linked 5	1063
0.202118839240844	0.114462206157917	0.222611391965173
	0.637057332022885	0.912904093515797
284406	ZFP82	ZFP82 zinc finger protein
2623	-0.134284725100905	1.3980371856277 0.222567869331424
	0.637090257952572	0.912904093515797
4720	NDUFS2	NADH:ubiquinone oxidoreductase core subunit S2	3658
	0.0608417792592198	6.28023713246944
0.222330038541813	0.637270252272842	0.912904093515797
5780	PTPN9	protein tyrosine phosphatase non-receptor type 9
	8632	0.0600716156807045	6.36248288250252
0.222189702830711	0.637376515897437	0.912904093515797
55153	SDAD1	SDA1 domain containing 1
3491	-0.0646213135519094	5.6413636820777 0.222176184302988
	0.637386754429404	0.912904093515797
1356	CP	ceruloplasmin	7296	-0.375448136828901
3.13345018205458	0.222060879319786	0.63747409847899
	0.912904093515797
81611	ANP32E	acidic nuclear phosphoprotein 32 family member E
	3605	-0.0728322581311807	4.95072053852058
0.22199116736066	0.637526919048168	0.912904093515797
9253	NUMBL	NUMB like endocytic adaptor protein

4300	-0.0970370231864296	4.14307548505299
0.221970250196513	0.637542769905431	0.912904093515797
81888	HYI	hydroxypyruvate isomerase (putative)
2740	-0.0882742279142616	5.25040591747253
0.221962182329335	0.637548883913601	0.912904093515797
54955	C1orf109	chromosome 1 open reading frame 109	3149
	0.0908684018030568	4.19397612773699
0.221909450147918	0.637588848871213	0.912904093515797
728621	CCDC30	coiled-coil domain containing 30	12416
0.153094726616965	1.36675420070245	0.221798925826687
	0.637672632484435	0.912904093515797
8846	ALKBH1	alkB homolog 1, histone H2A dioxygenase
2844	-0.0781135287082016	3.80144808367468
0.221631248162975	0.637799790237885	0.912904093515797
22848	AAK1	AP2 associated kinase 1 23608	-0.0842636929039568
	6.84036138922254	0.221522597267096
0.637882216631859	0.912904093515797
83480	PUS3	pseudouridine synthase 3
2010	-0.0922196870652911	3.25450485182631
0.221458869646801	0.637930574136955	0.912904093515797
163882	CNST	consortin, connexin sorting protein
5922	-0.0880163157800442	5.14137853887798
0.221445392126427	0.637940802176817	0.912904093515797
9904	RBM19	RNA binding motif protein 19
5908	-0.0667589858315984	4.91270257711386
0.221406569630735	0.6379702665487 0.912904093515797
100505876	CEBPZOS CEBPZ opposite strand	1761
0.0857960392743172	4.31977686770975	0.221375069954621
	0.637994175578017	0.912904093515797
55172	DNAAF2	dynein axonemal assembly factor 2	2963
0.0763888037661561	4.03601729524272	0.221374996757151
	0.637994231139118	0.912904093515797
51750	RTEL1	regulator of telomere elongation helicase 1	5995
	0.10256795288879	4.85397246445584
0.221371895238917	0.637996585380605	0.912904093515797
100527943	TGIF2-RAB5IF	TGIF2-RAB5IF readthrough	918
	-0.0867630788412219	6.13671965139826
0.221230564654714	0.638103885291445	0.912959332844903
146705	TEPSIN	TEPSIN adaptor related protein complex 4 accessory
protein 4762	0.0978058484545239	4.63993871956264
0.221098151800845	0.638204452736637	0.912959332844903
11217	NA	NA	7862	0.107655692519662
7.43855267260062	0.220989191500507	0.638287235571581
	0.912959332844903
539	ATP5PO	ATP synthase peripheral stalk subunit OSCP
3759	-0.0905318458994662	6.71431927289601
0.22097778204509	0.638295905372324	0.912959332844903
26024	PTCD1	pentatricopeptide repeat domain 1	5752
0.0752223661390166	4.47991499532938	0.220842262294353
	0.638398904833912	0.913013422504453
92140	MTDH	metadherin	6391	-0.0733426908960154
7.4662833212756 0.220602102180209	0.638581529217175
0.913142782553417
64864	RFX7	regulatory factor X7	10588	0.100539780910124

5.88057946531697 0.220472846768949
0.638679868831911 0.913142782553417
55684 RABL6 RAB, member RAS oncogene family like 6
10444 -0.06370589295526 6.91410872503693
0.220432252447999 0.638710760909825 0.913142782553417
57574 MARCHF4 membrane associated ring-CH-type finger 4
4447 -0.279353842895627 1.41181154537545
0.220380533561441 0.638750123756577 0.913142782553417
26258 BLOC1S6 biogenesis of lysosomal organelles complex 1 subunit
6 6812 -0.0637099407663774 6.47975076643433
0.220270648558849 0.63883377510122 0.913145741555646
23401 FRAT2 FRAT regulator of WNT signaling pathway 2 2213
0.108220243672334 3.1334207436761 0.220188170112442
0.638896579588465 0.913145741555646
27097 TAF5L TATA-box binding protein associated factor 5 like
6412 -0.0645315108870181 5.22469557322724
0.220102666883484 0.638961702496859 0.913145741555646
6170 RPL39 ribosomal protein L39 2141 0.126374353720407
6.36561181920035 0.220002853955492
0.639037743809397 0.913145741555646
121642 ALKBH2 alkB homolog 2, alpha-ketoglutarate dependent
dioxygenase 2290 0.122095092796562 2.21843719797969
0.219942263849276 0.639083913938946
0.913145741555646
10498 CARM1 coactivator associated arginine methyltransferase 1
4455 -0.0548055270602793 6.01796411417683
0.219864284534232 0.639143346199154 0.913145741555646
55205 ZNF532 zinc finger protein 532 11951 -0.0810549283419767
7.0655709287428 0.219769926037316 0.639215279114617
0.91315537119413
7311 UBA52 ubiquitin A-52 residue ribosomal protein fusion
product 1 3315 0.0879774866931079 7.56446478339186
0.219613752479326 0.639334377298572
0.913177611580516
4784 NFIX nuclear factor I X 6302 0.12042009003546
5.28873717835804 0.219468510109799
0.639445185702247 0.913177611580516
79894 ZNF672 zinc finger protein 672 3741 0.0839982753168274
4.5248691313618 0.219455961426798 0.639454761449967
0.913177611580516
9538 EI24 EI24 autophagy associated transmembrane protein 3558
0.0656665688464685 7.38142788224523
0.219407657057754 0.639491625044141 0.913177611580516
26505 CNNM3 cyclin and CBS domain divalent metal cation
transport mediator 3 4416 0.0995640632583784
4.44253188016417 0.219256626270095 0.639606916480951
0.91324914160537
9781 RNF144A ring finger protein 144A 6473
0.168340506612233 3.22801053378312 0.218872715984883
0.639900197799864 0.913363328123882
158960 CH17-340M24.3 uncharacterized protein BC009467 598
0.144629279171368 1.03594254463068
0.218834640040121 0.639929302314238 0.913363328123882
10229 COQ7 coenzyme Q7, hydroxylase

4120	-0.0789530848598778	4.08236243742101
0.218765116681867	0.639982452585025	0.913363328123882
5440	POLR2K	RNA polymerase II subunit K	1711
0.0789532443747799	5.37051991728823	0.218733631379573
	0.640006526332523	0.913363328123882
55779	CFAP44	cilia and flagella associated protein 44
12160	0.129115485515787	4.04992968435097
0.218625314594313	0.640089361771536	0.913363328123882
220992	ZNF485	zinc finger protein 485	2397	0.158320481822764
	0.730370272505464	0.218452368565285
0.640221674384335	0.913363328123882
5373	PMM2	phosphomannomutase 2	7192	-0.0963076847085732
	5.5797291299755	0.218451643133953	0.640222229511022
	0.913363328123882
79728	PALB2	partner and localizer of BRCA2	4747
0.092990293587158	4.77521082635193	0.218425587119707
	0.640242169274349	0.913363328123882
84262	PSMG3	proteasome assembly chaperone 3	1960
0.072238014668591	4.12494943725698	0.218343704274005
	0.640304840807911	0.913363328123882
27101	CACYBP	calcyclin binding protein	4511
0.0673519503968368	5.21581312381322	0.218243613967992
	0.640381467426283	0.913363328123882
7919	DDX39B	DExD-box helicase 39B	6708	-0.0569521954582813
	7.776598856248	0.218213961665569	0.640404172592607
	0.913363328123882
83690	CRISPLD1	cysteine rich secretory protein LCCL domain containing 1	4543	0.164862217434914	2.71250577253088
	0.218053193521832	0.640527307646856
0.913397673412638
84656	GLYR1	glyoxylate reductase 1 homolog
5470	-0.053922516176674	6.75494157709493
0.218012252448109	0.64055867394391	0.913397673412638
5294	PIK3CG	phosphatidylinositol-4,5-bisphosphate 3-kinase catalytic subunit gamma	5930	0.397446456523867
0.0513530962883716	0.217773748760575	0.64074147078192
	0.913565327895191
440603	BCL2L15	BCL2 like 15	4251	-0.267166490359156
2.39602486085133	0.217647196027329	0.640838514340124
	0.91361069418444
202333	CMYA5	cardiomyopathy associated 5
14179	-0.204610086712951	2.82798258548615
0.217261581020232	0.641134424811377	0.913742174640749
8974	P4HA2	prolyl 4-hydroxylase subunit alpha 2
5667	-0.111163388699685	7.53418775287419
0.217236866025644	0.64115340132383	0.913742174640749
81554	RCC1L	RCC1 like	4196	-0.0627305060047645
5.51118668440688	0.217188196659624	0.641190774173623
	0.913742174640749
8515	ITGA10	integrin subunit alpha 10	6320
0.187594998645971	1.01895576068742	0.217187019645895
	0.641191678056966	0.913742174640749
100506076	LOC100506076	uncharacterized LOC100506076
4575	-0.156912136515221	3.26443918607411

0.216916586537128	0.64139943524174	0.913860625475215
6711	SPTBN1	spectrin beta, non-erythrocytic 1
18504	-0.0840511796590291	10.357219302105 0.216908976666105
	0.641405283718629	0.913860625475215
643854	CTAGE9	CTAGE family member 9	2334	-0.193311470637138
	3.93378681821999	0.216790419477078
0.641496415579758	0.913897507427347
9831	ZNF623	zinc finger protein 623 7146	-0.0697362144245905
	5.31129683998401	0.216522208186174
0.641702694643074	0.913960286785426
5413	SEPTIN5 septin 5	3104	-0.160362358049801
3.3675316890022 0.216365143065417	0.641823564283307
0.913960286785426
7574	ZNF26	zinc finger protein 26	20541	0.111096883994728
	3.91884165527208	0.21634299828029
0.641840610119542	0.913960286785426
90407	TMEM41A transmembrane protein 41A
8030	-0.0539248926986375	6.06172276760758
0.216342437987713	0.64184104141568	0.913960286785426
9489	PGS1	phosphatidylglycerophosphate synthase 1 4965
0.0860465084662391	5.11344217631462	0.216306031396076
	0.641869067555867	0.913960286785426
283638	CEP170B centrosomal protein 170B
7199	-0.0734789181330877	7.54593990749751
0.216129356928469	0.642005113994069	0.913960286785426
11211	FZD10	frizzled class receptor 10
3282	-0.262914225574761	2.25729999300663
0.216096819748302	0.642030176303209	0.913960286785426
28990	ASTE1	asteroid homolog 1	3427	-0.109667356430069
	2.34440074532931	0.215917279623589
0.642168511375621	0.913960286785426
146059	CDAN1	codanin 1	4913	-0.0889162856616376
4.34886354776855	0.215897435127198	0.642183805786037
	0.913960286785426
115123	MARCHF3 membrane associated ring-CH-type finger 3
5179	-0.136346230388653	0.138490887482737
0.215885525477643	0.642192985117596	0.913960286785426
10542	LAMTOR5 late endosomal/lysosomal adaptor, MAPK and MTOR
activator 5	2388	0.0733845841569924	5.94357341500783
	0.215690767935612	0.64234313768769
0.913961188141345
708	C1QBP	complement C1q binding protein	2335
0.0819849554301573	5.60902537332373	0.21565340177299
	0.642371955366687	0.913961188141345
25771	TBC1D22A	TBC1 domain family member 22A
5315	-0.065250268293145	5.26426492938795
0.215359490260366	0.642598732849482	0.913961188141345
100216001	MANCR	mitotically associated long non coding RNA
	1920	0.226223172290321	3.99723289880499
0.215308858084484	0.642637818825086	0.913961188141345
10421	CD2BP2	CD2 cytoplasmic tail binding protein 2
3773	-0.0558277534674459	6.96717302923455
0.215288137283949	0.642653816048923	0.913961188141345
5527	PPP2R5C protein phosphatase 2 regulatory subunit B'gamma

13980 -0.0497350883306058 6.62135311328454
0.215278256799365 0.642661444477622 0.913961188141345
132014 IL17RE interleukin 17 receptor E
4676 -0.197070970041372 2.89357794674655
0.215141113240264 0.642767350923285 0.913961188141345
100287932 TIMM23 translocase of inner mitochondrial membrane
23 1222 0.0779898359495405 5.74883921801125
0.21510217269803 0.642797429476078 0.913961188141345
377677 CA13 carbonic anhydrase 13 4764 0.11461208660149
3.01375878633512 0.214975886131884
0.642894998834275 0.913961188141345
65220 NADK NAD kinase 5570 0.0626589459525733
6.35942184406703 0.214965831175945 0.642902768818074
0.913961188141345
23193 GANAB glucosidase II alpha subunit 5027
0.0625803536026316 8.13437105549338 0.214954698409201
0.642911371939507 0.913961188141345
84445 LZTS2 leucine zipper tumor suppressor 2 6547
0.0853047989178825 6.47526219054094 0.214756990261314
0.643064200916934 0.914085677110158
7360 UGP2 UDP-glucose pyrophosphorylase 2 6495
0.0654347601033022 6.98554785816076 0.214379324053329
0.643356375695302 0.914408194253079
55111 PLEKHJ1 pleckstrin homology domain containing J1 4655
0.0616409443232607 5.53484934968176
0.213804495318819 0.643801682214422 0.914721427079121
6497 SKI SKI proto-oncogene 6080 0.0587890453202443
6.30687467116866 0.21371822666052
0.643868575236885 0.914721427079121
10237 SLC35B1 solute carrier family 35 member B1 5919
0.0592945473509765 5.6258679023657 0.213700840339198
0.64388205863581 0.914721427079121
29999 FSCN3 fascin actin-bundling protein 3 3214
0.0715245740967704 5.17839243340452 0.213623193770314
0.643942283046345 0.914721427079121
10570 DPYSL4 dihydropyrimidinase like 4 3176
0.535257408499979 1.99995517802637 0.213529505111559
0.644014967745287 0.914721427079121
154007 SNRNP48 small nuclear ribonucleoprotein U11/U12 subunit 48
5836 0.0757846806688092 4.3120346539477
0.21351538229152 0.644025926063711 0.914721427079121
126070 ZNF440 zinc finger protein 440 4801 -0.0854315273108884
4.31320468450664 0.213505117386039
0.644033891186856 0.914721427079121
3183 HNRNPC heterogeneous nuclear ribonucleoprotein C 7309
0.0625338861723737 7.77036526855438
0.213245751413979 0.64423522512593 0.914826029667595
7542 ZFPL1 zinc finger protein like 1 2784
0.0698180702576537 4.4629746909836 0.213197147260285
0.644272970832303 0.914826029667595
64802 NMNAT1 nicotinamide nucleotide adenylyltransferase 1
5050 -0.0710136943974991 4.2155944252937 0.213157871988719
0.644303475595833 0.914826029667595
54777 CFAP46 cilia and flagella associated protein 46

10561	0.225786282875985	0.451576441575224
0.213052822878025	0.644385083089091	0.914849164689732
55143	CDCA8	cell division cycle associated 8	2469
0.127081413807177	1.85288472881026	0.212681959050472
	0.644673384291998	0.915156546124506
93663	ARHGAP18	Rho GTPase activating protein 18	4640
	-0.166034722305528	6.80069374952517
0.212397423263042	0.644894782734471	0.915156546124506
10632	ATP5MG	ATP synthase membrane subunit g 3382
0.0785336959840776	6.48017598714362	0.212279521535592
	0.644986575230875	0.915156546124506
29893	PSMC3IP PSMC3 interacting protein	2573
0.133704412081538	1.47830720165322	0.212179012703391
	0.645064850876674	0.915156546124506
57462	MYORG	myogenesis regulating glycosidase (putative)
6398	-0.0832585189351235	6.38467660043989
0.212169599614541	0.645072182881858	0.915156546124506
29911	HOOK2	hook microtubule tethering protein 2
5909	-0.0914011156211584	5.65850993925084
0.212158856119345	0.645080551402709	0.915156546124506
25832	NBPF14	NBPF member 14	11985	-0.0774308781187904
5.4544824887841 0.212141340861532	0.645094195260142
0.915156546124506
6147	RPL23A	ribosomal protein L23a	2824	0.0942186446523761
	6.45185117405036	0.212102727675983
0.64512427617858	0.915156546124506
10280	SIGMAR1 sigma non-opioid intracellular receptor 1	2196
	0.0780948373573386	5.37765248892503
0.211959327437722	0.645236018653035	0.915197218653345
10294	DNAJA2	DnaJ heat shock protein family (Hsp40) member A2
	5065	-0.0577598433952086	6.55983388274344
0.211898251949719	0.645283624717672	0.915197218653345
11179	ZNF277	zinc finger protein 277 4014	0.111332616254034
	3.93747556162979	0.21159399645834
0.645520904481848	0.915377308438298
63951	DMRTA1	DMRT like family A1	5569	-0.152666884645237
	2.41156355907152	0.211567850539958
0.645541304544407	0.915377308438298
9961	MVP	major vault protein	6141	-0.0624036660704058
	8.22340991323675	0.211410244989793
0.645664306885601	0.915406340483904
51112	TRAPPC12	trafficking protein particle complex 12
2838	-0.0652780744828295	4.79097864020001
0.211363292225339	0.645700961637182	0.915406340483904
126017	ZNF813	zinc finger protein 813 6244	-0.109541919469766
	3.16762059688038	0.211290447939394
0.645757838960623	0.915406340483904
5361	PLXNA1	plexin A1	10944	-0.0674608439054189
8.71465911779717	0.211097509713568	0.645908543871199
	0.915527319773718
7103	TSPAN8	tetraspanin 8	2519	0.173319106077691
1.92575818808825	0.210926962881947	0.646041828229887
	0.915623584743098
1431	CS	citrate synthase	6402	-0.0610362649334363

7.39997224542121 0.210688854721411
0.646228021748858 0.915651479023826
7752 ZNF200 zinc finger protein 200 4271 0.0884496357149869
3.49981009877231 0.210338064264844
0.646502561684747 0.915651479023826
27043 PELP1 proline, glutamate and leucine rich protein 1 6577
0.085969849549707 4.87783249875977
0.210253587528424 0.646568717299753 0.915651479023826
4247 MGAT2 alpha-1,6-mannosyl-glycoprotein 2-beta-N-acetylglucosaminyltransferase 2687 -0.0784213618553868
5.55749236465844 0.210218701326834 0.646596042154493
0.915651479023826
11216 AKAP10 A-kinase anchoring protein 10
4848 -0.0582859464897244 5.55593985236222
0.210148900789193 0.646650722138893 0.915651479023826
64121 RRAGC Ras related GTP binding C 6155
0.0757178143743694 6.85744802458385 0.209880441649943
0.646861128740079 0.915651479023826
57504 MTA3 metastasis associated 1 family member 3 10580
0.0626491653888037 5.13135401873477 0.209824021475704
0.646905369121332 0.915651479023826
64951 MRPS24 mitochondrial ribosomal protein S24 2613
0.0935098945314585 5.06294705016487 0.209622390012722
0.647063531913132 0.915651479023826
64770 CCDC14 coiled-coil domain containing 14 11423
0.122834993187115 4.9055201088181 0.209605981625359
0.647076406950218 0.915651479023826
6836 SURF4 surfeit 4 3446 0.0644967174265436
8.23045121624125 0.20953893180689 0.647129024726588
0.915651479023826
10767 HBS1L HBS1 like translational GTPase
11099 -0.0563863285971072 5.8576761054023
0.209454000813619 0.647195689489821 0.915651479023826
54807 ZNF586 zinc finger protein 586 3261 -0.107222844685561
2.5869812540772 0.209452034779712 0.647197232879577
0.915651479023826
79705 LRRK1 leucine rich repeat kinase 1 17600
0.125706519625939 4.02868001687254 0.209345095008658
0.64728119669182 0.915651479023826
2739 GLO1 glyoxalase I 2352 0.0740928504790717
7.10625437554345 0.209341040418821 0.647284380665227
0.915651479023826
55825 PECR peroxisomal trans-2-enoyl-CoA reductase 3182
0.127028397758849 1.76028603892223 0.209337916697298
0.647286833675214 0.915651479023826
57828 CATSPERG cation channel sperm associated auxiliary subunit gamma 9190 0.142985790500238 0.0192355925264531
0.20925094095349 0.64735514326514
0.915651479023826
100131827 ZNF717 zinc finger protein 717 6871
0.133260233627204 2.07340180365325 0.209205312119713
0.64739098640871 0.915651479023826
89978 DPH6 diphthamine biosynthesis 6
9345 -0.141385502060916 1.38469531802302

0.209171365140708	0.647417656087929	0.915651479023826
221477	C6orf89	chromosome 6 open reading frame 89	7340
0.050469963320788	6.76584620523883	0.209105545211945
	0.647469373488589	0.915651479023826
80254	CEP63	centrosomal protein 63	10116	-0.069692002811675
	4.44996876979093	0.209008093668862
0.647545963213304	0.915651479023826
3795	KHK	ketohexokinase	3060	0.142551276041693
1.41341779426347	0.208946720611131	0.647594208988939
	0.915651479023826
105369154	PKD1P6-NPIPP1	PKD1P6-NPIPP1 readthrough
3043	-0.0918996347468329	4.03772016838802
0.208894125708323	0.647635561013951	0.915651479023826
81543	LRRC3	leucine rich repeat containing 3
5869	-0.151919288838944	2.37932694902669
0.20887624413192	0.647649621593038	0.915651479023826
203068	TUBB	tubulin beta class I	3359	-0.0872738446166366
	9.88948913156712	0.208832186304278
0.647684268096108	0.915651479023826
57085	AGTRAP	angiotensin II receptor associated protein
2435	-0.0910799830486809	5.11180074287409
0.208817542784079	0.647695784548302	0.915651479023826
136853	SSC4D	scavenger receptor cysteine rich family member with 4 domains	2819	0.365981470797918	4.22844485701426
	0.208303179595681	0.648100617494049
0.916070958629276
57657	HCN3	hyperpolarization activated cyclic nucleotide gated potassium channel 3	4121	-0.111398430867291
3.45728300894205	0.20827437045208	0.648123309774836
	0.916070958629276
27069	GHITM	growth hormone inducible transmembrane protein
2438	-0.0599359541148865	7.74018144853414
0.208115999784809	0.648248088478868	0.91615487592811
11015	KDELR3	KDEL endoplasmic reticulum protein retention receptor 3	1929	0.184610961128137	3.4355086868759
0.207904855943653	0.648414535894271	0.916214986599103
10953	TOMM34	translocase of outer mitochondrial membrane 34
2062	-0.0858113338923453	5.42093103521441
0.207855493058787	0.648453464016643	0.916214986599103
23438	HARS2	histidyl-tRNA synthetase 2, mitochondrial
3408	-0.0667549358485885	5.47973875237909
0.207813157270628	0.648486854939923	0.916214986599103
100313843	MIR711	microRNA 711	76	-0.141563524341999
	2.464868287087	0.207602460743584	0.648653095835191
	0.916357429486992
51155	JPT1	Jupiter microtubule associated homolog 1
3414	-0.0871383358124619	8.09607373047804
0.207484440884173	0.648746258769618	0.916396616234977
346157	ZNF391	zinc finger protein 391	4232	0.128476845562189
	2.67350256030934	0.207157038474185
0.64900487243835	0.916669481521022
2665	GDI2	GDP dissociation inhibitor 2
4033	-0.0697399900307093	7.83689586070801
0.206942513235845	0.649174458754328	0.916816560170587

92935	MARS2	methionyl-tRNA synthetase 2, mitochondrial	3019
	0.108445311403328	2.99920188496324
0.206750397136241	0.649326420285839	0.916846578384847
163131	ZNF780B	zinc finger protein 780B
9217	-0.0955359442503166	3.6413774751693	0.20667395199652
	0.649386911171124	0.916846578384847
8520	HAT1	histone acetyltransferase 1	6790
0.0655740228127029	4.98005817648486	0.206667417126553
	0.649392082826877	0.916846578384847
54798	DCHS2	dachsous cadherin-related 2
14844	-0.306191231367914	1.90931660877628
0.206532093841389	0.649499199021683	0.916905390112549
100131801	PET100	PET100 cytochrome c oxidase chaperone	1204
	0.0934528532508836	3.42940593847642
0.20630114587269	0.649682105505782	0.91707117321571
144110	TMEM86A	transmembrane protein 86A
4243	-0.12712693128324	4.69737693573678
0.206152117513177	0.64980019875623	0.917145443770507
553103	MIR3936HG	MIR3936 host gene	2857
0.156620315437731	1.38278017444375	0.205916183451503
	0.64998726325629	0.917236920234312
168544	ZNF467	zinc finger protein 467	2734	0.198843152676322
	1.55016933890037	0.205745951713832
0.650122314782686	0.917236920234312
283788	LOC283788	FSHD region gene 1 pseudogene	372
0.181625096444293	1.25473599578258	0.205468846944818
	0.650342297035304	0.917236920234312
400961	PAIP2B	poly(A) binding protein interacting protein 2B	6279
	0.144360491819275	1.290070183136	0.205372743513912
	0.650418631418299	0.917236920234312
650669	GAS6-AS1	GAS6 antisense RNA 1	7169
0.203970730187338	0.752104053822768	0.205360010470803
	0.650428746813673	0.917236920234312
57706	DENND1A	DENN domain containing 1A	10256
0.0839977627668159	3.94327826256122	0.205358992490827
	0.650429555534544	0.917236920234312
26010	SPATS2L	spermatogenesis associated serine rich 2 like
10437	0.0622836600958007	7.79562893821822
0.205339769900114	0.65044482712378	0.917236920234312
6837	MED22	mediator complex subunit 22
7224	-0.0564652646474042	5.41335288840909
0.20533799369109	0.650446238295192	0.917236920234312
11129	CLASRP	CLK4 associating serine/arginine rich protein	4931
	0.09042512116711	4.56795801726701
0.205327761548244	0.650454367722383	0.917236920234312
29907	SNX15	sorting nexin 15	4915	-0.065710833249736
	4.65315158562105	0.205048429809358
0.650676390877702	0.917365319028868
126282	TNFAIP8L1	TNF alpha induced protein 8 like 1
3957	-0.160392888365846	1.62488855675454
0.205048408483171	0.650676407835497	0.917365319028868
100506581	C16orf95	chromosome 16 open reading frame 95
	1622	0.115028855432517	1.59476029871967
0.20491871476107	0.650779555142586	0.917418400652396

10969	EBNA1BP2	EBNA1 binding protein 2 2851
0.0774701650116574	5.67486771194678	0.204788626459532
	0.650883055781604	0.917471969642038
6625	SNRNP70 small nuclear ribonucleoprotein U1 subunit 70	4684
	0.0667450569950138	7.46590659797949
0.204530478724678	0.651088560189673	0.917570127451976
79966	SCD5	stearoyl-CoA desaturase 5	4292
0.0829484077262689	8.04979955093373	0.204469746730155
	0.651136929979493	0.917570127451976
51184	GPN3	GPN-loop GTPase 3	2595	0.0843317786481245
	3.70864918604157	0.204380071363893
0.651208367441438	0.917570127451976
23169	SLC35D1 solute carrier family 35 member D1
6162	-0.0638680635793103	5.21694222744841
0.204284570592856	0.651284466312389	0.917570127451976
4001	LMNB1	lamin B1	3633	0.161608354936906
1.04169615694885	0.204241026352917	0.6513191712317
0.917570127451976
10592	SMC2	structural maintenance of chromosomes 2 6468
0.0716364577715737	4.59259721052701	0.204182487132122
	0.651365834227155	0.917570127451976
6903	TBCC	tubulin folding cofactor C	1616
0.0631536120302978	4.74110649685673	0.204002261174926
	0.651509547186983	0.917570127451976
9219	MTA2	metastasis associated 1 family member 2 4236
0.0569054787133369	7.40039722278856	0.203859219448077
	0.651623663694661	0.917570127451976
27299	ADAMDEC1	ADAM like decysin 1	2438
0.472164057347291	0.650125981713476	0.203844736895514
	0.651635220340909	0.917570127451976
2079	ERH	ERH mRNA splicing and mitosis factor	1446
0.0707812558056473	5.97097759602379	0.203836029860733
	0.651642168533608	0.917570127451976
5049	PAFAH1B2	platelet activating factor acetylhydrolase
1b catalytic subunit 2	7104	-0.0579556073757056
7.46052495331805	0.203727853668511	0.651728507774516
	0.917570127451976
22875	ENPP4	ectonucleotide pyrophosphatase/phosphodiesterase 4
	4651	-0.0963862070724309	4.99522835416082
0.20371497615891	0.651738787607604	0.917570127451976
55669	MFN1	mitofusin 1	5957	0.07239286356444
6.41353866038009	0.203568556343157	0.65185569884433
	0.917642489820354
5550	PREP	prolyl endopeptidase	3762	-0.0858743280521024
	6.39984796021683	0.203382035667289
0.652004702565956	0.91766996283996
55142	HAUS2	HAUS augmin like complex subunit 2
4323	-0.0743959115551859	5.05274373247512
0.203380105724477	0.65200624474786	0.91766996283996
4323	MMP14	matrix metallopeptidase 14	4464
0.0638654788447122	9.76555688092824	0.20317136669928
	0.652173096279113	0.917695764706234
90321	ZNF766	zinc finger protein 766 7335	-0.0721782605551335
	4.15237543830503	0.203158617199788

0.652183290687954	0.917695764706234
1384	CRAT	carnitine O-acetyltransferase
3803	-0.116057558321542	5.07394140070751
0.20304562671862	0.652273653894376	0.917695764706234
5052	PRDX1	peroxiredoxin 1 1522	0.0870463525758799
8.31551528198534	0.203019354621063	0.652294669121977
	0.917695764706234
5217	PFN2	profilin 2	3440	-0.0755726248872672
6.39567954901574	0.202947490945627	0.65235216171771
	0.917695764706234
143244	EIF5AL1 eukaryotic translation initiation factor 5A like 1
	3840	-0.0823777165864827	5.92230248042795
0.202793135796526	0.652475690800736	0.91777736503775
760	CA2	carbonic anhydrase 2	2123	0.23622231635985
	0.458874034491943	0.202679833195392
0.652566401878506	0.917803830216467
115110	TNFRSF14-AS1	TNFRSF14 antisense RNA 1
6137	-0.211288456415898	1.78796060652035
0.202605968539281	0.65262555500507	0.917803830216467
115708	TRMT61A tRNA methyltransferase 61A	3229
0.0797510538323158	4.07573644116822	0.20240436729859
	0.652787069605691	0.917938511305327
11104	KATNA1	katanin catalytic subunit A1	2308
0.066665161842945	4.23599221747604	0.202194384183487
	0.652955402284295	0.917938511305327
54544	CRCT1	cysteine rich C-terminal 1
700	-0.221456660402702	4.33045676190605
0.202132006595576	0.653005427436356	0.917938511305327
10945	KDELR1	KDEL endoplasmic reticulum protein retention receptor 1	2251	0.0621830425420994	7.5300967502065
0.20212296829822	0.653012676677993	0.917938511305327
24146	CLDN15	claudin 15	2893	0.172294645091828
2.93924696883652	0.202021586609417	0.653094004058736
	0.917938511305327
7520	XRCC5	X-ray repair cross complementing 5
5463	-0.0578737091835941	8.65333219837671
0.201996005666075	0.653114528711993	0.917938511305327
51128	SAR1B	secretion associated Ras related GTPase 1B
10963	-0.0523656621886057	6.0507858836835 0.201190167878629
	0.653761887045827	0.918706328916637
5605	MAP2K2	mitogen-activated protein kinase kinase 2
5236	-0.080890051434298	7.07735747855514
0.201152717420911	0.653792010183554	0.918706328916637
1050	CEBPA	CCAAT enhancer binding protein alpha	2631
0.142484150688864	2.52520541626715	0.201024780981014
	0.653894940801495	0.918758795847782
55012	PPP2R3C protein phosphatase 2 regulatory subunit B''gamma
	3863	-0.0688309115368739	4.3776973028366
0.200854749994789	0.65403179927436	0.91880508999466
100526772	STIMATE-MUSTN1	STIMATE-MUSTN1 readthrough
2289	-0.0906226858247078	3.45289200079597
0.200804877991406	0.654071954599629	0.91880508999466
57479	PRR12	proline rich 12 7093	-0.0829761670449007
5.4585516926216 0.200667232716162	0.654182813204378

0.91880508999466
8228 PNPLA4 patatin like phospholipase domain containing 4 3441
0.104298302337131 3.0813381978777 0.200621663172629
0.654219524675085 0.91880508999466
728734 NPIPB8 nuclear pore complex interacting protein family
member B8 2338 -0.133031119635184 1.95981680597723
0.200545777685252 0.654280670210633
0.91880508999466
445328 ARHGEF35 Rho guanine nucleotide exchange factor 35
2431 -0.122139500896001 5.07861924398551
0.200495157246529 0.654321465888251 0.91880508999466
92342 METTL18 methyltransferase like 18
1578 -0.104236642535284 2.29390684042157
0.200171635834986 0.654582342180709 0.919031981677006
55298 RNF121 ring finger protein 121 3567 0.0572572805819724
5.75245156113671 0.200132062497443
0.654614270092917 0.919031981677006
2109 ETFB electron transfer flavoprotein subunit beta 4056
0.0907318386071706 4.26591533059847
0.199900774894665 0.654800949582389 0.91911476179713
284385 ZNF561-AS1 ZNF561 antisense RNA 1 (head to head) 2614
0.12501771382778 2.15621153392498
0.199896415088229 0.65480446976323 0.91911476179713
10055 SAE1 SUMO1 activating enzyme subunit 1 3396
0.0573723225054119 6.26935162901153 0.199735056965249
0.654934785429826 0.91920556508172
9783 RIMS3 regulating synaptic membrane exocytosis 3
7235 -0.150463795971951 2.40021164132674
0.199609460461566 0.655036263097346 0.919255879892248
5230 PGK1 phosphoglycerate kinase 1 5415
0.108948956711603 8.65316773307471 0.199480871384822
0.655140198346761 0.919274834888862
10013 HDAC6 histone deacetylase 6 4892 0.0612541784414531
5.66856923774681 0.19942576857083
0.655184748835418 0.919274834888862
6838 SURF6 surfeit 6 4626 -0.0684332643603037
5.18626401730515 0.1990814038531 0.655463334215561
0.919274834888862
79725 THAP9 THAP domain containing 9 4332
0.119954415299786 1.79607962566557 0.198773868655943
0.65571236944145 0.919274834888862
93649 MYOCD myocardin 8875 -0.292180241347358
0.908075666968129 0.198738361637419 0.65574113711099
0.919274834888862
1622 DBI diazepam binding inhibitor, acyl-CoA binding protein
2509 0.11583315090361 5.97765017979591
0.198679427233301 0.655788892371908 0.919274834888862
159091 FAM122C family with sequence similarity 122C 5894
0.142655867784013 0.626769149190321 0.198536012380428
0.655905138641379 0.919274834888862
3646 EIF3E eukaryotic translation initiation factor 3 subunit E
5418 -0.0643155052667248 6.4024249265368
0.198534285125419 0.655906538990955 0.919274834888862
93643 TJAP1 tight junction associated protein 1 6741

0.076763567454337	4.77304011605447	0.198527350867373
	0.655912160924136	0.919274834888862
100506668	NRAV	negative regulator of antiviral response
	1901	-0.0869163595242205	3.21061974836087
0.198519545281674	0.655918489424921	0.919274834888862
4258	MGST2	microsomal glutathione S-transferase 2	3182
0.0808082342878224	4.44284407999479	0.198489209591981
	0.65594308597615	0.919274834888862
23096	IQSEC2	IQ motif and Sec7 domain ArfGEF 2	6962
0.101956911108295	3.28692397930139	0.198458895384365
	0.655967667359816	0.919274834888862
5788	PTPRC	protein tyrosine phosphatase receptor type C	8371
	0.398812329700401	2.86715145113672
0.198408554508006	0.656008493071121	0.919274834888862
51491	NOP16	NOP16 nucleolar protein 2862	0.0797004109501859
	4.24290204278083	0.198373670697254
0.656036786970114	0.919274834888862
10603	SH2B2	SH2B adaptor protein 2	2384	0.116030507747777
	1.13272818002476	0.198280226621397
0.656112593187718	0.919274834888862
79027	ZNF655	zinc finger protein 655 8401	-0.0617757402910211
	6.37221917970803	0.198149036393581
0.656219056982929	0.919274834888862
8674	VAMP4	vesicle associated membrane protein 4
5159	-0.105795579092521	3.87840352960175
0.198041141111474	0.656306648060481	0.919274834888862
55295	KLHL26	kelch like family member 26	4797
0.11930355050693	2.45163584125114	0.197993448029433
	0.656345375161174	0.919274834888862
51807	TUBA8	tubulin alpha 8 4374	0.1111109669417
2.43701237236052	0.197917858268232	0.65640676599939
	0.919274834888862
10289	EIF1B	eukaryotic translation initiation factor 1B	1912
	0.0803460628491641	4.95619370156159
0.197855030210518	0.656457803004716	0.919274834888862
5252	PHF1	PHD finger protein 1	3596	0.106303327924739
	5.37640045364341	0.197691729762624
0.656590501986546	0.919274834888862
5609	MAP2K7	mitogen-activated protein kinase kinase 7	5145
	0.0614692283949883	5.41743236881064
0.197618201083538	0.656650273291434	0.919274834888862
84335	AKT1S1	AKT1 substrate 1	5737	0.0923616966007169
	6.35381177336473	0.197616153245583
0.656651938165098	0.919274834888862
56241	SUSD2	sushi domain containing 2	5279
0.240651693353271	2.35380894361558	0.197598147866223
	0.656666576819795	0.919274834888862
55013	MCUB	mitochondrial calcium uniporter dominant negative
beta subunit	3566	-0.0803715551417769	4.40651839773732
	0.197533011143044	0.656719540665594
0.919274834888862
26259	FBXW8	F-box and WD repeat domain containing 8 5169
0.0957189204110498	3.62232265617662	0.197472879053395
	0.656768444437355	0.919274834888862

1268	CNR1	cannabinoid receptor 1	6329	-0.296862226610634
	0.349736894132163	0.197398116885132
0.656829258884506	0.919274834888862
5859	QARS1	glutaminyl-tRNA synthetase 1	4607
0.0559123149809708	6.88026254596643	0.19732665609105
	0.656887400756952	0.919274834888862
25917	THUMPD3 THUMP domain containing 3
5365	-0.0683312382605863	5.03456757883309
0.197126662139981	0.657050186703156	0.919410785729382
79685	SAP30L	SAP30 like	6755	0.0891165756522292
5.28656925682281	0.197041492199928	0.657119541160441
	0.919415983721337
26519	TIMM10	translocase of inner mitochondrial membrane 10	777
	0.119921515358601	3.55139644522076
0.196829421739167	0.65729230956196	0.919534592272566
100131187	TSTD1	thiosulfate sulfurtransferase like domain
containing 1	1210	0.109544116778365	3.41175306478581
	0.196776294033784	0.657335608797401
0.919534592272566
64834	ELOVL1	ELOVL fatty acid elongase 1
2507	-0.0729010065864437	7.30441886468741
0.196387352081956	0.657652810884533	0.91977371344337
55287	TMEM40	transmembrane protein 40
3562	-0.0819393204571285	6.64525818789377
0.196343375717639	0.657688699503628	0.91977371344337
55676	SLC30A6 solute carrier family 30 member 6
5410	-0.0579433357674441	5.84652854123343
0.196325189993164	0.657703542075305	0.91977371344337
51102	MECR	mitochondrial trans-2-enoyl-CoA reductase	3922
	0.10101415845375	3.57542198390102
0.196108259912826	0.657880656498907	0.919929555313985
54923	LIME1	Lck interacting transmembrane adaptor 1
4224	-0.0648955227076164	5.17270981498939
0.196018177015759	0.657954239884006	0.919940610706256
123879	DCUN1D3 defective in cullin neddylation 1 domain containing
3	6248	-0.0843094402295513	4.45522398782921
0.195890566883534	0.658058511667309	0.919994567613933
439931	THAP7-AS1	THAP7 antisense RNA 1	2975
0.238606767219592	0.488900780590141	0.195731446817859
	0.658188587512138	0.920069523570475
23070	CMTR1	cap methyltransferase 1 5274	0.058423747678378
	6.01535837202475	0.195609375060791
0.658288420345995	0.920069523570475
56917	MEIS3	Meis homeobox 3 5122	0.229043845889396
0.00901171206637164	0.195583989604585	0.658309185780203
	0.920069523570475
84747	UNC119B unc-119 lipid binding chaperone B
4560	-0.0599699507690245	6.18589272830621
0.195427014731514	0.658437627834727	0.920092438651971
5901	RAN	RAN, member RAS oncogene family 4825
0.0882776530098324	8.01052531756983	0.195403397473456
	0.658456957596013	0.920092438651971
81575	APOLD1	apolipoprotein L domain containing 1	5610
0.0993230839378791	3.1155236464665 0.195065442543502

0.658733712987086	0.920290185624395
64766	S100PBP S100P binding protein	9194	0.0728093850855415
	5.07713815607906	0.195047062414233
0.658748772910467	0.920290185624395
9816	URB2	URB2 ribosome biogenesis homolog
6001	-0.0857450484931138	3.75539740483363
0.194866090492344	0.658897099171795	0.920290185624395
352954	CASTOR3 CASTOR family member 3	4860	0.117440611993274
	4.06908847838218	0.194807981879974
0.658944742982417	0.920290185624395
57799	RAB40C	RAB40C, member RAS oncogene family	4180
0.0662873087222436	4.75566255865441	0.194686293173886
	0.659044544236847	0.920290185624395
57480	PLEKHG1 pleckstrin homology and RhoGEF domain containing G1
	7398	-0.0998328666246539	5.72453964171402
0.194549716893615	0.659156599712313	0.920290185624395
64747	MFSD1	major facilitator superfamily domain containing 1
	5288	-0.0540598338122997	6.60358831983903
0.194398430836017	0.659280778843296	0.920290185624395
56951	C5orf15 chromosome 5 open reading frame 15	2422
0.0558322600935964	6.44120704530027	0.194339836747861
	0.659328889745001	0.920290185624395
79002	TRIR	telomerase RNA component interacting RNase	1402
	0.0535081897621805	6.0106108764693 0.19421430769424
	0.659431989307348	0.920290185624395
27254	CSDC2	cold shock domain containing C2
3250	-0.184810908370493	0.249524522315945
0.194188018915115	0.659453585854571	0.920290185624395
7062	TCHH	trichohyalin	6995	-0.296882491925428
1.26316986750513	0.194119192091154	0.659510136185062
	0.920290185624395
9453	GGPS1	geranylgeranyl diphosphate synthase 1	3809
0.05456305436029	5.3545847021205 0.194102736645396
0.659523658281323	0.920290185624395
3790	KCNS3	potassium voltage-gated channel modifier subfamily S
member 3	3131	-0.132383601199855	4.55581460271215
	0.194093419324147	0.65953131499939
0.920290185624395
79412	KREMEN2 kringle containing transmembrane protein 2	2614
	0.224954437256885	1.6624085470494 0.194078533323354
	0.659543548360374	0.920290185624395
22	ABCB7	ATP binding cassette subfamily B member 7
2970	-0.0775040397812702	3.95318635919988
0.193970483437117	0.659632360882214	0.920290185624395
100996579	LINC01806	long intergenic non-protein coding
RNA 1806	2377	-0.149080079393159	0.839028140454898
	0.193949382692818	0.659649708265076
0.920290185624395
101928578	RAP2C-AS1	RAP2C antisense RNA 1	3536
0.125874512902606	0.768709761098375	0.193748310672426
	0.659815070514478	0.920324843764661
23241	PACS2	phosphofurin acidic cluster sorting protein 2
12363	0.05595049782328	6.45526188934545
0.193671497530609	0.659878268918793	0.920324843764661

57523	NYNRIN	NYN domain and retroviral integrase containing
7955	-0.0786436248139965	7.41713628512995
0.193652305510767	0.659894061586826	0.920324843764661
56131	PCDHB4	protocadherin beta 4	3825	0.416670434498368
	0.403371533711463	0.193599681779331
0.65993736922769	0.920324843764661
9246	UBE2L6	ubiquitin conjugating enzyme E2 L6
2197	-0.139959945066103	4.43609549555353
0.193497517288616	0.660021467385688	0.920350492152447
10579	TACC2	transforming acidic coiled-coil containing protein 2
	18844	0.0535961932837254	5.93719655458425
0.193401578948935	0.660100464526417	0.920369023155636
388743	CAPN8	calpain 8	5192	-0.32792584699465
3.26750998882096	0.193206650410458	0.660261043805457
	0.920501288004681
5464	PPA1	inorganic pyrophosphatase 1	5171
0.0999872322737405	6.84758744547194	0.192995390930122
	0.66043518545461	0.920652432589841
7092	TLL1	tolloid like 1	7522	-0.245173946722366
1.19552179286543	0.192809080455635	0.660588855964879
	0.920775012986373
8125	ANP32A	acidic nuclear phosphoprotein 32 family member A
	7355	0.0589462113618701	5.7832745053829
0.192670447053411	0.660703259484629	0.920842841353353
151194	METTL21A	methyltransferase like 21A
6797	-0.110843487054279	2.76267131633813
0.192515854021355	0.660830891169043	0.920923916153919
353497	POLN	DNA polymerase nu	7721	-0.127375699787299
	1.71646342278318	0.192428568729294
0.660902980718406	0.920923916153919
1175	AP2S1	adaptor related protein complex 2 subunit sigma 1
	1704	0.07100966383516	5.88415256404371
0.192253241304066	0.661047843829949	0.920923916153919
6832	SUPV3L1	Suv3 like RNA helicase	4283	0.0688234977670098
	4.62440388346829	0.192224137944745
0.66107189790644	0.920923916153919
80161	ASMTL-AS1	ASMTL antisense RNA 1	2670
0.169614992846088	1.51979919984816	0.192202035000967
	0.661090167553912	0.920923916153919
100533107	RTEL1-TNFRSF6B	RTEL1-TNFRSF6B readthrough (NMD candidate)	8738	0.103927342477118	5.12163742062084
	0.192044672294799	0.661220275180959
0.921013563490422
79684	MSANTD2	Myb/SANT DNA binding domain containing 2
4335	-0.125034923614689	2.97045361103547
0.191888860347524	0.66134916326059	0.921091492254999
56674	TMEM9B	TMEM9 domain family member B
3064	-0.0603119201643756	5.93836969132357
0.191602976492774	0.661585809478704	0.921091492254999
25888	ZNF473	zinc finger protein 473	5295	-0.084042521313342
	3.60482754584115	0.191570744539622
0.661612503332092	0.921091492254999
132789	GNPDA2	glucosamine-6-phosphate deaminase 2	7150
0.0796550551999713	4.41909491841578	0.191386492507203

0.661765148538609	0.921091492254999
7469	NELFA	negative elongation factor complex member A
5935	-0.0724287503113467	4.77469174257398
0.191373203734576	0.661776161122448	0.921091492254999
202658	TRIM39-RPP21	TRIM39-RPP21 readthrough
1780	-0.0683999487452787	4.46605963879774
0.191199223982283	0.661920382512664	0.921091492254999
57593	EBF4	EBF family member 4	3717	0.179760341530052
3.81089114665437	0.191179187124796
0.661936997185063	0.921091492254999
102157401	PCBP2-OT1	PCBP2 overlapping transcript 1	590
0.0804831177477181	3.87854689439807
0.19094575114693	0.662130640050092	0.921091492254999
3423	IDS	iduronate 2-sulfatase	10489	-0.0658921068262487
8.61878334611008	0.190891826692308
0.662175392210413	0.921091492254999
124152	IQCK	IQ motif containing K	4993	-0.107957370996441
3.16675593612766	0.190706391291293
0.662329343402816	0.921091492254999
25957	PNISR	PNN interacting serine and arginine rich protein
5135	-0.0656189727047427	5.99735638465341
0.190617193215004	0.6624034287021 0.921091492254999
23093	TTLL5	tubulin tyrosine ligase like 5
12220	-0.0677903459202232	4.75836090916848
0.190561677778007	0.662449548611765	0.921091492254999
4758	NEU1	neuraminidase 1 4822	0.0924453483654991
6.55952442899143	0.190536191687556	0.66247072407151
0.921091492254999
113730	KLHDC7B kelch domain containing 7B
2991	-0.349294434841909	1.38012158509841
0.190488944588823	0.662509984417066	0.921091492254999
441531	PGAM4	phosphoglycerate mutase family member 4 765
0.0708165999491637	5.38647453678326	0.190443622850012
0.662547650316997	0.921091492254999
7965	AIMP2	aminoacyl tRNA synthetase complex interacting
multifunctional protein 2	1838	0.0783213926129096
4.66781426428947	0.190232886940664	0.662722858409659
0.921091492254999
1489	CTF1	cardiotrophin 1 1664	0.121544527219252
1.92584269953057	0.190223257783822	0.662730866952846
0.921091492254999
399844	LINC01002	long intergenic non-protein coding RNA 1002
2871	0.135511768108685	1.6797181691185
0.19021314699664	0.662739276326321	0.921091492254999
200014	CC2D1B	coiled-coil and C2 domain containing 1B
7495	-0.0547059419622397	6.38036187635668
0.190172515164697	0.662773073435136	0.921091492254999
256051	ZNF549	zinc finger protein 549 9279	-0.0986321425915808
3.31049368937075	0.190148244650453
0.662793263430737	0.921091492254999
79671	NLRX1	NLR family member X1	5320	0.0673867789247676
4.43961300496626	0.19011148344266
0.662823847022917	0.921091492254999
23207	PLEKHM2 pleckstrin homology and RUN domain containing M2

4122	0.0576630758728878	6.91503794233249
0.190082622736028	0.662847860236497	0.921091492254999
23291	FBXW11	F-box and WD repeat domain containing 11
5577	-0.0491195654098095	6.85866138603244
0.189996725741958	0.66291934266615	0.921091492254999
79586	CHPF	chondroitin polymerizing factor
3354	-0.0747355846542796	7.09582232914309
0.189916261504138	0.662986321467288	0.921091492254999
9706	ULK2	unc-51 like autophagy activating kinase 2	6985
0.0644209737474744	5.01492045083878
0.18949140321829	0.66334025565921	0.921091492254999
8493	PPM1D	protein phosphatase, Mg2+/Mn2+ dependent 1D
5044	-0.122491386619164	4.24599212465	0.189474397023659
0.663354432730085	0.921091492254999
283932	FBXL19-AS1	FBXL19 antisense RNA 1
3951	-0.0861656697229615	4.04060482286793
0.189460464184449	0.663366048288106	0.921091492254999
6427	SRSF2	serine and arginine rich splicing factor 2	3017
0.0508322001805837	6.76768378480871
0.189195985958335	0.663586635451457	0.921091492254999
6879	TAF7	TATA-box binding protein associated factor 7
2334	-0.0583485266891185	6.69304679768274
0.189057528650586	0.663702188422155	0.921091492254999
5678	PSG9	pregnancy specific beta-1-glycoprotein 9	3437
0.299251395457972	1.83066218855044
0.189016121646574	0.663736755434329	0.921091492254999
56893	UBQLN4	ubiquilin 4	3878	0.0499503221257385
6.03486544181285	0.188923020347882	0.663814493843518
0.921091492254999
90809	PIP4P1	phosphatidylinositol-4,5-bisphosphate 4-phosphatase
1	3803	0.0727351352792797	5.49465111524377
0.188906861925563	0.663827988241326	0.921091492254999
64857	PLEKHG2	pleckstrin homology and RhoGEF domain containing G2
9200	0.0750142784946953	6.18709099775658
0.188883121155897	0.663847816136438	0.921091492254999
1639	DCTN1	dynactin subunit 1	10439	0.0585846030115841
7.80979727584799	0.188880846276817
0.663849716155305	0.921091492254999
7027	TFDP1	transcription factor Dp-1	3741
0.0511039822629961	6.5008828511976	0.18884503864868
0.663879625106108	0.921091492254999
6202	RPS8	ribosomal protein S8	2847	0.0962778937067181
7.64704634662261	0.188824923724042
0.663896427929239	0.921091492254999
51010	EXOSC3	exosome component 3	3023	-0.0733998166671719
3.77471077153823	0.188810242504503
0.663908692426896	0.921091492254999
64772	ENGASE	endo-beta-N-acetylglucosaminidase	6173
0.110083248410316	4.31002025893539	0.188657607967697
0.66403623490523	0.921091492254999
124454	EARS2	glutamyl-tRNA synthetase 2, mitochondrial	5730
0.0861743199645292	4.04677548962032
0.188642886951703	0.664048539130681	0.921091492254999
119559	SFXN4	sideroflexin 4	1868	-0.129163852647945

2.84294131646554	0.188579653038893	0.664101398245611
	0.921091492254999
84135	UTP15	UTP15 small subunit processome component	6321
	0.0636844216300243	4.39833951620964
0.188567571749784	0.664111498554209	0.921091492254999
81556	INTS14	integrator complex subunit 14	4210
0.0548705355363442	5.35399931801306	0.188530782639127
	0.664142257688014	0.921091492254999
5979	RET	ret proto-oncogene	6661	0.188498403029754
	0.00953209254643427	0.188360938816975
0.664284309256361	0.921091492254999
84124	ZNF394	zinc finger protein 394 3505	-0.0629962603468541
	4.28908021255492	0.188138410805984
0.664470539305851	0.921091492254999
4707	NDUFB1	NADH:ubiquinone oxidoreductase subunit B1	1940
	0.103772742485837	4.24129322917249
0.188046426390056	0.664547557783378	0.921091492254999
51225	ABI3	ABI family member 3	2456	0.371244896368297
	-0.251281576427892	0.188008351303393
0.664579444571846	0.921091492254999
79716	NPEPL1	aminopeptidase like 1	7072	-0.083021126379645
	5.22364495890335	0.187983161191113
0.664600542673803	0.921091492254999
100874188	LINC00571	long intergenic non-protein coding
RNA 571 1183	0.167247984400939	0.450833753415995
0.187926513666177	0.66464799422108	0.921091492254999
55033	FKBP14	FKBP prolyl isomerase 14
5326	-0.0767692642194883	5.81505248395488
0.187896727569825	0.664672948346139	0.921091492254999
100128494	LOC100128494	uncharacterized LOC100128494	4233
	0.171466705415297	0.323596078098833
0.187786463125867	0.664765345883162	0.921091492254999
10376	TUBA1B	tubulin alpha 1b	3574	-0.109920595267854
	8.64968429705793	0.187765804445576
0.664782660678991	0.921091492254999
26059	ERC2	ELKS/RAB6-interacting/CAST family member 2	7933
	0.161668119532752	0.453626523201486
0.187734576615756	0.66480883601354	0.921091492254999
79612	NAA16	N(alpha)-acetyltransferase 16, NatA auxiliary
subunit 5867	-0.113147803735767	3.25901559945506
0.187657622915861	0.664873350056548	0.921091492254999
79145	CHCHD7	coiled-coil-helix-coiled-coil-helix domain
containing 7	5866	-0.0997624153615238	4.26343413941146
	0.187649612715393	0.664880066301308
0.921091492254999
57494	RIMKLB	ribosomal modification protein rimK like family
member B	8619	-0.0896769283551822	4.61575288046608
	0.187602549571558	0.664919530374862
0.921091492254999
9183	ZW10	zw10 kinetochore protein
2992	-0.0595097952510498	4.61559184016816
0.187482460726085	0.665020255669791	0.921091492254999
6629	SNRPB2	small nuclear ribonucleoprotein polypeptide B2
2607	-0.0726189939052158	5.04705364716108

0.187404426567619	0.665085727859473	0.921091492254999
1801	DPH1	diphthamide biosynthesis 1	5757
0.0567443466451796	5.64037459064938	0.187399018693156
	0.665090265771904	0.921091492254999
401427	OR2A7	olfactory receptor family 2 subfamily A member 7
	1116	-0.248397067878539	1.0864323340938
0.18726882307899	0.665199540358549	0.921120762686167
4927	NUP88	nucleoporin 88	4443	-0.0631499930621735
5.31352120772288	0.187217148469795	0.665242923918988
	0.921120762686167
79788	ZNF665	zinc finger protein 665	7095	-0.135424256874841
	0.547028661270088	0.187062512966786
0.665372791058179	0.921208943745917
100529239	RPS10-NUDT3	RPS10-NUDT3 readthrough	876
0.0809708517479026	5.17916164190879	0.186922261380289
	0.665490633304693	0.921208943745917
658	BMPR1B	bone morphogenetic protein receptor type 1B
6312	-0.086985787694746	4.60580162309043
0.186904597392269	0.665505478668295	0.921208943745917
100379251	NPPA-AS1	NPPA antisense RNA 1	1427
0.143541897709447	1.62447372843754	0.186828217589039
	0.665569680210514	0.921208943745917
100033431	SNORD116-20	small nucleolar RNA, C/D box 116-20
	1236	0.136944941962086	1.5503946950185
0.186604780143601	0.665757581484696	0.921255649932869
9167	COX7A2L	cytochrome c oxidase subunit 7A2 like
4701	-0.0741759971787433	5.87631139448223
0.186521587470793	0.665827577044856	0.921255649932869
387	RHOA	ras homolog family member A
2368	-0.0685344334799625	8.90506809066638
0.186517669995887	0.665830873535102	0.921255649932869
51006	SLC35C2	solute carrier family 35 member C2	7708
0.0619705231324529	5.21362636147526	0.186376132226958
	0.665950002784474	0.921255649932869
3590	IL11RA	interleukin 11 receptor subunit alpha	4271
0.128183109306133	2.13646284214907	0.186352903546847
	0.665969558983268	0.921255649932869
51013	EXOSC1	exosome component 1	3345	0.0708188832113732
	4.11273113827603	0.186261387345358
0.666046620440433	0.921255649932869
3553	IL1B	interleukin 1 beta	3096	-0.323922739561997
	4.79335890049226	0.186181301408396
0.666114075440908	0.921255649932869
26515	TIMM10B	translocase of inner mitochondrial membrane 10B
3513	-0.0601885372536808	5.32148311162163
0.186105584616463	0.666177866214213	0.921255649932869
25854	FAM149A	family with sequence similarity 149 member A	5218
	0.143084541150073	2.7245667326194	0.185963221914683
	0.666297847305074	0.921255649932869
201163	FLCN	folliculin	10318	0.0983430942231465
6.27678733958075	0.185936583331184	0.666320303941499
	0.921255649932869
5355	PLP2	proteolipid protein 2	1279	-0.0997389958051699
	6.9469195841739	0.185898290942063	0.666352588214375

0.921255649932869
7325 UBE2E2 ubiquitin conjugating enzyme E2 E2
3274 -0.0970202199255914 3.80335612758518
0.185796509082228 0.666438419567222 0.921255649932869
10856 RUVBL2 RuvB like AAA ATPase 2 2525 0.0601401545640307
5.14647766934568 0.185772761750837
0.666458449401711 0.921255649932869
387804 VSTM5 V-set and transmembrane domain containing 5
3849 -0.164931175903008 0.598635370817084
0.185609412739696 0.666596268292638 0.921355232876948
55775 TDP1 tyrosyl-DNA phosphodiesterase 1
6227 -0.0711280833067359 4.17768128771074
0.18551875987076 0.66667278388228 0.921370072406423
7775 ZNF232 zinc finger protein 232 2794 -0.123410889188497
3.70610948138799 0.185355236237676
0.666810862133735 0.921469982824311
283459 GATC glutamyl-tRNA amidotransferase subunit C 4597
0.0548820539218555 6.48331760855381
0.185254672036514 0.666895813722167 0.921490076281743
90226 UCN2 urocortin 2 1481 -0.1104615754928
5.66395618763903 0.185182286651361 0.666956978178362
0.921490076281743
2182 ACSL4 acyl-CoA synthetase long chain family member 4
6181 -0.0670238855257238 7.60204309012403
0.184991591486806 0.667118180301785 0.921621890865677
11187 PKP3 plakophilin 3 3787 -0.0621008904250338
7.12959280153731 0.184774638651049 0.667301698837281
0.921784506470789
6747 SSR3 signal sequence receptor subunit 3
4627 -0.096672273318981 7.97101334697613
0.184686254134817 0.6673764991325 0.921796925682765
55732 C1orf112 chromosome 1 open reading frame 112
5967 -0.0971732653398038 2.56919822327967
0.18454635049679 0.667494943697645 0.92183501854533
401232 LINC01011 long intergenic non-protein coding RNA 1011
2811 -0.126905928753717 0.350716047581453
0.184498221091637 0.66753570308455 0.92183501854533
1018 CDK3 cyclin dependent kinase 3
1729 -0.129222823728534 2.48357209204206
0.18408520495886 0.667885733528307 0.922227471365438
57819 LSM2 LSM2 homolog, U6 small nuclear RNA and mRNA
degradation associated 1077 0.108343507019109
3.32841001297752 0.183818588066679 0.668111937891822
0.922313287507933
132949 AASDH aminoadipate-semialdehyde dehydrogenase 4654
0.0850303927282552 3.34889104508491 0.183697597791443
0.668214653096709 0.922313287507933
56890 MDM1 Mdm1 nuclear protein 6215 0.130004956339087
2.69519193054687 0.183543705612998
0.66834535838036 0.922313287507933
84892 POMGNT2 protein O-linked mannose N-
acetylglucosaminyltransferase 2 (beta 1,4-) 2668
0.0890880984756482 3.98679840390312 0.183422390043106
0.668448441085 0.922313287507933

54800	KLHL24	kelch like family member 24	8650
0.120383192279784	6.84918778616289	0.183363010379963
	0.668498911114158	0.922313287507933
147138	TMC8	transmembrane channel like 8	5459
0.213270564512147	2.2340689590561 0.183351300885166
0.668508864796845	0.922313287507933
30836	DNTTIP2 deoxynucleotidyltransferase terminal interacting
protein 2	7519	0.0861014821843322	6.41474881139671
	0.183298066809428	0.668554121286395
0.922313287507933
57835	SLC4A5	solute carrier family 4 member 5	9275
0.0597424234801716	5.36419585508153	0.183215616192931
	0.66862423131756	0.922313287507933
79657	RPAP3	RNA polymerase II associated protein 3	5381
0.0639789749364879	5.10330903965474	0.183210915755437
	0.668628228791436	0.922313287507933
11077	HSF2BP	heat shock transcription factor 2 binding protein
	1908	0.10186551852748	3.17359107712127
0.183183868920722	0.668651231875212	0.922313287507933
94059	LENG9	leukocyte receptor cluster member 9
1916	-0.138283099321885	2.09563661793741
0.183127869940219	0.668698864879684	0.922313287507933
9276	COPB2	coatomer protein complex subunit beta 2
8177	-0.0547045557040199	8.00376823768249
0.183081820109578	0.668738041545697	0.922313287507933
55561	CDC42BPG	CDC42 binding protein kinase gamma
6112	-0.0873774930802129	7.06017620190405
0.182758380399896	0.669013370662695	0.922582880198741
125170	MIEF2	mitochondrial elongation factor 2	3969
0.0579042899953881	5.81086926206252	0.18255717611521
	0.669184791886524	0.922582880198741
26270	FBXO6	F-box protein 6 1811	-0.0875498231065531
3.69265826173423	0.182489003562624	0.669242898602645
	0.922582880198741
102465492	MIR6820 microRNA 6820	54538	0.0526875541274083
	5.54079816778296	0.18244124796761
0.669283610617226	0.922582880198741
84278	MFSD14C major facilitator superfamily domain containing 14C
	1313	-0.0816136410049862	3.35453690739744
0.182238347394645	0.669456655212718	0.922582880198741
129831	RBM45	RNA binding motif protein 45	4784
0.0867154141050754	3.40014024073709	0.182213968004384
	0.669477454939929	0.922582880198741
85444	LRRCC1	leucine rich repeat and coiled-coil centrosomal
protein 1	5065	0.127505720386676	2.12571141374268
	0.18220440274924	0.669485616163396
0.922582880198741
283970	PDXDC2P-NPIPB14P	nuclear pore complex-interacting
protein 4969	0.0927954239121918	5.32755011888188
0.182187205784786	0.669500289516126	0.922582880198741
314	AOC2	amine oxidase copper containing 2	2657
0.15136161166384	2.1387200190583 0.182126228909875
0.669552324785382	0.922582880198741
10972	TMED10	transmembrane p24 trafficking protein 10

4540	-0.0596556985250024	7.84986350461837
0.182079542033177	0.669592172492354	0.922582880198741
29116	MYLIP	myosin regulatory light chain interacting protein
	3033	0.155218203742561	2.44376576179581
0.181932968106462	0.669717314029586	0.922664543878471
8604	SLC25A12	solute carrier family 25 member 12	5169
	0.0964643421382442	3.69907679157068
0.181852435631162	0.669786096216736	0.92266855327575
4217	MAP3K5	mitogen-activated protein kinase kinase kinase 5
	5303	-0.0813551351831254	4.7302301622043
0.181697154548566	0.669918771462311	0.922760569561668
64783	RBM15	RNA binding motif protein 15
4248	-0.105808864671636	3.83022746554843
0.181424150398797	0.670152194161617	0.922991325823183
219736	STOX1	storkhead box 1 3568
0.263574809294005	-0.494641974734434	0.18121401391732
	0.670332005406672	0.923148205656401
6992	PPP1R11 protein phosphatase 1 regulatory inhibitor subunit
11	2538	-0.0591378377991781	5.95778272525795
0.181006561367468	0.670509640882541	0.923302058576656
51154	MRTO4	MRT4 homolog, ribosome maturation factor	2822
	0.0866290383738479	4.81767277396019
0.180914930284047	0.670588140144919	0.923319382582314
54463	RETREG1 reticulophagy regulator 1	4568
0.152983037166336	3.65218342661499	0.180446995408722
	0.67098938087559	0.92378103577004
149013	NBPF12	NBPF member 12	7272	-0.0841393480454098
5.51567799105406	0.180335511985326	0.671085065281916
	0.923821966526171
57698	SHTN1	shootin 1	8247	0.0910780948695213
4.69100023727936	0.180025747156517	0.671351114777775
	0.923961839837547
6322	SCML1	Scm polycomb group protein like 1	5497
0.0974325714095139	2.59273070389418	0.180016540099653
	0.671359026627657	0.923961839837547
7398	USP1	ubiquitin specific peptidase 1
4263	-0.0614182640092914	5.00181280693913
0.17998682254661	0.671384565279257	0.923961839837547
5650	KLK7	kallikrein related peptidase 7
2434	-0.235070886943083	8.78888120204591
0.17980649279707	0.671539590318719	0.924026120463883
80723	SLC35G2 solute carrier family 35 member G2	2060
0.1339288003207 2.77374993911553	0.179687264854947
0.671642138013688	0.924026120463883
60437	CDH26	cadherin 26	5748	0.140600489705641
2.47863285268229	0.179673356069801	0.67165410354358
	0.924026120463883
134265	AFAP1L1 actin filament associated protein 1 like 1
4721	-0.0981550308332686	4.80595393052915
0.179625648933182	0.6716951494655 0.924026120463883
19	ABCA1	ATP binding cassette subfamily A member 1
11350	-0.0894199287997401	6.16349571240783
0.179192465844352	0.67206814298383	0.924220615296558
6574	SLC20A1 solute carrier family 20 member 1

5577	-0.110805583175258	6.75638257320467
0.179173274200423	0.672084680296333	0.924220615296558
205428	DIPK2A	divergent protein kinase domain 2A
5487	-0.0949974514525421	3.37308832697378
0.179166574232791	0.672090453860019	0.924220615296558
55740	ENAH	ENAH actin regulator	15044	-0.0976102178328183
	7.38904486010459	0.179154959234754
0.672100463154904	0.924220615296558
6783	SULT1E1 sulfotransferase family 1E member 1	2304
0.33382144347408	2.64124188093215	0.17896721260594
	0.672262308048887	0.924286537619201
523	ATP6V1A ATPase H+ transporting V1 subunit A
5130	-0.0556467570319675	6.77133152147379
0.178890417086485	0.672328537614998	0.924286537619201
400027	LINC00938	long intergenic non-protein coding RNA 938
	9730	-0.10029003066564	2.52696909243868
0.178869749150861	0.672346364803458	0.924286537619201
91661	ZNF765	zinc finger protein 765 12344	-0.0798929785549158
	4.30275595010964	0.178682658304284
0.672507795829805	0.924377780729272
51619	UBE2D4	ubiquitin conjugating enzyme E2 D4 (putative)	4713
	0.0827036304700536	3.53344040028539
0.178354620242155	0.672791083544769	0.924377780729272
26086	GPSM1	G protein signaling modulator 1
4950	-0.0863910488411528	4.78262351384268
0.178307409108871	0.672831879486647	0.924377780729272
4595	MUTYH	mutY DNA glycosylase	3909	0.127111847739425
	1.9155503638878 0.178252329161038	0.672879483048859
	0.924377780729272
146050	ZSCAN29 zinc finger and SCAN domain containing 29
5880	-0.0659325224869882	5.77255687684515
0.178209857981869	0.67291619523608	0.924377780729272
2177	FANCD2	FA complementation group D2
8378	-0.111224464010454	2.27015258464572
0.17814772339335	0.672969913807917	0.924377780729272
6882	TAF11	TATA-box binding protein associated factor 11	1580
	0.0642243392120043	5.00201383898636
0.177994895221507	0.673102088657749	0.924377780729272
51319	RSRC1	arginine and serine rich coiled-coil 1
5117	-0.0673608394270262	4.24528289363549
0.177966899995347	0.673126307829626	0.924377780729272
10868	USP20	ubiquitin specific peptidase 20
5521	-0.0789019114250202	5.12686970036682
0.177928343057209	0.673159667801543	0.924377780729272
29781	NCAPH2	non-SMC condensin II complex subunit H2 3159
0.0715889995255421	4.04610538205004	0.17792424862622
	0.673163210607355	0.924377780729272
153364	MBLAC2	metallo-beta-lactamase domain containing 2	4280
	0.119121471991992	2.43607033492958
0.177831597158915	0.673243392378759	0.924377780729272
87178	PNPT1	polyribonucleotide nucleotidyltransferase 1
5134	-0.0709147589709226	4.64070071099385
0.177676947189112	0.673377283273402	0.924377780729272
100506866	TTN-AS1 TTN antisense RNA 1

14536	-0.221486210920915	2.96606394069135
0.177624668306187	0.673422560142773	0.924377780729272
6100	RP9	RP9 pre-mRNA splicing factor	1898
0.087773064586693	3.26469595508246	0.177536852049132
	0.673498632323216	0.924377780729272
63915	BLOC1S5 biogenesis of lysosomal organelles complex 1 subunit
5	3315	-0.0715935582374138	4.05838538852999
0.177491297537292	0.673538103345605	0.924377780729272
83547	RILP	Rab interacting lysosomal protein	2103
0.0933723912247533	2.79285538236628	0.177382631779334
	0.673632281632873	0.924377780729272
5279	PIGC	phosphatidylinositol glycan anchor biosynthesis
class C 3279	0.0650546934315855	4.22581294666608
0.177373896041029	0.673639854185161	0.924377780729272
5514	PPP1R10 protein phosphatase 1 regulatory subunit 10
5215	-0.0771764649910226	6.81441496700743
0.177362086839011	0.673650091310579	0.924377780729272
64743	WDR13	WD repeat domain 13	7884	0.0499650509710584
	5.5766360454481 0.177338422555575	0.673670606540931
	0.924377780729272
167691	LCA5	lebercilin LCA5 5964	0.0877837348245025
4.02217669318723	0.177266905131138	0.673732616796255
	0.924377780729272
57763	ANKRA2	ankyrin repeat family A member 2	2659
0.0675111581625033	4.33863199872928	0.177163751851785
	0.673822083337622	0.924409982498538
10300	KATNB1	katanin regulatory subunit B1	4591
0.0720709176531031	4.40976382370346	0.17705410248292
	0.67391721766827	0.924440409994967
317662	FAM149B1	family with sequence similarity 149 member
B1	6692	-0.0647450153229445	4.47535217634863
0.176978143811986	0.673983141501276	0.924440409994967
25895	EEF1AKMT3	EEF1A lysine methyltransferase 3	3172
	0.100383797038443	2.4195964998158 0.176644097187652
	0.674273255291311	0.924440409994967
284615	ANKRD34A	ankyrin repeat domain 34A	3677
0.209827435053282	0.251732701194297	0.176601497573201
	0.674310275528985	0.924440409994967
79091	METTL22 methyltransferase like 22	8254
0.0781054159553002	4.10808323118596	0.176425780420352
	0.674463034123827	0.924440409994967
84946	LTV1	LTV1 ribosome biogenesis factor 1847
0.0697626640846631	4.61809800958245	0.176417576408519
	0.674470168415957	0.924440409994967
91801	ALKBH8	alkB homolog 8, tRNA methyltransferase
4506	-0.0841328775031578	3.48561643235672
0.176410080709423	0.674476686922731	0.924440409994967
54522	ANKRD16 ankyrin repeat domain 16	3059
0.115792877581626	1.83942915021806	0.176321589862985
	0.674553653794341	0.924440409994967
22881	ANKRD6	ankyrin repeat domain 6 8329	0.0790059231905444
	4.4460000336104 0.176315583925599	0.674558878414516
	0.924440409994967
653857	ACTR3C	actin related protein 3C	6454

0.106310909911234	2.88126556665274	0.176261805278611
	0.674605665619782	0.924440409994967
6871	TADA2A	transcriptional adaptor 2A	5654
0.105799963869892	3.27143795304289	0.176215808204384
	0.674645689549977	0.924440409994967
5166	PDK4	pyruvate dehydrogenase kinase 4	4519
0.133249761662433	2.95384688698879	0.176153277087794
	0.674700110279897	0.924440409994967
595	CCND1	cyclin D1	4830	0.0846519544103861
10.4670415405194	0.176137051875841	0.67471423291199
	0.924440409994967
6251	RSU1	Ras suppressor protein 1	4191
0.0950545585845256	6.53158851032477	0.17605460145974
	0.674786010633182	0.924440409994967
55254	TMEM39A	transmembrane protein 39A
5333	-0.0701272393467361	4.97685684934499
0.175871093400938	0.674945835696895	0.924440409994967
51734	MSRB1	methionine sulfoxide reductase B1	2157
0.0747455847918549	4.88715521145188	0.17570356492628
	0.675091829088076	0.924440409994967
23240	TMEM131L	transmembrane 131 like	6393
0.122477432115168	3.83657064225832	0.175618681156372
	0.675165832641404	0.924440409994967
55909	BIN3	bridging integrator 3	5072	-0.0630493108575644
	5.07544258113553	0.175608814271026
0.675174436174848	0.924440409994967
191	AHCY	adenosylhomocysteinase	3372	0.0909029724274841
	6.41264196793386	0.175584060483033
0.675196021748811	0.924440409994967
653308	ASAH2B	N-acylsphingosine amidohydrolase 2B
632	-0.116392511477576	0.784027940681296
0.175545623365056	0.67522954288167	0.924440409994967
100287896	LOC100287896	uncharacterized LOC100287896
2493	-0.127516366039089	1.22502933013442
0.175544835280738	0.675230230217642	0.924440409994967
574407	OBSCN-AS1	OBSCN antisense RNA 1
3647	-0.13172532292037	0.571934311777178
0.175346254818749	0.67540348208075	0.924587233532551
107	ADCY1	adenylate cyclase 1	13530	0.312903791273758
	1.24913749373916	0.174981893622935
0.675721669743313	0.924932417482125
5928	RBBP4	RB binding protein 4, chromatin remodeling factor
	9470	0.052872838921125	7.00533200724119
0.174844931657219	0.675841375912867	0.925005877702905
25912	C1orf43	chromosome 1 open reading frame 43	2582
0.048068018929978	7.34078525642322	0.174586421291345
	0.676067466820593	0.925168835147745
100287015	LOC100287015	uncharacterized LOC100287015	2413
	0.126418914327665	0.67857096482037
0.174450563745807	0.676186365465139	0.925168835147745
64946	CENPH	centromere protein H	2244	0.121732714192319
	0.395830638862093	0.174325915035219
0.676295502296983	0.925168835147745
54960	GEMIN8	gem nuclear organelle associated protein 8	2424

0.106811143778969	2.66601314921541
0.174271652176949	0.676343026742169	0.925168835147745
378108	TRIM74	tripartite motif containing 74
6782	-0.112143269605234	2.22755955188638
0.174223089493089	0.676385566233711	0.925168835147745
9209	LRRFIP2 LRR binding FLII interacting protein 2
10644	-0.0532514621695938	6.48021104346855
0.174160305882921	0.676440573160285	0.925168835147745
11346	SYNPO	synaptopodin	10119	0.064118648648146
8.06070640708401	0.174026385785652	0.676557944173933
0.925168835147745
642658	SCX	scleraxis bHLH transcription factor	999
0.162335153246061	0.876533356631535	0.174006288090802
0.676575562885069	0.925168835147745
100132929	FAM25BP protein FAM25	239	0.0839299947004936
3.34430979545344	0.173885176430566
0.676681761121829	0.925168835147745
9189	ZBED1	zinc finger BED-type containing 1
9510	-0.0574448957158773	6.03491959855516
0.173861087198313	0.676702889235061	0.925168835147745
5558	PRIM2	DNA primase subunit 2	4667	0.0934584403704263
2.96654461682683	0.173856282358344
0.676707103655551	0.925168835147745
1471	CST3	cystatin C	3695	0.0584534274850757
6.06291514547844	0.173647738770594	0.676890087285873
0.925168835147745
2584	GALK1	galactokinase 1 2891	-0.116850074696052
2.17819567606874	0.173600114502172	0.676931892614404
0.925168835147745
26133	TRPC4AP transient receptor potential cation channel subfamily C member 4 associated protein 3226	0.0536116889376811
7.21803064100752	0.173533063720072
0.676990762249747	0.925168835147745
11336	EXOC3	exocyst complex component 3
8182	-0.0492018204767433	6.04694465404118
0.173453758194558	0.677060408607288	0.925168835147745
57559	STAMBPL1	STAM binding protein like 1	6034
0.151835399008531	1.18704871593559	0.173444382114774
0.677068643944466	0.925168835147745
137994	LETM2	leucine zipper and EF-hand containing transmembrane protein 2	6114	0.162827248045192	2.07766672266887
0.17335809062952	0.677144449025937
0.925168835147745
9554	SEC22B	SEC22 homolog B, vesicle trafficking protein (gene/pseudogene)	9034	-0.0490733039273725	6.37708337073824
0.173352516413111	0.677149346607745
0.925168835147745
54606	DDX56	DEAD-box helicase 56	4068	0.0482591016790464
6.07238299708612	0.173110425373764
0.677362140264698	0.925263336784042
2110	ETFDH	electron transfer flavoprotein dehydrogenase	4493
0.0846067706440691	4.44514024136978
0.173000235366551	0.677459053101346	0.925263336784042
98	ACYP2	acylphosphatase 2	3431	0.109933626939564

2.90992848687037	0.17295503155318
0.677498820625055	0.925263336784042
116828	N4BP2L2-IT2	N4BPL2 intronic transcript 2
4890	-0.275625158755281	0.870702330268438
0.172926977779021	0.677523503668742	0.925263336784042
100505854	APTR	Alu-mediated CDKN1A/p21 transcriptional
regulator	3100	0.107086839261682	1.70579760931682
0.172898229427699	0.677548800270859
0.925263336784042
64333	ARHGAP9 Rho GTPase activating protein 9 5127
0.35203997308072	1.26235366049931	0.172778248070557
0.677654402377143	0.925264202458932
28992	MACROD1 mono-ADP ribosylhydrolase 1	2089
0.126789064471161	0.321089255949784	0.172725165373002
0.677701137062392	0.925264202458932
4756	NEO1	neogenin 1	8117	0.062994636257645
5.99368013049543	0.172564721118221	0.677842445451363
0.925264202458932
79661	NEIL1	nei like DNA glycosylase 1	6938
0.106364042540668	4.68769865289386	0.172541711324676
0.67786271722761	0.925264202458932
10390	CEPT1	choline/ethanolamine phosphotransferase 1	8413
0.0618895883241192	5.24744173717083
0.172522412836422	0.677879720542126	0.925264202458932
440993	MIR570HG	MIR570 host gene
10458	-0.104886862428892	4.81737762984407
0.172252522258646	0.678117629896526	0.925498747146121
51763	INPP5K	inositol polyphosphate-5-phosphatase K	5325
0.0537380935775715	5.31033797368532	0.172169833045224
0.678190564408224	0.925508109476422
29103	DNAJC15 DnaJ heat shock protein family (Hsp40) member C15
8042	-0.151277575875253	5.99803889740497
0.171801672545877	0.678515543079437	0.925673839129814
57089	ENTPD7	ectonucleoside triphosphate diphosphohydrolase 7
3657	-0.101784175574417	5.86598237509674
0.17172818944141	0.678580456132218	0.925673839129814
222223	KIAA1324L	KIAA1324 like	8115	0.219557672005799
2.4275490549798 0.171487328409775	0.678793340624823
0.925673839129814
79803	HPS6	HPS6 biogenesis of lysosomal organelles complex 2
subunit 3	2649	-0.0599210460163164	5.20196412196526
0.171438908809051	0.678836157322508
0.925673839129814
10480	EIF3M	eukaryotic translation initiation factor 3 subunit M
7428	-0.0606597035186385	6.69365948766968
0.171433297233619	0.678841120009124	0.925673839129814
60626	RIC8A	RIC8 guanine nucleotide exchange factor A
4543	-0.0530994885408891	7.39561393729064
0.171365180766962	0.678901367486022	0.925673839129814
100526836	BLOC1S5-TXNDC5	BLOC1S5-TXNDC5 readthrough (NMD
candidate)	3030	0.0879419156726525	7.1953371125326
0.171240666439701	0.679011533860079	0.925673839129814
84955	NUDCD1	NudC domain containing 1
4439	-0.0748312650818175	5.03736954624297

0.171210890832242	0.679037885334283	0.925673839129814
5681	PSKH1	protein serine kinase H1	3977
0.0528391233530602	6.20270653291715	0.171200367163305
	0.679047199444941	0.925673839129814
100132159	CALML3-AS1	CALML3 antisense RNA 1
4757	-0.211292194744514	0.844140743158591
0.171103456910386	0.679132986894117	0.925673839129814
11100	HNRNPUL1	heterogeneous nuclear ribonucleoprotein U like 1	5443	-0.0518926679940394	7.74911830796766
0.171099263873501	0.679136699320318	0.925673839129814
55090	MED9	mediator complex subunit 9	2521
0.0805911899100304	3.69592389015972	0.170965433907614
	0.679255217537416	0.925673839129814
83744	ZNF484	zinc finger protein 484	4031	-0.0995633769077434
	2.86654960032622	0.170809068747379
0.679393761481019	0.925673839129814
9344	TAOK2	TAO kinase 2	9058	-0.0612464712513653
6.72614642943634	0.1707102386071	0.679481366071219
0.925673839129814
54940	OCIAD1	OCIA domain containing 1	4714
0.0508084801936993	7.04452674068855	0.170525202100183
	0.679645465198108	0.925673839129814
7334	UBE2N	ubiquitin conjugating enzyme E2 N
6252	-0.0689978184858634	6.41877724775072
0.170456047360801	0.679706821644359	0.925673839129814
283229	CRACR2B	calcium release activated channel regulator 2B	3911
	0.121421127510821	2.89475637725974
0.170344989380784	0.67980538656671	0.925673839129814
79884	MAP9	microtubule associated protein 9	8857
0.0708073088350485	5.24227833430905	0.170337196684848
	0.679812304063773	0.925673839129814
5211	PFKL	phosphofructokinase, liver type	8926
0.0552791418706972	6.46230613440889	0.170274063049427
	0.679868353979744	0.925673839129814
89866	SEC16B	SEC16 homolog B, endoplasmic reticulum export factor
	10192	0.135470375619235	1.58613478184244
0.170264634694869	0.679876725497736	0.925673839129814
139221	PWWP3B	PWWP domain containing 3B
4343	-0.279939332104743	3.90182214006938
0.170229405954629	0.679908007802967	0.925673839129814
5675	PSG6	pregnancy specific beta-1-glycoprotein 6
5140	-0.34270774170387	3.15345297792879
0.170176718858333	0.679954799793577	0.925673839129814
122704	MRPL52	mitochondrial ribosomal protein L52	3213
0.0701579994406738	4.69390294093453	0.170054683493376
	0.680063213307978	0.925673839129814
9939	RBM8A	RNA binding motif protein 8A	5585
0.0483116710041917	6.48618840903356	0.170042820032585
	0.680073754970864	0.925673839129814
100526833	SEPT5-GP1BB	SEPT5-GP1BB readthrough
5062	-0.144553456216455	3.46072037199692
0.170011641149756	0.680101462030004	0.925673839129814
57696	DDX55	DEAD-box helicase 55	5714	0.0667529628754649
	4.21791617041427	0.169869814891825

0.680227533211597	0.925673839129814
93953	GCNA	germ cell nuclear acidic peptidase
3621	-0.186061871591512	1.30633990307005
0.169817191600319	0.680274326391423	0.925673839129814
162966	ZNF600	zinc finger protein 600 3829	-0.082074269677663 7
	4.17557106300444	0.169810057479921
0.680280670778536	0.925673839129814
79624	ARMT1	acidic residue methyltransferase 1
2819	-0.05965544653679	5.4510416203220 8
0.169803203298148	0.680286766362534	0.925673839129814
2958	GTF2A2	general transcription factor IIA subunit 2
1961	-0.0730217785283161	5.14931253410947
0.16979439196454	0.680294602698815	0.925673839129814
84693	MCEE	methylmalonyl-CoA epimerase	1624
0.127180286852073	1.23107701835119	0.169637321824492
	0.680434332541196	0.925689552119055
59352	LGR6	leucine rich repeat containing G protein-coupled
receptor 6	4583	0.353958047563651	3.08439323420317
	0.169632834326023	0.680438325738402
0.925689552119055
1789	DNMT3B	DNA methyltransferase 3 beta	4630
0.142962331985182	0.940162381031457	0.169306700354392
	0.680728701360022	0.925838987719305
57823	SLAMF7	SLAM family member 7	4833	0.189356978795346
	6.39620225389733	0.169276652266831
0.680755471329704	0.925838987719305
11269	DDX19B	DEAD-box helicase 19B	4698	0.0584235654920167
	4.55927232609645	0.169265388295474
0.680765507165856	0.925838987719305
3417	IDH1	isocitrate dehydrogenase (NADP(+)) 1	4721
0.0881422630551132	7.36540070373526	0.169212579563066
	0.680812563254719	0.925838987719305
754	PTTG1IP PTTG1 interacting protein
3537	-0.0592557757846589	9.19589381163826
0.169090932132008	0.680920991872976	0.925859663447087
55958	KLHL9	kelch like family member 9
5710	-0.0726529994240653	5.45837388984993
0.168985613416863	0.68101490296835	0.925859663447087
221120	ALKBH3	alkB homolog 3, alpha-ketoglutaratedependent
dioxygenase	2372	0.0718722552571918	4.21504698725124
	0.168955754120745	0.681041534272433
0.925859663447087
26160	IFT172	intraflagellar transport 172	8401
0.0980547630667994	2.62484097780798	0.168798975584166
	0.681181409146833	0.925859663447087
147081	LINC02210	long intergenic non-protein coding RNA 2210
	2991	-0.108876734159369	2.70067433274295
0.168727998317977	0.681244758701413	0.925859663447087
5768	QSOX1	quiescin sulfhydryl oxidase 1
9311	-0.0762263668040354	9.1957798077055 0.168686918280741
	0.681281431106645	0.925859663447087
25904	CNOT10	CCR4-NOT transcription complex subunit 10
4846	-0.0667598488666559	4.31598232377792
0.168676798875794	0.681290465563584	0.925859663447087

116064	LRRC58	leucine rich repeat containing 58
7903	-0.0911712733536007	5.25604757072362
0.168431642066565	0.681509434827614	0.925966734736502
91978	TPGS1	tubulin polyglutamylase complex subunit 1	1114
	0.15841994916884	1.71290162311764
0.168349004863643	0.681583286718379	0.925966734736502
84864	RIOX2	ribosomal oxygenase 2	6880	-0.0915211546301434
	4.31461381378913	0.168210152403681
0.68170742522036	0.925966734736502
390927	ZNF793	zinc finger protein 793 12007	-0.145166930882339
	3.05162076912298	0.168191764922312
0.681723868699216	0.925966734736502
56984	PSMG2	proteasome assembly chaperone 2
3075	-0.058561338374407	5.11050324278835
0.168185055168724	0.681729869330687	0.925966734736502
9989	PPP4R1	protein phosphatase 4 regulatory subunit 1
8131	-0.0619637126436489	7.86883130038868
0.16814477195021	0.681765898146466	0.925966734736502
730051	ZNF814	zinc finger protein 814 7401	-0.116503799829889
	2.70900956067926	0.168005566407858
0.681890440699772	0.926046092968946
9853	RUSC2	RUN and SH3 domain containing 2
5682	-0.069315649223636	5.89174275583362
0.167829447402255	0.682048094881634	0.926100131931697
883	KYAT1	kynurenine aminotransferase 1	3624
0.100206444068355	3.37247988851346	0.167718202599789
	0.682147726245516	0.926100131931697
55157	DARS2	aspartyl-tRNA synthetase 2, mitochondrial	3689
	0.0778077975948045	4.54410307047099
0.16769212095805	0.68217109066867	0.926100131931697
127281	PRXL2B	peroxiredoxin like 2B	3371	0.0775366560389592
	5.22817237159892	0.167597025820644
0.68225629664839	0.926100131931697
128989	TANGO2	transport and golgi organization 2 homolog	5764
	0.0597796796281989	4.94760146285856
0.167591981893128	0.682260816834667	0.926100131931697
10531	PITRM1	pitrilysin metallopeptidase 1	6979
0.0506198139906127	7.18601393747271	0.167431562191908
	0.682404620646243	0.926205573778287
1984	EIF5A	eukaryotic translation initiation factor 5A
2815	-0.0762713101972494	7.75915560428952
0.1671908892381 0.682620516187867	0.92640883346996
81567	TXNDC5	thioredoxin domain containing 5 4387
0.0884957609993254	7.17913060527131	0.166961099674666
	0.682826818110794	0.926471130044391
54499	TMCO1	transmembrane and coiled-coil domains 1
5584	-0.053196921115792	6.78443153316101
0.166917451506578	0.682866023558944	0.926471130044391
55902	ACSS2	acyl-CoA synthetase short chain family member 2
5385	-0.108862301770772	5.17266409151264
0.16685291298824	0.682924003999811	0.926471130044391
64848	YTHDC2	YTH domain containing 2 10816	-0.0615933340499572
	5.45682209180298	0.166748394889442
0.683017929257827	0.926471130044391

203522	INTS6L	integrator complex subunit 6 like
7177	-0.103790428646547	2.97934048066479
0.166720811096113	0.683042723179653	0.926471130044391
22909	FAN1	FANCD2 and FANCI associated nuclease 1	6194
0.0614143126765183	4.93048827659747	0.16669794342918
	0.683063279786423	0.926471130044391
10617	STAMBP	STAM binding protein	7698	0.0474870841307503
	6.03132345880193	0.166617721403405
0.683135407396292	0.926479245875277
3700	ITIH4	inter-alpha-trypsin inhibitor heavy chain 4
7062	-0.13885329765939	1.59459702482128
0.166429626775799	0.683304602709219	0.926618992172334
9128	PRPF4	pre-mRNA processing factor 4
3215	-0.0572552021611973	4.83626615503292
0.166207263271332	0.683504767553714	0.92663455080393
50848	F11R	F11 receptor	5327	-0.0637435432800519
8.7632169084168 0.166171621378453	0.68353686581685
0.92663455080393
79081	LBHD1	LBH domain containing 1 3362	0.0670180785076992
	4.68133323044271	0.166055359771013
0.683641596239994	0.92663455080393
3661	IRF3	interferon regulatory factor 3
3794	-0.046404219262872	5.8484018816032 0.165967285984301
	0.683720963060657	0.92663455080393
285498	RNF212	ring finger protein 212 4965	0.231950575455287
	0.681746485826075	0.165847710581783
0.6838287565579 0.92663455080393
5223	PGAM1	phosphoglycerate mutase 1	2568
0.0669821395122604	6.86841313441937	0.165846733157252
	0.683829637862289	0.92663455080393
5575	PRKAR1B protein kinase cAMP-dependent type I regulatory subunit beta	3889	0.0841421991301718	4.07082134421638
	0.16583540812864	0.683839849405901
0.92663455080393
54567	DLL4	delta like canonical Notch ligand 4	4753
0.222209638323247	0.882183768137967	0.165635833880227
	0.684019868108533	0.92663455080393
64753	CCDC136 coiled-coil domain containing 136	6188
0.247267999123026	1.26275137041898	0.16554273681999
	0.684103886171674	0.92663455080393
9093	DNAJA3	DnaJ heat shock protein family (Hsp40) member A3
	7017	-0.0544223495765173	5.45823964667785
0.165431513564883	0.684204298837095	0.92663455080393
25938	HEATR5A HEAT repeat containing 5A
10662	-0.0594127592920543	5.48089945682262
0.165409662552435	0.684224030615341	0.92663455080393
51123	ZNF706	zinc finger protein 706 7393	-0.0678499891172446
	5.32985130905954	0.165397988972018
0.684234572648689	0.92663455080393
4968	OGG1	8-oxoguanine DNA glycosylase	4022
0.0824279842878083	3.04998529148027	0.165358580766129
	0.684270164128891	0.92663455080393
57683	ZDBF2	zinc finger DBF-type containing 2
10340	-0.0996153393282355	3.91296419590555

0.165310551426874	0.684313548459833	0.92663455080393
10311	VPS26C	VPS26 endosomal protein sorting factor C
8995	-0.0443282321500017	6.30815609156678
0.16524408905469	0.684373595245028	0.92663455080393
54663	WDR74	WD repeat domain 74	4010	0.0620871012415329
	4.38990644656942	0.165243024024162
0.684374557583113	0.92663455080393
79058	ASPSCR1 ASPSCR1 tether for SLC2A4, UBX domain containing
	12097	-0.101886971824452	3.47032293010373
0.164975160636367	0.684616707867954	0.926767949861507
23078	VWA8	von Willebrand factor A domain containing 8	7633
	0.0639746061787999	5.27798427847593
0.164958399022808	0.684631868089924	0.926767949861507
399664	MEX3D	mex-3 RNA binding family member D
2850	-0.059450656402407	5.3022681395798 0.164914502990278
	0.684671574581771	0.926767949861507
132321	C4orf33 chromosome 4 open reading frame 33
7015	-0.105353034226324	3.09869356677501
0.164673782384741	0.684889429759557	0.926859170936466
120526	DNAJC24 DnaJ heat shock protein family (Hsp40) member C24
	7592	0.0777243370007699	3.4414525574066
0.164654706814501	0.684906701319176	0.926859170936466
66005	CHID1	chitinase domain containing 1	5984
0.0690475060764923	5.13008231375345	0.164620716812117
	0.684937479714668	0.926859170936466
440295	GOLGA6L9	golgin A6 family like 9
6814	-0.0987244658898869	3.5538194074221 0.164430073643536
	0.685110178239053	0.927003310142428
10908	PNPLA6	patatin like phospholipase domain containing 6	7225
	0.0648213037540037	7.43685335797436
0.164324949661001	0.685205457099313	0.927018682712046
1609	DGKQ	diacylglycerol kinase theta
4970	-0.0786820277038726	5.45326091054719
0.164174201867411	0.68534214893394	0.927018682712046
2150	F2RL1	F2R like trypsin receptor 1
2958	-0.0754472852987463	7.14480705264156
0.164093341790476	0.685415499597857	0.927018682712046
10952	SEC61B	SEC61 translocon beta subunit
1190	-0.075973386653248	5.85207559732443
0.164039499139506	0.685464353592439	0.927018682712046
729262	NUTM2B	NUT family member 2B	5080	0.151024711594936
	0.977404395044244	0.164017622066876
0.685484206365969	0.927018682712046
23788	MTCH2	mitochondrial carrier 2 2897	0.0530983505109073
	6.38620586406313	0.163979687310558
0.685518634649202	0.927018682712046
165055	CCDC138 coiled-coil domain containing 138
3006	-0.117752062201117	1.23041088642014
0.16386716576659	0.68562078261539	0.927028450941133
9669	EIF5B	eukaryotic translation initiation factor 5B
7883	-0.0501039039816477	7.1541531739191 0.163825932657275
	0.685658224584146	0.927028450941133
84875	PARP10	poly(ADP-ribose) polymerase family member 10
5562	-0.0644870900787478	4.54473733903266

0.163612704480709	0.685851935244709	0.927200854837626
4782	NFIC	nuclear factor I C	9347	0.0735671029442857
	6.03939604120955	0.16347003000871
0.685981632260232	0.927224657495981
6173	RPL36A	ribosomal protein L36a	3964	0.102365211018832
	3.3777821309302 0.163447700601655	0.686001936576772
	0.927224657495981
114899	C1QTNF3 C1q and TNF related 3	3576
0.120384531842283	-0.046280509150004	0.163236813603814
	0.686193777518959	0.927394465622159
340706	VWA2	von Willebrand factor A domain containing 2
3301	-0.0891774220477294	4.21192484470521
0.162958590196624	0.686447094111718	0.927527732478083
9238	TBRG4	transforming growth factor beta regulator 4	5803
	0.0590178294151273	5.54341817652133
0.162914480752475	0.686487277929331	0.927527732478083
10657	KHDRBS1 KH RNA binding domain containing, signal
transduction associated 1	3588	-0.0491644602645346
7.08469300936108	0.162816259646142	0.6865767803439
0.927527732478083
2137	EXTL3	exostosin like glycosyltransferase 3	13220
0.0491015327378478	6.74052472876226	0.16280595327148
	0.686586173682889	0.927527732478083
5062	PAK2	p21 (RAC1) activated kinase 2	6270
0.059549086267411	8.03509027791529	0.16276502657206
	0.686623478122741	0.927527732478083
284001	CCDC57	coiled-coil domain containing 57
11583	-0.0855169860207265	4.32846437737739
0.162541024139102	0.686827751656297	0.927714206118587
9159	PCSK7	proprotein convertase subtilisin/kexin type 7
13149	-0.0513550678703214	6.17290934021386
0.162433640843819	0.686925735251175	0.927757089351384
55657	ZNF692	zinc finger protein 692 3929	0.0689168420880329
	4.24476580241335	0.162023923257649
0.687299934819073	0.927993885999964
148741	ANKRD35 ankyrin repeat domain 35
3363	-0.182952137318729	1.72491899133007
0.162021460988477	0.687302185297691	0.927993885999964
3299	HSF4	heat shock transcription factor 4	4507
0.171030097411401	1.81631919381903	0.161609153591466
	0.687679308924547	0.927993885999964
342977	NANOS3	nanos C2HC-type zinc finger 3	988
0.115533796994967	0.0801100102162501	0.161565993239308
	0.687718818537129	0.927993885999964
6141	RPL18	ribosomal protein L18	3521	0.0600222731236406
	7.29200901543022	0.161474615413624
0.687802487356348	0.927993885999964
114926	SMIM19	small integral membrane protein 19
4056	-0.0921501635463186	3.36595754650831
0.161406698722719	0.687864692132944	0.927993885999964
200185	KRTCAP2 keratinocyte associated protein 2	3874
0.0608222993533422	5.43161498249772	0.161385965777548
	0.687883684428778	0.927993885999964
255520	ELMOD2	ELMO domain containing 2

6627	-0.0600004865386314	5.79758640142909
0.161378600288828	0.687890431883665	0.927993885999964
4913	NTHL1	nth like DNA glycosylase 1	1658
0.0988047710442444	2.72503324971036	0.161326811463549
	0.687937880189036	0.927993885999964
256302	NATD1	N-acetyltransferase domain containing 1 4931
0.0919887925521305	5.09650702281468	0.161061154289456
	0.6881814112797 0.927993885999964
9751	SNPH	syntaphilin	5574	0.0783763283973536
4.11519439418986	0.161042376385426	0.688198634029395
	0.927993885999964
84668	FAM126A family with sequence similarity 126 member A
13961	-0.102991601509367	5.59307176540987
0.16097441338551	0.68826097819547	0.927993885999964
9837	GINS1	GINS complex subunit 1	4068	0.107894806314283
	2.0515054825041 0.160844527949314	0.688380167883269
	0.927993885999964
23560	GTPBP4	GTP binding protein 4	6540	0.0780359704082029
	6.49100999054387	0.160796449189732
0.688424301641104	0.927993885999964
91445	RNF185	ring finger protein 185 3447	-0.0520499364440235
	5.86220501263206	0.160788380594976
0.688431708934745	0.927993885999964
81844	TRIM56	tripartite motif containing 56
11173	-0.0926103859628584	6.35978009816948
0.160772497725134	0.688446290676712	0.927993885999964
993	CDC25A	cell division cycle 25A 4117	-0.114640137756528
	2.69229412690301	0.160572124803561
0.688630320856456	0.927993885999964
8451	CUL4A	cullin 4A	7869	-0.0504801293387866
6.59287455518985	0.160512879514386	0.688684759527712
	0.927993885999964
146227	BEAN1	brain expressed associated with NEDD4 1
6634	-0.141951830445582	2.60646084473322
0.160506284982397	0.688690819761103	0.927993885999964
2059	EPS8	epidermal growth factor receptor pathway substrate 8
	6200	0.0922540250045187	5.09098587047121
0.160493935347127	0.688702169199552	0.927993885999964
5281	PIGF	phosphatidylinositol glycan anchor biosynthesis
class F 4978	0.0685567803346108	3.88397545687813
0.160457458187345	0.688735695033708	0.927993885999964
388324	INCA1	inhibitor of CDK, cyclin A1 interacting protein 1
	1553	0.125031063422614	0.720151315402481
0.160413812161424	0.68877581552095	0.927993885999964
10597	TRAPPC2B	trafficking protein particle complex 2B
732	-0.0892287367812888	3.21399965271265
0.160391905471614	0.688795955069658	0.927993885999964
755	CFAP410 cilia and flagella associated protein 410
6063	-0.117218945794063	3.13949616758667
0.160369444659405	0.688816605698603	0.927993885999964
27102	EIF2AK1 eukaryotic translation initiation factor 2 alpha
kinase 1	5550	-0.0467295594703433	7.20435737946511
	0.160256277108463	0.688920678276713
0.927993885999964

151050	KANSL1L KAT8 regulatory NSL complex subunit 1 like
8491	-0.0971502641065954	3.55072353332217
0.160198310374987	0.688974002892152	0.927993885999964
65988	ZNF747	zinc finger protein 747 4065	0.0725942790600155
	4.22704639535266	0.160172031499251
0.688998180978547	0.927993885999964
7982	ST7	suppression of tumorigenicity 7 6293
0.0654833436703257	4.54355540757153	0.160166556648264
	0.689003218447911	0.927993885999964
26520	TIMM9	translocase of inner mitochondrial membrane 9	1500
	0.103468205537969	2.72146710652794
0.16014573720396	0.689022375555053	0.927993885999964
162972	ZNF550	zinc finger protein 550 5087	-0.0933645811513775
	3.02459781248332	0.159843752462149
0.689300410794252	0.92811428888417
4294	MAP3K10 mitogen-activated protein kinase kinase kinase 10
	4566	-0.0776471165210654	4.65190687935032
0.159811828058203	0.689329821168613	0.92811428888417
7114	TMSB4X	thymosin beta 4 X-linked	1705
0.067890094145646	10.5125935038644	0.159765304044868
	0.689372687542013	0.92811428888417
11040	PIM2	Pim-2 proto-oncogene, serine/threonine kinase	2761
	0.112833814959852	2.25176785356461
0.159760823746958	0.689376815988499	0.92811428888417
347344	ZNF81	zinc finger protein 81	8443	-0.112245695448933
	3.49337949707487	0.15964411593702
0.689484382039836	0.928165808082547
57805	CCAR2	cell cycle and apoptosis regulator 2
6534	-0.0535277770810066	6.77389909925686
0.159575535488839	0.689547611830298	0.928165808082547
54920	DUS2	dihydrouridine synthase 2
4472	-0.0728916019494564	3.45008050988518
0.159313106301028	0.689789711848019	0.92840246890124
51001	MTERF3	mitochondrial transcription termination factor 3
	1801	0.0879518153425053	2.72664033852912
0.159127296424729	0.689961267712567	0.928526709220845
10793	ZNF273	zinc finger protein 273 6039	0.106768755805733
	2.03689369360888	0.159069557451584
0.690014601005196	0.928526709220845
6932	TCF7	transcription factor 7	10063	-0.117509729363411
	3.09954034341329	0.158827327230526
0.690238470579073	0.928662215573899
1856	DVL2	dishevelled segment polarity protein 2	5350
0.095467653707356	5.21794752175013	0.158753345238928
	0.690306884293409	0.928662215573899
374659	HDDC3	HD domain containing 3	2606	0.0934617406011193
	2.66509888238645	0.158745436116924
0.690314199225776	0.928662215573899
1039	CDR2	cerebellar degeneration related protein 2
3289	-0.0916160821266275	6.12886897438259
0.158512273221746	0.690529940194953	0.928663634899814
441108	IRF1-AS1	IRF1 antisense RNA 1	7217
0.148109251758069	2.04030585282777	0.15849414170467
	0.690546724626904	0.928663634899814

3482	IGF2R	insulin like growth factor 2 receptor	14911
0.0554720254846499	9.21207731125995	0.158427030754883
	0.690608859232434	0.928663634899814
170685	NUDT10	nudix hydrolase 10	2054	0.161901857439568
	0.35384405118593	0.15838496925469
0.690647809601554	0.928663634899814
63922	CHTF18	chromosome transmission fidelity factor 18	5753
	0.141233464384657	2.18373794961884
0.158368346091756	0.690663204864362	0.928663634899814
53637	S1PR5	sphingosine-1-phosphate receptor 5	2806
0.121451714836823	2.35931137535981	0.158314527926368
	0.690713054071983	0.928663634899814
3182	HNRNPAB heterogeneous nuclear ribonucleoprotein A/B	2161
	0.053620763453656	7.27014354952905
0.1580965073166 0.690915096667892	0.928678065602992
55733	HHAT	hedgehog acyltransferase	6641
0.0894044138503651	4.77581009409178	0.158074708279327
	0.690935306995797	0.928678065602992
100131017	ZNF316	zinc finger protein 316 5811
0.0789836043398687	5.20138696331648	0.158063648248774
	0.690945561590703	0.928678065602992
57107	PDSS2	decaprenyl diphosphate synthase subunit 2
3926	-0.0591370296520352	4.45908563698741
0.157954808062444	0.691046497768683	0.928678065602992
158431	ZNF782	zinc finger protein 782 4622	0.127324622207645
	1.72322205108363	0.157945327026471
0.69105529219533	0.928678065602992
100506713	ALOX12-AS1	ALOX12 antisense RNA 1	3217
0.118651510517306	0.451429361995687	0.157860967167311
	0.691133556240379	0.928694140661837
9938	ARHGAP25	Rho GTPase activating protein 25	5452
	-0.106862471294011	1.94637881697415
0.157788365665598	0.691200930971033	0.928695581785253
26115	TANC2	tetratricopeptide repeat, ankyrin repeat and coiled-coil containing 2	15978	-0.0645305669993311
7.46718453340614	0.157612721163993	0.691364004628154
	0.928775273896187
8659	ALDH4A1 aldehyde dehydrogenase 4 family member A1	6541
	0.0921465465699926	6.58732024515616
0.157466435118295	0.691499901310858	0.928775273896187
5723	PSPH	phosphoserine phosphatase	2762
0.0691058147868358	4.35391959992359	0.157414168439131
	0.69154847368046	0.928775273896187
54973	INTS11	integrator complex subunit 11
8220	-0.0440844510041987	6.01920080576103
0.157379675078512	0.691580534096284	0.928775273896187
11186	RASSF1	Ras association domain family member 1
3683	-0.0700039548687448	5.2669733786388 0.157317315916963
	0.691638505156938	0.928775273896187
126119	JOSD2	Josephin domain containing 2	1514
0.0796274384990228	3.51521762981779	0.157296250656243
	0.691658091098103	0.928775273896187
4781	NFIB	nuclear factor I B	10777	0.0748783962988429
	5.79507752007973	0.15719361422600З

0.691753541547724	0.928777545631524
401320	GARS-DT GARS divergent transcript
21706	-0.104515518200253	3.24242609760814
0.157132106575856	0.691810760086112	0.928777545631524
9521	EEF1E1	eukaryotic translation elongation factor 1 epsilon 1
	2107	0.0834351427708345	3.7328596328805
0.157080576110907	0.691858707154945	0.928777545631524
6477	SIAH1	siah E3 ubiquitin protein ligase 1	4769
0.0637332391837715	4.58677123119721	0.156913694282984
	0.692014046608256	0.928877551003355
1009	CDH11	cadherin 11	9920	-0.160307535265158
5.76184384464997	0.156856156639471	0.692067626817146
	0.928877551003355
10910	SUGT1	SGT1 homolog, MIS12 kinetochore complex assembly
cochaperone	14512	0.0556127583071218	5.65073443427675
	0.156723186245969	0.692191495026961
0.928877551003355
29964	PRICKLE4	prickle planar cell polarity protein 4	4729
	0.0567309078027012	6.08590331859407
0.156651384252779	0.692258407253193	0.928877551003355
114987	WDR31	WD repeat domain 31	4933	-0.121094568890069
	2.09317425017659	0.156644548214921
0.692264778676663	0.928877551003355
79077	DCTPP1	dCTP pyrophosphatase 1	1581	0.0680270477868405
	5.13912410240329	0.156191097544635
0.692687769965792	0.928914673859672
3703	STT3A	STT3 oligosaccharyltransferase complex catalytic
subunit A	7345	-0.0554584587857944	7.25523539594468
	0.15617641926444	0.692701474126178
0.928914673859672
56985	ADPRM	ADP-ribose/CDP-alcohol diphosphatase, manganese
dependent	1620	0.11926771401646	2.3417716599821
0.156135886085661	0.692739321199638	0.928914673859672
51312	SLC25A37	solute carrier family 25 member 37	8147
	0.0684167591685172	5.79224170185528
0.156037552996203	0.692831161433066	0.928914673859672
144233	BCDIN3D BCDIN3 domain containing RNA methyltransferase
3721	-0.0988759552375326	2.71971670462393
0.15602867364262	0.692839456136616	0.928914673859672
100129387	GABPB1-AS1	GABPB1 antisense RNA 1	1076
0.11667189951805	1.21284627515999	0.1559990422347
0.692867138475864	0.928914673859672
89782	LMLN	leishmanolysin like peptidase	8692
0.0788518499787908	3.79566082699273	0.15587391823188
	0.692984065706803	0.928914673859672
538	ATP7A	ATPase copper transporting alpha	8529
0.0703892097134271	4.52513158976006	0.155817142142314
	0.693037140332847	0.928914673859672
23408	SIRT5	sirtuin 5	6023	0.0732367096528979
3.50097074487644	0.15581423173546	0.693039861299595
	0.928914673859672
51380	CSAD	cysteine sulfinic acid decarboxylase
4390	-0.100376880395222	3.14671074819029
0.155605712187215	0.69323488467293	0.928914673859672

55139 ANKZF1 ankyrin repeat and zinc finger peptidyl tRNA hydrolase 1 6119 0.0702642698230763 5.0743443581387 0.155592567079694 0.69324718403791 0.928914673859672
118924 FRA10AC1 FRA10A associated CGG repeat 1 8142 -0.0857505870895351 3.19259473182142 0.155556452279385 0.693280978340508 0.928914673859672
10169 SERF2 small EDRK-rich factor 2 4905 0.0731117334233382 7.95018024651999 0.155537589857005 0.693298630590776 0.928914673859672
9310 ZNF235 zinc finger protein 235 4617 0.113360748614297 2.28552264164912 0.15549677628201 1 0.693336829889212 0.928914673859672
79807 GSTCD glutathione S-transferase C-terminal domain containing 6295 0.0663535574233594 3.98537098275583 0.15538799300194 0.69343867345881 0.928914673859672
51389 RWDD1 RWD domain containing 1 5909 0.0683298418416025 5.10461296587163 0.155235067732992 0.693581912700626 0.928914673859672
9921 RNF10 ring finger protein 10 5769 -0.0492168254060346 7.75621782800784 0.155233228569838 0.69358363587023 0.928914673859672
4086 SMAD1 SMAD family member 1 6058 -0.0629336892996828 4.18160872798677 0.155154889087978 0.693657045537734 0.928914673859672
10548 TM9SF1 transmembrane 9 superfamily member 1 4180 0.0445915167978015 7.0116658965364 0.155014187444259 0.693788946759701 0.928914673859672
10714 POLD3 DNA polymerase delta 3, accessory subunit 5508 -0.0673591351210754 3.81526680784914 0.154982416063944 0.693818740513608 0.928914673859672
3070 HELLS helicase, lymphoid specific 10148 0.113886581445062 1.76676931172266 0.154867615425084 0.693926424733153 0.928914673859672
100506178 LOC100506178 uncharacterized LOC100506178 2432 0.125011664071819 1.93760931764891 0.154834370045673 0.693957617859003 0.928914673859672
11033 ADAP1 ArfGAP with dual PH domains 1 5296 -0.118202178877674 4.09257045559686 0.15480372836582 0.693986371438009 0.928914673859672
4188 MDFI MyoD family inhibitor 2523 0.0575441167379701 5.6564110230856 0.154741888835417 0.694044410506165 0.928914673859672
5917 RARS1 arginyl-tRNA synthetase 1 3421 -0.0491616433658023 6.39815796030206 0.154670665946156 0.694111272857793 0.928914673859672
389906 LOC389906 zinc finger protein 839 pseudogene 3054 -0.113657077113827 3.86734872447181 0.15458017287185 0.694196251245142 0.928914673859672
147700 KLC3 kinesin light chain 3 3654 -0.0843336247334475 3.37226336557315 0.154528362116826 0.694244917559307 0.928914673859672
55599 RNPC3 RNA binding region (RNP1, RRM) containing 3 6867 -0.0942013010147828 3.97935695801165

0.154517339219886	0.694255272681937	0.928914673859672
488	ATP2A2	ATPase sarcoplasmic/endoplasmic reticulum Ca2+ transporting 2	13745	-0.0576729905545059	9.44777460163995	0.154437830864822	0.694329977002674	0.928914673859672
2782	GNB1	G protein subunit beta 1	6362	-0.0574259325384582	9.5112320125711	0.154299367448751	0.694460127227608	0.928914673859672
390928	ACP7	acid phosphatase 7, tartrate resistant (putative)	3107	-0.173841161066123	1.49794622011539	0.154286882231672	0.694471866158609	0.928914673859672
117155	CATSPER2	cation channel sperm associated 2	8403	-0.178686484519103	1.78330823051297	0.154161692255606	0.694589603403202	0.928914673859672
26232	FBXO2	F-box protein 2	2333	0.128759671980995	6.77519397727571	0.154156170736783	0.694594797488656	0.928914673859672
347732	CATSPER3	cation channel sperm associated 3	2145	0.102633455442451	1.61420844653708	0.154132358923928	0.694617198459439	0.928914673859672
10076	PTPRU	protein tyrosine phosphatase receptor type U	6682	0.0689858370037422	6.64423168103207	0.154040065682295	0.694704042232866	0.928914673859672
57716	PRX	periaxin	5767	0.147929004101683	1.26659748495791	0.154023196376994	0.694719918732708	0.928914673859672
518	ATP5MC3	ATP synthase membrane subunit c locus 3	6780	0.0584606608704751	6.71968272952974	0.153982163725885	0.694758540687215	0.928914673859672
57336	ZNF287	zinc finger protein 287	3566	-0.0916679535999089	2.27588093254671	0.153798778512182	0.694931224507154	0.928914673859672
113402	SFT2D1	SFT2 domain containing 1	3104	-0.0602500255536905	5.37336805917683	0.153765655348273	0.694962427441507	0.928914673859672
56342	PPAN	peter pan homolog	2583	0.0679492681516922	4.07999248913094	0.153531950632423	0.695182694003089	0.928914673859672
8087	FXR1	FMR1 autosomal homolog 1	12054	0.0439143094115934	6.61210790379366	0.153399653826089	0.695307469398519	0.928914673859672
8509	NDST2	N-deacetylase and N-sulfotransferase 2	6615	-0.0616883903010144	5.33122087343774	0.153343114018014	0.695360813702964	0.928914673859672
3868	KRT16	keratin 16	1754	-0.145343036827283	8.03667941989458	0.15329157838039	0.695409446543429	0.928914673859672
1564	CYP2D7	cytochrome P450 family 2 subfamily D member 7 (gene/pseudogene)	1877	-0.14269758688435	0.192404439820071	0.153259363688339	0.69543985149915	0.928914673859672
55591	VEZT	vezatin, adherens junctions transmembrane protein	8757	0.0660579461954645	7.46942768687615	0.153219739192195	0.695477255050398	0.928914673859672

79882	ZC3H14	zinc finger CCCH-type containing 14	20977
0.0467854340116179	6.10573576136242	0.153204534539921
	0.695491608965747	0.928914673859672
8649	LAMTOR3	late endosomal/lysosomal adaptor, MAPK and MTOR
activator 3	4226	-0.0516588270355376	5.93696127064219
	0.153171875398272	0.695522443527002
0.928914673859672
7168	TPM1	tropomyosin 1	12492	0.0802036690687798
10.586568176958	0.153165834068204	0.695528147759297
0.928914673859672
10791	VAMP5	vesicle associated membrane protein 5	819
0.154442745641877	3.57499439217032	0.15299773949803
	0.69568691493528	0.928914673859672
10320	IKZF1	IKAROS family zinc finger 1	9784
0.212642206976591	1.88909446111463	0.152943527996435
	0.695738139741076	0.928914673859672
84364	ARFGAP2	ADP ribosylation factor GTPase activating protein 2
	5918	-0.0411602170712649	6.36326803081396
0.152943416900655	0.695738244726958	0.928914673859672
54101	RIPK4	receptor interacting serine/threonine kinase 4	4033
	0.0553803962739113	6.5433218007491	0.152940527247964
	0.695740975473822	0.928914673859672
55226	NAT10	N-acetyltransferase 10	5013	0.0553113982040878
	4.84647465165807	0.152862260135374
0.695814949889645	0.928924895454085
29946	SERTAD3	SERTA domain containing 3
2218	-0.0565092827449714	5.23099504276164
0.152682712518731	0.695984732432566	0.929063008403979
56122	PCDHB14	protocadherin beta 14	4828	-0.174634819259433
	1.97344615608233	0.152608919724085
0.696054545216533	0.929067659124069
1875	E2F5	E2F transcription factor 5
5928	-0.0863774465784879	2.95864120711246
0.152427700608758	0.696226072879452	0.929208061250356
23450	SF3B3	splicing factor 3b subunit 3
12417	-0.0433306143045227	6.98068529504304
0.152299411908409	0.696347571852094	0.929244771054207
8994	LIMD1	LIM domains containing 1
12673	-0.0703839249484852	5.7155728427495	0.152258573462441
	0.696386261280503	0.929244771054207
23464	GCAT	glycine C-acetyltransferase	2104
0.105600312088785	1.30790482248224	0.15190408709217
	0.696722345171455	0.929604676016057
23306	NEMP1	nuclear envelope integral membrane protein 1
5708	-0.10925230847517	4.43666441338693
0.151829612579053	0.696793010890505	0.929610410852777
23230	VPS13A	vacuolar protein sorting 13 homolog A
20196	-0.0899672496508853	6.5623404135904	0.151582995819581
	0.697027157715327	0.929834228392246
57620	STIM2	stromal interaction molecule 2
6602	-0.0693915654903167	4.62674045675231
0.151292503935046	0.697303242683551	0.930093500454139
408050	NOMO3	NODAL modulator 3	4534	-0.0619303169938318
	6.60893703704385	0.151238797803579

0.697354318681328	0.930093500454139
57120	GOPC	golgi associated PDZ and coiled-coil motif containing	5011	-0.0598453916607999	6.16214876167657	0.151016242144928	0.697566086536346	0.930261217263745
146547	PRSS36	serine protease 36	3685	-0.114264359374452	0.337182754432393	0.150967074194026	0.697612895419318	0.930261217263745
79072	FASTKD3	FAST kinase domains 3	2857	-0.0909150235886294	2.78786600228963	0.150895287945074	0.697681253137162	0.930263808918822
3178	HNRNPA1	heterogeneous nuclear ribonucleoprotein A1	5093	-0.0545780113674152	6.96508250665966	0.150702394080874	0.697865027086968	0.930334653045711
80816	ASXL3	ASXL transcriptional regulator 3	12546	-0.313102563403936	0.51563630131718	0.150650372561154	0.697914612227481	0.930334653045711
56990	CDC42SE2	CDC42 small effector 2	5414	-0.0455349744489049	6.42745085236285	0.150630409697365	0.697933642764641	0.930334653045711
1962	EHHADH	enoyl-CoA hydratase and 3-hydroxyacyl CoA dehydrogenase	4700	0.114248730502041	2.83576185110114	0.150266919686782	0.698280410715805	0.930708317908106
51130	ASB3	ankyrin repeat and SOCS box containing 3	2952	0.0776145750828031	3.83310744034619	0.150129277227567	0.698411846972525	0.930777835976065
729438	CASTOR2	cytosolic arginine sensor for mTORC1 subunit 2	1363	-0.091124754798319	2.242639362279	0.149901491770511	0.698629514060436	0.930777835976065
10346	TRIM22	tripartite motif containing 22	6020	0.0783060382093379	7.2215042521583	0.149883788603058	0.698646438804714	0.930777835976065
8220	ESS2	ess-2 splicing factor homolog	6003	0.0634665947967186	4.81647840712077	0.149874539381337	0.698655281787475	0.930777835976065
206358	SLC36A1	solute carrier family 36 member 1	6980	0.070954041879888	5.41230646067004	0.149701907793094	0.698820388761232	0.930777835976065
63916	ELMO2	engulfment and cell motility 2	6598	0.0595168737413597	6.08859326332319	0.149660070669768	0.69886041877561	0.930777835976065
91012	CERS5	ceramide synthase 5	6749	0.0639148714254477	5.50586647135561	0.149636083952885	0.698883372313688	0.930777835976065
100303755	PET117	PET117 cytochrome c oxidase chaperone	1528	-0.0781771507765467	3.50631003331203	0.149543359132637	0.698972123097959	0.930777835976065
1429	CRYZ	crystallin zeta	4085	-0.169821978425942	5.3152953719069	0.149408874428332	0.699100900227573	0.930777835976065
51463	GPR89B	G protein-coupled receptor 89B	5143	0.0713857450901607	3.88444009415355	0.149332799512155	0.69917377600149	0.930777835976065

7342	UBP1	upstream binding protein 1
9760	-0.0490389361284247	6.57597000175726
0.14930955203511	0.699196050119277	0.930777835976065
100532726	NDUFC2-KCTD14	NDUFC2-KCTD14 readthrough	960
	0.0528840963839982	5.37191898672786
0.14930948819377	0.699196111290359	0.930777835976065
1803	DPP4	dipeptidyl peptidase 4	5009	-0.133641318999579
	2.74209520131844	0.149266358459469
0.699237440495296	0.930777835976065
92912	UBE2Q2	ubiquitin conjugating enzyme E2 Q2
3852	-0.0555218542722898	6.47232359943884
0.149239813646663	0.699262880558016	0.930777835976065
26040	SETBP1	SET binding protein 1	11151	-0.105748270846949
	5.13567987918615	0.148963626697885
0.699527727471095	0.9308186753537
64118	DUS1L	dihydrouridine synthase 1 like	5999
0.0511042229189745	6.18431231827187	0.14894894812754
	0.699541811242193	0.9308186753537
54862	CC2D1A	coiled-coil and C2 domain containing 1A	5039
0.0584144677434089	6.14769574598093	0.148912101065449
	0.699577168715386	0.9308186753537
102288414	C11orf98	chromosome 11 open reading frame 98
	887	0.0708418390032539	4.44211264573352
0.148893395039147	0.699595120450415	0.9308186753537
22916	NCBP2	nuclear cap binding protein subunit 2
5110	-0.047754974287765	6.34861187151912
0.148861397283312	0.699625830950507	0.9308186753537
9863	MAGI2	membrane associated guanylate kinase, WW and PDZ domain containing 2	10218	0.123050182111005
0.716906962884754	0.148641031522257	0.699837434544958
	0.930918518560718
152992	TRMT44	tRNA methyltransferase 44 homolog	7128
0.0646442045592865	3.93938897617518	0.148588841290945
	0.699887575977312	0.930918518560718
55270	NUDT15	nudix hydrolase 15	2098	-0.0598368386497949
	4.49227528591843	0.148539234664078
0.699935244604636	0.930918518560718
1933	EEF1B2	eukaryotic translation elongation factor 1 beta 2
	1915	0.08550749438908	5.94760222452229
0.148459888600712	0.700011509837205	0.930918518560718
55971	BAIAP2L1	BAR/IMD domain containing adaptor protein 2 like 1	4798	-0.0488066522902837	7.40001086868426
0.14843734720796	0.700033180267013	0.930918518560718
55314	TMEM144	transmembrane protein 144	9243
0.0853967491655168	3.03901429105427	0.148272075748719
	0.700192123652116	0.931041491930434
10126	DNAL4	dynein axonemal light chain 4	2430
0.0675166066036675	3.7572943861268	0.148132440735917
0.700326491476764	0.93104861200013
9723	SEMA3E	semaphorin 3E	6748	0.20997014023743
1.6326977829184	0.148099073368947	0.700358610952277
0.93104861200013
81621	KAZALD1	Kazal type serine peptidase inhibitor domain 1
3102	-0.149825471792505	0.724243740422588

0.147981681107166	0.70047164594077	0.93104861200013
4714	NDUFB8	NADH:ubiquinone oxidoreductase subunit B8	2572
	0.0515354480009915	5.56104884963225
0.147899747001517	0.7005505693823 0.93104861200013
641371	ACOT1	acyl-CoA thioesterase 1 1713	0.105495362030615
	3.5939278722789 0.147861261389856	0.700587649504382
	0.93104861200013
92400	RBM18	RNA binding motif protein 18
5193	-0.054928834302424	5.32692146097403
0.147837784978666	0.70061027128017	0.93104861200013
3708	ITPR1	inositol 1,4,5-trisphosphate receptor type 1
12895	-0.129841051153886	3.40411114445793
0.147783311158008	0.700662769978823	0.93104861200013
7381	UQCRB	ubiquinol-cytochrome c reductase binding protein
	9715	0.0585688836510694	6.19706379934987
0.147638570690916	0.70080231639725	0.931145707244952
10134	BCAP31	B cell receptor associated protein 31	2624
0.0556927763180238	7.66879712771681	0.147467281032498
	0.700967560742179	0.931211868431081
27010	TPK1	thiamin pyrophosphokinase 1	8438
0.138215166583966	1.45667253686413	0.147449133072469
	0.700985074658193	0.931211868431081
126695	KDF1	keratinocyte differentiation factor 1	1793
0.0579096087665808	4.75415457491769	0.147150442910338
	0.701273507190367	0.931278666264991
25980	AAR2	AAR2 splicing factor	3101	-0.0474506858989791
	5.45140425063184	0.14711574420773
0.701307036033958	0.931278666264991
55192	DNAJC17 DnaJ heat shock protein family (Hsp40) member C17
	5634	-0.0688868766722801	3.7161285178367
0.147050282530993	0.701370303037221	0.931278666264991
7178	TPT1	tumor protein, translationally-controlled 1
7269	-0.0742831418964043	10.0190385825691
0.146973803688979	0.70144423830621	0.931278666264991
84734	FAM167B family with sequence similarity 167 member B
936	-0.141160045616243	0.582918235139554
0.146940282541221	0.701476651544948	0.931278666264991
10094	ARPC3	actin related protein 2/3 complex subunit 3
3025	-0.0670583736357736	7.55189809543436
0.146795078934957	0.701617105008423	0.931278666264991
79571	GCC1	GRIP and coiled-coil domain containing 1
4454	-0.0484112543332915	5.75980838858841
0.146785516160655	0.701626357744789	0.931278666264991
51230	PHF20	PHD finger protein 20	8384	-0.0488598799311847
	6.04256471662123	0.146769944731044
0.701641425067485	0.931278666264991
10954	PDIA5	protein disulfide isomerase family A member 5	3540
	0.12167349473218	3.40151419280124
0.146746808249596	0.701663814225078	0.931278666264991
55215	FANCI	FA complementation group I	7766
0.0853398258361566	4.09320380662364	0.146709187168078
	0.701700224442116	0.931278666264991
4077	NBR1	NBR1 autophagy cargo receptor
8405	-0.0519617508987975	7.60506562810331

0.146599728615257 0.701806190439638 0.9313310572703
157567 ANKRD46 ankyrin repeat domain 46
5535 -0.0775225512206251 4.08614667939696
0.146452065033548 0.701949214331356 0.931432611324299
23417 MLYCD malonyl-CoA decarboxylase
13653 -0.0656267344802879 3.85460022827076
0.146306076804819 0.702090696795618 0.931532100977192
79902 NUP85 nucleoporin 85 4980 -0.0567931444973259
4.92854997903419 0.146101992186004 0.702288618256723
0.93170644780469
26750 RPS6KC1 ribosomal protein S6 kinase C1 6190
0.0570470364040525 5.02626916645157 0.145592614670434
0.702783305234448 0.932274434739929
4343 MOV10 Mov10 RISC complex RNA helicase 5851
0.0587463562769536 5.94861588729784 0.145289117452137
0.703078520993594 0.932531578466544
1303 COL12A1 collagen type XII alpha 1 chain 12481
0.0877345632141756 7.449360051008 0.145209098554403
0.703156415253435 0.932531578466544
83394 PITPNM3 PITPNM family member 3 10609 0.051343420057256
5.02735669668337 0.145188081735435
0.703176878115488 0.932531578466544
37 ACADVL acyl-CoA dehydrogenase very long chain
4711 -0.0543693017640596 8.9695935007375 0.144957217387645
0.703401769116229 0.932577358078186
57645 POGK pogo transposable element derived with KRAB domain
8448 0.0578208535230807 6.63542316448511
0.144937948204984 0.703420549000144 0.932577358078186
25937 WWTR1 WW domain containing transcription regulator 1 8413
0.0507922002592609 6.7693618219204 0.144918748150143
0.703439262933267 0.932577358078186
121665 SPPL3 signal peptide peptidase like 3
6672 -0.0445330383033931 5.98309761814181
0.144864730018426 0.703491921007628 0.932577358078186
4597 MVD mevalonate diphosphate decarboxylase
4717 -0.0986440897291848 5.96754360777526
0.144811014539584 0.70354429519613 0.932577358078186
79144 PPDPF pancreatic progenitor cell differentiation and
proliferation factor 1094 -0.0816567764392098
6.37478442469816 0.144723929166513 0.703629229657263
0.932579221528139
9451 EIF2AK3 eukaryotic translation initiation factor 2 alpha
kinase 3 5434 -0.0467768418044857 6.02607842542395
0.14467305582286 0.703678860022196
0.932579221528139
3621 ING1 inhibitor of growth family member 1
6388 -0.0683045769442892 3.82044017761872
0.144546023017 0.703802832686337 0.932613323765984
7013 TERF1 telomeric repeat binding factor 1
5531 -0.058262105568997 4.35695586442937
0.144510249386824 0.703837755802829 0.932613323765984
654433 PAX8-AS1 PAX8 antisense RNA 1
16398 -0.263675010589605 6.79378022123243
0.144142347958677 0.704197197819511 0.93279106860708

55784	MCTP2	multiple C2 and transmembrane domain containing 2
	12978	-0.0701374083623858	3.90967751670304
0.14407446110585	0.70426358104452	0.93279106860708
10401	PIAS3	protein inhibitor of activated STAT 3
4805	-0.0505087445595009	6.27610151934209
0.144072915948659	0.704265092186227	0.93279106860708
6530	SLC6A2	solute carrier family 6 member 2	5463
0.242390964451771	4.3134483055726 0.144067384037442
0.704270502392548	0.93279106860708
114971	PTPMT1	protein tyrosine phosphatase mitochondrial 1	2740
	0.0531995257619731	5.25281127089532
0.143989079192262	0.704347097240458	0.93279106860708
90550	MCU	mitochondrial calcium uniporter
4533	-0.052000456214493	6.10563323359131
0.143964170888175	0.704371466599349	0.93279106860708
3921	RPSA	ribosomal protein SA	3936	0.0738689873530069
	6.76396536942319	0.143820990342647
0.704511595919768	0.932882278146207
1389	CREBL2	cAMP responsive element binding protein like 2	3922
	0.0498334578475777	5.67451320419263
0.143705150048012	0.704625026017291	0.932882278146207
55191	NADSYN1 NAD synthetase 1	14360	0.0511503795948066
	6.22464699367711	0.143689714443777
0.70464014441257	0.932882278146207
10524	KAT5	lysine acetyltransferase 5	3732
0.0473463629764173	5.31801748873724	0.143550668204895
	0.704776375068297	0.932974452847712
27338	UBE2S	ubiquitin conjugating enzyme E2 S
2635	-0.0799920400186449	2.98342052660826
0.14344978922413	0.704875258557725	0.932994047112599
283820	NOMO2	NODAL modulator 2	14012	-0.0567280732055885
	7.95495374488651	0.143399676288862
0.704924394987695	0.932994047112599
3363	HTR7	5-hydroxytryptamine receptor 7	3131
0.166924270640265	2.42309735760981	0.143230525455202
	0.705090322259919	0.933125486006678
22884	WDR37	WD repeat domain 37	5656	-0.0507834943088634
	5.53461995752609	0.14278307857364
0.705529782732079	0.933531055245561
3561	IL2RG	interleukin 2 receptor subunit gamma
2760	-0.319608185899122	1.40268012649558
0.142677857106783	0.705633240368857	0.933531055245561
643529	LINC00865	long intergenic non-protein coding RNA 865
	4272	0.116127108724759	3.48083373079924
0.14254557130635	0.705763370530072	0.933531055245561
100507577	NA	NA	2286	-0.104527944049908
1.65214824304794	0.142493450509052	0.705814660948037
	0.933531055245561
4802	NFYC	nuclear transcription factor Y subunit gamma	6068
	0.0646293575632452	5.15496727507095
0.142492332124077	0.705815761632541	0.933531055245561
10213	PSMD14	proteasome 26S subunit, non-ATPase 14
7698	-0.0567272264745572	6.72000826515593
0.142302370702332	0.706002788222746	0.933531055245561

574036	SERTAD4-AS1	SERTAD4 antisense RNA 1
1004	-0.122234033146251	1.44603963984977
0.142246075589122	0.706058240986801	0.933531055245561
23158	TBC1D9	TBC1 domain family member 9	5603
0.0445736630227716	6.86725012400287	0.142184443007096
	0.706118965729191	0.933531055245561
1111	CHEK1	checkpoint kinase 1	6558	0.106398553070111
	3.31790181366905	0.142045485183395
0.706255931897605	0.933531055245561
113026	PLCD3	phospholipase C delta 3 6740	0.0624071440464628
	6.45928814272684	0.141987395762889
0.706313211411349	0.933531055245561
100616408	MIR5047 microRNA 5047	100	-0.080895325979948
	3.26371027978794	0.141828620070632
0.706469841688106	0.933531055245561
29950	SERTAD1 SERTA domain containing 1
2114	-0.0636063499822122	4.92938190894236
0.141809712694908	0.706488500252074	0.933531055245561
29926	GMPPA	GDP-mannose pyrophosphorylase A
5332	-0.0703634658724938	4.64803150250285
0.141744002271322	0.70655335700614	0.933531055245561
147808	ZNF784	zinc finger protein 784 2058	-0.0932048216806633
	2.42554043402611	0.141700788729956
0.706596018499732	0.933531055245561
51503	CWC15	CWC15 spliceosome associated protein homolog	2758
	0.0569989242407338	5.08205577615953
0.141690976455058	0.706605706458625	0.933531055245561
64897	C12orf43	chromosome 12 open reading frame 43	6140
	0.0644706797249719	4.10751569721419
0.141669860689568	0.706626555998193	0.933531055245561
64147	KIF9	kinesin family member 9 5850	0.0987014051287236
	1.78463327011955	0.141645279921635
0.706650829088301	0.933531055245561
340385	ZNF517	zinc finger protein 517 5308	0.107088441558037
	2.16659061842227	0.141612470548292
0.706683231533106	0.933531055245561
5518	PPP2R1A protein phosphatase 2 scaffold subunit Aalpha	7437
	0.0524215169209491	8.25224627341212
0.141593373851535	0.706702093350072	0.933531055245561
1827	RCAN1	regulator of calcineurin 1	5531
0.0791039630169983	6.38259701366719	0.141475009501054
	0.706819034282365	0.933531055245561
23484	LEPROTL1	leptin receptor overlapping transcript like
1	5421	-0.0416015574705057	5.95586212494106
0.141438868321558	0.70685475196523	0.933531055245561
4350	MPG	N-methylpurine DNA glycosylase
1661	-0.0566473011480098	4.80037081265823
0.141280138957155	0.707011683069199	0.933531055245561
51163	DBR1	debranching RNA lariats 1
3160	-0.0583402327889986	4.17590708544302
0.141109426049836	0.707180574285931	0.933531055245561
8427	ZNF282	zinc finger protein 282 3832	0.0595442767673843
	5.55272088227353	0.141066967610362
0.70722597749201	0.933531055245561

55137	FIGN	fidgetin, microtubule severing factor
9942	-0.134517984670117	3.21547362965824
0.140989693316435	0.707299098909229	0.933531055245561
5430	POLR2A	RNA polymerase II subunit A
8357	-0.0514111941163743	8.27945028792432
0.140980373757756	0.70730832684145	0.933531055245561
55607	PPP1R9A	protein phosphatase 1 regulatory subunit 9A
11151	-0.15284194481405	3.80029132220759
0.140953521588742	0.707334916955898	0.933531055245561
400566	C17orf97	chromosome 17 open reading frame 97
2265	-0.176459892301645	1.6207067336371	0.140953118670697
0.707335315963807	0.933531055245561
342538	NACA2	nascent polypeptide associated complex subunit alpha
2	777	-0.0822429223448848	3.14714968358152
0.140884600882141	0.707403178284157	0.933531055245561
81610	FAM83D	family with sequence similarity 83 member D
2475	-0.178931318962424	-0.158509699132606
0.140806161254157	0.707480890645455	0.933531055245561
8997	KALRN	kalirin RhoGEF kinase	24490	-0.10705335018391
3.8505885430341	0.14058924606212	0.707695923315797
0.933531055245561
5692	PSMB4	proteasome 20S subunit beta 4	2158
0.0526838459445811	7.22893185180106	0.140572097708992
0.707712930913395	0.933531055245561
29990	PILRB	paired immunoglobin like type 2 receptor beta	1149
0.116673958784766	4.43624349635578
0.140491718610775	0.707792666026105	0.933531055245561
8577	TMEFF1	transmembrane protein with EGF like and two
follistatin like domains 1	2611	-0.0914046325028918
2.95887089682935	0.140472449146662	0.707811784971631
0.933531055245561
7386	UQCRFS1	ubiquinol-cytochrome c reductase, Rieske iron-sulfur
polypeptide 1	3475	0.0587681537330597	5.77974255367565
0.140421331174707	0.707862510895413
0.933531055245561
283596	SNHG10	small nucleolar RNA host gene 10	1945
0.112028077768727	0.206300514221562	0.140293507658107
0.707989400173299	0.933531055245561
8703	B4GALT3	beta-1,4-galactosyltransferase 3	3890
0.0508823615422393	5.52719976705303	0.140289270192871
0.707993607796281	0.933531055245561
22794	CASC3	CASC3 exon junction complex subunit
6380	-0.0579567031695289	6.2493717019674	0.140269553381358
0.708013186705627	0.933531055245561
53832	IL20RA	interleukin 20 receptor subunit alpha
4730	-0.14287155451727	3.13799488972615
0.140181353729119	0.708100788690699	0.933531055245561
85459	CEP295	centrosomal protein 295	9495	-0.0859947979041975
4.17533915248069	0.1401541779503	0.708127786650503
0.933531055245561
10989	IMMT	inner membrane mitochondrial protein	3986
0.0451672511310811	6.38788525961675	0.140100198946868
0.708181421305801	0.933531055245561
5886	RAD23A	RAD23 homolog A, nucleotide excision repair protein

2449    -0.048747240724276      6.0570680551581
0.140019423833664       0.708261703138092       0.933531055245561
56978   PRDM8   PR/SET domain 8 4588    0.151742764199403
1.64268624142954        0.139989819932617       0.708291132863265
0.933531055245561
54097   FAM3B   family with sequence similarity 3 member B
3599    -0.287326635724112      2.72037398632642
0.139951464529291       0.708329267876763       0.933531055245561
9842    PLEKHM1 pleckstrin homology and RUN domain containing M1
8734    0.0627295877384542      6.63046472493331
0.139794160896471       0.708485729988558       0.933594795957108
5803    PTPRZ1  protein tyrosine phosphatase receptor type Z1   9917
0.152650248236012       0.575685925022366
0.139768828711554       0.708510936033702       0.933594795957108
8733    GPAA1   glycosylphosphatidylinositol anchor attachment 1
3268    0.0510877743903955      6.04058397008982
0.139632890665599       0.708646241676001       0.933601012441316
8874    ARHGEF7 Rho guanine nucleotide exchange factor 7
11768   -0.0473656307282682     6.25520162167347
0.139544023750602       0.708734735746877       0.933601012441316
5191    PEX7    peroxisomal biogenesis factor 7 1835
0.107134527057261       3.00195608631453        0.139532980576484
0.708745734831832       0.933601012441316
26118   WSB1    WD repeat and SOCS box containing 1     8602
0.0602904876157464      6.71676457544974        0.139496308953881
0.708782263603981       0.933601012441316
9358    ITGBL1  integrin subunit beta like 1
6174    -0.271858019736035      2.22192029061204
0.13930899865295        0.708968929651571       0.933662143664905
6171    RPL41   ribosomal protein L41   1088    0.0757073469278576
7.52772284968442        0.139198269368649
0.709079345332978       0.933662143664905
7098    TLR3    toll like receptor 3    6724    0.125356365467383
4.07108633465608        0.139171276677517
0.709106269160479       0.933662143664905
10444   ZER1    zyg-11 related cell cycle regulator     4370
0.0564936012451292      6.17151715927785        0.139064285098083
0.70921301705316        0.933662143664905
80305   TRABD   TraB domain containing  3153    -0.0569342499939092
5.11306773327806        0.139040070889327
0.709237182601485       0.933662143664905
139322  APOOL   apolipoprotein O like   6548    -0.0635031966902764
5.2474261102019 0.138965948520038       0.709311170917197
0.933662143664905
643783  USP46-AS1       USP46 antisense RNA 1
1048    -0.119580360438956      0.928474454077586
0.138932015186976       0.709345050379889       0.933662143664905
284323  ZNF780A zinc finger protein 780A
8066    -0.0882928154176871     3.92658671754449
0.138915111963406       0.709361928527302       0.933662143664905
57659   ZBTB4   zinc finger and BTB domain containing 4
6126    -0.0510982855938066     7.71796616056704
0.138729504075272       0.709547338039091       0.933754035414895
22994   CEP131  centrosomal protein 131 3966    0.0957409460526665

3.31176028018058	0.138658743569396
0.709618060102255	0.933754035414895
149428	BNIPL	BCL2 interacting protein like
2892	-0.148886285383412	2.20227879886078
0.138612331532251	0.709664458083906	0.933754035414895
4863	NPAT	nuclear protein, coactivator of histone
transcription	6084	-0.0700615572462065	4.29989088480581
0.138578386919239	0.709698398017203
0.933754035414895
101060691	NUTM2B-AS1	NUTM2B antisense RNA 1	14755
0.127154154009355	4.86362057824504	0.138477328325481
0.709799470706992	0.933799303671724
29927	SEC61A1 SEC61 translocon alpha 1 subunit	4697
0.0500169336543413	9.25360463305799	0.138144819758338
0.710132322305076	0.933858167517032
10520	ZNF211	zinc finger protein 211 3300	-0.0752239841092659
4.31828281404479	0.138108624681806
0.710168582227988	0.933858167517032
100996541	FAM157C family with sequence similarity 157 member C
2041	-0.129682377669958	0.580522259349272
0.138069492848452	0.710207790253273	0.933858167517032
79842	ZBTB3	zinc finger and BTB domain containing 3 2978
0.0822763809630242	2.57342003188446	0.138019094548688
0.71025829599879	0.933858167517032
27245	AHDC1	AT-hook DNA binding motif containing 1
7157	-0.0785799848425285	6.22904187518048
0.137927300447153	0.710350312768715	0.933858167517032
728577	CNTNAP3B	contactin associated protein like 3B
13220	0.12376357249445	1.52862482023998
0.137921001094946	0.710356628678949	0.933858167517032
29777	ABT1	activator of basal transcription 1	2949
0.0584628147696193	4.87048954381421	0.137886625746042
0.710391097276328	0.933858167517032
727956	SDHAP2	succinate dehydrogenase complex flavoprotein subunit
A pseudogene 2	8449	0.116461145479782	4.456721587776
0.137873982738363	0.710403775809857	0.933858167517032
25983	NGDN	neuroguidin	4654	-0.0560461625833535
4.34546955610806	0.137833624713072	0.710444251664274
0.933858167517032
23293	SMG6	SMG6 nonsense mediated mRNA decay factor
9915	-0.0445258257066018	6.31326265711982
0.137764215605421	0.710513879190915	0.933862053657422
90799	CEP95	centrosomal protein 95	8048	-0.0688426131863202
4.41813215728664	0.13765090614637
0.710627588110275	0.933866656773079
135154	SDHAF4	succinate dehydrogenase complex assembly factor 4
1440	-0.101602790036884	2.73385130203851
0.137601952422489	0.710676730885781	0.933866656773079
84641	MFSD14B major facilitator superfamily domain containing 14B
3428	-0.043425583129878	7.08423583271004
0.137561451560174	0.710717395673665	0.933866656773079
66004	LYNX1	Ly6/neurotoxin 1	6039	-0.234843556808583
5.00190489451276	0.137436246735071
0.710843150342176	0.93394428354215

7568	ZNF20	zinc finger protein 20	4148	-0.0888228326251092
	3.11601252723077	0.137261505933162
0.711018767493994	0.933997022763441
10388	SYCP2	synaptonemal complex protein 2
6466	-0.161556313308906	1.64137660886839
0.137197634577201	0.711082990899538	0.933997022763441
205327	C2orf69	chromosome 2 open reading frame 69
4296	-0.0806379984654757	3.70679144458946
0.137132005254145	0.711148999672982	0.933997022763441
624	BDKRB2	bradykinin receptor B2	6845	0.114903731180068
	3.0684890794213	0.137059971733143	0.711221470350965
	0.933997022763441
375346	STIMATE	STIM activating enhancer	4939
0.0461573273596382	5.57274476585191	0.136912496225061
	0.711369908434396	0.933997022763441
2956	MSH6	mutS homolog 6	9196	0.0622010853568859
5.26552706083344	0.136910583843707	0.711371833894726
	0.933997022763441
55003	PAK1IP1	PAK1 interacting protein 1	1800
0.0712600540617959	4.02887947721417	0.13682857252563
	0.711454420487181	0.933997022763441
10217	CTDSPL	CTD small phosphatase like
5796	-0.0487298300812922	6.1900003461648	0.136815200395098
	0.711467889085195	0.933997022763441
55793	MINDY1	MINDY lysine 48 deubiquitinase 1
3792	-0.0782071716644039	4.89685146680104
0.136775269914866	0.711508112080419	0.933997022763441
143686	SESN3	sestrin 3	10573	0.0717112264397991
5.67627158036084	0.136457729054385	0.711828216957485
	0.933997022763441
115209	OMA1	OMA1 zinc metallopeptidase
3718	-0.0918413395291886	3.03850839014773
0.136354614949472	0.711932254699031	0.933997022763441
2067	ERCC1	ERCC excision repair 1, endonuclease non-catalytic
subunit	5576	-0.0535761330313123	5.14448552946246
0.136340470179228	0.711946529656184	0.933997022763441
643641	ZNF862	zinc finger protein 862	8430	0.0767296313008552
	4.35383677325757	0.136310273759563
0.711977006816181	0.933997022763441
79036	KXD1	KxDL motif containing 1	4112	0.0520830001959132
	5.58272606538314	0.136109618348902
0.71217962524747	0.933997022763441
57213	SPRYD7	SPRY domain containing 7	3340
0.0666476014246955	4.08551117601006	0.13607645659595
	0.71221312775898	0.933997022763441
162967	ZNF320	zinc finger protein 320	7590	-0.0647161197236453
	5.82609636607044	0.136056414409495
0.712233378140475	0.933997022763441
84709	MGARP	mitochondria localized glutamic acid rich protein
	1607	0.0793043347695748	3.41432474061787
0.135970897802653	0.712319802124858	0.933997022763441
199731	CADM4	cell adhesion molecule 4	2246
0.0915810654098855	5.55961001421045	0.135924316736606
	0.712366890430005	0.933997022763441

91942	NDUFAF2 NADH:ubiquinone oxidoreductase complex assembly factor 2	1017	0.106326252406711	2.75902622059826	0.135917056472597	0.712374230579368	0.933997022763441
54982	CLN6	CLN6 transmembrane ER protein	4225	0.0602466697880753	3.96296549848185	0.135861160019823	0.712430749524453	0.933997022763441
2589	GALNT1	polypeptide N-acetylgalactosaminyltransferase 1 4939	0.0694923657016905	7.28287907216772	0.135857294807458	0.712434658269999	0.933997022763441
23224	SYNE2	spectrin repeat containing nuclear envelope protein 2	30702	-0.18399157897502	9.12978461785799	0.135739345405206	0.712553966497052	0.933997022763441
57109	REXO4	REX4 homolog, 3'-5' exonuclease 2654	0.0603563125075592	4.95263275337025	0.135724796124876	0.712568687469357	0.933997022763441
9488	PIGB	phosphatidylinositol glycan anchor biosynthesis class B 4542	0.0715268670771431	3.84728898807654	0.135675495163897	0.712618576881127	0.933997022763441
54994	GID8	GID complex subunit 8 homolog	4622	0.0400609543024853	6.36748319296102	0.135645391870874	0.712649044545671	0.933997022763441
8409	UXT	ubiquitously expressed prefoldin like chaperone 1243	0.0615554800819995	4.11776396346584	0.135540543623614	0.712755191864182	0.933997022763441
90678	LRSAM1	leucine rich repeat and sterile alpha motif containing 1	4365	0.0602725329484553	4.49915833643008	0.135494032542141	0.712802294146278	0.933997022763441
339318	ZNF181	zinc finger protein 181 6266	0.0944575623413176	2.86219156908245	0.135493929750124	0.712802398255045	0.933997022763441
6196	RPS6KA2 ribosomal protein S6 kinase A2	8243	0.082897261691902	6.39521389407806	0.135479481099765	0.712817032436723	0.933997022763441
58488	PCTP	phosphatidylcholine transfer protein	5880	0.0677467969043686	4.13293128961572	0.135330378597978	0.712968101302731	0.934107585347241
114790	STK11IP serine/threonine kinase 11 interacting protein	5694	0.0567713241170948	5.01036017622011	0.13487657087223	0.713428476070345	0.934613808001238
128977	C22orf39	chromosome 22 open reading frame 39	4159	-0.0723178191773748	3.42997606812657	0.134810672089324	0.713495401526309	0.934613808001238
84296	GINS4	GINS complex subunit 4	5475	-0.111188228095484	1.38617492090945	0.134752342131752	0.713554655726796	0.934613808001238
440926	H3P6	H3 histone pseudogene 6 1629	0.0558323027882703	5.78020440332646	0.134657502868318	0.713651028800417	0.934614260647674
781	CACNA2D1	calcium voltage-gated channel auxiliary subunit alpha2delta 1	9010	0.176554960818832	3.06868519818455	0.134619748828005	0.713689404142587	0.934614260647674

88745 RRP36 ribosomal RNA processing 36 1326
0.0604521587323569 5.59984099267936 0.134544299014657
0.713766113897902 0.934614260647674
10948 STARD3 StAR related lipid transfer domain containing 3 5160
0.0464526673993559 5.5725293642376 0.134447024264407
0.713865049066527 0.934614260647674
440574 MICOS10 mitochondrial contact site and cristae organizing
system subunit 10 4552 0.0578038848946111
5.45013909033801 0.134423849589822 0.713888625306594
0.934614260647674
9732 DOCK4 dedicator of cytokinesis 4
11348 -0.106758245755735 4.17802645928741
0.134282801140222 0.714032167551828 0.93466577842863
55063 ZCWPW1 zinc finger CW-type and PWWP domain containing 1
3318 0.120790965736118 1.37533475493819
0.13425405422877 0.71406143320945 0.93466577842863
28969 BZW2 basic leucine zipper and W2 domains 2 3662
0.0965487685640063 4.89954171073509 0.134138423874873
0.714179186056493 0.934732560184386
26012 NSMF NMDA receptor synaptonuclear signaling and neuronal
migration factor 4915 0.0667723757204482
5.59431935439428 0.133882414287353 0.714440099871232
0.93485093002956
84164 ASCC2 activating signal cointegrator 1 complex subunit 2
5416 -0.0520284376301978 6.05774652863504
0.133836393282081 0.714487032461624 0.93485093002956
23568 ARL2BP ADP ribosylation factor like GTPase 2 binding
protein 2155 0.0546691072281006 5.67924747555366
0.133795434227068 0.714528810530064 0.93485093002956
10848 PPP1R13L protein phosphatase 1 regulatory subunit 13
like 5032 0.0483413034144413 6.82440049895624
0.133650436237573 0.714676766177554 0.93485093002956
55504 TNFRSF19 TNF receptor superfamily member 19
5113 -0.110654152347339 3.79821624850328
0.133615010063572 0.714712928801225 0.93485093002956
50810 HDGFL3 HDGF like 3 16253 -0.0609658504024654
6.04044370741833 0.133576993545397 0.714751741663563
0.93485093002956
85013 TMEM128 transmembrane protein 128
1737 -0.0759991224572019 3.70129826302717
0.133516106930932 0.714813916735054 0.93485093002956
51300 TIMMDC1 translocase of inner mitochondrial membrane domain
containing 1 2730 0.0481957047804703 5.76115703311787
0.133474792032238 0.714856115074364
0.93485093002956
25923 ATL3 atlastin GTPase 3 7600 -0.0554577408848116
7.56211327951848 0.133460904342476
0.71487030138835 0.93485093002956
55102 ATG2B autophagy related 2B 14778 -0.0663021947102002
5.72039396198206 0.132983946922735
0.715358023353433 0.9354014033898
203523 ZNF449 zinc finger protein 449 5330 0.0834768440641896
2.87107624899841 0.132725397564644
0.715622822233675 0.935544997705457

100130872	LOC100130872	uncharacterized LOC100130872	2796
	0.133354396709983	0.122079469779066
0.132686649980404	0.715662531570393	0.935544997705457
5208	PFKFB2	6-phosphofructo-2-kinase/fructose-2,6-biphosphatase
2	10036	-0.0925537399748863	6.34622710076249
0.132681107310148	0.715668212351965	0.935544997705457
84065	TMEM222	transmembrane protein 222	4476
0.0523445500964613	5.43673508835764	0.132544583581298
	0.715808180402164	0.93563867292553
6339	SCNN1D	sodium channel epithelial 1 delta subunit
4025	-0.10881268003231	1.28807518384394
0.132480930388503	0.715873467390798	0.93563867292553
8462	KLF11	Kruppel like factor 11	4589	0.0729749588398617
	4.62821474212584	0.132197021065548
0.71616488034947	0.935888246781714
53840	TRIM34	tripartite motif containing 34
2690	-0.0865986860181374	3.51607827418112
0.132164725214363	0.716198052206267	0.935888246781714
56255	TMX4	thioredoxin related transmembrane protein 4	6579
	0.0554778752683402	5.02125528635814
0.131934675807739	0.716434474425634	0.936083720695479
10396	ATP8A1	ATPase phospholipid transporting 8A1	11199
0.133856717172466	2.28601928919573	0.131889139305543
	0.716481300034	0.936083720695479
59286	UBL5	ubiquitin like 5	1931	0.0674458613419363
	6.48588451671528	0.131794257249165
0.716578897537052	0.93612391510926
11098	PRSS23	serine protease 23	11725	-0.0752160162260104
	8.20460081453155	0.13159346816235
0.716785564224233	0.936246968910865
23152	CIC	capicua transcriptional repressor	9494
0.0623391086033693	6.89432933519437	0.131541137194972
	0.716839456356738	0.936246968910865
56949	XAB2	XPA binding protein 2	4673	-0.0509626866898198
	5.16654635005684	0.13150797304095
0.716873616161136	0.936246968910865
56063	TMEM234	transmembrane protein 234	3571
0.0700443819022596	3.81265717899197	0.131418290944534
	0.716966015099378	0.936280344349896
54984	PINX1	PIN2 (TERF1) interacting telomerase inhibitor 1	2802
	0.0699429186012712	3.3848311472999	0.131339747596521
	0.717046967155032	0.936298766564793
388228	SBK1	SH3 domain binding kinase 1	4992
0.139718182620392	4.8159450184615	0.131270094694599
0.717118779032639	0.936305251483051
83871	RAB34	RAB34, member RAS oncogene family	2946
0.0477066813837478	6.34093747829898	0.131175203579744
	0.71721664610254	0.936345750951466
10465	PPIH	peptidylprolyl isomerase H	1444
0.0913976596751501	2.88613340855673	0.131083089055938
	0.717311687811144	0.936382554629143
6207	RPS13	ribosomal protein S13	2186	0.0695453628679457
	6.8158325473214	0.130934233125572	0.71746535390545
	0.936495872905939

92797	HELB	DNA helicase B	13796	-0.127980652153579
1.74330376495742	0.130826850849321	0.717576267502603
	0.936505920015767
126129	CPT1C	carnitine palmitoyltransferase 1C	5864
0.20162331110012	2.00240071507573	0.1307420265589
	0.717663917681405	0.936505920015767
100506070	RBFADN	RBFA downstream neighbor
7693	-0.102651794490124	0.511163373257654
0.13057274134526	0.717838938713003	0.936505920015767
54733	SLC35F2 solute carrier family 35 member F2	3717
0.0433299108903469	6.32751572941539	0.130525484599746
	0.717887819538842	0.936505920015767
55813	UTP6	UTP6 small subunit processome component
6450	-0.0619385404070413	5.3327825178864 0.130456211625269
	0.717959491312398	0.936505920015767
3615	IMPDH2	inosine monophosphate dehydrogenase 2
2773	-0.0677559455540269	5.48399989740521
0.130430547176418	0.717986049936232	0.936505920015767
10019	SH2B3	SH2B adaptor protein 3	5682	0.118629861830549
	3.80277199362446	0.130375471022932
0.718043054970342	0.936505920015767
9382	COG1	component of oligomeric golgi complex 1 5317
0.0633221687410999	4.9155831019804 0.130359981396284
0.718059089527809	0.936505920015767
9692	PRORP	protein only RNase P catalytic subunit
7981	-0.0524762439045519	7.20937703235729
0.130344815685547	0.718074789817187	0.936505920015767
23641	LDOC1	LDOC1 regulator of NFKB signaling
1381	-0.0986576041834689	4.00855742966018
0.130258411193733	0.718164259698263	0.936534321917039
64434	NOM1	nucleolar protein with MIF4G domain 1	7788
0.0550049831924016	4.68993629001525	0.130194663002463
	0.718230290999702	0.936534321917039
101929693	RARA-AS1	RARA antisense RNA 1
1790	-0.105827324110614	0.380616576849492
0.130004519862798	0.718427352608747	0.936654530214236
101927468	LOC101927468	uncharacterized LOC101927468
5944	-0.121509243761114	0.690869091860018
0.12996444300923	0.718468908457843	0.936654530214236
221785	ZSCAN25 zinc finger and SCAN domain containing 25
7094	-0.0641361044077428	4.31466298470495
0.129848202624538	0.718589479543617	0.936654530214236
11193	WBP4	WW domain binding protein 4
2623	-0.0626283371895158	4.69237396947971
0.129847738719713	0.718589960854014	0.936654530214236
6876	TAGLN	transgelin	3917	-0.110413359195453
10.4369437808934	0.129631367934962	0.718814555780773
	0.93672438166945
90525	SHF	Src homology 2 domain containing F
4894	-0.166205791508134	2.36128333561767
0.129585875924359	0.718861803853509	0.93672438166945
54386	TERF2IP TERF2 interacting protein
2248	-0.0465124519472078	6.00529259070402
0.129533095734867	0.718916633181191	0.93672438166945

54903	MKS1	MKS transition zone complex subunit 1	2541
0.0844321103344054	2.75005966040332	0.129486294372033
	0.718965262115714	0.93672438166945
10541	ANP32B	acidic nuclear phosphoprotein 32 family member B
	1850	-0.0523601245360163	5.7020071869924
0.129474106763134	0.718977927274096	0.93672438166945
3992	FADS1	fatty acid desaturase 1	10033	0.0865540980534499
	6.01962746533753	0.12939380212562
0.719061395342172	0.936745997447712
9513	FXR2	FMR1 autosomal homolog 2
4331	-0.048100659352333	6.59667750580624
0.129284297397177	0.719175260931492	0.936788091933265
283254	HARBI1	harbinger transposase derived 1
2662	-0.0831220115648411	2.53850861185944
0.129234107523622	0.719227467741153	0.936788091933265
6749	SSRP1	structure specific recognition protein 1	6323
	0.0398728501571598	6.30370481577767
0.129014148619769	0.719456401061141	0.936892301321506
84940	CORO6	coronin 6	4321	-0.188891794268851
3.87524500451048	0.128851496731095	0.71962583095858
	0.936892301321506
79738	BBS10	Bardet-Biedl syndrome 10
3595	-0.0734675832743231	3.7609081099438	0.128826315049794
	0.719652072797334	0.936892301321506
100302652	GPR75-ASB3	GPR75-ASB3 readthrough	4726
0.0730621934374049	3.71634528950353	0.128775244638168
	0.719705302182054	0.936892301321506
11152	WDR45	WD repeat domain 45	4541	-0.0573068617501702
	5.86822568361287	0.128734247562406
0.719748041007883	0.936892301321506
84787	KMT5C	lysine methyltransferase 5C	4775
0.0813901297651949	3.20967619527732	0.128703804032452
	0.719779782893411	0.936892301321506
51318	MRPL35	mitochondrial ribosomal protein L35
3829	-0.0598305063279235	5.04174253059987
0.128630317579791	0.719856420858876	0.936892301321506
340156	MYLK4	myosin light chain kinase family member 4	5806
	0.129774335187582	1.57206213450331
0.128540455694332	0.719950170114957	0.936892301321506
78990	OTUB2	OTU deubiquitinase, ubiquitin aldehyde binding 2
	4252	-0.0751930527517773	4.28666875383327
0.128524890602307	0.719966412300728	0.936892301321506
1659	DHX8	DEAH-box helicase 8	6656	-0.0435862356529964
	5.99257117579762	0.128496126010628
0.719996431095868	0.936892301321506
439990	LINC00857	long intergenic non-protein coding RNA 857
	2171	0.0933418993297832	3.68709740466183
0.128451282723855	0.720043237218963	0.936892301321506
3020	H3-3A	H3.3 histone A	2548	0.0535594546777936
5.80491099695656	0.128318103430599	0.720182300279161
	0.936968074989656
10988	METAP2	methionyl aminopeptidase 2
4491	-0.0503516204334809	6.3669203218231	0.128196519631672
	0.72030932668239	0.936968074989656

64849	SLC13A3	solute carrier family 13 member 3
6055	-0.110143164199241	0.297269665570176
0.128147422023311	0.720360641369254	0.936968074989656
168455	CCDC71L	coiled-coil domain containing 71 like	4232
0.0720086586278382	4.22014196580833	0.128139383426091
	0.720369044018249	0.936968074989656
78996	CYREN	cell cycle regulator of NHEJ	5321
0.0504572401538773	5.32963377838811	0.12776446091841
	0.720761276803565	0.93728398426041
11171	STRAP	serine/threonine kinase receptor associated protein
	2211	-0.0539343674808602	7.26028846986785
0.127715316234116	0.720812738605225	0.93728398426041
81855	SFXN3	sideroflexin 3	4631	-0.0432347661400324
6.81232069214241	0.127704087500767	0.72082449832789
	0.93728398426041
150962	PUS10	pseudouridine synthase 10	6366
0.0831903950624947	3.07751729442906	0.127651494534751
	0.720879586095338	0.93728398426041
6990	DYNLT3	dynein light chain Tctex-type 3
3186	-0.0681597383814125	6.04515834175399
0.127291683682259	0.721256808638039	0.937615160940042
80179	MYO19	myosin XIX	12821	-0.0478183701903906
6.07838860175719	0.127280848678382	0.721268177277715
	0.937615160940042
118424	UBE2J2	ubiquitin conjugating enzyme E2 J2	4712
0.0419877039943063	5.75085948449154	0.127006096144302
	0.721556644154986	0.937694258680072
2745	GLRX	glutaredoxin	6508	-0.099274376492182
5.9597484359221	0.126955693691002	0.721609600621736
0.937694258680072
1119	CHKA	choline kinase alpha	3877	0.0922217950340162
	4.01734271940157	0.12693041435886
0.721636165379332	0.937694258680072
541471	MIR4435-2HG	MIR4435-2 host gene	1516
0.109499973432732	4.02884498657935	0.126905215163717
	0.721662648892133	0.937694258680072
80318	GKAP1	G kinase anchoring protein 1	3381
0.116820561565834	0.710148872285684	0.126859538970609
	0.721710660600215	0.937694258680072
984	CDK11B	cyclin dependent kinase 11B
3488	-0.0527411541981186	5.66332593500218
0.126840484732117	0.72173069199899	0.937694258680072
23562	CLDN14	claudin 14	2853	0.149141432755755
1.62695803369936	0.126638717673448	0.721942910402673
	0.937829912200244
7738	ZNF184	zinc finger protein 184	3122	0.0704519580795146
	3.82396167248321	0.126253842877063
0.72234825014312	0.937829912200244
26121	PRPF31	pre-mRNA processing factor 31	2673
0.0480113130478301	5.41633588399513	0.126245210475308
	0.72235734952925	0.937829912200244
92610	TIFA	TRAF interacting protein with forkhead associated
domain	4250	-0.0886992401268116	2.85420942036224
0.126235908432463	0.722367155174043	0.937829912200244

22952	CYP2G1P cytochrome P450 family 2 subfamily G member 1, pseudogene	575	0.0990095880475312	1.34876277691426	0.126230855417968	0.722372481923503	0.937829912200244
55299	BRIX1	biogenesis of ribosomes BRX1	5011	-0.0639122737299231	4.14987101024859	0.126208621353264	0.722395921892109	0.937829912200244
148213	ZNF681	zinc finger protein 681	6497	-0.110520006712854	1.91291201785341	0.126145287927847	0.722462703066023	0.937829912200244
147685	C19orf18	chromosome 19 open reading frame 18	915	-0.125160056722598	0.655808881405156	0.126100880419488	0.722509539293105	0.937829912200244
56910	STARD7	StAR related lipid transfer domain containing 7	4414	0.0460761388805321	6.89007455067278	0.126053911336186	0.722559087298747	0.937829912200244
54540	FAM193B	family with sequence similarity 193 member B	5804	0.0696184725066484	5.6435884082574	0.125968683651003	0.7226490211141	0.937829912200244
55800	SCN3B	sodium voltage-gated channel beta subunit 3	9464	0.118922426888096	1.22317193547821	0.12596685901871	0.722650946874029	0.937829912200244
80726	IQCN	IQ motif containing N	4642	-0.156941891928817	0.917893919617161	0.125962652786334	0.722655386290652	0.937829912200244
1615	DARS1	aspartyl-tRNA synthetase 1	4014	-0.0484090446722405	6.00865488724343	0.12584168307793	0.722783098024395	0.937829912200244
8738	CRADD	CASP2 and RIPK1 domain containing adaptor with death domain	3820	0.0789241978558229	3.13250363716436	0.125818223761202	0.722807872802874	0.937829912200244
155054	ZNF425	zinc finger protein 425	3387	0.0789927608335345	3.48786002812147	0.125788356793487	0.722839418298982	0.937829912200244
2885	GRB2	growth factor receptor bound protein 2	4716	-0.0486810126553642	7.25503712138519	0.125699412463291	0.722933386295648	0.93786495710319
80308	FLAD1	flavin adenine dinucleotide synthetase 1	4019	-0.0544692441136956	5.07310774647427	0.125559517869466	0.723081258030678	0.937969918617865
5342	PLGLB2	plasminogen like B2	3065	0.115968929398329	0.160919125511567	0.12545913234035	0.723187424943014	0.938020766846634
163	AP2B1	adaptor related protein complex 2 subunit beta 1	7587	-0.0595768371164368	8.63161407602905	0.125328158698355	0.723326013488764	0.938113654716399
28955	DEXI	Dexi homolog	1722	0.0637996156458755	4.49228995235165	0.125199471971865	0.723462261548171	0.938188565491521
94081	SFXN1	sideroflexin 1	7572	-0.0419861427955813	5.6177004548891	0.125147100427668	0.723517732879232	0.938188565491521
9528	TMEM59	transmembrane protein 59	9149	0.0475493663858265	8.51329148595989	0.124948180773927

0.723728545349222	0.938375056438634
4731	NDUFV3	NADH:ubiquinone oxidoreductase subunit V3
6345	-0.0916544611154259	4.60822763284256
0.124702300253574	0.723989387386967	0.938573050255342
58480	RHOU	ras homolog family member U	4372
0.165419645253519	2.4951728192941 0.124677922388513
0.724015264368456	0.938573050255342
1837	DTNA	dystrobrevin alpha	12634	-0.0753717560996631
5.55133341042264	0.124532870718831
0.724169294850389	0.938636835139183
10438	C1D	C1D nuclear receptor corepressor
3581	-0.0802154103433921	3.6169152303457 0.124487584471844
0.724217405013877	0.938636835139183
440836	ODF3B	outer dense fiber of sperm tails 3B
2301	-0.167202704499315	2.08279564142208
0.124442318026681	0.724265503975462	0.938636835139183
83860	TAF3	TATA-box binding protein associated factor 3
4872	-0.0695372898637139	3.75831410446889
0.12419296635742	0.724530635146347	0.938757106524869
153090	DAB2IP	DAB2 interacting protein	8408
0.0594598507842589	8.0165032023019 0.124165206725451
0.724560169970819	0.938757106524869
2537	IFI6	interferon alpha inducible protein 6
865	-0.22672744346582	3.37285470771743
0.124100722918026	0.724628791761563	0.938757106524869
6102	RP2	RP2 activator of ARL3 GTPase
3803	-0.0498539881459964	6.24473708329015
0.124027360132999	0.724706886676133	0.938757106524869
9547	CXCL14	C-X-C motif chemokine ligand 14 1975
0.231184554931003	5.31513016540022	0.124008109398062
0.724727383443605	0.938757106524869
83930	STARD3NL	STARD3 N-terminal like	4348
0.0474495177636845	5.52120060435941	0.123949841522631
0.72478943369397	0.938757106524869
51663	ZFR	zinc finger RNA binding protein
6792	-0.0515146647299884	7.39176629945554
0.123871931135241	0.724872427139807	0.938757106524869
100533183	ZNF664-RFLNA	ZNF664-RFLNA readthrough	2998
0.103892756941373	1.80657700383205
0.123851241416516	0.724894471633682	0.938757106524869
91355	LRP5L	LDL receptor related protein 5 like	4555
0.109551675765673	1.5570807451524 0.123687555628218
0.725068948656591	0.938882402106867
8535	CBX4	chromobox 4	2880	-0.0720778769615547
4.10533605586808	0.123578919615376	0.72518481829651
0.938882402106867
2010	EMD	emerin	2020	0.0609206928912027
4.85517726095795	0.12357189597013	0.725192311586649
0.938882402106867
3619	INCENP	inner centromere protein	4291
0.0679121580200606	4.0169293092009 0.123213933809657
0.725574526622617	0.938926995952206
5985	RFC5	replication factor C subunit 5
3249	-0.0803593989892864	2.76813947348932

0.123173067152712	0.725618201747255	0.938926995952206
60436	TGIF2	TGFB induced factor homeobox 2	3784
0.0586774070792327	4.24572982584607	0.12312799126066
	0.725666384816296	0.938926995952206
64081	PBLD	phenazine biosynthesis like protein domain
containing	5172	0.0769852515922878	3.22041610388349
	0.123106841596695	0.725688995785774
0.938926995952206
79989	TTC26	tetratricopeptide repeat domain 26	4678
0.0955923016655464	2.28762852535103	0.12310379643985
	0.725692251522983	0.938926995952206
79629	OCEL1	occludin/ELL domain containing 1	1867
0.100508811552299	3.01316128171797	0.12308990017897
	0.725707109319865	0.938926995952206
6936	GCFC2	GC-rich sequence DNA-binding factor 2	5465
0.103425088321867	2.27109229814338	0.123081240546869
	0.725716368621761	0.938926995952206
65082	VPS33A	VPS33A core subunit of CORVET and HOPS complexes
	8681	-0.0382254882088391	6.4749226665015
0.122989219844866	0.725814784259775	0.938926995952206
55502	HES6	hes family bHLH transcription factor 6	2037
0.113671471338291	2.1416717398177	0.122974949395637
0.72583005012997	0.938926995952206
79087	ALG12	ALG12 alpha-1,6-mannosyltransferase
5350	-0.0700307396627412	5.2160073722867	0.122719549861557
	0.726103433170941	0.939019464518477
83608	C18orf21	chromosome 18 open reading frame 21	2158
	0.0673506357881303	3.53421365091831
0.122638127812741	0.726190655544108	0.939019464518477
56133	PCDHB2	protocadherin beta 2	4126	0.230162911083642
	1.48208159407231	0.122603505629193
0.726227753984056	0.939019464518477
9790	BMS1	BMS1 ribosome biogenesis factor
7753	-0.0426340777571313	6.07743527958724
0.1225786062262	0.726254437883673	0.939019464518477
83440	ADPGK	ADP dependent glucokinase	5931
0.0463565495552741	5.79412725712404	0.122553005609542
	0.726281876423984	0.939019464518477
63894	VIPAS39	VPS33B interacting protein, apical-basolateral
polarity regulator, spe-39 homolog	3695	0.0508506714423096
	4.63480733117817	0.122520554005253
0.726316662423231	0.939019464518477
54780	NSMCE4A	NSE4 homolog A, SMC5-SMC6 complex component	3546
	0.0662124758015012	3.57207971962402
0.122457016648672	0.726384785294808	0.939019464518477
574449	MIR492	microRNA 492
116	-0.119220801790686	-0.0315612750500356
0.122391252957057	0.726455316071991	0.939019464518477
84515	MCM8	minichromosome maintenance 8 homologous
recombination repair factor	7691	0.0946896383861821
3.52863437171998	0.122345045341135	0.726504885913239
	0.939019464518477
643246	MAP1LC3B2	microtubule associated protein 1 light chain
3 beta 2	1212	-0.0513656503275637	5.13849773331381

0.122234045837854 0.726624005069964
0.939086772376359
53916 RAB4B RAB4B, member RAS oncogene family 3931
0.0535810369386117 4.29364109584231 0.122130522131865
0.726735156286816 0.939143232275869
10072 DPP3 dipeptidyl peptidase 3 3977 -0.0469919991811846
6.65294910660427 0.121858496390445
0.727027476756976 0.939143232275869
284361 EMC10 ER membrane protein complex subunit 10 9976
0.0440508876474209 7.46617657142897 0.121827785984713
0.727060501327282 0.939143232275869
59338 PLEKHA1 pleckstrin homology domain containing A1
23986 -0.0542742354857328 7.22709499531744
0.121803784600459 0.727086314572673 0.939143232275869
9667 SAFB2 scaffold attachment factor B2 6222
0.0577348342175512 5.20813926051654 0.121775258534921
0.72711699777766 0.939143232275869
84935 MEDAG mesenteric estrogen dependent adipogenesis
2374 -0.132813614734813 2.85794967805029
0.121756765453333 0.727136891458999 0.939143232275869
10265 IRX5 iroquois homeobox 5 2845 0.0847528375154218
3.36168150235786 0.121681288856651
0.727218101960691 0.939143232275869
7290 HIRA histone cell cycle regulator
4421 -0.0440343584922897 5.59856077664727
0.121658041225958 0.727243121388659 0.939143232275869
5094 PCBP2 poly(rC) binding protein 2 8400
0.041826884775895 8.19853680945087 0.12163202418634
0.727271124473217 0.939143232275869
5152 PDE9A phosphodiesterase 9A 5266 0.140873590544839
2.24791666897579 0.121359908519143
0.727564213961911 0.93923039991881
9258 MFHAS1 malignant fibrous histiocytoma amplified sequence 1
6883 -0.058913749497532 6.37938851195042
0.121302590057496 0.727625997320673 0.93923039991881
79134 TMEM185B transmembrane protein 185B 2131
0.0590613303038502 4.69849931946329 0.121295894208794
0.727633215818969 0.93923039991881
10197 PSME3 proteasome activator subunit 3 4999
0.0565477635183672 7.46089044494856 0.12127824812714
0.727652240349274 0.93923039991881
8545 CGGBP1 CGG triplet repeat binding protein 1
5282 -0.0459754871279208 6.61217362713213
0.121258160759368 0.727673898759115 0.93923039991881
1742 DLG4 discs large MAGUK scaffold protein 4
4489 -0.0768431058142897 3.98790025730346
0.121059147506269 0.727888585694568 0.939420936130454
84708 LNX1 ligand of numb-protein X 1
5725 -0.126216781511745 3.02602398548904
0.120959600869845 0.727996046553324 0.939429359088837
11201 POLI DNA polymerase iota 9471 0.0672320308596202
4.09870860081205 0.120902851066361
0.728057330301757 0.939429359088837
84924 ZNF566 zinc finger protein 566 6481 0.100086956428312

2.54181023034283	0.120866755817747
0.728096317717457	0.939429359088837
23203	PMPCA	peptidase, mitochondrial processing alpha subunit
6590	0.0472400139130317	5.57714416957377
0.120744675261021	0.728228228474614	0.939442052387524
284273	ZADH2	zinc binding alcohol dehydrogenase domain containing
2	8157	0.058108834175702	4.51731865639092
0.120707410302003	0.728268508979493	0.939442052387524
5058	PAK1	p21 (RAC1) activated kinase 1	6299
0.0520330989149301	5.45836273595714	0.120667441208895
0.728311720187568	0.939442052387524
11198	SUPT16H SPT16 homolog, facilitates chromatin remodeling
subunit 7151	0.0387105423816823	6.63809761351345
0.120609446510702	0.72837443342104	0.939442052387524
103910	MYL12B	myosin light chain 12B	1761	-0.0661007417581682
9.59913736022808	0.120495530286234
0.72849766724694	0.939462119102532
8601	RGS20	regulator of G protein signaling 20	3192
0.0823146941913502	3.8196647910039 0.120352770823516
0.728652195688068	0.939462119102532
10780	ZNF234	zinc finger protein 234 5210	0.068602980864547
3.67645224608146	0.120323540642541
0.728683848244866	0.939462119102532
84171	LOXL4	lysyl oxidase like 4	3653	0.228361052134795
8.31521788507619	0.120311654528678
0.728696720621611	0.939462119102532
57589	RIC1	RIC1 homolog, RAB6A GEF complex partner 1
8591	-0.0481108099437213	6.06433429616269
0.120153896237262	0.728867636587498	0.939462119102532
22858	ICK	intestinal cell kinase	6259	-0.0605813925910093
4.82754457909692	0.120092462663006
0.728934227969331	0.939462119102532
51699	VPS29	VPS29 retromer complex component
3211	-0.0540694423838749	5.84452952428528
0.120068364822728	0.728960354212793	0.939462119102532
8284	KDM5D	lysine demethylase 5D	9463	-0.0651839675730136
5.50214751599211	0.1200401389071 0.728990959741285
0.939462119102532
23523	CABIN1	calcineurin binding protein 1	9388
0.0490418352895505	5.94921392864441	0.120037676630062
0.728993629794061	0.939462119102532
23185	LARP4B	La ribonucleoprotein 4B 12836	0.0515835986099489
6.99547920980534	0.119762485675437
0.729292235833871	0.939707281895068
84938	ATG4C	autophagy related 4C cysteine peptidase
3110	-0.0815576026699338	2.82354871647139
0.119721271463933	0.72933698989089	0.939707281895068
6605	SMARCE1 SWI/SNF related, matrix associated, actin dependent
regulator of chromatin, subfamily e, member 1	11022
0.0419803685433869	6.79159391936422	0.119676944413882
0.72938513377334	0.939707281895068
89801	PPP1R3F protein phosphatase 1 regulatory subunit 3F	4173
0.120048755811327	3.23056611078001
0.119594185612266	0.729475045343248	0.939716760512628

58516   SINHCAF SIN3-HDAC complex associated factor
5748    -0.0496480737340646     6.83546010975127
0.119546683199481       0.729526669080768       0.939716760512628
348654  GEN1    GEN1 Holliday junction 5' flap endonuclease
10235   0.0858187841406572      4.00623483662024
0.119383698987868       0.729703881192798       0.939756033340186
4356    MPP3    membrane palmitoylated protein 3
4491    -0.0881918419435169     2.73941638081383
0.119303671720971       0.729790944103625       0.939756033340186
92106   OXNAD1  oxidoreductase NAD binding domain containing 1
5659    -0.0748262894826069     2.98913177860239
0.119274147825622       0.729823071862496       0.939756033340186
100133315       LOC100133315    transient receptor potential cation
channel, subfamily C, member 2-like     3651    -0.124606654801531
        0.612723817020789       0.119115612180919
0.729995665735816       0.939756033340186
55786   ZNF415  zinc finger protein 415 3780    -0.0735507232884616
        2.90442734738455        0.119059526191043
0.730056755958188       0.939756033340186
84182   MINDY4  MINDY lysine 48 deubiquitinase 4        7336
0.089779467337132       1.53542317321383        0.119048599803861
        0.730068659118422       0.939756033340186
9570    GOSR2   golgi SNAP receptor complex member 2
2758    -0.053205520631119      4.59864181822799
0.119001695369734       0.730119763557073       0.939756033340186
6120    RPE     ribulose-5-phosphate-3-epimerase
3522    -0.0609505222793228     5.14626542913233
0.118965317405582       0.730159406705281       0.939756033340186
5977    DPF2    double PHD fingers 2    7715    -0.0460401772156869
        5.87868441036115        0.118963869631521
0.730160984567805       0.939756033340186
124093  CCDC78  coiled-coil domain containing 78
4147    -0.103588744626748      0.237514105615412
0.118879232484737       0.730253245389556       0.939788424124541
5564    PRKAB1  protein kinase AMP-activated non-catalytic subunit
beta 1  6076    0.0785081829307034      5.58486174867123
0.118804206301359       0.730335060272087       0.939807366948196
389125  MUSTN1  musculoskeletal, embryonic nuclear protein 1    6252
        0.0465507535323214      5.16479223784318
0.11842537587934        0.730748610976476       0.94022156836982
51247   PAIP2   poly(A) binding protein interacting protein 2   3158
        0.0441958760567537      5.76734673928724
0.118357603656417       0.730822672590644       0.94022156836982
152559  PAQR3   progestin and adipoQ receptor family member 3
12403   0.0804586561248259      4.21453307936169
0.118213194957507       0.730980561827127       0.94022156836982
9815    GIT2    GIT ArfGAP 2    13542   -0.0436033862879639
5.94023015532526        0.118170108512512       0.731027691273507
        0.94022156836982
203     AK1     adenylate kinase 1      3066    0.0503249435989376
        5.49193690986642        0.118145152880658
0.731054993002182       0.94022156836982
667     DST     dystonin        42341   0.0706568433511556
9.31710911142709        0.118085054861353       0.731120754113135

0.94022156836982
91574 C12orf65 chromosome 12 open reading frame 65 7090
0.100099752156115 2.06590169152689
0.118079515145435 0.731126816783328 0.94022156836982
338376 IFNE interferon epsilon 1475 0.16420325653881
1.21634365865914 0.117864036847571
0.731362759918904 0.940438646670714
100500842 MIR3652 microRNA 3652 131 -0.137018237842191
0.888570910281889 0.117749029149442
0.731488789049368 0.940460357620092
7355 SLC35A2 solute carrier family 35 member A2
3745 -0.0500919753027587 5.93361192925949
0.117726095702154 0.73151392848368 0.940460357620092
54847 SIDT1 SID1 transmembrane family member 1
5095 -0.136780367514011 3.30755979096193
0.117450240194183 0.731816533447918 0.940736117556612
286205 SCAI suppressor of cancer cell invasion
13014 -0.0947764070656628 3.27450792038301
0.117408145195578 0.73186274527125 0.940736117556612
80232 WDR26 WD repeat domain 26 10481 -0.0399342341815928
7.50183275719055 0.11731282926662
0.731968017274261 0.940785099831214
81 ACTN4 actinin alpha 4 5264 -0.0440739420620671
9.72156948871425 0.117199806149756 0.732091595938253
0.94085760017498
83932 SPRTN SprT-like N-terminal domain
5906 -0.0526927991887182 4.64945841999347
0.116820138160847 0.732509228518839 0.941157170889005
5834 PYGB glycogen phosphorylase B
4474 -0.0659553446661919 7.72744599860601
0.116799771462752 0.732531653175615 0.941157170889005
8992 ATP6V0E1 ATPase H+ transporting V0 subunit e1 1623
0.0577811536251381 8.04145323536207
0.116782304532337 0.732550886796236 0.941157170889005
101927768 HDAC2-AS2 HDAC2 and HS3ST5 antisense RNA 2
6637 0.112065983689354 0.463025959846583
0.116743644455156 0.732593462854489 0.941157170889005
112479 ERI2 ERI1 exoribonuclease family member 2 5832
0.0526404088704411 4.07606894612907 0.116387947454765
0.732985559293063 0.941425227512871
10443 N4BP2L2 NEDD4 binding protein 2 like 2
16960 -0.0673139772786604 6.17011657708028
0.116386998561552 0.732986606183103 0.941425227512871
54823 SWT1 SWT1 RNA endoribonuclease homolog
4054 -0.0892863997975845 2.86267560699972
0.116371459826107 0.733003750357348 0.941425227512871
6302 TSPAN31 tetraspanin 31 5385 -0.0449421457839924
5.56123648767931 0.1162127039903 0.733178981871829
0.941563949672569
9278 ZBTB22 zinc finger and BTB domain containing 22 2728
0.0519946777478758 4.413854199187 0.116127698695676
0.733272863938397 0.941594462869685
101180901 IPO11-LRRC70 IPO11-LRRC70 readthrough 1823
-0.076693536753270 7 3.3233870379065 0.116069476463176

0.733337188223416	0.941594462869685
84206	MEX3B	mex-3 RNA binding family member B	3528
0.115804129078443	3.00892514698883	0.115973900124042
0.733442820608909	0.941643775276718
22880	MORC2	MORC family CW-type zinc finger 2
6258	-0.042788699677491	6.02321512273303
0.115890415521572	0.733535128780577	0.941675973680648
55151	TMEM38B transmembrane protein 38B	4511
0.0993583289349223	2.81819206054483	0.115824921360682
0.733607571056753	0.941682665670083
10445	MCRS1	microspherule protein 1 3269	-0.0407121425491895
5.69865213361015	0.115635306956535
0.73381743013699	0.941829691021917
83693	HSDL1	hydroxysteroid dehydrogenase like 1
3896	-0.0603492582657477	5.57120246246159
0.115594994889761	0.733862070965479	0.941829691021917
9530	BAG4	BCL2 associated athanogene 4
4809	-0.0596066848192459	4.70880410646133
0.115539238072387	0.733923829335634	0.941829691021917
4245	MGAT1	alpha-1,3-mannosyl-glycoprotein 2-beta-N-acetylglucosaminyltransferase	11534	0.0413918312364755
7.18305610860562	0.115394188762664	0.734084569376676
0.941949666843083
526	ATP6V1B2	ATPase H+ transporting V1 subunit B2
7816	-0.0465399258147492	7.25757177525133
0.115253374206901	0.734240724399766	0.941991218350499
5672	PSG4	pregnancy specific beta-1-glycoprotein 4
6274	-0.277618053116612	5.48805601407483
0.115199382921631	0.734300625760567	0.941991218350499
388753	COA6	cytochrome c oxidase assembly factor 6	1257
0.0716190594931601	3.59909303019543	0.115117930883424
0.734391023454014	0.941991218350499
10623	POLR3C	RNA polymerase III subunit C	2793
0.0521086129795939	4.65133098304168	0.115111995725537
0.734397611847908	0.941991218350499
26136	TES	testin LIM domain protein	4280
0.0517503721690866	8.39570801164489	0.115061917981933
0.734453208796016	0.941991218350499
54820	NDE1	nudE neurodevelopment protein 1
5525	-0.0595602982298392	5.30328879978032
0.114884930423596	0.734649810801517	0.942110004700669
7107	GPR137B G protein-coupled receptor 137B 2309
0.0541672073622239	5.29059638533488	0.114857456875727
0.73468034420971	0.942110004700669
51455	REV1	REV1 DNA directed polymerase
5679	-0.0530029260480771	5.23647439551592
0.114792343291516	0.734752726067239	0.94211657213693
1410	CRYAB	crystallin alpha B	3034	0.15930851012884
2.44077499162145	0.114695174119223
0.734860784273228	0.942168879205044
440348	NPIPB15 nuclear pore complex interacting protein family member B15	1428	0.242971299330274	1.41685309120171
0.114422529010119	0.73516425541325
0.942259622440454

57804	POLD4	DNA polymerase delta 4, accessory subunit
3110	-0.0473581656853308	5.29319567329785
0.114409028583761	0.735179292701299	0.942259622440454
112703	FAM71E1 family with sequence similarity 71 member E1	1566
	0.132766650984173	0.654708993215503
0.114187336815843	0.735426362971432	0.942259622440454
57498	KIDINS220	kinase D interacting substrate 220
10771	-0.0568887977359608	7.64781924915129
0.114165341313822	0.735450891009479	0.942259622440454
642778	NPIPA3	nuclear pore complex interacting protein family
member A3	1155	-0.0922746322805049	2.21698434847423
	0.114121147613144	0.735500181080147
0.942259622440454
5873	RAB27A	RAB27A, member RAS oncogene family	5077
0.0556353269573879	4.7667956049767 0.114093568589836
0.735530945875136	0.942259622440454
54810	GIPC2	GIPC PDZ domain containing family member 2
3973	-0.166988632031898	-0.369067765274581
0.114068465079931	0.735558952801165	0.942259622440454
537	ATP6AP1 ATPase H+ transporting accessory protein 1
4884	-0.0464386188823368	7.42144707275221
0.114019251449202	0.735613868335636	0.942259622440454
84627	ZNF469	zinc finger protein 469 13203	0.109377438701445
	4.79005787165577	0.113897855115866
0.735749386154427	0.942259622440454
85369	STRIP1	striatin interacting protein 1	7352
0.0415938606400481	6.1156043847819 0.113858819763854
0.735792979403796	0.942259622440454
5612	THAP12	THAP domain containing 12
7727	-0.0480707646570228	4.90250512652787
0.113685225145028	0.73598694438942	0.942259622440454
23247	KIAA0556	KIAA0556	9784	-0.049927964436479
	5.64378067039817	0.113582581113219
0.736101710728863	0.942259622440454
50487	PLA2G3	phospholipase A2 group III
2697	-0.147203584037869	0.834608468387464
0.113540374177067	0.736148919078295	0.942259622440454
59345	GNB4	G protein subunit beta 4	7035
0.124200715816791	4.10239384402392	0.113514676506544
	0.736177666642238	0.942259622440454
5524	PTPA	protein phosphatase 2 phosphatase activator
5624	-0.0485477594705038	7.40320631462922
0.113454126737338	0.736245417009123	0.942259622440454
27315	PGAP2	post-GPI attachment to proteins 2
3781	-0.0548107962333858	4.28805726699505
0.113293614915428	0.736425114415526	0.942259622440454
6193	RPS5	ribosomal protein S5	2815	0.0531296557474019
	7.09524952976291	0.113291996531927
0.736426926961878	0.942259622440454
1965	EIF2S1	eukaryotic translation initiation factor 2 subunit
alpha	5651	-0.0441449569823407	6.72547499920772
0.11315809471597	0.736576943381029	0.942259622440454
11334	TUSC2	tumor suppressor 2, mitochondrial calcium regulator
	1897	0.055836350233723	4.86755043848616

0.113155230019116	0.736580153913122	0.942259622440454
23603	CORO1C	coronin 1C	6532	-0.0623104958909764
8.18755496132876	0.113142433874501	0.736594495401041
	0.942259622440454
1781	DYNC1I2 dynein cytoplasmic 1 intermediate chain 2
4695	-0.0482407695006761	7.33303277410157
0.113135593494526	0.736602162238815	0.942259622440454
9739	SETD1A	SET domain containing 1A, histone lysine
methyltransferase	6903	0.0691629987031843
5.39781625766612	0.113117666570231	0.736622256326362
	0.942259622440454
84221	SPATC1L spermatogenesis and centriole associated 1 like
2333	-0.135236016514418	1.17256023759213
0.113093163072888	0.73664972488772	0.942259622440454
26263	FBXO22	F-box protein 22	11837	0.0479470196784119
	5.27263973642902	0.113041785258034
0.736707330468948	0.942259622440454
3163	HMOX2	heme oxygenase 2	3793	0.0784428487807406
	6.73570448712857	0.112985630114508
0.736770309131659	0.942259622440454
6712	SPTBN2	spectrin beta, non-erythrocytic 2
9137	-0.0689350862542974	7.19379568626034
0.112812816484361	0.736964231033332	0.942259622440454
22995	CEP152	centrosomal protein 152 8168	-0.0886599497949816
	3.12665400224846	0.1127543233948 0.737029906183003
	0.942259622440454
79897	RPP21	ribonuclease P/MRP subunit p21	1606
0.0714442091657439	3.48052200999792	0.112685704587225
	0.737106974501061	0.942259622440454
55208	DCUN1D2 defective in cullin neddylation 1 domain containing
2	4266	0.0715678723726982	4.4521265125196
0.112626092897468	0.737173947788409	0.942259622440454
81853	TMEM14B transmembrane protein 14B	4815
0.0615283615682101	5.00781490609024	0.112537232162865
	0.737273818776776	0.942259622440454
79414	LRFN3	leucine rich repeat and fibronectin type III domain
containing 3	3689	0.0676584440869178	3.79279769073735
	0.112535403250743	0.737275874760172
0.942259622440454
100534592	URGCP-MRPS24	URGCP-MRPS24 readthrough	971
	0.0611561075479895	5.10256696376548
0.112512601574926	0.737301508972124	0.942259622440454
5833	PCYT2	phosphate cytidylyltransferase 2, ethanolamine
6897	-0.0569821794799787	4.92524655611705
0.112507029081574	0.737307774147365	0.942259622440454
29928	TIMM22	translocase of inner mitochondrial membrane 22
3194	-0.0508923489121432	4.8079772200517 0.112501718277969
	0.737313745264486	0.942259622440454
107303344	SETDB2-PHF11	SETDB2-PHF11 readthrough	2613
	0.0822321300185072	2.6339925114194 0.112484421694583
	0.737333193488433	0.942259622440454
55544	RBM38	RNA binding motif protein 38
2686	-0.0599473834217431	5.06596359406169
0.112466538130858	0.7373533034604 0.942259622440454

55737 VPS35 VPS35 retromer complex component
9733 -0.0394353528700623 7.86800936409261
0.112215825149718 0.737635416323344 0.94236142377535
63901 FAM111A family with sequence similarity 111 member A
7221 -0.0624752773551688 4.98540911263674
0.112207288576926 0.737645028204445 0.94236142377535
100507436 MICA MHC class I polypeptide-related sequence A
2386 -0.0759254129254495 5.48238503826735
0.112174360354777 0.73768210803598 0.94236142377535
4869 NPM1 nucleophosmin 1 3828 -0.064366121287939
7.2418231523686 0.111975657939183 0.737905991536217
0.94236142377535
91687 CENPL centromere protein L 4914 0.0589198182504171
3.49243498640872 0.111912059060579
0.737977696809042 0.94236142377535
100132891 MSC-AS1 MSC antisense RNA 1
6065 -0.175740740791141 -0.11049667292642
0.111777560598147 0.738129413267724 0.94236142377535
1876 E2F6 E2F transcription factor 6 4067
0.0558862637992573 3.77678555940459 0.111648254210155
0.738275368691711 0.94236142377535
4254 KITLG KIT ligand 6040 0.0635300333050295
4.74485948779484 0.111486442149101 0.738458147520099
0.94236142377535
22838 RNF44 ring finger protein 44 5660 -0.0527337040695713
6.64499257543176 0.111481622391739
0.738463594057083 0.94236142377535
80709 AKNA AT-hook transcription factor
10601 -0.10137599376182 3.94352290010778
0.111418688825779 0.738534723757192 0.94236142377535
817 CAMK2D calcium/calmodulin dependent protein kinase II delta
6673 -0.0713254468641556 4.63869906005156
0.111409519415294 0.738545089206439 0.94236142377535
8487 GEMIN2 gem nuclear organelle associated protein 2
2123 -0.0745093458178278 2.56683070459274
0.111370259041874 0.738589475982336 0.94236142377535
197342 EME2 essential meiotic structure-specific endonuclease subunit 2 6909 0.0632331469382227 5.37104283411518
0.1113142807811 0.738652778614451 0.94236142377535
646817 SETSIP SET like protein 909 -0.088469620202186
1.81920512294955 0.11128175826928
0.738689564597907 0.94236142377535
23585 TMEM50A transmembrane protein 50A 3410
0.0423887382227387 7.40676430675106 0.111198735996794
0.738783497600615 0.94236142377535
9045 RPL14 ribosomal protein L14 8626 0.0553422754524378
7.04978967918093 0.111198657725254
0.738783586176929 0.94236142377535
3092 HIP1 huntingtin interacting protein 1 8953
0.0955797343717492 4.11074956215116 0.111168983227458
0.738817169942389 0.94236142377535
100616387 MIR3064 microRNA 3064 67 -0.0962131943467007
2.28886303349181 0.111125281205345
0.738866638264112 0.94236142377535

8833	GMPS	guanine monophosphate synthase
8885	-0.043273138302475	5.17527351083509
0.111022934314691	0.738982531710703	0.94236142377535
143570	XRRA1	X-ray radiation resistance associated 1
9086	-0.159032587556038	3.47607473146843
0.111014243605581	0.738992375448743	0.94236142377535
9556	ATP5MPL ATP synthase membrane subunit 6.8PL
3415	-0.0691504076727424	5.09281305035721
0.110880741158471	0.739143643992547	0.94236142377535
401207	C5orf63 chromosome 5 open reading frame 63
7190	-0.0855445417468246	1.781027419651	0.110849696958017
	0.739178833955518	0.94236142377535
118490	MSS51	MSS51 mitochondrial translational activator
2423	-0.12297398412858	0.0848839756588758
0.110731312404202	0.739313078300881	0.94236142377535
8078	USP5	ubiquitin specific peptidase 5	3738
0.0441584590670701	6.77549679947513	0.110719773443375
	0.739326167380504	0.94236142377535
2521	FUS	FUS RNA binding protein 10283	-0.0426749693075785
	7.45166893537782	0.110718303250227
0.739327835130598	0.94236142377535
55808	ST6GALNAC1	ST6 N-acetylgalactosaminide alpha-2,6-
sialyltransferase 1	3448	-0.118381265608302
5.64657945779621	0.110611690199534	0.73944880708961
	0.94236142377535
10489	LRRC41	leucine rich repeat containing 41	8606
0.0380165012631459	7.17260802425026	0.110545808598935
	0.739523594157326	0.94236142377535
10051	SMC4	structural maintenance of chromosomes 4
8457	-0.0559729213101101	4.16874814729429
0.11052455393755	0.739547727172352	0.94236142377535
11218	DDX20	DEAD-box helicase 20	6019	-0.048375598233885
	4.85229270675247	0.110511103832237
0.739563000052262	0.94236142377535
23457	ABCB9	ATP binding cassette subfamily B member 9
13039	0.0962533275897602	2.66625069633438
0.110497978909413	0.739577904675589	0.94236142377535
26011	TENM4	teneurin transmembrane protein 4
15642	-0.118577309069461	3.8967967364331 0.11042723311256
	0.739658260326238	0.94236142377535
6830	SUPT6H	SPT6 homolog, histone chaperone and transcription
elongation factor	10620	-0.04515224784719
7.68174218265462	0.110352807406116	0.739742826609449
	0.94236142377535
399818	EEF1AKMT2	EEF1A lysine methyltransferase 2
3887	-0.071630566794275	3.07845837760055
0.11031471800293	0.739786117981339	0.94236142377535
57477	SHROOM4 shroom family member 4	10333	-0.0827933073763058
	4.72277284175968	0.110294490263676
0.739809111649127	0.94236142377535
78995	HROB	homologous recombination factor with OB-fold	2918
	0.112966342727584	0.251981629620706
0.110271703482264	0.739835017085122	0.94236142377535
136242	PRSS37	serine protease 37	2203	-0.440447621565191

2.55697018013084	0.110254153620131
0.739854970890093	0.94236142377535
55268	ECHDC2	enoyl-CoA hydratase domain containing 2	7916
0.0553404278032456	5.2980196680845	0.110102437147209
0.740027542715908	0.942495525624816
7633	ZNF79	zinc finger protein 79	2302	0.0768593452498262
3.12374451660711	0.109942918263606
0.740209131897773	0.942614034380683
54765	TRIM44	tripartite motif containing 44
13077	-0.0405917979628644	6.83875788163794
0.109902482727774	0.740255185135112	0.942614034380683
100129917	LOC100129917	uncharacterized LOC100129917	1575
0.105581513559793	-0.143737320465794
0.109547212738753	0.740660217567653	0.942805805866068
200895	DHFR2	dihydrofolate reductase 2	4659
0.0824315115143054	2.83127311183206	0.1094859704103
0.740730111677716	0.942805805866068
160857	CCDC122	coiled-coil domain containing 122
4773	-0.1181528117361	1.37687929605933
0.109343002771083	0.740893360993353	0.942805805866068
3692	EIF6	eukaryotic translation initiation factor 6
1882	-0.0511400832248398	7.57715493067913
0.109320046663882	0.740919584729666	0.942805805866068
9070	ASH2L	ASH2 like, histone lysine methyltransferase complex subunit	5910	0.038904062048667	5.77538520164538
0.109271316711095	0.740975261149629	0.942805805866068
25764	HYPK	huntingtin interacting protein K
6393	-0.0627139241632953	4.66652221162236
0.10924862797395	0.741001188870676	0.942805805866068
124975	GGT6	gamma-glutamyltransferase 6	3161
0.13284650023705	4.77569157497274	0.10913954030255
0.741125891325441	0.942805805866068
7419	VDAC3	voltage dependent anion channel 3
3141	-0.0457792056960262	6.41008508791964
0.109100779344651	0.741170217160958	0.942805805866068
1155	TBCB	tubulin folding cofactor B	3220
0.0419918856771606	5.99363747857946	0.10909290697295
0.741179220829256	0.942805805866068
9520	NPEPPS	aminopeptidase puromycin sensitive
11514	-0.0476777188917328	8.49429307609976
0.109069910665994	0.741205523876641	0.942805805866068
286187	PPP1R42	protein phosphatase 1 regulatory subunit 42	2239
0.0688223479791073	3.26487569438348
0.10905255735037	0.741225374538077	0.942805805866068
389362	PSMG4	proteasome assembly chaperone 4	8002
0.117567182787383	1.99835247347771	0.10902004328671
0.741262572473273	0.942805805866068
148046	CIRBP-AS1	CIRBP antisense RNA 1
994	-0.121260759150398	0.467627170815856
0.108984377832455	0.741303382853884	0.942805805866068
55234	SMU1	SMU1 DNA replication regulator and spliceosomal factor	7122	0.0419392923751495	5.99942364484994
0.108945286283912	0.741348122068171	0.942805805866068
339524	LINC01140	long intergenic non-protein coding RNA 1140

7562	-0.0790702858922735	3.03171601547867
0.10878919448601	0.741526853671286	0.942909538196829
51506	UFC1	ubiquitin-fold modifier conjugating enzyme 1	2274
	0.0447697232948927	6.30873678229399
0.108756487070401	0.741564323036786	0.942909538196829
8030	CCDC6	coiled-coil domain containing 6
7345	-0.0489245601147198	7.24840990035639
0.108676750236391	0.74165569513113	20.942914804063645
219670	ENKUR	enkurin, TRPC channel interacting protein
4076	-0.145334015378664	1.00085395890178
0.108635395299268	0.741703099248464	0.942914804063645
153918	ZC2HC1B zinc finger C2HC-type containing 1B	1711
0.0779790068394952	3.1866698327441 0.108387679970457
0.741987258685718	0.943116295265963
6601	SMARCC2 SWI/SNF related, matrix associated, actin dependent regulator of chromatin subfamily c member 2	8555
0.054500124317058	6.88475249307313	0.108348538641458
	0.74203219144108	0.943116295265963
8395	PIP5K1B phosphatidylinositol-4-phosphate 5-kinase type 1 beta	4379	-0.194894920831755	1.50510447499382
0.108321192656241	0.742063588928834	0.943116295265963
55147	RBM23	RNA binding motif protein 23	14022
0.0436633989750254	6.18628069542214	0.108230247750697
	0.742168039576651	0.943163466734726
100528019	HSPB2-C11orf52	HSPB2-C11orf52 readthrough (NMD candidate)	1662	0.0797094279306526	2.78027240245577
	0.108095368853572	0.742323038207521
0.943270431083548
91894	C11orf52	chromosome 11 open reading frame 52	1477
	0.0802549368175007	2.78625277415614
0.107923749286031	0.742520413066007	0.943270431083548
102465976	MIR6807 microRNA 6807	92	0.0852066652064476
	1.73634416737265	0.107912383245906
0.742533490967539	0.943270431083548
65999	LRRC61	leucine rich repeat containing 61	2735
0.1559362946304 4.60789369436031	0.107865250315754
0.74258773082435	0.943270431083548
51307	FAM53C	family with sequence similarity 53 member C	6615
	0.0501148290351573	6.48112657508809
0.107762641050527	0.742705857419392	0.943270431083548
55082	ARGLU1	arginine and glutamate rich 1
4074	-0.0460650649485803	5.92260002889517
0.10775577207927	0.742713767391609	0.943270431083548
8111	GPR68	G protein-coupled receptor 68
3278	-0.0875854696561352	3.95099248019148
0.107747226449806	0.742723608511491	0.943270431083548
55520	ELAC1	elaC ribonuclease Z 1	4162	-0.0729441093531695
	2.76022982115071	0.107685120996329
0.742795141946292	0.9432757527871
10980	COPS6	COP9 signalosome subunit 6	2807
0.0496400285183301	5.56372461342	0.107243596943079
0.743304351877161	0.943834598310329
10135	NAMPT	nicotinamide phosphoribosyltransferase
9049	-0.0647703804241359	6.45402358230493

0.107186766705297	0.743369978479299	0.943834598310329
201255	LRRC45	leucine rich repeat containing 45	3606
0.0792871195403411	3.23082780404879	0.10712509871782
	0.743441213428779	0.943839488488798
441733	PRKXP1	PRKX pseudogene 1	2492	-0.19721604682959
	1.13236226295009	0.106956772560057
0.743635768752904	0.943955406173613
80789	INTS5	integrator complex subunit 5	3556
0.0447506096013989	5.14376014031952	0.106845727465732
	0.743764210071563	0.943955406173613
64881	PCDH20	protocadherin 20	6548	-0.12009124259164
	2.24200618052947	0.106814190113165
0.743800701506319	0.943955406173613
100306951	PITPNA-AS1	PITPNA antisense RNA 1	542
0.105383239594011	0.758382783960622	0.10681299372094
	0.743802085952607	0.943955406173613
10081	PDCD7	programmed cell death 7	3263	-0.0571168809109609
	4.23594703545966	0.106362022426836
0.744324555386117	0.944232838451701
153768	PRELID2	PRELI domain containing 2
5160	-0.0884972519751998	1.78339515866194
0.10636040237756	0.744326434489207	0.944232838451701
92259	MRPS36	mitochondrial ribosomal protein S36
1530	-0.0567802324337902	4.24978527885443
0.106266547662724	0.744435324072777	0.944232838451701
84365	NIFK	nucleolar protein interacting with the FHA domain of MKI67	2447	-0.0575426886478847	4.92092623023372
0.105896764777619	0.744864862047203	0.944232838451701
55164	SHQ1	SHQ1, H/ACA ribonucleoprotein assembly factor	5841
	0.0524415243683335	3.91777147706016
0.105872664313068	0.744892885798611	0.944232838451701
84259	DCUN1D5	defective in cullin neddylation 1 domain containing 5	1700	-0.0635284402837475	4.80826845945852
0.105872657945696	0.744892893202987	0.944232838451701
56413	LTB4R2	leukotriene B4 receptor 2
1705	-0.0813544537949209	1.96935357151549
0.105814615235625	0.744960399101464	0.944232838451701
114879	OSBPL5	oxysterol binding protein like 5
7466	-0.0736593939768292	3.63917320822695
0.105795799342365	0.744982287101308	0.944232838451701
51399	TRAPPC4	trafficking protein particle complex 4	5140
0.0775669098067947	5.36834703407741	0.105691235539126
	0.745103962467483	0.944232838451701
7627	ZNF75A	zinc finger protein 75a	4758	-0.0576796280754426
	4.07426045414127	0.105664419534214
0.745135177554952	0.944232838451701
51067	YARS2	tyrosyl-tRNA synthetase 2	2152
0.0502453359177324	4.62424271254261	0.105656633473225
	0.745144241715518	0.944232838451701
51016	EMC9	ER membrane protein complex subunit 9
1350	-0.0894683036734011	1.58460511406633
0.105609203355014	0.745199465573018	0.944232838451701
79956	ERMP1	endoplasmic reticulum metallopeptidase 1
6528	-0.0544728666497451	6.23595764607299

0.105600715686176	0.745209349385779	0.944232838451701
54455	FBXO42	F-box protein 42	8984	-0.0404977852100465
	5.36816343130382	0.105564348945027
0.745251703095942	0.944232838451701
10686	CLDN16	claudin 16	3583	0.0863190801803582
2.17873536632536	0.105558789841473	0.745258178091983
	0.944232838451701
124961	ZFP3	ZFP3 zinc finger protein
5206	-0.102397742754357	2.77742713286965
0.105467282263437	0.745364789140134	0.944232838451701
84314	TMEM107 transmembrane protein 107
3163	-0.093757900541854	1.75241316643802
0.105413327947842	0.745427672662343	0.944232838451701
84108	PCGF6	polycomb group ring finger 6	2613
0.0702249932551169	2.80463466524621	0.105385270336587
	0.745460380718294	0.944232838451701
55330	BLOC1S4 biogenesis of lysosomal organelles complex 1 subunit
4	1617	0.0690885510901871	3.31136277121639
0.105325873198144	0.745529638594085	0.944232838451701
51520	LARS1	leucyl-tRNA synthetase 1
8118	-0.0530861581312867	6.6418597356076 0.105294619506139
	0.745566089493704	0.944232838451701
9252	RPS6KA5 ribosomal protein S6 kinase A5	27404
0.114130890825868	1.85497342961402	0.105231398163229
	0.745639842261963	0.944232838451701
65991	FUNDC2	FUN14 domain containing 2	10252
0.0487266369027343	4.94810489387099	0.105225438414614
	0.745646796049629	0.944232838451701
347918	EP400P1 EP400 pseudogene 1	12647	0.0733721959768195
	3.64574183364067	0.105221585881019
0.745651291271254	0.944232838451701
4552	MTRR	5-methyltetrahydrofolate-homocysteine
methyltransferase reductase	5788	-0.0497801758802843
4.26248554366519	0.105161110670195	0.745721867011139
	0.944232838451701
79822	ARHGAP28	Rho GTPase activating protein 28
10009	-0.111001254259628	1.11228674623265
0.10512850772076	0.745759924588057	0.944232838451701
7728	ZNF175	zinc finger protein 175 6587	-0.0655099767197641
	3.55085485448009	0.105116972481334
0.745773391289439	0.944232838451701
64506	CPEB1	cytoplasmic polyadenylation element binding protein
1	7878	0.0922231154975408	2.8447998887753
0.104951488548886	0.74596667382409	0.944310885509521
90204	ZSWIM1	zinc finger SWIM-type containing 1	2769
0.0644779046564596	3.32415613726645	0.104948754547992
	0.74596986850595	0.944310885509521
57456	KIAA1143	KIAA1143	5259	-0.0463332899595881
	4.78024101330532	0.104777489410454
0.746170083658813	0.944375679085262
1760	DMPK	DM1 protein kinase	5721	-0.0583160531732314
	5.69193949804288	0.104637465322739
0.746333911253722	0.944375679085262
9086	EIF1AY	eukaryotic translation initiation factor 1A Y-linked

4074	0.071357820286488	4.07750399910036
0.104632520055306	0.746339699409012	0.944375679085262
54039	PCBP3	poly(rC) binding protein 3
8077	-0.219736398581648	0.0730974524967165
0.104598047258275	0.746380052062775	0.944375679085262
221908	PPP1R35 protein phosphatase 1 regulatory subunit 35	1187
0.0660790010516441	3.15129423562546
0.104469958286252	0.74653005292912	0.944375679085262
147686	ZNF418	zinc finger protein 418 4501	-0.136650271966599
1.17361154873505	0.104445388465107
0.746558837462389	0.944375679085262
4016	LOXL1	lysyl oxidase like 1	3637	0.0853772749353037
4.94619380589717	0.104426005710049
0.746581547777775	0.944375679085262
1747	DLX3	distal-less homeobox 3	2715	0.147633741943191
1.16869162213392	0.104370571015531
0.746646512148958	0.944375679085262
54892	NCAPG2	non-SMC condensin II complex subunit G2 6988
0.0710399813464905	2.89349394082101	0.104364014358515
0.746654197208794	0.944375679085262
7767	ZNF224	zinc finger protein 224 4914	-0.0726927957205844
4.67972597779946	0.104223513805165
0.746818942060925	0.944375679085262
401546	C9orf152	chromosome 9 open reading frame 152
2947	-0.149772159945117	0.564942177991211
0.104221791621919	0.746820962178759	0.944375679085262
283104	SBF2-AS1	SBF2 antisense RNA 1	3858
0.0921611872778298	2.16780710998266	0.104213990325047
0.746830113316731	0.944375679085262
63920	ZBED8	zinc finger BED-type containing 8	2925
0.097845541777859	1.93835706946788	0.104115016166586
0.746946245606416	0.944437268659421
54927	CHCHD3	coiled-coil-helix-coiled-coil-helix domain
containing 3	2871	0.0502097226082158	4.99352048685855
0.103890377082823	0.747210054391711
0.944553942472782
8351	H3C4	H3 clustered histone 4	1637	0.103676359703573
2.73894375620678	0.103815096124098
0.747298532312255	0.944553942472782
284459	ZNF875	zinc finger protein 875 6275	-0.0538300059992456
4.81476242843954	0.103769965748111
0.74735159114694	0.944553942472782
84190	METTL25 methyltransferase like 25
3672	-0.0783668399353722	2.87397793417021
0.103761955467571	0.747361009996164	0.944553942472782
2235	FECH	ferrochelatase	4473	0.0470706471128647
4.86181265730103	0.103712437095704	0.747419244841591
0.944553942472782
25929	GEMIN5	gem nuclear organelle associated protein 5	5637
0.0560514252589399	4.11978946412169
0.10362880962442	0.747517627690708	0.944553942472782
10349	ABCA10	ATP binding cassette subfamily A member 10
8405	-0.135706249576659	0.689029844327346
0.103514032199964	0.74765272800003	0.944553942472782

10159	ATP6AP2 ATPase H+ transporting accessory protein 2	2573
	0.0393202927853626	6.99110407790544
0.103498723347521	0.747670753736749	0.944553942472782
51117	COQ4	coenzyme Q4	4814	0.0482164826159889
4.59938992756609	0.103440430222143	0.74773940569687
	0.944553942472782
104355426	LENG8-AS1	LENG8 antisense RNA 1	1248
0.109937390357288	0.166696235104087	0.103438836199417
	0.747741283281225	0.944553942472782
29082	CHMP4A	charged multivesicular body protein 4A
3191	-0.0578499615496462	4.80885041538056
0.10320531714931	0.748016515827797	0.944553942472782
8888	MCM3AP	minichromosome maintenance complex component 3
associated protein	8908	-0.0429528810570842
6.5525859866374 0.103199717983927	0.748023119386966
0.944553942472782
9095	TBX19	T-box transcription factor 19
3647	-0.0935485180095707	2.86573287266148
0.103053421337226	0.748195729219329	0.944553942472782
54108	CHRAC1	chromatin accessibility complex subunit 1
3301	-0.0471740836464697	4.49692031070206
0.103014034888393	0.748242222875181	0.944553942472782
6165	RPL35A	ribosomal protein L35a	2977	0.0620178533517605
	7.20698115499868	0.103008718521522
0.748248499320895	0.944553942472782
55171	TBCCD1	TBCC domain containing 1
3405	-0.0712235800220667	3.24315772529937
0.102943949564663	0.748324979207239	0.944553942472782
4621	MYH3	myosin heavy chain 3	6684	0.143321202270817
	0.135867265171496	0.102844040158818
0.748443005371769	0.944553942472782
5775	PTPN4	protein tyrosine phosphatase non-receptor type 4
	12486	-0.0761509420777382	4.21654863137921
0.102727827387669	0.748580370791318	0.944553942472782
7008	TEF	TEF transcription factor, PAR bZIP family member
	4715	0.0704003454614849	4.32042143521354
0.102645458533956	0.748677784033496	0.944553942472782
9638	FEZ1	fasciculation and elongation protein zeta 1	5937
	0.097842152397116	5.34508235978409
0.102578487124561	0.748757019311338	0.944553942472782
148189	LINC00662	long intergenic non-protein coding RNA 662
	5462	0.0748601726597484	2.70487663636999
0.102529835695008	0.748814597722673	0.944553942472782
79969	ATAT1	alpha tubulin acetyltransferase 1	4035
0.0609761625294003	3.81476539045106	0.102527254112101
	0.748817653417333	0.944553942472782
9605	VPS9D1	VPS9 domain containing 1
3428	-0.0660509725145862	4.7838884114877 0.102446971055366
	0.748912701760705	0.944553942472782
10752	CHL1	cell adhesion molecule L1 like
11861	-0.132047961441586	5.52075566361864
0.102423018020311	0.748941068078622	0.944553942472782
9191	DEDD	death effector domain containing
2769	-0.0420234594228146	5.37362878011587

0.102374694168955	0.748998306602758	0.944553942472782
6785	ELOVL4	ELOVL fatty acid elongase 4	3042
0.0650904279710724	3.92928447775467	0.10232226803487
	0.749060469891717	0.944553942472782
51805	COQ3	coenzyme Q3, methyltransferase	1458
0.0835677612967415	2.17323421298562	0.10230042347273
	0.749086307626307	0.944553942472782
29883	CNOT7	CCR4-NOT transcription complex subunit 7
8066	-0.0382385482915733	6.33625785999969
0.102255166870448	0.749139948024663	0.944553942472782
123720	WHAMM	WASP homolog associated with actin, golgi membranes
and microtubules	5123	-0.0570987236149728
4.9574972499955	0.102244847889303	0.749152180431135
0.944553942472782
1124	CHN2	chimerin 2	8803	0.133136305147817
0.105270272735437	0.102219129240678	0.749182670992006
	0.944553942472782
113115	MTFR2	mitochondrial fission regulator 2	2450
0.114119601507058	1.02336335025803	0.102198651198172
	0.749206951613095	0.944553942472782
57579	FAM135A	family with sequence similarity 135 member A
10589	0.0598656719237611	5.49702291061351
0.10218010497058	0.74922894401339	0.944553942472782
6184	RPN1	ribophorin I	3992	0.0462107043072999
8.00949987890966	0.102150666009948	0.749263857700798
	0.944553942472782
8726	EED	embryonic ectoderm development
7016	-0.0512857092771894	3.956593958992	0.101979357922134
	0.749467133835045	0.944661294489433
285605	DTWD2	DTW domain containing 2	4617	-0.0678740064034408
	2.93721183004788	0.101965236847286
0.749483898455134	0.944661294489433
5589	PRKCSH	protein kinase C substrate 80K-H	5061
0.0452943755673243	7.52833637998543	0.101614435120524
	0.749900783553647	0.945051539644329
55862	ECHDC1	ethylmalonyl-CoA decarboxylase 1	5678
0.048834482068976	5.77562435200188	0.101570412774294
	0.74995315469971	0.945051539644329
2271	FH	fumarate hydratase	2142	-0.0430869281068078
	5.26542437823383	0.101508776120864
0.750026501701653	0.945051539644329
10238	DCAF7	DDB1 and CUL4 associated factor 7	6575
0.0390969335844806	7.13198308535247	0.101477781330196
	0.75006339446177	0.945051539644329
649	BMP1	bone morphogenetic protein 1
7471	-0.0435262107532456	8.21918270298243
0.101274355385954	0.750305684305208	0.945149917852646
51637	RTRAF	RNA transcription, translation and transport factor
	8323	0.0490065664776683	6.30710303823074
0.101213020971443	0.75037878904384	0.945149917852646
1629	DBT	dihydrolipoamide branched chain transacylase E2
11303	-0.0468579389387794	5.11274221834842
0.10117975580583	0.750418448132868	0.945149917852646
201294	UNC13D	unc-13 homolog D	7752	0.0803792481376766

6.45381309123059	0.10116191268412
0.750439723853109	0.945149917852646
51283	BFAR	bifunctional apoptosis regulator	4086
0.0411936263052378	6.56039979850699	0.101129095578827
0.750478859595711	0.945149917852646
8076	MFAP5	microfibril associated protein 5	3876
0.154307739475848	1.60460473645607	0.10098783839701
0.750647394442861	0.945157023551133
4281	MID1	midline 1	9295	-0.0725750743333489
6.50852089073558	0.100975344068374	0.750662307751712
0.945157023551133
29101	SSU72	SSU72 homolog, RNA polymerase II CTD phosphatase
6925	0.0431749308834484	6.45507264001651
0.100898372147547	0.750754204331353	0.945157023551133
90529	STPG1	sperm tail PG-rich repeat containing 1	8511
0.0593688628254725	4.66331581938831	0.100825301390163
0.750841479033713	0.945157023551133
5432	POLR2C	RNA polymerase II subunit C	3940
0.0428956127727992	5.76615244568513	0.100794009023925
0.750878864850086	0.945157023551133
79156	PLEKHF1 pleckstrin homology and FYVE domain containing 1
2681	-0.123531303853234	5.11027650616095
0.100707091173561	0.750982741428626	0.945157023551133
9551	ATP5MF	ATP synthase membrane subunit f 2275
0.0666854627327797	6.16066000991021	0.100687462982471
0.751006206146903	0.945157023551133
148268	ZNF570	zinc finger protein 570 6019	-0.0619731799783842
3.58297097508623	0.100672310913034
0.751024321562369	0.945157023551133
283742	FAM98B	family with sequence similarity 98 member B
6851	-0.0433320021241131	5.02974104987928
0.100570255119927	0.751146375890252	0.945207760247571
166785	MMAA	metabolism of cobalamin associated A
7677	-0.0793823225178092	2.5816126730684 0.100525770190224
0.751199599277216	0.945207760247571
317781	DDX51	DEAD-box helicase 51	6365	-0.0583872901008808
4.14976039070137	0.100409466843679
0.751338810092402	0.945265523387315
81565	NDEL1	nudE neurodevelopment protein 1 like 1
4832	-0.0458806716360281	6.49784630692329
0.100264607920593	0.751512325002951	0.945265523387315
2953	GSTT2	glutathione S-transferase theta 2 (gene/pseudogene)
2283	0.188360768081358	2.29818740294804
0.100263090069294	0.751514143845428	0.945265523387315
8195	MKKS	McKusick-Kaufman syndrome	6953
0.0432204435800013	5.34682706723322	0.100168076880406
0.751628028394391	0.945265523387315
84902	CEP89	centrosomal protein 89	9233	0.0429443828914062
5.32487222392774	0.100008473306488
0.751819465984969	0.945265523387315
54557	SGTB	small glutamine rich tetratricopeptide repeat containing beta 5748	-0.0852219870963654	4.13374248272753
0.0999431186050828	0.75189790461562
0.945265523387315

27243	CHMP2A	charged multivesicular body protein 2A
2129	-0.0420509206851136	5.99042149897678
0.0998916608579954	0.751959683990895	0.945265523387315
145748	LYSMD4	LysM domain containing 4	5695
0.0742027138274163	3.54660932328645	0.099891602444746
	0.751959754130883	0.945265523387315
100885782	MYO16-AS1	MYO16 antisense RNA 1
580	-0.145698436385577	-0.228385597061599
0.0998374140652913	0.752024830804664	0.945265523387315
91409	CCDC74B	coiled-coil domain containing 74B
4487	-0.139549083008025	0.770499591689195
0.0997998628878776	0.752069938689591	0.945265523387315
3625	INHBB	inhibin subunit beta B	3204	0.116305757341219
	3.15337473371147	0.0997768390116072
0.75209760045038	0.945265523387315
1743	DLST	dihydrolipoamide S-succinyltransferase
4550	-0.0418976642511028	7.08185458375786
0.0997119923005698	0.752175528625463	0.945265523387315
130827	TMEM182	transmembrane protein 182	3593
0.0886744477847171	1.93788347654589	0.0996094242104659
	0.752298844593767	0.945265523387315
57038	RARS2	arginyl-tRNA synthetase 2, mitochondrial	2545
	0.055348304670942	5.13470566457825
0.099584680732816	0.752328603731094	0.945265523387315
105369340	PPP1R14B-AS1	PPP1R14B antisense RNA 1	597
	-0.0844913394981563	2.68256652984303
0.0995800243073965	0.752334204497932	0.945265523387315
10627	MYL12A	myosin light chain 12A	2510	-0.0565988000141954
	9.22649366222055	0.0994681605325241
0.752468797989568	0.945265523387315
8612	PLPP2	phospholipid phosphatase 2
7059	-0.0474201897801682	6.36679691456678
0.0994227376018486	0.752523474206594	0.945265523387315
54995	OXSM	3-oxoacyl-ACP synthase, mitochondrial	2064
0.0701877682601227	2.96714455473777	0.0993585481887265
	0.752600763319576	0.945265523387315
6566	SLC16A1	solute carrier family 16 member 1
5312	-0.0631354301033633	5.4615287138263	0.0993503506132285
	0.75261063582284	0.945265523387315
26230	TIAM2	TIAM Rac1 associated GEF 2	8936
0.0505768555507546	4.74516442426178	0.0993282709097212
	0.752637229076453	0.945265523387315
80772	CPTP	ceramide-1-phosphate transfer protein
2556	-0.0468672908067777	5.37012339779155
0.0993071289296097	0.752662695961928	0.945265523387315
92399	MRRF	mitochondrial ribosome recycling factor	11041
0.0483213990428787	4.2632619486616	0.0992121068098646
0.752777193076418	0.945324560105916
84881	RPUSD4	RNA pseudouridine synthase D4
4083	-0.0458869935638978	4.43608377884385
0.0991263492925771	0.752880578541468	0.945337464890843
9636	ISG15	ISG15 ubiquitin like modifier
942	-0.0902120587573762	3.92351159152523
0.0990768256369154	0.75294030445797	0.945337464890843

57020	VPS35L	VPS35 endosomal protein sorting factor like
9767	-0.0434749581951091	6.01024232782348
0.0990321481813154	0.752994199897217	0.945337464890843
84893	FBH1	F-box DNA helicase 1	6578	-0.0362198064214306
	6.2785570446904 0.0988955264413889	0.753159092782699
	0.945337464890843
89848	FCHSD1	FCH and double SH3 domains 1
6073	-0.0550820005594688	5.72585879579807
0.0987440161672524	0.753342101518502	0.945337464890843
9895	TECPR2	tectonin beta-propeller repeat containing 2	9384
	0.0492762866341356	5.59678563859605
0.0987433478906254	0.753342909069133	0.945337464890843
8624	PSMG1	proteasome assembly chaperone 1 2515
0.0841981006999475	3.63422802126642	0.098583445063019
	0.753536223258301	0.945337464890843
11138	TBC1D8	TBC1 domain family member 8
5620	-0.0683582000498067	4.79518280191819
0.0984919193209208	0.753646950627138	0.945337464890843
65123	INTS3	integrator complex subunit 3	10450
0.0379209845001058	6.03797625697725	0.0984027902304163
	0.753754832867176	0.945337464890843
2064	ERBB2	erb-b2 receptor tyrosine kinase 2	9917
0.0376956807423512	7.96130237506207	0.098299485440819
	0.753879940603685	0.945337464890843
10566	AKAP3	A-kinase anchoring protein 3	3668
0.151766898202964	1.50155319234687	0.0982767980402954
	0.753907425951922	0.945337464890843
27304	MOCS3	molybdenum cofactor synthesis 3
5106	-0.0526618163858709	3.91085985767271
0.0982635281204418	0.75392350381896	0.945337464890843
440894	LINC01123	long intergenic non-protein coding RNA 1123
	3295	-0.0894857160169405	3.71085731028023
0.0982127270483346	0.753985065542602	0.945337464890843
5883	RAD9A	RAD9 checkpoint clamp component A
3175	-0.0820796396310626	2.93828202202682
0.098121217120319	0.75409600317862	0.945337464890843
10068	IL18BP	interleukin 18 binding protein
4323	-0.106801103120906	3.042810306722	0.0981069931429328
	0.754113252030328	0.945337464890843
124402	UBALD1	UBA like domain containing 1	2243
0.061339920028951	3.76264882827816	0.0980549375099145
	0.754176389536671	0.945337464890843
161436	EML5	EMAP like 5	11000	-0.14704231347646
1.13282670990315	0.0979927275758143	0.754251867186489
	0.945337464890843
445577	C9orf129	chromosome 9 open reading frame 129
1154	-0.0852028093372733	1.845184445209	0.0978630511916445
	0.75440928468812	0.945337464890843
116028	RMI2	RecQ mediated genome instability 2	2208
0.119071061037616	0.750110047111606	0.0978185085647461
	0.754463382548224	0.945337464890843
27303	RBMS3	RNA binding motif single stranded interacting
protein 3	10350	0.0943578599323207	4.17409147458213
	0.0978136708070956	0.75446925891008

0.945337464890843
4521	NUDT1	nudix hydrolase 1	2093	0.0887632188824208
	1.35476848696091	0.0977509793299038
0.754545423872661	0.945337464890843
79774	GRTP1	growth hormone regulated TBC protein 1
2916	-0.114720573915209	2.28386490555633
0.0976465059923042	0.75467240993367	0.945337464890843
84923	FAM104A family with sequence similarity 104 member A
3162	-0.0395651373861087	5.24861660914943
0.0975761849930468	0.754757926237733	0.945337464890843
56061	UBFD1	ubiquitin family domain containing 1	8619
0.0336372758000181	6.44360076043908	0.0975504961036924
	0.754789174541931	0.945337464890843
8454	CUL1	cullin 1	3548	0.0360417219677213
6.47503669171719	0.0975185957533995	0.754827984835568
	0.945337464890843
11079	RER1	retention in endoplasmic reticulum sorting receptor
1	5691	0.0376147441388481	6.95820542341472
0.0975136297118375	0.754834027199378	0.945337464890843
79039	DDX54	DEAD-box helicase 54	5064	0.044777762051794
	5.56359101140519	0.097483775433779
0.754870355547292	0.945337464890843
7549	ZNF2	zinc finger protein 2	3996	0.0870928091047493
	2.62074889460249	0.0974531626403641
0.754907613239156	0.945337464890843
5688	PSMA7	proteasome 20S subunit alpha 7
1750	-0.0468676240903512	6.96331624079324
0.0973846619872916	0.754991006104982	0.945337464890843
51150	SDF4	stromal cell derived factor 4	4709
0.0366446109641795	7.13329553687073	0.0973657865374271
	0.755013990780695	0.945337464890843
7324	UBE2E1	ubiquitin conjugating enzyme E2 E1
3074	-0.048353941786017	6.51924688781158
0.0973077037306282	0.755084733679307	0.945337464890843
54535	CCHCR1	coiled-coil alpha-helical rod protein 1 5086
0.0821716797118226	2.90599441874186	0.0972416803194385
	0.755165176106898	0.945337464890843
60412	EXOC4	exocyst complex component 4	8456
0.0423135616287728	6.58800067712301	0.0972239590075037
	0.755186772720491	0.945337464890843
83549	UCK1	uridine-cytidine kinase 1	2688
0.0548667514131729	5.13858265464989	0.0971834680462687
	0.755236126362724	0.945337464890843
389741	GLIDR	glioblastoma down-regulated RNA 2606
0.0979732952966153	1.4275348201901 0.0971496850177203
0.755277312469682	0.945337464890843
339894	LINC00880	long intergenic non-protein coding RNA 880
	2018	0.150984118180886	1.15457878370776
0.0970001057749812	0.755459764381782	0.945337464890843
10038	PARP2	poly(ADP-ribose) polymerase 2
4096	-0.0550320602188783	4.18609358161979
0.0969983310380975	0.755461930074374	0.945337464890843
6390	SDHB	succinate dehydrogenase complex iron sulfur subunit
B	2462	0.0447344754108625	5.6103483254828

0.0969442129134848	0.755527980286746	0.945337464890843
10838	ZNF275	zinc finger protein 275 7661	0.0664591475261813
	3.92149373130424	0.0969106974484575
0.755568895458861	0.945337464890843
100526767	RNF103-CHMP3	RNF103-CHMP3 readthrough	1135
	-0.0382573159955099	6.35783294651979
0.0968834054733954	0.755602218821892	0.945337464890843
1070	CETN3	centrin 3	3106	0.0693906589638829
2.96277406737903	0.0968665617262872	0.755622787520862
	0.945337464890843
652968	CASTOR1 cytosolic arginine sensor for mTORC1 subunit 1
3541	-0.0534114980215651	4.87845200628392
0.0968467803515445	0.755646946001022	0.945337464890843
9695	EDEM1	ER degradation enhancing alpha-mannosidase like
protein 1	6758	-0.0556820151020072	6.72835416704483
	0.0968118969241232	0.755689554821358
0.945337464890843
285855	RPL7L1	ribosomal protein L7 like 1
7809	-0.0391434943618334	7.43747518472978
0.0967199482807075	0.755801907060222	0.945388524300588
1973	EIF4A1	eukaryotic translation initiation factor 4A1
4810	-0.0558794527103567	9.14690295276722
0.0966584355801388	0.755877102258563	0.945388524300588
23498	HAAO	3-hydroxyanthranilate 3,4-dioxygenase
2726	-0.147682433965909	0.817762566094187
0.0966128430691136	0.755932853013964	0.945388524300588
6631	SNRPC	small nuclear ribonucleoprotein polypeptide C	1277
	0.0454485048565042	5.50368213896845
0.0965152369870301	0.756052254747612	0.945434942039779
127018	LYPLAL1 lysophospholipase like 1	2966
0.0661709687633987	3.27911682495747	0.0964721671012585
	0.756104963284758	0.945434942039779
9025	RNF8	ring finger protein 8	6575	0.0424264064659609
	5.18584653265073	0.096258184807855
0.756367024338911	0.945678203152292
100287616	LOXL1-AS1	LOXL1 antisense RNA 1	4221
0.102370168170748	3.07798725276172	0.0961523584911674
	0.756496746334061	0.945746693240901
285671	RNF180	ring finger protein 180 5434	0.062797836444716
	3.32854050081683	0.0959565310015851
0.756736998378453	0.945746693240901
51116	MRPS2	mitochondrial ribosomal protein S2	2464
0.0456233812127859	4.55216245104745	0.0959107885836712
	0.756793156419581	0.945746693240901
23097	CDK19	cyclin dependent kinase 19	7492
0.0573542600175145	5.28247119872182	0.0958630865956724
	0.75685173586287	0.945746693240901
8189	SYMPK	symplekin	10628	0.0365741802247572
6.23341455727797	0.095823366733228	0.756900525209107
	0.945746693240901
199221	DZIP1L	DAZ interacting zinc finger protein 1 like	5045
	0.12630173814908	0.235390945656725
0.0957969530246432	0.75693297625919	0.945746693240901
55717	WDR11	WD repeat domain 11	9562	-0.0367246010664992

6.87285320954638 0.0956870549381676
0.757068046242682 0.945746693240901
1676 DFFA DNA fragmentation factor subunit alpha 6622
0.0357229831198664 6.27344112773273 0.0956639027449313
0.757096512227719 0.945746693240901
10874 NMU neuromedin U 975 -0.180446759107089
0.538454848343941 0.0954242006300028 0.757391451815216
0.945746693240901
7942 TFEB transcription factor EB 5648 -0.0720851719701549
3.57948568839385 0.0953677633745436
0.757460953597393 0.945746693240901
22934 RPIA ribose 5-phosphate isomerase A
1848 -0.0632356686509181 3.16102895858129
0.0953549244533916 0.757476767714978 0.945746693240901
493754 GS1-124K5.11 RAB guanine nucleotide exchange factor 1
pseudogene 7429 0.0737167175955377 3.4152405312879
0.0953544200794552 0.757477388992597 0.945746693240901
10576 CCT2 chaperonin containing TCP1 subunit 2 4322
0.0491149352381674 7.76020695032923 0.0953358668355975
0.75750024375423 0.945746693240901
83460 EMC6 ER membrane protein complex subunit 6
769 -0.047863053902111 4.02325262461118
0.0952893382650366 0.757557570560421 0.945746693240901
3385 ICAM3 intercellular adhesion molecule 3
2950 -0.080790709542523 2.41822808331091
0.0952178994389961 0.75764561857765 0.945746693240901
64601 VPS16 VPS16 core subunit of CORVET and HOPS complexes 3364
0.0472602829645896 5.11335657234898
0.0952053932326153 0.757661036138587 0.945746693240901
25792 CIZ1 CDKN1A interacting zinc finger protein 1 6993
0.0439775663868498 6.62626310861414
0.0951689312941766 0.757705992484261 0.945746693240901
137209 ZNF572 zinc finger protein 572 3275 -0.0965776483134099
0.262849478598382 0.0951567594396159
0.757721002071925 0.945746693240901
653598 PPIAL4C peptidylprolyl isomerase A like 4C 600
0.0659716415761229 4.34939864030651 0.0951199871604302
0.757766353787545 0.945746693240901
55337 SHFL shiftless antiviral inhibitor of ribosomal
frameshifting 3902 0.0520994425338346 5.1018658574039
0.0951152513618223 0.757772195205443 0.945746693240901
57178 ZMIZ1 zinc finger MIZ-type containing 1 11048
0.0650538137136953 6.63559021922249 0.0950310766335605
0.757876047943547 0.945746739710319
26778 SNORA70 small nucleolar RNA, H/ACA box 70
135 -0.0994911678566369 0.44584435289724
0.0950025922008324 0.757911202750696 0.945746739710319
84895 MIGA2 mitoguardin 2 6302 -0.0538466412387209
5.06937329002184 0.0949510811035026 0.757974791174987
0.945746739710319
285033 STARD7-AS1 STARD7 antisense RNA 1 2285
0.106417492762562 1.60036287580233 0.0948346211380757
0.758118626075039 0.945752523657763
10723 SLC12A7 solute carrier family 12 member 7

5984	-0.0580429222934227	6.87614254100462
0.0947773285530502	0.758189421269508	0.945752523657763
257218	SHPRH	SNF2 histone linker PHD RING helicase
17637	-0.056626595329932	4.67632941137752
0.0947753519600791	0.758191864120204	0.945752523657763
7343	UBTF	upstream binding transcription factor
6993	-0.0387039990391181	5.95649793563661
0.094599113455331	0.758409787653645	0.945752523657763
10797	MTHFD2	methylenetetrahydrofolate dehydrogenase (NADP+
dependent) 2, methenyltetrahydrofolate cyclohydrolase
4870	-0.0672156164104993	6.42306244261035
0.0945672240527529	0.758449243471188	0.945752523657763
55615	PRR5	proline rich 5	5247	-0.069458669401311
3.19806155081821	0.0945363951187428	0.758487394125287
	0.945752523657763
8891	EIF2B3	eukaryotic translation initiation factor 2B subunit
gamma	3411	0.0625601754521167	3.82973922010558
0.094499414568225	0.758533166357735	0.945752523657763
3014	H2AX	H2A.X variant histone	1614	0.0554889237525844
	4.10498442534653	0.0944943502512352
0.75853943541836	0.945752523657763
8725	URI1	URI1 prefoldin like chaperone	5432
0.044940482280782	5.68689349939146	0.0944084634826368
	0.758645781688589	0.945752523657763
84181	CHD6	chromodomain helicase DNA binding protein 6
14126	-0.0604000197823651	6.37924776674762
0.0944013220712066	0.758654626673708	0.945752523657763
57228	SMAGP	small cell adhesion glycoprotein
3558	-0.0578725919826515	4.83363972430856
0.0941757941157793	0.758934142584351	0.945856261452476
125893	ZNF816	zinc finger protein 816	3310	0.0535968192613878
	3.72838032144819	0.0940903325592783
0.759040158006019	0.945856261452476
100526832	PHOSPHO2-KLHL23	PHOSPHO2-KLHL23 readthrough	2180
	0.0930986782916311	2.34147910731379
0.0940525981030969	0.759086984522393	0.945856261452476
100507495	SDCBP2-AS1	SDCBP2 antisense RNA 1	4058
0.0999083085688316	1.22451222565894	0.0940062168602256
	0.759144555337103	0.945856261452476
55680	RUFY2	RUN and FYVE domain containing 2
8137	-0.0439813607367732	5.03866099153345
0.0939697833486131	0.759189789406617	0.945856261452476
8667	EIF3H	eukaryotic translation initiation factor 3 subunit H
	6434	-0.043929405368645	6.76838518664089
0.0938552919825781	0.759331998768596	0.945856261452476
3305	HSPA1L	heat shock protein family A (Hsp70) member 1 like
	2544	0.10100483910683	1.01803991830985
0.0938119554691812	0.759385851666679	0.945856261452476
10758	TRAF3IP2	TRAF3 interacting protein 2
6553	-0.0410318742660042	5.55417040929342
0.0937805522721025	0.759424883916766	0.945856261452476
118487	CHCHD1	coiled-coil-helix-coiled-coil-helix domain
containing 1	1483	-0.0553140857183303	4.71949167398547
	0.0937396864616904	0.759475688320356

0.945856261452476
143684	FAM76B	family with sequence similarity 76 member B
6042	-0.0743337431843796	3.76723624401374
0.0937173069250496	0.759503515708573	0.945856261452476
54465	ETAA1	ETAA1 activator of ATR kinase	3486
0.0489448057019422	4.04414471263914	0.0936819474834163
	0.75954749010824	0.945856261452476
223	ALDH9A1	aldehyde dehydrogenase 9 family member A1
3308	-0.0592116925516437	5.15297447472244
0.0936428056027587	0.759596179084551	0.945856261452476
145483	FAM161B	FAM161 centrosomal protein B
8199	-0.0673874998893666	3.17159794858205
0.0935670065667562	0.759690497914519	0.945856261452476
79132	DHX58	DExH-box helicase 58	3843	-0.0845789901032134
	2.89406534234488	0.0935506588816182
0.759710845266177	0.945856261452476
9868	TOMM70	translocase of outer mitochondrial membrane 70
5387	-0.0451242518993356	5.93448020565517
0.0935186003033408	0.759750753023618	0.945856261452476
55662	HIF1AN	hypoxia inducible factor 1 subunit alpha inhibitor
	13950	-0.0350254992802119	6.42649346457071
0.0933009861126948	0.76002184550477	0.946029625314381
2002	ELK1	ETS transcription factor ELK1	3398
0.0465074685762692	4.51158580849817	0.0932983868848496
	0.760025085575978	0.946029625314381
5571	PRKAG1	protein kinase AMP-activated non-catalytic subunit
gamma 1	4556	0.0388411546787034	5.50845024237939
0.0932011413642755	0.760146342594931	0.946096483092873
84461	NEURL4	neuralized E3 ubiquitin protein ligase 4
6170	-0.0588120444415105	5.34539609394761
0.0930751357635486	0.76030356392812	0.946142094697851
9185	REPS2	RALBP1 associated Eps domain containing 2
11311	0.0852008484047485	3.80978358823131
0.0930635024227655	0.760318085094525	0.946142094697851
55957	LIN37	lin-37 DREAM MuvB core complex component
3105	-0.048223344612785	4.96110710205243
0.0929233464470798	0.760493110999487	0.946195581600211
3281	HSBP1	heat shock factor binding protein 1	9525
0.0417188640609609	7.09514103572707	0.0928273708393448
	0.760613047969307	0.946195581600211
3280	HES1	hes family bHLH transcription factor 1	2062
0.05097734417588	5.02954490346426	0.09282283265888
	0.760619407601941	0.946195581600211
7988	ZNF212	zinc finger protein 212	3182	-0.0663633190442225
	4.15071365461761	0.0928127911506671
0.76063127356453	0.946195581600211
6495	SIX1	SIX homeobox 1	5270	0.070403915037998
4.8458378766901	0.0927365332493153	0.760726626588802
0.946210313926834
9633	TESMIN	testis expressed metallothionein like protein
5833	-0.100734912632716	0.144034726302892
0.0926952844722742	0.760778221992147	0.946210313926834
6595	SMARCA2	SWI/SNF related, matrix associated, actin dependent
regulator of chromatin, subfamily a, member 2	9299

0.0419287866932386	5.82289954524293	0.092546433419983
	0.760964514438693	0.946306910116213
83743	GRWD1	glutamate rich WD repeat containing 1
5763	-0.0464692703826122	5.13817334439301
0.092525275966608	0.760991007035698	0.946306910116213
100505696	SH3BP5-AS1	SH3BP5 antisense RNA 1	5733
0.0879577703485757	3.26193333509066	0.0924273201695183
	0.761113706886296	0.946314325804806
100129361	SMIM10L1	small integral membrane protein 10
like 1	4795	0.0594236347388029	4.09465732543543
0.0924005210019336	0.761147288011662	0.946314325804806
7984	ARHGEF5	Rho guanine nucleotide exchange factor 5
5818	-0.069320270383865	6.6738856402386	0.0923587417324327
	0.761199650806222	0.946314325804806
2188	FANCF	FA complementation group F	4269
0.0555080262799611	3.94108157586707	0.092262714364427
	0.761320052888814	0.946339940864727
115677	NOSTRIN	nitric oxide synthase trafficking
5248	-0.107462698996531	1.30983670615426
0.0922345501770039	0.761355378996545	0.946339940864727
9092	SART1	spliceosome associated factor 1, recruiter of U4/
U6.U5 tri-snRNP	4323	-0.0432152161073122	5.29184340614451
	0.0919539736280353	0.761707625968483
0.946590275255772
57464	STRIP2	striatin interacting protein 2	5876
0.149183347777072	2.45978229614683	0.0919207181781605
	0.761749415064148	0.946590275255772
83861	RSPH3	radial spoke head 3	6561	0.0490945044596082
	5.15909690583758	0.0918908717210272
0.761786927417965	0.946590275255772
284403	WDR62	WD repeat domain 62	9775	0.105451716757334
	0.636575923974022	0.0918181244321898
0.761878387318015	0.946590275255772
115572	TENT5B	terminal nucleotidyltransferase 5B
2368	-0.0782876286956158	4.51355249848453
0.0918051696269426	0.761894678605952	0.946590275255772
91607	SLFN11	schlafen family member 11	5982
0.117744533887836	2.21125197251126	0.091735020834177
	0.761982915875669	0.946595968281355
51062	ATL1	atlastin GTPase 1	4360	0.053387948710149
	5.11307364438175	0.0916940875014092
0.762034421242276	0.946595968281355
5871	MAP4K2	mitogen-activated protein kinase kinase kinase
kinase 2	7649	-0.0663010732187816	4.08140420651978
	0.0915679040544157	0.762193273670986
0.946709336198413
130340	AP1S3	adaptor related protein complex 1 subunit sigma 3
	4284	-0.0508259023538035	5.93257515419213
0.0913784681434695	0.762431979148821	0.946883681604765
114884	OSBPL10	oxysterol binding protein like 10
11976	-0.0553618932327055	5.10260811870917
0.0912654436169618	0.762574528377365	0.946883681604765
2071	ERCC3	ERCC excision repair 3, TFIIH core complex helicase
subunit	7401	-0.0364997283315798	6.46793872761423

0.0912174289642067	0.762635114753879	0.946883681604765
116159	CYYR1	cysteine and tyrosine rich 1	3266
0.162737247167574	1.97589557180257	0.0910920273076625
	0.762793432562129	0.946883681604765
114881	OSBPL7	oxysterol binding protein like 7
6572	-0.0714043188375782	5.08573435316345
0.0910345970780781	0.762865977106218	0.946883681604765
26751	SH3YL1	SH3 and SYLF domain containing 1
9688	-0.0631167387874228	5.1276405277841 0.0910332945426831
	0.762867622728449	0.946883681604765
219844	HYLS1	HYLS1 centriolar and ciliogenesis associated	2999
	0.0790225809098217	2.1054847960399 0.0910081259156166
	0.762899423275172	0.946883681604765
55713	ZNF334	zinc finger protein 334 6865	-0.0735844283106182
	2.82416567246291	0.0908817635697456
0.763059154583159	0.946883681604765
100529261	CHURC1-FNTB	CHURC1-FNTB readthrough 3907
0.0489423600240793	4.80150954335762	0.0908092968826582
	0.763150812470786	0.946883681604765
56848	SPHK2	sphingosine kinase 2	5557	0.0651371595348397
	5.46183044237609	0.0907937608957248
0.763170468013128	0.946883681604765
2145	EZH1	enhancer of zeste 1 polycomb repressive complex 2
subunit 5389	-0.048280446094898	4.5881693406111
0.0907542314753647	0.763220487411564	0.946883681604765
25988	HINFP	histone H4 transcription factor 9836
0.0548331344941301	4.7366675913981 0.0906380074713962
0.763367622805902	0.946883681604765
9908	G3BP2	G3BP stress granule assembly factor 2
7183	-0.0373267184546636	7.96046486747249
0.0906061926590809	0.763407917144638	0.946883681604765
84524	ZC3H8	zinc finger CCCH-type containing 8	6600
0.0681203430943438	2.88751756883929	0.0905956405860939
	0.763421283340405	0.946883681604765
347735	SERINC2 serine incorporator 2	3549	0.060672135593558
	9.07997820909915	0.0905286979370885
0.76350609864656	0.946883681604765
101928736	MIR762HG	MIR762 host gene	1815
0.0973360236191097	0.597628055586763	0.0905241272281998
	0.76351189091141	0.946883681604765
124245	ZC3H18	zinc finger CCCH-type containing 18
8648	-0.0440677528320618	5.49300592549365
0.09052217652183	0.763514363007125	0.946883681604765
441024	MTHFD2L methylenetetrahydrofolate dehydrogenase (NADP+
dependent) 2 like	9748	-0.08120528763096
2.45460944094729	0.0904445616382255	0.763612746557652
	0.946883681604765
80344	DCAF11	DDB1 and CUL4 associated factor 11	6904
0.0341472178510288	5.94443396715768	0.0904075131443598
	0.763659724945719	0.946883681604765
23463	ICMT	isoprenylcysteine carboxyl methyltransferase	4910
	0.0357949077418243	7.52993266454622
0.090387068551216	0.76368565367952	0.946883681604765
648	BMI1	BMI1 proto-oncogene, polycomb ring finger

3859	-0.0580412180630753	5.89658575626798
0.0902789238770652	0.763822860738165	0.946969977904008
23428	SLC7A8	solute carrier family 7 member 8	5991
0.110583726535586	3.8373908429687	0.0901338555171662
0.764007054994997	0.947114507417906
6119	RPA3	replication protein A3	3232	0.0776880562311911
	2.84491309247974	0.0900147196706875
0.764158443445868	0.947218346667812
6993	DYNLT1	dynein light chain Tctex-type 1	2949
0.0480818938908667	6.7327569384298	0.0897366432625972
0.764512226473671	0.94753525042198
10857	PGRMC1	progesterone receptor membrane component 1
1919	-0.0367788317065378	6.63652520773169
0.0896802656171545	0.764584025800919	0.94753525042198
6146	RPL22	ribosomal protein L22	4900	0.050445768466936
	6.75593022593828	0.0896302607815471
0.764647729666481	0.94753525042198
5360	PLTP	phospholipid transfer protein
2541	-0.149779435139284	4.38896731893335
0.0895254910101357	0.764781264359777	0.94753525042198
79819	WDR78	WD repeat domain 78	6704	-0.0875279919166053
	2.38323227980628	0.0895069650591083
0.764804885534346	0.94753525042198
100302174	MIR1307	microRNA 1307	149	0.149369918939148
	0.464745009547038	0.0894780596710483
0.764841746140204	0.94753525042198
81609	SNX27	sorting nexin 27	7467	-0.0520597328149372
	5.51468075810592	0.0894420831110381
0.764887633077842	0.94753525042198
10476	ATP5PD	ATP synthase peripheral stalk subunit d	3231
0.0466572445635791	5.99859541812869	0.0893383080159822
	0.765020051206792	0.947615480832467
144481	SOCS2-AS1	SOCS2 antisense RNA 1
3816	-0.109206523649519	1.96407226476373
0.089169688217245	0.765235390649487	0.94765457812183
2709	GJB5	gap junction protein beta 5
1355	-0.0497779326563265	5.35966258819763
0.0891095278266825	0.765312273404864	0.94765457812183
10420	TESK2	testis associated actin remodelling kinase 2	3322
	0.111979277307088	2.64592047192407
0.0890400748863698	0.765401066855262	0.94765457812183
56254	RNF20	ring finger protein 20	4694	-0.0384282149111642
	5.69890762241708	0.0890101327170467
0.765439358632997	0.94765457812183
7753	ZNF202	zinc finger protein 202	4808	-0.0527232593852064
	3.84938268573577	0.0889995465228104
0.765452898548618	0.94765457812183
50628	GEMIN4	gem nuclear organelle associated protein 4
4432	-0.0476669633054551	4.73742154148113
0.0889959102736704	0.765457549573098	0.94765457812183
55005	RMND1	required for meiotic nuclear division 1 homolog	2373
	0.0590163667914866	3.42143346234352
0.0888583557128872	0.765633567810505	0.947788721548541
10066	SCAMP2	secretory carrier membrane protein 2	5000

0.0401683308239873	7.20261485619645	0.0887899993404559
	0.765721093503733	0.947813304763767
1454	CSNK1E	casein kinase 1 epsilon 6246	-0.0362958921434125
	7.56986049176697	0.0887021218943147
0.765833668515969	0.94786888706399
23231	SEL1L3	SEL1L family member 3	7489	0.0532648179908344
	8.61194410742218	0.0884059567691793
0.766213517256366	0.948255233805435
257160	RNF214	ring finger protein 214 4398	-0.0545484152196973
	4.37779318997969	0.0882884835101265
0.766364375243688	0.948358141535324
92014	SLC25A51	solute carrier family 25 member 51	3397
	0.0509440522187415	4.02101692357465
0.088214700403197	0.766459182624992	0.948374080072586
7057	THBS1	thrombospondin 1	9158	0.0911699307673356
	11.3788948283925	0.0880886101977012
0.766621301776901	0.948374080072586
51759	C9orf78 chromosome 9 open reading frame 78	5099
0.038969644173551	5.69380668835154	0.0880823597392411
	0.766629341521345	0.948374080072586
26278	SACS	sacsin molecular chaperone	15909
0.115979149987398	5.53194574678828	0.0880159376480094
	0.766714797097861	0.948374080072586
6883	TAF12	TATA-box binding protein associated factor 12	2337
	0.0532078469331549	3.82233938347478
0.0879972444063295	0.766738853279792	0.948374080072586
2017	CTTN	cortactin	6949	0.0404336373424961
8.89237059032033	0.0879069291162722	0.766855118477662
	0.948374080072586
115196	ZNF554	zinc finger protein 554 4685	0.0728339750209007
	3.02356395573676	0.0878724565543134
0.766899513026953	0.948374080072586
302	ANXA2	annexin A2	8824	0.046805112557478
11.410331500792 0.0878573952209081	0.766918912328277
0.948374080072586
51603	EEF1AKNMT	eEF1A lysine and N-terminal
methyltransferase	3827	-0.0398536742285261
5.66897354047108	0.0877915145639294	0.767003789211975
	0.948395310336523
11156	PTP4A3	protein tyrosine phosphatase 4A3	2877
0.0954958412061253	1.02843690274409	0.0876846119548471
	0.767141590187446	0.948481971379253
1663	DDX11	DEAD/H-box helicase 11	8374	-0.154441946184561
	4.43926008317402	0.0875513302997142
0.767313522648407	0.948572879104672
148523	CIART	circadian associated repressor of transcription
2696	-0.0738936315547006	3.3647879547015 0.0874956768672721
	0.767385357333449	0.948572879104672
1506	CTRL	chymotrypsin like	5774	-0.0735899884846567
	3.48774385487094	0.087470175687919
0.767418281288937	0.948572879104672
1352	COX10	cytochrome c oxidase assembly factor heme
A:farnesyltransferase COX10	3172	-0.0551279977533992
3.77244418655067	0.0873137805845694	0.767620313469604

0.94869088912099
7711 ZNF155 zinc finger protein 155 3454 -0.0790513121704454
2.86438653217967 0.0872700693063373
0.767676815146859 0.94869088912099
4641 MYO1C myosin IC 8152 0.0407913060587213
8.3989418797053 0.0872390320840282 0.767716943541545
0.94869088912099
162417 NAGS N-acetylglutamate synthase 2822
0.0864944615384174 2.43990426825064 0.0871098315083412
0.767884071957215 0.948709861229325
7769 ZNF226 zinc finger protein 226 6922 0.05867322014076
3.87189098949174 0.0870617927676332
0.767946247206824 0.948709861229325
2961 GTF2E2 general transcription factor IIE subunit 2
2595 -0.04676246702656 5.24745736926358
0.0870178801616817 0.768003098426527 0.948709861229325
25786 DGCR11 DiGeorge syndrome critical region gene 11
2214 -0.0875383432080894 2.74885517967874
0.0869178002329711 0.768132724660389 0.948709861229325
84769 MPV17L2 MPV17 mitochondrial inner membrane protein like 2
2135 0.0588876095153879 3.83220809212187
0.0869088042842199 0.768144380429321 0.948709861229325
55084 SOBP sine oculis binding protein homolog
6904 -0.190653351036319 1.88640069436461
0.0868337759694384 0.768241617814524 0.948709861229325
10557 RPP38 ribonuclease P/MRP subunit p38
1769 -0.0449079938158873 4.09503804645234
0.0868243079491287 0.76825389169968 0.948709861229325
22897 CEP164 centrosomal protein 164 11851 0.0463150508387613
4.61001204630481 0.0867393024815382
0.76836412128426 0.948709861229325
10622 POLR3G RNA polymerase III subunit G
4019 -0.0897538420778733 0.812895266555791
0.0867005366059539 0.768414409849124 0.948709861229325
84988 PPP1R16A protein phosphatase 1 regulatory subunit 16A
4472 -0.0504395843967013 4.9301488187564
0.0866889093155279 0.768429495593173 0.948709861229325
79845 RNF122 ring finger protein 122 1868 0.0719862552449799
2.9445712943965 0.0866520398652857 0.768477338866847
0.948709861229325
9487 PIGL phosphatidylinositol glycan anchor biosynthesis
class L 4191 -0.0692908403609866 2.7672436540781
0.0862752461148486 0.768966916900335 0.949230598838723
11062 DUS4L dihydrouridine synthase 4 like 5439
0.0640297783278788 3.09242370344995 0.0861021690239578
0.769192189767457 0.949375541198657
139324 HDX highly divergent homeobox
7258 -0.0664035429269895 3.11295234743662
0.0859839188290432 0.769346242733969 0.949375541198657
6382 SDC1 syndecan 1 3595 -0.0485559421054625
9.33189926513741 0.0859261839245846 0.769421499910293
0.949375541198657
51626 DYNC2LI1 dynein cytoplasmic 2 light intermediate
chain 1 4618 0.0585938615488866 3.61716790563824

0.0859229670597905	0.769425693885815	0.949375541198657
26073	POLDIP2 DNA polymerase delta interacting protein 2	2121
	0.040414339554051	5.9884338958243 0.0858582771204794
	0.769510051255934	0.949375541198657
221981	THSD7A	thrombospondin type 1 domain containing 7A
10929	0.168756640659481	3.6508865597932 0.0857571925263656
	0.769641937325514	0.949375541198657
56981	PRDM11	PR/SET domain 11	13954	-0.0654223206565679
	3.68955588801592	0.0856989407289257
0.769717977354764	0.949375541198657
5588	PRKCQ	protein kinase C theta	3295	-0.0860604691135997
	0.47017906571497	0.0856497407959167
0.769782223229943	0.949375541198657
23385	NCSTN	nicastrin	4175	0.037626369142279
7.14368915340262	0.0856243550901183	0.769815380030504
	0.949375541198657
100271927	RASA4B	RAS p21 protein activator 4B
10265	-0.0923203803002896	4.96679903952718
0.0855250556311322	0.769945128412301	0.949375541198657
54768	HYDIN	HYDIN axonemal central pair apparatus protein
23408	-0.148154225965392	1.3296401326289 0.0854745031607251
	0.770011213567792	0.949375541198657
79591	ARMH3	armadillo like helical domain containing 3	4747
	0.0404486383182402	5.33919673149924
0.0854717743725928	0.770014781403116	0.949375541198657
81887	LAS1L	LAS1 like ribosome biogenesis factor
5589	-0.043093958554172	4.56705734599915
0.0854582118986329	0.770032514980934	0.949375541198657
197320	ZNF778	zinc finger protein 778 11273	-0.0621963200701726
	3.50019882090735	0.0854576612096363
0.770033235065178	0.949375541198657
1355	COX15	cytochrome c oxidase assembly homolog COX15	3992
	0.0428751077462866	4.43218857649125
0.0853346229127672	0.770194183865683	0.949435418914561
127707	KLHDC7A kelch domain containing 7A	5145
0.0714439187556802	1.95474390621994	0.0853169080469016
	0.770217367396741	0.949435418914561
4676	NAP1L4	nucleosome assembly protein 1 like 4	5200
0.0356366891647368	7.35537914752185	0.0852079317195802
	0.770360042741364	0.949469092646106
221496	LEMD2	LEM domain containing 2 6788	0.0394219782269112
	6.56522635550082	0.0851775478338723
0.770399839959944	0.949469092646106
23270	TSPYL4	TSPY like 4	4180	0.0577481280763544
4.23901253044957	0.0851407559542885	0.770448040766448
	0.949469092646106
6132	RPL8	ribosomal protein L8	2159	0.0584361539877409
	8.26774797830653	0.0849037306572882
0.770758837361061	0.94966030485168
100131213	ZNF503-AS2	ZNF503 antisense RNA 2	3348
0.0749604317923451	2.15914273994479	0.0847880082027943
	0.770910748049002	0.94966030485168
54951	COMMD8	COMM domain containing 8	1766
0.0581195784441239	3.98891955227165	0.084688764943273

0.771041115838882	0.94966030485168
81669	CCNL2	cyclin L2	6850	-0.0427650691708787
7.72197806835833	0.0846878516076117	0.771042315998251
0.94966030485168
57175	CORO1B	coronin 1B	5327	-0.0368237583029706
6.69210125662161	0.0846495558251981	0.771092644498033
0.94966030485168
10713	USP39	ubiquitin specific peptidase 39 3952
0.0379547211577649	5.48235381522086	0.08455624500902
0.771215325738612	0.94966030485168
81671	VMP1	vacuole membrane protein 1
4302	-0.0396444779782951	7.19932829767561
0.0845080294799914	0.771278746311634	0.94966030485168
1912	PHC2	polyhomeotic homolog 2	4975	-0.0379007559619728
6.97018932947027	0.0845019874872861
0.771286695064749	0.94966030485168
10016	PDCD6	programmed cell death 6 9493	-0.0332381775801684
6.4379378718269 0.0844990127939882	0.771290608638934
0.94966030485168
6612	SUMO3	small ubiquitin like modifier 3 2732
0.0400579124383037	6.28052208662656	0.0844941686385532
0.77129698187993	0.94966030485168
79581	SLC52A2 solute carrier family 52 member 2
2926	-0.044339085050276	6.15638299638324
0.0844546448722383	0.771348988955348	0.94966030485168
11160	ERLIN2	ER lipid raft associated 2
9464	-0.0333539741023938	6.04209310870807
0.0843139266818973	0.771534259228952	0.949767203893445
2926	GRSF1	G-rich RNA sequence binding factor 1	7785
0.04129550155239	6.92898838304016	0.0842393391957899
0.771632529469844	0.949767203893445
51029	DESI2	desumoylating isopeptidase 2	4483
0.0373010652279847	6.21080174628249	0.0842342501287376
0.771639236118184	0.949767203893445
2850	GPR27	G protein-coupled receptor 27
2447	-0.162067357230924	0.944249897240726
0.0839995164762684	0.771948819898152	0.950064767622654
57142	RTN4	reticulon 4	5275	-0.0448622233667341
9.52882873836771	0.0838679696691997	0.772122518425354
0.950136228828382
2936	GSR	glutathione-disulfide reductase
3237	-0.0441993084229418	5.41211496329749
0.0838428731378502	0.772155673439809	0.950136228828382
4007	PRICKLE3	prickle planar cell polarity protein 3
3194	-0.0568534412801976	3.46506170122441
0.0838014702025394	0.772210382593153	0.950136228828382
9724	UTP14C	UTP14C small subunit processome component
5529	-0.034478308765502	6.10561401475339
0.083736813586486	0.772295848080632	0.950157922184051
9674	KIAA0040	KIAA0040	4783	0.0724351793745611
4.70944927169344	0.0835787353569266
0.772504952384616	0.950313492166625
10043	TOM1	target of myb1 membrane trafficking protein
5871	-0.046337492362123	6.20527749942043

0.0835386674739027	0.772557987813333	0.950313492166625
9392	TGFBRAP1	transforming growth factor beta receptor associated protein 1	6005	0.0361459402217663	5.62577939437556	0.0833696873481493	0.772781808317108	0.950328305479207
57291	DANCR	differentiation antagonizing non-protein coding RNA	1641	-0.0598034706568783	3.50620896376064	0.0833688338797636	0.772782939391382	0.950328305479207
57623	ZFAT	zinc finger and AT-hook domain containing	8900	0.0570371775636824	3.83120081974785	0.0833628547665484	0.772790863494222	0.950328305479207
91647	ATPAF2	ATP synthase mitochondrial F1 complex assembly factor 2	5564	-0.048597024266132	3.993034594936	0.0832756464380704	0.77290647546942	0.950328305479207
55275	VPS53	VPS53 subunit of GARP complex	16295	0.0379465946676814	6.34927300757739	0.0832735438666159	0.772909263655253	0.950328305479207
4207	BORCS8-MEF2B	BORCS8-MEF2B readthrough	2364	0.0597603516668885	3.03245160953751	0.0831491569858107	0.773074278978699	0.950445111272127
1161	ERCC8	ERCC excision repair 8, CSA ubiquitin ligase complex subunit	4847	0.0690009332089902	3.29843607231392	0.082894205636908	0.773412923320869	0.950445111272127
26000	TBC1D10B	TBC1 domain family member 10B	6699	-0.0363063595342077	6.17865559849889	0.0828629116674287	0.773454529015516	0.950445111272127
10168	ZNF197	zinc finger protein 197	6767	0.0645577774103163	4.10670767884764	0.0828607801746557	0.77345736316927	0.950445111272127
5089	PBX2	PBX homeobox 2	4344	-0.0377358062601299	5.47088858594554	0.0828589859286577	0.773459748931172	0.950445111272127
23506	BICRAL	BRD4 interacting chromatin remodeling complex associated protein like	6843	-0.0562271512447258	4.09375492108405	0.0827981222256113	0.773540694375915	0.950445111272127
26156	RSL1D1	ribosomal L1 domain containing 1	6167	0.0499404283373152	6.10312703643932	0.0827662026143798	0.773583158616289	0.950445111272127
5442	POLRMT	RNA polymerase mitochondrial	5124	0.0480873105292045	4.57248328559591	0.0827606156284517	0.77359059217219	0.950445111272127
10257	ABCC4	ATP binding cassette subfamily C member 4	7819	-0.0492330310066908	3.70107563845401	0.0826449756920629	0.773744513626751	0.950445111272127
120534	ARL14EP	ADP ribosylation factor like GTPase 14 effector protein	5465	-0.0541955394914483	3.97665220478037	0.0825866403148883	0.77382204626668	0.950445111272127
2983	GUCY1B1	guanylate cyclase 1 soluble subunit beta 1	3724	0.0978656565009777	1.84394990083846	0.0825789650835036	0.773832428703541	0.950445111272127
3977	LIFR	LIF receptor subunit alpha	11864	0.109702896022722	2.98321701101706	0.0825075935486552	0.77392752644620	0.950445111272127

9374	PPT2	palmitoyl-protein thioesterase 2
2940	-0.104445578158428	4.64145911739947
0.0824878802636242	0.773953800911979	0.950445111272127
54855	TENT5C	terminal nucleotidyltransferase 5C
5751	-0.0917699610690436	0.485908102038184
0.0824875562284566	0.773954232824294	0.950445111272127
100130890	TSTD3	thiosulfate sulfurtransferase like domain
containing 3	962	-0.0925940060830533	0.53464802217385
	0.0823563601665338	0.77412918193382
0.950576615354346
375743	PTAR1	protein prenyltransferase alpha subunit repeat
containing 1	12200	-0.0448340313330315	6.09848018242775
	0.0820553319542796	0.774531172011224
0.95073647893842
100506136	NA	NA	2229	0.0864261339892637
0.193131414298504	0.0819988599254073	0.774606673034528
	0.95073647893842
10270	AKAP8	A-kinase anchoring protein 8
4296	-0.0364981353157667	5.58383692344744
0.0819598599700377	0.774658830961945	0.95073647893842
6391	SDHC	succinate dehydrogenase complex subunit C
14160	-0.0415895303440082	6.25365798324798
0.0818517870814013	0.774803436137234	0.95073647893842
51023	MRPS18C mitochondrial ribosomal protein S18C	4465
0.0511114685902449	3.92056218205968	0.0818353298129662
	0.774825465577607	0.95073647893842
7626	ZNF75D	zinc finger protein 75D 6819	0.052283858124455
	4.3628034636232 0.0818141224974394	0.774853856890172
	0.95073647893842
4257	MGST1	microsomal glutathione S-transferase 1	4159
0.0648091405137578	7.05213226938488	0.0818111360423757
	0.774857855329298	0.95073647893842
956	ENTPD3	ectonucleoside triphosphate diphosphohydrolase 3
	3365	0.0590585814739229	5.74010611570185
0.0818055782498561	0.774865296634222	0.95073647893842
23566	LPAR3	lysophosphatidic acid receptor 3	3543
0.111299491383293	1.27125657581054	0.0818018760102532
	0.774870253698936	0.95073647893842
388272	C16orf87	chromosome 16 open reading frame 87
7443	-0.0483126719615856	3.96640073184437
0.0816983793057169	0.775008878385092	0.950739705639358
100129842	ZNF737	zinc finger protein 737
3213	-0.0565448145177578	3.17896836583549
0.081664615755519	0.775054122220797	0.950739705639358
8233	ZRSR2	zinc finger CCCH-type, RNA binding motif and serine/
arginine rich 2 2077	-0.0601103214726649	3.05757395963316
	0.0816479102678678	0.775076511651019
0.950739705639358
80326	WNT10A	Wnt family member 10A	3021	0.0706369072712919
	4.60644984713446	0.0815115452985467
0.775259366712521	0.950821180308983
57666	FBRSL1	fibrosin like 1 9311	0.04416366252487
5.27695833658292	0.0814119304775947	0.77539304719946
	0.950821180308983

9604	RNF14	ring finger protein 14	4987	-0.041257873798895
	6.4095365671345	0.0813537732707914	0.775471133545135
	0.950821180308983
143879	KBTBD3	kelch repeat and BTB domain containing 3
5516	-0.0855366327073951	1.79331656803916
0.0813536225958753	0.775471335892077	0.950821180308983
50807	ASAP1	ArfGAP with SH3 domain, ankyrin repeat and PH domain
1	8280	0.0351975361974016	6.24878782850385
0.0813109280097488	0.775528680202881	0.950821180308983
64149	C17orf75	chromosome 17 open reading frame 75	5740
	0.0443969880091086	4.33006796549572
0.0812390327194481	0.775625281608183	0.950821180308983
374443	LOC374443	C-type lectin domain family 2 member D
pseudogene	615	0.0840621262840169	0.621423796366526
	0.0812247842337541	0.775644431933844
0.950821180308983
51131	PHF11	PHD finger protein 11	4841	-0.0439888136488316
	4.36235820212807	0.0811047153953286
0.775805879676725	0.950821180308983
23510	KCTD2	potassium channel tetramerization domain containing
2	5279	-0.0504840435024813	5.91813834880496
0.0810591944964472	0.775867122058315	0.950821180308983
220064	LTO1	LTO1 maturation factor of ABCE1	6094
0.0495388162045141	4.69093084483537	0.0810490951545617
	0.775880711913923	0.950821180308983
79154	DHRS11	dehydrogenase/reductase 11	2376
0.0797642498238971	2.06336540922266	0.0810424657129274
	0.775889633107138	0.950821180308983
8570	KHSRP	KH-type splicing regulatory protein
4563	-0.0397927464750238	6.80070563475422
0.0807632200224564	0.776265770920594	0.951184452647478
23081	KDM4C	lysine demethylase 4C	12811	-0.0406513326388746
	4.68468961840539	0.0806648893725139
0.77639838742618	0.951184452647478
147929	ZNF565	zinc finger protein 565	3953	-0.0737118971505226
	2.09827021648786	0.0806480115799857
0.776421158936278	0.951184452647478
284021	MILR1	mast cell immunoglobulin like receptor 1	1816
	0.193854061239739	0.420994250294279
0.0805160427266625	0.776599300101324	0.951184452647478
8877	SPHK1	sphingosine kinase 1	4416	-0.0933546613963198
	6.39313751213456	0.0804930166348186
0.776630398555687	0.951184452647478
89797	NAV2	neuron navigator 2	14535	-0.0632004476848887
	7.20402357246352	0.0803322907451616
0.776847604836001	0.951184452647478
83877	TM2D2	TM2 domain containing 2	4459	-0.0398059598567082
	6.71060893586052	0.0802995861501969
0.776891830828667	0.951184452647478
283377	SPRYD4	SPRY domain containing 4	10779
0.0575410269785387	3.20622861073566	0.0802735930712162
	0.77692698786656	0.951184452647478
27229	TUBGCP4	tubulin gamma complex associated protein 4
11171	-0.054879344646196	5.10647704459643

0.0801967362008575	0.777030976903498	0.951184452647478
4338	MOCS2	molybdenum cofactor synthesis 2 6010
0.055446682471288	4.39292911116339	0.0801818333685347
	0.777051146993192	0.951184452647478
1345	COX6C	cytochrome c oxidase subunit 6C
2537	-0.0487918137131829	6.17599064677076
0.0801150314158692	0.777141584317746	0.951184452647478
158880	USP51	ubiquitin specific peptidase 51 4407
0.0933033935451969	0.0386627566947864	0.0800616139386858
	0.777213930878125	0.951184452647478
9875	URB1	URB1 ribosome biogenesis homolog
11461	-0.0532215030742501	6.01084077056358
0.0800386219130722	0.77724507841089	0.951184452647478
26610	ELP4	elongator acetyltransferase complex subunit 4	2219
	0.070205797182056	2.62316674324447
0.0800051364524639	0.777290450151688	0.951184452647478
65250	CPLANE1 ciliogenesis and planar polarity effector 1
15146	0.0567028462316939	5.63553155704821
0.0800009640579873	0.777296104332771	0.951184452647478
114785	MBD6	methyl-CpG binding domain protein 6
5646	-0.048011858641168	5.81361849064258
0.0799984591164389	0.77729949895722	0.951184452647478
1396	CRIP1	cysteine rich protein 1 3036	0.0822202948701132
	2.95976200547625	0.0799682047898163
0.777340503280908	0.951184452647478
56897	WRNIP1	WRN helicase interacting protein 1
3877	-0.0333707519953948	6.04872939748993
0.0799110613023686	0.777417973907553	0.951196153085526
192683	SCAMP5	secretory carrier membrane protein 5
4873	-0.109289779642771	3.06452142430341
0.079799495043261	0.777569312886013	0.951200898695352
25778	DSTYK	dual serine/threonine and tyrosine protein kinase
	8339	0.0418982914141278	5.76657722396623
0.0797594111671085	0.777623714322245	0.951200898695352
52	ACP1	acid phosphatase 1	3655	0.0394554591669006
	5.97668364847925	0.0797580371237814
0.777625579421752	0.951200898695352
401152	C4orf3	chromosome 4 open reading frame 3	3148
0.0487404580984119	6.17441968114234	0.07967575727357
	0.777737296115936	0.951254480151582
221092	HNRNPUL2	heterogeneous nuclear ribonucleoprotein U like 2	5106	-0.0330038869998064	7.62527723814993
0.0796155526109459	0.777819079357987	0.951271443688756
131076	CCDC58	coiled-coil domain containing 58
1746	-0.0709683370615987	2.67397651051018
0.0794597897701745	0.778030825818023	0.951274503362887
10607	TBL3	transducin beta like 3	7139	0.0489378618990026
	4.90328017578545	0.0794382640352405
0.778060105855404	0.951274503362887
29071	C1GALT1C1	C1GALT1 specific chaperone 1
1785	-0.049176875402377	5.25686449317905
0.0794236869685268	0.778079936509644	0.951274503362887
3038	HAS3	hyaluronan synthase 3	5289	0.122314362389212
	6.49109062161099	0.0794139101329279

0.778093238025887	0.951274503362887
440498	HSBP1L1	heat shock factor binding protein 1 like 1
1659	-0.0437939768477613	4.18969284267882
0.0792724443062855	0.778285803182547	0.951426884724903
3437	IFIT3	interferon induced protein with tetratricopeptide
repeats 3	2640	0.119889942106361	4.32676658757175
	0.0791765385531065	0.778416456859786
0.951503561500569
4439	MSH5	mutS homolog 5	6234	0.099475837128327
1.23005148450173	0.079074786487201	0.778555168442179
	0.951590073679721
252884	ZNF396	zinc finger protein 396	2788	0.102350169789196
	1.12947746465007	0.0789496652665047
0.7787258696678	0.951674917991018
162963	ZNF610	zinc finger protein 610	3546	0.0982101413723397
	1.65288534801805	0.0788606637994445
0.778847382039	0.951674917991018
4717	NDUFC1	NADH:ubiquinone oxidoreductase subunit C1
1581	-0.0482217269768017	4.38237094366376
0.0788482523451446	0.778864333088222	0.951674917991018
55778	ZNF839	zinc finger protein 839	5590	-0.0585745583126265
	3.46624360942699	0.0787648947513548
0.778978216743744	0.951674917991018
283989	TSEN54	tRNA splicing endonuclease subunit 54	3440
0.0597777789402687	3.85261004850378	0.0787265987419374
	0.779030558780242	0.951674917991018
9534	ZNF254	zinc finger protein 254	6710	0.0478160829837534
	5.14710996121896	0.0787253675916304
0.779032241713787	0.951674917991018
6273	S100A2	S100 calcium binding protein A2	1567
0.0851172625644732	10.8512199437421	0.0786208084446116
	0.779175221742586	0.951719472128357
22999	RIMS1	regulating synaptic membrane exocytosis 1
12788	0.124732620883361	0.554665427479737
0.0785993299775054	0.779204605295067	0.951719472128357
10131	TRAP1	TNF receptor associated protein 1
5227	-0.0551977789298392	5.32898037324294
0.0784257186083295	0.779442272932755	0.951926751850126
5778	PTPN7	protein tyrosine phosphatase non-receptor type 7
	5906	-0.238040896048968	0.459134258283286
0.0782760537878371	0.77964738641695	0.952094240762182
192111	PGAM5	PGAM family member 5, mitochondrial serine/threonine
protein phosphatase	3952	-0.0518153972982001
4.40199721381302	0.077888791918288	0.780179106583815
	0.95260991059198
55035	NOL8	nucleolar protein 8	7795	-0.0404387507427165
	4.90568784181332	0.0778694685542529
0.780205675327422	0.95260991059198
4261	CIITA	class II major histocompatibility complex
transactivator	13913	0.198015834488585	1.67374255044903
	0.0778043804676436	0.780295194626553
0.952628433683229
100532731	COMMD3-BMI1	COMMD3-BMI1 readthrough
2192	-0.043297257385662	5.36867702848517

0.0777595558870283	0.780356867857598	0.952628433683229
51654	CDK5RAP1	CDK5 regulatory subunit associated protein 1
	4567	0.0436034550890467	4.46508834325551
0.077684646823112	0.780459976493731	0.952663258795955
7108	TM7SF2	transmembrane 7 superfamily member 2
3408	-0.0701227642186559	4.99236999718075
0.0776400247815268	0.78052142199917	0.952663258795955
6222	RPS18	ribosomal protein S18	1898	0.0581270593162937
	7.36033021895262	0.0775123564724254
0.780697329164553	0.952794936796018
57700	FAM160B1	family with sequence similarity 160 member
B1	6216	-0.0491059845031667	5.95623500622601
0.0774198351366309	0.780824906814312	0.952867613673817
58986	PGAP6	post-glycosylphosphatidylinositol attachment to
proteins 6	4447	-0.0415587442738027	6.94048700676653
	0.0773408237733406	0.780933920600572
0.952917625738498
91526	ANKRD44 ankyrin repeat domain 44
11665	-0.0794114452347504	2.05977686886673
0.0772792267304196	0.78101894907209	0.952920171777029
23338	JADE2	jade family PHD finger 2
9877	-0.068938976145322	5.28588220963866
0.077240757438652	0.781072070548445	0.952920171777029
6134	RPL10	ribosomal protein L10	4825	-0.0448220894820884
	7.95476185337922	0.0769980921560602
0.781407491119317	0.953064541731691
8602	NOP14	NOP14 nucleolar protein 3538	0.0397987447425029
	5.33343618114651	0.0769786704932045
0.78143436102046	0.953064541731691
6281	S100A10 S100 calcium binding protein A10	1543
0.0524141806577305	9.63129648710446	0.0769644989547373
	0.781453969670019	0.953064541731691
23423	TMED3	transmembrane p24 trafficking protein 3 19470
0.0380523943898303	6.58629066061579	0.0769582815186993
	0.781462573126899	0.953064541731691
100507419	MMP25-AS1	MMP25 antisense RNA 1
5377	-0.0684613811034669	2.75948766277091
0.0767080752495191	0.781809110351088	0.953404162344392
114134	SLC2A13 solute carrier family 2 member 13	9695
0.0840135385696401	1.93408166779562	0.0766293682311954
	0.78191824584756	0.953411797813612
57062	DDX24	DEAD-box helicase 24	8636	0.0364197550898727
	7.10761824635799	0.0765355161552588
0.78204846045281	0.953411797813612
56927	GPR108	G protein-coupled receptor 108	3213
0.0357780880868667	6.76435330570998	0.0765211769263896
	0.782068362913672	0.953411797813612
51256	TBC1D7	TBC1 domain family member 7	6078
0.0813299699303002	4.44723665470966	0.0765072904968704
	0.782087638814764	0.953411797813612
1515	CTSV	cathepsin V	4700	-0.0820766730234448
3.71159449846098	0.0763998843084135	0.782236794145416
	0.953422070117809
51537	MTFP1	mitochondrial fission process 1

2286	-0.0426097580628884	5.85134064642451
0.0763730678388868	0.782274051869183	0.953422070117809
9984	THOC1	THO complex 1	8102	0.0440763725748513
4.30056403860151	0.0763542017620971	0.782300267856356
	0.953422070117809
10978	CLP1	cleavage and polyadenylation factor I subunit 1	2793
	0.0451207845419148	4.37498678798601
0.076282940741935	0.782399322452293	0.95344257262089
63899	NSUN3	NOP2/Sun RNA methyltransferase 3	5378
0.047928581016976	3.67324361706237	0.0762275335939044
	0.782476374186392	0.95344257262089
58190	CTDSP1	CTD small phosphatase 1	4606	-0.0308588862676599
	5.88195943420578	0.0761837425720513
0.78253729332503	0.95344257262089
84726	PRRC2B	proline rich coiled-coil 2B
12293	-0.0460845536481609	8.40587634211261
0.0761320303026878	0.782609256241031	0.95344257262089
139596	UPRT	uracil phosphoribosyltransferase homolog	2825
	0.0503540591673326	3.81620158933799
0.0760974193861834	0.782657435567573	0.95344257262089
147912	SIX5	SIX homeobox 5	3556	0.064650157450688
3.47980486923114	0.0758147763872117	0.783051324262566
	0.953839455513155
693207	MIR622	microRNA 622	96	-0.0820745060208587
0.394895961744934	0.0756209007739876	0.783321964471527
	0.954053841440073
84542	KIAA1841	KIAA1841	7992	-0.0369751254983743
	5.66660384934375	0.0755911349982554
0.783363548968536	0.954053841440073
121268	RHEBL1	RHEB like 1	1290
0.0953675862490882	-0.0108709923803134	0.0753488256322239
	0.783702397180272	0.954106105685445
4799	NFX1	nuclear transcription factor, X-box binding 1	7147
	0.0357051209545448	5.5125656038819	0.0753297087482157
	0.783729155372049	0.954106105685445
255758	TCTEX1D2	Tctex1 domain containing 2	1567
0.0462635094608316	3.83140804617826	0.0753117069062674
	0.783754356160624	0.954106105685445
8106	PABPN1	poly(A) binding protein nuclear 1	3630
0.0342148881466676	6.75291424572976	0.0752650358764724
	0.783819706047705	0.954106105685445
55683	KANSL3	KAT8 regulatory NSL complex subunit 3
9015	-0.0434721834599994	6.56907761643313
0.0752363619918981	0.783859866708491	0.954106105685445
4849	CNOT3	CCR4-NOT transcription complex subunit 3
6676	-0.0415932376755697	5.59358582212958
0.0752180399920936	0.783885532821971	0.954106105685445
375484	SIMC1	SUMO interacting motifs containing 1
5536	-0.0591768112397217	3.23023338578442
0.0752060488423929	0.783902332273622	0.954106105685445
51239	ANKRD39	ankyrin repeat domain 39
976	-0.0655875532215605	2.57811238722206
0.0751533098231487	0.783976236083497	0.954106105685445
10869	USP19	ubiquitin specific peptidase 19

5731	-0.0360734488843448	5.9378869382871 0.0750724583544979
	0.784089588353329	0.954106105685445
5479	PPIB	peptidylprolyl isomerase B
1911	-0.0603680193513308	7.63495593857888
0.0750441014083805	0.784129359812379	0.954106105685445
55181	SMG8	SMG8 nonsense mediated mRNA decay factor	3840
	0.038736733077584	5.43592750847167
0.0750252936224669	0.784155742746544	0.954106105685445
51188	SS18L2	SS18 like 2	1301	-0.0572342509711335
3.26910619747215	0.074939222445928	0.784276525903391
	0.954140302542346
10598	AHSA1	activator of HSP90 ATPase activity 1	2400
0.0391938664041036	6.3683168219104 0.0748612622176665
0.784385991384696	0.954140302542346
57150	SMIM8	small integral membrane protein 8
2926	-0.0782639804092303	1.40972102041342
0.0748204534672241	0.784443316178704	0.954140302542346
23637	RABGAP1 RAB GTPase activating protein 1
7667	-0.0398008822563922	6.46729170980247
0.0748111953532486	0.784456323558054	0.954140302542346
1359	CPA3	carboxypeptidase A3	1848	-0.30326765768364
	0.290777019950422	0.0747410438264566
0.784554912524957	0.954177360400891
123803	NTAN1	N-terminal asparagine amidase	3540
0.0462331511517888	5.73848871118563	0.0746285242741749
	0.784713148210909	0.954217219677571
115	ADCY9	adenylate cyclase 9	8189	-0.085318609918276
	6.87637156395673	0.0746091148000403
0.784740456618607	0.954217219677571
104909134	LINC02210-CRHR1 LINC02210-CRHR1 readthrough	2771
	0.0787067451964718	1.65991703028013
0.0745724457553236	0.784792059019463	0.954217219677571
5884	RAD17	RAD17 checkpoint clamp loader component 4159
0.0382988960495844	5.33922287880313	0.0745152188062135
	0.784872618716095	0.954225335455846
101927599	ZNF529-AS1	ZNF529 antisense RNA 1
1868	-0.0786072713689546	0.626795432727084
0.0743816292916235	0.785060805213574	0.954225335455846
10884	MRPS30	mitochondrial ribosomal protein S30	2175
0.0445094377375733	4.31674144572702	0.0743694130002703
	0.785078023264896	0.954225335455846
4668	NAGA	alpha-N-acetylgalactosaminidase
3706	-0.0483591922142221	4.45140200931027
0.0743326616585711	0.785129831002726	0.954225335455846
64087	MCCC2	methylcrotonoyl-CoA carboxylase 2
6034	-0.0519053866684085	5.57656253345398
0.0743259042817925	0.785139358258629	0.954225335455846
54867	TMEM214 transmembrane protein 214
4910	-0.0384551087242406	6.26515228876929
0.0741930572163625	0.78532675450236	0.954273336353029
51248	PDZD11	PDZ domain containing 11
1393	-0.0427613207592933	4.68197107290822
0.0741563754880445	0.785378530094819	0.954273336353029
63970	TP53AIP1	tumor protein p53 regulated apoptosis

inducing protein 1	3445	0.10583560337969
1.90964922034165	0.074153040192261	0.785383238486308
	0.954273336353029
27090	ST6GALNAC4	ST6 N-acetylgalactosaminide alpha-2,6-sialyltransferase 4	2630	0.0590096527353026
3.0675491822748 0.0740847330483243	0.785479691777149
0.954307749390452
100528022	BLOC1S1-RDH5	BLOC1S1-RDH5 readthrough	3382
	0.0578263984800156	3.26427090560343
0.0740268039563263	0.785561528144554	0.95432439936867
125113	KRT222	keratin 222	4661	0.03679899683874
5.87292600451121	0.0739400212330317	0.785684190383056
	0.954390638686624
23335	WDR7	WD repeat domain 7	15694	-0.0467771069636861
	4.96723757412087	0.0738440679400156
0.785819904704094	0.954412230665221
4324	MMP15	matrix metallopeptidase 15
4419	-0.0424576306215652	6.39808086628136
0.0738311081320404	0.785838242040551	0.954412230665221
55074	OXR1	oxidation resistance 1	8075	0.0393556758758078
	5.53739094602822	0.0737818982778045
0.785907886722164	0.954414060110746
79139	DERL1	derlin 1	7081	-0.0310540730332219
6.59400643695487	0.0735910460556397	0.786178228201542
	0.954659596221846
84909	AOPEP	aminopeptidase O (putative)	10175
0.0515962981111927	6.48219790658663	0.0734714239033618
	0.786347864458366	0.954680911448599
606495	CYB5RL	cytochrome b5 reductase like	7949
0.0790286019106964	1.0164566551481 0.0734421211409635
0.786389441326765	0.954680911448599
23779	ARHGAP8 Rho GTPase activating protein 8
4990	-0.0498046935538042	5.31231944800216
0.0733031169965273	0.786586791809653	0.954680911448599
10615	SPAG5	sperm associated antigen 5	5472
0.0734379787462288	1.82807029726985	0.0732846119414226
	0.786613079418175	0.954680911448599
113230	MISP3	MISP family member 3	1665	-0.0849455681081447
	1.70222032417301	0.0732531410412811
0.786657794016413	0.954680911448599
150381	PRR34-AS1	PRR34 antisense RNA 1	1093
0.0728634773316583	2.0240439995583 0.0732264779986815
0.786695685564463	0.954680911448599
10616	RBCK1	RANBP2-type and C3HC4-type zinc finger containing 1
	4024	0.0404116857483533	6.03873276476233
0.0731615555810397	0.786787979484364	0.954680911448599
132001	TAMM41	TAM41 mitochondrial translocator assembly and maintenance homolog	5558	-0.0637144454468255
2.73778071636514	0.0730954632491649	0.78688198171032
	0.954680911448599
6624	FSCN1	fascin actin-bundling protein 1 3692
0.0557660246908369	7.55065613843349	0.0730875617409765
	0.786893222969621	0.954680911448599
2793	GNGT2	G protein subunit gamma transducin 2	1689

0.176530344028708	0.11857920304971	0.0730774468241648
	0.786907614136423	0.954680911448599
201254	CENPX	centromere protein X	1531	-0.0637102628512715
	4.29379693546264	0.0730399252389446
0.786961007936028	0.954680911448599
25845	PP7080	uncharacterized LOC25845
2766	-0.0632749069016859	2.68671143275614
0.0730029275540369	0.787013670628755	0.954680911448599
23173	METAP1	methionyl aminopeptidase 1	4117
0.036339512065898	6.27221517851339	0.0729543035055311
	0.787082904126482	0.954682216669521
223082	ZNRF2	zinc and ring finger 2	4044	-0.0528769628039603
	3.67289977516998	0.0728083375309332
0.787290887041212	0.954851801436163
57474	ZNF490	zinc finger protein 490 4921	-0.0710715569111834
	2.59715816241723	0.0726669747243411
0.787492524328524	0.955013661322046
79903	NAA60	N(alpha)-acetyltransferase 60, NatF catalytic
subunit 5933	0.0315730881187996	6.08136369269652
0.0724613199053508	0.787786242569264	0.955187389767859
598	BCL2L1	BCL2 like 1	3065	-0.0378108379495198
8.45437988233937	0.0724302715577494	0.787830624946675
	0.955187389767859
100507299	SMC5-AS1	SMC5 antisense RNA 1 (head to head)
	1818	-0.102957440223301	0.0889907658104411
0.0723373148135726	0.787963563939275	0.955187389767859
339201	ASB16-AS1	ASB16 antisense RNA 1	2460
0.0553361493413323	3.03784719841357	0.072297565687518
	0.788020437785478	0.955187389767859
123207	C15orf40	chromosome 15 open reading frame 40
13893	0.0448747163979741	4.14299096257052
0.072293427777069	0.78802635935405	0.955187389767859
100529067	SERF2-C15ORF63	SERF2-C15orf63 readthrough	2816
	0.0512677233936987	5.36801633466984
0.0722804446235727	0.788044940112613	0.955187389767859
10944	C11orf58	chromosome 11 open reading frame 58
8468	-0.0329214874516332	7.08139514082654
0.0721704906540683	0.788202371843109	0.955205336728381
339344	MYPOP	Myb related transcription factor, partner of
profilin	1890	-0.0706271089842243	2.80950842458773
	0.0721446309618057	0.788239416338451
0.955205336728381
22929	SEPHS1	selenophosphate synthetase 1	3888
0.0357782145146451	5.61949869951582	0.0720890204661764
	0.788319103538797	0.955205336728381
55216	NKAPD1	NKAP domain containing 1
5755	-0.0370165951498499	5.01075778897206
0.0720575874572749	0.788364160125602	0.955205336728381
3831	KLC1	kinesin light chain 1	19767	-0.0386086783312822
	7.68677733461658	0.0719884581140171
0.788463288329429	0.955205336728381
54962	TIPIN	TIMELESS interacting protein
2084	-0.0860480554276605	1.66226012355181
0.0719845347519663	0.788468915774509	0.955205336728381

54496	PRMT7	protein arginine methyltransferase 7
9458	-0.0450023572990389	3.86103120637902
0.0718583632200165	0.788649976696457	0.955310148797058
9869	SETDB1	SET domain bifurcated histone lysine
methyltransferase 1	7093	-0.0391390452832437
4.97429381282171	0.07182920993716	0.788691836987876
	0.955310148797058
5685	PSMA4	proteasome 20S subunit alpha 4
6996	-0.0399565608056648	6.8835664305066 0.071711990776862
	0.788860240301369	0.955431507643863
201895	SMIM14	small integral membrane protein 14	6721
0.0488277629752039	6.91775621375469	0.0715164322955957
	0.789141518707211	0.95561932715111
58513	EPS15L1 epidermal growth factor receptor pathway substrate
15 like 1	8431	-0.0345480687434517	5.51394981945668
	0.0714339802753545	0.789260235740023
0.95561932715111
400673	VMAC	vimentin type intermediate filament associated
coiled-coil protein	2254	0.0726110352656013
2.02170802850707	0.0714333379625702	0.789261160850419
	0.95561932715111
10801	SEPTIN9 septin 9	12844	-0.0411200007983042
8.58696177013145	0.071387800314028	0.789326759185554
	0.95561932715111
10236	HNRNPR	heterogeneous nuclear ribonucleoprotein R	8340
	0.0302202848056093	7.03445168997935
0.0713672033068677	0.78935643714845	0.95561932715111
51493	RTCB	RNA 2',3'-cyclic phosphate and 5'-OH ligase
2948	-0.0352538679301537	7.06536498397446
0.0713097845938933	0.789439195265968	0.955636920585119
197258	FCSK	fucose kinase	7405	-0.0530035120184687
3.92975180519026	0.0711584587986174	0.789657473667684
	0.955818547672247
79640	C22orf46	CTA-216E10.6	5341	-0.0472402618291714
	4.20176789868712	0.0709927143652873
0.789896835555045	0.955986795440578
692312	PPAN-P2RY11	PPAN-P2RY11 readthrough 2661
0.0449076365346105	3.95997586048908	0.0709565775289294
	0.789949062741762	0.955986795440578
2342	FNTB	farnesyltransferase, CAAX box, beta	3948
0.0417495013418719	4.4052536628335 0.0709204955181004
0.790001224903597	0.955986795440578
23584	VSIG2	V-set and immunoglobulin domain containing 2
1432	-0.0655192534027606	4.79911207961414
0.0707871576150012	0.790194108977124	0.956116437932166
7162	TPBG	trophoblast glycoprotein
6904	-0.0382788488367342	6.36831190640832
0.0707520842827023	0.79024487769977	0.956116437932166
27072	VPS41	VPS41 subunit of HOPS complex	7801
0.0426902588210079	6.84556783701651	0.0705056448803703
	0.790601979805053	0.956319294665764
80895	ILKAP	ILK associated serine/threonine phosphatase
4771	-0.0381969000591494	4.26556356950665
0.0704556741952409	0.790674471076398	0.956319294665764

10138 YAF2 YY1 associated factor 2 10053 -0.0541288621662257
3.783601892478 0.0703729237361941 0.790794575719336
0.956319294665764
26469 PTPN18 protein tyrosine phosphatase non-receptor type 18
5774 -0.0536303228849044 5.11123361217129
0.0702544359334913 0.790966681483859 0.956319294665764
8573 CASK calcium/calmodulin dependent serine protein kinase
10529 -0.0632453285567802 7.04440509735736
0.0702529817837103 0.790968794628285 0.956319294665764
10328 EMC8 ER membrane protein complex subunit 8 2773
0.0384677045271396 4.69875315940646 0.0702397106293047
0.790988081113926 0.956319294665764
100527978 TLCD4-RWDD3 TLCD4-RWDD3 readthrough 1562
0.0614036082759693 2.40729423176887 0.0701362806086629
0.791138459063316 0.956319294665764
10161 LPAR6 lysophosphatidic acid receptor 6 4062
0.0555731962926208 3.44113166985685 0.0701135923925413
0.7911714615558 0.956319294665764
63906 GPATCH3 G-patch domain containing 3 2225
0.0494204719355947 3.80753373576692 0.0701046885416083
0.791184414742306 0.956319294665764
203259 FAM219A family with sequence similarity 219 member A 3782
0.03886623582329 5.54024307198815
0.0700942618235647 0.791199584491652 0.956319294665764
55109 AGGF1 angiogenic factor with G-patch and FHA domains 1
5051 -0.0339560826140197 5.41116785862164
0.0700728064007734 0.791230803614817 0.956319294665764
25913 POT1 protection of telomeres 1 5897
0.0557454960767825 3.59313041777105 0.0700720979802512
0.791231834502859 0.956319294665764
55699 IARS2 isoleucyl-tRNA synthetase 2, mitochondrial
4165 -0.0315834576592153 6.85977140810359
0.0700108524868739 0.791320979558171 0.956344517745582
84239 ATP13A4 ATPase 13A4 6013 -0.131371140364932
5.08896244107849 0.0698847185448557 0.791504703657664
0.956484029344569
6045 RNF2 ring finger protein 2 4073 0.0457111969730028
3.98236013815478 0.0697217527293734
0.791742339147084 0.956636127288622
145282 MIPOL1 mirror-image polydactyly 1
18149 -0.0697823150828517 2.52151554389748
0.0696767027826515 0.79180808303376 0.956636127288622
51347 TAOK3 TAO kinase 3 10570 -0.0344469387960046
5.31353047536319 0.0696579491378531 0.791835457969892
0.956636127288622
25 ABL1 ABL proto-oncogene 1, non-receptor tyrosine kinase
6276 -0.0357631972341508 7.06296560550957
0.0695507995357616 0.79199194127029 0.956680279318685
10251 SPRY3 sprouty RTK signaling antagonist 3 18038
0.0722831728711275 2.0694065407641 0.0695393950426109
0.792008604196435 0.956680279318685
58 ACTA1 actin alpha 1, skeletal muscle
1495 -0.200011941120813 3.47191106752297
0.0694727643211905 0.79210598652527 0.956715405123828

284344  LOC284344       uncharacterized LOC284344       2830
0.154765881650913       0.873205997437306       0.069346410034175
        0.792290793457107       0.956834357366156
151056  PLB1    phospholipase B1        8238    0.114610854917896
        1.07136970436948        0.0692692284003442
0.792403768608646       0.956834357366156
9860    LRIG2   leucine rich repeats and immunoglobulin like domains
2       12456   0.0540926454547363      4.39014212574302
0.0692653785017292      0.792409405686257       0.956834357366156
29117   BRD7    bromodomain containing 7
8492    -0.0345712273647894     5.96730374580478
0.0691648151858715      0.792556711333924       0.956869993929869
51301   GCNT4   glucosaminyl (N-acetyl) transferase 4
5554    -0.122770314125992      3.36075726814907
0.0691202301605962      0.792622056335156       0.956869993929869
1025    CDK9    cyclin dependent kinase 9       3000
0.0341603730752483      6.07348578135449        0.0691053572587723
        0.792643859468  0.956869993929869
23310   NCAPD3  non-SMC condensin II complex subunit D3 9114
0.0399953507975356      4.32781005338473        0.0690126639203754
        0.792779801095697       0.956946501299999
91544   UBXN11  UBX domain protein 11   4704    -0.0691883697979354
        3.95787706574002        0.0689689979257082
0.792843874319018       0.956946501299999
1785    DNM2    dynamin 2       12802   -0.0312504627351585
7.73611922819218        0.0688916856871273      0.792957371502056
        0.9569782584385
55127   HEATR1  HEAT repeat containing 1
9049    -0.0366956070004914     5.53201345972931
0.0687876535925867      0.793110201887247       0.9569782584385
3611    ILK     integrin linked kinase  3484    0.0367341967843358
        7.37265641935413        0.0687665160561437
0.793141269494074       0.9569782584385
54959   ODAM    odontogenic, ameloblast associated      1562
0.220545545599583       0.286466240132088       0.0686961048140837
        0.793244795619301       0.9569782584385
64853   AIDA    axin interactor, dorsalization associated       4232
        0.0427314384140841      4.86704135480286
0.0686827724774783      0.793264404627611       0.9569782584385
25885   POLR1A  RNA polymerase I subunit A
13519   -0.0371325928546994     6.32955233298781
0.0686720924413748      0.79328011413789        0.9569782584385
22862   FNDC3A  fibronectin type III domain containing 3A
7698    -0.0448752368049811     6.25127834098938
0.0685981994536178      0.793388840810135       0.957026997349945
57692   MAGEE1  MAGE family member E1   3630    0.0767896896738502
        2.71432087460571        0.068497121331589
0.793537669276869       0.957040752398452
51317   PHF21A  PHD finger protein 21A  11190   0.0482008157774507
        5.20788571093592        0.0684336750272649
0.793631148214178       0.957040752398452
81532   MOB2    MOB kinase activator 2  4518    -0.0419594651221067
        4.98844328024192        0.0684183359738544
0.793653755036131       0.957040752398452

23283	CSTF2T	cleavage stimulation factor subunit 2 tau variant
	4105	-0.0402769637779922	4.69627459264764
0.0683843372874691	0.793703872230555	0.957040752398452
117608	ZNF354B	zinc finger protein 354B	3112
0.0580889048197678	3.11782701874141	0.0683160978583568
	0.793804503616265	0.957040752398452
23065	EMC1	ER membrane protein complex subunit 1
10650	-0.0399035008995711	6.48934206257246
0.0683122366416988	0.793810199283587	0.957040752398452
55133	SRBD1	S1 RNA binding domain 1 5045	0.056395994029941
	3.32175029145107	0.068180986552477
0.794003908201625	0.957191904826591
9270	ITGB1BP1	integrin subunit beta 1 binding protein 1
	6678	-0.0384474409853933	6.13908532269261
0.0680773100128675	0.794157062805279	0.957294146188773
22996	TTC39A	tetratricopeptide repeat domain 39A
9362	-0.112299533956239	3.17542827130823
0.0678785703121383	0.79445099665063	0.957566053703242
123355	LRRC28	leucine rich repeat containing 28	6072
0.0487086488950796	3.7258712305089 0.0678124225912069
0.794548930311698	0.957601692210975
9399	STOML1	stomatin like 1 10461	-0.0444440833434671
4.44112076356015	0.0677452470052131	0.794648437970225
	0.957615228697636
9518	GDF15	growth differentiation factor 15
1814	-0.116995069863228	7.80811374289921
0.0677125470363933	0.794696895722233	0.957615228697636
6635	SNRPE	small nuclear ribonucleoprotein polypeptide E
1474	-0.0621517691486022	3.02699971002862
0.0675194128577594	0.794983354425661	0.957712010099765
114883	OSBPL9	oxysterol binding protein like 9
9523	-0.0281873419381855	6.44227121393332
0.0675063377277398	0.795002763398207	0.957712010099765
112495	GTF3C6	general transcription factor IIIC subunit 6	1067
	0.0506052500930858	5.0357014242234 0.0674523580517405
	0.795082913108274	0.957712010099765
9205	ZMYM5	zinc finger MYM-type containing 5	3713
0.0705154107095367	4.58277669680975	0.0674347028274092
	0.795109135239255	0.957712010099765
116541	MRPL54	mitochondrial ribosomal protein L54	1003
0.0508913211419105	4.14312240587982	0.0674280071454234
	0.795119080848873	0.957712010099765
80010	RMI1	RecQ mediated genome instability 1
3507	-0.0671268066113235	2.93634294877686
0.067262762271346	0.795364698947023	0.957895705565736
100505549	LOC100505549	uncharacterized LOC100505549	3879
	0.0687687947295507	1.21242132387762
0.0672022942195387	0.795454658434876	0.957895705565736
23399	CTDNEP1	CTD nuclear envelope phosphatase 1
2584	-0.0324831412630702	5.9234770163212 0.0671715407940212
	0.795500427469707	0.957895705565736
10956	OS9	OS9 endoplasmic reticulum lectin	5169
0.0311910276430968	7.83836516457396	0.0671191103222526
	0.795578483332012	0.957895705565736

1628	DBP	D-box binding PAR bZIP transcription factor	6544
	0.12572988515527	1.61897419773847
0.066995358469816	0.795762848006221	0.957895705565736
7329	UBE2I	ubiquitin conjugating enzyme E2 I	8895
0.0324820516273782	6.66487563423909	0.066872310619857
	0.795946344087089	0.957895705565736
147949	ZNF583	zinc finger protein 583 5613	-0.0631535426022661
	2.85219172228396	0.0668359521084279
0.79600059847259	0.957895705565736
643707	GOLGA6L4	golgin A6 family like 4
6557	-0.0588766349158285	3.81312488449059
0.0668064848747214	0.796044581204114	0.957895705565736
201266	SLC39A11	solute carrier family 39 member 11	5583
	0.0640459650342617	4.63617501277104
0.066669860123076	0.79624864236159	0.957895705565736
11183	MAP4K5	mitogen-activated protein kinase kinase kinase
kinase 5	7620	0.0333694710390655	6.8405739227382
0.0666276532170116	0.796311727196321	0.957895705565736
2717	GLA	galactosidase alpha	2065	0.0484293337290893
	4.53162066381149	0.0665433823794608
0.796437746955894	0.957895705565736
7564	ZNF16	zinc finger protein 16	5237	-0.0569464062151068
	2.80450206739623	0.0665165836909587
0.796477839913998	0.957895705565736
51164	DCTN4	dynactin subunit 4	6417	0.0297776408528868
	6.42686900774079	0.0664735805540868
0.796542194014025	0.957895705565736
6049	RNF6	ring finger protein 6	4621	-0.0432780058041007
	6.96833198255178	0.0664436328824749
0.796587023753168	0.957895705565736
101927686	LINC02541	long intergenic non-protein coding
RNA 2541	637	0.125727523683842	0.411341186692759
	0.0664376775507494	0.796595939786129
0.957895705565736
414918	DENND6B	DENN domain containing 6B	5935
0.0804828786455758	2.22083430722694	0.0664144759047165
	0.796630680226855	0.957895705565736
6583	SLC22A4	solute carrier family 22 member 4
2688	-0.0699233541761631	1.73989794853527
0.0663546383086961	0.796720306525253	0.957895705565736
10272	FSTL3	follistatin like 3	3067	-0.042226619559012
	8.85300193210647	0.0663478836926288
0.796730426470224	0.957895705565736
84516	DCTN5	dynactin subunit 5	12191	0.042477088641615
	7.38442046081915	0.066324221770472
0.79676588172294	0.957895705565736
51009	DERL2	derlin 2	6386	-0.0367239252935887
5.50535627787869	0.0662958810762682	0.79680835657016
	0.957895705565736
56243	KIAA1217	KIAA1217	10442	-0.0341182264953171
	6.97851813571625	0.0662661848549391
0.796852873364436	0.957895705565736
79037	PVRIG	PVR related immunoglobulin domain containing	3058
	0.0875793179394446	2.7547854600778

0.0662648584911381	0.796854861928288	0.957895705565736
57017	COQ9	coenzyme Q9	4324	-0.0357491235501725
5.15435325070279	0.0661890974903585	0.796968482589599
	0.957895705565736
3609	ILF3	interleukin enhancer binding factor 3	11022
0.0301316344942859	7.78187096165543	0.0661723356143682
	0.796993630157965	0.957895705565736
147965	FAM98C	family with sequence similarity 98 member C	4598
	0.0575927134564048	3.0313165699983 0.066094063341966
	0.797111105743756	0.957895705565736
166336	PRICKLE2	prickle planar cell polarity protein 2	8726
	0.0506397274607139	5.25798388044858
0.0660646685877886	0.797155242256433	0.957895705565736
26128	KIFBP	kinesin family binding protein
3054	-0.0307390210309955	6.0421751145697 0.0660636536387074
	0.797156766399406	0.957895705565736
10534	ZNRD2	zinc ribbon domain containing 2 1428
0.0490168734510306	3.77891584550826	0.066006284754522
	0.797242937210016	0.957895705565736
91663	MYADM	myeloid associated differentiation marker	4230
	0.0673865825232269	8.00278623757843
0.0659972959709307	0.797256442410199	0.957895705565736
129685	TAF8	TATA-box binding protein associated factor 8	9286
	0.0455542029113909	4.83916363562044
0.0659528745579294	0.797323197771893	0.957895705565736
23246	BOP1	BOP1 ribosomal biogenesis factor	3716
0.0535023054299928	4.0853758546483 0.0657820437560162
0.797580141456916	0.958070024113826
23601	CLEC5A	C-type lectin domain containing 5A
4129	-0.278112485649906	1.15265867640094
0.0657335661940159	0.797653120483138	0.958070024113826
8498	RANBP3	RAN binding protein 3	7445	0.0353201627470415
	5.92736021820051	0.0657200365766926
0.797673493340147	0.958070024113826
51025	PAM16	presequence translocase associated motor 16
2383	-0.0631713615829331	3.05496287403952
0.0655906410688019	0.797868449706322	0.958156883575909
100289187	TMEM225B	transmembrane protein 225B	1666
	0.0692906122057856	2.23113584032508
0.0655188576401122	0.79797669204513	0.958156883575909
22827	PUF60	poly(U) binding splicing factor 60	5424
0.0315782713276407	6.78940673997144	0.0654861266312379
	0.798026068154226	0.958156883575909
58499	ZNF462	zinc finger protein 462 12259	-0.0527596647835552
	4.62039055079877	0.0654244802772475
0.798119099995242	0.958156883575909
326625	MMAB	metabolism of cobalamin associated B
6290	-0.0674153614676101	3.88282082388729
0.0653682202248511	0.798204044016368	0.958156883575909
101928069	LOC101928069	uncharacterized LOC101928069	1833
	0.0874000942132136	0.766216574044971
0.0651811935668438	0.798486705743346	0.958156883575909
55188	RIC8B	RIC8 guanine nucleotide exchange factor B	7795
	0.0494213155137444	3.86031278956198

0.0651732069092974	0.798498785946264	0.958156883575909
23595	ORC3	origin recognition complex subunit 3	3689
0.0416044626541476	4.16868200041045	0.0650014053421408
	0.79875883519786	0.958156883575909
6867	TACC1	transforming acidic coiled-coil containing protein 1
	12550	-0.0665103098527859	7.24577353444683
0.064951890956479	0.798833851096647	0.958156883575909
126308	MOB3A	MOB kinase activator 3A 4149	0.0414485807346003
	5.38214950180187	0.0649096852787601
0.798897818109667	0.958156883575909
348180	CTU2	cytosolic thiouridylase subunit 2
3532	-0.0704293904677506	1.84446330221676
0.0648926468963782	0.798923647792196	0.958156883575909
92960	PEX11G	peroxisomal biogenesis factor 11 gamma	1466
0.0798151652755243	0.412379928907737	0.0648281512594302
	0.799021453984344	0.958156883575909
55173	MRPS10	mitochondrial ribosomal protein S10
2110	-0.0432904649732351	5.31658222514953
0.0648197222696254	0.799034240186144	0.958156883575909
53	ACP2	acid phosphatase 2, lysosomal
3484	-0.0344554027988279	5.87228133178402
0.0648123477579521	0.799045427542704	0.958156883575909
7846	TUBA1A	tubulin alpha 1a	3421	0.0585335513259621
	9.82055814051004	0.0647251667546698
0.799177735149856	0.958156883575909
85236	H2BC12	H2B clustered histone 12	381
0.0546698433581588	4.27996722899453	0.0647231947275202
	0.799180729034049	0.958156883575909
11338	U2AF2	U2 small nuclear RNA auxiliary factor 2 4115
0.0323087202947602	7.35065214516722	0.0647192470554643
	0.799186722440738	0.958156883575909
5935	RBM3	RNA binding motif protein 3
3731	-0.0513280790248095	7.52111458129816
0.064700225053798	0.799215604617899	0.958156883575909
284366	KLK9	kallikrein related peptidase 9
1986	-0.116002201792613	2.01784632278854
0.0646288615558106	0.799324000149109	0.958156883575909
23091	ZC3H13	zinc finger CCCH-type containing 13
10265	-0.0322803454903285	6.22702747567857
0.0645618953413902	0.799425774513423	0.958156883575909
54704	PDP1	pyruvate dehyrogenase phosphatase catalytic subunit
1	5215	-0.0431585207303659	6.27565205312524
0.0644905347619513	0.799534289236332	0.958156883575909
25963	TMEM87A transmembrane protein 87A	5668
0.0364555445819501	7.32135746620158	0.0644802357055454
	0.799549955810176	0.958156883575909
79958	DENND1C DENN domain containing 1C	4657
0.128738206696601	0.46072228493698	0.0644233893632196
	0.799636452502869	0.958156883575909
55131	RBM28	RNA binding motif protein 28
15866	-0.0382367148244526	4.73953756091639
0.0643558329192686	0.799739298391468	0.958156883575909
285761	DCBLD1	discoidin, CUB and LCCL domain containing 1	6596
	0.0625160592594973	6.23391309118607

0.0642508520336662 0.799899232195931 0.958156883575909
7277 TUBA4A tubulin alpha 4a 4079 -0.0466734613270875
7.42377368333431 0.0642433347260294
0.79991068981999 0.958156883575909
26015 RPAP1 RNA polymerase II associated protein 1
7046 -0.0404542774441896 4.88783278452938
0.0642377482631371 0.799919204978395 0.958156883575909
3920 LAMP2 lysosomal associated membrane protein 2 5526
0.0325410121072241 7.85964710375745 0.0642162737242415
0.799951941184093 0.958156883575909
114876 OSBPL1A oxysterol binding protein like 1A 8022
0.0467305519982792 5.43226535788995 0.0641979150288847
0.799979932156131 0.958156883575909
29104 N6AMT1 N-6 adenine-specific DNA methyltransferase 1
4865 -0.0642220324403064 2.74882448925527
0.0640723408734418 0.800171505745904 0.958156883575909
9973 CCS copper chaperone for superoxide dismutase 2474
0.0442063065120029 3.86193145203264
0.0640455039165904 0.800212473608257 0.958156883575909
9933 PUM3 pumilio RNA binding family member 3 2707
0.0520229615879238 4.71810100300781 0.0640208614810334
0.800250099475967 0.958156883575909
342357 ZKSCAN2 zinc finger with KRAB and SCAN domains 2 7970
0.0547035584964904 3.51292370949206
0.0640103664658049 0.800266126370085 0.958156883575909
29090 TIMM21 translocase of inner mitochondrial membrane 21
5208 -0.0453018873230498 3.6480474701292 0.0639946318322906
0.800290157282814 0.958156883575909
340390 WDR97 WD repeat domain 97 8967 -0.0777919629137162
0.781576181233891 0.0639792306479912
0.800313681975451 0.958156883575909
91227 GGTLC2 gamma-glutamyltransferase light chain 2 1229
0.112425309865245 0.356264288076132 0.0639659466694891
0.800333975140938 0.958156883575909
55324 ABCF3 ATP binding cassette subfamily F member 3 5207
0.0312663470538018 6.03441918898395
0.0639217149416993 0.80040156155816 0.958156883575909
340527 NHSL2 NHS like 2 15788 -0.123581014870797
1.27308990708148 0.0639215851278818 0.800401759951271
0.958156883575909
339804 C2orf74 chromosome 2 open reading frame 74 2131
0.0621299223357611 3.25093953083983 0.0639138194651849
0.800413628523004 0.958156883575909
1606 DGKA diacylglycerol kinase alpha
10215 -0.0414757894623438 7.217272790273 0.0637775699963363
0.800621988942562 0.958324406009098
2527 FUT5 fucosyltransferase 5 2463 -0.0393929780138329
5.5903488676991 0.0636891738072336 0.800757296066075
0.958404464325174
23094 SIPA1L3 signal induced proliferation associated 1 like 3
10335 0.047901283943755 6.19250325683184
0.0636067664872968 0.800883526076988 0.958473645741659
6182 MRPL12 mitochondrial ribosomal protein L12 1028
0.043038460703861 4.44342349908983 0.0634241868122949

0.801163508405603	0.958671109331576
148789	B3GALNT2	beta-1,3-N-acetylgalactosaminyltransferase 2
5702	0.0386612819529113	4.4727344097525
0.0634099177913257	0.80118540771296	0.958671109331576
100499183	BAALC-AS1	BAALC antisense RNA 1	2027
0.0768633873363846	-0.139112550164062	0.0633202483009896
0.801323087516735	0.958713818834539
375790	AGRN	agrin	10758	0.0348194765528494
9.17449275682015	0.0632975262178377	0.801357991783735
0.958713818834539
338699	ANKRD42	ankyrin repeat domain 42
6724	-0.0400955490762483	4.44060022665779
0.0629547270413027	0.8018853892246	0.95926284455282
1736	DKC1	dyskerin pseudouridine synthase 1	4203
0.0395618716405788	5.05202770112272	0.0628197272899627
0.802093505521002	0.959429866083063
285755	PPIL6	peptidylprolyl isomerase like 6	4481
0.103512956922386	-0.00883416348354442	0.0626879351196798
0.802296906385352	0.959498675822601
100507412	LOC100507412	uncharacterized LOC100507412	4996
0.0856976554583155	-0.0399772751956147
0.0626785758432824	0.802311359620594	0.959498675822601
5318	PKP2	plakophilin 2	7043	-0.0428009823516613
6.46091779996251	0.0626493268588888	0.80235653529736
0.959498675822601
55734	ZFP64	ZFP64 zinc finger protein
6997	-0.0456961847335434	4.49469309411617
0.0624619149775754	0.802646263227054	0.959636469293424
84878	ZBTB45	zinc finger and BTB domain containing 45	3418
0.0411109102583748	4.20367360566302
0.0624517987696924	0.802661915428006	0.959636469293424
51226	COPZ2	coatomer protein complex subunit zeta 2
1851	-0.0810428597971735	1.48582198363356
0.0624188277350957	0.802712938880451	0.959636469293424
9650	MTFR1	mitochondrial fission regulator 1
4386	-0.0383264398028324	4.88915672234431
0.0623975938267254	0.802745806433287	0.959636469293424
51057	WDPCP	WD repeat containing planar cell polarity effector
11776	0.0705391956092334	1.57197220014704
0.0623251769507318	0.802857943809572	0.959688617421119
55672	NBPF1	NBPF member 1	11006	0.0479987339163418
5.22730459779906	0.062229959105963	0.80300549381219
0.959767697382943
29760	BLNK	B cell linker	3655	-0.092761421915661
1.15966539416357	0.0621940680888144	0.803061141809673
0.959767697382943
10092	ARPC5	actin related protein 2/3 complex subunit 5
8044	-0.0331126218430197	8.02668285207318
0.0620578924260826	0.803272433549393	0.959925430051341
257236	CCDC96	coiled-coil domain containing 96	2091
0.0767984512784861	1.72350377573174	0.0620083264408615
0.803349401819177	0.959925430051341
65055	REEP1	receptor accessory protein 1	5193
0.127950077826221	3.79814012360527	0.0619765801064993

0.803398716105652 0.959925430051341
23029 RBM34 RNA binding motif protein 34 3113
0.0399301859687042 5.4308180998679 0.0619269611516629
0.803475820329411 0.959935671729279
65125 WNK1 WNK lysine deficient protein kinase 1 14208
0.0398969878896067 8.44042693667063 0.0618377568265487
0.803614519860595 0.959940145454588
401265 KLHL31 kelch like family member 31 5743
0.080226639813953 0.614291472324055 0.0618363998000317
0.803616630656136 0.959940145454588
51125 GOLGA7 golgin A7 2474 -0.030423273203826
6.00677374167424 0.0617012845551272 0.80382691978514
0.959981665739911
7203 CCT3 chaperonin containing TCP1 subunit 3
4018 -0.0361060538356271 7.86837045965074
0.0616643862038018 0.803884389707533 0.959981665739911
388567 ZNF749 zinc finger protein 749 2587 0.0556783232991454
2.77401204047546 0.0616190834590957
0.803954974611679 0.959981665739911
286128 ZFP41 ZFP41 zinc finger protein 7516
0.0535279156168761 4.10251929286813 0.0616186396985103
0.803955666158502 0.959981665739911
100529145 TEN1-CDK3 TEN1-CDK3 readthrough (NMD candidate) 3898 0.0394128882271307 4.99803931084747
0.0615939997050052 0.803994068736696
0.959981665739911
100526740 ATP5MF-PTCD1 ATP5MF-PTCD1 readthrough 2877
0.0369711203076677 4.69824639118121
0.0615314620252088 0.804091573252394 0.95999048855483
220906 WAC-AS1 WAC antisense RNA 1 (head to head) 2943
0.0564932222208974 4.85298615705484 0.0615013562508473
0.804138530892684 0.95999048855483
10380 BPNT1 3'(2'), 5'-bisphosphate nucleotidase 1 3113
0.0485953058473473 5.88780188886469 0.0610964767477356
0.804771232859484 0.960591514607271
220594 USP32P2 ubiquitin specific peptidase 32 pseudogene 2
6980 -0.10756402524416 1.84996270348668
0.0610914253235357 0.804779140709383 0.960591514607271
51678 MPP6 membrane palmitoylated protein 6 9369
0.0549941340682598 2.72835516813212 0.0610355286312165
0.804866668437632 0.96061412958469
4000 LMNA lamin A/C 6585 -0.0405541897973051
8.08303218202657 0.0608964094039948 0.80508469778939
0.960679425488034
10827 FAM114A2 family with sequence similarity 114 member A2 6543 0.0486466467170101 3.85343201762057
0.0608395772677208 0.805173841816639 0.960679425488034
81037 CLPTM1L CLPTM1 like 6557 -0.0307512550201869
6.98101772207624 0.060834710918499 0.805181476979593
0.960679425488034
5295 PIK3R1 phosphoinositide-3-kinase regulatory subunit 1
10767 -0.0472399446737177 4.77836680803347
0.0608097216097754 0.805220689595342 0.960679425488034
8496 PPFIBP1 PPFIA binding protein 1 10330 0.0462111902506426

6.6391564340381 0.0607819322404675 0.805264306036625
0.960679425488034
126382 NR2C2AP nuclear receptor 2C2 associated protein 1794
0.0521506342161502 2.83495424581953 0.0606177603723113
0.805522195922243 0.960731738818335
10924 SMPDL3A sphingomyelin phosphodiesterase acid like 3A 2349
0.0661597241920159 3.2930539267317 0.0606067928636316
0.805539437458753 0.960731738818335
8019 BRD3 bromodomain containing 3
6876 -0.0456950327945631 5.77547235151489
0.0605555968522538 0.805619942346008 0.960731738818335
85371 KIAA1656 KIAA1656 protein
8141 -0.0611744781878055 3.63230061226898
0.060547060580781 0.805633369002916 0.960731738818335
166793 ZBTB49 zinc finger and BTB domain containing 49 2889
0.0738210229106067 2.20452375914911
0.0605357866063589 0.805651103316925 0.960731738818335
51363 CHST15 carbohydrate sulfotransferase 15 6554
0.0351954529910532 7.12834852292524 0.0603937271160646
0.805874717571834 0.960874995280864
10093 ARPC4 actin related protein 2/3 complex subunit 4
3827 -0.0431298198383843 7.52010399128285
0.0603139748895956 0.806000376902281 0.960874995280864
441908 NBPF18P NBPF member 18, pseudogene 4109
0.107680386063923 -0.324520929711259 0.0602952941399906
0.806029823426314 0.960874995280864
84945 ABHD13 abhydrolase domain containing 13
5338 -0.0374296012333893 5.03819691588683
0.0602484203176754 0.80610373207965 0.960874995280864
140831 ZSWIM3 zinc finger SWIM-type containing 3
2773 -0.0663092784793341 2.69267594881815
0.0602234091247054 0.806143181139507 0.960874995280864
163059 ZNF433 zinc finger protein 433 3079 -0.0698791228415573
2.12561479795739 0.0601734370931357
0.806222025857143 0.960874995280864
116983 ACAP3 ArfGAP with coiled-coil, ankyrin repeat and PH
domains 3 8809 -0.0468681975322007 6.31854312594433
0.0601548044960154 0.806251432821874
0.960874995280864
79165 LENG1 leukocyte receptor cluster member 1
1542 -0.0535209432618704 2.86931089609243
0.0600085307448617 0.806482457828268 0.960886819621833
8883 NAE1 NEDD8 activating enzyme E1 subunit 1
4810 -0.0345091597527561 5.73156789450082
0.0600048125771133 0.806488334196372 0.960886819621833
55972 SLC25A40 solute carrier family 25 member 40
5081 -0.0454268271613975 3.53381798671131
0.0599922838761537 0.806508136568597 0.960886819621833
26507 CNNM1 cyclin and CBS domain divalent metal cation
transport mediator 1 6413 0.169460786156155
3.53340351522584 0.0599321970605473 0.806603137903455
0.960886819621833
285973 ATG9B autophagy related 9B 6386 -0.137094195037863
3.11204300872171 0.0599314263617998

0.8066043567583 0.960886819621833
51691 LSM8 LSM8 homolog, U6 small nuclear RNA associated
6210 -0.0361675952356524 4.46279657416041
0.0598326183970892 0.806760689740424 0.960916176513882
2027 ENO3 enolase 3 4292 0.0785092668359626
2.29681266664121 0.0598291340292523 0.806766205169076
0.960916176513882
85437 ZCRB1 zinc finger CCHC-type and RNA binding motif
containing 1 2446 -0.0371168955024884 4.35226173581862
0.0597728830165067 0.806855268830705
0.960929276539392
83642 SELENOO selenoprotein O 2363 -0.0430171421705449
4.46917310996692 0.0597355455626456 0.806914410698675
0.960929276539392
100188893 TOMM6 translocase of outer mitochondrial membrane
6 637 0.0403178190977881 5.4850656953837
0.0594815012141652 0.807317333506874 0.961114048358953
23274 CLEC16A C-type lectin domain containing 16A 12387
0.0360164972274069 5.94238107441757 0.0594631107041508
0.807346536825337 0.961114048358953
9454 HOMER3 homer scaffold protein 3
3686 -0.0367430576665418 7.67377421264212
0.0594623112754675 0.807347806391316 0.961114048358953
6203 RPS9 ribosomal protein S9 856 0.0460191950908816
7.23984889800597 0.0594305702906581
0.807398221291051 0.961114048358953
10212 DDX39A DExD-box helicase 39A 3840 0.0352997914058135
5.31493465271198 0.0594214868645153
0.80741265132004 0.961114048358953
84272 YIPF4 Yip1 domain family member 4
12233 -0.0321090782165866 6.1983423835874 0.0593702574034403
0.807494056882615 0.961129270458428
8339 H2BC8 H2B clustered histone 8 1534 -0.0741292211488713
3.25114927535793 0.0593062558204664
0.807595810127588 0.961157491832912
4337 MOCS1 molybdenum cofactor synthesis 1
5174 -0.0854895730552023 4.63829011673076
0.059248305368377 0.807687993135276 0.961157491832912
3476 IGBP1 immunoglobulin binding protein 1 1862
0.0315196361225022 5.64879299396878 0.0592027059267028
0.807760562706246 0.961157491832912
55707 NECAP2 NECAP endocytosis associated 2 5559
0.0334347244252285 7.01162271416397 0.0591410139645063
0.80785878997048 0.961157491832912
55692 LUC7L LUC7 like 6176 0.0354582706112723
5.44165324413221 0.0590812071035209 0.807954067549895
0.961157491832912
22828 SCAF8 SR-related CTD associated factor 8
5304 -0.0275207813865471 6.03637007345855
0.0590127914266816 0.808063122379535 0.961157491832912
54850 FBXL12 F-box and leucine rich repeat protein 12 2913
0.0387141553560751 4.10413337752635
0.0590118921977814 0.808064556199193 0.961157491832912
23192 ATG4B autophagy related 4B cysteine peptidase 6638

0.0284419931956455	6.05535783799502	0.0590105317324827
	0.808066725481857	0.961157491832912
727866	FAM156B family with sequence similarity 156 member B	1701
	0.0666429187491391	1.41123143313058
0.0589115269232039	0.808224661303429	0.961185572651711
55689	YEATS2	YEATS domain containing 2
10212	-0.0347623248998314	6.43109470917508
0.0589096998323129	0.808227577260788	0.961185572651711
116844	LRG1	leucine rich alpha-2-glycoprotein 1
3133	-0.109988820610159	2.64088340255327
0.0588227722498909	0.808366365262749	0.961269010801879
6895	TARBP2	TARBP2 subunit of RISC loading complex	4090
0.0397046357031311	4.84969285958387	0.0586474067369114
	0.808646683531064	0.961481551617946
54806	AHI1	Abelson helper integration site 1
8705	-0.0556859535105637	3.783274075766	0.0585770337546476
	0.808759298072765	0.961481551617946
79770	TXNDC15 thioredoxin domain containing 15
5265	-0.0531125904716367	5.03234956576114
0.0585316942763541	0.80883189062071	0.961481551617946
79834	PEAK1	pseudopodium enriched atypical kinase 1
24184	-0.0453019725840658	5.20664121943862
0.0583734770934612	0.809085443843739	0.961481551617946
10500	SEMA6C	semaphorin 6C	5809	0.0825177334583677
3.31772919877906	0.0583591015821412	0.809108499540268
	0.961481551617946
54494	C11orf71	chromosome 11 open reading frame 71
2242	-0.0688436001357553	0.402687759229501
0.0583353795410804	0.809146551937203	0.961481551617946
129401	NUP35	nucleoporin 35	4046	-0.0543275515379737
2.69730255360272	0.0583225509506775	0.809167133620709
	0.961481551617946
54865	GPATCH4 G-patch domain containing 4
4437	-0.0430722821826442	4.42551988438241
0.0582747271294011	0.809243881388552	0.961481551617946
79723	SUV39H2 suppressor of variegation 3-9 homolog 2 3565
0.055855366412341	2.68680464908164	0.0582186408714587
	0.809333931205476	0.961481551617946
126231	ZNF573	zinc finger protein 573 3495	-0.0783558417512668
	1.04629125324272	0.0581994573883251
0.809364741960793	0.961481551617946
553158	PRR5-ARHGAP8	PRR5-ARHGAP8 readthrough	3264
0.0390943323280809	5.1035035186148 0.0581867421040059
0.809385167048282	0.961481551617946
4677	NARS1	asparaginyl-tRNA synthetase 1
4089	-0.0302592407003607	7.85782318804568
0.0581794295781606	0.809396914530817	0.961481551617946
53344	CHIC1	cysteine rich hydrophobic domain 1
6939	-0.0625462334793896	3.79359399215121
0.0581541967556491	0.809437456748684	0.961481551617946
54475	NLE1	notchless homolog 1	5463	0.0655259233450478
	2.76901233875901	0.0580080741100373
0.80967241893499	0.961559600071823
57489	ODF2L	outer dense fiber of sperm tails 2 like

10898	-0.0354061804138082	6.09016165948257
0.0579973193478054	0.809689724734229	0.961559600071823
8971	H1-10	H1.10 linker histone	1621	-0.0387074571373601
	5.51840695311686	0.0579852744730687
0.809709108506257	0.961559600071823
9968	MED12	mediator complex subunit 12
7425	-0.037873341588289	5.67717659084477
0.0578527397306026	0.809922536986318	0.961574001673542
102724316	SVIL-AS1	SVIL antisense RNA 1	4995
0.0481695858523958	5.17992652677793	0.0578289517612429
	0.80996087151617	0.961574001673542
8771	TNFRSF6B	TNF receptor superfamily member 6b	1124
	-0.101707973355097	1.05672323116263
0.0578144316822531	0.809984274855407	0.961574001673542
22835	ZFP30	ZFP30 zinc finger protein
6839	-0.0623266460126089	3.87685727556317
0.0578072608290725	0.809995833922048	0.961574001673542
171391	LOC171391	uncharacterized LOC171391	1059
0.0896864014743268	0.212420943332268	0.0577622200289269
	0.810068454849521	0.961578715854003
1400	CRMP1	collapsin response mediator protein 1	5823
0.247704983414121	0.954360920699784	0.0576676263062428
	0.81022106932335	0.961678376240332
23197	FAF2	Fas associated factor family member 2
5954	-0.0273714021938062	7.09561263280429
0.0575835443349284	0.810356835650527	0.961691774412753
284352	PPP1R37	protein phosphatase 1 regulatory subunit 37	7354
	0.0355504745923894	5.2089786051143	0.0575652209140332
	0.81038643620605	0.961691774412753
10313	RTN3	reticulon 3	5792	0.0295538819854205
7.09436664965134	0.057507321547078	0.810480002450123
	0.961691774412753
509	ATP5F1C	ATP synthase F1 subunit gamma
2326	-0.0408407768419083	6.62412180324707
0.0574127836795526	0.810632884185652	0.961691774412753
79710	MORC4	MORC family CW-type zinc finger 4	6055
0.0383880604092515	4.47721095928693	0.0573947080158099
	0.81066213036503	0.961691774412753
310	ANXA7	annexin A7	2862	0.0307982681155267
7.12093750043536	0.0573833164487088	0.810680564268055
	0.961691774412753
56137	PCDHA12	protocadherin alpha 12	5454	-0.154990673554807
	3.15664666857825	0.0573503063471517
0.810733992343491	0.961691774412753
7539	ZFP37	ZFP37 zinc finger protein	6354
0.100921130929026	0.881142987751072	0.0573208878930345
	0.810781620906704	0.961691774412753
84289	ING5	inhibitor of growth family member 5	8200
0.0450454911929117	3.75446950675166	0.0572739133607953
	0.810857699610916	0.961700575651999
56005	MYDGF	myeloid derived growth factor
1678	-0.0482925031693123	6.03190091989886
0.0571235860247423	0.811101387702108	0.961847544509177
64762	GAREM1	GRB2 associated regulator of MAPK1 subtype 1	7568

0.0632980961297598	4.91678918831644
0.05709157590735	0.811153321454307	0.961847544509177
55028	C17orf80	chromosome 17 open reading frame 80
4663	-0.038129619077848	4.43680770623237
0.0570573223644786	0.811208912026604	0.961847544509177
84890	ADO	2-aminoethanethiol dioxygenase
3627	-0.0366278047712118	4.93124196019944
0.0570281361806813	0.811256292627359	0.961847544509177
57332	CBX8	chromobox 8	4164	-0.0560017728247021
3.00820116041102	0.0569237970075775	0.81142578083184
0.961853121613566
5150	PDE7A	phosphodiesterase 7A	7425	0.0452210742833747
5.01565444381106	0.0569109760337385
0.811446618501647	0.961853121613566
84159	ARID5B	AT-rich interaction domain 5B
8164	-0.0726789273544591	5.53375437174761
0.0568766340190621	0.811502446109289	0.961853121613566
10572	SIVA1	SIVA1 apoptosis inducing factor 4776
0.0409264068357273	3.92502874726221	0.0568561990415288
0.811535674393455	0.961853121613566
199746	U2AF1L4 U2 small nuclear RNA auxiliary factor 1 like 4	2875
0.061511633763786	3.07033048819404
0.0567600658099141	0.81169207652326	0.961957095850859
340547	VSIG1	V-set and immunoglobulin domain containing 1	3769
0.0851796227847551	0.915279123557812
0.0566671025271503	0.811843454539421	0.961973004261191
23741	EID1	EP300 interacting inhibitor of differentiation 1
2298	0.0329577596570665	6.23662202352425
0.0566380643248863	0.811890766177375	0.961973004261191
100506302	LOC100506302	uncharacterized LOC100506302	1683
0.0689542506442991	0.178308307872849
0.0566253202726137	0.811911533974142	0.961973004261191
2339	FNTA	farnesyltransferase, CAAX box, alpha	9622
0.0354757807595258	5.79484664953779	0.0565823892615001
0.811981512826552	0.961974545391315
23549	DNPEP	aspartyl aminopeptidase 4734	0.0399192450133648
4.86913363203947	0.0563762829559469
0.812317863582921	0.962291636942318
9541	CIR1	corepressor interacting with RBPJ, 1	3181
0.0329511825998556	4.74713523055072	0.0562883171751434
0.812461615192648	0.962380536490776
284098	PIGW	phosphatidylinositol glycan anchor biosynthesis class W 2799	0.049818610039214	3.68259573986119
0.0562110987818798	0.812587901495231	0.962398474026633
401052	NA	NA	2533	0.0809258843399135
0.0814346210562213	0.0561458151361887	0.812694740684514
0.962398474026633
81550	TDRD3	tudor domain containing 3
6814	-0.041936434403223	3.6210849442753 0.0561034270632539
0.812764145520001	0.962398474026633
132158	GLYCTK	glycerate kinase	3013	0.0653245279185344
2.026583848854	0.0560506969869921	0.812850522670377
0.962398474026633
1198	CLK3	CDC like kinase 3	17606	-0.0299120319360297

5.83533976692094	0.0560341556059321
0.81287762795117	0.962398474026633
84193	SETD3	SET domain containing 3, actin histidine methyltransferase	5188	0.028795412669701
6.35578045089946	0.0560272110526654	0.812889008796251
0.962398474026633
8896	BUD31	BUD31 homolog	2381	-0.031957602183404
5.82675649303668	0.0558885040660773	0.813116480225082
0.962586421442928
80304	WDCP	WD repeat and coiled coil containing	3847
0.0553284980789838	3.17762222500407	0.0556938132683484
0.813436265215321	0.962667626104856
79664	ICE2	interactor of little elongation complex ELL subunit 2	16010	-0.0357873764123555	4.93074214699376
0.0555465462512839	0.813678547517722	0.962667626104856
586	BCAT1	branched chain amino acid transaminase 1
11775	-0.0918450564328754	5.12812611774661
0.0555172771041867	0.813726741212571	0.962667626104856
128869	PIGU	phosphatidylinositol glycan anchor biosynthesis class U	2705	0.0353841446112097	4.5982933394945
0.0555090481977445	0.81374029309657	0.962667626104856
151246	SGO2	shugoshin 2	6022	-0.0589546767678581
2.83957055341833	0.055500995181907	0.813753556339329
0.962667626104856
374395	TMEM179B	transmembrane protein 179B	1496
0.0331584733462737	4.93365740415933	0.0554401960815127
0.813853724692706	0.962667626104856
25979	DHRS7B	dehydrogenase/reductase 7B	3867
0.0346434655284488	4.41536641224397	0.0554112101621556
0.813901500276779	0.962667626104856
64801	ARV1	ARV1 homolog, fatty acid homeostasis modulator	1987
0.0431622590804217	3.71215069211127
0.0553172202658185	0.81405650842082	0.962667626104856
169841	ZNF169	zinc finger protein 169	5286	-0.0666199280271367
0.932031125596798	0.0553124403087062
0.814064395238675	0.962667626104856
113179	ADAT3	adenosine deaminase tRNA specific 3	2237
0.082225381146892	1.38875413108595	0.0552708034964082
0.814133110224073	0.962667626104856
26993	AKAP8L	A-kinase anchoring protein 8 like	5081
0.0348714612458075	6.16055554228157	0.0551959866848009
0.814256652227275	0.962667626104856
23218	NBEAL2	neurobeachin like 2	9819	-0.0489517941552937
7.97920824215459	0.0551806818806426
0.814281935320498	0.962667626104856
80778	ZNF34	zinc finger protein 34	2222	0.0702130687486167
0.969346203525536	0.0550987260886924
0.814417386907732	0.962667626104856
131870	NUDT16	nudix hydrolase 16	7029	0.040312002747585
4.84365517808522	0.0550369219499807
0.814519603372893	0.962667626104856
27258	LSM3	LSM3 homolog, U6 small nuclear RNA and mRNA degradation associated	3815	0.0383670787624642
4.4419555474756	0.0549893454111547	0.814598330386639

0.962667626104856
132320	SCLT1	sodium channel and clathrin linker 1	8292
0.0561255795972181	2.29244317002654	0.0549761030838973
	0.814620249445811	0.962667626104856
150274	HSCB	HscB mitochondrial iron-sulfur cluster cochaperone
	1462	-0.0558584382907608	2.62276684054418
0.0549467230708949	0.814668889985979	0.962667626104856
6810	STX4	syntaxin 4	4078	-0.0314679626825896
5.69937129521482	0.0549210071595354	0.814711475625482
	0.962667626104856
11313	LYPLA2	lysophospholipase 2	2196	0.0312622555733308
	6.48316098391899	0.0548372945189719
0.814850176944834	0.962667626104856
23314	SATB2	SATB homeobox 2	10958	-0.0613373389708679
3.20215216038396	0.054832245597936	0.814858545941831
	0.962667626104856
5511	PPP1R8	protein phosphatase 1 regulatory subunit 8	2819
	0.0290382707337978	5.76837276370474
0.05480688423697	0.814900590608598	0.962667626104856
85027	SMIM3	small integral membrane protein 3	2332
0.0497233763097843	4.25427161890544	0.0547533891546799
	0.814989309688229	0.962667626104856
1600	DAB1	DAB adaptor protein 1	7548	-0.075098910220443
	0.359339044958805	0.0547470451524252
0.814999833949303	0.962667626104856
728642	CDK11A	cyclin dependent kinase 11A
7332	-0.0469720383995851	5.45043219650199
0.0547320402484957	0.815024728607219	0.962667626104856
284358	MAMSTR	MEF2 activating motif and SAP domain containing
transcriptional regulator	2265	-0.0781366702701143
0.182928437413729	0.054636041777953	0.815184085052784
	0.962667626104856
283130	SLC25A45	solute carrier family 25 member 45
7855	-0.0668327465615433	1.41438021574078
0.0545963809466459	0.815249964711597	0.962667626104856
2628	GATM	glycine amidinotransferase
5186	-0.11935567588558	2.46569753394157
0.0545772686266801	0.815281720739886	0.962667626104856
63931	MRPS14	mitochondrial ribosomal protein S14	2207
0.044997986599849	3.78719349303387	0.0545722070593246
	0.815290131756747	0.962667626104856
79810	PTCD2	pentatricopeptide repeat domain 2
12115	-0.0475043308270605	3.19734291857528
0.0545544866872572	0.815319581676962	0.962667626104856
100507171	BOLA3-AS1	BOLA3 divergent transcript	1834
	0.072304794879872	1.05939009936717
0.0545461476140354	0.815333442332241	0.962667626104856
401884	NA	NA	12734	0.0482553446577808
4.18953228002438	0.0545319500010306	0.815357043235095
	0.962667626104856
57097	PARP11	poly(ADP-ribose) polymerase family member 11	6200
	0.0564999498463271	3.60435935941335
0.0544367260303673	0.815515419257913	0.962667626104856
79018	GID4	GID complex subunit 4 homolog

5210	-0.0397600547445448	4.43838883105048
0.0544309149957094	0.815525088868157	0.962667626104856
390637	GDPGP1	GDP-D-glucose phosphorylase 1
5390	-0.0595201764062237	1.93812470512147
0.0544083464439229	0.815562648289427	0.962667626104856
10772	SRSF10	serine and arginine rich splicing factor 10
10277	-0.0292253208402193	6.37889110378088
0.054382564095377	0.815605566283277	0.962667626104856
84125	LRRIQ1	leucine rich repeats and IQ motif containing 1
7560	-0.103438920603018	0.995123548903663
0.0543503118128736	0.81565926940904	0.962667626104856
8816	DCAF5	DDB1 and CUL4 associated factor 5
8295	-0.0316401154202898	7.09495425789145
0.0542667962678678	0.815798408879465	0.96275072147398
89887	ZNF628	zinc finger protein 628 3847	-0.0614285461761261
	2.72647604871086	0.0541992565236873
0.815911014748549	0.962802492298477
10296	MAEA	macrophage erythroblast attacher	9417
0.0277896558742496	6.37662969830507	0.0540170969289733
	0.816215090217702	0.963080175948055
29967	LRP12	LDL receptor related protein 12
9777	-0.0298596772221391	6.53434622523775
0.0538727227850302	0.816456475265286	0.963127905443801
6949	TCOF1	treacle ribosome biogenesis factor 1
11278	-0.0369544758722697	4.60557042727421
0.0537707902565243	0.816627105793048	0.963127905443801
7247	TSN	translin	4407	-0.0290116175428717
6.25141213034647	0.0537676848428577	0.816632306791747
	0.963127905443801
10923	SUB1	SUB1 regulator of transcription 9324
0.0360055759132518	6.13490261196949	0.053732445294628
	0.816691337667806	0.963127905443801
28991	COMMD5	COMM domain containing 5
2215	-0.0312642950120266	4.77384729590227
0.0537094403253269	0.81672988502063	0.963127905443801
57473	ZNF512B	zinc finger protein 512B	6022
0.0355633531540704	5.83555777318228	0.0536936496424261
	0.816756349079066	0.963127905443801
8763	CD164	CD164 molecule	6944	-0.0349783586677083
8.03907018477485	0.0536818708563942	0.816776092154595
	0.963127905443801
138241	C9orf85	chromosome 9 open reading frame 85
4973	-0.0486297119741579	3.04325309782253
0.0536642540276899	0.816805624956587	0.963127905443801
9412	MED21	mediator complex subunit 21
2551	-0.0341551913282512	5.20642350400837
0.0535597829592938	0.816980865008887	0.963253449173357
7391	USF1	upstream transcription factor 1
2336	-0.0364466175364499	5.62231264734367
0.0534682899558589	0.817134483576349	0.96328273702941
85461	TANC1	tetratricopeptide repeat, ankyrin repeat and coiled-coil containing 1	14921	0.035857007579872
6.8362974902996	0.0534630721935798	0.81714324847458
0.96328273702941

440288 UBL7-AS1 UBL7 antisense RNA 1 (head to head)
1710 -0.061027192867629 0.828824821000374
0.0533472169380498 0.817337980573154 0.963363401849934
26503 SLC17A5 solute carrier family 17 member 5 3604
0.0300497566746638 6.25991923941525 0.053340528044675
0.817349230212366 0.963363401849934
2036 EPB41L1 erythrocyte membrane protein band 4.1 like 1 7859
0.0305806679348587 6.2694248489004 0.0531207717640179
0.81771923882109 0.963718416337148
56906 THAP10 THAP domain containing 10 2399
0.0722692515923999 1.28404906391425 0.0529342731185865
0.818033883559105 0.963923197311256
55721 IQCC IQ motif containing C 2478 -0.0763523471520138
1.33609271149283 0.0528188766606688
0.81822886305849 0.963923197311256
4706 NDUFAB1 NADH:ubiquinone oxidoreductase subunit AB1
1600 -0.0404648473653118 5.16453039155521
0.0527620152584234 0.818325021337632 0.963923197311256
85364 ZCCHC3 zinc finger CCHC-type containing 3
3354 -0.0444900415555667 3.61376003233471
0.0526166083530479 0.818571167235706 0.963923197311256
4999 ORC2 origin recognition complex subunit 2
5835 -0.0353217078157375 4.73804591183588
0.0526154782044648 0.818573081760914 0.963923197311256
6877 TAF5 TATA-box binding protein associated factor 5
3268 -0.0613231888992933 2.00630608625203
0.0525997612954057 0.818599709192753 0.963923197311256
6194 RPS6 ribosomal protein S6 2714 0.0495013371373581
8.23211943614229 0.0525872588552829
0.818620893693626 0.963923197311256
80195 TMEM254 transmembrane protein 254
2996 -0.0527689764176366 3.87167318367823
0.052576499027186 0.81863912758398 0.963923197311256
91010 FMNL3 formin like 3 13424 0.0946165490773029
2.45038606094835 0.0525267927999664 0.81872338657862
0.963923197311256
2887 GRB10 growth factor receptor bound protein 10 8990
0.0388378338137203 6.57458763921856 0.0524738248911163
0.818813220773794 0.963923197311256
116985 ARAP1 ArfGAP with RhoGAP domain, ankyrin repeat and PH
domain 1 9056 -0.0294502783604919 7.03407288367424
0.0524403795191581 0.818869969097686
0.963923197311256
89796 NAV1 neuron navigator 1 22374 -0.0436515408758154
6.62223755021042 0.0524153373243053
0.818912471828617 0.963923197311256
90835 CCDC189 coiled-coil domain containing 189 3081
0.0753294965610421 1.06331517432411 0.0524138264296532
0.818915036528223 0.963923197311256
7761 ZNF214 zinc finger protein 214 2949 -0.065917311414923
1.11467693040596 0.052412357057162
0.818917530782486 0.963923197311256
9342 SNAP29 synaptosome associated protein 29 4953
0.030682260234308 5.83795434751055 0.0524078067842417

0.818925255087024 0.963923197311256
57544 TXNDC16 thioredoxin domain containing 16 4739
0.0715399225542316 1.92736112216346 0.05227012352311
0.819159146070214 0.963946642863039
84811 BUD13 BUD13 homolog 2196 0.0369284960510256
4.00681830966536 0.0521620085454959 0.819343034896017
0.963946642863039
7562 ZNF708 zinc finger protein 708 4956 -0.0499857357500022
3.52006747075105 0.0521257232203993
0.819404796217727 0.963946642863039
8878 SQSTM1 sequestosome 1 7654 -0.0457642006587307
9.52621224335358 0.0521161454144021 0.819421102392356
0.963946642863039
23076 RRP1B ribosomal RNA processing 1B
5762 -0.0280665853929624 5.7080192428284 0.0520854278775431
0.819473409508471 0.963946642863039
54879 ST7L suppression of tumorigenicity 7 like 8111
0.055254413487491 3.72419720890419 0.0520783409008168
0.819485479814757 0.963946642863039
147727 ILF3-DT ILF3 divergent transcript 1983
0.0681573842311366 1.37869949372498 0.0520210922884892
0.819583015470919 0.963946642863039
2962 GTF2F1 general transcription factor IIF subunit 1 3134
0.0274149344804501 6.35513315810496
0.0520151343998805 0.819593169293183 0.963946642863039
101927111 SUCLG2-AS1 SUCLG2 antisense RNA 1 (head to
head) 3032 0.076256987714162 0.12391941232869
0.051984207508168 0.8196458867513 0.963946642863039
133418 EMB embigin 5257 -0.0368469918585561
6.4690963877762 0.0519145533875474 0.819764678439373
0.963946642863039
4092 SMAD7 SMAD family member 7 3692 0.0486149126952404
5.6759915864292 0.0518920129862295 0.819803137940156
0.963946642863039
91523 PCED1B PC-esterase domain containing 1B 2618
0.0498032556976914 3.0045011022987 0.0518765077410022
0.819829598825283 0.963946642863039
128061 C1orf131 chromosome 1 open reading frame 131
5413 -0.0490737142760823 3.04249835039429
0.0518705192037801 0.819839819834896 0.963946642863039
146540 ZNF785 zinc finger protein 785 4800 0.0497707764007163
3.61559586091461 0.0518190623771524
0.819927669996096 0.963969017427843
22921 MSRB2 methionine sulfoxide reductase B2 2052
0.0501476057861825 3.5364307496866 0.0514979817466426
0.820476877195649 0.964341554856705
56947 MFF mitochondrial fission factor 6150
0.0303066273313761 5.64710720248623 0.0514802400933529
0.820507276646659 0.964341554856705
9531 BAG3 BCL2 associated athanogene 3 2747
0.0379314566292147 7.24599071640354 0.0514162775635896
0.820616919046162 0.964341554856705
9150 CTDP1 CTD phosphatase subunit 1
7415 -0.0325124295102053 5.33205247303805

0.0514125816049376	0.820623256723976	0.964341554856705
10025	MED16	mediator complex subunit 16	4222
0.0384743836603726	4.65006592917775	0.0513755358912533
	0.820686794425857	0.964341554856705
91862	MARVELD3	MARVEL domain containing 3	4798
0.041888021100353	6.08841503167939	0.0513561380562919
	0.820720073573427	0.964341554856705
55752	SEPTIN11	septin 11	9787	-0.0343962815237421
	8.11514748495436	0.0513524097093523
0.820726470725122	0.964341554856705
5664	PSEN2	presenilin 2	3752	-0.0490503910765169
4.89249986901131	0.0513040486086069	0.820809471535304
	0.964358183844573
30968	STOML2	stomatin like 2 1806	0.0330879446364023
5.44154883016287	0.0511970681748295	0.82099322530756
	0.964493173436467
79595	SAP130	Sin3A associated protein 130
4373	-0.0411809507336104	5.13453220247023
0.0511286133217226	0.821110911940876	0.964550531998981
64080	RBKS	ribokinase	3585	0.0769666272105525
0.664291155660941	0.0508922663719034	0.82151787412812
	0.964947661460175
118471	PRAP1	proline rich acidic protein 1	1478
0.0614857144253536	2.10747724768582	0.0507348745882439
	0.821789435738179	0.964949225910808
25950	RWDD3	RWD domain containing 3 2192	0.0444138878236177
	3.36072197617403	0.0507229840203811
0.821809969530602	0.964949225910808
26205	GMEB2	glucocorticoid modulatory element binding protein 2
	4707	-0.0333479031108517	4.8897643794772
0.050704808807799	0.821841361148142	0.964949225910808
9987	HNRNPDL heterogeneous nuclear ribonucleoprotein D like	5476
	0.0255105367432273	7.3101839375403 0.0506855660658161
	0.821874603009806	0.964949225910808
7322	UBE2D2	ubiquitin conjugating enzyme E2 D2
3556	-0.0300915222374987	6.57666893353695
0.0506094949607245	0.822006080896226	0.964949225910808
57590	WDFY1	WD repeat and FYVE domain containing 1	5315
0.0311925605175424	6.60402443528408	0.0505481736655753
	0.822112141462863	0.964949225910808
1181	CLCN2	chloride voltage-gated channel 2
4290	-0.0673472507761152	2.69478103145017
0.0505356514522726	0.822133808039186	0.964949225910808
80765	STARD5	StAR related lipid transfer domain containing 5 6339
	0.0675091902359219	3.27974132817305
0.0504995066988272	0.822196363369514	0.964949225910808
1603	DAD1	defender against cell death 1	808
0.0408118504597587	6.39579460146965	0.0504716080143908
	0.822244663401417	0.964949225910808
11083	DIDO1	death inducer-obliterator 1
13281	-0.0400558830660451	6.85226583530711
0.0504704068351316	0.822246743276555	0.964949225910808
79811	SLTM	SAFB like transcription modulator
8157	-0.0325775035237584	7.00799377229221

0.0504063315065579	0.822357729189717	0.964949225910808
1822	ATN1	atrophin 1	4762	0.0330740107633059
7.27780739019598	0.0503721839900493	0.822416906923131
	0.964949225910808
23677	SH3BP4	SH3 domain binding protein 4
6692	-0.0372841528007064	7.92087446378056
0.0503298608627603	0.822490282257587	0.964949225910808
54552	GNL3L	G protein nucleolar 3 like
2589	-0.0429851127194342	3.88239528584061
0.0502831165449344	0.822571360249936	0.964949225910808
79050	NOC4L	nucleolar complex associated 4 homolog	2029
0.0469073147188076	3.48646231417992	0.050279028782084
	0.82257845235393	0.964949225910808
100271849	MEF2B	myocyte enhancer factor 2B	1848
0.0682519759449953	1.1151997465495 0.05025424711621
0.822621454030196	0.964949225910808
6607	SMN2	survival of motor neuron 2, centromeric
3424	-0.0440088184109475	5.41471708377673
0.050170630224585	0.82276663007964	0.964976217975434
50865	HEBP1	heme binding protein 1	3442	-0.0361496290124641
	5.16912128634477	0.0501616381762524
0.822782249680917	0.964976217975434
84885	ZDHHC12 zinc finger DHHC-type containing 12	2597
0.0316475506711506	4.82104379128656	0.0500102971738476
	0.823045356882851	0.965203978052419
154796	AMOT	angiomotin	8502	-0.082763636376071
4.64873416532358	0.0498849197565931	0.823263643011577
	0.965244987874708
80728	ARHGAP39	Rho GTPase activating protein 39	4888
	-0.0565055873442081	3.51463652553632
0.0498752743882491	0.823280447833447	0.965244987874708
78998	RHPN1-AS1	RHPN1 antisense RNA 1 (head to head)
1918	-0.109978864545154	0.00650167595979535
0.0498714786599734	0.823287061479198	0.965244987874708
6634	SNRPD3	small nuclear ribonucleoprotein D3 polypeptide
2544	-0.0303405341565349	5.98912190287024
0.04979519024414	0.823420041853316	0.965247177618329
151613	TTC14	tetratricopeptide repeat domain 14
7783	-0.0376426138968311	5.02346586575071
0.049791341966654	0.823426752720884	0.965247177618329
51372	TMA7	translation machinery associated 7 homolog	1139
	0.0408375619588955	4.61735052972496
0.0497232567307719	0.823545529071651	0.965305625582144
389668	XKR9	XK related 9	3905	-0.143637925197735
1.32038815214752	0.0496690850675723	0.823640093848487
	0.965335686932956
26354	GNL3	G protein nucleolar 3	3318	-0.0407835503911367
	5.71651772571603	0.0495796120961991
0.823796400972395	0.965438101440791
222166	MTURN	maturin, neural progenitor differentiation regulator
homolog 6532	0.0365585476476317	4.3915343208266
0.0494162958633955	0.82408209265982	0.965692116781235
112476	PRRT2	proline rich transmembrane protein 2	4471
0.0427217650230881	3.59477829129177	0.0493671656336812

0.824168133862846	0.965712150829587
6206	RPS12	ribosomal protein S12	767	-0.051268011270152
7.4855205170129	0.0493042269717989	0.824278423282788
0.965753137463342
57801	HES4	hes family bHLH transcription factor 4
1118	-0.0565278373808284	3.37573025630568
0.0492152852048537	0.824434404791225	0.965753137463342
80196	RNF34	ring finger protein 34	4719	0.0327669580249678
5.3682848138527	0.0491915460515915	0.824476062298755
0.965753137463342
51726	DNAJB11	DnaJ heat shock protein family (Hsp40) member B11
5781	-0.0307937912589417	6.43229323402806
0.0491541517537257	0.824541703273766	0.965753137463342
9768	PCLAF	PCNA clamp associated factor
4053	-0.0569487677514566	1.99800780778072
0.0491506493927625	0.824547852568295	0.965753137463342
10479	SLC9A6	solute carrier family 9 member A6	5200
0.0358530575327519	5.03425736280658	0.0490167088088214
0.824783192389256	0.965873931715697
5786	PTPRA	protein tyrosine phosphatase receptor type A	4469
0.0252109868575467	6.50846321392992
0.0490088734247252	0.824796969959919	0.965873931715697
645	BLVRB	biliverdin reductase B	1665	-0.0420405920966643
4.6480387318915	0.0489321963675113	0.824931858264657
0.965873931715697
100529063	BCL2L2-PABPN1	BCL2L2-PABPN1 readthrough
2729	-0.0246240947154283	6.22302663859021
0.0489211583976275	0.82495128510256	0.965873931715697
100529264	RAB4B-EGLN2	RAB4B-EGLN2 readthrough (NMD candidate)	2589	0.0287760106262183	6.18613313553382
0.0488958890921478	0.824995767762305
0.965873931715697
23787	MTCH1	mitochondrial carrier 1	3126	0.0295970259203977
7.57091716448925	0.0487488131547593
0.825254911569211	0.966096577212699
54980	C2orf42	chromosome 2 open reading frame 42	3630
0.035550221985949	4.09428958984661	0.0486322595042274
0.825460567260117	0.966250773352824
147407	SLC25A52	solute carrier family 25 member 52	1319
0.0734523298916186	1.25744706199061
0.0485661060420934	0.825577408190906	0.966250773352824
57794	SUGP1	SURP and G-patch domain containing 1
5356	-0.0346842596034352	4.9393823559713	0.0484886923027013
0.825714243144368	0.966250773352824
126731	CCSAP	centriole, cilia and spindle associated protein	6263
0.0534655194030542	3.65241897395509
0.0484855210001847	0.825719851115404	0.966250773352824
23616	SH3BP1	SH3 domain binding protein 1
6170	-0.046878569494435	5.49754495760496
0.0484789082178381	0.825731545443943	0.966250773352824
166647	ADGRA3	adhesion G protein-coupled receptor A3	9250
0.0292683579875004	6.27447214311026	0.0483778080137824
0.825910439619944	0.966335720185129
647042	GOLGA6L10	golgin A6 family like 10

4447 -0.0616252733434839 2.93652888498317
0.048359917025512 0.825942117659482 0.966335720185129
55051 NRDE2 NRDE-2, necessary for RNA interference, domain
containing 15157 0.0357686073083191 4.89678945699922
0.0482587227999396 0.826121409349582
0.966421147712656
54344 DPM3 dolichyl-phosphate mannosyltransferase subunit 3,
regulatory 622 0.0608047158280573 2.24304124481865
0.0482408336650364 0.826153125059709
0.966421147712656
51593 SRRT serrate, RNA effector molecule
4588 -0.026351176808466 6.60536724588235
0.0480925421175806 0.826416269343623 0.966630619855024
126069 ZNF491 zinc finger protein 491 3032
0.0680484424480831 -0.0738123788818822 0.0480448019469986
0.826501075036052 0.966630619855024
8822 FGF17 fibroblast growth factor 17 3287
0.143513426082326 0.719953462375878 0.0479845606798648
0.826608150731886 0.966630619855024
6817 SULT1A1 sulfotransferase family 1A member 1
8909 -0.0673395529140159 1.97973530435299
0.0479845126041099 0.826608236212142 0.966630619855024
7043 TGFB3 transforming growth factor beta 3 4198
0.0792192458655989 0.971220322452592 0.0478566233942841
0.82683578643859 0.966695009857263
51162 EGFL7 EGF like domain multiple 7 2553
0.0861883871942412 0.763862845838986 0.0478343761747082
0.826875402905732 0.966695009857263
4600 MX2 MX dynamin like GTPase 2 13047
0.0825465169060741 3.6947508982805 0.0478263986427692
0.826889611150557 0.966695009857263
489 ATP2A3 ATPase sarcoplasmic/endoplasmic reticulum Ca2+
transporting 3 6893 0.138799800628605 1.98990872169919
0.0477984718461215 0.826939359470959
0.966695009857263
3300 DNAJB2 DnaJ heat shock protein family (Hsp40) member B2
5886 0.0300318460680953 7.13346613658776
0.0475628877214547 0.827359632462971 0.966903668040735
9553 MRPL33 mitochondrial ribosomal protein L33 1092
0.0412199343101645 5.53590941503091 0.047518834758173
0.827438342173575 0.966903668040735
27429 HTRA2 HtrA serine peptidase 2 2370 0.0330221863695769
4.814561780056 0.0475186927867668 0.827438595896562
0.966903668040735
11252 PACSIN2 protein kinase C and casein kinase substrate in
neurons 2 4263 -0.0238663128112982 7.09257606810037
0.0474148328920521 0.827624314721451
0.966903668040735
6122 RPL3 ribosomal protein L3 4787 0.0404071563660502
8.30240468722422 0.0474133318703025
0.827627000359857 0.966903668040735
130557 ZNF513 zinc finger protein 513 2365 -0.0366080895746928
4.86502200910678 0.0474097767221173
0.827633361432992 0.966903668040735

51018	RRP15	ribosomal RNA processing 15 homolog
7883	-0.0583400724519768	3.98849199331277
0.0473605992281492	0.827721378239674	0.966903668040735
5795	PTPRJ	protein tyrosine phosphatase receptor type J
9620	-0.0495014975580918	5.86780284672492
0.0473192278924444	0.827795460829666	0.966903668040735
9341	VAMP3	vesicle associated membrane protein 3
2760	-0.0294118828046506	7.42828208681657
0.0473014478821199	0.827827309458707	0.966903668040735
57862	ZNF410	zinc finger protein 410 7628	0.0266392984265327
	5.91274420525703	0.0472815220675358
0.827863009232309	0.966903668040735
5878	RAB5C	RAB5C, member RAS oncogene family
3166	-0.0273738422018708	7.43742854291262
0.0472484655966454	0.827922251720863	0.966903668040735
10899	JTB	jumping translocation breakpoint
1496	-0.0251702028469473	6.06463509007645
0.0472272196862598	0.827960339251732	0.966903668040735
83857	TMTC1	transmembrane O-mannosyltransferase targeting
cadherins 1	14636	-0.127721132707248	2.03790379438826
	0.0471631910012009	0.828075177729177
0.966903668040735
6433	SFSWAP	splicing factor SWAP	10157	-0.0333296684654219
	5.21696979910233	0.0471581222393773
0.828084272279336	0.966903668040735
106699570	LOC106699570	uncharacterized LOC106699570
2894	-0.0665314075266202	0.228266368422863
0.0470695197803961	0.828243328657971	0.967008777570409
29915	HCFC2	host cell factor C2	6773	-0.0474300905489142
	4.11305194663315	0.0469370600319152
0.828481408891627	0.967206125549676
4709	NDUFB3	NADH:ubiquinone oxidoreductase subunit B3	1103
	0.0362250832568078	4.64550189869938
0.0468924766575185	0.828561621162888	0.967219153898539
994	CDC25B	cell division cycle 25B 4064	-0.0605245963475804
	3.66336882731215	0.0468129343554153
0.828704829067228	0.967250813768198
8894	EIF2S2	eukaryotic translation initiation factor 2 subunit
beta	2656	-0.0307024469010022	6.89357914863741
0.0468007083740645	0.828726852004865	0.967250813768198
23483	TGDS	TDP-glucose 4,6-dehydratase
2589	-0.0440378115364652	3.06776977231676
0.0467557985538902	0.828807774939664	0.967264664520903
7620	ZNF69	zinc finger protein 69	1490	-0.0583127840181076
	3.41293667803036	0.0467145527020278
0.828882131621368	0.967270850422436
10084	PQBP1	polyglutamine binding protein 1
1903	-0.0354505793178759	5.18594718934092
0.0466727944123759	0.828957447119989	0.967278154099441
93973	ACTR8	actin related protein 8 4066	-0.0331958497211196
	5.53649154685047	0.0465063791102978
0.829257945595264	0.967311877604423
63929	XPNPEP3 X-prolyl aminopeptidase 3
9574	-0.0394225999201341	3.89531007289501

0.0464813594224189	0.829303172628161	0.967311877604423
57404	CYP20A1 cytochrome P450 family 20 subfamily A member 1
10961	0.0400390715774802	4.56829769888139
0.0464510565438729	0.829357966921717	0.967311877604423
468	ATF4	activating transcription factor 4	2514
0.0273996800049427	8.61362663156473	0.0464255637247355
	0.829404078053305	0.967311877604423
100526842	RPL17-C18orf32	RPL17-C18orf32 readthrough
1529	-0.0356490976687006	7.55070294652322
0.0464235087863472	0.829407795580242	0.967311877604423
7555	CNBP	CCHC-type zinc finger nucleic acid binding protein
	1927	-0.0295373271701459	7.34562459217757
0.0464175737322705	0.82941853299067	0.967311877604423
100533955	SENP3-EIF4A1	SENP3-EIF4A1 readthrough (NMD
candidate)	3519	-0.0369283841766316	9.0540071910322
0.0463511630828179	0.829538728920245	0.967311877604423
7678	ZNF124	zinc finger protein 124 4517	0.0565140272271646
	2.47386470820021	0.0463150941630843
0.829604047432637	0.967311877604423
285268	ZNF621	zinc finger protein 621 8354	-0.0290297643457594
	5.76032659977403	0.0462559630272956
0.829711187748131	0.967311877604423
91133	L3MBTL4 L3MBTL histone methyl-lysine binding protein 4	7501
	0.056183744127457	3.41383515031735
0.0462415454467795	0.829737321974211	0.967311877604423
122060	SLAIN1	SLAIN motif family member 1
4133	-0.06106035498473	2.76208039112097
0.0462367590185973	0.829745999101674	0.967311877604423
55967	NDUFA12 NADH:ubiquinone oxidoreductase subunit A12
1704	-0.0390595263162533	4.72570804007852
0.0461360250373986	0.829928724898531	0.967444378300077
55178	MRM3	mitochondrial rRNA methyltransferase 3	2065
0.0533490411416826	3.18275377388027	0.0460843839773464
	0.830022479804823	0.967473152586016
9776	ATG13	autophagy related 13	7001	-0.0239419606393667
	6.84322374282655	0.0459878813972363
0.830197828692965	0.967559541581844
197187	SNAI3-AS1	SNAI3 antisense RNA 1	2844
0.0820045579713638	0.500539714967948	0.0459206430873254
	0.830320117059503	0.967559541581844
118460	EXOSC6	exosome component 6	5153	0.0497645661131906
	4.66497759943502	0.0458183146331663
0.830506404774269	0.967559541581844
6613	SUMO2	small ubiquitin like modifier 2
3372	-0.0288731954989859	7.21555679418437
0.0456822820179354	0.830754387666449	0.967559541581844
63908	NAPB	NSF attachment protein beta	5234
0.0406609291409919	3.98351293510465	0.0456270055641177
	0.830855265094444	0.967559541581844
9170	LPAR2	lysophosphatidic acid receptor 2	2980
0.0341290542913374	4.77275769823945	0.0455837261177496
	0.830934293045774	0.967559541581844
677	ZFP36L1 ZFP36 ring finger protein like 1
5160	-0.0398685617349209	8.78281588051904

0.0455526472885595	0.830991066960665	0.967559541581844
2629	GBA	glucosylceramidase beta 3648	0.0385342853162306
	6.68036290985531	0.0455429612942737
0.831008765196257	0.967559541581844
81892	SLIRP	SRA stem-loop interacting RNA binding protein	2631
	0.0490818020663314	4.34163647555257
0.0455141358905422	0.831061446579388	0.967559541581844
23168	RTF1	RTF1 homolog, Paf1/RNA polymerase II complex
component	5770	0.0242607701186986	6.47639048830551
	0.0454977363830729	0.831091426146502
0.967559541581844
84522	JAGN1	jagunal homolog 1	1799	0.0347737011527198
	4.8604863698843 0.0454922114934604	0.831101527343777
	0.967559541581844
51000	SLC35B3 solute carrier family 35 member B3	2137
0.0326724388233974	4.83086611651147	0.0454529316309689
	0.831173361506484	0.967559541581844
100652772	NNT-AS1 NNT antisense RNA 1	6580
0.0488029993052245	3.60498592829695	0.045447717502384
	0.831182899431328	0.967559541581844
25949	SYF2	SYF2 pre-mRNA splicing factor
3370	-0.0281152508076894	5.6352910959055 0.0454412975312035
	0.8311946439249 0.967559541581844
28987	NOB1	NIN1 (RPN12) binding protein 1 homolog
2119	-0.038330628206109	4.78232917274544
0.0453891054029043	0.831290154768941	0.967559541581844
161142	FAM71D	family with sequence similarity 71 member D
4399	-0.058361898489267	0.830728348496447
0.0453837195906541	0.831300013994753	0.967559541581844
9016	SLC25A14	solute carrier family 25 member 14	3009
	0.0439028632933415	3.14867555980069
0.045373428281227	0.831318854886264	0.967559541581844
51263	MRPL30	mitochondrial ribosomal protein L30
3187	-0.0348581713744323	4.93040415706459
0.0453618913898879	0.831339978791853	0.967559541581844
84167	C19orf44	chromosome 19 open reading frame 44
4421	-0.066071068566741	2.22103469045924
0.0451530552211725	0.831722841787768	0.967910331183352
22920	KIFAP3	kinesin associated protein 3	4978
0.0412241811361575	4.71710529226282	0.0451221465950864
	0.831779585668167	0.967910331183352
84837	ARHGAP5-AS1	ARHGAP5 antisense RNA 1 (head to head)
1848	-0.0745450775478994	1.36383277939337
0.0450503131093161	0.831911540328141	0.967931518718684
56919	DHX33	DEAH-box helicase 33	5872	-0.0300143430158133
	5.1950962453515 0.0450370042496564	0.831936000132379
	0.967931518718684
10397	NDRG1	N-myc downstream regulated 1	7415
0.0353781843987408	7.01676111390287	0.0449356363083488
	0.832122424153999	0.967993881568345
5713	PSMD7	proteasome 26S subunit, non-ATPase 7
2740	-0.0304740671254884	6.62827746754813
0.044931210458202	0.832130568661888	0.967993881568345
56941	HMCES	5-hydroxymethylcytosine binding, ES cell specific

2783	-0.0306756287807963	5.22870569320356
0.0448951601533309	0.832196924526737	0.967993881568345
1819	DRG2	developmentally regulated GTP binding protein 2 7411
0.0302976995371017	4.93361894791871
0.0447703248310272	0.832426917570985	0.968117226016493
54602	NDFIP2	Nedd4 family interacting protein 2
4831	-0.032024374387331	7.51961407316818
0.0447359104539196	0.832490380530921	0.968117226016493
50809	HP1BP3	heterochromatin protein 1 binding protein 3
7442	-0.0280311964085481	7.35399288171505
0.0447251031428095	0.832510315368649	0.968117226016493
6746	SSR2	signal sequence receptor subunit 2	6651
0.0357831329496554	7.05802983250113	0.0446371767325218
0.832672595351926	0.968225555627962
84131	CEP78	centrosomal protein 78	14355	0.0331147291811362
4.13203117212251	0.0445642200671639
0.832807373452125	0.968229010856909
81854	ZNF205-AS1	ZNF205 antisense RNA 1
3439	-0.063746841253808	2.01430925507528
0.0445607341349508	0.832813816149357	0.968229010856909
161176	SYNE3	spectrin repeat containing nuclear envelope family
member 3	18164	-0.0720579690446036	2.14838150305364
0.0444723890907621	0.832977183346605
0.968338568108216
79817	MOB3B	MOB kinase activator 3B 7087	-0.0487586469603753
4.42951740628586	0.0444209979193246
0.833072293612504	0.968368764865589
22901	ARSG	arylsulfatase G 7084	-0.0811388126203934
0.369491818735757	0.0443061190208152	0.833285109384528
0.968535766919681
79798	ARMC5	armadillo repeat containing 5	5545
0.0368125951528281	4.03789087637521	0.0440648924116127
0.833732926841185	0.968888339917693
55052	MRPL20	mitochondrial ribosomal protein L20	2788
0.0324433014281895	5.2055582860708 0.0440505475737609
0.833759597128191	0.968888339917693
7112	TMPO	thymopoietin	8593	0.0326858533011468
5.22142387377814	0.0440309919808186	0.83379596268779
0.968888339917693
81857	MED25	mediator complex subunit 25
6120	-0.034624487453769	5.62216544408537
0.043864479138878	0.834105951927393	0.96889276216357
8705	B3GALT4 beta-1,3-galactosyltransferase 4
1694	-0.0459192217625134	3.24951744569136
0.0438468022354854	0.834138896235817	0.96889276216357
84996	URB1-AS1	URB1 antisense RNA 1 (head to head)	831
0.0650823364514224	0.461502832671762
0.0438266561094145	0.834176450868641	0.96889276216357
55719	SLF2	SMC5-SMC6 complex localization factor 2
9131	-0.0461972932762785	5.32757316860902
0.0438106689240811	0.83420625918295	0.96889276216357
7905	REEP5	receptor accessory protein 5
4230	-0.0308602129409242	7.78399520414021
0.0437843347534397	0.834255371965439	0.96889276216357

101241892	NPTN-IT1	NPTN intronic transcript 1	2283	-0.0776405350560139	0.906495239288383	0.0437632809896265	0.834294647968964	0.96889276216357
23252	OTUD3	OTU deubiquitinase 3	6767	-0.0418432836742229	4.55194836615901	0.0437581792449784	0.834304166807982	0.96889276216357
284086	NEK8	NIMA related kinase 8	4420	0.0535260897800431	2.49205023775185	0.0436605666822416	0.834486404074512	0.96889276216357
57126	CD177	CD177 molecule	2501	0.145418844068523	1.70165943285499	0.0436305761935891	0.834542437349572	0.96889276216357
729877	TBC1D3H	TBC1 domain family member 3H	7112	-0.0676937027185156	4.24374552185776	0.0436261440372192	0.834550719953949	0.96889276216357
9639	ARHGEF10	Rho guanine nucleotide exchange factor 10	10792	-0.0328735786669874	5.89211970158824	0.0435789210067075	0.834638995397717	0.96889276216357
8038	ADAM12	ADAM metallopeptidase domain 12	9860	-0.111367903862021	4.23152212915589	0.0435432484547675	0.834705712279792	0.96889276216357
64693	CTAGE1	cutaneous T cell lymphoma-associated antigen 1	4315	-0.0595666864517644	1.38362858194064	0.0435374927822392	0.834716479545008	0.96889276216357
2245	FGD1	FYVE, RhoGEF and PH domain containing 1	4275	-0.0696492188427971	2.95169949377307	0.0435098633433917	0.834768176896549	0.96889276216357
8836	GGH	gamma-glutamyl hydrolase	3130	0.0418771506215423	3.970960388376	0.0434588056100473	0.83486375594419	0.96892340952111
25865	PRKD2	protein kinase D2	5446	-0.0304750829788719	5.96750092532589	0.0433866871107398	0.834998860194502	0.968935463200112
55622	TTC27	tetratricopeptide repeat domain 27	3185	0.0441059798629231	3.51280888948926	0.0433794139064112	0.835012492060295	0.968935463200112
95	ACY1	aminoacylase 1	3285	-0.0420097651833398	3.57150651530703	0.0433393535974469	0.835087596818295	0.968942343326198
64755	C16orf58	chromosome 16 open reading frame 58	3630	0.0304522399912527	5.50508055232475	0.0431747486802916	0.835396577600549	0.969088782942087
5437	POLR2H	RNA polymerase II subunit H	3084	-0.0355685196898684	5.52987426164133	0.0431502756978155	0.835442568403514	0.969088782942087
55929	DMAP1	DNA methyltransferase 1 associated protein 1	5599	0.0379684049192263	5.40068474073964	0.043132207739248	0.835476531308654	0.969088782942087
11099	PTPN21	protein tyrosine phosphatase non-receptor type 21	7396	0.03282119148201	6.21067018807999	0.0431247507327104	0.835490550639583	0.969088782942087
9256	TSPOAP1	TSPO associated protein 1	8698	0.127098285724298	1.67776628097102	0.043047574643424	0.835635717648581	0.969176906276697

291	SLC25A4 solute carrier family 25 member 4	4844
0.0303402002517992	5.85378546457402	0.0427673939841231
	0.836163877088318	0.969483859850801
644096	SDHAF1	succinate dehydrogenase complex assembly factor 1
	1117	-0.048000534915246	2.52595089737671
0.0427195595423058	0.836254228354545	0.969483859850801
8567	MADD	MAP kinase activating death domain
8366	-0.0288136588271539	6.22936634649627
0.0426802725744437	0.836328474328665	0.969483859850801
6947	TCN1	transcobalamin 1	1586	-0.151009962224291
	1.72311154402356	0.042666682827349
0.836354165032063	0.969483859850801
84964	ALKBH6	alkB homolog 6	4347	-0.0425680826090479
3.49802392741484	0.0426398323498036	0.836404936993978
	0.969483859850801
57168	ASPHD2	aspartate beta-hydroxylase domain containing 2
3357	-0.0480229550878955	3.59714821545329
0.042609175362335	0.836462927124672	0.969483859850801
158747	MOSPD2	motile sperm domain containing 2
5656	-0.0310031529094689	4.17450738135249
0.0425914028037049	0.836496555293223	0.969483859850801
30827	CXXC1	CXXC finger protein 1	4319	0.0464496160276632
	5.81852898072738	0.042590035859007
0.836499142046845	0.969483859850801
573	BAG1	BCL2 associated athanogene 1	7114
0.0283477768838427	6.43902512535689	0.0425465374895122
	0.8365814793024 0.969483859850801
55069	TMEM248 transmembrane protein 248
5086	-0.0231828681122077	6.97304148664734
0.042540707018361	0.836592519027389	0.969483859850801
51250	MTRES1	mitochondrial transcription rescue factor 1	2036
	0.0426639126731381	3.30127783608381
0.0424055355211155	0.836848681390909	0.969668033032474
5336	PLCG2	phospholipase C gamma 2 13416	-0.0609560299097168
	2.76924408109657	0.042383805439794
0.836889901572755	0.969668033032474
162632	USP32P1 ubiquitin specific peptidase 32 pseudogene 1	3243
	0.177240229052592	1.22567034767628
0.0423389342785896	0.836975053354105	0.969686482409508
54797	MED18	mediator complex subunit 18	2019
0.0314959231005435	4.72456618001613	0.0421767407687752
	0.837283240085023	0.969821033652292
2196	FAT2	FAT atypical cadherin 2 14537	-0.0987797524023504
	6.11066644921835	0.0421136486768887
0.837403289287466	0.969821033652292
256329	LMNTD2	lamin tail domain containing 2
2934	-0.0722273624380619	1.59982606770085
0.0420971112068385	0.837434771647141	0.969821033652292
1877	E4F1	E4F transcription factor 1
6402	-0.0372367052011449	4.49338178321618
0.0420903701148454	0.837447606503712	0.969821033652292
23513	SCRIB	scribble planar cell polarity protein	6192
0.0351021025456526	6.79155971734407	0.0420830724093548
	0.837461502344734	0.969821033652292

10973	ASCC3	activating signal cointegrator 1 complex subunit 3
	11731	-0.0353884514764706	7.22788336401073
0.0420539342846951	0.837516997884429	0.969821033652292
90576	ZNF799	zinc finger protein 799 6840	-0.049727012400395
	2.71315771521982	0.0419657672860194
0.837685040058519	0.969821033652292
9497	SLC4A7	solute carrier family 4 member 7
9046	-0.0298697823017679	6.2503741187797 0.0418977090418977
	0.837814881715457	0.969821033652292
10921	RNPS1	RNA binding protein with serine rich domain 1
7324	-0.026719273831688	6.46579391997245
0.0418871469161473	0.837835042007058	0.969821033652292
51008	ASCC1	activating signal cointegrator 1 complex subunit 1
	5586	0.0286308969829524	4.78920037346823
0.0418826888151499	0.837843552132086	0.969821033652292
8764	TNFRSF14	TNF receptor superfamily member 14	6219
	0.0378169299644478	5.46764438618198
0.0418778385894951	0.837852811324794	0.969821033652292
10279	PRSS16	serine protease 16	5290	-0.0495278519400645
	2.82112882533933	0.0418002918429226
0.83800092578896	0.969912325857376
5601	MAPK9	mitogen-activated protein kinase 9
8955	-0.0271444040955046	5.8627055394118 0.0417331420705125
	0.838129297683306	0.969980754577384
80821	DDHD1	DDHD domain containing 1	17458
0.0438988331739876	3.79161106921822	0.0416602012170975
	0.838268862495917	0.970062124667904
3608	ILF2	interleukin enhancer binding factor 2
2281	-0.0264481688361958	7.414760032156	0.0416078230201116
	0.83836916130804	0.970074036904864
2222	FDFT1	farnesyl-diphosphate farnesyltransferase 1
5202	-0.0506118004450162	8.31155019030925
0.0415825035316857	0.838417669077625	0.970074036904864
4952	OCRL	OCRL inositol polyphosphate-5-phosphatase
5779	-0.0249905828651226	6.28224361648434
0.0414361625011406	0.838698335096005	0.970089276144772
177	AGER	advanced glycosylation end-product specific receptor
	2077	-0.0619351230462206	0.736537641533104
0.0413985640895143	0.838770528028884	0.970089276144772
5091	PC	pyruvate carboxylase	5364	-0.062584949825236
	4.72457631207594	0.0413920686492197
0.838783003420271	0.970089276144772
57513	CASKIN2 CASK interacting protein 2
5823	-0.0530102222169456	5.35530853979006
0.0413889976432813	0.838788902066794	0.970089276144772
4200	ME2	malic enzyme 2	9363	0.0352418262355182
5.33305101126144	0.0413870989726144	0.838792549059818
	0.970089276144772
7768	ZNF225	zinc finger protein 225 4871	0.0583710700156497
	2.79362279410786	0.0413422914042183
0.838878641360725	0.970089276144772
124930	ANKRD13B	ankyrin repeat domain 13B	3971
0.0531995254309028	3.11794365585124	0.0413230416640253
	0.838915642317529	0.970089276144772

494143	CHAC2	ChaC cation transport regulator homolog 2	1321
	0.0535114186203926	1.54295618523958
0.0412851674872985	0.838988468509353	0.970093402543381
6767	ST13	ST13 Hsp70 interacting protein
4147	-0.0255105720876929	6.64695825325017
0.0412409603288495	0.839073516055056	0.970111658223951
441027	TMEM150C	transmembrane protein 150C	3671
0.0759561994347577	2.78631657099308	0.0409654731412195
	0.839604582354072	0.970590923292675
5711	PSMD5	proteasome 26S subunit, non-ATPase 5	4432
0.0227007852841813	6.07524662384482	0.040781854675366
	0.839959583139338	0.970590923292675
57393	CLTRN	collectrin, amino acid transport regulator	1578
	0.0676738042263862	3.54344439320206
0.0407792722275673	0.839964581868648	0.970590923292675
57511	COG6	component of oligomeric golgi complex 6 8051
0.0248861386712356	5.50662270601941	0.040778377061548
	0.839966314640132	0.970590923292675
11319	ECD	ecdysoneless cell cycle regulator	3382
0.0276355087028587	5.10136853833732	0.040773947479078
	0.83997488926743	0.970590923292675
54069	MIS18A	MIS18 kinetochore protein A
1723	-0.0470303520082727	2.4792051392393 0.0407510448644293
	0.840019231064185	0.970590923292675
7424	VEGFC	vascular endothelial growth factor C
2420	-0.0923659309889042	4.71325999851934
0.0407495718846178	0.840022083347054	0.970590923292675
4683	NBN	nibrin	6156	-0.0346067045468545
5.34399145309168	0.040739085662004	0.840042390453137
	0.970590923292675
4050	LTB	lymphotoxin beta	1535	0.0506958371893693
	4.06966350731127	0.0406659747545337
0.840184049116469	0.970674527876474
100130967	C6orf99 chromosome 6 open reading frame 99
859	-0.0819319690327431	1.07529223415016
0.0405750458012903	0.840360416798951	0.970684211775055
728927	ZNF736	zinc finger protein 736 7909	-0.0605752640232012
	3.38550252835103	0.0405488243193517
0.840411314721835	0.970684211775055
54801	HAUS6	HAUS augmin like complex subunit 6	7932
0.0283588329553822	4.99951017509455	0.0404651880326243
	0.840573773734825	0.970684211775055
285908	LINC00174	long intergenic non-protein coding RNA 174
	4849	0.0743193412381143	1.28952907591765
0.0404233949645363	0.840655019990086	0.970684211775055
81579	PLA2G12A	phospholipase A2 group XIIA	5228
0.0319455657614547	4.86042046469598	0.040383148109393
	0.84073330170032	0.970684211775055
5939	RBMS2	RNA binding motif single stranded interacting
protein 2	8805	-0.0270456363421476	7.45397758786298
	0.0403825626538001	0.840734440733795
0.970684211775055
22913	RALY	RALY heterogeneous nuclear ribonucleoprotein
7513	-0.028958155921251	4	7.11171438977814

0.040356395439316	0.840785359148324	0.970684211775055
8892	EIF2B2	eukaryotic translation initiation factor 2B subunit beta	5446	-0.0285492067208025	5.23533381250214
0.0403256261756937	0.840845254600715	0.970684211775055
84216	TMEM117	transmembrane protein 117
3440	-0.0395527740381908	3.28371573489606
0.0403116926995715	0.840872385326858	0.970684211775055
400954	EML6	EMAP like 6	14253	-0.075544607267205
0.539121475859003	0.0403049939817759	0.840885430547478
0.970684211775055
421	ARVCF	ARVCF delta catenin family member	5621
0.0435752643805311	5.44154912386278	0.0401804975923739
0.841128082925494	0.970744918655602
54102	CLIC6	chloride intracellular channel 6
3860	-0.0902599309852568	6.42828039105422
0.0401748021054971	0.841139193184693	0.970744918655602
4157	MC1R	melanocortin 1 receptor 3099	0.0645148989079841
2.38575577997047	0.0401296544122678
0.841227292366996	0.970744918655602
6284	S100A13 S100 calcium binding protein A13
3271	-0.0326966610445859	5.77308711312594
0.04007596164017	0.841332133163499	0.970744918655602
91782	CHMP7	charged multivesicular body protein 7	4698
0.0229968423450133	5.98017379234869	0.0400699834006488
0.841343810824474	0.970744918655602
153733	CCDC112 coiled-coil domain containing 112	3289
0.0430661981866179	4.00967730051535	0.0400431210925376
0.841396293799526	0.970744918655602
84503	ZNF527	zinc finger protein 527 6436	0.0451583263611233
2.45123614104731	0.040010812040528
0.84145944275999	0.970744918655602
5827	PXMP2	peroxisomal membrane protein 2	1565
0.0500099905336521	1.87860673266008	0.0399939278507198
0.841492453938482	0.970744918655602
22820	COPG1	coatomer protein complex subunit gamma 1
4425	-0.0295580960705315	7.32022736048153
0.0398870131189142	0.841701657062439	0.97087967629954
81847	RNF146	ring finger protein 146 3508	0.034733818499782
5.65865761177461	0.0398634021093756
0.841747896693197	0.97087967629954
808	CALM3	calmodulin 3	3667	0.0392385783384727
6.61657216743804	0.0397609389216882	0.841948725029415
0.971031353765913
653121	ZBTB8A	zinc finger and BTB domain containing 8A
8740	-0.0337628335239959	4.0211301538766 0.0396974828298369
0.842073234207011	0.971047497549933
10987	COPS5	COP9 signalosome subunit 5	7740
0.0229869672504622	5.94387614103593	0.0395713148718997
0.842321100027005	0.971047497549933
4803	NGF	nerve growth factor	1047	-0.0783074818398448
2.15937802543749	0.0395329209149189
0.842396609153768	0.971047497549933
84614	ZBTB37	zinc finger and BTB domain containing 37
19464	-0.0634405274669268	3.526312485017	0.0395090750539033

0.842443525831599	0.971047497549933
84948	TIGD5	tigger transposable element derived 5	5390
0.0552214737734595	3.84596460361662	0.0394381701074931
0.842583118163901	0.971047497549933
875	CBS	cystathionine beta-synthase	9986
0.0621924742369978	5.06788882264918	0.0394341653006869
0.842591006416857	0.971047497549933
345757	FAM174A family with sequence similarity 174 member A	1416
0.0342736688635316	4.45307412199836
0.0393412941194207	0.842774051345704	0.971047497549933
51082	POLR1D	RNA polymerase I and III subunit D	5365
0.0270201050328968	6.08968432957267	0.0393373643548482
0.842781801684668	0.971047497549933
7408	VASP	vasodilator stimulated phosphoprotein
3601	-0.0298749564753599	7.46861075037645
0.0393338745259939	0.842788684713314	0.971047497549933
861	RUNX1	RUNX family transcription factor 1
10075	-0.040950340224127	7.72742833440145
0.0393289132751278	0.842798470390412	0.971047497549933
78991	PCYOX1L prenylcysteine oxidase 1 like
6069	-0.076688980830356	1.71898187758724
0.0393285281232005	0.842799230099126	0.971047497549933
100293516	ZNF587B zinc finger protein 587B
4576	-0.0490752194170859	3.06514741424217
0.0391123222112295	0.843226306229143	0.971047497549933
8861	LDB1	LIM domain binding 1	2790	0.0303501332336844
6.47306271587747	0.0390453755494065
0.843358795981337	0.971047497549933
100505633	LINC01133	long intergenic non-protein coding
RNA 1133	3155	0.0969579122621074	2.33623062395238
0.0390444083422299	0.843360710982196
0.971047497549933
9761	MLEC	malectin	6723	0.0275827102945336
7.40792829248421	0.0390113793805398	0.8434261207549
0.971047497549933
114049	BUD23	BUD23 rRNA methyltransferase and ribosome maturation
factor	7944	0.0302200127723195	4.94790459808947
0.0390051920080978	0.843438377279773	0.971047497549933
401024	FSIP2	fibrous sheath interacting protein 2	23761
0.18811271372945	0.903457578582318	0.0389257455328398
0.843595842106992	0.971047497549933
57102	C12orf4 chromosome 12 open reading frame 4
4939	-0.0347772820921249	4.58671362493754
0.0389230606457858	0.843601166532794	0.971047497549933
56931	DUS3L	dihydrouridine synthase 3 like	4004
0.0422037610036062	2.99450651368864	0.0389117633803253
0.843623572341102	0.971047497549933
4329	ALDH6A1 aldehyde dehydrogenase 6 family member A1	6680
0.0440186172449502	2.96700303690765
0.0388749225174929	0.843696662120773	0.971047497549933
51692	CPSF3	cleavage and polyadenylation specific factor 3
2345	-0.0277329950368272	5.15954706699625
0.0388629840115655	0.843720355027128	0.971047497549933
389792	IER5L	immediate early response 5 like

2711 -0.074794258105427 3.30533779760359
0.0388590284627757 0.843728205959395 0.971047497549933
64927 TTC23 tetratricopeptide repeat domain 23 6148
0.0339066939469346 4.41787226463419 0.0388413133465457
0.843763371832173 0.971047497549933
130355 C2orf76 chromosome 2 open reading frame 76
950 -0.0538862847313268 1.5012312584924З
0.0388105114189354 0.843824535876675 0.971047497549933
2898 GRIK2 glutamate ionotropic receptor kainate type subunit 2
7375 -0.0709166942413436 2.19256787465629
0.0386817072165808 0.844080578690795 0.971047497549933
22869 ZNF510 zinc finger protein 510 5921 -0.0412364453786087
4.15802038711459 0.0385892159774315
0.844264710197548 0.971047497549933
10393 ANAPC10 anaphase promoting complex subunit 10
3418 -0.0418896307620595 3.16998677783313
0.0385699297738133 0.844303134028376 0.971047497549933
10471 PFDN6 prefoldin subunit 6 2953 -0.0369510245445414
4.23205718671802 0.0385575402425857
0.844327822908347 0.971047497549933
5583 PRKCH protein kinase C eta 8395 -0.0319212874377686
5.42351642994122 0.0385437537935083
0.844355300298873 0.971047497549933
9111 NMI N-myc and STAT interactor
3480 -0.0448400885452318 4.04372791192604
0.0384996293434838 0.844443277843192 0.971047497549933
6902 TBCA tubulin folding cofactor A
3058 -0.0340928238709304 5.30507598111543
0.0384243786097827 0.844593437402681 0.971047497549933
84437 MSANTD4 Myb/SANT DNA binding domain containing 4 with
coiled-coils 4263 -0.0332036849683254 4.76730587809872
0.0383464806467053 0.844749040318807
0.971047497549933
3965 LGALS9 galectin 9 6150 -0.099308660221317
4.54474415245706 0.038342298029896 0.844757399829416
0.971047497549933
3431 SP110 SP110 nuclear body protein 6865
0.0380801351310152 4.49361341937023 0.0383073474329301
0.844827271715823 0.971047497549933
9415 FADS2 fatty acid desaturase 2 10977 -0.0571512523562383
7.76257836027208 0.0382920134200475
0.844857937328556 0.971047497549933
6502 SKP2 S-phase kinase associated protein 2 3837
0.0646439250090513 2.77245319559931 0.038231094958654
0.844979827680922 0.971047497549933
4841 NONO non-POU domain containing octamer binding 5292
0.0233892608437115 8.23561482431544
0.0382045303390157 0.845033011787532 0.971047497549933
81572 PDRG1 p53 and DNA damage regulated 1 1998
0.0303198454654052 4.88478607832745 0.0381814867002932
0.845079162210676 0.971047497549933
9883 POM121 POM121 transmembrane nucleoporin 9822
0.0255082502883369 6.17992798327499 0.0381349250074123
0.845172457330337 0.971047497549933

25992	SNED1	sushi, nidogen and EGF like domains 1	13592
0.0935136467121827	1.37920582109881	0.0381103853079887
	0.845221651032883	0.971047497549933
84274	COQ5	coenzyme Q5, methyltransferase
2523	-0.0384140662004488	3.71760578166969
0.0380847711847032	0.845273016128252	0.971047497549933
3570	IL6R	interleukin 6 receptor	6453	0.0498285034634333
	2.9220558472336	0.0380599949015878	0.84532271813033
	0.971047497549933
55135	WRAP53	WD repeat containing antisense to TP53	5087
0.0405565202398415	3.38541666606602	0.0380394993481197
	0.845363845547414	0.971047497549933
84337	ELOF1	elongation factor 1 homolog
4173	-0.0308674928764751	4.82472382085198
0.038030441568603	0.84538202501169	0.971047497549933
1072	CFL1	cofilin 1	4553	-0.0326196694802927
9.90839817691717	0.0380004155990719	0.84544230487004
	0.971047497549933
80006	TRAPPC13	trafficking protein particle complex 13	8671
	0.0286574973754181	4.96272188341331
0.0379983985633245	0.845446355138261	0.971047497549933
10478	SLC25A17	solute carrier family 25 member 17
3989	-0.0325189172904157	4.67584760848675
0.0379903113003053	0.845462595725441	0.971047497549933
9600	PITPNM1	phosphatidylinositol transfer protein membrane
associated 1	6449	-0.0235544647919672	6.26599111195658
	0.0379423174617184	0.84555901238356
0.971047497549933
197322	ACSF3	acyl-CoA synthetase family member 3
5427	-0.0325983426080947	3.92226510059698
0.0378704534343226	0.845703501193433	0.971047497549933
26589	MRPL46	mitochondrial ribosomal protein L46	6522
0.034682492347473	3.95878875590777	0.0378219141201477
	0.845801174183533	0.971047497549933
27237	ARHGEF16	Rho guanine nucleotide exchange factor 16
	5563	-0.0420496011581351	5.03516136239881
0.0378005656713345	0.845844153086791	0.971047497549933
8082	SSPN	sarcospan	5303	0.0726211547945544
2.41863396594402	0.0377581809682184	0.845929519722713
	0.971047497549933
148203	ZNF738	zinc finger protein 738	3905	-0.0476115759465983
	2.77896068367384	0.0377341909473206
0.845977859837798	0.971047497549933
79707	NOL9	nucleolar protein 9	7293	0.0299553683963324
	4.25426403369335	0.0376378867123961
0.846172074438884	0.971047497549933
10906	TRAFD1	TRAF-type zinc finger domain containing 1
3940	-0.0321266834524259	6.58062827203936
0.0375940390565859	0.846260586442238	0.971047497549933
7920	ABHD16A	abhydrolase domain containing 16A	4690
0.0252292983874063	5.37925725135458	0.0375924579803879
	0.846263779043409	0.971047497549933
163033	ZNF579	zinc finger protein 579	3321	0.047416740859351
	3.5521461411448	0.037579659822045	0.846289624394881

0.971047497549933
51056 LAP3 leucine aminopeptidase 3
3785 -0.032483211531371 5.04491591530994
0.0375744075329258 0.84630023249706 0.971047497549933
5251 PHEX phosphate regulating endopeptidase homolog X-linked
6384 0.0657380453877346 1.21898222819473
0.0375713087030816 0.846306491596295 0.971047497549933
6233 RPS27A ribosomal protein S27a 3130 -0.0427408022902933
7.10102054151638 0.0375650378284718
0.846319158495618 0.971047497549933
1635 DCTD dCMP deaminase 3466 0.0217506043076128
6.36880137455074 0.0375396388437235 0.846370474584685
0.971047497549933
81570 CLPB ClpB homolog, mitochondrial AAA ATPase chaperonin
4264 0.0316670928853215 3.9007694392088
0.0374906048022265 0.846469593900162 0.971047497549933
55425 GPALPP1 GPALPP motifs containing 1 10134
0.0313020664710679 4.60050995179734 0.0374722821456572
0.846506649288125 0.971047497549933
466 ATF1 activating transcription factor 1
2790 -0.0300791560186059 4.451656162883 0.037471871074783
0.846507480737735 0.971047497549933
64116 SLC39A8 solute carrier family 39 member 8
6480 -0.0622377042800666 3.53192779513234
0.0374586107360528 0.846534304206985 0.971047497549933
3707 ITPKB inositol-trisphosphate 3-kinase B
7036 -0.0412849842707077 4.73974092061756
0.0374566700261063 0.846538230354982 0.971047497549933
101060200 ZNF891 zinc finger protein 891
15455 -0.0600181769485378 2.32043546421754
0.0373894250571354 0.846674335336556 0.971096548335379
55112 WDR60 WD repeat domain 60 6667 0.0377937149334064
4.38387955059901 0.0373670502317935
0.846719650519025 0.971096548335379
93100 NAPRT nicotinate phosphoribosyltransferase 3079
0.0663769526598221 3.87971093265536 0.03719879410197
0.847060867305566 0.971336620581296
231 AKR1B1 aldo-keto reductase family 1 member B
3660 -0.127682040294649 8.40423528913874
0.0371954448251017 0.847067667623369 0.971336620581296
4061 LY6E lymphocyte antigen 6 family member E
2146 -0.041620726410908 5.48860077904005
0.0371243193942599 0.847212154597902 0.971387759068409
107080638 TBC1D7-LOC100130357 TBC1D7-LOC100130357
readthrough 3459 0.0581055934423109 4.10640230367534
0.0371052270396746 0.847250963985061
0.971387759068409
7295 TXN thioredoxin 1142 0.0429294032312659
7.32116915742892 0.0370507074153466 0.847361844086383
0.971435369955637
6347 CCL2 C-C motif chemokine ligand 2
1123 -0.0750230357610511 7.55919183927492
0.0369343678947871 0.847598734552993 0.971596800714987
7421 VDR vitamin D receptor 5674 -0.0274469327408759

5.9532361030945 0.0369134410643781 0.84764138678783
0.971596800714987
998 CDC42 cell division cycle 42 3598 -0.0320172316352001
8.43017613483548 0.0367803513477156
0.847912938840791 0.971611307113757
55831 EMC3 ER membrane protein complex subunit 3
4270 -0.0264484185378741 6.25014532407062
0.0367715038943857 0.847931008948359 0.971611307113757
100527960 MROH7-TTC4 MROH7-TTC4 readthrough (NMD candidate) 7090 0.0352327208873371 4.27750953021869
0.036764552501396 0.847945208108654
0.971611307113757
51121 RPL26L1 ribosomal protein L26 like 1 1164
0.0392452216813718 3.69243824099353 0.0367386621060541
0.847998104990975 0.971611307113757
9699 RIMS2 regulating synaptic membrane exocytosis 2
13621 -0.140709060301812 0.0215189896566576
0.036737307516173 0.848000873096714 0.971611307113757
257068 PLCXD2 phosphatidylinositol specific phospholipase C X domain containing 2 2578 0.0454628673336232
3.95938109211502 0.0366218449554054 0.848237015710118
0.971802378459972
145173 B3GLCT beta 3-glucosyltransferase 4566
0.040951889295361 4.71347138661862 0.0365048628500695
0.848476659951193 0.971976982831917
72 ACTG2 actin gamma 2, smooth muscle 3811
0.166424550445491 0.197391664465632 0.0364582316368178
0.848572297472493 0.971976982831917
28998 MRPL13 mitochondrial ribosomal protein L13 4209
0.0347715885817343 4.70753565452109 0.0364459018186913
0.848597595705826 0.971976982831917
9644 SH3PXD2A SH3 and PX domains 2A
11756 -0.0311253147983836 8.34524865415986
0.0363649549686826 0.848763792130394 0.972087852524156
132 ADK adenosine kinase 3184 0.0336924151506712
5.66957836256774 0.0362754976453887
0.848947685021014 0.972218970165101
490 ATP2B1 ATPase plasma membrane Ca2+ transporting 1 9315
0.0422902064069692 4.97549597109692
0.0361746313827096 0.849155312746585 0.972377245392529
51676 ASB2 ankyrin repeat and SOCS box containing 2 5040
0.0783489810592059 1.53830474162852
0.0361005289015797 0.849308039867195 0.972424425492824
167555 FAM151B family with sequence similarity 151 member B
3109 -0.0548071391194558 0.358937646826814
0.0360872804262249 0.849335362424445 0.972424425492824
55687 TRMU tRNA 5-methylaminomethyl-2-thiouridylate methyltransferase 7200 -0.0427495885649088
3.87435226617379 0.0360285088255203 0.849456630801984
0.972483778311678
5565 PRKAB2 protein kinase AMP-activated non-catalytic subunit beta 2 5460 0.0290541816218784 5.64790997304283
0.0359822785528081 0.849552093628757 0.972513580864498
84876 ORAI1 ORAI calcium release-activated calcium modulator 1

2262 -0.0384874965605649 3.61143025685582
0.0359464915351451 0.849626035374669 0.972518744585518
81620 CDT1 chromatin licensing and DNA replication factor 1
3336 0.0791398639593258 0.923496377628539
0.0358344053838895 0.849857870482596 0.972625595353009
79415 CYBC1 cytochrome b-245 chaperone 1 7234
0.0234418551285184 5.85081739348684 0.0358277794463397
0.849871587091539 0.972625595353009
9146 HGS hepatocyte growth factor-regulated tyrosine kinase
substrate 5218 0.0247992897368975 7.36380352987046
0.0358006804173172 0.849927699516016
0.972625595353009
148423 C1orf52 chromosome 1 open reading frame 52 3420
0.0305620166748374 4.55616879272259 0.0357472570678192
0.850038384684436 0.972672792604763
91120 ZNF682 zinc finger protein 682 4055 0.0510031274388308
1.67178702146244 0.0356497569515632
0.850240611360743 0.972758488501589
427 ASAH1 N-acylsphingosine amidohydrolase 1
5051 -0.0260360790487888 8.41980731584761
0.0355706392296042 0.850404921384142 0.972758488501589
51061 TXNDC11 thioredoxin domain containing 11
3576 -0.0283529806279086 5.81034088745032
0.0355492171252507 0.850449442908293 0.972758488501589
6124 RPL4 ribosomal protein L4 5508 0.0347550476959397
8.35407007923796 0.0355477457214874
0.850452501434085 0.972758488501589
84191 CIAO2A cytosolic iron-sulfur assembly component 2A 2427
0.028743655406009 4.56829551809848
0.0355301571456899 0.850489066911968 0.972758488501589
55296 TBC1D19 TBC1 domain family member 19
3434 -0.0342548837325858 4.4069309319865 0.0354677489481361
0.850618885080047 0.972758488501589
4361 MRE11 MRE11 homolog, double strand break repair nuclease
7689 0.0275661539664985 4.50888706114779
0.0354310562917171 0.850695266378074 0.972758488501589
6908 TBP TATA-box binding protein 2038
0.0329866663084614 3.8903251371415 0.035410497012144
0.850738081521097 0.972758488501589
27332 ZNF638 zinc finger protein 638 12474 -0.0246521797377985
6.80017130539583 0.0354103881396099
0.850738308284748 0.972758488501589
8637 EIF4EBP3 eukaryotic translation initiation factor 4E
binding protein 3 691 0.0458184863773954
2.72284554688506 0.0353373379051312 0.850890541408058
0.972853139621473
338440 ANO9 anoctamin 9 6207 -0.0294403565970371
5.63142829223681 0.0352678419872916 0.851035518795941
0.972912165642446
92806 CENPBD1 CENPB DNA-binding domain containing 1 2744
0.0372967597850554 4.03134845732451 0.0352460138630732
0.851081085572706 0.972912165642446
4719 NDUFS1 NADH:ubiquinone oxidoreductase core subunit S1
12268 0.0241978748401248 7.02627211016692

0.0352096857176818	0.851156953917509	0.972919491882043
2050	EPHB4	EPH receptor B4	6435	0.0326631856041648
7.59606882782117	0.0351184801461955	0.851347607994532
	0.973058012662538
404636	DENND10	DENN domain containing 10	6161
0.0338981626400206	4.90248018515573	0.0350738029472693
	0.851441093721249	0.973075325046077
643911	CRNDE	colorectal neoplasia differentially expressed	2250
	0.0416295381824126	2.85541623428874
0.0350130075443893	0.851568405363171	0.973075325046077
1209	CLPTM1	CLPTM1 regulator of GABA type A receptor forward trafficking	5229	0.0246317868163777	7.35851621578571
	0.0350022693020833	0.851590904199022
0.973075325046077
90957	DHX57	DExH-box helicase 57	7985	0.0252679454183044
	4.8482425062609	0.0349785386356647	0.851640637519801
	0.973075325046077
10283	CWC27	CWC27 spliceosome associated cyclophilin
3740	-0.0271834934636967	4.79168924797214
0.0349152930468897	0.851773269208082	0.973122818357463
55863	TMEM126B	transmembrane protein 126B
2416	-0.0345183421691975	4.71892117304286
0.0348924746340087	0.851821151986925	0.973122818357463
54623	PAF1	PAF1 homolog, Paf1/RNA polymerase II complex component	2668	0.0268405713428431	6.10618212987313
	0.0347203919904731	0.852182777940823
0.973405611959478
3773	KCNJ16	potassium inwardly rectifying channel subfamily J member 16	6340	0.149874293644735	0.71843193660818
	0.0347085567921752	0.852207683262588
0.973405611959478
7532	YWHAG	tyrosine 3-monooxygenase/tryptophan 5-monooxygenase activation protein gamma	3750	-0.029431488182021
8.97990187540313	0.0346733114429298	0.85228187761709
	0.973410980086643
283237	TTC9C	tetratricopeptide repeat domain 9C
2620	-0.0320087244205918	3.81316584519317
0.0346020453382609	0.852432017918	0.973441995300717
51185	CRBN	cereblon	7163	0.0266344954688051
4.96033753413642	0.0345696809786107	0.852500254546783
	0.973441995300717
11231	SEC63	SEC63 homolog, protein translocation regulator
8587	-0.0284834772494139	6.77977213239971
0.0345614927086046	0.852517523834789	0.973441995300717
54957	TXNL4B	thioredoxin like 4B	3170	0.0322472336386131
	3.67761390600754	0.0345249414770161
0.852594637215103	0.973450691512223
9617	MTRF1	mitochondrial translation release factor 1	3839
	0.0372207333883566	3.42522284162399
0.0344493651493818	0.852754217212466	0.973553534965769
81876	RAB1B	RAB1B, member RAS oncogene family
2007	-0.027712450399564	7.77896752179013
0.0343948437923629	0.852869451941043	0.973566326498692
9855	FARP2	FERM, ARH/RhoGEF and pleckstrin domain protein 2

14413 -0.0293355630940972 5.73643442297086
0.0343783019537796 0.852904432959624 0.973566326498692
151534 LBX2-AS1 LBX2 antisense RNA 1 2083
0.0746901403139149 1.31691186676748 0.0343090179292975
0.85305104252632 0.973588265650907
2034 EPAS1 endothelial PAS domain protein 1 7302
0.0396853355473705 7.34905546621026 0.0343035273274097
0.853062667547197 0.973588265650907
22928 SEPHS2 selenophosphate synthetase 2
1801 -0.0249572696866494 5.32205628061209
0.0341745974591099 0.853335922443227 0.973706220076824
6282 S100A11 S100 calcium binding protein A11
680 -0.0306256145617215 9.45008204516221
0.0341271795296905 0.853436554396793 0.973706220076824
11047 ADRM1 adhesion regulating molecule 1
2142 -0.0257089527412001 6.67941057386681
0.0341016050115925 0.853490859527959 0.973706220076824
50 ACO2 aconitase 2 4041 -0.0243409020655867
6.65715693194058 0.0340805409304679 0.85353560297841
0.973706220076824
9848 MFAP3L microfibril associated protein 3 like 8741
0.0648597630376305 2.60708325336631 0.0340721889753253
0.853553347814853 0.973706220076824
10922 FASTK Fas activated serine/threonine kinase 3805
0.0255956233060648 5.70670487666022 0.034047344594569
0.853606146296117 0.973706220076824
51265 CDKL3 cyclin dependent kinase like 3
3575 -0.0284030253715722 4.65621905367729
0.0340254795915929 0.853652629581166 0.973706220076824
7268 TTC4 tetratricopeptide repeat domain 4 3527
0.0327547788328342 4.33364577904585 0.0339654086432368
0.853780415270772 0.973741779836737
81558 FAM117A family with sequence similarity 117 member A 3836
0.0548885157784522 3.26077079840149
0.0339128349923783 0.853892348272071 0.973741779836737
55290 BRF2 BRF2 RNA polymerase III transcription initiation
factor subunit 2760 0.0330488824324776 4.28827311425759
0.0339128296465105 0.853892359658359
0.973741779836737
751580 LINC00106 long intergenic non-protein coding RNA 106
1184 0.0679366599820575 0.507071914209356
0.0338441685863042 0.85403867906267 0.973759156202903
27095 TRAPPC3 trafficking protein particle complex 3 3527
0.0238002498807089 6.32146360345958 0.0337915876876878
0.854150834792823 0.973759156202903
285521 COX18 cytochrome c oxidase assembly factor COX18 6976
0.0351557748170064 4.21364221084748
0.0337114522777249 0.854321938291427 0.973759156202903
6143 RPL19 ribosomal protein L19 2031 0.0322194921808268
7.91798969751733 0.0337095394049811
0.854326025185989 0.973759156202903
134353 LSM11 LSM11, U7 small nuclear RNA associated 6584
0.0454834434274506 2.89579951948065 0.0336674967654655
0.854415880494377 0.973759156202903

169981	SPIN3	spindlin family member 3	2222
0.0486495478118395	1.5136596956076 0.0336455867345968
0.854462730526298	0.973759156202903
343578	ARHGAP40	Rho GTPase activating protein 40	3195
	0.109020714812427	1.81231235237562
0.0334937083080842	0.854787925251189	0.973759156202903
1633	DCK	deoxycytidine kinase	2948	0.0401295034853351
	3.99627788719541	0.0334718035575499
0.854834889469332	0.973759156202903
388552	BLOC1S3 biogenesis of lysosomal organelles complex 1 subunit
3	2601	0.0416456897669141	3.56948685729181
0.0334547628118749	0.85487143614931	0.973759156202903
4942	OAT	ornithine aminotransferase
3388	-0.0365723876238573	9.35777160945067
0.0334257798476756	0.854933616979724	0.973759156202903
5256	PHKA2	phosphorylase kinase regulatory subunit alpha 2 9029
	0.0292666477383913	4.79646852034251
0.03340419298104	0.854979948127955	0.973759156202903
11154	AP4S1	adaptor related protein complex 4 subunit sigma 1
	7223	-0.046925916395476	2.51839540935404
0.0333729985779989	0.855046926939671	0.973759156202903
2049	EPHB3	EPH receptor B3 4532	-0.034908798858898
5.0145817256133 0.0333050971509152	0.8551928328709
0.973759156202903
8694	DGAT1	diacylglycerol O-acyltransferase 1	4131
0.0277069899071708	5.38111646894741	0.0332959515194169
	0.855212496655694	0.973759156202903
102724862	TBC1D3I TBC1 domain family member 3I
2065	-0.0786404369767294	2.87805600820277
0.0332905690728085	0.855224070618809	0.973759156202903
84498	FAM120B family with sequence similarity 120B
6029	-0.0219534544062011	5.58683118947901
0.0332576234527009	0.855294935204634	0.973759156202903
55326	AGPAT5	1-acylglycerol-3-phosphate O-acyltransferase 5
4684	-0.0296713828804458	4.2773990740351 0.0331898819513055
	0.855440758246203	0.973759156202903
163183	SYNE4	spectrin repeat containing nuclear envelope family
member 4	2528	0.0470410875150643	3.5231640012742
0.0331428586445419	0.855542072908579	0.973759156202903
2767	GNA11	G protein subunit alpha 11	6242
0.0251910553279005	6.73488243892156	0.0331327678429574
	0.855563823848363	0.973759156202903
79109	MAPKAP1 MAPK associated protein 1	6741
0.0230369090931471	6.61402851651784	0.033012513441971
	0.85582329863288	0.973759156202903
8638	OASL	2'-5'-oligoadenylate synthetase like	3517
0.113130334118886	0.661119876999663	0.0329925082631348
	0.855866511453374	0.973759156202903
55644	OSGEP	O-sialoglycoprotein endopeptidase	5037
0.0383466294871332	3.94087790336069	0.0329552848320134
	0.855946953138473	0.973759156202903
7046	TGFBR1	transforming growth factor beta receptor 1	6844
	0.0401327366857355	7.61582632447624
0.0329509565051964	0.855956309912863	0.973759156202903

8625	RFXANK	regulatory factor X associated ankyrin containing protein 2503	0.0289804079720018	4.96193045431694	0.0329243436656004	0.856013854259795	0.973759156202903
121273	C12orf54	chromosome 12 open reading frame 54	1857	0.0703938521330424	2.6121213004507	0.0329215805257554	0.85601983031155	0.973759156202903
126375	ZNF792	zinc finger protein 792	4234	0.0510567035831459	2.7341928159603	0.0329009371070939	0.856064485595548	0.973759156202903
3832	KIF11	kinesin family member 11	4860	0.0476580433340072	1.98701104968429	0.0328220144735143	0.856235342449061	0.973759156202903
8569	MKNK1	MAPK interacting serine/threonine kinase 1	8070	-0.024510755294331	5.77868739966677	0.0327869101907297	0.856311406653201	0.973759156202903
284307	ZIK1	zinc finger protein interacting with K protein 1	4547	-0.0641622759767896	2.31901394477073	0.0327450332435753	0.856402200943029	0.973759156202903
54741	LEPROT	leptin receptor overlapping transcript	5237	0.0233129587509929	8.49428650719829	0.0326030659867502	0.856710450141943	0.973759156202903
51490	SPOUT1	SPOUT domain containing methyltransferase 1	4593	-0.0237799271032756	5.1843400432331	0.0326021075587271	0.856712533500294	0.973759156202903
81034	SLC25A32	solute carrier family 25 member 32	3654	-0.0337439614867568	4.32115604038024	0.0325998800990042	0.856717375505924	0.973759156202903
400660	LINC00683	long intergenic non-protein coding RNA 683	4928	0.0617111185082885	-0.123576103382412	0.0325302059047154	0.856868918145424	0.973759156202903
27031	NPHP3	nephrocystin 3	9701	-0.04401013830213	4.86238388531484	0.0325244424767019	0.856881461206921	0.973759156202903
642799	NPIPA2	nuclear pore complex interacting protein family member A2	2891	-0.0460379490670439	2.65494041303373	0.0325040354294632	0.856925882691351	0.973759156202903
10056	FARSB	phenylalanyl-tRNA synthetase subunit beta	3369	-0.0453822402891706	4.20433094087429	0.0324921326088088	0.856951799063627	0.973759156202903
55450	CAMK2N1	calcium/calmodulin dependent protein kinase II inhibitor 1	2615	0.0376026516305362	6.47445768750836	0.032459893172188	0.857022019592214	0.973759156202903
26959	HBP1	HMG-box transcription factor 1	6167	0.0259095230113882	7.03515547243742	0.0324407570876168	0.857063716814753	0.973759156202903
23563	CHST5	carbohydrate sulfotransferase 5	3484	0.051555353979093	0.0457936556222782	0.0324048132913539	0.857142072109558	0.973759156202903
60370	AVPI1	arginine vasopressin induced 1	1457	-0.0454560712454694	4.68839067135938	0.0323852459149947	0.857184746663205	0.973759156202903
729970	LOC729970	hCG2028352-like	1012	-0.0716490727725923

1.79258471702204	0.0323828381121753
0.857189998767626	0.973759156202903
54436	SH3TC1	SH3 domain and tetratricopeptide repeats 1
10154	-0.033226728049314	4.43486946799963
0.0323770677218036	0.85720258645467	0.973759156202903
25799	ZNF324	zinc finger protein 324 5664	0.0322236094038486
4.38936685066004	0.0323207862856911
0.857325421094916	0.973759156202903
684	BST2	bone marrow stromal cell antigen 2	1119
0.138489444536525	2.76688569645763	0.0323158785651998
0.857336137461907	0.973759156202903
5201	PFDN1	prefoldin subunit 1	2156	0.0282960514029135
6.13988979992973	0.0322999795879007
0.8573708598127 0.973759156202903
405	ARNT	aryl hydrocarbon receptor nuclear translocator	6516
0.0249690257939696	5.67248545414088
0.032299219479782	0.85737252006121	0.973759156202903
404281	YY2	YY2 transcription factor
2741	-0.0551547466995	1.76132943532766
0.0322945994436088	0.857382611701785	0.973759156202903
3015	H2AZ1	H2A.Z variant histone 1 2303	-0.0266374929709524
6.59073001083686	0.0322536327538785
0.857472128707541	0.973759156202903
166348	KBTBD12 kelch repeat and BTB domain containing 12	5818
0.144092156356951	0.330250361256104
0.0322258148616754	0.857532947511137	0.973759156202903
84901	NFATC2IP	nuclear factor of activated T cells 2
interacting protein	5738	0.0225473884011609
6.39238493719239	0.0322161275992077	0.85755413332368
0.973759156202903
25794	FSCN2	fascin actin-bundling protein 2, retinal	2019
0.0635951216787672	0.0796543884674585
0.032144780296413	0.857710269562678	0.973759156202903
11146	GLMN	glomulin, FKBP associated protein	2910
0.0418986099988966	3.24640325360051	0.0321359631702407
0.857729577348901	0.973759156202903
137682	NDUFAF6 NADH:ubiquinone oxidoreductase complex assembly
factor 6	4598	-0.0424486373344123	2.60954959361963
0.0321289739493258	0.857744884321747
0.973759156202903
4173	MCM4	minichromosome maintenance complex component 4
6824	-0.0269775082975607	5.39486381442574
0.032027081769666	0.857968232067273	0.973759156202903
54107	POLE3	DNA polymerase epsilon 3, accessory subunit	2604
0.0243345854255721	5.78234867670239
0.0320183564835474	0.857987374934885	0.973759156202903
8446	DUSP11	dual specificity phosphatase 11 3544
0.026191705145111	5.21178263974603	0.0320028274233453
0.85802145162291	0.973759156202903
100506054	RNASEH1-AS1	RNASEH1 antisense RNA 1 1218
0.06327257183078	1.00127416178556	0.0319668042225869
0.858100533148744	0.973759156202903
26157	GIMAP2	GTPase, IMAP family member 2	2097
0.0674039521115374	0.153008520067205	0.0319169217780031

0.858210115753584	0.973759156202903
84942	WDR73	WD repeat domain 73	9626	0.0283863001579257
4.85181340150668	0.0318976955867463
0.858252375769679	0.973759156202903
84134	TOMM40L translocase of outer mitochondrial membrane 40 like
3289	-0.0304888187622679	4.68314420043153
0.0318964527056318	0.858255108129366	0.973759156202903
64965	MRPS9	mitochondrial ribosomal protein S9	1915
0.0295060291141318	4.32752235066037	0.0318638555887887
0.858326789512331	0.973759156202903
374882	TMEM205 transmembrane protein 205	1399
0.0312406142772807	4.21526192654751	0.0318551795284243
0.858345874629386	0.973759156202903
9812	DELE1	DAP3 binding cell death enhancer 1
6047	-0.025541519913318	5.48122570992349
0.0318171793080761	0.858429497022137	0.973759156202903
84844	PHF5A	PHD finger protein 5A	1235	0.030027079064025
4.49444294284503	0.0318060455704092
0.85845400742257	0.973759156202903
10486	CAP2	cyclase associated actin cytoskeleton regulatory
protein 2	3637	-0.0364456573790286	3.963659959462
0.0317651096625804	0.858544163977999	0.973759156202903
4771	NF2	neurofibromin 2 7170	0.027618365612427
6.14016777984762	0.0317545361413814	0.858567460662261
0.973759156202903
22949	PTGR1	prostaglandin reductase 1	3722
0.0371750819385291	5.81978386356938	0.0317439429615654
0.858590804673741	0.973759156202903
51257	MARCHF2 membrane associated ring-CH-type finger 2
2216	-0.0298105947122912	4.53024807347264
0.0316803681260822	0.858730987973304	0.973759156202903
27042	UTP25	UTP25 small subunit processor component
10376	-0.0247279830725368	5.47237584764201
0.0316678499279277	0.858758607850343	0.973759156202903
1788	DNMT3A	DNA methyltransferase 3 alpha
12020	-0.0252349590222377	5.6732256324957 0.0316608929907432
0.858773959918217	0.973759156202903
100526835	FPGT-TNNI3K	FPGT-TNNI3K readthrough
10714	-0.0606394820572617	0.634203602688822
0.0315611721788831	0.858994208144506	0.973863898783988
100287171	WASHC1	WASH complex subunit 1
1827	-0.0539434668864288	3.99982258104202
0.0315192620632132	0.859086879974536	0.973863898783988
26003	GORASP2 golgi reassembly stacking protein 2
5340	-0.0248879850311767	7.00104970520473
0.0314961527967483	0.859138006469454	0.973863898783988
81693	AMN	amnion associated transmembrane protein
4646	-0.0630192044208108	1.29308383456254
0.0314648267807591	0.859207342440621	0.973863898783988
57761	TRIB3	tribbles pseudokinase 3 3434	-0.0478949448316318
5.05762926418367	0.0314618338247499
0.859213968813631	0.973863898783988
387882	C12orf75	chromosome 12 open reading frame 75	2693
0.0467600393152376	5.25558178674408

0.0313264611683621	0.859514023058359	0.974125165545629
100048912	CDKN2B-AS1	CDKN2B antisense RNA 1	4685
0.0415192562941299	3.22296570967573	0.0312887539428957
	0.859597720341825	0.974126563526496
2923	PDIA3	protein disulfide isomerase family A member 3
4240	-0.0389279644238233	8.05145912829671
0.0312582577876128	0.859665449539181	0.974126563526496
440585	FAM183A	family with sequence similarity 183 member A	843
	0.0704279436746314	0.144075622451374
0.031227179349897	0.859734506995934	0.974126563526496
255403	ZNF718	zinc finger protein 718	4118	0.0725125981402592
	2.83382643273172	0.0311855031966459
0.859827168622434	0.974126563526496
84309	NUDT16L1	nudix hydrolase 16 like 1
2068	-0.0275387702404266	4.38638106909099
0.0311413277426862	0.85992545685832	0.974126563526496
9380	GRHPR	glyoxylate and hydroxypyruvate reductase	6908
	0.0373959864915951	5.45189163739632
0.0311077436475671	0.860000227943787	0.974126563526496
246184	CDC26	cell division cycle 26	1175	-0.0447927122852585
	1.78423476576998	0.031106913619825
0.860002076430974	0.974126563526496
64983	MRPL32	mitochondrial ribosomal protein L32	3534
0.0252177258323423	5.33284787439238	0.0309870770882821
	0.860269221981538	0.974350367291615
9616	RNF7	ring finger protein 7	3147	-0.0256750496844779
	6.14216406057431	0.0308385060204763
0.860601164504428	0.974585312210587
645553	IGFL2-AS1	IGFL2 antisense RNA 1
1751	-0.111775751778563	5.36509850906333
0.0308245698121681	0.860632343499996	0.974585312210587
23456	ABCB10	ATP binding cassette subfamily B member 10	4256
	0.0346558829529406	4.61940154265459
0.0308008651208809	0.860685393877701	0.974585312210587
1390	CREM	cAMP responsive element modulator
5699	-0.0308724994012602	3.55227532990625
0.0307276750275367	0.860849323999389	0.974629467202034
553115	PEF1	penta-EF-hand domain containing 1	2227
0.0251815278907642	6.12224686353077	0.0306344608973372
	0.86105839528969	0.974629467202034
9997	SCO2	SCO cytochrome c oxidase assembly protein 2	1361
	0.0306577397550407	4.22565606542149
0.0306091529126498	0.861115215493951	0.974629467202034
81873	ARPC5L	actin related protein 2/3 complex subunit 5 like
	3610	0.0232329259767193	6.25785451943492
0.030606656823096	0.861120820899062	0.974629467202034
81789	TIGD6	tigger transposable element derived 6	4640
0.040651066873532	2.76502700260776	0.0305690018182219
	0.861205410388396	0.974629467202034
80045	GPR157	G protein-coupled receptor 157
5372	-0.0354489038166186	8.18506718832103
0.0305533637840654	0.861240555981038	0.974629467202034
23786	BCL2L13	BCL2 like 13	7662	0.0197661608422604
6.40750946502091	0.030491844400526	0.861378907332556

0.974629467202034
51204 TACO1 translational activator of cytochrome c oxidase I
2315 -0.0273760566036085 3.92717570364092
0.0304494184633333 0.861474403116176 0.974629467202034
118980 SFXN2 sideroflexin 2 8034 0.0542970108438008
0.438686939370428 0.0304370112768995 0.861502343184232
0.974629467202034
64960 MRPS15 mitochondrial ribosomal protein S15
2916 -0.0282117221883654 5.34592495908624
0.0304297866368728 0.861518615243643 0.974629467202034
57002 YAE1 YAE1 maturation factor of ABCE1 2569
0.039203973033873 3.00796576599802 0.030366623306306
0.861660962890353 0.974629467202034
10139 ARFRP1 ADP ribosylation factor related protein 1 7627
0.0216710505241405 5.53149335690327
0.0303620268441307 0.861671327630328 0.974629467202034
7073 TIAL1 TIA1 cytotoxic granule associated RNA binding
protein like 1 7355 -0.0211165004545951 6.64637739071638
0.0302842571711039 0.861846816163299
0.974629467202034
200765 TIGD1 tigger transposable element derived 1
2625 -0.0476103059693368 1.16499820129313
0.0302841780444212 0.861846994832096 0.974629467202034
8492 PRSS12 serine protease 12 5725 0.0376314114441025
4.6960858469014 0.0302602369095286 0.861901065188986
0.974629467202034
9645 MICAL2 microtubule associated monooxygenase, calponin and
LIM domain containing 2 15160 -0.0530732065067638
7.27547385791401 0.0301611544016396 0.862125074810989
0.974629467202034
1455 CSNK1G2 casein kinase 1 gamma 2 3804 0.0232552501987237
6.19746961720694 0.0300513079275504
0.862373863533703 0.974629467202034
6809 STX3 syntaxin 3 6566 -0.0337816102645936
7.28633700031075 0.030037999788874 0.862404036615503
0.974629467202034
54922 RASIP1 Ras interacting protein 1
4149 -0.0898301410753656 1.97621698802157
0.030007094264402 0.862474134210587 0.974629467202034
100529097 RPL36A-HNRNPH2 RPL36A-HNRNPH2 readthrough 695
0.0265653121527303 4.72591333196919
0.0299517193841012 0.862599824420123 0.974629467202034
10575 CCT4 chaperonin containing TCP1 subunit 4 2830
0.0300197069199155 7.25078372312377 0.0298954641455111
0.862727635469246 0.974629467202034
661 POLR3D RNA polymerase III subunit D 5611
0.0312090570419254 5.22697224747073 0.0298852415051982
0.862750874446952 0.974629467202034
22829 NLGN4Y neuroligin 4 Y-linked 8206 -0.0425358335149656
4.42636123313086 0.0298839766704617
0.862753750061685 0.974629467202034
23184 MESD mesoderm development LRP chaperone
7510 -0.0220590752509719 5.95765610868662
0.029879160262665 0.86276470078792 0.974629467202034

23022	PALLD	palladin, cytoskeletal associated protein
10114	0.029988894793263	9.11219543274946
0.0298649977876551	0.862796906274211	0.974629467202034
145508	CEP128	centrosomal protein 128 8795	0.046704925188352
	1.1934709390745 0.0298624661839595	0.862802663972721
	0.974629467202034
9966	TNFSF15 TNF superfamily member 15
6687	-0.0774016338851902	4.77234773318618
0.0297798240336249	0.862990757583561	0.974629467202034
125228	FAM210A family with sequence similarity 210 member A
4681	-0.033767779926265	4.38733238634877
0.0297313863239594	0.863101126800407	0.974629467202034
84896	ATAD1	ATPase family AAA domain containing 1
4640	-0.0207974393229082	6.62434442006441
0.0297200350140301	0.863127005060302	0.974629467202034
23729	SHPK	sedoheptulokinase	5873	-0.0326959405087463
	4.08046303458879	0.0296968457349323
0.863179886867635	0.974629467202034
10755	GIPC1	GIPC PDZ domain containing family member 1	3156
	0.0223035382808737	8.4963026687805 0.0296614709247365
	0.863260598041947	0.974629467202034
219402	MTIF3	mitochondrial translational initiation factor 3
1603	-0.0287592363639629	4.45933292916258
0.0296533376824129	0.86327916185501	0.974629467202034
4882	NPR2	natriuretic peptide receptor 2
3865	-0.0525242039428915	2.89490077370391
0.0296389218277877	0.863312071931019	0.974629467202034
8683	SRSF9	serine and arginine rich splicing factor 9
2510	-0.0226229940005875	6.45520604396395
0.0295746806678565	0.863458828897305	0.974629467202034
55702	YJU2	YJU2 splicing factor homolog
1671	-0.0310394632760075	3.78460457385615
0.0295690162176512	0.86347177703021	0.974629467202034
2730	GCLM	glutamate-cysteine ligase modifier subunit
5420	-0.0365325190781012	5.50675947990029
0.029544587587722	0.863527632059936	0.974629467202034
27330	RPS6KA6 ribosomal protein S6 kinase A6	8502
0.0500079353462787	3.63572942129093	0.0295061644008925
	0.863615533178052	0.974629467202034
80021	TMEM62	transmembrane protein 62	4810
0.0321079325546026	5.35854714349776	0.0294619604889093
	0.863716731837242	0.974629467202034
51660	MPC1	mitochondrial pyruvate carrier 1	1546
0.0344755824691303	3.89073978482781	0.0294515839882736
	0.86374049870498	0.974629467202034
1371	CPOX	coproporphyrinogen oxidase	3236
0.0340871699946856	4.99908239450888	0.0293237546871321
	0.864033639808674	0.974629467202034
2073	ERCC5	ERCC excision repair 5, endonuclease
6670	-0.0217965547450424	6.14275795848372
0.0293092350456448	0.864066978146594	0.974629467202034
285051	STPG4	sperm-tail PG-rich repeat containing 4	2397
0.0533750202346187	2.03113570889529	0.0292765525487582
	0.864142051065494	0.974629467202034

24148	PRPF6	pre-mRNA processing factor 6
3044	-0.0214931932564163	6.30224486016821
0.0292562591081351	0.864188687564768	0.974629467202034
6602	SMARCD1 SWI/SNF related, matrix associated, actin dependent regulator of chromatin, subfamily d, member 1	5136
0.0206634976297154	6.01331178923943	0.0292291222448036
	0.864251077005184	0.974629467202034
219293	ATAD3C	ATPase family AAA domain containing 3C	3976
0.0384785724986617	2.79813939719676	0.0292176504604869
	0.864277460358505	0.974629467202034
6294	SAFB	scaffold attachment factor B
7036	-0.0243691362164174	6.37786906084115
0.029209602361675	0.86429597293955	0.974629467202034
57149	LYRM1	LYR motif containing 1	3593	-0.0390680810929614
	3.6288176961638 0.0291747027997644	0.864376280790708
	0.974629467202034
83544	DNAL1	dynein axonemal light chain 1
8884	-0.0245669263957485	4.69538967367296
0.0291633588676206	0.864402395110229	0.974629467202034
64841	GNPNAT1 glucosamine-phosphate N-acetyltransferase 1	4009
	0.0390322817467213	4.93372096596196
0.0291471622564252	0.864439689609376	0.974629467202034
9653	HS2ST1	heparan sulfate 2-O-sulfotransferase 1
7429	-0.0313013031743461	5.15539885333559
0.0291012030647888	0.864545573861305	0.974629467202034
5818	NECTIN1 nectin cell adhesion molecule 1
7179	-0.0329781002739721	7.1851842950909 0.0290982261301451
	0.86455243531408	0.974629467202034
1026	CDKN1A	cyclin dependent kinase inhibitor 1A
3050	-0.0244287138828392	10.2454984324732
0.0290965487774884	0.864556301556049	0.974629467202034
389524	GTF2IRD2B	GTF2I repeat domain containing 2B	5672
	0.0370863014953283	3.82795343695614
0.0290845690299317	0.864583917809982	0.974629467202034
7416	VDAC1	voltage dependent anion channel 1
3163	-0.0275793133442346	7.11926702309218
0.0290723272947844	0.864612144056478	0.974629467202034
64757	MARC1	mitochondrial amidoxime reducing component 1	9258
	0.0502471162628697	2.08692392420906
0.0290498380690316	0.864664014283547	0.974629467202034
6720	SREBF1	sterol regulatory element binding transcription factor 1	5239	0.0314687987296066	6.28381982130145
	0.0290198708489697	0.864733164233859
0.974629467202034
60401	EDA2R	ectodysplasin A2 receptor	3579
0.0622759157801217	4.19777981185303	0.0289369543499154
	0.864924687375794	0.974629467202034
10873	ME3	malic enzyme 3	3838	0.036437905038433
3.56987768661375	0.02892522652892	0.8649517994465
0.974629467202034
134510	UBLCP1	ubiquitin like domain containing CTD phosphatase 1
	2676	0.0243500380339942	5.23913673126177
0.0289176559567323	0.864969303901572	0.974629467202034
10098	TSPAN5	tetraspanin 5	7430	0.031023501441803

4.38566415910074	0.0288859951297411	0.865042534696374
	0.974629467202034
5082	PDCL	phosducin like	3147	0.026539137762782
5.19479143054859	0.0288849504156685	0.865044951800557
	0.974629467202034
4149	MAX	MYC associated factor X	4430	-0.0250498925609195
	5.81547493375197	0.0288668296504682
0.865086884091924	0.974629467202034
29105	CFAP20	cilia and flagella associated protein 20	2375
	0.0269815322069134	4.74109494752525
0.0288258678695854	0.865181721482017	0.974629467202034
23023	TMCC1	transmembrane and coiled-coil domain family 1
7561	-0.0249425054310322	6.97394239303428
0.0287721342143357	0.865306234328952	0.974629467202034
79589	RNF128	ring finger protein 128	3432	-0.0374975732092974
	3.51905836610688	0.028764315008833
0.865324363135853	0.974629467202034
5714	PSMD8	proteasome 26S subunit, non-ATPase 8
3105	-0.02382988580083	7.02007809323614
0.0287575424451916	0.865340067355183	0.974629467202034
6742	SSBP1	single stranded DNA binding protein 1
4988	-0.027342217321247	5.63223878411268
0.0287375850393801	0.865386355650153	0.974629467202034
57156	TMEM63C	transmembrane protein 63C	6603
0.055175326115578	2.19456597568262	0.028652561191457
	0.865583741592529	0.974634787282742
64499	TPSB2	tryptase beta 2 (gene/pseudogene)
1738	-0.155164429601722	-0.110515808602178
0.0286243526182055	0.865649295356221	0.974634787282742
6128	RPL6	ribosomal protein L6	4447	0.0329769761080793
	7.71142580573007	0.0285968430337107
0.865713256751755	0.974634787282742
55851	PSENEN	presenilin enhancer, gamma-secretase subunit
1845	-0.026500386106429	5.43623879287042
0.0285895623539432	0.865730190053979	0.974634787282742
84319	CMSS1	cms1 ribosomal small subunit homolog
4081	-0.0509625843137674	2.50679633078624
0.0285857793796991	0.865738989317618	0.974634787282742
79679	VTCN1	V-set domain containing T cell activation inhibitor
1	3027	-0.0672948890125106	5.23110386909256
0.0285164560312694	0.865900342793483	0.974733676687043
22933	SIRT2	sirtuin 2	6224	0.024027678606589
5.3697676808047	0.0284882686824659	0.865966007902732
0.974733676687043
80777	CYB5B	cytochrome b5 type B	6903	-0.0246565422912617
	6.49401095691912	0.0283496566681327
0.866289404956869	0.975019340717626
55593	OTUD5	OTU deubiquitinase 5	3592	-0.020404986647583
	5.76735240447793	0.0282750334537711
0.866463845827313	0.975137320303967
100500840	NA	NA	69	-0.0601750165884985
0.956280270195199	0.0282448866344343	0.866534384817115
	0.975138356883854
5546	PRCC	proline rich mitotic checkpoint control factor	4108

0.0244286651722149 6.15958402565757
0.028165205621093 0.866721012889182 0.97527002149251
90507 SCRN2 secernin 2 3043 -0.0396884944959067
4.11679592908903 0.02806736379540 1 0.866950548694778
0.975389580364618
146880 ARHGAP27P1 Rho GTPase activating protein 27 pseudogene
1 3038 -0.0345611601886078 5.18759037419444
0.0280605553978255 0.866966536413186 0.975389580364618
51138 COPS4 COP9 signalosome subunit 4 3566
0.0267340517395069 4.9747597037478 0.0279350756865471
0.867261550077532 0.975632404265804
7884 SLBP stem-loop binding protein
2026 -0.0224646122964245 5.26220500908793
0.0279095380096344 0.867321674728193 0.975632404265804
7913 DEK DEK proto-oncogene 4361 -0.0277212389590606
6.33549708652062 0.027861501465523
0.867434846268523 0.975681353222774
79986 ZNF702P zinc finger protein 702, pseudogene
553 -0.0514045455476513 1.98152353989245
0.0278063515894509 0.867564900163195 0.975749281884204
1112 FOXN3 forkhead box N3 11203 0.0310091152600942
6.06098355543057 0.0277311023454487 0.867742566202532
0.975870744664702
84859 LRCH3 leucine rich repeats and calponin homology domain
containing 3 12070 -0.0250775723642805 5.84918435384879
0.0276297100115812 0.86798234921829
0.975979018765688
6484 ST3GAL4 ST3 beta-galactoside alpha-2,3-sialyltransferase 4
4465 0.0397775155962136 4.57534312831314
0.0275864717206957 0.868084741073056 0.975979018765688
5218 CDK14 cyclin dependent kinase 14 6841
0.0439191796663515 6.16176734236623 0.0275656819751475
0.868134002267523 0.975979018765688
79838 TMC5 transmembrane channel like 5 6393
0.0474104751242362 4.73581198257411 0.0275491435654374
0.868173203577558 0.975979018765688
79672 FN3KRP fructosamine 3 kinase related protein 2399
0.0300794629414286 4.79614495905887 0.0275432259783095
0.86818723308492 0.975979018765688
90826 PRMT9 protein arginine methyltransferase 9
4019 -0.0331505695163359 3.26289533448731
0.027402985896682 0.868520170798199 0.976076034366732
51304 ZDHHC3 zinc finger DHHC-type containing 3 7063
0.0199358628992475 6.94671546088142 0.0273946591879337
0.8685399663637 0.976076034366732
83940 TATDN1 TatD DNase domain containing 1 4778
0.0417337724476907 3.25934105299671 0.0273903669017783
0.868550171870737 0.976076034366732
4478 MSN moesin 4180 -0.0301302496259624
9.23940866844048 0.0273894831074131 0.868552273316695
0.976076034366732
114780 PKD1L2 polycystin 1 like 2 (gene/pseudogene)
18637 -0.0970356577835363 0.256142424121667
0.0273375910492692 0.868675721009536 0.976105290327359

51643	TMBIM4	transmembrane BAX inhibitor motif containing 4
4457	-0.0225773930247783	6.05868137529992
0.0272813369146143	0.868809681833488	0.976105290327359
201516	ZSCAN4	zinc finger and SCAN domain containing 4	2249
	0.0584958381999437	-0.18445612056034
0.0272798421453508	0.868813243337555	0.976105290327359
79703	C11orf80	chromosome 11 open reading frame 80	3765
	0.0293034532964548	4.91161290019522
0.0272614580454871	0.868857054315807	0.976105290327359
284443	ZNF493	zinc finger protein 493	8176	0.0466413590170342
	2.8859068860557	0.0272140062270516	0.868970206488397
	0.976154116792283
58155	PTBP2	polypyrimidine tract binding protein 2	12937
0.0462934097541322	2.85491216610601	0.0271574082223633
	0.869105300975702	0.976227582258754
28989	NTMT1	N-terminal Xaa-Pro-Lys N-methyltransferase 1
3029	-0.0238050746791576	4.23886084432397
0.027104483328582	0.869231758954692	0.976252157995422
129285	PPP1R21	protein phosphatase 1 regulatory subunit 21	3995
	0.0304201615374677	4.85276426643403
0.027083860860607	0.869281068312277	0.976252157995422
80851	SH3BP5L	SH3 binding domain protein 5 like	658
0.039218035477858	2.75055008393301	0.0270607826502918
	0.869336272340751	0.976252157995422
6304	SATB1	SATB homeobox 1	8303	-0.0398449640130403
4.92416385598497	0.0269993771675843	0.869483274635668
	0.976267188899573
283131	NEAT1	nuclear paraspeckle assembly transcript 1
22743	-0.0537788656586528	9.65996783862453
0.0269775903539085	0.869535472661291	0.976267188899573
84961	FBXL20	F-box and leucine rich repeat protein 20
11038	0.0307011770093645	5.60281416265762
0.0269678764749273	0.869558752674453	0.976267188899573
25861	WHRN	whirlin	5542	-0.0727076384055312
2.69168119116945	0.0269053366389898	0.869708737121626
	0.976299003434223
1938	EEF2	eukaryotic translation elongation factor 2	4027
	0.0258578181687187	9.93344978892197
0.026878637037953	0.869772823227891	0.976299003434223
79791	FBXO31	F-box protein 31	6810	0.0242536396195396
	4.92301653591792	0.0268689044345329
0.869796192179315	0.976299003434223
14	AAMP	angio associated migratory cell protein
3140	-0.020294605829179	6.58158809318024
0.0267722937895769	0.870028400068151	0.976421093168474
100131691	MZF1-AS1	MZF1 antisense RNA 1	2968
0.0583734924874887	1.33900121357234	0.0267656508870573
	0.870044382368511	0.976421093168474
3083	HGFAC	HGF activator	2847
0.0844290224894954	-0.39490480576604	0.0267363229022628
	0.870114967543803	0.976422075820064
55970	GNG12	G protein subunit gamma 12
4724	-0.0248660351659374	8.35490369670249
0.026668131006276	0.870277598075626	0.97643980097168

79031	PDCL3	phosducin like 3	1831	0.0317836988436333
	4.4629310790289 0.0266425690805789	0.870340876733152
	0.976439800971684
51621	KLF13	Kruppel like factor 13	7694	0.028314406763054
	7.74857723835915	0.0266085449828424
0.870422962089557	0.976439800971684
118813	ZFYVE27 zinc finger FYVE-type containing 27	3899
0.0183514120720004	5.89276340649018	0.0265503082044063
	0.870563587247014	0.976439800971684
675	BRCA2	BRCA2 DNA repair associated	12273
0.0585374377462098	0.841835230247694	0.0265309058534484
	0.870610473527769	0.976439800971684
338692	ANKRD13D	ankyrin repeat domain 13D	6601
0.0211352342411439	5.57735570772872	0.0265196697175374
	0.870637633986875	0.976439800971684
25766	PRPF40B pre-mRNA processing factor 40 homolog B
6824	-0.0349206466629846	3.93238744909751
0.0265154631218998	0.870647803865839	0.976439800971684
10768	AHCYL1	adenosylhomocysteinase like 1	5113
0.0204918969445014	7.46947704738572	0.0264986541150449
	0.87068844964206	0.976439800971684
5510	PPP1R7	protein phosphatase 1 regulatory subunit 7
3781	-0.0240351874620501	5.89034299937623
0.0264493319365897	0.870807791799423	0.976484090330982
215	ABCD1	ATP binding cassette subfamily D member 1
4233	-0.0382540435589575	3.09795535348805
0.0263701946046009	0.87099951493934	0.976484090330982
2048	EPHB2	EPH receptor B2 12516	0.0311273219773802
8.08559156430931	0.0263053123058228	0.871156923317301
	0.976484090330982
10147	SUGP2	SURP and G-patch domain containing 2	7107
0.0217689724128906	6.65505815708293	0.0262623571359981
	0.871261244791866	0.976484090330982
896	CCND3	cyclin D3	5300	0.0242234996291717
6.14516658291206	0.0262596625745566	0.871267791755667
	0.976484090330982
64320	RNF25	ring finger protein 25	2323	0.0262950108482148
	4.90610062983079	0.0262573313381651
0.871273456228367	0.976484090330982
197131	UBR1	ubiquitin protein ligase E3 component n-recognin 1
	9928	-0.0222880869495617	5.99766048122428
0.0262481220056259	0.871295835724641	0.976484090330982
57209	ZNF248	zinc finger protein 248 7951	0.0336034545323587
	3.73618039590866	0.0262210928388029
0.871361542277224	0.976484090330982
341208	HEPHL1	hephaestin like 1	5345	0.170092614167529
	1.05897655902117	0.0261220229715402
0.871602673792626	0.976484090330982
338657	CCDC84	coiled-coil domain containing 84
5105	-0.0399747893352844	4.74473994133884
0.0260869970085622	0.871688037620206	0.976484090330982
56666	PANX2	pannexin 2	3102	-0.0559526784190708
1.1365513921524 0.0260859739557997	0.871690531845641
0.976484090330982

55690	PACS1	phosphofurin acidic cluster sorting protein 1	9037
	0.0236636819693661	6.97262050032188
0.0260821603478547	0.871699829949048	0.976484090330982
7332	UBE2L3	ubiquitin conjugating enzyme E2 L3
3838	-0.0225377957035815	7.21376377349235
0.0260778284444569	0.871710392574589	0.976484090330982
126321	MFSD12	major facilitator superfamily domain containing 12
	3453	-0.0219864939358836	5.6172884467385
0.0260653757635891	0.87174076137876	0.976484090330982
860	RUNX2	RUNX family transcription factor 2	8192
0.0633702162946197	3.10330910621642	0.0260518922812594
	0.871773652430137	0.976484090330982
51535	PPHLN1	periphilin 1	9018	0.0192775871832545
5.84672499137649	0.0260101382965061	0.871875560813668
	0.976488037028563
8848	TSC22D1	TSC22 domain family member 1
8398	-0.0318902285322044	6.95054266562063
0.0259873332150189	0.871931256304561	0.976488037028563
51538	ZCCHC17	zinc finger CCHC-type containing 17	1921
0.0230741457309693	5.29361457918399	0.0259315313599764
	0.872067643673627	0.976488037028563
56683	CFAP298	cilia and flagella associated protein 298	3579
	0.0223537713948636	5.11455275135503
0.0259153316975973	0.872107266056324	0.976488037028563
56616	DIABLO	diablo IAP-binding mitochondrial protein	5967
	0.021157920222531	5.68704851012179
0.0259077773699179	0.872125747356845	0.976488037028563
57600	FNIP2	folliculin interacting protein 2	10868
0.0306944684748934	4.75734131117264	0.025858183778027
	0.872247144402944	0.976545899740391
29761	USP25	ubiquitin specific peptidase 25	7823
0.0177295600127698	6.14521475345477	0.0257382163163165
	0.872541300004836	0.976707495719729
643904	RNF222	ring finger protein 222	3062
0.08146966951106	-0.256677107276801	0.0257368508558953
	0.872544652104207	0.976707495719729
25777	SUN2	Sad1 and UNC84 domain containing 2	6489
0.0232176046806988	7.87822115796783	0.0256933808366036
	0.872651415340297	0.976707495719729
80127	BBOF1	basal body orientation factor 1
4620	-0.0523155941230925	2.54282705391473
0.0256856544670061	0.872670401151739	0.976707495719729
9040	UBE2M	ubiquitin conjugating enzyme E2 M
1831	-0.0211250989302205	5.81404813774866
0.025651723716853	0.872753813014763	0.9767228075182
284611	FAM102B	family with sequence similarity 102 member B
9355	-0.0448697463804497	3.60465571101444
0.0255686923541631	0.872958167717908	0.976804712040402
29960	MRM2	mitochondrial rRNA methyltransferase 2	2843
0.0261512806954774	4.82140045763963	0.0255653208116353
	0.872966472857982	0.976804712040402
9836	LCMT2	leucine carboxyl methyltransferase 2
6957	-0.0273635750638192	3.81668632045291
0.0254428012471914	0.873268657941377	0.976945631781426

445372	TRIM6-TRIM34	TRIM6-TRIM34 readthrough
3391	-0.0362153325872913	3.75464362655311
0.0254253306906627	0.873311808455857	0.976945631781426
56034	PDGFC	platelet derived growth factor C
5593	-0.0281449191308052	5.41108068702048
0.0253969589283898	0.873381916252738	0.976945631781426
11202	KLK8	kallikrein related peptidase 8	1865
0.0840674801342521	2.47692996753893	0.0253753266644736
	0.873435397454597	0.976945631781426
27343	POLL	DNA polymerase lambda	5420	0.023602960269016
	4.51250928708041	0.0253555451062679
0.873484323641856	0.976945631781426
100505715	LOC100505715	uncharacterized LOC100505715	2388
	0.0558011861953145	0.0226958856122474
0.0253448056388157	0.873510894012321	0.976945631781426
6901	TAZ	tafazzin	4721	0.0353873926829102
4.28910661288805	0.0253052875742199	0.87360871484956
	0.976977027456909
100527964	LHX4-AS1	LHX4 antisense RNA 1	5189
0.0527021651119297	2.24335878130381	0.0252733260305433
	0.873687887989448	0.976987565628948
79898	ZNF613	zinc finger protein 613	3722	0.0309743442367162
	3.17042786852301	0.0251844207483281
0.873908388953831	0.977156127091587
152189	CMTM8	CKLF like MARVEL transmembrane domain containing 8
	1174	0.0384993583300892	2.73091083266886
0.0250936768991394	0.874133861748153	0.977230711364702
100526694	MSANTD3-TMEFF1	MSANTD3-TMEFF1 readthrough
2069	-0.0287012596332625	3.90398169818588
0.0250772308948699	0.874174770125332	0.977230711364702
55319	TMA16	translation machinery associated 16 homolog
3513	-0.03733215722986	2.99866071018707
0.0250513701816519	0.874239124806098	0.977230711364702
26013	L3MBTL1	L3MBTL histone methyl-lysine binding protein 1
10674	-0.0508662402860782	2.98888740514606
0.0250453295933006	0.874254161784185	0.977230711364702
54556	ING3	inhibitor of growth family member 3
5214	-0.0296317470495853	3.63736460128632
0.0250093493579477	0.874343766761493	0.977252883899476
58505	OSTC	oligosaccharyltransferase complex non-catalytic
subunit	1617	-0.0323724151759056	6.27216461891357
0.0249458896669239	0.874501967593852	0.977333690196529
7165	TPD52L2	TPD52 like 2	4694	-0.0232277105741671
7.59027915684271	0.0248991548565982	0.874618606313988
	0.977333690196529
57553	MICAL3	microtubule associated monooxygenase, calponin and
LIM domain containing 3	22573	0.0456665555641747
7.20023255463053	0.0248882374355617	0.874645869702872
	0.977333690196529
9588	PRDX6	peroxiredoxin 6	2367	0.0220138727285172
6.7598813352856	0.0248568463711649	0.874724294795479
0.977333690196529
51605	TRMT6	tRNA methyltransferase 6	3776
0.0266473088898256	4.56413649084854	0.0248191918377607

0.874818435053054	0.977333690196529
25973	PARS2	prolyl-tRNA synthetase 2, mitochondrial
2347	-0.0411244963614264	2.38755259510419
0.0248126843905103	0.87483471176441	0.977333690196529
10982	MAPRE2	microtubule associated protein RP/EB family member 2
6180	-0.0352549461312636	4.31944001888037
0.0246884293879219	0.875145924451694	0.977603394512711
55833	UBAP2	ubiquitin associated protein 2
10622	-0.0213768717623422	6.16071648029929
0.0246220913034065	0.875312405374628	0.977682349980137
23378	RRP8	ribosomal RNA processing 8	6922
0.0303177900264398	3.67778118344429	0.0246020151343433
0.875362833469017	0.977682349980137
51077	FCF1	FCF1 rRNA-processing protein
4747	-0.0208744413971992	5.60569574825697
0.0245421934866781	0.875513220913134	0.977682349980137
12	SERPINA3	serpin family A member 3
5979	-0.135820311533362	7.45934664801888
0.0245421215554416	0.875513401856207	0.977682349980137
6917	TCEA1	transcription elongation factor A1	4391
0.0190839868773216	6.00435728761667	0.0245213743614272
0.875565602780812	0.977682349980137
120114	FAT3	FAT atypical cadherin 3 19280	-0.0983567799849145
1.73126484704196	0.0244448031551876
0.875758455670531	0.977685581031527
8379	MAD1L1	mitotic arrest deficient 1 like 1	7230
0.0284676270659043	4.504559014622	0.0243814840732313
0.875918165508254	0.977685581031527
79800	CARF	calcium responsive transcription factor 7194
0.0392957427394547	3.19801832694576	0.0243486776363966
0.876000996796482	0.977685581031527
89894	TMEM116 transmembrane protein 116	5393
0.0417861705966926	2.37506515162392	0.0243462451698129
0.876007140682099	0.977685581031527
126432	RINL	Ras and Rab interactor like	4201
0.046517438161541	1.92614495570909	0.0243365013197692
0.876031754697031	0.977685581031527
51645	PPIL1	peptidylprolyl isomerase like 1
1810	-0.0332494605287545	4.73820157154525
0.0243167752217719	0.876081600402864	0.977685581031527
27351	DESI1	desumoylating isopeptidase 1
4972	-0.0326380522698687	5.7310098721683 0.0242654791957193
0.87621131691514	0.977685581031527
7733	ZNF180	zinc finger protein 180 4783	0.0289459206532189
3.84156761839465	0.0242538860711354
0.876240652860716	0.977685581031527
79918	SETD6	SET domain containing 6, protein lysine
methyltransferase	5078	0.0313235951270411
3.42836224915349	0.0242438758113863	0.87626598920704
0.977685581031527
7587	ZNF37A	zinc finger protein 37A 9626	-0.0264121759451358
5.07926187451257	0.0242303121222829
0.876300327966804	0.977685581031527
124801	LSM12	LSM12 homolog	2811	-0.0238586465570519

4.37306731512482	0.0242161098396538	0.876336293985209
	0.977685581031527
10668	CGRRF1	cell growth regulator with ring finger domain 1 4564
	0.0279533576230084	4.29723814349537
0.0241010903728744	0.87662796943247	0.977889113604925
51497	NELFCD	negative elongation factor complex member C/D	6298
	0.0181517387564488	5.79552349787733
0.0240891236459735	0.876658356502966	0.977889113604925
4696	NDUFA3	NADH:ubiquinone oxidoreductase subunit A3
2197	-0.0294672817065345	4.37188791857939
0.0240020047407263	0.876879810611774	0.978011255187843
84674	CARD6	caspase recruitment domain family member 6
4399	-0.0319948842168884	5.28865600034046
0.0239180923642941	0.877093503244814	0.978011255187843
10436	EMG1	EMG1 N1-specific pseudouridine methyltransferase
	6268	0.0247655578057045	4.68334295685905
0.0238801322302571	0.87719029922603	0.978011255187843
54894	RNF43	ring finger protein 43	7492	-0.0266522265745088
	4.80208753722226	0.0238468000002596
0.877275359311477	0.978011255187843
79609	VCPKMT	valosin containing protein lysine methyltransferase
	2055	-0.0361384588553925	2.98046593503639
0.0237401334404748	0.877547970128235	0.978011255187843
11278	KLF12	Kruppel like factor 12	11090	-0.0367858762187999
	4.76612758479353	0.0236835009768974
0.877692962189479	0.978011255187843
54662	TBC1D13	TBC1 domain family member 13	3982
0.0262153520259687	5.64284433475448	0.0236733752661422
	0.877718905019626	0.978011255187843
57592	ZNF687	zinc finger protein 687 6349	-0.0220614501080686
	5.45399897546957	0.0236553928513672
0.877764991317897	0.978011255187843
57721	METTL14	methyltransferase like 14
6862	-0.0206369651053409	4.89828555549043
0.0236175684090687	0.877861988391461	0.978011255187843
11328	FKBP9	FKBP prolyl isomerase 9 5170	0.0190469058173494
	6.8916918310181 0.0236099082141852	0.877881641889277
	0.978011255187843
9586	CREB5	cAMP responsive element binding protein 5
10724	-0.0650790470432777	3.87949115303014
0.0236004786083015	0.877905839585778	0.978011255187843
55039	TRMT12	tRNA methyltransferase 12 homolog
2207	-0.0291874928968934	3.51518029282538
0.0235941299743887	0.877922133861673	0.978011255187843
92595	ZNF764	zinc finger protein 764 3197	-0.0379318756912805
	2.85467251369835	0.0235799068180738
0.877958646874947	0.978011255187843
2975	GTF3C1	general transcription factor IIIC subunit 1
9628	-0.0195724718059545	7.30566643490426
0.0235490677760595	0.878037854138057	0.978011255187843
2161	F12	coagulation factor XII	2769	0.0462815534276472
	1.32902719977645	0.0235291101270096
0.878089141833373	0.978011255187843
10059	DNM1L	dynamin 1 like	7282	-0.0206145863191909

6.8658707760528 0.0234990712535694	0.878166378516507
0.978011255187843
2597	GAPDH	glyceraldehyde-3-phosphate dehydrogenase	2981
	0.0304334054468073	10.5507639053068
0.0234811601085312	0.878212456135881	0.978011255187843
4331	MNAT1	MNAT1 component of CDK activating kinase
3943	-0.0245735065349735	4.93612644781893
0.0234731869646012	0.878232973367577	0.978011255187843
29093	MRPL22	mitochondrial ribosomal protein L22	5267
0.0284203272330344	4.13911469190246	0.0234686291957047
	0.878244703442266	0.978011255187843
23332	CLASP1	cytoplasmic linker associated protein 1 11052
0.0222556966765165	7.32948864131315	0.0234535699962652
	0.878283468752317	0.978011255187843
83862	TMEM120A	transmembrane protein 120A	1997
0.0288024657117037	4.63655265996461	0.0234332017478422
	0.878335920857679	0.978011255187843
2618	GART	phosphoribosylglycinamide formyltransferase,
phosphoribosylglycinamide synthetase, phosphoribosylaminoimidazole
synthetase	7751	-0.0233286950154166	5.92716390421473
	0.0234090308975858	0.878398195663497
0.978011255187843
26301	GBGT1	globoside alpha-1,3-N-
acetylgalactosaminyltransferase 1 (FORS blood group)	3713
0.0651538955022224	1.27175394695502	0.0233935680699489
	0.878438052008706	0.978011255187843
5438	POLR2I	RNA polymerase II subunit I	1269
0.0346929718555595	3.74465770391965	0.0233582240256922
	0.878529204007149	0.978011255187843
2139	EYA2	EYA transcriptional coactivator and phosphatase 2
	3618	0.025910907528643	5.58187653384616
0.0233047326484002	0.878667292161047	0.978011255187843
5680	PSG11	pregnancy specific beta-1-glycoprotein 11	5032
	0.0844234325746223	-0.214574509641514
0.0232967092516079	0.878688018565625	0.978011255187843
54700	RRN3	RRN3 homolog, RNA polymerase I transcription factor
	4156	-0.0255068368733944	5.69556770038812
0.0232840993675438	0.878720600375313	0.978011255187843
6921	ELOC	elongin C	4934	-0.0258913702954082
5.90831696028384	0.02326570133269	0.878768153987994
	0.978011255187843
5694	PSMB6	proteasome 20S subunit beta 6
1448	-0.020028559979706	6.54704830189532
0.0231859849368448	0.878974420493622	0.978011255187843
27034	ACAD8	acyl-CoA dehydrogenase family member 8	7174
0.0323451186792297	5.38143462447332	0.0231663138467333
	0.87902537530824	0.978011255187843
80263	TRIM45	tripartite motif containing 45
4368	-0.0427997102711453	2.12226215758148
0.0231596910513145	0.879042535581451	0.978011255187843
201973	PRIMPOL	primase and DNA directed polymerase	2843
0.0366365503294537	2.31879638316332	0.0231172335153613
	0.879152606653974	0.978011255187843
8572	PDLIM4	PDZ and LIM domain 4	3192	0.0243298535882951

6.52545963494358	0.0230852888964108
0.879235491203937	0.978011255187843
3315	HSPB1	heat shock protein family B (small) member 1
1634	-0.0349561607368049	8.90782544137039
0.0230824316486746	0.879242907569837	0.978011255187843
91433	RCCD1	RCC1 domain containing 1	4969
0.0361758122581184	2.52951182515821	0.0230144335308626
0.879419544486672	0.978011255187843
11091	WDR5	WD repeat domain 5	3629	0.022107497592174
5.21873432246763	0.0229893285943668
0.879484826379421	0.978011255187843
7592	ZNF41	zinc finger protein 41	4784	-0.0248082982079256
4.12004601077242	0.0229819631713539
0.879503986054342	0.978011255187843
57171	DOLPP1	dolichyldiphosphatase 1 2249	0.0218747815346668
4.8590514285713 0.0229760487908308	0.879519373406134
0.978011255187843
57798	GATAD1	GATA zinc finger domain containing 1	4791
0.0255065388481503	5.34299891678864	0.0229688492707991
0.879538107019782	0.978011255187843
10732	TCFL5	transcription factor like 5	3436
0.0342633778491814	2.66876024188829	0.0229601391069707
0.879560775440941	0.978011255187843
5828	PEX2	peroxisomal biogenesis factor 2 4969
0.0202637898209359	5.31625401076015	0.0228732784231749
0.879787073633301	0.97818523101934
11054	OGFR	opioid growth factor receptor
4702	-0.02347355267387	5.53305700178033
0.022836388643384	0.879883315551257	0.978214588532936
55723	ASF1B	anti-silencing function 1B histone chaperone	2304
0.0532984683957185	-0.194691776815207
0.022696676264875	0.880248534529377	0.978359713674608
92703	TMEM183A	transmembrane protein 183A
4026	-0.0200837795852677	5.4587614233454 0.0226845310816648
0.880280337223571	0.978359713674608
10447	FAM3C	family with sequence similarity 3 member C	3422
0.0387766885543394	7.23151175742492
0.0226774881255665	0.880298783501104	0.978359713674608
148206	ZNF714	zinc finger protein 714 8033	0.0306382893337928
3.80748659627935	0.0226629749375817
0.880336804378613	0.978359713674608
5693	PSMB5	proteasome 20S subunit beta 5
1765	-0.0287737072364188	7.31512226795074
0.0226526436047152	0.880363877440993	0.978359713674608
79675	FASTKD1 FAST kinase domains 1	3203	0.0233278854559317
4.52546840036495	0.022582080342918
0.880548956198713	0.978359713674608
159	ADSS2	adenylosuccinate synthase 2	3666
0.0200055543923439	5.80589163326181	0.0225788786409886
0.880557360877016	0.978359713674608
100533106	ZHX1-C8orf76	ZHX1-C8orf76 readthrough	1325
-0.0251773439589994	4.35897342244405
0.0225678275984524	0.880586375273447	0.978359713674608
11102	RPP14	ribonuclease P/MRP subunit p14	8608

0.0187897210057817	5.46267438161495	0.0224775343443859
	0.880823711925303	0.978359713674608
7638	ZNF221	zinc finger protein 221 2717	0.0461196291706133
	1.05684157687319	0.0224315837057252
0.880944680750968	0.978359713674608
4935	GPR143	G protein-coupled receptor 143	2486
0.0395303921132005	2.43837487059936	0.0224164978000836
	0.880984423252848	0.978359713674608
5965	RECQL	RecQ like helicase	3778	-0.0184387987099751
	5.75835635040208	0.0224037977951141
0.881017890909047	0.978359713674608
100529257	SYNJ2BP-COX16	SYNJ2BP-COX16 readthrough
1028	-0.0249734366513245	5.14987871575433
0.0223832684563696	0.881072011313631	0.978359713674608
79171	RBM42	RNA binding motif protein 42	2206
0.0214776249692981	5.58057734527561	0.0223727062081274
	0.8810998658933 0.978359713674608
353088	ZNF429	zinc finger protein 429 3960	0.0358164140835203
	2.60221717171604	0.0223235415989791
0.881229610512197	0.978359713674608
375248	ANKRD36 ankyrin repeat domain 36
9670	-0.0657825158819009	3.04569653353117
0.0223118816201264	0.881260402432369	0.978359713674608
150678	COPS9	COP9 signalosome subunit 9	2039
0.0267576646480444	4.89575228493737	0.0222765843202239
	0.881353666464274	0.978359713674608
8449	DHX16	DEAH-box helicase 16	4008	-0.018916611426291
	5.88192371275386	0.0222716955066709
0.881366589849837	0.978359713674608
26224	FBXL3	F-box and leucine rich repeat protein 3 7082
0.0209799542530915	6.0886731737878 0.0222405400730779
0.881448982094054	0.978359713674608
84905	ZNF341	zinc finger protein 341 5457	0.0337332618950354
	2.67061014692461	0.0222326625378684
0.881469824008038	0.978359713674608
5700	PSMC1	proteasome 26S subunit, ATPase 1
6161	-0.0203502043311712	5.89339585444834
0.0221873542607796	0.881589771305565	0.978359713674608
10113	PREB	prolactin regulatory element binding
3221	-0.0182469757967018	5.64550925123117
0.0221335116571737	0.8817324749295 0.978359713674608
4528	MTIF2	mitochondrial translational initiation factor 2 3095
	0.020771935724514	4.7212528626903
0.0221079820530683	0.881800200209969	0.978359713674608
137695	TMEM68	transmembrane protein 68
8574	-0.0236164723758395	4.75437687956174
0.0221065154075859	0.881804092160737	0.978359713674608
29925	GMPPB	GDP-mannose pyrophosphorylase B
6742	-0.022692757785789	5.26022122664262
0.0220926252378995	0.881840958231031	0.978359713674608
100133161	LINC01001	long intergenic non-protein coding
RNA 1001	2553	-0.0511832307841507	2.4609416950741
0.022063481113678	0.881918348529807	0.978359713674608
57129	MRPL47	mitochondrial ribosomal protein L47	1370

0.0233559616704858	4.85959669037291	0.0219999695448081
	0.882087180480549	0.978359713674608
27030	MLH3	mutL homolog 3	8363	0.0305706915664312
4.15822429873692	0.0219679970665894	0.882172266741828
	0.978359713674608
25847	ANAPC13	anaphase promoting complex subunit 13	2359
0.0306112889121637	5.33300186455354	0.0219663963273078
	0.882176528347216	0.978359713674608
55768	NGLY1	N-glycanase 1	8326	-0.0298127014663513
4.38948801229066	0.021962849768471	0.882185970821284
	0.978359713674608
57092	PCNP	PEST proteolytic signal containing nuclear protein
	3102	-0.0224479944937891	6.92506633148674
0.0219030363994648	0.88234533741832	0.978359713674608
55341	LSG1	large 60S subunit nuclear export GTPase 1	4818
	0.0252626686505399	5.5471595959433 0.0219020283838054
	0.882348025076939	0.978359713674608
100130311	C17orf107	chromosome 17 open reading frame 107
	3408	-0.0469689464660317	0.351818024412762
0.0218965717768063	0.88236257505212	0.978359713674608
521	ATP5ME	ATP synthase membrane subunit e 940
0.0291084422422449	5.16059589481931	0.0218849449338521
	0.882393584065959	0.978359713674608
1857	DVL3	dishevelled segment polarity protein 3	6090
0.0215834090411888	7.275942429587	0.0218606038438338
0.882458529489898	0.978359713674608
7813	EVI5	ecotropic viral integration site 5	8276
0.0249051172433717	6.19082955225328	0.0218071012198457
	0.882601411938256	0.978440678830166
4163	MCC	MCC regulator of WNT signaling pathway
10746	-0.0284987467565146	6.50761484099043
0.0216811634313814	0.88293844605941	0.97867879783922
87	ACTN1	actinin alpha 1 10626	0.0269505215332612
9.66768662278979	0.0216746336254232	0.882955948332564
	0.97867879783922
9443	MED7	mediator complex subunit 7	2913
0.0278779343760138	3.55537332930617	0.0215860260519656
	0.883193715893586	0.978864881984606
137886	UBXN2B	UBX domain protein 2B	5390	0.0235039507077772
	5.0132674901232 0.0215059753182736	0.883408951234904
	0.978992303612277
51195	RAPGEFL1	Rap guanine nucleotide exchange factor like
1	5711	0.0373496535029816	3.40473468023579
0.0214745024326159	0.883493685582842	0.978992303612277
549	AUH	AU RNA binding methylglutaconyl-CoA hydratase	2258
	0.0327099927129939	3.13867866121188
0.0214234725082076	0.883631208185381	0.978992303612277
57708	MIER1	MIER1 transcriptional regulator
8273	-0.0248529096776116	5.93182159480106
0.0214192274389688	0.883642655927542	0.978992303612277
1196	CLK2	CDC like kinase 2	3670	0.0230521770842377
	4.8585116502285 0.0213832269333629	0.883739785650254
	0.978992303612277
10153	CEBPZ	CCAAT enhancer binding protein zeta	3629

0.019969136182939	5.7384413487389 0.0213830695786896
0.883740210377964	0.978992303612277
6319	SCD	stearoyl-CoA desaturase 5362	-0.0538563704551977
	8.65165920436998	0.021334289083967
0.883871954474068	0.978992303612277
5629	PROX1	prospero homeobox 1	8928	0.0661712914515391
	1.80594101502908	0.0212664112017009
0.884055532124148	0.978992303612277
5836	PYGL	glycogen phosphorylase L
4798	-0.037248749864096	6.96371238219141
0.021239934386557	0.884127220522249	0.978992303612277
9584	RBM39	RNA binding motif protein 39
9339	-0.0178501689837914	7.67115647341709
0.0212321561562945	0.88414828946259	0.978992303612277
119391	GSTO2	glutathione S-transferase omega 2
9921	-0.030295484597665	3.09675398188704
0.0211782932405953	0.884294296543434	0.978992303612277
9397	NMT2	N-myristoyltransferase 2	4700
0.0410112349273424	4.16568359661717	0.021167894068963
	0.884322507575661	0.978992303612277
7594	ZNF43	zinc finger protein 43	6876	-0.0307854711979317
	3.85802552661248	0.0211533027154935
0.884362103155995	0.978992303612277
23657	SLC7A11 solute carrier family 7 member 11	9645
0.0437172721333782	4.64765592676811	0.0211385794953181
	0.884402070711548	0.978992303612277
23608	MKRN1	makorin ring finger protein 1	5869
0.0166472420868003	6.77371748238837	0.0211322452428959
	0.884419270003712	0.978992303612277
6009	RHEB	Ras homolog, mTORC1 binding	3535
0.0189464958072034	5.91469447318808	0.0211294086778793
	0.884426972935657	0.978992303612277
147011	PROCA1	protein interacting with cyclin A1	4829
0.0513717070802516	0.438899473515788	0.020977379729036
	0.884840595600847	0.979233545343834
387787	LIPT2	lipoyl(octanoyl) transferase 2
1187	-0.0474316359206416	-0.0320033423452604
0.0209551920009829	0.884901089195635	0.979233545343834
1497	CTNS	cystinosin, lysosomal cystine transporter	3398
	0.0280375411493531	5.32979447224109
0.02095134712558	0.884911575355928	0.979233545343834
23255	MTCL1	microtubule crosslinking factor 1	10981
0.0500403283651447	4.69619686628441	0.0209465891357823
	0.884924553220693	0.979233545343834
2954	GSTZ1	glutathione S-transferase zeta 1	8477
0.0343440355944893	3.44248418236802	0.0207854229519651
	0.885365041289396	0.979643584275264
729967	MORN2	MORN repeat containing 2	1364
0.0343494349498773	3.05458013081418	0.0207198858780071
	0.885544661020493	0.979689945222896
134218	DNAJC21 DnaJ heat shock protein family (Hsp40) member C21
	9137	-0.0183578205375591	5.66066391763121
0.0207190964736137	0.885546826334482	0.979689945222896
6157	RPL27A	ribosomal protein L27a	7117	0.0275334051360979

7.51724461982993	0.0206878016619498
0.885632701065854	0.97970756940684
122769	LRR1	leucine rich repeat protein 1	2359
0.0299599645153688	3.52294224365236	0.0206351905184867
0.885777218561297	0.979790057678913
51272	BET1L	Bet1 golgi vesicular membrane trafficking protein like	3868	0.0200957799203817	5.87194224395635
0.0205650354117353	0.885970220404474	0.979890498550071
8924	HERC2	HECT and RLD domain containing E3 ubiquitin protein ligase 2	20027	0.0317315214953162	7.32629834085959
0.0205513401216848	0.886007936327312
0.979890498550071
57482	CRACD	capping protein inhibiting regulator of actin dynamics	6323	0.0744082414044859	0.603445305575137
0.0204843491602418	0.886192610074452
0.980017360596309
85476	GFM1	G elongation factor mitochondrial 1	9032
0.0198378193444742	5.65041353965206	0.0204075578513567
0.886404679892199	0.980068061375199
10365	KLF2	Kruppel like factor 2	2018	0.0523917946467902
1.05170442951161	0.0203746188958265
0.886495770133725	0.980068061375199
79895	ATP8B4	ATPase phospholipid transporting 8B4 (putative)
10861	-0.101640766384434	-0.222859690290178
0.0203690782267696	0.886511099823536	0.980068061375199
9922	IQSEC1	IQ motif and Sec7 domain ArfGEF 1
6882	-0.0358762035210978	4.98526118417653
0.0203664629586271	0.886518336376367	0.980068061375199
147923	ZNF420	zinc finger protein 420 4020	-0.0361031888345909
2.84735767632175	0.0202205219673739
0.886922915058937	0.980425576018452
83992	CTTNBP2 cortactin binding protein 2	7035
0.0778864162511898	2.99181586101263	0.0201933563737899
0.886998387976975	0.980425576018452
50514	DEC1	deleted in esophageal cancer 1	4707
0.0904565050859153	4.37435015788978	0.0201607339376277
0.88708908986328	0.980425576018452
7697	ZNF138	zinc finger protein 138 2850	-0.0362587736675211
1.98945364601191	0.0201418736295589
0.887141562330526	0.980425576018452
8208	CHAF1B	chromatin assembly factor 1 subunit B	5118
0.0339722670598905	2.20071949594303	0.0201238600599796
0.887191702436916	0.980425576018452
64921	CASD1	CAS1 domain containing 1
5271	-0.0229855626414214	4.64476799502254
0.0200005947729256	0.887535422865488	0.980728042606255
1408	CRY2	cryptochrome circadian regulator 2	6200
0.0302467308107773	4.49609877088942	0.0199277405283595
0.887739082209351	0.980827587864936
9260	PDLIM7	PDZ and LIM domain 7	4407	0.0229813014314514
7.55955732323769	0.0199182797128401
0.887765557146545	0.980827587864936
23590	PDSS1	decaprenyl diphosphate synthase subunit 1
1872	-0.0474410580375689	1.62331115704812

0.0198710891339893	0.887897710091202	0.980847582860566
83541	FAM110A family with sequence similarity 110 member A	2007
	0.0295885564403789	3.76186012751563
0.0198618129384016	0.887923706052115	0.980847582860566
22944	KIN	Kin17 DNA and RNA binding protein
6844	-0.0238779563313928	3.83344584425885
0.019820241809559	0.888040282667308	0.980865287798745
3189	HNRNPH3 heterogeneous nuclear ribonucleoprotein H3
4270	-0.0190472637925448	6.52128021093858
0.0198061645856171	0.888079787239501	0.980865287798745
25897	RNF19A	ring finger protein 19A, RBR E3 ubiquitin protein
ligase	6049	-0.0322449333983944	8.01328133283868
0.0197340154890639	0.888282482597155	0.98095268743331
84218	TBC1D3F TBC1 domain family member 3F
2117	-0.0629348056293178	0.198180710383609
0.0197185098728117	0.888326093311072	0.98095268743331
768213	MIR671	microRNA 671	118	0.0369771058469441
1.12977953281326	0.0196212546431205	0.88860003064523
	0.98095268743331
9184	BUB3	BUB3 mitotic checkpoint protein 8300
0.0171560022069781	6.59821972915428	0.0196101072968133
	0.88863147338474	0.98095268743331
130612	TMEM198 transmembrane protein 198	2578
0.0423770347803987	1.95600676039166	0.0196054585968448
	0.888644588423021	0.98095268743331
54896	SLC66A1 solute carrier family 66 member 1
2686	-0.0278872285362801	4.83346853397302
0.0195867523426685	0.88869737904576	0.98095268743331
401388	NA	NA	3095	0.0272226656962855
4.3060557546314 0.0195762393684475	0.888727058825179
0.98095268743331
96764	TGS1	trimethylguanosine synthase 1
3869	-0.0213346221908907	4.1533828603436 0.019514031234829
	0.888902848702905	0.98095268743331
283991	UBALD2	UBA like domain containing 2
1797	-0.0212260567068551	4.65546693963413
0.0195069662058911	0.888922831357257	0.98095268743331
900	CCNG1	cyclin G1	2848	0.0230083776766918
5.51091346026349	0.0195031312330372	0.888933679699386
	0.98095268743331
51406	NOL7	nucleolar protein 7	1865	-0.0229581901537872
	5.16792109087985	0.0194577077108704
0.889062256221925	0.98095268743331
85453	TSPYL5	TSPY like 5	4491	-0.0646328811496118
4.0585032044785 0.0194496471714274	0.889085088487534
0.98095268743331
56993	TOMM22	translocase of outer mitochondrial membrane 22	2191
	0.0256499741313979	5.35611008491306
0.0194345767481519	0.889127789876677	0.98095268743331
54861	SNRK	SNF related kinase	6981	-0.0230023806694329
	5.17430284969614	0.0194197926759063
0.889169696305781	0.98095268743331
5824	PEX19	peroxisomal biogenesis factor 19
4888	-0.016280424566523	6.38713890824721

0.0193944205813916	0.88924165310125	0.98095268743331
101929709	LOC101929709	uncharacterized LOC101929709
3344	-0.05926599357433	2.02644435362463
0.0193795012964522	0.889283987499378	0.98095268743331
56902	PNO1	partner of NOB1 homolog 2674	-0.0223165945582178
	4.77333264370824	0.0193081571060532
0.889486661100099	0.98095268743331
79576	NKAP	NFKB activating protein 7894	-0.0204613342932026
	4.20611332476968	0.019288814290022
0.889541484644136	0.98095268743331
57510	XPO5	exportin 5	8522	-0.0194400724823044
5.84703975797626	0.0192831169664984	0.889557885250757
	0.98095268743331
84186	ZCCHC7	zinc finger CCHC-type containing 7
3992	-0.0289309264860772	3.73447234581338
0.0192553445035841	0.889636936373236	0.98095268743331
80817	CEP44	centrosomal protein 44	7132	0.0293734694411485
	3.55143560511268	0.0192446226977019
0.889667470306174	0.98095268743331
57179	KIAA1191	KIAA1191	3973	-0.0199788718741506
	8.07705635724875	0.0192333276998653
0.88969964597364	0.98095268743331
27094	KCNMB3	potassium calcium-activated channel subfamily M regulatory beta subunit 3	8979	-0.0336476471703385
2.93581141735311	0.0191613794774907	0.889904828792888
	0.981024559093019
4724	NDUFS4	NADH:ubiquinone oxidoreductase subunit S4	1176
	0.0247428431524227	5.11795642040834
0.0191613516927021	0.889904908105654	0.981024559093019
3833	KIFC1	kinesin family member C1
3345	-0.0433399388380159	0.992573143858752
0.0190987291732636	0.89008381569745	0.981104280361624
25796	PGLS	6-phosphogluconolactonase
1587	-0.0221225119849997	5.36187479782344
0.0190113111079455	0.890334062821299	0.981104280361624
10992	SF3B2	splicing factor 3b subunit 2
8992	-0.016650446349383	7.72283084609626
0.0189919340546965	0.890389611830758	0.981104280361624
79724	ZNF768	zinc finger protein 768 2395	-0.022762868231581
	5.08526799364286	0.0189886692386469
0.890398974057601	0.981104280361624
1347	COX7A2	cytochrome c oxidase subunit 7A2
1922	-0.0310359066955805	6.45082801826052
0.0189821919141764	0.890417550941243	0.981104280361624
23307	FKBP15	FKBP prolyl isomerase 15
11060	-0.0154644504349108	5.90356373947807
0.0189555290018522	0.890494053862857	0.981104280361624
54329	GPR85	G protein-coupled receptor 85	7396
0.0672639604937935	1.00781186561678	0.0189348294264873
	0.890553484209888	0.981104280361624
1678	TIMM8A	translocase of inner mitochondrial membrane 8A
2629	-0.0352693148113566	2.07631116539052
0.0189290518789162	0.89057007797779	0.981104280361624
967	CD63	CD63 molecule	2633	-0.0214009542568784

9.08822545993241	0.0188971978819978	0.89066161261415
	0.981104280361624
55037	PTCD3	pentatricopeptide repeat domain 3
10164	-0.017471151044103	5.60090618558698
0.0188916150887337	0.890677663245406	0.981104280361624
144699	FBXL14	F-box and leucine rich repeat protein 14	2111
	0.0300374658163795	2.63100631598846
0.018780499424165	0.890997626888751	0.981287450454805
8050	PDHX	pyruvate dehydrogenase complex component X
3760	-0.0210449903479593	4.88243892800197
0.0187749463924214	0.891013642415581	0.981287450454805
90488	TMEM263 transmembrane protein 263	4125
0.0253610362110973	5.61633917134582	0.0187609084767715
	0.891054140001976	0.981287450454805
5333	PLCD1	phospholipase C delta 1 4314	0.0374854017223527
	6.4013669467384 0.0187366432073155	0.891124178609632
	0.981287450454805
6790	AURKA	aurora kinase A 2928	-0.0331612497000748
3.3080933303444 0.0187055013513069	0.891214133340171
0.981309359774842
4733	DRG1	developmentally regulated GTP binding protein 1
1940	-0.0201120696734018	5.28357648167862
0.0186467989887404	0.891383905560646	0.98141914519635
23150	FRMD4B	FERM domain containing 4B
9639	-0.0276831383697139	5.48746339082648
0.0185994849711779	0.891520939940968	0.981492871630366
101060389	TBC1D3D TBC1 domain family member 3D
2785	-0.0618413406376307	2.57416031054051
0.0185416321495353	0.891688739075269	0.981582898397909
56940	DUSP22	dual specificity phosphatase 22 5686
0.0182263699581008	5.2555323716419 0.0184564673192718
0.891936241190347	0.981582898397909
3683	ITGAL	integrin subunit alpha L
6472	-0.120491240966754	1.35964557486236
0.0184346766024461	0.891999661759289	0.981582898397909
3684	ITGAM	integrin subunit alpha M	5665
0.0621520446963328	1.57849033731143	0.0183764120655363
	0.89216942492151	0.981582898397909
654841	MFF-DT	MFF divergent transcript
3652	-0.0418788200569629	-0.125654764573193
0.0183759228472185	0.892170851497702	0.981582898397909
100533496	TVP23C-CDRT4	TVP23C-CDRT4 readthrough	3097
	0.0216859272213145	4.68831874936539
0.0183515479805889	0.892241953886244	0.981582898397909
7507	XPA	XPA, DNA damage recognition and repair factor	1833
	0.0327833143588501	2.63319654374326
0.0183448587295487	0.892261475087026	0.981582898397909
10130	PDIA6	protein disulfide isomerase family A member 6
4101	-0.0286040588640214	7.71328187444499
0.0183053730935683	0.892376779692328	0.981582898397909
54331	GNG2	G protein subunit gamma 2
6379	-0.0963984722236096	1.03724604247691
0.0182925300099193	0.892414310915037	0.981582898397909
1439	CSF2RB	colony stimulating factor 2 receptor beta common

subunit 5000	0.127521279802089	1.1534033388671
0.0182829509056468	0.892442312577304	0.981582898397909
117583	PARD3B	par-3 family cell polarity regulator beta
10382	0.0248200553250939	4.58676778281807
0.018258826201162	0.892512867088048	0.981582898397909
54443	ANLN	anillin actin binding protein
6012	-0.0447569349768807	3.35202776840281
0.0182550712434377	0.892523853011643	0.981582898397909
79959	CEP76	centrosomal protein 76	6158	0.0249174420867514
	3.56116719735946	0.0182505228321936
0.892537161892411	0.981582898397909
170679	PSORS1C1	psoriasis susceptibility 1 candidate 1	5781
	0.0564437399243034	2.17885583964951
0.018211428830357	0.892651622617895	0.981582898397909
144577	C12orf66	chromosome 12 open reading frame 66
5898	-0.0368662780605855	1.70398718088255
0.0181944003383769	0.892701518314318	0.981582898397909
116412	ZNF837	zinc finger protein 837	2076	0.0437560605985472
	1.02138770263282	0.018168702249961
0.892776862088388	0.981582898397909
57216	VANGL2	VANGL planar cell polarity protein 2	5340
0.0282120632999318	6.0923553568137	0.0181587003720267
0.892806201084295	0.981582898397909
6130	RPL7A	ribosomal protein L7a	2242	0.0211301907886676
	7.18942942618248	0.0181118160054439
0.892943839073059	0.981582898397909
10897	YIF1A	Yip1 interacting factor homolog A, membrane
trafficking protein	2375	-0.0273342394706555
5.0518867150932	0.0180793712484615	0.893039193049781
0.981582898397909
4780	NFE2L2	nuclear factor, erythroid 2 like 2	5979
0.0172046759460908	7.84040092409832	0.0180522829865701
	0.893118871222698	0.981582898397909
57104	PNPLA2	patatin like phospholipase domain containing 2
3843	-0.0211087461072835	6.58072973521255
0.0180375180494377	0.893162326850718	0.981582898397909
7791	ZYX	zyxin	3482	-0.025694786536693
8.24540878368959	0.0180143211977111	0.893230635569749
	0.981582898397909
642517	AGAP9	ArfGAP with GTPase domain, ankyrin repeat and PH
domain 9	2387	0.0521600529242448	2.39386659238354
	0.0179822042487672	0.893325285614644
0.981582898397909
84791	LINC00467	long intergenic non-protein coding RNA 467
	4448	0.0425794284942398	1.38587316392506
0.017980732812755	0.893329624061099	0.981582898397909
1825	DSC3	desmocollin 3	7056	0.0282524832816695
8.65249333016907	0.01795657166673	0.893400887705256
	0.981582898397909
10128	LRPPRC	leucine rich pentatricopeptide repeat containing
	9760	0.0209846417771698	6.56381038023271
0.0179484875736264	0.893424742748264	0.981582898397909
503637	DUXAP8	double homeobox A pseudogene 8	5464
0.132247131132462	0.941566526298659	0.0178279390752358

0.893781115619165 0.981710589999371
284565 NBPF15 NBPF member 15 5306 -0.0249231749398913
4.61003925637033 0.0178254809009211 0.893788395363414
0.981710589999371
54583 EGLN1 egl-9 family hypoxia inducible factor 1 7265
0.0200181369198795 5.41822125479932 0.017795986377382
0.893875781599208 0.981710589999371
728558 ENTPD1-AS1 ENTPD1 antisense RNA 1 12353
0.0362166084609565 1.74054624080887 0.0177937237726191
0.893882488276839 0.981710589999371
5710 PSMD4 proteasome 26S subunit, non-ATPase 4
3027 -0.0161656803624928 6.34153701272379
0.017762246637858 0.893975835970345 0.981710589999371
91050 CCDC149 coiled-coil domain containing 149
9582 -0.0209887114554659 4.59317736120023
0.0177462900228078 0.894023188606858 0.981710589999371
9125 CNOT9 CCR4-NOT transcription complex subunit 9
5289 -0.0166213734131541 5.5742507810992 0.0177434644247647
0.894031576071391 0.981710589999371
55858 TMEM165 transmembrane protein 165
7045 -0.0195874620864942 7.61714546920325
0.0176866598708898 0.89420033829671 0.981818933802777
8226 PUDP pseudouridine 5'-phosphatase 3168
0.0245827133288191 3.76868795777561 0.017608532863985
0.894432899150723 0.981997305095169
7979 SEM1 SEM1 26S proteasome complex subunit 7393
0.0214456926998555 5.5125547762007 0.0175472029569832
0.894615828106508 0.9821211618691
3084 NRG1 neuregulin 1 9456 -0.0214434647933763
5.8010543883902 0.017403179830759 0.895046689490441
0.982517161417916
102 ADAM10 ADAM metallopeptidase domain 10 16168
0.0200128983781269 7.76106612193145 0.0173228672642836
0.895287741554342 0.98264270589524
10523 CHERP calcium homeostasis endoplasmic reticulum protein
6113 -0.0183859521279558 5.55878875160377
0.0173183340761547 0.895301364505965 0.98264270589524
93134 ZNF561 zinc finger protein 561 6467 -0.0172116522965535
6.7158602933665 0.0172715517752664 0.895442058846575
0.982720122092296
5190 PEX6 peroxisomal biogenesis factor 6 3499
0.0445839609758512 4.85387469181423 0.0172333120830217
0.895557205956582 0.982769491055613
11264 PXMP4 peroxisomal membrane protein 4 5723
0.0275284944322935 3.44829046180322 0.0171897467414794
0.895688548102941 0.982826839992867
2230 FDX1 ferredoxin 1 3206 0.0206197212889136
4.38773975250509 0.0171553625640919 0.895792330173976
0.982826839992867
1388 ATF6B activating transcription factor 6 beta
3660 -0.0164744893679357 6.78760016871333
0.0171462123473418 0.895819966173266 0.982826839992867
26147 PHF19 PHD finger protein 19 6733 0.0363409536282215
2.02048465995451 0.0170604625773438

0.896079317813986	0.982895018162232
5439	POLR2J	RNA polymerase II subunit J	1355
0.0201301300193704	4.67737197316726	0.0170420577910502
	0.896135069754236	0.982895018162232
90441	ZNF622	zinc finger protein 622 1699	0.0189400031798013
	5.5284256628797 0.0170326361915203	0.896163621595024
	0.982895018162232
9308	CD83	CD83 molecule	2515	-0.0632865539728711
2.14627478886767	0.0170041936620331	0.896249864477879
	0.982895018162232
5686	PSMA5	proteasome 20S subunit alpha 5
5724	-0.0198491720582847	6.39953966814368
0.0169958761424596	0.896275098558339	0.982895018162232
5566	PRKACA	protein kinase cAMP-activated catalytic subunit
alpha	4880	-0.016146108201512	5.74111098929128
0.0169866360559752	0.896303138929548	0.982895018162232
29079	MED4	mediator complex subunit 4
2358	-0.0176791348832003	5.12275685095871
0.0168489237573866	0.896721969494891	0.983199321194541
54476	RNF216	ring finger protein 216 7619	-0.0193844866136499
	5.76137828709806	0.01684036660705 6
0.89674805207375	0.983199321194541
83933	HDAC10	histone deacetylase 10	8436	-0.0213175364479517
	4.29413015989534	0.0168066353720064
0.896850932083817	0.983199321194541
84100	ARL6	ADP ribosylation factor like GTPase 6	4574
0.0296339804040372	2.89420546978817	0.0167700395789536
	0.896962667771836	0.983199321194541
51205	ACP6	acid phosphatase 6, lysophosphatidic	11960
0.0259656672922557	4.56354041156403	0.0167647614582904
	0.896978793358278	0.983199321194541
79840	NHEJ1	non-homologous end joining factor 1
5725	-0.028273708335825	3.77911640007897
0.0167572345718341	0.897001793784897	0.983199321194541
51003	MED31	mediator complex subunit 31
2103	-0.0304413530610863	3.41351142581587
0.0167209485503328	0.897112749248887	0.98324399677067
23164	MPRIP	myosin phosphatase Rho interacting protein
17536	0.0271942999951014	8.98023197908069
0.0166806881744037	0.897236000846778	0.983302140979643
86	ACTL6A	actin like 6A	3675	-0.0169029772527357
5.59076456444348	0.016640356544805	0.897359622293828
	0.983360681385514
6523	SLC5A1	solute carrier family 5 member 1	5310
0.0886286119734196	1.39653756112953	0.0165993888429785
	0.897485349407411	0.983421519920958
5298	PI4KB	phosphatidylinositol 4-kinase beta
5456	-0.0148600798608498	6.53043872649365
0.0164942936681811	0.897808602542124	0.983426981306394
55534	MAML3	mastermind like transcriptional coactivator 3	7012
	0.030620357208686	2.5091829703412 0.0164875510763434
	0.897829377182121	0.983426981306394
387032	ZKSCAN4 zinc finger with KRAB and SCAN domains 4
2375	-0.0395219517905237	2.6872151392213 0.0164813856544441

0.897848377280028	0.983426981306394
3151	HMGN2	high mobility group nucleosomal binding domain 2
2509	0.0324298514389863	3.96487833903161
0.0164567284080377	0.897924400120931	0.983426981306394
6432	SRSF7	serine and arginine rich splicing factor 7
4045	-0.0179819650979946	5.8070851821828 0.0164504686527209
0.897943709313949	0.983426981306394
140823	ROMO1	reactive oxygen species modulator 1
702	-0.022589544503285	4.93415591843533
0.0164378048049798	0.897982784352555	0.983426981306394
55788	LMBRD1	LMBR1 domain containing 1
2402	-0.0172591841306865	6.11653020242449
0.0164136901813272	0.898057233753394	0.983426981306394
55239	OGFOD1	2-oxoglutarate and iron dependent oxygenase domain containing 1	5806	-0.020169748849813	5.82752703440569
0.0163710697649435	0.898188952406319
0.983426981306394
6632	SNRPD1	small nuclear ribonucleoprotein D1 polypeptide
4909	-0.0235199412836625	4.82175292079486
0.0163574367295212	0.898231122162509	0.983426981306394
8676	STX11	syntaxin 11	1926	0.0520478464094152
2.22965129106883	0.0163519477200609	0.898248105830156
0.983426981306394
414777	HCG18	HLA complex group 18	5832	0.0196839957138347
5.10334371292229	0.0163288929629437
0.89831947173148	0.983426981306394
133308	SLC9B2	solute carrier family 9 member B2
4977	-0.0381692994545478	3.2367108093234 0.0163245730478652
0.898332849704812	0.983426981306394
7476	WNT7A	Wnt family member 7A	4579	-0.0221704399237604
7.36946796527271	0.0162932622581735
0.898429867216385	0.983456326203494
145864	HAPLN3	hyaluronan and proteoglycan link protein 3	3524
0.0323683068789601	5.09723485866602
0.0162414169442968	0.898590720169568	0.98350489894061
131965	METTL6	methyltransferase like 6
5517	-0.0225096001180157	3.52129799076882
0.0162332014381943	0.898616233166377	0.98350489894061
90527	DUOXA1	dual oxidase maturation factor 1
3721	-0.0257647944466201	6.2962009918102 0.0162015212831932
0.898714676375733	0.98350489894061
79760	GEMIN7	gem nuclear organelle associated protein 7	1915
0.0239591064791635	3.78613052576238
0.0161737203558943	0.898801145842275	0.98350489894061
730087	ZNF726	zinc finger protein 726 6289	-0.0546617789352275
0.503309788441291	0.0161657840758922
0.898825843976783	0.98350489894061
8897	MTMR3	myotubularin related protein 3	9903
0.016485417837564	7.24881768059733	0.0161434018243654
0.898895531965281	0.98350489894061
9377	COX5A	cytochrome c oxidase subunit 5A
1411	-0.019378838718428	5.58293974272888
0.0161205000504481	0.8989668883374 0.983506147382798
55854	ZC3H15	zinc finger CCCH-type containing 15

3125	-0.0187271816593168	6.0151497124249	0.0160433905214177
	0.899207522338124	0.983692577740557
8677	STX10	syntaxin 10	1796	-0.0191418391810603
5.30930452497739	0.0160042787578081	0.899329802157402
	0.983706109954698
1728	NQO1	NAD(P)H quinone dehydrogenase 1	3254
0.0327542149739582	5.34124613274221	0.0159837372594041
	0.899394084323004	0.983706109954698
3861	KRT14	keratin 14	2281	-0.0869820384616022
8.41155903795952	0.0159720805872041	0.899430581151554
	0.983706109954698
100128782	LINC00476	long intergenic non-protein coding
RNA 476	5100	-0.0518616885044115	0.924038496693115
0.0159254427890598	0.899576738786739	0.983789145860857
83875	BCO2	beta-carotene oxygenase 2	7220
0.0296143876215288	2.65883209690459	0.0158833381049011
	0.899708877011195	0.983856838183762
2967	GTF2H3	general transcription factor IIH subunit 3	3996
	0.0194637546424546	4.61551570001481
0.0158233074080059	0.899897580961427	0.98398637131356
9926	LPGAT1	lysophosphatidylglycerol acyltransferase 1
8799	-0.018983658592926	6.46882414722684
0.0157953441991765	0.899985606256277	0.984005806481279
25790	CFAP45	cilia and flagella associated protein 45
2625	-0.0430777631459399	0.155925898462601
0.0157607693339363	0.900094553866686	0.984010575719279
85004	RERG	RAS like estrogen regulated growth inhibitor	3926
	0.108584284898168	0.59292827007012
0.015744749381426	0.900145074910713	0.984010575719279
84310	C7orf50	chromosome 7 open reading frame 50	4961
0.0288908210632494	4.14451862951845	0.0157271134679977
	0.900200722300766	0.984010575719279
23421	ITGB3BP	integrin subunit beta 3 binding protein	5459
0.0487815275285722	1.85842767449454	0.0156279610509973
	0.900514174026293	0.984064634631121
23299	BICD2	BICD cargo adaptor 2	6427	-0.014045187673628
	7.32614647593989	0.0156091572161259
0.900573732629546	0.984064634631121
9223	MAGI1	membrane associated guanylate kinase, WW and PDZ
domain containing 1	11649	0.0256122436756136
5.54837195685725	0.0156051911060331	0.900586299398744
	0.984064634631121
440452	TBC1D3P2	TBC1 domain family member 3 pseudogene 2
	397	-0.0337879188230961	0.399310521135256
0.0156024817296037	0.900594885092433	0.984064634631121
9498	SLC4A8	solute carrier family 4 member 8
17980	-0.0536229655280525	0.400244342796098
0.0156004092834481	0.900601452940411	0.984064634631121
51760	SYT17	synaptotagmin 17	7538	0.0311910712137387
	4.32327735004229	0.0155385182167294
0.900797798261775	0.984202399395685
55818	KDM3A	lysine demethylase 3A	13423	0.0173390233384787
	6.47063637821305	0.0154499294639976
0.901079532520662	0.984373656829182

7771	ZNF112	zinc finger protein 112	3947	-0.0315088070320541
	2.90208437766297	0.0154210786632571
0.901171462211526	0.984373656829182
60343	FAM3A	family with sequence similarity 3 member A
5098	-0.0291863454383842	4.36505173418994
0.0154113863026311	0.90120236537349	0.984373656829182
130888	FBXO36	F-box protein 36	3945	-0.0340812283045852
	2.05585384989982	0.0154009499147678
0.901235651829616	0.984373656829182
149478	BTBD19	BTB domain containing 19	3520
0.0355588193657258	3.06900429486005	0.0152264478358015
	0.901793925405891	0.984897495484634
54808	DYM	dymeclin	6766	0.0166592366059422
5.20962368983717	0.0152071453928018	0.901855877567353
	0.984897495484634
23412	COMMD3	COMM domain containing 3
3408	-0.0188856183561285	5.23040266283216
0.0150497061355068	0.902362684582634	0.985121124423048
285203	EOGT	EGF domain specific O-linked N-acetylglucosamine transferase	5645	-0.0152572883217099	6.44107119484711
	0.0150273082864416	0.902435002927066
0.985121124423048
5771	PTPN2	protein tyrosine phosphatase non-receptor type 2
	5024	-0.0172497563804926	4.4912168208581
0.015006579822153	0.902501979905368	0.985121124423048
54969	HPF1	histone PARylation factor 1	1436
0.0271361699389943	3.08795443963527	0.014973266780764
	0.902609718099572	0.985121124423048
54954	FAM120C	family with sequence similarity 120C
8799	-0.0431362181319414	2.75407572792366
0.0149661813561952	0.90263264885783	0.985121124423048
1052	CEBPD	CCAAT enhancer binding protein delta	2178
0.0289735612521673	3.14848311192652	0.0149621189680467
	0.902645798563455	0.985121124423048
2625	GATA3	GATA binding protein 3	3253	0.0552006438185582
	2.09322710690783	0.0149576897901893
0.902660137610062	0.985121124423048
57117	INTS12	integrator complex subunit 12	3897
0.0213392491960327	4.18239443667223	0.0149412296326616
	0.90271344471379	0.985121124423048
8100	IFT88	intraflagellar transport 88	6111
0.0312560666677083	2.97351160669281	0.0149192463375627
	0.902784685296023	0.985121124423048
6748	SSR4	signal sequence receptor subunit 4
4590	-0.0188435495375664	6.04222032560179
0.0148657432707324	0.902958293932735	0.985121124423048
126374	WTIP	WT1 interacting protein	13733	-0.024903444741726
	4.50096771652923	0.0148355254239574
0.903056485990289	0.985121124423048
5473	PPBP	pro-platelet basic protein
715	-0.105651266203144	4.16357538938113
0.0148105867342991	0.903137600037999	0.985121124423048
554313	H4C15	H4 clustered histone 15	1715	0.0314622317807337
	2.80585204602039	0.0148006999985966

0.903169776208149	0.985121124423048
83886	PRSS27	serine protease 27	3152	0.044488018837663
	1.13670467175497	0.0147840614002455
0.903223950801366	0.985121124423048
388559	ZNF888	zinc finger protein 888 551	-0.0313048080447857
	1.39067016835474	0.0147605902528622
0.903300424520574	0.985121124423048
9753	ZSCAN12 zinc finger and SCAN domain containing 12	5495
	0.0354513910780969	3.12233932185917
0.0147477161924421	0.903342397002518	0.985121124423048
2529	FUT7	fucosyltransferase 7	2761	-0.0398239504778207
	2.94527243613709	0.014712927481753
0.903455909507896	0.985121124423048
54852	PAQR5	progestin and adipoQ receptor family member 5
6224	-0.0822283125404519	2.31711288375881
0.0146887159812366	0.903534989890231	0.985121124423048
80745	THUMPD2 THUMP domain containing 2	2091
0.0268827805563973	2.91230683205432	0.0146770448603064
	0.903573134103823	0.985121124423048
22948	CCT5	chaperonin containing TCP1 subunit 5
5839	-0.0199795953116385	7.98609494092484
0.0146612630237826	0.903624737670853	0.985121124423048
23256	SCFD1	sec1 family domain containing 1
6571	-0.020108941602135	5.67440455904524
0.0146562227114941	0.903641224458486	0.985121124423048
219854	TMEM218 transmembrane protein 218	5424
0.0328447842149036	2.68363284387161	0.0146485076061196
	0.903666466027341	0.985121124423048
54534	MRPL50	mitochondrial ribosomal protein L50
3332	-0.017568314836702	4.70652074719408
0.0145620622552549	0.903949752393135	0.985121124423048
9643	MORF4L2 mortality factor 4 like 2	2648
0.0221268353506749	7.95496284942167	0.0145329089764346
	0.904045481633454	0.985121124423048
126526	C19orf47	chromosome 19 open reading frame 47	4875
	0.021710334664034	3.85258376789085
0.0144606542089534	0.904283161929626	0.985121124423048
203286	ANKS6	ankyrin repeat and sterile alpha motif domain
containing 6	8846	0.0177602445113725	5.39366222458753
	0.0144537653598711	0.904305854053975
0.985121124423048
146434	ZNF597	zinc finger protein 597 5513	0.0304279536523141
	3.7735206205328 0.0144295738496414	0.904385585274888
	0.985121124423048
200539	ANKRD23 ankyrin repeat domain 23
6055	-0.027302633160085	3.06783356589414
0.0144169668274046	0.90442716281105	0.985121124423048
55664	CDC37L1 cell division cycle 37 like 1
3949	-0.0184419407714332	4.43219502050575
0.0144148051714401	0.904434293731673	0.985121124423048
51433	ANAPC5	anaphase promoting complex subunit 5
12304	-0.0139724712427021	6.59102489761915
0.0142956276463906	0.904828282013324	0.985121124423048
129790	NA	NA	2292	-0.0363780423987081

1.19125014668536	0.0142946371161017	0.904831563567562
	0.985121124423048
7174	TPP2	tripeptidyl peptidase 2 6752	-0.0162890988872305
	5.48335128908791	0.0142776697865941
0.904887793003246	0.985121124423048
7327	UBE2G2	ubiquitin conjugating enzyme E2 G2
7388	-0.0145863631796263	6.20797801981642
0.0142707072134218	0.904910876655701	0.985121124423048
79717	PPCS	phosphopantothenoylcysteine synthetase	3391
0.01734055187304	5.16594552447591	0.0142561577817801
	0.904959132155475	0.985121124423048
80790	CMIP	c-Maf inducing protein	11318	-0.0154376650826546
	7.15415285809364	0.0142541598911805
0.904965760426693	0.985121124423048
100528021	ST20-MTHFS	ST20-MTHFS readthrough
982	-0.0296647777775844	3.69932322494563
0.0142537221646624	0.904967212706306	0.985121124423048
79879	CCDC134 coiled-coil domain containing 134	1270
0.0335337046853376	2.24587217804631	0.0142151904449057
	0.905095141158231	0.985121124423048
84951	TNS4	tensin 4	4847	0.0190882828159625
8.85304050808023	0.0141770312698668	0.905222006152624
	0.985121124423048
112858	TP53RK	TP53 regulating kinase	3520	0.016702876585855
	4.84560438739649	0.0141604700030848
0.905277120053922	0.985121124423048
55071	C9orf40 chromosome 9 open reading frame 40	2327
0.0307958810337009	1.72868831960585	0.0141237456659802
	0.905399450942843	0.985121124423048
27433	TOR2A	torsin family 2 member A
3794	-0.0245323868882427	3.10344815882236
0.0141183546355967	0.905417422346645	0.985121124423048
55219	MACO1	macoilin 1	3979	-0.0160240657707576
5.88599316083786	0.0141153552483217	0.905427422535433
	0.985121124423048
8467	SMARCA5 SWI/SNF related, matrix associated, actin dependent regulator of chromatin, subfamily a, member 5
8385	-0.01426193611961	7.44924237251728
0.014109927378172	0.905445522213257	0.985121124423048
2736	GLI2	GLI family zinc finger 2	7406
0.0791977012484276	2.9612794045149 0.0140999851585342
0.905478684517655	0.985121124423048
140809	SRXN1	sulfiredoxin 1	3120	0.022752920475175
6.08326993945322	0.0140963342979319	0.905490864950673
	0.985121124423048
5987	TRIM27	tripartite motif containing 27	7986
0.0183076724802565	5.87101206750386	0.0140832158733346
	0.90553464539602	0.985121124423048
3984	LIMK1	LIM domain kinase 1	6902	-0.0185141212187677
	6.11762321235933	0.014078083244641
0.905551780273208	0.985121124423048
9620	CELSR1	cadherin EGF LAG seven-pass G-type receptor 1
12806	0.0289328153151915	8.89512432831301
0.0140606430813932	0.905610026566044	0.985121124423048

79365	BHLHE41 basic helix-loop-helix family member e41
3948	-0.0555889363318406	2.26711524522643
0.0140566827737061	0.905623258221592	0.985121124423048
10469	TIMM44	translocase of inner mitochondrial membrane 44
3701	-0.017767200982598	4.55810024092892
0.014030489877058	0.905710818090495	0.985121124423048
79191	IRX3	iroquois homeobox 3	2911	0.0295250234036545
2.86770885434958	0.0139942700643125
0.905832033450503	0.985121124423048
80724	ACAD10	acyl-CoA dehydrogenase family member 10
8931	-0.0210141382984593	3.91927967557257
0.0139905519022463	0.905844485868566	0.985121124423048
57577	CCDC191 coiled-coil domain containing 191	5303
0.0304556272846021	2.50154518475848	0.0139679570817943
0.905920193809394	0.985121124423048
51031	GLOD4	glyoxalase domain containing 4	5378
0.0149331073327288	5.61060842844434	0.0139655240519225
0.905928349807853	0.985121124423048
5110	PCMT1	protein-L-isoaspartate (D-aspartate) O-
methyltransferase	2832	0.0171074622405319
6.26660114704121	0.0139653790678693	0.905928835845883
0.985121124423048
11130	ZWINT	ZW10 interacting kinetochore protein	2378
0.0230013525105605	3.20709919436225	0.0138563230803834
0.906295157733822	0.985254991279941
283643	TEDC1	tubulin epsilon and delta complex 1	7068
0.0298375855292584	2.93430959764888	0.0138379176234977
0.906357126107025	0.985254991279941
283624	LINC00641	long intergenic non-protein coding RNA 641
5715	-0.0377686755530418	3.01291782922561
0.013829972557442	0.90638388884585	0.985254991279941
93621	MRFAP1	Morf4 family associated protein 1
2184	-0.0150770318643865	8.17047853117971
0.0138263207142018	0.90639619259481	0.985254991279941
26984	SEC22A	SEC22 homolog A, vesicle trafficking protein	4930
0.0205759792691339	4.24875531682804
0.0138172244904116	0.906426846652946	0.985254991279941
55846	ITFG2	integrin alpha FG-GAP repeat containing 2
4281	-0.0225622386715116	3.74374043753569
0.0138032434193889	0.906473982481945	0.985254991279941
55869	HDAC8	histone deacetylase 8	5379	-0.0193927885829679
4.13003047333518	0.0137456662224338
0.906668353956347	0.985389791578721
3603	IL16	interleukin 16	11668	0.0691078200706828
0.535928845485604	0.0137113511415352	0.906784392532014
0.98543944341655
5709	PSMD3	proteasome 26S subunit, non-ATPase 3	4964
0.0138382751568485	7.0174990014812 0.0136771939522191
0.906900043459707	0.985447130648635
3856	KRT8	keratin 8	6881	0.0247135252612055
10.6002081530998	0.0136432952952248	0.907014963857328
0.985447130648635
6430	SRSF5	serine and arginine rich splicing factor 5	5357
0.0175216668810005	7.18252453153442

0.0136406935826408	0.90702378994279	0.985447130648635
100131755	ARMCX4	armadillo repeat containing X-linked 4
13351	-0.02998767210193	3.21322595288165
0.0136262271000192	0.907072881805729	0.985447130648635
57647	DHX37	DEAH-box helicase 37	6618	0.0177262607973555
	4.89188036111469	0.013542356901219
0.907358016434696	0.985505925183466
196483	EEF2KMT eukaryotic elongation factor 2 lysine
methyltransferase	3128	-0.0315522536919917
2.40821663321437	0.0135319103924849	0.907393594172991
	0.985505925183466
51324	SPG21	SPG21 abhydrolase domain containing, maspardin	3723
	0.0191266515585 5.63559190273311	0.0135164343750205
	0.907446326533108	0.985505925183466
8314	BAP1	BRCA1 associated protein 1	5055
0.0157648023084165	6.93978266500718	0.0134909960772518
	0.907533070490454	0.985505925183466
1880	GPR183	G protein-coupled receptor 183	1700
0.0701538466578726	1.48420915315558	0.0134164312146634
	0.907787813445082	0.985505925183466
64130	LIN7B	lin-7 homolog B, crumbs cell polarity complex
component	1753	0.0313259166950174	0.854513036025234
	0.0134149709017739	0.907792809602527
0.985505925183466
672	BRCA1	BRCA1 DNA repair associated
8922	-0.0343882012136348	2.10996260828475
0.013408520998329	0.907814879905727	0.985505925183466
4093	SMAD9	SMAD family member 9	5723	0.050636096583503
	0.236208752195385	0.0133292027286114
0.908086732542811	0.985505925183466
93349	SP140L	SP140 nuclear body protein like
5650	-0.0237963435335056	4.33559353093613
0.0132764313005449	0.908268053769954	0.985505925183466
55604	CARMIL1 capping protein regulator and myosin 1 linker 1
6799	-0.0197466628434837	4.96950403421876
0.0132508148622459	0.908356202816115	0.985505925183466
56267	KYAT3	kynurenine aminotransferase 3	2306
0.0312760206521441	4.81522770324095	0.0132463187243772
	0.908371683436027	0.985505925183466
440944	THUMPD3-AS1	THUMPD3 antisense RNA 1
6275	-0.0296253691718369	3.41490540306046
0.013232458424314	0.9084194224948 0.985505925183466
6533	SLC6A6	solute carrier family 6 member 6	8254
0.0283672738549474	6.71910282998667	0.0132287472016976
	0.908432209365339	0.985505925183466
55217	TMLHE	trimethyllysine hydroxylase, epsilon
3922	-0.029706547210443	2.58645477382341
0.0132252259560257	0.908444343358138	0.985505925183466
403150	FLJ31356	uncharacterized protein FLJ31356	3001
	0.0532705019035938	0.302944086337508
0.0131968181620303	0.908542294768412	0.985505925183466
282969	FUOM	fucose mutarotase	1284	0.0593275697272737
	0.59944005441974	0.0131561233386304
0.908682798733093	0.985505925183466

9709 HERPUD1 homocysteine inducible ER protein with ubiquitin like domain 1 5155 0.0192903130858787 6.33857586476625 0.0131269846669113 0.908783538947206 0.985505925183466
22981 NINL ninein like 6008 0.0238145500233946 4.46431261228207 0.0131236551511762 0.908795057185593 0.985505925183466
7181 NR2C1 nuclear receptor subfamily 2 group C member 1 6529 -0.0185439411957641 4.75531737373713 0.0131147600568102 0.90882583643536 0.985505925183466
221178 SPATA13 spermatogenesis associated 13 11704 -0.0355389796897832 2.68216610535411 0.0131006544406951 0.908874667081718 0.985505925183466
79869 CPSF7 cleavage and polyadenylation specific factor 7 6036 -0.0140802438213571 6.52922635474424 0.0130977307425226 0.908884791635356 0.985505925183466
7093 TLL2 tolloid like 2 6853 -0.0399109178422087 3.88230701133015 0.0130895076538451 0.908913273731529 0.985505925183466
11333 PDAP1 PDGFA associated protein 1 2854 -0.0151032988374381 6.35060948235739 0.0130776798508805 0.90895425728079 0.985505925183466
65005 MRPL9 mitochondrial ribosomal protein L9 1812 -0.0174698298957781 4.99526479203193 0.0130398390067175 0.909085502762965 0.985505925183466
145553 MDP1 magnesium dependent phosphatase 1 2380 -0.0266494668983637 2.27227024570788 0.0130302295928648 0.909118862326521 0.985505925183466
138428 PTRH1 peptidyl-tRNA hydrolase 1 homolog 2730 0.033939088783348 2.30521752163899 0.0130275408279288 0.909128198740214 0.985505925183466
9240 PNMA1 PNMA family member 1 2590 0.0134124765513687 6.01617306083124 0.0130236949036 0.909141554947741 0.985505925183466
51142 CHCHD2 coiled-coil-helix-coiled-coil-helix domain containing 2 1132 -0.018856133441494 6.7335075406437 0.0129964919955867 0.909236082877092 0.985505925183466
81628 TSC22D4 TSC22 domain family member 4 3715 0.0165625191120713 6.06293511465469 0.0129753079384489 0.909309765209082 0.985505925183466
11034 DSTN destrin, actin depolymerizing factor 3851 -0.0196021010033622 8.66847817553072 0.0129590150371541 0.909366476596324 0.985505925183466
100131998 RRN3P3 RRN3 homolog, RNA polymerase I transcription factor pseudogene 3 1120 0.0234185835933928 3.65231474021827 0.0129353308487348 0.909448979588882 0.985505925183466
388561 ZNF761 zinc finger protein 761 4714 0.0242305571556896 4.65166664649242 0.0129168831661559 0.90951329442526 0.985505925183466
3269 HRH1 histamine receptor H1 5546 0.0304556261407174 4.97420809234019 0.0128955879018946 0.909587594763385 0.985505925183466
26502 NARF nuclear prelamin A recognition factor

7962	-0.0174955477193629	5.81897975077491
0.0128842598357011	0.90962714432436	0.985505925183466
10785	WDR4	WD repeat domain 4	2738	0.0258504927240147
	3.31784910864374	0.0128687075008926
0.909681470727156	0.985505925183466
116092	DNTTIP1 deoxynucleotidyltransferase terminal interacting protein 1	1883	0.019703451369625	4.968357463672
0.0128184758038854	0.909857164322637	0.985505925183466
125476	INO80C	INO80 complex subunit C 5249	0.0274044724810063
	4.07489720421404	0.0128010797835572
0.909918091013608	0.985505925183466
8893	EIF2B5	eukaryotic translation initiation factor 2B subunit epsilon 5975	-0.013363978614993	5.62044498027736
0.0127939881296522	0.909942940390784	0.985505925183466
9391	CIAO1	cytosolic iron-sulfur assembly component 1
4031	-0.0131481734674803	6.3186673346264 0.0127910537270211
	0.909953224669576	0.985505925183466
10440	TIMM17A translocase of inner mitochondrial membrane 17A 3513
	0.0204425735108481	6.4589123740018 0.0127879248437353
	0.909964191866117	0.985505925183466
23220	DTX4	deltex E3 ubiquitin ligase 4
6072	-0.0280540795699188	6.32065927462146
0.0127486430657484	0.910101996070587	0.985505925183466
5738	PTGFRN	prostaglandin F2 receptor inhibitor
6352	-0.0209294058637364	8.56457810000296
0.0127057539315274	0.91025270098542	0.985505925183466
4047	LSS	lanosterol synthase	7238	-0.0234640474469558
	7.44604274734979	0.0126619972883191
0.910406719868871	0.985505925183466
1786	DNMT1	DNA methyltransferase 1 9796	0.0156434642553747
	6.06119661779447	0.0126593990105448
0.910415874009538	0.985505925183466
79447	PAGR1	PAXIP1 associated glutamate rich protein 1	1856
	0.0164026370931743	4.28308542340831
0.0126572999531853	0.910423270013004	0.985505925183466
100507331	ZSWIM8-AS1	ZSWIM8 antisense RNA 1
1632	-0.0330584289902158	1.61670880232063
0.0126530299955991	0.910438317074257	0.985505925183466
23759	PPIL2	peptidylprolyl isomerase like 2
8437	-0.0134509322409073	5.60212146874781
0.0126407904799963	0.910481462608327	0.985505925183466
79603	CERS4	ceramide synthase 4	4059	0.0525208888059016
	2.1013991236342 0.0126343458236438	0.910504189176786
	0.985505925183466
64427	TTC31	tetratricopeptide repeat domain 31	5094
0.016022775328962	4.41145058289407	0.0125541435412933
	0.910787508685109	0.985702373186257
147015	DHRS13	dehydrogenase/reductase 13
2944	-0.0298451589084462	2.12718356631644
0.0125431456451386	0.910826430795297	0.985702373186257
129563	DIS3L2	DIS3 like 3'-5' exoribonuclease 2	11337
0.0154293328712118	4.85317023886067	0.0124548416539454
	0.911139571436572	0.985713442629109
146754	DNAH2	dynein axonemal heavy chain 2

17231   -0.0476916868538885     0.209967706829023
0.0124351397317604      0.911209590938328       0.985713442629109
7177    TPSAB1  tryptase alpha/beta 1   1512    0.111218007031215
        0.788265551699531       0.0124219174853266
0.911256613539512       0.985713442629109
200316  APOBEC3F        apolipoprotein B mRNA editing enzyme
catalytic subunit 3F    3242    -0.034597903199205
2.2408607502529 0.0123980489808879      0.911341561886656
0.985713442629109
135398  C6orf141        chromosome 6 open reading frame 141
1450    -0.0305667960726265     2.56337052875679
0.012397788167128       0.91134249058357        0.985713442629109
29115   SAP30BP SAP30 binding protein   7936    0.0128755378355408
        6.35187557959284        0.0123736495828766
0.911428485301971       0.985713442629109
113     ADCY7   adenylate cyclase 7     8630    -0.0225368121315419
        4.8212070109495 0.0123640478810474      0.911462715372096
        0.985713442629109
57834   CYP4F11 cytochrome P450 family 4 subfamily F member 11  4027
        0.0964835073816386      0.880549747051402
0.0123321233844322      0.911576623062532       0.985713442629109
10090   UST     uronyl 2-sulfotransferase
4563    -0.0374928662784103     3.73664343651859
0.0123147636381722      0.911638625880022       0.985713442629109
6235    RPS29   ribosomal protein S29   8384    0.0270882396516889
        6.01580848927187        0.0123108196092971
0.91165271870958        0.985713442629109
80818   ZNF436  zinc finger protein 436 4392    0.0193081868945768
        4.91869370316627        0.0122875881382924
0.911735775966945       0.985713442629109
1211    CLTA    clathrin light chain A  2515    -0.0149567108543374
        7.36663309581405        0.0122811358105537
0.911758858457715       0.985713442629109
100507420       LINC01605       long intergenic non-protein coding
RNA 1605        2032    0.0640969355370937      -0.0221385563803211
        0.012256638478422       0.911846550900872
0.985713442629109
51022   GLRX2   glutaredoxin 2  1299    -0.026458764730559
2.99115321611674        0.0122510381880403      0.911866610564886
        0.985713442629109
5349    FXYD3   FXYD domain containing ion transport regulator 3
        3334    -0.0311718521134537     7.02782914998877
0.0122298406278958      0.911942580105664       0.985713442629109
349114  LINC00265       long intergenic non-protein coding RNA 265
        3115    0.0394564910064691      2.69533628201472
0.0122242503653531      0.911962626046307       0.985713442629109
171392  ZNF675  zinc finger protein 675 4458    -0.0224407149857547
        3.64687943459935        0.0121950414620748
0.912067440836448       0.98571356750983
79133   NDUFAF5 NADH:ubiquinone oxidoreductase complex assembly
factor 5        7217    0.0200291538236638      3.71456078429575
        0.0121850044718679      0.91210348743449
0.98571356750983
64412   GZF1    GDNF inducible zinc finger protein 1    5463

0.0168358641503394	4.31950390354282	0.0121177522858995
	0.912345403928394	0.98589894096783
2074	ERCC6	ERCC excision repair 6, chromatin remodeling factor
	14404	0.0212222903391416	5.12656454021666
0.012030632860327	0.912659798366643	0.985980293694092
126068	ZNF441	zinc finger protein 441 4748	0.0312470259554943
	1.97204237765229	0.0119842741026375
0.912827566401429	0.985980293694092
51650	MRPS33	mitochondrial ribosomal protein S33
5262	-0.0217515918622752	4.31620262087243
0.0119776699942893	0.912851492759463	0.985980293694092
60528	ELAC2	elaC ribonuclease Z 2	6345	0.0126428017805309
	6.25656059259725	0.0119762497850218
0.912856638979447	0.985980293694092
60558	GUF1	GUF1 homolog, GTPase	4454	0.0154280990465661
	5.01162843535087	0.0119533505526022
0.91293965849463	0.985980293694092
10875	FGL2	fibrinogen like 2	4261	0.11166933346847
	0.669454816374104	0.0119325831125465
0.913015018960986	0.985980293694092
115201	ATG4A	autophagy related 4A cysteine peptidase 2582
0.019840354392665	4.86596492854383	0.0119231364175612
	0.913049320894703	0.985980293694092
10519	CIB1	calcium and integrin binding 1	1367
0.0176383138083336	6.16475872416086	0.0119167751136464
	0.91307242719999	0.985980293694092
259307	IL4I1	interleukin 4 induced 1 3274	-0.0337529733409313
	1.67574170265636	0.011906388548486
0.913110167968534	0.985980293694092
93627	TBCK	TBC1 domain containing kinase	11112
0.0194317434737705	4.66232766313413	0.0119024160308285
	0.913124606967927	0.985980293694092
23596	OPN3	opsin 3 3442	0.0186685604051904
5.85798818383248	0.0117527612786716	0.913670347655463
	0.986477483431522
2969	GTF2I	general transcription factor IIi	10438
0.0165368249570417	8.16634570530809	0.0117375775754169
	0.913725913474732	0.986477483431522
9738	CCP110	centriolar coiled-coil protein 110
10142	-0.0224169629040861	4.13167800032356
0.0116876482138579	0.91390889029264	0.986501169162673
23326	USP22	ubiquitin specific peptidase 22 6808
0.0130557606491861	8.5772419954666 0.0116410210258593
0.914080122969521	0.986501169162673
6337	SCNN1A	sodium channel epithelial 1 alpha subunit	5874
	0.0214322198519772	7.27638058777793
0.0116305346875603	0.914118680530451	0.986501169162673
100506365	OTUD6B-AS1	OTUD6B antisense RNA 1 (head to
head)	6009	-0.0209256194048584	3.96494634258808
0.0115957670711193	0.914246644678124	0.986501169162673
102724594	U2AF1L5 U2 small nuclear RNA auxiliary factor 1 like
5	8389	-0.0151580313422566	5.80492874961919
0.0115902638514171	0.914266917365413	0.986501169162673
90353	CTU1	cytosolic thiouridylase subunit 1	2087

0.0375534888992235	1.15951341139381	0.011564536980309
	0.914361754314987	0.986501169162673
7531	YWHAE	tyrosine 3-monooxygenase/tryptophan 5-monooxygenase activation protein epsilon	2856	-0.0152028864151874
8.94981821430359	0.0115428043395696	0.914441950512249
	0.986501169162673
51244	CCDC174 coiled-coil domain containing 174	2942
0.0163330225731786	4.65857669677765	0.0115406836537524
	0.914449780199622	0.986501169162673
55670	PEX26	peroxisomal biogenesis factor 26	18696
0.0189163519901811	6.3312083274684 0.0115360807917568
0.914466776720551	0.986501169162673
10574	CCT7	chaperonin containing TCP1 subunit 7	3591
0.015380765618709	7.63536033331961	0.011514130316769
	0.914547878229971	0.986501169162673
284058	KANSL1	KAT8 regulatory NSL complex subunit 1
13318	-0.01511416873761	6.29956433130715
0.0115058752907906	0.914578398715421	0.986501169162673
118491	CFAP70	cilia and flagella associated protein 70	7428
	0.0402845670011444	1.56611241539482
0.0115019264555709	0.914593002266472	0.986501169162673
58506	SCAF1	SR-related CTD associated factor 1
4548	-0.0171512622134108	6.59640568133975
0.011481317385531	0.914669259749451	0.986507455248368
3059	HCLS1	hematopoietic cell-specific Lyn substrate 1	5093
	0.0780141583564322	2.1092121855291 0.0114096874623044
	0.914934843987605	0.986694096396111
9066	SYT7	synaptotagmin 7 5953	-0.0492988986551523
2.20015027619328	0.0113966711396838	0.914983195408659
	0.986694096396111
5203	PFDN4	prefoldin subunit 4	1975	0.0201928574510477
	4.26813520364659	0.0113448982876569
0.915175791773623	0.986768160473148
9215	LARGE1	LARGE xylosyl- and glucuronyltransferase 1
10537	0.0213206317849561	4.85866188980588
0.0113403391675391	0.915192773092501	0.986768160473148
578	BAK1	BCL2 antagonist/killer 1
2212	-0.0156253499284917	5.75214193267578
0.0112681049157093	0.915462285511918	0.986905961797866
158572	USP27X-AS1	USP27X antisense RNA 1 (head to head)
2175	-0.0325019499717041	0.135798642413702
0.0112390699064036	0.91557086378916	0.986905961797866
9100	USP10	ubiquitin specific peptidase 10
5083	-0.0153020069217253	5.75150533083681
0.0112158354172323	0.915657852846838	0.986905961797866
9737	GPRASP1 G protein-coupled receptor associated sorting protein 1	6340	-0.0336202183586808	0.471166697212123
	0.0112130542119244	0.915668271678985
0.986905961797866
128637	TBC1D20 TBC1 domain family member 20
5831	-0.0126860670009695	6.29409455351764
0.011210476812046	0.915677928190512	0.986905961797866
221154	MICU2	mitochondrial calcium uptake 2	3645
0.0152005813716549	5.46227484832094	0.0111930291532651

0.915743327299698 0.986905961797866
10916 MAGED2 MAGE family member D2 2940 0.0138376139657842
7.42230583437179 0.0111368949421617
0.915954085736494 0.986906568938262
51255 RNF181 ring finger protein 181 1220 0.0164943804582894
5.56368720486774 0.0111288586370284
0.915984302317074 0.986906568938262
1460 CSNK2B casein kinase 2 beta 2933 0.0149940991987429
6.58710670461292 0.0111247173622786
0.91599987785498 0.986906568938262
2029 ENSA endosulfine alpha 5477 -0.0144633492383994
6.94835872254551 0.0111137706599607
0.916041063061078 0.986906568938262
3939 LDHA lactate dehydrogenase A 6210 -0.0287462565519981
9.81825548160413 0.0110991292328455
0.916096181147661 0.986906568938262
10488 CREB3 cAMP responsive element binding protein 3 1921
0.0160907400919247 6.23910369119983
0.0110516035326498 0.916275346820719 0.986958108201917
5499 PPP1CA protein phosphatase 1 catalytic subunit alpha
2711 -0.0128790981104617 6.86563323182312
0.0110081499009738 0.916439502502972 0.986958108201917
56144 PCDHA4 protocadherin alpha 4 13423 -0.0733363656609103
3.31478949411031 0.0109840557966905
0.916530664482521 0.986958108201917
7356 SCGB1A1 secretoglobin family 1A member 1 582
0.091359472278578 5.55622285953065 0.0109420656483508
0.916689779498618 0.986958108201917
5297 PI4KA phosphatidylinositol 4-kinase alpha 11407
0.0160584397722925 6.37795547596814 0.010921828791993
0.916766574076302 0.986958108201917
388685 LINC01138 long intergenic non-protein coding RNA 1138
3938 0.022579734646243 2.71833784473091
0.0108496109039062 0.917041213621221 0.986958108201917
7318 UBA7 ubiquitin like modifier activating enzyme 7
4545 -0.0284065313002701 4.89480118669325
0.0108304593393456 0.917114200558339 0.986958108201917
3550 IK IK cytokine 4095 -0.0129383906963757
6.00111405719577 0.0108286732199616 0.917121010813871
0.986958108201917
10907 TXNL4A thioredoxin like 4A 5231 0.0168157628742699
5.3354696001912 0.0108259133024049 0.917131535159664
0.986958108201917
8458 TTF2 transcription termination factor 2
10181 -0.0244226287731379 4.28320612061146
0.0108033197293196 0.917217741878932 0.986958108201917
440567 UQCRHL ubiquinol-cytochrome c reductase hinge protein like
2180 0.0199032528175829 4.9873165975399
0.0107971925048247 0.91724113626162 0.986958108201917
4494 MT1F metallothionein 1F 734 0.0357050063294787
0.909578972002733 0.0107946979307663
0.917250662727635 0.986958108201917
83858 ATAD3B ATPase family AAA domain containing 3B 7682
0.0166426589418924 5.07366203277506 0.0107924003296187

0.91725943796313	0.986958108201917
51315	KRCC1	lysine rich coiled-coil 1
1753	-0.0160410143576992	5.41771258488515
0.0107922286291711	0.917260093777119	0.986958108201917
55954	ZMAT5	zinc finger matrin-type 5	3200
0.0212327307273989	3.70212651116272	0.0107834357791337
0.917293685331611	0.986958108201917
100126314	MIR877	microRNA 877	114	0.0334688192993395
0.745061619453159	0.0107670550574763
0.91735630196706	0.986958108201917
4594	MMUT	methylmalonyl-CoA mutase	3749
0.0214635745152875	5.115930072797	0.0107662094029228
0.917359535856543	0.986958108201917
3838	KPNA2	karyopherin subunit alpha 2	3140
0.0209227057456957	6.14716125403284	0.0107524074391137
0.917412334460553	0.986958108201917
90410	IFT20	intraflagellar transport 20	2666
0.0174679323692967	4.08770253982753	0.0107242552688973
0.917520135224857	0.986959961614517
3104	ZBTB48	zinc finger and BTB domain containing 48
4503	-0.0191505522940709	3.77551803888268
0.0106769508152289	0.917701596780993	0.986959961614517
7407	VARS1	valyl-tRNA synthetase 1 6081	0.0206404711543826
5.79421508824489	0.0106742346756192
0.917712028310267	0.986959961614517
8503	PIK3R3	phosphoinositide-3-kinase regulatory subunit 3	7815
0.0251894520314435	5.69940602449111
0.0106654323352613	0.917745843551207	0.986959961614517
100507321	ERVK13-1	endogenous retrovirus group K13
member 1	7023	0.0300202450933667	2.9830944646659
0.0106600917766713	0.917766366818668	0.986959961614517
64419	MTMR14	myotubularin related protein 14 3657
0.014768297794952	5.26716038354338	0.0106124382339701
0.917949725434905	0.987081360676087
29894	CPSF1	cleavage and polyadenylation specific factor 1	6072
0.0261401582125115	6.72247472518103
0.0105733223493782	0.918100544471186	0.987167753620013
100132273	NDUFA6-DT	NDUFA6 divergent transcript	5125
0.0380980149224916	1.36351772202425
0.0105530170065578	0.918178946906239	0.987176274893743
84328	LZIC	leucine zipper and CTNNBIP1 domain containing
2794	-0.0149021365658929	4.66845237540698
0.010519311015118	0.918309260028864	0.987210994716614
9168	TMSB10	thymosin beta 10	496	-0.0240312903519271
9.97948623167786	0.0105082164382111
0.918352199697114	0.987210994716614
56134	PCDHAC2	protocadherin alpha subfamily C, 2
6112	-0.0666380908103432	2.97124663541264
0.0104420631092559	0.918608711736448	0.987362034419248
2632	GBE1	1,4-alpha-glucan branching enzyme 1
5171	-0.0152500566203671	5.79547083372052
0.0104003015703977	0.918771066617698	0.987362034419248
23098	SARM1	sterile alpha and TIR motif containing 1
11137	0.027399107673338	2.94675364397132

0.0103996791197369	0.918773488986146	0.987362034419248
10245	TIMM17B translocase of inner mitochondrial membrane 17B
2562	-0.0156797460559777	4.36125896756182
0.0103840447392951	0.918834356774857	0.987362034419248
84649	DGAT2	diacylglycerol O-acyltransferase 2	7905
0.0538602027368404	1.50942524290997	0.0103619449941998
	0.918920474558389	0.987362034419248
9235	IL32	interleukin 32	3209	-0.0353614334172761
8.48681278838666	0.0103472385154078	0.918977833857103
	0.987362034419248
5983	RFC3	replication factor C subunit 3
2874	-0.0256650473376197	2.27052878754408
0.0103278877728581	0.919053369840505	0.987362034419248
10926	DBF4	DBF4 zinc finger	4583	0.0267496956358271
	2.32682280735485	0.0103270532129525
0.919056629168141	0.987362034419248
51686	OAZ3	ornithine decarboxylase antizyme 3	4633
0.0302896951473031	0.0563421005930988	0.0102940504750544
	0.919185626376891	0.987391112161883
54531	MIER2	MIER family member 2	7931	0.0154314519651629
	5.04763085468657	0.010272741783389
0.919269026293385	0.987391112161883
51478	HSD17B7 hydroxysteroid 17-beta dehydrogenase 7
4124	-0.0229639683342614	3.881446285842	0.010242138532
0.919388957136405	0.987391112161883
10124	ARL4A	ADP ribosylation factor like GTPase 4A	3561
0.0244861699838894	3.00944790219811	0.010220980942532
	0.919471977309244	0.987391112161883
60673	ATG101	autophagy related 101	3199	0.0131208882357562
	5.51291815757175	0.0102206966491885
0.919473093437392	0.987391112161883
100505538	RBM26-AS1	RBM26 antisense RNA 1	3747
0.0445678753375771	0.563582909591637	0.0102121508193669
	0.919506651462811	0.987391112161883
1479	CSTF3	cleavage stimulation factor subunit 3
4517	-0.0157123322044073	4.65629936660852
0.0101856410221686	0.919610841290904	0.987407390716406
100131193	CCDC183-AS1	CCDC183 antisense RNA 1
2386	-0.0320030173170785	1.70089450436218
0.0101720882824825	0.919664159702704	0.987407390716406
8813	DPM1	dolichyl-phosphate mannosyltransferase subunit 1,
catalytic	1207	-0.0144264136355789	5.12569992270243
	0.0101046806912635	0.91992988541327
0.987407390716406
6513	SLC2A1	solute carrier family 2 member 1
4841	-0.0161904850217699	8.3691299366073 0.0100942716445047
	0.91997099835399	0.987407390716406
56339	METTL3	methyltransferase like 3	5894
0.0163412997451923	4.93351655668671	0.0100850571441189
	0.920007411027456	0.987407390716406
54788	DNAJB12 DnaJ heat shock protein family (Hsp40) member B12
	5649	0.0110401665342936	6.12667182704238
0.0100805020521797	0.920025417458035	0.987407390716406
3241	HPCAL1	hippocalcin like 1	6237	0.0253975321956563

4.00414796783446	0.0100334550725734
0.920211636678645	0.987407390716406
9479	MAPK8IP1	mitogen-activated protein kinase 8
interacting protein 1	3498	0.0300927198977811
3.42741987923529	0.00998043342475796	0.920422033875625
0.987407390716406
7597	ZBTB25	zinc finger and BTB domain containing 25
12229	0.0221153124359465	3.23252554175664
0.00996948921519625	0.920465532231013	0.987407390716406
728229	TMEM191B	transmembrane protein 191B	2067
0.0358351122753749	1.14632944584362	0.00996891338491857
0.920467821567317	0.987407390716406
6029	RN7SL1	RNA component of signal recognition particle 7SL1
304	-0.0550133021642491	5.59827064504347
0.00990659939239436	0.920715960009438	0.987407390716406
125488	TTC39C	tetratricopeptide repeat domain 39C
7188	-0.0174504371076429	3.76522815170438
0.00989689816082784	0.920754661782415	0.987407390716406
6117	RPA1	replication protein A1	5492	0.0118903211517747
5.86774464424229	0.00988200167925513
0.920814126623965	0.987407390716406
27124	INPP5J	inositol polyphosphate-5-phosphatase J	4822
0.0351820577505463	2.35546132916968	0.00986566946774481
0.920879374755152	0.987407390716406
493753	COA5	cytochrome c oxidase assembly factor 5	2760
0.0173662936290218	3.58844611301072	0.00984751292821096
0.920951975234461	0.987407390716406
5216	PFN1	profilin 1	1987	-0.0156643164115658
9.11232609049861	0.00983573283019723	0.920999115122685
0.987407390716406
25989	ULK3	unc-51 like kinase 3	3516	0.0148338013656368
4.86854417003187	0.00981337731893106
0.921088652614052	0.987407390716406
7156	TOP3A	DNA topoisomerase III alpha
5731	-0.015775972030655	4.75892489270884
0.00980519213765874	0.921121461361296	0.987407390716406
9978	RBX1	ring-box 1	3092	0.0164625374350637
5.48364761636679	0.00974767026588097	0.921352417999382
0.987407390716406
53349	ZFYVE1	zinc finger FYVE-type containing 1	5306
0.0130240091821105	6.02782365769238	0.00973845582877431
0.921389478855926	0.987407390716406
6935	ZEB1	zinc finger E-box binding homeobox 1
8129	-0.0322346095427796	1.61298396379955
0.00973695875104585	0.921395501837597	0.987407390716406
93081	TEX30	testis expressed 30	2154	0.0237292969366756
2.06237899761592	0.00972889455214421
0.921427953436744	0.987407390716406
4637	MYL6	myosin light chain 6	3638	-0.0175345830679137
10.4704181686491	0.00972256641688318
0.921453428354718	0.987407390716406
55154	MSTO1	misato mitochondrial distribution and morphology
regulator 1	3683	0.0180300919257388	4.34487711564656
0.00971611112489601	0.921479423788896

0.987407390716406
84173	ELMOD3	ELMO domain containing 3	7259
0.0173589026107026	4.64893626234656	0.00971308265606297
	0.92149162242527	0.987407390716406
63971	KIF13A	kinesin family member 13A
11006	-0.0193380281072284	7.53924454868547
0.00970062307260378	0.921541829691195	0.987407390716406
57515	SERINC1	serine incorporator 1	3137	0.0124915579240221
	9.0395940846737	0.00969979785872965	0.921545156130736
	0.987407390716406
1140	CHRNB1	cholinergic receptor nicotinic beta 1 subunit	4329
	0.0181738869898893	7.23616586770114
0.00969485622505317	0.921565078860692	0.987407390716406
6621	SNAPC4	small nuclear RNA activating complex polypeptide 4
	5010	0.0140880574201577	5.14356965163312
0.00969459489969005	0.921566132564831	0.987407390716406
80108	ZFP2	ZFP2 zinc finger protein	2641
0.0336237195319305	0.540583549535236	0.0096253684864549
	0.921845770964305	0.98763146044799
8728	ADAM19	ADAM metallopeptidase domain 19
7704	-0.0549332317895122	6.46497569403923
0.00958565730624628	0.922006641683482	0.987678876861142
101929767	LOC101929767	uncharacterized LOC101929767
8246	-0.0320738323604972	2.07635998116626
0.00957963797132066	0.922031055460577	0.987678876861142
8736	MYOM1	myomesin 1	6574	-0.0544770873042049
0.732879243757786	0.00949481020937792	0.922375934801441
	0.98793509866227
100131564	CCDC18-AS1	CCDC18 antisense RNA 1
4452	-0.0255822296221413	1.93835235446481
0.00948613071343907	0.922411310081042	0.98793509866227
58478	ENOPH1	enolase-phosphatase 1	2025	-0.0150449240008295
	5.68316096908714	0.00946270682705119
0.922506861132402	0.987961893407871
10439	OLFM1	olfactomedin 1	6302	-0.0439404241983268
0.158752250723398	0.00944490836476142	0.922579544718921
	0.987964195938679
23157	SEPTIN6	septin 6	6245	-0.0229619954055988
4.52652195361569	0.00940088880318513	0.922759604891368
	0.98806741024135
100505761	RPARP-AS1	RPARP antisense RNA 1	2481
0.0346141406644677	1.21681145103713	0.00936991729317516
	0.922886547703851	0.98806741024135
8636	SSNA1	SS nuclear autoantigen 1	1183
0.0142310054577109	5.11545714518317	0.00935483644640556
	0.922948436175054	0.98806741024135
148254	ZNF555	zinc finger protein 555	8557	0.0239858765606036
	3.229354024248	0.00935248462519134	0.922958092068096
	0.98806741024135
57407	NMRAL1	NmrA like redox sensor 1	3078
0.0194703973463035	4.15618885081131	0.00931303679688256
	0.923120236395852	0.988075398536065
8872	CDC123	cell division cycle 123	2404	0.0129563766232075
	5.85737721172192	0.00929128085851971

0.923209809195157	0.988075398536065
100507487	LINC02615	long intergenic non-protein coding RNA 2615	3283	0.0460888324277183	1.01022816874498	0.00928751444070741	0.923225326904716	0.988075398536065
286	ANK1	ankyrin 1	10799	-0.0786718753734236	5.04498508152513	0.00927444818415779	0.923279184751364	0.988075398536065
54584	GNB1L	G protein subunit beta 1 like	6988	-0.0265064817649035	2.48695470708763	0.00926497368201495	0.923318261686196	0.988075398536065
36	ACADSB	acyl-CoA dehydrogenase short/branched chain	6255	-0.0201188499459595	3.58513058384642	0.00917750346791379	0.923679983323848	0.988203468995772
56965	PARP6	poly(ADP-ribose) polymerase family member 6	7696	0.0146552225857072	6.04486365552035	0.00914842868164101	0.923800603228859	0.988203468995772
5092	PCBD1	pterin-4 alpha-carbinolamine dehydratase 1	1614	-0.0249008977376401	4.81717669886601	0.00912650459081554	0.923891685669455	0.988203468995772
374920	ZSWIM9	zinc finger SWIM-type containing 9	3751	0.0216387227228249	3.07314872666408	0.00911719801564281	0.923930382704485	0.988203468995772
8905	AP1S2	adaptor related protein complex 1 subunit sigma 2	5454	0.0214624356229074	3.17647053510694	0.00911052636540388	0.923958135904446	0.988203468995772
54816	ZNF280D	zinc finger protein 280D	13858	-0.0160101066996468	4.38625356504627	0.00910867756049072	0.923965828507579	0.988203468995772
100500888	MIR3940	microRNA 3940	102	-0.0249777194694262	1.82696141280353	0.00907916916149976	0.924088715380038	0.988203468995772
27336	HTATSF1	HIV-1 Tat specific factor 1	3136	-0.0113357403967631	6.19832290245265	0.00907757758301742	0.924095349185955	0.988203468995772
1453	CSNK1D	casein kinase 1 delta	10927	0.0112605563318556	7.47238220519084	0.00907746339013804	0.924095825172101	0.988203468995772
7637	ZNF84	zinc finger protein 84	9482	0.0180877424597627	5.3303614038376	0.00906186632556683	0.924160866276049	0.988203468995772
93611	FBXO44	F-box protein 44	3527	0.0231493111252462	4.86599655263592	0.00904573088340044	0.924228211927454	0.988203468995772
4580	MTX1	metaxin 1	3278	0.0184828820016924	3.77587780257955	0.00903224466873986	0.924284546820419	0.988203468995772
9098	USP6	ubiquitin specific peptidase 6	11270	0.0377418977432353	0.784507175470957	0.00900428081863502	0.924401493374536	0.988253069584577
29066	ZC3H7A	zinc finger CCCH-type containing 7A	7076	-0.0114133020783322	5.9847876974527	0.00897683508816072	0.924516451424938	0.988300536908273
57212	TP73-AS1	TP73 antisense RNA 1	6822

0.0201308937628805	4.77966302731816	0.00891203556039954
	0.924788573578809	0.9885159913094
10140	TOB1	transducer of ERBB2, 1	2521	-0.0245815337897619
	6.1269903903944 0.00885485488766946	0.92502953091977
	0.988649663747194
548596	CKMT1A	creatine kinase, mitochondrial 1A
1779	-0.0419334175008147	1.78874457377878
0.00884887066477624	0.925054793536854	0.988649663747194
1059	CENPB	centromere protein B	2840	0.0103164527066584
	6.36141015131719	0.00882452051029947
0.925157677593286	0.988684183264604
8818	DPM2	dolichyl-phosphate mannosyltransferase subunit 2,
regulatory	3133	-0.0143022229455459	4.83741141664097
	0.0087679370436673	0.925397307957124
0.988864822440909
25896	INTS7	integrator complex subunit 7
5311	-0.0138743162737647	4.69586599295145
0.00874180730222962	0.925508230367567	0.98888095620971
23191	CYFIP1	cytoplasmic FMR1 interacting protein 1
10414	-0.0134273867897252	7.56198622483322
0.00872432356376862	0.925582543352102	0.98888095620971
10484	SEC23A	Sec23 homolog A, coat complex II component
8945	-0.0165833415359682	7.05862687241101
0.00870952227200661	0.925645513580149	0.98888095620971
7267	TTC3	tetratricopeptide repeat domain 3	15883
0.012113610138827	7.66898421583352	0.00869794565925019
	0.925694802443187	0.98888095620971
119587	CPXM2	carboxypeptidase X, M14 family member 2
4859	-0.0625780887070706	2.25166603252029
0.00864645308972989	0.925914440372193	0.988984385869201
8420	SNHG3	small nucleolar RNA host gene 3
4088	-0.015493331164732	4.51344275669381
0.0086390890040926	0.925945905220906	0.988984385869201
23478	SEC11A	SEC11 homolog A, signal peptidase complex subunit
	5093	-0.0129540594823487	6.66850239226234
0.0086016282820367	0.92610617510148	0.988984385869201
102157402	AK6	adenylate kinase 6
2162	-0.0171602208894188	3.86489088890836
0.00859551678977155	0.926132355545219	0.988984385869201
6911	TBX6	T-box transcription factor 6	3137
0.0687495020865146	2.01998726228581	0.00857375686228323
	0.926225647137078	0.988984385869201
7673	ZNF222	zinc finger protein 222 2003	0.0261057415192063
	1.849576054996	0.00855687939971972	0.926298088329082
	0.988984385869201
7049	TGFBR3	transforming growth factor beta receptor 3	8264
	0.0419935985005432	4.2486062991002 0.008551442550635
	0.926321439632537	0.988984385869201
996	CDC27	cell division cycle 27	6824	0.0109985694076088
	6.23062115211464	0.00854028749424529
0.92636937412502	0.988984385869201
84263	HSDL2	hydroxysteroid dehydrogenase like 2	3455
0.014592580390423	4.79368597042058	0.00850663121521933
	0.926514190471036	0.988984385869201

55344	PLCXD1	phosphatidylinositol specific phospholipase C X domain containing 1	12382	-0.0273202250453931	3.81949284513459	0.0085001008066854	0.926542322982596	0.988984385869201
146057	TTBK2	tau tubulin kinase 2	12917	0.0192677607342981	5.4177884914125	0.00847302665481209	0.926659072822347	0.988984385869201
51646	YPEL5	yippee like 5	3421	0.0130460137321878	7.81783991364384	0.00846388106684515	0.926698553150053	0.988984385869201
10203	CALCRL	calcitonin receptor like receptor	6144	0.028237519798341	0.266782651406531	0.00846134449864477	0.926709506998877	0.988984385869201
4204	MECP2	methyl-CpG binding protein 2	15888	-0.0141367337971204	5.90867714341926	0.00842192080845905	0.926879966547758	0.989090940989978
80347	COASY	Coenzyme A synthase	3594	0.0128611381668017	6.09864675970071	0.00840419480917465	0.926956741056289	0.989097514434024
90933	TRIM41	tripartite motif containing 41	12803	-0.0150043077739833	5.3897063256443	0.0083622518153561	0.927138729546698	0.989216345578961
7033	TFF3	trefoil factor 3	1168	0.0342476787554084	0.466592055669799	0.00832935628217246	0.927281783824003	0.989293620688766
91369	ANKRD40	ankyrin repeat domain 40	4849	-0.0126545202707158	5.88368113434393	0.00831111704307474	0.927361224383035	0.98930302132012
348262	MCRIP1	MAPK regulated corepressor interacting protein 1	3726	-0.0152657151327616	4.7862992078763	0.00829266026318631	0.927441701958112	0.989313526717484
64718	UNKL	unk like zinc finger	7494	-0.020171714239319	4.15026039106479	0.00821746728037454	0.927770504927182	0.989570658331506
10928	RALBP1	ralA binding protein 1	4979	-0.0101982030301673	7.04782082892496	0.00818767629700545	0.927901193886312	0.989570658331506
353	APRT	adenine phosphoribosyltransferase	1803	-0.0182004294517797	5.15346291925977	0.0081598984998692	0.928023267398873	0.989570658331506
10653	SPINT2	serine peptidase inhibitor, Kunitz type 2	5199	-0.0119354568513805	9.5388088048859	0.00815601949622646	0.928040330895029	0.989570658331506
56135	PCDHAC1	protocadherin alpha subfamily C, 1	6943	-0.0552973758792576	3.05846199452551	0.00813466049397782	0.928134361184816	0.989570658331506
5799	PTPRN2	protein tyrosine phosphatase receptor type N2	4833	0.0464486586354269	1.3473213534948	0.00813256585458966	0.928143589265519	0.989570658331506
23642	SNHG1	small nucleolar RNA host gene 1	3756	-0.0151234383557852	4.43437636166532	0.00811660429739636	0.928213948410035	0.989570658331506
4144	MAT2A	methionine adenosyltransferase 2A	4487	0.011163296964065 9	7.21697148594673	0.0080940426218276 7

0.928313520196099 0.989570658331506
23395 LARS2 leucyl-tRNA synthetase 2, mitochondrial 5393
0.0131468942716082 4.79542545811028 0.00809289535789048
0.928318587168986 0.989570658331506
9967 THRAP3 thyroid hormone receptor associated protein 3 4604
0.0100589177812578 7.27421150385323
0.00803580647998281 0.92857118312663 0.98976459645801
283489 CHAMP1 chromosome alignment maintaining phosphoprotein 1
4171 0.0151402245542512 4.29899503964635
0.00801073622071513 0.928682394973457 0.989807815126557
130617 ZFAND2B zinc finger AN1-type containing 2B 3396
0.014758933041086 4.48729062620223 0.00797001860809576
0.928863393063233 0.989888296713168
27252 KLHL20 kelch like family member 20
4254 -0.0127706046894761 4.97563392651798
0.00796008288214622 0.928907630079728 0.989888296713168
8701 DNAH11 dynein axonemal heavy chain 11 15998
0.0492236888870192 2.04582144157692 0.00793480523583412
0.929020299829896 0.989888296713168
64429 ZDHHC6 zinc finger DHHC-type containing 6
3748 -0.0152948392421093 5.46069774663971
0.00793025733633357 0.929040590318505 0.989888296713168
80022 MYO15B myosin XVB 12761 0.0378084814780026
4.48239279910626 0.00791267080063562 0.929119108020169
0.9898966567307
79731 NARS2 asparaginyl-tRNA synthetase 2, mitochondrial
3153 -0.0174898074764434 3.76323997259919
0.00788192951165056 0.92925656859218 0.989905893507228
80013 MINDY3 MINDY lysine 48 deubiquitinase 3 4094
0.0121261831813268 5.18155823504945 0.00786120267473933
0.929349401793791 0.989905893507228
7442 TRPV1 transient receptor potential cation channel
subfamily V member 1 5237 -0.0255367600382581
2.94491148262354 0.00785116167827393 0.929394418651946
0.989905893507228
114659 LRRC37B leucine rich repeat containing 37B 5534
0.0209788457941422 3.49874194484337 0.00784758375443584
0.929410466589102 0.989905893507228
51306 FAM13B family with sequence similarity 13 member B
6808 -0.0115002822929017 6.32646906620837
0.00776946441131443 0.929761776124358 0.990158631313729
221545 C6orf136 chromosome 6 open reading frame 136
2118 -0.0166563367703721 3.61239823517535
0.0077633962437087 0.929789139549473 0.990158631313729
84934 RITA1 RBPJ interacting and tubulin associated 1
2053 -0.0167605534943249 4.02798118724954
0.00774288674028867 0.929881703712466 0.990181923665844
103611081 LOC103611081 uncharacterized LOC103611081
3994 -0.0224662141972178 2.93799530467789
0.0077244614271379 0.929964966844257 0.990195309052642
7593 MZF1 myeloid zinc finger 1 5581 -0.0144983487129494
4.06162469431098 0.00764307924134378
0.93033393191055 0.990258351769625
85315 PAQR8 progestin and adipoQ receptor family member 8 5057

0.0449953196472223 2.9160522516938 0.00761758632014065 0.930449916736838 0.990258351769625
6392 SDHD succinate dehydrogenase complex subunit D 2844 -0.0129217506132606 5.09969736061909 0.00760873013255292 0.930490255380082 0.990258351769625
54941 RNF125 ring finger protein 125 6534 -0.0274531559387948 1.51209321730329 0.00757827795452481 0.930629141428821 0.990258351769625
79913 ACTR5 actin related protein 5 2565 0.0187762327177875 2.94210642692247 0.00756484957507197 0.930690474791672 0.990258351769625
102723099 GHET1 gastric carcinoma proliferation enhancing transcript 1 1906 -0.028472798433205 0.043779344616652 0.00753871018379293 0.930810022481888 0.990258351769625
119032 BORCS7 BLOC-1 related complex subunit 7 2708 0.0165059509916426 4.72223292841357 0.0075353232051576 0.930825527995078 0.990258351769625
5362 PLXNA2 plexin A2 13781 0.0150558670070614 7.29295585371632 0.00753406932427936 0.930831269124432 0.990258351769625
966 CD59 CD59 molecule (CD59 blood group) 4942 -0.0178598026504767 9.94414392604949 0.00752471856503689 0.930874098521902 0.990258351769625
130940 CCDC148 coiled-coil domain containing 148 3790 -0.027813648210884 1.00164237711235 0.00752101153663309 0.930891085290779 0.990258351769625
55703 POLR3B RNA polymerase III subunit B 4858 0.0130657340740452 4.21054025386052 0.00750043870292361 0.930985433229589 0.990258351769625
5831 PYCR1 pyrroline-5-carboxylate reductase 1 2886 0.0270040430715606 3.53400285226716 0.00749438870640518 0.931013203583778 0.990258351769625
8842 PROM1 prominin 1 7119 0.0825875647667719 4.73627893498392 0.00746754206532074 0.931136569987455 0.990258351769625
11244 ZHX1 zinc fingers and homeoboxes 1 5848 -0.0114657984697952 5.38843057667038 0.00746423376405403 0.931151787833634 0.990258351769625
105221694 BISPR BST2 interferon stimulated positive regulator 2809 -0.0524470612711852 0.329471368650378 0.00742414956102344 0.931336441721333 0.990258351769625
60685 ZFAND3 zinc finger AN1-type containing 3 4036 -0.0100551039781913 6.81302721353995 0.0073838450518362 0.931522617518982 0.990258351769625
85014 TMEM141 transmembrane protein 141 1443 -0.0175540925411501 3.90213642726647 0.00733792212427176 0.93173537105785 0.990258351769625
4691 NCL nucleolin 6929 0.0116796599809284 8.4422477913754 0.00731790878830907 0.931828299400513 0.990258351769625
2585 GALK2 galactokinase 2 7531 -0.0162002068372192 3.42582897115823 0.007316414086322 0.93183524491559

0.990258351769625
2189 FANCG FA complementation group G 3614
0.0216013840295543 2.96918230879511 0.00731470652183042
0.93184318042536 0.990258351769625
94240 EPSTI1 epithelial stromal interaction 1
4924 -0.0241238161110859 4.11718898948985
0.00730765211183382 0.931875974052551 0.990258351769625
11064 CNTRL centriolin 9417 -0.0208516867559954
2.91745183811725 0.00730631338760901 0.931882199139994
0.990258351769625
9950 GOLGA5 golgin A5 3295 -0.0104301717861096
6.27546704303413 0.007300810958073 0.931907791553817
0.990258351769625
6139 RPL17 ribosomal protein L17 2466 -0.0168054725070221
7.10584875467959 0.0072932071826699
0.931943173554993 0.990258351769625
170394 PWWP2B PWWP domain containing 2B 2651
0.0153013062513458 4.40136627479589 0.00728222813609136
0.931994294214396 0.990258351769625
100528062 ARMCX5-GPRASP2 ARMCX5-GPRASP2 readthrough
1523 -0.0217040131980575 2.00323981364057
0.00727586644574174 0.932023933293584 0.990258351769625
284106 CISD3 CDGSH iron sulfur domain 3
2663 -0.0188940435464207 3.55408703214466
0.0072680511033445 0.932060362801199 0.990258351769625
142940 TRUB1 TruB pseudouridine synthase family member 1
3601 -0.0113213274914785 5.18236293268975
0.00725744945531659 0.932109811610043 0.990258351769625
64799 IQCH IQ motif containing H 6740 -0.0276044637091941
0.682244058584779 0.0072475641303158
0.932155952090175 0.990258351769625
220042 DDIAS DNA damage induced apoptosis suppressor 5339
0.0276105439565014 -0.00883676610622602 0.00724672762952761
0.932159857973096 0.990258351769625
29924 EPN1 epsin 1 17616 0.0117593172524188
6.3942138338403 0.00722405716373231 0.932265800042768
0.990258351769625
11140 CDC37 cell division cycle 37 3041 0.0117767352380303
7.64933669945603 0.00720798841848591
0.932340992834837 0.990258351769625
6510 SLC1A5 solute carrier family 1 member 5 3332
0.0165534526208105 6.28100348768542 0.00719212611134878
0.932415302454573 0.990258351769625
1613 DAPK3 death associated protein kinase 3
3549 -0.0115907105392402 6.92658477594286
0.00718745760688932 0.932437188557994 0.990258351769625
7866 IFRD2 interferon related developmental regulator 2 3712
0.0147768734494926 5.2114231310462 0.00717436964470153
0.932498583554034 0.990258351769625
5634 PRPS2 phosphoribosyl pyrophosphate synthetase 2 2758
0.0116315237979385 4.64114549707755
0.00714203422927895 0.932650509668962 0.990322985504453
81576 CCDC130 coiled-coil domain containing 130
3158 -0.0142445453781594 4.34006353938176

0.00713133591029624	0.932700851343953	0.990322985504453
3397	ID1	inhibitor of DNA binding 1, HLH protein
1233	-0.0150320083298998	5.25466390138321
0.00710352977927187	0.932831873161235	0.990387027012008
203245	NAIF1	nuclear apoptosis inducing factor 1	3621
0.0157086610466119	3.58431023618532	0.00705976220187887
	0.933038629212319	0.99047910964050б
159371	SLC35G1 solute carrier family 35 member G1	5895
0.0208530048658933	2.5234697980434 0.00705523953782006
0.933060030820766	0.990479109640506
280	AMY2B	amylase alpha 2B	7491	0.0250218961781013
	2.75710155751632	0.00695933532729409
0.933515494463612	0.990722628184908
283848	CES4A	carboxylesterase 4A	4340	0.0216310631730556
	2.46328470744226	0.00694880460140102
0.933565698521382	0.990722628184908
149466	C1orf210	chromosome 1 open reading frame 210
1516	-0.0199054347203691	2.53036082359689
0.00694309192507747	0.933592949093676	0.990722628184908
100526760	ABHD14A-ACY1	ABHD14A-ACY1 readthrough	2164
	-0.0174494637396829	3.5919252516240З
0.00689496944028711	0.933822952195317	0.990722628184908
163051	ZNF709	zinc finger protein 709 5386	0.0258361665283328
	1.75355223758167	0.00689355246516854
0.933829736906454	0.990722628184908
50507	NOX4	NADPH oxidase 4 6327	-0.0316497082300814
3.09452361658913	0.00689062161876031	0.933843772511972
	0.990722628184908
259230	SGMS1	sphingomyelin synthase 1	5131
0.0120818434660711	4.99195479414662	0.00688177617696084
	0.933886150908207	0.990722628184908
23543	RBFOX2	RNA binding fox-1 homolog 2	10014
0.0104407502783863	7.73752670916969	0.00687776174205057
	0.933905393060228	0.990722628184908
10581	IFITM2	interferon induced transmembrane protein 2
1192	-0.0397024443143173	3.70885968333754
0.00686954415055041	0.933944799594276	0.990722628184908
3748	KCNC3	potassium voltage-gated channel subfamily C member 3
	6277	0.0226951780396498	4.26869722810565
0.0068575896462626	0.934002168470027	0.990722628184908
221443	OARD1	O-acyl-ADP-ribose deacylase 1	5423
0.0156548356329375	3.89954622656936	0.00684122519857278
	0.93408078206663	0.990722628184908
11194	ABCB8	ATP binding cassette subfamily B member 8	9336
	0.0136836019432518	5.43645165976534
0.00681866276450904	0.934189325733834	0.990722628184908
727910	TLCD2	TLC domain containing 2 5971	0.0269597609923316
	4.20961240700293	0.00681459186323252
0.934208929325785	0.990722628184908
63877	FAM204A family with sequence similarity 204 member A
15018	0.010809658273766	5.17261065379399
0.00677799711737137	0.934385418188869	0.990755373248302
728743	LOC728743	zinc finger protein pseudogene	4584
0.0255344470163702	1.12044759337953	0.0067621637722648

0.9344619277656 0.990755373248302
389072 PLEKHM3 pleckstrin homology domain containing M3
4656 -0.0219659726533551 2.70987391503157
0.00674012148502445 0.934568590382607 0.990755373248302
51236 HGH1 HGH1 homolog 2814 -0.0132836766646668
4.06822191783534 0.00673591558436071 0.934588962672128
0.990755373248302
202915 TMEM184A transmembrane protein 184A
7805 -0.0196974821010746 5.78406787998313
0.00673498503256553 0.934593470888114 0.990755373248302
5531 PPP4C protein phosphatase 4 catalytic subunit
2734 -0.0105361467528638 6.74044697349354
0.0067068506942789 0.934729920808651 0.990825034112818
6728 SRP19 signal recognition particle 19 5480
0.0112647492275868 5.51336588744544 0.00665105129529309
0.935001399717062 0.9908886246181
5716 PSMD10 proteasome 26S subunit, non-ATPase 10 2459
0.0111770357850969 5.356045878217 0.00664001792503655
0.935055215575743 0.9908886246181
5054 SERPINE1 serpin family E member 1
3190 -0.0186225317285737 10.6978223613349
0.00663583168034165 0.935075645987063 0.9908886246181
90416 CCDC32 coiled-coil domain containing 32 8648
0.0146774897301319 4.21066337271375 0.00662667692126195
0.93512034719011 0.9908886246181
64216 TFB2M transcription factor B2, mitochondrial
1848 -0.0166792219608854 3.49569064314837
0.00661094085576863 0.935197256536493 0.9908886246181
8491 MAP4K3 mitogen-activated protein kinase kinase kinase
kinase 3 6067 -0.0124753948853858 5.77197183796213
0.00659407599868356 0.935279785163775
0.9908886246181
100526737 RBM14-RBM4 RBM14-RBM4 readthrough
2826 -0.00972727363074504 6.5674118284383 0.00658321890796554
0.935332970845209 0.9908886246181
55558 PLXNA3 plexin A3 11710 0.0127480840933164
7.57004732017805 0.00657480394232834 0.935374223651786
0.9908886246181
284013 VMO1 vitelline membrane outer layer 1 homolog
803 -0.0432342022888625 0.14625633043802
0.00655188953654307 0.935486692162599 0.9908886246181
100132815 IPO5P1 importin 5 pseudogene 1 4062
0.0276596510273996 2.31233548472501 0.00654972323458125
0.935497335043175 0.9908886246181
3383 ICAM1 intercellular adhesion molecule 1 3451
0.0258502852832855 9.63114637339651 0.00653267233656152
0.935581166762208 0.990902487927277
126074 SWSAP1 SWIM-type zinc finger 7 associated protein 1 1684
0.0208875514629074 1.80288442553372
0.00648491082911651 0.935816576962331 0.990987449852644
58497 PRUNE1 prune exopolyphosphatase 1 3063
0.0124971326155884 4.41745931784843 0.00646865689398712
0.935896889353383 0.990987449852644
91392 ZNF502 zinc finger protein 502 3273 0.0255517603783976

2.0809161293115 0.00645933588523118 0.935942991252763
0.990987449852644
7319 UBE2A ubiquitin conjugating enzyme E2 A
5276 -0.0107079680397606 6.77848847853184
0.00645905458264195 0.935944383101352 0.990987449852644
100529209 RNASEK-C17orf49 RNASEK-C17orf49 readthrough
1947 -0.012432349191104 7.08544628929535
0.00644350379047864 0.936021373907345 0.990994057771744
51096 UTP18 UTP18 small subunit processome component 2721
0.0138574091736386 4.30638162965485
0.00641520585585909 0.936161715129717 0.991067730619148
54626 HES2 hes family bHLH transcription factor 2
5407 -0.0140290439362528 8.40066025268754
0.00636825424021481 0.936395256693445 0.991186448852025
101752399 STAG3L5P-PVRIG2P-PILRB STAG3L5P-PVRIG2P-PILRB
readthrough 6539 0.0272319877954821 6.39884647679906
0.00636314992769726 0.936420698144741
0.991186448852025
26054 SENP6 SUMO specific peptidase 6
11414 -0.0089982414348826 6.17076427310434
0.00632631088755886 0.936604620493254 0.991186448852025
90806 ANGEL2 angel homolog 2 7279 0.0133143630500544
4.51063490095011 0.00629486143881763 0.936762061647439
0.991186448852025
64375 IKZF4 IKAROS family zinc finger 4 6628
0.0219653440816912 2.93785352489657 0.00629034826845043
0.936784687770691 0.991186448852025
161779 PGBD4 piggyBac transposable element derived 4
6612 -0.0207613892783627 2.37353662908077
0.00628228437327571 0.936825135280627 0.991186448852025
10307 APBB3 amyloid beta precursor protein binding family B
member 3 4409 0.0190393421683338 3.19584304485326
0.00627922192166208 0.936840503006114
0.991186448852025
84219 WDR24 WD repeat domain 24 3570 0.019929591726575
3.42055398851276 0.00626315132699062
0.936921208940397 0.991186448852025
970 CD70 CD70 molecule 1532 -0.0260577803698183
0.403404585757825 0.00623510056218857 0.937062328894965
0.991186448852025
6175 RPLP0 ribosomal protein lateral stalk subunit P0 3614
0.017093161373203 8.74350908312611
0.00620231301763852 0.937227684500912 0.991186448852025
4515 MTCP1 mature T cell proliferation 1 2552
0.0212231389900863 1.29901579738751 0.00615799433331654
0.937451895020565 0.991186448852025
6835 SURF2 surfeit 2 1008 -0.0171910049333956
3.28796451290574 0.00614340363818755 0.937525887619454
0.991186448852025
5557 PRIM1 DNA primase subunit 1 2437 -0.0277493137704909
1.11258465662171 0.00613837344205415
0.937551417331104 0.991186448852025
9814 SFI1 SFI1 centrin binding protein
12339 -0.0121471618098355 4.82772614273779

0.00613199248593332	0.937583817691521	0.991186448852025
6606	SMN1	survival of motor neuron 1, telomeric
5145	-0.0148144386532046	5.20790253636899
0.00610299718529461	0.93773126022519	0.991186448852025
515	ATP5PB	ATP synthase peripheral stalk-membrane subunit b
	3383	0.0107766671903021	6.51209823234546
0.00610231825960739	0.937734716813247	0.991186448852025
5757	PTMA	prothymosin alpha	3378	0.00961598114641942
	8.61871296977186	0.00607916232699779
0.937852725753802	0.991186448852025
1407	CRY1	cryptochrome circadian regulator 1
5062	-0.0108548386700052	4.97340614632151
0.00606738066722556	0.937912855095898	0.991186448852025
10299	MARCHF6	membrane associated ring-CH-type finger 6
11336	-0.0115550273806796	7.36544764432877
0.0060422475672226	0.938041321972788	0.991186448852025
230	ALDOC	aldolase, fructose-bisphosphate C
2928	-0.0303298632382202	2.501363994918	0.00604150310829965
	0.938045131340961	0.991186448852025
11068	CYB561D2	cytochrome b561 family member D2	3288
	-0.014987044493288	3.83750322292986
0.00603364573673382	0.938085351620861	0.991186448852025
84305	PYM1	PYM homolog 1, exon junction complex associated
factor	2425	-0.0129063434125226	4.72040510769116
0.00603322590539079	0.938087501393007	0.991186448852025
5428	POLG	DNA polymerase gamma, catalytic subunit	5640
0.00866424491896374	6.46413705858461	0.0060305792164943
	0.93810105565869	0.991186448852025
10898	CPSF4	cleavage and polyadenylation specific factor 4	4250
	0.0138524255623815	4.31732193643487
0.00600659735414011	0.938224008607949	0.991186448852025
92017	SNX29	sorting nexin 29	11298	0.0137390484462263
	4.6438685991871	0.00600403817894701	0.93823714384912
	0.991186448852025
55750	AGK	acylglycerol kinase	6547	-0.0136554999443177
	4.25967714729474	0.00599280431198057
0.938294836217632	0.991186448852025
29083	GTPBP8	GTP binding protein 8 (putative)	3828
0.0135382941270587	3.96459778238102	0.00599031459654498
	0.938307629700527	0.991186448852025
644019	CBWD6	COBW domain containing 6	10393
0.0118198865862157	5.06114868604456	0.00598146486054407
	0.938353126016922	0.991186448852025
100506314	GPRC5D-AS1	GPRC5D and HEBP1 antisense RNA 1
	2407	-0.0280545865633743	1.91425091578325
0.00598103191884469	0.938355352629985	0.991186448852025
51281	ANKMY1	ankyrin repeat and MYND domain containing 1
11477	0.0176178965289239	3.01162040516617
0.0059681116889081	0.93842183854008	0.991186448852025
64005	MYO1G	myosin IG	8653	0.0676518677833937
0.584561273271844	0.00595924071074627	0.938467529426398
	0.991186448852025
51141	INSIG2	insulin induced gene 2	3559	-0.0108104188499997
	6.07152009100384	0.00592284903650864

0.938655327492396	0.991301506031999
9727	RAB11FIP3	RAB11 family interacting protein 3	6317
	0.0106373936953718	5.78805174645129
0.00591072718449936	0.938718010709534	0.991301506031999
5501	PPP1CC	protein phosphatase 1 catalytic subunit gamma
6906	-0.00907354267583191	6.65514378862495
0.00588739806207528	0.938838829980997	0.991354352924525
11143	KAT7	lysine acetyltransferase 7
11749	-0.00998757719042694	5.3620786348456 0.0058626690200409б
	0.938967162487734	0.991363541933917
93183	PIGM	phosphatidylinositol glycan anchor biosynthesis
class M 4319	-0.0121734460207147	4.02586208523829
0.00585844993293527	0.938989084803118	0.991363541933917
93594	TBC1D31 TBC1 domain family member 31
8611	-0.0199906002785254	1.94794012507383
0.00584294324397749	0.939069725632585	0.991373955904102
5896	RAG1	recombination activating 1	7468
0.0245480943187641	2.43145752404551	0.00577906952021756
	0.939403034162011	0.991651088967314
80270	HSD3B7	hydroxy-delta-5-steroid dehydrogenase, 3 beta- and
steroid delta-isomerase 7	2495	-0.0343730660964932
3.86952624114813	0.00574284563853666	0.939592882708469
	0.991656618293821
84315	MON1A	MON1 homolog A, secretory trafficking associated
	4532	0.0155953345015362	3.70881292944958
0.00569298718476574	0.939855177324784	0.991656618293821
56145	PCDHA3	protocadherin alpha 3	7460	-0.046567215812908
	2.97145635068445	0.00567538818300228
0.939948037547492	0.991656618293821
56242	ZNF253	zinc finger protein 253 3626	-0.0177904492592532
	3.05838434699759	0.0056730385022874
0.939960446466514	0.991656618293821
84872	ZC3H10	zinc finger CCCH-type containing 10	8464
0.0170352519576157	3.15411413234573	0.00566844888090756
	0.939984692211846	0.991656618293821
28952	CCDC22	coiled-coil domain containing 22
2516	-0.0139722971430733	3.76282396779369
0.00565143843085281	0.940074640084825	0.991656618293821
340485	ACER2	alkaline ceramidase 2	2238	0.0188131800588006
	3.9598151923521 0.00564107562507132	0.940129503277773
	0.991656618293821
56139	PCDHA10 protocadherin alpha 10	15224	0.0510070202776381
	3.2398447882158 0.00564063664941372	0.940131828439126
	0.991656618293821
149951	COMMD7	COMM domain containing 7	2584
0.0120479520380408	5.27239438319431	0.00562378413638953
	0.940221161539232	0.991656618293821
27430	MAT2B	methionine adenosyltransferase 2B	5153
0.00868885725375223	6.11446479034601	0.00560957549628682
	0.940296584432257	0.991656618293821
9752	PCDHA9	protocadherin alpha 9	10075	-0.0491437711838939
	2.93986149608136	0.00559111914268584
0.940394698783176	0.991656618293821
26164	MTG2	mitochondrial ribosome associated GTPase 2	4648

0.0133846854555841 4.32799413493891
0.0055860619645296 0.940421611258039 0.991656618293821
729440 CCDC61 coiled-coil domain containing 61 2351
0.0180120468210123 2.19638038560112 0.00558585681180979
0.94042270326519 0.991656618293821
84250 SLF1 SMC5-SMC6 complex localization factor 1
7185 -0.0186042238039041 2.70164034843677
0.00556564135244697 0.940530407180175 0.991656618293821
130814 SLC66A3 solute carrier family 66 member 3
2275 -0.0187520049389999 3.28737616899259
0.00554259344325203 0.940653442058964 0.991656618293821
64073 C19orf33 chromosome 19 open reading frame 33
859 -0.00881089885650549 8.13156982264565
0.00552964112706888 0.940722697322182 0.991656618293821
8930 MBD4 methyl-CpG binding domain 4, DNA glycosylase
3356 -0.0121963056127006 4.89343673183117
0.0055161776397199 0.940794772292135 0.991656618293821
23329 TBC1D30 TBC1 domain family member 30
8332 -0.0299543161061122 1.33428122637742
0.00550282083164166 0.94086636361633 0.991656618293821
1950 EGF epidermal growth factor 6127 0.0315374198971249
4.41590108364099 0.0054972230170236
0.940896393390547 0.991656618293821
64943 NT5DC2 5'-nucleotidase domain containing 2 4719
0.0108354608854 7.54672822483246 0.00549678800095421
0.940898727701029 0.991656618293821
753 LDLRAD4 low density lipoprotein receptor class A domain containing 4 13058 -0.0369011849864883 3.99429370947738
0.00548712181057454 0.940950620776286
0.991656618293821
10929 SRSF8 serine and arginine rich splicing factor 8 4028
0.010092520172582 5.51514216697968
0.00547241643603513 0.941029655002736 0.991656618293821
643072 LOC643072 uncharacterized LOC643072 1494
0.0246137332916343 0.627614177069494 0.00547112365944447
0.941036608150315 0.991656618293821
101927204 FAM111A-DT FAM111A divergent transcript
2645 -0.0231127408109746 2.75460961250281
0.00539113683229431 0.941468432399179 0.992037036983021
25880 TMEM186 transmembrane protein 186 1978
0.0159189843183341 2.748239365789 0.00533215766848549
0.941788909452738 0.992123623543096
1525 CXADR CXADR Ig-like cell adhesion molecule 6682
0.0103947133299561 6.26554157904854 0.00532773319370961
0.941813022552028 0.992123623543096
51808 PHAX phosphorylated adaptor for RNA export 4319
0.00916626250717402 6.46721971331285 0.00532242431339114
0.941841968892801 0.992123623543096
58498 MYL7 myosin light chain 7 1458 -0.0473951836880461
0.941730444657854 0.00532035585363211
0.941853250970102 0.992123623543096
51241 COX16 cytochrome c oxidase assembly factor COX16
1863 -0.0123543446652717 5.14391551713749
0.00531091766472169 0.9419047580407 0.992123623543096

2130	EWSR1	EWS RNA binding protein 1
10326	-0.00957353865037498	7.84221467655951
0.00525405111045529	0.942216074706059	0.992252915094959
100616414	MIR4761 microRNA 4761	82	-0.0362794382714821
	0.372090192752655	0.00525345742290284
0.942219333765588	0.992252915094959
100288142	NBPF20	NBPF member 20	18995	0.0171421850170572
	4.17806400803145	0.0052236599228852
0.942383146209797	0.992252915094959
5393	EXOSC9	exosome component 9	4711	0.011678788697104
	4.04740855620278	0.00521975730348245
0.942404635698209	0.992252915094959
58527	ABRACL	ABRA C-terminal like	1200	0.0138842248445533
	6.16548992780272	0.00517724658568852
0.942639242969836	0.992252915094959
80832	APOL4	apolipoprotein L4	4377	0.0498521363923512
	-0.411121201846666	0.00517316568327253
0.942661815418604	0.992252915094959
79955	PDZD7	PDZ domain containing 7 5821	0.0290599903444898
	1.34695538558518	0.00514212753531318
0.942833788549672	0.992252915094959
11014	KDELR2	KDEL endoplasmic reticulum protein retention receptor 2	3990	0.0134079930079615	7.62105499793726
	0.00513080230329876	0.942896668140591
0.992252915094959
84553	FAXC	failed axon connections homolog, metaxin like GST domain containing	11662	-0.0155795720287873
4.65422444635585	0.00512597292548023	0.942923502870249
	0.992252915094959
170082	TCEANC	transcription elongation factor A N-terminal and central domain containing	3200	0.020425608945456
0.718148497224558	0.00512428082926686	0.942932908110701
	0.992252915094959
100507246	SNHG16	small nucleolar RNA host gene 16
3972	-0.0208697321632381	5.18020645600911
0.0051160825718739	0.942978498902006	0.992252915094959
97	ACYP1	acylphosphatase 1	1841	0.0208494719167926
	1.7021828273292 0.00509585211527686	0.943091158501057
	0.992252915094959
100529144	CORO7-PAM16	CORO7-PAM16 readthrough 5760
0.0108115563806775	4.92367394302757	0.00508355422467943
	0.943159753006299	0.992252915094959
64236	PDLIM2	PDZ and LIM domain 2	7909	-0.0115502565938495
	4.73069840664423	0.00503357001682758
0.943439414203565	0.992252915094959
10347	ABCA7	ATP binding cassette subfamily A member 7	8367
	0.0181377181970859	6.20070813423303
0.00501727946405595	0.943530860976889	0.992252915094959
29889	GNL2	G protein nucleolar 2	5745	-0.00925589651761563
	5.85451440950731	0.00500506159227143
0.943599543771431	0.992252915094959
91107	TRIM47	tripartite motif containing 47	3142
0.0107786257679215	6.25039578199641	0.00500213093472546
	0.943616030993847	0.992252915094959

83937	RASSF4	Ras association domain family member 4
9481	-0.0363553213905175	4.33392918300649
0.00497857063629681	0.943748752706854	0.992252915094959
150472	CBWD2	COBW domain containing 2
8396	-0.0112031250681884	4.3678829139267 0.00497771369576228
	0.943753586033147	0.992252915094959
284348	LYPD5	LY6/PLAUR domain containing 5
3226	-0.0259025985069764	0.634883319084325
0.00497416830323072	0.943773587243405	0.992252915094959
100131378	C11orf91	chromosome 11 open reading frame 91
	891	0.0338730402825935	2.41192468651145
0.00495243281196167	0.943896364163009	0.992252915094959
10039	PARP3	poly(ADP-ribose) polymerase family member 3
2928	-0.0157688218992608	4.21492091012373
0.00492910799826518	0.944028420224873	0.992252915094959
51427	ZNF107	zinc finger protein 107 7162	-0.025135415582812
	3.42406161369042	0.0049261176061286
0.944045373355483	0.992252915094959
9204	ZMYM6	zinc finger MYM-type containing 6
10389	-0.00881769839463681	5.78873861965381
0.00489242801636181	0.944236724829176	0.992252915094959
55505	NOP10	NOP10 ribonucleoprotein 554	0.0133205061938247
	6.00270191665575	0.00486977364072239
0.944365770562373	0.992252915094959
389203	SMIM20	small integral membrane protein 20	1612
0.0134628541314074	3.511350313879	0.00486567160427276
0.944389169151242	0.992252915094959
285753	CEP57L1	centrosomal protein 57 like 1	6432
0.0186139080683036	2.70736543395835	0.00485985410915646
	0.944422369957608	0.992252915094959
100288152	SLC9A3-AS1	SLC9A3 antisense RNA 1
2808	-0.0245987922361688	3.90058807579596
0.00484499278161366	0.944507275250224	0.992252915094959
5989	RFX1	regulatory factor X1	5175	-0.0152092355060598
	3.8643507325419 0.0048000594686437	0.94476478590061
	0.992252915094959
166379	BBS12	Bardet-Biedl syndrome 12
3432	-0.0195303589592577	2.72507872958188
0.00477783668679654	0.944892591184961	0.992252915094959
80146	UXS1	UDP-glucuronate decarboxylase 1
7023	-0.00959551365670264	6.23747388244049
0.00476659922710354	0.944957332554028	0.992252915094959
7095	SEC62	SEC62 homolog, preprotein translocation factor	7928
	0.00902560683701404	7.3014664270068 0.00475123133007571
	0.945045994503812	0.992252915094959
51253	MRPL37	mitochondrial ribosomal protein L37
2549	-0.011870403991462	5.36300948033769
0.00471565395747575	0.945251805366737	0.992252915094959
83700	JAM3	junctional adhesion molecule 3	5451
0.0402399852888612	2.26171817241866	0.00470459722303218
	0.945315926026958	0.992252915094959
161742	SPRED1	sprouty related EVH1 domain containing 1	8260
	0.0109041407663777	5.40973199262409
0.00470338556394534	0.945322957329618	0.992252915094959

83787 ARMC10 armadillo repeat containing 10 4580
0.0120729241250012 4.2153292096926 0.00470140135955033
0.945334473706825 0.992252915094959
256586 LYSMD2 LysM domain containing 2 2074
0.0194465524213237 1.65532315687243 0.00469159614471204
0.945391419323653 0.992252915094959
8672 EIF4G3 eukaryotic translation initiation factor 4 gamma 3
9287 -0.00908087127584184 7.20764287736248
0.00465992088933831 0.945575788981912 0.992252915094959
51642 MRPL48 mitochondrial ribosomal protein L48
4386 -0.013468260430066 3.92784545754812
0.00465966000493268 0.945577310100704 0.992252915094959
55345 ZGRF1 zinc finger GRF-type containing 1 9103
0.0212505025528256 1.4382830599262 0.00465874893702223
0.94558262253043 0.992252915094959
54512 EXOSC4 exosome component 4 1407 0.013958264671892
3.9301087292448 0.00465415979607897 0.945609389713137
0.992252915094959
100289678 ZNF783 zinc finger family member 783 4452
0.0156886039771951 3.62304149194739 0.0046344882695637
0.945724278756737 0.992252915094959
2800 GOLGA1 golgin A1 5229 0.00946274534988675
4.95367969518701 0.00463007336777466 0.94575009706599
0.992252915094959
27153 ZNF777 zinc finger protein 777 3233 -0.0110878711177188
4.57742901536844 0.00459991713982433
0.945926782155856 0.992252915094959
5789 PTPRD protein tyrosine phosphatase receptor type D
11045 0.0248664681722471 1.29380921167316
0.00458382712723093 0.946021291470018 0.992252915094959
150209 AIFM3 apoptosis inducing factor mitochondria associated 3
5193 -0.0219421346551726 -0.00188694940245114
0.00458260792006682 0.946028459620349 0.992252915094959
9807 IP6K1 inositol hexakisphosphate kinase 1
5200 -0.00905575699059333 6.24781760019799
0.00457189986033768 0.946091457290188 0.992252915094959
11311 VPS45 vacuolar protein sorting 45 homolog 6262
0.00962954645473542 5.2679239907161 0.00456837772978647
0.946112194887427 0.992252915094959
56143 PCDHA5 protocadherin alpha 5 7114 -0.0445321260887912
2.93391077116845 0.00455949584760162
0.946164525313267 0.992252915094959
49855 SCAPER S-phase cyclin A associated protein in the ER
11610 0.0160268056331972 3.52709174255523
0.00454836585701646 0.946230173518089 0.992252915094959
54885 TBC1D8B TBC1 domain family member 8B 9503
0.0143068411534304 4.12338361177704 0.0045417180818994
0.946269422692162 0.992252915094959
22943 DKK1 dickkopf WNT signaling pathway inhibitor 1 2206
0.0274184008507415 4.80673539007783
0.0045329576740869 0.946321189169141 0.992252915094959
131474 CHCHD4 coiled-coil-helix-coiled-coil-helix domain
containing 4 1772 -0.013843955006852 3.03379193211682
0.0045189882778341 0.946403840348112

0.992252915094959
23301	EHBP1	EH domain binding protein 1	8219
0.0122981227475331	6.44845373517025	0.00451519066322703
	0.946426331450308	0.992252915094959
81552	VOPP1	VOPP1 WW domain binding protein
4641	-0.0107589501062063	7.55877097527661
0.00451359796841189	0.946435766899932	0.992252915094959
57654	UVSSA	UV stimulated scaffold protein A	7830
0.0221989280848627	4.278324769218	0.00451176581383805
0.946446623026616	0.992252915094959
6764	DENND2B	DENN domain containing 2B	14067
0.0134873801568522	5.92342905163729	0.00451006647853092
	0.946456694134276	0.992252915094959
56140	PCDHA8	protocadherin alpha 8	5482	-0.044147246725146
	2.93595889444455	0.00450896048619498
0.946463249819779	0.992252915094959
91750	LIN52	lin-52 DREAM MuvB core complex component	3692
	0.0148702527469768	3.34700054323244
0.00450472707375127	0.946488350515493	0.992252915094959
100861437	NA	NA	1308	0.013435734453521
3.67452007273261	0.00449421294318331	0.946550742065912
	0.992252915094959
57685	CACHD1	cache domain containing 1	6345
0.0134907996307428	4.50203477957817	0.00449278690082444
	0.94655920993963	0.992252915094959
51136	RNFT1	ring finger protein, transmembrane 1	3180
0.0146436409675423	3.78604881707882	0.00448238900765752
	0.946620993728213	0.992252915094959
10174	SORBS3	sorbin and SH3 domain containing 3
6097	-0.0103073642550222	6.18540273332564
0.00446654473961416	0.9467152781953	0.992252915094959
54795	TRPM4	transient receptor potential cation channel
subfamily M member 4	4557	-0.0106491339620324
6.19362758264196	0.00444620180371569	0.946836579510739
	0.992252915094959
57035	RSRP1	arginine and serine rich protein 1	3018
0.0195462370099178	4.99808295426377	0.00444289277965026
	0.946856336970769	0.992252915094959
8347	H2BC4	H2B clustered histone 4	571	0.0256102557588256
	1.83377257318335	0.00444228329176255
0.946859976895712	0.992252915094959
253639	ZNF620	zinc finger protein 620	3457	-0.0187075988543441
	1.20658913641409	0.00443629584227878
0.946895747908478	0.992252915094959
387338	NSUN4	NOP2/Sun RNA methyltransferase 4
8468	-0.00934710247389634	4.95321078293388
0.00442852812174088	0.946942191003497	0.992252915094959
729852	UMAD1	UBAP1-MVB12-associated (UMA) domain containing 1
	3549	-0.0149754282368131	3.50199848200537
0.00441856211297065	0.947001837591941	0.992252915094959
246243	RNASEH1	ribonuclease H1	3192	0.00971546843670296
4.72342755511048	0.00441392909455107	0.947029589223629
	0.992252915094959
107282092	BOLA2-SMG1P6	BOLA2-SMG1P6 readthrough	3591

0.0178370032934598	3.20664446554986
0.0044093336669766	0.947057130142394	0.992252915094959
5306	PITPNA	phosphatidylinositol transfer protein alpha
4784	-0.00881787535012297	6.82290479912696
0.00439658157807443	0.947133630447796	0.992258845301591
80185	TTI2	TELO2 interacting protein 2	4222
0.0132999922010648	3.47715747461346	0.00436257055785605
0.947338210599545	0.992348445943735
22800	RRAS2	RAS related 2	3453	-0.00900179745505592
6.59339077273363	0.00435381711332772	0.947390993039684
0.992348445943735
23318	TUT4	terminal uridylyl transferase 4
16199	-0.0121218264486125	5.33558708800277
0.00434707293961267	0.947431696123765	0.992348445943735
572	BAD	BCL2 associated agonist of cell death	1708
0.0105606583506599	5.41998305180121	0.00428323353395044
0.947818567630614	0.992432418701484
7064	THOP1	thimet oligopeptidase 1	7464	0.0104582779829061
4.78006822894464	0.0042570791786769
0.947977900528056	0.992432418701484
51397	COMMD10	COMM domain containing 10	2026
0.0121824682175181	4.23399522352698	0.00425569410434745
0.947986352116666	0.992432418701484
21	ABCA3	ATP binding cassette subfamily A member 3
7804	-0.0179506915074534	4.96682826835278
0.00423821551177728	0.948093123709337	0.992432418701484
114822	RHPN1	rhophilin Rho GTPase binding protein 1	4204
0.0170730051383283	4.5027400504846	0.00423056191365845
0.948139946890577	0.992432418701484
3777	KCNK3	potassium two pore domain channel subfamily K member
3	6223	-0.0343613801612737	1.86456450775588
0.00421328895433248	0.948245775961919	0.992432418701484
143	PARP4	poly(ADP-ribose) polymerase family member 4	6080
0.0116938239554021	7.35167648854106
0.00420074910357471	0.948322742468299	0.992432418701484
23328	SASH1	SAM and SH3 domain containing 1
9476	-0.017297189178035	4.85376188774599
0.00418941070134693	0.948392434151874	0.992432418701484
91543	RSAD2	radical S-adenosyl methionine domain containing 2
4283	-0.020800334420672	3.23308868057335
0.00415869627398724	0.948581698074234	0.992432418701484
11019	LIAS	lipoic acid synthetase	2464	0.0155104112944092
2.43434539792145	0.00415116993368603
0.948628182739847	0.992432418701484
90417	KNSTRN	kinetochore localized astrin (SPAG5) binding protein
3654	0.0180433080650122	3.92587913361422
0.00413727458611834	0.948714115390316	0.992432418701484
7726	TRIM26	tripartite motif containing 26	4377
0.00728164073549397	6.20688429415471	0.00412219716773166
0.948807522371098	0.992432418701484
55968	NSFL1C	NSFL1 cofactor	6531	-0.00835586927427303
6.71858457956957	0.00412203097265218	0.948808552930711
0.992432418701484
25939	SAMHD1	SAM and HD domain containing deoxynucleoside

triphosphate triphosphohydrolase 1 5038 -0.0163717716083856
4.98209122352413 0.0041197523530343
0.94882268453485 0.992432418701484
79980 DSN1 DSN1 component of MIS12 kinetochore complex
2739 -0.0125755051308168 3.43987545410818
0.00411625306735175 0.948844394141192 0.992432418701484
10181 RBM5 RNA binding motif protein 5 10303
0.00981369770239097 6.82584582581136 0.00411357153682701
0.94886103665718 0.992432418701484
56141 PCDHA7 protocadherin alpha 7 5788 -0.0416130057767414
2.98278127097222 0.00410906554694179
0.948889014676759 0.992432418701484
3017 H2BC5 H2B clustered histone 5 871 -0.0125625756515505
4.65580177552871 0.00410286387499781
0.948927546510786 0.992432418701484
149986 LSM14B LSM family member 14B 3381 0.00997505822630254
5.04330728364091 0.00409906469749188
0.94895116576367 0.992432418701484
204 AK2 adenylate kinase 2 5727 -0.0109895702009287
5.64296508649159 0.00408920179432926
0.949012534177406 0.992432418701484
26511 CHIC2 cysteine rich hydrophobic domain 2
1318 -0.0122608893277451 4.16538592681477
0.00408036595288053 0.949067575201113 0.992432418701484
2197 FAU FAU ubiquitin like and ribosomal protein S30 fusion
1752 -0.0115991391599127 6.705231270809
0.00404453326871135 0.94929140315413 0.992432418701484
112398 EGLN2 egl-9 family hypoxia inducible factor 2 6577
0.00926341588122026 6.30278121162196 0.00404127598991977
0.949311798935705 0.992432418701484
124641 OVCA2 OVCA2 serine hydrolase domain containing 1359
0.00933190183319504 5.26475397668909
0.00404014923646656 0.949318856137784 0.992432418701484
7307 U2AF1 U2 small nuclear RNA auxiliary factor 1
10245 -0.00976850770093107 5.64681287111755
0.00402335944335164 0.949424133048217 0.992432418701484
10109 ARPC2 actin related protein 2/3 complex subunit 2
6562 -0.0113456866458648 8.89988712372985
0.00401135945890019 0.949499511665417 0.992432418701484
120103 SLC36A4 solute carrier family 36 member 4
9685 -0.0140203472770581 4.26430102572358
0.00400587898660376 0.949533975231612 0.992432418701484
84916 UTP4 UTP4 small subunit processome component 5681
0.00972581526544742 5.28110340865708 0.00400502001390635
0.949539378966372 0.992432418701484
9194 SLC16A7 solute carrier family 16 member 7
13665 -0.0141125933843537 3.85051489671988
0.00399906377792547 0.949576865229042 0.992432418701484
84266 ALKBH7 alkB homolog 7 1455 0.0145002630684063
2.715349032426 0.00398944741586238 0.949637446123794
0.992432418701484
4302 MLLT6 MLLT6, PHD finger containing
12371 -0.0106408419706524 6.77774923422572
0.00397462234082013 0.949730984523833 0.992456125054113

101060376	TBC1D3L TBC1 domain family member 3L	4767
0.0218210155388955	1.93601438584297	0.00395783592392362
	0.94983710963509	0.992472857305124
3421	IDH3G	isocitrate dehydrogenase (NAD(+)) 3 non-catalytic
subunit gamma	2717	-0.00976783166757432	5.06371612126197
	0.00393511548183767	0.949981110669342
0.992472857305124
81555	YIPF5	Yip1 domain family member 5	3701
0.0109781194098591	6.78033513741648	0.00393311971680887
	0.949993779645709	0.992472857305124
6144	RPL21	ribosomal protein L21	2249	-0.0120860924085347
	5.69036478121517	0.00392354772765202
0.950054586863119	0.992472857305124
1030	CDKN2B	cyclin dependent kinase inhibitor 2B
4034	-0.00933689190445636	9.25373329770458
0.00391588826187217	0.950103298197955	0.992472857305124
100505641	FGD5-AS1	FGD5 antisense RNA 1
5131	-0.0118763280373738	6.47538808092852
0.00389703978211742	0.950223371517617	0.992472857305124
51174	TUBD1	tubulin delta 1 3330	-0.0147782225586132
3.10262119503775	0.00389396531545572	0.950242984887486
	0.992472857305124
339210	C17orf67	chromosome 17 open reading frame 67
2410	-0.0226803393484456	0.111920701550223
0.00388249131903073	0.950316251261237	0.992475375142868
222950	NYAP1	neuronal tyrosine phosphorylated phosphoinositide-3-
kinase adaptor 1	3586	-0.0290853003302361
1.97456884806828	0.00386872213163514	0.950404317027945
	0.992493347395096
389813	AJM1	apical junction component 1 homolog
4387	-0.0253462108514905	0.172692821106398
0.00385518434002208	0.950491056333326	0.992509931866773
9249	DHRS3	dehydrogenase/reductase 3
3036	-0.0108363668312679	7.05017818890975
0.00382725503051251	0.95067048932611	0.992564329889782
64786	TBC1D15 TBC1 domain family member 15	8227
0.0131011193136437	4.92090906986335	0.00382502020560427
	0.950684875405249	0.992564329889782
552900	BOLA2	bolA family member 2	2377	-0.0121302283458629
	4.68889392142943	0.00376459961830733
0.951075427896	0.992898078448183
25920	NELFB	negative elongation factor complex member B	2699
	0.00773746897189415	5.94550890248842
0.0037268579834091	0.951320983933371	0.993057249340189
10984	KCNQ1OT1	KCNQ1 opposite strand/antisense transcript 1
	91667	-0.0341036224135673	2.86069522900452
0.00369295154388283	0.951542653674246	0.993057249340189
100129583	FAM47E	family with sequence similarity 47 member E
	7651	0.0229309626266917	1.47805864675963
0.00367969914849464	0.951629571192962	0.993057249340189
56136	PCDHA13 protocadherin alpha 13	5354	-0.039923396130556
	2.93551727510335	0.00367290997269265
0.951674159720011	0.993057249340189
2783	GNB2	G protein subunit beta 2	2857

0.00820613424884988	7.64290833276011	0.00366831242389765
	0.951704378030464	0.993057249340189
79568	MAIP1	matrix AAA peptidase interacting protein 1
1624	-0.0106253452147154	3.63033868583935
0.00365601309238173	0.951785311410344	0.993057249340189
22824	HSPA4L	heat shock protein family A (Hsp70) member 4 like
	11621	-0.0115894979806079	5.19092999790853
0.00364114770277979	0.951883312825768	0.993057249340189
1427	CRYGS	crystallin gamma S	2151	0.0183259775074781
	0.798090353979556	0.00362332700290136
0.952001062390141	0.993057249340189
201965	RWDD4	RWD domain containing 4 2776	0.00989441017159811
	4.47482730282975	0.00361961167709524
0.9520256478415 0.993057249340189
56138	PCDHA11 protocadherin alpha 11	6284	0.0411200368093157
	3.06635038300816	0.00360924118196948
0.952094339675568	0.993057249340189
100130418	CECR7	cat eye syndrome chromosome region,
candidate 7	4354	0.0276553361375015	1.26972087148203
	0.00359553857992001	0.95218525470879
0.993057249340189
51330	TNFRSF12A	TNF receptor superfamily member 12A	1848
	0.00766491685017098	8.45824446215061
0.00359524886084905	0.95218717883339	0.993057249340189
8673	VAMP8	vesicle associated membrane protein 8
990	-0.00949976594685208	7.20329958553475
0.00359417844302357	0.952194288523427	0.993057249340189
9329	GTF3C4	general transcription factor IIIC subunit 4
9043	-0.00834495183369875	6.07281557124255
0.00359024022981203	0.952220455192981	0.993057249340189
51028	VPS36	vacuolar protein sorting 36 homolog
5614	-0.0108237852134208	5.35757105049092
0.00356267235163799	0.952404029178371	0.993132877037949
90313	TP53I13 tumor protein p53 inducible protein 13	2813
0.0105462386186043	4.70727443650883	0.00355766563684767
	0.952437445185691	0.993132877037949
4296	MAP3K11 mitogen-activated protein kinase kinase kinase 11
	6842	-0.00803645256539644	6.04686369832389
0.00354745288825598	0.952505680740187	0.993132877037949
1994	ELAVL1	ELAV like RNA binding protein 1
7063	-0.00683755598464187	6.25542242397367
0.00349362242475948	0.952866984427295	0.993297052777545
10558	SPTLC1	serine palmitoyltransferase long chain base subunit
1	3658	0.00708290759449417	6.60196538862971
0.00349184932158764	0.952878932635055	0.993297052777545
23089	PEG10	paternally expressed 10 6689	-0.0360661306927673
	5.36124096623245	0.00348869470801105
0.952900197797495	0.993297052777545
90665	TBL1Y	transducin beta like 1 Y-linked 2407
0.0214211084171423	1.35657978938483	0.00347962449207095
	0.952961393678113	0.993297052777545
55617	TASP1	taspase 1	2905	-0.0121996656978033
3.11407137852534	0.0034712876742482	0.953017712020948
	0.993297052777545

7916	PRRC2A	proline rich coiled-coil 2A	9115
0.00822383775349001	8.21489592182935	0.00345021793158651
	0.953160349291176	0.993321498573797
1176	AP3S1	adaptor related protein complex 3 subunit sigma 1
	3198	-0.0103019939889458	5.43478825784414
0.00343627657472112	0.953254969420343	0.993321498573797
953	ENTPD1	ectonucleoside triphosphate diphosphohydrolase 1
	6975	0.0486571701121782	0.369402062971575
0.0034000672059733	0.953501626366449	0.993321498573797
100132287	LOC100132287	uncharacterized LOC100132287	3362
	0.0253630416530987	0.19546143984285
0.00338525815833535	0.953602884614011	0.993321498573797
642826	BMS1P2	BMS1 pseudogene 2	1625	-0.0273259764379821
	1.60306034633262	0.00337027055673556
0.953705590262063	0.993321498573797
9681	DEPDC5	DEP domain containing 5, GATOR1 subcomplex subunit
	8196	0.00970094560989206	3.91107830635639
0.00337020793345033	0.95370601988129	0.993321498573797
5345	SERPINF2	serpin family F member 2
2348	-0.0181094005101922	2.16388584853169
0.00336965692833679	0.953709800155708	0.993321498573797
84851	TRIM52	tripartite motif containing 52	4681
0.0149358795901572	2.95866855132078	0.00332824358426365
	0.953994817233331	0.993321498573797
6340	SCNN1G	sodium channel epithelial 1 gamma subunit	3507
	0.0272933185235806	0.775017263672217
0.00332344909625348	0.954027928747599	0.993321498573797
51127	TRIM17	tripartite motif containing 17	3548
0.0172595132794717	2.5985766199235 0.00330180685208514
0.954177692694948	0.993321498573797
9612	NCOR2	nuclear receptor corepressor 2	13975
0.0113574899978674	7.41908425153432	0.00329284958820608
	0.954239820847465	0.993321498573797
10733	PLK4	polo like kinase 4	4482	-0.0166806992437287
	1.40911094057526	0.00328862709123046
0.954269137756878	0.993321498573797
26017	FAM32A	family with sequence similarity 32 member A	1921
	0.00732656568711329	6.89693105260085
0.00327830644131311	0.95434087388605	0.993321498573797
29095	ORMDL2	ORMDL sphingolipid biosynthesis regulator 2
2810	-0.0108337058611607	5.27464744566811
0.00327668255312119	0.954352171416187	0.993321498573797
92211	CDHR1	cadherin related family member 1
7633	-0.0225643309982203	1.76849003030244
0.00326329145823134	0.954445441700221	0.993321498573797
51567	TDP2	tyrosyl-DNA phosphodiesterase 2
3082	-0.00797918570394024	5.71840544059033
0.00326203228997279	0.954454221797409	0.993321498573797
54462	CCSER2	coiled-coil serine rich protein 2
8424	-0.00834366933325638	6.79743248488294
0.00324911706991493	0.954544377109292	0.993321498573797
5916	RARG	retinoic acid receptor gamma	5127
0.0115234641202454	5.91480747232719	0.0032427240601578
	0.954589070334864	0.993321498573797

10226	PLIN3	perilipin 3	2813	-0.00829987437688958
6.71854961344526	0.00322457314700841	0.954716203946079
	0.993321498573797
6166	RPL36AL	ribosomal protein L36a like
746	-0.0100695017933895	6.50101735306068
0.0032082582174553	0.954830784547471	0.993321498573797
8662	EIF3B	eukaryotic translation initiation factor 3 subunit B
	8998	0.00920891615653059	7.38651172782275
0.00320727770866824	0.954837680011962	0.993321498573797
2286	FKBP2	FKBP prolyl isomerase 2	1608	-0.00922499398584777
	4.63345776335409	0.0032027197720054
0.954869747762098	0.993321498573797
59341	TRPV4	transient receptor potential cation channel
subfamily V member 4	3759	0.0175858214845047
4.68070086238787	0.00319811020064975	0.95490220208309
	0.993321498573797
23420	NOMO1	NODAL modulator 1	5819	-0.00696432791332112
	7.7883097472856	0.00319548769573696	0.954920676661965
	0.993321498573797
79175	ZNF343	zinc finger protein 343	4818	-0.00798498374533334
	5.24402301209356	0.0031784748088195
0.955040711322936	0.993321498573797
4116	MAGOH	mago homolog, exon junction complex subunit	1463
	0.0104113043172617	3.92682558689099
0.00317284478386171	0.955080504980411	0.993321498573797
5810	RAD1	RAD1 checkpoint DNA exonuclease
6766	-0.00928839525529174	4.67425819103283
0.00317133620170296	0.955091173826286	0.993321498573797
10160	FARP1	FERM, ARH/RhoGEF and pleckstrin domain protein 1
	15748	-0.0111580556357448	6.73373514201015
0.00316087026713774	0.955165260173604	0.993321498573797
8175	SF3A2	splicing factor 3a subunit 2
3134	-0.0105200595815989	4.99208696023816
0.00315891358035358	0.955179124829854	0.993321498573797
10220	GDF11	growth differentiation factor 11
8657	-0.0137292237015671	4.56598158981748
0.00315440558912883	0.95521108387687	0.993321498573797
57060	PCBP4	poly(rC) binding protein 4	3738
0.0139058767192147	5.37207038019372	0.00313273186338847
	0.955365058868362	0.993321498573797
80339	PNPLA3	patatin like phospholipase domain containing 3	3232
	0.013459024249964	3.22339149827676
0.00313270551548328	0.955365246374375	0.993321498573797
55611	OTUB1	OTU deubiquitinase, ubiquitin aldehyde binding 1
	5170	-0.00694900392293707	6.75403210940519
0.00313043647442868	0.955381397071907	0.993321498573797
5028	P2RY1	purinergic receptor P2Y1
6747	-0.0205708644762186	1.75818078522778
0.0031009915214248	0.955591516777937	0.993363428538655
117246	FTSJ3	FtsJ RNA 2'-O-methyltransferase 3	4858
0.00903356940452153	6.04511201310442	0.00309658351442366
	0.95562305836755	0.993363428538655
178	AGL	amylo-alpha-1, 6-glucosidase, 4-alpha-
glucanotransferase	8434	0.00894195302725659

5.78903783199296	0.00309498771070338	0.955634482727092
	0.993363428538655
388730	TMEM81	transmembrane protein 81
1332	-0.0157957447242871	1.25139092009087
0.00305955711348638	0.955888895638911	0.993448442991934
7133	TNFRSF1B	TNF receptor superfamily member 1B	3766
	0.0444300624858063	2.11281046336406
0.00305796981341189	0.955900327895335	0.993448442991934
7019	TFAM	transcription factor A, mitochondrial	5606
0.00832741535690308	4.96630477519439	0.00305389717064486
	0.955929674027616	0.993448442991934
10473	HMGN4	high mobility group nucleosomal binding domain 4
	1274	-0.0086080738952178	5.20459124918135
0.00304374368417371	0.956002922310021	0.993448442991934
85379	KIAA1671	KIAA1671	15818	-0.0102123945064702
	7.26532808269631	0.00303061841993113
0.956097791134431	0.993448442991934
50626	CYHR1	cysteine and histidine rich 1	9787
0.00992106563464805	5.59263262034789	0.00301491976759394
	0.95621153131981	0.993448442991934
401491	VLDLR-AS1	VLDLR antisense RNA 1	4504
0.017427218658976	0.332566767497124	0.00301475257971617
	0.956212744229118	0.993448442991934
104968399	LOC104968399	uncharacterized LOC104968399
1063	-0.0158516816103998	0.342561835635258
0.00298758045688174	0.956410321966816	0.993509489602913
9522	SCAMP1	secretory carrier membrane protein 1	5106
0.00839068302012296	6.07786622826691	0.00298342024980425
	0.95644065167167	0.993509489602913
222643	UNC5CL	unc-5 family C-terminal like	3634
0.019646189738288	1.45350999495494	0.00297743948261076
	0.956484291232618	0.993509489602913
56146	PCDHA2	protocadherin alpha 2	6054	-0.0355706906393886
	2.93328761771914	0.00295928749551777
0.956617009784972	0.993573665731729
9410	SNRNP40	small nuclear ribonucleoprotein U5 subunit 40
3881	-0.00834189955138725	4.36847858019853
0.00292266941512587	0.956885991774111	0.993766858160623
8312	AXIN1	axin 1	7487	0.00790654230326331
5.48773230708562	0.00291211266586799	0.956963850918062
	0.993766858160623
6455	SH3GL1	SH3 domain containing GRB2 like 1, endophilin A2
	3685	0.00774312498808415	7.32443239432414
0.00289019842739435	0.957125928048899	0.993766858160623
57459	GATAD2B	GATA zinc finger domain containing 2B
8113	-0.00827135969074694	5.41159591215025
0.00287535036697051	0.957236094367794	0.993766858160623
117143	TADA1	transcriptional adaptor 1	2963
0.0124190044738431	3.24766113939903	0.00287044280297977
	0.957272569110627	0.993766858160623
4839	NOP2	NOP2 nucleolar protein	4687	-0.00681206188949427
	5.47640971800343	0.00284877742227252
0.957433968840195	0.993766858160623
9688	NUP93	nucleoporin 93	11136	0.00715993337519196

5.51724853616892	0.00284855211559787	0.957435650526052
	0.993766858160623
388789	SMIM26	small integral membrane protein 26
1098	-0.00981251290449135	4.23278418806896
0.00283729560428014	0.957519754040487	0.993766858160623
378706	RN7SL2	RNA component of signal recognition particle 7SL2
	303	-0.0309454467630896	5.47926058467216
0.00283365650783907	0.957546979482878	0.993766858160623
5096	PCCB	propionyl-CoA carboxylase subunit beta	7532
0.0112840712142582	4.27523736707519	0.00281841629166379
	0.957661188145738	0.993766858160623
101929639	B4GALT1-AS1	B4GALT1 antisense RNA 1 843
0.0185629552748641	-0.159631371682914	0.00280948141189352
	0.957728289266648	0.993766858160623
63897	HEATR6	HEAT repeat containing 6
4868	-0.00874915832505616	4.22009397779972
0.00280463044203927	0.957764764964118	0.993766858160623
1327	COX4I1	cytochrome c oxidase subunit 4I1
3034	-0.00806079976725516	7.61171032419611
0.00279837378742798	0.95781185711198	0.993766858160623
51588	PIAS4	protein inhibitor of activated STAT 4	3646
0.00878737411014529	4.78888574980894	0.00278691485600646
	0.957898242479817	0.993766858160623
55613	MTMR8	myotubularin related protein 8	2782
0.0173043494222623	-0.0448361788683742	0.00277089267744568
	0.958019327913133	0.993766858160623
1605	DAG1	dystroglycan 1	6923	-0.00823867952491702
8.12279315149721	0.00276666077962773	0.958051368471077
	0.993766858160623
65078	RTN4R	reticulon 4 receptor	3339	-0.0142638228962659
	2.75151270746589	0.00276565386981176
0.958058995607955	0.993766858160623
100506599	PPP1R26-AS1	PPP1R26 antisense RNA 1
3024	-0.0152321568233129	0.122840339180857
0.00274635548949909	0.958205446883679	0.993766858160623
58496	LY6G5B	lymphocyte antigen 6 family member G5B	3477
0.00668507771307912	6.25410855348199	0.00273664414047259
	0.958279339387009	0.993766858160623
9720	CCDC144A	coiled-coil domain containing 144A
19303	-0.0339091017476723	3.14844637977324
0.00272637830258304	0.958357594013908	0.993766858160623
729020	RPEL1	ribulose-5-phosphate-3-epimerase like 1 2130
0.0134532753941583	1.93585274663191	0.00270540651257978
	0.958517918681386	0.993766858160623
84450	ZNF512	zinc finger protein 512 4751	-0.0105261080697797
	4.59745615471071	0.00270484900268109
0.958522189207164	0.993766858160623
51107	APH1A	aph-1 homolog A, gamma-secretase subunit	3289
	0.00623981731189886	7.55224198823877
0.00270318657040036	0.958534926058867	0.993766858160623
56147	PCDHA1	protocadherin alpha 1	5801	-0.0339799562279057
	2.93215107268232	0.00268874895660076
0.958645706504225	0.993766858160623
79446	WDR25	WD repeat domain 25	4218	-0.011532267281910

2.76469749503311 0.00268198630269634
0.958697699197993 0.993766858160623
84292 WDR83 WD repeat domain 83 2360 -0.0118626643648921
3.21476848572927 0.00267511173626955
0.958750619709016 0.993766858160623
7832 BTG2 BTG anti-proliferation factor 2
2757 -0.00940357872282326 6.23503411906868
0.00264810342540045 0.958959193106932 0.993766858160623
286046 XKR6 XK related 6 5209 -0.0159165026122329
1.62072608576132 0.00264779299422013 0.958961596615442
0.993766858160623
84992 PIGY phosphatidylinositol glycan anchor biosynthesis class Y 1361 0.0083008893727979 5.26554100824065
0.00264056191986128 0.959017623118958 0.993766858160623
119504 ANAPC16 anaphase promoting complex subunit 16 4324
0.00605988624399627 5.88195384389503 0.00263823971912824
0.959035631887733 0.993766858160623
10556 RPP30 ribonuclease P/MRP subunit p30
4653 -0.00887764201328829 4.01655502719026
0.00261468753029792 0.959218731071879 0.993766858160623
51434 ANAPC7 anaphase promoting complex subunit 7 6272
0.00688819652439811 6.08881779021505 0.00261275957890916
0.959233755874102 0.993766858160623
84148 KAT8 lysine acetyltransferase 8
4334 -0.00781167562695027 4.32487012836133
0.00261074414250118 0.959249468402748 0.993766858160623
4191 MDH2 malate dehydrogenase 2 3641 0.00802763539441915
6.85999066385263 0.0026099313805048
0.959255806490896 0.993766858160623
6925 TCF4 transcription factor 4 24564 0.0184453177426632
4.41820808270233 0.00259929077683907
0.959338875596414 0.993766858160623
5585 PKN1 protein kinase N1 4606 -0.0104129424594416
5.8538063977325 0.00259147307099283 0.959400015577035
0.993766858160623
3069 HDLBP high density lipoprotein binding protein
12182 -0.00695409136253021 9.24409785858543
0.00258788757764705 0.959428087592355 0.993766858160623
8450 CUL4B cullin 4B 7031 -0.00791817982614627
6.35318016260098 0.00257611564447302 0.959520391313056
0.993788976717094
475 ATOX1 antioxidant 1 copper chaperone 5239
0.00890174959746832 4.56437399370036 0.00255779251049759
0.959664484703096 0.993864727337247
26235 FBXL4 F-box and leucine rich repeat protein 4 8115
0.00821060009973508 4.75863281193297 0.00252014414987656
0.959962184921325 0.993932024494202
389384 BNIP5 BCL2 interacting protein 5 3736
0.0351239220422685 1.41069342302166 0.00251901751372507
0.959971127905936 0.993932024494202
84440 RAB11FIP4 RAB11 family interacting protein 4
15383 -0.0168050334008066 6.6303349476789 0.00250642390625444
0.960071229921007 0.993932024494202
11228 RASSF8 Ras association domain family member 8

7583	-0.01031981123486	4.20361976774511
0.0025014061273394	0.960111184644736	0.993932024494202
6637	SNRPG	small nuclear ribonucleoprotein polypeptide G	1795
	0.0101627275317052	3.65616117866939
0.00248180981921386	0.960267608960691	0.993932024494202
11161	ERG28	ergosterol biosynthesis 28 homolog
2633	-0.0107280224927801	5.29037656680917
0.00247089034067116	0.960355040570841	0.993932024494202
27347	STK39	serine/threonine kinase 39
4247	-0.0100307648973391	5.61874636460509
0.00246592351520203	0.960394873719237	0.993932024494202
404093	CUEDC1	CUE domain containing 1 5577	0.00673556492178422
	6.6235805915677 0.0024582561462978	0.960456443835499
	0.993932024494202
55781	RIOK2	RIO kinase 2	5502	-0.00852511989811857
4.25166374598345	0.00244407755325682	0.960570554164291
	0.993932024494202
79668	PARP8	poly(ADP-ribose) polymerase family member 8
10056	-0.00787203563498471	5.18474412649253
0.00242255656685586	0.960744392503263	0.993932024494202
147798	TMC4	transmembrane channel like 4
3062	-0.0122417808280795	5.68073140759309
0.00240953213872075	0.960849975026329	0.993932024494202
54913	RPP25	ribonuclease P and MRP subunit p25	3049
0.0132762279383285	3.53157878133056	0.00240891418059164
	0.9608549916029 0.993932024494202
677778	SCARNA15	small Cajal body-specific RNA 15
4325	-0.0163911884979946	0.433590574386068
0.00240461050698881	0.960889946675614	0.993932024494202
4610	MYCL	MYCL proto-oncogene, bHLH transcription factor	5063
	0.0141650071096018	2.37837276544768
0.00239479749757976	0.960969766833294	0.993932024494202
9512	PMPCB	peptidase, mitochondrial processing beta subunit
	4835	0.00669391716353666	5.47842331606319
0.00238293646310694	0.96106646500201	0.993932024494202
9319	TRIP13	thyroid hormone receptor interactor 13
3560	-0.0192533465851793	1.05257530154419
0.00238188027285879	0.961075087374596	0.993932024494202
5570	PKIB	cAMP-dependent protein kinase inhibitor beta
9408	-0.0161947434995477	4.21464845623306
0.00236602372973493	0.961204765797833	0.993932024494202
100861548	PINK1-AS	PINK1 antisense RNA	4443
0.00744413861094774	5.46960813382254	0.002365972337941
	0.961205186798597	0.993932024494202
64837	KLC2	kinesin light chain 2	5479	-0.00632677610208773
	6.48600545920052	0.00234843672875229
0.961349106808991	0.993932024494202
30	ACAA1	acetyl-CoA acyltransferase 1	6329
0.00577381544942365	6.34428617272611	0.00234775414977495
	0.961354719811293	0.993932024494202
10960	LMAN2	lectin, mannose binding 2	2357
0.00580306209520148	6.76087871271064	0.00234251725625301
	0.961397811215681	0.993932024494202
316	AOX1	aldehyde oxidase 1	5556	0.0295425508906641

1.26988455414483	0.00234041686171338
0.961415107725913	0.993932024494202
7582	ZNF33B	zinc finger protein 33B	8307	0.00822045824991547
4.69690779233942	0.00233771973513885
0.961437329669374	0.993932024494202
9924	PAN2	poly(A) specific ribonuclease subunit PAN2
7288	-0.0108275520323843	4.03044408964172
0.00232944449561501	0.961505590651907	0.993932024494202
118	ADD1	adducin 1	13106	-0.00568962277397755
8.30864016197913	0.00232238645438088	0.961563907272907
0.993932024494202
9236	CCPG1	cell cycle progression 1
8655	-0.00778681440977899	7.60523175938449
0.00231503161145952	0.961624770780344	0.993932024494202
79585	CORO7	coronin 7	6855	-0.00814681262673458
4.94930555010932	0.00231254416976867	0.961645377021876
0.993932024494202
2752	GLUL	glutamate-ammonia ligase	9032
0.00839712407766583	8.34328917614059	0.00229943232500318
0.961754181061612	0.99397113658452
8697	CDC23	cell division cycle 23	4618	-0.00580942307247632
5.38120783012575	0.00227865021543305
0.961927273041686	0.994076679466939
375341	C3orf62	chromosome 3 open reading frame 62	4813
0.00926444367517881	3.88843355767843	0.00225968994577386
0.96208588256363	0.994093013447221
379025	PSMA3-AS1	PSMA3 antisense RNA 1	7409
0.00943311987311292	4.85127856146916	0.00225807894473462
0.962099389858331	0.994093013447221
51095	TRNT1	tRNA nucleotidyl transferase 1
5518	-0.00772454669005064	3.96779781784367
0.00225133429903934	0.962155992239879	0.994093013447221
284408	ZNF790-AS1	ZNF790 antisense RNA 1	1506
0.0152977700290633	0.105829257827842	0.00224059651828856
0.962246281412185	0.994112971215554
285989	ZNF789	zinc finger protein 789	6683	-0.0156670419434193
3.0041042984697	0.00221956956161762	0.962423717894975
0.994222952766054
10943	MSL3	MSL complex subunit 3	6496	-0.00610098013588014
5.29843276783148	0.00220190997773795
0.962573390573557	0.99425271658762
64077	LHPP	phospholysine phosphohistidine inorganic
pyrophosphate phosphatase	4064	0.0138392698908638
2.69805812308021	0.0021994256098381	0.962594494870044
0.99425271658762
414236	C10orf55	chromosome 10 open reading frame 55	2514
0.0157066427745568	1.5625270612703	0.00212468032320245
0.963235125168952	0.994802907214121
208	AKT2	AKT serine/threonine kinase 2	12366
0.00544280977013651	6.68223194501569	0.00212073971882987
0.96326921043372	0.994802907214121
51091	SEPSECS	Sep (O-phosphoserine) tRNA:Sec (selenocysteine) tRNA
synthase	6232	-0.00793266383917042	4.0737553038495
0.00210798440495141	0.963379758946358	0.994811017188673

121053	C12orf45	chromosome 12 open reading frame 45
1131	-0.014090284577194	1.77284892656841
0.00210345351498198	0.963419108171939	0.994811017188673
253714	MMS22L	MMS22 like, DNA repair protein
21793	-0.00981719501584749	3.18350755609225
0.00208122634341912	0.963612761664471	0.994893042270351
23370	ARHGEF18	Rho/Rac guanine nucleotide exchange factor
18	6438	-0.00770903768969754	7.83900743631952
0.00207773867637684	0.96364324173247	0.994893042270351
54762	GRAMD1C	GRAM domain containing 1C
7814	-0.0173622045639652	0.947971644834616
0.00206992407861328	0.963711629722576	0.994893042270351
10023	FRAT1	FRAT regulator of WNT signaling pathway 1
2649	-0.0127874324612629	1.72912043929007
0.0020615715021215	0.963784868912097	0.994895324552417
84936	ZFYVE19	zinc finger FYVE-type containing 19
4779	-0.00767799646958732	5.05879420008974
0.00203086747513126	0.964055381840449	0.995101233120056
54461	FBXW5	F-box and WD repeat domain containing 5	3252
0.00624896527886077	6.4499885213181	0.00200238319337132
0.96430817629118	0.995288823617313
22898	DENND3	DENN domain containing 3
14670	-0.00818463847753617	6.24720153892626
0.00199033621881028	0.964415633523873	0.995311819704268
23234	DNAJC9	DnaJ heat shock protein family (Hsp40) member C9
	3247	0.0061662221108254	5.26758591890899
0.00198396753537011	0.964472572916829	0.995311819704268
344148	NCKAP5	NCK associated protein 5	9552
0.024250364283881	1.53455282820241	0.00195067401276106
	0.964771736632522	0.995442731183119
54815	GATAD2A	GATA zinc finger domain containing 2A
7296	-0.00587060626641358	6.70755670924787
0.00193874978398867	0.964879504912605	0.995442731183119
56986	DTWD1	DTW domain containing 1	15053	0.00947553189969952
	2.88492643072447	0.00193000801008214
0.964958722041022	0.995442731183119
25850	ZNF345	zinc finger protein 345	4738	0.011830850984753
	1.65308339732472	0.00191958079406973
0.965053447931795	0.995442731183119
10309	CCNO	cyclin O	1809	-0.0175649622552209
2.63608368314372	0.00191958043013329	0.965053451242463
	0.995442731183119
27013	CNPPD1	cyclin Pas1/PHO80 domain containing 1
2141	-0.00644838788007456	5.76438476639115
0.00191645054228751	0.965081934937753	0.995442731183119
145482	PTGR2	prostaglandin reductase 2	8072
0.0113249462728046	3.05145719242428	0.00189269971401229
	0.965298844306852	0.995442731183119
8850	KAT2B	lysine acetyltransferase 2B
5634	-0.0126934284369237	4.08309744615257
0.00189260015865855	0.965299756377658	0.995442731183119
7436	VLDLR	very low density lipoprotein receptor
9163	-0.00643046761784039	5.67620795642836
0.00188570441709857	0.965362989920475	0.995442731183119

154807	VKORC1L1	vitamin K epoxide reductase complex subunit 1 like 1	5899	0.00598394603511596	6.29106583508573	0.00188432395864879	0.965375662551063	0.995442731183119
85015	USP45	ubiquitin specific peptidase 45	8051	-0.0105075514604249	3.80070399924414	0.00188212619609729	0.965395847646255	0.995442731183119
8045	RASSF7	Ras association domain family member 7	2492	0.00635401635395188	5.82373440747457	0.00186718005164721	0.965533433115434	0.995442731183119
9678	PHF14	PHD finger protein 14	8734	-0.00591014401193727	5.06295799042765	0.00186516534386882	0.965552021472011	0.995442731183119
5894	RAF1	Raf-1 proto-oncogene, serine/threonine kinase	7847	-0.00531659912338683	7.13428816530303	0.00185826370144326	0.965615774658711	0.995442731183119
810	CALML3	calmodulin like 3	2763	0.0183011759852679	3.09979614911721	0.00184179223332226	0.965768409112781	0.995442731183119
261726	TIPRL	TOR signaling pathway regulator	3220	0.00705468338192258	5.54691488819246	0.0018393671574275	0.965790938969718	0.995442731183119
1185	CLCN6	chloride voltage-gated channel 6	6938	-0.0081803918837557	4.87065144730825	0.00182759214886818	0.965900545309564	0.995442731183119
8372	HYAL3	hyaluronidase 3	2030	-0.0107509658660219	3.41396113498702	0.00182481111783162	0.965926483818958	0.995442731183119
84669	USP32	ubiquitin specific peptidase 32	7411	0.00532474913895027	5.86531060685189	0.00182232258374881	0.965949711018845	0.995442731183119
440400	RNASEK	ribonuclease K	1789	-0.00732458070110439	6.98924897742419	0.00181250658519616	0.966041485787179	0.995464069110646
26234	FBXL5	F-box and leucine rich repeat protein 5	4513	-0.00534660150524128	6.17672356216886	0.00180265697825144	0.966133825364052	0.995485985866873
9909	DENND4B	DENN domain containing 4B	7584	0.00729005739708854	5.4020597393781	0.00178439948339815	0.96630565936117	0.995536078125864
93655	ST7-OT3	ST7 overlapping transcript 3	1189	-0.0107338338366426	2.01322266376208	0.00177677028392775	0.966377723847453	0.995536078125864
7737	RNF113A	ring finger protein 113A	1353	-0.00788602481687721	3.57331438676046	0.00177487365781559	0.966395663188218	0.995536078125864
90637	ZFAND2A	zinc finger AN1-type containing 2A	1918	-0.00930152910182117	4.31245704332189	0.00176622413005845	0.966477597225437	0.995547264622496
5936	RBM4	RNA binding motif protein 4	4787	-0.00488907569247189	6.63272250766749	0.00171646004843451	0.966952951037511	0.995601089916844
747	DAGLA	diacylglycerol lipase alpha	5757	0.0126324896402236	3.28563358332704	0.0017154145196443

0.9669630116113 0.995601089916844
5863 RGL2 ral guanine nucleotide dissociation stimulator like 2 4541 -0.00759557163114605 6.53463198135608 0.00170696050275865 0.967044473247293 0.995601089916844
65268 WNK2 WNK lysine deficient protein kinase 2 13198 -0.0201697354941231 4.93216593257796 0.00169434354766196 0.967166425084931 0.995601089916844
5925 RB1 RB transcriptional corepressor 1 6169 0.00503724851451396 6.12218122219651 0.00168619663235603 0.967245412607452 0.995601089916844
4710 NDUFB4 NADH:ubiquinone oxidoreductase subunit B4 1889 0.00618824394041328 5.85379481430302 0.0016736969930804 0.967366974019482 0.995601089916844
79726 WDR59 WD repeat domain 59 9422 -0.00556960543513494 5.06021175600966 0.00167137860569255 0.967389570715312 0.995601089916844
55803 ADAP2 ArfGAP with dual PH domains 2 3449 0.017782004406406 1.49542587647087 0.00166510448752888 0.967450801698258 0.995601089916844
114908 TMEM123 transmembrane protein 123 4647 0.00699555573582883 8.22118720287686 0.00166358019441759 0.96746569516589 0.995601089916844
126567 C2CD4C C2 calcium dependent domain containing 4C 3129 -0.0240926781750729 0.461302993027715 0.0016301121963096 0.967794441777012 0.995601089916844
55722 CEP72 centrosomal protein 72 2595 -0.0142012326921428 2.40571486441669 0.00162158026355108 0.967878787891655 0.995601089916844
55634 KRBOX4 KRAB box domain containing 4 4120 -0.00895833657162273 3.39752121005953 0.00161687217385298 0.967925426924394 0.995601089916844
1351 COX8A cytochrome c oxidase subunit 8A 507 0.00614762404496205 5.94676457658701 0.00161493719816441 0.967944614797544 0.995601089916844
3875 KRT18 keratin 18 3218 0.00822439030175133 9.16613677529035 0.00161396919469325 0.967954218167985 0.995601089916844
23406 COTL1 coactosin like F-actin binding protein 1 2786 0.00871571170491252 6.76198500979162 0.00161308810135097 0.967962961828273 0.995601089916844
7561 ZNF14 zinc finger protein 14 2949 -0.0103103492765404 2.03513065204232 0.00161115608238305 0.967982142880976 0.995601089916844
10962 MLLT11 MLLT11 transcription factor 7 cofactor 4075 -0.0112333501295142 6.41332508251613 0.00161098652323233 0.967983826811145 0.995601089916844
10399 RACK1 receptor for activated C kinase 1 5990 -0.0065904059026316 8.63393789154737 0.00159937529019771 0.968099352499474 0.995601089916844
135 ADORA2A adenosine A2a receptor 4193 0.0158596556798178 0.553833766794123 0.0015988178754025 0.968104909033425 0.995601089916844
54751 FBLIM1 filamin binding LIM protein 1 4716 0.00572567726579523 8.82468990789133 0.00159097189893487

0.968183224048508 0.995601089916844
221960 CCZ1B CCZ1 homolog B, vacuolar protein trafficking and biogenesis associated 6769 0.0060615036418599
6.56500624725666 0.00157907883246367 0.968302305024272
0.995601089916844
83696 TRAPPC9 trafficking protein particle complex 9 4598
0.00629340673173109 4.72403653368397 0.00157615350049856
0.968331664075738 0.995601089916844
9463 PICK1 protein interacting with PRKCA 1
4371 -0.00506206236902181 5.59403252232947
0.00157312960920564 0.968362040979633 0.995601089916844
101927720 ZNF793-AS1 ZNF793 antisense RNA 1 (head to head) 1321 0.0143057539825137 0.632579343766339
0.00156339015014062 0.968460079301101 0.995601089916844
7695 ZNF136 zinc finger protein 136 5425 -0.00752152012575449
3.3748858555852 0.00155275748904238 0.96856745854054
0.995601089916844
55652 SLC48A1 solute carrier family 48 member 1
4348 -0.00756825978548268 3.45869286552244
0.00154489485164788 0.968647100328432 0.995601089916844
119016 AGAP4 ArfGAP with GTPase domain, ankyrin repeat and PH domain 4 3474 0.0106887569499563 2.56484570638859
0.00153811852124619 0.968715901757854
0.995601089916844
79020 C7orf25 chromosome 7 open reading frame 25
4029 -0.00563728133992232 6.84634045346714
0.00153372711948663 0.96876056961623 0.995601089916844
79940 LINC00472 long intergenic non-protein coding RNA 472
3908 -0.0171859721493725 2.12612237232634
0.0015228045736162 0.968871948618834 0.995601089916844
257415 FAM133B family with sequence similarity 133 member B
4462 -0.00862211565214843 3.57856995611847
0.00152263071964498 0.968873724665354 0.995601089916844
8322 FZD4 frizzled class receptor 4 7383
0.00980908205067164 2.27864542188487 0.00151481176728296
0.968953706281057 0.995601089916844
11258 DCTN3 dynactin subunit 3 3740 0.00593172070096645
5.18492212995978 0.00151245111194953
0.968977894502212 0.995601089916844
342945 ZSCAN22 zinc finger and SCAN domain containing 22 4629
0.00969810153769462 2.70654283130757
0.00151174296967582 0.968985154097506 0.995601089916844
51531 TRMO tRNA methyltransferase O 4203
0.00688895442143449 3.62083407630156 0.00150320901035639
0.969072775161662 0.995601089916844
3396 MRPL58 mitochondrial ribosomal protein L58 1108
0.00783051811015804 3.93241013437741 0.00149770523183168
0.969129416510024 0.995601089916844
127002 ATXN7L2 ataxin 7 like 2 5232 -0.0112573008229622
2.7621189570789 0.00148878377317008 0.969221452291029
0.995601089916844
9866 TRIM66 tripartite motif containing 66
10816 -0.0100411727586725 3.71967239958361
0.0014742327306827 0.969372158707189 0.995601089916844

10775	POP4	POP4 homolog, ribonuclease P/MRP subunit	6574
	0.00577995188778927	4.58252556130999
0.0014730608924296	0.969384327879554	0.995601089916844
9725	TMEM63A transmembrane protein 63A
10339	-0.00526634724385646	6.7431665864231 0.00146735904351392
	0.969443609087174	0.995601089916844
9145	SYNGR1	synaptogyrin 1	6953	0.0181692304670479
4.0334261189438 0.00146380259866419	0.969480643297685
0.995601089916844
6934	TCF7L2	transcription factor 7 like 2
4935	-0.0058413911844122 6	4.41412399196832
0.00145881706121109	0.969532634963787	0.995601089916844
54543	TOMM7	translocase of outer mitochondrial membrane 7
1751	-0.00857900910141223	5.09605303770838
0.00145497879529444	0.969572722955082	0.995601089916844
57558	USP35	ubiquitin specific peptidase 35
5233	-0.00568847332370851	4.5012908353048 0.00145368570696469
	0.969586240276695	0.995601089916844
3107	HLA-C	major histocompatibility complex, class I, C	2507
	0.011312540244172	9.27350686483929
0.00143180163308898	0.969815925728008	0.995705425696215
8987	STBD1	starch binding domain 1 3420	-0.00568816690101495
	6.13912513142955	0.00143046394430257
0.969830022329797	0.995705425696215
89122	TRIM4	tripartite motif containing 4	4891
0.00629764249149493	5.44258040306481	0.00137506976835766
	0.970419677354253	0.995962001531513
55559	HAUS7	HAUS augmin like complex subunit 7	4847
0.0100563978013599	1.59576509033289	0.00137214988793399
	0.970451085711585	0.995962001531513
89781	HPS4	HPS4 biogenesis of lysosomal organelles complex 3
subunit 2	10782	0.00496295329624222	5.53015106541733
	0.00136570892574994	0.970520487926395
0.995962001531513
113452	TMEM54	transmembrane protein 54	1417
0.00633736860205157	5.14192174740169	0.00135294692437071
	0.970658485915096	0.995962001531513
285266	ENTPD3-AS1	ENTPD3 antisense RNA 1	2961
0.0117982908183599	0.0480590467440954	0.00134271610969705
	0.97076958502629	0.995962001531513
8859	STK19	serine/threonine kinase 19
3954	-0.0068994114889751	3.83690709846057
0.00134042423383107	0.970794531156321	0.995962001531513
1756	DMD	dystrophin	18861	-0.0106117729170892
3.09886329672268	0.00133098239610518	0.970897527600251
	0.995962001531513
284359	IZUMO1	izumo sperm-egg fusion 1	1621
0.0206530810213695	0.226228658041745	0.00132195388570899
	0.970996357953328	0.995962001531513
8518	ELP1	elongator complex protein 1	6048
0.00419655188533561	5.91666157983391	0.00131928171843754
	0.971025673528519	0.995962001531513
259217	HSPA12A heat shock protein family A (Hsp70) member 12A	6754
	0.011856185745559	2.27216301942416

0.00131388449714365	0.971084975685144	0.995962001531513
1159	CKMT1B	creatine kinase, mitochondrial 1B	1943
0.0115503239374629	4.23046174657865	0.00130602455833539
	0.971171555600449	0.995962001531513
252983	STXBP4	syntaxin binding protein 4
18055	-0.00724955960475918	3.32682419448487
0.00129791433262483	0.971261166413675	0.995962001531513
6293	VPS52	VPS52 subunit of GARP complex	4503
0.00481530168343224	5.34872961568914	0.00129420894772458
	0.97130220092252	0.995962001531513
11020	IFT27	intraflagellar transport 27
8009	-0.00807289500820008	3.58277573180081
0.00129336617382592	0.971311542252642	0.995962001531513
51340	CRNKL1	crooked neck pre-mRNA splicing factor 1
4567	-0.00477206799475386	5.10256601202165
0.00129297799955097	0.9713158458125 0.995962001531513
7536	SF1	splicing factor 1	5972	0.00441488834112327
	7.35983525446371	0.00128390569510373
0.971416612309815	0.995962001531513
84933	C8orf76 chromosome 8 open reading frame 76
2408	-0.00556940405673281	4.66145835045504
0.00127713702893928	0.971492024588645	0.995962001531513
286148	DPY19L4 dpy-19 like 4	7136	0.0046458452133887
5.77115357622537	0.00127192344712057	0.971550247598695
	0.995962001531513
79017	GGCT	gamma-glutamylcyclotransferase	1843
0.00620842454075501	4.62993622805149	0.00127079972692457
	0.971562812458502	0.995962001531513
4053	LTBP2	latent transforming growth factor beta binding
protein 2	9091	-0.00618006042355641	8.3004033911466
0.00126283167257313	0.971652067156941	0.995962001531513
3376	IARS1	isoleucyl-tRNA synthetase 1
7401	-0.00441418335550787	7.26177799419828
0.00125960585884854	0.97168828147886	0.995962001531513
4817	NIT1	nitrilase 1	3793	-0.00579646546181019
4.28246733006234	0.00125272667545628	0.971765665452037
	0.995962001531513
152404	IGSF11	immunoglobulin superfamily member 11	4899
0.0132539699724331	0.17802732537262	0.00124167557416044
	0.971890426099172	0.995962001531513
55313	CPPED1	calcineurin like phosphoesterase domain containing 1
	6484	-0.00645897982123011	5.46454939749673
0.00122909448766251	0.972033138001409	0.995962001531513
57231	SNX14	sorting nexin 14	7448	-0.00454772941286756
	6.80887768752736	0.00122698453273529
0.972057143484414	0.995962001531513
79979	TRMT2B	tRNA methyltransferase 2 homolog B
4055	-0.00571982603743906	4.54957760215184
0.00121836492979988	0.972155426173911	0.995962001531513
3622	ING2	inhibitor of growth family member 2	1426
0.00723323246015328	3.42721504262752	0.00121463776194419
	0.972198031970391	0.995962001531513
631	BFSP1	beaded filament structural protein 1
3226	-0.0136409284499545	1.23773720654482

0.00120915592816839	0.972260814703617	0.995962001531513
5187	PER1	period circadian regulator 1	7002
0.00622689489289231	4.49403467851726	0.00120394915270339
	0.972320579308102	0.995962001531513
200879	LIPH	lipase H	4122	-0.00864608452632326
7.81286200531743	0.00120113451202997	0.972352940303697
	0.995962001531513
58491	ZNF71	zinc finger protein 71	6170	0.00528610699467938
	4.42437046065828	0.00119003021445963
0.972480982454762	0.995962001531513
9798	IST1	IST1 factor associated with ESCRT-III	7806
0.00396077071886391	7.61135690730215	0.0011867759064188
	0.972518620658407	0.995962001531513
8192	CLPP	caseinolytic mitochondrial matrix peptidase
proteolytic subunit	2276	0.00676816887698538
4.81083275726163	0.00118574059473531	0.972530605546287
	0.995962001531513
5719	PSMD13	proteasome 26S subunit, non-ATPase 13	3626
0.00562118905586317	6.7594360216716 0.00118034759380237
0.972593120437934	0.995962001531513
79659	DYNC2H1 dynein cytoplasmic 2 heavy chain 1	14284
0.0118097102936725	4.0500141330748 0.00117155718614548
0.972695324843449	0.995962001531513
84456	L3MBTL3 L3MBTL histone methyl-lysine binding protein 3
5113	-0.00527406199287242	4.99717371974231
0.0011699444049853	0.972714117971679	0.995962001531513
7773	ZNF230	zinc finger protein 230 4653	0.00852135243702179
	2.1533124639352 0.00116116280325507	0.972816674981753
	0.995962001531513
100302692	FTX	FTX transcript, XIST regulator	6132
0.0110884419861508	2.82377837040392	0.00115884059195182
	0.972843860064003	0.995962001531513
57568	SIPA1L2 signal induced proliferation associated 1 like 2
	7641	-0.00833739664891253	5.31745384214239
0.00114950951206083	0.972953370723436	0.995962001531513
55758	RCOR3	REST corepressor 3	9092	-0.0055822936125608
	5.23922683392302	0.00114734678881234
0.97297881615201	0.995962001531513
55224	ETNK2	ethanolamine kinase 2	4472	-0.00775538418735283
	4.249916446847	0.00113508139993446	0.973123580487815
	0.995962001531513
9686	VGLL4	vestigial like family member 4	7013
0.00394863262142337	6.53490891724717	0.00113492904392842
	0.973125383606244	0.995962001531513
506	ATP5F1B ATP synthase F1 subunit beta	2322
0.00513394212257969	8.80558377682651	0.00112908039625381
	0.973194693615661	0.995962001531513
9064	MAP3K6	mitogen-activated protein kinase kinase kinase 6
	4951	-0.0071622631344605	4.85037719555008
0.00112791442764504	0.973208532516729	0.995962001531513
7345	UCHL1	ubiquitin C-terminal hydrolase L1
2282	-0.014858984396067	4.29394821262641
0.00112148393715827	0.97328498510535	0.995967474165009
134359	POC5	POC5 centriolar protein 3448	-0.0068226239260227

3.49473951029431	0.00110784341272563
0.973447888148461	0.996061404726094
388962	BOLA3	bolA family member 3	3156	0.00776386450075268
3.05129598195725	0.00108415618833391
0.973733178676831	0.996196497478218
55823	VPS11	VPS11 core subunit of CORVET and HOPS complexes 5907
0.00449045707248337	5.54318189243098
0.00108216922668802	0.973757250974229	0.996196497478218
2354	FOSB	FosB proto-oncogene, AP-1 transcription factor subunit 5553	0.0150730778097721	2.42926892754185
0.00107919892189301	0.97379327789475	0.996196497478218
5877	RABIF	RAB interacting factor	2430	0.00450041361648031
4.94360803511082	0.00104299810872632
0.974236412772353	0.996457591887545
51629	SLC25A39	solute carrier family 25 member 39
3448	-0.00408410719214224	6.90363612141366
0.00103055642826888	0.974390484587127	0.996457591887545
220004	PPP1R32 protein phosphatase 1 regulatory subunit 32	2791
0.00572783141886221	3.50582686448322
0.00102976929246523	0.974400263331641	0.996457591887545
84823	LMNB2	lamin B2	5432	0.00392103384690336
5.86407631272805	0.00102959858497265	0.974402384558352
0.996457591887545
29803	REPIN1	replication initiator 1 4649	0.00542083091479709
5.66618942937592	0.00102352222343427
0.974478004959792	0.996457591887545
6421	SFPQ	splicing factor proline and glutamine rich	3973
0.00393211220885825	7.47853052064223
0.0010229363633556	0.974485307858922	0.996457591887545
55893	ZNF395	zinc finger protein 395 6527	-0.00557044537148585
4.92047837196956	0.00101803551341817
0.974546480421752	0.996457591887545
22911	WDR47	WD repeat domain 47	4423	0.00375215493244462
6.10292175528437	0.00100167799788409
0.974751729774076	0.996535376603299
11094	CACFD1	calcium channel flower domain containing 1	3520
0.00634140249088261	5.00443791205941
0.00100063721497712	0.974764845780096	0.996535376603299
8357	H3C10	H3 clustered histone 10
1237	-0.0133585859422347	-0.12741246274501
0.000976380449204228	0.975072488193399	0.996713021189026
1809	DPYSL3	dihydropyrimidinase like 3	7164
0.00819799682690168	7.80647852198569	0.000962675236451105
0.975248000627055	0.996713021189026
729013	ZBED5-AS1	ZBED5 antisense RNA 1
1995	-0.00910054041868407	1.76773080212136
0.000961528738031792	0.975262739493101	0.996713021189026
125050	RN7SK	RNA component of 7SK nuclear ribonucleoprotein	331
0.0151314194987738	1.45480197384125
0.000946368291913302	0.975458468801545	0.996713021189026
8464	SUPT3H	SPT3 homolog, SAGA and STAGA complex component	5205
0.00899849269256001	1.78755490695524
0.000936628210767232	0.975585047286065	0.996713021189026
25759	SHC2	SHC adaptor protein 2	4131	0.0108168322312061

4.80368535500142	0.000934532756232898
0.9756123650181 0.996713021189026
11222	MRPL3	mitochondrial ribosomal protein L3	2715
0.00729112825687509	5.30062668036065	0.000928799298662852
0.975687267247544	0.996713021189026
79957	PAQR6	progestin and adipoQ receptor family member 6
2439	-0.00984043715033988	1.31807778837555
0.000922053233558984	0.975775695175939	0.996713021189026
10588	MTHFS	methenyltetrahydrofolate synthetase	1208
0.00785349848542593	3.75693547995554	0.000921980865193461
0.975776645538109	0.996713021189026
84920	ALG10	ALG10 alpha-1,2-glucosyltransferase	4389
0.00928061506632551	1.92713619431803	0.000916278026057427
0.975851654608597	0.996713021189026
23670	CEMIP2	cell migration inducing hyaluronidase 2 9625
0.00648668169617159	8.1085429476479 0.00091203969691378
0.975907552534594	0.996713021189026
3337	DNAJB1	DnaJ heat shock protein family (Hsp40) member B1
3672	-0.00509646252874836	8.08560315610768
0.000908922129411849	0.975948752130776	0.996713021189026
8724	SNX3	sorting nexin 3 1842	-0.00462762069090199
6.74749013756889	0.000891259481349493	0.976183516668743
0.996713021189026
79178	THTPA	thiamine triphosphatase 6414	-0.00768046740130946
3.10881280572297	0.000888594575583213
0.976219138939505	0.996713021189026
23446	SLC44A1 solute carrier family 44 member 1
12118	-0.00433187622401352	6.60866352737279
0.000885652015394811	0.976258534831832	0.996713021189026
80164	PRR36	proline rich 36 4456	0.0110814049007051
2.5633291287301 0.000878734657266733	0.976351405315876
0.996713021189026
84869	CBR4	carbonyl reductase 4	5110	-0.00389262785064633
5.28102906909608	0.000875834587159119
0.976390449643901	0.996713021189026
56142	PCDHA6	protocadherin alpha 6	6133	-0.0195943630974295
2.95904697583987	0.000873551140205819
0.976421237777084	0.996713021189026
4698	NDUFA5	NADH:ubiquinone oxidoreductase subunit A5
10261	-0.00404459748163112	5.18260457919298
0.000870131270922947	0.976467423927723	0.996713021189026
256471	MFSD8	major facilitator superfamily domain containing 8
5023	-0.00540433961743483	4.28881656861727
0.000866117568733671	0.976521745959368	0.996713021189026
5176	SERPINF1	serpin family F member 1	3069
0.0157836986292635	4.88698979000923	0.000846228510695823
0.976792805638341	0.996713021189026
132299	OCIAD2	OCIA domain containing 2
2483	-0.00579341297594016	7.88340688940448
0.00084434117593446	0.976818692176782	0.996713021189026
90780	PYGO2	pygopus family PHD finger 2	3630
0.00337833431763717	6.18879144308384	0.000842337309393315
0.97684620876931	0.996713021189026
2261	FGFR3	fibroblast growth factor receptor 3

4834 -0.0111922142532919 3.6116508075054 0.000839347259301348
0.976887328365713 0.996713021189026
11022 TDRKH tudor and KH domain containing 5092
0.00678060644243442 2.80039476104785 0.00083455463796156
0.976953390390974 0.996713021189026
734 OSGIN2 oxidative stress induced growth inhibitor family
member 2 4681 -0.00390249278153497 5.70394948536588
0.000832828415164855 0.976977231323804
0.996713021189026
51249 TMEM69 transmembrane protein 69 2439
0.00442954106900342 4.47156450466612 0.000831307513491808
0.976998257057855 0.996713021189026
53905 DUOX1 dual oxidase 1 7572 0.00405163858360473
6.80502843846516 0.000831161739256459 0.977000273327096
0.996713021189026
64784 CRTC3 CREB regulated transcription coactivator 3
10783 -0.00450979790503593 5.07805762088989
0.000831022632621803 0.977002197538754 0.996713021189026
55658 RNF126 ring finger protein 126 3496 -0.0046748332237826
5.18849303104288 0.000817212500557218
0.977194037099 0.99683612829697
57226 LYRM2 LYR motif containing 2 8679 -0.00381182705291685
5.408605419889 0.000785971876304359 0.977634084637186
0.997065400450625
55745 AP5M1 adaptor related protein complex 5 subunit mu 1
13034 -0.00392682064943336 5.81720792097671
0.000785763065311418 0.977637055058962 0.997065400450625
3798 KIF5A kinesin family member 5A
6261 -0.00942179928843498 0.215714136187293
0.000773361270463457 0.977814189653803 0.997065400450625
79009 DDX50 DExD-box helicase 50 2692 -0.00425662457996385
4.81629921531121 0.000772262596663964
0.97782995030426 0.997065400450625
6150 MRPL23 mitochondrial ribosomal protein L23
3715 -0.00615146154593223 3.2350895701285 0.00076684919598291
0.977907770750893 0.997065400450625
64343 AZI2 5-azacytidine induced 2 6349 0.00364870766999019
5.5899643122803 0.000764371274698306 0.977943483854906
0.997065400450625
201725 C4orf46 chromosome 4 open reading frame 46 3730
0.00498946100269262 3.53960783381968 0.000757952816359264
0.978036260339259 0.997065400450625
9398 CD101 CD101 molecule 3828 -0.0122421410125269
1.92869839753508 0.000756714753970655 0.978054201251165
0.997065400450625
1650 DDOST dolichyl-diphosphooligosaccharide--protein
glycosyltransferase non-catalytic subunit
2721 -0.00403898976295409 7.72567179021369
0.000756353302051593 0.978059441864181 0.997065400450625
196513 DCP1B decapping mRNA 1B 4609 -0.0062252478417373
3.2380718029317 0.000731513193493072 0.978422645484054
0.997363073669686
57521 RPTOR regulatory associated protein of MTOR complex 1
9389 -0.00372399752994933 5.45668294658257

0.00072291938180058	0.9785497345195 0.997420035760052
116143	WDR92	WD repeat domain 92	5194	0.00366861226996995
	4.79322733281986	0.000707948361887256
0.978772951534646	0.997434985700113
6646	SOAT1	sterol O-acyltransferase 1	7005
0.00419729687583324	6.67315603271481	0.000696537986613066
	0.978944670129212	0.997434985700113
114788	CSMD3	CUB and Sushi multiple domains 3
20567	-0.0137381829202074	0.130523925873541
0.00069236405094486	0.979007836321732	0.997434985700113
79142	PHF23	PHD finger protein 23	2783	-0.00388378824456633
	6.39773466969394	0.000692356485849643
0.979007950980706	0.997434985700113
83636	C19orf12	chromosome 19 open reading frame 12
5378	-0.00352298892598674	5.12728374902358
0.000682253846150616	0.979161633267264	0.997434985700113
1850	DUSP8	dual specificity phosphatase 8	4616
0.00663047997894563	4.85695814859006	0.000679899580713794
	0.979197609836405	0.997434985700113
54849	DEF8	differentially expressed in FDCP 8 homolog	6565
	0.00435785477335385	5.95497485524835
0.000678477486093954	0.979219371693908	0.997434985700113
150737	TTC30B	tetratricopeptide repeat domain 30B
4017	-0.00707422007089523	2.99301946637827
0.000670086093819577	0.979348249774203	0.997434985700113
274	BIN1	bridging integrator 1	6929	-0.00841550024443304
	3.50098091610747	0.000667941746009149
0.979381312798974	0.997434985700113
6793	STK10	serine/threonine kinase 10	9083
0.00327436109218105	6.58782796961297	0.000665877670684267
	0.979413188327945	0.997434985700113
93664	CADPS2	calcium dependent secretion activator 2
6351	-0.00874458466729576	3.0981734377935 0.000658507420795651
	0.979527412501268	0.997434985700113
100533467	BIVM-ERCC5	BIVM-ERCC5 readthrough	4155
0.00317269040326742	5.85422191671175	0.000645176972930983
	0.979735644613242	0.997434985700113
6628	SNRPB	small nuclear ribonucleoprotein polypeptides B and
B1	1278	0.00374545181807125	6.89275962704865
0.000639667939324795	0.979822328151273	0.997434985700113
1944	EFNA3	ephrin A3	2098	0.00697052801890436
1.98501207670426	0.000638151642973206	0.979846252257549
	0.997434985700113
27044	SND1	staphylococcal nuclease and tudor domain containing
1	6500	-0.00494976861311784	7.4262902122461
0.000637317399053927	0.979859427077641	0.997434985700113
57493	HEG1	heart development protein with EGF like domains 1
	9925	0.00623049601369221	7.36518785299034
0.000636083689585121	0.979878926293475	0.997434985700113
5705	PSMC5	proteasome 26S subunit, ATPase 5	4490
0.00332297085699052	6.92180010288174	0.000635274677268427
	0.979891723294078	0.997434985700113
65057	ACD	ACD shelterin complex subunit and telomerase
recruitment factor	3223	-0.00536639744221344

3.32739464226935	0.000623562502656938	0.98007790879018
	0.997434985700113
3772	KCNJ15	potassium inwardly rectifying channel subfamily J
member 15	9728	-0.00656343213888118	5.67964346917857
	0.000621698491485745	0.980107701383014
0.997434985700113
100132406	NBPF10	NBPF member 10	14379	-0.00421584867732155
	5.11605181207605	0.000617605197064108
0.980173281956915	0.997434985700113
29945	ANAPC4	anaphase promoting complex subunit 4	7494
0.00362404058905023	4.55121901609475	0.000617247675820387
	0.980179020283215	0.997434985700113
27292	DIMT1	DIMT1 rRNA methyltransferase and ribosome maturation
factor	3736	0.00395692067827801	4.41520711132759
0.000615493506346709	0.980207199370817	0.997434985700113
84513	PLPP5	phospholipid phosphatase 5
3895	-0.00374077786230583	4.99085931649353
0.000605720360368878	0.980364936696185	0.997434985700113
26151	NAT9	N-acetyltransferase 9 (putative)
4049	-0.00377991111080609	4.76341243872256
0.000600509382984171	0.980449561813601	0.997434985700113
8517	IKBKG	inhibitor of nuclear factor kappa B kinase
regulatory subunit gamma	3473	-0.00380255278419677
3.94783137042559	0.000597203635212651	0.980503436997569
	0.997434985700113
55756	INTS9	integrator complex subunit 9	4905
0.00381906267732944	4.05494496973348	0.000595036173798746
	0.980538842059845	0.997434985700113
2483	FRG1	FSHD region gene 1	1707	0.00423635462779465
	3.82272017589043	0.000593239324913286
0.98056824218342	0.997434985700113
55749	CCAR1	cell division cycle and apoptosis regulator 1
6059	-0.00311135619879505	6.91749171500592
0.000582741120521035	0.980740912102387	0.997434985700113
10243	GPHN	gephyrin	6007	-0.00545032872660358
2.6292472031806	0.000579917787416395	0.980787614062496
0.997434985700113
27085	MTBP	MDM2 binding protein	8124	0.00723587689532867
	1.24514728127584	0.00056269144862986
0.981075060857265	0.997434985700113
6636	SNRPF	small nuclear ribonucleoprotein polypeptide F	3043
	0.00477019256762792	4.2858715952327	0.000561944578617357
	0.981087622371891	0.997434985700113
8407	TAGLN2	transgelin 2	2487	-0.00359841180894901
9.77332450824581	0.000559820948694778	0.981123385150596
	0.997434985700113
11262	SP140	SP140 nuclear body protein	5427
0.00852475540460887	0.789402793445485	0.000555442585727484
	0.981197333387368	0.997434985700113
84632	AFAP1L2	actin filament associated protein 1 like 2	5789
	0.00331928470247956	7.42557336744478
0.000552234653298456	0.981251698945451	0.997434985700113
7037	TFRC	transferrin receptor	7887	0.00541406999426827
	8.41425603605614	0.000550296709434406

0.98128461829476        0.997434985700113
51647   CIAO2B  cytosolic iron-sulfur assembly component 2B
1070    -0.0036871040964562     5.21078392697172
0.000549482552452574    0.981298465466382       0.997434985700113
3150    HMGN1   high mobility group nucleosome binding domain 1 5102
        0.00328458775805501     7.00308537907043
0.00054857829615429     0.981313857082351       0.997434985700113
29074   MRPL18  mitochondrial ribosomal protein L18     1375
0.00353077551769534     5.14976034142294        0.000547497344939529
        0.981332272939707       0.997434985700113
51277   DNAJC27 DnaJ heat shock protein family (Hsp40) member C27
        5624    -0.00626162989548705    2.8576244891487
0.000546951343864954    0.981341581918559       0.997434985700113
64210   MMS19   MMS19 homolog, cytosolic iron-sulfur assembly
component       5849    0.00264033271650214     6.03946558545558
        0.000538697424373513    0.981482876623356
0.997485610014729
10687   PNMA2   PNMA family member 2    5579    0.0130548400510517
        3.09298019358009        0.000533247236234047
0.981576770094384       0.997485610014729
166378  SPATA5  spermatogenesis associated 5
8247    -0.00517907480799302    2.75719342167333
0.000531612303574125    0.981605029516886       0.997485610014729
126208  ZNF787  zinc finger protein 787 3671    0.0042729865238916
        3.84528882565562        0.000515538916019409
0.981885202904814       0.997633834568509
79023   NUP37   nucleoporin 37  3492    0.00484011172585743
3.48752522224879        0.000513951449885752    0.981913109492955
        0.997633834568509
23047   PDS5B   PDS5 cohesin associated factor B        13212
0.00398832563479498     4.79454392428725        0.00051046427274315
        0.981974563521821       0.997633834568509
23548   TTC33   tetratricopeptide repeat domain 33      2612
0.00438360977934584     3.7585665753946 0.000507001816594688
0.982035790035739       0.997633834568509
28      ABO     ABO, alpha 1-3-N-acetylgalactosaminyltransferase and
alpha 1-3-galactosyltransferase 6354    0.00593793872839604
4.86902611087638        0.00050249401222402     0.982115815709222
        0.997636591521385
27238   GPKOW   G-patch domain and KOW motifs
1742    -0.00355096970317945    4.37320871947773
0.000496082842644796    0.982230252428713       0.997636591521385
64091   POPDC2  popeye domain containing 2      2666
0.00539011979954631     2.91594766910504        0.000491623665006102
        0.982310283705882       0.997636591521385
6002    RGS12   regulator of G protein signaling 12
13536   -0.00329802355459241    6.94075719349908
0.000490894747542825    0.982323400461408       0.997636591521385
1298    COL9A2  collagen type IX alpha 2 chain  4151
0.00921318631522897     4.37359030541035        0.000482821386128762
        0.982469336075086       0.997637382111722
81631   MAP1LC3B        microtubule associated protein 1 light chain
3 beta  4591    0.003319040382568       7.85631536839176
0.0004799767514284      0.982520665795377       0.997637382111722

8623	ASMTL	acetylserotonin O-methyltransferase like	7012
	0.00581642808048249	2.54741713455779
0.000477194908587109	0.982571764379606	0.997637382111722
65260	COA7	cytochrome c oxidase assembly factor 7 (putative)
	4165	-0.0039668633270819	4.02888092450057
0.000473578951137199	0.982637910947491	0.997637382111722
636	BICD1	BICD cargo adaptor 1	11933	0.00495965024974757
	5.00829899514849	0.000471268956076187
0.982680299921142	0.997637382111722
162962	ZNF836	zinc finger protein 836 3794	0.00638491302996629
	3.15672308968436	0.000465851859132727
0.982780114492737	0.997666405544265
9782	MATR3	matrin 3	13304	0.00255586225205959
8.00179499198378	0.000456109696358453	0.982961094250326
	0.997777813404182
23015	GOLGA8A golgin A8 family member A	6750
0.00558882962610285	6.28704825888598	0.000449550646862917
	0.983084032723064	0.997819846493786
55835	CENPJ	centromere protein J	5812	0.00438788365492362
	3.02175396743587	0.000446316716461226
0.98314497761553	0.997819846493786
54826	GIN1	gypsy retrotransposon integrase 1
4182	-0.00451524864866934	2.80980115975119
0.000432103412952856	0.983415490402718	0.997884715787137
10221	TRIB1	tribbles pseudokinase 1 4138	0.0040567332199992
	5.5281408871601 0.000432048017831477	0.983416553340468
	0.997884715787137
91442	FAAP24	FA core complex associated protein 24
2370	-0.00524982835560981	1.46071772377064
0.000431732001569762	0.983422618456888	0.997884715787137
1773	DNASE1	deoxyribonuclease 1	13801	-0.00470475827427287
	4.95288971620819	0.000416539260882587
0.983716870260042	0.998046816805069
3915	LAMC1	laminin subunit gamma 1 8406	-0.00278075255945195
	8.0428329595742 0.000412543375034602	0.983795150197835
	0.998046816805069
54870	QRICH1	glutamine rich 1	5499	-0.00230348861172061
	6.65053915004572	0.000408933121367028
0.983866202322407	0.998046816805069
318	NUDT2	nudix hydrolase 2	1075	-0.00555853224112964
	3.46533643117255	0.000408873217434547
0.983867383911216	0.998046816805069
54093	SETD4	SET domain containing 4 7312	-0.00344692328623179
	4.25625543262405	0.000396541490992774
0.984112495842352	0.998164802466981
91368	CDKN2AIPNL	CDKN2A interacting protein N-terminal like
	1757	-0.00407136418666482	3.14257101524068
0.000393136866414068	0.98418083733297	0.998164802466981
2822	GPLD1	glycosylphosphatidylinositol specific phospholipase
D1	4919	-0.00686600919188392	-0.162094596474972
0.000392169673190779	0.984200305879093	0.998164802466981
91653	BOC	BOC cell adhesion associated, oncogene regulated
	10330	-0.0119148295963818	2.25823238132518
0.000388779300198649	0.984268740749193	0.998164802466981

60491	NIF3L1	NGG1 interacting factor 3 like 1
1985	-0.00311791941174228	4.21667542767666
0.000378918459155386	0.984469497192467	0.998273090749873
378884	NHLRC1	NHL repeat containing E3 ubiquitin protein ligase 1
	1188	-0.00534733373324671	1.12042125330499
0.000376552131559293	0.984518060656725	0.998273090749873
112611	RWDD2A	RWD domain containing 2A	3744
0.00396975030258599	3.27187896684498	0.000371964543383285
	0.984612647235916	0.998296731801192
160897	GPR180	G protein-coupled receptor 180
8822	-0.00409530722692449	5.11797600406401
0.00036849710387532	0.984684526474601	0.998297348170942
203062	TSNARE1	t-SNARE domain containing 1	4070
0.00480782924188918	2.94490427290268	0.000361814506305791
	0.984824015976556	0.998315486863472
253782	CERS6	ceramide synthase 6	7241	-0.00394096810951056
	6.93980432712225	0.000357046382362114
0.984924332970918	0.998315486863472
5639	PRRG2	proline rich and Gla domain 2	1701
0.00329140472775467	4.304643197374	0.000346942260430083
0.985139153394014	0.998315486863472
51371	POMP	proteasome maturation protein	1428
0.00324361074964267	6.36387747369923	0.000344727594146832
	0.98518665512033	0.998315486863472
285352	KIF9-AS1	KIF9 antisense RNA 1
4898	-0.00563992532432947	2.52415039322185
0.000343189865660065	0.985219727248242	0.998315486863472
9779	TBC1D5	TBC1 domain family member 5	11189
0.0032916316588598	5.40932031889437	0.000342376850561266
	0.985237242825477	0.998315486863472
56652	TWNK	twinkle mtDNA helicase	3508	-0.00356414144186982
	3.39397213180017	0.000335608396904163
0.985383877338732	0.998315486863472
286319	TUSC1	tumor suppressor candidate 1	2052
0.00462772477728689	3.46288502144998	0.000335114441273987
	0.985394636258355	0.998315486863472
8574	AKR7A2	aldo-keto reductase family 7 member A2	1781
0.00321122879143437	4.15846496436153	0.000333350380046227
	0.985433124426427	0.998315486863472
729092	AGAP5	ArfGAP with GTPase domain, ankyrin repeat and PH
domain 5	4595	-0.00482039472005088	2.53377450278315
	0.00033153613512571	0.985472813908548
0.998315486863472
200030	NBPF11	NBPF member 11	5947	0.00345909672077005
4.4311449050195	0.000326414549739695	0.985585446652289
0.998315486863472
1869	E2F1	E2F transcription factor 1
2503	-0.00557086864833063	1.43848299866346
0.000324733984167658	0.985622597622211	0.998315486863472
24144	TFIP11	tuftelin interacting protein 11
5436	-0.00212200020066281	5.78338223802513
0.000324446529096356	0.985628961793029	0.998315486863472
49856	WRAP73	WD repeat containing, antisense to TP73	7604
0.00268798518786639	4.51512403660657	0.000315576420984343

0.985826748519962	0.998322200907023
3835	KIF22	kinesin family member 22	4312
0.00338823260098548	3.72993835190089	0.000312830012795473
0.985888550536565	0.998322200907023
51200	CPA4	carboxypeptidase A4	3958	-0.0113336332093154
4.84261827602446	0.000310995429913419
0.985929985179196	0.998322200907023
100616256	MIR4721 microRNA 4721	89	-0.00412002976555334
2.37983415756072	0.000309772984534495
0.985957662403452	0.998322200907023
80144	FRAS1	Fraser extracellular matrix complex subunit 1
20379	-0.00415979898254974	6.05701523532056
0.000308261649174746	0.985991955975423	0.998322200907023
6201	RPS7	ribosomal protein S7	5583	0.00349351069016841
6.69666904105926	0.000304825025693845
0.986070250468991	0.998329309600228
3306	HSPA2	heat shock protein family A (Hsp70) member 2
2767	-0.00750893061062308	3.66314870654999
0.000300253583198185	0.986175086320907	0.998333017482618
165215	FAM171B family with sequence similarity 171 member B	5826
0.00595277752725083	5.26823542131109
0.000297428608121209	0.986240270296842	0.998333017482618
120425	JAML	junction adhesion molecule like 8587
0.0168541452899017	1.35454895129106	0.000294795138021442
0.986301314906446	0.998333017482618
81603	TRIM8	tripartite motif containing 8	2916
0.00225169139718692	6.62744528678686	0.000289077302600305
0.986434802080892	0.998333017482618
23265	EXOC7	exocyst complex component 7	10086
0.00218397004911908	7.75417808840567	0.000286535935515531
0.9864945558961 0.998333017482618
10244	RABEPK	Rab9 effector protein with kelch motifs
3401	-0.00489550207002065	2.92409549931502
0.00028205058045927	0.986600668081285	0.998333017482618
100533483	DNAAF4-CCPG1	DNAAF4-CCPG1 readthrough (NMD
candidate)	4589	0.00272848082436148	7.28707521727627
0.000282028250541799	0.986601198453832
0.998333017482618
57211	ADGRG6	adhesion G protein-coupled receptor G6	8473
0.00525815775688895	6.30031345251302	0.000279721206140948
0.986656108222076	0.998333017482618
55108	BSDC1	BSD domain containing 1 6271	-0.0019003752481771
7.18667711068771	0.000275507293133526
0.986756991131558	0.998333017482618
657	BMPR1A	bone morphogenetic protein receptor type 1A
11360	-0.00233269132187606	5.6726814408605 0.000273704445003631
0.986800387686802	0.998333017482618
25821	MTO1	mitochondrial tRNA translation optimization 1
12071	0.00254648341882142	4.47226869036456
0.000267518359301278	0.986950391149227	0.998333017482618
6396	SEC13	SEC13 homolog, nuclear pore and COPII coat complex
component	5446	-0.00268450372647367	6.96296009587129
0.000266535002424462	0.9869743952862 0.998333017482618
5864	RAB3A	RAB3A, member RAS oncogene family

1818    -0.00563391114466205    1.12332632537518
0.000262246292813018    0.987079605965029       0.998333017482618
1975    EIF4B   eukaryotic translation initiation factor 4B
6146    -0.00188227421114974    7.16664821468229
0.000257166125130937    0.987205352418896       0.998333017482618
1032    CDKN2D  cyclin dependent kinase inhibitor 2D
1503    -0.00340089882323832    3.07629136355279
0.000256156445936462    0.987230491984394       0.998333017482618
100529262       MIA-RAB4B       MIA-RAB4B readthrough (NMD
candidate)      1229    -0.00355502757732047    2.8780870977169
0.000254084758466888    0.987282229674567       0.998333017482618
64771   ILRUN   inflammation and lipid regulator with UBA-like and
NBR1-like domains       4551    -0.00218907294345503
8.04195572931501        0.000252479553068952    0.987322462814342
        0.998333017482618
6618    SNAPC2  small nuclear RNA activating complex polypeptide 2
        2000    0.00311537562624407     3.51699749230825
0.000248186929692196    0.987430686724987       0.998333017482618
90268   OTULIN  OTU deubiquitinase with linear linkage specificity
        8614    0.00257579608583049     5.86928069424481
0.000243985166719085    0.987537530356252       0.998333017482618
440689  H2BC18  H2B clustered histone 18        3196
0.00448546927724889     1.01015143822159        0.000241600378571682
        0.987598581046827       0.998333017482618
102466515       MIR1199 microRNA 1199   2187    0.00481289201183972
        2.08270544050035        0.00023983734272548
0.987643908843121       0.998333017482618
55742   PARVA   parvin alpha    8294    -0.00201962832892395
7.04278928850125        0.000232487938006898    0.987834682540434
        0.998333017482618
148137  PROSER3 proline and serine rich 3       5943
0.00308120602551854     4.30606422157701        0.000232380339934224
        0.987837497772047       0.998333017482618
23480   SEC61G  SEC61 translocon gamma subunit
1842    -0.00352706401261247    5.62506281857491
0.000229022212579366    0.987925690953542       0.998333017482618
80237   ELL3    elongation factor for RNA polymerase II 3
3648    -0.00643006576326967    3.35166876657234
0.000217065661792581    0.988245073625869       0.998333017482618
64395   GMCL1   germ cell-less 1, spermatogenesis associated    4922
        0.00280541909466387     5.03778576656921
0.000213299396762068    0.988347491291229       0.998333017482618
26290   GALNT8  polypeptide N-acetylgalactosaminyltransferase 8
2921    -0.00296364950432977    3.55516117898141
0.000211945440114381    0.988384530753914       0.998333017482618
100507459       NRSN2-AS1       NRSN2 antisense RNA 1   875
0.00582997638943902     -0.00858950312144115    0.00021161201238673
        0.988393670300671       0.998333017482618
83640   RAMAC   RNA guanine-7 methyltransferase activating subunit
        1568    -0.00272678329173826    3.52561844272364
0.000208198518780733    0.98848765464203        0.998333017482618
1653    DDX1    DEAD-box helicase 1     3833    0.00191609711406868
        6.92665525645529        0.000208027800255195
0.988492375226395       0.998333017482618

57570	TRMT5	tRNA methyltransferase 5
5692	-0.0024966847651573	3.88034162650375
0.000207624987666577	0.988503521191304	0.998333017482618
64092	SAMSN1	SAM domain, SH3 domain and nuclear localization
signals 1	3169	0.0117335073974582	0.75648103655842
	0.000203296584642132	0.988623978926645
0.998333017482618
2166	FAAH	fatty acid amide hydrolase
2776	-0.00289505030545423	4.10557351264836
0.000202095821293824	0.988657622424614	0.998333017482618
56123	PCDHB13	protocadherin beta 13	5061	0.00703879168772547
	1.89366606059424	0.000201400209181024
0.988677158121966	0.998333017482618
100885776	UGDH-AS1	UGDH antisense RNA 1
2787	-0.00622991746474023	0.0861683928706007
0.000201078734329840	0.988686197880324	0.998333017482618
22932	POMZP3	POM121 and ZP3 fusion	1861	0.00293750174767966
	3.20369064925273	0.000200534856351409
0.988701508008715	0.998333017482618
128338	DRAM2	DNA damage regulated autophagy modulator 2
3138	-0.00197458913475072	5.21918769846977
0.000200196407668329	0.988711045799877	0.998333017482618
100463486	MTRNR2L8	MT-RNR2 like 8	1303
0.016175330461444	4.80834989021955	0.000196462297651578
	0.988816816510705	0.998367847535893
401466	RBIS	ribosomal biogenesis factor	2514
0.00301319687596788	3.99973610341257	0.000188599113755572
	0.989042884636413	0.99843580878927
729096	BMS1P4	BMS1 pseudogene 4	2883	-0.00404235901250135
	1.76460433158594	0.000187592136802195
0.989072173316068	0.99843580878927
100506639	LOC100506639	uncharacterized LOC100506639	2393
	0.0033733675403172	2.12072662084948
0.00018670740996507	0.989097971211531	0.99843580878927
115098	CCDC124	coiled-coil domain containing 124
1660	-0.00220493557058012	5.04784772712425
0.000181116117346392	0.989262442518508	0.998529871900032
100130691	LOC100130691	uncharacterized LOC100130691	7171
	0.00491757935360193	0.634731703236855
0.000177828055990759	0.989360350276746	0.998556739179015
92840	REEP6	receptor accessory protein 6
2449	-0.00491445983995586	1.2107999513335	0.000172427634544725
	0.989523143003168	0.998649085960471
57232	ZNF630	zinc finger protein 630	3676	0.00305014751281488
	2.62973648186883	0.000169412698028282
0.989615136831619	0.998657185068097
170487	ACTL10	actin like 10	2028	-0.0044838648778521
0.917358410875776	0.000167505232115062	0.989673762113961
	0.998657185068097
79443	FYCO1	FYVE and coiled-coil domain containing 1	8895
	0.00246414097164107	6.51351129362608
0.000160945517222899	0.989877964539698	0.998700400972824
100861541	DOCK9-DT	DOCK9 divergent transcript	807
	-0.00475580942944339	0.00593963650872101

0.000156785397674941    0.990009631239101       0.998700400972824
5683    PSMA2   proteasome 20S subunit alpha 2
6157    -0.00178382778327749    6.74042970164167
0.000156314945805036    0.990024630319008       0.998700400972824
84833   ATP5MD  ATP synthase membrane subunit DAPIT
898     -0.0023032393041688     6.07072272111467
0.000155045981017565    0.99006520075503        0.998700400972824
79411   GLB1L   galactosidase beta 1 like
3411    -0.00381425918736912    1.72306170119372
0.000151284153110609    0.990186456983455       0.998700400972824
158584  FAAH2   fatty acid amide hydrolase 2    2019
0.00315735467414478     2.39081751905488        0.000151267748446315
        0.99018698904224        0.998700400972824
8668    EIF3I   eukaryotic translation initiation factor 3 subunit I
        1725    0.0020452616410664      7.14956070393343
0.000150142703773781    0.990223547149948       0.998700400972824
10240   MRPS31  mitochondrial ribosomal protein S31     2124
0.00243595947296649     3.41784924750446        0.000146563056855342
        0.990340787729001       0.998700400972824
123096  SLC25A29        solute carrier family 25 member 29      7380
        0.00191632257308913     5.68736913903667
0.000144675151858564    0.990403197370033       0.998700400972824
6500    SKP1    S-phase kinase associated protein 1
12771   -0.00176086117959652    7.65532872532963
0.000143880445762079    0.990429590198722       0.998700400972824
1353    COX11   cytochrome c oxidase copper chaperone COX11
3502    -0.0023368479603317     4.62932058300962
0.000136016972973607    0.990694777351168       0.99884232319915
81605   URM1    ubiquitin related modifier 1    4410
0.00179370240342932     5.40647303712229        0.000135485978773886
        0.990712957535932       0.99884232319915
226     ALDOA   aldolase, fructose-bisphosphate A       6153
0.00187565607829591     9.36541366978124        0.00012700817828204
        0.99100819848266        0.999068075150901
55056   GABPB1-IT1      GABPB1 intronic transcript      6266
0.00306832010974152     2.1410871657195 0.000124180703855004
0.991108846019215       0.99909763268738
100631383       FAM47E-STBD1    FAM47E-STBD1 readthrough        3242
        0.00163160263622611     6.08361235225874
0.000120413645063877    0.991244736981736       0.999115603386373
100129405       MSTO2P  misato family member 2, pseudogene
1863    -0.00237526931575256    3.35258941158833
0.000119738030061711    0.991269332495211       0.999115603386373
7100    TLR5    toll like receptor 5    4608    -0.00474162649051814
        2.3580176396196 0.000117121415105892    0.991365250566725
        0.999140384565269
124872  B4GALNT2        beta-1,4-N-acetyl-galactosaminyltransferase
2       8961    -0.00645461995064169    4.72871657250818
0.000114981873736042    0.991444479629387       0.999148343490095
10099   TSPAN3  tetraspanin 3   8506    0.00184000361377258
8.11140013067167        0.000106665483201596    0.991759675897787
        0.999225041865627
4122    MAN2A2  mannosidase alpha class 2A member 2     6709
0.00143033021489387     6.16347357356112        0.000106288564413859

0.991774247487975	0.999225041865627
56987	BBX	BBX high mobility group box domain containing
11877	0.0017080724917872	6.75495345209879
0.000105498762195566	0.991804865091095	0.999225041865627
2132	EXT2	exostosin glycosyltransferase 2 5286
0.00121434938886797	7.21417299908358	0.00010244798842507
0.991924222149509	0.999225041865627
122970	ACOT4	acyl-CoA thioesterase 4 2362	0.0040979187522205
1.00864347666461	0.000101171830827695
0.991974676587734	0.999225041865627
83989	FAM172A family with sequence similarity 172 member A	5071
0.00143427136255688	4.7292871879562 9.89135904960747e-05
0.992064745038383	0.999225041865627
2778	GNAS	GNAS complex locus	13320	-0.00164706117247613
9.84506974976281	9.45044732816314e-05
0.992243614174582	0.999225041865627
369	ARAF	A-Raf proto-oncogene, serine/threonine kinase	3535
0.0011656340645459	5.84443637292219
9.11863145196534e-05	0.992380994399256	0.999225041865627
1107	CHD3	chromodomain helicase DNA binding protein 3
9758	-0.00125186872582955	7.78803945366836
9.10403160077067e-05	0.992387096056677	0.999225041865627
10410	IFITM3	interferon induced transmembrane protein 3	1348
0.0022033207627637	5.93338444276269
8.94466442673547e-05	0.992454020637039	0.999225041865627
5422	POLA1	DNA polymerase alpha 1, catalytic subunit
6336	-0.00227279043928162	3.11259441506978
8.91762941286345e-05	0.992465432689945	0.999225041865627
100101267	POM121C POM121 transmembrane nucleoporin C
8767	-0.00110850848546944	6.3392879801553 8.42865174490726e-05
0.992674909145952	0.999225041865627
498	ATP5F1A ATP synthase F1 subunit alpha	7672
0.00154601484468873	8.15945436611814	8.04135668204253e-05
0.992845176841466	0.999225041865627
23396	PIP5K1C phosphatidylinositol-4-phosphate 5-kinase type 1
gamma	6694	0.00138803321144907	6.21931979146222
7.97194326480621e-05	0.992876123353258	0.999225041865627
100009676	ZBTB11-AS1	ZBTB11 antisense RNA 1
2743	-0.00255992995952574	2.25050755300202
7.87427434145727e-05	0.992919896071212	0.999225041865627
5689	PSMB1	proteasome 20S subunit beta 1	2256
0.00151142215072346	6.58114553322938	7.68744153560874e-05
0.993004392895878	0.999225041865627
7837	PXDN	peroxidasin	11257	0.00135627445886671
9.15074575079736	7.6155420174473e-05	0.993037183360567
0.999225041865627
23149	FCHO1	FCH and mu domain containing endocytic adaptor 1
5694	0.00481893826487192	0.407442047428132
7.60598528053436e-05	0.993041553437409	0.999225041865627
2650	GCNT1	glucosaminyl (N-acetyl) transferase 1	6463
0.00219006472950298	4.51694221372028	7.35497244228611e-05
0.993157335248985	0.999225041865627
23014	FBXO21	F-box protein 21
6949	-0.000999999185962338	5.9025859668077 7.33796579268642e-05

0.993165250662735	0.999225041865627
8800	PEX11A	peroxisomal biogenesis factor 11 alpha	3171
0.00230825518098086	2.69543767979189	7.331472116201e-05
0.993168275437623	0.999225041865627
11157	LSM6	LSM6 homolog, U6 small nuclear RNA and mRNA
degradation associated	2605	0.0023156233803756
2.40737377009268	7.25317768228706e-05	0.993204851224557
0.999225041865627
7867	MAPKAPK3	MAPK activated protein kinase 3
3921	-0.00115450586754864	6.32249324607347
7.21031787924709e-05	0.993224957117902	0.999225041865627
10011	SRA1	steroid receptor RNA activator 1
3454	-0.00165157615551811	5.06519032082143
7.19387218310885e-05	0.99323268779147	0.999225041865627
55760	DHX32	DEAH-box helicase 32 (putative)	3948
0.00130362336871375	6.6669859403917	6.99707949110007e-05
0.993325889377546	0.999247036233215
8480	RAE1	ribonucleic acid export 1	5855
0.00116251019750728	5.24562097067474	6.64083486938694e-05
0.993498005536942	0.999320550928023
9777	TM9SF4	transmembrane 9 superfamily member 4
5415	-0.000923105716468876	6.8888519720582	6.55196471228692e-05
0.993541657187381	0.999320550928023
79140	CCDC28B	coiled-coil domain containing 28B
2818	-0.00199736616049946	1.93974229166129
6.02141881147489e-05	0.993808653350522	0.99938290053889
8828	NRP2	neuropilin 2	13896	-0.00185786849414426
4.7221211490542	5.93920219813526e-05	0.993851066191244
0.99938290053889
83463	MXD3	MAX dimerization protein 3	4130
0.00252610235886128	1.15404946004271	5.88942870338727e-05
0.993876885468248	0.99938290053889
4175	MCM6	minichromosome maintenance complex component 6	4118
0.00191121817625389	2.92690052712359
5.86606722405847e-05	0.993889041515438	0.99938290053889
161823	ADAL	adenosine deaminase like
5718	-0.00184791382291658	2.59979225756083
5.21952831604722e-05	0.994235629161235	0.999539855387015
283337	ZNF740	zinc finger protein 740	9063	-0.00122978164468619
4.99849440837628	5.10949462082522e-05
0.994296711560471	0.999539855387015
29882	ANAPC2	anaphase promoting complex subunit 2
5447	-0.00108630523356057	5.2747946360454	4.84486824774422e-05
0.994446362353489	0.999539855387015
51669	SARAF	store-operated calcium entry associated regulatory
factor	2990	0.00117562314121491	8.22126572806718
4.74385977042857e-05	0.99450455900558	0.999539855387015
3276	PRMT1	protein arginine methyltransferase 1
4796	-0.000993512316532254	6.50395746446297
4.63310997531607e-05	0.994569084914414	0.999539855387015
51074	APIP	APAF1 interacting protein
5337	-0.00199299895018947	2.87605552369107
4.63111319906062e-05	0.994570255329489	0.999539855387015
7869	SEMA3B	semaphorin 3B	5768	-0.00179760948743611

4.20936714993591	4.50904581796863e-05	0.994642290995292	0.999539855387015
116988	AGAP3	ArfGAP with GTPase domain, ankyrin repeat and PH domain 3	11032	0.000913032619670894	7.35690158684765	4.45525987231576e-05	0.994674340975446	0.999539855387015
79862	ZNF669	zinc finger protein 669 1980	-0.00143350716603609	3.13085153112578	4.41146056235908e-05	0.994700583313277	0.999539855387015
27434	POLM	DNA polymerase mu	4487	0.00144918592292772	3.36971164539507	4.00687989241533e-05	0.994949430828728	0.999539855387015
126823	KLHDC9	kelch domain containing 9	1720	-0.00177219219345997	0.858182331104461	3.96139803404694e-05	0.994978176611916	0.999539855387015
27113	BBC3	BCL2 binding component 3	2528	0.0016067137501811	3.2644813167864 3.66484983018722e-05	0.995169795694623	0.999539855387015
22818	COPZ1	coatomer protein complex subunit zeta 1	3901	-0.00104299884289728	6.33939499515532	3.60975516855433e-05	0.99520623961517	0.999539855387015
100652748	TIMM23B translocase of inner mitochondrial membrane 23 homolog B	3636	-0.00136651831564582	3.29643580772447	3.60744311862504e-05	0.995207775045528	0.999539855387015
10933	MORF4L1 mortality factor 4 like 1	6584	-0.000785610221670325	7.65104313036788	3.55828434521754e-05	0.995240538579003	0.999539855387015
54758	KLHDC4	kelch domain containing 4	9270	0.00112903378845199	4.32956515776502	3.46308568879294e-05	0.995304636996741	0.999539855387015
162239	ZFP1	ZFP1 zinc finger protein	3761	0.00134159257880719	3.21787447319571	3.44249682182607e-05	0.995318615167342	0.999539855387015
10456	HAX1	HCLS1 associated protein X-1	2155	-0.000875209422877408	5.63613922534769	3.42633950438653e-05	0.995329613974304	0.999539855387015
51129	ANGPTL4 angiopoietin like 4	2032	-0.00368377161359084	5.63105180717098	3.11832275201596e-05	0.995544480629744	0.999553340504714
5598	MAPK7	mitogen-activated protein kinase 7	5196	-0.00097108697990276	5.09884528502178	2.94585833557903e-05	0.995669442013216	0.999553340504714
81602	CDADC1	cytidine and dCMP deaminase domain containing 1 3863	0.00125332453204556	3.05676564191472	2.90584095425928e-05	0.995698956070416	0.999553340504714
64645	MFSD14A major facilitator superfamily domain containing 14A	2943	0.000610914281567259	6.47297614797484	2.81754064488382e-05	0.995764807869768	0.999553340504714
968	CD68	CD68 molecule	4954	-0.00102641803220361	9.20545929797129	2.7821067305922e-05	0.995791523163188	0.999553340504714
5636	PRPSAP2 phosphoribosyl pyrophosphate synthetase associated protein 2	4734	0.000801741991605352	4.72164689617763

2.63570468952423e-05 0.995903749302471
0.999553340504714
728537 C2orf92 chromosome 2 open reading frame 92 6234
0.00212634782528978 0.796031561154339 2.54023048533725e-05
0.995978623046502 0.999553340504714
10482 NXF1 nuclear RNA export factor 1 6194
0.000878949248054741 6.26993447054171 2.48184479190172e-05
0.996025105793145 0.999553340504714
153222 CREBRF CREB3 regulatory factor 12079 0.00168119965533187
4.53867956782448 2.46866291320202e-05
0.996035675732952 0.999553340504714
9284 NPIPA1 nuclear pore complex interacting protein family
member A1 8801 -0.000872492176231683 5.63579866648236
2.38606159399524e-05 0.996102562499083
0.999553340504714
23341 DNAJC16 DnaJ heat shock protein family (Hsp40) member C16
8563 0.000675733880404644 5.17739655336623
2.31532902716935e-05 0.996160764578158 0.999553340504714
100132403 FAM157B family with sequence similarity 157 member B
2233 0.00206459812316075 1.2743117708788
2.26899428472649e-05 0.996199374130492 0.999553340504714
400451 FAM174B family with sequence similarity 174 member B
3810 -0.000762993671856756 5.16561455933131
2.09905450985204e-05 0.996344469594697 0.999627317929441
157574 FBXO16 F-box protein 16 1919 0.00122919135219561
1.30128733120447 1.88361893966871e-05
0.9965371376991 0.999725706132217
7458 EIF4H eukaryotic translation initiation factor 4H
3191 -0.000535746294821349 8.26471704515052
1.8316074545055e-05 0.996585281268233 0.999725706132217
5372 PMM1 phosphomannomutase 1
5082 -0.000942477079631387 4.40557269734277
1.73911849472574e-05 0.996672612433154 0.9997417131446
401548 SNX30 sorting nexin family member 30 11578
0.00088683004590701 5.5790075626688 1.55279930069696e-05
0.996855898187612 0.999793097811188
9898 UBAP2L ubiquitin associated protein 2 like
7503 -0.000459825729178094 7.24931897774507
1.54225084969539e-05 0.99686659556158 0.999793097811188
4798 NFRKB nuclear factor related to kappaB binding protein
6335 0.000543035925027416 5.3013294009801
1.46559267193425e-05 0.996945461221249 0.999800606810771
10312 TCIRG1 T cell immune regulator 1, ATPase H+ transporting V0
subunit a3 4384 -0.000432831396834744 6.52965574300648
1.18886992481748e-05 0.997248900035318
0.999833968381908
283663 LINC00926 long intergenic non-protein coding RNA 926
3188 -0.0010481155499829 0.108594196709658
1.13542956308166e-05 0.997311442442075 0.999833968381908
9140 ATG12 autophagy related 12 8888 0.000476116601324188
5.76969756939566 1.0097868518244e-05
0.997464555045071 0.999833968381908
1983 EIF5 eukaryotic translation initiation factor 5
8505 -0.000428696071933997 8.37802006086805

8.91233366928645e-06    0.997618036319266       0.999833968381908
55093   WDYHV1  WDYHV motif containing 1
2132    -0.000720894315255321   2.75517445814084
8.23975053076964e-06    0.997709678254077       0.999833968381908
57646   USP28   ubiquitin specific peptidase 28
5719    -0.000428560642960223   4.51907792869052
6.2322459628561e-06     0.99800812581624        0.999833968381908
5607    MAP2K5  mitogen-activated protein kinase kinase 5       5031
        0.000506138546629867    3.47487670423789
6.1781078239953e-06     0.998016796153476       0.999833968381908
159013  CXorf38 chromosome X open reading frame 38      6043
0.000366891939419084    4.68486255203305        6.16053721458343e-06
        0.998019618282878       0.999833968381908
83606   GUCD1   guanylyl cyclase domain containing 1
4529    -0.000322838726305689   6.28832331613146
5.99309518634072e-06    0.998046716796916       0.999833968381908
29121   CLEC2D  C-type lectin domain family 2 member D  6957
0.000490286353098212    3.95517505982599        5.93193984599338e-06
        0.998056708275282       0.999833968381908
10949   HNRNPA0 heterogeneous nuclear ribonucleoprotein A0
8726    -0.000310898233712856   7.05422901041556
5.83510591489755e-06    0.998072634810425       0.999833968381908
113444  SMIM12  small integral membrane protein 12      8091
0.000380346727744723    5.40539769692558        5.61881915217555e-06
        0.998108692338608       0.999833968381908
55027   HEATR3  HEAT repeat containing 3
4183    -0.000487147430547017   4.23749275790487
5.46283271418702e-06    0.998135129766924       0.999833968381908
89944   GLB1L2  galactosidase beta 1 like 2     7280
0.000722003822424705    1.96129721356732        4.86513266650945e-06
        0.998240103801962       0.999833968381908
253152  EPHX4   epoxide hydrolase 4     2251    0.0006670436590388
        3.33103921688601        4.51807872181575e-06
0.998304035992232       0.999833968381908
55609   ZNF280C zinc finger protein 280C        4634
0.000418390248740189    1.3989474222447 4.01847111142217e-06
0.998400551739665       0.999833968381908
23386   NUDCD3  NudC domain containing 3        10113
0.000253939829215737    6.69270558587773        3.91048524761572e-06
        0.998422188546389       0.999833968381908
84071   ARMC2   armadillo repeat containing 2   4250
0.000857178419675389    0.772260250758903       3.80569761693206e-06
        0.998443472044217       0.999833968381908
10184   LHFPL2  LHFPL tetraspan subfamily member 2      7977
0.000386976665985073    6.35492580582368        3.41500765443925e-06
        0.998525530969877       0.999833968381908
140461  ASB8    ankyrin repeat and SOCS box containing 8        4087
        0.000264858543177685    4.81634114161556
3.17359960799735e-06    0.998578601349817       0.999833968381908
7411    VBP1    VHL binding protein 1   2018    0.000235206221981904
        5.55422997322926        2.22379460268485e-06
0.998810163639681       0.999833968381908
90580   TIMM29  translocase of inner mitochondrial membrane 29
1425    -0.000237774265358117   3.64754963998034

1.87426386233369e-06    0.99890766640263        0.999833968381908
25879   DCAF13  DDB1 and CUL4 associated factor 13
6186    -0.000211915668589746   5.14904418590952
1.85086067006068e-06    0.998914507590305       0.999833968381908
9884    LRRC37A leucine rich repeat containing 37A      6506
0.000356732658541654    2.69541876199037        1.70752925221507e-06
        0.998957384891182       0.999833968381908
100128553       CTAGE4  CTAGE family member 4
5176    -0.000478962117485344   4.07897991292992
1.47930977334454e-06    0.999029558148685       0.999833968381908
653319  KIAA0895L       KIAA0895 like
4778    -0.000222244852064017   4.30287148411405
1.27770150015749e-06    0.999098107716103       0.999833968381908
84275   SLC25A33        solute carrier family 25 member 33      3854
        0.000252654202490503    2.51792151546084
1.13906049392654e-06    0.999148443685346       0.999833968381908
8831    SYNGAP1 synaptic Ras GTPase activating protein 1
11004   0.00017663109735088     4.82626964225726
1.06192834792296e-06    0.999177780841976       0.999833968381908
23492   CBX7    chromobox 7     7461    -0.000218033679244401
2.42822527805168        9.81048968284881e-07    0.999209712098896
        0.999833968381908
5160    PDHA1   pyruvate dehydrogenase E1 alpha 1 subunit
4493    -0.000162153139466568   5.66745976681953
8.90619517690538e-07    0.999247015387743       0.999833968381908
1491    CTH     cystathionine gamma-lyase       2414
0.000256686143799687    1.40416729114429        5.82998355369568e-07
        0.999390780656775       0.999833968381908
285962  WEE2-AS1        WEE2 antisense RNA 1
3504    -0.000414845965885698   0.700647698424957
4.29843090898885e-07    0.999476887611049       0.999833968381908
5436    POLR2G  RNA polymerase II subunit G     1923
9.35689868239121e-05    4.98932290965156        3.89066526551574e-07
        0.999502317961499       0.999833968381908
4700    NDUFA6  NADH:ubiquinone oxidoreductase subunit A6       1416
        9.90909141487666e-05    5.23495322614667
3.7442825906453e-07     0.999511770132721       0.999833968381908
10004   NAALADL1        N-acetylated alpha-linked acidic dipeptidase
like 1  4044    -6.20869843193742e-05   1.16063043184882
2.60864167600516e-07    0.999592481567833       0.999833968381908
54472   TOLLIP  toll interacting protein
5983    -7.0388434766863e-05    6.6950524605076 2.4219147576332e-07
        0.99960733746076        0.999833968381908
6169    RPL38   ribosomal protein L38   3829    9.44094383601305e-05
        6.57589290543883        2.27031591748528e-07
0.999619825317359       0.999833968381908
92558   BICDL1  BICD family like cargo adaptor 1
5199    -3.89758453763667e-05   2.92155411396266
2.80997936030758e-08    0.999866250667143       0.99990127760834
78987   CRELD1  cysteine rich with EGF like domains 1   3778
1.62672734051249e-05    4.30014747325513        1.55923438605043e-08
        0.99990036870799        0.99990127760834
9603    NFE2L3  nuclear factor, erythroid 2 like 3      3776
2.17672630784136e-05    5.15332479417643        1.53091548327211e-08

| | |
|---|---|
| 0.99990127760834 | 0.99990127760834 |
